# Tools and applications for integrative analysis of DNA methylation in social insects

**DOI:** 10.1101/2021.08.19.457008

**Authors:** Claire Morandin, Volker P. Brendel

**Affiliations:** Department of Ecology and Evolution, Biophore, University of Lausanne, 1015, Lausanne, Switzerland; Departments of Biology and Computer Science, Indiana University, Bloomington IN 47405, U.S.A

**Keywords:** DNA methylation, BS-seq, arthropods, workflow, reproducibility

## Abstract

DNA methylation is a common epigenetic signaling tool and an important biological process which is widely studied in a large array of species. The presence, level, and function of DNA methylation varies greatly across species. In insects, DNA methylation systems are reduced, and methylation rates are often low. Low methylation levels probed by whole genome bisulfite sequencing require great care with respect to data quality control and interpretation. Here we introduce `BWASP/R`, a complete workflow that allows efficient, scalable, and entirely reproducible analyses of raw DNA methylation sequencing data. Consistent application of quality control filters and analysis parameters provides fair comparisons among different studies and an integrated view of all experiments on one species. We describe the capabilities of the `BWASP/R` workflow by re-analyzing several publicly available social insect WGBS data sets, comprising 70 samples and cumulatively 147 replicates from four different species. We show that the CpG methylome comprises only about 1.5% of CpG sites in the honeybee genome and that the cumulative data are consistent with genetic signatures of site accessibility and physiological control of methylation levels.

**Significance Statement:** DNA methylation in the honeybee genome occurs almost entirely at CpG sites. Methylation rates are small compared to rates in mammalian or plant genomes. De novo analysis of all published honeybee methylation studies and statistical modeling suggests that the CpG methylome consists of about only 300,000 sites. The development of a fully reproducible, scalable, portable workflow allows for easy accessible updates of integrative views of all current experiments. The integrated results for the honeybee are consistent with genetic determination of methylation site accessibility by yet uncharacterized sequence features and physiological control of methylation levels at those sites.

---

DNA methylation is a heritable, reversible biological process and a common epigenetic signaling tool that can alter the activity of a gene, via regulating its expression, without changing its nucleotide sequence. DNA methylation is found across a wide array of species, including mammals, plants, insects, bacteria, and fungi (1, 2). However, its functions, biological characteristics, and genomic distribution are distinct for different taxonomic lineages (3).

In insects, the presence and levels of DNA methylation vary greatly (4). DNA methylation systems are reduced in some insect lineages. For example, the *Drosophila melanogaster* genome is missing most of the methylation machinery and, as a result, lacks any detectable DNA methylation patterns (5). The role of DNA methylation in social insects remains enigmatic, even after more than a decade of studies since the initial discovery of a full complement of vertebrate-like DNA methyltransferase genes in the genome of the honeybee *Apis mellifera* (6), including genes encoding the CpG-specific Dnmt1 and Dnmt3. Most strikingly, even within social insect species, DNA methylation is not always present. The gene Dnmt3 seems to have been lost in the genus *Polistes*, thus *Polistes dominula* and *P. canadensis* have greatly reduced genome wide methylation compared to other Hymenoptera species (7).

In insect species where DNA methylation is present, DNA methylation is largely confined to genic regions and elevated in coding regions (1, 2, 8–10). Gene body methylation has been suggested to affect gene expression and function via alternative splicing (8, 9, 11–14), nucleosome stability (15), or regulation of transcription elongation (16–20). However, its precise function remains unclear(21, 22). It is nonetheless evident that DNA methylation in insects is involved in a wide range of biological processes, such as nutritional control of reproductive status (23), development (9, 24–26), embryogenesis (27), alternative splicing (8, 11–14), host-parasite evolution (28), memory processing (29–31), age-related changes in worker behavior (32), modulation of context-dependent gene expression (33), maternal care (34), defense against territorial intrusion (35), longevity(36, 37), or caste determination in social insects (23, 26, 32, 38, 39).

In honeybee, a standard social insect model organism, the methylome size is reduced and DNA methylation occurs at much lower levels than in vertebrates, with fewer than 1% of CpG dinucleotides methylated (9). The CpG methylation system was demonstrated to be functional, and identification of an active methyl-DNA binding domain encoding gene was suggestive of pathways for molecular recognition of DNA methylation marks (40). The difference of the honeybee genome-based findings compared to the observed scant DNA methylation in the model insect *Drosophila melanogaster* (which lacks Dnmt1 and Dnmt3 genes) (41) raised the possibility of involvement of DNA methylation in the determination of the complex social phenotypes in the honeybee.

DNA methylation has been largely studied by whole-genome bisulfite-sequencing (BS-seq) for different samples, probing a wide variety of developmental and environmental conditions (2, 9–11, 14, 32, 35, 42–52). Although all studies have reported low levels of DNA methylation, restricted almost exclusively to cytosines in CpG context, it has been difficult to compare studies even on the same species due to several factors: (1) potential differences in data quality control; (2) use of different computational methods and detection thresholds; (3) mapping of BS-seq reads to different genome assemblies and annotation versions (e.g., (53)).

Here we present a reproducible, scalable workflow for BS-seq data analysis, tailored to, but not exclusive to, studies of species with low levels of DNA methylation. The workflow provides turnkey computation, starting with just a few edits of configuration files. The implementation works on any UNIX/LINUX system and produces a complete analysis, from data quality control, read mapping, methylation site calling, to statistical analysis of methylation patterns relative to genome annotation. We demonstrate the power of the approach by re-analysis of publicly available BS-seq data for social insects, showing (1) the validity of the workflow by comparison with trusted data sets; (2) critical re-evaluation of published results; (3) re-analysis of published BS-seq data mapped to updated genome versions; (4) evaluation of published BS-seq data relative to an only recently made available genome annotation; and (5) integration and comparative analysis of a large set of published experiments for *A. mellifera.* The integrative analysis provides a solid estimate of the size of the honeybee CpG methylome, and the consistent application of strict data quality control measures suggests alternative interpretations of some puzzling results in the literature.

## Materials and Methods

The following subsections describe the essential elements of the data processing and analyses in this study. Technical aspects are given in full detail in *SI Text.*

### Data sets

Within the scope of this paper, we collected publicly available BS-seq data from studies on four arthropod species: the paper wasp *Polistes canadensis*, the raider ant *Ooceraea biroi*, the African social spider *Stegodyphus dumicola* (one study each; Dataset S1); and the honeybee *Apis mellifera* (17 different studies; Dataset S2). Another large data set for *Apis mellifera* (nine samples and a total of 30 replicates) by Yagound *et al.* (52) became available after completion of our study but is being discussed as an independent data set to test our conclusions (see section ‘Validation’).

### Read quality control, mapping, and methylation status analysis

After standard read quality control (see *SI Text*, ‘Read quality control and trimming.’), `Bismark` (54) was used for BS-seq read mapping (using the `Bowtie2` (55) option) and methylation calling. Non-converted reads were filtered using an iterative process (described in *SI Text*, ‘Removal of non-converted reads’). A standard binomial test with correction for multiple applications is used to derive reliable sets of methylation sites and levels for analysis (*SI Text*, ‘Determination of significant methylation sites’). Only sites that have sufficient coverage of reads to be detectable as statistically solid methylation sites (calls unlikely to have resulted from failed BS-conversion) enter further analysis. These scd sites are distinguished as highly significantly methylated (`hsm`) sites (with methylation levels unlikely to have resulted from failed BS-conversion) or (otherwise) not significantly methylated (`nsm`) sites (Fig. 1).

**Fig. 1.**
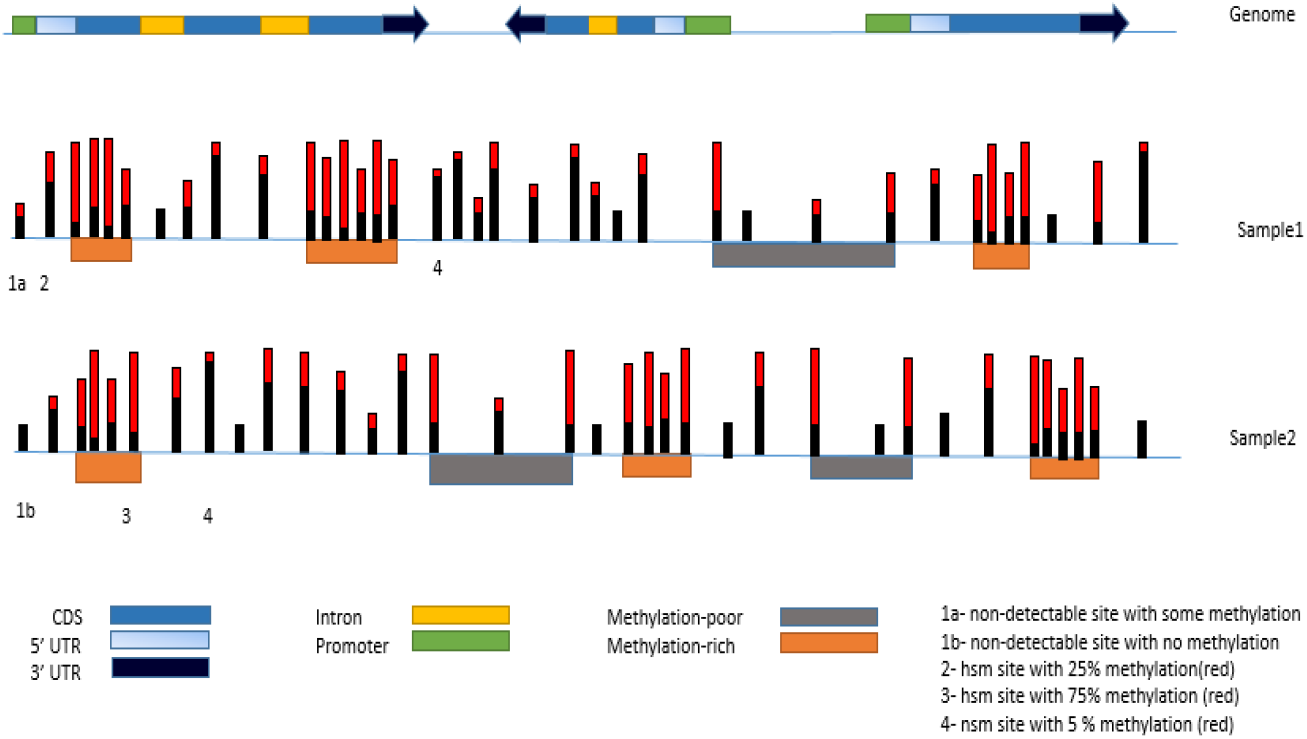
Schematic distribution of methylation sites. Track 1 shows a hypothetical genome annotation, displaying three genes with one to three exons each. Transcriptional direction is indicated by the arrow of the last exon for each gene. Putative promoter regions were defined as 500 bp upstream of the gene annotation start. Tracks 2 and 3 show methylation sites (vertical bars) and methylation-rich and -poor regions (orange and gray horizontal bars, respectively). Methylation percentages are indicated by the relative length of the red portions of the vertical bars. Labeled sites illustrate different classifications, depending on depth of read coverage and the proportion of reads indicating non-converted (presumed methylated) Cs: 1 corresponds to sites with coverage too low to establish methylation status; 2 and 3 indicate highly supported methylation (hsm) sites with methylation levels of 25% and 75%, respectively; 4 labels not significantly methylated (nsm) sites. Thresholds for site classification are discussed in the text.

### BWASP workflow

The entire workflow for obtaining the methylation site data is defined using `GNU Make` (56) and requires as input only the relevant genome assembly file in multi-FASTA format, and the BS-seq reads, provided either as fastq files or specified for download by `NCBI SRA` identifiers. To facilitate portability along with reproducibility, the newly developed code is distributed via github (https://github.com/BrendelGroup/BWASP), and the entire workflow is encapsulated in a Singularity container (57) that produces identical results in any compatible compute environment (https://BrendelGroup.org/SingularityHub/bwasp.sif).

### BWASPR – R scripts for statistical analysis

Statistical analysis of single base resolution methylation levels calculated by `BWASP` was done using a suite of `R` scripts called `BWASPR`, described in detail in *SI Text.* The code is available on github (https://github.com/BrendelGroup/BWASPR) and bundled with all dependencies in another Singularity container (https://BrendelGroup.org/SingularityHub/bwaspr.sif).

### Statistical assessment of overlap between different sets of sites

When comparing two sets of sites from the same genome, assessment of the statistical significance of their overlap amounts to determination whether the overlap is or is not consistent with the two sets being random samples from a shared pool of sites. Comparing scd CpG sets, the candidate shared pool of sites is the set of all genomic CpG sites. Comparing `hsm` CpG sites, the candidate shared pool of sites is the set of common scd sites. In each case, the size of the shared pool of sites for sampling was estimated using a mark-and-capture approach as described in the legend to Fig. 2. As a scale-invariant measure of overlap we use *ω*, defined as follows. Let *S*_1_, *S*_2_, and *S*_12_ be the number of sites unique to sample S1, unique to sample S2, and common between both samples, respectively (Fig. 2), where *S*_1_ < *S*_2_. Then

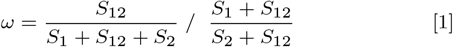

gives a value of 0 if *S*_12_ =0 (no overlap between samples S1 and S2) and a value of 1 if *S*_1_ = 0 (sample S1 is completely contained in sample S2).

**Fig. 2.**
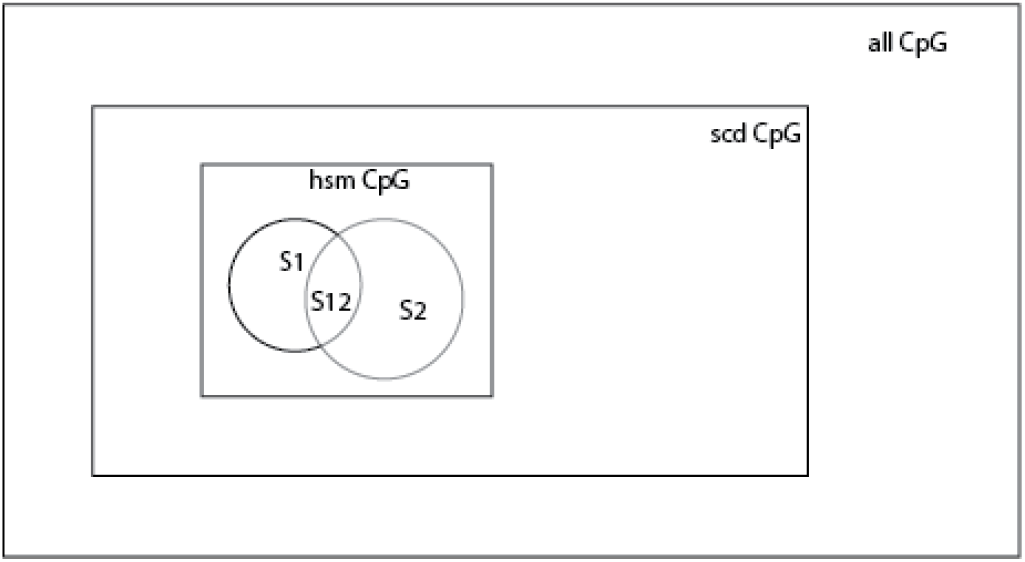
Estimating the number of methylated genomic CpGs. The hsmCpG rectangle represents the set of methylated genomic CpGs that are detectable as a subset of the sufficiently covered, detectable (scd) CpG set, which in turn is a subset of all genomic CpGs. The overlap of two independent random samples S_1_ and S_2_ of the hsmCpG set provides the mark-and-capture population size estimate for hsmCpG from the identity *S*_12_/*S*_2_ = *S*_1_/*hsmCpG*, where S_12_ denotes the overlap between S_1_ and S_2_, and italicized labels denote numbers of sites in the respective sets. Multiplying *hsmCpG* by the ratio of the number of all CpGs in the genome over the number of scd sites gives the estimated size of the CpG methylome.

### Distribution of methylation sites: co-occurrence with annotated genome annotation

To correlate methylation site occurrence with transcriptional activity, site density was calculated per genome feature type, including genic (exon and intron) and intergenic (promoter and other). For the purpose of these statistics, promoter simply refers to the region 500 nucleotides upstream of the annotated 5′-end of a gene, or shorter to avoid overlap with an upstream gene annotation. Site density was defined as number of sites within a feature divided by feature length and is reported as values normalized per 10kb. For exons, statistics were also calculated for the subcategories 5′-UTR, CDS, 3′-UTR, and Other (non-coding genes). Expected values were derived under the assumption of random positioning of sites relative to annotated genome features.

### Comparing ranked lists

The `BWASPR` workflow generates lists of genome regions (genes and promoters) ranked by extent of methylation (density of `hsm` sites or overall methylation level), presented as tab-delimited files in output directory `RNK`. It is of interest to compare such lists between different samples to determine whether the same regions are relatively hypermethylated under different conditions. List comparisons can be made with the included script `xcmprnks` which calculates the rank-biased overlap (`rbo`) measure (58) for all pairwise comparisons of specified lists. Parameters to the script include the number of ranks to consider (default: 40), the rbo parameter *p* (default: 0.95), and, for statistical evaluation, the number of ranks to shuffle (default: 40), the number of permutations to generate (default: 100), and the significance level *t* for evaluation (default: 5%). The permutation test is performed by randomly shuffling the association of identifier (e.g., gene name) and the corresponding ranked score. Observed `rbo` values in the top *t* % of values generated by shuffling are starred as significant, indicating more than expected overlap of the ranked lists.

## Results

The `BWASP/R` workflow proceeds through a large number of steps, from data download and quality control to read mapping, statistical and visual annotation of results, and testing for significant methylation differences between groups of biological samples. Our aim is to offer researchers a reliable, stress-free, and reproducible method to analyze whole-genome bisulfite sequencing data sets, especially targeting data sets with low methylation rates, as observed in insects. In order to demonstrate the many ways our workflow can be beneficial, short read data generated from bisulfite-treated genomes were obtained from `NCBI SRA` as referred to in the published studies listed in Datasets S1 and S2. In total, we re-analyzed publicly available BS-seq read sets of 70 samples and cumulatively 147 replicates from four different species and 20 studies (*SI Text*, Table S1). These re-analyses illustrate how the `BWASP/R` workflow can be used to rapidly re-evaluate published results, easily re-analyze data in the context of a new genome assembly and/or genome annotation, and, perhaps most importantly, compare and integrate multiple studies in an efficient, consistent manner.

### Description and validation of workflow

To demonstrate and validate the functionality of the `BWASP/R` workflows, we chose a medium-sized study with multiple samples and biological replicates. The study compared queen and worker samples (three biological replicates each) from *Polistes canadensis* (59). Our goal was to provide scripts that download the published BS-seq data sets from `NCBI SRA` and, with minimal initial setup configuration by the user, execute a complete analysis of the data for comparison with the published results. Our implementation is accessible as described in the **Materials and Methods** section.

For the `BWASP` workflow, three configuration files are needed. In the first file, parameters for the different programs are set to values appropriate for the available computational resources (mostly, number of cores and amount of memory to use; examples are provided in https://github.com/BrendelGroup/BWASP/data/machines.cfgdir/). The second file specifies the genome assembly and annotation files. For *Polistes*, this amounts to specifying the `NCBI` download site for the FASTA genome sequence file and the `GFF` annotation file (see https://github.com/BrendelGroup/BWASP/data/species.cfgdir/). The third configuration file specifies the design and data source of the study:

~~~
SPECIES=Pcan
GENOME=Pcan.gdna
STUDY=Patalano2015
SAMPLES=( Queen Worker )
NREPS=( 3 3 )
PORS=( p p )
SRAID=( SRR1519132 SRR1519133 SRR1519134 \
 SRR1519135 SRR1519136 SRR1519137 \
 )
~~~

Here, `SAMPLES` provides labels for the two samples, `NREPS` indicates that each sample has three replicates, `PORS` says that the sequence data are paired-end, and `SRAID` gives the `NCBI` accession numbers for the data sets. After these preparatory steps,

~~~
xsetup -m mycpu -s Pcan Patalano2015
~~~

will generate a complete directory structure populated with the data files and workflow makefiles, ready for execution (`mycpu, Pcan`, and `Patalano2015` refer to the three configuration files, respectively). In general, the user can of course alternatively specify local files instead of download sites, for example in case pre-publication genome, annotation, or BS-seq data are to be analyzed.

A complete script for running the `BWASP` workflow for this example is given in *SI Text*, ‘`BWASP` workflow: design and output.’ The entire process, including download of all required code (bundled as a Singularity image), involves fewer than a dozen lines of commands and creates 266 Gb of output files (in about 12h on our old 32-processor Linux server). Obviously, running times will vary with available resource allocation. For comparison, we also ran the worker samples on the Indiana University Carbonate cluster (https://kb.iu.edu/d/aolp), allocating one node with 16 threads, which produced identical results in five hours.

The point we wish to make here is that the user time to set up the workflow is marginal, while the computational execution is roughly overnight without any additional user intervention. Moreover, the workflow takes advantage of the `GNU Make` environment (56) that allows seamless integration of different parts of the computation run separately. For example, downloading the raw data from `NCBI SRA` can be run independently first to make sure that there is no interference by network problems. For large data sets, each replicate can be run separately, followed by the cumulative sample analysis. Depending on available resources and queuing times on a computing resource, such strategies can enhance real-time performance. It is these workflow design features that enable the large-scale, multi-study integration of experimental data being presented here.

The dependencies of the workflow are depicted in Figs. S1-S3, and details are discussed in *SI Text*, ‘`BWASP` workflow: design and output.’ Briefly, the workflow takes the raw BS-seq input data, subjects the data to several quality filtering steps, and ultimately derives `*.mcalls` files that for each C context record position, read coverage, and percent methylation for every sufficiently covered (`scd`) site in the genome. Data are derived initially for each replicate and then cumulated for each sample. Given sufficient coverage, between-replicate comparisons can probe the robustness of between-sample comparisons. Per sample mapping statistics for the *Polistes canadensis* study are shown in Dataset S3. Cumulative read numbers are seen to be in excess of 90 million per sample, with a mapping efficiency of about 83%. Two to three million of the mapped reads per sample were identified as PCR duplicates and removed.

While the starting point of the analysis presented here is the same as what is presented in (59) and in each case the `Bismark` (54) software was used to make methylation calls, different quality control setting choices may impact result details. Here, our mapping efficiency is about 10% higher than reported in (59) due to our slightly more error-tolerant `bowtie2 min_score` setting. More importantly, inclusion of removal of low quality reads, PCR duplicates, and likely unconverted reads as done in the `BWASP` workflow is not discussed in (59) (although a procedure for elimination of false positive methylation regions is described). Our point here is not to argue for particular choices of quality control but to emphasize that the workflow-enabled approach allows transparent re-analysis, either with original or with modified workflow steps and parameter sets.

The final genome coverage was estimated as 35.8 for the queen sample and as 31.7 for the worker sample (Dataset-S3). Five to eleven 5′-positions and one to three 3′-positions of the mapped reads were significantly biased for methylation calls and conservatively ignored for summary results (see *SI Text*, ‘`BWASP` workflow: design and output.’) Overall, more than 80% of CpG sites were covered by at least one read, and 21.32% and 24.73% of CpG sites were covered by at least 20 reads in queen and worker, respectively (Dataset S5). There were 13,840 and 12,036 `hsm` CpG sites identified in the queen and worker samples, respectively, corresponding to the fractions 0.16% and 0.15% of `scd` CpG sites (Dataset S7). Overall CpG methylation levels were calculated as 0.99% for queen and 0.98% for worker (Dataset S9). Patalano *et al.* (59) report the global CpG methylation level as 2.79%. Beside the aforementioned differences in data quality control, their estimate was based on averaging over values in 1-kb windows that excluded regions with fewer than 20 methylation calls and thus should be higher than `BWASP` reported overall level.

Further analyses of the methylation calls (`*.mcalls` output files from `BWASP`) were done via the `BWASPR` workflow, again facilitated by user-supplied configuration parameters. Settings and output for the *Polistes* study are shown in Dataset S15. The workflow compiles a large number of statistics concerning the extent and distribution of methylation (for details, see *SI Text*, ‘`BWASPR - R` scripts for statistical analysis.’). Here we highlight the design aspects of the analysis that are tailored to the problem of producing reliable estimates in view of very low overall methylation rates (and, typically, less than statistically desirable read coverage for replicate samples).

Key to our strategy is to discard all methylation call data at sites that are covered by fewer than *t* reads, where *t* is calculated as the minimal coverage required for a binomial test to detect statistically significant methylation at the site (as opposed to chance events due to incomplete BS-conversion). For the *Polistes* study, *t* = 4 (*SI Text*, ‘Determination of significant methylation sites’). Thus, low coverage sites (examples labeled “1” in Fig. 1) are ignored in subsequent analyses.

About 70% of the `hsm` CpG sites are in annotated genes, with about 3.5-fold over-representation of within-exon sites (Dataset S11). Dataset S13 provides more detail and shows that more than 90% of these exonic sites occur in coding sequences. These data would seem to nuance the Patalano *et al.* (59) observation of “relatively little gene body-specific methylation targeting,” which was based on overlap with highly methylated regions rather than analysis of methylation sites. `BWASPR` combines site-based and region-based analyses as shown in Fig. 1. In particular, methylation-rich and -poor regions are determined as clustering or overdispersion of sites (*SI Text*, ‘Methylation-rich and -poor regions (MRPR)’) as an empirical proxy for statistical *r*-scan analysis (following ideas reviewed in (60)). For the *Polistes* data, it is clear that there is great variation in the dispersion of methylation sites, ranging from tight clustering to stretches of multiple hundred kb devoid of any sites (see Dataset S15, pp. 170-187).

Fig. 3 illustrates the complementarity of the site density versus regional methylation measures. Genes of lengths 500 to 5000 bp with at least five `hsm` sites were measured by overall percent methylation and `hsm` site density. A wide scatter is seen, indicating both genes with relatively high density of sites but relatively low overall methylation level, and vice versa.

**Fig. 3.**
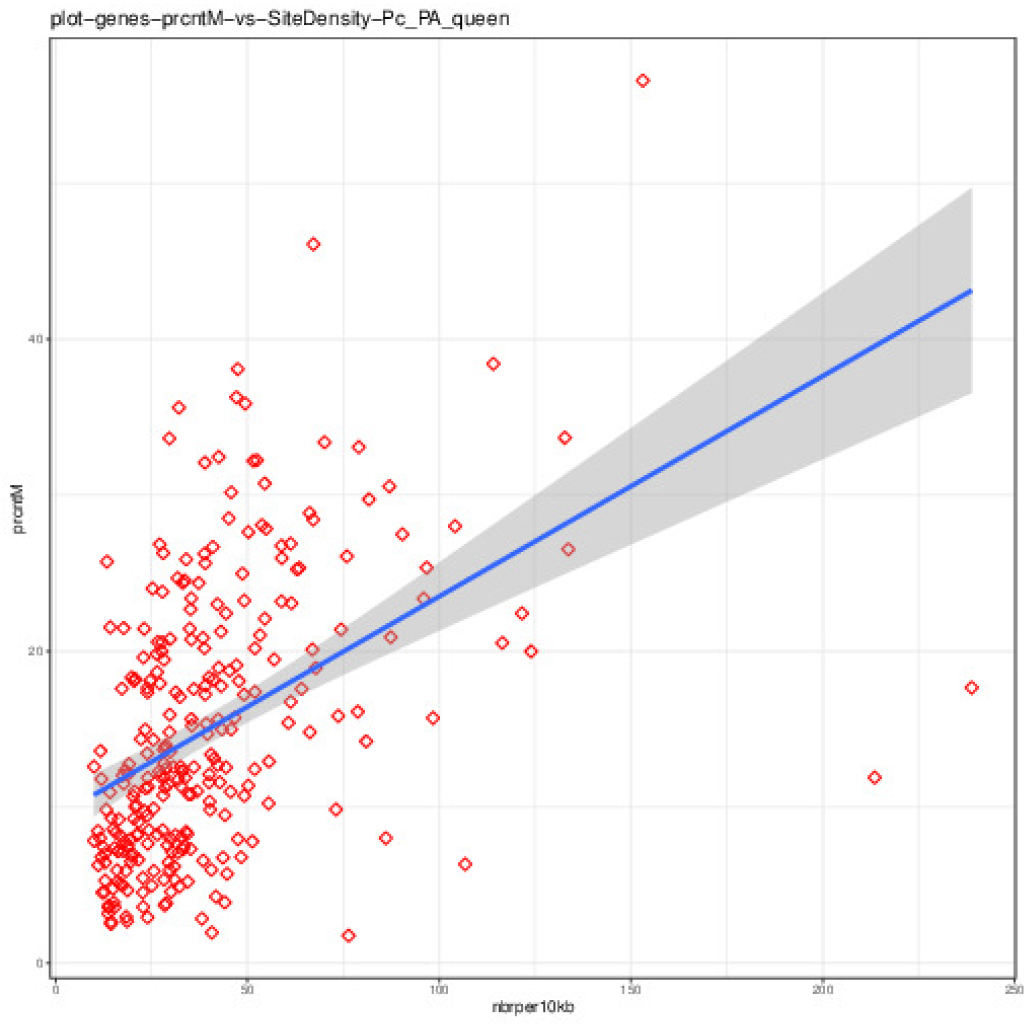
Correlation of two methylation measures for genes. The y-axis score is the overall methylation percentage for a gene. The x-axis score is the number of hsm sites in the gene, normalized to 10kb length.

In summary, this case study demonstrates the validity and capabilities of the `BWASP/R` workflow as well as the difficulty of detailed comparison with previous computational results for which precise reproduction is rendered impractical or even impossible without the original analysis scripts. The latter theme is taken up also in the next case study, which shows that an easy-to-use workflow enables individualized re-evaluation of published results - and thereby scientific discussion beyond the point of initial peer review. For practical considerations of how to adapt the workflow to analysis of very large genomes with limited computing resources please see *SI Text*, ‘Case study of a large genome.’

### Using workflow-enabled re-analysis to evaluate published results

One motivation for our work was to significantly lower the burden of computational reproducibility. A frequently encountered problem is that a computational analysis depends on a large number of software and parameter choices. If the analysis yields surprising results, one has to evaluate whether the findings reflect biologically significant features or whether differences in software and parameter choices provide the explanation. If the analysis can easily be re-run with changed parameters, for example, then this would provide more robust evaluation. In current practice, publications rarely provide enough details to completely reproduce a genome-wide computational analysis, and even if one succeeded in such effort, it would likely be a highly time-consuming task.

For DNA methylation studies, `BWASP/R` offers a solution that allows authors and readers to easily re-evaluate and fairly compare results. For illustration, we discuss `BWASP/R` results for the data reported in (45). The paper analyzed differential DNA methylation patterns in *Apis mellifera capensis* female embryos produced either sexually by fertilization of eggs with sperm or asexually from two maternal genomes via a process called thelytokous parthenogenesis. The central goal of the study was to probe parent-of-origin effects on DNA methylation, and the authors’ thorough analysis showed patterns of differential methylation between the two types of embryos, consistent with genomic imprinting. However, effects of *cis*-mediated allele-specific methylation were also demonstrated and shown to confound interpretation of the genome-wide analysis.

As a surprising collateral result of the study, the authors reported a high level of non-CpG methylation at over 50,000 sites, compared to the 114,156 and 99,923 methylated cytosines in the CG context detected in fertilized and thelytokous embryos, respectively. Only few studies on honeybees have revealed significant non-CpG methylation (11, 42). For a consistency check, we set up a `BWASP/R` re-analysis of the deposited BS-seq data, following the analogous procedure described in the previous section. Our analysis gave consistent results with previous studies, tallying only 13 and 30 CHG and 54 and 104 CHH sites in fertilized and thelytokous embryos, respectively (Dataset S8). It would seem, therefore, that the surprising numbers of non-CpG methylation sites reported in (45) reflect software and parameter choices.

Our concern here is not to investigate all possible causes for the widely discrepant results but rather to demonstrate how `BWASP/R` allows nearly effortless re-analysis of published BS-seq studies, with complete documentation and reproducibility, as well as the flexibility to change parameter settings for robustness of results analyses. For this example, several differences in the respective methods stand out: 1) Mapping efficiency and quality control. `BWASP` yields mapping efficiencies of 39.3% and 32.9% for the two data sets; after mapping, 1.76% and 2.31% of the reads were identified as PCR duplicates and removed (Dataset S4). No mapping efficiencies nor PCR duplicate removal are discussed in (45). It is likely that the more stringent quality control choices in `BWASP` (including also removal of non-converted reads; see *SI Text*) explain the lower coverage reported in Dataset S6 relative to (45). 2) Correction for potential incomplete BS-conversion by a binomial test with false discovery rate adjustment (*SI Text*, ‘Determination of significant methylation sites’) yielded a minimum per site read coverage of four for reliable methylation site detection in our workflow, compared to a coverage of two in (45). The very low number of common non-CpG methylation sites detected by the authors (only 561, compared to about 75,000 postulated common CpG sites) suggests to us that their threshold setting is too liberal and that our more conservative approach correctly shows lack of significant non-CpG methylation, consistent with other studies (discussed in more detail below in the section ‘Non-CpG methylation’).

### Workflow-enabled exploration of grouping and aggregation statistics

Replication and aggregation statistics are essential to any typical large-scale multi-sample data analysis, and BS-Seq studies are no exception. Clearly, the per site methylation percentages are aggregate statistics over multiple DNA molecules in the respective sample. In some studies, it may not be clear a priori how the input data should be partitioned into homogeneous samples when several biological criteria can be used to group samples before comparing methylation patterns. For example, pooling different cell types may be acceptable for some questions about DNA methylation status but not when cell type specific methylation is being probed. A random partitioning of data may be the most desirable option to serve as a control for differences between biologically motivated groupings of samples.

Here, we illustrate how the `BWASPR` workflow can easily be run to provide the requisite analyses for different data partitionings, with only minimal editing of the workflow configuration files. We chose to replicate a study by Libbrecht *et al.* (13), in which the authors compared reproductive (R phase) and brood care (BC phase) samples from the the clonal raider ant *Ooceraea biroi* (known as *Cerapachys biroi* at the time).

The authors discussed analysis of three data partitionings: (1) R phase versus BC phase with four replicates each, derived from two clonal lineages (A and B) and two batches of library preparation and sequencing (1 and 2); (2) Four separate comparisons of R phase versus BC phase with two replicates each (i.e., A1 samples of R phase versus A1 samples of BC phase; and the same for A2, B1, and B2 samples); (3) Comparison of the eight individual samples.

Any of these partitionings (and more, e.g., grouping by line or batch) are easily analyzed with the `BWASPR` workflow. All that is required to change the data partitioning are minor, intuitive edits to the configuration file, specifying the design and data source of the study. Samples to be compared are indicated in the `samplelist` variable. For example, comparing the phases with four replicates each, samples are specified as `BCphase` and `Rphase` (Dataset S17, page 1), whereas the global comparisons of the eight individual samples is indicated by labels `BCphaseA1`, `BCphaseA2`, etc. (Dataset S18, page 1). The corresponding `.dat` file includes columns with the species name, the name of the study, the samples to compare (matching the labels in the configuration file), the replicate number detailing whether the samples should be aggregated or analyzed individually, and the locations of the respective `*.mcalls` files (Dataset S17, pages 4-6). After these preparatory steps, `BWASPR` can be executed with the generic script as before (*SI Text*, ‘`BWASPR` – `R` scripts for statistical analysis’).

#### Similarity of DNA methylation patterns

Similarity of DNA methylation patterns across individuals, or groups of individuals, can be assessed on different levels. The `BWASPR` package implements functions and scripts to calculate all of the following measures.

(1) To what extent do two sets of methylation sites overlap, i.e., are the two corresponding samples methylated at the same sites? The output directory `PWC` shows common and unique sites comparing two samples (or groups of samples) and calculates an overlap index indicating the degree of congruence between them (see Methods, ‘Statistical assessment of overlap between different sets of sites’). The overlap index is normalized to a value between 0 and 1, with 1 indicating that one set is contained in the other, and 0 indicating absence of any overlap. Results for the 28 pairwise sample comparisons are shown in Table S2. Values range between 0.75 and 0.91, indicative of high pairwise overlap of `hsm` CpG sites.

The overlap index measure is easily extended to multiple sample comparisons: How large are the sets of methylation sites shared between biologically motivated groups of samples relative to groups of samples generated by random assortment? `BWASP` includes the script `hsmsetcmp.pl` which counts the `hsm` CpG sites shared between particular subsets of a set of `*CpGhsm.mcalls` files. This script is a generalization of a clever approach by Libbrecht *et al.* (13) who analyzed the separation of two sets of four samples by a simple criterion: how many sites are consistently methylated in one set versus consistently unmethylated in the complementary set? For their data, one would hypothesize that if differential DNA methylation were associated with behavioral phase, then the set of brood care phase samples compared to reproductive phase samples should be significantly more consistent then other partitions of the eight samples, including partitions by line or by sequencing batch. Their data (figure S3 of (13)) did not support that hypothesis and in fact showed a surprising outlier point for the partitioning by sequencing batch. The generalized `BWASP` results are available as Dataset S19, showing results for all 2^8^ = 256 partitionings of the eight samples. The homogeneity of the samples is indeed striking. The second largest site count occurs for the partitioning of all samples sharing `hsm` status (following the expected, overwhelming count of all samples being `nsm`), tallying a count of 136,608 sites. Fig. S5 shows the equivalent of figure S3 of (13), based on `BWASP` determined `hsm` and `nsm` sites. While our data are consistent in not showing any evidence of differential DNA methylation by phase, there is no outlier behavior for the partitioning by sequencing batch. It is conceivable that the corresponding data point reported by (13) is an artifact of their data processing, in particular the apparent lack of PCR duplicate removal (their data processing was done using a protocol for reduced representation bisulfite sequencing, which relies on PCR quantification, while their experimental protocol was BS-seq, which relies on PCR duplicate removal); according to Dataset S7, sequencing batch 2 had on average 29.3% PCR duplicates, much higher than the 19.3% for batch 1.

(2) How well do the methylation percentages at the sites common to the two samples correlate? Groups or individuals may show similar methylome size and location, but perhaps may differ largely in the level of methylation present at these sites. `BWASPR` calculates the correlation in methylation percentage across all common sites between groups or individuals. The correlations are shown in the output directory `CRL` (determined as described in *SI Text*, ‘Correlations between aggregate samples’) and summarized here in Table S2. The high values ranging between 0.90 and 0.93 show that not only are the methylation sites highly conserved between the samples but so are the methylation levels at the common sites. These results are in accord with the observation by Libbrecht *et al.* (13) that DNA methylation was not associated with reproduction and behavior in the context of colony cycles in *O. biroi*.

(3) How similar are respective lists of genome features (genes, promoters) ranked by degree of methylation? Ranked lists of genes and promoters are given in output directory RNK and compared by the rank-biased overlap measure (58) implemented via the `BWASPR` script `xcmprnks` (see **Materials and Methods**). Results for the comparison of the sample gene lists ranked by site density are shown in Table S3. Six of the 28 pairwise comparisons show significant congruence of the ranked lists, however there is no pattern of similarity by grouping, neither by phase, nor line, nor batch.

#### Re-analysis by read mapping to a novel genome assembly and annotation

As sequencing technologies advance or new resources become available, the reference genome and annotation of a species of interest may have been updated for a current BS-seq study relative to previously published work. In order for the current work to be comparable to the previous work, one would ideally have the analyses done with respect to the same reference genome and annotation. Thus, one option would be to re-run the previous analyses as published, but now on the new genome. This is not a small task, and ultimately it would be nearly impossible for any reader to repeat the authors’ steps with the new genome - unless that work was done with a reproducible workflow. A second option is to apply the current workflow to all data sets and, potentially, to both the previous and current reference genome and annotation. This can be done very easily with `BWASP/R`, as reviewed in *SI Text*, ‘Re-analysis by read mapping to a novel genome assembly and annotation’.

### Integrative analysis of multiple *Apis mellifera* BS-seq studies

Of all arthropod species, DNA methylation has been studied the most in the honeybee *Apis mellifera*, going back to the initial demonstration of a functional CpG methylation system (40) that accompanied the genome paper (6). However, 15 years of research have not clarified the precise role of DNA methylation in diverse processes like development, caste differentiation, or gene regulation. An integrated picture of the various experimental studies has been difficult to obtain because of a number of complicating factors. First, since the initial genome release, multiple updates to the genome assembly and annotation make comparisons between older and more current studies cumbersome. Second, studies are highly variable in terms of computational data processing, relying on distinct data quality control, software, and parameter choices, thus obscuring differences in reported results due to technical rather than biological factors. Third, documentation of computational details and interim results (rather than interpreted summaries) has been lacking, making precise comparisons across studies impossible.

Our re-analysis of published BS-seq data sets overcomes the aforementioned problems. First, all data were mapped to the most current `NCBI` *Apis mellifera* reference genome assembly (63), deposited as version HAv3.1 (64) and annotation release 104 (65). Second, all data sets were analyzed with the same `BWASP/R` approach, ensuring consistent data quality control. Third, the `BWASP/R` workflow guarantees complete reproducibility as all required software is available with this publication, together with all interim and final results.

Our integration over multiple studies addresses the following questions: To what extent are the same genomic CpG sites methylated in different samples? Conversely, what sites have rarely or never been seen methylated? The answers to the first two questions describes the observed and estimated CpG methylome as a subset of all CpG sites in the genome. Another question addresses the evidence for differential methylation between biologically distinct samples, evaluated not only within each study but also across different studies. Lastly, we probe the extent of non-CpG methylation, based on consistent computational assessment of all available data sets.

#### Summary statistics

At the time of completion of this work, we analyzed 17 BS-seq studies on *Apis mellifera* (Dataset S2), comprising 58 samples, a total of 126 replicates, and overall more than 13.7 billion processed reads (*SI Text*, Table S1). We observed a wide range of mapping efficiency, PCR duplicates, and genome coverage, but on average 94% and 38% of CpG sites were covered by at least one read and by 20 or more reads, respectively (Datasets S4 and S6; *SI Text*, Table S1). Thus, in aggregate, a large part of the CpG methylome has been probed under some conditions, supporting our goal of providing an integrated view of DNA methylation of the honeybee genome.

#### Determination of the CpG methylome

Coverage of the 19, 687, 378 genomic CpG sites in the 58 samples range such that between 4.79% and 98.53% of sites were sufficiently covered for `hsm` detection (Dataset S8). Numbers of `hsm` sites detected in each sample range from 8, 540 to 187, 243 (Dataset S8), totaling 6, 634, 422 identifications. The `hsm` fraction of `scd` sites ranges between 0.18% and 1.15%. Overall CpG methylation rates were determined in the range 0.38% – 1.98%, with an average of 0.99% (Dataset S10).

Dataset S12 data show that on average 94.5% of all `hsm` CpG sites are found in annotated genic regions, which however is only 1.15 times more than expected if genomic sites were picked randomly irrespective of annotation (Dataset S12). Further inspection shows that on average 83.78% of all `hsm` CpG sites are in exons, which is 4.83 times higher than expected. But note that on average only 3.98% of exonic `scd` sites are identified as `hsm`. Within exons, the vast majority of the `hsm` CpG sites are within protein coding sequences, about 2-fold higher than expected based on random selection of sites anywhere in the exon regions, the balance being in untranslated mRNA regions and non-coding RNAs (Dataset S14).

Fig. 4 shows the cumulative number of *Apis mellifera* CpG sites identified as `hsm` in any of the 17 analyzed studies, ordered by time of publication. The total number of sites discovered stands at 287,455 (calculated over all 58 samples). The graphic shows that rapid initial discovery of sites has slowed to apparently asymptotic increases now.

**Fig. 4.**
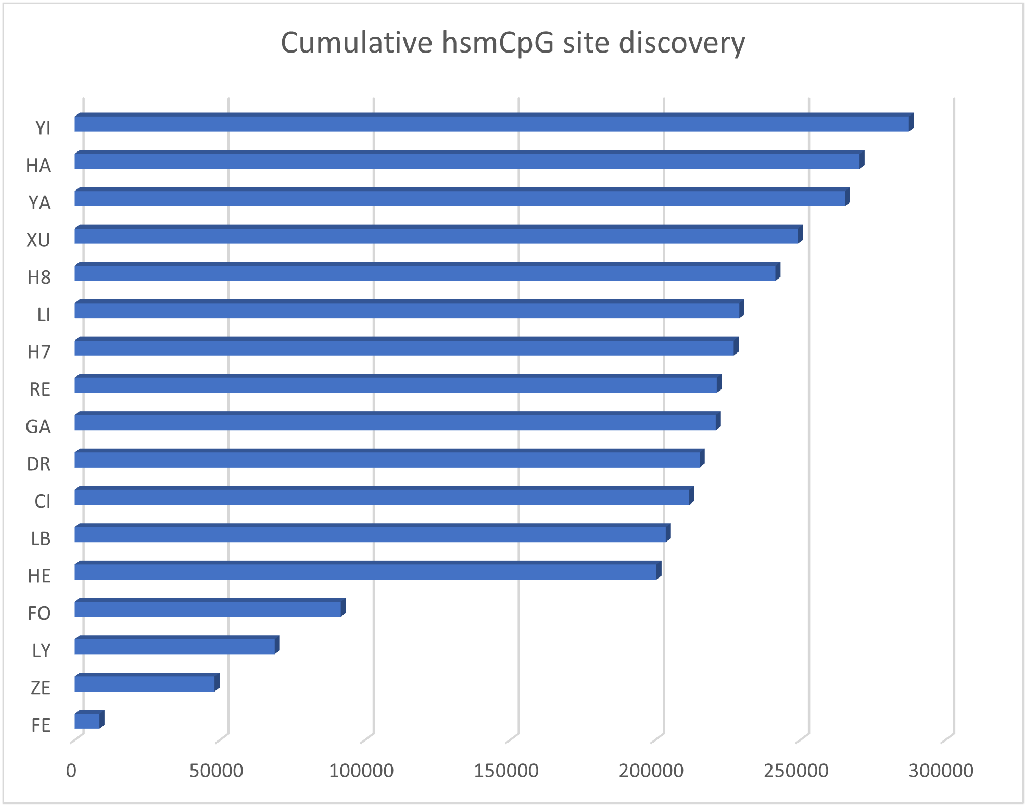
Novel hsm CpG site identification by successive experiments. The row labels refer to the *Apis mellifera* studies listed in Dataset 2, from earliest (bottom) to latest (top). The horizontal bars show the cumulative number of sites discovered from earliest to current study.

To probe whether the increases in number of sites (or lack thereof) is entirely explained by the genome coverage values of the respective BS-Seq experiments, we plotted the number of sites that are unique to each sample relative to the genome coverage of the sample (Fig. 5). It is seen that there is no clear correlation, with some high coverage samples contributing few novel sites and some low coverage sites contributing large numbers. The two outlier samples with more than 6000 unique sites each are from the YA (49) study on sperm. As sperm has been relatively under-sampled compared to other tissues (Dataset S2), this result may reflect caste- and tissue-specific DNA methylation.

**Fig. 5.**
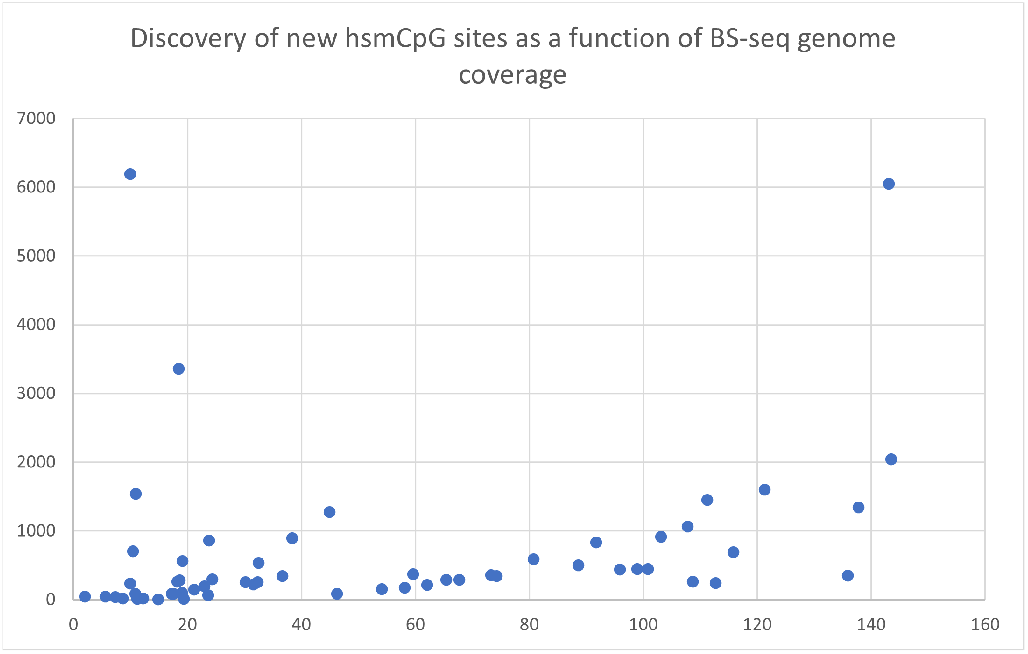
The graph shows the number of novel `hsm` CpG sites discovered in each of the 58 *Apis mellifera* studies shown in Dataset S4, ordered by genome coverage of the respective BS-seq data set.

The relative lack of unique sites in high-coverage samples could result from high-coverage studies on multiple similar samples, thus with few sites that are unique to a single sample. To explore this possibility further, Fig. 6 shows a Venn diagram of site overlap between all previous studies and the two most recent data sets available at `NCBI SRA`. It is seen that these two large experiments contribute 22,022 novel sites (16,979 unique to YI; 2,940 unique to HA; and 2,103 observed in both but not in earlier studies).

**Fig. 6.**
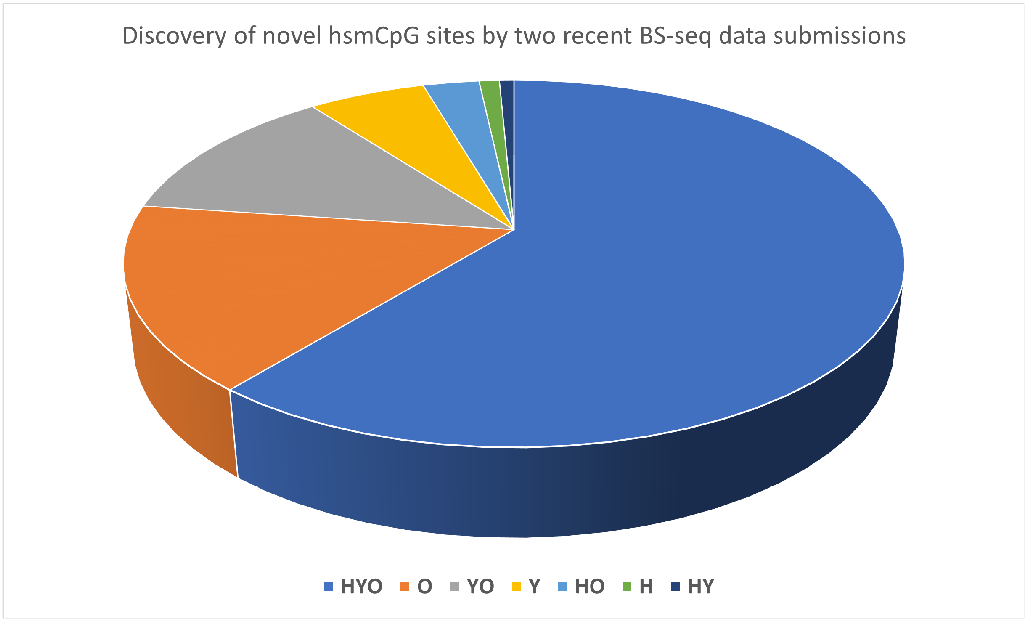
Novel `hsm` CpG sites discovered by the most recent *Apis mellifera* BS-seq studies. Abbreviations used: H = H7 (50); Y = YI (51); O = all other studies (see Dataset S2). Overlap sets are labeled by the respective combination of letters. The numbers of sites in each segment are: HYO, 178162; O, 47600; YO, 36603; Y, 16979; HO, 8111; H, 2940; HY, 2103.

Lastly, Fig. 7 records the numbers of `hsm` CpG sites shared by several samples in the set of experiments analyzed. The figures gives further evidence to considerable overlap between sets of sites shown as methylated in different samples. We cannot exclude the possibility that others sites are going to be found methylated under different physiological conditions from those that have been used in current experiments. However, a consistent, straightforward interpretation of the integrative analysis over all studies would seem to be that the *Apis mellifera* CpG methylome is far smaller than the set of all genomic CpGs and closer in size to the currently observed 287,455 value.

**Fig. 7.**
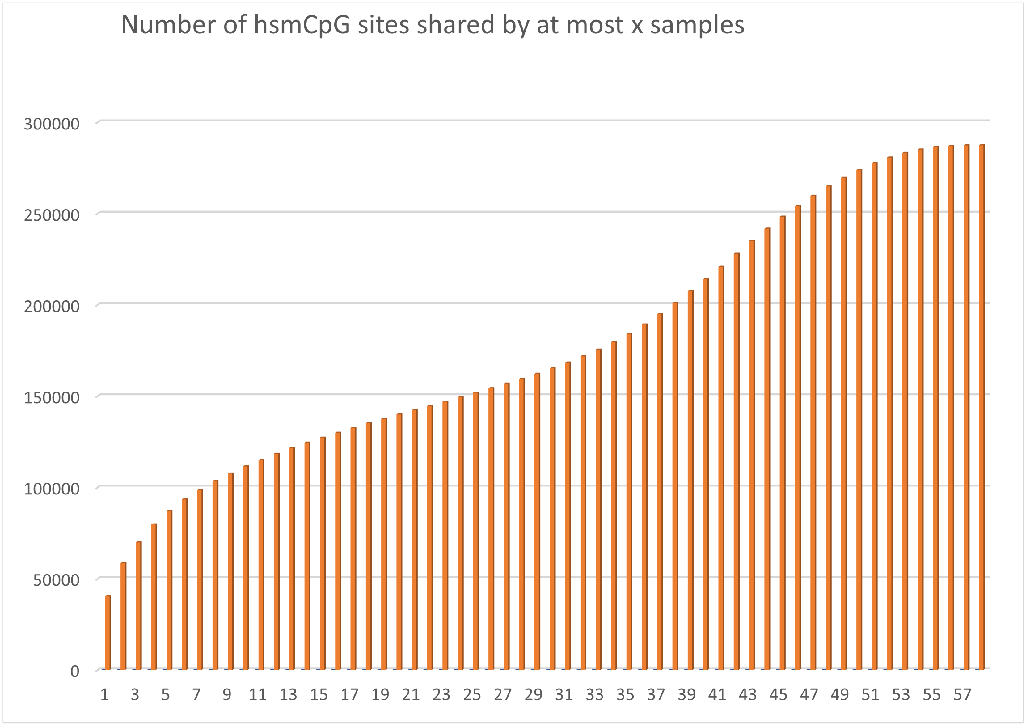
The graph shows the number of `hsm` CpG sites shared by the x-axis indicated number of experimental *Apis mellifera* samples shown in Dataset S2. For example, there are 40,545 sites unique to one experiment and 58,607 sites seen in only two samples.

#### Features of methylated CpG sites

To further explore characteristics of CpG methylation sites, we applied the `BWASP` script `hsmsetexplore.pl` to the set of `*.mcalls` files of the 58 *Apis mellifera* samples. The script output shows that of the 19,687,378 CpGs in the *Apis mellifera* genome, 19,544,535 (99.27%) were identified as `scd` in at least one sample, but only 287,455 (1.46%) as `hsm`. 91,043 sites were identified as `scd` in all 58 samples, but only 57 sites as `hsm`. Thus, the CpG methylome is small relative to the entirety of all genomic CpGs and to a large extent conserved under different conditions (as shown in the previous section), but still modulated in sample-specific manner. On average, 0.69% of CpG sites that are `scd` for the Cs on both forward and reverse strand are `hsm` in both positions, with 0.06% each being `hsm` on only forward or reverse strand, respectively. In the following we focus on CpG sites methylated on both strands.

We used the `hsmsetexplore.pl` script to pull out representative highly methylated sites (criteria: read depth between 33 and 330; at least 90% methylation on both strands; shared by at least 15 samples) and strongly non-methylated sites (criteria: read depth between 33 and 330; at most 4% methylation on both strands; shared by at least 15 samples), which generated 1,183 and 782,849 sites, respectively. We further restricted these sets to unique sites (no overlap of +/-25 base segments around the CpG with other sites in the set) and differentiated sites completely within coding regions from sites with no overlap with coding regions (again including 25 nucleotide flanks) to end up with four sets: `hhCDS` (553 highly methylated sites in coding sequences); `hhNCS` (213 highly methylated sites in non-coding sequences); `nnCDS` (858 non-methylated sites in coding sequences); and `nnNCS` (950 non-methylated sites in non-coding sequences). (The `nn` sets have the additional constraint to include only sites shared by at least 17 or 25 samples, respectively, to generate set sizes similar to those for the hh sets.) There is no rationale for the parameters used other than to generate sets of several hundred sites each that are representative to the most consistently methylated and unmethylated sites in coding and non-coding regions. Standard sequence logos for each set (reflecting the frequency distribution of nucleotides in the positions around the CpG) are shown in Fig. 8. No obvious diagnostic motifs are found that correlate with consistently high methylation. We note, however, the strong 3-periodic signature unique to the `hhCDS` set. It is therefore conceivable that a methylation code is hidden in the third codon positions of the coding sequences surrounding strongly methylated CpGs.

**Fig. 8.**
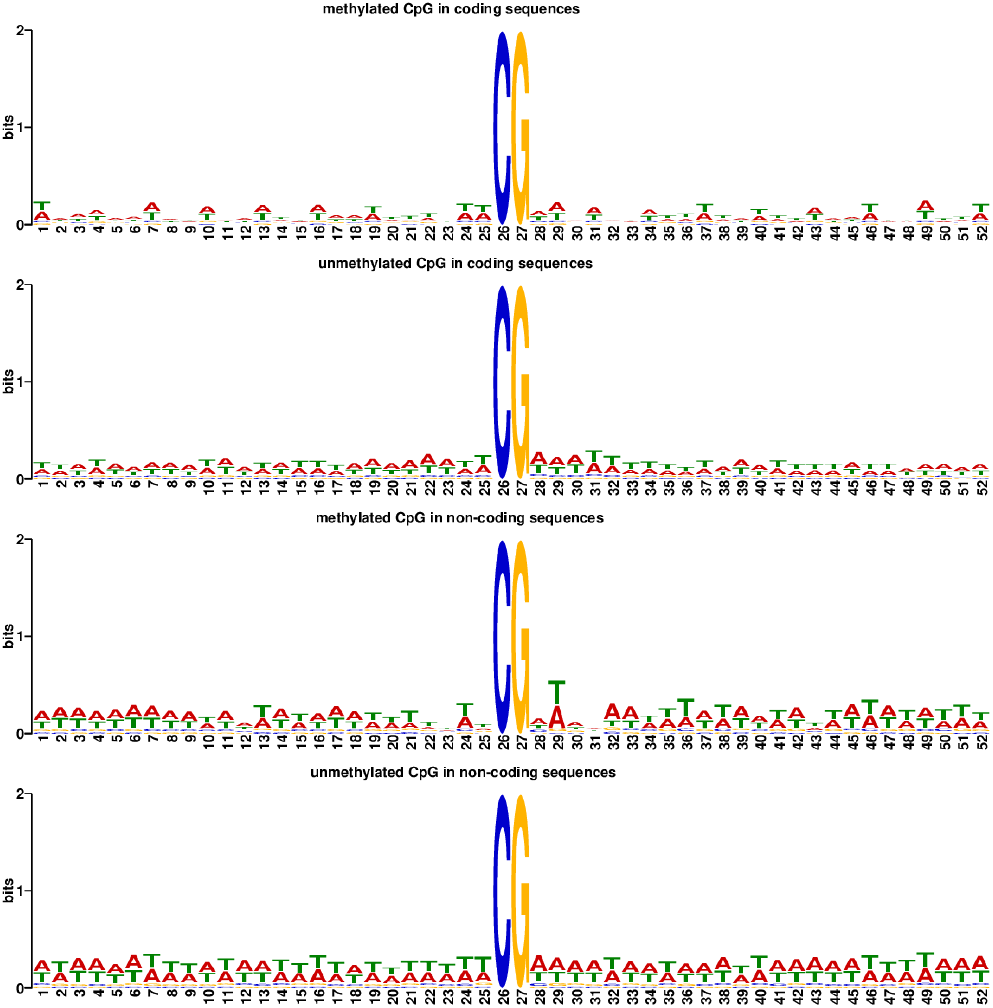
Sequence logos produced by the MEME software for CpG isites from four distinct sets, in order top to bottom: highly methylated sites in coding sequences; strongly non-methylated sites in coding sequences; highly methylated sites in non-coding sequences; and strongly ‘non-methylated sites in non-coding sequences.

#### Non-CpG methylation

The extent and role of cytosine methylation in non-CpG context has not been settled in social insect genomics. Cingolani *et al.* (42) discuss experimental and computational caveats that might lead to under-detection of non-CpG methylation and report significant levels of non-CpG methylation in honeybee introns, with a potential role in the regulation of alternative splicing. While our study cannot address any experimental biases against non-CpG methylation site detection, our consistent workflow application can eliminate biases in the computational and statistical treatment of the data. Our re-evaluation of the Remnant *et al.* (45) study (section ‘Using workflow-enabled re-analysis to evaluate published results’ above) points to the importance of analysis parameter choices. A similar argument can be made concerning the data of He *et al.* (46) who report the average context of methylation sites as 77.55% CpG, 20.5% CHH, and 1.95% CHG, which is in stark contrast to our analysis results that found only a few hundred potential `hsm`CHH sites (Dataset S8). The criteria for site definition are not discussed in (46), but lenient treatment of the statistical problem of multiple comparisons would be one explanation of the results. For example, calling a site methylated on the evidence of one read would give context proportions of 26.21%, 63.52%, and 9.67% for CpG, CHH, and CHG, respectively, for the aggregate QWE sample. A threshold of two reads would change the proportions to 77.84%, 19.93%, and 2.24%, respectively; and a threshold of three reads would lead to proportions 95.63%, 3.96%, and 0.41%. Thus, evidence for non-CpG methylation sites seems to largely be at the statistical noise level, and the substantial number of reported sites is accounted for by the large number of genomic CHH sites.

The overall methylation levels per C-context are shown in Dataset S10 for all 58 *Apis mellifera* samples. CpG methylation was observed between 0.38% and 1.98%, with a mean of 0.99%; CHG methylation was observed between 0.06% and 0.94%, with a mean of 0.34%; and CHH methylation was observed between 0.06% and 4.72%, with a mean of 0.58%. However, CHH methylation in excess of 1.0% was observed in only five samples, and without those samples, the average is 0.35%. Thus, significant CHH methylation levels remain outliers. The five samples are: queen and worker samples of Foret *et al.* (11); the data of Cingolani *et al.* (42) on European and Africanized bees; and the low count sperm sample of Yagound *et al.* (49). Looking at the fraction of scdCHH sites that are `hsm`CHH (Dataset S8), the 2.5% value of the low count sperm sample of Yagound *et al.* (49) is the only outlier (noted also in (49), but nor pursued by the authors); without that data point, the average is 0.05% (compared to 0.69% for CpG sites). Thus, our global analysis confirms the impression that DNA methylation in honeybee is overwhelmingly in CpG context, consistent with observations in other social insect species (e.g., (66)). Obviously, this computational result does not preclude the possibility of biologically significant non-CpG methylation at specific sites. A recent study by Harris *et al.* (50) postulates that low level but elevated CpA methylation in CpG-methylated genes in honeybee head tissues may be involved in the regulation of gene expression during development.

#### Validation

After completion of the data analyses described above, Yagound *et al.* published another DNA methylation study on *Apis mellifera*, comparing methylation sites and levels between drones (samples of DNA taken from thorax labeled as “drone” and samples from semen labeled as “semen”) and their daughters derived from instrumental insemination of queens (“worker” samples) (52). The goal of the study was to investigate the existence of epigenetic inheritance, and the authors concluded that there is no DNA methylation reprogramming in bees and that epigenetic information is transferred from fathers to daughters within patrilines. The author’s interpretation has been critically discussed (67, 68). Here, we have pursued two major aims with our re-analysis of the study’s BS-seq data via our `BWASP/R` workflows: (1) We wanted to know whether the new data are consistent with our earlier findings on the size of the CpG methylome and the lack of non-CpG methylation; (2) We wanted to probe the robustness of the published similarities of DNA methylation patterns within patrilines for different methylation site selection criteria and with respect to the different `BWASPR`-implemented similarity measures.

Summary statistics for the aggregate samples from colonies B1 and B2 (four replicates each for drone, semen, and worker) and colony B4 (two replicates for drone, semen, and worker) have been added to Datasets S4, S6, S8, S10, S12, and S14. All samples give high genome coverage (97X or higher) except for B4D and B4S that were sequenced about half as deeply (Datasets S4 and S6). Overall methylation levels are similar to values observed in the previous studies (Dataset S10), and the mapping of methylation sites to genomic feature regions also gave consistent results (Datasets S12 and S14). Non-CpG methylation levels are marginal, as for most studies as discussed in the previous section.

The tally of new `hsm`CpG sites was 12,282, thus increasing the total observed CpG methylome to 299,737 sites (update to Fig. 4). Unique sites (update to Fig. 5) were predominantly seen in the semen samples (B1S, 2107 sites; B2S, 1788 sites; B4S, 1067 sites) compared to worker (512, 632, and 910 sites) and drone (269, 235, and 208 sites). Thus, this additional large data set does not greatly change our estimate of the size of the CpG methylome, nor the overall summary statistics of methylation levels.

To probe the similarity of the replicate samples, we used the same approach as introduced in the subsection ‘Similarity of DNA methylation patterns’ above. The equivalent of Table S2 is shown in Dataset S23. We excluded B4 samples because of the large difference in sequencing depth relative to the other samples. We then transformed the correlation and overlap values to distances by subtracting them from 1 and generated dendograms based on hierarchical clustering of the distance matrices (using standard `R` functions and functions from the `ggdendro` and `ggplot2` libraries; clustering by Ward.D2 method). Fig. 9 shows the clustering based on the correlation of methylation levels, and Fig. 10 shows the clustering based on the overlap index distances.

**Fig. 9.**
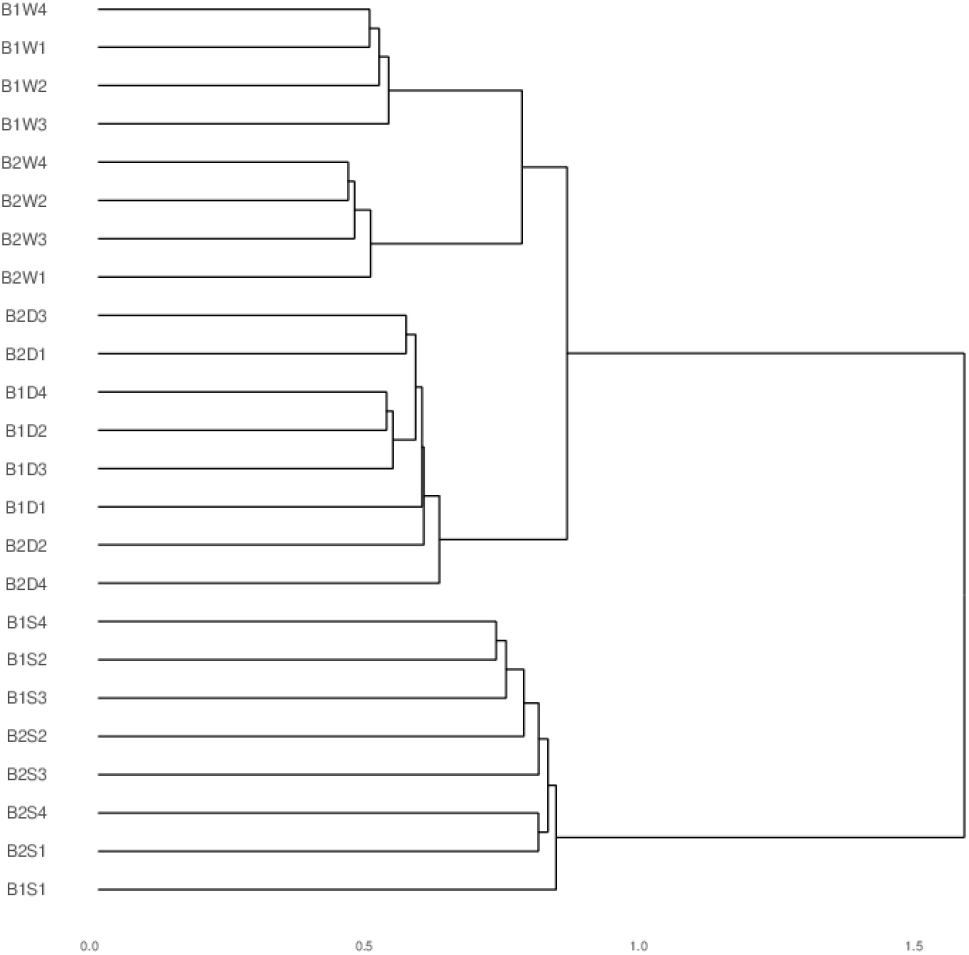
Dendrogram based on correlation distances between methylation levels of the indicated samples from Yagound *et al.* (52).

**Fig. 10.**
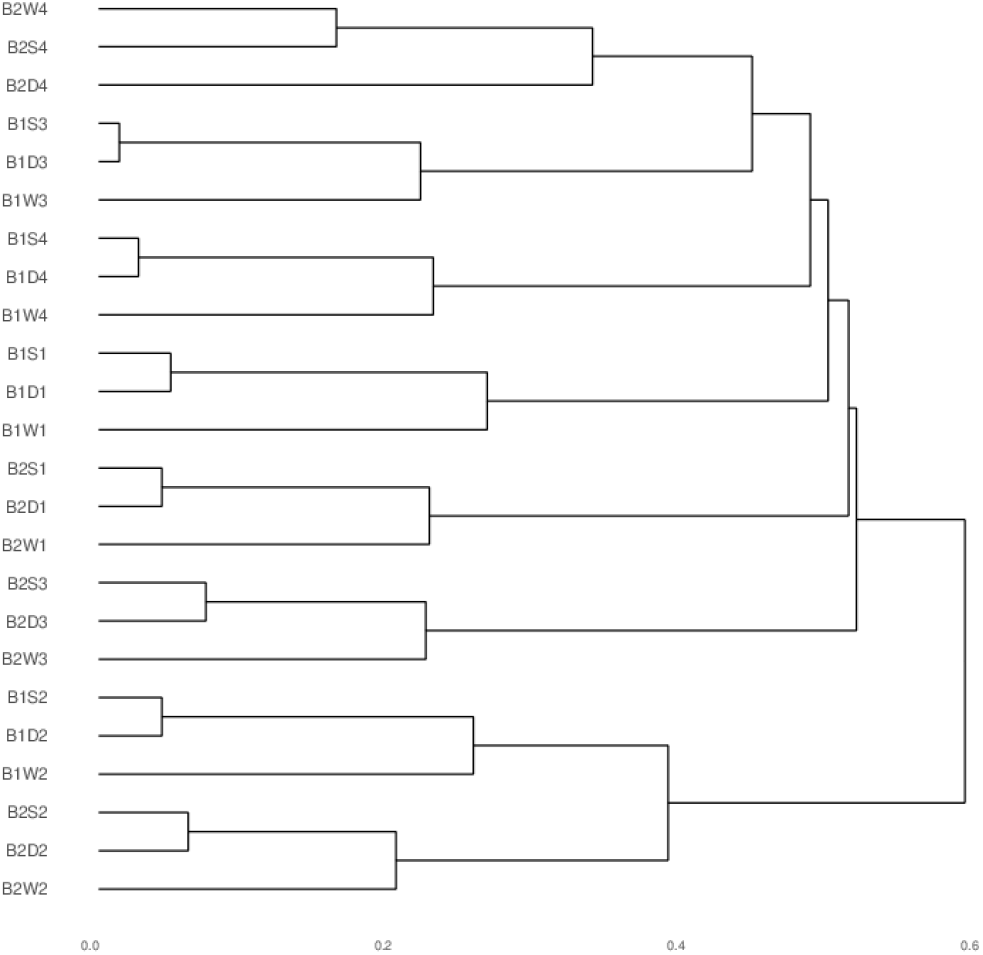
Dendrogram based on overlap index distances between hsmCpG sets of the indicated samples from Yagound *et al.* (52).

The correlation-based clustering shows clear grouping by type: the drone samples form a subcluster, as do the workers, and these clusters are separated from the semen cluster. In contrast, the overlap-based clusters shows consistent clustering by patriline (related drone, semen, and worker samples forming subclusters). What was initially puzzling is that figures 2 and S3 of (52) showed correlation-based clustering along patrilines like our overlap-based clustering. However, the explanation is easy enough: Yagound *et al.* build the methylation level matrix from all sites that were covered by at least 10 reads in all samples and classified as `hsm`CpG in at least one sample. The correlations derived from that matrix are heavily biased by the zero values at sites that are sufficiently covered but not methylated (`nsm` sites in our notation). Thus the authors’ approach convolutes the two factors separated in our analysis: the overlap index measures how similar two sets of `hsm`CpG sites are, whereas the correlation measures how similar the methylation levels are at shared `hsm` sites. The first measure reflects genetic distance (e.g., drone thorax and semen from the same individual represent the same haploid genome), whereas the second measure reflects the activity of methylation and de-methylation enzymes in the respective samples. In this purview, the data of (52) would seem to be perfectly consistent with classical genetic inheritance with subsequent methylation level control based on environmental (sample) conditions (see Discussion).

We should note that the clustering patterns shown in Figs. 9 and 10 are unchanged for different coverage thresholds or sample subset choices. Also, clustering the methylation level matrix from Yagound *et al.* (kindly provided by the authors) after deletion of rows with zero values (which reduces the number of sites from 20523 to 5680) similarly shifted the pattern to what we show in Fig. 9. Thus, we are confident that the two components of genetic and physiological difference are properly reflected in the overlap and correlation distance measures, respectively.

## Discussion

DNA methylation systems have not been found universally in all social insects, and, where present, methylation levels are very low compared to vertebrates and plants, with typical observations at around 1% of CpGs being methylated in honeybee tissues. The low levels complicate statistical analysis of whole genome bisulfite-sequencing experiments, as different approaches to data quality control can significantly alter reported methylation levels. We have presented a conceptual framework for analyzing such data in a consistent manner and implemented the computational steps in completely and easily reproducible workflows. The framework (summarized in Fig. 1) centers on the identification of sites that are sufficiently read-covered (`scd` sites) to allow statistically solid determination of methylation levels at accessible (`hsm`) sites. We have shown that re-analysis of published studies in this way with conservative data quality control (including unique mapping of reads; removal of likely unconverted BS-seq reads and of PCR duplicates; and culling of technically biased read positions from further analyses) may suggest alternative interpretations of the data. Most importantly, different studies can be fairly compared in this way, with the original data equally quality-controlled and mapped to the most current genome and annotation.

### Identification of the Apis mellifera CpG methylome

Although low levels of cytosine methylation in largely CpG context, predominantly in exon regions of the genome, have been consistently reported for *Apis mellifera*, only by means of an integrative study as presented here can we assess the extent to which different studies have identified the same methylation sites. This assessment is particularly challenging for the honeybee because there have been multiple genome and annotation versions in use over the period of more than ten years of methylation studies. Our results show that the set of all CpGs identified as methylated in at least one experiment numbers just below 300,000 sites currently, or about 1.5% of all genomic CpGs (Fig. 4). By the same method, we find no evidence for significant levels of non-CpG methylation.

Our workflow implements three measures of evaluating similarity of DNA methylation patterns between samples. The overlap index measures the extent to which two sets of methylation sites overlap. Correlation analysis of methylation levels at common sites measures whether the shared sites are methylated in similar proportions in samples being compared. A flexible script applies rank-biased overlap statistics to compared lists of genes (or other regions) ordered by methylation level. The power of these complementary measures was demonstrated in our re-analysis of the recent Yagound *et al.* (52) study comparing methylation levels from honeybee drone thorax and semen samples and worker daughters derived from artificially inseminated queens. We show that the `hsm`CpG sets are closely related within patrilines (Fig. 10), whereas methylation levels at common sites cluster by sample type (Fig. 9). This refined analysis offers a much simpler explanation for the data then the epigenetic inheritance model proposed by Yagound *et al.* that was based on a correlation analysis that convoluted these two measures of similarity. It seems to us that the most parsimonious model for the data involves sequence-determined sets of methylation-accessible CpGs in combination with dynamically determined methylation levels by physiologically regulated methylation/de-methylation activity. As the genetic background of drone thorax and semen is identical and semi-conserved in the daughter workers, `hsm`CpG sets are most similar within a patriline, but methylation level similarity goes with cellular type; see also the discussion in (67) and (68).

An open question remains concerning the sequence characteristics that distinguish the 1.5% of methylable CpGs from the other 98.5%. A preliminary search for characteristic sequence motifs did not result in the identification of clear signatures (Fig. 8). However, our analysis should provide good data sets for more sophisticated machine learning approaches.

### Standards of reproducibility

Motivated by open questions concerning the extent and role of DNA methylation in social insects, we have implemented easily accessible computational tools for the analysis of whole genome bisulfite sequencing data. Beyond the introduction of several new concepts and measures for the presentation and statistical treatment of the data, a significant component of our work has been the emphasis on complete reproducibility of all steps in the data processing, from download of the raw sequence data from a public repository to the generation of summary tables and figures. Our general philosophy with respect to computational reproducibility has been previously discussed (69). In brief, we argue that, for the most part, computational reproducibility should encapsulate the ability of bitwise regeneration of published results. Obviously, there can be changes like time stamps or expected fluctuations in stochastic models, but the practical requirement should be that every data point in a publication can be reproduced without ambiguity.

It could be debated whether such standard of reproducibility is realistic. Initially, there may seem to be few apparent incentives to put a premium on providing complete workflows. Workflows often are the result of a lot of trial and error and keeping track of what worked and what did not requires diligent documentation - often pieced together at the manuscript writing stage, rather than incrementally put into an executable script that retraces everything done and validated up to that point. A standard accepted by most reviewers and journals is that sufficient detail is provided in the publication to reproduce the work “in principle.” In practice, this would typically require much effort and additional communication with the original authors, with obstacles for resolving any differences in outcome.

We would like to argue that our large-scale data work with the `BWASP/R` software has demonstrated that: (1) There are now software solutions that do allow complete reproducibility of even very complex workflows. (2) Implementation of reproducible workflows is feasible and provides no particular technical difficulties beyond the implementation of the original data analyses in a study. (3) The adherence to such workflow standard for dissemination of scientific work enhances peer review, democratizes science, and accelerates discussion and community efforts. (4) The workflow approach opens new possibilities for integrative studies that incorporate raw data from multiple original sources.

The workflow approach demonstrated here has the advantage of capturing what researchers are already doing, whatever software and scripts in any programming language they are using. The only add-on is the demonstrated and verified reproducibility of the entire data analysis, with the discussed benefits of scalability and re-usability. For more narrowly defined bioinformatics workflows there are now a number of alternative workflow managements systems, adoption of which will greatly help the cause of reproducibility (e.g., (70, 71)).

## ACKNOWLEDGMENTS

The authors wish to thank Dr. Murat Öztürk for contributions to the implementation of the `BWASP` workflow; students Saranya Sankaranarayanan, Shengyao Chen, and Kate Mortensen for contributions in the early stages of the code development as well as presentation and testing of the workflow; and Dr. Amy Toth for numerous patient discussions on the biology of social insects. We are obliged to colleagues at `NCBI` who quickly followed up on our request for an annotation of the *Stegodyphus dumicola* genome. Thanks also to Dr. Boris Yagound for additional explanation and data for the published study (52). The authors are grateful for use of the Extreme Science and Engineering Discovery Environment (XSEDE) Jetstream resource at Indiana University and the Texas Advanced Computing Center through allocation TG-BIO160012 (Computational Genomics) to V.B.. XSEDE is supported by National Science Foundation grant number ACI-1548562. Use of the IU Carbonate system was supported in part by Lilly Endowment, Inc., through its support for the Indiana University Pervasive Technology Institute.

## References

1. Suzuki MM, Bird A (2008) DNA methylation landscapes: provocative insights from epigenomics. Nature Reviews Genetics 9(6):465–476.

2. Feng S, et al. (2010) Conservation and divergence of methylation patterning in plants and animals. Proc. Natl. Acad. Sci. U.S.A. 107(19):8689–8694.

3. Colot V, Rossignol JL (1999) Eukaryotic DNA methylation as an evolutionary device. BioEssays 21(5):402–411.

4. Glastad KM, et al. (2017) Variation in DNA methylation is not consistently reflected by sociality in hymenoptera. Genome Biol Evol 9(6):1687–1698.

5. Raddatz G, et al. (2013) Dnmt2-dependent methylomes lack defined DNA methylation patterns. Proc. Natl. Acad. Sci. U.S.A. 110(21):8627–8631.

6. Consortium, The Honeybee Genome Sequencing (2006) Insights into social insects from the genome of the honeybee *Apis mellifera*. Nature 443(7114):931–949.

7. Standage DS, et al. (2016) Genome, transcriptome and methylome sequencing of a primitively eusocial wasp reveal a greatly reduced DNA methylation system in a social insect. Molecular Ecology 25(8):1769–1784.

8. Bonasio R, et al. (2012) Genome-wide and caste-specific DNA methylomes of the ants Camponotus floridanus and Harpegnathos saltator. Curr. Biol. 22(19):1755–64.

9. Lyko F, et al. (2010) The honey bee epigenomes: Differential methylation of brain DNA in queens and workers. PLOS Biology 8(11):e1000506.

10. Zemach A, McDaniel IE, Silva P, Zilberman D (2010) Genome-wide evolutionary analysis of eukaryotic DNA methylation. Science 328(5980):916–919.

11. Foret S, et al. (2012) DNA methylation dynamics, metabolic fluxes, gene splicing, and alternative phenotypes in honey bees. Proc. Natl. Acad. Sci. U.S.A. 109(13):4968–73.

12. Flores K, et al. (2012) Genome-wide association between DNA methylation and alternative splicing in an invertebrate. BMC Genomics 13:480.

13. Libbrecht R, Oxley PR, Keller L, Kronauer DJC (2016) Robust DNA methylation in the clonal raider ant brain. Curr. Biol. 26(3):391–5.

14. Li-Byarlay H, et al. (2013) RNA interference knockdown of DNA methyl-transferase 3 affects gene alternative splicing in the honey bee. Proc. Natl. Acad. Sci. U.S.A. 110(31):12750–5.

15. Hunt BG, Glastad KM, Yi SV, Goodisman MAD (2013) Patterning and regulatory associations of dna methylation are mirrored by histone modifications in insects. Genome Biology and Evolution 5(3):591–598.

16. Bird A (2002) DNA methylation patterns and epigenetic memory. Genes & Development 16(1):6–21.

17. Lorincz MC, Dickerson DR, Schmitt M, Groudine M (2004) Intragenic DNA methylation alters chromatin structure and elongation efficiency in mammalian cells. Nature Structural & Molecular Biology 11(11):1068–1075.

18. Zilberman D, Henikoff S (2007) Genome-wide analysis of DNA methylation patterns. Development 134(22):3959–3965.

19. Luco RF, et al. (2010) Regulation of alternative splicing by histone modifications. Science 327(5968):996–1000.

20. Maunakea AK, et al. (2010) Conserved role of intragenic DNA methylation in regulating alternative promoters. Nature 466(7303):253–257.

21. Zilberman D (2017) An evolutionary case for functional gene body methylation in plants and animals. Genome Biology 18(1):87.

22. Dyson CJ, Goodisman MAD (2020) Gene duplication in the honeybee: Patterns of DNA methylation, gene expression, and genomic environment. Mol. Biol. Evol. 37(8):2322–2331.

23. Kucharski R, Maleszka J, Foret S, Maleszka R (2008) Nutritional control of reproductive status in honeybees via DNA methylation. Science 319(5871):1827–1830.

24. Shi YY, et al. (2013) Genomewide analysis indicates that queen larvae have lower methylation levels in the honey bee (Apis mellifera). Naturwissenschaften 100(2):193–7.

25. Yang SX, Guo C, Zhao XT, Sun JT, Hong XY (2018) Divergent methylation pattern in adult stage between two forms of Tetranychus urticae (Acari: Tetranychidae). Insect Science 25(4):667–678.

26. Elango N, Hunt BG, Goodisman MAD, Yi SV (2009) DNA methylation is widespread and associated with differential gene expression in castes of the honeybee, Apis mellifera. Proc. Natl. Acad. Sci. U.S.A. 106(27):11206–11.

27. Kay S, Skowronski D, Hunt BG (2018) Developmental DNA methyltransferase expression in the fire ant Solenopsis invicta. Insect Science 25(1):57–65.

28. Vilcinskas A (2016) The role of epigenetics in host-parasite coevolution: lessons from the model host insects Galleria mellonella and Tribolium castaneum. Zoology (Jena) 119(4):273–280.

29. Biergans SD, Jones JC, Treiber N, Galizia CG, Szyszka P (2012) DNA methylation mediates the discriminatory power of associative long-term memory in honeybees. PlOS One 7(6):e39349.

30. Lockett GA, Helliwell P, Maleszka R (2010) Involvement of DNA methylation in memory processing in the honeybee. Neuroreport 21(12):812–816.

31. Gong Z, Tan K, Nieh JC (2018) First demonstration of olfactory learning and long-term memory in honey bee queens. J. Exp. Biol. 221(Pt 14).

32. Herb BR, et al. (2012) Reversible switching between epigenetic states in honeybee behavioral subcastes. Nature Neuroscience 15(10):1371–1373.

33. Wedd L, Kucharski R, Maleszka R (2016) Differentially methylated obligatory epialleles modulate context-dependent LAM gene expression in the honeybee Apis mellifera. Epigenetics 11(1):1–10.

34. Arsenault SV, Hunt BG, Rehan SM (2018) The effect of maternal care on gene expression and DNA methylation in a subsocial bee. Nature Communications 9(1):3468.

35. Herb BR, Shook MS, Fields CJ, Robinson GE (2018) Defense against territorial intrusion is associated with DNA methylation changes in the honey bee brain. BMC Genomics 19(1):216.

36. Cardoso-Júnior CAM, Guidugli-Lazzarini KR, Hartfelder K (2018) DNA methylation affects the lifespan of honey bee (Apis mellifera L.) workers - Evidence for a regulatory module that involves vitellogenin expression but is independent of juvenile hormone function. Insect Biochem. Mol. Biol. 92:21–29.

37. Yan H, et al. (2015) DNA methylation in social insects: how epigenetics can control behavior and longevity. Annual Review of Entomology 60:435–452.

38. Morandin C, Brendel VP, Sundström L, Helanterä H, Mikheyev AS (2019) Changes in gene DNA methylation and expression networks accompany caste specialization and age-related physiological changes in a social insect. Mol. Ecol. 28(8):1975–1993.

39. Marshall H, Lonsdale ZN, Mallon EB (2019) Methylation and gene expression differences between reproductive and sterile bumblebee workers. Evol Lett 3(5):485–499.

40. Wang Y, et al. (2006) Functional CpG methylation system in a social insect. Science 314(5799):645–7.

41. Bewick AJ, Vogel KJ, Moore AJ, Schmitz RJ (2017) Evolution of DNA methylation across insects. Mol. Biol. Evol. 34(3):654–665.

42. Cingolani P, et al. (2013) Intronic non-cg dna hydroxymethylation and alternative mrna splicing in honey bees. BMC Genomics 14(1):666.

43. Drewell RA, et al. (2014) The dynamic DNA methylation cycle from egg to sperm in the honey bee Apis mellifera. Development 141(13):2702–11.

44. Galbraith DA, Yang X, Niño EL, Yi S, Grozinger C (2015) Parallel epigenomic and transcriptomic responses to viral infection in honey bees (*Apis mellifera*). PLOS Pathogens 11(3):1–24.

45. Remnant EJ, et al. (2016) Parent-of-origin effects on genome-wide DNA methylation in the Cape honey bee (Apis mellifera capensis) may be confounded by allele-specific methylation. BMC Genomics 17:226.

46. He XJ, et al. (2017) Making a queen: an epigenetic analysis of the robustness of the honeybee *(Apis mellifera)* queen developmental pathway. Molecular Ecology 26(6):1598–1607.

47. Li Y, et al. (2017) Genome-wide DNA methylation changes associated with olfactory learning and memory in *Apis mellifera*. Scientific Reports 7(1):17017.

48. Xu X, et al. (2019) Evolutionary transition between invertebrates and vertebrates via methylation reprogramming in embryogenesis. National Science Review 6(5):993–1003.

49. Yagound B, Smith NMA, Buchmann G, Oldroyd BP, Remnant EJ (2019) Unique DNA methylation profiles are associated with cis-variation in honey bees. Genome Biol Evol 11(9):2517–2530.

50. Harris KD, Lloyd JPB, Domb K, Zilberman D, Zemach A (2019) DNA methylation is maintained with high fidelity in the honey bee germline and exhibits global non-functional fluctuations during somatic development. Epigenetics & Chromatin 12(1):62.

51. Yi Y, et al. (2020) Transgenerational accumulation of methylome changes discovered in commercially reared honey bee *(Apis mellifera)* queens. Insect Biochem. Mol. Biol. 127:103476.

52. Yagound B, Remnant EJ, Buchmann G, Oldroyd BP (2020) Intergenerational transfer of DNA methylation marks in the honey bee. Proceedings of the National Academy of Sciences 117(51):32519–32527.

53. Elsik CG, et al. (2014) Finding the missing honey bee genes: lessons learned from a genome upgrade. BMC Genomics 15:86.

54. Krueger F, Andrews SR (2011) Bismark: a flexible aligner and methylation caller for bisulfite-seq applications. Bioinformatics 27(11):1571–1572.

55. Langmead B, Salzberg SL (2012) Fast gapped-read alignment with Bowtie 2. Nature Methods 9(4):357–359.

56. Free Software Foundation (2020) GNU Make. https://www.gnu.org/software/make/.

57. Sochat V (2017) Singularity registry: open source registry for singularity images. Journal of Open Source Software 2. 18:426.

58. Webber W, Moffat A, Zobel J (2010) A similarity measure for indefinite rankings. ACM Trans. Inf. Syst. 28(4).

59. Patalano S, Hore TA, Reik W, Sumner S (2012) Shifting behaviour: epigenetic reprogramming in eusocial insects. Curr. Opin. Cell Biol. 24(3):367–73.

60. Karlin S, Brendel V (1992) Chance and statistical significance in protein and DNA sequence analysis. Science 257(5066):39–49.

61. McKenzie SK, Kronauer DJ (2018) The genomic architecture and molecular evolution of ant odorant receptors. Genome Research 28(11):1757–1765.

62. Liu S, Aageaard A, Bechsgaard J, Bilde T (2019) DNA methylation patterns in the social spider, *Stegodyphus dumicola*. Genes 10(2).

63. Wallberg A, et al. (2019) A hybrid de novo genome assembly of the honeybee, *Apis mellifera*, with chromosome-length scaffolds. BMC Genomics 20(1):275.

64. NCBI (2018) Apis mellifera genome assembly HAv3.1; https://www.ncbi.nlm.nih.gov/assembly/GCF_003254395.2/.

65. NCBI (2018) Apis mellifera Annotation Release 104; https://www.ncbi.nlm.nih.gov/genome/annotation_euk/Apis_mellifera/104/.

66. Bebane PSA, et al. (2019) The effects of the neonicotinoid imidacloprid on gene expression and DNA methylation in the buff-tailed bumblebee Bombus terrestris. Proc. Biol. Sci. 286(1905):20190718.

67. Soley FG (2021) Still no evidence for transgenerational inheritance or absence of epigenetic reprogramming in the honey bee. Proceedings of the National Academy of Sciences 118(28).

68. Yagound B, Remnant EJ, Buchmann G, Oldroyd BP (2021) Reply to Soley: DNA methylation marks are stably transferred across generations in honey bees. Proceedings of the National Academy of Sciences 118(28).

69. Brendel VP (2018) From small RNA discoveries to a new paradigm in computational genomics? New Phytologist 220(3):659–660.

70. Baichoo S, et al. (2018) Developing reproducible bioinformatics analysis workflows for heterogeneous computing environments to support African genomics. BMC Bioinformatics 19(1):457.

71. Jackson M, Kavoussanakis K, Wallace E (2021) Using prototyping to choose a bioinformatics workflow management system. PLOS Computational Biology 17:e1008622.

