## Supplemental Information for "Tools and applications for integrative analysis of DNA methylation in social insects"

#### This PDF file includes:

Supplementary text
Figs. S1 to S5
Tables S1 to S3
Legends for Datasets S1 to S23
SI References

#### Other supplementary materials for this manuscript include the following:

Datasets S1 to S23

### Supporting Information Text

### Methods

**Read quality control and trimming.** Quality of the sequenced reads was assessed with `FastQC` (1). Quality and adapter trimming was done using `Trim Galore!` (2). `Trim Galore!` first trims low quality bases from the 3′-end of the reads and then invokes `Cutadapt` (3) to remove adapter sequences. The `-phred` flag was set according to the `FastQC`-reported encoding method (Illumina 1.5 or 1.9+). Otherwise, default parameters were used.

**Removal of non-converted reads.** An implicit assumption in bisulfite sequencing is that all genomic DNA is equally exposed to the treatment, resulting in unbiased methylation calls in the BS-seq reads. Reads for which every C is found unconverted might represent methylation hot spots, or alternatively, and in contrast to the aforementioned assumption of equal exposure, genomic DNA segments that were inaccessible to the treatment. As overall methylation rates in the samples analyzed in this study are very low, the latter interpretation appears more prudent, particularly when Cs in all contexts are found unconverted. Conservative interpretation of the data would require removal of these reads from consideration.

To formalize this approach, we inserted an iterative filter (script `xfilterMsam.pl`) into our `BWASP` workflow. The script first calculates the overall frequencies of the six C-methylation calls (z, Z, x, X, h, H); this is using the conventional nomenclature that equates z with unmethylated C in CpG context; Z with methylated C in CpG context; x with unmethylated C in CHG context; X with methylated C in CHG context; h with unmethylated C in CHH context; and H with methylated C in CHH context.

Using these frequencies as probabilities, the probability of a particular read methylation call string is calculated, assuming independence of all positions. Reads with probability less than 0.01 divided by the number of reads analyzed (Bonferroni adjustment for multiple tests) are candidates for elimination. To safeguard against removal of reads from real, short methylation hot spots detected within a read, the script also requires that the fraction of non-methylation calls (z, x, h) be less than 1/4 of all Cs in the read. For paired-end read samples, both read-ends are eliminated if one of the reads fails the acceptance threshold. After removal of reads, frequencies of C-methylation calls are re-calculated, and the procedure is repeated until no more reads fail the acceptance threshold. For the data in this study, the script terminated after 1-4 rounds of elimination and removed between thousands and hundred thousands of reads, depending on sample (**Datasets S3** and **S4**, rows "rejected reads.")

**Determination of significant methylation sites.** During the sequencing process, C-to-T conversion at unmethylated sites fails with a small probability $p$. For a given C-position in the genome covered by $n$ mapped reads of which $x$ are C and $n - x$ are T, the null hypothesis that this position is unmethylated can be rejected if the binomial probability of observing $x$ or more Cs is below a significance threshold $t$. For an individual site, we would choose $t = 0.01$, but because many sites are evaluated simultaneously, $t$ needs to be adjusted downward. A false-discovery rate approach is common, but because our goal is to identify only the most prominent methylation sites, we use the more conservative Bonferroni correction of a threshold $t^* = t/S$, where $S$ is the number of sites tested. For example, if we have $S = 10^6$ sites with coverage $n = 20$, then at a given site $x$ would have to be 6 or larger for the site to be deemed a methylation site at the 1% significance level, assuming $p = 0.005$. For further analysis, we considered only genomic sites with read coverage high enough that statistically significant methylation could be detected. These `s`ufficiently `c`overed `d`etectable (`scd`) sites were classified as either `h`ighly `s`upported `m`ethylation (`hsm`) sites (if the null hypothesis that the site is unmethylated was rejected) or as `n`ot `s`ignificantly `m`ethylated (`nsm`) sites (otherwise).

We should note that the workflow can be used to estimate $p$ if the experiment included bacteriophage lambda DNA (for example) as unmethylated DNA control. In that case, the reads should be mapped to the lambda DNA by the same procedure. The percent methylation calls on lambda DNA CpGs provides the estimate for $p$, as the only explanation for such calls would be conversion failure of the known unmethylated Cs in the lambda DNA (see (4)).

As a complementary measure of genome methylation, the workflow also determines the overall fraction of mapped read calls supporting methylation (unconverted Cs) versus read calls supporting unmethylated status (converted Cs), independent of particular genomic sites (specifically, no coverage thresholds are imposed); output is recorded in the `*.mstats` output files.

### `BWASP` workflow: design and output

The workflow is organized into distinct stages. The first stage, resulting in the mapped BS-seq reads, is depicted in **Fig. S1**. In the example, a user chose NCBI SRA Accession `SRR1519132` as `sample`, with `label Pcan-21Q` (this is a *Polistes canadensis* queen sample (5); we use the generic `sample` and `label` below to indicate file names). While the reference genome for the species of interest is being processed using the `bismark-genome_preparation` script, the specified `sample` data are simultaneously being downloaded in `fastq` format using the SRA Toolkit `fasterq-dump` utility, producing files for the left and right reads in files `sample_1.fastq` and `sample_2.fastq`. The `fastq` reads are then subjected to quality assessment and trimming, as described above. The resulting validated read files are labeled `sample_1_val_1.fq` and `sample_2_val_2.fq` and enter the mapping process.

Read mapping with the `bismark` script (`bowtie2` option) results in the output file `sample_1_val_1.fq_bismark_bt2_pe.sam`. The alignment option `-score_min L,0,-0.6` was used throughout, roughly allowing up to 10 mismatches per alignment for a read of length 100. Bismark uses only uniquely mapped alignments in subsequent analysis steps to reduce misalignment errors. The alignment file is piped into `deduplicate_bismark` to remove likely PCR artifacts that over-sequenced particular reads. This step produces a non-redundant `SAM` file, which gets renamed to `label.sam` and serves as input to the next step of the workflow. Other scripts derive genome and (before and after trimming) read statistics reported in files `genome.stats` (`genome` refers to the label used for the genome file, here `Pcan.gdna`) and `sample.stats`, respectively (**Fig. S2**).

As the first step in the remaining part of the workflow (**Fig. S3**) our customized `bismark_methylation_extractor` script (described below) is invoked with just the `-mbias_only` parameter to determine the number of bases to be removed from the read ends due to technical methylation bias. The resulting file is `label.M-bias.txt` which shows the methylation proportion in all positions in the read and is parsed by a Python script `eval_prmbias.py` to flag the positions that needs to be culled from the reads. Precisely, first the mean and the standard deviation of the methylation percentage are calculated from the central half of the reads, and then consecutive end bases that show deviation of methylation percentage of more than three standard deviations from the mean are marked for removal. In detail, the script returns the four values `R1-fp, R2-fp, R1-tp`, and `R2-tp`, which indicate the number of bases to be ignored at the 5′- (fp) and 3′-ends (tp) of left (`R1`) and right (`R2`) reads, respectively (our customized script returns `R1-tp` and `R2-tp` as the desired maximal read lengths, rather than number of bases to be trimmed, so that only the biased positions of full-length reads are eliminated). These parameters are provided to the `bismark_methyl_extractor` script, which is then run again to produce the coverage file for the sample, `label.cov`, from which the genome-wide cytosine report (`label.Creport`) is generated via the `coverage2cytosine` script.

The `label.Creport` file lists all cytosine positions in the genome, their sequence context, and read counts supporting methylation or not in that position. If there are more than one data sets for a particular sample (for example, if data from different lanes on a sequencer were kept as separate `.fastq` files or stored at `NCBI SRA` under distinct accession numbers), the individual `Creports` are merged and then used for downstream processing.

The `label.Creport` has the methylation data for all the three contexts (CpG, CHG, and CHH). In order to analyze the contexts separately, the script `Creport2CXreport.py` is used to separate the `label.Creport` file into `label.CpGreport, label.CHGreport`, and `label.CHHreport`. The script also creates `label.HSMthresholds`, reporting the minimum coverages required to detect significant methylation based on the respective sample sizes. Script `CXreport2hsm.py` is used to derive the `scd = hsm + nsm` calls in files `label.CXscd.mcalls, label.CXnsm.mcalls`, and `label.CXhsm.mcalls`, respectively (where `X` is one of the three distinct contexts). These resulting files report the site positions, read coverage, and percent methylation and serve as input into the `mstats.sh` script to produce methylation summary statistics in file `label.mstats`.

Depending on the scientific questions being pursued, pooling of replicate or sample data may be desired prior to statistical analysis. For example, sequencing depth may be insufficient to rely on per-replicate statistics. But it may be appropriate to view the replicates as multiple sources of genetic material for each sample (in much the same way as DNA from an individual typically averages over many cells and possibly tissues). `BWASP` includes the `Makefile_merge_template` recipe to merge replicate `*.Creport` files and generate aggregate sample statistics. For example, for the *Polistes canadensis* study (5), there are three replicates of queen and worker castes, respectively. The `BWASPR` analysis scripts described below will analyze and compare both aggregate samples and individual replicates.

```
mkdir BWASPonPatalano2015
cd BWASPonPatalano2015
singularity pull http://BrendelGroup.org/SingularityHub/bwasp.sif
git clone https://github.com/BrendelGroup/BWASP

cd BWASP/data
xsetup -m bggnomic -s Pcan Patalano2015
mv Pcan ../../
mv xdoitPatalano2015 xgetdPatalano2015 ../..
cd ../..

time singularity exec -e -B ${PWD} bwasp.sif  ./xgetdPatalano2015 >& errd
#real   14m6.323s
#user   23m59.166s
#sys    5m11.965s
# ... 59G downloaded

time singularity exec -e -B ${PWD} bwasp.sif  ./xdoitPatalano2015 >& err

#real   738m32.074s
#user   5826m55.671s
#sys    745m9.883s
# ... 266G output saved on disk
```

Simple `bash` and `Perl` scripts were written to scrape data from the `BWASP` output files to provide summary tables of statistics from multiple experiments (available from the corresponding author upon request). Quality control and mapping statistics (**Datasets S3** and **S4**) record number and lengths of raw and trimmed reads; mapping efficiency; number and percentage of removed PCR duplicates; number and percentage of filtered reads; genome coverage; and length of discarded read ends removed due to bias. Coverage statistics (**Datasets S5** and **S6**) record number and percentage of CpG, CHG, and CHH sites covered by zero and above selected threshold numbers of reads. Numbers of `scd` and `hsm` sites analyzed are shown in (**Datasets S7** and **S8**). Overall methylation levels (**Datasets S9** and **S10**) are given as percent reads showing methylation per site (C, CpG, CHG, or CHH).

The script displayed above illustrates the simplicity of running a typical `BWASP` workflow. Any novice user could execute these few lines of script essentially as displayed. The only adjustment would be to edit/replace the references "bggnomic.conf" machine configuration file, appropriate to the user's facilities. The precise performance statistics depend on download speeds and numbers of threads available, but, roughly, the promise of the workflow is minimal set-up effort, overnight execution, and reliable generation of hundreds of Gb of meaningful output for summary statistics and further processing.

### `BWASPR` – `R` scripts for statistical analysis

The next phase of a BS-seq study evaluation involves statistical analysis of genome-wide methylation patterns. Starting points are the `BWASP`-derived `*.mcalls` files. These files are simple tab-delimited tables with the following columns:

```
SeqID.Pos  SequenceID   Position  Strand Coverage Prcnt_Meth  Prcnt_Unmeth
```

These data are read into `Bioconductor` (6) `methylKit` (7) package `methylRaw(List)` objects: `studymk`, storing aggregate sample `hsm` data; `studymc`, storing `scd` data; `studyhc`, storing `scd` data restricted to sites with coverage greater or equal to the parameter `highcoverage`; and `mkrd`, storing `hsm` replicate data, when applicable. As a record of all the data work in this study and as a template for analyses on new data sets in the future, all our statistical analyses were run within `R` (8) using scripts and functions collected in the Brendel Group github repository `BWASPR` (https://github.com/BrendelGroup/BWASPR). The `BWASPR` functions invoke `methylKit` package functions (7) and provide additional capabilities tailored to the data analysis needs in this study. The following paragraphs give details.

All `R` analyses were scripted and run through the `Rscript` front-end for `R` as documented in the `BWASPR` demo directory. The package provides a generic `Rscript.BWASPR` file that the user can adjust and individualize through settings in a configuration file that is supplied to `Rscript.BWASPR` as a command-line argument. Different sections of the script derive coverage and methylation statistics for specified aggregate or replicate samples; pairwise sample correlations and comparisons; annotation of conserved sites and overlay of methylation sites with genome annotation; and differentially methylated sites and genes. `Rscript.BWASPR` automatically generates a host of summary statistics, tables, and figures with minimal configuration effort and thus provides a fast way of surveying relevant outcomes of the user's experiment. With the supplied code and documentation, the user can easily modify and expand the `R` code to facilitate additional analyses. Output of the `BWASPR` workflow is collected in several output sub-directories, as described next.

**Coverage and methylation statistics (`CMS`).** Output directory `CMS` contains files `cms-*.txt` and `cms-*.pdf` that record read depth (coverage) and methylation levels for CpG `hsm` sites for the specified aggregate samples. The `cms-*.pdf` files show histograms of coverage and methylation levels for different ranges of coverage. The first range is all sites of minimal coverage ($\geq t$), determined as described in section "**Determination of significant methylation sites**" above. The other ranges are all sites above threshold levels given in the `covlist` array in the configuration file, and lastly all sites with coverage in the range `locount` to `hicount`, also set in the configuration file.

The coverage histograms are easily interpreted and reflect the sequencing depth. The methylation level histograms must be more cautiously evaluated because of the bias to high methylation levels for low coverage sites (`hsm` sites are selected to have statistically significant methylation levels, which translates to high methylation percentages for minimally covered sites). The same bias at high coverage would suggest that the sample includes a preponderance of consistently methylated genomic sites.

**Pairwise comparisons between all samples (PWC).** Output directory `PWC` contains files `pwc-*.txt` and `pwc-*.pdf` that record common and distinct methylation sites in pairwise comparisons of any two samples. The output is generated by the `BWASPR::cmpSites()` function, which also calculates the overlap index between the samples and estimates the size of the common pool of potential methylation sites as described in the **Materials and Methods** section of the main text. Input consists of the `studymk` and `studymc` objects, as well as the parameter `nbrpms`. The `CpGscd` data input (`studymc`) is necessary to determine sites that are detectable in both samples, and `nbrpms` (scraped from the `*.par` file specified in the `*.conf` configuration file as argument to `TOTALNBRPMSITES`) provides the total number of potential methylation sites (typically, all `CpG` sites), a necessary parameter for the estimations.

The `*.pdf` output files display side-by-side histograms of methylation levels comparing any two samples as well as scatter plots of respective methylation levels at common sites. If more than two samples are specified in the input, then pairwise comparisons are computed in parallel. If input consists of just two samples, then a Venn diagram is displayed to visualize the sizes of the common and distinct `hsm` sets.

**Correlations between aggregate samples (CRL).** Output directory `CRL` contains files `crl-*.txt` and `crl-*.pdf`, generated by `BWASPR::cmpSamples()`, which uses `methylKit::unite()` to determine the `hsm` sites common to all samples and then applies `methylKit::getCorrelation()` and `methylKit::PCASamples()` to the intersection to determine the correlation between sample methylation levels. `methylKit::PCASamples()` performs standard principal component analysis on the percent methylation matrix.

The `*.txt` files show the correlations in a table, and the `*.pdf` files show graphics. The user can set `destrand` to `TRUE` in the configuration file to merge methylation counts on both strands of a `CpG` site (*i.e.*, counts for a `CpG` in positions $i, i+1$ in a sequence are added up and and displayed for position $i$). Beware that the number of common sites (shown at the bottom of the `*.txt` files) may be small, which may make the correlations less informative.

**Coverage and methylation statistics for replicate samples (REPCMS).** Output directory `REPCMS` contains files `repcms-*.txt` and `repcms-*.pdf` that record read depth (coverage) and methylation levels for CpG `hsm` sites for each replicate of the specified aggregate samples. Parameters `replocount` and `rephicount` are the equivalents of parameters `locount` and `hicount` for aggregate samples; otherwise the descriptions given for output directory `CMS` apply with replicate instead of aggregate sample data sets.

**Correlations between replicates (REPCRL).** Output directory `REPCRL` contains files `repcrl-*.txt` and `repcrl-*.pdf`, analogous to output in directory `CRL` but applied to replicate instead of aggregate sample data sets.

**Mapping of methylation sites on genome annotation (MMP).** Output directory `MMP` contains files `mmp-*.txt` that summarize genome annotation in terms of genic versus intergenic regions, exons, introns, untranslated transcript regions (`UTR`s) and coding sequences (`CDS`) and the mapping of methylation sites onto annotated genome features. Initially, the `BWASPR::get_genome_annotation()` function reads data from the `BWASP genome/GFF3DIR` directory, which should have been generated with the `BWASP Makefile_parse_GFF3` recipe from the input genome `FASTA` sequence and `GFF3` annotation files, as deposited in the `genome` directory or, for large-scale data processing, as specified in the species configuration file https://github.com/BrendelGroup/BWASP/blob/master/data/README.md. The summary output is generated by the `BWASPR::map_methylome()` function and gives an accounting for every sample as to where the `CpGhsm` and `CpGscd` (control) sites reside relative to the genome annotation. Also shown are `O/E` ratios, representing, observed over expected values, where the expected percentages are based on the respective feature proportions in the genome, as annotated.

Overlapping feature regions can complicate assignment of sites to features. `BWASPR::map_methylome()` uses the following conventions and approximations:

- the intergenic region size is calculated as genome size minus aggregate width of annotated genes
- promoter regions are defined are 500 nucleotides upstream of 5′-ends of annotated genes (adjustable in `BWASP::gene2promoter.py`)
- sites in regions overlapping genes and promoters are counted as genic
- sites in intergenic and promoters are counted as in promoters
- sites in 5′-`UTR`s are counted as within-`CDS` in case of overlap with the `CDS` region of another gene
- sites in 3′-`UTR`s are only counted for this region if they overlap `CDS` or 5′-`UTR` regions of other genes

For whole genome statistics, these conventions should give close approximations to other ways of accounting, unless there is a great amount of gene overlap. Site to annotation mapping can easily be analyzed in greater detail for specific regions of interest (see next section). Simple `bash` and `Perl` scripts were written to scrape data from the `mmp-*.txt` output files to provide summary tables of sites in genes (**Datasets S11** and **S12**) and exons (**Datasets S13** and **S14**).

**Annotation of conserved methylation sites (ACS).** Output directory `ACS` contains the file `acs-*.txt`, generated by the `BWASPR::annotate_methylome()` function, which uses `methylKit::unite()` to determine the sites common to all specified experiments and then applies `genomation::annotateWithFeature()`. The output is a table showing the common sites with methylation levels in all samples and columns with booleans indicating whether the site fits a specific annotation feature. The template `Rscript.BWASPR` applies the function to `studymk`, but of course the input can easily be changed to get the annotation for other data sets.

**Genomic feature regions ranked by CpGhsm statistics (RNK).** Output directory `RNK` contains the files `ranked-genes-*.txt`, `sites-in-genes-*.txt`, `ranked-promoters-*.txt`, `sites-in-promoters-*.txt`, `plot-genes-prcntM-vs-Sitedensity-*.pdf`, and `plot-promoters-prcntM-vs-Sitedensity-*.pdf`, generated by the `BWASPR::rank_rbm()` function. The files `ranked-genes-*.txt` provide ranked lists of genes based on overall methylation percentage (`ranked-genes-byPrcntM-*.txt` files) or the occurrence of `CpGhms` sites within the annotated gene bounds, measured as number of sites per 10kb (`ranked-genes-bySiteDensity-*.txt` files). The output columns include: `region_ID`; `rwidth` (= region width); `nbrsites` (= number of `CpGhsm` sites in the region); `nbrper10kb` (= number of `CpGhsm` sites in the region normalized to 10kb width = `SiteDensity`); `pmrpersite` (= average % methylation per `CpGhsm` site in the region); `pmrpernucl` (= average % `CpGhsm` methylation per nucleotide in the region); `prcntM` (= overall % `CpGhsm` methylation in the region). The `ranked-promoters-*.txt` files provide analogous tables for promoter regions. The files `sites-*.txt` provide details for the `CpGhsm` sites in the respective regions.

The files `plot-*.pdf` explore concordance of the two methylation measures. The optional parameters `minrwidth`, `maxrwidth`, and `minnbrsites` to the `BWASPR::rank_rbm()` function specify the length range and minimal number of methylation sites for regions to be included in the plot.

**Methylation-rich and -poor regions (MRPR).** Output directory `MRPR` contains the text output files `dst-*.txt` and `mdr-*.bed` and the plots `1ds-*.pdf` and `5ds-*.pdf` generated by the `BWASPR` function `det_mrpr()`, as well as files `rmp-*.txt`, `gwp-*.txt`, and `gwr-*.txt` generated by the `BWASPR` function `map_mrpr()`.

Methylation-rich and -poor regions are determined based on the spacing between neighboring `CpGhsm` sites. If sites occur at positions $a, b, c, d, e$, and $f$ (and not in between), then $b-a, c-b, \cdots, f-e$ are 1-distances, and $f-a$ is a 5-distance. The file `dst-*.txt` shows the empirical distribution of $d$-distances for the sample. Methylation-poor regions are determined as long 1-distances (in the top *nbrxtrms*; merged if adjacent 1-distances are both in the top *nbrxtrms*), and methylation-rich regions are determined as short 5-distances (in the low *nbrxtrms*; merged if adjacent 5-distances are both in the low *nbrxtrms*); see **Fig. 1** for illustration. *nbrxtrms* is a parameter to the functions, set to 100 by default in `Rscript.BWASPR`.

Algorithmically, the $d$-distances are first ordered by size (decreasing to determine methylation-poor regions, and increasing to determine methylation-rich regions). Then the distance value of the *nbrxtrms*-th entry is used as a threshold, and the regions above the threshold (for methylation-poor regions) or below the threshold (for methylation-rich) regions are selected for display. The merging takes into account any overlap of selected regions. For example, if the next site at position $g$ in the above example made the distance $g-b$ also a top *nbrxtrms* 5-distance, then the region from $a$ to $g$ would be reported as a single methylation-rich region (assuming there is no further merging by these criteria). In consequence, the actual number of regions displayed may be less than *nbrxtrms* (because of merging) or larger than *nbrxtrms* (because of ties).

Parts of the distribution are plotted in the `1ds-*.pdf` and `5ds-*.pdf` files. The methylation-rich and -poor regions are listed in files `dst-*.txt` and `mdr-*.bed`, the latter in `BED` format for potential display in genome browsers.

The overlap of methylation regions with genome features is summarized in files `rmp-*.txt`. As in the `MMP` directory, for the summary statistics gene annotations take precedence over other feature annotations in case of overlaps. Files `gwr-*.txt` show genes overlapping with methylation-rich regions, ordered by site density in the methylation-rich region. Files `gwp-*.txt` show genes overlapping with methylation-poor regions, ordered by site density in the methylation-poor region.

**Differentially methylated tiles (DMT).** This and the following two sections assume that the user supplies at least two data samples for comparison. If the input consists of more than two samples, then all pairwise comparisons are performed. Output directory `DMT` contains the `BWASPR::det_dmt()`-generated output files `dmt-*.txt` and `dmg-*.txt` that show differentially methylated tiles and genes overlapping such tiles, respectively.

Tiles refer to sliding windows (of size `wsize`) along the genome (offset by `stepsize`) within which methylation calls are cumulated by `methylKit::tileMethylCounts()`. Differentially methylated tiles are determined by the `methylKit::getMethylDiff()` function with parameters `difference=threshold` and `qvalue`, which are read from the configuration file by `Rscript.BWASPR` (as are `wsize` and `stepsize`).

A positive `meth.diff` value in the comparison labeled `A.vs.B` means that the `B` methylation percentage in that tile is higher than the `A` methylation percentage.

**Differentially methylated sites and genes (DMSG).** Output directory `DMSG` further explores differential methylation, in this case based on sites and annotated genes. Only “high coverage" sites are evaluated (otherwise, %-methylation differences become statistically meaningless: for example, is $4/4 = 100\%$ indicative of higher methylation than $4/5 = 80\%$ or $6/8 = 75\%$?). The threshold for high-coverage is set in the configuration script as `highcoverage` (suggested value: 20).

Output tables `dms-*.txt` and `dmg-*.txt` are generated by `BWASPR::det_dmsg()` and show the differentially methylated sites and genes as determined by `methylKit::getMethylDiff()` with parameters `difference=threshold` and `qvalue`, read from the configuration file as in the previous section. The table of differentially methylated genes contains all genes with at least one differentially methylated site.

The files `dmg-*details.txt`, generated by `BWASPR::show_dmsg()`, show all `CpGscd` sites in the differentially methylated genes, with coverage numbers and methylation percentages.

The files `dmg-*heatmaps.pdf`, also generated by `BWASPR::show_dmsg()`, give heatmap displays for genes meeting the following criteria: 1) there are between `minNsites` and `maxNsites` common `CpGscd` sites; 2) at least `minPdmsites` % of these sites are differentially methylated sites.

**Ordered gene lists (OGL [and OGLall]).** Output directory `OGL` shows ordered gene lists based on methylation and differential methylation levels. The output derives from `BWASPR::explore_dmsg()` and `BWASPR::rank_dmg()`. `DMSG` output is required, as well as the genome annotation object; `maxgwidth` and `minnbrdmsites` are optional parameters.

Output files `ogl-<study>_<sample>.txt` give tables for each sample with columns

```
gene\_ID gwidth \#Sites \#per10Kb \%perSite \%pNucl
```

ordered by `%pNucl`. If available, a link to an NCBI entry of `gene_ID` is inserted as second column. Abbreviations used: `%perSite`, percent methylation per site; `%perNucl`, percent methylation per nucleotide of the gene.

Output files `ogl-<study>_<sample1>.vs.<sample2>.txt` give tables for each comparison with columns

```
gene\_ID gwidth \#Sites \#per10Kb \#dmSites \#dmsp10kb \%dmSites \%pSite1 \%pSite2
```

and (continuing)

```
DMpSite ADMpSite DMpNucl ADMpNucl
```

ordered by DMpSite. As before, links to NCBI records for the gene_ID entries are inserted, if available. Abbreviations used: %dmSites, percent sites that are differentially methylated; %pSite1, average per site % methylation for sample1; %pSite2, average per site % methylation for sample2; DMpSite, average per site difference in % methylation between sample1 and sample2; ADMpSite, absolute value of DMpSite; DMpNucl, average per nucleotide difference in % methylation between sample1 and sample2; ADMpNucl, absolute value of DMpNucl. Output files `rnk-dmg-<study>_<sample1>.vs.<sample2>.txt` are equivalent to files `ogl-<study>_<sample1>.vs.<sample2>.txt` but ordered by ADMpNucl.

Output files `rnk-dmg-<study>_<sample1>.vs.<sample2>.pdf` provide visualization of the distribution of ADMpNucl values. Output file `wrt-<study>.txt` gives results of the Wilcoxon signed rank test comparing the %pSite1 and %pSite2 vectors.

In output directory OGL, tables are restricted to genes with gwidth $\leq$ maxgwidth and, for pairwise comparisons, #dmSites $\geq$ minnbrdmsites. If either maxgwidth or minnbrdmsites is set, then output directory OGLall will additionally show the full tables with all genes, for reference.

### Results

***Re-analysis by read mapping to a novel genome assembly and annotation.*** The Libbrecht *et al.* (9) study provides an example as a new genome assembly has since been released, with updated species name *Ooceraea biroi* (? ). The BWASP/R workflow makes it easy to run all analyses reported in the main text from scratch with respect to the current assembly, as this requires only a small change in the configuration files (as described in the main text section 'Description and validation of workflow'). The output of BWASP on *O. biroi* is shown side-by-side with the *C. biroi* data in Datasets S3, S5, S7, S9, S11, and S13, and the BWASPR output is shown in Datasets S18 and S20, respectively. We shall refer to our workflow output derived on the original *C. biroi* genome assembly with the label Cbir and to our workflow output derived on the current *O. biroi* genome assembly with the label Obir.

Comparing the assembly statistics between Cbir and Obir, it is clear that the current long-read assembly has filled many previous gaps in the genome: the overall genome size increased from 212.8 Mb to 223.9 Mb, the number of scaffolds decreased from 4579 to 139, and the N50 and N90 measures increased from 1.35 Mb to 16.9 Mb and from 104 Kb to 13.6 Mb, respectively (BWASP *.gdna.stats output). Based on these statistics, one might expect significant changes for the BS-seq read mapping as well. However, this expectation is not born out: average read mapping efficiency dropped from 80.6% for Cbir to 79.3% for Obir, and average genome coverage dropped from 36.6X to 34.7X (Dataset S3). These data are likely a consequence of the gap-filling of assemblies largely targeting repetitive rather than unique genome regions, which will result in some reads that previously mapped uniquely now being recognized as matching multiple genome regions (and therefore being discarded in the BS-seq analysis).

BS-seq coverage statistics are similar. For example, the average percentage of CpG sites covered by at least five reads was calculated as 72.4% for Cbir compared to 68.8% for Obir, and coverage with at least 20 reads were 27.6% versus 25.9% (Dataset S5). The Cbir replicates yielded on average 218763 hsmCpG sites, compared to 218021 for Obir (Dataset S7). The average overall CpG methylation level was calculated as 1.82% for both assemblies (Dataset S9).

Datasets S11 and S13 summarize differences of methylation site mapping to genome features. The Obir annotation includes 166.8 Mb regions in genes, compared to 152.7 Mb in Cbir. Correlated measures are the fraction of the genome that is annotated as exon (20.1% in Obir, compared to 18.7% in Cbir) and hsmCpG site density in exons (44.3 per 10kb for Cbir, compared to 39.2 for Obir; 86.7 versus 81.0 in coding regions). By the method explained in Fig. 2 applied to all pairwise comparisons between samples, the Cbir CpG methylome is estimated between 354661 to 401372 sites for Cbir (Dataset S18, pages 105-133), compared to the range 373342 to 422785 sites for Obir (Dataset S20, pages 105-133).

While the above statistics point to subtle global differences in methylation statistics in wake of improved genome assemblies and annotation, more critical may be differences for specific gene models that have to be re-assessed based on current knowledge. Without workflow-enabled re-analysis such re-assessments would be very difficult, as the factors of a changed genome assembly/annotation and possible changes in methods from original to current analysis are entangled. To illustrate these points, we compare detection of potential differentially methylated genes in Cbir versus Obir. The BWASPR workflow casts a wide net by default, listing all genes that harbor at least one differentially methylated hsmCpG site determined per specified workflow parameters (see *SI Text*, 'Differentially methylated sites and genes (DMSG).'). Results are summarized in Fig. S1. Comparing the reproductive and brood care phases, BWASPR tagged 1533 genes as differentially methylated for Cbir (Dataset S17), compared to 1618 genes for Obir (Dataset S21), with an overlap of 1281 gene models sharing the same gene model identifier. Thus, only about 80% of the tagged gene models are conserved between the analyses of the original and current genome assembly/annotation pairs. Because the default BWASPR settings tag any gene with at least one differentially methylated site as of interest, most of the differences are explained by just one site passing or failing the significance threshold in one or the other set. For example, of the 285 differentially methylated genes specific to Obir, 251 have just one differentially methylated site; and of the 197 differentially methylated genes specific to Cbir, 185 have just one site. Differences involving multiple sites are due to refined annotation of gene models, involving resolution of repetitive sequences and novel assignment of exons. Scripts in https://github.com/BrendelGroup/BWASP/data/CbirObir-Explore provide the summary statistics and facilitate review of particular gene models. Our point here is to reiterate that only the workflow-enabled approach allows for facile re-evaluation of prior sequence data in the context of a novel genome assembly and annotation.

**Case study of a large genome.** A recent publication explored DNA methylation patterns in the social spider, *Stegodyphus dumicola* (11). The underlying genome assembly is just over $2.5 \times 10^9$ nucleotides, which is an order of magnitude larger than

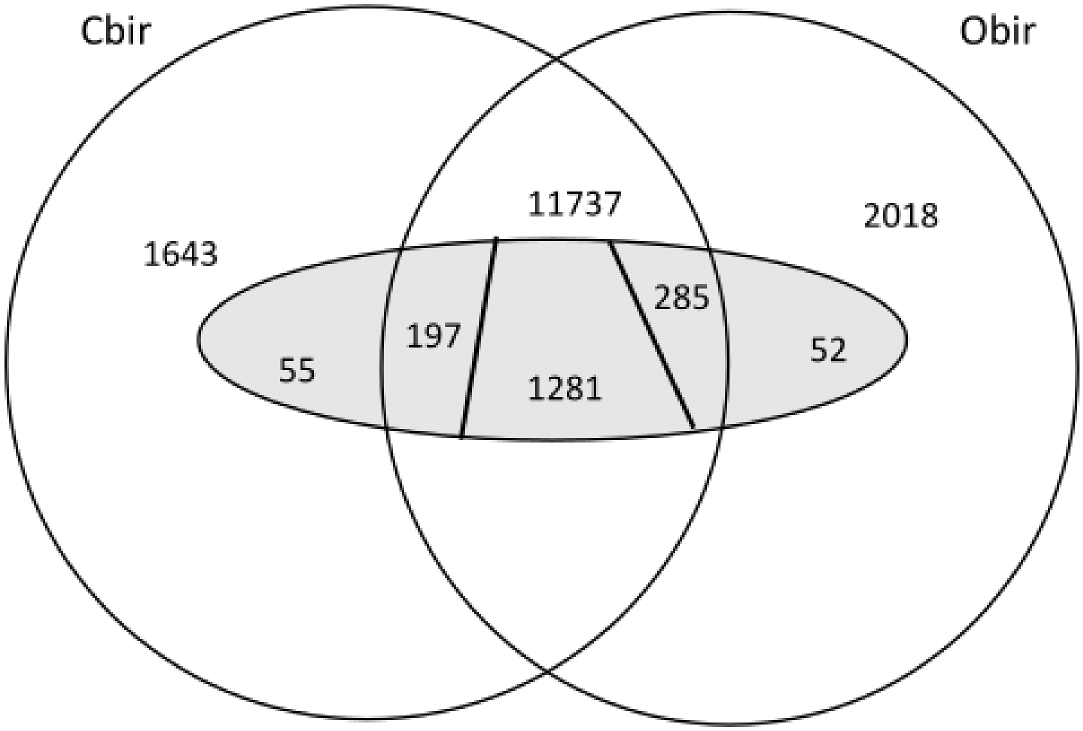


**Fig. S1.** Overlap of gene models and genes identified as differentially methylated comparing two genome assembly/annotation versions of *Cerachys biroi* (`Cbir`)/*Oochera biroi* (`Obir`). From the original `Cbir` assembly 13350 gene models, 11737 transferred to the current `Obir` assembly (although possibly modified). Thus there are 1643 and 2018 gene models unique to `Cbir` and `Obir`, respectively. The inner oval represents genes with differentially methylated CpG sites. Of the 1533 and 1618 genes in `Cbir` and `Obir`, respectively, 1281 are shared. The 252 `Cbir` unique genes comprise 55 gene models unique to `Cbir` and 197 gene models also present in `Obir` but not harboring differentially methylated sites by the `Obir` analysis. The 337 `Obir` unique genes comprise 52 gene models unique to `Obir` and 285 gene models also present in `Cbir` but not harboring differentially methylated sites by the `Cbir` analysis.

the insect genomes we analyzed (`SI Text`, Table S1). The authors evaluated BS-seq data from one individual each from two nests sampled in climatically different locations in Southern Africa (Betta and Karasburg, respectively), for a total of 200 Gb of sequencing data. The purpose of the study was to probe the extent of DNA methylation in chelicerate species, as well as its potential role in the regulation of gene expression.

Our first aim here was to prove that the `BWASP/R` workflows can be applied to such large data sets. While we did not have the computational resources to do the methylation calling on each of the downloaded samples in one workflow submission, the replicate merging functionality of `BWASP` could be easily applied to get the job done. First, we split each downloaded BS-seq read data file into eight chunks and distributed these smaller files as “pseudo-replicates” into distinct replicate directories. Then, each replicate was run separately, and the usual sample-level merging of `*.mcalls` data provided the desired output. In summary, the `BWASP` workflow is trivially parallelizable, which thus also allows to run independent parts in serial, adjusted to available computational resources. The subsequent `BWASPR` workflow took about nine times as long as the analysis of the *Polistes canadensis* data and proportionally more memory for data ingest, but otherwise also worked flawlessly.

A fine point of the pseudo-replicate approach is that it may miss PCR duplicates that are split between different replicate alignment sets. Here, about 15% of alignments per replicate were removed as likely PCR duplicates (*SI Text*, Table S1). Combining the replicated `*.bam` alignment files and running `deduplicate_bismark` on the aggregate identified an additional 3.2% and 4.8% of alignments as duplicates for the Betta and Karasburg samples, respectively. The most conservative approach to data scrubbing might be to also remove these reads before the next workflow steps. However, as a control, we also checked for between-replicate duplicates for the *P. canadensis* study and found a similar fraction of 4.3% of alignments flagged by `deduplicate_bismark`. Thus, it may be difficult to correctly separate all PCR duplicates from independently generated equivalent data points. Given large enough samples, statistical analysis of the results should be robust to precise choices.

Our second aim with this case study was to underscore the value of workflow-based analysis for any in-flux data. At the time of publication, Liu *et al.* (11) did not provide a genome annotation linked to public databases to go along with their study. As part of our efforts, we petitioned `NCBI` colleagues to run their standard genome annotation pipeline for *Stegodyphus dumicola*, and they graciously agreed. Without a reproducible workflow, re-analysis of the published data relative to the current reference genome annotation would be at minimum cumbersome and hugely time-consuming. Within the `BWASP/R` framework, specification of the relevant `GFF3` annotation file is all that is needed.

Although details of initial read processing and mapping differ between our re-analysis and the published study, overall methylation rates were very close: our approach reports 14.6% `hsm` CpG sites (Dataset S7) compared to the "about 15%" stated by Liu *et al.* (11), and in both workflows CHG and CHH methylation were each found to be less than 0.02%. Moreover, the stated "about a third of the CpGs in genes were methylated, while only 5% of the intergenic CpGs were methylated" and "exons and introns were methylated to more or less the same extent" (11) are reproduced with the `BWASPR` statistics shown in Dataset S11, reporting about 29% CpG methylation in exons and introns, and 5.2% CpG methylation in intergenic regions.

The `BWASPR` output directories `RNK`, `DMSG`, and `OGL` provide ranked gene lists based on overall and differential methylation levels (Dataset S22). Comparison with the published results by Liu *et al.* (11) would require detailed comparison of individual gene structure models and is beyond the scope of our consideration here. What we should like to emphasize is that our workflow readily enables incorporation of the now available `NCBI` genome annotation, which then provides a documented, publicly available reference set for transparent communication of annotation-based analyses.

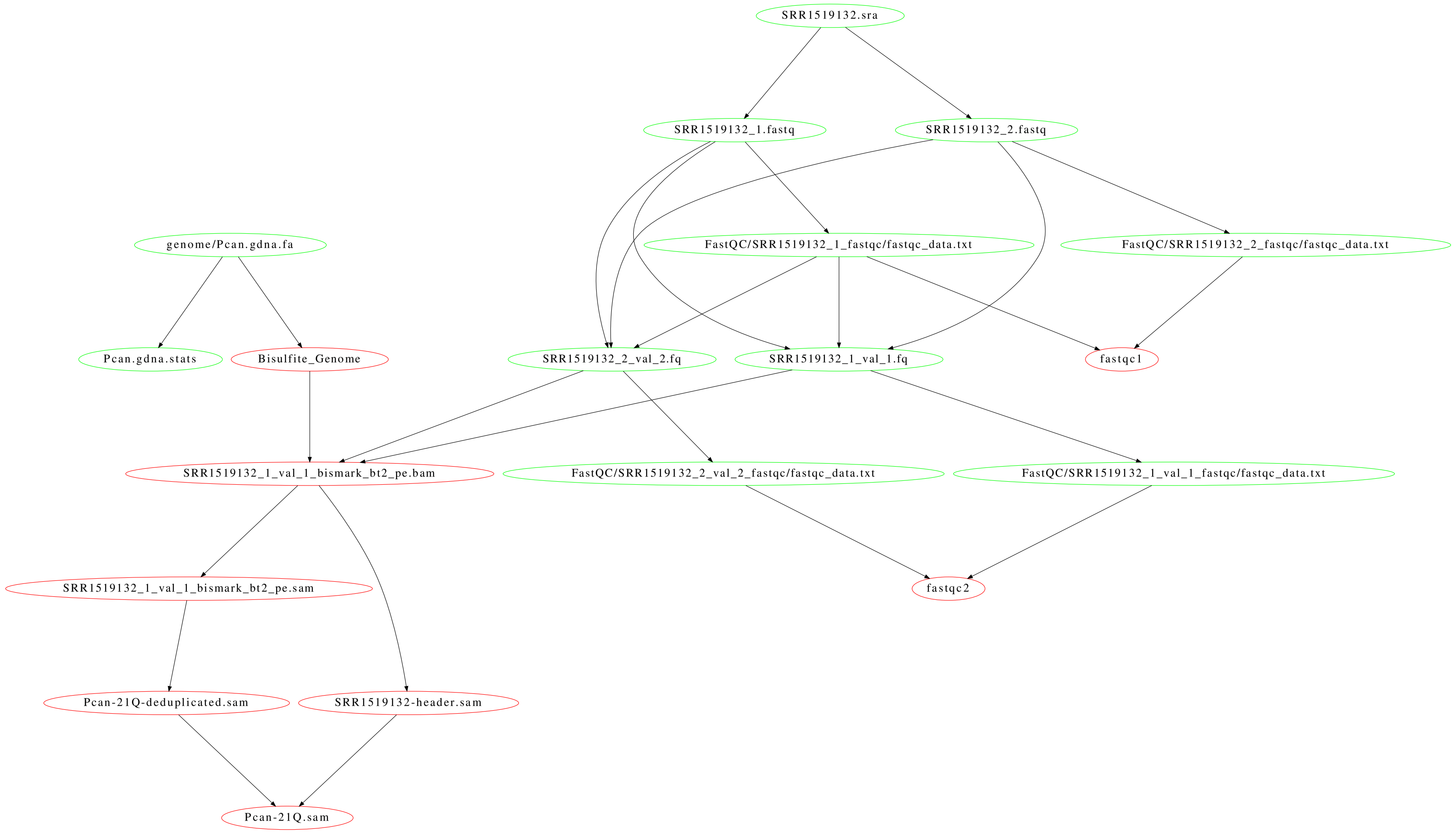


**Fig. S2.** `BWASP` workflow to generate the BS-seq read alignments.

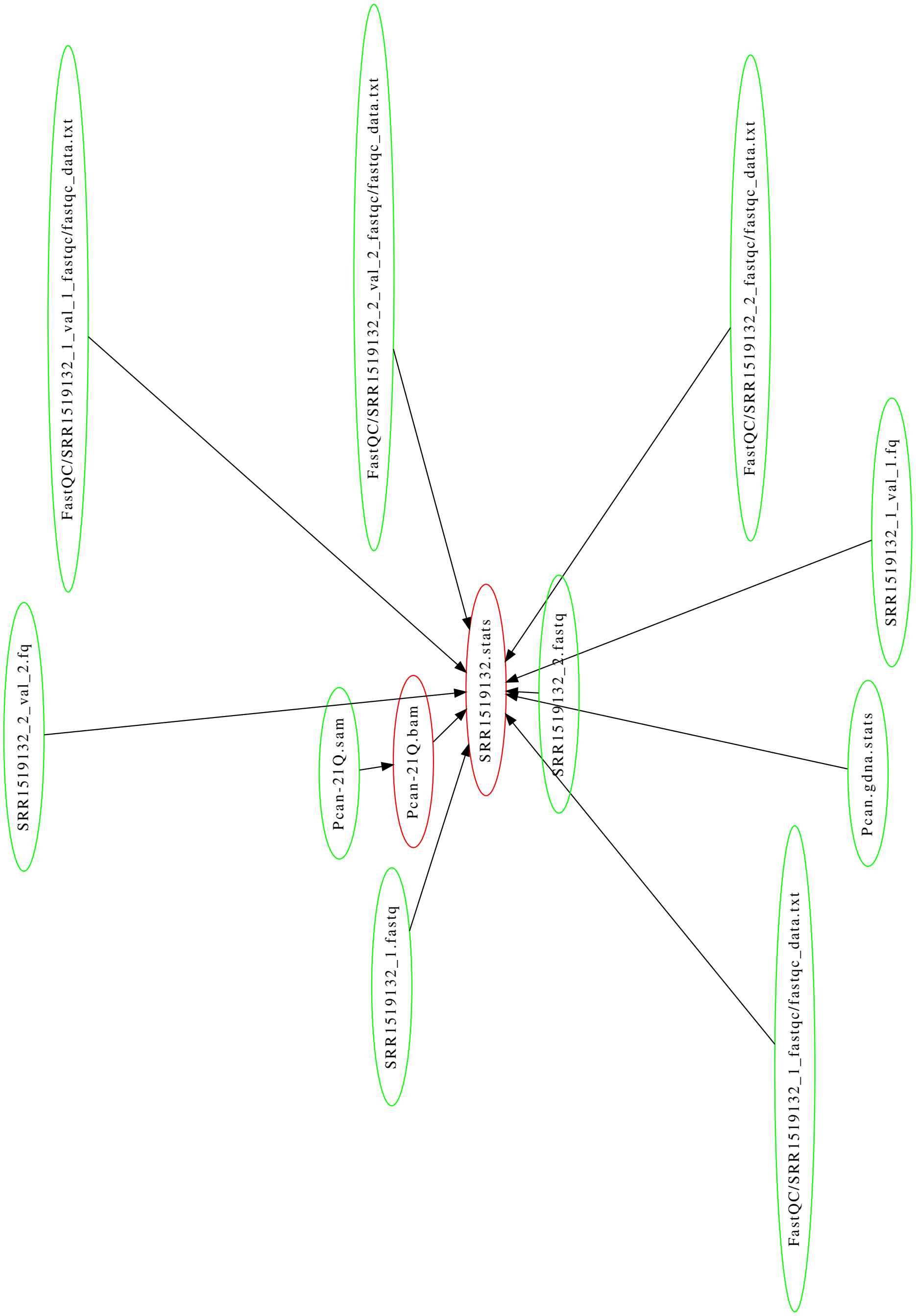


**Fig. S3.** BWASP workflow to derive sample statistics.

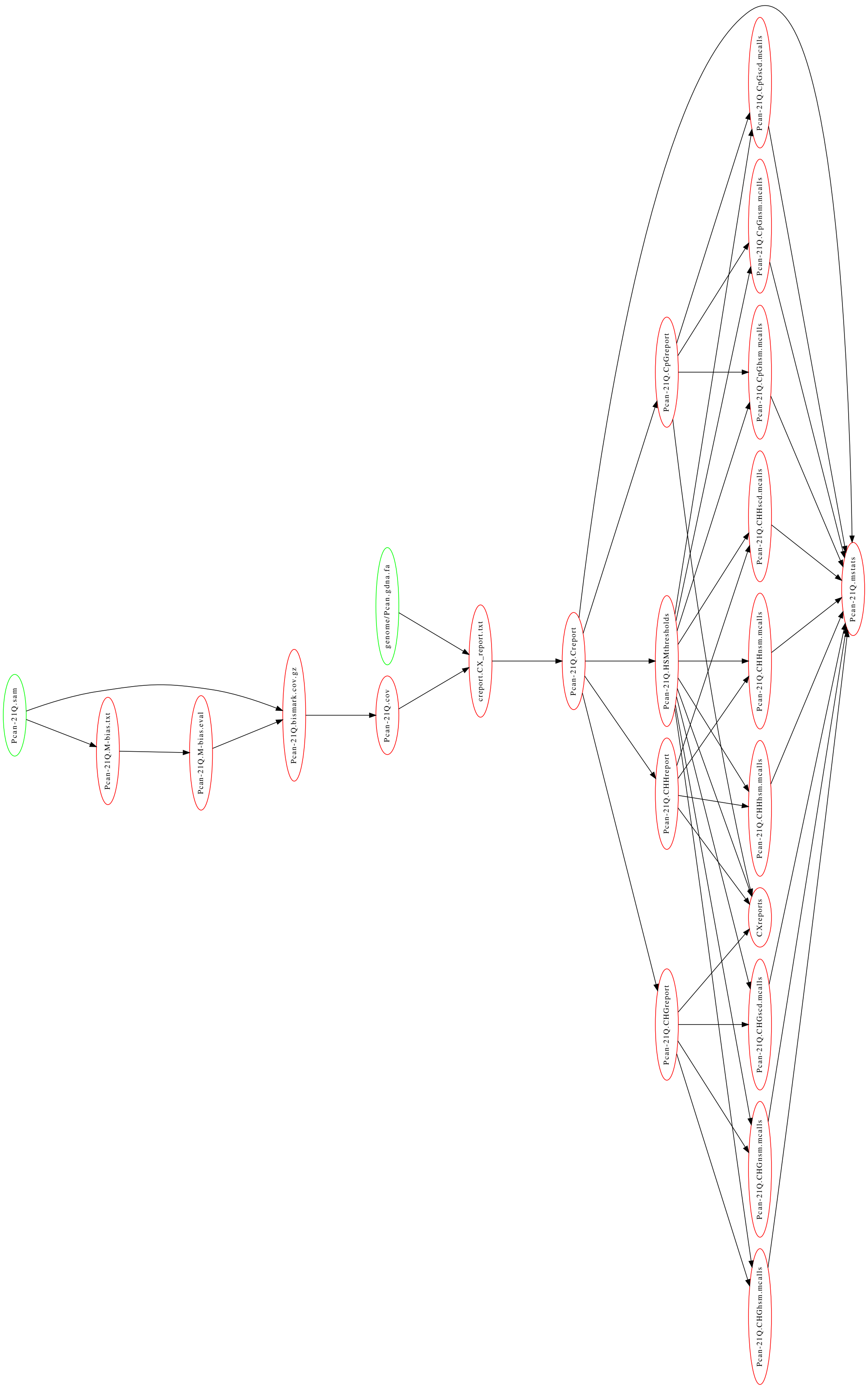


**Fig. S4.** `BWASP` workflow to derive methylation calls and statistics.

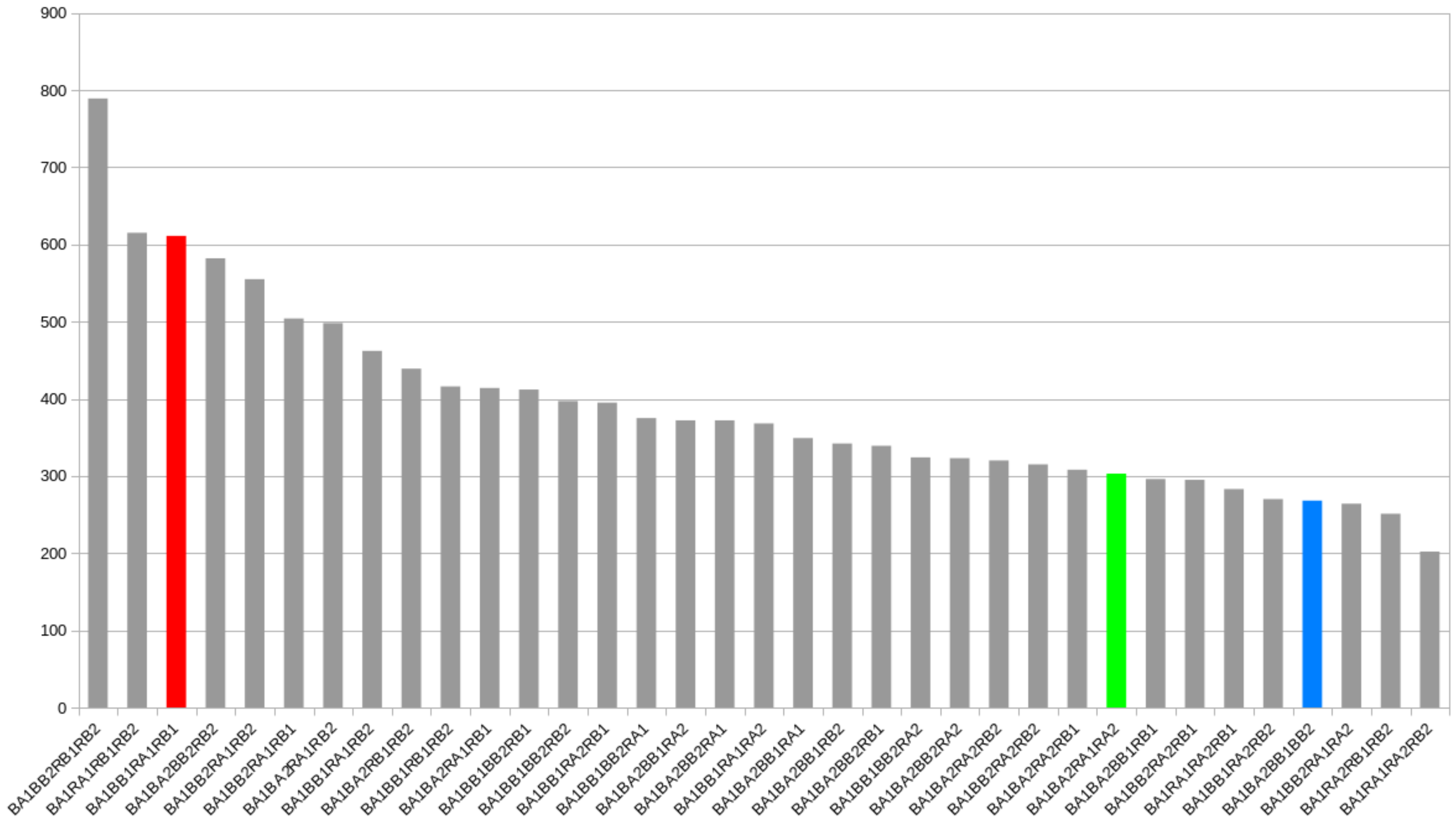


**Fig. S5.** Comparison of the extent of `hsm` site conservation between balanced sets of samples from the Libbrecht *et al.* (9) *Cerapachys biroi* study. The displayed counts are from Dataset S19 (rows "4-4" and column "cum. # sites") and refer to the numbers of CpG sites that are consistently `hsm` in one set versus consistently `nsm` in the complementary set. The plot is analogous to Figure S3 of Libbrecht *et al.* (9); the blue bar compares brood care versus reproductive phase; the green bar compares line A and B; the blue bar compares sequencing batches 1 and 2; the gray bar are the remaining set partitionings. Abbreviations used: B (in first position), brood care phase; R, reproductive phase; A, line A; B (in second position), line B; 1, sequencing batch 1; 2, sequencing batch 2.

**Table S1. Average statistics for datasets analyzed.**

| | ***Apis mellifera*** | ***Polistes canadensis*** | ***Cerapachys biroi*** | ***Ooceraea biroi*** | ***Stegodyphus dumicola*** |
|---|---|---|---|---|---|
| Genome size (Mp) | 225.25 | 211.20 | 212.83 | 223.88 | 2551.87 |
| Number of studies | 17 | 1 | 1 | 1 | 1 |
| Number of samples | 58 | 2 | 4 | 8 | 2 |
| Number of replicates | 148 | 6 | 8 | 8 | 2 |
| Number of reads aligned (in millions) | 12868.83 | 189.19 | 1142.48 | 1142.48 | 1331.81 |
| Mapping efficiency (%) | 68.30 | 83.50 | 80.61 | 79.31 | 67.57 |
| range (%) | 26.42 - 88.17 | 83.27 - 83.72 | 78.20 - 81.80 | 77.00 - 80.40 | 67.30 - 67.83 |
| Percent reads removed[a] | 19.29 | 7.01 | 24.29 | 23.40 | 15.38 |
| range (%) | 0.55 - 73.85 | 5.36 - 8.66 | 15.69 - 30.63 | 14.57 - 29.91 | 14.16 - 16.6 |
| Percent reads rejected[b] | 0.20 | 1.21 | 0.03 | 0.02 | 0.00 |
| range (%) | 0 - 2.87 | 1.03 - 1.38 | 0.02 - 0.04 | 0.01 - 0.04 | 0.00 - 0.00 |
| Genome coverage[c] | 53.16 | 33.75 | 36.59 | 34.67 | 21.91 |
| range | 2.01 - 143.48 | 31.74 - 35.76 | 29.95 - 49.83 | 28.35 - 47.25 | 21.28 - 22.54 |
| Percent CpG sites: | | | | | |
| covered by $\geq$ 1 reads | 93.91 | 82.66 | 91.96 | 88.67 | 76.67 |
| range | 31.71 - 99.21 | 81.03 - 84.29 | 91.04 - 92.92 | 88.67 - 89.85 | 76.24 - 77.09 |
| covered by $\geq$ 20 reads | 37.60 | 23.03 | 27.64 | 25.91 | 2.58 |
| range | 0.00 - 94.08 | 21.32 - 24.73 | 21.07 - 37.22 | 19.62 - 34.97 | 2.43 - 2.73 |

Averages and ranges were calculated from the respective rows of Datasets S3-S6.
[a] Values refer to percentages of reads removed as likely PCR duplicates; see *SI Text*, '`BWASP` workflow: design and output.'
[b] Values refer to percentages of reads removed as likely BS-seq failures; see *SI Text*, 'Removal of unconverted reads.'
[c] Genome coverage was calculated as the ratio of total length of all accepted reads over genome size.

**Table S2. Comparison of methylation sites and levels between all samples of the Libbrecht *et al.* (9) study.**

| | BphA1 | BphA2 | BphB1 | BphB2 | RphA1 | RphA2 | RphB1 | RphB2 |
|---|---|---|---|---|---|---|---|---|
| BphA1 | | 0.905 | 0.908 | 0.913 | 0.905 | 0.915 | 0.914 | 0.909 |
| BphA2 | 0.802 | | 0.922 | 0.913 | 0.918 | 0.934 | 0.912 | 0.910 |
| BphB1 | 0.826 | 0.782 | | 0.914 | 0.923 | 0.933 | 0.915 | 0.911 |
| BphB2 | 0.795 | 0.774 | 0.788 | | 0.910 | 0.923 | 0.919 | 0.919 |
| RphA1 | 0.791 | 0.773 | 0.801 | 0.758 | | 0.929 | 0.912 | 0.908 |
| RphA2 | 0.900 | 0.890 | 0.856 | 0.888 | 0.882 | | 0.922 | 0.920 |
| RphB1 | 0.809 | 0.754 | 0.779 | 0.769 | 0.774 | 0.866 | | 0.916 |
| RphB2 | 0.762 | 0.806 | 0.817 | 0.798 | 0.780 | 0.908 | 0.799 | |

Entries below the diagonal are overlap index values. Entries above the diagonal are correlation coefficients. Samples are abbreviated as follows: B (in first position), brood care phase; R, reproductive phase; A, line A; B (in second position), line B; 1, sequencing batch 1; 2, sequencing batch 2.

**Table S3. Comparison of genes ranked by methylation level between all samples of the Libbrecht *et al.* (9) study.**

| | BphA1 | BphA2 | BphB1 | BphB2 | RphA1 | RphA2 | RphB1 | RphB2 |
|---|---|---|---|---|---|---|---|---|
| BphA1 | | 0.36 | 0.35 | 0.37 | *0.47 | 0.41 | 0.44 | 0.36 |
| BphA2 | | | *0.56 | 0.34 | *0.50 | 0.46 | 0.43 | 0.37 |
| BphB1 | | | | *0.48 | *0.51 | 0.37 | *0.47 | 0.35 |
| BphB2 | | | | | 0.37 | 0.28 | 0.45 | 0.43 |
| RphA1 | | | | | | 0.40 | 0.39 | 0.35 |
| RphA2 | | | | | | | 0.41 | 0.39 |
| RphB1 | | | | | | | | 0.39 |

Table entries are rank-biased overlap values. Significantly high values are starred (see text for details). Samples are abbreviated as follows: B (in first position), brood care phase; R, reproductive phase; A, line A; B (in second position), line B; 1, sequencing batch 1; 2, sequencing batch 2.

**Additional Dataset S1** (**Dataset_S01-STUDIES-PcanObirSdum.xlsx**)

**Published studies of BS-seq experiments on *Polistes canadensis*, *Ooceraea biroi*, and *Stegodyphus dumicola*.** Listing of the BS-seq experiments on *Polistes canadensis* (1 study), *Ooceraea biroi* (1 study, 2 different genome assemblies), and *Stegodyphus dumicola* (1 study) re-analyzed in this paper, including details about the authors, publication year, samples analyzed, life-stage of the samples, tissues analyzed, and the accession numbers of the publicly available datasets.

**Additional Dataset S2** (**Dataset_S02-STUDIES-Amel.xlsx**)

**Published studies of BS-seq experiments on *Apis mellifera*.** Listing of the BS-seq experiments on *Apis mellifera* (16 studies) re-analyzed in this paper, including details about the authors, publication year, samples analyzed, life-stage of the samples, tissues analyzed, and the accession numbers of the publicly available datasets.

**Additional Dataset S3** (**Dataset_S03-tableQCM-PcanObirSdum.xlsx**)

**Quality control and mapping statistics for BS-seq experiments on *Polistes canadensis*, *Ooceraea biroi*, and *Stegodyphus dumicola*.** Quality control and mapping statistics for BS-seq experiments on *Polistes canadensis*, *Ooceraea biroi*, and *Stegodyphus dumicola*. The table includes the number of reads for each sample, the read length range, the number of trimmed reads after quality control and median length of the trimmed reads, total sequence set size (bp), mapping efficiency, the number and percentage of PCR read duplicates removed, the number and percentage of rejected reads (non-converted reads), genome coverage, and the numbers of left and right read biased positions being ignored. Details are given in *SI Text, Methods*.

**Additional Dataset S4** (**Dataset_S04-tableQCM-Amel.xlsx**)

**Quality control and mapping statistics for BS-seq experiments on *Apis mellifera*.** Quality control and mapping statistics for BS-seq experiments on *Apis mellifera*. The table includes the number of reads for each sample, the read length range, the number of trimmed reads after quality control and median length of the trimmed reads, total sequence set size (bp), mapping efficiency, the number and percentage of PCR read duplicates removed, the number and percentage of rejected reads (non-converted reads), genome coverage, and the numbers of left and right read biased positions being ignored. Details are given in *SI Text, Methods*.

**Additional Dataset S5** (**Dataset_S05-tableCOV-PcanObirSdum.xlsx**)

**Coverage statistics for BS-seq experiments on *Polistes canadensis*, *Ooceraea biroi*, and *Stegodyphus dumicola*.** Coverage statistics of sites at different levels of minimum coverage for BS-seq experiments on *Polistes canadensis*, *Ooceraea biroi*, and *Stegodyphus dumicola*. The table includes the number and percentage of CpG, CHG, and CHH sites covered by zero or at least one, two, five, ten, or twenty reads.

**Additional Dataset S6** (**Dataset_S06-tableCOV-Amel.xlsx**)

**Coverage statistics for BS-seq experiments on *Apis mellifera*.** Coverage statistics of sites at different levels of minimum coverage for BS-seq experiments on *Apis mellifera*. The table includes the number and percentage of CpG, CHG, and CHH sites covered by zero or at least one, two, five, ten, or twenty reads.

**Additional Dataset S7** (**Dataset_S07-tableNBS-PcanObirSdum.xlsx**)

**Methylation site statistics for BS-seq experiments on *Polistes canadensis*, *Ooceraea biroi*, and *Stegodyphus dumicola***. Methylation site statistics for BS-seq experiments on *Polistes canadensis*, *Ooceraea biroi*, and *Stegodyphus dumicola*. The table includes the number of sufficiently covered detectable (`scd`) and the highly significantly methylated (`hsm`) CpG, CHG and CHH sites.

**Additional Dataset S8** (**Dataset_S08-tableNBS-Amel.xlsx**)

**Methylation site statistics for BS-seq experiments on *Apis mellifera***. The table includes the number of sufficiently covered detectable (`scd`) and the highly significantly methylated (`hsm`) CpG, CHG and CHH sites.

**Additional Dataset S9** (**Dataset_S09-tableOML-PcanObirSdum.xlsx**)

**Overall methylation levels deduced from BS-seq experiments on *Polistes canadensis*, *Ooceraea biroi*, and *Stegodyphus dumicola***. Overall methylation levels deduced from BS-seq experiments on *Polistes canadensis*, *Ooceraea biroi*, and *Stegodyphus dumicola*, determined the overall percent of mapped read calls supporting methylation (unconverted Cs) versus read calls supporting unmethylated status (converted Cs).

**Additional Dataset S10** (**Dataset_S10-tableOML-Amel.xlsx**)

**Overall methylation levels deduced from BS-seq experiments on *Apis mellifera***. Overall methylation levels deduced from BS-seq experiments on *Apis mellifera*, determined the overall percent of mapped read calls supporting methylation (unconverted Cs) versus read calls supporting unmethylated status (converted Cs).

**Additional Dataset S11** (**Dataset_S11-tableMSG-PcanObirSdum.xlsx**)

**Methylation site distribution in genes for BS-seq experiments on *Polistes canadensis*, *Ooceraea biroi*, and *Stegodyphus dumicola*.** Methylation site distribution in genomic regions for BS-seq experiments on *Polistes canadensis*, *Ooceraea biroi*, and *Stegodyphus dumicola*.

**Additional Dataset S12 (Dataset_S12-tableMSG-Amel.xlsx)**

**Methylation site distribution in genes for BS-seq experiments on *Apis mellifera*.** Methylation site distribution in genomic regions for BS-seq experiments on *Apis mellifera*.

**Additional Dataset S13 (Dataset_S13-tableMSE-PcanObirSdum.xlsx)**

**Methylation site distribution in exons for BS-seq experiments on *Polistes canadensis*, *Ooceraea biroi*, and *Stegodyphus dumicola*.** Methylation site distribution in exons for BS-seq experiments on *Polistes canadensis*, *Ooceraea biroi*, and *Stegodyphus dumicola*.

**Additional Dataset S14 (Dataset_S14-tableMSE-Amel.xlsx)**

**Methylation site distribution in exons for BS-seq experiments on *Apis mellifera*.** Methylation site distribution in exons for BS-seq experiments on *Apis mellifera*.

**Additional Dataset S15 (Dataset_S15-Pc_PA-BWASPR-output.pdf)**

**Configuration files, `Rscript` commands, and `BWASPR` output for the Patalano *et al.* (5) *Polistes canadensis* study.** Pages 1-2: `Pc_PA.conf`. Page 3: `Pc.par`. Page 4: `Pc.dat`. Pages 5-18: `Rscript.BWASPR`. Page 19: `0README` in output directory. Pages 20-44: `CMS` output directory: coverage and methylation statistics. Pages 45-62: `PWC` output directory: pairwise comparisons. Pages 63-66: `CRL` output directory: between sample correlations. Pages 67-139: `REPCMS` output directory: coverage and methylation statistics for replicates. Pages 140-146 `REPCRL` output directory: between replicate correlations. Pages 147-149: `MMP` output directory: mapping of methylation sites on genome annotation. Pages 150-151: `ACS` output directory: annotation of conserved methylation sites. Pages 152-170: `RNK` output directory: genome feature regions ranked by methylation density. Pages 171-187: `MRPR` output directory: methylation-rich and -poor regions. Pages 188-190: `DMT` output directory: differentially methylated tiles. Pages 191-194: `DMSG` output directory: differentially methylated sites and genes. Pages 195-202: `OGL` output directory: ordered gene lists. Pages 203-210: `OGLall` output directory: unfiltered ordered gene lists.

**Additional Dataset S16 (Dataset_S16-Am_RE-BWASPR-output.pdf)**

**Configuration files and `BWASPR` output for the Remnant *et al.* (10) *Apis mellifera* study.** Pages 1-2: `Am_RE.conf`. Page 3: `Am.par`. Page 4-10: `Am.dat`. Page 11: `0README` in output directory. Pages 12-36: `CMS` output directory: coverage and methylation statistics. Pages 37-54: `PWC` output directory: pairwise comparisons. Pages 55-58: `CRL` output directory: between sample correlations. Pages 59-61: `MMP` output directory: mapping of methylation sites on genome annotation. Pages 62-63: `ACS` output directory: annotation of conserved methylation sites. Pages 64-88: `RNK` output directory: genome feature regions ranked by methylation density. Pages 89-107: `MRPR` output directory: methylation-rich and -poor regions. Pages 108-110: `DMT` output directory: differentially methylated tiles. Pages 111-117: `DMSG` output directory: differentially methylated sites and genes. Pages 118-125: `OGL` output directory: ordered gene lists. Pages 126-133: `OGLall` output directory: unfiltered ordered gene lists.

**Additional Dataset S17 (Dataset_S17-Cb_LIp-BWASPR-output.pdf)**

**Configuration files and `BWASPR` output for the Libbrecht *et al.* (9) *Cerapachys biroi* study, comparing samples by phase.** Pages 1-2: `Cb_LIp.conf`. Page 3: `Cb.par`. Page 4-6: `Cb.dat`. Page 7: `0README` in output directory. Pages 8-32: `CMS` output directory: coverage and methylation statistics. Pages 33-50: `PWC` output directory: pairwise comparisons. Pages 51-54: `CRL` output directory: between sample correlations. Pages 55-151: `REPCMS` output directory: coverage and methylation statistics for replicates. Pages 152-158 `REPCRL` output directory: between replicate correlations. Pages 159-161: `MMP` output directory: mapping of methylation sites on genome annotation. Pages 162-163: `ACS` output directory: annotation of conserved methylation sites. Pages 164-180: `RNK` output directory: genome feature regions ranked by methylation density. Pages 181-202: `MRPR` output directory: methylation-rich and -poor regions. Pages 203-205: `DMT` output directory: differentially methylated tiles. Pages 206-219: `DMSG` output directory: differentially methylated sites and genes. Pages 220-227: `OGL` output directory: ordered gene lists. Pages 228-235: `OGLall` output directory: unfiltered ordered gene lists.

**Additional Dataset S18 (Dataset_S18-Cb_LI-BWASPR-output.pdf)**

**Configuration files and `BWASPR` output for the Libbrecht *et al.* (9) *Cerapachys biroi* study, comparing all samples.** Pages 1-2: `Cb_LI.conf`. Page 3: `Cb.par`. Page 4-6: `Cb.dat`. Page 7: `0README` in output directory. Pages 8-104: `CMS` output directory: coverage and methylation statistics. Pages 105-133: `PWC` output directory: pairwise comparisons. Pages 134-137: `CRL` output directory: between sample correlations. Pages 138-146: `MMP` output directory: mapping of methylation sites on genome annotation. Pages 147-148: `ACS` output directory: annotation of conserved methylation sites. Pages 149-213: `RNK` output directory: genome feature regions ranked by methylation density. Pages 214-282: `MRPR` output directory: methylation-rich and -poor regions. Pages 283-285: `DMT` output directory: differentially methylated tiles. Pages 286-1590: `DMSG` output directory: differentially methylated sites and genes. Pages 1591-1780: `OGL` output directory: ordered gene lists. Pages 1781-1960: `OGLall` output directory: unfiltered ordered gene lists.

**Additional Dataset S19 (Dataset_S19-Cbir-hsmSetsOrdered-byCount.xlsx)**

**Comparison of sets of samples with common methylation sites for the Libbrecht *et al.* (9) *Cerapachys biroi* study.**

**Additional Dataset S20 (Dataset_S20-Ob_LI-BWASPR-output.pdf)**

**Configuration files and `BWASPR` output for the Libbrecht *et al.* (9) data mapped to the current genome assembly of *Ooceraea biroi*.** Pages 1-2: `Ob_LI.conf`. Page 3: `Ob.par`. Page 4: `Ob.dat`. Page 7: `OREADME` in output directory. Pages 8-104: `CMS` output directory: coverage and methylation statistics. Pages 105-133: `PWC` output directory: pairwise comparisons. Pages 134-137: `CRL` output directory: between sample correlations. Pages 138-146: `MMP` output directory: mapping of methylation sites on genome annotation. Pages 147-148: `ACS` output directory: annotation of conserved methylation sites. Pages 149-213: `RNK` output directory: genome feature regions ranked by methylation density. Pages 214-287: `MRPR` output directory: methylation-rich and -poor regions. Pages 288-290: `DMT` output directory: differentially methylated tiles. Pages 291-1564: `DMSG` output directory: differentially methylated sites and genes. Pages 1565-1742: `OGL` output directory: ordered gene lists. Pages 1743-1924: `OGLall` output directory: unfiltered ordered gene lists.

**Additional Dataset S21 (Dataset_S21-Ob_LIp-BWASPR-output.pdf)**

**Configuration files and `BWASPR` output for the Libbrecht *et al.* (9) data mapped to the current genome assembly of *Ooceraea biroi*, comparing samples by phase.** Pages 1-2: `Ob_LIp.conf`. Page 3: `Ob.par`. Page 4-6: `Ob.dat`. Page 7: `OREADME` in output directory. Pages 8-32: `CMS` output directory: coverage and methylation statistics. Pages 33-50: `PWC` output directory: pairwise comparisons. Pages 51-54: `CRL` output directory: between sample correlations. Pages 55-151: `REPCMS` output directory: coverage and methylation statistics for replicates. Pages 152-158 `REPCRL` output directory: between replicate correlations. Pages 159-161: `MMP` output directory: mapping of methylation sites on genome annotation. Pages 162-163: `ACS` output directory: annotation of conserved methylation sites. Pages 164-180: `RNK` output directory: genome feature regions ranked by methylation density. Pages 181-202: `MRPR` output directory: methylation-rich and -poor regions. Pages 203-205: `DMT` output directory: differentially methylated tiles. Pages 206-218: `DMSG` output directory: differentially methylated sites and genes. Pages 219-226: `OGL` output directory: ordered gene lists. Pages 227-234: `OGLall` output directory: unfiltered ordered gene lists.

**Additional Dataset S22 (Dataset_S22-Sd_LI-BWASPR-output.pdf)**

**Configuration files and `BWASPR` output for the Liu *et al.* (11) *Stegodyphus dumicola* study.** Pages 1-2: `Sd_LI.conf`. Page 3: `Sd.par`. Page 4: `Sd.dat`. Page 5: `OREADME` in output directory. Pages 6-30: `CMS` output directory: coverage and methylation statistics. Pages 31-48: `PWC` output directory: pairwise comparisons. Pages 49-52: `CRL` output directory: between sample correlations. Pages 53-245: `REPCMS` output directory: coverage and methylation statistics for replicates. Pages 246-252 `REPCRL` output directory: between replicate correlations. Pages 253-255: `MMP` output directory: mapping of methylation sites on genome annotation. Pages 256-257: `ACS` output directory: annotation of conserved methylation sites. Pages 258-274: `RNK` output directory: genome feature regions ranked by methylation density. Pages 275-292: `MRPR` output directory: methylation-rich and -poor regions. Pages 293-295: `DMT` output directory: differentially methylated tiles. Pages 296-370: `DMSG` output directory: differentially methylated sites and genes. Pages 371-378: `OGL` output directory: ordered gene lists. Pages 379-386: `OGLall` output directory: unfiltered ordered gene lists.

**Additional Dataset S23 (Dataset_S23-cor-ovi-Y0.xlsx)**

**Comparison of methylation sites and levels for samples from the Yagound *et al.* (12) *Apis mellifera* study.**

### References


1. Andrews A (2010) FastQC: A quality control tool for high throughput sequence data. http://www.bioinformatics.babraham.ac.uk/projects/fastqc/.
2. Krueger F (2012) Trim galore! http://www.bioinformatics.babraham.ac.uk/projects/trim_galore/.
3. Martin M (2011) Cutadapt removes adapter sequences from high-throughput sequencing reads. *EMBnet.journal* 17(1):10–12.
4. Morandin C, Brendel VP, Sundström L, Helanterä H, Mikheyev AS (2019) Changes in gene DNA methylation and expression networks accompany caste specialization and age-related physiological changes in a social insect. *Mol. Ecol.* 28(8):1975–1993.
5. Patalano S, Hore TA, Reik W, Sumner S (2012) Shifting behaviour: epigenetic reprogramming in eusocial insects. *Curr. Opin. Cell Biol.* 24(3):367–73.
6. Gentleman RC, et al. (2004) Bioconductor: open software development for computational biology and bioinformatics. *Genome Biology* 5(10):R80.
7. Akalin A, et al. (2012) methylKit: a comprehensive R package for the analysis of genome-wide DNA methylation profiles. *Genome Biology* 13(10):R87.
8. The R Foundation (2020) The R Project for Statistical Computing. https://www.r-project.org.
9. Libbrecht R, Oxley PR, Keller L, Kronauer DJC (2016) Robust DNA methylation in the clonal raider ant brain. *Curr. Biol.* 26(3):391–5.

10. Remnant EJ, et al. (2016) Parent-of-origin effects on genome-wide DNA methylation in the Cape honey bee (Apis mellifera capensis) may be confounded by allele-specific methylation. *BMC Genomics* 17:226.
11. Liu S, Aageaard A, Bechsgaard J, Bilde T (2019) DNA methylation patterns in the social spider, *Stegodyphus dumicola*. *Genes* 10(2).
12. Yagound B, Remnant EJ, Buchmann G, Oldroyd BP (2020) Intergenerational transfer of DNA methylation marks in the honey bee. *Proceedings of the National Academy of Sciences* 117(51):32519–32527.
