## Supplementary material for "Tools and applications for integrative analysis of DNA methylation in social insects": Dataset S15

Pc_PA.conf

```
#Customize variables here:#####################################################
###############################################################################

#Load the files:
#
infiles       <- setup_BWASPR(datafile="./RWORK/Pcan/Pc.dat",
                              parfile="./RWORK/Pcan/Pc.par")

#Set the study and samples in the study:
#
species       <- "Pc"
study         <- "PA"
samplelist    <- list("queen","worker")
## The following two variables are used for output file labeling:
studyLabel    <- "PA"
sampleLabels  <- list("Pc_PA_qn","Pc_PA_wr")

hasreplicates <- TRUE
type          <- "CpG"
destrand      <- TRUE
covlist       <- c(6,10,15)
locount       <- 100
hicount       <- 1000
repcovlist    <- c(6,10,15)
replocount    <- 10
rephicount    <- 100
hheight       <- 0.10
nbrpnts       <- 5000

## The following four variables are sent to rank_rbm() and determine what data
#  points get plotted:
minnbrmsprm   <- 5      # minimum number of methylation sites per promoter
mingenewidth  <- 500    # minimum gene width
maxgenewidth  <- 5000   # maximum gene width
minnbrmsgene  <- 5      # minimum number of methylation sites per gene

## Other parameters (see Rscript.BWASPR for usage notes):
#
highcoverage  <- 20     # high read coverage threshold for studyhc methylRawList object
threshold     <- 25.0   # "difference" threshold for getMethylDiff(), called by det_dmsg()
qvalue        <- 0.01   # "qvalue" setting for getMethylDiff(), called by det_dmsg()
wsize         <- 1000   # "win.size" parameter for tileMethylCounts() in det_dmt()
stepsize      <- 1000   # "step.size" parameter for tileMethylCounts() in det_dmt()

minNsites     <- 10     # minimum number of hc sites in a gene to be heatmapped in show_dmsg()
maxNsites     <- 60     # maximum number of hc sites in a gene to be heatmapped in show_dmsg()
minPdmsites   <- 10     # minimum ratio of dm/hc (in %) in a gene to be heatmapped in show_dmsg()

maxgwidth     <- 20000  # maximal gene width for a gene to be considered by explore_dmsg()
minnbrdmsites <- 2      # minimum number of differentially methylated sites for a gene to be
                        #  considered in sample comparisons by explore_dmsg()
glink         <- "NCBIgene"   # URL to show in explore_dmsg(); options: "" or "NCBIgene"

#Set the number of processors to use:
#
numprc <- 6

#Determine what analyses to run:
#
RUNload    <- FALSE
```

```
RUNcms     <- TRUE
RUNpwc     <- TRUE
RUNcrl     <- TRUE

RUNrepcms  <- TRUE
RUNrepcrl  <- TRUE

RUNmmp     <- TRUE
RUNacs     <- TRUE
RUNrnk     <- TRUE
RUNmrpr    <- TRUE

RUNdmt     <- TRUE
RUNdmsg    <- TRUE
RUNdmgdtls <- TRUE
RUNogl     <- TRUE

RUNsave    <- TRUE


mymessage <- sprintf("\nAnalyzing %s study %s for type %s\n\n",species,studyLabel,type)
message(mymessage)

################################################################################
#End of typical customization.##################################################
```

Pc.par

```
SPECIESNAME Polistes canadensis
TOTALNBRPMSITES   14357872
ASSEMBLYVERSION   ASM131383v1
GENOMESIZE  211202212
SPECIESGFF3DIR    ./MCALLS/Pcan/genome/GFF3DIR
GENELISTGFF3      Pcan.gene.gff3
EXONLISTGFF3      Pcan.exon.gff3
PCGEXNLISTGFF3    Pcan.pcg-exon.gff3
PROMOTRLISTGFF3   Pcan.promoter.gff3
CDSLISTGFF3 Pcan.pcg-CDS.gff3
UTRFLAGSET  1
5UTRLISTGFF3      Pcan.pcg-5pUTR.gff3
3UTRLISTGFF3      Pcan.pcg-3pUTR.gff3
```

Pc.dat

```
# Samples from Patalano et al. (2015) PNAS:
# qn = queen; wr = worker
#
Pc	PA	queen	0	CpGhsm	./MCALLS/Pcan/Patalano2015/Queen/Queen.CpGhsm.mcalls
Pc	PA	queen	0	CpGscd	./MCALLS/Pcan/Patalano2015/Queen/Queen.CpGscd.mcalls
Pc	PA	worker	0	CpGhsm	./MCALLS/Pcan/Patalano2015/Worker/Worker.CpGhsm.mcalls
Pc	PA	worker	0	CpGscd	./MCALLS/Pcan/Patalano2015/Worker/Worker.CpGscd.mcalls

# ... continuing: replicates
#
Pc	PA	queen	1	CpGhsm	./MCALLS/Pcan/Patalano2015/Queen/replicate1/Queen1.CpGhsm.mcalls
Pc	PA	queen	1	CpGscd	./MCALLS/Pcan/Patalano2015/Queen/replicate1/Queen1.CpGscd.mcalls
Pc	PA	queen	2	CpGhsm	./MCALLS/Pcan/Patalano2015/Queen/replicate2/Queen2.CpGhsm.mcalls
Pc	PA	queen	2	CpGscd	./MCALLS/Pcan/Patalano2015/Queen/replicate2/Queen2.CpGscd.mcalls
Pc	PA	queen	3	CpGhsm	./MCALLS/Pcan/Patalano2015/Queen/replicate3/Queen3.CpGhsm.mcalls
Pc	PA	queen	3	CpGscd	./MCALLS/Pcan/Patalano2015/Queen/replicate3/Queen3.CpGscd.mcalls
Pc	PA	worker	1	CpGhsm	./MCALLS/Pcan/Patalano2015/Worker/replicate1/Worker1.CpGhsm.mcalls
Pc	PA	worker	1	CpGscd	./MCALLS/Pcan/Patalano2015/Worker/replicate1/Worker1.CpGscd.mcalls
Pc	PA	worker	2	CpGhsm	./MCALLS/Pcan/Patalano2015/Worker/replicate2/Worker2.CpGhsm.mcalls
Pc	PA	worker	2	CpGscd	./MCALLS/Pcan/Patalano2015/Worker/replicate2/Worker2.CpGscd.mcalls
Pc	PA	worker	3	CpGhsm	./MCALLS/Pcan/Patalano2015/Worker/replicate3/Worker3.CpGhsm.mcalls
Pc	PA	worker	3	CpGscd	./MCALLS/Pcan/Patalano2015/Worker/replicate3/Worker3.CpGscd.mcalls
```

```
#Rscript.BWASPR
#Version:   February 12, 2021
#Contact:   Volker Brendel
#Documentation:   https://github.com/brendelgroup/BWASPR

#Load required libraries and set options:
#
library("BWASPR")
library("parallel")

options(width=240)
options(max.print=1000000)
options(digits=3)
options(warn=-1)


#Read the required BWASPR .conf file to set parameters.
#Usage example: Rscript Rscript.BWASPR Am_HE
#  would read the configuration files Am_HE.conf
#
args       <- commandArgs(trailingOnly = TRUE)
if (length(args) == 0) {
  exit
} else {
  configfile <- paste(args[1],"conf",sep=".")
  outputdir <- paste("Routput",args[1],sep="-")
  if (length(args) > 1) {
    outputdir <- args[2]
  }
}
source(configfile)

message("Rscript.BWASPR is being run with .conf file ",configfile," and output directory ",outputdir,"\n\n")
cmd <- paste("mkdir -p",outputdir,sep=" ")
message(".. now executing: ",cmd)
system(cmd)
readme <- paste("Rscript.BWASPR output for .conf file ",configfile,".\n\n",
                "Data input are the specified *.mcalls files and parameters set in ",configfile,".\n",
                "The *.mcalls input data are saved in the following data structures:\n\n",
                "studymk - a methylKit methylRaw(List) object storing the CpGhsm site data for all samples\n",
                "studymc - a methylKit methylRaw(List) object storing the CpGscd site data for all samples\n",
                "studyhc - a methylKit methylRaw(List) object storing the CpGscd site data for all samples,\n",
                 "           restricted to sites with high coverage (here set to ",highcoverage,")\n",
                "mkrd    - a methylKit methylRawList object storing the CpGhsm site data for all\n",
                 "           replicates of a given sample\n\n",
                "Output of the analysis is stored in the following subdirectories (if the corresponding\n",
                 " function calls were specified in ",configfile,"; please consult the 0README\n",
                 " files in the subdirectories for details on the output files):\n\n",
                "CMS - Coverage and methylation statistics for aggregate samples\n",
                "PWC - Pairwise comparisons between all samples\n",
                "CRL - Correlations between aggregate samples\n",
                "REPCMS - Coverage and methylation statistics for replicate samples\n",
                "REPCRL - Correlations between replicates\n",
                "MMP - Mapping of methylation sites on genome annotation\n",
                "ACS - Annotation of conserved methylation sites\n",
                "RNK - Genomic feature regions ranked by CpGhsm statistics\n",
                "MRPR - Methylation-rich and -poor regions\n",
                "DMSG - Differentially methylateds sites and genes\n",
                "OGL - Ordered gene lists\n",
                sep="")
readmefile <- paste(outputdir,"0README",sep="/")
```

```
sink(readmefile)
cat(sprintf("%s",readme))
sink()

#Based on the configuration file, set some basic parameters ...
#
speciesstudy <- paste(species,studyLabel,sep="_")
stype <- paste(type,"hsm",sep="")
ctype <- paste(type,"scd",sep="")

#... and pull out the assembly version, number of potential (CpG or RBBS) methylation
#  sites, genome size, and status of UTR annotation (from the parameter file):
#
asmblv     <- infiles$parameters[infiles$parameters$Variable == "ASSEMBLYVERSION",2]
nbrpms     <- as.numeric(infiles$parameters[infiles$parameters$Variable == "TOTALNBRPMSITES",2])
gnmsize    <- as.numeric(infiles$parameters[infiles$parameters$Variable == "GENOMESIZE",2])
UTRFLAGSET <- as.numeric(infiles$parameters[infiles$parameters$Variable == "UTRFLAGSET",2])


#The RUN* logicals in the configuration file determine which parts of the
#  workflow should be run.
#


### Section: Load necessary data to proceed.
##
#
if (RUNload) {
#Either read data from a previously saved workflow run or call mcalls2mkobj() to
#  produce the required methylRawList objects. studymk contains the data for the
#  hsm sites, while studymc contains the data for all scd sites.
#
  rdatafile <- paste(speciesstudy,"RData",sep=".")
  message(".. loading previously stored image ", rdatafile, " ...")
  load(rdatafile)
#Read the configuration file again so that the load command does not overwrite
#  intended current settings.
  source(configfile)
  message(".. done ...")
} else {
#Create the methylKit raw objects:
#
  studymk <- mcalls2mkobj(infiles$datafiles,species=species,study=study,sample=samplelist,
                          replicate=c(0),type=stype,mincov=1,assembly=asmblv)
  studymc <- mcalls2mkobj(infiles$datafiles,species=species,study=study,sample=samplelist,
                          replicate=c(0),type=ctype,mincov=1,assembly=asmblv)
}
################################################################################



### Section: Coverage and methylation statistics.
##
#
if (RUNcms) {
#Produce cms-*txt and cms-*.pdf files reporting on coverage and methylation statistics.
#
  CMSdir <- paste(outputdir,"CMS",sep="/")
  cmd <- paste("mkdir -p ",CMSdir,sep="")
  message("\n.. now executing: ",cmd)
  system(cmd)
  plotfile <- paste("cms",species,sep="-")
  plotfile <- paste(plotfile,studyLabel,sep="_")
  message("\n.. coverage and methylation statistics for aggregate samples ...")
```

```
  tmpN <- lapply(studymk,function(x) {plotfile <- paste(plotfile,,sep="_");
                                       outfile  <- paste(plotfile,"txt",sep=".");
                                       cmStats(x,stype,covlist,locount,hicount,outfile=outfile,plotfile=plotfile)
                                       cmd <- paste("mv ",plotfile,".pdf ",outfile," ",CMSdir,sep="")
                                       message("\n.. now executing: ",cmd)
                                       system(cmd)
                                      }
                )
  readme <- paste("CMS - Coverage and methylation statistics for aggregate samples.\n\n",
                  "Input: studymk, covlist, locount, hicount\n\n",
                  "Output: files cms-*.txt and cms-*.pdf\n\n",
                  "Notes: The statistics are given for CpGhsm sites restricted to minimum or\n",
                  "        higher level coverage.  At the minimum level, the number of sites will be\n",
                  "        the number of lines of the corresponding *CpGhsm.mcalls file minus one\n",
                  "        (consistency check).  The distribution of methylation levels is really only\n",
                  "        interesting for high coverage because at low coverage there will be a strong\n",
                  "        bias towards high methylation levels (by definition of hsm sites).\n",
                  "        The same bias at high coverage would suggest that the sample includes a\n",
                  "        preponderance of consistently methylated genomic sites.\n\n",
                  "        locount and hicount set bounds on the coverage to exclude sites with too few\n",
                  "        or too many covering reads to provide statistics on a typical range.\n",
                  sep="")
  readmefile <- paste(CMSdir,"0READMEcms",sep="/")
  sink(readmefile)
  cat(sprintf("%s",readme))
  sink()
  message(".. done ...")
}
################################################################################



### Section: Comparing aggregate data among samples.
##
#
if (RUNpwc) {
  l <- length(studymk@treatment)
  if (l > 1) {
    PWCdir <- paste(outputdir,"PWC",sep="/")
    cmd <- paste("mkdir -p",PWCdir,sep=" ")
    message("\n.. now executing: ",cmd)
    system(cmd)
  }
  if (l < 2) {
    message("\n.. no pairs to compare ....")
  } else if (l == 2) {
#First, produce pairwise sample site comparisons and save the output in file
#  pwc-*.txt and plots in pwc-*.pdf.
#
    sample1hsm <- methylKit::getData(studymk[[1]])
    sample1scd <- methylKit::getData(studymc[[1]])
    s1l <- sampleLabels[[1]]
    sample2hsm <- methylKit::getData(studymk[[2]])
    sample2scd <- methylKit::getData(studymc[[2]])
    s2l <- sampleLabels[[2]]
    outf1 <- "pwc"
    outf2<- paste(s1l,s2l,sep=".vs.")
    outfile <- paste(outf1,outf2,sep="-")
    plotfile <- paste(outfile,"pdf",sep=".")
    outfile <- paste(outfile,"txt",sep=".")
    sink(outfile)
    message("... comparing ", s1l, " versus ", s2l, " ...")
    cuSlist <- cmpSites(sample1hsm,sample1scd,s1l,sample2hsm,sample2scd,s2l,
                        nbrpms,plotfile,covlist,hheight,nbrpnts)
```

```
    sink()
    cmd <- paste("mv",outfile,plotfile,PWCdir,sep=" ")
    message("\n.. now executing: ",cmd)
    system(cmd)
  } else {
#First, produce pairwise sample site comparisons and save the output in files
#  pwc-* (done in parallel).
#  Note: no plots are generated - for unclear reasons, venneuler stalls when invoked on a cluster ...
#
  v <- c(1:length(studymk@treatment))
  numprcINcl <- dim(combn(v,2))[2]
  if (numprcINcl > numprc) {numprcINcl <- numprc}
  cl <- makeCluster(numprcINcl,type="PSOCK",useXDR=FALSE,outfile="")
  clusterExport(cl=cl, varlist=c("studymk", "studymc", "sampleLabels",
                              "nbrpms", "covlist", "PWCdir"), envir=environment())
  clusterEvalQ(cl=cl,library(BWASPR))
  message("\n.. running pairwise comparisons ...")

  cusData <- parApply(cl, combn(v,2), 2, function (x) {
      sample1hsm <- methylKit::getData(studymk[[x[1]]])
      sample1scd <- methylKit::getData(studymc[[x[1]]])
      s1l <- sampleLabels[[x[1]]]
      sample2hsm <- methylKit::getData(studymk[[x[2]]])
      sample2scd <- methylKit::getData(studymc[[x[2]]])
      s2l <- sampleLabels[[x[2]]]
      outf1 <- "pwc"
      outf2<- paste(s1l,s2l,sep=".vs.")
      outfile <- paste(outf1,outf2,sep="-")
      outfile <- paste(outfile,"txt",sep=".")
      sink(outfile)
      message("... comparing ", s1l, " versus ", s2l, " ...")
      cuSlist <- cmpSites(sample1hsm,sample1scd,s1l,sample2hsm,sample2scd,s2l,
                          nbrpms,plotfile="",covlist,hheight,nbrpnts)
      sink()
      cmd <- paste("mv",outfile,PWCdir,sep=" ")
      message("\n.. now executing: ",cmd)
      system(cmd)
      return(cuSlist)
    })
  stopCluster(cl)
  #cusData is a list of lists.  The outer list contains data for each pairwise
  #  comparison.  These data are in the form of lists of data frames as returned
  #  by cmpSites(): for each comparions, the data frames are
  #    commonHSM, unique1HSM, unique2HSM,
  #    commonSCD, unique1SCD, unique2SCD,
  #    hsm1cSCD, unique1HSMn2SCD,
  #    hsm2cSCD, unique2HSMn1SCD,
  #    unique1HSM2SCD, unique2HSM1SCD.
  #  For details, see the *.txt output files of the above code section.
  }
  readme <- paste("PWC - Pairwise comparisons between all samples.\n\n",
                  "Input: studymk, studymc, nbrpms, hheight, nbrpnts\n\n",
                  "Output: files pwc-*.txt pwc-.*pdf\n\n",
                  "Notes: The output is generated by BWASPR::cmpSites(), which determines for each pairwise\n",
                  "        comparison the numbers of common and distinct sites.  The function also calculates\n",
                  "        the overlap index between the samples and estimates the size of the common pool of\n",
                  "        potential methylation sites.\n\n",
                  "       The CpGscd data input (studymc) is necessary to determine sites that are detectable\n",
                  "        in both samples, and nbrpms (listed in the *.par file specified in the ",configfile,"\n",
                  "        configuration file as argument to TOTALNBRPMSITES) provides the total number of\n",
                  "        potential methylation sites (typically, all CpG sites), a necessary parameter for\n",
                  "        the estimations.\n",
                  "       hheight (default: 0.10) specifies the y-axis limit in the histogram plots.\n",
                  "       nbrpnts (default: 5000) specifies the number of common sites to be plotted in the\n",
```

```
                  "        methylation levels scatter plot.\n\n",
                  sep="")
  readmefile <- paste(PWCdir,"0READMEpwc",sep="/")
  sink(readmefile)
  cat(sprintf("%s",readme))
  sink()
  message(".. done ...")
}


if (RUNcrl) {
#Second, produce correlations between methylation levels in common sites of
#  aggregate samples.
#  Output will be in crl-*txt and crl-*pdf files.
#
  CRLdir <- paste(outputdir,"CRL",sep="/")
  cmd <- paste("mkdir -p ",CRLdir,sep="")
  message("\n.. now executing: ",cmd)
  system(cmd)
  plotfile <- paste("crl",speciesstudy,sep="-")
  outfile <- paste(plotfile,"txt",sep=".")
  sink(outfile)
  message("\n.. running aggregate sample pairwise correlations ...")
  mymessage <- sprintf("\n... comparing aggregate type %s data for all %s samples (destrand=%s) ...",type,studyLabel,destrand)
  message(mymessage)
  studyData <- cmpSamples(studymk,destrand=destrand,plotfile=plotfile)
  cat(sprintf("\nThe number of conserved sites is %6d.\n\n",dim(studyData)[1]))
  sink()

  cmd <- paste("mv ",outfile," ",plotfile,".pdf ",CRLdir,sep="")
  message("\n.. now executing: ",cmd)
  system(cmd)
  readme <- paste("CRL - Correlations between aggregate samples.\n\n",
                  "Input: studymk\n\n",
                  "Output: files crl-*.txt crl-*.pdf\n\n",
                  "Notes: The output is generated by BWASPR::cmpSamples(), which uses methylKit::unite()\n",
                  "        to determine the sites common to all samples and applies methylKit::getCorrelation()\n",
                  "        and methylKit::PCASamples().  The *.txt files show the correlations in a table, and\n",
                  "        the *.pdf files show graphics.\n\n",
                  "       Beware that the number of common sites (shown at the bottom of the *.txt files) may be\n",
                  "        small, which may make the correlations less informative.\n\n",
                  sep="")
  readmefile <- paste(CRLdir,"0READMEcrl",sep="/")
  sink(readmefile)
  cat(sprintf("%s",readme))
  sink()
  message(".. done ...")
}
################################################################################



### Section: Comparisons between replicate samples.
##
#
if (hasreplicates & (RUNrepcms | RUNrepcrl)) {
#Compare replicates for all samples.
#
  mymessage <- sprintf("\n\n  Comparison of replicates for all samples in %s study %s.\n",species,studyLabel)
  message(mymessage)

  if (RUNrepcms) {
      REPCMSdir <- paste(outputdir,"REPCMS",sep="/")
      cmd <- paste("mkdir -p",REPCMSdir,sep=" ")
```

```
      message("\n.. now executing: ",cmd)
      system(cmd)
  }
  if (RUNrepcrl) {
      REPCRLdir <- paste(outputdir,"REPCRL",sep="/")
      cmd <- paste("mkdir -p",REPCRLdir,sep=" ")
      message("\n.. now executing: ",cmd)
      system(cmd)
  }

  csR <- lapply(samplelist,function(x) {
           mkrd <- mcalls2mkobj(infiles$datafiles,species,study,sample=x,
                                replicate=c(1:10),type=stype,mincov=1,assembly=asmblv
                               )
           if (is.null(mkrd)) {    # ... return if there are no replicates for a particular sample
               return(NULL)
           }


if (RUNrepcms) {
#Produce repcms-*txt and repcms-*.pdf files reporting on coverage and methylation statistics for
#  all sample replicates separately.
#
           rplotfile <- paste("repcms",speciesstudy,sep="-")
           message("\n.. coverage and methylation statistics for replicates ...")
           tmpN <- lapply(mkrd,function(y) {rplotfile <- paste(rplotfile,,sep="_")
                     routfile <- paste(rplotfile,"txt",sep=".")
                     cmStats(y,stype,repcovlist,replocount,rephicount,outfile=routfile,plotfile=rplotfile)
                   cmd <- paste("mv ",routfile," ",rplotfile,".pdf ",REPCMSdir,sep="")
                   message("\n.. now executing: ",cmd)
                   system(cmd)
                   }
                   )
           message(".. done ...")
}


if (RUNrepcrl) {
#Produce correlations between methylation levels in common sites of replicates.
#  Output will be in repcrl-*txt and repcrl-*pdf files.
#
           outfile <- paste("repcrl",speciesstudy,sep="-")
           outfile <- paste(outfile,x,sep="_")
           outfile <- paste(outfile,"txt",sep=".")
           sink(outfile)
           mymessage <- sprintf("\n.. comparing replicates of sample %s ...\n",x)
           message(mymessage)
           plotfile <- paste("repcrl",species,sep="-")
           plotfile <- paste(plotfile,studyLabel,x,sep="_")
           mkurd <- cmpSamples(mkrd,destrand=destrand,plotfile=plotfile)
           cat(sprintf("\nThe number of conserved sites is %6d.\n\n",dim(mkurd)[1]))
           sink()
         cmd <- paste("mv ",outfile," ",plotfile,".pdf ",REPCRLdir,sep="")
         message("\n.. now executing: ",cmd)
         system(cmd)
           message(".. done ...")

           return(mkurd)
} else {
           return(mkrd)
}

           }
         ) # end lapply
```

```
if (RUNrepcms) {
  readme <- paste("REPCMS - Coverage and methylation statistics for replicate samples.\n\n",
                  "Input: studymk, repcovlist, replocount, rephicount\n\n",
                  "Output: files repcms-*txt and repcms-*pdf\n\n",
                  "Notes: The statistics are given for CpGhsm sites restricted to minimum or\n",
                  "        higher level coverage.  At the minimum level, the number of sites will be\n",
                  "        the number of lines of the corresponding *CpGhsm.mcalls file minus one\n",
                  "        (consistency check).  The distribution of methylation levels is really only\n",
                  "        interesting for high coverage because at low coverage there will be a strong\n",
                  "        bias towards high methylation levels (by definition of hsm sites).\n",
                  "        The same bias at high coverage would suggest that the sample includes a\n",
                  "        preponderance of consistently methylated genomic sites.\n\n",
                  "        replocount and rephicount set bounds on the coverage to exclude sites with too few\n",
                  "        or too many covering reads to provide statistics on a typical range.\n",
                  sep="")
  readmefile <- paste(REPCMSdir,"0READMErepcms",sep="/")
  sink(readmefile)
  cat(sprintf("%s",readme))
  sink()
  message(".. done ...")
}


if (RUNrepcrl) {
  readme <- paste("REPCRL - Correlations between replicates.\n\n",
                  "Input: mkrd (methylKit methylRawList object of replicate data)\n\n",
                  "Output: files repcrl-*.txt repcrl-*.pdf\n\n",
                  "Notes: The output is generated by BWASPR::cmpSamples(), which uses methylKit::unite()\n",
                  "        to determine the sites common to all replicates and applies methylKit::getCorrelation()\n",
                  "        and methylKit::PCASamples().  The *.txt files show the correlations in a table, and\n",
                  "        the *.pdf files show graphics.\n\n",
                  "       Beware that the number of common sites (shown at the bottom of the *.txt files) may be\n",
                  "        small, which may make the correlations less informative.\n\n",
                  sep="")
  readmefile <- paste(REPCRLdir,"0READMErepcrl",sep="/")
  sink(readmefile)
  cat(sprintf("%s",readme))
  sink()
  message(".. done ...")
}

#csR is a list of methylRaw objects, each representing the methylKit::unite() set of
#  sites common to all replicates of a particular sample in case RUNreprcl is set;
#  otherwise the list has the individual (not united) replicate data.
#  For details, see the *.txt output files of the above code section.
} ### end if (hasreplicates & (RUNrepcms | RUNrepcrl))
################################################################################



### Section: Connecting methylation patterns with genome annotation.
##
#
if (RUNmmp) {
#Determine where the hsm sites fall relative to the genome annotation features
#  (genes, exons, etc.).
#  Output will be in mmp-* files.
#
  MMPdir <- paste(outputdir,"MMP",sep="/")
  cmd <- paste("mkdir -p",MMPdir,sep=" ")
  message("\n.. now executing: ",cmd)
  system(cmd)
```

```
  message("\n.. generate methylation to annotation maps ...")
  if (!exists("genome_ann")) {
    genome_ann <- get_genome_annotation(infiles$parameters)
  }
  tmpN <- lapply(c(1:length(sampleLabels)),function(i) {
    sampleL  <- paste(sampleLabels[i],"hsm",sep="_")
    controlL <- paste(sampleLabels[i],"scd",sep="_")
    outfile  <- paste("mmp",sampleLabels[i],sep="-")
    outfile  <- paste(outfile,"txt",sep=".")
    tmp <- map_methylome(studymk[[i]],sampleL,studymc[[i]],controlL,genome_ann,
                         species,gnmsize,UTRflag=UTRFLAGSET,outfile)
    cmd <- paste("mv",outfile,MMPdir,sep=" ")
    message("\n.. now executing: ",cmd)
    system(cmd)
    }
    )
  readme <- paste("MMP - Mapping of methylation sites on genome annotation.\n\n",
                 "Input: studymk, studymc, genome_ann (a list of GRanges objects providing annotated region labels\n",
                 "        and boundaries; output of BWASPR::get_genome_annotation())\n\n",
                 "Output: files mmp-*.txt\n\n",
                 "Notes: The output is generated by BWASPR::map_methylome() and gives an accounting for every sample\n",
                 "        as to where the CpGhsm and CpGscd (control) sites reside relative to the genome annotation.\n",
                 "       Abbreviations used: O/E, observed over expected\n",
                 "                           (expected percentages are based on the respective feature proportions\n",
                 "                            in the genome, as annotated).\n\n",
                 sep="")
  readmefile <- paste(MMPdir,"0READMEmmp",sep="/")
  sink(readmefile)
  cat(sprintf("%s",readme))
  sink()
  message(".. done ...")
}


if (RUNacs) {
#Show methylation / annotation connections for sites common to all studymk
#  samples.
#  Output will be in the acs-*.txt file.
#
  library(methods)
  library(utils)
  ACSdir <- paste(outputdir,"ACS",sep="/")
  cmd <- paste("mkdir -p",ACSdir,sep=" ")
  message("\n.. now executing: ",cmd)
  system(cmd)
  message("\n.. annotating conserved sites ...")
  if (!exists("genome_ann")) {
    genome_ann <- get_genome_annotation(infiles$parameters)
  }
  outfile <- paste("acs",speciesstudy,sep="-")
  outfile <- paste(outfile,"txt",sep=".")
  methylome_ann <- annotate_methylome(studymk,genome_ann,destrand=destrand,
                                      outfile=outfile)
  cmd <- paste("mv",outfile,ACSdir,sep=" ")
  message("\n.. now executing: ",cmd)
  system(cmd)
  readme <- paste("ACS - Annotation of conserved methylation sites.\n\n",
                 "Input: studymk, genome_ann (a list of GRanges objects providing annotated region labels and bounds;\n",
                 "                            output of BWASPR::get_genome_annotation())\n\n",
                 "Output: file acs-*.txt\n\n",
                 "Notes: The output is generated by BWASPR::annotate_methylome(), which uses methylKit::unite()\n",
                 "        to determine the sites common to all experiments and applies genomation::annotateWithFeature().\n",
                 "        The acs-*.txt files show the common sites with methylation levels in all samples\n",
                 "        and columns with booleans indicating whether the site fits a specific annotation feature.\n\n",
```

```
                  "       Abbreviations used: pc, protein coding; nc, non-coding; fp, five prime; tp, three prime.\n",
                  "                           UTR, untranslated region; unique, not overlapping with other categories.\n\n",
                  sep="")
  readmefile <- paste(ACSdir,"0READMEacs",sep="/")
  sink(readmefile)
  cat(sprintf("%s",readme))
  sink()
  message(".. done ...")
}
################################################################################



### Section: Determination of methylation site rich and poor regions.
##
#
if (RUNrnk) {
#Produce ranked*.txt, sites*.txt, and plot*.pdf files showing methylation-percentage
# and site density ranked annotated regions and exploratory plots comparing the
# measures.
#
  RNKdir <- paste(outputdir,"RNK",sep="/")
  cmd <- paste("mkdir -p",RNKdir,sep=" ")
  message("\n.. now executing: ",cmd)
  system(cmd)
  if (!exists("genome_ann")) {
    genome_ann <- get_genome_annotation(infiles$parameters)
  }
  message("\n.. ranked genes for each sample ...")
  rgL <- rank_rbm(studymc,studymk,region.gr=genome_ann$gene,rlabel="genes",
                  minrwidth=mingenewidth, maxrwidth=maxgenewidth,
                  minnbrsites=minnbrmsgene,
                  withglink=glink,
                  outflabel=speciesstudy)
  message(".. done ...")
  message("\n.. ranked promoters for each sample ...")
  rpL <- rank_rbm(studymc,studymk,region.gr=genome_ann$promoter,
                  rlabel="promoters",
                  minnbrsites=minnbrmsprm,
                  withglink=glink,
                  outflabel=speciesstudy)
  cmd <- paste("mv ranked-*-",speciesstudy,"_*.txt sites-*",speciesstudy,"_*.txt plot-*",speciesstudy,"_*.pdf ",RNKdir,sep="")
  message("\n.. now executing: ",cmd)
  system(cmd)
  readme <- paste("RNK - Genomic feature regions ranked by CpG methylation and hsm statistics.\n\n",
                  "Input: studymc, studymk, region.gr (GRanges object providing annotated region labels and bounds)\n\n",
                  "Output: files ranked-genes-*.txt sites-in-genes-*.txt\n",
                  "              ranked-promoters-*.txt sites-in-promoters-*.txt\n",
                  "              plot*.pdf\n\n",
                  "Notes: The output is generated by BWASPR::rank_rbm().\n",
                  "       The files ranked-genes-*.txt provide ranked lists of genes based on\n",
                  "        overall methylation percentage or the occurrence of CpGhms sites\n",
                  "        within the annotated gene bounds.  The output columns include\n",
                  "          region_ID    (= region/gene name)\n",
                  "          rwidth       (= region/gene width)\n",
                  "          nbrsites     (= number of CpGhsm sites in the region/gene)\n",
                  "          nbrper10kb   (= number of CpGhsm sites in the region/gene normalized to 10kb width)\n",
                  "          pmrpersite   (= average % methylation per CpGhsm site in the region/gene)\n",
                  "          pmrpernucl   (= average % CpGhsm methylation per nucleotide in the region/gene)\n",
                  "          prcntM       (= overall % CpGhsm methylation in the region/gene)\n\n",
                  "       The tables are sorted by prcntM (*byPrcntM* files) or by nbper10kb\n",
                  "         (*bySiteDensity* files) in descending order.\n\n",
                  "       The ranked-promoters-*.txt files provide analogous tables for promoter regions.\n",
                  "        If the promoter annotation was derived with BWASP, then these regions are simply\n",
```

```
                 "        defined here as 500 nucleotides upstream of the 5'-end of the gene (shorter if the\n",
                 "        scaffold ends before).\n\n",
                 "       The files sites-*.txt provide details for the CpGhsm sites in the respective\n",
                 "        regions.\n",
                 "       The files plot-*.pdf explore concordance of the two methylation measures.\n",
                 "       The optional parameters minrwidth, maxrwidth, and minnbrsites specify\n",
                 "       the length range and minimal number of methylation sites for regions\n",
                 "       to be included in the plot.\n\n",
                 sep="")
  readmefile <- paste(RNKdir,"0READMErnk",sep="/")
  sink(readmefile)
  cat(sprintf("%s",readme))
  sink()
  message(".. done ...")
}


if (RUNmrpr) {
#Produces the dst-*txt and mdr-*.bed files reporting on methylation-rich and -poor regions
# based on spacings between neighboring methylation sites.
#Also: 1ds-*.pdf 5ds-*.pdf (distance histograms)
#      gwr-*.txt (genes with methylation-dense regions)
#      gwp-*.txt (genes in methylation-poor regions)
#      rmp-*.txt (regions mapped to genome features)
#
  MRPRdir <- paste(outputdir,"MRPR",sep="/")
  cmd <- paste("mkdir -p",MRPRdir,sep=" ")
  message("\n.. now executing: ",cmd)
  system(cmd)
  ddset <- c(1,5)
  nbrxtrms <- 100L
  doplots <- TRUE
  message("\n.. methylation-rich and -poor regions for aggregate samples ...")
  hsmrList <- lapply(c(1:length(sampleLabels)),function(i) {
    outfile  <- paste("dst",sampleLabels[i],sep="-")
    outfile  <- paste(outfile,"txt",sep=".")
    hsmrL <- det_mrpr(studymk[[i]],sampleLabels[i],ddset,nbrxtrms,outfile,doplots)
    cmd <- paste("mv ",outfile," m*r-",sampleLabels[i],".* ",MRPRdir,sep="")
    message("\n.. now executing: ",cmd)
    system(cmd)
    if (doplots) {
# The following cmd will get rid of empty first pages in the PDF files on a
# Fedora Linux system.  Can be omitted or replaced as needed:
      cmd <- paste("for fn in 1ds-",sampleLabels[i],".pdf 5ds-",sampleLabels[i],".pdf; do if [ `pdfinfo $fn | grep Pages | awk '{print $2}'` -gt 1 ]; then pdfseparate -f 2 $fn
tmp-$fn; \\mv tmp-$fn $fn; fi; mv $fn ",MRPRdir,"; done",sep="")
      message("\n.. now executing: ",cmd)
      system(cmd)
      }
    return(hsmrL)
    })
  message(".. done ...")

  message("\n.. mapping of methylation-rich and -poor regions to genome features ...")
  if (!exists("genome_ann")) {
    genome_ann <- get_genome_annotation(infiles$parameters)
  }
  gwrList <- lapply(c(1:length(sampleLabels)),function(i) {
     outfile   <- paste("rmp",sampleLabels[i],sep="-")
     outfile   <- paste(outfile,"txt",sep=".")
     gwithrdfL <- map_mrpr(hsmrList[[i]],species=species,sampleLabels[i],
                           genome_ann,gnmsize,UTRflag=UTRFLAGSET,outfile)
     cmd <- paste("mv ",outfile," gw*-",sampleLabels[i],".* ",MRPRdir,sep="")
     message("\n.. now executing: ",cmd)
     system(cmd)
```

```
    })
  readme <- paste("MRPR - Methylation-rich and -poor regions.\n\n",
                  "Input: studymk, genome_ann (a list of GRanges objects providing annotated region labels and bounds;\n",
                  "                            output of BWASPR::get_genome_annotation())\n",
                  "       nbrxtrms (specifies the number of richest and poorest regions to be displayed; default: 100)\n\n",
                  "Output: files dst-*.txt 1ds-*.pdf 5ds-*.pdf   mdr-*.bed   rmp-* gwp-* gwr-*\n\n",
                  "Notes: The output is generated by BWASPR functions det_mrpr() and map_mrpr().\n",
                  "       Methylation-rich and -poor regions are determined based on the spacing between\n",
                  "        neighboring CpGhsm sites.  If sites occur at positions a, b, c, d, e, and f\n",
                  "        (and not in between), then b-a, c-b, ... are 1-distances, and f-a is a 5-distance.\n",
                  "        The file dst-*.txt shows the empirical distribution of d-distances for the sample.\n",
                  "        Methylation-poor regions are determined as long 1-distances (in the top <nbrxtrms>; merged\n",
                  "        if adjacent 1-distances are both in the top <nbrxtrms>).  Methylation-rich regions are determined\n",
                  "        as short 5-distances (in the low <nbrxtrms>; merged if adjacent 5-distances are both in the low\n",
                  "        <nbrxtrms>).  Parts of the distribution are plotted in the 1ds-*.pdf and 5ds-*.pdf files.\n\n",
                  "       The methylation-rich and -poor regions are listed in files dst-*.txt and mdr-*.bed,\n",
                  "        the latter in BED format for potential display in genome browsers.\n\n",
                  "       The overlap of methylation regions with genome features is summarized in files rmp-*.txt.\n",
                  "        Files gwr-*.txt show genes overlapping with methylation-rich regions, ordered by site\n",
                  "        density in the methylation-rich region.\n",
                  "        Files gwp-*.txt show genes overlapping with methylation-poor regions, ordered by site\n",
                  "        density in the methylation-poor region.\n",
                  sep="")
  readmefile <- paste(MRPRdir,"0READMEmrpr",sep="/")
  sink(readmefile)
  cat(sprintf("%s",readme))
  sink()
  message(".. done ...")
}
################################################################################



### Section: Evaluating differential methylation patterns.
##
#
if (RUNdmt) {
#Determine differentially methylated sites and genes.
#  Output will be in dmt-* and dmg-* files.
#
  DMTdir  <- paste(outputdir,"DMT",sep="/")
  cmd <- paste("mkdir -p",DMTdir,sep=" ")
  message("\n.. now executing: ",cmd)
  system(cmd)
  message("\n.. determining differentially methylated tiles and genes ...")
  if (!exists("genome_ann")) {
    genome_ann <- get_genome_annotation(infiles$parameters)
  }

  outfile1 <- paste("dmt",speciesstudy,sep="-")
  outfile1 <- paste(outfile1,"txt",sep=".")
  outfile2 <- paste("dmg",speciesstudy,sep="-")
  outfile2 <- paste(outfile2,"txt",sep=".")
  mtList   <- det_dmt(studymc,genome_ann,wsize=wsize,stepsize=stepsize,
                      threshold,qvalue,mc.cores=numprc,
                      destrand,outfile1,outfile2)
  cmd <- paste("mv",outfile1,outfile2,DMTdir,sep=" ")
  message("\n.. now executing: ",cmd)
  system(cmd)
  readme <- paste("DMT - Differentially methylated tiles.\n\n",
                  "Input: studymc wsize stepsize threshold qvalue\n",
                  "Output: files dmt-*.txt dmg-*.txt\n\n",
                  "Notes: Output files dmt-*.txt and dmg-*.txt are generated by BWASPR::det_dmt() and show the\n",
                  "        differentially methylated tiles and genes as determined by methylKit::getMethylDiff with\n",
```

```
                  "        parameters difference=threshold and qvalue, as provided in the ",configfile,"\n",
                  "        configuration file.  Tiles refer to sliding windows along the genome within which\n",
                  "        methylation calls are cumulated by methylKit::tileMethylCounts.\n",
                  "        The dmg-*.txt files show all genes with at least one differentially methylated tile.\n\n",
                  "        A positive meth.diff value in comparison A.vs.B means that the B methylation percentage\n",
                  "        in that tile is higher than the A methylation percentage.\n\n",
                  sep="")
  readmefile <- paste(DMTdir,"0READMEdmt",sep="/")
  sink(readmefile)
  cat(sprintf("%s",readme))
  sink()
  message(".. done ...")
}


if (RUNdmsg) {
#Determine differentially methylated sites and genes.
#  Output will be in dms-* and dmg-* files.
#
  DMSGdir <- paste(outputdir,"DMSG",sep="/")
  cmd <- paste("mkdir -p",DMSGdir,sep=" ")
  message("\n.. now executing: ",cmd)
  system(cmd)
  message("\n.. determining differentially methylated sites and genes ...")
  if (!exists("genome_ann")) {
    genome_ann <- get_genome_annotation(infiles$parameters)
  }
  studyhc <- mcalls2mkobj(infiles$datafiles,species,study,
                          sample=samplelist,replicate=c(0),type=ctype,
                          mincov=highcoverage,assembly=asmblv)

  outfile1 <- paste("dms",speciesstudy,sep="-")
  outfile1 <- paste(outfile1,"txt",sep=".")
  outfile2 <- paste("dmg",speciesstudy,sep="-")
  outfile2 <- paste(outfile2,"txt",sep=".")
  dmsgList <- det_dmsg(studyhc,genome_ann,threshold,qvalue,mc.cores=numprc,
                       destrand,outfile1,outfile2)
  cmd <- paste("mv",outfile1,outfile2,DMSGdir,sep=" ")
  message("\n.. now executing: ",cmd)
  system(cmd)
  readme <- paste("DMSG - Differentially methylated sites and genes.\n\n",
                  "Input: studyhc threshold qvalue\n",
                  "       The studyhc methylKit raw object contains all CpGscd sites with coverage at least\n",
                  "        ",highcoverage," reads.\n",
                  "Output: files dms-*.txt dmg-*.txt dmg-*details.txt dmg-*heatmaps.pdf\n\n",
                  "Notes: Output files dms-*.txt and dmg-*.txt are generated by BWASPR::det_dmsg() and show the\n",
                  "        differentially methylated sites and genes as determined by methylKit::getMethylDiff with\n",
                  "        parameters difference=threshold and qvalue, as provided in the ",configfile,"\n",
                  "        configuration file.  The table of differentially methylated genes contains all genes\n",
                  "        with at least one differentially methylated site.\n\n",
                  "       The files dmg-*details.txt, generated by BWASPR::show_dmsg(), show all CpGscd sites in\n",
                  "        the differentially methylated genes, with coverage numbers and methylation percentages.\n\n",
                  "       The files dmg-*heatmaps.pdf, generated by BWASPR::show_dmsg(), give heatmap displays for\n",
                  "        genes meeting the following criteria: 1) there are between minNsites and maxNsites common\n",
                  "        CpGscd sites; 2) at least minPdmsites % of these sites are differentially methylated sites.\n",
                  "        The parameters are set in the ",configfile," configuration file.\n\n",
                  sep="")
  readmefile <- paste(DMSGdir,"0READMEdmsg",sep="/")
  sink(readmefile)
  cat(sprintf("%s",readme))
  sink()
  message(".. done ...")
}
```

```
if (RUNdmgdtls) {
#Provide details for differentially methylated genes.
#  Output will be in file dmg-*details.txt.
#
  if (!RUNload & !RUNdmsg) {
    message("ERROR: missing input.  Set RUNload=TRUE or RUNdmsg=TRUE.")
    q()
  }
  outflabel <- paste(species,studyLabel,sep="_")
  message("\n.. deriving details for differentially methylated genes ...")
  dmgprp <- show_dmsg(studyhc,dmsgList,destrand,minNsites,maxNsites,minPdmsites,
                      mc.cores=numprc,outflabel)
  cmd <- paste("mv dmg-",outflabel,"_*.vs.* ",DMSGdir,sep="")
  message("\n.. now executing: ",cmd)
  system(cmd)
  message(".. done ...")
}


if (RUNogl) {
#Print out ordered gene lists with tabulated methylation properties (files ogl-*).
#
  if (!RUNload & !(RUNdmsg & RUNdmgdtls)) {
    message("ERROR: missing input.  Set RUNload=TRUE or RUNdmsg=RUNdmgdtls=TRUE.")
    q()
  }
  OGLalldir <- paste(outputdir,"OGLall",sep="/")
  cmd <- paste("mkdir -p",OGLalldir,sep=" ")
  message("\n.. now executing: ",cmd)
  system(cmd)
  outflabel <- paste(species,studyLabel,sep="_")
  message("\n.. printing ordered gene lists ...")
  if (maxgwidth > 0  |  minnbrdmsites > 1) {
    summaries <- explore_dmsg(studyhc,genome_ann,dmgprp,-1,
                              1,withglink=glink,outflabel)
    rnk_summaries <- rank_dmg(summaries,outflabel)
    cmd <- paste("mv ogl-",speciesstudy,"_* rnk-dmg-",speciesstudy,"*.vs.* wrt-",speciesstudy,".txt ",OGLalldir,sep="")
    message("\n.. now executing: ",cmd)
    system(cmd)
  }

  summaries <- explore_dmsg(studyhc,genome_ann,dmgprp,maxgwidth,
                            minnbrdmsites,withglink=glink,outflabel)
  rnk_summaries <- rank_dmg(summaries,outflabel)
  OGLdir <- paste(outputdir,"OGL",sep="/")
  cmd <- paste("mkdir -p ",OGLdir,"; mv ogl-",outflabel,"_* rnk-dmg-",outflabel,"*.vs.* wrt-",outflabel,".txt ",OGLdir,sep="")
  message("\n.. now executing: ",cmd)
  system(cmd)

  readme <- paste("OGL - Ordered gene lists.\n\n",
                  "Input: studyhc annotation   maxgwidth minnbrdmsites\n",
                  "       dmgprp (= output of BWASPR:show_dmsg())\n\n",
                  "Output: files ogl-*.txt rnk-dmg-*.txt\n",
                  "         ogl-<study>_<sample>.txt ogl-<study>_<sample1>.vs.<sample2>.txt\n",
                  "         rnk-dmg--<study>_<sample1>.vs.<sample2>.txt rnk-dmg--<study>_<sample1>.vs.<sample2>.pdf\n",
                  "         wrt-<study>.txt\n\n",
                  "Notes: Output files ogl-<study>_<sample>.txt give tables for each sample with columns\n\n",
                  "        gene_ID gwidth #Sites #per10Kb %perSite %pNucl\n\n",
                  "       ordered by %pNucl.  If available, a link to an NCBI entry of gene_ID is inserted as second column.\n",
                  "       Abbreviations used: %perSite, percent methylation per site\n",
                  "                           %perNucl, percent methylation per nucleotide of the gene\n\n",
                  "       Output files ogl-<study>_<sample1>.vs.<sample2>.txt give tables for each comparison with columns\n\n",
                  "        gene_ID gwidth #Sites #per10Kb #dmSites #dmsp10kb %dmSites %pSite1 %pSite2 DMpSite ADMpSite DMpNucl ADMpNucl\n\n",
```

```
                  "       ordered by DMpSite.  If available, a link to an NCBI entry of gene_ID is inserted as second column.\n",
                  "       Abbreviations used: %dmSites, percent sites that are differentially methylated\n",
                  "                           %pSite1, average per site % methylation for sample1\n",
                  "                           %pSite2, average per site % methylation for sample2\n",
                  "                           DMpSite, average per site difference in % methylation between sample1 and sample2\n",
                  "                           ADMpSite, absolute value of DMpSite\n",
                  "                           DMpNucl, average per nucleotide difference in % methylation between sample1 and sample2\n",
                  "                           ADMpNucl, absolute value of DMpNucl\n\n",
                  "       Output files rnk-dmg-<study>_<sample1>.vs.<sample2>.txt are equivalent to files\n",
                  "        ogl-<study>_<sample1>.vs.<sample2>.txt but ordered by ADMpNucl.\n",
                  "       Output files rnk-dmg-<study>_<sample1>.vs.<sample2>.pdf provide visualization of the distribution\n",
                  "        of ADMpNucl values.\n\n",
                  "       Output file wrt-<study>.txt gives results of the Wilcoxon signed rank test comparing\n",
                  "        the %pSite1 and %pSite2 vectors\n\n",
                  "       In output directory OGL, tables are restricted to genes with gwidth <= maxgwidth and, for\n",
                  "        pairwise comparisons, #dmSites >= minnbrdmsites\n\n",
                  "       If either maxgwidth or minnbrdmsites is set, then output directory OGLall will show the full tables\n",
                  "        with all genes (for reference).\n\n",
                  sep="")
if (maxgwidth > 0  |  minnbrdmsites > 1) {
  readmefile <- paste(OGLalldir,"0READMEogl",sep="/")
  sink(readmefile)
  cat(sprintf("%s",readme))
  sink()
}
  readmefile <- paste(OGLdir,"0READMEogl",sep="/")
  sink(readmefile)
  cat(sprintf("%s",readme))
  sink()
  message(".. done ...")
}
################################################################################



if (RUNsave) {
# Save everything ... (for later restart with the load command).
#
  outflabel <- paste(species,studyLabel,sep="_")
  rdatafile <- paste(outflabel,"RData",sep=".")
  save.image(file=rdatafile)
}
```

```
0README


Rscript.BWASPR output for .conf file Pc_PA.conf.

Data input are the specified *.mcalls files and parameters set in Pc_PA.conf.
The *.mcalls input data are saved in the following data structures:

studymk - a methylKit methylRaw(List) object storing the CpGhsm site data for all samples
studymc - a methylKit methylRaw(List) object storing the CpGscd site data for all samples
studyhc - a methylKit methylRaw(List) object storing the CpGscd site data for all samples,
           restricted to sites with high coverage (here set to 20)
mkrd    - a methylKit methylRawList object storing the CpGhsm site data for all
           replicates of a given sample

Output of the analysis is stored in the following subdirectories (if the corresponding
 function calls were specified in Pc_PA.conf; please consult the 0README
 files in the subdirectories for details on the output files):

CMS - Coverage and methylation statistics for aggregate samples
PWC - Pairwise comparisons between all samples
CRL - Correlations between aggregate samples
REPCMS - Coverage and methylation statistics for replicate samples
REPCRL - Correlations between replicates
MMP - Mapping of methylation sites on genome annotation
ACS - Annotation of conserved methylation sites
RNK - Genomic feature regions ranked by CpGhsm statistics
MRPR - Methylation-rich and -poor regions
DMSG - Differentially methylateds sites and genes
OGL - Ordered gene lists
```

```
Directory: CMS File: 0READMEcms


CMS - Coverage and methylation statistics for aggregate samples.

Input: studymk, covlist, locount, hicount

Output: files cms-*.txt and cms-*.pdf

Notes: The statistics are given for CpGhsm sites restricted to minimum or
        higher level coverage.  At the minimum level, the number of sites will be
        the number of lines of the corresponding *CpGhsm.mcalls file minus one
        (consistency check).  The distribution of methylation levels is really only
        interesting for high coverage because at low coverage there will be a strong
        bias towards high methylation levels (by definition of hsm sites).
        The same bias at high coverage would suggest that the sample includes a
        preponderance of consistently methylated genomic sites.

        locount and hicount set bounds on the coverage to exclude sites with too few
        or too many covering reads to provide statistics on a typical range.
```

```
Directory: CMS File: cms-Pc_PA_queen.txt


Number of "queen" CpGhsm-sites with minimal and higher level coverage:

  number of "queen" CpGhsm-sites with coverage >=  4:  13840
  number of "queen" CpGhsm-sites with coverage >=  6:  13635
  number of "queen" CpGhsm-sites with coverage >= 10:  12768
  number of "queen" CpGhsm-sites with coverage >= 15:  11607


Coverage and methylation statistics for "queen" CpGhsm-sites at different levels of minimum coverage:


methylKit::getCoverageStats output for "queen" CpGhsm-sites (#: 13840) at minimum coverage 4   - read coverage statistics per base
summary:
   Min. 1st Qu.  Median    Mean 3rd Qu.    Max. 
      4      20      45     486     104   28934 
percentiles:
   0%   10%   20%   30%   40%   50%   60%   70%   80%   90%   95%   99% 99.5% 99.9%  100% 
    4    11    17    24    33    45    60    86   127   451  1822 13486 17555 25031 28934 

methylKit::getMethylationStats output for "queen" CpGhsm-sites (#: 13840) at minimum coverage 4      - methylation statistics per base
summary:
   Min. 1st Qu.  Median    Mean 3rd Qu.    Max. 
    0.8    13.9    31.8    37.4    57.1   100.0 
percentiles:
     0%     10%     20%     30%     40%     50%     60%     70%     80%     90%     95%     99%   99.5%   99.9%    100% 
  0.819   5.348  11.111  16.981  23.913  31.818  41.667  52.174  62.857  77.778  87.500 100.000 100.000 100.000 100.000 



methylKit::getCoverageStats output for "queen" CpGhsm-sites (#: 13635) at minimum coverage 6   - read coverage statistics per base
summary:
   Min. 1st Qu.  Median    Mean 3rd Qu.    Max. 
      6      21      45     493     106   28934 
percentiles:
     0%     10%     20%     30%     40%     50%     60%     70%     80%     90%     95%     99%   99.5%   99.9%    100% 
    6.0    11.0    18.0    24.2    34.0    45.0    61.0    87.0   129.0   469.6  1862.0 13576.6 17681.0 25103.2 28934.0 

methylKit::getMethylationStats output for "queen" CpGhsm-sites (#: 13635) at minimum coverage 6      - methylation statistics per base
summary:
   Min. 1st Qu.  Median    Mean 3rd Qu.    Max. 
    0.8    13.7    31.1    36.4    55.6   100.0 
percentiles:
     0%     10%     20%     30%     40%     50%     60%     70%     80%     90%     95%     99%   99.5%   99.9%    100% 
  0.819   5.211  10.976  16.667  23.333  31.111  40.856  50.000  61.912  75.000  85.000 100.000 100.000 100.000 100.000 



methylKit::getCoverageStats output for "queen" CpGhsm-sites (#: 12768) at minimum coverage 10  - read coverage statistics per base
summary:
   Min. 1st Qu.  Median    Mean 3rd Qu.    Max. 
     10      24      50     526     112   28934 
percentiles:
   0%   10%   20%   30%   40%   50%   60%   70%   80%   90%   95%   99% 99.5% 99.9%  100% 
   10    15    21    28    38    50    67    93   138   549  2042 13959 17944 25334 28934 

methylKit::getMethylationStats output for "queen" CpGhsm-sites (#: 12768) at minimum coverage 10     - methylation statistics per base
summary:
   Min. 1st Qu.  Median    Mean 3rd Qu.    Max. 
    0.8    12.9    28.6    33.6    50.0   100.0 
percentiles:
     0%     10%     20%     30%     40%     50%     60%     70%     80%     90%     95%     99%   99.5%   99.9%    100% 
```

```
  0.819   4.762  10.377  15.385  21.519  28.571  36.842  46.512  56.250  70.000  80.000  92.593  95.141 100.000 100.000
```

```
methylKit::getCoverageStats output for "queen" CpGhsm-sites (#: 11607) at minimum coverage 15  - read coverage statistics per base
summary:
   Min. 1st Qu.  Median    Mean 3rd Qu.    Max.
     15      30      57     577     122   28934
percentiles:
   0%   10%   20%   30%   40%   50%   60%   70%   80%   90%   95%   99% 99.5% 99.9%  100%
   15    20    26    34    44    57    76   103   154   658  2406 14467 18452 25387 28934

methylKit::getMethylationStats output for "queen" CpGhsm-sites (#: 11607) at minimum coverage 15     - methylation statistics per base
summary:
   Min. 1st Qu.  Median    Mean 3rd Qu.    Max.
    0.8    11.6    25.5    30.4    45.0   100.0
percentiles:
     0%     10%     20%     30%     40%     50%     60%     70%     80%     90%     95%     99%   99.5%   99.9%    100%
  0.819   4.281   9.687  13.924  19.298  25.455  32.000  40.000  50.000  65.000  75.542  89.474  93.750  96.951 100.000
```

```
methylKit::getCoverageStats output for "queen" CpGhsm-sites (#: 2665) in coverage range [100-1000]  - read coverage statistics per base
summary:
   Min. 1st Qu.  Median    Mean 3rd Qu.    Max.
    100     120     156     256     287     999
percentiles:
   0%   10%   20%   30%   40%   50%   60%   70%   80%   90%   95%   99% 99.5% 99.9%  100%
  100   108   116   125   138   156   185   240   371   609   772   966   981   995   999

methylKit::getMethylationStats output for "queen" CpGhsm-sites (#: 2665) in coverage range [100-1000]     - methylation statistics per base
summary:
   Min. 1st Qu.  Median    Mean 3rd Qu.    Max.
    2.4     6.5     9.4    14.4    14.4    92.8
percentiles:
   0%   10%   20%   30%   40%   50%   60%   70%   80%   90%   95%   99% 99.5% 99.9%  100%
 2.45  4.19  5.68  7.22  8.47  9.43 10.65 12.76 17.57 33.51 47.26 72.52 80.92 87.82 92.81
```

```
Directory: CMS File: cms-Pc_PA_worker.txt


Number of "worker" CpGhsm-sites with minimal and higher level coverage:

  number of "worker" CpGhsm-sites with coverage >=  4:  12036
  number of "worker" CpGhsm-sites with coverage >=  6:  11800
  number of "worker" CpGhsm-sites with coverage >= 10:  10941
  number of "worker" CpGhsm-sites with coverage >= 15:   9904


Coverage and methylation statistics for "worker" CpGhsm-sites at different levels of minimum coverage:


methylKit::getCoverageStats output for "worker" CpGhsm-sites (#: 12036) at minimum coverage 4  - read coverage statistics per base
summary:
   Min. 1st Qu.  Median    Mean 3rd Qu.    Max.
      4      19      43     446     102   27503
percentiles:
   0%   10%   20%   30%   40%   50%   60%   70%   80%   90%   95%   99% 99.5% 99.9%  100%
    4    10    16    22    31    43    59    85   125   326  1662 12000 15495 22905 27503

methylKit::getMethylationStats output for "worker" CpGhsm-sites (#: 12036) at minimum coverage 4     - methylation statistics per base
summary:
   Min. 1st Qu.  Median    Mean 3rd Qu.    Max.
    0.8    14.2    33.3    38.3    59.1   100.0
percentiles:
     0%     10%     20%     30%     40%     50%     60%     70%     80%     90%     95%     99%   99.5%   99.9%    100%
  0.808   6.267  11.607  17.500  25.000  33.333  43.182  53.846  64.286  77.778  87.969 100.000 100.000 100.000 100.000



methylKit::getCoverageStats output for "worker" CpGhsm-sites (#: 11800) at minimum coverage 6  - read coverage statistics per base
summary:
   Min. 1st Qu.  Median    Mean 3rd Qu.    Max.
      6      20      44     455     104   27503
percentiles:
     0%     10%     20%     30%     40%     50%     60%     70%     80%     90%     95%     99%   99.5%   99.9%    100%
    6.0    11.0    17.0    23.7    33.0    44.0    61.0    86.0   127.0   345.3  1716.5 12345.7 15694.7 22965.5 27503.0

methylKit::getMethylationStats output for "worker" CpGhsm-sites (#: 11800) at minimum coverage 6     - methylation statistics per base
summary:
   Min. 1st Qu.  Median    Mean 3rd Qu.    Max.
    0.8    14.0    32.4    37.1    57.1   100.0
percentiles:
     0%     10%     20%     30%     40%     50%     60%     70%     80%     90%     95%     99%   99.5%   99.9%    100%
  0.808   6.136  11.392  17.183  24.242  32.432  42.105  52.185  62.500  75.000  84.615 100.000 100.000 100.000 100.000



methylKit::getCoverageStats output for "worker" CpGhsm-sites (#: 10941) at minimum coverage 10 - read coverage statistics per base
summary:
   Min. 1st Qu.  Median    Mean 3rd Qu.    Max.
     10      24      50     490     112   27503
percentiles:
   0%   10%   20%   30%   40%   50%   60%   70%   80%   90%   95%   99% 99.5% 99.9%  100%
   10    15    21    28    37    50    67    93   137   425  2012 12738 16329 23214 27503

methylKit::getMethylationStats output for "worker" CpGhsm-sites (#: 10941) at minimum coverage 10    - methylation statistics per base
summary:
   Min. 1st Qu.  Median    Mean 3rd Qu.    Max.
    0.8    13.0    29.3    34.0    51.4   100.0
percentiles:
     0%     10%     20%     30%     40%     50%     60%     70%     80%     90%     95%     99%   99.5%   99.9%    100%
```

```
  0.808   5.447  10.656  15.625  21.875  29.268  37.500  46.875  56.977  70.000  80.000  92.308  95.271 100.000 100.000


methylKit::getCoverageStats output for "worker" CpGhsm-sites (#: 9904) at minimum coverage 15  - read coverage statistics per base
summary:
   Min. 1st Qu.  Median    Mean 3rd Qu.    Max. 
     15      30      57     540     122   27503 
percentiles:
   0%   10%   20%   30%   40%   50%   60%   70%   80%   90%   95%   99% 99.5% 99.9%  100% 
   15    20    26    34    44    57    76   104   151   531  2378 13106 16857 23448 27503 

methylKit::getMethylationStats output for "worker" CpGhsm-sites (#: 9904) at minimum coverage 15     - methylation statistics per base
summary:
   Min. 1st Qu.  Median    Mean 3rd Qu.    Max. 
    0.8    11.9    25.8    30.7    45.5   100.0 
percentiles:
     0%     10%     20%     30%     40%     50%     60%     70%     80%     90%     95%     99%   99.5%   99.9%    100% 
  0.808   4.798   9.804  14.044  19.298  25.806  33.333  40.541  50.000  65.000  75.000  88.889  93.333 100.000 100.000 



methylKit::getCoverageStats output for "worker" CpGhsm-sites (#: 2295) in coverage range [100-1000]  - read coverage statistics per base
summary:
   Min. 1st Qu.  Median    Mean 3rd Qu.    Max. 
    100     121     156     232     245     996 
percentiles:
   0%   10%   20%   30%   40%   50%   60%   70%   80%   90%   95%   99% 99.5% 99.9%  100% 
  100   108   116   126   139   156   175   211   303   508   671   904   952   992   996 

methylKit::getMethylationStats output for "worker" CpGhsm-sites (#: 2295) in coverage range [100-1000]     - methylation statistics per base
summary:
   Min. 1st Qu.  Median    Mean 3rd Qu.    Max. 
    2.4     7.0     9.6    15.2    15.7    89.8 
percentiles:
   0%   10%   20%   30%   40%   50%   60%   70%   80%   90%   95%   99% 99.5% 99.9%  100% 
 2.45  4.62  6.38  7.56  8.52  9.57 11.04 13.69 19.30 37.09 50.10 73.11 82.93 88.17 89.83 
```

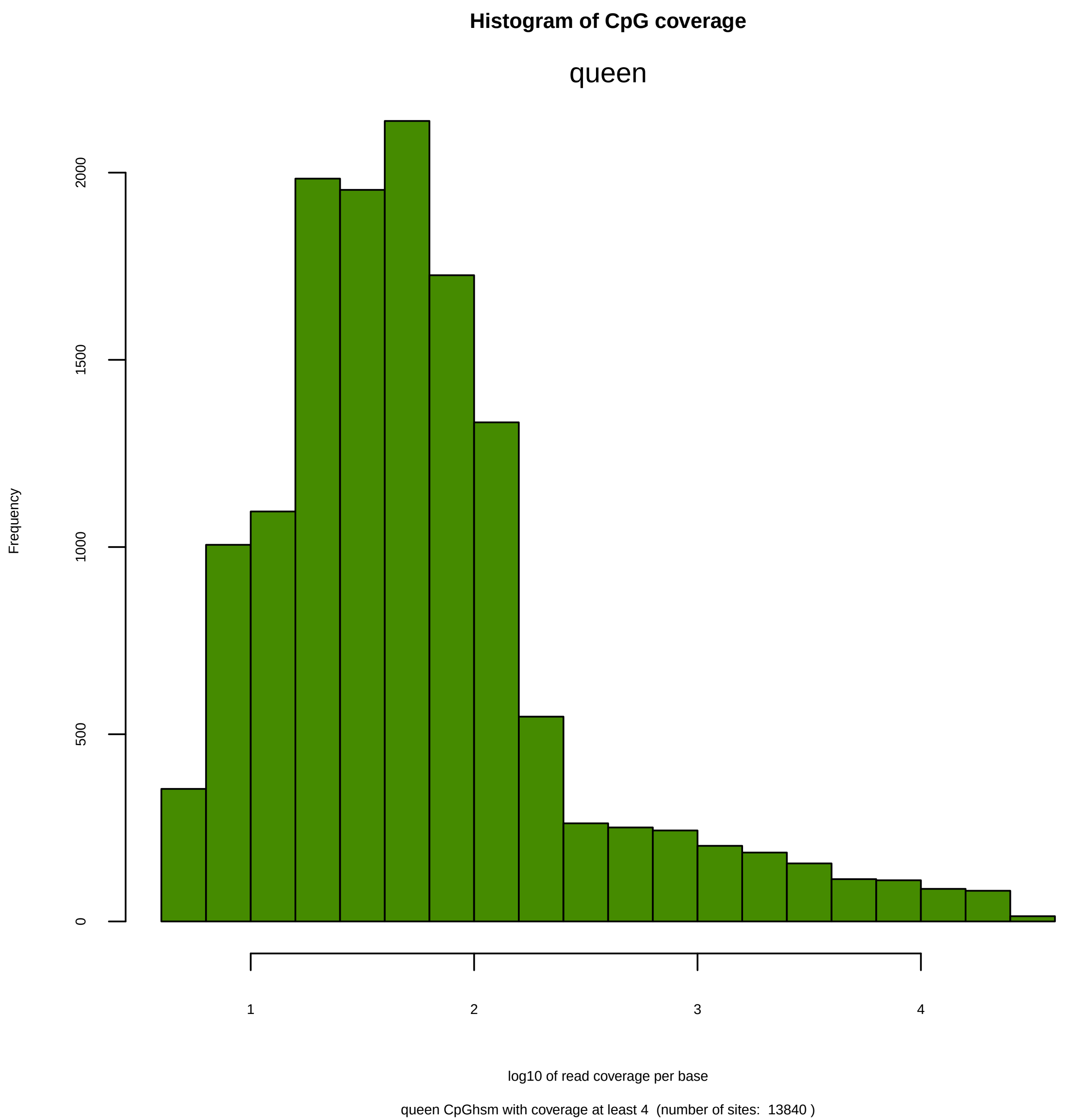
Histogram of CpG coverage
queen
Frequency
0
500
1000
1500
2000
1
2
3
4
log10 of read coverage per base
queen CpGhsm with coverage at least 4  (number of sites:  13840 )

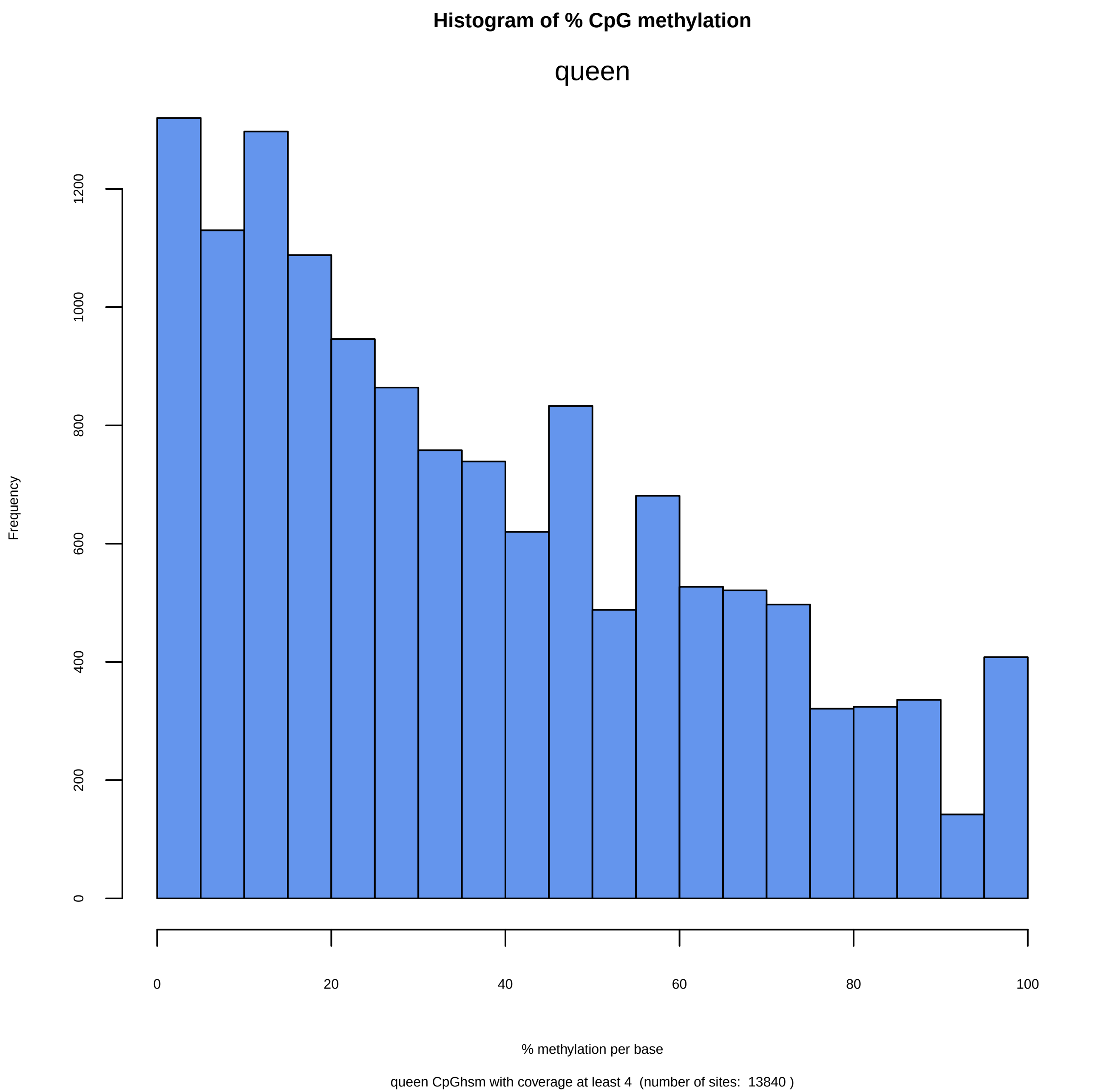
Histogram of % CpG methylation
queen
Frequency
0
200
400
600
800
1000
1200
0
20
40
60
80
100
% methylation per base
queen CpGhsm with coverage at least 4 (number of sites: 13840 )

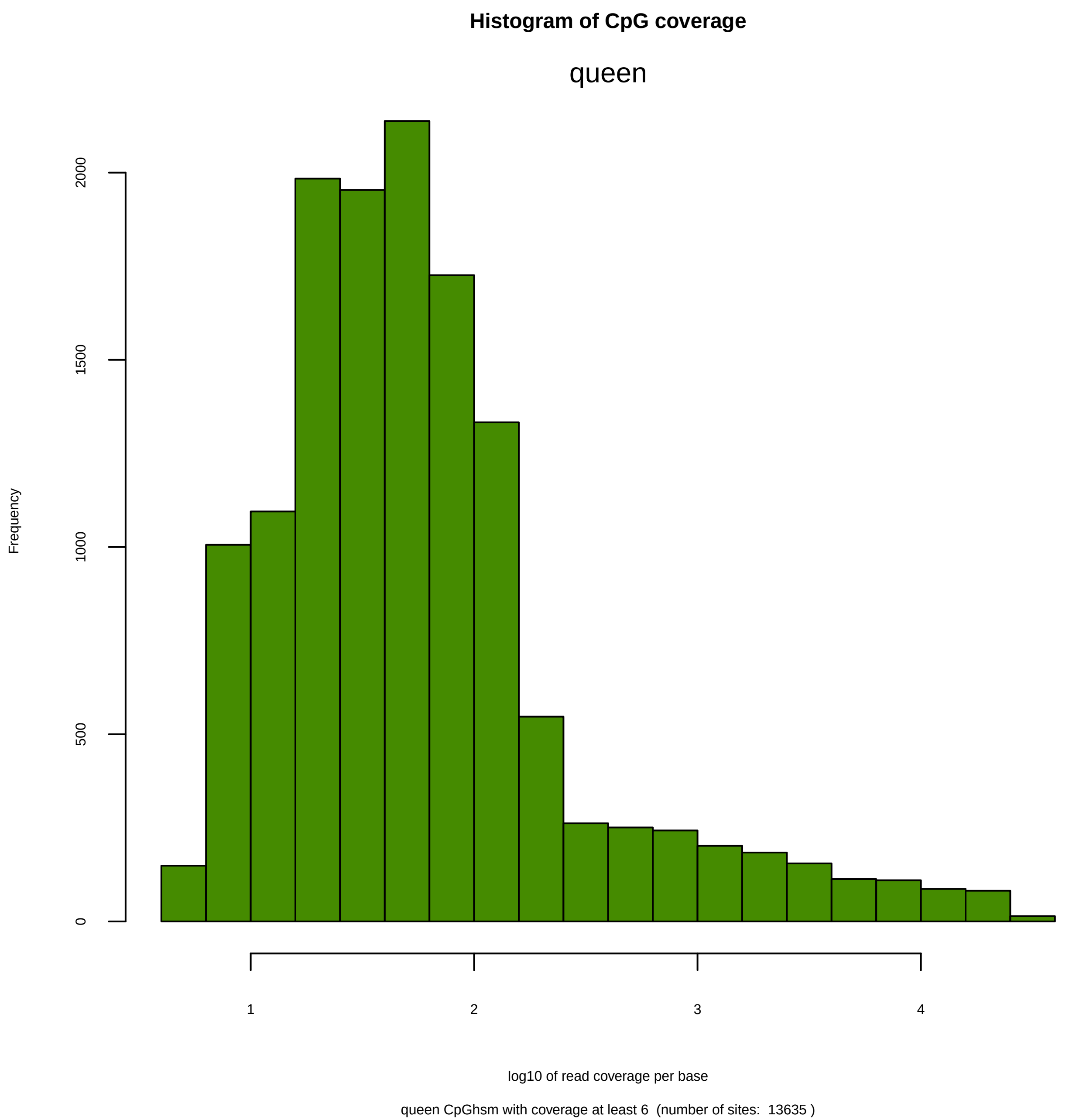

Histogram of CpG coverage
queen
Frequency
0
500
1000
1500
2000
1
2
3
4
log10 of read coverage per base
queen CpGhsm with coverage at least 6 (number of sites: 13635 )

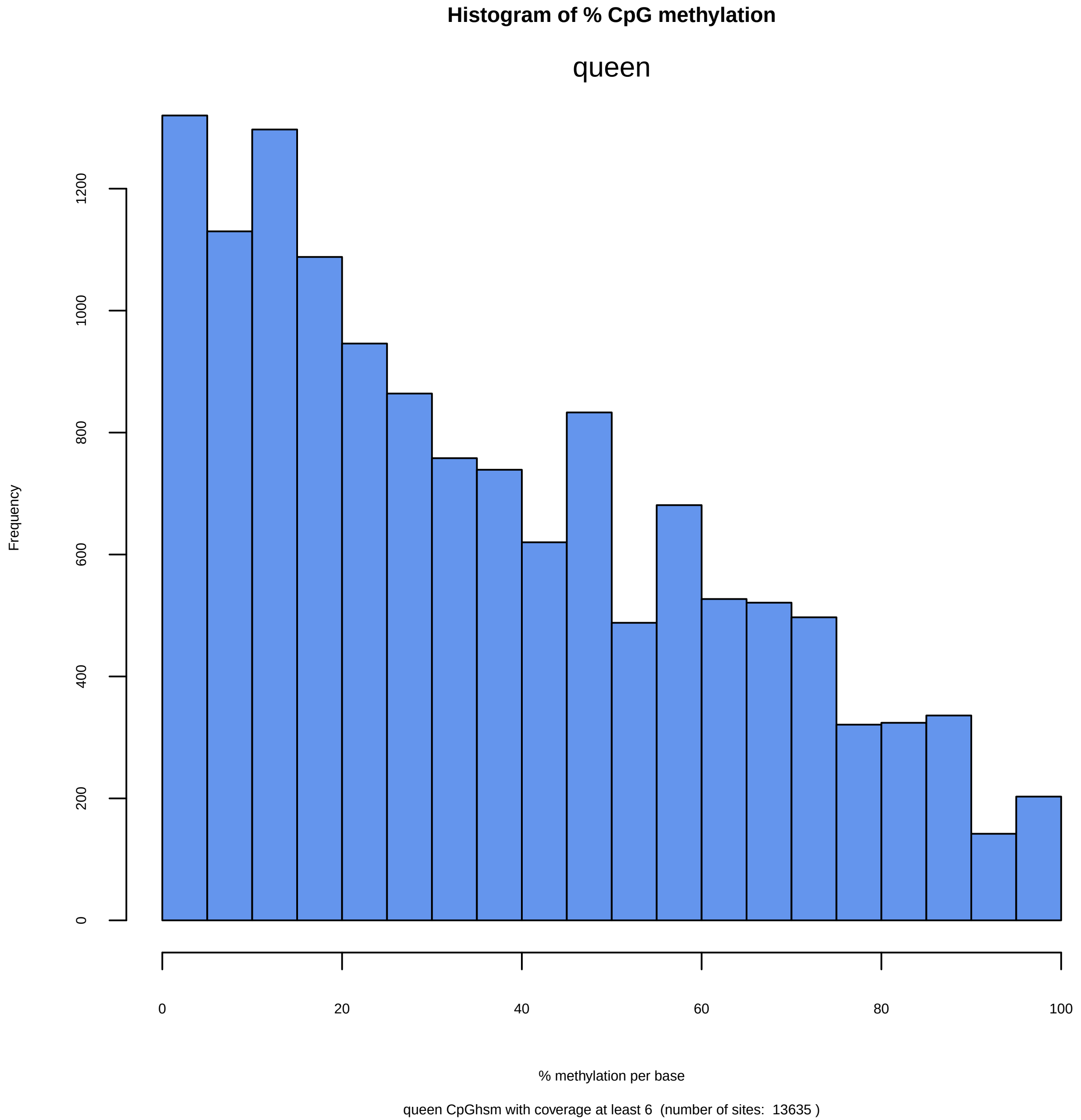
Histogram of % CpG methylation
queen
Frequency
0
200
400
600
800
1000
1200
0
20
40
60
80
100
% methylation per base
queen CpGhsm with coverage at least 6  (number of sites:  13635 )

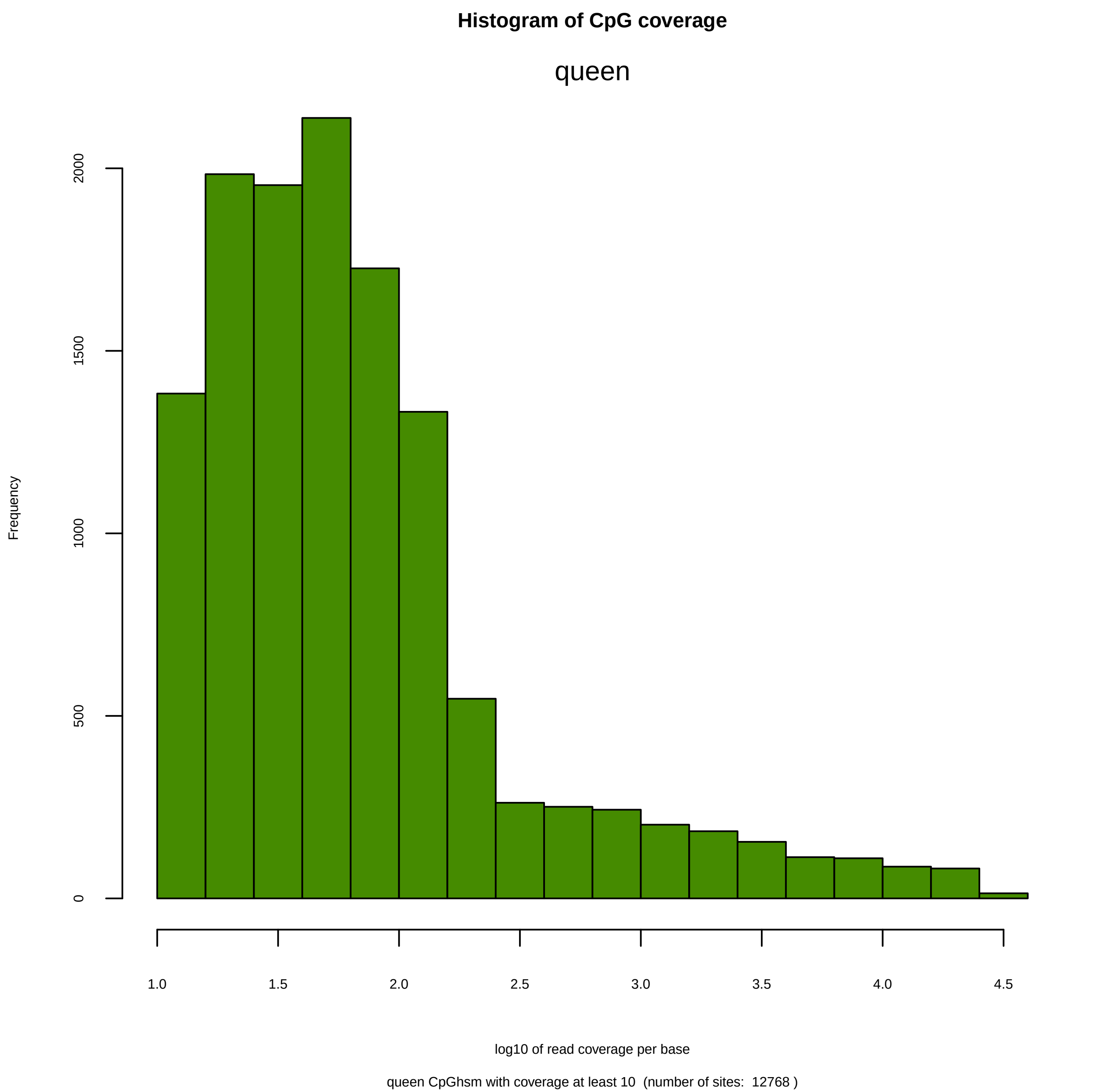

Histogram of CpG coverage
queen
Frequency
0
500
1000
1500
2000
1.0
1.5
2.0
2.5
3.0
3.5
4.0
4.5
log10 of read coverage per base
queen CpGhsm with coverage at least 10  (number of sites:  12768 )

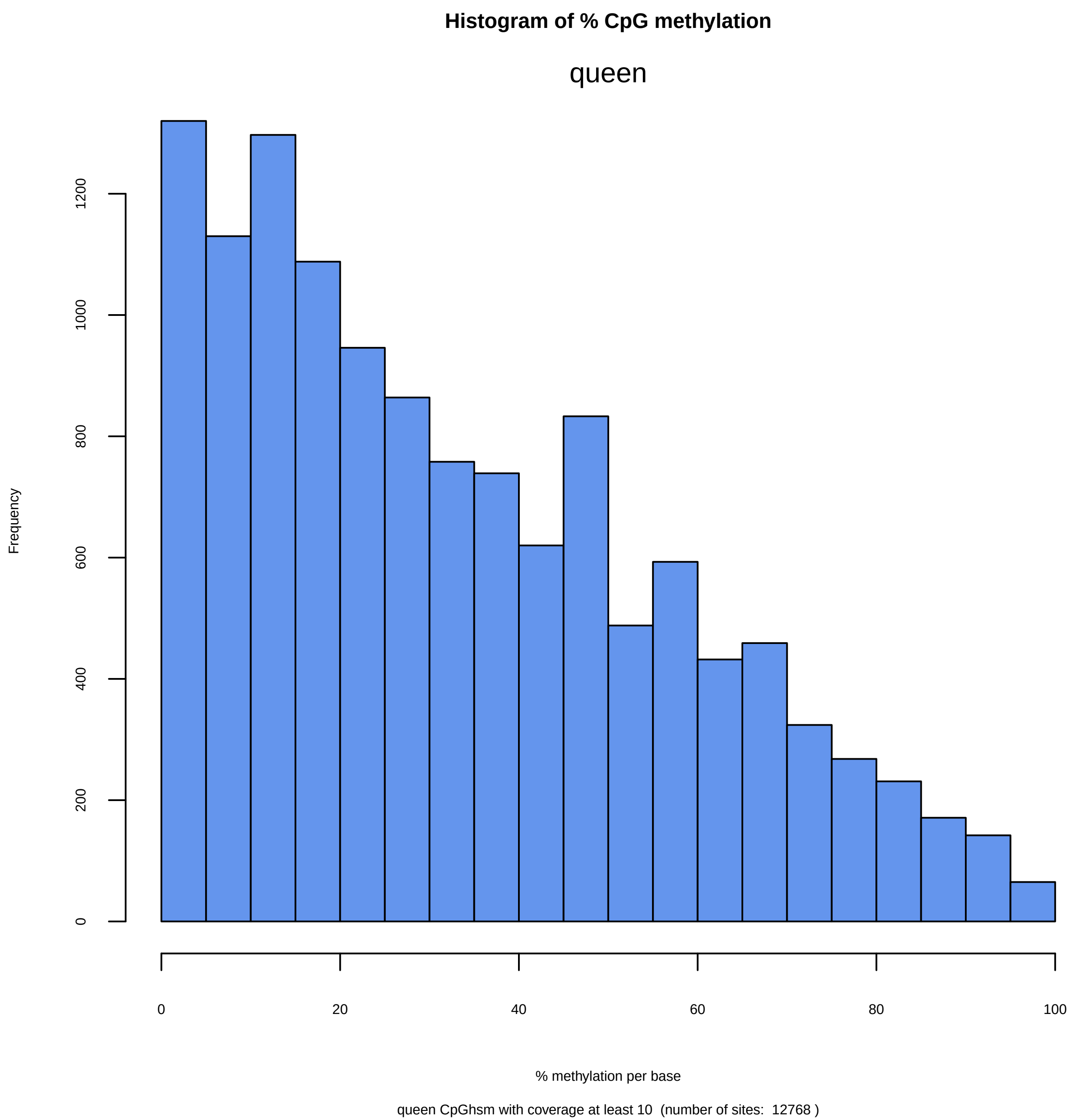

Histogram of % CpG methylation
queen
Frequency
0
200
400
600
800
1000
1200
0
20
40
60
80
100
% methylation per base
queen CpGhsm with coverage at least 10  (number of sites:  12768 )

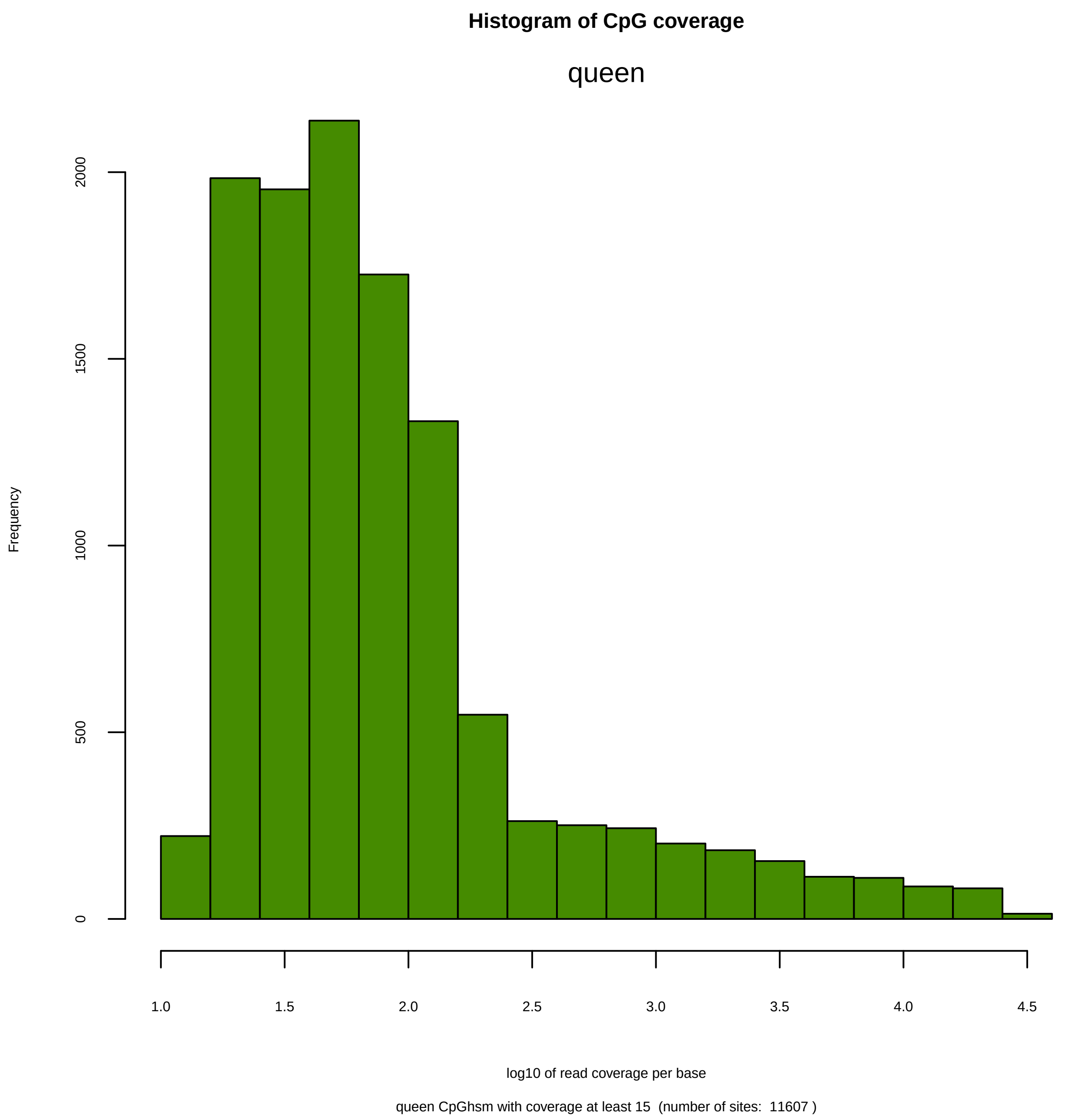

Histogram of CpG coverage
queen
Frequency
0
500
1000
1500
2000
1.0
1.5
2.0
2.5
3.0
3.5
4.0
4.5
log10 of read coverage per base
queen CpGhsm with coverage at least 15  (number of sites:  11607 )

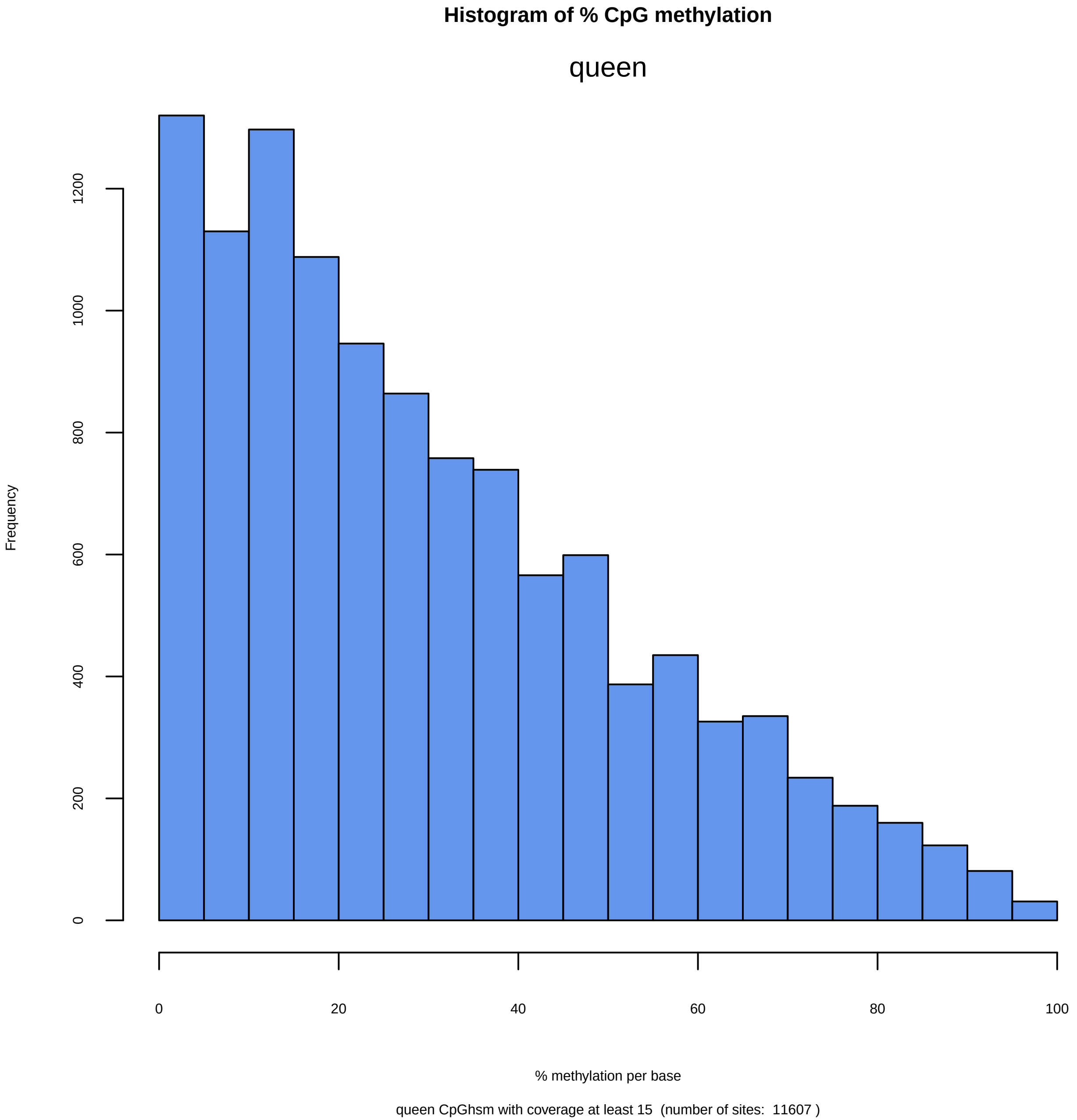
Histogram of % CpG methylation
queen
Frequency
0
200
400
600
800
1000
1200
0
20
40
60
80
100
% methylation per base
queen CpGhsm with coverage at least 15 (number of sites: 11607 )

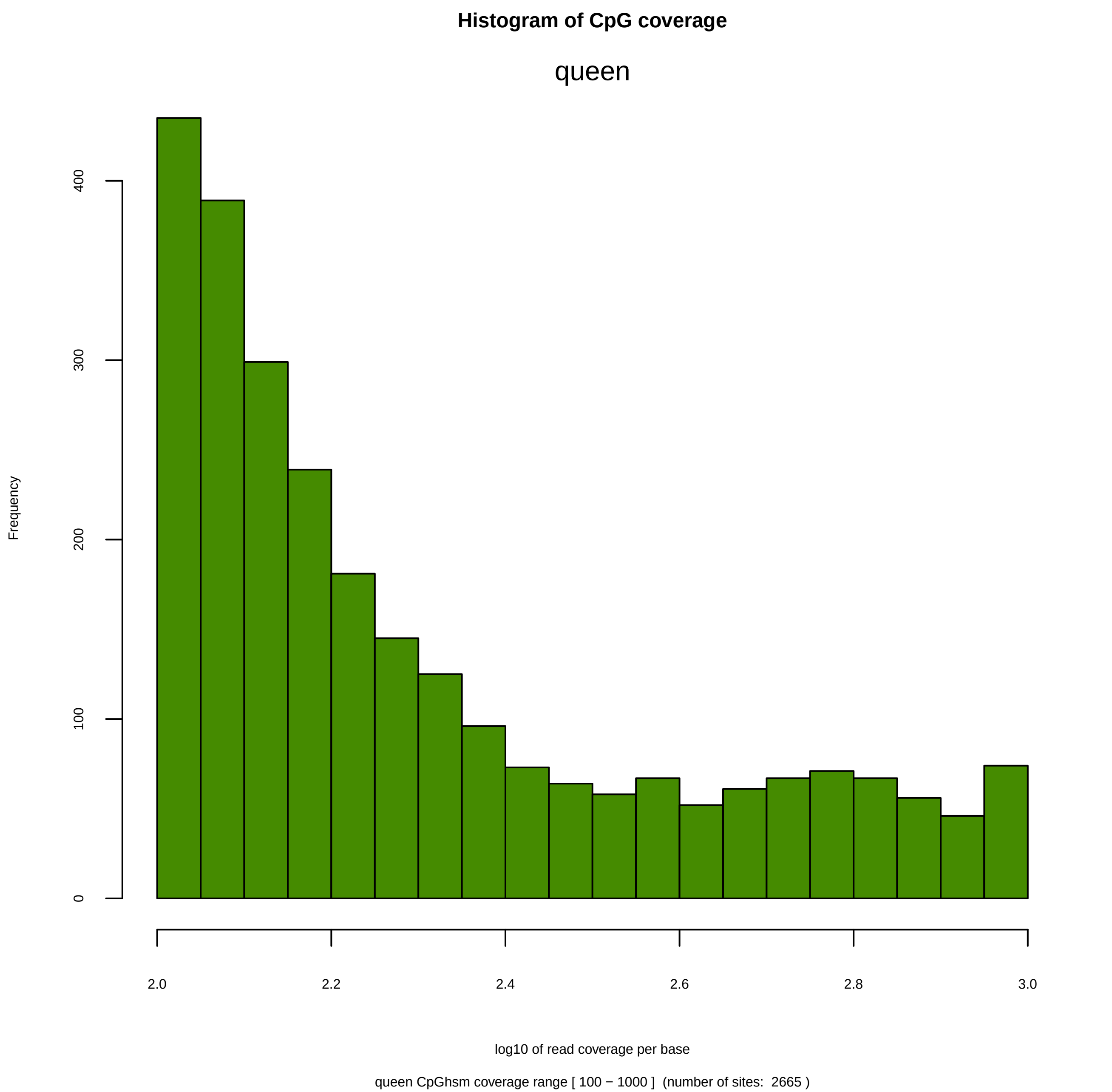
Histogram of CpG coverage
queen
Frequency
0
100
200
300
400
2.0
2.2
2.4
2.6
2.8
3.0
log10 of read coverage per base
queen CpGhsm coverage range [ 100 − 1000 ]  (number of sites:  2665 )

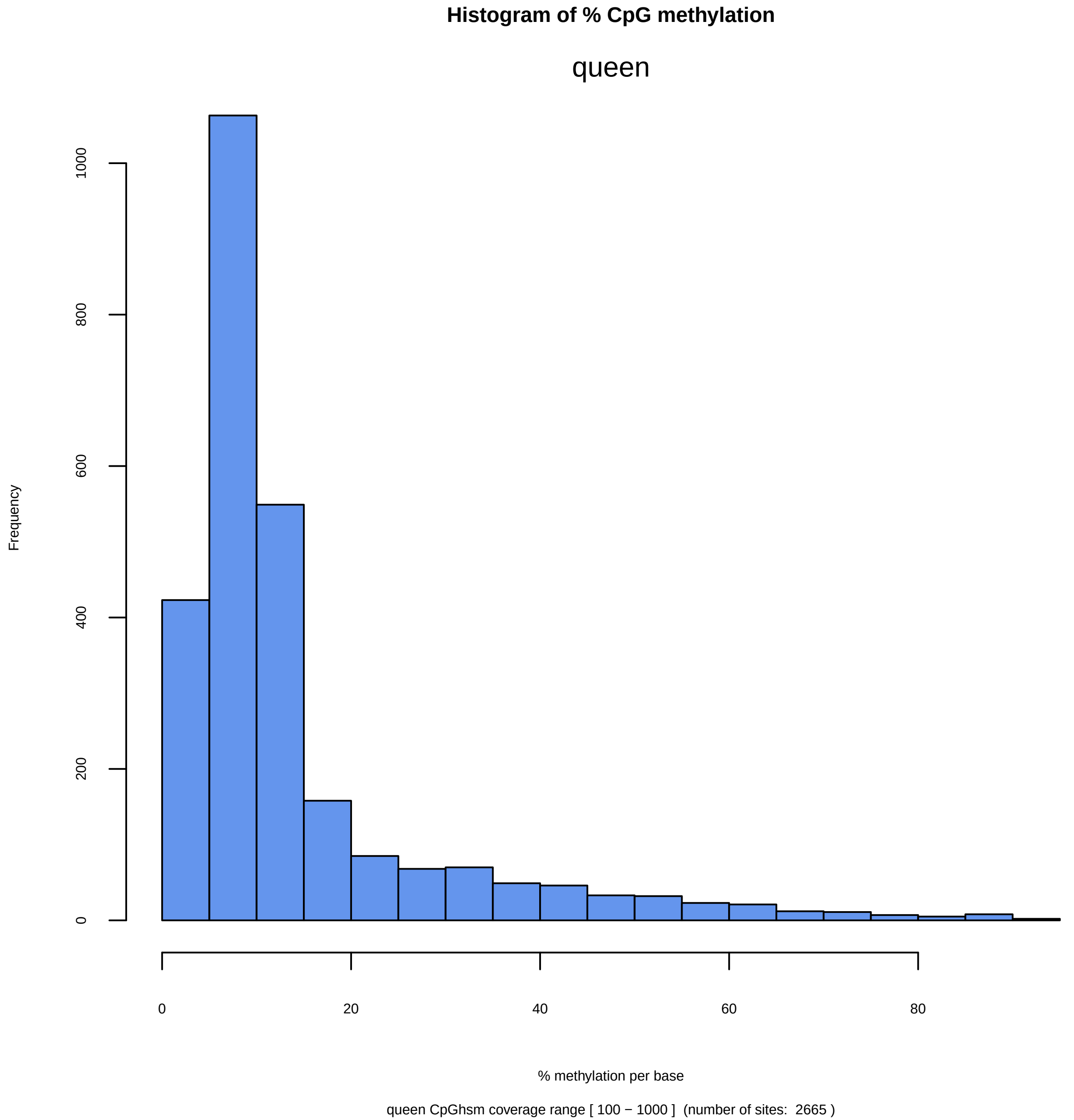
Histogram of % CpG methylation
queen
Frequency
0
200
400
600
800
1000
0
20
40
60
80
% methylation per base
queen CpGhsm coverage range [ 100 − 1000 ]  (number of sites:  2665 )

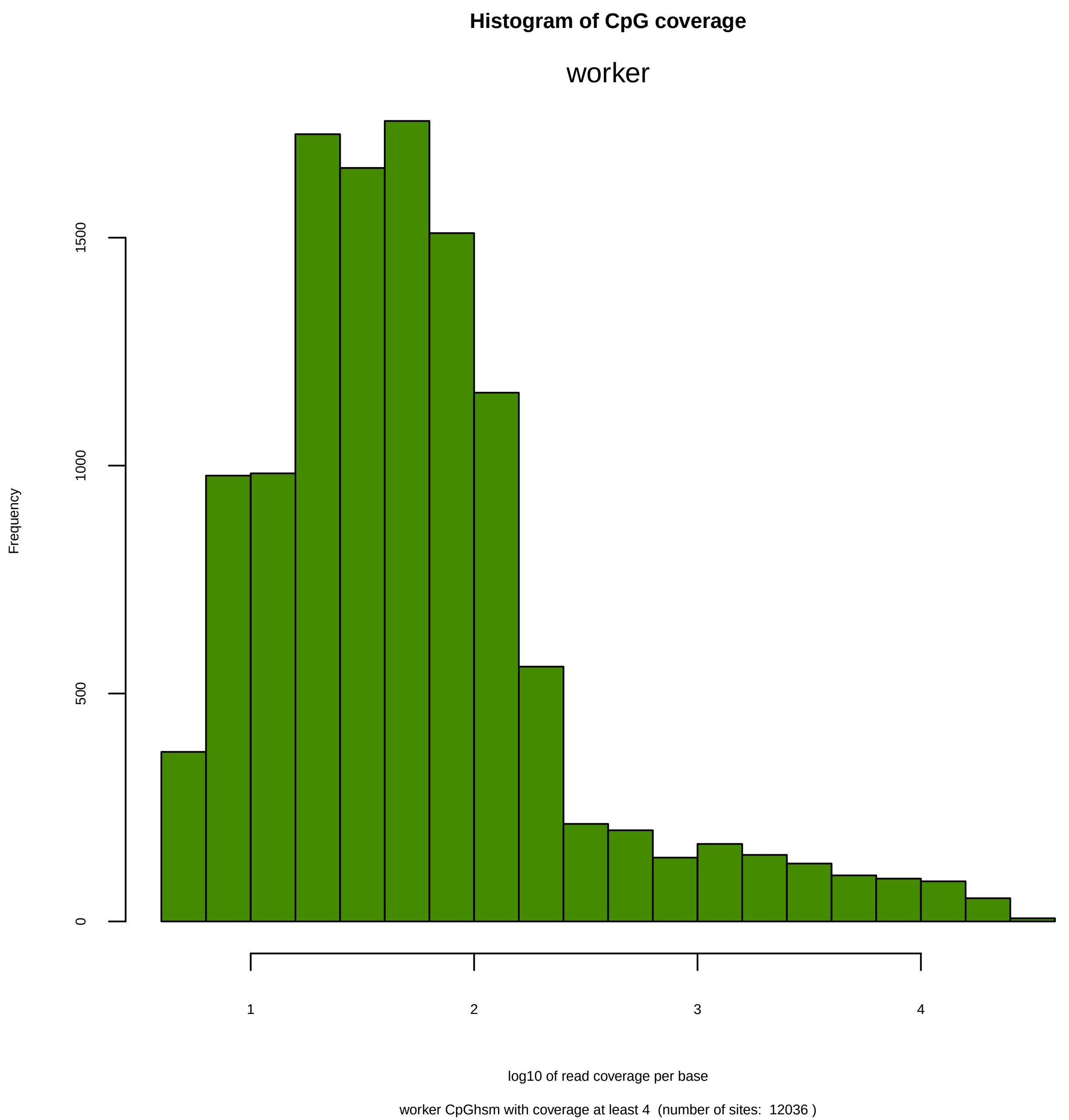
Histogram of CpG coverage
worker
Frequency
0
500
1000
1500
1
2
3
4
log10 of read coverage per base
worker CpGhsm with coverage at least 4  (number of sites:  12036 )

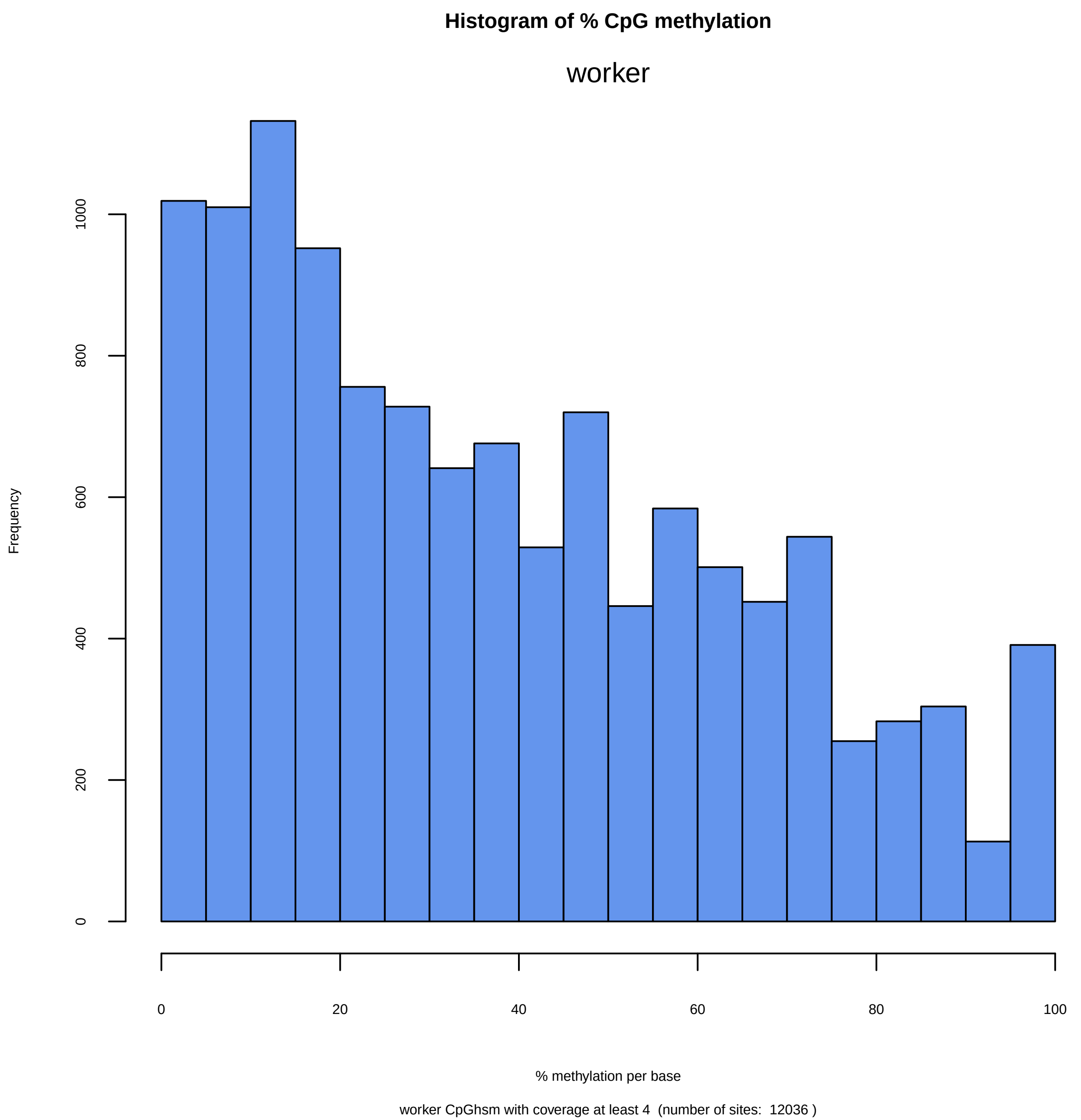
Histogram of % CpG methylation
worker
Frequency
0
200
400
600
800
1000
0
20
40
60
80
100
% methylation per base
worker CpGhsm with coverage at least 4 (number of sites: 12036 )

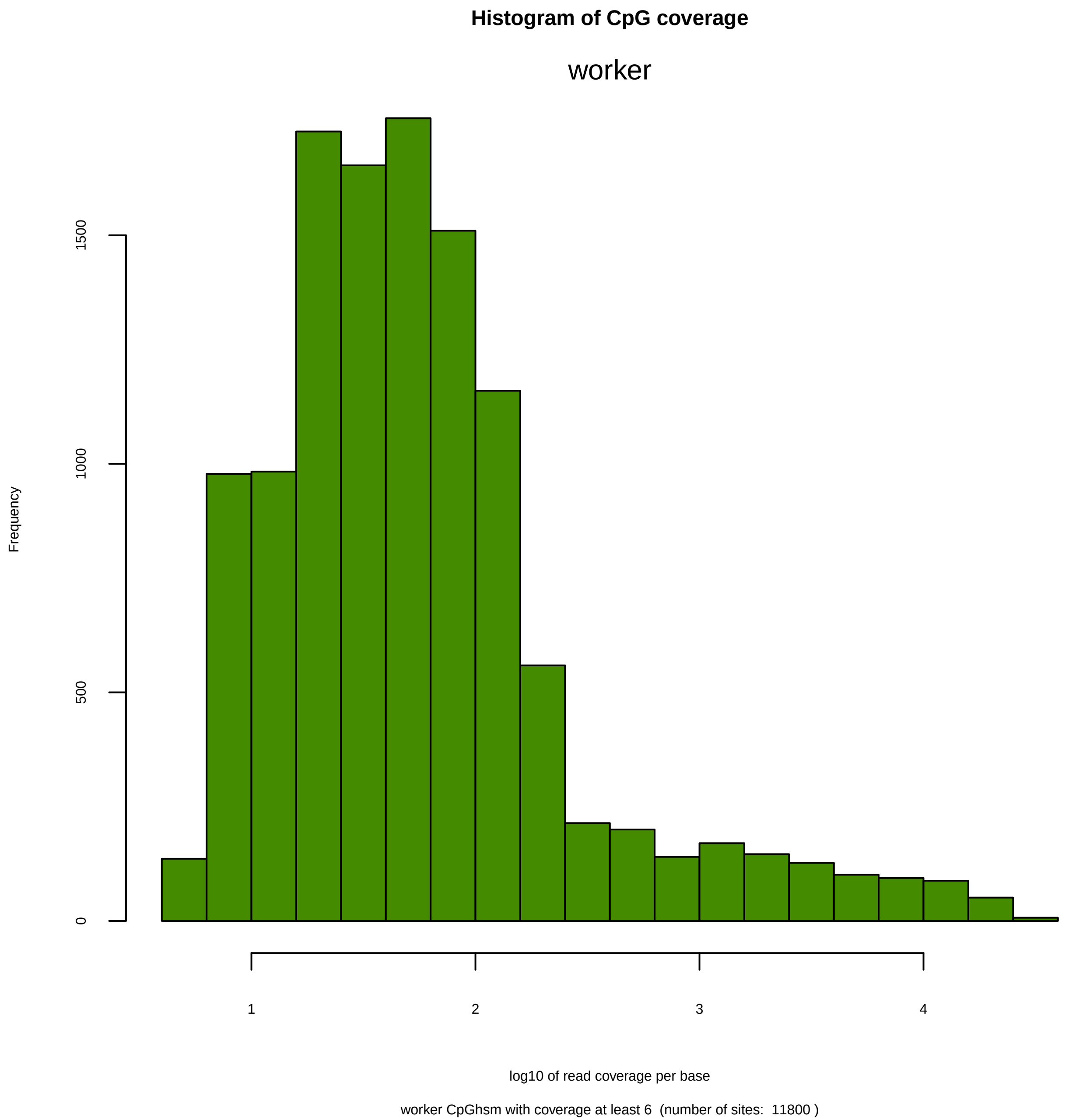

Histogram of CpG coverage
worker
Frequency
0
500
1000
1500
1
2
3
4
log10 of read coverage per base
worker CpGhsm with coverage at least 6 (number of sites: 11800 )

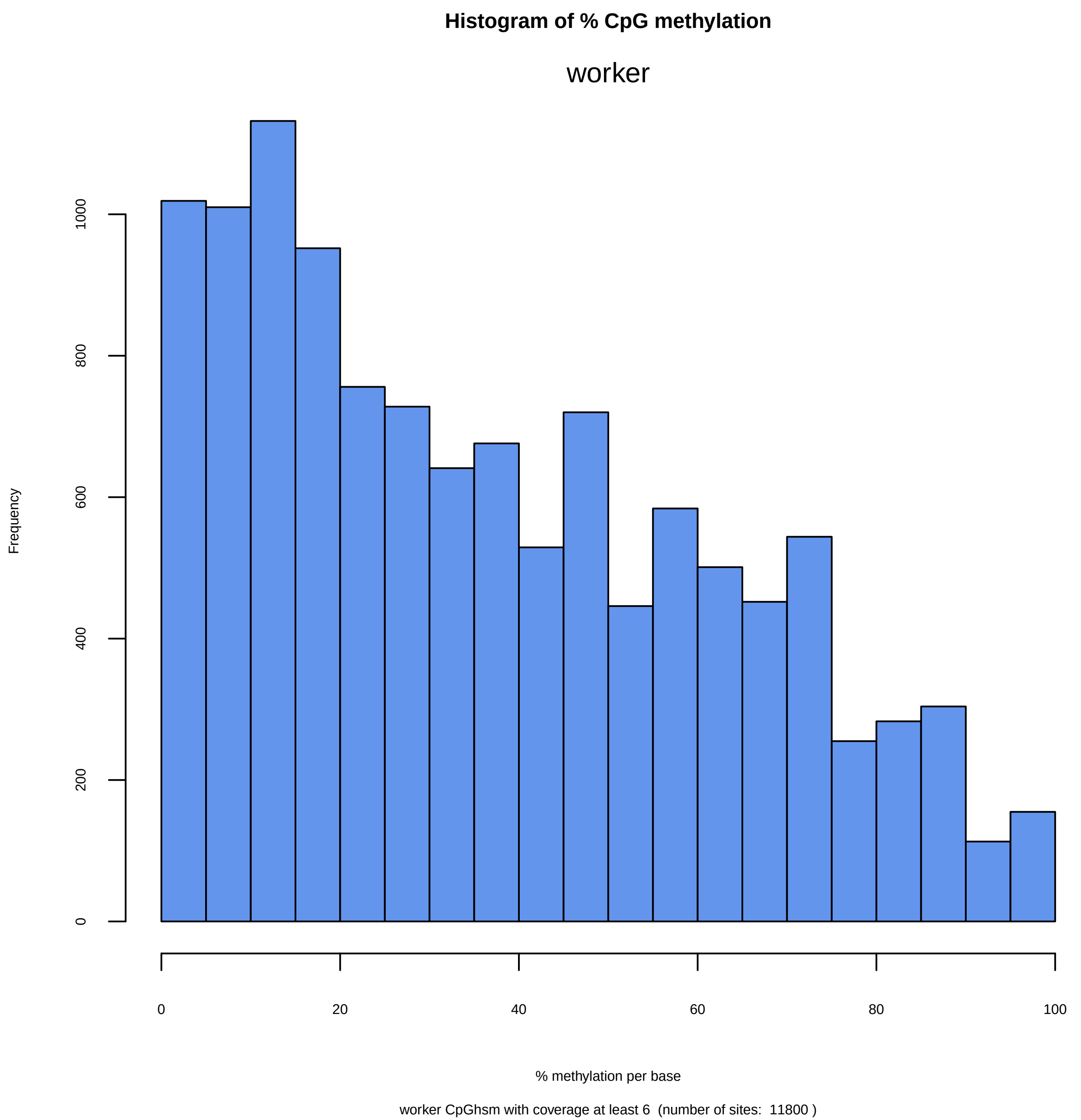
Histogram of % CpG methylation
worker
Frequency
0
200
400
600
800
1000
0
20
40
60
80
100
% methylation per base
worker CpGhsm with coverage at least 6 (number of sites: 11800 )

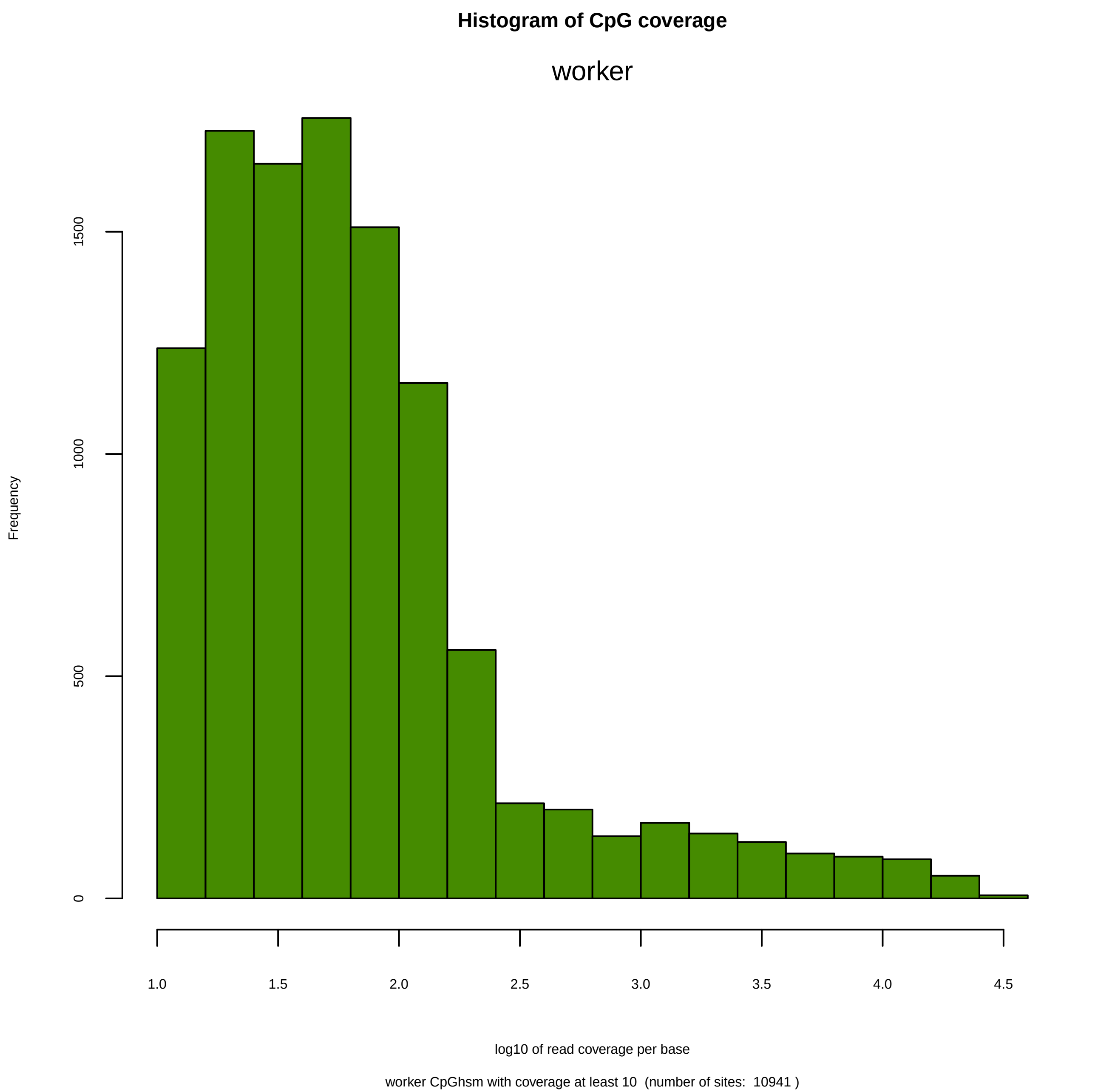
Histogram of CpG coverage
worker
Frequency
0
500
1000
1500
1.0
1.5
2.0
2.5
3.0
3.5
4.0
4.5
log10 of read coverage per base
worker CpGhsm with coverage at least 10 (number of sites: 10941 )

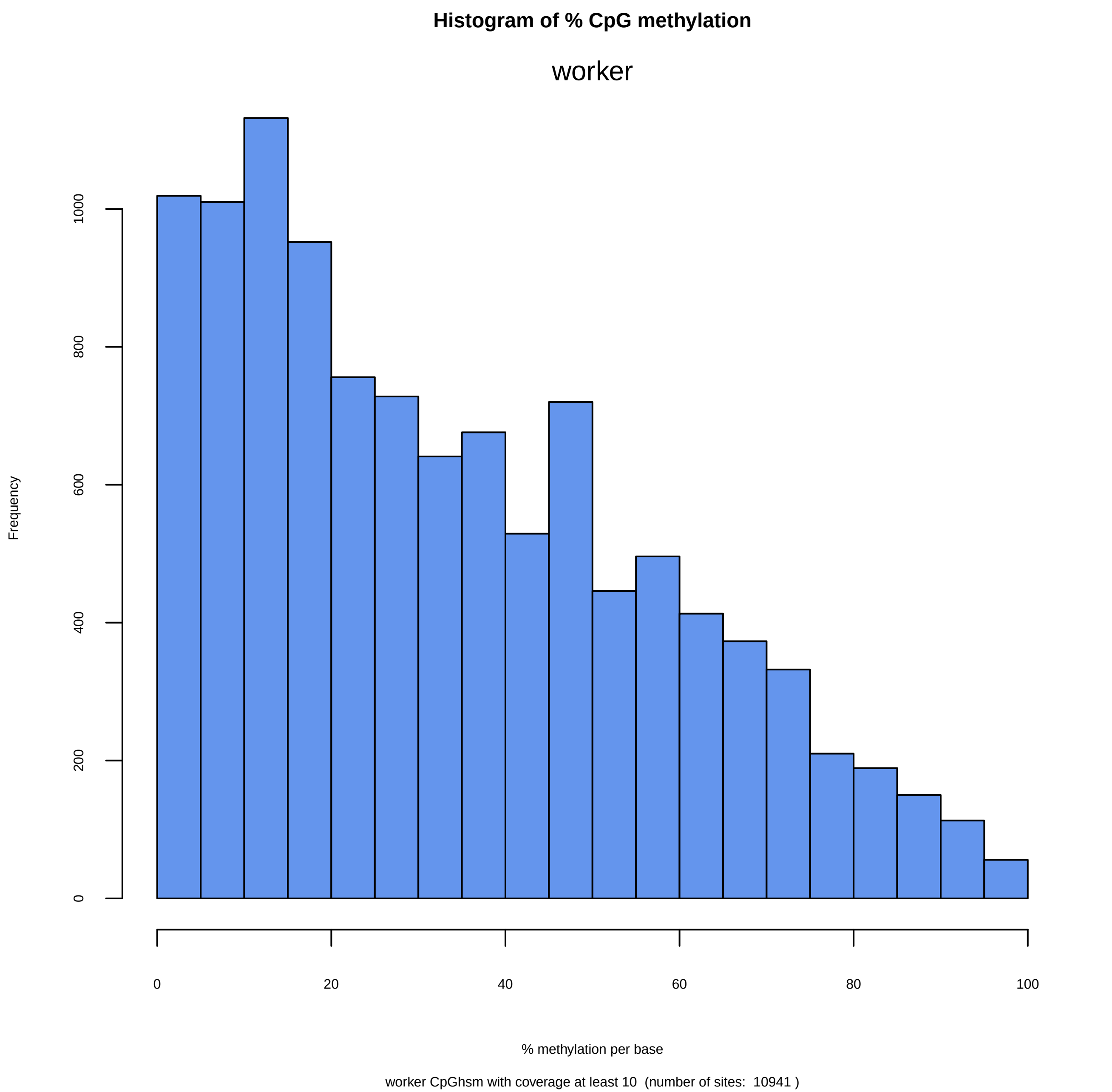
Histogram of % CpG methylation
worker
Frequency
0
200
400
600
800
1000
0
20
40
60
80
100
% methylation per base
worker CpGhsm with coverage at least 10  (number of sites:  10941 )

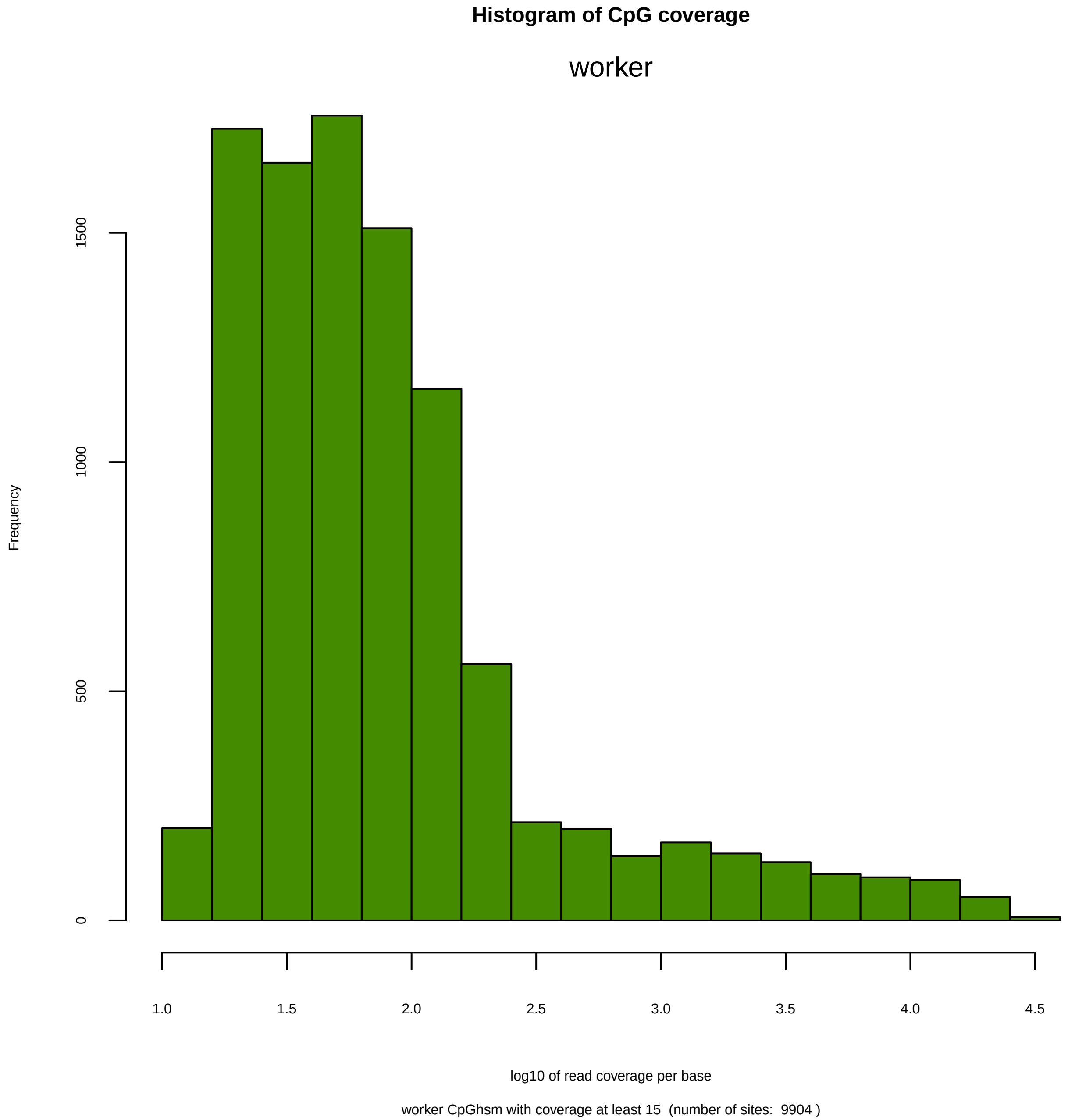
Histogram of CpG coverage
worker
Frequency
0
500
1000
1500
1.0
1.5
2.0
2.5
3.0
3.5
4.0
4.5
log10 of read coverage per base
worker CpGhsm with coverage at least 15  (number of sites:  9904 )

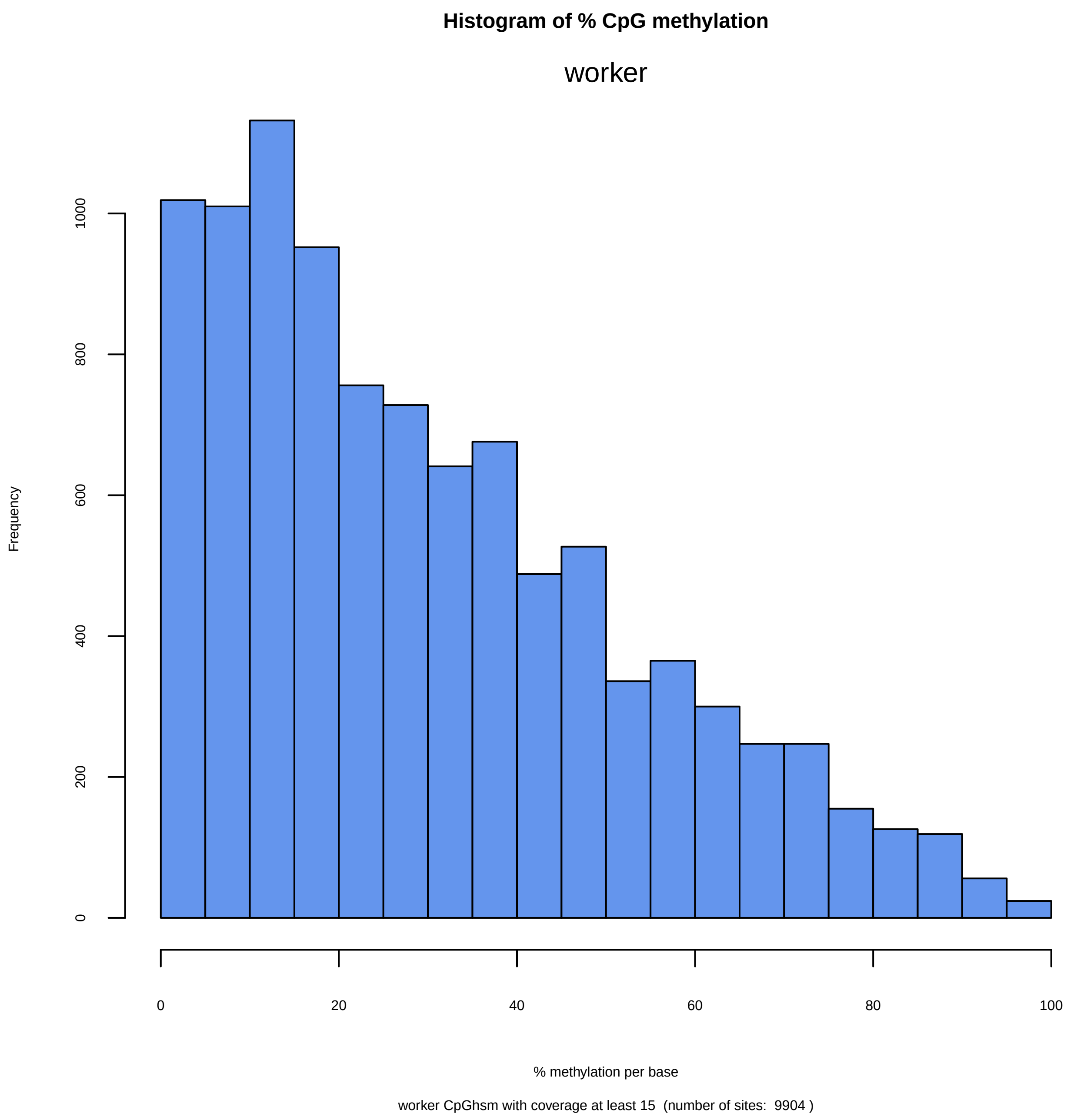
Histogram of % CpG methylation
worker
Frequency
0
200
400
600
800
1000
0
20
40
60
80
100
% methylation per base
worker CpGhsm with coverage at least 15 (number of sites: 9904 )

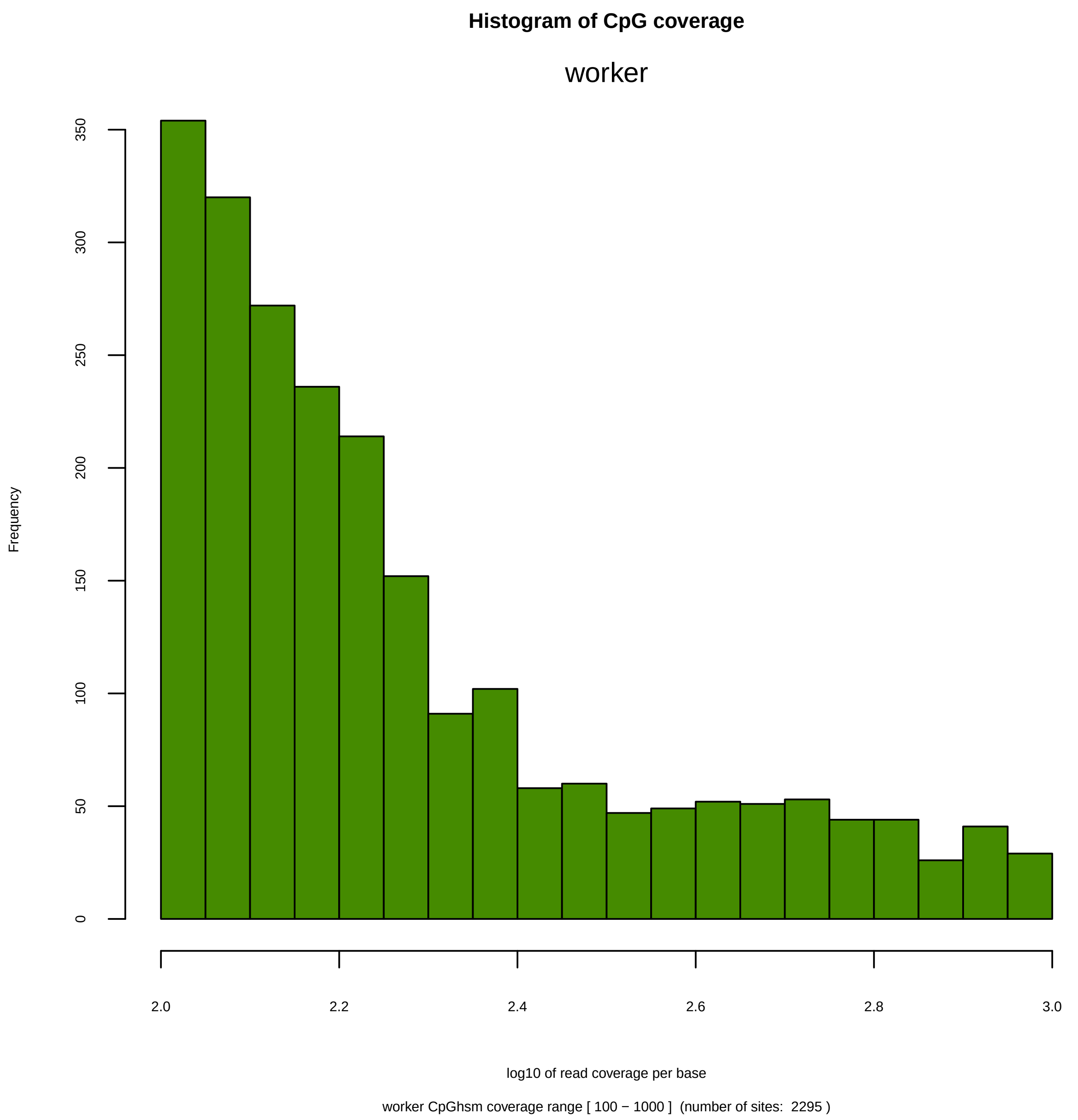
Histogram of CpG coverage
worker
Frequency
0
50
100
150
200
250
300
350
2.0
2.2
2.4
2.6
2.8
3.0
log10 of read coverage per base
worker CpGhsm coverage range [ 100 − 1000 ]  (number of sites:  2295 )

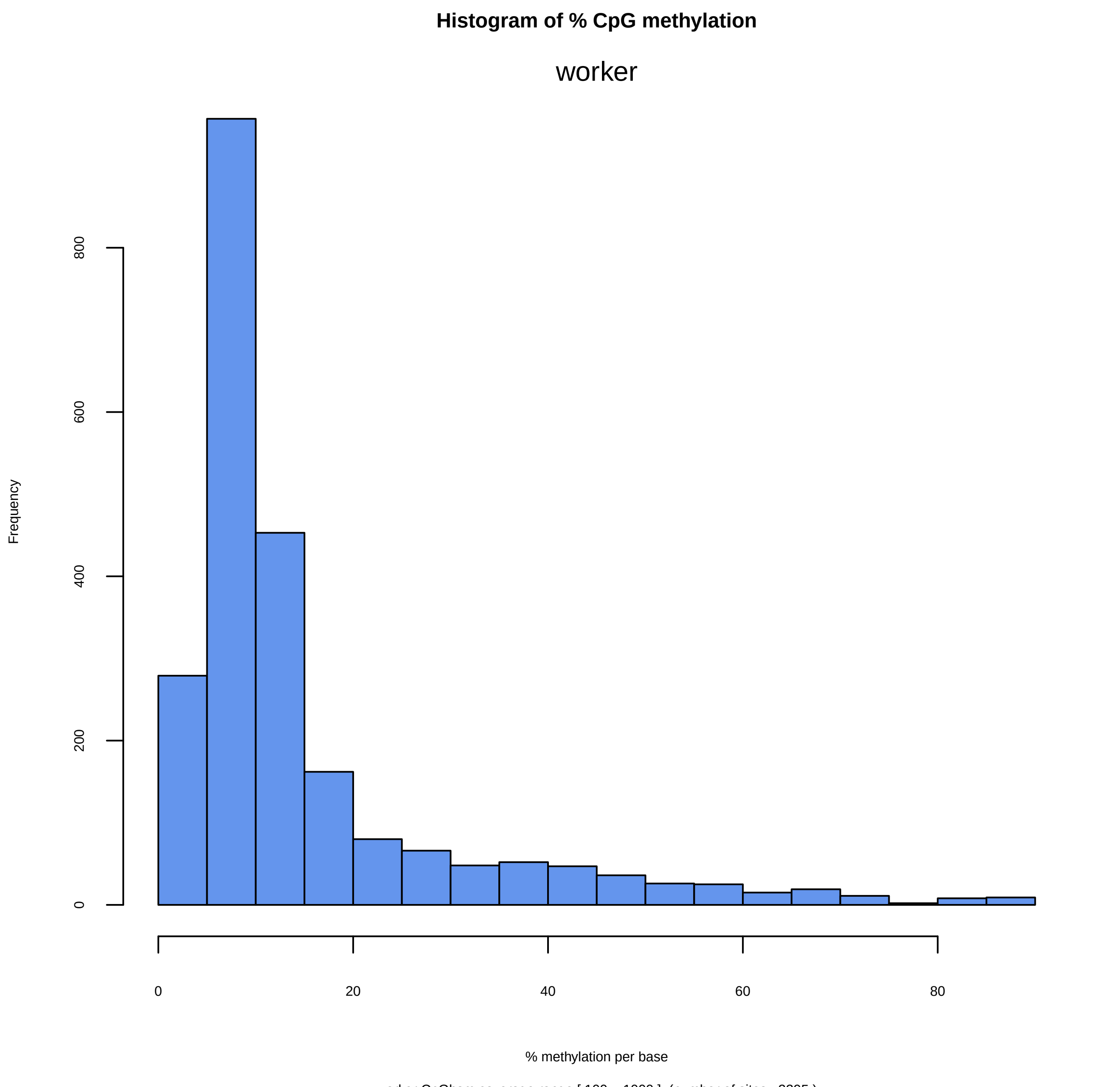
Histogram of % CpG methylation
worker
Frequency
0
200
400
600
800
0
20
40
60
80
% methylation per base
worker CpGhsm coverage range [ 100 − 1000 ]  (number of sites:  2295 )

```
Directory: PWC File: 0READMEpwc


PWC - Pairwise comparisons between all samples.

Input: studymk, studymc, nbrpms, hheight, nbrpnts

Output: files pwc-*.txt pwc-.*pdf

Notes: The output is generated by BWASPR::cmpSites(), which determines for each pairwise
        comparison the numbers of common and distinct sites.  The function also calculates
        the overlap index between the samples and estimates the size of the common pool of
        potential methylation sites.

       The CpGscd data input (studymc) is necessary to determine sites that are detectable
        in both samples, and nbrpms (listed in the *.par file specified in the Pc_PA.conf
        configuration file as argument to TOTALNBRPMSITES) provides the total number of
        potential methylation sites (typically, all CpG sites), a necessary parameter for
        the estimations.
       hheight (default: 0.10) specifies the y-axis limit in the histogram plots.
       nbrpnts (default: 5000) specifies the number of common sites to be plotted in the
        methylation levels scatter plot.
```

```
Directory: PWC File: pwc-Pc_PA_qn.vs.Pc_PA_wr.txt


Numbers of common and distinct sites comparing Pc_PA_qn versus Pc_PA_wr
================================================================================

  total number of potential sites:  14357872

  number of "Pc_PA_qn_hsm"         sites: 13840
  number of "Pc_PA_wr_hsm"         sites: 12036
    number of "Pc_PA_qn_hsm"-unique sites:       5330
    number of common sites:          8510
    number of "Pc_PA_wr_hsm"-unique sites:       3526
  total number of "Pc_PA_qn_hsm+Pc_PA_wr_hsm"-sites observed:     17366

  number of "Pc_PA_qn_scd"        sites:   8534313 ( 59.44% of total)
  number of "Pc_PA_wr_scd"        sites:   7865066 ( 54.78% of total)
    number of "Pc_PA_qn_scd"-unique sites:       1205641
    number of sites in common:       7328672   (Expected:   4674992; O/E:       1.6)
    number of "Pc_PA_wr_scd"-unique sites:        536394
  total number of "Pc_PA_qn_scd+Pc_PA_wr_scd"-sites observed:      9070707

  number of sites in "Pc_PA_qn_hsm" that are not  detectable in "Pc_PA_wr_hsm":       365
  number of sites in "Pc_PA_qn_hsm" that are also detectable in "Pc_PA_wr_hsm":     13475
    number of sites unique to "Pc_PA_qn_hsm" although detectable in "Pc_PA_wr_hsm":             4965
    number of "Pc_PA_qn_hsm" / "Pc_PA_wr_hsm" common sites                        :             8510      (Expected:     22; O/E:     386.8)

  number of sites in "Pc_PA_wr_hsm" that are not  detectable in "Pc_PA_qn_hsm":       153
  number of sites in "Pc_PA_wr_hsm" that are also detectable in "Pc_PA_qn_hsm":     11883
    number of sites unique to "Pc_PA_wr_hsm" although detectable in "Pc_PA_qn_hsm":             3373
    number of "Pc_PA_wr_hsm" / "Pc_PA_qn_hsm" common sites                        :             8510      (Expected:     22; O/E:     386.8)

  Overlap index of "Pc_PA_qn_hsm" with "Pc_PA_wr_hsm":           0.573
  Estimated number of "Pc_PA_qn_hsm" = "Pc_PA_wr_hsm" sites (assuming sampling from one population):          18816
  Adjusted population size of "Pc_PA_qn_hsm" = "Pc_PA_wr_hsm" sites (assuming all sites detectable):          36863 (  2.12x of observed)


  number of "Pc_PA_qn_hsm" sites with coverage >=  4:  13840
  number of "Pc_PA_wr_hsm" sites with coverage >=  4:  12036
  number of common sites with coverage >=  4:  8510
  number of "Pc_PA_qn_hsm" sites with coverage >=  6:  13635
  number of "Pc_PA_wr_hsm" sites with coverage >=  6:  11800
  number of common sites with coverage >=  6:  8365
  number of "Pc_PA_qn_hsm" sites with coverage >= 10:  12768
  number of "Pc_PA_wr_hsm" sites with coverage >= 10:  10941
  number of common sites with coverage >= 10:  7782
  number of "Pc_PA_qn_hsm" sites with coverage >= 15:  11607
  number of "Pc_PA_wr_hsm" sites with coverage >= 15:   9904
  number of common sites with coverage >= 15:  7061
```

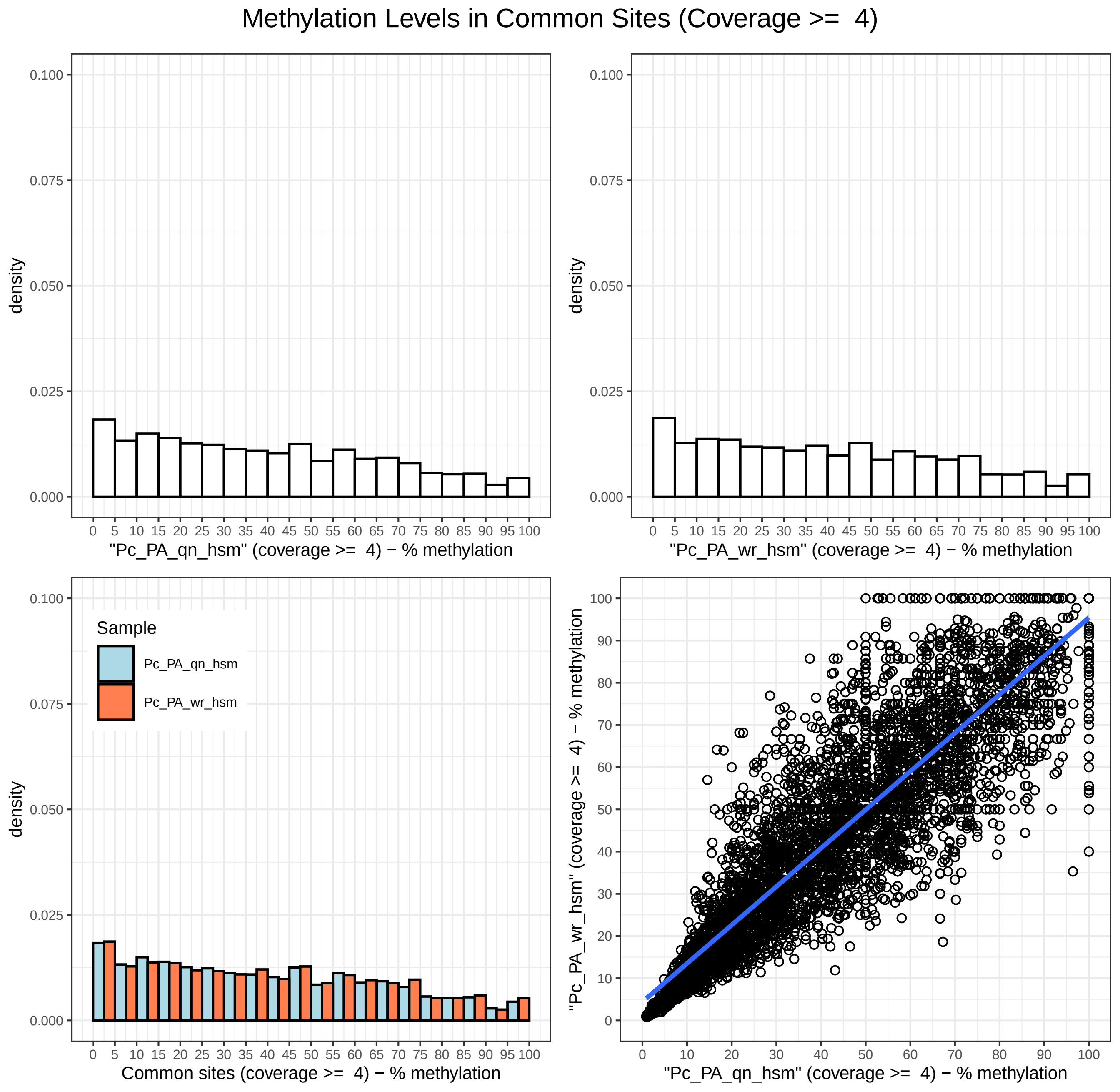
Methylation Levels in Common Sites (Coverage >= 4)
density
"Pc_PA_qn_hsm" (coverage >= 4) − % methylation
"Pc_PA_wr_hsm" (coverage >= 4) − % methylation
Sample
Pc_PA_qn_hsm
Pc_PA_wr_hsm
Common sites (coverage >= 4) − % methylation
"Pc_PA_wr_hsm" (coverage >= 4) − % methylation
"Pc_PA_qn_hsm" (coverage >= 4) − % methylation

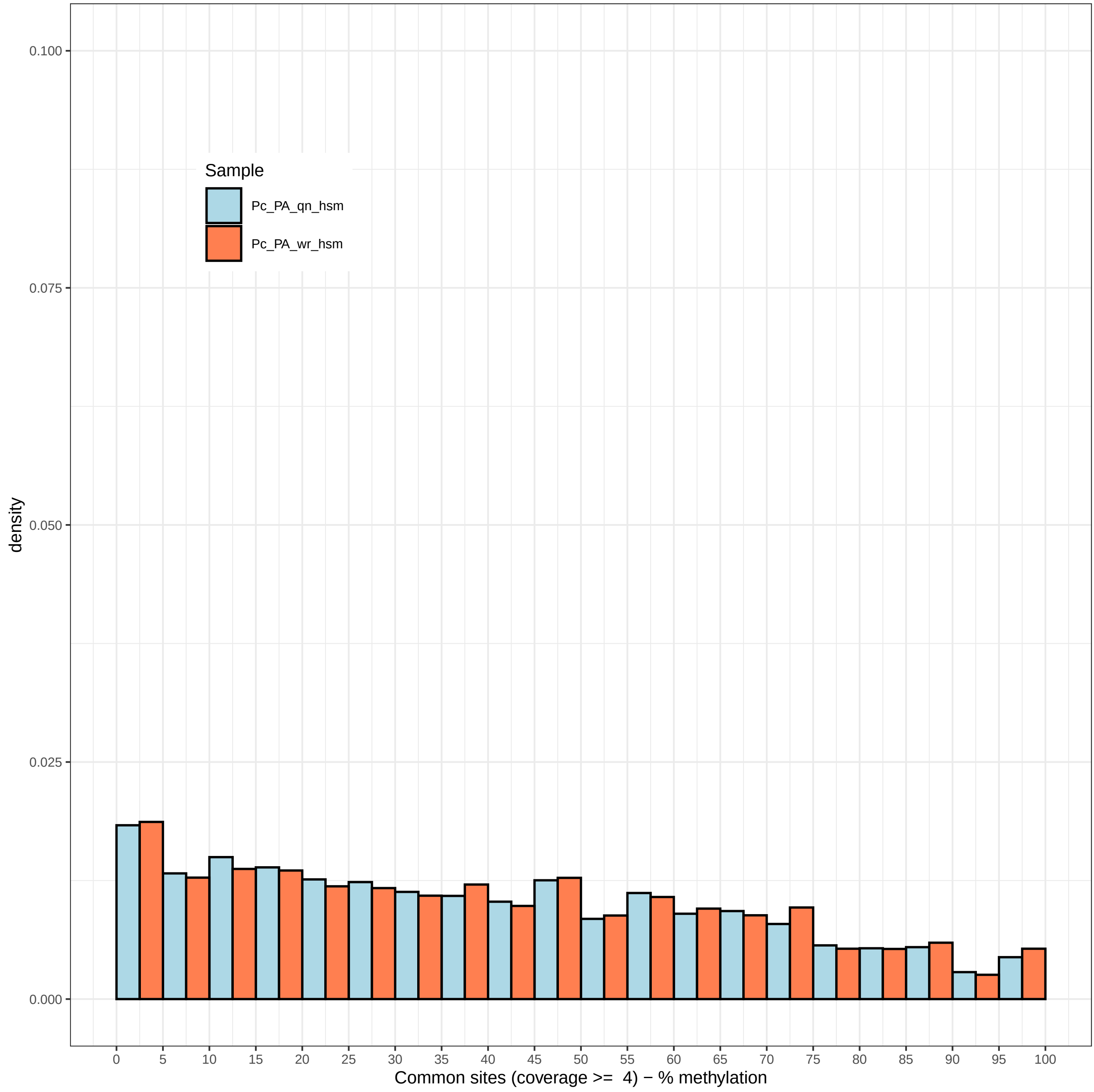

Sample
Pc_PA_qn_hsm
Pc_PA_wr_hsm
density
0.100
0.075
0.050
0.025
0.000
0 5 10 15 20 25 30 35 40 45 50 55 60 65 70 75 80 85 90 95 100
Common sites (coverage >= 4) − % methylation

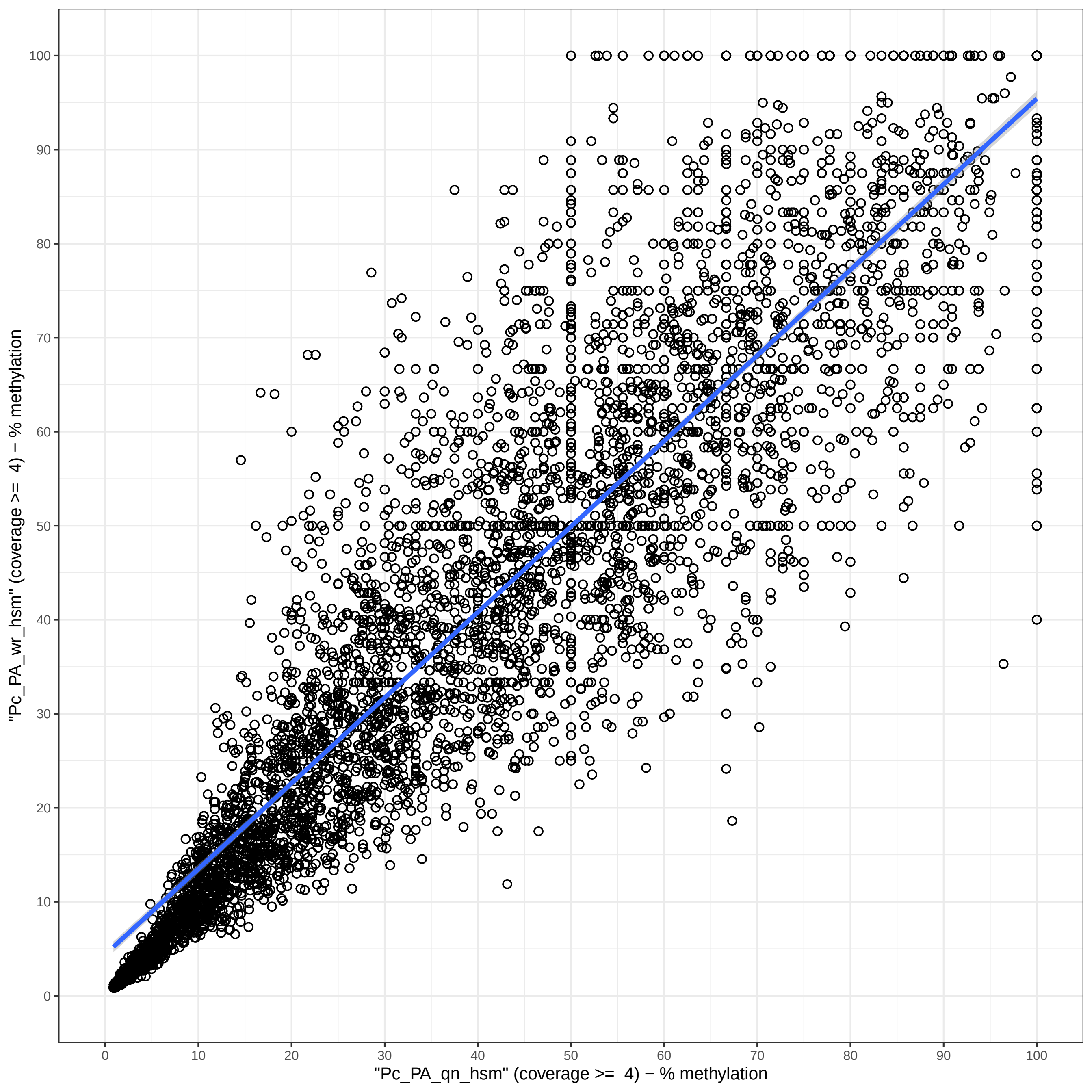

"Pc_PA_wr_hsm" (coverage >= 4) − % methylation
"Pc_PA_qn_hsm" (coverage >= 4) − % methylation
0
10
20
30
40
50
60
70
80
90
100

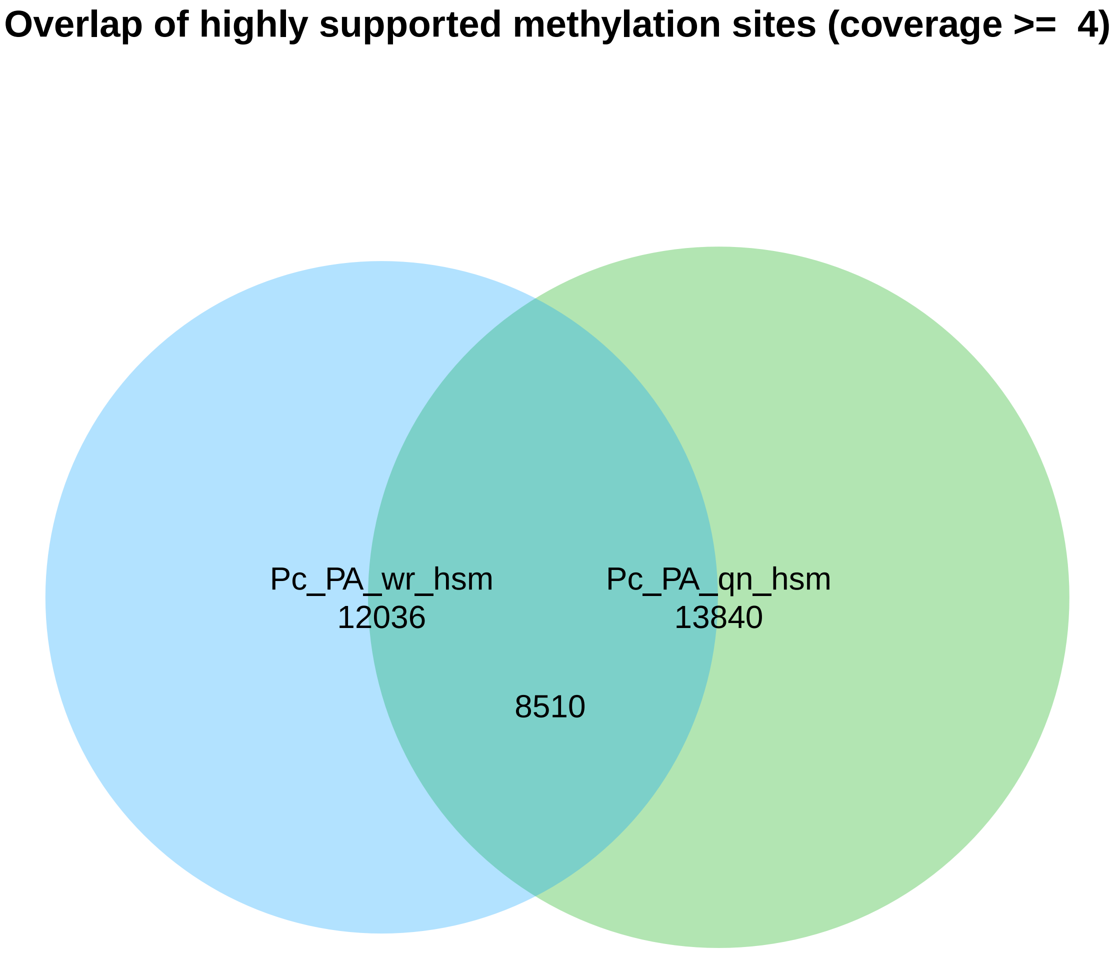
Overlap of highly supported methylation sites (coverage >= 4)
Pc_PA_wr_hsm
12036
Pc_PA_qn_hsm
13840
8510

Methylation Levels in Common Sites (Coverage >=  6)

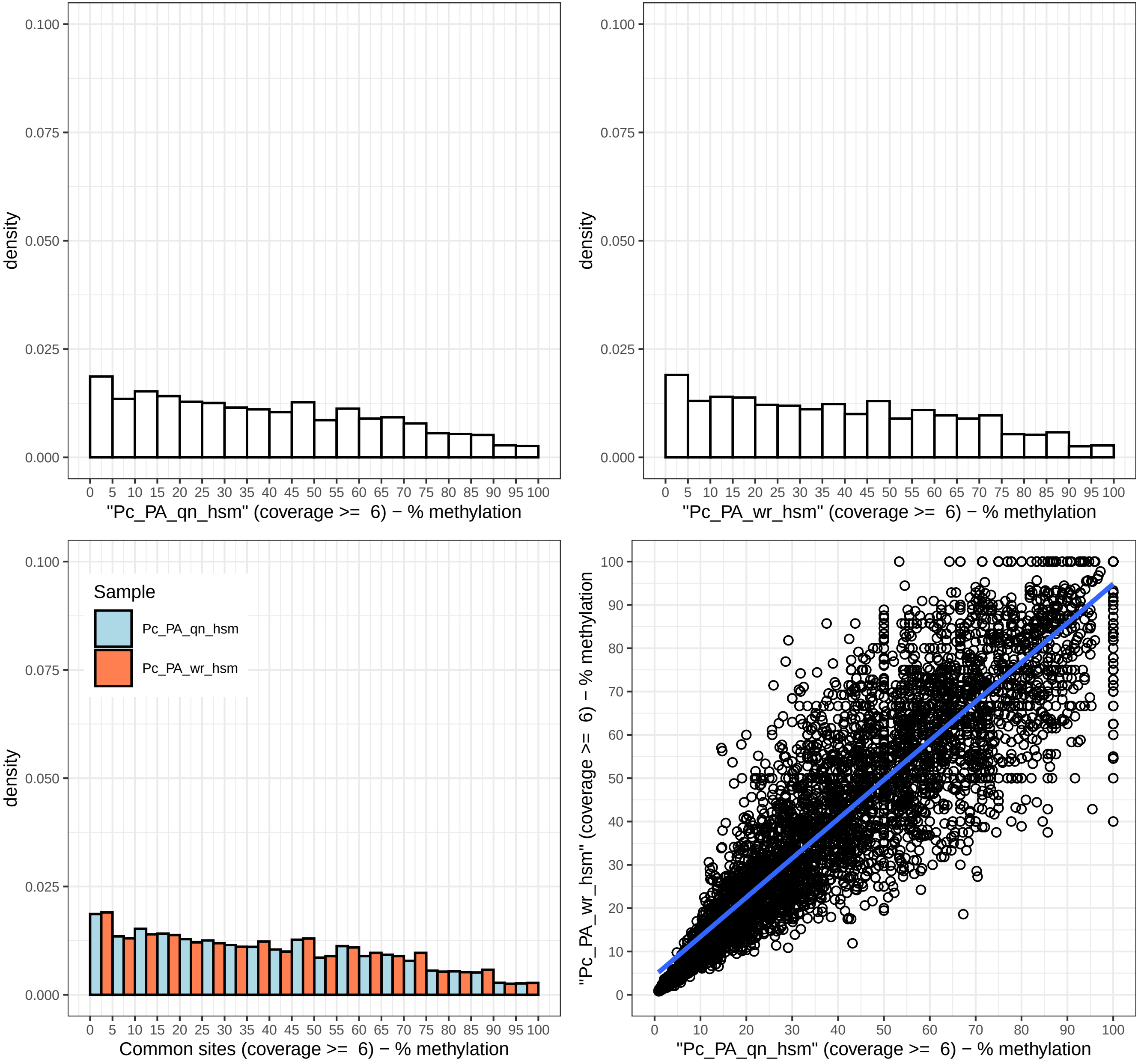

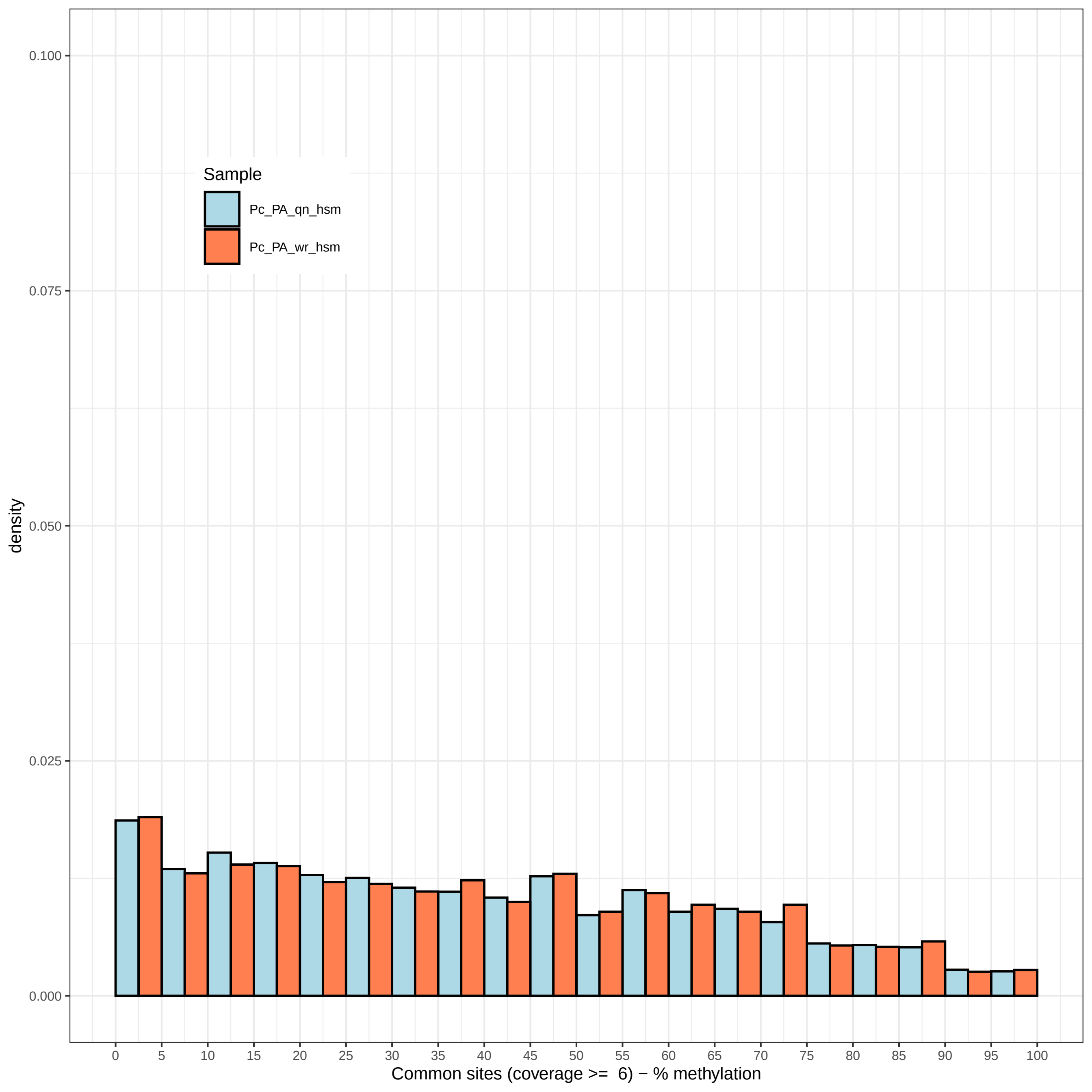

Sample
Pc_PA_qn_hsm
Pc_PA_wr_hsm
density
0.000
0.025
0.050
0.075
0.100
0 5 10 15 20 25 30 35 40 45 50 55 60 65 70 75 80 85 90 95 100
Common sites (coverage >= 6) − % methylation

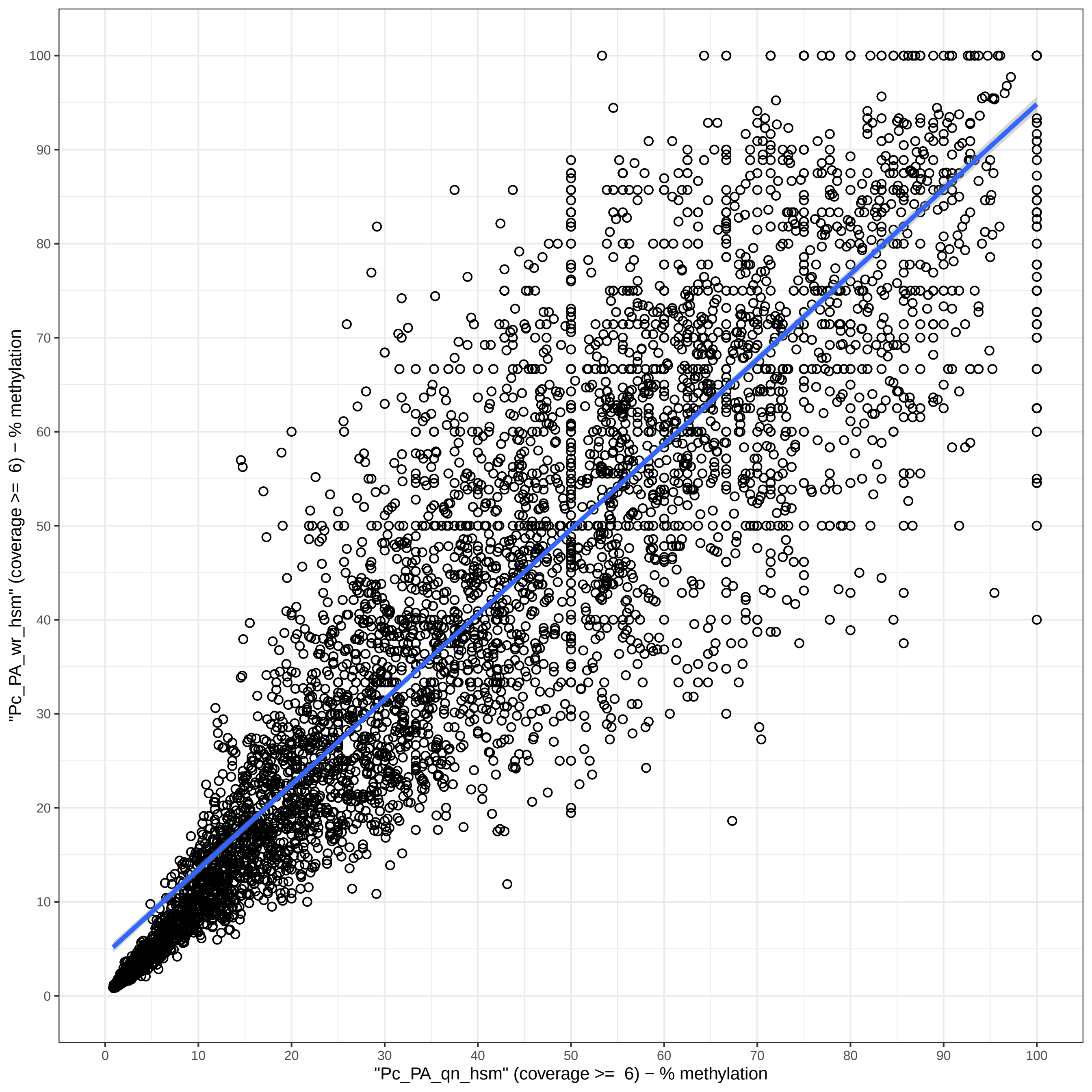
"Pc_PA_wr_hsm" (coverage >= 6) − % methylation
"Pc_PA_qn_hsm" (coverage >= 6) − % methylation
0
10
20
30
40
50
60
70
80
90
100

#### Overlap of highly supported methylation sites (coverage >= 6)

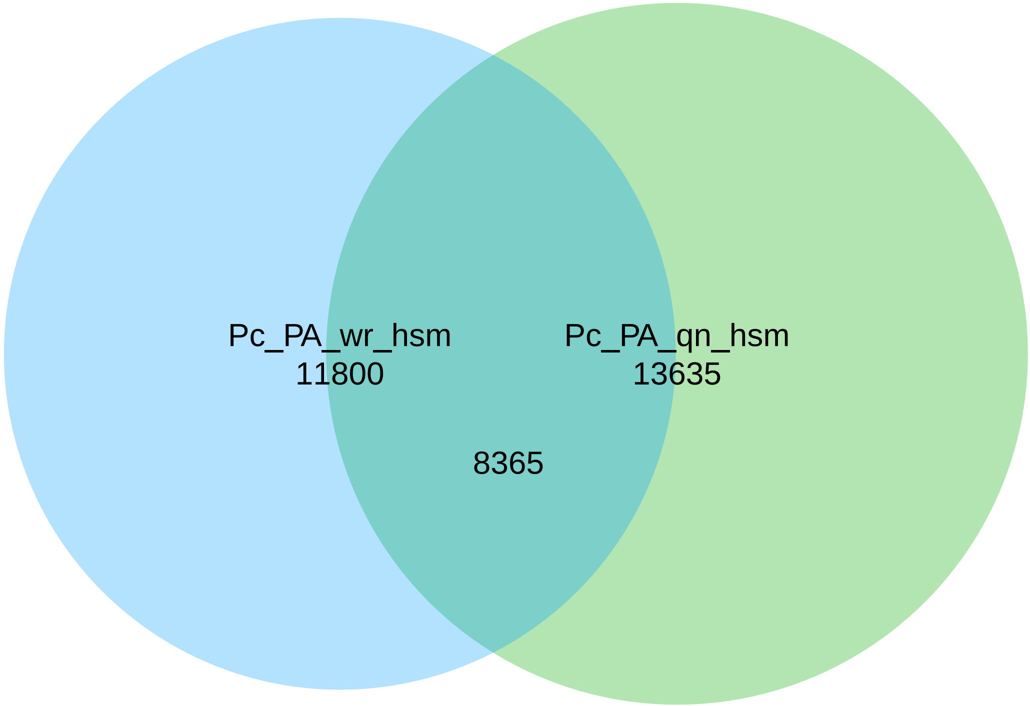

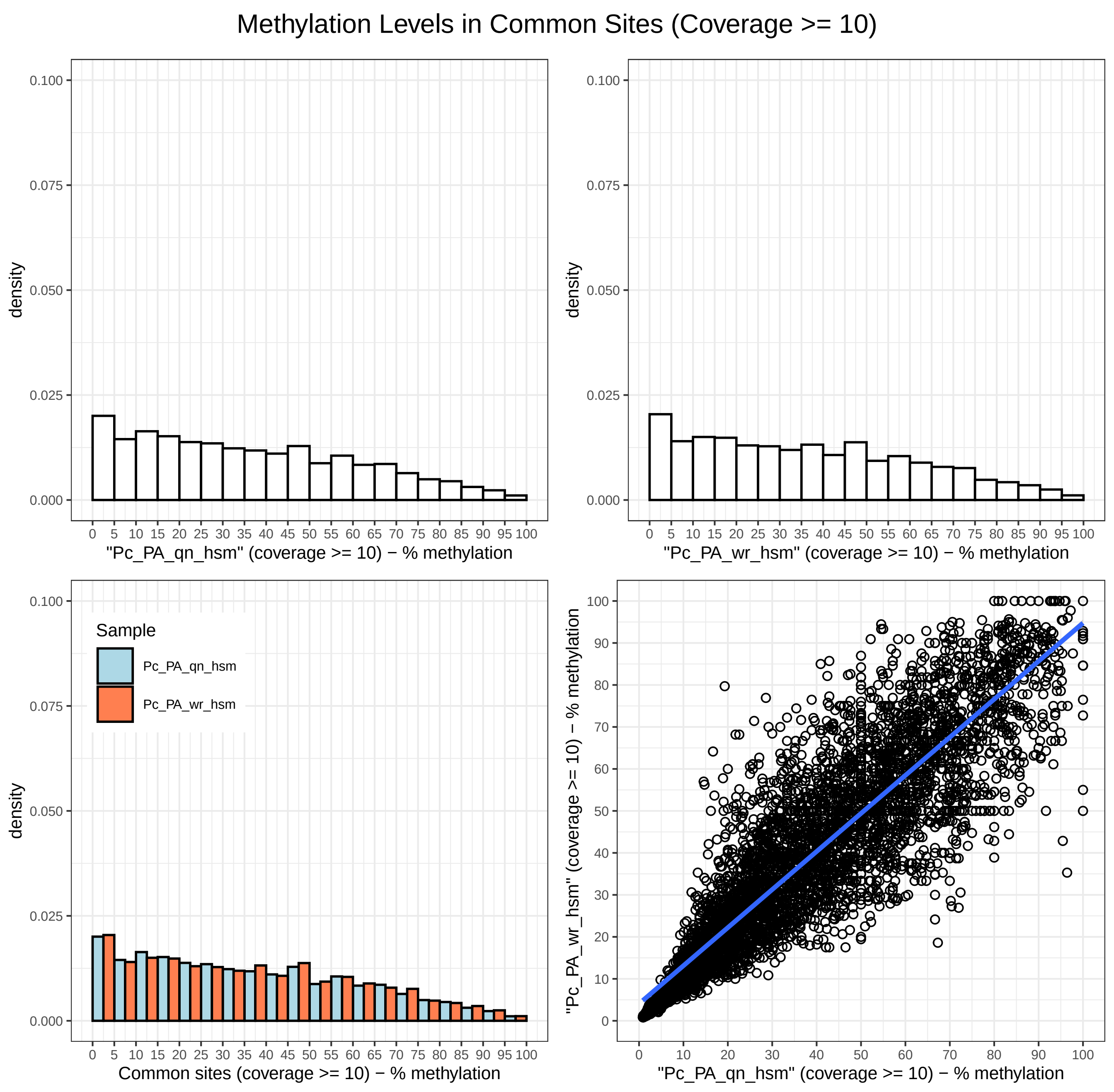

Methylation Levels in Common Sites (Coverage >= 10)
density
"Pc_PA_qn_hsm" (coverage >= 10) − % methylation
density
"Pc_PA_wr_hsm" (coverage >= 10) − % methylation
Sample
Pc_PA_qn_hsm
Pc_PA_wr_hsm
density
Common sites (coverage >= 10) − % methylation
"Pc_PA_wr_hsm" (coverage >= 10) − % methylation
"Pc_PA_qn_hsm" (coverage >= 10) − % methylation

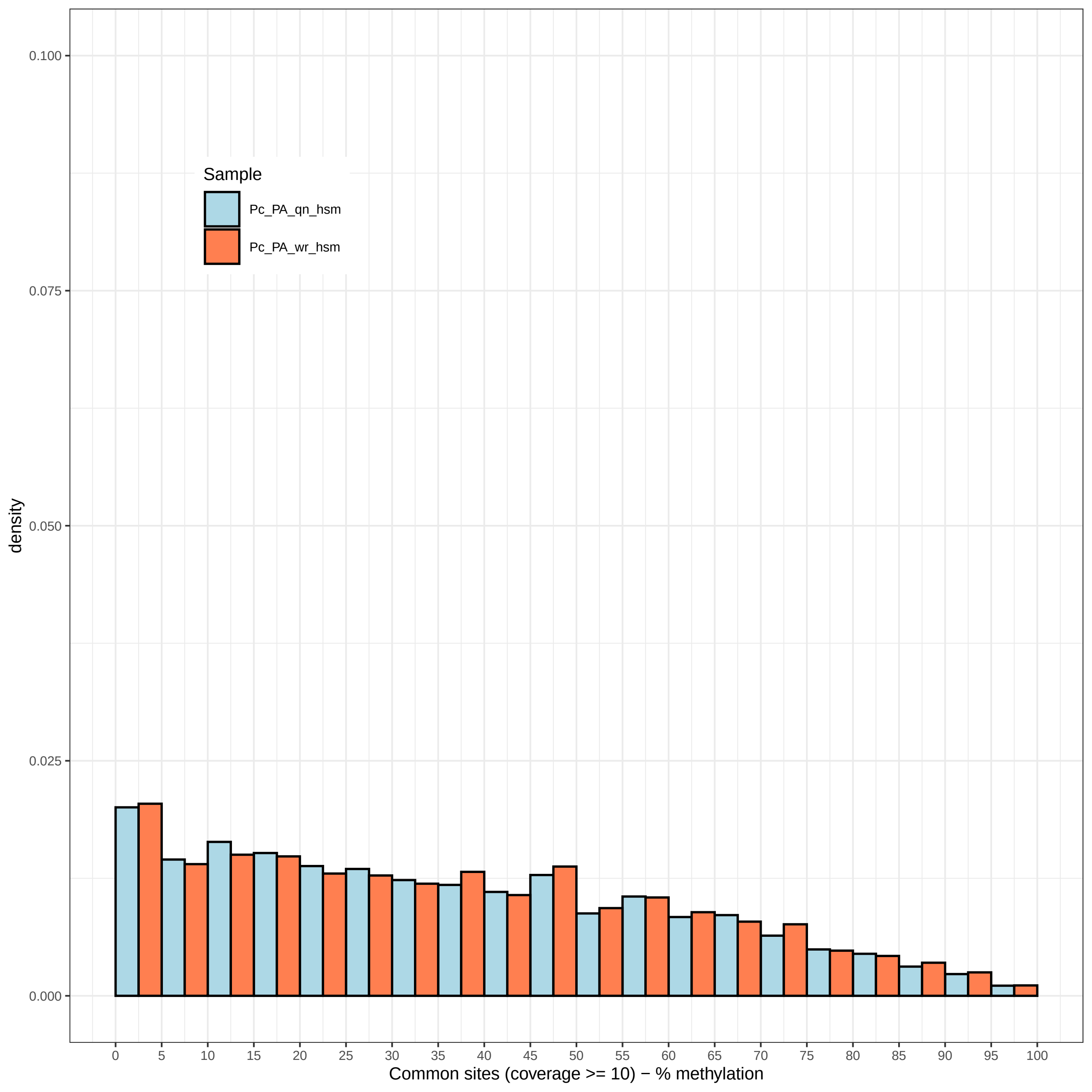
Sample
Pc_PA_qn_hsm
Pc_PA_wr_hsm
density
0.100
0.075
0.050
0.025
0.000
0 5 10 15 20 25 30 35 40 45 50 55 60 65 70 75 80 85 90 95 100
Common sites (coverage >= 10) − % methylation

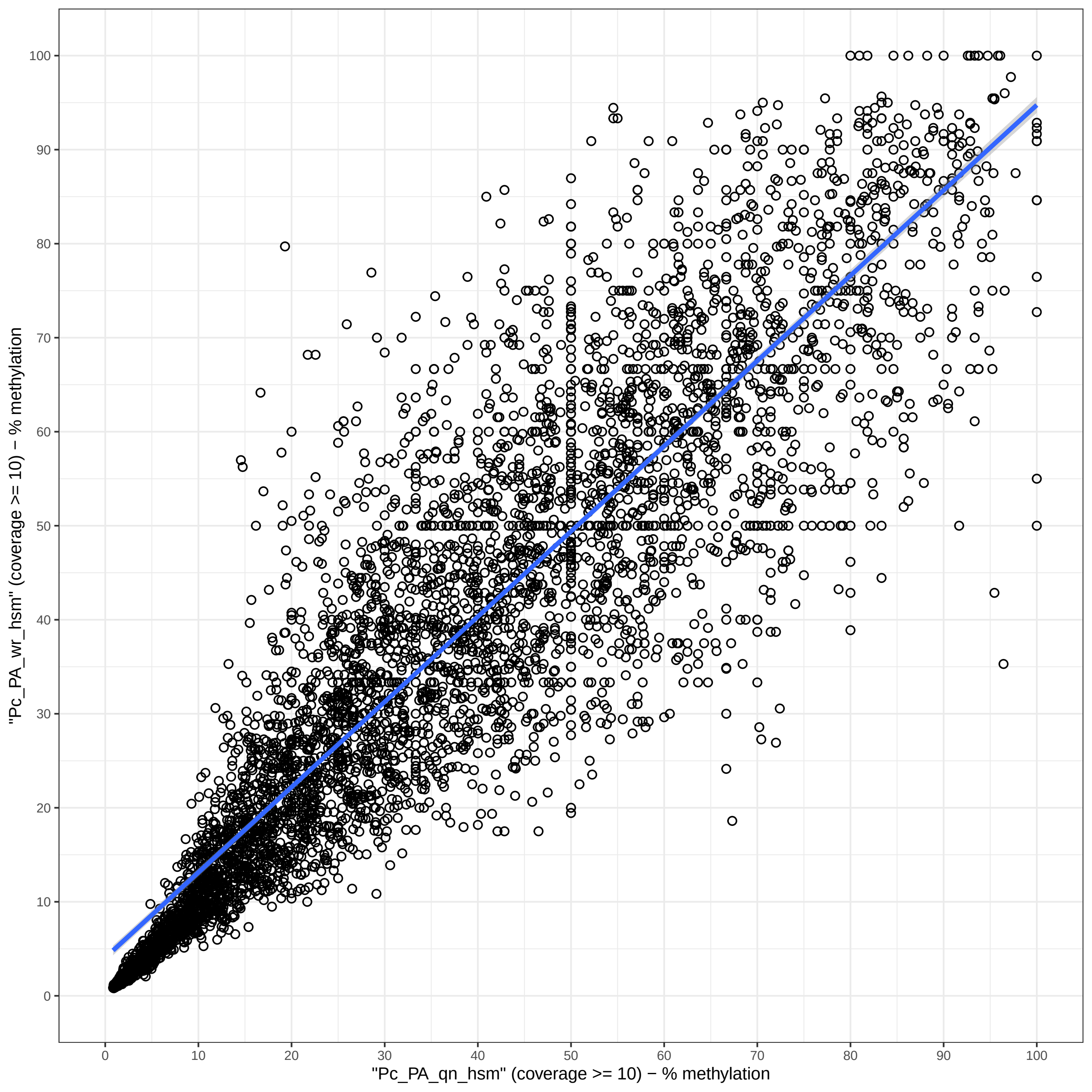

"Pc_PA_wr_hsm" (coverage >= 10) − % methylation
"Pc_PA_qn_hsm" (coverage >= 10) − % methylation
0
10
20
30
40
50
60
70
80
90
100

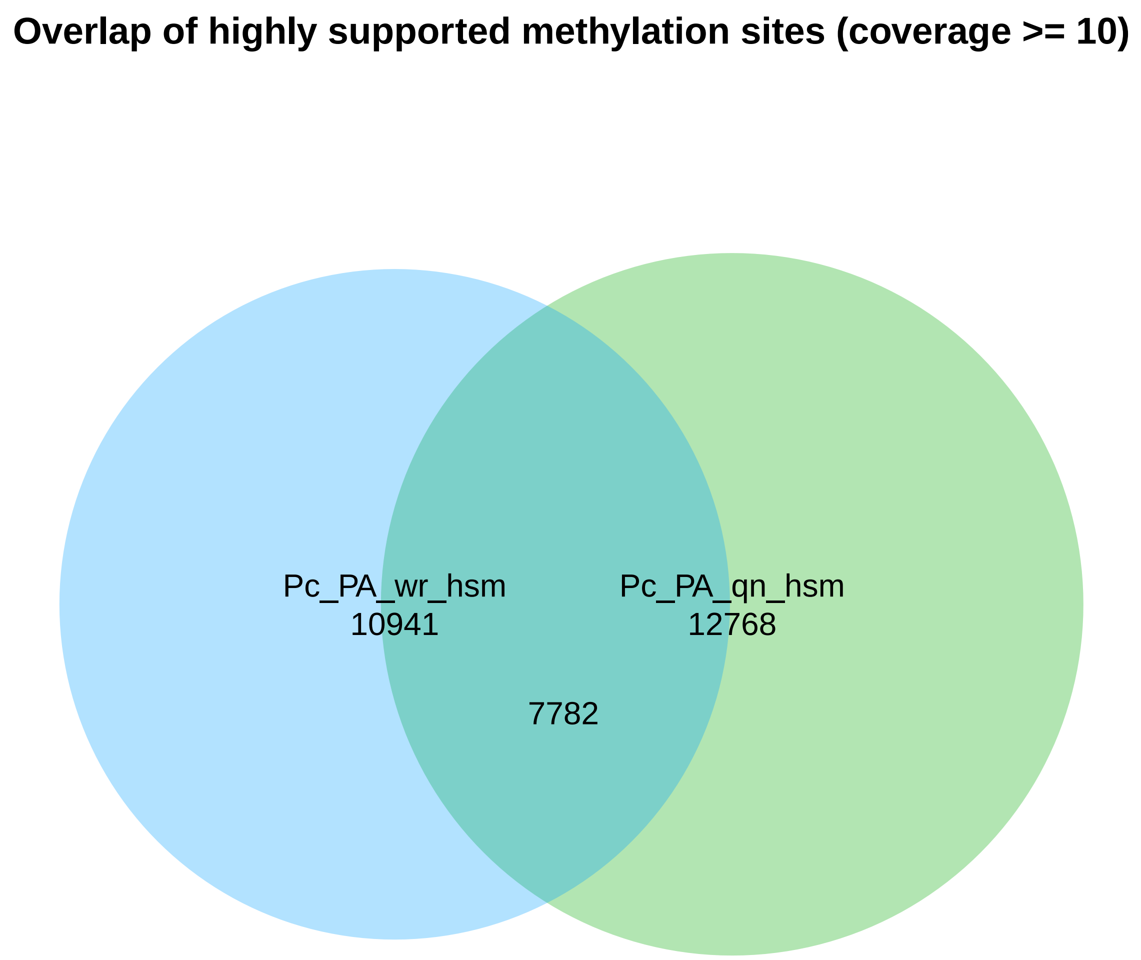
Overlap of highly supported methylation sites (coverage >= 10)
Pc_PA_wr_hsm
10941
Pc_PA_qn_hsm
12768
7782

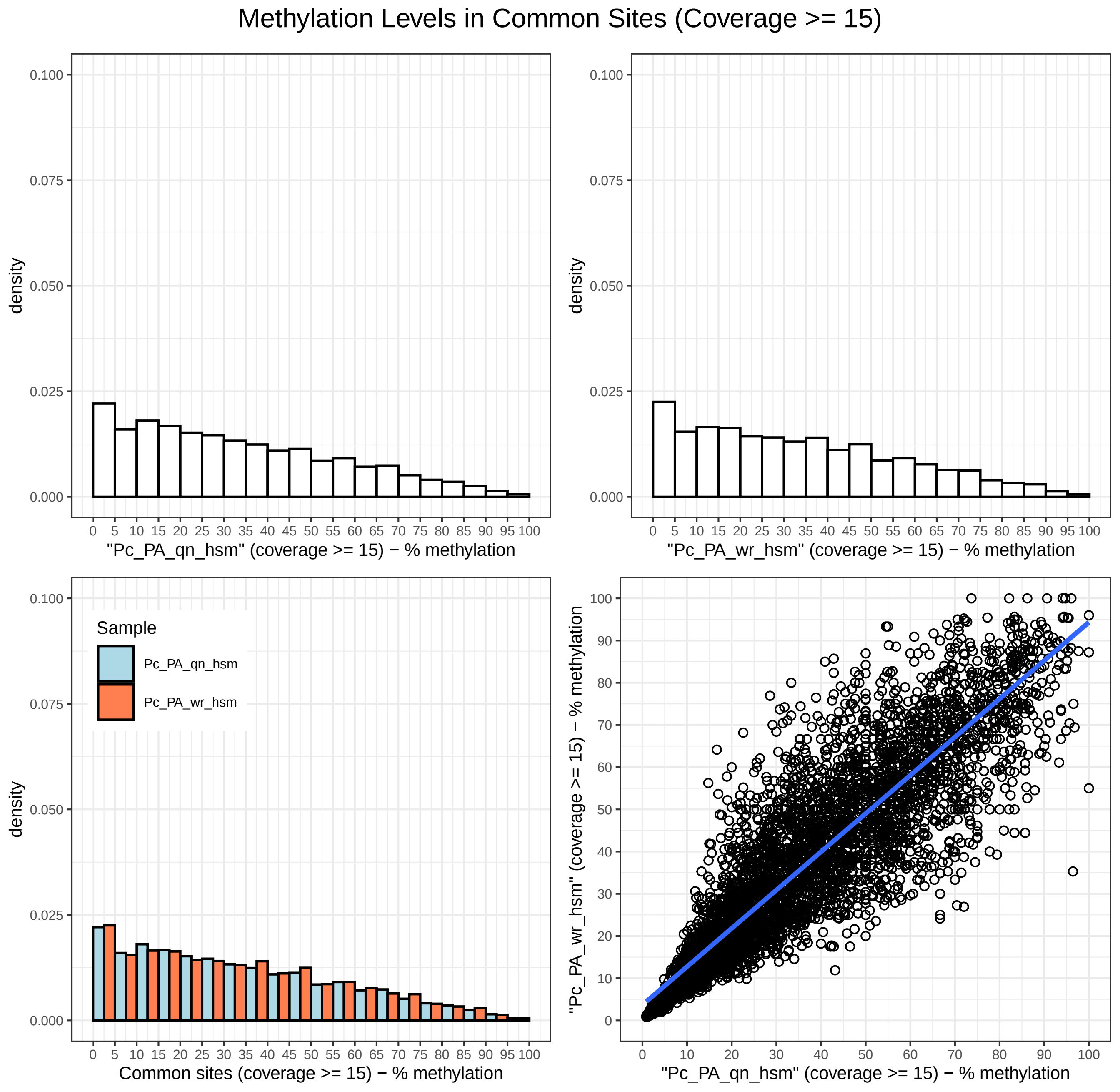
Methylation Levels in Common Sites (Coverage >= 15)
density
"Pc_PA_qn_hsm" (coverage >= 15) − % methylation
"Pc_PA_wr_hsm" (coverage >= 15) − % methylation
Sample
Pc_PA_qn_hsm
Pc_PA_wr_hsm
Common sites (coverage >= 15) − % methylation
"Pc_PA_wr_hsm" (coverage >= 15) − % methylation
"Pc_PA_qn_hsm" (coverage >= 15) − % methylation

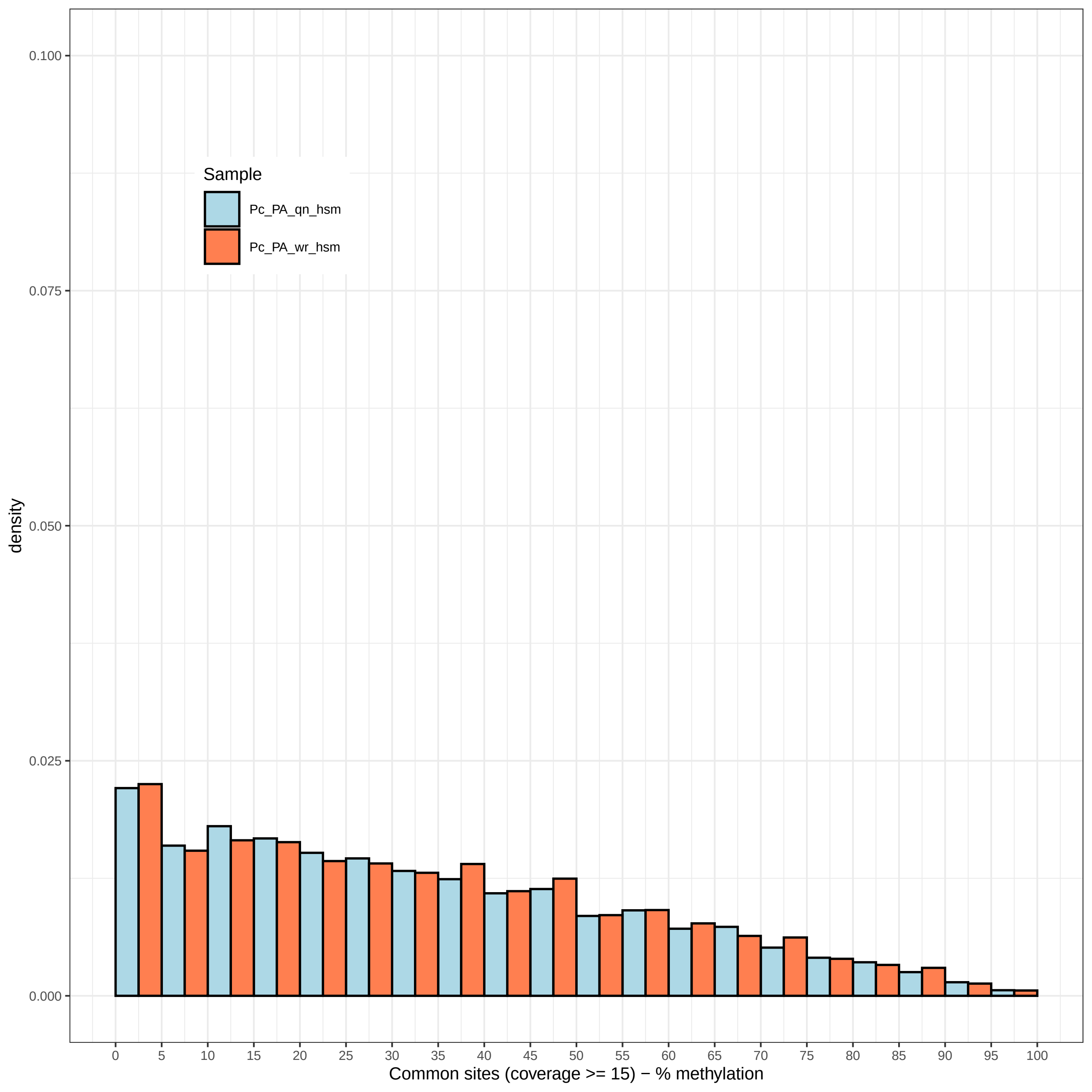
Sample
Pc_PA_qn_hsm
Pc_PA_wr_hsm
density
0.100
0.075
0.050
0.025
0.000
0 5 10 15 20 25 30 35 40 45 50 55 60 65 70 75 80 85 90 95 100
Common sites (coverage >= 15) − % methylation

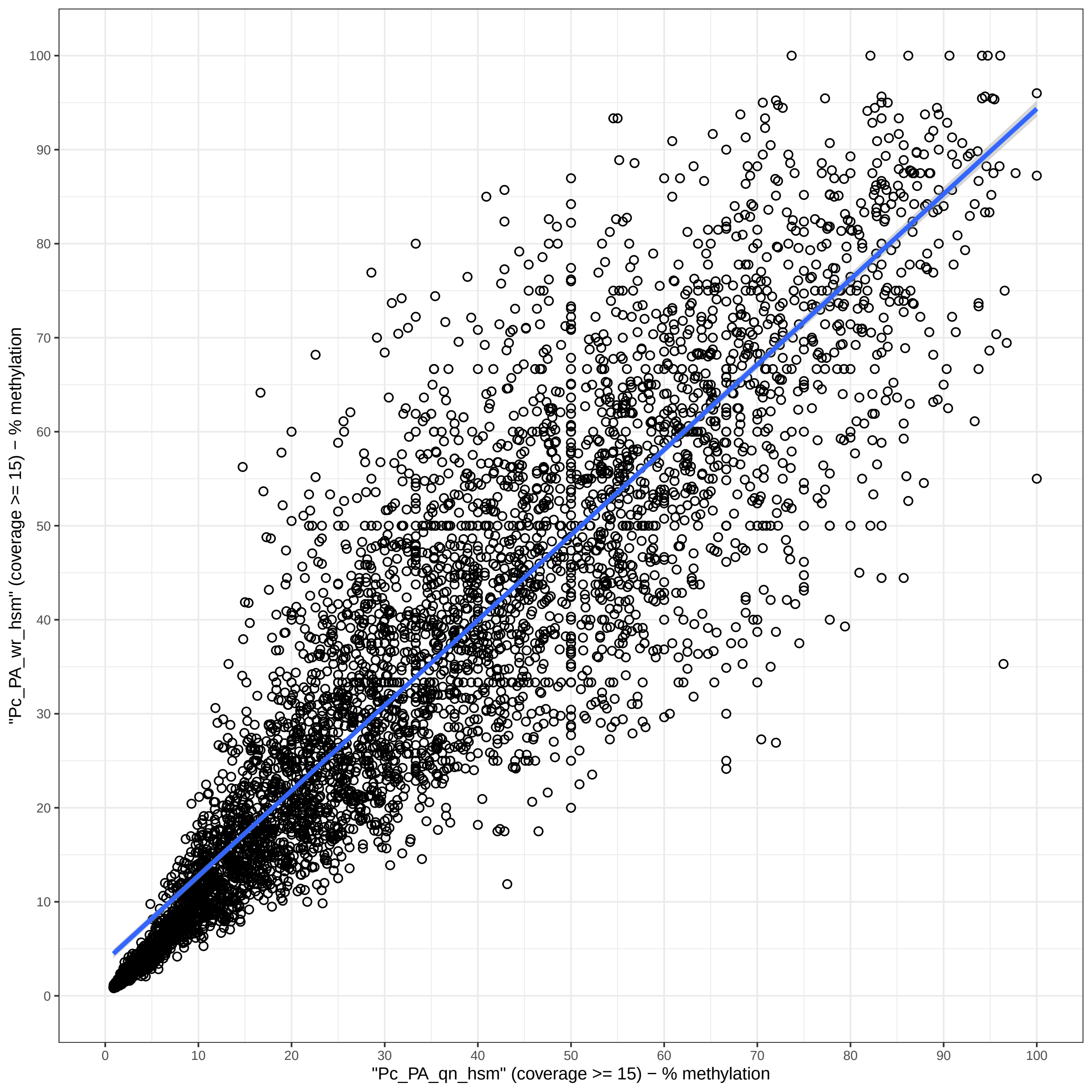

"Pc_PA_wr_hsm" (coverage >= 15) − % methylation
"Pc_PA_qn_hsm" (coverage >= 15) − % methylation
0
10
20
30
40
50
60
70
80
90
100

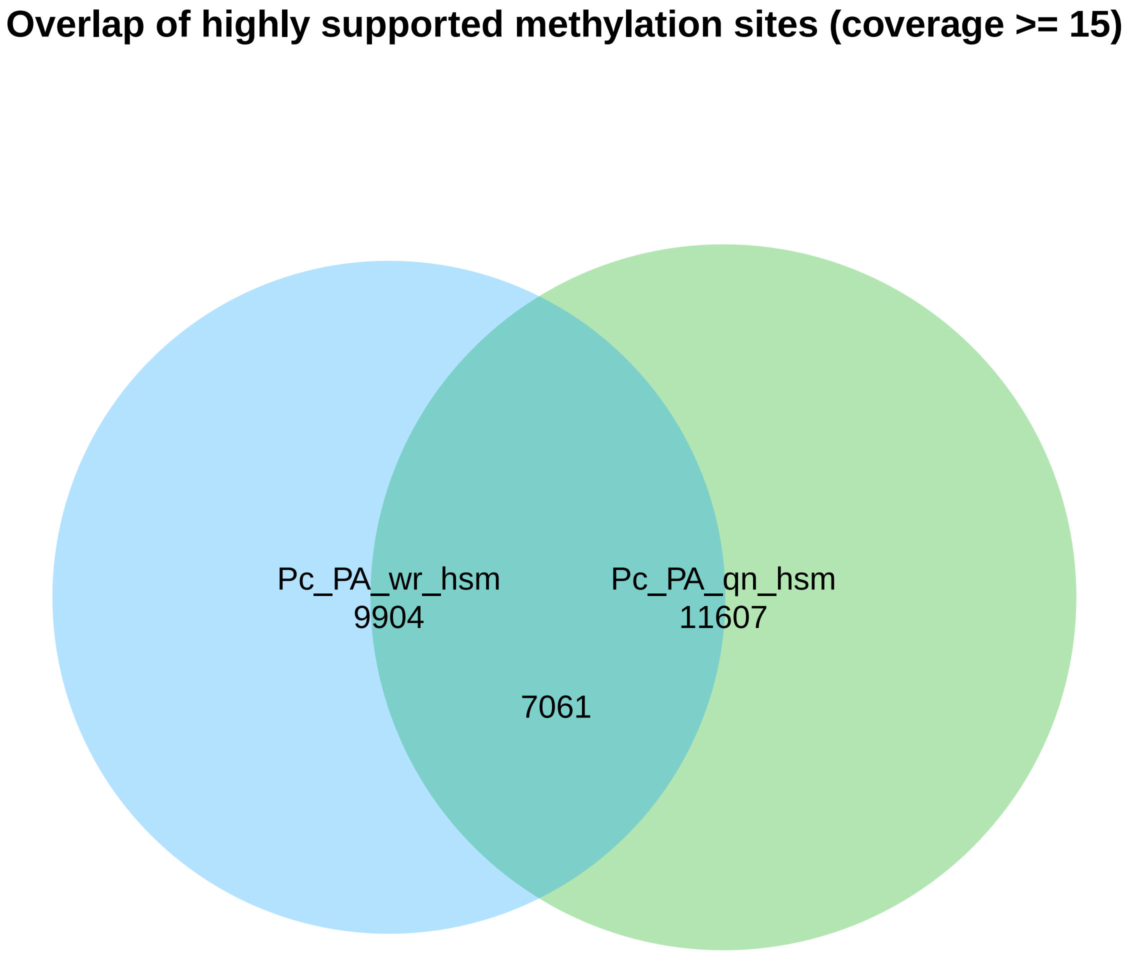
Overlap of highly supported methylation sites (coverage >= 15)
Pc_PA_wr_hsm
9904
Pc_PA_qn_hsm
11607
7061

```
Directory: CRL File: 0READMEcrl


CRL - Correlations between aggregate samples.

Input: studymk

Output: files crl-*.txt crl-*.pdf

Notes: The output is generated by BWASPR::cmpSamples(), which uses methylKit::unite()
        to determine the sites common to all samples and applies methylKit::getCorrelation()
        and methylKit::PCASamples().  The *.txt files show the correlations in a table, and
        the *.pdf files show graphics.

       Beware that the number of common sites (shown at the bottom of the *.txt files) may be
        small, which may make the correlations less informative.
```

Directory: CRL File: crl-Pc_PA.txt

| | queen | worker |
|---|---|---|
| queen | 1.000 | 0.906 |
| worker | 0.906 | 1.000 |

The number of conserved sites is 8061.

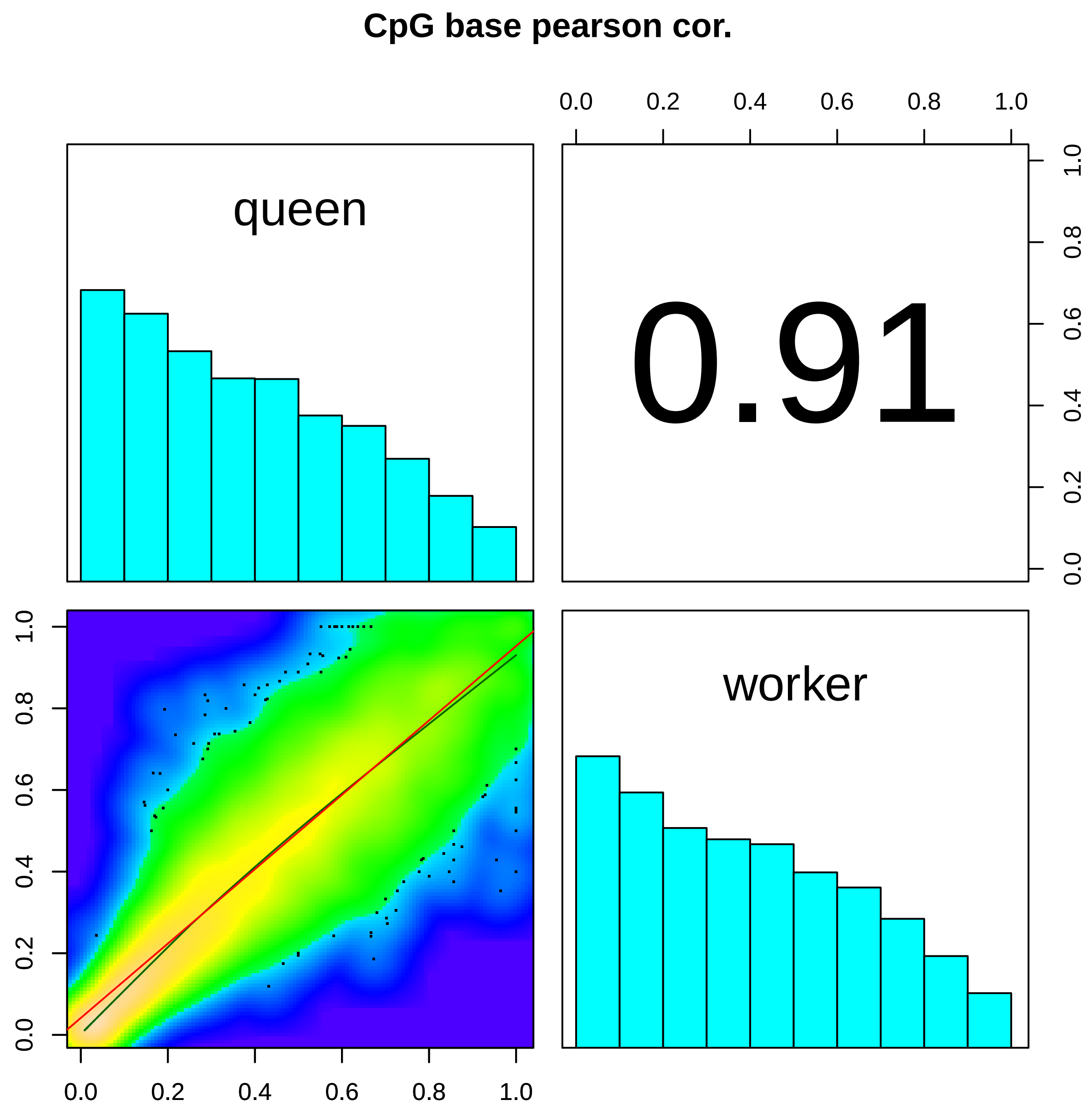

CpG base pearson cor.
queen
0.91
worker
0.0
0.2
0.4
0.6
0.8
1.0

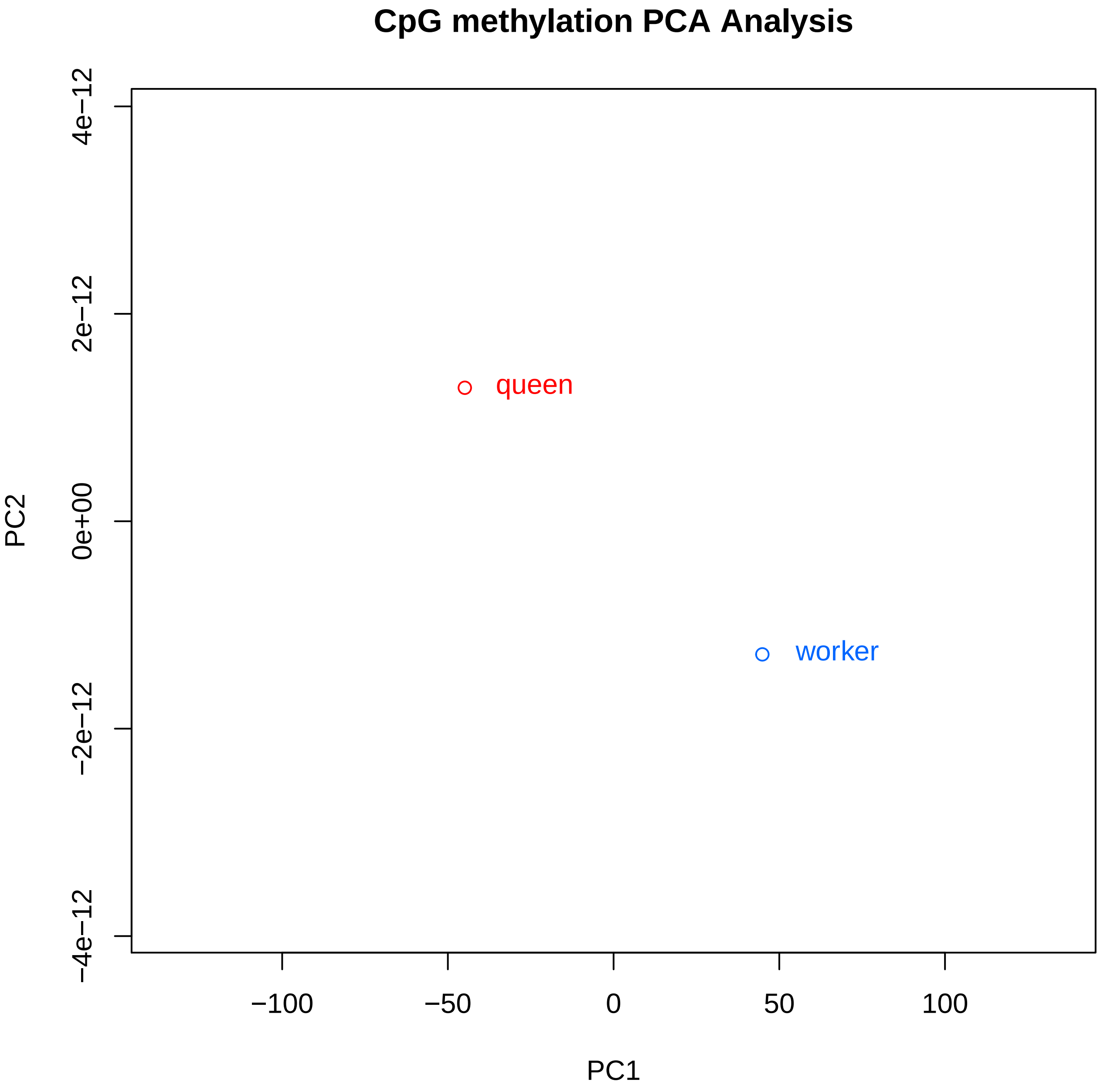
CpG methylation PCA Analysis
4e−12
2e−12
0e+00
−2e−12
−4e−12
PC2
queen
worker
−100
−50
0
50
100
PC1

```
Directory: REPCMS File: 0READMErepcms


REPCMS - Coverage and methylation statistics for replicate samples.

Input: studymk, repcovlist, replocount, rephicount

Output: files repcms-*txt and repcms-*pdf

Notes: The statistics are given for CpGhsm sites restricted to minimum or
        higher level coverage.  At the minimum level, the number of sites will be
        the number of lines of the corresponding *CpGhsm.mcalls file minus one
        (consistency check).  The distribution of methylation levels is really only
        interesting for high coverage because at low coverage there will be a strong
        bias towards high methylation levels (by definition of hsm sites).
        The same bias at high coverage would suggest that the sample includes a
        preponderance of consistently methylated genomic sites.

        replocount and rephicount set bounds on the coverage to exclude sites with too few
        or too many covering reads to provide statistics on a typical range.
```

```
Directory: REPCMS File: repcms-Pc_PA_queen_1.txt


Number of "queen_1" CpGhsm-sites with minimal and higher level coverage:

  number of "queen_1" CpGhsm-sites with coverage >=  4:   3988
  number of "queen_1" CpGhsm-sites with coverage >=  6:   3735
  number of "queen_1" CpGhsm-sites with coverage >= 10:   2916
  number of "queen_1" CpGhsm-sites with coverage >= 15:   2117


Coverage and methylation statistics for "queen_1" CpGhsm-sites at different levels of minimum coverage:


methylKit::getCoverageStats output for "queen_1" CpGhsm-sites (#: 3988) at minimum coverage 4  - read coverage statistics per base
summary:
   Min. 1st Qu.  Median    Mean 3rd Qu.    Max.
      4       9      16     113      28    7031
percentiles:
   0%   10%   20%   30%   40%   50%   60%   70%   80%   90%   95%   99% 99.5% 99.9%  100%
    4     6     8    10    12    16    19    24    33    56   295  3429  4273  6446  7031

methylKit::getMethylationStats output for "queen_1" CpGhsm-sites (#: 3988) at minimum coverage 4     - methylation statistics per base
summary:
   Min. 1st Qu.  Median    Mean 3rd Qu.    Max.
    1.1    36.8    55.6    56.3    76.9   100.0
percentiles:
    0%    10%    20%    30%    40%    50%    60%    70%    80%    90%    95%    99%  99.5%  99.9%   100%
  1.12  20.00  31.82  41.38  50.00  55.56  63.64  71.43  83.33 100.00 100.00 100.00 100.00 100.00 100.00



methylKit::getCoverageStats output for "queen_1" CpGhsm-sites (#: 3735) at minimum coverage 6  - read coverage statistics per base
summary:
   Min. 1st Qu.  Median    Mean 3rd Qu.    Max.
      6      10      17     120      30    7031
percentiles:
   0%   10%   20%   30%   40%   50%   60%   70%   80%   90%   95%   99% 99.5% 99.9%  100%
    6     7     9    11    13    17    20    26    35    59   339  3546  4345  6471  7031

methylKit::getMethylationStats output for "queen_1" CpGhsm-sites (#: 3735) at minimum coverage 6     - methylation statistics per base
summary:
   Min. 1st Qu.  Median    Mean 3rd Qu.    Max.
    1.1    35.3    53.8    53.4    71.4   100.0
percentiles:
    0%    10%    20%    30%    40%    50%    60%    70%    80%    90%    95%    99%  99.5%  99.9%   100%
  1.12  19.05  30.56  40.00  47.06  53.85  61.54  69.23  76.92  85.71  91.67 100.00 100.00 100.00 100.00



methylKit::getCoverageStats output for "queen_1" CpGhsm-sites (#: 2916) at minimum coverage 10 - read coverage statistics per base
summary:
   Min. 1st Qu.  Median    Mean 3rd Qu.    Max.
     10      14      20     152      35    7031
percentiles:
   0%   10%   20%   30%   40%   50%   60%   70%   80%   90%   95%   99% 99.5% 99.9%  100%
   10    11    13    15    18    20    25    31    41    89   589  3900  4620  6578  7031

methylKit::getMethylationStats output for "queen_1" CpGhsm-sites (#: 2916) at minimum coverage 10    - methylation statistics per base
summary:
   Min. 1st Qu.  Median    Mean 3rd Qu.    Max.
    1.1    30.0    46.2    46.4    61.5   100.0
percentiles:
    0%    10%    20%    30%    40%    50%    60%    70%    80%    90%    95%    99%  99.5%  99.9%   100%
```

```
  1.12  14.63  26.92  33.33  41.03  46.15  50.00  58.33  66.67  78.57  85.32 100.00 100.00 100.00 100.00


methylKit::getCoverageStats output for "queen_1" CpGhsm-sites (#: 2117) at minimum coverage 15 - read coverage statistics per base
summary:
   Min. 1st Qu.  Median    Mean 3rd Qu.    Max.
     15      19      27     205      43    7031
percentiles:
   0%   10%   20%   30%   40%   50%   60%   70%   80%   90%   95%   99% 99.5% 99.9%  100%
   15    16    18    20    23    27    32    39    53   257   938  4171  5243  6902  7031

methylKit::getMethylationStats output for "queen_1" CpGhsm-sites (#: 2117) at minimum coverage 15   - methylation statistics per base
summary:
   Min. 1st Qu.  Median    Mean 3rd Qu.    Max.
    1.1    25.0    38.5    40.1    53.8   100.0
percentiles:
    0%    10%    20%    30%    40%    50%    60%    70%    80%    90%    95%    99%  99.5%  99.9%   100%
  1.12   8.38  20.95  28.12  33.33  38.46  44.07  50.00  59.07  72.73  82.35  93.33  95.24 100.00 100.00



methylKit::getCoverageStats output for "queen_1" CpGhsm-sites (#: 2634) in coverage range [10-100]  - read coverage statistics per base
summary:
   Min. 1st Qu.  Median    Mean 3rd Qu.    Max.
   10.0    13.0    19.0    23.7    30.0   100.0
percentiles:
   0%   10%   20%   30%   40%   50%   60%   70%   80%   90%   95%   99% 99.5% 99.9%  100%
 10.0  11.0  12.0  14.0  17.0  19.0  22.0  27.0  33.0  42.0  54.0  76.0  83.8  94.7 100.0

methylKit::getMethylationStats output for "queen_1" CpGhsm-sites (#: 2634) in coverage range [10-100]     - methylation statistics per base
summary:
   Min. 1st Qu.  Median    Mean 3rd Qu.    Max.
    9.6    35.3    50.0    50.5    63.6   100.0
percentiles:
    0%    10%    20%    30%    40%    50%    60%    70%    80%    90%    95%    99%  99.5%  99.9%   100%
  9.57  25.71  31.82  38.71  44.44  50.00  53.85  60.00  68.75  80.00  86.36 100.00 100.00 100.00 100.00
```

Directory: REPCMS File: repcms-Pc_PA_queen_2.txt

Number of "queen_2" CpGhsm-sites with minimal and higher level coverage:

```
  number of "queen_2" CpGhsm-sites with coverage >=  4:   4835
  number of "queen_2" CpGhsm-sites with coverage >=  6:   4599
  number of "queen_2" CpGhsm-sites with coverage >= 10:   3780
  number of "queen_2" CpGhsm-sites with coverage >= 15:   2878
```

Coverage and methylation statistics for "queen_2" CpGhsm-sites at different levels of minimum coverage:

```
methylKit::getCoverageStats output for "queen_2" CpGhsm-sites (#: 4835) at minimum coverage 4  - read coverage statistics per base
summary:
   Min. 1st Qu.  Median    Mean 3rd Qu.    Max. 
      4      10      18     165      32    9729 
percentiles:
   0%   10%   20%   30%   40%   50%   60%   70%   80%   90%   95%   99% 99.5% 99.9%  100% 
    4     7     9    11    14    18    22    28    37    63   348  5462  7011  8890  9729 

methylKit::getMethylationStats output for "queen_2" CpGhsm-sites (#: 4835) at minimum coverage 4     - methylation statistics per base
summary:
   Min. 1st Qu.  Median    Mean 3rd Qu.    Max. 
    1.0    33.3    53.3    53.5    73.3   100.0 
percentiles:
    0%    10%    20%    30%    40%    50%    60%    70%    80%    90%    95%    99%  99.5%  99.9%   100% 
  1.04  18.85  29.03  37.50  45.45  53.33  60.00  69.54  78.57  90.91 100.00 100.00 100.00 100.00 100.00 



methylKit::getCoverageStats output for "queen_2" CpGhsm-sites (#: 4599) at minimum coverage 6  - read coverage statistics per base
summary:
   Min. 1st Qu.  Median    Mean 3rd Qu.    Max. 
      6      11      19     173      33    9729 
percentiles:
   0%   10%   20%   30%   40%   50%   60%   70%   80%   90%   95%   99% 99.5% 99.9%  100% 
    6     8    10    12    15    19    23    29    38    66   386  5602  7226  8897  9729 

methylKit::getMethylationStats output for "queen_2" CpGhsm-sites (#: 4599) at minimum coverage 6     - methylation statistics per base
summary:
   Min. 1st Qu.  Median    Mean 3rd Qu.    Max. 
    1.0    32.0    50.0    51.1    70.6   100.0 
percentiles:
    0%    10%    20%    30%    40%    50%    60%    70%    80%    90%    95%    99%  99.5%  99.9%   100% 
  1.04  18.18  28.08  35.76  43.75  50.00  58.33  66.67  75.00  85.71  90.95 100.00 100.00 100.00 100.00 



methylKit::getCoverageStats output for "queen_2" CpGhsm-sites (#: 3780) at minimum coverage 10 - read coverage statistics per base
summary:
   Min. 1st Qu.  Median    Mean 3rd Qu.    Max. 
     10      15      22     209      38    9729 
percentiles:
    0%    10%    20%    30%    40%    50%    60%    70%    80%    90%    95%    99%  99.5%  99.9%   100% 
  10.0   11.0   14.0   16.0   19.0   22.0   27.0   33.0   44.0   88.1  610.3 5884.2 7601.9 8949.8 9729.0 

methylKit::getMethylationStats output for "queen_2" CpGhsm-sites (#: 3780) at minimum coverage 10    - methylation statistics per base
summary:
   Min. 1st Qu.  Median    Mean 3rd Qu.    Max. 
    1.0    28.6    45.0    45.2    61.1   100.0 
percentiles:
    0%    10%    20%    30%    40%    50%    60%    70%    80%    90%    95%    99%  99.5%  99.9%   100% 
```

```
  1.04  14.81  25.48  31.82  38.10  45.00  50.00  57.14  66.67  76.92  85.71  94.51 100.00 100.00 100.00


methylKit::getCoverageStats output for "queen_2" CpGhsm-sites (#: 2878) at minimum coverage 15 - read coverage statistics per base
summary:
   Min. 1st Qu.  Median    Mean 3rd Qu.    Max.
     15      20      28     271      45    9729
percentiles:
   0%   10%   20%   30%   40%   50%   60%   70%   80%   90%   95%   99% 99.5% 99.9%  100%
   15    16    19    21    24    28    33    40    53   190  1124  6582  7792  9128  9729

methylKit::getMethylationStats output for "queen_2" CpGhsm-sites (#: 2878) at minimum coverage 15   - methylation statistics per base
summary:
   Min. 1st Qu.  Median    Mean 3rd Qu.    Max.
    1.0    25.0    37.0    39.4    53.3   100.0
percentiles:
    0%    10%    20%    30%    40%    50%    60%    70%    80%    90%    95%    99%  99.5%  99.9%   100%
  1.04   9.98  21.21  27.27  32.00  37.04  41.95  50.00  58.33  72.22  80.00  93.33  94.12 100.00 100.00


methylKit::getCoverageStats output for "queen_2" CpGhsm-sites (#: 3425) in coverage range [10-100]  - read coverage statistics per base
summary:
   Min. 1st Qu.  Median    Mean 3rd Qu.    Max.
   10.0    14.0    20.0    25.5    32.0    99.0
percentiles:
   0%   10%   20%   30%   40%   50%   60%   70%   80%   90%   95%   99% 99.5% 99.9%  100%
 10.0  11.0  13.0  15.0  18.0  20.0  24.0  29.0  35.0  46.0  57.0  84.0  90.0  96.6  99.0

methylKit::getMethylationStats output for "queen_2" CpGhsm-sites (#: 3425) in coverage range [10-100]     - methylation statistics per base
summary:
   Min. 1st Qu.  Median    Mean 3rd Qu.    Max.
    9.6    32.7    47.1    49.0    63.2   100.0
percentiles:
    0%    10%    20%    30%    40%    50%    60%    70%    80%    90%    95%    99%  99.5%  99.9%   100%
  9.57  23.53  30.00  35.33  41.67  47.06  52.94  60.00  68.00  78.45  86.20  95.24 100.00 100.00 100.00
```

```
Directory: REPCMS File: repcms-Pc_PA_queen_3.txt


Number of "queen_3" CpGhsm-sites with minimal and higher level coverage:

  number of "queen_3" CpGhsm-sites with coverage >=  4:   5147
  number of "queen_3" CpGhsm-sites with coverage >=  6:   4867
  number of "queen_3" CpGhsm-sites with coverage >= 10:   4131
  number of "queen_3" CpGhsm-sites with coverage >= 15:   3311


Coverage and methylation statistics for "queen_3" CpGhsm-sites at different levels of minimum coverage:


methylKit::getCoverageStats output for "queen_3" CpGhsm-sites (#: 5147) at minimum coverage 4  - read coverage statistics per base
summary:
   Min. 1st Qu.  Median    Mean 3rd Qu.    Max.
      4      11      21     289      53   12407
percentiles:
     0%     10%     20%     30%     40%     50%     60%     70%     80%     90%     95%     99%   99.5%   99.9%    100%
    4.0     7.0    10.0    12.0    16.0    21.0    29.0    41.0    88.8   415.2  1096.7  7271.2  8916.6 11219.5 12407.0

methylKit::getMethylationStats output for "queen_3" CpGhsm-sites (#: 5147) at minimum coverage 4     - methylation statistics per base
summary:
   Min. 1st Qu.  Median    Mean 3rd Qu.    Max.
    0.9    20.6    47.4    47.7    71.4   100.0
percentiles:
     0%     10%     20%     30%     40%     50%     60%     70%     80%     90%     95%     99%   99.5%   99.9%    100%
  0.944   5.506  13.924  26.923  37.500  47.368  56.522  66.667  77.778  90.909 100.000 100.000 100.000 100.000 100.000



methylKit::getCoverageStats output for "queen_3" CpGhsm-sites (#: 4867) at minimum coverage 6  - read coverage statistics per base
summary:
   Min. 1st Qu.  Median    Mean 3rd Qu.    Max.
      6      12      23     305      59   12407
percentiles:
   0%   10%   20%   30%   40%   50%   60%   70%   80%   90%   95%   99% 99.5% 99.9%  100%
    6     8    11    14    18    23    32    44   106   454  1165  7473  9059 11256 12407

methylKit::getMethylationStats output for "queen_3" CpGhsm-sites (#: 4867) at minimum coverage 6     - methylation statistics per base
summary:
   Min. 1st Qu.  Median    Mean 3rd Qu.    Max.
    0.9    18.9    45.5    44.7    67.3   100.0
percentiles:
     0%     10%     20%     30%     40%     50%     60%     70%     80%     90%     95%     99%   99.5%   99.9%    100%
  0.944   5.189  12.321  25.000  35.000  45.455  54.545  62.500  72.727  83.333  90.000 100.000 100.000 100.000 100.000



methylKit::getCoverageStats output for "queen_3" CpGhsm-sites (#: 4131) at minimum coverage 10 - read coverage statistics per base
summary:
   Min. 1st Qu.  Median    Mean 3rd Qu.    Max.
     10      16      29     358      88   12407
percentiles:
   0%   10%   20%   30%   40%   50%   60%   70%   80%   90%   95%   99% 99.5% 99.9%  100%
   10    11    15    18    22    29    39    57   173   551  1456  7824  9210 11374 12407

methylKit::getMethylationStats output for "queen_3" CpGhsm-sites (#: 4131) at minimum coverage 10    - methylation statistics per base
summary:
   Min. 1st Qu.  Median    Mean 3rd Qu.    Max.
    0.9    14.0    37.5    38.6    58.3   100.0
percentiles:
     0%     10%     20%     30%     40%     50%     60%     70%     80%     90%     95%     99%   99.5%   99.9%    100%
```

```
  0.944   4.607   9.016  19.512  29.167  37.500  45.455  53.846  63.158  76.923  84.211  95.238 100.000 100.000 100.000
```

```
methylKit::getCoverageStats output for "queen_3" CpGhsm-sites (#: 3311) at minimum coverage 15 - read coverage statistics per base
summary:
   Min. 1st Qu.  Median    Mean 3rd Qu.    Max.
     15      22      39     444     172   12407
percentiles:
   0%   10%   20%   30%   40%   50%   60%   70%   80%   90%   95%   99% 99.5% 99.9%  100%
   15    17    20    24    31    39    51    97   281   817  2109  8399  9624 11554 12407

methylKit::getMethylationStats output for "queen_3" CpGhsm-sites (#: 3311) at minimum coverage 15   - methylation statistics per base
summary:
   Min. 1st Qu.  Median    Mean 3rd Qu.    Max.
    0.9     9.0    29.2    32.4    48.6   100.0
percentiles:
     0%     10%     20%     30%     40%     50%     60%     70%     80%     90%     95%     99%   99.5%   99.9%    100%
  0.944   3.939   6.780  12.903  21.622  29.167  35.484  43.750  54.545  69.231  80.000  94.118 100.000 100.000 100.000
```

```
methylKit::getCoverageStats output for "queen_3" CpGhsm-sites (#: 3146) in coverage range [10-100]  - read coverage statistics per base
summary:
   Min. 1st Qu.  Median    Mean 3rd Qu.    Max.
   10.0    14.0    21.0    27.6    36.0   100.0
percentiles:
   0%   10%   20%   30%   40%   50%   60%   70%   80%   90%   95%   99% 99.5% 99.9%  100%
 10.0  11.0  13.0  15.5  18.0  21.0  26.0  32.0  39.0  53.0  67.0  91.0  95.0  98.9 100.0

methylKit::getMethylationStats output for "queen_3" CpGhsm-sites (#: 3146) in coverage range [10-100]     - methylation statistics per base
summary:
   Min. 1st Qu.  Median    Mean 3rd Qu.    Max.
    9.3    31.1    46.7    48.7    63.6   100.0
percentiles:
    0%    10%    20%    30%    40%    50%    60%    70%    80%    90%    95%    99%  99.5%  99.9%   100%
  9.28  20.51  28.00  34.48  41.67  46.67  53.33  60.00  69.23  80.00  87.50 100.00 100.00 100.00 100.00
```

```
Directory: REPCMS File: repcms-Pc_PA_worker_1.txt


Number of "worker_1" CpGhsm-sites with minimal and higher level coverage:

  number of "worker_1" CpGhsm-sites with coverage >=  4:   4476
  number of "worker_1" CpGhsm-sites with coverage >=  6:   4282
  number of "worker_1" CpGhsm-sites with coverage >= 10:   3684
  number of "worker_1" CpGhsm-sites with coverage >= 15:   3003


Coverage and methylation statistics for "worker_1" CpGhsm-sites at different levels of minimum coverage:


methylKit::getCoverageStats output for "worker_1" CpGhsm-sites (#: 4476) at minimum coverage 4 - read coverage statistics per base
summary:
   Min. 1st Qu.  Median    Mean 3rd Qu.    Max.
      4      11      22     226      46    9741
percentiles:
    0%    10%    20%    30%    40%    50%    60%    70%    80%    90%    95%    99%  99.5%  99.9%   100%
   4.0    8.0   10.0   13.0   18.0   22.5   30.0   40.0   58.0  207.5 1130.2 5143.5 6634.6 9028.4 9741.0

methylKit::getMethylationStats output for "worker_1" CpGhsm-sites (#: 4476) at minimum coverage 4   - methylation statistics per base
summary:
   Min. 1st Qu.  Median    Mean 3rd Qu.    Max.
    1.0    25.0    46.4    47.4    66.7   100.0
percentiles:
    0%    10%    20%    30%    40%    50%    60%    70%    80%    90%    95%    99%  99.5%  99.9%   100%
  1.04   8.89  20.00  30.00  38.89  46.43  54.55  62.50  72.73  85.71 100.00 100.00 100.00 100.00 100.00



methylKit::getCoverageStats output for "worker_1" CpGhsm-sites (#: 4282) at minimum coverage 6 - read coverage statistics per base
summary:
   Min. 1st Qu.  Median    Mean 3rd Qu.    Max.
      6      13      24     236      48    9741
percentiles:
   0%   10%   20%   30%   40%   50%   60%   70%   80%   90%   95%   99% 99.5% 99.9%  100%
    6     9    11    15    19    24    32    42    61   228  1222  5271  6714  9043  9741

methylKit::getMethylationStats output for "worker_1" CpGhsm-sites (#: 4282) at minimum coverage 6   - methylation statistics per base
summary:
   Min. 1st Qu.  Median    Mean 3rd Qu.    Max.
    1.0    24.1    45.5    45.0    63.8   100.0
percentiles:
    0%    10%    20%    30%    40%    50%    60%    70%    80%    90%    95%    99%  99.5%  99.9%   100%
  1.04   8.40  19.12  28.57  37.50  45.45  52.51  60.00  70.00  81.48  86.67 100.00 100.00 100.00 100.00



methylKit::getCoverageStats output for "worker_1" CpGhsm-sites (#: 3684) at minimum coverage 10     - read coverage statistics per base
summary:
   Min. 1st Qu.  Median    Mean 3rd Qu.    Max.
     10      17      29     273      57    9741
percentiles:
    0%    10%    20%    30%    40%    50%    60%    70%    80%    90%    95%    99%  99.5%  99.9%   100%
  10.0   12.0   15.0   19.0   23.0   29.0   37.0   47.0   68.4  379.7 1485.4 5382.2 6869.7 9248.5 9741.0

methylKit::getMethylationStats output for "worker_1" CpGhsm-sites (#: 3684) at minimum coverage 10  - methylation statistics per base
summary:
   Min. 1st Qu.  Median    Mean 3rd Qu.    Max.
    1.0    20.6    39.2    39.8    56.5   100.0
percentiles:
    0%    10%    20%    30%    40%    50%    60%    70%    80%    90%    95%    99%  99.5%  99.9%   100%
```

```
  1.04   6.34  17.01  25.00  32.50  39.17  46.15  52.78  60.71  72.73  80.95  91.67  96.85 100.00 100.00


methylKit::getCoverageStats output for "worker_1" CpGhsm-sites (#: 3003) at minimum coverage 15     - read coverage statistics per base
summary:
   Min. 1st Qu.  Median    Mean 3rd Qu.    Max.
     15      22      36     332      67    9741
percentiles:
   0%   10%   20%   30%   40%   50%   60%   70%   80%   90%   95%   99% 99.5% 99.9%  100%
   15    17    21    24    30    36    44    58    85   606  1974  5864  7173  9645  9741

methylKit::getMethylationStats output for "worker_1" CpGhsm-sites (#: 3003) at minimum coverage 15  - methylation statistics per base
summary:
   Min. 1st Qu.  Median    Mean 3rd Qu.    Max.
    1.0    17.3    33.3    34.9    50.0   100.0
percentiles:
    0%    10%    20%    30%    40%    50%    60%    70%    80%    90%    95%    99%  99.5%  99.9%   100%
  1.04   4.42  13.56  20.00  26.97  33.33  38.89  45.63  54.66  68.15  77.78  88.88  93.48 100.00 100.00


methylKit::getCoverageStats output for "worker_1" CpGhsm-sites (#: 3140) in coverage range [10-100] - read coverage statistics per base
summary:
   Min. 1st Qu.  Median    Mean 3rd Qu.    Max.
   10.0    16.0    24.0    30.6    40.0   100.0
percentiles:
   0%   10%   20%   30%   40%   50%   60%   70%   80%   90%   95%   99% 99.5% 99.9%  100%
 10.0  11.0  14.0  17.0  20.0  24.0  30.0  37.0  45.0  61.0  72.0  91.0  93.3  97.9 100.0

methylKit::getMethylationStats output for "worker_1" CpGhsm-sites (#: 3140) in coverage range [10-100]    - methylation statistics per base
summary:
   Min. 1st Qu.  Median    Mean 3rd Qu.    Max.
    9.3    29.4    44.4    45.6    60.0   100.0
percentiles:
    0%    10%    20%    30%    40%    50%    60%    70%    80%    90%    95%    99%  99.5%  99.9%   100%
  9.28  18.46  25.85  32.50  38.86  44.44  50.00  56.25  63.64  75.00  81.82  93.33 100.00 100.00 100.00
```

```
Directory: REPCMS File: repcms-Pc_PA_worker_2.txt


Number of "worker_2" CpGhsm-sites with minimal and higher level coverage:

  number of "worker_2" CpGhsm-sites with coverage >=  4:   3377
  number of "worker_2" CpGhsm-sites with coverage >=  6:   3150
  number of "worker_2" CpGhsm-sites with coverage >= 10:   2401
  number of "worker_2" CpGhsm-sites with coverage >= 15:   1678


Coverage and methylation statistics for "worker_2" CpGhsm-sites at different levels of minimum coverage:


methylKit::getCoverageStats output for "worker_2" CpGhsm-sites (#: 3377) at minimum coverage 4 - read coverage statistics per base
summary:
   Min. 1st Qu.  Median    Mean 3rd Qu.    Max.
      4       9      14     143      26    8394
percentiles:
   0%   10%   20%   30%   40%   50%   60%   70%   80%   90%   95%   99% 99.5% 99.9%  100%
    4     6     8    10    11    14    18    22    30    48   266  4453  5715  7507  8394

methylKit::getMethylationStats output for "worker_2" CpGhsm-sites (#: 3377) at minimum coverage 4    - methylation statistics per base
summary:
   Min. 1st Qu.  Median    Mean 3rd Qu.    Max.
    1.0    38.9    58.3    58.0    78.3   100.0
percentiles:
    0%    10%    20%    30%    40%    50%    60%    70%    80%    90%    95%    99%  99.5%  99.9%   100%
  1.04  22.22  33.33  43.75  50.00  58.33  66.67  75.00  83.33 100.00 100.00 100.00 100.00 100.00 100.00



methylKit::getCoverageStats output for "worker_2" CpGhsm-sites (#: 3150) at minimum coverage 6 - read coverage statistics per base
summary:
   Min. 1st Qu.  Median    Mean 3rd Qu.    Max.
      6      10      16     153      27    8394
percentiles:
   0%   10%   20%   30%   40%   50%   60%   70%   80%   90%   95%   99% 99.5% 99.9%  100%
    6     7     9    10    12    16    19    24    31    53   333  4547  5970  7546  8394

methylKit::getMethylationStats output for "worker_2" CpGhsm-sites (#: 3150) at minimum coverage 6    - methylation statistics per base
summary:
   Min. 1st Qu.  Median    Mean 3rd Qu.    Max.
    1.0    37.5    55.6    55.0    75.0   100.0
percentiles:
    0%    10%    20%    30%    40%    50%    60%    70%    80%    90%    95%    99%  99.5%  99.9%   100%
  1.04  20.59  33.33  42.11  50.00  55.56  62.50  71.01  77.78  85.71  93.33 100.00 100.00 100.00 100.00



methylKit::getCoverageStats output for "worker_2" CpGhsm-sites (#: 2401) at minimum coverage 10      - read coverage statistics per base
summary:
   Min. 1st Qu.  Median    Mean 3rd Qu.    Max.
     10      13      20     198      32    8394
percentiles:
   0%   10%   20%   30%   40%   50%   60%   70%   80%   90%   95%   99% 99.5% 99.9%  100%
   10    11    12    14    17    20    23    29    37    85   669  5023  6241  7892  8394

methylKit::getMethylationStats output for "worker_2" CpGhsm-sites (#: 2401) at minimum coverage 10   - methylation statistics per base
summary:
   Min. 1st Qu.  Median    Mean 3rd Qu.    Max.
    1.0    31.8    48.1    47.9    64.0   100.0
percentiles:
    0%    10%    20%    30%    40%    50%    60%    70%    80%    90%    95%    99%  99.5%  99.9%   100%
```

```
  1.04  16.33  28.57  35.29  42.86  48.15  53.85  60.00  69.23  80.00  87.50  96.30 100.00 100.00 100.00
```

```
methylKit::getCoverageStats output for "worker_2" CpGhsm-sites (#: 1678) at minimum coverage 15      - read coverage statistics per base
summary:
   Min. 1st Qu.  Median    Mean 3rd Qu.    Max.
     15      19      26     278      40    8394
percentiles:
   0%   10%   20%   30%   40%   50%   60%   70%   80%   90%   95%   99% 99.5% 99.9%  100%
   15    16    18    20    22    26    30    36    49   279  1574  5742  6594  8151  8394

methylKit::getMethylationStats output for "worker_2" CpGhsm-sites (#: 1678) at minimum coverage 15   - methylation statistics per base
summary:
   Min. 1st Qu.  Median    Mean 3rd Qu.    Max.
    1.0    25.8    38.8    41.2    56.9   100.0
percentiles:
    0%    10%    20%    30%    40%    50%    60%    70%    80%    90%    95%    99%  99.5%  99.9%   100%
  1.04   8.52  21.98  29.03  33.33  38.85  44.44  52.94  61.85  75.00  83.33  93.43  94.12 100.00 100.00
```

```
methylKit::getCoverageStats output for "worker_2" CpGhsm-sites (#: 2171) in coverage range [10-100]  - read coverage statistics per base
summary:
   Min. 1st Qu.  Median    Mean 3rd Qu.    Max.
   10.0    13.0    18.0    22.3    27.0    99.0
percentiles:
   0%   10%   20%   30%   40%   50%   60%   70%   80%   90%   95%   99% 99.5% 99.9%  100%
 10.0  11.0  12.0  14.0  16.0  18.0  21.0  25.0  30.0  39.0  48.0  74.3  84.2  97.7  99.0

methylKit::getMethylationStats output for "worker_2" CpGhsm-sites (#: 2171) in coverage range [10-100]     - methylation statistics per base
summary:
   Min. 1st Qu.  Median    Mean 3rd Qu.    Max.
   10.4    36.7    50.0    52.2    66.7   100.0
percentiles:
   0%   10%   20%   30%   40%   50%   60%   70%   80%   90%   95%   99% 99.5% 99.9%  100%
 10.4  26.1  33.3  40.0  45.5  50.0  55.6  62.5  70.0  80.0  89.3 100.0 100.0 100.0 100.0
```

```
Directory: REPCMS File: repcms-Pc_PA_worker_3.txt


Number of "worker_3" CpGhsm-sites with minimal and higher level coverage:

  number of "worker_3" CpGhsm-sites with coverage >=  4:   4032
  number of "worker_3" CpGhsm-sites with coverage >=  6:   3750
  number of "worker_3" CpGhsm-sites with coverage >= 10:   2998
  number of "worker_3" CpGhsm-sites with coverage >= 15:   2243


Coverage and methylation statistics for "worker_3" CpGhsm-sites at different levels of minimum coverage:


methylKit::getCoverageStats output for "worker_3" CpGhsm-sites (#: 4032) at minimum coverage 4 - read coverage statistics per base
summary:
   Min. 1st Qu.  Median    Mean 3rd Qu.    Max.
      4       9      17     126      32    9451
percentiles:
    0%    10%    20%    30%    40%    50%    60%    70%    80%    90%    95%    99%  99.5%  99.9%   100%
   4.0    6.0    8.0   10.0   13.0   17.0   21.0   28.0   39.8   86.0  388.4 3280.4 5157.3 8075.8 9451.0

methylKit::getMethylationStats output for "worker_3" CpGhsm-sites (#: 4032) at minimum coverage 4    - methylation statistics per base
summary:
   Min. 1st Qu.  Median    Mean 3rd Qu.    Max.
    1.0    33.3    56.2    56.1    80.0   100.0
percentiles:
    0%    10%    20%    30%    40%    50%    60%    70%    80%    90%    95%    99%  99.5%  99.9%   100%
  1.04  15.37  28.57  38.89  48.82  56.25  66.67  75.00  84.62 100.00 100.00 100.00 100.00 100.00 100.00



methylKit::getCoverageStats output for "worker_3" CpGhsm-sites (#: 3750) at minimum coverage 6 - read coverage statistics per base
summary:
   Min. 1st Qu.  Median    Mean 3rd Qu.    Max.
      6      10      18     136      35    9451
percentiles:
   0%   10%   20%   30%   40%   50%   60%   70%   80%   90%   95%   99% 99.5% 99.9%  100%
    6     8     9    11    14    18    22    29    41    96   438  3385  5193  8226  9451

methylKit::getMethylationStats output for "worker_3" CpGhsm-sites (#: 3750) at minimum coverage 6    - methylation statistics per base
summary:
   Min. 1st Qu.  Median    Mean 3rd Qu.    Max.
    1.0    31.8    53.8    52.8    75.0   100.0
percentiles:
    0%    10%    20%    30%    40%    50%    60%    70%    80%    90%    95%    99%  99.5%  99.9%   100%
  1.04  14.02  27.27  36.84  46.15  53.85  62.33  70.59  78.95  87.50 100.00 100.00 100.00 100.00 100.00



methylKit::getCoverageStats output for "worker_3" CpGhsm-sites (#: 2998) at minimum coverage 10      - read coverage statistics per base
summary:
   Min. 1st Qu.  Median    Mean 3rd Qu.    Max.
     10      14      22     168      41    9451
percentiles:
   0%   10%   20%   30%   40%   50%   60%   70%   80%   90%   95%   99% 99.5% 99.9%  100%
   10    11    13    16    19    22    28    36    50   174   584  4571  5927  8666  9451

methylKit::getMethylationStats output for "worker_3" CpGhsm-sites (#: 2998) at minimum coverage 10   - methylation statistics per base
summary:
   Min. 1st Qu.  Median    Mean 3rd Qu.    Max.
    1.0    27.3    46.0    45.9    64.0   100.0
percentiles:
    0%    10%    20%    30%    40%    50%    60%    70%    80%    90%    95%    99%  99.5%  99.9%   100%
```

```
  1.04   9.45  22.22  31.25  38.85  45.99  52.17  60.00  69.64  80.00  90.00 100.00 100.00 100.00 100.00



methylKit::getCoverageStats output for "worker_3" CpGhsm-sites (#: 2243) at minimum coverage 15      - read coverage statistics per base
summary:
   Min. 1st Qu.  Median    Mean 3rd Qu.    Max.
     15      20      30     220      53    9451
percentiles:
    0%    10%    20%    30%    40%    50%    60%    70%    80%    90%    95%    99%  99.5%  99.9%   100%
  15.0   17.0   19.0   21.0   25.0   30.0   36.0   45.0   68.6  332.2  878.7 4946.3 6250.5 8845.4 9451.0

methylKit::getMethylationStats output for "worker_3" CpGhsm-sites (#: 2243) at minimum coverage 15  - methylation statistics per base
summary:
   Min. 1st Qu.  Median    Mean 3rd Qu.    Max.
    1.0    20.7    36.8    39.2    56.5   100.0
percentiles:
    0%    10%    20%    30%    40%    50%    60%    70%    80%    90%    95%    99%  99.5%  99.9%   100%
  1.04   6.16  16.73  25.00  31.03  36.84  42.42  51.11  61.90  75.00  82.59  94.12  99.30 100.00 100.00



methylKit::getCoverageStats output for "worker_3" CpGhsm-sites (#: 2631) in coverage range [10-100] - read coverage statistics per base
summary:
   Min. 1st Qu.  Median    Mean 3rd Qu.    Max.
   10.0    14.0    20.0    25.6    32.0   100.0
percentiles:
   0%   10%   20%   30%   40%   50%   60%   70%   80%   90%   95%   99% 99.5% 99.9%  100%
 10.0  11.0  13.0  15.0  17.0  20.0  23.0  29.0  36.0  48.0  60.0  89.0  94.0  98.1 100.0

methylKit::getMethylationStats output for "worker_3" CpGhsm-sites (#: 2631) in coverage range [10-100]    - methylation statistics per base
summary:
   Min. 1st Qu.  Median    Mean 3rd Qu.    Max.
    9.8    34.3    50.0    51.3    66.7   100.0
percentiles:
    0%    10%    20%    30%    40%    50%    60%    70%    80%    90%    95%    99%  99.5%  99.9%   100%
  9.78  22.73  30.61  37.50  43.75  50.00  55.81  62.50  71.43  81.82  90.00 100.00 100.00 100.00 100.00
```

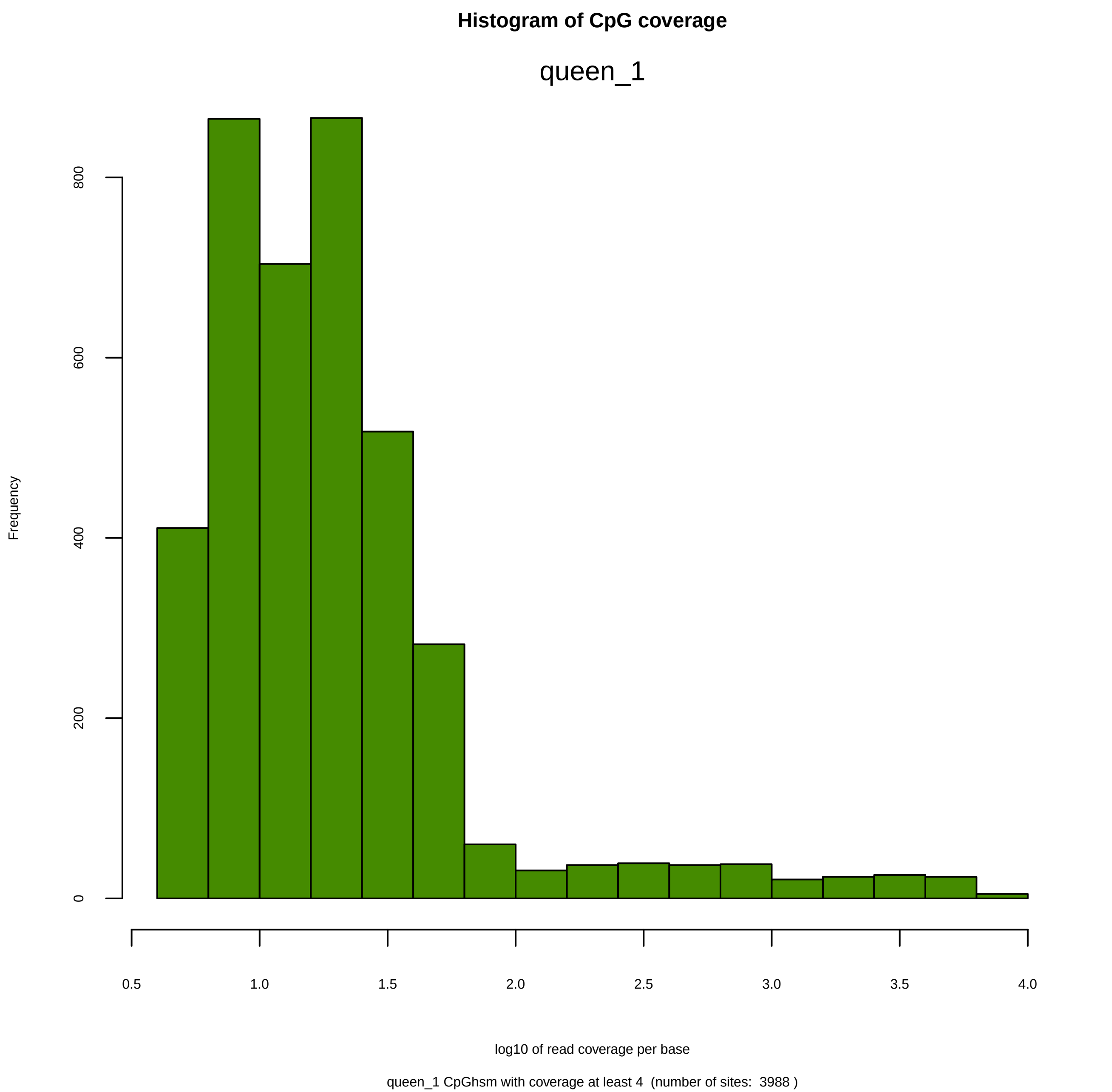

Histogram of CpG coverage
queen_1
Frequency
0
200
400
600
800
0.5
1.0
1.5
2.0
2.5
3.0
3.5
4.0
log10 of read coverage per base
queen_1 CpGhsm with coverage at least 4 (number of sites: 3988 )

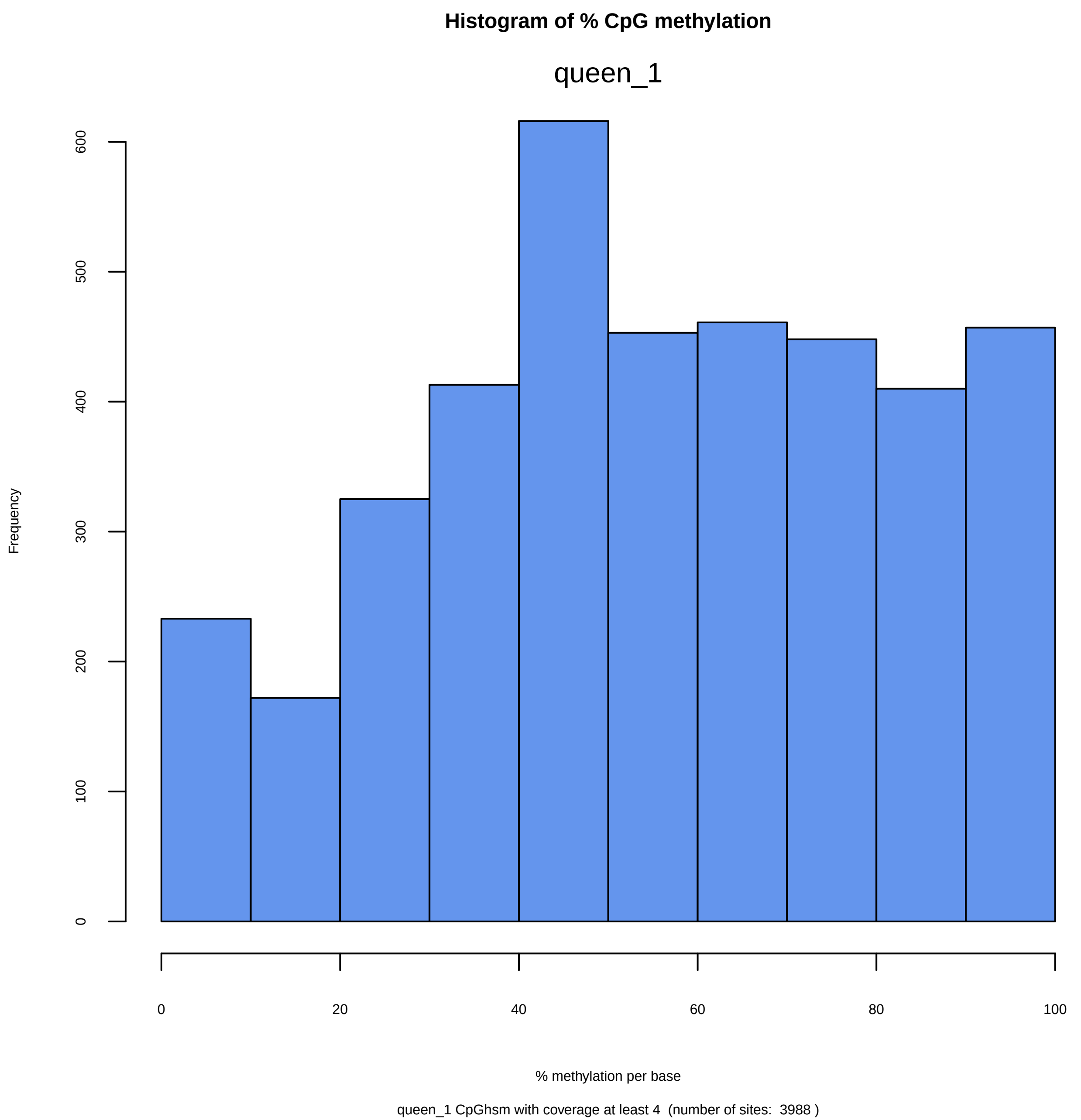

Histogram of % CpG methylation
queen_1
Frequency
0
100
200
300
400
500
600
0
20
40
60
80
100
% methylation per base
queen_1 CpGhsm with coverage at least 4 (number of sites: 3988 )

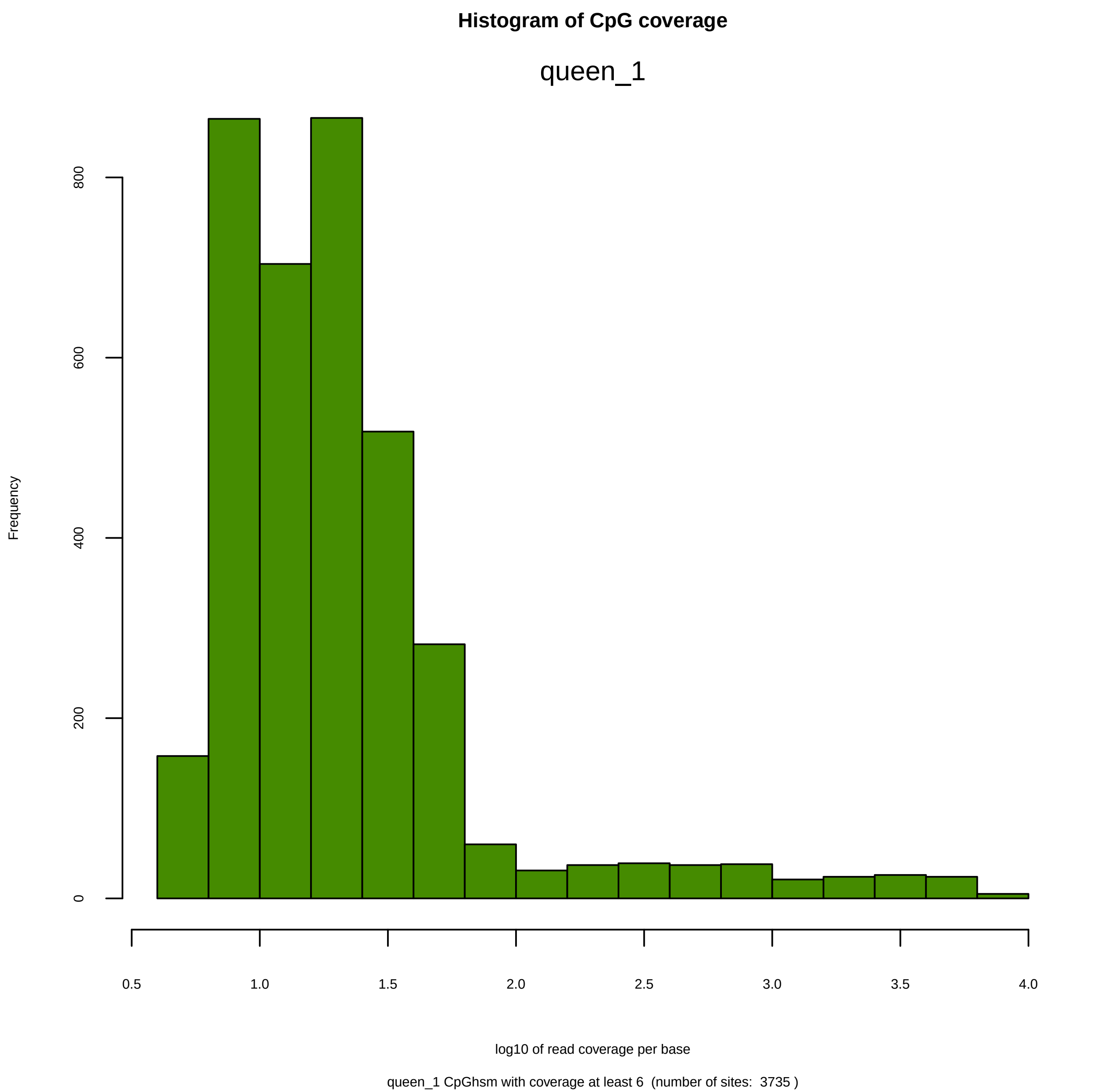
Histogram of CpG coverage
queen_1
Frequency
0
200
400
600
800
0.5
1.0
1.5
2.0
2.5
3.0
3.5
4.0
log10 of read coverage per base
queen_1 CpGhsm with coverage at least 6 (number of sites: 3735 )

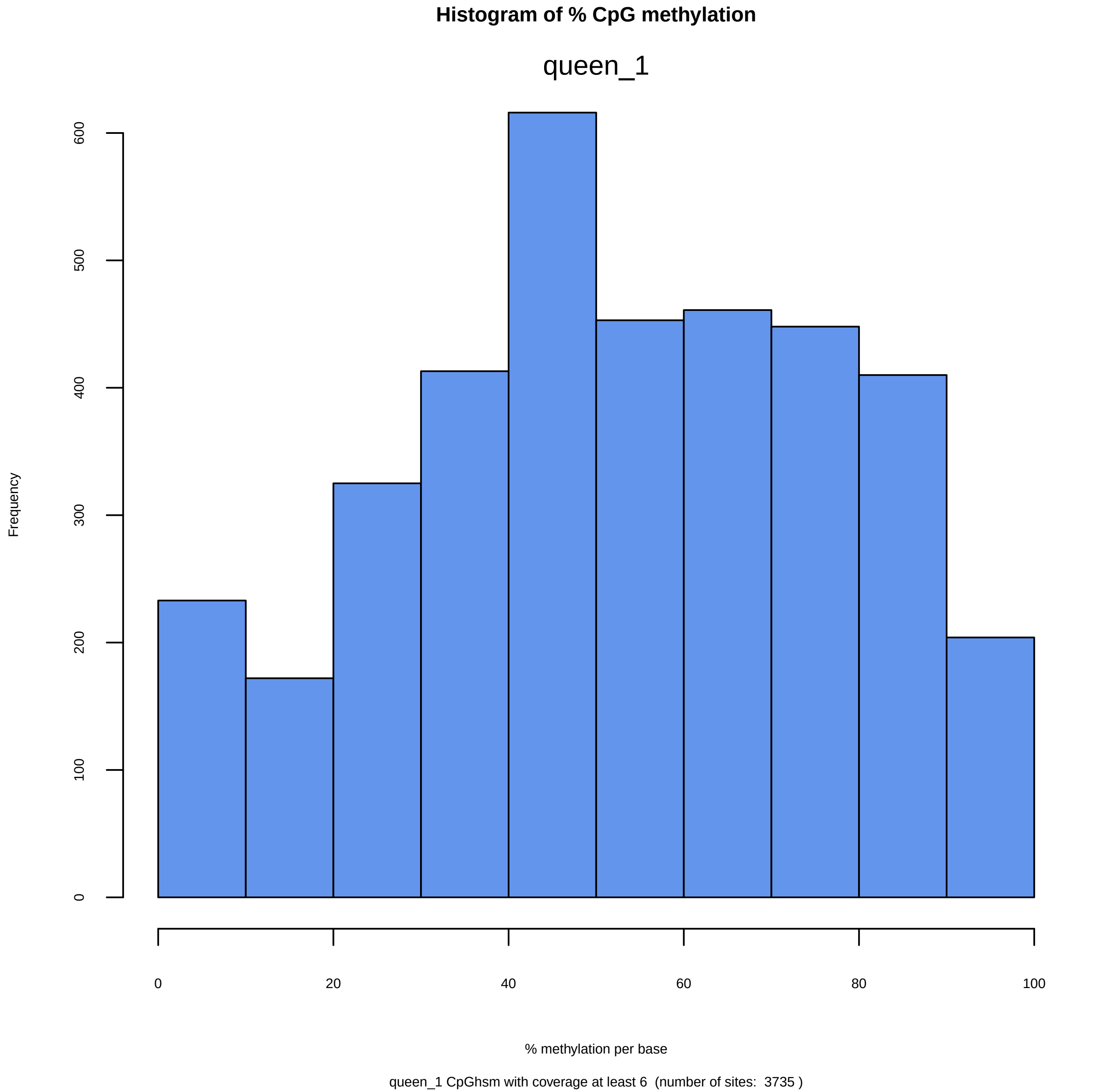
Histogram of % CpG methylation
queen_1
Frequency
0
100
200
300
400
500
600
0
20
40
60
80
100
% methylation per base
queen_1 CpGhsm with coverage at least 6  (number of sites:  3735 )

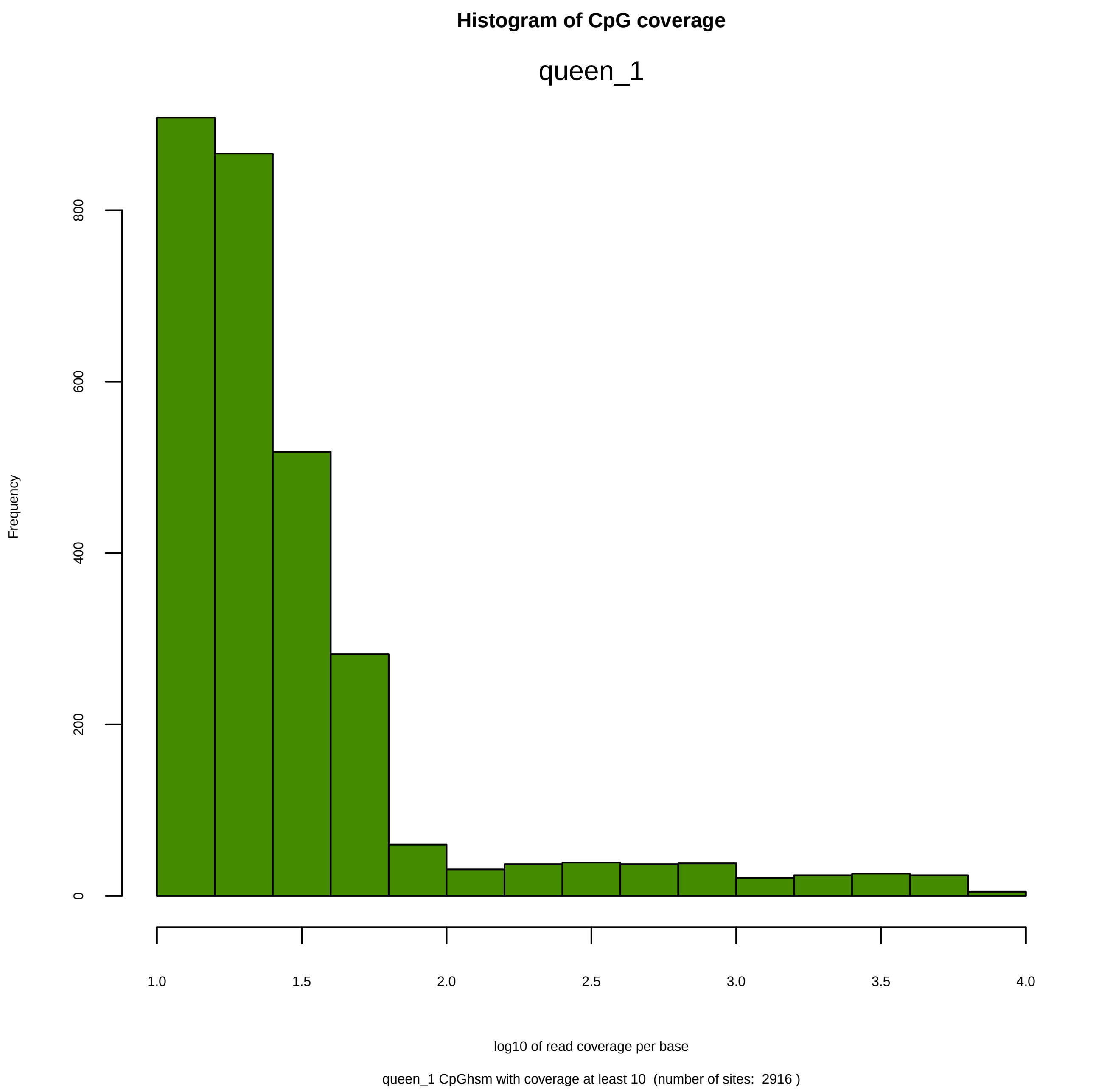
Histogram of CpG coverage
queen_1
Frequency
0
200
400
600
800
1.0
1.5
2.0
2.5
3.0
3.5
4.0
log10 of read coverage per base
queen_1 CpGhsm with coverage at least 10  (number of sites:  2916 )

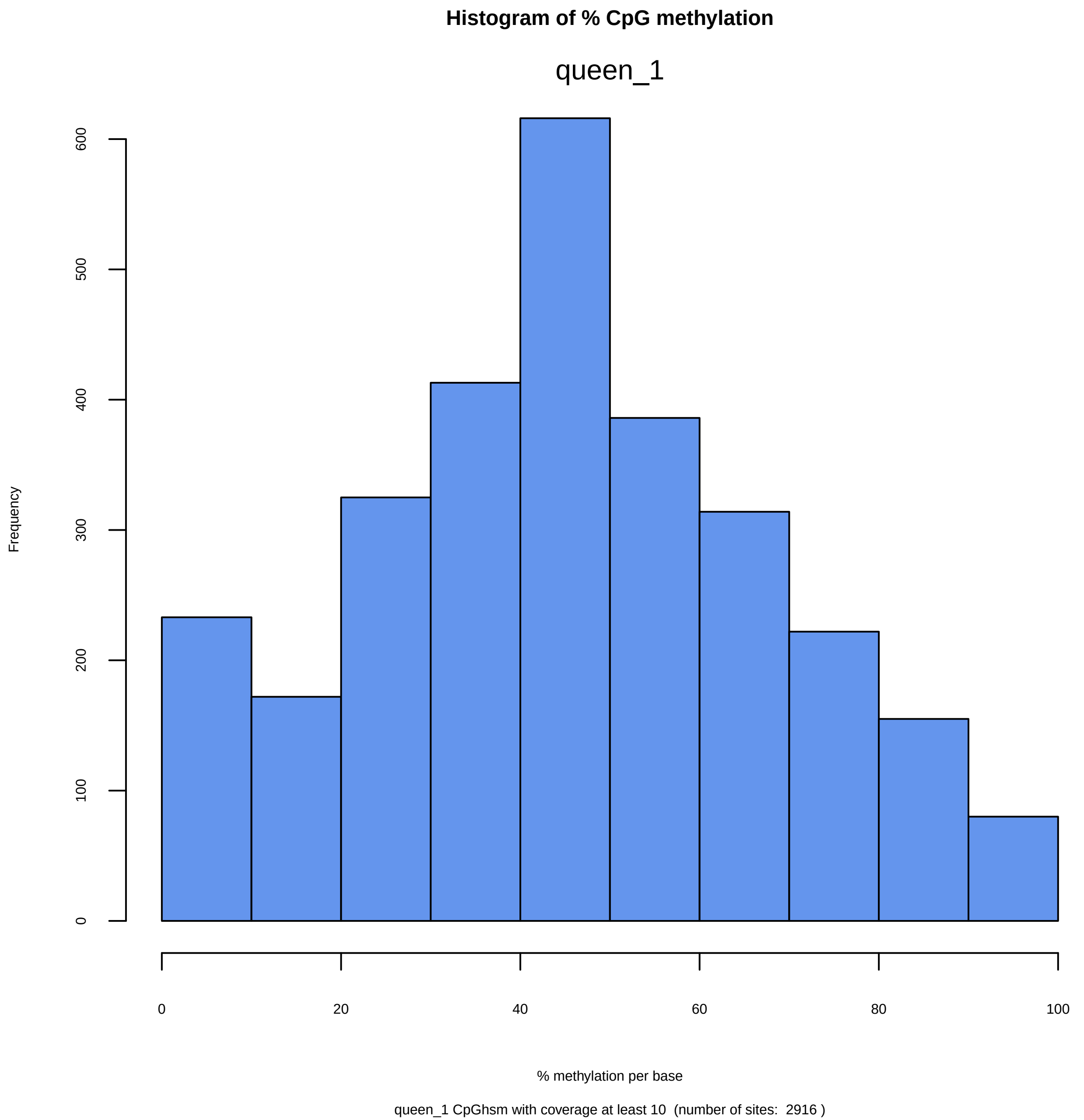

Histogram of % CpG methylation
queen_1
Frequency
0
100
200
300
400
500
600
0
20
40
60
80
100
% methylation per base
queen_1 CpGhsm with coverage at least 10  (number of sites:  2916 )

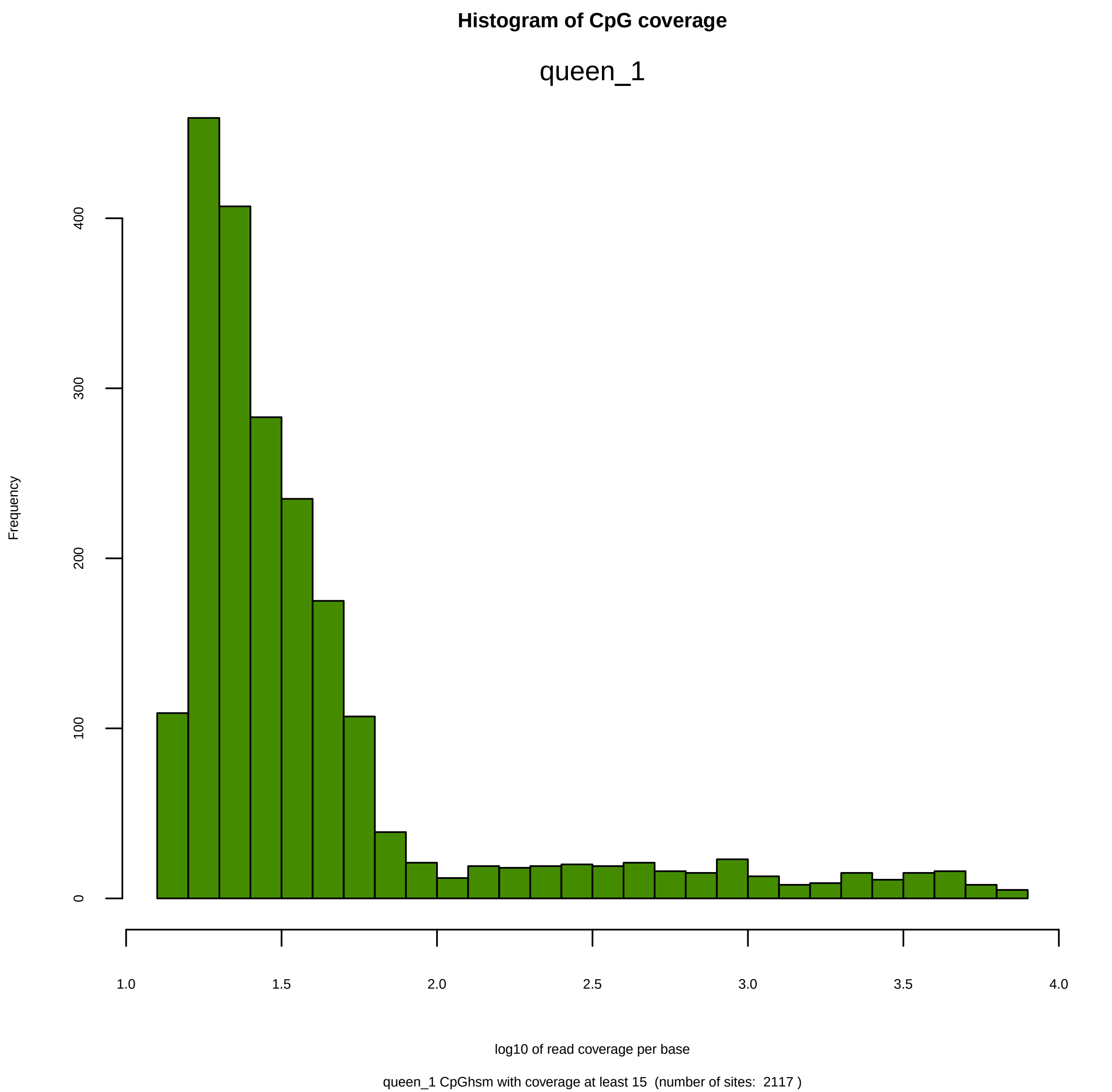

Histogram of CpG coverage
queen_1
Frequency
0
100
200
300
400
1.0
1.5
2.0
2.5
3.0
3.5
4.0
log10 of read coverage per base
queen_1 CpGhsm with coverage at least 15  (number of sites:  2117 )

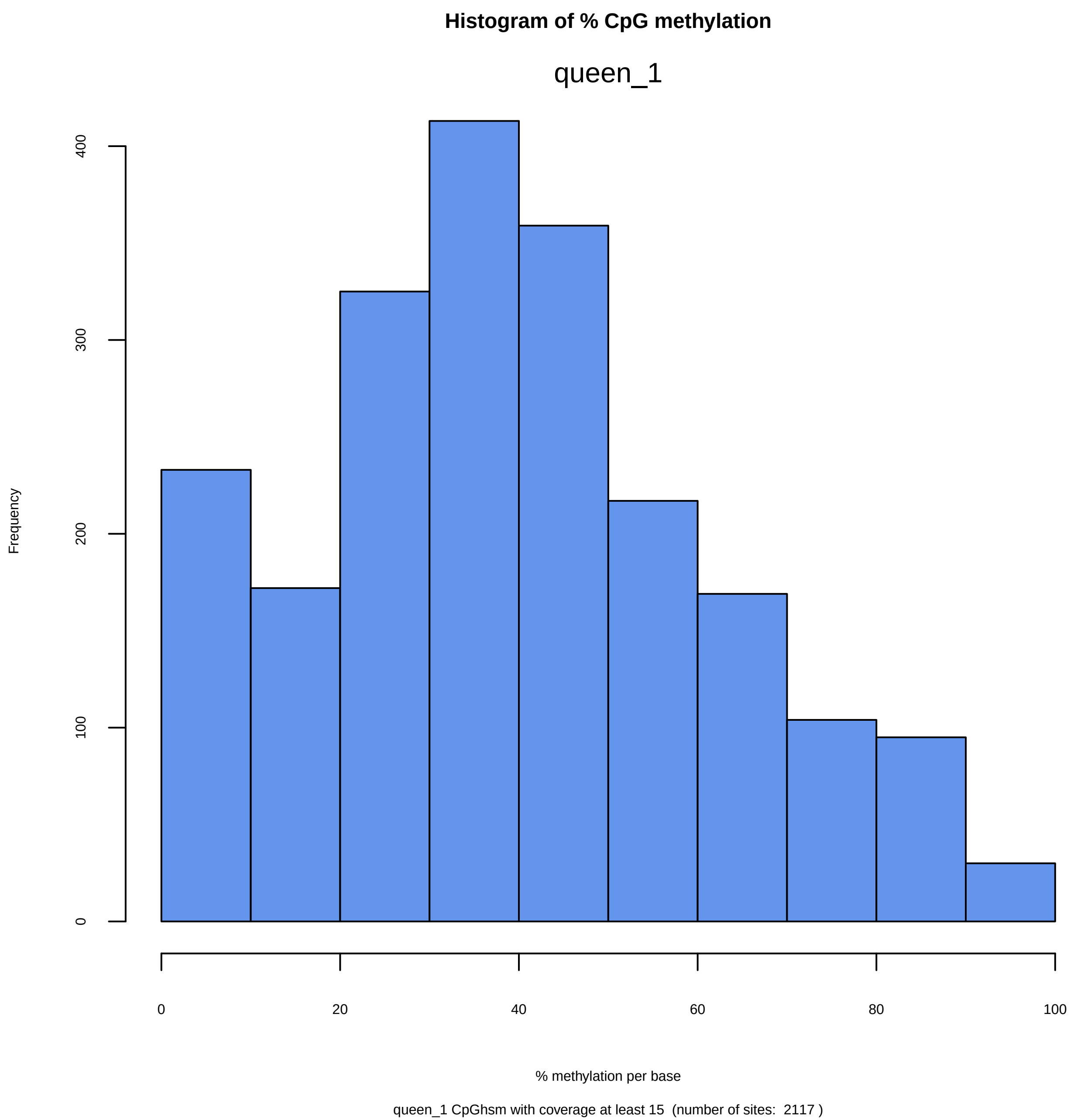
Histogram of % CpG methylation
queen_1
Frequency
0
100
200
300
400
0
20
40
60
80
100
% methylation per base
queen_1 CpGhsm with coverage at least 15  (number of sites:  2117 )

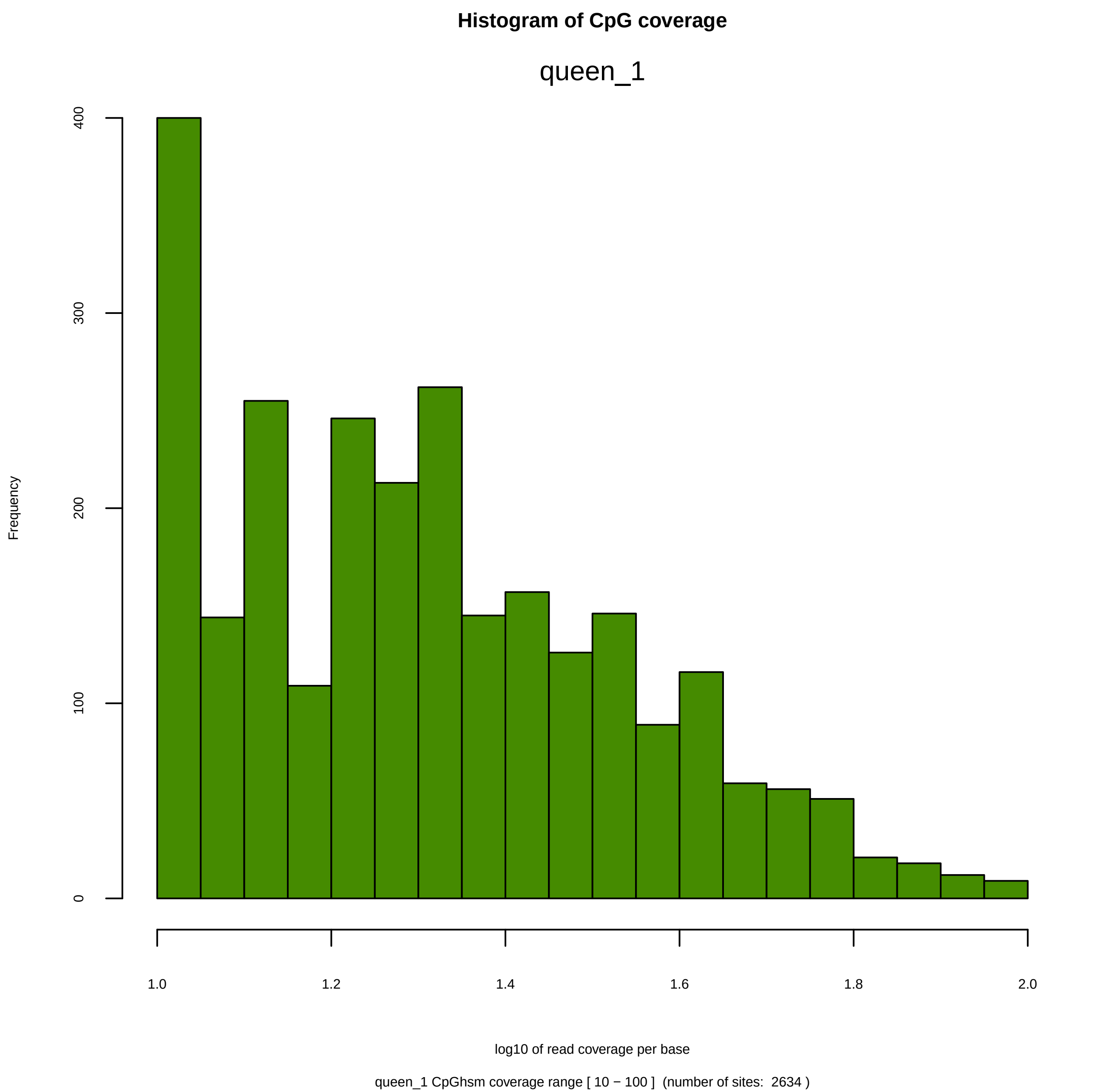
Histogram of CpG coverage
queen_1
Frequency
0
100
200
300
400
1.0
1.2
1.4
1.6
1.8
2.0
log10 of read coverage per base
queen_1 CpGhsm coverage range [ 10 − 100 ]  (number of sites:  2634 )

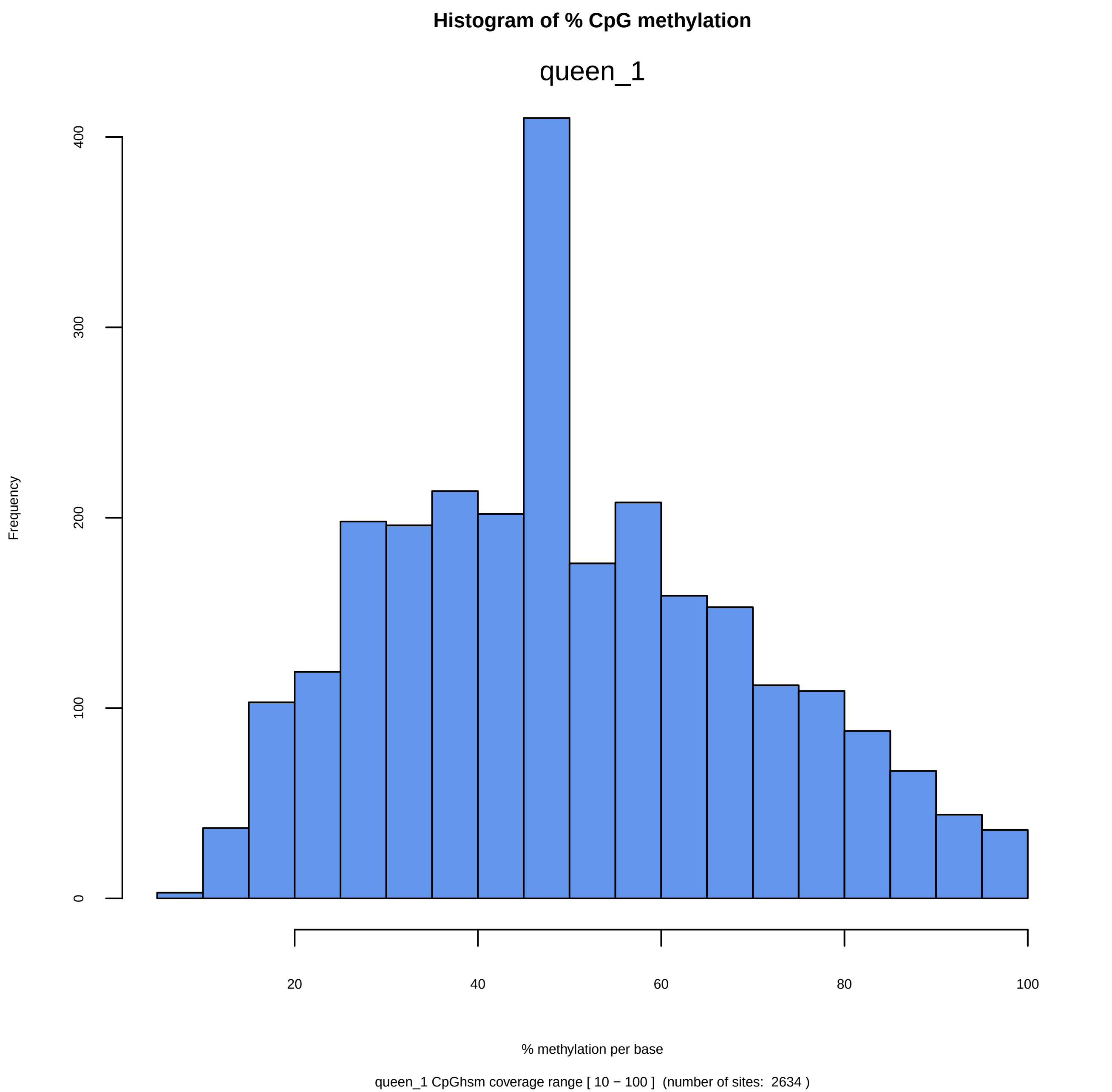
Histogram of % CpG methylation
queen_1
Frequency
0
100
200
300
400
20
40
60
80
100
% methylation per base
queen_1 CpGhsm coverage range [ 10 − 100 ]  (number of sites:  2634 )

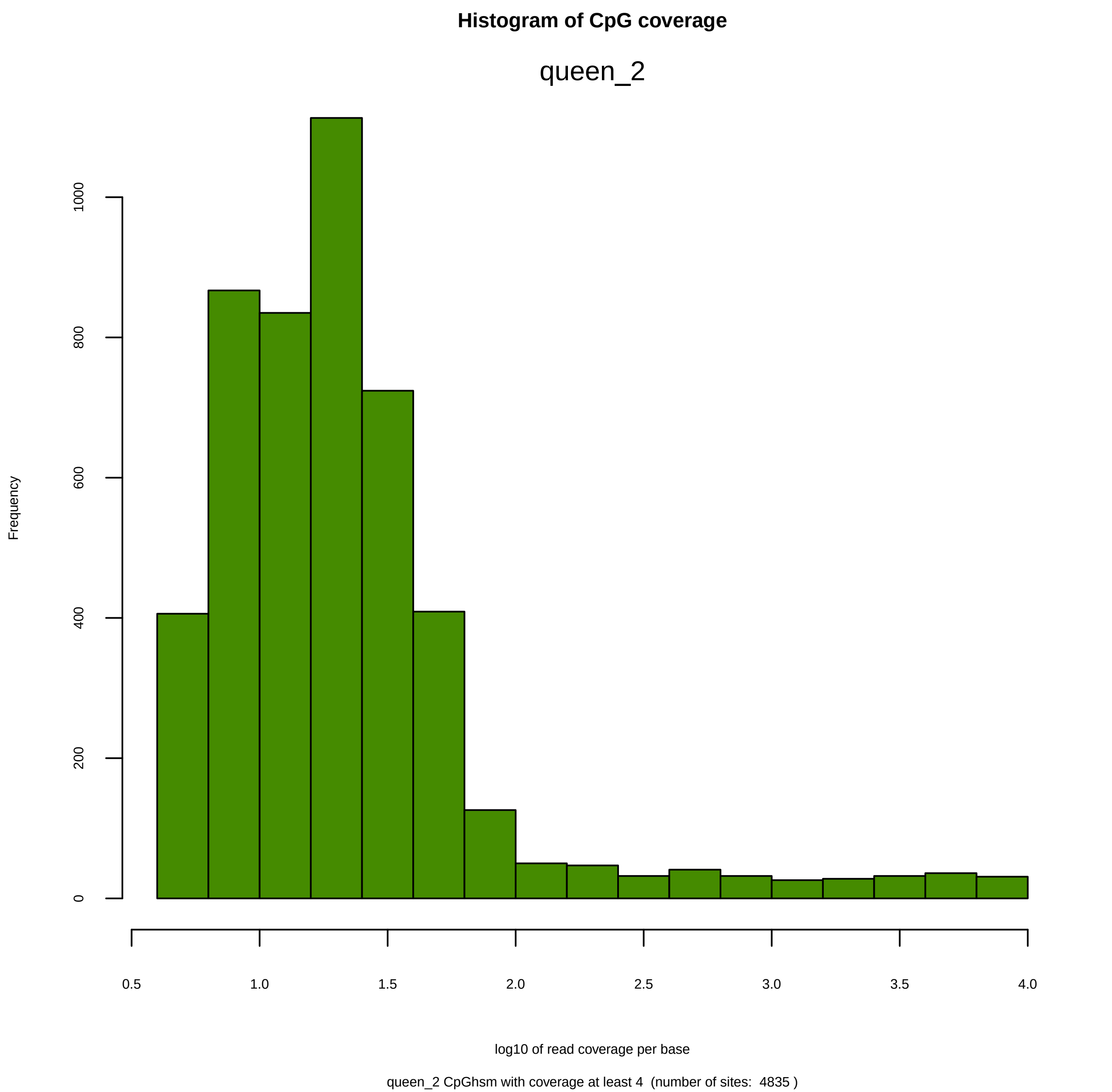

Histogram of CpG coverage
queen_2
Frequency
0
200
400
600
800
1000
0.5
1.0
1.5
2.0
2.5
3.0
3.5
4.0
log10 of read coverage per base
queen_2 CpGhsm with coverage at least 4 (number of sites: 4835 )

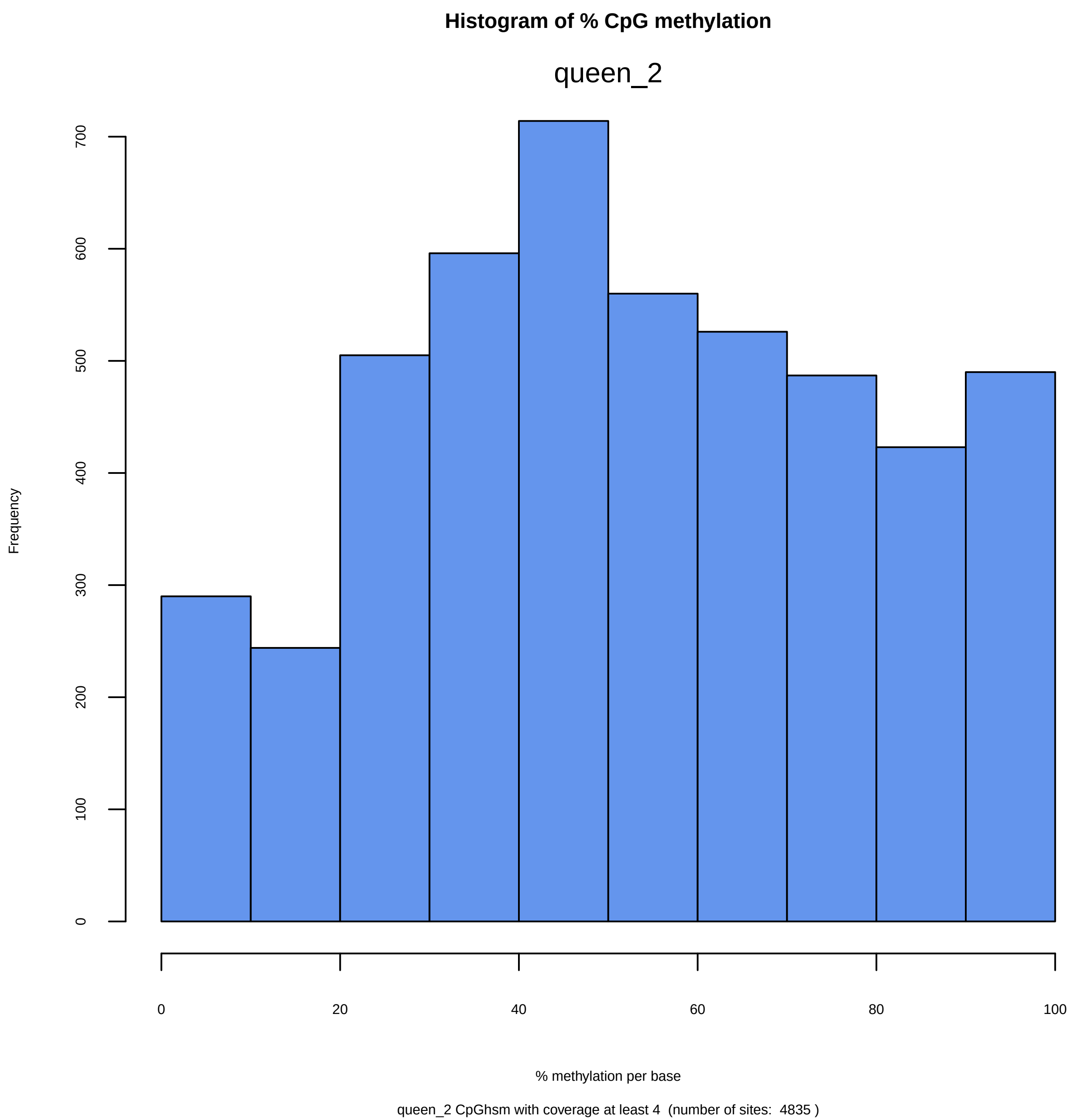
Histogram of % CpG methylation
queen_2
Frequency
0
100
200
300
400
500
600
700
0
20
40
60
80
100
% methylation per base
queen_2 CpGhsm with coverage at least 4 (number of sites: 4835 )

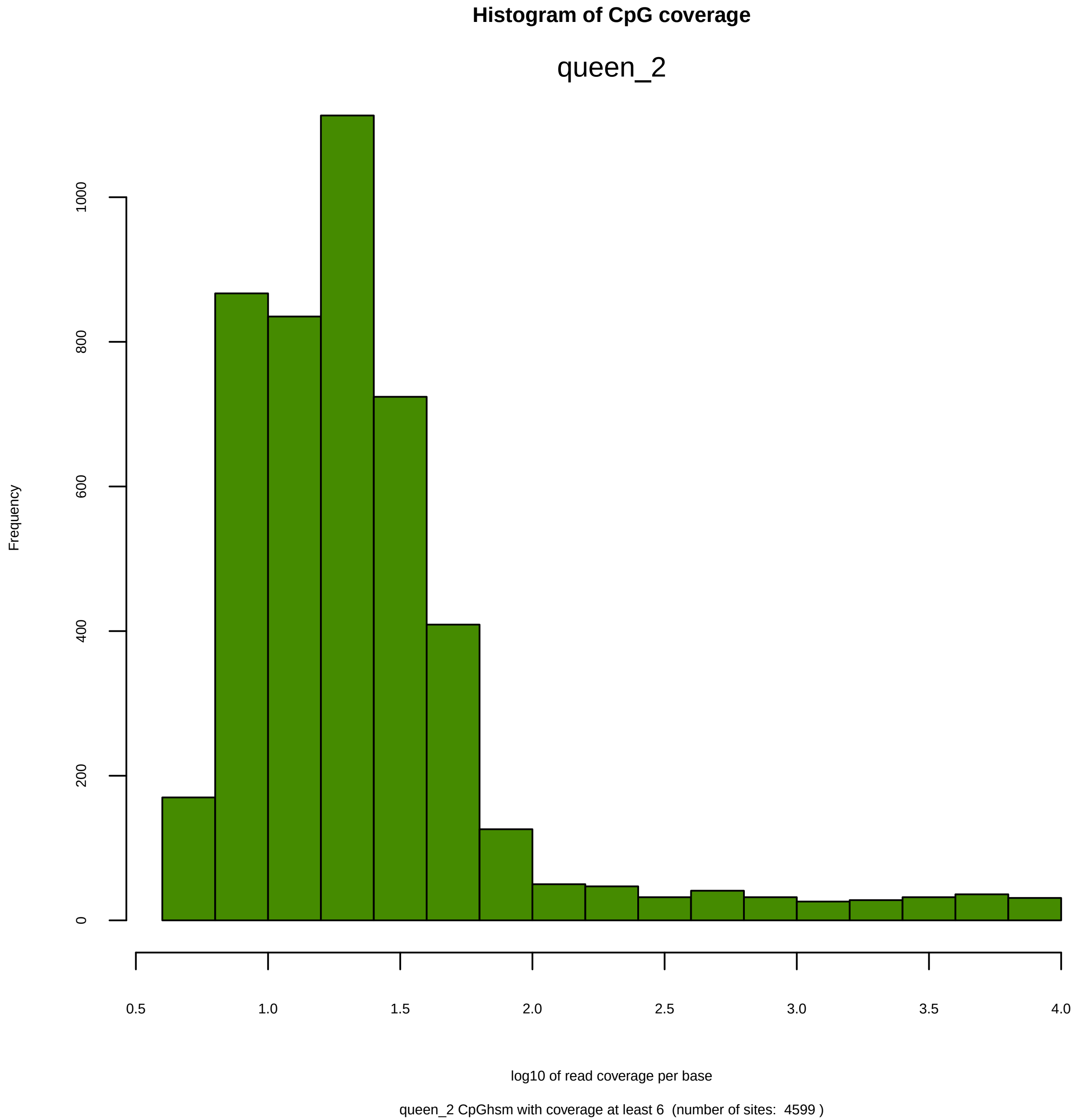
Histogram of CpG coverage
queen_2
Frequency
0
200
400
600
800
1000
0.5
1.0
1.5
2.0
2.5
3.0
3.5
4.0
log10 of read coverage per base
queen_2 CpGhsm with coverage at least 6 (number of sites: 4599 )

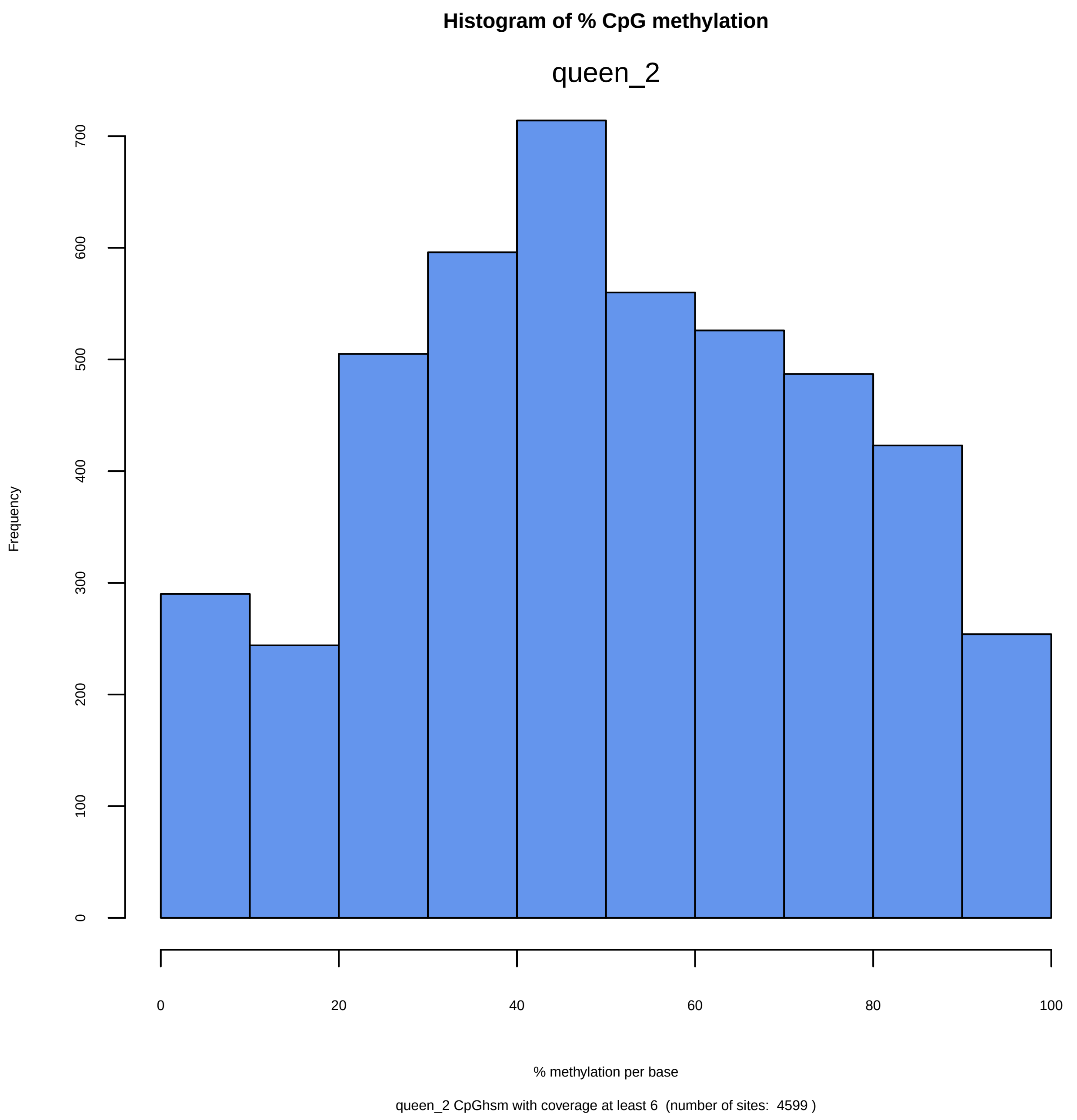
Histogram of % CpG methylation
queen_2
Frequency
0
100
200
300
400
500
600
700
0
20
40
60
80
100
% methylation per base
queen_2 CpGhsm with coverage at least 6 (number of sites: 4599 )

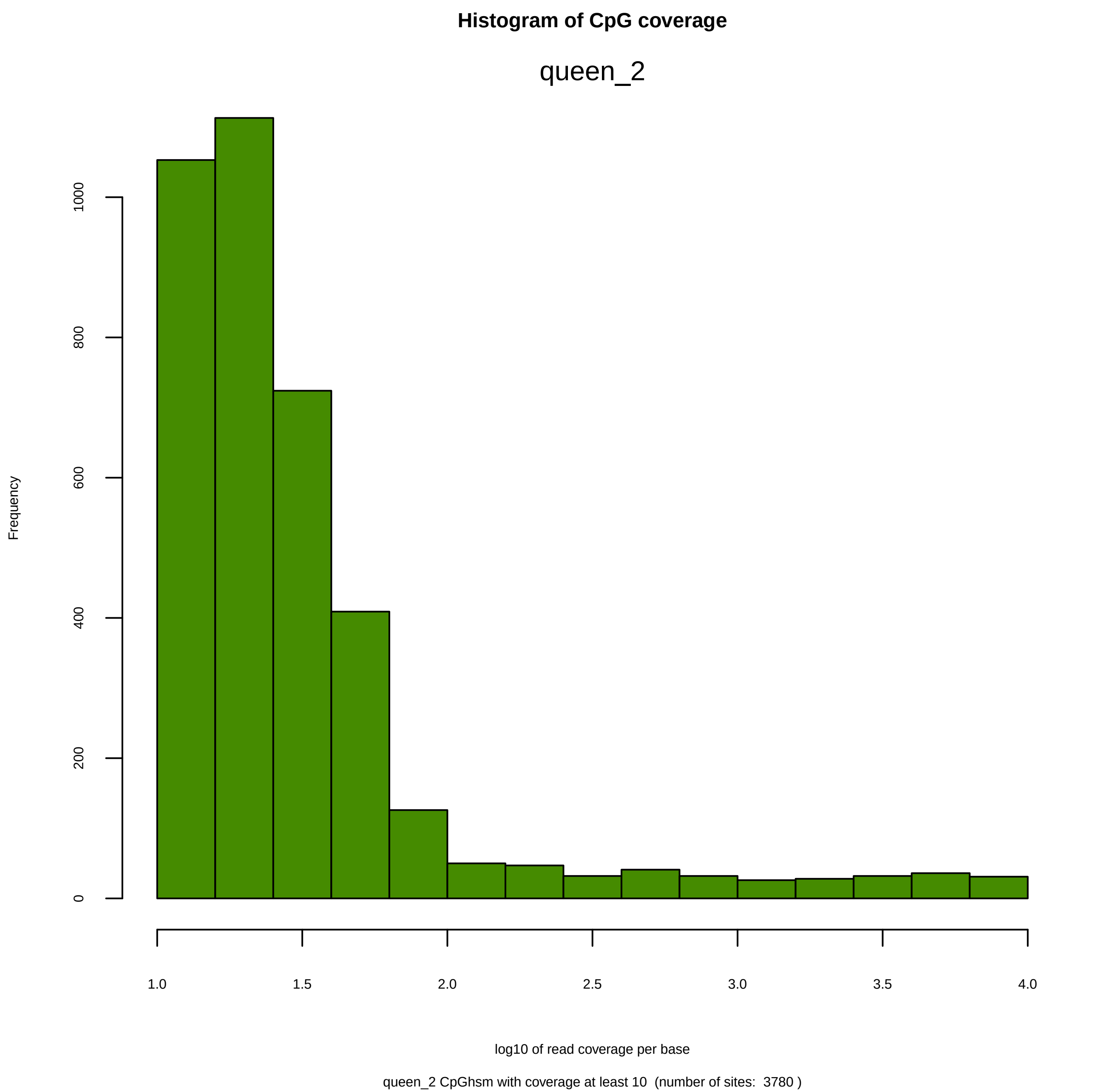

Histogram of CpG coverage
queen_2
Frequency
0
200
400
600
800
1000
1.0
1.5
2.0
2.5
3.0
3.5
4.0
log10 of read coverage per base
queen_2 CpGhsm with coverage at least 10  (number of sites:  3780 )

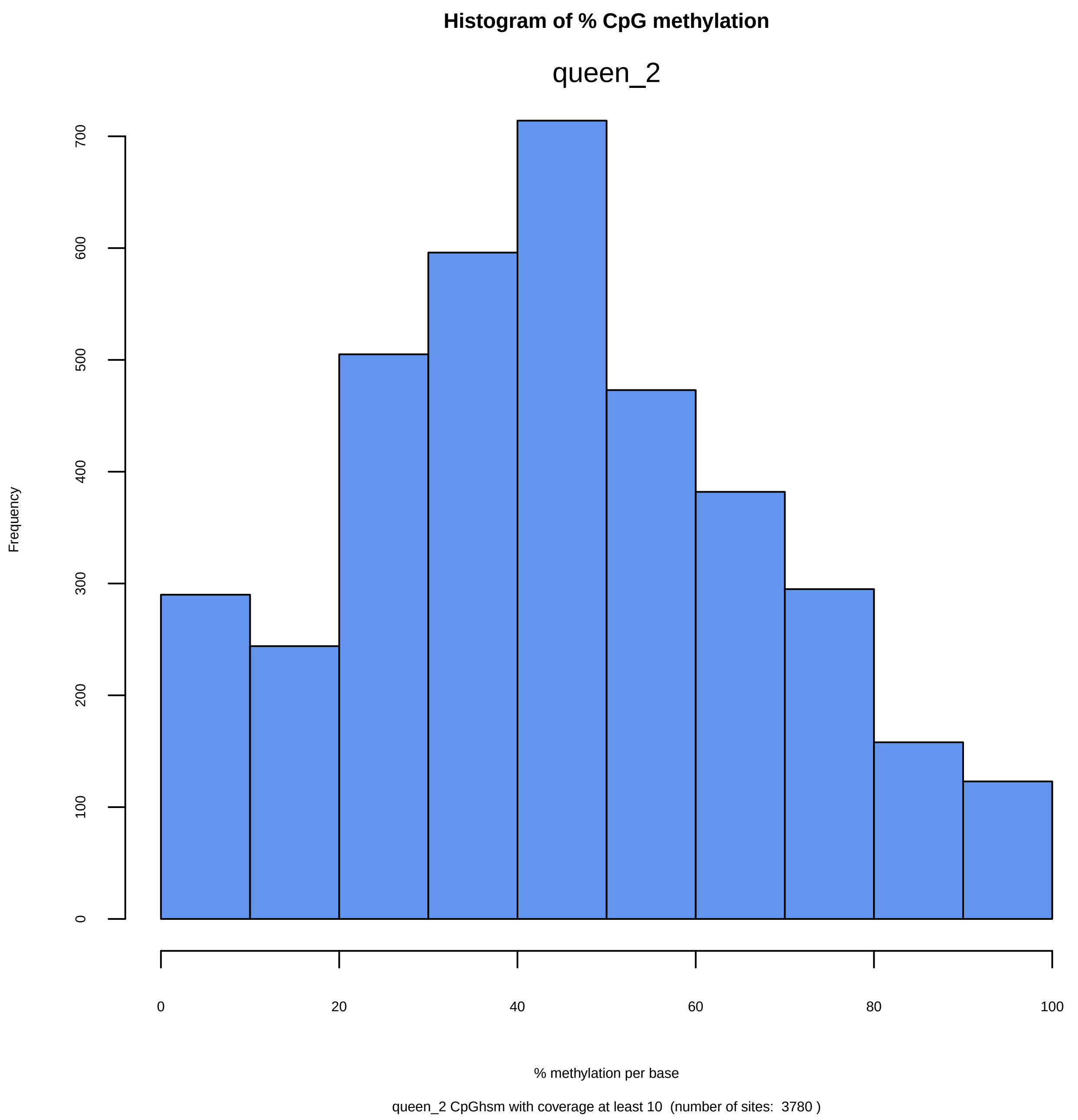
Histogram of % CpG methylation
queen_2
Frequency
0
100
200
300
400
500
600
700
0
20
40
60
80
100
% methylation per base
queen_2 CpGhsm with coverage at least 10 (number of sites: 3780 )

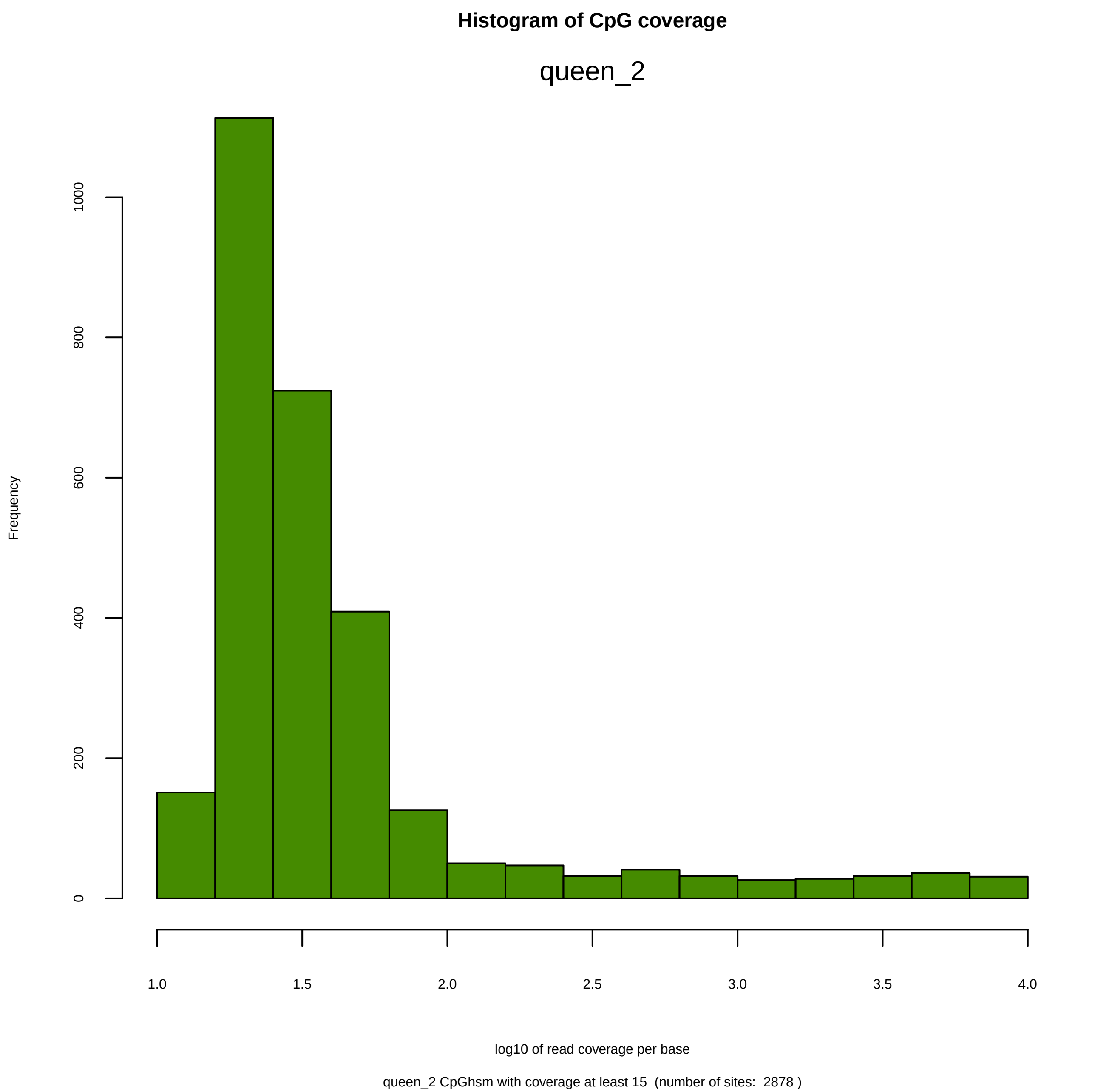
Histogram of CpG coverage
queen_2
Frequency
0
200
400
600
800
1000
1.0
1.5
2.0
2.5
3.0
3.5
4.0
log10 of read coverage per base
queen_2 CpGhsm with coverage at least 15 (number of sites: 2878 )

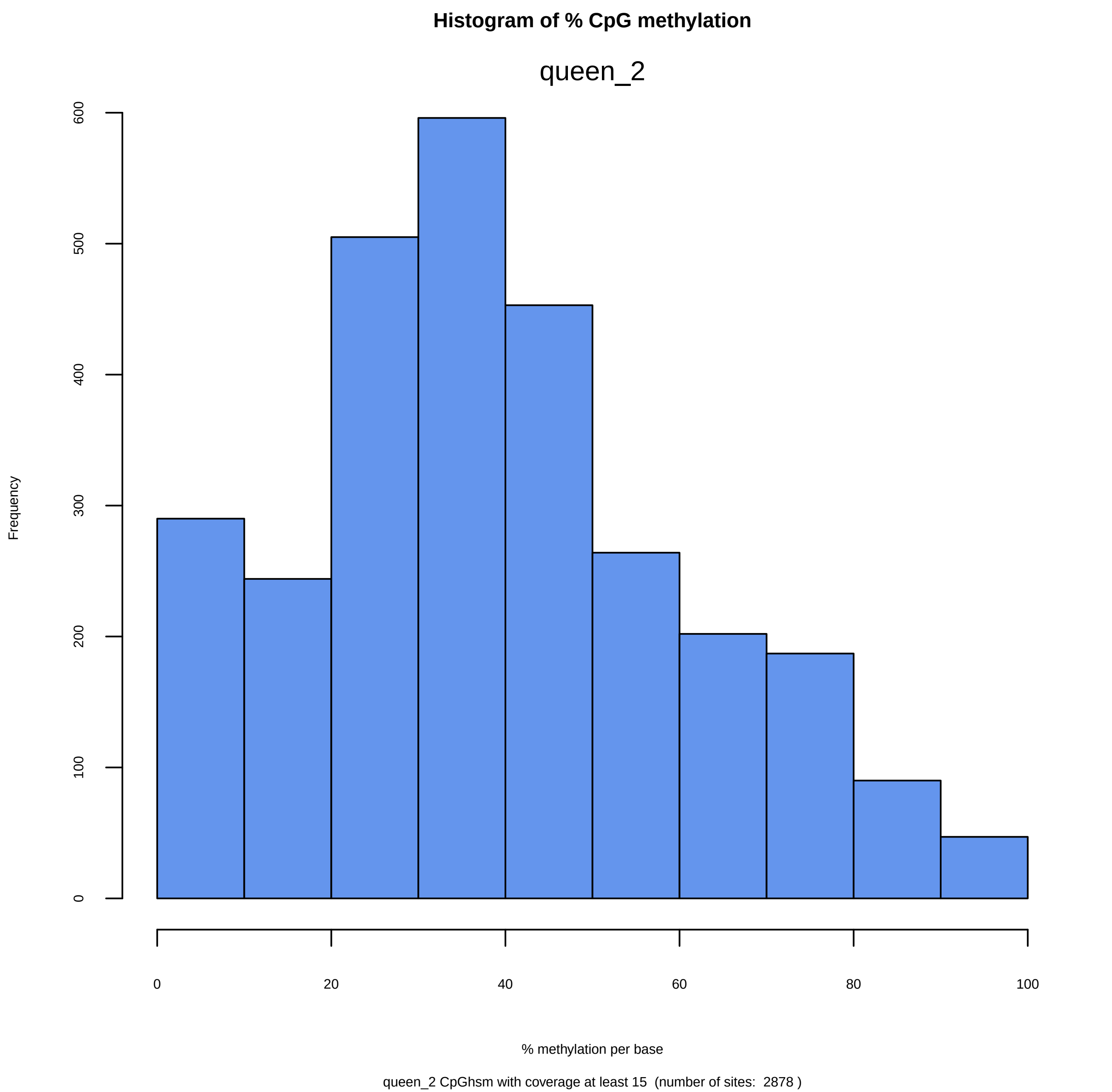
Histogram of % CpG methylation
queen_2
Frequency
0
100
200
300
400
500
600
0
20
40
60
80
100
% methylation per base
queen_2 CpGhsm with coverage at least 15  (number of sites:  2878 )

### Histogram of CpG coverage

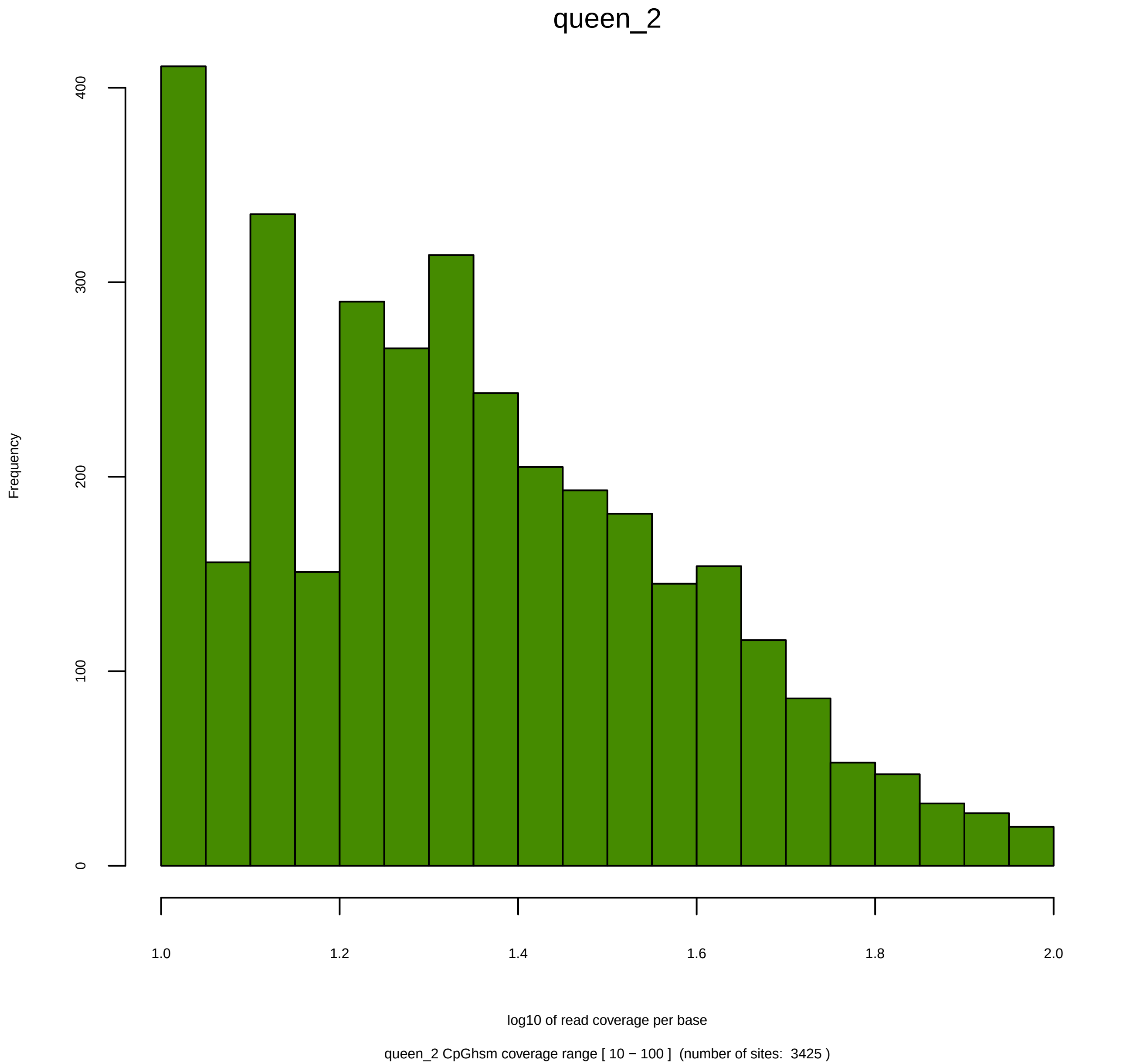

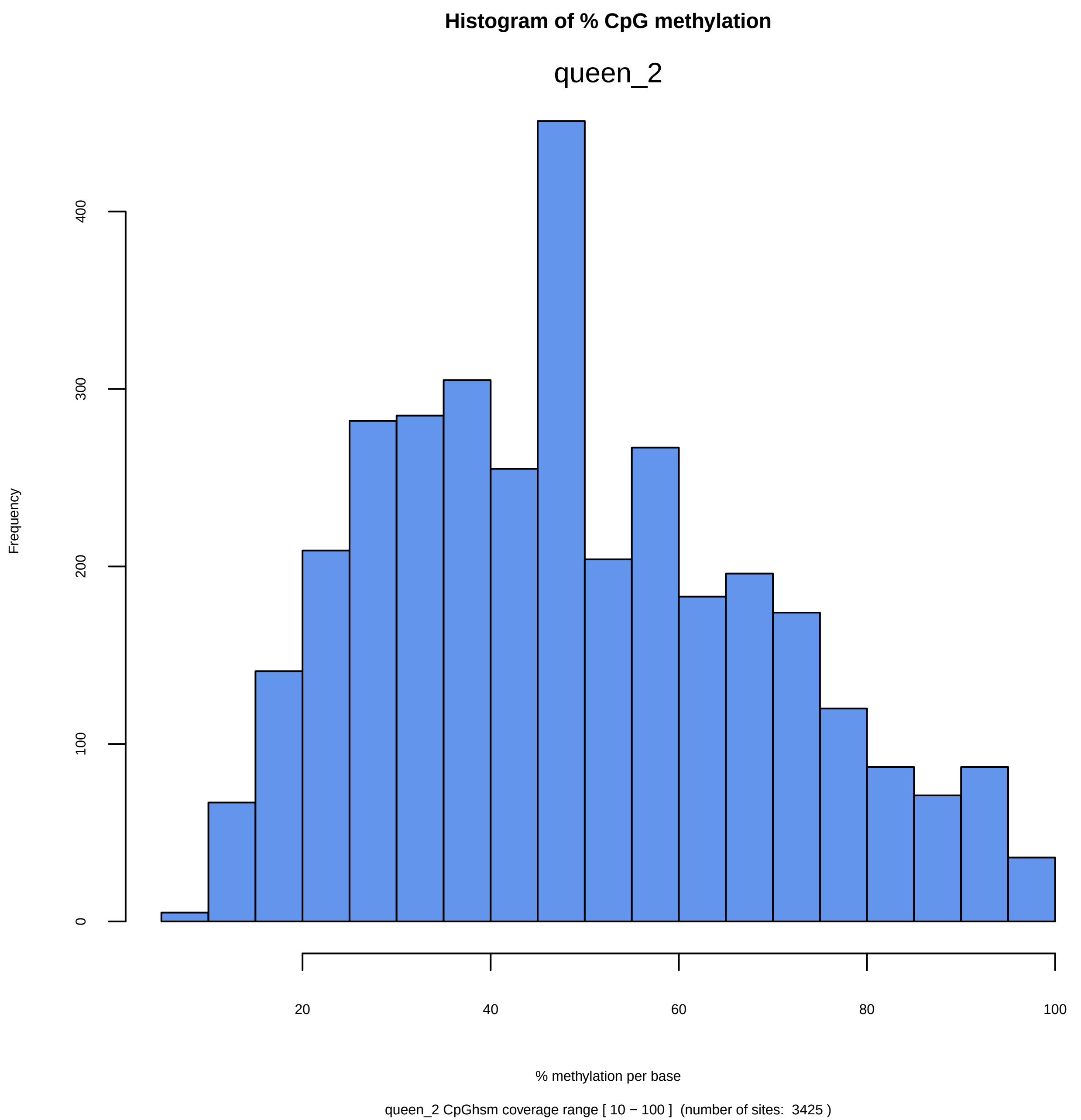

Histogram of % CpG methylation
queen_2
Frequency
0
100
200
300
400
20
40
60
80
100
% methylation per base
queen_2 CpGhsm coverage range [ 10 − 100 ]  (number of sites:  3425 )

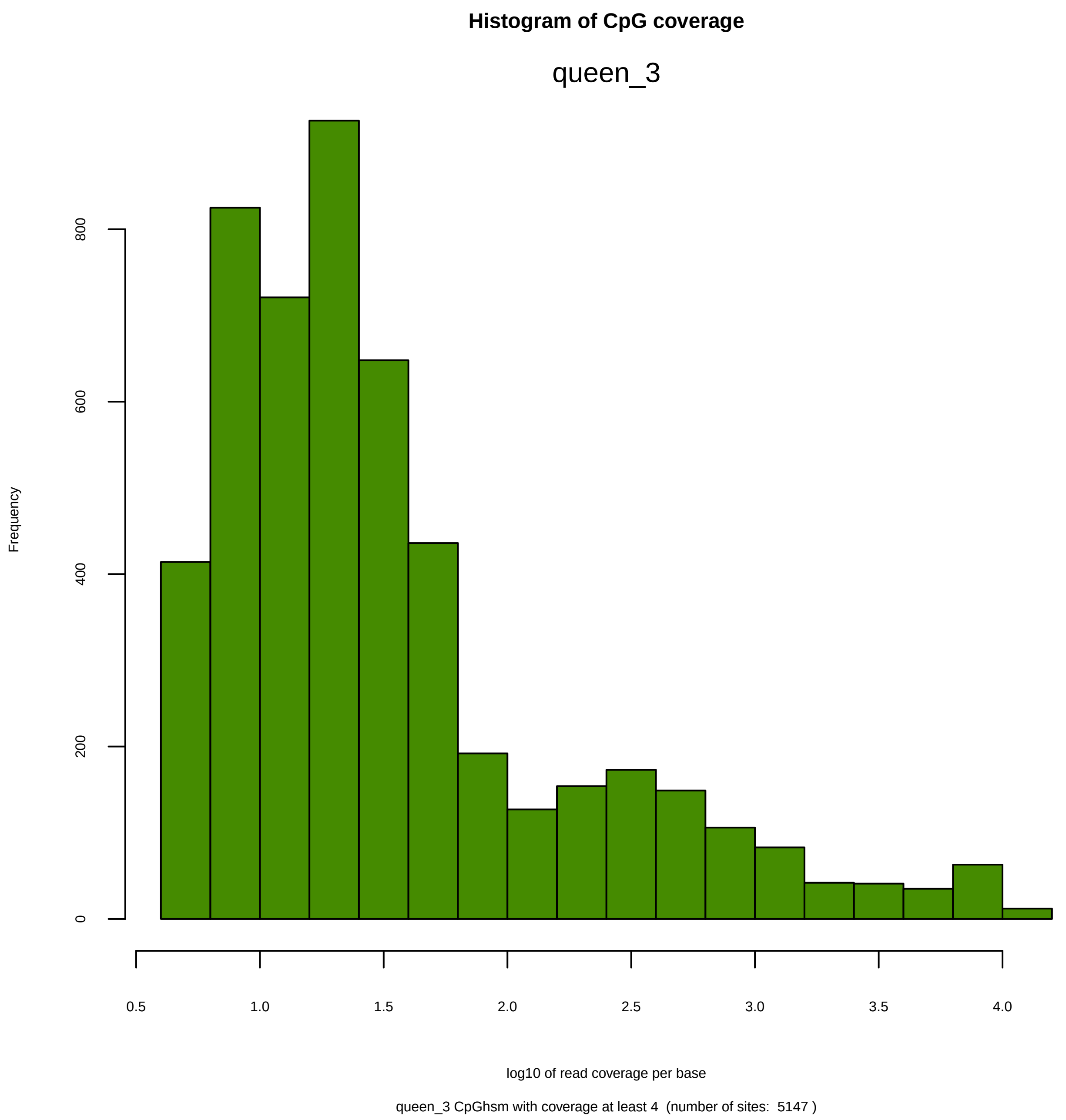
Histogram of CpG coverage
queen_3
Frequency
0
200
400
600
800
0.5
1.0
1.5
2.0
2.5
3.0
3.5
4.0
log10 of read coverage per base
queen_3 CpGhsm with coverage at least 4 (number of sites: 5147 )

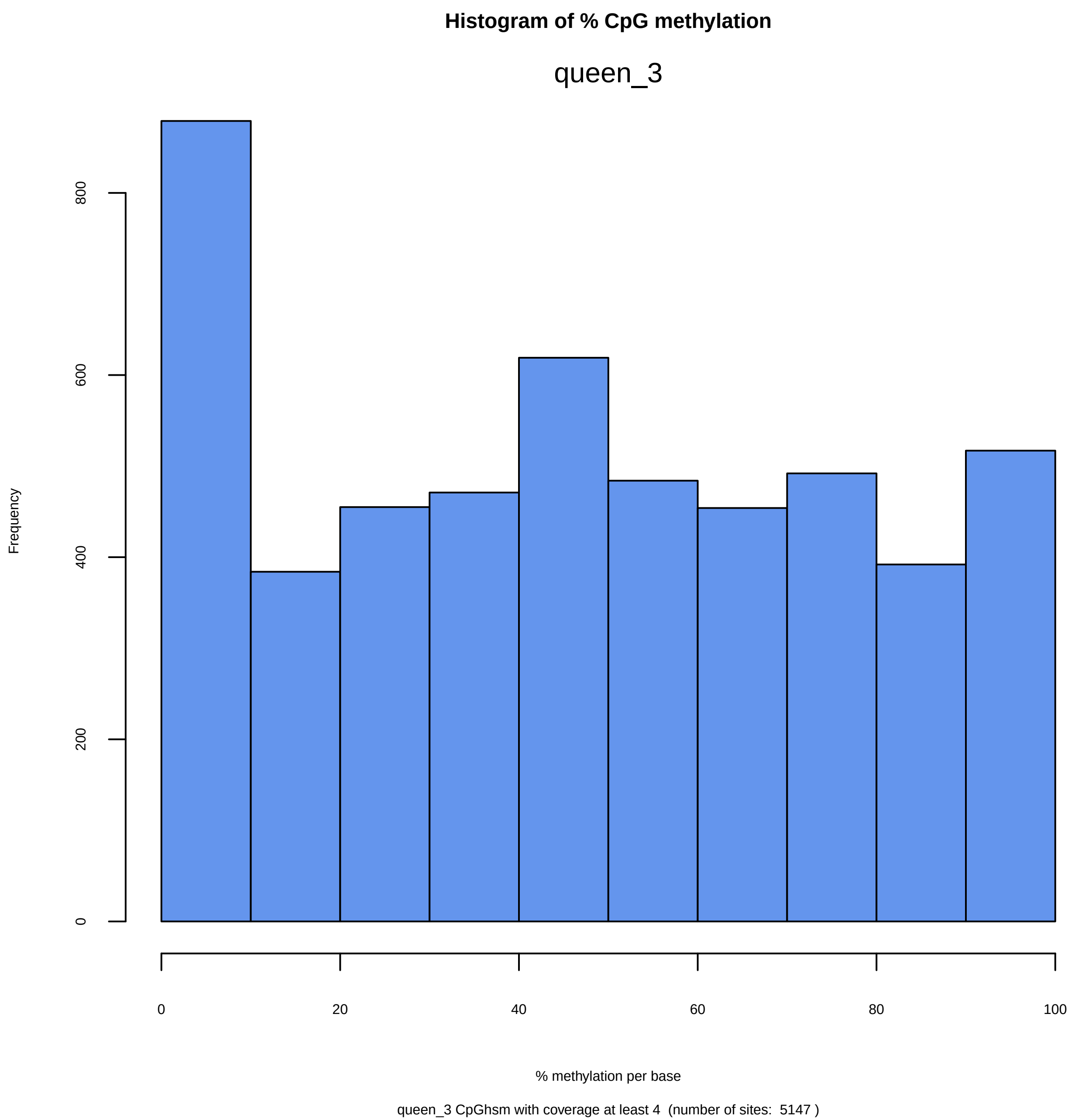

Histogram of % CpG methylation
queen_3
Frequency
0
200
400
600
800
0
20
40
60
80
100
% methylation per base
queen_3 CpGhsm with coverage at least 4  (number of sites:  5147 )

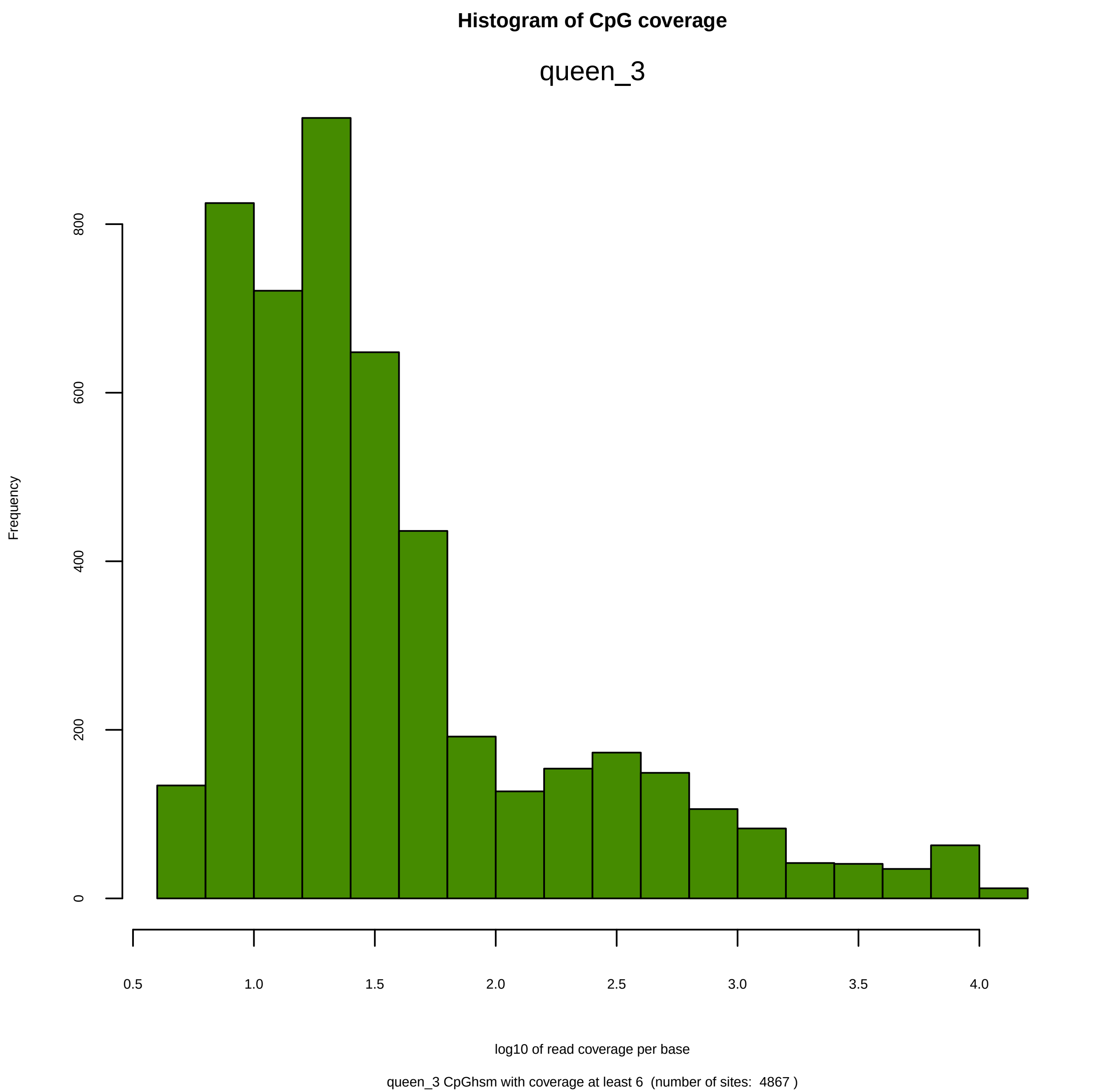
Histogram of CpG coverage
queen_3
Frequency
0
200
400
600
800
0.5
1.0
1.5
2.0
2.5
3.0
3.5
4.0
log10 of read coverage per base
queen_3 CpGhsm with coverage at least 6  (number of sites:  4867 )

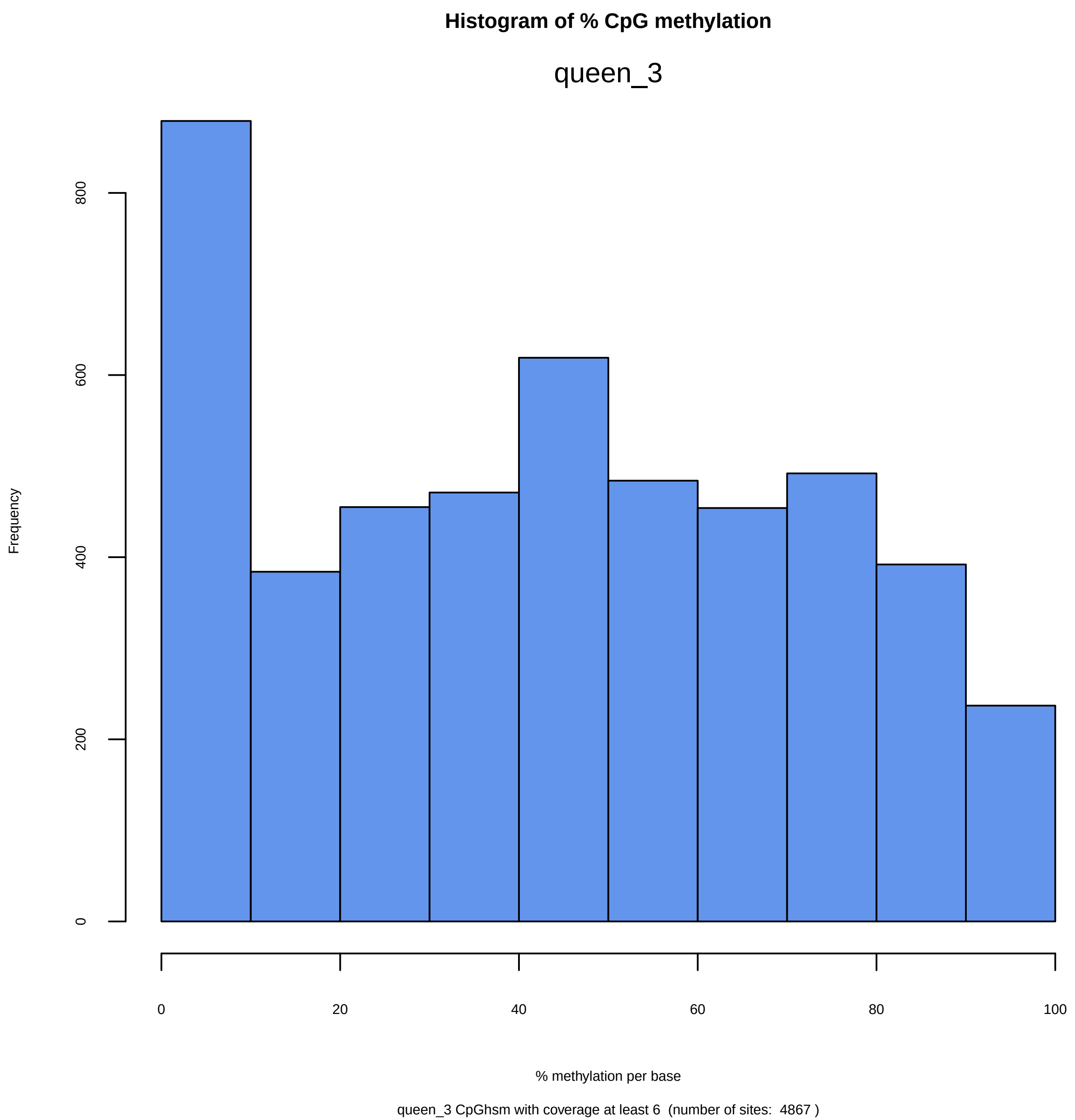
Histogram of % CpG methylation
queen_3
Frequency
0
200
400
600
800
0
20
40
60
80
100
% methylation per base
queen_3 CpGhsm with coverage at least 6  (number of sites:  4867 )

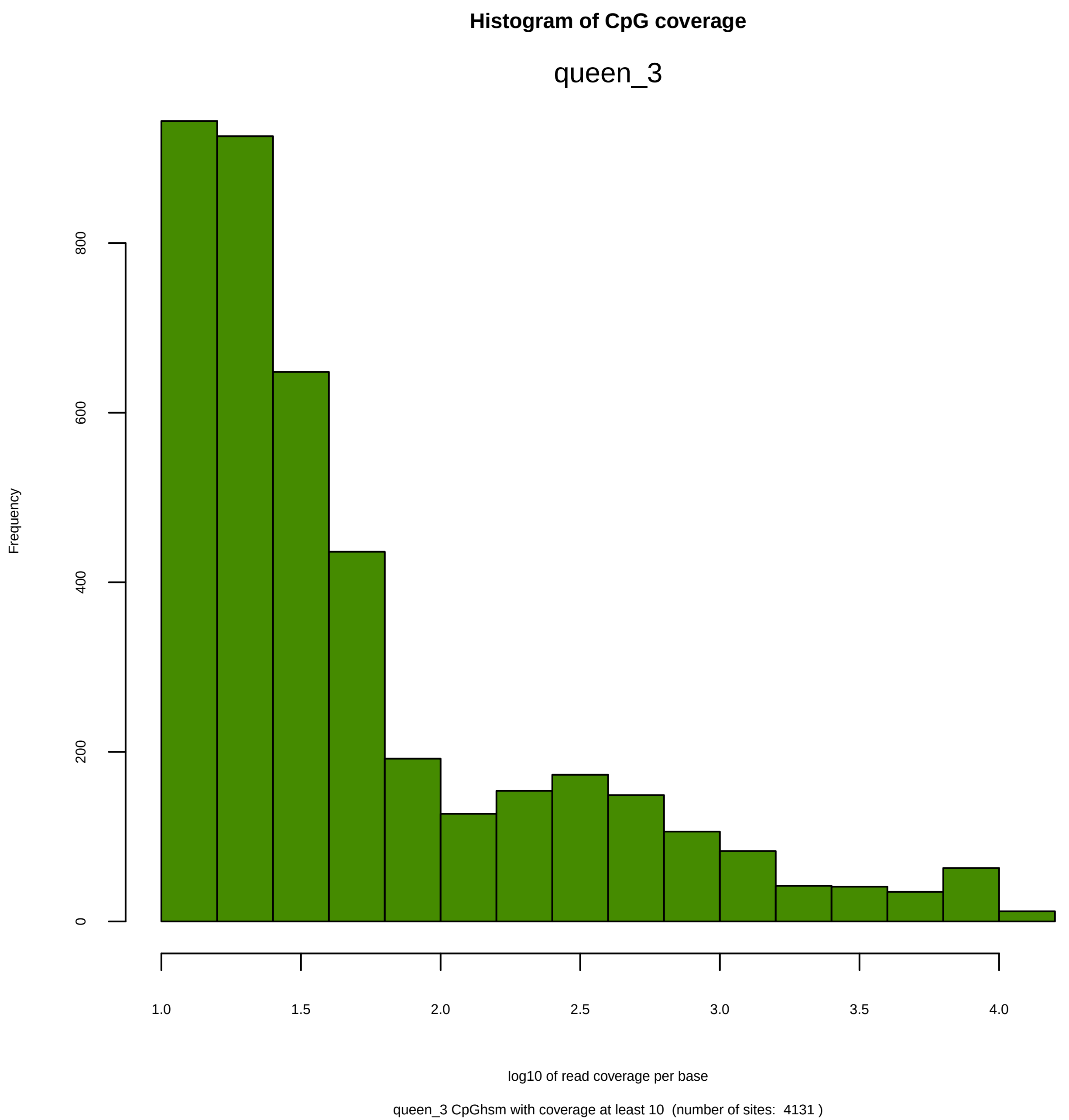
Histogram of CpG coverage
queen_3
Frequency
0
200
400
600
800
1.0
1.5
2.0
2.5
3.0
3.5
4.0
log10 of read coverage per base
queen_3 CpGhsm with coverage at least 10  (number of sites:  4131 )

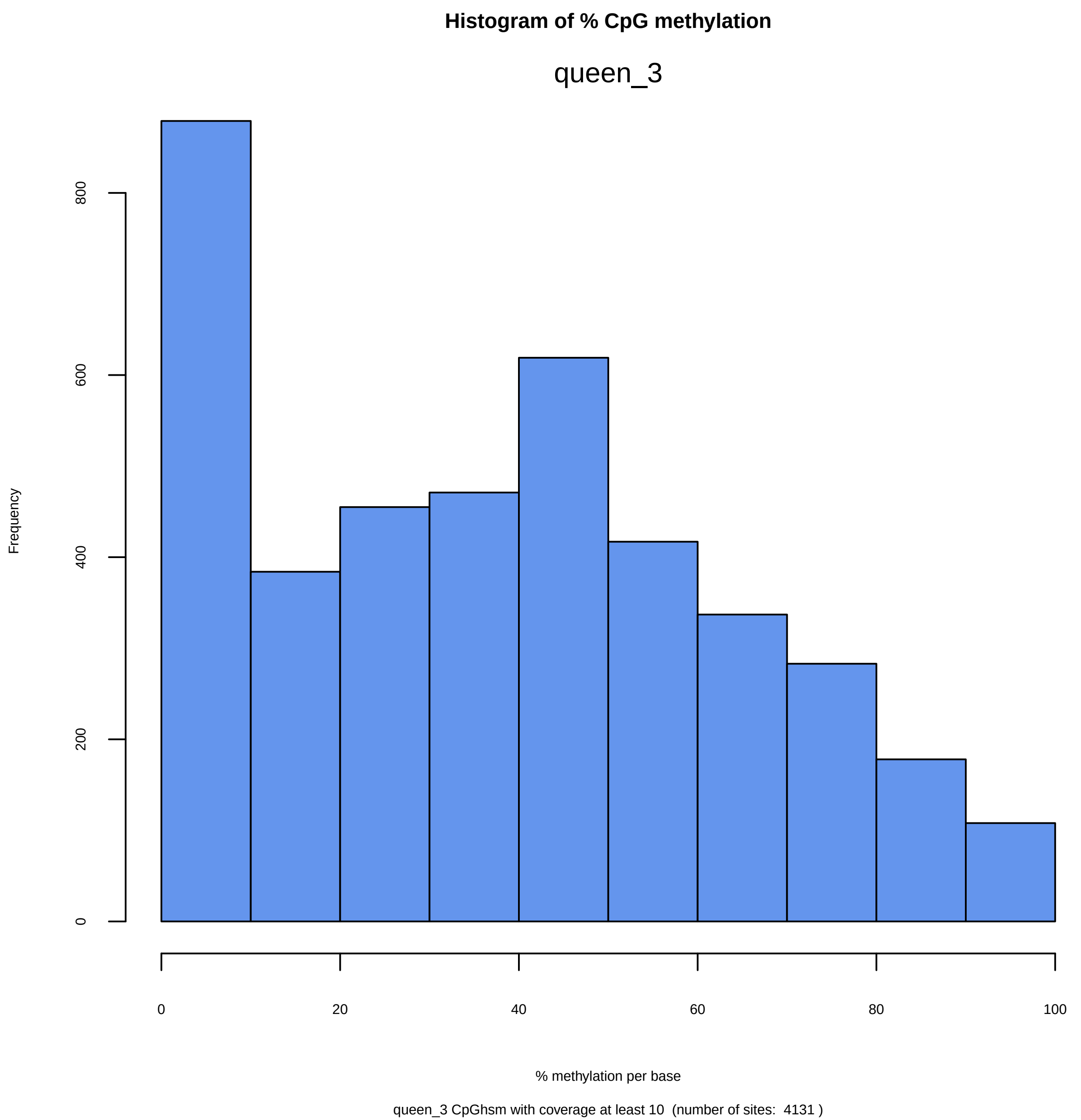

Histogram of % CpG methylation
queen_3
Frequency
0
200
400
600
800
0
20
40
60
80
100
% methylation per base
queen_3 CpGhsm with coverage at least 10 (number of sites: 4131 )

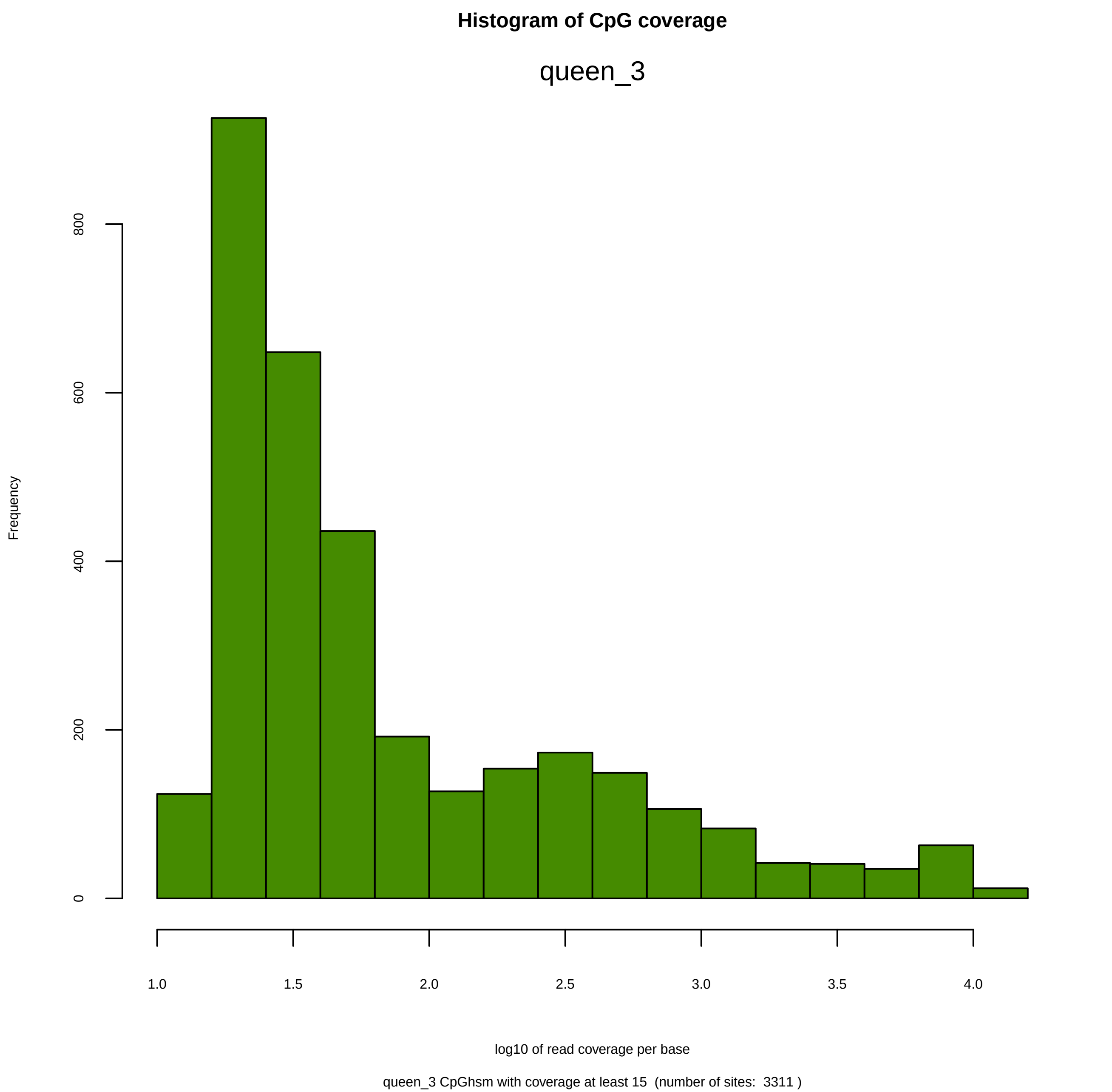
Histogram of CpG coverage
queen_3
Frequency
0
200
400
600
800
1.0
1.5
2.0
2.5
3.0
3.5
4.0
log10 of read coverage per base
queen_3 CpGhsm with coverage at least 15  (number of sites:  3311 )

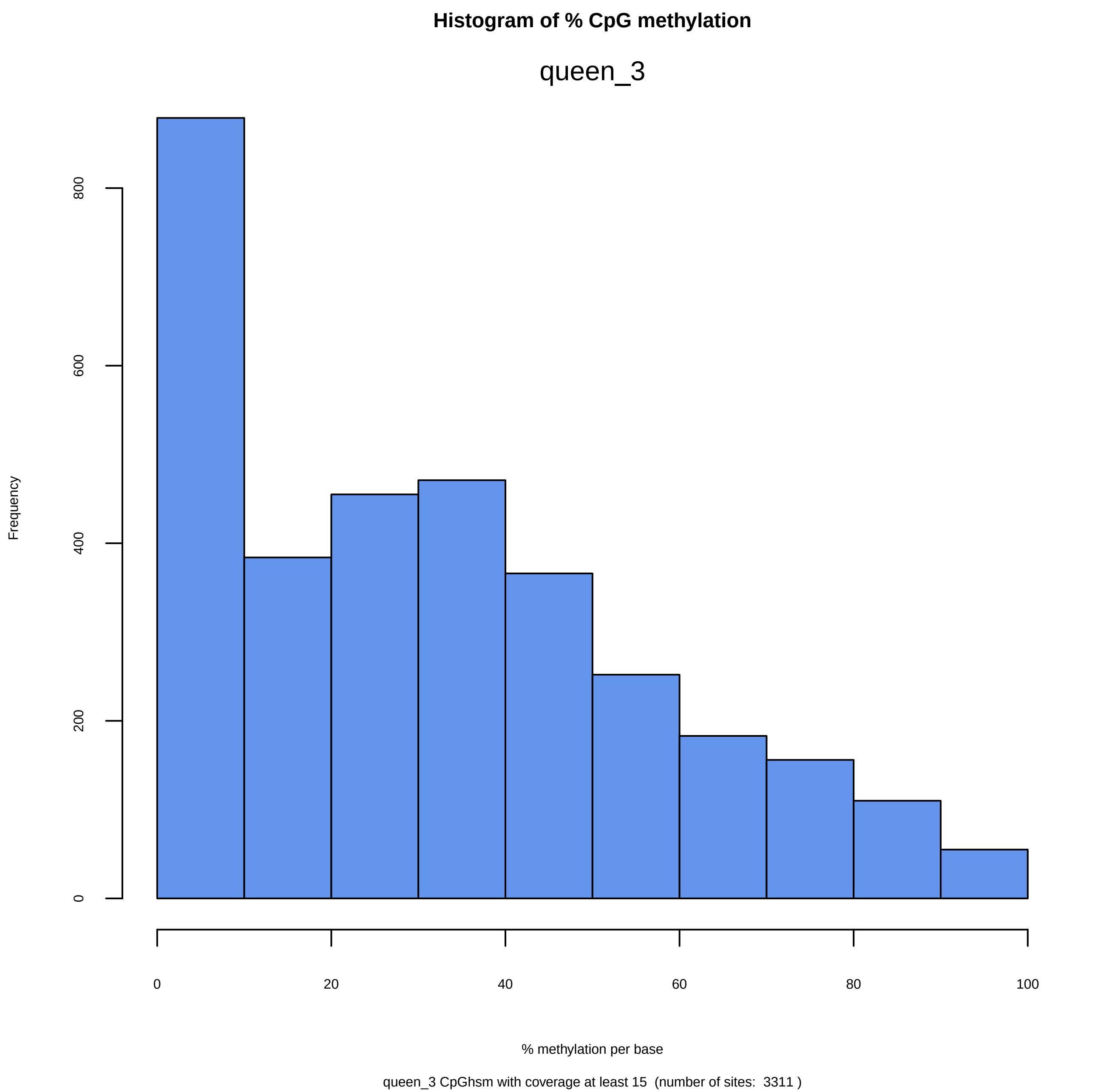
Histogram of % CpG methylation
queen_3
Frequency
0
200
400
600
800
0
20
40
60
80
100
% methylation per base
queen_3 CpGhsm with coverage at least 15  (number of sites:  3311 )

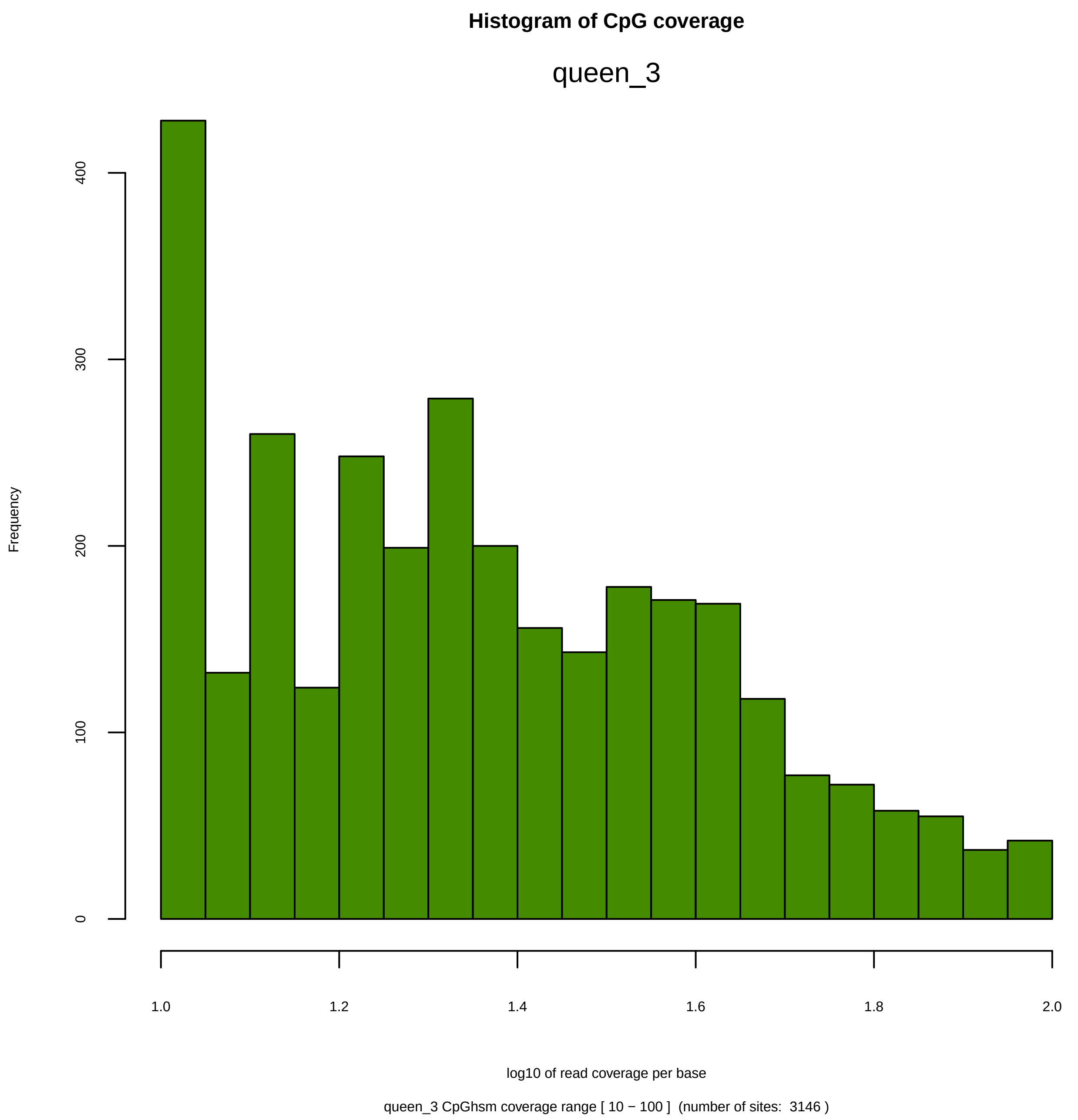
Histogram of CpG coverage
queen_3
Frequency
0
100
200
300
400
1.0
1.2
1.4
1.6
1.8
2.0
log10 of read coverage per base
queen_3 CpGhsm coverage range [ 10 − 100 ]  (number of sites:  3146 )

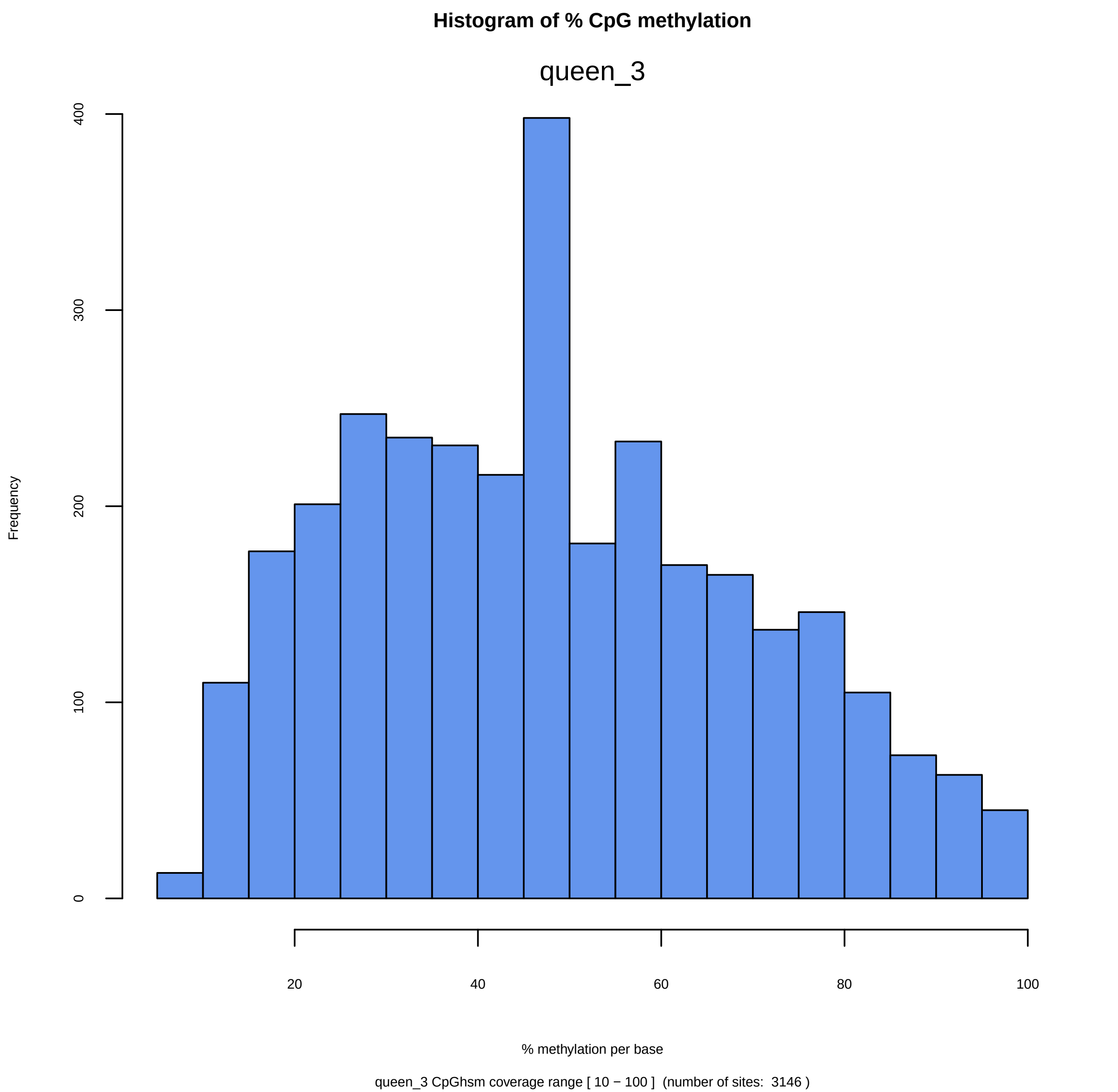

Histogram of % CpG methylation
queen_3
Frequency
0
100
200
300
400
20
40
60
80
100
% methylation per base
queen_3 CpGhsm coverage range [ 10 − 100 ]  (number of sites:  3146 )

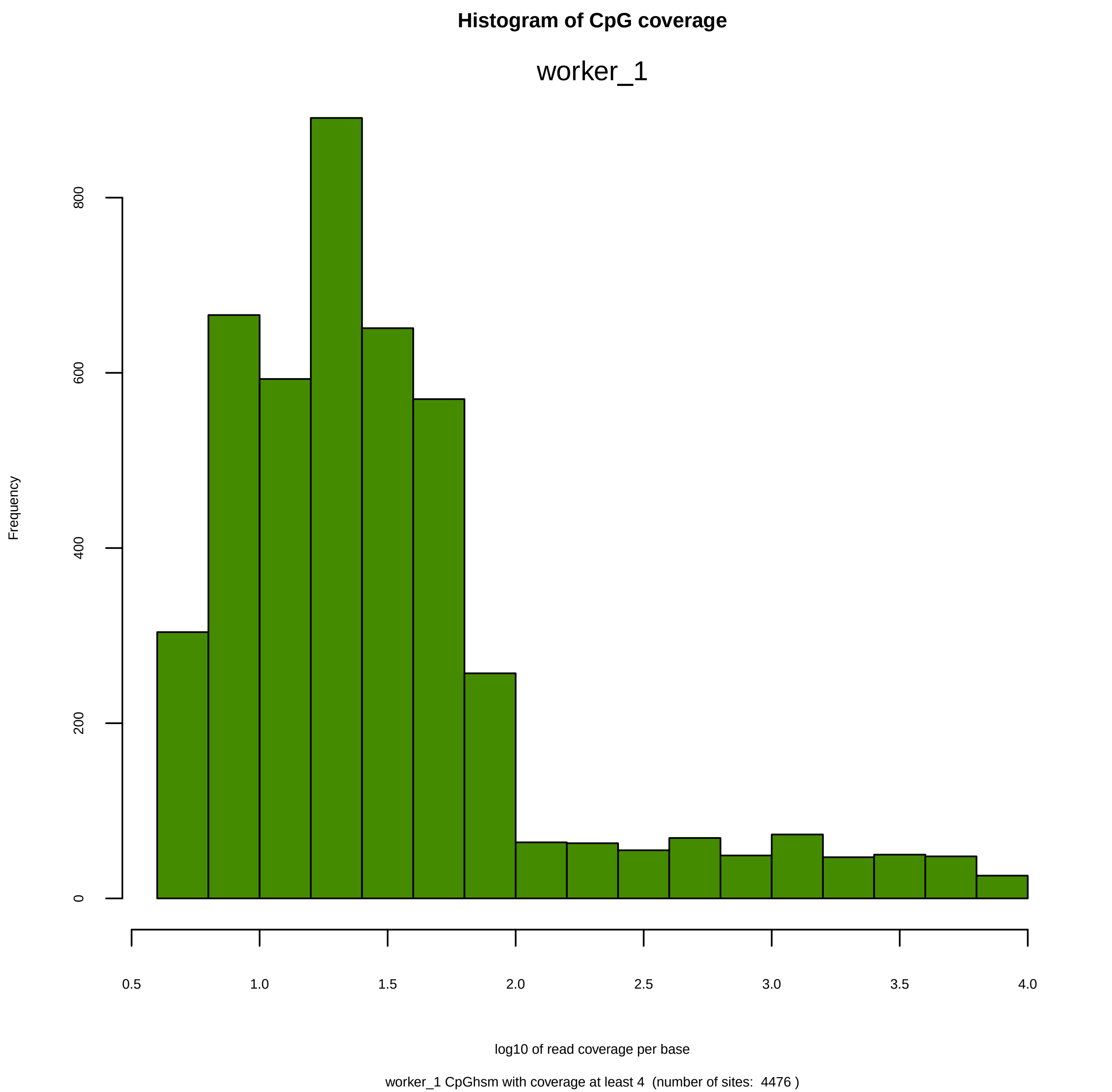
Histogram of CpG coverage
worker_1
Frequency
0
200
400
600
800
0.5
1.0
1.5
2.0
2.5
3.0
3.5
4.0
log10 of read coverage per base
worker_1 CpGhsm with coverage at least 4  (number of sites:  4476 )

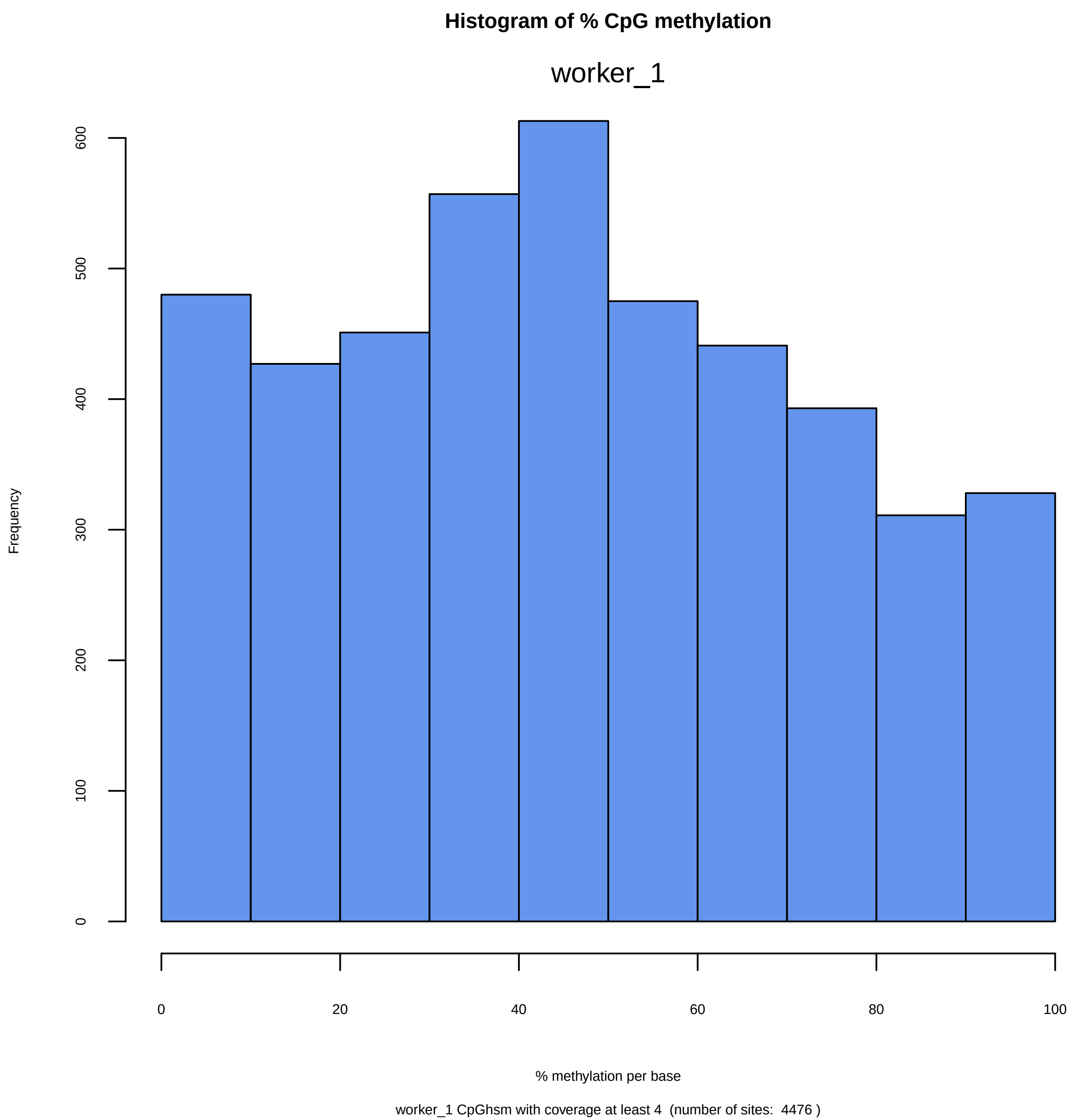
Histogram of % CpG methylation
worker_1
Frequency
0
100
200
300
400
500
600
0
20
40
60
80
100
% methylation per base
worker_1 CpGhsm with coverage at least 4  (number of sites:  4476 )

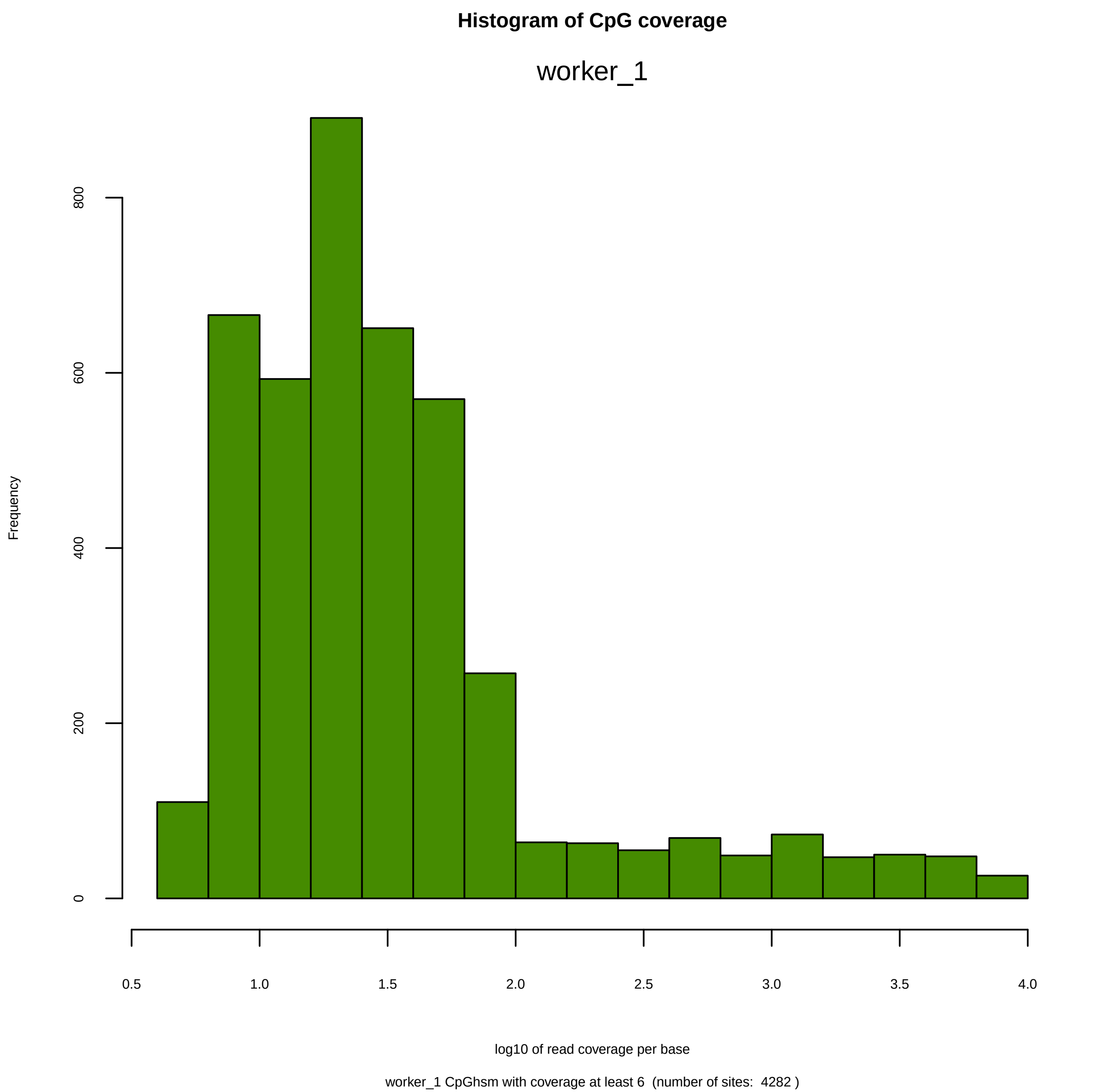
Histogram of CpG coverage
worker_1
Frequency
0
200
400
600
800
0.5
1.0
1.5
2.0
2.5
3.0
3.5
4.0
log10 of read coverage per base
worker_1 CpGhsm with coverage at least 6  (number of sites:  4282 )

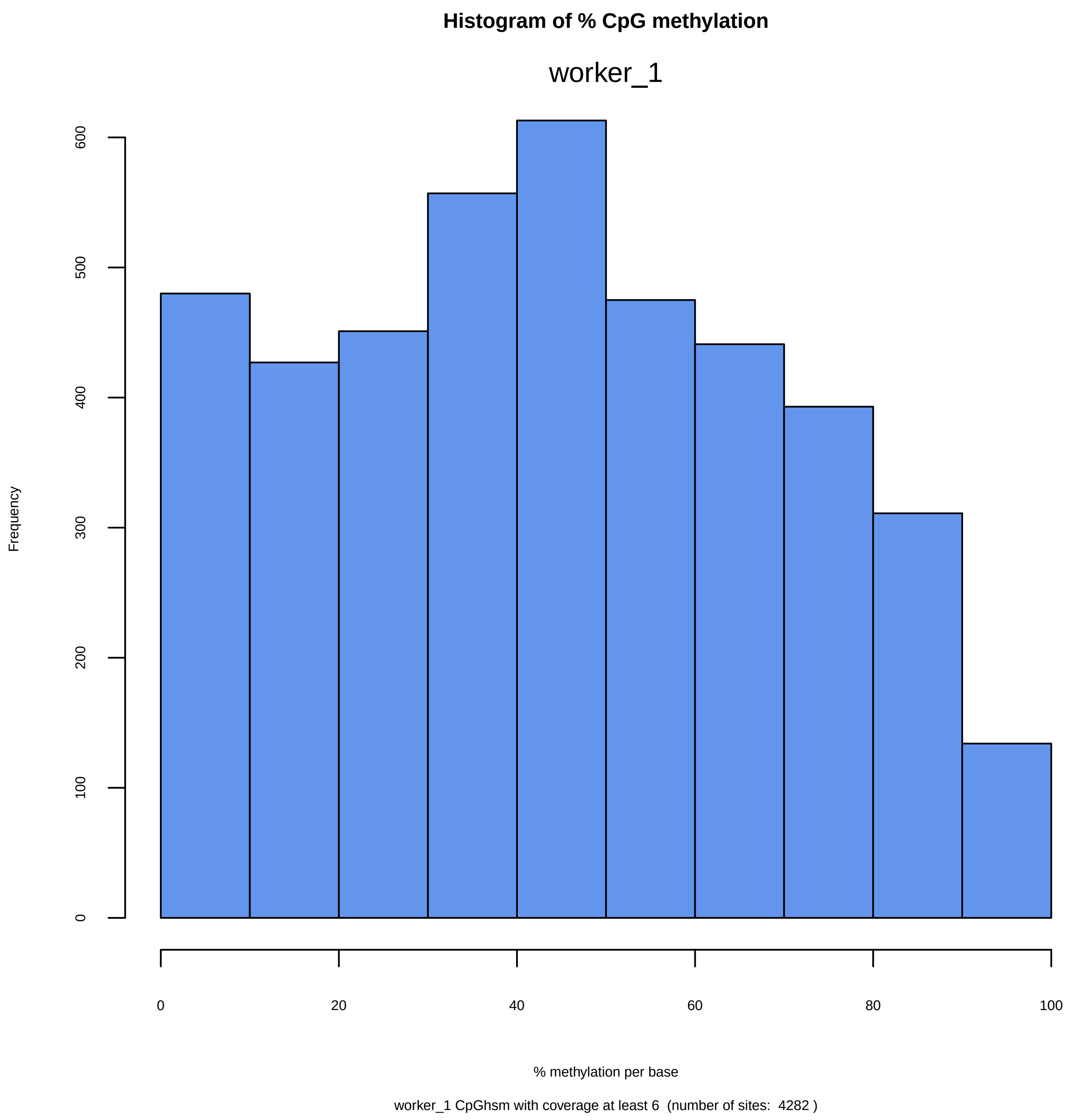

Histogram of % CpG methylation
worker_1
Frequency
0
100
200
300
400
500
600
0
20
40
60
80
100
% methylation per base
worker_1 CpGhsm with coverage at least 6 (number of sites: 4282 )

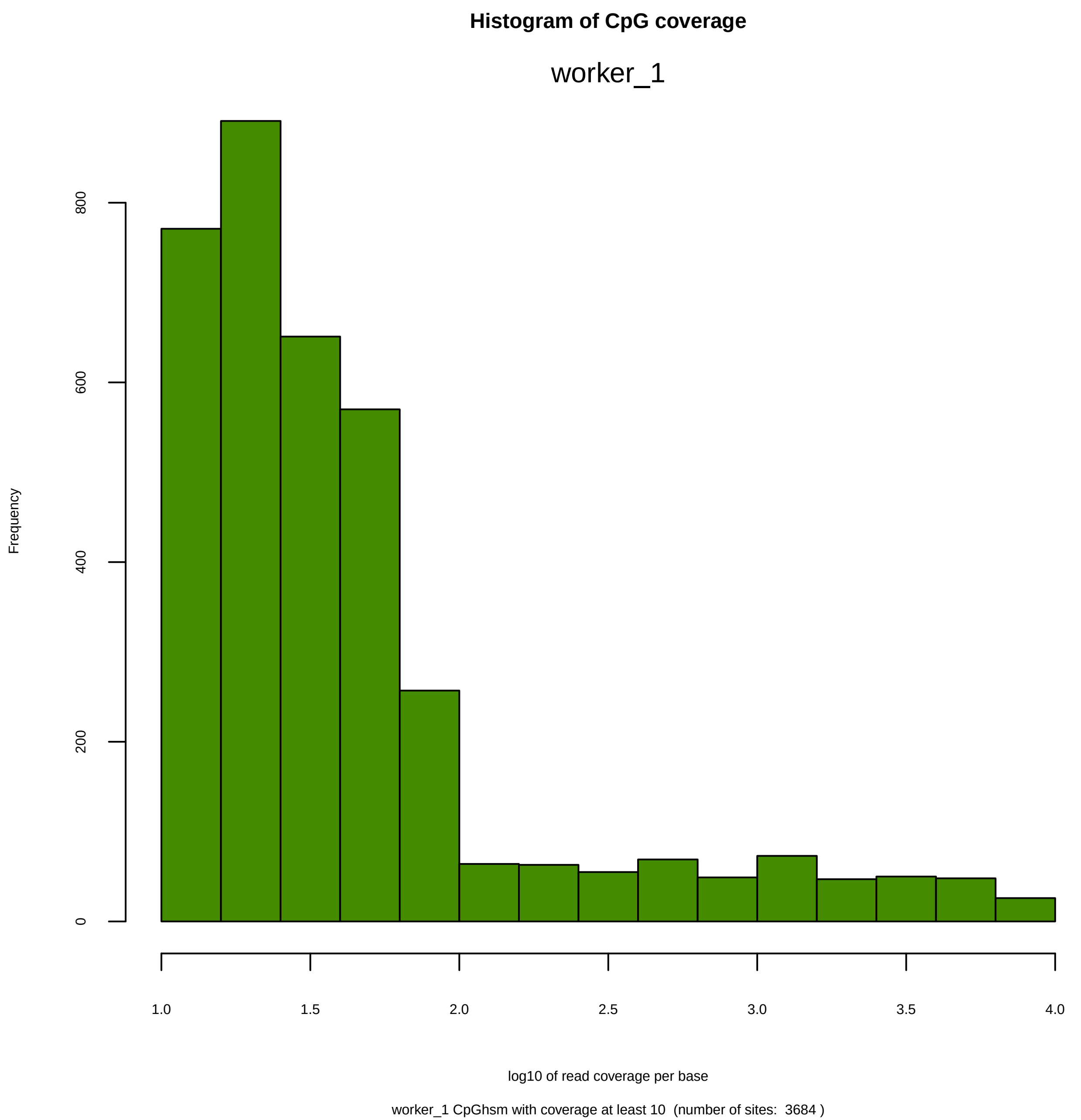
Histogram of CpG coverage
worker_1
Frequency
0
200
400
600
800
1.0
1.5
2.0
2.5
3.0
3.5
4.0
log10 of read coverage per base
worker_1 CpGhsm with coverage at least 10  (number of sites:  3684 )

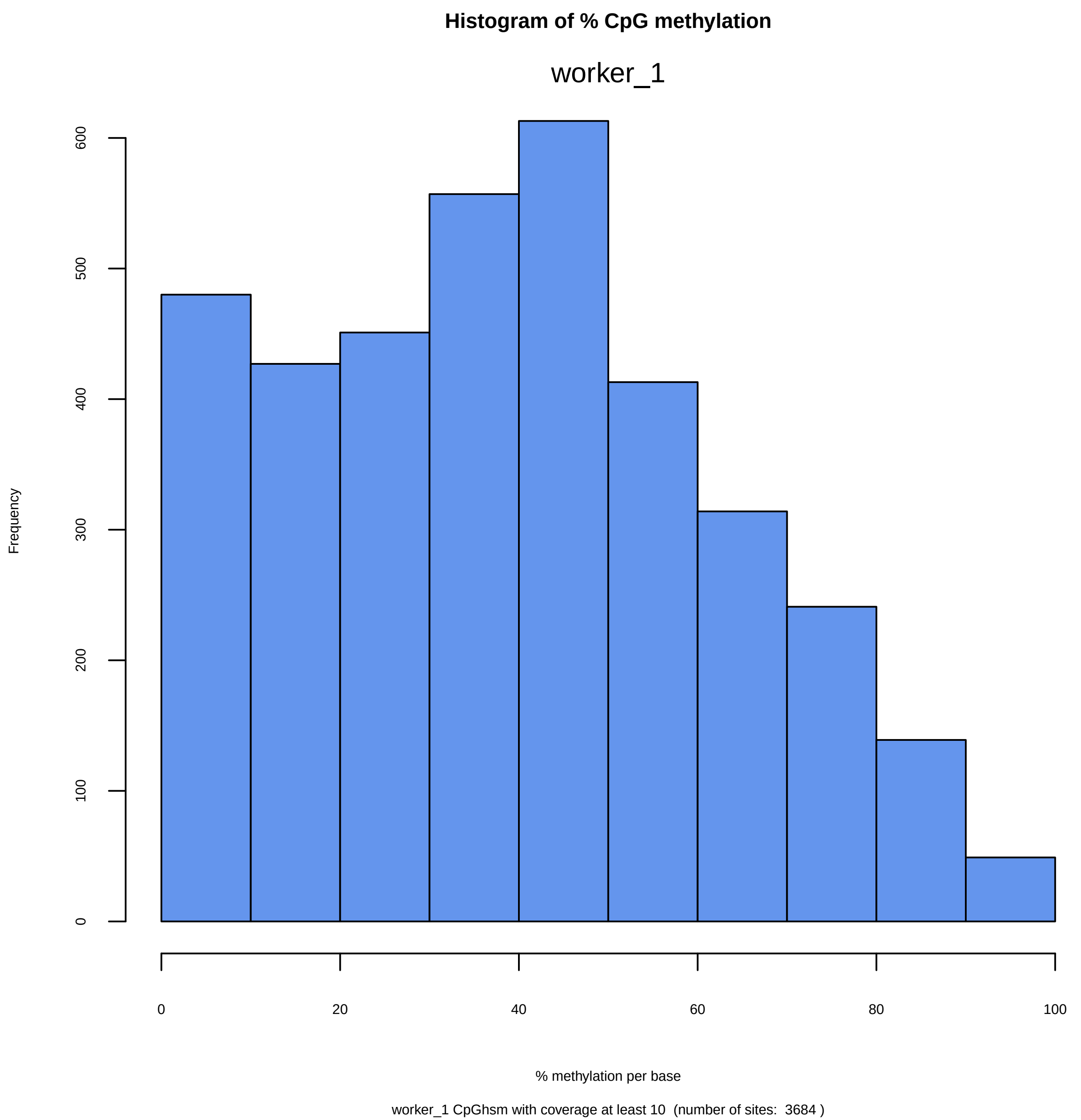
Histogram of % CpG methylation
worker_1
Frequency
0
100
200
300
400
500
600
0
20
40
60
80
100
% methylation per base
worker_1 CpGhsm with coverage at least 10  (number of sites:  3684 )

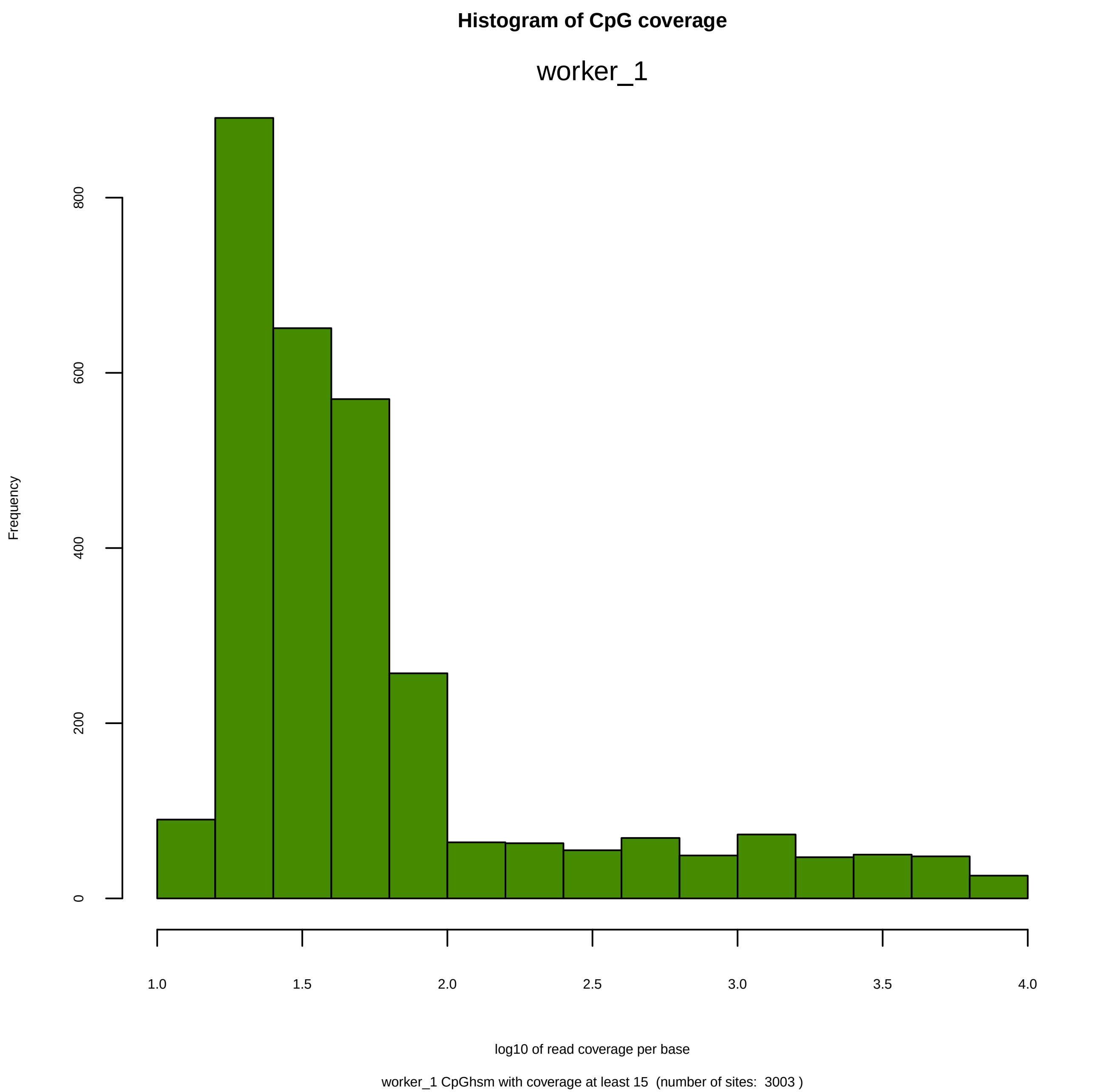
Histogram of CpG coverage
worker_1
Frequency
0
200
400
600
800
1.0
1.5
2.0
2.5
3.0
3.5
4.0
log10 of read coverage per base
worker_1 CpGhsm with coverage at least 15  (number of sites:  3003 )

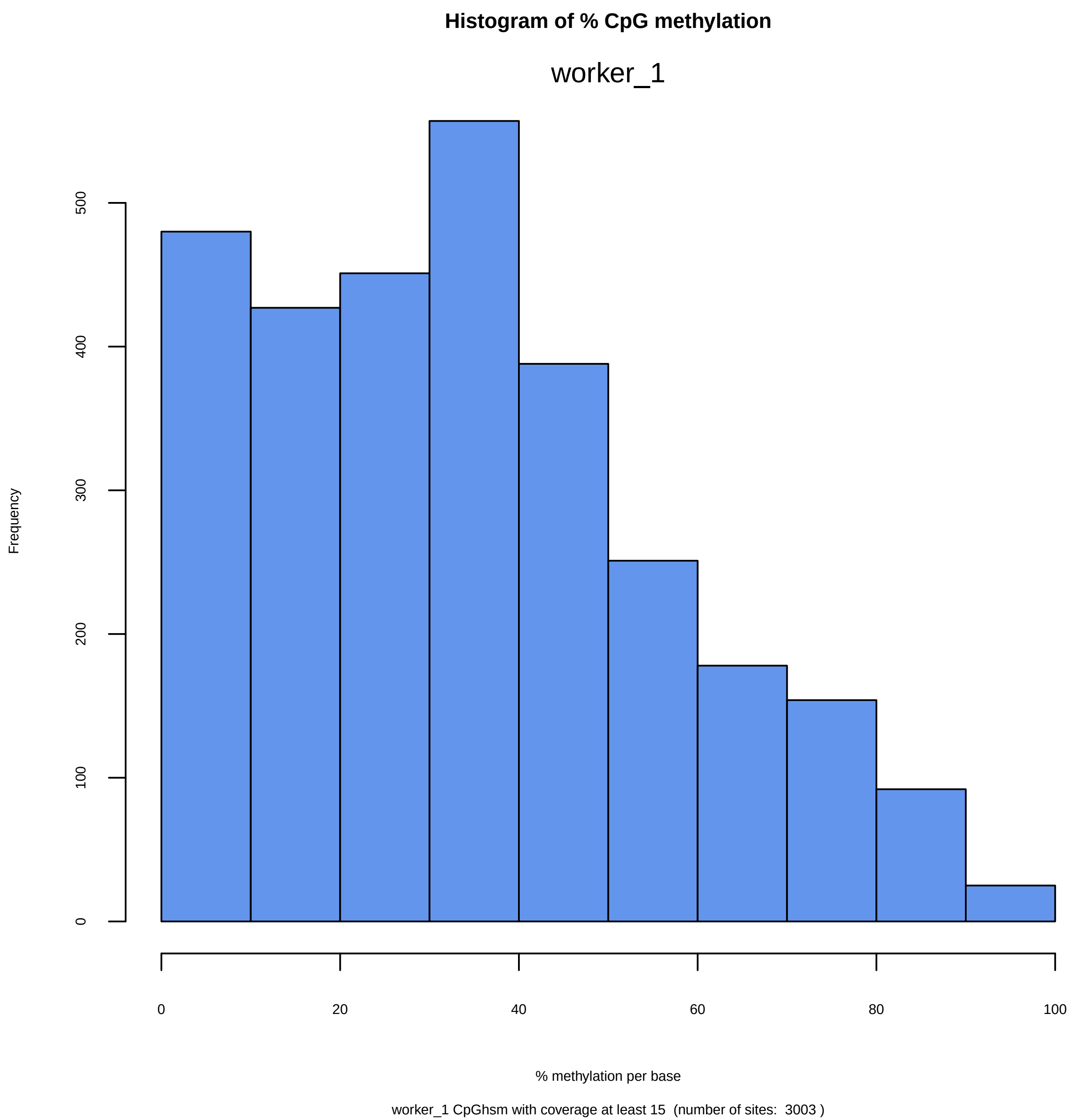
Histogram of % CpG methylation
worker_1
Frequency
0
100
200
300
400
500
0
20
40
60
80
100
% methylation per base
worker_1 CpGhsm with coverage at least 15 (number of sites: 3003 )

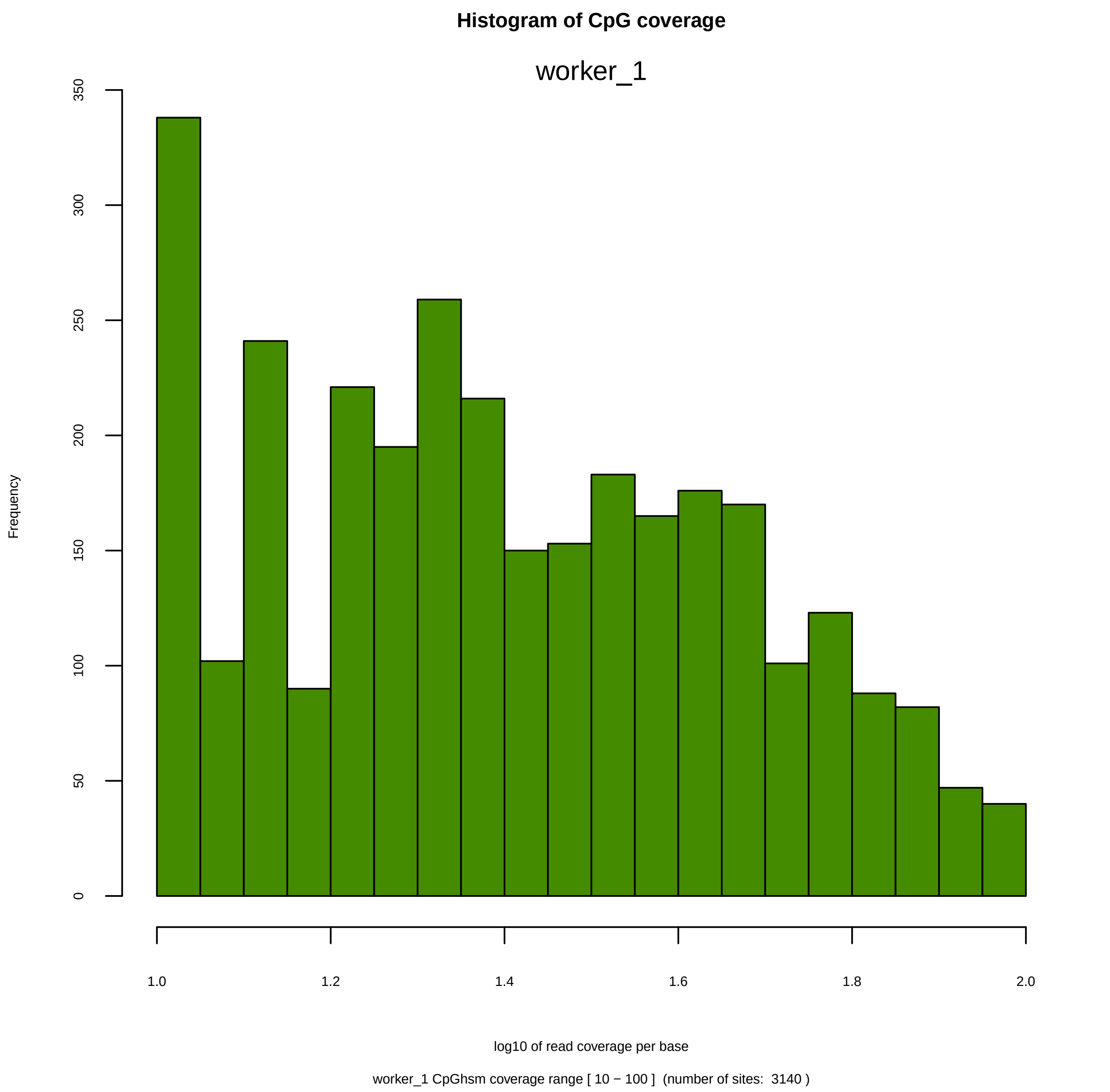

Histogram of CpG coverage
worker_1
Frequency
0
50
100
150
200
250
300
350
1.0
1.2
1.4
1.6
1.8
2.0
log10 of read coverage per base
worker_1 CpGhsm coverage range [ 10 − 100 ]  (number of sites:  3140 )

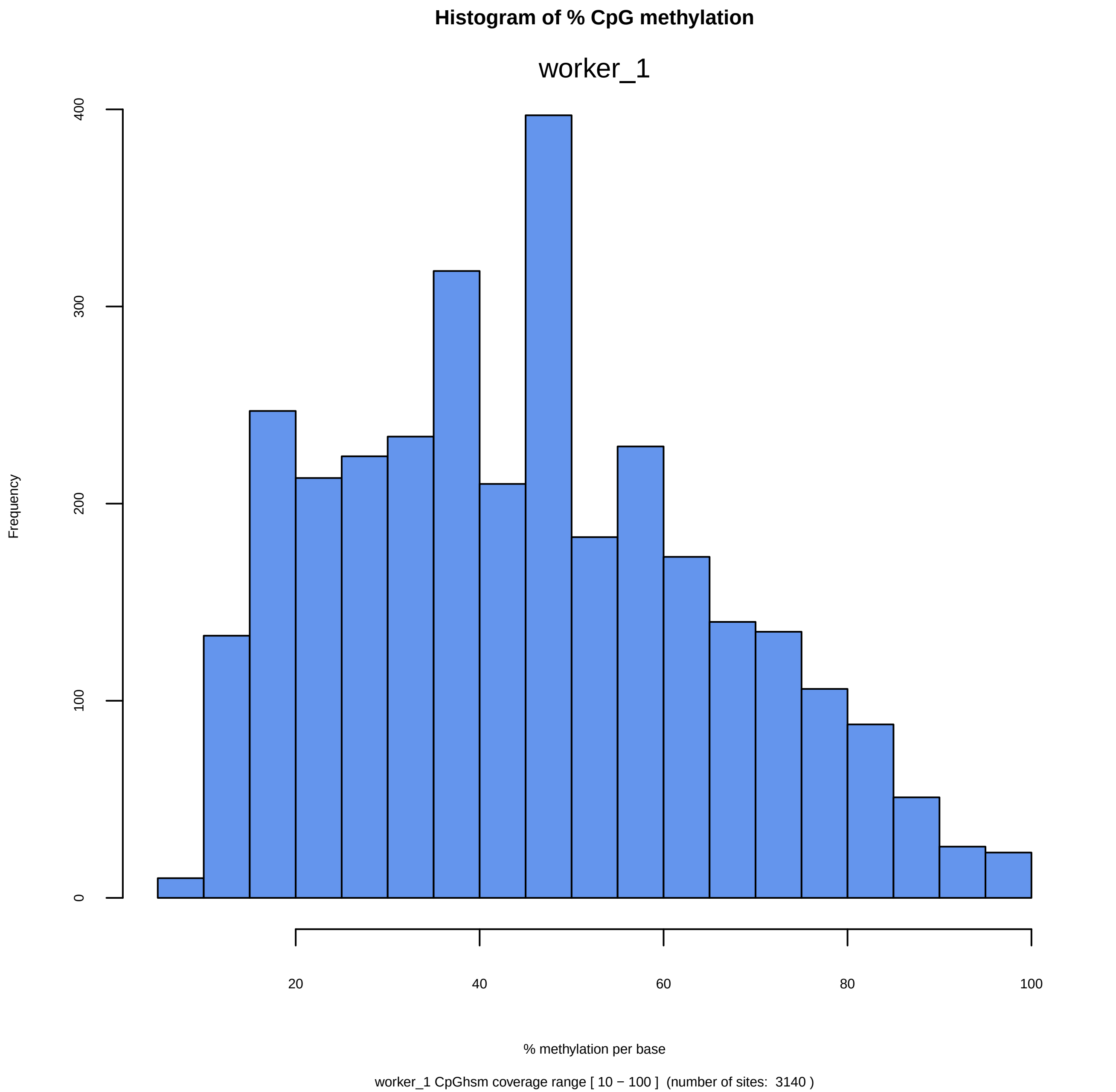
Histogram of % CpG methylation
worker_1
Frequency
0
100
200
300
400
20
40
60
80
100
% methylation per base
worker_1 CpGhsm coverage range [ 10 − 100 ]  (number of sites:  3140 )

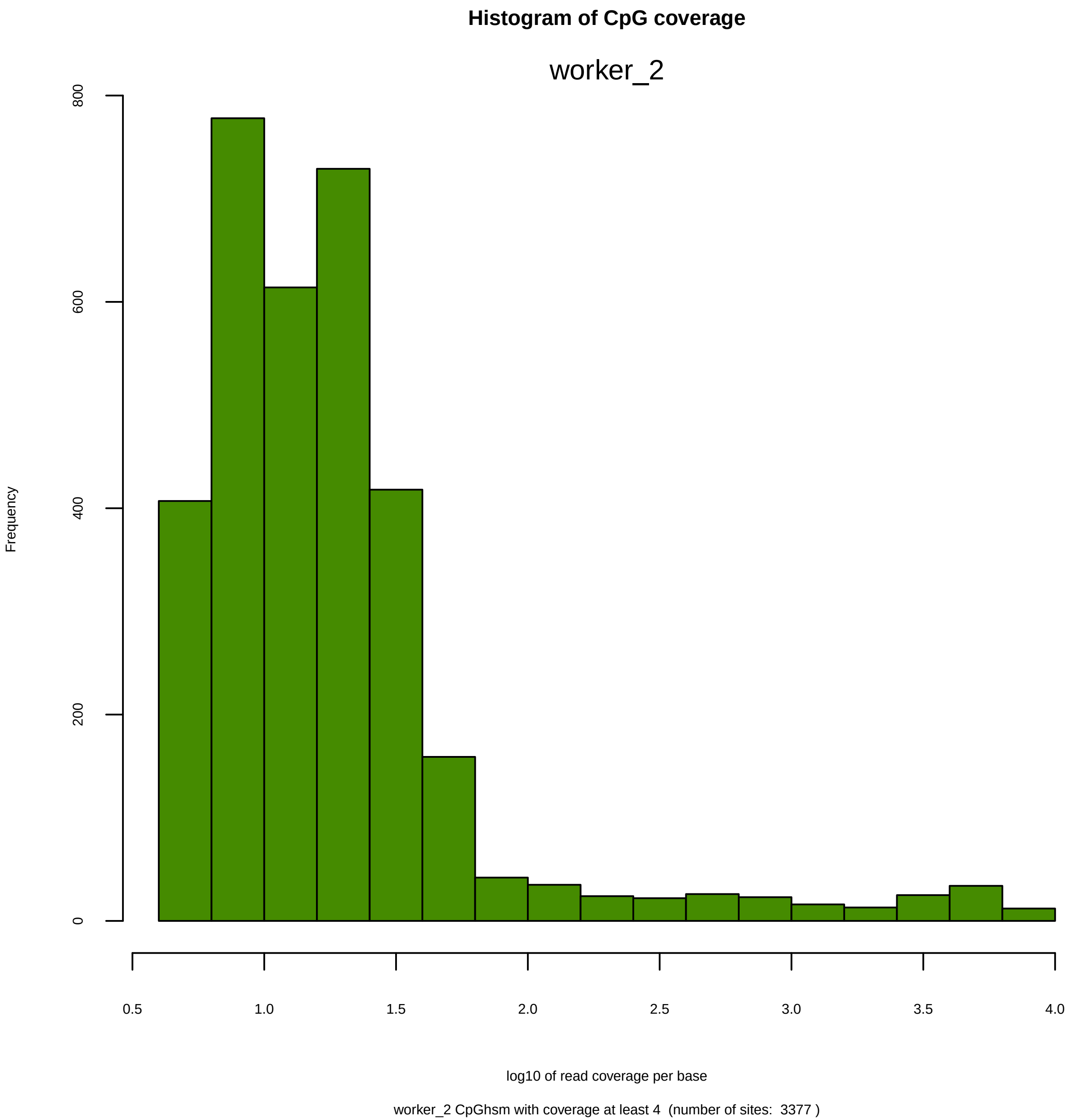
Histogram of CpG coverage
worker_2
Frequency
0
200
400
600
800
0.5
1.0
1.5
2.0
2.5
3.0
3.5
4.0
log10 of read coverage per base
worker_2 CpGhsm with coverage at least 4 (number of sites: 3377 )

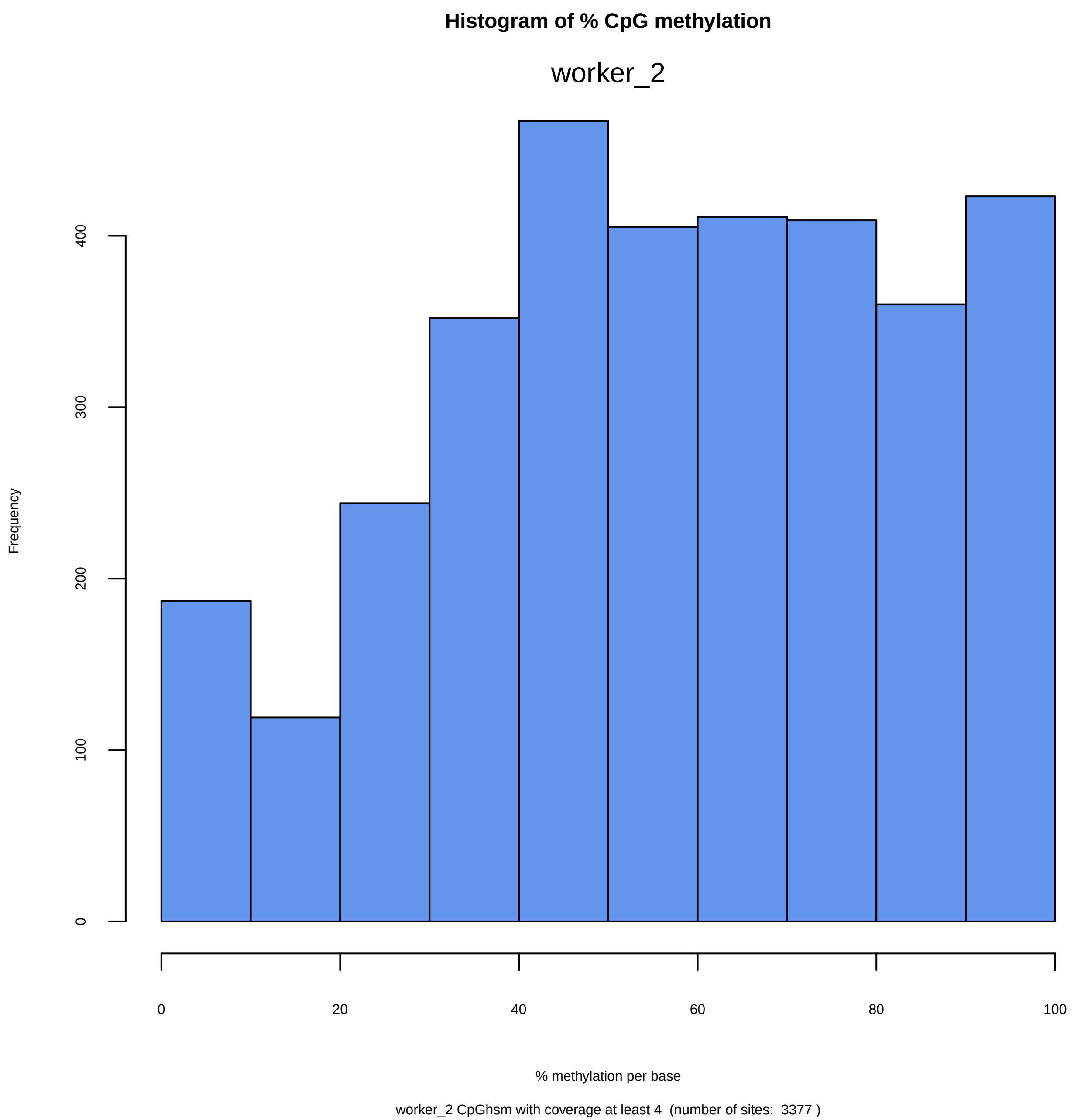
Histogram of % CpG methylation
worker_2
Frequency
0
100
200
300
400
0
20
40
60
80
100
% methylation per base
worker_2 CpGhsm with coverage at least 4  (number of sites:  3377 )

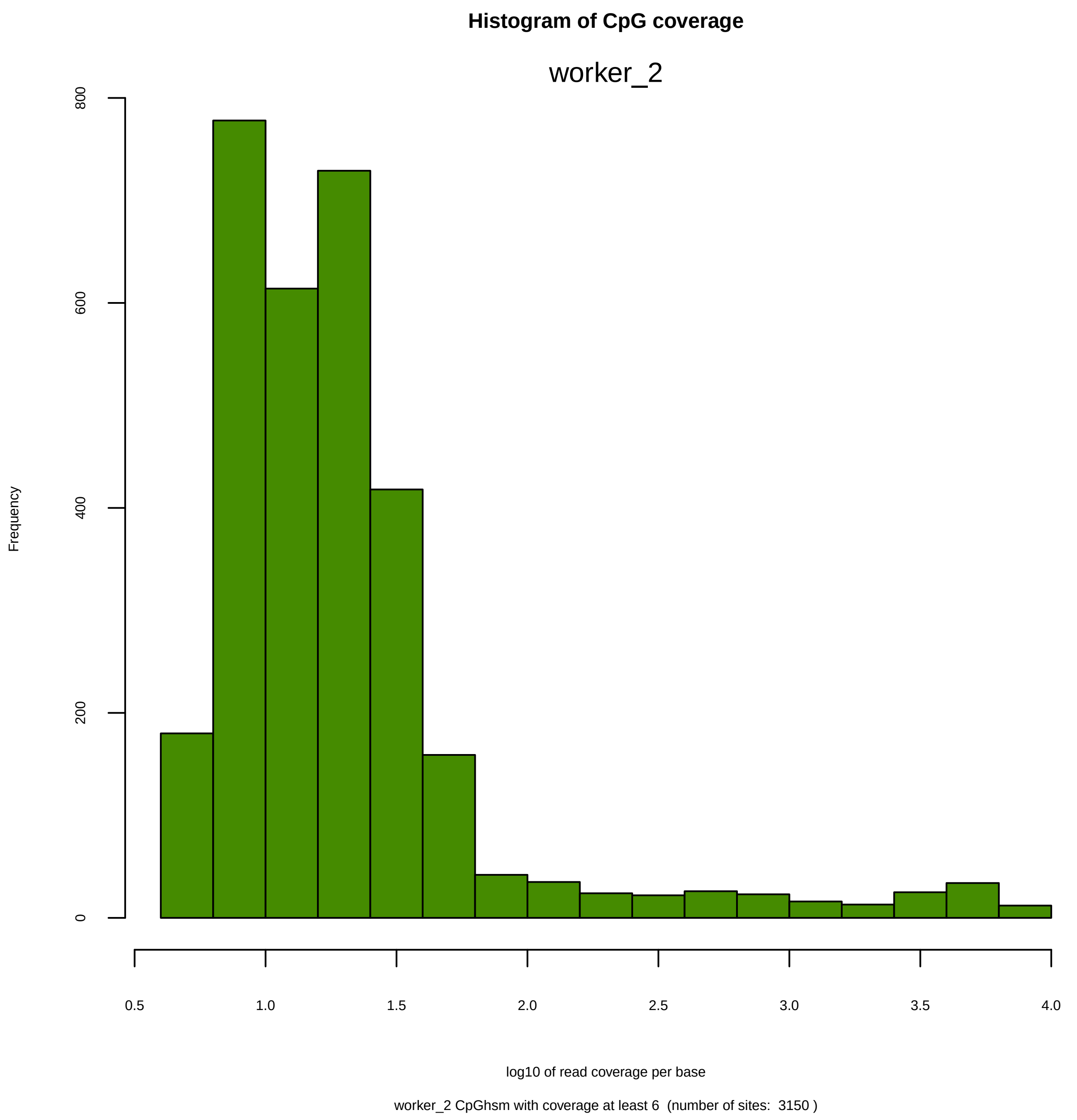
Histogram of CpG coverage
worker_2
Frequency
0
200
400
600
800
0.5
1.0
1.5
2.0
2.5
3.0
3.5
4.0
log10 of read coverage per base
worker_2 CpGhsm with coverage at least 6  (number of sites:  3150 )

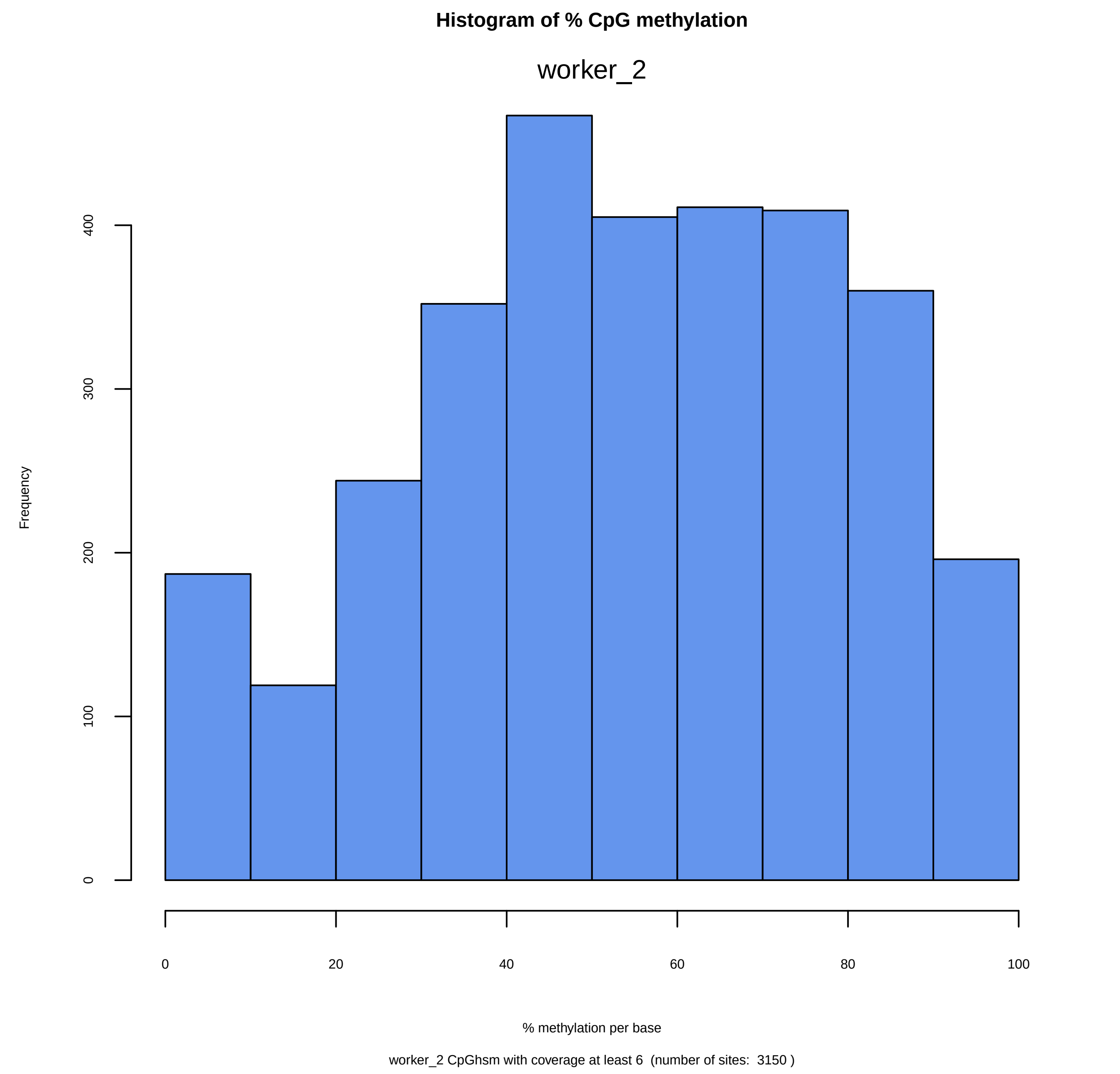
Histogram of % CpG methylation
worker_2
Frequency
0
100
200
300
400
0
20
40
60
80
100
% methylation per base
worker_2 CpGhsm with coverage at least 6  (number of sites:  3150 )

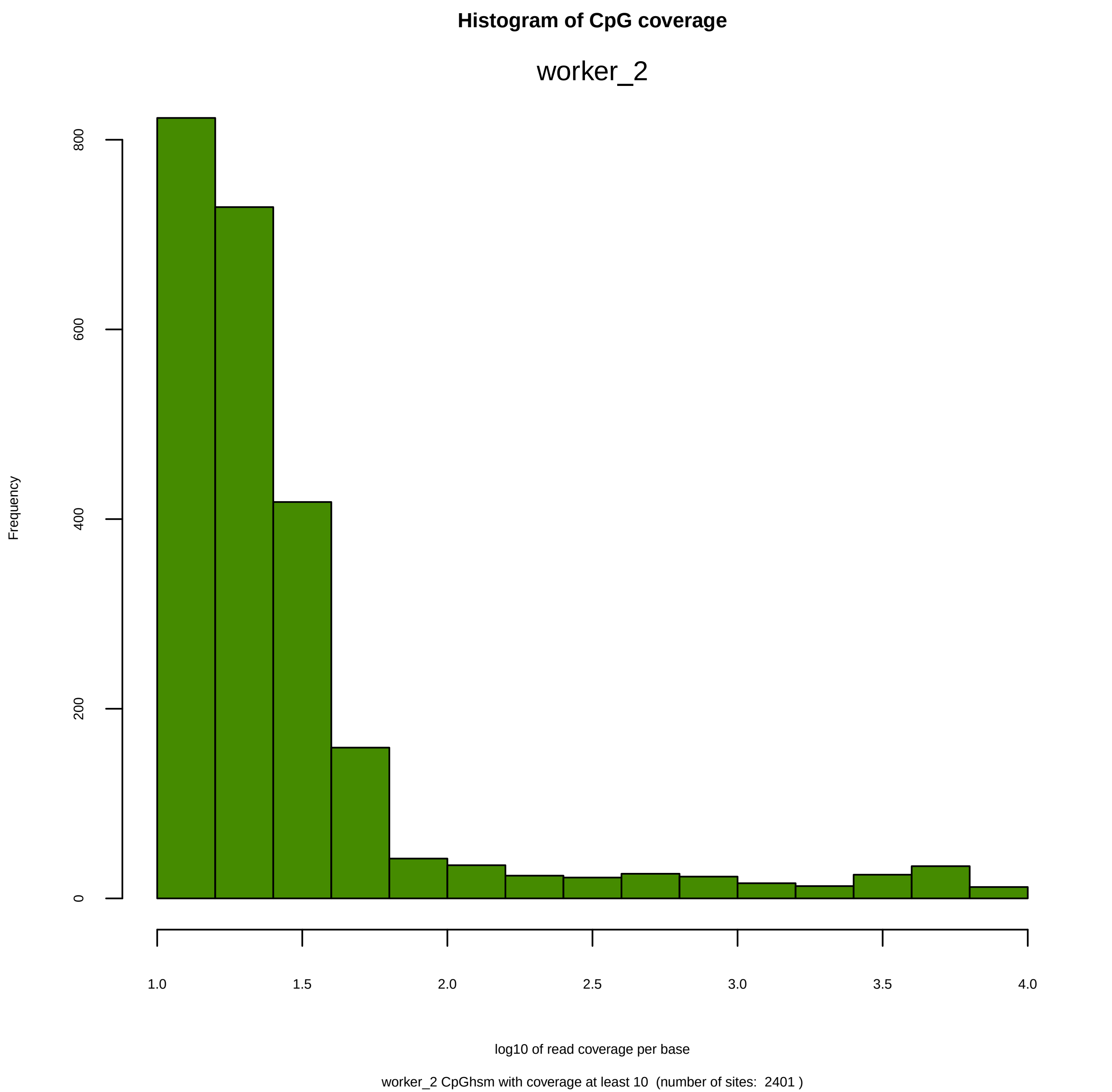
Histogram of CpG coverage
worker_2
Frequency
0
200
400
600
800
1.0
1.5
2.0
2.5
3.0
3.5
4.0
log10 of read coverage per base
worker_2 CpGhsm with coverage at least 10  (number of sites:  2401 )

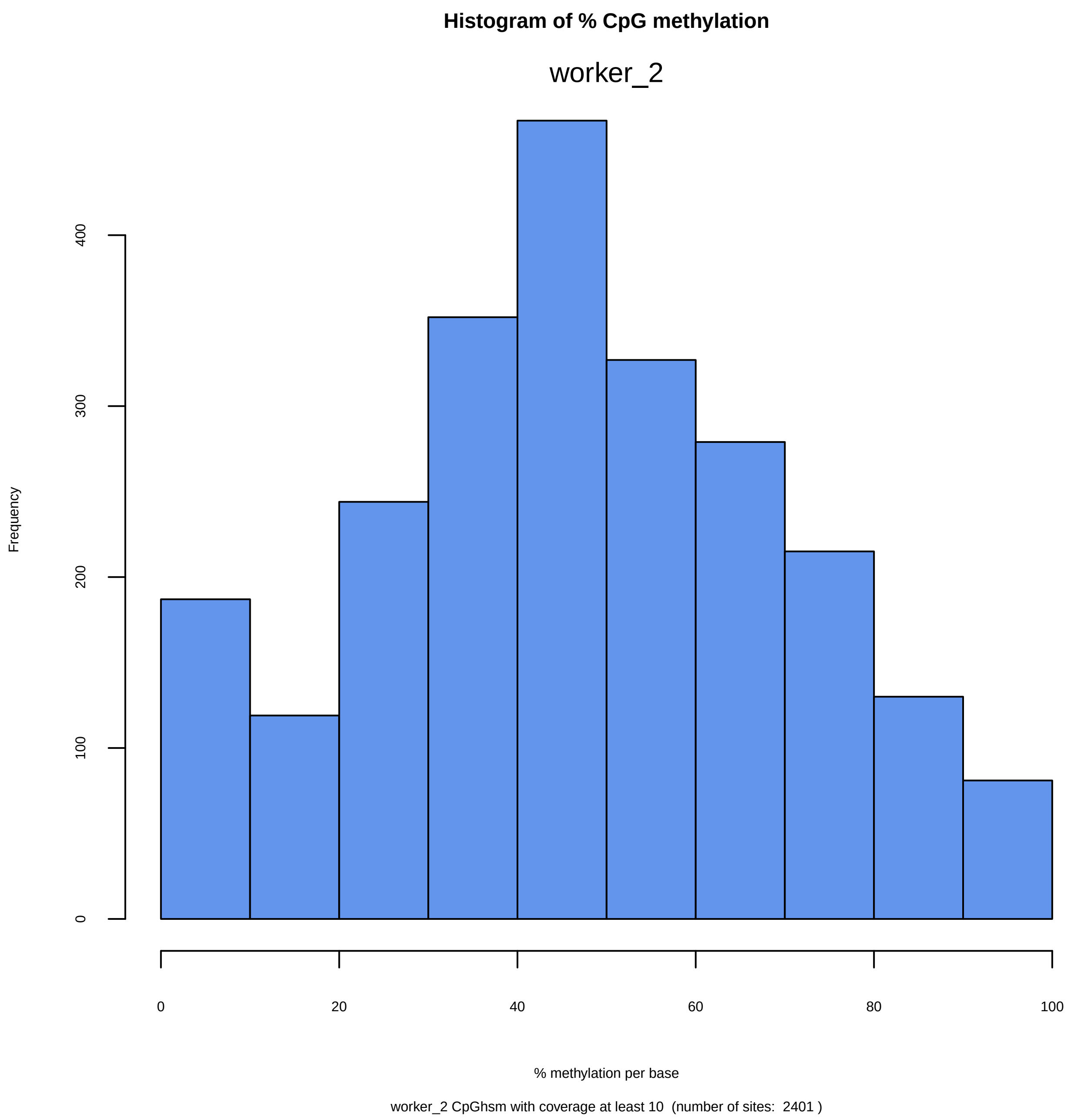

Histogram of % CpG methylation
worker_2
Frequency
0
100
200
300
400
0
20
40
60
80
100
% methylation per base
worker_2 CpGhsm with coverage at least 10  (number of sites:  2401 )

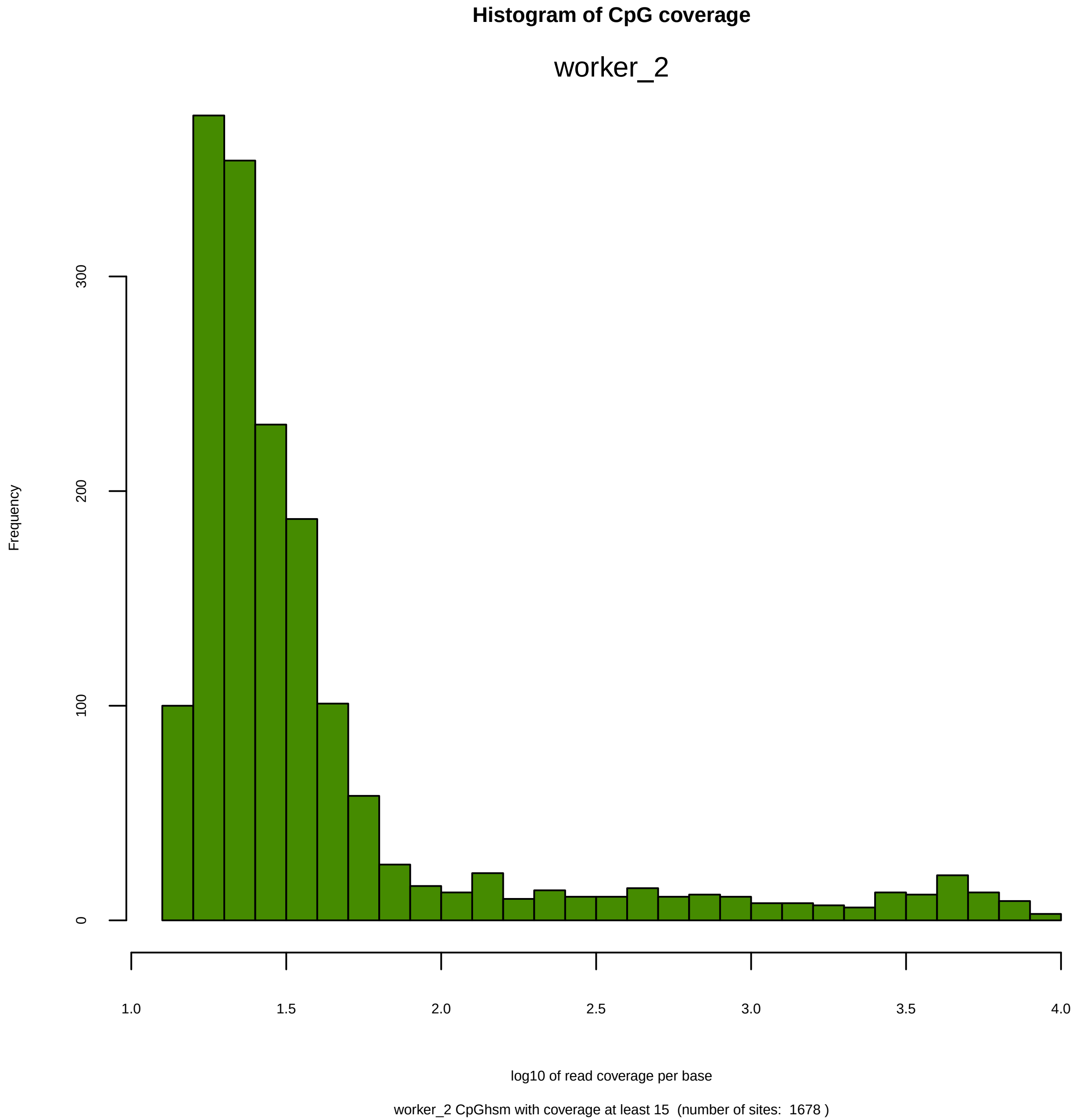
Histogram of CpG coverage
worker_2
Frequency
0
100
200
300
1.0
1.5
2.0
2.5
3.0
3.5
4.0
log10 of read coverage per base
worker_2 CpGhsm with coverage at least 15  (number of sites:  1678 )

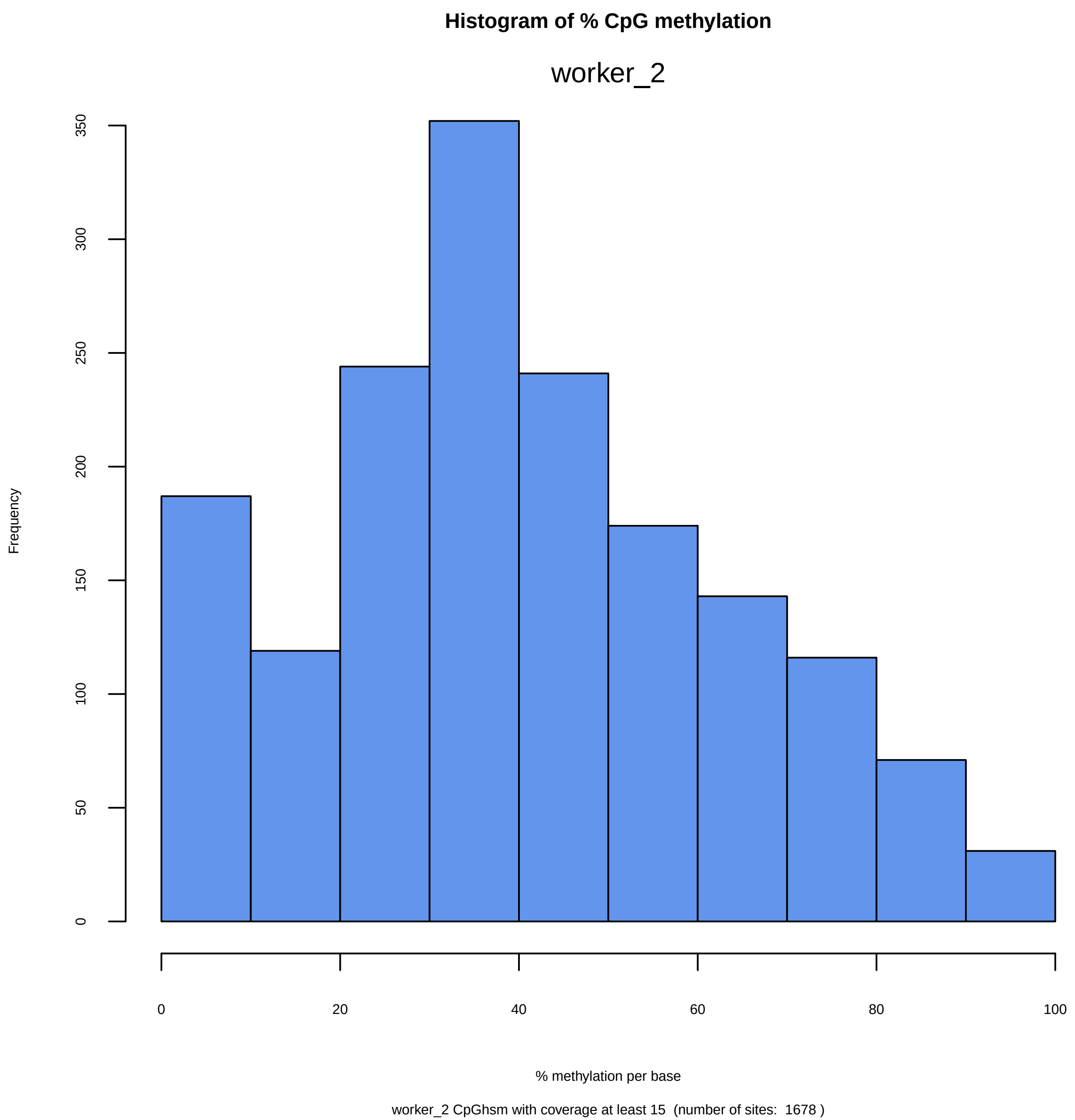
Histogram of % CpG methylation
worker_2
Frequency
0
50
100
150
200
250
300
350
0
20
40
60
80
100
% methylation per base
worker_2 CpGhsm with coverage at least 15  (number of sites:  1678 )

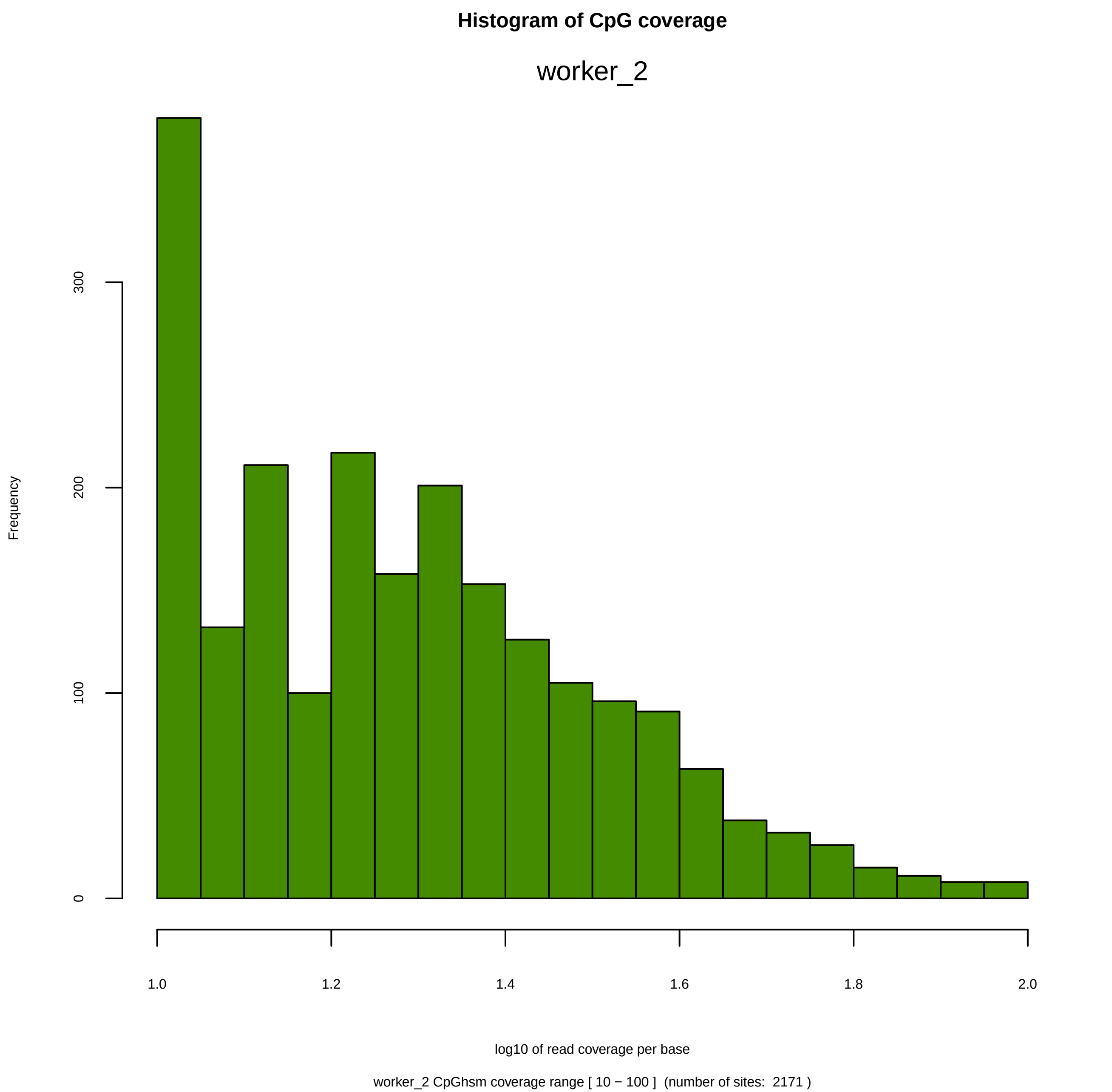

Histogram of CpG coverage
worker_2
Frequency
0
100
200
300
1.0
1.2
1.4
1.6
1.8
2.0
log10 of read coverage per base
worker_2 CpGhsm coverage range [ 10 − 100 ]  (number of sites:  2171 )

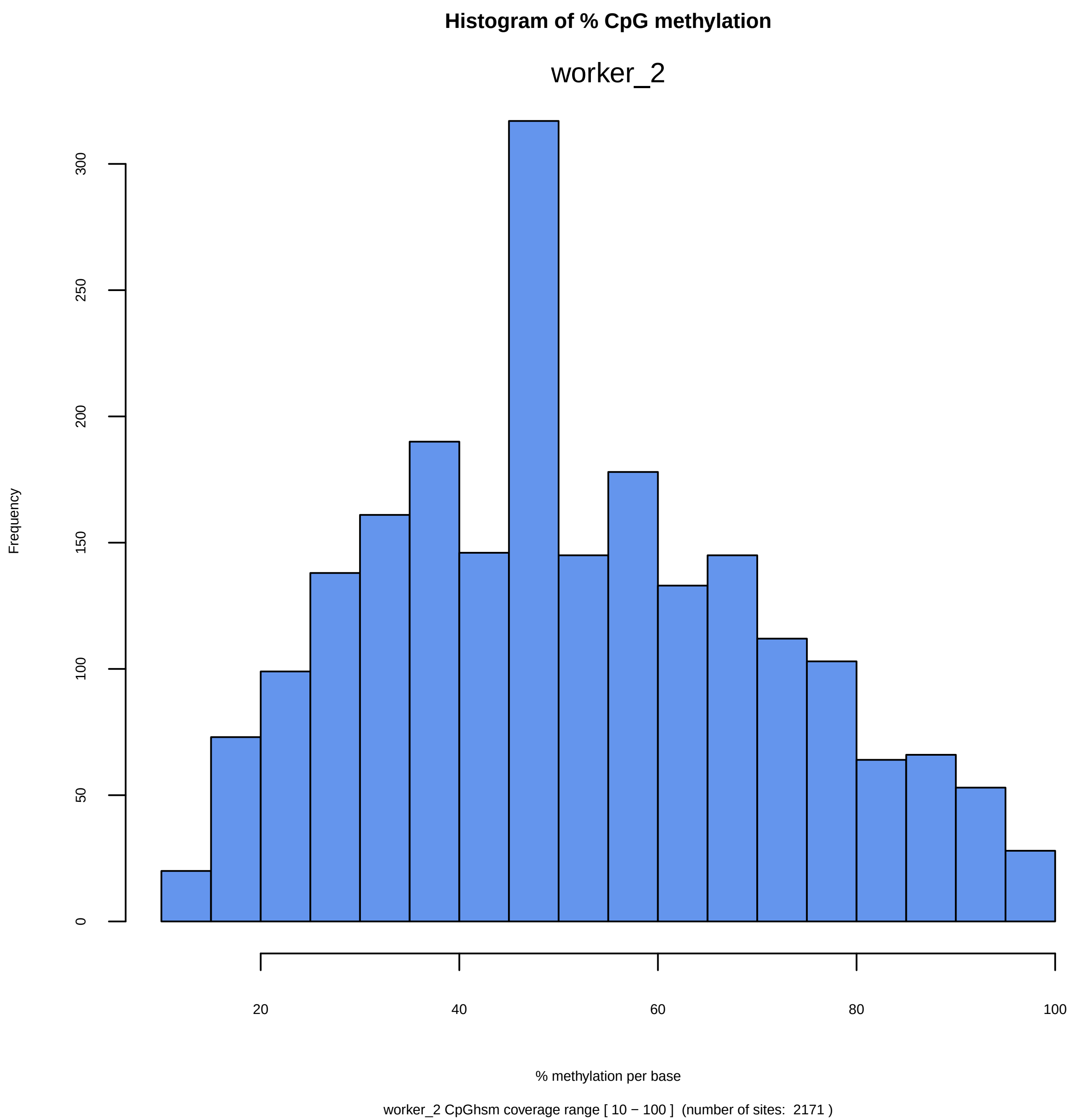

Histogram of % CpG methylation
worker_2
Frequency
0
50
100
150
200
250
300
20
40
60
80
100
% methylation per base
worker_2 CpGhsm coverage range [ 10 − 100 ]  (number of sites:  2171 )

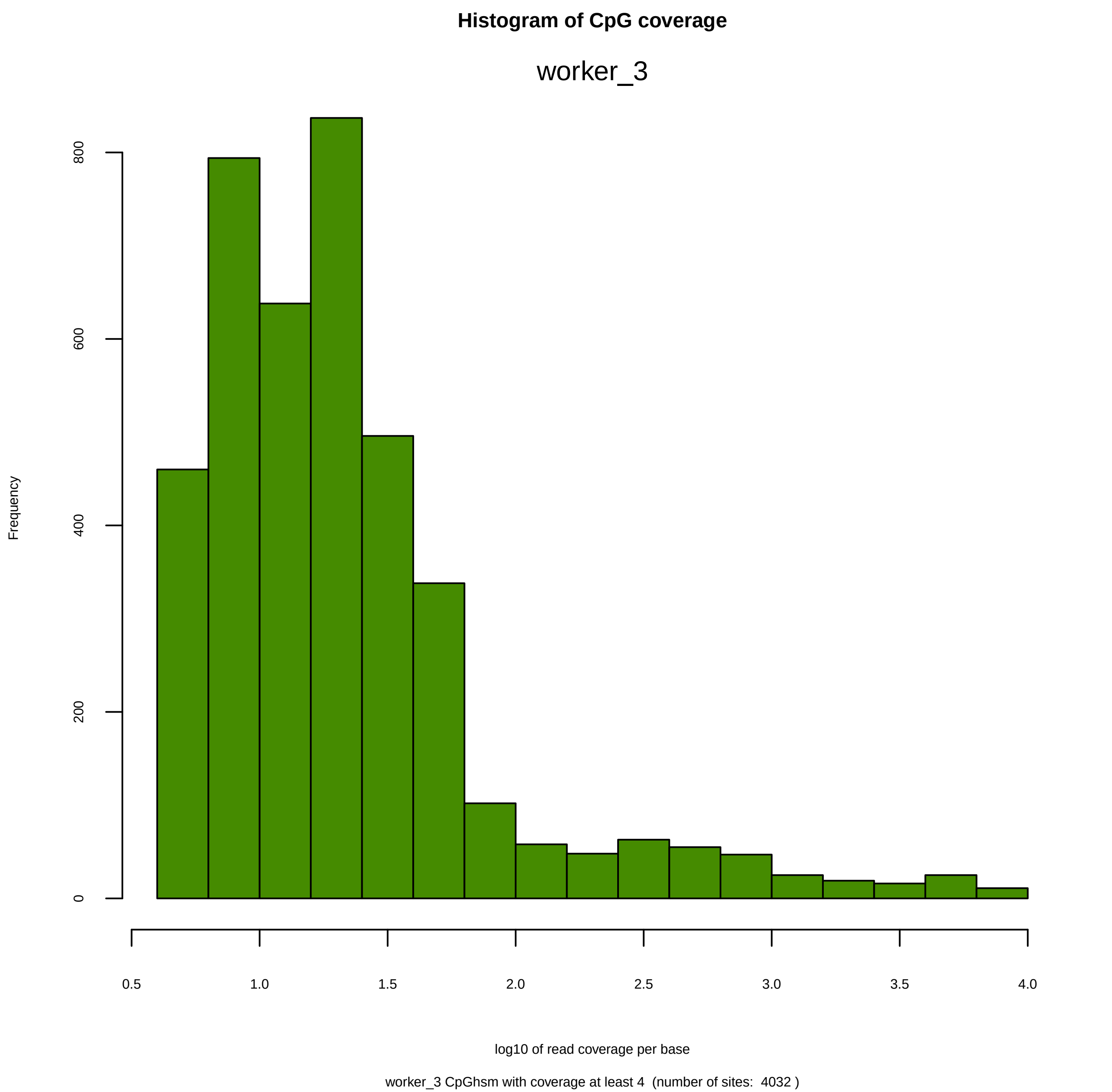

Histogram of CpG coverage
worker_3
Frequency
0
200
400
600
800
0.5
1.0
1.5
2.0
2.5
3.0
3.5
4.0
log10 of read coverage per base
worker_3 CpGhsm with coverage at least 4  (number of sites:  4032 )

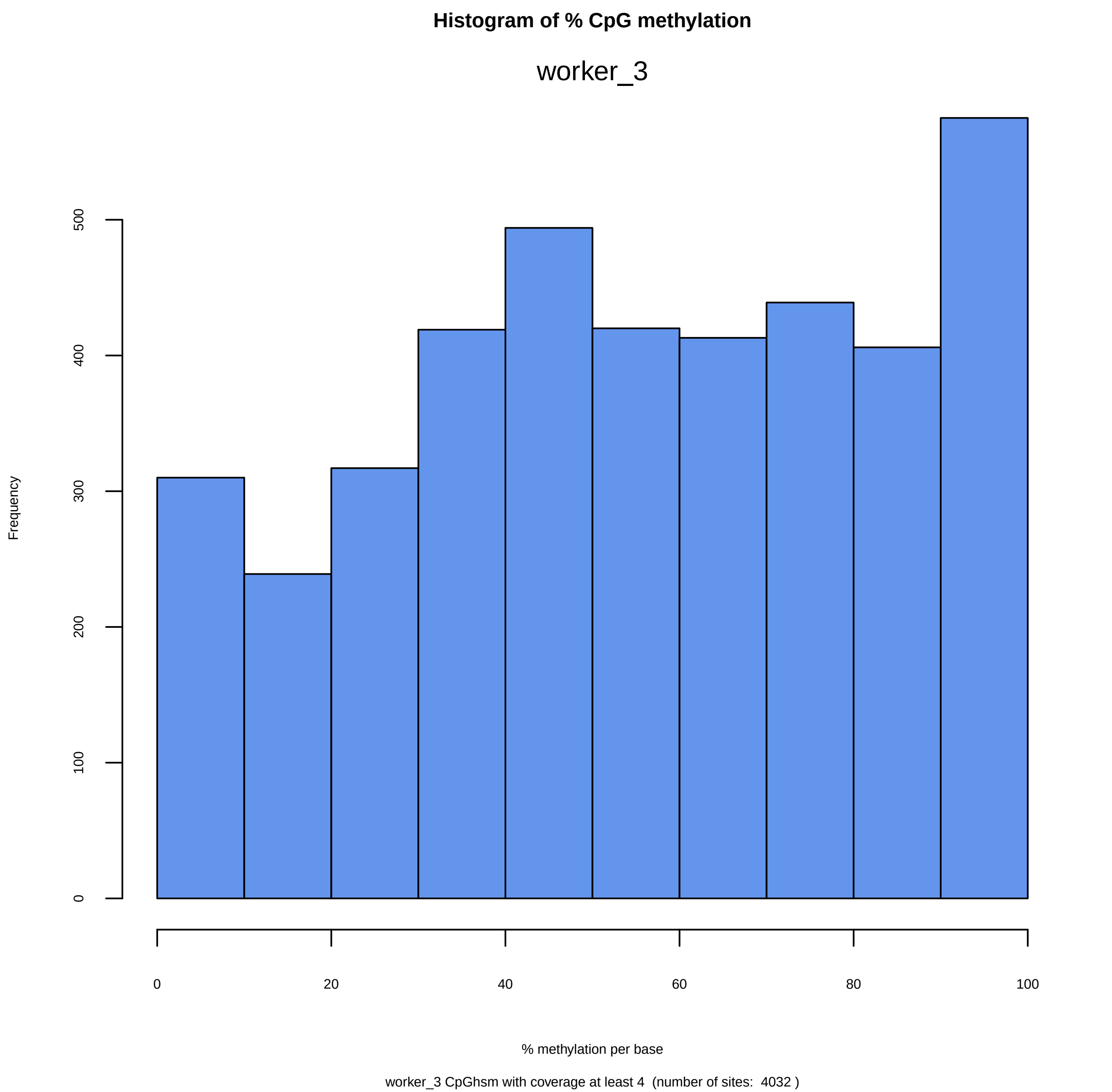
Histogram of % CpG methylation
worker_3
Frequency
0
100
200
300
400
500
0
20
40
60
80
100
% methylation per base
worker_3 CpGhsm with coverage at least 4 (number of sites: 4032 )

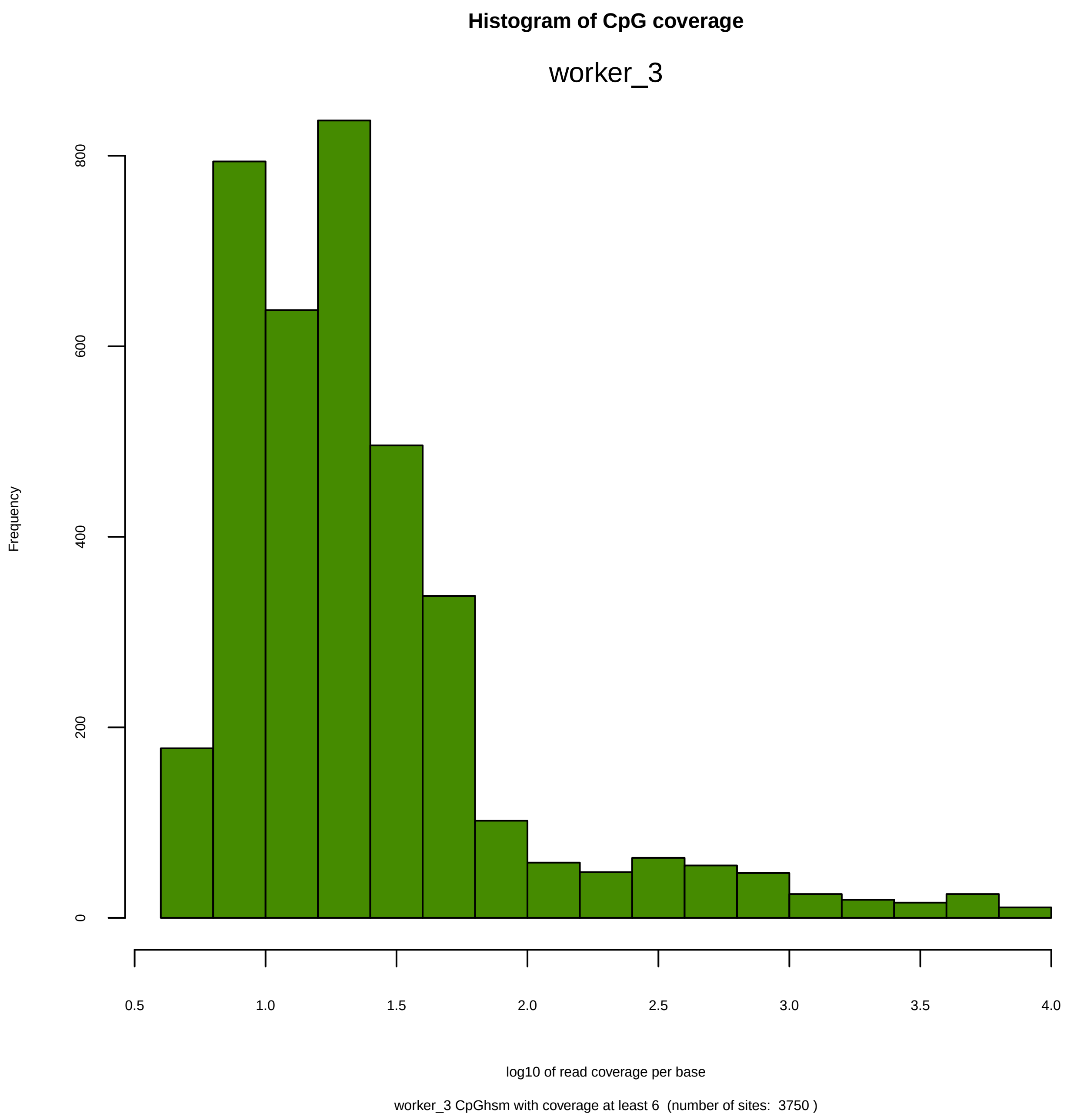
Histogram of CpG coverage
worker_3
Frequency
0
200
400
600
800
0.5
1.0
1.5
2.0
2.5
3.0
3.5
4.0
log10 of read coverage per base
worker_3 CpGhsm with coverage at least 6  (number of sites:  3750 )

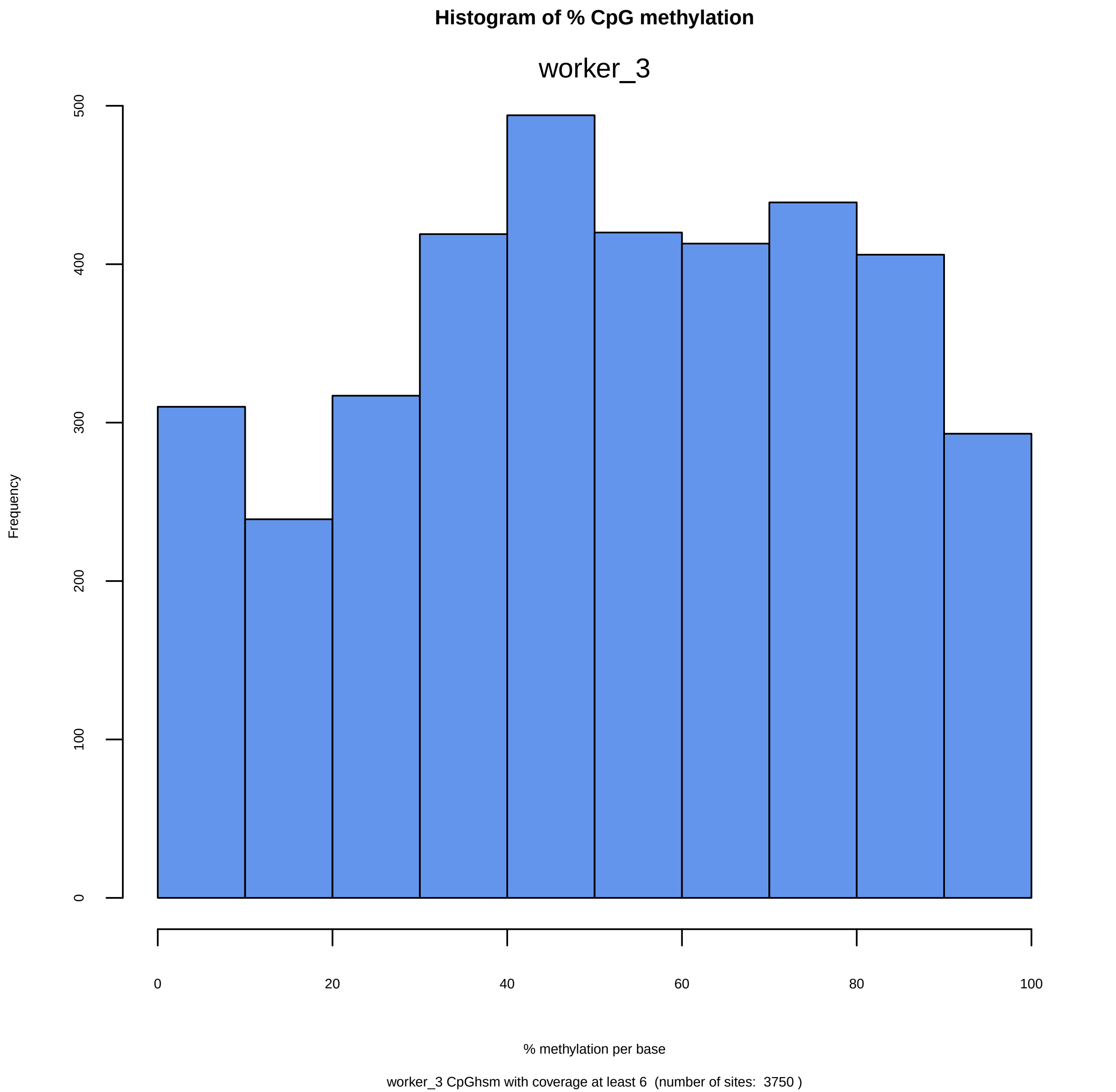
Histogram of % CpG methylation
worker_3
Frequency
0
100
200
300
400
500
0
20
40
60
80
100
% methylation per base
worker_3 CpGhsm with coverage at least 6  (number of sites:  3750 )

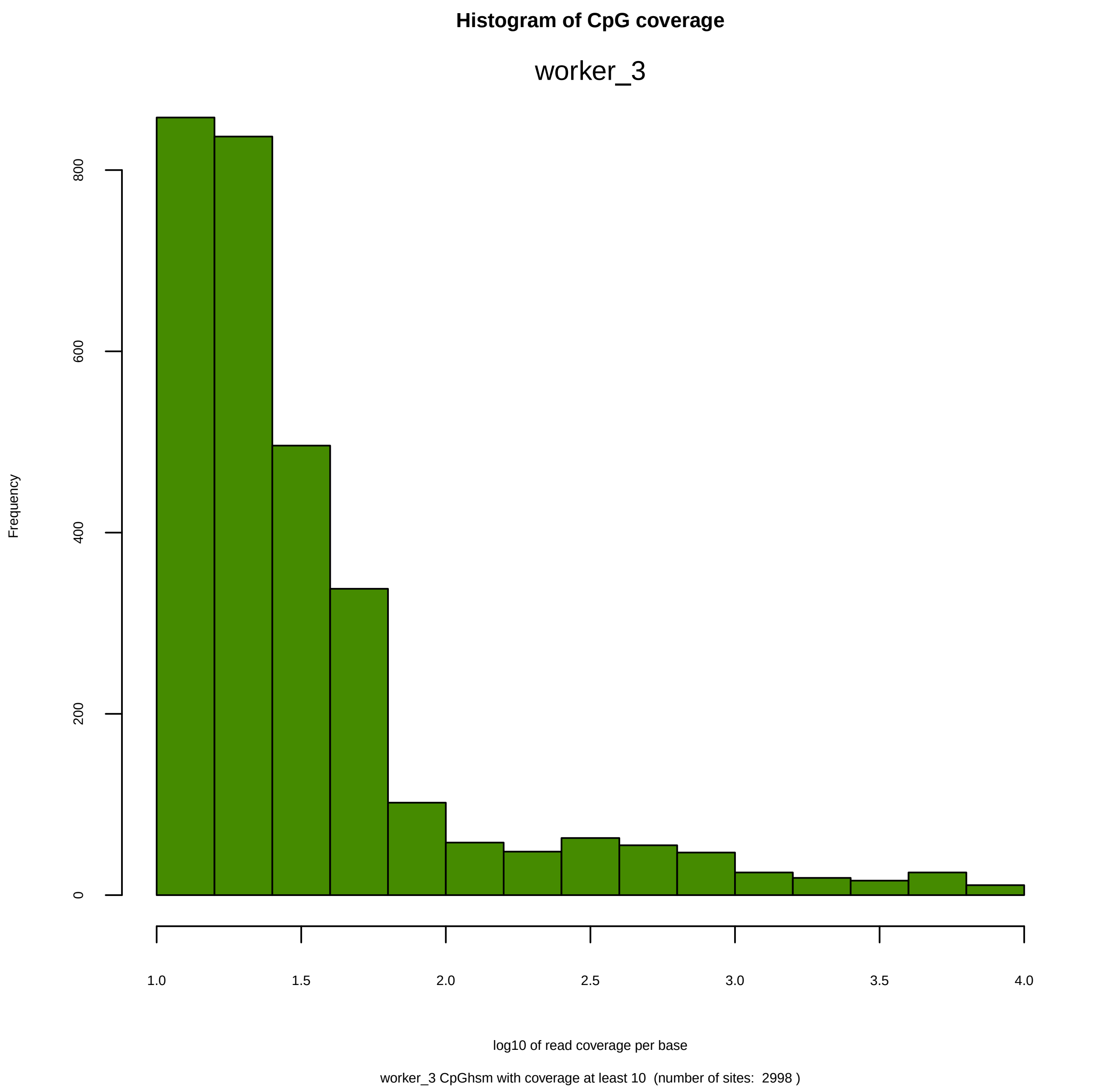
Histogram of CpG coverage
worker_3
Frequency
0
200
400
600
800
1.0
1.5
2.0
2.5
3.0
3.5
4.0
log10 of read coverage per base
worker_3 CpGhsm with coverage at least 10  (number of sites:  2998 )

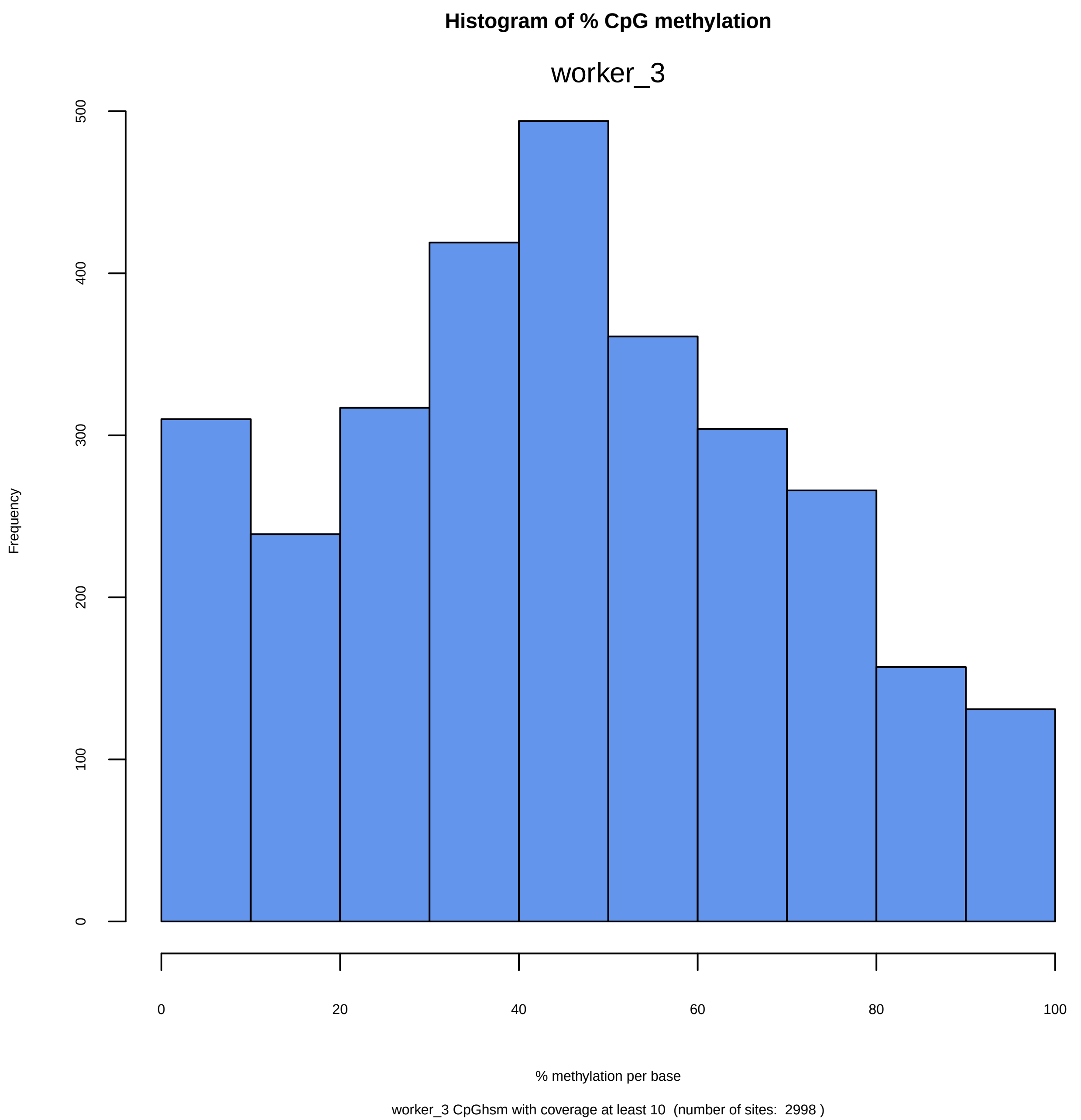
Histogram of % CpG methylation
worker_3
Frequency
0
100
200
300
400
500
0
20
40
60
80
100
% methylation per base
worker_3 CpGhsm with coverage at least 10  (number of sites:  2998 )

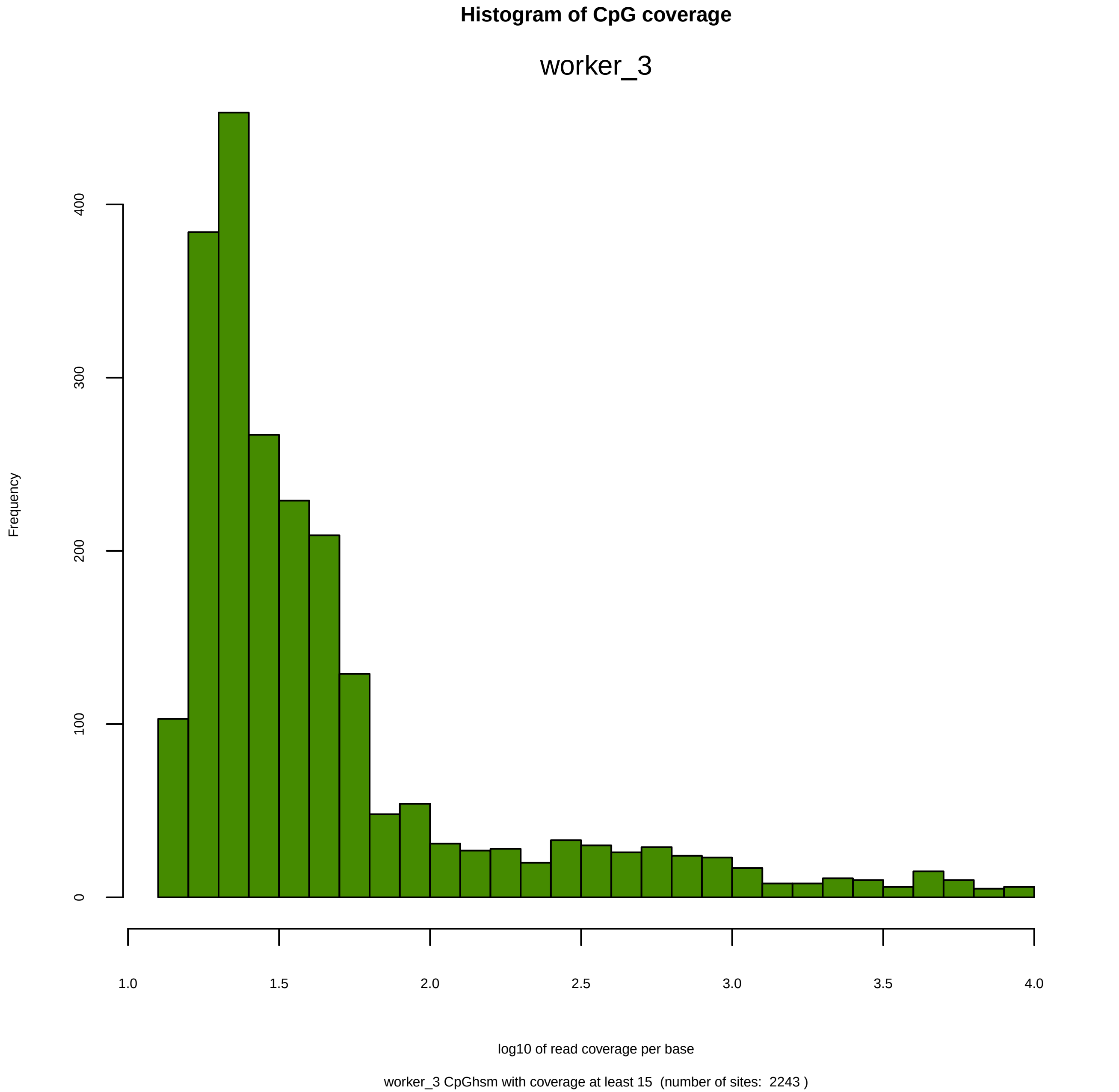
Histogram of CpG coverage
worker_3
Frequency
0
100
200
300
400
1.0
1.5
2.0
2.5
3.0
3.5
4.0
log10 of read coverage per base
worker_3 CpGhsm with coverage at least 15  (number of sites:  2243 )

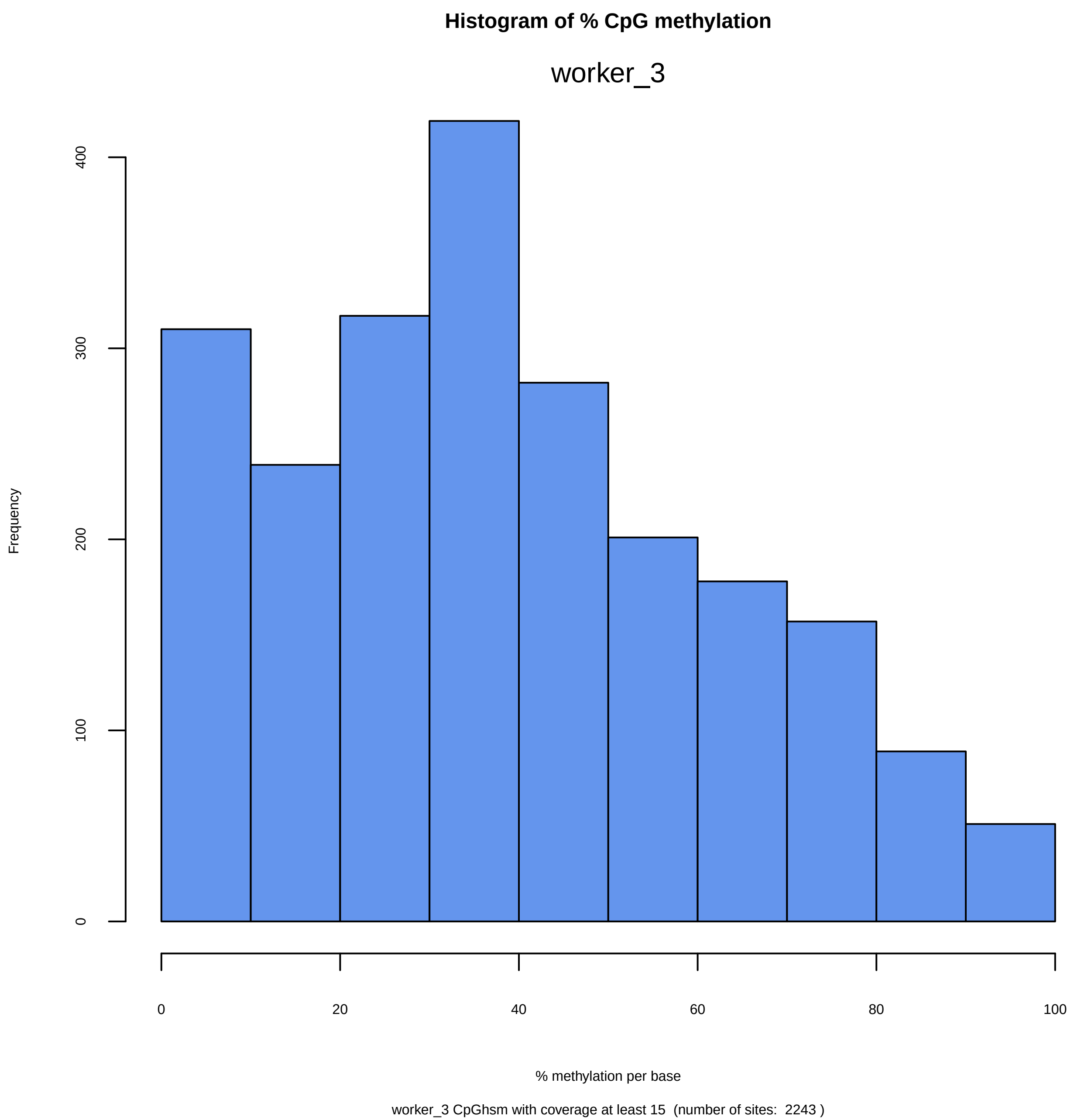
Histogram of % CpG methylation
worker_3
Frequency
0
100
200
300
400
0
20
40
60
80
100
% methylation per base
worker_3 CpGhsm with coverage at least 15  (number of sites:  2243 )

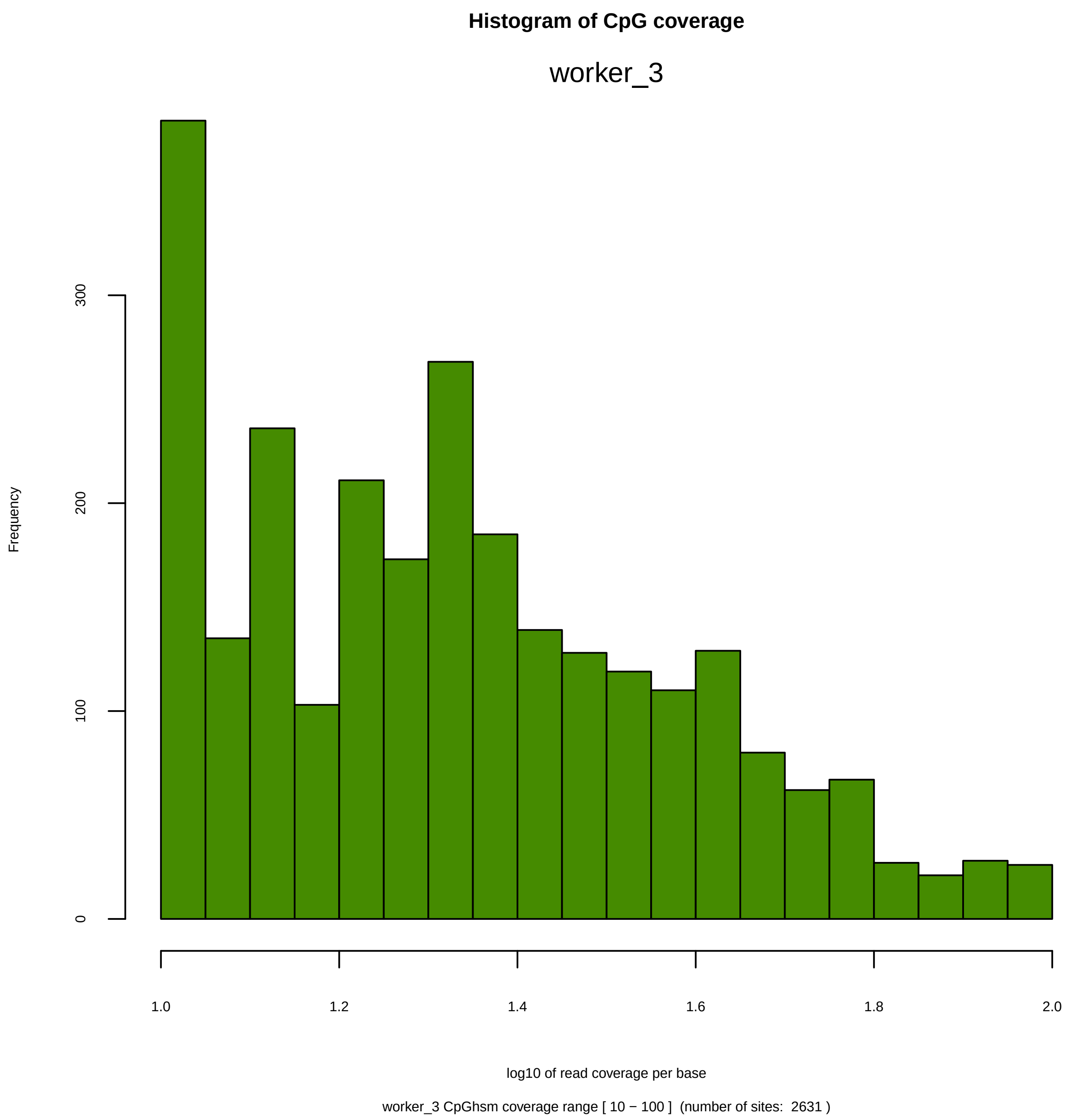

Histogram of CpG coverage
worker_3
Frequency
0
100
200
300
1.0
1.2
1.4
1.6
1.8
2.0
log10 of read coverage per base
worker_3 CpGhsm coverage range [ 10 − 100 ]  (number of sites:  2631 )

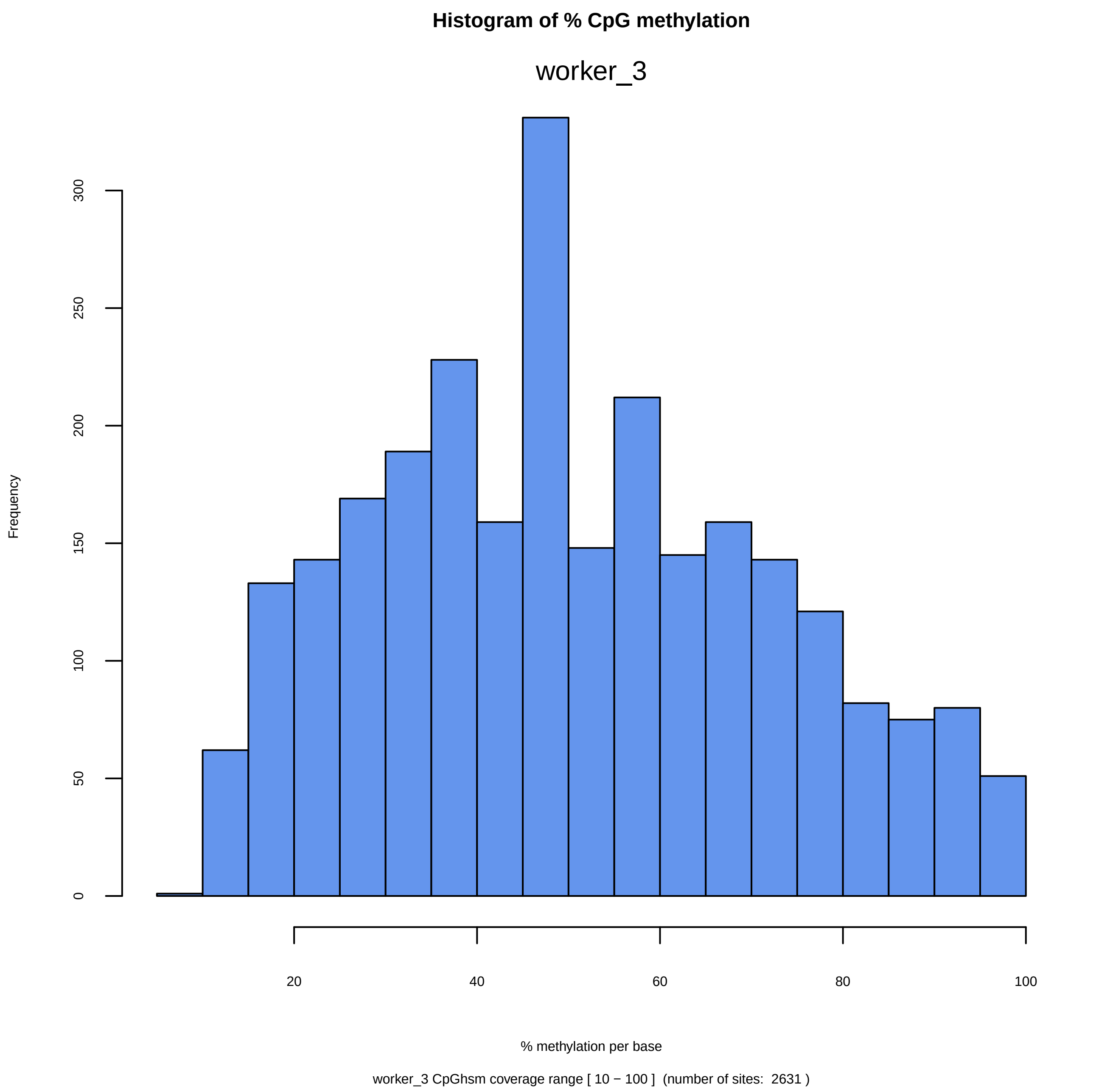

Histogram of % CpG methylation
worker_3
Frequency
0
50
100
150
200
250
300
20
40
60
80
100
% methylation per base
worker_3 CpGhsm coverage range [ 10 − 100 ] (number of sites: 2631 )

```
Directory: REPCRL File: 0READMErepcrl


REPCRL - Correlations between replicates.

Input: mkrd (methylKit methylRawList object of replicate data)

Output: files repcrl-*.txt repcrl-*.pdf

Notes: The output is generated by BWASPR::cmpSamples(), which uses methylKit::unite()
        to determine the sites common to all replicates and applies methylKit::getCorrelation()
        and methylKit::PCASamples().  The *.txt files show the correlations in a table, and
        the *.pdf files show graphics.

       Beware that the number of common sites (shown at the bottom of the *.txt files) may be
        small, which may make the correlations less informative.
```

Directory: REPCRL File: repcrl-Pc_PA_queen.txt

| | queen_1 | queen_2 | queen_3 |
|---|---|---|---|
| queen_1 | 1.000 | 0.823 | 0.830 |
| queen_2 | 0.823 | 1.000 | 0.831 |
| queen_3 | 0.830 | 0.831 | 1.000 |

The number of conserved sites is 1673.

Directory: REPCRL File: repcrl-Pc_PA_worker.txt

| | worker_1 | worker_2 | worker_3 |
|---|---|---|---|
| worker_1 | 1.000 | 0.830 | 0.821 |
| worker_2 | 0.830 | 1.000 | 0.833 |
| worker_3 | 0.821 | 0.833 | 1.000 |

The number of conserved sites is 1455.

#### CpG base pearson cor.

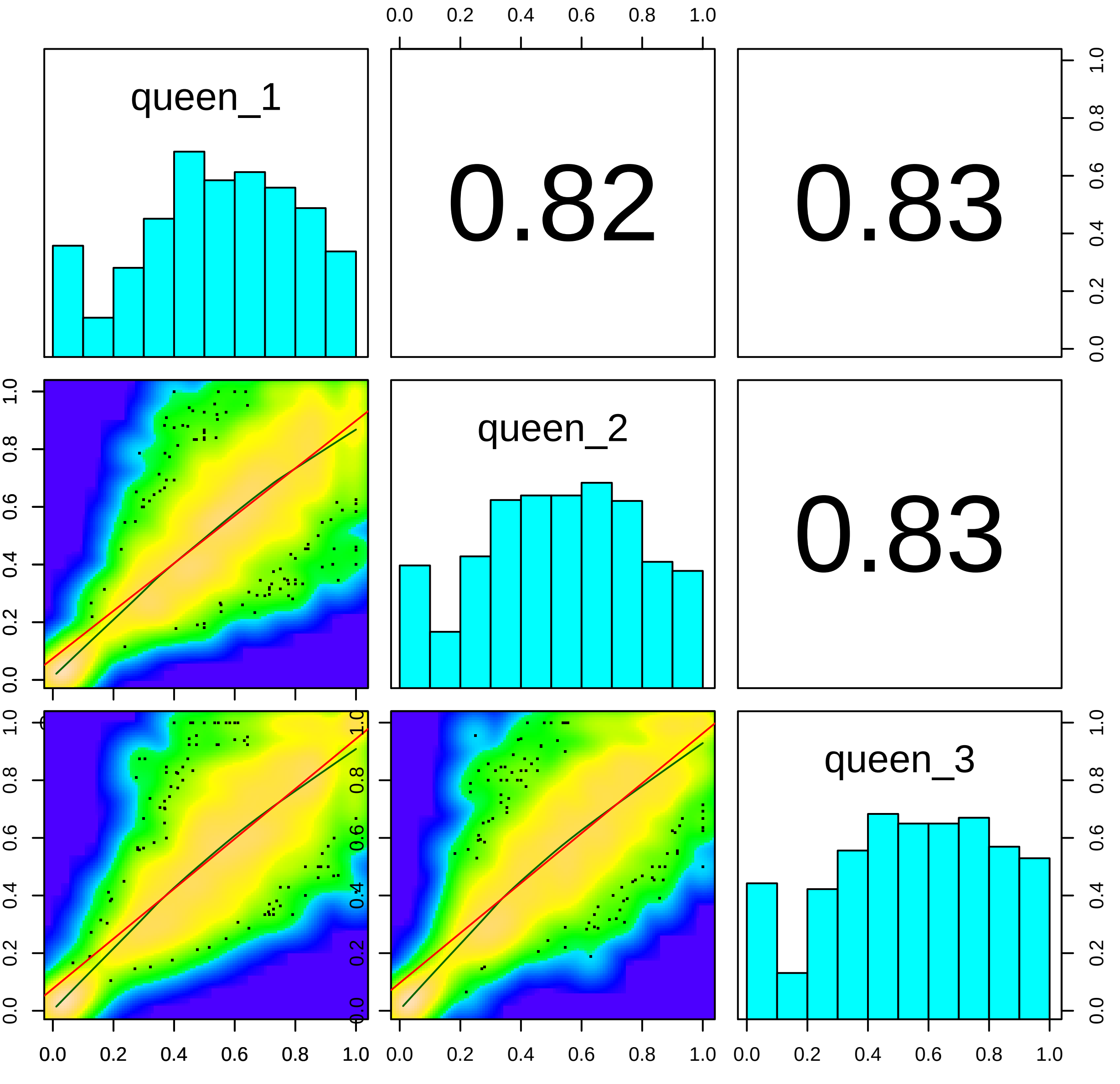

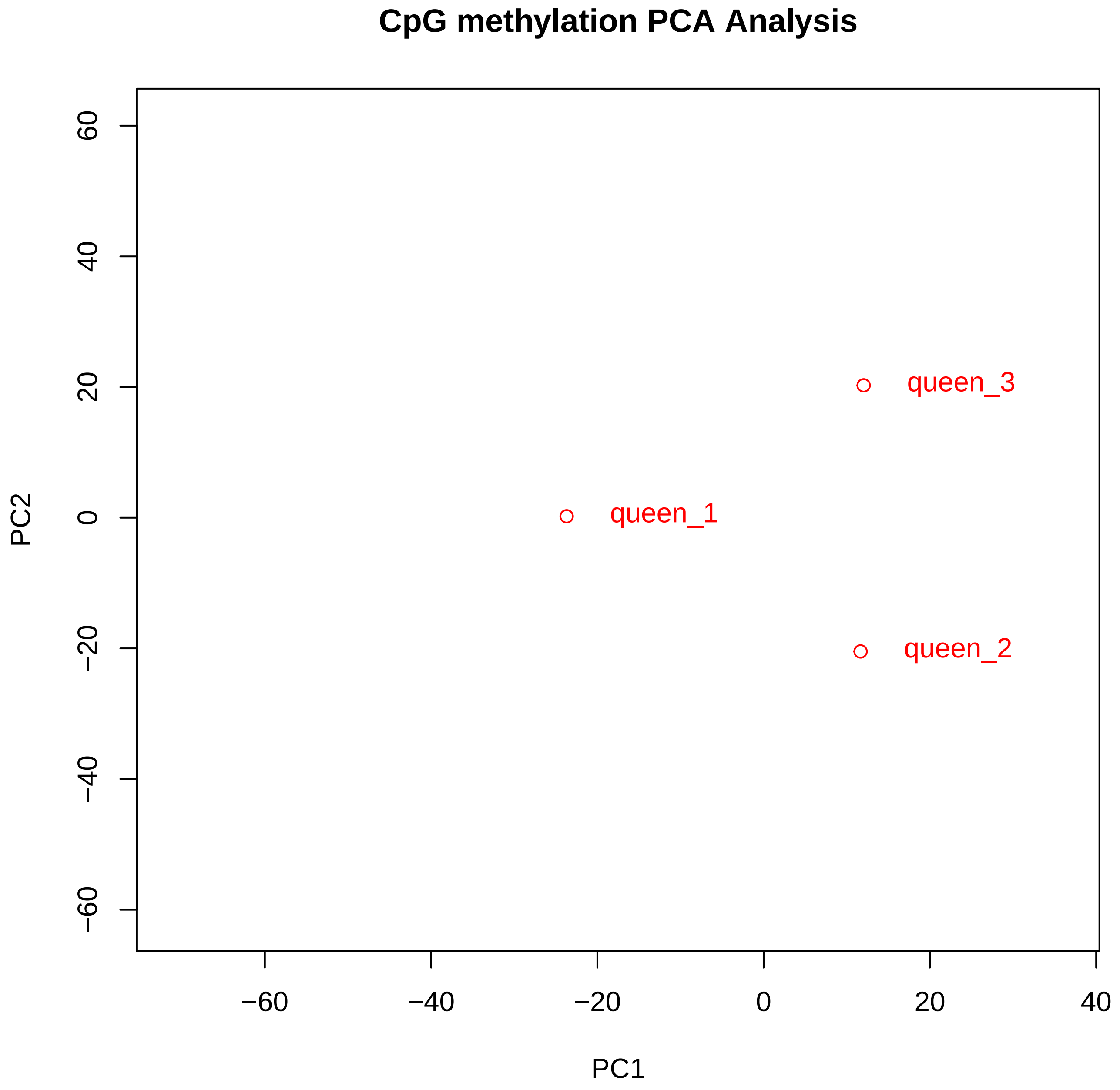
CpG methylation PCA Analysis
PC1
PC2
−60
−40
−20
0
20
40
60
queen_1
queen_2
queen_3

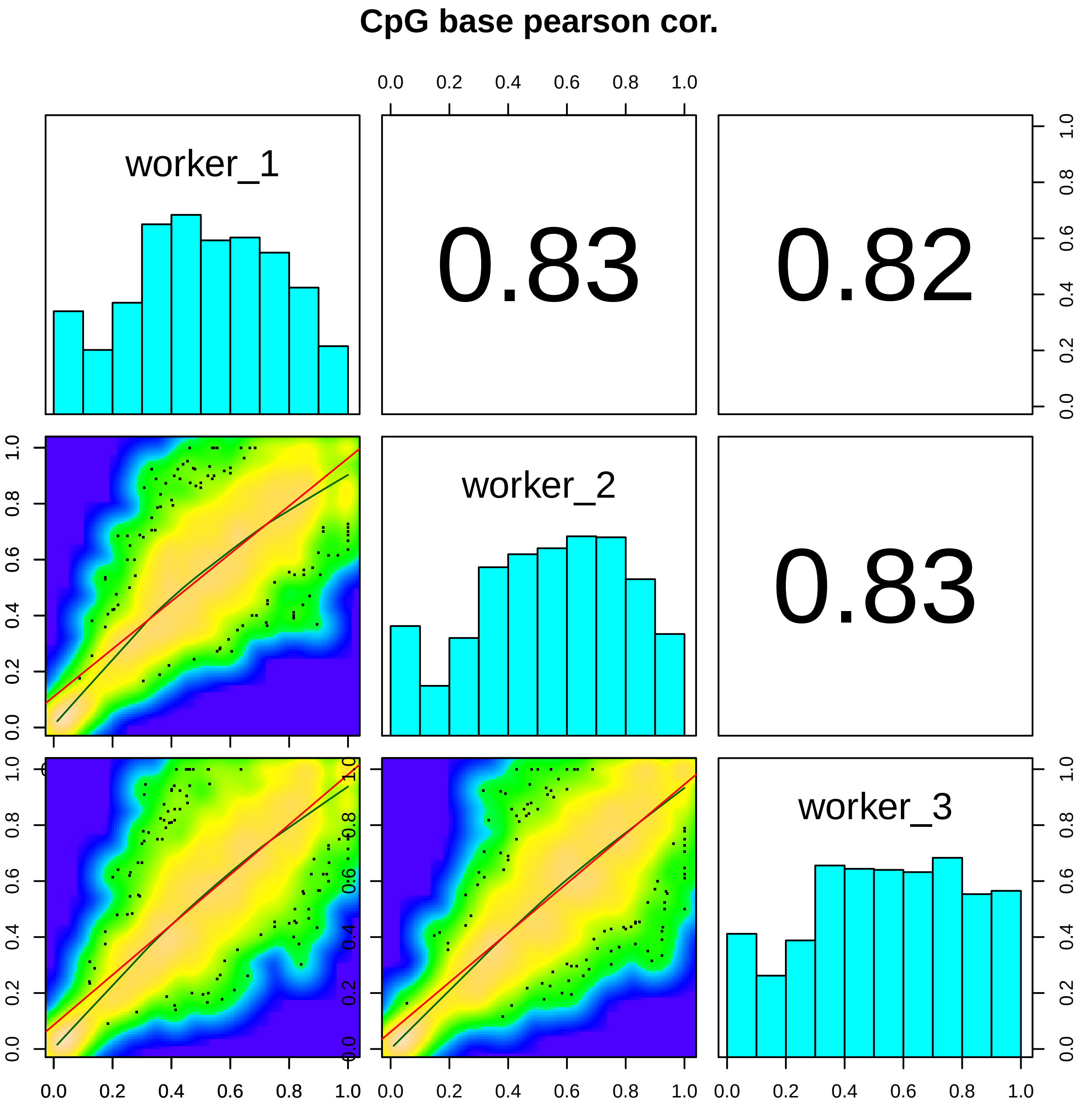

CpG base pearson cor.
worker_1
0.83
0.82
worker_2
0.83
worker_3
0.0
0.2
0.4
0.6
0.8
1.0

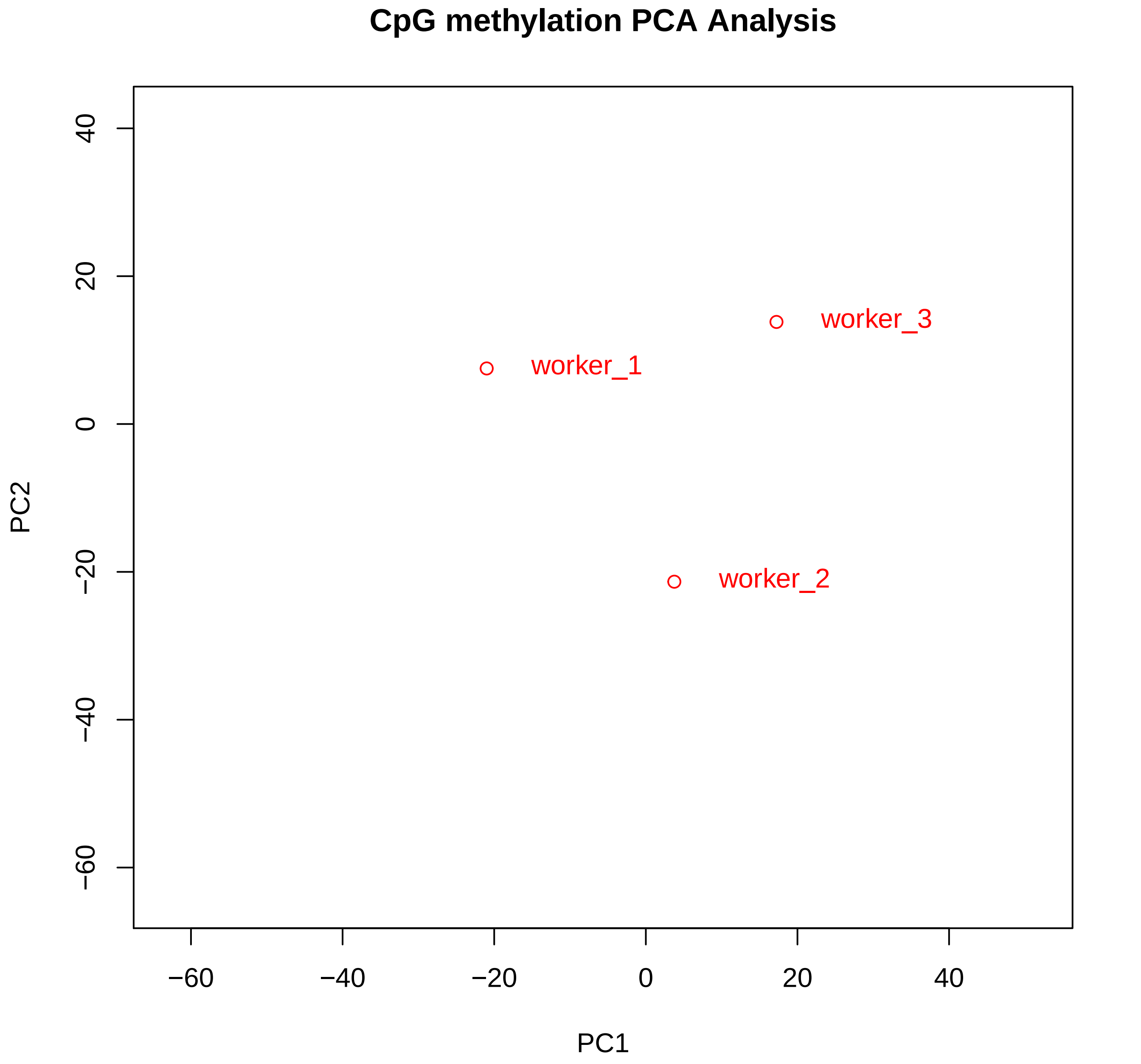
CpG methylation PCA Analysis
worker_1
worker_2
worker_3
PC1
PC2
−60
−40
−20
0
20
40

```
Directory: MMP File: 0READMEmmp


MMP - Mapping of methylation sites on genome annotation.

Input: studymk, studymc, genome_ann (a list of GRanges objects providing annotated region labels
        and boundaries; output of BWASPR::get_genome_annotation())

Output: files mmp-*.txt

Notes: The output is generated by BWASPR::map_methylome() and gives an accounting for every sample
        as to where the CpGhsm and CpGscd (control) sites reside relative to the genome annotation.
       Abbreviations used: O/E, observed over expected
                           (expected percentages are based on the respective feature proportions
                            in the genome, as annotated).
```

Directory: MMP File: mmp-Pc_PA_qn.txt

================================================================================

#### Genomic composition in terms of feature regions

================================================================================

| Region | Size | Percentage |
|---|---|---|
| Pc genome size: | 211202212 bp | |
| Pc genic region size: | 125747138 bp | ( 59.5%) |
| Pc exon region size: | 27615710 bp | ( 13.1%) |
| Pc CDS region size: | 17712034 bp | ( 64.1%) |
| Pc five-prime UTR region size: | 3011712 bp | ( 10.9%) |
| Pc three-prime UTR region size: | 4732072 bp | ( 17.1%) |
| Pc other exon region size: | 2159892 bp | ( 7.8%) |
| Pc intron region size: | 98131428 bp | ( 46.5%) |
| Pc intergenic region size: | 85455074 bp | ( 40.5%) |
| Pc promoter region size: | 5302351 bp | ( 2.5%) |
| Pc other intergenic region size: | 80152723 bp | ( 38.0%) |

================================================================================

#### Methylation sites in genomic feature regions

================================================================================

Number and density (per 10kb) of sites in Pc_PA_qn_scd

| | | | | |
|---|---|---|---|---|
| Number of sites identified in Pc Pc_PA_qn_scd: | 8534313 | ( 404.08) | | |
| Number of sites identified in Pc Pc_PA_qn_scd genic regions: | 5244698 | ( 417.08) | 61.45% | ( 1.03 O/E) |
| Number of sites identified in Pc Pc_PA_qn_scd exon regions: | 1301866 | ( 471.42) | 15.25% | ( 1.17 O/E) |
| Number of sites identified in Pc Pc_PA_qn_scd CDS regions: | 957804 | ( 540.76) | 73.57% | ( 1.15 O/E) |
| Number of sites identified in Pc Pc_PA_qn_scd five-prime UTR regions: | 180989 | ( 600.95) | 13.90% | ( 1.27 O/E) |
| Number of sites identified in Pc Pc_PA_qn_scd three-prime UTR regions: | 114133 | ( 241.19) | 8.77% | ( 0.51 O/E) |
| Number of sites identified in Pc Pc_PA_qn_scd other exon regions: | 48940 | ( 226.59) | 3.76% | ( 0.48 O/E) |
| Number of sites identified in Pc Pc_PA_qn_scd intron regions: | 3942832 | ( 401.79) | 46.20% | ( 0.99 O/E) |
| Number of sites identified in Pc Pc_PA_qn_scd intergenic regions: | 3289615 | ( 384.95) | 38.55% | ( 0.95 O/E) |
| Number of sites identified in Pc Pc_PA_qn_scd promoter regions: | 121113 | ( 228.41) | 1.42% | ( 0.57 O/E) |
| Number of sites identified in Pc Pc_PA_qn_scd other intergenic regions: | 3168502 | ( 395.31) | 37.13% | ( 0.98 O/E) |

Number and density (per 10kb) of sites in Pc_PA_qn_hsm

| | | | | | |
|---|---|---|---|---|---|
| Number of sites identified in Pc Pc_PA_qn_hsm: | 13840 | ( 0.66) | ( 0.16% of control) | | |
| Number of sites identified in Pc Pc_PA_qn_hsm genic regions: | 9694 | ( 0.77) | ( 0.18% of control) | 70.04% | ( 1.18 O/E) |
| Number of sites identified in Pc Pc_PA_qn_hsm exon regions: | 6575 | ( 2.38) | ( 0.51% of control) | 47.51% | ( 3.63 O/E) |
| Number of sites identified in Pc Pc_PA_qn_hsm CDS regions: | 5994 | ( 3.38) | ( 0.63% of control) | 91.16% | ( 1.42 O/E) |
| Number of sites identified in Pc Pc_PA_qn_hsm five-prime UTR regions: | 292 | ( 0.97) | ( 0.16% of control) | 4.44% | ( 0.41 O/E) |
| Number of sites identified in Pc Pc_PA_qn_hsm three-prime UTR regions: | 145 | ( 0.31) | ( 0.13% of control) | 2.21% | ( 0.13 O/E) |
| Number of sites identified in Pc Pc_PA_qn_hsm other exon regions: | 144 | ( 0.67) | ( 0.29% of control) | 2.19% | ( 0.28 O/E) |
| Number of sites identified in Pc Pc_PA_qn_hsm intron regions: | 3119 | ( 0.32) | ( 0.08% of control) | 22.54% | ( 0.49 O/E) |
| Number of sites identified in Pc Pc_PA_qn_hsm intergenic regions: | 4146 | ( 0.49) | ( 0.13% of control) | 29.96% | ( 0.74 O/E) |
| Number of sites identified in Pc Pc_PA_qn_hsm promoter regions: | 188 | ( 0.35) | ( 0.16% of control) | 1.36% | ( 0.54 O/E) |
| Number of sites identified in Pc Pc_PA_qn_hsm other intergenic regions: | 3958 | ( 0.49) | ( 0.12% of control) | 28.60% | ( 0.75 O/E) |

Directory: MMP File: mmp-Pc_PA_wr.txt

#### Genomic composition in terms of feature regions

| Region | Size | Percent |
|---|---|---|
| Pc genome size: | 211202212 bp | |
| Pc genic region size: | 125747138 bp | ( 59.5%) |
| Pc exon region size: | 27615710 bp | ( 13.1%) |
| Pc CDS region size: | 17712034 bp | ( 64.1%) |
| Pc five-prime UTR region size: | 3011712 bp | ( 10.9%) |
| Pc three-prime UTR region size: | 4732072 bp | ( 17.1%) |
| Pc other exon region size: | 2159892 bp | ( 7.8%) |
| Pc intron region size: | 98131428 bp | ( 46.5%) |
| Pc intergenic region size: | 85455074 bp | ( 40.5%) |
| Pc promoter region size: | 5302351 bp | ( 2.5%) |
| Pc other intergenic region size: | 80152723 bp | ( 38.0%) |

#### Methylation sites in genomic feature regions

Number and density (per 10kb) of sites in Pc_PA_wr_scd

| | | | | |
|---|---|---|---|---|
| Number of sites identified in Pc Pc_PA_wr_scd: | 7865066 | ( 372.40) | | |
| Number of sites identified in Pc Pc_PA_wr_scd genic regions: | 4816298 | ( 383.01) | 61.24% | ( 1.03 O/E) |
| Number of sites identified in Pc Pc_PA_wr_scd exon regions: | 1197709 | ( 433.71) | 15.23% | ( 1.16 O/E) |
| Number of sites identified in Pc Pc_PA_wr_scd CDS regions: | 884230 | ( 499.23) | 73.83% | ( 1.15 O/E) |
| Number of sites identified in Pc Pc_PA_wr_scd five-prime UTR regions: | 168001 | ( 557.83) | 14.03% | ( 1.29 O/E) |
| Number of sites identified in Pc Pc_PA_wr_scd three-prime UTR regions: | 99761 | ( 210.82) | 8.33% | ( 0.49 O/E) |
| Number of sites identified in Pc Pc_PA_wr_scd other exon regions: | 45717 | ( 211.66) | 3.82% | ( 0.49 O/E) |
| Number of sites identified in Pc Pc_PA_wr_scd intron regions: | 3618589 | ( 368.75) | 46.01% | ( 0.99 O/E) |
| Number of sites identified in Pc Pc_PA_wr_scd intergenic regions: | 3048768 | ( 356.77) | 38.76% | ( 0.96 O/E) |
| Number of sites identified in Pc Pc_PA_wr_scd promoter regions: | 109902 | ( 207.27) | 1.40% | ( 0.56 O/E) |
| Number of sites identified in Pc Pc_PA_wr_scd other intergenic regions: | 2938866 | ( 366.66) | 37.37% | ( 0.98 O/E) |

Number and density (per 10kb) of sites in Pc_PA_wr_hsm

| | | | | | |
|---|---|---|---|---|---|
| Number of sites identified in Pc Pc_PA_wr_hsm: | 12036 | ( 0.57) | ( 0.15% of control) | | |
| Number of sites identified in Pc Pc_PA_wr_hsm genic regions: | 8318 | ( 0.66) | ( 0.17% of control) | 69.11% | ( 1.16 O/E) |
| Number of sites identified in Pc Pc_PA_wr_hsm exon regions: | 5555 | ( 2.01) | ( 0.46% of control) | 46.15% | ( 3.53 O/E) |
| Number of sites identified in Pc Pc_PA_wr_hsm CDS regions: | 5006 | ( 2.83) | ( 0.57% of control) | 90.12% | ( 1.41 O/E) |
| Number of sites identified in Pc Pc_PA_wr_hsm five-prime UTR regions: | 286 | ( 0.95) | ( 0.17% of control) | 5.15% | ( 0.47 O/E) |
| Number of sites identified in Pc Pc_PA_wr_hsm three-prime UTR regions: | 149 | ( 0.31) | ( 0.15% of control) | 2.68% | ( 0.16 O/E) |
| Number of sites identified in Pc Pc_PA_wr_hsm other exon regions: | 114 | ( 0.53) | ( 0.25% of control) | 2.05% | ( 0.26 O/E) |
| Number of sites identified in Pc Pc_PA_wr_hsm intron regions: | 2763 | ( 0.28) | ( 0.08% of control) | 22.96% | ( 0.49 O/E) |
| Number of sites identified in Pc Pc_PA_wr_hsm intergenic regions: | 3718 | ( 0.44) | ( 0.12% of control) | 30.89% | ( 0.76 O/E) |
| Number of sites identified in Pc Pc_PA_wr_hsm promoter regions: | 171 | ( 0.32) | ( 0.16% of control) | 1.42% | ( 0.57 O/E) |
| Number of sites identified in Pc Pc_PA_wr_hsm other intergenic regions: | 3547 | ( 0.44) | ( 0.12% of control) | 29.47% | ( 0.78 O/E) |

```
Directory: ACS File: 0READMEacs


ACS - Annotation of conserved methylation sites.

Input: studymk, genome_ann (a list of GRanges objects providing annotated region labels and bounds;
                            output of BWASPR::get_genome_annotation())

Output: file acs-*.txt

Notes: The output is generated by BWASPR::annotate_methylome(), which uses methylKit::unite()
        to determine the sites common to all experiments and applies genomation::annotateWithFeature().
        The acs-*.txt files show the common sites with methylation levels in all samples
        and columns with booleans indicating whether the site fits a specific annotation feature.

       Abbreviations used: pc, protein coding; nc, non-coding; fp, five prime; tp, three prime.
                           UTR, untranslated region; unique, not overlapping with other categories.
```

Directory: ACS File: acs-Pc_PA.txt

| queen | worker | chr | start | end | strand | gene | exon | pcexon | promoter | CDS | fpUTR | tpUTR | fpUTRnotCDS | tpUTRnotCDS | tpUTRunique | ncexon |
|---|---|---|---|---|---|---|---|---|---|---|---|---|---|---|---|---|
| 16.67 | 24.56 | NW_014569547.1 | 32395 | 32395 | + | 0 | 0 | 0 | 0 | 0 | 0 | 0 | 0 | 0 | 0 | 0 |
| 15.07 | 21.43 | NW_014569547.1 | 99867 | 99867 | + | 1 | 0 | 0 | 0 | 0 | 0 | 0 | 0 | 0 | 0 | 0 |
| 44.12 | 41.67 | NW_014569547.1 | 127777 | 127777 | + | 1 | 0 | 0 | 0 | 0 | 0 | 0 | 0 | 0 | 0 | 0 |
| 55.88 | 46.15 | NW_014569547.1 | 127781 | 127781 | + | 1 | 0 | 0 | 0 | 0 | 0 | 0 | 0 | 0 | 0 | 0 |
| 31.11 | 32.43 | NW_014569547.1 | 127799 | 127799 | + | 1 | 0 | 0 | 0 | 0 | 0 | 0 | 0 | 0 | 0 | 0 |
| 12.62 | 16.33 | NW_014569547.1 | 205368 | 205368 | + | 0 | 0 | 0 | 0 | 0 | 0 | 0 | 0 | 0 | 0 | 0 |
| 20 | 25.58 | NW_014569547.1 | 228148 | 228148 | + | 1 | 1 | 1 | 0 | 1 | 0 | 0 | 0 | 0 | 0 | 0 |
| 17.89 | 9.48 | NW_014569547.1 | 247617 | 247617 | + | 1 | 0 | 0 | 0 | 0 | 0 | 0 | 0 | 0 | 0 | 0 |
| 35.82 | 47.76 | NW_014569547.1 | 282304 | 282304 | + | 1 | 0 | 0 | 0 | 0 | 0 | 0 | 0 | 0 | 0 | 0 |
| 62.04 | 73.1 | NW_014569547.1 | 282316 | 282316 | + | 1 | 0 | 0 | 0 | 0 | 0 | 0 | 0 | 0 | 0 | 0 |
| 58.39 | 56.08 | NW_014569547.1 | 282334 | 282334 | + | 1 | 0 | 0 | 0 | 0 | 0 | 0 | 0 | 0 | 0 | 0 |
| 70.07 | 70.83 | NW_014569547.1 | 282340 | 282340 | + | 1 | 0 | 0 | 0 | 0 | 0 | 0 | 0 | 0 | 0 | 0 |
| 65.44 | 75.35 | NW_014569547.1 | 282343 | 282343 | + | 1 | 0 | 0 | 0 | 0 | 0 | 0 | 0 | 0 | 0 | 0 |
| 57.14 | 68.06 | NW_014569547.1 | 282346 | 282346 | + | 1 | 0 | 0 | 0 | 0 | 0 | 0 | 0 | 0 | 0 | 0 |
| 32.58 | 44.44 | NW_014569547.1 | 362588 | 362588 | + | 1 | 0 | 0 | 0 | 0 | 0 | 0 | 0 | 0 | 0 | 0 |
| 27.37 | 37.23 | NW_014569547.1 | 362604 | 362604 | + | 1 | 0 | 0 | 0 | 0 | 0 | 0 | 0 | 0 | 0 | 0 |
| 13.1 | 11.58 | NW_014569547.1 | 460148 | 460148 | + | 0 | 0 | 0 | 0 | 0 | 0 | 0 | 0 | 0 | 0 | 0 |
| 5.31 | 6.58 | NW_014569547.1 | 467359 | 467359 | + | 0 | 0 | 0 | 0 | 0 | 0 | 0 | 0 | 0 | 0 | 0 |
| 42.11 | 32.76 | NW_014569547.1 | 519866 | 519866 | + | 1 | 0 | 0 | 0 | 0 | 0 | 0 | 0 | 0 | 0 | 0 |
| 38.82 | 28.57 | NW_014569547.1 | 519882 | 519882 | + | 1 | 0 | 0 | 0 | 0 | 0 | 0 | 0 | 0 | 0 | 0 |
| 16.5 | 12.5 | NW_014569547.1 | 617105 | 617105 | + | 0 | 0 | 0 | 0 | 0 | 0 | 0 | 0 | 0 | 0 | 0 |
| 30.61 | 23.81 | NW_014569547.1 | 762717 | 762717 | + | 1 | 0 | 0 | 0 | 0 | 0 | 0 | 0 | 0 | 0 | 0 |
| 26 | 20.93 | NW_014569547.1 | 762721 | 762721 | + | 1 | 0 | 0 | 0 | 0 | 0 | 0 | 0 | 0 | 0 | 0 |
| 25 | 28.89 | NW_014569547.1 | 762727 | 762727 | + | 1 | 0 | 0 | 0 | 0 | 0 | 0 | 0 | 0 | 0 | 0 |

...

... only first 25 lines shown ...

```
Directory: RNK File: 0READMErnk


RNK - Genomic feature regions ranked by CpG methylation and hsm statistics.

Input: studymc, studymk, region.gr (GRanges object providing annotated region labels and bounds)

Output: files ranked-genes-*.txt sites-in-genes-*.txt
              ranked-promoters-*.txt sites-in-promoters-*.txt
              plot*.pdf

Notes: The output is generated by BWASPR::rank_rbm().
       The files ranked-genes-*.txt provide ranked lists of genes based on
        overall methylation percentage or the occurrence of CpGhms sites
        within the annotated gene bounds.  The output columns include
          region_ID    (= region/gene name)
          rwidth       (= region/gene width)
          nbrsites     (= number of CpGhsm sites in the region/gene)
          nbrper10kb   (= number of CpGhsm sites in the region/gene normalized to 10kb width)
          pmrpersite   (= average % methylation per CpGhsm site in the region/gene)
          pmrpernucl   (= average % CpGhsm methylation per nucleotide in the region/gene)
          prcntM       (= overall % CpGhsm methylation in the region/gene)

       The tables are sorted by prcntM (*byPrcntM* files) or by nbper10kb
         (*bySiteDensity* files) in descending order.

       The ranked-promoters-*.txt files provide analogous tables for promoter regions.
        If the promoter annotation was derived with BWASP, then these regions are simply
        defined here as 500 nucleotides upstream of the 5'-end of the gene (shorter if the
        scaffold ends before).

       The files sites-*.txt provide details for the CpGhsm sites in the respective
        regions.
       The files plot-*.pdf explore concordance of the two methylation measures.
       The optional parameters minrwidth, maxrwidth, and minnbrsites specify
       the length range and minimal number of methylation sites for regions
       to be included in the plot.
```

Directory: RNK File: ranked-genes-byPrcntM-Pc_PA_queen.txt

| region_ID | seqnames | start | end | width | strand | coverage | numCs | numTs | prcntM |
|---|---|---|---|---|---|---|---|---|---|
| gene-LOC106792910 | NW_014571011.1 | 245 | 1289 | 1045 | + | 394 | 223 | 171 | 56.6 |
| gene-LOC106790095 | NW_014569761.1 | 20476 | 21813 | 1338 | + | 180 | 83 | 97 | 46.11 |
| gene-LOC106792704 | NW_014570116.1 | 10909 | 11269 | 361 | + | 52 | 20 | 32 | 38.46 |
| gene-LOC106792957 | NW_014572754.1 | 173 | 1048 | 876 | - | 445 | 171 | 274 | 38.43 |
| gene-LOC106784739 | NW_014569586.1 | 366853 | 368115 | 1263 | - | 336 | 128 | 208 | 38.1 |
| gene-LOC106793822 | NW_014569566.1 | 834325 | 837505 | 3181 | - | 532 | 193 | 339 | 36.28 |
| gene-LOC106792777 | NW_014569557.1 | 1535000 | 1537224 | 2225 | - | 652 | 234 | 418 | 35.89 |
| gene-LOC106787554 | NW_014569652.1 | 318974 | 320837 | 1864 | - | 160 | 57 | 103 | 35.62 |
| gene-LOC106789903 | NW_014569753.1 | 199603 | 200355 | 753 | - | 377 | 127 | 250 | 33.69 |
| gene-LOC106786039 | NW_014569610.1 | 688308 | 691004 | 2697 | + | 318 | 107 | 211 | 33.65 |
| gene-LOC106789759 | NW_014569553.1 | 1171515 | 1173654 | 2140 | + | 473 | 158 | 315 | 33.4 |
| gene-LOC106788699 | NW_014569697.1 | 365860 | 366871 | 1012 | - | 420 | 139 | 281 | 33.1 |
| gene-LOC106785449 | NW_014569597.1 | 452539 | 456537 | 3999 | + | 1288 | 418 | 870 | 32.45 |
| gene-LOC106784090 | NW_014569577.1 | 921282 | 924136 | 2855 | + | 1098 | 354 | 744 | 32.24 |
| gene-LOC106792139 | NW_014569930.1 | 101128 | 102680 | 1553 | - | 267 | 86 | 181 | 32.21 |
| gene-LOC106788173 | NW_014569675.1 | 321032 | 323374 | 2343 | + | 137 | 44 | 93 | 32.12 |
| gene-LOC106791088 | NW_014569832.1 | 123300 | 124584 | 1285 | - | 349 | 112 | 237 | 32.09 |
| gene-LOC106790592 | NW_014569554.1 | 49877 | 51710 | 1834 | - | 400 | 123 | 277 | 30.75 |
| gene-LOC106792739 | NW_014570148.1 | 17546 | 19959 | 2414 | - | 2349 | 718 | 1631 | 30.57 |
| gene-LOC106792807 | NW_014570273.1 | 5117 | 8568 | 3452 | + | 158 | 48 | 110 | 30.38 |
| gene-LOC106786868 | NW_014569633.1 | 259239 | 260986 | 1748 | - | 560 | 169 | 391 | 30.18 |
| gene-LOC106792948 | NW_014569557.1 | 1393260 | 1395217 | 1958 | + | 750 | 223 | 527 | 29.73 |
| gene-LOC106784622 | NW_014569585.1 | 114990 | 116348 | 1359 | + | 402 | 116 | 286 | 28.86 |
| gene-LOC106790391 | NW_014569781.1 | 227977 | 229306 | 1330 | - | 256 | 73 | 183 | 28.52 |

...
... only first 25 lines shown ...

Directory: RNK File: ranked-genes-byPrcntM-Pc_PA_worker.txt

| region_ID | seqnames | start | end | width | strand | coverage | numCs | numTs | prcntM |
|---|---|---|---|---|---|---|---|---|---|
| gene-LOC106792910 | NW_014571011.1 | 245 | 1289 | 1045 | + | 306 | 169 | 137 | 55.23 |
| gene-LOC106792807 | NW_014570273.1 | 5117 | 8568 | 3452 | + | 174 | 78 | 96 | 44.83 |
| gene-LOC106786039 | NW_014569610.1 | 688308 | 691004 | 2697 | + | 272 | 109 | 163 | 40.07 |
| gene-LOC106791088 | NW_014569832.1 | 123300 | 124584 | 1285 | - | 317 | 121 | 196 | 38.17 |
| gene-LOC106792957 | NW_014572754.1 | 173 | 1048 | 876 | - | 536 | 202 | 334 | 37.69 |
| gene-LOC106785145 | NW_014569592.1 | 307477 | 309238 | 1762 | - | 271 | 101 | 170 | 37.27 |
| gene-LOC106784090 | NW_014569577.1 | 921282 | 924136 | 2855 | + | 937 | 347 | 590 | 37.03 |
| gene-LOC106793822 | NW_014569566.1 | 834325 | 837505 | 3181 | - | 477 | 174 | 303 | 36.48 |
| gene-LOC106784604 | NW_014569584.1 | 645507 | 646846 | 1340 | - | 126 | 43 | 83 | 34.13 |
| gene-LOC106789759 | NW_014569553.1 | 1171515 | 1173654 | 2140 | + | 417 | 141 | 276 | 33.81 |
| gene-LOC106785790 | NW_014569550.1 | 2110 | 3323 | 1214 | + | 36 | 12 | 24 | 33.33 |
| gene-LOC106790592 | NW_014569554.1 | 49877 | 51710 | 1834 | - | 516 | 170 | 346 | 32.95 |
| gene-LOC106790095 | NW_014569761.1 | 20476 | 21813 | 1338 | + | 113 | 36 | 77 | 31.86 |
| gene-LOC106784739 | NW_014569586.1 | 366853 | 368115 | 1263 | - | 289 | 91 | 198 | 31.49 |
| gene-LOC106790223 | NW_014569772.1 | 259797 | 262084 | 2288 | - | 697 | 217 | 480 | 31.13 |
| gene-LOC106792739 | NW_014570148.1 | 17546 | 19959 | 2414 | - | 1979 | 614 | 1365 | 31.03 |
| gene-LOC106793843 | NW_014569566.1 | 820919 | 822584 | 1666 | - | 189 | 58 | 131 | 30.69 |
| gene-LOC106785449 | NW_014569597.1 | 452539 | 456537 | 3999 | + | 1180 | 362 | 818 | 30.68 |
| gene-LOC106792777 | NW_014569557.1 | 1535000 | 1537224 | 2225 | - | 597 | 174 | 423 | 29.15 |
| gene-LOC106793792 | NW_014569566.1 | 274063 | 276096 | 2034 | - | 446 | 129 | 317 | 28.92 |
| gene-LOC106790121 | NW_014569762.1 | 235821 | 237897 | 2077 | - | 607 | 175 | 432 | 28.83 |
| gene-LOC106793570 | NW_014569562.1 | 197070 | 198642 | 1573 | + | 159 | 45 | 114 | 28.3 |
| gene-LOC106791105 | NW_014569832.1 | 155770 | 157298 | 1529 | - | 71 | 20 | 51 | 28.17 |
| gene-LOC106785630 | NW_014569600.1 | 104316 | 106828 | 2513 | - | 253 | 71 | 182 | 28.06 |

...
... only first 25 lines shown ...

Directory: RNK File: ranked-genes-bySiteDensity-Pc_PA_queen.txt

| region_ID | rwidth | nbrsites | nbrper10kb | pmpersite | pmpernucl | pglink | seqnames | start | end | strand | coverage | numCs | numTs | prcntM |
|---|---|---|---|---|---|---|---|---|---|---|---|---|---|---|
| gene-LOC106792344 | 369 | 15 | 406.5 | 1.27 | 0.05 | https://www.ncbi.nlm.nih.gov/gene/?term=LOC106792344 | NW_014569955.1 | 55529 | 55897 | - | 764589 | 6093 | 758496 | 0.8 |
| gene-LOC106786260 | 2638 | 63 | 238.82 | 52 | 1.24 | https://www.ncbi.nlm.nih.gov/gene/?term=LOC106786260 | NW_014569617.1 | 65215 | 67852 | + | 5886 | 1039 | 4847 | 17.65 |
| gene-LOC106792553 | 2482 | 53 | 213.54 | 25.99 | 0.55 | https://www.ncbi.nlm.nih.gov/gene/?term=LOC106792553 | NW_014570016.1 | 51933 | 54414 | - | 56416 | 6703 | 49713 | 11.88 |
| gene-LOC106792910 | 1045 | 16 | 153.11 | 75.03 | 1.15 | https://www.ncbi.nlm.nih.gov/gene/?term=LOC106792910 | NW_014571011.1 | 245 | 1289 | + | 394 | 223 | 171 | 56.6 |
| gene-LOC106786231 | 2170 | 29 | 133.64 | 59.48 | 0.79 | https://www.ncbi.nlm.nih.gov/gene/?term=LOC106786231 | NW_014569616.1 | 377938 | 380107 | + | 1549 | 411 | 1138 | 26.53 |
| gene-LOC106789903 | 753 | 10 | 132.8 | 55.17 | 0.73 | https://www.ncbi.nlm.nih.gov/gene/?term=LOC106789903 | NW_014569753.1 | 199603 | 200355 | - | 377 | 127 | 250 | 33.69 |
| gene-LOC106793481 | 3790 | 47 | 124.01 | 51.13 | 0.63 | https://www.ncbi.nlm.nih.gov/gene/?term=LOC106793481 | NW_014569561.1 | 707023 | 710812 | + | 8533 | 1705 | 6828 | 19.98 |
| gene-LOC106793930 | 2962 | 36 | 121.54 | 57.12 | 0.69 | https://www.ncbi.nlm.nih.gov/gene/?term=LOC106793930 | NW_014569568.1 | 363121 | 366082 | + | 3038 | 681 | 2357 | 22.42 |
| gene-LOC106793786 | 1202 | 14 | 116.47 | 57.66 | 0.67 | https://www.ncbi.nlm.nih.gov/gene/?term=LOC106793786 | NW_014569566.1 | 796582 | 797783 | - | 950 | 195 | 755 | 20.53 |
| gene-LOC106792957 | 876 | 10 | 114.16 | 48.5 | 0.55 | https://www.ncbi.nlm.nih.gov/gene/?term=LOC106792957 | NW_014572754.1 | 173 | 1048 | - | 445 | 171 | 274 | 38.43 |
| gene-LOC106786090 | 2903 | 31 | 106.79 | 63.91 | 0.68 | https://www.ncbi.nlm.nih.gov/gene/?term=LOC106786090 | NW_014569612.1 | 510700 | 513602 | - | 11993 | 759 | 11234 | 6.33 |
| gene-LOC106784969 | 2112 | 22 | 104.17 | 55.51 | 0.58 | https://www.ncbi.nlm.nih.gov/gene/?term=LOC106784969 | NW_014569589.1 | 334267 | 336378 | - | 1642 | 460 | 1182 | 28.01 |
| gene-LOC106787127 | 1220 | 12 | 98.36 | 57.07 | 0.56 | https://www.ncbi.nlm.nih.gov/gene/?term=LOC106787127 | NW_014569639.1 | 57160 | 58379 | - | 1356 | 213 | 1143 | 15.71 |
| gene-LOC106788343 | 1965 | 19 | 96.69 | 55.44 | 0.54 | https://www.ncbi.nlm.nih.gov/gene/?term=LOC106788343 | NW_014569682.1 | 57478 | 59442 | - | 927 | 235 | 692 | 25.35 |
| gene-LOC106789767 | 938 | 9 | 95.95 | 68.39 | 0.66 | https://www.ncbi.nlm.nih.gov/gene/?term=LOC106789767 | NW_014569553.1 | 1173704 | 1174641 | - | 646 | 151 | 495 | 23.37 |
| gene-LOC106789851 | 1327 | 12 | 90.43 | 61.33 | 0.55 | https://www.ncbi.nlm.nih.gov/gene/?term=LOC106789851 | NW_014569752.1 | 20642 | 21968 | + | 869 | 239 | 630 | 27.5 |
| gene-LOC106790223 | 2288 | 20 | 87.41 | 64.72 | 0.57 | https://www.ncbi.nlm.nih.gov/gene/?term=LOC106790223 | NW_014569772.1 | 259797 | 262084 | - | 1077 | 225 | 852 | 20.89 |
| gene-LOC106792739 | 2414 | 21 | 86.99 | 51.2 | 0.45 | https://www.ncbi.nlm.nih.gov/gene/?term=LOC106792739 | NW_014570148.1 | 17546 | 19959 | - | 2349 | 718 | 1631 | 30.57 |
| gene-LOC106793024 | 4066 | 35 | 86.08 | 62.34 | 0.54 | https://www.ncbi.nlm.nih.gov/gene/?term=LOC106793024 | NW_014569557.1 | 100847 | 104912 | - | 7489 | 599 | 6890 | 8 |
| gene-LOC106792948 | 1958 | 16 | 81.72 | 52.34 | 0.43 | https://www.ncbi.nlm.nih.gov/gene/?term=LOC106792948 | NW_014569557.1 | 1393260 | 1395217 | + | 750 | 223 | 527 | 29.73 |
| gene-LOC106791200 | 1235 | 10 | 80.97 | 55.61 | 0.45 | https://www.ncbi.nlm.nih.gov/gene/?term=LOC106791200 | NW_014569838.1 | 189455 | 190689 | - | 1479 | 210 | 1269 | 14.2 |
| gene-LOC106788699 | 1012 | 8 | 79.05 | 61.48 | 0.49 | https://www.ncbi.nlm.nih.gov/gene/?term=LOC106788699 | NW_014569697.1 | 365860 | 366871 | - | 420 | 139 | 281 | 33.1 |
| gene-LOC106787164 | 1270 | 10 | 78.74 | 37.55 | 0.3 | https://www.ncbi.nlm.nih.gov/gene/?term=LOC106787164 | NW_014569641.1 | 133532 | 134801 | - | 1316 | 212 | 1104 | 16.11 |
| gene-LOC106792397 | 1701 | 13 | 76.43 | 2.4 | 0.02 | https://www.ncbi.nlm.nih.gov/gene/?term=LOC106792397 | NW_014569967.1 | 73881 | 75581 | + | 57256 | 1001 | 56255 | 1.75 |

...
... only first 25 lines shown ...

Directory: RNK File: ranked-genes-bySiteDensity-Pc_PA_worker.txt

| region_ID | rwidth | nbrsites | nbrper10kb | pmpersite | pmpernucl | pglink | seqnames | start | end | strand | coverage | numCs | numTs | prcntM |
|---|---|---|---|---|---|---|---|---|---|---|---|---|---|---|
| gene-LOC106792344 | 369 | 15 | 406.5 | 1.27 | 0.05 | https://www.ncbi.nlm.nih.gov/gene/?term=LOC106792344 | NW_014569955.1 | 55529 | 55897 | - | 679823 | 5242 | 674581 | 0.77 |
| gene-LOC106786260 | 2638 | 59 | 223.65 | 59.69 | 1.34 | https://www.ncbi.nlm.nih.gov/gene/?term=LOC106786260 | NW_014569617.1 | 65215 | 67852 | + | 4754 | 977 | 3777 | 20.55 |
| gene-LOC106792553 | 2482 | 48 | 193.39 | 26.16 | 0.51 | https://www.ncbi.nlm.nih.gov/gene/?term=LOC106792553 | NW_014570016.1 | 51933 | 54414 | - | 43652 | 4751 | 38901 | 10.88 |
| gene-LOC106792910 | 1045 | 15 | 143.54 | 74.99 | 1.08 | https://www.ncbi.nlm.nih.gov/gene/?term=LOC106792910 | NW_014571011.1 | 245 | 1289 | + | 306 | 169 | 137 | 55.23 |
| gene-LOC106793481 | 3790 | 49 | 129.29 | 54.37 | 0.7 | https://www.ncbi.nlm.nih.gov/gene/?term=LOC106793481 | NW_014569561.1 | 707023 | 710812 | + | 7216 | 1618 | 5598 | 22.42 |
| gene-LOC106784969 | 2112 | 21 | 99.43 | 56.95 | 0.57 | https://www.ncbi.nlm.nih.gov/gene/?term=LOC106784969 | NW_014569589.1 | 334267 | 336378 | - | 1347 | 377 | 970 | 27.99 |
| gene-LOC106793930 | 2962 | 29 | 97.91 | 58.58 | 0.57 | https://www.ncbi.nlm.nih.gov/gene/?term=LOC106793930 | NW_014569568.1 | 363121 | 366082 | + | 2694 | 619 | 2075 | 22.98 |
| gene-LOC106786090 | 2903 | 28 | 96.45 | 70.61 | 0.68 | https://www.ncbi.nlm.nih.gov/gene/?term=LOC106786090 | NW_014569612.1 | 510700 | 513602 | - | 10118 | 661 | 9457 | 6.53 |
| gene-LOC106786231 | 2170 | 20 | 92.17 | 59.28 | 0.55 | https://www.ncbi.nlm.nih.gov/gene/?term=LOC106786231 | NW_014569616.1 | 377938 | 380107 | + | 1357 | 323 | 1034 | 23.8 |
| gene-LOC106792957 | 876 | 8 | 91.32 | 53.18 | 0.49 | https://www.ncbi.nlm.nih.gov/gene/?term=LOC106792957 | NW_014572754.1 | 173 | 1048 | - | 536 | 202 | 334 | 37.69 |
| gene-LOC106787127 | 1220 | 11 | 90.16 | 63.91 | 0.58 | https://www.ncbi.nlm.nih.gov/gene/?term=LOC106787127 | NW_014569639.1 | 57160 | 58379 | - | 1140 | 150 | 990 | 13.16 |
| gene-LOC106790223 | 2288 | 20 | 87.41 | 73.72 | 0.64 | https://www.ncbi.nlm.nih.gov/gene/?term=LOC106790223 | NW_014569772.1 | 259797 | 262084 | - | 697 | 217 | 480 | 31.13 |
| gene-LOC106792739 | 2414 | 21 | 86.99 | 47.09 | 0.41 | https://www.ncbi.nlm.nih.gov/gene/?term=LOC106792739 | NW_014570148.1 | 17546 | 19959 | - | 1979 | 614 | 1365 | 31.03 |
| gene-LOC106789767 | 938 | 8 | 85.29 | 64.74 | 0.55 | https://www.ncbi.nlm.nih.gov/gene/?term=LOC106789767 | NW_014569553.1 | 1173704 | 1174641 | - | 382 | 77 | 305 | 20.16 |
| gene-LOC106791087 | 2016 | 17 | 84.33 | 61.58 | 0.52 | https://www.ncbi.nlm.nih.gov/gene/?term=LOC106791087 | NW_014569832.1 | 124408 | 126423 | - | 1445 | 293 | 1152 | 20.28 |
| gene-LOC106788200 | 2717 | 22 | 80.97 | 52.42 | 0.42 | https://www.ncbi.nlm.nih.gov/gene/?term=LOC106788200 | NW_014569676.1 | 375648 | 378364 | + | 2098 | 348 | 1750 | 16.59 |
| gene-LOC106791200 | 1235 | 10 | 80.97 | 60.15 | 0.49 | https://www.ncbi.nlm.nih.gov/gene/?term=LOC106791200 | NW_014569838.1 | 189455 | 190689 | - | 1304 | 202 | 1102 | 15.49 |
| gene-LOC106787164 | 1270 | 9 | 70.87 | 40.38 | 0.29 | https://www.ncbi.nlm.nih.gov/gene/?term=LOC106787164 | NW_014569641.1 | 133532 | 134801 | - | 995 | 181 | 814 | 18.19 |
| gene-LOC106793792 | 2034 | 14 | 68.83 | 61.05 | 0.42 | https://www.ncbi.nlm.nih.gov/gene/?term=LOC106793792 | NW_014569566.1 | 274063 | 276096 | - | 446 | 129 | 317 | 28.92 |
| gene-LOC106790592 | 1834 | 12 | 65.43 | 70.89 | 0.46 | https://www.ncbi.nlm.nih.gov/gene/?term=LOC106790592 | NW_014569554.1 | 49877 | 51710 | - | 516 | 170 | 346 | 32.95 |
| gene-LOC106792397 | 1701 | 11 | 64.67 | 2.39 | 0.02 | https://www.ncbi.nlm.nih.gov/gene/?term=LOC106792397 | NW_014569967.1 | 73881 | 75581 | + | 54746 | 886 | 53860 | 1.62 |
| gene-LOC106787992 | 3099 | 20 | 64.54 | 48.33 | 0.31 | https://www.ncbi.nlm.nih.gov/gene/?term=LOC106787992 | NW_014569668.1 | 209651 | 212749 | + | 1694 | 293 | 1401 | 17.3 |
| gene-LOC106790088 | 3286 | 21 | 63.91 | 53.45 | 0.34 | https://www.ncbi.nlm.nih.gov/gene/?term=LOC106790088 | NW_014569554.1 | 103426 | 106711 | - | 2447 | 547 | 1900 | 22.35 |
| gene-LOC106785145 | 1762 | 11 | 62.43 | 55.66 | 0.35 | https://www.ncbi.nlm.nih.gov/gene/?term=LOC106785145 | NW_014569592.1 | 307477 | 309238 | - | 271 | 101 | 170 | 37.27 |

...
... only first 25 lines shown ...

Directory: RNK File: ranked-promoters-byPrcntM-Pc_PA_queen.txt

| region_ID | seqnames | start | end | width | strand | coverage | numCs | numTs | prcntM |
|---|---|---|---|---|---|---|---|---|---|
| gene-LOC106785085 | NW_014569549.1 | 2123847 | 2124346 | 500 | + | 85 | 56 | 29 | 65.88 |
| gene-LOC106787597 | NW_014569654.1 | 22088 | 22587 | 500 | - | 4 | 2 | 2 | 50 |
| gene-LOC106788339 | NW_014569682.1 | 411042 | 411541 | 500 | - | 44 | 22 | 22 | 50 |
| gene-LOC106786867 | NW_014569633.1 | 260124 | 260623 | 500 | + | 246 | 122 | 124 | 49.59 |
| gene-LOC106788639 | NW_014569552.1 | 2074962 | 2075461 | 500 | + | 25 | 12 | 13 | 48 |
| gene-LOC106793571 | NW_014569562.1 | 198130 | 198629 | 500 | + | 94 | 38 | 56 | 40.43 |
| gene-LOC106790059 | NW_014569758.1 | 4912 | 5411 | 500 | - | 11 | 4 | 7 | 36.36 |
| gene-LOC106789852 | NW_014569752.1 | 21487 | 21986 | 500 | + | 20 | 7 | 13 | 35 |
| gene-LOC106792704 | NW_014570116.1 | 10409 | 10908 | 500 | + | 158 | 47 | 111 | 29.75 |
| gene-LOC106792709 | NW_014570116.1 | 11498 | 11997 | 500 | - | 99 | 28 | 71 | 28.28 |
| gene-LOC106793851 | NW_014569566.1 | 796259 | 796758 | 500 | - | 65 | 17 | 48 | 26.15 |
| gene-LOC106787052 | NW_014569550.1 | 1013576 | 1014075 | 500 | - | 4 | 1 | 3 | 25 |
| gene-LOC106784836 | NW_014569588.1 | 182287 | 182786 | 500 | - | 4 | 1 | 3 | 25 |
| gene-LOC106784992 | NW_014569590.1 | 421864 | 422363 | 500 | + | 4 | 1 | 3 | 25 |
| gene-LOC106785498 | NW_014569598.1 | 779078 | 779577 | 500 | + | 4 | 1 | 3 | 25 |
| gene-LOC106786027 | NW_014569610.1 | 662740 | 663239 | 500 | - | 4 | 1 | 3 | 25 |
| gene-LOC106788259 | NW_014569677.1 | 15719 | 16218 | 500 | - | 12 | 3 | 9 | 25 |
| gene-LOC106789361 | NW_014569728.1 | 101038 | 101537 | 500 | - | 4 | 1 | 3 | 25 |
| gene-LOC106790357 | NW_014569779.1 | 232218 | 232717 | 500 | + | 4 | 1 | 3 | 25 |
| gene-LOC106790862 | NW_014569813.1 | 125485 | 125984 | 500 | + | 8 | 2 | 6 | 25 |
| gene-LOC106788490 | NW_014569687.1 | 411960 | 412459 | 500 | - | 1299 | 323 | 976 | 24.87 |
| gene-LOC106785314 | NW_014569595.1 | 551303 | 551802 | 500 | + | 585 | 135 | 450 | 23.08 |
| gene-LOC106793309 | NW_014569560.1 | 636478 | 636977 | 500 | - | 131 | 27 | 104 | 20.61 |
| gene-LOC106785137 | NW_014569592.1 | 434079 | 434578 | 500 | - | 5 | 1 | 4 | 20 |

...
... only first 25 lines shown ...

Directory: RNK File: ranked-promoters-byPrcntM-Pc_PA_worker.txt

| region_ID | seqnames | start | end | width | strand | coverage | numCs | numTs | prcntM |
|---|---|---|---|---|---|---|---|---|---|
| gene-LOC106785085 | NW_014569549.1 | 2123847 | 2124346 | 500 | + | 63 | 37 | 26 | 58.73 |
| gene-LOC106788339 | NW_014569682.1 | 411042 | 411541 | 500 | - | 56 | 27 | 29 | 48.21 |
| gene-LOC106789852 | NW_014569752.1 | 21487 | 21986 | 500 | + | 18 | 7 | 11 | 38.89 |
| gene-LOC106793571 | NW_014569562.1 | 198130 | 198629 | 500 | + | 96 | 37 | 59 | 38.54 |
| gene-LOC106786867 | NW_014569633.1 | 260124 | 260623 | 500 | + | 215 | 72 | 143 | 33.49 |
| gene-LOC106792262 | NW_014569945.1 | 91969 | 92468 | 500 | - | 61 | 20 | 41 | 32.79 |
| gene-LOC106788490 | NW_014569687.1 | 411960 | 412459 | 500 | - | 1368 | 379 | 989 | 27.7 |
| gene-LOC106790208 | NW_014569769.1 | 184168 | 184667 | 500 | - | 34 | 9 | 25 | 26.47 |
| gene-LOC106785426 | NW_014569597.1 | 241882 | 242381 | 500 | - | 20 | 5 | 15 | 25 |
| gene-LOC106785746 | NW_014569602.1 | 829015 | 829514 | 500 | + | 4 | 1 | 3 | 25 |
| gene-LOC106786368 | NW_014569619.1 | 411768 | 412267 | 500 | + | 4 | 1 | 3 | 25 |
| gene-LOC106787231 | NW_014569642.1 | 413647 | 414146 | 500 | - | 8 | 2 | 6 | 25 |
| gene-LOC106790518 | NW_014569790.1 | 6512 | 7011 | 500 | + | 4 | 1 | 3 | 25 |
| gene-LOC106790628 | NW_014569798.1 | 25003 | 25502 | 500 | - | 4 | 1 | 3 | 25 |
| gene-LOC106791192 | NW_014569838.1 | 102271 | 102770 | 500 | + | 4 | 1 | 3 | 25 |
| gene-LOC106792549 | NW_014570014.1 | 60680 | 61179 | 500 | - | 4 | 1 | 3 | 25 |
| gene-LOC106792952 | NW_014572278.1 | 1 | 96 | 96 | + | 4 | 1 | 3 | 25 |
| gene-LOC106793851 | NW_014569566.1 | 796259 | 796758 | 500 | - | 41 | 10 | 31 | 24.39 |
| gene-LOC106792704 | NW_014570116.1 | 10409 | 10908 | 500 | + | 128 | 27 | 101 | 21.09 |
| gene-LOC106788779 | NW_014569700.1 | 219692 | 220191 | 500 | + | 115 | 24 | 91 | 20.87 |
| gene-LOC106786486 | NW_014569622.1 | 432832 | 433331 | 500 | - | 5 | 1 | 4 | 20 |
| gene-LOC106791040 | NW_014569827.1 | 43361 | 43860 | 500 | - | 98 | 19 | 79 | 19.39 |
| gene-LOC106791724 | NW_014569883.1 | 90156 | 90655 | 500 | - | 220 | 41 | 179 | 18.64 |
| gene-LOC106793700 | NW_014569564.1 | 879896 | 880395 | 500 | - | 243 | 43 | 200 | 17.7 |

...

... only first 25 lines shown ...

Directory: RNK File: ranked-promoters-bySiteDensity-Pc_PA_queen.txt

| region_ID | rwidth | nbrsites | nbrper10kb | pmpersite | pmpernucl | pglink | seqnames | start | end | strand | coverage | numCs | numTs | prcntM |
|---|---|---|---|---|---|---|---|---|---|---|---|---|---|---|
| gene-LOC106792344 | 500 | 23 | 460 | 1.14 | 0.05 | https://www.ncbi.nlm.nih.gov/gene/?term=LOC106792344 | NW_014569955.1 | 55898 | 56397 | - | 1215478 | 9780 | 1205698 | 0.8 |
| gene-LOC106785314 | 500 | 11 | 220 | 74.38 | 1.64 | https://www.ncbi.nlm.nih.gov/gene/?term=LOC106785314 | NW_014569595.1 | 551303 | 551802 | + | 585 | 135 | 450 | 23.08 |
| gene-LOC106787867 | 500 | 11 | 220 | 19.49 | 0.43 | https://www.ncbi.nlm.nih.gov/gene/?term=LOC106787867 | NW_014569664.1 | 337446 | 337945 | + | 3721 | 171 | 3550 | 4.6 |
| gene-LOC106791550 | 500 | 8 | 160 | 25.52 | 0.41 | https://www.ncbi.nlm.nih.gov/gene/?term=LOC106791550 | NW_014569865.1 | 85298 | 85797 | - | 6530 | 319 | 6211 | 4.89 |
| gene-LOC106790993 | 500 | 7 | 140 | 65.49 | 0.92 | https://www.ncbi.nlm.nih.gov/gene/?term=LOC106790993 | NW_014569822.1 | 88866 | 89365 | + | 2009 | 361 | 1648 | 17.97 |
| gene-LOC106786867 | 500 | 6 | 120 | 75.82 | 0.91 | https://www.ncbi.nlm.nih.gov/gene/?term=LOC106786867 | NW_014569633.1 | 260124 | 260623 | + | 246 | 122 | 124 | 49.59 |
| gene-LOC106787307 | 500 | 6 | 120 | 45.62 | 0.55 | https://www.ncbi.nlm.nih.gov/gene/?term=LOC106787307 | NW_014569645.1 | 320505 | 321004 | + | 3064 | 321 | 2743 | 10.48 |
| gene-LOC106793172 | 500 | 6 | 120 | 9.12 | 0.11 | https://www.ncbi.nlm.nih.gov/gene/?term=LOC106793172 | NW_014569559.1 | 1151932 | 1152431 | + | 5149 | 155 | 4994 | 3.01 |
| gene-LOC106786771 | 500 | 5 | 100 | 16.46 | 0.16 | https://www.ncbi.nlm.nih.gov/gene/?term=LOC106786771 | NW_014569628.1 | 539233 | 539732 | + | 2117 | 115 | 2002 | 5.43 |
| gene-LOC106787745 | 500 | 5 | 100 | 43.75 | 0.44 | https://www.ncbi.nlm.nih.gov/gene/?term=LOC106787745 | NW_014569551.1 | 2222289 | 2222788 | + | 881 | 92 | 789 | 10.44 |
| gene-LOC106788490 | 500 | 5 | 100 | 48.01 | 0.48 | https://www.ncbi.nlm.nih.gov/gene/?term=LOC106788490 | NW_014569687.1 | 411960 | 412459 | - | 1299 | 323 | 976 | 24.87 |
| gene-LOC106793074 | 500 | 5 | 100 | 20.83 | 0.21 | https://www.ncbi.nlm.nih.gov/gene/?term=LOC106793074 | NW_014569558.1 | 934812 | 935311 | - | 1706 | 124 | 1582 | 7.27 |
| gene-LOC106785085 | 500 | 4 | 80 | 81.7 | 0.65 | https://www.ncbi.nlm.nih.gov/gene/?term=LOC106785085 | NW_014569549.1 | 2123847 | 2124346 | + | 85 | 56 | 29 | 65.88 |
| gene-LOC106785672 | 500 | 4 | 80 | 14.63 | 0.12 | https://www.ncbi.nlm.nih.gov/gene/?term=LOC106785672 | NW_014569602.1 | 734501 | 735000 | + | 3574 | 106 | 3468 | 2.97 |
| gene-LOC106788202 | 500 | 4 | 80 | 82.12 | 0.66 | https://www.ncbi.nlm.nih.gov/gene/?term=LOC106788202 | NW_014569676.1 | 375634 | 376133 | - | 459 | 66 | 393 | 14.38 |
| gene-LOC106791306 | 500 | 4 | 80 | 26.55 | 0.21 | https://www.ncbi.nlm.nih.gov/gene/?term=LOC106791306 | NW_014569842.1 | 53546 | 54045 | - | 2057 | 198 | 1859 | 9.63 |
| gene-LOC106791724 | 500 | 4 | 80 | 54.46 | 0.44 | https://www.ncbi.nlm.nih.gov/gene/?term=LOC106791724 | NW_014569883.1 | 90156 | 90655 | - | 231 | 37 | 194 | 16.02 |
| gene-LOC106791733 | 500 | 4 | 80 | 54.46 | 0.44 | https://www.ncbi.nlm.nih.gov/gene/?term=LOC106791733 | NW_014569883.1 | 90350 | 90849 | + | 242 | 37 | 205 | 15.29 |
| gene-LOC106791842 | 500 | 4 | 80 | 26.02 | 0.21 | https://www.ncbi.nlm.nih.gov/gene/?term=LOC106791842 | NW_014569894.1 | 75828 | 76327 | + | 6190 | 182 | 6008 | 2.94 |
| gene-LOC106792704 | 500 | 4 | 80 | 73.94 | 0.59 | https://www.ncbi.nlm.nih.gov/gene/?term=LOC106792704 | NW_014570116.1 | 10409 | 10908 | + | 158 | 47 | 111 | 29.75 |
| gene-LOC106784134 | 500 | 3 | 60 | 15.57 | 0.09 | https://www.ncbi.nlm.nih.gov/gene/?term=LOC106784134 | NW_014569578.1 | 222786 | 223285 | + | 3459 | 78 | 3381 | 2.25 |
| gene-LOC106786084 | 500 | 3 | 60 | 55.77 | 0.33 | https://www.ncbi.nlm.nih.gov/gene/?term=LOC106786084 | NW_014569550.1 | 1117335 | 1117834 | - | 642 | 94 | 548 | 14.64 |
| gene-LOC106788339 | 500 | 3 | 60 | 63.61 | 0.38 | https://www.ncbi.nlm.nih.gov/gene/?term=LOC106788339 | NW_014569682.1 | 411042 | 411541 | - | 44 | 22 | 22 | 50 |
| gene-LOC106790057 | 500 | 3 | 60 | 26.83 | 0.16 | https://www.ncbi.nlm.nih.gov/gene/?term=LOC106790057 | NW_014569758.1 | 16736 | 17235 | - | 235 | 35 | 200 | 14.89 |
| gene-LOC106791088 | 500 | 3 | 60 | 70.38 | 0.42 | https://www.ncbi.nlm.nih.gov/gene/?term=LOC106791088 | NW_014569832.1 | 124585 | 125084 | - | 572 | 73 | 499 | 12.76 |
| gene-LOC106793571 | 500 | 3 | 60 | 86.31 | 0.52 | https://www.ncbi.nlm.nih.gov/gene/?term=LOC106793571 | NW_014569562.1 | 198130 | 198629 | + | 94 | 38 | 56 | 40.43 |
| gene-Trnai-aau-2 | 500 | 3 | 60 | 68.91 | 0.41 | https://www.ncbi.nlm.nih.gov/gene/?term=Trnai-aau | NW_014569564.1 | 1136734 | 1137233 | + | 546 | 52 | 494 | 9.52 |
| gene-LOC106784544 | 500 | 2 | 40 | 29.28 | 0.12 | https://www.ncbi.nlm.nih.gov/gene/?term=LOC106784544 | NW_014569584.1 | 993361 | 993860 | - | 1276 | 34 | 1242 | 2.66 |
| gene-LOC106784545 | 500 | 2 | 40 | 29.28 | 0.12 | https://www.ncbi.nlm.nih.gov/gene/?term=LOC106784545 | NW_014569584.1 | 992939 | 993438 | + | 231 | 26 | 205 | 11.26 |
| gene-LOC106784958 | 500 | 2 | 40 | 5.83 | 0.02 | https://www.ncbi.nlm.nih.gov/gene/?term=LOC106784958 | NW_014569589.1 | 409835 | 410334 | - | 4255 | 74 | 4181 | 1.74 |
| gene-LOC106786691 | 500 | 2 | 40 | 62.5 | 0.25 | https://www.ncbi.nlm.nih.gov/gene/?term=LOC106786691 | NW_014569627.1 | 208195 | 208694 | - | 527 | 17 | 510 | 3.23 |
| gene-LOC106786692 | 500 | 2 | 40 | 38.75 | 0.16 | https://www.ncbi.nlm.nih.gov/gene/?term=LOC106786692 | NW_014569627.1 | 207776 | 208275 | + | 176 | 18 | 158 | 10.23 |
| gene-LOC106786756 | 500 | 2 | 40 | 51.98 | 0.21 | https://www.ncbi.nlm.nih.gov/gene/?term=LOC106786756 | NW_014569628.1 | 69713 | 70212 | - | 391 | 17 | 374 | 4.35 |
| gene-LOC106787461 | 500 | 2 | 40 | 19.12 | 0.08 | https://www.ncbi.nlm.nih.gov/gene/?term=LOC106787461 | NW_014569648.1 | 505358 | 505857 | + | 1548 | 53 | 1495 | 3.42 |
| gene-LOC106789356 | 500 | 2 | 40 | 16.04 | 0.06 | https://www.ncbi.nlm.nih.gov/gene/?term=LOC106789356 | NW_014569727.1 | 280749 | 281248 | - | 1101 | 48 | 1053 | 4.36 |
| gene-LOC106789732 | 500 | 2 | 40 | 41.56 | 0.17 | https://www.ncbi.nlm.nih.gov/gene/?term=LOC106789732 | NW_014569746.1 | 142611 | 143110 | + | 187 | 32 | 155 | 17.11 |
| gene-LOC106790092 | 500 | 2 | 40 | 20.41 | 0.08 | https://www.ncbi.nlm.nih.gov/gene/?term=LOC106790092 | NW_014569761.1 | 88919 | 89418 | - | 4383 | 60 | 4323 | 1.37 |
| gene-LOC106791040 | 500 | 2 | 40 | 33.03 | 0.13 | https://www.ncbi.nlm.nih.gov/gene/?term=LOC106791040 | NW_014569827.1 | 43361 | 43860 | - | 133 | 24 | 109 | 18.05 |
| gene-LOC106791092 | 500 | 2 | 40 | 50.35 | 0.2 | https://www.ncbi.nlm.nih.gov/gene/?term=LOC106791092 | NW_014569832.1 | 60504 | 61003 | + | 842 | 33 | 809 | 3.92 |
| gene-LOC106791482 | 500 | 2 | 40 | 8.97 | 0.04 | https://www.ncbi.nlm.nih.gov/gene/?term=LOC106791482 | NW_014569859.1 | 88265 | 88764 | + | 2119 | 50 | 2069 | 2.36 |
| gene-LOC106792709 | 500 | 2 | 40 | 67.64 | 0.27 | https://www.ncbi.nlm.nih.gov/gene/?term=LOC106792709 | NW_014570116.1 | 11498 | 11997 | - | 99 | 28 | 71 | 28.28 |
| gene-LOC106793309 | 500 | 2 | 40 | 48.08 | 0.19 | https://www.ncbi.nlm.nih.gov/gene/?term=LOC106793309 | NW_014569560.1 | 636478 | 636977 | - | 131 | 27 | 104 | 20.61 |
| gene-LOC106793700 | 500 | 2 | 40 | 20.25 | 0.08 | https://www.ncbi.nlm.nih.gov/gene/?term=LOC106793700 | NW_014569564.1 | 879896 | 880395 | - | 286 | 32 | 254 | 11.19 |
| gene-LOC106783580 | 500 | 1 | 20 | 17.02 | 0.03 | https://www.ncbi.nlm.nih.gov/gene/?term=LOC106783580 | NW_014569570.1 | 1050869 | 1051368 | - | 1095 | 23 | 1072 | 2.1 |
| gene-LOC106783733 | 500 | 1 | 20 | 13.64 | 0.03 | https://www.ncbi.nlm.nih.gov/gene/?term=LOC106783733 | NW_014569572.1 | 822511 | 823010 | - | 1910 | 32 | 1878 | 1.68 |
| gene-LOC106783887 | 500 | 1 | 20 | 39.42 | 0.08 | https://www.ncbi.nlm.nih.gov/gene/?term=LOC106783887 | NW_014569574.1 | 274326 | 274825 | + | 2951 | 69 | 2882 | 2.34 |
| gene-LOC106783888 | 500 | 1 | 20 | 39.42 | 0.08 | https://www.ncbi.nlm.nih.gov/gene/?term=LOC106783888 | NW_014569574.1 | 274388 | 274887 | - | 3252 | 73 | 3179 | 2.24 |
| gene-LOC106784279 | 500 | 1 | 20 | 26.92 | 0.05 | https://www.ncbi.nlm.nih.gov/gene/?term=LOC106784279 | NW_014569580.1 | 841658 | 842157 | + | 632 | 11 | 621 | 1.74 |
| gene-LOC106784302 | 500 | 1 | 20 | 11.39 | 0.02 | https://www.ncbi.nlm.nih.gov/gene/?term=LOC106784302 | NW_014569580.1 | 520579 | 521078 | + | 869 | 19 | 850 | 2.19 |
| gene-LOC106784378 | 500 | 1 | 20 | 28.21 | 0.06 | https://www.ncbi.nlm.nih.gov/gene/?term=LOC106784378 | NW_014569581.1 | 763831 | 764330 | + | 2221 | 38 | 2183 | 1.71 |
| gene-LOC106784677 | 500 | 1 | 20 | 35.29 | 0.07 | https://www.ncbi.nlm.nih.gov/gene/?term=LOC106784677 | NW_014569585.1 | 716168 | 716667 | + | 276 | 19 | 257 | 6.88 |
| gene-LOC106784703 | 500 | 1 | 20 | 10.71 | 0.02 | https://www.ncbi.nlm.nih.gov/gene/?term=LOC106784703 | NW_014569586.1 | 371307 | 371806 | + | 2272 | 31 | 2241 | 1.36 |
| gene-LOC106784967 | 500 | 1 | 20 | 15 | 0.03 | https://www.ncbi.nlm.nih.gov/gene/?term=LOC106784967 | NW_014569589.1 | 371818 | 372317 | + | 1066 | 27 | 1039 | 2.53 |
| gene-LOC106785018 | 500 | 1 | 20 | 8.93 | 0.02 | https://www.ncbi.nlm.nih.gov/gene/?term=LOC106785018 | NW_014569590.1 | 120087 | 120586 | + | 1281 | 24 | 1257 | 1.87 |
| gene-LOC106785021 | 500 | 1 | 20 | 16.22 | 0.03 | https://www.ncbi.nlm.nih.gov/gene/?term=LOC106785021 | NW_014569549.1 | 603372 | 603871 | + | 916 | 28 | 888 | 3.06 |
| gene-LOC106785315 | 500 | 1 | 20 | 8.94 | 0.02 | https://www.ncbi.nlm.nih.gov/gene/?term=LOC106785315 | NW_014569549.1 | 370564 | 371063 | + | 2004 | 31 | 1973 | 1.55 |
| gene-LOC106785612 | 500 | 1 | 20 | 10.23 | 0.02 | https://www.ncbi.nlm.nih.gov/gene/?term=LOC106785612 | NW_014569600.1 | 732075 | 732574 | + | 1140 | 25 | 1115 | 2.19 |
| gene-LOC106785729 | 500 | 1 | 20 | 75 | 0.15 | https://www.ncbi.nlm.nih.gov/gene/?term=LOC106785729 | NW_014569602.1 | 287206 | 287705 | - | 422 | 11 | 411 | 2.61 |
| gene-LOC106785832 | 500 | 1 | 20 | 28.57 | 0.06 | https://www.ncbi.nlm.nih.gov/gene/?term=LOC106785832 | NW_014569604.1 | 212685 | 213184 | - | 259 | 21 | 238 | 8.11 |
| gene-LOC106786025 | 500 | 1 | 20 | 11.11 | 0.02 | https://www.ncbi.nlm.nih.gov/gene/?term=LOC106786025 | NW_014569610.1 | 561518 | 562017 | - | 4795 | 92 | 4703 | 1.92 |

| | | | | | | | | | | | | | | |
|---|---|---|---|---|---|---|---|---|---|---|---|---|---|---|
| gene-LOC106786031 | 500 | 1 | 20 | 10.34 | 0.02 | https://www.ncbi.nlm.nih.gov/gene/?term=LOC106786031 | NW_014569610.1 | 691791 | 692290 | + | 759 | 10 | 749 | 1.32 |
| gene-LOC106786164 | 500 | 1 | 20 | 9.82 | 0.02 | https://www.ncbi.nlm.nih.gov/gene/?term=LOC106786164 | NW_014569613.1 | 416879 | 417378 | + | 2751 | 68 | 2683 | 2.47 |
| gene-LOC106786213 | 500 | 1 | 20 | 9.74 | 0.02 | https://www.ncbi.nlm.nih.gov/gene/?term=LOC106786213 | NW_014569615.1 | 500122 | 500621 | - | 2377 | 58 | 2319 | 2.44 |
| gene-LOC106786223 | 500 | 1 | 20 | 7.05 | 0.01 | https://www.ncbi.nlm.nih.gov/gene/?term=LOC106786223 | NW_014569615.1 | 533811 | 534310 | - | 1911 | 22 | 1889 | 1.15 |
| gene-LOC106786379 | 500 | 1 | 20 | 21.62 | 0.04 | https://www.ncbi.nlm.nih.gov/gene/?term=LOC106786379 | NW_014569550.1 | 1942801 | 1943300 | - | 1807 | 26 | 1781 | 1.44 |
| gene-LOC106786576 | 500 | 1 | 20 | 7.38 | 0.01 | https://www.ncbi.nlm.nih.gov/gene/?term=LOC106786576 | NW_014569624.1 | 160695 | 161194 | + | 2248 | 34 | 2214 | 1.51 |
| gene-LOC106786610 | 500 | 1 | 20 | 20 | 0.04 | https://www.ncbi.nlm.nih.gov/gene/?term=LOC106786610 | NW_014569625.1 | 10936 | 11435 | + | 712 | 25 | 687 | 3.51 |
| gene-LOC106786701 | 500 | 1 | 20 | 25.81 | 0.05 | https://www.ncbi.nlm.nih.gov/gene/?term=LOC106786701 | NW_014569627.1 | 117103 | 117602 | + | 153 | 14 | 139 | 9.15 |
| gene-LOC106786761 | 500 | 1 | 20 | 8.28 | 0.02 | https://www.ncbi.nlm.nih.gov/gene/?term=LOC106786761 | NW_014569628.1 | 186031 | 186530 | - | 2324 | 55 | 2269 | 2.37 |
| gene-LOC106786924 | 500 | 1 | 20 | 12.86 | 0.03 | https://www.ncbi.nlm.nih.gov/gene/?term=LOC106786924 | NW_014569635.1 | 536212 | 536711 | + | 4365 | 74 | 4291 | 1.7 |
| gene-LOC106787161 | 500 | 1 | 20 | 8.12 | 0.02 | https://www.ncbi.nlm.nih.gov/gene/?term=LOC106787161 | NW_014569641.1 | 17313 | 17812 | + | 3085 | 56 | 3029 | 1.82 |
| gene-LOC106787219 | 500 | 1 | 20 | 18.42 | 0.04 | https://www.ncbi.nlm.nih.gov/gene/?term=LOC106787219 | NW_014569642.1 | 291517 | 292016 | + | 1036 | 18 | 1018 | 1.74 |
| gene-LOC106787338 | 500 | 1 | 20 | 18.42 | 0.04 | https://www.ncbi.nlm.nih.gov/gene/?term=LOC106787338 | NW_014569645.1 | 85968 | 86467 | - | 868 | 14 | 854 | 1.61 |
| gene-LOC106787384 | 500 | 1 | 20 | 8.55 | 0.02 | https://www.ncbi.nlm.nih.gov/gene/?term=LOC106787384 | NW_014569551.1 | 1621220 | 1621719 | - | 2239 | 32 | 2207 | 1.43 |
| gene-LOC106787574 | 500 | 1 | 20 | 9.71 | 0.02 | https://www.ncbi.nlm.nih.gov/gene/?term=LOC106787574 | NW_014569653.1 | 281290 | 281789 | - | 3574 | 56 | 3518 | 1.57 |
| gene-LOC106787736 | 500 | 1 | 20 | 21.62 | 0.04 | https://www.ncbi.nlm.nih.gov/gene/?term=LOC106787736 | NW_014569660.1 | 497436 | 497935 | + | 280 | 10 | 270 | 3.57 |
| gene-LOC106787743 | 500 | 1 | 20 | 19.35 | 0.04 | https://www.ncbi.nlm.nih.gov/gene/?term=LOC106787743 | NW_014569660.1 | 69931 | 70430 | - | 1524 | 36 | 1488 | 2.36 |
| gene-LOC106787879 | 500 | 1 | 20 | 21.82 | 0.04 | https://www.ncbi.nlm.nih.gov/gene/?term=LOC106787879 | NW_014569665.1 | 437872 | 438371 | - | 2554 | 46 | 2508 | 1.8 |
| gene-LOC106787963 | 500 | 1 | 20 | 19.3 | 0.04 | https://www.ncbi.nlm.nih.gov/gene/?term=LOC106787963 | NW_014569668.1 | 413196 | 413695 | - | 4388 | 68 | 4320 | 1.55 |
| gene-LOC106788418 | 500 | 1 | 20 | 20.59 | 0.04 | https://www.ncbi.nlm.nih.gov/gene/?term=LOC106788418 | NW_014569552.1 | 513121 | 513620 | + | 1878 | 35 | 1843 | 1.86 |
| gene-LOC106788674 | 500 | 1 | 20 | 12.16 | 0.02 | https://www.ncbi.nlm.nih.gov/gene/?term=LOC106788674 | NW_014569695.1 | 76070 | 76569 | - | 1171 | 24 | 1147 | 2.05 |
| gene-LOC106788754 | 500 | 1 | 20 | 14.67 | 0.03 | https://www.ncbi.nlm.nih.gov/gene/?term=LOC106788754 | NW_014569700.1 | 205383 | 205882 | - | 3082 | 61 | 3021 | 1.98 |
| gene-LOC106788896 | 500 | 1 | 20 | 18.52 | 0.04 | https://www.ncbi.nlm.nih.gov/gene/?term=LOC106788896 | NW_014569703.1 | 368439 | 368938 | - | 1181 | 26 | 1155 | 2.2 |
| gene-LOC106788960 | 500 | 1 | 20 | 13.41 | 0.03 | https://www.ncbi.nlm.nih.gov/gene/?term=LOC106788960 | NW_014569705.1 | 195563 | 196062 | - | 1881 | 32 | 1849 | 1.7 |
| gene-LOC106789287 | 500 | 1 | 20 | 13.7 | 0.03 | https://www.ncbi.nlm.nih.gov/gene/?term=LOC106789287 | NW_014569721.1 | 144570 | 145069 | - | 1529 | 31 | 1498 | 2.03 |
| gene-LOC106789368 | 500 | 1 | 20 | 44 | 0.09 | https://www.ncbi.nlm.nih.gov/gene/?term=LOC106789368 | NW_014569728.1 | 76522 | 77021 | + | 132 | 11 | 121 | 8.33 |
| gene-LOC106789541 | 500 | 1 | 20 | 9.52 | 0.02 | https://www.ncbi.nlm.nih.gov/gene/?term=LOC106789541 | NW_014569736.1 | 128805 | 129304 | + | 1205 | 26 | 1179 | 2.16 |
| gene-LOC106789584 | 500 | 1 | 20 | 10.48 | 0.02 | https://www.ncbi.nlm.nih.gov/gene/?term=LOC106789584 | NW_014569739.1 | 103551 | 104050 | - | 3330 | 51 | 3279 | 1.53 |
| gene-LOC106789631 | 500 | 1 | 20 | 25.93 | 0.05 | https://www.ncbi.nlm.nih.gov/gene/?term=LOC106789631 | NW_014569553.1 | 1499503 | 1500002 | + | 708 | 12 | 696 | 1.69 |
| gene-LOC106789765 | 500 | 1 | 20 | 26 | 0.05 | https://www.ncbi.nlm.nih.gov/gene/?term=LOC106789765 | NW_014569749.1 | 71315 | 71814 | + | 1048 | 20 | 1028 | 1.91 |
| gene-LOC106789768 | 500 | 1 | 20 | 26 | 0.05 | https://www.ncbi.nlm.nih.gov/gene/?term=LOC106789768 | NW_014569749.1 | 70928 | 71427 | - | 905 | 21 | 884 | 2.32 |
| gene-LOC106789966 | 500 | 1 | 20 | 28.57 | 0.06 | https://www.ncbi.nlm.nih.gov/gene/?term=LOC106789966 | NW_014569756.1 | 48261 | 48760 | - | 105 | 16 | 89 | 15.24 |
| gene-LOC106790323 | 500 | 1 | 20 | 16.67 | 0.03 | https://www.ncbi.nlm.nih.gov/gene/?term=LOC106790323 | NW_014569777.1 | 65272 | 65771 | - | 2337 | 45 | 2292 | 1.93 |
| gene-LOC106790436 | 500 | 1 | 20 | 12.5 | 0.03 | https://www.ncbi.nlm.nih.gov/gene/?term=LOC106790436 | NW_014569786.1 | 208666 | 209165 | + | 1376 | 30 | 1346 | 2.18 |
| gene-LOC106790654 | 500 | 1 | 20 | 1.82 | 0 | https://www.ncbi.nlm.nih.gov/gene/?term=LOC106790654 | NW_014569800.1 | 4731 | 5230 | - | 34870 | 272 | 34598 | 0.78 |
| gene-LOC106790793 | 500 | 1 | 20 | 35.29 | 0.07 | https://www.ncbi.nlm.nih.gov/gene/?term=LOC106790793 | NW_014569807.1 | 153999 | 154498 | - | 146 | 8 | 138 | 5.48 |
| gene-LOC106790827 | 500 | 1 | 20 | 16.45 | 0.03 | https://www.ncbi.nlm.nih.gov/gene/?term=LOC106790827 | NW_014569554.1 | 889524 | 890023 | - | 1576 | 38 | 1538 | 2.41 |
| gene-LOC106790837 | 500 | 1 | 20 | 14.58 | 0.03 | https://www.ncbi.nlm.nih.gov/gene/?term=LOC106790837 | NW_014569812.1 | 14291 | 14790 | + | 3241 | 52 | 3189 | 1.6 |
| gene-LOC106791239 | 500 | 1 | 20 | 14.08 | 0.03 | https://www.ncbi.nlm.nih.gov/gene/?term=LOC106791239 | NW_014569840.1 | 47190 | 47689 | + | 1322 | 22 | 1300 | 1.66 |
| gene-LOC106791253 | 500 | 1 | 20 | 76.92 | 0.15 | https://www.ncbi.nlm.nih.gov/gene/?term=LOC106791253 | NW_014569840.1 | 104895 | 105394 | - | 77 | 10 | 67 | 12.99 |
| gene-LOC106791261 | 500 | 1 | 20 | 9.26 | 0.02 | https://www.ncbi.nlm.nih.gov/gene/?term=LOC106791261 | NW_014569840.1 | 27593 | 28092 | - | 4206 | 67 | 4139 | 1.59 |
| gene-LOC106791400 | 500 | 1 | 20 | 7.48 | 0.01 | https://www.ncbi.nlm.nih.gov/gene/?term=LOC106791400 | NW_014569850.1 | 102609 | 103108 | + | 997 | 18 | 979 | 1.81 |
| gene-LOC106791479 | 500 | 1 | 20 | 18.92 | 0.04 | https://www.ncbi.nlm.nih.gov/gene/?term=LOC106791479 | NW_014569858.1 | 106487 | 106986 | + | 2118 | 29 | 2089 | 1.37 |
| gene-LOC106791625 | 500 | 1 | 20 | 15.62 | 0.03 | https://www.ncbi.nlm.nih.gov/gene/?term=LOC106791625 | NW_014569873.1 | 25604 | 26103 | - | 1872 | 28 | 1844 | 1.5 |
| gene-LOC106791782 | 500 | 1 | 20 | 26.32 | 0.05 | https://www.ncbi.nlm.nih.gov/gene/?term=LOC106791782 | NW_014569889.1 | 13934 | 14433 | + | 1418 | 41 | 1377 | 2.89 |
| gene-LOC106791783 | 500 | 1 | 20 | 26.32 | 0.05 | https://www.ncbi.nlm.nih.gov/gene/?term=LOC106791783 | NW_014569889.1 | 13665 | 14164 | - | 835 | 40 | 795 | 4.79 |
| gene-LOC106792003 | 500 | 1 | 20 | 7.59 | 0.02 | https://www.ncbi.nlm.nih.gov/gene/?term=LOC106792003 | NW_014569555.1 | 830071 | 830570 | + | 1939 | 37 | 1902 | 1.91 |
| gene-LOC106792461 | 500 | 1 | 20 | 2.28 | 0 | https://www.ncbi.nlm.nih.gov/gene/?term=LOC106792461 | NW_014569983.1 | 71397 | 71896 | - | 59675 | 454 | 59221 | 0.76 |
| gene-LOC106792882 | 500 | 1 | 20 | 2.26 | 0 | https://www.ncbi.nlm.nih.gov/gene/?term=LOC106792882 | NW_014570595.1 | 5669 | 6168 | - | 7012 | 87 | 6925 | 1.24 |
| gene-LOC106793168 | 500 | 1 | 20 | 15.19 | 0.03 | https://www.ncbi.nlm.nih.gov/gene/?term=LOC106793168 | NW_014569559.1 | 321867 | 322366 | + | 2108 | 37 | 2071 | 1.76 |
| gene-LOC106793249 | 500 | 1 | 20 | 22.5 | 0.04 | https://www.ncbi.nlm.nih.gov/gene/?term=LOC106793249 | NW_014569559.1 | 376190 | 376689 | + | 2189 | 35 | 2154 | 1.6 |
| gene-LOC106793276 | 500 | 1 | 20 | 42.86 | 0.09 | https://www.ncbi.nlm.nih.gov/gene/?term=LOC106793276 | NW_014569559.1 | 590930 | 591429 | + | 438 | 18 | 420 | 4.11 |
| gene-LOC106793387 | 500 | 1 | 20 | 9.24 | 0.02 | https://www.ncbi.nlm.nih.gov/gene/?term=LOC106793387 | NW_014569560.1 | 357805 | 358304 | - | 3337 | 51 | 3286 | 1.53 |
| gene-LOC106793622 | 500 | 1 | 20 | 13.4 | 0.03 | https://www.ncbi.nlm.nih.gov/gene/?term=LOC106793622 | NW_014569548.1 | 1791422 | 1791921 | + | 3755 | 55 | 3700 | 1.46 |
| gene-LOC106793851 | 500 | 1 | 20 | 46.67 | 0.09 | https://www.ncbi.nlm.nih.gov/gene/?term=LOC106793851 | NW_014569566.1 | 796259 | 796758 | - | 65 | 17 | 48 | 26.15 |
| gene-LOC106793963 | 500 | 1 | 20 | 23.33 | 0.05 | https://www.ncbi.nlm.nih.gov/gene/?term=LOC106793963 | NW_014569568.1 | 1081208 | 1081707 | + | 2690 | 26 | 2664 | 0.97 |
| gene-Trnad-guc-5 | 500 | 1 | 20 | 2.8 | 0.01 | https://www.ncbi.nlm.nih.gov/gene/?term=Trnad-guc | NW_014569595.1 | 862960 | 863459 | - | 4922 | 69 | 4853 | 1.4 |

Directory: RNK File: ranked-promoters-bySiteDensity-Pc_PA_worker.txt

| region_ID | rwidth | nbrsites | nbrper10kb | pmpersite | pmpernucl | pglink | seqnames | start | end | strand | coverage | numCs | numTs | prcntM |
|---|---|---|---|---|---|---|---|---|---|---|---|---|---|---|
| gene-LOC106792344 | 500 | 21 | 420 | 1.13 | 0.05 | https://www.ncbi.nlm.nih.gov/gene/?term=LOC106792344 | NW_014569955.1 | 55898 | 56397 | - | 1053463 | 8291 | 1045172 | 0.79 |
| gene-LOC106791550 | 500 | 10 | 200 | 28.23 | 0.56 | https://www.ncbi.nlm.nih.gov/gene/?term=LOC106791550 | NW_014569865.1 | 85298 | 85797 | - | 6341 | 390 | 5951 | 6.15 |
| gene-LOC106787745 | 500 | 8 | 160 | 42.79 | 0.68 | https://www.ncbi.nlm.nih.gov/gene/?term=LOC106787745 | NW_014569551.1 | 2222289 | 2222788 | + | 731 | 119 | 612 | 16.28 |
| gene-LOC106790993 | 500 | 7 | 140 | 64.72 | 0.91 | https://www.ncbi.nlm.nih.gov/gene/?term=LOC106790993 | NW_014569822.1 | 88866 | 89365 | + | 1860 | 325 | 1535 | 17.47 |
| gene-LOC106793172 | 500 | 7 | 140 | 10.39 | 0.15 | https://www.ncbi.nlm.nih.gov/gene/?term=LOC106793172 | NW_014569559.1 | 1151932 | 1152431 | + | 4962 | 175 | 4787 | 3.53 |
| gene-LOC106786771 | 500 | 6 | 120 | 14.94 | 0.18 | https://www.ncbi.nlm.nih.gov/gene/?term=LOC106786771 | NW_014569628.1 | 539233 | 539732 | + | 2156 | 100 | 2056 | 4.64 |
| gene-LOC106787307 | 500 | 6 | 120 | 47.72 | 0.57 | https://www.ncbi.nlm.nih.gov/gene/?term=LOC106787307 | NW_014569645.1 | 320505 | 321004 | + | 3133 | 363 | 2770 | 11.59 |
| gene-LOC106785314 | 500 | 5 | 100 | 62.86 | 0.63 | https://www.ncbi.nlm.nih.gov/gene/?term=LOC106785314 | NW_014569595.1 | 551303 | 551802 | + | 482 | 57 | 425 | 11.83 |
| gene-LOC106786867 | 500 | 5 | 100 | 72.45 | 0.72 | https://www.ncbi.nlm.nih.gov/gene/?term=LOC106786867 | NW_014569633.1 | 260124 | 260623 | + | 215 | 72 | 143 | 33.49 |
| gene-LOC106788490 | 500 | 5 | 100 | 50.45 | 0.5 | https://www.ncbi.nlm.nih.gov/gene/?term=LOC106788490 | NW_014569687.1 | 411960 | 412459 | - | 1368 | 379 | 989 | 27.7 |
| gene-LOC106791724 | 500 | 5 | 100 | 43.18 | 0.43 | https://www.ncbi.nlm.nih.gov/gene/?term=LOC106791724 | NW_014569883.1 | 90156 | 90655 | - | 220 | 41 | 179 | 18.64 |
| gene-LOC106791733 | 500 | 5 | 100 | 43.18 | 0.43 | https://www.ncbi.nlm.nih.gov/gene/?term=LOC106791733 | NW_014569883.1 | 90350 | 90849 | + | 240 | 41 | 199 | 17.08 |
| gene-LOC106785085 | 500 | 4 | 80 | 66.92 | 0.54 | https://www.ncbi.nlm.nih.gov/gene/?term=LOC106785085 | NW_014569549.1 | 2123847 | 2124346 | + | 63 | 37 | 26 | 58.73 |
| gene-LOC106788202 | 500 | 4 | 80 | 81.94 | 0.66 | https://www.ncbi.nlm.nih.gov/gene/?term=LOC106788202 | NW_014569676.1 | 375634 | 376133 | - | 395 | 49 | 346 | 12.41 |
| gene-LOC106791306 | 500 | 4 | 80 | 28.46 | 0.23 | https://www.ncbi.nlm.nih.gov/gene/?term=LOC106791306 | NW_014569842.1 | 53546 | 54045 | - | 2080 | 194 | 1886 | 9.33 |
| gene-LOC106791842 | 500 | 4 | 80 | 37.22 | 0.3 | https://www.ncbi.nlm.nih.gov/gene/?term=LOC106791842 | NW_014569894.1 | 75828 | 76327 | + | 6283 | 244 | 6039 | 3.88 |
| gene-LOC106793074 | 500 | 4 | 80 | 28.46 | 0.23 | https://www.ncbi.nlm.nih.gov/gene/?term=LOC106793074 | NW_014569558.1 | 934812 | 935311 | - | 1570 | 131 | 1439 | 8.34 |
| gene-LOC106785672 | 500 | 3 | 60 | 10.74 | 0.06 | https://www.ncbi.nlm.nih.gov/gene/?term=LOC106785672 | NW_014569602.1 | 734501 | 735000 | + | 3187 | 92 | 3095 | 2.89 |
| gene-LOC106786084 | 500 | 3 | 60 | 53.52 | 0.32 | https://www.ncbi.nlm.nih.gov/gene/?term=LOC106786084 | NW_014569550.1 | 1117335 | 1117834 | - | 479 | 69 | 410 | 14.41 |
| gene-LOC106788339 | 500 | 3 | 60 | 67.68 | 0.41 | https://www.ncbi.nlm.nih.gov/gene/?term=LOC106788339 | NW_014569682.1 | 411042 | 411541 | - | 56 | 27 | 29 | 48.21 |
| gene-LOC106792704 | 500 | 3 | 60 | 62.96 | 0.38 | https://www.ncbi.nlm.nih.gov/gene/?term=LOC106792704 | NW_014570116.1 | 10409 | 10908 | + | 128 | 27 | 101 | 21.09 |
| gene-LOC106793571 | 500 | 3 | 60 | 68.24 | 0.41 | https://www.ncbi.nlm.nih.gov/gene/?term=LOC106793571 | NW_014569562.1 | 198130 | 198629 | + | 96 | 37 | 59 | 38.54 |
| gene-LOC106793700 | 500 | 3 | 60 | 32.38 | 0.19 | https://www.ncbi.nlm.nih.gov/gene/?term=LOC106793700 | NW_014569564.1 | 879896 | 880395 | - | 243 | 43 | 200 | 17.7 |
| gene-Trnai-aau-2 | 500 | 3 | 60 | 61.9 | 0.37 | https://www.ncbi.nlm.nih.gov/gene/?term=Trnai-aau | NW_014569564.1 | 1136734 | 1137233 | + | 520 | 45 | 475 | 8.65 |
| gene-LOC106783822 | 500 | 2 | 40 | 14.84 | 0.06 | https://www.ncbi.nlm.nih.gov/gene/?term=LOC106783822 | NW_014569573.1 | 915537 | 916036 | + | 3216 | 98 | 3118 | 3.05 |
| gene-LOC106784544 | 500 | 2 | 40 | 43.72 | 0.17 | https://www.ncbi.nlm.nih.gov/gene/?term=LOC106784544 | NW_014569584.1 | 993361 | 993860 | - | 1032 | 34 | 998 | 3.29 |
| gene-LOC106784545 | 500 | 2 | 40 | 43.72 | 0.17 | https://www.ncbi.nlm.nih.gov/gene/?term=LOC106784545 | NW_014569584.1 | 992939 | 993438 | + | 212 | 24 | 188 | 11.32 |
| gene-LOC106786332 | 500 | 2 | 40 | 10.21 | 0.04 | https://www.ncbi.nlm.nih.gov/gene/?term=LOC106786332 | NW_014569619.1 | 591461 | 591960 | - | 1203 | 42 | 1161 | 3.49 |
| gene-LOC106786756 | 500 | 2 | 40 | 54.62 | 0.22 | https://www.ncbi.nlm.nih.gov/gene/?term=LOC106786756 | NW_014569628.1 | 69713 | 70212 | - | 251 | 15 | 236 | 5.98 |
| gene-LOC106786908 | 500 | 2 | 40 | 21.36 | 0.09 | https://www.ncbi.nlm.nih.gov/gene/?term=LOC106786908 | NW_014569635.1 | 197580 | 198079 | - | 2080 | 39 | 2041 | 1.88 |
| gene-LOC106787461 | 500 | 2 | 40 | 15.7 | 0.06 | https://www.ncbi.nlm.nih.gov/gene/?term=LOC106787461 | NW_014569648.1 | 505358 | 505857 | + | 1423 | 38 | 1385 | 2.67 |
| gene-LOC106787743 | 500 | 2 | 40 | 15.12 | 0.06 | https://www.ncbi.nlm.nih.gov/gene/?term=LOC106787743 | NW_014569660.1 | 69931 | 70430 | - | 1517 | 54 | 1463 | 3.56 |
| gene-LOC106790590 | 500 | 2 | 40 | 35.15 | 0.14 | https://www.ncbi.nlm.nih.gov/gene/?term=LOC106790590 | NW_014569796.1 | 107031 | 107530 | + | 156 | 23 | 133 | 14.74 |
| gene-LOC106791142 | 500 | 2 | 40 | 11.38 | 0.05 | https://www.ncbi.nlm.nih.gov/gene/?term=LOC106791142 | NW_014569834.1 | 58898 | 59397 | + | 6419 | 89 | 6330 | 1.39 |
| gene-LOC106792690 | 500 | 2 | 40 | 64.58 | 0.26 | https://www.ncbi.nlm.nih.gov/gene/?term=LOC106792690 | NW_014569557.1 | 1529494 | 1529993 | - | 2065 | 33 | 2032 | 1.6 |
| gene-LOC106792768 | 500 | 2 | 40 | 19.78 | 0.08 | https://www.ncbi.nlm.nih.gov/gene/?term=LOC106792768 | NW_014570192.1 | 8982 | 9481 | + | 1653 | 32 | 1621 | 1.94 |
| gene-LOC106783887 | 500 | 1 | 20 | 35.78 | 0.07 | https://www.ncbi.nlm.nih.gov/gene/?term=LOC106783887 | NW_014569574.1 | 274326 | 274825 | + | 2516 | 58 | 2458 | 2.31 |
| gene-LOC106783888 | 500 | 1 | 20 | 35.78 | 0.07 | https://www.ncbi.nlm.nih.gov/gene/?term=LOC106783888 | NW_014569574.1 | 274388 | 274887 | - | 2739 | 59 | 2680 | 2.15 |
| gene-LOC106783974 | 500 | 1 | 20 | 30 | 0.06 | https://www.ncbi.nlm.nih.gov/gene/?term=LOC106783974 | NW_014569575.1 | 976069 | 976568 | + | 138 | 7 | 131 | 5.07 |
| gene-LOC106784052 | 500 | 1 | 20 | 25.93 | 0.05 | https://www.ncbi.nlm.nih.gov/gene/?term=LOC106784052 | NW_014569576.1 | 37636 | 38135 | - | 262 | 12 | 250 | 4.58 |
| gene-LOC106784134 | 500 | 1 | 20 | 16 | 0.03 | https://www.ncbi.nlm.nih.gov/gene/?term=LOC106784134 | NW_014569578.1 | 222786 | 223285 | + | 2484 | 53 | 2431 | 2.13 |
| gene-LOC106784378 | 500 | 1 | 20 | 32.43 | 0.06 | https://www.ncbi.nlm.nih.gov/gene/?term=LOC106784378 | NW_014569581.1 | 763831 | 764330 | + | 1712 | 38 | 1674 | 2.22 |
| gene-LOC106784646 | 500 | 1 | 20 | 7.89 | 0.02 | https://www.ncbi.nlm.nih.gov/gene/?term=LOC106784646 | NW_014569549.1 | 319635 | 320134 | - | 1244 | 30 | 1214 | 2.41 |
| gene-LOC106784780 | 500 | 1 | 20 | 17.19 | 0.03 | https://www.ncbi.nlm.nih.gov/gene/?term=LOC106784780 | NW_014569587.1 | 576862 | 577361 | - | 2881 | 47 | 2834 | 1.63 |
| gene-LOC106784946 | 500 | 1 | 20 | 15.7 | 0.03 | https://www.ncbi.nlm.nih.gov/gene/?term=LOC106784946 | NW_014569549.1 | 1284120 | 1284619 | - | 1904 | 29 | 1875 | 1.52 |
| gene-LOC106784958 | 500 | 1 | 20 | 6.03 | 0.01 | https://www.ncbi.nlm.nih.gov/gene/?term=LOC106784958 | NW_014569589.1 | 409835 | 410334 | - | 4063 | 61 | 4002 | 1.5 |
| gene-LOC106784967 | 500 | 1 | 20 | 12.61 | 0.03 | https://www.ncbi.nlm.nih.gov/gene/?term=LOC106784967 | NW_014569589.1 | 371818 | 372317 | + | 1190 | 24 | 1166 | 2.02 |
| gene-LOC106785021 | 500 | 1 | 20 | 16.83 | 0.03 | https://www.ncbi.nlm.nih.gov/gene/?term=LOC106785021 | NW_014569549.1 | 603372 | 603871 | + | 814 | 20 | 794 | 2.46 |
| gene-LOC106785026 | 500 | 1 | 20 | 18.92 | 0.04 | https://www.ncbi.nlm.nih.gov/gene/?term=LOC106785026 | NW_014569591.1 | 916047 | 916546 | - | 2383 | 22 | 2361 | 0.92 |
| gene-LOC106785315 | 500 | 1 | 20 | 7.73 | 0.02 | https://www.ncbi.nlm.nih.gov/gene/?term=LOC106785315 | NW_014569549.1 | 370564 | 371063 | + | 2036 | 31 | 2005 | 1.52 |
| gene-LOC106785612 | 500 | 1 | 20 | 12.15 | 0.02 | https://www.ncbi.nlm.nih.gov/gene/?term=LOC106785612 | NW_014569600.1 | 732075 | 732574 | + | 1323 | 24 | 1299 | 1.81 |
| gene-LOC106785729 | 500 | 1 | 20 | 66.67 | 0.13 | https://www.ncbi.nlm.nih.gov/gene/?term=LOC106785729 | NW_014569602.1 | 287206 | 287705 | - | 422 | 13 | 409 | 3.08 |
| gene-LOC106785832 | 500 | 1 | 20 | 37.5 | 0.07 | https://www.ncbi.nlm.nih.gov/gene/?term=LOC106785832 | NW_014569604.1 | 212685 | 213184 | - | 172 | 21 | 151 | 12.21 |
| gene-LOC106786213 | 500 | 1 | 20 | 11.03 | 0.02 | https://www.ncbi.nlm.nih.gov/gene/?term=LOC106786213 | NW_014569615.1 | 500122 | 500621 | - | 2084 | 42 | 2042 | 2.02 |
| gene-LOC106786233 | 500 | 1 | 20 | 22.22 | 0.04 | https://www.ncbi.nlm.nih.gov/gene/?term=LOC106786233 | NW_014569616.1 | 545242 | 545741 | + | 2339 | 43 | 2296 | 1.84 |
| gene-LOC106786379 | 500 | 1 | 20 | 32.5 | 0.06 | https://www.ncbi.nlm.nih.gov/gene/?term=LOC106786379 | NW_014569550.1 | 1942801 | 1943300 | - | 1656 | 30 | 1626 | 1.81 |
| gene-LOC106786586 | 500 | 1 | 20 | 8.33 | 0.02 | https://www.ncbi.nlm.nih.gov/gene/?term=LOC106786586 | NW_014569550.1 | 751495 | 751994 | - | 1816 | 25 | 1791 | 1.38 |
| gene-LOC106786610 | 500 | 1 | 20 | 26.76 | 0.05 | https://www.ncbi.nlm.nih.gov/gene/?term=LOC106786610 | NW_014569625.1 | 10936 | 11435 | + | 649 | 31 | 618 | 4.78 |
| gene-LOC106786761 | 500 | 1 | 20 | 13.43 | 0.03 | https://www.ncbi.nlm.nih.gov/gene/?term=LOC106786761 | NW_014569628.1 | 186031 | 186530 | - | 2412 | 79 | 2333 | 3.28 |
| gene-LOC106786858 | 500 | 1 | 20 | 12.94 | 0.03 | https://www.ncbi.nlm.nih.gov/gene/?term=LOC106786858 | NW_014569633.1 | 449768 | 450267 | + | 1996 | 67 | 1929 | 3.36 |

| | | | | | | | | | | | | | | |
|---|---|---|---|---|---|---|---|---|---|---|---|---|---|---|
| gene-LOC106786924 | 500 | 1 | 20 | 12.61 | 0.03 | https://www.ncbi.nlm.nih.gov/gene/?term=LOC106786924 | NW_014569635.1 | 536212 | 536711 | + | 3149 | 51 | 3098 | 1.62 |
| gene-LOC106786937 | 500 | 1 | 20 | 12 | 0.02 | https://www.ncbi.nlm.nih.gov/gene/?term=LOC106786937 | NW_014569635.1 | 418119 | 418618 | - | 979 | 27 | 952 | 2.76 |
| gene-LOC106787059 | 500 | 1 | 20 | 7.51 | 0.02 | https://www.ncbi.nlm.nih.gov/gene/?term=LOC106787059 | NW_014569638.1 | 406493 | 406992 | + | 1914 | 29 | 1885 | 1.52 |
| gene-LOC106787064 | 500 | 1 | 20 | 7.51 | 0.02 | https://www.ncbi.nlm.nih.gov/gene/?term=LOC106787064 | NW_014569638.1 | 406832 | 407331 | - | 1273 | 23 | 1250 | 1.81 |
| gene-LOC106787134 | 500 | 1 | 20 | 17.5 | 0.04 | https://www.ncbi.nlm.nih.gov/gene/?term=LOC106787134 | NW_014569640.1 | 288626 | 289125 | + | 2012 | 32 | 1980 | 1.59 |
| gene-LOC106787154 | 500 | 1 | 20 | 10.2 | 0.02 | https://www.ncbi.nlm.nih.gov/gene/?term=LOC106787154 | NW_014569551.1 | 2278087 | 2278586 | - | 2889 | 62 | 2827 | 2.15 |
| gene-LOC106787338 | 500 | 1 | 20 | 15.69 | 0.03 | https://www.ncbi.nlm.nih.gov/gene/?term=LOC106787338 | NW_014569645.1 | 85968 | 86467 | - | 1029 | 18 | 1011 | 1.75 |
| gene-LOC106787770 | 500 | 1 | 20 | 8.72 | 0.02 | https://www.ncbi.nlm.nih.gov/gene/?term=LOC106787770 | NW_014569551.1 | 2088589 | 2089088 | + | 3440 | 52 | 3388 | 1.51 |
| gene-LOC106787879 | 500 | 1 | 20 | 18.52 | 0.04 | https://www.ncbi.nlm.nih.gov/gene/?term=LOC106787879 | NW_014569665.1 | 437872 | 438371 | - | 2682 | 36 | 2646 | 1.34 |
| gene-LOC106788091 | 500 | 1 | 20 | 8.18 | 0.02 | https://www.ncbi.nlm.nih.gov/gene/?term=LOC106788091 | NW_014569672.1 | 64535 | 65034 | + | 769 | 19 | 750 | 2.47 |
| gene-LOC106788165 | 500 | 1 | 20 | 9.78 | 0.02 | https://www.ncbi.nlm.nih.gov/gene/?term=LOC106788165 | NW_014569675.1 | 239927 | 240426 | + | 1438 | 26 | 1412 | 1.81 |
| gene-LOC106788209 | 500 | 1 | 20 | 11.5 | 0.02 | https://www.ncbi.nlm.nih.gov/gene/?term=LOC106788209 | NW_014569676.1 | 35906 | 36405 | - | 2742 | 42 | 2700 | 1.53 |
| gene-LOC106788418 | 500 | 1 | 20 | 19.44 | 0.04 | https://www.ncbi.nlm.nih.gov/gene/?term=LOC106788418 | NW_014569552.1 | 513121 | 513620 | + | 1770 | 29 | 1741 | 1.64 |
| gene-LOC106788514 | 500 | 1 | 20 | 15.79 | 0.03 | https://www.ncbi.nlm.nih.gov/gene/?term=LOC106788514 | NW_014569689.1 | 63633 | 64132 | - | 996 | 24 | 972 | 2.41 |
| gene-LOC106788713 | 500 | 1 | 20 | 25 | 0.05 | https://www.ncbi.nlm.nih.gov/gene/?term=LOC106788713 | NW_014569698.1 | 12926 | 13425 | + | 2720 | 40 | 2680 | 1.47 |
| gene-LOC106788816 | 500 | 1 | 20 | 15.49 | 0.03 | https://www.ncbi.nlm.nih.gov/gene/?term=LOC106788816 | NW_014569702.1 | 185679 | 186178 | - | 2494 | 42 | 2452 | 1.68 |
| gene-LOC106788838 | 500 | 1 | 20 | 15.87 | 0.03 | https://www.ncbi.nlm.nih.gov/gene/?term=LOC106788838 | NW_014569702.1 | 308593 | 309092 | - | 3930 | 55 | 3875 | 1.4 |
| gene-LOC106788896 | 500 | 1 | 20 | 27.27 | 0.05 | https://www.ncbi.nlm.nih.gov/gene/?term=LOC106788896 | NW_014569703.1 | 368439 | 368938 | - | 1041 | 26 | 1015 | 2.5 |
| gene-LOC106789541 | 500 | 1 | 20 | 14.78 | 0.03 | https://www.ncbi.nlm.nih.gov/gene/?term=LOC106789541 | NW_014569736.1 | 128805 | 129304 | + | 1161 | 27 | 1134 | 2.33 |
| gene-LOC106789584 | 500 | 1 | 20 | 9.73 | 0.02 | https://www.ncbi.nlm.nih.gov/gene/?term=LOC106789584 | NW_014569739.1 | 103551 | 104050 | - | 3303 | 64 | 3239 | 1.94 |
| gene-LOC106789631 | 500 | 1 | 20 | 33.33 | 0.07 | https://www.ncbi.nlm.nih.gov/gene/?term=LOC106789631 | NW_014569553.1 | 1499503 | 1500002 | + | 690 | 18 | 672 | 2.61 |
| gene-LOC106789732 | 500 | 1 | 20 | 36.84 | 0.07 | https://www.ncbi.nlm.nih.gov/gene/?term=LOC106789732 | NW_014569746.1 | 142611 | 143110 | + | 113 | 12 | 101 | 10.62 |
| gene-LOC106789765 | 500 | 1 | 20 | 23.53 | 0.05 | https://www.ncbi.nlm.nih.gov/gene/?term=LOC106789765 | NW_014569749.1 | 71315 | 71814 | + | 904 | 20 | 884 | 2.21 |
| gene-LOC106789768 | 500 | 1 | 20 | 23.53 | 0.05 | https://www.ncbi.nlm.nih.gov/gene/?term=LOC106789768 | NW_014569749.1 | 70928 | 71427 | - | 905 | 18 | 887 | 1.99 |
| gene-LOC106789852 | 500 | 1 | 20 | 38.89 | 0.08 | https://www.ncbi.nlm.nih.gov/gene/?term=LOC106789852 | NW_014569752.1 | 21487 | 21986 | + | 18 | 7 | 11 | 38.89 |
| gene-LOC106789956 | 500 | 1 | 20 | 12.5 | 0.03 | https://www.ncbi.nlm.nih.gov/gene/?term=LOC106789956 | NW_014569756.1 | 197277 | 197776 | + | 5607 | 80 | 5527 | 1.43 |
| gene-LOC106790323 | 500 | 1 | 20 | 13 | 0.03 | https://www.ncbi.nlm.nih.gov/gene/?term=LOC106790323 | NW_014569777.1 | 65272 | 65771 | - | 2143 | 41 | 2102 | 1.91 |
| gene-LOC106790433 | 500 | 1 | 20 | 14.77 | 0.03 | https://www.ncbi.nlm.nih.gov/gene/?term=LOC106790433 | NW_014569786.1 | 181182 | 181681 | - | 1613 | 28 | 1585 | 1.74 |
| gene-LOC106790436 | 500 | 1 | 20 | 10.13 | 0.02 | https://www.ncbi.nlm.nih.gov/gene/?term=LOC106790436 | NW_014569786.1 | 208666 | 209165 | + | 1310 | 28 | 1282 | 2.14 |
| gene-LOC106790450 | 500 | 1 | 20 | 40 | 0.08 | https://www.ncbi.nlm.nih.gov/gene/?term=LOC106790450 | NW_014569787.1 | 171155 | 171654 | + | 77 | 13 | 64 | 16.88 |
| gene-LOC106790512 | 500 | 1 | 20 | 11.46 | 0.02 | https://www.ncbi.nlm.nih.gov/gene/?term=LOC106790512 | NW_014569790.1 | 207053 | 207552 | - | 460 | 12 | 448 | 2.61 |
| gene-LOC106790654 | 500 | 1 | 20 | 2.44 | 0 | https://www.ncbi.nlm.nih.gov/gene/?term=LOC106790654 | NW_014569800.1 | 4731 | 5230 | - | 35812 | 307 | 35505 | 0.86 |
| gene-LOC106790827 | 500 | 1 | 20 | 10.64 | 0.02 | https://www.ncbi.nlm.nih.gov/gene/?term=LOC106790827 | NW_014569554.1 | 889524 | 890023 | - | 1184 | 28 | 1156 | 2.36 |
| gene-LOC106790923 | 500 | 1 | 20 | 12.68 | 0.03 | https://www.ncbi.nlm.nih.gov/gene/?term=LOC106790923 | NW_014569816.1 | 7283 | 7782 | + | 2046 | 35 | 2011 | 1.71 |
| gene-LOC106791040 | 500 | 1 | 20 | 47.83 | 0.1 | https://www.ncbi.nlm.nih.gov/gene/?term=LOC106791040 | NW_014569827.1 | 43361 | 43860 | - | 98 | 19 | 79 | 19.39 |
| gene-LOC106791088 | 500 | 1 | 20 | 43.59 | 0.09 | https://www.ncbi.nlm.nih.gov/gene/?term=LOC106791088 | NW_014569832.1 | 124585 | 125084 | - | 527 | 37 | 490 | 7.02 |
| gene-LOC106791253 | 500 | 1 | 20 | 58.33 | 0.12 | https://www.ncbi.nlm.nih.gov/gene/?term=LOC106791253 | NW_014569840.1 | 104895 | 105394 | - | 104 | 8 | 96 | 7.69 |
| gene-LOC106791400 | 500 | 1 | 20 | 10 | 0.02 | https://www.ncbi.nlm.nih.gov/gene/?term=LOC106791400 | NW_014569850.1 | 102609 | 103108 | + | 782 | 13 | 769 | 1.66 |
| gene-LOC106791426 | 500 | 1 | 20 | 17.91 | 0.04 | https://www.ncbi.nlm.nih.gov/gene/?term=LOC106791426 | NW_014569854.1 | 158389 | 158888 | - | 1427 | 24 | 1403 | 1.68 |
| gene-LOC106791625 | 500 | 1 | 20 | 13.22 | 0.03 | https://www.ncbi.nlm.nih.gov/gene/?term=LOC106791625 | NW_014569873.1 | 25604 | 26103 | - | 1715 | 28 | 1687 | 1.63 |
| gene-LOC106792003 | 500 | 1 | 20 | 10.38 | 0.02 | https://www.ncbi.nlm.nih.gov/gene/?term=LOC106792003 | NW_014569555.1 | 830071 | 830570 | + | 1734 | 49 | 1685 | 2.83 |
| gene-LOC106792179 | 500 | 1 | 20 | 8.47 | 0.02 | https://www.ncbi.nlm.nih.gov/gene/?term=LOC106792179 | NW_014569934.1 | 77912 | 78411 | - | 1632 | 41 | 1591 | 2.51 |
| gene-LOC106792262 | 500 | 1 | 20 | 42.86 | 0.09 | https://www.ncbi.nlm.nih.gov/gene/?term=LOC106792262 | NW_014569945.1 | 91969 | 92468 | - | 61 | 20 | 41 | 32.79 |
| gene-LOC106792351 | 500 | 1 | 20 | 11.7 | 0.02 | https://www.ncbi.nlm.nih.gov/gene/?term=LOC106792351 | NW_014569958.1 | 62671 | 63170 | - | 2717 | 47 | 2670 | 1.73 |
| gene-LOC106792453 | 500 | 1 | 20 | 1.94 | 0 | https://www.ncbi.nlm.nih.gov/gene/?term=LOC106792453 | NW_014569980.1 | 20924 | 21423 | + | 32191 | 253 | 31938 | 0.79 |
| gene-LOC106792589 | 500 | 1 | 20 | 18.92 | 0.04 | https://www.ncbi.nlm.nih.gov/gene/?term=LOC106792589 | NW_014569547.1 | 1301789 | 1302288 | + | 1246 | 35 | 1211 | 2.81 |
| gene-LOC106792709 | 500 | 1 | 20 | 75 | 0.15 | https://www.ncbi.nlm.nih.gov/gene/?term=LOC106792709 | NW_014570116.1 | 11498 | 11997 | - | 141 | 20 | 121 | 14.18 |
| gene-LOC106792754 | 500 | 1 | 20 | 7.69 | 0.02 | https://www.ncbi.nlm.nih.gov/gene/?term=LOC106792754 | NW_014570169.1 | 21714 | 22213 | + | 3977 | 30 | 3947 | 0.75 |
| gene-LOC106793249 | 500 | 1 | 20 | 21.74 | 0.04 | https://www.ncbi.nlm.nih.gov/gene/?term=LOC106793249 | NW_014569559.1 | 376190 | 376689 | + | 2151 | 28 | 2123 | 1.3 |
| gene-LOC106793276 | 500 | 1 | 20 | 27.27 | 0.05 | https://www.ncbi.nlm.nih.gov/gene/?term=LOC106793276 | NW_014569559.1 | 590930 | 591429 | + | 384 | 10 | 374 | 2.6 |
| gene-LOC106793304 | 500 | 1 | 20 | 9.38 | 0.02 | https://www.ncbi.nlm.nih.gov/gene/?term=LOC106793304 | NW_014569560.1 | 94052 | 94551 | + | 2722 | 35 | 2687 | 1.29 |
| gene-LOC106793309 | 500 | 1 | 20 | 33.33 | 0.07 | https://www.ncbi.nlm.nih.gov/gene/?term=LOC106793309 | NW_014569560.1 | 636478 | 636977 | - | 113 | 8 | 105 | 7.08 |
| gene-LOC106793622 | 500 | 1 | 20 | 15.53 | 0.03 | https://www.ncbi.nlm.nih.gov/gene/?term=LOC106793622 | NW_014569548.1 | 1791422 | 1791921 | + | 3292 | 62 | 3230 | 1.88 |
| gene-LOC106793763 | 500 | 1 | 20 | 17.5 | 0.04 | https://www.ncbi.nlm.nih.gov/gene/?term=LOC106793763 | NW_014569565.1 | 732641 | 733140 | - | 1778 | 41 | 1737 | 2.31 |
| gene-LOC106793923 | 500 | 1 | 20 | 17.02 | 0.03 | https://www.ncbi.nlm.nih.gov/gene/?term=LOC106793923 | NW_014569568.1 | 685086 | 685585 | + | 1342 | 17 | 1325 | 1.27 |
| gene-LOC106794090 | 500 | 1 | 20 | 7.27 | 0.01 | https://www.ncbi.nlm.nih.gov/gene/?term=LOC106794090 | NW_014569570.1 | 758160 | 758659 | + | 1363 | 36 | 1327 | 2.64 |

Directory: RNK File: sites-in-genes-Pc_PA_queen.txt

| seqnames | start | end | width | strand | coverage | numCs | numTs | perc_meth | region_seqnames | region_start | region_end | region_width | region_strand | region_source | region_type | region_score | region_phase | region_ID | region_Dbxref | region_Name | region_gbkey | region_gene | region_gene_biotype | region_partial | region_start_range | region_end_range |
|---|---|---|---|---|---|---|---|---|---|---|---|---|---|---|---|---|---|---|---|---|---|---|---|---|---|---|
| NW_014569547.1 | 99868 | 99868 | 1 | - | 73 | 11 | 62 | 15.0684931506849 | NW_014569547.1 | 57733 | 165446 | 107714 | + | Gnomon | gene | NA | NA | gene-LOC106783879 | GeneID:106783879 | LOC106783879 | Gene | LOC106783879 | protein_coding | NA | character(0) | character(0) |
| NW_014569547.1 | 127778 | 127778 | 1 | - | 34 | 15 | 19 | 44.1176470588235 | NW_014569547.1 | 57733 | 165446 | 107714 | + | Gnomon | gene | NA | NA | gene-LOC106783879 | GeneID:106783879 | LOC106783879 | Gene | LOC106783879 | protein_coding | NA | character(0) | character(0) |
| NW_014569547.1 | 127782 | 127782 | 1 | - | 34 | 19 | 15 | 55.8823529411765 | NW_014569547.1 | 57733 | 165446 | 107714 | + | Gnomon | gene | NA | NA | gene-LOC106783879 | GeneID:106783879 | LOC106783879 | Gene | LOC106783879 | protein_coding | NA | character(0) | character(0) |
| NW_014569547.1 | 127800 | 127800 | 1 | - | 45 | 14 | 31 | 31.1111111111111 | NW_014569547.1 | 57733 | 165446 | 107714 | + | Gnomon | gene | NA | NA | gene-LOC106783879 | GeneID:106783879 | LOC106783879 | Gene | LOC106783879 | protein_coding | NA | character(0) | character(0) |
| NW_014569547.1 | 145879 | 145879 | 1 | - | 31 | 9 | 22 | 29.0322580645161 | NW_014569547.1 | 57733 | 165446 | 107714 | + | Gnomon | gene | NA | NA | gene-LOC106783879 | GeneID:106783879 | LOC106783879 | Gene | LOC106783879 | protein_coding | NA | character(0) | character(0) |
| NW_014569547.1 | 145882 | 145882 | 1 | - | 30 | 7 | 23 | 23.3333333333333 | NW_014569547.1 | 57733 | 165446 | 107714 | + | Gnomon | gene | NA | NA | gene-LOC106783879 | GeneID:106783879 | LOC106783879 | Gene | LOC106783879 | protein_coding | NA | character(0) | character(0) |
| NW_014569547.1 | 145885 | 145885 | 1 | - | 26 | 8 | 18 | 30.7692307692308 | NW_014569547.1 | 57733 | 165446 | 107714 | + | Gnomon | gene | NA | NA | gene-LOC106783879 | GeneID:106783879 | LOC106783879 | Gene | LOC106783879 | protein_coding | NA | character(0) | character(0) |
| NW_014569547.1 | 228148 | 228148 | 1 | + | 50 | 10 | 40 | 20 | NW_014569547.1 | 226579 | 292274 | 65696 | + | Gnomon | gene | NA | NA | gene-LOC106793296 | GeneID:106793296 | LOC106793296 | Gene | LOC106793296 | protein_coding | NA | character(0) | character(0) |
| NW_014569547.1 | 232915 | 232915 | 1 | + | 168 | 13 | 155 | 7.73809523809524 | NW_014569547.1 | 226579 | 292274 | 65696 | + | Gnomon | gene | NA | NA | gene-LOC106793296 | GeneID:106793296 | LOC106793296 | Gene | LOC106793296 | protein_coding | NA | character(0) | character(0) |
| NW_014569547.1 | 247617 | 247617 | 1 | + | 95 | 17 | 78 | 17.8947368421053 | NW_014569547.1 | 226579 | 292274 | 65696 | + | Gnomon | gene | NA | NA | gene-LOC106793296 | GeneID:106793296 | LOC106793296 | Gene | LOC106793296 | protein_coding | NA | character(0) | character(0) |
| NW_014569547.1 | 282304 | 282304 | 1 | + | 134 | 48 | 86 | 35.8208955223881 | NW_014569547.1 | 226579 | 292274 | 65696 | + | Gnomon | gene | NA | NA | gene-LOC106793296 | GeneID:106793296 | LOC106793296 | Gene | LOC106793296 | protein_coding | NA | character(0) | character(0) |
| NW_014569547.1 | 282316 | 282316 | 1 | + | 137 | 85 | 52 | 62.043795620438 | NW_014569547.1 | 226579 | 292274 | 65696 | + | Gnomon | gene | NA | NA | gene-LOC106793296 | GeneID:106793296 | LOC106793296 | Gene | LOC106793296 | protein_coding | NA | character(0) | character(0) |
| NW_014569547.1 | 282334 | 282334 | 1 | + | 137 | 80 | 57 | 58.3941605839416 | NW_014569547.1 | 226579 | 292274 | 65696 | + | Gnomon | gene | NA | NA | gene-LOC106793296 | GeneID:106793296 | LOC106793296 | Gene | LOC106793296 | protein_coding | NA | character(0) | character(0) |
| NW_014569547.1 | 282340 | 282340 | 1 | + | 137 | 96 | 41 | 70.0729927007299 | NW_014569547.1 | 226579 | 292274 | 65696 | + | Gnomon | gene | NA | NA | gene-LOC106793296 | GeneID:106793296 | LOC106793296 | Gene | LOC106793296 | protein_coding | NA | character(0) | character(0) |
| NW_014569547.1 | 282343 | 282343 | 1 | + | 136 | 89 | 47 | 65.4411764705882 | NW_014569547.1 | 226579 | 292274 | 65696 | + | Gnomon | gene | NA | NA | gene-LOC106793296 | GeneID:106793296 | LOC106793296 | Gene | LOC106793296 | protein_coding | NA | character(0) | character(0) |
| NW_014569547.1 | 282346 | 282346 | 1 | + | 133 | 76 | 57 | 57.1428571428571 | NW_014569547.1 | 226579 | 292274 | 65696 | + | Gnomon | gene | NA | NA | gene-LOC106793296 | GeneID:106793296 | LOC106793296 | Gene | LOC106793296 | protein_coding | NA | character(0) | character(0) |
| NW_014569547.1 | 362588 | 362588 | 1 | + | 89 | 29 | 60 | 32.5842696629214 | NW_014569547.1 | 307772 | 417909 | 110138 | - | Gnomon | gene | NA | NA | gene-LOC106785363 | GeneID:106785363 | LOC106785363 | Gene | LOC106785363 | protein_coding | NA | character(0) | character(0) |
| NW_014569547.1 | 362604 | 362604 | 1 | + | 95 | 26 | 69 | 27.3684210526316 | NW_014569547.1 | 307772 | 417909 | 110138 | - | Gnomon | gene | NA | NA | gene-LOC106785363 | GeneID:106785363 | LOC106785363 | Gene | LOC106785363 | protein_coding | NA | character(0) | character(0) |
| NW_014569547.1 | 370455 | 370455 | 1 | + | 93 | 12 | 81 | 12.9032258064516 | NW_014569547.1 | 307772 | 417909 | 110138 | - | Gnomon | gene | NA | NA | gene-LOC106785363 | GeneID:106785363 | LOC106785363 | Gene | LOC106785363 | protein_coding | NA | character(0) | character(0) |
| NW_014569547.1 | 398989 | 398989 | 1 | + | 14 | 7 | 7 | 50 | NW_014569547.1 | 307772 | 417909 | 110138 | - | Gnomon | gene | NA | NA | gene-LOC106785363 | GeneID:106785363 | LOC106785363 | Gene | LOC106785363 | protein_coding | NA | character(0) | character(0) |
| NW_014569547.1 | 519866 | 519866 | 1 | + | 76 | 32 | 44 | 42.1052631578947 | NW_014569547.1 | 517588 | 600078 | 82491 | - | Gnomon | gene | NA | NA | gene-LOC106789119 | GeneID:106789119 | LOC106789119 | Gene | LOC106789119 | protein_coding | NA | character(0) | character(0) |
| NW_014569547.1 | 519882 | 519882 | 1 | + | 85 | 33 | 52 | 38.8235294117647 | NW_014569547.1 | 517588 | 600078 | 82491 | - | Gnomon | gene | NA | NA | gene-LOC106789119 | GeneID:106789119 | LOC106789119 | Gene | LOC106789119 | protein_coding | NA | character(0) | character(0) |
| NW_014569547.1 | 560337 | 560337 | 1 | - | 70 | 11 | 59 | 15.7142857142857 | NW_014569547.1 | 517588 | 600078 | 82491 | - | Gnomon | gene | NA | NA | gene-LOC106789119 | GeneID:106789119 | LOC106789119 | Gene | LOC106789119 | protein_coding | NA | character(0) | character(0) |
| NW_014569547.1 | 762718 | 762718 | 1 | - | 49 | 15 | 34 | 30.6122448979592 | NW_014569547.1 | 692122 | 841696 | 149575 | + | Gnomon | gene | NA | NA | gene-LOC106790448 | GeneID:106790448 | LOC106790448 | Gene | LOC106790448 | protein_coding | NA | character(0) | character(0) |

...
... only first 25 lines shown ...

Directory: RNK File: sites-in-genes-Pc_PA_worker.txt

```
seqnames	start	end	width	strand	coverage	numCs	numTs	perc_meth	region_seqnames	region_start	region_end	region_width	region_strand	region_source
	region_type	region_score	region_phase	region_ID	region_Dbxref	region_Name	region_gbkey	region_gene	region_gene_biotype	region_partial
	region_start_range	region_end_range
NW_014569547.1	99868	99868	1	-	56	12	44	21.4285714285714	NW_014569547.1	57733	165446	107714	+	Gnomon	gene	NA	NA	gene-LOC106783879
	GeneID:106783879	LOC106783879	Gene	LOC106783879	protein_coding	NA	character(0)	character(0)
NW_014569547.1	127778	127778	1	-	24	10	14	41.6666666666667	NW_014569547.1	57733	165446	107714	+	Gnomon	gene	NA	NA	gene-
LOC106783879	GeneID:106783879	LOC106783879	Gene	LOC106783879	protein_coding	NA	character(0)	character(0)
NW_014569547.1	127782	127782	1	-	26	12	14	46.1538461538462	NW_014569547.1	57733	165446	107714	+	Gnomon	gene	NA	NA	gene-
LOC106783879	GeneID:106783879	LOC106783879	Gene	LOC106783879	protein_coding	NA	character(0)	character(0)
NW_014569547.1	127800	127800	1	-	37	12	25	32.4324324324324	NW_014569547.1	57733	165446	107714	+	Gnomon	gene	NA	NA	gene-
LOC106783879	GeneID:106783879	LOC106783879	Gene	LOC106783879	protein_coding	NA	character(0)	character(0)
NW_014569547.1	220042	220042	1	-	64	11	53	17.1875	NW_014569547.1	214575	226272	11698	+	Gnomon	gene	NA	NA	gene-
LOC106793447	GeneID:106793447	LOC106793447	Gene	LOC106793447	lncRNA	NA	character(0)	character(0)
NW_014569547.1	228148	228148	1	+	43	11	32	25.5813953488372	NW_014569547.1	226579	292274	65696	+	Gnomon	gene	NA	NA	gene-
LOC106793296	GeneID:106793296	LOC106793296	Gene	LOC106793296	protein_coding	NA	character(0)	character(0)
NW_014569547.1	247617	247617	1	+	116	11	105	9.48275862068965	NW_014569547.1	226579	292274	65696	+	Gnomon	gene	NA	NA	gene-
LOC106793296	GeneID:106793296	LOC106793296	Gene	LOC106793296	protein_coding	NA	character(0)	character(0)
NW_014569547.1	282304	282304	1	+	134	64	70	47.7611940298507	NW_014569547.1	226579	292274	65696	+	Gnomon	gene	NA	NA	gene-
LOC106793296	GeneID:106793296	LOC106793296	Gene	LOC106793296	protein_coding	NA	character(0)	character(0)
NW_014569547.1	282316	282316	1	+	145	106	39	73.1034482758621	NW_014569547.1	226579	292274	65696	+	Gnomon	gene	NA	NA	gene-
LOC106793296	GeneID:106793296	LOC106793296	Gene	LOC106793296	protein_coding	NA	character(0)	character(0)
NW_014569547.1	282328	282328	1	+	151	17	134	11.2582781456954	NW_014569547.1	226579	292274	65696	+	Gnomon	gene	NA	NA	gene-
LOC106793296	GeneID:106793296	LOC106793296	Gene	LOC106793296	protein_coding	NA	character(0)	character(0)
NW_014569547.1	282334	282334	1	+	148	83	65	56.0810810810811	NW_014569547.1	226579	292274	65696	+	Gnomon	gene	NA	NA	gene-
LOC106793296	GeneID:106793296	LOC106793296	Gene	LOC106793296	protein_coding	NA	character(0)	character(0)
NW_014569547.1	282340	282340	1	+	144	102	42	70.8333333333333	NW_014569547.1	226579	292274	65696	+	Gnomon	gene	NA	NA	gene-
LOC106793296	GeneID:106793296	LOC106793296	Gene	LOC106793296	protein_coding	NA	character(0)	character(0)
NW_014569547.1	282343	282343	1	+	142	107	35	75.3521126760563	NW_014569547.1	226579	292274	65696	+	Gnomon	gene	NA	NA	gene-
LOC106793296	GeneID:106793296	LOC106793296	Gene	LOC106793296	protein_coding	NA	character(0)	character(0)
NW_014569547.1	282346	282346	1	+	144	98	46	68.0555555555556	NW_014569547.1	226579	292274	65696	+	Gnomon	gene	NA	NA	gene-
LOC106793296	GeneID:106793296	LOC106793296	Gene	LOC106793296	protein_coding	NA	character(0)	character(0)
NW_014569547.1	362588	362588	1	+	90	40	50	44.4444444444444	NW_014569547.1	307772	417909	110138	-	Gnomon	gene	NA	NA
	gene-LOC106785363	GeneID:106785363	LOC106785363	Gene	LOC106785363	protein_coding	NA	character(0)	character(0)
NW_014569547.1	362604	362604	1	+	94	35	59	37.2340425531915	NW_014569547.1	307772	417909	110138	-	Gnomon	gene	NA	NA
	gene-LOC106785363	GeneID:106785363	LOC106785363	Gene	LOC106785363	protein_coding	NA	character(0)	character(0)
NW_014569547.1	382379	382379	1	-	86	11	75	12.7906976744186	NW_014569547.1	307772	417909	110138	-	Gnomon	gene	NA	NA
	gene-LOC106785363	GeneID:106785363	LOC106785363	Gene	LOC106785363	protein_coding	NA	character(0)	character(0)
NW_014569547.1	519866	519866	1	+	58	19	39	32.7586206896552	NW_014569547.1	517588	600078	82491	-	Gnomon	gene	NA	NA	gene-
LOC106789119	GeneID:106789119	LOC106789119	Gene	LOC106789119	protein_coding	NA	character(0)	character(0)
NW_014569547.1	519882	519882	1	+	70	20	50	28.5714285714286	NW_014569547.1	517588	600078	82491	-	Gnomon	gene	NA	NA	gene-
LOC106789119	GeneID:106789119	LOC106789119	Gene	LOC106789119	protein_coding	NA	character(0)	character(0)
NW_014569547.1	560245	560245	1	-	80	9	71	11.25	NW_014569547.1	517588	600078	82491	-	Gnomon	gene	NA	NA	gene-LOC106789119
	GeneID:106789119	LOC106789119	Gene	LOC106789119	protein_coding	NA	character(0)	character(0)
NW_014569547.1	762718	762718	1	-	42	10	32	23.8095238095238	NW_014569547.1	692122	841696	149575	+	Gnomon	gene	NA	NA
	gene-LOC106790448	GeneID:106790448	LOC106790448	Gene	LOC106790448	protein_coding	NA	character(0)	character(0)
NW_014569547.1	762722	762722	1	-	43	9	34	20.9302325581395	NW_014569547.1	692122	841696	149575	+	Gnomon	gene	NA	NA
	gene-LOC106790448	GeneID:106790448	LOC106790448	Gene	LOC106790448	protein_coding	NA	character(0)	character(0)
NW_014569547.1	762728	762728	1	-	45	13	32	28.8888888888889	NW_014569547.1	692122	841696	149575	+	Gnomon	gene	NA	NA
	gene-LOC106790448	GeneID:106790448	LOC106790448	Gene	LOC106790448	protein_coding	NA	character(0)	character(0)
NW_014569547.1	762732	762732	1	-	48	11	37	22.9166666666667	NW_014569547.1	692122	841696	149575	+	Gnomon	gene	NA	NA
	gene-LOC106790448	GeneID:106790448	LOC106790448	Gene	LOC106790448	protein_coding	NA	character(0)	character(0)
...
... only first 25 lines shown ...
```

Directory: RNK File: sites-in-promoters-Pc_PA_queen.txt

| seqnames | start | end | width | strand | coverage | numCs | numTs | perc_meth | region_seqnames | region_start | region_end | region_width | region_strand | region_source | region_type | region_score | region_phase | region_ID | region_Dbxref | region_Name | region_gbkey | region_gene | region_gene_biotype | region_partial | region_start_range | region_end_range |
|---|---|---|---|---|---|---|---|---|---|---|---|---|---|---|---|---|---|---|---|---|---|---|---|---|---|---|
| NW_014569548.1 | 1791722 | 1791722 | 1 | + | 97 | 13 | 84 | 13.4020618556701 | NW_014569548.1 | 1791422 | 1791921 | 500 | + | Gnomon | promoter | NA | NA | gene-LOC106793622 | GeneID:106793622 | LOC106793622 | Gene | LOC106793622 | protein_coding | NA | character(0) | character(0) |
| NW_014569549.1 | 371056 | 371056 | 1 | + | 235 | 21 | 214 | 8.93617021276596 | NW_014569549.1 | 370564 | 371063 | 500 | + | Gnomon | promoter | NA | NA | gene-LOC106785315 | GeneID:106785315 | LOC106785315 | Gene | LOC106785315 | protein_coding | true | c(".", "371064") | character(0) |
| NW_014569549.1 | 603869 | 603869 | 1 | + | 111 | 18 | 93 | 16.2162162162162 | NW_014569549.1 | 603372 | 603871 | 500 | + | Gnomon | promoter | NA | NA | gene-LOC106785021 | GeneID:106785021 | LOC106785021 | Gene | LOC106785021 | protein_coding | NA | character(0) | character(0) |
| NW_014569549.1 | 2124086 | 2124086 | 1 | + | 11 | 9 | 2 | 81.8181818181818 | NW_014569549.1 | 2123847 | 2124346 | 500 | + | Gnomon | promoter | NA | NA | gene-LOC106785085 | GeneID:106785085 | LOC106785085 | Gene | LOC106785085 | protein_coding | NA | character(0) | character(0) |
| NW_014569549.1 | 2124105 | 2124105 | 1 | + | 15 | 9 | 6 | 60 | NW_014569549.1 | 2123847 | 2124346 | 500 | + | Gnomon | promoter | NA | NA | gene-LOC106785085 | GeneID:106785085 | LOC106785085 | Gene | LOC106785085 | protein_coding | NA | character(0) | character(0) |
| NW_014569549.1 | 2124138 | 2124138 | 1 | + | 18 | 18 | 0 | 100 | NW_014569549.1 | 2123847 | 2124346 | 500 | + | Gnomon | promoter | NA | NA | gene-LOC106785085 | GeneID:106785085 | LOC106785085 | Gene | LOC106785085 | protein_coding | NA | character(0) | character(0) |
| NW_014569549.1 | 2124160 | 2124160 | 1 | + | 20 | 17 | 3 | 85 | NW_014569549.1 | 2123847 | 2124346 | 500 | + | Gnomon | promoter | NA | NA | gene-LOC106785085 | GeneID:106785085 | LOC106785085 | Gene | LOC106785085 | protein_coding | NA | character(0) | character(0) |
| NW_014569550.1 | 1117758 | 1117758 | 1 | - | 52 | 35 | 17 | 67.3076923076923 | NW_014569550.1 | 1117335 | 1117834 | 500 | - | Gnomon | promoter | NA | NA | gene-LOC106786084 | GeneID:106786084 | LOC106786084 | Gene | LOC106786084 | protein_coding | NA | character(0) | character(0) |
| NW_014569550.1 | 1117764 | 1117764 | 1 | - | 54 | 35 | 19 | 64.8148148148148 | NW_014569550.1 | 1117335 | 1117834 | 500 | - | Gnomon | promoter | NA | NA | gene-LOC106786084 | GeneID:106786084 | LOC106786084 | Gene | LOC106786084 | protein_coding | NA | character(0) | character(0) |
| NW_014569550.1 | 1117768 | 1117768 | 1 | - | 54 | 19 | 35 | 35.1851851851852 | NW_014569550.1 | 1117335 | 1117834 | 500 | - | Gnomon | promoter | NA | NA | gene-LOC106786084 | GeneID:106786084 | LOC106786084 | Gene | LOC106786084 | protein_coding | NA | character(0) | character(0) |
| NW_014569550.1 | 1942811 | 1942811 | 1 | - | 37 | 8 | 29 | 21.6216216216216 | NW_014569550.1 | 1942801 | 1943300 | 500 | - | Gnomon | promoter | NA | NA | gene-LOC106786379 | GeneID:106786379 | LOC106786379 | Gene | LOC106786379 | protein_coding | NA | character(0) | character(0) |
| NW_014569551.1 | 1621261 | 1621261 | 1 | - | 117 | 10 | 107 | 8.54700854700855 | NW_014569551.1 | 1621220 | 1621719 | 500 | - | Gnomon | promoter | NA | NA | gene-LOC106787384 | GeneID:106787384 | LOC106787384 | Gene | LOC106787384 | lncRNA | NA | character(0) | character(0) |
| NW_014569551.1 | 2222568 | 2222568 | 1 | + | 44 | 19 | 25 | 43.1818181818182 | NW_014569551.1 | 2222289 | 2222788 | 500 | + | Gnomon | promoter | NA | NA | gene-LOC106787745 | GeneID:106787745 | LOC106787745 | Gene | LOC106787745 | protein_coding | NA | character(0) | character(0) |
| NW_014569551.1 | 2222569 | 2222569 | 1 | - | 20 | 9 | 11 | 45 | NW_014569551.1 | 2222289 | 2222788 | 500 | + | Gnomon | promoter | NA | NA | gene-LOC106787745 | GeneID:106787745 | LOC106787745 | Gene | LOC106787745 | protein_coding | NA | character(0) | character(0) |
| NW_014569551.1 | 2222622 | 2222622 | 1 | + | 33 | 18 | 15 | 54.5454545454545 | NW_014569551.1 | 2222289 | 2222788 | 500 | + | Gnomon | promoter | NA | NA | gene-LOC106787745 | GeneID:106787745 | LOC106787745 | Gene | LOC106787745 | protein_coding | NA | character(0) | character(0) |
| NW_014569551.1 | 2222623 | 2222623 | 1 | - | 25 | 10 | 15 | 40 | NW_014569551.1 | 2222289 | 2222788 | 500 | + | Gnomon | promoter | NA | NA | gene-LOC106787745 | GeneID:106787745 | LOC106787745 | Gene | LOC106787745 | protein_coding | NA | character(0) | character(0) |
| NW_014569551.1 | 2222642 | 2222642 | 1 | + | 25 | 9 | 16 | 36 | NW_014569551.1 | 2222289 | 2222788 | 500 | + | Gnomon | promoter | NA | NA | gene-LOC106787745 | GeneID:106787745 | LOC106787745 | Gene | LOC106787745 | protein_coding | NA | character(0) | character(0) |
| NW_014569552.1 | 513346 | 513346 | 1 | - | 34 | 7 | 27 | 20.5882352941176 | NW_014569552.1 | 513121 | 513620 | 500 | + | Gnomon | promoter | NA | NA | gene-LOC106788418 | GeneID:106788418 | LOC106788418 | Gene | LOC106788418 | protein_coding | NA | character(0) | character(0) |
| NW_014569553.1 | 1499839 | 1499839 | 1 | - | 27 | 7 | 20 | 25.9259259259259 | NW_014569553.1 | 1499503 | 1500002 | 500 | + | Gnomon | promoter | NA | NA | gene-LOC106789631 | GeneID:106789631 | LOC106789631 | Gene | LOC106789631 | protein_coding | NA | character(0) | character(0) |
| NW_014569554.1 | 889559 | 889559 | 1 | - | 152 | 25 | 127 | 16.4473684210526 | NW_014569554.1 | 889524 | 890023 | 500 | - | Gnomon | promoter | NA | NA | gene-LOC106790827 | GeneID:106790827 | LOC106790827 | Gene | LOC106790827 | protein_coding | NA | character(0) | character(0) |
| NW_014569555.1 | 830345 | 830345 | 1 | + | 158 | 12 | 146 | 7.59493670886076 | NW_014569555.1 | 830071 | 830570 | 500 | + | Gnomon | promoter | NA | NA | gene-LOC106792003 | GeneID:106792003 | LOC106792003 | Gene | LOC106792003 | lncRNA | NA | character(0) | character(0) |
| NW_014569558.1 | 935120 | 935120 | 1 | - | 99 | 13 | 86 | 13.1313131313131 | NW_014569558.1 | 934812 | 935311 | 500 | - | Gnomon | promoter | NA | NA | gene-LOC106793074 | GeneID:106793074 | LOC106793074 | Gene | LOC106793074 | protein_coding | NA | character(0) | character(0) |
| NW_014569558.1 | 935132 | 935132 | 1 | - | 100 | 22 | 78 | 22 | NW_014569558.1 | 934812 | 935311 | 500 | - | Gnomon | promoter | NA | NA | gene-LOC106793074 | GeneID:106793074 | LOC106793074 | Gene | LOC106793074 | protein_coding | NA | character(0) | character(0) |
| NW_014569558.1 | 935136 | 935136 | 1 | - | 101 | 22 | 79 | 21.7821782178218 | NW_014569558.1 | 934812 | 935311 | 500 | - | Gnomon | promoter | NA | NA | gene-LOC106793074 | GeneID:106793074 | LOC106793074 | Gene | LOC106793074 | protein_coding | NA | character(0) | character(0) |

...

... only first 25 lines shown ...

Directory: RNK File: sites-in-promoters-Pc_PA_worker.txt

| seqnames | start | end | width | strand | coverage | numCs | numTs | perc_meth | region_seqnames | region_start | region_end | region_width | region_strand | region_source | region_type | region_score | region_phase | region_ID | region_Dbxref | region_Name | region_gbkey | region_gene | region_gene_biotype | region_partial | region_start_range | region_end_range |
|---|---|---|---|---|---|---|---|---|---|---|---|---|---|---|---|---|---|---|---|---|---|---|---|---|---|---|
| NW_014569547.1 | 1301946 | 1301946 | 1 | + | 37 | 7 | 30 | 18.9189189189189 | NW_014569547.1 | 1301789 | 1302288 | 500 | + | Gnomon | promoter | NA | NA | gene-LOC106792589 | GeneID:106792589 | LOC106792589 | Gene | LOC106792589 | protein_coding | NA | character(0) | character(0) |
| NW_014569548.1 | 1791711 | 1791711 | 1 | + | 103 | 16 | 87 | 15.5339805825243 | NW_014569548.1 | 1791422 | 1791921 | 500 | + | Gnomon | promoter | NA | NA | gene-LOC106793622 | GeneID:106793622 | LOC106793622 | Gene | LOC106793622 | protein_coding | NA | character(0) | character(0) |
| NW_014569549.1 | 320105 | 320105 | 1 | - | 190 | 15 | 175 | 7.89473684210526 | NW_014569549.1 | 319635 | 320134 | 500 | - | Gnomon | promoter | NA | NA | gene-LOC106784646 | GeneID:106784646 | LOC106784646 | Gene | LOC106784646 | protein_coding | NA | character(0) | character(0) |
| NW_014569549.1 | 371056 | 371056 | 1 | + | 233 | 18 | 215 | 7.72532188841202 | NW_014569549.1 | 370564 | 371063 | 500 | + | Gnomon | promoter | NA | NA | gene-LOC106785315 | GeneID:106785315 | LOC106785315 | Gene | LOC106785315 | protein_coding | true | c(".", "371064") | character(0) |
| NW_014569549.1 | 603869 | 603869 | 1 | + | 101 | 17 | 84 | 16.8316831683168 | NW_014569549.1 | 603372 | 603871 | 500 | + | Gnomon | promoter | NA | NA | gene-LOC106785021 | GeneID:106785021 | LOC106785021 | Gene | LOC106785021 | protein_coding | NA | character(0) | character(0) |
| NW_014569549.1 | 1284586 | 1284586 | 1 | - | 121 | 19 | 102 | 15.702479338843 | NW_014569549.1 | 1284120 | 1284619 | 500 | - | Gnomon | promoter | NA | NA | gene-LOC106784946 | GeneID:106784946 | LOC106784946 | Gene | LOC106784946 | protein_coding | NA | character(0) | character(0) |
| NW_014569549.1 | 2124086 | 2124086 | 1 | + | 10 | 6 | 4 | 60 | NW_014569549.1 | 2123847 | 2124346 | 500 | + | Gnomon | promoter | NA | NA | gene-LOC106785085 | GeneID:106785085 | LOC106785085 | Gene | LOC106785085 | protein_coding | NA | character(0) | character(0) |
| NW_014569549.1 | 2124105 | 2124105 | 1 | + | 13 | 6 | 7 | 46.1538461538462 | NW_014569549.1 | 2123847 | 2124346 | 500 | + | Gnomon | promoter | NA | NA | gene-LOC106785085 | GeneID:106785085 | LOC106785085 | Gene | LOC106785085 | protein_coding | NA | character(0) | character(0) |
| NW_014569549.1 | 2124138 | 2124138 | 1 | + | 13 | 12 | 1 | 92.3076923076923 | NW_014569549.1 | 2123847 | 2124346 | 500 | + | Gnomon | promoter | NA | NA | gene-LOC106785085 | GeneID:106785085 | LOC106785085 | Gene | LOC106785085 | protein_coding | NA | character(0) | character(0) |
| NW_014569549.1 | 2124160 | 2124160 | 1 | + | 13 | 9 | 4 | 69.2307692307692 | NW_014569549.1 | 2123847 | 2124346 | 500 | + | Gnomon | promoter | NA | NA | gene-LOC106785085 | GeneID:106785085 | LOC106785085 | Gene | LOC106785085 | protein_coding | NA | character(0) | character(0) |
| NW_014569550.1 | 751585 | 751585 | 1 | - | 144 | 12 | 132 | 8.33333333333333 | NW_014569550.1 | 751495 | 751994 | 500 | - | Gnomon | promoter | NA | NA | gene-LOC106786586 | GeneID:106786586 | LOC106786586 | Gene | LOC106786586 | protein_coding | NA | character(0) | character(0) |
| NW_014569550.1 | 1117758 | 1117758 | 1 | - | 39 | 25 | 14 | 64.1025641025641 | NW_014569550.1 | 1117335 | 1117834 | 500 | - | Gnomon | promoter | NA | NA | gene-LOC106786084 | GeneID:106786084 | LOC106786084 | Gene | LOC106786084 | protein_coding | NA | character(0) | character(0) |
| NW_014569550.1 | 1117764 | 1117764 | 1 | - | 40 | 22 | 18 | 55 | NW_014569550.1 | 1117335 | 1117834 | 500 | - | Gnomon | promoter | NA | NA | gene-LOC106786084 | GeneID:106786084 | LOC106786084 | Gene | LOC106786084 | protein_coding | NA | character(0) | character(0) |
| NW_014569550.1 | 1117768 | 1117768 | 1 | - | 41 | 17 | 24 | 41.4634146341463 | NW_014569550.1 | 1117335 | 1117834 | 500 | - | Gnomon | promoter | NA | NA | gene-LOC106786084 | GeneID:106786084 | LOC106786084 | Gene | LOC106786084 | protein_coding | NA | character(0) | character(0) |
| NW_014569550.1 | 1942811 | 1942811 | 1 | - | 40 | 13 | 27 | 32.5 | NW_014569550.1 | 1942801 | 1943300 | 500 | - | Gnomon | promoter | NA | NA | gene-LOC106786379 | GeneID:106786379 | LOC106786379 | Gene | LOC106786379 | protein_coding | NA | character(0) | character(0) |
| NW_014569551.1 | 2089009 | 2089009 | 1 | + | 172 | 15 | 157 | 8.72093023255814 | NW_014569551.1 | 2088589 | 2089088 | 500 | + | Gnomon | promoter | NA | NA | gene-LOC106787770 | GeneID:106787770 | LOC106787770 | Gene | LOC106787770 | protein_coding | NA | character(0) | character(0) |
| NW_014569551.1 | 2222544 | 2222544 | 1 | + | 26 | 7 | 19 | 26.9230769230769 | NW_014569551.1 | 2222289 | 2222788 | 500 | + | Gnomon | promoter | NA | NA | gene-LOC106787745 | GeneID:106787745 | LOC106787745 | Gene | LOC106787745 | protein_coding | NA | character(0) | character(0) |
| NW_014569551.1 | 2222548 | 2222548 | 1 | + | 26 | 8 | 18 | 30.7692307692308 | NW_014569551.1 | 2222289 | 2222788 | 500 | + | Gnomon | promoter | NA | NA | gene-LOC106787745 | GeneID:106787745 | LOC106787745 | Gene | LOC106787745 | protein_coding | NA | character(0) | character(0) |
| NW_014569551.1 | 2222568 | 2222568 | 1 | + | 32 | 14 | 18 | 43.75 | NW_014569551.1 | 2222289 | 2222788 | 500 | + | Gnomon | promoter | NA | NA | gene-LOC106787745 | GeneID:106787745 | LOC106787745 | Gene | LOC106787745 | protein_coding | NA | character(0) | character(0) |
| NW_014569551.1 | 2222569 | 2222569 | 1 | - | 19 | 10 | 9 | 52.6315789473684 | NW_014569551.1 | 2222289 | 2222788 | 500 | + | Gnomon | promoter | NA | NA | gene-LOC106787745 | GeneID:106787745 | LOC106787745 | Gene | LOC106787745 | protein_coding | NA | character(0) | character(0) |
| NW_014569551.1 | 2222622 | 2222622 | 1 | + | 26 | 12 | 14 | 46.1538461538462 | NW_014569551.1 | 2222289 | 2222788 | 500 | + | Gnomon | promoter | NA | NA | gene-LOC106787745 | GeneID:106787745 | LOC106787745 | Gene | LOC106787745 | protein_coding | NA | character(0) | character(0) |
| NW_014569551.1 | 2222623 | 2222623 | 1 | - | 24 | 12 | 12 | 50 | NW_014569551.1 | 2222289 | 2222788 | 500 | + | Gnomon | promoter | NA | NA | gene-LOC106787745 | GeneID:106787745 | LOC106787745 | Gene | LOC106787745 | protein_coding | NA | character(0) | character(0) |
| NW_014569551.1 | 2222642 | 2222642 | 1 | + | 21 | 10 | 11 | 47.6190476190476 | NW_014569551.1 | 2222289 | 2222788 | 500 | + | Gnomon | promoter | NA | NA | gene-LOC106787745 | GeneID:106787745 | LOC106787745 | Gene | LOC106787745 | protein_coding | NA | character(0) | character(0) |
| NW_014569551.1 | 2222646 | 2222646 | 1 | + | 18 | 8 | 10 | 44.4444444444444 | NW_014569551.1 | 2222289 | 2222788 | 500 | + | Gnomon | promoter | NA | NA | gene-LOC106787745 | GeneID:106787745 | LOC106787745 | Gene | LOC106787745 | protein_coding | NA | character(0) | character(0) |

...
... only first 25 lines shown ...

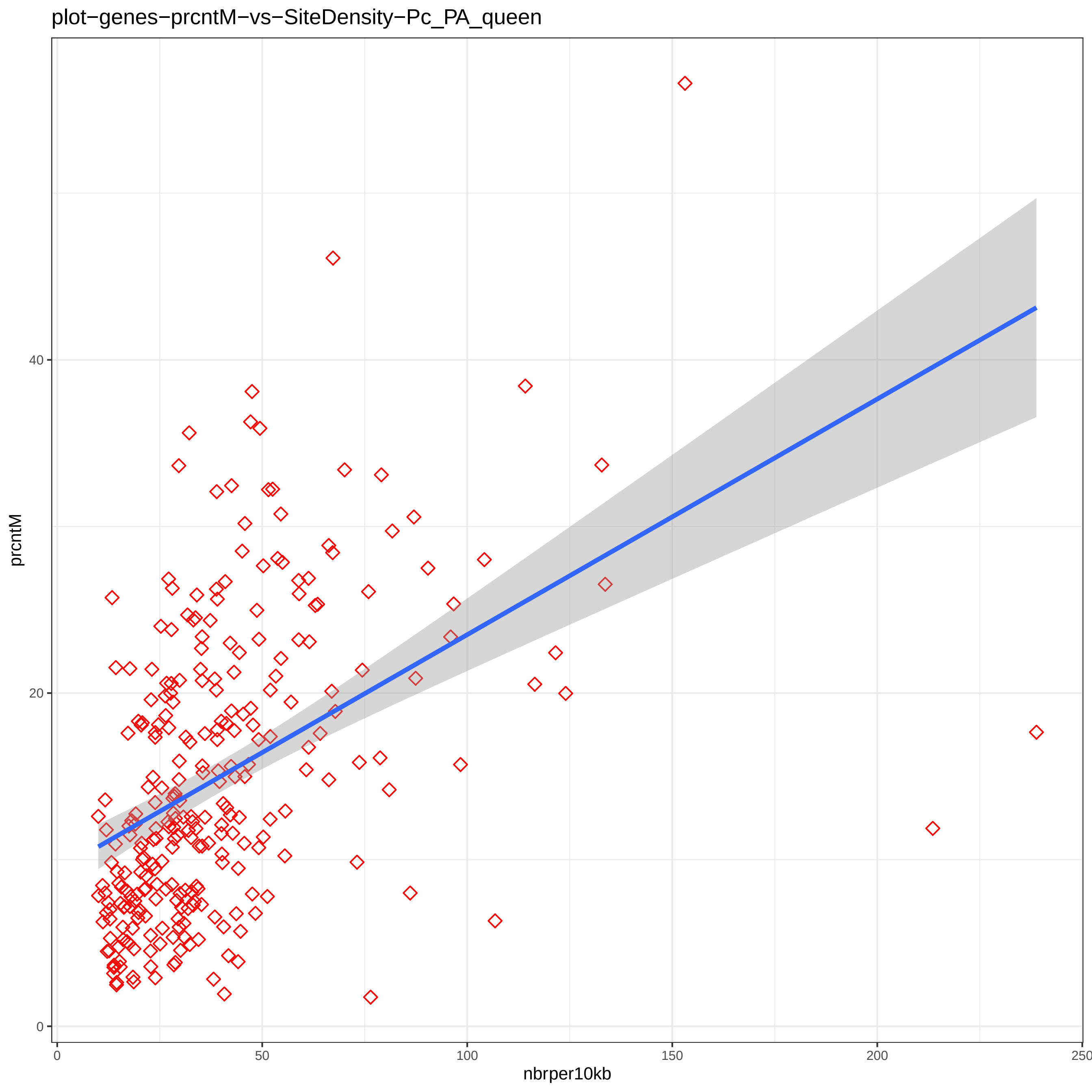
plot−genes−prcntM−vs−SiteDensity−Pc_PA_queen
prcntM
0
20
40
nbrper10kb
0
50
100
150
200
250

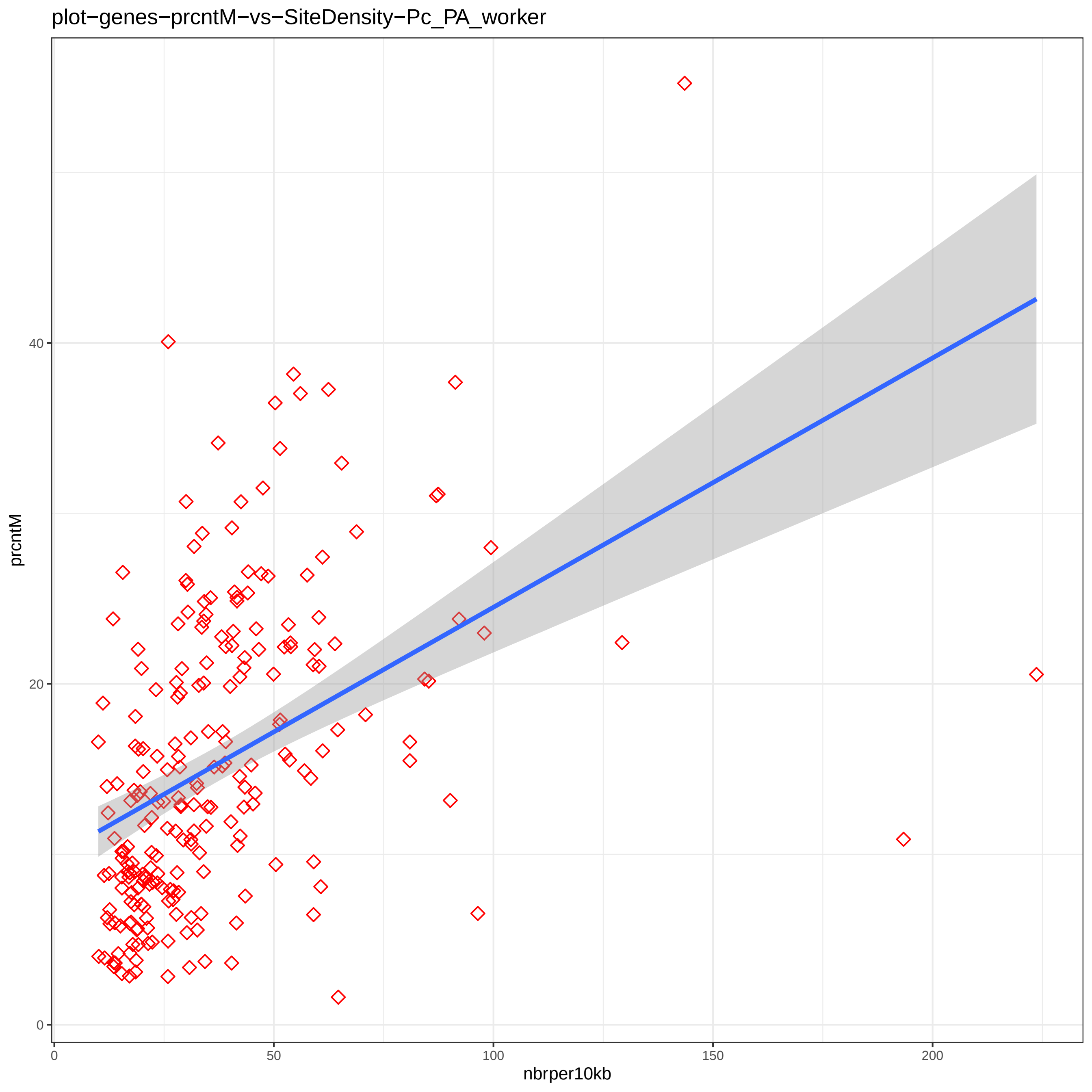
plot−genes−prcntM−vs−SiteDensity−Pc_PA_worker
prcntM
nbrper10kb
0
20
40
0
50
100
150
200

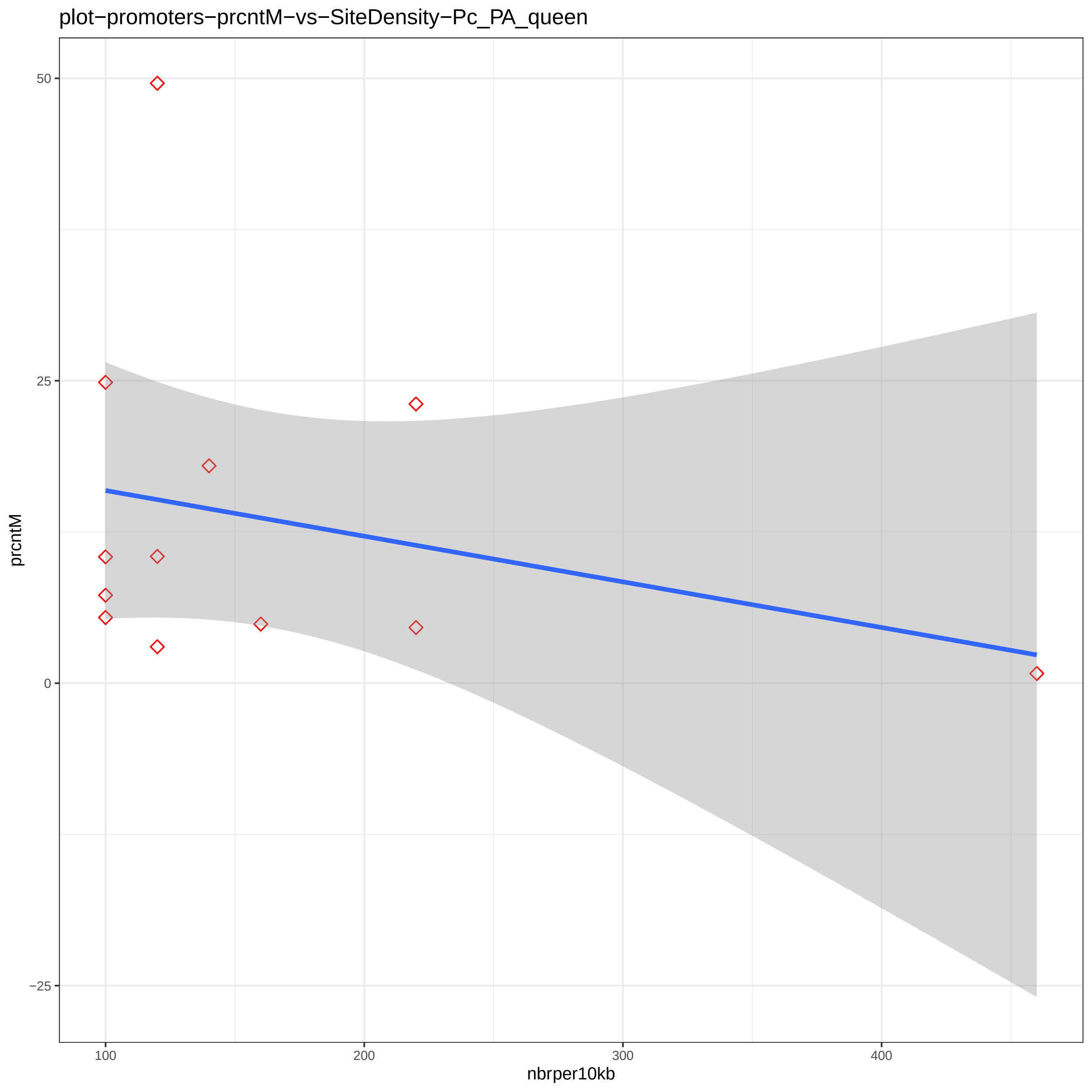

plot−promoters−prcntM−vs−SiteDensity−Pc_PA_queen
prcntM
50
25
0
−25
100
200
300
400
nbrper10kb

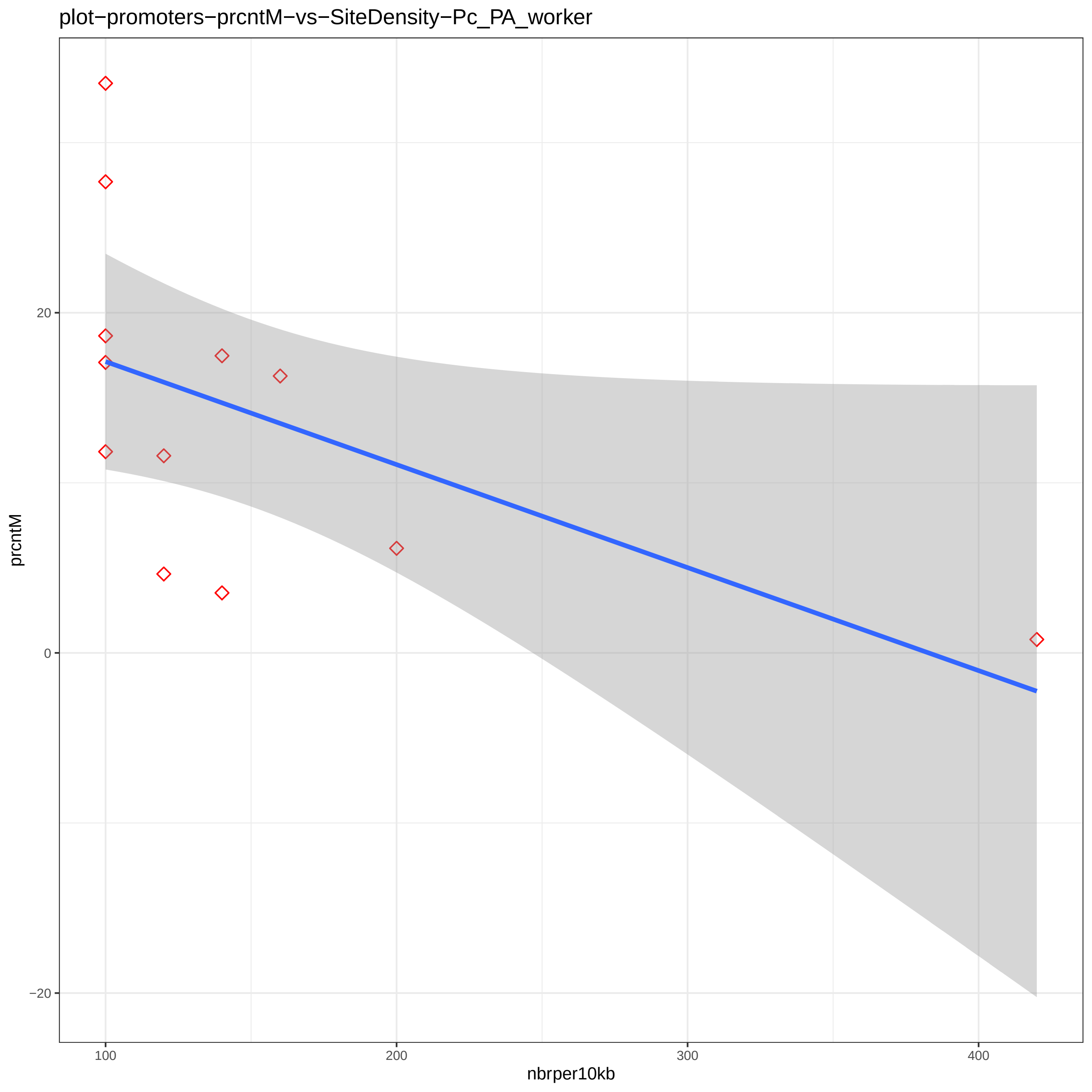
plot−promoters−prcntM−vs−SiteDensity−Pc_PA_worker
prcntM
20
0
−20
100
200
300
400
nbrper10kb

```
Directory: MRPR File: 0READMEmrpr


MRPR - Methylation-rich and -poor regions.

Input: studymk, genome_ann (a list of GRanges objects providing annotated region labels and bounds;
                             output of BWASPR::get_genome_annotation())
       nbrxtrms (specifies the number of richest and poorest regions to be displayed; default: 100)

Output: files dst-*.txt 1ds-*.pdf 5ds-*.pdf    mdr-*.bed   rmp-* gwp-* gwr-*

Notes: The output is generated by BWASPR functions det_mrpr() and map_mrpr().
       Methylation-rich and -poor regions are determined based on the spacing between
        neighboring CpGhsm sites.  If sites occur at positions a, b, c, d, e, and f
        (and not in between), then b-a, c-b, ... are 1-distances, and f-a is a 5-distance.
        The file dst-*.txt shows the empirical distribution of d-distances for the sample.
        Methylation-poor regions are determined as long 1-distances (in the top <nbrxtrms>; merged
        if adjacent 1-distances are both in the top <nbrxtrms>).  Methylation-rich regions are determined
        as short 5-distances (in the low <nbrxtrms>; merged if adjacent 5-distances are both in the low
        <nbrxtrms>).  Parts of the distribution are plotted in the 1ds-*.pdf and 5ds-*.pdf files.

       The methylation-rich and -poor regions are listed in files dst-*.txt and mdr-*.bed,
        the latter in BED format for potential display in genome browsers.

       The overlap of methylation regions with genome features is summarized in files rmp-*.txt.
        Files gwr-*.txt show genes overlapping with methylation-rich regions, ordered by site
        density in the methylation-rich region.
        Files gwp-*.txt show genes overlapping with methylation-poor regions, ordered by site
        density in the methylation-poor region.
```

Directory: MRPR File: dst-Pc_PA_qn.txt

Analysis of 1-distances for sample "Pc_PA_qn" (13840 sites):

Total number of 1-distances: 12898
Median: 36.0 Mean: 10631.2 Std: 29821.3
Quantiles:

| 0% | 5% | 10% | 15% | 20% | 25% | 30% | 35% | 40% | 45% | 50% | 55% | 60% | 65% | 70% | 75% | 80% | 85% | 90% | 95% | 100% |
|---|---|---|---|---|---|---|---|---|---|---|---|---|---|---|---|---|---|---|---|---|
| 1 | 1 | 2 | 3 | 5 | 7 | 11 | 16 | 22 | 28 | 36 | 57 | 101 | 177 | 486 | 2954 | 8831 | 19165 | 35099 | 66084 | 458452 |

Low density region distance cutoff (lowest 100): 156518
High density region distance cutoff (highest 100): 1

Ordered lists of methylation-poor regions:

Top 100 (out of 89) methylation-poor regions in "Pc_PA_qn":
(Sdnsty = sites per 1kb)

| | Rtype | Sample | SeqID | From | To | Rlgth | NbrSites | Sdnsty |
|---|---|---|---|---|---|---|---|---|
| 1 | Poor | queen | NW_014569596.1 | 107147 | 565599 | 458453 | 2 | 0.00436 |
| 2 | Poor | queen | NW_014569579.1 | 83024 | 523197 | 440174 | 2 | 0.00454 |
| 3 | Poor | queen | NW_014569611.1 | 283276 | 700510 | 417235 | 2 | 0.00479 |
| 4 | Poor | queen | NW_014569571.1 | 725749 | 1142276 | 416528 | 2 | 0.00480 |
| 5 | Poor | queen | NW_014569549.1 | 1460342 | 2057028 | 596687 | 3 | 0.00503 |
| 6 | Poor | queen | NW_014569554.1 | 1423594 | 1801461 | 377868 | 2 | 0.00529 |
| 7 | Poor | queen | NW_014569599.1 | 363927 | 730170 | 366244 | 2 | 0.00546 |
| 8 | Poor | queen | NW_014569590.1 | 476592 | 837225 | 360634 | 2 | 0.00555 |
| 9 | Poor | queen | NW_014569671.1 | 4141 | 335853 | 331713 | 2 | 0.00603 |
| 10 | Poor | queen | NW_014569553.1 | 1547908 | 1873709 | 325802 | 2 | 0.00614 |
| 11 | Poor | queen | NW_014569553.1 | 495395 | 815513 | 320119 | 2 | 0.00625 |
| 12 | Poor | queen | NW_014569612.1 | 172316 | 489724 | 317409 | 2 | 0.00630 |
| 13 | Poor | queen | NW_014569585.1 | 124812 | 578186 | 453375 | 3 | 0.00662 |
| 14 | Poor | queen | NW_014569573.1 | 412311 | 1012953 | 600643 | 4 | 0.00666 |
| 15 | Poor | queen | NW_014569552.1 | 1628763 | 2077304 | 448542 | 3 | 0.00669 |
| 16 | Poor | queen | NW_014569639.1 | 179565 | 476011 | 296447 | 2 | 0.00675 |
| 17 | Poor | queen | NW_014569563.1 | 807504 | 1228194 | 420691 | 3 | 0.00713 |
| 18 | Poor | queen | NW_014569735.1 | 795 | 269998 | 269204 | 2 | 0.00743 |
| 19 | Poor | queen | NW_014569551.1 | 478719 | 746954 | 268236 | 2 | 0.00746 |
| 20 | Poor | queen | NW_014569549.1 | 1072665 | 1333083 | 260419 | 2 | 0.00768 |
| 21 | Poor | queen | NW_014569651.1 | 35133 | 421453 | 386321 | 3 | 0.00777 |
| 22 | Poor | queen | NW_014569550.1 | 732921 | 1117758 | 384838 | 3 | 0.00780 |
| 23 | Poor | queen | NW_014569596.1 | 595137 | 851082 | 255946 | 2 | 0.00781 |
| 24 | Poor | queen | NW_014569594.1 | 59065 | 314253 | 255189 | 2 | 0.00784 |
| 25 | Poor | queen | NW_014569572.1 | 173801 | 427777 | 253977 | 2 | 0.00787 |
| 26 | Poor | queen | NW_014569562.1 | 546593 | 918826 | 372234 | 3 | 0.00806 |
| 27 | Poor | queen | NW_014569587.1 | 386650 | 758674 | 372025 | 3 | 0.00806 |
| 28 | Poor | queen | NW_014569620.1 | 963 | 243732 | 242770 | 2 | 0.00824 |
| 29 | Poor | queen | NW_014569600.1 | 493239 | 732502 | 239264 | 2 | 0.00836 |
| 30 | Poor | queen | NW_014569569.1 | 477917 | 714611 | 236695 | 2 | 0.00845 |
| 31 | Poor | queen | NW_014569549.1 | 2247823 | 2483498 | 235676 | 2 | 0.00849 |
| 32 | Poor | queen | NW_014569551.1 | 213157 | 447988 | 234832 | 2 | 0.00852 |
| 33 | Poor | queen | NW_014569591.1 | 721675 | 953637 | 231963 | 2 | 0.00862 |
| 34 | Poor | queen | NW_014569635.1 | 120005 | 351824 | 231820 | 2 | 0.00863 |
| 35 | Poor | queen | NW_014569551.1 | 791909 | 1133085 | 341177 | 3 | 0.00879 |
| 36 | Poor | queen | NW_014569557.1 | 362770 | 586992 | 224223 | 2 | 0.00892 |
| 37 | Poor | queen | NW_014569602.1 | 510697 | 734891 | 224195 | 2 | 0.00892 |
| 38 | Poor | queen | NW_014569550.1 | 1967141 | 2190487 | 223347 | 2 | 0.00895 |
| 39 | Poor | queen | NW_014569570.1 | 826534 | 1048535 | 222002 | 2 | 0.00901 |
| 40 | Poor | queen | NW_014569550.1 | 1622584 | 1842753 | 220170 | 2 | 0.00908 |
| 41 | Poor | queen | NW_014569642.1 | 24223 | 244267 | 220045 | 2 | 0.00909 |
| 42 | Poor | queen | NW_014569683.1 | 152245 | 372052 | 219808 | 2 | 0.00910 |

| | | | | | | | | |
|---|---|---|---|---|---|---|---|---|
| 43 | Poor | queen | NW_014569578.1 | 779177 | 997550 | 218374 | 2 | 0.00916 |
| 44 | Poor | queen | NW_014569552.1 | 162090 | 380083 | 217994 | 2 | 0.00917 |
| 45 | Poor | queen | NW_014569552.1 | 1114773 | 1330803 | 216031 | 2 | 0.00926 |
| 46 | Poor | queen | NW_014569769.1 | 36282 | 250910 | 214629 | 2 | 0.00932 |
| 47 | Poor | queen | NW_014569580.1 | 260297 | 474843 | 214547 | 2 | 0.00932 |
| 48 | Poor | queen | NW_014569547.1 | 1140640 | 1353767 | 213128 | 2 | 0.00938 |
| 49 | Poor | queen | NW_014569554.1 | 1139666 | 1347273 | 207608 | 2 | 0.00963 |
| 50 | Poor | queen | NW_014569688.1 | 175756 | 383360 | 207605 | 2 | 0.00963 |
| 51 | Poor | queen | NW_014569658.1 | 195409 | 402148 | 206740 | 2 | 0.00967 |
| 52 | Poor | queen | NW_014569547.1 | 2955179 | 3159928 | 204750 | 2 | 0.00977 |
| 53 | Poor | queen | NW_014569618.1 | 74621 | 275430 | 200810 | 2 | 0.00996 |
| 54 | Poor | queen | NW_014569663.1 | 111049 | 310644 | 199596 | 2 | 0.01002 |
| 55 | Poor | queen | NW_014569549.1 | 371056 | 566984 | 195929 | 2 | 0.01021 |
| 56 | Poor | queen | NW_014569630.1 | 271732 | 467606 | 195875 | 2 | 0.01021 |
| 57 | Poor | queen | NW_014569579.1 | 556514 | 749198 | 192685 | 2 | 0.01038 |
| 58 | Poor | queen | NW_014569559.1 | 902429 | 1093064 | 190636 | 2 | 0.01049 |
| 59 | Poor | queen | NW_014569619.1 | 405507 | 592004 | 186498 | 2 | 0.01072 |
| 60 | Poor | queen | NW_014569556.1 | 465259 | 648486 | 183228 | 2 | 0.01092 |
| 61 | Poor | queen | NW_014569774.1 | 8134 | 187368 | 179235 | 2 | 0.01116 |
| 62 | Poor | queen | NW_014569708.1 | 119942 | 298704 | 178763 | 2 | 0.01119 |
| 63 | Poor | queen | NW_014569560.1 | 927083 | 1105788 | 178706 | 2 | 0.01119 |
| 64 | Poor | queen | NW_014569547.1 | 2017386 | 2194102 | 176717 | 2 | 0.01132 |
| 65 | Poor | queen | NW_014569744.1 | 70149 | 246501 | 176353 | 2 | 0.01134 |
| 66 | Poor | queen | NW_014569704.1 | 86812 | 262413 | 175602 | 2 | 0.01139 |
| 67 | Poor | queen | NW_014569682.1 | 236525 | 411380 | 174856 | 2 | 0.01144 |
| 68 | Poor | queen | NW_014569556.1 | 1392398 | 1566904 | 174507 | 2 | 0.01146 |
| 69 | Poor | queen | NW_014569552.1 | 635498 | 808511 | 173014 | 2 | 0.01156 |
| 70 | Poor | queen | NW_014569594.1 | 372668 | 545378 | 172711 | 2 | 0.01158 |
| 71 | Poor | queen | NW_014569575.1 | 580775 | 751922 | 171148 | 2 | 0.01169 |
| 72 | Poor | queen | NW_014569698.1 | 176106 | 346124 | 170019 | 2 | 0.01176 |
| 73 | Poor | queen | NW_014569684.1 | 97683 | 267286 | 169604 | 2 | 0.01179 |
| 74 | Poor | queen | NW_014569588.1 | 540699 | 709688 | 168990 | 2 | 0.01184 |
| 75 | Poor | queen | NW_014569564.1 | 636653 | 805008 | 168356 | 2 | 0.01188 |
| 76 | Poor | queen | NW_014569551.1 | 1913625 | 2081883 | 168259 | 2 | 0.01189 |
| 77 | Poor | queen | NW_014569614.1 | 285567 | 452448 | 166882 | 2 | 0.01198 |
| 78 | Poor | queen | NW_014569558.1 | 1345918 | 1511285 | 165368 | 2 | 0.01209 |
| 79 | Poor | queen | NW_014569561.1 | 948225 | 1113385 | 165161 | 2 | 0.01211 |
| 80 | Poor | queen | NW_014569558.1 | 358602 | 522951 | 164350 | 2 | 0.01217 |
| 81 | Poor | queen | NW_014569564.1 | 109305 | 271171 | 161867 | 2 | 0.01236 |
| 82 | Poor | queen | NW_014569591.1 | 60471 | 221478 | 161008 | 2 | 0.01242 |
| 83 | Poor | queen | NW_014569603.1 | 64580 | 225354 | 160775 | 2 | 0.01244 |
| 84 | Poor | queen | NW_014569607.1 | 54847 | 215264 | 160418 | 2 | 0.01247 |
| 85 | Poor | queen | NW_014569566.1 | 277465 | 437601 | 160137 | 2 | 0.01249 |
| 86 | Poor | queen | NW_014569591.1 | 473855 | 633900 | 160046 | 2 | 0.01250 |
| 87 | Poor | queen | NW_014569789.1 | 25396 | 184648 | 159253 | 2 | 0.01256 |
| 88 | Poor | queen | NW_014569552.1 | 1399182 | 1555741 | 156560 | 2 | 0.01277 |
| 89 | Poor | queen | NW_014569636.1 | 396375 | 552924 | 156550 | 2 | 0.01278 |

Analysis of 5-distances for sample "Pc_PA_qn" (13840 sites):

Total number of 5-distances: 10624
Median: 13467.5 Mean: 50693.3 Std: 80426.9
Quantiles:

| 0% | 5% | 10% | 15% | 20% | 25% | 30% | 35% | 40% | 45% | 50% | 55% | 60% | 65% | 70% | 75% | 80% | 85% | 90% | 95% | 100% |
|---|---|---|---|---|---|---|---|---|---|---|---|---|---|---|---|---|---|---|---|---|
| 7 | 42 | 82 | 148 | 211 | 321 | 503 | 1078 | 3398 | 7429 | 13468 | 20621 | 32381 | 43401 | 56922 | 72498 | 90199 | 115188 | 150605 | 207185 | 743935 |

Low density region distance cutoff (lowest 100): 388213
High density region distance cutoff (highest 100): 15

Ordered lists of methylation-rich regions:

Top 100 (out of 46) methylation-rich regions in "Pc_PA_qn":
(Sdnsty = sites per 1kb)

| | Rtype | Sample | SeqID | From | To | Rlgth | NbrSites | Sdnsty |
|---|---|---|---|---|---|---|---|---|
| 1 | Rich | queen | NW_014569572.1 | 1094647 | 1094662 | 16 | 10 | 625 |
| 2 | Rich | queen | NW_014569572.1 | 1094710 | 1094724 | 15 | 9 | 600 |
| 3 | Rich | queen | NW_014569595.1 | 551488 | 551497 | 10 | 6 | 600 |
| 4 | Rich | queen | NW_014569624.1 | 326546 | 326555 | 10 | 6 | 600 |
| 5 | Rich | queen | NW_014569597.1 | 285114 | 285127 | 14 | 8 | 571 |
| 6 | Rich | queen | NW_014569597.1 | 285188 | 285201 | 14 | 8 | 571 |
| 7 | Rich | queen | NW_014569601.1 | 298560 | 298573 | 14 | 7 | 500 |
| 8 | Rich | queen | NW_014569616.1 | 379670 | 379681 | 12 | 6 | 500 |
| 9 | Rich | queen | NW_014569559.1 | 279940 | 279952 | 13 | 6 | 462 |
| 10 | Rich | queen | NW_014569601.1 | 298540 | 298552 | 13 | 6 | 462 |
| 11 | Rich | queen | NW_014569615.1 | 136009 | 136021 | 13 | 6 | 462 |
| 12 | Rich | queen | NW_014569571.1 | 510294 | 510315 | 22 | 10 | 455 |
| 13 | Rich | queen | NW_014569617.1 | 66084 | 66098 | 15 | 6 | 400 |
| 14 | Rich | queen | NW_014569644.1 | 512191 | 512205 | 15 | 6 | 400 |
| 15 | Rich | queen | NW_014569553.1 | 1451862 | 1451877 | 16 | 6 | 375 |
| 16 | Rich | queen | NW_014569559.1 | 1151733 | 1151748 | 16 | 6 | 375 |
| 17 | Rich | queen | NW_014569560.1 | 576448 | 576463 | 16 | 6 | 375 |
| 18 | Rich | queen | NW_014569574.1 | 963687 | 963702 | 16 | 6 | 375 |
| 19 | Rich | queen | NW_014569586.1 | 604958 | 604973 | 16 | 6 | 375 |
| 20 | Rich | queen | NW_014569598.1 | 680231 | 680246 | 16 | 6 | 375 |
| 21 | Rich | queen | NW_014569602.1 | 823342 | 823357 | 16 | 6 | 375 |
| 22 | Rich | queen | NW_014569627.1 | 494217 | 494232 | 16 | 6 | 375 |
| 23 | Rich | queen | NW_014569645.1 | 320940 | 320955 | 16 | 6 | 375 |
| 24 | Rich | queen | NW_014569664.1 | 434571 | 434586 | 16 | 6 | 375 |
| 25 | Rich | queen | NW_014569910.1 | 14533 | 14548 | 16 | 6 | 375 |
| 26 | Rich | queen | NW_014570016.1 | 53123 | 53138 | 16 | 6 | 375 |
| 27 | Rich | queen | NW_014569565.1 | 1315126 | 1315144 | 19 | 7 | 368 |
| 28 | Rich | queen | NW_014569574.1 | 620870 | 620888 | 19 | 7 | 368 |
| 29 | Rich | queen | NW_014569591.1 | 439928 | 439946 | 19 | 7 | 368 |
| 30 | Rich | queen | NW_014569609.1 | 272078 | 272096 | 19 | 7 | 368 |
| 31 | Rich | queen | NW_014569619.1 | 50206 | 50224 | 19 | 7 | 368 |
| 32 | Rich | queen | NW_014569629.1 | 370658 | 370676 | 19 | 7 | 368 |
| 33 | Rich | queen | NW_014569662.1 | 381625 | 381643 | 19 | 7 | 368 |
| 34 | Rich | queen | NW_014569672.1 | 276958 | 276976 | 19 | 7 | 368 |
| 35 | Rich | queen | NW_014569865.1 | 85306 | 85324 | 19 | 7 | 368 |
| 36 | Rich | queen | NW_014569975.1 | 5349 | 5367 | 19 | 7 | 368 |
| 37 | Rich | queen | NW_014569571.1 | 530528 | 530549 | 22 | 8 | 364 |
| 38 | Rich | queen | NW_014569593.1 | 458070 | 458091 | 22 | 8 | 364 |
| 39 | Rich | queen | NW_014569601.1 | 446509 | 446530 | 22 | 8 | 364 |
| 40 | Rich | queen | NW_014569606.1 | 96573 | 96594 | 22 | 8 | 364 |
| 41 | Rich | queen | NW_014569568.1 | 911614 | 911638 | 25 | 9 | 360 |
| 42 | Rich | queen | NW_014569768.1 | 159776 | 159800 | 25 | 9 | 360 |
| 43 | Rich | queen | NW_014569569.1 | 888237 | 888264 | 28 | 10 | 357 |
| 44 | Rich | queen | NW_014569548.1 | 2472708 | 2472738 | 31 | 11 | 355 |
| 45 | Rich | queen | NW_014569664.1 | 337878 | 337908 | 31 | 11 | 355 |
| 46 | Rich | queen | NW_014569559.1 | 117897 | 117933 | 37 | 13 | 351 |

Directory: MRPR File: dst-Pc_PA_wr.txt

Analysis of 1-distances for sample "Pc_PA_wr" (12036 sites):

Total number of 1-distances: 11167
Median: 40.0 Mean: 12189.4 Std: 33060.4
Quantiles:

| 0% | 5% | 10% | 15% | 20% | 25% | 30% | 35% | 40% | 45% | 50% | 55% | 60% | 65% | 70% | 75% | 80% | 85% | 90% | 95% | 100% |
|---|---|---|---|---|---|---|---|---|---|---|---|---|---|---|---|---|---|---|---|---|
| 1 | 1 | 3 | 4 | 6 | 8 | 12 | 17 | 23 | 30 | 40 | 66 | 122 | 214 | 766 | 3904 | 11525 | 23401 | 42689 | 74907 | 662769 |

Low density region distance cutoff (lowest 100): 167405
High density region distance cutoff (highest 100): 1

Ordered lists of methylation-poor regions:

Top 100 (out of 97) methylation-poor regions in "Pc_PA_wr":
(Sdnsty = sites per 1kb)

| | Rtype | Sample | SeqID | From | To | Rlgth | NbrSites | Sdnsty |
|---|---|---|---|---|---|---|---|---|
| 1 | Poor | worker | NW_014569549.1 | 1394259 | 2057028 | 662770 | 2 | 0.00302 |
| 2 | Poor | worker | NW_014569596.1 | 107205 | 565599 | 458395 | 2 | 0.00436 |
| 3 | Poor | worker | NW_014569590.1 | 411702 | 869309 | 457608 | 2 | 0.00437 |
| 4 | Poor | worker | NW_014569599.1 | 319335 | 730170 | 410836 | 2 | 0.00487 |
| 5 | Poor | worker | NW_014569583.1 | 181964 | 575444 | 393481 | 2 | 0.00508 |
| 6 | Poor | worker | NW_014569548.1 | 1009569 | 1378263 | 368695 | 2 | 0.00542 |
| 7 | Poor | worker | NW_014569572.1 | 173801 | 507263 | 333463 | 2 | 0.00600 |
| 8 | Poor | worker | NW_014569547.1 | 1652854 | 1978021 | 325168 | 2 | 0.00615 |
| 9 | Poor | worker | NW_014569611.1 | 288166 | 608498 | 320333 | 2 | 0.00624 |
| 10 | Poor | worker | NW_014569613.1 | 47760 | 367879 | 320120 | 2 | 0.00625 |
| 11 | Poor | worker | NW_014569569.1 | 463643 | 775235 | 311593 | 2 | 0.00642 |
| 12 | Poor | worker | NW_014569552.1 | 1702945 | 1997741 | 294797 | 2 | 0.00678 |
| 13 | Poor | worker | NW_014569579.1 | 83024 | 524431 | 441408 | 3 | 0.00680 |
| 14 | Poor | worker | NW_014569594.1 | 191023 | 480013 | 288991 | 2 | 0.00692 |
| 15 | Poor | worker | NW_014569554.1 | 1516545 | 1801461 | 284917 | 2 | 0.00702 |
| 16 | Poor | worker | NW_014569593.1 | 549787 | 818644 | 268858 | 2 | 0.00744 |
| 17 | Poor | worker | NW_014569553.1 | 1605486 | 1873709 | 268224 | 2 | 0.00746 |
| 18 | Poor | worker | NW_014569551.1 | 527883 | 791885 | 264003 | 2 | 0.00758 |
| 19 | Poor | worker | NW_014569579.1 | 556514 | 946448 | 389935 | 3 | 0.00769 |
| 20 | Poor | worker | NW_014569630.1 | 2297 | 259254 | 256958 | 2 | 0.00778 |
| 21 | Poor | worker | NW_014569671.1 | 4141 | 255750 | 251610 | 2 | 0.00795 |
| 22 | Poor | worker | NW_014569550.1 | 1942811 | 2192438 | 249628 | 2 | 0.00801 |
| 23 | Poor | worker | NW_014569680.1 | 124805 | 372297 | 247493 | 2 | 0.00808 |
| 24 | Poor | worker | NW_014569573.1 | 672608 | 915926 | 243319 | 2 | 0.00822 |
| 25 | Poor | worker | NW_014569600.1 | 493418 | 732502 | 239085 | 2 | 0.00837 |
| 26 | Poor | worker | NW_014569547.1 | 2081921 | 2319957 | 238037 | 2 | 0.00840 |
| 27 | Poor | worker | NW_014569552.1 | 274471 | 511408 | 236938 | 2 | 0.00844 |
| 28 | Poor | worker | NW_014569698.1 | 13279 | 360380 | 347102 | 3 | 0.00864 |
| 29 | Poor | worker | NW_014569647.1 | 85651 | 312828 | 227178 | 2 | 0.00880 |
| 30 | Poor | worker | NW_014569551.1 | 213157 | 440041 | 226885 | 2 | 0.00882 |
| 31 | Poor | worker | NW_014569595.1 | 60797 | 285751 | 224955 | 2 | 0.00889 |
| 32 | Poor | worker | NW_014569575.1 | 527034 | 751922 | 224889 | 2 | 0.00889 |
| 33 | Poor | worker | NW_014569557.1 | 362770 | 587012 | 224243 | 2 | 0.00892 |
| 34 | Poor | worker | NW_014569602.1 | 510697 | 734897 | 224201 | 2 | 0.00892 |
| 35 | Poor | worker | NW_014569573.1 | 353694 | 577792 | 224099 | 2 | 0.00892 |
| 36 | Poor | worker | NW_014569549.1 | 848989 | 1072659 | 223671 | 2 | 0.00894 |
| 37 | Poor | worker | NW_014569612.1 | 99942 | 322495 | 222554 | 2 | 0.00899 |
| 38 | Poor | worker | NW_014569622.1 | 207709 | 429966 | 222258 | 2 | 0.00900 |
| 39 | Poor | worker | NW_014569554.1 | 565378 | 785823 | 220446 | 2 | 0.00907 |
| 40 | Poor | worker | NW_014569550.1 | 1622584 | 1842753 | 220170 | 2 | 0.00908 |
| 41 | Poor | worker | NW_014569683.1 | 152245 | 372052 | 219808 | 2 | 0.00910 |
| 42 | Poor | worker | NW_014569730.1 | 34043 | 252713 | 218671 | 2 | 0.00915 |

| | | | | | | | |
|---|---|---|---|---|---|---|---|
| 43 | Poor worker | NW_014569609.1 | 486592 | 704809 | 218218 | 2 | 0.00917 |
| 44 | Poor worker | NW_014569635.1 | 197647 | 413895 | 216249 | 2 | 0.00925 |
| 45 | Poor worker | NW_014569607.1 | 130843 | 345877 | 215035 | 2 | 0.00930 |
| 46 | Poor worker | NW_014569566.1 | 491270 | 706151 | 214882 | 2 | 0.00931 |
| 47 | Poor worker | NW_014569771.1 | 22714 | 237000 | 214287 | 2 | 0.00933 |
| 48 | Poor worker | NW_014569549.1 | 371056 | 583728 | 212673 | 2 | 0.00940 |
| 49 | Poor worker | NW_014569549.1 | 1072665 | 1284586 | 211922 | 2 | 0.00944 |
| 50 | Poor worker | NW_014569659.1 | 125619 | 333431 | 207813 | 2 | 0.00962 |
| 51 | Poor worker | NW_014569565.1 | 958204 | 1165122 | 206919 | 2 | 0.00967 |
| 52 | Poor worker | NW_014569591.1 | 710727 | 916307 | 205581 | 2 | 0.00973 |
| 53 | Poor worker | NW_014569630.1 | 262337 | 467606 | 205270 | 2 | 0.00974 |
| 54 | Poor worker | NW_014569574.1 | 620888 | 824918 | 204031 | 2 | 0.00980 |
| 55 | Poor worker | NW_014569669.1 | 25846 | 229786 | 203941 | 2 | 0.00981 |
| 56 | Poor worker | NW_014569585.1 | 775841 | 977878 | 202038 | 2 | 0.00990 |
| 57 | Poor worker | NW_014569554.1 | 1139813 | 1341302 | 201490 | 2 | 0.00993 |
| 58 | Poor worker | NW_014569575.1 | 774954 | 975924 | 200971 | 2 | 0.00995 |
| 59 | Poor worker | NW_014569584.1 | 280990 | 481463 | 200474 | 2 | 0.00998 |
| 60 | Poor worker | NW_014569603.1 | 550187 | 750221 | 200035 | 2 | 0.01000 |
| 61 | Poor worker | NW_014569637.1 | 311063 | 511084 | 200022 | 2 | 0.01000 |
| 62 | Poor worker | NW_014569584.1 | 777757 | 977474 | 199718 | 2 | 0.01001 |
| 63 | Poor worker | NW_014569581.1 | 258620 | 457287 | 198668 | 2 | 0.01007 |
| 64 | Poor worker | NW_014569585.1 | 407040 | 603603 | 196564 | 2 | 0.01017 |
| 65 | Poor worker | NW_014569587.1 | 577046 | 773328 | 196283 | 2 | 0.01019 |
| 66 | Poor worker | NW_014569550.1 | 332449 | 528361 | 195913 | 2 | 0.01021 |
| 67 | Poor worker | NW_014569645.1 | 115507 | 311259 | 195753 | 2 | 0.01022 |
| 68 | Poor worker | NW_014569576.1 | 37800 | 233275 | 195476 | 2 | 0.01023 |
| 69 | Poor worker | NW_014569658.1 | 207162 | 402148 | 194987 | 2 | 0.01026 |
| 70 | Poor worker | NW_014569555.1 | 1211977 | 1406167 | 194191 | 2 | 0.01030 |
| 71 | Poor worker | NW_014569586.1 | 172833 | 366192 | 193360 | 2 | 0.01034 |
| 72 | Poor worker | NW_014569571.1 | 725749 | 918324 | 192576 | 2 | 0.01039 |
| 73 | Poor worker | NW_014569547.1 | 1110705 | 1301946 | 191242 | 2 | 0.01046 |
| 74 | Poor worker | NW_014569570.1 | 221293 | 410891 | 189599 | 2 | 0.01055 |
| 75 | Poor worker | NW_014569550.1 | 928377 | 1117758 | 189382 | 2 | 0.01056 |
| 76 | Poor worker | NW_014569556.1 | 649066 | 836250 | 187185 | 2 | 0.01068 |
| 77 | Poor worker | NW_014569619.1 | 405316 | 591548 | 186233 | 2 | 0.01074 |
| 78 | Poor worker | NW_014569655.1 | 26424 | 211707 | 185284 | 2 | 0.01079 |
| 79 | Poor worker | NW_014569704.1 | 86812 | 270919 | 184108 | 2 | 0.01086 |
| 80 | Poor worker | NW_014569587.1 | 61554 | 245321 | 183768 | 2 | 0.01088 |
| 81 | Poor worker | NW_014569636.1 | 369375 | 552924 | 183550 | 2 | 0.01090 |
| 82 | Poor worker | NW_014569708.1 | 119942 | 298629 | 178688 | 2 | 0.01119 |
| 83 | Poor worker | NW_014569632.1 | 97581 | 275438 | 177858 | 2 | 0.01124 |
| 84 | Poor worker | NW_014569757.1 | 38384 | 215254 | 176871 | 2 | 0.01131 |
| 85 | Poor worker | NW_014569548.1 | 1987032 | 2163879 | 176848 | 2 | 0.01131 |
| 86 | Poor worker | NW_014569619.1 | 50224 | 226975 | 176752 | 2 | 0.01132 |
| 87 | Poor worker | NW_014569555.1 | 1499618 | 1675986 | 176369 | 2 | 0.01134 |
| 88 | Poor worker | NW_014569553.1 | 652760 | 828603 | 175844 | 2 | 0.01137 |
| 89 | Poor worker | NW_014569585.1 | 124946 | 300234 | 175289 | 2 | 0.01141 |
| 90 | Poor worker | NW_014569562.1 | 744009 | 919099 | 175091 | 2 | 0.01142 |
| 91 | Poor worker | NW_014569602.1 | 336871 | 510689 | 173819 | 2 | 0.01151 |
| 92 | Poor worker | NW_014569558.1 | 168867 | 339911 | 171045 | 2 | 0.01169 |
| 93 | Poor worker | NW_014569598.1 | 477994 | 649010 | 171017 | 2 | 0.01169 |
| 94 | Poor worker | NW_014569654.1 | 71150 | 241282 | 170133 | 2 | 0.01176 |
| 95 | Poor worker | NW_014569588.1 | 540699 | 709688 | 168990 | 2 | 0.01184 |
| 96 | Poor worker | NW_014569551.1 | 1913795 | 2081877 | 168083 | 2 | 0.01190 |
| 97 | Poor worker | NW_014569575.1 | 157240 | 324645 | 167406 | 2 | 0.01195 |

Analysis of 5-distances for sample "Pc_PA_wr" (12036 sites):

Total number of 5-distances: 9081
Median: 17880.0 Mean: 58446.7 Std: 87567.6
Quantiles:

| 0% | 5% | 10% | 15% | 20% | 25% | 30% | 35% | 40% | 45% | 50% | 55% | 60% | 65% | 70% | 75% | 80% | 85% | 90% | 95% | 100% |
|---|---|---|---|---|---|---|---|---|---|---|---|---|---|---|---|---|---|---|---|---|

| | | | | | | | | | | | | | | | | | | | | |
|---|---|---|---|---|---|---|---|---|---|---|---|---|---|---|---|---|---|---|---|---|
| 5 | 42 | 97 | 171 | 262 | 418 | 827 | 2362 | 5855 | 10522 | 17880 | 28307 | 40521 | 54049 | 68658 | 84902 | 105548 | 130518 | 173384 | 234356 | 984363 |

Low density region distance cutoff (lowest 100): 396076
High density region distance cutoff (highest 100): 15

Ordered lists of methylation-rich regions:

Top 100 (out of 41) methylation-rich regions in "Pc_PA_wr":
(Sdnsty = sites per 1kb)

| | Rtype | Sample | SeqID | From | To | Rlgth | NbrSites | Sdnsty |
|---|---|---|---|---|---|---|---|---|
| 1 | Rich | worker | NW_014569756.1 | 80429 | 80434 | 6 | 6 | 1000 |
| 2 | Rich | worker | NW_014569597.1 | 285188 | 285201 | 14 | 8 | 571 |
| 3 | Rich | worker | NW_014569597.1 | 285114 | 285126 | 13 | 7 | 538 |
| 4 | Rich | worker | NW_014569613.1 | 571082 | 571094 | 13 | 6 | 462 |
| 5 | Rich | worker | NW_014569553.1 | 1172276 | 1172293 | 18 | 8 | 444 |
| 6 | Rich | worker | NW_014569601.1 | 298540 | 298571 | 32 | 13 | 406 |
| 7 | Rich | worker | NW_014569617.1 | 66084 | 66098 | 15 | 6 | 400 |
| 8 | Rich | worker | NW_014569644.1 | 512191 | 512205 | 15 | 6 | 400 |
| 9 | Rich | worker | NW_014569667.1 | 255801 | 255815 | 15 | 6 | 400 |
| 10 | Rich | worker | NW_014569959.1 | 66990 | 67004 | 15 | 6 | 400 |
| 11 | Rich | worker | NW_014569559.1 | 1151733 | 1151748 | 16 | 6 | 375 |
| 12 | Rich | worker | NW_014569586.1 | 604958 | 604973 | 16 | 6 | 375 |
| 13 | Rich | worker | NW_014569598.1 | 680231 | 680246 | 16 | 6 | 375 |
| 14 | Rich | worker | NW_014569602.1 | 823342 | 823357 | 16 | 6 | 375 |
| 15 | Rich | worker | NW_014569609.1 | 272078 | 272093 | 16 | 6 | 375 |
| 16 | Rich | worker | NW_014569627.1 | 494217 | 494232 | 16 | 6 | 375 |
| 17 | Rich | worker | NW_014569632.1 | 372328 | 372343 | 16 | 6 | 375 |
| 18 | Rich | worker | NW_014569645.1 | 320940 | 320955 | 16 | 6 | 375 |
| 19 | Rich | worker | NW_014569872.1 | 112887 | 112902 | 16 | 6 | 375 |
| 20 | Rich | worker | NW_014569910.1 | 14533 | 14548 | 16 | 6 | 375 |
| 21 | Rich | worker | NW_014570016.1 | 53123 | 53138 | 16 | 6 | 375 |
| 22 | Rich | worker | NW_014569565.1 | 1315126 | 1315144 | 19 | 7 | 368 |
| 23 | Rich | worker | NW_014569574.1 | 620870 | 620888 | 19 | 7 | 368 |
| 24 | Rich | worker | NW_014569591.1 | 439928 | 439946 | 19 | 7 | 368 |
| 25 | Rich | worker | NW_014569619.1 | 50206 | 50224 | 19 | 7 | 368 |
| 26 | Rich | worker | NW_014569662.1 | 381625 | 381643 | 19 | 7 | 368 |
| 27 | Rich | worker | NW_014569672.1 | 276958 | 276976 | 19 | 7 | 368 |
| 28 | Rich | worker | NW_014569975.1 | 5349 | 5367 | 19 | 7 | 368 |
| 29 | Rich | worker | NW_014569553.1 | 1451862 | 1451883 | 22 | 8 | 364 |
| 30 | Rich | worker | NW_014569571.1 | 530528 | 530549 | 22 | 8 | 364 |
| 31 | Rich | worker | NW_014569593.1 | 458070 | 458091 | 22 | 8 | 364 |
| 32 | Rich | worker | NW_014569606.1 | 96573 | 96594 | 22 | 8 | 364 |
| 33 | Rich | worker | NW_014569653.1 | 214404 | 214425 | 22 | 8 | 364 |
| 34 | Rich | worker | NW_014569664.1 | 434571 | 434595 | 25 | 9 | 360 |
| 35 | Rich | worker | NW_014569865.1 | 85303 | 85327 | 25 | 9 | 360 |
| 36 | Rich | worker | NW_014569568.1 | 911614 | 911641 | 28 | 10 | 357 |
| 37 | Rich | worker | NW_014569569.1 | 888237 | 888264 | 28 | 10 | 357 |
| 38 | Rich | worker | NW_014569601.1 | 446503 | 446530 | 28 | 10 | 357 |
| 39 | Rich | worker | NW_014569768.1 | 159770 | 159800 | 31 | 11 | 355 |
| 40 | Rich | worker | NW_014569548.1 | 2472708 | 2472741 | 34 | 12 | 353 |
| 41 | Rich | worker | NW_014569559.1 | 117894 | 117933 | 40 | 14 | 350 |

Directory: MRPR File: gwp-Pc_PA_qn.txt

| seqnames | start | end | width | strand | source | type | score | phase | ID | Dbxref | Name | gbkey | gene | gene_biotype | partial | start_range | end_range | seqnames | start | end | width | strand | NbrSites | Sdnsty |
|---|---|---|---|---|---|---|---|---|---|---|---|---|---|---|---|---|---|---|---|---|---|---|---|---|
| NW_014569596.1 | 104166 | 108409 | 4244 | - | Gnomon | gene | NA | NA | gene-LOC106785379 | GeneID:106785379 | LOC106785379 | Gene | LOC106785379 | protein_coding | NA | character(0) | character(0) | NW_014569596.1 | 107147 | 565599 | 458453 | * | 2 | 0.00436249735523598 |
| NW_014569596.1 | 146177 | 148054 | 1878 | - | Gnomon | gene | NA | NA | gene-LOC106785392 | GeneID:106785392 | LOC106785392 | Gene | LOC106785392 | lncRNA | NA | character(0) | character(0) | NW_014569596.1 | 107147 | 565599 | 458453 | * | 2 | 0.00436249735523598 |
| NW_014569596.1 | 147348 | 151903 | 4556 | + | Gnomon | gene | NA | NA | gene-LOC106785390 | GeneID:106785390 | LOC106785390 | Gene | LOC106785390 | protein_coding | NA | character(0) | character(0) | NW_014569596.1 | 107147 | 565599 | 458453 | * | 2 | 0.00436249735523598 |
| NW_014569596.1 | 151875 | 155826 | 3952 | - | Gnomon | gene | NA | NA | gene-LOC106785389 | GeneID:106785389 | LOC106785389 | Gene | LOC106785389 | protein_coding | NA | character(0) | character(0) | NW_014569596.1 | 107147 | 565599 | 458453 | * | 2 | 0.00436249735523598 |
| NW_014569596.1 | 156178 | 163510 | 7333 | + | Gnomon | gene | NA | NA | gene-LOC106785388 | GeneID:106785388 | LOC106785388 | Gene | LOC106785388 | protein_coding | NA | character(0) | character(0) | NW_014569596.1 | 107147 | 565599 | 458453 | * | 2 | 0.00436249735523598 |
| NW_014569596.1 | 163640 | 204917 | 41278 | - | Gnomon | gene | NA | NA | gene-LOC106785387 | GeneID:106785387 | LOC106785387 | Gene | LOC106785387 | protein_coding | NA | character(0) | character(0) | NW_014569596.1 | 107147 | 565599 | 458453 | * | 2 | 0.00436249735523598 |
| NW_014569596.1 | 214502 | 217044 | 2543 | + | Gnomon | gene | NA | NA | gene-LOC106785399 | GeneID:106785399 | LOC106785399 | Gene | LOC106785399 | protein_coding | NA | character(0) | character(0) | NW_014569596.1 | 107147 | 565599 | 458453 | * | 2 | 0.00436249735523598 |
| NW_014569596.1 | 224623 | 257819 | 33197 | + | Gnomon | gene | NA | NA | gene-LOC106785404 | GeneID:106785404 | LOC106785404 | Gene | LOC106785404 | protein_coding | NA | character(0) | character(0) | NW_014569596.1 | 107147 | 565599 | 458453 | * | 2 | 0.00436249735523598 |
| NW_014569596.1 | 260991 | 261809 | 819 | - | Gnomon | gene | NA | NA | gene-LOC106785370 | GeneID:106785370 | LOC106785370 | Gene | LOC106785370 | protein_coding | NA | character(0) | character(0) | NW_014569596.1 | 107147 | 565599 | 458453 | * | 2 | 0.00436249735523598 |
| NW_014569596.1 | 295243 | 297337 | 2095 | - | Gnomon | gene | NA | NA | gene-LOC106785401 | GeneID:106785401 | LOC106785401 | Gene | LOC106785401 | protein_coding | NA | character(0) | character(0) | NW_014569596.1 | 107147 | 565599 | 458453 | * | 2 | 0.00436249735523598 |
| NW_014569596.1 | 299057 | 311144 | 12088 | - | Gnomon | gene | NA | NA | gene-LOC106785386 | GeneID:106785386 | LOC106785386 | Gene | LOC106785386 | protein_coding | NA | character(0) | character(0) | NW_014569596.1 | 107147 | 565599 | 458453 | * | 2 | 0.00436249735523598 |
| NW_014569596.1 | 347702 | 531555 | 183854 | - | Gnomon | gene | NA | NA | gene-LOC106785371 | GeneID:106785371 | LOC106785371 | Gene | LOC106785371 | protein_coding | NA | character(0) | character(0) | NW_014569596.1 | 107147 | 565599 | 458453 | * | 2 | 0.00436249735523598 |
| NW_014569596.1 | 385266 | 387266 | 2001 | + | Gnomon | gene | NA | NA | gene-LOC106785373 | GeneID:106785373 | LOC106785373 | Gene | LOC106785373 | protein_coding | NA | character(0) | character(0) | NW_014569596.1 | 107147 | 565599 | 458453 | * | 2 | 0.00436249735523598 |
| NW_014569596.1 | 395858 | 402796 | 6939 | + | Gnomon | gene | NA | NA | gene-LOC106785372 | GeneID:106785372 | LOC106785372 | Gene | LOC106785372 | protein_coding | NA | character(0) | character(0) | NW_014569596.1 | 107147 | 565599 | 458453 | * | 2 | 0.00436249735523598 |
| NW_014569596.1 | 541747 | 547209 | 5463 | + | Gnomon | gene | NA | NA | gene-LOC106785384 | GeneID:106785384 | LOC106785384 | Gene | LOC106785384 | protein_coding | NA | character(0) | character(0) | NW_014569596.1 | 107147 | 565599 | 458453 | * | 2 | 0.00436249735523598 |
| NW_014569596.1 | 558081 | 692185 | 134105 | - | Gnomon | gene | NA | NA | gene-LOC106785374 | GeneID:106785374 | LOC106785374 | Gene | LOC106785374 | protein_coding | NA | character(0) | character(0) | NW_014569596.1 | 107147 | 565599 | 458453 | * | 2 | 0.00436249735523598 |
| NW_014569579.1 | 3229 | 277131 | 273903 | - | Gnomon | gene | NA | NA | gene-LOC106784216 | GeneID:106784216 | LOC106784216 | Gene | LOC106784216 | protein_coding | NA | character(0) | character(0) | NW_014569579.1 | 83024 | 523197 | 440174 | * | 2 | 0.00454365773535011 |
| NW_014569579.1 | 275942 | 276034 | 93 | - | tRNAscan-SE | gene | NA | NA | gene-Trnay-gua | GeneID:106783543 | Trnay-gua | Gene | Trnay-gua | tRNA | NA | character(0) | character(0) | NW_014569579.1 | 83024 | 523197 | 440174 | * | 2 | 0.00454365773535011 |
| NW_014569579.1 | 278459 | 280080 | 1622 | + | Gnomon | gene | NA | NA | gene-LOC106784202 | GeneID:106784202 | LOC106784202 | Gene | LOC106784202 | protein_coding | NA | character(0) | character(0) | NW_014569579.1 | 83024 | 523197 | 440174 | * | 2 | 0.00454365773535011 |
| NW_014569579.1 | 280437 | 286185 | 5749 | - | Gnomon | gene | NA | NA | gene-LOC106784250 | GeneID:106784250 | LOC106784250 | Gene | LOC106784250 | protein_coding | NA | character(0) | character(0) | NW_014569579.1 | 83024 | 523197 | 440174 | * | 2 | 0.00454365773535011 |
| NW_014569579.1 | 298909 | 342575 | 43667 | - | Gnomon | gene | NA | NA | gene-LOC106784222 | GeneID:106784222 | LOC106784222 | Gene | LOC106784222 | protein_coding | NA | character(0) | character(0) | NW_014569579.1 | 83024 | 523197 | 440174 | * | 2 | 0.00454365773535011 |
| NW_014569579.1 | 346312 | 352165 | 5854 | - | Gnomon | gene | NA | NA | gene-LOC106784220 | GeneID:106784220 | LOC106784220 | Gene | LOC106784220 | protein_coding | NA | character(0) | character(0) | NW_014569579.1 | 83024 | 523197 | 440174 | * | 2 | 0.00454365773535011 |
| NW_014569579.1 | 352375 | 356805 | 4431 | - | Gnomon | gene | NA | NA | gene-LOC106784219 | GeneID:106784219 | LOC106784219 | Gene | LOC106784219 | protein_coding | NA | character(0) | character(0) | NW_014569579.1 | 83024 | 523197 | 440174 | * | 2 | 0.00454365773535011 |
| NW_014569579.1 | 357122 | 359672 | 2551 | + | Gnomon | gene | NA | NA | gene-LOC106784221 | GeneID:106784221 | LOC106784221 | Gene | LOC106784221 | protein_coding | NA | character(0) | character(0) | NW_014569579.1 | 83024 | 523197 | 440174 | * | 2 | 0.00454365773535011 |

...
... only first 25 lines shown ...

Directory: MRPR File: gwp-Pc_PA_wr.txt

| seqnames | start | end | width | strand | source | type | score | phase | ID | Dbxref | Name | gbkey | gene | gene_biotype | partial | start_range | end_range | seqnames | start | end | width | strand | NbrSites | Sdnsty |
|---|---|---|---|---|---|---|---|---|---|---|---|---|---|---|---|---|---|---|---|---|---|---|---|---|
| NW_014569549.1 | 1400338 | 1401329 | 992 | - | Gnomon | gene | NA | NA | gene-LOC106784707 | GeneID:106784707 | LOC106784707 | Gene | LOC106784707 | lncRNA | NA | character(0) | character(0) | NW_014569549.1 | 1394259 | 2057028 | 662770 | * | 2 | 0.00301763809466331 |
| NW_014569549.1 | 1402110 | 1403799 | 1690 | - | Gnomon | gene | NA | NA | gene-LOC106785487 | GeneID:106785487 | LOC106785487 | Gene | LOC106785487 | lncRNA | NA | character(0) | character(0) | NW_014569549.1 | 1394259 | 2057028 | 662770 | * | 2 | 0.00301763809466331 |
| NW_014569549.1 | 1404754 | 1408640 | 3887 | + | Gnomon | gene | NA | NA | gene-LOC106785454 | GeneID:106785454 | LOC106785454 | Gene | LOC106785454 | protein_coding | NA | character(0) | character(0) | NW_014569549.1 | 1394259 | 2057028 | 662770 | * | 2 | 0.00301763809466331 |
| NW_014569549.1 | 1409463 | 1415266 | 5804 | + | Gnomon | gene | NA | NA | gene-LOC106785447 | GeneID:106785447 | LOC106785447 | Gene | LOC106785447 | protein_coding | NA | character(0) | character(0) | NW_014569549.1 | 1394259 | 2057028 | 662770 | * | 2 | 0.00301763809466331 |
| NW_014569549.1 | 1416939 | 1421496 | 4558 | + | Gnomon | gene | NA | NA | gene-LOC106785529 | GeneID:106785529 | LOC106785529 | Gene | LOC106785529 | protein_coding | NA | character(0) | character(0) | NW_014569549.1 | 1394259 | 2057028 | 662770 | * | 2 | 0.00301763809466331 |
| NW_014569549.1 | 1427704 | 1665158 | 237455 | - | Gnomon | gene | NA | NA | gene-LOC106785193 | GeneID:106785193 | LOC106785193 | Gene | LOC106785193 | protein_coding | NA | character(0) | character(0) | NW_014569549.1 | 1394259 | 2057028 | 662770 | * | 2 | 0.00301763809466331 |
| NW_014569549.1 | 1675128 | 1706347 | 31220 | - | Gnomon | gene | NA | NA | gene-LOC106785349 | GeneID:106785349 | LOC106785349 | Gene | LOC106785349 | protein_coding | NA | character(0) | character(0) | NW_014569549.1 | 1394259 | 2057028 | 662770 | * | 2 | 0.00301763809466331 |
| NW_014569549.1 | 1722619 | 1738182 | 15564 | + | Gnomon | gene | NA | NA | gene-LOC106785246 | GeneID:106785246 | LOC106785246 | Gene | LOC106785246 | protein_coding | NA | character(0) | character(0) | NW_014569549.1 | 1394259 | 2057028 | 662770 | * | 2 | 0.00301763809466331 |
| NW_014569549.1 | 1733835 | 1737907 | 4073 | - | Gnomon | gene | NA | NA | gene-LOC106785260 | GeneID:106785260 | LOC106785260 | Gene | LOC106785260 | protein_coding | NA | character(0) | character(0) | NW_014569549.1 | 1394259 | 2057028 | 662770 | * | 2 | 0.00301763809466331 |
| NW_014569549.1 | 1740625 | 1768132 | 27508 | + | Gnomon | gene | NA | NA | gene-LOC106785692 | GeneID:106785692 | LOC106785692 | Gene | LOC106785692 | protein_coding | NA | character(0) | character(0) | NW_014569549.1 | 1394259 | 2057028 | 662770 | * | 2 | 0.00301763809466331 |
| NW_014569549.1 | 1773020 | 1804458 | 31439 | + | Gnomon | gene | NA | NA | gene-LOC106785270 | GeneID:106785270 | LOC106785270 | Gene | LOC106785270 | protein_coding | NA | character(0) | character(0) | NW_014569549.1 | 1394259 | 2057028 | 662770 | * | 2 | 0.00301763809466331 |
| NW_014569549.1 | 1809819 | 2003488 | 193670 | - | Gnomon | gene | NA | NA | gene-LOC106784764 | GeneID:106784764 | LOC106784764 | Gene | LOC106784764 | protein_coding | NA | character(0) | character(0) | NW_014569549.1 | 1394259 | 2057028 | 662770 | * | 2 | 0.00301763809466331 |
| NW_014569549.1 | 1837449 | 1839708 | 2260 | + | Gnomon | gene | NA | NA | gene-LOC106784838 | GeneID:106784838 | LOC106784838 | Gene | LOC106784838 | protein_coding | NA | character(0) | character(0) | NW_014569549.1 | 1394259 | 2057028 | 662770 | * | 2 | 0.00301763809466331 |
| NW_014569549.1 | 1841069 | 1842598 | 1530 | + | Gnomon | gene | NA | NA | gene-LOC106784834 | GeneID:106784834 | LOC106784834 | Gene | LOC106784834 | protein_coding | NA | character(0) | character(0) | NW_014569549.1 | 1394259 | 2057028 | 662770 | * | 2 | 0.00301763809466331 |
| NW_014569549.1 | 1981449 | 2107584 | 126136 | - | Gnomon | gene | NA | NA | gene-LOC106784849 | GeneID:106784849 | LOC106784849 | Gene | LOC106784849 | protein_coding | NA | character(0) | character(0) | NW_014569549.1 | 1394259 | 2057028 | 662770 | * | 2 | 0.00301763809466331 |
| NW_014569596.1 | 104166 | 108409 | 4244 | - | Gnomon | gene | NA | NA | gene-LOC106785379 | GeneID:106785379 | LOC106785379 | Gene | LOC106785379 | protein_coding | NA | character(0) | character(0) | NW_014569596.1 | 107205 | 565599 | 458395 | * | 2 | 0.00436304933518036 |
| NW_014569596.1 | 146177 | 148054 | 1878 | - | Gnomon | gene | NA | NA | gene-LOC106785392 | GeneID:106785392 | LOC106785392 | Gene | LOC106785392 | lncRNA | NA | character(0) | character(0) | NW_014569596.1 | 107205 | 565599 | 458395 | * | 2 | 0.00436304933518036 |
| NW_014569596.1 | 147348 | 151903 | 4556 | + | Gnomon | gene | NA | NA | gene-LOC106785390 | GeneID:106785390 | LOC106785390 | Gene | LOC106785390 | protein_coding | NA | character(0) | character(0) | NW_014569596.1 | 107205 | 565599 | 458395 | * | 2 | 0.00436304933518036 |
| NW_014569596.1 | 151875 | 155826 | 3952 | - | Gnomon | gene | NA | NA | gene-LOC106785389 | GeneID:106785389 | LOC106785389 | Gene | LOC106785389 | protein_coding | NA | character(0) | character(0) | NW_014569596.1 | 107205 | 565599 | 458395 | * | 2 | 0.00436304933518036 |
| NW_014569596.1 | 156178 | 163510 | 7333 | + | Gnomon | gene | NA | NA | gene-LOC106785388 | GeneID:106785388 | LOC106785388 | Gene | LOC106785388 | protein_coding | NA | character(0) | character(0) | NW_014569596.1 | 107205 | 565599 | 458395 | * | 2 | 0.00436304933518036 |
| NW_014569596.1 | 163640 | 204917 | 41278 | - | Gnomon | gene | NA | NA | gene-LOC106785387 | GeneID:106785387 | LOC106785387 | Gene | LOC106785387 | protein_coding | NA | character(0) | character(0) | NW_014569596.1 | 107205 | 565599 | 458395 | * | 2 | 0.00436304933518036 |
| NW_014569596.1 | 214502 | 217044 | 2543 | + | Gnomon | gene | NA | NA | gene-LOC106785399 | GeneID:106785399 | LOC106785399 | Gene | LOC106785399 | protein_coding | NA | character(0) | character(0) | NW_014569596.1 | 107205 | 565599 | 458395 | * | 2 | 0.00436304933518036 |
| NW_014569596.1 | 224623 | 257819 | 33197 | + | Gnomon | gene | NA | NA | gene-LOC106785404 | GeneID:106785404 | LOC106785404 | Gene | LOC106785404 | protein_coding | NA | character(0) | character(0) | NW_014569596.1 | 107205 | 565599 | 458395 | * | 2 | 0.00436304933518036 |
| NW_014569596.1 | 260991 | 261809 | 819 | - | Gnomon | gene | NA | NA | gene-LOC106785370 | GeneID:106785370 | LOC106785370 | Gene | LOC106785370 | protein_coding | NA | character(0) | character(0) | NW_014569596.1 | 107205 | 565599 | 458395 | * | 2 | 0.00436304933518036 |

...

... only first 25 lines shown ...

Directory: MRPR File: gwr-Pc_PA_qn.txt

| seqnames | start | end | width | strand | source | type | score | phase | ID | Dbxref | Name | gbkey | gene | gene_biotype | partial | start_range | end_range | seqnames | start | end | width | strand | NbrSites | Sdnsty |
|---|---|---|---|---|---|---|---|---|---|---|---|---|---|---|---|---|---|---|---|---|---|---|---|---|
| NW_014569572.1 | 1083329 | 1143708 | 60380 | - | Gnomon | gene | NA | NA | gene-LOC106783780 | GeneID:106783780 | LOC106783780 | Gene | LOC106783780 | protein_coding | NA | character(0) | character(0) | NW_014569572.1 | 1094647 | 1094662 | 16 | * | 10 | 625 |
| NW_014569572.1 | 1083329 | 1143708 | 60380 | - | Gnomon | gene | NA | NA | gene-LOC106783780 | GeneID:106783780 | LOC106783780 | Gene | LOC106783780 | protein_coding | NA | character(0) | character(0) | NW_014569572.1 | 1094710 | 1094724 | 15 | * | 9 | 600 |
| NW_014569601.1 | 163289 | 500575 | 337287 | + | Gnomon | gene | NA | NA | gene-LOC106785642 | GeneID:106785642 | LOC106785642 | Gene | LOC106785642 | protein_coding | NA | character(0) | character(0) | NW_014569601.1 | 298560 | 298573 | 14 | * | 7 | 500 |
| NW_014569616.1 | 377938 | 380107 | 2170 | + | Gnomon | gene | NA | NA | gene-LOC106786231 | GeneID:106786231 | LOC106786231 | Gene | LOC106786231 | protein_coding | NA | character(0) | character(0) | NW_014569616.1 | 379670 | 379681 | 12 | * | 6 | 500 |
| NW_014569559.1 | 278931 | 280191 | 1261 | + | Gnomon | gene | NA | NA | gene-LOC106793216 | GeneID:106793216 | LOC106793216 | Gene | LOC106793216 | protein_coding | NA | character(0) | character(0) | NW_014569559.1 | 279940 | 279952 | 13 | * | 6 | 461.538461538462 |
| NW_014569601.1 | 163289 | 500575 | 337287 | + | Gnomon | gene | NA | NA | gene-LOC106785642 | GeneID:106785642 | LOC106785642 | Gene | LOC106785642 | protein_coding | NA | character(0) | character(0) | NW_014569601.1 | 298540 | 298552 | 13 | * | 6 | 461.538461538462 |
| NW_014569615.1 | 134518 | 137701 | 3184 | + | Gnomon | gene | NA | NA | gene-LOC106786225 | GeneID:106786225 | LOC106786225 | Gene | LOC106786225 | protein_coding | NA | character(0) | character(0) | NW_014569615.1 | 136009 | 136021 | 13 | * | 6 | 461.538461538462 |
| NW_014569617.1 | 65215 | 67852 | 2638 | + | Gnomon | gene | NA | NA | gene-LOC106786260 | GeneID:106786260 | LOC106786260 | Gene | LOC106786260 | protein_coding | NA | character(0) | character(0) | NW_014569617.1 | 66084 | 66098 | 15 | * | 6 | 400 |
| NW_014569644.1 | 511063 | 516728 | 5666 | + | Gnomon | gene | NA | NA | gene-LOC106787290 | GeneID:106787290 | LOC106787290 | Gene | LOC106787290 | protein_coding | NA | character(0) | character(0) | NW_014569644.1 | 512191 | 512205 | 15 | * | 6 | 400 |
| NW_014569553.1 | 1432863 | 1452812 | 19950 | - | Gnomon | gene | NA | NA | gene-LOC106788969 | GeneID:106788969 | LOC106788969 | Gene | LOC106788969 | protein_coding | NA | character(0) | character(0) | NW_014569553.1 | 1451862 | 1451877 | 16 | * | 6 | 375 |
| NW_014569574.1 | 962557 | 968466 | 5910 | + | Gnomon | gene | NA | NA | gene-LOC106783913 | GeneID:106783913 | LOC106783913 | Gene | LOC106783913 | protein_coding | NA | character(0) | character(0) | NW_014569574.1 | 963687 | 963702 | 16 | * | 6 | 375 |
| NW_014569586.1 | 526658 | 648722 | 122065 | - | Gnomon | gene | NA | NA | gene-LOC106784713 | GeneID:106784713 | LOC106784713 | Gene | LOC106784713 | protein_coding | NA | character(0) | character(0) | NW_014569586.1 | 604958 | 604973 | 16 | * | 6 | 375 |
| NW_014569602.1 | 821699 | 826707 | 5009 | - | Gnomon | gene | NA | NA | gene-LOC106785716 | GeneID:106785716 | LOC106785716 | Gene | LOC106785716 | protein_coding | NA | character(0) | character(0) | NW_014569602.1 | 823342 | 823357 | 16 | * | 6 | 375 |
| NW_014569664.1 | 430677 | 458086 | 27410 | + | Gnomon | gene | NA | NA | gene-LOC106787876 | GeneID:106787876 | LOC106787876 | Gene | LOC106787876 | lncRNA | NA | character(0) | character(0) | NW_014569664.1 | 434571 | 434586 | 16 | * | 6 | 375 |
| NW_014569910.1 | 2611 | 26228 | 23618 | + | Gnomon | gene | NA | NA | gene-LOC106791980 | GeneID:106791980 | LOC106791980 | Gene | LOC106791980 | protein_coding | NA | character(0) | character(0) | NW_014569910.1 | 14533 | 14548 | 16 | * | 6 | 375 |
| NW_014570016.1 | 51933 | 54414 | 2482 | - | Gnomon | gene | NA | NA | gene-LOC106792553 | GeneID:106792553 | LOC106792553 | Gene | LOC106792553 | protein_coding | NA | character(0) | character(0) | NW_014570016.1 | 53123 | 53138 | 16 | * | 6 | 375 |
| NW_014569574.1 | 586111 | 631586 | 45476 | + | Gnomon | gene | NA | NA | gene-LOC106783850 | GeneID:106783850 | LOC106783850 | Gene | LOC106783850 | protein_coding | true | c(".", "586111") | character(0) | NW_014569574.1 | 620870 | 620888 | 19 | * | 7 | 368.421052631579 |
| NW_014569591.1 | 259959 | 509338 | 249380 | - | Gnomon | gene | NA | NA | gene-LOC106785034 | GeneID:106785034 | LOC106785034 | Gene | LOC106785034 | protein_coding | NA | character(0) | character(0) | NW_014569591.1 | 439928 | 439946 | 19 | * | 7 | 368.421052631579 |
| NW_014569619.1 | 14836 | 60089 | 45254 | + | Gnomon | gene | NA | NA | gene-LOC106786381 | GeneID:106786381 | LOC106786381 | Gene | LOC106786381 | protein_coding | NA | character(0) | character(0) | NW_014569619.1 | 50206 | 50224 | 19 | * | 7 | 368.421052631579 |
| NW_014569662.1 | 233378 | 455905 | 222528 | + | Gnomon | gene | NA | NA | gene-LOC106787804 | GeneID:106787804 | LOC106787804 | Gene | LOC106787804 | protein_coding | NA | character(0) | character(0) | NW_014569662.1 | 381625 | 381643 | 19 | * | 7 | 368.421052631579 |
| NW_014569672.1 | 65035 | 344355 | 279321 | + | Gnomon | gene | NA | NA | gene-LOC106788091 | GeneID:106788091 | LOC106788091 | Gene | LOC106788091 | protein_coding | NA | character(0) | character(0) | NW_014569672.1 | 276958 | 276976 | 19 | * | 7 | 368.421052631579 |
| NW_014569593.1 | 417287 | 487336 | 70050 | - | Gnomon | gene | NA | NA | gene-LOC106785173 | GeneID:106785173 | LOC106785173 | Gene | LOC106785173 | protein_coding | NA | character(0) | character(0) | NW_014569593.1 | 458070 | 458091 | 22 | * | 8 | 363.636363636364 |
| NW_014569601.1 | 163289 | 500575 | 337287 | + | Gnomon | gene | NA | NA | gene-LOC106785642 | GeneID:106785642 | LOC106785642 | Gene | LOC106785642 | protein_coding | NA | character(0) | character(0) | NW_014569601.1 | 446509 | 446530 | 22 | * | 8 | 363.636363636364 |
| NW_014569569.1 | 753050 | 992491 | 239442 | - | Gnomon | gene | NA | NA | gene-LOC106794051 | GeneID:106794051 | LOC106794051 | Gene | LOC106794051 | protein_coding | NA | character(0) | character(0) | NW_014569569.1 | 888237 | 888264 | 28 | * | 10 | 357.142857142857 |
| NW_014569548.1 | 2459355 | 2479961 | 20607 | + | Gnomon | gene | NA | NA | gene-LOC106784077 | GeneID:106784077 | LOC106784077 | Gene | LOC106784077 | protein_coding | NA | character(0) | character(0) | NW_014569548.1 | 2472708 | 2472738 | 31 | * | 11 | 354.838709677419 |
| NW_014569559.1 | 39048 | 193247 | 154200 | - | Gnomon | gene | NA | NA | gene-LOC106793193 | GeneID:106793193 | LOC106793193 | Gene | LOC106793193 | protein_coding | NA | character(0) | character(0) | NW_014569559.1 | 117897 | 117933 | 37 | * | 13 | 351.351351351351 |

Directory: MRPR File: gwr-Pc_PA_wr.txt

| seqnames | start | end | width | strand | source | type | score | phase | ID | Dbxref | Name | gbkey | gene | gene_biotype | partial | start_range | end_range | seqnames | start | end | width | strand | NbrSites | Sdnsty |
|---|---|---|---|---|---|---|---|---|---|---|---|---|---|---|---|---|---|---|---|---|---|---|---|---|
| NW_014569756.1 | 78660 | 80705 | 2046 | + | Gnomon | gene | NA | NA | gene-LOC106789972 | GeneID:106789972 | LOC106789972 | Gene | LOC106789972 | protein_coding | NA | character(0) | character(0) | NW_014569756.1 | 80429 | 80434 | 6 | * | 6 | 1000 |
| NW_014569613.1 | 554468 | 575845 | 21378 | - | Gnomon | gene | NA | NA | gene-LOC106786120 | GeneID:106786120 | LOC106786120 | Gene | LOC106786120 | protein_coding | NA | character(0) | character(0) | NW_014569613.1 | 571082 | 571094 | 13 | * | 6 | 461.538461538462 |
| NW_014569553.1 | 1171515 | 1173654 | 2140 | + | Gnomon | gene | NA | NA | gene-LOC106789759 | GeneID:106789759 | LOC106789759 | Gene | LOC106789759 | protein_coding | NA | character(0) | character(0) | NW_014569553.1 | 1172276 | 1172293 | 18 | * | 8 | 444.444444444444 |
| NW_014569601.1 | 163289 | 500575 | 337287 | + | Gnomon | gene | NA | NA | gene-LOC106785642 | GeneID:106785642 | LOC106785642 | Gene | LOC106785642 | protein_coding | NA | character(0) | character(0) | NW_014569601.1 | 298540 | 298571 | 32 | * | 13 | 406.25 |
| NW_014569617.1 | 65215 | 67852 | 2638 | + | Gnomon | gene | NA | NA | gene-LOC106786260 | GeneID:106786260 | LOC106786260 | Gene | LOC106786260 | protein_coding | NA | character(0) | character(0) | NW_014569617.1 | 66084 | 66098 | 15 | * | 6 | 400 |
| NW_014569644.1 | 511063 | 516728 | 5666 | + | Gnomon | gene | NA | NA | gene-LOC106787290 | GeneID:106787290 | LOC106787290 | Gene | LOC106787290 | protein_coding | NA | character(0) | character(0) | NW_014569644.1 | 512191 | 512205 | 15 | * | 6 | 400 |
| NW_014569667.1 | 254947 | 258077 | 3131 | - | Gnomon | gene | NA | NA | gene-LOC106787918 | GeneID:106787918 | LOC106787918 | Gene | LOC106787918 | protein_coding | NA | character(0) | character(0) | NW_014569667.1 | 255801 | 255815 | 15 | * | 6 | 400 |
| NW_014569959.1 | 65227 | 68675 | 3449 | - | Gnomon | gene | NA | NA | gene-LOC106792366 | GeneID:106792366 | LOC106792366 | Gene | LOC106792366 | protein_coding | NA | character(0) | character(0) | NW_014569959.1 | 66990 | 67004 | 15 | * | 6 | 400 |
| NW_014569586.1 | 526658 | 648722 | 122065 | - | Gnomon | gene | NA | NA | gene-LOC106784713 | GeneID:106784713 | LOC106784713 | Gene | LOC106784713 | protein_coding | NA | character(0) | character(0) | NW_014569586.1 | 604958 | 604973 | 16 | * | 6 | 375 |
| NW_014569602.1 | 821699 | 826707 | 5009 | - | Gnomon | gene | NA | NA | gene-LOC106785716 | GeneID:106785716 | LOC106785716 | Gene | LOC106785716 | protein_coding | NA | character(0) | character(0) | NW_014569602.1 | 823342 | 823357 | 16 | * | 6 | 375 |
| NW_014569632.1 | 352527 | 484228 | 131702 | + | Gnomon | gene | NA | NA | gene-LOC106786852 | GeneID:106786852 | LOC106786852 | Gene | LOC106786852 | protein_coding | NA | character(0) | character(0) | NW_014569632.1 | 372328 | 372343 | 16 | * | 6 | 375 |
| NW_014569872.1 | 112597 | 143816 | 31220 | + | Gnomon | gene | NA | NA | gene-LOC106791611 | GeneID:106791611 | LOC106791611 | Gene | LOC106791611 | protein_coding | NA | character(0) | character(0) | NW_014569872.1 | 112887 | 112902 | 16 | * | 6 | 375 |
| NW_014569910.1 | 2611 | 26228 | 23618 | + | Gnomon | gene | NA | NA | gene-LOC106791980 | GeneID:106791980 | LOC106791980 | Gene | LOC106791980 | protein_coding | NA | character(0) | character(0) | NW_014569910.1 | 14533 | 14548 | 16 | * | 6 | 375 |
| NW_014570016.1 | 51933 | 54414 | 2482 | - | Gnomon | gene | NA | NA | gene-LOC106792553 | GeneID:106792553 | LOC106792553 | Gene | LOC106792553 | protein_coding | NA | character(0) | character(0) | NW_014570016.1 | 53123 | 53138 | 16 | * | 6 | 375 |
| NW_014569574.1 | 586111 | 631586 | 45476 | + | Gnomon | gene | NA | NA | gene-LOC106783850 | GeneID:106783850 | LOC106783850 | Gene | LOC106783850 | protein_coding | true | c(".", "586111") | character(0) | NW_014569574.1 | 620870 | 620888 | 19 | * | 7 | 368.421052631579 |
| NW_014569591.1 | 259959 | 509338 | 249380 | - | Gnomon | gene | NA | NA | gene-LOC106785034 | GeneID:106785034 | LOC106785034 | Gene | LOC106785034 | protein_coding | NA | character(0) | character(0) | NW_014569591.1 | 439928 | 439946 | 19 | * | 7 | 368.421052631579 |
| NW_014569619.1 | 14836 | 60089 | 45254 | + | Gnomon | gene | NA | NA | gene-LOC106786381 | GeneID:106786381 | LOC106786381 | Gene | LOC106786381 | protein_coding | NA | character(0) | character(0) | NW_014569619.1 | 50206 | 50224 | 19 | * | 7 | 368.421052631579 |
| NW_014569662.1 | 233378 | 455905 | 222528 | + | Gnomon | gene | NA | NA | gene-LOC106787804 | GeneID:106787804 | LOC106787804 | Gene | LOC106787804 | protein_coding | NA | character(0) | character(0) | NW_014569662.1 | 381625 | 381643 | 19 | * | 7 | 368.421052631579 |
| NW_014569672.1 | 65035 | 344355 | 279321 | + | Gnomon | gene | NA | NA | gene-LOC106788091 | GeneID:106788091 | LOC106788091 | Gene | LOC106788091 | protein_coding | NA | character(0) | character(0) | NW_014569672.1 | 276958 | 276976 | 19 | * | 7 | 368.421052631579 |
| NW_014569553.1 | 1432863 | 1452812 | 19950 | - | Gnomon | gene | NA | NA | gene-LOC106788969 | GeneID:106788969 | LOC106788969 | Gene | LOC106788969 | protein_coding | NA | character(0) | character(0) | NW_014569553.1 | 1451862 | 1451883 | 22 | * | 8 | 363.636363636364 |
| NW_014569593.1 | 417287 | 487336 | 70050 | - | Gnomon | gene | NA | NA | gene-LOC106785173 | GeneID:106785173 | LOC106785173 | Gene | LOC106785173 | protein_coding | NA | character(0) | character(0) | NW_014569593.1 | 458070 | 458091 | 22 | * | 8 | 363.636363636364 |
| NW_014569653.1 | 46258 | 281289 | 235032 | - | Gnomon | gene | NA | NA | gene-LOC106787574 | GeneID:106787574 | LOC106787574 | Gene | LOC106787574 | protein_coding | NA | character(0) | character(0) | NW_014569653.1 | 214404 | 214425 | 22 | * | 8 | 363.636363636364 |
| NW_014569664.1 | 430677 | 458086 | 27410 | + | Gnomon | gene | NA | NA | gene-LOC106787876 | GeneID:106787876 | LOC106787876 | Gene | LOC106787876 | lncRNA | NA | character(0) | character(0) | NW_014569664.1 | 434571 | 434595 | 25 | * | 9 | 360 |
| NW_014569569.1 | 753050 | 992491 | 239442 | - | Gnomon | gene | NA | NA | gene-LOC106794051 | GeneID:106794051 | LOC106794051 | Gene | LOC106794051 | protein_coding | NA | character(0) | character(0) | NW_014569569.1 | 888237 | 888264 | 28 | * | 10 | 357.142857142857 |
| NW_014569601.1 | 163289 | 500575 | 337287 | + | Gnomon | gene | NA | NA | gene-LOC106785642 | GeneID:106785642 | LOC106785642 | Gene | LOC106785642 | protein_coding | NA | character(0) | character(0) | NW_014569601.1 | 446503 | 446530 | 28 | * | 10 | 357.142857142857 |
| NW_014569548.1 | 2459355 | 2479961 | 20607 | + | Gnomon | gene | NA | NA | gene-LOC106784077 | GeneID:106784077 | LOC106784077 | Gene | LOC106784077 | protein_coding | NA | character(0) | character(0) | NW_014569548.1 | 2472708 | 2472741 | 34 | * | 12 | 352.941176470588 |
| NW_014569559.1 | 39048 | 193247 | 154200 | - | Gnomon | gene | NA | NA | gene-LOC106793193 | GeneID:106793193 | LOC106793193 | Gene | LOC106793193 | protein_coding | NA | character(0) | character(0) | NW_014569559.1 | 117894 | 117933 | 40 | * | 14 | 350 |

```
Directory: MRPR File: rmp-Pc_PA_qn.txt


Overlap of methylation-rich regions with genome features for Pc sample Pc_PA_qn

Total size of the methylation-rich regions of Pc_PA_qn:                                         843 bp (  0.0% of genome)
Size of overlap of methylation-rich regions of Pc Pc_PA_qn with genic regions:                     473 bp ( 56.1% of total size)   ( 0.94 O/E)
Size of overlap of methylation-rich regions of Pc Pc_PA_qn with   exon regions:                      151 bp ( 17.9% of total size) ( 2.92 O/E)
Size of overlap of methylation-rich regions of Pc Pc_PA_qn with   intron regions:                    322 bp ( 38.2% of total size) ( 0.82 O/E)
Size of overlap of methylation-rich regions of Pc Pc_PA_qn with intergenic regions:                370 bp ( 43.9% of total size)   ( 1.39 O/E)
Size of overlap of methylation-rich regions of Pc Pc_PA_qn with   promoter regions:                   76 bp (  9.0% of total size) ( 3.59 O/E)
Size of overlap of methylation-rich regions of Pc Pc_PA_qn with   other intergenic regions:          294 bp ( 34.9% of total size) ( 0.92 O/E)


Overlap of methylation-poor regions with genome features for Pc sample Pc_PA_qn

Total size of the methylation-poor regions of Pc_PA_qn:                                    22116818 bp ( 10.5% of genome)
Size of overlap of methylation-poor regions of Pc Pc_PA_qn with genic regions:               16463545 bp ( 74.4% of total size)   ( 1.25 O/E)
Size of overlap of methylation-poor regions of Pc Pc_PA_qn with   exon regions:                 2695453 bp ( 12.2% of total size) ( 4.76 O/E)
Size of overlap of methylation-poor regions of Pc Pc_PA_qn with   intron regions:              13768092 bp ( 62.2% of total size) ( 1.34 O/E)
Size of overlap of methylation-poor regions of Pc Pc_PA_qn with intergenic regions:           5653273 bp ( 25.6% of total size)   ( 1.84 O/E)
Size of overlap of methylation-poor regions of Pc Pc_PA_qn with   promoter regions:              333472 bp (  1.5% of total size) ( 0.60 O/E)
Size of overlap of methylation-poor regions of Pc Pc_PA_qn with   other intergenic regions:     5319801 bp ( 24.1% of total size) ( 0.63 O/E)
```

Directory: MRPR File: rmp-Pc_PA_wr.txt

```
Overlap of methylation-rich regions with genome features for Pc sample Pc_PA_wr

Total size of the methylation-rich regions of Pc_PA_wr:                                        814 bp (  0.0% of genome)
Size of overlap of methylation-rich regions of Pc Pc_PA_wr with genic regions:                    541 bp ( 66.5% of total size)   ( 1.12 O/E)
Size of overlap of methylation-rich regions of Pc Pc_PA_wr with   exon regions:                   160 bp ( 19.7% of total size) ( 3.58 O/E)
Size of overlap of methylation-rich regions of Pc Pc_PA_wr with   intron regions:                 381 bp ( 46.8% of total size) ( 1.01 O/E)
Size of overlap of methylation-rich regions of Pc Pc_PA_wr with intergenic regions:               273 bp ( 33.5% of total size)   ( 1.64 O/E)
Size of overlap of methylation-rich regions of Pc Pc_PA_wr with   promoter regions:                41 bp (  5.0% of total size) ( 2.01 O/E)
Size of overlap of methylation-rich regions of Pc Pc_PA_wr with   other intergenic regions:       232 bp ( 28.5% of total size) ( 0.75 O/E)


Overlap of methylation-poor regions with genome features for Pc sample Pc_PA_wr

Total size of the methylation-poor regions of Pc_PA_wr:                                   22853587 bp ( 10.8% of genome)
Size of overlap of methylation-poor regions of Pc Pc_PA_wr with genic regions:               17391493 bp ( 76.1% of total size)   ( 1.28 O/E)
Size of overlap of methylation-poor regions of Pc Pc_PA_wr with   exon regions:               2784904 bp ( 12.2% of total size) ( 4.89 O/E)
Size of overlap of methylation-poor regions of Pc Pc_PA_wr with   intron regions:            14606589 bp ( 63.9% of total size) ( 1.38 O/E)
Size of overlap of methylation-poor regions of Pc Pc_PA_wr with intergenic regions:           5462094 bp ( 23.9% of total size)   ( 1.88 O/E)
Size of overlap of methylation-poor regions of Pc Pc_PA_wr with   promoter regions:            365392 bp (  1.6% of total size) ( 0.64 O/E)
Size of overlap of methylation-poor regions of Pc Pc_PA_wr with   other intergenic regions:   5096702 bp ( 22.3% of total size) ( 0.59 O/E)
```

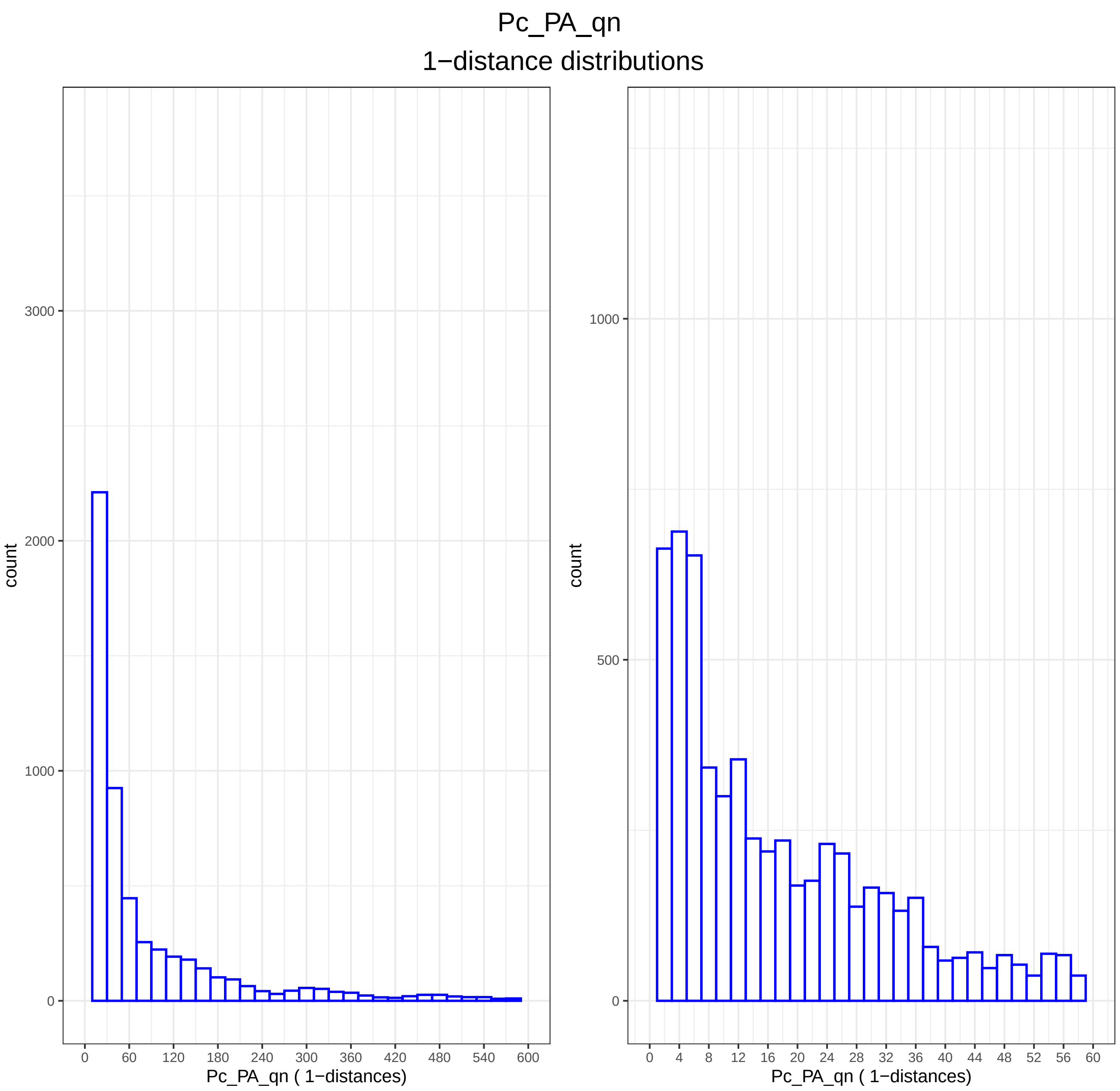
Pc_PA_qn
1−distance distributions
count
0
1000
2000
3000
0
60
120
180
240
300
360
420
480
540
600
Pc_PA_qn ( 1−distances)
count
0
500
1000
0
4
8
12
16
20
24
28
32
36
40
44
48
52
56
60
Pc_PA_qn ( 1−distances)

Pc_PA_wr
1−distance distributions

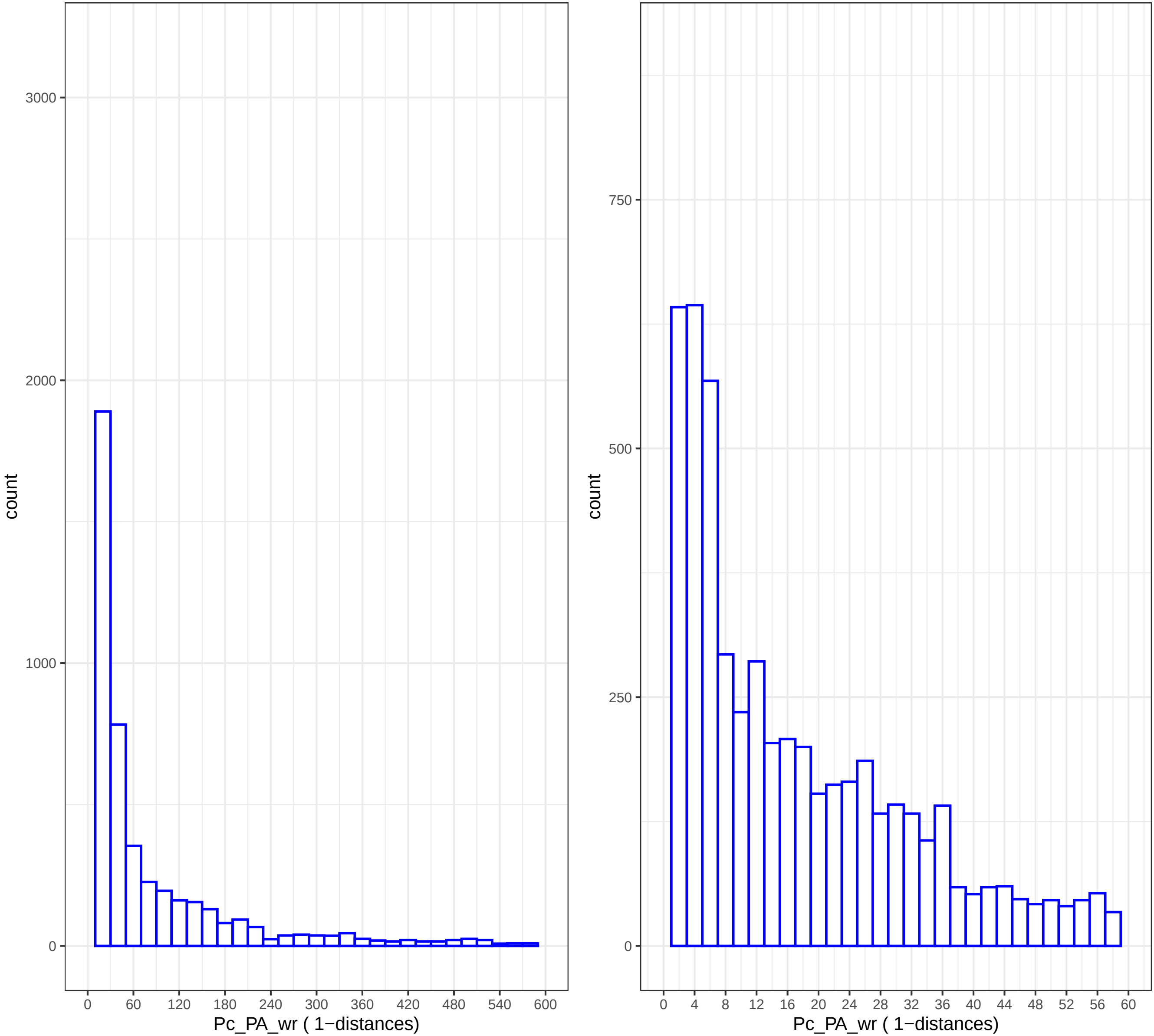

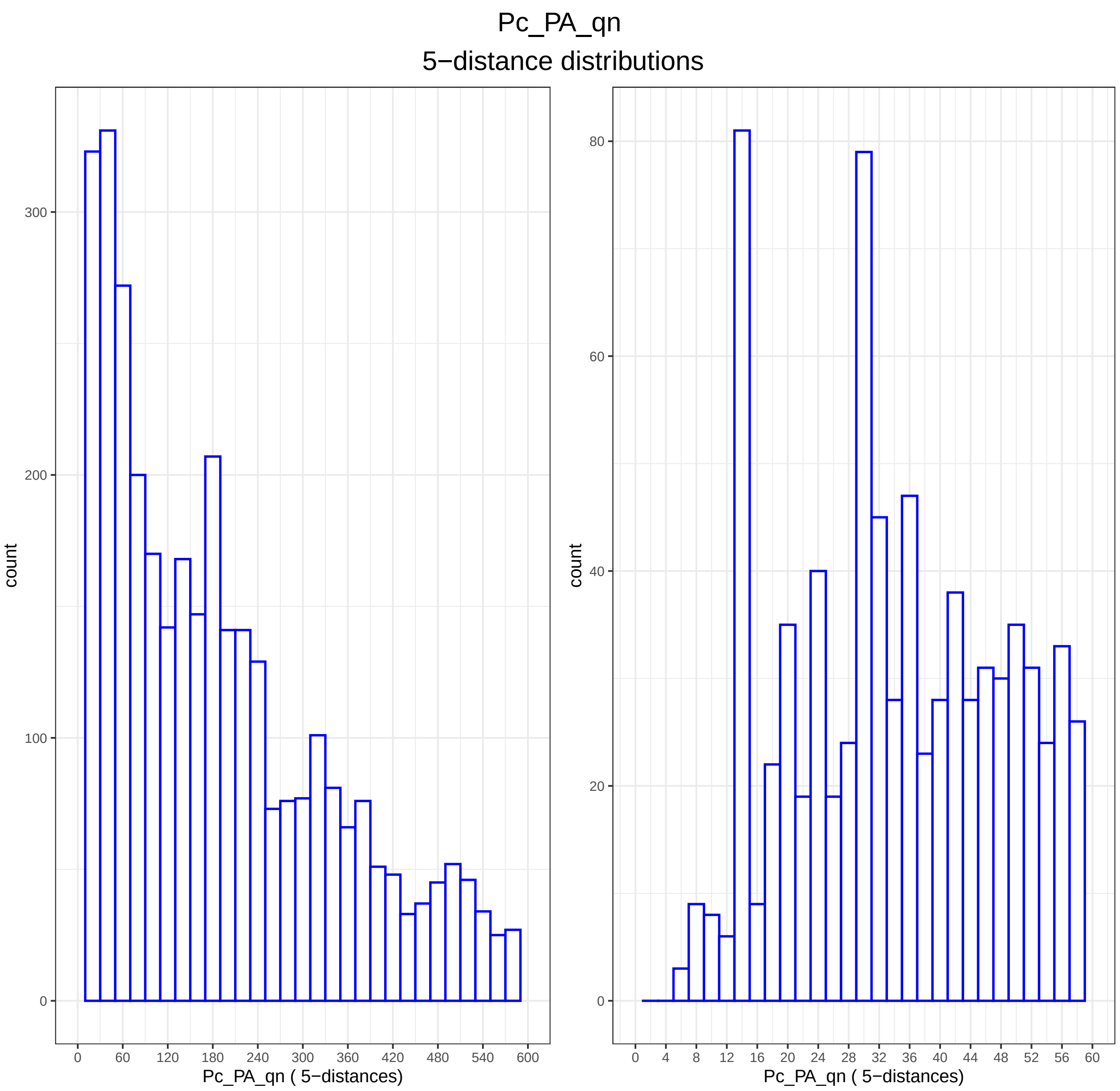

Pc_PA_qn
5−distance distributions
count
300
200
100
0
0 60 120 180 240 300 360 420 480 540 600
Pc_PA_qn ( 5−distances)
count
80
60
40
20
0
0 4 8 12 16 20 24 28 32 36 40 44 48 52 56 60
Pc_PA_qn ( 5−distances)

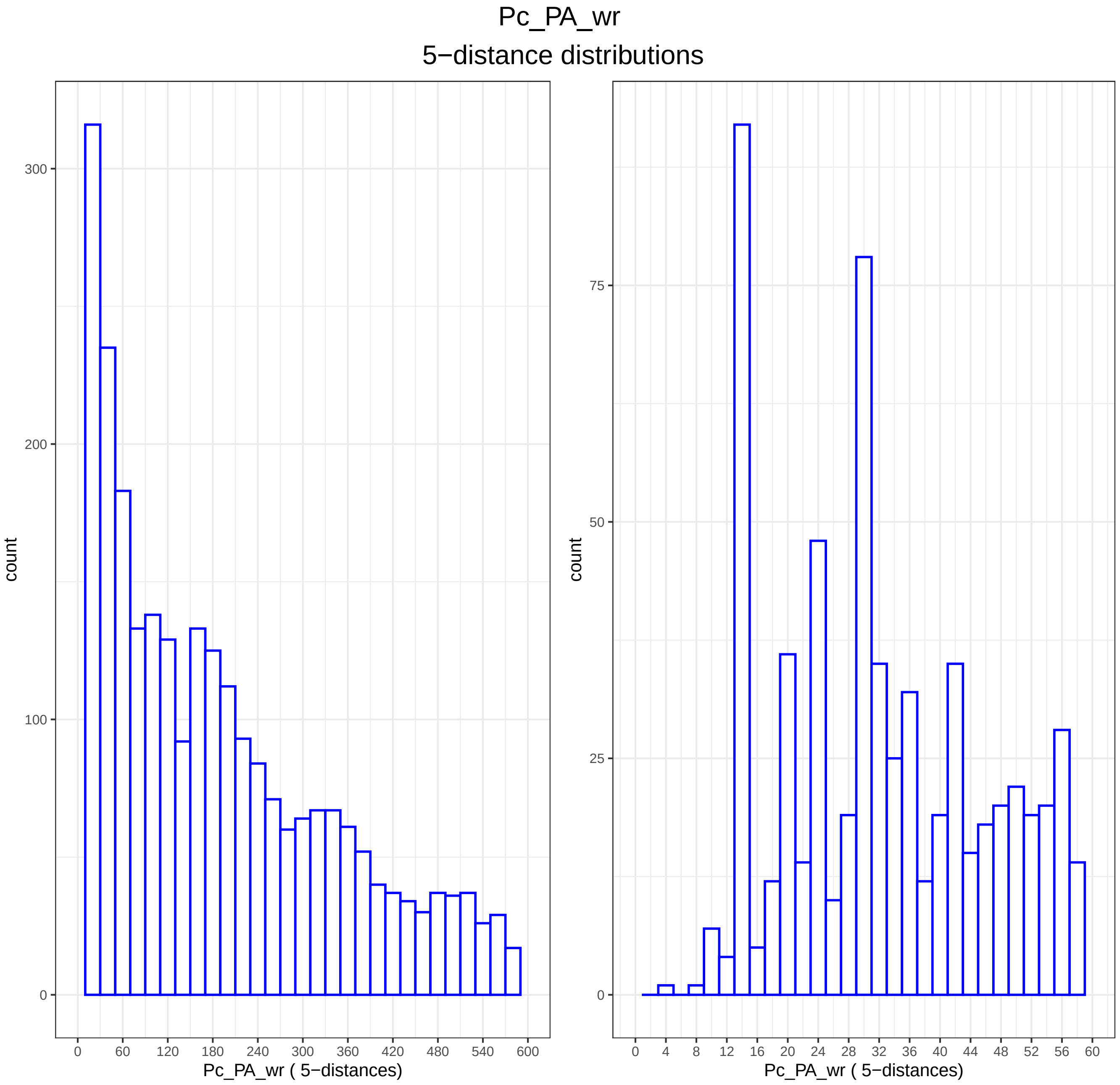

Pc_PA_wr
5−distance distributions
count
300
200
100
0
0 60 120 180 240 300 360 420 480 540 600
Pc_PA_wr ( 5−distances)
count
75
50
25
0
0 4 8 12 16 20 24 28 32 36 40 44 48 52 56 60
Pc_PA_wr ( 5−distances)

```
Directory: DMT File: 0READMEdmt


DMT - Differentially methylated tiles.

Input: studymc wsize stepsize threshold qvalue
Output: files dmt-*.txt dmg-*.txt

Notes: Output files dmt-*.txt and dmg-*.txt are generated by BWASPR::det_dmt() and show the
        differentially methylated tiles and genes as determined by methylKit::getMethylDiff with
        parameters difference=threshold and qvalue, as provided in the Pc_PA.conf
        configuration file.  Tiles refer to sliding windows along the genome within which
        methylation calls are cumulated by methylKit::tileMethylCounts.
        The dmg-*.txt files show all genes with at least one differentially methylated tile.

        A positive meth.diff value in comparison A.vs.B means that the B methylation percentage
        in that tile is higher than the A methylation percentage.
```

Directory: DMT File: dmg-Pc_PA.txt

| seqnames | start | end | width | strand | source | type | score | phase | ID | Dbxref | Name | gbkey | gene | gene_biotype | partial | start_range | end_range | pvalue | qvalue | meth.diff | comparison |
|---|---|---|---|---|---|---|---|---|---|---|---|---|---|---|---|---|---|---|---|---|---|
| NW_014569547.1 | 1307608 | 1312253 | 4646 | - | Gnomon | gene | NA | NA | gene-LOC106792988 | GeneID:106792988 | LOC106792988 | Gene | LOC106792988 | protein_coding | NA | character(0) | character(0) | 6.77875590409025e-09 | 1.88686082955098e-05 | 26.5775881428251 | queen.vs.worker |
| NW_014569589.1 | 825211 | 829866 | 4656 | + | Gnomon | gene | NA | NA | gene-LOC106784950 | GeneID:106784950 | LOC106784950 | Gene | LOC106784950 | protein_coding | NA | character(0) | character(0) | 3.89392027386584e-15 | 4.5387028305882e-11 | 31.2830547916226 | queen.vs.worker |
| NW_014569762.1 | 235821 | 237897 | 2077 | - | Gnomon | gene | NA | NA | gene-LOC106790121 | GeneID:106790121 | LOC106790121 | Gene | LOC106790121 | protein_coding | NA | character(0) | character(0) | 2.27771734989549e-11 | 1.14805523079309e-07 | 25.0655361677726 | queen.vs.worker |
| NW_014569841.1 | 191242 | 197243 | 6002 | + | Gnomon | gene | NA | NA | gene-LOC106791291 | GeneID:106791291 | LOC106791291 | Gene | LOC106791291 | protein_coding | NA | character(0) | character(0) | 9.88176657335068e-06 | 0.00777590454347625 | -25.7142857142857 | queen.vs.worker |
| NW_014569870.1 | 11096 | 71850 | 60755 | - | Gnomon | gene | NA | NA | gene-LOC106791597 | GeneID:106791597 | LOC106791597 | Gene | LOC106791597 | protein_coding | NA | character(0) | character(0) | 9.22397708850916e-09 | 2.3891883857952e-05 | 34.2790516906335 | queen.vs.worker |
| NW_014569870.1 | 49833 | 54006 | 4174 | + | Gnomon | gene | NA | NA | gene-LOC106791596 | GeneID:106791596 | LOC106791596 | Gene | LOC106791596 | protein_coding | NA | character(0) | character(0) | 9.22397708850916e-09 | 2.3891883857952e-05 | 34.2790516906335 | queen.vs.worker |

Directory: DMT File: dmt-Pc_PA.txt

| seqnames | start | end | width | strand | pvalue | qvalue | meth.diff | comparison |
|---|---|---|---|---|---|---|---|---|
| NW_014569547.1 | 1310001 | 1311000 | 1000 | * | 6.77875590409025e-09 | 0 | 26.58 | queen.vs.worker |
| NW_014569589.1 | 826001 | 827000 | 1000 | * | 3.89392027386584e-15 | 0 | 31.28 | queen.vs.worker |
| NW_014569762.1 | 236001 | 237000 | 1000 | * | 2.27771734989549e-11 | 0 | 25.07 | queen.vs.worker |
| NW_014569841.1 | 192001 | 193000 | 1000 | * | 9.88176657335068e-06 | 0.008 | -25.71 | queen.vs.worker |
| NW_014569870.1 | 51001 | 52000 | 1000 | * | 9.22397708850916e-09 | 0 | 34.28 | queen.vs.worker |

```
Directory: DMSG File: 0READMEdmsg


DMSG - Differentially methylated sites and genes.

Input: studyhc threshold qvalue
       The studyhc methylKit raw object contains all CpGscd sites with coverage at least
        20 reads.
Output: files dms-*.txt dmg-*.txt  dmg-*details.txt dmg-*heatmaps.pdf

Notes: Output files dms-*.txt and dmg-*.txt are generated by BWASPR::det_dmsg() and show the
        differentially methylated sites and genes as determined by methylKit::getMethylDiff with
        parameters difference=threshold and qvalue, as provided in the Pc_PA.conf
        configuration file.  The table of differentially methylated genes contains all genes
        with at least one differentially methylated site.

       The files dmg-*details.txt, generated by BWASPR::show_dmsg(), show all CpGscd sites in
        the differentially methylated genes, with coverage numbers and methylation percentages.

       The files dmg-*heatmaps.pdf, generated by BWASPR::show_dmsg(), give heatmap displays for
        genes meeting the following criteria: 1) there are between minNsites and maxNsites common
        CpGscd sites; 2) at least minPdmsites % of these sites are differentially methylated sites.
        The parameters are set in the Pc_PA.conf configuration file.
```

Directory: DMSG File: dmg-Pc_PA.txt

| seqnames | start | end | width | strand | source | type | score | phase | ID | Dbxref | Name | gbkey | gene | gene_biotype | partial | start_range | end_range | comparison |
|---|---|---|---|---|---|---|---|---|---|---|---|---|---|---|---|---|---|---|
| NW_014569550.1 | 2330862 | 2338113 | 7252 | - | Gnomon | gene | NA | NA | gene-LOC106786709 | GeneID:106786709 | LOC106786709 | Gene | LOC106786709 | protein_coding | NA | character(0) | character(0) | queen.vs.worker |
| NW_014569553.1 | 827076 | 853561 | 26486 | + | Gnomon | gene | NA | NA | gene-LOC106789019 | GeneID:106789019 | LOC106789019 | Gene | LOC106789019 | protein_coding | NA | character(0) | character(0) | queen.vs.worker |
| NW_014569564.1 | 840784 | 888336 | 47553 | - | Gnomon | gene | NA | NA | gene-LOC106793677 | GeneID:106793677 | LOC106793677 | Gene | LOC106793677 | protein_coding | NA | character(0) | character(0) | queen.vs.worker |
| NW_014569564.1 | 860813 | 883639 | 22827 | - | Gnomon | gene | NA | NA | gene-LOC106793698 | GeneID:106793698 | LOC106793698 | Gene | LOC106793698 | protein_coding | NA | character(0) | character(0) | queen.vs.worker |
| NW_014569636.1 | 549495 | 554919 | 5425 | + | Gnomon | gene | NA | NA | gene-LOC106786951 | GeneID:106786951 | LOC106786951 | Gene | LOC106786951 | protein_coding | NA | character(0) | character(0) | queen.vs.worker |
| NW_014569637.1 | 218086 | 224301 | 6216 | - | Gnomon | gene | NA | NA | gene-LOC106787037 | GeneID:106787037 | LOC106787037 | Gene | LOC106787037 | protein_coding | NA | character(0) | character(0) | queen.vs.worker |

Directory: DMSG File: dmg-Pc_PA_queen.vs.worker_details.txt

| seqnames | start | end | width | strand | coverage1 | numCs1 | numTs1 | coverage2 | numCs2 | numTs2 | queen | worker | gene_seqnames | gene_start | gene_end | gene_width | gene_strand | gene_source | gene_type | gene_ID | gene_Dbxref | gene_Name | gene_gbkey | gene_gene | gene_gene_biotype | gene_start_range | gene_end_range | gene_comparison | is.dm |
|---|---|---|---|---|---|---|---|---|---|---|---|---|---|---|---|---|---|---|---|---|---|---|---|---|---|---|---|---|---|
| NW_014569550.1 | 2330950 | 2330950 | 1 | + | 43 | 1 | 42 | 36 | 0 | 36 | 2.33 | 0 | NW_014569550.1 | 2330862 | 2338113 | 7252 | - | Gnomon | gene | gene-LOC106786709 | GeneID:106786709 | LOC106786709 | Gene | LOC106786709 | protein_coding | character(0) | character(0) | queen.vs.worker | FALSE |
| NW_014569550.1 | 2331005 | 2331005 | 1 | + | 49 | 0 | 49 | 36 | 0 | 36 | 0 | 0 | NW_014569550.1 | 2330862 | 2338113 | 7252 | - | Gnomon | gene | gene-LOC106786709 | GeneID:106786709 | LOC106786709 | Gene | LOC106786709 | protein_coding | character(0) | character(0) | queen.vs.worker | FALSE |
| NW_014569550.1 | 2331013 | 2331013 | 1 | + | 48 | 2 | 46 | 34 | 0 | 34 | 4.17 | 0 | NW_014569550.1 | 2330862 | 2338113 | 7252 | - | Gnomon | gene | gene-LOC106786709 | GeneID:106786709 | LOC106786709 | Gene | LOC106786709 | protein_coding | character(0) | character(0) | queen.vs.worker | FALSE |
| NW_014569550.1 | 2331020 | 2331020 | 1 | + | 47 | 0 | 47 | 33 | 0 | 33 | 0 | 0 | NW_014569550.1 | 2330862 | 2338113 | 7252 | - | Gnomon | gene | gene-LOC106786709 | GeneID:106786709 | LOC106786709 | Gene | LOC106786709 | protein_coding | character(0) | character(0) | queen.vs.worker | FALSE |
| NW_014569550.1 | 2331030 | 2331030 | 1 | + | 46 | 1 | 45 | 29 | 0 | 29 | 2.17 | 0 | NW_014569550.1 | 2330862 | 2338113 | 7252 | - | Gnomon | gene | gene-LOC106786709 | GeneID:106786709 | LOC106786709 | Gene | LOC106786709 | protein_coding | character(0) | character(0) | queen.vs.worker | FALSE |
| NW_014569550.1 | 2331037 | 2331037 | 1 | + | 46 | 2 | 44 | 25 | 0 | 25 | 4.35 | 0 | NW_014569550.1 | 2330862 | 2338113 | 7252 | - | Gnomon | gene | gene-LOC106786709 | GeneID:106786709 | LOC106786709 | Gene | LOC106786709 | protein_coding | character(0) | character(0) | queen.vs.worker | FALSE |
| NW_014569550.1 | 2333787 | 2333787 | 1 | + | 26 | 1 | 25 | 28 | 1 | 27 | 3.85 | 3.57 | NW_014569550.1 | 2330862 | 2338113 | 7252 | - | Gnomon | gene | gene-LOC106786709 | GeneID:106786709 | LOC106786709 | Gene | LOC106786709 | protein_coding | character(0) | character(0) | queen.vs.worker | FALSE |
| NW_014569550.1 | 2333789 | 2333789 | 1 | + | 26 | 0 | 26 | 29 | 1 | 28 | 0 | 3.45 | NW_014569550.1 | 2330862 | 2338113 | 7252 | - | Gnomon | gene | gene-LOC106786709 | GeneID:106786709 | LOC106786709 | Gene | LOC106786709 | protein_coding | character(0) | character(0) | queen.vs.worker | FALSE |
| NW_014569550.1 | 2333799 | 2333799 | 1 | + | 27 | 0 | 27 | 30 | 0 | 30 | 0 | 0 | NW_014569550.1 | 2330862 | 2338113 | 7252 | - | Gnomon | gene | gene-LOC106786709 | GeneID:106786709 | LOC106786709 | Gene | LOC106786709 | protein_coding | character(0) | character(0) | queen.vs.worker | FALSE |
| NW_014569550.1 | 2333802 | 2333802 | 1 | + | 26 | 0 | 26 | 29 | 2 | 27 | 0 | 6.9 | NW_014569550.1 | 2330862 | 2338113 | 7252 | - | Gnomon | gene | gene-LOC106786709 | GeneID:106786709 | LOC106786709 | Gene | LOC106786709 | protein_coding | character(0) | character(0) | queen.vs.worker | FALSE |
| NW_014569550.1 | 2333839 | 2333839 | 1 | + | 29 | 1 | 28 | 26 | 0 | 26 | 3.45 | 0 | NW_014569550.1 | 2330862 | 2338113 | 7252 | - | Gnomon | gene | gene-LOC106786709 | GeneID:106786709 | LOC106786709 | Gene | LOC106786709 | protein_coding | character(0) | character(0) | queen.vs.worker | FALSE |
| NW_014569550.1 | 2333844 | 2333844 | 1 | + | 29 | 1 | 28 | 29 | 1 | 28 | 3.45 | 3.45 | NW_014569550.1 | 2330862 | 2338113 | 7252 | - | Gnomon | gene | gene-LOC106786709 | GeneID:106786709 | LOC106786709 | Gene | LOC106786709 | protein_coding | character(0) | character(0) | queen.vs.worker | FALSE |
| NW_014569550.1 | 2333872 | 2333872 | 1 | + | 36 | 0 | 36 | 33 | 0 | 33 | 0 | 0 | NW_014569550.1 | 2330862 | 2338113 | 7252 | - | Gnomon | gene | gene-LOC106786709 | GeneID:106786709 | LOC106786709 | Gene | LOC106786709 | protein_coding | character(0) | character(0) | queen.vs.worker | FALSE |
| NW_014569550.1 | 2333879 | 2333879 | 1 | + | 34 | 0 | 34 | 34 | 0 | 34 | 0 | 0 | NW_014569550.1 | 2330862 | 2338113 | 7252 | - | Gnomon | gene | gene-LOC106786709 | GeneID:106786709 | LOC106786709 | Gene | LOC106786709 | protein_coding | character(0) | character(0) | queen.vs.worker | FALSE |
| NW_014569550.1 | 2333890 | 2333890 | 1 | + | 36 | 1 | 35 | 32 | 1 | 31 | 2.78 | 3.12 | NW_014569550.1 | 2330862 | 2338113 | 7252 | - | Gnomon | gene | gene-LOC106786709 | GeneID:106786709 | LOC106786709 | Gene | LOC106786709 | protein_coding | character(0) | character(0) | queen.vs.worker | FALSE |
| NW_014569550.1 | 2333930 | 2333930 | 1 | + | 31 | 0 | 31 | 29 | 0 | 29 | 0 | 0 | NW_014569550.1 | 2330862 | 2338113 | 7252 | - | Gnomon | gene | gene-LOC106786709 | GeneID:106786709 | LOC106786709 | Gene | LOC106786709 | protein_coding | character(0) | character(0) | queen.vs.worker | FALSE |
| NW_014569550.1 | 2333944 | 2333944 | 1 | + | 30 | 0 | 30 | 29 | 0 | 29 | 0 | 0 | NW_014569550.1 | 2330862 | 2338113 | 7252 | - | Gnomon | gene | gene-LOC106786709 | GeneID:106786709 | LOC106786709 | Gene | LOC106786709 | protein_coding | character(0) | character(0) | queen.vs.worker | FALSE |
| NW_014569550.1 | 2333958 | 2333958 | 1 | + | 34 | 0 | 34 | 33 | 0 | 33 | 0 | 0 | NW_014569550.1 | 2330862 | 2338113 | 7252 | - | Gnomon | gene | gene-LOC106786709 | GeneID:106786709 | LOC106786709 | Gene | LOC106786709 | protein_coding | character(0) | character(0) | queen.vs.worker | FALSE |
| NW_014569550.1 | 2333966 | 2333966 | 1 | + | 36 | 0 | 36 | 36 | 1 | 35 | 0 | 2.78 | NW_014569550.1 | 2330862 | 2338113 | 7252 | - | Gnomon | gene | gene-LOC106786709 | GeneID:106786709 | LOC106786709 | Gene | LOC106786709 | protein_coding | character(0) | character(0) | queen.vs.worker | FALSE |
| NW_014569550.1 | 2333996 | 2333996 | 1 | + | 54 | 0 | 54 | 52 | 0 | 52 | 0 | 0 | NW_014569550.1 | 2330862 | 2338113 | 7252 | - | Gnomon | gene | gene-LOC106786709 | GeneID:106786709 | LOC106786709 | Gene | LOC106786709 | protein_coding | character(0) | character(0) | queen.vs.worker | FALSE |
| NW_014569550.1 | 2334043 | 2334043 | 1 | + | 69 | 1 | 68 | 76 | 0 | 76 | 1.45 | 0 | NW_014569550.1 | 2330862 | 2338113 | 7252 | - | Gnomon | gene | gene-LOC106786709 | GeneID:106786709 | LOC106786709 | Gene | LOC106786709 | protein_coding | character(0) | character(0) | queen.vs.worker | FALSE |
| NW_014569550.1 | 2334074 | 2334074 | 1 | + | 74 | 0 | 74 | 77 | 0 | 77 | 0 | 0 | NW_014569550.1 | 2330862 | 2338113 | 7252 | - | Gnomon | gene | gene-LOC106786709 | GeneID:106786709 | LOC106786709 | Gene | LOC106786709 | protein_coding | character(0) | character(0) | queen.vs.worker | FALSE |
| NW_014569550.1 | 2334078 | 2334078 | 1 | + | 73 | 3 | 70 | 77 | 1 | 76 | 4.11 | 1.3 | NW_014569550.1 | 2330862 | 2338113 | 7252 | - | Gnomon | gene | gene-LOC106786709 | GeneID:106786709 | LOC106786709 | Gene | LOC106786709 | protein_coding | character(0) | character(0) | queen.vs.worker | FALSE |
| NW_014569550.1 | 2334110 | 2334110 | 1 | + | 80 | 0 | 80 | 74 | 1 | 73 | 0 | 1.35 | NW_014569550.1 | 2330862 | 2338113 | 7252 | - | Gnomon | gene | gene-LOC106786709 | GeneID:106786709 | LOC106786709 | Gene | LOC106786709 | protein_coding | character(0) | character(0) | queen.vs.worker | FALSE |

...

... only first 25 lines shown ...

Directory: DMSG File: dms-Pc_PA.txt

| seqnames | start | end | width | strand | pvalue | qvalue | meth.diff | comparison |
|---|---|---|---|---|---|---|---|---|
| NW_014569550.1 | 2336628 | 2336628 | 1 | + | 7.76379632746078e-12 | 0 | 60.41 | queen.vs.worker |
| NW_014569553.1 | 830837 | 830837 | 1 | + | 1.33072553065564e-08 | 0.006 | 40.5 | queen.vs.worker |
| NW_014569564.1 | 880765 | 880765 | 1 | + | 9.80191488184107e-09 | 0.005 | 29.41 | queen.vs.worker |
| NW_014569636.1 | 554174 | 554174 | 1 | + | 1.51392932012244e-11 | 0 | 28.51 | queen.vs.worker |
| NW_014569637.1 | 219972 | 219972 | 1 | + | 2.76347915661357e-08 | 0.009 | 36.48 | queen.vs.worker |
| NW_014569637.1 | 219986 | 219986 | 1 | + | 7.81498957500295e-10 | 0.001 | 42.41 | queen.vs.worker |

```
Directory: OGL File: 0READMEogl


OGL - Ordered gene lists.

Input: studyhc annotation   maxgwidth minnbrdmsites
       dmgprp (= output of BWASPR:show_dmsg())

Output: files ogl-*.txt rnk-dmg-*.txt
         ogl-<study>_<sample>.txt ogl-<study>_<sample1>.vs.<sample2>.txt
         rnk-dmg--<study>_<sample1>.vs.<sample2>.txt rnk-dmg--<study>_<sample1>.vs.<sample2>.pdf
         wrt-<study>.txt

Notes: Output files ogl-<study>_<sample>.txt give tables for each sample with columns

        gene_ID gwidth #Sites #per10Kb %perSite %pNucl

       ordered by %pNucl.  If available, a link to an NCBI entry of gene_ID is inserted as second column.
       Abbreviations used: %perSite, percent methylation per site
                           %perNucl, percent methylation per nucleotide of the gene

       Output files ogl-<study>_<sample1>.vs.<sample2>.txt give tables for each comparison with columns

        gene_ID gwidth #Sites #per10Kb #dmSites #dmsp10kb %dmSites %pSite1 %pSite2 DMpSite ADMpSite DMpNucl ADMpNucl

       ordered by DMpSite.  If available, a link to an NCBI entry of gene_ID is inserted as second column.
       Abbreviations used: %dmSites, percent sites that are differentially methylated
                           %pSite1, average per site % methylation for sample1
                           %pSite2, average per site % methylation for sample2
                           DMpSite, average per site difference in % methylation between sample1 and sample2
                           ADMpSite, absolute value of DMpSite
                           DMpNucl, average per nucleotide difference in % methylation between sample1 and sample2
                           ADMpNucl, absolute value of DMpNucl

       Output files rnk-dmg-<study>_<sample1>.vs.<sample2>.txt are equivalent to files
        ogl-<study>_<sample1>.vs.<sample2>.txt but ordered by ADMpNucl.
       Output files rnk-dmg-<study>_<sample1>.vs.<sample2>.pdf provide visualization of the distribution
        of ADMpNucl values.

       Output file wrt-<study>.txt gives results of the Wilcoxon signed rank test comparing
        the %pSite1 and %pSite2 vectors

       In output directory OGL, tables are restricted to genes with gwidth <= maxgwidth and, for
        pairwise comparisons, #dmSites >= minnbrdmsites

       If either maxgwidth or minnbrdmsites is set, then output directory OGLall will show the full tables
        with all genes (for reference).
```

Directory: OGL File: ogl-Pc_PA_queen.txt

| gene_ID | gene_link | gwidth | #Sites | #per10Kb | %perSite | %pNucl |
|---|---|---|---|---|---|---|
| gene-LOC106786260 | https://www.ncbi.nlm.nih.gov/gene/?term=LOC106786260 | 2638 | 113 | 428.35 | 17.94 | 0.77 |
| gene-LOC106792553 | https://www.ncbi.nlm.nih.gov/gene/?term=LOC106792553 | 2482 | 122 | 491.54 | 11.46 | 0.56 |
| gene-LOC106786090 | https://www.ncbi.nlm.nih.gov/gene/?term=LOC106786090 | 2903 | 232 | 799.17 | 6.78 | 0.54 |
| gene-LOC106793786 | https://www.ncbi.nlm.nih.gov/gene/?term=LOC106793786 | 1202 | 22 | 183.03 | 28.32 | 0.52 |
| gene-LOC106793481 | https://www.ncbi.nlm.nih.gov/gene/?term=LOC106793481 | 3790 | 119 | 313.98 | 16.1 | 0.51 |
| gene-LOC106792739 | https://www.ncbi.nlm.nih.gov/gene/?term=LOC106792739 | 2414 | 34 | 140.85 | 33.06 | 0.47 |
| gene-LOC106792957 | https://www.ncbi.nlm.nih.gov/gene/?term=LOC106792957 | 876 | 10 | 114.16 | 39.39 | 0.45 |
| gene-LOC106784969 | https://www.ncbi.nlm.nih.gov/gene/?term=LOC106784969 | 2112 | 32 | 151.52 | 28.91 | 0.44 |
| gene-LOC106791200 | https://www.ncbi.nlm.nih.gov/gene/?term=LOC106791200 | 1235 | 33 | 267.21 | 15.71 | 0.42 |
| gene-LOC106789851 | https://www.ncbi.nlm.nih.gov/gene/?term=LOC106789851 | 1327 | 21 | 158.25 | 26.09 | 0.41 |
| gene-LOC106792910 | https://www.ncbi.nlm.nih.gov/gene/?term=LOC106792910 | 1045 | 7 | 66.99 | 60.6 | 0.41 |
| gene-LOC106789903 | https://www.ncbi.nlm.nih.gov/gene/?term=LOC106789903 | 753 | 8 | 106.24 | 37.71 | 0.4 |
| gene-LOC106791087 | https://www.ncbi.nlm.nih.gov/gene/?term=LOC106791087 | 2016 | 31 | 153.77 | 24.25 | 0.37 |
| gene-LOC106793930 | https://www.ncbi.nlm.nih.gov/gene/?term=LOC106793930 | 2962 | 53 | 178.93 | 20.62 | 0.37 |
| gene-LOC106788699 | https://www.ncbi.nlm.nih.gov/gene/?term=LOC106788699 | 1012 | 8 | 79.05 | 45.13 | 0.36 |
| gene-LOC106789767 | https://www.ncbi.nlm.nih.gov/gene/?term=LOC106789767 | 938 | 10 | 106.61 | 31.18 | 0.33 |
| gene-LOC106786499 | https://www.ncbi.nlm.nih.gov/gene/?term=LOC106786499 | 2765 | 35 | 126.58 | 25.25 | 0.32 |
| gene-LOC106792948 | https://www.ncbi.nlm.nih.gov/gene/?term=LOC106792948 | 1958 | 18 | 91.93 | 33.66 | 0.31 |
| gene-LOC106787127 | https://www.ncbi.nlm.nih.gov/gene/?term=LOC106787127 | 1220 | 25 | 204.92 | 13.95 | 0.29 |
| gene-LOC106792784 | https://www.ncbi.nlm.nih.gov/gene/?term=LOC106792784 | 2084 | 16 | 76.78 | 37.73 | 0.29 |
| gene-LOC106786231 | https://www.ncbi.nlm.nih.gov/gene/?term=LOC106786231 | 2170 | 29 | 133.64 | 20.78 | 0.28 |
| gene-LOC106786868 | https://www.ncbi.nlm.nih.gov/gene/?term=LOC106786868 | 1748 | 10 | 57.21 | 48.25 | 0.28 |
| gene-LOC106785716 | https://www.ncbi.nlm.nih.gov/gene/?term=LOC106785716 | 5009 | 86 | 171.69 | 15.93 | 0.27 |
| gene-LOC106787695 | https://www.ncbi.nlm.nih.gov/gene/?term=LOC106787695 | 1712 | 13 | 75.93 | 35.92 | 0.27 |

...

... only first 25 lines shown ...

Directory: OGL File: ogl-Pc_PA_queen.vs.worker.txt

| gene_ID | gene_link | gwidth | #Sites | #per10Kb | #dmSites | #dmsp10kb | %dmSites | %pSite1 | %pSite2 | DMpSite | ADMpSite | DMpNucl | ADMpNucl |
|---|---|---|---|---|---|---|---|---|---|---|---|---|---|
| gene-LOC106787037 | https://www.ncbi.nlm.nih.gov/gene/?term=LOC106787037 | 6216 | 57 | 91.7 | 2 | 3.22 | 3.51 | 1.8 | 4.5 | -2.7 | 2.7 | -0.02 | 0.02 |

Directory: OGL File: ogl-Pc_PA_worker.txt

| gene_ID | gene_link | gwidth | #Sites | #per10Kb | %perSite | %pNucl |
|---|---|---|---|---|---|---|
| gene-LOC106786260 | https://www.ncbi.nlm.nih.gov/gene/?term=LOC106786260 | 2638 | 93 | 352.54 | 19.44 | 0.69 |
| gene-LOC106793481 | https://www.ncbi.nlm.nih.gov/gene/?term=LOC106793481 | 3790 | 108 | 284.96 | 18.86 | 0.54 |
| gene-LOC106792553 | https://www.ncbi.nlm.nih.gov/gene/?term=LOC106792553 | 2482 | 120 | 483.48 | 10.63 | 0.51 |
| gene-LOC106792957 | https://www.ncbi.nlm.nih.gov/gene/?term=LOC106792957 | 876 | 11 | 125.57 | 40.23 | 0.51 |
| gene-LOC106786090 | https://www.ncbi.nlm.nih.gov/gene/?term=LOC106786090 | 2903 | 203 | 699.28 | 6.89 | 0.48 |
| gene-LOC106792739 | https://www.ncbi.nlm.nih.gov/gene/?term=LOC106792739 | 2414 | 33 | 136.7 | 31.32 | 0.43 |
| gene-LOC106793930 | https://www.ncbi.nlm.nih.gov/gene/?term=LOC106793930 | 2962 | 55 | 185.69 | 20.97 | 0.39 |
| gene-LOC106791200 | https://www.ncbi.nlm.nih.gov/gene/?term=LOC106791200 | 1235 | 15 | 121.46 | 31.03 | 0.38 |
| gene-LOC106791087 | https://www.ncbi.nlm.nih.gov/gene/?term=LOC106791087 | 2016 | 30 | 148.81 | 20.52 | 0.31 |
| gene-LOC106787918 | https://www.ncbi.nlm.nih.gov/gene/?term=LOC106787918 | 3131 | 71 | 226.76 | 13 | 0.29 |
| gene-LOC106787164 | https://www.ncbi.nlm.nih.gov/gene/?term=LOC106787164 | 1270 | 20 | 157.48 | 17.66 | 0.28 |
| gene-LOC106788200 | https://www.ncbi.nlm.nih.gov/gene/?term=LOC106788200 | 2717 | 35 | 128.82 | 21.51 | 0.28 |
| gene-LOC106784969 | https://www.ncbi.nlm.nih.gov/gene/?term=LOC106784969 | 2112 | 24 | 113.64 | 23.25 | 0.26 |
| gene-LOC106792910 | https://www.ncbi.nlm.nih.gov/gene/?term=LOC106792910 | 1045 | 5 | 47.85 | 53.91 | 0.26 |
| gene-LOC106784090 | https://www.ncbi.nlm.nih.gov/gene/?term=LOC106784090 | 2855 | 14 | 49.04 | 50.03 | 0.25 |
| gene-LOC106786231 | https://www.ncbi.nlm.nih.gov/gene/?term=LOC106786231 | 2170 | 25 | 115.21 | 21.38 | 0.25 |
| gene-LOC106784739 | https://www.ncbi.nlm.nih.gov/gene/?term=LOC106784739 | 1263 | 8 | 63.34 | 37.96 | 0.24 |
| gene-LOC106785716 | https://www.ncbi.nlm.nih.gov/gene/?term=LOC106785716 | 5009 | 78 | 155.72 | 15.35 | 0.24 |
| gene-LOC106790989 | https://www.ncbi.nlm.nih.gov/gene/?term=LOC106790989 | 2986 | 50 | 167.45 | 14.07 | 0.24 |
| gene-LOC106792948 | https://www.ncbi.nlm.nih.gov/gene/?term=LOC106792948 | 1958 | 15 | 76.61 | 30.81 | 0.24 |
| gene-LOC106786682 | https://www.ncbi.nlm.nih.gov/gene/?term=LOC106786682 | 1658 | 28 | 168.88 | 13.9 | 0.23 |
| gene-LOC106788528 | https://www.ncbi.nlm.nih.gov/gene/?term=LOC106788528 | 7476 | 177 | 236.76 | 9.86 | 0.23 |
| gene-LOC106788699 | https://www.ncbi.nlm.nih.gov/gene/?term=LOC106788699 | 1012 | 8 | 79.05 | 29.35 | 0.23 |
| gene-LOC106789851 | https://www.ncbi.nlm.nih.gov/gene/?term=LOC106789851 | 1327 | 12 | 90.43 | 25.17 | 0.23 |

...

... only first 25 lines shown ...

Directory: OGL File: rnk-dmg-Pc_PA_queen.vs.worker.txt

| gene_ID | gene_link | gwidth | #Sites | #per10Kb | #dmSites | #dmsp10kb | %dmSites | %pSite1 | %pSite2 | DMpSite | ADMpSite | DMpNucl | ADMpNucl |
|---|---|---|---|---|---|---|---|---|---|---|---|---|---|
| gene-LOC106787037 | https://www.ncbi.nlm.nih.gov/gene/?term=LOC106787037 | 6216 | 57 | 91.7 | 2 | 3.22 | 3.51 | 1.8 | 4.5 | -2.7 | 2.7 | -0.02 | 0.02 |

```
Directory: OGL File: wrt-Pc_PA.txt


... comparing  queen.vs.worker ...

Number of genes <=  20000  and with #dmsites >=  2 :  1
Only 1 gene, so there is nothing to test here ...
```

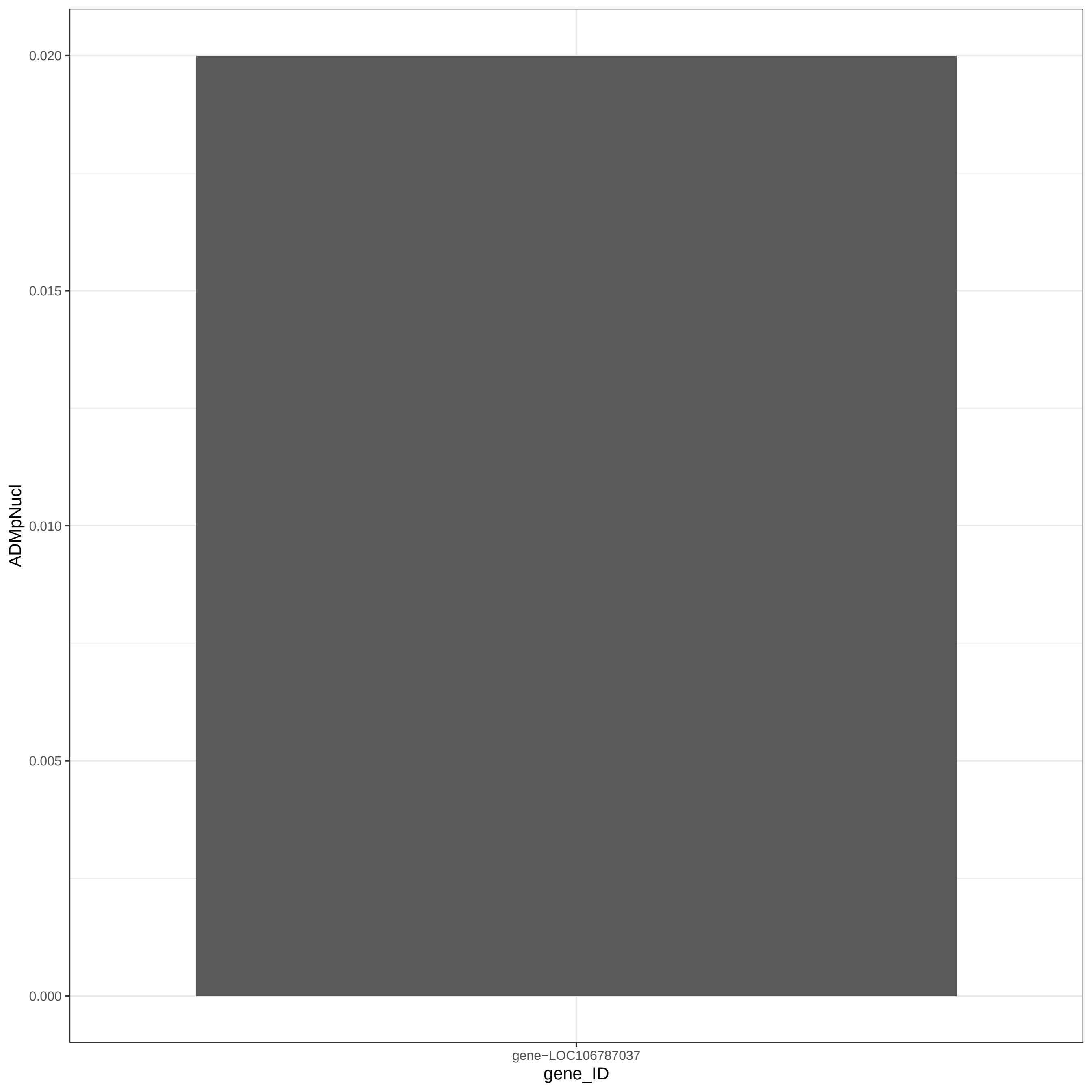
0.020
0.015
0.010
0.005
0.000
ADMpNucl
gene−LOC106787037
gene_ID

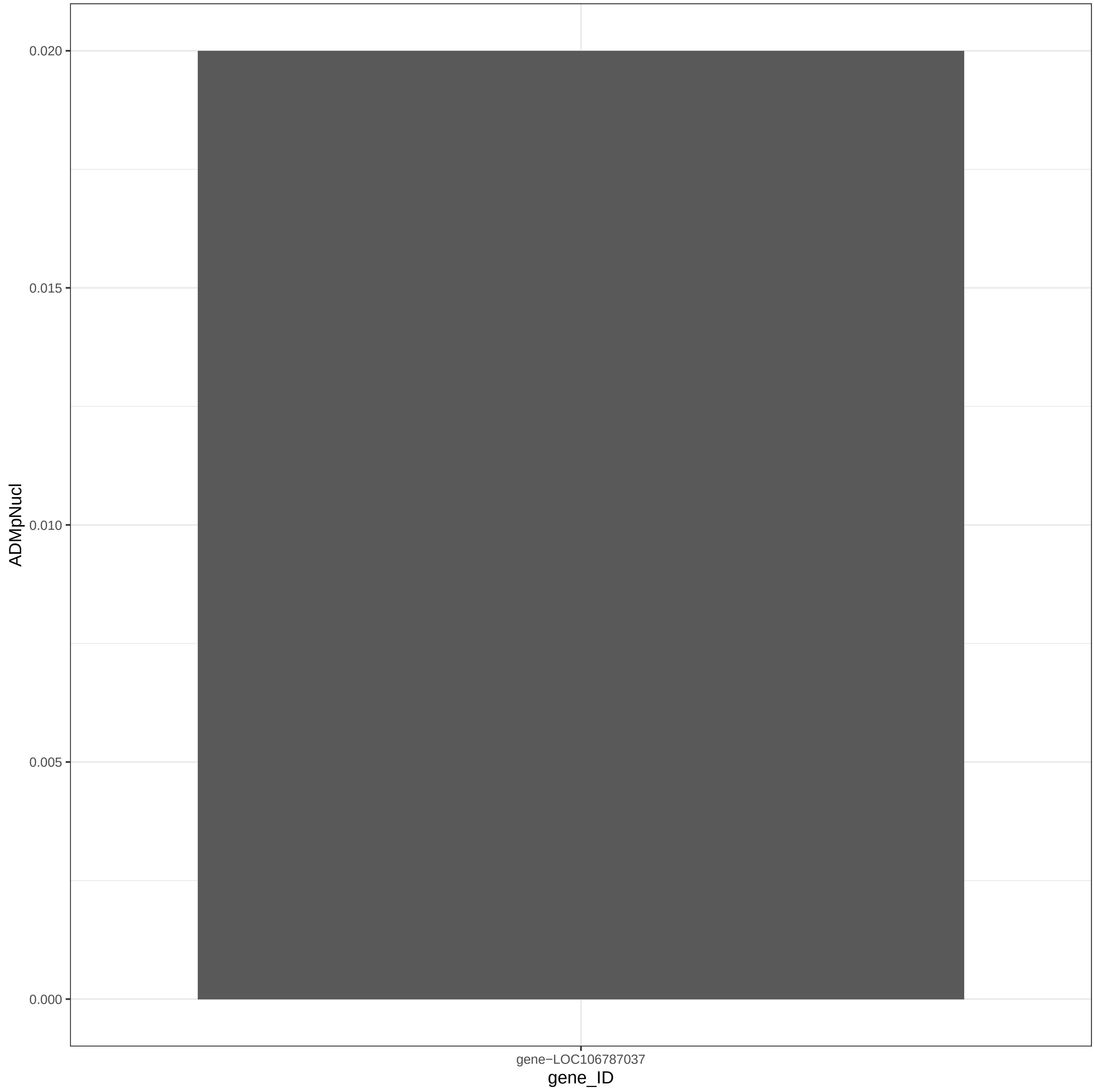
0.020
0.015
0.010
0.005
0.000
ADMpNucl
gene−LOC106787037
gene_ID

```
Directory: OGLall File: 0READMEogl


OGL - Ordered gene lists.

Input: studyhc annotation   maxgwidth minnbrdmsites
       dmgprp (= output of BWASPR:show_dmsg())

Output: files ogl-*.txt rnk-dmg-*.txt
         ogl-<study>_<sample>.txt ogl-<study>_<sample1>.vs.<sample2>.txt
         rnk-dmg--<study>_<sample1>.vs.<sample2>.txt rnk-dmg--<study>_<sample1>.vs.<sample2>.pdf
         wrt-<study>.txt

Notes: Output files ogl-<study>_<sample>.txt give tables for each sample with columns

        gene_ID gwidth #Sites #per10Kb %perSite %pNucl

       ordered by %pNucl.  If available, a link to an NCBI entry of gene_ID is inserted as second column.
       Abbreviations used: %perSite, percent methylation per site
                           %perNucl, percent methylation per nucleotide of the gene

       Output files ogl-<study>_<sample1>.vs.<sample2>.txt give tables for each comparison with columns

        gene_ID gwidth #Sites #per10Kb #dmSites #dmsp10kb %dmSites %pSite1 %pSite2 DMpSite ADMpSite DMpNucl ADMpNucl

       ordered by DMpSite.  If available, a link to an NCBI entry of gene_ID is inserted as second column.
       Abbreviations used: %dmSites, percent sites that are differentially methylated
                           %pSite1, average per site % methylation for sample1
                           %pSite2, average per site % methylation for sample2
                           DMpSite, average per site difference in % methylation between sample1 and sample2
                           ADMpSite, absolute value of DMpSite
                           DMpNucl, average per nucleotide difference in % methylation between sample1 and sample2
                           ADMpNucl, absolute value of DMpNucl

       Output files rnk-dmg-<study>_<sample1>.vs.<sample2>.txt are equivalent to files
        ogl-<study>_<sample1>.vs.<sample2>.txt but ordered by ADMpNucl.
       Output files rnk-dmg-<study>_<sample1>.vs.<sample2>.pdf provide visualization of the distribution
        of ADMpNucl values.

       Output file wrt-<study>.txt gives results of the Wilcoxon signed rank test comparing
        the %pSite1 and %pSite2 vectors

       In output directory OGL, tables are restricted to genes with gwidth <= maxgwidth and, for
        pairwise comparisons, #dmSites >= minnbrdmsites

       If either maxgwidth or minnbrdmsites is set, then output directory OGLall will show the full tables
        with all genes (for reference).
```

Directory: OGLall File: ogl-Pc_PA_queen.txt

| gene_ID | gene_link | gwidth | #Sites | #per10Kb | %perSite | %pNucl |
|---|---|---|---|---|---|---|
| gene-LOC106786260 | https://www.ncbi.nlm.nih.gov/gene/?term=LOC106786260 | 2638 | 113 | 428.35 | 17.94 | 0.77 |
| gene-LOC106792553 | https://www.ncbi.nlm.nih.gov/gene/?term=LOC106792553 | 2482 | 122 | 491.54 | 11.46 | 0.56 |
| gene-LOC106786090 | https://www.ncbi.nlm.nih.gov/gene/?term=LOC106786090 | 2903 | 232 | 799.17 | 6.78 | 0.54 |
| gene-LOC106793786 | https://www.ncbi.nlm.nih.gov/gene/?term=LOC106793786 | 1202 | 22 | 183.03 | 28.32 | 0.52 |
| gene-LOC106793481 | https://www.ncbi.nlm.nih.gov/gene/?term=LOC106793481 | 3790 | 119 | 313.98 | 16.1 | 0.51 |
| gene-LOC106792739 | https://www.ncbi.nlm.nih.gov/gene/?term=LOC106792739 | 2414 | 34 | 140.85 | 33.06 | 0.47 |
| gene-LOC106792957 | https://www.ncbi.nlm.nih.gov/gene/?term=LOC106792957 | 876 | 10 | 114.16 | 39.39 | 0.45 |
| gene-LOC106784969 | https://www.ncbi.nlm.nih.gov/gene/?term=LOC106784969 | 2112 | 32 | 151.52 | 28.91 | 0.44 |
| gene-LOC106791200 | https://www.ncbi.nlm.nih.gov/gene/?term=LOC106791200 | 1235 | 33 | 267.21 | 15.71 | 0.42 |
| gene-LOC106789851 | https://www.ncbi.nlm.nih.gov/gene/?term=LOC106789851 | 1327 | 21 | 158.25 | 26.09 | 0.41 |
| gene-LOC106792910 | https://www.ncbi.nlm.nih.gov/gene/?term=LOC106792910 | 1045 | 7 | 66.99 | 60.6 | 0.41 |
| gene-LOC106789903 | https://www.ncbi.nlm.nih.gov/gene/?term=LOC106789903 | 753 | 8 | 106.24 | 37.71 | 0.4 |
| gene-LOC106791087 | https://www.ncbi.nlm.nih.gov/gene/?term=LOC106791087 | 2016 | 31 | 153.77 | 24.25 | 0.37 |
| gene-LOC106793930 | https://www.ncbi.nlm.nih.gov/gene/?term=LOC106793930 | 2962 | 53 | 178.93 | 20.62 | 0.37 |
| gene-LOC106788699 | https://www.ncbi.nlm.nih.gov/gene/?term=LOC106788699 | 1012 | 8 | 79.05 | 45.13 | 0.36 |
| gene-LOC106789767 | https://www.ncbi.nlm.nih.gov/gene/?term=LOC106789767 | 938 | 10 | 106.61 | 31.18 | 0.33 |
| gene-LOC106786499 | https://www.ncbi.nlm.nih.gov/gene/?term=LOC106786499 | 2765 | 35 | 126.58 | 25.25 | 0.32 |
| gene-LOC106792948 | https://www.ncbi.nlm.nih.gov/gene/?term=LOC106792948 | 1958 | 18 | 91.93 | 33.66 | 0.31 |
| gene-LOC106787127 | https://www.ncbi.nlm.nih.gov/gene/?term=LOC106787127 | 1220 | 25 | 204.92 | 13.95 | 0.29 |
| gene-LOC106792784 | https://www.ncbi.nlm.nih.gov/gene/?term=LOC106792784 | 2084 | 16 | 76.78 | 37.73 | 0.29 |
| gene-LOC106786231 | https://www.ncbi.nlm.nih.gov/gene/?term=LOC106786231 | 2170 | 29 | 133.64 | 20.78 | 0.28 |
| gene-LOC106786868 | https://www.ncbi.nlm.nih.gov/gene/?term=LOC106786868 | 1748 | 10 | 57.21 | 48.25 | 0.28 |
| gene-LOC106785716 | https://www.ncbi.nlm.nih.gov/gene/?term=LOC106785716 | 5009 | 86 | 171.69 | 15.93 | 0.27 |
| gene-LOC106787695 | https://www.ncbi.nlm.nih.gov/gene/?term=LOC106787695 | 1712 | 13 | 75.93 | 35.92 | 0.27 |

...

... only first 25 lines shown ...

Directory: OGLall File: ogl-Pc_PA_queen.vs.worker.txt

| gene_ID | gene_link | gwidth | #Sites | #per10Kb | #dmSites | #dmsp10kb | %dmSites | %pSite1 | %pSite2 | DMpSite | ADMpSite | DMpNucl | ADMpNucl |
|---|---|---|---|---|---|---|---|---|---|---|---|---|---|
| gene-LOC106789019 | https://www.ncbi.nlm.nih.gov/gene/?term=LOC106789019 | 26486 | 304 | 114.78 | 1 | 0.38 | 0.33 | 3.5 | 4.26 | -0.76 | 0.76 | -0.01 | 0.01 |
| gene-LOC106793698 | https://www.ncbi.nlm.nih.gov/gene/?term=LOC106793698 | 22827 | 216 | 94.62 | 1 | 0.44 | 0.46 | 1.27 | 2.14 | -0.87 | 0.87 | -0.01 | 0.01 |
| gene-LOC106793677 | https://www.ncbi.nlm.nih.gov/gene/?term=LOC106793677 | 47553 | 327 | 68.77 | 1 | 0.21 | 0.31 | 1.03 | 1.92 | -0.89 | 0.89 | -0.01 | 0.01 |
| gene-LOC106786951 | https://www.ncbi.nlm.nih.gov/gene/?term=LOC106786951 | 5425 | 33 | 60.83 | 1 | 1.84 | 3.03 | 11.94 | 14.05 | -2.11 | 2.11 | -0.01 | 0.01 |
| gene-LOC106786709 | https://www.ncbi.nlm.nih.gov/gene/?term=LOC106786709 | 7252 | 66 | 91.01 | 1 | 1.38 | 1.52 | 2.34 | 4.87 | -2.53 | 2.53 | -0.02 | 0.02 |
| gene-LOC106787037 | https://www.ncbi.nlm.nih.gov/gene/?term=LOC106787037 | 6216 | 57 | 91.7 | 2 | 3.22 | 3.51 | 1.8 | 4.5 | -2.7 | 2.7 | -0.02 | 0.02 |

Directory: OGLall File: ogl-Pc_PA_worker.txt

| gene_ID | gene_link | gwidth | #Sites | #per10Kb | %perSite | %pNucl |
|---|---|---|---|---|---|---|
| gene-LOC106786260 | https://www.ncbi.nlm.nih.gov/gene/?term=LOC106786260 | 2638 | 93 | 352.54 | 19.44 | 0.69 |
| gene-LOC106793481 | https://www.ncbi.nlm.nih.gov/gene/?term=LOC106793481 | 3790 | 108 | 284.96 | 18.86 | 0.54 |
| gene-LOC106792553 | https://www.ncbi.nlm.nih.gov/gene/?term=LOC106792553 | 2482 | 120 | 483.48 | 10.63 | 0.51 |
| gene-LOC106792957 | https://www.ncbi.nlm.nih.gov/gene/?term=LOC106792957 | 876 | 11 | 125.57 | 40.23 | 0.51 |
| gene-LOC106786090 | https://www.ncbi.nlm.nih.gov/gene/?term=LOC106786090 | 2903 | 203 | 699.28 | 6.89 | 0.48 |
| gene-LOC106792739 | https://www.ncbi.nlm.nih.gov/gene/?term=LOC106792739 | 2414 | 33 | 136.7 | 31.32 | 0.43 |
| gene-LOC106793930 | https://www.ncbi.nlm.nih.gov/gene/?term=LOC106793930 | 2962 | 55 | 185.69 | 20.97 | 0.39 |
| gene-LOC106791200 | https://www.ncbi.nlm.nih.gov/gene/?term=LOC106791200 | 1235 | 15 | 121.46 | 31.03 | 0.38 |
| gene-LOC106791087 | https://www.ncbi.nlm.nih.gov/gene/?term=LOC106791087 | 2016 | 30 | 148.81 | 20.52 | 0.31 |
| gene-LOC106787918 | https://www.ncbi.nlm.nih.gov/gene/?term=LOC106787918 | 3131 | 71 | 226.76 | 13 | 0.29 |
| gene-LOC106787164 | https://www.ncbi.nlm.nih.gov/gene/?term=LOC106787164 | 1270 | 20 | 157.48 | 17.66 | 0.28 |
| gene-LOC106788200 | https://www.ncbi.nlm.nih.gov/gene/?term=LOC106788200 | 2717 | 35 | 128.82 | 21.51 | 0.28 |
| gene-LOC106784969 | https://www.ncbi.nlm.nih.gov/gene/?term=LOC106784969 | 2112 | 24 | 113.64 | 23.25 | 0.26 |
| gene-LOC106792910 | https://www.ncbi.nlm.nih.gov/gene/?term=LOC106792910 | 1045 | 5 | 47.85 | 53.91 | 0.26 |
| gene-LOC106784090 | https://www.ncbi.nlm.nih.gov/gene/?term=LOC106784090 | 2855 | 14 | 49.04 | 50.03 | 0.25 |
| gene-LOC106786231 | https://www.ncbi.nlm.nih.gov/gene/?term=LOC106786231 | 2170 | 25 | 115.21 | 21.38 | 0.25 |
| gene-LOC106784739 | https://www.ncbi.nlm.nih.gov/gene/?term=LOC106784739 | 1263 | 8 | 63.34 | 37.96 | 0.24 |
| gene-LOC106785716 | https://www.ncbi.nlm.nih.gov/gene/?term=LOC106785716 | 5009 | 78 | 155.72 | 15.35 | 0.24 |
| gene-LOC106790989 | https://www.ncbi.nlm.nih.gov/gene/?term=LOC106790989 | 2986 | 50 | 167.45 | 14.07 | 0.24 |
| gene-LOC106792948 | https://www.ncbi.nlm.nih.gov/gene/?term=LOC106792948 | 1958 | 15 | 76.61 | 30.81 | 0.24 |
| gene-LOC106786682 | https://www.ncbi.nlm.nih.gov/gene/?term=LOC106786682 | 1658 | 28 | 168.88 | 13.9 | 0.23 |
| gene-LOC106788528 | https://www.ncbi.nlm.nih.gov/gene/?term=LOC106788528 | 7476 | 177 | 236.76 | 9.86 | 0.23 |
| gene-LOC106788699 | https://www.ncbi.nlm.nih.gov/gene/?term=LOC106788699 | 1012 | 8 | 79.05 | 29.35 | 0.23 |
| gene-LOC106789851 | https://www.ncbi.nlm.nih.gov/gene/?term=LOC106789851 | 1327 | 12 | 90.43 | 25.17 | 0.23 |

...

... only first 25 lines shown ...

Directory: OGLall File: rnk-dmg-Pc_PA_queen.vs.worker.txt

| gene_ID | gene_link | gwidth | #Sites | #per10Kb | #dmSites | #dmsp10kb | %dmSites | %pSite1 | %pSite2 | DMpSite | ADMpSite | DMpNucl | ADMpNucl |
|---|---|---|---|---|---|---|---|---|---|---|---|---|---|
| gene-LOC106786709 | https://www.ncbi.nlm.nih.gov/gene/?term=LOC106786709 | 7252 | 66 | 91.01 | 1 | 1.38 | 1.52 | 2.34 | 4.87 | -2.53 | 2.53 | -0.02 | 0.02 |
| gene-LOC106787037 | https://www.ncbi.nlm.nih.gov/gene/?term=LOC106787037 | 6216 | 57 | 91.7 | 2 | 3.22 | 3.51 | 1.8 | 4.5 | -2.7 | 2.7 | -0.02 | 0.02 |
| gene-LOC106789019 | https://www.ncbi.nlm.nih.gov/gene/?term=LOC106789019 | 26486 | 304 | 114.78 | 1 | 0.38 | 0.33 | 3.5 | 4.26 | -0.76 | 0.76 | -0.01 | 0.01 |
| gene-LOC106793698 | https://www.ncbi.nlm.nih.gov/gene/?term=LOC106793698 | 22827 | 216 | 94.62 | 1 | 0.44 | 0.46 | 1.27 | 2.14 | -0.87 | 0.87 | -0.01 | 0.01 |
| gene-LOC106793677 | https://www.ncbi.nlm.nih.gov/gene/?term=LOC106793677 | 47553 | 327 | 68.77 | 1 | 0.21 | 0.31 | 1.03 | 1.92 | -0.89 | 0.89 | -0.01 | 0.01 |
| gene-LOC106786951 | https://www.ncbi.nlm.nih.gov/gene/?term=LOC106786951 | 5425 | 33 | 60.83 | 1 | 1.84 | 3.03 | 11.94 | 14.05 | -2.11 | 2.11 | -0.01 | 0.01 |

```
Directory: OGLall File: wrt-Pc_PA.txt


... comparing  queen.vs.worker ...

Number of genes with #dmsites >=  1 :  6

sample1     sum of percent methylation:    21.88     average:     3.65
sample2     sum of percent methylation:    31.74     average:     5.29

sum difference:    -9.86     average difference:  -1.6433

      Wilcoxon signed rank test with continuity correction

data:  pw_summary$`%pSite1` and pw_summary$`%pSite2`
V = 0, p-value = 0.04
alternative hypothesis: true location shift is not equal to 0
```

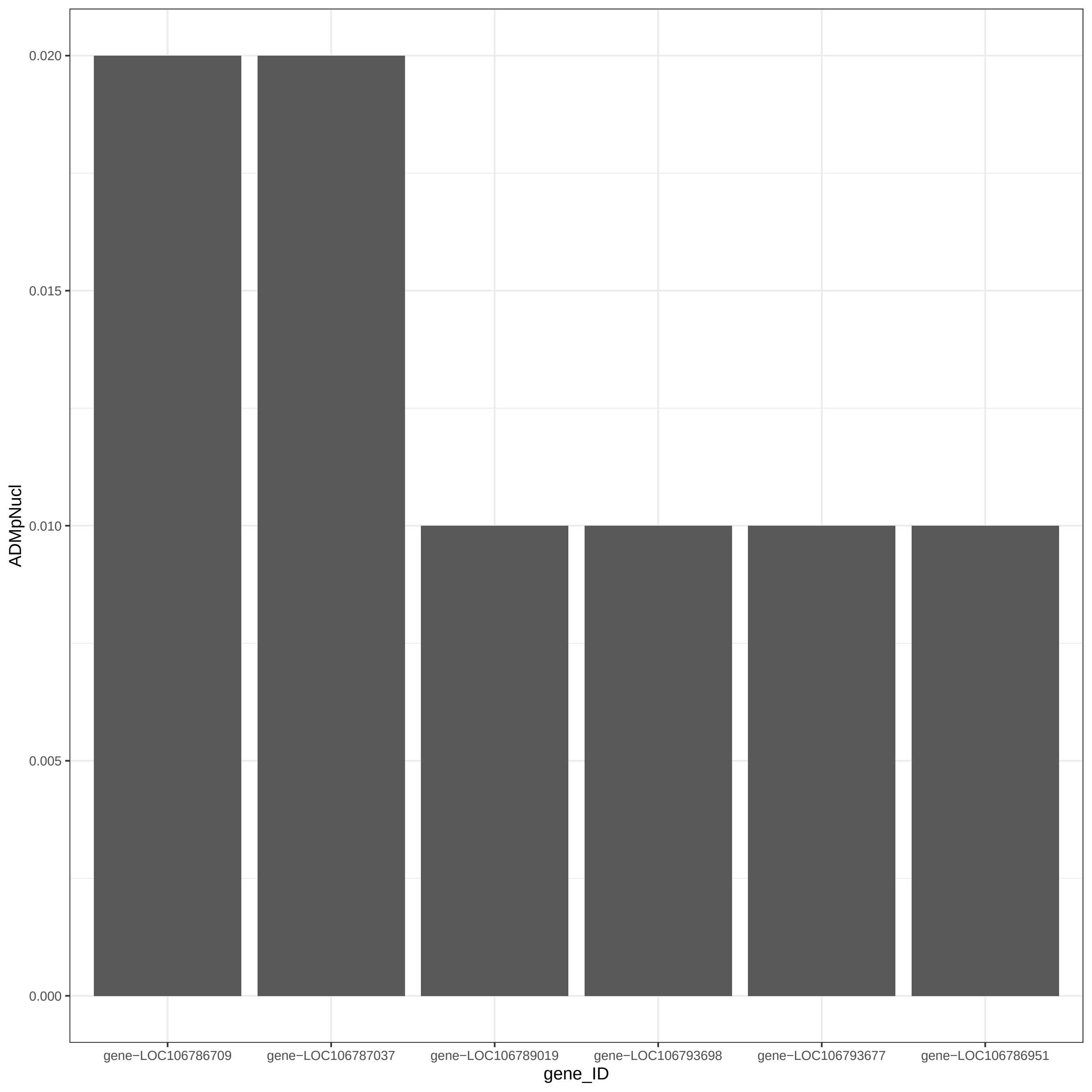

ADMpNucl
0.000
0.005
0.010
0.015
0.020
gene−LOC106786709
gene−LOC106787037
gene−LOC106789019
gene−LOC106793698
gene−LOC106793677
gene−LOC106786951
gene_ID

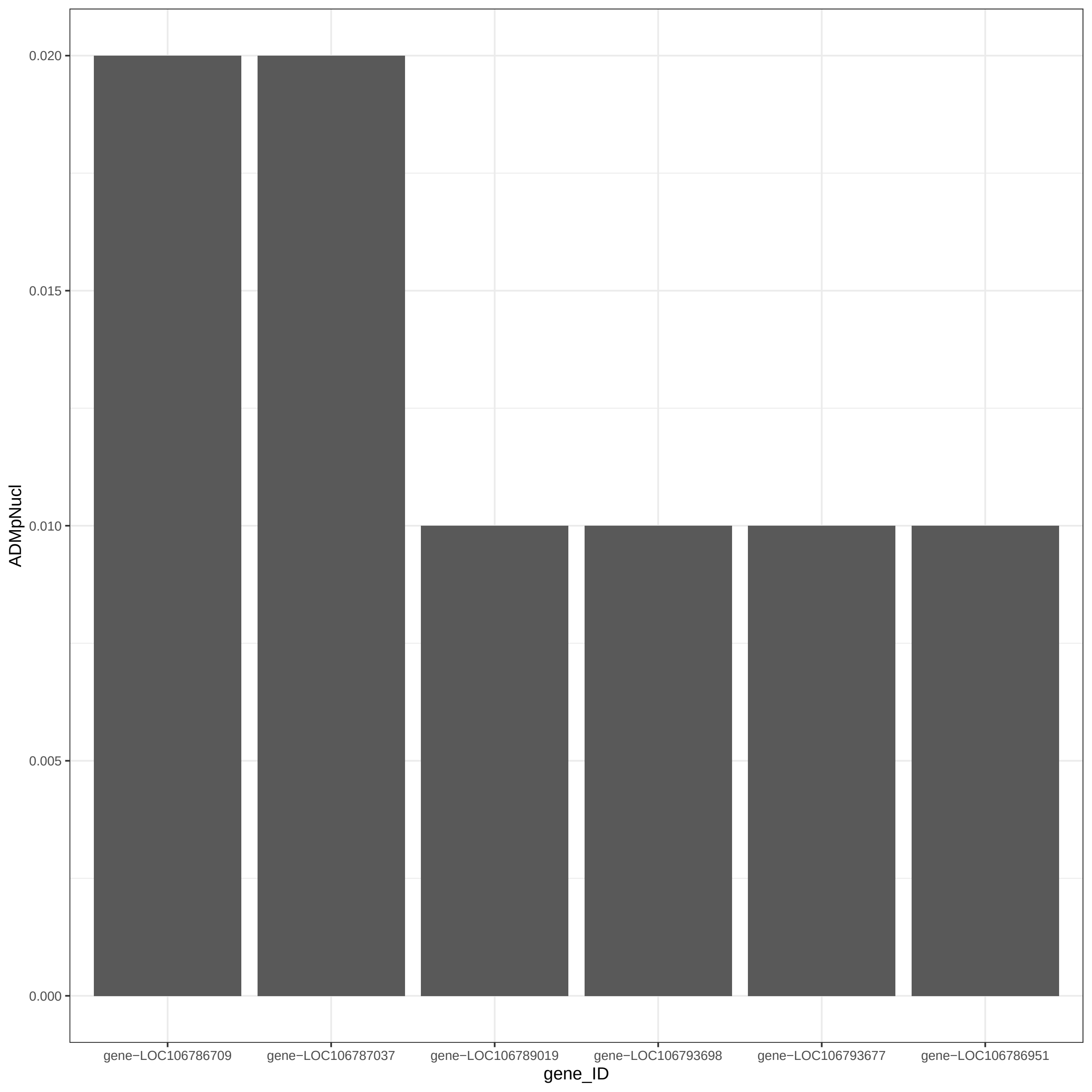
0.020
0.015
0.010
0.005
0.000
ADMpNucl
gene−LOC106786709
gene−LOC106787037
gene−LOC106789019
gene−LOC106793698
gene−LOC106793677
gene−LOC106786951
gene_ID
