## Supplementary material for "Tools and applications for integrative analysis of DNA methylation in social insects": Dataset S16

Am_RE.conf

```
#Customize variables here:#####################################################
###############################################################################

#Load the files:
#
infiles       <- setup_BWASPR(datafile="./RWORK/Amel/Am.dat",
                              parfile="./RWORK/Amel/Am.par")

#Set the study and samples in the study:
#
species       <- "Am"
study         <- "RE"
samplelist    <- list("fe","te")
## The following two variables are used for output file labeling:
studyLabel    <- "RE"
sampleLabels  <- list("Am_RE_fe","Am_RE_te")

hasreplicates <- FALSE
type          <- "CpG"
destrand      <- TRUE
covlist       <- c(4,10,15)
locount       <- 4
hicount       <- 12
repcovlist    <- c(4,10,15)
replocount    <- 10
rephicount    <- 100
hheight       <- 0.10
nbrpnts       <- 5000

## The following four variables are sent to rank_rbm() and determine what data
#  points get plotted:
minnbrmsprm   <- 5      # minimum number of methylation sites per promoter
mingenewidth  <- 500    # minimum gene width
maxgenewidth  <- 5000   # maximum gene width
minnbrmsgene  <- 5      # minimum number of methylation sites per gene

## Other parameters (see Rscript.BWASPR for usage notes):
#
highcoverage  <- 10     # high read coverage threshold for studyhc methylRawList object
threshold     <- 25.0   # "difference" threshold for getMethylDiff(), called by det_dmsg()
qvalue        <- 0.01   # "qvalue" setting for getMethylDiff(), called by det_dmsg()
wsize         <- 1000   # "win.size" parameter for tileMethylCounts() in det_dmt()
stepsize      <- 1000   # "step.size" parameter for tileMethylCounts() in det_dmt()

minNsites     <- 10     # minimum number of hc sites in a gene to be heatmapped in show_dmsg()
maxNsites     <- 60     # maximum number of hc sites in a gene to be heatmapped in show_dmsg()
minPdmsites   <- 10     # minimum ratio of dm/hc (in %) in a gene to be heatmapped in show_dmsg()

maxgwidth     <- 20000  # maximal gene width for a gene to be considered by explore_dmsg()
minnbrdmsites <- 2      # minimum number of differentially methylated sites for a gene to be
                        #  considered in sample comparisons by explore_dmsg()
glink         <- "NCBIgene"   # URL to show in explore_dmsg(); options: "" or "NCBIgene"

#Set the number of processors to use:
#
numprc <- 6

#Determine what analyses to run:
#
RUNload    <- FALSE
```

```
RUNcms     <- TRUE
RUNpwc     <- TRUE
RUNcrl     <- TRUE

RUNrepcms  <- FALSE
RUNrepcrl  <- FALSE

RUNmmp     <- TRUE
RUNacs     <- TRUE
RUNrnk     <- TRUE
RUNmrpr    <- TRUE

RUNdmt     <- TRUE
RUNdmsg    <- TRUE
RUNdmgdtls <- TRUE
RUNogl     <- TRUE

RUNsave    <- TRUE


mymessage <- sprintf("\nAnalyzing %s study %s for type %s\n\n",species,studyLabel,type)
message(mymessage)

################################################################################
#End of typical customization.##################################################
```

Am.par

```
SPECIESNAME Apis mellifera
TOTALNBRPMSITES   19687378
ASSEMBLYVERSION   Amel_HAv3.1
GENOMESIZE  225250884
SPECIESGFF3DIR    ./MCALLS/Amel/genome/GFF3DIR
GENELISTGFF3      Amel.gene.gff3
EXONLISTGFF3      Amel.exon.gff3
PCGEXNLISTGFF3    Amel.pcg-exon.gff3
PROMOTRLISTGFF3   Amel.promoter.gff3
CDSLISTGFF3 Amel.pcg-CDS.gff3
UTRFLAGSET  1
5UTRLISTGFF3      Amel.pcg-5pUTR.gff3
3UTRLISTGFF3      Amel.pcg-3pUTR.gff3
```

Am.dat

```
# Samples from Feng et al. (2010) PNAS:
# im = immature male
#
Am    FE    immature_male     0     CpGhsm      ./MCALLS/Amel/Feng2010/immature_male/immature_male.CpGhsm.mcalls
Am    FE    immature_male     0     CpGscd      ./MCALLS/Amel/Feng2010/immature_male/immature_male.CpGscd.mcalls


# Samples from Zemach et al. (2010) Science:
# wr = whole adult worker
#
Am    ZE    worker      0     CpGhsm      ./MCALLS/Amel/Zemach2010/worker/worker.CpGhsm.mcalls
Am    ZE    worker      0     CpGscd      ./MCALLS/Amel/Zemach2010/worker/worker.CpGscd.mcalls

# ... continuing: replicates
#
Am    ZE    worker      1     CpGhsm      ./MCALLS/Amel/Zemach2010/worker/replicate1/worker1.CpGhsm.mcalls
Am    ZE    worker      1     CpGscd      ./MCALLS/Amel/Zemach2010/worker/replicate1/worker1.CpGscd.mcalls
Am    ZE    worker      2     CpGhsm      ./MCALLS/Amel/Zemach2010/worker/replicate2/worker2.CpGhsm.mcalls
Am    ZE    worker      2     CpGscd      ./MCALLS/Amel/Zemach2010/worker/replicate2/worker2.CpGscd.mcalls
Am    ZE    worker      3     CpGhsm      ./MCALLS/Amel/Zemach2010/worker/replicate3/worker3.CpGhsm.mcalls
Am    ZE    worker      3     CpGscd      ./MCALLS/Amel/Zemach2010/worker/replicate3/worker3.CpGscd.mcalls
Am    ZE    worker      4     CpGhsm      ./MCALLS/Amel/Zemach2010/worker/replicate4/worker4.CpGhsm.mcalls
Am    ZE    worker      4     CpGscd      ./MCALLS/Amel/Zemach2010/worker/replicate4/worker4.CpGscd.mcalls


# Samples from Lyko et al. (2010) PLoS Biology:
# qb = queen brain; wb = worker brain
#
Am    LY    queen 0     CpGhsm      ./MCALLS/Amel/Lyko2010/queen/queen.CpGhsm.mcalls
Am    LY    queen 0     CpGscd      ./MCALLS/Amel/Lyko2010/queen/queen.CpGscd.mcalls
Am    LY    worker      0     CpGhsm      ./MCALLS/Amel/Lyko2010/worker/worker.CpGhsm.mcalls
Am    LY    worker      0     CpGscd      ./MCALLS/Amel/Lyko2010/worker/worker.CpGscd.mcalls


# Samples from Foret et al. (2012) PNAS:
# ql = queen larvae; wl = worker larvae
#
Am    FO    queen 0     CpGhsm      ./MCALLS/Amel/Foret2012/queen/queen.CpGhsm.mcalls
Am    FO    queen 0     CpGscd      ./MCALLS/Amel/Foret2012/queen/queen.CpGscd.mcalls
Am    FO    worker      0     CpGhsm      ./MCALLS/Amel/Foret2012/worker/worker.CpGhsm.mcalls
Am    FO    worker      0     CpGscd      ./MCALLS/Amel/Foret2012/worker/worker.CpGscd.mcalls

# ... continuing: replicates
#
Am    FO    queen 1     CpGhsm      ./MCALLS/Amel/Foret2012/queen/replicate1/queen1.CpGhsm.mcalls
Am    FO    queen 1     CpGscd      ./MCALLS/Amel/Foret2012/queen/replicate1/queen1.CpGscd.mcalls
Am    FO    queen 2     CpGhsm      ./MCALLS/Amel/Foret2012/queen/replicate2/queen2.CpGhsm.mcalls
Am    FO    queen 2     CpGscd      ./MCALLS/Amel/Foret2012/queen/replicate2/queen2.CpGscd.mcalls
Am    FO    worker      1     CpGhsm      ./MCALLS/Amel/Foret2012/worker/replicate1/worker1.CpGhsm.mcalls
Am    FO    worker      1     CpGscd      ./MCALLS/Amel/Foret2012/worker/replicate1/worker1.CpGscd.mcalls
Am    FO    worker      2     CpGhsm      ./MCALLS/Amel/Foret2012/worker/replicate2/worker2.CpGhsm.mcalls
Am    FO    worker      2     CpGscd      ./MCALLS/Amel/Foret2012/worker/replicate2/worker2.CpGscd.mcalls


# Samples from Herb et al. (2012) Nature Neuroscience:
# fr = forager; qn = queen; rn = reverted nurse; wr = worker
#
Am    HE    fr    0     CpGhsm      ./MCALLS/Amel/Herb2012/forager/forager.CpGhsm.mcalls
Am    HE    fr    0     CpGscd      ./MCALLS/Amel/Herb2012/forager/forager.CpGscd.mcalls
Am    HE    qn    0     CpGhsm      ./MCALLS/Amel/Herb2012/queen/queen.CpGhsm.mcalls
Am    HE    qn    0     CpGscd      ./MCALLS/Amel/Herb2012/queen/queen.CpGscd.mcalls
```

```
Am    HE    rn    0     CpGhsm      ./MCALLS/Amel/Herb2012/reverted_nurse/reverted_nurse.CpGhsm.mcalls
Am    HE    rn    0     CpGscd      ./MCALLS/Amel/Herb2012/reverted_nurse/reverted_nurse.CpGscd.mcalls
Am    HE    wr    0     CpGhsm      ./MCALLS/Amel/Herb2012/worker/worker.CpGhsm.mcalls
Am    HE    wr    0     CpGscd      ./MCALLS/Amel/Herb2012/worker/worker.CpGscd.mcalls

# ... continuing: replicates
#
Am    HE    fr    1     CpGhsm      ./MCALLS/Amel/Herb2012/forager/replicate1/forager1.CpGhsm.mcalls
Am    HE    fr    1     CpGscd      ./MCALLS/Amel/Herb2012/forager/replicate1/forager1.CpGscd.mcalls
Am    HE    fr    2     CpGhsm      ./MCALLS/Amel/Herb2012/forager/replicate2/forager2.CpGhsm.mcalls
Am    HE    fr    2     CpGscd      ./MCALLS/Amel/Herb2012/forager/replicate2/forager2.CpGscd.mcalls
Am    HE    fr    3     CpGhsm      ./MCALLS/Amel/Herb2012/forager/replicate3/forager3.CpGhsm.mcalls
Am    HE    fr    3     CpGscd      ./MCALLS/Amel/Herb2012/forager/replicate3/forager3.CpGscd.mcalls
Am    HE    fr    4     CpGhsm      ./MCALLS/Amel/Herb2012/forager/replicate4/forager4.CpGhsm.mcalls
Am    HE    fr    4     CpGscd      ./MCALLS/Amel/Herb2012/forager/replicate4/forager4.CpGscd.mcalls
Am    HE    fr    5     CpGhsm      ./MCALLS/Amel/Herb2012/forager/replicate5/forager5.CpGhsm.mcalls
Am    HE    fr    5     CpGscd      ./MCALLS/Amel/Herb2012/forager/replicate5/forager5.CpGscd.mcalls
Am    HE    fr    6     CpGhsm      ./MCALLS/Amel/Herb2012/forager/replicate6/forager6.CpGhsm.mcalls
Am    HE    fr    6     CpGscd      ./MCALLS/Amel/Herb2012/forager/replicate6/forager6.CpGscd.mcalls

Am    HE    qn    1     CpGhsm      ./MCALLS/Amel/Herb2012/queen/replicate1/queen1.CpGhsm.mcalls
Am    HE    qn    1     CpGscd      ./MCALLS/Amel/Herb2012/queen/replicate1/queen1.CpGscd.mcalls
Am    HE    qn    2     CpGhsm      ./MCALLS/Amel/Herb2012/queen/replicate2/queen2.CpGhsm.mcalls
Am    HE    qn    2     CpGscd      ./MCALLS/Amel/Herb2012/queen/replicate2/queen2.CpGscd.mcalls
Am    HE    qn    3     CpGhsm      ./MCALLS/Amel/Herb2012/queen/replicate3/queen3.CpGhsm.mcalls
Am    HE    qn    3     CpGscd      ./MCALLS/Amel/Herb2012/queen/replicate3/queen3.CpGscd.mcalls
Am    HE    qn    4     CpGhsm      ./MCALLS/Amel/Herb2012/queen/replicate4/queen4.CpGhsm.mcalls
Am    HE    qn    4     CpGscd      ./MCALLS/Amel/Herb2012/queen/replicate4/queen4.CpGscd.mcalls
Am    HE    qn    5     CpGhsm      ./MCALLS/Amel/Herb2012/queen/replicate5/queen5.CpGhsm.mcalls
Am    HE    qn    5     CpGscd      ./MCALLS/Amel/Herb2012/queen/replicate5/queen5.CpGscd.mcalls

Am    HE    rn    1     CpGhsm      ./MCALLS/Amel/Herb2012/reverted_nurse/replicate1/reverted_nurse1.CpGhsm.mcalls
Am    HE    rn    1     CpGscd      ./MCALLS/Amel/Herb2012/reverted_nurse/replicate1/reverted_nurse1.CpGscd.mcalls
Am    HE    rn    2     CpGhsm      ./MCALLS/Amel/Herb2012/reverted_nurse/replicate2/reverted_nurse2.CpGhsm.mcalls
Am    HE    rn    2     CpGscd      ./MCALLS/Amel/Herb2012/reverted_nurse/replicate2/reverted_nurse2.CpGscd.mcalls
Am    HE    rn    3     CpGhsm      ./MCALLS/Amel/Herb2012/reverted_nurse/replicate3/reverted_nurse3.CpGhsm.mcalls
Am    HE    rn    3     CpGscd      ./MCALLS/Amel/Herb2012/reverted_nurse/replicate3/reverted_nurse3.CpGscd.mcalls
Am    HE    rn    4     CpGhsm      ./MCALLS/Amel/Herb2012/reverted_nurse/replicate4/reverted_nurse4.CpGhsm.mcalls
Am    HE    rn    4     CpGscd      ./MCALLS/Amel/Herb2012/reverted_nurse/replicate4/reverted_nurse4.CpGscd.mcalls
Am    HE    rn    5     CpGhsm      ./MCALLS/Amel/Herb2012/reverted_nurse/replicate5/reverted_nurse5.CpGhsm.mcalls
Am    HE    rn    5     CpGscd      ./MCALLS/Amel/Herb2012/reverted_nurse/replicate5/reverted_nurse5.CpGscd.mcalls
Am    HE    rn    6     CpGhsm      ./MCALLS/Amel/Herb2012/reverted_nurse/replicate6/reverted_nurse6.CpGhsm.mcalls
Am    HE    rn    6     CpGscd      ./MCALLS/Amel/Herb2012/reverted_nurse/replicate6/reverted_nurse6.CpGscd.mcalls

Am    HE    wr    1     CpGhsm      ./MCALLS/Amel/Herb2012/worker/replicate1/worker1.CpGhsm.mcalls
Am    HE    wr    1     CpGscd      ./MCALLS/Amel/Herb2012/worker/replicate1/worker1.CpGscd.mcalls
Am    HE    wr    2     CpGhsm      ./MCALLS/Amel/Herb2012/worker/replicate2/worker2.CpGhsm.mcalls
Am    HE    wr    2     CpGscd      ./MCALLS/Amel/Herb2012/worker/replicate2/worker2.CpGscd.mcalls
Am    HE    wr    3     CpGhsm      ./MCALLS/Amel/Herb2012/worker/replicate3/worker3.CpGhsm.mcalls
Am    HE    wr    3     CpGscd      ./MCALLS/Amel/Herb2012/worker/replicate3/worker3.CpGscd.mcalls
Am    HE    wr    4     CpGhsm      ./MCALLS/Amel/Herb2012/worker/replicate4/worker4.CpGhsm.mcalls
Am    HE    wr    4     CpGscd      ./MCALLS/Amel/Herb2012/worker/replicate4/worker4.CpGscd.mcalls
Am    HE    wr    5     CpGhsm      ./MCALLS/Amel/Herb2012/worker/replicate5/worker5.CpGhsm.mcalls
Am    HE    wr    5     CpGscd      ./MCALLS/Amel/Herb2012/worker/replicate5/worker5.CpGscd.mcalls


# Samples from Li-Byarlay et al. (2013) PNAS:
# ac = abdominal fat bodies (workers), control; ak = abdominal fat bodies (workers), Dmnt3 knockdown
#
Am    LB    ac    0     CpGhsm      ./MCALLS/Amel/Li-Byarlay2013/control/control.CpGhsm.mcalls
Am    LB    ac    0     CpGscd      ./MCALLS/Amel/Li-Byarlay2013/control/control.CpGscd.mcalls
Am    LB    ak    0     CpGhsm      ./MCALLS/Amel/Li-Byarlay2013/knockdown/knockdown.CpGhsm.mcalls
Am    LB    ak    0     CpGscd      ./MCALLS/Amel/Li-Byarlay2013/knockdown/knockdown.CpGscd.mcalls
```

```
# Samples from Cingolani et al (2013) BMC Genomics:
# ahb = africanized honey bee, ehb = european honey bee
#
Am    CI    ahb   0     CpGhsm      ./MCALLS/Amel/Cingolani2013/africanized/africanized.CpGhsm.mcalls
Am    CI    ahb   0     CpGscd      ./MCALLS/Amel/Cingolani2013/africanized/africanized.CpGscd.mcalls
Am    CI    ehb   0     CpGhsm      ./MCALLS/Amel/Cingolani2013/european/european.CpGhsm.mcalls
Am    CI    ehb   0     CpGscd      ./MCALLS/Amel/Cingolani2013/european/european.CpGscd.mcalls


# Samples from Drewell et al (2014) Development:
# dr = drone, he = haploid_egg, sp = sperm
#
Am    DR    dr    0     CpGhsm      ./MCALLS/Amel/Drewell2014/drone/drone.CpGhsm.mcalls
Am    DR    dr    0     CpGscd      ./MCALLS/Amel/Drewell2014/drone/drone.CpGscd.mcalls
Am    DR    he    0     CpGhsm      ./MCALLS/Amel/Drewell2014/haploid_egg/haploid_egg.CpGhsm.mcalls
Am    DR    he    0     CpGscd      ./MCALLS/Amel/Drewell2014/haploid_egg/haploid_egg.CpGscd.mcalls
Am    DR    sp    0     CpGhsm      ./MCALLS/Amel/Drewell2014/sperm/sperm.CpGhsm.mcalls
Am    DR    sp    0     CpGscd      ./MCALLS/Amel/Drewell2014/sperm/sperm.CpGscd.mcalls


# Samples from Galbraith et al (2015) PLoS Pathogen:
# ctrl = control, infd = infected
#
Am    GA    ctrl  0     CpGhsm      ./MCALLS/Amel/Galbraith2015/control/control.CpGhsm.mcalls
Am    GA    ctrl  0     CpGscd      ./MCALLS/Amel/Galbraith2015/control/control.CpGscd.mcalls
Am    GA    infd  0     CpGhsm      ./MCALLS/Amel/Galbraith2015/infected/infected.CpGhsm.mcalls
Am    GA    infd  0     CpGscd      ./MCALLS/Amel/Galbraith2015/infected/infected.CpGscd.mcalls


# Samples from Remnant et al. (2016) BMC Genomics:
# fe = female embryos; te = thelytokous embryos
#
Am    RE    fe    0     CpGhsm      ./MCALLS/Amel/Remnant2016/fembryo/fembryo.CpGhsm.mcalls
Am    RE    fe    0     CpGscd      ./MCALLS/Amel/Remnant2016/fembryo/fembryo.CpGscd.mcalls
Am    RE    te    0     CpGhsm      ./MCALLS/Amel/Remnant2016/tembryo/tembryo.CpGhsm.mcalls
Am    RE    te    0     CpGscd      ./MCALLS/Amel/Remnant2016/tembryo/tembryo.CpGscd.mcalls


# Samples from Li et al. [Wang last author] (2017) Scientific Reports:
# tr = trained; nt = not trained
#
Am    LI    tr    0     CpGhsm      ./MCALLS/Amel/Li2017/trained/trained.CpGhsm.mcalls
Am    LI    tr    0     CpGscd      ./MCALLS/Amel/Li2017/trained/trained.CpGscd.mcalls
Am    LI    nt    0     CpGhsm      ./MCALLS/Amel/Li2017/control/control.CpGhsm.mcalls
Am    LI    nt    0     CpGscd      ./MCALLS/Amel/Li2017/control/control.CpGscd.mcalls


# Samples from Herb et al. (2018) BMC Genomics:
# A,C : aggressive, control; 5m, 2h: 5 minute, 120 minute
#
Am    H8    a2h   0     CpGhsm      ./MCALLS/Amel/Herb2018/A_120min/A_120min.CpGhsm.mcalls
Am    H8    a2h   0     CpGscd      ./MCALLS/Amel/Herb2018/A_120min/A_120min.CpGscd.mcalls
Am    H8    a5m   0     CpGhsm      ./MCALLS/Amel/Herb2018/A_5min/A_5min.CpGhsm.mcalls
Am    H8    a5m   0     CpGscd      ./MCALLS/Amel/Herb2018/A_5min/A_5min.CpGscd.mcalls
Am    H8    c2h   0     CpGhsm      ./MCALLS/Amel/Herb2018/C_120min/C_120min.CpGhsm.mcalls
Am    H8    c2h   0     CpGscd      ./MCALLS/Amel/Herb2018/C_120min/C_120min.CpGscd.mcalls
Am    H8    c5m   0     CpGhsm      ./MCALLS/Amel/Herb2018/C_5min/C_5min.CpGhsm.mcalls
Am    H8    c5m   0     CpGscd      ./MCALLS/Amel/Herb2018/C_5min/C_5min.CpGscd.mcalls

# ... continuing: replicates
#
Am    H8    a2h   1     CpGhsm      ./MCALLS/Amel/Herb2018/A_120min/replicate1/A_120min1.CpGhsm.mcalls
Am    H8    a2h   1     CpGscd      ./MCALLS/Amel/Herb2018/A_120min/replicate1/A_120min1.CpGscd.mcalls
```

```
Am    H8    a2h   2     CpGhsm      ./MCALLS/Amel/Herb2018/A_120min/replicate2/A_120min2.CpGhsm.mcalls
Am    H8    a2h   2     CpGscd      ./MCALLS/Amel/Herb2018/A_120min/replicate2/A_120min2.CpGscd.mcalls
Am    H8    a2h   3     CpGhsm      ./MCALLS/Amel/Herb2018/A_120min/replicate3/A_120min3.CpGhsm.mcalls
Am    H8    a2h   3     CpGscd      ./MCALLS/Amel/Herb2018/A_120min/replicate3/A_120min3.CpGscd.mcalls
Am    H8    a2h   4     CpGhsm      ./MCALLS/Amel/Herb2018/A_120min/replicate4/A_120min4.CpGhsm.mcalls
Am    H8    a2h   4     CpGscd      ./MCALLS/Amel/Herb2018/A_120min/replicate4/A_120min4.CpGscd.mcalls
Am    H8    a2h   5     CpGhsm      ./MCALLS/Amel/Herb2018/A_120min/replicate5/A_120min5.CpGhsm.mcalls
Am    H8    a2h   5     CpGscd      ./MCALLS/Amel/Herb2018/A_120min/replicate5/A_120min5.CpGscd.mcalls
Am    H8    a2h   6     CpGhsm      ./MCALLS/Amel/Herb2018/A_120min/replicate6/A_120min6.CpGhsm.mcalls
Am    H8    a2h   6     CpGscd      ./MCALLS/Amel/Herb2018/A_120min/replicate6/A_120min6.CpGscd.mcalls

Am    H8    a5m   1     CpGhsm      ./MCALLS/Amel/Herb2018/A_5min/replicate1/A_5min1.CpGhsm.mcalls
Am    H8    a5m   1     CpGscd      ./MCALLS/Amel/Herb2018/A_5min/replicate1/A_5min1.CpGscd.mcalls
Am    H8    a5m   2     CpGhsm      ./MCALLS/Amel/Herb2018/A_5min/replicate2/A_5min2.CpGhsm.mcalls
Am    H8    a5m   2     CpGscd      ./MCALLS/Amel/Herb2018/A_5min/replicate2/A_5min2.CpGscd.mcalls
Am    H8    a5m   3     CpGhsm      ./MCALLS/Amel/Herb2018/A_5min/replicate3/A_5min3.CpGhsm.mcalls
Am    H8    a5m   3     CpGscd      ./MCALLS/Amel/Herb2018/A_5min/replicate3/A_5min3.CpGscd.mcalls
Am    H8    a5m   4     CpGhsm      ./MCALLS/Amel/Herb2018/A_5min/replicate4/A_5min4.CpGhsm.mcalls
Am    H8    a5m   4     CpGscd      ./MCALLS/Amel/Herb2018/A_5min/replicate4/A_5min4.CpGscd.mcalls
Am    H8    a5m   5     CpGhsm      ./MCALLS/Amel/Herb2018/A_5min/replicate5/A_5min5.CpGhsm.mcalls
Am    H8    a5m   5     CpGscd      ./MCALLS/Amel/Herb2018/A_5min/replicate5/A_5min5.CpGscd.mcalls
Am    H8    a5m   6     CpGhsm      ./MCALLS/Amel/Herb2018/A_5min/replicate6/A_5min6.CpGhsm.mcalls
Am    H8    a5m   6     CpGscd      ./MCALLS/Amel/Herb2018/A_5min/replicate6/A_5min6.CpGscd.mcalls

Am    H8    c2h   1     CpGhsm      ./MCALLS/Amel/Herb2018/C_120min/replicate1/C_120min1.CpGhsm.mcalls
Am    H8    c2h   1     CpGscd      ./MCALLS/Amel/Herb2018/C_120min/replicate1/C_120min1.CpGscd.mcalls
Am    H8    c2h   2     CpGhsm      ./MCALLS/Amel/Herb2018/C_120min/replicate2/C_120min2.CpGhsm.mcalls
Am    H8    c2h   2     CpGscd      ./MCALLS/Amel/Herb2018/C_120min/replicate2/C_120min2.CpGscd.mcalls
Am    H8    c2h   3     CpGhsm      ./MCALLS/Amel/Herb2018/C_120min/replicate3/C_120min3.CpGhsm.mcalls
Am    H8    c2h   3     CpGscd      ./MCALLS/Amel/Herb2018/C_120min/replicate3/C_120min3.CpGscd.mcalls
Am    H8    c2h   4     CpGhsm      ./MCALLS/Amel/Herb2018/C_120min/replicate4/C_120min4.CpGhsm.mcalls
Am    H8    c2h   4     CpGscd      ./MCALLS/Amel/Herb2018/C_120min/replicate4/C_120min4.CpGscd.mcalls
Am    H8    c2h   5     CpGhsm      ./MCALLS/Amel/Herb2018/C_120min/replicate5/C_120min5.CpGhsm.mcalls
Am    H8    c2h   5     CpGscd      ./MCALLS/Amel/Herb2018/C_120min/replicate5/C_120min5.CpGscd.mcalls
Am    H8    c2h   6     CpGhsm      ./MCALLS/Amel/Herb2018/C_120min/replicate6/C_120min6.CpGhsm.mcalls
Am    H8    c2h   6     CpGscd      ./MCALLS/Amel/Herb2018/C_120min/replicate6/C_120min6.CpGscd.mcalls

Am    H8    c5m   1     CpGhsm      ./MCALLS/Amel/Herb2018/C_5min/replicate1/C_5min1.CpGhsm.mcalls
Am    H8    c5m   1     CpGscd      ./MCALLS/Amel/Herb2018/C_5min/replicate1/C_5min1.CpGscd.mcalls
Am    H8    c5m   2     CpGhsm      ./MCALLS/Amel/Herb2018/C_5min/replicate2/C_5min2.CpGhsm.mcalls
Am    H8    c5m   2     CpGscd      ./MCALLS/Amel/Herb2018/C_5min/replicate2/C_5min2.CpGscd.mcalls
Am    H8    c5m   3     CpGhsm      ./MCALLS/Amel/Herb2018/C_5min/replicate3/C_5min3.CpGhsm.mcalls
Am    H8    c5m   3     CpGscd      ./MCALLS/Amel/Herb2018/C_5min/replicate3/C_5min3.CpGscd.mcalls
Am    H8    c5m   4     CpGhsm      ./MCALLS/Amel/Herb2018/C_5min/replicate4/C_5min4.CpGhsm.mcalls
Am    H8    c5m   4     CpGscd      ./MCALLS/Amel/Herb2018/C_5min/replicate4/C_5min4.CpGscd.mcalls
Am    H8    c5m   5     CpGhsm      ./MCALLS/Amel/Herb2018/C_5min/replicate5/C_5min5.CpGhsm.mcalls
Am    H8    c5m   5     CpGscd      ./MCALLS/Amel/Herb2018/C_5min/replicate5/C_5min5.CpGscd.mcalls
Am    H8    c5m   6     CpGhsm      ./MCALLS/Amel/Herb2018/C_5min/replicate6/C_5min6.CpGhsm.mcalls
Am    H8    c5m   6     CpGscd      ./MCALLS/Amel/Herb2018/C_5min/replicate6/C_5min6.CpGscd.mcalls


# Samples from Yagound et al. (2019) Genome Biology and Evolution:
# sl = sperm_lc; sh = sperm_hc
#
Am    YA    sperm_lc    0     CpGhsm      ./MCALLS/Amel/Yagound2019/sperm_lc/sperm_lc.CpGhsm.mcalls
Am    YA    sperm_lc    0     CpGscd      ./MCALLS/Amel/Yagound2019/sperm_lc/sperm_lc.CpGscd.mcalls
Am    YA    sperm_hc    0     CpGhsm      ./MCALLS/Amel/Yagound2019/sperm_hc/sperm_hc.CpGhsm.mcalls
Am    YA    sperm_hc    0     CpGscd      ./MCALLS/Amel/Yagound2019/sperm_hc/sperm_hc.CpGscd.mcalls

# ... continuing: replicates
#
Am    YA    sperm_lc    1     CpGhsm      ./MCALLS/Amel/Yagound2019/sperm_lc/replicate1/sperm_lc1.CpGhsm.mcalls
Am    YA    sperm_lc    1     CpGscd      ./MCALLS/Amel/Yagound2019/sperm_lc/replicate1/sperm_lc1.CpGscd.mcalls
Am    YA    sperm_lc    2     CpGhsm      ./MCALLS/Amel/Yagound2019/sperm_lc/replicate2/sperm_lc2.CpGhsm.mcalls
```

```
Am    YA    sperm_lc    2     CpGscd      ./MCALLS/Amel/Yagound2019/sperm_lc/replicate2/sperm_lc2.CpGscd.mcalls
Am    YA    sperm_lc    3     CpGhsm      ./MCALLS/Amel/Yagound2019/sperm_lc/replicate3/sperm_lc3.CpGhsm.mcalls
Am    YA    sperm_lc    3     CpGscd      ./MCALLS/Amel/Yagound2019/sperm_lc/replicate3/sperm_lc3.CpGscd.mcalls
Am    YA    sperm_hc    1     CpGhsm      ./MCALLS/Amel/Yagound2019/sperm_hc/replicate1/sperm_hc1.CpGhsm.mcalls
Am    YA    sperm_hc    1     CpGscd      ./MCALLS/Amel/Yagound2019/sperm_hc/replicate1/sperm_hc1.CpGscd.mcalls
Am    YA    sperm_hc    2     CpGhsm      ./MCALLS/Amel/Yagound2019/sperm_hc/replicate2/sperm_hc2.CpGhsm.mcalls
Am    YA    sperm_hc    2     CpGscd      ./MCALLS/Amel/Yagound2019/sperm_hc/replicate2/sperm_hc2.CpGscd.mcalls
Am    YA    sperm_hc    3     CpGhsm      ./MCALLS/Amel/Yagound2019/sperm_hc/replicate3/sperm_hc3.CpGhsm.mcalls
Am    YA    sperm_hc    3     CpGscd      ./MCALLS/Amel/Yagound2019/sperm_hc/replicate3/sperm_hc3.CpGscd.mcalls
Am    YA    sperm_hc    4     CpGhsm      ./MCALLS/Amel/Yagound2019/sperm_hc/replicate4/sperm_hc4.CpGhsm.mcalls
Am    YA    sperm_hc    4     CpGscd      ./MCALLS/Amel/Yagound2019/sperm_hc/replicate4/sperm_hc4.CpGscd.mcalls


# Samples from Harris et al. (2019) Epigenetics & Chromatin:
# dh = drone_head; dl = drone_larva; ds = drone_sperm; qh = queen_head; we = worker_embryo; wh = worker_head; wp = worker_pupa
#
Am    HA    dh    0     CpGhsm      ./MCALLS/Amel/Harris2019/drone_head/drone_head.CpGhsm.mcalls
Am    HA    dh    0     CpGscd      ./MCALLS/Amel/Harris2019/drone_head/drone_head.CpGscd.mcalls
Am    HA    dl    0     CpGhsm      ./MCALLS/Amel/Harris2019/drone_larva/drone_larva.CpGhsm.mcalls
Am    HA    dl    0     CpGscd      ./MCALLS/Amel/Harris2019/drone_larva/drone_larva.CpGscd.mcalls
Am    HA    ds    0     CpGhsm      ./MCALLS/Amel/Harris2019/drone_sperm/drone_sperm.CpGhsm.mcalls
Am    HA    ds    0     CpGscd      ./MCALLS/Amel/Harris2019/drone_sperm/drone_sperm.CpGscd.mcalls
Am    HA    qh    0     CpGhsm      ./MCALLS/Amel/Harris2019/queen_head/queen_head.CpGhsm.mcalls
Am    HA    qh    0     CpGscd      ./MCALLS/Amel/Harris2019/queen_head/queen_head.CpGscd.mcalls
Am    HA    we    0     CpGhsm      ./MCALLS/Amel/Harris2019/worker_embryo/worker_embryo.CpGhsm.mcalls
Am    HA    we    0     CpGscd      ./MCALLS/Amel/Harris2019/worker_embryo/worker_embryo.CpGscd.mcalls
Am    HA    wh    0     CpGhsm      ./MCALLS/Amel/Harris2019/worker_head/worker_head.CpGhsm.mcalls
Am    HA    wh    0     CpGscd      ./MCALLS/Amel/Harris2019/worker_head/worker_head.CpGscd.mcalls
Am    HA    wp    0     CpGhsm      ./MCALLS/Amel/Harris2019/worker_pupa/worker_pupa.CpGhsm.mcalls
Am    HA    wp    0     CpGscd      ./MCALLS/Amel/Harris2019/worker_pupa/worker_pupa.CpGscd.mcalls

# ... continuing: replicates
#
Am    HA    dh    1     CpGhsm      ./MCALLS/Amel/Harris2019/drone_head/replicate1/drone_head1.CpGhsm.mcalls
Am    HA    dh    1     CpGscd      ./MCALLS/Amel/Harris2019/drone_head/replicate1/drone_head1.CpGscd.mcalls
Am    HA    dh    2     CpGhsm      ./MCALLS/Amel/Harris2019/drone_head/replicate2/drone_head2.CpGhsm.mcalls
Am    HA    dh    2     CpGscd      ./MCALLS/Amel/Harris2019/drone_head/replicate2/drone_head2.CpGscd.mcalls
Am    HA    dl    1     CpGhsm      ./MCALLS/Amel/Harris2019/drone_larva/replicate1/drone_larva1.CpGhsm.mcalls
Am    HA    dl    1     CpGscd      ./MCALLS/Amel/Harris2019/drone_larva/replicate1/drone_larva1.CpGscd.mcalls
Am    HA    dl    2     CpGhsm      ./MCALLS/Amel/Harris2019/drone_larva/replicate2/drone_larva2.CpGhsm.mcalls
Am    HA    dl    2     CpGscd      ./MCALLS/Amel/Harris2019/drone_larva/replicate2/drone_larva2.CpGscd.mcalls
Am    HA    ds    1     CpGhsm      ./MCALLS/Amel/Harris2019/drone_sperm/replicate1/drone_sperm1.CpGhsm.mcalls
Am    HA    ds    1     CpGscd      ./MCALLS/Amel/Harris2019/drone_sperm/replicate1/drone_sperm1.CpGscd.mcalls
Am    HA    ds    2     CpGhsm      ./MCALLS/Amel/Harris2019/drone_sperm/replicate2/drone_sperm2.CpGhsm.mcalls
Am    HA    ds    2     CpGscd      ./MCALLS/Amel/Harris2019/drone_sperm/replicate2/drone_sperm2.CpGscd.mcalls
Am    HA    qh    1     CpGhsm      ./MCALLS/Amel/Harris2019/queen_head/replicate1/queen_head1.CpGhsm.mcalls
Am    HA    qh    1     CpGscd      ./MCALLS/Amel/Harris2019/queen_head/replicate1/queen_head1.CpGscd.mcalls
Am    HA    qh    2     CpGhsm      ./MCALLS/Amel/Harris2019/queen_head/replicate2/queen_head2.CpGhsm.mcalls
Am    HA    qh    2     CpGscd      ./MCALLS/Amel/Harris2019/queen_head/replicate2/queen_head2.CpGscd.mcalls
Am    HA    we    1     CpGhsm      ./MCALLS/Amel/Harris2019/worker_embryo/replicate1/worker_embryo1.CpGhsm.mcalls
Am    HA    we    1     CpGscd      ./MCALLS/Amel/Harris2019/worker_embryo/replicate1/worker_embryo1.CpGscd.mcalls
Am    HA    we    2     CpGhsm      ./MCALLS/Amel/Harris2019/worker_embryo/replicate2/worker_embryo2.CpGhsm.mcalls
Am    HA    we    2     CpGscd      ./MCALLS/Amel/Harris2019/worker_embryo/replicate2/worker_embryo2.CpGscd.mcalls
Am    HA    wh    1     CpGhsm      ./MCALLS/Amel/Harris2019/worker_head/replicate1/worker_head1.CpGhsm.mcalls
Am    HA    wh    1     CpGscd      ./MCALLS/Amel/Harris2019/worker_head/replicate1/worker_head1.CpGscd.mcalls
Am    HA    wh    2     CpGhsm      ./MCALLS/Amel/Harris2019/worker_head/replicate2/worker_head2.CpGhsm.mcalls
Am    HA    wh    2     CpGscd      ./MCALLS/Amel/Harris2019/worker_head/replicate2/worker_head2.CpGscd.mcalls
Am    HA    wp    1     CpGhsm      ./MCALLS/Amel/Harris2019/worker_pupa/replicate1/worker_pupa1.CpGhsm.mcalls
Am    HA    wp    1     CpGscd      ./MCALLS/Amel/Harris2019/worker_pupa/replicate1/worker_pupa1.CpGscd.mcalls
Am    HA    wp    2     CpGhsm      ./MCALLS/Amel/Harris2019/worker_pupa/replicate2/worker_pupa2.CpGhsm.mcalls
Am    HA    wp    2     CpGscd      ./MCALLS/Amel/Harris2019/worker_pupa/replicate2/worker_pupa2.CpGscd.mcalls


# Samples from NCBI SRA BGI (2020) ???:
```

```
# sp = sperm; oo = oocyte; wb = worker_blastoderm; wg = worker_gastrula; db = drone_blastoderm; dg = drone_gastrula
#
Am    BG    sp    0     CpGhsm      ./MCALLS/Amel/BGI2020/sperm/sperm.CpGhsm.mcalls
Am    BG    sp    0     CpGscd      ./MCALLS/Amel/BGI2020/sperm/sperm.CpGscd.mcalls
Am    BG    oo    0     CpGhsm      ./MCALLS/Amel/BGI2020/oocyte/oocyte.CpGhsm.mcalls
Am    BG    oo    0     CpGscd      ./MCALLS/Amel/BGI2020/oocyte/oocyte.CpGscd.mcalls
Am    BG    wb    0     CpGhsm      ./MCALLS/Amel/BGI2020/worker_blastoderm/worker_blastoderm.CpGhsm.mcalls
Am    BG    wb    0     CpGscd      ./MCALLS/Amel/BGI2020/worker_blastoderm/worker_blastoderm.CpGscd.mcalls
Am    BG    wg    0     CpGhsm      ./MCALLS/Amel/BGI2020/worker_gastrula/worker_gastrula.CpGhsm.mcalls
Am    BG    wg    0     CpGscd      ./MCALLS/Amel/BGI2020/worker_gastrula/worker_gastrula.CpGscd.mcalls
Am    BG    db    0     CpGhsm      ./MCALLS/Amel/BGI2020/drone_blastoderm/drone_blastoderm.CpGhsm.mcalls
Am    BG    db    0     CpGscd      ./MCALLS/Amel/BGI2020/drone_blastoderm/drone_blastoderm.CpGscd.mcalls
Am    BG    dg    0     CpGhsm      ./MCALLS/Amel/BGI2020/drone_gastrula/drone_gastrula.CpGhsm.mcalls
Am    BG    dg    0     CpGscd      ./MCALLS/Amel/BGI2020/drone_gastrula/drone_gastrula.CpGscd.mcalls


# Samples from NCBI SRA Jiangxi (2020) ???:
# G = generation; E = eggs; L = larvae
#
Am    JI    G1E   0     CpGhsm      ./MCALLS/Amel/Jiangxi2020/G1E/G1E.CpGhsm.mcalls
Am    JI    G1E   0     CpGscd      ./MCALLS/Amel/Jiangxi2020/G1E/G1E.CpGscd.mcalls
Am    JI    G1L1  0     CpGhsm      ./MCALLS/Amel/Jiangxi2020/G1L1/G1L1.CpGhsm.mcalls
Am    JI    G1L1  0     CpGscd      ./MCALLS/Amel/Jiangxi2020/G1L1/G1L1.CpGscd.mcalls
Am    JI    G1L2  0     CpGhsm      ./MCALLS/Amel/Jiangxi2020/G1L2/G1L2.CpGhsm.mcalls
Am    JI    G1L2  0     CpGscd      ./MCALLS/Amel/Jiangxi2020/G1L2/G1L2.CpGscd.mcalls
Am    JI    G2E   0     CpGhsm      ./MCALLS/Amel/Jiangxi2020/G2E/G2E.CpGhsm.mcalls
Am    JI    G2E   0     CpGscd      ./MCALLS/Amel/Jiangxi2020/G2E/G2E.CpGscd.mcalls
Am    JI    G2L1  0     CpGhsm      ./MCALLS/Amel/Jiangxi2020/G2L1/G2L1.CpGhsm.mcalls
Am    JI    G2L1  0     CpGscd      ./MCALLS/Amel/Jiangxi2020/G2L1/G2L1.CpGscd.mcalls
Am    JI    G2L2  0     CpGhsm      ./MCALLS/Amel/Jiangxi2020/G2L2/G2L2.CpGhsm.mcalls
Am    JI    G2L2  0     CpGscd      ./MCALLS/Amel/Jiangxi2020/G2L2/G2L2.CpGscd.mcalls
Am    JI    G3E   0     CpGhsm      ./MCALLS/Amel/Jiangxi2020/G3E/G3E.CpGhsm.mcalls
Am    JI    G3E   0     CpGscd      ./MCALLS/Amel/Jiangxi2020/G3E/G3E.CpGscd.mcalls
Am    JI    G3L1  0     CpGhsm      ./MCALLS/Amel/Jiangxi2020/G3L1/G3L1.CpGhsm.mcalls
Am    JI    G3L1  0     CpGscd      ./MCALLS/Amel/Jiangxi2020/G3L1/G3L1.CpGscd.mcalls
Am    JI    G3L2  0     CpGhsm      ./MCALLS/Amel/Jiangxi2020/G3L2/G3L2.CpGhsm.mcalls
Am    JI    G3L2  0     CpGscd      ./MCALLS/Amel/Jiangxi2020/G3L2/G3L2.CpGscd.mcalls
Am    JI    G4E   0     CpGhsm      ./MCALLS/Amel/Jiangxi2020/G4E/G4E.CpGhsm.mcalls
Am    JI    G4E   0     CpGscd      ./MCALLS/Amel/Jiangxi2020/G4E/G4E.CpGscd.mcalls
Am    JI    G4L1  0     CpGhsm      ./MCALLS/Amel/Jiangxi2020/G4L1/G4L1.CpGhsm.mcalls
Am    JI    G4L1  0     CpGscd      ./MCALLS/Amel/Jiangxi2020/G4L1/G4L1.CpGscd.mcalls
Am    JI    G4L2  0     CpGhsm      ./MCALLS/Amel/Jiangxi2020/G4L2/G4L2.CpGhsm.mcalls
Am    JI    G4L2  0     CpGscd      ./MCALLS/Amel/Jiangxi2020/G4L2/G4L2.CpGscd.mcalls

# ... continuing: replicates
#
Am    JI    G1E   1     CpGhsm      ./MCALLS/Amel/Jiangxi2020/G1E/replicate1/G1E1.CpGhsm.mcalls
Am    JI    G1E   1     CpGscd      ./MCALLS/Amel/Jiangxi2020/G1E/replicate1/G1E1.CpGscd.mcalls
Am    JI    G1E   2     CpGhsm      ./MCALLS/Amel/Jiangxi2020/G1E/replicate2/G1E2.CpGhsm.mcalls
Am    JI    G1E   2     CpGscd      ./MCALLS/Amel/Jiangxi2020/G1E/replicate2/G1E2.CpGscd.mcalls
Am    JI    G1E   3     CpGhsm      ./MCALLS/Amel/Jiangxi2020/G1E/replicate3/G1E3.CpGhsm.mcalls
Am    JI    G1E   3     CpGscd      ./MCALLS/Amel/Jiangxi2020/G1E/replicate3/G1E3.CpGscd.mcalls
Am    JI    G1L1  1     CpGhsm      ./MCALLS/Amel/Jiangxi2020/G1L1/replicate1/G1L11.CpGhsm.mcalls
Am    JI    G1L1  1     CpGscd      ./MCALLS/Amel/Jiangxi2020/G1L1/replicate1/G1L11.CpGscd.mcalls
Am    JI    G1L1  2     CpGhsm      ./MCALLS/Amel/Jiangxi2020/G1L1/replicate2/G1L12.CpGhsm.mcalls
Am    JI    G1L1  2     CpGscd      ./MCALLS/Amel/Jiangxi2020/G1L1/replicate2/G1L12.CpGscd.mcalls
Am    JI    G1L1  3     CpGhsm      ./MCALLS/Amel/Jiangxi2020/G1L1/replicate3/G1L13.CpGhsm.mcalls
Am    JI    G1L1  3     CpGscd      ./MCALLS/Amel/Jiangxi2020/G1L1/replicate3/G1L13.CpGscd.mcalls
Am    JI    G1L2  1     CpGhsm      ./MCALLS/Amel/Jiangxi2020/G1L2/replicate1/G1L21.CpGhsm.mcalls
Am    JI    G1L2  1     CpGscd      ./MCALLS/Amel/Jiangxi2020/G1L2/replicate1/G1L21.CpGscd.mcalls
Am    JI    G1L2  2     CpGhsm      ./MCALLS/Amel/Jiangxi2020/G1L2/replicate2/G1L22.CpGhsm.mcalls
Am    JI    G1L2  2     CpGscd      ./MCALLS/Amel/Jiangxi2020/G1L2/replicate2/G1L22.CpGscd.mcalls
Am    JI    G1L2  3     CpGhsm      ./MCALLS/Amel/Jiangxi2020/G1L2/replicate3/G1L23.CpGhsm.mcalls
Am    JI    G1L2  3     CpGscd      ./MCALLS/Amel/Jiangxi2020/G1L2/replicate3/G1L23.CpGscd.mcalls
```

| | | | | | |
|---|---|---|---|---|---|
| Am | JI | G1L2 | 4 | CpGhsm | ./MCALLS/Amel/Jiangxi2020/G1L2/replicate4/G1L24.CpGhsm.mcalls |
| Am | JI | G1L2 | 4 | CpGscd | ./MCALLS/Amel/Jiangxi2020/G1L2/replicate4/G1L24.CpGscd.mcalls |
| Am | JI | G2E | 1 | CpGhsm | ./MCALLS/Amel/Jiangxi2020/G2E/replicate1/G2E1.CpGhsm.mcalls |
| Am | JI | G2E | 1 | CpGscd | ./MCALLS/Amel/Jiangxi2020/G2E/replicate1/G2E1.CpGscd.mcalls |
| Am | JI | G2E | 2 | CpGhsm | ./MCALLS/Amel/Jiangxi2020/G2E/replicate2/G2E2.CpGhsm.mcalls |
| Am | JI | G2E | 2 | CpGscd | ./MCALLS/Amel/Jiangxi2020/G2E/replicate2/G2E2.CpGscd.mcalls |
| Am | JI | G2E | 3 | CpGhsm | ./MCALLS/Amel/Jiangxi2020/G2E/replicate3/G2E3.CpGhsm.mcalls |
| Am | JI | G2E | 3 | CpGscd | ./MCALLS/Amel/Jiangxi2020/G2E/replicate3/G2E3.CpGscd.mcalls |
| Am | JI | G2L1 | 1 | CpGhsm | ./MCALLS/Amel/Jiangxi2020/G2L1/replicate1/G2L11.CpGhsm.mcalls |
| Am | JI | G2L1 | 1 | CpGscd | ./MCALLS/Amel/Jiangxi2020/G2L1/replicate1/G2L11.CpGscd.mcalls |
| Am | JI | G2L1 | 2 | CpGhsm | ./MCALLS/Amel/Jiangxi2020/G2L1/replicate2/G2L12.CpGhsm.mcalls |
| Am | JI | G2L1 | 2 | CpGscd | ./MCALLS/Amel/Jiangxi2020/G2L1/replicate2/G2L12.CpGscd.mcalls |
| Am | JI | G2L1 | 3 | CpGhsm | ./MCALLS/Amel/Jiangxi2020/G2L1/replicate3/G2L13.CpGhsm.mcalls |
| Am | JI | G2L1 | 3 | CpGscd | ./MCALLS/Amel/Jiangxi2020/G2L1/replicate3/G2L13.CpGscd.mcalls |
| Am | JI | G2L2 | 1 | CpGhsm | ./MCALLS/Amel/Jiangxi2020/G2L2/replicate1/G2L21.CpGhsm.mcalls |
| Am | JI | G2L2 | 1 | CpGscd | ./MCALLS/Amel/Jiangxi2020/G2L2/replicate1/G2L21.CpGscd.mcalls |
| Am | JI | G2L2 | 2 | CpGhsm | ./MCALLS/Amel/Jiangxi2020/G2L2/replicate2/G2L22.CpGhsm.mcalls |
| Am | JI | G2L2 | 2 | CpGscd | ./MCALLS/Amel/Jiangxi2020/G2L2/replicate2/G2L22.CpGscd.mcalls |
| Am | JI | G3E | 1 | CpGhsm | ./MCALLS/Amel/Jiangxi2020/G3E/replicate1/G3E1.CpGhsm.mcalls |
| Am | JI | G3E | 1 | CpGscd | ./MCALLS/Amel/Jiangxi2020/G3E/replicate1/G3E1.CpGscd.mcalls |
| Am | JI | G3E | 2 | CpGhsm | ./MCALLS/Amel/Jiangxi2020/G3E/replicate2/G3E2.CpGhsm.mcalls |
| Am | JI | G3E | 2 | CpGscd | ./MCALLS/Amel/Jiangxi2020/G3E/replicate2/G3E2.CpGscd.mcalls |
| Am | JI | G3E | 3 | CpGhsm | ./MCALLS/Amel/Jiangxi2020/G3E/replicate3/G3E3.CpGhsm.mcalls |
| Am | JI | G3E | 3 | CpGscd | ./MCALLS/Amel/Jiangxi2020/G3E/replicate3/G3E3.CpGscd.mcalls |
| Am | JI | G3L1 | 1 | CpGhsm | ./MCALLS/Amel/Jiangxi2020/G3L1/replicate1/G3L11.CpGhsm.mcalls |
| Am | JI | G3L1 | 1 | CpGscd | ./MCALLS/Amel/Jiangxi2020/G3L1/replicate1/G3L11.CpGscd.mcalls |
| Am | JI | G3L1 | 2 | CpGhsm | ./MCALLS/Amel/Jiangxi2020/G3L1/replicate2/G3L12.CpGhsm.mcalls |
| Am | JI | G3L1 | 2 | CpGscd | ./MCALLS/Amel/Jiangxi2020/G3L1/replicate2/G3L12.CpGscd.mcalls |
| Am | JI | G3L1 | 3 | CpGhsm | ./MCALLS/Amel/Jiangxi2020/G3L1/replicate3/G3L13.CpGhsm.mcalls |
| Am | JI | G3L1 | 3 | CpGscd | ./MCALLS/Amel/Jiangxi2020/G3L1/replicate3/G3L13.CpGscd.mcalls |
| Am | JI | G3L2 | 1 | CpGhsm | ./MCALLS/Amel/Jiangxi2020/G3L2/replicate1/G3L21.CpGhsm.mcalls |
| Am | JI | G3L2 | 1 | CpGscd | ./MCALLS/Amel/Jiangxi2020/G3L2/replicate1/G3L21.CpGscd.mcalls |
| Am | JI | G3L2 | 2 | CpGhsm | ./MCALLS/Amel/Jiangxi2020/G3L2/replicate2/G3L22.CpGhsm.mcalls |
| Am | JI | G3L2 | 2 | CpGscd | ./MCALLS/Amel/Jiangxi2020/G3L2/replicate2/G3L22.CpGscd.mcalls |
| Am | JI | G3L2 | 3 | CpGhsm | ./MCALLS/Amel/Jiangxi2020/G3L2/replicate3/G3L23.CpGhsm.mcalls |
| Am | JI | G3L2 | 3 | CpGscd | ./MCALLS/Amel/Jiangxi2020/G3L2/replicate3/G3L23.CpGscd.mcalls |
| Am | JI | G4E | 1 | CpGhsm | ./MCALLS/Amel/Jiangxi2020/G4E/replicate1/G4E1.CpGhsm.mcalls |
| Am | JI | G4E | 1 | CpGscd | ./MCALLS/Amel/Jiangxi2020/G4E/replicate1/G4E1.CpGscd.mcalls |
| Am | JI | G4E | 2 | CpGhsm | ./MCALLS/Amel/Jiangxi2020/G4E/replicate2/G4E2.CpGhsm.mcalls |
| Am | JI | G4E | 2 | CpGscd | ./MCALLS/Amel/Jiangxi2020/G4E/replicate2/G4E2.CpGscd.mcalls |
| Am | JI | G4E | 3 | CpGhsm | ./MCALLS/Amel/Jiangxi2020/G4E/replicate3/G4E3.CpGhsm.mcalls |
| Am | JI | G4E | 3 | CpGscd | ./MCALLS/Amel/Jiangxi2020/G4E/replicate3/G4E3.CpGscd.mcalls |
| Am | JI | G4E | 4 | CpGhsm | ./MCALLS/Amel/Jiangxi2020/G4E/replicate4/G4E4.CpGhsm.mcalls |
| Am | JI | G4E | 4 | CpGscd | ./MCALLS/Amel/Jiangxi2020/G4E/replicate4/G4E4.CpGscd.mcalls |
| Am | JI | G4L1 | 1 | CpGhsm | ./MCALLS/Amel/Jiangxi2020/G4L1/replicate1/G4L11.CpGhsm.mcalls |
| Am | JI | G4L1 | 1 | CpGscd | ./MCALLS/Amel/Jiangxi2020/G4L1/replicate1/G4L11.CpGscd.mcalls |
| Am | JI | G4L1 | 2 | CpGhsm | ./MCALLS/Amel/Jiangxi2020/G4L1/replicate2/G4L12.CpGhsm.mcalls |
| Am | JI | G4L1 | 2 | CpGscd | ./MCALLS/Amel/Jiangxi2020/G4L1/replicate2/G4L12.CpGscd.mcalls |
| Am | JI | G4L1 | 3 | CpGhsm | ./MCALLS/Amel/Jiangxi2020/G4L1/replicate3/G4L13.CpGhsm.mcalls |
| Am | JI | G4L1 | 3 | CpGscd | ./MCALLS/Amel/Jiangxi2020/G4L1/replicate3/G4L13.CpGscd.mcalls |
| Am | JI | G4L1 | 4 | CpGhsm | ./MCALLS/Amel/Jiangxi2020/G4L1/replicate4/G4L14.CpGhsm.mcalls |
| Am | JI | G4L1 | 4 | CpGscd | ./MCALLS/Amel/Jiangxi2020/G4L1/replicate4/G4L14.CpGscd.mcalls |
| Am | JI | G4L2 | 1 | CpGhsm | ./MCALLS/Amel/Jiangxi2020/G4L2/replicate1/G4L21.CpGhsm.mcalls |
| Am | JI | G4L2 | 1 | CpGscd | ./MCALLS/Amel/Jiangxi2020/G4L2/replicate1/G4L21.CpGscd.mcalls |
| Am | JI | G4L2 | 2 | CpGhsm | ./MCALLS/Amel/Jiangxi2020/G4L2/replicate2/G4L22.CpGhsm.mcalls |
| Am | JI | G4L2 | 2 | CpGscd | ./MCALLS/Amel/Jiangxi2020/G4L2/replicate2/G4L22.CpGscd.mcalls |
| Am | JI | G4L2 | 3 | CpGhsm | ./MCALLS/Amel/Jiangxi2020/G4L2/replicate3/G4L23.CpGhsm.mcalls |
| Am | JI | G4L2 | 3 | CpGscd | ./MCALLS/Amel/Jiangxi2020/G4L2/replicate3/G4L23.CpGscd.mcalls |
| Am | JI | G4L2 | 4 | CpGhsm | ./MCALLS/Amel/Jiangxi2020/G4L2/replicate4/G4L24.CpGhsm.mcalls |
| Am | JI | G4L2 | 4 | CpGscd | ./MCALLS/Amel/Jiangxi2020/G4L2/replicate4/G4L24.CpGscd.mcalls |

```
0README


Rscript.BWASPR output for .conf file Am_RE.conf.

Data input are the specified *.mcalls files and parameters set in Am_RE.conf.
The *.mcalls input data are saved in the following data structures:

studymk - a methylKit methylRaw(List) object storing the CpGhsm site data for all samples
studymc - a methylKit methylRaw(List) object storing the CpGscd site data for all samples
studyhc - a methylKit methylRaw(List) object storing the CpGscd site data for all samples,
           restricted to sites with high coverage (here set to 10)
mkrd    - a methylKit methylRawList object storing the CpGhsm site data for all
           replicates of a given sample

Output of the analysis is stored in the following subdirectories (if the corresponding
 function calls were specified in Am_RE.conf; please consult the 0README
 files in the subdirectories for details on the output files):

CMS - Coverage and methylation statistics for aggregate samples
PWC - Pairwise comparisons between all samples
CRL - Correlations between aggregate samples
REPCMS - Coverage and methylation statistics for replicate samples
REPCRL - Correlations between replicates
MMP - Mapping of methylation sites on genome annotation
ACS - Annotation of conserved methylation sites
RNK - Genomic feature regions ranked by CpGhsm statistics
MRPR - Methylation-rich and -poor regions
DMSG - Differentially methylateds sites and genes
OGL - Ordered gene lists
```

```
Directory: CMS File: 0READMEcms


CMS - Coverage and methylation statistics for aggregate samples.

Input: studymk, covlist, locount, hicount

Output: files cms-*.txt and cms-*.pdf

Notes: The statistics are given for CpGhsm sites restricted to minimum or
        higher level coverage.  At the minimum level, the number of sites will be
        the number of lines of the corresponding *CpGhsm.mcalls file minus one
        (consistency check).  The distribution of methylation levels is really only
        interesting for high coverage because at low coverage there will be a strong
        bias towards high methylation levels (by definition of hsm sites).
        The same bias at high coverage would suggest that the sample includes a
        preponderance of consistently methylated genomic sites.

        locount and hicount set bounds on the coverage to exclude sites with too few
        or too many covering reads to provide statistics on a typical range.
```

```
Directory: CMS File: cms-Am_RE_fe.txt


Number of "fe" CpGhsm-sites with minimal and higher level coverage:

  number of "fe" CpGhsm-sites with coverage >=  4:  18314
  number of "fe" CpGhsm-sites with coverage >=  4:  18314
  number of "fe" CpGhsm-sites with coverage >= 10:   1684
  number of "fe" CpGhsm-sites with coverage >= 15:    405


Coverage and methylation statistics for "fe" CpGhsm-sites at different levels of minimum coverage:


methylKit::getCoverageStats output for "fe" CpGhsm-sites (#: 18314) at minimum coverage 4      - read coverage statistics per base
summary:
   Min. 1st Qu.  Median    Mean 3rd Qu.    Max.
    4.0     4.0     5.0     6.2     7.0   122.0
percentiles:
   0%   10%   20%   30%   40%   50%   60%   70%   80%   90%   95%   99% 99.5% 99.9%  100%
    4     4     4     4     5     5     6     6     7     9    11    19    26    43   122

methylKit::getMethylationStats output for "fe" CpGhsm-sites (#: 18314) at minimum coverage 4   - methylation statistics per base
summary:
   Min. 1st Qu.  Median    Mean 3rd Qu.    Max.
   15.9   100.0   100.0    95.5   100.0   100.0
percentiles:
   0%   10%   20%   30%   40%   50%   60%   70%   80%   90%   95%   99% 99.5% 99.9%  100%
 15.9  83.3  90.9 100.0 100.0 100.0 100.0 100.0 100.0 100.0 100.0 100.0 100.0 100.0 100.0



methylKit::getCoverageStats output for "fe" CpGhsm-sites (#: 18314) at minimum coverage 4      - read coverage statistics per base
summary:
   Min. 1st Qu.  Median    Mean 3rd Qu.    Max.
    4.0     4.0     5.0     6.2     7.0   122.0
percentiles:
   0%   10%   20%   30%   40%   50%   60%   70%   80%   90%   95%   99% 99.5% 99.9%  100%
    4     4     4     4     5     5     6     6     7     9    11    19    26    43   122

methylKit::getMethylationStats output for "fe" CpGhsm-sites (#: 18314) at minimum coverage 4   - methylation statistics per base
summary:
   Min. 1st Qu.  Median    Mean 3rd Qu.    Max.
   15.9   100.0   100.0    95.5   100.0   100.0
percentiles:
   0%   10%   20%   30%   40%   50%   60%   70%   80%   90%   95%   99% 99.5% 99.9%  100%
 15.9  83.3  90.9 100.0 100.0 100.0 100.0 100.0 100.0 100.0 100.0 100.0 100.0 100.0 100.0



methylKit::getCoverageStats output for "fe" CpGhsm-sites (#: 1684) at minimum coverage 10      - read coverage statistics per base
summary:
   Min. 1st Qu.  Median    Mean 3rd Qu.    Max.
   10.0    10.0    12.0    14.1    14.0   122.0
percentiles:
   0%   10%   20%   30%   40%   50%   60%   70%   80%   90%   95%   99% 99.5% 99.9%  100%
 10.0  10.0  10.0  11.0  11.0  12.0  13.0  14.0  15.0  20.0  27.0  46.3  54.6  70.9 122.0

methylKit::getMethylationStats output for "fe" CpGhsm-sites (#: 1684) at minimum coverage 10   - methylation statistics per base
summary:
   Min. 1st Qu.  Median    Mean 3rd Qu.    Max.
   15.9    70.0    90.0    81.6   100.0   100.0
percentiles:
   0%   10%   20%   30%   40%   50%   60%   70%   80%   90%   95%   99% 99.5% 99.9%  100%
```

```
 15.9  50.0  63.6  72.7  81.8  90.0  90.9  94.4 100.0 100.0 100.0 100.0 100.0 100.0 100.0



methylKit::getCoverageStats output for "fe" CpGhsm-sites (#: 405) at minimum coverage 15 - read coverage statistics per base
summary:
   Min. 1st Qu.  Median    Mean 3rd Qu.    Max. 
   15.0    16.0    19.0    22.6    25.0   122.0 
percentiles:
   0%   10%   20%   30%   40%   50%   60%   70%   80%   90%   95%   99% 99.5% 99.9%  100% 
 15.0  15.0  16.0  17.0  17.0  19.0  21.0  23.0  27.0  35.0  41.0  65.9  69.0 103.0 122.0 

methylKit::getMethylationStats output for "fe" CpGhsm-sites (#: 405) at minimum coverage 15    - methylation statistics per base
summary:
   Min. 1st Qu.  Median    Mean 3rd Qu.    Max. 
   15.9    50.0    73.3    70.4    93.3   100.0 
percentiles:
   0%   10%   20%   30%   40%   50%   60%   70%   80%   90%   95%   99% 99.5% 99.9%  100% 
 15.9  37.7  45.4  55.6  65.0  73.3  82.4  88.0  93.8 100.0 100.0 100.0 100.0 100.0 100.0 



methylKit::getCoverageStats output for "fe" CpGhsm-sites (#: 17623) in coverage range [4-12]   - read coverage statistics per base
summary:
   Min. 1st Qu.  Median    Mean 3rd Qu.    Max. 
   4.00    4.00    5.00    5.66    7.00   12.00 
percentiles:
   0%   10%   20%   30%   40%   50%   60%   70%   80%   90%   95%   99% 99.5% 99.9%  100% 
    4     4     4     4     5     5     6     6     7     8    10    12    12    12    12 

methylKit::getMethylationStats output for "fe" CpGhsm-sites (#: 17623) in coverage range [4-12]      - methylation statistics per base
summary:
   Min. 1st Qu.  Median    Mean 3rd Qu.    Max. 
   50.0   100.0   100.0    96.3   100.0   100.0 
percentiles:
   0%   10%   20%   30%   40%   50%   60%   70%   80%   90%   95%   99% 99.5% 99.9%  100% 
 50.0  83.3 100.0 100.0 100.0 100.0 100.0 100.0 100.0 100.0 100.0 100.0 100.0 100.0 100.0 
```

```
Directory: CMS File: cms-Am_RE_te.txt


Number of "te" CpGhsm-sites with minimal and higher level coverage:

  number of "te" CpGhsm-sites with coverage >=  4:  27814
  number of "te" CpGhsm-sites with coverage >=  4:  27814
  number of "te" CpGhsm-sites with coverage >= 10:   2246
  number of "te" CpGhsm-sites with coverage >= 15:    407


Coverage and methylation statistics for "te" CpGhsm-sites at different levels of minimum coverage:


methylKit::getCoverageStats output for "te" CpGhsm-sites (#: 27814) at minimum coverage 4      - read coverage statistics per base
summary:
   Min. 1st Qu.  Median    Mean 3rd Qu.    Max.
    4.0     4.0     5.0     6.2     7.0    99.0
percentiles:
   0%   10%   20%   30%   40%   50%   60%   70%   80%   90%   95%   99% 99.5% 99.9%  100%
  4.0   4.0   4.0   4.0   5.0   5.0   6.0   7.0   8.0   9.0  11.0  16.0  24.0  51.2  99.0

methylKit::getMethylationStats output for "te" CpGhsm-sites (#: 27814) at minimum coverage 4   - methylation statistics per base
summary:
   Min. 1st Qu.  Median    Mean 3rd Qu.    Max.
   16.7   100.0   100.0    95.6   100.0   100.0
percentiles:
   0%   10%   20%   30%   40%   50%   60%   70%   80%   90%   95%   99% 99.5% 99.9%  100%
 16.7  83.3  91.7 100.0 100.0 100.0 100.0 100.0 100.0 100.0 100.0 100.0 100.0 100.0 100.0
```


```
methylKit::getCoverageStats output for "te" CpGhsm-sites (#: 27814) at minimum coverage 4      - read coverage statistics per base
summary:
   Min. 1st Qu.  Median    Mean 3rd Qu.    Max.
    4.0     4.0     5.0     6.2     7.0    99.0
percentiles:
   0%   10%   20%   30%   40%   50%   60%   70%   80%   90%   95%   99% 99.5% 99.9%  100%
  4.0   4.0   4.0   4.0   5.0   5.0   6.0   7.0   8.0   9.0  11.0  16.0  24.0  51.2  99.0

methylKit::getMethylationStats output for "te" CpGhsm-sites (#: 27814) at minimum coverage 4   - methylation statistics per base
summary:
   Min. 1st Qu.  Median    Mean 3rd Qu.    Max.
   16.7   100.0   100.0    95.6   100.0   100.0
percentiles:
   0%   10%   20%   30%   40%   50%   60%   70%   80%   90%   95%   99% 99.5% 99.9%  100%
 16.7  83.3  91.7 100.0 100.0 100.0 100.0 100.0 100.0 100.0 100.0 100.0 100.0 100.0 100.0
```


```
methylKit::getCoverageStats output for "te" CpGhsm-sites (#: 2246) at minimum coverage 10      - read coverage statistics per base
summary:
   Min. 1st Qu.  Median    Mean 3rd Qu.    Max.
   10.0    10.0    11.0    13.6    13.0    99.0
percentiles:
   0%   10%   20%   30%   40%   50%   60%   70%   80%   90%   95%   99% 99.5% 99.9%  100%
 10.0  10.0  10.0  10.0  11.0  11.0  12.0  12.0  14.0  18.0  28.0  54.0  59.8  87.0  99.0

methylKit::getMethylationStats output for "te" CpGhsm-sites (#: 2246) at minimum coverage 10   - methylation statistics per base
summary:
   Min. 1st Qu.  Median    Mean 3rd Qu.    Max.
   16.7    70.4    90.0    83.2   100.0   100.0
percentiles:
   0%   10%   20%   30%   40%   50%   60%   70%   80%   90%   95%   99% 99.5% 99.9%  100%
```

```
 16.7  50.0  63.6  78.9  84.6  90.0  92.9 100.0 100.0 100.0 100.0 100.0 100.0 100.0 100.0
```

```
methylKit::getCoverageStats output for "te" CpGhsm-sites (#: 407) at minimum coverage 15 - read coverage statistics per base
summary:
   Min. 1st Qu.  Median    Mean 3rd Qu.    Max.
   15.0    16.0    19.0    25.4    30.0    99.0
percentiles:
   0%   10%   20%   30%   40%   50%   60%   70%   80%   90%   95%   99% 99.5% 99.9%  100%
 15.0  15.0  16.0  16.0  17.0  19.0  21.0  26.2  35.0  46.4  55.0  72.9  87.9  95.8  99.0

methylKit::getMethylationStats output for "te" CpGhsm-sites (#: 407) at minimum coverage 15    - methylation statistics per base
summary:
   Min. 1st Qu.  Median    Mean 3rd Qu.    Max.
   16.7    46.7    66.7    66.2    87.5   100.0
percentiles:
   0%   10%   20%   30%   40%   50%   60%   70%   80%   90%   95%   99% 99.5% 99.9%  100%
 16.7  33.3  42.6  50.0  58.3  66.7  75.7  84.5  93.2 100.0 100.0 100.0 100.0 100.0 100.0
```

```
methylKit::getCoverageStats output for "te" CpGhsm-sites (#: 27153) in coverage range [4-12]   - read coverage statistics per base
summary:
   Min. 1st Qu.  Median    Mean 3rd Qu.    Max.
   4.00    4.00    5.00    5.81    7.00   12.00
percentiles:
   0%   10%   20%   30%   40%   50%   60%   70%   80%   90%   95%   99% 99.5% 99.9%  100%
    4     4     4     4     5     5     6     6     7     9    10    12    12    12    12

methylKit::getMethylationStats output for "te" CpGhsm-sites (#: 27153) in coverage range [4-12]      - methylation statistics per base
summary:
   Min. 1st Qu.  Median    Mean 3rd Qu.    Max.
   50.0   100.0   100.0    96.2   100.0   100.0
percentiles:
   0%   10%   20%   30%   40%   50%   60%   70%   80%   90%   95%   99% 99.5% 99.9%  100%
 50.0  83.3 100.0 100.0 100.0 100.0 100.0 100.0 100.0 100.0 100.0 100.0 100.0 100.0 100.0
```

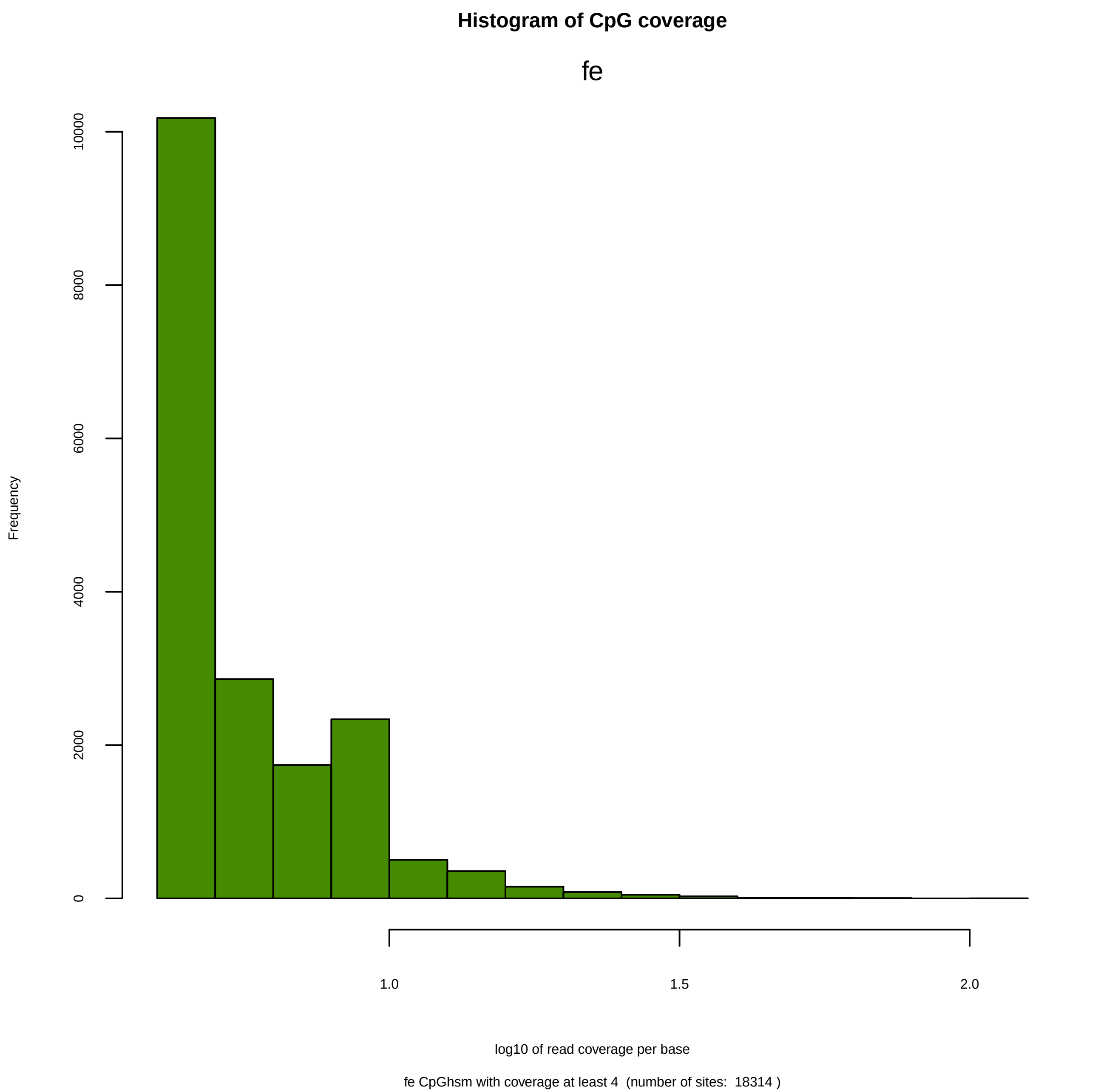

Histogram of CpG coverage
fe
Frequency
0
2000
4000
6000
8000
10000
1.0
1.5
2.0
log10 of read coverage per base
fe CpGhsm with coverage at least 4 (number of sites: 18314 )

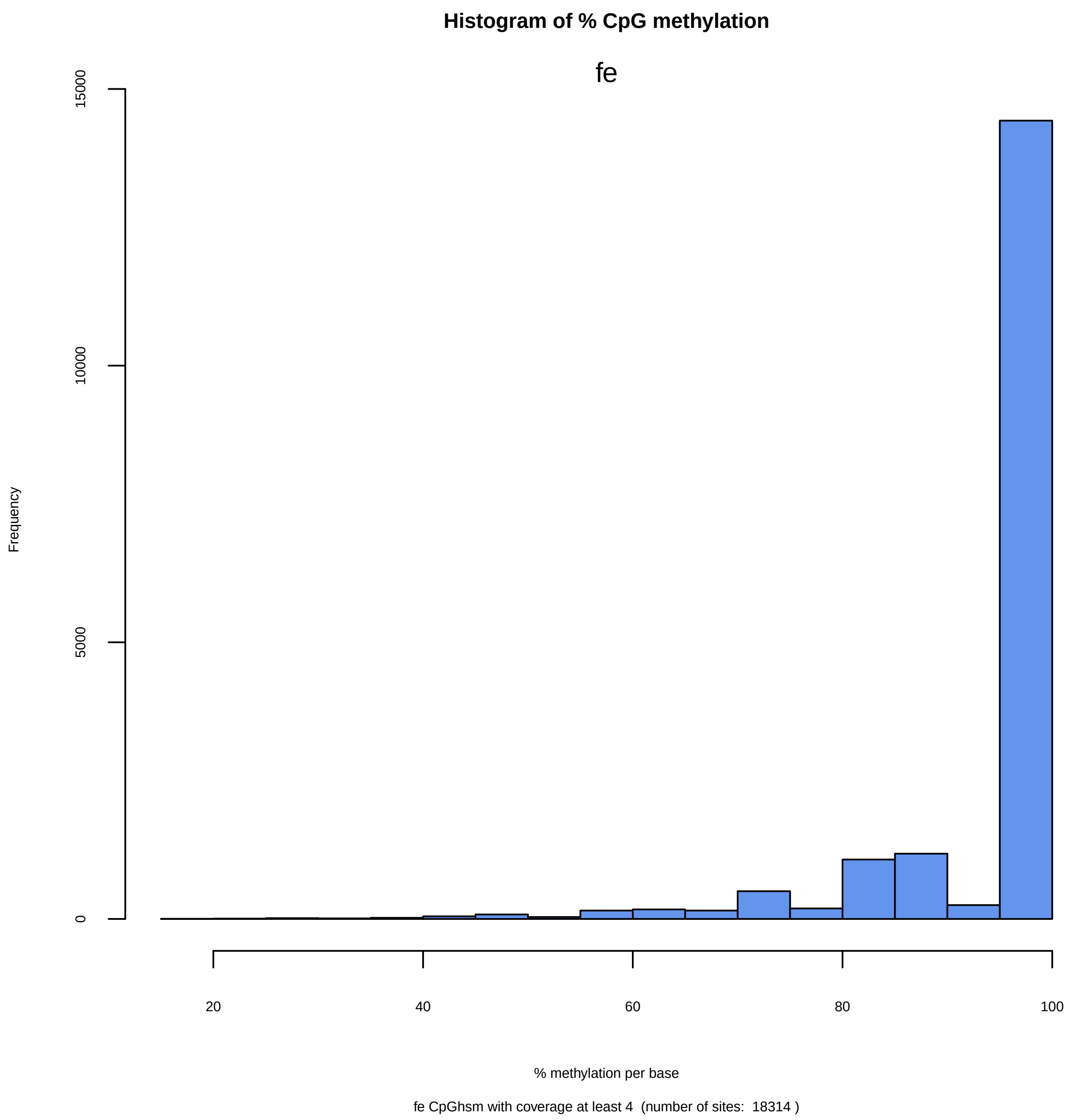
Histogram of % CpG methylation
fe
Frequency
15000
10000
5000
0
20
40
60
80
100
% methylation per base
fe CpGhsm with coverage at least 4 (number of sites: 18314 )

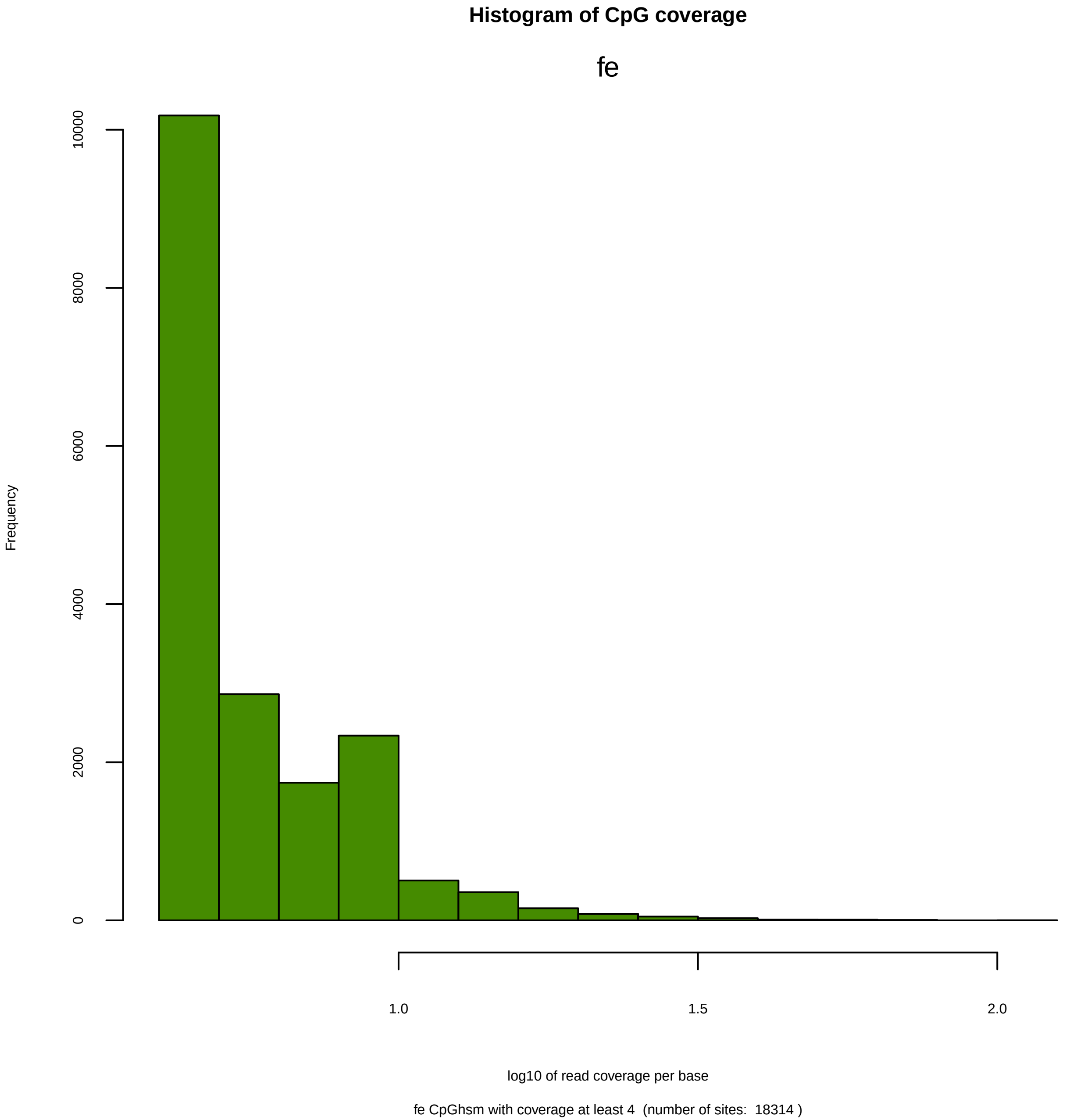
Histogram of CpG coverage
fe
Frequency
0
2000
4000
6000
8000
10000
1.0
1.5
2.0
log10 of read coverage per base
fe CpGhsm with coverage at least 4 (number of sites: 18314 )

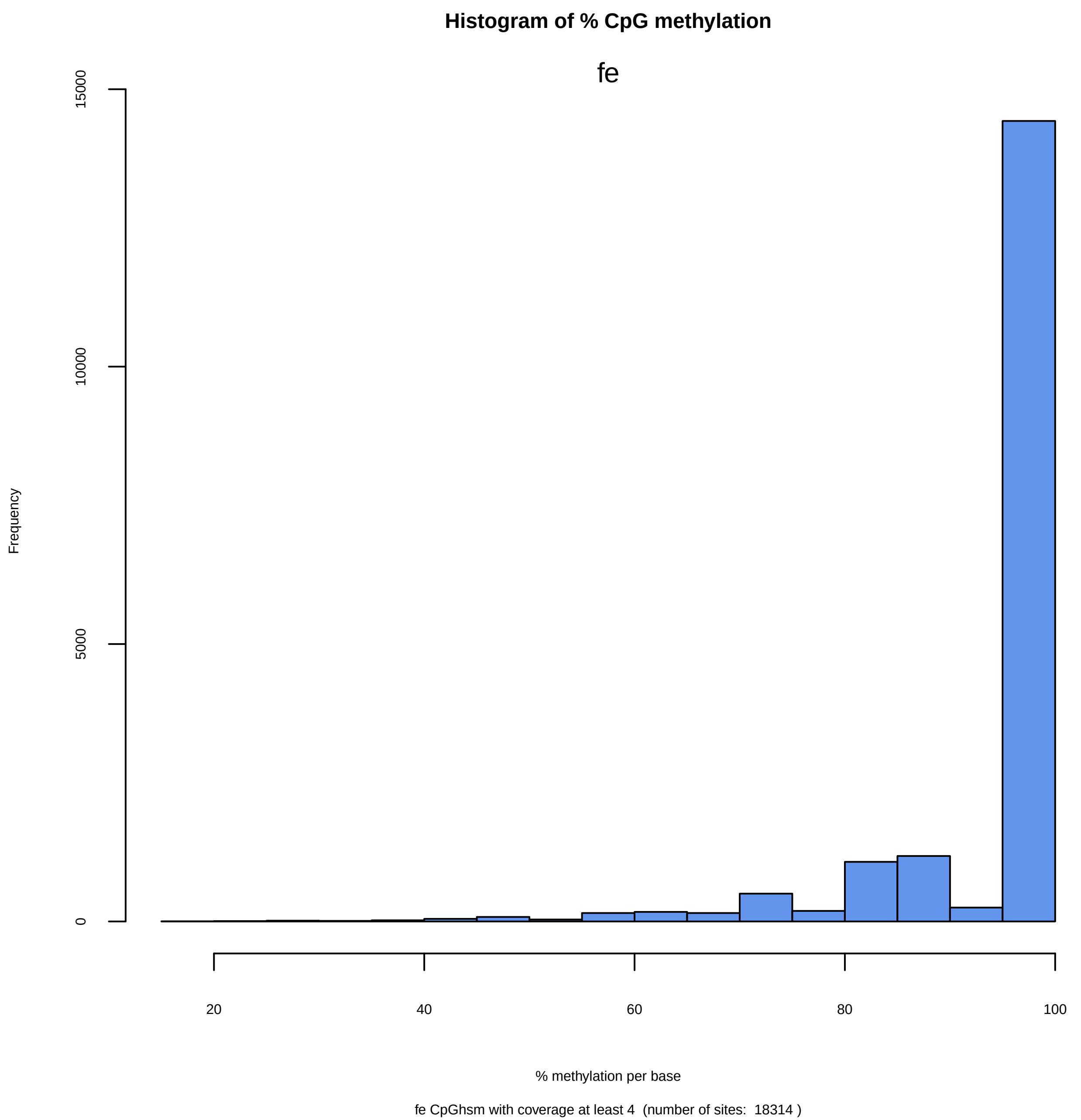

Histogram of % CpG methylation
fe
Frequency
0
5000
10000
15000
20
40
60
80
100
% methylation per base
fe CpGhsm with coverage at least 4 (number of sites: 18314 )

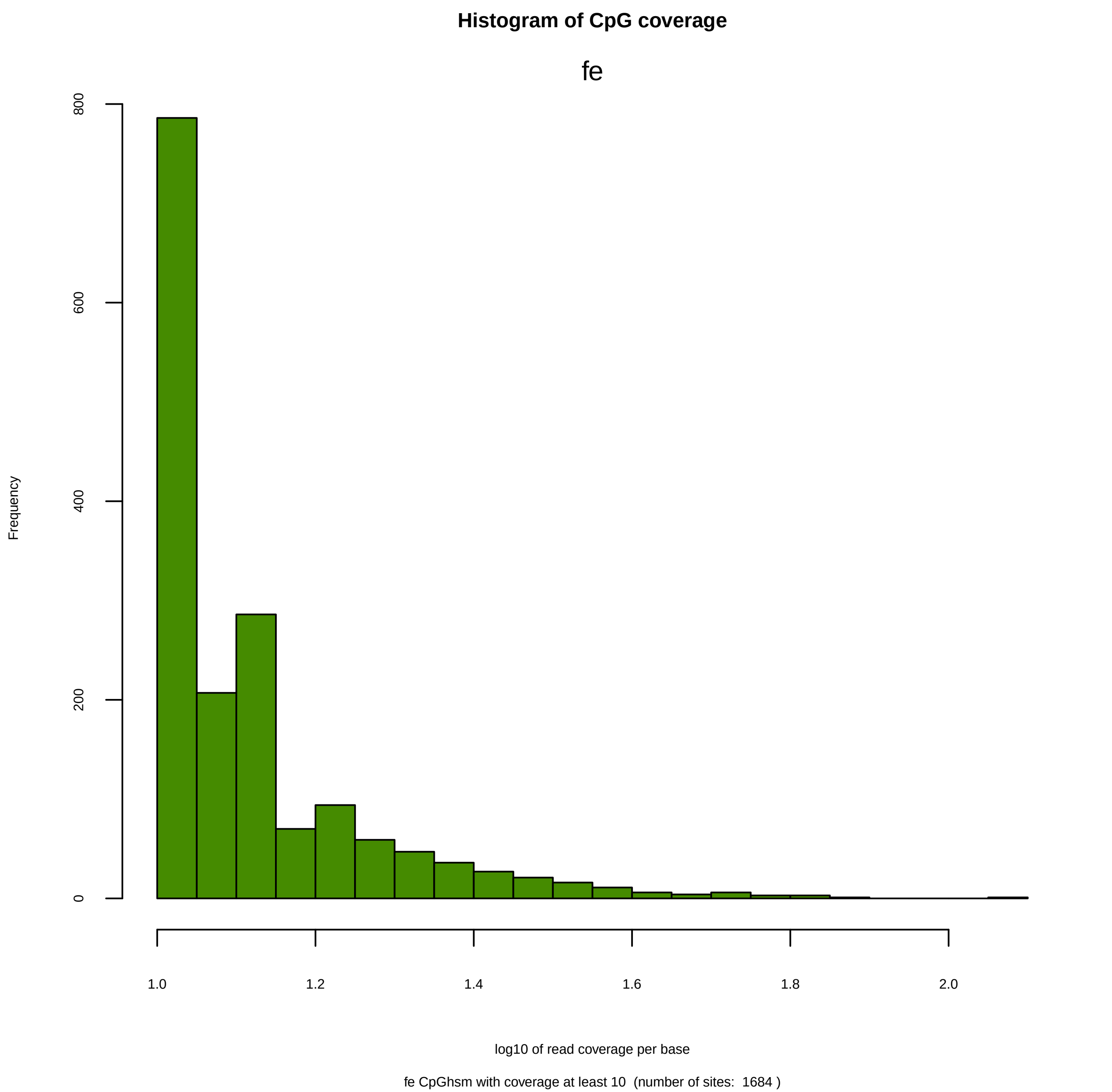
Histogram of CpG coverage
fe
Frequency
0
200
400
600
800
1.0
1.2
1.4
1.6
1.8
2.0
log10 of read coverage per base
fe CpGhsm with coverage at least 10 (number of sites: 1684 )

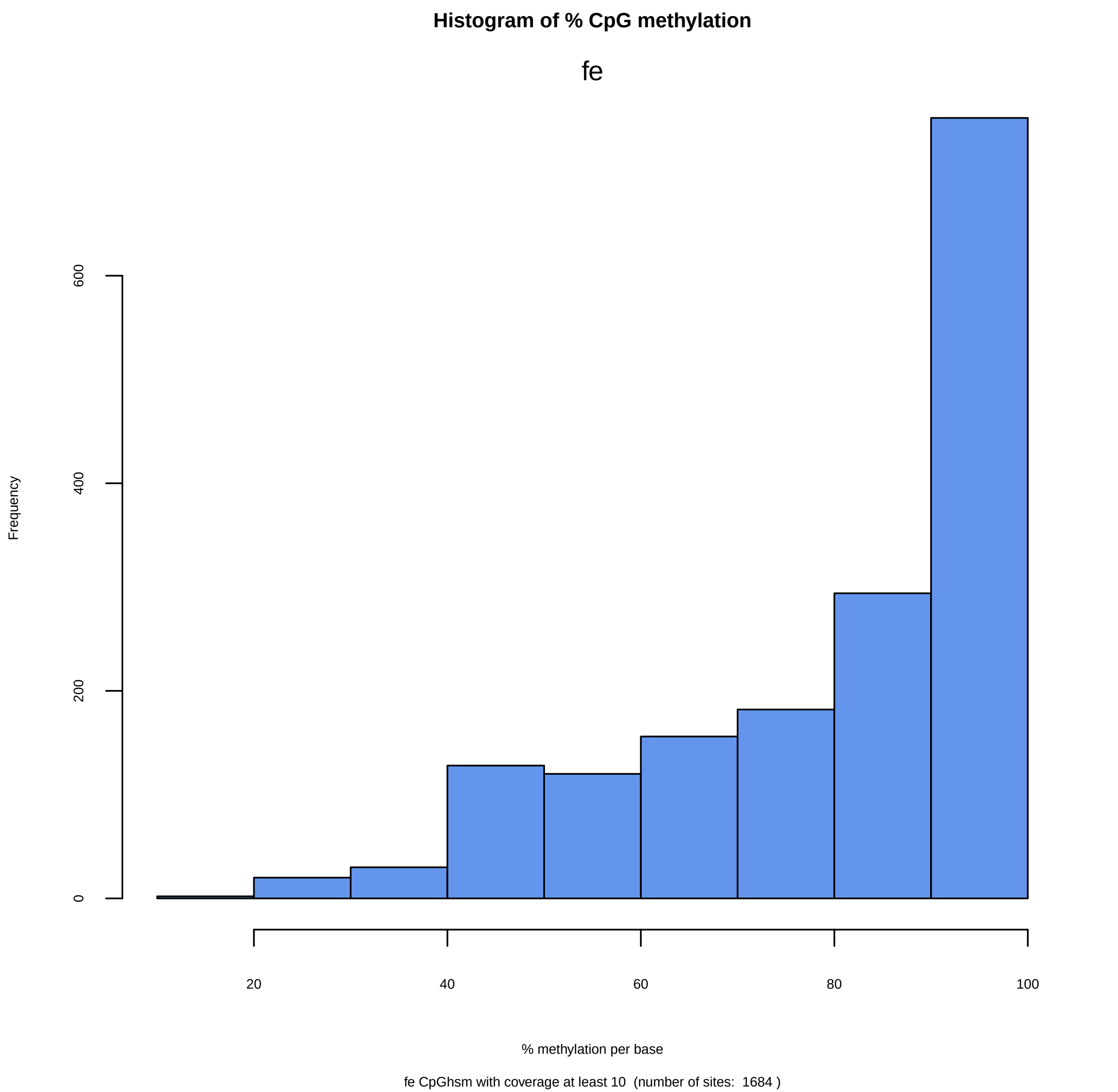

Histogram of % CpG methylation
fe
Frequency
0
200
400
600
20
40
60
80
100
% methylation per base
fe CpGhsm with coverage at least 10 (number of sites: 1684 )

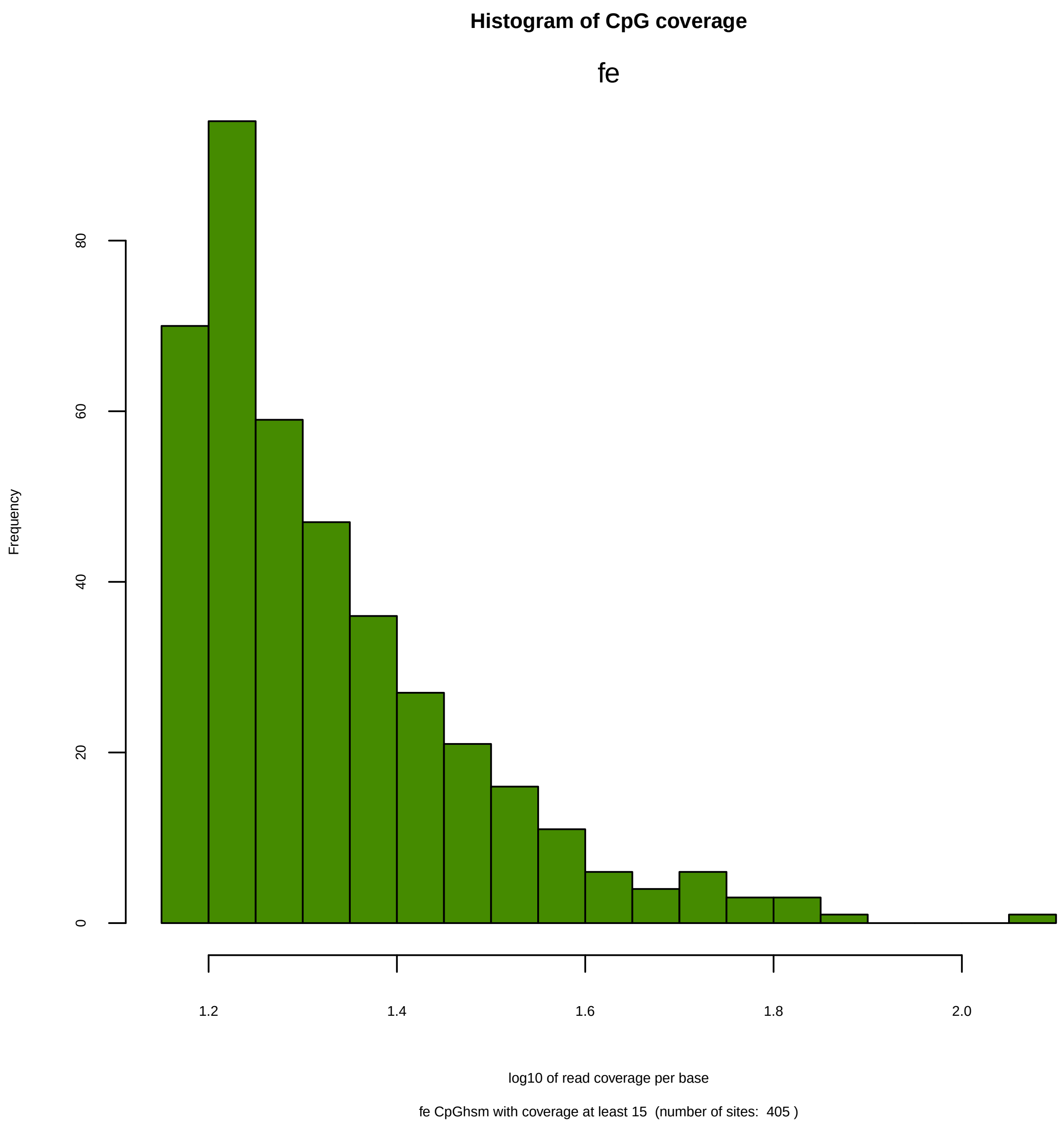

Histogram of CpG coverage
fe
Frequency
0
20
40
60
80
1.2
1.4
1.6
1.8
2.0
log10 of read coverage per base
fe CpGhsm with coverage at least 15 (number of sites: 405 )

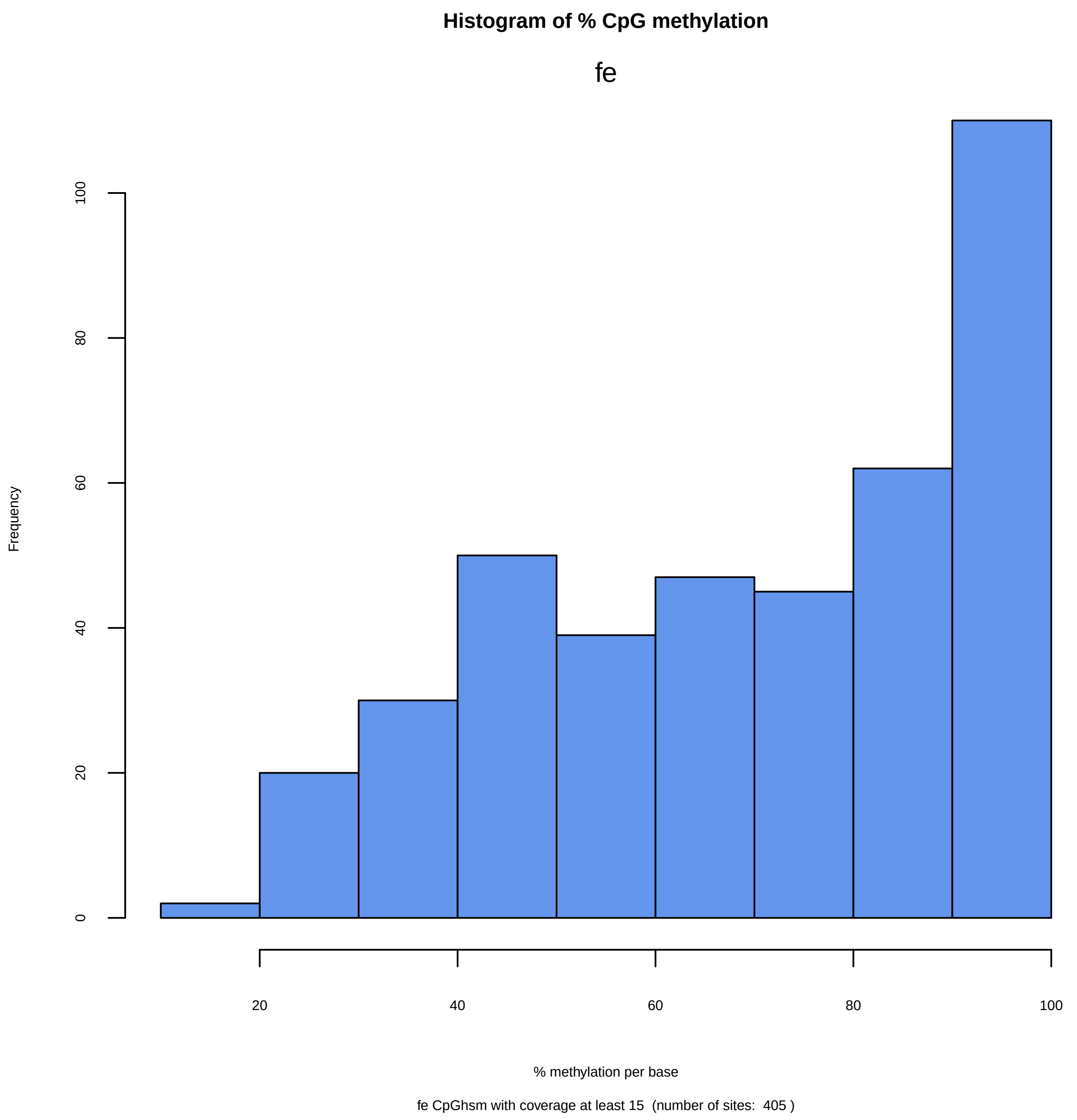

Histogram of % CpG methylation
fe
Frequency
0
20
40
60
80
100
20
40
60
80
100
% methylation per base
fe CpGhsm with coverage at least 15 (number of sites: 405 )

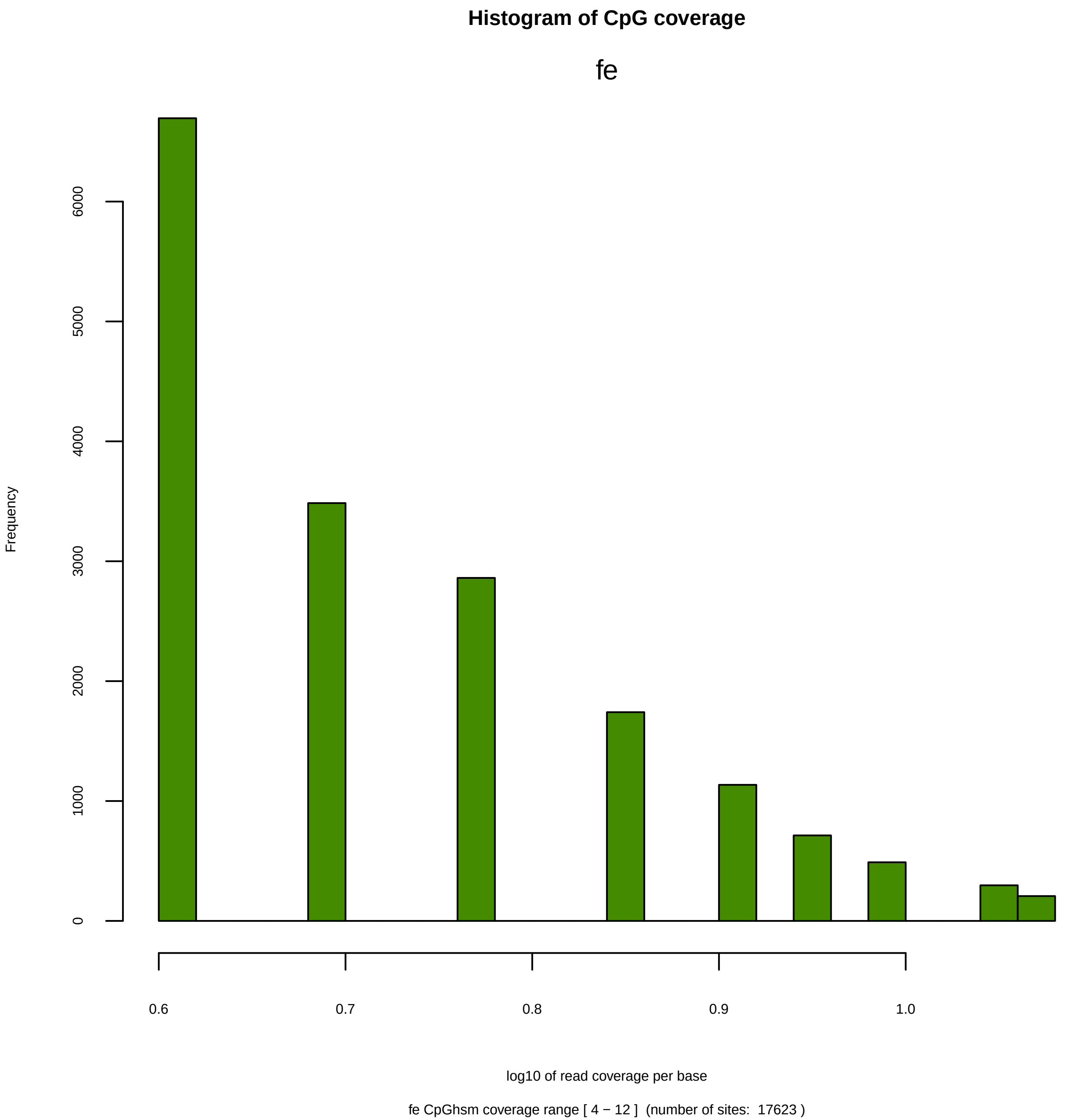

Histogram of CpG coverage
fe
Frequency
0
1000
2000
3000
4000
5000
6000
0.6
0.7
0.8
0.9
1.0
log10 of read coverage per base
fe CpGhsm coverage range [ 4 − 12 ]  (number of sites:  17623 )

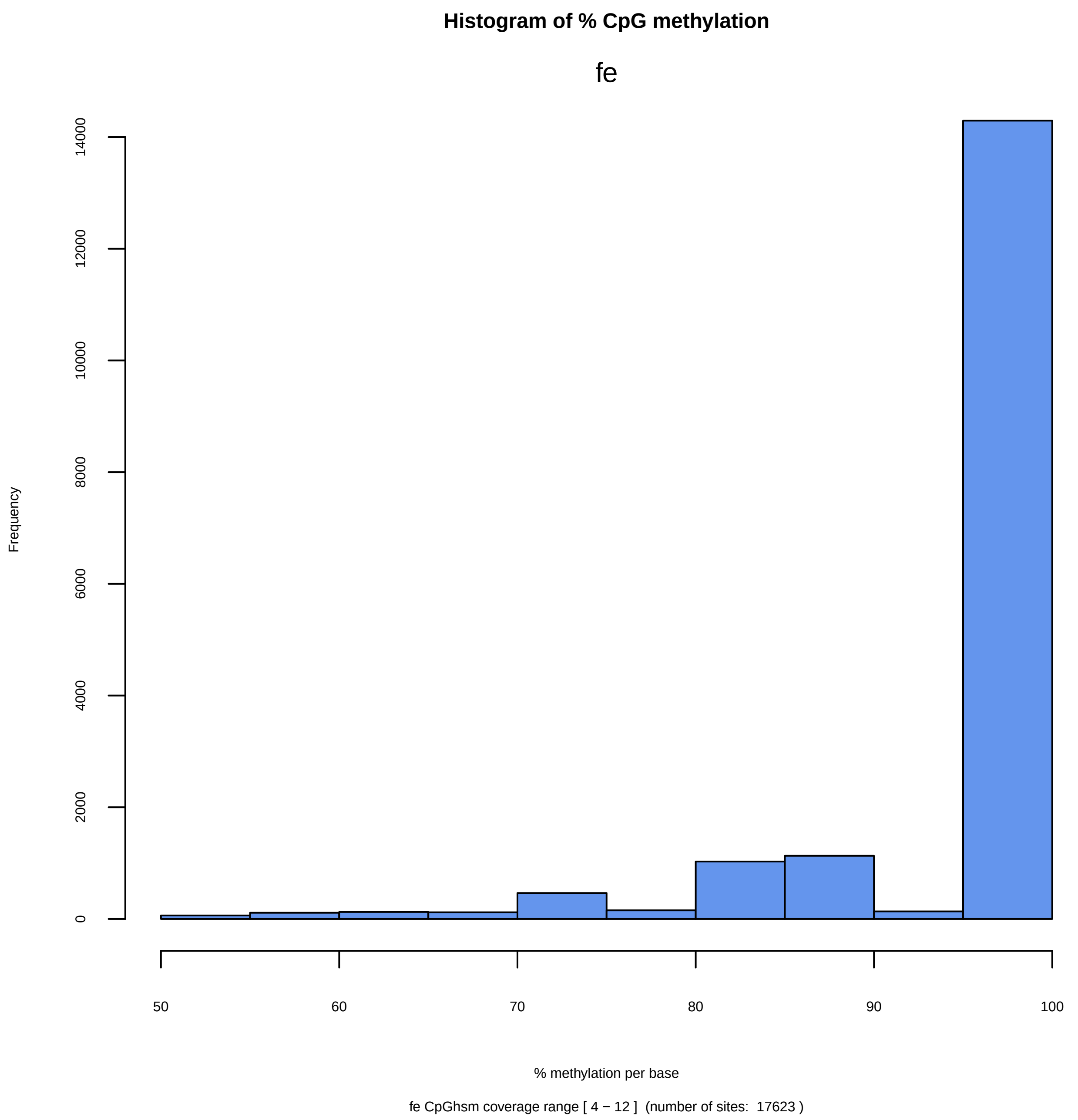

Histogram of % CpG methylation
fe
Frequency
0
2000
4000
6000
8000
10000
12000
14000
50
60
70
80
90
100
% methylation per base
fe CpGhsm coverage range [ 4 − 12 ] (number of sites: 17623 )

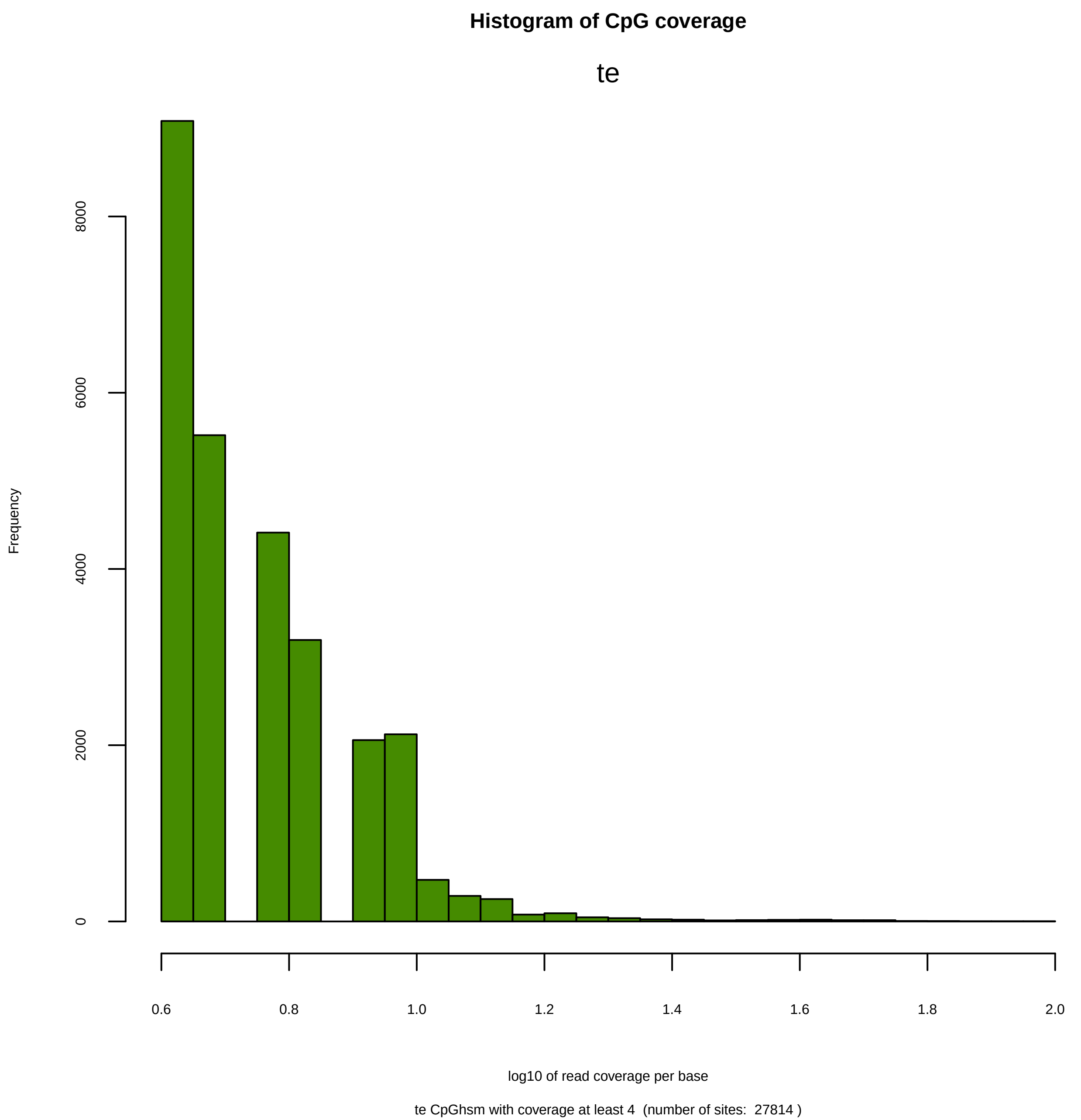
Histogram of CpG coverage
te
Frequency
0
2000
4000
6000
8000
0.6
0.8
1.0
1.2
1.4
1.6
1.8
2.0
log10 of read coverage per base
te CpGhsm with coverage at least 4  (number of sites:  27814 )

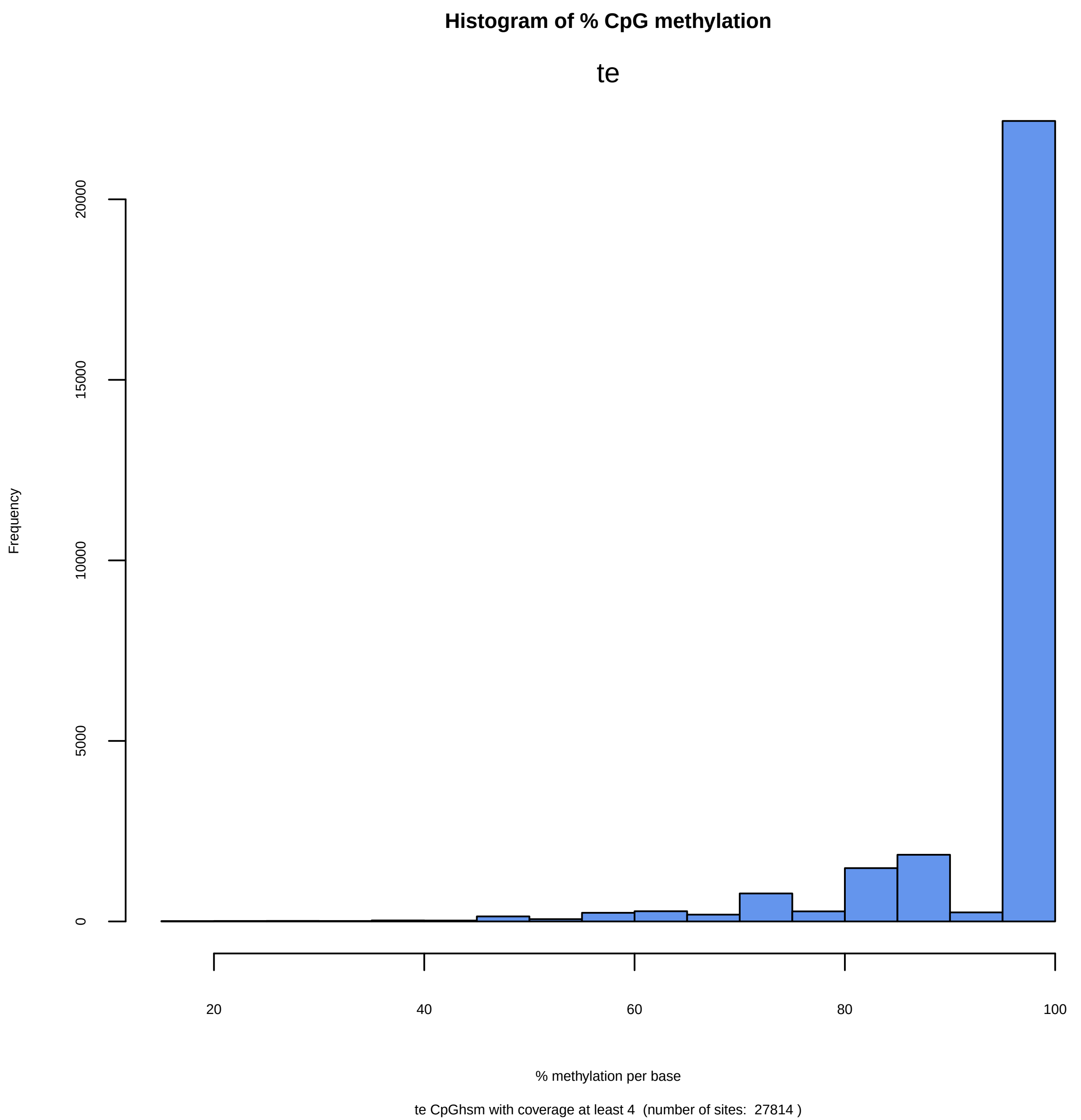
Histogram of % CpG methylation
te
Frequency
0
5000
10000
15000
20000
20
40
60
80
100
% methylation per base
te CpGhsm with coverage at least 4 (number of sites: 27814 )

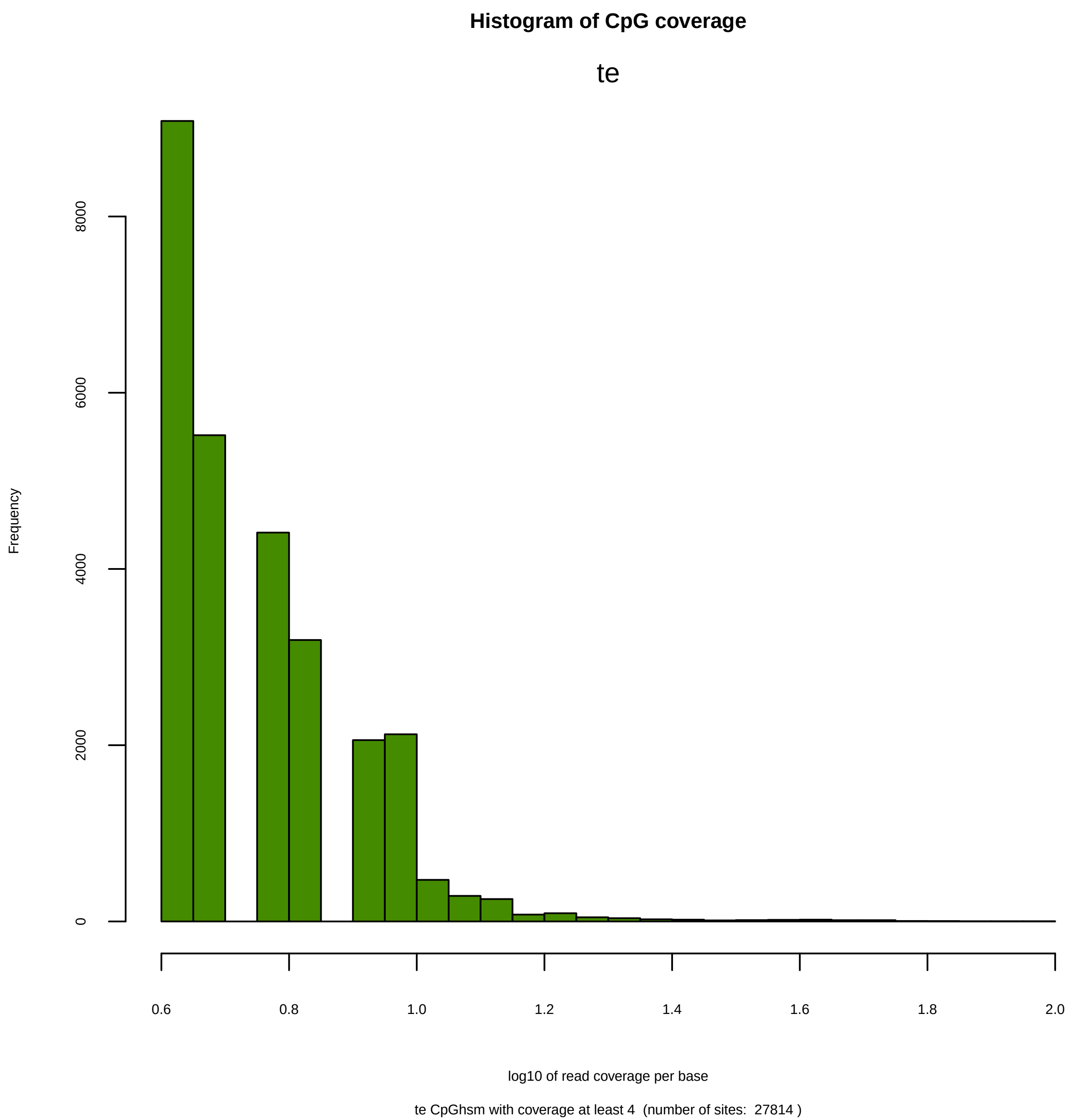
Histogram of CpG coverage
te
Frequency
0
2000
4000
6000
8000
0.6
0.8
1.0
1.2
1.4
1.6
1.8
2.0
log10 of read coverage per base
te CpGhsm with coverage at least 4 (number of sites: 27814 )

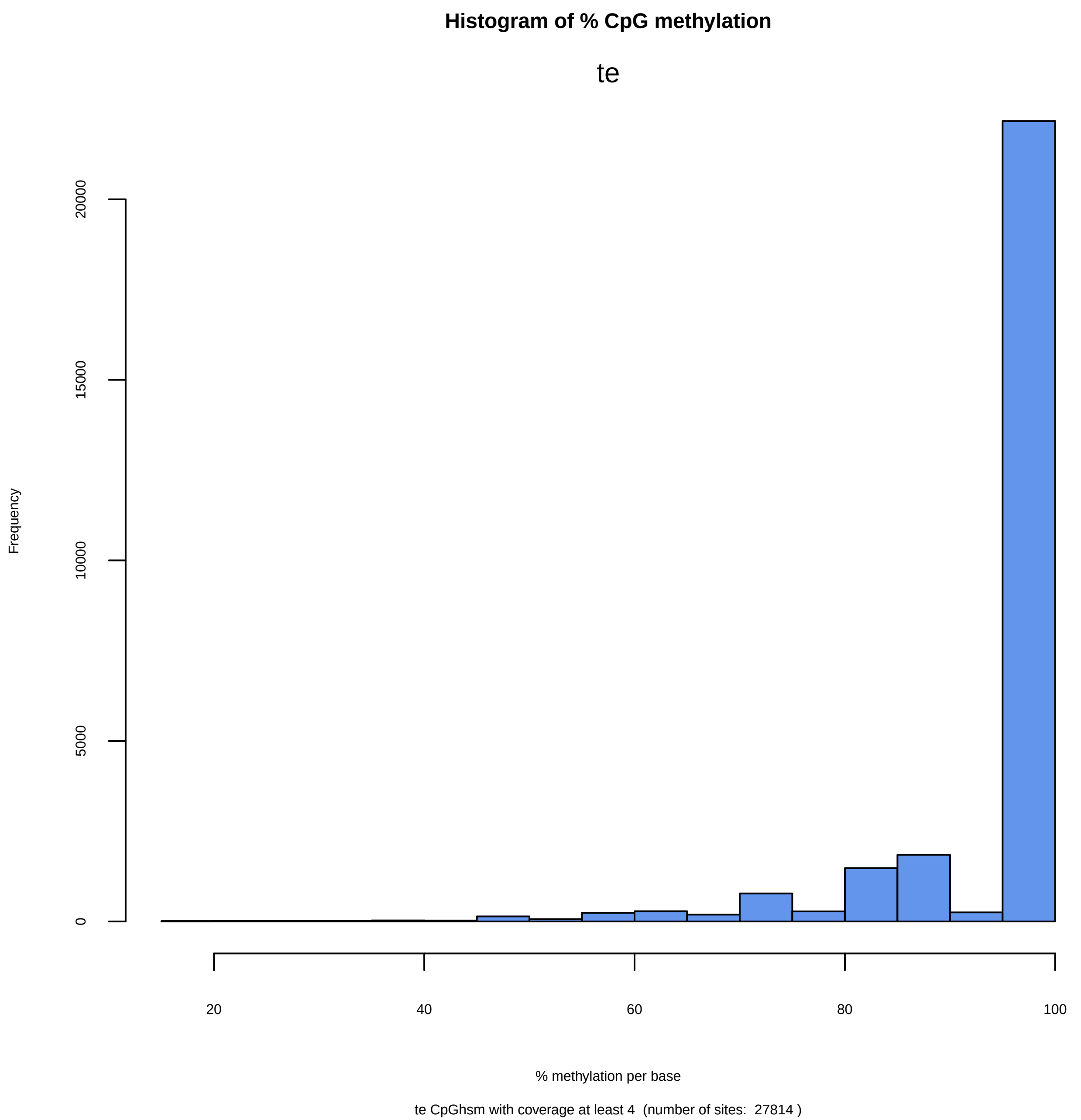

Histogram of % CpG methylation
te
Frequency
0
5000
10000
15000
20000
20
40
60
80
100
% methylation per base
te CpGhsm with coverage at least 4  (number of sites:  27814 )

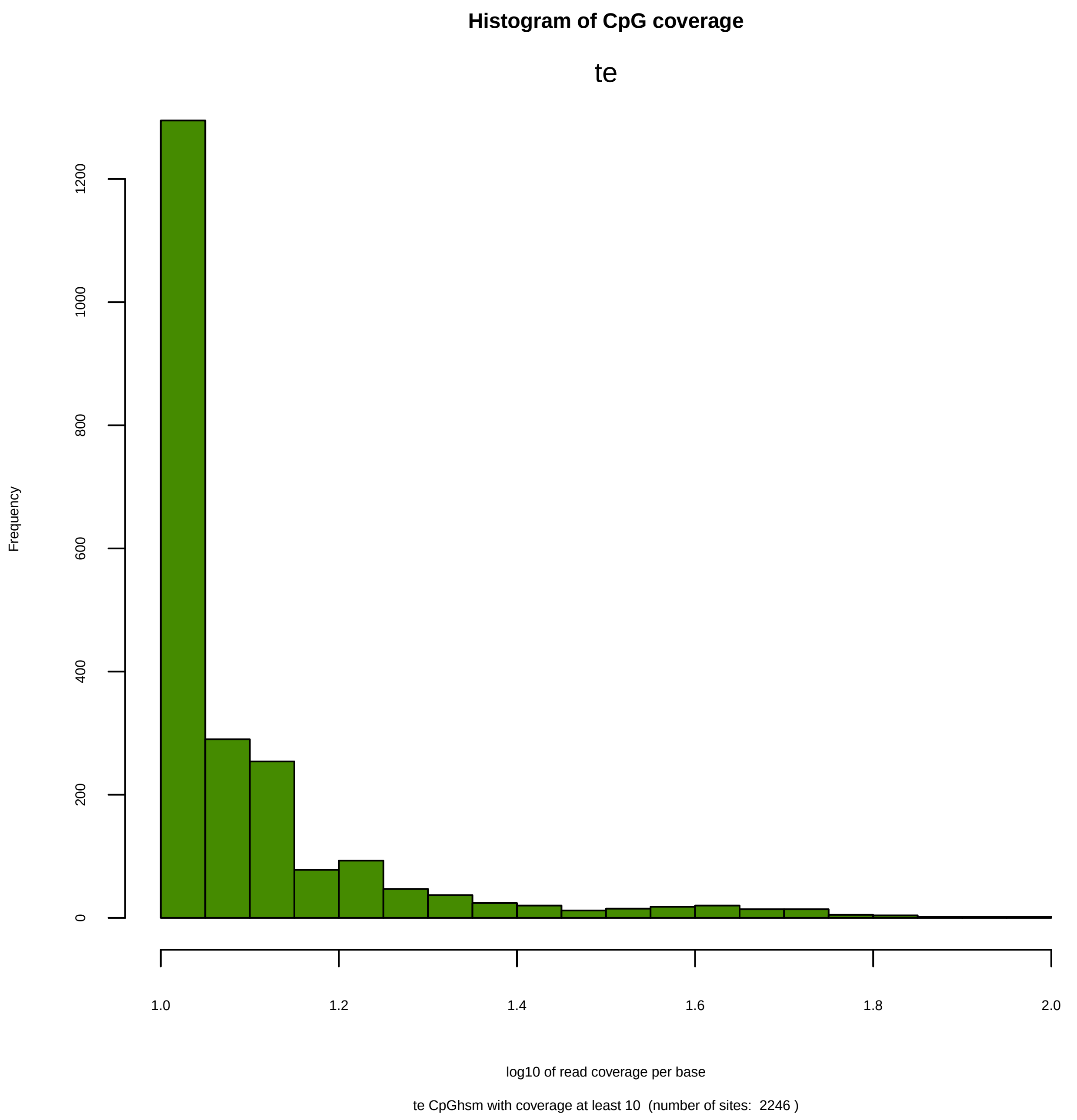
Histogram of CpG coverage
te
Frequency
0
200
400
600
800
1000
1200
1.0
1.2
1.4
1.6
1.8
2.0
log10 of read coverage per base
te CpGhsm with coverage at least 10  (number of sites:  2246 )

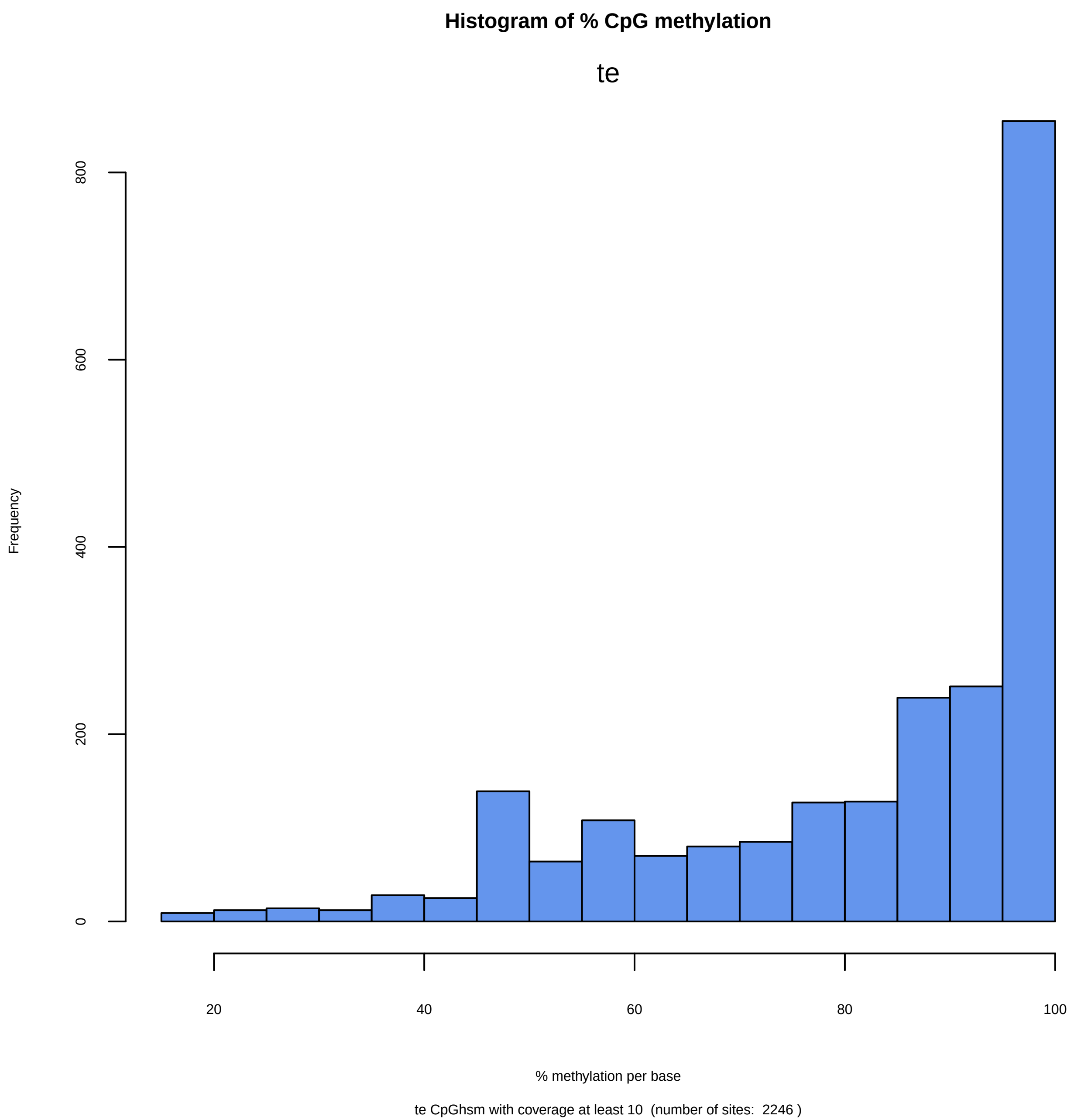
Histogram of % CpG methylation
te
Frequency
0
200
400
600
800
20
40
60
80
100
% methylation per base
te CpGhsm with coverage at least 10  (number of sites:  2246 )

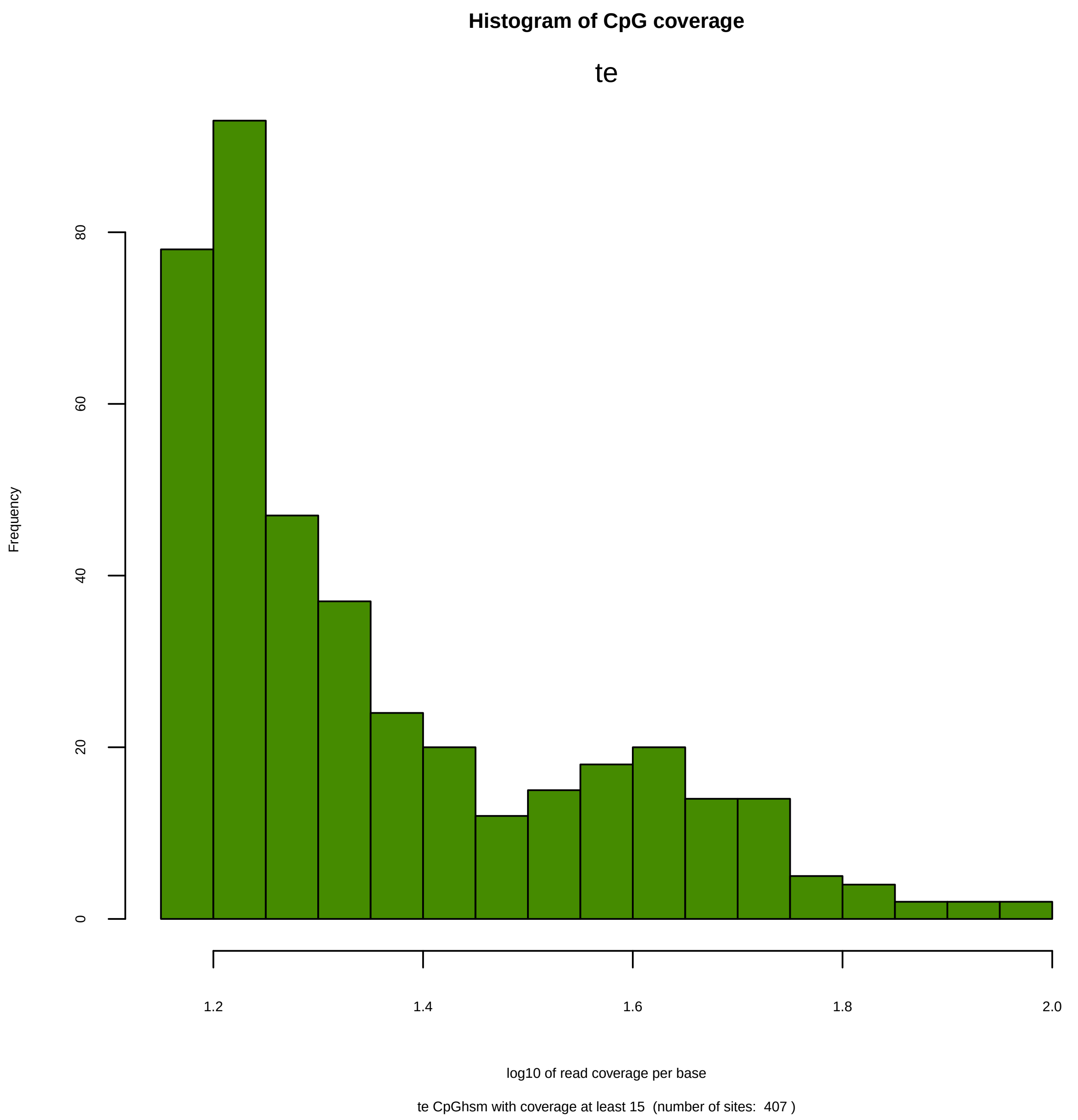

Histogram of CpG coverage
te
Frequency
0
20
40
60
80
1.2
1.4
1.6
1.8
2.0
log10 of read coverage per base
te CpGhsm with coverage at least 15 (number of sites: 407 )

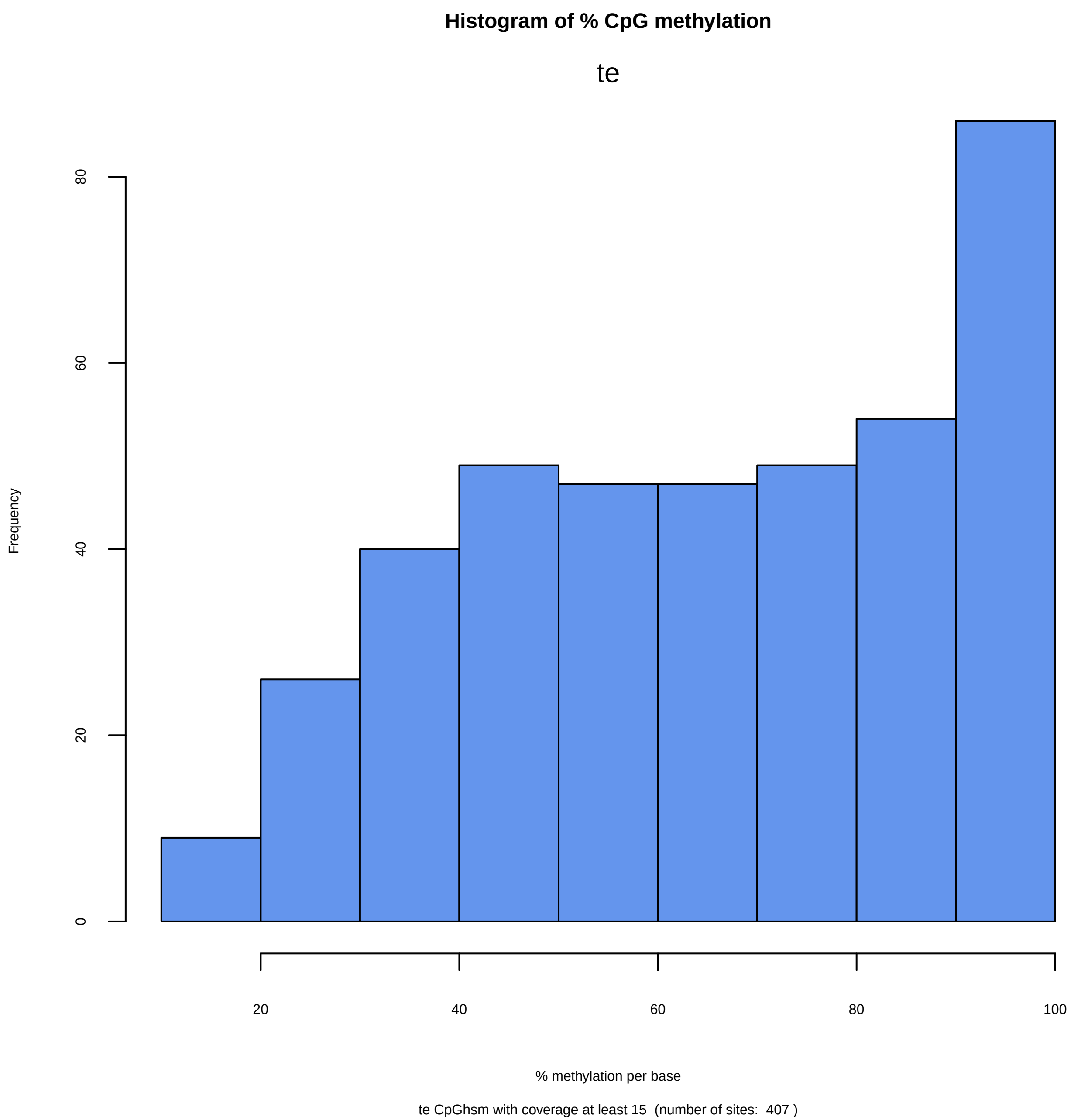

Histogram of % CpG methylation
te
Frequency
0
20
40
60
80
20
40
60
80
100
% methylation per base
te CpGhsm with coverage at least 15 (number of sites: 407 )

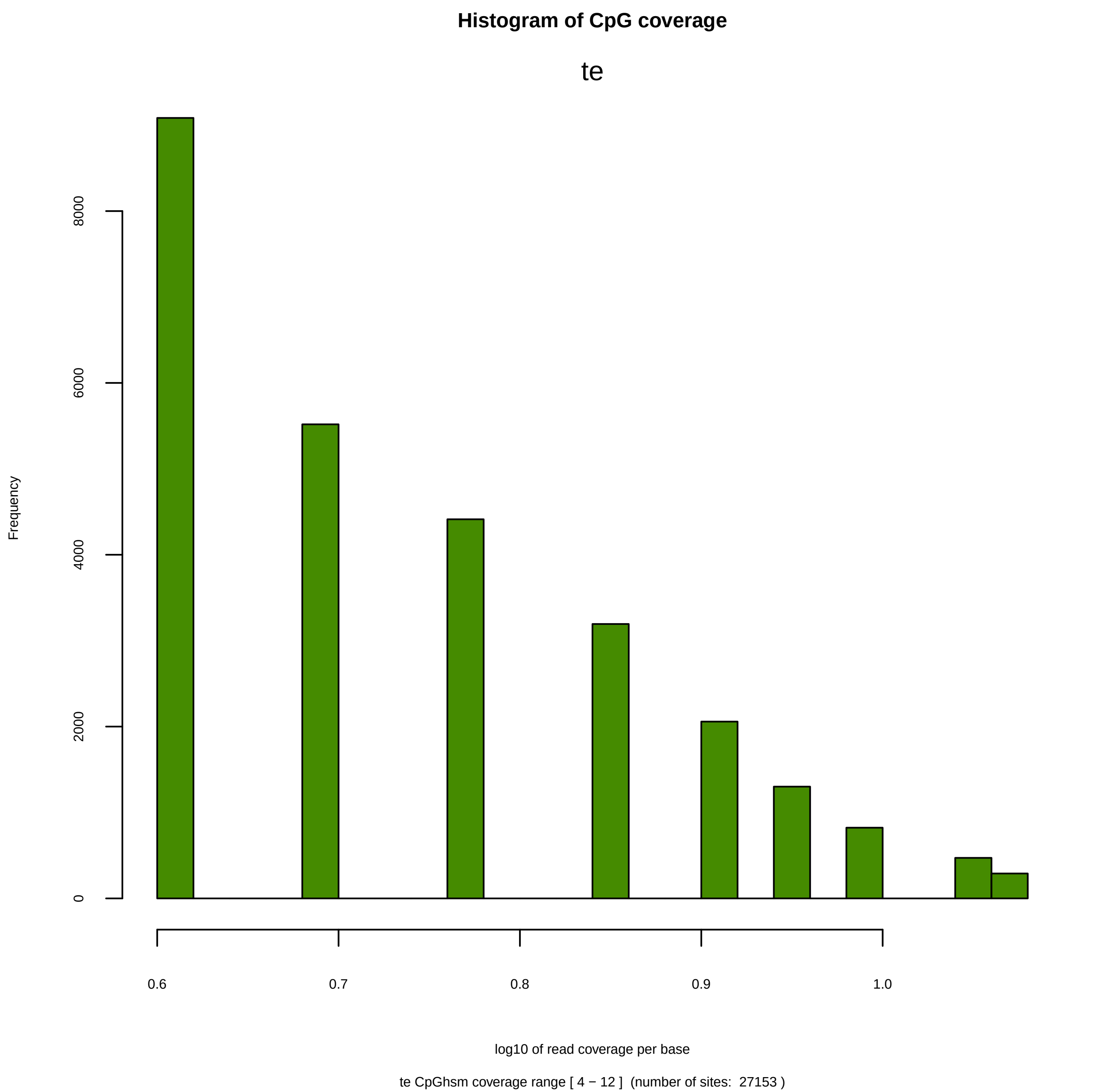
Histogram of CpG coverage
te
Frequency
0
2000
4000
6000
8000
0.6
0.7
0.8
0.9
1.0
log10 of read coverage per base
te CpGhsm coverage range [ 4 − 12 ]  (number of sites:  27153 )

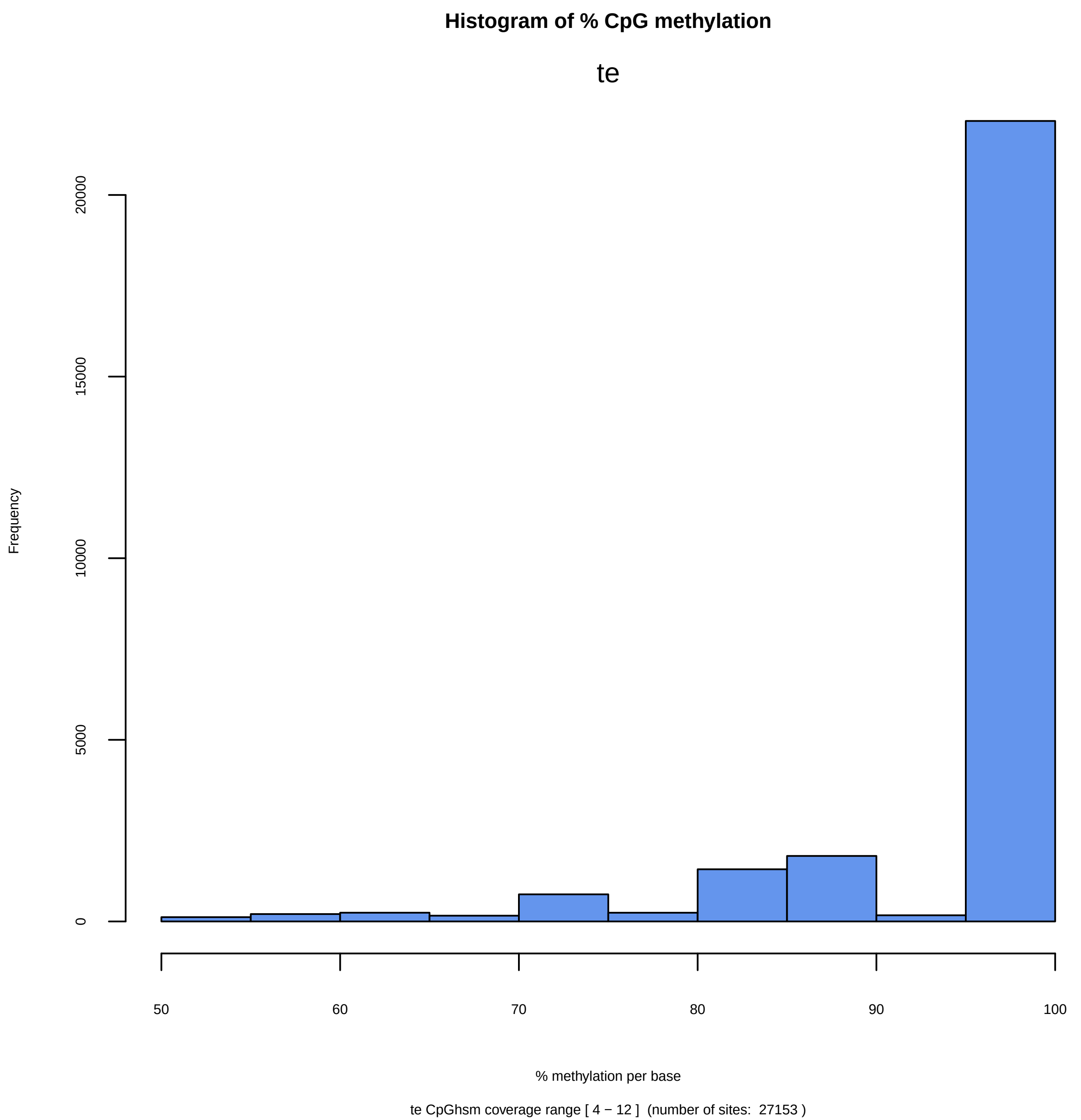

Histogram of % CpG methylation
te
Frequency
0
5000
10000
15000
20000
50
60
70
80
90
100
% methylation per base
te CpGhsm coverage range [ 4 − 12 ] (number of sites: 27153 )

```
Directory: PWC File: 0READMEpwc


PWC - Pairwise comparisons between all samples.

Input: studymk, studymc, nbrpms, hheight, nbrpnts

Output: files pwc-*.txt pwc-.*pdf

Notes: The output is generated by BWASPR::cmpSites(), which determines for each pairwise
        comparison the numbers of common and distinct sites.  The function also calculates
        the overlap index between the samples and estimates the size of the common pool of
        potential methylation sites.

       The CpGscd data input (studymc) is necessary to determine sites that are detectable
        in both samples, and nbrpms (listed in the *.par file specified in the Am_RE.conf
        configuration file as argument to TOTALNBRPMSITES) provides the total number of
        potential methylation sites (typically, all CpG sites), a necessary parameter for
        the estimations.
       hheight (default: 0.10) specifies the y-axis limit in the histogram plots.
       nbrpnts (default: 5000) specifies the number of common sites to be plotted in the
        methylation levels scatter plot.
```

```
Directory: PWC File: pwc-Am_RE_fe.vs.Am_RE_te.txt


Numbers of common and distinct sites comparing Am_RE_fe versus Am_RE_te
================================================================================

  total number of potential sites:  19687378

  number of "Am_RE_fe_hsm"         sites: 18314
  number of "Am_RE_te_hsm"         sites: 27814
    number of "Am_RE_fe_hsm"-unique sites:       9952
    number of common sites:          8362
    number of "Am_RE_te_hsm"-unique sites:      19452
  total number of "Am_RE_fe_hsm+Am_RE_te_hsm"-sites observed:     37766

  number of "Am_RE_fe_scd"        sites:   7682417 ( 39.02% of total)
  number of "Am_RE_te_scd"        sites:   8517171 ( 43.26% of total)
    number of "Am_RE_fe_scd"-unique sites:       2436175
    number of sites in common:       5246242   (Expected:   3323574; O/E:       1.6)
    number of "Am_RE_te_scd"-unique sites:       3270929
  total number of "Am_RE_fe_scd+Am_RE_te_scd"-sites observed:     10953346

  number of sites in "Am_RE_fe_hsm" that are not  detectable in "Am_RE_te_hsm":      8340
  number of sites in "Am_RE_fe_hsm" that are also detectable in "Am_RE_te_hsm":      9974
    number of sites unique to "Am_RE_fe_hsm" although detectable in "Am_RE_te_hsm":              1612
    number of "Am_RE_fe_hsm" / "Am_RE_te_hsm" common sites                        :              8362      (Expected:     20; O/E:     418.1)

  number of sites in "Am_RE_te_hsm" that are not  detectable in "Am_RE_fe_hsm":     17113
  number of sites in "Am_RE_te_hsm" that are also detectable in "Am_RE_fe_hsm":     10701
    number of sites unique to "Am_RE_te_hsm" although detectable in "Am_RE_fe_hsm":              2339
    number of "Am_RE_te_hsm" / "Am_RE_fe_hsm" common sites                        :              8362      (Expected:     20; O/E:     418.1)

  Overlap index of "Am_RE_fe_hsm" with "Am_RE_te_hsm":           0.729
  Estimated number of "Am_RE_fe_hsm" = "Am_RE_te_hsm" sites (assuming sampling from one population):          12764
  Adjusted population size of "Am_RE_fe_hsm" = "Am_RE_te_hsm" sites (assuming all sites detectable):          47899 (  1.27x of observed)


  number of "Am_RE_fe_hsm" sites with coverage >=  4:  18314
  number of "Am_RE_te_hsm" sites with coverage >=  4:  27814
  number of common sites with coverage >=  4:  8362
  number of "Am_RE_fe_hsm" sites with coverage >=  4:  18314
  number of "Am_RE_te_hsm" sites with coverage >=  4:  27814
  number of common sites with coverage >=  4:  8362
  number of "Am_RE_fe_hsm" sites with coverage >= 10:   1684
  number of "Am_RE_te_hsm" sites with coverage >= 10:   2246
  number of common sites with coverage >= 10:   525
  number of "Am_RE_fe_hsm" sites with coverage >= 15:    405
  number of "Am_RE_te_hsm" sites with coverage >= 15:    407
  number of common sites with coverage >= 15:   160
```

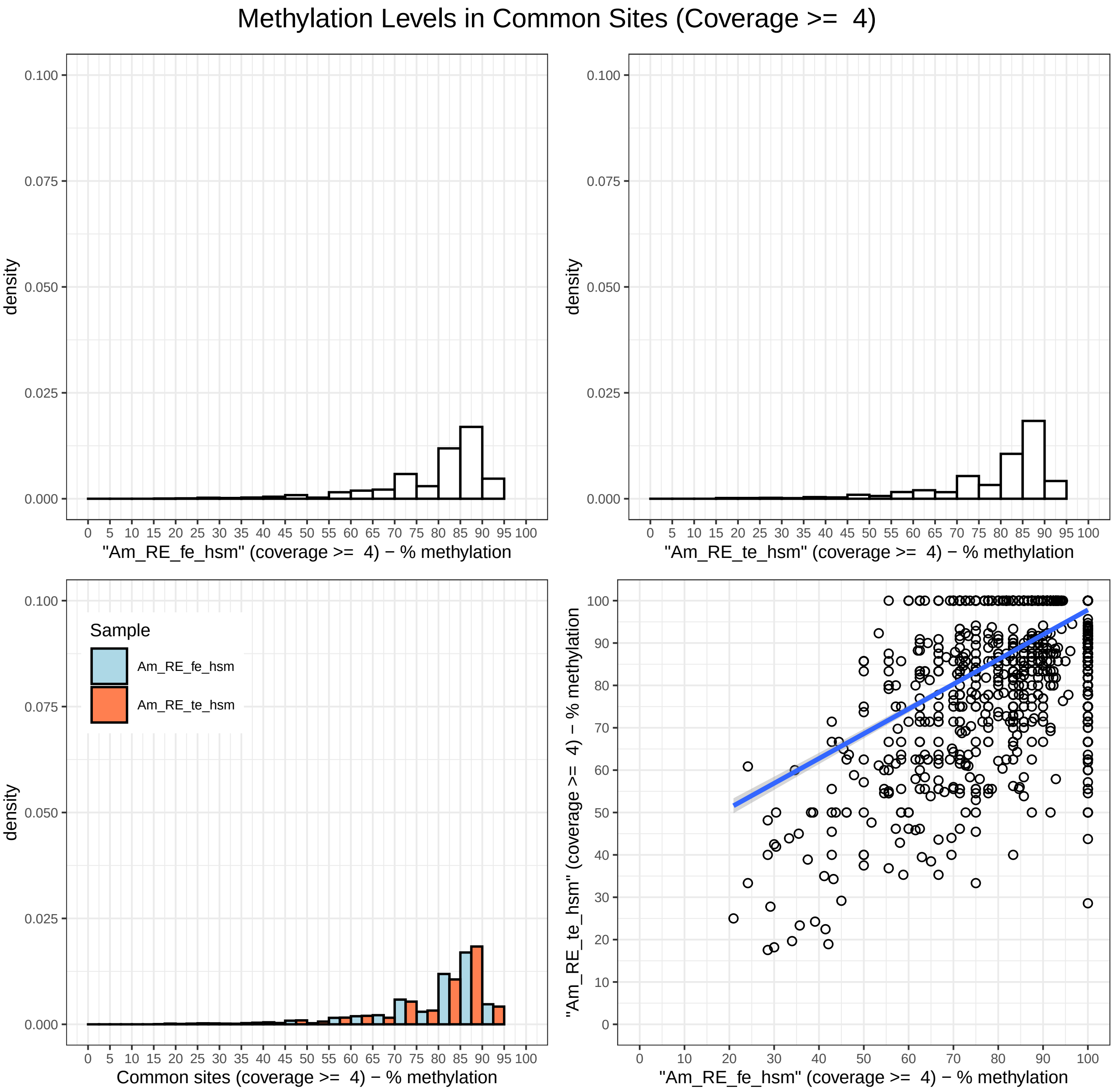

Methylation Levels in Common Sites (Coverage >=  4)
density
"Am_RE_fe_hsm" (coverage >=  4) − % methylation
"Am_RE_te_hsm" (coverage >=  4) − % methylation
Sample
Am_RE_fe_hsm
Am_RE_te_hsm
Common sites (coverage >=  4) − % methylation
"Am_RE_te_hsm" (coverage >=  4) − % methylation
"Am_RE_fe_hsm" (coverage >=  4) − % methylation

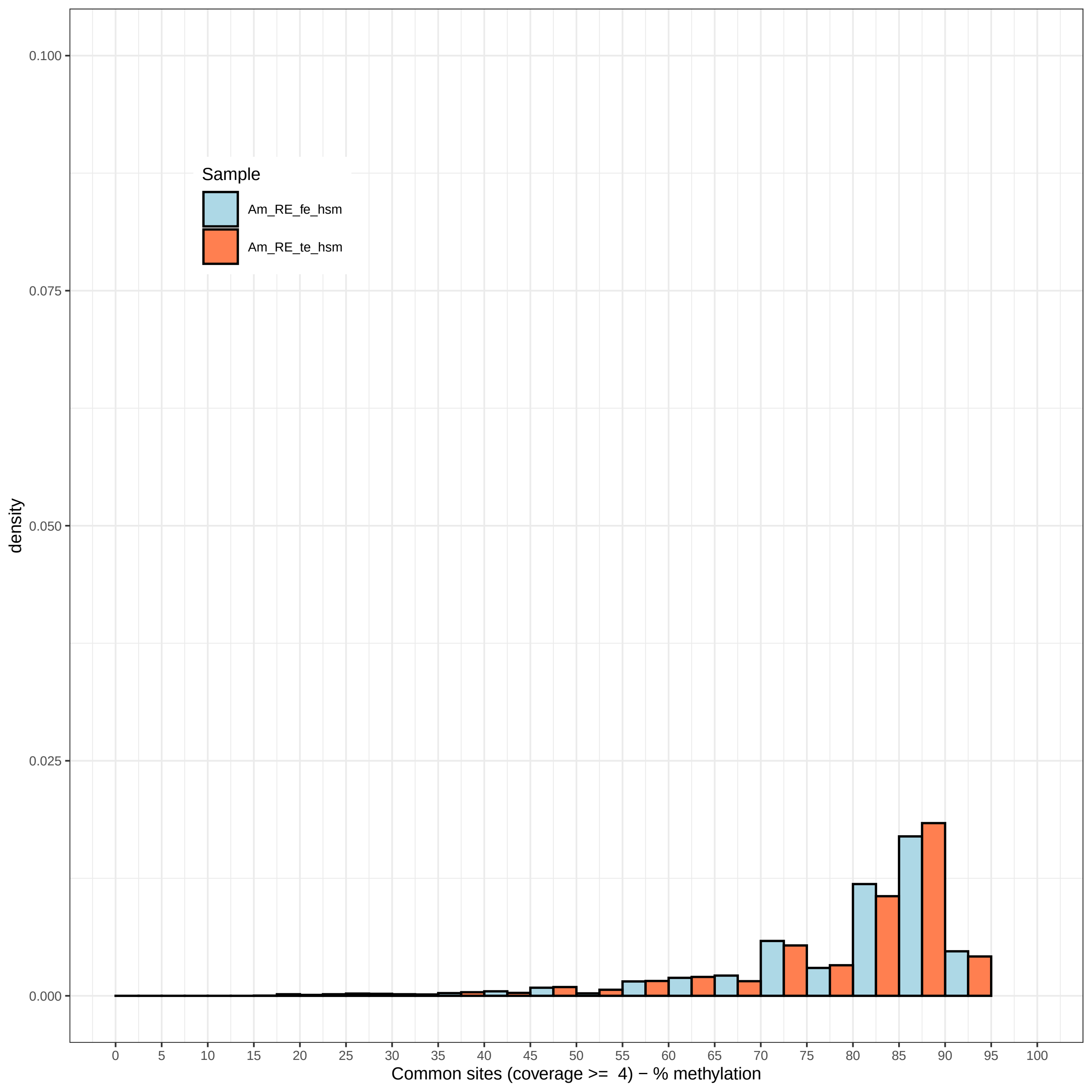
Sample
Am_RE_fe_hsm
Am_RE_te_hsm
density
0.100
0.075
0.050
0.025
0.000
0
5
10
15
20
25
30
35
40
45
50
55
60
65
70
75
80
85
90
95
100
Common sites (coverage >= 4) − % methylation

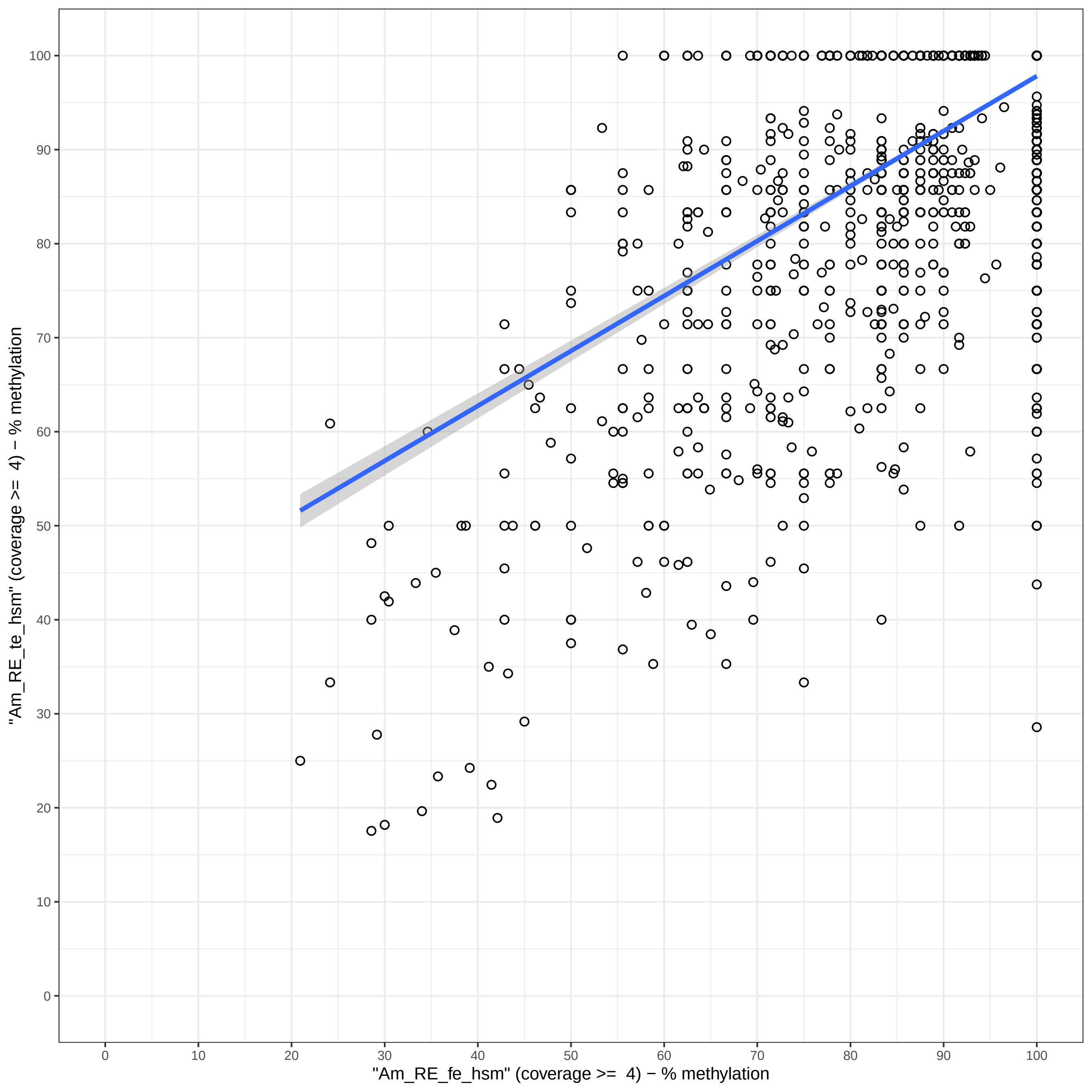
"Am_RE_te_hsm" (coverage >= 4) − % methylation
"Am_RE_fe_hsm" (coverage >= 4) − % methylation
0
10
20
30
40
50
60
70
80
90
100

#### Overlap of highly supported methylation sites (coverage >= 4)

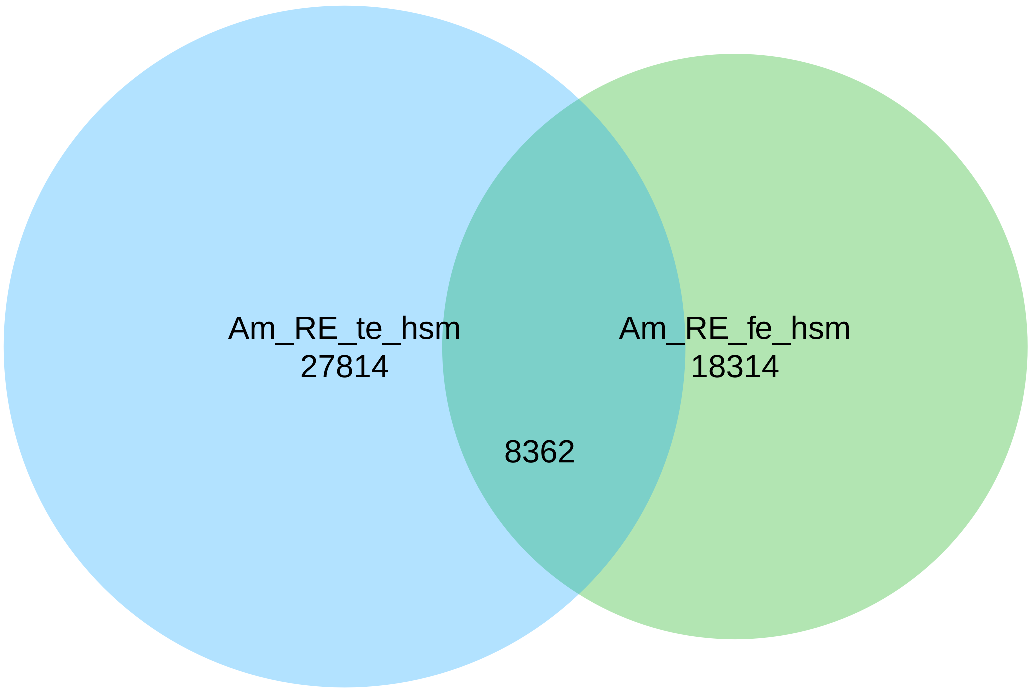

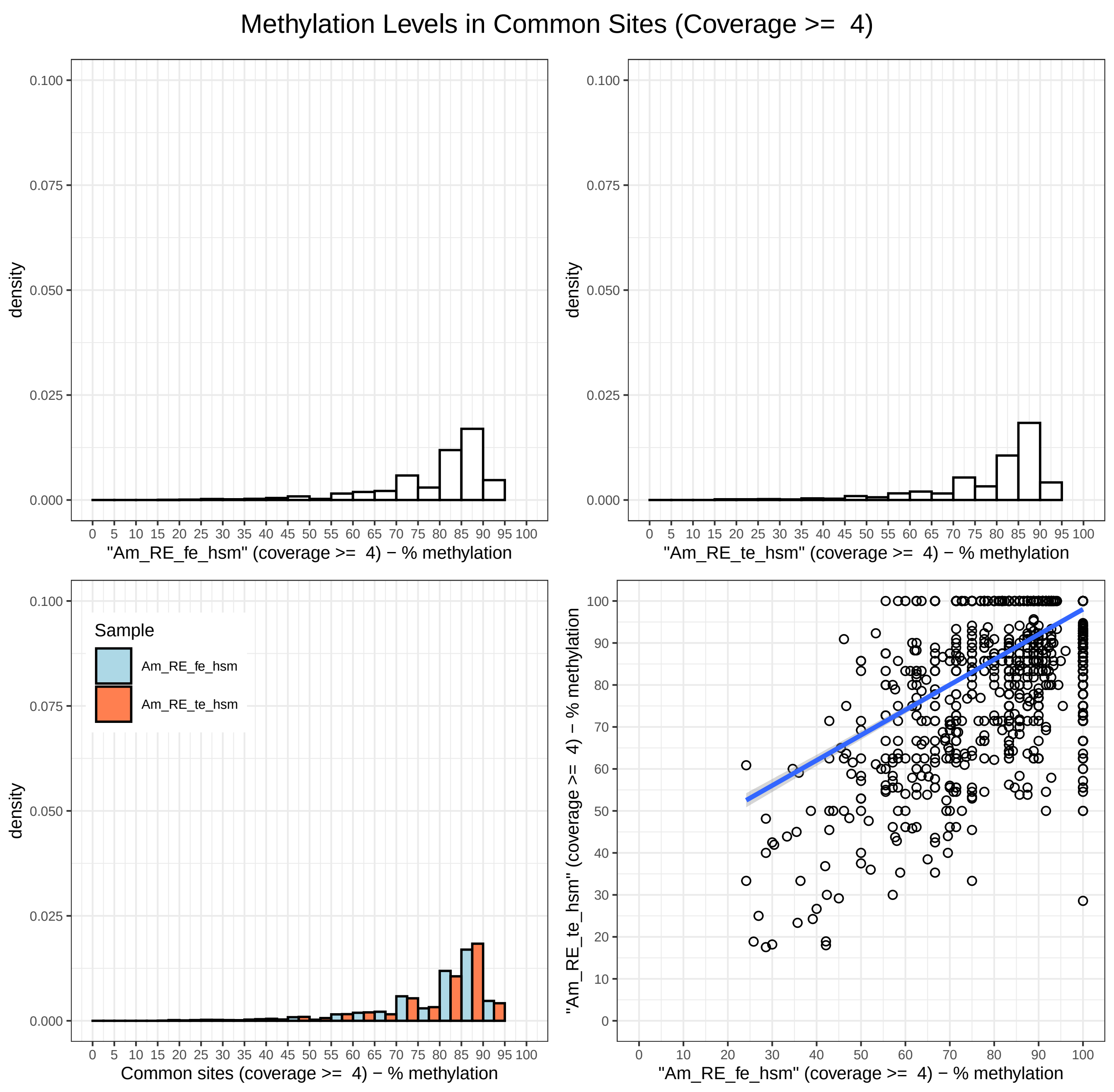
Methylation Levels in Common Sites (Coverage >=  4)
density
"Am_RE_fe_hsm" (coverage >=  4) − % methylation
"Am_RE_te_hsm" (coverage >=  4) − % methylation
Sample
Am_RE_fe_hsm
Am_RE_te_hsm
Common sites (coverage >=  4) − % methylation
"Am_RE_te_hsm" (coverage >=  4) − % methylation
"Am_RE_fe_hsm" (coverage >=  4) − % methylation

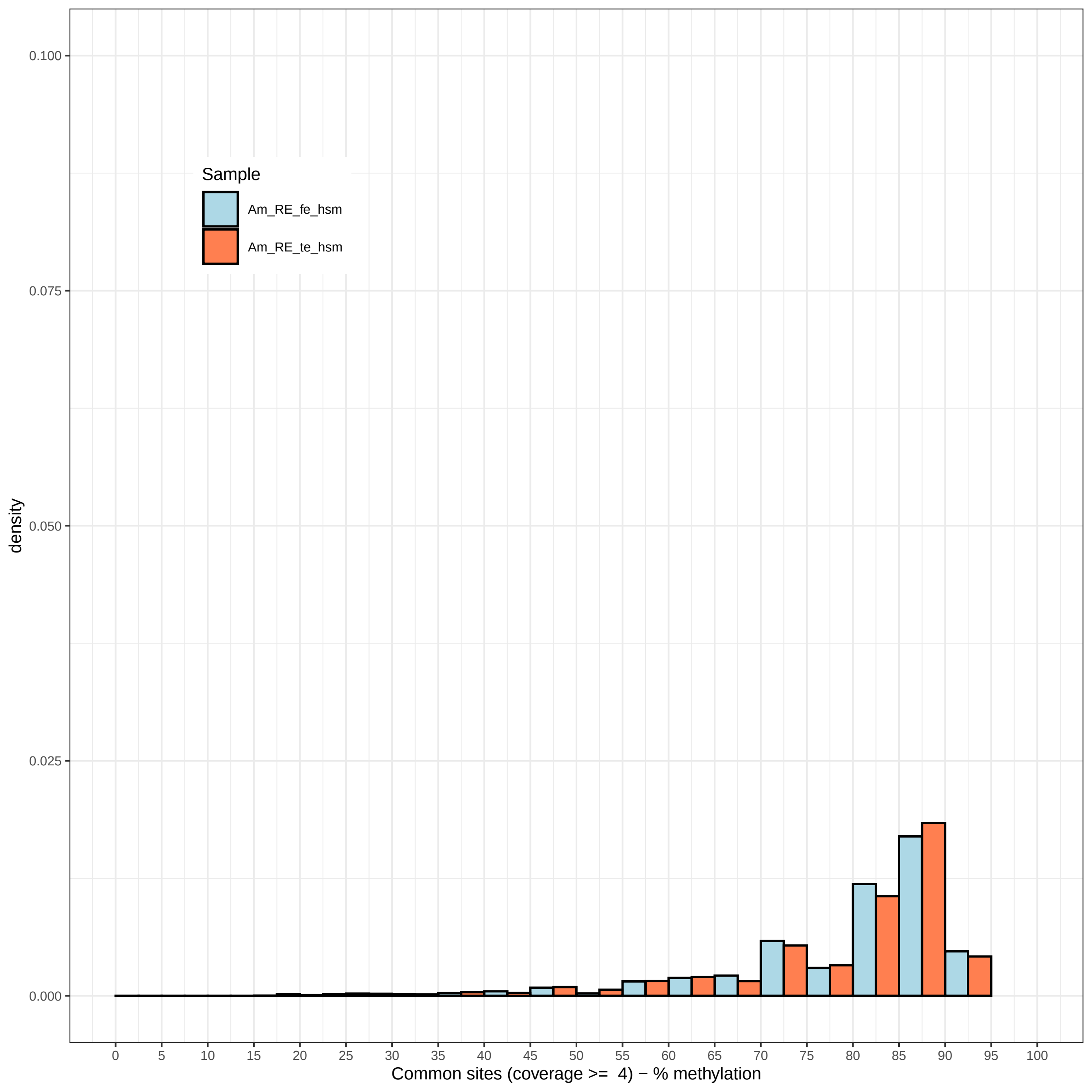

Sample
Am_RE_fe_hsm
Am_RE_te_hsm
density
0.100
0.075
0.050
0.025
0.000
0
5
10
15
20
25
30
35
40
45
50
55
60
65
70
75
80
85
90
95
100
Common sites (coverage >= 4) − % methylation

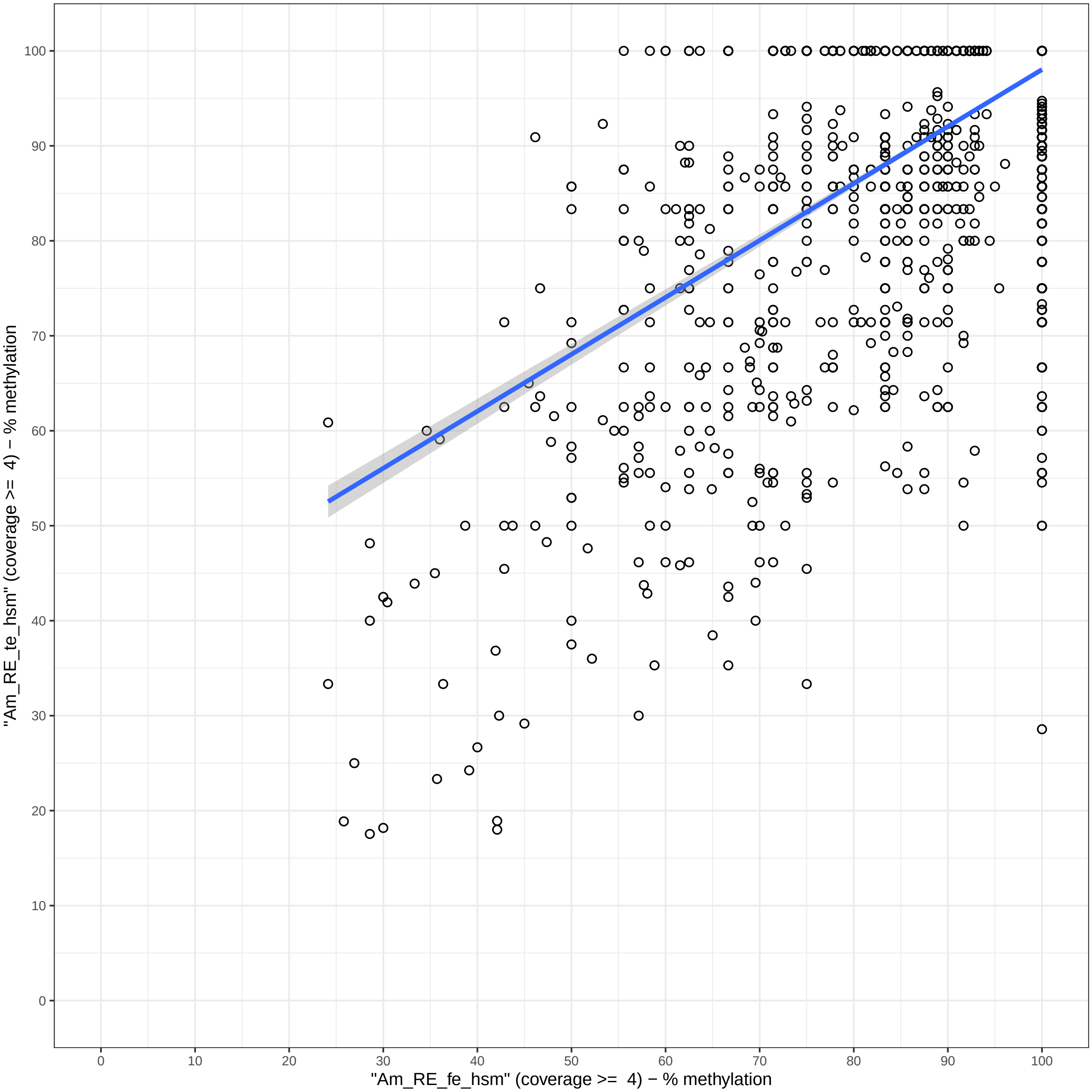

"Am_RE_te_hsm" (coverage >= 4) − % methylation
"Am_RE_fe_hsm" (coverage >= 4) − % methylation
0
10
20
30
40
50
60
70
80
90
100

### Overlap of highly supported methylation sites (coverage >= 4)

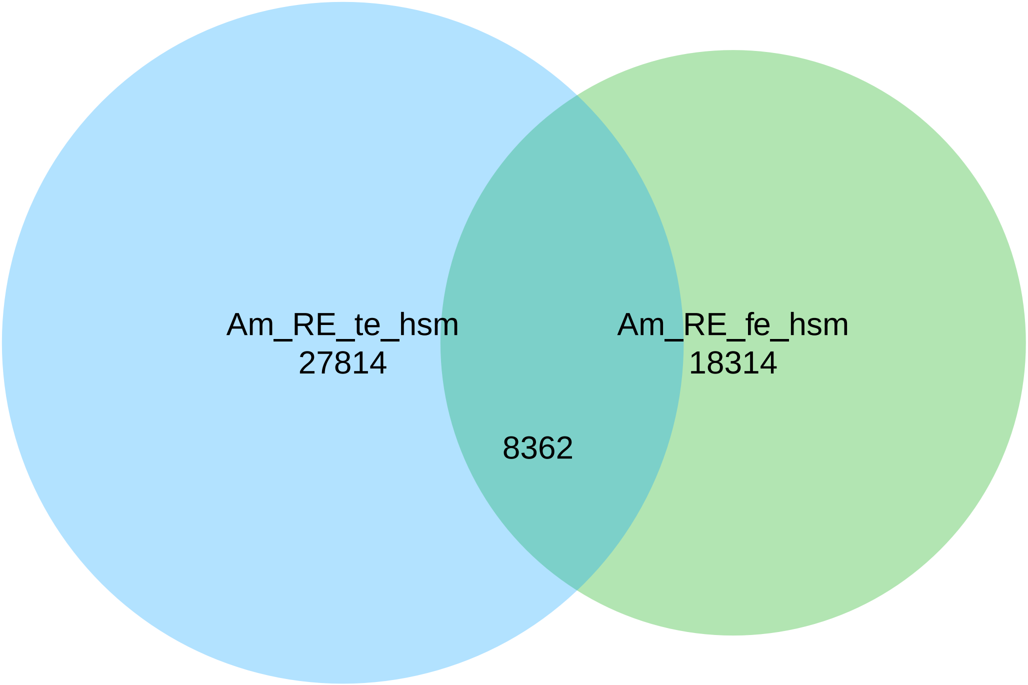

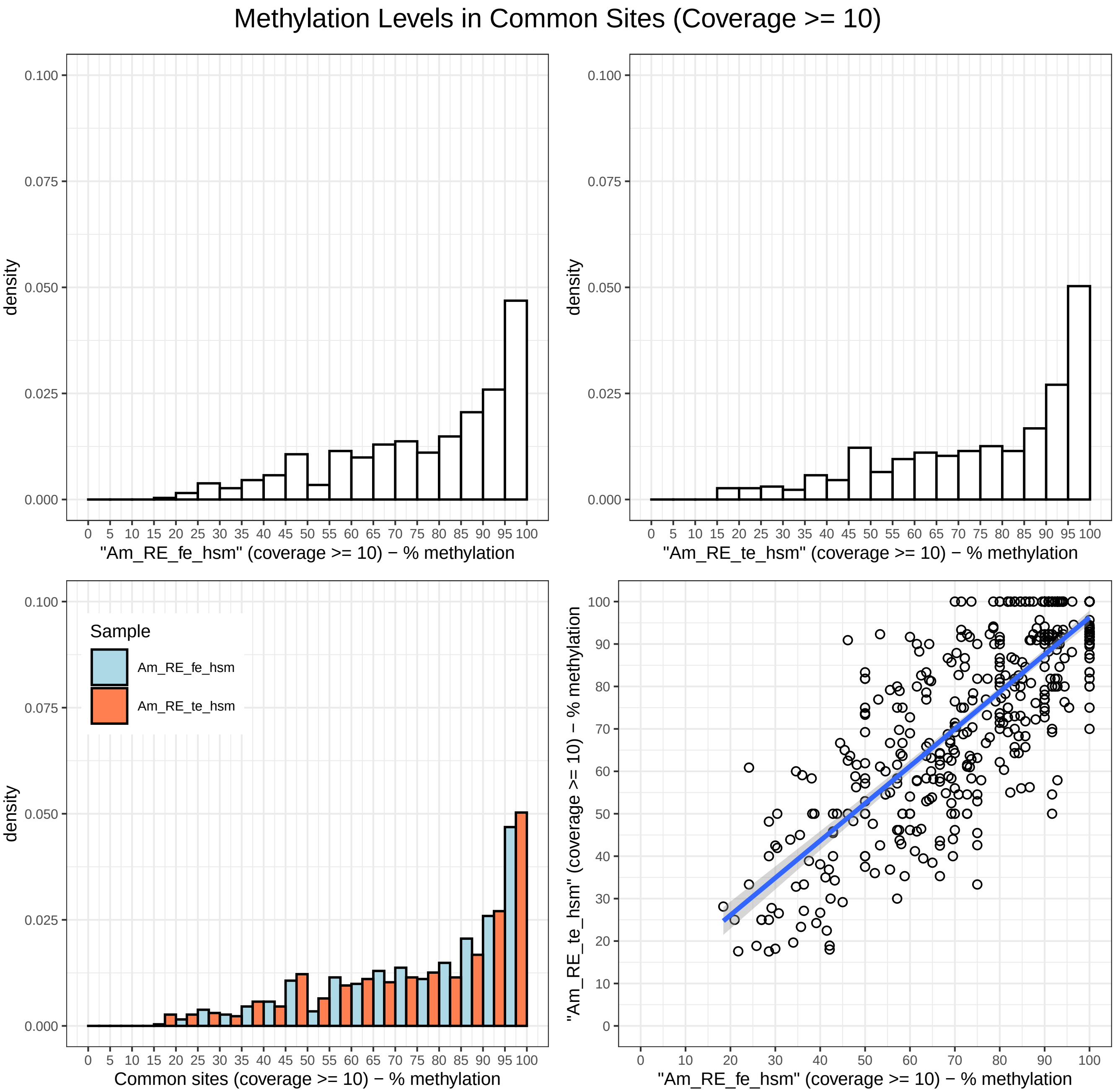

Methylation Levels in Common Sites (Coverage >= 10)
density
0.000
0.025
0.050
0.075
0.100
0 5 10 15 20 25 30 35 40 45 50 55 60 65 70 75 80 85 90 95 100
"Am_RE_fe_hsm" (coverage >= 10) − % methylation
"Am_RE_te_hsm" (coverage >= 10) − % methylation
Sample
Am_RE_fe_hsm
Am_RE_te_hsm
Common sites (coverage >= 10) − % methylation
"Am_RE_te_hsm" (coverage >= 10) − % methylation
0 10 20 30 40 50 60 70 80 90 100
"Am_RE_fe_hsm" (coverage >= 10) − % methylation

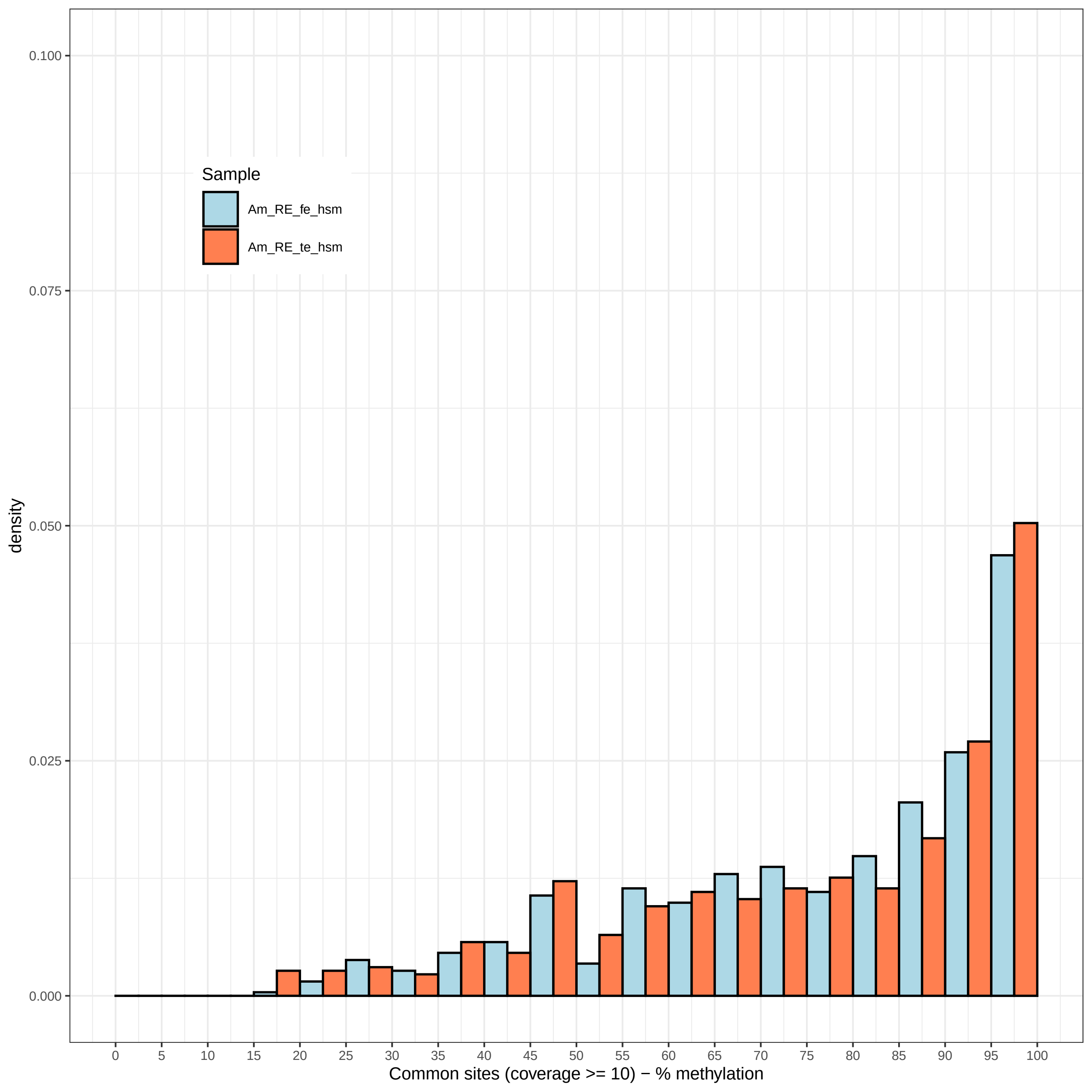

Sample
Am_RE_fe_hsm
Am_RE_te_hsm
density
0.000
0.025
0.050
0.075
0.100
0 5 10 15 20 25 30 35 40 45 50 55 60 65 70 75 80 85 90 95 100
Common sites (coverage >= 10) − % methylation

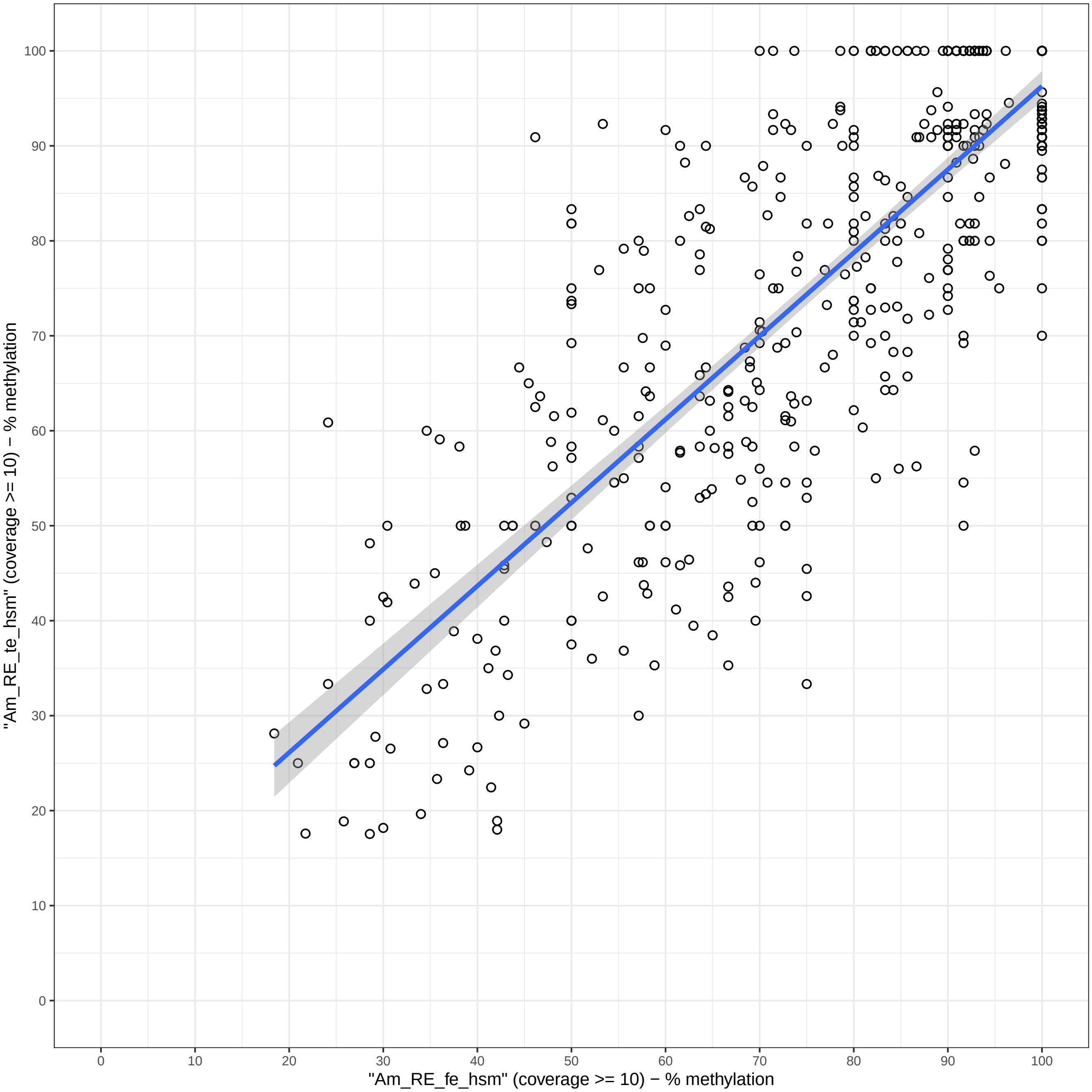

0
10
20
30
40
50
60
70
80
90
100
"Am_RE_te_hsm" (coverage >= 10) − % methylation
0
10
20
30
40
50
60
70
80
90
100
"Am_RE_fe_hsm" (coverage >= 10) − % methylation

### Overlap of highly supported methylation sites (coverage >= 10)

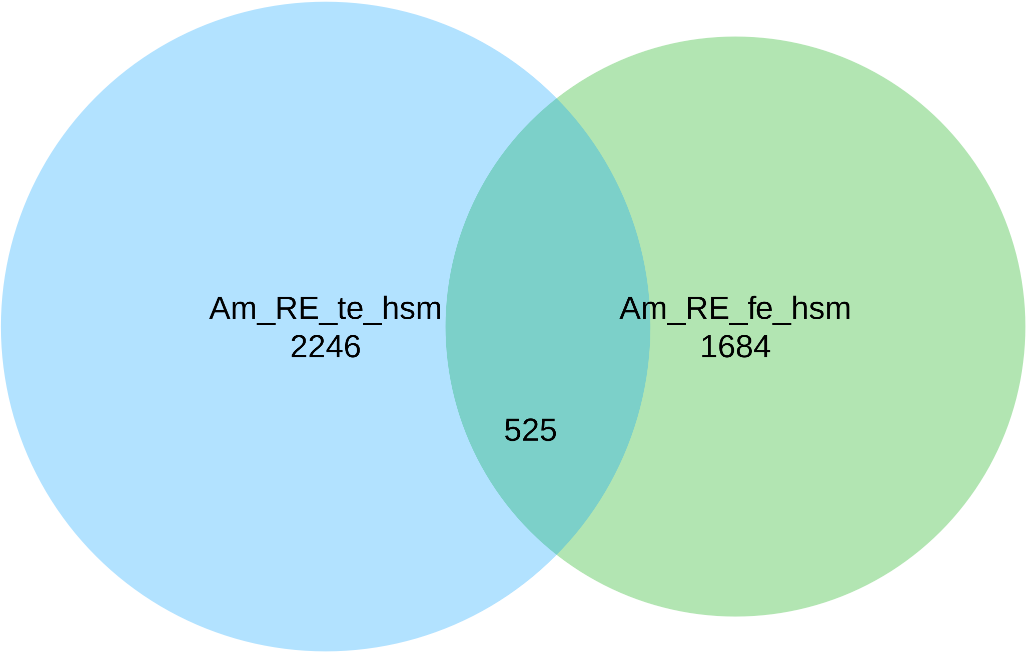

### Methylation Levels in Common Sites (Coverage >= 15)

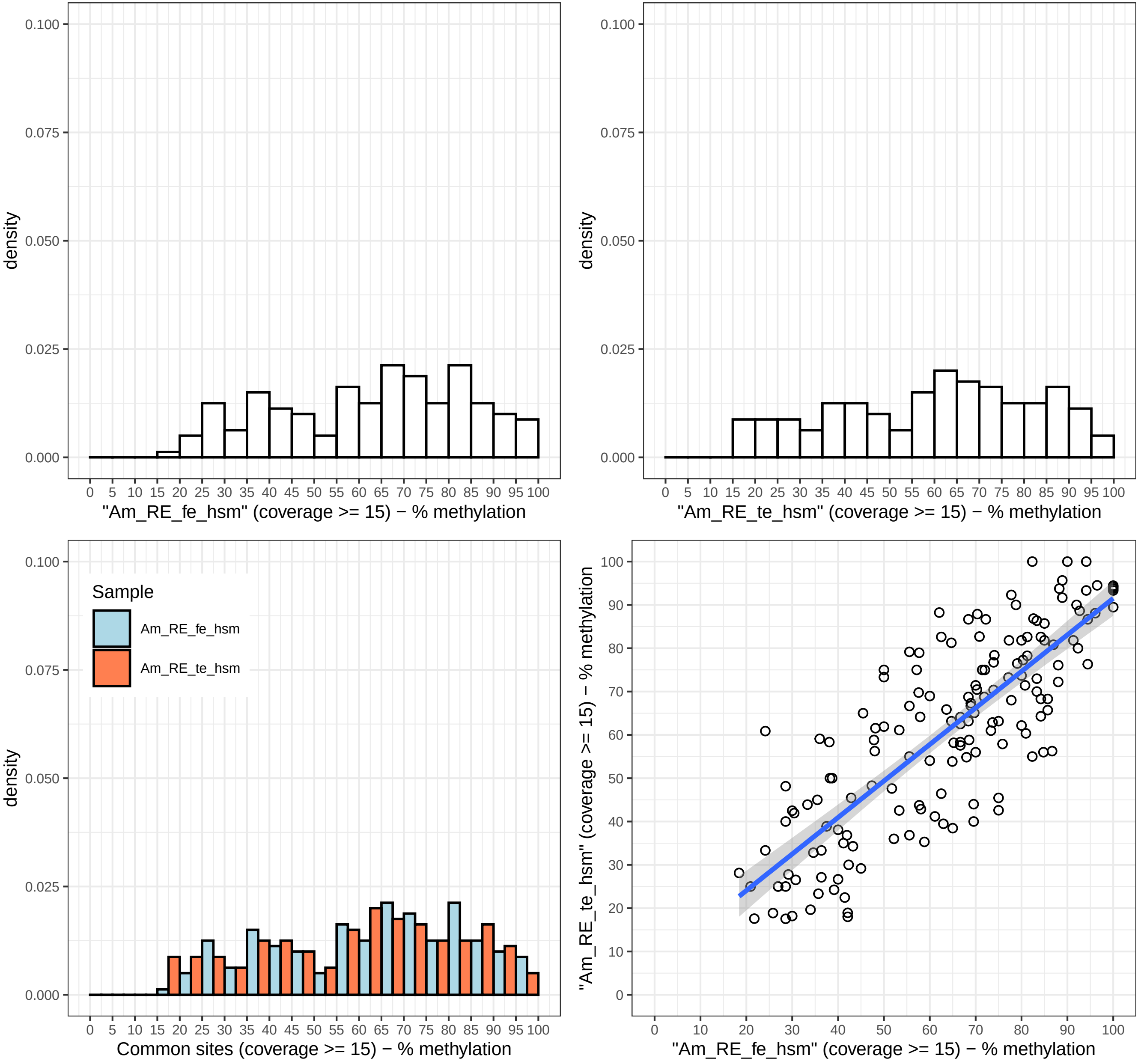

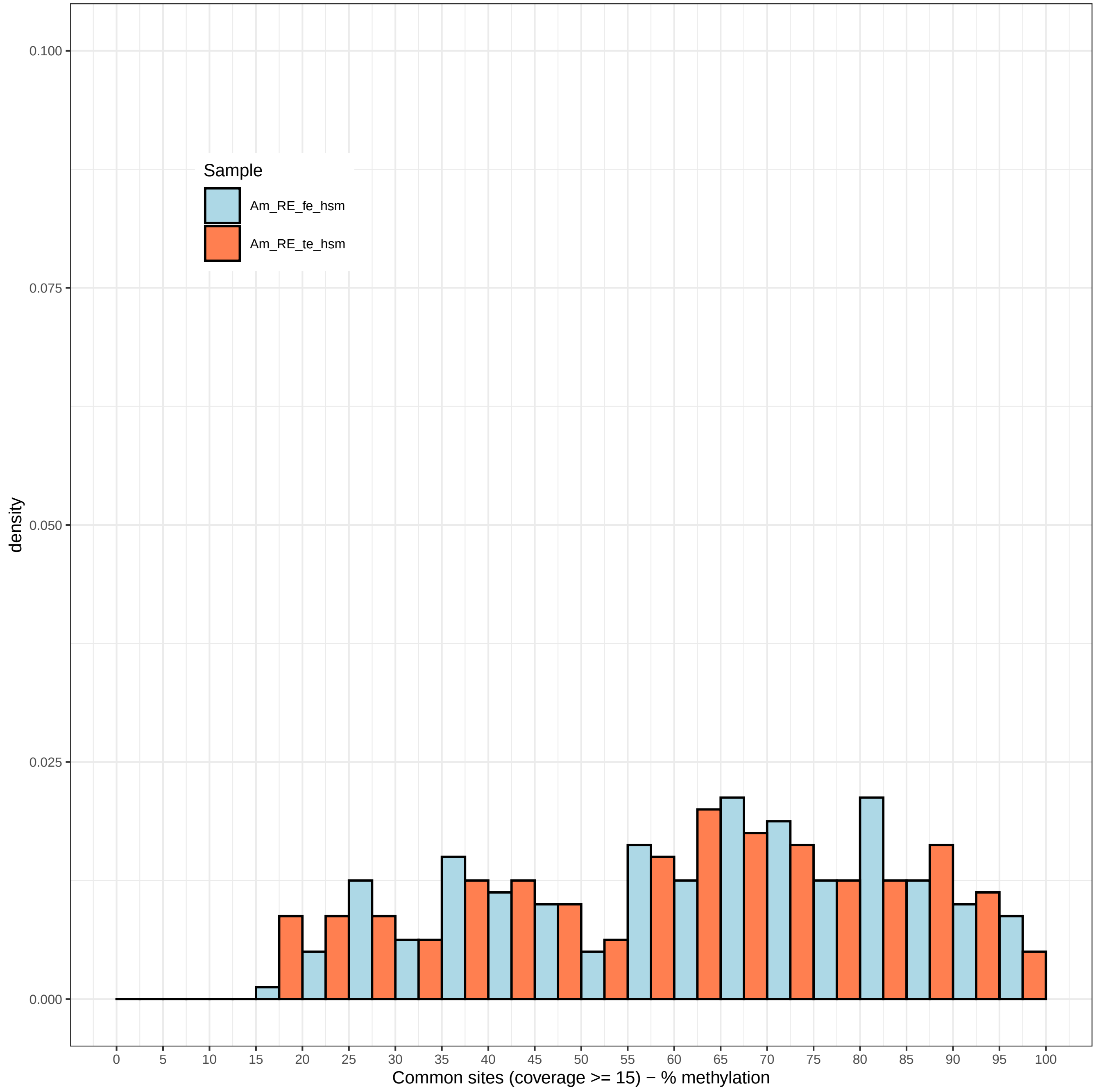

Sample
Am_RE_fe_hsm
Am_RE_te_hsm
density
0.100
0.075
0.050
0.025
0.000
0
5
10
15
20
25
30
35
40
45
50
55
60
65
70
75
80
85
90
95
100
Common sites (coverage >= 15) − % methylation

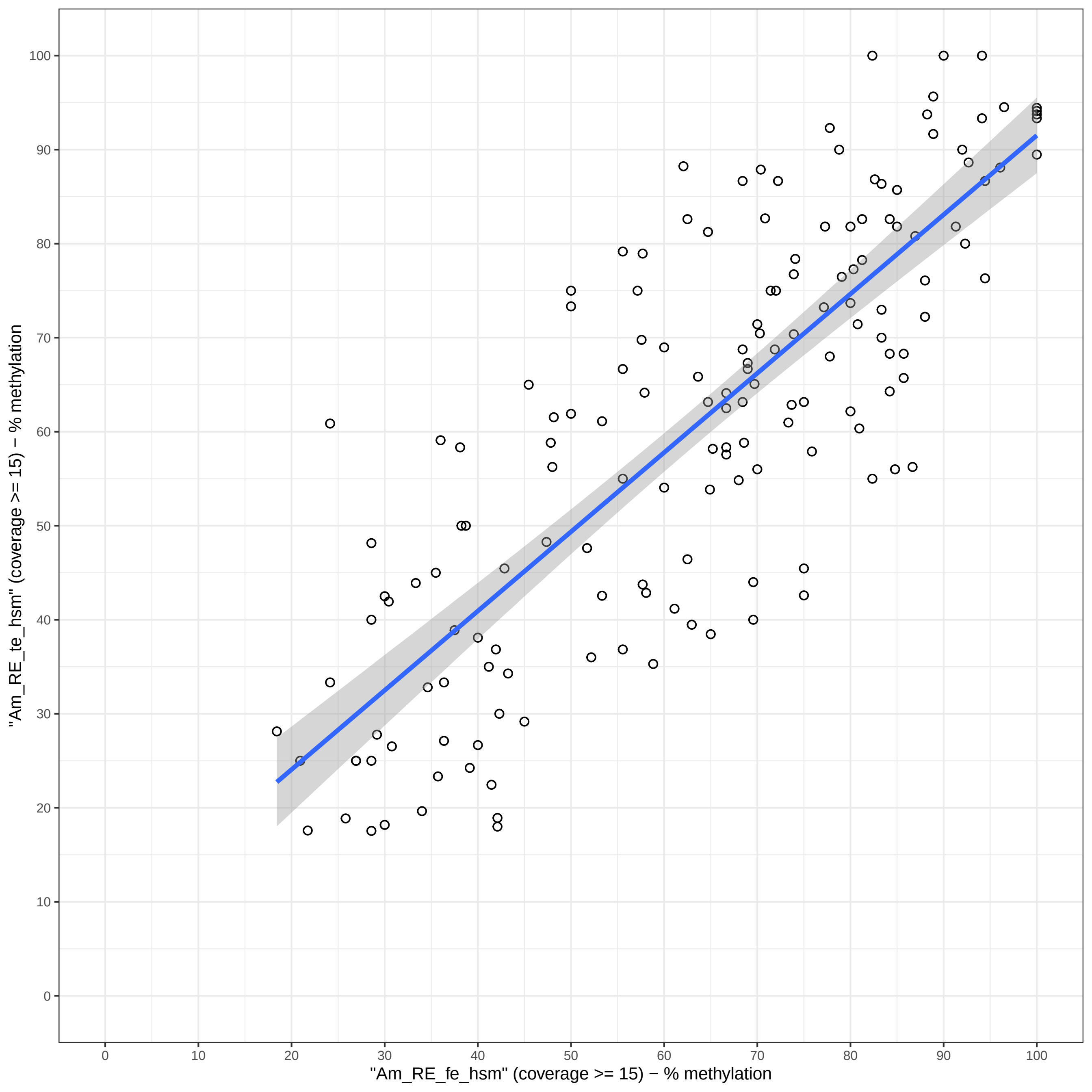

"Am_RE_te_hsm" (coverage >= 15) − % methylation
"Am_RE_fe_hsm" (coverage >= 15) − % methylation
0
10
20
30
40
50
60
70
80
90
100

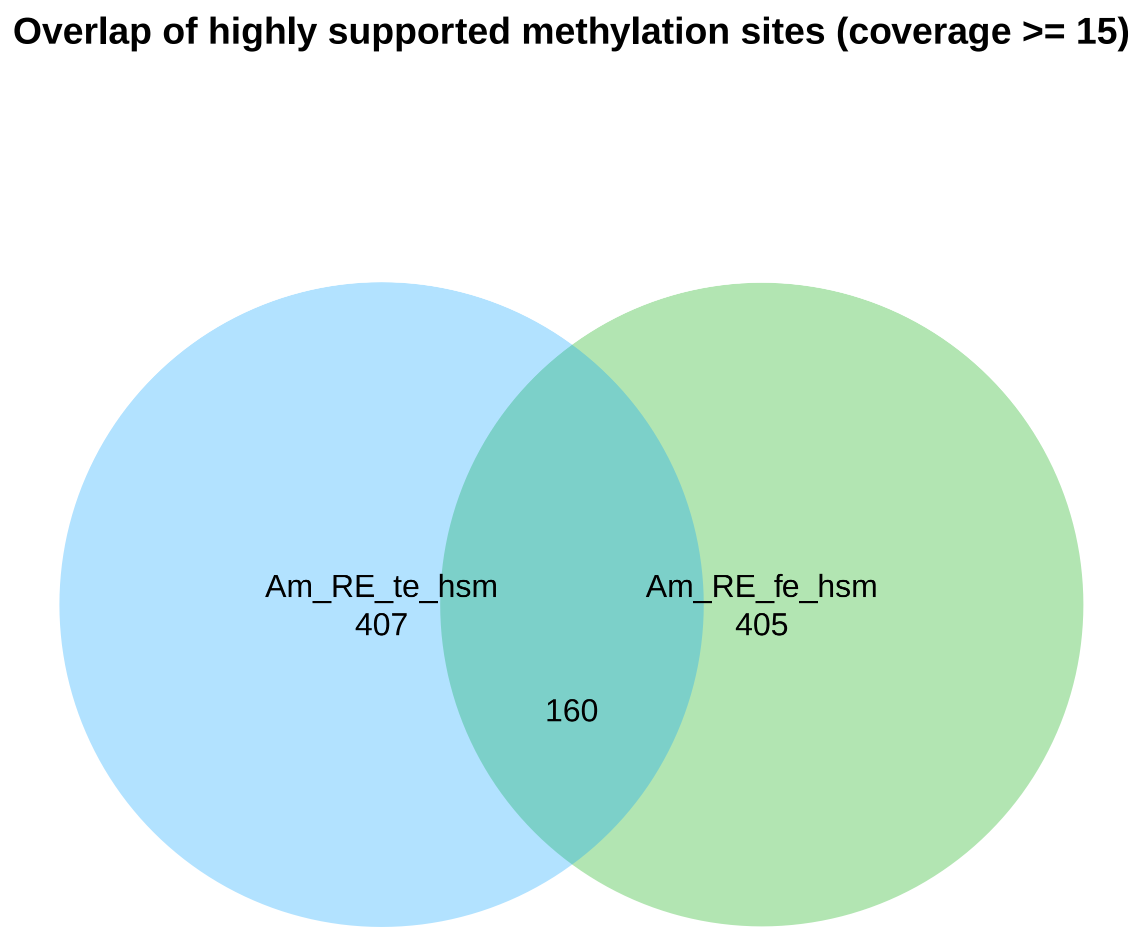
Overlap of highly supported methylation sites (coverage >= 15)
Am_RE_te_hsm
407
Am_RE_fe_hsm
405
160

```
Directory: CRL File: 0READMEcrl


CRL - Correlations between aggregate samples.

Input: studymk

Output: files crl-*.txt crl-*.pdf

Notes: The output is generated by BWASPR::cmpSamples(), which uses methylKit::unite()
        to determine the sites common to all samples and applies methylKit::getCorrelation()
        and methylKit::PCASamples().  The *.txt files show the correlations in a table, and
        the *.pdf files show graphics.

       Beware that the number of common sites (shown at the bottom of the *.txt files) may be
        small, which may make the correlations less informative.
```

Directory: CRL File: crl-Am_RE.txt

| | fe | te |
|---|---|---|
| fe | 1.000 | 0.543 |
| te | 0.543 | 1.000 |

The number of conserved sites is 9320.

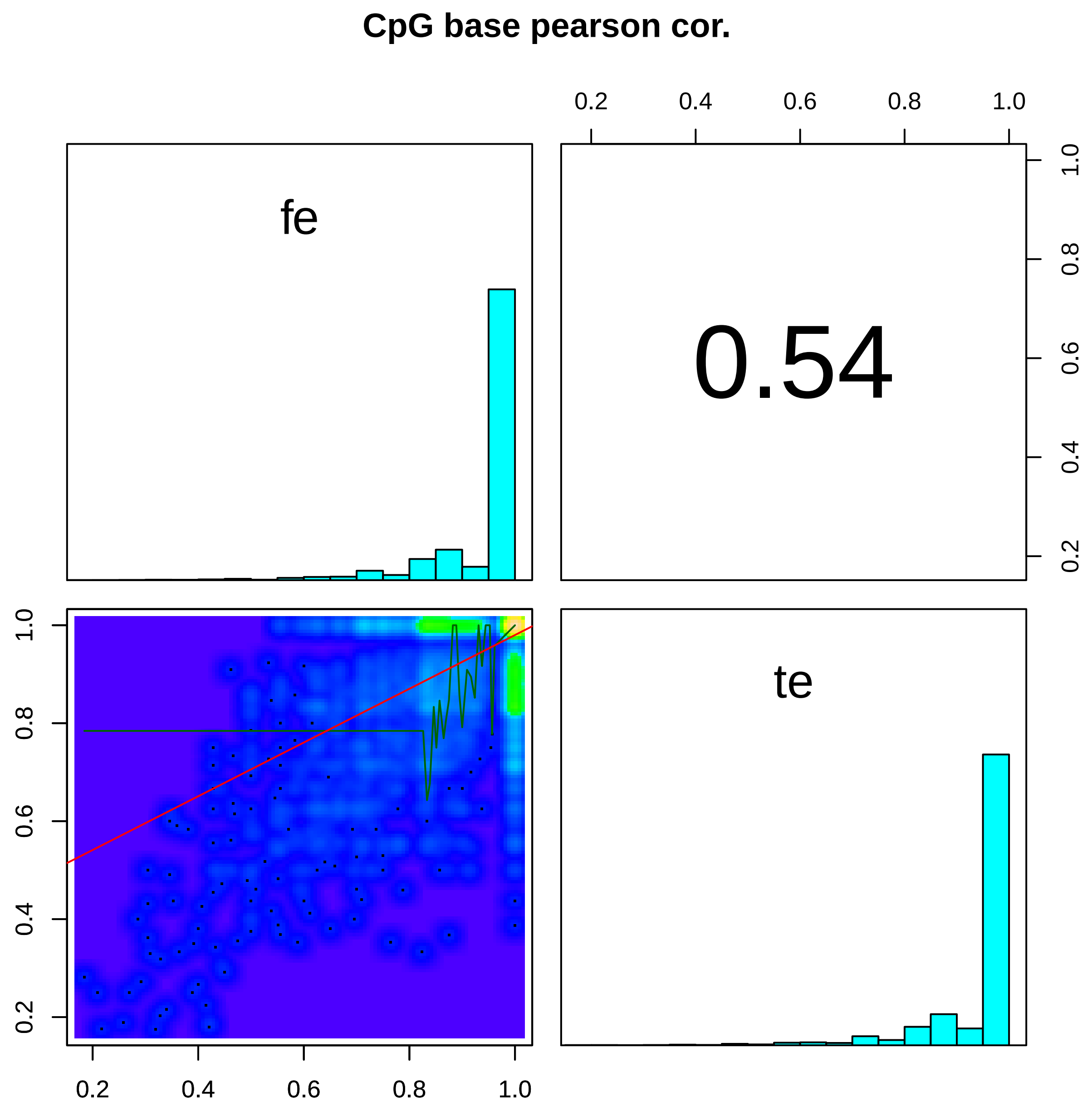

CpG base pearson cor.
fe
0.54
te
0.2
0.4
0.6
0.8
1.0

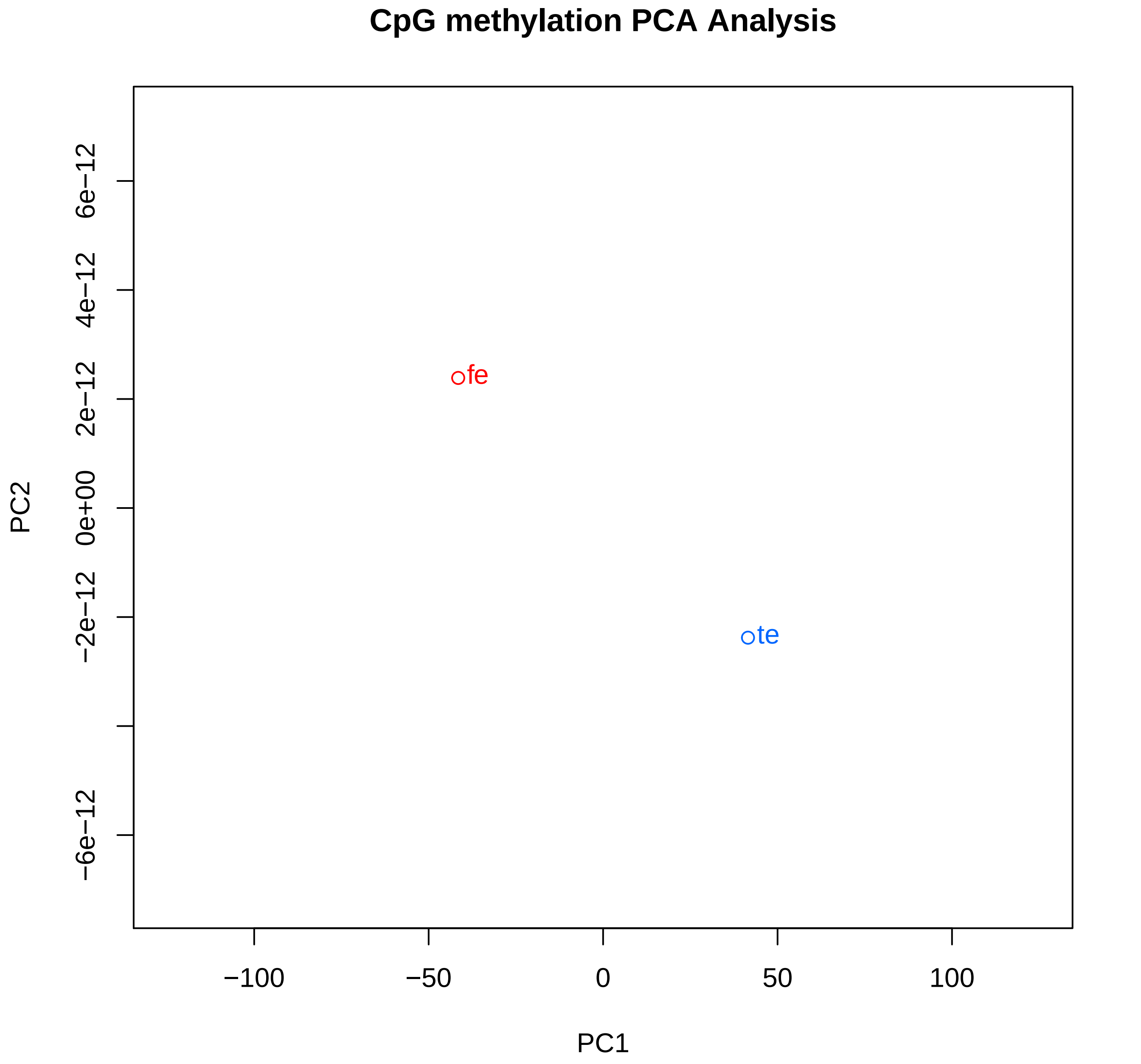
CpG methylation PCA Analysis
PC2
6e−12
4e−12
2e−12
0e+00
−2e−12
−6e−12
fe
te
−100
−50
0
50
100
PC1

Directory: MMP File: 0READMEmmp

MMP - Mapping of methylation sites on genome annotation.

Input: studymk, studymc, genome_ann (a list of GRanges objects providing annotated region labels
and boundaries; output of BWASPR::get_genome_annotation())

Output: files mmp-*.txt

Notes: The output is generated by BWASPR::map_methylome() and gives an accounting for every sample
as to where the CpGhsm and CpGscd (control) sites reside relative to the genome annotation.
Abbreviations used: O/E, observed over expected
(expected percentages are based on the respective feature proportions
in the genome, as annotated).

Directory: MMP File: mmp-Am_RE_fe.txt

==============================================================================

#### Genomic composition in terms of feature regions

==============================================================================

| | | |
|---|---|---|
| Am genome size: | 225250884 bp | |
| Am genic region size: | 185146880 bp | ( 82.2%) |
| Am exon region size: | 39060230 bp | ( 17.3%) |
| Am CDS region size: | 18098288 bp | ( 46.3%) |
| Am five-prime UTR region size: | 3360721 bp | ( 8.6%) |
| Am three-prime UTR region size: | 5581464 bp | ( 14.3%) |
| Am other exon region size: | 12019757 bp | ( 30.8%) |
| Am intron region size: | 146086650 bp | ( 64.9%) |
| Am intergenic region size: | 40104004 bp | ( 17.8%) |
| Am promoter region size: | 6164949 bp | ( 2.7%) |
| Am other intergenic region size: | 33939055 bp | ( 15.1%) |

==============================================================================

#### Methylation sites in genomic feature regions

==============================================================================

Number and density (per 10kb) of sites in Am_RE_fe_scd

| | | | | |
|---|---|---|---|---|
| Number of sites identified in Am Am_RE_fe_scd: | 7682417 | ( 341.06) | | |
| Number of sites identified in Am Am_RE_fe_scd genic regions: | 6086957 | ( 328.76) | 79.23% | ( 0.96 O/E) |
| Number of sites identified in Am Am_RE_fe_scd exon regions: | 1252192 | ( 320.58) | 16.30% | ( 0.94 O/E) |
| Number of sites identified in Am Am_RE_fe_scd CDS regions: | 740450 | ( 409.13) | 59.13% | ( 1.28 O/E) |
| Number of sites identified in Am Am_RE_fe_scd five-prime UTR regions: | 171876 | ( 511.43) | 13.73% | ( 1.60 O/E) |
| Number of sites identified in Am Am_RE_fe_scd three-prime UTR regions: | 130984 | ( 234.68) | 10.46% | ( 0.73 O/E) |
| Number of sites identified in Am Am_RE_fe_scd other exon regions: | 208882 | ( 173.78) | 16.68% | ( 0.54 O/E) |
| Number of sites identified in Am Am_RE_fe_scd intron regions: | 4834765 | ( 330.95) | 62.93% | ( 0.97 O/E) |
| Number of sites identified in Am Am_RE_fe_scd intergenic regions: | 1595460 | ( 397.83) | 20.77% | ( 1.17 O/E) |
| Number of sites identified in Am Am_RE_fe_scd promoter regions: | 60925 | ( 98.82) | 0.79% | ( 0.29 O/E) |
| Number of sites identified in Am Am_RE_fe_scd other intergenic regions: | 1534535 | ( 452.14) | 19.97% | ( 1.33 O/E) |

Number and density (per 10kb) of sites in Am_RE_fe_hsm

| | | | | | |
|---|---|---|---|---|---|
| Number of sites identified in Am Am_RE_fe_hsm: | 18314 | ( 0.81) | ( 0.24% of control) | | |
| Number of sites identified in Am Am_RE_fe_hsm genic regions: | 17885 | ( 0.97) | ( 0.29% of control) | 97.66% | ( 1.19 O/E) |
| Number of sites identified in Am Am_RE_fe_hsm exon regions: | 16668 | ( 4.27) | ( 1.33% of control) | 91.01% | ( 5.25 O/E) |
| Number of sites identified in Am Am_RE_fe_hsm CDS regions: | 15825 | ( 8.74) | ( 2.14% of control) | 94.94% | ( 2.05 O/E) |
| Number of sites identified in Am Am_RE_fe_hsm five-prime UTR regions: | 143 | ( 0.43) | ( 0.08% of control) | 0.86% | ( 0.10 O/E) |
| Number of sites identified in Am Am_RE_fe_hsm three-prime UTR regions: | 452 | ( 0.81) | ( 0.35% of control) | 2.71% | ( 0.19 O/E) |
| Number of sites identified in Am Am_RE_fe_hsm other exon regions: | 248 | ( 0.21) | ( 0.12% of control) | 1.49% | ( 0.05 O/E) |
| Number of sites identified in Am Am_RE_fe_hsm intron regions: | 1217 | ( 0.08) | ( 0.03% of control) | 6.65% | ( 0.10 O/E) |
| Number of sites identified in Am Am_RE_fe_hsm intergenic regions: | 429 | ( 0.11) | ( 0.03% of control) | 2.34% | ( 0.13 O/E) |
| Number of sites identified in Am Am_RE_fe_hsm promoter regions: | 25 | ( 0.04) | ( 0.04% of control) | 0.14% | ( 0.05 O/E) |
| Number of sites identified in Am Am_RE_fe_hsm other intergenic regions: | 404 | ( 0.12) | ( 0.03% of control) | 2.21% | ( 0.15 O/E) |

```
Directory: MMP File: mmp-Am_RE_te.txt


================================================================================
Genomic composition in terms of feature regions
================================================================================

Am genome size:                        225250884 bp
  Am genic region size:                  185146880 bp ( 82.2%)
    Am exon region size:                    39060230 bp ( 17.3%)
      Am CDS region size:                      18098288 bp ( 46.3%)
      Am five-prime UTR region size:            3360721 bp (  8.6%)
      Am three-prime UTR region size:           5581464 bp ( 14.3%)
      Am other exon region size:               12019757 bp ( 30.8%)
    Am intron region size:                 146086650 bp ( 64.9%)
  Am intergenic region size:              40104004 bp ( 17.8%)
    Am promoter region size:                 6164949 bp (  2.7%)
    Am other intergenic region size:        33939055 bp ( 15.1%)


================================================================================
Methylation sites in genomic feature regions
================================================================================

Number and density (per 10kb) of sites in Am_RE_te_scd

Number of sites identified in Am Am_RE_te_scd:                              8517171     (  378.12)
Number of sites identified in Am Am_RE_te_scd genic regions:                6770476     (  365.68)                          79.49%     ( 0.97 O/E)
Number of sites identified in Am Am_RE_te_scd   exon regions:                 1331210   (  340.81)                            15.63%   ( 0.90 O/E)
Number of sites identified in Am Am_RE_te_scd     CDS regions:                   753499 (  416.34)                              56.60% ( 1.22 O/E)
Number of sites identified in Am Am_RE_te_scd     five-prime UTR regions:        171246 (  509.55)                              12.86% ( 1.50 O/E)
Number of sites identified in Am Am_RE_te_scd     three-prime UTR regions:       166253 (  297.87)                              12.49% ( 0.87 O/E)
Number of sites identified in Am Am_RE_te_scd     other exon regions:            240212 (  199.85)                              18.04% ( 0.59 O/E)
Number of sites identified in Am Am_RE_te_scd   intron regions:               5439266   (  372.33)                            63.86%   ( 0.98 O/E)
Number of sites identified in Am Am_RE_te_scd intergenic regions:           1746695     (  435.54)                          20.51%     ( 1.15 O/E)
Number of sites identified in Am Am_RE_te_scd   promoter regions:               63791   (  103.47)                             0.75%   ( 0.27 O/E)
Number of sites identified in Am Am_RE_te_scd   other intergenic regions:     1682904   (  495.86)                            19.76%   ( 1.31 O/E)


Number and density (per 10kb) of sites in Am_RE_te_hsm

Number of sites identified in Am Am_RE_te_hsm:                                27814     (    1.23) (  0.33% of control)
Number of sites identified in Am Am_RE_te_hsm genic regions:                  27309     (    1.47) (  0.40% of control)  98.18%     ( 1.19 O/E)
Number of sites identified in Am Am_RE_te_hsm   exon regions:                   25521   (    6.53) (  1.92% of control)    91.76%   ( 5.29 O/E)
Number of sites identified in Am Am_RE_te_hsm     CDS regions:                    24228 (   13.39) (  3.22% of control)      94.93% ( 2.05 O/E)
Number of sites identified in Am Am_RE_te_hsm     five-prime UTR regions:           197 (    0.59) (  0.12% of control)       0.77% ( 0.09 O/E)
Number of sites identified in Am Am_RE_te_hsm     three-prime UTR regions:          770 (    1.38) (  0.46% of control)       3.02% ( 0.21 O/E)
Number of sites identified in Am Am_RE_te_hsm     other exon regions:               326 (    0.27) (  0.14% of control)       1.28% ( 0.04 O/E)
Number of sites identified in Am Am_RE_te_hsm   intron regions:                  1788   (    0.12) (  0.03% of control)     6.43%   ( 0.10 O/E)
Number of sites identified in Am Am_RE_te_hsm intergenic regions:               505     (    0.13) (  0.03% of control)   1.82%     ( 0.10 O/E)
Number of sites identified in Am Am_RE_te_hsm   promoter regions:                  23   (    0.04) (  0.04% of control)     0.08%   ( 0.03 O/E)
Number of sites identified in Am Am_RE_te_hsm   other intergenic regions:         482   (    0.14) (  0.03% of control)     1.73%   ( 0.12 O/E)
```

```
Directory: ACS File: 0READMEacs


ACS - Annotation of conserved methylation sites.

Input: studymk, genome_ann (a list of GRanges objects providing annotated region labels and bounds;
                            output of BWASPR::get_genome_annotation())

Output: file acs-*.txt

Notes: The output is generated by BWASPR::annotate_methylome(), which uses methylKit::unite()
        to determine the sites common to all experiments and applies genomation::annotateWithFeature().
        The acs-*.txt files show the common sites with methylation levels in all samples
        and columns with booleans indicating whether the site fits a specific annotation feature.

       Abbreviations used: pc, protein coding; nc, non-coding; fp, five prime; tp, three prime.
                           UTR, untranslated region; unique, not overlapping with other categories.
```

Directory: ACS File: acs-Am_RE.txt

| fe | te | chr | start | end | strand | gene | exon | pcexon | promoter | CDS | fpUTR | tpUTR | fpUTRnotCDS | tpUTRnotCDS | tpUTRunique | ncexon |
|---|---|---|---|---|---|---|---|---|---|---|---|---|---|---|---|---|
| 80 | 90.91 | NC_037638.1 | 13129 | 13129 | + | 1 | 1 | 1 | 0 | 1 | 0 | 0 | 0 | 0 | 0 | 0 |
| 89.47 | 100 | NC_037638.1 | 13165 | 13165 | + | 1 | 1 | 1 | 0 | 1 | 0 | 0 | 0 | 0 | 0 | 0 |
| 92.86 | 93.33 | NC_037638.1 | 13208 | 13208 | + | 1 | 1 | 1 | 0 | 1 | 0 | 0 | 0 | 0 | 0 | 0 |
| 90.91 | 90.91 | NC_037638.1 | 13244 | 13244 | + | 1 | 1 | 1 | 0 | 1 | 0 | 0 | 0 | 0 | 0 | 0 |
| 100 | 100 | NC_037638.1 | 13253 | 13253 | + | 1 | 1 | 1 | 0 | 1 | 0 | 0 | 0 | 0 | 0 | 0 |
| 100 | 100 | NC_037638.1 | 13273 | 13273 | + | 1 | 1 | 1 | 0 | 1 | 0 | 0 | 0 | 0 | 0 | 0 |
| 100 | 100 | NC_037638.1 | 13291 | 13291 | + | 1 | 1 | 1 | 0 | 1 | 0 | 0 | 0 | 0 | 0 | 0 |
| 100 | 90.91 | NC_037638.1 | 13329 | 13329 | + | 1 | 1 | 1 | 0 | 1 | 0 | 0 | 0 | 0 | 0 | 0 |
| 93.75 | 100 | NC_037638.1 | 13372 | 13372 | + | 1 | 1 | 1 | 0 | 1 | 0 | 0 | 0 | 0 | 0 | 0 |
| 100 | 100 | NC_037638.1 | 13392 | 13392 | + | 1 | 1 | 1 | 0 | 1 | 0 | 0 | 0 | 0 | 0 | 0 |
| 100 | 100 | NC_037638.1 | 13443 | 13443 | + | 1 | 1 | 1 | 0 | 1 | 0 | 0 | 0 | 0 | 0 | 0 |
| 85.71 | 100 | NC_037638.1 | 13459 | 13459 | + | 1 | 1 | 1 | 0 | 1 | 0 | 0 | 0 | 0 | 0 | 0 |
| 100 | 100 | NC_037638.1 | 13479 | 13479 | + | 1 | 1 | 1 | 0 | 1 | 0 | 0 | 0 | 0 | 0 | 0 |
| 100 | 100 | NC_037638.1 | 13498 | 13498 | + | 1 | 1 | 1 | 0 | 1 | 0 | 0 | 0 | 0 | 0 | 0 |
| 100 | 100 | NC_037638.1 | 13514 | 13514 | + | 1 | 1 | 1 | 0 | 1 | 0 | 0 | 0 | 0 | 0 | 0 |
| 100 | 100 | NC_037638.1 | 24663 | 24663 | + | 1 | 1 | 1 | 0 | 1 | 0 | 0 | 0 | 0 | 0 | 0 |
| 95 | 100 | NC_037638.1 | 24701 | 24701 | + | 1 | 1 | 1 | 0 | 1 | 0 | 0 | 0 | 0 | 0 | 0 |
| 100 | 100 | NC_037638.1 | 24743 | 24743 | + | 1 | 1 | 1 | 0 | 1 | 0 | 0 | 0 | 0 | 0 | 0 |
| 94.12 | 93.33 | NC_037638.1 | 24781 | 24781 | + | 1 | 1 | 1 | 0 | 1 | 0 | 0 | 0 | 0 | 0 | 0 |
| 88.24 | 93.75 | NC_037638.1 | 24796 | 24796 | + | 1 | 1 | 1 | 0 | 1 | 0 | 0 | 0 | 0 | 0 | 0 |
| 88.89 | 100 | NC_037638.1 | 24938 | 24938 | + | 1 | 1 | 1 | 0 | 1 | 0 | 0 | 0 | 0 | 0 | 0 |
| 77.78 | 75 | NC_037638.1 | 24950 | 24950 | + | 1 | 1 | 1 | 0 | 1 | 0 | 0 | 0 | 0 | 0 | 0 |
| 100 | 100 | NC_037638.1 | 25318 | 25318 | + | 1 | 1 | 1 | 0 | 1 | 0 | 0 | 0 | 0 | 0 | 0 |
| 100 | 100 | NC_037638.1 | 25361 | 25361 | + | 1 | 1 | 1 | 0 | 1 | 0 | 0 | 0 | 0 | 0 | 0 |

...
... only first 25 lines shown ...

```
Directory: RNK File: 0READMErnk


RNK - Genomic feature regions ranked by CpG methylation and hsm statistics.

Input: studymc, studymk, region.gr (GRanges object providing annotated region labels and bounds)

Output: files ranked-genes-*.txt sites-in-genes-*.txt
              ranked-promoters-*.txt sites-in-promoters-*.txt
              plot*.pdf

Notes: The output is generated by BWASPR::rank_rbm().
       The files ranked-genes-*.txt provide ranked lists of genes based on
        overall methylation percentage or the occurrence of CpGhms sites
        within the annotated gene bounds.  The output columns include
          region_ID    (= region/gene name)
          rwidth       (= region/gene width)
          nbrsites     (= number of CpGhsm sites in the region/gene)
          nbrper10kb   (= number of CpGhsm sites in the region/gene normalized to 10kb width)
          pmrpersite   (= average % methylation per CpGhsm site in the region/gene)
          pmrpernucl   (= average % CpGhsm methylation per nucleotide in the region/gene)
          prcntM       (= overall % CpGhsm methylation in the region/gene)

       The tables are sorted by prcntM (*byPrcntM* files) or by nbper10kb
         (*bySiteDensity* files) in descending order.

       The ranked-promoters-*.txt files provide analogous tables for promoter regions.
        If the promoter annotation was derived with BWASP, then these regions are simply
        defined here as 500 nucleotides upstream of the 5'-end of the gene (shorter if the
        scaffold ends before).

       The files sites-*.txt provide details for the CpGhsm sites in the respective
        regions.
       The files plot-*.pdf explore concordance of the two methylation measures.
       The optional parameters minrwidth, maxrwidth, and minnbrsites specify
       the length range and minimal number of methylation sites for regions
       to be included in the plot.
```

Directory: RNK File: ranked-genes-byPrcntM-Am_RE_fe.txt

| region_ID | seqnames | start | end | width | strand | coverage | numCs | numTs | prcntM |
|---|---|---|---|---|---|---|---|---|---|
| gene-LOC408625 | NC_037638.1 | 1696663 | 1699582 | 2920 | - | 4 | 4 | 0 | 100 |
| gene-LOC727419 | NC_037638.1 | 2241599 | 2242631 | 1033 | - | 10 | 10 | 0 | 100 |
| gene-LOC726316 | NC_037638.1 | 2492814 | 2493891 | 1078 | - | 4 | 4 | 0 | 100 |
| gene-LOC102655778 | NC_037638.1 | 2505276 | 2507841 | 2566 | - | 10 | 10 | 0 | 100 |
| gene-LOC724233 | NC_037638.1 | 4375803 | 4376815 | 1013 | + | 6 | 6 | 0 | 100 |
| gene-LOC413437 | NC_037638.1 | 4599212 | 4600692 | 1481 | + | 4 | 4 | 0 | 100 |
| gene-LOC412141 | NC_037638.1 | 4763774 | 4766067 | 2294 | + | 4 | 4 | 0 | 100 |
| gene-LOC552038 | NC_037638.1 | 6019689 | 6021000 | 1312 | + | 5 | 5 | 0 | 100 |
| gene-LOC413868 | NC_037638.1 | 6569130 | 6571369 | 2240 | + | 5 | 5 | 0 | 100 |
| gene-LOC100576770 | NC_037638.1 | 7968300 | 7970147 | 1848 | - | 4 | 4 | 0 | 100 |
| gene-LOC102654365 | NC_037638.1 | 9116104 | 9119332 | 3229 | + | 9 | 9 | 0 | 100 |
| gene-LOC413470 | NC_037638.1 | 9273481 | 9275481 | 2001 | + | 14 | 14 | 0 | 100 |
| gene-LOC725817 | NC_037638.1 | 11071285 | 11073537 | 2253 | + | 5 | 5 | 0 | 100 |
| gene-LOC107964507 | NC_037638.1 | 11074328 | 11075115 | 788 | + | 4 | 4 | 0 | 100 |
| gene-LOC551354 | NC_037638.1 | 11485404 | 11486356 | 953 | - | 4 | 4 | 0 | 100 |
| gene-LOC724330 | NC_037638.1 | 12198744 | 12200333 | 1590 | - | 8 | 8 | 0 | 100 |
| gene-LOC409298 | NC_037638.1 | 14064075 | 14065563 | 1489 | + | 17 | 17 | 0 | 100 |
| gene-LOC551978 | NC_037638.1 | 15101259 | 15102679 | 1421 | + | 5 | 5 | 0 | 100 |
| gene-LOC550996 | NC_037638.1 | 18545078 | 18546497 | 1420 | - | 4 | 4 | 0 | 100 |
| gene-LOC413499 | NC_037638.1 | 19777725 | 19780145 | 2421 | - | 4 | 4 | 0 | 100 |
| gene-LOC102656360 | NC_037638.1 | 20629670 | 20635127 | 5458 | - | 4 | 4 | 0 | 100 |
| gene-LOC551606 | NC_037638.1 | 20643921 | 20646310 | 2390 | + | 10 | 10 | 0 | 100 |
| gene-LOC100578271 | NC_037638.1 | 20648774 | 20654433 | 5660 | + | 10 | 10 | 0 | 100 |
| gene-LOC412362 | NC_037638.1 | 20655228 | 20658636 | 3409 | - | 4 | 4 | 0 | 100 |

...

... only first 25 lines shown ...

Directory: RNK File: ranked-genes-byPrcntM-Am_RE_te.txt

| region_ID | seqnames | start | end | width | strand | coverage | numCs | numTs | prcntM |
|---|---|---|---|---|---|---|---|---|---|
| gene-LOC100576088 | NC_037638.1 | 1682530 | 1683635 | 1106 | + | 4 | 4 | 0 | 100 |
| gene-LOC113219401 | NC_037638.1 | 1812159 | 1812431 | 273 | - | 25 | 25 | 0 | 100 |
| gene-LOC102655618 | NC_037638.1 | 2205034 | 2207747 | 2714 | - | 4 | 4 | 0 | 100 |
| gene-egh | NC_037638.1 | 2223306 | 2226752 | 3447 | - | 21 | 21 | 0 | 100 |
| gene-LOC727419 | NC_037638.1 | 2241599 | 2242631 | 1033 | - | 8 | 8 | 0 | 100 |
| gene-LOC412000 | NC_037638.1 | 2479748 | 2483349 | 3602 | - | 4 | 4 | 0 | 100 |
| gene-LOC100577369 | NC_037638.1 | 4556117 | 4561037 | 4921 | + | 17 | 17 | 0 | 100 |
| gene-LOC414038 | NC_037638.1 | 4600849 | 4602360 | 1512 | - | 15 | 15 | 0 | 100 |
| gene-LOC726449 | NC_037638.1 | 5312656 | 5313472 | 817 | - | 8 | 8 | 0 | 100 |
| gene-LOC411408 | NC_037638.1 | 5647180 | 5649218 | 2039 | + | 6 | 6 | 0 | 100 |
| gene-LOC724585 | NC_037638.1 | 6533013 | 6535592 | 2580 | - | 10 | 10 | 0 | 100 |
| gene-LOC724455 | NC_037638.1 | 6550365 | 6554904 | 4540 | - | 9 | 9 | 0 | 100 |
| gene-LOC551974 | NC_037638.1 | 6571641 | 6572750 | 1110 | - | 4 | 4 | 0 | 100 |
| gene-LOC727538 | NC_037638.1 | 6611382 | 6614040 | 2659 | - | 5 | 5 | 0 | 100 |
| gene-LOC100577194 | NC_037638.1 | 7060376 | 7064019 | 3644 | - | 7 | 7 | 0 | 100 |
| gene-LOC551790 | NC_037638.1 | 10792754 | 10796244 | 3491 | - | 20 | 20 | 0 | 100 |
| gene-LOC107964502 | NC_037638.1 | 11124497 | 11125070 | 574 | + | 4 | 4 | 0 | 100 |
| gene-LOC409079 | NC_037638.1 | 11646550 | 11648373 | 1824 | + | 4 | 4 | 0 | 100 |
| gene-LOC102654495 | NC_037638.1 | 11678171 | 11679543 | 1373 | - | 5 | 5 | 0 | 100 |
| gene-LOC100577499 | NC_037638.1 | 12051755 | 12053930 | 2176 | - | 10 | 10 | 0 | 100 |
| gene-LOC551730 | NC_037638.1 | 12171035 | 12172877 | 1843 | + | 18 | 18 | 0 | 100 |
| gene-LOC724330 | NC_037638.1 | 12198744 | 12200333 | 1590 | - | 11 | 11 | 0 | 100 |
| gene-LOC552091 | NC_037638.1 | 14352996 | 14355214 | 2219 | - | 12 | 12 | 0 | 100 |
| gene-LOC409832 | NC_037638.1 | 14846347 | 14847647 | 1301 | - | 4 | 4 | 0 | 100 |

...

... only first 25 lines shown ...

Directory: RNK File: ranked-genes-bySiteDensity-Am_RE_fe.txt

| region_ID | rwidth | nbrsites | nbrper10kb | pmpersite | pmpernucl | pglink | seqnames | start | end | strand | coverage | numCs | numTs | prcntM |
|---|---|---|---|---|---|---|---|---|---|---|---|---|---|---|
| gene-LOC113219401 | 273 | 7 | 256.41 | 100 | 2.56 | https://www.ncbi.nlm.nih.gov/gene/?term=LOC113219401 | NC_037638.1 | 1812159 | 1812431 | - | 48 | 47 | 1 | 97.92 |
| gene-LOC113218557 | 2000 | 38 | 190 | 67.25 | 1.28 | https://www.ncbi.nlm.nih.gov/gene/?term=LOC113218557 | NC_037641.1 | 1779220 | 1781219 | - | 1126 | 594 | 532 | 52.75 |
| gene-LOC726419 | 1386 | 26 | 187.59 | 91.75 | 1.72 | https://www.ncbi.nlm.nih.gov/gene/?term=LOC726419 | NC_037639.1 | 15973484 | 15974869 | - | 308 | 202 | 106 | 65.58 |
| gene-LOC408391 | 1806 | 27 | 149.5 | 93.7 | 1.4 | https://www.ncbi.nlm.nih.gov/gene/?term=LOC408391 | NC_037646.1 | 7351184 | 7352989 | - | 298 | 239 | 59 | 80.2 |
| gene-LOC102656390 | 1760 | 24 | 136.36 | 59.41 | 0.81 | https://www.ncbi.nlm.nih.gov/gene/?term=LOC102656390 | NW_020555869.1 | 40445 | 42204 | - | 1370 | 281 | 1089 | 20.51 |
| gene-LOC725372 | 1843 | 25 | 135.65 | 97.17 | 1.32 | https://www.ncbi.nlm.nih.gov/gene/?term=LOC725372 | NC_037646.1 | 7764886 | 7766728 | - | 172 | 155 | 17 | 90.12 |
| gene-LOC102656501 | 3344 | 43 | 128.59 | 97.15 | 1.25 | https://www.ncbi.nlm.nih.gov/gene/?term=LOC102656501 | NC_037642.1 | 9865790 | 9869133 | - | 487 | 273 | 214 | 56.06 |
| gene-LOC100577929 | 2006 | 24 | 119.64 | 61.19 | 0.73 | https://www.ncbi.nlm.nih.gov/gene/?term=LOC100577929 | NW_020555870.1 | 33316 | 35321 | - | 1939 | 388 | 1551 | 20.01 |
| gene-LOC726270 | 1018 | 12 | 117.88 | 91.67 | 1.08 | https://www.ncbi.nlm.nih.gov/gene/?term=LOC726270 | NC_037647.1 | 5752275 | 5753292 | + | 78 | 68 | 10 | 87.18 |
| gene-LOC113219373 | 2604 | 30 | 115.21 | 61.93 | 0.71 | https://www.ncbi.nlm.nih.gov/gene/?term=LOC113219373 | NW_020555894.1 | 7811 | 10414 | - | 689 | 311 | 378 | 45.14 |
| gene-LOC408986 | 1391 | 16 | 115.03 | 98.61 | 1.13 | https://www.ncbi.nlm.nih.gov/gene/?term=LOC408986 | NC_037646.1 | 11559280 | 11560670 | + | 166 | 96 | 70 | 57.83 |
| gene-LOC102654498 | 849 | 8 | 94.23 | 100 | 0.94 | https://www.ncbi.nlm.nih.gov/gene/?term=LOC102654498 | NC_037646.1 | 1562620 | 1563468 | + | 42 | 42 | 0 | 100 |
| gene-LOC410280 | 2829 | 26 | 91.91 | 90.99 | 0.84 | https://www.ncbi.nlm.nih.gov/gene/?term=LOC410280 | NC_037648.1 | 16196034 | 16198862 | + | 566 | 207 | 359 | 36.57 |
| gene-LOC100576404 | 4488 | 40 | 89.13 | 95.51 | 0.85 | https://www.ncbi.nlm.nih.gov/gene/?term=LOC100576404 | NC_037642.1 | 12528666 | 12533153 | - | 2816 | 405 | 2411 | 14.38 |
| gene-LOC726957 | 2819 | 25 | 88.68 | 94.92 | 0.84 | https://www.ncbi.nlm.nih.gov/gene/?term=LOC726957 | NC_037642.1 | 13819098 | 13821916 | - | 330 | 207 | 123 | 62.73 |
| gene-LOC413271 | 2177 | 19 | 87.28 | 95.11 | 0.83 | https://www.ncbi.nlm.nih.gov/gene/?term=LOC413271 | NC_037647.1 | 3303140 | 3305316 | + | 210 | 109 | 101 | 51.9 |
| gene-LOC551550 | 2868 | 25 | 87.17 | 97.12 | 0.85 | https://www.ncbi.nlm.nih.gov/gene/?term=LOC551550 | NC_037642.1 | 479262 | 482129 | - | 142 | 136 | 6 | 95.77 |
| gene-LOC102656249 | 1514 | 13 | 85.87 | 96.52 | 0.83 | https://www.ncbi.nlm.nih.gov/gene/?term=LOC102656249 | NC_037642.1 | 9504339 | 9505852 | + | 225 | 68 | 157 | 30.22 |
| gene-LOC408853 | 1908 | 16 | 83.86 | 95.11 | 0.8 | https://www.ncbi.nlm.nih.gov/gene/?term=LOC408853 | NC_037642.1 | 11251428 | 11253335 | - | 312 | 147 | 165 | 47.12 |
| gene-LOC726028 | 1200 | 10 | 83.33 | 94.17 | 0.78 | https://www.ncbi.nlm.nih.gov/gene/?term=LOC726028 | NC_037645.1 | 10544803 | 10546002 | + | 89 | 70 | 19 | 78.65 |
| gene-LOC726706 | 2427 | 20 | 82.41 | 94.75 | 0.78 | https://www.ncbi.nlm.nih.gov/gene/?term=LOC726706 | NC_037652.1 | 3751884 | 3754310 | - | 184 | 158 | 26 | 85.87 |
| gene-LOC409500 | 1222 | 10 | 81.83 | 94.22 | 0.77 | https://www.ncbi.nlm.nih.gov/gene/?term=LOC409500 | NC_037642.1 | 13888315 | 13889536 | + | 439 | 138 | 301 | 31.44 |
| gene-LOC102654390 | 1226 | 10 | 81.57 | 92.93 | 0.76 | https://www.ncbi.nlm.nih.gov/gene/?term=LOC102654390 | NW_020555859.1 | 11520 | 12745 | + | 116 | 67 | 49 | 57.76 |
| gene-LOC408901 | 1641 | 13 | 79.22 | 91.92 | 0.73 | https://www.ncbi.nlm.nih.gov/gene/?term=LOC408901 | NC_037644.1 | 13783829 | 13785469 | - | 145 | 85 | 60 | 58.62 |

...

... only first 25 lines shown ...

Directory: RNK File: ranked-genes-bySiteDensity-Am_RE_te.txt

| region_ID | rwidth | nbrsites | nbrper10kb | pmpersite | pmpernucl | pglink | seqnames | start | end | strand | coverage | numCs | numTs | prcntM |
|---|---|---|---|---|---|---|---|---|---|---|---|---|---|---|
| gene-LOC726419 | 1386 | 31 | 223.67 | 96.35 | 2.16 | https://www.ncbi.nlm.nih.gov/gene/?term=LOC726419 | NC_037639.1 | 15973484 | 15974869 | - | 562 | 280 | 282 | 49.82 |
| gene-LOC113219401 | 273 | 6 | 219.78 | 100 | 2.2 | https://www.ncbi.nlm.nih.gov/gene/?term=LOC113219401 | NC_037638.1 | 1812159 | 1812431 | - | 25 | 25 | 0 | 100 |
| gene-LOC113218557 | 2000 | 42 | 210 | 61.62 | 1.29 | https://www.ncbi.nlm.nih.gov/gene/?term=LOC113218557 | NC_037641.1 | 1779220 | 1781219 | - | 1705 | 886 | 819 | 51.96 |
| gene-LOC102656390 | 1760 | 29 | 164.77 | 63.2 | 1.04 | https://www.ncbi.nlm.nih.gov/gene/?term=LOC102656390 | NW_020555869.1 | 40445 | 42204 | - | 1590 | 382 | 1208 | 24.03 |
| gene-LOC408391 | 1806 | 29 | 160.58 | 97.35 | 1.56 | https://www.ncbi.nlm.nih.gov/gene/?term=LOC408391 | NC_037646.1 | 7351184 | 7352989 | - | 258 | 195 | 63 | 75.58 |
| gene-LOC100577929 | 2006 | 28 | 139.58 | 63.74 | 0.89 | https://www.ncbi.nlm.nih.gov/gene/?term=LOC100577929 | NW_020555870.1 | 33316 | 35321 | - | 1966 | 449 | 1517 | 22.84 |
| gene-LOC726028 | 1200 | 16 | 133.33 | 94.98 | 1.27 | https://www.ncbi.nlm.nih.gov/gene/?term=LOC726028 | NC_037645.1 | 10544803 | 10546002 | + | 156 | 113 | 43 | 72.44 |
| gene-LOC725607 | 612 | 8 | 130.72 | 93.18 | 1.22 | https://www.ncbi.nlm.nih.gov/gene/?term=LOC725607 | NC_037651.1 | 6474420 | 6475031 | + | 59 | 53 | 6 | 89.83 |
| gene-LOC113219373 | 2604 | 34 | 130.57 | 48.51 | 0.63 | https://www.ncbi.nlm.nih.gov/gene/?term=LOC113219373 | NW_020555894.1 | 7811 | 10414 | - | 1258 | 469 | 789 | 37.28 |
| gene-Mir3758 | 80 | 1 | 125 | 90.91 | 1.14 | https://www.ncbi.nlm.nih.gov/gene/?term=Mir3758 | NC_037650.1 | 796222 | 796301 | - | 15 | 13 | 2 | 86.67 |
| gene-LOC408853 | 1908 | 21 | 110.06 | 87.24 | 0.96 | https://www.ncbi.nlm.nih.gov/gene/?term=LOC408853 | NC_037642.1 | 11251428 | 11253335 | - | 494 | 192 | 302 | 38.87 |
| gene-LOC410280 | 2829 | 31 | 109.58 | 94.74 | 1.04 | https://www.ncbi.nlm.nih.gov/gene/?term=LOC410280 | NC_037648.1 | 16196034 | 16198862 | + | 1082 | 247 | 835 | 22.83 |
| gene-LOC408986 | 1391 | 15 | 107.84 | 98.89 | 1.07 | https://www.ncbi.nlm.nih.gov/gene/?term=LOC408986 | NC_037646.1 | 11559280 | 11560670 | + | 331 | 110 | 221 | 33.23 |
| gene-LOC100576404 | 4488 | 48 | 106.95 | 93.47 | 1 | https://www.ncbi.nlm.nih.gov/gene/?term=LOC100576404 | NC_037642.1 | 12528666 | 12533153 | - | 2369 | 399 | 1970 | 16.84 |
| gene-Mir9865 | 94 | 1 | 106.38 | 100 | 1.06 | https://www.ncbi.nlm.nih.gov/gene/?term=Mir9865 | NC_037648.1 | 1463541 | 1463634 | - | 4 | 4 | 0 | 100 |
| gene-PDHB | 1983 | 21 | 105.9 | 97.46 | 1.03 | https://www.ncbi.nlm.nih.gov/gene/?term=PDHB | NC_037639.1 | 15991467 | 15993449 | + | 425 | 151 | 274 | 35.53 |
| gene-LOC102654884 | 1889 | 20 | 105.88 | 86.25 | 0.91 | https://www.ncbi.nlm.nih.gov/gene/?term=LOC102654884 | NW_020555846.1 | 19139 | 21027 | + | 324 | 147 | 177 | 45.37 |
| gene-LOC413271 | 2177 | 23 | 105.65 | 98.73 | 1.04 | https://www.ncbi.nlm.nih.gov/gene/?term=LOC413271 | NC_037647.1 | 3303140 | 3305316 | + | 297 | 145 | 152 | 48.82 |
| gene-LOC725102 | 1152 | 12 | 104.17 | 98.61 | 1.03 | https://www.ncbi.nlm.nih.gov/gene/?term=LOC725102 | NC_037641.1 | 11819133 | 11820284 | - | 337 | 74 | 263 | 21.96 |
| gene-LOC102656501 | 3344 | 33 | 98.68 | 97.83 | 0.97 | https://www.ncbi.nlm.nih.gov/gene/?term=LOC102656501 | NC_037642.1 | 9865790 | 9869133 | - | 558 | 247 | 311 | 44.27 |
| gene-LOC409500 | 1222 | 12 | 98.2 | 92.99 | 0.91 | https://www.ncbi.nlm.nih.gov/gene/?term=LOC409500 | NC_037642.1 | 13888315 | 13889536 | + | 490 | 100 | 390 | 20.41 |
| gene-LOC102655284 | 1637 | 16 | 97.74 | 96.61 | 0.94 | https://www.ncbi.nlm.nih.gov/gene/?term=LOC102655284 | NC_037643.1 | 15555577 | 15557213 | - | 158 | 115 | 43 | 72.78 |
| gene-LOC726706 | 2427 | 23 | 94.77 | 95.1 | 0.9 | https://www.ncbi.nlm.nih.gov/gene/?term=LOC726706 | NC_037652.1 | 3751884 | 3754310 | - | 165 | 144 | 21 | 87.27 |
| gene-LOC726337 | 1588 | 15 | 94.46 | 99.17 | 0.94 | https://www.ncbi.nlm.nih.gov/gene/?term=LOC726337 | NC_037642.1 | 11271301 | 11272888 | + | 216 | 102 | 114 | 47.22 |

...

... only first 25 lines shown ...

Directory: RNK File: ranked-promoters-byPrcntM-Am_RE_fe.txt

| region_ID | seqnames | start | end | width | strand | coverage | numCs | numTs | prcntM |
|---|---|---|---|---|---|---|---|---|---|
| gene-LOC551448 | NC_037638.1 | 29440 | 29939 | 500 | + | 4 | 4 | 0 | 100 |
| gene-LOC102654014 | NC_037638.1 | 10630803 | 10631302 | 500 | - | 4 | 4 | 0 | 100 |
| gene-LOC551785 | NC_037638.1 | 11731008 | 11731507 | 500 | - | 8 | 8 | 0 | 100 |
| gene-LOC724495 | NC_037638.1 | 12199970 | 12200469 | 500 | + | 4 | 4 | 0 | 100 |
| gene-LOC113219399 | NC_037638.1 | 14979923 | 14980422 | 500 | + | 4 | 4 | 0 | 100 |
| gene-LOC102655034 | NC_037638.1 | 20135801 | 20136300 | 500 | - | 8 | 8 | 0 | 100 |
| gene-LOC724850 | NC_037639.1 | 9678412 | 9678911 | 500 | + | 4 | 4 | 0 | 100 |
| gene-LOC551008 | NC_037639.1 | 11351106 | 11351605 | 500 | - | 9 | 9 | 0 | 100 |
| gene-LOC100576818 | NC_037639.1 | 14428536 | 14429035 | 500 | + | 4 | 4 | 0 | 100 |
| gene-LOC113218626 | NC_037640.1 | 8174128 | 8174627 | 500 | - | 4 | 4 | 0 | 100 |
| gene-LOC726826 | NC_037641.1 | 3345469 | 3345968 | 500 | - | 4 | 4 | 0 | 100 |
| gene-LOC552194 | NC_037641.1 | 12755131 | 12755630 | 500 | - | 5 | 5 | 0 | 100 |
| gene-LOC725267 | NC_037642.1 | 382084 | 382583 | 500 | - | 4 | 4 | 0 | 100 |
| gene-LOC113218789 | NC_037642.1 | 2997817 | 2998316 | 500 | + | 4 | 4 | 0 | 100 |
| gene-LOC726568 | NC_037642.1 | 6572032 | 6572531 | 500 | - | 4 | 4 | 0 | 100 |
| gene-LOC552607 | NC_037642.1 | 9535192 | 9535691 | 500 | + | 4 | 4 | 0 | 100 |
| gene-LOC100576101 | NC_037642.1 | 9552222 | 9552721 | 500 | + | 4 | 4 | 0 | 100 |
| gene-LOC113218816 | NC_037642.1 | 12958242 | 12958741 | 500 | - | 12 | 12 | 0 | 100 |
| gene-LOC408866 | NC_037642.1 | 13475230 | 13475729 | 500 | + | 6 | 6 | 0 | 100 |
| gene-Imd | NC_037642.1 | 13682786 | 13683285 | 500 | - | 7 | 7 | 0 | 100 |
| gene-LOC727501 | NC_037643.1 | 13312355 | 13312854 | 500 | - | 8 | 8 | 0 | 100 |
| gene-LOC551193 | NC_037644.1 | 1350072 | 1350571 | 500 | + | 9 | 9 | 0 | 100 |
| gene-LOC100578363 | NC_037645.1 | 1963287 | 1963786 | 500 | + | 9 | 9 | 0 | 100 |
| gene-LOC552596 | NC_037645.1 | 2246110 | 2246609 | 500 | + | 8 | 8 | 0 | 100 |

...
... only first 25 lines shown ...

Directory: RNK File: ranked-promoters-byPrcntM-Am_RE_te.txt

| region_ID | seqnames | start | end | width | strand | coverage | numCs | numTs | prcntM |
|---|---|---|---|---|---|---|---|---|---|
| gene-LOC724235 | NC_037638.1 | 12172654 | 12173153 | 500 | + | 5 | 5 | 0 | 100 |
| gene-LOC726591 | NC_037638.1 | 20564061 | 20564560 | 500 | - | 5 | 5 | 0 | 100 |
| gene-LOC725042 | NC_037638.1 | 21488758 | 21489257 | 500 | + | 4 | 4 | 0 | 100 |
| gene-LOC408708 | NC_037639.1 | 6615024 | 6615523 | 500 | - | 5 | 5 | 0 | 100 |
| gene-LOC113218590 | NC_037639.1 | 8162800 | 8163299 | 500 | - | 4 | 4 | 0 | 100 |
| gene-LOC724850 | NC_037639.1 | 9678412 | 9678911 | 500 | + | 5 | 5 | 0 | 100 |
| gene-LOC551008 | NC_037639.1 | 11351106 | 11351605 | 500 | - | 4 | 4 | 0 | 100 |
| gene-LOC100578691 | NC_037639.1 | 11351287 | 11351786 | 500 | + | 4 | 4 | 0 | 100 |
| gene-LOC107964045 | NC_037639.1 | 14436780 | 14437279 | 500 | + | 5 | 5 | 0 | 100 |
| gene-LOC102653600 | NC_037639.1 | 15540044 | 15540543 | 500 | - | 19 | 19 | 0 | 100 |
| gene-LOC410091 | NC_037640.1 | 4537987 | 4538486 | 500 | + | 4 | 4 | 0 | 100 |
| gene-LOC409346 | NC_037640.1 | 4903861 | 4904360 | 500 | - | 4 | 4 | 0 | 100 |
| gene-LOC411519 | NC_037641.1 | 3679148 | 3679647 | 500 | - | 5 | 5 | 0 | 100 |
| gene-LOC102655796 | NC_037641.1 | 5686954 | 5687453 | 500 | - | 4 | 4 | 0 | 100 |
| gene-LOC409596 | NC_037641.1 | 8880354 | 8880853 | 500 | - | 12 | 12 | 0 | 100 |
| gene-LOC725062 | NC_037641.1 | 11568622 | 11569121 | 500 | - | 4 | 4 | 0 | 100 |
| gene-LOC552731 | NC_037641.1 | 12732330 | 12732829 | 500 | - | 8 | 8 | 0 | 100 |
| gene-LOC551956 | NC_037642.1 | 797283 | 797782 | 500 | - | 10 | 10 | 0 | 100 |
| gene-LOC725170 | NC_037642.1 | 5471902 | 5472401 | 500 | - | 27 | 27 | 0 | 100 |
| gene-LOC408831 | NC_037642.1 | 6871391 | 6871890 | 500 | - | 5 | 5 | 0 | 100 |
| gene-LOC100578355 | NC_037642.1 | 7465773 | 7466272 | 500 | - | 4 | 4 | 0 | 100 |
| gene-LOC551548 | NC_037642.1 | 8518784 | 8519283 | 500 | - | 8 | 8 | 0 | 100 |
| gene-Coq7 | NC_037642.1 | 9904290 | 9904789 | 500 | + | 36 | 36 | 0 | 100 |
| gene-LOC412232 | NC_037642.1 | 11440583 | 11441082 | 500 | + | 6 | 6 | 0 | 100 |

...
... only first 25 lines shown ...

Directory: RNK File: ranked-promoters-bySiteDensity-Am_RE_fe.txt

| region_ID | rwidth | nbrsites | nbrper10kb | pmpersite | pmpernucl | pglink | seqnames | start | end | strand | coverage | numCs | numTs | prcntM |
|---|---|---|---|---|---|---|---|---|---|---|---|---|---|---|
| gene-LOC113219401 | 500 | 7 | 140 | 95.58 | 1.34 | https://www.ncbi.nlm.nih.gov/gene/?term=LOC113219401 | NC_037638.1 | 1812432 | 1812931 | - | 41 | 39 | 2 | 95.12 |
| gene-LOC408500 | 500 | 7 | 140 | 92.74 | 1.3 | https://www.ncbi.nlm.nih.gov/gene/?term=LOC408500 | NC_037651.1 | 10488842 | 10489341 | + | 130 | 41 | 89 | 31.54 |
| gene-LOC412289 | 500 | 7 | 140 | 94.05 | 1.32 | https://www.ncbi.nlm.nih.gov/gene/?term=LOC412289 | NC_037646.1 | 12339177 | 12339676 | - | 94 | 59 | 35 | 62.77 |
| gene-LOC410566 | 500 | 6 | 120 | 85 | 1.02 | https://www.ncbi.nlm.nih.gov/gene/?term=LOC410566 | NC_037651.1 | 10651890 | 10652389 | - | 87 | 37 | 50 | 42.53 |
| gene-LOC413683 | 500 | 5 | 100 | 93.33 | 0.93 | https://www.ncbi.nlm.nih.gov/gene/?term=LOC413683 | NC_037647.1 | 5753080 | 5753579 | + | 27 | 25 | 2 | 92.59 |
| gene-LOC102654261 | 500 | 4 | 80 | 96.43 | 0.77 | https://www.ncbi.nlm.nih.gov/gene/?term=LOC102654261 | NC_037652.1 | 5098331 | 5098830 | - | 23 | 22 | 1 | 95.65 |
| gene-LOC102656321 | 500 | 4 | 80 | 83.71 | 0.67 | https://www.ncbi.nlm.nih.gov/gene/?term=LOC102656321 | NW_020555869.1 | 7786 | 8285 | + | 1453 | 142 | 1311 | 9.77 |
| gene-LOC551338 | 500 | 4 | 80 | 97.22 | 0.78 | https://www.ncbi.nlm.nih.gov/gene/?term=LOC551338 | NC_037645.1 | 6576093 | 6576592 | + | 28 | 27 | 1 | 96.43 |
| gene-LOC113219036 | 500 | 3 | 60 | 82.21 | 0.49 | https://www.ncbi.nlm.nih.gov/gene/?term=LOC113219036 | NC_037647.1 | 15047 | 15546 | - | 54 | 46 | 8 | 85.19 |
| gene-LOC412907 | 500 | 3 | 60 | 81.94 | 0.49 | https://www.ncbi.nlm.nih.gov/gene/?term=LOC412907 | NC_037652.1 | 5989340 | 5989839 | - | 25 | 20 | 5 | 80 |
| gene-LOC551451 | 500 | 3 | 60 | 100 | 0.6 | https://www.ncbi.nlm.nih.gov/gene/?term=LOC551451 | NC_037638.1 | 20102989 | 20103488 | - | 18 | 17 | 1 | 94.44 |
| gene-LOC726550 | 500 | 3 | 60 | 87.18 | 0.52 | https://www.ncbi.nlm.nih.gov/gene/?term=LOC726550 | NC_037641.1 | 12159286 | 12159785 | + | 90 | 17 | 73 | 18.89 |
| gene-LOC727151 | 500 | 3 | 60 | 100 | 0.6 | https://www.ncbi.nlm.nih.gov/gene/?term=LOC727151 | NC_037651.1 | 9854645 | 9855144 | + | 28 | 28 | 0 | 100 |
| gene-Mir9865 | 500 | 3 | 60 | 100 | 0.6 | https://www.ncbi.nlm.nih.gov/gene/?term=Mir9865 | NC_037648.1 | 1463635 | 1464134 | - | 12 | 12 | 0 | 100 |
| gene-Coq7 | 500 | 2 | 40 | 100 | 0.4 | https://www.ncbi.nlm.nih.gov/gene/?term=Coq7 | NC_037642.1 | 9904290 | 9904789 | + | 16 | 15 | 1 | 93.75 |
| gene-LOC100577699 | 500 | 2 | 40 | 100 | 0.4 | https://www.ncbi.nlm.nih.gov/gene/?term=LOC100577699 | NC_037648.1 | 10530337 | 10530836 | + | 8 | 8 | 0 | 100 |
| gene-LOC100578363 | 500 | 2 | 40 | 100 | 0.4 | https://www.ncbi.nlm.nih.gov/gene/?term=LOC100578363 | NC_037645.1 | 1963287 | 1963786 | + | 9 | 9 | 0 | 100 |
| gene-LOC100578691 | 500 | 2 | 40 | 100 | 0.4 | https://www.ncbi.nlm.nih.gov/gene/?term=LOC100578691 | NC_037639.1 | 11351287 | 11351786 | + | 13 | 9 | 4 | 69.23 |
| gene-LOC102655034 | 500 | 2 | 40 | 100 | 0.4 | https://www.ncbi.nlm.nih.gov/gene/?term=LOC102655034 | NC_037638.1 | 20135801 | 20136300 | - | 8 | 8 | 0 | 100 |
| gene-LOC102655138 | 500 | 2 | 40 | 100 | 0.4 | https://www.ncbi.nlm.nih.gov/gene/?term=LOC102655138 | NC_037645.1 | 6985360 | 6985859 | + | 8 | 8 | 0 | 100 |
| gene-LOC102655648 | 500 | 2 | 40 | 100 | 0.4 | https://www.ncbi.nlm.nih.gov/gene/?term=LOC102655648 | NC_037647.1 | 354951 | 355450 | - | 10 | 10 | 0 | 100 |
| gene-LOC113218816 | 500 | 2 | 40 | 100 | 0.4 | https://www.ncbi.nlm.nih.gov/gene/?term=LOC113218816 | NC_037642.1 | 12958242 | 12958741 | - | 12 | 12 | 0 | 100 |
| gene-LOC113219178 | 500 | 2 | 40 | 100 | 0.4 | https://www.ncbi.nlm.nih.gov/gene/?term=LOC113219178 | NC_037649.1 | 5229514 | 5230013 | - | 8 | 8 | 0 | 100 |
| gene-LOC113219199 | 500 | 2 | 40 | 100 | 0.4 | https://www.ncbi.nlm.nih.gov/gene/?term=LOC113219199 | NC_037650.1 | 5955044 | 5955543 | - | 12 | 12 | 0 | 100 |
| gene-LOC409088 | 500 | 2 | 40 | 100 | 0.4 | https://www.ncbi.nlm.nih.gov/gene/?term=LOC409088 | NC_037639.1 | 15260436 | 15260935 | - | 15 | 12 | 3 | 80 |
| gene-LOC411093 | 500 | 2 | 40 | 91.67 | 0.37 | https://www.ncbi.nlm.nih.gov/gene/?term=LOC411093 | NC_037642.1 | 8842778 | 8843277 | + | 90 | 9 | 81 | 10 |
| gene-LOC412851 | 500 | 2 | 40 | 100 | 0.4 | https://www.ncbi.nlm.nih.gov/gene/?term=LOC412851 | NC_037640.1 | 4598201 | 4598700 | - | 121 | 8 | 113 | 6.61 |
| gene-LOC551008 | 500 | 2 | 40 | 100 | 0.4 | https://www.ncbi.nlm.nih.gov/gene/?term=LOC551008 | NC_037639.1 | 11351106 | 11351605 | - | 9 | 9 | 0 | 100 |
| gene-LOC551193 | 500 | 2 | 40 | 100 | 0.4 | https://www.ncbi.nlm.nih.gov/gene/?term=LOC551193 | NC_037644.1 | 1350072 | 1350571 | + | 9 | 9 | 0 | 100 |
| gene-LOC551785 | 500 | 2 | 40 | 100 | 0.4 | https://www.ncbi.nlm.nih.gov/gene/?term=LOC551785 | NC_037638.1 | 11731008 | 11731507 | - | 8 | 8 | 0 | 100 |
| gene-LOC552508 | 500 | 2 | 40 | 100 | 0.4 | https://www.ncbi.nlm.nih.gov/gene/?term=LOC552508 | NC_037651.1 | 6846579 | 6847078 | - | 22 | 16 | 6 | 72.73 |
| gene-LOC552574 | 500 | 2 | 40 | 100 | 0.4 | https://www.ncbi.nlm.nih.gov/gene/?term=LOC552574 | NC_037642.1 | 7404503 | 7405002 | - | 14 | 12 | 2 | 85.71 |
| gene-LOC552603 | 500 | 2 | 40 | 100 | 0.4 | https://www.ncbi.nlm.nih.gov/gene/?term=LOC552603 | NC_037640.1 | 4469685 | 4470184 | + | 12 | 10 | 2 | 83.33 |
| gene-LOC724388 | 500 | 2 | 40 | 100 | 0.4 | https://www.ncbi.nlm.nih.gov/gene/?term=LOC724388 | NC_037642.1 | 11379858 | 11380357 | - | 21 | 18 | 3 | 85.71 |
| gene-LOC725170 | 500 | 2 | 40 | 91.67 | 0.37 | https://www.ncbi.nlm.nih.gov/gene/?term=LOC725170 | NC_037642.1 | 5471902 | 5472401 | - | 10 | 9 | 1 | 90 |
| gene-LOC726521 | 500 | 2 | 40 | 100 | 0.4 | https://www.ncbi.nlm.nih.gov/gene/?term=LOC726521 | NC_037653.1 | 1302284 | 1302783 | - | 12 | 11 | 1 | 91.67 |
| gene-LOC727501 | 500 | 2 | 40 | 100 | 0.4 | https://www.ncbi.nlm.nih.gov/gene/?term=LOC727501 | NC_037643.1 | 13312355 | 13312854 | - | 8 | 8 | 0 | 100 |
| gene-Mir3758 | 500 | 2 | 40 | 87.5 | 0.35 | https://www.ncbi.nlm.nih.gov/gene/?term=Mir3758 | NC_037650.1 | 796302 | 796801 | - | 23 | 17 | 6 | 73.91 |
| gene-Imd | 500 | 1 | 20 | 100 | 0.2 | https://www.ncbi.nlm.nih.gov/gene/?term=Imd | NC_037642.1 | 13682786 | 13683285 | - | 7 | 7 | 0 | 100 |
| gene-LOC100576101 | 500 | 1 | 20 | 100 | 0.2 | https://www.ncbi.nlm.nih.gov/gene/?term=LOC100576101 | NC_037642.1 | 9552222 | 9552721 | + | 4 | 4 | 0 | 100 |
| gene-LOC100576155 | 500 | 1 | 20 | 100 | 0.2 | https://www.ncbi.nlm.nih.gov/gene/?term=LOC100576155 | NC_037642.1 | 544461 | 544960 | + | 12 | 7 | 5 | 58.33 |
| gene-LOC100576400 | 500 | 1 | 20 | 87.5 | 0.17 | https://www.ncbi.nlm.nih.gov/gene/?term=LOC100576400 | NC_037643.1 | 13196766 | 13197265 | + | 16 | 11 | 5 | 68.75 |
| gene-LOC100576608 | 500 | 1 | 20 | 85.71 | 0.17 | https://www.ncbi.nlm.nih.gov/gene/?term=LOC100576608 | NC_037651.1 | 10476098 | 10476597 | + | 7 | 6 | 1 | 85.71 |
| gene-LOC100576818 | 500 | 1 | 20 | 100 | 0.2 | https://www.ncbi.nlm.nih.gov/gene/?term=LOC100576818 | NC_037639.1 | 14428536 | 14429035 | + | 4 | 4 | 0 | 100 |
| gene-LOC100577702 | 500 | 1 | 20 | 83.33 | 0.17 | https://www.ncbi.nlm.nih.gov/gene/?term=LOC100577702 | NC_037642.1 | 11233386 | 11233885 | - | 20 | 12 | 8 | 60 |
| gene-LOC100578212 | 500 | 1 | 20 | 100 | 0.2 | https://www.ncbi.nlm.nih.gov/gene/?term=LOC100578212 | NC_037646.1 | 12334987 | 12335486 | + | 7 | 7 | 0 | 100 |
| gene-LOC102653824 | 500 | 1 | 20 | 100 | 0.2 | https://www.ncbi.nlm.nih.gov/gene/?term=LOC102653824 | NC_037648.1 | 14282631 | 14283130 | - | 22 | 4 | 18 | 18.18 |
| gene-LOC102654014 | 500 | 1 | 20 | 100 | 0.2 | https://www.ncbi.nlm.nih.gov/gene/?term=LOC102654014 | NC_037638.1 | 10630803 | 10631302 | - | 4 | 4 | 0 | 100 |
| gene-LOC102654994 | 500 | 1 | 20 | 100 | 0.2 | https://www.ncbi.nlm.nih.gov/gene/?term=LOC102654994 | NC_037642.1 | 13877729 | 13878228 | - | 55 | 18 | 37 | 32.73 |
| gene-LOC102655461 | 500 | 1 | 20 | 100 | 0.2 | https://www.ncbi.nlm.nih.gov/gene/?term=LOC102655461 | NC_037652.1 | 6577021 | 6577520 | - | 5 | 5 | 0 | 100 |
| gene-LOC102656914 | 500 | 1 | 20 | 100 | 0.2 | https://www.ncbi.nlm.nih.gov/gene/?term=LOC102656914 | NC_037648.1 | 5027280 | 5027779 | - | 4 | 4 | 0 | 100 |
| gene-LOC107964222 | 500 | 1 | 20 | 100 | 0.2 | https://www.ncbi.nlm.nih.gov/gene/?term=LOC107964222 | NC_037640.1 | 6085859 | 6086358 | - | 12 | 10 | 2 | 83.33 |
| gene-LOC107964260 | 500 | 1 | 20 | 75 | 0.15 | https://www.ncbi.nlm.nih.gov/gene/?term=LOC107964260 | NC_037641.1 | 11557856 | 11558355 | + | 8 | 6 | 2 | 75 |
| gene-LOC107964531 | 500 | 1 | 20 | 83.33 | 0.17 | https://www.ncbi.nlm.nih.gov/gene/?term=LOC107964531 | NC_037643.1 | 16276897 | 16277396 | - | 6 | 5 | 1 | 83.33 |
| gene-LOC107965260 | 500 | 1 | 20 | 100 | 0.2 | https://www.ncbi.nlm.nih.gov/gene/?term=LOC107965260 | NC_037648.1 | 16099189 | 16099688 | + | 4 | 4 | 0 | 100 |
| gene-LOC107965368 | 500 | 1 | 20 | 71.43 | 0.14 | https://www.ncbi.nlm.nih.gov/gene/?term=LOC107965368 | NC_037649.1 | 2396713 | 2397212 | + | 59 | 6 | 53 | 10.17 |
| gene-LOC113218553 | 500 | 1 | 20 | 90.91 | 0.18 | https://www.ncbi.nlm.nih.gov/gene/?term=LOC113218553 | NC_037639.1 | 12452945 | 12453444 | - | 25 | 16 | 9 | 64 |
| gene-LOC113218626 | 500 | 1 | 20 | 100 | 0.2 | https://www.ncbi.nlm.nih.gov/gene/?term=LOC113218626 | NC_037640.1 | 8174128 | 8174627 | - | 4 | 4 | 0 | 100 |
| gene-LOC113218704 | 500 | 1 | 20 | 92.31 | 0.18 | https://www.ncbi.nlm.nih.gov/gene/?term=LOC113218704 | NC_037640.1 | 6890661 | 6891160 | - | 40 | 16 | 24 | 40 |
| gene-LOC113218789 | 500 | 1 | 20 | 100 | 0.2 | https://www.ncbi.nlm.nih.gov/gene/?term=LOC113218789 | NC_037642.1 | 2997817 | 2998316 | + | 4 | 4 | 0 | 100 |

| | | | | | | | | | | | | | | |
|---|---|---|---|---|---|---|---|---|---|---|---|---|---|---|
| gene-LOC113218977 | 500 | 1 | 20 | 100 | 0.2 | https://www.ncbi.nlm.nih.gov/gene/?term=LOC113218977 | NC_037646.1 | 1694180 | 1694679 | - | 8 | 6 | 2 | 75 |
| gene-LOC113219080 | 500 | 1 | 20 | 100 | 0.2 | https://www.ncbi.nlm.nih.gov/gene/?term=LOC113219080 | NC_037648.1 | 15378785 | 15379284 | - | 5 | 5 | 0 | 100 |
| gene-LOC113219395 | 500 | 1 | 20 | 100 | 0.2 | https://www.ncbi.nlm.nih.gov/gene/?term=LOC113219395 | NC_037638.1 | 1680961 | 1681460 | - | 8 | 5 | 3 | 62.5 |
| gene-LOC113219399 | 500 | 1 | 20 | 100 | 0.2 | https://www.ncbi.nlm.nih.gov/gene/?term=LOC113219399 | NC_037638.1 | 14979923 | 14980422 | + | 4 | 4 | 0 | 100 |
| gene-LOC408860 | 500 | 1 | 20 | 100 | 0.2 | https://www.ncbi.nlm.nih.gov/gene/?term=LOC408860 | NC_037642.1 | 12216212 | 12216711 | - | 12 | 4 | 8 | 33.33 |
| gene-LOC408866 | 500 | 1 | 20 | 100 | 0.2 | https://www.ncbi.nlm.nih.gov/gene/?term=LOC408866 | NC_037642.1 | 13475230 | 13475729 | + | 6 | 6 | 0 | 100 |
| gene-LOC409017 | 500 | 1 | 20 | 100 | 0.2 | https://www.ncbi.nlm.nih.gov/gene/?term=LOC409017 | NC_037646.1 | 8278485 | 8278984 | + | 4 | 4 | 0 | 100 |
| gene-LOC409056 | 500 | 1 | 20 | 100 | 0.2 | https://www.ncbi.nlm.nih.gov/gene/?term=LOC409056 | NC_037647.1 | 4836722 | 4837221 | + | 4 | 4 | 0 | 100 |
| gene-LOC409153 | 500 | 1 | 20 | 100 | 0.2 | https://www.ncbi.nlm.nih.gov/gene/?term=LOC409153 | NC_037642.1 | 12636357 | 12636856 | + | 12 | 4 | 8 | 33.33 |
| gene-LOC409443 | 500 | 1 | 20 | 100 | 0.2 | https://www.ncbi.nlm.nih.gov/gene/?term=LOC409443 | NC_037640.1 | 4424618 | 4425117 | - | 9 | 5 | 4 | 55.56 |
| gene-LOC409610 | 500 | 1 | 20 | 100 | 0.2 | https://www.ncbi.nlm.nih.gov/gene/?term=LOC409610 | NC_037652.1 | 7880975 | 7881474 | + | 14 | 7 | 7 | 50 |
| gene-LOC409614 | 500 | 1 | 20 | 71.43 | 0.14 | https://www.ncbi.nlm.nih.gov/gene/?term=LOC409614 | NC_037651.1 | 10573988 | 10574487 | + | 7 | 5 | 2 | 71.43 |
| gene-LOC409983 | 500 | 1 | 20 | 100 | 0.2 | https://www.ncbi.nlm.nih.gov/gene/?term=LOC409983 | NC_037641.1 | 12676653 | 12677152 | + | 18 | 5 | 13 | 27.78 |
| gene-LOC410402 | 500 | 1 | 20 | 100 | 0.2 | https://www.ncbi.nlm.nih.gov/gene/?term=LOC410402 | NC_037650.1 | 2451545 | 2452044 | + | 4 | 4 | 0 | 100 |
| gene-LOC410617 | 500 | 1 | 20 | 100 | 0.2 | https://www.ncbi.nlm.nih.gov/gene/?term=LOC410617 | NC_037652.1 | 6229095 | 6229594 | - | 13 | 4 | 9 | 30.77 |
| gene-LOC411640 | 500 | 1 | 20 | 83.33 | 0.17 | https://www.ncbi.nlm.nih.gov/gene/?term=LOC411640 | NC_037641.1 | 4817094 | 4817593 | - | 13 | 6 | 7 | 46.15 |
| gene-LOC412065 | 500 | 1 | 20 | 65 | 0.13 | https://www.ncbi.nlm.nih.gov/gene/?term=LOC412065 | NC_037649.1 | 3084955 | 3085454 | + | 204 | 24 | 180 | 11.76 |
| gene-LOC412256 | 500 | 1 | 20 | 100 | 0.2 | https://www.ncbi.nlm.nih.gov/gene/?term=LOC412256 | NC_037640.1 | 10222330 | 10222829 | + | 16 | 4 | 12 | 25 |
| gene-LOC412827 | 500 | 1 | 20 | 100 | 0.2 | https://www.ncbi.nlm.nih.gov/gene/?term=LOC412827 | NC_037645.1 | 4209689 | 4210188 | - | 5 | 5 | 0 | 100 |
| gene-LOC412972 | 500 | 1 | 20 | 100 | 0.2 | https://www.ncbi.nlm.nih.gov/gene/?term=LOC412972 | NC_037648.1 | 8043010 | 8043509 | + | 4 | 4 | 0 | 100 |
| gene-LOC413813 | 500 | 1 | 20 | 100 | 0.2 | https://www.ncbi.nlm.nih.gov/gene/?term=LOC413813 | NC_037638.1 | 7052487 | 7052986 | + | 8 | 4 | 4 | 50 |
| gene-LOC494508 | 500 | 1 | 20 | 100 | 0.2 | https://www.ncbi.nlm.nih.gov/gene/?term=LOC494508 | NC_037648.1 | 10532625 | 10533124 | - | 5 | 5 | 0 | 100 |
| gene-LOC551448 | 500 | 1 | 20 | 100 | 0.2 | https://www.ncbi.nlm.nih.gov/gene/?term=LOC551448 | NC_037638.1 | 29440 | 29939 | + | 4 | 4 | 0 | 100 |
| gene-LOC551814 | 500 | 1 | 20 | 75 | 0.15 | https://www.ncbi.nlm.nih.gov/gene/?term=LOC551814 | NC_037647.1 | 4839082 | 4839581 | + | 8 | 6 | 2 | 75 |
| gene-LOC552046 | 500 | 1 | 20 | 100 | 0.2 | https://www.ncbi.nlm.nih.gov/gene/?term=LOC552046 | NC_037643.1 | 5154685 | 5155184 | - | 31 | 7 | 24 | 22.58 |
| gene-LOC552194 | 500 | 1 | 20 | 100 | 0.2 | https://www.ncbi.nlm.nih.gov/gene/?term=LOC552194 | NC_037641.1 | 12755131 | 12755630 | - | 5 | 5 | 0 | 100 |
| gene-LOC552347 | 500 | 1 | 20 | 100 | 0.2 | https://www.ncbi.nlm.nih.gov/gene/?term=LOC552347 | NC_037649.1 | 7696058 | 7696557 | + | 15 | 7 | 8 | 46.67 |
| gene-LOC552494 | 500 | 1 | 20 | 100 | 0.2 | https://www.ncbi.nlm.nih.gov/gene/?term=LOC552494 | NC_037645.1 | 2615624 | 2616123 | + | 10 | 4 | 6 | 40 |
| gene-LOC552596 | 500 | 1 | 20 | 100 | 0.2 | https://www.ncbi.nlm.nih.gov/gene/?term=LOC552596 | NC_037645.1 | 2246110 | 2246609 | + | 8 | 8 | 0 | 100 |
| gene-LOC552607 | 500 | 1 | 20 | 100 | 0.2 | https://www.ncbi.nlm.nih.gov/gene/?term=LOC552607 | NC_037642.1 | 9535192 | 9535691 | + | 4 | 4 | 0 | 100 |
| gene-LOC724423 | 500 | 1 | 20 | 83.33 | 0.17 | https://www.ncbi.nlm.nih.gov/gene/?term=LOC724423 | NC_037653.1 | 6399796 | 6400295 | - | 6 | 5 | 1 | 83.33 |
| gene-LOC724495 | 500 | 1 | 20 | 100 | 0.2 | https://www.ncbi.nlm.nih.gov/gene/?term=LOC724495 | NC_037638.1 | 12199970 | 12200469 | + | 4 | 4 | 0 | 100 |
| gene-LOC724850 | 500 | 1 | 20 | 100 | 0.2 | https://www.ncbi.nlm.nih.gov/gene/?term=LOC724850 | NC_037639.1 | 9678412 | 9678911 | + | 4 | 4 | 0 | 100 |
| gene-LOC724878 | 500 | 1 | 20 | 100 | 0.2 | https://www.ncbi.nlm.nih.gov/gene/?term=LOC724878 | NC_037648.1 | 16005777 | 16006276 | + | 4 | 4 | 0 | 100 |
| gene-LOC725093 | 500 | 1 | 20 | 100 | 0.2 | https://www.ncbi.nlm.nih.gov/gene/?term=LOC725093 | NC_037642.1 | 2944866 | 2945365 | - | 47 | 6 | 41 | 12.77 |
| gene-LOC725267 | 500 | 1 | 20 | 100 | 0.2 | https://www.ncbi.nlm.nih.gov/gene/?term=LOC725267 | NC_037642.1 | 382084 | 382583 | - | 4 | 4 | 0 | 100 |
| gene-LOC725948 | 500 | 1 | 20 | 100 | 0.2 | https://www.ncbi.nlm.nih.gov/gene/?term=LOC725948 | NC_037647.1 | 4904215 | 4904714 | - | 9 | 4 | 5 | 44.44 |
| gene-LOC726063 | 500 | 1 | 20 | 85.71 | 0.17 | https://www.ncbi.nlm.nih.gov/gene/?term=LOC726063 | NC_037647.1 | 6316824 | 6317323 | + | 7 | 6 | 1 | 85.71 |
| gene-LOC726568 | 500 | 1 | 20 | 100 | 0.2 | https://www.ncbi.nlm.nih.gov/gene/?term=LOC726568 | NC_037642.1 | 6572032 | 6572531 | - | 4 | 4 | 0 | 100 |
| gene-LOC726826 | 500 | 1 | 20 | 100 | 0.2 | https://www.ncbi.nlm.nih.gov/gene/?term=LOC726826 | NC_037641.1 | 3345469 | 3345968 | - | 4 | 4 | 0 | 100 |
| gene-LOC726879 | 500 | 1 | 20 | 100 | 0.2 | https://www.ncbi.nlm.nih.gov/gene/?term=LOC726879 | NC_037642.1 | 13725907 | 13726406 | + | 95 | 6 | 89 | 6.32 |
| gene-LOC727014 | 500 | 1 | 20 | 100 | 0.2 | https://www.ncbi.nlm.nih.gov/gene/?term=LOC727014 | NC_037642.1 | 13860538 | 13861037 | + | 8 | 4 | 4 | 50 |
| gene-LOC727030 | 500 | 1 | 20 | 85.71 | 0.17 | https://www.ncbi.nlm.nih.gov/gene/?term=LOC727030 | NC_037645.1 | 1961022 | 1961521 | - | 7 | 6 | 1 | 85.71 |
| gene-LOC727254 | 500 | 1 | 20 | 100 | 0.2 | https://www.ncbi.nlm.nih.gov/gene/?term=LOC727254 | NC_037651.1 | 10057690 | 10058189 | + | 4 | 4 | 0 | 100 |
| gene-Lop2 | 500 | 1 | 20 | 100 | 0.2 | https://www.ncbi.nlm.nih.gov/gene/?term=Lop2 | NC_037652.1 | 7419330 | 7419829 | - | 5 | 5 | 0 | 100 |
| gene-TRNAI-AAU-3 | 500 | 1 | 20 | 100 | 0.2 | https://www.ncbi.nlm.nih.gov/gene/?term=TRNAI-AAU | NC_037648.1 | 15654947 | 15655446 | - | 4 | 4 | 0 | 100 |

Directory: RNK File: ranked-promoters-bySiteDensity-Am_RE_te.txt

| region_ID | rwidth | nbrsites | nbrper10kb | pmpersite | pmpernucl | pglink | seqnames | start | end | strand | coverage | numCs | numTs | prcntM |
|---|---|---|---|---|---|---|---|---|---|---|---|---|---|---|
| gene-LOC412289 | 500 | 12 | 240 | 98.96 | 2.38 | https://www.ncbi.nlm.nih.gov/gene/?term=LOC412289 | NC_037646.1 | 12339177 | 12339676 | - | 121 | 84 | 37 | 69.42 |
| gene-LOC408382 | 500 | 8 | 160 | 95.04 | 1.52 | https://www.ncbi.nlm.nih.gov/gene/?term=LOC408382 | NC_037648.1 | 664817 | 665316 | - | 64 | 60 | 4 | 93.75 |
| gene-LOC410566 | 500 | 7 | 140 | 78.48 | 1.1 | https://www.ncbi.nlm.nih.gov/gene/?term=LOC410566 | NC_037651.1 | 10651890 | 10652389 | - | 99 | 45 | 54 | 45.45 |
| gene-Coq7 | 500 | 6 | 120 | 100 | 1.2 | https://www.ncbi.nlm.nih.gov/gene/?term=Coq7 | NC_037642.1 | 9904290 | 9904789 | + | 36 | 36 | 0 | 100 |
| gene-LOC113219280 | 500 | 6 | 120 | 100 | 1.2 | https://www.ncbi.nlm.nih.gov/gene/?term=LOC113219280 | NC_037652.1 | 2284312 | 2284811 | + | 37 | 34 | 3 | 91.89 |
| gene-LOC408500 | 500 | 6 | 120 | 100 | 1.2 | https://www.ncbi.nlm.nih.gov/gene/?term=LOC408500 | NC_037651.1 | 10488842 | 10489341 | + | 53 | 33 | 20 | 62.26 |
| gene-LOC411093 | 500 | 6 | 120 | 100 | 1.2 | https://www.ncbi.nlm.nih.gov/gene/?term=LOC411093 | NC_037642.1 | 8842778 | 8843277 | + | 88 | 30 | 58 | 34.09 |
| gene-LOC551338 | 500 | 6 | 120 | 95.14 | 1.14 | https://www.ncbi.nlm.nih.gov/gene/?term=LOC551338 | NC_037645.1 | 6576093 | 6576592 | + | 52 | 45 | 7 | 86.54 |
| gene-LOC727151 | 500 | 5 | 100 | 95 | 0.95 | https://www.ncbi.nlm.nih.gov/gene/?term=LOC727151 | NC_037651.1 | 9854645 | 9855144 | + | 37 | 35 | 2 | 94.59 |
| gene-LOC102653600 | 500 | 4 | 80 | 100 | 0.8 | https://www.ncbi.nlm.nih.gov/gene/?term=LOC102653600 | NC_037639.1 | 15540044 | 15540543 | - | 19 | 19 | 0 | 100 |
| gene-LOC102656321 | 500 | 4 | 80 | 79.41 | 0.64 | https://www.ncbi.nlm.nih.gov/gene/?term=LOC102656321 | NW_020555869.1 | 7786 | 8285 | + | 1948 | 131 | 1817 | 6.72 |
| gene-LOC113219022 | 500 | 4 | 80 | 83.84 | 0.67 | https://www.ncbi.nlm.nih.gov/gene/?term=LOC113219022 | NC_037646.1 | 7394724 | 7395223 | - | 36 | 30 | 6 | 83.33 |
| gene-LOC113219401 | 500 | 4 | 80 | 100 | 0.8 | https://www.ncbi.nlm.nih.gov/gene/?term=LOC113219401 | NC_037638.1 | 1812432 | 1812931 | - | 22 | 21 | 1 | 95.45 |
| gene-LOC724731 | 500 | 4 | 80 | 100 | 0.8 | https://www.ncbi.nlm.nih.gov/gene/?term=LOC724731 | NC_037651.1 | 8201763 | 8202262 | + | 19 | 19 | 0 | 100 |
| gene-LOC725170 | 500 | 4 | 80 | 100 | 0.8 | https://www.ncbi.nlm.nih.gov/gene/?term=LOC725170 | NC_037642.1 | 5471902 | 5472401 | - | 27 | 27 | 0 | 100 |
| gene-Mir9865 | 500 | 4 | 80 | 100 | 0.8 | https://www.ncbi.nlm.nih.gov/gene/?term=Mir9865 | NC_037648.1 | 1463635 | 1464134 | - | 30 | 28 | 2 | 93.33 |
| gene-LOC100576400 | 500 | 3 | 60 | 88.89 | 0.53 | https://www.ncbi.nlm.nih.gov/gene/?term=LOC100576400 | NC_037643.1 | 13196766 | 13197265 | + | 30 | 23 | 7 | 76.67 |
| gene-LOC100577516 | 500 | 3 | 60 | 100 | 0.6 | https://www.ncbi.nlm.nih.gov/gene/?term=LOC100577516 | NC_037653.1 | 6389020 | 6389519 | - | 17 | 17 | 0 | 100 |
| gene-LOC100577687 | 500 | 3 | 60 | 94.44 | 0.57 | https://www.ncbi.nlm.nih.gov/gene/?term=LOC100577687 | NC_037648.1 | 14172650 | 14173149 | + | 29 | 15 | 14 | 51.72 |
| gene-LOC102656914 | 500 | 3 | 60 | 100 | 0.6 | https://www.ncbi.nlm.nih.gov/gene/?term=LOC102656914 | NC_037648.1 | 5027280 | 5027779 | - | 18 | 18 | 0 | 100 |
| gene-LOC113219319 | 500 | 3 | 60 | 100 | 0.6 | https://www.ncbi.nlm.nih.gov/gene/?term=LOC113219319 | NC_037638.1 | 12309993 | 12310492 | + | 16 | 14 | 2 | 87.5 |
| gene-LOC409596 | 500 | 3 | 60 | 100 | 0.6 | https://www.ncbi.nlm.nih.gov/gene/?term=LOC409596 | NC_037641.1 | 8880354 | 8880853 | - | 12 | 12 | 0 | 100 |
| gene-LOC412907 | 500 | 3 | 60 | 88.5 | 0.53 | https://www.ncbi.nlm.nih.gov/gene/?term=LOC412907 | NC_037652.1 | 5989340 | 5989839 | - | 27 | 24 | 3 | 88.89 |
| gene-LOC413683 | 500 | 3 | 60 | 100 | 0.6 | https://www.ncbi.nlm.nih.gov/gene/?term=LOC413683 | NC_037647.1 | 5753080 | 5753579 | + | 13 | 13 | 0 | 100 |
| gene-LOC552046 | 500 | 3 | 60 | 100 | 0.6 | https://www.ncbi.nlm.nih.gov/gene/?term=LOC552046 | NC_037643.1 | 5154685 | 5155184 | - | 28 | 18 | 10 | 64.29 |
| gene-LOC724791 | 500 | 3 | 60 | 100 | 0.6 | https://www.ncbi.nlm.nih.gov/gene/?term=LOC724791 | NC_037648.1 | 13462249 | 13462748 | + | 23 | 17 | 6 | 73.91 |
| gene-LOC726815 | 500 | 3 | 60 | 95.24 | 0.57 | https://www.ncbi.nlm.nih.gov/gene/?term=LOC726815 | NC_037650.1 | 6855686 | 6856185 | + | 23 | 14 | 9 | 60.87 |
| gene-LOC726879 | 500 | 3 | 60 | 100 | 0.6 | https://www.ncbi.nlm.nih.gov/gene/?term=LOC726879 | NC_037642.1 | 13725907 | 13726406 | + | 148 | 16 | 132 | 10.81 |
| gene-LOC100576818 | 500 | 2 | 40 | 100 | 0.4 | https://www.ncbi.nlm.nih.gov/gene/?term=LOC100576818 | NC_037639.1 | 14428536 | 14429035 | + | 18 | 17 | 1 | 94.44 |
| gene-LOC100577699 | 500 | 2 | 40 | 100 | 0.4 | https://www.ncbi.nlm.nih.gov/gene/?term=LOC100577699 | NC_037648.1 | 10530337 | 10530836 | + | 8 | 8 | 0 | 100 |
| gene-LOC100578363 | 500 | 2 | 40 | 100 | 0.4 | https://www.ncbi.nlm.nih.gov/gene/?term=LOC100578363 | NC_037645.1 | 1963287 | 1963786 | + | 25 | 8 | 17 | 32 |
| gene-LOC102653824 | 500 | 2 | 40 | 100 | 0.4 | https://www.ncbi.nlm.nih.gov/gene/?term=LOC102653824 | NC_037648.1 | 14282631 | 14283130 | - | 70 | 12 | 58 | 17.14 |
| gene-LOC102654090 | 500 | 2 | 40 | 100 | 0.4 | https://www.ncbi.nlm.nih.gov/gene/?term=LOC102654090 | NC_037648.1 | 14289244 | 14289743 | + | 18 | 13 | 5 | 72.22 |
| gene-LOC102654261 | 500 | 2 | 40 | 100 | 0.4 | https://www.ncbi.nlm.nih.gov/gene/?term=LOC102654261 | NC_037652.1 | 5098331 | 5098830 | - | 14 | 14 | 0 | 100 |
| gene-LOC107965260 | 500 | 2 | 40 | 91.67 | 0.37 | https://www.ncbi.nlm.nih.gov/gene/?term=LOC107965260 | NC_037648.1 | 16099189 | 16099688 | + | 57 | 26 | 31 | 45.61 |
| gene-LOC107965626 | 500 | 2 | 40 | 100 | 0.4 | https://www.ncbi.nlm.nih.gov/gene/?term=LOC107965626 | NC_037652.1 | 3773928 | 3774427 | + | 9 | 9 | 0 | 100 |
| gene-LOC113218553 | 500 | 2 | 40 | 100 | 0.4 | https://www.ncbi.nlm.nih.gov/gene/?term=LOC113218553 | NC_037639.1 | 12452945 | 12453444 | - | 12 | 8 | 4 | 66.67 |
| gene-LOC113218658 | 500 | 2 | 40 | 100 | 0.4 | https://www.ncbi.nlm.nih.gov/gene/?term=LOC113218658 | NC_037640.1 | 8288137 | 8288636 | + | 18 | 16 | 2 | 88.89 |
| gene-LOC113218789 | 500 | 2 | 40 | 92.86 | 0.37 | https://www.ncbi.nlm.nih.gov/gene/?term=LOC113218789 | NC_037642.1 | 2997817 | 2998316 | + | 14 | 13 | 1 | 92.86 |
| gene-LOC113218816 | 500 | 2 | 40 | 100 | 0.4 | https://www.ncbi.nlm.nih.gov/gene/?term=LOC113218816 | NC_037642.1 | 12958242 | 12958741 | - | 8 | 8 | 0 | 100 |
| gene-LOC113218872 | 500 | 2 | 40 | 100 | 0.4 | https://www.ncbi.nlm.nih.gov/gene/?term=LOC113218872 | NC_037643.1 | 54099 | 54598 | + | 10 | 10 | 0 | 100 |
| gene-LOC113219064 | 500 | 2 | 40 | 100 | 0.4 | https://www.ncbi.nlm.nih.gov/gene/?term=LOC113219064 | NC_037647.1 | 9422072 | 9422571 | + | 17 | 10 | 7 | 58.82 |
| gene-LOC408522 | 500 | 2 | 40 | 100 | 0.4 | https://www.ncbi.nlm.nih.gov/gene/?term=LOC408522 | NC_037652.1 | 6983350 | 6983849 | - | 11 | 11 | 0 | 100 |
| gene-LOC408860 | 500 | 2 | 40 | 100 | 0.4 | https://www.ncbi.nlm.nih.gov/gene/?term=LOC408860 | NC_037642.1 | 12216212 | 12216711 | - | 36 | 15 | 21 | 41.67 |
| gene-LOC408866 | 500 | 2 | 40 | 100 | 0.4 | https://www.ncbi.nlm.nih.gov/gene/?term=LOC408866 | NC_037642.1 | 13475230 | 13475729 | + | 36 | 17 | 19 | 47.22 |
| gene-LOC409600 | 500 | 2 | 40 | 100 | 0.4 | https://www.ncbi.nlm.nih.gov/gene/?term=LOC409600 | NC_037646.1 | 7031214 | 7031713 | - | 8 | 8 | 0 | 100 |
| gene-LOC410451 | 500 | 2 | 40 | 100 | 0.4 | https://www.ncbi.nlm.nih.gov/gene/?term=LOC410451 | NC_037650.1 | 5957782 | 5958281 | - | 14 | 14 | 0 | 100 |
| gene-LOC410875 | 500 | 2 | 40 | 100 | 0.4 | https://www.ncbi.nlm.nih.gov/gene/?term=LOC410875 | NC_037639.1 | 10379035 | 10379534 | + | 22 | 17 | 5 | 77.27 |
| gene-LOC411335 | 500 | 2 | 40 | 100 | 0.4 | https://www.ncbi.nlm.nih.gov/gene/?term=LOC411335 | NC_037646.1 | 8362817 | 8363316 | + | 11 | 11 | 0 | 100 |
| gene-LOC411371 | 500 | 2 | 40 | 88.89 | 0.36 | https://www.ncbi.nlm.nih.gov/gene/?term=LOC411371 | NC_037646.1 | 6743274 | 6743773 | + | 19 | 12 | 7 | 63.16 |
| gene-LOC413793 | 500 | 2 | 40 | 100 | 0.4 | https://www.ncbi.nlm.nih.gov/gene/?term=LOC413793 | NC_037651.1 | 8245796 | 8246295 | - | 14 | 10 | 4 | 71.43 |
| gene-LOC551548 | 500 | 2 | 40 | 100 | 0.4 | https://www.ncbi.nlm.nih.gov/gene/?term=LOC551548 | NC_037642.1 | 8518784 | 8519283 | - | 8 | 8 | 0 | 100 |
| gene-LOC551956 | 500 | 2 | 40 | 100 | 0.4 | https://www.ncbi.nlm.nih.gov/gene/?term=LOC551956 | NC_037642.1 | 797283 | 797782 | - | 10 | 10 | 0 | 100 |
| gene-LOC552731 | 500 | 2 | 40 | 100 | 0.4 | https://www.ncbi.nlm.nih.gov/gene/?term=LOC552731 | NC_037641.1 | 12732330 | 12732829 | - | 8 | 8 | 0 | 100 |
| gene-LOC724123 | 500 | 2 | 40 | 100 | 0.4 | https://www.ncbi.nlm.nih.gov/gene/?term=LOC724123 | NC_037652.1 | 2475870 | 2476369 | - | 12 | 10 | 2 | 83.33 |
| gene-LOC724388 | 500 | 2 | 40 | 100 | 0.4 | https://www.ncbi.nlm.nih.gov/gene/?term=LOC724388 | NC_037642.1 | 11379858 | 11380357 | - | 24 | 17 | 7 | 70.83 |
| gene-LOC724451 | 500 | 2 | 40 | 86.67 | 0.35 | https://www.ncbi.nlm.nih.gov/gene/?term=LOC724451 | NC_037646.1 | 7466366 | 7466865 | - | 16 | 14 | 2 | 87.5 |
| gene-LOC725502 | 500 | 2 | 40 | 100 | 0.4 | https://www.ncbi.nlm.nih.gov/gene/?term=LOC725502 | NC_037651.1 | 8359474 | 8359973 | - | 20 | 18 | 2 | 90 |
| gene-LOC726550 | 500 | 2 | 40 | 100 | 0.4 | https://www.ncbi.nlm.nih.gov/gene/?term=LOC726550 | NC_037641.1 | 12159286 | 12159785 | + | 57 | 15 | 42 | 26.32 |
| gene-LOC727254 | 500 | 2 | 40 | 100 | 0.4 | https://www.ncbi.nlm.nih.gov/gene/?term=LOC727254 | NC_037651.1 | 10057690 | 10058189 | + | 8 | 8 | 0 | 100 |

| | | | | | | | | | | | | | | |
|---|---|---|---|---|---|---|---|---|---|---|---|---|---|---|
| gene-Mir9b | 500 | 2 | 40 | 92.86 | 0.37 | https://www.ncbi.nlm.nih.gov/gene/?term=Mir9b | NC_037652.1 | 7427028 | 7427527 | - | 64 | 16 | 48 | 25 |
| gene-H | 500 | 1 | 20 | 100 | 0.2 | https://www.ncbi.nlm.nih.gov/gene/?term=H | NC_037642.1 | 9576253 | 9576752 | - | 140 | 9 | 131 | 6.43 |
| gene-LOC100576100 | 500 | 1 | 20 | 100 | 0.2 | https://www.ncbi.nlm.nih.gov/gene/?term=LOC100576100 | NC_037641.1 | 11262817 | 11263316 | + | 17 | 5 | 12 | 29.41 |
| gene-LOC100576155 | 500 | 1 | 20 | 100 | 0.2 | https://www.ncbi.nlm.nih.gov/gene/?term=LOC100576155 | NC_037642.1 | 544461 | 544960 | + | 13 | 4 | 9 | 30.77 |
| gene-LOC100577128 | 500 | 1 | 20 | 100 | 0.2 | https://www.ncbi.nlm.nih.gov/gene/?term=LOC100577128 | NC_037642.1 | 12083316 | 12083815 | + | 4 | 4 | 0 | 100 |
| gene-LOC100578212 | 500 | 1 | 20 | 100 | 0.2 | https://www.ncbi.nlm.nih.gov/gene/?term=LOC100578212 | NC_037646.1 | 12334987 | 12335486 | + | 5 | 5 | 0 | 100 |
| gene-LOC100578259 | 500 | 1 | 20 | 100 | 0.2 | https://www.ncbi.nlm.nih.gov/gene/?term=LOC100578259 | NC_037645.1 | 515132 | 515631 | + | 4 | 4 | 0 | 100 |
| gene-LOC100578355 | 500 | 1 | 20 | 100 | 0.2 | https://www.ncbi.nlm.nih.gov/gene/?term=LOC100578355 | NC_037642.1 | 7465773 | 7466272 | - | 4 | 4 | 0 | 100 |
| gene-LOC100578691 | 500 | 1 | 20 | 100 | 0.2 | https://www.ncbi.nlm.nih.gov/gene/?term=LOC100578691 | NC_037639.1 | 11351287 | 11351786 | + | 4 | 4 | 0 | 100 |
| gene-LOC102653702 | 500 | 1 | 20 | 83.33 | 0.17 | https://www.ncbi.nlm.nih.gov/gene/?term=LOC102653702 | NC_037648.1 | 8302625 | 8303124 | - | 36 | 6 | 30 | 16.67 |
| gene-LOC102654498 | 500 | 1 | 20 | 100 | 0.2 | https://www.ncbi.nlm.nih.gov/gene/?term=LOC102654498 | NC_037646.1 | 1562120 | 1562619 | + | 5 | 5 | 0 | 100 |
| gene-LOC102655138 | 500 | 1 | 20 | 100 | 0.2 | https://www.ncbi.nlm.nih.gov/gene/?term=LOC102655138 | NC_037645.1 | 6985360 | 6985859 | + | 4 | 4 | 0 | 100 |
| gene-LOC102655796 | 500 | 1 | 20 | 100 | 0.2 | https://www.ncbi.nlm.nih.gov/gene/?term=LOC102655796 | NC_037641.1 | 5686954 | 5687453 | - | 4 | 4 | 0 | 100 |
| gene-LOC102655798 | 500 | 1 | 20 | 62.5 | 0.12 | https://www.ncbi.nlm.nih.gov/gene/?term=LOC102655798 | NC_037652.1 | 6900217 | 6900716 | + | 8 | 5 | 3 | 62.5 |
| gene-LOC107963972 | 500 | 1 | 20 | 100 | 0.2 | https://www.ncbi.nlm.nih.gov/gene/?term=LOC107963972 | NC_037651.1 | 9088238 | 9088737 | - | 8 | 4 | 4 | 50 |
| gene-LOC107964045 | 500 | 1 | 20 | 100 | 0.2 | https://www.ncbi.nlm.nih.gov/gene/?term=LOC107964045 | NC_037639.1 | 14436780 | 14437279 | + | 5 | 5 | 0 | 100 |
| gene-LOC107964211 | 500 | 1 | 20 | 58.33 | 0.12 | https://www.ncbi.nlm.nih.gov/gene/?term=LOC107964211 | NC_037640.1 | 4695677 | 4696176 | + | 196 | 15 | 181 | 7.65 |
| gene-LOC107964413 | 500 | 1 | 20 | 100 | 0.2 | https://www.ncbi.nlm.nih.gov/gene/?term=LOC107964413 | NC_037642.1 | 9898088 | 9898587 | + | 70 | 6 | 64 | 8.57 |
| gene-LOC107964531 | 500 | 1 | 20 | 100 | 0.2 | https://www.ncbi.nlm.nih.gov/gene/?term=LOC107964531 | NC_037643.1 | 16276897 | 16277396 | - | 4 | 4 | 0 | 100 |
| gene-LOC113218590 | 500 | 1 | 20 | 100 | 0.2 | https://www.ncbi.nlm.nih.gov/gene/?term=LOC113218590 | NC_037639.1 | 8162800 | 8163299 | - | 4 | 4 | 0 | 100 |
| gene-LOC113218704 | 500 | 1 | 20 | 100 | 0.2 | https://www.ncbi.nlm.nih.gov/gene/?term=LOC113218704 | NC_037640.1 | 6890661 | 6891160 | - | 85 | 11 | 74 | 12.94 |
| gene-LOC113219178 | 500 | 1 | 20 | 100 | 0.2 | https://www.ncbi.nlm.nih.gov/gene/?term=LOC113219178 | NC_037649.1 | 5229514 | 5230013 | - | 11 | 9 | 2 | 81.82 |
| gene-LOC113219199 | 500 | 1 | 20 | 100 | 0.2 | https://www.ncbi.nlm.nih.gov/gene/?term=LOC113219199 | NC_037650.1 | 5955044 | 5955543 | - | 7 | 7 | 0 | 100 |
| gene-LOC113219399 | 500 | 1 | 20 | 100 | 0.2 | https://www.ncbi.nlm.nih.gov/gene/?term=LOC113219399 | NC_037638.1 | 14979923 | 14980422 | + | 12 | 10 | 2 | 83.33 |
| gene-LOC408271 | 500 | 1 | 20 | 62.5 | 0.12 | https://www.ncbi.nlm.nih.gov/gene/?term=LOC408271 | NC_037647.1 | 11824356 | 11824855 | - | 50 | 14 | 36 | 28 |
| gene-LOC408708 | 500 | 1 | 20 | 100 | 0.2 | https://www.ncbi.nlm.nih.gov/gene/?term=LOC408708 | NC_037639.1 | 6615024 | 6615523 | - | 5 | 5 | 0 | 100 |
| gene-LOC408831 | 500 | 1 | 20 | 100 | 0.2 | https://www.ncbi.nlm.nih.gov/gene/?term=LOC408831 | NC_037642.1 | 6871391 | 6871890 | - | 5 | 5 | 0 | 100 |
| gene-LOC408839 | 500 | 1 | 20 | 100 | 0.2 | https://www.ncbi.nlm.nih.gov/gene/?term=LOC408839 | NC_037642.1 | 9838392 | 9838891 | - | 87 | 10 | 77 | 11.49 |
| gene-LOC409063 | 500 | 1 | 20 | 100 | 0.2 | https://www.ncbi.nlm.nih.gov/gene/?term=LOC409063 | NC_037642.1 | 3632694 | 3633193 | + | 81 | 8 | 73 | 9.88 |
| gene-LOC409346 | 500 | 1 | 20 | 100 | 0.2 | https://www.ncbi.nlm.nih.gov/gene/?term=LOC409346 | NC_037640.1 | 4903861 | 4904360 | - | 4 | 4 | 0 | 100 |
| gene-LOC409610 | 500 | 1 | 20 | 100 | 0.2 | https://www.ncbi.nlm.nih.gov/gene/?term=LOC409610 | NC_037652.1 | 7880975 | 7881474 | + | 23 | 12 | 11 | 52.17 |
| gene-LOC409614 | 500 | 1 | 20 | 83.33 | 0.17 | https://www.ncbi.nlm.nih.gov/gene/?term=LOC409614 | NC_037651.1 | 10573988 | 10574487 | + | 12 | 10 | 2 | 83.33 |
| gene-LOC410091 | 500 | 1 | 20 | 100 | 0.2 | https://www.ncbi.nlm.nih.gov/gene/?term=LOC410091 | NC_037640.1 | 4537987 | 4538486 | + | 4 | 4 | 0 | 100 |
| gene-LOC411192 | 500 | 1 | 20 | 100 | 0.2 | https://www.ncbi.nlm.nih.gov/gene/?term=LOC411192 | NC_037644.1 | 13772019 | 13772518 | - | 4 | 4 | 0 | 100 |
| gene-LOC411519 | 500 | 1 | 20 | 100 | 0.2 | https://www.ncbi.nlm.nih.gov/gene/?term=LOC411519 | NC_037641.1 | 3679148 | 3679647 | - | 5 | 5 | 0 | 100 |
| gene-LOC412232 | 500 | 1 | 20 | 100 | 0.2 | https://www.ncbi.nlm.nih.gov/gene/?term=LOC412232 | NC_037642.1 | 11440583 | 11441082 | + | 6 | 6 | 0 | 100 |
| gene-LOC412837 | 500 | 1 | 20 | 100 | 0.2 | https://www.ncbi.nlm.nih.gov/gene/?term=LOC412837 | NC_037645.1 | 6744571 | 6745070 | - | 31 | 4 | 27 | 12.9 |
| gene-LOC412972 | 500 | 1 | 20 | 100 | 0.2 | https://www.ncbi.nlm.nih.gov/gene/?term=LOC412972 | NC_037648.1 | 8043010 | 8043509 | + | 21 | 11 | 10 | 52.38 |
| gene-LOC414014 | 500 | 1 | 20 | 100 | 0.2 | https://www.ncbi.nlm.nih.gov/gene/?term=LOC414014 | NC_037644.1 | 12110431 | 12110930 | + | 8 | 4 | 4 | 50 |
| gene-LOC551008 | 500 | 1 | 20 | 100 | 0.2 | https://www.ncbi.nlm.nih.gov/gene/?term=LOC551008 | NC_037639.1 | 11351106 | 11351605 | - | 4 | 4 | 0 | 100 |
| gene-LOC551451 | 500 | 1 | 20 | 100 | 0.2 | https://www.ncbi.nlm.nih.gov/gene/?term=LOC551451 | NC_037638.1 | 20102989 | 20103488 | - | 38 | 20 | 18 | 52.63 |
| gene-LOC551785 | 500 | 1 | 20 | 100 | 0.2 | https://www.ncbi.nlm.nih.gov/gene/?term=LOC551785 | NC_037638.1 | 11731008 | 11731507 | - | 11 | 10 | 1 | 90.91 |
| gene-LOC551811 | 500 | 1 | 20 | 100 | 0.2 | https://www.ncbi.nlm.nih.gov/gene/?term=LOC551811 | NC_037643.1 | 16599155 | 16599654 | - | 10 | 9 | 1 | 90 |
| gene-LOC551814 | 500 | 1 | 20 | 50 | 0.1 | https://www.ncbi.nlm.nih.gov/gene/?term=LOC551814 | NC_037647.1 | 4839082 | 4839581 | + | 12 | 6 | 6 | 50 |
| gene-LOC552194 | 500 | 1 | 20 | 100 | 0.2 | https://www.ncbi.nlm.nih.gov/gene/?term=LOC552194 | NC_037641.1 | 12755131 | 12755630 | - | 14 | 8 | 6 | 57.14 |
| gene-LOC552345 | 500 | 1 | 20 | 100 | 0.2 | https://www.ncbi.nlm.nih.gov/gene/?term=LOC552345 | NC_037646.1 | 7838082 | 7838581 | - | 4 | 4 | 0 | 100 |
| gene-LOC552400 | 500 | 1 | 20 | 100 | 0.2 | https://www.ncbi.nlm.nih.gov/gene/?term=LOC552400 | NC_037645.1 | 2629443 | 2629942 | + | 20 | 4 | 16 | 20 |
| gene-LOC552640 | 500 | 1 | 20 | 100 | 0.2 | https://www.ncbi.nlm.nih.gov/gene/?term=LOC552640 | NC_037651.1 | 8274345 | 8274844 | - | 85 | 15 | 70 | 17.65 |
| gene-LOC552652 | 500 | 1 | 20 | 100 | 0.2 | https://www.ncbi.nlm.nih.gov/gene/?term=LOC552652 | NC_037651.1 | 8268657 | 8269156 | + | 8 | 4 | 4 | 50 |
| gene-LOC552845 | 500 | 1 | 20 | 100 | 0.2 | https://www.ncbi.nlm.nih.gov/gene/?term=LOC552845 | NC_037652.1 | 6573840 | 6574339 | - | 17 | 5 | 12 | 29.41 |
| gene-LOC724235 | 500 | 1 | 20 | 100 | 0.2 | https://www.ncbi.nlm.nih.gov/gene/?term=LOC724235 | NC_037638.1 | 12172654 | 12173153 | + | 5 | 5 | 0 | 100 |
| gene-LOC724850 | 500 | 1 | 20 | 100 | 0.2 | https://www.ncbi.nlm.nih.gov/gene/?term=LOC724850 | NC_037639.1 | 9678412 | 9678911 | + | 5 | 5 | 0 | 100 |
| gene-LOC724959 | 500 | 1 | 20 | 100 | 0.2 | https://www.ncbi.nlm.nih.gov/gene/?term=LOC724959 | NC_037649.1 | 9968146 | 9968645 | - | 8 | 4 | 4 | 50 |
| gene-LOC725042 | 500 | 1 | 20 | 100 | 0.2 | https://www.ncbi.nlm.nih.gov/gene/?term=LOC725042 | NC_037638.1 | 21488758 | 21489257 | + | 4 | 4 | 0 | 100 |
| gene-LOC725062 | 500 | 1 | 20 | 100 | 0.2 | https://www.ncbi.nlm.nih.gov/gene/?term=LOC725062 | NC_037641.1 | 11568622 | 11569121 | - | 4 | 4 | 0 | 100 |
| gene-LOC725065 | 500 | 1 | 20 | 100 | 0.2 | https://www.ncbi.nlm.nih.gov/gene/?term=LOC725065 | NC_037645.1 | 1441538 | 1442037 | + | 4 | 4 | 0 | 100 |
| gene-LOC725093 | 500 | 1 | 20 | 100 | 0.2 | https://www.ncbi.nlm.nih.gov/gene/?term=LOC725093 | NC_037642.1 | 2944866 | 2945365 | - | 19 | 4 | 15 | 21.05 |
| gene-LOC725178 | 500 | 1 | 20 | 100 | 0.2 | https://www.ncbi.nlm.nih.gov/gene/?term=LOC725178 | NC_037646.1 | 7546323 | 7546822 | + | 4 | 4 | 0 | 100 |
| gene-LOC725360 | 500 | 1 | 20 | 100 | 0.2 | https://www.ncbi.nlm.nih.gov/gene/?term=LOC725360 | NC_037641.1 | 12871976 | 12872475 | + | 36 | 4 | 32 | 11.11 |
| gene-LOC725565 | 500 | 1 | 20 | 100 | 0.2 | https://www.ncbi.nlm.nih.gov/gene/?term=LOC725565 | NC_037648.1 | 1074794 | 1075293 | + | 9 | 4 | 5 | 44.44 |
| gene-LOC725603 | 500 | 1 | 20 | 100 | 0.2 | https://www.ncbi.nlm.nih.gov/gene/?term=LOC725603 | NC_037647.1 | 6434923 | 6435422 | - | 5 | 5 | 0 | 100 |
| gene-LOC725980 | 500 | 1 | 20 | 100 | 0.2 | https://www.ncbi.nlm.nih.gov/gene/?term=LOC725980 | NC_037638.1 | 20112501 | 20113000 | - | 11 | 9 | 2 | 81.82 |
| gene-LOC726063 | 500 | 1 | 20 | 100 | 0.2 | https://www.ncbi.nlm.nih.gov/gene/?term=LOC726063 | NC_037647.1 | 6316824 | 6317323 | + | 6 | 6 | 0 | 100 |
| gene-LOC726132 | 500 | 1 | 20 | 100 | 0.2 | https://www.ncbi.nlm.nih.gov/gene/?term=LOC726132 | NC_037641.1 | 11118027 | 11118526 | - | 19 | 10 | 9 | 52.63 |

| | | | | | | | | | | | | | | |
|---|---|---|---|---|---|---|---|---|---|---|---|---|---|---|
| gene-LOC726504 | 500 | 1 | 20 | 100 | 0.2 | https://www.ncbi.nlm.nih.gov/gene/?term=LOC726504 | NC_037645.1 | 1901517 | 1902016 | + | 5 | 5 | 0 | 100 |
| gene-LOC726591 | 500 | 1 | 20 | 100 | 0.2 | https://www.ncbi.nlm.nih.gov/gene/?term=LOC726591 | NC_037638.1 | 20564061 | 20564560 | - | 5 | 5 | 0 | 100 |
| gene-LOC726888 | 500 | 1 | 20 | 83.33 | 0.17 | https://www.ncbi.nlm.nih.gov/gene/?term=LOC726888 | NC_037653.1 | 1473424 | 1473923 | - | 6 | 5 | 1 | 83.33 |
| gene-LOC726989 | 500 | 1 | 20 | 100 | 0.2 | https://www.ncbi.nlm.nih.gov/gene/?term=LOC726989 | NC_037641.1 | 3493187 | 3493686 | - | 33 | 4 | 29 | 12.12 |
| gene-LOC727014 | 500 | 1 | 20 | 100 | 0.2 | https://www.ncbi.nlm.nih.gov/gene/?term=LOC727014 | NC_037642.1 | 13860538 | 13861037 | + | 28 | 4 | 24 | 14.29 |
| gene-LOC727030 | 500 | 1 | 20 | 100 | 0.2 | https://www.ncbi.nlm.nih.gov/gene/?term=LOC727030 | NC_037645.1 | 1961022 | 1961521 | - | 10 | 8 | 2 | 80 |
| gene-LOC727501 | 500 | 1 | 20 | 100 | 0.2 | https://www.ncbi.nlm.nih.gov/gene/?term=LOC727501 | NC_037643.1 | 13312355 | 13312854 | - | 4 | 4 | 0 | 100 |
| gene-Mir3758 | 500 | 1 | 20 | 71.43 | 0.14 | https://www.ncbi.nlm.nih.gov/gene/?term=Mir3758 | NC_037650.1 | 796302 | 796801 | - | 15 | 11 | 4 | 73.33 |
| gene-Mir6037 | 500 | 1 | 20 | 100 | 0.2 | https://www.ncbi.nlm.nih.gov/gene/?term=Mir6037 | NC_037639.1 | 9360768 | 9361267 | - | 13 | 4 | 9 | 30.77 |
| gene-Ndufs5 | 500 | 1 | 20 | 85.71 | 0.17 | https://www.ncbi.nlm.nih.gov/gene/?term=Ndufs5 | NC_037642.1 | 13859323 | 13859822 | - | 13 | 10 | 3 | 76.92 |
| gene-Nrk1 | 500 | 1 | 20 | 100 | 0.2 | https://www.ncbi.nlm.nih.gov/gene/?term=Nrk1 | NC_037646.1 | 11436629 | 11437128 | - | 6 | 6 | 0 | 100 |
| gene-TRNAR-ACG-4 | 500 | 1 | 20 | 100 | 0.2 | https://www.ncbi.nlm.nih.gov/gene/?term=TRNAR-ACG | NC_037645.1 | 3931986 | 3932485 | - | 7 | 7 | 0 | 100 |
| gene-Tg | 500 | 1 | 20 | 100 | 0.2 | https://www.ncbi.nlm.nih.gov/gene/?term=Tg | NC_037642.1 | 13518767 | 13519266 | + | 89 | 4 | 85 | 4.49 |
| gene-c-mos | 500 | 1 | 20 | 100 | 0.2 | https://www.ncbi.nlm.nih.gov/gene/?term=c-mos | NC_037648.1 | 2801260 | 2801759 | + | 6 | 6 | 0 | 100 |

Directory: RNK File: sites-in-genes-Am_RE_fe.txt

```
seqnames    start end   width strand      coverage    numCs numTs perc_meth   region_seqnames   region_start      region_end  region_width      region_strand     region_source
      region_type region_score      region_phase      region_ID   region_Dbxref     region_Name region_gbkey      region_gene region_gene_biotype     region_description
      region_gene_synonym     region_end_range  region_partial    region_start_range      region_exception
NC_037638.1 13072 13072 1     +     11    7     4     63.6363636363636  NC_037638.1 10792 17180 6389  +     Gnomon      gene  NA    NA    gene-LOC551555    c("BEEBASE:GB42138",
"GeneID:551555")  LOC551555   Gene  LOC551555   protein_coding    NA    character(0)      character(0)      NA    character(0)      NA
NC_037638.1 13129 13129 1     +     15    12    3     80    NC_037638.1 10792 17180 6389  +     Gnomon      gene  NA    NA    gene-LOC551555    c("BEEBASE:GB42138", "GeneID:551555")
      LOC551555   Gene  LOC551555   protein_coding    NA    character(0)      character(0)      NA    character(0)      NA
NC_037638.1 13165 13165 1     +     19    17    2     89.4736842105263  NC_037638.1 10792 17180 6389  +     Gnomon      gene  NA    NA    gene-LOC551555    c("BEEBASE:GB42138",
"GeneID:551555")  LOC551555   Gene  LOC551555   protein_coding    NA    character(0)      character(0)      NA    character(0)      NA
NC_037638.1 13208 13208 1     +     14    13    1     92.8571428571429  NC_037638.1 10792 17180 6389  +     Gnomon      gene  NA    NA    gene-LOC551555    c("BEEBASE:GB42138",
"GeneID:551555")  LOC551555   Gene  LOC551555   protein_coding    NA    character(0)      character(0)      NA    character(0)      NA
NC_037638.1 13244 13244 1     +     11    10    1     90.9090909090909  NC_037638.1 10792 17180 6389  +     Gnomon      gene  NA    NA    gene-LOC551555    c("BEEBASE:GB42138",
"GeneID:551555")  LOC551555   Gene  LOC551555   protein_coding    NA    character(0)      character(0)      NA    character(0)      NA
NC_037638.1 13253 13253 1     +     10    10    0     100   NC_037638.1 10792 17180 6389  +     Gnomon      gene  NA    NA    gene-LOC551555    c("BEEBASE:GB42138", "GeneID:551555")
      LOC551555   Gene  LOC551555   protein_coding    NA    character(0)      character(0)      NA    character(0)      NA
NC_037638.1 13273 13273 1     +     8     8     0     100   NC_037638.1 10792 17180 6389  +     Gnomon      gene  NA    NA    gene-LOC551555    c("BEEBASE:GB42138", "GeneID:551555")
      LOC551555   Gene  LOC551555   protein_coding    NA    character(0)      character(0)      NA    character(0)      NA
NC_037638.1 13291 13291 1     +     10    10    0     100   NC_037638.1 10792 17180 6389  +     Gnomon      gene  NA    NA    gene-LOC551555    c("BEEBASE:GB42138", "GeneID:551555")
      LOC551555   Gene  LOC551555   protein_coding    NA    character(0)      character(0)      NA    character(0)      NA
NC_037638.1 13329 13329 1     +     16    16    0     100   NC_037638.1 10792 17180 6389  +     Gnomon      gene  NA    NA    gene-LOC551555    c("BEEBASE:GB42138", "GeneID:551555")
      LOC551555   Gene  LOC551555   protein_coding    NA    character(0)      character(0)      NA    character(0)      NA
NC_037638.1 13372 13372 1     +     11    10    1     90.9090909090909  NC_037638.1 10792 17180 6389  +     Gnomon      gene  NA    NA    gene-LOC551555    c("BEEBASE:GB42138",
"GeneID:551555")  LOC551555   Gene  LOC551555   protein_coding    NA    character(0)      character(0)      NA    character(0)      NA
NC_037638.1 13373 13373 1     -     5     5     0     100   NC_037638.1 10792 17180 6389  +     Gnomon      gene  NA    NA    gene-LOC551555    c("BEEBASE:GB42138", "GeneID:551555")
      LOC551555   Gene  LOC551555   protein_coding    NA    character(0)      character(0)      NA    character(0)      NA
NC_037638.1 13392 13392 1     +     7     7     0     100   NC_037638.1 10792 17180 6389  +     Gnomon      gene  NA    NA    gene-LOC551555    c("BEEBASE:GB42138", "GeneID:551555")
      LOC551555   Gene  LOC551555   protein_coding    NA    character(0)      character(0)      NA    character(0)      NA
NC_037638.1 13393 13393 1     -     4     4     0     100   NC_037638.1 10792 17180 6389  +     Gnomon      gene  NA    NA    gene-LOC551555    c("BEEBASE:GB42138", "GeneID:551555")
      LOC551555   Gene  LOC551555   protein_coding    NA    character(0)      character(0)      NA    character(0)      NA
NC_037638.1 13443 13443 1     +     6     6     0     100   NC_037638.1 10792 17180 6389  +     Gnomon      gene  NA    NA    gene-LOC551555    c("BEEBASE:GB42138", "GeneID:551555")
      LOC551555   Gene  LOC551555   protein_coding    NA    character(0)      character(0)      NA    character(0)      NA
NC_037638.1 13459 13459 1     +     7     6     1     85.7142857142857  NC_037638.1 10792 17180 6389  +     Gnomon      gene  NA    NA    gene-LOC551555    c("BEEBASE:GB42138",
"GeneID:551555")  LOC551555   Gene  LOC551555   protein_coding    NA    character(0)      character(0)      NA    character(0)      NA
NC_037638.1 13479 13479 1     +     11    11    0     100   NC_037638.1 10792 17180 6389  +     Gnomon      gene  NA    NA    gene-LOC551555    c("BEEBASE:GB42138", "GeneID:551555")
      LOC551555   Gene  LOC551555   protein_coding    NA    character(0)      character(0)      NA    character(0)      NA
NC_037638.1 13498 13498 1     +     11    11    0     100   NC_037638.1 10792 17180 6389  +     Gnomon      gene  NA    NA    gene-LOC551555    c("BEEBASE:GB42138", "GeneID:551555")
      LOC551555   Gene  LOC551555   protein_coding    NA    character(0)      character(0)      NA    character(0)      NA
NC_037638.1 13514 13514 1     +     9     9     0     100   NC_037638.1 10792 17180 6389  +     Gnomon      gene  NA    NA    gene-LOC551555    c("BEEBASE:GB42138", "GeneID:551555")
      LOC551555   Gene  LOC551555   protein_coding    NA    character(0)      character(0)      NA    character(0)      NA
NC_037638.1 13757 13757 1     +     4     4     0     100   NC_037638.1 10792 17180 6389  +     Gnomon      gene  NA    NA    gene-LOC551555    c("BEEBASE:GB42138", "GeneID:551555")
      LOC551555   Gene  LOC551555   protein_coding    NA    character(0)      character(0)      NA    character(0)      NA
NC_037638.1 24663 24663 1     +     9     9     0     100   NC_037638.1 23613 26208 2596  +     BestRefSeq%2CGnomon     gene  NA    NA    gene-Rfwd3  c("BEEBASE:GB42139",
"GeneID:409940")  Rfwd3 Gene  Rfwd3 protein_coding    ring finger and WD repeat domain 3  character(0)      character(0)      NA    character(0)      NA
NC_037638.1 24701 24701 1     +     15    14    1     93.3333333333333  NC_037638.1 23613 26208 2596  +     BestRefSeq%2CGnomon     gene  NA    NA    gene-Rfwd3  c("BEEBASE:GB42139",
"GeneID:409940")  Rfwd3 Gene  Rfwd3 protein_coding    ring finger and WD repeat domain 3  character(0)      character(0)      NA    character(0)      NA
NC_037638.1 24702 24702 1     -     5     5     0     100   NC_037638.1 23613 26208 2596  +     BestRefSeq%2CGnomon     gene  NA    NA    gene-Rfwd3  c("BEEBASE:GB42139",
"GeneID:409940")  Rfwd3 Gene  Rfwd3 protein_coding    ring finger and WD repeat domain 3  character(0)      character(0)      NA    character(0)      NA
NC_037638.1 24743 24743 1     +     16    16    0     100   NC_037638.1 23613 26208 2596  +     BestRefSeq%2CGnomon     gene  NA    NA    gene-Rfwd3  c("BEEBASE:GB42139",
"GeneID:409940")  Rfwd3 Gene  Rfwd3 protein_coding    ring finger and WD repeat domain 3  character(0)      character(0)      NA    character(0)      NA
NC_037638.1 24781 24781 1     +     17    16    1     94.1176470588235  NC_037638.1 23613 26208 2596  +     BestRefSeq%2CGnomon     gene  NA    NA    gene-Rfwd3  c("BEEBASE:GB42139",
"GeneID:409940")  Rfwd3 Gene  Rfwd3 protein_coding    ring finger and WD repeat domain 3  character(0)      character(0)      NA    character(0)      NA
...
... only first 25 lines shown ...
```

Directory: RNK File: sites-in-genes-Am_RE_te.txt

```
seqnames    start end   width strand      coverage    numCs numTs perc_meth   region_seqnames   region_start      region_end  region_width      region_strand     region_source
      region_type region_score      region_phase      region_ID   region_Dbxref     region_Name region_gbkey      region_gene region_gene_biotype     region_description
      region_gene_synonym     region_end_range  region_partial    region_start_range      region_exception
NC_037638.1 13129 13129 1     +     11    10    1     90.9090909090909  NC_037638.1 10792 17180 6389  +     Gnomon      gene  NA    NA    gene-LOC551555    c("BEEBASE:GB42138",
"GeneID:551555")  LOC551555   Gene  LOC551555   protein_coding    NA    character(0)      character(0)      NA    character(0)      NA
NC_037638.1 13165 13165 1     +     14    14    0     100   NC_037638.1 10792 17180 6389  +     Gnomon      gene  NA    NA    gene-LOC551555    c("BEEBASE:GB42138", "GeneID:551555")
      LOC551555   Gene  LOC551555   protein_coding    NA    character(0)      character(0)      NA    character(0)      NA
NC_037638.1 13208 13208 1     +     15    14    1     93.3333333333333  NC_037638.1 10792 17180 6389  +     Gnomon      gene  NA    NA    gene-LOC551555    c("BEEBASE:GB42138",
"GeneID:551555")  LOC551555   Gene  LOC551555   protein_coding    NA    character(0)      character(0)      NA    character(0)      NA
NC_037638.1 13244 13244 1     +     11    10    1     90.9090909090909  NC_037638.1 10792 17180 6389  +     Gnomon      gene  NA    NA    gene-LOC551555    c("BEEBASE:GB42138",
"GeneID:551555")  LOC551555   Gene  LOC551555   protein_coding    NA    character(0)      character(0)      NA    character(0)      NA
NC_037638.1 13253 13253 1     +     9     9     0     100   NC_037638.1 10792 17180 6389  +     Gnomon      gene  NA    NA    gene-LOC551555    c("BEEBASE:GB42138", "GeneID:551555")
      LOC551555   Gene  LOC551555   protein_coding    NA    character(0)      character(0)      NA    character(0)      NA
NC_037638.1 13273 13273 1     +     9     9     0     100   NC_037638.1 10792 17180 6389  +     Gnomon      gene  NA    NA    gene-LOC551555    c("BEEBASE:GB42138", "GeneID:551555")
      LOC551555   Gene  LOC551555   protein_coding    NA    character(0)      character(0)      NA    character(0)      NA
NC_037638.1 13291 13291 1     +     7     7     0     100   NC_037638.1 10792 17180 6389  +     Gnomon      gene  NA    NA    gene-LOC551555    c("BEEBASE:GB42138", "GeneID:551555")
      LOC551555   Gene  LOC551555   protein_coding    NA    character(0)      character(0)      NA    character(0)      NA
NC_037638.1 13329 13329 1     +     11    10    1     90.9090909090909  NC_037638.1 10792 17180 6389  +     Gnomon      gene  NA    NA    gene-LOC551555    c("BEEBASE:GB42138",
"GeneID:551555")  LOC551555   Gene  LOC551555   protein_coding    NA    character(0)      character(0)      NA    character(0)      NA
NC_037638.1 13372 13372 1     +     8     8     0     100   NC_037638.1 10792 17180 6389  +     Gnomon      gene  NA    NA    gene-LOC551555    c("BEEBASE:GB42138", "GeneID:551555")
      LOC551555   Gene  LOC551555   protein_coding    NA    character(0)      character(0)      NA    character(0)      NA
NC_037638.1 13373 13373 1     -     4     4     0     100   NC_037638.1 10792 17180 6389  +     Gnomon      gene  NA    NA    gene-LOC551555    c("BEEBASE:GB42138", "GeneID:551555")
      LOC551555   Gene  LOC551555   protein_coding    NA    character(0)      character(0)      NA    character(0)      NA
NC_037638.1 13392 13392 1     +     5     5     0     100   NC_037638.1 10792 17180 6389  +     Gnomon      gene  NA    NA    gene-LOC551555    c("BEEBASE:GB42138", "GeneID:551555")
      LOC551555   Gene  LOC551555   protein_coding    NA    character(0)      character(0)      NA    character(0)      NA
NC_037638.1 13443 13443 1     +     4     4     0     100   NC_037638.1 10792 17180 6389  +     Gnomon      gene  NA    NA    gene-LOC551555    c("BEEBASE:GB42138", "GeneID:551555")
      LOC551555   Gene  LOC551555   protein_coding    NA    character(0)      character(0)      NA    character(0)      NA
NC_037638.1 13459 13459 1     +     5     5     0     100   NC_037638.1 10792 17180 6389  +     Gnomon      gene  NA    NA    gene-LOC551555    c("BEEBASE:GB42138", "GeneID:551555")
      LOC551555   Gene  LOC551555   protein_coding    NA    character(0)      character(0)      NA    character(0)      NA
NC_037638.1 13460 13460 1     -     4     4     0     100   NC_037638.1 10792 17180 6389  +     Gnomon      gene  NA    NA    gene-LOC551555    c("BEEBASE:GB42138", "GeneID:551555")
      LOC551555   Gene  LOC551555   protein_coding    NA    character(0)      character(0)      NA    character(0)      NA
NC_037638.1 13479 13479 1     +     5     5     0     100   NC_037638.1 10792 17180 6389  +     Gnomon      gene  NA    NA    gene-LOC551555    c("BEEBASE:GB42138", "GeneID:551555")
      LOC551555   Gene  LOC551555   protein_coding    NA    character(0)      character(0)      NA    character(0)      NA
NC_037638.1 13480 13480 1     -     5     5     0     100   NC_037638.1 10792 17180 6389  +     Gnomon      gene  NA    NA    gene-LOC551555    c("BEEBASE:GB42138", "GeneID:551555")
      LOC551555   Gene  LOC551555   protein_coding    NA    character(0)      character(0)      NA    character(0)      NA
NC_037638.1 13498 13498 1     +     7     7     0     100   NC_037638.1 10792 17180 6389  +     Gnomon      gene  NA    NA    gene-LOC551555    c("BEEBASE:GB42138", "GeneID:551555")
      LOC551555   Gene  LOC551555   protein_coding    NA    character(0)      character(0)      NA    character(0)      NA
NC_037638.1 13499 13499 1     -     5     5     0     100   NC_037638.1 10792 17180 6389  +     Gnomon      gene  NA    NA    gene-LOC551555    c("BEEBASE:GB42138", "GeneID:551555")
      LOC551555   Gene  LOC551555   protein_coding    NA    character(0)      character(0)      NA    character(0)      NA
NC_037638.1 13514 13514 1     +     6     6     0     100   NC_037638.1 10792 17180 6389  +     Gnomon      gene  NA    NA    gene-LOC551555    c("BEEBASE:GB42138", "GeneID:551555")
      LOC551555   Gene  LOC551555   protein_coding    NA    character(0)      character(0)      NA    character(0)      NA
NC_037638.1 13515 13515 1     -     6     6     0     100   NC_037638.1 10792 17180 6389  +     Gnomon      gene  NA    NA    gene-LOC551555    c("BEEBASE:GB42138", "GeneID:551555")
      LOC551555   Gene  LOC551555   protein_coding    NA    character(0)      character(0)      NA    character(0)      NA
NC_037638.1 13570 13570 1     +     9     8     1     88.8888888888889  NC_037638.1 10792 17180 6389  +     Gnomon      gene  NA    NA    gene-LOC551555    c("BEEBASE:GB42138",
"GeneID:551555")  LOC551555   Gene  LOC551555   protein_coding    NA    character(0)      character(0)      NA    character(0)      NA
NC_037638.1 13581 13581 1     +     10    9     1     90    NC_037638.1 10792 17180 6389  +     Gnomon      gene  NA    NA    gene-LOC551555    c("BEEBASE:GB42138", "GeneID:551555")
      LOC551555   Gene  LOC551555   protein_coding    NA    character(0)      character(0)      NA    character(0)      NA
NC_037638.1 13603 13603 1     +     9     9     0     100   NC_037638.1 10792 17180 6389  +     Gnomon      gene  NA    NA    gene-LOC551555    c("BEEBASE:GB42138", "GeneID:551555")
      LOC551555   Gene  LOC551555   protein_coding    NA    character(0)      character(0)      NA    character(0)      NA
NC_037638.1 24663 24663 1     +     4     4     0     100   NC_037638.1 23613 26208 2596  +     BestRefSeq%2CGnomon     gene  NA    NA    gene-Rfwd3  c("BEEBASE:GB42139",
"GeneID:409940")  Rfwd3 Gene  Rfwd3 protein_coding    ring finger and WD repeat domain 3  character(0)      character(0)      NA    character(0)      NA
...
... only first 25 lines shown ...
```

Directory: RNK File: sites-in-promoters-Am_RE_fe.txt

seqnames start end width strand coverage numCs numTs perc_meth region_seqnames region_start region_end region_width region_strand region_source
region_type region_score region_phase region_ID region_Dbxref region_Name region_gbkey region_gene region_gene_biotype region_description
region_gene_synonym region_end_range region_partial region_start_range region_exception
NC_037638.1 29442 29442 1 + 4 4 0 100 NC_037638.1 29440 29939 500 + Gnomon promoter NA NA gene-LOC551448 c("BEEBASE:GB42140",
"GeneID:551448") LOC551448 Gene LOC551448 protein_coding NA character(0) character(0) NA character(0) NA
NC_037638.1 1681374 1681374 1 - 4 4 0 100 NC_037638.1 1680961 1681460 500 - Gnomon promoter NA NA gene-LOC113219395
GeneID:113219395 LOC113219395 Gene LOC113219395 protein_coding NA character(0) character(0) NA character(0) NA
NC_037638.1 1812432 1812432 1 - 5 5 0 100 NC_037638.1 1812432 1812931 500 - Gnomon promoter NA NA gene-LOC113219401
GeneID:113219401 LOC113219401 Gene LOC113219401 lncRNA NA character(0) character(0) NA character(0) NA
NC_037638.1 1812440 1812440 1 + 8 8 0 100 NC_037638.1 1812432 1812931 500 - Gnomon promoter NA NA gene-LOC113219401
GeneID:113219401 LOC113219401 Gene LOC113219401 lncRNA NA character(0) character(0) NA character(0) NA
NC_037638.1 1812441 1812441 1 - 5 5 0 100 NC_037638.1 1812432 1812931 500 - Gnomon promoter NA NA gene-LOC113219401
GeneID:113219401 LOC113219401 Gene LOC113219401 lncRNA NA character(0) character(0) NA character(0) NA
NC_037638.1 1812498 1812498 1 - 5 5 0 100 NC_037638.1 1812432 1812931 500 - Gnomon promoter NA NA gene-LOC113219401
GeneID:113219401 LOC113219401 Gene LOC113219401 lncRNA NA character(0) character(0) NA character(0) NA
NC_037638.1 1812516 1812516 1 - 6 5 1 83.3333333333333 NC_037638.1 1812432 1812931 500 - Gnomon promoter NA NA gene-
LOC113219401 GeneID:113219401 LOC113219401 Gene LOC113219401 lncRNA NA character(0) character(0) NA character(0) NA
NC_037638.1 1812518 1812518 1 - 5 5 0 100 NC_037638.1 1812432 1812931 500 - Gnomon promoter NA NA gene-LOC113219401
GeneID:113219401 LOC113219401 Gene LOC113219401 lncRNA NA character(0) character(0) NA character(0) NA
NC_037638.1 1812549 1812549 1 - 7 6 1 85.7142857142857 NC_037638.1 1812432 1812931 500 - Gnomon promoter NA NA gene-
LOC113219401 GeneID:113219401 LOC113219401 Gene LOC113219401 lncRNA NA character(0) character(0) NA character(0) NA
NC_037638.1 7052688 7052688 1 + 4 4 0 100 NC_037638.1 7052487 7052986 500 + Gnomon promoter NA NA gene-LOC413813
c("BEEBASE:GB51867", "GeneID:413813") LOC413813 Gene LOC413813 protein_coding NA character(0) character(0) NA character(0) NA
NC_037638.1 10631079 10631079 1 - 4 4 0 100 NC_037638.1 10630803 10631302 500 - Gnomon promoter NA NA gene-LOC102654014
GeneID:102654014 LOC102654014 Gene LOC102654014 protein_coding NA character(0) character(0) NA character(0) NA
NC_037638.1 11731482 11731482 1 - 4 4 0 100 NC_037638.1 11731008 11731507 500 - Gnomon promoter NA NA gene-LOC551785
c("BEEBASE:GB46462", "GeneID:551785") LOC551785 Gene LOC551785 protein_coding NA character(0) character(0) NA character(0) NA
NC_037638.1 11731506 11731506 1 - 4 4 0 100 NC_037638.1 11731008 11731507 500 - Gnomon promoter NA NA gene-LOC551785
c("BEEBASE:GB46462", "GeneID:551785") LOC551785 Gene LOC551785 protein_coding NA character(0) character(0) NA character(0) NA
NC_037638.1 12200029 12200029 1 - 4 4 0 100 NC_037638.1 12199970 12200469 500 + Gnomon promoter NA NA gene-LOC724495
c("BEEBASE:GB46500", "GeneID:724495") LOC724495 Gene LOC724495 protein_coding NA character(0) character(0) NA character(0) NA
NC_037638.1 14979994 14979994 1 - 4 4 0 100 NC_037638.1 14979923 14980422 500 + Gnomon promoter NA NA gene-LOC113219399
GeneID:113219399 LOC113219399 Gene LOC113219399 lncRNA NA character(0) character(0) NA character(0) NA
NC_037638.1 20103161 20103161 1 - 5 5 0 100 NC_037638.1 20102989 20103488 500 - Gnomon promoter NA NA gene-LOC551451
c("BEEBASE:GB45498", "GeneID:551451") LOC551451 Gene LOC551451 protein_coding NA character(0) character(0) NA character(0) NA
NC_037638.1 20103165 20103165 1 - 5 5 0 100 NC_037638.1 20102989 20103488 500 - Gnomon promoter NA NA gene-LOC551451
c("BEEBASE:GB45498", "GeneID:551451") LOC551451 Gene LOC551451 protein_coding NA character(0) character(0) NA character(0) NA
NC_037638.1 20103390 20103390 1 - 4 4 0 100 NC_037638.1 20102989 20103488 500 - Gnomon promoter NA NA gene-LOC551451
c("BEEBASE:GB45498", "GeneID:551451") LOC551451 Gene LOC551451 protein_coding NA character(0) character(0) NA character(0) NA
NC_037638.1 20136253 20136253 1 + 4 4 0 100 NC_037638.1 20135801 20136300 500 - Gnomon promoter NA NA gene-LOC102655034
GeneID:102655034 LOC102655034 Gene LOC102655034 protein_coding NA character(0) character(0) NA character(0) NA
NC_037638.1 20136254 20136254 1 - 4 4 0 100 NC_037638.1 20135801 20136300 500 - Gnomon promoter NA NA gene-LOC102655034
GeneID:102655034 LOC102655034 Gene LOC102655034 protein_coding NA character(0) character(0) NA character(0) NA
NC_037639.1 9678421 9678421 1 - 4 4 0 100 NC_037639.1 9678412 9678911 500 + Gnomon promoter NA NA gene-LOC724850
c("BEEBASE:GB52350", "GeneID:724850") LOC724850 Gene LOC724850 protein_coding NA character(0) character(0) NA character(0) NA
NC_037639.1 11351287 11351287 1 + 5 5 0 100 NC_037639.1 11351106 11351605 500 - Gnomon promoter NA NA gene-LOC551008
c("BEEBASE:GB49807", "GeneID:551008") LOC551008 Gene LOC551008 protein_coding NA character(0) character(0) NA character(0) NA
NC_037639.1 11351287 11351287 1 + 5 5 0 100 NC_037639.1 11351287 11351786 500 + Gnomon promoter NA NA gene-LOC100578691
c("BEEBASE:GB49758", "GeneID:100578691") LOC100578691 Gene LOC100578691 protein_coding NA character(0) character(0) NA character(0) NA
NC_037639.1 11351327 11351327 1 + 4 4 0 100 NC_037639.1 11351106 11351605 500 - Gnomon promoter NA NA gene-LOC551008
c("BEEBASE:GB49807", "GeneID:551008") LOC551008 Gene LOC551008 protein_coding NA character(0) character(0) NA character(0) NA
NC_037639.1 11351327 11351327 1 + 4 4 0 100 NC_037639.1 11351287 11351786 500 + Gnomon promoter NA NA gene-LOC100578691
c("BEEBASE:GB49758", "GeneID:100578691") LOC100578691 Gene LOC100578691 protein_coding NA character(0) character(0) NA character(0) NA
NC_037639.1 12453314 12453314 1 + 11 10 1 90.9090909090909 NC_037639.1 12452945 12453444 500 - Gnomon promoter NA NA gene-
LOC113218553 GeneID:113218553 LOC113218553 Gene LOC113218553 lncRNA NA character(0) character(0) NA character(0) NA
NC_037639.1 14428831 14428831 1 - 4 4 0 100 NC_037639.1 14428536 14429035 500 + Gnomon promoter NA NA gene-LOC100576818
c("BEEBASE:GB55419", "GeneID:100576818") LOC100576818 Gene LOC100576818 protein_coding NA character(0) character(0) NA character(0) NA
NC_037639.1 15260689 15260689 1 - 4 4 0 100 NC_037639.1 15260436 15260935 500 - Gnomon promoter NA NA gene-LOC409088
c("BEEBASE:GB55558", "GeneID:409088") LOC409088 Gene LOC409088 protein_coding NA character(0) character(0) NA character(0) NA
NC_037639.1 15260717 15260717 1 - 6 6 0 100 NC_037639.1 15260436 15260935 500 - Gnomon promoter NA NA gene-LOC409088
c("BEEBASE:GB55558", "GeneID:409088") LOC409088 Gene LOC409088 protein_coding NA character(0) character(0) NA character(0) NA

NC_037640.1 4424974 4424974 1 - 4 4 0 100 NC_037640.1 4424618 4425117 500 - Gnomon promoter NA NA gene-LOC409443
c("BEEBASE:GB46915", "GeneID:409443") LOC409443 Gene LOC409443 protein_coding NA character(0) character(0) NA character(0) NA
NC_037640.1 4469691 4469691 1 - 4 4 0 100 NC_037640.1 4469685 4470184 500 + Gnomon promoter NA NA gene-LOC552603
c("BEEBASE:GB46895", "GeneID:552603") LOC552603 Gene LOC552603 protein_coding NA character(0) character(0) NA character(0) NA
NC_037640.1 4469730 4469730 1 - 4 4 0 100 NC_037640.1 4469685 4470184 500 + Gnomon promoter NA NA gene-LOC552603
c("BEEBASE:GB46895", "GeneID:552603") LOC552603 Gene LOC552603 protein_coding NA character(0) character(0) NA character(0) NA
NC_037640.1 4598646 4598646 1 + 4 4 0 100 NC_037640.1 4598201 4598700 500 - Gnomon promoter NA NA gene-LOC412851
c("BEEBASE:GB51524", "GeneID:412851") LOC412851 Gene LOC412851 protein_coding NA character(0) character(0) NA character(0) NA
NC_037640.1 4598675 4598675 1 + 4 4 0 100 NC_037640.1 4598201 4598700 500 - Gnomon promoter NA NA gene-LOC412851
c("BEEBASE:GB51524", "GeneID:412851") LOC412851 Gene LOC412851 protein_coding NA character(0) character(0) NA character(0) NA
NC_037640.1 6086317 6086317 1 + 7 7 0 100 NC_037640.1 6085859 6086358 500 - Gnomon promoter NA NA gene-LOC107964222
GeneID:107964222 LOC107964222 Gene LOC107964222 lncRNA NA character(0) character(0) NA character(0) NA
NC_037640.1 6891129 6891129 1 + 13 12 1 92.3076923076923 NC_037640.1 6890661 6891160 500 - cmsearch promoter NA NA gene-
LOC113218704 GeneID:113218704 LOC113218704 Gene LOC113218704 snoRNA NA character(0) character(0) NA character(0) NA
NC_037640.1 8174327 8174327 1 + 4 4 0 100 NC_037640.1 8174128 8174627 500 - Gnomon promoter NA NA gene-LOC113218626
GeneID:113218626 LOC113218626 Gene LOC113218626 lncRNA NA character(0) character(0) NA character(0) NA
NC_037640.1 10222397 10222397 1 - 4 4 0 100 NC_037640.1 10222330 10222829 500 + Gnomon promoter NA NA gene-LOC412256
c("BEEBASE:GB49172", "GeneID:412256") LOC412256 Gene LOC412256 protein_coding NA character(0) character(0) NA character(0) NA
NC_037641.1 3345495 3345495 1 - 4 4 0 100 NC_037641.1 3345469 3345968 500 - Gnomon promoter NA NA gene-LOC726826
c("BEEBASE:GB53154", "GeneID:726826") LOC726826 Gene LOC726826 protein_coding NA character(0) character(0) NA character(0) NA
NC_037641.1 4817398 4817398 1 - 6 5 1 83.3333333333333 NC_037641.1 4817094 4817593 500 - Gnomon promoter NA NA gene-LOC411640
c("BEEBASE:GB49556", "GeneID:411640") LOC411640 Gene LOC411640 protein_coding NA character(0) character(0) NA character(0) NA
NC_037641.1 11557869 11557869 1 + 8 6 2 75 NC_037641.1 11557856 11558355 500 + Gnomon promoter NA NA gene-LOC107964260
GeneID:107964260 LOC107964260 Gene LOC107964260 protein_coding NA character(0) character(0) NA character(0) NA
NC_037641.1 12159345 12159345 1 - 13 8 5 61.5384615384615 NC_037641.1 12159286 12159785 500 + Gnomon promoter NA NA gene-LOC726550
c("BEEBASE:GB44084", "GeneID:726550") LOC726550 Gene LOC726550 protein_coding NA character(0) character(0) NA character(0) NA
NC_037641.1 12159397 12159397 1 + 4 4 0 100 NC_037641.1 12159286 12159785 500 + Gnomon promoter NA NA gene-LOC726550
c("BEEBASE:GB44084", "GeneID:726550") LOC726550 Gene LOC726550 protein_coding NA character(0) character(0) NA character(0) NA
NC_037641.1 12159398 12159398 1 - 5 5 0 100 NC_037641.1 12159286 12159785 500 + Gnomon promoter NA NA gene-LOC726550
c("BEEBASE:GB44084", "GeneID:726550") LOC726550 Gene LOC726550 protein_coding NA character(0) character(0) NA character(0) NA
NC_037641.1 12676747 12676747 1 + 5 5 0 100 NC_037641.1 12676653 12677152 500 + Gnomon promoter NA NA gene-LOC409983
c("BEEBASE:GB52985", "GeneID:409983") LOC409983 Gene LOC409983 protein_coding NA character(0) character(0) NA character(0) NA
NC_037641.1 12755450 12755450 1 - 5 5 0 100 NC_037641.1 12755131 12755630 500 - Gnomon promoter NA NA gene-LOC552194
c("BEEBASE:GB53030", "GeneID:552194") LOC552194 Gene LOC552194 protein_coding NA character(0) character(0) NA character(0) NA
NC_037642.1 382379 382379 1 - 4 4 0 100 NC_037642.1 382084 382583 500 - Gnomon promoter NA NA gene-LOC725267
c("BEEBASE:GB55106", "GeneID:725267") LOC725267 Gene LOC725267 protein_coding NA character(0) character(0) NA character(0) NA
NC_037642.1 544735 544735 1 - 4 4 0 100 NC_037642.1 544461 544960 500 + Gnomon promoter NA NA gene-LOC100576155
GeneID:100576155 LOC100576155 Gene LOC100576155 protein_coding NA character(0) character(0) NA character(0) NA
NC_037642.1 2945365 2945365 1 - 6 6 0 100 NC_037642.1 2944866 2945365 500 - Gnomon promoter NA NA gene-LOC725093
c("BEEBASE:GB48889", "GeneID:725093") LOC725093 Gene LOC725093 protein_coding NA character(0) character(0) NA character(0) NA
NC_037642.1 2998069 2998069 1 + 4 4 0 100 NC_037642.1 2997817 2998316 500 + Gnomon promoter NA NA gene-LOC113218789
GeneID:113218789 LOC113218789 Gene LOC113218789 protein_coding NA character(0) character(0) NA character(0) NA
NC_037642.1 5472330 5472330 1 + 6 5 1 83.3333333333333 NC_037642.1 5471902 5472401 500 - Gnomon promoter NA NA gene-LOC725170
c("BEEBASE:GB46777", "GeneID:725170") LOC725170 Gene LOC725170 protein_coding NA character(0) character(0) NA character(0) NA
NC_037642.1 5472393 5472393 1 + 4 4 0 100 NC_037642.1 5471902 5472401 500 - Gnomon promoter NA NA gene-LOC725170
c("BEEBASE:GB46777", "GeneID:725170") LOC725170 Gene LOC725170 protein_coding NA character(0) character(0) NA character(0) NA
NC_037642.1 6572144 6572144 1 - 4 4 0 100 NC_037642.1 6572032 6572531 500 - Gnomon promoter NA NA gene-LOC726568
c("BEEBASE:GB46745", "GeneID:726568") LOC726568 Gene LOC726568 protein_coding NA character(0) character(0) NA character(0) NA
NC_037642.1 7404760 7404760 1 - 6 6 0 100 NC_037642.1 7404503 7405002 500 - Gnomon promoter NA NA gene-LOC552574
c("BEEBASE:GB42020", "GeneID:552574") LOC552574 Gene LOC552574 protein_coding NA character(0) character(0) NA character(0) NA
NC_037642.1 7404775 7404775 1 - 4 4 0 100 NC_037642.1 7404503 7405002 500 - Gnomon promoter NA NA gene-LOC552574
c("BEEBASE:GB42020", "GeneID:552574") LOC552574 Gene LOC552574 protein_coding NA character(0) character(0) NA character(0) NA
NC_037642.1 8842869 8842869 1 - 6 5 1 83.3333333333333 NC_037642.1 8842778 8843277 500 + Gnomon promoter NA NA gene-LOC411093
c("BEEBASE:GB50297", "GeneID:411093") LOC411093 Gene LOC411093 protein_coding NA character(0) character(0) NA character(0) NA
NC_037642.1 8842886 8842886 1 - 4 4 0 100 NC_037642.1 8842778 8843277 500 + Gnomon promoter NA NA gene-LOC411093
c("BEEBASE:GB50297", "GeneID:411093") LOC411093 Gene LOC411093 protein_coding NA character(0) character(0) NA character(0) NA
NC_037642.1 9535232 9535232 1 + 4 4 0 100 NC_037642.1 9535192 9535691 500 + Gnomon promoter NA NA gene-LOC552607
c("BEEBASE:GB44432", "GeneID:552607") LOC552607 Gene LOC552607 protein_coding NA character(0) character(0) NA character(0) NA
NC_037642.1 9552505 9552505 1 - 4 4 0 100 NC_037642.1 9552222 9552721 500 + Gnomon promoter NA NA gene-LOC100576101
c("BEEBASE:GB44435", "GeneID:100576101") LOC100576101 Gene LOC100576101 protein_coding NA character(0) character(0) NA character(0) NA
NC_037642.1 9904440 9904440 1 - 5 5 0 100 NC_037642.1 9904290 9904789 500 + BestRefSeq promoter NA NA gene-Coq7
c("BEEBASE:GB44464", "GeneID:409224") Coq7 Gene Coq7 protein_coding ubiquinone biosynthesis protein COQ7 Clk-1 character(0) true c(".", "9904790") NA
NC_037642.1 9904459 9904459 1 - 6 6 0 100 NC_037642.1 9904290 9904789 500 + BestRefSeq promoter NA NA gene-Coq7
c("BEEBASE:GB44464", "GeneID:409224") Coq7 Gene Coq7 protein_coding ubiquinone biosynthesis protein COQ7 Clk-1 character(0) true c(".", "9904790") NA

| | | | | | | | | | | | | | | | | | | | | | | | | | | | | | |
|---|---|---|---|---|---|---|---|---|---|---|---|---|---|---|---|---|---|---|---|---|---|---|---|---|---|---|---|---|---|
| NC_037642.1 | 11233802 | 11233802 | 1 | + | 6 | 5 | 1 | 83.3333333333333 | NC_037642.1 | 11233386 | 11233885 | 500 | - | Gnomon | promoter | NA | NA | gene-LOC100577702 | c("BEEBASE:GB44586", "GeneID:100577702") | LOC100577702 | Gene | LOC100577702 | protein_coding | NA | character(0) | character(0) | NA | character(0) | NA |
| NC_037642.1 | 11379884 | 11379884 | 1 | + | 9 | 9 | 0 | 100 | NC_037642.1 | 11379858 | 11380357 | 500 | - | Gnomon | promoter | NA | NA | gene-LOC724388 | c("BEEBASE:GB44577", "GeneID:724388") | LOC724388 | Gene | LOC724388 | protein_coding | NA | character(0) | character(0) | NA | character(0) | NA |
| NC_037642.1 | 11380028 | 11380028 | 1 | + | 7 | 7 | 0 | 100 | NC_037642.1 | 11379858 | 11380357 | 500 | - | Gnomon | promoter | NA | NA | gene-LOC724388 | c("BEEBASE:GB44577", "GeneID:724388") | LOC724388 | Gene | LOC724388 | protein_coding | NA | character(0) | character(0) | NA | character(0) | NA |
| NC_037642.1 | 12216670 | 12216670 | 1 | + | 4 | 4 | 0 | 100 | NC_037642.1 | 12216212 | 12216711 | 500 | - | Gnomon | promoter | NA | NA | gene-LOC408860 | c("BEEBASE:GB47815", "GeneID:408860") | LOC408860 | Gene | LOC408860 | protein_coding | NA | character(0) | character(0) | NA | character(0) | NA |
| NC_037642.1 | 12636366 | 12636366 | 1 | - | 4 | 4 | 0 | 100 | NC_037642.1 | 12636357 | 12636856 | 500 | + | Gnomon | promoter | NA | NA | gene-LOC409153 | c("BEEBASE:GB41288", "GeneID:409153") | LOC409153 | Gene | LOC409153 | protein_coding | NA | character(0) | character(0) | NA | character(0) | NA |
| NC_037642.1 | 12958689 | 12958689 | 1 | - | 6 | 6 | 0 | 100 | NC_037642.1 | 12958242 | 12958741 | 500 | - | cmsearch | promoter | NA | NA | gene-LOC113218816 | GeneID:113218816 | LOC113218816 | Gene | LOC113218816 | guide_RNA | NA | character(0) | character(0) | NA | character(0) | NA |
| NC_037642.1 | 12958692 | 12958692 | 1 | - | 6 | 6 | 0 | 100 | NC_037642.1 | 12958242 | 12958741 | 500 | - | cmsearch | promoter | NA | NA | gene-LOC113218816 | GeneID:113218816 | LOC113218816 | Gene | LOC113218816 | guide_RNA | NA | character(0) | character(0) | NA | character(0) | NA |
| NC_037642.1 | 13475250 | 13475250 | 1 | + | 6 | 6 | 0 | 100 | NC_037642.1 | 13475230 | 13475729 | 500 | + | Gnomon | promoter | NA | NA | gene-LOC408866 | c("BEEBASE:GB45703", "GeneID:408866") | LOC408866 | Gene | LOC408866 | protein_coding | NA | character(0) | character(0) | NA | character(0) | NA |
| NC_037642.1 | 13683256 | 13683256 | 1 | - | 7 | 7 | 0 | 100 | NC_037642.1 | 13682786 | 13683285 | 500 | - | BestRefSeq%2CGnomon | promoter | NA | NA | gene-Imd | GeneID:100302584 | Imd | Gene | Imd | protein_coding | immune deficiency | character(0) | character(0) | NA | character(0) | NA |
| NC_037642.1 | 13725928 | 13725928 | 1 | + | 6 | 6 | 0 | 100 | NC_037642.1 | 13725907 | 13726406 | 500 | + | Gnomon | promoter | NA | NA | gene-LOC726879 | c("BEEBASE:GB45732", "GeneID:726879") | LOC726879 | Gene | LOC726879 | protein_coding | NA | character(0) | character(0) | NA | character(0) | NA |
| NC_037642.1 | 13860586 | 13860586 | 1 | - | 4 | 4 | 0 | 100 | NC_037642.1 | 13860538 | 13861037 | 500 | + | Gnomon | promoter | NA | NA | gene-LOC727014 | c("BEEBASE:GB45740", "GeneID:727014") | LOC727014 | Gene | LOC727014 | protein_coding | NA | character(0) | character(0) | NA | character(0) | NA |
| NC_037642.1 | 13878080 | 13878080 | 1 | - | 4 | 4 | 0 | 100 | NC_037642.1 | 13877729 | 13878228 | 500 | - | Gnomon | promoter | NA | NA | gene-LOC102654994 | GeneID:102654994 | LOC102654994 | Gene | LOC102654994 | protein_coding | NA | character(0) | character(0) | NA | character(0) | NA |
| NC_037643.1 | 5155083 | 5155083 | 1 | - | 4 | 4 | 0 | 100 | NC_037643.1 | 5154685 | 5155184 | 500 | - | Gnomon | promoter | NA | NA | gene-LOC552046 | c("BEEBASE:GB52211", "GeneID:552046") | LOC552046 | Gene | LOC552046 | protein_coding | NA | character(0) | character(0) | NA | character(0) | NA |
| NC_037643.1 | 13196903 | 13196903 | 1 | + | 8 | 7 | 1 | 87.5 | NC_037643.1 | 13196766 | 13197265 | 500 | + | Gnomon | promoter | NA | NA | gene-LOC100576400 | c("BEEBASE:GB46060", "GeneID:100576400") | LOC100576400 | Gene | LOC100576400 | protein_coding | NA | character(0) | character(0) | NA | character(0) | NA |
| NC_037643.1 | 13312803 | 13312803 | 1 | + | 4 | 4 | 0 | 100 | NC_037643.1 | 13312355 | 13312854 | 500 | - | Gnomon | promoter | NA | NA | gene-LOC727501 | c("BEEBASE:GB14598", "GeneID:727501") | LOC727501 | Gene | LOC727501 | protein_coding | NA | character(0) | character(0) | NA | character(0) | NA |
| NC_037643.1 | 13312836 | 13312836 | 1 | + | 4 | 4 | 0 | 100 | NC_037643.1 | 13312355 | 13312854 | 500 | - | Gnomon | promoter | NA | NA | gene-LOC727501 | c("BEEBASE:GB14598", "GeneID:727501") | LOC727501 | Gene | LOC727501 | protein_coding | NA | character(0) | character(0) | NA | character(0) | NA |
| NC_037643.1 | 16277354 | 16277354 | 1 | + | 6 | 5 | 1 | 83.3333333333333 | NC_037643.1 | 16276897 | 16277396 | 500 | - | Gnomon | promoter | NA | NA | gene-LOC107964531 | GeneID:107964531 | LOC107964531 | Gene | LOC107964531 | protein_coding | NA | character(0) | character(0) | NA | character(0) | NA |
| NC_037644.1 | 1350179 | 1350179 | 1 | + | 5 | 5 | 0 | 100 | NC_037644.1 | 1350072 | 1350571 | 500 | + | Gnomon | promoter | NA | NA | gene-LOC551193 | c("BEEBASE:GB46241", "GeneID:551193") | LOC551193 | Gene | LOC551193 | protein_coding | NA | character(0) | character(0) | NA | character(0) | NA |
| NC_037644.1 | 1350180 | 1350180 | 1 | - | 4 | 4 | 0 | 100 | NC_037644.1 | 1350072 | 1350571 | 500 | + | Gnomon | promoter | NA | NA | gene-LOC551193 | c("BEEBASE:GB46241", "GeneID:551193") | LOC551193 | Gene | LOC551193 | protein_coding | NA | character(0) | character(0) | NA | character(0) | NA |
| NC_037645.1 | 1961511 | 1961511 | 1 | + | 7 | 6 | 1 | 85.7142857142857 | NC_037645.1 | 1961022 | 1961521 | 500 | - | Gnomon | promoter | NA | NA | gene-LOC727030 | c("BEEBASE:GB41164", "GeneID:727030") | LOC727030 | Gene | LOC727030 | protein_coding | NA | character(0) | character(0) | NA | character(0) | NA |
| NC_037645.1 | 1963375 | 1963375 | 1 | + | 5 | 5 | 0 | 100 | NC_037645.1 | 1963287 | 1963786 | 500 | + | Gnomon | promoter | NA | NA | gene-LOC100578363 | GeneID:100578363 | LOC100578363 | Gene | LOC100578363 | protein_coding | NA | character(0) | character(0) | NA | character(0) | NA |
| NC_037645.1 | 1963621 | 1963621 | 1 | - | 4 | 4 | 0 | 100 | NC_037645.1 | 1963287 | 1963786 | 500 | + | Gnomon | promoter | NA | NA | gene-LOC100578363 | GeneID:100578363 | LOC100578363 | Gene | LOC100578363 | protein_coding | NA | character(0) | character(0) | NA | character(0) | NA |
| NC_037645.1 | 2246116 | 2246116 | 1 | - | 8 | 8 | 0 | 100 | NC_037645.1 | 2246110 | 2246609 | 500 | + | Gnomon | promoter | NA | NA | gene-LOC552596 | c("BEEBASE:GB40459", "GeneID:552596") | LOC552596 | Gene | LOC552596 | protein_coding | NA | character(0) | character(0) | NA | character(0) | NA |
| NC_037645.1 | 2615979 | 2615979 | 1 | + | 4 | 4 | 0 | 100 | NC_037645.1 | 2615624 | 2616123 | 500 | + | Gnomon | promoter | NA | NA | gene-LOC552494 | c("BEEBASE:GB40492", "GeneID:552494") | LOC552494 | Gene | LOC552494 | protein_coding | NA | character(0) | character(0) | NA | character(0) | NA |
| NC_037645.1 | 4210110 | 4210110 | 1 | - | 5 | 5 | 0 | 100 | NC_037645.1 | 4209689 | 4210188 | 500 | - | Gnomon | promoter | NA | NA | gene-LOC412827 | c("BEEBASE:GB54522", "GeneID:412827") | LOC412827 | Gene | LOC412827 | protein_coding | NA | character(0) | character(0) | NA | character(0) | NA |
| NC_037645.1 | 6576115 | 6576115 | 1 | - | 9 | 8 | 1 | 88.8888888888889 | NC_037645.1 | 6576093 | 6576592 | 500 | + | Gnomon | promoter | NA | NA | gene-LOC551338 | c("BEEBASE:GB43840", "GeneID:551338") | LOC551338 | Gene | LOC551338 | protein_coding | NA | character(0) | character(0) | NA | character(0) | NA |
| NC_037645.1 | 6576168 | 6576168 | 1 | - | 7 | 7 | 0 | 100 | NC_037645.1 | 6576093 | 6576592 | 500 | + | Gnomon | promoter | NA | NA | gene-LOC551338 | c("BEEBASE:GB43840", "GeneID:551338") | LOC551338 | Gene | LOC551338 | protein_coding | NA | character(0) | character(0) | NA | character(0) | NA |
| NC_037645.1 | 6576215 | 6576215 | 1 | - | 8 | 8 | 0 | 100 | NC_037645.1 | 6576093 | 6576592 | 500 | + | Gnomon | promoter | NA | NA | gene-LOC551338 | c("BEEBASE:GB43840", "GeneID:551338") | LOC551338 | Gene | LOC551338 | protein_coding | NA | character(0) | character(0) | NA | character(0) | NA |
| NC_037645.1 | 6576517 | 6576517 | 1 | - | 4 | 4 | 0 | 100 | NC_037645.1 | 6576093 | 6576592 | 500 | + | Gnomon | promoter | NA | NA | gene-LOC551338 | c("BEEBASE:GB43840", "GeneID:551338") | LOC551338 | Gene | LOC551338 | protein_coding | NA | character(0) | character(0) | NA | character(0) | NA |
| NC_037645.1 | 6985383 | 6985383 | 1 | - | 4 | 4 | 0 | 100 | NC_037645.1 | 6985360 | 6985859 | 500 | + | Gnomon | promoter | NA | NA | gene-LOC102655138 | GeneID:102655138 | LOC102655138 | Gene | LOC102655138 | protein_coding | NA | character(0) | character(0) | NA | character(0) | NA |
| NC_037645.1 | 6985571 | 6985571 | 1 | + | 4 | 4 | 0 | 100 | NC_037645.1 | 6985360 | 6985859 | 500 | + | Gnomon | promoter | NA | NA | gene-LOC102655138 | GeneID:102655138 | LOC102655138 | Gene | LOC102655138 | protein_coding | NA | character(0) | character(0) | NA | character(0) | NA |
| NC_037646.1 | 1694184 | 1694184 | 1 | + | 4 | 4 | 0 | 100 | NC_037646.1 | 1694180 | 1694679 | 500 | - | Gnomon | promoter | NA | NA | gene-LOC113218977 | | | | | | | | | | | |

GeneID:113218977 LOC113218977 Gene LOC113218977 lncRNA NA character(0) character(0) NA character(0) NA
NC_037646.1 8278668 8278668 1 - 4 4 0 100 NC_037646.1 8278485 8278984 500 + Gnomon promoter NA NA gene-LOC409017
c("BEEBASE:GB42856", "GeneID:409017") LOC409017 Gene LOC409017 protein_coding NA character(0) character(0) NA character(0) NA
NC_037646.1 12335076 12335076 1 - 7 7 0 100 NC_037646.1 12334987 12335486 500 + Gnomon promoter NA NA gene-LOC100578212
c("BEEBASE:GB53377", "GeneID:100578212") LOC100578212 Gene LOC100578212 protein_coding NA character(0) character(0) NA character(0) NA
NC_037646.1 12339207 12339207 1 - 6 6 0 100 NC_037646.1 12339177 12339676 500 - Gnomon promoter NA NA gene-LOC412289
c("BEEBASE:GB53382", "GeneID:412289") LOC412289 Gene LOC412289 protein_coding NA character(0) character(0) NA character(0) NA
NC_037646.1 12339219 12339219 1 - 7 7 0 100 NC_037646.1 12339177 12339676 500 - Gnomon promoter NA NA gene-LOC412289
c("BEEBASE:GB53382", "GeneID:412289") LOC412289 Gene LOC412289 protein_coding NA character(0) character(0) NA character(0) NA
NC_037646.1 12339239 12339239 1 - 6 6 0 100 NC_037646.1 12339177 12339676 500 - Gnomon promoter NA NA gene-LOC412289
c("BEEBASE:GB53382", "GeneID:412289") LOC412289 Gene LOC412289 protein_coding NA character(0) character(0) NA character(0) NA
NC_037646.1 12339292 12339292 1 + 4 4 0 100 NC_037646.1 12339177 12339676 500 - Gnomon promoter NA NA gene-LOC412289
c("BEEBASE:GB53382", "GeneID:412289") LOC412289 Gene LOC412289 protein_coding NA character(0) character(0) NA character(0) NA
NC_037646.1 12339293 12339293 1 - 6 5 1 83.3333333333333 NC_037646.1 12339177 12339676 500 - Gnomon promoter NA NA gene-LOC412289
c("BEEBASE:GB53382", "GeneID:412289") LOC412289 Gene LOC412289 protein_coding NA character(0) character(0) NA character(0) NA
NC_037646.1 12339431 12339431 1 + 4 4 0 100 NC_037646.1 12339177 12339676 500 - Gnomon promoter NA NA gene-LOC412289
c("BEEBASE:GB53382", "GeneID:412289") LOC412289 Gene LOC412289 protein_coding NA character(0) character(0) NA character(0) NA
NC_037646.1 12339481 12339481 1 + 8 6 2 75 NC_037646.1 12339177 12339676 500 - Gnomon promoter NA NA gene-LOC412289
c("BEEBASE:GB53382", "GeneID:412289") LOC412289 Gene LOC412289 protein_coding NA character(0) character(0) NA character(0) NA
NC_037647.1 15101 15101 1 + 14 13 1 92.8571428571429 NC_037647.1 15047 15546 500 - Gnomon promoter NA NA gene-LOC113219036 GeneID:113219036
LOC113219036 Gene LOC113219036 lncRNA NA character(0) character(0) NA character(0) NA
NC_037647.1 15102 15102 1 - 9 6 3 66.6666666666667 NC_037647.1 15047 15546 500 - Gnomon promoter NA NA gene-LOC113219036 GeneID:113219036
LOC113219036 Gene LOC113219036 lncRNA NA character(0) character(0) NA character(0) NA
NC_037647.1 15137 15137 1 + 31 27 4 87.0967741935484 NC_037647.1 15047 15546 500 - Gnomon promoter NA NA gene-LOC113219036 GeneID:113219036
LOC113219036 Gene LOC113219036 lncRNA NA character(0) character(0) NA character(0) NA
NC_037647.1 355423 355423 1 - 5 5 0 100 NC_037647.1 354951 355450 500 - Gnomon promoter NA NA gene-LOC102655648
GeneID:102655648 LOC102655648 Gene LOC102655648 protein_coding NA character(0) character(0) NA character(0) NA
NC_037647.1 355448 355448 1 - 5 5 0 100 NC_037647.1 354951 355450 500 - Gnomon promoter NA NA gene-LOC102655648
GeneID:102655648 LOC102655648 Gene LOC102655648 protein_coding NA character(0) character(0) NA character(0) NA
NC_037647.1 4837100 4837100 1 - 4 4 0 100 NC_037647.1 4836722 4837221 500 + Gnomon promoter NA NA gene-LOC409056
c("BEEBASE:GB54598", "GeneID:409056") LOC409056 Gene LOC409056 protein_coding NA character(0) character(0) NA character(0) NA
NC_037647.1 4839188 4839188 1 + 8 6 2 75 NC_037647.1 4839082 4839581 500 + Gnomon promoter NA NA gene-LOC551814
c("BEEBASE:GB54599", "GeneID:551814") LOC551814 Gene LOC551814 protein_coding NA character(0) character(0) NA character(0) NA
NC_037647.1 4904656 4904656 1 - 4 4 0 100 NC_037647.1 4904215 4904714 500 - Gnomon promoter NA NA gene-LOC725948
c("BEEBASE:GB54583", "GeneID:725948") LOC725948 Gene LOC725948 protein_coding NA character(0) character(0) NA character(0) NA
NC_037647.1 5753084 5753084 1 + 6 5 1 83.3333333333333 NC_037647.1 5753080 5753579 500 + Gnomon promoter NA NA gene-LOC413683
c("BEEBASE:GB48336", "GeneID:413683") LOC413683 Gene LOC413683 protein_coding NA character(0) character(0) NA character(0) NA
NC_037647.1 5753085 5753085 1 - 6 6 0 100 NC_037647.1 5753080 5753579 500 + Gnomon promoter NA NA gene-LOC413683
c("BEEBASE:GB48336", "GeneID:413683") LOC413683 Gene LOC413683 protein_coding NA character(0) character(0) NA character(0) NA
NC_037647.1 5753145 5753145 1 - 4 4 0 100 NC_037647.1 5753080 5753579 500 + Gnomon promoter NA NA gene-LOC413683
c("BEEBASE:GB48336", "GeneID:413683") LOC413683 Gene LOC413683 protein_coding NA character(0) character(0) NA character(0) NA
NC_037647.1 5753166 5753166 1 - 6 5 1 83.3333333333333 NC_037647.1 5753080 5753579 500 + Gnomon promoter NA NA gene-LOC413683
c("BEEBASE:GB48336", "GeneID:413683") LOC413683 Gene LOC413683 protein_coding NA character(0) character(0) NA character(0) NA
NC_037647.1 5753181 5753181 1 - 5 5 0 100 NC_037647.1 5753080 5753579 500 + Gnomon promoter NA NA gene-LOC413683
c("BEEBASE:GB48336", "GeneID:413683") LOC413683 Gene LOC413683 protein_coding NA character(0) character(0) NA character(0) NA
NC_037647.1 6316923 6316923 1 + 7 6 1 85.7142857142857 NC_037647.1 6316824 6317323 500 + Gnomon promoter NA NA gene-LOC726063
c("BEEBASE:GB48360", "GeneID:726063") LOC726063 Gene LOC726063 protein_coding NA character(0) character(0) NA character(0) NA
NC_037648.1 1464051 1464051 1 - 4 4 0 100 NC_037648.1 1463635 1464134 500 - BestRefSeq promoter NA NA gene-Mir9865
c("GeneID:104796158", "miRBase:MI0031791") Mir9865 Gene Mir9865 miRNA microRNA 9865 ame-mir-9865 character(0) NA character(0) NA
NC_037648.1 1464075 1464075 1 - 4 4 0 100 NC_037648.1 1463635 1464134 500 - BestRefSeq promoter NA NA gene-Mir9865
c("GeneID:104796158", "miRBase:MI0031791") Mir9865 Gene Mir9865 miRNA microRNA 9865 ame-mir-9865 character(0) NA character(0) NA
NC_037648.1 1464129 1464129 1 - 4 4 0 100 NC_037648.1 1463635 1464134 500 - BestRefSeq promoter NA NA gene-Mir9865
c("GeneID:104796158", "miRBase:MI0031791") Mir9865 Gene Mir9865 miRNA microRNA 9865 ame-mir-9865 character(0) NA character(0) NA
NC_037648.1 5027314 5027314 1 - 4 4 0 100 NC_037648.1 5027280 5027779 500 - Gnomon promoter NA NA gene-LOC102656914
GeneID:102656914 LOC102656914 Gene LOC102656914 protein_coding NA character(0) character(0) NA character(0) NA
NC_037648.1 8043122 8043122 1 - 4 4 0 100 NC_037648.1 8043010 8043509 500 + Gnomon promoter NA NA gene-LOC412972
c("BEEBASE:GB49837", "GeneID:412972") LOC412972 Gene LOC412972 protein_coding NA character(0) character(0) NA character(0) NA
NC_037648.1 10530339 10530339 1 + 4 4 0 100 NC_037648.1 10530337 10530836 500 + Gnomon promoter NA NA gene-LOC100577699
c("BEEBASE:GB47304", "GeneID:100577699") LOC100577699 Gene LOC100577699 protein_coding NA character(0) character(0) NA character(0) NA
NC_037648.1 10530340 10530340 1 - 4 4 0 100 NC_037648.1 10530337 10530836 500 + Gnomon promoter NA NA gene-LOC100577699
c("BEEBASE:GB47304", "GeneID:100577699") LOC100577699 Gene LOC100577699 protein_coding NA character(0) character(0) NA character(0) NA
NC_037648.1 10532654 10532654 1 - 5 5 0 100 NC_037648.1 10532625 10533124 500 - Gnomon promoter NA NA gene-LOC494508
GeneID:494508 LOC494508 Gene LOC494508 misc_RNA NA character(0) character(0) NA character(0) NA
NC_037648.1 14282732 14282732 1 + 4 4 0 100 NC_037648.1 14282631 14283130 500 - Gnomon promoter NA NA gene-LOC102653824

GeneID:102653824 LOC102653824 Gene LOC102653824 protein_coding NA character(0) character(0) NA character(0) NA
NC_037648.1 15379123 15379123 1 + 5 5 0 100 NC_037648.1 15378785 15379284 500 - Gnomon promoter NA NA gene-LOC113219080
GeneID:113219080 LOC113219080 Gene LOC113219080 lncRNA NA character(0) character(0) NA character(0) NA
NC_037648.1 15655275 15655275 1 - 4 4 0 100 NC_037648.1 15654947 15655446 500 - tRNAscan-SE promoter NA NA gene-TRNAI-AAU-3
GeneID:107965304 TRNAI-AAU Gene TRNAI-AAU tRNA NA character(0) character(0) NA character(0) NA
NC_037648.1 16005779 16005779 1 + 4 4 0 100 NC_037648.1 16005777 16006276 500 + Gnomon promoter NA NA gene-LOC724878
c("BEEBASE:GB43211", "GeneID:724878") LOC724878 Gene LOC724878 protein_coding NA character(0) character(0) NA character(0) NA
NC_037648.1 16099282 16099282 1 - 4 4 0 100 NC_037648.1 16099189 16099688 500 + Gnomon promoter NA NA gene-LOC107965260
GeneID:107965260 LOC107965260 Gene LOC107965260 lncRNA NA character(0) character(0) NA character(0) NA
NC_037649.1 2396758 2396758 1 - 7 5 2 71.4285714285714 NC_037649.1 2396713 2397212 500 + Gnomon promoter NA NA gene-
LOC107965368 GeneID:107965368 LOC107965368 Gene LOC107965368 protein_coding NA character(0) character(0) NA character(0) NA
NC_037649.1 3084966 3084966 1 - 20 13 7 65 NC_037649.1 3084955 3085454 500 + Gnomon promoter NA NA gene-LOC412065
c("BEEBASE:GB48307", "GeneID:412065") LOC412065 Gene LOC412065 protein_coding NA character(0) character(0) NA character(0) NA
NC_037649.1 5229663 5229663 1 + 4 4 0 100 NC_037649.1 5229514 5230013 500 - cmsearch promoter NA NA gene-LOC113219178
GeneID:113219178 LOC113219178 Gene LOC113219178 snoRNA NA character(0) character(0) NA character(0) NA
NC_037649.1 5229705 5229705 1 + 4 4 0 100 NC_037649.1 5229514 5230013 500 - cmsearch promoter NA NA gene-LOC113219178
GeneID:113219178 LOC113219178 Gene LOC113219178 snoRNA NA character(0) character(0) NA character(0) NA
NC_037649.1 7696161 7696161 1 - 7 7 0 100 NC_037649.1 7696058 7696557 500 + Gnomon promoter NA NA gene-LOC552347
c("BEEBASE:GB49358", "GeneID:552347") LOC552347 Gene LOC552347 protein_coding NA character(0) character(0) NA character(0) NA
NC_037650.1 796370 796370 1 + 8 6 2 75 NC_037650.1 796302 796801 500 - BestRefSeq promoter NA NA gene-Mir3758
c("GeneID:100629098", "miRBase:MI0016160") Mir3758 Gene Mir3758 miRNA microRNA 3758 ame-mir-3758 character(0) NA character(0) NA
NC_037650.1 796371 796371 1 - 7 7 0 100 NC_037650.1 796302 796801 500 - BestRefSeq promoter NA NA gene-Mir3758
c("GeneID:100629098", "miRBase:MI0016160") Mir3758 Gene Mir3758 miRNA microRNA 3758 ame-mir-3758 character(0) NA character(0) NA
NC_037650.1 2451643 2451643 1 - 4 4 0 100 NC_037650.1 2451545 2452044 500 + Gnomon promoter NA NA gene-LOC410402
c("BEEBASE:GB54762", "GeneID:410402") LOC410402 Gene LOC410402 protein_coding NA character(0) character(0) NA character(0) NA
NC_037650.1 5955079 5955079 1 + 8 8 0 100 NC_037650.1 5955044 5955543 500 - Gnomon promoter NA NA gene-LOC113219199
GeneID:113219199 LOC113219199 Gene LOC113219199 lncRNA NA character(0) character(0) NA character(0) NA
NC_037650.1 5955310 5955310 1 - 4 4 0 100 NC_037650.1 5955044 5955543 500 - Gnomon promoter NA NA gene-LOC113219199
GeneID:113219199 LOC113219199 Gene LOC113219199 lncRNA NA character(0) character(0) NA character(0) NA
NC_037651.1 6847001 6847001 1 - 4 4 0 100 NC_037651.1 6846579 6847078 500 - Gnomon promoter NA NA gene-LOC552508
c("BEEBASE:GB54193", "GeneID:552508") LOC552508 Gene LOC552508 protein_coding NA character(0) character(0) NA character(0) NA
NC_037651.1 6847049 6847049 1 - 6 6 0 100 NC_037651.1 6846579 6847078 500 - Gnomon promoter NA NA gene-LOC552508
c("BEEBASE:GB54193", "GeneID:552508") LOC552508 Gene LOC552508 protein_coding NA character(0) character(0) NA character(0) NA
NC_037651.1 9854796 9854796 1 + 11 11 0 100 NC_037651.1 9854645 9855144 500 + Gnomon promoter NA NA gene-LOC727151
c("BEEBASE:GB41627", "GeneID:727151") LOC727151 Gene LOC727151 protein_coding NA character(0) character(0) NA character(0) NA
NC_037651.1 9854839 9854839 1 + 11 11 0 100 NC_037651.1 9854645 9855144 500 + Gnomon promoter NA NA gene-LOC727151
c("BEEBASE:GB41627", "GeneID:727151") LOC727151 Gene LOC727151 protein_coding NA character(0) character(0) NA character(0) NA
NC_037651.1 9854970 9854970 1 + 6 6 0 100 NC_037651.1 9854645 9855144 500 + Gnomon promoter NA NA gene-LOC727151
c("BEEBASE:GB41627", "GeneID:727151") LOC727151 Gene LOC727151 protein_coding NA character(0) character(0) NA character(0) NA
NC_037651.1 10058086 10058086 1 - 4 4 0 100 NC_037651.1 10057690 10058189 500 + Gnomon promoter NA NA gene-LOC727254
c("BEEBASE:GB41641", "GeneID:727254") LOC727254 Gene LOC727254 protein_coding NA character(0) character(0) NA character(0) NA
NC_037651.1 10476129 10476129 1 - 7 6 1 85.7142857142857 NC_037651.1 10476098 10476597 500 + Gnomon promoter NA NA gene-
LOC100576608 c("BEEBASE:GB41656", "GeneID:100576608") LOC100576608 Gene LOC100576608 protein_coding NA character(0) character(0) NA character(0)
NA
NC_037651.1 10489076 10489076 1 + 4 4 0 100 NC_037651.1 10488842 10489341 500 + Gnomon promoter NA NA gene-LOC408500
c("BEEBASE:GB41659", "GeneID:408500") LOC408500 Gene LOC408500 protein_coding NA character(0) character(0) NA character(0) NA
NC_037651.1 10489142 10489142 1 + 7 5 2 71.4285714285714 NC_037651.1 10488842 10489341 500 + Gnomon promoter NA NA gene-LOC408500
c("BEEBASE:GB41659", "GeneID:408500") LOC408500 Gene LOC408500 protein_coding NA character(0) character(0) NA character(0) NA
NC_037651.1 10489143 10489143 1 - 9 7 2 77.7777777777778 NC_037651.1 10488842 10489341 500 + Gnomon promoter NA NA gene-LOC408500
c("BEEBASE:GB41659", "GeneID:408500") LOC408500 Gene LOC408500 protein_coding NA character(0) character(0) NA character(0) NA
NC_037651.1 10489160 10489160 1 + 6 6 0 100 NC_037651.1 10488842 10489341 500 + Gnomon promoter NA NA gene-LOC408500
c("BEEBASE:GB41659", "GeneID:408500") LOC408500 Gene LOC408500 protein_coding NA character(0) character(0) NA character(0) NA
NC_037651.1 10489161 10489161 1 - 5 5 0 100 NC_037651.1 10488842 10489341 500 + Gnomon promoter NA NA gene-LOC408500
c("BEEBASE:GB41659", "GeneID:408500") LOC408500 Gene LOC408500 protein_coding NA character(0) character(0) NA character(0) NA
NC_037651.1 10489170 10489170 1 + 4 4 0 100 NC_037651.1 10488842 10489341 500 + Gnomon promoter NA NA gene-LOC408500
c("BEEBASE:GB41659", "GeneID:408500") LOC408500 Gene LOC408500 protein_coding NA character(0) character(0) NA character(0) NA
NC_037651.1 10489171 10489171 1 - 5 5 0 100 NC_037651.1 10488842 10489341 500 + Gnomon promoter NA NA gene-LOC408500
c("BEEBASE:GB41659", "GeneID:408500") LOC408500 Gene LOC408500 protein_coding NA character(0) character(0) NA character(0) NA
NC_037651.1 10574043 10574043 1 + 7 5 2 71.4285714285714 NC_037651.1 10573988 10574487 500 + Gnomon promoter NA NA gene-LOC409614
c("BEEBASE:GB41664", "GeneID:409614") LOC409614 Gene LOC409614 protein_coding NA character(0) character(0) NA character(0) NA
NC_037651.1 10652153 10652153 1 - 4 4 0 100 NC_037651.1 10651890 10652389 500 - Gnomon promoter NA NA gene-LOC410566
c("BEEBASE:GB41670", "GeneID:410566") LOC410566 Gene LOC410566 protein_coding NA character(0) character(0) NA character(0) NA
NC_037651.1 10652170 10652170 1 - 6 5 1 83.3333333333333 NC_037651.1 10651890 10652389 500 - Gnomon promoter NA NA gene-LOC410566
c("BEEBASE:GB41670", "GeneID:410566") LOC410566 Gene LOC410566 protein_coding NA character(0) character(0) NA character(0) NA

NC_037651.1 10652196 10652196 1 - 9 7 2 77.7777777777778 NC_037651.1 10651890 10652389 500 - Gnomon promoter NA NA gene-LOC410566
c("BEEBASE:GB41670", "GeneID:410566") LOC410566 Gene LOC410566 protein_coding NA character(0) character(0) NA character(0) NA
NC_037651.1 10652241 10652241 1 - 4 4 0 100 NC_037651.1 10651890 10652389 500 - Gnomon promoter NA NA gene-LOC410566
c("BEEBASE:GB41670", "GeneID:410566") LOC410566 Gene LOC410566 protein_coding NA character(0) character(0) NA character(0) NA
NC_037651.1 10652319 10652319 1 - 9 8 1 88.8888888888889 NC_037651.1 10651890 10652389 500 - Gnomon promoter NA NA gene-LOC410566
c("BEEBASE:GB41670", "GeneID:410566") LOC410566 Gene LOC410566 protein_coding NA character(0) character(0) NA character(0) NA
NC_037651.1 10652328 10652328 1 - 10 6 4 60 NC_037651.1 10651890 10652389 500 - Gnomon promoter NA NA gene-LOC410566
c("BEEBASE:GB41670", "GeneID:410566") LOC410566 Gene LOC410566 protein_coding NA character(0) character(0) NA character(0) NA
NC_037652.1 5098826 5098826 1 + 4 4 0 100 NC_037652.1 5098331 5098830 500 - Gnomon promoter NA NA gene-LOC102654261
GeneID:102654261 LOC102654261 Gene LOC102654261 protein_coding NA character(0) character(0) NA character(0) NA
NC_037652.1 5098827 5098827 1 - 7 6 1 85.7142857142857 NC_037652.1 5098331 5098830 500 - Gnomon promoter NA NA gene-
LOC102654261 GeneID:102654261 LOC102654261 Gene LOC102654261 protein_coding NA character(0) character(0) NA character(0) NA
NC_037652.1 5098829 5098829 1 + 5 5 0 100 NC_037652.1 5098331 5098830 500 - Gnomon promoter NA NA gene-LOC102654261
GeneID:102654261 LOC102654261 Gene LOC102654261 protein_coding NA character(0) character(0) NA character(0) NA
NC_037652.1 5098830 5098830 1 - 7 7 0 100 NC_037652.1 5098331 5098830 500 - Gnomon promoter NA NA gene-LOC102654261
GeneID:102654261 LOC102654261 Gene LOC102654261 protein_coding NA character(0) character(0) NA character(0) NA
NC_037652.1 5989440 5989440 1 - 6 6 0 100 NC_037652.1 5989340 5989839 500 - Gnomon promoter NA NA gene-LOC412907
c("BEEBASE:GB16006", "GeneID:412907") LOC412907 Gene LOC412907 protein_coding NA character(0) character(0) NA character(0) NA
NC_037652.1 5989494 5989494 1 + 8 5 3 62.5 NC_037652.1 5989340 5989839 500 - Gnomon promoter NA NA gene-LOC412907
c("BEEBASE:GB16006", "GeneID:412907") LOC412907 Gene LOC412907 protein_coding NA character(0) character(0) NA character(0) NA
NC_037652.1 5989572 5989572 1 + 6 5 1 83.3333333333333 NC_037652.1 5989340 5989839 500 - Gnomon promoter NA NA gene-LOC412907
c("BEEBASE:GB16006", "GeneID:412907") LOC412907 Gene LOC412907 protein_coding NA character(0) character(0) NA character(0) NA
NC_037652.1 6229444 6229444 1 + 4 4 0 100 NC_037652.1 6229095 6229594 500 - Gnomon promoter NA NA gene-LOC410617
c("BEEBASE:GB50093", "GeneID:410617") LOC410617 Gene LOC410617 protein_coding NA character(0) character(0) NA character(0) NA
NC_037652.1 6577421 6577421 1 - 5 5 0 100 NC_037652.1 6577021 6577520 500 - Gnomon promoter NA NA gene-LOC102655461
GeneID:102655461 LOC102655461 Gene LOC102655461 protein_coding NA character(0) character(0) NA character(0) NA
NC_037652.1 7419435 7419435 1 - 5 5 0 100 NC_037652.1 7419330 7419829 500 - BestRefSeq promoter NA NA gene-Lop2
c("BEEBASE:GB50034", "GeneID:768250") Lop2 Gene Lop2 protein_coding long wavelength sensitive opsin 2 character(0) character(0) NA character(0) NA
NC_037652.1 7881114 7881114 1 + 5 5 0 100 NC_037652.1 7880975 7881474 500 + Gnomon promoter NA NA gene-LOC409610
c("BEEBASE:GB50243", "GeneID:409610") LOC409610 Gene LOC409610 protein_coding NA character(0) character(0) NA character(0) NA
NC_037653.1 1302381 1302381 1 + 4 4 0 100 NC_037653.1 1302284 1302783 500 - Gnomon promoter NA NA gene-LOC726521
c("BEEBASE:GB54336", "GeneID:726521") LOC726521 Gene LOC726521 protein_coding NA character(0) character(0) NA character(0) NA
NC_037653.1 1302418 1302418 1 + 4 4 0 100 NC_037653.1 1302284 1302783 500 - Gnomon promoter NA NA gene-LOC726521
c("BEEBASE:GB54336", "GeneID:726521") LOC726521 Gene LOC726521 protein_coding NA character(0) character(0) NA character(0) NA
NC_037653.1 6399828 6399828 1 - 6 5 1 83.3333333333333 NC_037653.1 6399796 6400295 500 - Gnomon promoter NA NA gene-LOC724423
c("BEEBASE:GB45982", "GeneID:724423") LOC724423 Gene LOC724423 protein_coding NA character(0) character(0) NA character(0) NA
NW_020555869.1 8251 8251 1 + 23 19 4 82.6086956521739 NW_020555869.1 7786 8285 500 + Gnomon promoter NA NA gene-LOC102656321
GeneID:102656321 LOC102656321 Gene LOC102656321 protein_coding NA character(0) character(0) NA character(0) NA
NW_020555869.1 8252 8252 1 - 54 51 3 94.4444444444444 NW_020555869.1 7786 8285 500 + Gnomon promoter NA NA gene-LOC102656321
GeneID:102656321 LOC102656321 Gene LOC102656321 protein_coding NA character(0) character(0) NA character(0) NA
NW_020555869.1 8255 8255 1 + 27 21 6 77.7777777777778 NW_020555869.1 7786 8285 500 + Gnomon promoter NA NA gene-LOC102656321
GeneID:102656321 LOC102656321 Gene LOC102656321 protein_coding NA character(0) character(0) NA character(0) NA
NW_020555869.1 8256 8256 1 - 55 44 11 80 NW_020555869.1 7786 8285 500 + Gnomon promoter NA NA gene-LOC102656321 GeneID:102656321
LOC102656321 Gene LOC102656321 protein_coding NA character(0) character(0) NA character(0) NA

Directory: RNK File: sites-in-promoters-Am_RE_te.txt

```
seqnames	start	end	width	strand	coverage	numCs	numTs	perc_meth	region_seqnames	region_start	region_end	region_width	region_strand	region_source
	region_type	region_score	region_phase	region_ID	region_Dbxref	region_Name	region_gbkey	region_gene	region_gene_biotype	region_description
	region_gene_synonym	region_end_range	region_partial	region_start_range	region_exception
NC_037638.1	1812432	1812432	1	-	5	5	0	100	NC_037638.1	1812432	1812931	500	-	Gnomon	promoter	NA	NA	gene-LOC113219401
	GeneID:113219401	LOC113219401	Gene	LOC113219401	lncRNA	NA	character(0)	character(0)	NA	character(0)	NA
NC_037638.1	1812497	1812497	1	+	4	4	0	100	NC_037638.1	1812432	1812931	500	-	Gnomon	promoter	NA	NA	gene-LOC113219401
	GeneID:113219401	LOC113219401	Gene	LOC113219401	lncRNA	NA	character(0)	character(0)	NA	character(0)	NA
NC_037638.1	1812515	1812515	1	+	4	4	0	100	NC_037638.1	1812432	1812931	500	-	Gnomon	promoter	NA	NA	gene-LOC113219401
	GeneID:113219401	LOC113219401	Gene	LOC113219401	lncRNA	NA	character(0)	character(0)	NA	character(0)	NA
NC_037638.1	1812548	1812548	1	+	4	4	0	100	NC_037638.1	1812432	1812931	500	-	Gnomon	promoter	NA	NA	gene-LOC113219401
	GeneID:113219401	LOC113219401	Gene	LOC113219401	lncRNA	NA	character(0)	character(0)	NA	character(0)	NA
NC_037638.1	11731482	11731482	1	-	6	6	0	100	NC_037638.1	11731008	11731507	500	-	Gnomon	promoter	NA	NA	gene-LOC551785
	c("BEEBASE:GB46462", "GeneID:551785")	LOC551785	Gene	LOC551785	protein_coding	NA	character(0)	character(0)	NA	character(0)	NA
NC_037638.1	12172749	12172749	1	-	5	5	0	100	NC_037638.1	12172654	12173153	500	+	Gnomon	promoter	NA	NA	gene-LOC724235
	c("BEEBASE:GB46496", "GeneID:724235")	LOC724235	Gene	LOC724235	protein_coding	NA	character(0)	character(0)	NA	character(0)	NA
NC_037638.1	12310026	12310026	1	+	4	4	0	100	NC_037638.1	12309993	12310492	500	+	Gnomon	promoter	NA	NA	gene-LOC113219319
	GeneID:113219319	LOC113219319	Gene	LOC113219319	protein_coding	NA	character(0)	character(0)	NA	character(0)	NA
NC_037638.1	12310035	12310035	1	-	4	4	0	100	NC_037638.1	12309993	12310492	500	+	Gnomon	promoter	NA	NA	gene-LOC113219319
	GeneID:113219319	LOC113219319	Gene	LOC113219319	protein_coding	NA	character(0)	character(0)	NA	character(0)	NA
NC_037638.1	12310234	12310234	1	-	4	4	0	100	NC_037638.1	12309993	12310492	500	+	Gnomon	promoter	NA	NA	gene-LOC113219319
	GeneID:113219319	LOC113219319	Gene	LOC113219319	protein_coding	NA	character(0)	character(0)	NA	character(0)	NA
NC_037638.1	14979993	14979993	1	+	4	4	0	100	NC_037638.1	14979923	14980422	500	+	Gnomon	promoter	NA	NA	gene-LOC113219399
	GeneID:113219399	LOC113219399	Gene	LOC113219399	lncRNA	NA	character(0)	character(0)	NA	character(0)	NA
NC_037638.1	20103160	20103160	1	+	6	6	0	100	NC_037638.1	20102989	20103488	500	-	Gnomon	promoter	NA	NA	gene-LOC551451
	c("BEEBASE:GB45498", "GeneID:551451")	LOC551451	Gene	LOC551451	protein_coding	NA	character(0)	character(0)	NA	character(0)	NA
NC_037638.1	20112973	20112973	1	+	7	7	0	100	NC_037638.1	20112501	20113000	500	-	Gnomon	promoter	NA	NA	gene-LOC725980
	c("BEEBASE:GB45496", "GeneID:725980")	LOC725980	Gene	LOC725980	protein_coding	NA	character(0)	character(0)	NA	character(0)	NA
NC_037638.1	20564380	20564380	1	-	5	5	0	100	NC_037638.1	20564061	20564560	500	-	Gnomon	promoter	NA	NA	gene-LOC726591
	c("BEEBASE:GB48930", "GeneID:726591")	LOC726591	Gene	LOC726591	protein_coding	NA	character(0)	character(0)	NA	character(0)	NA
NC_037638.1	21488858	21488858	1	-	4	4	0	100	NC_037638.1	21488758	21489257	500	+	Gnomon	promoter	NA	NA	gene-LOC725042
	c("BEEBASE:GB51553", "GeneID:725042")	LOC725042	Gene	LOC725042	protein_coding	NA	character(0)	character(0)	NA	character(0)	NA
NC_037639.1	6615176	6615176	1	-	5	5	0	100	NC_037639.1	6615024	6615523	500	-	Gnomon	promoter	NA	NA	gene-LOC408708
	c("BEEBASE:GB50740", "GeneID:408708")	LOC408708	Gene	LOC408708	protein_coding	NA	character(0)	character(0)	NA	character(0)	NA
NC_037639.1	8163195	8163195	1	-	4	4	0	100	NC_037639.1	8162800	8163299	500	-	Gnomon	promoter	NA	NA	gene-LOC113218590
	GeneID:113218590	LOC113218590	Gene	LOC113218590	protein_coding	NA	character(0)	character(0)	NA	character(0)	NA
NC_037639.1	9361235	9361235	1	-	4	4	0	100	NC_037639.1	9360768	9361267	500	-	BestRefSeq	promoter	NA	NA	gene-Mir6037
	c("GeneID:104794335", "miRBase:MI0020304")	Mir6037	Gene	Mir6037	miRNA	microRNA 6037	ame-mir-6037	character(0)	NA	character(0)	NA
NC_037639.1	9678421	9678421	1	-	5	5	0	100	NC_037639.1	9678412	9678911	500	+	Gnomon	promoter	NA	NA	gene-LOC724850
	c("BEEBASE:GB52350", "GeneID:724850")	LOC724850	Gene	LOC724850	protein_coding	NA	character(0)	character(0)	NA	character(0)	NA
NC_037639.1	10379059	10379059	1	+	8	8	0	100	NC_037639.1	10379035	10379534	500	+	Gnomon	promoter	NA	NA	gene-LOC410875
	c("BEEBASE:GB52434", "GeneID:410875")	LOC410875	Gene	LOC410875	protein_coding	NA	character(0)	character(0)	NA	character(0)	NA
NC_037639.1	10379068	10379068	1	+	9	9	0	100	NC_037639.1	10379035	10379534	500	+	Gnomon	promoter	NA	NA	gene-LOC410875
	c("BEEBASE:GB52434", "GeneID:410875")	LOC410875	Gene	LOC410875	protein_coding	NA	character(0)	character(0)	NA	character(0)	NA
NC_037639.1	11351328	11351328	1	-	4	4	0	100	NC_037639.1	11351106	11351605	500	-	Gnomon	promoter	NA	NA	gene-LOC551008
	c("BEEBASE:GB49807", "GeneID:551008")	LOC551008	Gene	LOC551008	protein_coding	NA	character(0)	character(0)	NA	character(0)	NA
NC_037639.1	11351328	11351328	1	-	4	4	0	100	NC_037639.1	11351287	11351786	500	+	Gnomon	promoter	NA	NA	gene-LOC100578691	NA
	c("BEEBASE:GB49758", "GeneID:100578691")	LOC100578691	Gene	LOC100578691	protein_coding	NA	character(0)	character(0)	NA	character(0)	NA
NC_037639.1	12453240	12453240	1	-	4	4	0	100	NC_037639.1	12452945	12453444	500	-	Gnomon	promoter	NA	NA	gene-LOC113218553
	GeneID:113218553	LOC113218553	Gene	LOC113218553	lncRNA	NA	character(0)	character(0)	NA	character(0)	NA
NC_037639.1	12453314	12453314	1	+	4	4	0	100	NC_037639.1	12452945	12453444	500	-	Gnomon	promoter	NA	NA	gene-LOC113218553
	GeneID:113218553	LOC113218553	Gene	LOC113218553	lncRNA	NA	character(0)	character(0)	NA	character(0)	NA
...
... only first 25 lines shown ...
```

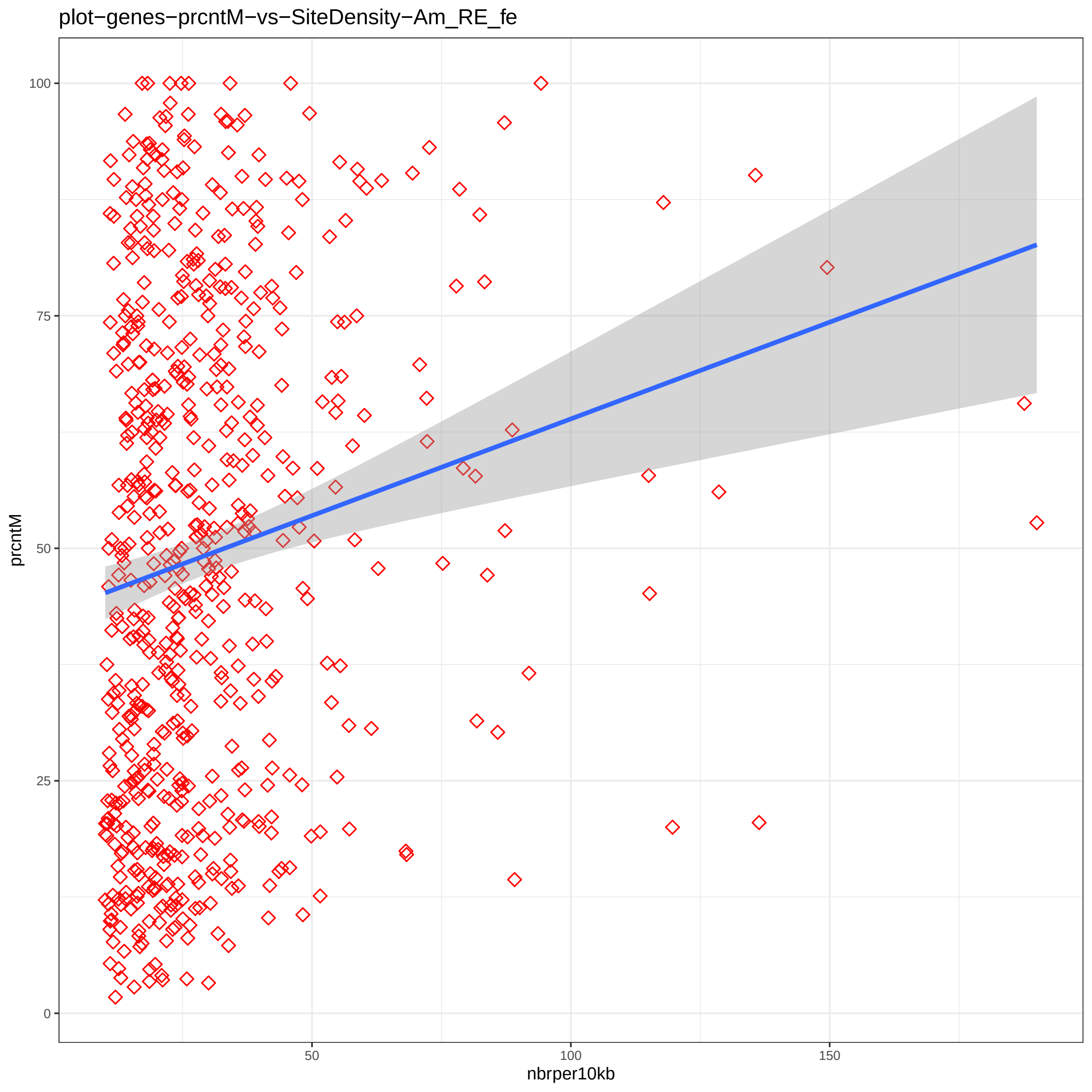
plot−genes−prcntM−vs−SiteDensity−Am_RE_fe
prcntM
nbrper10kb
0
25
50
75
100
50
100
150

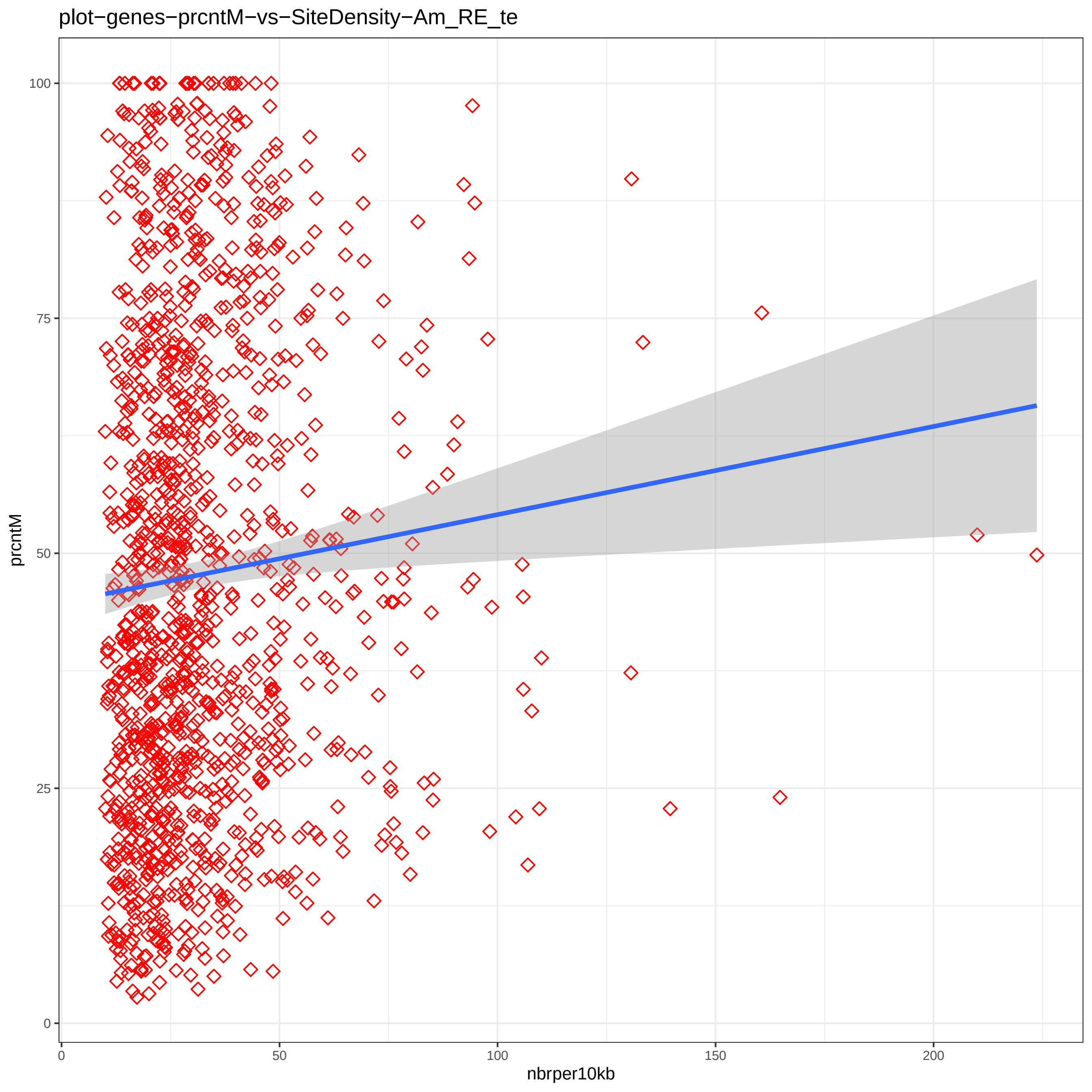
plot−genes−prcntM−vs−SiteDensity−Am_RE_te
prcntM
0
25
50
75
100
nbrper10kb
0
50
100
150
200

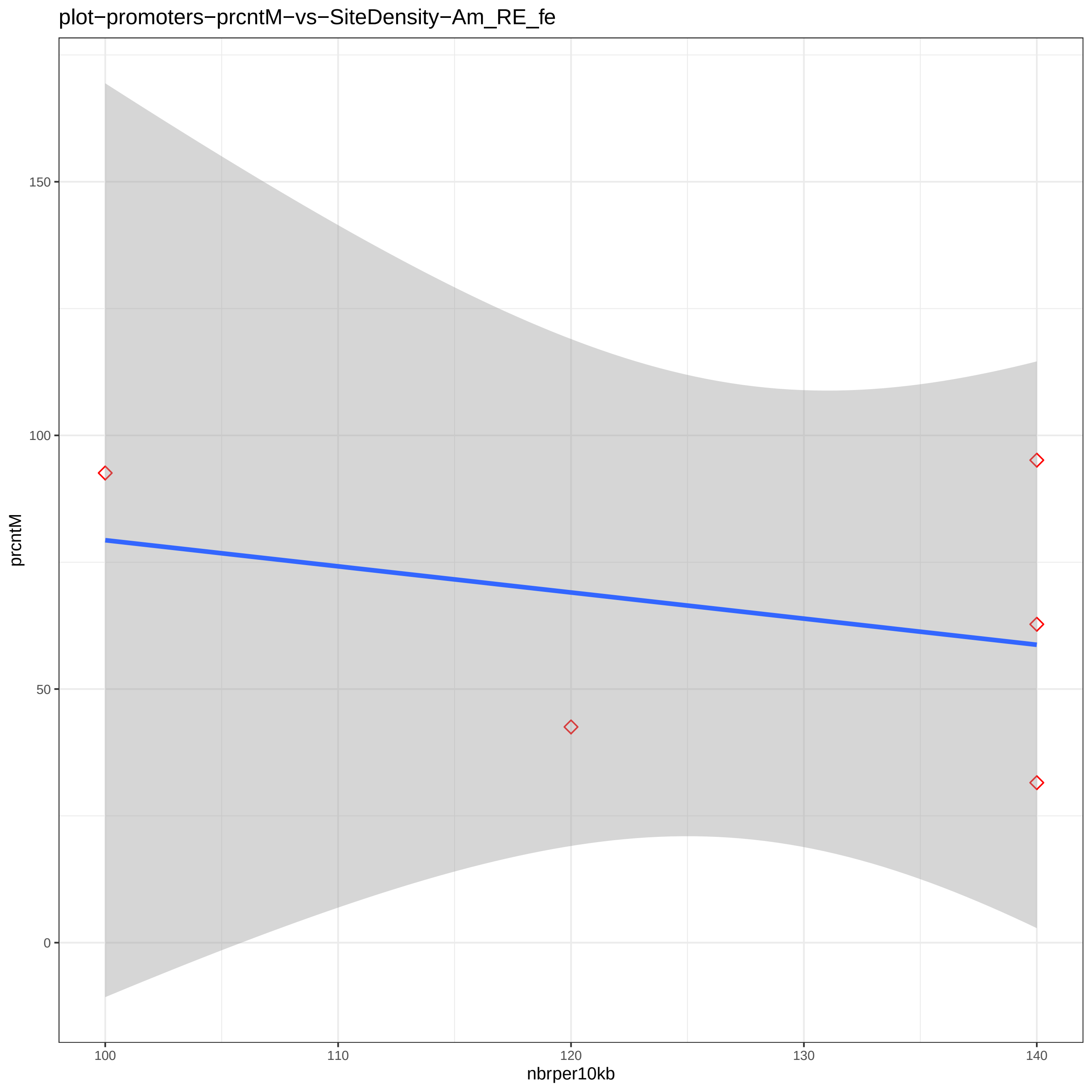

plot−promoters−prcntM−vs−SiteDensity−Am_RE_fe
prcntM
nbrper10kb
150
100
50
0
100
110
120
130
140

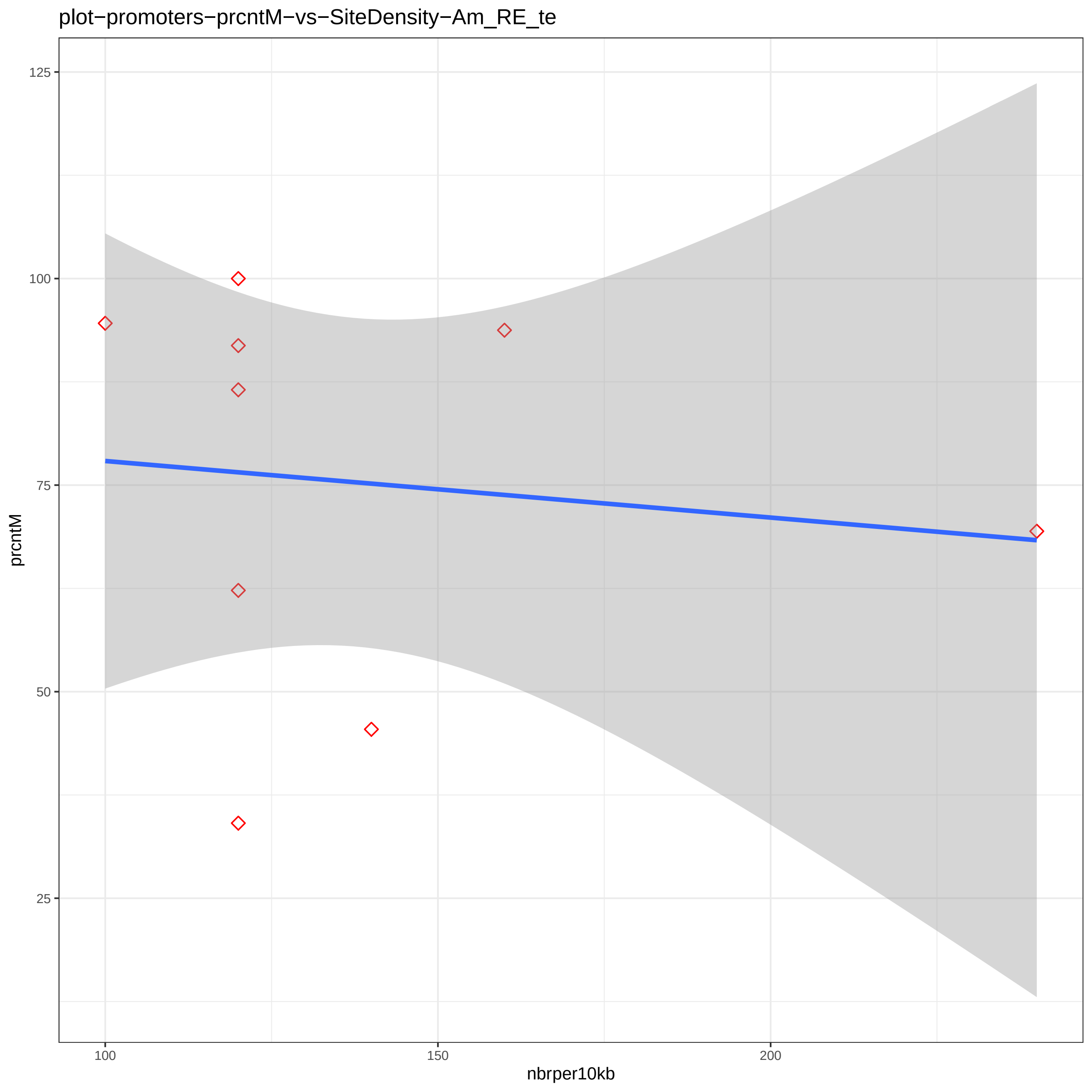
plot−promoters−prcntM−vs−SiteDensity−Am_RE_te
prcntM
nbrper10kb
125
100
75
50
25
100
150
200

```
Directory: MRPR File: 0READMEmrpr


MRPR - Methylation-rich and -poor regions.

Input: studymk, genome_ann (a list of GRanges objects providing annotated region labels and bounds;
                            output of BWASPR::get_genome_annotation())
       nbrxtrms (specifies the number of richest and poorest regions to be displayed; default: 100)

Output: files dst-*.txt 1ds-*.pdf 5ds-*.pdf    mdr-*.bed   rmp-* gwp-* gwr-*

Notes: The output is generated by BWASPR functions det_mrpr() and map_mrpr().
       Methylation-rich and -poor regions are determined based on the spacing between
        neighboring CpGhsm sites.  If sites occur at positions a, b, c, d, e, and f
        (and not in between), then b-a, c-b, ... are 1-distances, and f-a is a 5-distance.
        The file dst-*.txt shows the empirical distribution of d-distances for the sample.
        Methylation-poor regions are determined as long 1-distances (in the top <nbrxtrms>; merged
        if adjacent 1-distances are both in the top <nbrxtrms>).  Methylation-rich regions are determined
        as short 5-distances (in the low <nbrxtrms>; merged if adjacent 5-distances are both in the low
        <nbrxtrms>).  Parts of the distribution are plotted in the 1ds-*.pdf and 5ds-*.pdf files.

       The methylation-rich and -poor regions are listed in files dst-*.txt and mdr-*.bed,
        the latter in BED format for potential display in genome browsers.

       The overlap of methylation regions with genome features is summarized in files rmp-*.txt.
        Files gwr-*.txt show genes overlapping with methylation-rich regions, ordered by site
        density in the methylation-rich region.
        Files gwp-*.txt show genes overlapping with methylation-poor regions, ordered by site
        density in the methylation-poor region.
```

Directory: MRPR File: dst-Am_RE_fe.txt

Analysis of 1-distances for sample "Am_RE_fe" (18314 sites):

Total number of 1-distances: 18281
Median: 174.0 Mean: 11937.4 Std: 67350.7
Quantiles:

| 0% | 5% | 10% | 15% | 20% | 25% | 30% | 35% | 40% | 45% | 50% | 55% | 60% | 65% | 70% | 75% | 80% | 85% | 90% | 95% | 100% |
|---|---|---|---|---|---|---|---|---|---|---|---|---|---|---|---|---|---|---|---|---|
| 1 | 1 | 6 | 12 | 19 | 27 | 37 | 51 | 74 | 119 | 174 | 273 | 415 | 628 | 1006 | 1748 | 2844 | 4503 | 8580 | 35809 | 2054824 |

Low density region distance cutoff (lowest 100): 431566
High density region distance cutoff (highest 100): 1

Ordered lists of methylation-poor regions:

Top 100 (out of 100) methylation-poor regions in "Am_RE_fe":
(Sdnsty = sites per 1kb)

| | Rtype | Sample | SeqID | From | To | Rlgth | NbrSites | Sdnsty |
|---|---|---|---|---|---|---|---|---|
| 1 | Poor | fe | NC_037646.1 | 8838034 | 10892858 | 2054825 | 2 | 0.000973 |
| 2 | Poor | fe | NC_037644.1 | 5877852 | 7501133 | 1623282 | 2 | 0.001232 |
| 3 | Poor | fe | NC_037642.1 | 2998069 | 4544984 | 1546916 | 2 | 0.001293 |
| 4 | Poor | fe | NC_037650.1 | 7383993 | 8723904 | 1339912 | 2 | 0.001493 |
| 5 | Poor | fe | NC_037653.1 | 3186021 | 4513856 | 1327836 | 2 | 0.001506 |
| 6 | Poor | fe | NC_037649.1 | 6083838 | 7318003 | 1234166 | 2 | 0.001621 |
| 7 | Poor | fe | NC_037647.1 | 1069740 | 2258984 | 1189245 | 2 | 0.001682 |
| 8 | Poor | fe | NC_037651.1 | 3455944 | 4594019 | 1138076 | 2 | 0.001757 |
| 9 | Poor | fe | NC_037648.1 | 5031399 | 6163202 | 1131804 | 2 | 0.001767 |
| 10 | Poor | fe | NC_037648.1 | 8317397 | 9436732 | 1119336 | 2 | 0.001787 |
| 11 | Poor | fe | NC_037643.1 | 9763148 | 10865513 | 1102366 | 2 | 0.001814 |
| 12 | Poor | fe | NC_037643.1 | 606515 | 1649376 | 1042862 | 2 | 0.001918 |
| 13 | Poor | fe | NC_037644.1 | 11060801 | 12061035 | 1000235 | 2 | 0.002000 |
| 14 | Poor | fe | NC_037645.1 | 10545948 | 11545216 | 999269 | 2 | 0.002001 |
| 15 | Poor | fe | NC_037638.1 | 13030149 | 14020785 | 990637 | 2 | 0.002019 |
| 16 | Poor | fe | NC_037643.1 | 3800824 | 4791295 | 990472 | 2 | 0.002019 |
| 17 | Poor | fe | NC_037643.1 | 10866185 | 11851233 | 985049 | 2 | 0.002030 |
| 18 | Poor | fe | NC_037649.1 | 4255169 | 5205913 | 950745 | 2 | 0.002104 |
| 19 | Poor | fe | NC_037641.1 | 6967501 | 7857039 | 889539 | 2 | 0.002248 |
| 20 | Poor | fe | NC_037638.1 | 17009505 | 17888909 | 879405 | 2 | 0.002274 |
| 21 | Poor | fe | NC_037639.1 | 8165986 | 9031116 | 865131 | 2 | 0.002312 |
| 22 | Poor | fe | NC_037644.1 | 3800295 | 4657968 | 857674 | 2 | 0.002332 |
| 23 | Poor | fe | NC_037639.1 | 3022299 | 3870294 | 847996 | 2 | 0.002359 |
| 24 | Poor | fe | NC_037644.1 | 9656620 | 10486209 | 829590 | 2 | 0.002411 |
| 25 | Poor | fe | NC_037649.1 | 5254277 | 6076213 | 821937 | 2 | 0.002433 |
| 26 | Poor | fe | NC_037638.1 | 3281090 | 4102987 | 821898 | 2 | 0.002433 |
| 27 | Poor | fe | NC_037648.1 | 1601211 | 2414333 | 813123 | 2 | 0.002460 |
| 28 | Poor | fe | NC_037639.1 | 6991804 | 7803611 | 811808 | 2 | 0.002464 |
| 29 | Poor | fe | NC_037638.1 | 8246243 | 9057910 | 811668 | 2 | 0.002464 |
| 30 | Poor | fe | NC_037648.1 | 7240503 | 8042913 | 802411 | 2 | 0.002492 |
| 31 | Poor | fe | NC_037643.1 | 5667647 | 6465169 | 797523 | 2 | 0.002508 |
| 32 | Poor | fe | NC_037648.1 | 3462988 | 4260283 | 797296 | 2 | 0.002508 |
| 33 | Poor | fe | NC_037638.1 | 21787714 | 22583166 | 795453 | 2 | 0.002514 |
| 34 | Poor | fe | NC_037641.1 | 4010774 | 4793285 | 782512 | 2 | 0.002556 |
| 35 | Poor | fe | NC_037638.1 | 7147893 | 7926290 | 778398 | 2 | 0.002569 |
| 36 | Poor | fe | NC_037650.1 | 8738884 | 9496755 | 757872 | 2 | 0.002639 |
| 37 | Poor | fe | NC_037642.1 | 1016984 | 1761709 | 744726 | 2 | 0.002686 |
| 38 | Poor | fe | NC_037643.1 | 13392893 | 14129751 | 736859 | 2 | 0.002714 |
| 39 | Poor | fe | NC_037641.1 | 1018454 | 1752694 | 734241 | 2 | 0.002724 |
| 40 | Poor | fe | NC_037638.1 | 15874696 | 16591941 | 717246 | 2 | 0.002788 |
| 41 | Poor | fe | NC_037643.1 | 2979811 | 3696360 | 716550 | 2 | 0.002791 |
| 42 | Poor | fe | NC_037653.1 | 1996672 | 2696838 | 700167 | 2 | 0.002856 |

| | | | | | | | | |
|---|---|---|---|---|---|---|---|---|
| 43 | Poor | fe | NC_037650.1 | 9591 | 708402 | 698812 | 2 | 0.002862 |
| 44 | Poor | fe | NC_037643.1 | 12227477 | 12922276 | 694800 | 2 | 0.002879 |
| 45 | Poor | fe | NC_037643.1 | 7790129 | 8476118 | 685990 | 2 | 0.002915 |
| 46 | Poor | fe | NC_037649.1 | 10717773 | 11382988 | 665216 | 2 | 0.003007 |
| 47 | Poor | fe | NC_037649.1 | 1388680 | 2052185 | 663506 | 2 | 0.003014 |
| 48 | Poor | fe | NC_037643.1 | 17093624 | 17751684 | 658061 | 2 | 0.003039 |
| 49 | Poor | fe | NC_037638.1 | 26191020 | 26832973 | 641954 | 2 | 0.003115 |
| 50 | Poor | fe | NC_037641.1 | 5928690 | 6565365 | 636676 | 2 | 0.003141 |
| 51 | Poor | fe | NC_037645.1 | 9835111 | 10471468 | 636358 | 2 | 0.003143 |
| 52 | Poor | fe | NC_037652.1 | 4366627 | 5002947 | 636321 | 2 | 0.003143 |
| 53 | Poor | fe | NC_037651.1 | 1769175 | 2391989 | 622815 | 2 | 0.003211 |
| 54 | Poor | fe | NC_037642.1 | 7610999 | 8230196 | 619198 | 2 | 0.003230 |
| 55 | Poor | fe | NC_037638.1 | 26851028 | 27466838 | 615811 | 2 | 0.003248 |
| 56 | Poor | fe | NC_037646.1 | 1945917 | 2559144 | 613228 | 2 | 0.003261 |
| 57 | Poor | fe | NC_037642.1 | 5930821 | 6536663 | 605843 | 2 | 0.003301 |
| 58 | Poor | fe | NC_037645.1 | 5274338 | 5875939 | 601602 | 2 | 0.003324 |
| 59 | Poor | fe | NC_037645.1 | 4483870 | 5077208 | 593339 | 2 | 0.003371 |
| 60 | Poor | fe | NC_037647.1 | 2259073 | 2847510 | 588438 | 2 | 0.003399 |
| 61 | Poor | fe | NC_037649.1 | 10096825 | 10683401 | 586577 | 2 | 0.003410 |
| 62 | Poor | fe | NC_037653.1 | 5284669 | 5861118 | 576450 | 2 | 0.003470 |
| 63 | Poor | fe | NC_037638.1 | 248644 | 817455 | 568812 | 2 | 0.003516 |
| 64 | Poor | fe | NC_037641.1 | 2255580 | 2802874 | 547295 | 2 | 0.003654 |
| 65 | Poor | fe | NC_037652.1 | 818127 | 1362311 | 544185 | 2 | 0.003675 |
| 66 | Poor | fe | NC_037643.1 | 2410981 | 2951266 | 540286 | 2 | 0.003702 |
| 67 | Poor | fe | NC_037638.1 | 2648332 | 3188266 | 539935 | 2 | 0.003704 |
| 68 | Poor | fe | NC_037643.1 | 14511759 | 15041350 | 529592 | 2 | 0.003776 |
| 69 | Poor | fe | NC_037638.1 | 17902222 | 18431677 | 529456 | 2 | 0.003777 |
| 70 | Poor | fe | NC_037647.1 | 421823 | 945786 | 523964 | 2 | 0.003817 |
| 71 | Poor | fe | NC_037646.1 | 4166903 | 4689346 | 522444 | 2 | 0.003828 |
| 72 | Poor | fe | NC_037639.1 | 10532597 | 11051241 | 518645 | 2 | 0.003856 |
| 73 | Poor | fe | NC_037643.1 | 9162210 | 9674562 | 512353 | 2 | 0.003904 |
| 74 | Poor | fe | NC_037642.1 | 11443055 | 11955384 | 512330 | 2 | 0.003904 |
| 75 | Poor | fe | NC_037639.1 | 12731740 | 13243040 | 511301 | 2 | 0.003912 |
| 76 | Poor | fe | NC_037643.1 | 54911 | 562936 | 508026 | 2 | 0.003937 |
| 77 | Poor | fe | NC_037652.1 | 5227090 | 5734366 | 507277 | 2 | 0.003943 |
| 78 | Poor | fe | NC_037639.1 | 1133144 | 1636171 | 503028 | 2 | 0.003976 |
| 79 | Poor | fe | NC_037638.1 | 9727833 | 10228427 | 500595 | 2 | 0.003995 |
| 80 | Poor | fe | NC_037640.1 | 7102975 | 7601454 | 498480 | 2 | 0.004012 |
| 81 | Poor | fe | NC_037639.1 | 4821326 | 5318834 | 497509 | 2 | 0.004020 |
| 82 | Poor | fe | NC_037641.1 | 2837360 | 3333824 | 496465 | 2 | 0.004028 |
| 83 | Poor | fe | NC_037638.1 | 1157176 | 1652819 | 495644 | 2 | 0.004035 |
| 84 | Poor | fe | NC_037652.1 | 46924 | 540142 | 493219 | 2 | 0.004055 |
| 85 | Poor | fe | NC_037649.1 | 3349591 | 3839794 | 490204 | 2 | 0.004080 |
| 86 | Poor | fe | NC_037640.1 | 1437761 | 1925533 | 487773 | 2 | 0.004100 |
| 87 | Poor | fe | NC_037643.1 | 7305343 | 7786882 | 481540 | 2 | 0.004153 |
| 88 | Poor | fe | NC_037653.1 | 6650384 | 7131200 | 480817 | 2 | 0.004160 |
| 89 | Poor | fe | NC_037647.1 | 3828907 | 4304038 | 475132 | 2 | 0.004209 |
| 90 | Poor | fe | NC_037642.1 | 2288941 | 2764047 | 475107 | 2 | 0.004210 |
| 91 | Poor | fe | NC_037648.1 | 11442633 | 11913006 | 470374 | 2 | 0.004252 |
| 92 | Poor | fe | NC_037647.1 | 6491464 | 6961486 | 470023 | 2 | 0.004255 |
| 93 | Poor | fe | NC_037652.1 | 8350518 | 8815125 | 464608 | 2 | 0.004305 |
| 94 | Poor | fe | NC_037641.1 | 10606281 | 11068746 | 462466 | 2 | 0.004325 |
| 95 | Poor | fe | NC_037641.1 | 9059656 | 9512470 | 452815 | 2 | 0.004417 |
| 96 | Poor | fe | NC_037647.1 | 7068261 | 7516198 | 447938 | 2 | 0.004465 |
| 97 | Poor | fe | NC_037651.1 | 589291 | 1027173 | 437883 | 2 | 0.004567 |
| 98 | Poor | fe | NC_037642.1 | 8887078 | 9321991 | 434914 | 2 | 0.004599 |
| 99 | Poor | fe | NC_037645.1 | 2801114 | 3233156 | 432043 | 2 | 0.004629 |
| 100 | Poor | fe | NC_037638.1 | 22828702 | 23260268 | 431567 | 2 | 0.004634 |

Analysis of 5-distances for sample "Am_RE_fe" (18314 sites):

Total number of 5-distances: 18172

Median:  8018.5 Mean: 59616.2 Std: 156115.1
Quantiles:

| 0% | 5% | 10% | 15% | 20% | 25% | 30% | 35% | 40% | 45% | 50% | 55% | 60% | 65% | 70% | 75% | 80% | 85% | 90% | 95% | 100% |
|---|---|---|---|---|---|---|---|---|---|---|---|---|---|---|---|---|---|---|---|---|
| 9 | 144 | 341 | 638 | 1115 | 2017 | 3055 | 4027 | 5131 | 6472 | 8018 | 10155 | 12898 | 16882 | 22213 | 32292 | 50907 | 86974 | 159723 | 319699 | 2175714 |

Low  density region distance cutoff (lowest  100):  998004
High density region distance cutoff (highest 100):      38

Ordered lists of methylation-rich regions:

Top 100 (out of    55) methylation-rich regions in "Am_RE_fe":
(Sdnsty = sites per 1kb)

| | Rtype | Sample | SeqID | From | To | Rlgth | NbrSites | Sdnsty |
|---|---|---|---|---|---|---|---|---|
| 1 | Rich | fe | NC_037647.1 | 6387427 | 6387442 | 16 | 6 | 375 |
| 2 | Rich | fe | NC_037642.1 | 12532230 | 12532248 | 19 | 7 | 368 |
| 3 | Rich | fe | NC_037638.1 | 2214300 | 2214317 | 18 | 6 | 333 |
| 4 | Rich | fe | NC_037645.1 | 2212217 | 2212254 | 38 | 10 | 263 |
| 5 | Rich | fe | NC_037644.1 | 12524163 | 12524185 | 23 | 6 | 261 |
| 6 | Rich | fe | NC_037645.1 | 2688050 | 2688072 | 23 | 6 | 261 |
| 7 | Rich | fe | NC_037647.1 | 5753063 | 5753085 | 23 | 6 | 261 |
| 8 | Rich | fe | NC_037640.1 | 11065219 | 11065242 | 24 | 6 | 250 |
| 9 | Rich | fe | NC_037649.1 | 10710779 | 10710802 | 24 | 6 | 250 |
| 10 | Rich | fe | NC_037644.1 | 12764403 | 12764437 | 35 | 8 | 229 |
| 11 | Rich | fe | NC_037645.1 | 12285770 | 12285796 | 27 | 6 | 222 |
| 12 | Rich | fe | NC_037646.1 | 11560054 | 11560080 | 27 | 6 | 222 |
| 13 | Rich | fe | NC_037639.1 | 2517093 | 2517124 | 32 | 7 | 219 |
| 14 | Rich | fe | NC_037638.1 | 11747156 | 11747185 | 30 | 6 | 200 |
| 15 | Rich | fe | NC_037638.1 | 25095007 | 25095046 | 40 | 8 | 200 |
| 16 | Rich | fe | NC_037651.1 | 10489142 | 10489171 | 30 | 6 | 200 |
| 17 | Rich | fe | NC_037647.1 | 7525901 | 7525941 | 41 | 8 | 195 |
| 18 | Rich | fe | NC_037645.1 | 1950793 | 1950823 | 31 | 6 | 194 |
| 19 | Rich | fe | NC_037639.1 | 10191356 | 10191397 | 42 | 8 | 190 |
| 20 | Rich | fe | NC_037642.1 | 13820439 | 13820480 | 42 | 8 | 190 |
| 21 | Rich | fe | NC_037646.1 | 7758193 | 7758229 | 37 | 7 | 189 |
| 22 | Rich | fe | NC_037638.1 | 1812410 | 1812441 | 32 | 6 | 188 |
| 23 | Rich | fe | NC_037639.1 | 16058424 | 16058455 | 32 | 6 | 188 |
| 24 | Rich | fe | NC_037642.1 | 8472589 | 8472652 | 64 | 12 | 188 |
| 25 | Rich | fe | NC_037646.1 | 7023557 | 7023588 | 32 | 6 | 188 |
| 26 | Rich | fe | NC_037652.1 | 2790699 | 2790730 | 32 | 6 | 188 |
| 27 | Rich | fe | NC_037645.1 | 12289475 | 12289507 | 33 | 6 | 182 |
| 28 | Rich | fe | NC_037639.1 | 16050608 | 16050646 | 39 | 7 | 179 |
| 29 | Rich | fe | NC_037641.1 | 12681077 | 12681115 | 39 | 7 | 179 |
| 30 | Rich | fe | NW_020555859.1 | 12467 | 12505 | 39 | 7 | 179 |
| 31 | Rich | fe | NC_037638.1 | 20108766 | 20108810 | 45 | 8 | 178 |
| 32 | Rich | fe | NC_037641.1 | 1779461 | 1779494 | 34 | 6 | 176 |
| 33 | Rich | fe | NC_037648.1 | 10007513 | 10007546 | 34 | 6 | 176 |
| 34 | Rich | fe | NW_020555894.1 | 8349 | 8382 | 34 | 6 | 176 |
| 35 | Rich | fe | NC_037640.1 | 12174016 | 12174055 | 40 | 7 | 175 |
| 36 | Rich | fe | NC_037643.1 | 7301061 | 7301100 | 40 | 7 | 175 |
| 37 | Rich | fe | NC_037641.1 | 1779739 | 1779773 | 35 | 6 | 171 |
| 38 | Rich | fe | NC_037650.1 | 3497094 | 3497128 | 35 | 6 | 171 |
| 39 | Rich | fe | NC_037652.1 | 5098826 | 5098860 | 35 | 6 | 171 |
| 40 | Rich | fe | NC_037647.1 | 265987 | 266045 | 59 | 10 | 169 |
| 41 | Rich | fe | NC_037642.1 | 11252265 | 11252300 | 36 | 6 | 167 |
| 42 | Rich | fe | NC_037640.1 | 237176 | 237212 | 37 | 6 | 162 |
| 43 | Rich | fe | NC_037640.1 | 11950567 | 11950603 | 37 | 6 | 162 |
| 44 | Rich | fe | NC_037642.1 | 13758842 | 13758878 | 37 | 6 | 162 |
| 45 | Rich | fe | NC_037644.1 | 10937848 | 10937884 | 37 | 6 | 162 |
| 46 | Rich | fe | NC_037645.1 | 2212373 | 2212409 | 37 | 6 | 162 |
| 47 | Rich | fe | NC_037644.1 | 8429547 | 8429609 | 63 | 10 | 159 |
| 48 | Rich | fe | NC_037645.1 | 6582331 | 6582368 | 38 | 6 | 158 |
| 49 | Rich | fe | NC_037648.1 | 15962246 | 15962283 | 38 | 6 | 158 |

| | | | | | | | | |
|---|---|---|---|---|---|---|---|---|
| 50 | Rich | fe | NC_037648.1 | 15962171 | 15962209 | 39 | 6 | 154 |
| 51 | Rich | fe | NC_037650.1 | 3086594 | 3086646 | 53 | 8 | 151 |
| 52 | Rich | fe | NC_037646.1 | 7765421 | 7765527 | 107 | 16 | 150 |
| 53 | Rich | fe | NC_037639.1 | 15974371 | 15974432 | 62 | 9 | 145 |
| 54 | Rich | fe | NC_037652.1 | 1799916 | 1799984 | 69 | 10 | 145 |
| 55 | Rich | fe | NC_037642.1 | 12531818 | 12531932 | 115 | 16 | 139 |

Directory: MRPR File: dst-Am_RE_te.txt

Analysis of 1-distances for sample "Am_RE_te" (27814 sites):

Total number of 1-distances: 27781
Median: 119.0 Mean: 7844.9 Std: 53568.4
Quantiles:

| 0% | 5% | 10% | 15% | 20% | 25% | 30% | 35% | 40% | 45% | 50% | 55% | 60% | 65% | 70% | 75% | 80% | 85% | 90% | 95% | 100% |
|---|---|---|---|---|---|---|---|---|---|---|---|---|---|---|---|---|---|---|---|---|
| 1 | 1 | 1 | 5 | 13 | 22 | 32 | 44 | 62 | 87 | 119 | 161 | 230 | 330 | 480 | 741 | 1321 | 2410 | 4366 | 15195 | 1623393 |

Low density region distance cutoff (lowest 100): 446129
High density region distance cutoff (highest 100): 1

Ordered lists of methylation-poor regions:

Top 100 (out of 99) methylation-poor regions in "Am_RE_te":
(Sdnsty = sites per 1kb)

| | Rtype | Sample | SeqID | From | To | Rlgth | NbrSites | Sdnsty |
|---|---|---|---|---|---|---|---|---|
| 1 | Poor | te | NC_037644.1 | 5877978 | 7501371 | 1623394 | 2 | 0.00123 |
| 2 | Poor | te | NC_037646.1 | 9472317 | 10859195 | 1386879 | 2 | 0.00144 |
| 3 | Poor | te | NC_037653.1 | 3181830 | 4513371 | 1331542 | 2 | 0.00150 |
| 4 | Poor | te | NC_037650.1 | 7383994 | 8688940 | 1304947 | 2 | 0.00153 |
| 5 | Poor | te | NC_037647.1 | 1069917 | 2257658 | 1187742 | 2 | 0.00168 |
| 6 | Poor | te | NC_037651.1 | 3455587 | 4594644 | 1139058 | 2 | 0.00176 |

...
... only first 25 lines shown ...

Directory: MRPR File: gwp-Am_RE_fe.txt

```
seqnames	start	end	width	strand	source	type	score	phase	ID	Dbxref	Name	gbkey	gene	gene_biotype	description	gene_synonym	end_range	partial
	start_range	exception	seqnames	start	end	width	strand	NbrSites	Sdnsty
NC_037646.1	8835312	8843959	8648	+	Gnomon	gene	NA	NA	gene-LOC409004	c("BEEBASE:GB42881", "GeneID:409004")	LOC409004	Gene	LOC409004	protein_coding
	NA	character(0)	character(0)	NA	character(0)	NA	NC_037646.1	8838034	10892858	2054825	*	2	0.000973318895769713
NC_037646.1	8846893	9096263	249371	-	BestRefSeq%2CGnomon	gene	NA	NA	gene-NLG-3	c("BEEBASE:GB42603", "GeneID:409003")	NLG-3	Gene	NLG-3	protein_coding
	neuroligin 3	Nlgn3	character(0)	NA	character(0)	NA	NC_037646.1	8838034	10892858	2054825	*	2	0.000973318895769713
NC_037646.1	8868033	8885238	17206	+	Gnomon	gene	NA	NA	gene-LOC100578354	GeneID:100578354	LOC100578354	Gene	LOC100578354	lncRNA	NA
	character(0)	character(0)	NA	character(0)	NA	NC_037646.1	8838034	10892858	2054825	*	2	0.000973318895769713
NC_037646.1	9096466	9168877	72412	+	Gnomon	gene	NA	NA	gene-LOC724358	c("BEEBASE:GB42884", "GeneID:724358")	LOC724358	Gene	LOC724358	protein_coding
	NA	character(0)	character(0)	NA	character(0)	NA	NC_037646.1	8838034	10892858	2054825	*	2	0.000973318895769713
NC_037646.1	9180431	9465219	284789	+	BestRefSeq%2CGnomon	gene	NA	NA	gene-NLG-4	c("BEEBASE:GB42885", "GeneID:411321")	NLG-4	Gene	NLG-4	protein_coding
	neuroligin 4	Nlgn4	character(0)	NA	character(0)	NA	NC_037646.1	8838034	10892858	2054825	*	2	0.000973318895769713
NC_037646.1	9370774	9373667	2894	-	Gnomon	gene	NA	NA	gene-LOC100578400	c("BEEBASE:GB42602", "GeneID:100578400")	LOC100578400	Gene	LOC100578400
	protein_coding	NA	character(0)	character(0)	NA	character(0)	NA	NC_037646.1	8838034	10892858	2054825	*	2	0.000973318895769713
NC_037646.1	9469190	9470870	1681	+	Gnomon	gene	NA	NA	gene-LOC551922	c("BEEBASE:GB42887", "GeneID:551922")	LOC551922	Gene	LOC551922	protein_coding
	NA	character(0)	character(0)	NA	character(0)	NA	NC_037646.1	8838034	10892858	2054825	*	2	0.000973318895769713
NC_037646.1	9471354	9472837	1484	-	Gnomon	gene	NA	NA	gene-LOC411320	c("BEEBASE:GB42600", "GeneID:411320")	LOC411320	Gene	LOC411320	protein_coding
	NA	character(0)	character(0)	NA	character(0)	NA	NC_037646.1	8838034	10892858	2054825	*	2	0.000973318895769713
NC_037646.1	9480780	9483694	2915	+	Gnomon	gene	NA	NA	gene-LOC724521	c("BEEBASE:GB42888", "GeneID:724521")	LOC724521	Gene	LOC724521	protein_coding
	NA	character(0)	character(0)	NA	character(0)	NA	NC_037646.1	8838034	10892858	2054825	*	2	0.000973318895769713
NC_037646.1	9486764	9491937	5174	+	Gnomon	gene	NA	NA	gene-LOC724616	c("BEEBASE:GB42889", "GeneID:724616")	LOC724616	Gene	LOC724616	protein_coding
	NA	character(0)	character(0)	NA	character(0)	NA	NC_037646.1	8838034	10892858	2054825	*	2	0.000973318895769713
NC_037646.1	9492074	9494500	2427	-	Gnomon	gene	NA	NA	gene-LOC724664	c("BEEBASE:GB42599", "GeneID:724664")	LOC724664	Gene	LOC724664	protein_coding
	NA	character(0)	character(0)	NA	character(0)	NA	NC_037646.1	8838034	10892858	2054825	*	2	0.000973318895769713
NC_037646.1	9499193	9502351	3159	+	Gnomon	gene	NA	NA	gene-LOC100578860	c("BEEBASE:GB42890", "GeneID:100578860")	LOC100578860	Gene	LOC100578860
	protein_coding	NA	character(0)	character(0)	NA	character(0)	NA	NC_037646.1	8838034	10892858	2054825	*	2	0.000973318895769713
NC_037646.1	9503099	9505051	1953	-	Gnomon	gene	NA	NA	gene-LOC100578625	c("BEEBASE:GB42598", "GeneID:100578625")	LOC100578625	Gene	LOC100578625
	protein_coding	NA	character(0)	character(0)	NA	character(0)	NA	NC_037646.1	8838034	10892858	2054825	*	2	0.000973318895769713
NC_037646.1	9511443	9512568	1126	-	BestRefSeq	gene	NA	NA	gene-LOC409002	c("BEEBASE:GB42597", "GeneID:409002")	LOC409002	Gene	LOC409002	protein_coding
	cuticular protein	character(0)	character(0)	NA	character(0)	NA	NC_037646.1	8838034	10892858	2054825	*	2	0.000973318895769713
NC_037646.1	9527800	9531025	3226	-	BestRefSeq	gene	NA	NA	gene-LOC411317	c("BEEBASE:GB42596", "GeneID:411317")	LOC411317	Gene	LOC411317	protein_coding
	CG1567-like	character(0)	character(0)	NA	character(0)	NA	NC_037646.1	8838034	10892858	2054825	*	2	0.000973318895769713
NC_037646.1	9531281	9537447	6167	-	BestRefSeq	gene	NA	NA	gene-LOC100642173	GeneID:100642173	LOC100642173	Gene	LOC100642173	protein_coding
	uncharacterized LOC100642173	character(0)	character(0)	NA	character(0)	NA	NC_037646.1	8838034	10892858	2054825	*	2	0.000973318895769713
NC_037646.1	9549680	9559425	9746	-	Gnomon	gene	NA	NA	gene-LOC100578664	c("BEEBASE:GB42594", "GeneID:100578664")	LOC100578664	Gene	LOC100578664
	protein_coding	NA	character(0)	character(0)	NA	character(0)	NA	NC_037646.1	8838034	10892858	2054825	*	2	0.000973318895769713
NC_037646.1	9560992	9566506	5515	+	Gnomon	gene	NA	NA	gene-LOC102655319	GeneID:102655319	LOC102655319	Gene	LOC102655319	protein_coding	NA
	character(0)	character(0)	NA	character(0)	NA	NC_037646.1	8838034	10892858	2054825	*	2	0.000973318895769713
NC_037646.1	9566779	9575015	8237	+	Gnomon	gene	NA	NA	gene-LOC100578699	c("BEEBASE:GB42892", "GeneID:100578699")	LOC100578699	Gene	LOC100578699
	protein_coding	NA	character(0)	character(0)	NA	character(0)	NA	NC_037646.1	8838034	10892858	2054825	*	2	0.000973318895769713
NC_037646.1	9574934	9581055	6122	-	Gnomon	gene	NA	NA	gene-LOC411316	c("BEEBASE:GB42593", "GeneID:411316")	LOC411316	Gene	LOC411316	protein_coding
	NA	character(0)	character(0)	NA	character(0)	NA	NC_037646.1	8838034	10892858	2054825	*	2	0.000973318895769713
NC_037646.1	9581343	9628800	47458	+	Gnomon	gene	NA	NA	gene-LOC409001	c("BEEBASE:GB42894", "GeneID:409001")	LOC409001	Gene	LOC409001	protein_coding
	NA	character(0)	character(0)	NA	character(0)	NA	NC_037646.1	8838034	10892858	2054825	*	2	0.000973318895769713
NC_037646.1	9609953	9614134	4182	-	Gnomon	gene	NA	NA	gene-LOC102656114	GeneID:102656114	LOC102656114	Gene	LOC102656114	lncRNA	NA
	character(0)	character(0)	NA	character(0)	NA	NC_037646.1	8838034	10892858	2054825	*	2	0.000973318895769713
NC_037646.1	9630658	9684832	54175	-	Gnomon	gene	NA	NA	gene-LOC411315	c("BEEBASE:GB42592", "GeneID:411315")	LOC411315	Gene	LOC411315	protein_coding
	NA	character(0)	character(0)	NA	character(0)	NA	NC_037646.1	8838034	10892858	2054825	*	2	0.000973318895769713
NC_037646.1	9692542	9692649	108	-	BestRefSeq	gene	NA	NA	gene-Mir3752	c("GeneID:100629002", "miRBase:MI0016152")	Mir3752	Gene	Mir3752	miRNA
	microRNA 3752	ame-mir-3752	character(0)	NA	character(0)	NA	NC_037646.1	8838034	10892858	2054825	*	2	0.000973318895769713
...
... only first 25 lines shown ...
```

Directory: MRPR File: gwp-Am_RE_te.txt

```
seqnames    start end   width strand      source      type  score phase ID    Dbxref      Name  gbkey gene  gene_biotype      description gene_synonym      end_range   partial
      start_range exception    seqnames    start end   width strand      NbrSites    Sdnsty
NC_037644.1 5875772     5879187     3416  +     Gnomon      gene  NA    NA    gene-LOC409386    c("BEEBASE:GB45752", "GeneID:409386")    LOC409386   Gene  LOC409386   protein_coding
      NA    character(0)      character(0)      NA    character(0)      NA    NC_037644.1 5877978     7501371     1623394     *     2     0.00123198681281316
NC_037644.1 5897044     6049869     152826      +     Gnomon      gene  NA    NA    gene-LOC726401    c("BEEBASE:GB51953", "GeneID:726401")    LOC726401   Gene  LOC726401
      protein_coding    NA    character(0)      character(0)      NA    character(0)      NA    NC_037644.1 5877978     7501371     1623394     *     2     0.00123198681281316
NC_037644.1 6070348     6070420     73    -     tRNAscan-SE gene  NA    NA    gene-TRNAV-UAC-2  GeneID:113218916  TRNAV-UAC   Gene  TRNAV-UAC   tRNA  NA    character(0)
      character(0)      NA    character(0)      NA    NC_037644.1 5877978     7501371     1623394     *     2     0.00123198681281316
NC_037644.1 6106098     6110611     4514  +     Gnomon      gene  NA    NA    gene-LOC107965780 GeneID:107965780  LOC107965780      Gene  LOC107965780      lncRNA      NA
      character(0)      character(0)      NA    character(0)      NA    NC_037644.1 5877978     7501371     1623394     *     2     0.00123198681281316
NC_037644.1 6167158     6485529     318372      -     Gnomon      gene  NA    NA    gene-LOC724823    c("BEEBASE:GB54299", "GeneID:724823")    LOC724823   Gene  LOC724823
      protein_coding    NA    character(0)      character(0)      NA    character(0)      NA    NC_037644.1 5877978     7501371     1623394     *     2     0.00123198681281316
NC_037644.1 6750017     6764283     14267 +     Gnomon      gene  NA    NA    gene-LOC102656403 GeneID:102656403  LOC102656403      Gene  LOC102656403      protein_coding    NA
      character(0)      character(0)      NA    character(0)      NA    NC_037644.1 5877978     7501371     1623394     *     2     0.00123198681281316
NC_037644.1 7130504     7502881     372378      -     Gnomon      gene  NA    NA    gene-LOC113218567 GeneID:113218567  LOC113218567      Gene  LOC113218567      protein_coding    NA
      character(0)      character(0)      NA    character(0)      NA    NC_037644.1 5877978     7501371     1623394     *     2     0.00123198681281316
NC_037644.1 7333530     7334715     1186  +     Gnomon      gene  NA    NA    gene-LOC113218901 GeneID:113218901  LOC113218901      Gene  LOC113218901      lncRNA      NA
      character(0)      character(0)      NA    character(0)      NA    NC_037644.1 5877978     7501371     1623394     *     2     0.00123198681281316
NC_037644.1 7407877     7411198     3322  -     Gnomon      gene  NA    NA    gene-LOC107964762 GeneID:107964762  LOC107964762      Gene  LOC107964762      protein_coding    NA
      character(0)      character(0)      NA    character(0)      NA    NC_037644.1 5877978     7501371     1623394     *     2     0.00123198681281316
NC_037644.1 7424450     7427073     2624  +     Gnomon      gene  NA    NA    gene-LOC113218900 GeneID:113218900  LOC113218900      Gene  LOC113218900      protein_coding    NA
      character(0)      character(0)      NA    character(0)      NA    NC_037644.1 5877978     7501371     1623394     *     2     0.00123198681281316
NC_037646.1 9471354     9472837     1484  -     Gnomon      gene  NA    NA    gene-LOC411320    c("BEEBASE:GB42600", "GeneID:411320")    LOC411320   Gene  LOC411320   protein_coding
      NA    character(0)      character(0)      NA    character(0)      NA    NC_037646.1 9472317     10859195    1386879     *     2     0.0014420868727553
NC_037646.1 9480780     9483694     2915  +     Gnomon      gene  NA    NA    gene-LOC724521    c("BEEBASE:GB42888", "GeneID:724521")    LOC724521   Gene  LOC724521   protein_coding
      NA    character(0)      character(0)      NA    character(0)      NA    NC_037646.1 9472317     10859195    1386879     *     2     0.0014420868727553
NC_037646.1 9486764     9491937     5174  +     Gnomon      gene  NA    NA    gene-LOC724616    c("BEEBASE:GB42889", "GeneID:724616")    LOC724616   Gene  LOC724616   protein_coding
      NA    character(0)      character(0)      NA    character(0)      NA    NC_037646.1 9472317     10859195    1386879     *     2     0.0014420868727553
NC_037646.1 9492074     9494500     2427  -     Gnomon      gene  NA    NA    gene-LOC724664    c("BEEBASE:GB42599", "GeneID:724664")    LOC724664   Gene  LOC724664   protein_coding
      NA    character(0)      character(0)      NA    character(0)      NA    NC_037646.1 9472317     10859195    1386879     *     2     0.0014420868727553
NC_037646.1 9499193     9502351     3159  +     Gnomon      gene  NA    NA    gene-LOC100578860 c("BEEBASE:GB42890", "GeneID:100578860") LOC100578860      Gene  LOC100578860
      protein_coding    NA    character(0)      character(0)      NA    character(0)      NA    NC_037646.1 9472317     10859195    1386879     *     2     0.0014420868727553
NC_037646.1 9503099     9505051     1953  -     Gnomon      gene  NA    NA    gene-LOC100578625 c("BEEBASE:GB42598", "GeneID:100578625") LOC100578625      Gene  LOC100578625
      protein_coding    NA    character(0)      character(0)      NA    character(0)      NA    NC_037646.1 9472317     10859195    1386879     *     2     0.0014420868727553
NC_037646.1 9511443     9512568     1126  -     BestRefSeq  gene  NA    NA    gene-LOC409002    c("BEEBASE:GB42597", "GeneID:409002")    LOC409002   Gene  LOC409002   protein_coding
      cuticular protein character(0)      character(0)      NA    character(0)      NA    NC_037646.1 9472317     10859195    1386879     *     2     0.0014420868727553
NC_037646.1 9527800     9531025     3226  -     BestRefSeq  gene  NA    NA    gene-LOC411317    c("BEEBASE:GB42596", "GeneID:411317")    LOC411317   Gene  LOC411317   protein_coding
      CG1567-like character(0)      character(0)      NA    character(0)      NA    NC_037646.1 9472317     10859195    1386879     *     2     0.0014420868727553
NC_037646.1 9531281     9537447     6167  -     BestRefSeq  gene  NA    NA    gene-LOC100642173 GeneID:100642173  LOC100642173      Gene  LOC100642173      protein_coding
      uncharacterized LOC100642173  character(0)      character(0)      NA    character(0)      NA    NC_037646.1 9472317     10859195    1386879     *     2     0.0014420868727553
NC_037646.1 9549680     9559425     9746  -     Gnomon      gene  NA    NA    gene-LOC100578664 c("BEEBASE:GB42594", "GeneID:100578664") LOC100578664      Gene  LOC100578664
      protein_coding    NA    character(0)      character(0)      NA    character(0)      NA    NC_037646.1 9472317     10859195    1386879     *     2     0.0014420868727553
NC_037646.1 9560992     9566506     5515  +     Gnomon      gene  NA    NA    gene-LOC102655319 GeneID:102655319  LOC102655319      Gene  LOC102655319      protein_coding    NA
      character(0)      character(0)      NA    character(0)      NA    NC_037646.1 9472317     10859195    1386879     *     2     0.0014420868727553
NC_037646.1 9566779     9575015     8237  +     Gnomon      gene  NA    NA    gene-LOC100578699 c("BEEBASE:GB42892", "GeneID:100578699") LOC100578699      Gene  LOC100578699
      protein_coding    NA    character(0)      character(0)      NA    character(0)      NA    NC_037646.1 9472317     10859195    1386879     *     2     0.0014420868727553
NC_037646.1 9574934     9581055     6122  -     Gnomon      gene  NA    NA    gene-LOC411316    c("BEEBASE:GB42593", "GeneID:411316")    LOC411316   Gene  LOC411316   protein_coding
      NA    character(0)      character(0)      NA    character(0)      NA    NC_037646.1 9472317     10859195    1386879     *     2     0.0014420868727553
NC_037646.1 9581343     9628800     47458 +     Gnomon      gene  NA    NA    gene-LOC409001    c("BEEBASE:GB42894", "GeneID:409001")    LOC409001   Gene  LOC409001   protein_coding
      NA    character(0)      character(0)      NA    character(0)      NA    NC_037646.1 9472317     10859195    1386879     *     2     0.0014420868727553
...
... only first 25 lines shown ...
```

Directory: MRPR File: gwr-Am_RE_fe.txt

```
seqnames	start	end	width	strand	source	type	score	phase	ID	Dbxref	Name	gbkey	gene	gene_biotype	description	gene_synonym	end_range	partial
	start_range	exception	seqnames	start	end	width	strand	NbrSites	Sdnsty
NC_037647.1	6385866	6388080	2215	-	Gnomon	gene	NA	NA	gene-LOC410225	c("BEEBASE:GB48417", "GeneID:410225")	LOC410225	Gene	LOC410225	protein_coding
	NA	character(0)	character(0)	NA	character(0)	NA	NC_037647.1	6387427	6387442	16	*	6	375
NC_037642.1	12233711	12557089	323379	+	Gnomon	gene	NA	NA	gene-LOC408862	c("BEEBASE:GB41280", "GeneID:408862")	LOC408862	Gene	LOC408862
	protein_coding	NA	character(0)	character(0)	NA	character(0)	NA	NC_037642.1	12532230	12532248	19	*	7	368.421052631579
NC_037642.1	12528666	12533153	4488	-	Gnomon	gene	NA	NA	gene-LOC100576404	c("BEEBASE:GB41378", "GeneID:100576404")	LOC100576404	Gene	LOC100576404
	protein_coding	NA	character(0)	character(0)	NA	character(0)	NA	NC_037642.1	12532230	12532248	19	*	7	368.421052631579
NC_037638.1	2209902	2216661	6760	-	Gnomon	gene	NA	NA	gene-LOC550706	c("BEEBASE:GB50345", "GeneID:550706")	LOC550706	Gene	LOC550706	protein_coding
	NA	character(0)	character(0)	NA	character(0)	NA	NC_037638.1	2214300	2214317	18	*	6	333.333333333333
NC_037645.1	2209633	2219106	9474	-	Gnomon	gene	NA	NA	gene-LOC410085	c("BEEBASE:GB11704", "GeneID:410085")	LOC410085	Gene	LOC410085	protein_coding
	NA	character(0)	character(0)	NA	character(0)	NA	NC_037645.1	2212217	2212254	38	*	10	263.157894736842
NC_037644.1	12521310	12529896	8587	+	Gnomon	gene	NA	NA	gene-LOC411539	c("BEEBASE:GB44321", "GeneID:411539")	LOC411539	Gene	LOC411539	protein_coding
	NA	character(0)	character(0)	NA	character(0)	NA	NC_037644.1	12524163	12524185	23	*	6	260.869565217391
NC_037645.1	2686046	2691314	5269	+	Gnomon	gene	NA	NA	gene-LOC412115	c("BEEBASE:GB40500", "GeneID:412115")	LOC412115	Gene	LOC412115	protein_coding
	NA	character(0)	character(0)	NA	character(0)	NA	NC_037645.1	2688050	2688072	23	*	6	260.869565217391
NC_037647.1	5752275	5753292	1018	+	Gnomon	gene	NA	NA	gene-LOC726270	c("BEEBASE:GB50709", "GeneID:726270")	LOC726270	Gene	LOC726270	protein_coding
	NA	character(0)	character(0)	NA	character(0)	NA	NC_037647.1	5753063	5753085	23	*	6	260.869565217391
NC_037640.1	11060572	11083852	23281	+	Gnomon	gene	NA	NA	gene-LOC551826	c("BEEBASE:GB49193", "GeneID:551826")	LOC551826	Gene	LOC551826	protein_coding
	NA	character(0)	character(0)	NA	character(0)	NA	NC_037640.1	11065219	11065242	24	*	6	250
NC_037649.1	10710464	10712252	1789	-	Gnomon	gene	NA	NA	gene-LOC410395	c("BEEBASE:GB52058", "GeneID:410395")	LOC410395	Gene	LOC410395	protein_coding
	NA	character(0)	character(0)	NA	character(0)	NA	NC_037649.1	10710779	10710802	24	*	6	250
NC_037644.1	12756037	12765457	9421	-	Gnomon	gene	NA	NA	gene-LOC409045	c("BEEBASE:GB44331", "GeneID:409045")	LOC409045	Gene	LOC409045	protein_coding
	NA	character(0)	character(0)	NA	character(0)	NA	NC_037644.1	12764403	12764437	35	*	8	228.571428571429
NC_037645.1	12263746	12291782	28037	-	Gnomon	gene	NA	NA	gene-LOC412161	c("BEEBASE:GB51667", "GeneID:412161")	LOC412161	Gene	LOC412161	protein_coding
	NA	character(0)	character(0)	NA	character(0)	NA	NC_037645.1	12285770	12285796	27	*	6	222.222222222222
NC_037646.1	11559280	11560670	1391	+	Gnomon	gene	NA	NA	gene-LOC408986	c("BEEBASE:GB53349", "GeneID:408986")	LOC408986	Gene	LOC408986	protein_coding
	NA	character(0)	character(0)	NA	character(0)	NA	NC_037646.1	11560054	11560080	27	*	6	222.222222222222
NC_037638.1	11745931	11749466	3536	+	Gnomon	gene	NA	NA	gene-LOC726694	c("BEEBASE:GB46467", "GeneID:726694")	LOC726694	Gene	LOC726694	protein_coding
	NA	character(0)	character(0)	NA	character(0)	NA	NC_037638.1	11747156	11747185	30	*	6	200
NC_037638.1	25093307	25096555	3249	-	Gnomon	gene	NA	NA	gene-LOC412384	c("BEEBASE:GB54971", "GeneID:412384")	LOC412384	Gene	LOC412384	protein_coding
	NA	character(0)	character(0)	NA	character(0)	NA	NC_037638.1	25095007	25095046	40	*	8	200
NC_037651.1	10487864	10492899	5036	+	Gnomon	gene	NA	NA	gene-LOC100576461	c("BEEBASE:GB41658", "GeneID:100576461")	LOC100576461	Gene	LOC100576461
	protein_coding	NA	character(0)	character(0)	NA	character(0)	NA	NC_037651.1	10489142	10489171	30	*	6	200
NC_037647.1	7524180	7527616	3437	-	Gnomon	gene	NA	NA	gene-LOC411754	c("BEEBASE:GB54291", "GeneID:411754")	LOC411754	Gene	LOC411754	protein_coding
	NA	character(0)	character(0)	NA	character(0)	NA	NC_037647.1	7525901	7525941	41	*	8	195.121951219512
NC_037645.1	1947413	1952102	4690	-	Gnomon	gene	NA	NA	gene-LOC552550	c("BEEBASE:GB41165", "GeneID:552550")	LOC552550	Gene	LOC552550	protein_coding
	NA	character(0)	character(0)	NA	character(0)	NA	NC_037645.1	1950793	1950823	31	*	6	193.548387096774
NC_037639.1	10190225	10193905	3681	+	Gnomon	gene	NA	NA	gene-LOC552365	c("BEEBASE:GB52421", "GeneID:552365")	LOC552365	Gene	LOC552365	protein_coding
	NA	character(0)	character(0)	NA	character(0)	NA	NC_037639.1	10191356	10191397	42	*	8	190.47619047619
NC_037642.1	13819098	13821916	2819	-	Gnomon	gene	NA	NA	gene-LOC726957	c("BEEBASE:GB18386", "GeneID:726957")	LOC726957	Gene	LOC726957	protein_coding
	NA	character(0)	character(0)	NA	character(0)	NA	NC_037642.1	13820439	13820480	42	*	8	190.47619047619
NC_037646.1	7757020	7760724	3705	-	Gnomon	gene	NA	NA	gene-LOC411450	c("BEEBASE:GB42664", "GeneID:411450")	LOC411450	Gene	LOC411450	protein_coding
	NA	character(0)	character(0)	NA	character(0)	NA	NC_037646.1	7758193	7758229	37	*	7	189.189189189189
NC_037638.1	1810517	1813545	3029	+	Gnomon	gene	NA	NA	gene-LOC100577325	c("BEEBASE:GB47734", "GeneID:100577325")	LOC100577325	Gene	LOC100577325
	protein_coding	NA	character(0)	character(0)	NA	character(0)	NA	NC_037638.1	1812410	1812441	32	*	6	187.5
NC_037638.1	1812159	1812431	273	-	Gnomon	gene	NA	NA	gene-LOC113219401	GeneID:113219401	LOC113219401	Gene	LOC113219401	lncRNA	NA
	character(0)	character(0)	NA	character(0)	NA	NC_037638.1	1812410	1812441	32	*	6	187.5
NC_037639.1	16052535	16065479	12945	-	Gnomon	gene	NA	NA	gene-LOC410804	c("BEEBASE:GB55507", "GeneID:410804")	LOC410804	Gene	LOC410804	protein_coding
	NA	character(0)	character(0)	NA	character(0)	NA	NC_037639.1	16058424	16058455	32	*	6	187.5
NC_037639.1	16056665	16060549	3885	+	Gnomon	gene	NA	NA	gene-LOC408657	c("BEEBASE:GB55504", "GeneID:408657")	LOC408657	Gene	LOC408657	protein_coding
	NA	character(0)	character(0)	NA	character(0)	NA	NC_037639.1	16058424	16058455	32	*	6	187.5
NC_037642.1	8457126	8486687	29562	+	Gnomon	gene	NA	NA	gene-LOC414032	c("BEEBASE:GB48675", "GeneID:414032")	LOC414032	Gene	LOC414032	protein_coding
	NA	character(0)	character(0)	NA	character(0)	NA	NC_037642.1	8472589	8472652	64	*	12	187.5
NC_037642.1	8470184	8476085	5902	-	Gnomon	gene	NA	NA	gene-LOC100578106	GeneID:100578106	LOC100578106	Gene	LOC100578106	protein_coding	NA
	character(0)	character(0)	NA	character(0)	NA	NC_037642.1	8472589	8472652	64	*	12	187.5
NC_037646.1	7022176	7026424	4249	+	Gnomon	gene	NA	NA	gene-LOC412762	c("BEEBASE:GB42805", "GeneID:412762")	LOC412762	Gene	LOC412762	protein_coding
	NA	character(0)	character(0)	NA	character(0)	NA	NC_037646.1	7023557	7023588	32	*	6	187.5
NC_037652.1	2787901	2793072	5172	-	Gnomon	gene	NA	NA	gene-LOC409447	c("BEEBASE:GB54657", "GeneID:409447")	LOC409447	Gene	LOC409447	protein_coding
	NA	character(0)	character(0)	NA	character(0)	NA	NC_037652.1	2790699	2790730	32	*	6	187.5
NC_037645.1	12263746	12291782	28037	-	Gnomon	gene	NA	NA	gene-LOC412161	c("BEEBASE:GB51667", "GeneID:412161")	LOC412161	Gene	LOC412161	protein_coding
```

NA character(0) character(0) NA character(0) NA NC_037645.1 12289475 12289507 33 * 6 181.818181818182
NC_037639.1 16046083 16051632 5550 - Gnomon gene NA NA gene-LOC408656 c("BEEBASE:GB55508", "GeneID:408656") LOC408656 Gene LOC408656 protein_coding
NA character(0) character(0) NA character(0) NA NC_037639.1 16050608 16050646 39 * 7 179.487179487179
NC_037641.1 12677153 12694681 17529 + Gnomon gene NA NA gene-LOC409983 c("BEEBASE:GB52985", "GeneID:409983") LOC409983 Gene LOC409983 protein_coding
NA character(0) character(0) NA character(0) NA NC_037641.1 12681077 12681115 39 * 7 179.487179487179
NW_020555859.1 11520 12745 1226 + Gnomon gene NA NA gene-LOC102654390 GeneID:102654390 LOC102654390 Gene LOC102654390 lncRNA NA character(0)
character(0) NA character(0) NA NW_020555859.1 12467 12505 39 * 7 179.487179487179
NC_037638.1 20106642 20111054 4413 - Gnomon gene NA NA gene-LOC551494 c("BEEBASE:GB45497", "GeneID:551494") LOC551494 Gene LOC551494 protein_coding
NA character(0) character(0) NA character(0) NA NC_037638.1 20108766 20108810 45 * 8 177.777777777778
NC_037641.1 1779220 1781219 2000 - Gnomon gene NA NA gene-LOC113218557 GeneID:113218557 LOC113218557 Gene LOC113218557 lncRNA NA
character(0) character(0) NA character(0) NA NC_037641.1 1779461 1779494 34 * 6 176.470588235294
NC_037648.1 9987853 10015291 27439 + Gnomon gene NA NA gene-LOC724501 c("BEEBASE:GB47277", "GeneID:724501") LOC724501 Gene LOC724501 protein_coding
NA character(0) character(0) NA character(0) NA NC_037648.1 10007513 10007546 34 * 6 176.470588235294
NW_020555894.1 7811 10414 2604 - Gnomon gene NA NA gene-LOC113219373 GeneID:113219373 LOC113219373 Gene LOC113219373 lncRNA NA character(0)
character(0) NA character(0) NA NW_020555894.1 8349 8382 34 * 6 176.470588235294
NC_037640.1 12172999 12188461 15463 + Gnomon gene NA NA gene-LOC107964206 GeneID:107964206 LOC107964206 Gene LOC107964206 protein_coding NA
character(0) character(0) NA character(0) NA NC_037640.1 12174016 12174055 40 * 7 175
NC_037643.1 7294289 7312184 17896 - Gnomon gene NA NA gene-LOC411177 c("BEEBASE:GB48524", "GeneID:411177") LOC411177 Gene LOC411177 protein_coding
NA character(0) character(0) NA character(0) NA NC_037643.1 7301061 7301100 40 * 7 175
NC_037641.1 1779220 1781219 2000 - Gnomon gene NA NA gene-LOC113218557 GeneID:113218557 LOC113218557 Gene LOC113218557 lncRNA NA
character(0) character(0) NA character(0) NA NC_037641.1 1779739 1779773 35 * 6 171.428571428571
NC_037650.1 3491942 3503730 11789 - Gnomon gene NA NA gene-LOC410413 c("BEEBASE:GB48263", "GeneID:410413") LOC410413 Gene LOC410413 protein_coding
NA character(0) character(0) NA character(0) NA NC_037650.1 3497094 3497128 35 * 6 171.428571428571
NC_037652.1 5098612 5099750 1139 - Gnomon gene NA NA gene-LOC412554 c("BEEBASE:GB50519", "GeneID:412554") LOC412554 Gene LOC412554 protein_coding
NA character(0) character(0) NA character(0) NA NC_037652.1 5098826 5098860 35 * 6 171.428571428571
NC_037647.1 264211 268026 3816 - Gnomon gene NA NA gene-LOC411655 c("BEEBASE:GB46983", "GeneID:411655") LOC411655 Gene LOC411655 protein_coding
NA character(0) character(0) NA character(0) NA NC_037647.1 265987 266045 59 * 10 169.491525423729
NC_037642.1 11251428 11253335 1908 - Gnomon gene NA NA gene-LOC408853 c("BEEBASE:GB44585", "GeneID:408853") LOC408853 Gene LOC408853 protein_coding
NA character(0) character(0) NA character(0) NA NC_037642.1 11252265 11252300 36 * 6 166.666666666667
NC_037640.1 230965 238878 7914 - Gnomon gene NA NA gene-LOC412045 c("BEEBASE:GB47114", "GeneID:412045") LOC412045 Gene LOC412045 protein_coding
NA character(0) character(0) NA character(0) NA NC_037640.1 237176 237212 37 * 6 162.162162162162
NC_037640.1 11946155 11987911 41757 - Gnomon gene NA NA gene-LOC410922 c("BEEBASE:GB47014", "GeneID:410922") LOC410922 Gene LOC410922 protein_coding
NA character(0) character(0) NA character(0) NA NC_037640.1 11950567 11950603 37 * 6 162.162162162162
NC_037642.1 13757504 13760286 2783 - Gnomon gene NA NA gene-LOC409726 c("BEEBASE:GB45640", "GeneID:409726") LOC409726 Gene LOC409726 protein_coding
NA character(0) character(0) NA character(0) NA NC_037642.1 13758842 13758878 37 * 6 162.162162162162
NC_037644.1 10937513 10940303 2791 - BestRefSeq gene NA NA gene-Ant c("BEEBASE:GB42422", "GeneID:406075") Ant Gene Ant protein_coding ADP/ATP
translocase character(0) character(0) NA character(0) NA NC_037644.1 10937848 10937884 37 * 6 162.162162162162
NC_037645.1 2209633 2219106 9474 - Gnomon gene NA NA gene-LOC410085 c("BEEBASE:GB11704", "GeneID:410085") LOC410085 Gene LOC410085 protein_coding
NA character(0) character(0) NA character(0) NA NC_037645.1 2212373 2212409 37 * 6 162.162162162162
NC_037644.1 8420234 8431677 11444 - Gnomon gene NA NA gene-LOC408936 c("BEEBASE:GB43300", "GeneID:408936") LOC408936 Gene LOC408936 protein_coding
NA character(0) character(0) NA character(0) NA NC_037644.1 8429547 8429609 63 * 10 158.730158730159
NC_037645.1 6579241 6596535 17295 + Gnomon gene NA NA gene-LOC725679 c("BEEBASE:GB43841", "GeneID:725679") LOC725679 Gene LOC725679 protein_coding
NA character(0) character(0) NA character(0) NA NC_037645.1 6582331 6582368 38 * 6 157.894736842105
NC_037648.1 15961101 15965948 4848 + Gnomon gene NA NA gene-LOC552446 c("BEEBASE:GB43208", "GeneID:552446") LOC552446 Gene LOC552446 protein_coding
NA character(0) character(0) NA character(0) NA NC_037648.1 15962246 15962283 38 * 6 157.894736842105
NC_037648.1 15961101 15965948 4848 + Gnomon gene NA NA gene-LOC552446 c("BEEBASE:GB43208", "GeneID:552446") LOC552446 Gene LOC552446 protein_coding
NA character(0) character(0) NA character(0) NA NC_037648.1 15962171 15962209 39 * 6 153.846153846154
NC_037650.1 3084179 3087946 3768 - Gnomon gene NA NA gene-LOC413267 c("BEEBASE:GB54715", "GeneID:413267") LOC413267 Gene LOC413267 protein_coding
NA character(0) character(0) NA character(0) NA NC_037650.1 3086594 3086646 53 * 8 150.943396226415
NC_037646.1 7762983 7766728 3746 - BestRefSeq gene NA NA gene-Cox6b1 GeneID:100359409 Cox6b1 Gene Cox6b1 protein_coding cytochrome c oxidase
subunit VIb polypeptide 1 character(0) character(0) NA character(0) NA NC_037646.1 7765421 7765527 107 * 16 149.532710280374
NC_037646.1 7764886 7766728 1843 - BestRefSeq gene NA NA gene-LOC725372 c("BEEBASE:GB42662", "GeneID:725372") LOC725372 Gene LOC725372 protein_coding
uncharacterized LOC725372 character(0) character(0) NA character(0) NA NC_037646.1 7765421 7765527 107 * 16 149.532710280374
NC_037639.1 15973484 15974869 1386 - BestRefSeq gene NA NA gene-LOC726419 c("BEEBASE:GB55513", "GeneID:726419") LOC726419 Gene LOC726419 protein_coding
beta-1,3-galactosyltransferase B3GalT1 character(0) NA character(0) NA NC_037639.1 15974371 15974432 62 * 9 145.161290322581
NC_037652.1 1798228 1804590 6363 + Gnomon gene NA NA gene-LOC409573 c("BEEBASE:GB46211", "GeneID:409573") LOC409573 Gene LOC409573 protein_coding
NA character(0) character(0) NA character(0) NA NC_037652.1 1799916 1799984 69 * 10 144.927536231884
NC_037642.1 12233711 12557089 323379 + Gnomon gene NA NA gene-LOC408862 c("BEEBASE:GB41280", "GeneID:408862") LOC408862 Gene LOC408862
protein_coding NA character(0) character(0) NA character(0) NA NC_037642.1 12531818 12531932 115 * 16 139.130434782609
NC_037642.1 12528666 12533153 4488 - Gnomon gene NA NA gene-LOC100576404 c("BEEBASE:GB41378", "GeneID:100576404") LOC100576404 Gene LOC100576404
protein_coding NA character(0) character(0) NA character(0) NA NC_037642.1 12531818 12531932 115 * 16 139.130434782609

Directory: MRPR File: gwr-Am_RE_te.txt

```
seqnames	start	end	width	strand	source	type	score	phase	ID	Dbxref	Name	gbkey	gene	gene_biotype	description	gene_synonym	end_range	partial
	start_range	exception	seqnames	start	end	width	strand	NbrSites	Sdnsty
NC_037638.1	10853409	10856681	3273	+	Gnomon	gene	NA	NA	gene-LOC412343	c("BEEBASE:GB45877", "GeneID:412343")	LOC412343	Gene	LOC412343	protein_coding
	NA	character(0)	character(0)	NA	character(0)	NA	NC_037638.1	10855649	10855659	11	*	8	727.272727272727
NC_037646.1	5540266	5547250	6985	-	Gnomon	gene	NA	NA	gene-LOC409022	c("BEEBASE:GB17617", "GeneID:409022")	LOC409022	Gene	LOC409022	protein_coding
	NA	character(0)	character(0)	NA	character(0)	NA	NC_037646.1	5543629	5543644	16	*	6	375
NC_037642.1	12233711	12557089	323379	+	Gnomon	gene	NA	NA	gene-LOC408862	c("BEEBASE:GB41280", "GeneID:408862")	LOC408862	Gene	LOC408862
	protein_coding	NA	character(0)	character(0)	NA	character(0)	NA	NC_037642.1	12532230	12532248	19	*	7	368.421052631579
NC_037642.1	12528666	12533153	4488	-	Gnomon	gene	NA	NA	gene-LOC100576404	c("BEEBASE:GB41378", "GeneID:100576404")	LOC100576404	Gene	LOC100576404
	protein_coding	NA	character(0)	character(0)	NA	character(0)	NA	NC_037642.1	12532230	12532248	19	*	7	368.421052631579
NC_037652.1	1798228	1804590	6363	+	Gnomon	gene	NA	NA	gene-LOC409573	c("BEEBASE:GB46211", "GeneID:409573")	LOC409573	Gene	LOC409573	protein_coding
	NA	character(0)	character(0)	NA	character(0)	NA	NC_037652.1	1799208	1799225	18	*	6	333.333333333333
NW_020555846.1	19139	21027	1889	+	Gnomon	gene	NA	NA	gene-LOC102654884	GeneID:102654884	LOC102654884	Gene	LOC102654884	lncRNA	NA	character(0)
	character(0)	NA	character(0)	NA	NW_020555846.1	20957	20974	18	*	6	333.333333333333
NC_037642.1	12233711	12557089	323379	+	Gnomon	gene	NA	NA	gene-LOC408862	c("BEEBASE:GB41280", "GeneID:408862")	LOC408862	Gene	LOC408862
	protein_coding	NA	character(0)	character(0)	NA	character(0)	NA	NC_037642.1	12532356	12532374	19	*	6	315.789473684211
NC_037642.1	12528666	12533153	4488	-	Gnomon	gene	NA	NA	gene-LOC100576404	c("BEEBASE:GB41378", "GeneID:100576404")	LOC100576404	Gene	LOC100576404
	protein_coding	NA	character(0)	character(0)	NA	character(0)	NA	NC_037642.1	12532356	12532374	19	*	6	315.789473684211
NC_037643.1	17069372	17074875	5504	+	Gnomon	gene	NA	NA	gene-LOC726133	c("BEEBASE:GB55700", "GeneID:726133")	LOC726133	Gene	LOC726133	protein_coding
	NA	character(0)	character(0)	NA	character(0)	NA	NC_037643.1	17070228	17070246	19	*	6	315.789473684211
NC_037646.1	6559799	6565038	5240	+	Gnomon	gene	NA	NA	gene-LOC411879	c("BEEBASE:GB13929", "GeneID:411879")	LOC411879	Gene	LOC411879	protein_coding
	NA	character(0)	character(0)	NA	character(0)	NA	NC_037646.1	6561385	6561404	20	*	6	300
NC_037642.1	9903880	9908251	4372	-	Gnomon	gene	NA	NA	gene-LOC552805	c("BEEBASE:GB44651", "GeneID:552805")	LOC552805	Gene	LOC552805	protein_coding
	NA	character(0)	character(0)	NA	character(0)	NA	NC_037642.1	9904439	9904459	21	*	6	285.714285714286
NC_037646.1	1541638	1547588	5951	+	Gnomon	gene	NA	NA	gene-LOC552467	c("BEEBASE:GB45555", "GeneID:552467")	LOC552467	Gene	LOC552467	protein_coding
	NA	character(0)	character(0)	NA	character(0)	NA	NC_037646.1	1542811	1542831	21	*	6	285.714285714286
NC_037647.1	392556	396136	3581	-	Gnomon	gene	NA	NA	gene-LOC551953	c("BEEBASE:GB55979", "GeneID:551953")	LOC551953	Gene	LOC551953	protein_coding
	NA	character(0)	character(0)	NA	character(0)	NA	NC_037647.1	395331	395351	21	*	6	285.714285714286
NC_037648.1	14247574	14258092	10519	+	Gnomon	gene	NA	NA	gene-LOC107965250	GeneID:107965250	LOC107965250	Gene	LOC107965250	protein_coding	NA
	character(0)	character(0)	NA	character(0)	NA	NC_037648.1	14249408	14249430	23	*	6	260.869565217391
NC_037648.1	16196034	16198862	2829	+	Gnomon	gene	NA	NA	gene-LOC410280	c("BEEBASE:GB43229", "GeneID:410280")	LOC410280	Gene	LOC410280	protein_coding
	NA	character(0)	character(0)	NA	character(0)	NA	NC_037648.1	16196829	16196855	27	*	7	259.259259259259
NC_037638.1	2494572	2501772	7201	+	Gnomon	gene	NA	NA	gene-LOC411985	c("BEEBASE:GB50370", "GeneID:411985")	LOC411985	Gene	LOC411985	protein_coding
	NA	character(0)	character(0)	NA	character(0)	NA	NC_037638.1	2497442	2497466	25	*	6	240
NC_037648.1	16269838	16272912	3075	-	Gnomon	gene	NA	NA	gene-LOC100576660	c("BEEBASE:GB43081", "GeneID:100576660")	LOC100576660	Gene	LOC100576660
	protein_coding	NA	character(0)	character(0)	NA	character(0)	NA	NC_037648.1	16272169	16272193	25	*	6	240
NC_037647.1	7524180	7527616	3437	-	Gnomon	gene	NA	NA	gene-LOC411754	c("BEEBASE:GB54291", "GeneID:411754")	LOC411754	Gene	LOC411754	protein_coding
	NA	character(0)	character(0)	NA	character(0)	NA	NC_037647.1	7525901	7525926	26	*	6	230.769230769231
NC_037644.1	12756037	12765457	9421	-	Gnomon	gene	NA	NA	gene-LOC409045	c("BEEBASE:GB44331", "GeneID:409045")	LOC409045	Gene	LOC409045	protein_coding
	NA	character(0)	character(0)	NA	character(0)	NA	NC_037644.1	12764403	12764437	35	*	8	228.571428571429
NC_037641.1	11819133	11820284	1152	-	Gnomon	gene	NA	NA	gene-LOC725102	c("BEEBASE:GB44160", "GeneID:725102")	LOC725102	Gene	LOC725102	protein_coding
	NA	character(0)	character(0)	NA	character(0)	NA	NC_037641.1	11819439	11819465	27	*	6	222.222222222222
NC_037644.1	3614281	3615740	1460	+	Gnomon	gene	NA	NA	gene-LOC408939	c("BEEBASE:GB49244", "GeneID:408939")	LOC408939	Gene	LOC408939	protein_coding
	NA	character(0)	character(0)	NA	character(0)	NA	NC_037644.1	3615052	3615078	27	*	6	222.222222222222
NC_037642.1	8841745	8842956	1212	+	Gnomon	gene	NA	NA	gene-LOC724369	c("BEEBASE:GB50296", "GeneID:724369")	LOC724369	Gene	LOC724369	protein_coding
	NA	character(0)	character(0)	NA	character(0)	NA	NC_037642.1	8842885	8842912	28	*	6	214.285714285714
NC_037643.1	9758308	9763764	5457	-	Gnomon	gene	NA	NA	gene-LOC408882	c("BEEBASE:GB41469", "GeneID:408882")	LOC408882	Gene	LOC408882	protein_coding
	NA	character(0)	character(0)	NA	character(0)	NA	NC_037643.1	9762927	9762955	29	*	6	206.896551724138
NC_037641.1	12677153	12694681	17529	+	Gnomon	gene	NA	NA	gene-LOC409983	c("BEEBASE:GB52985", "GeneID:409983")	LOC409983	Gene	LOC409983	protein_coding
	NA	character(0)	character(0)	NA	character(0)	NA	NC_037641.1	12681077	12681115	39	*	8	205.128205128205
NC_037650.1	3084179	3087946	3768	-	Gnomon	gene	NA	NA	gene-LOC413267	c("BEEBASE:GB54715", "GeneID:413267")	LOC413267	Gene	LOC413267	protein_coding
	NA	character(0)	character(0)	NA	character(0)	NA	NC_037650.1	3086593	3086646	54	*	11	203.703703703704
NC_037638.1	10937632	10940839	3208	+	Gnomon	gene	NA	NA	gene-LOC725494	c("BEEBASE:GB45885", "GeneID:725494")	LOC725494	Gene	LOC725494	protein_coding
	NA	character(0)	character(0)	NA	character(0)	NA	NC_037638.1	10938810	10938844	35	*	7	200
NC_037642.1	9966263	9969780	3518	-	Gnomon	gene	NA	NA	gene-LOC100577295	c("BEEBASE:GB44646", "GeneID:100577295")	LOC100577295	Gene	LOC100577295
	protein_coding	NA	character(0)	character(0)	NA	character(0)	NA	NC_037642.1	9968634	9968663	30	*	6	200
NC_037648.1	14803730	14807137	3408	+	Gnomon	gene	NA	NA	gene-LOC724685	c("BEEBASE:GB45247", "GeneID:724685")	LOC724685	Gene	LOC724685	protein_coding
	NA	character(0)	character(0)	NA	character(0)	NA	NC_037648.1	14805590	14805624	35	*	7	200
NC_037642.1	9865790	9869133	3344	-	Gnomon	gene	NA	NA	gene-LOC102656501	GeneID:102656501	LOC102656501	Gene	LOC102656501	protein_coding	NA
	character(0)	character(0)	NA	character(0)	NA	NC_037642.1	9867238	9867273	36	*	7	194.444444444444
NC_037641.1	5501404	5505516	4113	-	Gnomon	gene	NA	NA	gene-LOC552614	c("BEEBASE:GB49519", "GeneID:552614")	LOC552614	Gene	LOC552614	protein_coding
```

NA character(0) character(0) NA character(0) NA NC_037641.1 5504951 5504982 32 * 6 187.5
NC_037646.1 7022176 7026424 4249 + Gnomon gene NA NA gene-LOC412762 c("BEEBASE:GB42805", "GeneID:412762") LOC412762 Gene LOC412762 protein_coding
NA character(0) character(0) NA character(0) NA NC_037646.1 7023557 7023588 32 * 6 187.5
NC_037648.1 16317895 16326171 8277 - Gnomon gene NA NA gene-LOC551500 c("BEEBASE:GB43074", "GeneID:551500") LOC551500 Gene LOC551500 protein_coding
NA character(0) character(0) NA character(0) NA NC_037648.1 16325204 16325251 48 * 9 187.5
NW_020555869.1 40445 42204 1760 - Gnomon gene NA NA gene-LOC102656390 GeneID:102656390 LOC102656390 Gene LOC102656390 lncRNA NA character(0)
character(0) NA character(0) NA NW_020555869.1 41083 41114 32 * 6 187.5
NC_037639.1 16046083 16051632 5550 - Gnomon gene NA NA gene-LOC408656 c("BEEBASE:GB55508", "GeneID:408656") LOC408656 Gene LOC408656 protein_coding
NA character(0) character(0) NA character(0) NA NC_037639.1 16050608 16050645 38 * 7 184.210526315789
NC_037640.1 23922 222272 198351 + Gnomon gene NA NA gene-LOC409422 GeneID:409422 LOC409422 Gene LOC409422 protein_coding NA character(0)
character(0) NA character(0) NA NC_037640.1 38622 38654 33 * 6 181.818181818182
NC_037646.1 11559280 11560670 1391 + Gnomon gene NA NA gene-LOC408986 c("BEEBASE:GB53349", "GeneID:408986") LOC408986 Gene LOC408986 protein_coding
NA character(0) character(0) NA character(0) NA NC_037646.1 11560037 11560080 44 * 8 181.818181818182
NC_037645.1 2798395 2805007 6613 + Gnomon gene NA NA gene-LOC413663 c("BEEBASE:GB40515", "GeneID:413663") LOC413663 Gene LOC413663 protein_coding
NA character(0) character(0) NA character(0) NA NC_037645.1 2800470 2800508 39 * 7 179.487179487179
NC_037644.1 2315532 2329396 13865 + Gnomon gene NA NA gene-LOC408952 c("BEEBASE:GB42559", "GeneID:408952") LOC408952 Gene LOC408952 protein_coding
NA character(0) character(0) NA character(0) NA NC_037644.1 2325294 2325338 45 * 8 177.777777777778
NC_037641.1 1779220 1781219 2000 - Gnomon gene NA NA gene-LOC113218557 GeneID:113218557 LOC113218557 Gene LOC113218557 lncRNA NA
character(0) character(0) NA character(0) NA NC_037641.1 1779461 1779494 34 * 6 176.470588235294
NC_037642.1 9839996 9843502 3507 + Gnomon gene NA NA gene-LOC411100 c("BEEBASE:GB44457", "GeneID:411100") LOC411100 Gene LOC411100 protein_coding
NA character(0) character(0) NA character(0) NA NC_037642.1 9841425 9841458 34 * 6 176.470588235294
NC_037645.1 6972337 6976482 4146 + Gnomon gene NA NA gene-LOC409992 c("BEEBASE:GB43857", "GeneID:409992") LOC409992 Gene LOC409992 protein_coding
NA character(0) character(0) NA character(0) NA NC_037645.1 6974327 6974360 34 * 6 176.470588235294
NW_020555894.1 7811 10414 2604 - Gnomon gene NA NA gene-LOC113219373 GeneID:113219373 LOC113219373 Gene LOC113219373 lncRNA NA character(0)
character(0) NA character(0) NA NW_020555894.1 8349 8382 34 * 6 176.470588235294
NC_037639.1 6948022 6956944 8923 + Gnomon gene NA NA gene-LOC413955 c("BEEBASE:GB50852", "GeneID:413955") LOC413955 Gene LOC413955 protein_coding
NA character(0) character(0) NA character(0) NA NC_037639.1 6949461 6949495 35 * 6 171.428571428571
NC_037641.1 1779220 1781219 2000 - Gnomon gene NA NA gene-LOC113218557 GeneID:113218557 LOC113218557 Gene LOC113218557 lncRNA NA
character(0) character(0) NA character(0) NA NC_037641.1 1779739 1779773 35 * 6 171.428571428571
NC_037647.1 264211 268026 3816 - Gnomon gene NA NA gene-LOC411655 c("BEEBASE:GB46983", "GeneID:411655") LOC411655 Gene LOC411655 protein_coding
NA character(0) character(0) NA character(0) NA NC_037647.1 265987 266045 59 * 10 169.491525423729
NC_037638.1 816289 820618 4330 + Gnomon gene NA NA gene-LOC410769 c("BEEBASE:GB42152", "GeneID:410769") LOC410769 Gene LOC410769 protein_coding
NA character(0) character(0) NA character(0) NA NC_037638.1 817573 817620 48 * 8 166.666666666667
NC_037643.1 6464125 6475423 11299 - Gnomon gene NA NA gene-LOC100577552 c("BEEBASE:GB48567", "GeneID:100577552") LOC100577552 Gene LOC100577552
protein_coding NA character(0) character(0) NA character(0) NA NC_037643.1 6465414 6465462 49 * 8 163.265306122449
NC_037638.1 10792 17180 6389 + Gnomon gene NA NA gene-LOC551555 c("BEEBASE:GB42138", "GeneID:551555") LOC551555 Gene LOC551555 protein_coding NA
character(0) character(0) NA character(0) NA NC_037638.1 13479 13515 37 * 6 162.162162162162
NC_037639.1 10170652 10173751 3100 + Gnomon gene NA NA gene-LOC550664 c("BEEBASE:GB52420", "GeneID:550664") LOC550664 Gene LOC550664 protein_coding
NA character(0) character(0) NA character(0) NA NC_037639.1 10172744 10172780 37 * 6 162.162162162162
NC_037644.1 10937513 10940303 2791 - BestRefSeq gene NA NA gene-Ant c("BEEBASE:GB42422", "GeneID:406075") Ant Gene Ant protein_coding ADP/ATP
translocase character(0) character(0) NA character(0) NA NC_037644.1 10937848 10937884 37 * 6 162.162162162162
NC_037648.1 16196034 16198862 2829 + Gnomon gene NA NA gene-LOC410280 c("BEEBASE:GB43229", "GeneID:410280") LOC410280 Gene LOC410280 protein_coding
NA character(0) character(0) NA character(0) NA NC_037648.1 16197344 16197380 37 * 6 162.162162162162
NC_037652.1 9083940 9090572 6633 + Gnomon gene NA NA gene-LOC409377 c("BEEBASE:GB50274", "GeneID:409377") LOC409377 Gene LOC409377 protein_coding
NA character(0) character(0) NA character(0) NA NC_037652.1 9085449 9085485 37 * 6 162.162162162162
NC_037639.1 15973484 15974869 1386 - BestRefSeq gene NA NA gene-LOC726419 c("BEEBASE:GB55513", "GeneID:726419") LOC726419 Gene LOC726419 protein_coding
beta-1,3-galactosyltransferase B3GalT1 character(0) NA character(0) NA NC_037639.1 15974371 15974433 63 * 10 158.730158730159
NC_037638.1 14288984 14290951 1968 - Gnomon gene NA NA gene-LOC551676 c("BEEBASE:GB55303", "GeneID:551676") LOC551676 Gene LOC551676 protein_coding
NA character(0) character(0) NA character(0) NA NC_037638.1 14289916 14289972 57 * 9 157.894736842105
NC_037641.1 1779220 1781219 2000 - Gnomon gene NA NA gene-LOC113218557 GeneID:113218557 LOC113218557 Gene LOC113218557 lncRNA NA
character(0) character(0) NA character(0) NA NC_037641.1 1779294 1779331 38 * 6 157.894736842105
NC_037641.1 12498995 12551707 52713 + Gnomon gene NA NA gene-LOC409658 c("BEEBASE:GB52975", "GeneID:409658") LOC409658 Gene LOC409658 protein_coding
NA character(0) character(0) NA character(0) NA NC_037641.1 12500779 12500816 38 * 6 157.894736842105
NC_037650.1 2776546 2783951 7406 + BestRefSeq%2CGnomon gene NA NA gene-arm c("BEEBASE:GB54774", "GeneID:408399") arm Gene arm protein_coding
armadillo segment polarity protein character(0) character(0) NA character(0) NA NC_037650.1 2777929 2777966 38 * 6 157.894736842105
NC_037651.1 6470760 6474112 3353 + Gnomon gene NA NA gene-LOC725575 GeneID:725575 LOC725575 Gene LOC725575 protein_coding NA character(0)
character(0) NA character(0) NA NC_037651.1 6473287 6473324 38 * 6 157.894736842105
NC_037639.1 14432742 14434626 1885 + Gnomon gene NA NA gene-LOC408672 c("BEEBASE:GB55420", "GeneID:408672") LOC408672 Gene LOC408672 protein_coding
NA character(0) character(0) NA character(0) NA NC_037639.1 14434062 14434100 39 * 6 153.846153846154
NC_037644.1 8420234 8431677 11444 - Gnomon gene NA NA gene-LOC408936 c("BEEBASE:GB43300", "GeneID:408936") LOC408936 Gene LOC408936 protein_coding
NA character(0) character(0) NA character(0) NA NC_037644.1 8429579 8429643 65 * 10 153.846153846154
NC_037646.1 1574656 1577281 2626 + Gnomon gene NA NA gene-LOC551554 c("BEEBASE:GB45558", "GeneID:551554") LOC551554 Gene LOC551554 protein_coding
NA character(0) character(0) NA character(0) NA NC_037646.1 1576158 1576196 39 * 6 153.846153846154
NC_037651.1 5394286 5398617 4332 - Gnomon gene NA NA gene-LOC411972 c("BEEBASE:GB51037", "GeneID:411972") LOC411972 Gene LOC411972 protein_coding

NA character(0) character(0) NA character(0) NA NC_037651.1 5396360 5396398 39 * 6 153.846153846154
NC_037642.1 12233711 12557089 323379 + Gnomon gene NA NA gene-LOC408862 c("BEEBASE:GB41280", "GeneID:408862") LOC408862 Gene LOC408862
protein_coding NA character(0) character(0) NA character(0) NA NC_037642.1 12531818 12531932 115 * 16 139.130434782609
NC_037642.1 12528666 12533153 4488 - Gnomon gene NA NA gene-LOC100576404 c("BEEBASE:GB41378", "GeneID:100576404") LOC100576404 Gene LOC100576404
protein_coding NA character(0) character(0) NA character(0) NA NC_037642.1 12531818 12531932 115 * 16 139.130434782609

```
Directory: MRPR File: rmp-Am_RE_fe.txt


Overlap of methylation-rich regions with genome features for Am sample Am_RE_fe

Total size of the methylation-rich regions of Am_RE_fe:                                       2160 bp (  0.0% of genome)
Size of overlap of methylation-rich regions of Am Am_RE_fe with genic regions:                   2128 bp ( 98.5% of total size)   ( 1.20 O/E)
Size of overlap of methylation-rich regions of Am Am_RE_fe with   exon regions:                    2004 bp ( 92.8% of total size) ( 0.33 O/E)
Size of overlap of methylation-rich regions of Am Am_RE_fe with   intron regions:                   124 bp (  5.7% of total size) ( 0.09 O/E)
Size of overlap of methylation-rich regions of Am Am_RE_fe with intergenic regions:                32 bp (  1.5% of total size)   ( 5.53 O/E)
Size of overlap of methylation-rich regions of Am Am_RE_fe with   promoter regions:                    0 bp (  0.0% of total size) ( 0.00 O/E)
Size of overlap of methylation-rich regions of Am Am_RE_fe with   other intergenic regions:          32 bp (  1.5% of total size) ( 0.10 O/E)


Overlap of methylation-poor regions with genome features for Am sample Am_RE_fe

Total size of the methylation-poor regions of Am_RE_fe:                                   72315096 bp ( 32.1% of genome)
Size of overlap of methylation-poor regions of Am Am_RE_fe with genic regions:               51571089 bp ( 71.3% of total size)   ( 0.87 O/E)
Size of overlap of methylation-poor regions of Am Am_RE_fe with   exon regions:                 4148195 bp (  5.7% of total size) ( 3.78 O/E)
Size of overlap of methylation-poor regions of Am Am_RE_fe with   intron regions:              47422894 bp ( 65.6% of total size) ( 1.01 O/E)
Size of overlap of methylation-poor regions of Am Am_RE_fe with intergenic regions:          20744007 bp ( 28.7% of total size)   ( 4.01 O/E)
Size of overlap of methylation-poor regions of Am Am_RE_fe with   promoter regions:              434150 bp (  0.6% of total size) ( 0.22 O/E)
Size of overlap of methylation-poor regions of Am Am_RE_fe with   other intergenic regions:    20309857 bp ( 28.1% of total size) ( 1.86 O/E)
```

```
Directory: MRPR File: rmp-Am_RE_te.txt


Overlap of methylation-rich regions with genome features for Am sample Am_RE_te

Total size of the methylation-rich regions of Am_RE_te:                                      2210 bp (  0.0% of genome)
Size of overlap of methylation-rich regions of Am Am_RE_te with genic regions:                  2163 bp ( 97.9% of total size)   ( 1.19 O/E)
Size of overlap of methylation-rich regions of Am Am_RE_te with   exon regions:                  2075 bp ( 93.9% of total size) ( 0.23 O/E)
Size of overlap of methylation-rich regions of Am Am_RE_te with   intron regions:                  88 bp (  4.0% of total size) ( 0.06 O/E)
Size of overlap of methylation-rich regions of Am Am_RE_te with intergenic regions:               47 bp (  2.1% of total size)   ( 5.50 O/E)
Size of overlap of methylation-rich regions of Am Am_RE_te with   promoter regions:                  0 bp (  0.0% of total size) ( 0.00 O/E)
Size of overlap of methylation-rich regions of Am Am_RE_te with   other intergenic regions:         47 bp (  2.1% of total size) ( 0.14 O/E)


Overlap of methylation-poor regions with genome features for Am sample Am_RE_te

Total size of the methylation-poor regions of Am_RE_te:                                  71620334 bp ( 31.8% of genome)
Size of overlap of methylation-poor regions of Am Am_RE_te with genic regions:              51142592 bp ( 71.4% of total size)   ( 0.87 O/E)
Size of overlap of methylation-poor regions of Am Am_RE_te with   exon regions:               3830926 bp (  5.3% of total size) ( 3.81 O/E)
Size of overlap of methylation-poor regions of Am Am_RE_te with   intron regions:            47311666 bp ( 66.1% of total size) ( 1.02 O/E)
Size of overlap of methylation-poor regions of Am Am_RE_te with intergenic regions:         20477742 bp ( 28.6% of total size)   ( 4.01 O/E)
Size of overlap of methylation-poor regions of Am Am_RE_te with   promoter regions:            398096 bp (  0.6% of total size) ( 0.20 O/E)
Size of overlap of methylation-poor regions of Am Am_RE_te with   other intergenic regions:   20079646 bp ( 28.0% of total size) ( 1.86 O/E)
```

Am_RE_fe
1−distance distributions

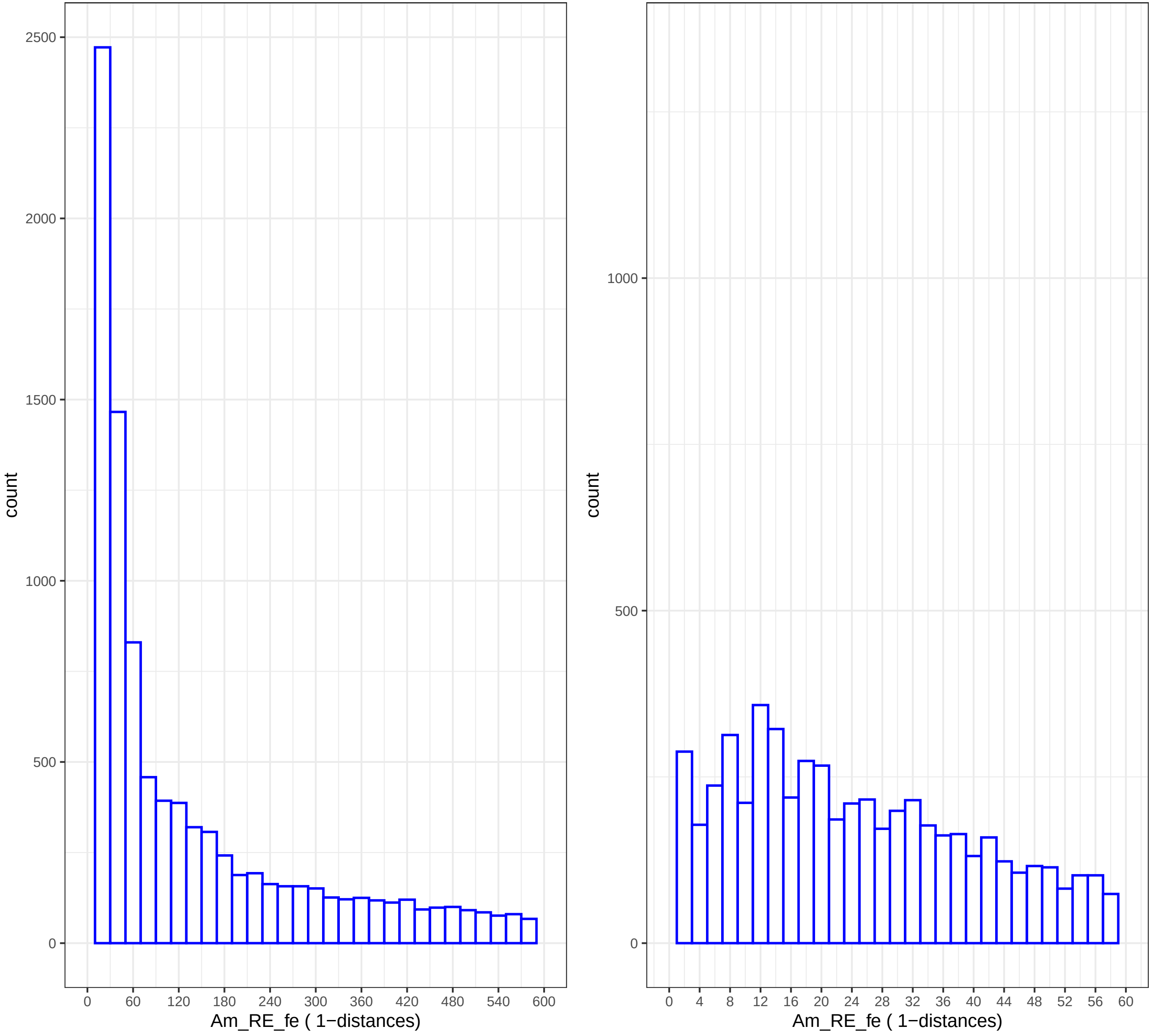

Am_RE_te
1−distance distributions

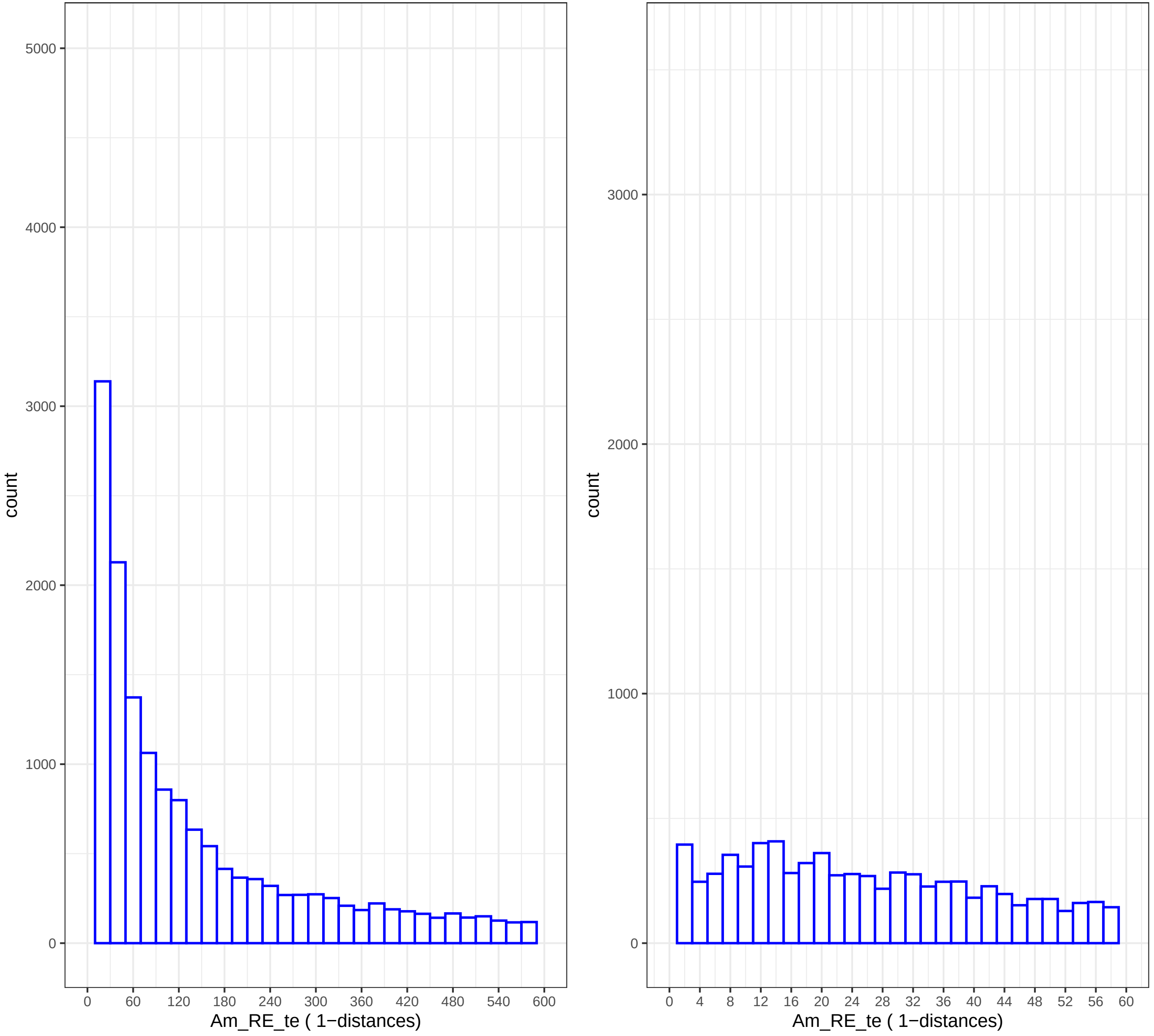

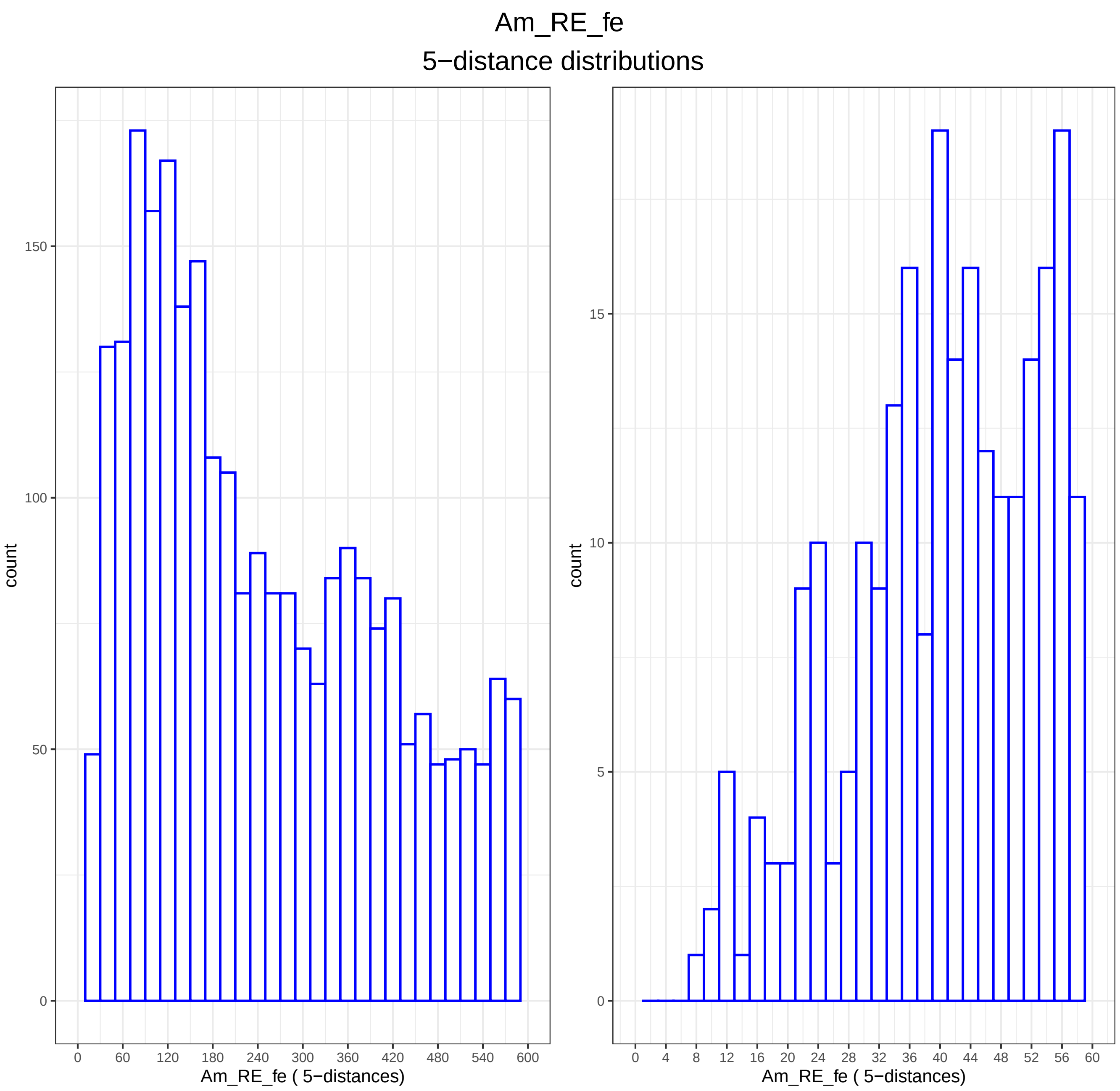

Am_RE_fe
5−distance distributions
count
Am_RE_fe ( 5−distances)
0
50
100
150
0
60
120
180
240
300
360
420
480
540
600
count
Am_RE_fe ( 5−distances)
0
5
10
15
0
4
8
12
16
20
24
28
32
36
40
44
48
52
56
60

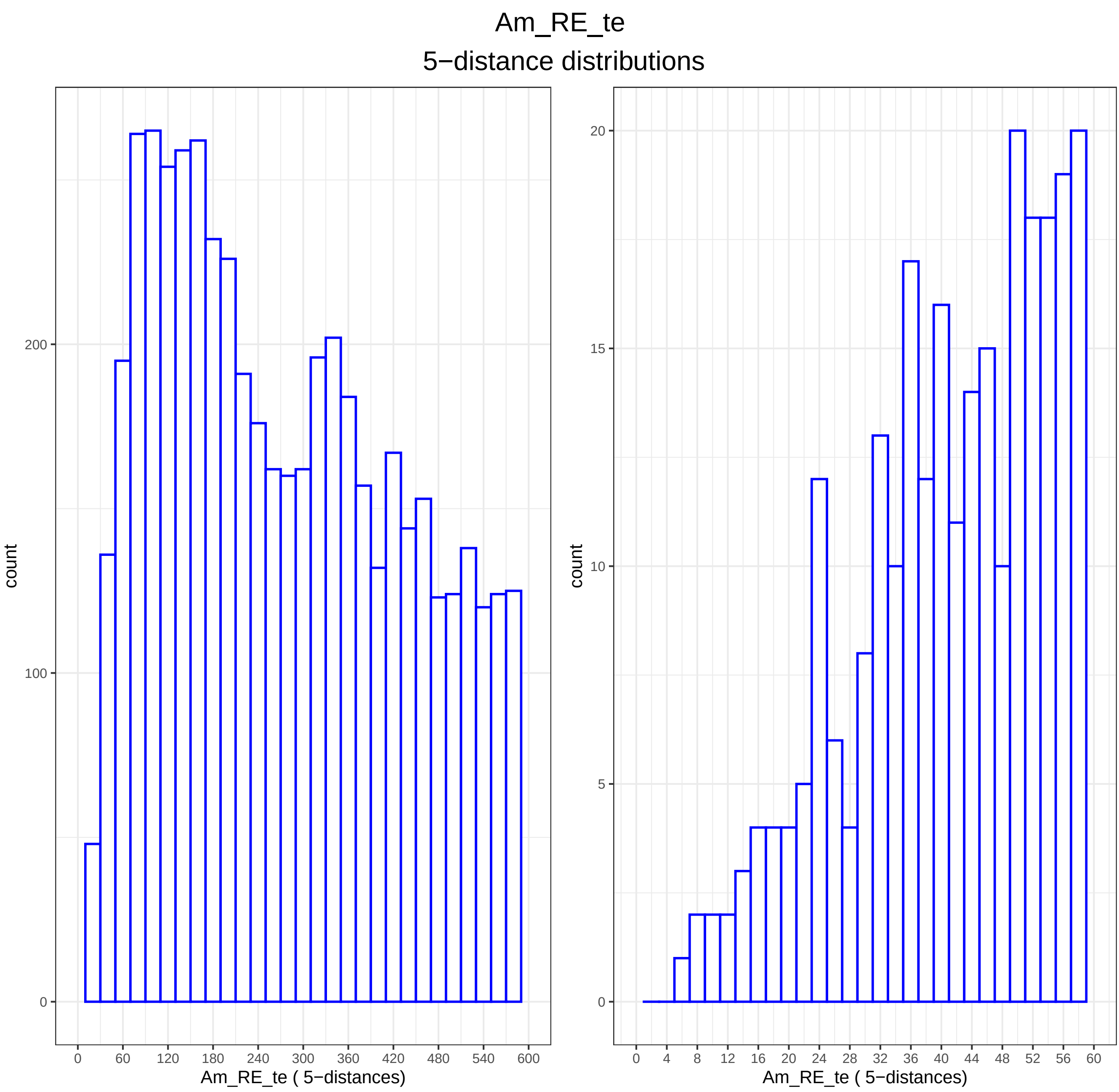

Am_RE_te
5−distance distributions
count
200
100
0
0
60
120
180
240
300
360
420
480
540
600
Am_RE_te ( 5−distances)
count
20
15
10
5
0
0
4
8
12
16
20
24
28
32
36
40
44
48
52
56
60
Am_RE_te ( 5−distances)

```
Directory: DMT File: 0READMEdmt


DMT - Differentially methylated tiles.

Input: studymc wsize stepsize threshold qvalue
Output: files dmt-*.txt dmg-*.txt

Notes: Output files dmt-*.txt and dmg-*.txt are generated by BWASPR::det_dmt() and show the
        differentially methylated tiles and genes as determined by methylKit::getMethylDiff with
        parameters difference=threshold and qvalue, as provided in the Am_RE.conf
        configuration file.  Tiles refer to sliding windows along the genome within which
        methylation calls are cumulated by methylKit::tileMethylCounts.
        The dmg-*.txt files show all genes with at least one differentially methylated tile.

        A positive meth.diff value in comparison A.vs.B means that the B methylation percentage
        in that tile is higher than the A methylation percentage.
```

Directory: DMT File: dmg-Am_RE.txt

```
seqnames    start end   width strand      source      type  score phase ID    Dbxref      Name  gbkey gene  gene_biotype      description gene_synonym      end_range   partial
      start_range exception   pvalue      qvalue      meth.diff   comparison
NC_037638.1 241926      245999      4074  -     Gnomon      gene  NA    NA    gene-LOC410772    c("BEEBASE:GB42186", "GeneID:410772")    LOC410772   Gene  LOC410772   protein_coding
      NA    character(0)      character(0)      NA    character(0)      NA    1.30605997461983e-06    0.00155101489327884     -42.4573953292657 fe.vs.te
NC_037638.1 2209902     2216661     6760  -     Gnomon      gene  NA    NA    gene-LOC550706    c("BEEBASE:GB50345", "GeneID:550706")    LOC550706   Gene  LOC550706   protein_coding
      NA    character(0)      character(0)      NA    character(0)      NA    1.22370562537476e-05    0.00859696829612803     88.8888888888889  fe.vs.te
NC_037638.1 2449902     2452507     2606  +     Gnomon      gene  NA    NA    gene-LOC724331    c("BEEBASE:GB50364", "GeneID:724331")    LOC724331   Gene  LOC724331   protein_coding
      NA    character(0)      character(0)      NA    character(0)      NA    6.02802929035427e-18    2.61288963498896e-13    76.7045454545455  fe.vs.te
NC_037638.1 2504095     2505287     1193  +     Gnomon      gene  NA    NA    gene-LOC412578    c("BEEBASE:GB50371", "GeneID:412578")    LOC412578   Gene  LOC412578   protein_coding
      NA    character(0)      character(0)      NA    character(0)      NA    3.66117307646542e-07    0.000561755741687899    -95   fe.vs.te
NC_037638.1 3187619     3194351     6733  +     Gnomon      gene  NA    NA    gene-LOC725982    c("BEEBASE:GB48031", "GeneID:725982")    LOC725982   Gene  LOC725982   protein_coding
      NA    character(0)      character(0)      NA    character(0)      NA    1.03078252101546e-05    0.00747781726606672     -55.9440559440559 fe.vs.te
NC_037638.1 4500967     4510818     9852  +     Gnomon      gene  NA    NA    gene-LOC414017    c("BEEBASE:GB53231", "GeneID:414017")    LOC414017   Gene  LOC414017   protein_coding
      NA    character(0)      character(0)      NA    character(0)      NA    2.28552421047006e-06    0.00227291803558675     -80.7692307692308 fe.vs.te
NC_037638.1 4563504     4567058     3555  +     Gnomon      gene  NA    NA    gene-LOC551842    c("BEEBASE:GB53240", "GeneID:551842")    LOC551842   Gene  LOC551842   protein_coding
      NA    character(0)      character(0)      NA    character(0)      NA    5.99414265269937e-09    2.16516770427567e-05    -67.8620689655172 fe.vs.te
NC_037638.1 4722480     4727235     4756  -     Gnomon      gene  NA    NA    gene-LOC412511    c("BEEBASE:GB53180", "GeneID:412511")    LOC412511   Gene  LOC412511   protein_coding
      NA    character(0)      character(0)      NA    character(0)      NA    3.81438201949993e-07    0.000575085012830779    -34.8635498201819 fe.vs.te
NC_037638.1 6598452     6601077     2626  -     Gnomon      gene  NA    NA    gene-LOC727087    c("BEEBASE:GB40798", "GeneID:727087")    LOC727087   Gene  LOC727087   protein_coding
      NA    character(0)      character(0)      NA    character(0)      NA    1.12958555640211e-06    0.00137922934793735     -47.2972972972973 fe.vs.te
NC_037638.1 7971326     7976718     5393  +     Gnomon      gene  NA    NA    gene-LOC413754    c("BEEBASE:GB16496", "GeneID:413754")    LOC413754   Gene  LOC413754   protein_coding
      NA    character(0)      character(0)      NA    character(0)      NA    2.49903835810243e-10    1.49410330990655e-06    52.3809523809524  fe.vs.te
NC_037638.1 10733614    10736662    3049  -     Gnomon      gene  NA    NA    gene-LOC410116    c("BEEBASE:GB45837", "GeneID:410116")    LOC410116   Gene  LOC410116   protein_coding
      NA    character(0)      character(0)      NA    character(0)      NA    2.69215237582448e-06    0.00260766807366988     91.6666666666667  fe.vs.te
NC_037638.1 10766913    10768883    1971  +     Gnomon      gene  NA    NA    gene-LOC413080    c("BEEBASE:GB45867", "GeneID:413080")    LOC413080   Gene  LOC413080   protein_coding
      NA    character(0)      character(0)      NA    character(0)      NA    3.05997724828698e-06    0.00286782187111452     -38.5167464114833 fe.vs.te
NC_037638.1 10768895    10772534    3640  -     Gnomon      gene  NA    NA    gene-LOC409756    c("BEEBASE:GB45829", "GeneID:409756")    LOC409756   Gene  LOC409756   protein_coding
      NA    character(0)      character(0)      NA    character(0)      NA    3.05997724828698e-06    0.00286782187111452     -38.5167464114833 fe.vs.te
NC_037638.1 10940386    10964255    23870 -     Gnomon      gene  NA    NA    gene-LOC408594    c("BEEBASE:GB45812", "GeneID:408594")    LOC408594   Gene  LOC408594   protein_coding
      NA    character(0)      character(0)      NA    character(0)      NA    1.93838271698062e-06    0.00201246697048085     -44.1471571906354 fe.vs.te
NC_037638.1 11500283    11506109    5827  -     Gnomon      gene  NA    NA    gene-LOC551573    c("BEEBASE:GB44779", "GeneID:551573")    LOC551573   Gene  LOC551573   protein_coding
      NA    character(0)      character(0)      NA    character(0)      NA    4.00160064025610e-06    0.0035996706732241      100   fe.vs.te
NC_037638.1 11667936    11671958    4023  -     Gnomon      gene  NA    NA    gene-LOC550977    c("BEEBASE:GB44768", "GeneID:550977")    LOC550977   Gene  LOC550977   protein_coding
      NA    character(0)      character(0)      NA    character(0)      NA    4.43919125495146e-06    0.00392693297177116     -71.4285714285714 fe.vs.te
NC_037638.1 11703719    11715797    12079 +     Gnomon      gene  NA    NA    gene-LOC552497    c("BEEBASE:GB44762", "GeneID:552497")    LOC552497   Gene  LOC552497   protein_coding
      NA    character(0)      character(0)      NA    character(0)      NA    1.90771886906619e-06    0.00199256267493314     -29.5081967213115 fe.vs.te
NC_037638.1 12531198    12532170    973   +     Gnomon      gene  NA    NA    gene-LOC102656491 GeneID:102656491  LOC102656491      Gene  LOC102656491      protein_coding    NA
      character(0)      character(0)      NA    character(0)      NA    8.45064424701539e-06    0.00639823283824847     55.5555555555556  fe.vs.te
NC_037638.1 14752856    14772072    19217 +     Gnomon      gene  NA    NA    gene-LOC725068    c("BEEBASE:GB47452", "GeneID:725068")    LOC725068   Gene  LOC725068   protein_coding
      NA    character(0)      character(0)      NA    character(0)      NA    1.09667142218585e-09    5.2817729642555e-06     -32.6751946607341 fe.vs.te
NC_037638.1 15090601    15097846    7246  -     Gnomon      gene  NA    NA    gene-LOC409282    c("BEEBASE:GB47418", "GeneID:409282")    LOC409282   Gene  LOC409282   protein_coding
      NA    character(0)      character(0)      NA    character(0)      NA    4.28133459218609e-07    0.000629075637060403    -60.4395604395604 fe.vs.te
NC_037638.1 19297850    19313191    15342 -     Gnomon      gene  NA    NA    gene-LOC551731    c("BEEBASE:GB42246", "GeneID:551731")    LOC551731   Gene  LOC551731   protein_coding
      NA    character(0)      character(0)      NA    character(0)      NA    4.71974173573221e-06    0.00412768573511023     -100  fe.vs.te
NC_037638.1 22790384    22793791    3408  -     Gnomon      gene  NA    NA    gene-LOC409312    c("BEEBASE:GB51599", "GeneID:409312")    LOC409312   Gene  LOC409312   protein_coding
      NA    character(0)      character(0)      NA    character(0)      NA    5.33491178913597e-06    0.00446850862956441     47.1869328493648  fe.vs.te
NC_037638.1 24225576    24234141    8566  +     Gnomon      gene  NA    NA    gene-LOC100576132 c("BEEBASE:GB54923", "GeneID:100576132") LOC100576132      Gene  LOC100576132
      protein_coding    NA    character(0)      character(0)      NA    character(0)      NA    1.87586445229613e-12    1.9131905881908e-08     -71.3489409141583 fe.vs.te
NC_037638.1 24271485    24277741    6257  -     Gnomon      gene  NA    NA    gene-LOC100578883 c("BEEBASE:GB55002", "GeneID:100578883") LOC100578883      Gene  LOC100578883
      protein_coding    NA    character(0)      character(0)      NA    character(0)      NA    2.26127523948216e-09    9.56258424219236e-06    62.737799834574   fe.vs.te
...
... only first 25 lines shown ...
```

Directory: DMT File: dmt-Am_RE.txt

| seqnames | start | end | width | strand | pvalue | qvalue | meth.diff | comparison |
|---|---|---|---|---|---|---|---|---|
| NC_037638.1 | 242001 | 243000 | 1000 | * | 1.30605997461983e-06 | 0.002 | -42.46 | fe.vs.te |
| NC_037638.1 | 2212001 | 2213000 | 1000 | * | 1.22370562537476e-05 | 0.009 | 88.89 | fe.vs.te |
| NC_037638.1 | 2450001 | 2451000 | 1000 | * | 6.02802929035427e-18 | 0 | 76.7 | fe.vs.te |
| NC_037638.1 | 2504001 | 2505000 | 1000 | * | 3.66117307646542e-07 | 0.001 | -95 | fe.vs.te |
| NC_037638.1 | 3189001 | 3190000 | 1000 | * | 1.03078252101546e-05 | 0.007 | -55.94 | fe.vs.te |
| NC_037638.1 | 4505001 | 4506000 | 1000 | * | 2.28552421047006e-06 | 0.002 | -80.77 | fe.vs.te |
| NC_037638.1 | 4566001 | 4567000 | 1000 | * | 5.99414265269937e-09 | 0 | -67.86 | fe.vs.te |
| NC_037638.1 | 4723001 | 4724000 | 1000 | * | 3.81438201949993e-07 | 0.001 | -34.86 | fe.vs.te |
| NC_037638.1 | 6600001 | 6601000 | 1000 | * | 1.12958555640211e-06 | 0.001 | -47.3 | fe.vs.te |
| NC_037638.1 | 7972001 | 7973000 | 1000 | * | 2.49903835810243e-10 | 0 | 52.38 | fe.vs.te |
| NC_037638.1 | 10734001 | 10735000 | 1000 | * | 2.69215237582448e-06 | 0.003 | 91.67 | fe.vs.te |
| NC_037638.1 | 10768001 | 10769000 | 1000 | * | 3.05997724828698e-06 | 0.003 | -38.52 | fe.vs.te |
| NC_037638.1 | 10963001 | 10964000 | 1000 | * | 1.93838271698062e-06 | 0.002 | -44.15 | fe.vs.te |
| NC_037638.1 | 11501001 | 11502000 | 1000 | * | 4.0016006402561e-06 | 0.004 | 100 | fe.vs.te |
| NC_037638.1 | 11669001 | 11670000 | 1000 | * | 4.43919125495146e-06 | 0.004 | -71.43 | fe.vs.te |
| NC_037638.1 | 11704001 | 11705000 | 1000 | * | 1.90771886906619e-06 | 0.002 | -29.51 | fe.vs.te |
| NC_037638.1 | 12531001 | 12532000 | 1000 | * | 8.45064424701539e-06 | 0.006 | 55.56 | fe.vs.te |
| NC_037638.1 | 14766001 | 14767000 | 1000 | * | 1.09667142218585e-09 | 0 | -32.68 | fe.vs.te |
| NC_037638.1 | 15096001 | 15097000 | 1000 | * | 4.28133459218609e-07 | 0.001 | -60.44 | fe.vs.te |
| NC_037638.1 | 19305001 | 19306000 | 1000 | * | 4.71974173573221e-06 | 0.004 | -100 | fe.vs.te |
| NC_037638.1 | 22791001 | 22792000 | 1000 | * | 5.33491178913597e-06 | 0.004 | 47.19 | fe.vs.te |
| NC_037638.1 | 24233001 | 24234000 | 1000 | * | 1.87586445229613e-12 | 0 | -71.35 | fe.vs.te |
| NC_037638.1 | 24277001 | 24278000 | 1000 | * | 2.26127523948216e-09 | 0 | 62.74 | fe.vs.te |
| NC_037638.1 | 24623001 | 24624000 | 1000 | * | 1.39954912478038e-05 | 0.009 | -78.74 | fe.vs.te |

...
... only first 25 lines shown ...

```
Directory: DMSG File: 0READMEdmsg


DMSG - Differentially methylated sites and genes.

Input: studyhc threshold qvalue
       The studyhc methylKit raw object contains all CpGscd sites with coverage at least
        10 reads.
Output: files dms-*.txt dmg-*.txt  dmg-*details.txt dmg-*heatmaps.pdf

Notes: Output files dms-*.txt and dmg-*.txt are generated by BWASPR::det_dmsg() and show the
        differentially methylated sites and genes as determined by methylKit::getMethylDiff with
        parameters difference=threshold and qvalue, as provided in the Am_RE.conf
        configuration file.  The table of differentially methylated genes contains all genes
        with at least one differentially methylated site.

       The files dmg-*details.txt, generated by BWASPR::show_dmsg(), show all CpGscd sites in
        the differentially methylated genes, with coverage numbers and methylation percentages.

       The files dmg-*heatmaps.pdf, generated by BWASPR::show_dmsg(), give heatmap displays for
        genes meeting the following criteria: 1) there are between minNsites and maxNsites common
        CpGscd sites; 2) at least minPdmsites % of these sites are differentially methylated sites.
        The parameters are set in the Am_RE.conf configuration file.
```

Directory: DMSG File: dmg-Am_RE.txt

| seqnames | start | end | width | strand | source | type | score | phase | ID | Dbxref | Name | gbkey | gene | gene_biotype | description | gene_synonym | end_range | partial | start_range | exception | comparison |
|---|---|---|---|---|---|---|---|---|---|---|---|---|---|---|---|---|---|---|---|---|---|
| NC_037638.1 | 867359 | 873957 | 6599 | + | Gnomon | gene | NA | NA | gene-LOC726756 | c("BEEBASE:GB42155", "GeneID:726756") | LOC726756 | Gene | LOC726756 | protein_coding | NA | character(0) | character(0) | NA | character(0) | NA | fe.vs.te |
| NC_037638.1 | 11352891 | 11356524 | 3634 | + | Gnomon | gene | NA | NA | gene-LOC100577140 | c("BEEBASE:GB44735", "GeneID:100577140") | LOC100577140 | Gene | LOC100577140 | protein_coding | NA | character(0) | character(0) | NA | character(0) | NA | fe.vs.te |
| NC_037639.1 | 14748246 | 14750873 | 2628 | + | Gnomon | gene | NA | NA | gene-LOC552747 | c("BEEBASE:GB55432", "GeneID:552747") | LOC552747 | Gene | LOC552747 | protein_coding | NA | character(0) | character(0) | NA | character(0) | NA | fe.vs.te |
| NC_037639.1 | 15049440 | 15067954 | 18515 | - | Gnomon | gene | NA | NA | gene-LOC409192 | c("BEEBASE:GB55574", "GeneID:409192") | LOC409192 | Gene | LOC409192 | protein_coding | NA | character(0) | character(0) | NA | character(0) | NA | fe.vs.te |
| NC_037639.1 | 16052535 | 16065479 | 12945 | - | Gnomon | gene | NA | NA | gene-LOC410804 | c("BEEBASE:GB55507", "GeneID:410804") | LOC410804 | Gene | LOC410804 | protein_coding | NA | character(0) | character(0) | NA | character(0) | NA | fe.vs.te |
| NC_037640.1 | 2104 | 38014 | 35911 | - | Gnomon | gene | NA | NA | gene-LOC100577216 | GeneID:100577216 | LOC100577216 | Gene | LOC100577216 | lncRNA | NA | character(0) | character(0) | NA | character(0) | NA | fe.vs.te |
| NC_037640.1 | 23922 | 222272 | 198351 | + | Gnomon | gene | NA | NA | gene-LOC409422 | GeneID:409422 | LOC409422 | Gene | LOC409422 | protein_coding | NA | character(0) | character(0) | NA | character(0) | NA | fe.vs.te |
| NC_037640.1 | 1955196 | 1958206 | 3011 | - | Gnomon | gene | NA | NA | gene-LOC413484 | c("BEEBASE:GB46542", "GeneID:413484") | LOC413484 | Gene | LOC413484 | protein_coding | NA | character(0) | character(0) | NA | character(0) | NA | fe.vs.te |
| NC_037640.1 | 3318321 | 3326801 | 8481 | - | Gnomon | gene | NA | NA | gene-LOC551291 | c("BEEBASE:GB49106", "GeneID:551291") | LOC551291 | Gene | LOC551291 | protein_coding | NA | character(0) | character(0) | NA | character(0) | NA | fe.vs.te |
| NC_037640.1 | 5372136 | 5376972 | 4837 | - | Gnomon | gene | NA | NA | gene-LOC410958 | c("BEEBASE:GB55847", "GeneID:410958") | LOC410958 | Gene | LOC410958 | protein_coding | NA | character(0) | character(0) | NA | character(0) | NA | fe.vs.te |
| NC_037640.1 | 8087735 | 8108431 | 20697 | - | Gnomon | gene | NA | NA | gene-LOC726727 | c("BEEBASE:GB49746", "GeneID:726727") | LOC726727 | Gene | LOC726727 | protein_coding | NA | character(0) | character(0) | NA | character(0) | NA | fe.vs.te |
| NC_037641.1 | 1779220 | 1781219 | 2000 | - | Gnomon | gene | NA | NA | gene-LOC113218557 | GeneID:113218557 | LOC113218557 | Gene | LOC113218557 | lncRNA | NA | character(0) | character(0) | NA | character(0) | NA | fe.vs.te |
| NC_037641.1 | 12660423 | 12666039 | 5617 | - | Gnomon | gene | NA | NA | gene-LOC411678 | c("BEEBASE:GB53036", "GeneID:411678") | LOC411678 | Gene | LOC411678 | protein_coding | NA | character(0) | character(0) | NA | character(0) | NA | fe.vs.te |
| NC_037642.1 | 13881140 | 13888076 | 6937 | - | Gnomon | gene | NA | NA | gene-LOC412468 | c("BEEBASE:GB45623", "GeneID:412468") | LOC412468 | Gene | LOC412468 | protein_coding | NA | character(0) | character(0) | NA | character(0) | NA | fe.vs.te |
| NC_037643.1 | 5146437 | 5154684 | 8248 | - | Gnomon | gene | NA | NA | gene-LOC552046 | c("BEEBASE:GB52211", "GeneID:552046") | LOC552046 | Gene | LOC552046 | protein_coding | NA | character(0) | character(0) | NA | character(0) | NA | fe.vs.te |
| NC_037643.1 | 6464125 | 6475423 | 11299 | - | Gnomon | gene | NA | NA | gene-LOC100577552 | c("BEEBASE:GB48567", "GeneID:100577552") | LOC100577552 | Gene | LOC100577552 | protein_coding | NA | character(0) | character(0) | NA | character(0) | NA | fe.vs.te |
| NC_037643.1 | 16784738 | 16794537 | 9800 | + | Gnomon | gene | NA | NA | gene-LOC409271 | c("BEEBASE:GB45373", "GeneID:409271") | LOC409271 | Gene | LOC409271 | protein_coding | NA | character(0) | character(0) | NA | character(0) | NA | fe.vs.te |
| NC_037643.1 | 17751535 | 17754492 | 2958 | - | Gnomon | gene | NA | NA | gene-LOC100578257 | c("BEEBASE:GB55711", "GeneID:100578257") | LOC100578257 | Gene | LOC100578257 | protein_coding | NA | character(0) | character(0) | NA | character(0) | NA | fe.vs.te |
| NC_037644.1 | 1303882 | 1309641 | 5760 | - | Gnomon | gene | NA | NA | gene-LOC413152 | c("BEEBASE:GB49680", "GeneID:413152") | LOC413152 | Gene | LOC413152 | protein_coding | NA | character(0) | character(0) | NA | character(0) | NA | fe.vs.te |
| NC_037644.1 | 8420234 | 8431677 | 11444 | - | Gnomon | gene | NA | NA | gene-LOC408936 | c("BEEBASE:GB43300", "GeneID:408936") | LOC408936 | Gene | LOC408936 | protein_coding | NA | character(0) | character(0) | NA | character(0) | NA | fe.vs.te |
| NC_037644.1 | 12076838 | 12087876 | 11039 | + | Gnomon | gene | NA | NA | gene-LOC409107 | c("BEEBASE:GB44285", "GeneID:409107") | LOC409107 | Gene | LOC409107 | protein_coding | NA | character(0) | character(0) | NA | character(0) | NA | fe.vs.te |
| NC_037644.1 | 12353582 | 12361285 | 7704 | - | Gnomon | gene | NA | NA | gene-LOC413433 | c("BEEBASE:GB44353", "GeneID:413433") | LOC413433 | Gene | LOC413433 | protein_coding | NA | character(0) | character(0) | NA | character(0) | NA | fe.vs.te |
| NC_037645.1 | 2219022 | 2224075 | 5054 | + | Gnomon | gene | NA | NA | gene-LOC411881 | c("BEEBASE:GB40453", "GeneID:411881") | LOC411881 | Gene | LOC411881 | protein_coding | NA | character(0) | character(0) | NA | character(0) | NA | fe.vs.te |
| NC_037645.1 | 7162825 | 7164882 | 2058 | - | Gnomon | gene | NA | NA | gene-LOC107964797 | GeneID:107964797 | LOC107964797 | Gene | LOC107964797 | lncRNA | NA | character(0) | character(0) | NA | character(0) | NA | fe.vs.te |
| NC_037645.1 | 7627700 | 7642782 | 15083 | + | Gnomon | gene | NA | NA | gene-LOC411270 | c("BEEBASE:GB41860", "GeneID:411270") | LOC411270 | Gene | LOC411270 | protein_coding | NA | character(0) | character(0) | NA | character(0) | NA | fe.vs.te |
| NC_037645.1 | 7637705 | 7642247 | 4543 | - | Gnomon | gene | NA | NA | gene-LOC113218927 | GeneID:113218927 | LOC113218927 | Gene | LOC113218927 | lncRNA | NA | character(0) | character(0) | NA | character(0) | NA | fe.vs.te |
| NC_037645.1 | 7723628 | 7731596 | 7969 | + | Gnomon | gene | NA | NA | gene-LOC413912 | c("BEEBASE:GB41869", "GeneID:413912") | LOC413912 | Gene | LOC413912 | protein_coding | NA | character(0) | character(0) | NA | character(0) | NA | fe.vs.te |
| NC_037645.1 | 8249532 | 8272000 | 22469 | + | Gnomon | gene | NA | NA | gene-LOC100577280 | c("BEEBASE:GB41888", "GeneID:100577280") | LOC100577280 | Gene | LOC100577280 | protein_coding | NA | character(0) | character(0) | NA | character(0) | NA | fe.vs.te |
| NC_037646.1 | 12264427 | 12267413 | 2987 | - | Gnomon | gene | NA | NA | gene-LOC411542 | c("BEEBASE:GB53389", "GeneID:411542") | LOC411542 | Gene | LOC411542 | protein_coding | NA | character(0) | character(0) | NA | character(0) | NA | fe.vs.te |
| NC_037648.1 | 6563300 | 6582126 | 18827 | - | Gnomon | gene | NA | NA | gene-LOC724670 | c("BEEBASE:GB48828", "GeneID:724670") | LOC724670 | Gene | LOC724670 | misc_RNA | NA | | | | | | |

character(0) character(0) NA character(0) NA fe.vs.te
NC_037648.1 9917400 9924509 7110 - Gnomon gene NA NA gene-LOC413344 c("BEEBASE:GB47231", "GeneID:413344") LOC413344 Gene LOC413344 protein_coding
NA character(0) character(0) NA character(0) NA fe.vs.te
NC_037648.1 15352724 15355797 3074 - Gnomon gene NA NA gene-LOC726432 c("BEEBASE:GB44865", "GeneID:726432") LOC726432 Gene LOC726432 protein_coding
NA character(0) character(0) NA character(0) NA fe.vs.te
NC_037648.1 15572654 15582091 9438 + Gnomon gene NA NA gene-LOC413428 c("BEEBASE:GB43141", "GeneID:413428") LOC413428 Gene LOC413428 protein_coding
NA character(0) character(0) NA character(0) NA fe.vs.te
NC_037648.1 15627088 15629378 2291 + Gnomon gene NA NA gene-LOC413429 c("BEEBASE:GB43184", "GeneID:413429") LOC413429 Gene LOC413429 protein_coding
NA character(0) character(0) NA character(0) NA fe.vs.te
NC_037649.1 3077569 3085347 7779 + Gnomon gene NA NA gene-LOC550829 c("BEEBASE:GB48305", "GeneID:550829") LOC550829 Gene LOC550829 protein_coding
NA character(0) character(0) NA character(0) NA fe.vs.te
NC_037651.1 7712396 7716924 4529 + Gnomon gene NA NA gene-LOC410510 c("BEEBASE:GB54140", "GeneID:410510") LOC410510 Gene LOC410510 protein_coding
NA character(0) character(0) NA character(0) NA fe.vs.te
NC_037651.1 8079571 8084767 5197 - Gnomon gene NA NA gene-LOC726791 c("BEEBASE:GB54158", "GeneID:726791") LOC726791 Gene LOC726791 protein_coding
NA character(0) character(0) NA character(0) NA fe.vs.te
NC_037651.1 9406063 9410790 4728 + Gnomon gene NA NA gene-LOC727008 c("BEEBASE:GB41609", "GeneID:727008") LOC727008 Gene LOC727008 protein_coding
NA character(0) character(0) NA character(0) NA fe.vs.te
NC_037652.1 628194 659504 31311 + Gnomon gene NA NA gene-LOC411466 c("BEEBASE:GB41196", "GeneID:411466") LOC411466 Gene LOC411466 protein_coding
NA character(0) character(0) NA character(0) NA fe.vs.te
NC_037652.1 2830254 2847042 16789 - Gnomon gene NA NA gene-LOC725503 c("BEEBASE:GB54649", "GeneID:725503") LOC725503 Gene LOC725503 protein_coding
NA character(0) character(0) NA character(0) NA fe.vs.te
NC_037653.1 1087555 1093753 6199 - Gnomon gene NA NA gene-LOC726354 c("BEEBASE:GB54354", "GeneID:726354") LOC726354 Gene LOC726354 protein_coding
NA character(0) character(0) NA character(0) NA fe.vs.te
NC_037653.1 3060190 3082831 22642 + Gnomon gene NA NA gene-LOC551848 c("BEEBASE:GB51276", "GeneID:551848") LOC551848 Gene LOC551848 protein_coding
NA character(0) character(0) NA character(0) NA fe.vs.te
NW_020555870.1 4446 46811 42366 + Gnomon gene NA NA gene-LOC113219367 GeneID:113219367 LOC113219367 Gene LOC113219367 protein_coding NA
character(0) character(0) NA character(0) NA fe.vs.te
NW_020555870.1 33316 35321 2006 - Gnomon gene NA NA gene-LOC100577929 GeneID:100577929 LOC100577929 Gene LOC100577929 lncRNA NA character(0)
character(0) NA character(0) NA fe.vs.te
NW_020555894.1 7811 10414 2604 - Gnomon gene NA NA gene-LOC113219373 GeneID:113219373 LOC113219373 Gene LOC113219373 lncRNA NA character(0)
character(0) NA character(0) NA fe.vs.te

Directory: DMSG File: dmg-Am_RE_fe.vs.te_details.txt

| seqnames | start | end | width | strand | coverage1 | numCs1 | numTs1 | coverage2 | numCs2 | numTs2 | fe | te | gene_seqnames | gene_start | gene_end | gene_width | gene_strand | gene_source | gene_type | gene_ID | gene_Dbxref | gene_Name | gene_gbkey | gene_gene | gene_gene_biotype | gene_gene_synonym | gene_end_range | gene_start_range | gene_comparison | is.dm |
|---|---|---|---|---|---|---|---|---|---|---|---|---|---|---|---|---|---|---|---|---|---|---|---|---|---|---|---|---|---|---|
| NC_037638.1 | 868756 | 868756 | 1 | + | 15 | 15 | 0 | 12 | 9 | 3 | 100 | 75 | NC_037638.1 | 867359 | 873957 | 6599 | + | Gnomon | gene | gene-LOC726756 | c("BEEBASE:GB42155", "GeneID:726756") | LOC726756 | Gene | LOC726756 | protein_coding | character(0) | character(0) | character(0) | fe.vs.te | FALSE |
| NC_037638.1 | 868808 | 868808 | 1 | + | 17 | 3 | 14 | 16 | 7 | 9 | 17.65 | 43.75 | NC_037638.1 | 867359 | 873957 | 6599 | + | Gnomon | gene | gene-LOC726756 | c("BEEBASE:GB42155", "GeneID:726756") | LOC726756 | Gene | LOC726756 | protein_coding | character(0) | character(0) | character(0) | fe.vs.te | TRUE |
| NC_037638.1 | 868820 | 868820 | 1 | + | 15 | 8 | 7 | 13 | 12 | 1 | 53.33 | 92.31 | NC_037638.1 | 867359 | 873957 | 6599 | + | Gnomon | gene | gene-LOC726756 | c("BEEBASE:GB42155", "GeneID:726756") | LOC726756 | Gene | LOC726756 | protein_coding | character(0) | character(0) | character(0) | fe.vs.te | TRUE |
| NC_037638.1 | 868921 | 868921 | 1 | + | 10 | 9 | 1 | 10 | 9 | 1 | 90 | 90 | NC_037638.1 | 867359 | 873957 | 6599 | + | Gnomon | gene | gene-LOC726756 | c("BEEBASE:GB42155", "GeneID:726756") | LOC726756 | Gene | LOC726756 | protein_coding | character(0) | character(0) | character(0) | fe.vs.te | FALSE |
| NC_037638.1 | 872523 | 872523 | 1 | + | 13 | 0 | 13 | 10 | 0 | 10 | 0 | 0 | NC_037638.1 | 867359 | 873957 | 6599 | + | Gnomon | gene | gene-LOC726756 | c("BEEBASE:GB42155", "GeneID:726756") | LOC726756 | Gene | LOC726756 | protein_coding | character(0) | character(0) | character(0) | fe.vs.te | FALSE |
| NC_037638.1 | 872527 | 872527 | 1 | + | 14 | 0 | 14 | 13 | 0 | 13 | 0 | 0 | NC_037638.1 | 867359 | 873957 | 6599 | + | Gnomon | gene | gene-LOC726756 | c("BEEBASE:GB42155", "GeneID:726756") | LOC726756 | Gene | LOC726756 | protein_coding | character(0) | character(0) | character(0) | fe.vs.te | FALSE |
| NC_037638.1 | 872531 | 872531 | 1 | + | 14 | 0 | 14 | 13 | 0 | 13 | 0 | 0 | NC_037638.1 | 867359 | 873957 | 6599 | + | Gnomon | gene | gene-LOC726756 | c("BEEBASE:GB42155", "GeneID:726756") | LOC726756 | Gene | LOC726756 | protein_coding | character(0) | character(0) | character(0) | fe.vs.te | FALSE |
| NC_037638.1 | 872535 | 872535 | 1 | + | 14 | 0 | 14 | 13 | 0 | 13 | 0 | 0 | NC_037638.1 | 867359 | 873957 | 6599 | + | Gnomon | gene | gene-LOC726756 | c("BEEBASE:GB42155", "GeneID:726756") | LOC726756 | Gene | LOC726756 | protein_coding | character(0) | character(0) | character(0) | fe.vs.te | FALSE |
| NC_037638.1 | 872543 | 872543 | 1 | + | 16 | 0 | 16 | 12 | 1 | 11 | 0 | 8.33 | NC_037638.1 | 867359 | 873957 | 6599 | + | Gnomon | gene | gene-LOC726756 | c("BEEBASE:GB42155", "GeneID:726756") | LOC726756 | Gene | LOC726756 | protein_coding | character(0) | character(0) | character(0) | fe.vs.te | FALSE |
| NC_037638.1 | 872549 | 872549 | 1 | + | 18 | 0 | 18 | 14 | 0 | 14 | 0 | 0 | NC_037638.1 | 867359 | 873957 | 6599 | + | Gnomon | gene | gene-LOC726756 | c("BEEBASE:GB42155", "GeneID:726756") | LOC726756 | Gene | LOC726756 | protein_coding | character(0) | character(0) | character(0) | fe.vs.te | FALSE |
| NC_037638.1 | 872553 | 872553 | 1 | + | 18 | 1 | 17 | 14 | 0 | 14 | 5.56 | 0 | NC_037638.1 | 867359 | 873957 | 6599 | + | Gnomon | gene | gene-LOC726756 | c("BEEBASE:GB42155", "GeneID:726756") | LOC726756 | Gene | LOC726756 | protein_coding | character(0) | character(0) | character(0) | fe.vs.te | FALSE |
| NC_037638.1 | 872557 | 872557 | 1 | + | 17 | 0 | 17 | 15 | 0 | 15 | 0 | 0 | NC_037638.1 | 867359 | 873957 | 6599 | + | Gnomon | gene | gene-LOC726756 | c("BEEBASE:GB42155", "GeneID:726756") | LOC726756 | Gene | LOC726756 | protein_coding | character(0) | character(0) | character(0) | fe.vs.te | FALSE |
| NC_037638.1 | 872570 | 872570 | 1 | + | 16 | 0 | 16 | 14 | 0 | 14 | 0 | 0 | NC_037638.1 | 867359 | 873957 | 6599 | + | Gnomon | gene | gene-LOC726756 | c("BEEBASE:GB42155", "GeneID:726756") | LOC726756 | Gene | LOC726756 | protein_coding | character(0) | character(0) | character(0) | fe.vs.te | FALSE |
| NC_037638.1 | 872581 | 872581 | 1 | + | 12 | 0 | 12 | 13 | 0 | 13 | 0 | 0 | NC_037638.1 | 867359 | 873957 | 6599 | + | Gnomon | gene | gene-LOC726756 | c("BEEBASE:GB42155", "GeneID:726756") | LOC726756 | Gene | LOC726756 | protein_coding | character(0) | character(0) | character(0) | fe.vs.te | FALSE |
| NC_037638.1 | 872592 | 872592 | 1 | + | 10 | 0 | 10 | 13 | 0 | 13 | 0 | 0 | NC_037638.1 | 867359 | 873957 | 6599 | + | Gnomon | gene | gene-LOC726756 | c("BEEBASE:GB42155", "GeneID:726756") | LOC726756 | Gene | LOC726756 | protein_coding | character(0) | character(0) | character(0) | fe.vs.te | FALSE |
| NC_037638.1 | 872668 | 872668 | 1 | + | 10 | 0 | 10 | 12 | 0 | 12 | 0 | 0 | NC_037638.1 | 867359 | 873957 | 6599 | + | Gnomon | gene | gene-LOC726756 | c("BEEBASE:GB42155", "GeneID:726756") | LOC726756 | Gene | LOC726756 | protein_coding | character(0) | character(0) | character(0) | fe.vs.te | FALSE |
| NC_037638.1 | 872672 | 872672 | 1 | + | 11 | 0 | 11 | 13 | 0 | 13 | 0 | 0 | NC_037638.1 | 867359 | 873957 | 6599 | + | Gnomon | gene | gene-LOC726756 | c("BEEBASE:GB42155", "GeneID:726756") | LOC726756 | Gene | LOC726756 | protein_coding | character(0) | character(0) | character(0) | fe.vs.te | FALSE |
| NC_037638.1 | 872682 | 872682 | 1 | + | 10 | 0 | 10 | 14 | 0 | 14 | 0 | 0 | NC_037638.1 | 867359 | 873957 | 6599 | + | Gnomon | gene | gene-LOC726756 | c("BEEBASE:GB42155", "GeneID:726756") | LOC726756 | Gene | LOC726756 | protein_coding | character(0) | character(0) | character(0) | fe.vs.te | FALSE |
| NC_037638.1 | 872685 | 872685 | 1 | + | 10 | 0 | 10 | 14 | 0 | 14 | 0 | 0 | NC_037638.1 | 867359 | 873957 | 6599 | + | Gnomon | gene | gene-LOC726756 | c("BEEBASE:GB42155", "GeneID:726756") | LOC726756 | Gene | LOC726756 | protein_coding | character(0) | character(0) | character(0) | fe.vs.te | FALSE |
| NC_037638.1 | 872690 | 872690 | 1 | + | 10 | 0 | 10 | 15 | 0 | 15 | 0 | 0 | NC_037638.1 | 867359 | 873957 | 6599 | + | Gnomon | gene | gene-LOC726756 | c("BEEBASE:GB42155", "GeneID:726756") | LOC726756 | Gene | LOC726756 | protein_coding | character(0) | character(0) | character(0) | fe.vs.te | FALSE |
| NC_037638.1 | 872727 | 872727 | 1 | + | 13 | 0 | 13 | 11 | 0 | 11 | 0 | 0 | NC_037638.1 | 867359 | 873957 | 6599 | + | Gnomon | gene | gene-LOC726756 | c("BEEBASE:GB42155", "GeneID:726756") | LOC726756 | Gene | LOC726756 | protein_coding | character(0) | character(0) | character(0) | fe.vs.te | FALSE |
| NC_037638.1 | 872729 | 872729 | 1 | + | 11 | 0 | 11 | 11 | 0 | 11 | 0 | 0 | NC_037638.1 | 867359 | 873957 | 6599 | + | Gnomon | gene | gene-LOC726756 | c("BEEBASE:GB42155", "GeneID:726756") | LOC726756 | Gene | LOC726756 | protein_coding | character(0) | character(0) | character(0) | fe.vs.te | FALSE |
| NC_037638.1 | 872756 | 872756 | 1 | + | 11 | 0 | 11 | 10 | 0 | 10 | 0 | 0 | NC_037638.1 | 867359 | 873957 | 6599 | + | Gnomon | gene | gene-LOC726756 | c("BEEBASE:GB42155", "GeneID:726756") | LOC726756 | Gene | LOC726756 | protein_coding | character(0) | character(0) | character(0) | fe.vs.te | FALSE |
| NC_037638.1 | 11356005 | 11356005 | 1 | + | 12 | 8 | 4 | 15 | 1 | 14 | 66.67 | 6.67 | NC_037638.1 | 11352891 | 11356524 | 3634 | + | Gnomon | gene | gene-LOC100577140 | c("BEEBASE:GB44735", "GeneID:100577140") | LOC100577140 | Gene | LOC100577140 | protein_coding | character(0) | character(0) | character(0) | fe.vs.te | TRUE |

...

... only first 25 lines shown ...

Directory: DMSG File: dms-Am_RE.txt

| seqnames | start | end | width | strand | pvalue | qvalue | meth.diff | comparison |
|---|---|---|---|---|---|---|---|---|
| NC_037638.1 | 868808 | 868808 | 1 | + | 0.141070325052527 | 0 | 26.1 | fe.vs.te |
| NC_037638.1 | 868820 | 868820 | 1 | + | 0.0376811594202899 | 0 | 38.97 | fe.vs.te |
| NC_037638.1 | 11356005 | 11356005 | 1 | + | 0.00269905533063428 | 0 | -60 | fe.vs.te |
| NC_037639.1 | 367900 | 367900 | 1 | + | 0.068649885583524 | 0 | -41.67 | fe.vs.te |
| NC_037639.1 | 14748887 | 14748887 | 1 | + | 0.193027325346615 | 0 | 33.33 | fe.vs.te |
| NC_037639.1 | 15051800 | 15051800 | 1 | + | 0.239668865257773 | 0 | -27.14 | fe.vs.te |
| NC_037639.1 | 16055459 | 16055459 | 1 | + | 0.340805075930934 | 0 | 25.71 | fe.vs.te |
| NC_037640.1 | 10865 | 10865 | 1 | + | 0.0652288016025502 | 0 | -30.34 | fe.vs.te |
| NC_037640.1 | 10879 | 10879 | 1 | + | 0.0335717774056555 | 0 | -34.62 | fe.vs.te |
| NC_037640.1 | 11940 | 11940 | 1 | + | 0.0109982384995441 | 0 | -28.78 | fe.vs.te |
| NC_037640.1 | 14889 | 14889 | 1 | + | 0.0169317154779532 | 0 | -51.92 | fe.vs.te |
| NC_037640.1 | 43864 | 43864 | 1 | + | 0.0289139352326091 | 0 | -36.54 | fe.vs.te |
| NC_037640.1 | 1955434 | 1955434 | 1 | + | 0.0918023960156145 | 0 | -38.57 | fe.vs.te |
| NC_037640.1 | 3325748 | 3325748 | 1 | + | 0.213756898640463 | 0 | -29.55 | fe.vs.te |
| NC_037640.1 | 5374969 | 5374969 | 1 | + | 0.232857719746938 | 0 | -29.76 | fe.vs.te |
| NC_037640.1 | 8096744 | 8096744 | 1 | + | 0.028042438111088 | 0 | 46.67 | fe.vs.te |
| NC_037641.1 | 1780111 | 1780111 | 1 | + | 0.0484814174669146 | 0 | -25.57 | fe.vs.te |
| NC_037641.1 | 1783232 | 1783232 | 1 | + | 0.000945276192807774 | 0 | -40.44 | fe.vs.te |
| NC_037641.1 | 12663940 | 12663940 | 1 | + | 0.239668865257773 | 0 | -27.14 | fe.vs.te |
| NC_037642.1 | 13887258 | 13887258 | 1 | + | 0.0743034055727555 | 0 | -43.33 | fe.vs.te |
| NC_037643.1 | 5153435 | 5153435 | 1 | + | 0.068649885583524 | 0 | -37.12 | fe.vs.te |
| NC_037643.1 | 6472890 | 6472890 | 1 | + | 0.128836680560819 | 0 | 26.32 | fe.vs.te |
| NC_037643.1 | 16787727 | 16787727 | 1 | + | 0.0804953560371517 | 0 | -41.82 | fe.vs.te |
| NC_037643.1 | 17753328 | 17753328 | 1 | + | 0.0461538461538461 | 0 | 30 | fe.vs.te |
| NC_037644.1 | 1305736 | 1305736 | 1 | + | 0.0725806451612903 | 0 | -25.65 | fe.vs.te |
| NC_037644.1 | 8429549 | 8429549 | 1 | + | 0.0902255639097746 | 0 | -30 | fe.vs.te |
| NC_037644.1 | 12079651 | 12079651 | 1 | + | 0.113965744400527 | 0 | 28.57 | fe.vs.te |
| NC_037644.1 | 12354845 | 12354845 | 1 | + | 0.135338345864662 | 0 | 31.67 | fe.vs.te |
| NC_037644.1 | 12354847 | 12354847 | 1 | + | 0.18266253869969 | 0 | 31.82 | fe.vs.te |
| NC_037645.1 | 2222930 | 2222930 | 1 | + | 0.394855918075732 | 0 | 27.27 | fe.vs.te |
| NC_037645.1 | 7163075 | 7163075 | 1 | + | 0.214285714285714 | 0 | -27.27 | fe.vs.te |
| NC_037645.1 | 7640570 | 7640570 | 1 | + | 0.227157087091129 | 0 | -29.87 | fe.vs.te |
| NC_037645.1 | 7726149 | 7726149 | 1 | + | 0.179024636690541 | 0 | 28.46 | fe.vs.te |
| NC_037645.1 | 7726161 | 7726161 | 1 | + | 0.0334724278727509 | 0 | 44.76 | fe.vs.te |
| NC_037645.1 | 8260627 | 8260627 | 1 | + | 0.0804953560371517 | 0 | -43.64 | fe.vs.te |
| NC_037645.1 | 8264690 | 8264690 | 1 | + | 0.169565217391304 | 0 | 27.38 | fe.vs.te |
| NC_037646.1 | 12264940 | 12264940 | 1 | + | 0.0505670345941581 | 0 | 41.82 | fe.vs.te |
| NC_037648.1 | 3435300 | 3435300 | 1 | + | 0.0349249860936683 | 0 | 26.76 | fe.vs.te |
| NC_037648.1 | 4953189 | 4953189 | 1 | + | 0.0385162490425648 | 0 | -29.47 | fe.vs.te |
| NC_037648.1 | 6564727 | 6564727 | 1 | + | 0.171826625386997 | 0 | 33.33 | fe.vs.te |
| NC_037648.1 | 9921203 | 9921203 | 1 | + | 0.213756898640463 | 0 | 31.54 | fe.vs.te |
| NC_037648.1 | 15353700 | 15353700 | 1 | + | 0.18266253869969 | 0 | 31.82 | fe.vs.te |
| NC_037648.1 | 15578819 | 15578819 | 1 | + | 0.110701305693902 | 0 | -35.26 | fe.vs.te |
| NC_037648.1 | 15628728 | 15628728 | 1 | + | 0.237743976309059 | 0 | 28.21 | fe.vs.te |
| NC_037649.1 | 3084665 | 3084665 | 1 | + | 0.0112497909805376 | 0 | -46.91 | fe.vs.te |
| NC_037649.1 | 3084716 | 3084716 | 1 | + | 0.26004904439276 | 0 | 25.38 | fe.vs.te |
| NC_037650.1 | 5640 | 5640 | 1 | + | 0.0986628452701867 | 0 | -29.57 | fe.vs.te |
| NC_037651.1 | 7716233 | 7716233 | 1 | + | 0.113144433974963 | 0 | -38.1 | fe.vs.te |
| NC_037651.1 | 8081991 | 8081991 | 1 | + | 0.0302059496567505 | 0 | -42.31 | fe.vs.te |
| NC_037651.1 | 9407621 | 9407621 | 1 | + | 0.0386484177266206 | 0 | 36.81 | fe.vs.te |
| NC_037652.1 | 648831 | 648831 | 1 | + | 0.192063657166632 | 0 | 25.71 | fe.vs.te |
| NC_037652.1 | 2834628 | 2834628 | 1 | + | 0.107093821510298 | 0 | -36.67 | fe.vs.te |
| NC_037653.1 | 1089030 | 1089030 | 1 | + | 0.0902255639097745 | 0 | -36.36 | fe.vs.te |
| NC_037653.1 | 1090399 | 1090399 | 1 | + | 0.227157087091129 | 0 | -29.87 | fe.vs.te |
| NC_037653.1 | 3078353 | 3078353 | 1 | + | 0.139419763802309 | 0 | -31.37 | fe.vs.te |
| NC_037653.1 | 3078365 | 3078365 | 1 | + | 0.0604329414240249 | 0 | -41.67 | fe.vs.te |
| NW_020555860.1 | 288491 | 288491 | 1 | + | 0.0662338185745837 | 0 | 35.71 | fe.vs.te |
| NW_020555870.1 | 34219 | 34219 | 1 | + | 0.0496216872817558 | 0 | 28.57 | fe.vs.te |
| NW_020555879.1 | 19480 | 19480 | 1 | + | 0.0419144274016838 | 0 | 38.76 | fe.vs.te |
| NW_020555894.1 | 5735 | 5735 | 1 | + | 0.0114993365767359 | 0 | 56.67 | fe.vs.te |

| | | | | | | | | |
|---|---|---|---|---|---|---|---|---|
| NW_020555894.1 | 5756 | 5756 | 1 | + | 0.0903024825490838 | 0 | 41.54 | fe.vs.te |
| NW_020555894.1 | 8202 | 8202 | 1 | + | 0.00164377545256409 | 0 | -32.84 | fe.vs.te |
| NW_020555894.1 | 8753 | 8753 | 1 | + | 0.11338524286244 | 0 | -30.42 | fe.vs.te |

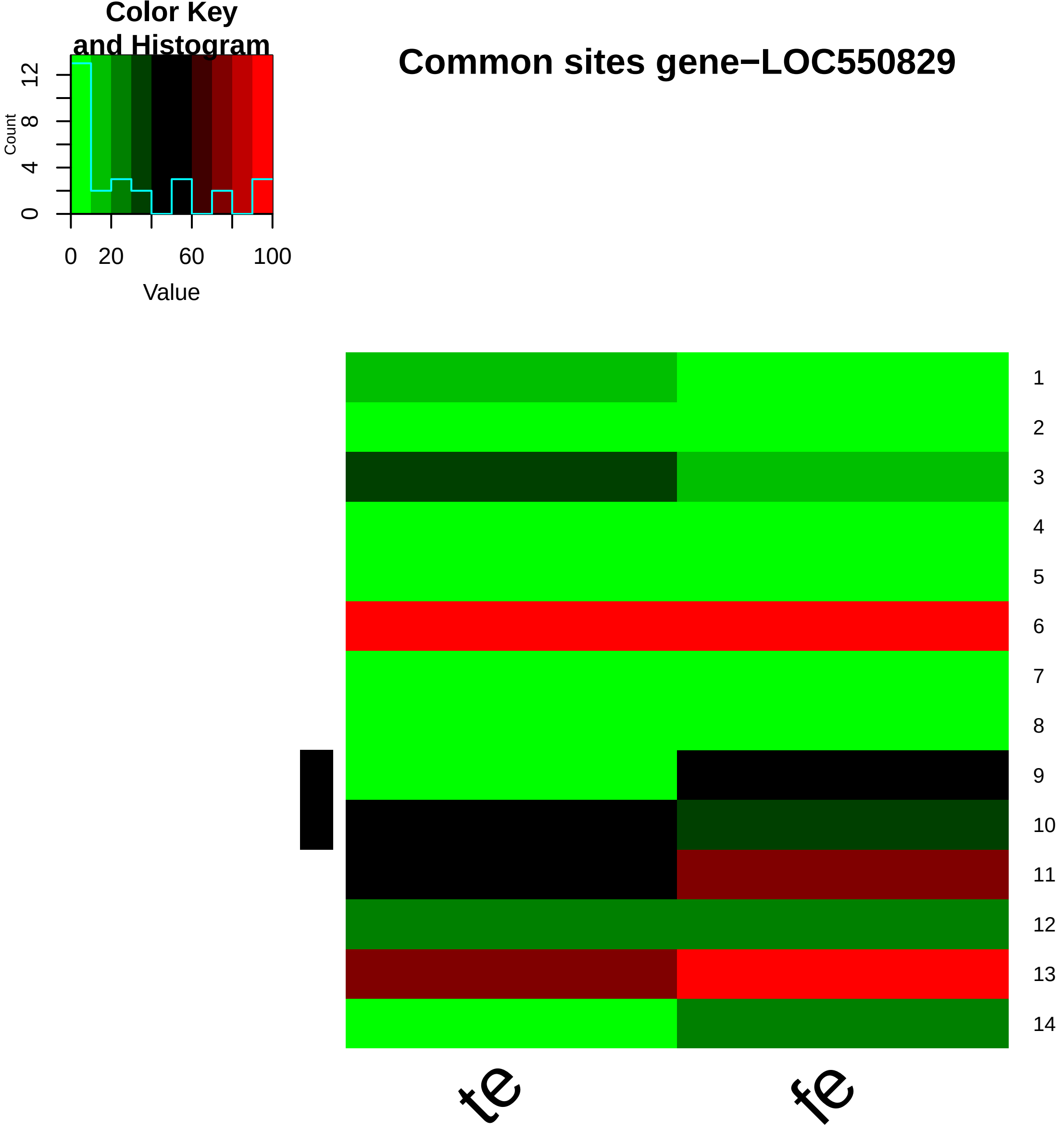
Color Key
and Histogram
Common sites gene−LOC550829
Count
0
4
8
12
0
20
60
100
Value
1
2
3
4
5
6
7
8
9
10
11
12
13
14
te
fe

```
Directory: OGL File: 0READMEogl


OGL - Ordered gene lists.

Input: studyhc annotation   maxgwidth minnbrdmsites
       dmgprp (= output of BWASPR:show_dmsg())

Output: files ogl-*.txt rnk-dmg-*.txt
         ogl-<study>_<sample>.txt ogl-<study>_<sample1>.vs.<sample2>.txt
         rnk-dmg--<study>_<sample1>.vs.<sample2>.txt rnk-dmg--<study>_<sample1>.vs.<sample2>.pdf
         wrt-<study>.txt

Notes: Output files ogl-<study>_<sample>.txt give tables for each sample with columns

        gene_ID gwidth #Sites #per10Kb %perSite %pNucl

       ordered by %pNucl.  If available, a link to an NCBI entry of gene_ID is inserted as second column.
       Abbreviations used: %perSite, percent methylation per site
                           %perNucl, percent methylation per nucleotide of the gene

       Output files ogl-<study>_<sample1>.vs.<sample2>.txt give tables for each comparison with columns

        gene_ID gwidth #Sites #per10Kb #dmSites #dmsp10kb %dmSites %pSite1 %pSite2 DMpSite ADMpSite DMpNucl ADMpNucl

       ordered by DMpSite.  If available, a link to an NCBI entry of gene_ID is inserted as second column.
       Abbreviations used: %dmSites, percent sites that are differentially methylated
                           %pSite1, average per site % methylation for sample1
                           %pSite2, average per site % methylation for sample2
                           DMpSite, average per site difference in % methylation between sample1 and sample2
                           ADMpSite, absolute value of DMpSite
                           DMpNucl, average per nucleotide difference in % methylation between sample1 and sample2
                           ADMpNucl, absolute value of DMpNucl

       Output files rnk-dmg-<study>_<sample1>.vs.<sample2>.txt are equivalent to files
        ogl-<study>_<sample1>.vs.<sample2>.txt but ordered by ADMpNucl.
       Output files rnk-dmg-<study>_<sample1>.vs.<sample2>.pdf provide visualization of the distribution
        of ADMpNucl values.

       Output file wrt-<study>.txt gives results of the Wilcoxon signed rank test comparing
        the %pSite1 and %pSite2 vectors

       In output directory OGL, tables are restricted to genes with gwidth <= maxgwidth and, for
        pairwise comparisons, #dmSites >= minnbrdmsites

       If either maxgwidth or minnbrdmsites is set, then output directory OGLall will show the full tables
        with all genes (for reference).
```

Directory: OGL File: ogl-Am_RE_fe.txt

| gene_ID | gene_link | gwidth | #Sites | #per10Kb | %perSite | %pNucl |
|---|---|---|---|---|---|---|
| gene-LOC113218557 | https://www.ncbi.nlm.nih.gov/gene/?term=LOC113218557 | 2000 | 49 | 245 | 51.47 | 1.26 |
| gene-LOC102656390 | https://www.ncbi.nlm.nih.gov/gene/?term=LOC102656390 | 1760 | 53 | 301.14 | 24.5 | 0.74 |
| gene-LOC100577929 | https://www.ncbi.nlm.nih.gov/gene/?term=LOC100577929 | 2006 | 57 | 284.15 | 22.5 | 0.64 |
| gene-LOC113219373 | https://www.ncbi.nlm.nih.gov/gene/?term=LOC113219373 | 2604 | 34 | 130.57 | 48.67 | 0.64 |
| gene-LOC408391 | https://www.ncbi.nlm.nih.gov/gene/?term=LOC408391 | 1806 | 13 | 71.98 | 87.97 | 0.63 |
| gene-LOC409500 | https://www.ncbi.nlm.nih.gov/gene/?term=LOC409500 | 1222 | 11 | 90.02 | 48.43 | 0.44 |
| gene-LOC408367 | https://www.ncbi.nlm.nih.gov/gene/?term=LOC408367 | 2232 | 11 | 49.28 | 79.27 | 0.39 |
| gene-LOC726800 | https://www.ncbi.nlm.nih.gov/gene/?term=LOC726800 | 1284 | 5 | 38.94 | 96.92 | 0.38 |
| gene-LOC724294 | https://www.ncbi.nlm.nih.gov/gene/?term=LOC724294 | 4301 | 18 | 41.85 | 82.6 | 0.35 |
| gene-LOC100576404 | https://www.ncbi.nlm.nih.gov/gene/?term=LOC100576404 | 4488 | 102 | 227.27 | 13.62 | 0.31 |
| gene-LOC725753 | https://www.ncbi.nlm.nih.gov/gene/?term=LOC725753 | 1556 | 6 | 38.56 | 80.43 | 0.31 |
| gene-LOC408939 | https://www.ncbi.nlm.nih.gov/gene/?term=LOC408939 | 1460 | 4 | 27.4 | 97.5 | 0.27 |
| gene-LOC412468 | https://www.ncbi.nlm.nih.gov/gene/?term=LOC412468 | 6937 | 68 | 98.03 | 27.3 | 0.27 |
| gene-LOC724992 | https://www.ncbi.nlm.nih.gov/gene/?term=LOC724992 | 1021 | 3 | 29.38 | 93.33 | 0.27 |
| gene-LOC726419 | https://www.ncbi.nlm.nih.gov/gene/?term=LOC726419 | 1386 | 4 | 28.86 | 93.67 | 0.27 |
| gene-LOC726706 | https://www.ncbi.nlm.nih.gov/gene/?term=LOC726706 | 2427 | 7 | 28.84 | 94.76 | 0.27 |
| gene-LOC411299 | https://www.ncbi.nlm.nih.gov/gene/?term=LOC411299 | 2514 | 7 | 27.84 | 93.51 | 0.26 |
| gene-LOC409726 | https://www.ncbi.nlm.nih.gov/gene/?term=LOC409726 | 2783 | 35 | 125.76 | 20 | 0.25 |
| gene-LOC413891 | https://www.ncbi.nlm.nih.gov/gene/?term=LOC413891 | 2259 | 10 | 44.27 | 55.63 | 0.25 |
| gene-LOC552747 | https://www.ncbi.nlm.nih.gov/gene/?term=LOC552747 | 2628 | 22 | 83.71 | 29.32 | 0.25 |
| gene-LOC412824 | https://www.ncbi.nlm.nih.gov/gene/?term=LOC412824 | 2823 | 8 | 28.34 | 82.94 | 0.24 |
| gene-LOC552446 | https://www.ncbi.nlm.nih.gov/gene/?term=LOC552446 | 4848 | 44 | 90.76 | 26.39 | 0.24 |
| gene-LOC410280 | https://www.ncbi.nlm.nih.gov/gene/?term=LOC410280 | 2829 | 9 | 31.81 | 72.32 | 0.23 |
| gene-LOC107964413 | https://www.ncbi.nlm.nih.gov/gene/?term=LOC107964413 | 3139 | 7 | 22.3 | 94.27 | 0.21 |

...

... only first 25 lines shown ...

Directory: OGL File: ogl-Am_RE_fe.vs.te.txt

| gene_ID | gene_link | gwidth | #Sites | #per10Kb | #dmSites | #dmsp10kb | %dmSites | %pSite1 | %pSite2 | DMpSite | ADMpSite | DMpNucl | ADMpNucl |
|---|---|---|---|---|---|---|---|---|---|---|---|---|---|
| gene-LOC113219373 | https://www.ncbi.nlm.nih.gov/gene/?term=LOC113219373 | 2604 | 21 | 80.65 | 2 | 7.68 | 9.52 | 50.62 | 42.31 | 8.31 | 8.31 | 0.07 | 0.07 |
| gene-LOC550829 | https://www.ncbi.nlm.nih.gov/gene/?term=LOC550829 | 7779 | 14 | 18 | 2 | 2.57 | 14.29 | 31.38 | 28.71 | 2.67 | 2.67 | 0 | 0 |
| gene-LOC726354 | https://www.ncbi.nlm.nih.gov/gene/?term=LOC726354 | 6199 | 28 | 45.17 | 2 | 3.23 | 7.14 | 16.54 | 15.09 | 1.45 | 1.45 | 0.01 | 0.01 |
| gene-LOC726756 | https://www.ncbi.nlm.nih.gov/gene/?term=LOC726756 | 6599 | 23 | 34.85 | 2 | 3.03 | 8.7 | 11.59 | 13.45 | -1.86 | 1.86 | -0.01 | 0.01 |
| gene-LOC413433 | https://www.ncbi.nlm.nih.gov/gene/?term=LOC413433 | 7704 | 6 | 7.79 | 2 | 2.6 | 33.33 | 67.41 | 75.58 | -8.18 | 8.18 | -0.01 | 0.01 |
| gene-LOC413912 | https://www.ncbi.nlm.nih.gov/gene/?term=LOC413912 | 7969 | 3 | 3.76 | 2 | 2.51 | 66.67 | 35.9 | 63.64 | -27.74 | 27.74 | -0.01 | 0.01 |

Directory: OGL File: ogl-Am_RE_te.txt

| gene_ID | gene_link | gwidth | #Sites | #per10Kb | %perSite | %pNucl |
|---|---|---|---|---|---|---|
| gene-LOC113218557 | https://www.ncbi.nlm.nih.gov/gene/?term=LOC113218557 | 2000 | 59 | 295 | 44.96 | 1.33 |
| gene-Mir3758 | https://www.ncbi.nlm.nih.gov/gene/?term=Mir3758 | 80 | 1 | 125 | 90.91 | 1.14 |
| gene-LOC100577929 | https://www.ncbi.nlm.nih.gov/gene/?term=LOC100577929 | 2006 | 71 | 353.94 | 27.47 | 0.97 |
| gene-LOC102656390 | https://www.ncbi.nlm.nih.gov/gene/?term=LOC102656390 | 1760 | 61 | 346.59 | 24.97 | 0.87 |
| gene-LOC726419 | https://www.ncbi.nlm.nih.gov/gene/?term=LOC726419 | 1386 | 15 | 108.23 | 59.01 | 0.64 |
| gene-LOC113219373 | https://www.ncbi.nlm.nih.gov/gene/?term=LOC113219373 | 2604 | 42 | 161.29 | 37.94 | 0.61 |
| gene-LOC726800 | https://www.ncbi.nlm.nih.gov/gene/?term=LOC726800 | 1284 | 8 | 62.31 | 93.86 | 0.58 |
| gene-LOC725753 | https://www.ncbi.nlm.nih.gov/gene/?term=LOC725753 | 1556 | 11 | 70.69 | 60.64 | 0.43 |
| gene-LOC408853 | https://www.ncbi.nlm.nih.gov/gene/?term=LOC408853 | 1908 | 19 | 99.58 | 41.91 | 0.42 |
| gene-LOC102654390 | https://www.ncbi.nlm.nih.gov/gene/?term=LOC102654390 | 1226 | 10 | 81.57 | 46.56 | 0.38 |
| gene-LOC102656302 | https://www.ncbi.nlm.nih.gov/gene/?term=LOC102656302 | 1400 | 9 | 64.29 | 55.51 | 0.36 |
| gene-LOC100577394 | https://www.ncbi.nlm.nih.gov/gene/?term=LOC100577394 | 2807 | 10 | 35.63 | 97.24 | 0.35 |
| gene-LOC725352 | https://www.ncbi.nlm.nih.gov/gene/?term=LOC725352 | 859 | 3 | 34.92 | 90.91 | 0.32 |
| gene-LOC100576404 | https://www.ncbi.nlm.nih.gov/gene/?term=LOC100576404 | 4488 | 76 | 169.34 | 17.63 | 0.3 |
| gene-LOC410518 | https://www.ncbi.nlm.nih.gov/gene/?term=LOC410518 | 1850 | 7 | 37.84 | 78.38 | 0.3 |
| gene-LOC113219365 | https://www.ncbi.nlm.nih.gov/gene/?term=LOC113219365 | 2702 | 15 | 55.51 | 53.07 | 0.29 |
| gene-LOC724992 | https://www.ncbi.nlm.nih.gov/gene/?term=LOC724992 | 1021 | 3 | 29.38 | 100 | 0.29 |
| gene-LOC410280 | https://www.ncbi.nlm.nih.gov/gene/?term=LOC410280 | 2829 | 32 | 113.11 | 24.24 | 0.27 |
| gene-LOC408328 | https://www.ncbi.nlm.nih.gov/gene/?term=LOC408328 | 2745 | 16 | 58.29 | 43.78 | 0.26 |
| gene-LOC102656905 | https://www.ncbi.nlm.nih.gov/gene/?term=LOC102656905 | 1931 | 17 | 88.04 | 26.72 | 0.24 |
| gene-LOC411655 | https://www.ncbi.nlm.nih.gov/gene/?term=LOC411655 | 3816 | 25 | 65.51 | 35.49 | 0.23 |
| gene-LOC725311 | https://www.ncbi.nlm.nih.gov/gene/?term=LOC725311 | 3076 | 38 | 123.54 | 18.23 | 0.23 |
| gene-LOC102655454 | https://www.ncbi.nlm.nih.gov/gene/?term=LOC102655454 | 684 | 4 | 58.48 | 37.41 | 0.22 |
| gene-LOC102655679 | https://www.ncbi.nlm.nih.gov/gene/?term=LOC102655679 | 876 | 4 | 45.66 | 48.94 | 0.22 |

...

... only first 25 lines shown ...

Directory: OGL File: rnk-dmg-Am_RE_fe.vs.te.txt

| gene_ID | gene_link | gwidth | #Sites | #per10Kb | #dmSites | #dmsp10kb | %dmSites | %pSite1 | %pSite2 | DMpSite | ADMpSite | DMpNucl | ADMpNucl |
|---|---|---|---|---|---|---|---|---|---|---|---|---|---|
| gene-LOC113219373 | https://www.ncbi.nlm.nih.gov/gene/?term=LOC113219373 | 2604 | 21 | 80.65 | 2 | 7.68 | 9.52 | 50.62 | 42.31 | 8.31 | 8.31 | 0.07 | 0.07 |
| gene-LOC726354 | https://www.ncbi.nlm.nih.gov/gene/?term=LOC726354 | 6199 | 28 | 45.17 | 2 | 3.23 | 7.14 | 16.54 | 15.09 | 1.45 | 1.45 | 0.01 | 0.01 |
| gene-LOC726756 | https://www.ncbi.nlm.nih.gov/gene/?term=LOC726756 | 6599 | 23 | 34.85 | 2 | 3.03 | 8.7 | 11.59 | 13.45 | -1.86 | 1.86 | -0.01 | 0.01 |
| gene-LOC413433 | https://www.ncbi.nlm.nih.gov/gene/?term=LOC413433 | 7704 | 6 | 7.79 | 2 | 2.6 | 33.33 | 67.41 | 75.58 | -8.18 | 8.18 | -0.01 | 0.01 |
| gene-LOC413912 | https://www.ncbi.nlm.nih.gov/gene/?term=LOC413912 | 7969 | 3 | 3.76 | 2 | 2.51 | 66.67 | 35.9 | 63.64 | -27.74 | 27.74 | -0.01 | 0.01 |
| gene-LOC550829 | https://www.ncbi.nlm.nih.gov/gene/?term=LOC550829 | 7779 | 14 | 18 | 2 | 2.57 | 14.29 | 31.38 | 28.71 | 2.67 | 2.67 | 0 | 0 |

```
Directory: OGL File: wrt-Am_RE.txt


... comparing  fe.vs.te ...

Number of genes <=  20000  and with #dmsites >=  2 :  6
sample1     sum of percent methylation:   213.44     average:    35.57
sample2     sum of percent methylation:   238.78     average:    39.80

sum difference:   -25.35     average difference:  -4.2250

      Wilcoxon signed rank test with continuity correction

data:  pw_summary$`%pSite1` and pw_summary$`%pSite2`
V = 9, p-value = 0.8
alternative hypothesis: true location shift is not equal to 0
```

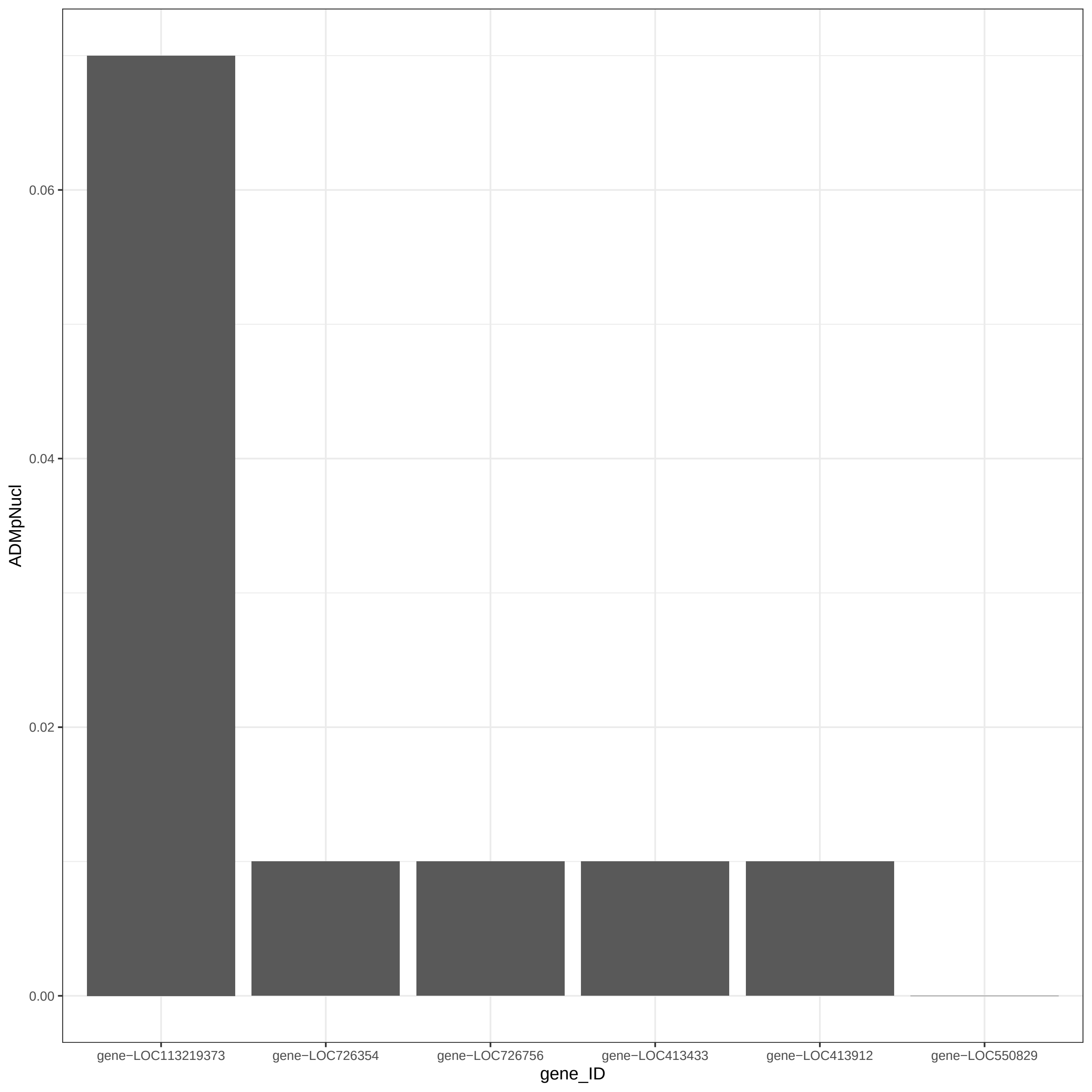

0.06
0.04
0.02
0.00
ADMpNucl
gene−LOC113219373
gene−LOC726354
gene−LOC726756
gene−LOC413433
gene−LOC413912
gene−LOC550829
gene_ID

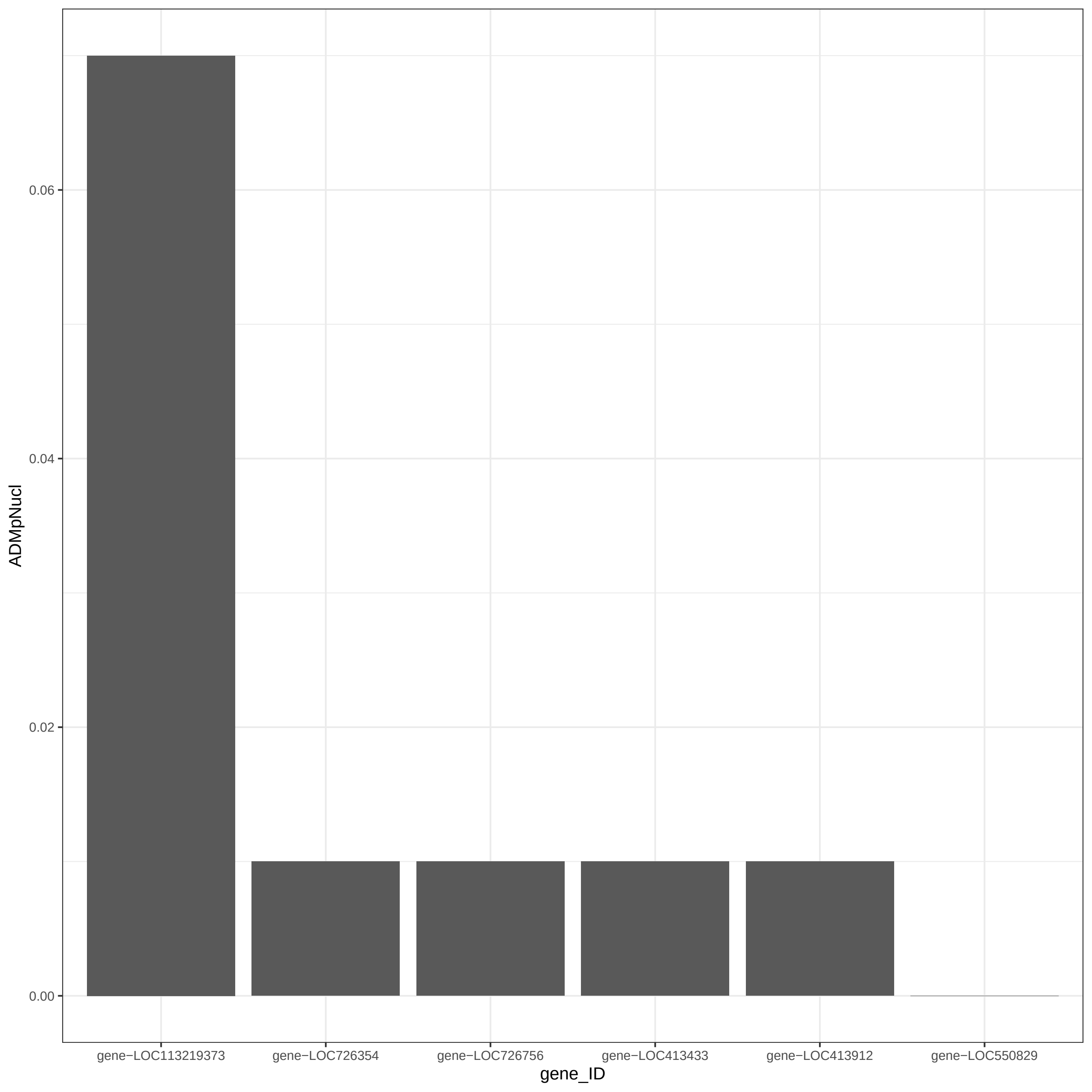

ADMpNucl
0.06
0.04
0.02
0.00
gene−LOC113219373
gene−LOC726354
gene−LOC726756
gene−LOC413433
gene−LOC413912
gene−LOC550829
gene_ID

```
Directory: OGLall File: 0READMEogl


OGL - Ordered gene lists.

Input: studyhc annotation   maxgwidth minnbrdmsites
       dmgprp (= output of BWASPR:show_dmsg())

Output: files ogl-*.txt rnk-dmg-*.txt
         ogl-<study>_<sample>.txt ogl-<study>_<sample1>.vs.<sample2>.txt
         rnk-dmg--<study>_<sample1>.vs.<sample2>.txt rnk-dmg--<study>_<sample1>.vs.<sample2>.pdf
         wrt-<study>.txt

Notes: Output files ogl-<study>_<sample>.txt give tables for each sample with columns

        gene_ID gwidth #Sites #per10Kb %perSite %pNucl

       ordered by %pNucl.  If available, a link to an NCBI entry of gene_ID is inserted as second column.
       Abbreviations used: %perSite, percent methylation per site
                           %perNucl, percent methylation per nucleotide of the gene

       Output files ogl-<study>_<sample1>.vs.<sample2>.txt give tables for each comparison with columns

        gene_ID gwidth #Sites #per10Kb #dmSites #dmsp10kb %dmSites %pSite1 %pSite2 DMpSite ADMpSite DMpNucl ADMpNucl

       ordered by DMpSite.  If available, a link to an NCBI entry of gene_ID is inserted as second column.
       Abbreviations used: %dmSites, percent sites that are differentially methylated
                           %pSite1, average per site % methylation for sample1
                           %pSite2, average per site % methylation for sample2
                           DMpSite, average per site difference in % methylation between sample1 and sample2
                           ADMpSite, absolute value of DMpSite
                           DMpNucl, average per nucleotide difference in % methylation between sample1 and sample2
                           ADMpNucl, absolute value of DMpNucl

       Output files rnk-dmg-<study>_<sample1>.vs.<sample2>.txt are equivalent to files
        ogl-<study>_<sample1>.vs.<sample2>.txt but ordered by ADMpNucl.
       Output files rnk-dmg-<study>_<sample1>.vs.<sample2>.pdf provide visualization of the distribution
        of ADMpNucl values.

       Output file wrt-<study>.txt gives results of the Wilcoxon signed rank test comparing
        the %pSite1 and %pSite2 vectors

       In output directory OGL, tables are restricted to genes with gwidth <= maxgwidth and, for
        pairwise comparisons, #dmSites >= minnbrdmsites

       If either maxgwidth or minnbrdmsites is set, then output directory OGLall will show the full tables
        with all genes (for reference).
```

Directory: OGLall File: ogl-Am_RE_fe.txt

| gene_ID | gene_link | gwidth | #Sites | #per10Kb | %perSite | %pNucl |
|---|---|---|---|---|---|---|
| gene-LOC113218557 | https://www.ncbi.nlm.nih.gov/gene/?term=LOC113218557 | 2000 | 49 | 245 | 51.47 | 1.26 |
| gene-LOC102656390 | https://www.ncbi.nlm.nih.gov/gene/?term=LOC102656390 | 1760 | 53 | 301.14 | 24.5 | 0.74 |
| gene-LOC100577929 | https://www.ncbi.nlm.nih.gov/gene/?term=LOC100577929 | 2006 | 57 | 284.15 | 22.5 | 0.64 |
| gene-LOC113219373 | https://www.ncbi.nlm.nih.gov/gene/?term=LOC113219373 | 2604 | 34 | 130.57 | 48.67 | 0.64 |
| gene-LOC408391 | https://www.ncbi.nlm.nih.gov/gene/?term=LOC408391 | 1806 | 13 | 71.98 | 87.97 | 0.63 |
| gene-LOC409500 | https://www.ncbi.nlm.nih.gov/gene/?term=LOC409500 | 1222 | 11 | 90.02 | 48.43 | 0.44 |
| gene-LOC408367 | https://www.ncbi.nlm.nih.gov/gene/?term=LOC408367 | 2232 | 11 | 49.28 | 79.27 | 0.39 |
| gene-LOC726800 | https://www.ncbi.nlm.nih.gov/gene/?term=LOC726800 | 1284 | 5 | 38.94 | 96.92 | 0.38 |
| gene-LOC724294 | https://www.ncbi.nlm.nih.gov/gene/?term=LOC724294 | 4301 | 18 | 41.85 | 82.6 | 0.35 |
| gene-LOC100576404 | https://www.ncbi.nlm.nih.gov/gene/?term=LOC100576404 | 4488 | 102 | 227.27 | 13.62 | 0.31 |
| gene-LOC725753 | https://www.ncbi.nlm.nih.gov/gene/?term=LOC725753 | 1556 | 6 | 38.56 | 80.43 | 0.31 |
| gene-LOC408939 | https://www.ncbi.nlm.nih.gov/gene/?term=LOC408939 | 1460 | 4 | 27.4 | 97.5 | 0.27 |
| gene-LOC412468 | https://www.ncbi.nlm.nih.gov/gene/?term=LOC412468 | 6937 | 68 | 98.03 | 27.3 | 0.27 |
| gene-LOC724992 | https://www.ncbi.nlm.nih.gov/gene/?term=LOC724992 | 1021 | 3 | 29.38 | 93.33 | 0.27 |
| gene-LOC726419 | https://www.ncbi.nlm.nih.gov/gene/?term=LOC726419 | 1386 | 4 | 28.86 | 93.67 | 0.27 |
| gene-LOC726706 | https://www.ncbi.nlm.nih.gov/gene/?term=LOC726706 | 2427 | 7 | 28.84 | 94.76 | 0.27 |
| gene-LOC411299 | https://www.ncbi.nlm.nih.gov/gene/?term=LOC411299 | 2514 | 7 | 27.84 | 93.51 | 0.26 |
| gene-LOC409726 | https://www.ncbi.nlm.nih.gov/gene/?term=LOC409726 | 2783 | 35 | 125.76 | 20 | 0.25 |
| gene-LOC413891 | https://www.ncbi.nlm.nih.gov/gene/?term=LOC413891 | 2259 | 10 | 44.27 | 55.63 | 0.25 |
| gene-LOC552747 | https://www.ncbi.nlm.nih.gov/gene/?term=LOC552747 | 2628 | 22 | 83.71 | 29.32 | 0.25 |
| gene-LOC412824 | https://www.ncbi.nlm.nih.gov/gene/?term=LOC412824 | 2823 | 8 | 28.34 | 82.94 | 0.24 |
| gene-LOC552446 | https://www.ncbi.nlm.nih.gov/gene/?term=LOC552446 | 4848 | 44 | 90.76 | 26.39 | 0.24 |
| gene-LOC410280 | https://www.ncbi.nlm.nih.gov/gene/?term=LOC410280 | 2829 | 9 | 31.81 | 72.32 | 0.23 |
| gene-LOC107964413 | https://www.ncbi.nlm.nih.gov/gene/?term=LOC107964413 | 3139 | 7 | 22.3 | 94.27 | 0.21 |

...

... only first 25 lines shown ...

Directory: OGLall File: ogl-Am_RE_fe.vs.te.txt

| gene_ID | gene_link | gwidth | #Sites | #per10Kb | #dmSites | #dmsp10kb | %dmSites | %pSite1 | %pSite2 | DMpSite | ADMpSite | DMpNucl | ADMpNucl |
|---|---|---|---|---|---|---|---|---|---|---|---|---|---|
| gene-LOC100577140 | https://www.ncbi.nlm.nih.gov/gene/?term=LOC100577140 | 3634 | 1 | 2.75 | 1 | 2.75 | 100 | 66.67 | 6.67 | 60 | 60 | 0.02 | 0.02 |
| gene-LOC726791 | https://www.ncbi.nlm.nih.gov/gene/?term=LOC726791 | 5197 | 1 | 1.92 | 1 | 1.92 | 100 | 50 | 7.69 | 42.31 | 42.31 | 0.01 | 0.01 |
| gene-LOC413484 | https://www.ncbi.nlm.nih.gov/gene/?term=LOC413484 | 3011 | 1 | 3.32 | 1 | 3.32 | 100 | 78.57 | 40 | 38.57 | 38.57 | 0.01 | 0.01 |
| gene-LOC408936 | https://www.ncbi.nlm.nih.gov/gene/?term=LOC408936 | 11444 | 1 | 0.87 | 1 | 0.87 | 100 | 100 | 70 | 30 | 30 | 0 | 0 |
| gene-LOC409271 | https://www.ncbi.nlm.nih.gov/gene/?term=LOC409271 | 9800 | 2 | 2.04 | 1 | 1.02 | 50 | 86.75 | 65 | 21.75 | 21.75 | 0 | 0 |
| gene-LOC107964797 | https://www.ncbi.nlm.nih.gov/gene/?term=LOC107964797 | 2058 | 2 | 9.72 | 1 | 4.86 | 50 | 13.63 | 0 | 13.63 | 13.63 | 0.01 | 0.01 |
| gene-LOC552046 | https://www.ncbi.nlm.nih.gov/gene/?term=LOC552046 | 8248 | 3 | 3.64 | 1 | 1.21 | 33.33 | 30.56 | 18.18 | 12.37 | 12.37 | 0 | 0 |
| gene-LOC410958 | https://www.ncbi.nlm.nih.gov/gene/?term=LOC410958 | 4837 | 5 | 10.34 | 1 | 2.07 | 20 | 52.95 | 41.92 | 11.03 | 11.03 | 0.01 | 0.01 |
| gene-LOC411678 | https://www.ncbi.nlm.nih.gov/gene/?term=LOC411678 | 5617 | 9 | 16.02 | 1 | 1.78 | 11.11 | 25.13 | 14.86 | 10.26 | 10.26 | 0.02 | 0.02 |
| gene-LOC410510 | https://www.ncbi.nlm.nih.gov/gene/?term=LOC410510 | 4529 | 6 | 13.25 | 1 | 2.21 | 16.67 | 24.21 | 15.59 | 8.61 | 8.61 | 0.01 | 0.01 |
| gene-LOC413152 | https://www.ncbi.nlm.nih.gov/gene/?term=LOC413152 | 5760 | 3 | 5.21 | 1 | 1.74 | 33.33 | 10 | 1.45 | 8.55 | 8.55 | 0 | 0 |
| gene-LOC113219373 | https://www.ncbi.nlm.nih.gov/gene/?term=LOC113219373 | 2604 | 21 | 80.65 | 2 | 7.68 | 9.52 | 50.62 | 42.31 | 8.31 | 8.31 | 0.07 | 0.07 |
| gene-LOC100577216 | https://www.ncbi.nlm.nih.gov/gene/?term=LOC100577216 | 35911 | 59 | 16.43 | 4 | 1.11 | 6.78 | 20.5 | 15.59 | 4.92 | 4.92 | 0.01 | 0.01 |
| gene-LOC412468 | https://www.ncbi.nlm.nih.gov/gene/?term=LOC412468 | 6937 | 11 | 15.86 | 1 | 1.44 | 9.09 | 14.37 | 10 | 4.37 | 4.37 | 0.01 | 0.01 |
| gene-LOC409192 | https://www.ncbi.nlm.nih.gov/gene/?term=LOC409192 | 18515 | 13 | 7.02 | 1 | 0.54 | 7.69 | 12.31 | 8.11 | 4.2 | 4.2 | 0 | 0 |
| gene-LOC725503 | https://www.ncbi.nlm.nih.gov/gene/?term=LOC725503 | 16789 | 8 | 4.77 | 1 | 0.6 | 12.5 | 44.92 | 40.86 | 4.06 | 4.06 | 0 | 0 |
| gene-LOC550829 | https://www.ncbi.nlm.nih.gov/gene/?term=LOC550829 | 7779 | 14 | 18 | 2 | 2.57 | 14.29 | 31.38 | 28.71 | 2.67 | 2.67 | 0 | 0 |
| gene-LOC113218927 | https://www.ncbi.nlm.nih.gov/gene/?term=LOC113218927 | 4543 | 11 | 24.21 | 1 | 2.2 | 9.09 | 5.84 | 3.24 | 2.61 | 2.61 | 0.01 | 0.01 |
| gene-LOC409422 | https://www.ncbi.nlm.nih.gov/gene/?term=LOC409422 | 198351 | 22 | 1.11 | 1 | 0.05 | 4.55 | 22.66 | 20.6 | 2.07 | 2.07 | 0 | 0 |
| gene-LOC113218557 | https://www.ncbi.nlm.nih.gov/gene/?term=LOC113218557 | 2000 | 30 | 150 | 1 | 5 | 3.33 | 46.43 | 44.36 | 2.06 | 2.06 | 0.03 | 0.03 |
| gene-LOC726354 | https://www.ncbi.nlm.nih.gov/gene/?term=LOC726354 | 6199 | 28 | 45.17 | 2 | 3.23 | 7.14 | 16.54 | 15.09 | 1.45 | 1.45 | 0.01 | 0.01 |
| gene-LOC100577280 | https://www.ncbi.nlm.nih.gov/gene/?term=LOC100577280 | 22469 | 33 | 14.69 | 2 | 0.89 | 6.06 | 5.49 | 4.14 | 1.34 | 1.34 | 0 | 0 |
| gene-LOC551291 | https://www.ncbi.nlm.nih.gov/gene/?term=LOC551291 | 8481 | 19 | 22.4 | 1 | 1.18 | 5.26 | 3.13 | 1.84 | 1.29 | 1.29 | 0 | 0 |
| gene-LOC551848 | https://www.ncbi.nlm.nih.gov/gene/?term=LOC551848 | 22642 | 83 | 36.66 | 2 | 0.88 | 2.41 | 3.71 | 2.8 | 0.91 | 0.91 | 0 | 0 |
| gene-LOC413428 | https://www.ncbi.nlm.nih.gov/gene/?term=LOC413428 | 9438 | 97 | 102.78 | 1 | 1.06 | 1.03 | 3.47 | 2.81 | 0.66 | 0.66 | 0.01 | 0.01 |
| gene-LOC411270 | https://www.ncbi.nlm.nih.gov/gene/?term=LOC411270 | 15083 | 54 | 35.8 | 1 | 0.66 | 1.85 | 1.19 | 0.77 | 0.42 | 0.42 | 0 | 0 |
| gene-LOC727008 | https://www.ncbi.nlm.nih.gov/gene/?term=LOC727008 | 4728 | 64 | 135.36 | 1 | 2.12 | 1.56 | 5.2 | 5.56 | -0.36 | 0.36 | 0 | 0 |
| gene-LOC113219367 | https://www.ncbi.nlm.nih.gov/gene/?term=LOC113219367 | 42366 | 367 | 86.63 | 1 | 0.24 | 0.27 | 5.35 | 5.85 | -0.49 | 0.49 | 0 | 0 |
| gene-LOC411466 | https://www.ncbi.nlm.nih.gov/gene/?term=LOC411466 | 31311 | 26 | 8.3 | 1 | 0.32 | 3.85 | 18.42 | 19.52 | -1.1 | 1.1 | 0 | 0 |
| gene-LOC724670 | https://www.ncbi.nlm.nih.gov/gene/?term=LOC724670 | 18827 | 15 | 7.97 | 1 | 0.53 | 6.67 | 14.44 | 16.15 | -1.71 | 1.71 | 0 | 0 |
| gene-LOC100577929 | https://www.ncbi.nlm.nih.gov/gene/?term=LOC100577929 | 2006 | 37 | 184.45 | 1 | 4.99 | 2.7 | 24.15 | 25.96 | -1.81 | 1.81 | -0.03 | 0.03 |
| gene-LOC726756 | https://www.ncbi.nlm.nih.gov/gene/?term=LOC726756 | 6599 | 23 | 34.85 | 2 | 3.03 | 8.7 | 11.59 | 13.45 | -1.86 | 1.86 | -0.01 | 0.01 |
| gene-LOC411542 | https://www.ncbi.nlm.nih.gov/gene/?term=LOC411542 | 2987 | 12 | 40.17 | 1 | 3.35 | 8.33 | 1.51 | 5 | -3.48 | 3.48 | -0.01 | 0.01 |
| gene-LOC410804 | https://www.ncbi.nlm.nih.gov/gene/?term=LOC410804 | 12945 | 11 | 8.5 | 1 | 0.77 | 9.09 | 19.83 | 23.55 | -3.72 | 3.72 | 0 | 0 |
| gene-LOC409107 | https://www.ncbi.nlm.nih.gov/gene/?term=LOC409107 | 11039 | 4 | 3.62 | 1 | 0.91 | 25 | 42.86 | 48.44 | -5.58 | 5.58 | 0 | 0 |
| gene-LOC100578257 | https://www.ncbi.nlm.nih.gov/gene/?term=LOC100578257 | 2958 | 4 | 13.52 | 1 | 3.38 | 25 | 36.82 | 42.47 | -5.65 | 5.65 | -0.01 | 0.01 |
| gene-LOC552747 | https://www.ncbi.nlm.nih.gov/gene/?term=LOC552747 | 2628 | 9 | 34.25 | 1 | 3.81 | 11.11 | 19.41 | 25.93 | -6.52 | 6.52 | -0.02 | 0.02 |
| gene-LOC726727 | https://www.ncbi.nlm.nih.gov/gene/?term=LOC726727 | 20697 | 6 | 2.9 | 1 | 0.48 | 16.67 | 6.67 | 14.45 | -7.78 | 7.78 | 0 | 0 |
| gene-LOC413433 | https://www.ncbi.nlm.nih.gov/gene/?term=LOC413433 | 7704 | 6 | 7.79 | 2 | 2.6 | 33.33 | 67.41 | 75.58 | -8.18 | 8.18 | -0.01 | 0.01 |
| gene-LOC413344 | https://www.ncbi.nlm.nih.gov/gene/?term=LOC413344 | 7110 | 4 | 5.63 | 1 | 1.41 | 25 | 18.54 | 27.5 | -8.96 | 8.96 | -0.01 | 0.01 |
| gene-LOC100577552 | https://www.ncbi.nlm.nih.gov/gene/?term=LOC100577552 | 11299 | 3 | 2.66 | 1 | 0.89 | 33.33 | 63.37 | 75.79 | -12.42 | 12.42 | 0 | 0 |
| gene-LOC413429 | https://www.ncbi.nlm.nih.gov/gene/?term=LOC413429 | 2291 | 2 | 8.73 | 1 | 4.36 | 50 | 19.23 | 33.34 | -14.11 | 14.11 | -0.01 | 0.01 |
| gene-LOC411881 | https://www.ncbi.nlm.nih.gov/gene/?term=LOC411881 | 5054 | 1 | 1.98 | 1 | 1.98 | 100 | 36.36 | 63.64 | -27.28 | 27.28 | -0.01 | 0.01 |
| gene-LOC413912 | https://www.ncbi.nlm.nih.gov/gene/?term=LOC413912 | 7969 | 3 | 3.76 | 2 | 2.51 | 66.67 | 35.9 | 63.64 | -27.74 | 27.74 | -0.01 | 0.01 |
| gene-LOC726432 | https://www.ncbi.nlm.nih.gov/gene/?term=LOC726432 | 3074 | 1 | 3.25 | 1 | 3.25 | 100 | 50 | 81.82 | -31.82 | 31.82 | -0.01 | 0.01 |

Directory: OGLall File: ogl-Am_RE_te.txt

| gene_ID | gene_link | gwidth | #Sites | #per10Kb | %perSite | %pNucl |
|---|---|---|---|---|---|---|
| gene-LOC113218557 | https://www.ncbi.nlm.nih.gov/gene/?term=LOC113218557 | 2000 | 59 | 295 | 44.96 | 1.33 |
| gene-Mir3758 | https://www.ncbi.nlm.nih.gov/gene/?term=Mir3758 | 80 | 1 | 125 | 90.91 | 1.14 |
| gene-LOC100577929 | https://www.ncbi.nlm.nih.gov/gene/?term=LOC100577929 | 2006 | 71 | 353.94 | 27.47 | 0.97 |
| gene-LOC102656390 | https://www.ncbi.nlm.nih.gov/gene/?term=LOC102656390 | 1760 | 61 | 346.59 | 24.97 | 0.87 |
| gene-LOC726419 | https://www.ncbi.nlm.nih.gov/gene/?term=LOC726419 | 1386 | 15 | 108.23 | 59.01 | 0.64 |
| gene-LOC113219373 | https://www.ncbi.nlm.nih.gov/gene/?term=LOC113219373 | 2604 | 42 | 161.29 | 37.94 | 0.61 |
| gene-LOC726800 | https://www.ncbi.nlm.nih.gov/gene/?term=LOC726800 | 1284 | 8 | 62.31 | 93.86 | 0.58 |
| gene-LOC725753 | https://www.ncbi.nlm.nih.gov/gene/?term=LOC725753 | 1556 | 11 | 70.69 | 60.64 | 0.43 |
| gene-LOC408853 | https://www.ncbi.nlm.nih.gov/gene/?term=LOC408853 | 1908 | 19 | 99.58 | 41.91 | 0.42 |
| gene-LOC102654390 | https://www.ncbi.nlm.nih.gov/gene/?term=LOC102654390 | 1226 | 10 | 81.57 | 46.56 | 0.38 |
| gene-LOC102656302 | https://www.ncbi.nlm.nih.gov/gene/?term=LOC102656302 | 1400 | 9 | 64.29 | 55.51 | 0.36 |
| gene-LOC100577394 | https://www.ncbi.nlm.nih.gov/gene/?term=LOC100577394 | 2807 | 10 | 35.63 | 97.24 | 0.35 |
| gene-LOC725352 | https://www.ncbi.nlm.nih.gov/gene/?term=LOC725352 | 859 | 3 | 34.92 | 90.91 | 0.32 |
| gene-LOC100576404 | https://www.ncbi.nlm.nih.gov/gene/?term=LOC100576404 | 4488 | 76 | 169.34 | 17.63 | 0.3 |
| gene-LOC410518 | https://www.ncbi.nlm.nih.gov/gene/?term=LOC410518 | 1850 | 7 | 37.84 | 78.38 | 0.3 |
| gene-LOC113219365 | https://www.ncbi.nlm.nih.gov/gene/?term=LOC113219365 | 2702 | 15 | 55.51 | 53.07 | 0.29 |
| gene-LOC724992 | https://www.ncbi.nlm.nih.gov/gene/?term=LOC724992 | 1021 | 3 | 29.38 | 100 | 0.29 |
| gene-LOC410280 | https://www.ncbi.nlm.nih.gov/gene/?term=LOC410280 | 2829 | 32 | 113.11 | 24.24 | 0.27 |
| gene-LOC408328 | https://www.ncbi.nlm.nih.gov/gene/?term=LOC408328 | 2745 | 16 | 58.29 | 43.78 | 0.26 |
| gene-LOC102656905 | https://www.ncbi.nlm.nih.gov/gene/?term=LOC102656905 | 1931 | 17 | 88.04 | 26.72 | 0.24 |
| gene-LOC411655 | https://www.ncbi.nlm.nih.gov/gene/?term=LOC411655 | 3816 | 25 | 65.51 | 35.49 | 0.23 |
| gene-LOC725311 | https://www.ncbi.nlm.nih.gov/gene/?term=LOC725311 | 3076 | 38 | 123.54 | 18.23 | 0.23 |
| gene-LOC102655454 | https://www.ncbi.nlm.nih.gov/gene/?term=LOC102655454 | 684 | 4 | 58.48 | 37.41 | 0.22 |
| gene-LOC102655679 | https://www.ncbi.nlm.nih.gov/gene/?term=LOC102655679 | 876 | 4 | 45.66 | 48.94 | 0.22 |

...
... only first 25 lines shown ...

Directory: OGLall File: rnk-dmg-Am_RE_fe.vs.te.txt

| gene_ID | gene_link | gwidth | #Sites | #per10Kb | #dmSites | #dmsp10kb | %dmSites | %pSite1 | %pSite2 | DMpSite | ADMpSite | DMpNucl | ADMpNucl |
|---|---|---|---|---|---|---|---|---|---|---|---|---|---|
| gene-LOC113219373 | https://www.ncbi.nlm.nih.gov/gene/?term=LOC113219373 | 2604 | 21 | 80.65 | 2 | 7.68 | 9.52 | 50.62 | 42.31 | 8.31 | 8.31 | 0.07 | 0.07 |
| gene-LOC113218557 | https://www.ncbi.nlm.nih.gov/gene/?term=LOC113218557 | 2000 | 30 | 150 | 1 | 5 | 3.33 | 46.43 | 44.36 | 2.06 | 2.06 | 0.03 | 0.03 |
| gene-LOC100577929 | https://www.ncbi.nlm.nih.gov/gene/?term=LOC100577929 | 2006 | 37 | 184.45 | 1 | 4.99 | 2.7 | 24.15 | 25.96 | -1.81 | 1.81 | -0.03 | 0.03 |
| gene-LOC100577140 | https://www.ncbi.nlm.nih.gov/gene/?term=LOC100577140 | 3634 | 1 | 2.75 | 1 | 2.75 | 100 | 66.67 | 6.67 | 60 | 60 | 0.02 | 0.02 |
| gene-LOC411678 | https://www.ncbi.nlm.nih.gov/gene/?term=LOC411678 | 5617 | 9 | 16.02 | 1 | 1.78 | 11.11 | 25.13 | 14.86 | 10.26 | 10.26 | 0.02 | 0.02 |
| gene-LOC552747 | https://www.ncbi.nlm.nih.gov/gene/?term=LOC552747 | 2628 | 9 | 34.25 | 1 | 3.81 | 11.11 | 19.41 | 25.93 | -6.52 | 6.52 | -0.02 | 0.02 |
| gene-LOC726791 | https://www.ncbi.nlm.nih.gov/gene/?term=LOC726791 | 5197 | 1 | 1.92 | 1 | 1.92 | 100 | 50 | 7.69 | 42.31 | 42.31 | 0.01 | 0.01 |
| gene-LOC413484 | https://www.ncbi.nlm.nih.gov/gene/?term=LOC413484 | 3011 | 1 | 3.32 | 1 | 3.32 | 100 | 78.57 | 40 | 38.57 | 38.57 | 0.01 | 0.01 |
| gene-LOC107964797 | https://www.ncbi.nlm.nih.gov/gene/?term=LOC107964797 | 2058 | 2 | 9.72 | 1 | 4.86 | 50 | 13.63 | 0 | 13.63 | 13.63 | 0.01 | 0.01 |
| gene-LOC410958 | https://www.ncbi.nlm.nih.gov/gene/?term=LOC410958 | 4837 | 5 | 10.34 | 1 | 2.07 | 20 | 52.95 | 41.92 | 11.03 | 11.03 | 0.01 | 0.01 |
| gene-LOC410510 | https://www.ncbi.nlm.nih.gov/gene/?term=LOC410510 | 4529 | 6 | 13.25 | 1 | 2.21 | 16.67 | 24.21 | 15.59 | 8.61 | 8.61 | 0.01 | 0.01 |
| gene-LOC100577216 | https://www.ncbi.nlm.nih.gov/gene/?term=LOC100577216 | 35911 | 59 | 16.43 | 4 | 1.11 | 6.78 | 20.5 | 15.59 | 4.92 | 4.92 | 0.01 | 0.01 |
| gene-LOC412468 | https://www.ncbi.nlm.nih.gov/gene/?term=LOC412468 | 6937 | 11 | 15.86 | 1 | 1.44 | 9.09 | 14.37 | 10 | 4.37 | 4.37 | 0.01 | 0.01 |
| gene-LOC113218927 | https://www.ncbi.nlm.nih.gov/gene/?term=LOC113218927 | 4543 | 11 | 24.21 | 1 | 2.2 | 9.09 | 5.84 | 3.24 | 2.61 | 2.61 | 0.01 | 0.01 |
| gene-LOC726354 | https://www.ncbi.nlm.nih.gov/gene/?term=LOC726354 | 6199 | 28 | 45.17 | 2 | 3.23 | 7.14 | 16.54 | 15.09 | 1.45 | 1.45 | 0.01 | 0.01 |
| gene-LOC413428 | https://www.ncbi.nlm.nih.gov/gene/?term=LOC413428 | 9438 | 97 | 102.78 | 1 | 1.06 | 1.03 | 3.47 | 2.81 | 0.66 | 0.66 | 0.01 | 0.01 |
| gene-LOC726756 | https://www.ncbi.nlm.nih.gov/gene/?term=LOC726756 | 6599 | 23 | 34.85 | 2 | 3.03 | 8.7 | 11.59 | 13.45 | -1.86 | 1.86 | -0.01 | 0.01 |
| gene-LOC411542 | https://www.ncbi.nlm.nih.gov/gene/?term=LOC411542 | 2987 | 12 | 40.17 | 1 | 3.35 | 8.33 | 1.51 | 5 | -3.48 | 3.48 | -0.01 | 0.01 |
| gene-LOC100578257 | https://www.ncbi.nlm.nih.gov/gene/?term=LOC100578257 | 2958 | 4 | 13.52 | 1 | 3.38 | 25 | 36.82 | 42.47 | -5.65 | 5.65 | -0.01 | 0.01 |
| gene-LOC413433 | https://www.ncbi.nlm.nih.gov/gene/?term=LOC413433 | 7704 | 6 | 7.79 | 2 | 2.6 | 33.33 | 67.41 | 75.58 | -8.18 | 8.18 | -0.01 | 0.01 |
| gene-LOC413344 | https://www.ncbi.nlm.nih.gov/gene/?term=LOC413344 | 7110 | 4 | 5.63 | 1 | 1.41 | 25 | 18.54 | 27.5 | -8.96 | 8.96 | -0.01 | 0.01 |
| gene-LOC413429 | https://www.ncbi.nlm.nih.gov/gene/?term=LOC413429 | 2291 | 2 | 8.73 | 1 | 4.36 | 50 | 19.23 | 33.34 | -14.11 | 14.11 | -0.01 | 0.01 |
| gene-LOC411881 | https://www.ncbi.nlm.nih.gov/gene/?term=LOC411881 | 5054 | 1 | 1.98 | 1 | 1.98 | 100 | 36.36 | 63.64 | -27.28 | 27.28 | -0.01 | 0.01 |
| gene-LOC413912 | https://www.ncbi.nlm.nih.gov/gene/?term=LOC413912 | 7969 | 3 | 3.76 | 2 | 2.51 | 66.67 | 35.9 | 63.64 | -27.74 | 27.74 | -0.01 | 0.01 |
| gene-LOC726432 | https://www.ncbi.nlm.nih.gov/gene/?term=LOC726432 | 3074 | 1 | 3.25 | 1 | 3.25 | 100 | 50 | 81.82 | -31.82 | 31.82 | -0.01 | 0.01 |
| gene-LOC408936 | https://www.ncbi.nlm.nih.gov/gene/?term=LOC408936 | 11444 | 1 | 0.87 | 1 | 0.87 | 100 | 100 | 70 | 30 | 30 | 0 | 0 |
| gene-LOC409271 | https://www.ncbi.nlm.nih.gov/gene/?term=LOC409271 | 9800 | 2 | 2.04 | 1 | 1.02 | 50 | 86.75 | 65 | 21.75 | 21.75 | 0 | 0 |
| gene-LOC552046 | https://www.ncbi.nlm.nih.gov/gene/?term=LOC552046 | 8248 | 3 | 3.64 | 1 | 1.21 | 33.33 | 30.56 | 18.18 | 12.37 | 12.37 | 0 | 0 |
| gene-LOC413152 | https://www.ncbi.nlm.nih.gov/gene/?term=LOC413152 | 5760 | 3 | 5.21 | 1 | 1.74 | 33.33 | 10 | 1.45 | 8.55 | 8.55 | 0 | 0 |
| gene-LOC409192 | https://www.ncbi.nlm.nih.gov/gene/?term=LOC409192 | 18515 | 13 | 7.02 | 1 | 0.54 | 7.69 | 12.31 | 8.11 | 4.2 | 4.2 | 0 | 0 |
| gene-LOC725503 | https://www.ncbi.nlm.nih.gov/gene/?term=LOC725503 | 16789 | 8 | 4.77 | 1 | 0.6 | 12.5 | 44.92 | 40.86 | 4.06 | 4.06 | 0 | 0 |
| gene-LOC550829 | https://www.ncbi.nlm.nih.gov/gene/?term=LOC550829 | 7779 | 14 | 18 | 2 | 2.57 | 14.29 | 31.38 | 28.71 | 2.67 | 2.67 | 0 | 0 |
| gene-LOC409422 | https://www.ncbi.nlm.nih.gov/gene/?term=LOC409422 | 198351 | 22 | 1.11 | 1 | 0.05 | 4.55 | 22.66 | 20.6 | 2.07 | 2.07 | 0 | 0 |
| gene-LOC100577280 | https://www.ncbi.nlm.nih.gov/gene/?term=LOC100577280 | 22469 | 33 | 14.69 | 2 | 0.89 | 6.06 | 5.49 | 4.14 | 1.34 | 1.34 | 0 | 0 |
| gene-LOC551291 | https://www.ncbi.nlm.nih.gov/gene/?term=LOC551291 | 8481 | 19 | 22.4 | 1 | 1.18 | 5.26 | 3.13 | 1.84 | 1.29 | 1.29 | 0 | 0 |
| gene-LOC551848 | https://www.ncbi.nlm.nih.gov/gene/?term=LOC551848 | 22642 | 83 | 36.66 | 2 | 0.88 | 2.41 | 3.71 | 2.8 | 0.91 | 0.91 | 0 | 0 |
| gene-LOC411270 | https://www.ncbi.nlm.nih.gov/gene/?term=LOC411270 | 15083 | 54 | 35.8 | 1 | 0.66 | 1.85 | 1.19 | 0.77 | 0.42 | 0.42 | 0 | 0 |
| gene-LOC727008 | https://www.ncbi.nlm.nih.gov/gene/?term=LOC727008 | 4728 | 64 | 135.36 | 1 | 2.12 | 1.56 | 5.2 | 5.56 | -0.36 | 0.36 | 0 | 0 |
| gene-LOC113219367 | https://www.ncbi.nlm.nih.gov/gene/?term=LOC113219367 | 42366 | 367 | 86.63 | 1 | 0.24 | 0.27 | 5.35 | 5.85 | -0.49 | 0.49 | 0 | 0 |
| gene-LOC411466 | https://www.ncbi.nlm.nih.gov/gene/?term=LOC411466 | 31311 | 26 | 8.3 | 1 | 0.32 | 3.85 | 18.42 | 19.52 | -1.1 | 1.1 | 0 | 0 |
| gene-LOC724670 | https://www.ncbi.nlm.nih.gov/gene/?term=LOC724670 | 18827 | 15 | 7.97 | 1 | 0.53 | 6.67 | 14.44 | 16.15 | -1.71 | 1.71 | 0 | 0 |
| gene-LOC410804 | https://www.ncbi.nlm.nih.gov/gene/?term=LOC410804 | 12945 | 11 | 8.5 | 1 | 0.77 | 9.09 | 19.83 | 23.55 | -3.72 | 3.72 | 0 | 0 |
| gene-LOC409107 | https://www.ncbi.nlm.nih.gov/gene/?term=LOC409107 | 11039 | 4 | 3.62 | 1 | 0.91 | 25 | 42.86 | 48.44 | -5.58 | 5.58 | 0 | 0 |
| gene-LOC726727 | https://www.ncbi.nlm.nih.gov/gene/?term=LOC726727 | 20697 | 6 | 2.9 | 1 | 0.48 | 16.67 | 6.67 | 14.45 | -7.78 | 7.78 | 0 | 0 |
| gene-LOC100577552 | https://www.ncbi.nlm.nih.gov/gene/?term=LOC100577552 | 11299 | 3 | 2.66 | 1 | 0.89 | 33.33 | 63.37 | 75.79 | -12.42 | 12.42 | 0 | 0 |

```
Directory: OGLall File: wrt-Am_RE.txt


... comparing  fe.vs.te ...

Number of genes with #dmsites >=  1 :  45

sample1     sum of percent methylation:  1318.09     average:    29.29
sample2     sum of percent methylation:  1190.23     average:    26.45

sum difference:   127.85     average difference:   2.8411

      Wilcoxon signed rank test with continuity correction

data:  pw_summary$`%pSite1` and pw_summary$`%pSite2`
V = 608, p-value = 0.3
alternative hypothesis: true location shift is not equal to 0
```

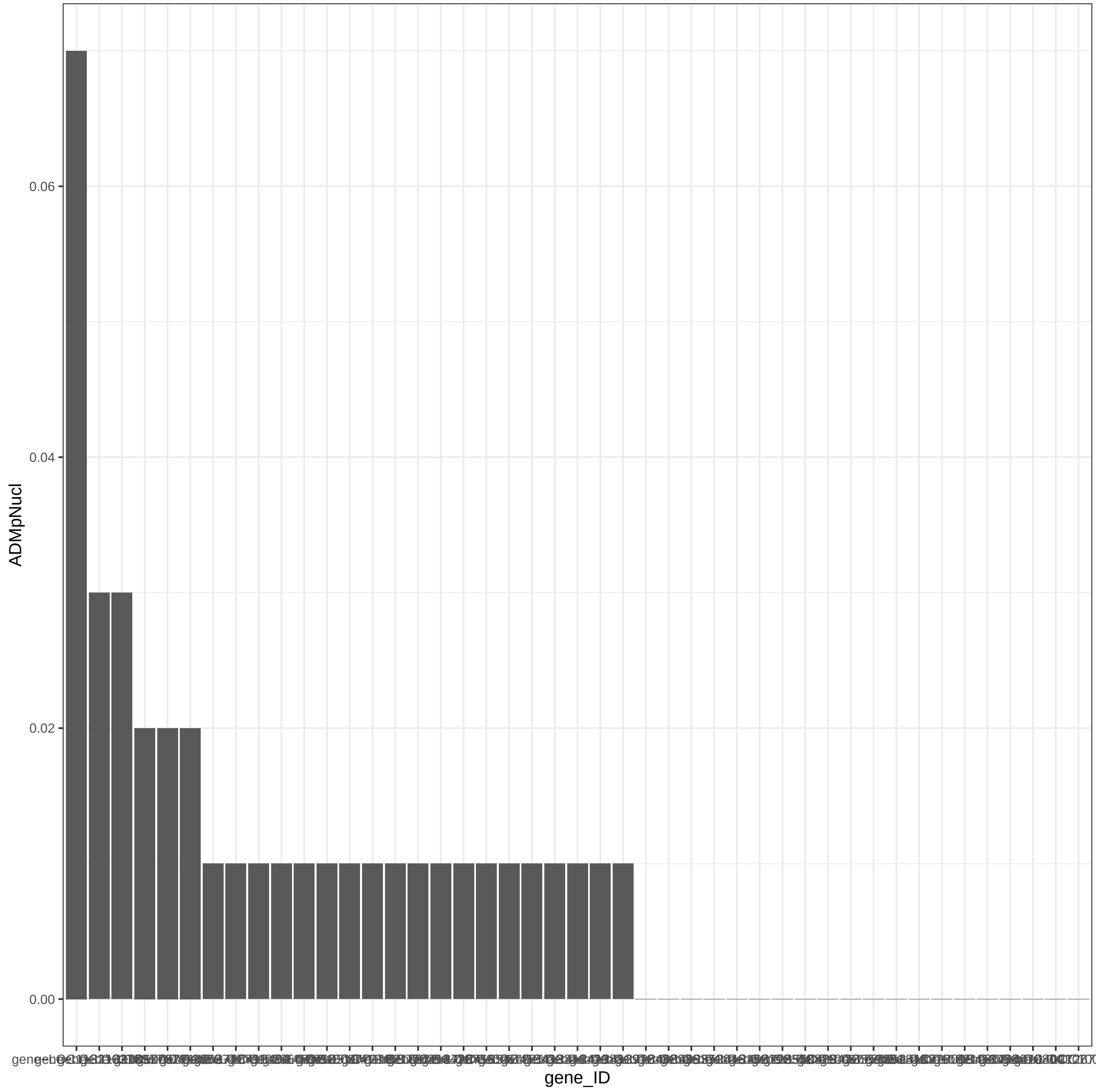

ADMpNucl
0.06
0.04
0.02
0.00
gene_ID

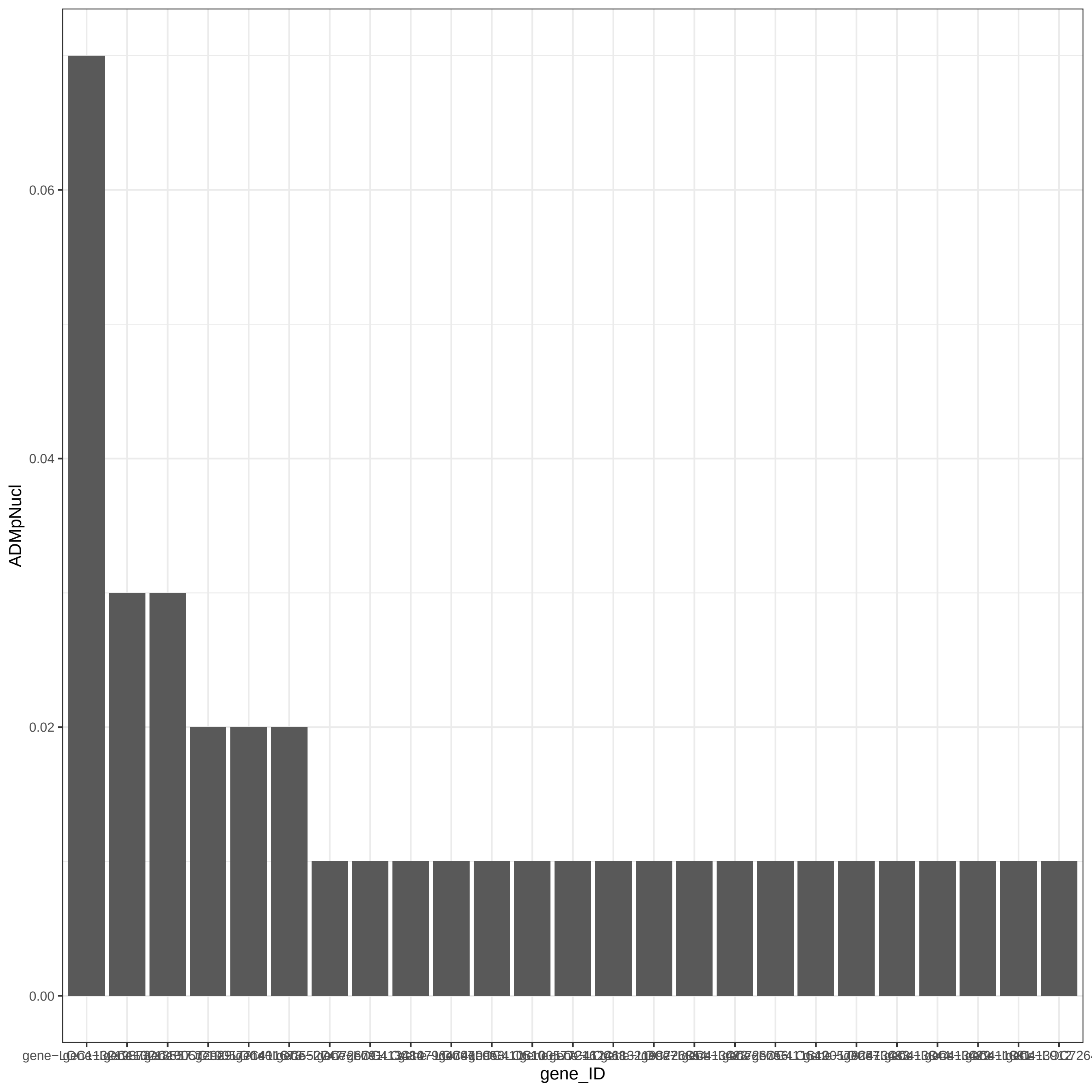

0.06
0.04
0.02
0.00
ADMpNucl
gene_ID
