## Supplementary material for "Tools and applications for integrative analysis of DNA methylation in social insects": Dataset S17

Cb_LIp.conf

```
#Customize variables here:######################################################
################################################################################

#Load the files:
#
infiles       <- setup_BWASPR(datafile="./Cb.dat",
                              parfile="./Cb.par")

#Set the study and samples in the study:
#
species       <- "Cb"
study         <- "LI"
samplelist    <- list("BCphase", "Rphase")
## The following two variables are used for output file labeling:
studyLabel    <- "LIp"
sampleLabels  <- list("Cb_LI_BCphase", "Cb_LI_Rphase")

hasreplicates <- TRUE
type          <- "CpG"
destrand      <- TRUE
covlist       <- c(6,20,50)
locount       <- 100
hicount       <- 1000
repcovlist    <- c(6,10,15)
replocount    <- 10
rephicount    <- 100
hheight       <- 0.035
nbrpnts       <- 5000

## The following four variables are sent to rank_rbm() and determine what data
#  points get plotted:
minnbrmsprm   <- 5      # minimum number of methylation sites per promoter
mingenewidth  <- 500    # minimum gene width
maxgenewidth  <- 5000   # maximum gene width
minnbrmsgene  <- 5      # minimum number of methylation sites per gene

## Other parameters (see Rscript.BWASPR for usage notes):
#
highcoverage  <- 20     # high read coverage threshold for studyhc methylRawList object
threshold     <- 20.0   # "difference" threshold for getMethylDiff(), called by det_dmsg()
qvalue        <- 0.05   # "qvalue" setting for getMethylDiff(), called by det_dmsg()
wsize         <- 500    # "win.size" parameter for tileMethylCounts() in det_dmt()
stepsize      <- 500    # "step.size" parameter for tileMethylCounts() in det_dmt()

minNsites     <-  1     # minimum number of hc sites in a gene to be heatmapped in show_dmsg()
maxNsites     <- 60     # maximum number of hc sites in a gene to be heatmapped in show_dmsg()
minPdmsites   <- 10     # minimum ratio of dm/hc (in %) in a gene to be heatmapped in show_dmsg()

maxgwidth     <- 20000  # maximal gene width for a gene to be considered by explore_dmsg()
minnbrdmsites <- 1      # minimum number of differentially methylated sites for a gene to be
                        #  considered in sample comparisons by explore_dmsg()
glink         <- "NCBIgene"   # URL to show in explore_dmsg(); options: "" or "NCBIgene"

#Set the number of processors to use:
#
numprc <- 6

#Determine what analyses to run:
#
RUNload    <- FALSE
```

```
RUNcms     <- TRUE
RUNpwc     <- TRUE
RUNcrl     <- TRUE

RUNrepcms  <- TRUE
RUNrepcrl  <- TRUE

RUNmmp     <- TRUE
RUNacs     <- TRUE
RUNrnk     <- TRUE
RUNmrpr    <- TRUE

RUNdmt     <- TRUE
RUNdmsg    <- TRUE
RUNdmgdtls <- TRUE
RUNogl     <- TRUE

RUNsave    <- TRUE

mymessage <- sprintf("\nAnalyzing %s study %s for type %s\n\n",species,studyLabel,type)
message(mymessage)

################################################################################
#End of typical customization.##################################################
```

Cb.par

```
SPECIESNAME Cerapachys biroi
TOTALNBRPMSITES   26684764
ASSEMBLYVERSION   CerBir1.0
GENOMESIZE  212825769
SPECIESGFF3DIR    ./MCALLS/Cbir/genome/GFF3DIR
GENELISTGFF3      Cbir.gene.gff3
EXONLISTGFF3      Cbir.exon.gff3
PCGEXNLISTGFF3    Cbir.pcg-exon.gff3
PROMOTRLISTGFF3   Cbir.promoter.gff3
CDSLISTGFF3 Cbir.pcg-CDS.gff3
UTRFLAGSET  1
5UTRLISTGFF3      Cbir.pcg-5pUTR.gff3
3UTRLISTGFF3      Cbir.pcg-3pUTR.gff3
```

Cb.dat

```
# Samples from Libbrecht et al. (2016) Current Biology:
#

### 1. Comparisons between all 8 data sets (no replicates) ###
#

Cb    LI    BCphaseA1   0     CpGhsm      ./MCALLS/Cbir/Libbrecht2016/BCphaseA/replicate1/BCphaseA1.CpGhsm.mcalls
Cb    LI    BCphaseA1   0     CpGscd      ./MCALLS/Cbir/Libbrecht2016/BCphaseA/replicate1/BCphaseA1.CpGscd.mcalls
Cb    LI    BCphaseA2   0     CpGhsm      ./MCALLS/Cbir/Libbrecht2016/BCphaseA/replicate2/BCphaseA2.CpGhsm.mcalls
Cb    LI    BCphaseA2   0     CpGscd      ./MCALLS/Cbir/Libbrecht2016/BCphaseA/replicate2/BCphaseA2.CpGscd.mcalls

Cb    LI    BCphaseB1   0     CpGhsm      ./MCALLS/Cbir/Libbrecht2016/BCphaseB/replicate1/BCphaseB1.CpGhsm.mcalls
Cb    LI    BCphaseB1   0     CpGscd      ./MCALLS/Cbir/Libbrecht2016/BCphaseB/replicate1/BCphaseB1.CpGscd.mcalls
Cb    LI    BCphaseB2   0     CpGhsm      ./MCALLS/Cbir/Libbrecht2016/BCphaseB/replicate2/BCphaseB2.CpGhsm.mcalls
Cb    LI    BCphaseB2   0     CpGscd      ./MCALLS/Cbir/Libbrecht2016/BCphaseB/replicate2/BCphaseB2.CpGscd.mcalls

Cb    LI    RphaseA1    0     CpGhsm      ./MCALLS/Cbir/Libbrecht2016/RphaseA/replicate1/RphaseA1.CpGhsm.mcalls
Cb    LI    RphaseA1    0     CpGscd      ./MCALLS/Cbir/Libbrecht2016/RphaseA/replicate1/RphaseA1.CpGscd.mcalls
Cb    LI    RphaseA2    0     CpGhsm      ./MCALLS/Cbir/Libbrecht2016/RphaseA/replicate2/RphaseA2.CpGhsm.mcalls
Cb    LI    RphaseA2    0     CpGscd      ./MCALLS/Cbir/Libbrecht2016/RphaseA/replicate2/RphaseA2.CpGscd.mcalls

Cb    LI    RphaseB1    0     CpGhsm      ./MCALLS/Cbir/Libbrecht2016/RphaseB/replicate1/RphaseB1.CpGhsm.mcalls
Cb    LI    RphaseB1    0     CpGscd      ./MCALLS/Cbir/Libbrecht2016/RphaseB/replicate1/RphaseB1.CpGscd.mcalls
Cb    LI    RphaseB2    0     CpGhsm      ./MCALLS/Cbir/Libbrecht2016/RphaseB/replicate2/RphaseB2.CpGhsm.mcalls
Cb    LI    RphaseB2    0     CpGscd      ./MCALLS/Cbir/Libbrecht2016/RphaseB/replicate2/RphaseB2.CpGscd.mcalls

### 2. Pairwise omparisons between data sets split accorinding to phase, or line, or batch (2 samples, each with 4 replicates) ###
### 2.1 Comparing BCphase versus Rphase ###
#
# BCphase
Cb    LI    BCphase     0     CpGhsm      ./MCALLS/Cbir/Libbrecht2016/PhaseBCvsR/PhaseBC/PhaseBC.CpGhsm.mcalls
Cb    LI    BCphase     0     CpGscd      ./MCALLS/Cbir/Libbrecht2016/PhaseBCvsR/PhaseBC/PhaseBC.CpGscd.mcalls

# ...with replicates
Cb    LI    BCphase     1     CpGhsm      ./MCALLS/Cbir/Libbrecht2016/BCphaseA/replicate1/BCphaseA1.CpGhsm.mcalls
Cb    LI    BCphase     1     CpGscd      ./MCALLS/Cbir/Libbrecht2016/BCphaseA/replicate1/BCphaseA1.CpGscd.mcalls
Cb    LI    BCphase     2     CpGhsm      ./MCALLS/Cbir/Libbrecht2016/BCphaseA/replicate2/BCphaseA2.CpGhsm.mcalls
Cb    LI    BCphase     2     CpGscd      ./MCALLS/Cbir/Libbrecht2016/BCphaseA/replicate2/BCphaseA2.CpGscd.mcalls
Cb    LI    BCphase     3     CpGhsm      ./MCALLS/Cbir/Libbrecht2016/BCphaseB/replicate1/BCphaseB1.CpGhsm.mcalls
Cb    LI    BCphase     3     CpGscd      ./MCALLS/Cbir/Libbrecht2016/BCphaseB/replicate1/BCphaseB1.CpGscd.mcalls
Cb    LI    BCphase     4     CpGhsm      ./MCALLS/Cbir/Libbrecht2016/BCphaseB/replicate2/BCphaseB2.CpGhsm.mcalls
Cb    LI    BCphase     4     CpGscd      ./MCALLS/Cbir/Libbrecht2016/BCphaseB/replicate2/BCphaseB2.CpGscd.mcalls

# Rphase
Cb    LI    Rphase      0     CpGhsm      ./MCALLS/Cbir/Libbrecht2016/PhaseBCvsR/PhaseR/PhaseR.CpGhsm.mcalls
Cb    LI    Rphase      0     CpGscd      ./MCALLS/Cbir/Libbrecht2016/PhaseBCvsR/PhaseR/PhaseR.CpGscd.mcalls

# ...with replicates
Cb    LI    Rphase      1     CpGhsm      ./MCALLS/Cbir/Libbrecht2016/RphaseA/replicate1/RphaseA1.CpGhsm.mcalls
Cb    LI    Rphase      1     CpGscd      ./MCALLS/Cbir/Libbrecht2016/RphaseA/replicate1/RphaseA1.CpGscd.mcalls
Cb    LI    Rphase      2     CpGhsm      ./MCALLS/Cbir/Libbrecht2016/RphaseA/replicate2/RphaseA2.CpGhsm.mcalls
Cb    LI    Rphase      2     CpGscd      ./MCALLS/Cbir/Libbrecht2016/RphaseA/replicate2/RphaseA2.CpGscd.mcalls
Cb    LI    Rphase      3     CpGhsm      ./MCALLS/Cbir/Libbrecht2016/RphaseB/replicate1/RphaseB1.CpGhsm.mcalls
Cb    LI    Rphase      3     CpGscd      ./MCALLS/Cbir/Libbrecht2016/RphaseB/replicate1/RphaseB1.CpGscd.mcalls
Cb    LI    Rphase      4     CpGhsm      ./MCALLS/Cbir/Libbrecht2016/RphaseB/replicate2/RphaseB2.CpGhsm.mcalls
Cb    LI    Rphase      4     CpGscd      ./MCALLS/Cbir/Libbrecht2016/RphaseB/replicate2/RphaseB2.CpGscd.mcalls

### 2.2 Comparing LineA versus LineB ###
```

```
#
# LineA
Cb    LI    LineA 0     CpGhsm      ./MCALLS/Cbir/Libbrecht2016/LineAvsB/LineA/LineA.CpGhsm.mcalls
Cb    LI    LineA 0     CpGscd      ./MCALLS/Cbir/Libbrecht2016/LineAvsB/LineA/LineA.CpGscd.mcalls

# ...with replicates
Cb    LI    LineA 1     CpGhsm      ./MCALLS/Cbir/Libbrecht2016/BCphaseA/replicate1/BCphaseA1.CpGhsm.mcalls
Cb    LI    LineA 1     CpGscd      ./MCALLS/Cbir/Libbrecht2016/BCphaseA/replicate1/BCphaseA1.CpGscd.mcalls
Cb    LI    LineA 2     CpGhsm      ./MCALLS/Cbir/Libbrecht2016/BCphaseA/replicate2/BCphaseA2.CpGhsm.mcalls
Cb    LI    LineA 2     CpGscd      ./MCALLS/Cbir/Libbrecht2016/BCphaseA/replicate2/BCphaseA2.CpGscd.mcalls
Cb    LI    LineA 3     CpGhsm      ./MCALLS/Cbir/Libbrecht2016/RphaseA/replicate1/RphaseA1.CpGhsm.mcalls
Cb    LI    LineA 3     CpGscd      ./MCALLS/Cbir/Libbrecht2016/RphaseA/replicate1/RphaseA1.CpGscd.mcalls
Cb    LI    LineA 4     CpGhsm      ./MCALLS/Cbir/Libbrecht2016/RphaseA/replicate2/RphaseA2.CpGhsm.mcalls
Cb    LI    LineA 4     CpGscd      ./MCALLS/Cbir/Libbrecht2016/RphaseA/replicate2/RphaseA2.CpGscd.mcalls

# LineB
Cb    LI    LineB 0     CpGhsm      ./MCALLS/Cbir/Libbrecht2016/LineAvsB/LineB/LineB.CpGhsm.mcalls
Cb    LI    LineB 0     CpGscd      ./MCALLS/Cbir/Libbrecht2016/LineAvsB/LineB/LineB.CpGscd.mcalls

# ...with replicates
Cb    LI    LineB 1     CpGhsm      ./MCALLS/Cbir/Libbrecht2016/BCphaseB/replicate1/BCphaseB1.CpGhsm.mcalls
Cb    LI    LineB 1     CpGscd      ./MCALLS/Cbir/Libbrecht2016/BCphaseB/replicate1/BCphaseB1.CpGscd.mcalls
Cb    LI    LineB 2     CpGhsm      ./MCALLS/Cbir/Libbrecht2016/BCphaseB/replicate2/BCphaseB2.CpGhsm.mcalls
Cb    LI    LineB 2     CpGscd      ./MCALLS/Cbir/Libbrecht2016/BCphaseB/replicate2/BCphaseB2.CpGscd.mcalls
Cb    LI    LineB 3     CpGhsm      ./MCALLS/Cbir/Libbrecht2016/RphaseB/replicate1/RphaseB1.CpGhsm.mcalls
Cb    LI    LineB 3     CpGscd      ./MCALLS/Cbir/Libbrecht2016/RphaseB/replicate1/RphaseB1.CpGscd.mcalls
Cb    LI    LineB 4     CpGhsm      ./MCALLS/Cbir/Libbrecht2016/RphaseB/replicate2/RphaseB2.CpGhsm.mcalls
Cb    LI    LineB 4     CpGscd      ./MCALLS/Cbir/Libbrecht2016/RphaseB/replicate2/RphaseB2.CpGscd.mcalls

### 2.2 2.2 Comparing Batch1 versus Batch2 ###
#
# Batch1
Cb    LI    Batch1      0     CpGhsm      ./MCALLS/Cbir/Libbrecht2016/Batch1vs2/Batch1/Batch1.CpGhsm.mcalls
Cb    LI    Batch1      0     CpGscd      ./MCALLS/Cbir/Libbrecht2016/Batch1vs2/Batch1/Batch1.CpGscd.mcalls

# ...with replicates
Cb    LI    Batch1      1     CpGhsm      ./MCALLS/Cbir/Libbrecht2016/BCphaseA/replicate1/BCphaseA1.CpGhsm.mcalls
Cb    LI    Batch1      1     CpGscd      ./MCALLS/Cbir/Libbrecht2016/BCphaseA/replicate1/BCphaseA1.CpGscd.mcalls
Cb    LI    Batch1      2     CpGhsm      ./MCALLS/Cbir/Libbrecht2016/BCphaseB/replicate1/BCphaseB1.CpGhsm.mcalls
Cb    LI    Batch1      2     CpGscd      ./MCALLS/Cbir/Libbrecht2016/BCphaseB/replicate1/BCphaseB1.CpGscd.mcalls
Cb    LI    Batch1      3     CpGhsm      ./MCALLS/Cbir/Libbrecht2016/RphaseA/replicate1/RphaseA1.CpGhsm.mcalls
Cb    LI    Batch1      3     CpGscd      ./MCALLS/Cbir/Libbrecht2016/RphaseA/replicate1/RphaseA1.CpGscd.mcalls
Cb    LI    Batch1      4     CpGhsm      ./MCALLS/Cbir/Libbrecht2016/RphaseB/replicate1/RphaseB1.CpGhsm.mcalls
Cb    LI    Batch1      4     CpGscd      ./MCALLS/Cbir/Libbrecht2016/RphaseB/replicate1/RphaseB1.CpGscd.mcalls

# Batch2
Cb    LI    Batch2      0     CpGhsm      ./MCALLS/Cbir/Libbrecht2016/Batch1vs2/Batch2/Batch2.CpGhsm.mcalls
Cb    LI    Batch2      0     CpGscd      ./MCALLS/Cbir/Libbrecht2016/Batch1vs2/Batch2/Batch2.CpGscd.mcalls

# ...with replicates
Cb    LI    Batch2      1     CpGhsm      ./MCALLS/Cbir/Libbrecht2016/BCphaseA/replicate2/BCphaseA2.CpGhsm.mcalls
Cb    LI    Batch2      1     CpGscd      ./MCALLS/Cbir/Libbrecht2016/BCphaseA/replicate2/BCphaseA2.CpGscd.mcalls
Cb    LI    Batch2      2     CpGhsm      ./MCALLS/Cbir/Libbrecht2016/BCphaseB/replicate2/BCphaseB2.CpGhsm.mcalls
Cb    LI    Batch2      2     CpGscd      ./MCALLS/Cbir/Libbrecht2016/BCphaseB/replicate2/BCphaseB2.CpGscd.mcalls
Cb    LI    Batch2      3     CpGhsm      ./MCALLS/Cbir/Libbrecht2016/RphaseA/replicate2/RphaseA2.CpGhsm.mcalls
Cb    LI    Batch2      3     CpGscd      ./MCALLS/Cbir/Libbrecht2016/RphaseA/replicate2/RphaseA2.CpGscd.mcalls
Cb    LI    Batch2      4     CpGhsm      ./MCALLS/Cbir/Libbrecht2016/RphaseB/replicate2/RphaseB2.CpGhsm.mcalls
Cb    LI    Batch2      4     CpGscd      ./MCALLS/Cbir/Libbrecht2016/RphaseB/replicate2/RphaseB2.CpGscd.mcalls

### 3. Pairwise comparisons between BCphaseA, BCphaseB, RphaseA, and RphaseB (2 replicates each, corresponding to batches 1 and 2) ###
#
```

```
# BCphaseA
Cb    LI    BCphaseA    0     CpGhsm      ./MCALLS/Cbir/Libbrecht2016/BCphaseA/BCphaseA.CpGhsm.mcalls
Cb    LI    BCphaseA    0     CpGscd      ./MCALLS/Cbir/Libbrecht2016/BCphaseA/BCphaseA.CpGscd.mcalls

# ... with replicates:
Cb    LI    BCphaseA    1     CpGhsm      ./MCALLS/Cbir/Libbrecht2016/BCphaseA/replicate1/BCphaseA1.CpGhsm.mcalls
Cb    LI    BCphaseA    1     CpGscd      ./MCALLS/Cbir/Libbrecht2016/BCphaseA/replicate1/BCphaseA1.CpGscd.mcalls
Cb    LI    BCphaseA    2     CpGhsm      ./MCALLS/Cbir/Libbrecht2016/BCphaseA/replicate2/BCphaseA2.CpGhsm.mcalls
Cb    LI    BCphaseA    2     CpGscd      ./MCALLS/Cbir/Libbrecht2016/BCphaseA/replicate2/BCphaseA2.CpGscd.mcalls

# BCphaseB
Cb    LI    BCphaseB    0     CpGhsm      ./MCALLS/Cbir/Libbrecht2016/BCphaseB/BCphaseB.CpGhsm.mcalls
Cb    LI    BCphaseB    0     CpGscd      ./MCALLS/Cbir/Libbrecht2016/BCphaseB/BCphaseB.CpGscd.mcalls

# ... with replicates:
Cb    LI    BCphaseB    1     CpGhsm      ./MCALLS/Cbir/Libbrecht2016/BCphaseB/replicate1/BCphaseB1.CpGhsm.mcalls
Cb    LI    BCphaseB    1     CpGscd      ./MCALLS/Cbir/Libbrecht2016/BCphaseB/replicate1/BCphaseB1.CpGscd.mcalls
Cb    LI    BCphaseB    2     CpGhsm      ./MCALLS/Cbir/Libbrecht2016/BCphaseB/replicate2/BCphaseB2.CpGhsm.mcalls
Cb    LI    BCphaseB    2     CpGscd      ./MCALLS/Cbir/Libbrecht2016/BCphaseB/replicate2/BCphaseB2.CpGscd.mcalls

# RphaseA
Cb    LI    RphaseA     0     CpGhsm      ./MCALLS/Cbir/Libbrecht2016/RphaseA/RphaseA.CpGhsm.mcalls
Cb    LI    RphaseA     0     CpGscd      ./MCALLS/Cbir/Libbrecht2016/RphaseA/RphaseA.CpGscd.mcalls

# ... with replicates:
Cb    LI    RphaseA     1     CpGhsm      ./MCALLS/Cbir/Libbrecht2016/RphaseA/replicate1/RphaseA1.CpGhsm.mcalls
Cb    LI    RphaseA     1     CpGscd      ./MCALLS/Cbir/Libbrecht2016/RphaseA/replicate1/RphaseA1.CpGscd.mcalls
Cb    LI    RphaseA     2     CpGhsm      ./MCALLS/Cbir/Libbrecht2016/RphaseA/replicate2/RphaseA2.CpGhsm.mcalls
Cb    LI    RphaseA     2     CpGscd      ./MCALLS/Cbir/Libbrecht2016/RphaseA/replicate2/RphaseA2.CpGscd.mcalls

# RphaseB
Cb    LI    RphaseB     0     CpGhsm      ./MCALLS/Cbir/Libbrecht2016/RphaseB/RphaseB.CpGhsm.mcalls
Cb    LI    RphaseB     0     CpGscd      ./MCALLS/Cbir/Libbrecht2016/RphaseB/RphaseB.CpGscd.mcalls

# ... with replicates:
Cb    LI    RphaseB     1     CpGhsm      ./MCALLS/Cbir/Libbrecht2016/RphaseB/replicate1/RphaseB1.CpGhsm.mcalls
Cb    LI    RphaseB     1     CpGscd      ./MCALLS/Cbir/Libbrecht2016/RphaseB/replicate1/RphaseB1.CpGscd.mcalls
Cb    LI    RphaseB     2     CpGhsm      ./MCALLS/Cbir/Libbrecht2016/RphaseB/replicate2/RphaseB2.CpGhsm.mcalls
Cb    LI    RphaseB     2     CpGscd      ./MCALLS/Cbir/Libbrecht2016/RphaseB/replicate2/RphaseB2.CpGscd.mcalls
```

```
0README

Rscript.BWASPR output for .conf file Cb_LIp.conf.

Data input are the specified *.mcalls files and parameters set in Cb_LIp.conf.
The *.mcalls input data are saved in the following data structures:

studymk - a methylKit methylRaw(List) object storing the CpGhsm site data for all samples
studymc - a methylKit methylRaw(List) object storing the CpGscd site data for all samples
studyhc - a methylKit methylRaw(List) object storing the CpGscd site data for all samples,
           restricted to sites with high coverage (here set to 20)
mkrd    - a methylKit methylRawList object storing the CpGhsm site data for all
           replicates of a given sample

Output of the analysis is stored in the following subdirectories (if the corresponding
 function calls were specified in Cb_LIp.conf; please consult the 0README
 files in the subdirectories for details on the output files):

CMS - Coverage and methylation statistics for aggregate samples
PWC - Pairwise comparisons between all samples
CRL - Correlations between aggregate samples
REPCMS - Coverage and methylation statistics for replicate samples
REPCRL - Correlations between replicates
MMP - Mapping of methylation sites on genome annotation
ACS - Annotation of conserved methylation sites
RNK - Genomic feature regions ranked by CpGhsm statistics
MRPR - Methylation-rich and -poor regions
DMSG - Differentially methylateds sites and genes
OGL - Ordered gene lists
```

Directory: CMS File: 0READMEcms

CMS - Coverage and methylation statistics for aggregate samples.

Input: studymk, covlist, locount, hicount

Output: files cms-*.txt and cms-*.pdf

Notes: The statistics are given for CpGhsm sites restricted to minimum or higher level coverage. At the minimum level, the number of sites will be the number of lines of the corresponding *CpGhsm.mcalls file minus one (consistency check). The distribution of methylation levels is really only interesting for high coverage because at low coverage there will be a strong bias towards high methylation levels (by definition of hsm sites). The same bias at high coverage would suggest that the sample includes a preponderance of consistently methylated genomic sites.

locount and hicount set bounds on the coverage to exclude sites with too few or too many covering reads to provide statistics on a typical range.

```
Directory: CMS File: cms-Cb_LIp_BCphase.txt

Number of "BCphase" CpGhsm-sites with minimal and higher level coverage:

  number of "BCphase" CpGhsm-sites with coverage >=  5: 417310
  number of "BCphase" CpGhsm-sites with coverage >=  6: 416306
  number of "BCphase" CpGhsm-sites with coverage >= 20: 384326
  number of "BCphase" CpGhsm-sites with coverage >= 50: 294489

Coverage and methylation statistics for "BCphase" CpGhsm-sites at different levels of minimum coverage:

methylKit::getCoverageStats output for "BCphase" CpGhsm-sites (#: 417310) at minimum coverage 5     - read coverage statistics per base
summary:
   Min. 1st Qu.  Median    Mean 3rd Qu.    Max.
      5      44      80      95     132    6323
percentiles:
   0%   10%   20%   30%   40%   50%   60%   70%   80%   90%   95%   99% 99.5% 99.9%  100%
    5    23    37    50    64    80    98   120   147   186   220   288   317   407  6323

methylKit::getMethylationStats output for "BCphase" CpGhsm-sites (#: 417310) at minimum coverage 5  - methylation statistics per base
summary:
   Min. 1st Qu.  Median    Mean 3rd Qu.    Max.
    1.3    20.0    41.2    48.1    77.1   100.0
percentiles:
    0%    10%    20%    30%    40%    50%    60%    70%    80%    90%    95%    99%  99.5%  99.9%   100%
  1.34  12.64  17.37  23.08  30.77  41.18  54.55  69.77  83.87  94.08  97.37 100.00 100.00 100.00 100.00

methylKit::getCoverageStats output for "BCphase" CpGhsm-sites (#: 416306) at minimum coverage 6     - read coverage statistics per base
summary:
   Min. 1st Qu.  Median    Mean 3rd Qu.    Max.
      6      44      80      95     133    6323
percentiles:
   0%   10%   20%   30%   40%   50%   60%   70%   80%   90%   95%   99% 99.5% 99.9%  100%
    6    23    37    50    65    80    99   120   147   186   220   288   318   407  6323

methylKit::getMethylationStats output for "BCphase" CpGhsm-sites (#: 416306) at minimum coverage 6  - methylation statistics per base
summary:
   Min. 1st Qu.  Median    Mean 3rd Qu.    Max.
    1.3    20.0    41.0    48.0    76.9   100.0
percentiles:
    0%    10%    20%    30%    40%    50%    60%    70%    80%    90%    95%    99%  99.5%  99.9%   100%
  1.34  12.63  17.33  23.02  30.77  41.03  54.38  69.49  83.54  93.88  97.22 100.00 100.00 100.00 100.00

methylKit::getCoverageStats output for "BCphase" CpGhsm-sites (#: 384326) at minimum coverage 20    - read coverage statistics per base
summary:
   Min. 1st Qu.  Median    Mean 3rd Qu.    Max.
     20      51      87     102     138    6323
percentiles:
   0%   10%   20%   30%   40%   50%   60%   70%   80%   90%   95%   99% 99.5% 99.9%  100%
   20    33    45    58    72    87   105   126   152   190   223   291   321   415  6323

methylKit::getMethylationStats output for "BCphase" CpGhsm-sites (#: 384326) at minimum coverage 20 - methylation statistics per base
summary:
   Min. 1st Qu.  Median    Mean 3rd Qu.    Max.
    1.3    19.0    37.0    45.8    73.2   100.0
percentiles:
    0%    10%    20%    30%    40%    50%    60%    70%    80%    90%    95%    99%  99.5%  99.9%   100%
```

```
  1.34  12.26  16.67  21.62  28.07  37.04  49.54  65.12  81.03  92.98  96.46 100.00 100.00 100.00 100.00

methylKit::getCoverageStats output for "BCphase" CpGhsm-sites (#: 294489) at minimum coverage 50    - read coverage statistics per base
summary:
   Min. 1st Qu.  Median    Mean 3rd Qu.    Max. 
     50      75     108     123     154    6323 
percentiles:
   0%   10%   20%   30%   40%   50%   60%   70%   80%   90%   95%   99% 99.5% 99.9%  100% 
   50    59    70    81    94   108   124   143   167   203   235   302   334   450  6323 

methylKit::getMethylationStats output for "BCphase" CpGhsm-sites (#: 294489) at minimum coverage 50  - methylation statistics per base
summary:
   Min. 1st Qu.  Median    Mean 3rd Qu.    Max. 
    1.3    16.2    32.7    43.1    70.0   100.0 
percentiles:
    0%    10%    20%    30%    40%    50%    60%    70%    80%    90%    95%    99%  99.5%  99.9%   100% 
  1.34  11.25  14.58  18.18  23.95  32.69  45.12  61.26  78.57  92.16  96.07  98.65  99.27 100.00 100.00 

methylKit::getCoverageStats output for "BCphase" CpGhsm-sites (#: 164371) in coverage range [100-1000]    - read coverage statistics per base
summary:
   Min. 1st Qu.  Median    Mean 3rd Qu.    Max. 
    100     121     147     160     187     997 
percentiles:
   0%   10%   20%   30%   40%   50%   60%   70%   80%   90%   95%   99% 99.5% 99.9%  100% 
  100   108   116   126   136   147   161   177   198   230   259   325   357   453   997 

methylKit::getMethylationStats output for "BCphase" CpGhsm-sites (#: 164371) in coverage range [100-1000] - methylation statistics per base
summary:
   Min. 1st Qu.  Median    Mean 3rd Qu.    Max. 
    2.6    14.4    32.6    42.5    70.4   100.0 
percentiles:
    0%    10%    20%    30%    40%    50%    60%    70%    80%    90%    95%    99%  99.5%  99.9%   100% 
  2.58   9.62  12.44  16.74  23.21  32.60  45.45  61.72  78.72  92.11  95.98  98.38  99.01  99.59 100.00 
```

```
Directory: CMS File: cms-Cb_LIp_Rphase.txt

Number of "Rphase" CpGhsm-sites with minimal and higher level coverage:

  number of "Rphase" CpGhsm-sites with coverage >=  5: 445317
  number of "Rphase" CpGhsm-sites with coverage >=  6: 444427
  number of "Rphase" CpGhsm-sites with coverage >= 20: 414586
  number of "Rphase" CpGhsm-sites with coverage >= 50: 328505

Coverage and methylation statistics for "Rphase" CpGhsm-sites at different levels of minimum coverage:

methylKit::getCoverageStats output for "Rphase" CpGhsm-sites (#: 445317) at minimum coverage 5 - read coverage statistics per base
summary:
   Min. 1st Qu.  Median    Mean 3rd Qu.    Max.
      5      48      87     104     144    6895
percentiles:
   0%   10%   20%   30%   40%   50%   60%   70%   80%   90%   95%   99% 99.5% 99.9%  100%
    5    25    40    55    70    87   107   130   160   203   241   317   350   447  6895

methylKit::getMethylationStats output for "Rphase" CpGhsm-sites (#: 445317) at minimum coverage 5    - methylation statistics per base
summary:
   Min. 1st Qu.  Median    Mean 3rd Qu.    Max.
    1.1    18.4    38.2    46.4    75.0   100.0
percentiles:
    0%    10%    20%    30%    40%    50%    60%    70%    80%    90%    95%    99%  99.5%  99.9%   100%
  1.13  11.76  16.00  21.21  28.18  38.17  51.54  67.07  82.71  93.62  97.14 100.00 100.00 100.00 100.00

methylKit::getCoverageStats output for "Rphase" CpGhsm-sites (#: 444427) at minimum coverage 6 - read coverage statistics per base
summary:
   Min. 1st Qu.  Median    Mean 3rd Qu.    Max.
      6      48      87     104     144    6895
percentiles:
   0%   10%   20%   30%   40%   50%   60%   70%   80%   90%   95%   99% 99.5% 99.9%  100%
    6    25    41    55    71    87   107   130   160   203   241   317   350   447  6895

methylKit::getMethylationStats output for "Rphase" CpGhsm-sites (#: 444427) at minimum coverage 6    - methylation statistics per base
summary:
   Min. 1st Qu.  Median    Mean 3rd Qu.    Max.
    1.1    18.4    38.1    46.3    75.0   100.0
percentiles:
    0%    10%    20%    30%    40%    50%    60%    70%    80%    90%    95%    99%  99.5%  99.9%   100%
  1.13  11.76  16.00  21.17  28.12  38.10  51.35  66.67  82.46  93.48  97.00 100.00 100.00 100.00 100.00

methylKit::getCoverageStats output for "Rphase" CpGhsm-sites (#: 414586) at minimum coverage 20      - read coverage statistics per base
summary:
   Min. 1st Qu.  Median    Mean 3rd Qu.    Max.
     20      55      93     110     149    6895
percentiles:
   0%   10%   20%   30%   40%   50%   60%   70%   80%   90%   95%   99% 99.5% 99.9%  100%
   20    34    48    62    77    93   113   136   165   207   244   320   354   454  6895

methylKit::getMethylationStats output for "Rphase" CpGhsm-sites (#: 414586) at minimum coverage 20   - methylation statistics per base
summary:
   Min. 1st Qu.  Median    Mean 3rd Qu.    Max.
    1.1    17.5    34.6    44.2    71.4   100.0
percentiles:
    0%    10%    20%    30%    40%    50%    60%    70%    80%    90%    95%    99%  99.5%  99.9%   100%
```

```
  1.13  11.50  15.38  20.00  25.93  34.57  46.85  62.71  79.69  92.59  96.35 100.00 100.00 100.00 100.00

methylKit::getCoverageStats output for "Rphase" CpGhsm-sites (#: 328505) at minimum coverage 50     - read coverage statistics per base
summary:
   Min. 1st Qu.  Median    Mean 3rd Qu.    Max.
     50      78     113     130     165    6895
percentiles:
   0%   10%   20%   30%   40%   50%   60%   70%   80%   90%   95%   99% 99.5% 99.9%  100%
   50    60    72    84    98   113   131   153   180   220   256   331   365   480  6895

methylKit::getMethylationStats output for "Rphase" CpGhsm-sites (#: 328505) at minimum coverage 50  - methylation statistics per base
summary:
   Min. 1st Qu.  Median    Mean 3rd Qu.    Max.
    1.1    15.4    30.2    41.7    68.0   100.0
percentiles:
    0%    10%    20%    30%    40%    50%    60%    70%    80%    90%    95%    99%  99.5%  99.9%   100%
  1.13  10.69  13.79  17.12  22.11  30.19  42.42  58.74  76.98  91.71  95.97  98.58  99.15 100.00 100.00

methylKit::getCoverageStats output for "Rphase" CpGhsm-sites (#: 193649) in coverage range [100-1000]     - read coverage statistics per base
summary:
   Min. 1st Qu.  Median    Mean 3rd Qu.    Max.
    100     124     154     168     198     997
percentiles:
   0%   10%   20%   30%   40%   50%   60%   70%   80%   90%   95%   99% 99.5% 99.9%  100%
  100   108   118   129   141   154   169   187   211   247   280   354   386   480   997

methylKit::getMethylationStats output for "Rphase" CpGhsm-sites (#: 193649) in coverage range [100-1000]  - methylation statistics per base
summary:
   Min. 1st Qu.  Median    Mean 3rd Qu.    Max.
    2.5    13.5    30.6    41.3    68.7   100.0
percentiles:
    0%    10%    20%    30%    40%    50%    60%    70%    80%    90%    95%    99%  99.5%  99.9%   100%
  2.51   9.25  11.73  15.57  21.57  30.56  43.30  59.67  77.44  91.80  95.91  98.33  98.94  99.56 100.00
```

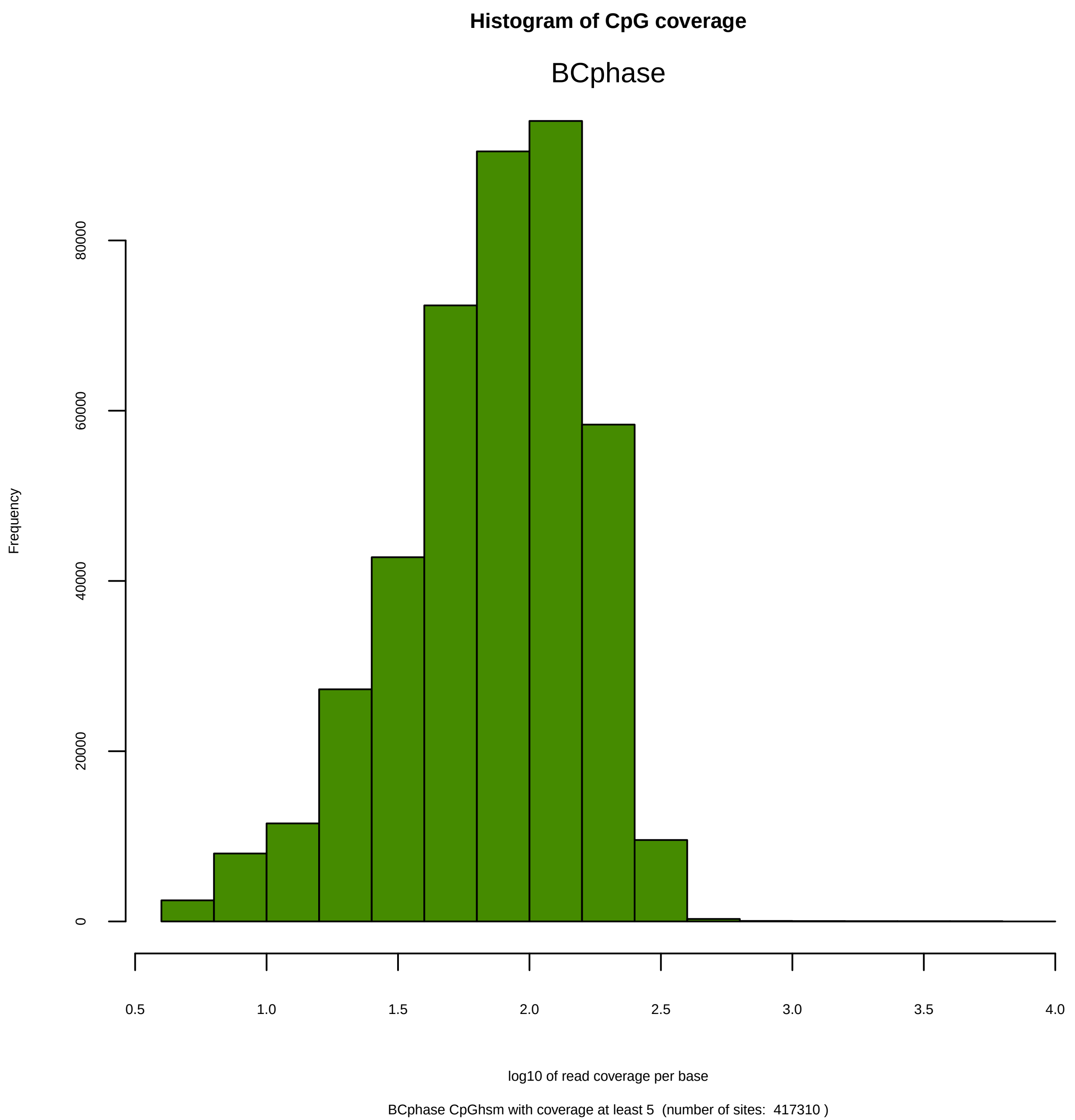
Histogram of CpG coverage
BCphase
Frequency
0
20000
40000
60000
80000
0.5
1.0
1.5
2.0
2.5
3.0
3.5
4.0
log10 of read coverage per base
BCphase CpGhsm with coverage at least 5  (number of sites:  417310 )

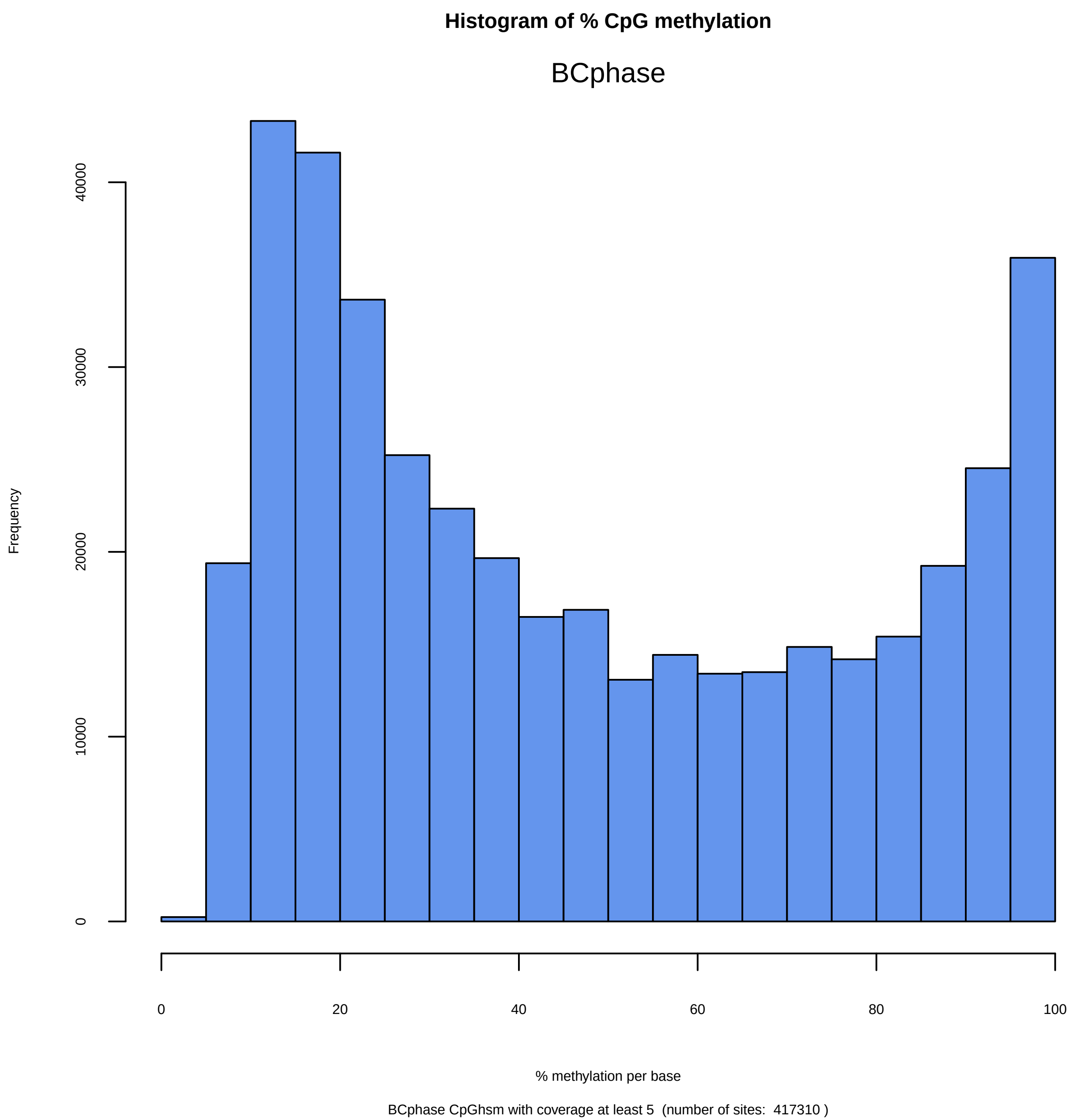
Histogram of % CpG methylation
BCphase
Frequency
0
10000
20000
30000
40000
0
20
40
60
80
100
% methylation per base
BCphase CpGhsm with coverage at least 5  (number of sites:  417310 )

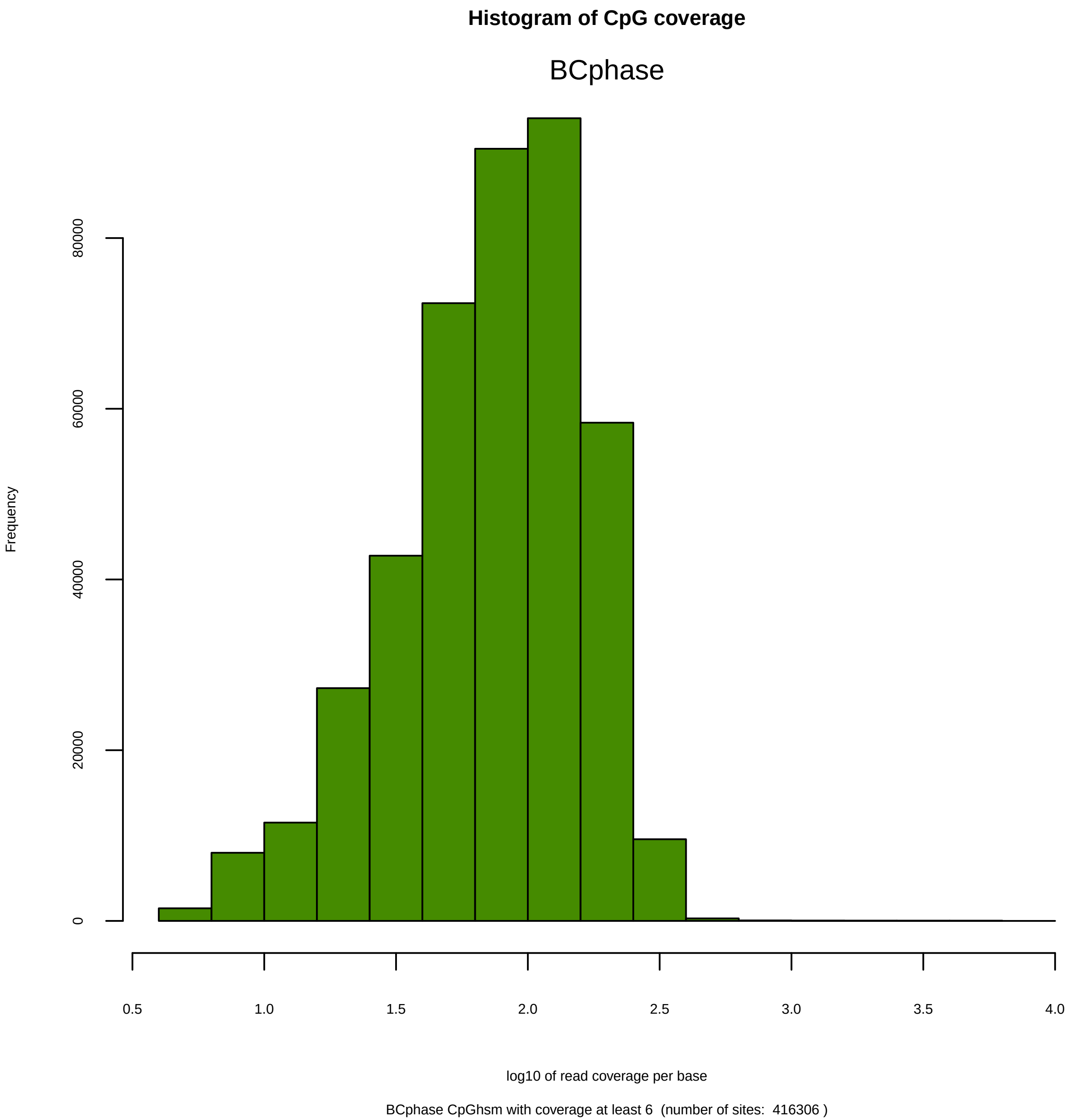
Histogram of CpG coverage
BCphase
Frequency
0
20000
40000
60000
80000
0.5
1.0
1.5
2.0
2.5
3.0
3.5
4.0
log10 of read coverage per base
BCphase CpGhsm with coverage at least 6  (number of sites:  416306 )

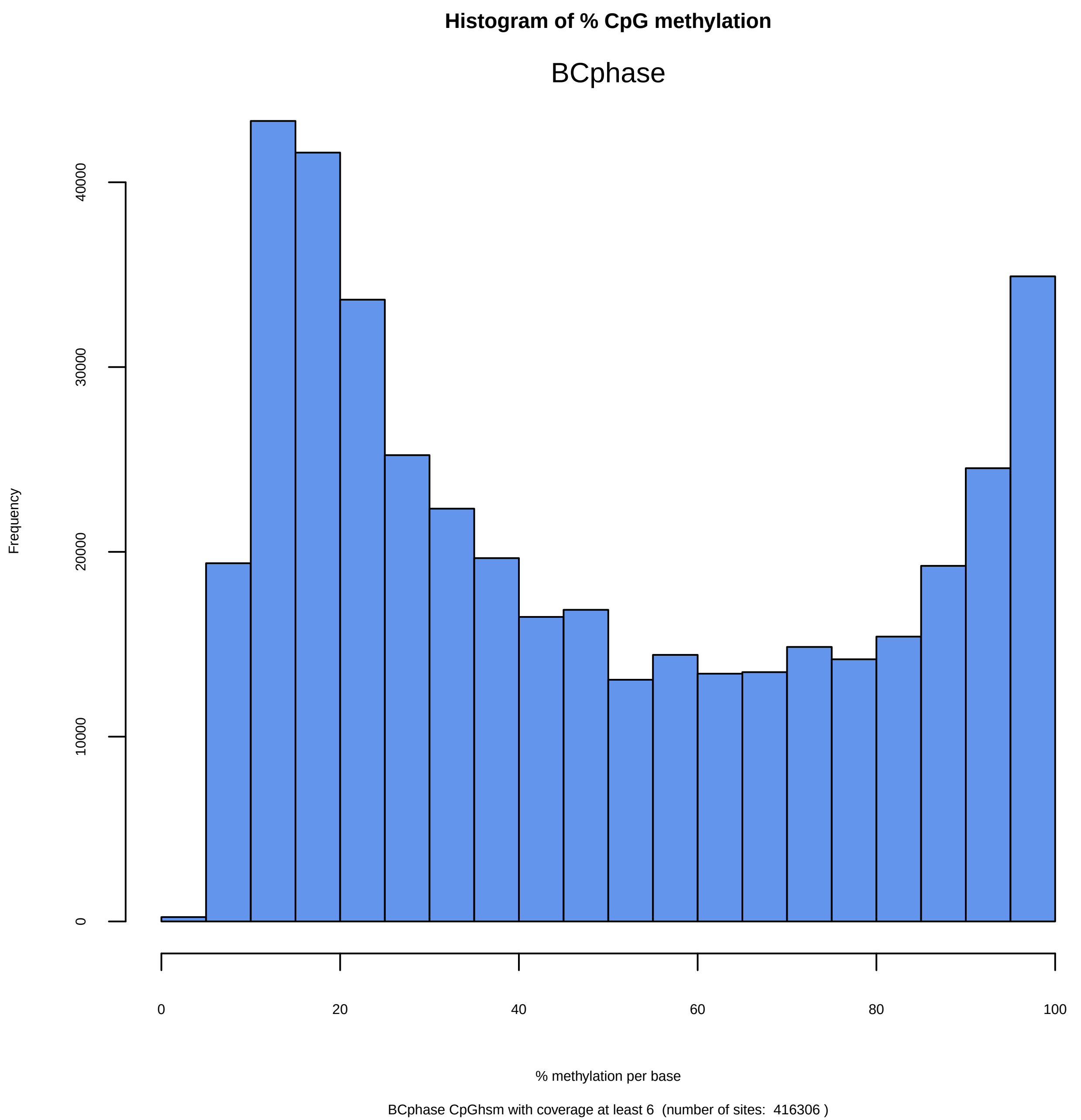

Histogram of % CpG methylation
BCphase
Frequency
0
10000
20000
30000
40000
0
20
40
60
80
100
% methylation per base
BCphase CpGhsm with coverage at least 6 (number of sites: 416306 )

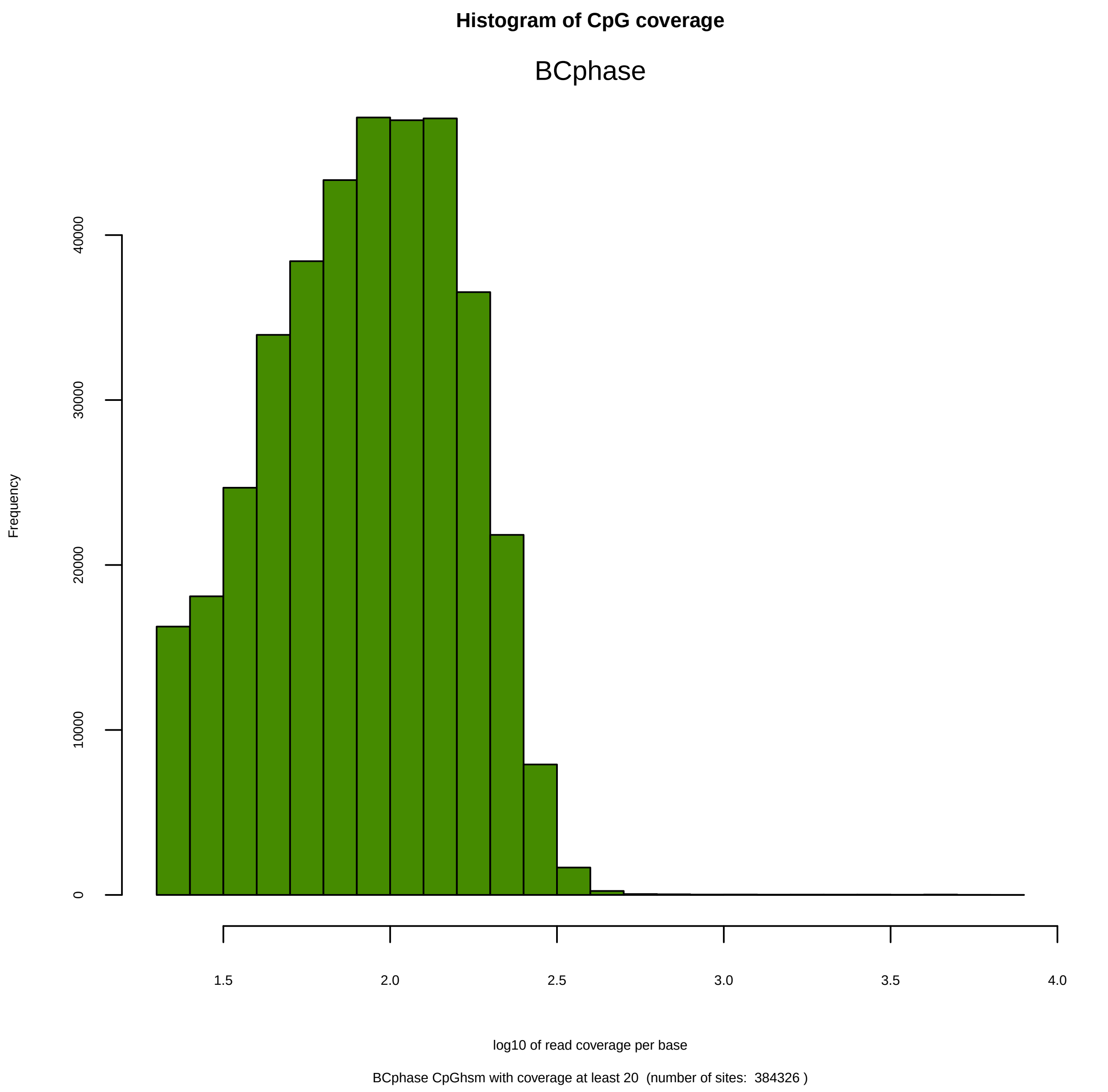

Histogram of CpG coverage
BCphase
Frequency
0
10000
20000
30000
40000
1.5
2.0
2.5
3.0
3.5
4.0
log10 of read coverage per base
BCphase CpGhsm with coverage at least 20  (number of sites:  384326 )

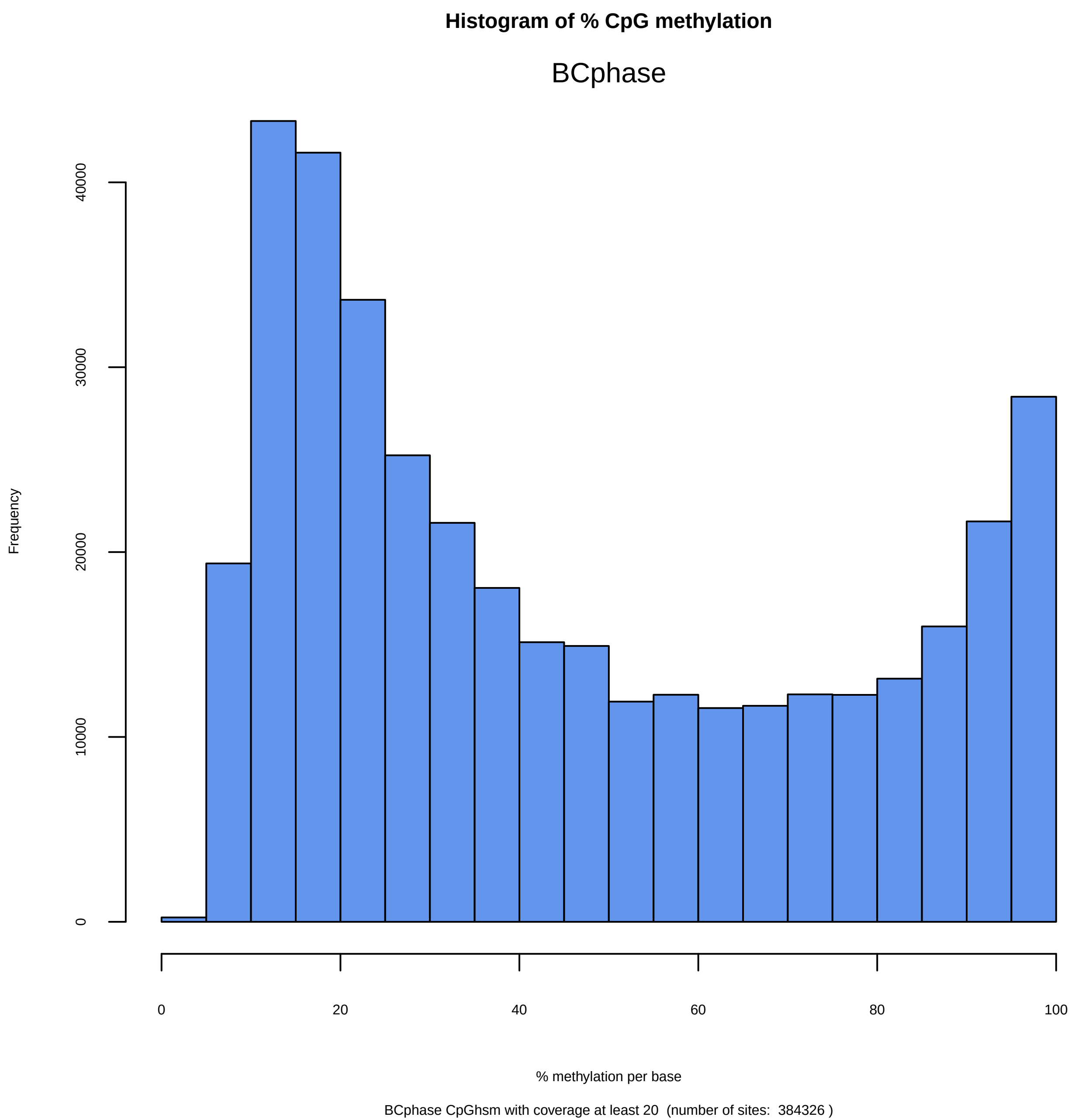
Histogram of % CpG methylation
BCphase
Frequency
0
10000
20000
30000
40000
0
20
40
60
80
100
% methylation per base
BCphase CpGhsm with coverage at least 20  (number of sites:  384326 )

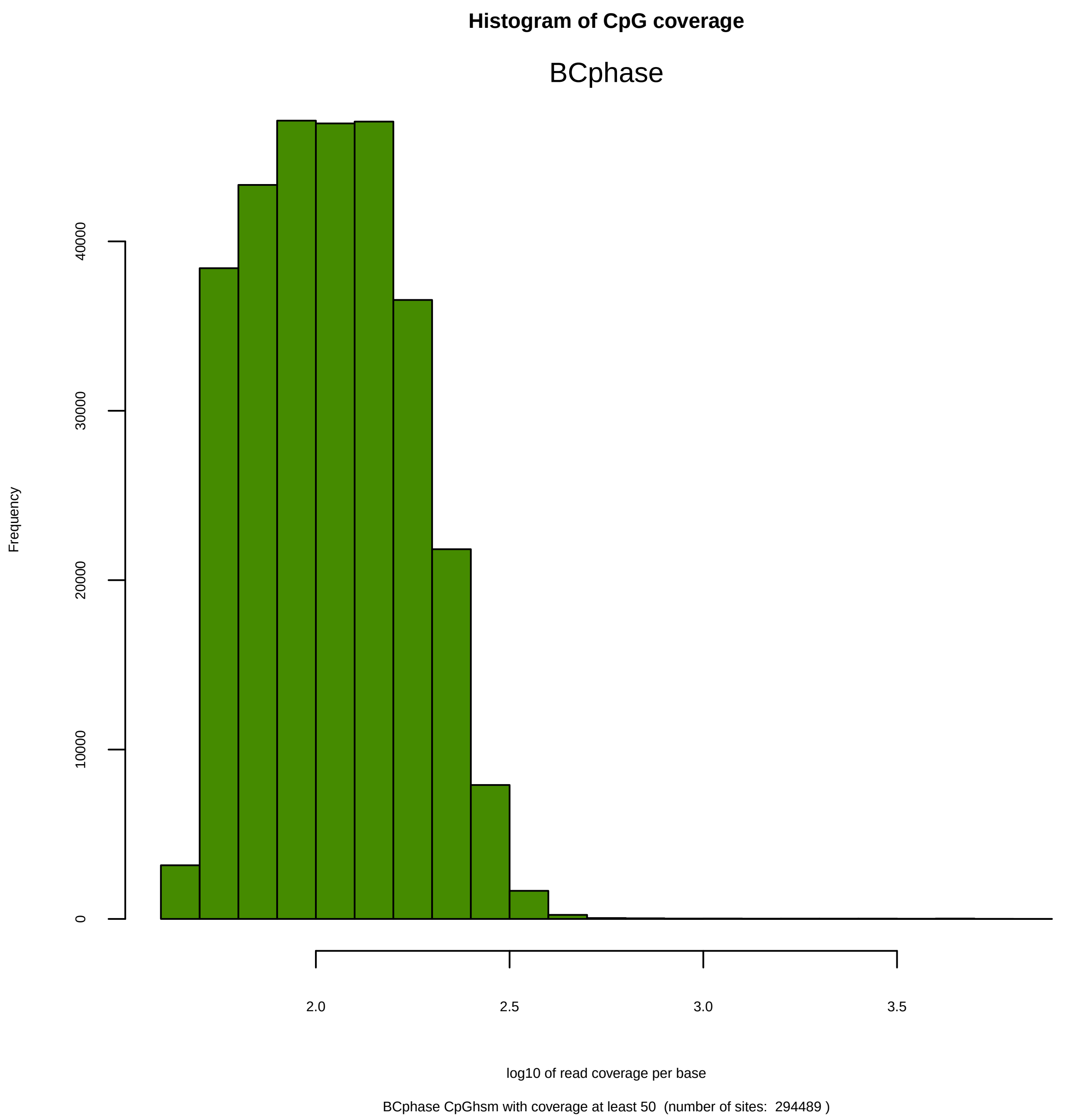
Histogram of CpG coverage
BCphase
Frequency
0
10000
20000
30000
40000
2.0
2.5
3.0
3.5
log10 of read coverage per base
BCphase CpGhsm with coverage at least 50  (number of sites:  294489 )

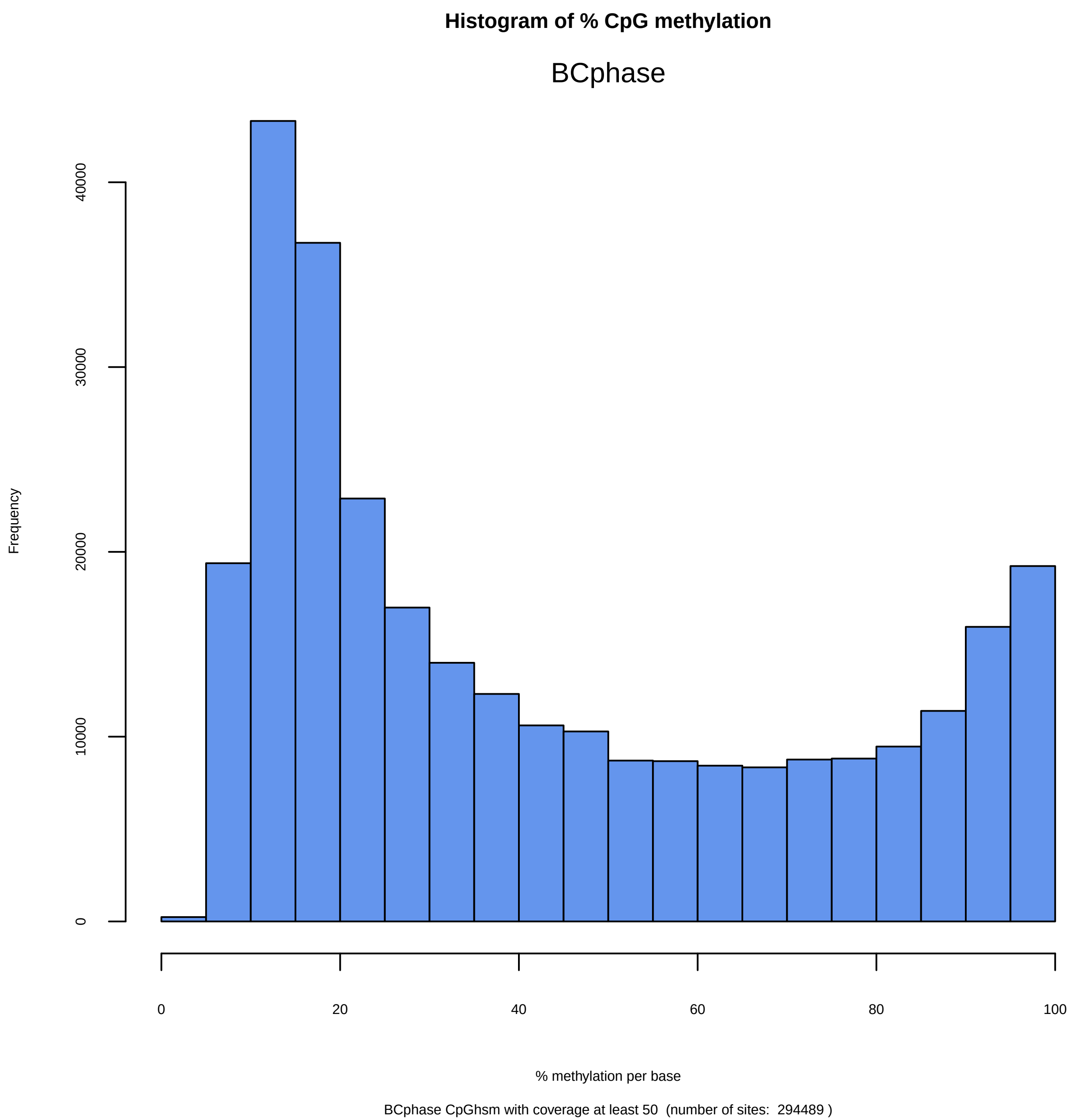

Histogram of % CpG methylation
BCphase
Frequency
0
10000
20000
30000
40000
0
20
40
60
80
100
% methylation per base
BCphase CpGhsm with coverage at least 50 (number of sites: 294489 )

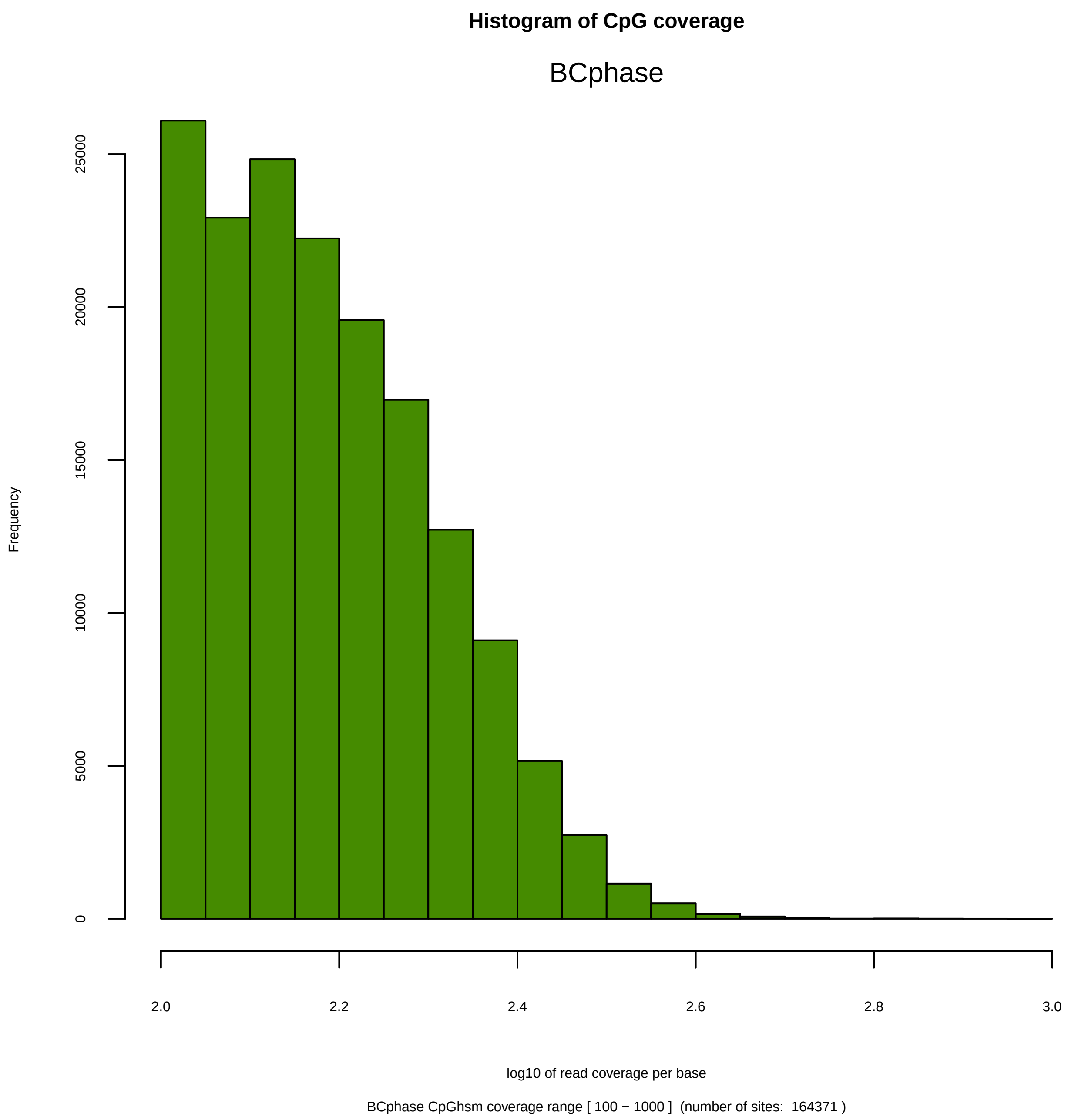

Histogram of CpG coverage
BCphase
Frequency
0
5000
10000
15000
20000
25000
2.0
2.2
2.4
2.6
2.8
3.0
log10 of read coverage per base
BCphase CpGhsm coverage range [ 100 − 1000 ]  (number of sites:  164371 )

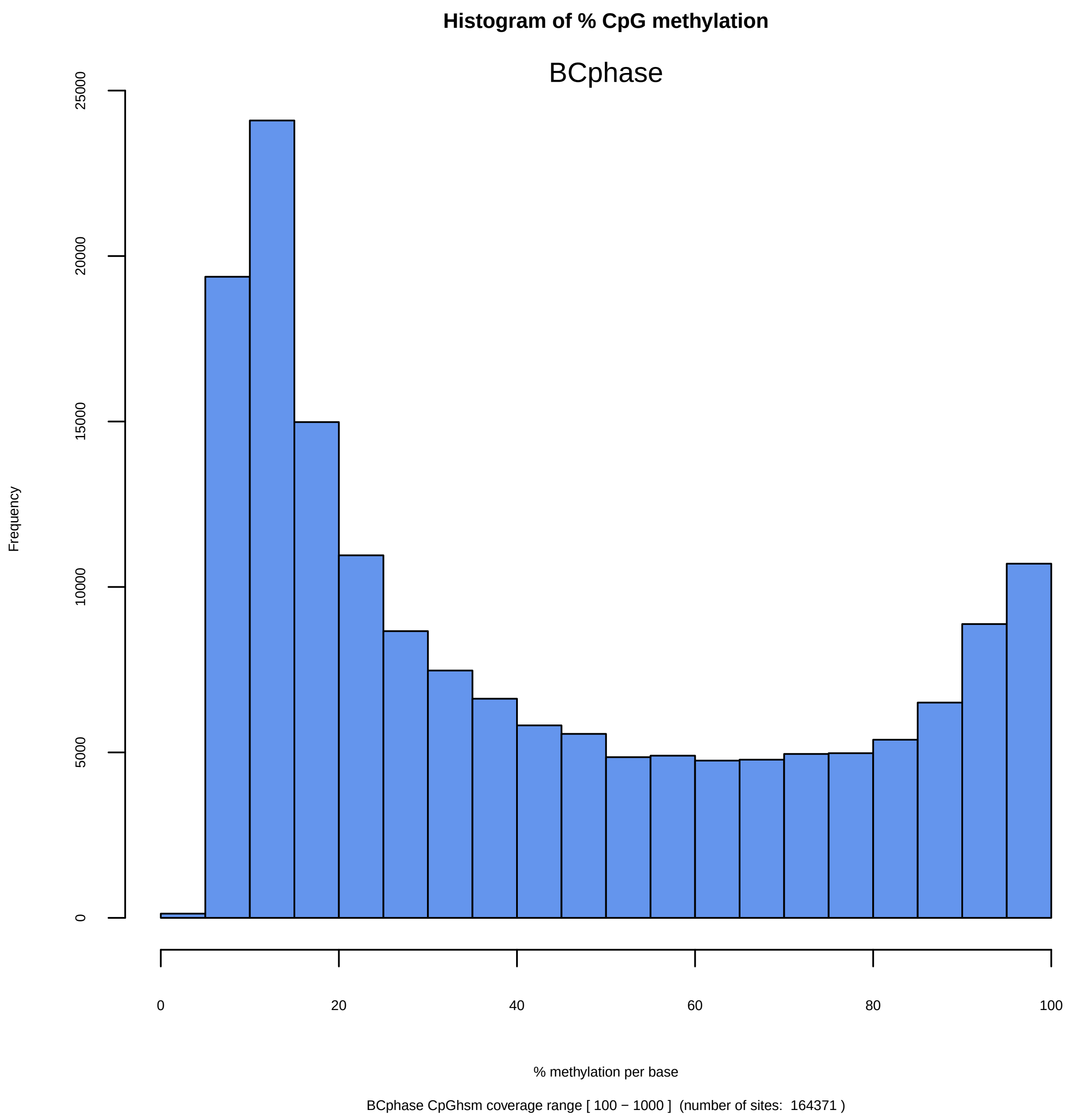
Histogram of % CpG methylation
BCphase
Frequency
0
5000
10000
15000
20000
25000
0
20
40
60
80
100
% methylation per base
BCphase CpGhsm coverage range [ 100 − 1000 ]  (number of sites:  164371 )

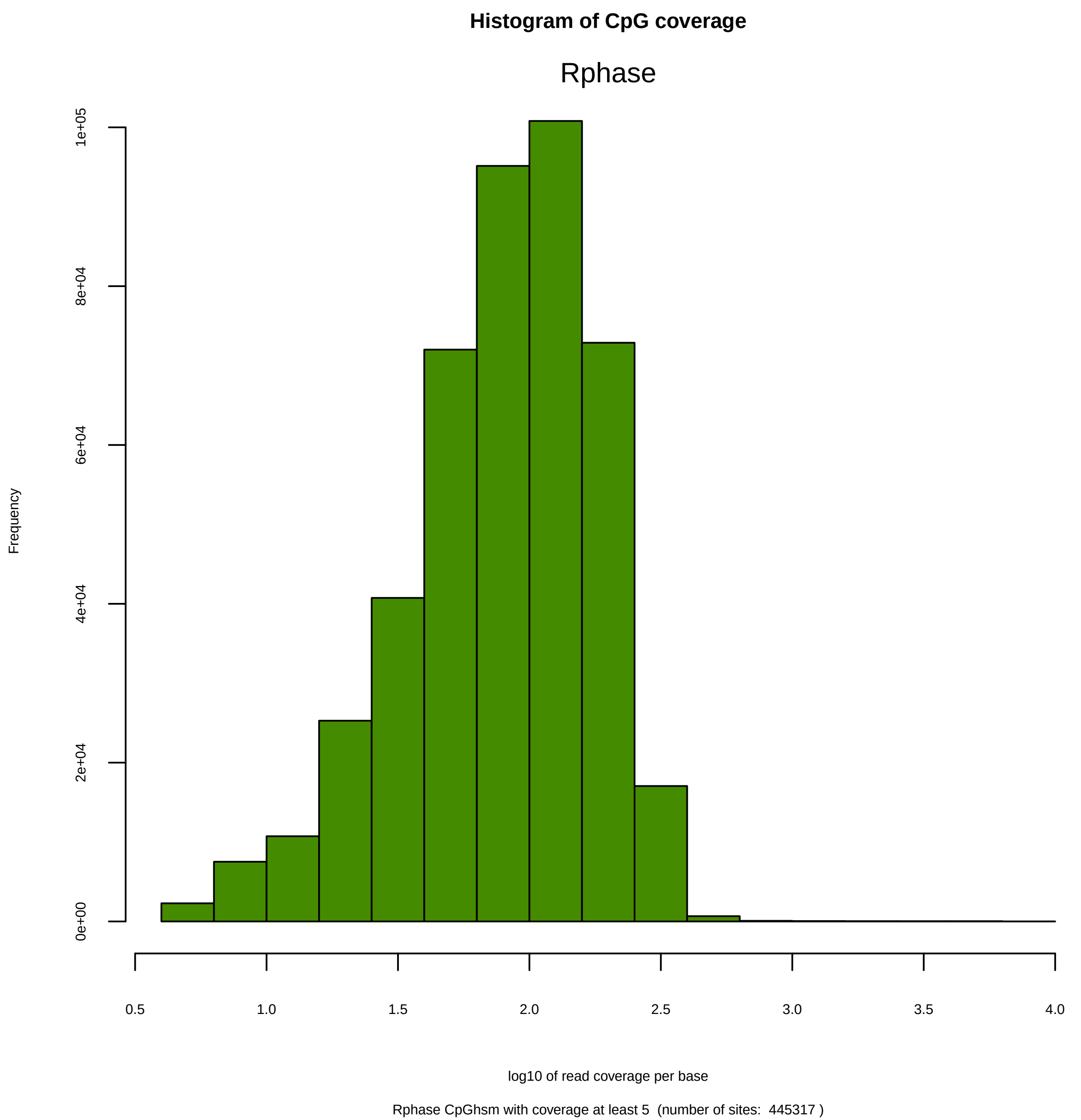

Histogram of CpG coverage
Rphase
Frequency
0e+00
2e+04
4e+04
6e+04
8e+04
1e+05
0.5
1.0
1.5
2.0
2.5
3.0
3.5
4.0
log10 of read coverage per base
Rphase CpGhsm with coverage at least 5 (number of sites: 445317 )

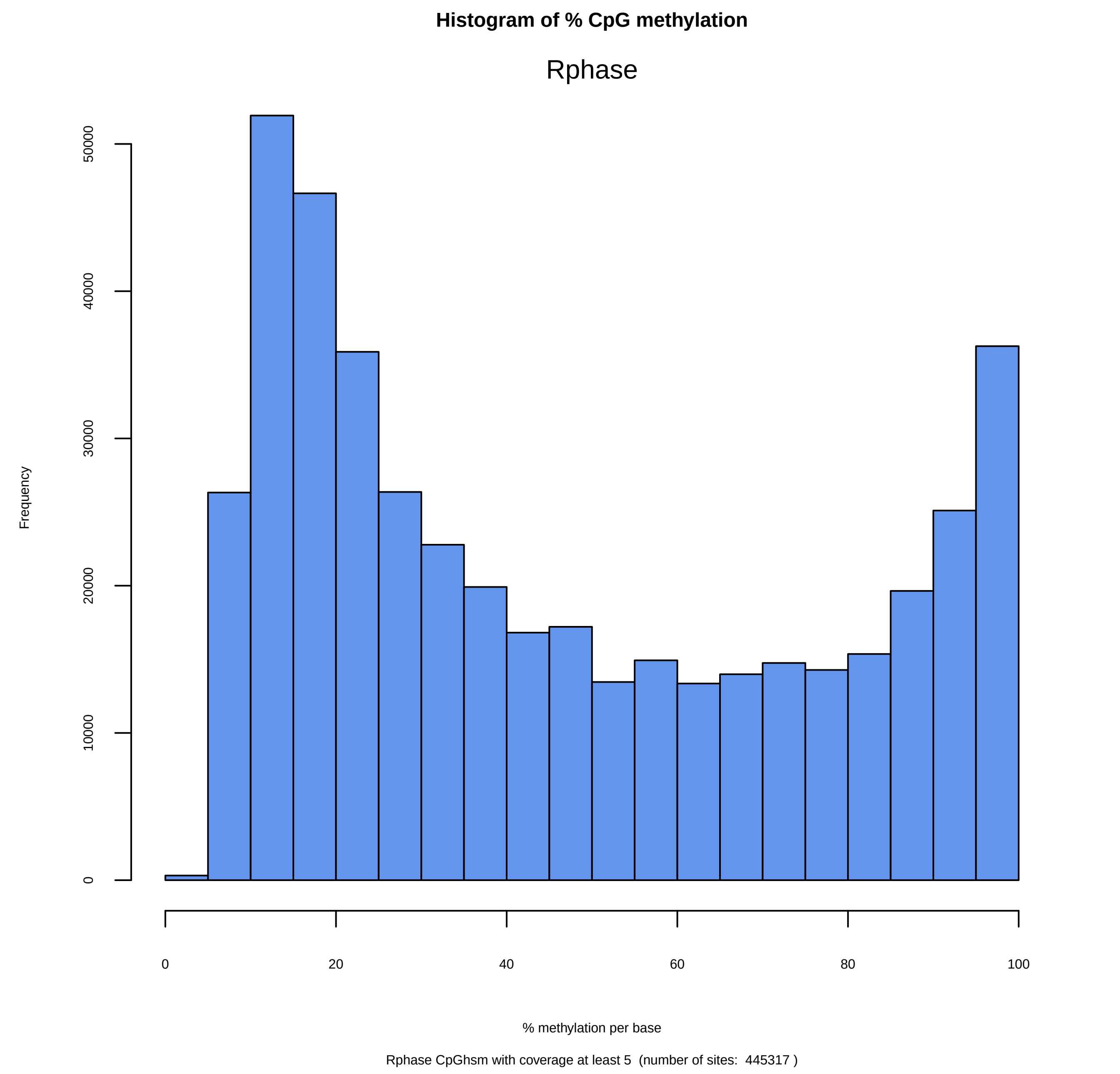
Histogram of % CpG methylation
Rphase
Frequency
0
10000
20000
30000
40000
50000
0
20
40
60
80
100
% methylation per base
Rphase CpGhsm with coverage at least 5 (number of sites: 445317 )

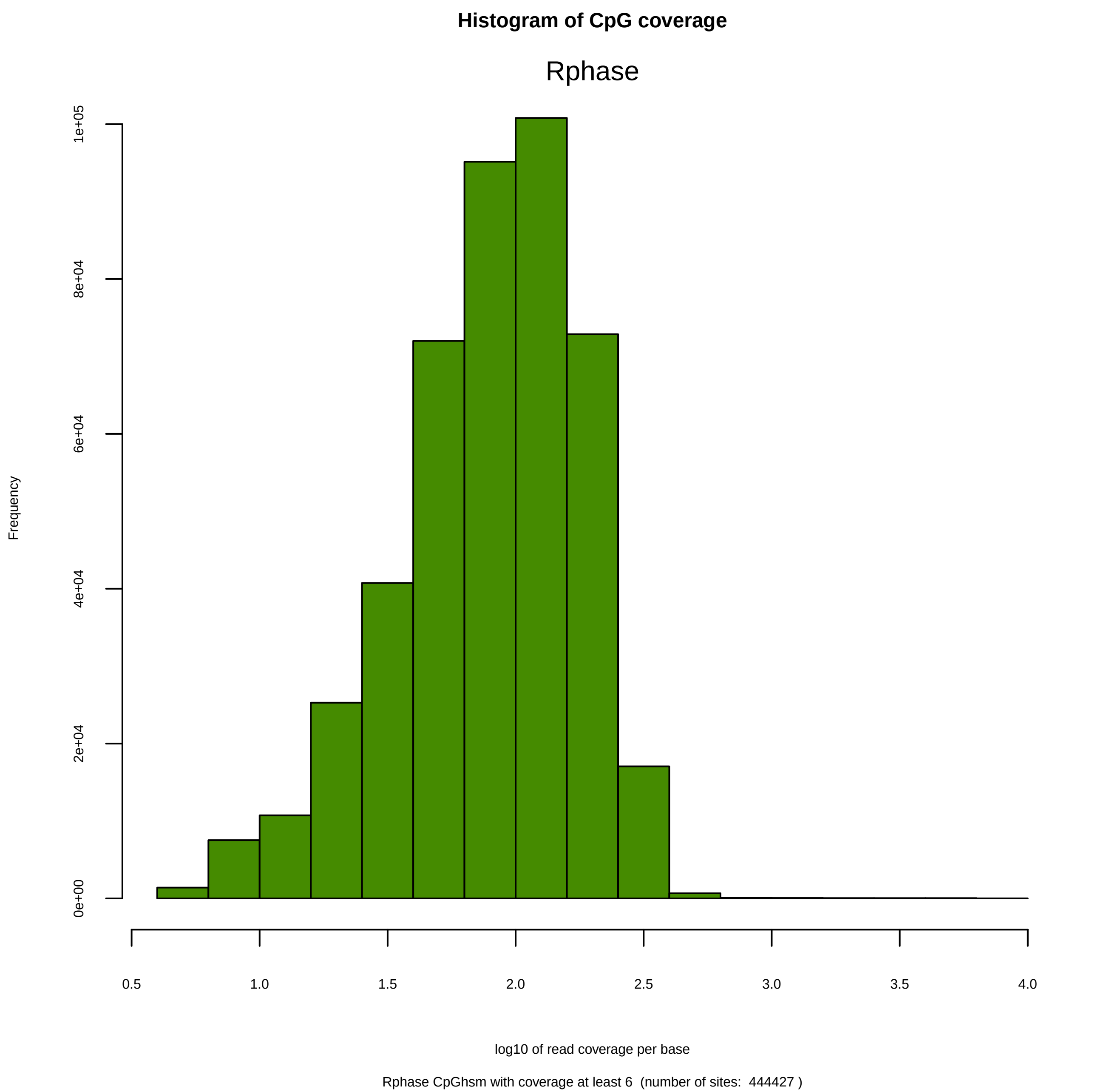
Histogram of CpG coverage
Rphase
Frequency
0e+00
2e+04
4e+04
6e+04
8e+04
1e+05
0.5
1.0
1.5
2.0
2.5
3.0
3.5
4.0
log10 of read coverage per base
Rphase CpGhsm with coverage at least 6 (number of sites: 444427 )

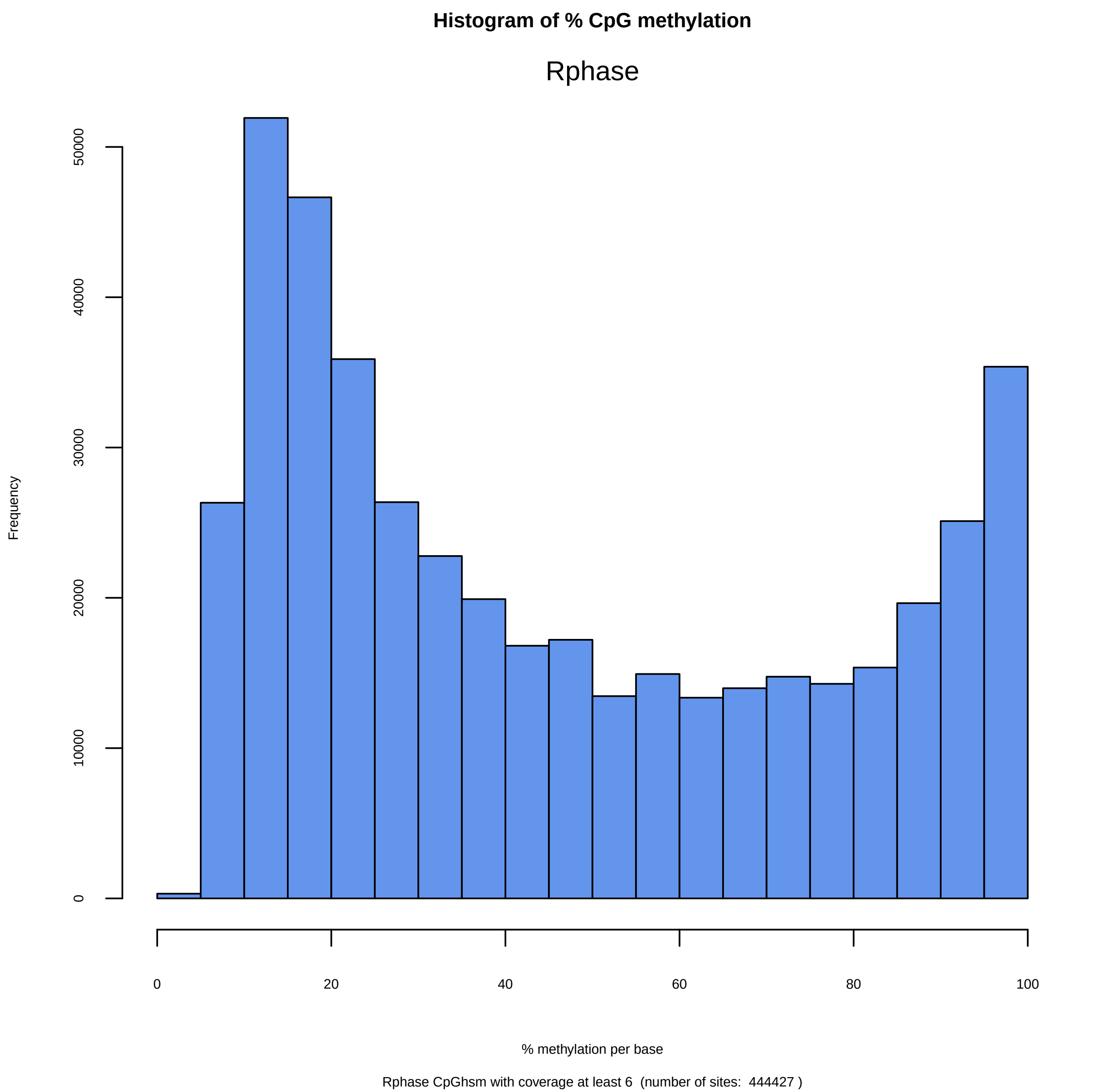

Histogram of % CpG methylation
Rphase
Frequency
0
10000
20000
30000
40000
50000
0
20
40
60
80
100
% methylation per base
Rphase CpGhsm with coverage at least 6 (number of sites: 444427 )

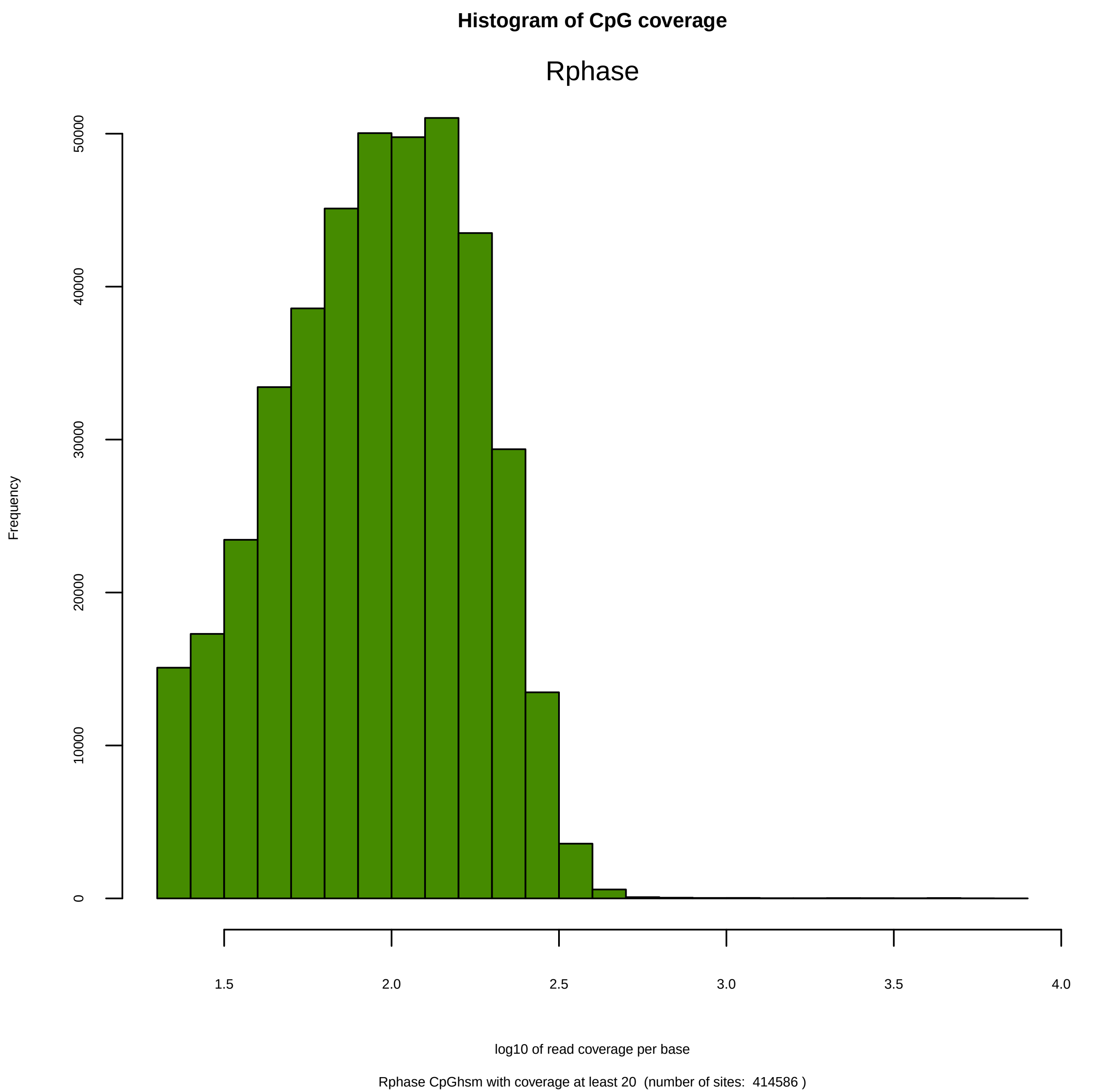
Histogram of CpG coverage
Rphase
Frequency
0
10000
20000
30000
40000
50000
1.5
2.0
2.5
3.0
3.5
4.0
log10 of read coverage per base
Rphase CpGhsm with coverage at least 20  (number of sites:  414586 )

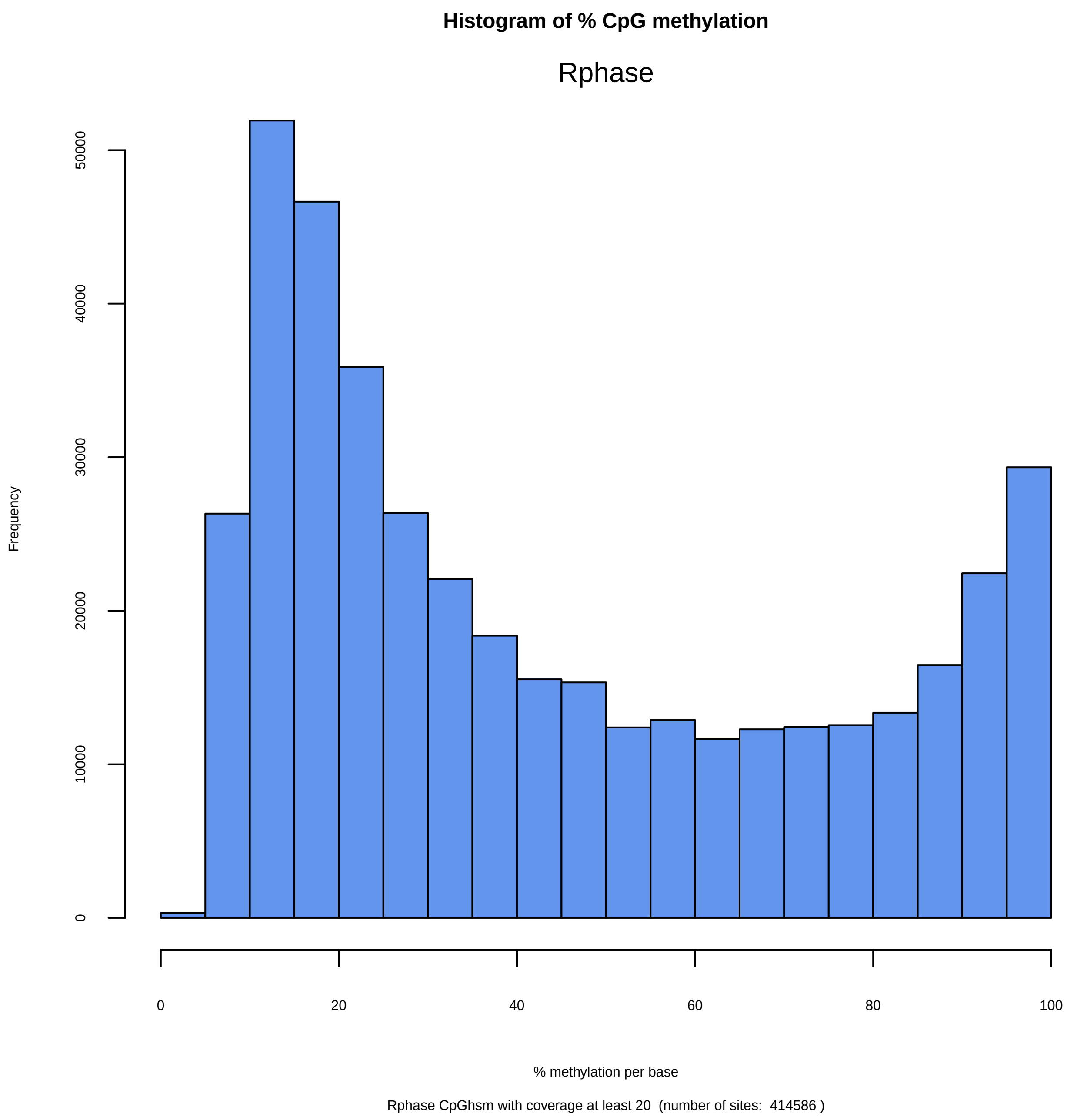
Histogram of % CpG methylation
Rphase
Frequency
0
10000
20000
30000
40000
50000
0
20
40
60
80
100
% methylation per base
Rphase CpGhsm with coverage at least 20 (number of sites: 414586 )

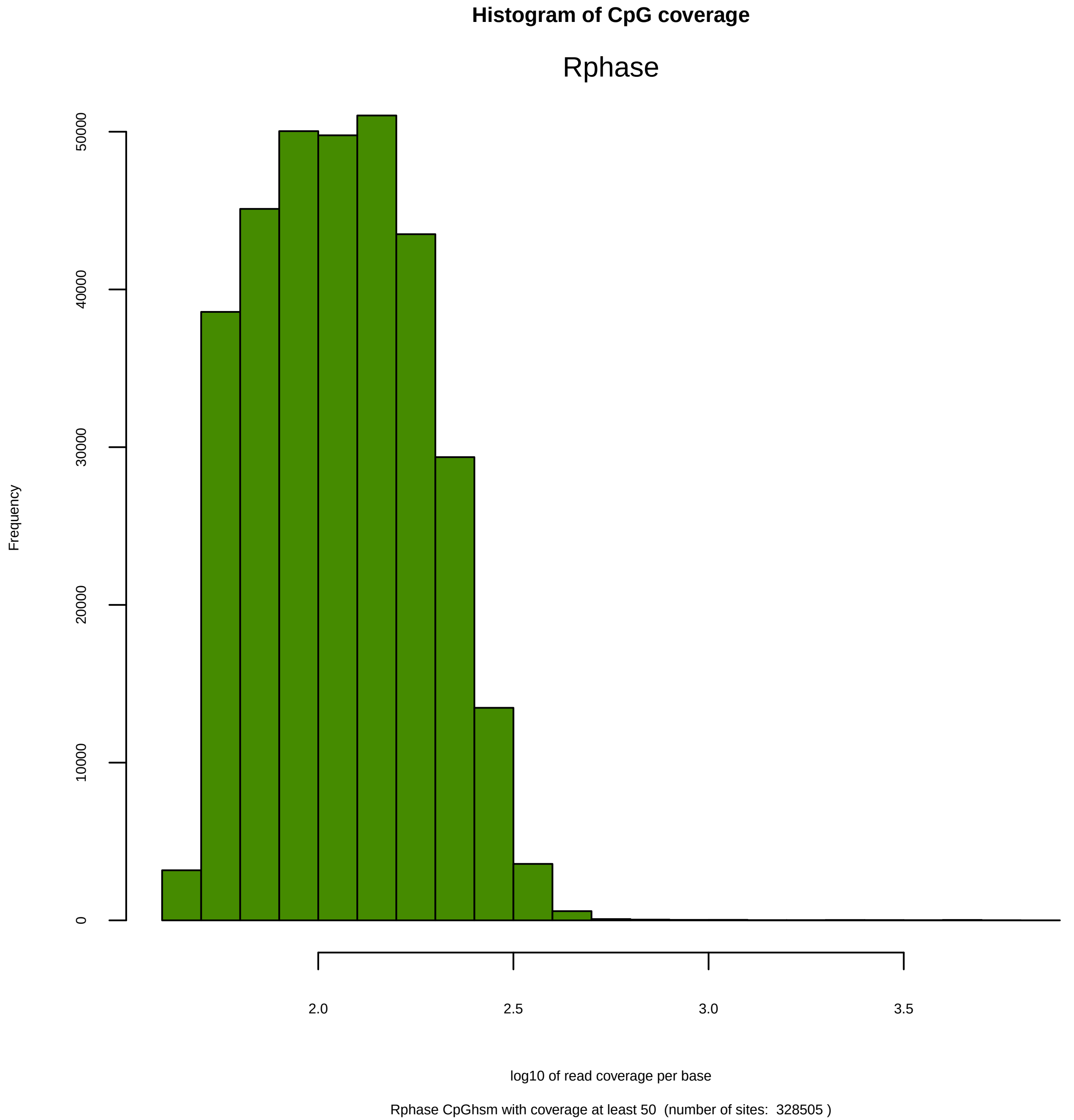
Histogram of CpG coverage
Rphase
Frequency
0
10000
20000
30000
40000
50000
2.0
2.5
3.0
3.5
log10 of read coverage per base
Rphase CpGhsm with coverage at least 50  (number of sites:  328505 )

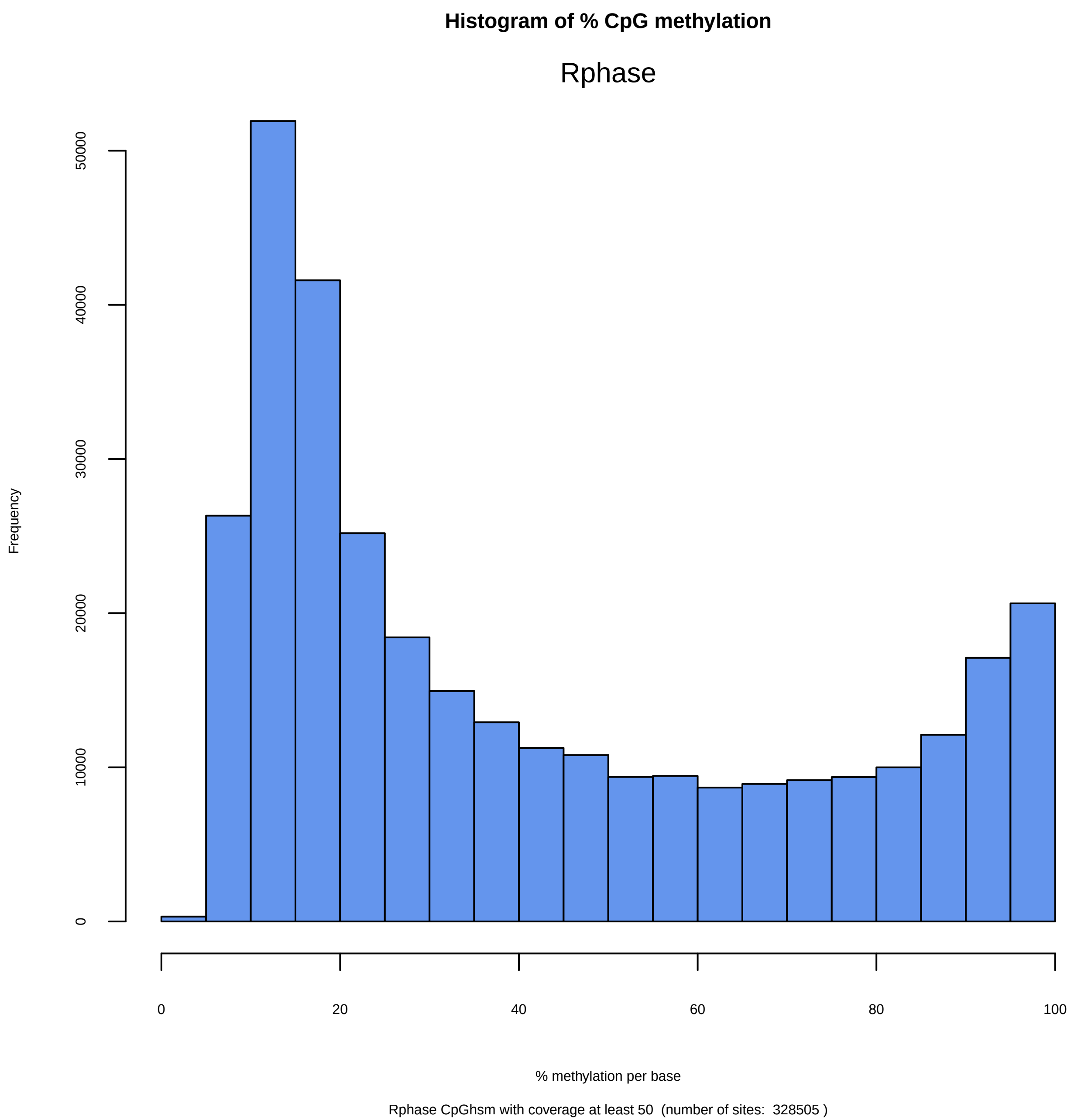

Histogram of % CpG methylation
Rphase
Frequency
0
10000
20000
30000
40000
50000
0
20
40
60
80
100
% methylation per base
Rphase CpGhsm with coverage at least 50 (number of sites: 328505 )

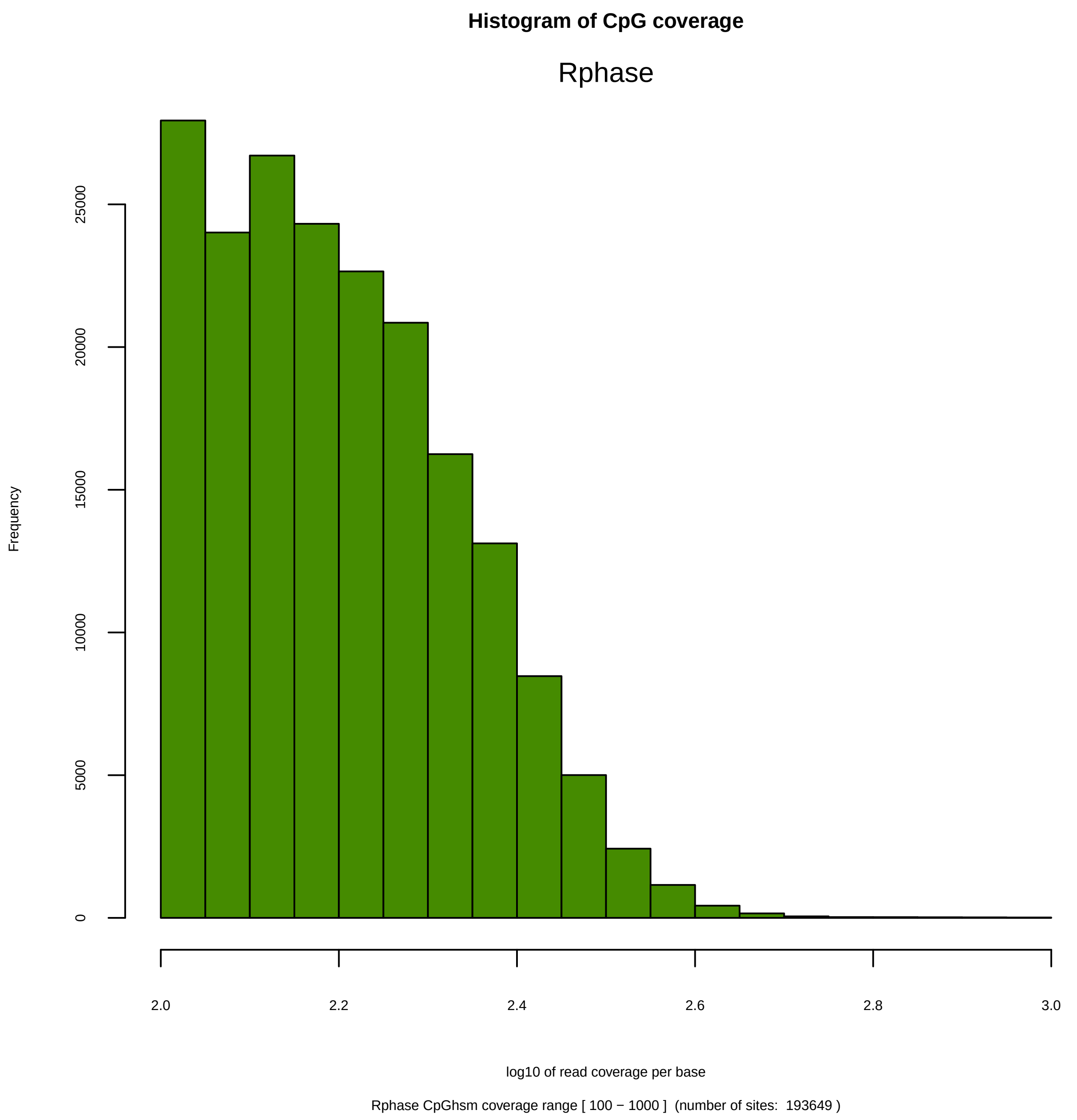
Histogram of CpG coverage
Rphase
Frequency
0
5000
10000
15000
20000
25000
2.0
2.2
2.4
2.6
2.8
3.0
log10 of read coverage per base
Rphase CpGhsm coverage range [ 100 − 1000 ]  (number of sites:  193649 )

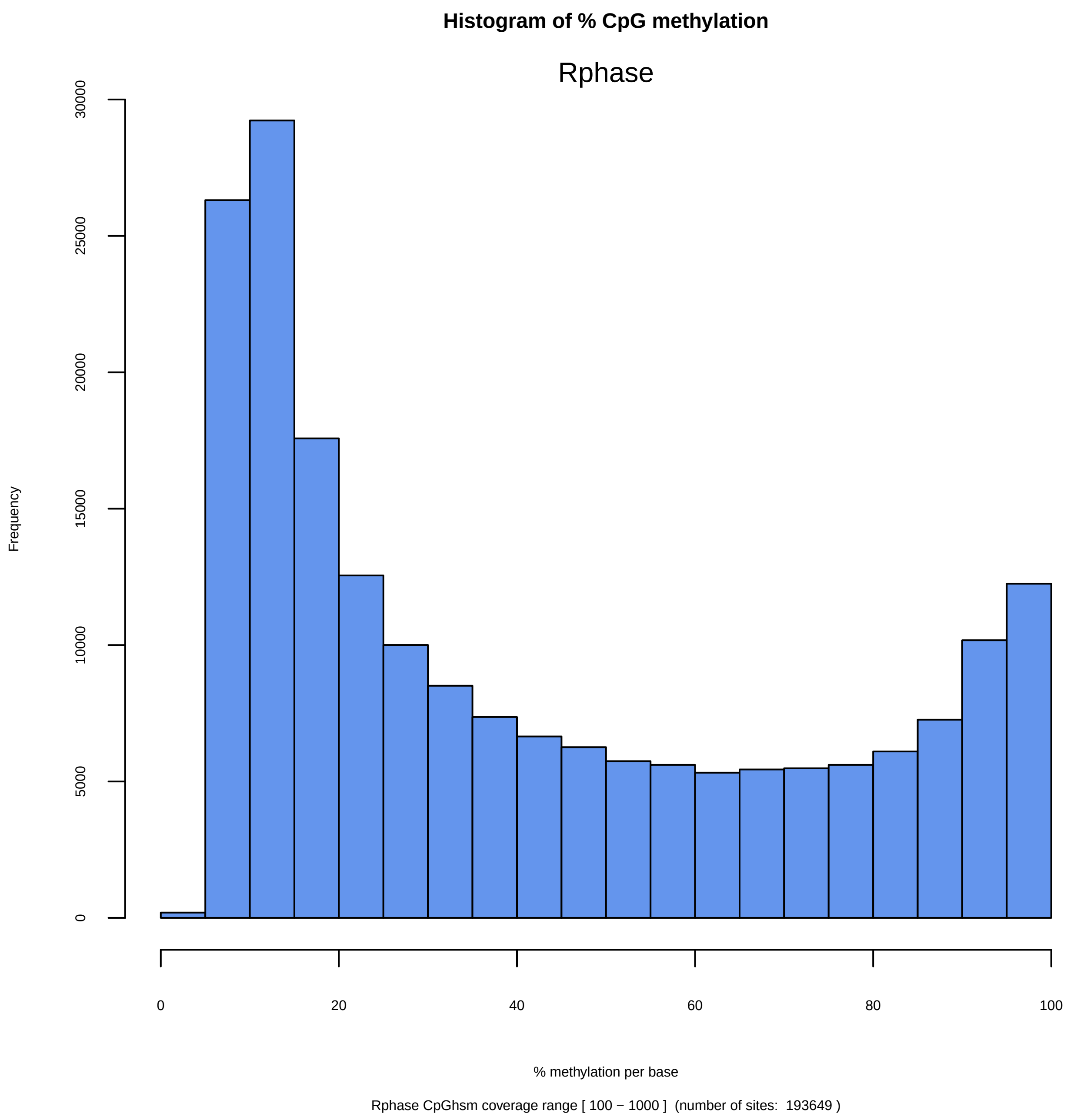

Histogram of % CpG methylation
Rphase
Frequency
0
5000
10000
15000
20000
25000
30000
0
20
40
60
80
100
% methylation per base
Rphase CpGhsm coverage range [ 100 − 1000 ]  (number of sites:  193649 )

```
Directory: PWC File: 0READMEpwc

PWC - Pairwise comparisons between all samples.

Input: studymk, studymc, nbrpms, hheight, nbrpnts

Output: files pwc-*.txt pwc-.*pdf

Notes: The output is generated by BWASPR::cmpSites(), which determines for each pairwise
        comparison the numbers of common and distinct sites.  The function also calculates
        the overlap index between the samples and estimates the size of the common pool of
        potential methylation sites.

       The CpGscd data input (studymc) is necessary to determine sites that are detectable
        in both samples, and nbrpms (listed in the *.par file specified in the Cb_LIp.conf
        configuration file as argument to TOTALNBRPMSITES) provides the total number of
        potential methylation sites (typically, all CpG sites), a necessary parameter for
        the estimations.
       hheight (default: 0.10) specifies the y-axis limit in the histogram plots.
       nbrpnts (default: 5000) specifies the number of common sites to be plotted in the
        methylation levels scatter plot.
```

```
Directory: PWC File: pwc-Cb_LI_BCphase.vs.Cb_LI_Rphase.txt

Numbers of common and distinct sites comparing Cb_LI_BCphase versus Cb_LI_Rphase
================================================================================

  total number of potential sites:  26684764

  number of "Cb_LI_BCphase_hsm"        sites: 417310
  number of "Cb_LI_Rphase_hsm"        sites:  445317
    number of "Cb_LI_BCphase_hsm"-unique sites: 48135
    number of common sites:        369175
    number of "Cb_LI_Rphase_hsm"-unique sites:  76142
  total number of "Cb_LI_BCphase_hsm+Cb_LI_Rphase_hsm"-sites observed: 493452

  number of "Cb_LI_BCphase_scd"       sites:   24098022 ( 90.31% of total)
  number of "Cb_LI_Rphase_scd"       sites:    24215006 ( 90.74% of total)
    number of "Cb_LI_BCphase_scd"-unique sites:   376760
    number of sites in common:      23721262   (Expected:  21867675; O/E:      1.1)
    number of "Cb_LI_Rphase_scd"-unique sites:    493744
  total number of "Cb_LI_BCphase_scd+Cb_LI_Rphase_scd"-sites observed:  24591766

  number of sites in "Cb_LI_BCphase_hsm" that are not  detectable in "Cb_LI_Rphase_hsm":   1478
  number of sites in "Cb_LI_BCphase_hsm" that are also detectable in "Cb_LI_Rphase_hsm": 415832
    number of sites unique to "Cb_LI_BCphase_hsm" although detectable in "Cb_LI_Rphase_hsm":         46657
    number of "Cb_LI_BCphase_hsm" / "Cb_LI_Rphase_hsm" common sites                        :        369175      (Expected:   7760; O/E:      47.6)

  number of sites in "Cb_LI_Rphase_hsm" that are not  detectable in "Cb_LI_BCphase_hsm":   2619
  number of sites in "Cb_LI_Rphase_hsm" that are also detectable in "Cb_LI_BCphase_hsm": 442698
    number of sites unique to "Cb_LI_Rphase_hsm" although detectable in "Cb_LI_BCphase_hsm":         73523
    number of "Cb_LI_Rphase_hsm" / "Cb_LI_BCphase_hsm" common sites                        :        369175      (Expected:   7760; O/E:      47.6)

  Overlap index of "Cb_LI_BCphase_hsm" with "Cb_LI_Rphase_hsm":        0.803
  Estimated number of "Cb_LI_BCphase_hsm" = "Cb_LI_Rphase_hsm" sites (assuming sampling from one population):            498647
  Adjusted population size of "Cb_LI_BCphase_hsm" = "Cb_LI_Rphase_hsm" sites (assuming all sites detectable):            560943 (  1.14x of observed)

  number of "Cb_LI_BCphase_hsm" sites with coverage >=  5: 417310
  number of "Cb_LI_Rphase_hsm" sites with coverage >=  5: 445317
  number of common sites with coverage >=  5: 369175
  number of "Cb_LI_BCphase_hsm" sites with coverage >=  6: 416306
  number of "Cb_LI_Rphase_hsm" sites with coverage >=  6: 444427
  number of common sites with coverage >=  6: 368280
  number of "Cb_LI_BCphase_hsm" sites with coverage >= 20: 384326
  number of "Cb_LI_Rphase_hsm" sites with coverage >= 20: 414586
  number of common sites with coverage >= 20: 339459
  number of "Cb_LI_BCphase_hsm" sites with coverage >= 50: 294489
  number of "Cb_LI_Rphase_hsm" sites with coverage >= 50: 328505
  number of common sites with coverage >= 50: 261901
```

### Methylation Levels in Common Sites (Coverage >=  5)

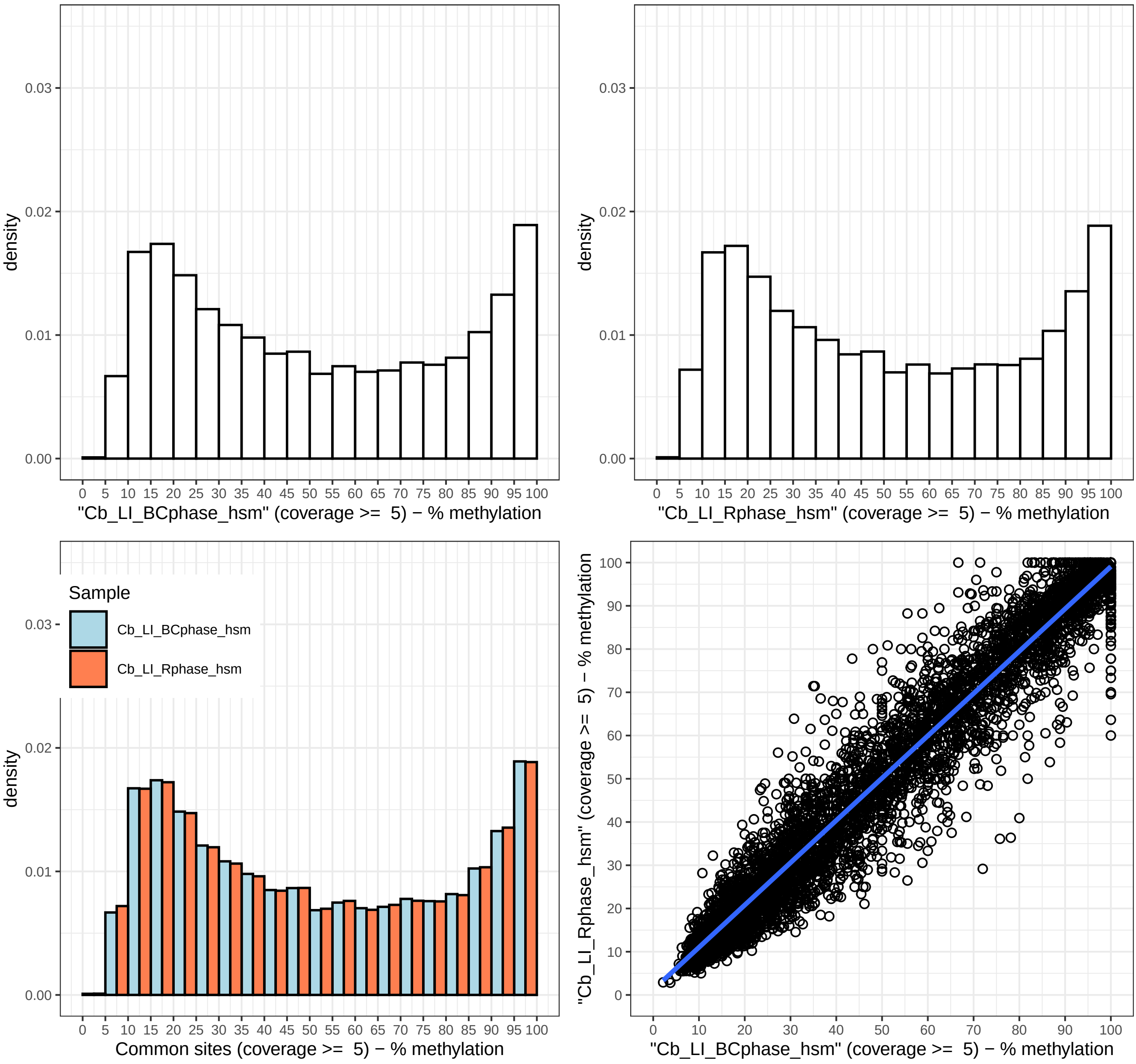

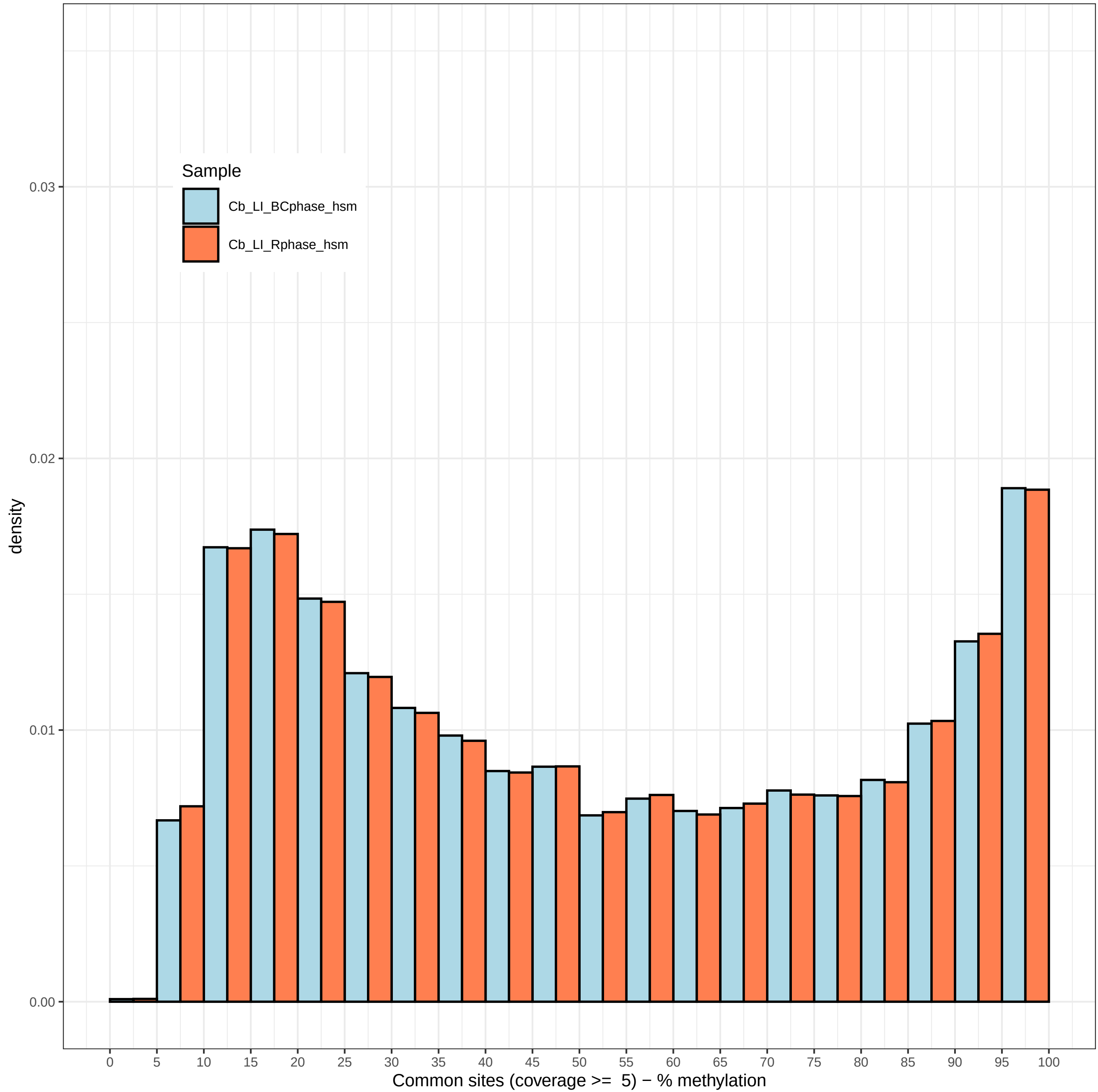
Sample
Cb_LI_BCphase_hsm
Cb_LI_Rphase_hsm
density
0.00
0.01
0.02
0.03
0
5
10
15
20
25
30
35
40
45
50
55
60
65
70
75
80
85
90
95
100
Common sites (coverage >= 5) − % methylation

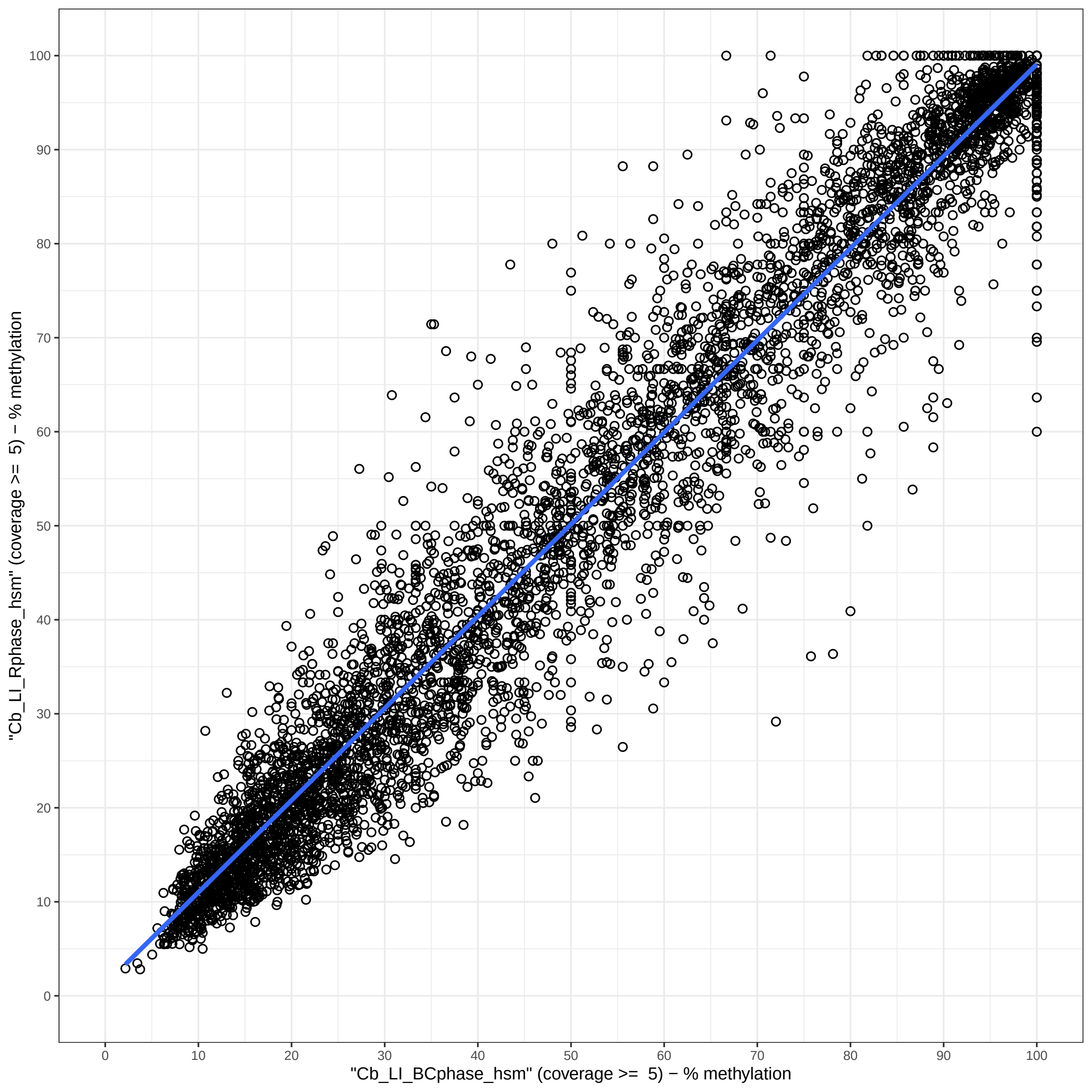

"Cb_LI_Rphase_hsm" (coverage >= 5) − % methylation
"Cb_LI_BCphase_hsm" (coverage >= 5) − % methylation
0
10
20
30
40
50
60
70
80
90
100

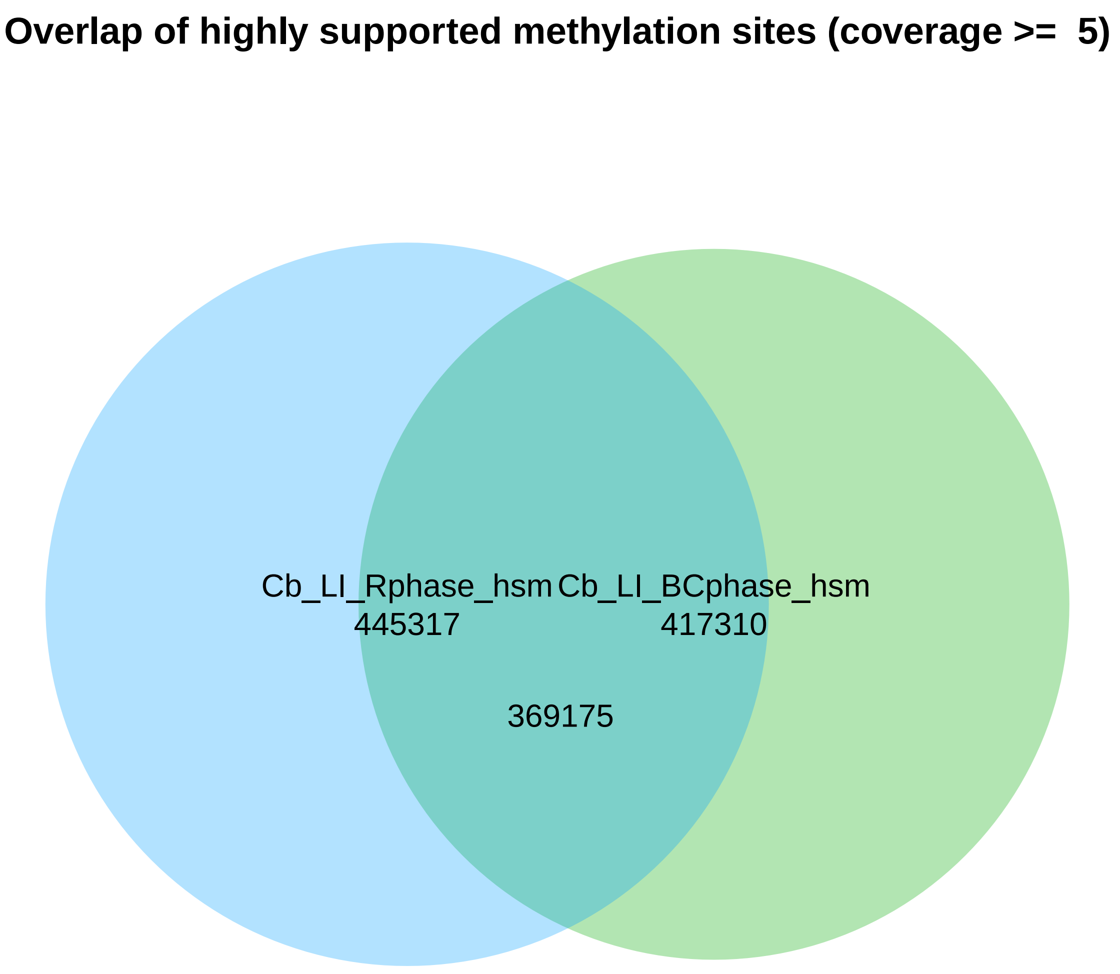
Overlap of highly supported methylation sites (coverage >= 5)
Cb_LI_Rphase_hsm
445317
Cb_LI_BCphase_hsm
417310
369175

Methylation Levels in Common Sites (Coverage >=  6)

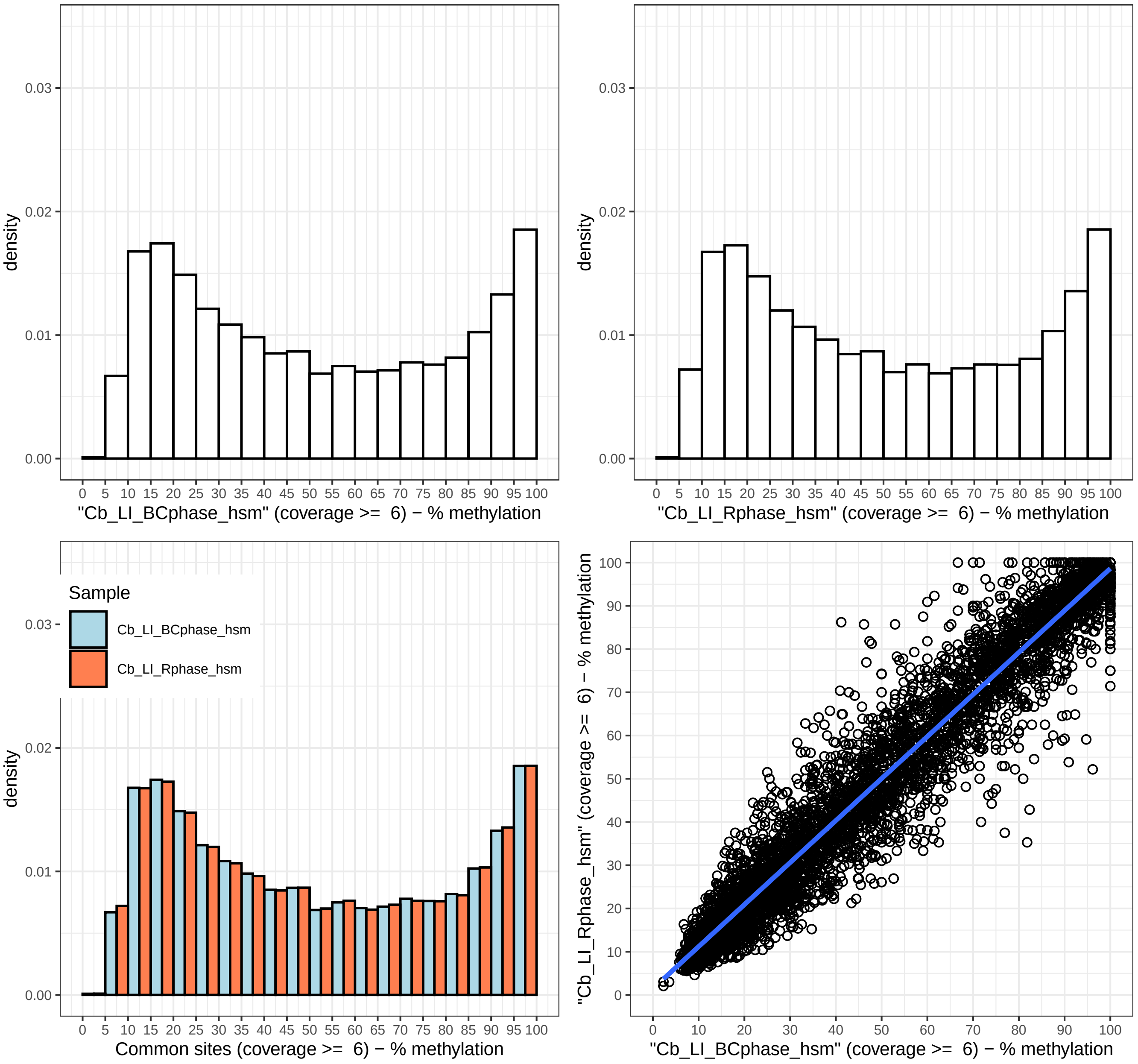

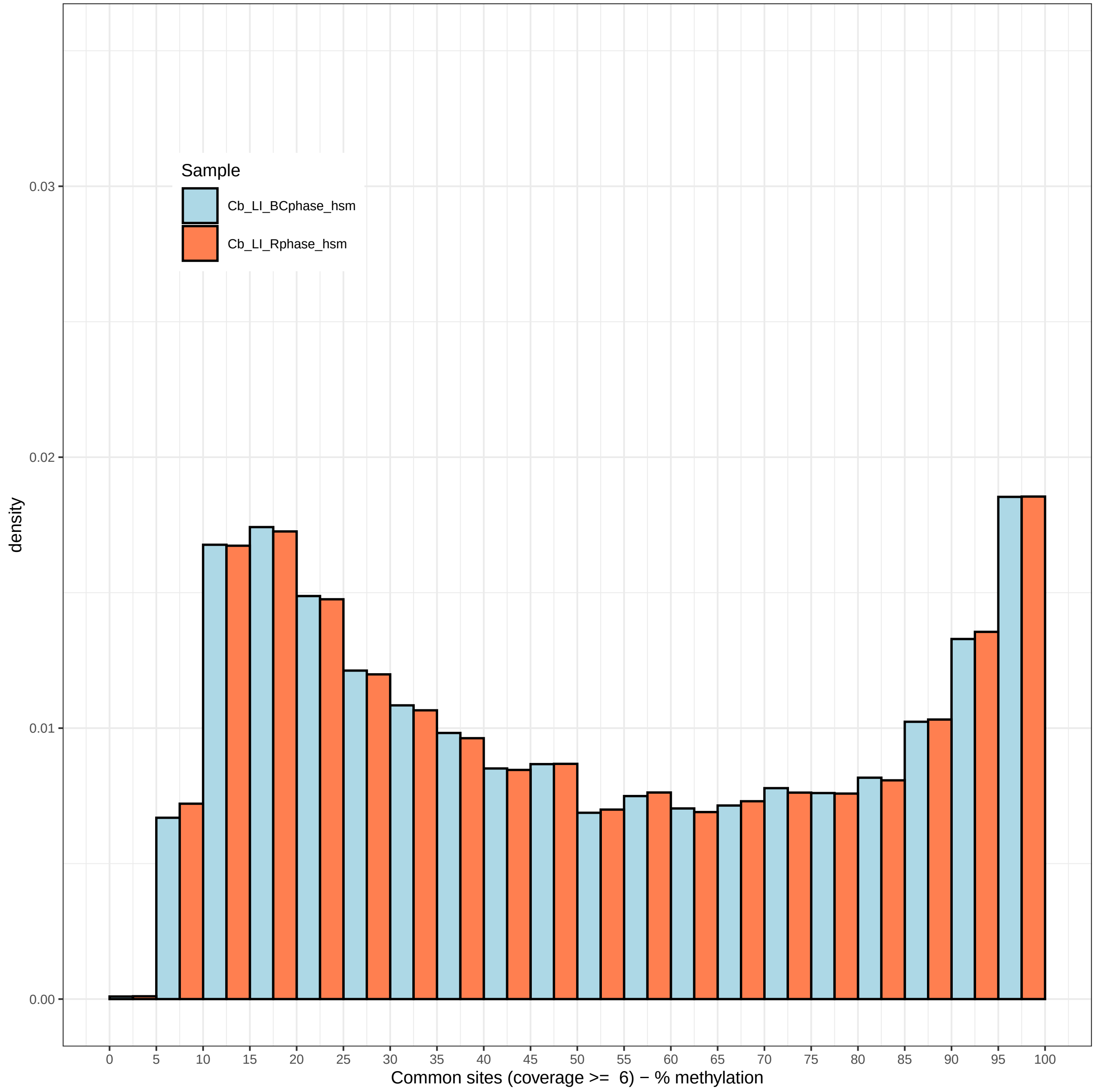
Sample
Cb_LI_BCphase_hsm
Cb_LI_Rphase_hsm
density
0.00
0.01
0.02
0.03
0 5 10 15 20 25 30 35 40 45 50 55 60 65 70 75 80 85 90 95 100
Common sites (coverage >= 6) − % methylation

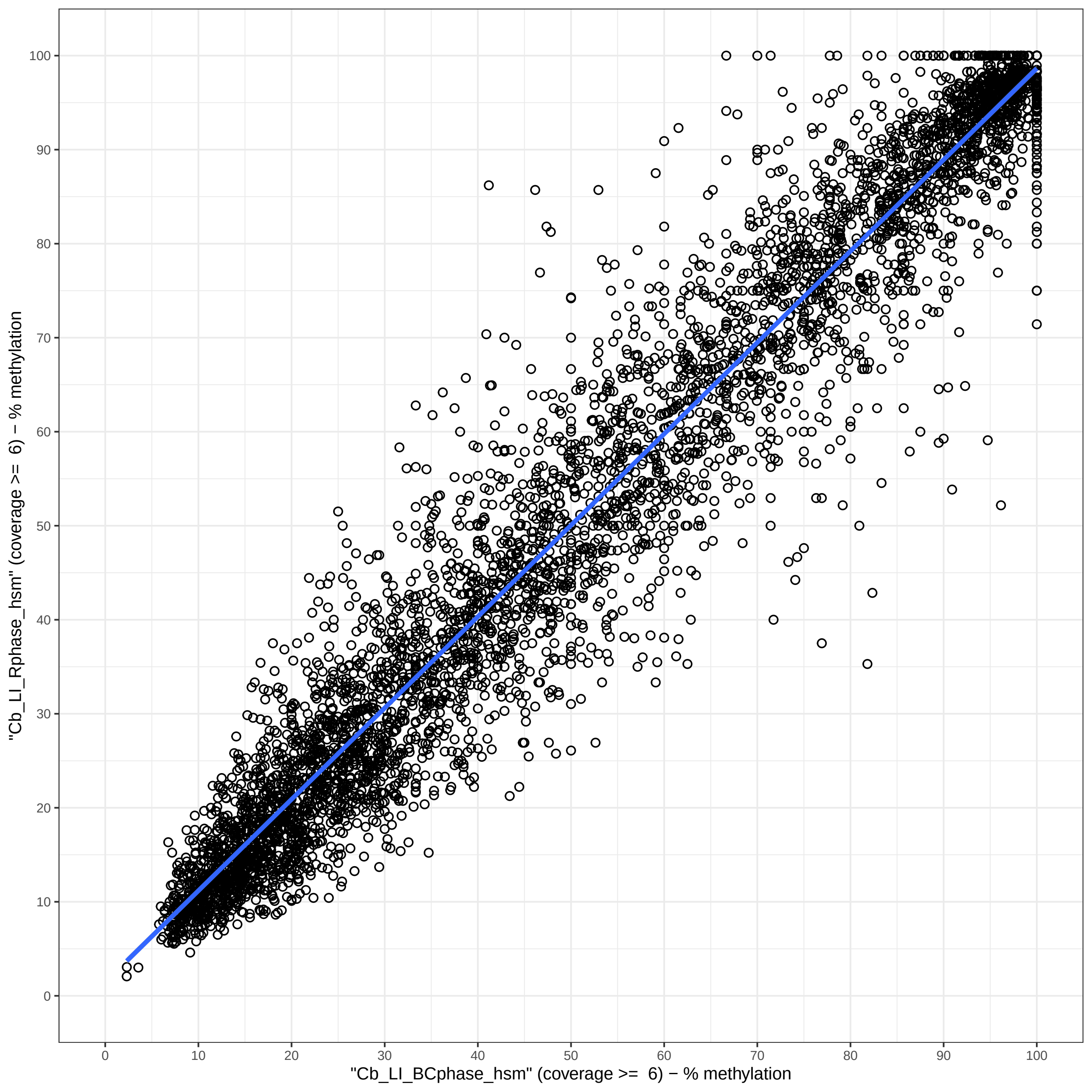

"Cb_LI_Rphase_hsm" (coverage >= 6) − % methylation
"Cb_LI_BCphase_hsm" (coverage >= 6) − % methylation
0
10
20
30
40
50
60
70
80
90
100

### Overlap of highly supported methylation sites (coverage >=  6)

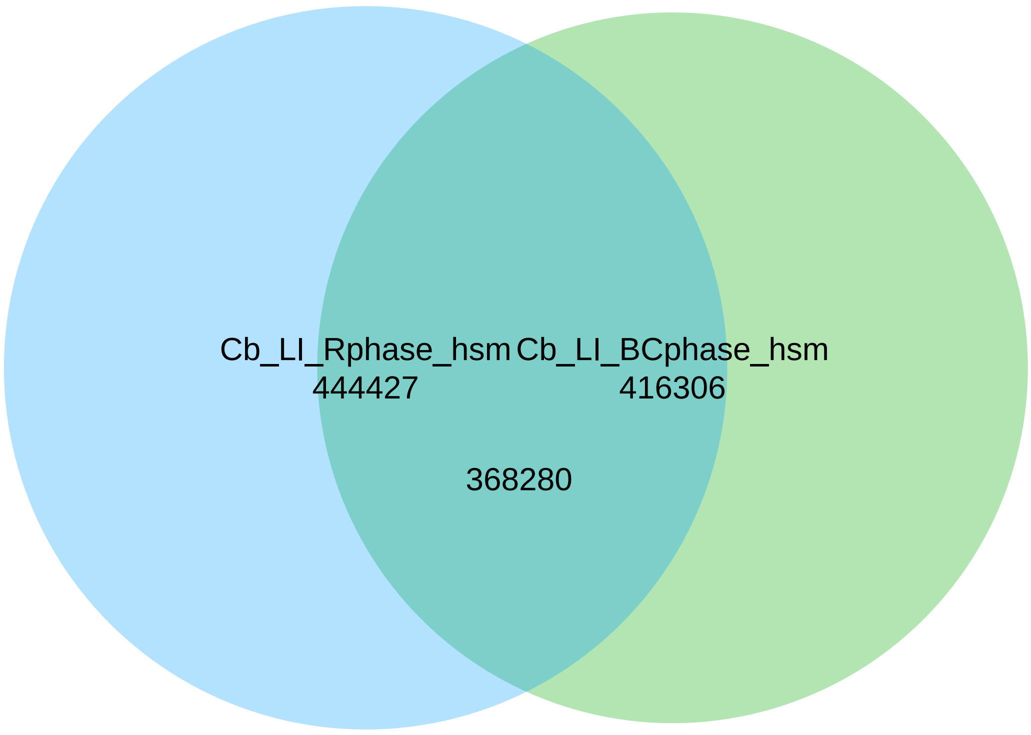

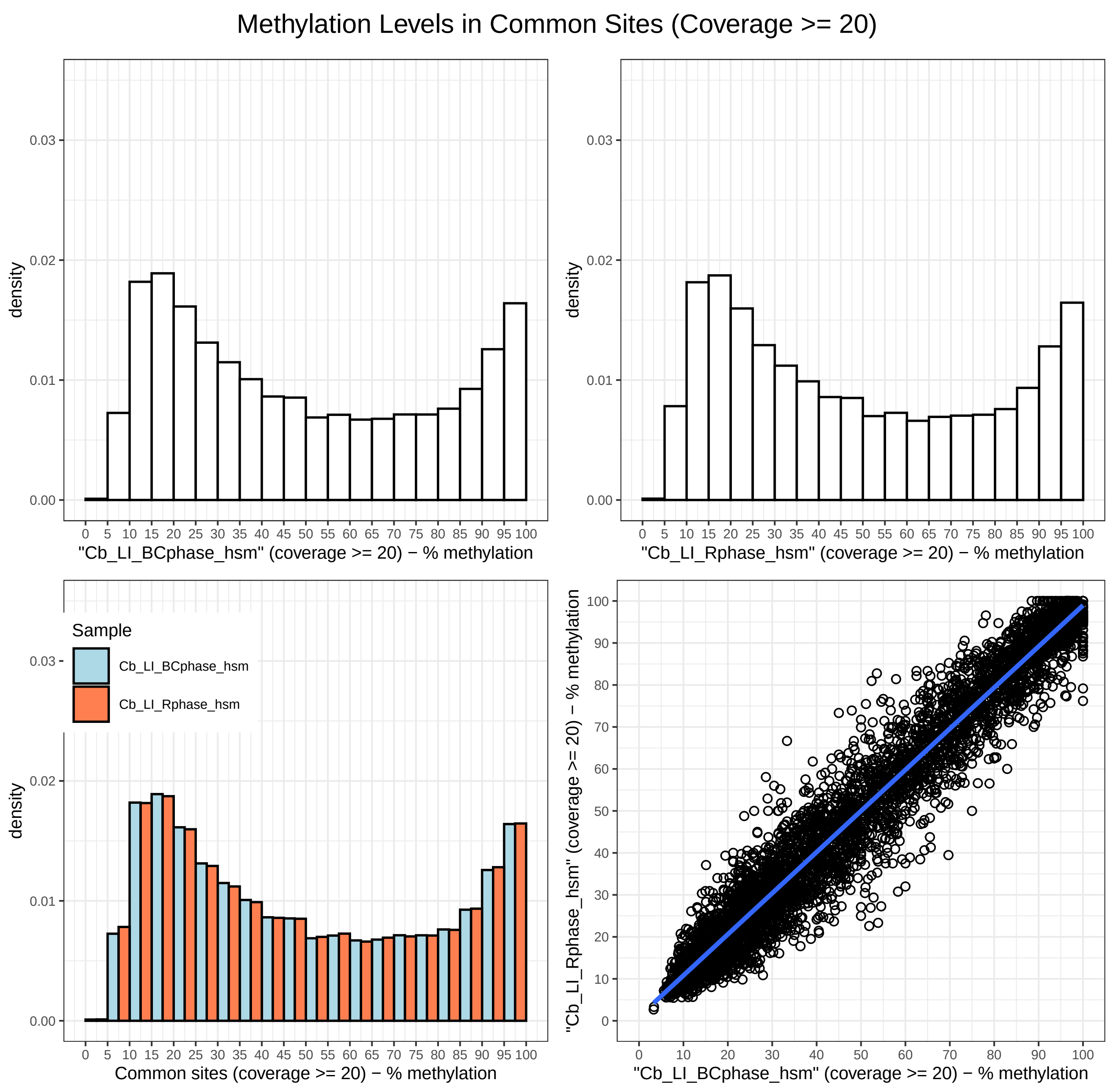
Methylation Levels in Common Sites (Coverage >= 20)
density
"Cb_LI_BCphase_hsm" (coverage >= 20) − % methylation
"Cb_LI_Rphase_hsm" (coverage >= 20) − % methylation
Sample
Cb_LI_BCphase_hsm
Cb_LI_Rphase_hsm
Common sites (coverage >= 20) − % methylation
"Cb_LI_Rphase_hsm" (coverage >= 20) − % methylation
"Cb_LI_BCphase_hsm" (coverage >= 20) − % methylation

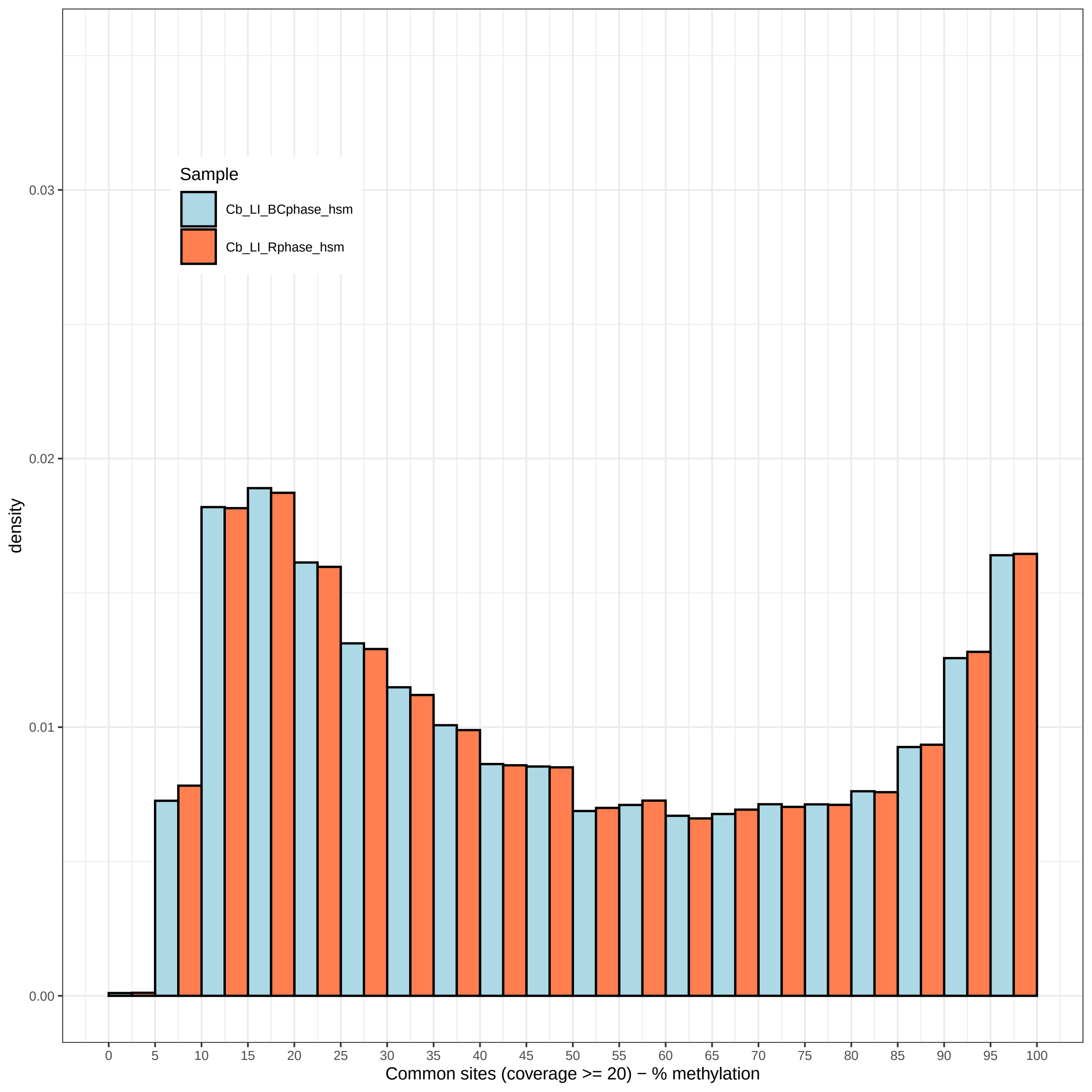

Sample
Cb_LI_BCphase_hsm
Cb_LI_Rphase_hsm
density
0.00
0.01
0.02
0.03
0
5
10
15
20
25
30
35
40
45
50
55
60
65
70
75
80
85
90
95
100
Common sites (coverage >= 20) − % methylation

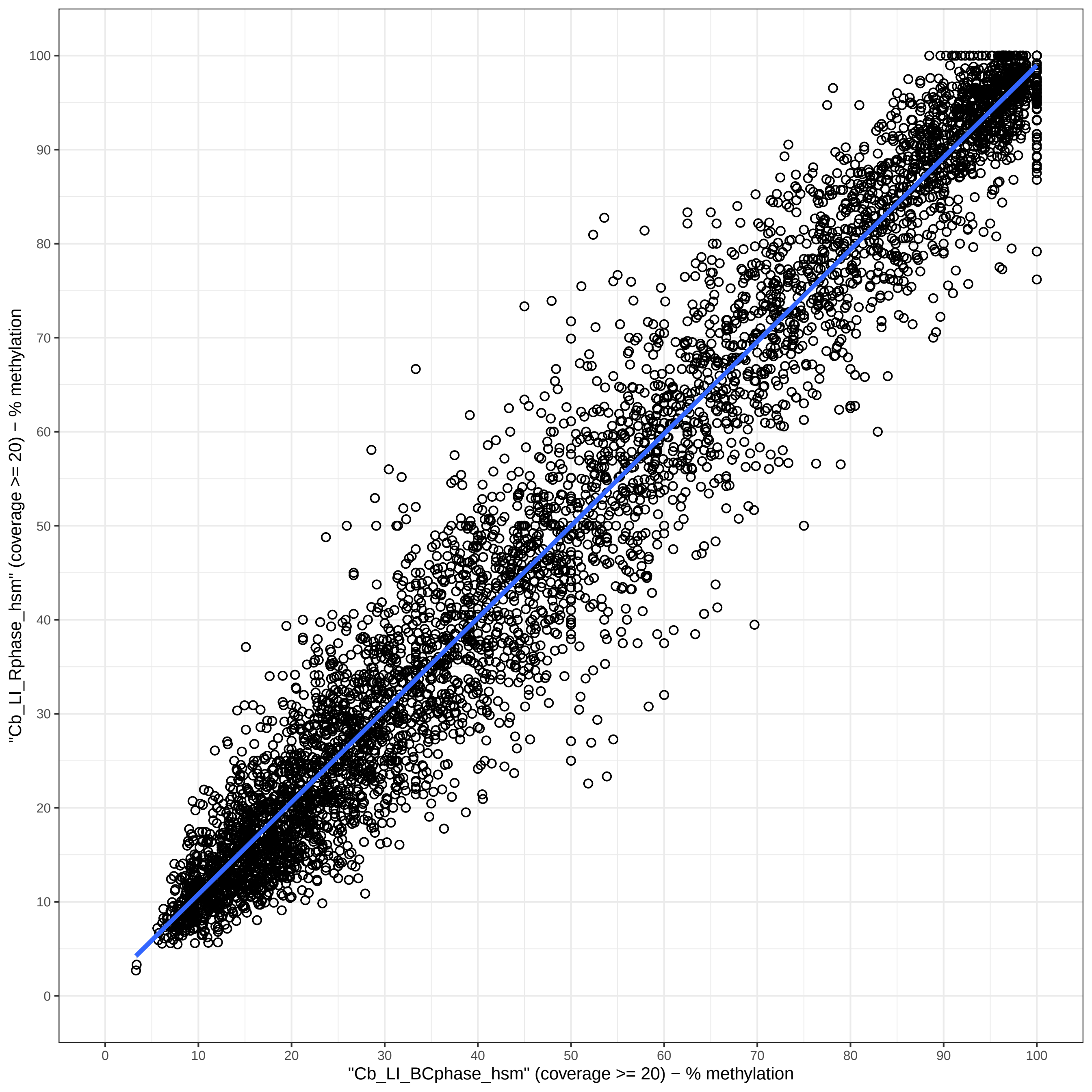

0
10
20
30
40
50
60
70
80
90
100
"Cb_LI_Rphase_hsm" (coverage >= 20) − % methylation
"Cb_LI_BCphase_hsm" (coverage >= 20) − % methylation

#### Overlap of highly supported methylation sites (coverage >= 20)

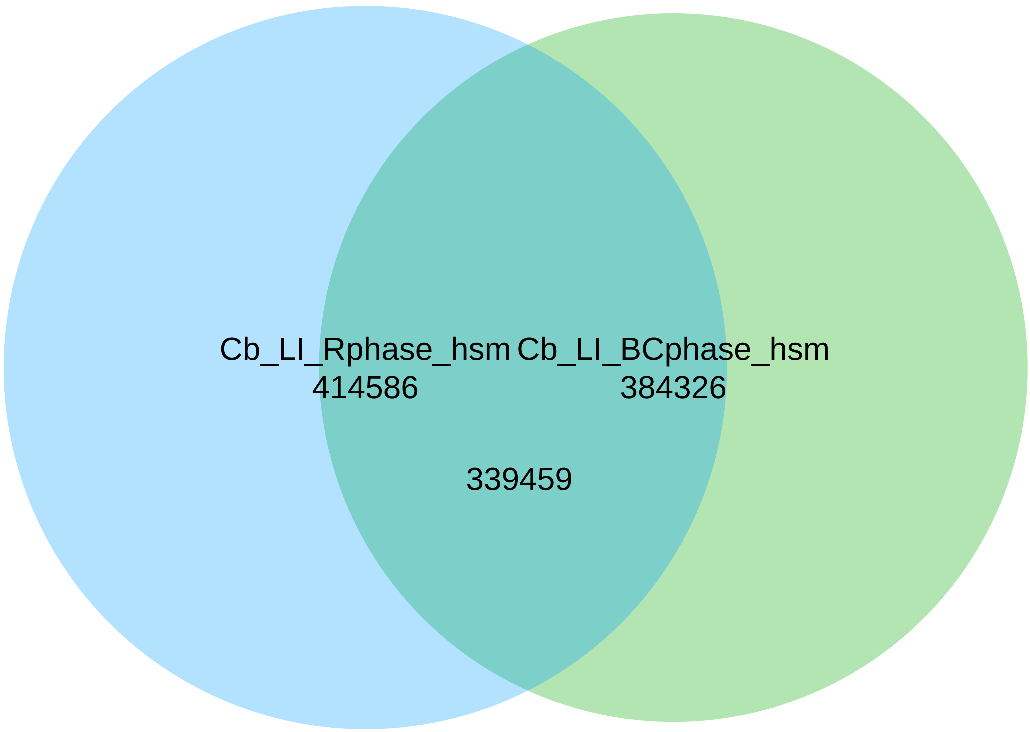

### Methylation Levels in Common Sites (Coverage >= 50)

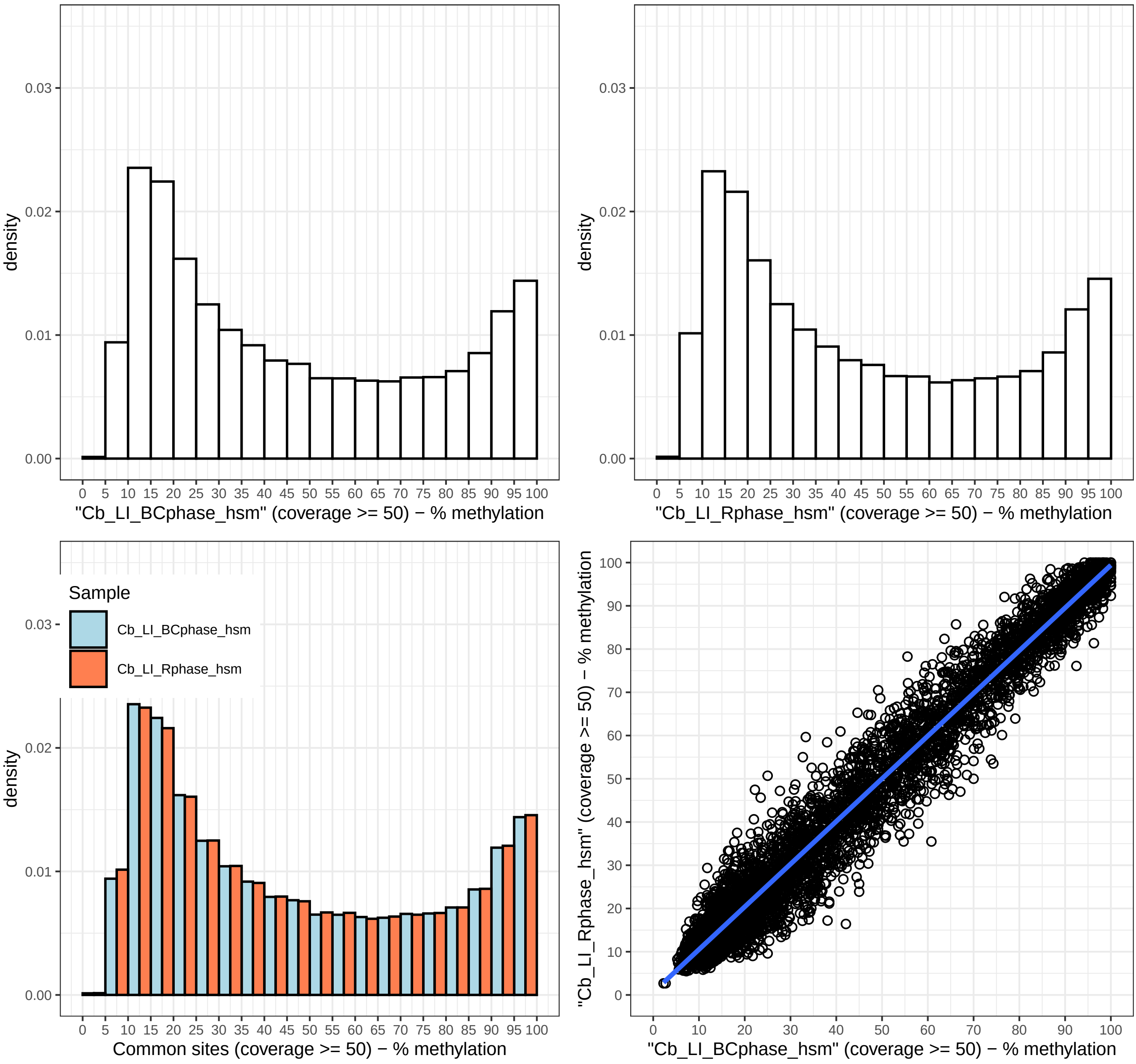

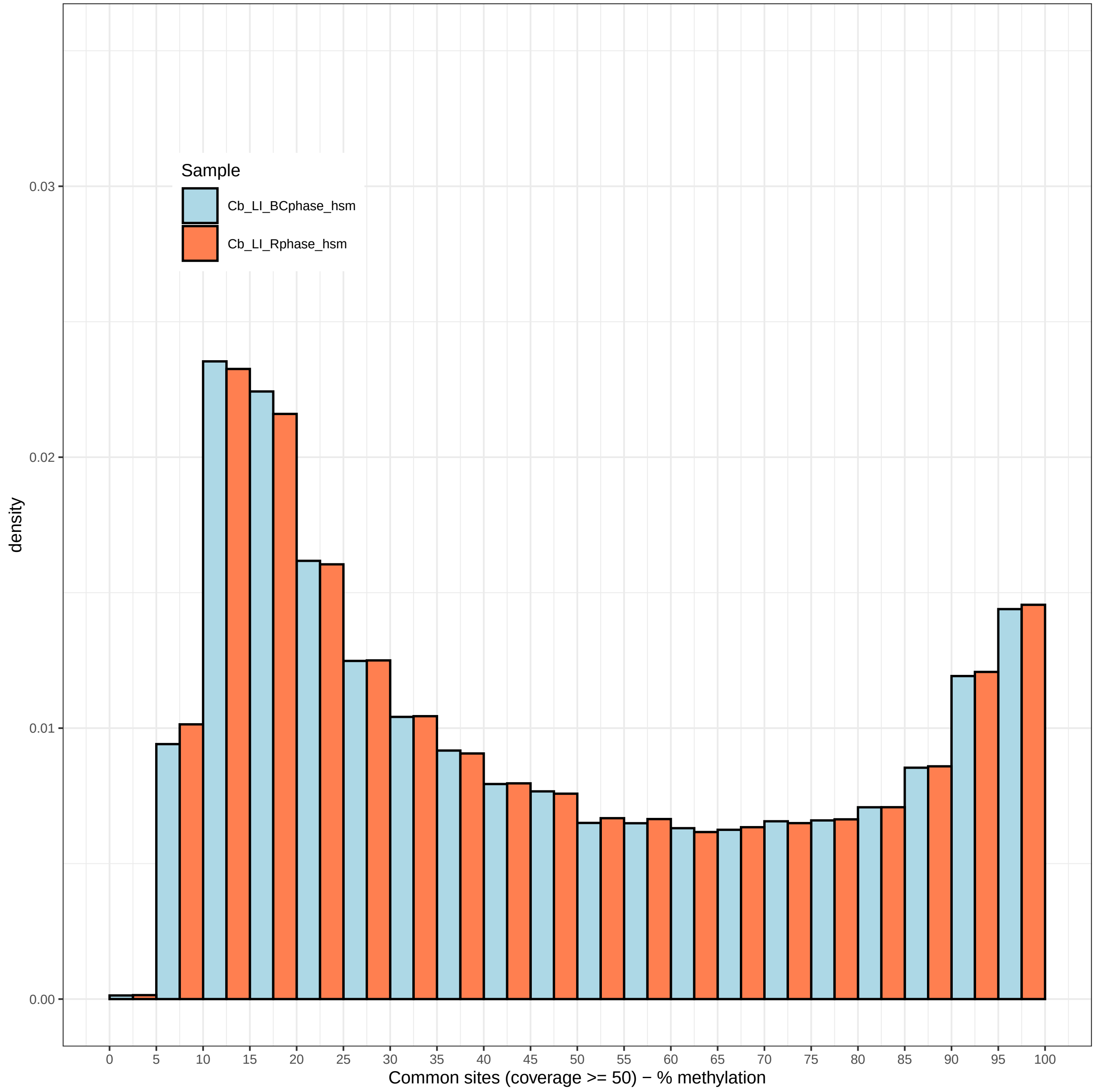

Sample
Cb_LI_BCphase_hsm
Cb_LI_Rphase_hsm
density
0.00
0.01
0.02
0.03
0
5
10
15
20
25
30
35
40
45
50
55
60
65
70
75
80
85
90
95
100
Common sites (coverage >= 50) − % methylation

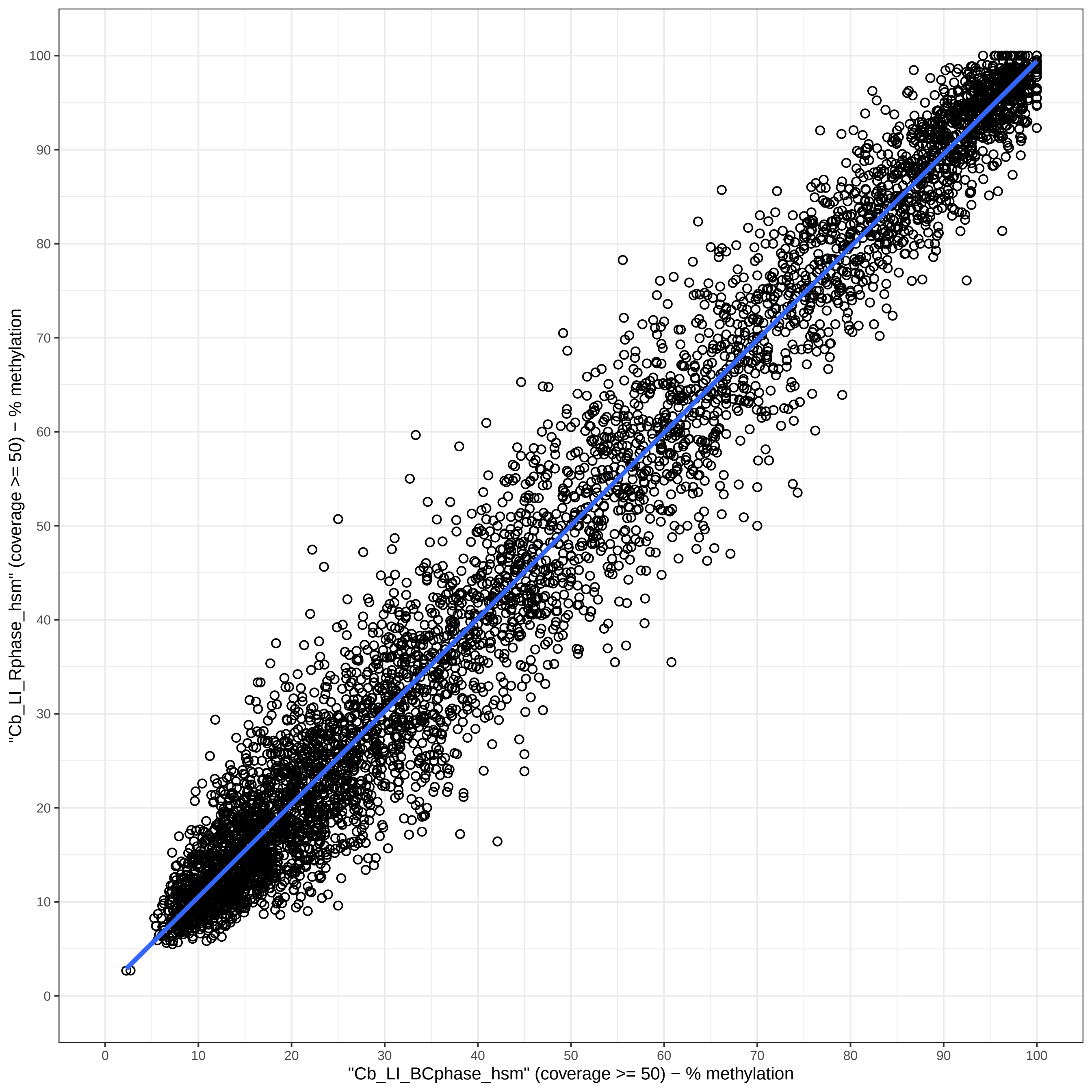

"Cb_LI_Rphase_hsm" (coverage >= 50) − % methylation
"Cb_LI_BCphase_hsm" (coverage >= 50) − % methylation
0
10
20
30
40
50
60
70
80
90
100

**Overlap of highly supported methylation sites (coverage >= 50)**

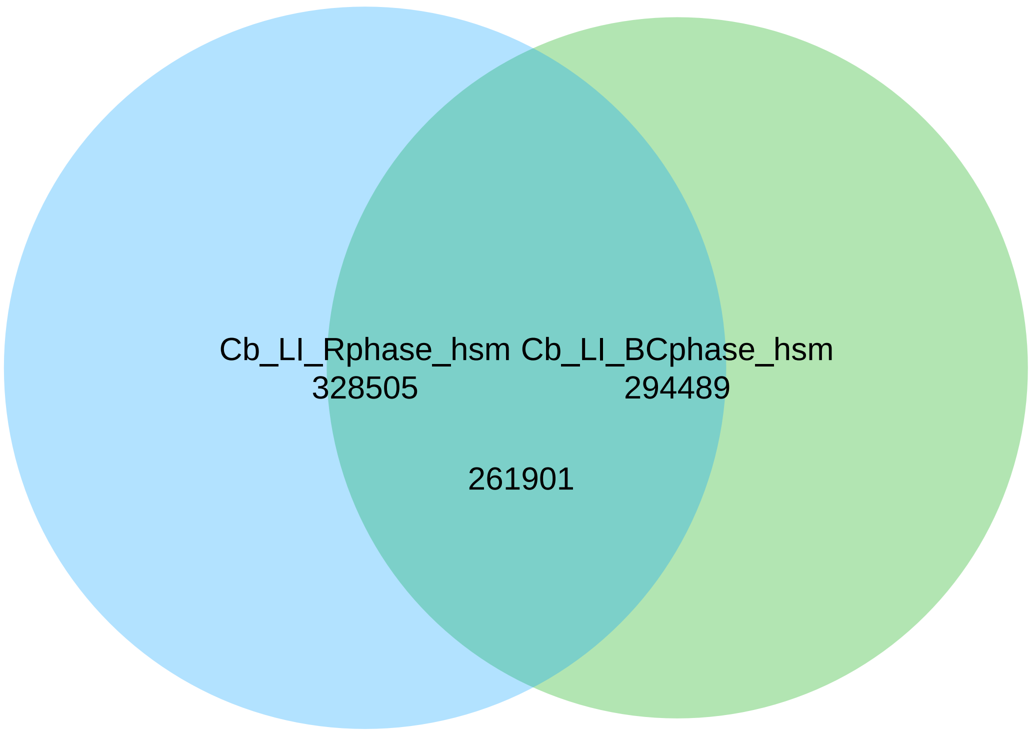

```
Directory: CRL File: 0READMEcrl

CRL - Correlations between aggregate samples.

Input: studymk

Output: files crl-*.txt crl-*.pdf

Notes: The output is generated by BWASPR::cmpSamples(), which uses methylKit::unite()
        to determine the sites common to all samples and applies methylKit::getCorrelation()
        and methylKit::PCASamples().  The *.txt files show the correlations in a table, and
        the *.pdf files show graphics.

       Beware that the number of common sites (shown at the bottom of the *.txt files) may be
        small, which may make the correlations less informative.
```

Directory: CRL File: crl-Cb_LIp.txt

| | BCphase | Rphase |
|---|---|---|
| BCphase | 1.000 | 0.983 |
| Rphase | 0.983 | 1.000 |

The number of conserved sites is 238539.

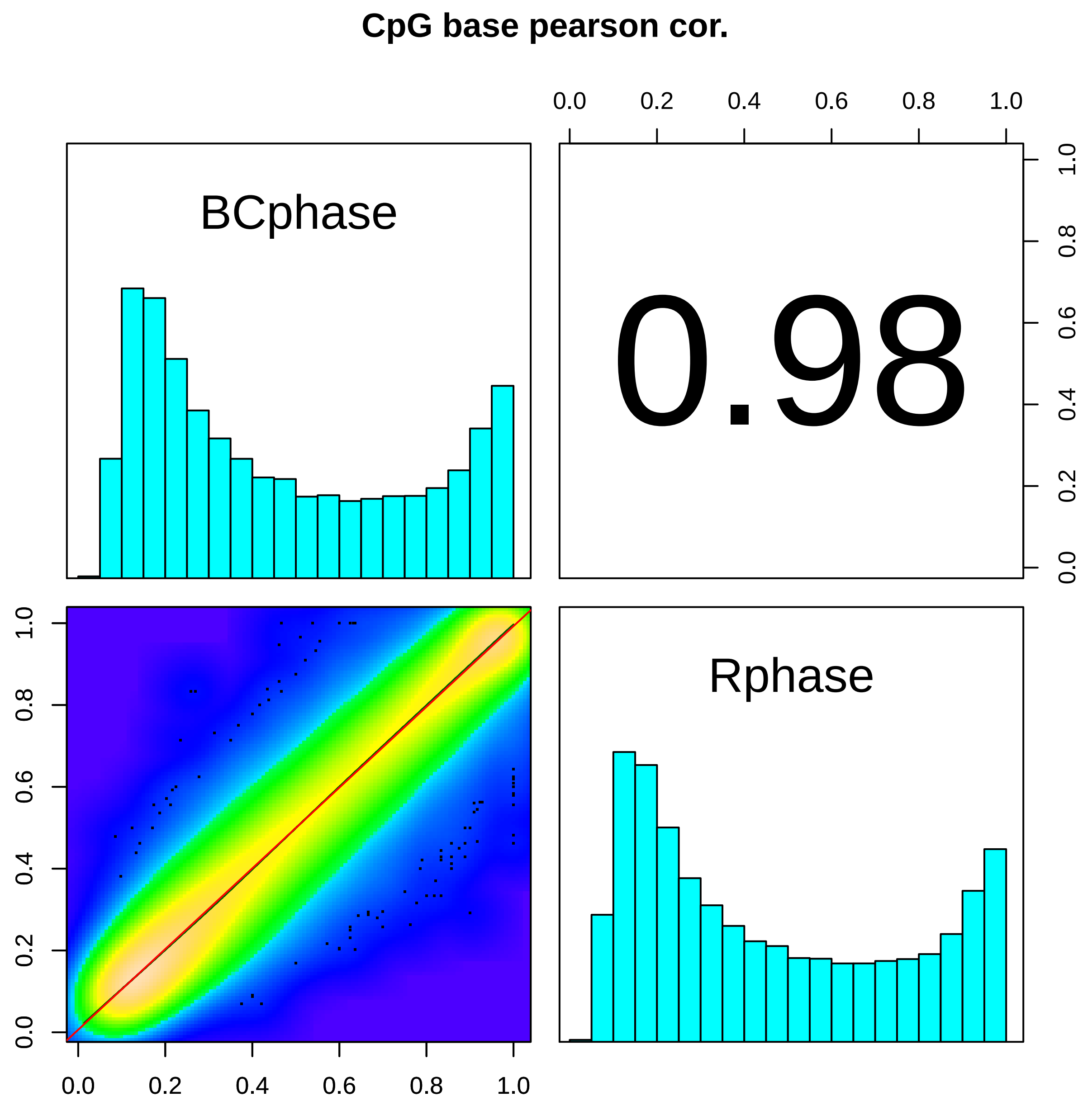

CpG base pearson cor.
BCphase
0.98
Rphase
0.0
0.2
0.4
0.6
0.8
1.0

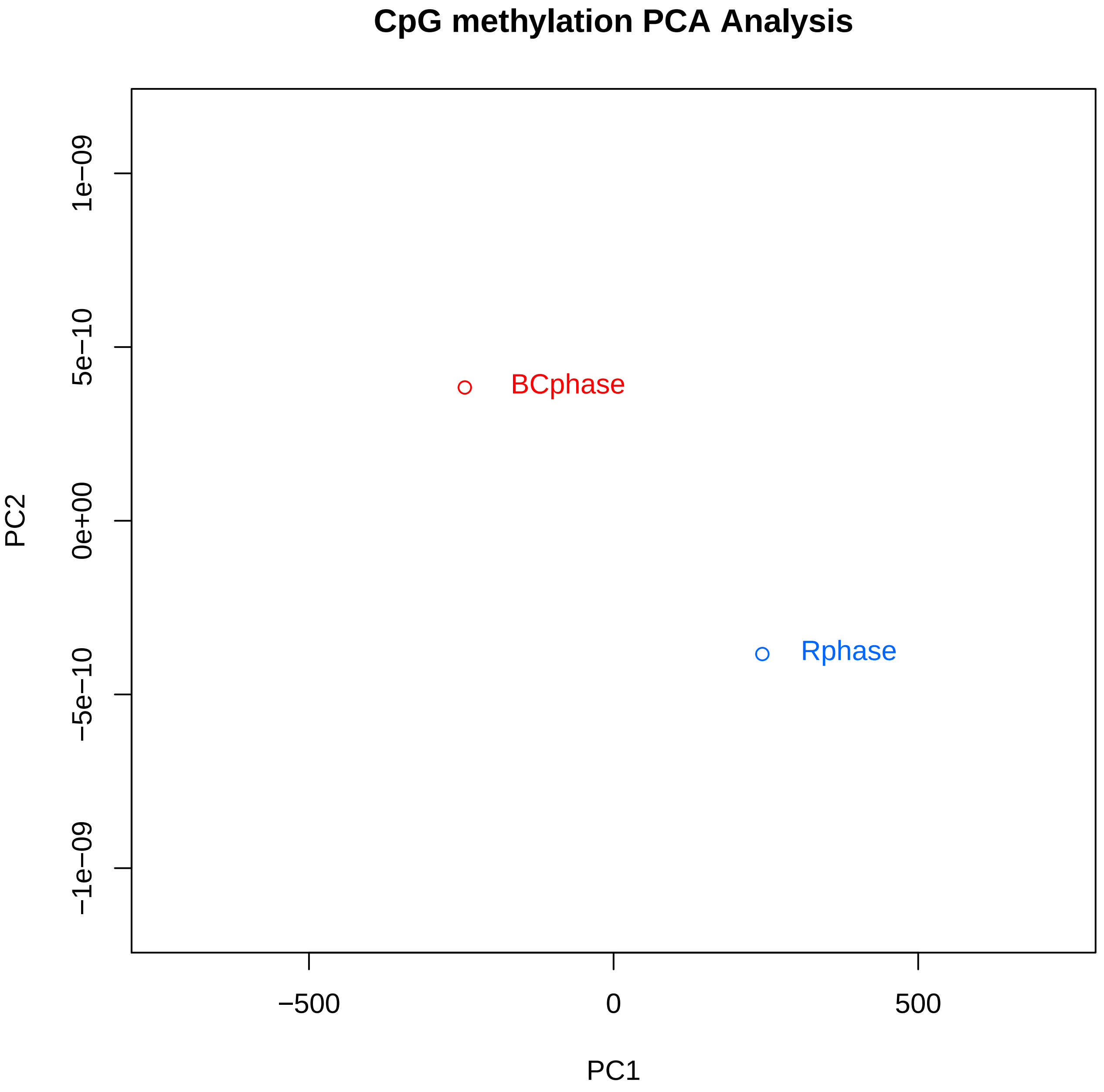
CpG methylation PCA Analysis
BCphase
Rphase
PC1
PC2
−500
0
500
1e−09
5e−10
0e+00
−5e−10
−1e−09

```
Directory: REPCMS File: 0READMErepcms

REPCMS - Coverage and methylation statistics for replicate samples.

Input: studymk, repcovlist, replocount, rephicount

Output: files repcms-*txt and repcms-*pdf

Notes: The statistics are given for CpGhsm sites restricted to minimum or
        higher level coverage.  At the minimum level, the number of sites will be
        the number of lines of the corresponding *CpGhsm.mcalls file minus one
        (consistency check).  The distribution of methylation levels is really only
        interesting for high coverage because at low coverage there will be a strong
        bias towards high methylation levels (by definition of hsm sites).
        The same bias at high coverage would suggest that the sample includes a
        preponderance of consistently methylated genomic sites.

        replocount and rephicount set bounds on the coverage to exclude sites with too few
        or too many covering reads to provide statistics on a typical range.
```

Directory: REPCMS File: repcms-Cb_LIp_BCphase_1.txt

Number of "BCphase_1" CpGhsm-sites with minimal and higher level coverage:

```
  number of "BCphase_1" CpGhsm-sites with coverage >=  5: 198408
  number of "BCphase_1" CpGhsm-sites with coverage >=  6: 195355
  number of "BCphase_1" CpGhsm-sites with coverage >= 10: 172578
  number of "BCphase_1" CpGhsm-sites with coverage >= 15: 143541
```

Coverage and methylation statistics for "BCphase_1" CpGhsm-sites at different levels of minimum coverage:

```
methylKit::getCoverageStats output for "BCphase_1" CpGhsm-sites (#: 198408) at minimum coverage 5    - read coverage statistics per base
summary:
   Min. 1st Qu.  Median    Mean 3rd Qu.    Max.
      5      14      22      25      33    1266
percentiles:
   0%   10%   20%   30%   40%   50%   60%   70%   80%   90%   95%   99% 99.5% 99.9%  100%
    5     9    12    15    19    22    26    31    36    44    50    64    69    84  1266

methylKit::getMethylationStats output for "BCphase_1" CpGhsm-sites (#: 198408) at minimum coverage 5 - methylation statistics per base
summary:
   Min. 1st Qu.  Median    Mean 3rd Qu.    Max.
    2.2    46.7    71.4    67.8    90.5   100.0
percentiles:
    0%    10%    20%    30%    40%    50%    60%    70%    80%    90%    95%    99%  99.5%  99.9%   100%
  2.21  31.71  41.40  52.00  61.54  71.43  80.00  87.50  93.55 100.00 100.00 100.00 100.00 100.00 100.00

methylKit::getCoverageStats output for "BCphase_1" CpGhsm-sites (#: 195355) at minimum coverage 6    - read coverage statistics per base
summary:
   Min. 1st Qu.  Median    Mean 3rd Qu.    Max.
      6      14      23      25      33    1266
percentiles:
   0%   10%   20%   30%   40%   50%   60%   70%   80%   90%   95%   99% 99.5% 99.9%  100%
    6     9    12    16    19    23    27    31    36    44    51    64    69    84  1266

methylKit::getMethylationStats output for "BCphase_1" CpGhsm-sites (#: 195355) at minimum coverage 6 - methylation statistics per base
summary:
   Min. 1st Qu.  Median    Mean 3rd Qu.    Max.
    2.2    46.2    70.4    67.3    90.0   100.0
percentiles:
    0%    10%    20%    30%    40%    50%    60%    70%    80%    90%    95%    99%  99.5%  99.9%   100%
  2.21  31.58  41.18  51.22  60.96  70.37  79.31  86.67  92.86 100.00 100.00 100.00 100.00 100.00 100.00

methylKit::getCoverageStats output for "BCphase_1" CpGhsm-sites (#: 172578) at minimum coverage 10   - read coverage statistics per base
summary:
   Min. 1st Qu.  Median    Mean 3rd Qu.    Max.
     10      17      25      27      35    1266
percentiles:
   0%   10%   20%   30%   40%   50%   60%   70%   80%   90%   95%   99% 99.5% 99.9%  100%
   10    13    15    18    21    25    29    33    38    45    52    65    70    85  1266

methylKit::getMethylationStats output for "BCphase_1" CpGhsm-sites (#: 172578) at minimum coverage 10      - methylation statistics per base
summary:
   Min. 1st Qu.  Median    Mean 3rd Qu.    Max.
    2.2    43.1    66.7    65.0    88.6   100.0
percentiles:
    0%    10%    20%    30%    40%    50%    60%    70%    80%    90%    95%    99%  99.5%  99.9%   100%
```

```
  2.21  30.43  38.89  47.50  57.14  66.67  75.86  84.62  91.67  96.97 100.00 100.00 100.00 100.00 100.00

methylKit::getCoverageStats output for "BCphase_1" CpGhsm-sites (#: 143541) at minimum coverage 15  - read coverage statistics per base
summary:
   Min. 1st Qu.  Median    Mean 3rd Qu.    Max. 
     15      21      28      31      37    1266 
percentiles:
   0%   10%   20%   30%   40%   50%   60%   70%   80%   90%   95%   99% 99.5% 99.9%  100% 
   15    17    19    22    25    28    31    35    40    47    53    66    72    88  1266 

methylKit::getMethylationStats output for "BCphase_1" CpGhsm-sites (#: 143541) at minimum coverage 15     - methylation statistics per base
summary:
   Min. 1st Qu.  Median    Mean 3rd Qu.    Max. 
    2.2    40.0    62.2    62.4    86.7   100.0 
percentiles:
    0%    10%    20%    30%    40%    50%    60%    70%    80%    90%    95%    99%  99.5%  99.9%   100% 
  2.21  28.57  36.00  43.59  52.38  62.16  72.55  82.35  90.32  95.92 100.00 100.00 100.00 100.00 100.00

methylKit::getCoverageStats output for "BCphase_1" CpGhsm-sites (#: 172489) in coverage range [10-100]    - read coverage statistics per base
summary:
   Min. 1st Qu.  Median    Mean 3rd Qu.    Max. 
   10.0    17.0    25.0    27.2    35.0   100.0 
percentiles:
   0%   10%   20%   30%   40%   50%   60%   70%   80%   90%   95%   99% 99.5% 99.9%  100% 
   10    13    15    18    21    25    29    33    38    45    52    64    69    81   100 

methylKit::getMethylationStats output for "BCphase_1" CpGhsm-sites (#: 172489) in coverage range [10-100] - methylation statistics per base
summary:
   Min. 1st Qu.  Median    Mean 3rd Qu.    Max. 
   10.7    43.2    66.7    65.0    88.6   100.0 
percentiles:
   0%   10%   20%   30%   40%   50%   60%   70%   80%   90%   95%   99% 99.5% 99.9%  100% 
 10.7  30.4  38.9  47.6  57.1  66.7  75.9  84.6  91.7  97.0 100.0 100.0 100.0 100.0 100.0 
```

```
Directory: REPCMS File: repcms-Cb_LIp_BCphase_2.txt

Number of "BCphase_2" CpGhsm-sites with minimal and higher level coverage:

  number of "BCphase_2" CpGhsm-sites with coverage >=  5: 213448
  number of "BCphase_2" CpGhsm-sites with coverage >=  6: 210593
  number of "BCphase_2" CpGhsm-sites with coverage >= 10: 190394
  number of "BCphase_2" CpGhsm-sites with coverage >= 15: 164338

Coverage and methylation statistics for "BCphase_2" CpGhsm-sites at different levels of minimum coverage:

methylKit::getCoverageStats output for "BCphase_2" CpGhsm-sites (#: 213448) at minimum coverage 5    - read coverage statistics per base
summary:
   Min. 1st Qu.  Median    Mean 3rd Qu.    Max.
      5      15      27      32      43    1272
percentiles:
   0%   10%   20%   30%   40%   50%   60%   70%   80%   90%   95%   99% 99.5% 99.9%  100%
    5     9    13    17    22    27    32    39    48    61    72    96   106   131  1272

methylKit::getMethylationStats output for "BCphase_2" CpGhsm-sites (#: 213448) at minimum coverage 5 - methylation statistics per base
summary:
   Min. 1st Qu.  Median    Mean 3rd Qu.    Max.
    2.5    42.9    68.8    65.6    90.0   100.0
percentiles:
    0%    10%    20%    30%    40%    50%    60%    70%    80%    90%    95%    99%  99.5%  99.9%   100%
  2.52  27.59  37.50  47.83  58.33  68.75  78.26  86.49  92.86 100.00 100.00 100.00 100.00 100.00 100.00

methylKit::getCoverageStats output for "BCphase_2" CpGhsm-sites (#: 210593) at minimum coverage 6    - read coverage statistics per base
summary:
   Min. 1st Qu.  Median    Mean 3rd Qu.    Max.
      6      16      27      32      43    1272
percentiles:
   0%   10%   20%   30%   40%   50%   60%   70%   80%   90%   95%   99% 99.5% 99.9%  100%
    6    10    14    18    22    27    33    39    48    61    72    96   106   131  1272

methylKit::getMethylationStats output for "BCphase_2" CpGhsm-sites (#: 210593) at minimum coverage 6 - methylation statistics per base
summary:
   Min. 1st Qu.  Median    Mean 3rd Qu.    Max.
    2.5    42.4    67.9    65.1    89.2   100.0
percentiles:
    0%    10%    20%    30%    40%    50%    60%    70%    80%    90%    95%    99%  99.5%  99.9%   100%
  2.52  27.50  37.50  47.37  57.89  67.92  77.78  85.71  92.31 100.00 100.00 100.00 100.00 100.00 100.00

methylKit::getCoverageStats output for "BCphase_2" CpGhsm-sites (#: 190394) at minimum coverage 10   - read coverage statistics per base
summary:
   Min. 1st Qu.  Median    Mean 3rd Qu.    Max.
     10      19      30      35      46    1272
percentiles:
   0%   10%   20%   30%   40%   50%   60%   70%   80%   90%   95%   99% 99.5% 99.9%  100%
   10    13    17    20    25    30    35    42    50    63    74    98   108   133  1272

methylKit::getMethylationStats output for "BCphase_2" CpGhsm-sites (#: 190394) at minimum coverage 10      - methylation statistics per base
summary:
   Min. 1st Qu.  Median    Mean 3rd Qu.    Max.
    2.5    40.0    63.9    62.9    88.0   100.0
percentiles:
    0%    10%    20%    30%    40%    50%    60%    70%    80%    90%    95%    99%  99.5%  99.9%   100%
```

```
  2.52  26.47  35.29  44.44  54.12  63.89  74.19  83.33  91.30  96.61 100.00 100.00 100.00 100.00 100.00

methylKit::getCoverageStats output for "BCphase_2" CpGhsm-sites (#: 164338) at minimum coverage 15  - read coverage statistics per base
summary:
   Min. 1st Qu.  Median    Mean 3rd Qu.    Max.
     15      23      33      38      49    1272
percentiles:
   0%   10%   20%   30%   40%   50%   60%   70%   80%   90%   95%   99% 99.5% 99.9%  100%
   15    18    21    25    29    33    38    45    53    65    76   100   110   135  1272

methylKit::getMethylationStats output for "BCphase_2" CpGhsm-sites (#: 164338) at minimum coverage 15     - methylation statistics per base
summary:
   Min. 1st Qu.  Median    Mean 3rd Qu.    Max.
    2.5    36.8    60.0    60.6    86.4   100.0
percentiles:
    0%    10%    20%    30%    40%    50%    60%    70%    80%    90%    95%    99%  99.5%  99.9%   100%
  2.52  25.00  33.33  40.74  50.00  60.00  70.97  81.36  90.00  95.83 100.00 100.00 100.00 100.00 100.00

methylKit::getCoverageStats output for "BCphase_2" CpGhsm-sites (#: 188845) in coverage range [10-100]    - read coverage statistics per base
summary:
   Min. 1st Qu.  Median    Mean 3rd Qu.    Max.
   10.0    18.0    29.0    33.8    45.0   100.0
percentiles:
   0%   10%   20%   30%   40%   50%   60%   70%   80%   90%   95%   99% 99.5% 99.9%  100%
   10    13    17    20    25    29    35    41    49    62    72    89    93    99   100

methylKit::getMethylationStats output for "BCphase_2" CpGhsm-sites (#: 188845) in coverage range [10-100] - methylation statistics per base
summary:
   Min. 1st Qu.  Median    Mean 3rd Qu.    Max.
     10      40      64      63      88     100
percentiles:
   0%   10%   20%   30%   40%   50%   60%   70%   80%   90%   95%   99% 99.5% 99.9%  100%
 10.0  26.5  35.4  44.4  54.2  64.0  74.3  83.3  91.3  96.7 100.0 100.0 100.0 100.0 100.0
```

```
Directory: REPCMS File: repcms-Cb_LIp_BCphase_3.txt

Number of "BCphase_3" CpGhsm-sites with minimal and higher level coverage:

  number of "BCphase_3" CpGhsm-sites with coverage >=  5: 226815
  number of "BCphase_3" CpGhsm-sites with coverage >=  6: 224272
  number of "BCphase_3" CpGhsm-sites with coverage >= 10: 205046
  number of "BCphase_3" CpGhsm-sites with coverage >= 15: 178656

Coverage and methylation statistics for "BCphase_3" CpGhsm-sites at different levels of minimum coverage:

methylKit::getCoverageStats output for "BCphase_3" CpGhsm-sites (#: 226815) at minimum coverage 5    - read coverage statistics per base
summary:
   Min. 1st Qu.  Median    Mean 3rd Qu.    Max.
      5      16      27      30      40    1524
percentiles:
   0%   10%   20%   30%   40%   50%   60%   70%   80%   90%   95%   99% 99.5% 99.9%  100%
    5    10    14    18    22    27    32    37    44    53    61    77    83   102  1524

methylKit::getMethylationStats output for "BCphase_3" CpGhsm-sites (#: 226815) at minimum coverage 5 - methylation statistics per base
summary:
   Min. 1st Qu.  Median    Mean 3rd Qu.    Max.
    2.4    41.9    66.7    65.0    89.3   100.0
percentiles:
    0%    10%    20%    30%    40%    50%    60%    70%    80%    90%    95%    99%  99.5%  99.9%   100%
  2.37  27.87  37.18  46.67  57.14  66.67  77.19  85.71  92.50 100.00 100.00 100.00 100.00 100.00 100.00

methylKit::getCoverageStats output for "BCphase_3" CpGhsm-sites (#: 224272) at minimum coverage 6    - read coverage statistics per base
summary:
   Min. 1st Qu.  Median    Mean 3rd Qu.    Max.
      6      16      27      30      40    1524
percentiles:
   0%   10%   20%   30%   40%   50%   60%   70%   80%   90%   95%   99% 99.5% 99.9%  100%
    6    10    14    18    23    27    32    37    44    53    61    77    83   102  1524

methylKit::getMethylationStats output for "BCphase_3" CpGhsm-sites (#: 224272) at minimum coverage 6 - methylation statistics per base
summary:
   Min. 1st Qu.  Median    Mean 3rd Qu.    Max.
    2.4    41.7    66.7    64.6    88.9   100.0
percentiles:
    0%    10%    20%    30%    40%    50%    60%    70%    80%    90%    95%    99%  99.5%  99.9%   100%
  2.37  27.78  36.96  46.67  56.41  66.67  76.47  85.37  91.94 100.00 100.00 100.00 100.00 100.00 100.00

methylKit::getCoverageStats output for "BCphase_3" CpGhsm-sites (#: 205046) at minimum coverage 10   - read coverage statistics per base
summary:
   Min. 1st Qu.  Median    Mean 3rd Qu.    Max.
     10      19      29      32      42    1524
percentiles:
   0%   10%   20%   30%   40%   50%   60%   70%   80%   90%   95%   99% 99.5% 99.9%  100%
   10    13    17    21    25    29    34    39    45    54    62    78    84   103  1524

methylKit::getMethylationStats output for "BCphase_3" CpGhsm-sites (#: 205046) at minimum coverage 10      - methylation statistics per base
summary:
   Min. 1st Qu.  Median    Mean 3rd Qu.    Max.
    2.4    39.6    63.2    62.6    87.5   100.0
percentiles:
    0%    10%    20%    30%    40%    50%    60%    70%    80%    90%    95%    99%  99.5%  99.9%   100%
```

```
  2.37  26.92  35.29  43.75  53.33  63.24  73.33  83.33  90.91  96.43 100.00 100.00 100.00 100.00 100.00

methylKit::getCoverageStats output for "BCphase_3" CpGhsm-sites (#: 178656) at minimum coverage 15  - read coverage statistics per base
summary:
   Min. 1st Qu.  Median    Mean 3rd Qu.    Max.
     15      23      32      35      44    1524
percentiles:
   0%   10%   20%   30%   40%   50%   60%   70%   80%   90%   95%   99% 99.5% 99.9%  100%
   15    18    21    25    28    32    36    41    47    56    63    79    86   105  1524

methylKit::getMethylationStats output for "BCphase_3" CpGhsm-sites (#: 178656) at minimum coverage 15     - methylation statistics per base
summary:
   Min. 1st Qu.  Median    Mean 3rd Qu.    Max.
    2.4    36.8    59.5    60.5    85.7   100.0
percentiles:
    0%    10%    20%    30%    40%    50%    60%    70%    80%    90%    95%    99%  99.5%  99.9%   100%
  2.37  25.71  33.33  40.54  50.00  59.46  70.59  81.08  89.74  95.65 100.00 100.00 100.00 100.00 100.00

methylKit::getCoverageStats output for "BCphase_3" CpGhsm-sites (#: 204793) in coverage range [10-100]    - read coverage statistics per base
summary:
   Min. 1st Qu.  Median    Mean 3rd Qu.    Max.
   10.0    19.0    29.0    31.8    42.0   100.0
percentiles:
   0%   10%   20%   30%   40%   50%   60%   70%   80%   90%   95%   99% 99.5% 99.9%  100%
   10    13    17    21    25    29    34    39    45    54    61    77    82    94   100

methylKit::getMethylationStats output for "BCphase_3" CpGhsm-sites (#: 204793) in coverage range [10-100] - methylation statistics per base
summary:
   Min. 1st Qu.  Median    Mean 3rd Qu.    Max.
   10.0    40.0    63.3    62.7    87.5   100.0
percentiles:
   0%   10%   20%   30%   40%   50%   60%   70%   80%   90%   95%   99% 99.5% 99.9%  100%
 10.0  26.9  35.3  43.9  53.3  63.3  73.3  83.3  90.9  96.4 100.0 100.0 100.0 100.0 100.0
```

```
Directory: REPCMS File: repcms-Cb_LIp_BCphase_4.txt

Number of "BCphase_4" CpGhsm-sites with minimal and higher level coverage:

  number of "BCphase_4" CpGhsm-sites with coverage >=  5: 212569
  number of "BCphase_4" CpGhsm-sites with coverage >=  6: 209742
  number of "BCphase_4" CpGhsm-sites with coverage >= 10: 190104
  number of "BCphase_4" CpGhsm-sites with coverage >= 15: 164033

Coverage and methylation statistics for "BCphase_4" CpGhsm-sites at different levels of minimum coverage:

methylKit::getCoverageStats output for "BCphase_4" CpGhsm-sites (#: 212569) at minimum coverage 5    - read coverage statistics per base
summary:
   Min. 1st Qu.  Median    Mean 3rd Qu.    Max.
      5      15      26      30      41    1287
percentiles:
   0%   10%   20%   30%   40%   50%   60%   70%   80%   90%   95%   99% 99.5% 99.9%  100%
    5     9    13    17    21    26    31    37    45    56    66    86    94   115  1287

methylKit::getMethylationStats output for "BCphase_4" CpGhsm-sites (#: 212569) at minimum coverage 5 - methylation statistics per base
summary:
   Min. 1st Qu.  Median    Mean 3rd Qu.    Max.
    2.4    42.9    68.4    65.6    89.7   100.0
percentiles:
    0%    10%    20%    30%    40%    50%    60%    70%    80%    90%    95%    99%  99.5%  99.9%   100%
  2.37  28.21  37.93  48.00  58.33  68.42  77.78  86.11  92.86 100.00 100.00 100.00 100.00 100.00 100.00

methylKit::getCoverageStats output for "BCphase_4" CpGhsm-sites (#: 209742) at minimum coverage 6    - read coverage statistics per base
summary:
   Min. 1st Qu.  Median    Mean 3rd Qu.    Max.
      6      16      27      30      41    1287
percentiles:
   0%   10%   20%   30%   40%   50%   60%   70%   80%   90%   95%   99% 99.5% 99.9%  100%
    6    10    14    18    22    27    32    38    45    56    66    86    94   115  1287

methylKit::getMethylationStats output for "BCphase_4" CpGhsm-sites (#: 209742) at minimum coverage 6 - methylation statistics per base
summary:
   Min. 1st Qu.  Median    Mean 3rd Qu.    Max.
    2.4    42.9    67.6    65.2    88.9   100.0
percentiles:
    0%    10%    20%    30%    40%    50%    60%    70%    80%    90%    95%    99%  99.5%  99.9%   100%
  2.37  28.07  37.50  47.62  57.89  67.65  77.27  85.71  92.11 100.00 100.00 100.00 100.00 100.00 100.00

methylKit::getCoverageStats output for "BCphase_4" CpGhsm-sites (#: 190104) at minimum coverage 10   - read coverage statistics per base
summary:
   Min. 1st Qu.  Median    Mean 3rd Qu.    Max.
     10      18      29      33      43    1287
percentiles:
   0%   10%   20%   30%   40%   50%   60%   70%   80%   90%   95%   99% 99.5% 99.9%  100%
   10    13    17    20    24    29    34    39    47    58    67    87    96   116  1287

methylKit::getMethylationStats output for "BCphase_4" CpGhsm-sites (#: 190104) at minimum coverage 10      - methylation statistics per base
summary:
   Min. 1st Qu.  Median    Mean 3rd Qu.    Max.
    2.4    40.0    63.8    63.1    87.5   100.0
percentiles:
    0%    10%    20%    30%    40%    50%    60%    70%    80%    90%    95%    99%  99.5%  99.9%   100%
```

```
  2.37  27.03  36.00  44.83  54.29  63.83  73.91  83.33  90.91  96.43 100.00 100.00 100.00 100.00 100.00

methylKit::getCoverageStats output for "BCphase_4" CpGhsm-sites (#: 164033) at minimum coverage 15   - read coverage statistics per base
summary:
   Min. 1st Qu.  Median    Mean 3rd Qu.    Max. 
     15      22      32      36      46    1287 
percentiles:
   0%   10%   20%   30%   40%   50%   60%   70%   80%   90%   95%   99% 99.5% 99.9%  100% 
   15    17    21    24    28    32    37    42    49    60    69    89    97   118  1287 

methylKit::getMethylationStats output for "BCphase_4" CpGhsm-sites (#: 164033) at minimum coverage 15      - methylation statistics per base
summary:
   Min. 1st Qu.  Median    Mean 3rd Qu.    Max. 
    2.4    37.5    60.0    60.8    86.2   100.0 
percentiles:
    0%    10%    20%    30%    40%    50%    60%    70%    80%    90%    95%    99%  99.5%  99.9%   100% 
  2.37  25.81  33.33  41.18  50.00  60.00  70.83  81.25  89.80  95.65 100.00 100.00 100.00 100.00 100.00

methylKit::getCoverageStats output for "BCphase_4" CpGhsm-sites (#: 189480) in coverage range [10-100]    - read coverage statistics per base
summary:
   Min. 1st Qu.  Median    Mean 3rd Qu.    Max. 
   10.0    18.0    29.0    32.5    43.0   100.0 
percentiles:
   0%   10%   20%   30%   40%   50%   60%   70%   80%   90%   95%   99% 99.5% 99.9%  100% 
   10    13    17    20    24    29    34    39    47    57    67    84    89    97   100 

methylKit::getMethylationStats output for "BCphase_4" CpGhsm-sites (#: 189480) in coverage range [10-100] - methylation statistics per base
summary:
   Min. 1st Qu.  Median    Mean 3rd Qu.    Max. 
   10.0    40.0    63.9    63.1    87.5   100.0 
percentiles:
   0%   10%   20%   30%   40%   50%   60%   70%   80%   90%   95%   99% 99.5% 99.9%  100% 
 10.0  27.1  36.0  44.9  54.3  63.9  73.9  83.3  90.9  96.5 100.0 100.0 100.0 100.0 100.0 
```

```
Directory: REPCMS File: repcms-Cb_LIp_Rphase_1.txt

Number of "Rphase_1" CpGhsm-sites with minimal and higher level coverage:

  number of "Rphase_1" CpGhsm-sites with coverage >=  5: 213610
  number of "Rphase_1" CpGhsm-sites with coverage >=  6: 210877
  number of "Rphase_1" CpGhsm-sites with coverage >= 10: 190451
  number of "Rphase_1" CpGhsm-sites with coverage >= 15: 162696

Coverage and methylation statistics for "Rphase_1" CpGhsm-sites at different levels of minimum coverage:

methylKit::getCoverageStats output for "Rphase_1" CpGhsm-sites (#: 213610) at minimum coverage 5    - read coverage statistics per base
summary:
   Min. 1st Qu.  Median    Mean 3rd Qu.    Max.
      5      15      25      27      37    1360
percentiles:
   0%   10%   20%   30%   40%   50%   60%   70%   80%   90%   95%   99% 99.5% 99.9%  100%
    5     9    13    17    21    25    29    34    40    48    54    69    74    91  1360

methylKit::getMethylationStats output for "Rphase_1" CpGhsm-sites (#: 213610) at minimum coverage 5  - methylation statistics per base
summary:
   Min. 1st Qu.  Median    Mean 3rd Qu.    Max.
    2.5    44.4    69.6    66.5    90.0   100.0
percentiles:
    0%    10%    20%    30%    40%    50%    60%    70%    80%    90%    95%    99%  99.5%  99.9%   100%
  2.48  29.73  39.39  50.00  59.52  69.57  78.79  86.84  93.10 100.00 100.00 100.00 100.00 100.00 100.00

methylKit::getCoverageStats output for "Rphase_1" CpGhsm-sites (#: 210877) at minimum coverage 6    - read coverage statistics per base
summary:
   Min. 1st Qu.  Median    Mean 3rd Qu.    Max.
      6      15      25      27      37    1360
percentiles:
   0%   10%   20%   30%   40%   50%   60%   70%   80%   90%   95%   99% 99.5% 99.9%  100%
    6    10    13    17    21    25    30    34    40    48    55    69    74    91  1360

methylKit::getMethylationStats output for "Rphase_1" CpGhsm-sites (#: 210877) at minimum coverage 6  - methylation statistics per base
summary:
   Min. 1st Qu.  Median    Mean 3rd Qu.    Max.
    2.5    43.8    68.8    66.1    89.5   100.0
percentiles:
    0%    10%    20%    30%    40%    50%    60%    70%    80%    90%    95%    99%  99.5%  99.9%   100%
  2.48  29.63  39.13  50.00  58.82  68.75  77.78  86.05  92.59 100.00 100.00 100.00 100.00 100.00 100.00

methylKit::getCoverageStats output for "Rphase_1" CpGhsm-sites (#: 190451) at minimum coverage 10   - read coverage statistics per base
summary:
   Min. 1st Qu.  Median    Mean 3rd Qu.    Max.
     10      18      27      30      38    1360
percentiles:
   0%   10%   20%   30%   40%   50%   60%   70%   80%   90%   95%   99% 99.5% 99.9%  100%
   10    13    16    20    23    27    31    36    41    49    55    70    75    92  1360

methylKit::getMethylationStats output for "Rphase_1" CpGhsm-sites (#: 190451) at minimum coverage 10 - methylation statistics per base
summary:
   Min. 1st Qu.  Median    Mean 3rd Qu.    Max.
    2.5    41.7    65.0    64.0    88.2   100.0
percentiles:
    0%    10%    20%    30%    40%    50%    60%    70%    80%    90%    95%    99%  99.5%  99.9%   100%
```

```
  2.48  28.57  37.50  46.15  55.17  65.00  75.00  84.21  91.67  96.88 100.00 100.00 100.00 100.00 100.00

methylKit::getCoverageStats output for "Rphase_1" CpGhsm-sites (#: 162696) at minimum coverage 15   - read coverage statistics per base
summary:
   Min. 1st Qu.  Median    Mean 3rd Qu.    Max.
     15      22      30      33      41    1360
percentiles:
   0%   10%   20%   30%   40%   50%   60%   70%   80%   90%   95%   99% 99.5% 99.9%  100%
   15    17    20    23    27    30    34    38    43    51    57    71    77    94  1360

methylKit::getMethylationStats output for "Rphase_1" CpGhsm-sites (#: 162696) at minimum coverage 15 - methylation statistics per base
summary:
   Min. 1st Qu.  Median    Mean 3rd Qu.    Max.
    2.5    38.5    61.1    61.7    86.7   100.0
percentiles:
    0%    10%    20%    30%    40%    50%    60%    70%    80%    90%    95%    99%  99.5%  99.9%   100%
  2.48  27.03  34.78  42.11  51.22  61.11  72.00  82.05  90.24  95.85 100.00 100.00 100.00 100.00 100.00

methylKit::getCoverageStats output for "Rphase_1" CpGhsm-sites (#: 190348) in coverage range [10-100]     - read coverage statistics per base
summary:
   Min. 1st Qu.  Median    Mean 3rd Qu.    Max.
   10.0    18.0    27.0    29.5    38.0   100.0
percentiles:
   0%   10%   20%   30%   40%   50%   60%   70%   80%   90%   95%   99% 99.5% 99.9%  100%
   10    13    16    20    23    27    31    36    41    49    55    69    74    87   100

methylKit::getMethylationStats output for "Rphase_1" CpGhsm-sites (#: 190348) in coverage range [10-100]  - methylation statistics per base
summary:
   Min. 1st Qu.  Median    Mean 3rd Qu.    Max.
   10.4    41.7    65.0    64.0    88.2   100.0
percentiles:
   0%   10%   20%   30%   40%   50%   60%   70%   80%   90%   95%   99% 99.5% 99.9%  100%
 10.4  28.6  37.5  46.2  55.2  65.0  75.0  84.2  91.7  96.9 100.0 100.0 100.0 100.0 100.0
```

Directory: REPCMS File: repcms-Cb_LIp_Rphase_2.txt

Number of "Rphase_2" CpGhsm-sites with minimal and higher level coverage:

```
  number of "Rphase_2" CpGhsm-sites with coverage >=  5: 265952
  number of "Rphase_2" CpGhsm-sites with coverage >=  6: 263685
  number of "Rphase_2" CpGhsm-sites with coverage >= 10: 247358
  number of "Rphase_2" CpGhsm-sites with coverage >= 15: 225302
```

Coverage and methylation statistics for "Rphase_2" CpGhsm-sites at different levels of minimum coverage:

```
methylKit::getCoverageStats output for "Rphase_2" CpGhsm-sites (#: 265952) at minimum coverage 5    - read coverage statistics per base
summary:
   Min. 1st Qu.  Median    Mean 3rd Qu.    Max. 
      5      20      36      42      58    1345 
percentiles:
   0%   10%   20%   30%   40%   50%   60%   70%   80%   90%   95%   99% 99.5% 99.9%  100% 
    5    11    17    23    29    36    44    53    65    83    98   130   143   175  1345 

methylKit::getMethylationStats output for "Rphase_2" CpGhsm-sites (#: 265952) at minimum coverage 5  - methylation statistics per base
summary:
   Min. 1st Qu.  Median    Mean 3rd Qu.    Max. 
    2.7    35.4    62.1    60.9    87.5   100.0 
percentiles:
    0%    10%    20%    30%    40%    50%    60%    70%    80%    90%    95%    99%  99.5%  99.9%   100% 
  2.68  22.45  31.25  40.00  50.67  62.07  73.08  83.33  91.23  97.37 100.00 100.00 100.00 100.00 100.00 
```

```
methylKit::getCoverageStats output for "Rphase_2" CpGhsm-sites (#: 263685) at minimum coverage 6    - read coverage statistics per base
summary:
   Min. 1st Qu.  Median    Mean 3rd Qu.    Max. 
      6      20      36      43      59    1345 
percentiles:
   0%   10%   20%   30%   40%   50%   60%   70%   80%   90%   95%   99% 99.5% 99.9%  100% 
    6    12    18    23    29    36    44    53    65    83    99   130   143   175  1345 

methylKit::getMethylationStats output for "Rphase_2" CpGhsm-sites (#: 263685) at minimum coverage 6  - methylation statistics per base
summary:
   Min. 1st Qu.  Median    Mean 3rd Qu.    Max. 
    2.7    35.3    61.5    60.6    87.2   100.0 
percentiles:
    0%    10%    20%    30%    40%    50%    60%    70%    80%    90%    95%    99%  99.5%  99.9%   100% 
  2.68  22.35  31.03  40.00  50.00  61.54  72.55  83.33  90.91  96.88 100.00 100.00 100.00 100.00 100.00 
```

```
methylKit::getCoverageStats output for "Rphase_2" CpGhsm-sites (#: 247358) at minimum coverage 10   - read coverage statistics per base
summary:
   Min. 1st Qu.  Median    Mean 3rd Qu.    Max. 
     10      23      38      45      60    1345 
percentiles:
   0%   10%   20%   30%   40%   50%   60%   70%   80%   90%   95%   99% 99.5% 99.9%  100% 
   10    15    20    26    32    38    46    55    67    85   100   131   144   177  1345 

methylKit::getMethylationStats output for "Rphase_2" CpGhsm-sites (#: 247358) at minimum coverage 10 - methylation statistics per base
summary:
   Min. 1st Qu.  Median    Mean 3rd Qu.    Max. 
    2.7    33.9    58.3    59.0    85.7   100.0 
percentiles:
    0%    10%    20%    30%    40%    50%    60%    70%    80%    90%    95%    99%  99.5%  99.9%   100% 
```

```
  2.68  21.88  29.85  38.24  47.73  58.33  69.77  80.77  90.00  95.83 100.00 100.00 100.00 100.00 100.00
```

```
methylKit::getCoverageStats output for "Rphase_2" CpGhsm-sites (#: 225302) at minimum coverage 15    - read coverage statistics per base
summary:
   Min. 1st Qu.  Median    Mean 3rd Qu.    Max. 
     15      27      42      48      63    1345 
percentiles:
   0%   10%   20%   30%   40%   50%   60%   70%   80%   90%   95%   99% 99.5% 99.9%  100% 
   15    19    24    30    35    42    49    58    69    87   102   133   146   180  1345 

methylKit::getMethylationStats output for "Rphase_2" CpGhsm-sites (#: 225302) at minimum coverage 15 - methylation statistics per base
summary:
   Min. 1st Qu.  Median    Mean 3rd Qu.    Max. 
    2.7    32.1    55.0    57.1    84.1   100.0 
percentiles:
    0%    10%    20%    30%    40%    50%    60%    70%    80%    90%    95%    99%  99.5%  99.9%   100% 
  2.68  21.21  28.30  35.96  44.44  55.00  66.67  78.79  88.68  95.29  97.96 100.00 100.00 100.00 100.00
```

```
methylKit::getCoverageStats output for "Rphase_2" CpGhsm-sites (#: 235297) in coverage range [10-100]      - read coverage statistics per base
summary:
   Min. 1st Qu.  Median    Mean 3rd Qu.    Max. 
   10.0    22.0    37.0    41.2    56.0   100.0 
percentiles:
   0%   10%   20%   30%   40%   50%   60%   70%   80%   90%   95%   99% 99.5% 99.9%  100% 
   10    15    20    25    31    37    44    52    62    76    85    97    99   100   100 

methylKit::getMethylationStats output for "Rphase_2" CpGhsm-sites (#: 235297) in coverage range [10-100]   - methylation statistics per base
summary:
   Min. 1st Qu.  Median    Mean 3rd Qu.    Max. 
   10.0    34.4    58.8    59.2    85.7   100.0 
percentiles:
   0%   10%   20%   30%   40%   50%   60%   70%   80%   90%   95%   99% 99.5% 99.9%  100% 
 10.0  22.2  30.4  38.7  48.1  58.8  70.0  81.0  90.0  95.8 100.0 100.0 100.0 100.0 100.0
```

```
Directory: REPCMS File: repcms-Cb_LIp_Rphase_3.txt

Number of "Rphase_3" CpGhsm-sites with minimal and higher level coverage:

  number of "Rphase_3" CpGhsm-sites with coverage >=  5: 218593
  number of "Rphase_3" CpGhsm-sites with coverage >=  6: 215760
  number of "Rphase_3" CpGhsm-sites with coverage >= 10: 195508
  number of "Rphase_3" CpGhsm-sites with coverage >= 15: 168399

Coverage and methylation statistics for "Rphase_3" CpGhsm-sites at different levels of minimum coverage:

methylKit::getCoverageStats output for "Rphase_3" CpGhsm-sites (#: 218593) at minimum coverage 5    - read coverage statistics per base
summary:
   Min. 1st Qu.  Median    Mean 3rd Qu.    Max.
      5      15      26      28      38    1983
percentiles:
   0%   10%   20%   30%   40%   50%   60%   70%   80%   90%   95%   99% 99.5% 99.9%  100%
    5     9    13    17    21    26    30    35    41    50    58    73    80    98  1983

methylKit::getMethylationStats output for "Rphase_3" CpGhsm-sites (#: 218593) at minimum coverage 5  - methylation statistics per base
summary:
   Min. 1st Qu.  Median    Mean 3rd Qu.    Max.
    2.0    43.3    68.4    65.9    89.7   100.0
percentiles:
   0%   10%   20%   30%   40%   50%   60%   70%   80%   90%   95%   99% 99.5% 99.9%  100%
  2.0  29.0  38.5  48.4  58.3  68.4  77.8  86.2  92.9 100.0 100.0 100.0 100.0 100.0 100.0

methylKit::getCoverageStats output for "Rphase_3" CpGhsm-sites (#: 215760) at minimum coverage 6    - read coverage statistics per base
summary:
   Min. 1st Qu.  Median    Mean 3rd Qu.    Max.
      6      16      26      29      38    1983
percentiles:
   0%   10%   20%   30%   40%   50%   60%   70%   80%   90%   95%   99% 99.5% 99.9%  100%
    6    10    14    17    21    26    30    35    42    50    58    73    80    98  1983

methylKit::getMethylationStats output for "Rphase_3" CpGhsm-sites (#: 215760) at minimum coverage 6  - methylation statistics per base
summary:
   Min. 1st Qu.  Median    Mean 3rd Qu.    Max.
    2.0    42.9    67.6    65.4    88.9   100.0
percentiles:
   0%   10%   20%   30%   40%   50%   60%   70%   80%   90%   95%   99% 99.5% 99.9%  100%
  2.0  28.9  38.1  47.8  57.9  67.6  77.3  85.7  92.3 100.0 100.0 100.0 100.0 100.0 100.0

methylKit::getCoverageStats output for "Rphase_3" CpGhsm-sites (#: 195508) at minimum coverage 10   - read coverage statistics per base
summary:
   Min. 1st Qu.  Median    Mean 3rd Qu.    Max.
     10      18      28      31      40    1983
percentiles:
   0%   10%   20%   30%   40%   50%   60%   70%   80%   90%   95%   99% 99.5% 99.9%  100%
   10    13    17    20    24    28    32    37    43    51    59    74    81    99  1983

methylKit::getMethylationStats output for "Rphase_3" CpGhsm-sites (#: 195508) at minimum coverage 10 - methylation statistics per base
summary:
   Min. 1st Qu.  Median    Mean 3rd Qu.    Max.
    2.0    40.7    63.9    63.4    87.8   100.0
percentiles:
   0%   10%   20%   30%   40%   50%   60%   70%   80%   90%   95%   99% 99.5% 99.9%  100%
```

```
  2.0  28.0  36.6  45.2  54.5  63.9  74.0  83.3  91.2  96.7 100.0 100.0 100.0 100.0 100.0

methylKit::getCoverageStats output for "Rphase_3" CpGhsm-sites (#: 168399) at minimum coverage 15    - read coverage statistics per base
summary:
   Min. 1st Qu.  Median    Mean 3rd Qu.    Max.
     15      22      31      34      42    1983
percentiles:
   0%   10%   20%   30%   40%   50%   60%   70%   80%   90%   95%   99% 99.5% 99.9%  100%
   15    17    20    24    27    31    35    39    45    53    60    76    82   103  1983

methylKit::getMethylationStats output for "Rphase_3" CpGhsm-sites (#: 168399) at minimum coverage 15 - methylation statistics per base
summary:
   Min. 1st Qu.  Median    Mean 3rd Qu.    Max.
    2.0    37.5    60.0    61.1    86.4   100.0
percentiles:
   0%   10%   20%   30%   40%   50%   60%   70%   80%   90%   95%   99% 99.5% 99.9%  100%
  2.0  26.7  34.3  41.5  50.0  60.0  71.2  81.4  90.0  95.8 100.0 100.0 100.0 100.0 100.0

methylKit::getCoverageStats output for "Rphase_3" CpGhsm-sites (#: 195324) in coverage range [10-100]     - read coverage statistics per base
summary:
   Min. 1st Qu.  Median    Mean 3rd Qu.    Max.
   10.0    18.0    28.0    30.5    40.0   100.0
percentiles:
   0%   10%   20%   30%   40%   50%   60%   70%   80%   90%   95%   99% 99.5% 99.9%  100%
   10    13    17    20    24    28    32    37    43    51    59    73    79    91   100

methylKit::getMethylationStats output for "Rphase_3" CpGhsm-sites (#: 195324) in coverage range [10-100]  - methylation statistics per base
summary:
   Min. 1st Qu.  Median    Mean 3rd Qu.    Max.
   10.2    40.7    64.0    63.4    87.8   100.0
percentiles:
   0%   10%   20%   30%   40%   50%   60%   70%   80%   90%   95%   99% 99.5% 99.9%  100%
 10.2  28.0  36.7  45.3  54.5  64.0  74.1  83.3  91.2  96.7 100.0 100.0 100.0 100.0 100.0
```

```
Directory: REPCMS File: repcms-Cb_LIp_Rphase_4.txt

Number of "Rphase_4" CpGhsm-sites with minimal and higher level coverage:

  number of "Rphase_4" CpGhsm-sites with coverage >=  5: 200707
  number of "Rphase_4" CpGhsm-sites with coverage >=  6: 197722
  number of "Rphase_4" CpGhsm-sites with coverage >= 10: 176448
  number of "Rphase_4" CpGhsm-sites with coverage >= 15: 149206

Coverage and methylation statistics for "Rphase_4" CpGhsm-sites at different levels of minimum coverage:

methylKit::getCoverageStats output for "Rphase_4" CpGhsm-sites (#: 200707) at minimum coverage 5     - read coverage statistics per base
summary:
   Min. 1st Qu.  Median    Mean 3rd Qu.    Max.
      5      14      24      28      38    1405
percentiles:
   0%   10%   20%   30%   40%   50%   60%   70%   80%   90%   95%   99% 99.5% 99.9%  100%
    5     9    12    16    20    24    29    35    42    53    63    82    90   110  1405

methylKit::getMethylationStats output for "Rphase_4" CpGhsm-sites (#: 200707) at minimum coverage 5  - methylation statistics per base
summary:
   Min. 1st Qu.  Median    Mean 3rd Qu.    Max.
    2.1    44.9    70.0    66.8    90.0   100.0
percentiles:
    0%    10%    20%    30%    40%    50%    60%    70%    80%    90%    95%    99%  99.5%  99.9%   100%
  2.06  29.79  40.00  50.00  60.00  70.00  79.31  87.10  93.33 100.00 100.00 100.00 100.00 100.00 100.00

methylKit::getCoverageStats output for "Rphase_4" CpGhsm-sites (#: 197722) at minimum coverage 6     - read coverage statistics per base
summary:
   Min. 1st Qu.  Median    Mean 3rd Qu.    Max.
      6      15      25      29      38    1405
percentiles:
   0%   10%   20%   30%   40%   50%   60%   70%   80%   90%   95%   99% 99.5% 99.9%  100%
    6     9    13    16    20    25    29    35    42    53    63    82    90   110  1405

methylKit::getMethylationStats output for "Rphase_4" CpGhsm-sites (#: 197722) at minimum coverage 6  - methylation statistics per base
summary:
   Min. 1st Qu.  Median    Mean 3rd Qu.    Max.
    2.1    44.4    69.4    66.3    89.5   100.0
percentiles:
    0%    10%    20%    30%    40%    50%    60%    70%    80%    90%    95%    99%  99.5%  99.9%   100%
  2.06  29.63  39.58  50.00  60.00  69.44  78.43  86.11  92.45 100.00 100.00 100.00 100.00 100.00 100.00

methylKit::getCoverageStats output for "Rphase_4" CpGhsm-sites (#: 176448) at minimum coverage 10    - read coverage statistics per base
summary:
   Min. 1st Qu.  Median    Mean 3rd Qu.    Max.
     10      18      27      31      40    1405
percentiles:
   0%   10%   20%   30%   40%   50%   60%   70%   80%   90%   95%   99% 99.5% 99.9%  100%
   10    13    16    19    23    27    32    37    44    55    64    84    91   112  1405

methylKit::getMethylationStats output for "Rphase_4" CpGhsm-sites (#: 176448) at minimum coverage 10 - methylation statistics per base
summary:
   Min. 1st Qu.  Median    Mean 3rd Qu.    Max.
    2.1    41.9    65.1    64.0    88.2   100.0
percentiles:
    0%    10%    20%    30%    40%    50%    60%    70%    80%    90%    95%    99%  99.5%  99.9%   100%
```

```
  2.06  28.57  37.50  46.32  55.56  65.12  75.00  84.09  91.43  96.77 100.00 100.00 100.00 100.00 100.00
```

```
methylKit::getCoverageStats output for "Rphase_4" CpGhsm-sites (#: 149206) at minimum coverage 15    - read coverage statistics per base
summary:
   Min. 1st Qu.  Median    Mean 3rd Qu.    Max.
     15      22      30      35      43    1405
percentiles:
   0%   10%   20%   30%   40%   50%   60%   70%   80%   90%   95%   99% 99.5% 99.9%  100%
   15    17    20    23    27    30    35    40    47    57    66    86    93   115  1405

methylKit::getMethylationStats output for "Rphase_4" CpGhsm-sites (#: 149206) at minimum coverage 15 - methylation statistics per base
summary:
   Min. 1st Qu.  Median    Mean 3rd Qu.    Max.
    2.1    38.5    61.1    61.5    86.7   100.0
percentiles:
    0%    10%    20%    30%    40%    50%    60%    70%    80%    90%    95%    99%  99.5%  99.9%   100%
  2.06  26.83  34.78  42.22  51.28  61.11  72.00  81.82  90.00  95.83 100.00 100.00 100.00 100.00 100.00
```

```
methylKit::getCoverageStats output for "Rphase_4" CpGhsm-sites (#: 176027) in coverage range [10-100]     - read coverage statistics per base
summary:
   Min. 1st Qu.  Median    Mean 3rd Qu.    Max.
   10.0    18.0    27.0    30.8    40.0   100.0
percentiles:
   0%   10%   20%   30%   40%   50%   60%   70%   80%   90%   95%   99% 99.5% 99.9%  100%
   10    13    16    19    23    27    32    37    44    55    64    81    87    96   100

methylKit::getMethylationStats output for "Rphase_4" CpGhsm-sites (#: 176027) in coverage range [10-100]  - methylation statistics per base
summary:
   Min. 1st Qu.  Median    Mean 3rd Qu.    Max.
   10.2    41.9    65.1    64.0    88.2   100.0
percentiles:
   0%   10%   20%   30%   40%   50%   60%   70%   80%   90%   95%   99% 99.5% 99.9%  100%
 10.2  28.6  37.5  46.3  55.6  65.1  75.0  84.1  91.4  96.8 100.0 100.0 100.0 100.0 100.0
```

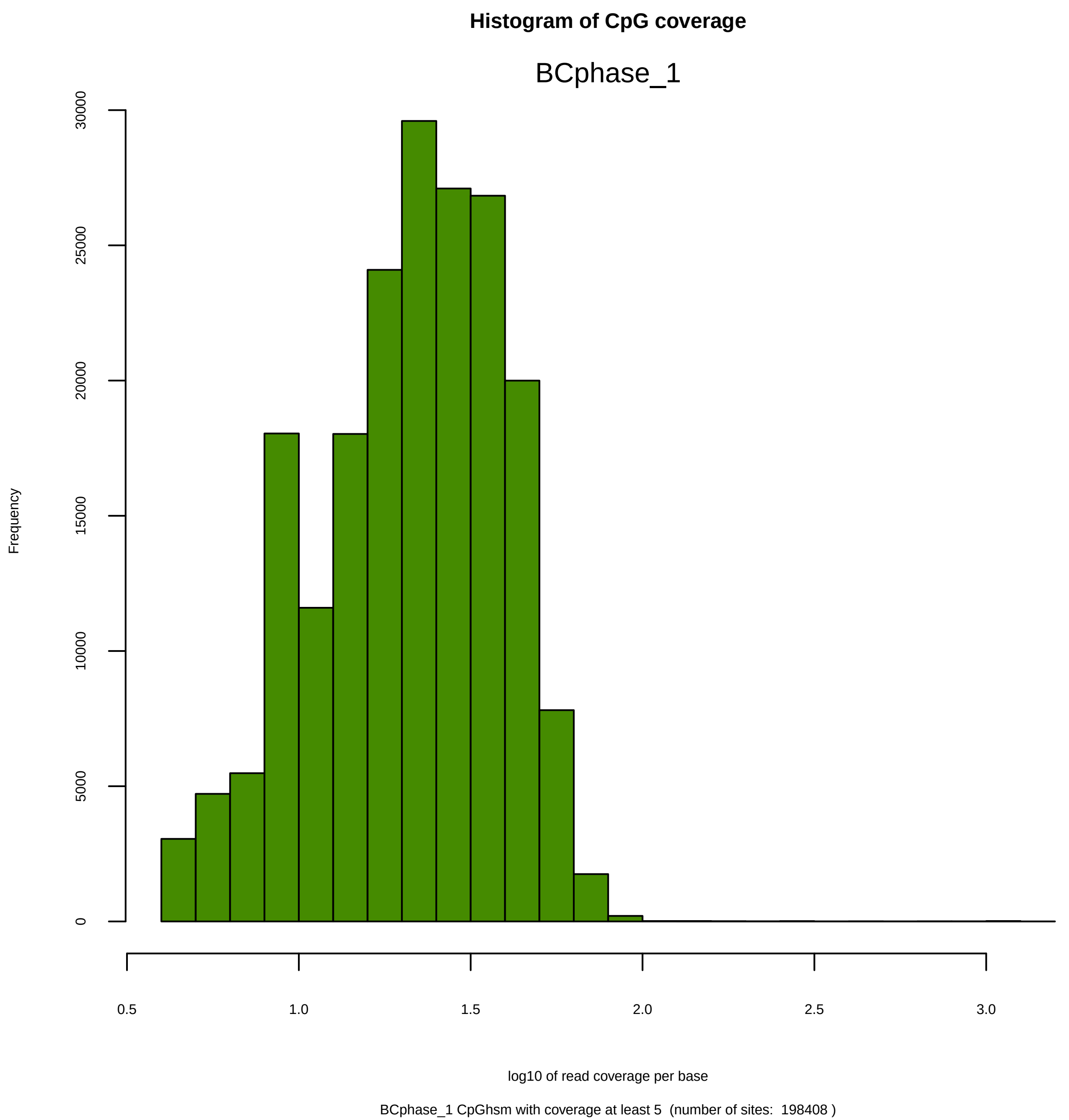

Histogram of CpG coverage
BCphase_1
Frequency
0
5000
10000
15000
20000
25000
30000
0.5
1.0
1.5
2.0
2.5
3.0
log10 of read coverage per base
BCphase_1 CpGhsm with coverage at least 5 (number of sites: 198408 )

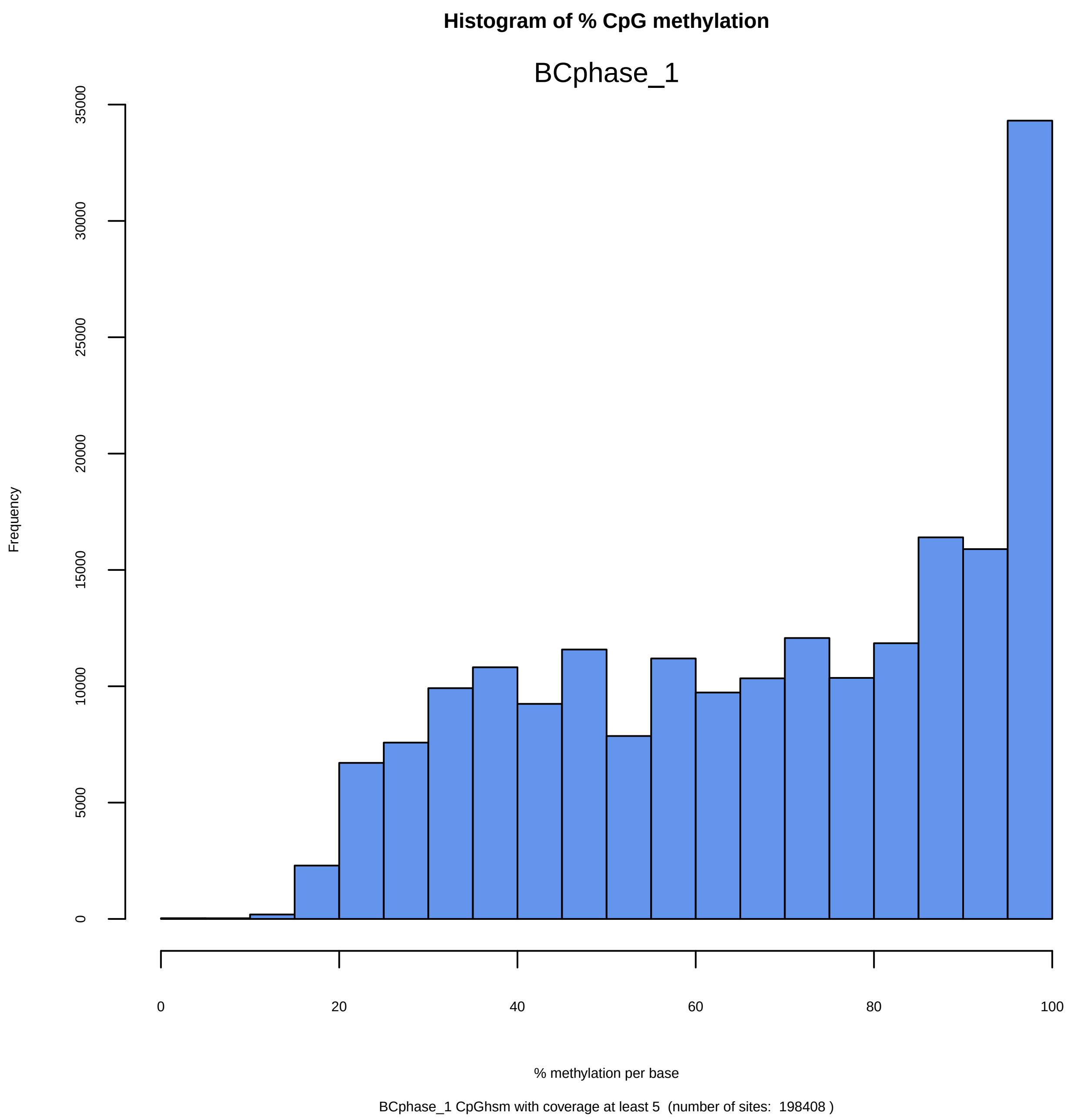
Histogram of % CpG methylation
BCphase_1
Frequency
0
5000
10000
15000
20000
25000
30000
35000
0
20
40
60
80
100
% methylation per base
BCphase_1 CpGhsm with coverage at least 5 (number of sites: 198408 )

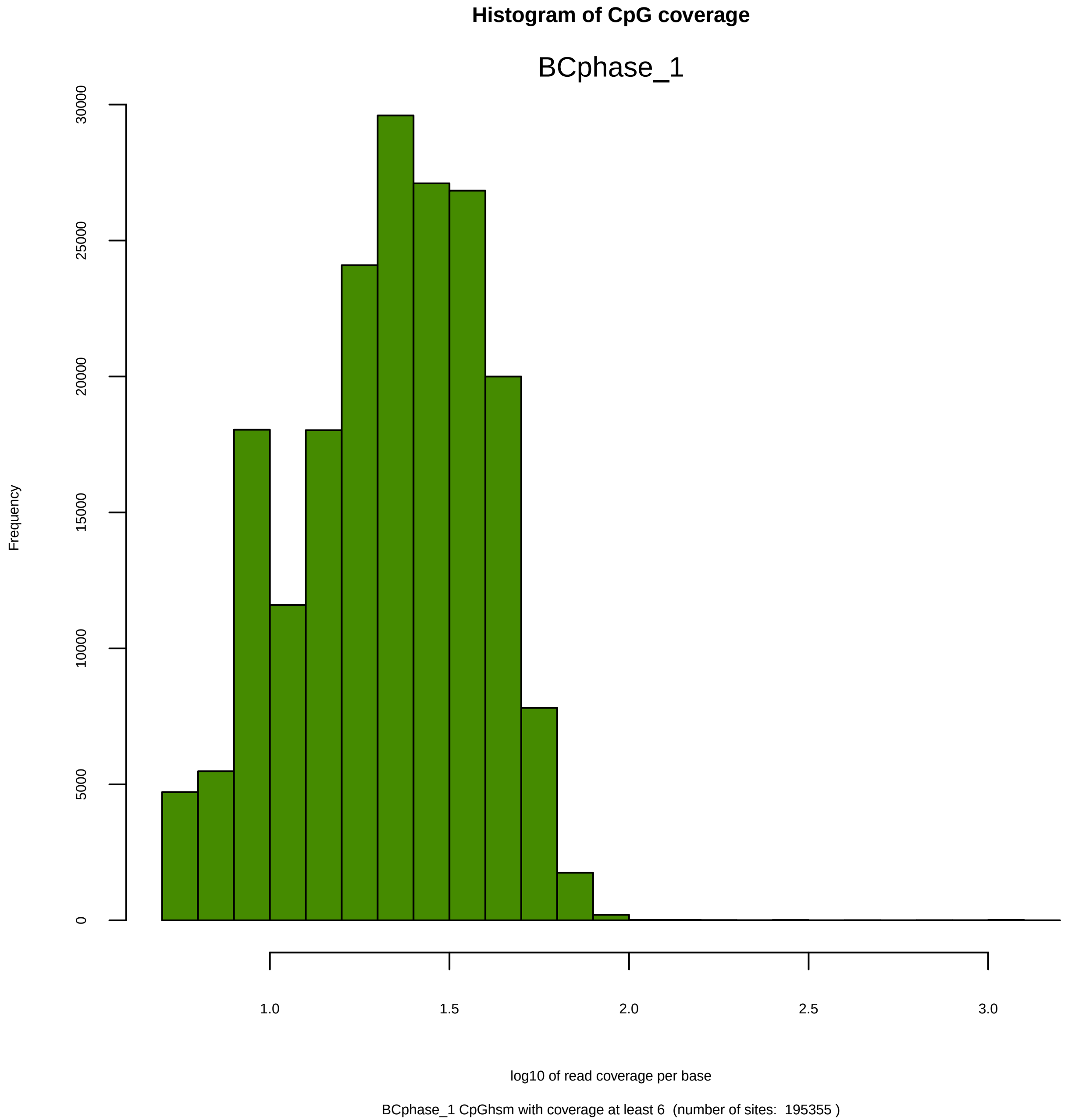
Histogram of CpG coverage
BCphase_1
Frequency
0
5000
10000
15000
20000
25000
30000
1.0
1.5
2.0
2.5
3.0
log10 of read coverage per base
BCphase_1 CpGhsm with coverage at least 6  (number of sites:  195355 )

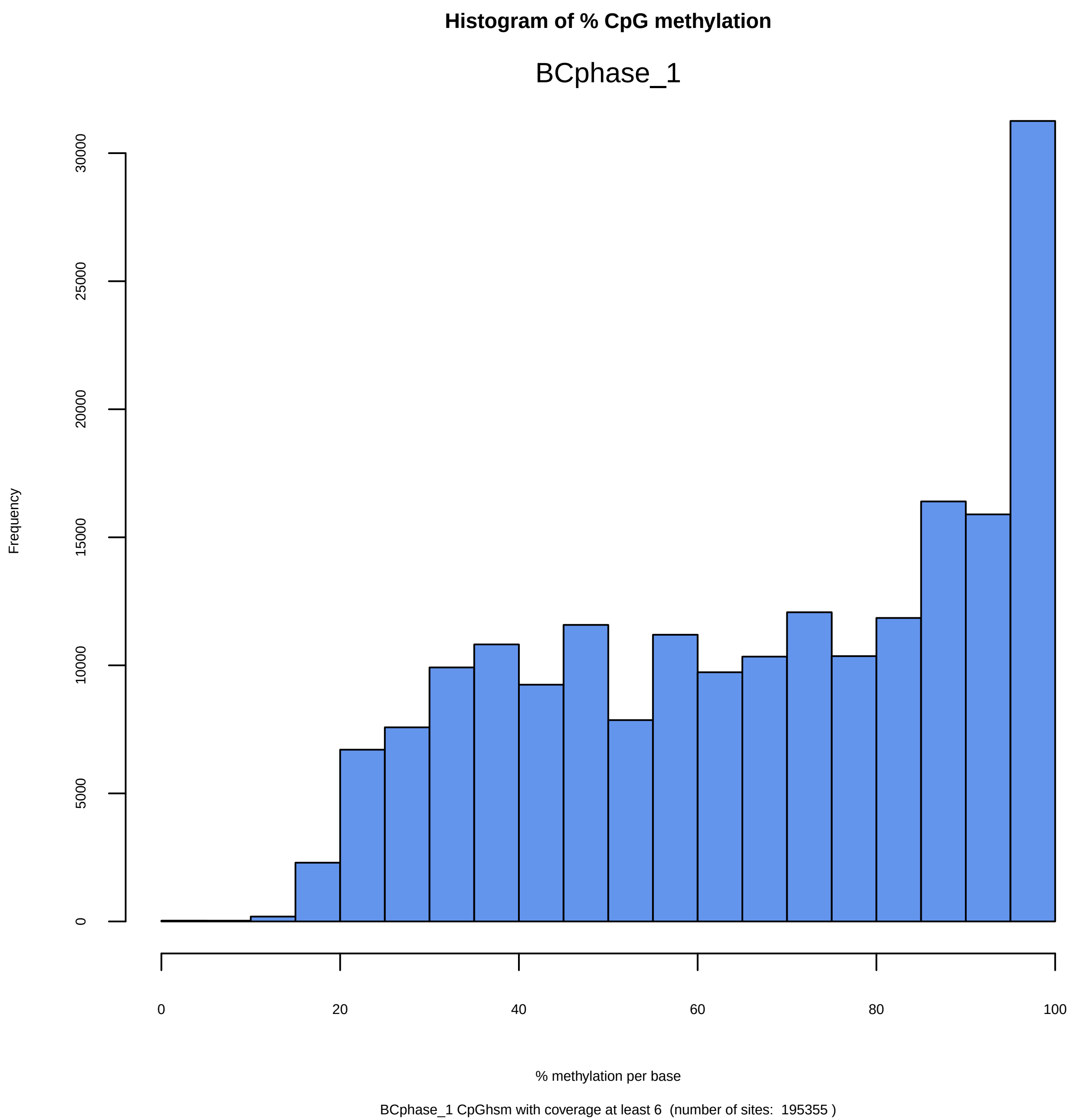

Histogram of % CpG methylation
BCphase_1
Frequency
0
5000
10000
15000
20000
25000
30000
0
20
40
60
80
100
% methylation per base
BCphase_1 CpGhsm with coverage at least 6 (number of sites: 195355 )

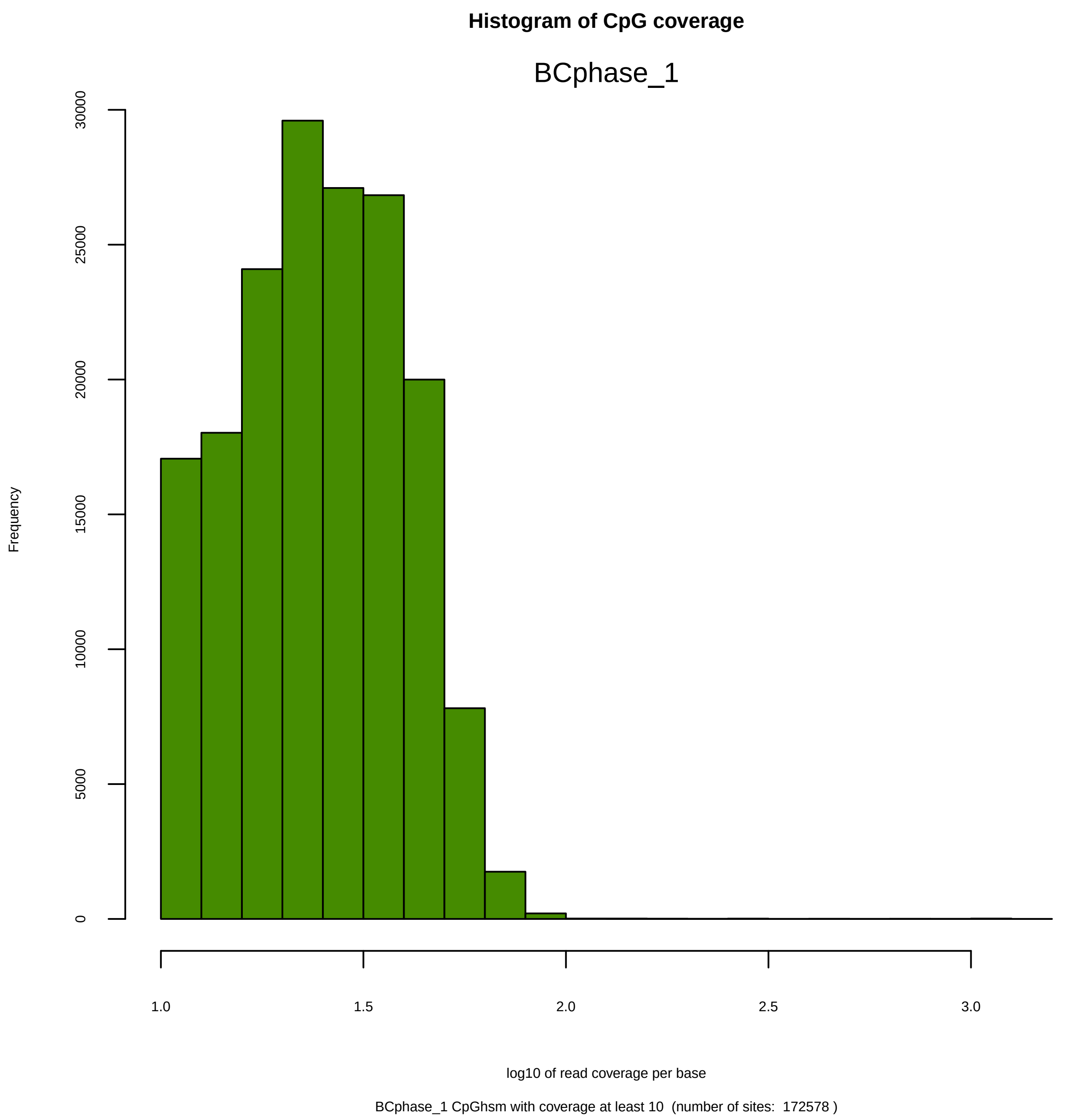

Histogram of CpG coverage
BCphase_1
Frequency
30000
25000
20000
15000
10000
5000
0
1.0
1.5
2.0
2.5
3.0
log10 of read coverage per base
BCphase_1 CpGhsm with coverage at least 10  (number of sites:  172578 )

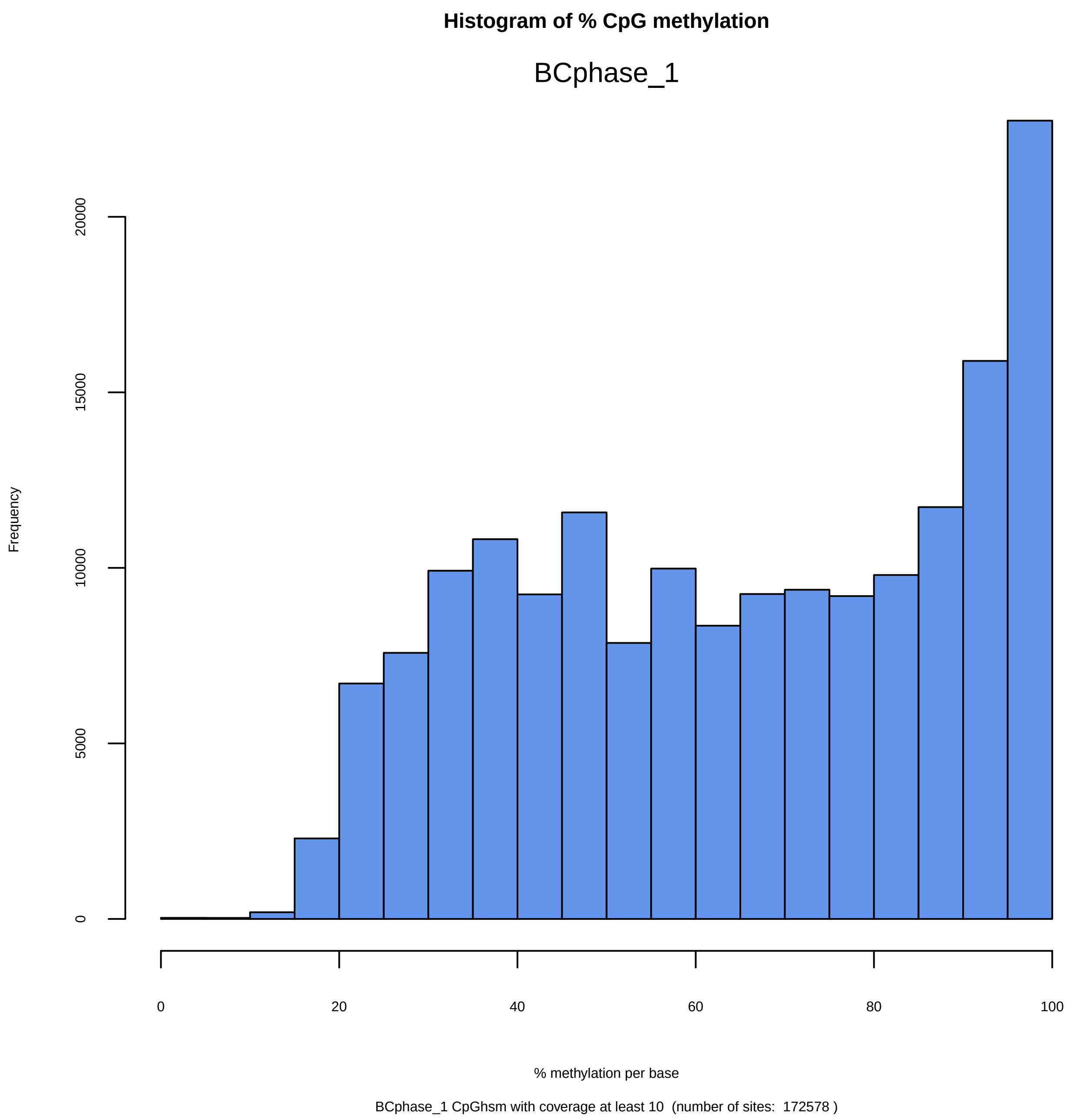

Histogram of % CpG methylation
BCphase_1
Frequency
0
5000
10000
15000
20000
0
20
40
60
80
100
% methylation per base
BCphase_1 CpGhsm with coverage at least 10 (number of sites: 172578 )

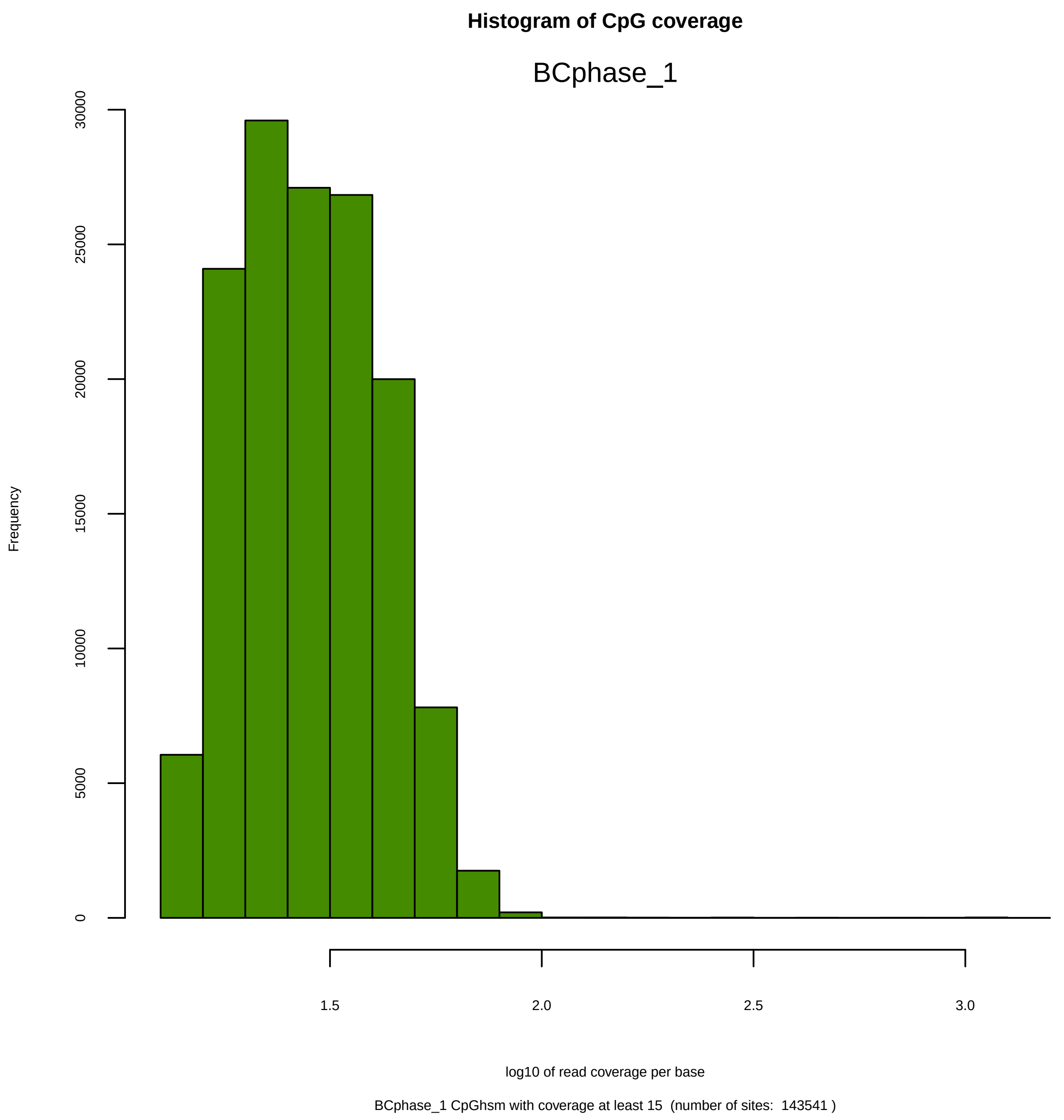

Histogram of CpG coverage
BCphase_1
Frequency
0
5000
10000
15000
20000
25000
30000
1.5
2.0
2.5
3.0
log10 of read coverage per base
BCphase_1 CpGhsm with coverage at least 15  (number of sites:  143541 )

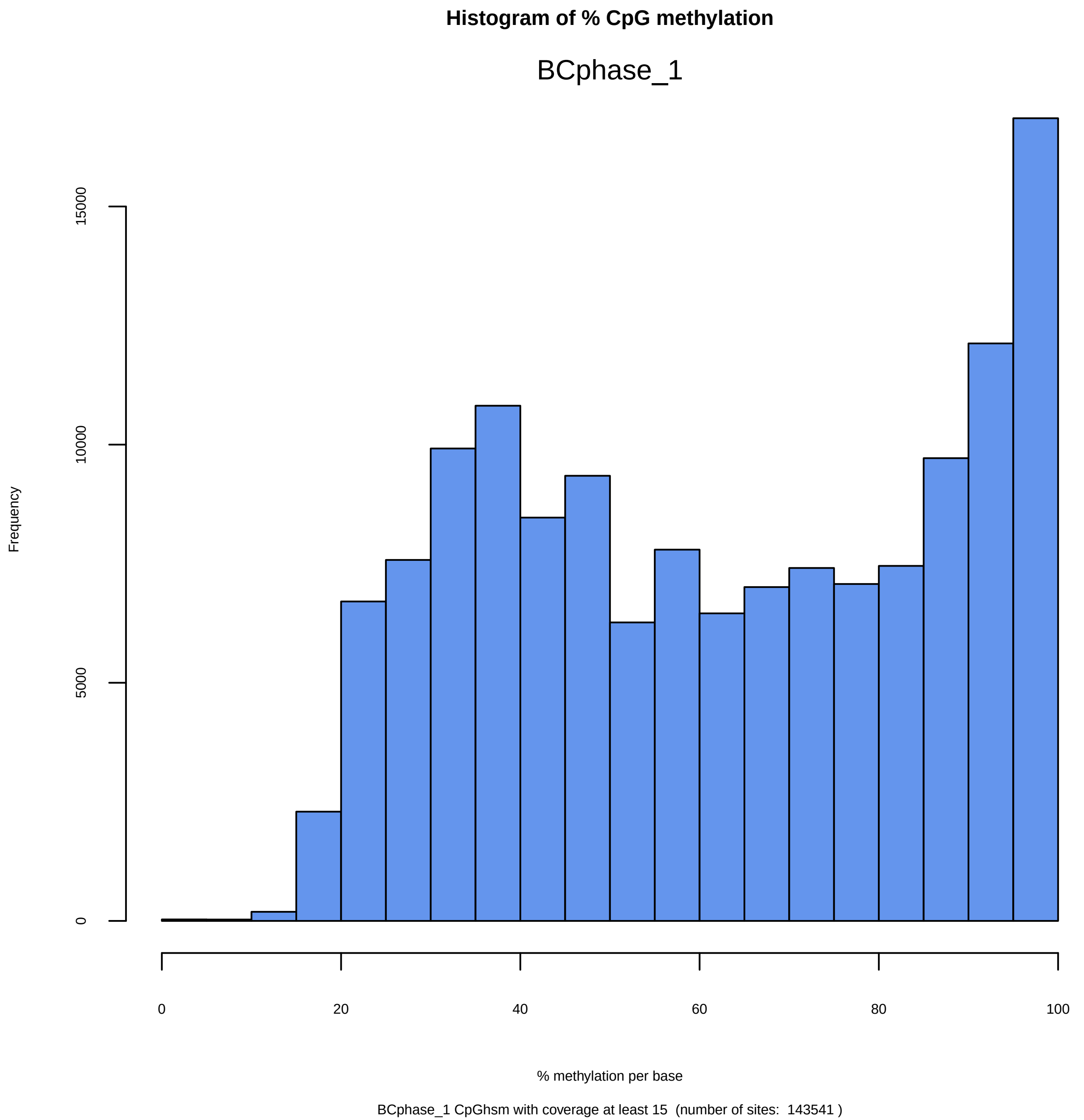

Histogram of % CpG methylation
BCphase_1
Frequency
0
5000
10000
15000
0
20
40
60
80
100
% methylation per base
BCphase_1 CpGhsm with coverage at least 15 (number of sites: 143541 )

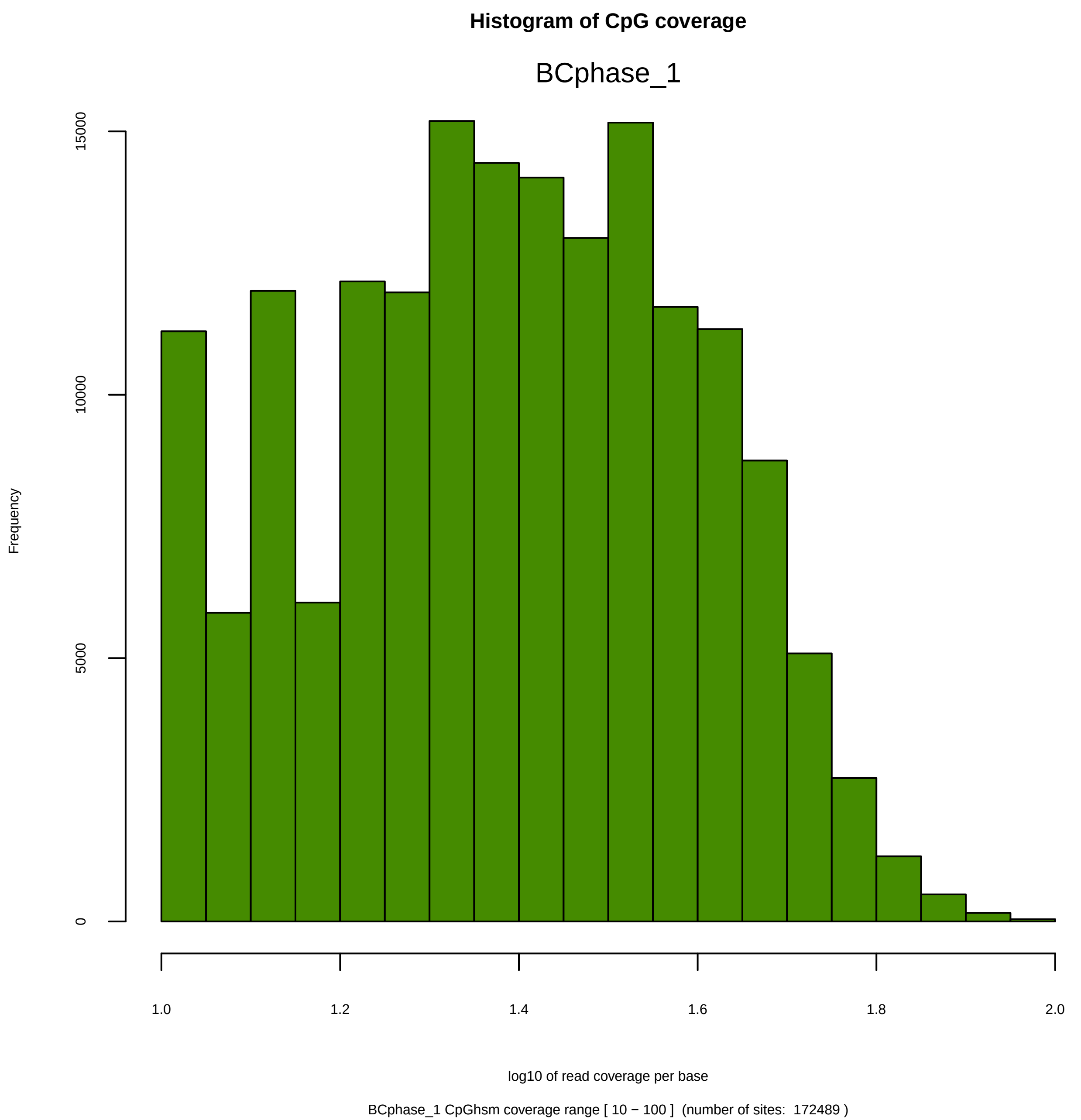
Histogram of CpG coverage
BCphase_1
Frequency
0
5000
10000
15000
1.0
1.2
1.4
1.6
1.8
2.0
log10 of read coverage per base
BCphase_1 CpGhsm coverage range [ 10 − 100 ]  (number of sites:  172489 )

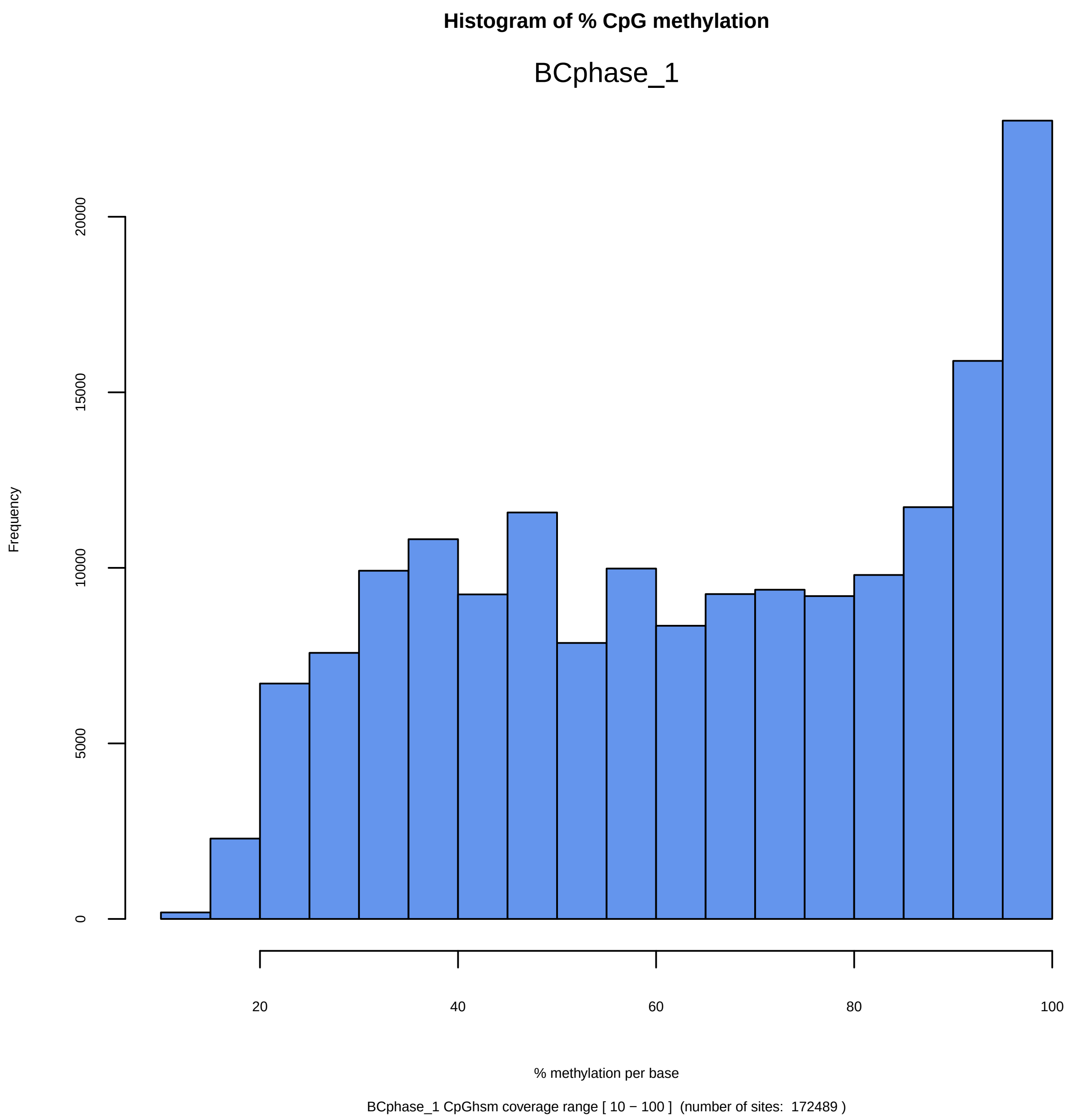
Histogram of % CpG methylation
BCphase_1
Frequency
0
5000
10000
15000
20000
20
40
60
80
100
% methylation per base
BCphase_1 CpGhsm coverage range [ 10 − 100 ]  (number of sites:  172489 )

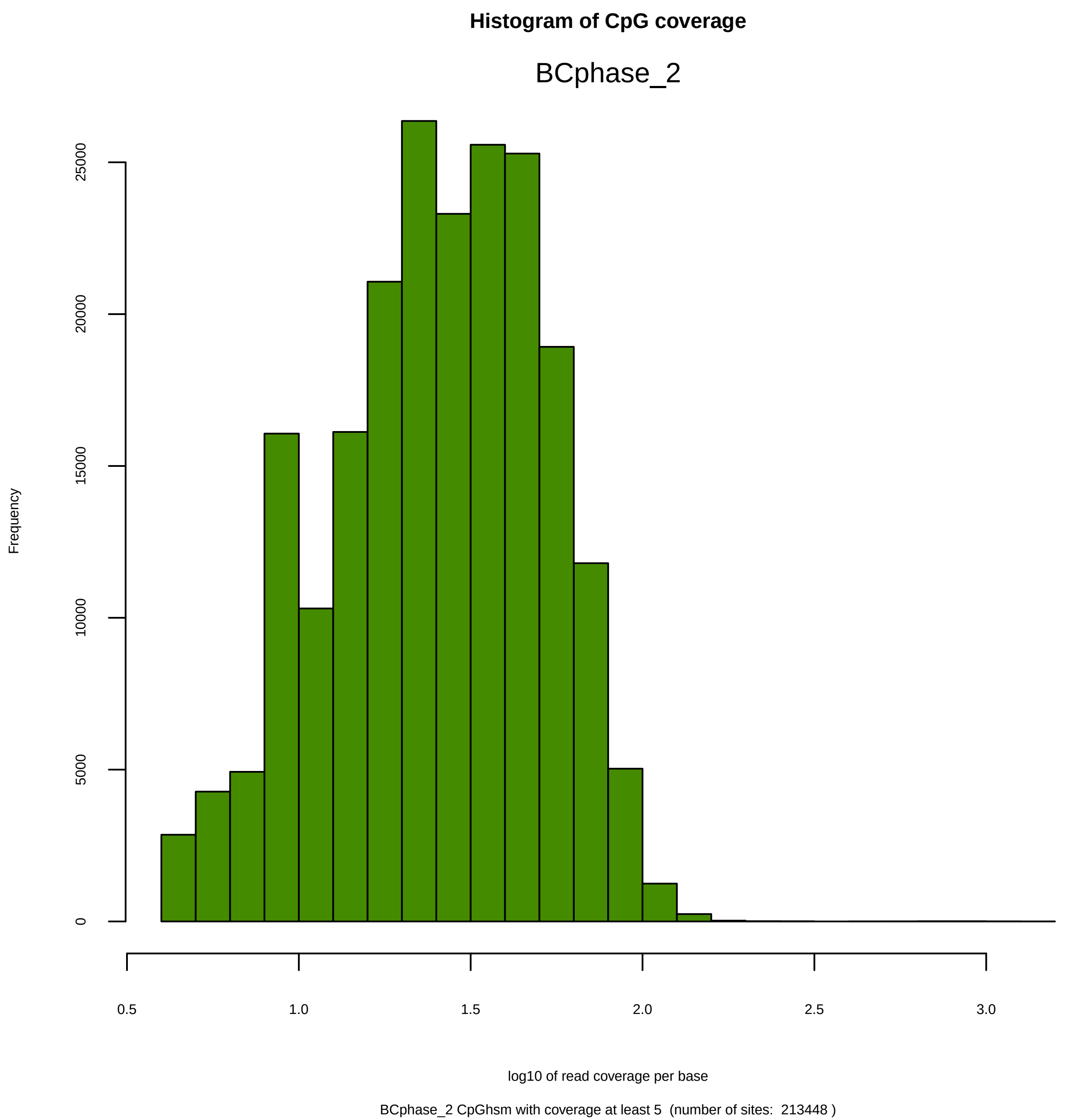
Histogram of CpG coverage
BCphase_2
Frequency
0
5000
10000
15000
20000
25000
0.5
1.0
1.5
2.0
2.5
3.0
log10 of read coverage per base
BCphase_2 CpGhsm with coverage at least 5 (number of sites: 213448 )

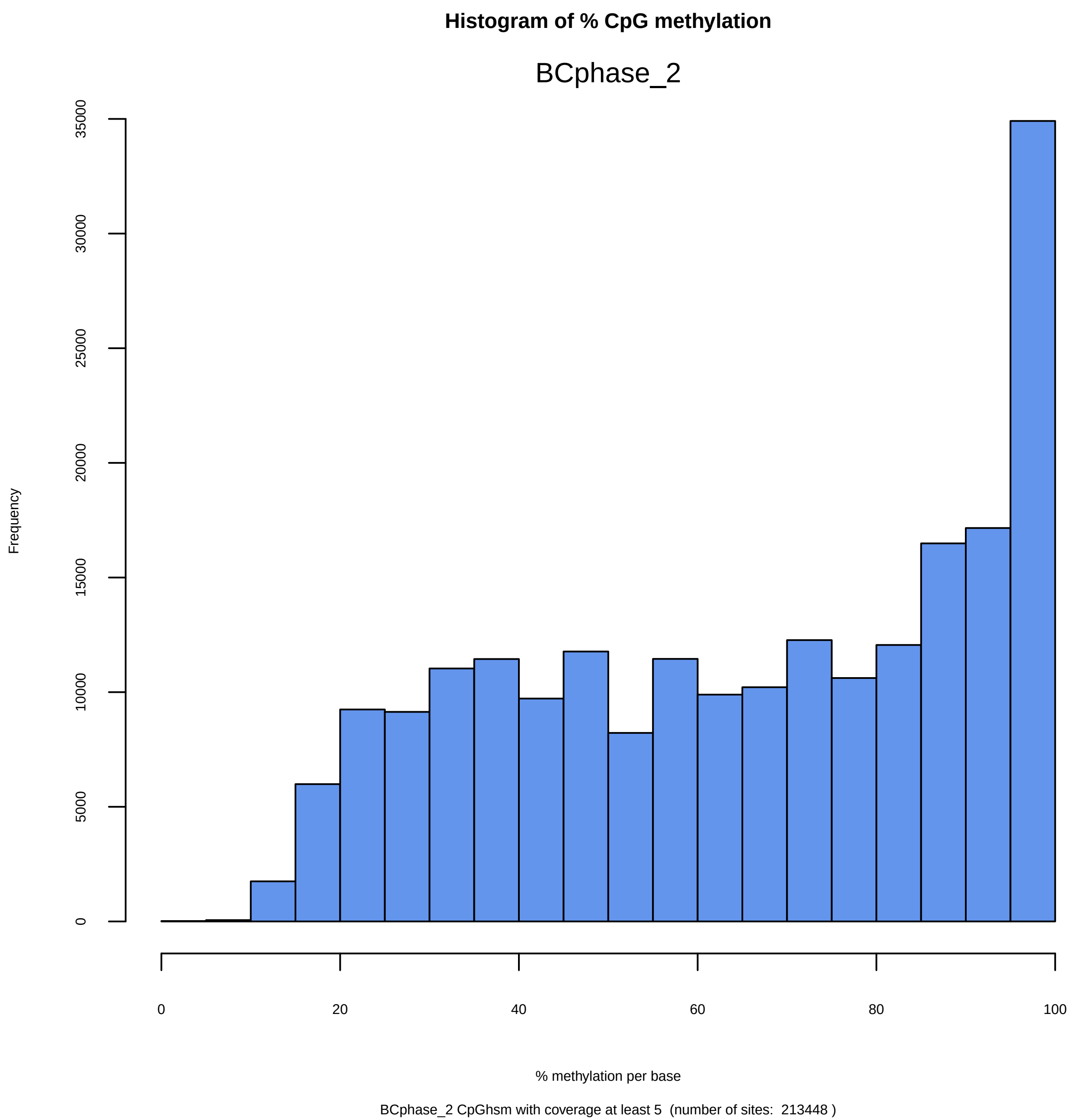
Histogram of % CpG methylation
BCphase_2
Frequency
0
5000
10000
15000
20000
25000
30000
35000
0
20
40
60
80
100
% methylation per base
BCphase_2 CpGhsm with coverage at least 5  (number of sites:  213448 )

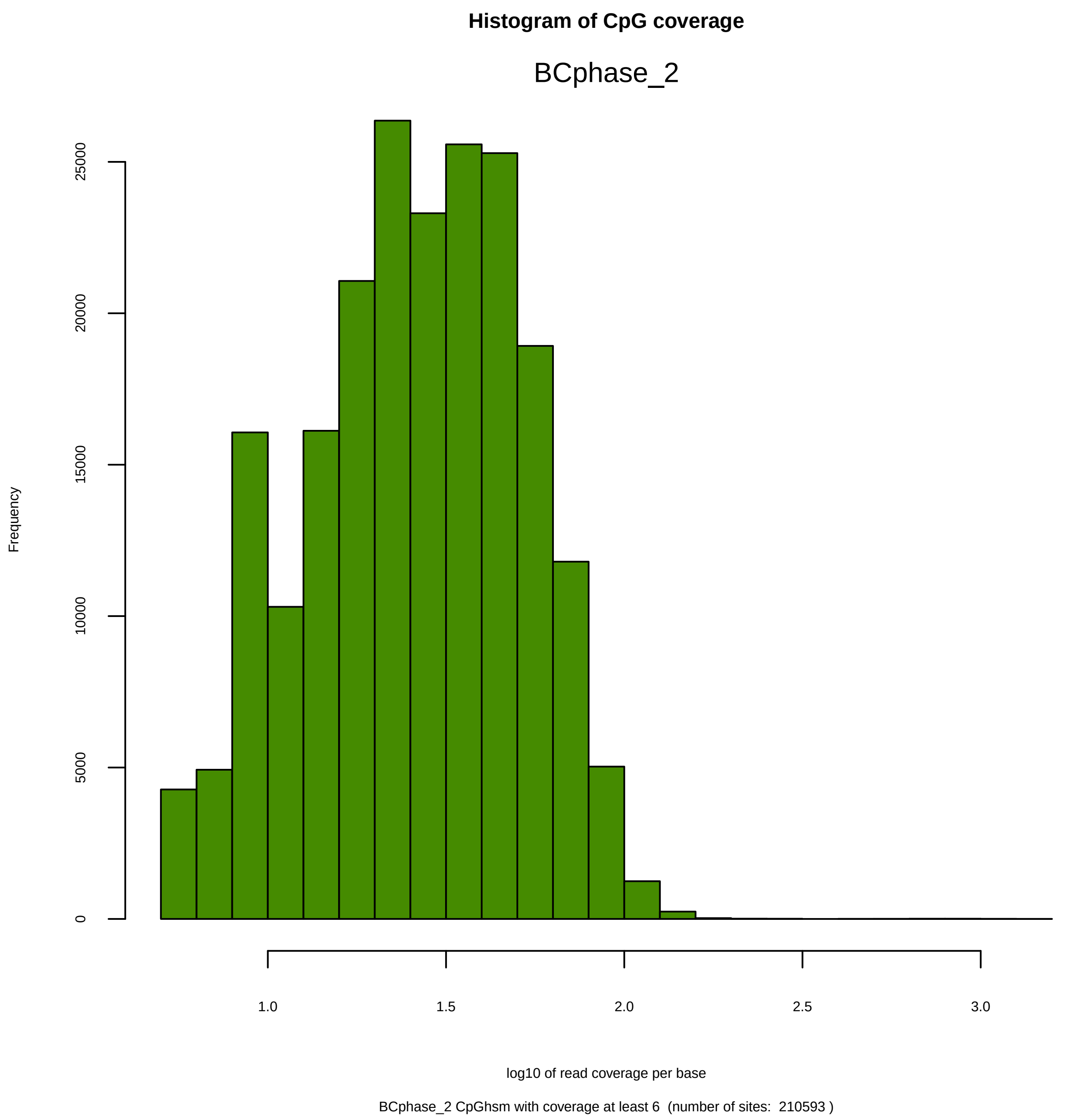

Histogram of CpG coverage
BCphase_2
Frequency
0
5000
10000
15000
20000
25000
1.0
1.5
2.0
2.5
3.0
log10 of read coverage per base
BCphase_2 CpGhsm with coverage at least 6  (number of sites:  210593 )

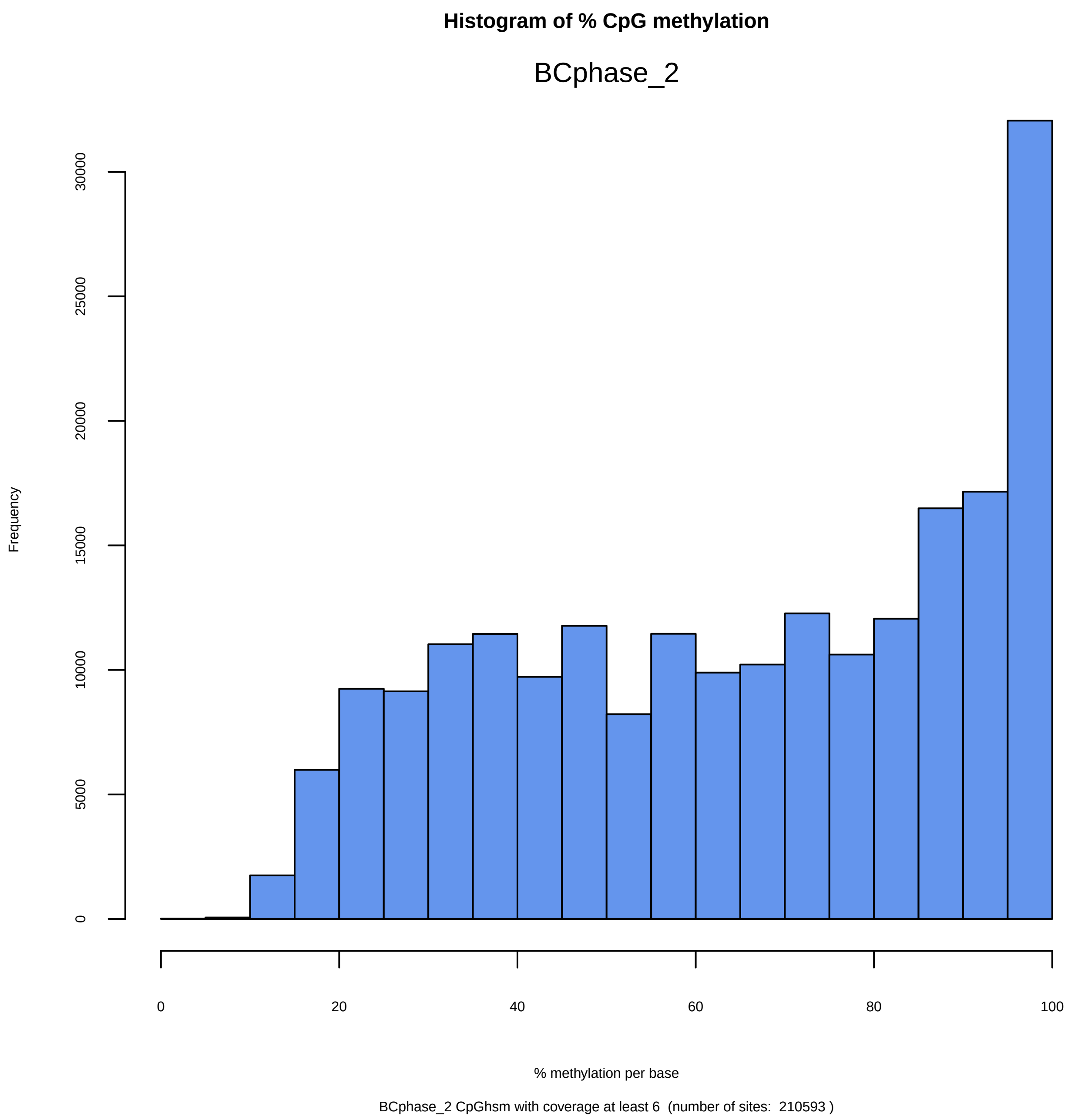

Histogram of % CpG methylation
BCphase_2
Frequency
0
5000
10000
15000
20000
25000
30000
0
20
40
60
80
100
% methylation per base
BCphase_2 CpGhsm with coverage at least 6 (number of sites: 210593 )

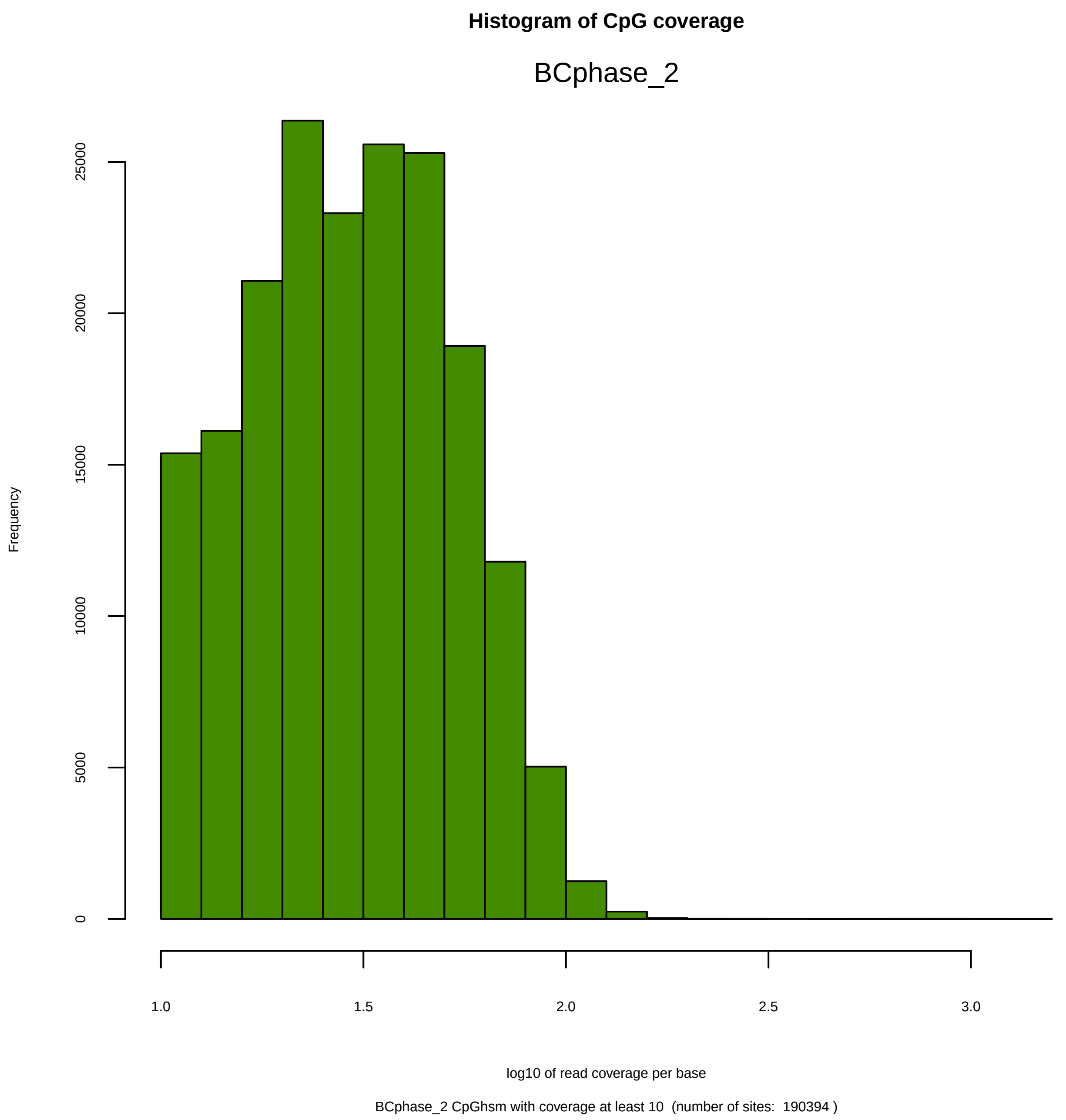

Histogram of CpG coverage
BCphase_2
Frequency
0
5000
10000
15000
20000
25000
1.0
1.5
2.0
2.5
3.0
log10 of read coverage per base
BCphase_2 CpGhsm with coverage at least 10  (number of sites:  190394 )

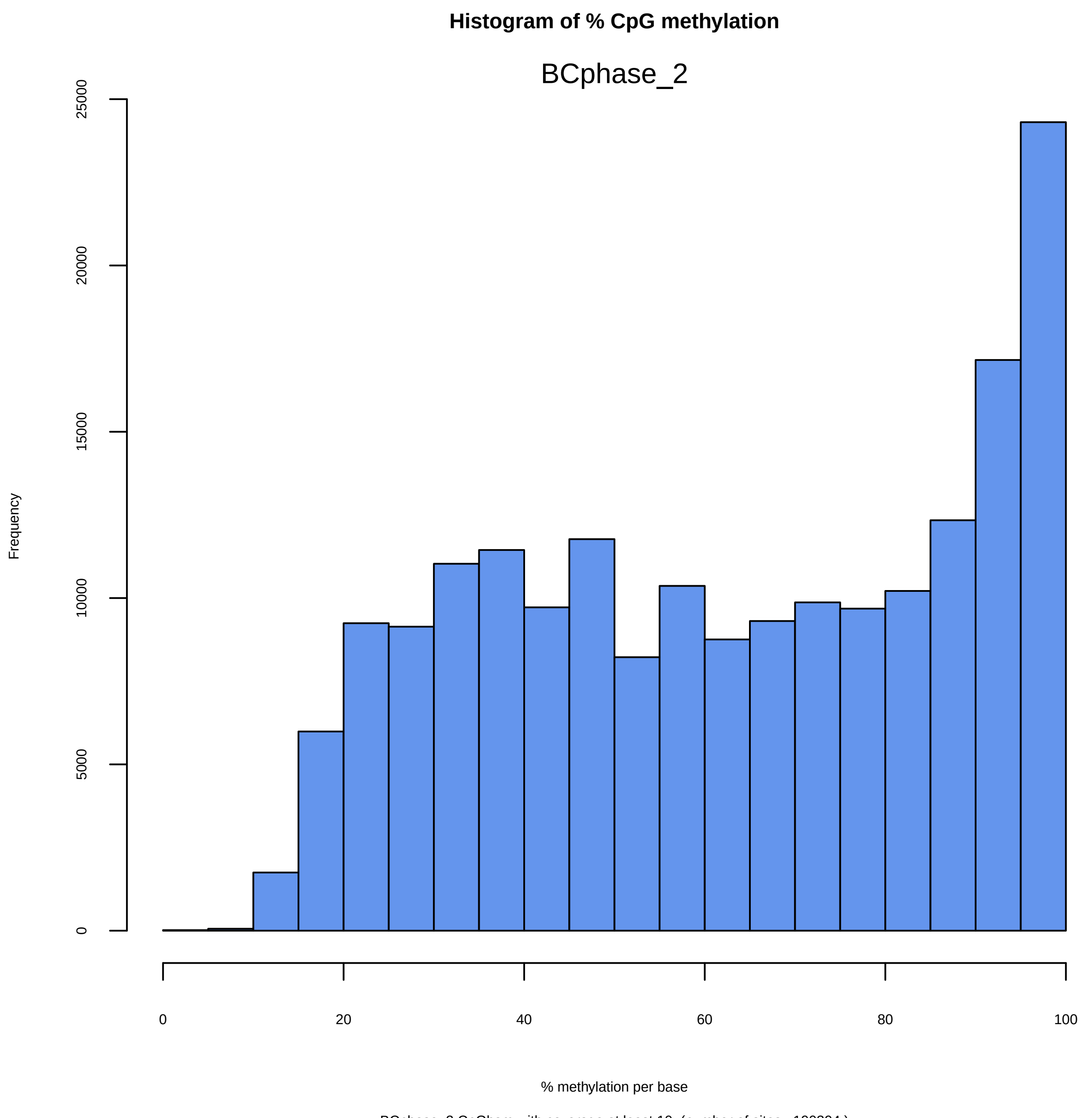

Histogram of % CpG methylation
BCphase_2
Frequency
0
5000
10000
15000
20000
25000
0
20
40
60
80
100
% methylation per base
BCphase_2 CpGhsm with coverage at least 10 (number of sites: 190394 )

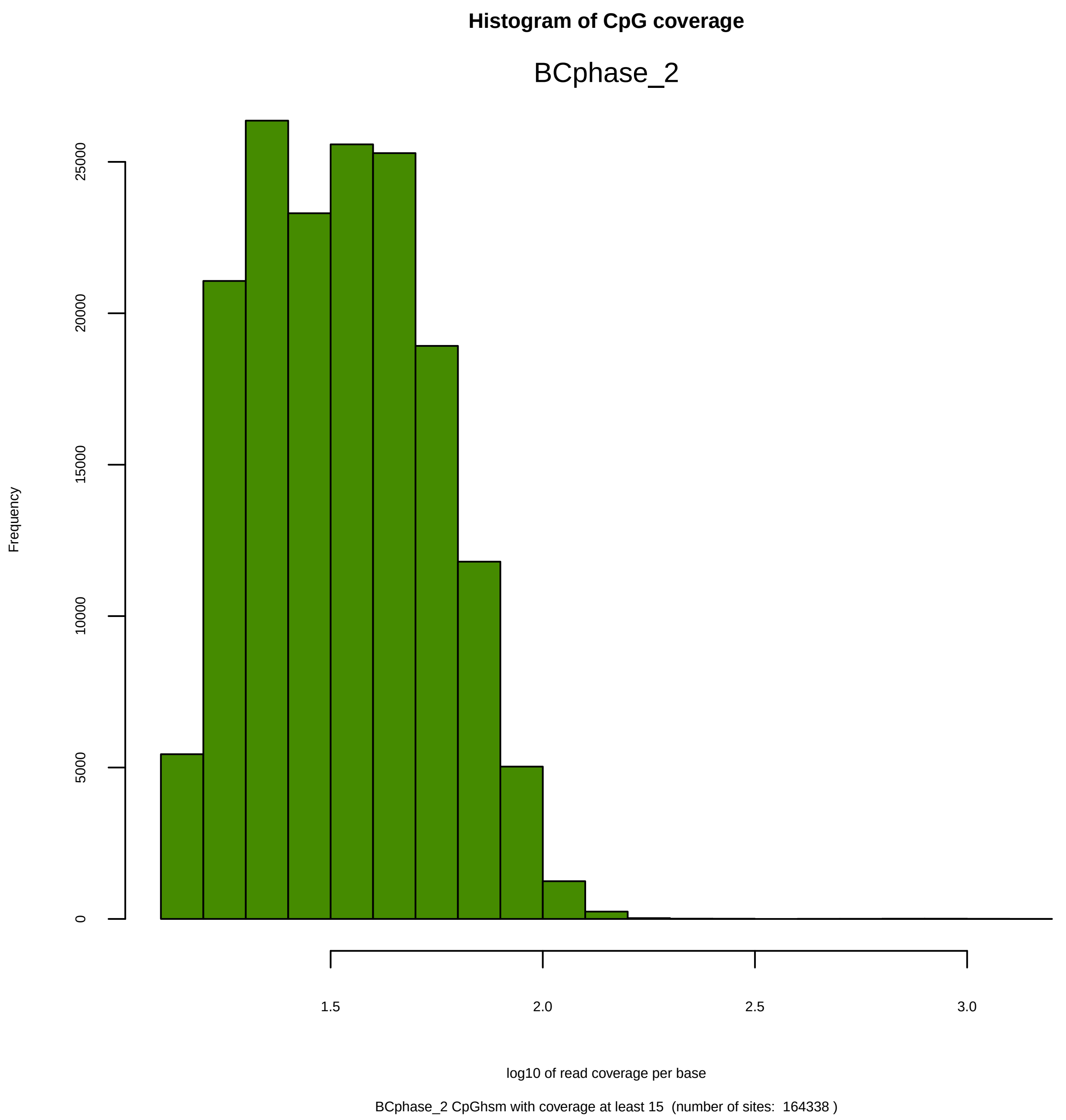

Histogram of CpG coverage
BCphase_2
Frequency
0
5000
10000
15000
20000
25000
1.5
2.0
2.5
3.0
log10 of read coverage per base
BCphase_2 CpGhsm with coverage at least 15 (number of sites: 164338 )

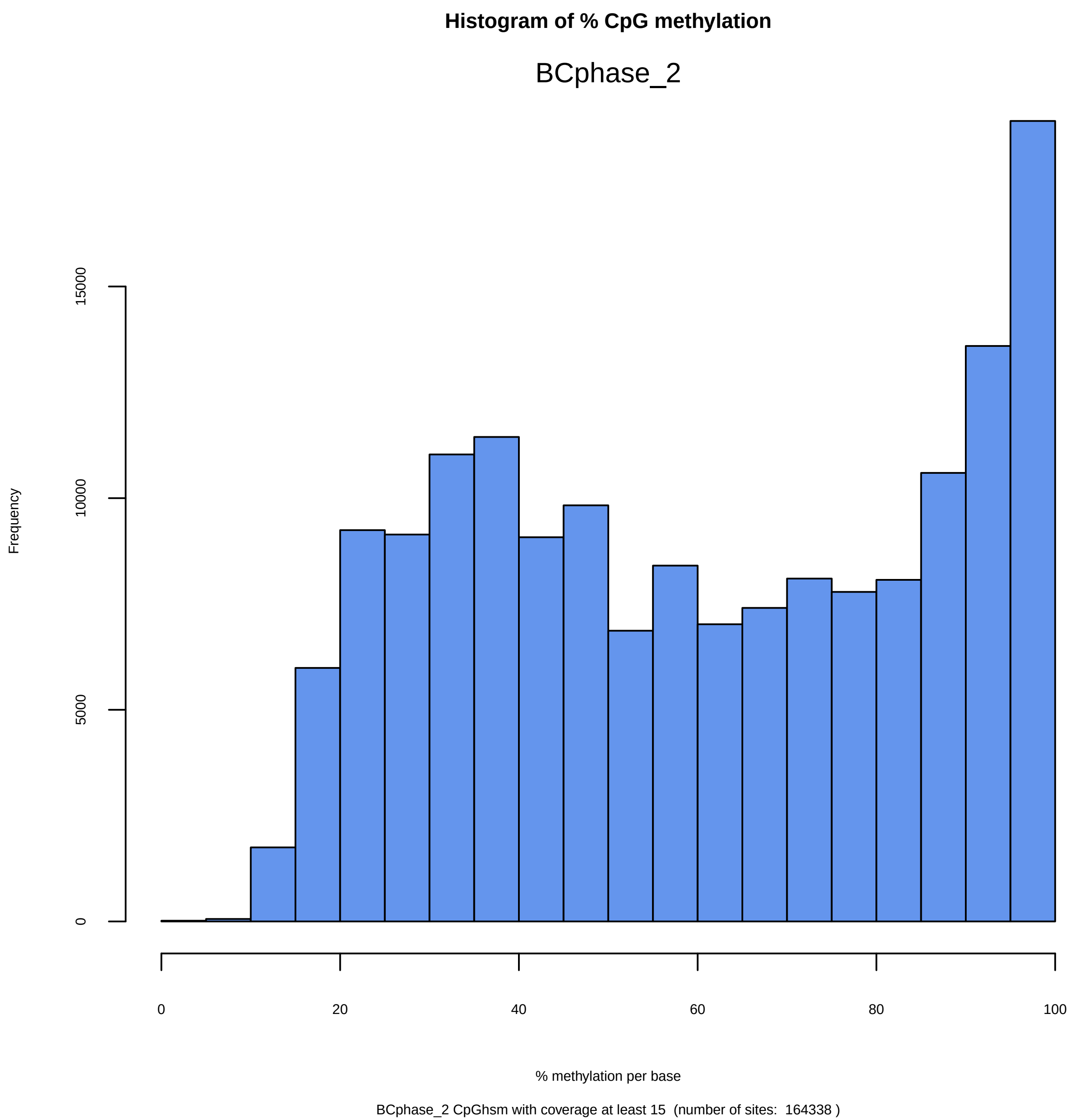
Histogram of % CpG methylation
BCphase_2
Frequency
0
5000
10000
15000
0
20
40
60
80
100
% methylation per base
BCphase_2 CpGhsm with coverage at least 15  (number of sites:  164338 )

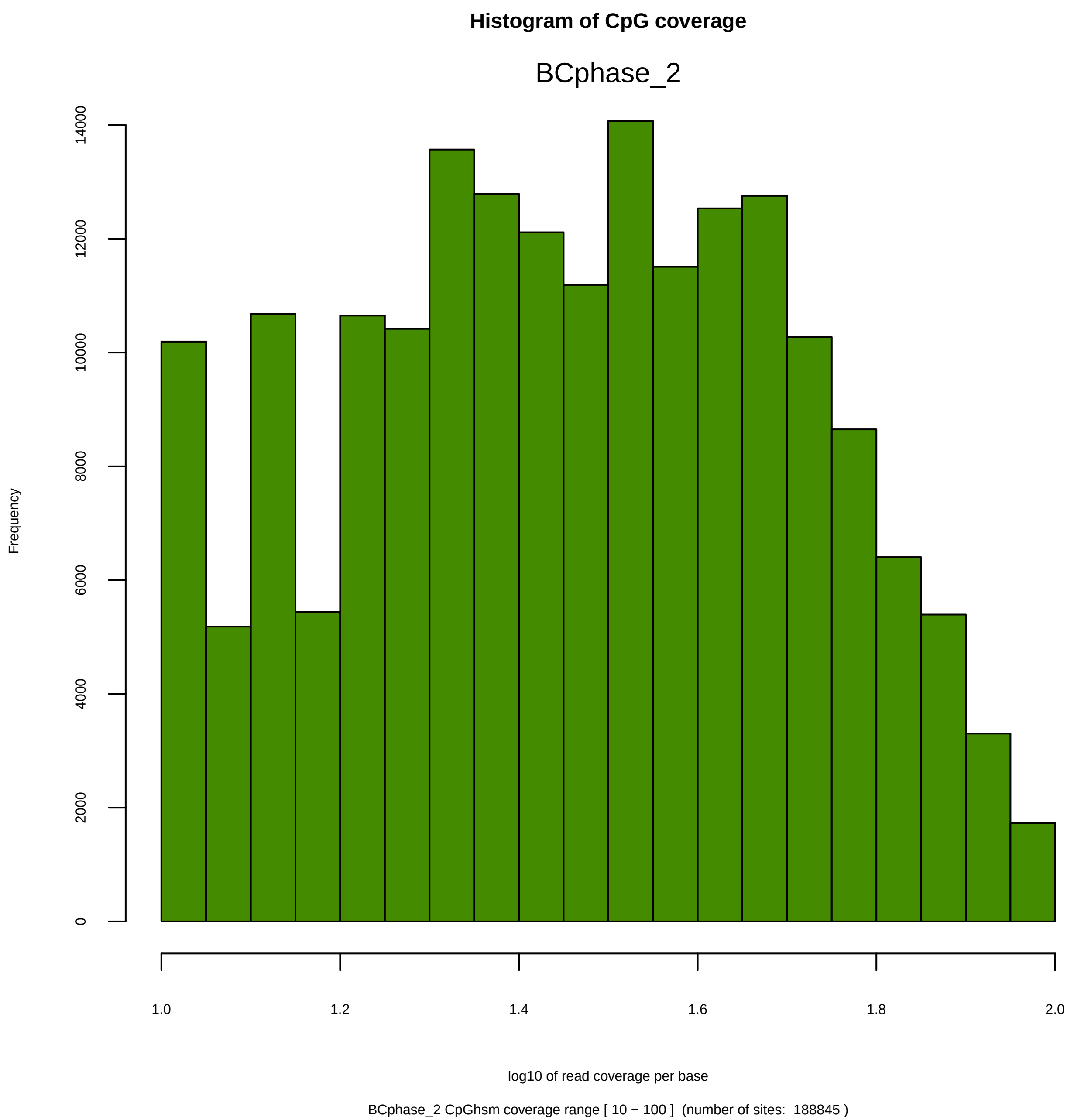

Histogram of CpG coverage
BCphase_2
Frequency
0
2000
4000
6000
8000
10000
12000
14000
1.0
1.2
1.4
1.6
1.8
2.0
log10 of read coverage per base
BCphase_2 CpGhsm coverage range [ 10 − 100 ] (number of sites: 188845 )

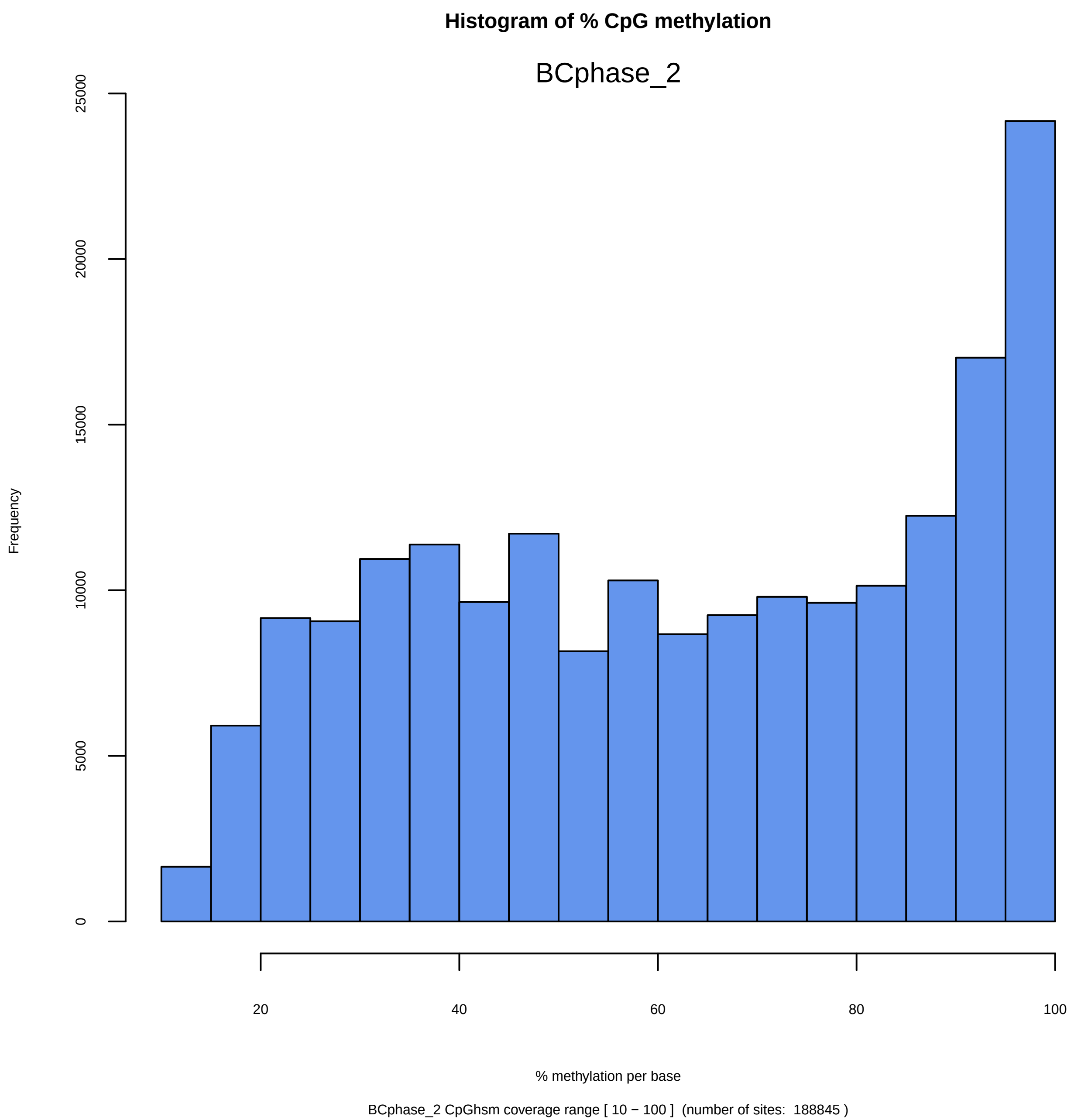

Histogram of % CpG methylation
BCphase_2
Frequency
0
5000
10000
15000
20000
25000
20
40
60
80
100
% methylation per base
BCphase_2 CpGhsm coverage range [ 10 − 100 ]  (number of sites:  188845 )

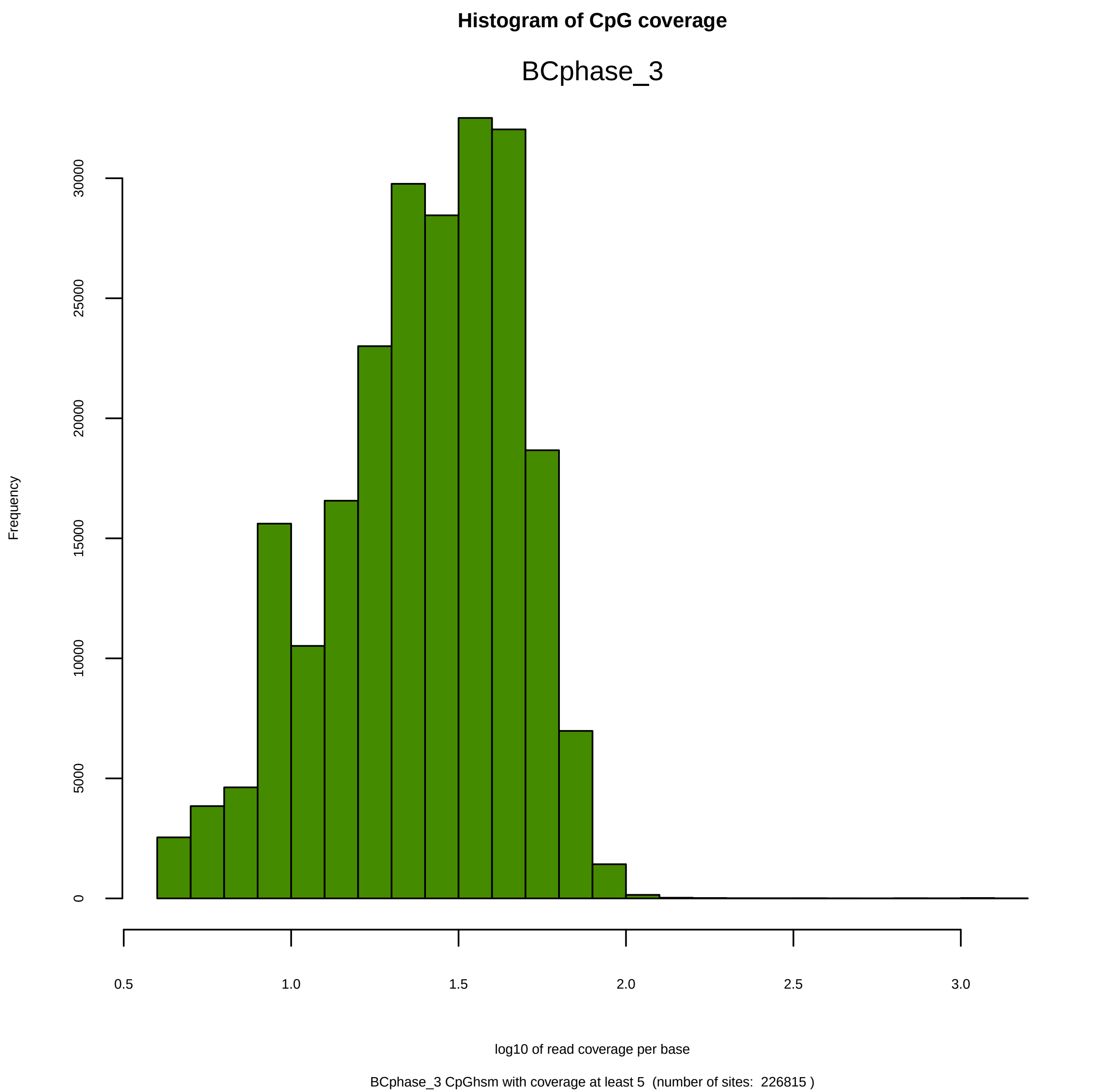
Histogram of CpG coverage
BCphase_3
Frequency
0
5000
10000
15000
20000
25000
30000
0.5
1.0
1.5
2.0
2.5
3.0
log10 of read coverage per base
BCphase_3 CpGhsm with coverage at least 5 (number of sites: 226815 )

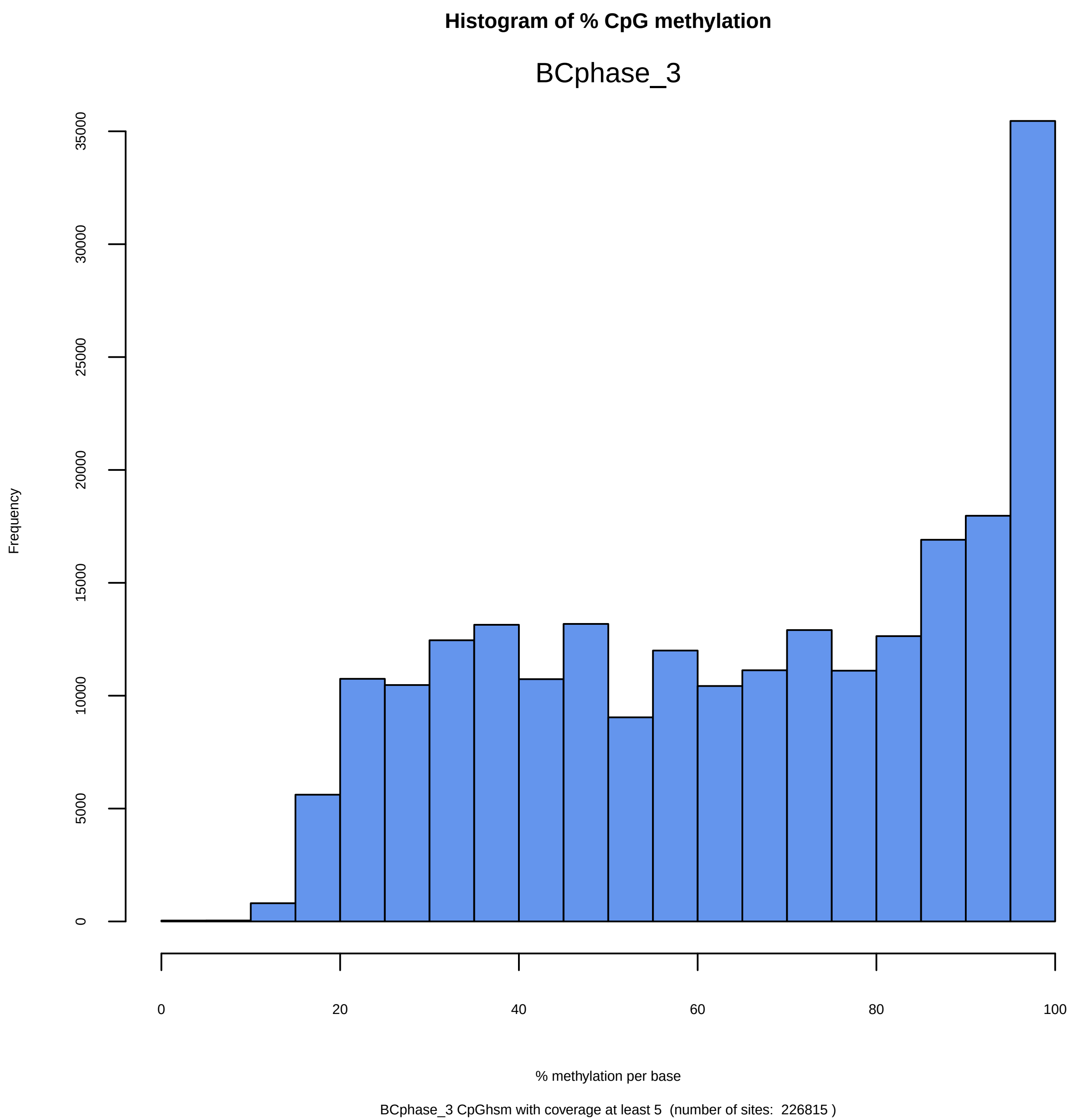

Histogram of % CpG methylation
BCphase_3
Frequency
0
5000
10000
15000
20000
25000
30000
35000
0
20
40
60
80
100
% methylation per base
BCphase_3 CpGhsm with coverage at least 5 (number of sites: 226815 )

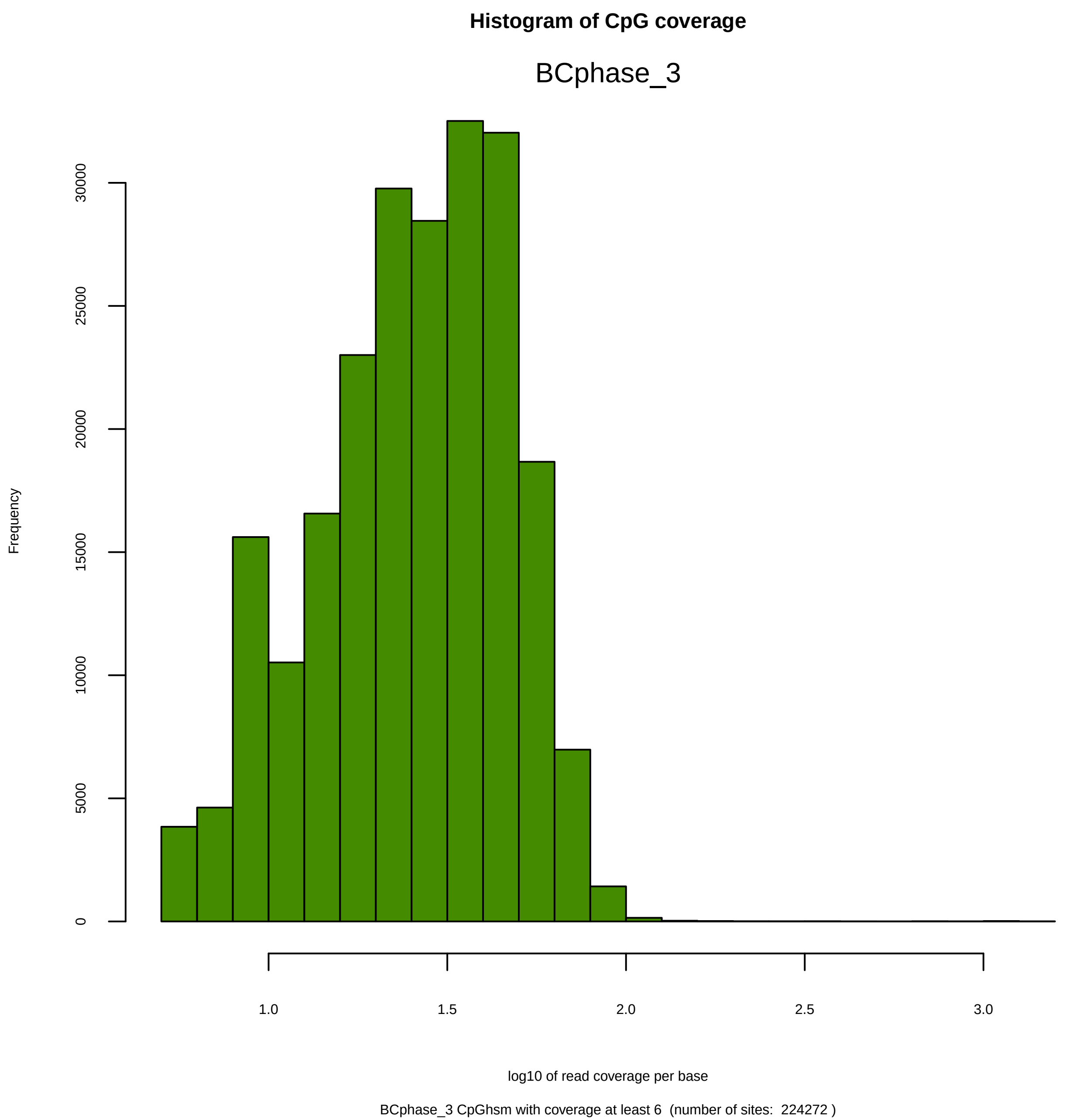
Histogram of CpG coverage
BCphase_3
Frequency
0
5000
10000
15000
20000
25000
30000
1.0
1.5
2.0
2.5
3.0
log10 of read coverage per base
BCphase_3 CpGhsm with coverage at least 6 (number of sites: 224272 )

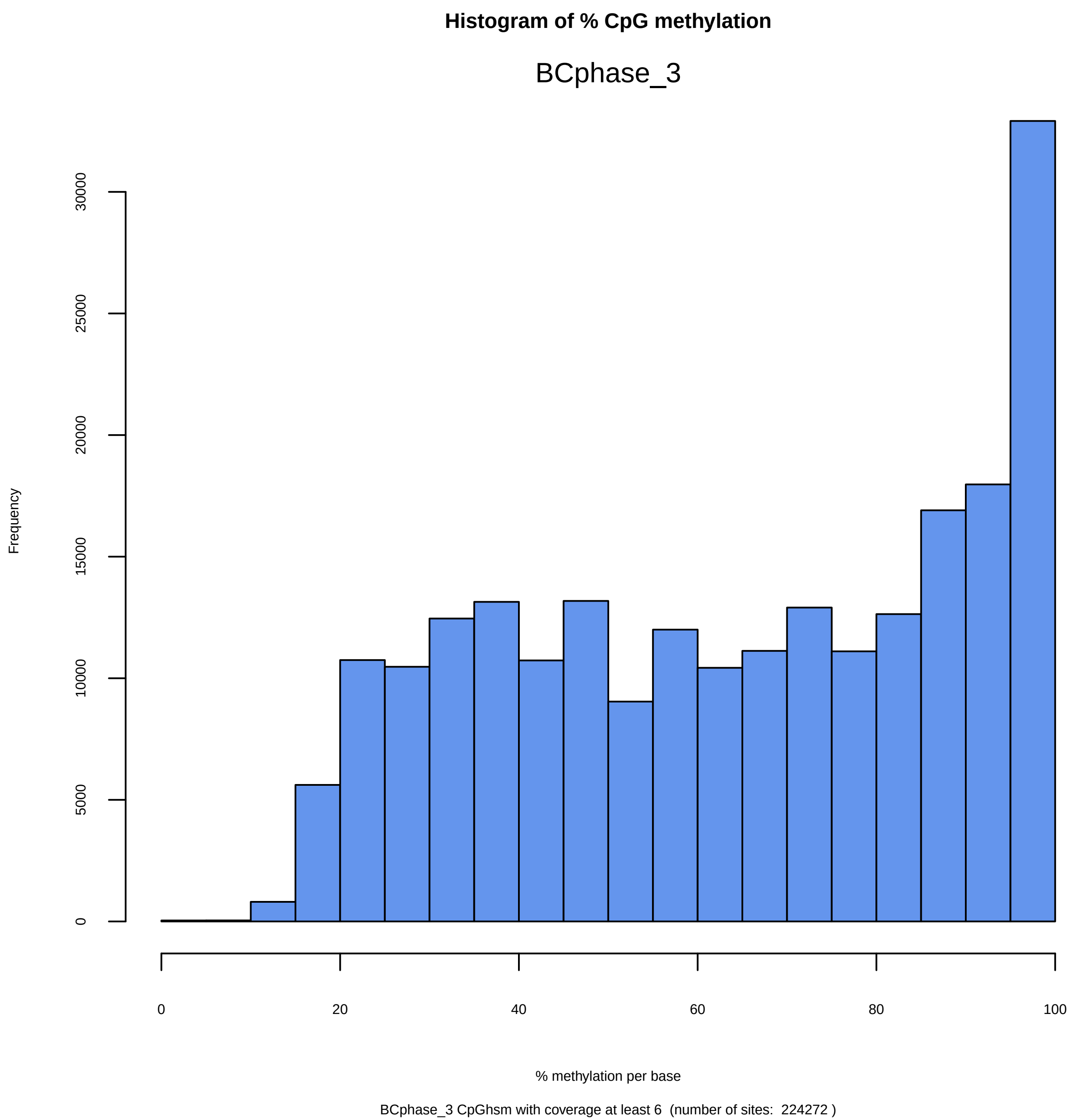

Histogram of % CpG methylation
BCphase_3
Frequency
0
5000
10000
15000
20000
25000
30000
0
20
40
60
80
100
% methylation per base
BCphase_3 CpGhsm with coverage at least 6 (number of sites: 224272 )

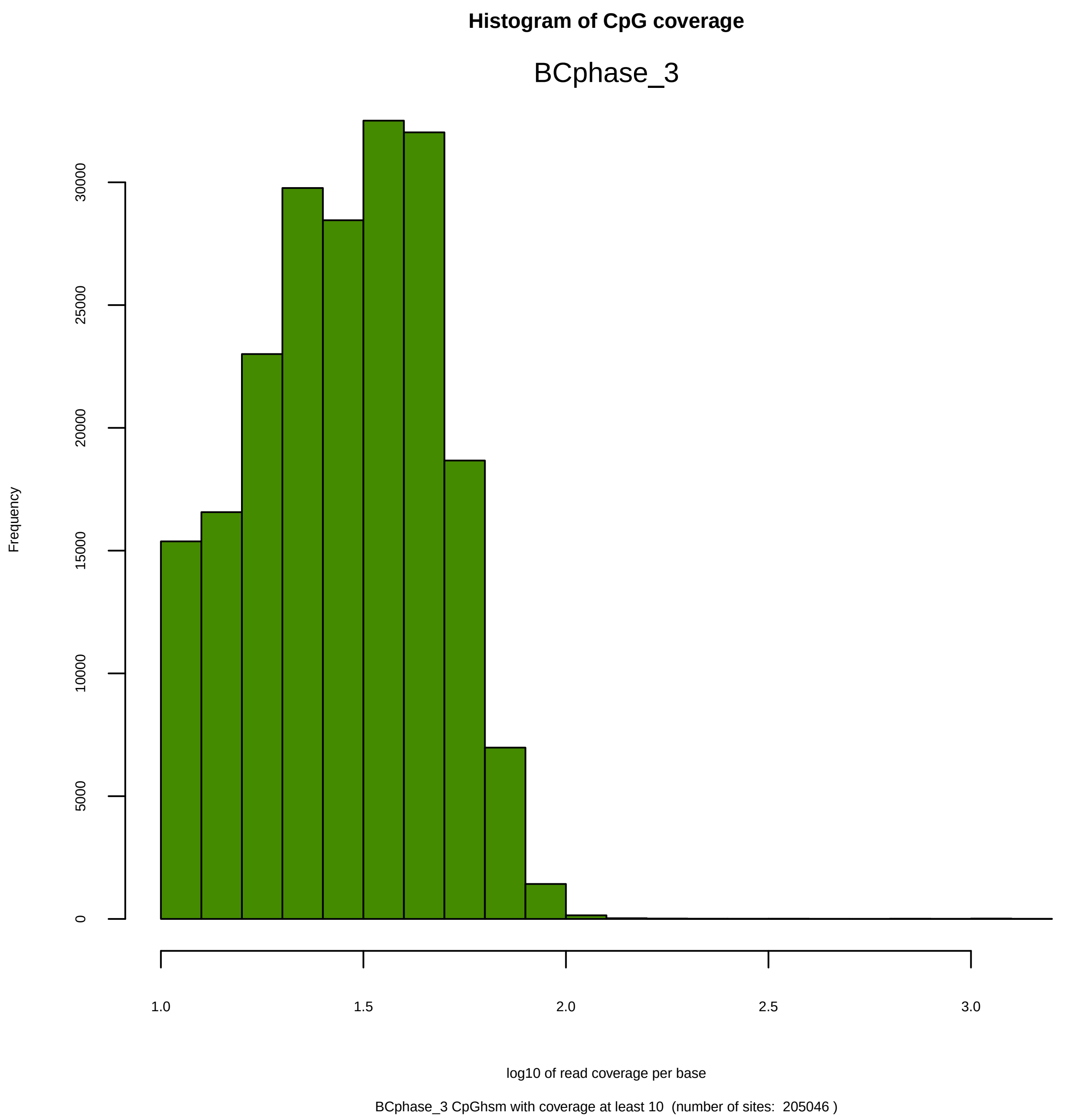
Histogram of CpG coverage
BCphase_3
Frequency
0
5000
10000
15000
20000
25000
30000
1.0
1.5
2.0
2.5
3.0
log10 of read coverage per base
BCphase_3 CpGhsm with coverage at least 10  (number of sites:  205046 )

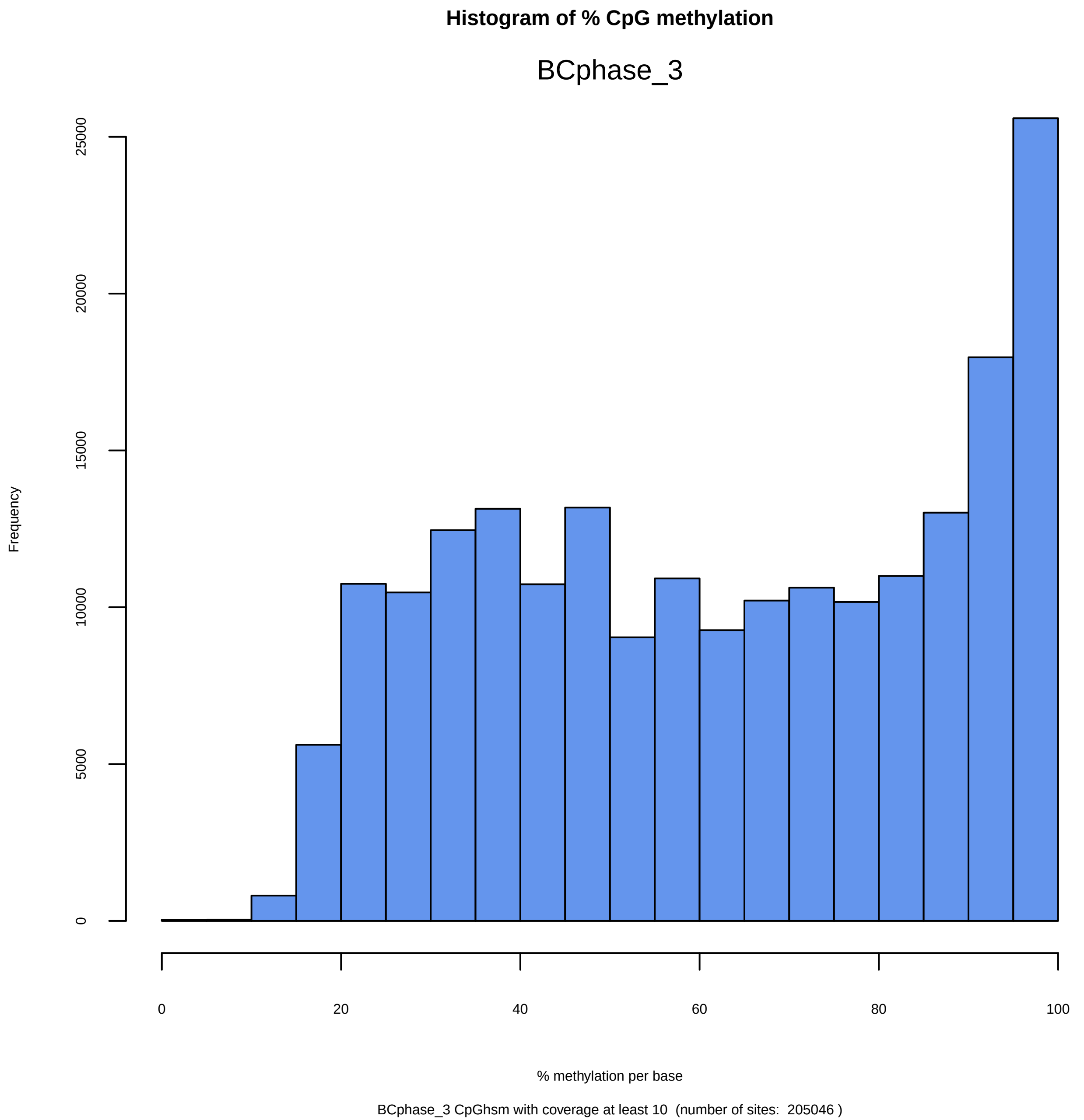
Histogram of % CpG methylation
BCphase_3
Frequency
0
5000
10000
15000
20000
25000
0
20
40
60
80
100
% methylation per base
BCphase_3 CpGhsm with coverage at least 10 (number of sites: 205046 )

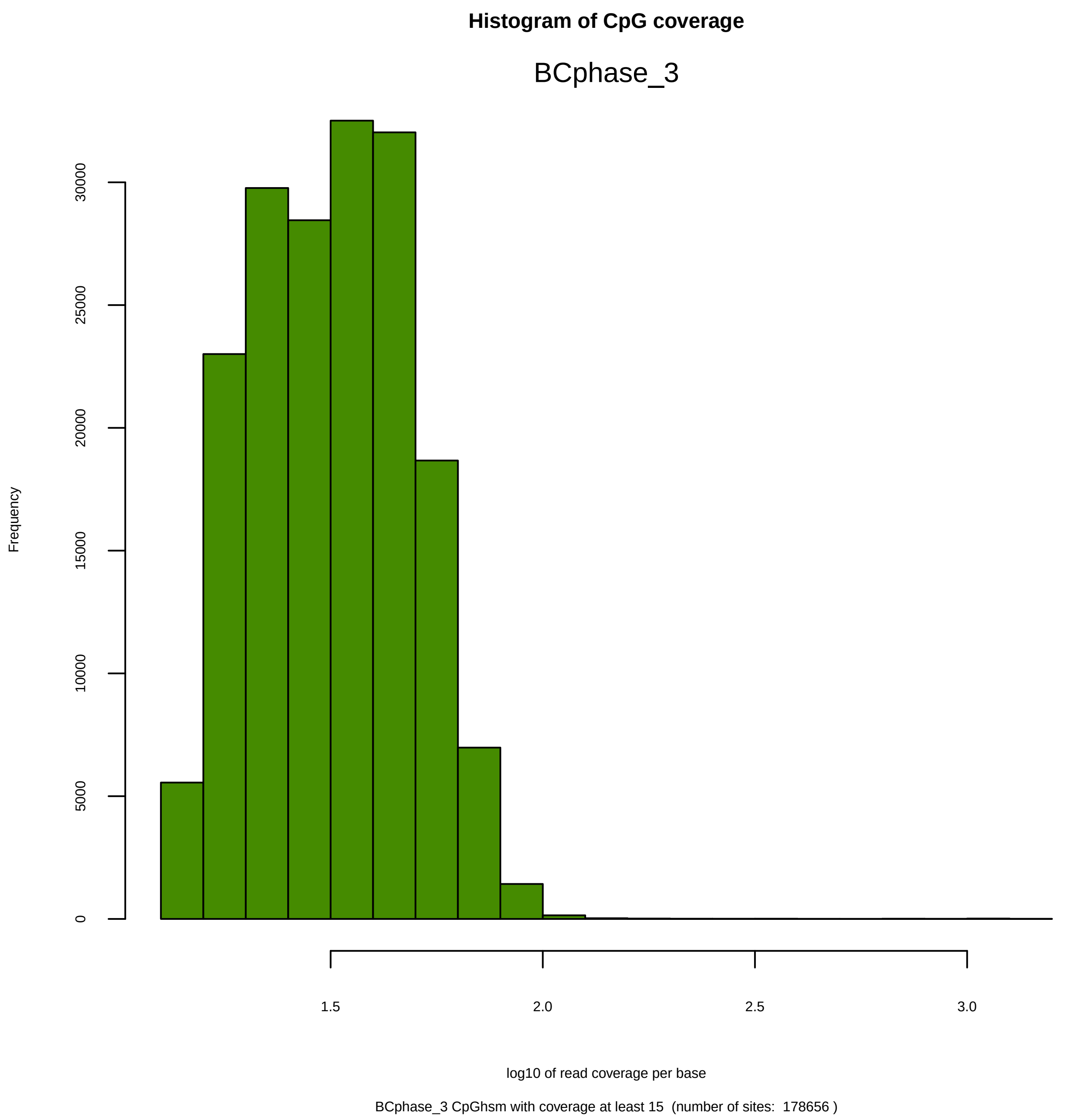
Histogram of CpG coverage
BCphase_3
Frequency
0
5000
10000
15000
20000
25000
30000
1.5
2.0
2.5
3.0
log10 of read coverage per base
BCphase_3 CpGhsm with coverage at least 15 (number of sites: 178656 )

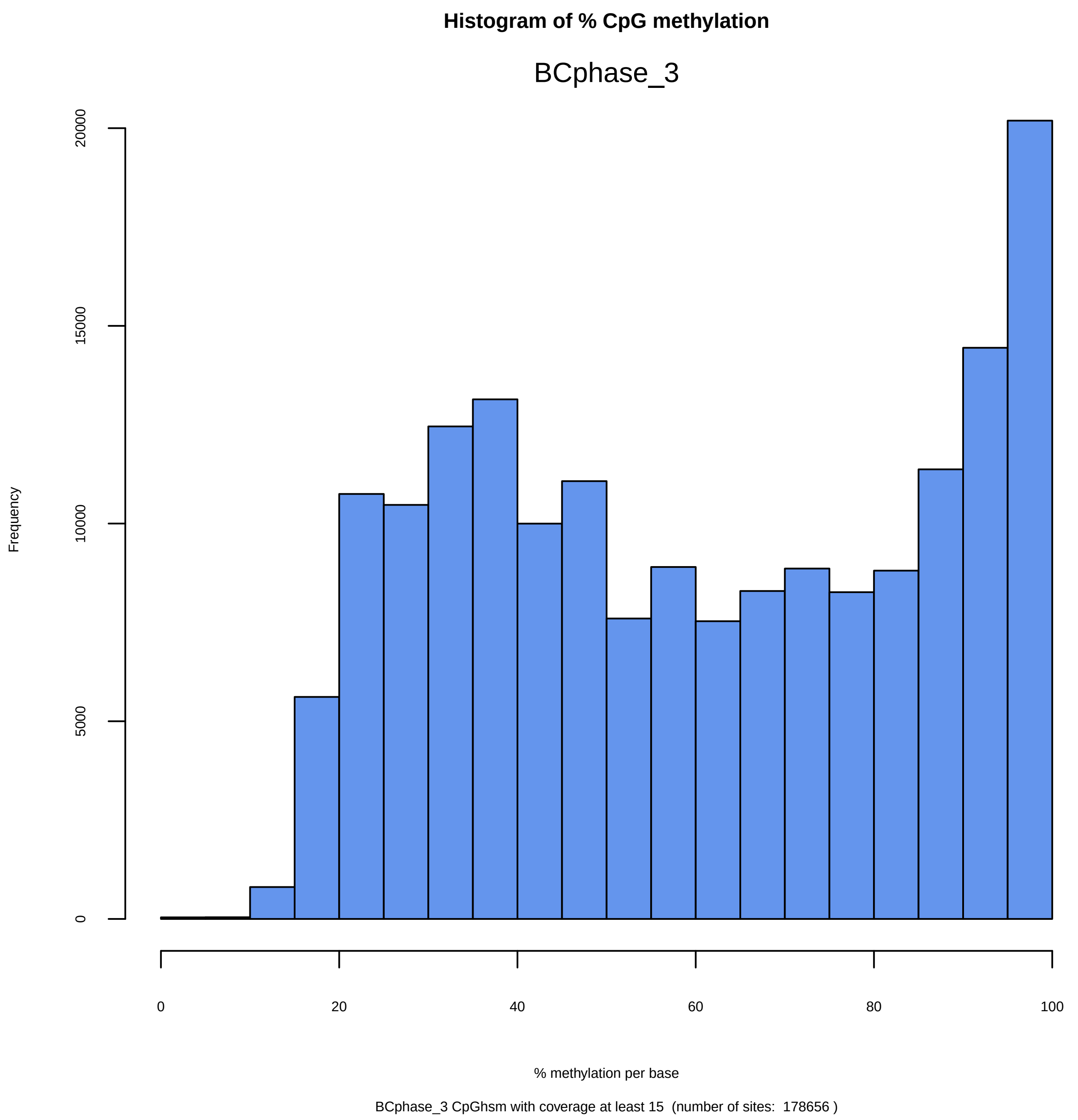
Histogram of % CpG methylation
BCphase_3
Frequency
0
5000
10000
15000
20000
0
20
40
60
80
100
% methylation per base
BCphase_3 CpGhsm with coverage at least 15 (number of sites: 178656 )

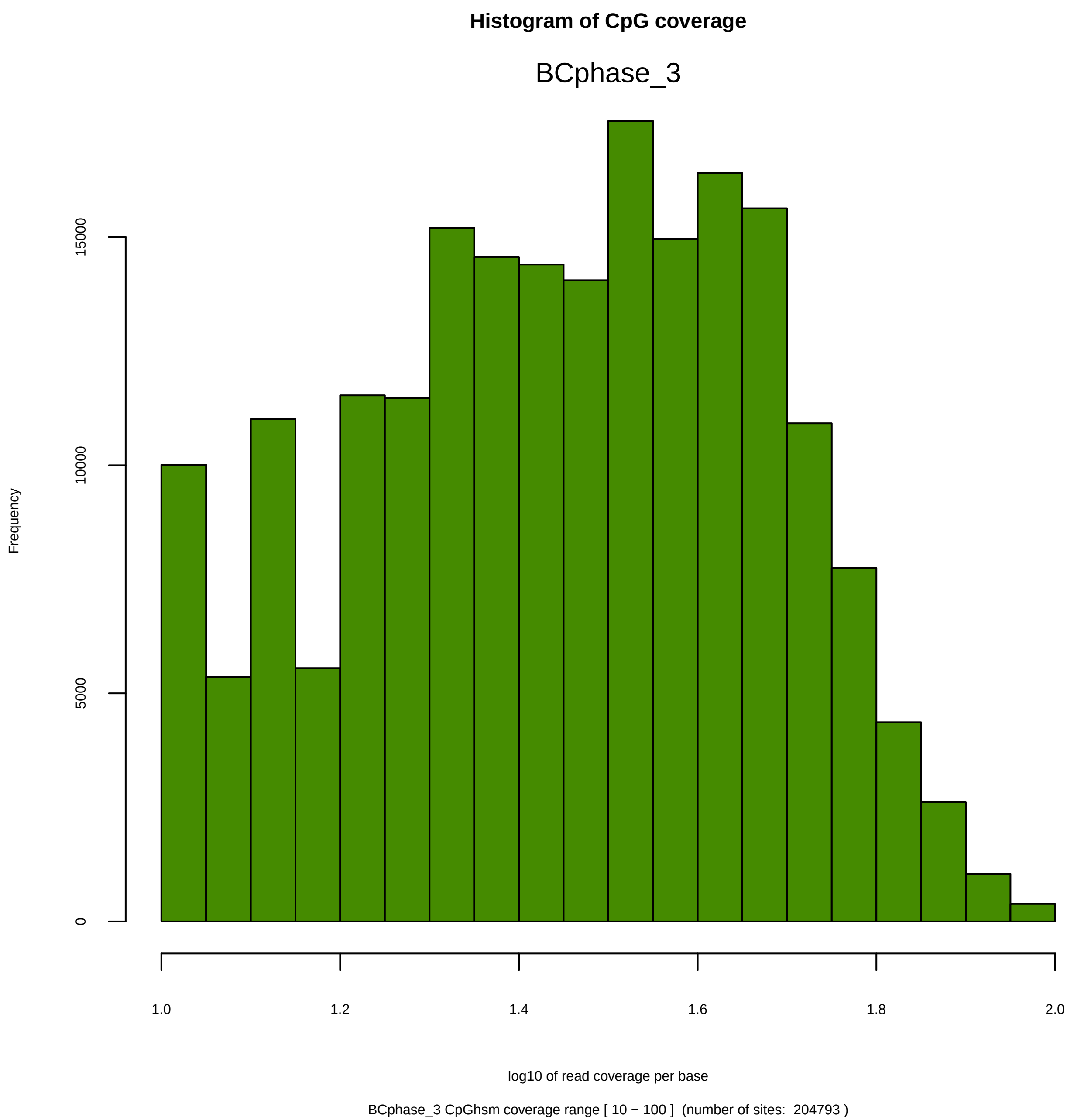

Histogram of CpG coverage
BCphase_3
Frequency
0
5000
10000
15000
1.0
1.2
1.4
1.6
1.8
2.0
log10 of read coverage per base
BCphase_3 CpGhsm coverage range [ 10 − 100 ]  (number of sites:  204793 )

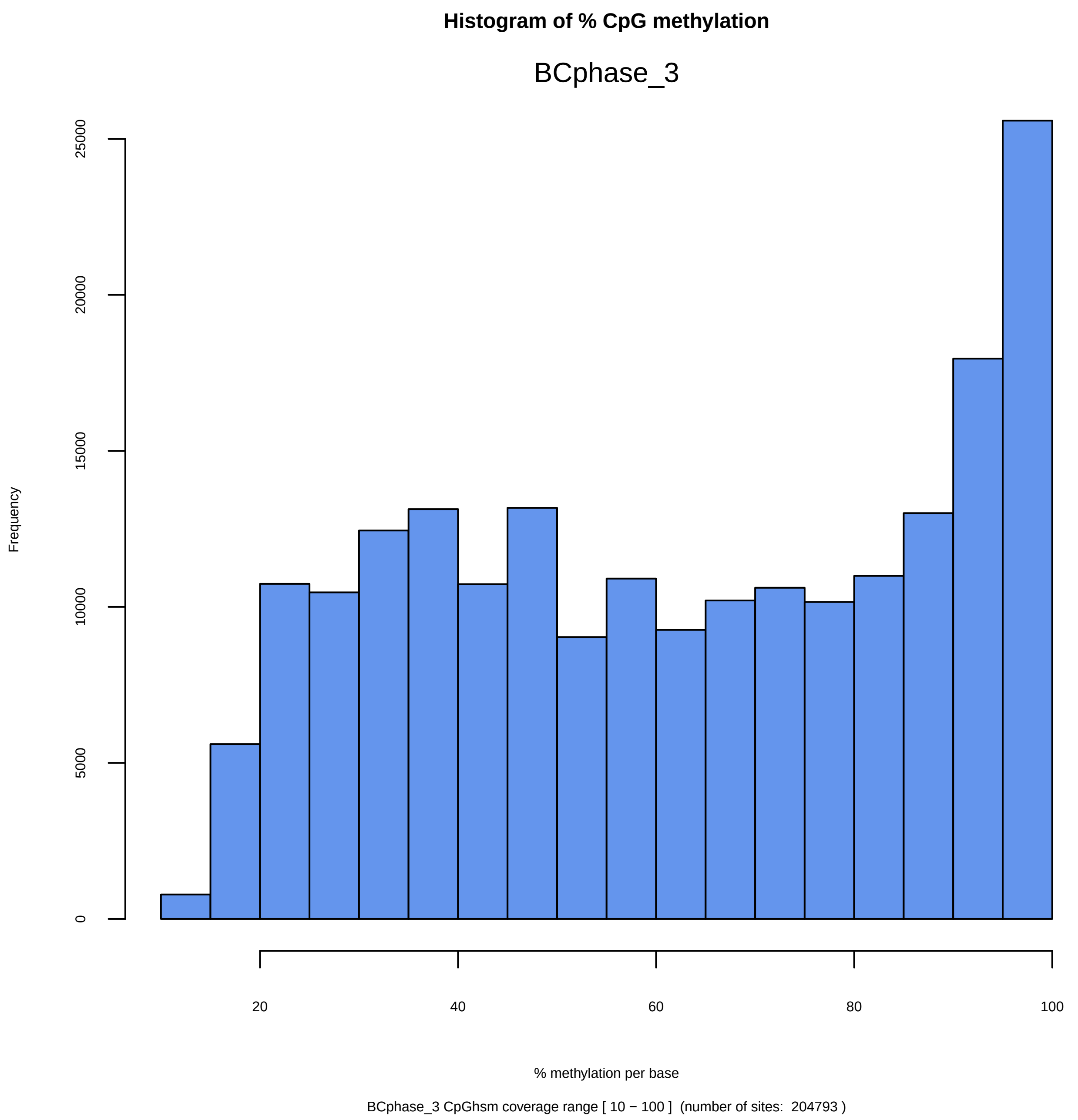
Histogram of % CpG methylation
BCphase_3
Frequency
0
5000
10000
15000
20000
25000
20
40
60
80
100
% methylation per base
BCphase_3 CpGhsm coverage range [ 10 − 100 ]  (number of sites:  204793 )

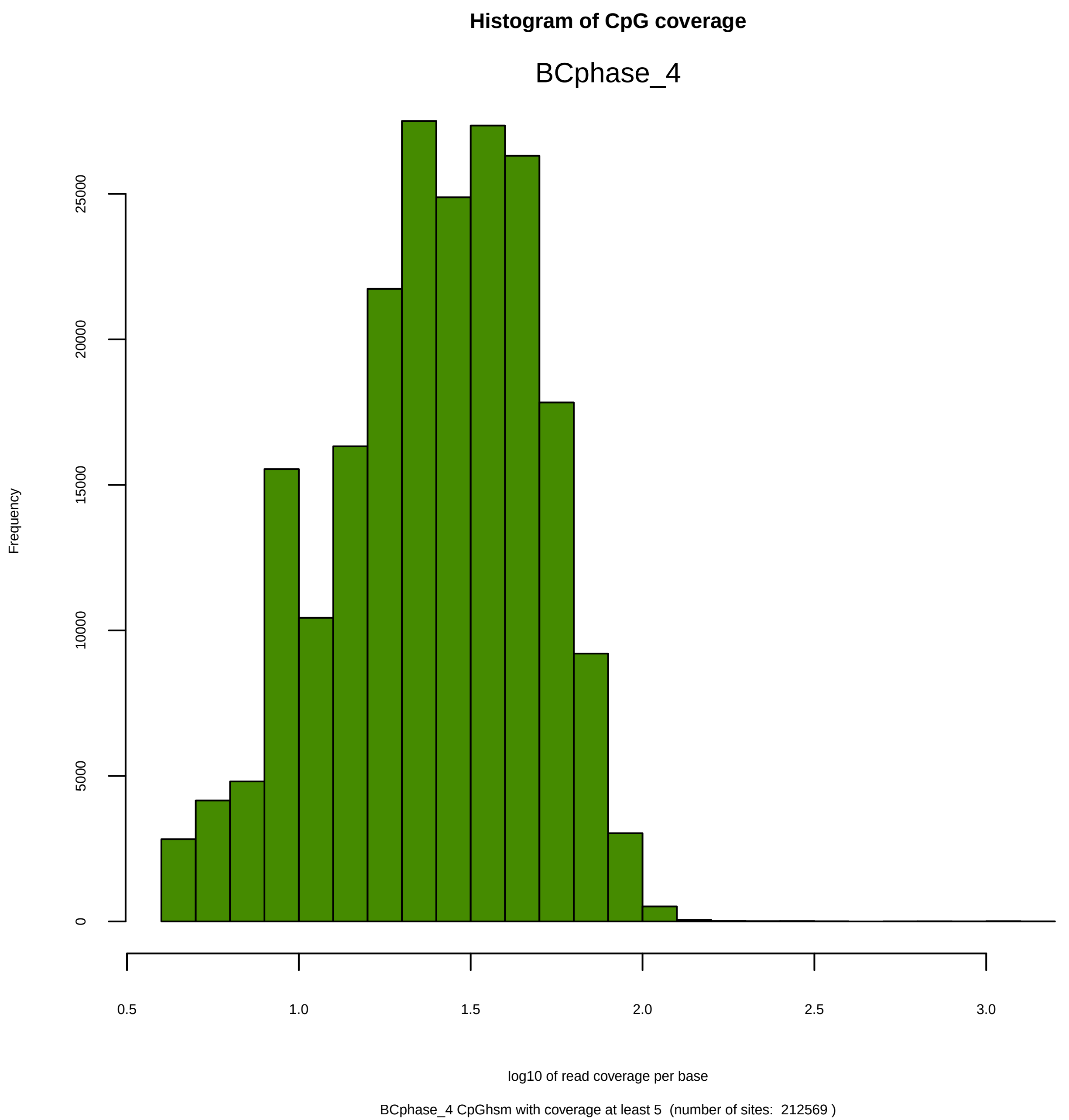
Histogram of CpG coverage
BCphase_4
Frequency
0
5000
10000
15000
20000
25000
0.5
1.0
1.5
2.0
2.5
3.0
log10 of read coverage per base
BCphase_4 CpGhsm with coverage at least 5  (number of sites:  212569 )

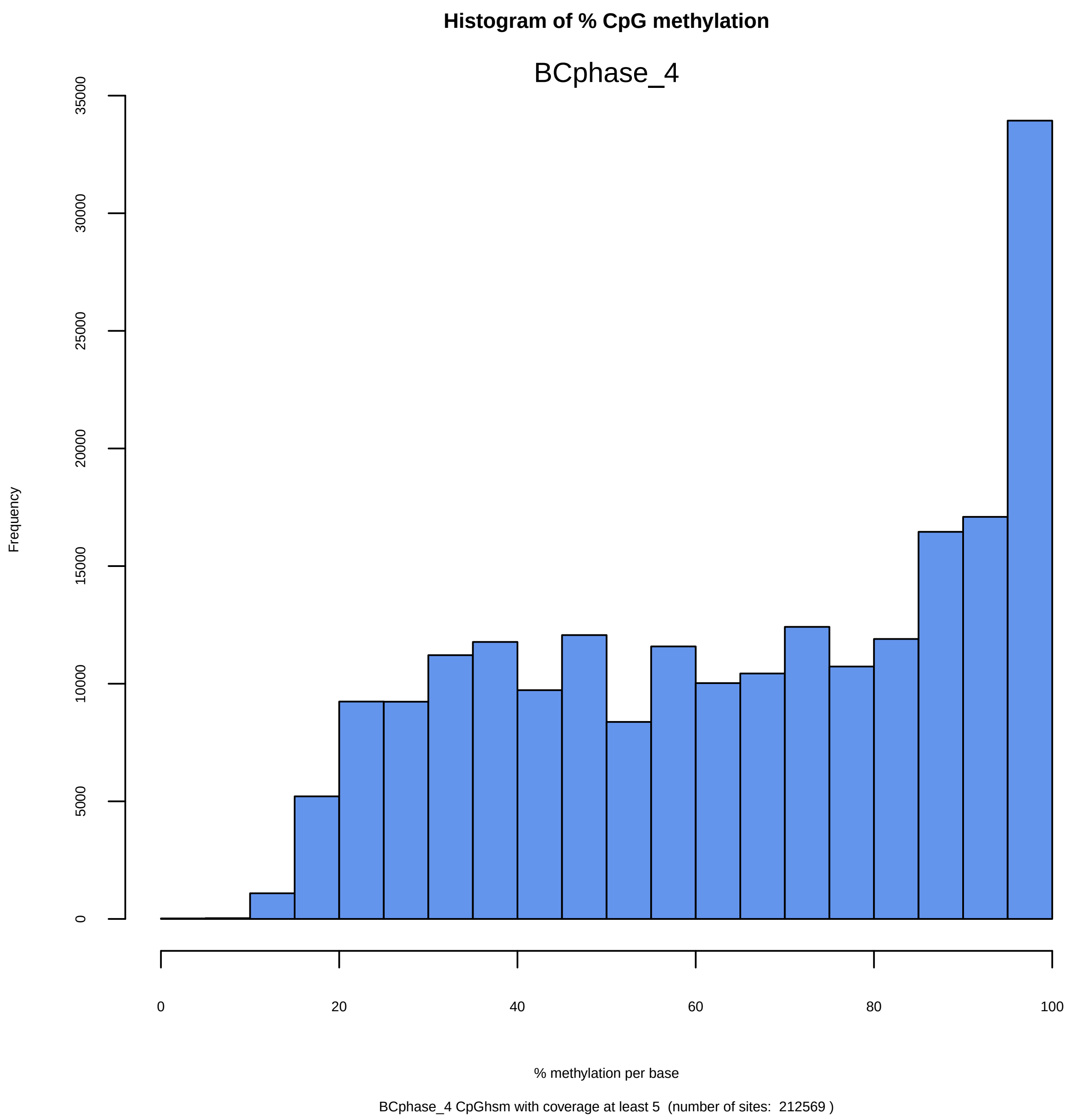

Histogram of % CpG methylation
BCphase_4
Frequency
0
5000
10000
15000
20000
25000
30000
35000
0
20
40
60
80
100
% methylation per base
BCphase_4 CpGhsm with coverage at least 5  (number of sites:  212569 )

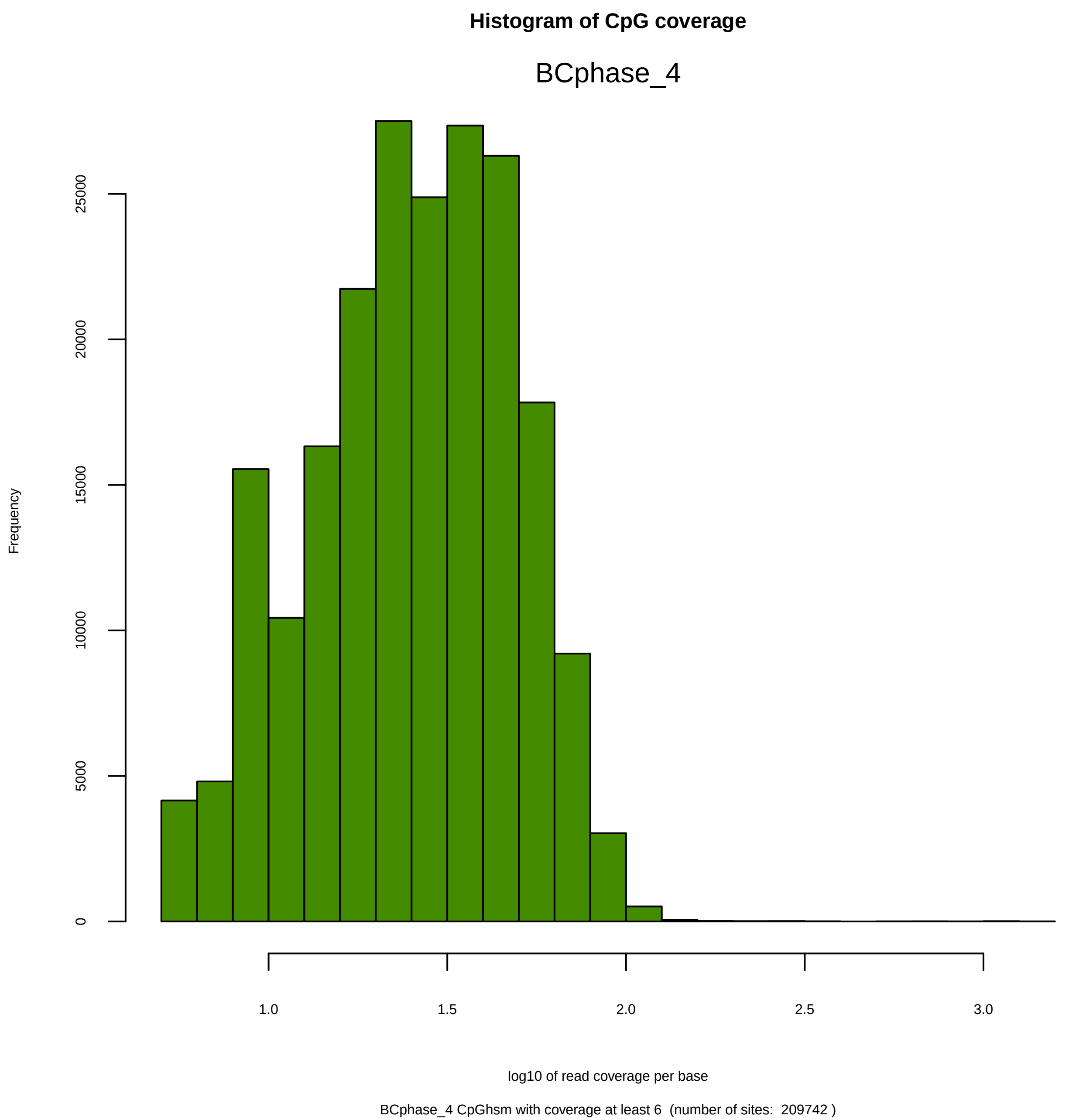
Histogram of CpG coverage
BCphase_4
Frequency
0
5000
10000
15000
20000
25000
1.0
1.5
2.0
2.5
3.0
log10 of read coverage per base
BCphase_4 CpGhsm with coverage at least 6 (number of sites: 209742 )

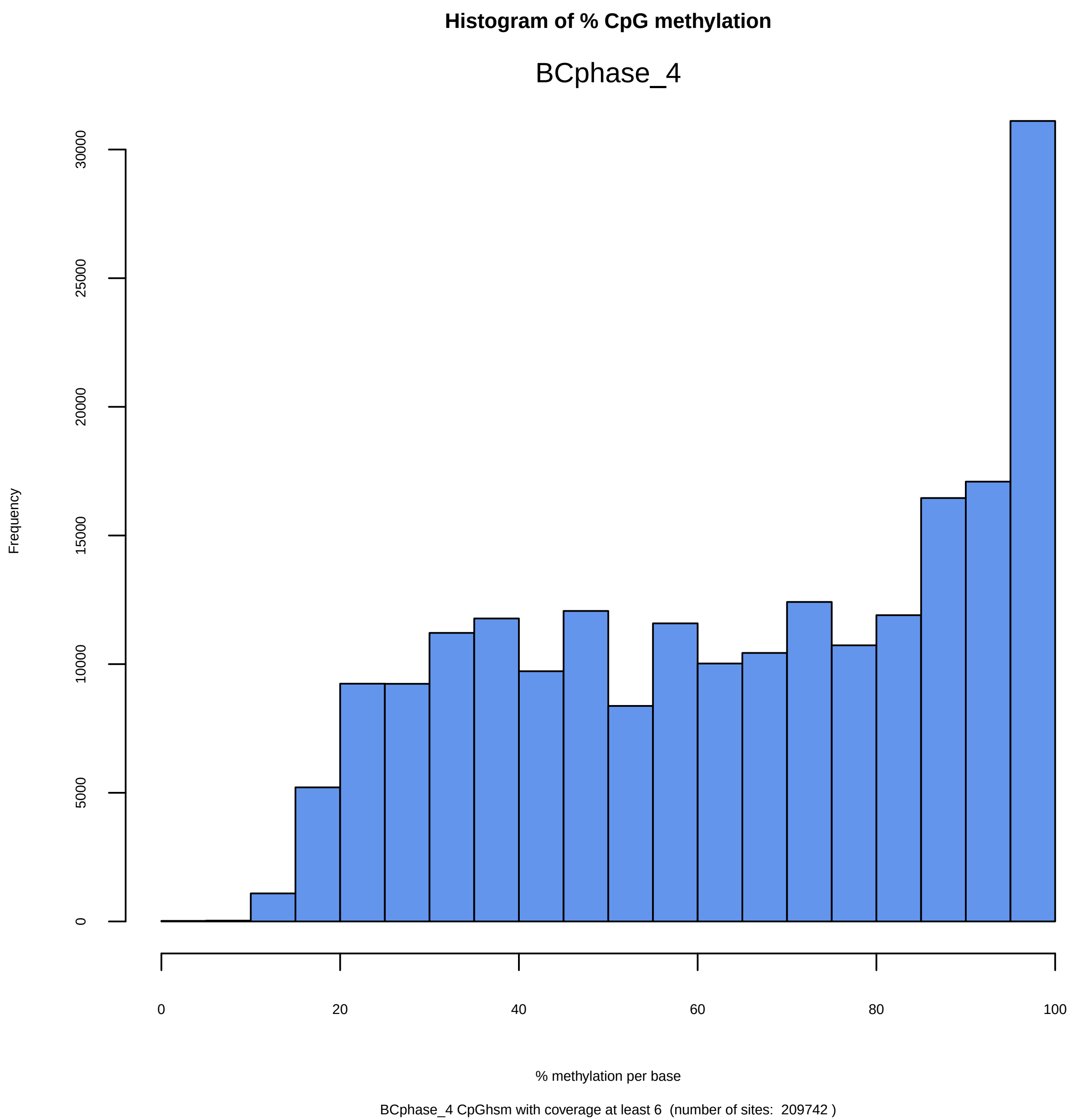

Histogram of % CpG methylation
BCphase_4
Frequency
0
5000
10000
15000
20000
25000
30000
0
20
40
60
80
100
% methylation per base
BCphase_4 CpGhsm with coverage at least 6 (number of sites: 209742 )

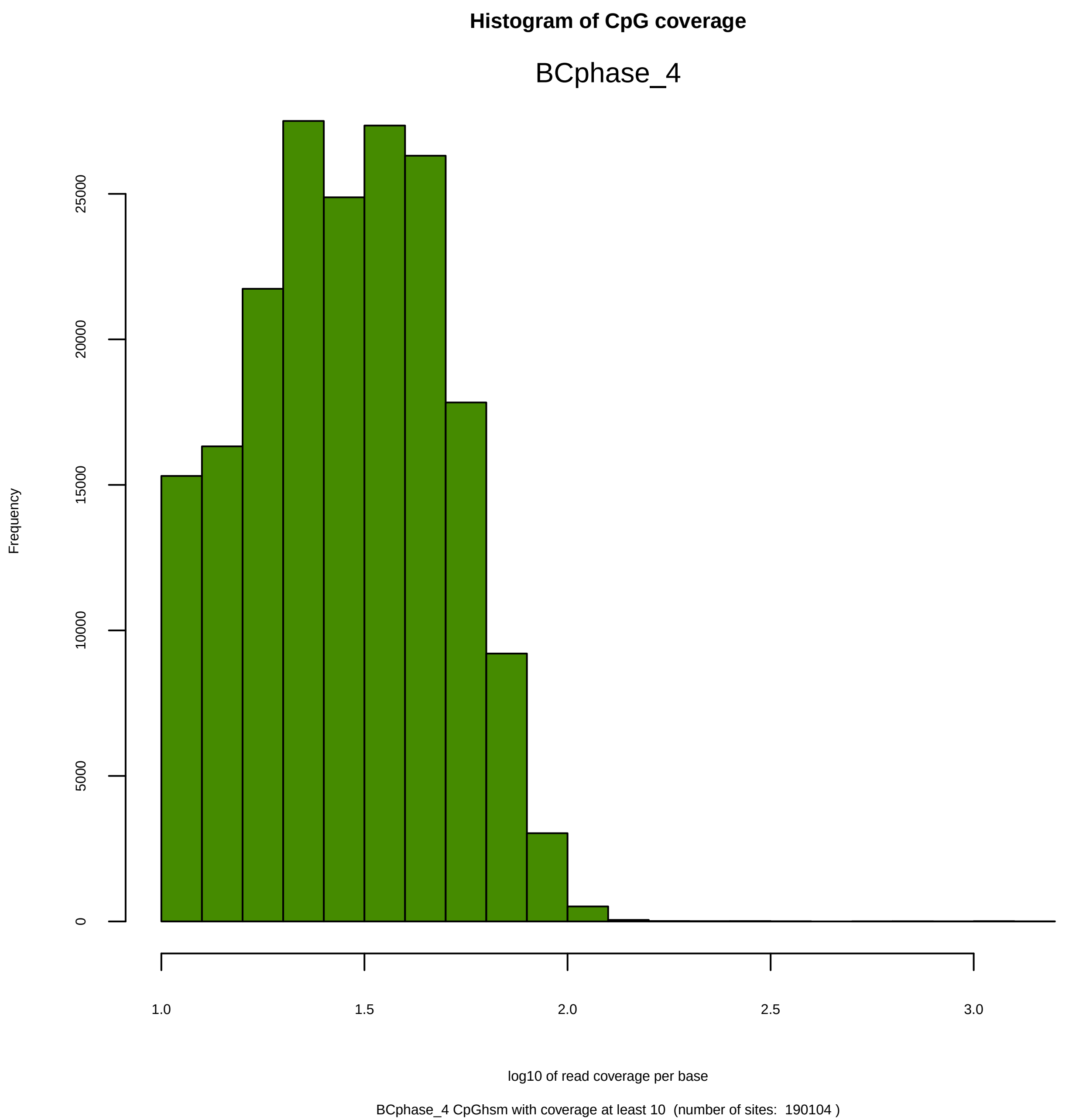

Histogram of CpG coverage
BCphase_4
Frequency
0
5000
10000
15000
20000
25000
1.0
1.5
2.0
2.5
3.0
log10 of read coverage per base
BCphase_4 CpGhsm with coverage at least 10  (number of sites:  190104 )

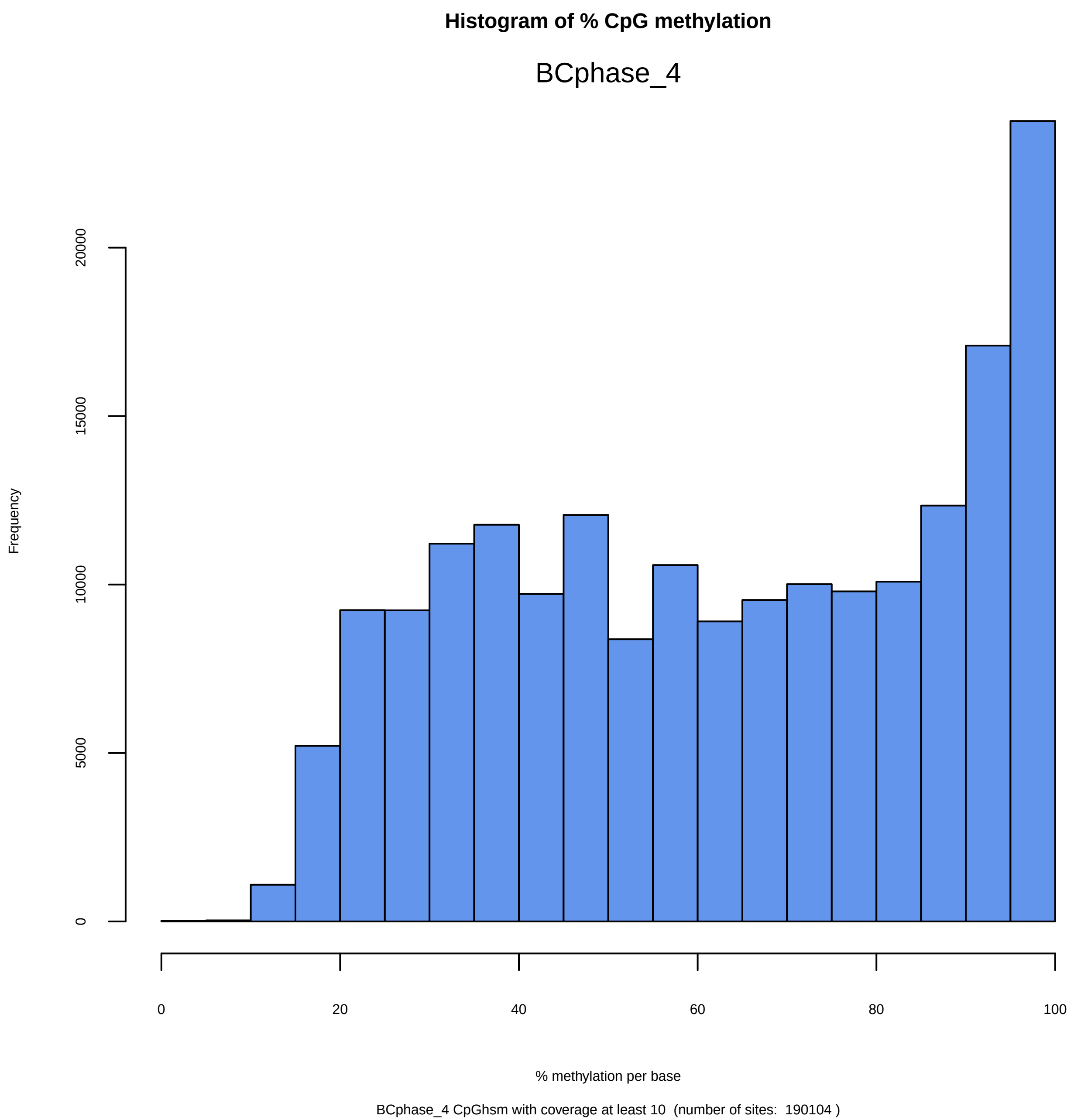

Histogram of % CpG methylation
BCphase_4
Frequency
0
5000
10000
15000
20000
0
20
40
60
80
100
% methylation per base
BCphase_4 CpGhsm with coverage at least 10 (number of sites: 190104 )

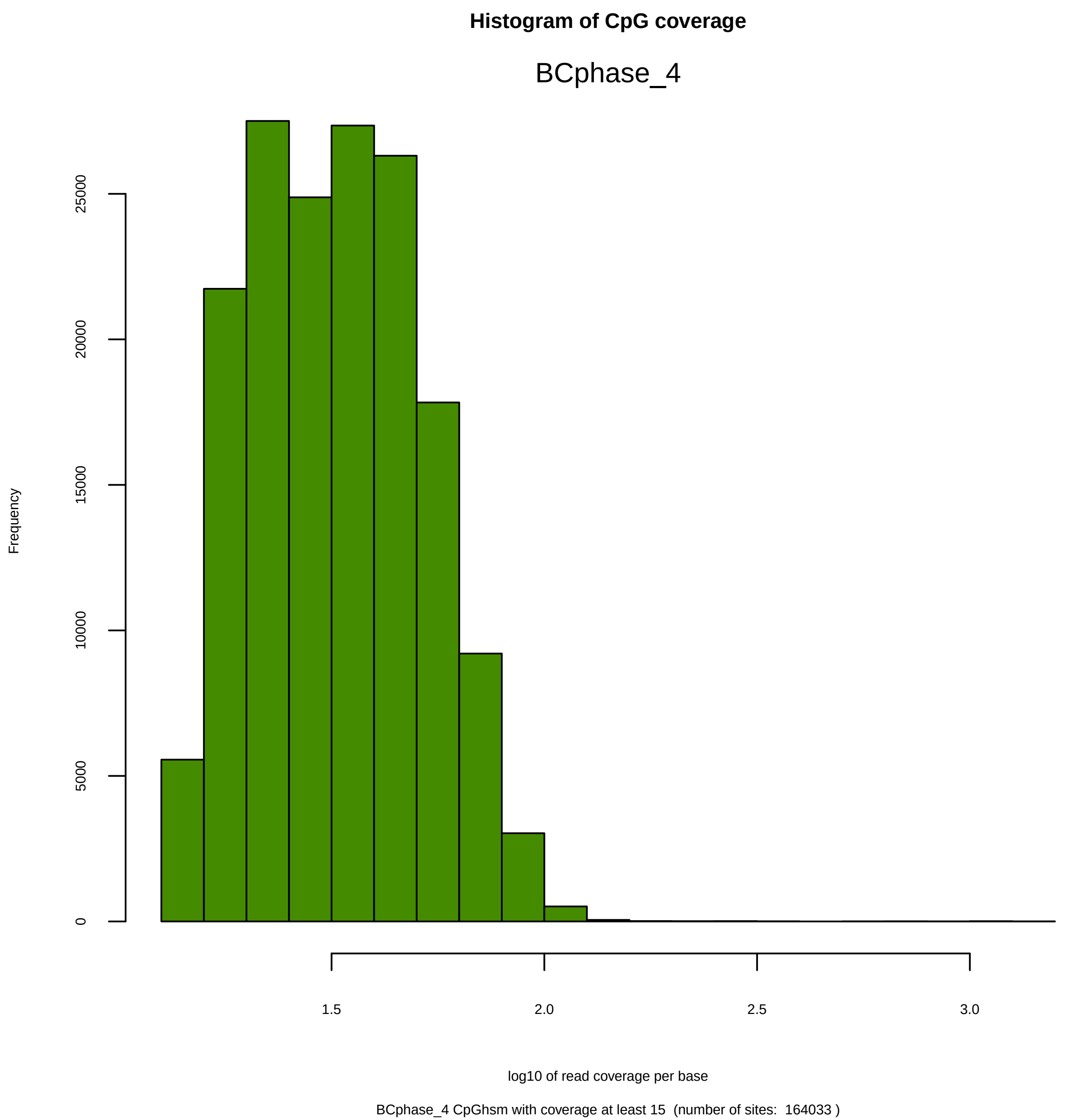

Histogram of CpG coverage
BCphase_4
Frequency
0
5000
10000
15000
20000
25000
1.5
2.0
2.5
3.0
log10 of read coverage per base
BCphase_4 CpGhsm with coverage at least 15  (number of sites:  164033 )

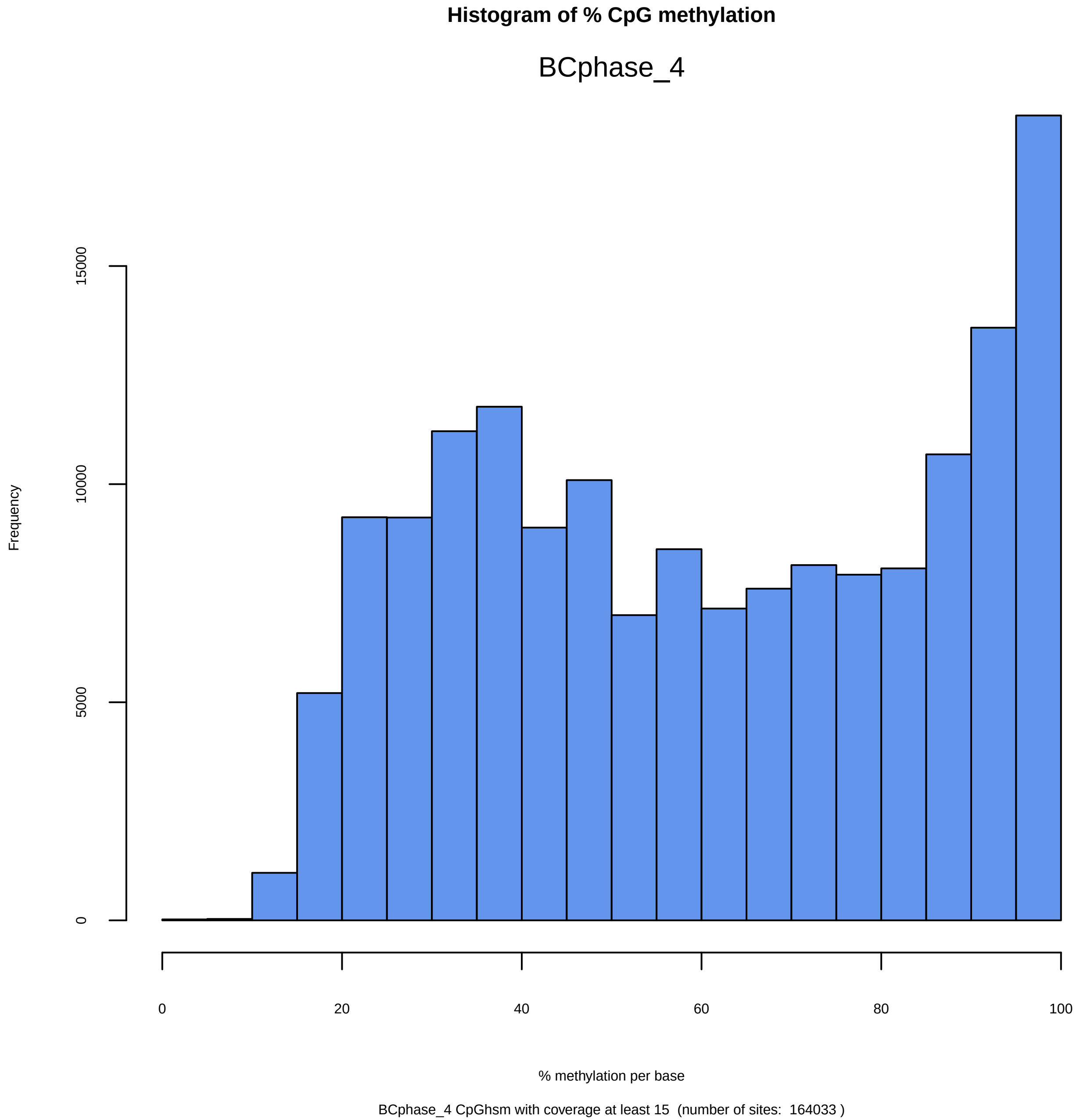

Histogram of % CpG methylation
BCphase_4
Frequency
0
5000
10000
15000
0
20
40
60
80
100
% methylation per base
BCphase_4 CpGhsm with coverage at least 15 (number of sites: 164033 )

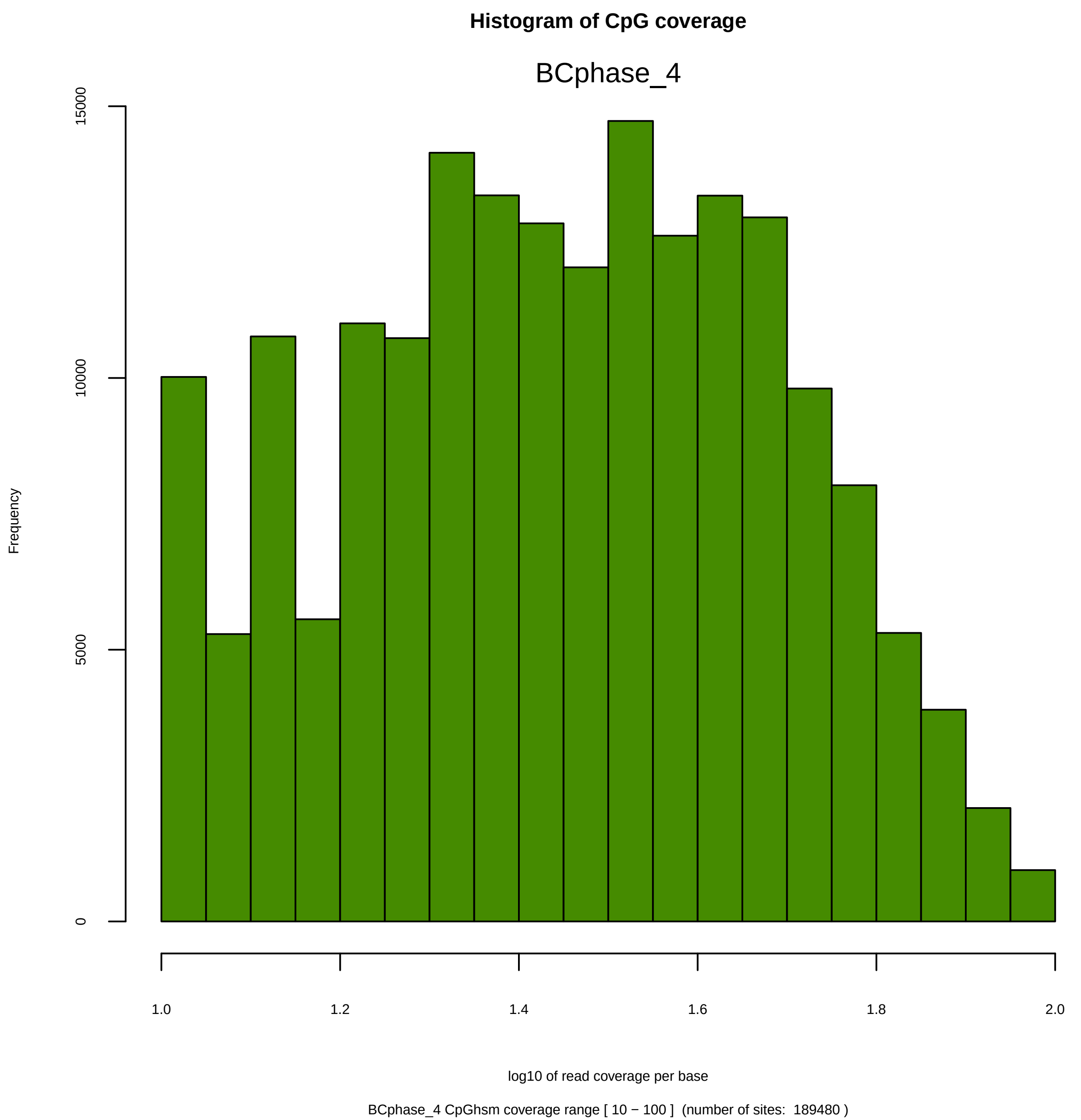

Histogram of CpG coverage
BCphase_4
Frequency
0
5000
10000
15000
1.0
1.2
1.4
1.6
1.8
2.0
log10 of read coverage per base
BCphase_4 CpGhsm coverage range [ 10 − 100 ]  (number of sites:  189480 )

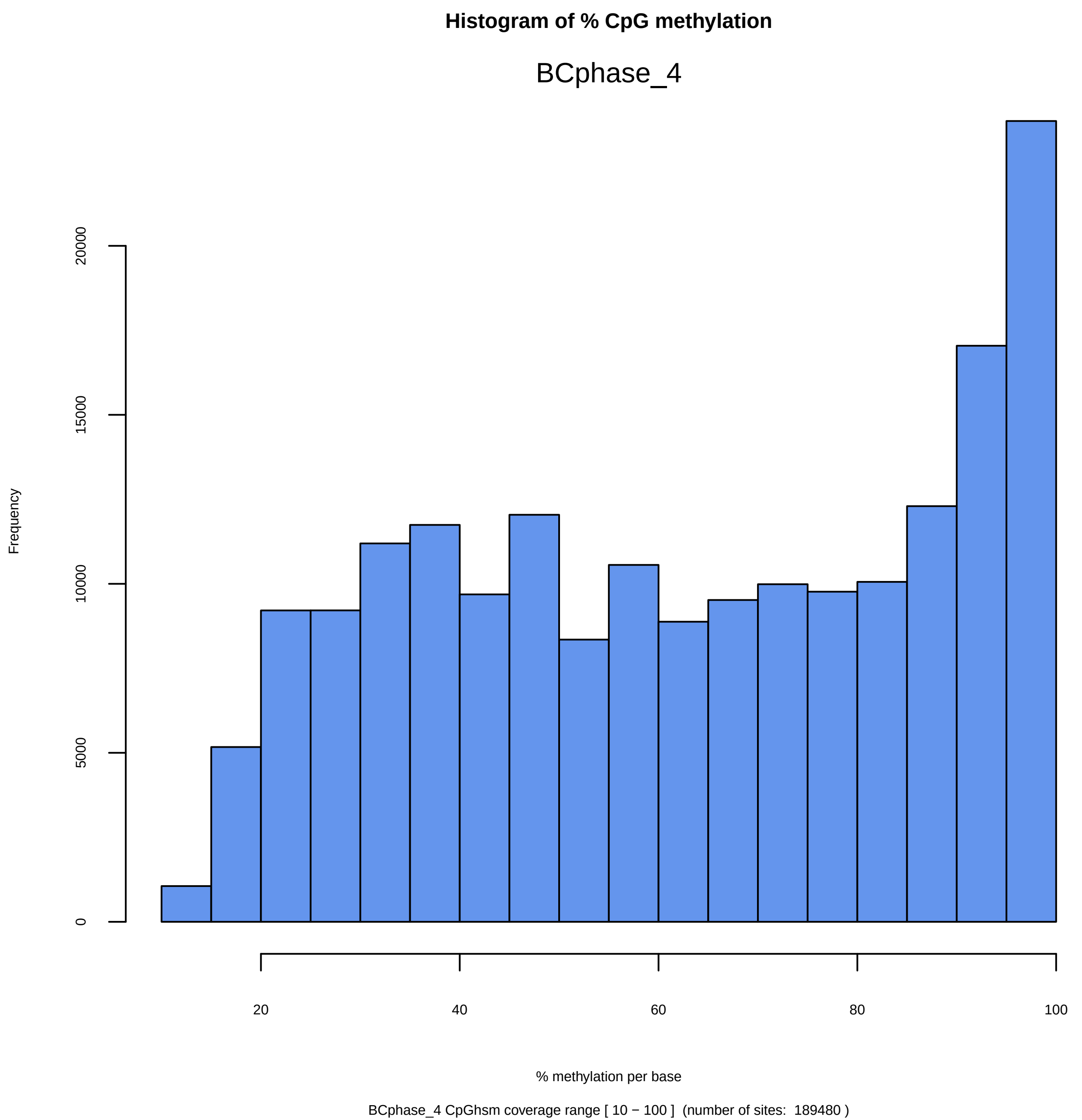

Histogram of % CpG methylation
BCphase_4
Frequency
0
5000
10000
15000
20000
20
40
60
80
100
% methylation per base
BCphase_4 CpGhsm coverage range [ 10 − 100 ]  (number of sites:  189480 )

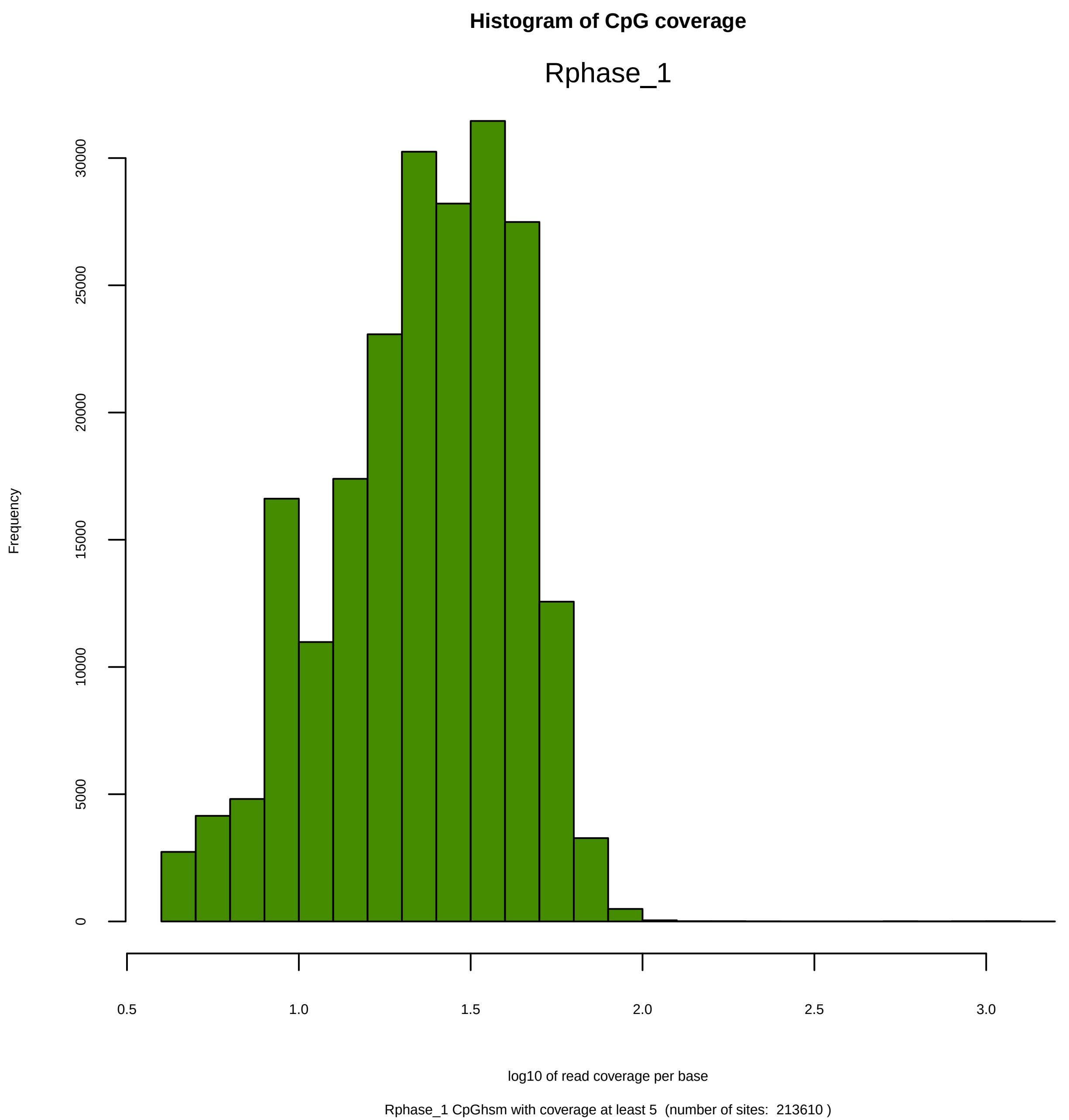

Histogram of CpG coverage
Rphase_1
Frequency
0
5000
10000
15000
20000
25000
30000
0.5
1.0
1.5
2.0
2.5
3.0
log10 of read coverage per base
Rphase_1 CpGhsm with coverage at least 5 (number of sites: 213610 )

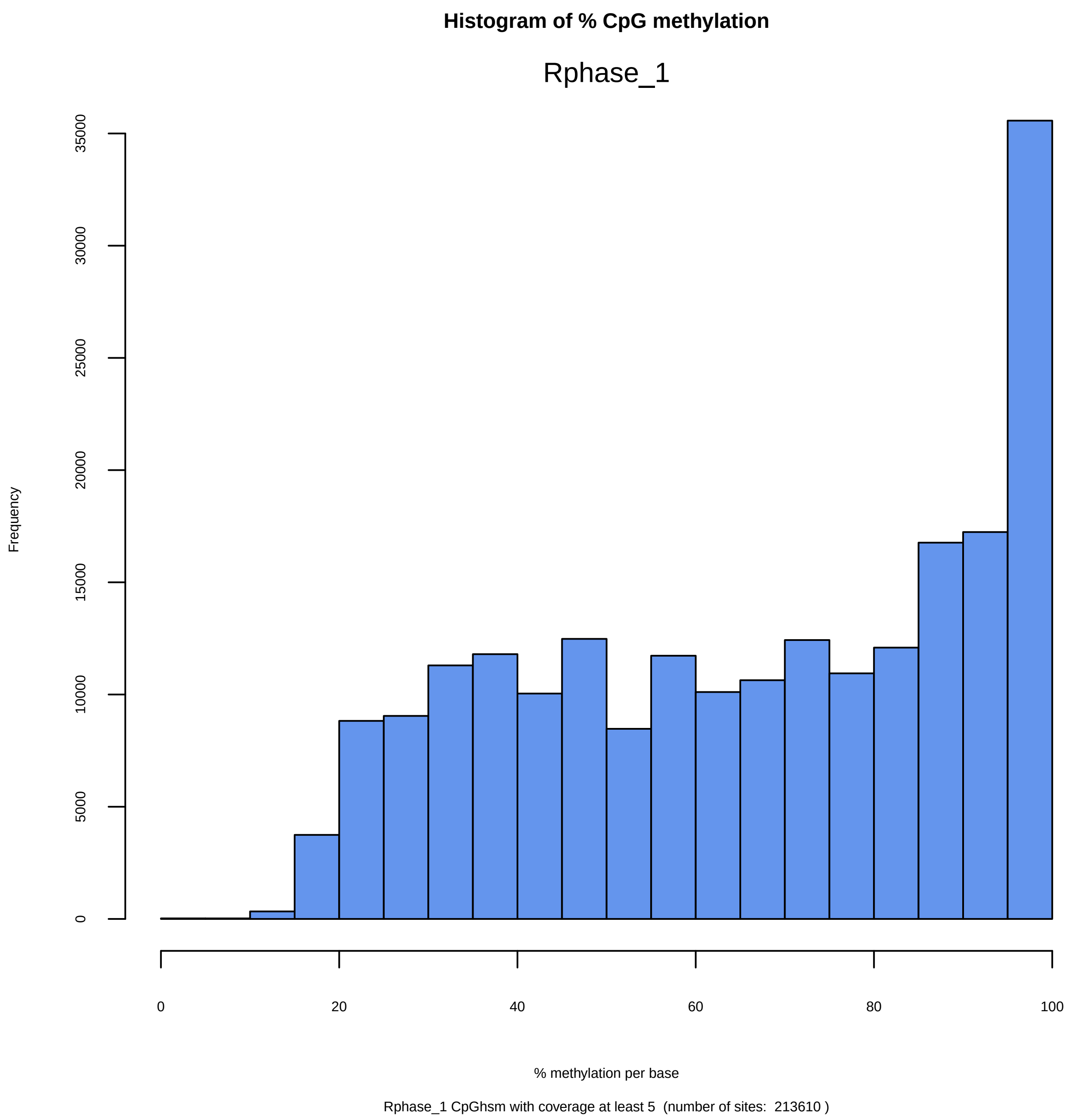
Histogram of % CpG methylation
Rphase_1
Frequency
0
5000
10000
15000
20000
25000
30000
35000
0
20
40
60
80
100
% methylation per base
Rphase_1 CpGhsm with coverage at least 5 (number of sites: 213610 )

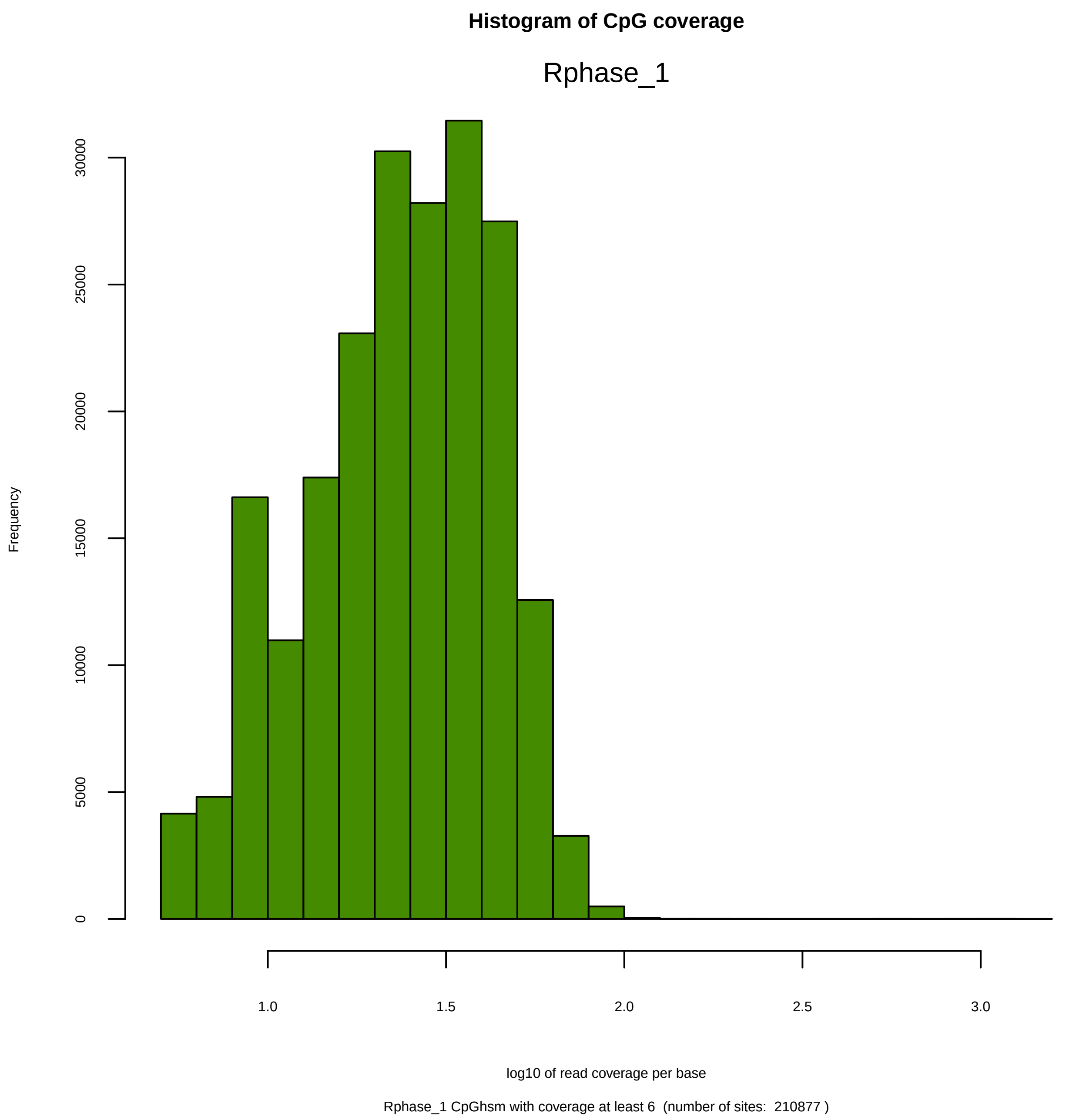

Histogram of CpG coverage
Rphase_1
Frequency
0
5000
10000
15000
20000
25000
30000
1.0
1.5
2.0
2.5
3.0
log10 of read coverage per base
Rphase_1 CpGhsm with coverage at least 6  (number of sites:  210877 )

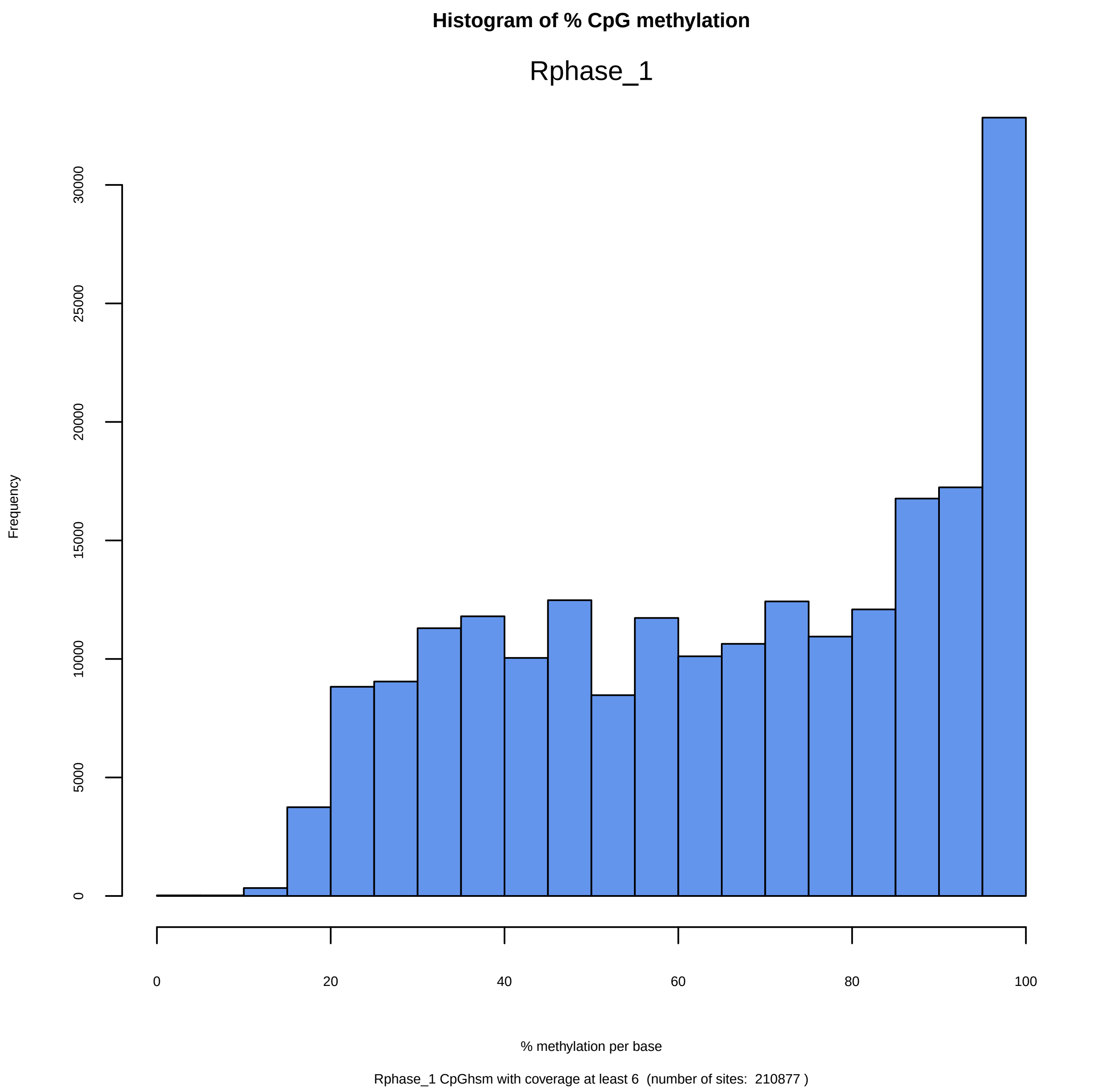
Histogram of % CpG methylation
Rphase_1
Frequency
0
5000
10000
15000
20000
25000
30000
0
20
40
60
80
100
% methylation per base
Rphase_1 CpGhsm with coverage at least 6 (number of sites: 210877 )

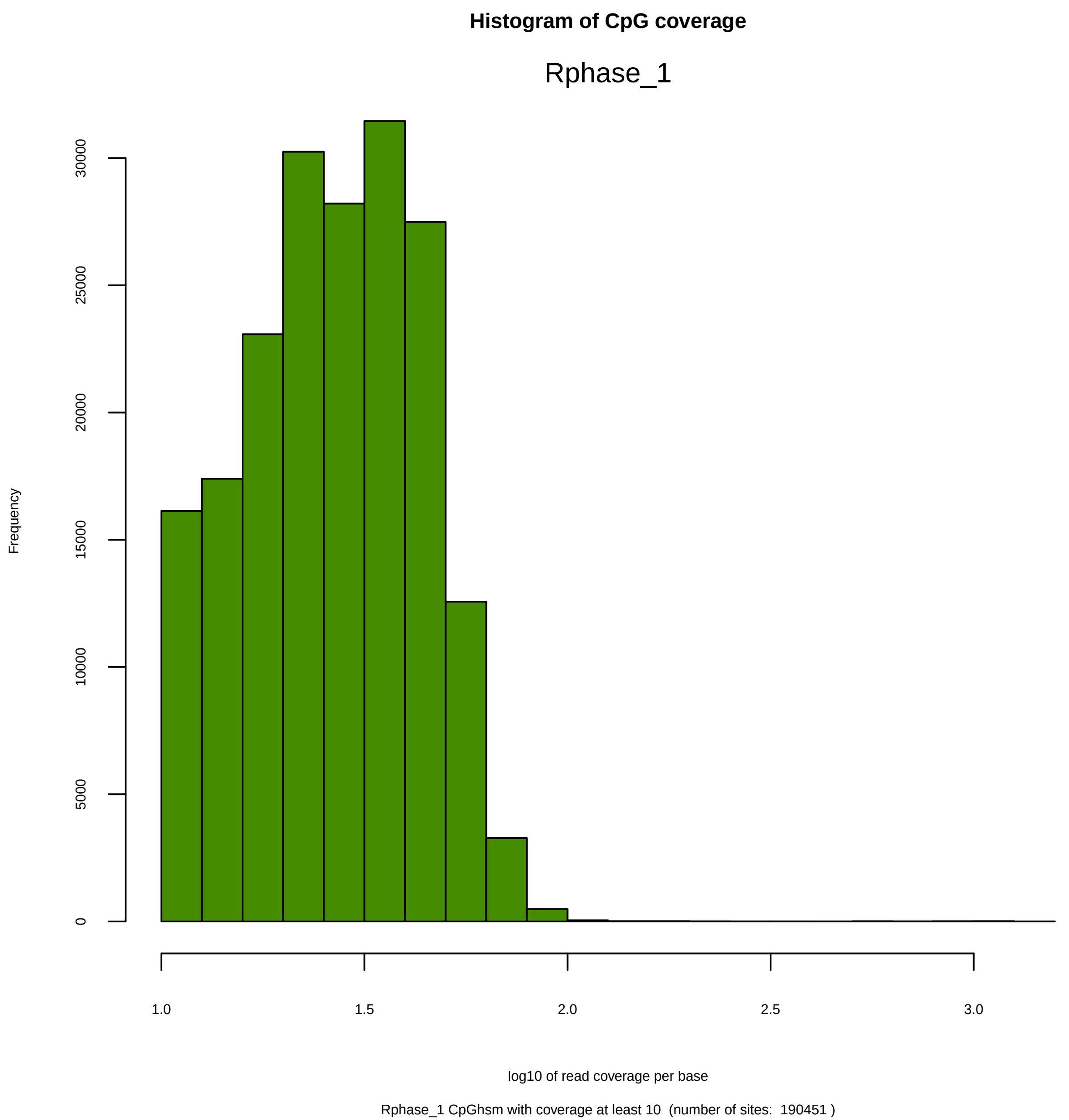

Histogram of CpG coverage
Rphase_1
Frequency
0
5000
10000
15000
20000
25000
30000
1.0
1.5
2.0
2.5
3.0
log10 of read coverage per base
Rphase_1 CpGhsm with coverage at least 10 (number of sites: 190451 )

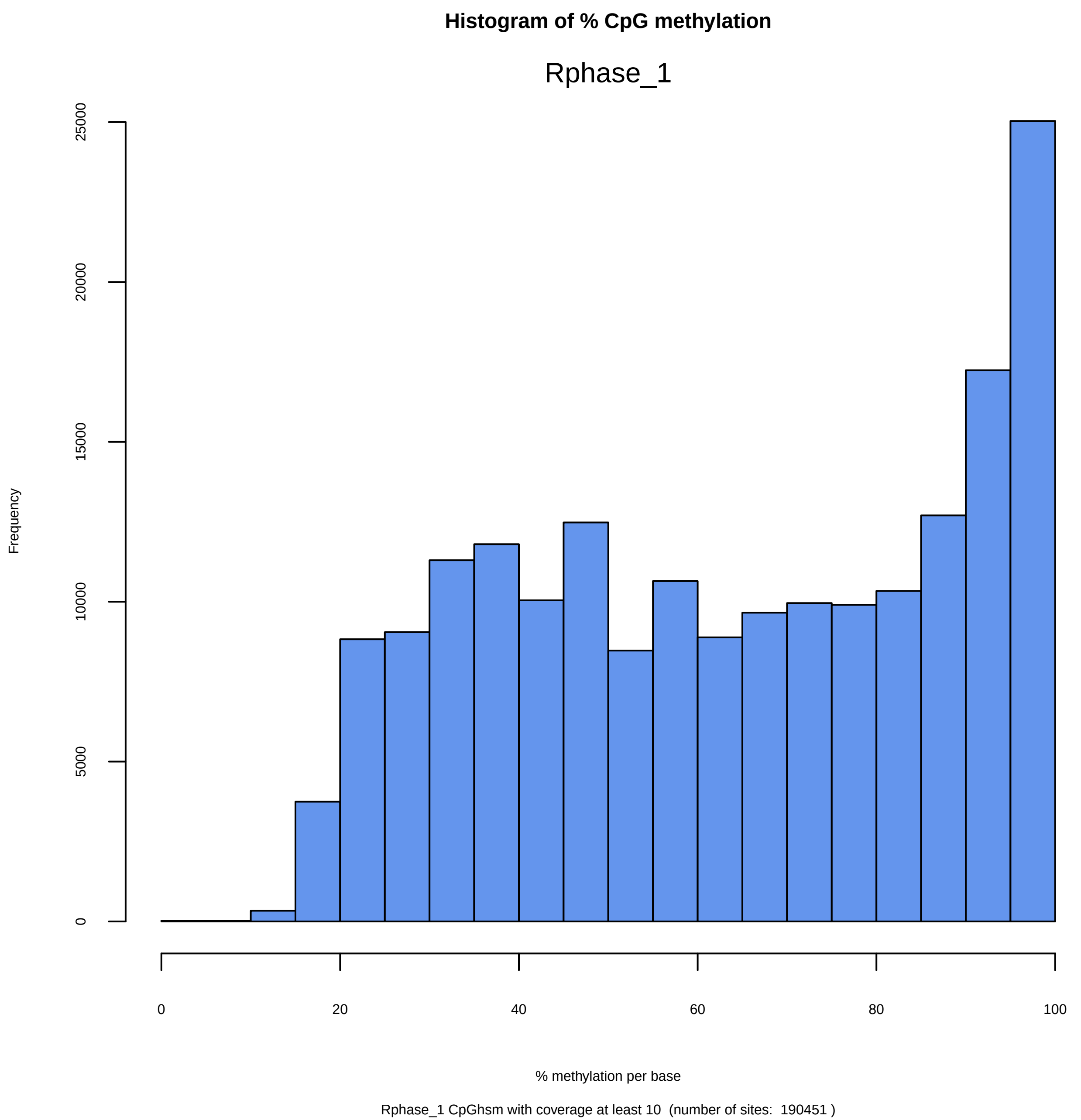

Histogram of % CpG methylation
Rphase_1
Frequency
0
5000
10000
15000
20000
25000
0
20
40
60
80
100
% methylation per base
Rphase_1 CpGhsm with coverage at least 10 (number of sites: 190451 )

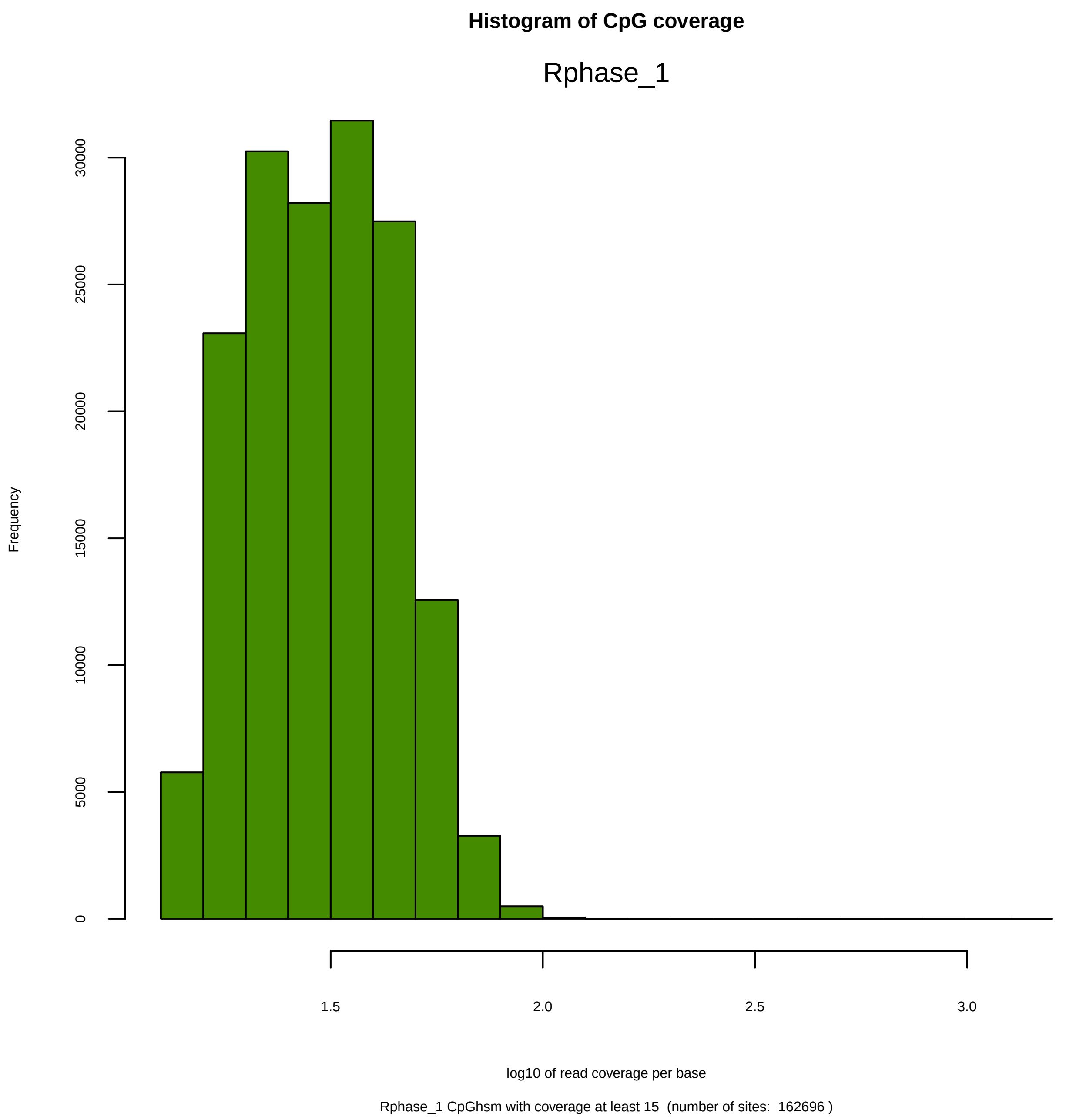

Histogram of CpG coverage
Rphase_1
Frequency
0
5000
10000
15000
20000
25000
30000
1.5
2.0
2.5
3.0
log10 of read coverage per base
Rphase_1 CpGhsm with coverage at least 15  (number of sites:  162696 )

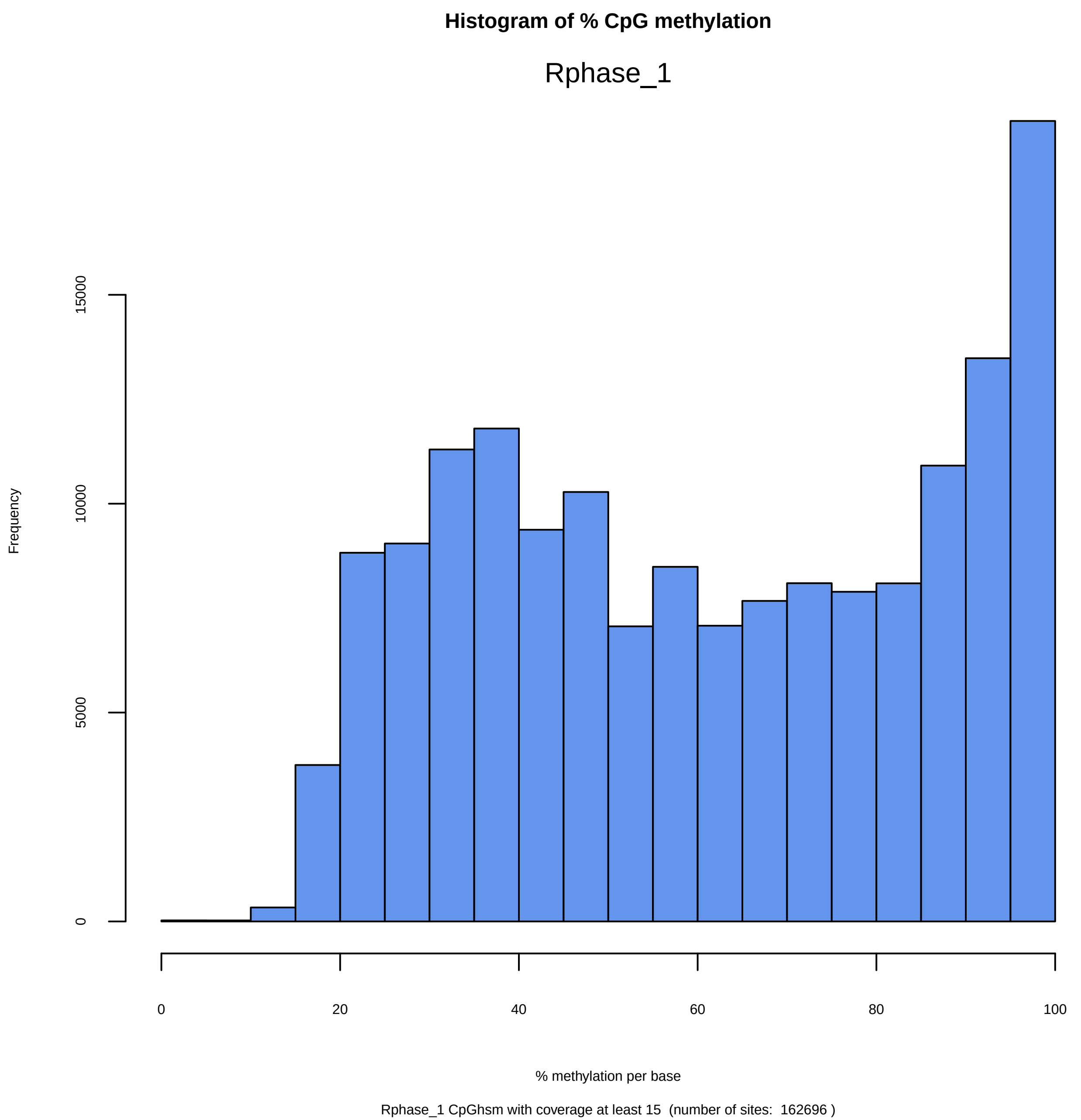
Histogram of % CpG methylation
Rphase_1
Frequency
0
5000
10000
15000
0
20
40
60
80
100
% methylation per base
Rphase_1 CpGhsm with coverage at least 15  (number of sites:  162696 )

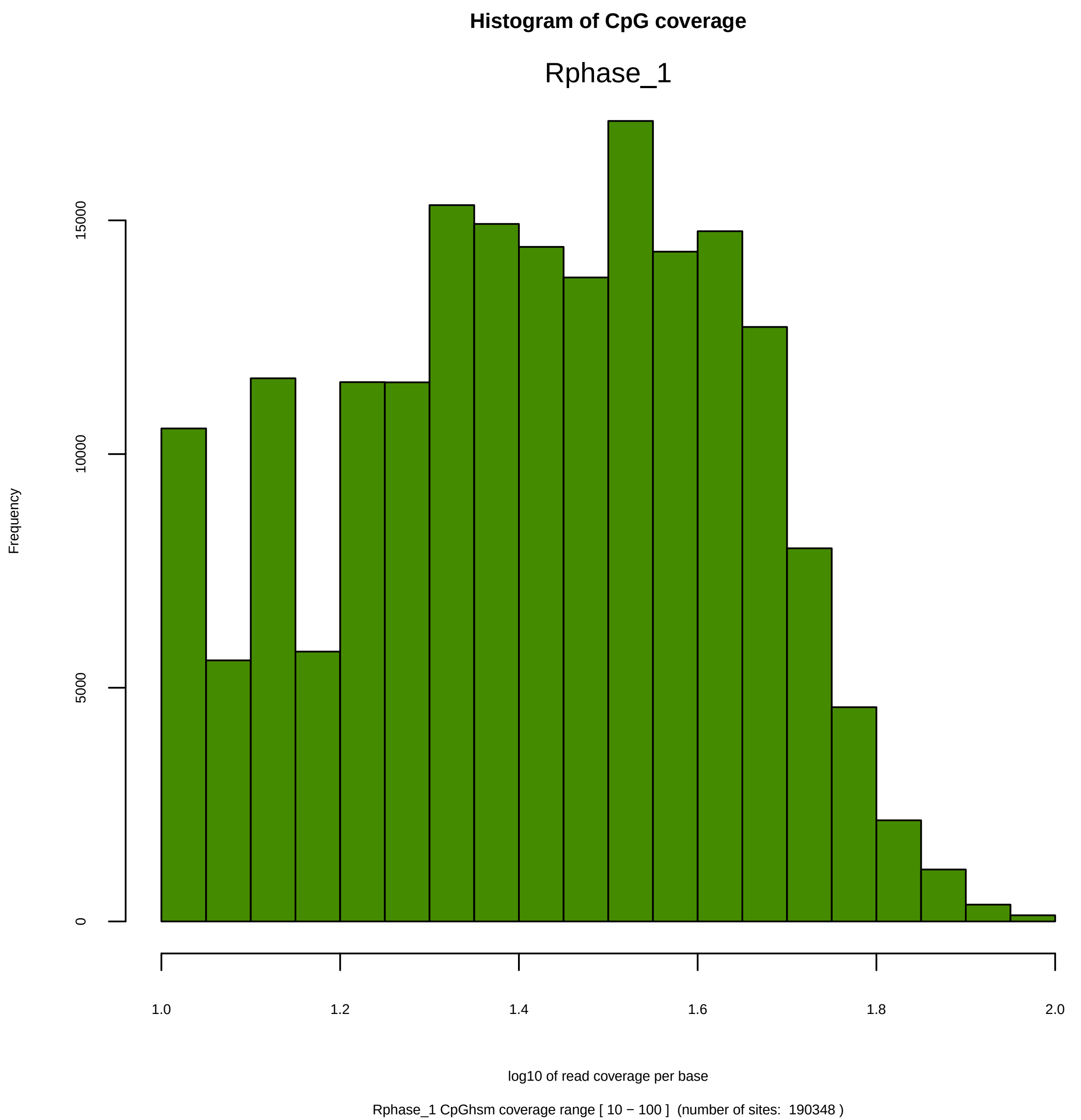
Histogram of CpG coverage
Rphase_1
Frequency
0
5000
10000
15000
1.0
1.2
1.4
1.6
1.8
2.0
log10 of read coverage per base
Rphase_1 CpGhsm coverage range [ 10 − 100 ]  (number of sites:  190348 )

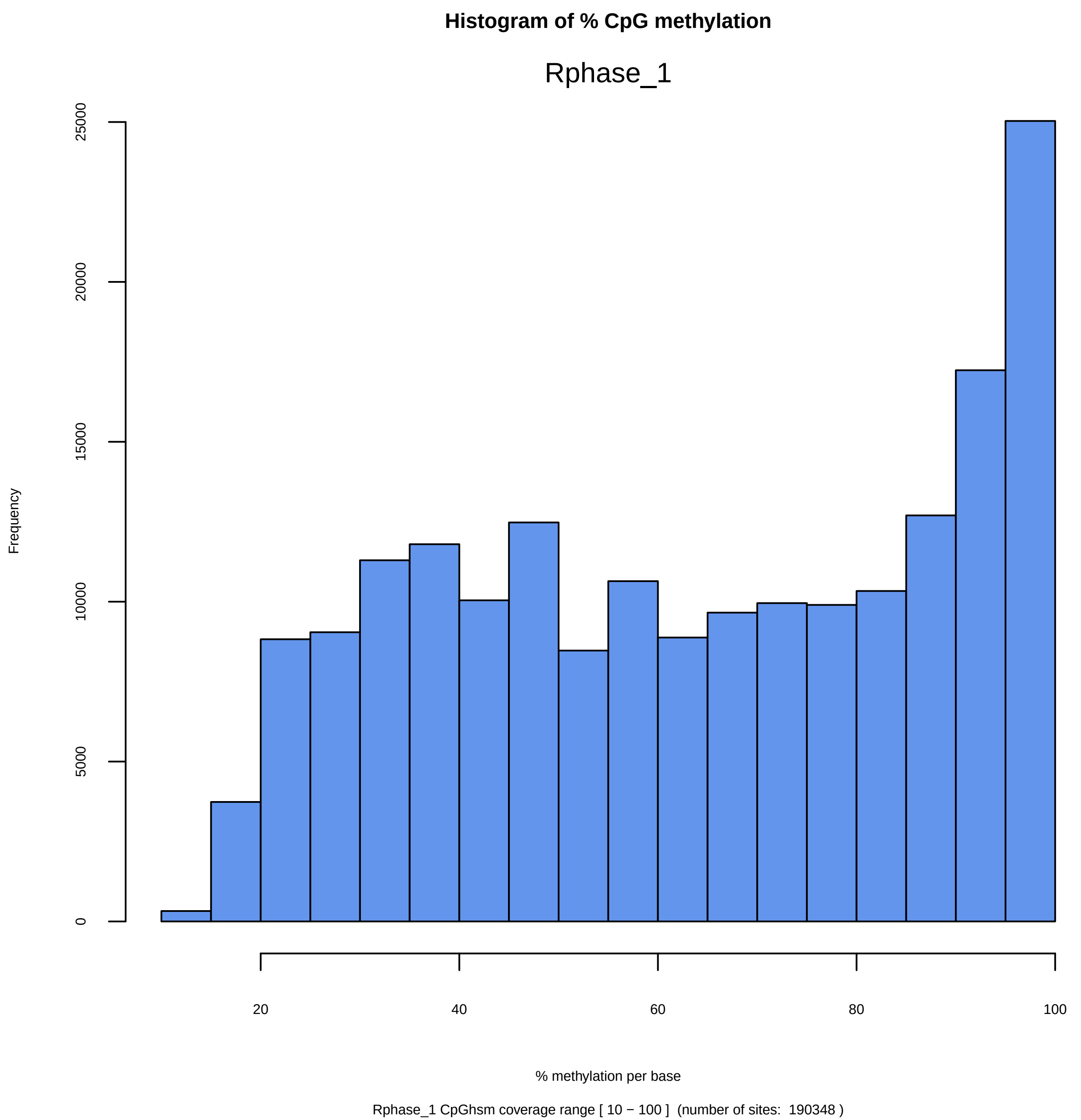
Histogram of % CpG methylation
Rphase_1
Frequency
0
5000
10000
15000
20000
25000
20
40
60
80
100
% methylation per base
Rphase_1 CpGhsm coverage range [ 10 − 100 ] (number of sites: 190348 )

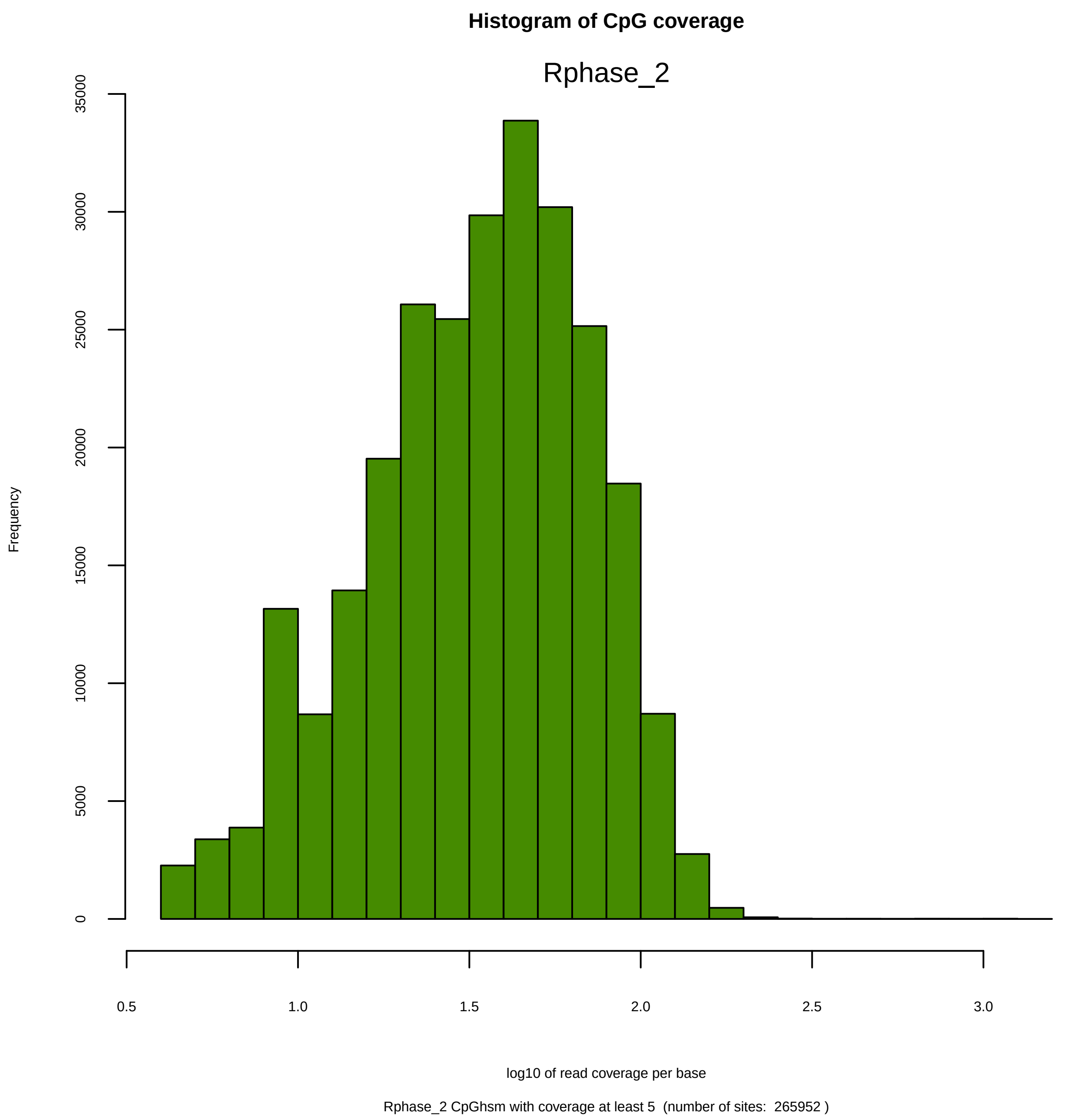

Histogram of CpG coverage
Rphase_2
Frequency
0
5000
10000
15000
20000
25000
30000
35000
0.5
1.0
1.5
2.0
2.5
3.0
log10 of read coverage per base
Rphase_2 CpGhsm with coverage at least 5 (number of sites: 265952 )

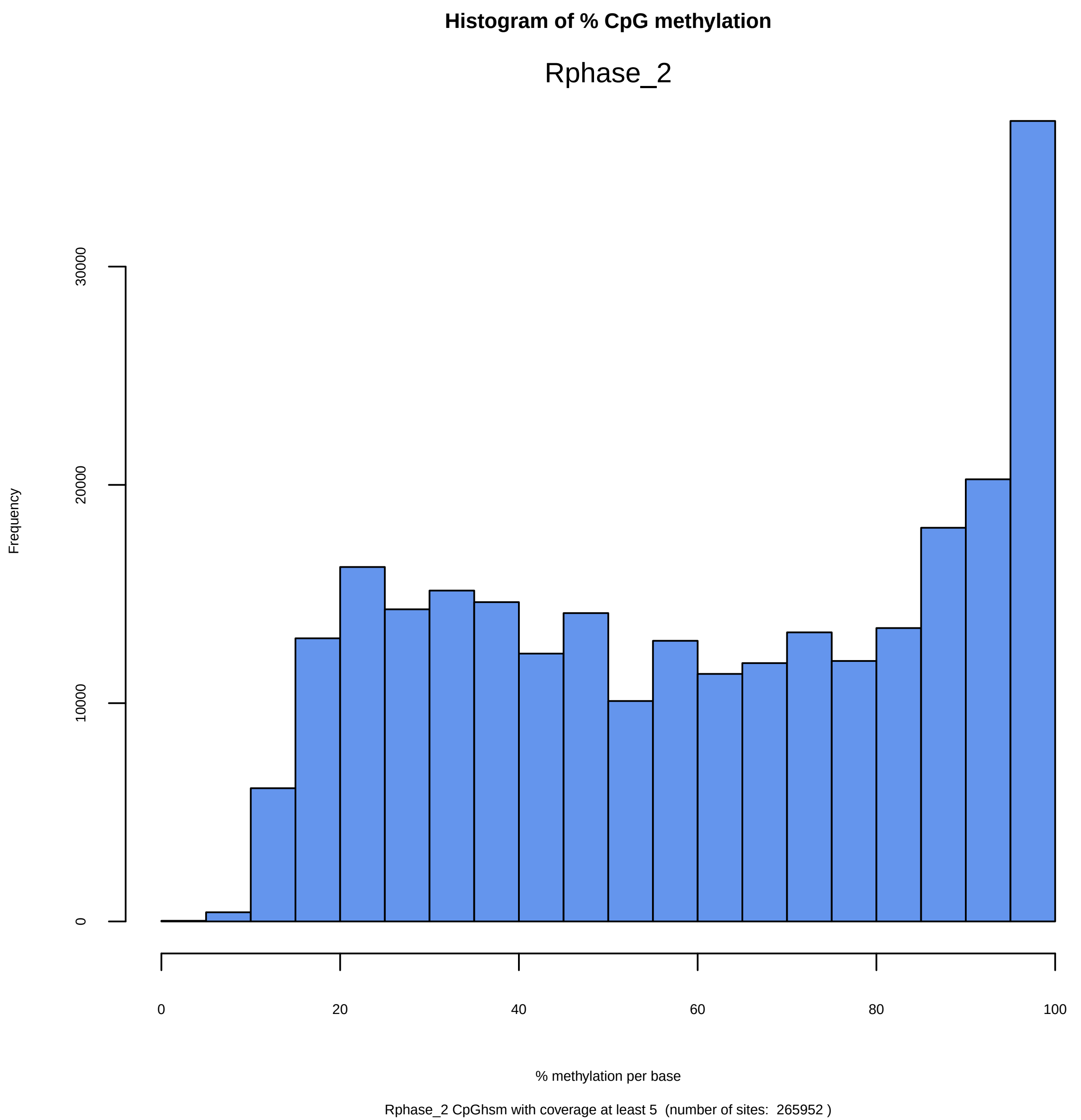
Histogram of % CpG methylation
Rphase_2
Frequency
0
10000
20000
30000
0
20
40
60
80
100
% methylation per base
Rphase_2 CpGhsm with coverage at least 5 (number of sites: 265952 )

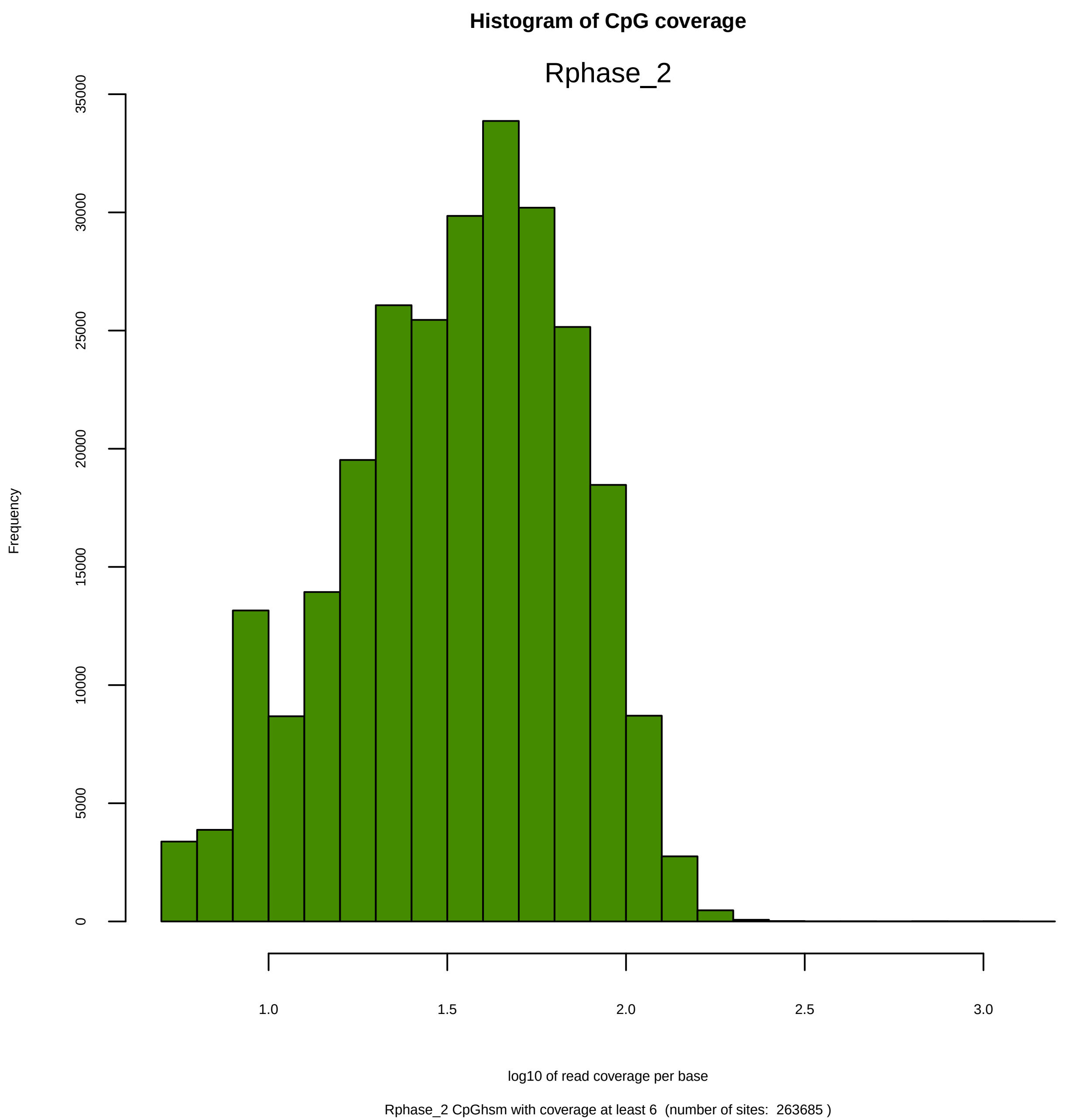
Histogram of CpG coverage
Rphase_2
Frequency
0
5000
10000
15000
20000
25000
30000
35000
1.0
1.5
2.0
2.5
3.0
log10 of read coverage per base
Rphase_2 CpGhsm with coverage at least 6 (number of sites: 263685 )

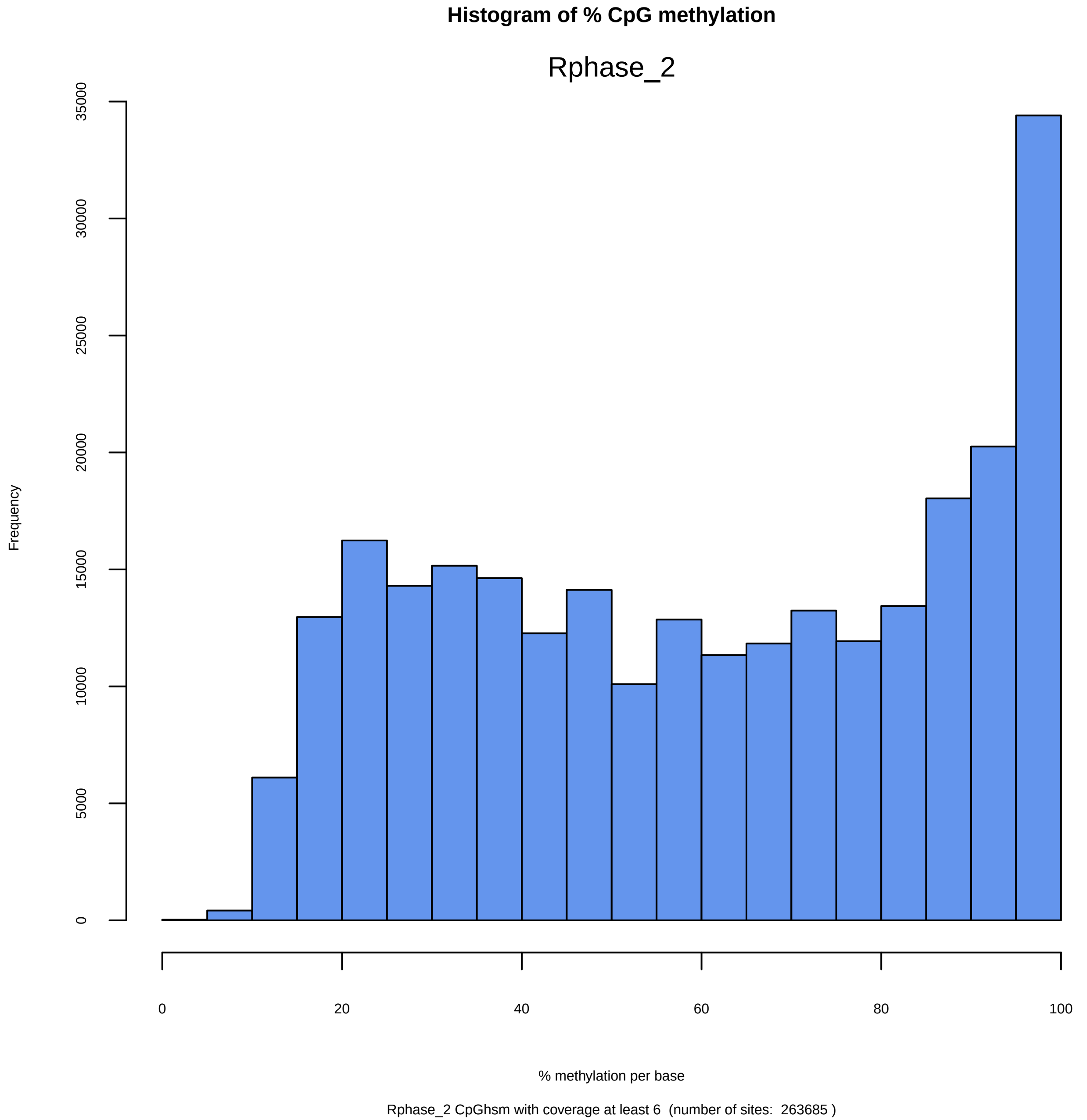
Histogram of % CpG methylation
Rphase_2
Frequency
0
5000
10000
15000
20000
25000
30000
35000
0
20
40
60
80
100
% methylation per base
Rphase_2 CpGhsm with coverage at least 6  (number of sites:  263685 )

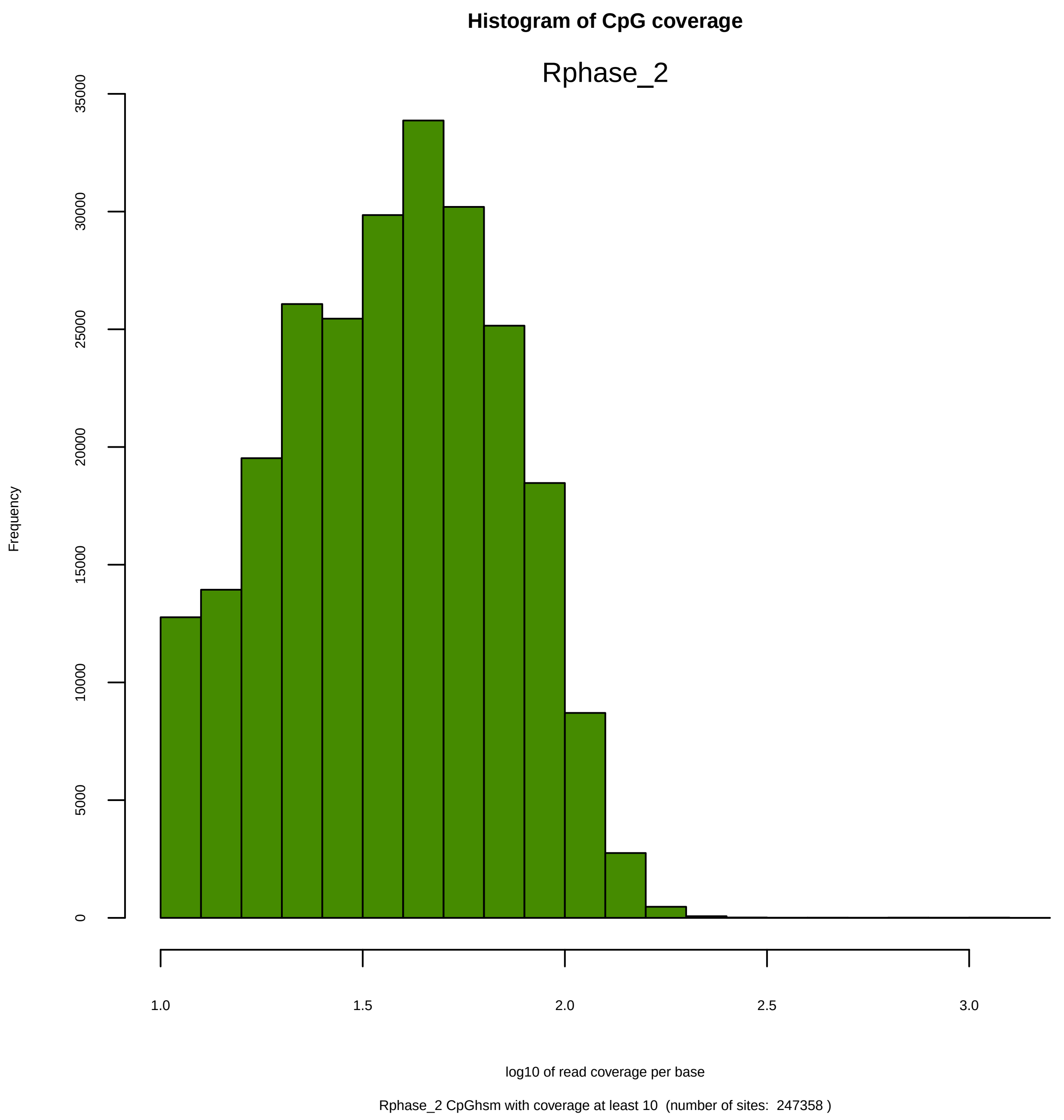
Histogram of CpG coverage
Rphase_2
Frequency
0
5000
10000
15000
20000
25000
30000
35000
1.0
1.5
2.0
2.5
3.0
log10 of read coverage per base
Rphase_2 CpGhsm with coverage at least 10  (number of sites:  247358 )

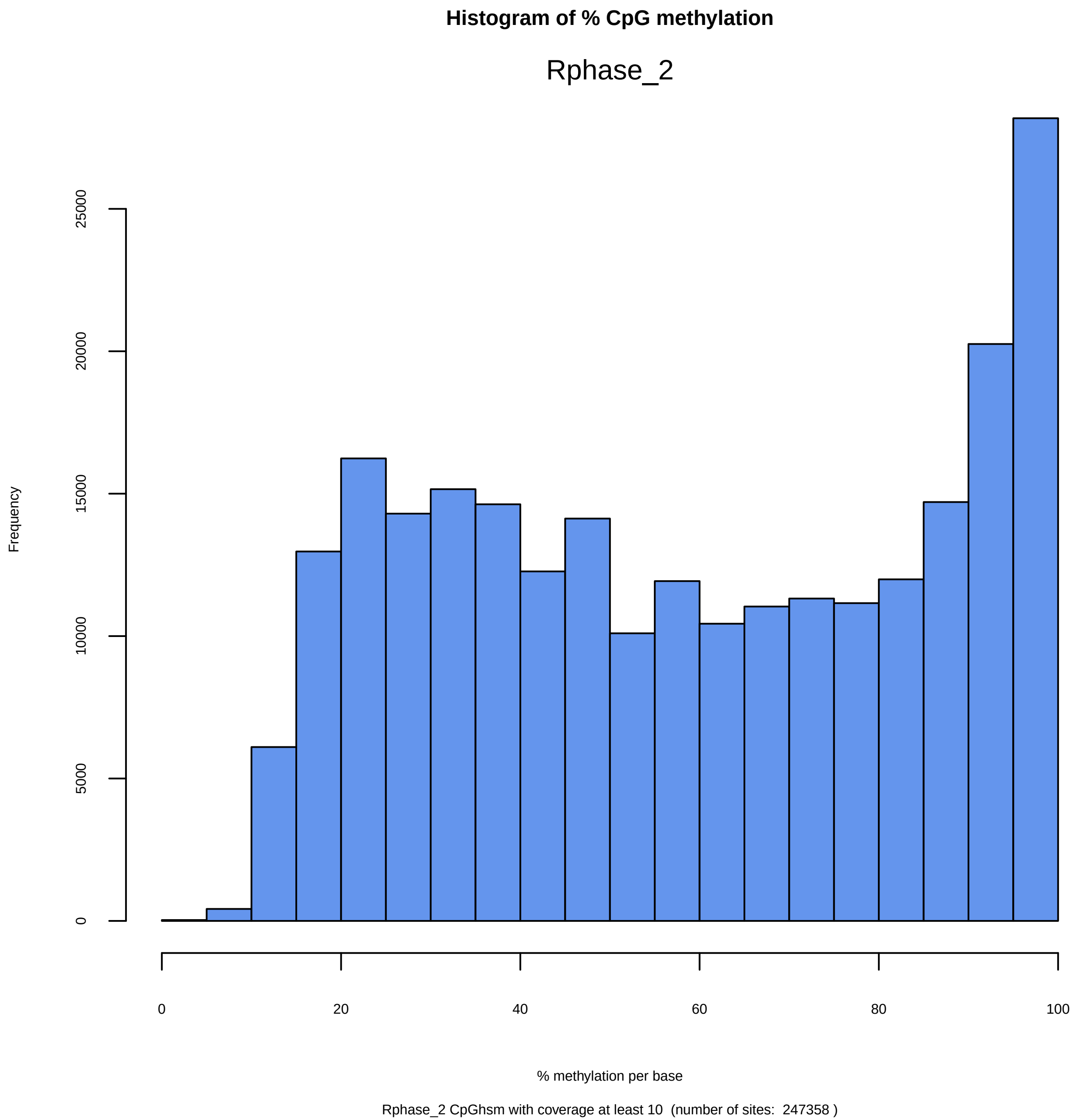
Histogram of % CpG methylation
Rphase_2
Frequency
0
5000
10000
15000
20000
25000
0
20
40
60
80
100
% methylation per base
Rphase_2 CpGhsm with coverage at least 10  (number of sites:  247358 )

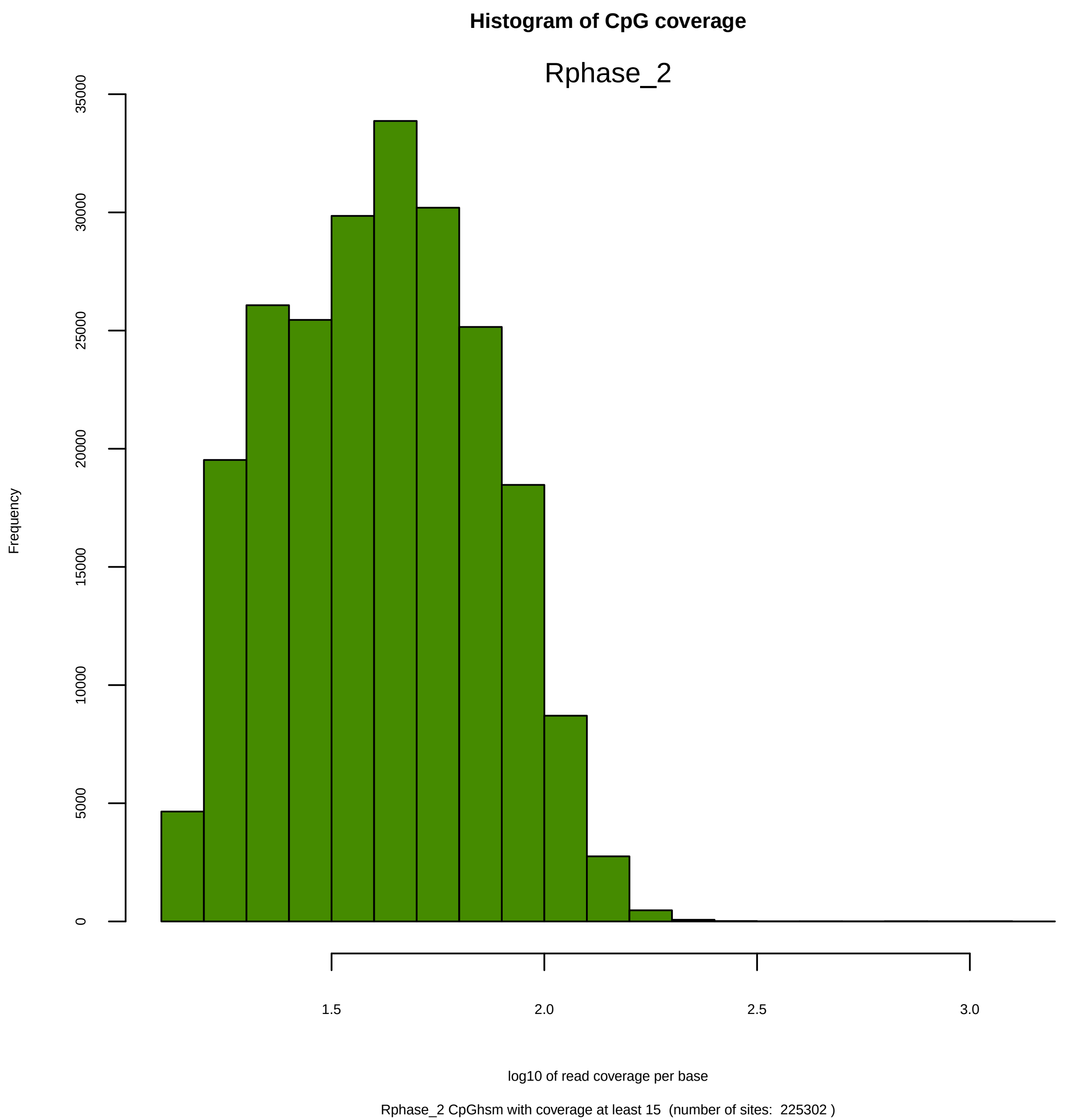
Histogram of CpG coverage
Rphase_2
Frequency
0
5000
10000
15000
20000
25000
30000
35000
1.5
2.0
2.5
3.0
log10 of read coverage per base
Rphase_2 CpGhsm with coverage at least 15 (number of sites: 225302 )

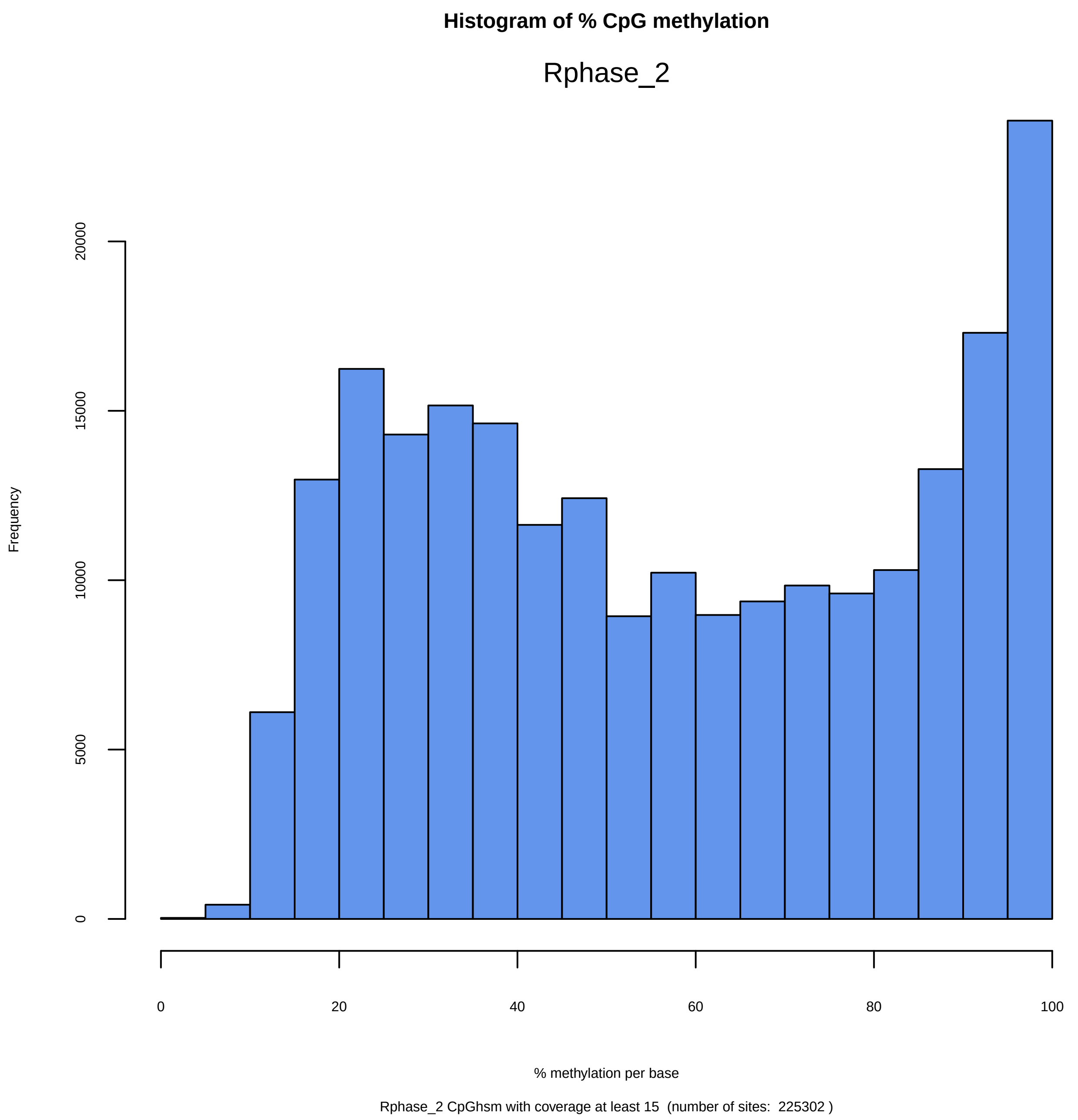
Histogram of % CpG methylation
Rphase_2
Frequency
0
5000
10000
15000
20000
0
20
40
60
80
100
% methylation per base
Rphase_2 CpGhsm with coverage at least 15 (number of sites: 225302 )

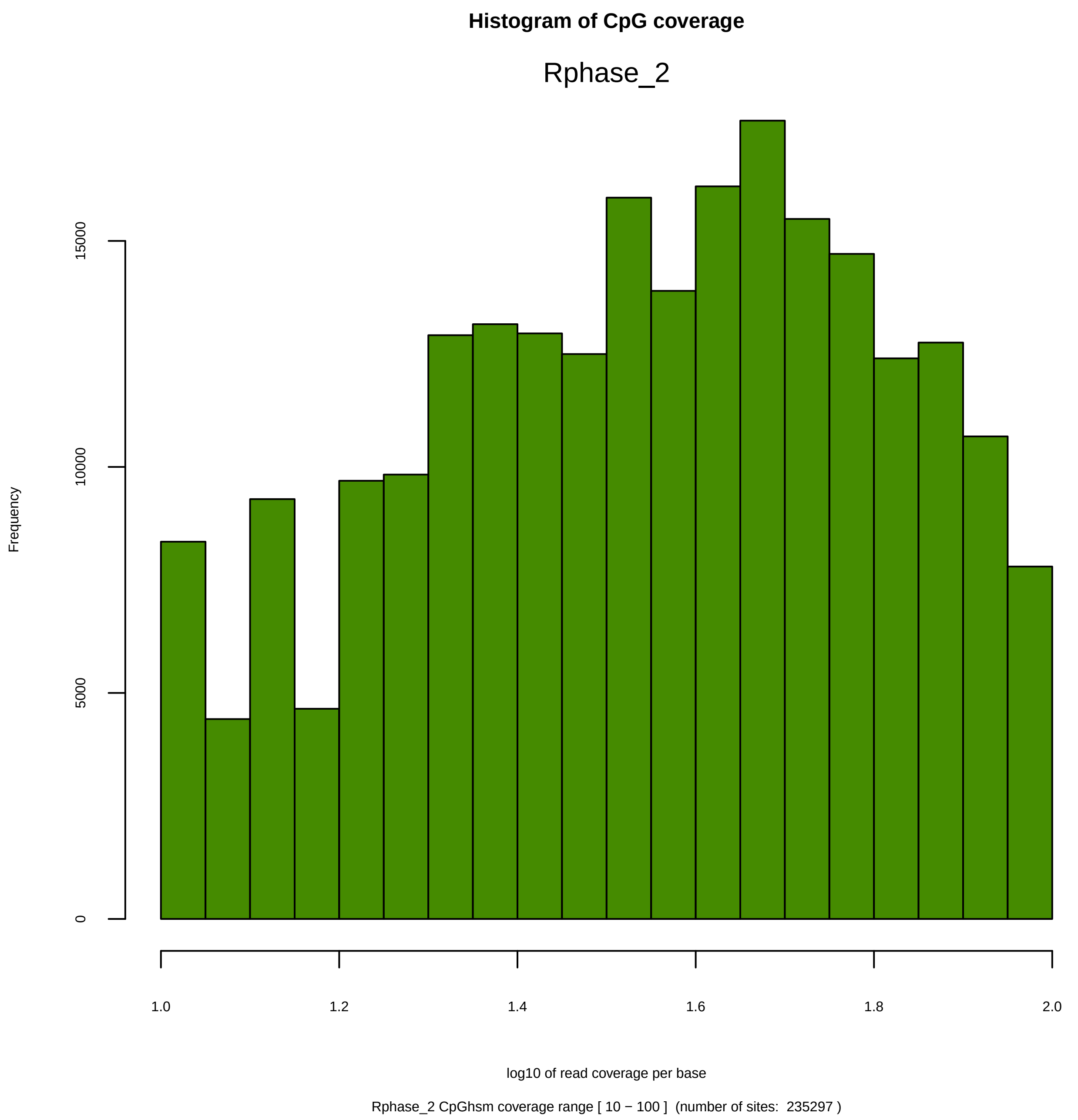

Histogram of CpG coverage
Rphase_2
Frequency
0
5000
10000
15000
1.0
1.2
1.4
1.6
1.8
2.0
log10 of read coverage per base
Rphase_2 CpGhsm coverage range [ 10 − 100 ]  (number of sites:  235297 )

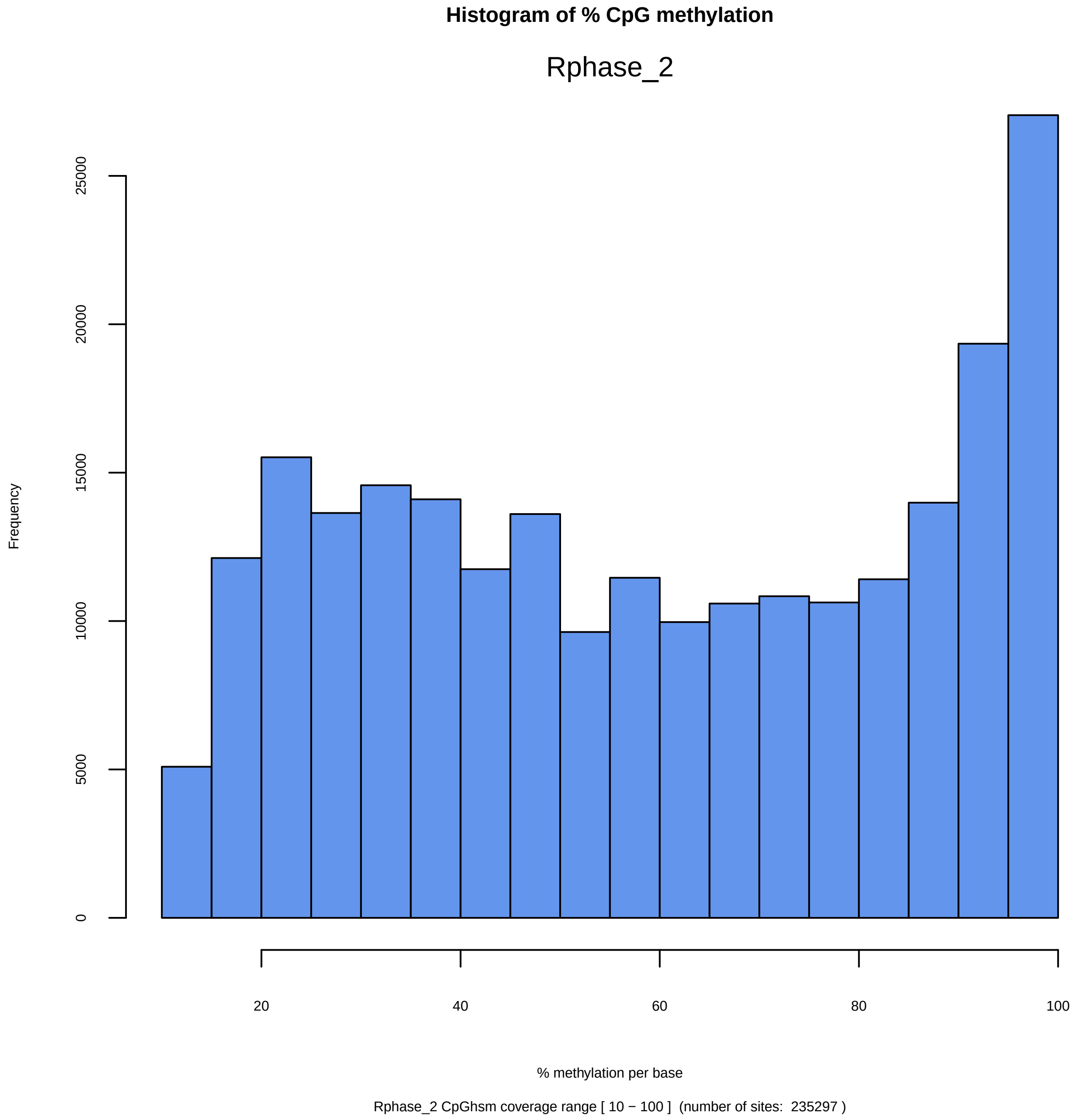
Histogram of % CpG methylation
Rphase_2
Frequency
0
5000
10000
15000
20000
25000
20
40
60
80
100
% methylation per base
Rphase_2 CpGhsm coverage range [ 10 − 100 ] (number of sites: 235297 )

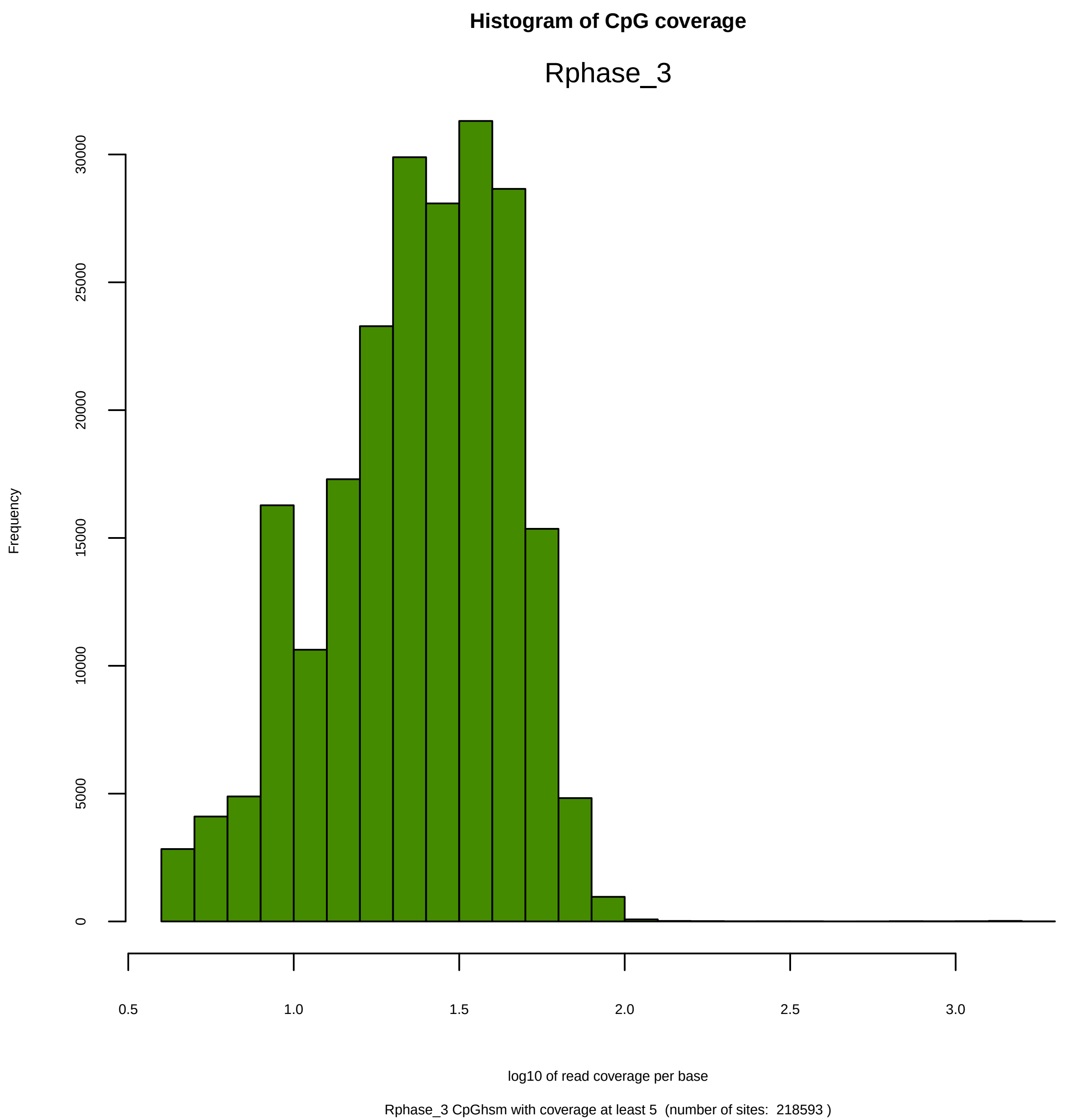
Histogram of CpG coverage
Rphase_3
Frequency
0
5000
10000
15000
20000
25000
30000
0.5
1.0
1.5
2.0
2.5
3.0
log10 of read coverage per base
Rphase_3 CpGhsm with coverage at least 5 (number of sites: 218593 )

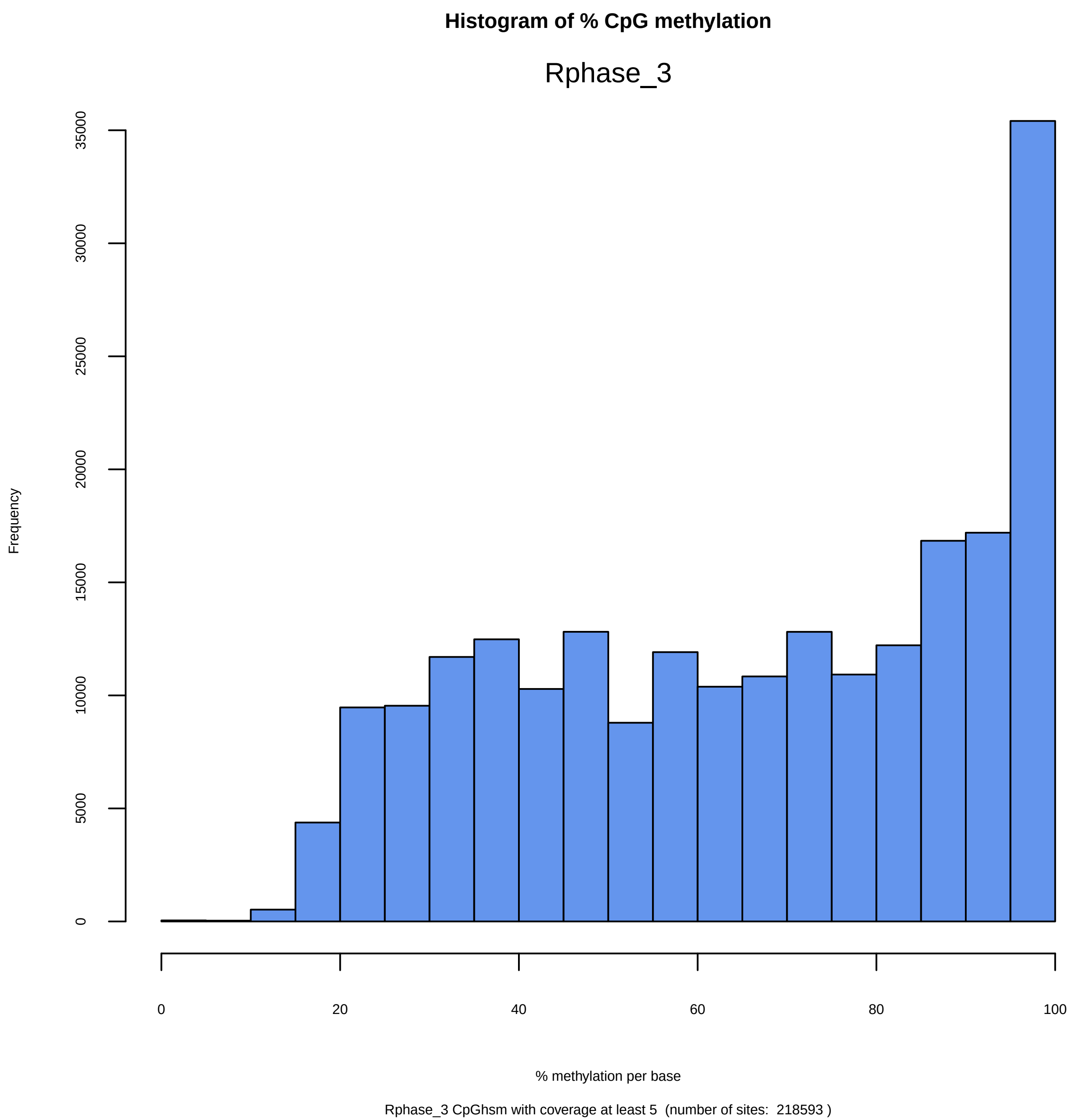
Histogram of % CpG methylation
Rphase_3
Frequency
0
5000
10000
15000
20000
25000
30000
35000
0
20
40
60
80
100
% methylation per base
Rphase_3 CpGhsm with coverage at least 5 (number of sites: 218593 )

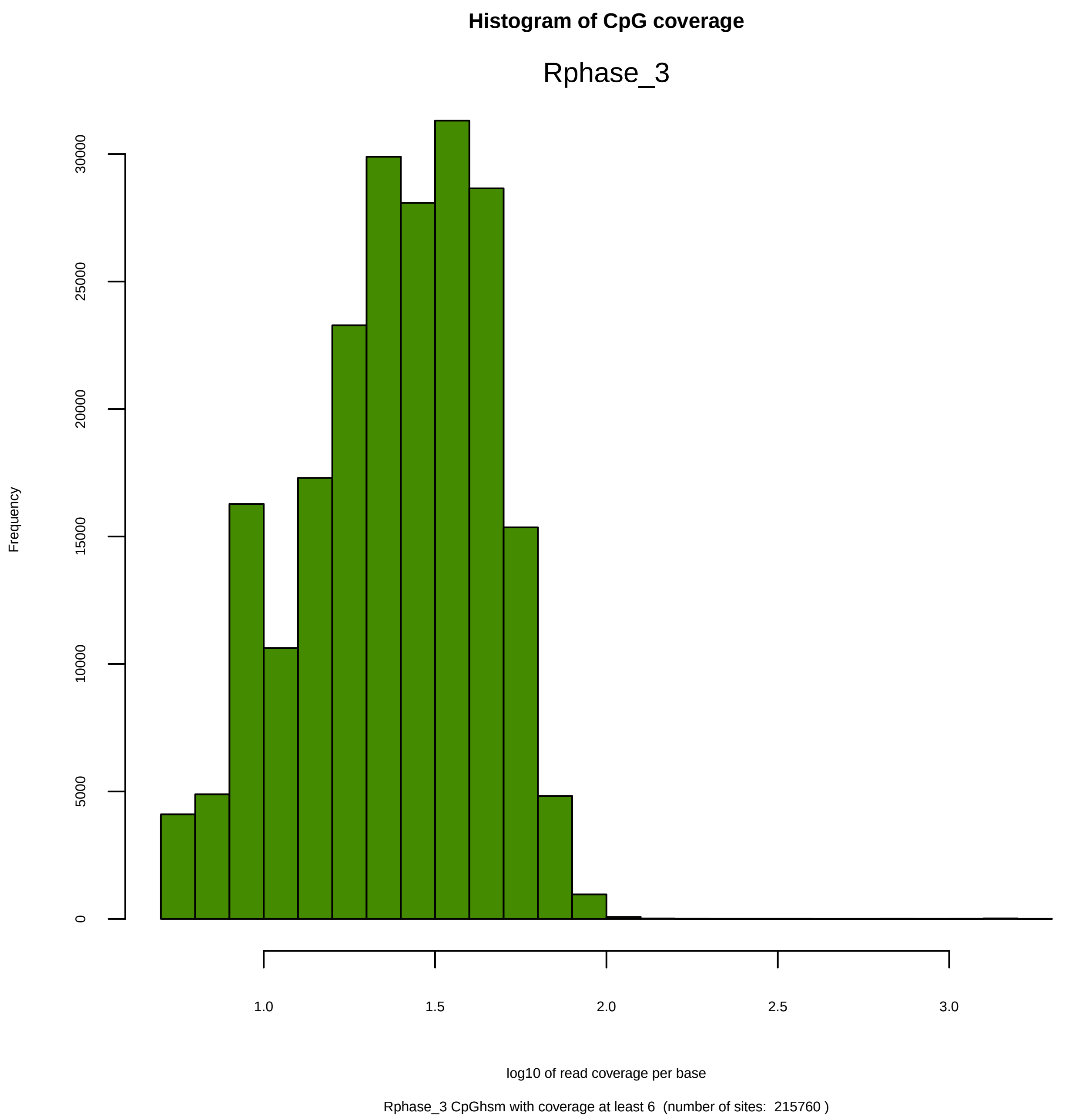
Histogram of CpG coverage
Rphase_3
Frequency
0
5000
10000
15000
20000
25000
30000
1.0
1.5
2.0
2.5
3.0
log10 of read coverage per base
Rphase_3 CpGhsm with coverage at least 6 (number of sites: 215760 )

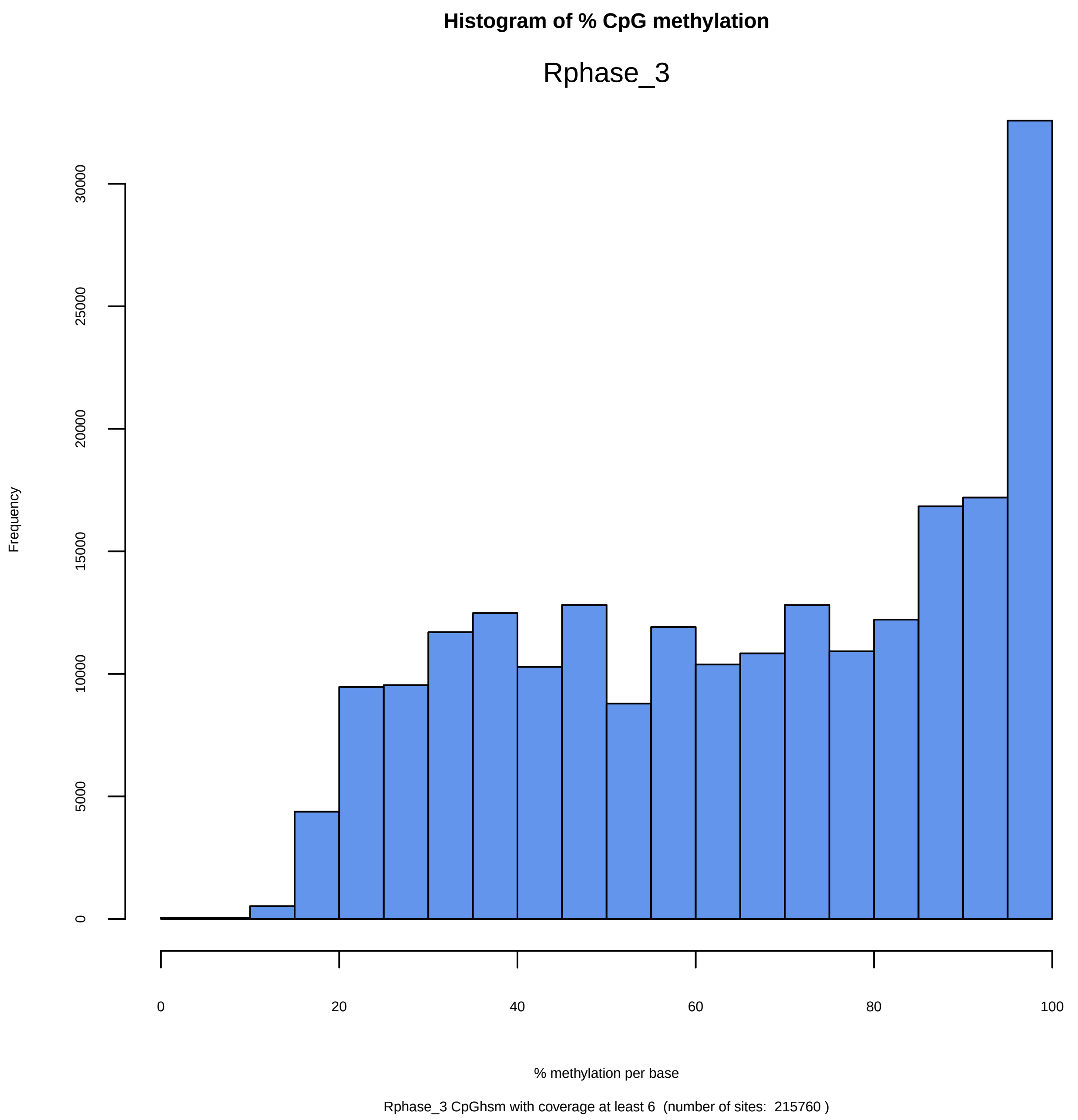

Histogram of % CpG methylation
Rphase_3
Frequency
0
5000
10000
15000
20000
25000
30000
0
20
40
60
80
100
% methylation per base
Rphase_3 CpGhsm with coverage at least 6  (number of sites:  215760 )

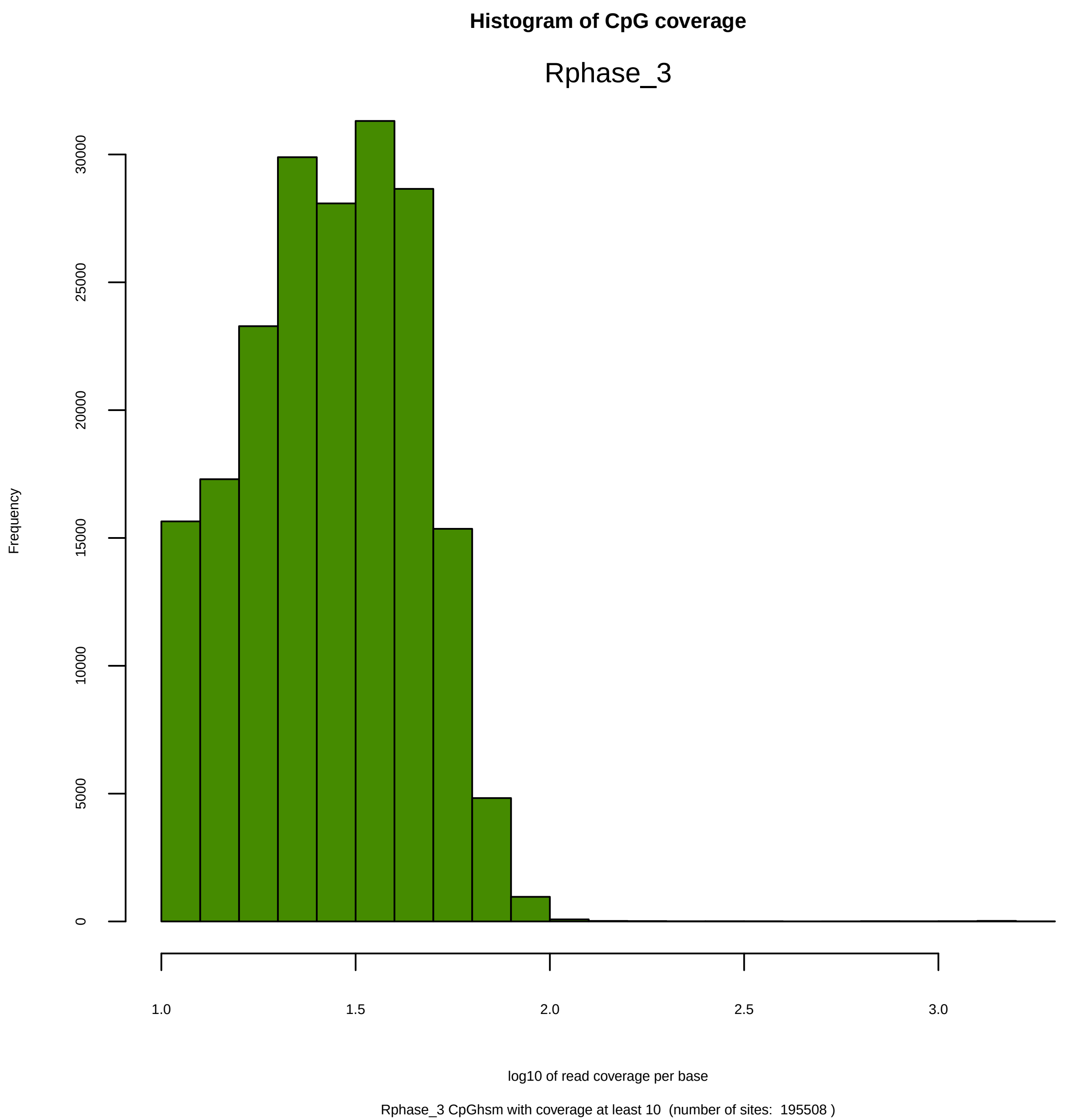
Histogram of CpG coverage
Rphase_3
Frequency
0
5000
10000
15000
20000
25000
30000
1.0
1.5
2.0
2.5
3.0
log10 of read coverage per base
Rphase_3 CpGhsm with coverage at least 10  (number of sites:  195508 )

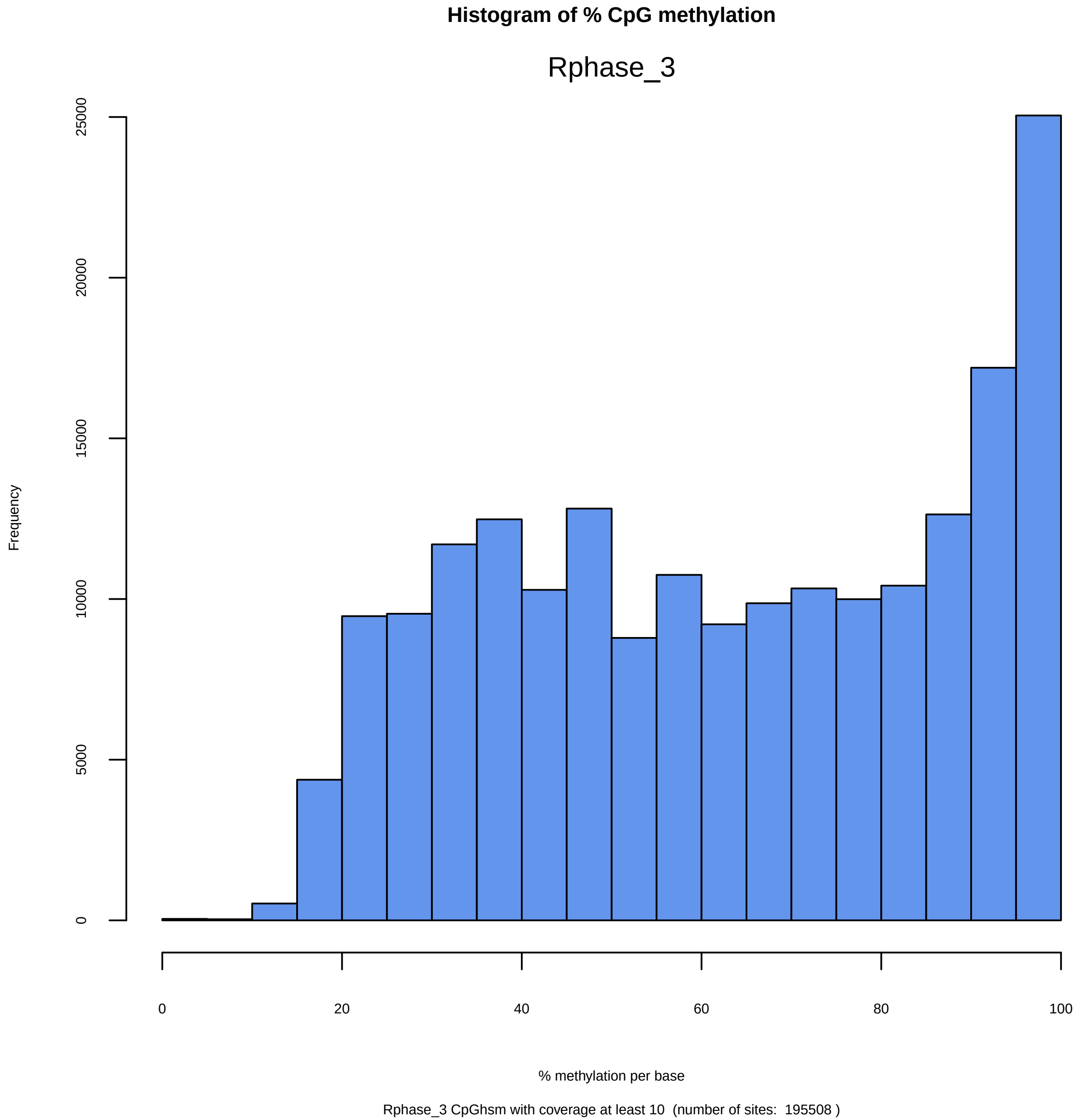
Histogram of % CpG methylation
Rphase_3
Frequency
0
5000
10000
15000
20000
25000
0
20
40
60
80
100
% methylation per base
Rphase_3 CpGhsm with coverage at least 10 (number of sites: 195508 )

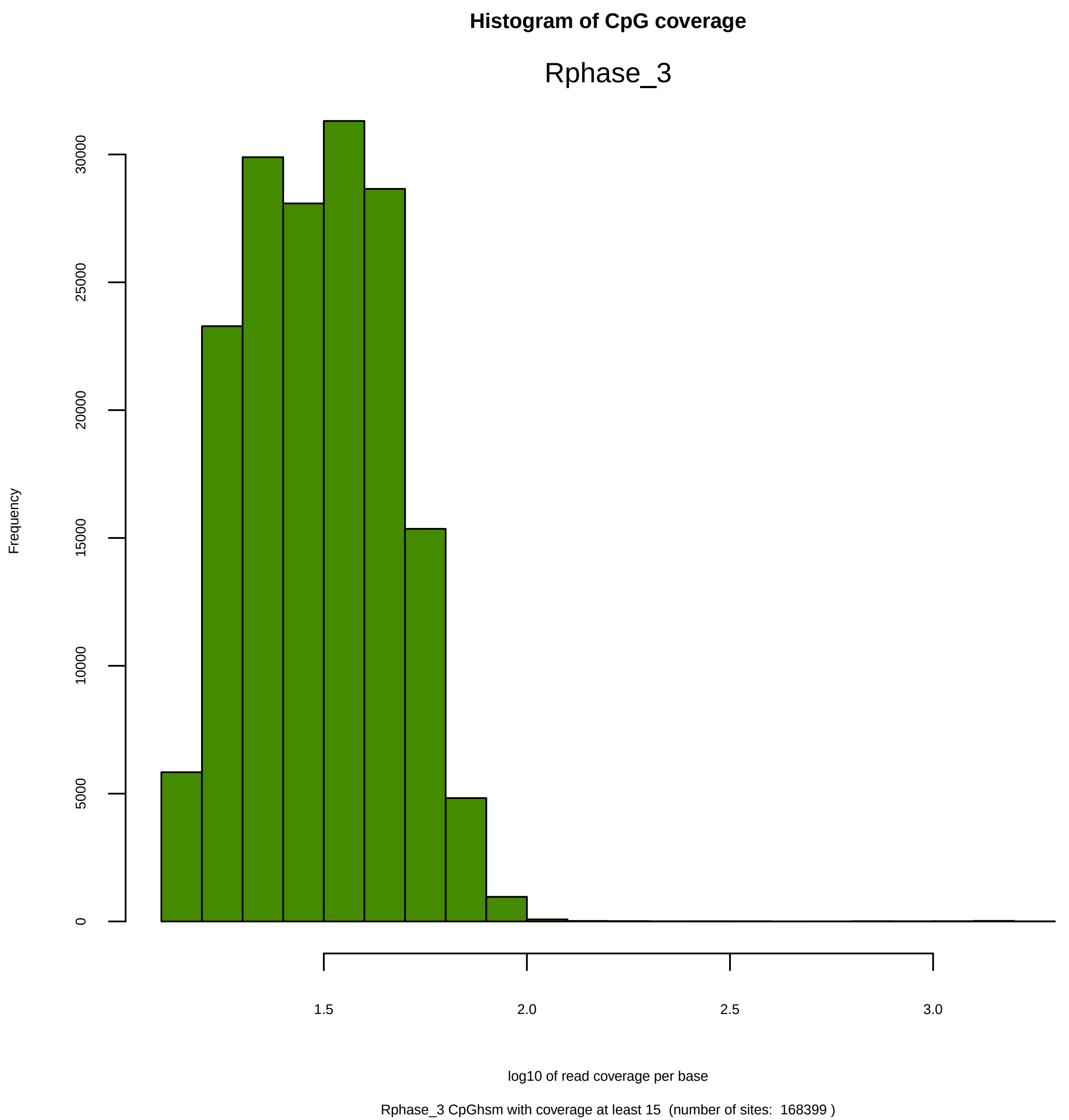
Histogram of CpG coverage
Rphase_3
Frequency
0
5000
10000
15000
20000
25000
30000
1.5
2.0
2.5
3.0
log10 of read coverage per base
Rphase_3 CpGhsm with coverage at least 15 (number of sites: 168399 )

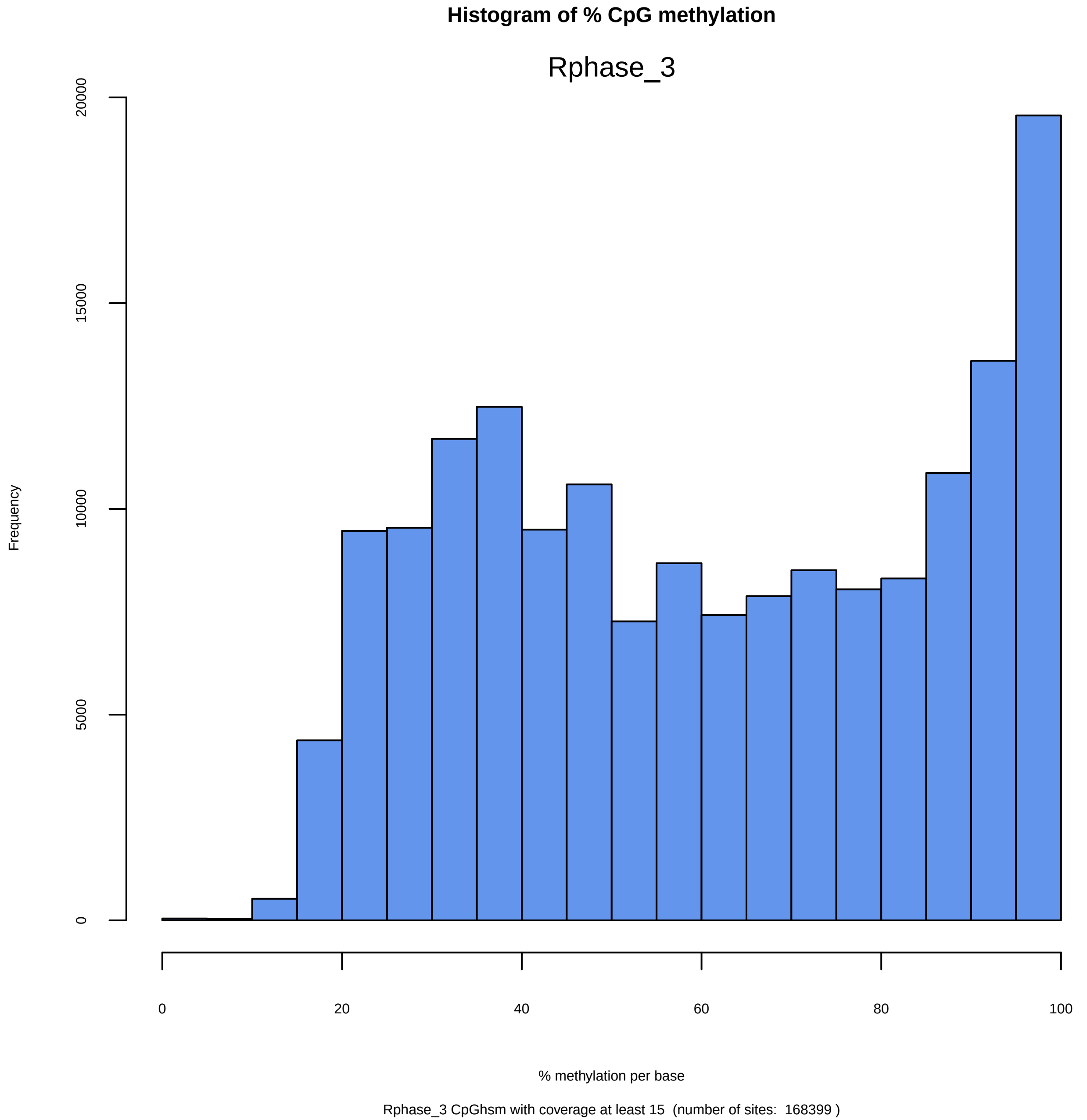

Histogram of % CpG methylation
Rphase_3
Frequency
0
5000
10000
15000
20000
0
20
40
60
80
100
% methylation per base
Rphase_3 CpGhsm with coverage at least 15 (number of sites: 168399 )

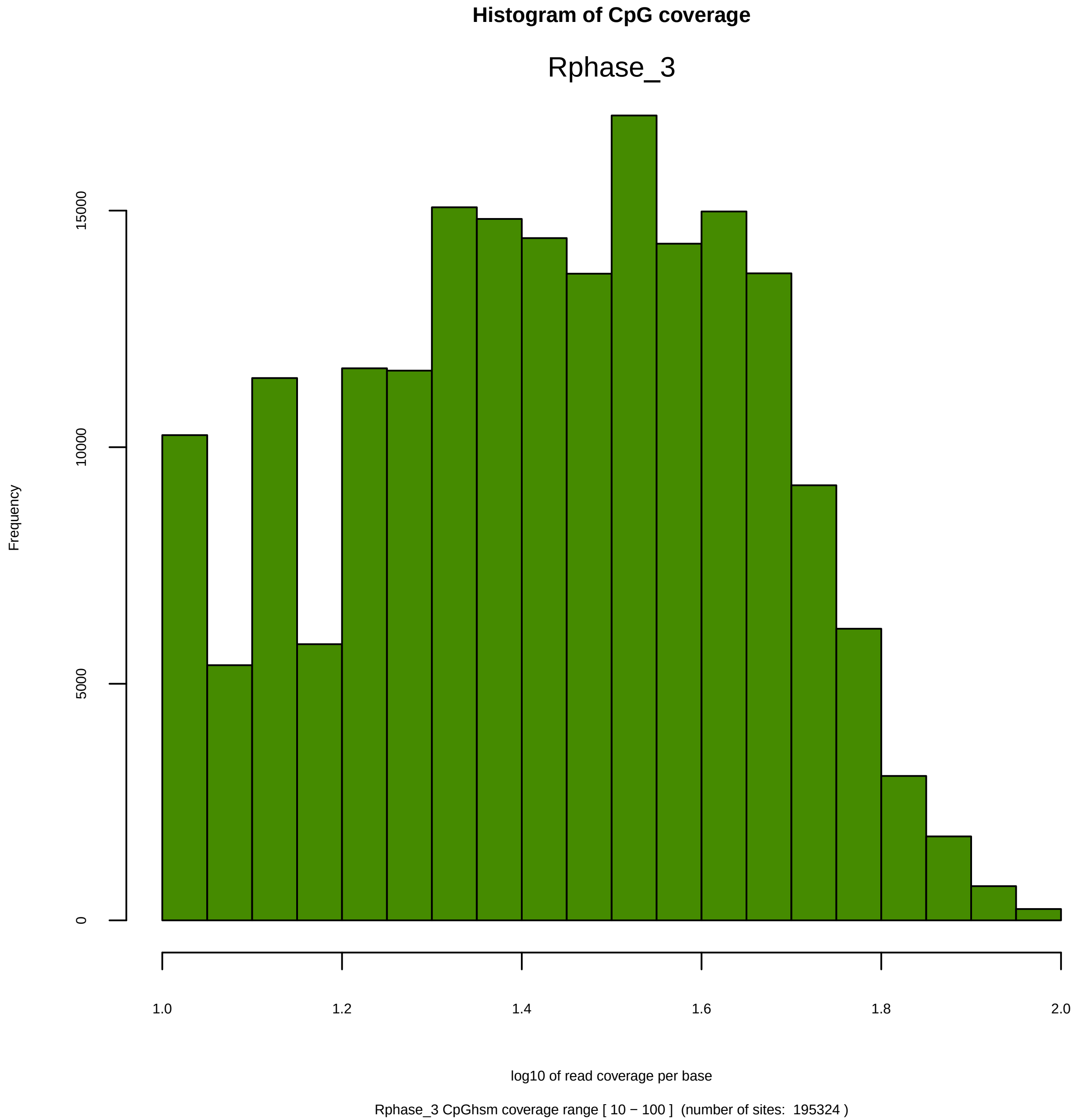
Histogram of CpG coverage
Rphase_3
Frequency
0
5000
10000
15000
1.0
1.2
1.4
1.6
1.8
2.0
log10 of read coverage per base
Rphase_3 CpGhsm coverage range [ 10 − 100 ]  (number of sites:  195324 )

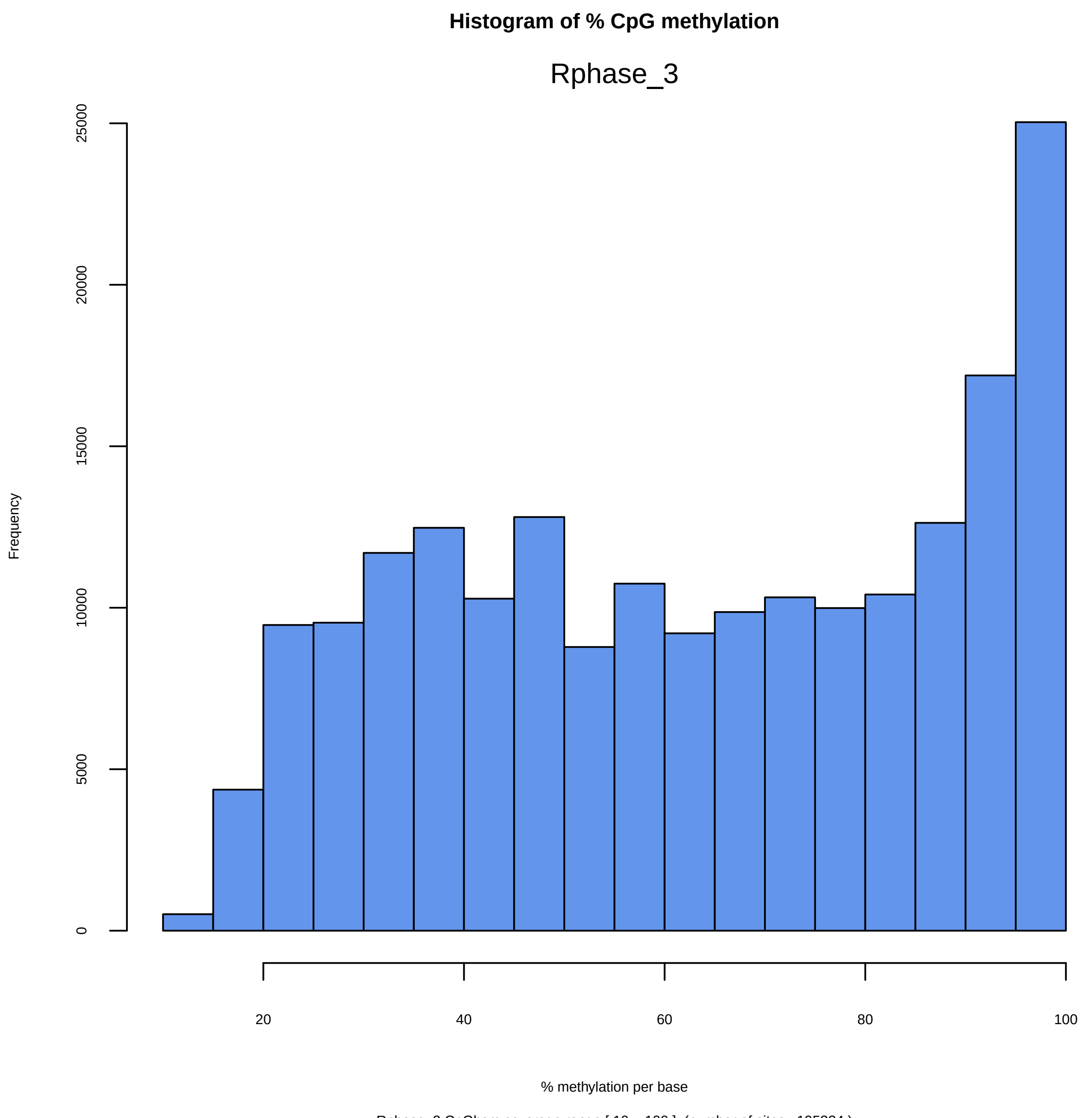

Histogram of % CpG methylation
Rphase_3
Frequency
0
5000
10000
15000
20000
25000
20
40
60
80
100
% methylation per base
Rphase_3 CpGhsm coverage range [ 10 − 100 ]  (number of sites:  195324 )

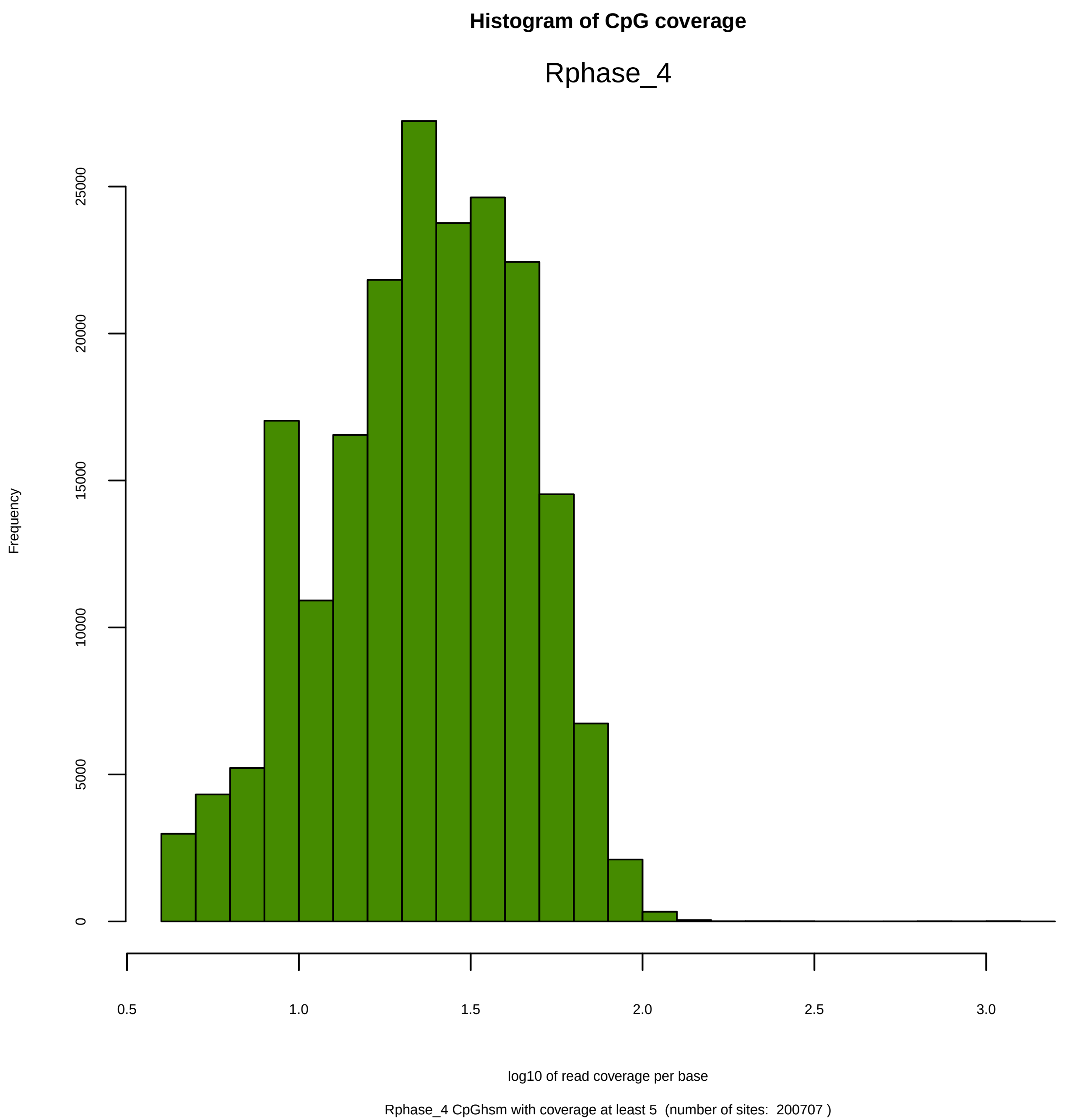
Histogram of CpG coverage
Rphase_4
Frequency
0
5000
10000
15000
20000
25000
0.5
1.0
1.5
2.0
2.5
3.0
log10 of read coverage per base
Rphase_4 CpGhsm with coverage at least 5 (number of sites: 200707 )

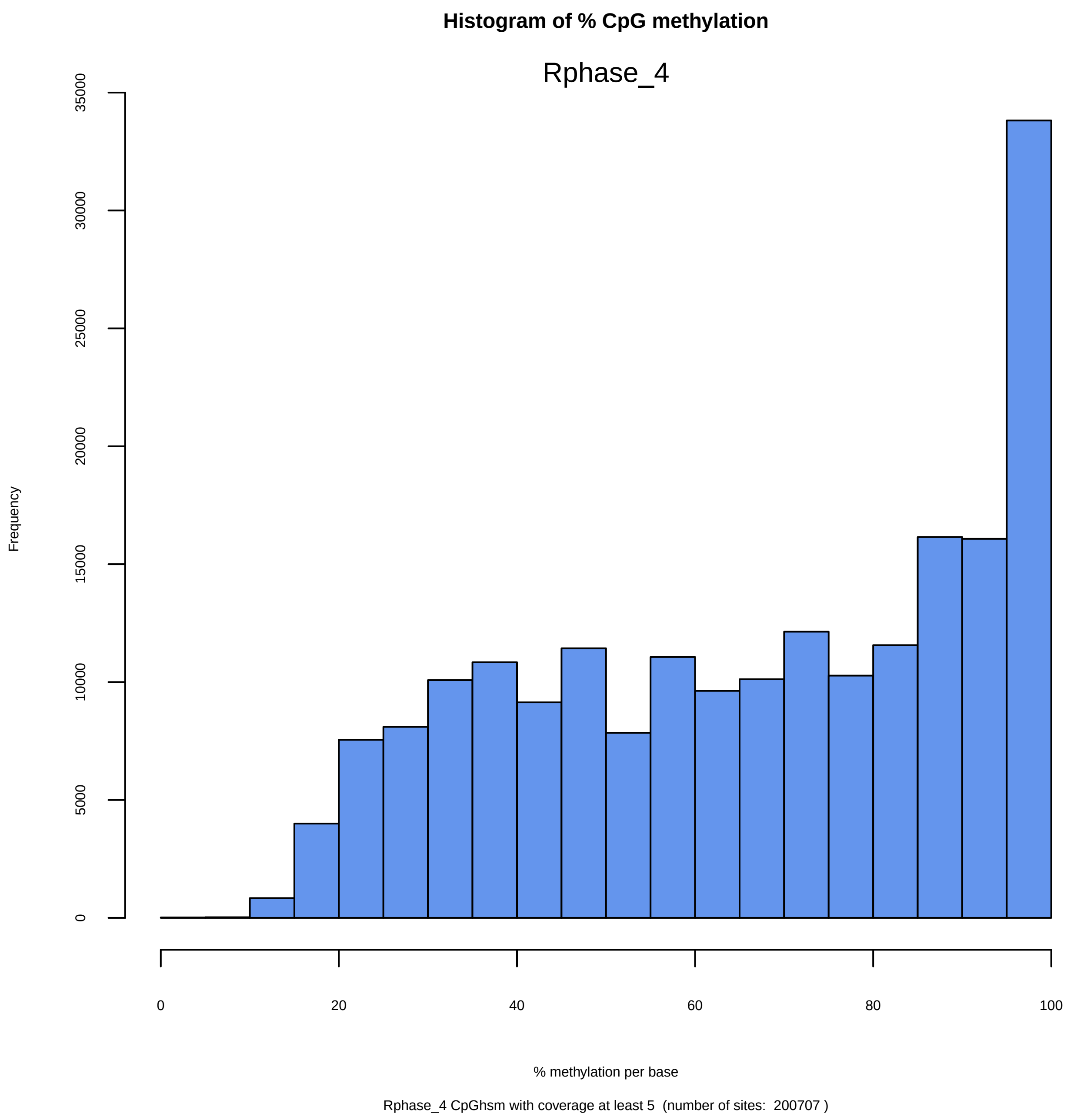
Histogram of % CpG methylation
Rphase_4
Frequency
0
5000
10000
15000
20000
25000
30000
35000
0
20
40
60
80
100
% methylation per base
Rphase_4 CpGhsm with coverage at least 5  (number of sites:  200707 )

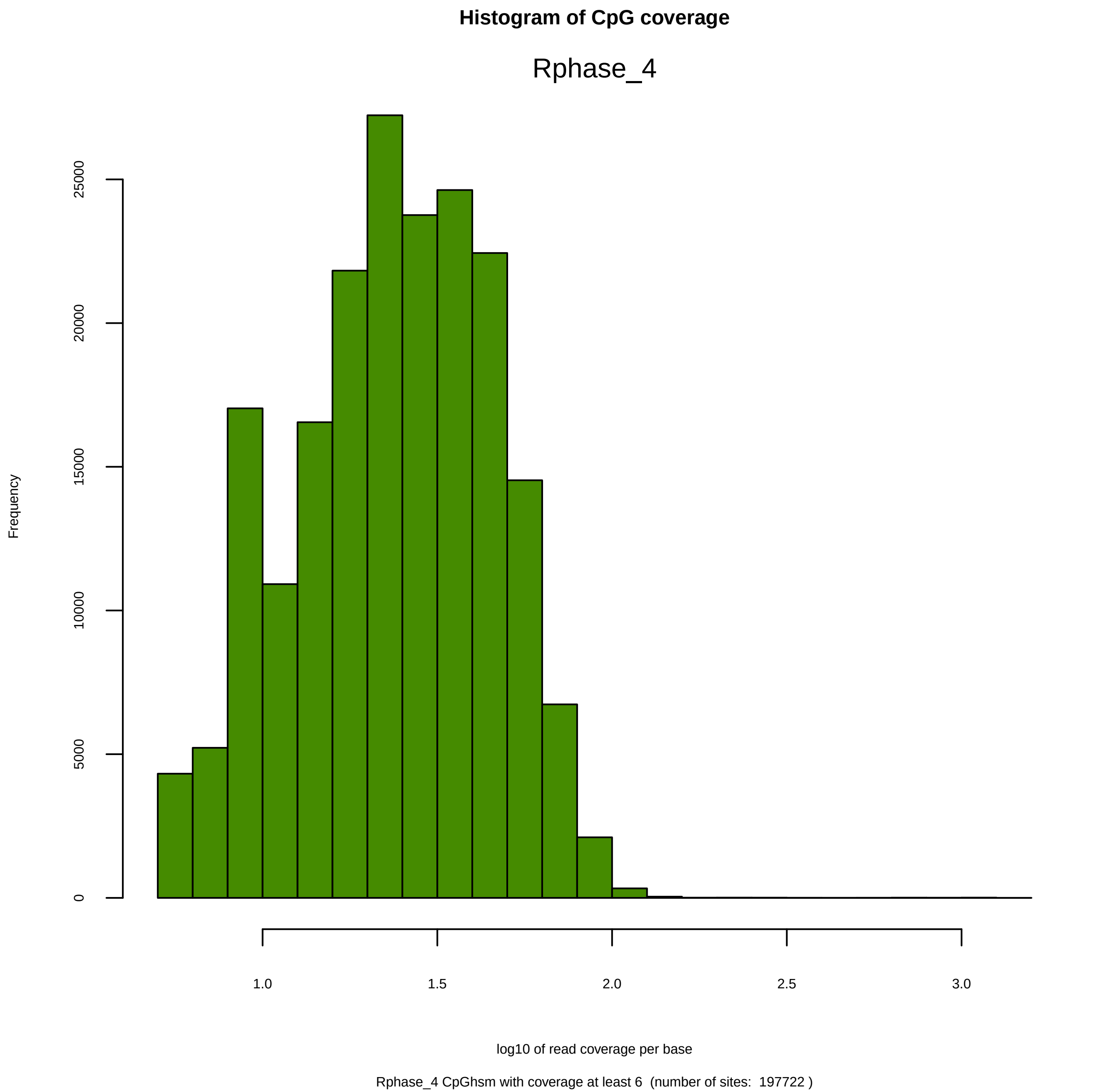

Histogram of CpG coverage
Rphase_4
Frequency
0
5000
10000
15000
20000
25000
1.0
1.5
2.0
2.5
3.0
log10 of read coverage per base
Rphase_4 CpGhsm with coverage at least 6 (number of sites: 197722 )

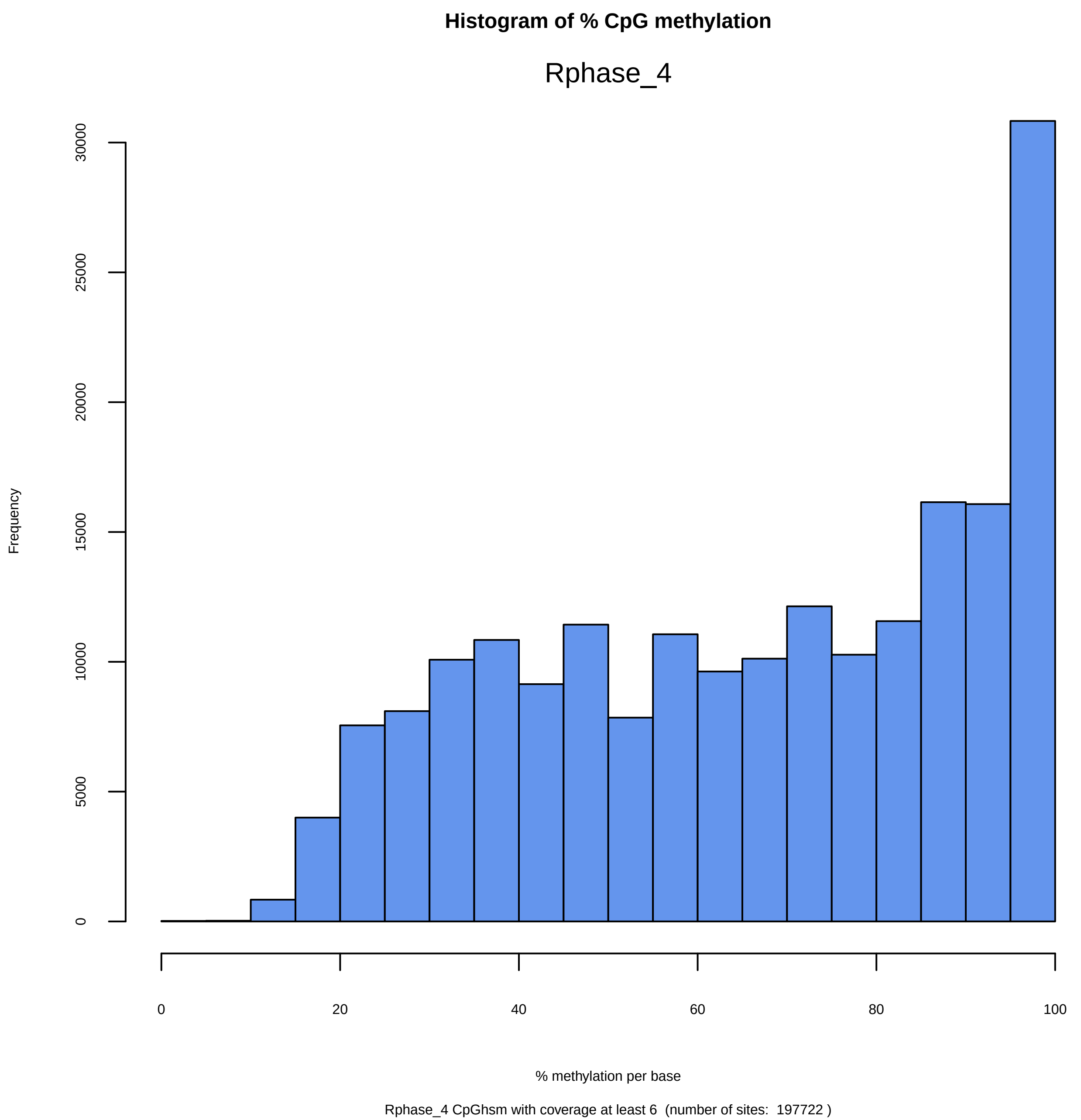
Histogram of % CpG methylation
Rphase_4
Frequency
0
5000
10000
15000
20000
25000
30000
0
20
40
60
80
100
% methylation per base
Rphase_4 CpGhsm with coverage at least 6 (number of sites: 197722 )

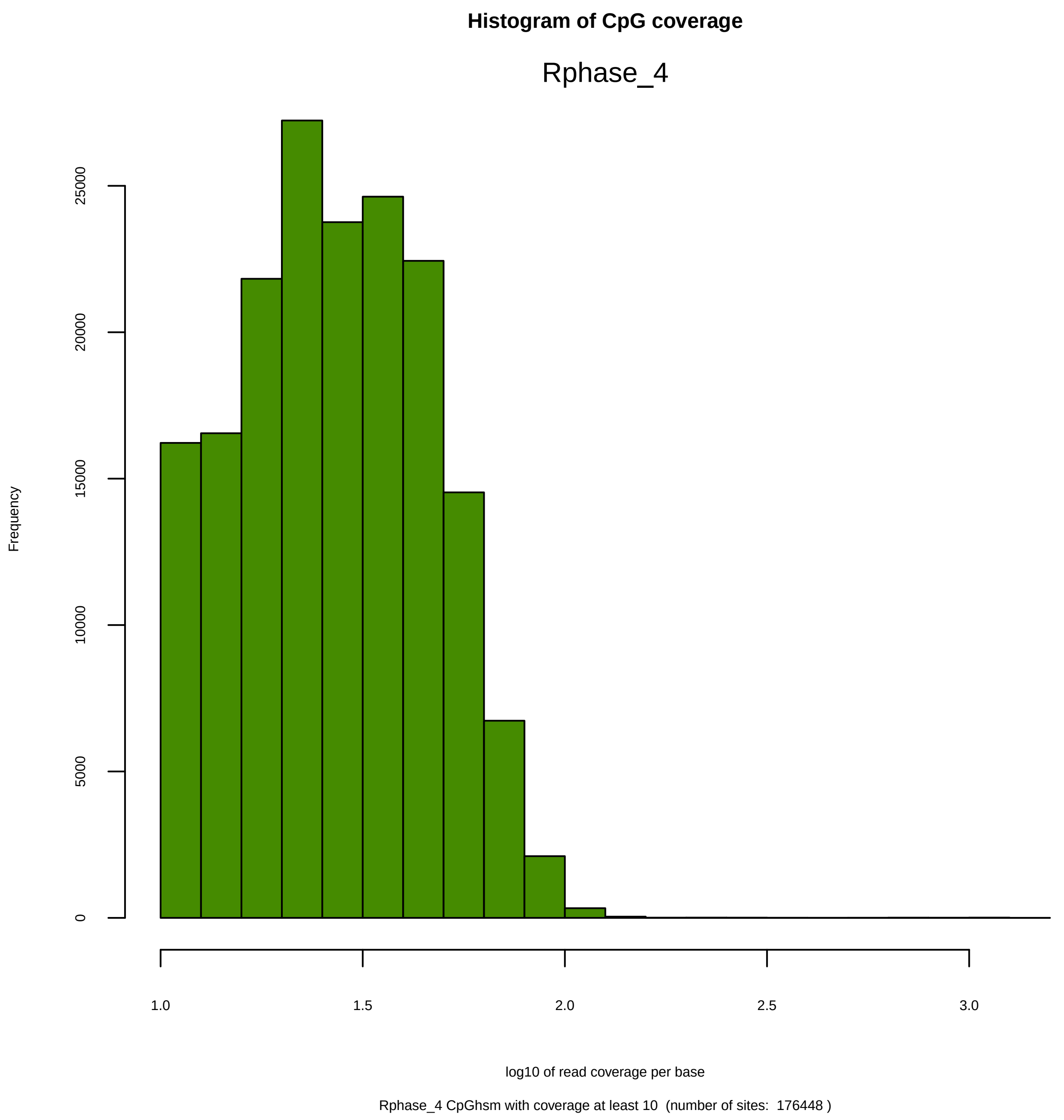

Histogram of CpG coverage
Rphase_4
Frequency
0
5000
10000
15000
20000
25000
1.0
1.5
2.0
2.5
3.0
log10 of read coverage per base
Rphase_4 CpGhsm with coverage at least 10 (number of sites: 176448 )

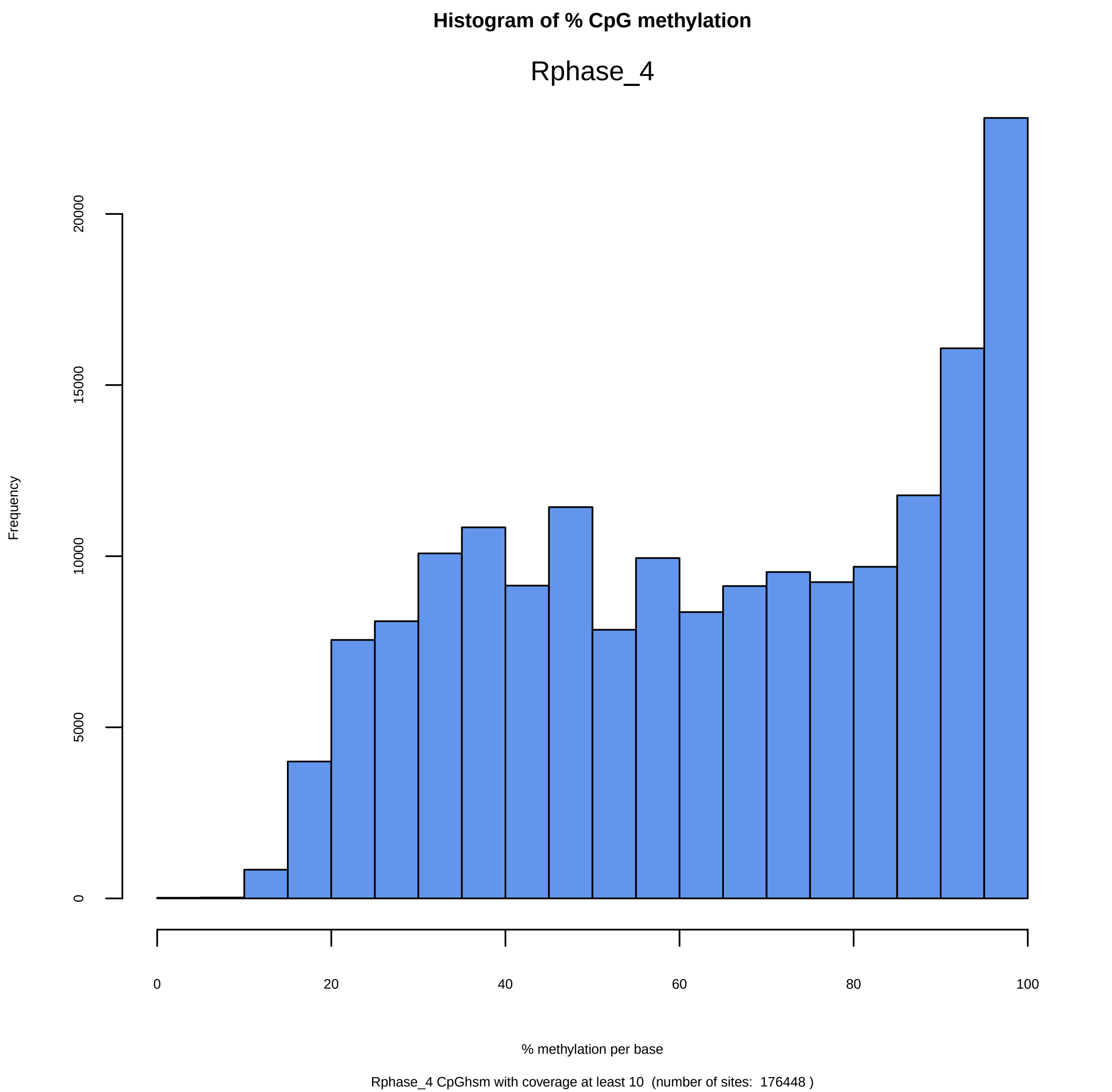

Histogram of % CpG methylation
Rphase_4
Frequency
0
5000
10000
15000
20000
0
20
40
60
80
100
% methylation per base
Rphase_4 CpGhsm with coverage at least 10 (number of sites: 176448 )

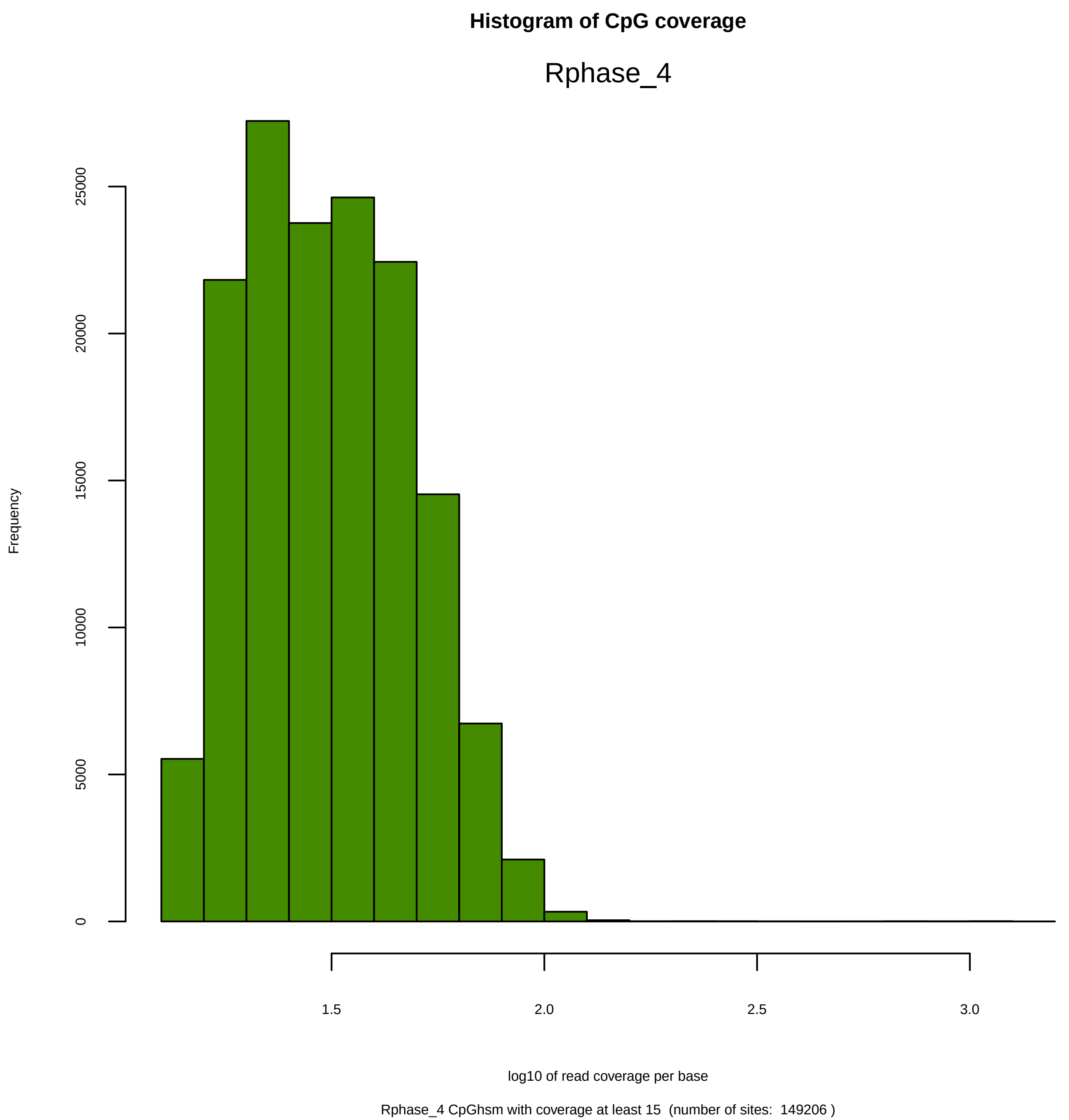

Histogram of CpG coverage
Rphase_4
Frequency
0
5000
10000
15000
20000
25000
1.5
2.0
2.5
3.0
log10 of read coverage per base
Rphase_4 CpGhsm with coverage at least 15 (number of sites: 149206 )

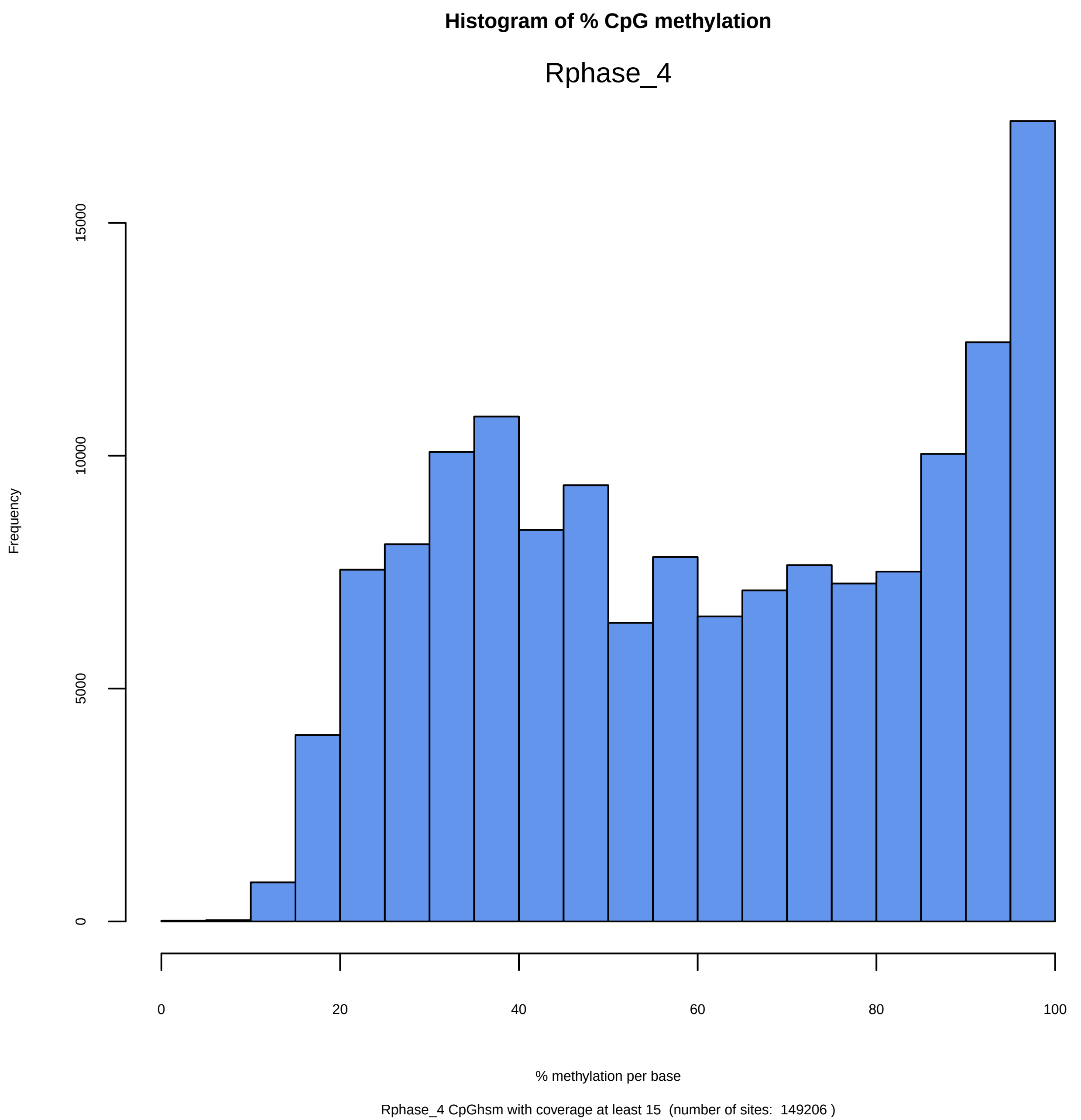
Histogram of % CpG methylation
Rphase_4
Frequency
0
5000
10000
15000
0
20
40
60
80
100
% methylation per base
Rphase_4 CpGhsm with coverage at least 15  (number of sites:  149206 )

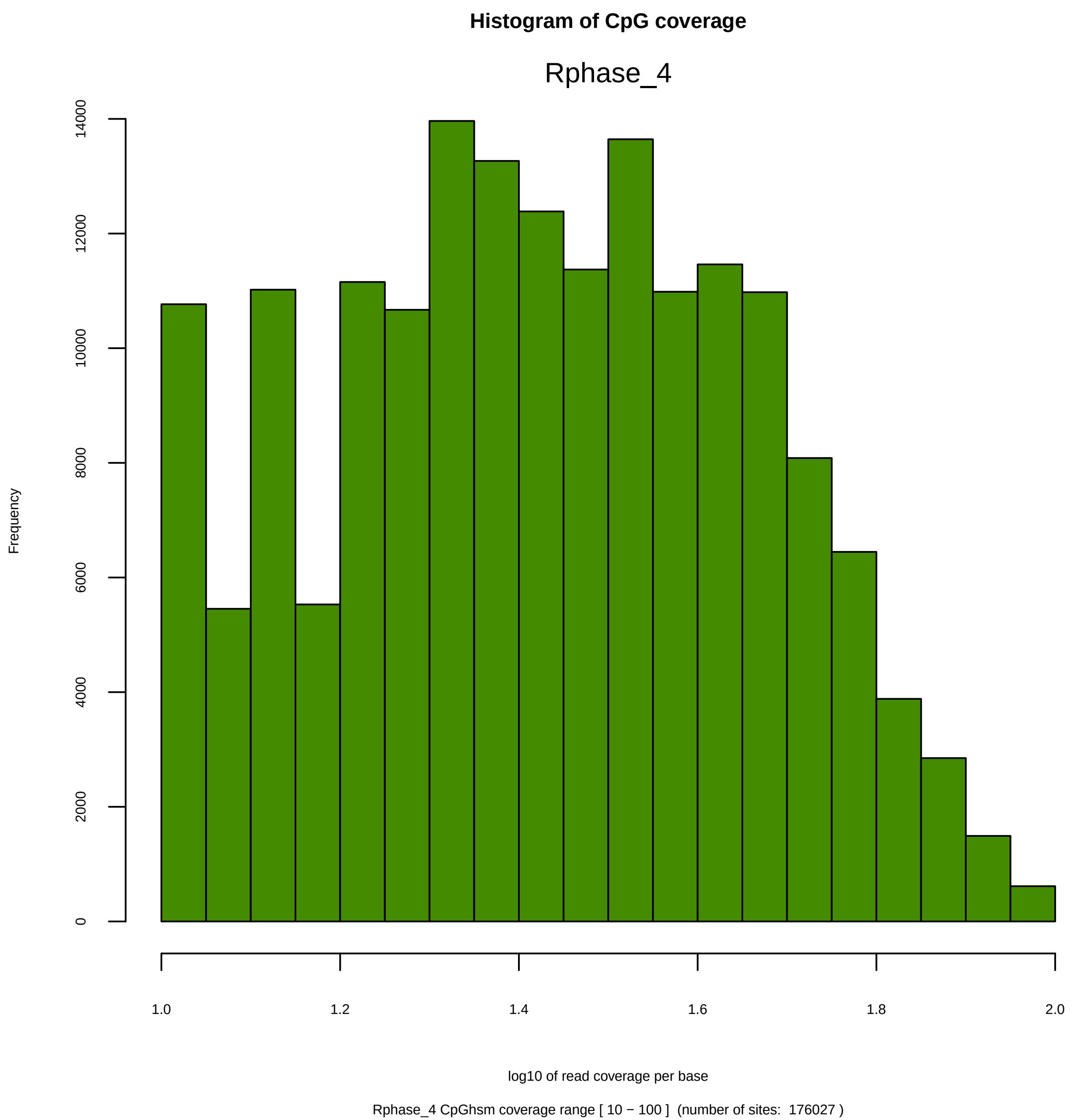

Histogram of CpG coverage
Rphase_4
Frequency
0
2000
4000
6000
8000
10000
12000
14000
1.0
1.2
1.4
1.6
1.8
2.0
log10 of read coverage per base
Rphase_4 CpGhsm coverage range [ 10 − 100 ]  (number of sites:  176027 )

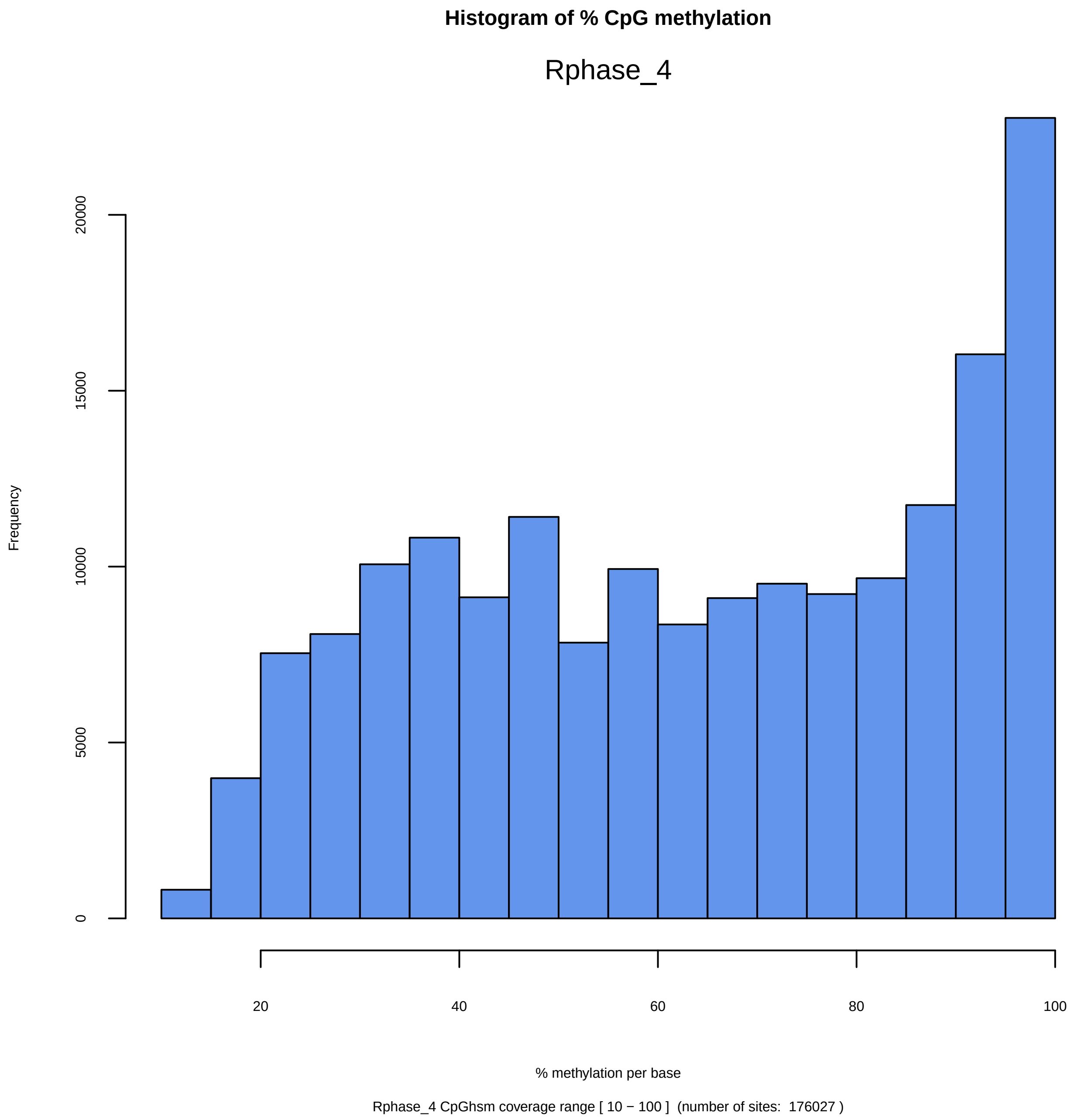
Histogram of % CpG methylation
Rphase_4
Frequency
0
5000
10000
15000
20000
20
40
60
80
100
% methylation per base
Rphase_4 CpGhsm coverage range [ 10 − 100 ]  (number of sites:  176027 )

```
Directory: REPCRL File: 0READMErepcrl

REPCRL - Correlations between replicates.

Input: mkrd (methylKit methylRawList object of replicate data)

Output: files repcrl-*.txt repcrl-*.pdf

Notes: The output is generated by BWASPR::cmpSamples(), which uses methylKit::unite()
        to determine the sites common to all replicates and applies methylKit::getCorrelation()
        and methylKit::PCASamples().  The *.txt files show the correlations in a table, and
        the *.pdf files show graphics.

       Beware that the number of common sites (shown at the bottom of the *.txt files) may be
        small, which may make the correlations less informative.
```

Directory: REPCRL File: repcrl-Cb_LIp_BCphase.txt

| | BCphase_1 | BCphase_2 | BCphase_3 | BCphase_4 |
|---|---|---|---|---|
| BCphase_1 | 1.000 | 0.908 | 0.911 | 0.916 |
| BCphase_2 | 0.908 | 1.000 | 0.924 | 0.916 |
| BCphase_3 | 0.911 | 0.924 | 1.000 | 0.918 |
| BCphase_4 | 0.916 | 0.916 | 0.918 | 1.000 |

The number of conserved sites is 113391.

Directory: REPCRL File: repcrl-Cb_LIp_Rphase.txt

| | Rphase_1 | Rphase_2 | Rphase_3 | Rphase_4 |
|---|---|---|---|---|
| Rphase_1 | 1.000 | 0.930 | 0.914 | 0.910 |
| Rphase_2 | 0.930 | 1.000 | 0.924 | 0.921 |
| Rphase_3 | 0.914 | 0.924 | 1.000 | 0.918 |
| Rphase_4 | 0.910 | 0.921 | 0.918 | 1.000 |

The number of conserved sites is 116489.

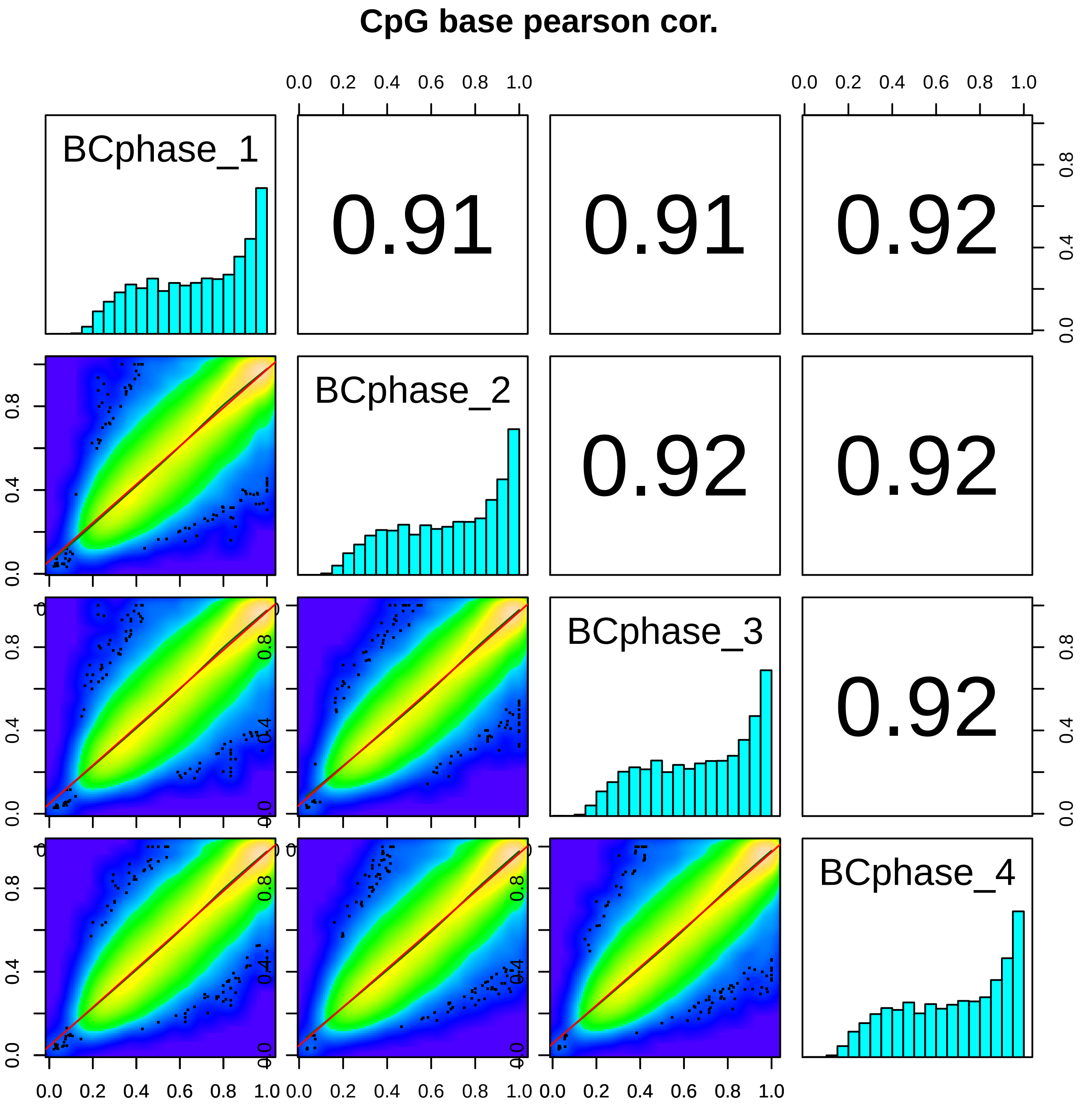

CpG base pearson cor.
BCphase_1
BCphase_2
BCphase_3
BCphase_4
0.91
0.91
0.92
0.92
0.92
0.92
0.0
0.2
0.4
0.6
0.8
1.0

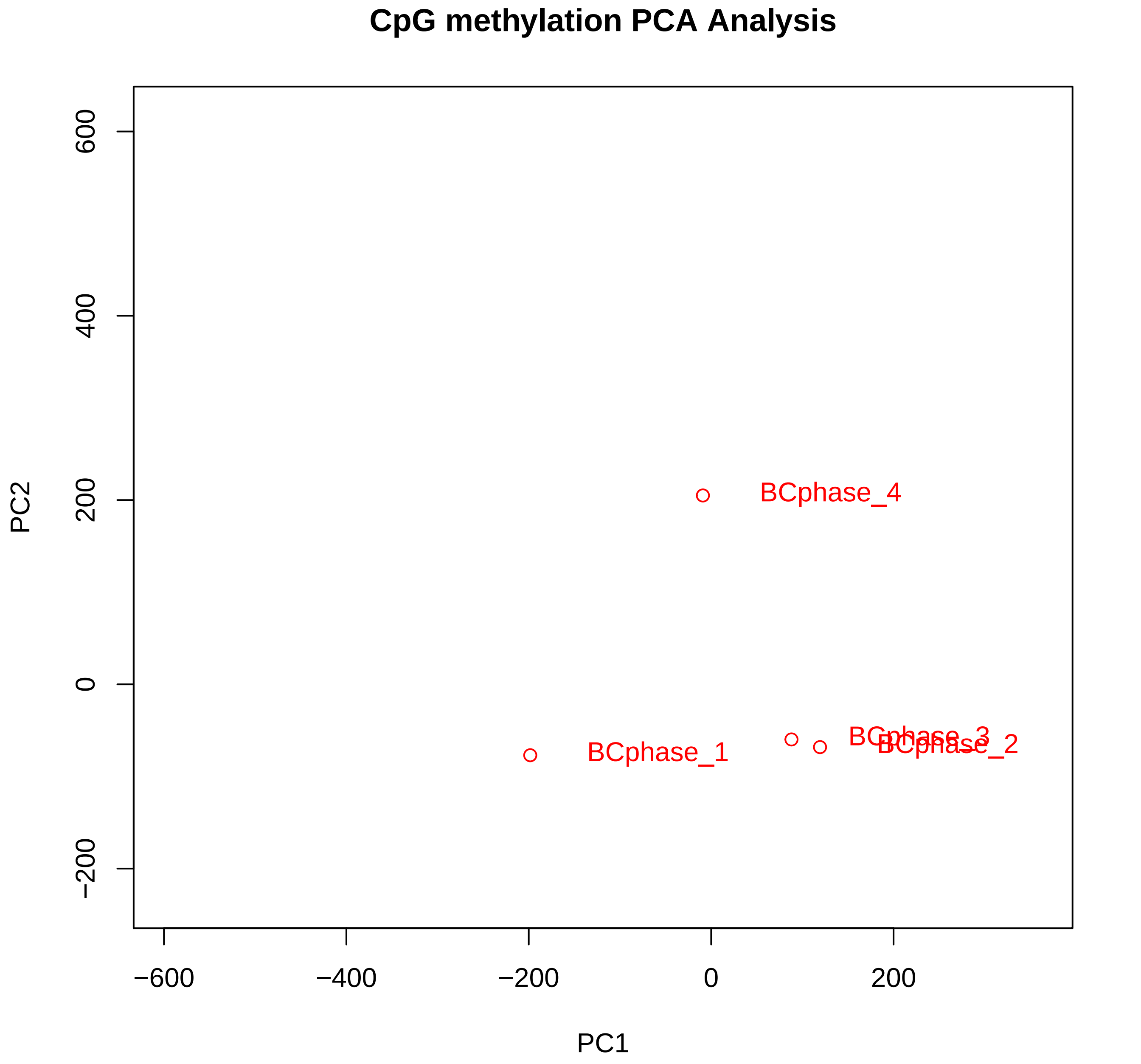

CpG methylation PCA Analysis
BCphase_4
BCphase_1
BCphase_3
BCphase_2
PC1
PC2
−600
−400
−200
0
200
400
600

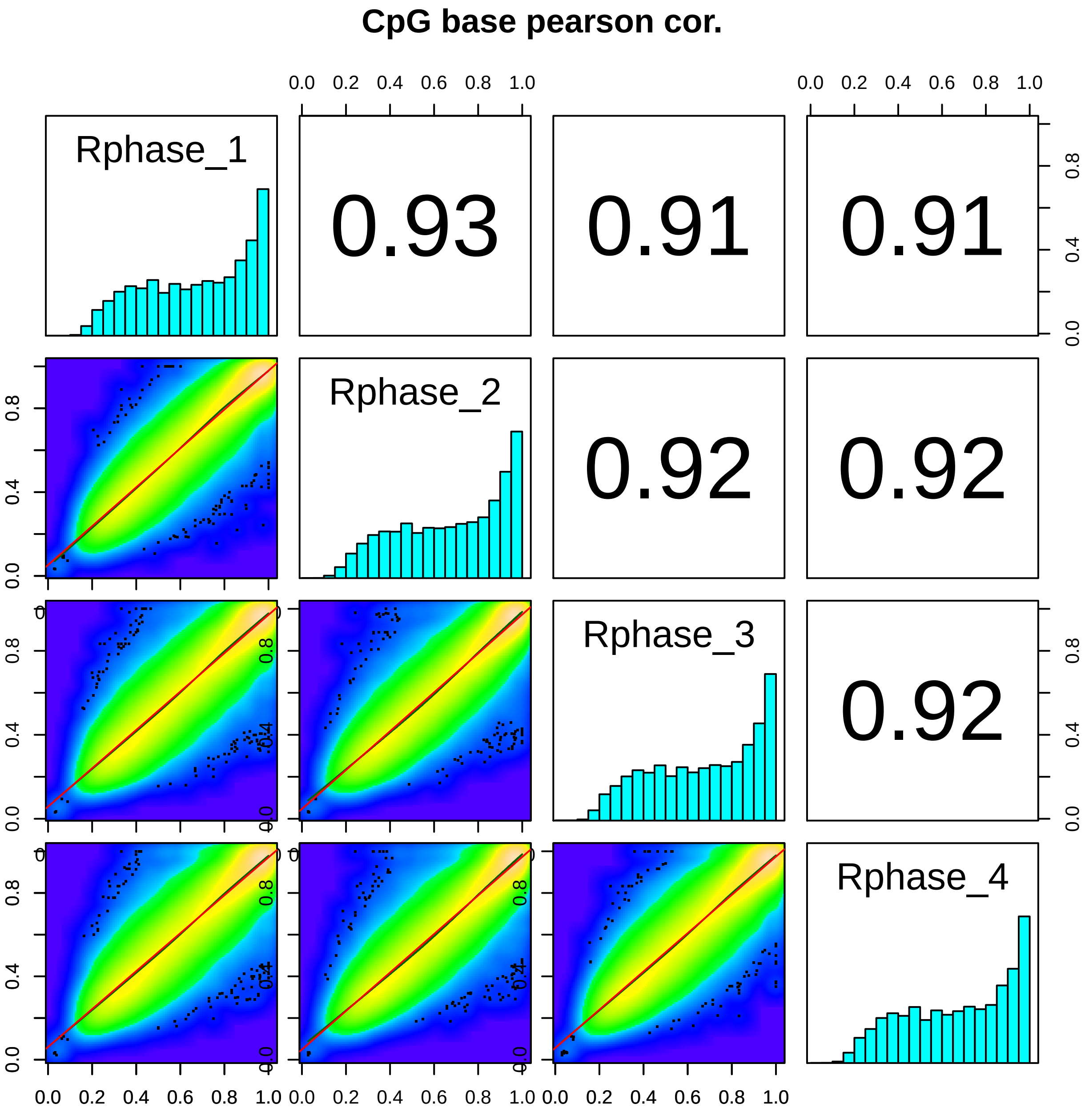
CpG base pearson cor.
Rphase_1
Rphase_2
Rphase_3
Rphase_4
0.93
0.91
0.91
0.92
0.92
0.92

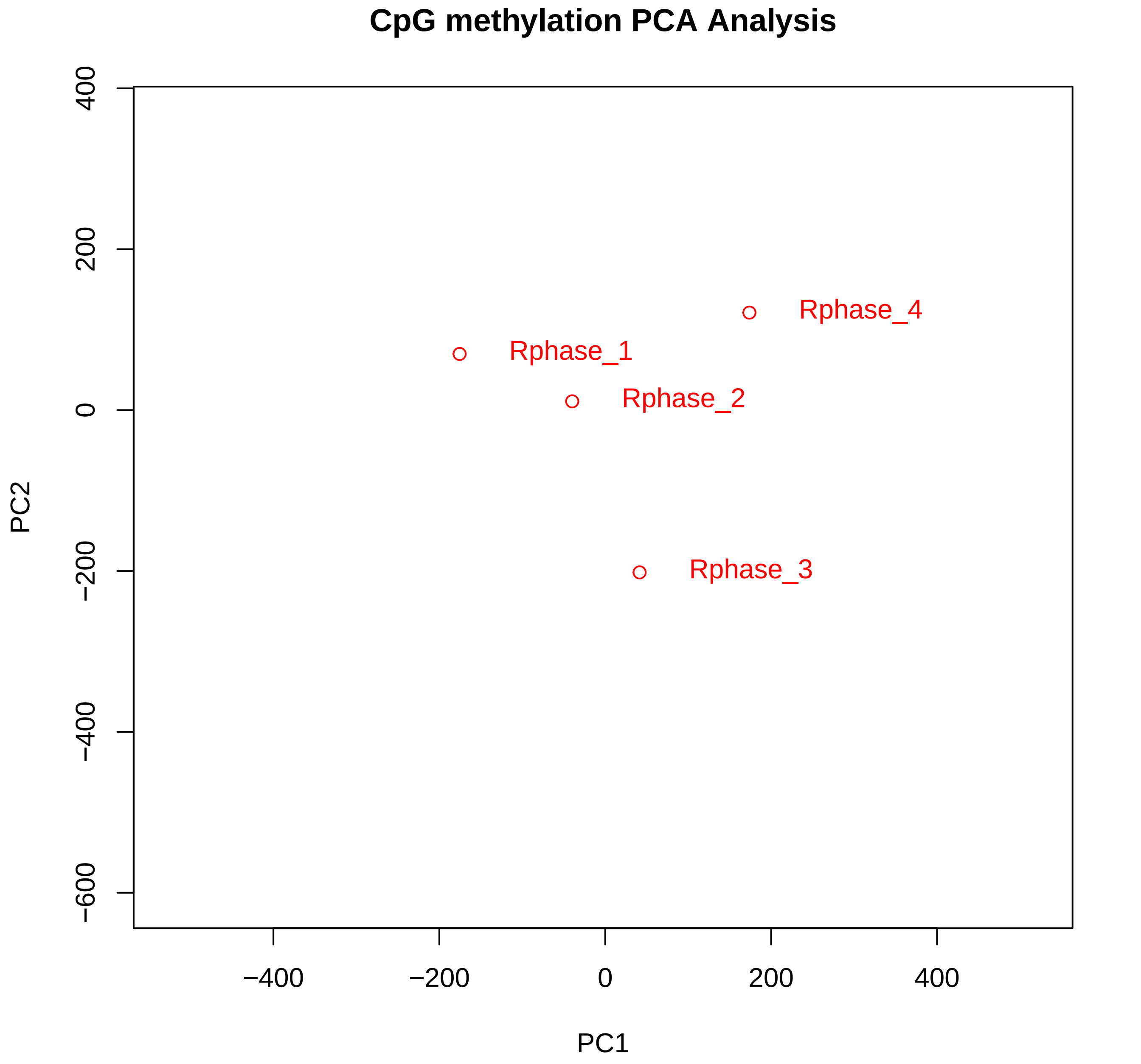
CpG methylation PCA Analysis
PC2
PC1
400
200
0
−200
−400
−600
−400
−200
0
200
400
Rphase_1
Rphase_2
Rphase_3
Rphase_4

```
Directory: MMP File: 0READMEmmp

MMP - Mapping of methylation sites on genome annotation.

Input: studymk, studymc, genome_ann (a list of GRanges objects providing annotated region labels
        and boundaries; output of BWASPR::get_genome_annotation())

Output: files mmp-*.txt

Notes: The output is generated by BWASPR::map_methylome() and gives an accounting for every sample
        as to where the CpGhsm and CpGscd (control) sites reside relative to the genome annotation.
       Abbreviations used: O/E, observed over expected
                           (expected percentages are based on the respective feature proportions
                            in the genome, as annotated).
```

```
Directory: MMP File: mmp-Cb_LI_BCphase.txt

================================================================================
Genomic composition in terms of feature regions
================================================================================

Cb genome size:                        212825769 bp
  Cb genic region size:                  152693430 bp ( 71.7%)
    Cb exon region size:                    39755998 bp ( 18.7%)
      Cb CDS region size:                     19364035 bp ( 48.7%)
      Cb five-prime UTR region size:           5122247 bp ( 12.9%)
      Cb three-prime UTR region size:          8663897 bp ( 21.8%)
      Cb other exon region size:               6605819 bp ( 16.6%)
    Cb intron region size:                 112937432 bp ( 53.1%)
  Cb intergenic region size:              60132339 bp ( 28.3%)
    Cb promoter region size:                 6533556 bp (  3.1%)
    Cb other intergenic region size:        53598783 bp ( 25.2%)

================================================================================
Methylation sites in genomic feature regions
================================================================================

Number and density (per 10kb) of sites in Cb_LI_BCphase_scd

Number of sites identified in Cb Cb_LI_BCphase_scd:                             24098022     ( 1132.29)
Number of sites identified in Cb Cb_LI_BCphase_scd genic regions:               16764357     ( 1097.91)                       69.57%     ( 0.97 O/E)
Number of sites identified in Cb Cb_LI_BCphase_scd   exon regions:                 4522126   ( 1137.47)                         18.77%   ( 1.00 O/E)
Number of sites identified in Cb Cb_LI_BCphase_scd     CDS regions:                  2548047 ( 1315.87)                           56.35% ( 1.16 O/E)
Number of sites identified in Cb Cb_LI_BCphase_scd     five-prime UTR regions:        727722 ( 1420.71)                           16.09% ( 1.25 O/E)
Number of sites identified in Cb Cb_LI_BCphase_scd     three-prime UTR regions:       788945 (  910.61)                           17.45% ( 0.80 O/E)
Number of sites identified in Cb Cb_LI_BCphase_scd     other exon regions:            457412 (  692.44)                           10.11% ( 0.61 O/E)
Number of sites identified in Cb Cb_LI_BCphase_scd   intron regions:              12242231   ( 1083.98)                         50.80%   ( 0.96 O/E)
Number of sites identified in Cb Cb_LI_BCphase_scd intergenic regions:           7333665     ( 1219.59)                       30.43%     ( 1.08 O/E)
Number of sites identified in Cb Cb_LI_BCphase_scd   promoter regions:              390780   (  598.11)                          1.62%   ( 0.53 O/E)
Number of sites identified in Cb Cb_LI_BCphase_scd   other intergenic regions:     6942885   ( 1295.34)                         28.81%   ( 1.14 O/E)

Number and density (per 10kb) of sites in Cb_LI_BCphase_hsm

Number of sites identified in Cb Cb_LI_BCphase_hsm:                               417310     (   19.61) (  1.73% of control)
Number of sites identified in Cb Cb_LI_BCphase_hsm genic regions:                 400176     (   26.21) (  2.39% of control)  95.89%     ( 1.34 O/E)
Number of sites identified in Cb Cb_LI_BCphase_hsm   exon regions:                  311652   (   78.39) (  6.89% of control)    74.68%   ( 4.00 O/E)
Number of sites identified in Cb Cb_LI_BCphase_hsm     CDS regions:                   289327 (  149.41) ( 11.35% of control)      92.84% ( 1.91 O/E)
Number of sites identified in Cb Cb_LI_BCphase_hsm     five-prime UTR regions:          5835 (   11.39) (  0.80% of control)       1.87% ( 0.15 O/E)
Number of sites identified in Cb Cb_LI_BCphase_hsm     three-prime UTR regions:        13818 (   15.95) (  1.75% of control)       4.43% ( 0.20 O/E)
Number of sites identified in Cb Cb_LI_BCphase_hsm     other exon regions:              2672 (    4.04) (  0.58% of control)       0.86% ( 0.05 O/E)
Number of sites identified in Cb Cb_LI_BCphase_hsm   intron regions:                 88524   (    7.84) (  0.72% of control)    21.21%   ( 0.40 O/E)
Number of sites identified in Cb Cb_LI_BCphase_hsm intergenic regions:             17134     (    2.85) (  0.23% of control)   4.11%     ( 0.15 O/E)
Number of sites identified in Cb Cb_LI_BCphase_hsm   promoter regions:                1691   (    2.59) (  0.43% of control)     0.41%   ( 0.13 O/E)
Number of sites identified in Cb Cb_LI_BCphase_hsm   other intergenic regions:       15443   (    2.88) (  0.22% of control)     3.70%   ( 0.15 O/E)
```

Directory: MMP File: mmp-Cb_LI_Rphase.txt

================================================================================
#### Genomic composition in terms of feature regions
================================================================================

| | | |
|---|---|---|
| Cb genome size: | 212825769 bp | |
| Cb genic region size: | 152693430 bp | ( 71.7%) |
| Cb exon region size: | 39755998 bp | ( 18.7%) |
| Cb CDS region size: | 19364035 bp | ( 48.7%) |
| Cb five-prime UTR region size: | 5122247 bp | ( 12.9%) |
| Cb three-prime UTR region size: | 8663897 bp | ( 21.8%) |
| Cb other exon region size: | 6605819 bp | ( 16.6%) |
| Cb intron region size: | 112937432 bp | ( 53.1%) |
| Cb intergenic region size: | 60132339 bp | ( 28.3%) |
| Cb promoter region size: | 6533556 bp | ( 3.1%) |
| Cb other intergenic region size: | 53598783 bp | ( 25.2%) |

================================================================================
#### Methylation sites in genomic feature regions
================================================================================

Number and density (per 10kb) of sites in Cb_LI_Rphase_scd

| | | | | |
|---|---|---|---|---|
| Number of sites identified in Cb Cb_LI_Rphase_scd: | 24215006 | ( 1137.79) | | |
| Number of sites identified in Cb Cb_LI_Rphase_scd genic regions: | 16847816 | ( 1103.38) | 69.58% | ( 0.97 O/E) |
| Number of sites identified in Cb Cb_LI_Rphase_scd exon regions: | 4545026 | ( 1143.23) | 18.77% | ( 1.00 O/E) |
| Number of sites identified in Cb Cb_LI_Rphase_scd CDS regions: | 2557897 | ( 1320.95) | 56.28% | ( 1.16 O/E) |
| Number of sites identified in Cb Cb_LI_Rphase_scd five-prime UTR regions: | 731210 | ( 1427.52) | 16.09% | ( 1.25 O/E) |
| Number of sites identified in Cb Cb_LI_Rphase_scd three-prime UTR regions: | 796019 | ( 918.78) | 17.51% | ( 0.80 O/E) |
| Number of sites identified in Cb Cb_LI_Rphase_scd other exon regions: | 459900 | ( 696.20) | 10.12% | ( 0.61 O/E) |
| Number of sites identified in Cb Cb_LI_Rphase_scd intron regions: | 12302790 | ( 1089.35) | 50.81% | ( 0.96 O/E) |
| Number of sites identified in Cb Cb_LI_Rphase_scd intergenic regions: | 7367190 | ( 1225.16) | 30.42% | ( 1.08 O/E) |
| Number of sites identified in Cb Cb_LI_Rphase_scd promoter regions: | 393158 | ( 601.75) | 1.62% | ( 0.53 O/E) |
| Number of sites identified in Cb Cb_LI_Rphase_scd other intergenic regions: | 6974032 | ( 1301.15) | 28.80% | ( 1.14 O/E) |

Number and density (per 10kb) of sites in Cb_LI_Rphase_hsm

| | | | | | |
|---|---|---|---|---|---|
| Number of sites identified in Cb Cb_LI_Rphase_hsm: | 445317 | ( 20.92) | ( 1.84% of control) | | |
| Number of sites identified in Cb Cb_LI_Rphase_hsm genic regions: | 424367 | ( 27.79) | ( 2.52% of control) | 95.30% | ( 1.33 O/E) |
| Number of sites identified in Cb Cb_LI_Rphase_hsm exon regions: | 326541 | ( 82.14) | ( 7.18% of control) | 73.33% | ( 3.93 O/E) |
| Number of sites identified in Cb Cb_LI_Rphase_hsm CDS regions: | 301663 | ( 155.79) | ( 11.79% of control) | 92.38% | ( 1.90 O/E) |
| Number of sites identified in Cb Cb_LI_Rphase_hsm five-prime UTR regions: | 6430 | ( 12.55) | ( 0.88% of control) | 1.97% | ( 0.15 O/E) |
| Number of sites identified in Cb Cb_LI_Rphase_hsm three-prime UTR regions: | 15428 | ( 17.81) | ( 1.94% of control) | 4.72% | ( 0.22 O/E) |
| Number of sites identified in Cb Cb_LI_Rphase_hsm other exon regions: | 3020 | ( 4.57) | ( 0.66% of control) | 0.92% | ( 0.06 O/E) |
| Number of sites identified in Cb Cb_LI_Rphase_hsm intron regions: | 97826 | ( 8.66) | ( 0.80% of control) | 21.97% | ( 0.41 O/E) |
| Number of sites identified in Cb Cb_LI_Rphase_hsm intergenic regions: | 20950 | ( 3.48) | ( 0.28% of control) | 4.70% | ( 0.17 O/E) |
| Number of sites identified in Cb Cb_LI_Rphase_hsm promoter regions: | 1986 | ( 3.04) | ( 0.51% of control) | 0.45% | ( 0.15 O/E) |
| Number of sites identified in Cb Cb_LI_Rphase_hsm other intergenic regions: | 18964 | ( 3.54) | ( 0.27% of control) | 4.26% | ( 0.17 O/E) |

```
Directory: ACS File: 0READMEacs

ACS - Annotation of conserved methylation sites.

Input: studymk, genome_ann (a list of GRanges objects providing annotated region labels and bounds;
                            output of BWASPR::get_genome_annotation())

Output: file acs-*.txt

Notes: The output is generated by BWASPR::annotate_methylome(), which uses methylKit::unite()
        to determine the sites common to all experiments and applies genomation::annotateWithFeature().
        The acs-*.txt files show the common sites with methylation levels in all samples
        and columns with booleans indicating whether the site fits a specific annotation feature.

       Abbreviations used: pc, protein coding; nc, non-coding; fp, five prime; tp, three prime.
                           UTR, untranslated region; unique, not overlapping with other categories.
```

Directory: ACS File: acs-Cb_LIp.txt

| BCphase | Rphase | chr | start | end | strand | gene | exon | pcexon | promoter | CDS | fpUTR | tpUTR | fpUTRnotCDS | tpUTRnotCDS | tpUTRunique | ncexon |
|---|---|---|---|---|---|---|---|---|---|---|---|---|---|---|---|---|
| 18.27 | 23.73 | NW_011924879.1 | 2162 | 2162 | + | 1 | 1 | 1 | 0 | 1 | 0 | 0 | 0 | 0 | 0 | 0 |
| 18.48 | 20 | NW_011924880.1 | 22922 | 22922 | + | 0 | 0 | 0 | 0 | 0 | 0 | 0 | 0 | 0 | 0 | 0 |
| 14.79 | 14.04 | NW_011924880.1 | 67566 | 67566 | + | 0 | 0 | 0 | 0 | 0 | 0 | 0 | 0 | 0 | 0 | 0 |
| 16.47 | 22.78 | NW_011924880.1 | 91058 | 91058 | + | 0 | 0 | 0 | 0 | 0 | 0 | 0 | 0 | 0 | 0 | 0 |
| 14.81 | 16.33 | NW_011924880.1 | 175720 | 175720 | + | 0 | 0 | 0 | 0 | 0 | 0 | 0 | 0 | 0 | 0 | 0 |
| 5.54 | 5.67 | NW_011924881.1 | 3405 | 3405 | + | 0 | 0 | 0 | 0 | 0 | 0 | 0 | 0 | 0 | 0 | 0 |
| 9.37 | 8.38 | NW_011924881.1 | 6387 | 6387 | + | 0 | 0 | 0 | 0 | 0 | 0 | 0 | 0 | 0 | 0 | 0 |
| 5.87 | 3.75 | NW_011924881.1 | 14174 | 14174 | + | 0 | 0 | 0 | 0 | 0 | 0 | 0 | 0 | 0 | 0 | 0 |
| 12.94 | 12.22 | NW_011924881.1 | 28467 | 28467 | + | 0 | 0 | 0 | 0 | 0 | 0 | 0 | 0 | 0 | 0 | 0 |
| 21.57 | 20.34 | NW_011924881.1 | 33583 | 33583 | + | 0 | 0 | 0 | 0 | 0 | 0 | 0 | 0 | 0 | 0 | 0 |
| 20 | 25.53 | NW_011924881.1 | 33588 | 33588 | + | 0 | 0 | 0 | 0 | 0 | 0 | 0 | 0 | 0 | 0 | 0 |
| 31.58 | 36.96 | NW_011924881.1 | 39844 | 39844 | + | 1 | 1 | 1 | 0 | 0 | 0 | 1 | 0 | 1 | 1 | 0 |
| 77.59 | 66.67 | NW_011924881.1 | 39967 | 39967 | + | 1 | 1 | 1 | 0 | 0 | 0 | 1 | 0 | 1 | 1 | 0 |
| 52.24 | 59.09 | NW_011924881.1 | 39987 | 39987 | + | 1 | 1 | 1 | 0 | 0 | 0 | 1 | 0 | 1 | 1 | 0 |
| 18.29 | 16.67 | NW_011924881.1 | 40035 | 40035 | + | 1 | 1 | 1 | 0 | 0 | 0 | 1 | 0 | 1 | 1 | 0 |
| 73.21 | 91.84 | NW_011924881.1 | 40061 | 40061 | + | 1 | 1 | 1 | 0 | 0 | 0 | 1 | 0 | 1 | 1 | 0 |
| 92.16 | 98.04 | NW_011924881.1 | 40072 | 40072 | + | 1 | 1 | 1 | 0 | 0 | 0 | 1 | 0 | 1 | 1 | 0 |
| 50 | 54.93 | NW_011924881.1 | 40095 | 40095 | + | 1 | 1 | 1 | 0 | 0 | 0 | 1 | 0 | 1 | 1 | 0 |
| 73.61 | 64.65 | NW_011924881.1 | 40120 | 40120 | + | 1 | 1 | 1 | 0 | 0 | 0 | 1 | 0 | 1 | 1 | 0 |
| 57.14 | 64.79 | NW_011924881.1 | 40159 | 40159 | + | 1 | 1 | 1 | 0 | 1 | 0 | 0 | 0 | 0 | 0 | 0 |
| 50 | 66.67 | NW_011924881.1 | 40202 | 40202 | + | 1 | 1 | 1 | 0 | 1 | 0 | 0 | 0 | 0 | 0 | 0 |
| 83.17 | 86.87 | NW_011924881.1 | 40256 | 40256 | + | 1 | 1 | 1 | 0 | 1 | 0 | 0 | 0 | 0 | 0 | 0 |
| 91 | 93.26 | NW_011924881.1 | 40259 | 40259 | + | 1 | 1 | 1 | 0 | 1 | 0 | 0 | 0 | 0 | 0 | 0 |
| 87 | 88.89 | NW_011924881.1 | 40270 | 40270 | + | 1 | 1 | 1 | 0 | 1 | 0 | 0 | 0 | 0 | 0 | 0 |

...

... only first 25 lines shown ...

```
Directory: RNK File: 0READMErnk

RNK - Genomic feature regions ranked by CpG methylation and hsm statistics.

Input: studymc, studymk, region.gr (GRanges object providing annotated region labels and bounds)

Output: files ranked-genes-*.txt sites-in-genes-*.txt
              ranked-promoters-*.txt sites-in-promoters-*.txt
              plot*.pdf

Notes: The output is generated by BWASPR::rank_rbm().
       The files ranked-genes-*.txt provide ranked lists of genes based on
        overall methylation percentage or the occurrence of CpGhms sites
        within the annotated gene bounds.  The output columns include
          region_ID    (= region/gene name)
          rwidth       (= region/gene width)
          nbrsites     (= number of CpGhsm sites in the region/gene)
          nbrper10kb   (= number of CpGhsm sites in the region/gene normalized to 10kb width)
          pmrpersite   (= average % methylation per CpGhsm site in the region/gene)
          pmrpernucl   (= average % CpGhsm methylation per nucleotide in the region/gene)
          prcntM       (= overall % CpGhsm methylation in the region/gene)

       The tables are sorted by prcntM (*byPrcntM* files) or by nbper10kb
         (*bySiteDensity* files) in descending order.

       The ranked-promoters-*.txt files provide analogous tables for promoter regions.
        If the promoter annotation was derived with BWASP, then these regions are simply
        defined here as 500 nucleotides upstream of the 5'-end of the gene (shorter if the
        scaffold ends before).

       The files sites-*.txt provide details for the CpGhsm sites in the respective
        regions.
       The files plot-*.pdf explore concordance of the two methylation measures.
       The optional parameters minrwidth, maxrwidth, and minnbrsites specify
       the length range and minimal number of methylation sites for regions
       to be included in the plot.
```

Directory: RNK File: ranked-genes-byPrcntM-Cb_LIp_BCphase.txt

| region_ID | seqnames | start | end | width | strand | coverage | numCs | numTs | prcntM |
|---|---|---|---|---|---|---|---|---|---|
| gene13336 | NW_011929072.1 | 30 | 517 | 488 | + | 459 | 428 | 31 | 93.25 |
| gene13213 | NW_011926558.1 | 4 | 1118 | 1115 | - | 145 | 128 | 17 | 88.28 |
| gene1643 | NW_011924932.1 | 342040 | 343422 | 1383 | - | 76 | 67 | 9 | 88.16 |
| gene10364 | NW_011925342.1 | 111577 | 112136 | 560 | + | 1491 | 1282 | 209 | 85.98 |
| gene8895 | NW_011925229.1 | 67030 | 70140 | 3111 | - | 246 | 210 | 36 | 85.37 |
| gene13380 | NW_011929316.1 | 135 | 1735 | 1601 | - | 217 | 180 | 37 | 82.95 |
| gene10653 | NW_011925362.1 | 80044 | 81082 | 1039 | - | 666 | 537 | 129 | 80.63 |
| gene10196 | NW_011925327.1 | 165617 | 166924 | 1308 | + | 1237 | 950 | 287 | 76.8 |
| gene13247 | NW_011926712.1 | 6 | 783 | 778 | + | 94 | 72 | 22 | 76.6 |
| gene5000 | NW_011925022.1 | 386851 | 387976 | 1126 | - | 1853 | 1402 | 451 | 75.66 |
| gene4579 | NW_011925016.1 | 246765 | 248208 | 1444 | - | 2754 | 2049 | 705 | 74.4 |
| gene3015 | NW_011924977.1 | 732939 | 733749 | 811 | - | 2809 | 2088 | 721 | 74.33 |
| gene8203 | NW_011925182.1 | 19245 | 20253 | 1009 | + | 1807 | 1330 | 477 | 73.6 |
| gene2956 | NW_011924977.1 | 323781 | 325133 | 1353 | + | 2431 | 1772 | 659 | 72.89 |
| gene903 | NW_011924914.1 | 17973 | 18954 | 982 | - | 638 | 455 | 183 | 71.32 |
| gene405 | NW_011924888.1 | 1707554 | 1708995 | 1442 | - | 4990 | 3522 | 1468 | 70.58 |
| gene2249 | NW_011924946.1 | 3301258 | 3302569 | 1312 | - | 1811 | 1270 | 541 | 70.13 |
| gene8315 | NW_011925189.1 | 490017 | 490870 | 854 | + | 2455 | 1717 | 738 | 69.94 |
| gene9525 | NW_011925273.1 | 781582 | 783179 | 1598 | + | 3746 | 2609 | 1137 | 69.65 |
| gene9952 | NW_011925321.1 | 87267 | 88468 | 1202 | + | 2407 | 1668 | 739 | 69.3 |
| gene4880 | NW_011925020.1 | 2602046 | 2603046 | 1001 | + | 2413 | 1661 | 752 | 68.84 |
| gene6173 | NW_011925075.1 | 824766 | 825641 | 876 | + | 930 | 630 | 300 | 67.74 |
| gene6356 | NW_011925092.1 | 13079 | 14878 | 1800 | + | 752 | 506 | 246 | 67.29 |
| gene2274 | NW_011924947.1 | 109214 | 110680 | 1467 | - | 8375 | 5628 | 2747 | 67.2 |

...

... only first 25 lines shown ...

Directory: RNK File: ranked-genes-byPrcntM-Cb_LIp_Rphase.txt

| region_ID | seqnames | start | end | width | strand | coverage | numCs | numTs | prcntM |
|---|---|---|---|---|---|---|---|---|---|
| gene13336 | NW_011929072.1 | 30 | 517 | 488 | + | 544 | 505 | 39 | 92.83 |
| gene1643 | NW_011924932.1 | 342040 | 343422 | 1383 | - | 121 | 108 | 13 | 89.26 |
| gene13213 | NW_011926558.1 | 4 | 1118 | 1115 | - | 182 | 157 | 25 | 86.26 |
| gene8895 | NW_011925229.1 | 67030 | 70140 | 3111 | - | 164 | 140 | 24 | 85.37 |
| gene10364 | NW_011925342.1 | 111577 | 112136 | 560 | + | 1763 | 1492 | 271 | 84.63 |
| gene10653 | NW_011925362.1 | 80044 | 81082 | 1039 | - | 712 | 589 | 123 | 82.72 |
| gene10196 | NW_011925327.1 | 165617 | 166924 | 1308 | + | 1446 | 1103 | 343 | 76.28 |
| gene5000 | NW_011925022.1 | 386851 | 387976 | 1126 | - | 1926 | 1428 | 498 | 74.14 |
| gene3015 | NW_011924977.1 | 732939 | 733749 | 811 | - | 2951 | 2187 | 764 | 74.11 |
| gene8203 | NW_011925182.1 | 19245 | 20253 | 1009 | + | 2094 | 1532 | 562 | 73.16 |
| gene2956 | NW_011924977.1 | 323781 | 325133 | 1353 | + | 2713 | 1981 | 732 | 73.02 |
| gene4579 | NW_011925016.1 | 246765 | 248208 | 1444 | - | 3257 | 2370 | 887 | 72.77 |
| gene903 | NW_011924914.1 | 17973 | 18954 | 982 | - | 713 | 516 | 197 | 72.37 |
| gene4880 | NW_011925020.1 | 2602046 | 2603046 | 1001 | + | 2640 | 1903 | 737 | 72.08 |
| gene2249 | NW_011924946.1 | 3301258 | 3302569 | 1312 | - | 1862 | 1329 | 533 | 71.37 |
| gene13380 | NW_011929316.1 | 135 | 1735 | 1601 | - | 250 | 177 | 73 | 70.8 |
| gene13247 | NW_011926712.1 | 6 | 783 | 778 | + | 124 | 87 | 37 | 70.16 |
| gene405 | NW_011924888.1 | 1707554 | 1708995 | 1442 | - | 5750 | 4033 | 1717 | 70.14 |
| gene9525 | NW_011925273.1 | 781582 | 783179 | 1598 | + | 3806 | 2666 | 1140 | 70.05 |
| gene8315 | NW_011925189.1 | 490017 | 490870 | 854 | + | 2834 | 1975 | 859 | 69.69 |
| gene4906 | NW_011925020.1 | 2960255 | 2961622 | 1368 | - | 4977 | 3376 | 1601 | 67.83 |
| gene2274 | NW_011924947.1 | 109214 | 110680 | 1467 | - | 9537 | 6456 | 3081 | 67.69 |
| gene4879 | NW_011925020.1 | 2601782 | 2603159 | 1378 | - | 3518 | 2371 | 1147 | 67.4 |
| gene6356 | NW_011925092.1 | 13079 | 14878 | 1800 | + | 1014 | 680 | 334 | 67.06 |

...

... only first 25 lines shown ...

Directory: RNK File: ranked-genes-bySiteDensity-Cb_LIp_BCphase.txt

| region_ID | rwidth | nbrsites | nbrper10kb | pmpersite | pmpernucl | pglink | seqnames | start | end | strand | coverage | numCs | numTs | prcntM |
|---|---|---|---|---|---|---|---|---|---|---|---|---|---|---|
| gene5917 | 823 | 118 | 1433.78 | 31.08 | 4.46 | https://www.ncbi.nlm.nih.gov/gene/?term=LOC105278849 | NW_011925053.1 | 1371807 | 1372629 | + | 23014 | 5499 | 17515 | 23.89 |
| gene13384 | 1877 | 215 | 1145.44 | 40.57 | 4.65 | https://www.ncbi.nlm.nih.gov/gene/?term=LOC105287447 | NW_011929339.1 | 7 | 1883 | + | 43736 | 13247 | 30489 | 30.29 |
| gene10219 | 235 | 24 | 1021.28 | 47.44 | 4.85 | https://www.ncbi.nlm.nih.gov/gene/?term=LOC105283830 | NW_011925327.1 | 293521 | 293755 | + | 2888 | 1028 | 1860 | 35.6 |
| gene5937 | 1902 | 182 | 956.89 | 31.96 | 3.06 | https://www.ncbi.nlm.nih.gov/gene/?term=LOC105278873 | NW_011925053.1 | 1492486 | 1494387 | - | 41204 | 8594 | 32610 | 20.86 |
| gene5309 | 1704 | 160 | 938.97 | 45.45 | 4.27 | https://www.ncbi.nlm.nih.gov/gene/?term=LOC105278160 | NW_011925031.1 | 150501 | 152204 | - | 28899 | 11318 | 17581 | 39.16 |
| gene5412 | 1868 | 175 | 936.83 | 41.04 | 3.84 | https://www.ncbi.nlm.nih.gov/gene/?term=LOC105278266 | NW_011925035.1 | 1477936 | 1479803 | + | 26811 | 8054 | 18757 | 30.04 |
| gene12538 | 372 | 33 | 887.1 | 49.96 | 4.43 | https://www.ncbi.nlm.nih.gov/gene/?term=LOC105286531 | NW_011925715.1 | 412527 | 412898 | - | 6631 | 2397 | 4234 | 36.15 |
| gene5301 | 1881 | 163 | 866.56 | 39.06 | 3.38 | https://www.ncbi.nlm.nih.gov/gene/?term=LOC105278148 | NW_011925031.1 | 114108 | 115988 | + | 32114 | 8671 | 23443 | 27 |
| gene11566 | 1603 | 134 | 835.93 | 46.74 | 3.91 | https://www.ncbi.nlm.nih.gov/gene/?term=LOC105285392 | NW_011925459.1 | 162939 | 164541 | + | 22113 | 7913 | 14200 | 35.78 |
| gene1993 | 1449 | 121 | 835.06 | 42.83 | 3.58 | https://www.ncbi.nlm.nih.gov/gene/?term=LOC105288193 | NW_011924946.1 | 775753 | 777201 | - | 20670 | 5912 | 14758 | 28.6 |
| gene11569 | 1311 | 108 | 823.8 | 72.61 | 5.98 | https://www.ncbi.nlm.nih.gov/gene/?term=LOC105285394 | NW_011925459.1 | 186244 | 187554 | - | 13404 | 8120 | 5284 | 60.58 |
| gene10943 | 1759 | 143 | 812.96 | 55.68 | 4.53 | https://www.ncbi.nlm.nih.gov/gene/?term=LOC105284659 | NW_011925387.1 | 118203 | 119961 | - | 18654 | 8542 | 10112 | 45.79 |
| gene1894 | 445 | 36 | 808.99 | 56.38 | 4.56 | https://www.ncbi.nlm.nih.gov/gene/?term=LOC105288083 | NW_011924945.1 | 117665 | 118109 | + | 2943 | 1631 | 1312 | 55.42 |
| gene6057 | 2105 | 166 | 788.6 | 44.15 | 3.48 | https://www.ncbi.nlm.nih.gov/gene/?term=LOC105279035 | NW_011925067.1 | 1035801 | 1037905 | - | 40667 | 7834 | 32833 | 19.26 |
| gene3850 | 2578 | 202 | 783.55 | 33.84 | 2.65 | https://www.ncbi.nlm.nih.gov/gene/?term=LOC105276490 | NW_011924996.1 | 1609008 | 1611585 | - | 53878 | 11425 | 42453 | 21.21 |
| gene10232 | 1591 | 124 | 779.38 | 40.17 | 3.13 | https://www.ncbi.nlm.nih.gov/gene/?term=LOC105283847 | NW_011925327.1 | 546966 | 548556 | + | 22494 | 7551 | 14943 | 33.57 |
| gene3719 | 1471 | 114 | 774.98 | 57.61 | 4.46 | https://www.ncbi.nlm.nih.gov/gene/?term=LOC105276360 | NW_011924994.1 | 94735 | 96205 | - | 26367 | 8359 | 18008 | 31.7 |
| gene1562 | 1491 | 115 | 771.29 | 49.25 | 3.8 | https://www.ncbi.nlm.nih.gov/gene/?term=LOC105287692 | NW_011924931.1 | 3966137 | 3967627 | - | 23250 | 7529 | 15721 | 32.38 |
| gene560 | 1584 | 122 | 770.2 | 30.66 | 2.36 | https://www.ncbi.nlm.nih.gov/gene/?term=LOC109610897 | NW_011924892.1 | 1109843 | 1111426 | + | 31762 | 4870 | 26892 | 15.33 |
| gene12364 | 1403 | 108 | 769.78 | 57.33 | 4.41 | https://www.ncbi.nlm.nih.gov/gene/?term=LOC105286269 | NW_011925664.1 | 291441 | 292843 | - | 16020 | 7747 | 8273 | 48.36 |
| gene4635 | 2067 | 157 | 759.55 | 52.8 | 4.01 | https://www.ncbi.nlm.nih.gov/gene/?term=LOC105277390 | NW_011925019.1 | 5988 | 8054 | - | 24700 | 10335 | 14365 | 41.84 |
| gene570 | 1673 | 127 | 759.12 | 52.15 | 3.96 | https://www.ncbi.nlm.nih.gov/gene/?term=LOC105278549 | NW_011924892.1 | 1160484 | 1162156 | + | 20893 | 8299 | 12594 | 39.72 |
| gene211 | 1280 | 97 | 757.81 | 54.83 | 4.16 | https://www.ncbi.nlm.nih.gov/gene/?term=LOC105274534 | NW_011924887.1 | 1565391 | 1566670 | + | 17371 | 7360 | 10011 | 42.37 |
| gene1237 | 2543 | 192 | 755.01 | 38.01 | 2.87 | https://www.ncbi.nlm.nih.gov/gene/?term=LOC105285861 | NW_011924922.1 | 2617834 | 2620376 | - | 26032 | 6833 | 19199 | 26.25 |

...

... only first 25 lines shown ...

Directory: RNK File: ranked-genes-bySiteDensity-Cb_LIp_Rphase.txt

| region_ID | rwidth | nbrsites | nbrper10kb | pmpersite | pmpernucl | pglink | seqnames | start | end | strand | coverage | numCs | numTs | prcntM |
|---|---|---|---|---|---|---|---|---|---|---|---|---|---|---|
| gene5917 | 823 | 124 | 1506.68 | 29.72 | 4.48 | https://www.ncbi.nlm.nih.gov/gene/?term=LOC105278849 | NW_011925053.1 | 1371807 | 1372629 | + | 25406 | 5922 | 19484 | 23.31 |
| gene13384 | 1877 | 216 | 1150.77 | 40.18 | 4.62 | https://www.ncbi.nlm.nih.gov/gene/?term=LOC105287447 | NW_011929339.1 | 7 | 1883 | + | 49418 | 14750 | 34668 | 29.85 |
| gene10219 | 235 | 23 | 978.72 | 50.11 | 4.9 | https://www.ncbi.nlm.nih.gov/gene/?term=LOC105283830 | NW_011925327.1 | 293521 | 293755 | + | 2789 | 1049 | 1740 | 37.61 |
| gene5937 | 1902 | 185 | 972.66 | 31.95 | 3.11 | https://www.ncbi.nlm.nih.gov/gene/?term=LOC105278873 | NW_011925053.1 | 1492486 | 1494387 | - | 45153 | 9534 | 35619 | 21.11 |
| gene5412 | 1868 | 180 | 963.6 | 41.08 | 3.96 | https://www.ncbi.nlm.nih.gov/gene/?term=LOC105278266 | NW_011925035.1 | 1477936 | 1479803 | + | 31190 | 9505 | 21685 | 30.47 |
| gene5309 | 1704 | 161 | 944.84 | 44.08 | 4.16 | https://www.ncbi.nlm.nih.gov/gene/?term=LOC105278160 | NW_011925031.1 | 150501 | 152204 | - | 33018 | 12554 | 20464 | 38.02 |
| gene1993 | 1449 | 131 | 904.07 | 42.78 | 3.87 | https://www.ncbi.nlm.nih.gov/gene/?term=LOC105288193 | NW_011924946.1 | 775753 | 777201 | - | 23349 | 6678 | 16671 | 28.6 |
| gene5301 | 1881 | 168 | 893.14 | 38.02 | 3.4 | https://www.ncbi.nlm.nih.gov/gene/?term=LOC105278148 | NW_011925031.1 | 114108 | 115988 | + | 35855 | 9709 | 26146 | 27.08 |
| gene12538 | 372 | 32 | 860.22 | 50.64 | 4.36 | https://www.ncbi.nlm.nih.gov/gene/?term=LOC105286531 | NW_011925715.1 | 412527 | 412898 | - | 7038 | 2470 | 4568 | 35.1 |
| gene11566 | 1603 | 136 | 848.41 | 44.79 | 3.8 | https://www.ncbi.nlm.nih.gov/gene/?term=LOC105285392 | NW_011925459.1 | 162939 | 164541 | + | 25389 | 9231 | 16158 | 36.36 |
| gene6057 | 2105 | 178 | 845.61 | 41.82 | 3.54 | https://www.ncbi.nlm.nih.gov/gene/?term=LOC105279035 | NW_011925067.1 | 1035801 | 1037905 | - | 44375 | 9093 | 35282 | 20.49 |
| gene10943 | 1759 | 148 | 841.39 | 54.51 | 4.59 | https://www.ncbi.nlm.nih.gov/gene/?term=LOC105284659 | NW_011925387.1 | 118203 | 119961 | - | 21022 | 9760 | 11262 | 46.43 |
| gene11569 | 1311 | 110 | 839.05 | 71.56 | 6 | https://www.ncbi.nlm.nih.gov/gene/?term=LOC105285394 | NW_011925459.1 | 186244 | 187554 | - | 14973 | 9268 | 5705 | 61.9 |
| gene11402 | 335 | 28 | 835.82 | 23.88 | 2 | https://www.ncbi.nlm.nih.gov/gene/?term=LOC105285201 | NW_011925438.1 | 138018 | 138352 | - | 6166 | 1275 | 4891 | 20.68 |
| gene3850 | 2578 | 210 | 814.58 | 33.8 | 2.75 | https://www.ncbi.nlm.nih.gov/gene/?term=LOC105276490 | NW_011924996.1 | 1609008 | 1611585 | - | 60131 | 12558 | 47573 | 20.88 |
| gene2130 | 1254 | 102 | 813.4 | 40.98 | 3.33 | https://www.ncbi.nlm.nih.gov/gene/?term=LOC105274542 | NW_011924946.1 | 2578206 | 2579459 | - | 15155 | 4345 | 10810 | 28.67 |
| gene1894 | 445 | 36 | 808.99 | 52.4 | 4.24 | https://www.ncbi.nlm.nih.gov/gene/?term=LOC105288083 | NW_011924945.1 | 117665 | 118109 | + | 3268 | 1698 | 1570 | 51.96 |
| gene10232 | 1591 | 128 | 804.53 | 38.64 | 3.11 | https://www.ncbi.nlm.nih.gov/gene/?term=LOC105283847 | NW_011925327.1 | 546966 | 548556 | + | 23890 | 8045 | 15845 | 33.68 |
| gene560 | 1584 | 127 | 801.77 | 29.91 | 2.4 | https://www.ncbi.nlm.nih.gov/gene/?term=LOC109610897 | NW_011924892.1 | 1109843 | 1111426 | + | 34360 | 5421 | 28939 | 15.78 |
| gene4635 | 2067 | 163 | 788.58 | 51.05 | 4.03 | https://www.ncbi.nlm.nih.gov/gene/?term=LOC105277390 | NW_011925019.1 | 5988 | 8054 | - | 27155 | 11249 | 15906 | 41.43 |
| gene12238 | 2081 | 163 | 783.28 | 28.13 | 2.2 | https://www.ncbi.nlm.nih.gov/gene/?term=LOC105286135 | NW_011925638.1 | 328060 | 330140 | + | 31144 | 6116 | 25028 | 19.64 |
| gene3719 | 1471 | 115 | 781.78 | 57.2 | 4.47 | https://www.ncbi.nlm.nih.gov/gene/?term=LOC105276360 | NW_011924994.1 | 94735 | 96205 | - | 29667 | 9522 | 20145 | 32.1 |
| gene12746 | 2688 | 208 | 773.81 | 40.55 | 3.14 | https://www.ncbi.nlm.nih.gov/gene/?term=LOC105286737 | NW_011925789.1 | 653088 | 655775 | + | 67190 | 18169 | 49021 | 27.04 |
| gene570 | 1673 | 129 | 771.07 | 51.35 | 3.96 | https://www.ncbi.nlm.nih.gov/gene/?term=LOC105278549 | NW_011924892.1 | 1160484 | 1162156 | + | 23272 | 9257 | 14015 | 39.78 |

...

... only first 25 lines shown ...

Directory: RNK File: ranked-promoters-byPrcntM-Cb_LIp_BCphase.txt

```
region_ID   seqnames    start end   width strand      coverage    numCs numTs prcntM
gene8748    NW_011925216.1    183604      184103      500   -     16    16    0     100
gene4040    NW_011925003.1    277741      278240      500   +     145   138   7     95.17
gene3546    NW_011924987.1    1804967     1805466     500   -     248   229   19    92.34
gene4016    NW_011925003.1    66624 67123 500   +     3461  3147  314   90.93
gene5713    NW_011925050.1    7072  7571  500   -     1754  1587  167   90.48
gene11598   NW_011925467.1    116917      117416      500   +     1359  1210  149   89.04
gene5601    NW_011925036.1    147325      147824      500   -     1567  1372  195   87.56
gene13313   NW_011928871.1    1     61    61    +     8     7     1     87.5
gene2337    NW_011924951.1    247713      248212      500   +     450   383   67    85.11
gene8331    NW_011925189.1    573460      573959      500   +     747   624   123   83.53
gene13190   NW_011926458.1    36701 37200 500   -     145   119   26    82.07
gene3373    NW_011924987.1    206378      206877      500   -     540   437   103   80.93
gene12609   NW_011925739.1    39939 40438 500   +     47    38    9     80.85
gene772     NW_011924904.1    1310349     1310848     500   +     2791  2249  542   80.58
gene9313    NW_011925258.1    34883 35382 500   +     813   654   159   80.44
gene12926   NW_011925919.1    43401 43900 500   -     3068  2466  602   80.38
gene3478    NW_011924987.1    1323515     1324014     500   -     1031  828   203   80.31
gene9622    NW_011925287.1    414625      415124      500   -     3131  2505  626   80.01
gene6765    NW_011925106.1    2070681     2071180     500   -     427   339   88    79.39
gene4521    NW_011925013.1    44114 44613 500   +     955   739   216   77.38
gene779     NW_011924904.1    1346112     1346611     500   -     267   205   62    76.78
gene617     NW_011924894.1    393331      393830      500   -     407   309   98    75.92
gene1675    NW_011924932.1    638769      639268      500   -     1285  973   312   75.72
gene5463    NW_011925035.1    1933485     1933984     500   -     1845  1394  451   75.56
...
... only first 25 lines shown ...
```

Directory: RNK File: ranked-promoters-byPrcntM-Cb_LIp_Rphase.txt

| region_ID | seqnames | start | end | width | strand | coverage | numCs | numTs | prcntM |
|---|---|---|---|---|---|---|---|---|---|
| gene8748 | NW_011925216.1 | 183604 | 184103 | 500 | - | 20 | 20 | 0 | 100 |
| gene13313 | NW_011928871.1 | 1 | 61 | 61 | + | 9 | 9 | 0 | 100 |
| gene4040 | NW_011925003.1 | 277741 | 278240 | 500 | + | 179 | 168 | 11 | 93.85 |
| gene4016 | NW_011925003.1 | 66624 | 67123 | 500 | + | 4066 | 3740 | 326 | 91.98 |
| gene3546 | NW_011924987.1 | 1804967 | 1805466 | 500 | - | 236 | 217 | 19 | 91.95 |
| gene5713 | NW_011925050.1 | 7072 | 7571 | 500 | - | 1837 | 1653 | 184 | 89.98 |
| gene11598 | NW_011925467.1 | 116917 | 117416 | 500 | + | 1436 | 1283 | 153 | 89.35 |
| gene5601 | NW_011925036.1 | 147325 | 147824 | 500 | - | 1830 | 1584 | 246 | 86.56 |
| gene2337 | NW_011924951.1 | 247713 | 248212 | 500 | + | 516 | 431 | 85 | 83.53 |
| gene13190 | NW_011926458.1 | 36701 | 37200 | 500 | - | 135 | 112 | 23 | 82.96 |
| gene12609 | NW_011925739.1 | 39939 | 40438 | 500 | + | 82 | 68 | 14 | 82.93 |
| gene3373 | NW_011924987.1 | 206378 | 206877 | 500 | - | 552 | 453 | 99 | 82.07 |
| gene617 | NW_011924894.1 | 393331 | 393830 | 500 | - | 509 | 416 | 93 | 81.73 |
| gene12926 | NW_011925919.1 | 43401 | 43900 | 500 | - | 3393 | 2739 | 654 | 80.73 |
| gene3478 | NW_011924987.1 | 1323515 | 1324014 | 500 | - | 1124 | 906 | 218 | 80.6 |
| gene8331 | NW_011925189.1 | 573460 | 573959 | 500 | + | 912 | 734 | 178 | 80.48 |
| gene9313 | NW_011925258.1 | 34883 | 35382 | 500 | + | 992 | 797 | 195 | 80.34 |
| gene8259 | NW_011925188.1 | 207457 | 207956 | 500 | - | 151 | 119 | 32 | 78.81 |
| gene6595 | NW_011925106.1 | 124931 | 125430 | 500 | + | 794 | 623 | 171 | 78.46 |
| gene772 | NW_011924904.1 | 1310349 | 1310848 | 500 | + | 3112 | 2436 | 676 | 78.28 |
| gene9622 | NW_011925287.1 | 414625 | 415124 | 500 | - | 3223 | 2484 | 739 | 77.07 |
| gene6765 | NW_011925106.1 | 2070681 | 2071180 | 500 | - | 391 | 299 | 92 | 76.47 |
| gene2532 | NW_011924955.1 | 1998697 | 1999196 | 500 | - | 913 | 697 | 216 | 76.34 |
| gene5463 | NW_011925035.1 | 1933485 | 1933984 | 500 | - | 2061 | 1569 | 492 | 76.13 |

...

... only first 25 lines shown ...

Directory: RNK File: ranked-promoters-bySiteDensity-Cb_LIp_BCphase.txt

| region_ID | rwidth | nbrsites | nbrper10kb | pmpersite | pmpernucl | pglink | seqnames | start | end | strand | coverage | numCs | numTs | prcntM |
|---|---|---|---|---|---|---|---|---|---|---|---|---|---|---|
| gene1399 | 500 | 61 | 1220 | 29.04 | 3.54 | https://www.ncbi.nlm.nih.gov/gene/?term=LOC105287518 | NW_011924931.1 | 442908 | 443407 | - | 8213 | 2166 | 6047 | 26.37 |
| gene12925 | 500 | 57 | 1140 | 36.64 | 4.18 | https://www.ncbi.nlm.nih.gov/gene/?term=LOC109611487 | NW_011925919.1 | 41730 | 42229 | - | 9987 | 2543 | 7444 | 25.46 |
| gene2406 | 500 | 55 | 1100 | 30.16 | 3.32 | https://www.ncbi.nlm.nih.gov/gene/?term=LOC105274855 | NW_011924953.1 | 426755 | 427254 | - | 7406 | 1672 | 5734 | 22.58 |
| gene7634 | 500 | 54 | 1080 | 60.76 | 6.56 | https://www.ncbi.nlm.nih.gov/gene/?term=LOC105280861 | NW_011925151.1 | 38041 | 38540 | - | 5864 | 2919 | 2945 | 49.78 |
| gene505 | 500 | 52 | 1040 | 45.52 | 4.73 | https://www.ncbi.nlm.nih.gov/gene/?term=LOC105277808 | NW_011924889.1 | 477908 | 478407 | + | 10835 | 3365 | 7470 | 31.06 |
| gene4515 | 500 | 50 | 1000 | 67.55 | 6.76 | https://www.ncbi.nlm.nih.gov/gene/?term=LOC109610839 | NW_011925013.1 | 19534 | 20033 | + | 5933 | 3852 | 2081 | 64.92 |
| gene10441 | 500 | 48 | 960 | 28.13 | 2.7 | https://www.ncbi.nlm.nih.gov/gene/?term=LOC105284094 | NW_011925353.1 | 595996 | 596495 | - | 8411 | 1692 | 6719 | 20.12 |
| gene10397 | 500 | 46 | 920 | 52.45 | 4.83 | https://www.ncbi.nlm.nih.gov/gene/?term=LOC105284046 | NW_011925353.1 | 2217 | 2716 | + | 4386 | 1625 | 2761 | 37.05 |
| gene8964 | 500 | 46 | 920 | 47.72 | 4.39 | https://www.ncbi.nlm.nih.gov/gene/?term=LOC105282404 | NW_011925232.1 | 562782 | 563281 | - | 6249 | 2622 | 3627 | 41.96 |
| gene12331 | 500 | 45 | 900 | 40.72 | 3.66 | https://www.ncbi.nlm.nih.gov/gene/?term=LOC105286234 | NW_011925664.1 | 139161 | 139660 | + | 7238 | 2224 | 5014 | 30.73 |
| gene12926 | 500 | 45 | 900 | 79.31 | 7.14 | https://www.ncbi.nlm.nih.gov/gene/?term=LOC105286944 | NW_011925919.1 | 43401 | 43900 | - | 3068 | 2466 | 602 | 80.38 |
| gene465 | 500 | 45 | 900 | 59.61 | 5.37 | https://www.ncbi.nlm.nih.gov/gene/?term=LOC105277304 | NW_011924889.1 | 171845 | 172344 | + | 5284 | 2592 | 2692 | 49.05 |
| gene10220 | 500 | 44 | 880 | 51.63 | 4.54 | https://www.ncbi.nlm.nih.gov/gene/?term=LOC105283833 | NW_011925327.1 | 293294 | 293793 | + | 5224 | 2062 | 3162 | 39.47 |
| gene128 | 500 | 43 | 860 | 52.1 | 4.48 | https://www.ncbi.nlm.nih.gov/gene/?term=LOC105286681 | NW_011924887.1 | 225822 | 226321 | + | 4381 | 1926 | 2455 | 43.96 |
| gene12924 | 500 | 43 | 860 | 33.71 | 2.9 | https://www.ncbi.nlm.nih.gov/gene/?term=LOC105286945 | NW_011925919.1 | 41354 | 41853 | - | 8228 | 1781 | 6447 | 21.65 |
| gene1179 | 500 | 42 | 840 | 43.21 | 3.63 | https://www.ncbi.nlm.nih.gov/gene/?term=LOC105285207 | NW_011924922.1 | 2259192 | 2259691 | - | 4895 | 1562 | 3333 | 31.91 |
| gene4634 | 500 | 41 | 820 | 34.32 | 2.81 | https://www.ncbi.nlm.nih.gov/gene/?term=LOC105277388 | NW_011925019.1 | 5897 | 6396 | - | 6769 | 1306 | 5463 | 19.29 |
| gene5753 | 500 | 41 | 820 | 74.03 | 6.07 | https://www.ncbi.nlm.nih.gov/gene/?term=LOC105278673 | NW_011925050.1 | 215094 | 215593 | + | 4322 | 2990 | 1332 | 69.18 |
| gene10085 | 500 | 40 | 800 | 48.01 | 3.84 | https://www.ncbi.nlm.nih.gov/gene/?term=LOC105283684 | NW_011925323.1 | 361110 | 361609 | + | 5077 | 2051 | 3026 | 40.4 |
| gene12539 | 500 | 40 | 800 | 52.8 | 4.22 | https://www.ncbi.nlm.nih.gov/gene/?term=LOC105286478 | NW_011925715.1 | 412371 | 412870 | + | 6419 | 2584 | 3835 | 40.26 |
| gene525 | 500 | 40 | 800 | 41.46 | 3.32 | https://www.ncbi.nlm.nih.gov/gene/?term=LOC105278116 | NW_011924889.1 | 635990 | 636489 | + | 5152 | 1612 | 3540 | 31.29 |
| gene3929 | 500 | 37 | 740 | 75.83 | 5.61 | https://www.ncbi.nlm.nih.gov/gene/?term=LOC105276585 | NW_011924999.1 | 140417 | 140916 | - | 2712 | 1941 | 771 | 71.57 |
| gene4878 | 500 | 37 | 740 | 69.54 | 5.15 | https://www.ncbi.nlm.nih.gov/gene/?term=LOC105277648 | NW_011925020.1 | 2601822 | 2602321 | - | 2082 | 1387 | 695 | 66.62 |
| gene772 | 500 | 37 | 740 | 77.82 | 5.76 | https://www.ncbi.nlm.nih.gov/gene/?term=LOC105280935 | NW_011924904.1 | 1310349 | 1310848 | + | 2791 | 2249 | 542 | 80.58 |

...
... only first 25 lines shown ...

Directory: RNK File: ranked-promoters-bySiteDensity-Cb_LIp_Rphase.txt

| region_ID | rwidth | nbrsites | nbrper10kb | pmpersite | pmpernucl | pglink | seqnames | start | end | strand | coverage | numCs | numTs | prcntM |
|---|---|---|---|---|---|---|---|---|---|---|---|---|---|---|
| gene12925 | 500 | 61 | 1220 | 37 | 4.51 | https://www.ncbi.nlm.nih.gov/gene/?term=LOC109611487 | NW_011925919.1 | 41730 | 42229 | - | 11803 | 3169 | 8634 | 26.85 |
| gene1399 | 500 | 58 | 1160 | 30.58 | 3.55 | https://www.ncbi.nlm.nih.gov/gene/?term=LOC105287518 | NW_011924931.1 | 442908 | 443407 | - | 8770 | 2414 | 6356 | 27.53 |
| gene505 | 500 | 58 | 1160 | 43.67 | 5.07 | https://www.ncbi.nlm.nih.gov/gene/?term=LOC105277808 | NW_011924889.1 | 477908 | 478407 | + | 12397 | 4022 | 8375 | 32.44 |
| gene4515 | 500 | 53 | 1060 | 63.95 | 6.78 | https://www.ncbi.nlm.nih.gov/gene/?term=LOC109610839 | NW_011925013.1 | 19534 | 20033 | + | 6724 | 4392 | 2332 | 65.32 |
| gene7634 | 500 | 53 | 1060 | 61.26 | 6.49 | https://www.ncbi.nlm.nih.gov/gene/?term=LOC105280861 | NW_011925151.1 | 38041 | 38540 | - | 6266 | 2951 | 3315 | 47.1 |
| gene2406 | 500 | 49 | 980 | 32.54 | 3.19 | https://www.ncbi.nlm.nih.gov/gene/?term=LOC105274855 | NW_011924953.1 | 426755 | 427254 | - | 8612 | 2056 | 6556 | 23.87 |
| gene8964 | 500 | 49 | 980 | 44.95 | 4.41 | https://www.ncbi.nlm.nih.gov/gene/?term=LOC105282404 | NW_011925232.1 | 562782 | 563281 | - | 6617 | 2648 | 3969 | 40.02 |
| gene10397 | 500 | 46 | 920 | 53.04 | 4.88 | https://www.ncbi.nlm.nih.gov/gene/?term=LOC105284046 | NW_011925353.1 | 2217 | 2716 | + | 4305 | 1577 | 2728 | 36.63 |
| gene128 | 500 | 45 | 900 | 50.41 | 4.54 | https://www.ncbi.nlm.nih.gov/gene/?term=LOC105286681 | NW_011924887.1 | 225822 | 226321 | + | 4774 | 2045 | 2729 | 42.84 |
| gene2830 | 500 | 45 | 900 | 27.68 | 2.49 | https://www.ncbi.nlm.nih.gov/gene/?term=LOC105275346 | NW_011924973.1 | 974217 | 974716 | + | 10399 | 1617 | 8782 | 15.55 |
| gene4634 | 500 | 45 | 900 | 29.77 | 2.68 | https://www.ncbi.nlm.nih.gov/gene/?term=LOC105277388 | NW_011925019.1 | 5897 | 6396 | - | 8073 | 1545 | 6528 | 19.14 |
| gene465 | 500 | 45 | 900 | 59.37 | 5.34 | https://www.ncbi.nlm.nih.gov/gene/?term=LOC105277304 | NW_011924889.1 | 171845 | 172344 | + | 6046 | 2990 | 3056 | 49.45 |
| gene10220 | 500 | 44 | 880 | 51.68 | 4.55 | https://www.ncbi.nlm.nih.gov/gene/?term=LOC105283833 | NW_011925327.1 | 293294 | 293793 | + | 5408 | 2269 | 3139 | 41.96 |
| gene12331 | 500 | 44 | 880 | 39.68 | 3.49 | https://www.ncbi.nlm.nih.gov/gene/?term=LOC105286234 | NW_011925664.1 | 139161 | 139660 | + | 7694 | 2396 | 5298 | 31.14 |
| gene10441 | 500 | 43 | 860 | 27.86 | 2.4 | https://www.ncbi.nlm.nih.gov/gene/?term=LOC105284094 | NW_011925353.1 | 595996 | 596495 | - | 9413 | 1849 | 7564 | 19.64 |
| gene12924 | 500 | 43 | 860 | 35.95 | 3.09 | https://www.ncbi.nlm.nih.gov/gene/?term=LOC105286945 | NW_011925919.1 | 41354 | 41853 | - | 9280 | 2140 | 7140 | 23.06 |
| gene12926 | 500 | 43 | 860 | 81.26 | 6.99 | https://www.ncbi.nlm.nih.gov/gene/?term=LOC105286944 | NW_011925919.1 | 43401 | 43900 | - | 3393 | 2739 | 654 | 80.73 |
| gene1179 | 500 | 42 | 840 | 44.35 | 3.73 | https://www.ncbi.nlm.nih.gov/gene/?term=LOC105285207 | NW_011924922.1 | 2259192 | 2259691 | - | 5520 | 1749 | 3771 | 31.68 |
| gene525 | 500 | 42 | 840 | 41.5 | 3.49 | https://www.ncbi.nlm.nih.gov/gene/?term=LOC105278116 | NW_011924889.1 | 635990 | 636489 | + | 5696 | 1852 | 3844 | 32.51 |
| gene5753 | 500 | 41 | 820 | 72.26 | 5.93 | https://www.ncbi.nlm.nih.gov/gene/?term=LOC105278673 | NW_011925050.1 | 215094 | 215593 | + | 4533 | 3062 | 1471 | 67.55 |
| gene12539 | 500 | 40 | 800 | 52.6 | 4.21 | https://www.ncbi.nlm.nih.gov/gene/?term=LOC105286478 | NW_011925715.1 | 412371 | 412870 | + | 6901 | 2693 | 4208 | 39.02 |
| gene10085 | 500 | 37 | 740 | 50.77 | 3.76 | https://www.ncbi.nlm.nih.gov/gene/?term=LOC105283684 | NW_011925323.1 | 361110 | 361609 | + | 5317 | 2138 | 3179 | 40.21 |
| gene3929 | 500 | 37 | 740 | 78.6 | 5.82 | https://www.ncbi.nlm.nih.gov/gene/?term=LOC105276585 | NW_011924999.1 | 140417 | 140916 | - | 3140 | 2378 | 762 | 75.73 |
| gene1010 | 500 | 36 | 720 | 63.13 | 4.55 | https://www.ncbi.nlm.nih.gov/gene/?term=LOC105283498 | NW_011924914.1 | 831409 | 831908 | - | 6643 | 3347 | 3296 | 50.38 |

...
... only first 25 lines shown ...

Directory: RNK File: sites-in-genes-Cb_LIp_BCphase.txt

| seqnames | start | end | width | strand | coverage | numCs | numTs | perc_meth | region_seqnames | region_start | region_end | region_width | region_strand | region_source | region_type | region_score | region_phase | region_ID | region_Dbxref | region_Name | region_gbkey | region_gene | region_gene_biotype | region_partial | region_start_range | region_end_range |
|---|---|---|---|---|---|---|---|---|---|---|---|---|---|---|---|---|---|---|---|---|---|---|---|---|---|---|
| NW_011924877.1 | 19784 | 19784 | 1 | + | 74 | 11 | 63 | 14.8648648648649 | NW_011924877.1 | 17682 | 20567 | 2886 | - | Gnomon | gene | NA | NA | gene0 | GeneID:105274440 | LOC105274440 | Gene | LOC105274440 | protein_coding | NA | character(0) | character(0) |
| NW_011924879.1 | 2162 | 2162 | 1 | + | 104 | 19 | 85 | 18.2692307692308 | NW_011924879.1 | 7 | 3079 | 3073 | + | Gnomon | gene | NA | NA | gene5 | GeneID:105275124 | LOC105275124 | Gene | LOC105275124 | protein_coding | NA | character(0) | character(0) |
| NW_011924880.1 | 61992 | 61992 | 1 | - | 91 | 13 | 78 | 14.2857142857143 | NW_011924880.1 | 60904 | 63136 | 2233 | - | Gnomon | gene | NA | NA | gene8 | GeneID:105278554 | LOC105278554 | Gene | LOC105278554 | protein_coding | NA | character(0) | character(0) |
| NW_011924881.1 | 39844 | 39844 | 1 | + | 19 | 6 | 13 | 31.5789473684211 | NW_011924881.1 | 39621 | 40962 | 1342 | - | Gnomon | gene | NA | NA | gene17 | GeneID:105286636 | LOC105286636 | Gene | LOC105286636 | protein_coding | NA | character(0) | character(0) |
| NW_011924881.1 | 39967 | 39967 | 1 | + | 48 | 38 | 10 | 79.1666666666667 | NW_011924881.1 | 39621 | 40962 | 1342 | - | Gnomon | gene | NA | NA | gene17 | GeneID:105286636 | LOC105286636 | Gene | LOC105286636 | protein_coding | NA | character(0) | character(0) |
| NW_011924881.1 | 39968 | 39968 | 1 | - | 10 | 7 | 3 | 70 | NW_011924881.1 | 39621 | 40962 | 1342 | - | Gnomon | gene | NA | NA | gene17 | GeneID:105286636 | LOC105286636 | Gene | LOC105286636 | protein_coding | NA | character(0) | character(0) |
| NW_011924881.1 | 39987 | 39987 | 1 | + | 58 | 30 | 28 | 51.7241379310345 | NW_011924881.1 | 39621 | 40962 | 1342 | - | Gnomon | gene | NA | NA | gene17 | GeneID:105286636 | LOC105286636 | Gene | LOC105286636 | protein_coding | NA | character(0) | character(0) |
| NW_011924881.1 | 39988 | 39988 | 1 | - | 9 | 5 | 4 | 55.5555555555556 | NW_011924881.1 | 39621 | 40962 | 1342 | - | Gnomon | gene | NA | NA | gene17 | GeneID:105286636 | LOC105286636 | Gene | LOC105286636 | protein_coding | NA | character(0) | character(0) |
| NW_011924881.1 | 40021 | 40021 | 1 | + | 70 | 9 | 61 | 12.8571428571429 | NW_011924881.1 | 39621 | 40962 | 1342 | - | Gnomon | gene | NA | NA | gene17 | GeneID:105286636 | LOC105286636 | Gene | LOC105286636 | protein_coding | NA | character(0) | character(0) |
| NW_011924881.1 | 40035 | 40035 | 1 | + | 82 | 15 | 67 | 18.2926829268293 | NW_011924881.1 | 39621 | 40962 | 1342 | - | Gnomon | gene | NA | NA | gene17 | GeneID:105286636 | LOC105286636 | Gene | LOC105286636 | protein_coding | NA | character(0) | character(0) |
| NW_011924881.1 | 40061 | 40061 | 1 | + | 49 | 34 | 15 | 69.3877551020408 | NW_011924881.1 | 39621 | 40962 | 1342 | - | Gnomon | gene | NA | NA | gene17 | GeneID:105286636 | LOC105286636 | Gene | LOC105286636 | protein_coding | NA | character(0) | character(0) |
| NW_011924881.1 | 40062 | 40062 | 1 | - | 7 | 7 | 0 | 100 | NW_011924881.1 | 39621 | 40962 | 1342 | - | Gnomon | gene | NA | NA | gene17 | GeneID:105286636 | LOC105286636 | Gene | LOC105286636 | protein_coding | NA | character(0) | character(0) |
| NW_011924881.1 | 40072 | 40072 | 1 | + | 45 | 42 | 3 | 93.3333333333333 | NW_011924881.1 | 39621 | 40962 | 1342 | - | Gnomon | gene | NA | NA | gene17 | GeneID:105286636 | LOC105286636 | Gene | LOC105286636 | protein_coding | NA | character(0) | character(0) |
| NW_011924881.1 | 40073 | 40073 | 1 | - | 6 | 5 | 1 | 83.3333333333333 | NW_011924881.1 | 39621 | 40962 | 1342 | - | Gnomon | gene | NA | NA | gene17 | GeneID:105286636 | LOC105286636 | Gene | LOC105286636 | protein_coding | NA | character(0) | character(0) |
| NW_011924881.1 | 40095 | 40095 | 1 | + | 60 | 30 | 30 | 50 | NW_011924881.1 | 39621 | 40962 | 1342 | - | Gnomon | gene | NA | NA | gene17 | GeneID:105286636 | LOC105286636 | Gene | LOC105286636 | protein_coding | NA | character(0) | character(0) |
| NW_011924881.1 | 40120 | 40120 | 1 | + | 58 | 44 | 14 | 75.8620689655172 | NW_011924881.1 | 39621 | 40962 | 1342 | - | Gnomon | gene | NA | NA | gene17 | GeneID:105286636 | LOC105286636 | Gene | LOC105286636 | protein_coding | NA | character(0) | character(0) |
| NW_011924881.1 | 40121 | 40121 | 1 | - | 14 | 9 | 5 | 64.2857142857143 | NW_011924881.1 | 39621 | 40962 | 1342 | - | Gnomon | gene | NA | NA | gene17 | GeneID:105286636 | LOC105286636 | Gene | LOC105286636 | protein_coding | NA | character(0) | character(0) |
| NW_011924881.1 | 40159 | 40159 | 1 | + | 48 | 27 | 21 | 56.25 | NW_011924881.1 | 39621 | 40962 | 1342 | - | Gnomon | gene | NA | NA | gene17 | GeneID:105286636 | LOC105286636 | Gene | LOC105286636 | protein_coding | NA | character(0) | character(0) |
| NW_011924881.1 | 40160 | 40160 | 1 | - | 8 | 5 | 3 | 62.5 | NW_011924881.1 | 39621 | 40962 | 1342 | - | Gnomon | gene | NA | NA | gene17 | GeneID:105286636 | LOC105286636 | Gene | LOC105286636 | protein_coding | NA | character(0) | character(0) |
| NW_011924881.1 | 40203 | 40203 | 1 | - | 12 | 6 | 6 | 50 | NW_011924881.1 | 39621 | 40962 | 1342 | - | Gnomon | gene | NA | NA | gene17 | GeneID:105286636 | LOC105286636 | Gene | LOC105286636 | protein_coding | NA | character(0) | character(0) |
| NW_011924881.1 | 40256 | 40256 | 1 | + | 9 | 8 | 1 | 88.8888888888889 | NW_011924881.1 | 39621 | 40962 | 1342 | - | Gnomon | gene | NA | NA | gene17 | GeneID:105286636 | LOC105286636 | Gene | LOC105286636 | protein_coding | NA | character(0) | character(0) |
| NW_011924881.1 | 40257 | 40257 | 1 | - | 92 | 76 | 16 | 82.6086956521739 | NW_011924881.1 | 39621 | 40962 | 1342 | - | Gnomon | gene | NA | NA | gene17 | GeneID:105286636 | LOC105286636 | Gene | LOC105286636 | protein_coding | NA | character(0) | character(0) |
| NW_011924881.1 | 40259 | 40259 | 1 | + | 11 | 11 | 0 | 100 | NW_011924881.1 | 39621 | 40962 | 1342 | - | Gnomon | gene | NA | NA | gene17 | GeneID:105286636 | LOC105286636 | Gene | LOC105286636 | protein_coding | NA | character(0) | character(0) |
| NW_011924881.1 | 40260 | 40260 | 1 | - | 89 | 80 | 9 | 89.8876404494382 | NW_011924881.1 | 39621 | 40962 | 1342 | - | Gnomon | gene | NA | NA | gene17 | GeneID:105286636 | LOC105286636 | Gene | LOC105286636 | protein_coding | NA | character(0) | character(0) |

...
... only first 25 lines shown ...

Directory: RNK File: sites-in-genes-Cb_LIp_Rphase.txt

| seqnames | start | end | width | strand | coverage | numCs | numTs | perc_meth | region_seqnames | region_start | region_end | region_width | region_strand | region_source | region_type | region_score | region_phase | region_ID | region_Dbxref | region_Name | region_gbkey | region_gene | region_gene_biotype | region_partial | region_start_range | region_end_range |
|---|---|---|---|---|---|---|---|---|---|---|---|---|---|---|---|---|---|---|---|---|---|---|---|---|---|---|
| NW_011924877.1 | 25848 | 25848 | 1 | - | 77 | 9 | 68 | 11.6883116883117 | NW_011924877.1 | 25368 | 26282 | 915 | - | Gnomon | gene | NA | NA | gene1 | GeneID:105281289 | LOC105281289 | Gene | LOC105281289 | protein_coding | NA | character(0) | character(0) |
| NW_011924879.1 | 2162 | 2162 | 1 | + | 118 | 28 | 90 | 23.728813559322 | NW_011924879.1 | 7 | 3079 | 3073 | + | Gnomon | gene | NA | NA | gene5 | GeneID:105275124 | LOC105275124 | Gene | LOC105275124 | protein_coding | NA | character(0) | character(0) |
| NW_011924880.1 | 69822 | 69822 | 1 | - | 34 | 10 | 24 | 29.4117647058824 | NW_011924880.1 | 69730 | 71017 | 1288 | - | Gnomon | gene | NA | NA | gene9 | GeneID:105283924 | LOC105283924 | Gene | LOC105283924 | protein_coding | NA | character(0) | character(0) |
| NW_011924881.1 | 39844 | 39844 | 1 | + | 11 | 6 | 5 | 54.5454545454545 | NW_011924881.1 | 39621 | 40962 | 1342 | - | Gnomon | gene | NA | NA | gene17 | GeneID:105286636 | LOC105286636 | Gene | LOC105286636 | protein_coding | NA | character(0) | character(0) |
| NW_011924881.1 | 39845 | 39845 | 1 | - | 35 | 11 | 24 | 31.4285714285714 | NW_011924881.1 | 39621 | 40962 | 1342 | - | Gnomon | gene | NA | NA | gene17 | GeneID:105286636 | LOC105286636 | Gene | LOC105286636 | protein_coding | NA | character(0) | character(0) |
| NW_011924881.1 | 39967 | 39967 | 1 | + | 40 | 28 | 12 | 70 | NW_011924881.1 | 39621 | 40962 | 1342 | - | Gnomon | gene | NA | NA | gene17 | GeneID:105286636 | LOC105286636 | Gene | LOC105286636 | protein_coding | NA | character(0) | character(0) |
| NW_011924881.1 | 39968 | 39968 | 1 | - | 14 | 8 | 6 | 57.1428571428571 | NW_011924881.1 | 39621 | 40962 | 1342 | - | Gnomon | gene | NA | NA | gene17 | GeneID:105286636 | LOC105286636 | Gene | LOC105286636 | protein_coding | NA | character(0) | character(0) |
| NW_011924881.1 | 39987 | 39987 | 1 | + | 55 | 31 | 24 | 56.3636363636364 | NW_011924881.1 | 39621 | 40962 | 1342 | - | Gnomon | gene | NA | NA | gene17 | GeneID:105286636 | LOC105286636 | Gene | LOC105286636 | protein_coding | NA | character(0) | character(0) |
| NW_011924881.1 | 39988 | 39988 | 1 | - | 11 | 8 | 3 | 72.7272727272727 | NW_011924881.1 | 39621 | 40962 | 1342 | - | Gnomon | gene | NA | NA | gene17 | GeneID:105286636 | LOC105286636 | Gene | LOC105286636 | protein_coding | NA | character(0) | character(0) |
| NW_011924881.1 | 40035 | 40035 | 1 | + | 78 | 13 | 65 | 16.6666666666667 | NW_011924881.1 | 39621 | 40962 | 1342 | - | Gnomon | gene | NA | NA | gene17 | GeneID:105286636 | LOC105286636 | Gene | LOC105286636 | protein_coding | NA | character(0) | character(0) |
| NW_011924881.1 | 40061 | 40061 | 1 | + | 49 | 45 | 4 | 91.8367346938776 | NW_011924881.1 | 39621 | 40962 | 1342 | - | Gnomon | gene | NA | NA | gene17 | GeneID:105286636 | LOC105286636 | Gene | LOC105286636 | protein_coding | NA | character(0) | character(0) |
| NW_011924881.1 | 40072 | 40072 | 1 | + | 51 | 50 | 1 | 98.0392156862745 | NW_011924881.1 | 39621 | 40962 | 1342 | - | Gnomon | gene | NA | NA | gene17 | GeneID:105286636 | LOC105286636 | Gene | LOC105286636 | protein_coding | NA | character(0) | character(0) |
| NW_011924881.1 | 40095 | 40095 | 1 | + | 71 | 39 | 32 | 54.9295774647887 | NW_011924881.1 | 39621 | 40962 | 1342 | - | Gnomon | gene | NA | NA | gene17 | GeneID:105286636 | LOC105286636 | Gene | LOC105286636 | protein_coding | NA | character(0) | character(0) |
| NW_011924881.1 | 40120 | 40120 | 1 | + | 75 | 46 | 29 | 61.3333333333333 | NW_011924881.1 | 39621 | 40962 | 1342 | - | Gnomon | gene | NA | NA | gene17 | GeneID:105286636 | LOC105286636 | Gene | LOC105286636 | protein_coding | NA | character(0) | character(0) |
| NW_011924881.1 | 40121 | 40121 | 1 | - | 24 | 18 | 6 | 75 | NW_011924881.1 | 39621 | 40962 | 1342 | - | Gnomon | gene | NA | NA | gene17 | GeneID:105286636 | LOC105286636 | Gene | LOC105286636 | protein_coding | NA | character(0) | character(0) |
| NW_011924881.1 | 40159 | 40159 | 1 | + | 47 | 29 | 18 | 61.7021276595745 | NW_011924881.1 | 39621 | 40962 | 1342 | - | Gnomon | gene | NA | NA | gene17 | GeneID:105286636 | LOC105286636 | Gene | LOC105286636 | protein_coding | NA | character(0) | character(0) |
| NW_011924881.1 | 40160 | 40160 | 1 | - | 24 | 17 | 7 | 70.8333333333333 | NW_011924881.1 | 39621 | 40962 | 1342 | - | Gnomon | gene | NA | NA | gene17 | GeneID:105286636 | LOC105286636 | Gene | LOC105286636 | protein_coding | NA | character(0) | character(0) |
| NW_011924881.1 | 40166 | 40166 | 1 | + | 48 | 8 | 40 | 16.6666666666667 | NW_011924881.1 | 39621 | 40962 | 1342 | - | Gnomon | gene | NA | NA | gene17 | GeneID:105286636 | LOC105286636 | Gene | LOC105286636 | protein_coding | NA | character(0) | character(0) |
| NW_011924881.1 | 40203 | 40203 | 1 | - | 15 | 10 | 5 | 66.6666666666667 | NW_011924881.1 | 39621 | 40962 | 1342 | - | Gnomon | gene | NA | NA | gene17 | GeneID:105286636 | LOC105286636 | Gene | LOC105286636 | protein_coding | NA | character(0) | character(0) |
| NW_011924881.1 | 40256 | 40256 | 1 | + | 13 | 12 | 1 | 92.3076923076923 | NW_011924881.1 | 39621 | 40962 | 1342 | - | Gnomon | gene | NA | NA | gene17 | GeneID:105286636 | LOC105286636 | Gene | LOC105286636 | protein_coding | NA | character(0) | character(0) |
| NW_011924881.1 | 40257 | 40257 | 1 | - | 86 | 74 | 12 | 86.046511627907 | NW_011924881.1 | 39621 | 40962 | 1342 | - | Gnomon | gene | NA | NA | gene17 | GeneID:105286636 | LOC105286636 | Gene | LOC105286636 | protein_coding | NA | character(0) | character(0) |
| NW_011924881.1 | 40259 | 40259 | 1 | + | 13 | 12 | 1 | 92.3076923076923 | NW_011924881.1 | 39621 | 40962 | 1342 | - | Gnomon | gene | NA | NA | gene17 | GeneID:105286636 | LOC105286636 | Gene | LOC105286636 | protein_coding | NA | character(0) | character(0) |
| NW_011924881.1 | 40260 | 40260 | 1 | - | 76 | 71 | 5 | 93.4210526315789 | NW_011924881.1 | 39621 | 40962 | 1342 | - | Gnomon | gene | NA | NA | gene17 | GeneID:105286636 | LOC105286636 | Gene | LOC105286636 | protein_coding | NA | character(0) | character(0) |
| NW_011924881.1 | 40270 | 40270 | 1 | + | 16 | 14 | 2 | 87.5 | NW_011924881.1 | 39621 | 40962 | 1342 | - | Gnomon | gene | NA | NA | gene17 | GeneID:105286636 | LOC105286636 | Gene | LOC105286636 | protein_coding | NA | character(0) | character(0) |

...
... only first 25 lines shown ...

Directory: RNK File: sites-in-promoters-Cb_LIp_BCphase.txt

| seqnames | start | end | width | strand | coverage | numCs | numTs | perc_meth | region_seqnames | region_start | region_end | region_width | region_strand | region_source | region_type | region_score | region_phase | region_ID | region_Dbxref | region_Name | region_gbkey | region_gene | region_gene_biotype | region_partial | region_start_range | region_end_range |
|---|---|---|---|---|---|---|---|---|---|---|---|---|---|---|---|---|---|---|---|---|---|---|---|---|---|---|
| NW_011924881.1 | 30121 | 30121 | 1 | + | 96 | 11 | 85 | 11.4583333333333 | NW_011924881.1 | 30079 | 30578 | 500 | - | Gnomon | promoter | NA | NA | gene16 | GeneID:105278149 | LOC105278149 | Gene | LOC105278149 | protein_coding | NA | character(0) | character(0) |
| NW_011924881.1 | 186433 | 186433 | 1 | + | 74 | 13 | 61 | 17.5675675675676 | NW_011924881.1 | 186350 | 186849 | 500 | - | Gnomon | promoter | NA | NA | gene21 | GeneID:105287749 | LOC105287749 | Gene | LOC105287749 | protein_coding | NA | character(0) | character(0) |
| NW_011924881.1 | 731753 | 731753 | 1 | - | 70 | 10 | 60 | 14.2857142857143 | NW_011924881.1 | 731568 | 732067 | 500 | - | Gnomon | promoter | NA | NA | gene34 | GeneID:105278416 | LOC105278416 | Gene | LOC105278416 | protein_coding | NA | character(0) | character(0) |
| NW_011924881.1 | 744660 | 744660 | 1 | - | 101 | 10 | 91 | 9.9009900990099 | NW_011924881.1 | 744598 | 745097 | 500 | - | Gnomon | promoter | NA | NA | gene40 | GeneID:105275462 | LOC105275462 | Gene | LOC105275462 | protein_coding | NA | character(0) | character(0) |
| NW_011924881.1 | 744706 | 744706 | 1 | - | 85 | 15 | 70 | 17.6470588235294 | NW_011924881.1 | 744598 | 745097 | 500 | - | Gnomon | promoter | NA | NA | gene40 | GeneID:105275462 | LOC105275462 | Gene | LOC105275462 | protein_coding | NA | character(0) | character(0) |
| NW_011924881.1 | 744708 | 744708 | 1 | - | 81 | 14 | 67 | 17.2839506172839 | NW_011924881.1 | 744598 | 745097 | 500 | - | Gnomon | promoter | NA | NA | gene40 | GeneID:105275462 | LOC105275462 | Gene | LOC105275462 | protein_coding | NA | character(0) | character(0) |
| NW_011924881.1 | 745918 | 745918 | 1 | + | 136 | 106 | 30 | 77.9411764705882 | NW_011924881.1 | 745918 | 746417 | 500 | + | Gnomon | promoter | NA | NA | gene42 | GeneID:105275579 | LOC105275579 | Gene | LOC105275579 | protein_coding | NA | character(0) | character(0) |
| NW_011924881.1 | 745919 | 745919 | 1 | - | 18 | 15 | 3 | 83.3333333333333 | NW_011924881.1 | 745918 | 746417 | 500 | + | Gnomon | promoter | NA | NA | gene42 | GeneID:105275579 | LOC105275579 | Gene | LOC105275579 | protein_coding | NA | character(0) | character(0) |
| NW_011924881.1 | 746036 | 746036 | 1 | - | 96 | 24 | 72 | 25 | NW_011924881.1 | 745918 | 746417 | 500 | + | Gnomon | promoter | NA | NA | gene42 | GeneID:105275579 | LOC105275579 | Gene | LOC105275579 | protein_coding | NA | character(0) | character(0) |
| NW_011924881.1 | 746261 | 746261 | 1 | - | 116 | 14 | 102 | 12.0689655172414 | NW_011924881.1 | 745918 | 746417 | 500 | + | Gnomon | promoter | NA | NA | gene42 | GeneID:105275579 | LOC105275579 | Gene | LOC105275579 | protein_coding | NA | character(0) | character(0) |
| NW_011924881.1 | 746617 | 746617 | 1 | - | 46 | 10 | 36 | 21.7391304347826 | NW_011924881.1 | 746408 | 746907 | 500 | - | Gnomon | promoter | NA | NA | gene41 | GeneID:105275643 | LOC105275643 | Gene | LOC105275643 | protein_coding | NA | character(0) | character(0) |
| NW_011924881.1 | 850306 | 850306 | 1 | - | 69 | 14 | 55 | 20.2898550724638 | NW_011924881.1 | 850004 | 850503 | 500 | - | Gnomon | promoter | NA | NA | gene57 | GeneID:109611563 | LOC109611563 | Gene | LOC109611563 | protein_coding | NA | character(0) | character(0) |
| NW_011924881.1 | 850348 | 850348 | 1 | - | 153 | 13 | 140 | 8.49673202614379 | NW_011924881.1 | 850004 | 850503 | 500 | - | Gnomon | promoter | NA | NA | gene57 | GeneID:109611563 | LOC109611563 | Gene | LOC109611563 | protein_coding | NA | character(0) | character(0) |
| NW_011924881.1 | 989261 | 989261 | 1 | + | 9 | 6 | 3 | 66.6666666666667 | NW_011924881.1 | 989253 | 989752 | 500 | + | Gnomon | promoter | NA | NA | gene61 | GeneID:105277580 | LOC105277580 | Gene | LOC105277580 | protein_coding | NA | character(0) | character(0) |
| NW_011924881.1 | 989262 | 989262 | 1 | - | 16 | 15 | 1 | 93.75 | NW_011924881.1 | 989253 | 989752 | 500 | + | Gnomon | promoter | NA | NA | gene61 | GeneID:105277580 | LOC105277580 | Gene | LOC105277580 | protein_coding | NA | character(0) | character(0) |
| NW_011924881.1 | 1012572 | 1012572 | 1 | + | 92 | 17 | 75 | 18.4782608695652 | NW_011924881.1 | 1012159 | 1012658 | 500 | + | Gnomon | promoter | NA | NA | gene65 | GeneID:105277639 | LOC105277639 | Gene | LOC105277639 | protein_coding | NA | character(0) | character(0) |
| NW_011924887.1 | 135338 | 135338 | 1 | + | 40 | 11 | 29 | 27.5 | NW_011924887.1 | 134949 | 135448 | 500 | - | Gnomon | promoter | NA | NA | gene114 | GeneID:105285264 | LOC105285264 | Gene | LOC105285264 | protein_coding | NA | character(0) | character(0) |
| NW_011924887.1 | 221081 | 221081 | 1 | + | 37 | 11 | 26 | 29.7297297297297 | NW_011924887.1 | 220963 | 221462 | 500 | + | Gnomon | promoter | NA | NA | gene126 | GeneID:105286831 | LOC105286831 | Gene | LOC105286831 | protein_coding | NA | character(0) | character(0) |
| NW_011924887.1 | 221082 | 221082 | 1 | - | 91 | 29 | 62 | 31.8681318681319 | NW_011924887.1 | 220963 | 221462 | 500 | + | Gnomon | promoter | NA | NA | gene126 | GeneID:105286831 | LOC105286831 | Gene | LOC105286831 | protein_coding | NA | character(0) | character(0) |
| NW_011924887.1 | 221097 | 221097 | 1 | + | 23 | 17 | 6 | 73.9130434782609 | NW_011924887.1 | 220963 | 221462 | 500 | + | Gnomon | promoter | NA | NA | gene126 | GeneID:105286831 | LOC105286831 | Gene | LOC105286831 | protein_coding | NA | character(0) | character(0) |
| NW_011924887.1 | 221098 | 221098 | 1 | - | 119 | 71 | 48 | 59.6638655462185 | NW_011924887.1 | 220963 | 221462 | 500 | + | Gnomon | promoter | NA | NA | gene126 | GeneID:105286831 | LOC105286831 | Gene | LOC105286831 | protein_coding | NA | character(0) | character(0) |
| NW_011924887.1 | 221105 | 221105 | 1 | - | 118 | 24 | 94 | 20.3389830508475 | NW_011924887.1 | 220963 | 221462 | 500 | + | Gnomon | promoter | NA | NA | gene126 | GeneID:105286831 | LOC105286831 | Gene | LOC105286831 | protein_coding | NA | character(0) | character(0) |
| NW_011924887.1 | 221135 | 221135 | 1 | - | 97 | 13 | 84 | 13.4020618556701 | NW_011924887.1 | 220963 | 221462 | 500 | + | Gnomon | promoter | NA | NA | gene126 | GeneID:105286831 | LOC105286831 | Gene | LOC105286831 | protein_coding | NA | character(0) | character(0) |
| NW_011924887.1 | 221214 | 221214 | 1 | - | 54 | 13 | 41 | 24.0740740740741 | NW_011924887.1 | 220963 | 221462 | 500 | + | Gnomon | promoter | NA | NA | gene126 | GeneID:105286831 | LOC105286831 | Gene | LOC105286831 | protein_coding | NA | character(0) | character(0) |

...
... only first 25 lines shown ...

Directory: RNK File: sites-in-promoters-Cb_LIp_Rphase.txt

```
seqnames    start end   width strand      coverage    numCs numTs perc_meth   region_seqnames   region_start      region_end  region_width      region_strand     region_source
      region_type region_score      region_phase      region_ID   region_Dbxref     region_Name region_gbkey      region_gene region_gene_biotype     region_partial
      region_start_range      region_end_range
NW_011924881.1    186433      186433      1     +     79    11    68    13.9240506329114  NW_011924881.1    186350      186849      500   -     Gnomon      promoter    NA    NA
      gene21      GeneID:105287749  LOC105287749      Gene  LOC105287749      protein_coding    NA    character(0)      character(0)
NW_011924881.1    446700      446700      1     +     48    10    38    20.8333333333333  NW_011924881.1    446689      447188      500   +     Gnomon      promoter    NA    NA
      gene25      GeneID:105274524  LOC105274524      Gene  LOC105274524      protein_coding    NA    character(0)      character(0)
NW_011924881.1    731753      731753      1     -     63    10    53    15.8730158730159  NW_011924881.1    731568      732067      500   -     Gnomon      promoter    NA    NA
      gene34      GeneID:105278416  LOC105278416      Gene  LOC105278416      protein_coding    NA    character(0)      character(0)
NW_011924881.1    744706      744706      1     -     72    13    59    18.0555555555556  NW_011924881.1    744598      745097      500   -     Gnomon      promoter    NA    NA
      gene40      GeneID:105275462  LOC105275462      Gene  LOC105275462      protein_coding    NA    character(0)      character(0)
NW_011924881.1    744707      744707      1     +     44    13    31    29.5454545454545  NW_011924881.1    744598      745097      500   -     Gnomon      promoter    NA    NA
      gene40      GeneID:105275462  LOC105275462      Gene  LOC105275462      protein_coding    NA    character(0)      character(0)
NW_011924881.1    744708      744708      1     -     71    12    59    16.9014084507042  NW_011924881.1    744598      745097      500   -     Gnomon      promoter    NA    NA
      gene40      GeneID:105275462  LOC105275462      Gene  LOC105275462      protein_coding    NA    character(0)      character(0)
NW_011924881.1    745031      745031      1     -     6     5     1     83.3333333333333  NW_011924881.1    744598      745097      500   -     Gnomon      promoter    NA    NA
      gene40      GeneID:105275462  LOC105275462      Gene  LOC105275462      protein_coding    NA    character(0)      character(0)
NW_011924881.1    745918      745918      1     +     154   122   32    79.2207792207792  NW_011924881.1    745918      746417      500   +     Gnomon      promoter    NA    NA
      gene42      GeneID:105275579  LOC105275579      Gene  LOC105275579      protein_coding    NA    character(0)      character(0)
NW_011924881.1    745919      745919      1     -     14    11    3     78.5714285714286  NW_011924881.1    745918      746417      500   +     Gnomon      promoter    NA    NA
      gene42      GeneID:105275579  LOC105275579      Gene  LOC105275579      protein_coding    NA    character(0)      character(0)
NW_011924881.1    746035      746035      1     +     24    7     17    29.1666666666667  NW_011924881.1    745918      746417      500   +     Gnomon      promoter    NA    NA
      gene42      GeneID:105275579  LOC105275579      Gene  LOC105275579      protein_coding    NA    character(0)      character(0)
NW_011924881.1    746036      746036      1     -     106   31    75    29.2452830188679  NW_011924881.1    745918      746417      500   +     Gnomon      promoter    NA    NA
      gene42      GeneID:105275579  LOC105275579      Gene  LOC105275579      protein_coding    NA    character(0)      character(0)
NW_011924881.1    746261      746261      1     -     154   24    130   15.5844155844156  NW_011924881.1    745918      746417      500   +     Gnomon      promoter    NA    NA
      gene42      GeneID:105275579  LOC105275579      Gene  LOC105275579      protein_coding    NA    character(0)      character(0)
NW_011924881.1    807838      807838      1     +     53    9     44    16.9811320754717  NW_011924881.1    807523      808022      500   -     Gnomon      promoter    NA    NA
      gene52      GeneID:105276184  LOC105276184      Gene  LOC105276184      protein_coding    NA    character(0)      character(0)
NW_011924881.1    850306      850306      1     -     47    13    34    27.6595744680851  NW_011924881.1    850004      850503      500   -     Gnomon      promoter    NA    NA
      gene57      GeneID:109611563  LOC109611563      Gene  LOC109611563      protein_coding    NA    character(0)      character(0)
NW_011924881.1    989261      989261      1     +     7     7     0     100   NW_011924881.1    989253      989752      500   +     Gnomon      promoter    NA    NA    gene61
      GeneID:105277580  LOC105277580      Gene  LOC105277580      protein_coding    NA    character(0)      character(0)
NW_011924881.1    989262      989262      1     -     24    22    2     91.6666666666667  NW_011924881.1    989253      989752      500   +     Gnomon      promoter    NA    NA
      gene61      GeneID:105277580  LOC105277580      Gene  LOC105277580      protein_coding    NA    character(0)      character(0)
NW_011924881.1    1012572     1012572     1     +     134   19    115   14.1791044776119  NW_011924881.1    1012159     1012658     500   +     Gnomon      promoter    NA    NA
      gene65      GeneID:105277639  LOC105277639      Gene  LOC105277639      protein_coding    NA    character(0)      character(0)
NW_011924885.1    58568 58568 1     +     38    8     30    21.0526315789474  NW_011924885.1    58397 58896 500   -     Gnomon      promoter    NA    NA    gene97
      GeneID:105282700  LOC105282700      Gene  LOC105282700      lncRNA      NA    character(0)      character(0)
NW_011924887.1    25536 25536 1     +     46    8     38    17.3913043478261  NW_011924887.1    25400 25899 500   +     Gnomon      promoter    NA    NA    gene106
      GeneID:105284438  LOC105284438      Gene  LOC105284438      protein_coding    NA    character(0)      character(0)
NW_011924887.1    135338      135338      1     +     55    12    43    21.8181818181818  NW_011924887.1    134949      135448      500   -     Gnomon      promoter    NA    NA
      gene114     GeneID:105285264  LOC105285264      Gene  LOC105285264      protein_coding    NA    character(0)      character(0)
NW_011924887.1    221081      221081      1     +     51    10    41    19.6078431372549  NW_011924887.1    220963      221462      500   +     Gnomon      promoter    NA    NA
      gene126     GeneID:105286831  LOC105286831      Gene  LOC105286831      protein_coding    NA    character(0)      character(0)
NW_011924887.1    221082      221082      1     -     92    23    69    25    NW_011924887.1    220963      221462      500   +     Gnomon      promoter    NA    NA    gene126
      GeneID:105286831  LOC105286831      Gene  LOC105286831      protein_coding    NA    character(0)      character(0)
NW_011924887.1    221097      221097      1     +     30    23    7     76.6666666666667  NW_011924887.1    220963      221462      500   +     Gnomon      promoter    NA    NA
      gene126     GeneID:105286831  LOC105286831      Gene  LOC105286831      protein_coding    NA    character(0)      character(0)
NW_011924887.1    221098      221098      1     -     125   66    59    52.8  NW_011924887.1    220963      221462      500   +     Gnomon      promoter    NA    NA    gene126
      GeneID:105286831  LOC105286831      Gene  LOC105286831      protein_coding    NA    character(0)      character(0)
...
... only first 25 lines shown ...
```

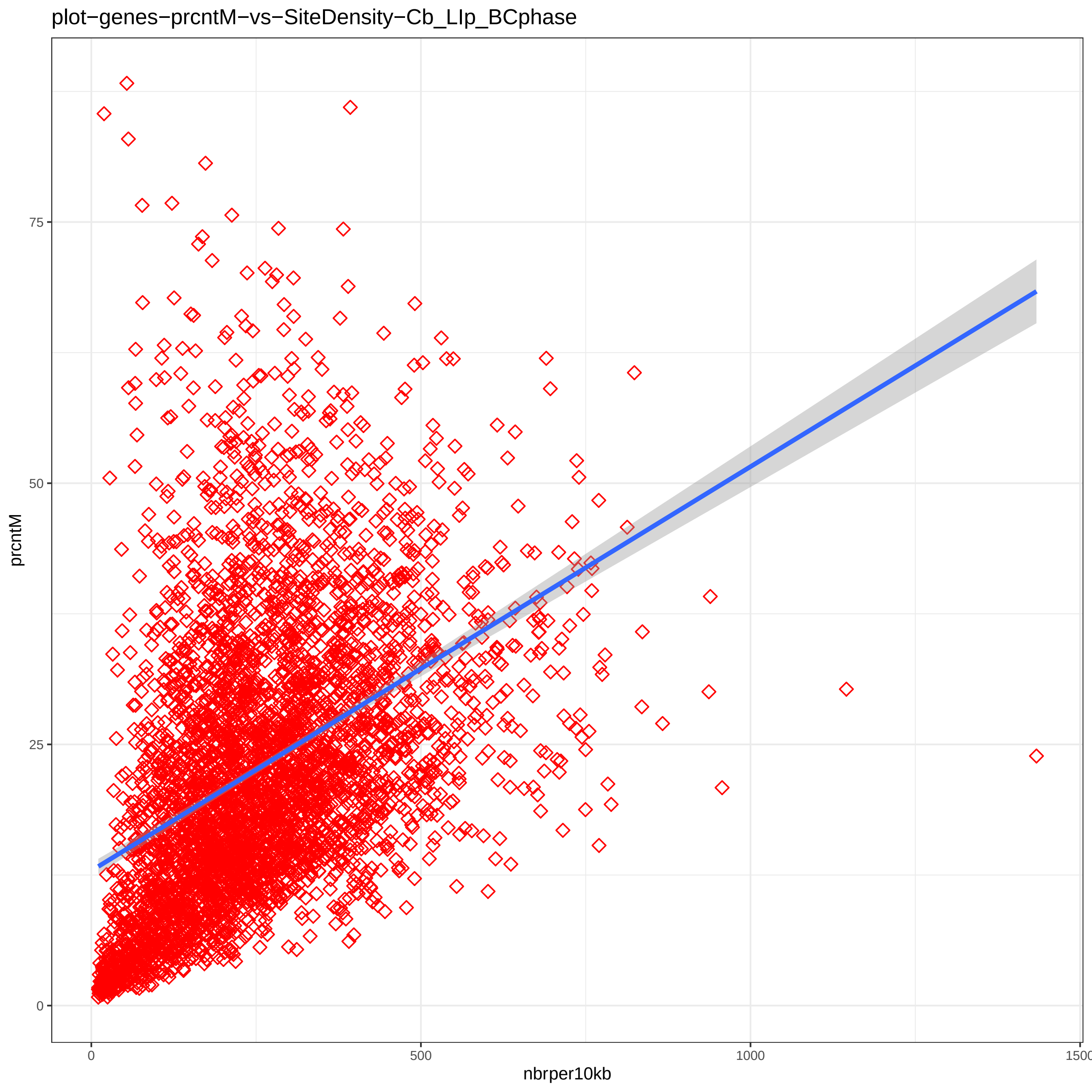
plot−genes−prcntM−vs−SiteDensity−Cb_LIp_BCphase
prcntM
nbrper10kb
0
25
50
75
0
500
1000
1500

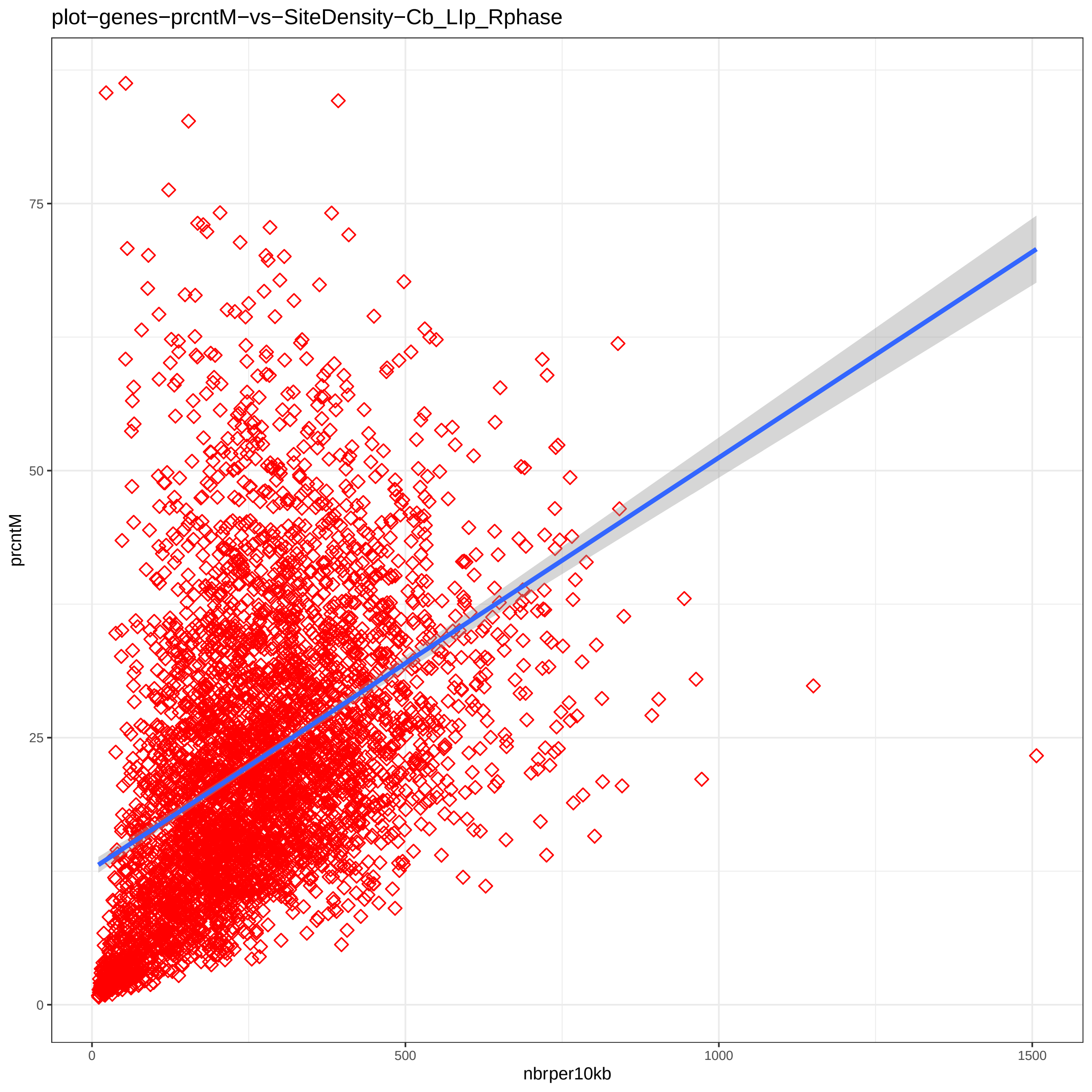

plot−genes−prcntM−vs−SiteDensity−Cb_LIp_Rphase
prcntM
75
50
25
0
0
500
1000
1500
nbrper10kb

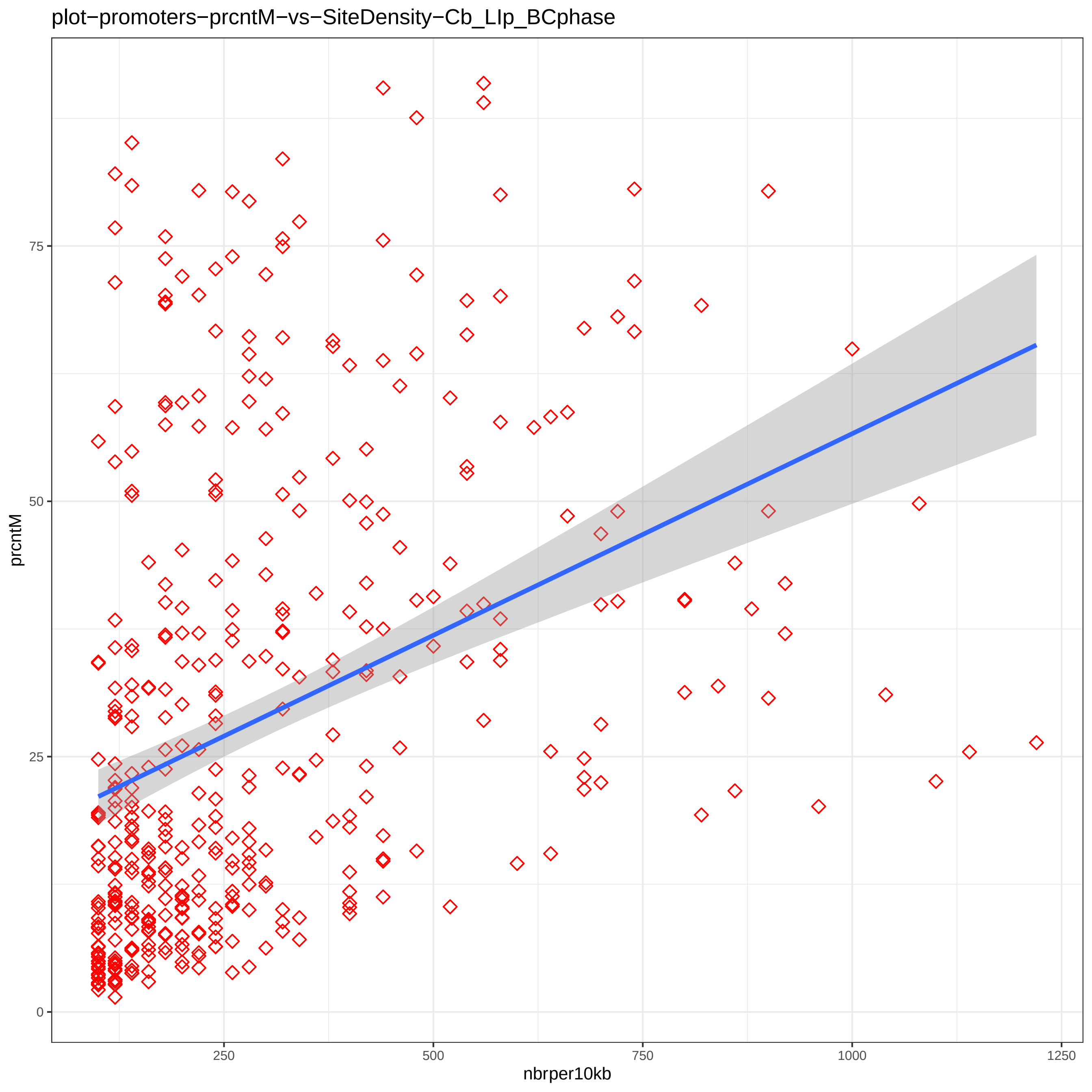
plot−promoters−prcntM−vs−SiteDensity−Cb_LIp_BCphase
prcntM
nbrper10kb
0
25
50
75
250
500
750
1000
1250

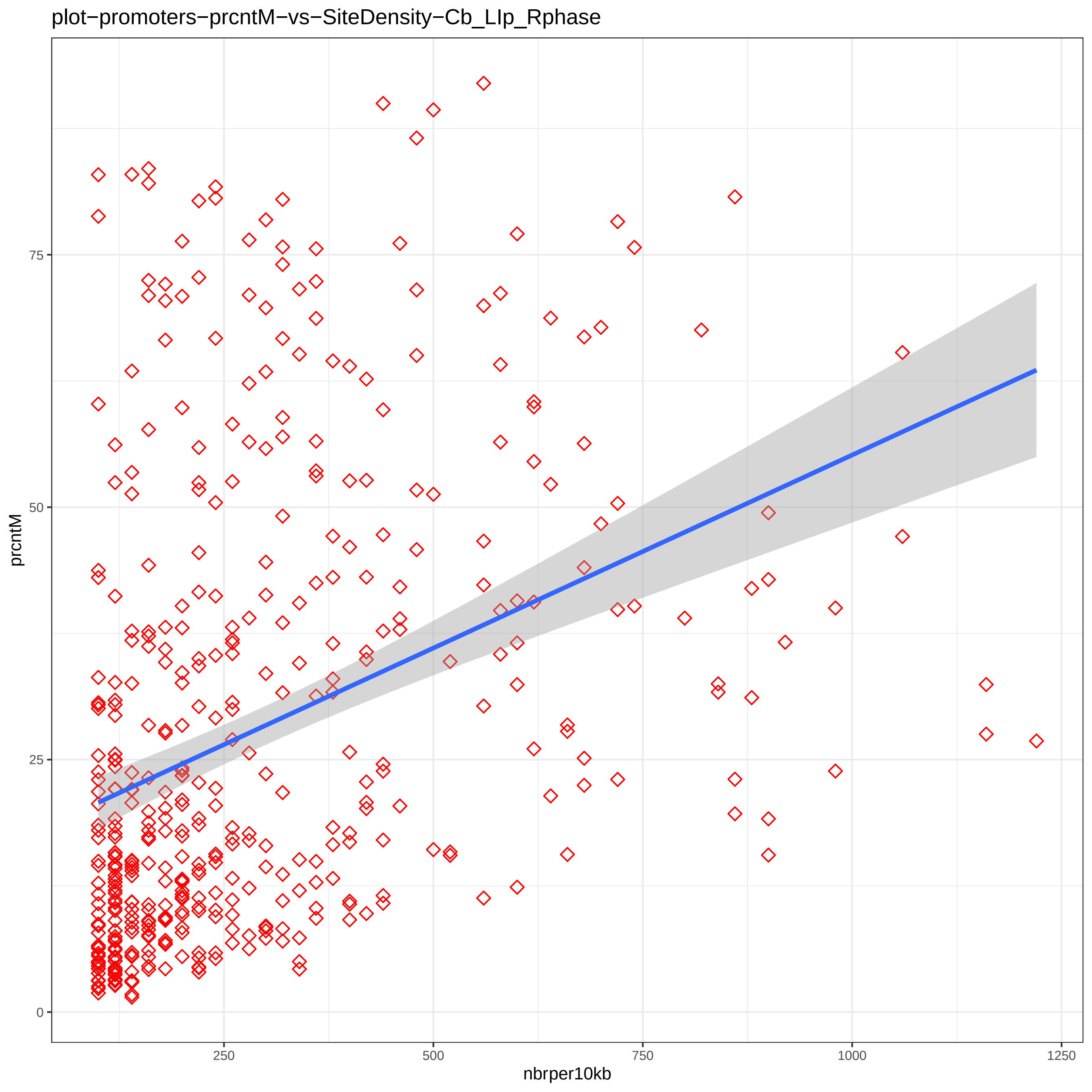

plot−promoters−prcntM−vs−SiteDensity−Cb_LIp_Rphase
prcntM
75
50
25
0
250
500
750
1000
1250
nbrper10kb

Directory: MRPR File: 0READMEmrpr

```
MRPR - Methylation-rich and -poor regions.

Input: studymk, genome_ann (a list of GRanges objects providing annotated region labels and bounds;
                             output of BWASPR::get_genome_annotation())
       nbrxtrms (specifies the number of richest and poorest regions to be displayed; default: 100)

Output: files dst-*.txt 1ds-*.pdf 5ds-*.pdf    mdr-*.bed   rmp-* gwp-* gwr-*

Notes: The output is generated by BWASPR functions det_mrpr() and map_mrpr().
       Methylation-rich and -poor regions are determined based on the spacing between
        neighboring CpGhsm sites.  If sites occur at positions a, b, c, d, e, and f
        (and not in between), then b-a, c-b, ... are 1-distances, and f-a is a 5-distance.
        The file dst-*.txt shows the empirical distribution of d-distances for the sample.
        Methylation-poor regions are determined as long 1-distances (in the top <nbrxtrms>; merged
        if adjacent 1-distances are both in the top <nbrxtrms>).  Methylation-rich regions are determined
        as short 5-distances (in the low <nbrxtrms>; merged if adjacent 5-distances are both in the low
        <nbrxtrms>).  Parts of the distribution are plotted in the 1ds-*.pdf and 5ds-*.pdf files.

       The methylation-rich and -poor regions are listed in files dst-*.txt and mdr-*.bed,
        the latter in BED format for potential display in genome browsers.

       The overlap of methylation regions with genome features is summarized in files rmp-*.txt.
        Files gwr-*.txt show genes overlapping with methylation-rich regions, ordered by site
        density in the methylation-rich region.
        Files gwp-*.txt show genes overlapping with methylation-poor regions, ordered by site
        density in the methylation-poor region.
```

```
Directory: MRPR File: dst-Cb_LI_BCphase.txt

Analysis of  1-distances for sample "Cb_LI_BCphase" (417310 sites):

  Total number of  1-distances:     416277
  Median:     9.0 Mean:   474.9 Std:  2245.5
  Quantiles:
   0%    5%   10%   15%   20%   25%   30%   35%   40%   45%   50%   55%   60%   65%   70%   75%   80%   85%   90%   95%  100%
    1     1     1     1     1     1     1     1     4     6     9    13    17    22    29    39    56    99   269  2399 82283

  Low  density region distance cutoff (lowest  100):   36645
  High density region distance cutoff (highest 100):       1

Ordered lists of methylation-poor regions:

 Top 100 (out of    99) methylation-poor regions in "Cb_LI_BCphase":
 (Sdnsty = sites per 1kb)

   Rtype  Sample          SeqID    From      To  Rlgth NbrSites Sdnsty
1   Poor BCphase NW_011925012.1 1218182 1300465  82284        2 0.0243
2   Poor BCphase NW_011924931.1 4498339 4575396  77058        2 0.0260
3   Poor BCphase NW_011925138.1   22282  128133 105852        3 0.0283
4   Poor BCphase NW_011925029.1  114432  184163  69732        2 0.0287
5   Poor BCphase NW_011925434.1  215433  285112  69680        2 0.0287
6   Poor BCphase NW_011924888.1 1808554 1878222  69669        2 0.0287
...
... only first 25 lines shown ...
```

Directory: MRPR File: dst-Cb_LI_Rphase.txt

Analysis of 1-distances for sample "Cb_LI_Rphase" (445317 sites):

Total number of 1-distances: 444180
Median: 10.0 Mean: 448.3 Std: 1906.9
Quantiles:

| 0% | 5% | 10% | 15% | 20% | 25% | 30% | 35% | 40% | 45% | 50% | 55% | 60% | 65% | 70% | 75% | 80% | 85% | 90% | 95% | 100% |
|---|---|---|---|---|---|---|---|---|---|---|---|---|---|---|---|---|---|---|---|---|
| 1 | 1 | 1 | 1 | 1 | 1 | 1 | 1 | 4 | 7 | 10 | 13 | 18 | 23 | 31 | 42 | 65 | 126 | 481 | 2669 | 57981 |

Low density region distance cutoff (lowest 100): 30653
High density region distance cutoff (highest 100): 1

Ordered lists of methylation-poor regions:

Top 100 (out of 99) methylation-poor regions in "Cb_LI_Rphase":
(Sdnsty = sites per 1kb)

| | Rtype | Sample | SeqID | From | To | Rlgth | NbrSites | Sdnsty |
|---|---|---|---|---|---|---|---|---|
| 1 | Poor | Rphase | NW_011925147.1 | 9998 | 67979 | 57982 | 2 | 0.0345 |
| 2 | Poor | Rphase | NW_011925234.1 | 35111 | 87222 | 52112 | 2 | 0.0384 |
| 3 | Poor | Rphase | NW_011924931.1 | 4497412 | 4548764 | 51353 | 2 | 0.0389 |
| 4 | Poor | Rphase | NW_011925300.1 | 473373 | 523294 | 49922 | 2 | 0.0401 |
| 5 | Poor | Rphase | NW_011925192.1 | 432387 | 482001 | 49615 | 2 | 0.0403 |
| 6 | Poor | Rphase | NW_011925431.1 | 33704 | 82028 | 48325 | 2 | 0.0414 |

...
... only first 25 lines shown ...

Directory: MRPR File: gwp-Cb_LI_BCphase.txt

| seqnames | start | end | width | strand | source | type | score | phase | ID | Dbxref | Name | gbkey | gene | gene_biotype | partial | start_range | end_range | seqnames | start | end | width | strand | NbrSites | Sdnsty |
|---|---|---|---|---|---|---|---|---|---|---|---|---|---|---|---|---|---|---|---|---|---|---|---|---|
| NW_011925012.1 | 1201498 | 1303795 | 102298 | + | Gnomon | gene | NA | NA | gene4489 | GeneID:105277228 | LOC105277228 | Gene | LOC105277228 | protein_coding | NA | character(0) | character(0) | NW_011925012.1 | 1218182 | 1300465 | 82284 | * | 2 | 0.0243060619318458 |
| NW_011924931.1 | 4493233 | 4499991 | 6759 | - | Gnomon | gene | NA | NA | gene1621 | GeneID:105287755 | LOC105287755 | Gene | LOC105287755 | protein_coding | NA | character(0) | character(0) | NW_011924931.1 | 4498339 | 4575396 | 77058 | * | 2 | 0.0259544758493602 |
| NW_011924931.1 | 4500817 | 4527583 | 26767 | + | Gnomon | gene | NA | NA | gene1622 | GeneID:105287758 | LOC105287758 | Gene | LOC105287758 | protein_coding | NA | character(0) | character(0) | NW_011924931.1 | 4498339 | 4575396 | 77058 | * | 2 | 0.0259544758493602 |
| NW_011924931.1 | 4519537 | 4523946 | 4410 | + | Gnomon | gene | NA | NA | gene1623 | GeneID:105287786 | LOC105287786 | Gene | LOC105287786 | protein_coding | NA | character(0) | character(0) | NW_011924931.1 | 4498339 | 4575396 | 77058 | * | 2 | 0.0259544758493602 |
| NW_011924931.1 | 4521493 | 4522142 | 650 | - | Gnomon | gene | NA | NA | gene1624 | GeneID:105287757 | LOC105287757 | Gene | LOC105287757 | protein_coding | NA | character(0) | character(0) | NW_011924931.1 | 4498339 | 4575396 | 77058 | * | 2 | 0.0259544758493602 |
| NW_011924931.1 | 4528303 | 4533712 | 5410 | + | Gnomon | gene | NA | NA | gene1625 | GeneID:105287787 | LOC105287787 | Gene | LOC105287787 | protein_coding | NA | character(0) | character(0) | NW_011924931.1 | 4498339 | 4575396 | 77058 | * | 2 | 0.0259544758493602 |
| NW_011924931.1 | 4533855 | 4534825 | 971 | + | Gnomon | gene | NA | NA | gene1626 | GeneID:105287760 | LOC105287760 | Gene | LOC105287760 | protein_coding | NA | character(0) | character(0) | NW_011924931.1 | 4498339 | 4575396 | 77058 | * | 2 | 0.0259544758493602 |
| NW_011924931.1 | 4535643 | 4542185 | 6543 | + | Gnomon | gene | NA | NA | gene1627 | GeneID:105287761 | LOC105287761 | Gene | LOC105287761 | protein_coding | NA | character(0) | character(0) | NW_011924931.1 | 4498339 | 4575396 | 77058 | * | 2 | 0.0259544758493602 |
| NW_011924931.1 | 4543478 | 4549354 | 5877 | + | Gnomon | gene | NA | NA | gene1628 | GeneID:105287788 | LOC105287788 | Gene | LOC105287788 | protein_coding | NA | character(0) | character(0) | NW_011924931.1 | 4498339 | 4575396 | 77058 | * | 2 | 0.0259544758493602 |
| NW_011924931.1 | 4550567 | 4556405 | 5839 | + | Gnomon | gene | NA | NA | gene1629 | GeneID:105287789 | LOC105287789 | Gene | LOC105287789 | protein_coding | NA | character(0) | character(0) | NW_011924931.1 | 4498339 | 4575396 | 77058 | * | 2 | 0.0259544758493602 |
| NW_011924931.1 | 4556498 | 4557472 | 975 | + | Gnomon | gene | NA | NA | gene1630 | GeneID:105287762 | LOC105287762 | Gene | LOC105287762 | protein_coding | NA | character(0) | character(0) | NW_011924931.1 | 4498339 | 4575396 | 77058 | * | 2 | 0.0259544758493602 |
| NW_011924931.1 | 4557517 | 4564469 | 6953 | + | Gnomon | gene | NA | NA | gene1631 | GeneID:105287763 | LOC105287763 | Gene | LOC105287763 | protein_coding | NA | character(0) | character(0) | NW_011924931.1 | 4498339 | 4575396 | 77058 | * | 2 | 0.0259544758493602 |
| NW_011924931.1 | 4565798 | 4583858 | 18061 | + | Gnomon | gene | NA | NA | gene1632 | GeneID:105287764 | LOC105287764 | Gene | LOC105287764 | protein_coding | NA | character(0) | character(0) | NW_011924931.1 | 4498339 | 4575396 | 77058 | * | 2 | 0.0259544758493602 |
| NW_011925138.1 | 39216 | 42889 | 3674 | + | Gnomon | gene | NA | NA | gene7431 | GeneID:105280632 | LOC105280632 | Gene | LOC105280632 | protein_coding | NA | character(0) | character(0) | NW_011925138.1 | 22282 | 128133 | 105852 | * | 3 | 0.0283414578845936 |
| NW_011925138.1 | 44652 | 47912 | 3261 | - | Gnomon | gene | NA | NA | gene7432 | GeneID:105280633 | LOC105280633 | Gene | LOC105280633 | protein_coding | NA | character(0) | character(0) | NW_011925138.1 | 22282 | 128133 | 105852 | * | 3 | 0.0283414578845936 |
| NW_011925138.1 | 66228 | 97549 | 31322 | - | Gnomon | gene | NA | NA | gene7433 | GeneID:105280634 | LOC105280634 | Gene | LOC105280634 | protein_coding | NA | character(0) | character(0) | NW_011925138.1 | 22282 | 128133 | 105852 | * | 3 | 0.0283414578845936 |
| NW_011925029.1 | 125886 | 127524 | 1639 | - | Gnomon | gene | NA | NA | gene5254 | GeneID:105278091 | LOC105278091 | Gene | LOC105278091 | lncRNA | NA | character(0) | character(0) | NW_011925029.1 | 114432 | 184163 | 69732 | * | 2 | 0.028681236734928 |
| NW_011925029.1 | 179417 | 181021 | 1605 | + | Gnomon | gene | NA | NA | gene5255 | GeneID:105278090 | LOC105278090 | Gene | LOC105278090 | protein_coding | NA | character(0) | character(0) | NW_011925029.1 | 114432 | 184163 | 69732 | * | 2 | 0.028681236734928 |
| NW_011925029.1 | 181274 | 183011 | 1738 | + | Gnomon | gene | NA | NA | gene5256 | GeneID:105278135 | LOC105278135 | Gene | LOC105278135 | protein_coding | NA | character(0) | character(0) | NW_011925029.1 | 114432 | 184163 | 69732 | * | 2 | 0.028681236734928 |
| NW_011925434.1 | 134845 | 291030 | 156186 | + | Gnomon | gene | NA | NA | gene11345 | GeneID:105285138 | LOC105285138 | Gene | LOC105285138 | protein_coding | NA | character(0) | character(0) | NW_011925434.1 | 215433 | 285112 | 69680 | * | 2 | 0.0287026406429392 |
| NW_011925434.1 | 213404 | 218149 | 4746 | - | Gnomon | gene | NA | NA | gene11347 | GeneID:105285137 | LOC105285137 | Gene | LOC105285137 | lncRNA | NA | character(0) | character(0) | NW_011925434.1 | 215433 | 285112 | 69680 | * | 2 | 0.0287026406429392 |
| NW_011925434.1 | 222347 | 227256 | 4910 | + | Gnomon | gene | NA | NA | gene11348 | GeneID:105285136 | LOC105285136 | Gene | LOC105285136 | lncRNA | NA | character(0) | character(0) | NW_011925434.1 | 215433 | 285112 | 69680 | * | 2 | 0.0287026406429392 |
| NW_011924888.1 | 1806051 | 1809677 | 3627 | + | Gnomon | gene | NA | NA | gene419 | GeneID:105276833 | LOC105276833 | Gene | LOC105276833 | protein_coding | NA | character(0) | character(0) | NW_011924888.1 | 1808554 | 1878222 | 69669 | * | 2 | 0.0287071724870459 |
| NW_011924888.1 | 1822902 | 1859000 | 36099 | + | Gnomon | gene | NA | NA | gene420 | GeneID:105276865 | LOC105276865 | Gene | LOC105276865 | protein_coding | NA | character(0) | character(0) | NW_011924888.1 | 1808554 | 1878222 | 69669 | * | 2 | 0.0287071724870459 |

...

... only first 25 lines shown ...

Directory: MRPR File: gwp-Cb_LI_Rphase.txt

| seqnames | start | end | width | strand | source | type | score | phase | ID | Dbxref | Name | gbkey | gene | gene_biotype | partial | start_range | end_range | seqnames | start | end | width | strand | NbrSites | Sdnsty |
|---|---|---|---|---|---|---|---|---|---|---|---|---|---|---|---|---|---|---|---|---|---|---|---|---|
| NW_011925147.1 | 25059 | 87139 | 62081 | + | Gnomon | gene | NA | NA | gene7608 | GeneID:105280833 | LOC105280833 | Gene | LOC105280833 | protein_coding | NA | character(0) | character(0) | NW_011925147.1 | 9998 | 67979 | 57982 | * | 2 | 0.0344934634886689 |
| NW_011925147.1 | 37666 | 37739 | 74 | + | tRNAscan-SE | gene | NA | NA | gene7609 | GeneID:109611062 | Trnas-gga | Gene | Trnas-gga | tRNA | NA | character(0) | character(0) | NW_011925147.1 | 9998 | 67979 | 57982 | * | 2 | 0.0344934634886689 |
| NW_011925234.1 | 4706 | 110846 | 106141 | - | Gnomon | gene | NA | NA | gene8986 | GeneID:105282438 | LOC105282438 | Gene | LOC105282438 | protein_coding | NA | character(0) | character(0) | NW_011925234.1 | 35111 | 87222 | 52112 | * | 2 | 0.0383788762665029 |
| NW_011925234.1 | 82911 | 84234 | 1324 | - | Gnomon | gene | NA | NA | gene8987 | GeneID:105282436 | LOC105282436 | Gene | LOC105282436 | protein_coding | NA | character(0) | character(0) | NW_011925234.1 | 35111 | 87222 | 52112 | * | 2 | 0.0383788762665029 |
| NW_011924931.1 | 4493233 | 4499991 | 6759 | - | Gnomon | gene | NA | NA | gene1621 | GeneID:105287755 | LOC105287755 | Gene | LOC105287755 | protein_coding | NA | character(0) | character(0) | NW_011924931.1 | 4497412 | 4548764 | 51353 | * | 2 | 0.0389461180456838 |
| NW_011924931.1 | 4500817 | 4527583 | 26767 | + | Gnomon | gene | NA | NA | gene1622 | GeneID:105287758 | LOC105287758 | Gene | LOC105287758 | protein_coding | NA | character(0) | character(0) | NW_011924931.1 | 4497412 | 4548764 | 51353 | * | 2 | 0.0389461180456838 |
| NW_011924931.1 | 4519537 | 4523946 | 4410 | + | Gnomon | gene | NA | NA | gene1623 | GeneID:105287786 | LOC105287786 | Gene | LOC105287786 | protein_coding | NA | character(0) | character(0) | NW_011924931.1 | 4497412 | 4548764 | 51353 | * | 2 | 0.0389461180456838 |
| NW_011924931.1 | 4521493 | 4522142 | 650 | - | Gnomon | gene | NA | NA | gene1624 | GeneID:105287757 | LOC105287757 | Gene | LOC105287757 | protein_coding | NA | character(0) | character(0) | NW_011924931.1 | 4497412 | 4548764 | 51353 | * | 2 | 0.0389461180456838 |
| NW_011924931.1 | 4528303 | 4533712 | 5410 | + | Gnomon | gene | NA | NA | gene1625 | GeneID:105287787 | LOC105287787 | Gene | LOC105287787 | protein_coding | NA | character(0) | character(0) | NW_011924931.1 | 4497412 | 4548764 | 51353 | * | 2 | 0.0389461180456838 |
| NW_011924931.1 | 4533855 | 4534825 | 971 | + | Gnomon | gene | NA | NA | gene1626 | GeneID:105287760 | LOC105287760 | Gene | LOC105287760 | protein_coding | NA | character(0) | character(0) | NW_011924931.1 | 4497412 | 4548764 | 51353 | * | 2 | 0.0389461180456838 |
| NW_011924931.1 | 4535643 | 4542185 | 6543 | + | Gnomon | gene | NA | NA | gene1627 | GeneID:105287761 | LOC105287761 | Gene | LOC105287761 | protein_coding | NA | character(0) | character(0) | NW_011924931.1 | 4497412 | 4548764 | 51353 | * | 2 | 0.0389461180456838 |
| NW_011924931.1 | 4543478 | 4549354 | 5877 | + | Gnomon | gene | NA | NA | gene1628 | GeneID:105287788 | LOC105287788 | Gene | LOC105287788 | protein_coding | NA | character(0) | character(0) | NW_011924931.1 | 4497412 | 4548764 | 51353 | * | 2 | 0.0389461180456838 |
| NW_011925300.1 | 465508 | 523096 | 57589 | - | Gnomon | gene | NA | NA | gene9783 | GeneID:105283340 | LOC105283340 | Gene | LOC105283340 | protein_coding | NA | character(0) | character(0) | NW_011925300.1 | 473373 | 523294 | 49922 | * | 2 | 0.0400624974960939 |
| NW_011925300.1 | 475032 | 478336 | 3305 | + | Gnomon | gene | NA | NA | gene9784 | GeneID:105283339 | LOC105283339 | Gene | LOC105283339 | protein_coding | NA | character(0) | character(0) | NW_011925300.1 | 473373 | 523294 | 49922 | * | 2 | 0.0400624974960939 |
| NW_011925300.1 | 480061 | 483598 | 3538 | + | Gnomon | gene | NA | NA | gene9785 | GeneID:105283338 | LOC105283338 | Gene | LOC105283338 | protein_coding | NA | character(0) | character(0) | NW_011925300.1 | 473373 | 523294 | 49922 | * | 2 | 0.0400624974960939 |
| NW_011925300.1 | 484353 | 488426 | 4074 | + | Gnomon | gene | NA | NA | gene9786 | GeneID:105283337 | LOC105283337 | Gene | LOC105283337 | protein_coding | NA | character(0) | character(0) | NW_011925300.1 | 473373 | 523294 | 49922 | * | 2 | 0.0400624974960939 |
| NW_011925300.1 | 490088 | 494811 | 4724 | + | Gnomon | gene | NA | NA | gene9787 | GeneID:105283336 | LOC105283336 | Gene | LOC105283336 | protein_coding | NA | character(0) | character(0) | NW_011925300.1 | 473373 | 523294 | 49922 | * | 2 | 0.0400624974960939 |
| NW_011925300.1 | 497928 | 499639 | 1712 | + | Gnomon | gene | NA | NA | gene9788 | GeneID:105283334 | LOC105283334 | Gene | LOC105283334 | protein_coding | NA | character(0) | character(0) | NW_011925300.1 | 473373 | 523294 | 49922 | * | 2 | 0.0400624974960939 |
| NW_011925300.1 | 501657 | 506794 | 5138 | + | Gnomon | gene | NA | NA | gene9789 | GeneID:105283333 | LOC105283333 | Gene | LOC105283333 | protein_coding | NA | character(0) | character(0) | NW_011925300.1 | 473373 | 523294 | 49922 | * | 2 | 0.0400624974960939 |
| NW_011925300.1 | 507501 | 511308 | 3808 | + | Gnomon | gene | NA | NA | gene9790 | GeneID:109610650 | LOC109610650 | Gene | LOC109610650 | protein_coding | NA | character(0) | character(0) | NW_011925300.1 | 473373 | 523294 | 49922 | * | 2 | 0.0400624974960939 |
| NW_011925300.1 | 511417 | 516212 | 4796 | + | Gnomon | gene | NA | NA | gene9791 | GeneID:105283332 | LOC105283332 | Gene | LOC105283332 | protein_coding | NA | character(0) | character(0) | NW_011925300.1 | 473373 | 523294 | 49922 | * | 2 | 0.0400624974960939 |
| NW_011925300.1 | 517463 | 519329 | 1867 | + | Gnomon | gene | NA | NA | gene9792 | GeneID:105283331 | LOC105283331 | Gene | LOC105283331 | protein_coding | NA | character(0) | character(0) | NW_011925300.1 | 473373 | 523294 | 49922 | * | 2 | 0.0400624974960939 |
| NW_011925300.1 | 519339 | 521536 | 2198 | + | Gnomon | gene | NA | NA | gene9793 | GeneID:105283330 | LOC105283330 | Gene | LOC105283330 | protein_coding | NA | character(0) | character(0) | NW_011925300.1 | 473373 | 523294 | 49922 | * | 2 | 0.0400624974960939 |
| NW_011925300.1 | 523179 | 526871 | 3693 | + | Gnomon | gene | NA | NA | gene9794 | GeneID:105283328 | LOC105283328 | Gene | LOC105283328 | protein_coding | NA | character(0) | character(0) | NW_011925300.1 | 473373 | 523294 | 49922 | * | 2 | 0.0400624974960939 |

...

... only first 25 lines shown ...

Directory: MRPR File: gwr-Cb_LI_BCphase.txt

| seqnames | start | end | width | strand | source | type | score | phase | ID | Dbxref | Name | gbkey | gene | gene_biotype | partial | start_range | end_range | seqnames | start | end | width | strand | NbrSites | Sdnsty |
|---|---|---|---|---|---|---|---|---|---|---|---|---|---|---|---|---|---|---|---|---|---|---|---|---|
| NW_011924887.1 | 1546715 | 1553169 | 6455 | + | Gnomon | gene | NA | NA | gene209 | GeneID:105274475 | LOC105274475 | Gene | LOC105274475 | protein_coding | NA | character(0) | character(0) | NW_011924887.1 | 1550201 | 1550206 | 6 | * | 6 | 1000 |
| NW_011924887.1 | 1817915 | 1820166 | 2252 | - | Gnomon | gene | NA | NA | gene234 | GeneID:105274775 | LOC105274775 | Gene | LOC105274775 | protein_coding | NA | character(0) | character(0) | NW_011924887.1 | 1819270 | 1819275 | 6 | * | 6 | 1000 |
| NW_011924888.1 | 66076 | 68170 | 2095 | + | Gnomon | gene | NA | NA | gene272 | GeneID:105275200 | LOC105275200 | Gene | LOC105275200 | protein_coding | NA | character(0) | character(0) | NW_011924888.1 | 67012 | 67017 | 6 | * | 6 | 1000 |
| NW_011924888.1 | 77334 | 78632 | 1299 | - | Gnomon | gene | NA | NA | gene276 | GeneID:105275214 | LOC105275214 | Gene | LOC105275214 | protein_coding | NA | character(0) | character(0) | NW_011924888.1 | 77560 | 77565 | 6 | * | 6 | 1000 |
| NW_011924888.1 | 368049 | 370372 | 2324 | - | Gnomon | gene | NA | NA | gene315 | GeneID:105275731 | LOC105275731 | Gene | LOC105275731 | protein_coding | NA | character(0) | character(0) | NW_011924888.1 | 369704 | 369709 | 6 | * | 6 | 1000 |
| NW_011924888.1 | 389413 | 407214 | 17802 | - | Gnomon | gene | NA | NA | gene320 | GeneID:105275770 | LOC105275770 | Gene | LOC105275770 | protein_coding | NA | character(0) | character(0) | NW_011924888.1 | 405832 | 405837 | 6 | * | 6 | 1000 |
| NW_011924888.1 | 1237213 | 1240061 | 2849 | + | Gnomon | gene | NA | NA | gene363 | GeneID:105276248 | LOC105276248 | Gene | LOC105276248 | protein_coding | NA | character(0) | character(0) | NW_011924888.1 | 1237974 | 1237981 | 8 | * | 8 | 1000 |
| NW_011924889.1 | 138414 | 140469 | 2056 | - | Gnomon | gene | NA | NA | gene455 | GeneID:105277237 | LOC105277237 | Gene | LOC105277237 | protein_coding | NA | character(0) | character(0) | NW_011924889.1 | 138461 | 138466 | 6 | * | 6 | 1000 |
| NW_011924889.1 | 237492 | 240474 | 2983 | + | Gnomon | gene | NA | NA | gene478 | GeneID:105277523 | LOC105277523 | Gene | LOC105277523 | protein_coding | NA | character(0) | character(0) | NW_011924889.1 | 238756 | 238761 | 6 | * | 6 | 1000 |
| NW_011924889.1 | 476236 | 478868 | 2633 | - | Gnomon | gene | NA | NA | gene504 | GeneID:105277817 | LOC105277817 | Gene | LOC105277817 | protein_coding | NA | character(0) | character(0) | NW_011924889.1 | 477660 | 477665 | 6 | * | 6 | 1000 |
| NW_011924892.1 | 1158089 | 1160303 | 2215 | - | Gnomon | gene | NA | NA | gene569 | GeneID:105278527 | LOC105278527 | Gene | LOC105278527 | protein_coding | NA | character(0) | character(0) | NW_011924892.1 | 1159381 | 1159386 | 6 | * | 6 | 1000 |
| NW_011924904.1 | 855071 | 858056 | 2986 | - | Gnomon | gene | NA | NA | gene755 | GeneID:105280619 | LOC105280619 | Gene | LOC105280619 | protein_coding | NA | character(0) | character(0) | NW_011924904.1 | 856949 | 856954 | 6 | * | 6 | 1000 |
| NW_011924905.1 | 521528 | 526485 | 4958 | - | Gnomon | gene | NA | NA | gene796 | GeneID:105281111 | LOC105281111 | Gene | LOC105281111 | protein_coding | NA | character(0) | character(0) | NW_011924905.1 | 525144 | 525149 | 6 | * | 6 | 1000 |
| NW_011924914.1 | 259948 | 262322 | 2375 | - | Gnomon | gene | NA | NA | gene952 | GeneID:105283016 | LOC105283016 | Gene | LOC105283016 | protein_coding | NA | character(0) | character(0) | NW_011924914.1 | 261398 | 261403 | 6 | * | 6 | 1000 |
| NW_011924931.1 | 214221 | 216344 | 2124 | + | Gnomon | gene | NA | NA | gene1375 | GeneID:105287333 | LOC105287333 | Gene | LOC105287333 | protein_coding | NA | character(0) | character(0) | NW_011924931.1 | 215617 | 215622 | 6 | * | 6 | 1000 |
| NW_011924931.1 | 3778005 | 3781577 | 3573 | - | Gnomon | gene | NA | NA | gene1536 | GeneID:105287663 | LOC105287663 | Gene | LOC105287663 | protein_coding | NA | character(0) | character(0) | NW_011924931.1 | 3780672 | 3780677 | 6 | * | 6 | 1000 |
| NW_011924931.1 | 3888281 | 3891433 | 3153 | + | Gnomon | gene | NA | NA | gene1547 | GeneID:105287677 | LOC105287677 | Gene | LOC105287677 | protein_coding | NA | character(0) | character(0) | NW_011924931.1 | 3889175 | 3889180 | 6 | * | 6 | 1000 |
| NW_011924931.1 | 3952621 | 3973270 | 20650 | + | Gnomon | gene | NA | NA | gene1560 | GeneID:105287694 | LOC105287694 | Gene | LOC105287694 | protein_coding | NA | character(0) | character(0) | NW_011924931.1 | 3966620 | 3966625 | 6 | * | 6 | 1000 |
| NW_011924931.1 | 3966137 | 3967627 | 1491 | - | Gnomon | gene | NA | NA | gene1562 | GeneID:105287692 | LOC105287692 | Gene | LOC105287692 | protein_coding | NA | character(0) | character(0) | NW_011924931.1 | 3966620 | 3966625 | 6 | * | 6 | 1000 |
| NW_011924931.1 | 4065100 | 4067558 | 2459 | - | Gnomon | gene | NA | NA | gene1584 | GeneID:105287715 | LOC105287715 | Gene | LOC105287715 | protein_coding | NA | character(0) | character(0) | NW_011924931.1 | 4066201 | 4066206 | 6 | * | 6 | 1000 |
| NW_011924931.1 | 4488185 | 4490767 | 2583 | + | Gnomon | gene | NA | NA | gene1620 | GeneID:105287754 | LOC105287754 | Gene | LOC105287754 | protein_coding | NA | character(0) | character(0) | NW_011924931.1 | 4489292 | 4489297 | 6 | * | 6 | 1000 |
| NW_011924932.1 | 543289 | 544100 | 812 | + | Gnomon | gene | NA | NA | gene1656 | GeneID:105287814 | LOC105287814 | Gene | LOC105287814 | protein_coding | NA | character(0) | character(0) | NW_011924932.1 | 543633 | 543638 | 6 | * | 6 | 1000 |
| NW_011924940.1 | 334573 | 349050 | 14478 | - | Gnomon | gene | NA | NA | gene1863 | GeneID:105288044 | LOC105288044 | Gene | LOC105288044 | protein_coding | NA | character(0) | character(0) | NW_011924940.1 | 347385 | 347390 | 6 | * | 6 | 1000 |
| NW_011924945.1 | 16609 | 22779 | 6171 | - | Gnomon | gene | NA | NA | gene1878 | GeneID:105288065 | LOC105288065 | Gene | LOC105288065 | protein_coding | NA | character(0) | character(0) | NW_011924945.1 | 20640 | 20645 | 6 | * | 6 | 1000 |
| NW_011924946.1 | 775753 | 777201 | 1449 | - | Gnomon | gene | NA | NA | gene1993 | GeneID:105288193 | LOC105288193 | Gene | LOC105288193 | protein_coding | NA | character(0) | character(0) | NW_011924946.1 | 776368 | 776373 | 6 | * | 6 | 1000 |
| NW_011924946.1 | 786370 | 788739 | 2370 | - | Gnomon | gene | NA | NA | gene1995 | GeneID:105288196 | LOC105288196 | Gene | LOC105288196 | protein_coding | NA | character(0) | character(0) | NW_011924946.1 | 787444 | 787449 | 6 | * | 6 | 1000 |
| NW_011924946.1 | 1411689 | 1414880 | 3192 | - | Gnomon | gene | NA | NA | gene2041 | GeneID:105288248 | LOC105288248 | Gene | LOC105288248 | protein_coding | NA | character(0) | character(0) | NW_011924946.1 | 1413935 | 1413940 | 6 | * | 6 | 1000 |
| NW_011924946.1 | 1502005 | 1507040 | 5036 | - | Gnomon | gene | NA | NA | gene2047 | GeneID:105274446 | LOC105274446 | Gene | LOC105274446 | protein_coding | NA | character(0) | character(0) | NW_011924946.1 | 1502183 | 1502188 | 6 | * | 6 | 1000 |
| NW_011924946.1 | 1607845 | 1608869 | 1025 | - | Gnomon | gene | NA | NA | gene2061 | GeneID:105274458 | LOC105274458 | Gene | LOC105274458 | protein_coding | NA | character(0) | character(0) | NW_011924946.1 | 1608077 | 1608082 | 6 | * | 6 | 1000 |
| NW_011924946.1 | 2832086 | 2841975 | 9890 | - | Gnomon | gene | NA | NA | gene2162 | GeneID:105274578 | LOC105274578 | Gene | LOC105274578 | protein_coding | NA | | | | | | | | | |

character(0)	character(0)	NW_011924946.1	2839967	2839972	6	*	6	1000
NW_011924951.1	64945	71389	6445	-	Gnomon	gene	NA	NA	gene2323	GeneID:105274768	LOC105274768	Gene	LOC105274768	protein_coding	NA	character(0)
character(0)	NW_011924951.1	69822	69827	6	*	6	1000
NW_011924955.1	1492124	1497652	5529	-	Gnomon	gene	NA	NA	gene2505	GeneID:105274952	LOC105274952	Gene	LOC105274952	protein_coding	NA
character(0)	character(0)	NW_011924955.1	1495077	1495082	6	*	6	1000
NW_011924956.1	717757	722032	4276	+	Gnomon	gene	NA	NA	gene2570	GeneID:105275036	LOC105275036	Gene	LOC105275036	protein_coding	NA
character(0)	character(0)	NW_011924956.1	719335	719344	10	*	10	1000
NW_011924956.1	789246	793927	4682	+	Gnomon	gene	NA	NA	gene2578	GeneID:105275047	LOC105275047	Gene	LOC105275047	protein_coding	NA
character(0)	character(0)	NW_011924956.1	790656	790661	6	*	6	1000
NW_011924973.1	236871	239530	2660	-	Gnomon	gene	NA	NA	gene2751	GeneID:105275264	LOC105275264	Gene	LOC105275264	protein_coding	NA
character(0)	character(0)	NW_011924973.1	238411	238416	6	*	6	1000
NW_011924973.1	244133	245537	1405	+	Gnomon	gene	NA	NA	gene2754	GeneID:105275270	LOC105275270	Gene	LOC105275270	protein_coding	NA
character(0)	character(0)	NW_011924973.1	244951	244956	6	*	6	1000
NW_011924973.1	828831	835120	6290	+	Gnomon	gene	NA	NA	gene2799	GeneID:105275316	LOC105275316	Gene	LOC105275316	protein_coding	NA
character(0)	character(0)	NW_011924973.1	829674	829679	6	*	6	1000
NW_011924973.1	935440	945867	10428	+	Gnomon	gene	NA	NA	gene2820	GeneID:105275342	LOC105275342	Gene	LOC105275342	protein_coding	NA
character(0)	character(0)	NW_011924973.1	936776	936781	6	*	6	1000
NW_011924973.1	997316	999807	2492	+	Gnomon	gene	NA	NA	gene2835	GeneID:105275357	LOC105275357	Gene	LOC105275357	protein_coding	NA
character(0)	character(0)	NW_011924973.1	997954	997959	6	*	6	1000
NW_011924973.1	1001098	1003969	2872	+	Gnomon	gene	NA	NA	gene2837	GeneID:105275355	LOC105275355	Gene	LOC105275355	protein_coding	NA
character(0)	character(0)	NW_011924973.1	1002105	1002110	6	*	6	1000
NW_011924977.1	31430	34619	3190	+	Gnomon	gene	NA	NA	gene2916	GeneID:109610741	LOC109610741	Gene	LOC109610741	protein_coding	NA	character(0)
character(0)	NW_011924977.1	32395	32400	6	*	6	1000
NW_011924977.1	516991	520258	3268	+	Gnomon	gene	NA	NA	gene2981	GeneID:105275526	LOC105275526	Gene	LOC105275526	protein_coding	NA
character(0)	character(0)	NW_011924977.1	519073	519078	6	*	6	1000
NW_011924977.1	662427	665182	2756	+	Gnomon	gene	NA	NA	gene3008	GeneID:105275550	LOC105275550	Gene	LOC105275550	protein_coding	NA
character(0)	character(0)	NW_011924977.1	663534	663539	6	*	6	1000
NW_011924977.1	3614512	3642583	28072	-	Gnomon	gene	NA	NA	gene3195	GeneID:105275750	LOC105275750	Gene	LOC105275750	protein_coding	NA
character(0)	character(0)	NW_011924977.1	3633490	3633495	6	*	6	1000
NW_011924977.1	3632271	3635543	3273	+	Gnomon	gene	NA	NA	gene3198	GeneID:105275746	LOC105275746	Gene	LOC105275746	protein_coding	NA
character(0)	character(0)	NW_011924977.1	3633490	3633495	6	*	6	1000
NW_011924977.1	3723831	3727852	4022	-	Gnomon	gene	NA	NA	gene3212	GeneID:105275760	LOC105275760	Gene	LOC105275760	protein_coding	NA
character(0)	character(0)	NW_011924977.1	3727104	3727109	6	*	6	1000
NW_011924977.1	3728770	3731404	2635	+	Gnomon	gene	NA	NA	gene3213	GeneID:105275761	LOC105275761	Gene	LOC105275761	protein_coding	NA
character(0)	character(0)	NW_011924977.1	3729892	3729897	6	*	6	1000
NW_011924979.1	16258	20038	3781	+	Gnomon	gene	NA	NA	gene3218	GeneID:105275781	LOC105275781	Gene	LOC105275781	protein_coding	NA	character(0)
character(0)	NW_011924979.1	17774	17779	6	*	6	1000
NW_011924979.1	1593634	1596428	2795	+	Gnomon	gene	NA	NA	gene3297	GeneID:105275861	LOC105275861	Gene	LOC105275861	protein_coding	NA
character(0)	character(0)	NW_011924979.1	1594463	1594468	6	*	6	1000
NW_011924979.1	1988584	1990897	2314	+	Gnomon	gene	NA	NA	gene3314	GeneID:105275878	LOC105275878	Gene	LOC105275878	protein_coding	NA
character(0)	character(0)	NW_011924979.1	1989444	1989449	6	*	6	1000
NW_011924979.1	2744383	2746071	1689	+	Gnomon	gene	NA	NA	gene3334	GeneID:105275897	LOC105275897	Gene	LOC105275897	protein_coding	NA
character(0)	character(0)	NW_011924979.1	2745453	2745458	6	*	6	1000
NW_011924996.1	75277	77259	1983	+	Gnomon	gene	NA	NA	gene3743	GeneID:105276382	LOC105276382	Gene	LOC105276382	protein_coding	NA	character(0)
character(0)	NW_011924996.1	76074	76079	6	*	6	1000
NW_011924996.1	1822758	1830685	7928	-	Gnomon	gene	NA	NA	gene3885	GeneID:105276527	LOC105276527	Gene	LOC105276527	protein_coding	NA
character(0)	character(0)	NW_011924996.1	1829379	1829384	6	*	6	1000
NW_011925007.1	926256	929694	3439	+	Gnomon	gene	NA	NA	gene4194	GeneID:105276881	LOC105276881	Gene	LOC105276881	protein_coding	NA
character(0)	character(0)	NW_011925007.1	927466	927471	6	*	6	1000
NW_011925007.1	1370227	1386638	16412	-	Gnomon	gene	NA	NA	gene4246	GeneID:105276941	LOC105276941	Gene	LOC105276941	protein_coding	NA
character(0)	character(0)	NW_011925007.1	1385156	1385161	6	*	6	1000
NW_011925007.1	1454403	1457778	3376	+	Gnomon	gene	NA	NA	gene4253	GeneID:105276950	LOC105276950	Gene	LOC105276950	protein_coding	NA
character(0)	character(0)	NW_011925007.1	1455553	1455558	6	*	6	1000
NW_011925012.1	1356105	1358935	2831	+	Gnomon	gene	NA	NA	gene4498	GeneID:105277239	LOC105277239	Gene	LOC105277239	protein_coding	NA
character(0)	character(0)	NW_011925012.1	1357276	1357281	6	*	6	1000
NW_011925016.1	800369	803160	2792	-	Gnomon	gene	NA	NA	gene4610	GeneID:105277357	LOC105277357	Gene	LOC105277357	protein_coding	NA
character(0)	character(0)	NW_011925016.1	801483	801490	8	*	8	1000
NW_011925020.1	469190	474352	5163	+	Gnomon	gene	NA	NA	gene4780	GeneID:105277552	LOC105277552	Gene	LOC105277552	protein_coding	NA
character(0)	character(0)	NW_011925020.1	470416	470421	6	*	6	1000
NW_011925022.1	386851	387976	1126	-	Gnomon	gene	NA	NA	gene5000	GeneID:105277804	LOC105277804	Gene	LOC105277804	protein_coding	NA
character(0)	character(0)	NW_011925022.1	387077	387082	6	*	6	1000
NW_011925035.1	818580	820886	2307	-	Gnomon	gene	NA	NA	gene5379	GeneID:105278234	LOC105278234	Gene	LOC105278234	protein_coding	NA
character(0)	character(0)	NW_011925035.1	820260	820265	6	*	6	1000
NW_011925050.1	692537	695061	2525	-	Gnomon	gene	NA	NA	gene5769	GeneID:105278683	LOC105278683	Gene	LOC105278683	protein_coding	NA

character(0)	character(0)	NW_011925050.1	693967	693972	6	*	6	1000
NW_011925051.1	754067	756870	2804	-	Gnomon	gene	NA	NA	gene5860	GeneID:105278789	LOC105278789	Gene	LOC105278789	protein_coding	NA
character(0)	character(0)	NW_011925051.1	755643	755648	6	*	6	1000
NW_011925053.1	649052	656133	7082	-	Gnomon	gene	NA	NA	gene5882	GeneID:105278812	LOC105278812	Gene	LOC105278812	protein_coding	NA
character(0)	character(0)	NW_011925053.1	655139	655144	6	*	6	1000
NW_011925053.1	1149732	1155700	5969	-	Gnomon	gene	NA	NA	gene5895	GeneID:105278828	LOC105278828	Gene	LOC105278828	protein_coding	NA
character(0)	character(0)	NW_011925053.1	1154241	1154246	6	*	6	1000
NW_011925053.1	1149732	1155700	5969	-	Gnomon	gene	NA	NA	gene5895	GeneID:105278828	LOC105278828	Gene	LOC105278828	protein_coding	NA
character(0)	character(0)	NW_011925053.1	1154686	1154691	6	*	6	1000
NW_011925053.1	1481875	1483457	1583	-	Gnomon	gene	NA	NA	gene5932	GeneID:105278862	LOC105278862	Gene	LOC105278862	protein_coding	NA
character(0)	character(0)	NW_011925053.1	1482901	1482906	6	*	6	1000
NW_011925067.1	201362	204847	3486	-	Gnomon	gene	NA	NA	gene6034	GeneID:105279004	LOC105279004	Gene	LOC105279004	protein_coding	NA
character(0)	character(0)	NW_011925067.1	203691	203696	6	*	6	1000
NW_011925075.1	793827	800147	6321	+	Gnomon	gene	NA	NA	gene6165	GeneID:105279155	LOC105279155	Gene	LOC105279155	protein_coding	NA
character(0)	character(0)	NW_011925075.1	794882	794887	6	*	6	1000
NW_011925086.1	128430	135139	6710	+	Gnomon	gene	NA	NA	gene6311	GeneID:105279333	LOC105279333	Gene	LOC105279333	protein_coding	NA
character(0)	character(0)	NW_011925086.1	129637	129644	8	*	8	1000
NW_011925086.1	250638	255287	4650	-	Gnomon	gene	NA	NA	gene6324	GeneID:105279346	LOC105279346	Gene	LOC105279346	protein_coding	NA
character(0)	character(0)	NW_011925086.1	253858	253863	6	*	6	1000
NW_011925086.1	604599	609266	4668	+	Gnomon	gene	NA	NA	gene6342	GeneID:105279367	LOC105279367	Gene	LOC105279367	protein_coding	NA
character(0)	character(0)	NW_011925086.1	606157	606162	6	*	6	1000
NW_011925099.1	12321	14914	2594	+	Gnomon	gene	NA	NA	gene6401	GeneID:105279437	LOC105279437	Gene	LOC105279437	protein_coding	NA	character(0)
character(0)	NW_011925099.1	14885	14890	6	*	6	1000
NW_011925106.1	528	7971	7444	-	Gnomon	gene	NA	NA	gene6583	GeneID:105279649	LOC105279649	Gene	LOC105279649	protein_coding	NA	character(0)
character(0)	NW_011925106.1	6714	6719	6	*	6	1000
NW_011925106.1	487858	493874	6017	+	Gnomon	gene	NA	NA	gene6649	GeneID:105279717	LOC105279717	Gene	LOC105279717	protein_coding	NA
character(0)	character(0)	NW_011925106.1	489460	489465	6	*	6	1000
NW_011925106.1	1997159	1999737	2579	-	Gnomon	gene	NA	NA	gene6753	GeneID:105279838	LOC105279838	Gene	LOC105279838	protein_coding	NA
character(0)	character(0)	NW_011925106.1	1998173	1998178	6	*	6	1000
NW_011925106.1	2086959	2096661	9703	+	Gnomon	gene	NA	NA	gene6771	GeneID:105279854	LOC105279854	Gene	LOC105279854	protein_coding	NA
character(0)	character(0)	NW_011925106.1	2088114	2088121	8	*	8	1000
NW_011925106.1	3275310	3277500	2191	-	Gnomon	gene	NA	NA	gene6816	GeneID:105279900	LOC105279900	Gene	LOC105279900	protein_coding	NA
character(0)	character(0)	NW_011925106.1	3276041	3276046	6	*	6	1000
NW_011925118.1	771350	783967	12618	-	Gnomon	gene	NA	NA	gene6991	GeneID:105280128	LOC105280128	Gene	LOC105280128	protein_coding	NA
character(0)	character(0)	NW_011925118.1	782879	782884	6	*	6	1000
NW_011925118.1	771350	783967	12618	-	Gnomon	gene	NA	NA	gene6991	GeneID:105280128	LOC105280128	Gene	LOC105280128	protein_coding	NA
character(0)	character(0)	NW_011925118.1	783338	783343	6	*	6	1000
NW_011925128.1	1738966	1741716	2751	-	Gnomon	gene	NA	NA	gene7206	GeneID:105280373	LOC105280373	Gene	LOC105280373	protein_coding	NA
character(0)	character(0)	NW_011925128.1	1740548	1740553	6	*	6	1000
NW_011925128.1	1875336	1880845	5510	+	Gnomon	gene	NA	NA	gene7225	GeneID:105280395	LOC105280395	Gene	LOC105280395	protein_coding	NA
character(0)	character(0)	NW_011925128.1	1876634	1876639	6	*	6	1000
NW_011925142.1	743409	745933	2525	-	Gnomon	gene	NA	NA	gene7568	GeneID:105280786	LOC105280786	Gene	LOC105280786	protein_coding	NA
character(0)	character(0)	NW_011925142.1	744366	744371	6	*	6	1000
NW_011925142.1	777887	780161	2275	-	Gnomon	gene	NA	NA	gene7574	GeneID:105280791	LOC105280791	Gene	LOC105280791	protein_coding	NA
character(0)	character(0)	NW_011925142.1	779002	779007	6	*	6	1000
NW_011925150.1	76140	79045	2906	-	Gnomon	gene	NA	NA	gene7626	GeneID:105280856	LOC105280856	Gene	LOC105280856	protein_coding	NA	character(0)
character(0)	NW_011925150.1	78377	78382	6	*	6	1000
NW_011925151.1	38146	39177	1032	-	Gnomon	gene	NA	NA	gene7635	GeneID:105280860	LOC105280860	Gene	LOC105280860	protein_coding	NA	character(0)
character(0)	NW_011925151.1	38257	38262	6	*	6	1000
NW_011925161.1	145622	148482	2861	-	Gnomon	gene	NA	NA	gene7710	GeneID:105280955	LOC105280955	Gene	LOC105280955	protein_coding	NA
character(0)	character(0)	NW_011925161.1	147171	147176	6	*	6	1000
NW_011925161.1	1702331	1707999	5669	-	Gnomon	gene	NA	NA	gene7821	GeneID:105281067	LOC105281067	Gene	LOC105281067	protein_coding	NA
character(0)	character(0)	NW_011925161.1	1707165	1707170	6	*	6	1000
NW_011925161.1	1705906	1731780	25875	+	Gnomon	gene	NA	NA	gene7822	GeneID:105281066	LOC105281066	Gene	LOC105281066	protein_coding	NA
character(0)	character(0)	NW_011925161.1	1707165	1707170	6	*	6	1000
NW_011925161.1	2209807	2215262	5456	-	Gnomon	gene	NA	NA	gene7866	GeneID:105281115	LOC105281115	Gene	LOC105281115	protein_coding	NA
character(0)	character(0)	NW_011925161.1	2212367	2212372	6	*	6	1000
NW_011925163.1	470647	473680	3034	+	Gnomon	gene	NA	NA	gene8040	GeneID:105281341	LOC105281341	Gene	LOC105281341	protein_coding	NA
character(0)	character(0)	NW_011925163.1	471213	471218	6	*	6	1000
NW_011925182.1	57229	59555	2327	-	Gnomon	gene	NA	NA	gene8209	GeneID:105281549	LOC105281549	Gene	LOC105281549	protein_coding	NA	character(0)
character(0)	NW_011925182.1	58922	58927	6	*	6	1000
NW_011925182.1	162285	170775	8491	+	Gnomon	gene	NA	NA	gene8218	GeneID:105281562	LOC105281562	Gene	LOC105281562	protein_coding	NA
character(0)	character(0)	NW_011925182.1	163572	163577	6	*	6	1000
NW_011925183.1	39511	42630	3120	+	Gnomon	gene	NA	NA	gene8236	GeneID:105281577	LOC105281577	Gene	LOC105281577	protein_coding	NA	character(0)

character(0) NW_011925183.1 40051 40056 6 * 6 1000
NW_011925188.1 196247 198859 2613 - Gnomon gene NA NA gene8256 GeneID:105281604 LOC105281604 Gene LOC105281604 protein_coding NA
character(0) character(0) NW_011925188.1 197361 197366 6 * 6 1000
NW_011925188.1 199014 202632 3619 - Gnomon gene NA NA gene8257 GeneID:105281603 LOC105281603 Gene LOC105281603 protein_coding NA
character(0) character(0) NW_011925188.1 201626 201631 6 * 6 1000
NW_011925189.1 165659 175663 10005 - Gnomon gene NA NA gene8294 GeneID:105281645 LOC105281645 Gene LOC105281645 protein_coding NA
character(0) character(0) NW_011925189.1 174276 174281 6 * 6 1000
NW_011925189.1 575609 577709 2101 - Gnomon gene NA NA gene8332 GeneID:105281690 LOC105281690 Gene LOC105281690 protein_coding NA
character(0) character(0) NW_011925189.1 577082 577087 6 * 6 1000
NW_011925203.1 63038 97839 34802 + Gnomon gene NA NA gene8521 GeneID:105281906 LOC105281906 Gene LOC105281906 protein_coding NA character(0)
character(0) NW_011925203.1 64603 64608 6 * 6 1000
NW_011925216.1 136610 140476 3867 - Gnomon gene NA NA gene8735 GeneID:105282144 LOC105282144 Gene LOC105282144 protein_coding NA
character(0) character(0) NW_011925216.1 137522 137527 6 * 6 1000
NW_011925216.1 136795 145548 8754 + Gnomon gene NA NA gene8736 GeneID:105282142 LOC105282142 Gene LOC105282142 protein_coding NA
character(0) character(0) NW_011925216.1 137522 137527 6 * 6 1000
NW_011925216.1 156691 162086 5396 + Gnomon gene NA NA gene8741 GeneID:105282150 LOC105282150 Gene LOC105282150 protein_coding NA
character(0) character(0) NW_011925216.1 157541 157546 6 * 6 1000
NW_011925224.1 305105 307887 2783 - Gnomon gene NA NA gene8866 GeneID:105282289 LOC105282289 Gene LOC105282289 protein_coding NA
character(0) character(0) NW_011925224.1 306866 306871 6 * 6 1000
NW_011925232.1 152153 159881 7729 + Gnomon gene NA NA gene8919 GeneID:105282356 LOC105282356 Gene LOC105282356 protein_coding NA
character(0) character(0) NW_011925232.1 153164 153169 6 * 6 1000
NW_011925239.1 3979 7808 3830 + Gnomon gene NA NA gene9001 GeneID:105282461 LOC105282461 Gene LOC105282461 protein_coding NA character(0)
character(0) NW_011925239.1 4877 4882 6 * 6 1000
NW_011925240.1 757237 760222 2986 - Gnomon gene NA NA gene9059 GeneID:105282525 LOC105282525 Gene LOC105282525 protein_coding NA
character(0) character(0) NW_011925240.1 758844 758849 6 * 6 1000
NW_011925240.1 2490948 2493602 2655 - Gnomon gene NA NA gene9134 GeneID:105282589 LOC105282589 Gene LOC105282589 protein_coding NA
character(0) character(0) NW_011925240.1 2492742 2492747 6 * 6 1000
NW_011925241.1 428008 431512 3505 + Gnomon gene NA NA gene9196 GeneID:105282662 LOC105282662 Gene LOC105282662 protein_coding NA
character(0) character(0) NW_011925241.1 428574 428579 6 * 6 1000
NW_011925241.1 428008 431512 3505 + Gnomon gene NA NA gene9196 GeneID:105282662 LOC105282662 Gene LOC105282662 protein_coding NA
character(0) character(0) NW_011925241.1 428600 428605 6 * 6 1000
NW_011925258.1 22829 32812 9984 + Gnomon gene NA NA gene9308 GeneID:105282804 LOC105282804 Gene LOC105282804 protein_coding NA character(0)
character(0) NW_011925258.1 28153 28158 6 * 6 1000
NW_011925258.1 404301 408788 4488 - Gnomon gene NA NA gene9347 GeneID:105282847 LOC105282847 Gene LOC105282847 protein_coding NA
character(0) character(0) NW_011925258.1 407771 407776 6 * 6 1000
NW_011925304.1 378599 380691 2093 - Gnomon gene NA NA gene9814 GeneID:105283373 LOC105283373 Gene LOC105283373 protein_coding NA
character(0) character(0) NW_011925304.1 379611 379616 6 * 6 1000
NW_011925304.1 468966 476624 7659 - Gnomon gene NA NA gene9824 GeneID:105283383 LOC105283383 Gene LOC105283383 protein_coding NA
character(0) character(0) NW_011925304.1 475523 475528 6 * 6 1000
NW_011925321.1 975185 978704 3520 - Gnomon gene NA NA gene10031 GeneID:105283620 LOC105283620 Gene LOC105283620 protein_coding NA
character(0) character(0) NW_011925321.1 977422 977427 6 * 6 1000
NW_011925321.1 1107715 1109901 2187 - Gnomon gene NA NA gene10043 GeneID:105283634 LOC105283634 Gene LOC105283634 protein_coding NA
character(0) character(0) NW_011925321.1 1108690 1108695 6 * 6 1000
NW_011925323.1 2107588 2112140 4553 - Gnomon gene NA NA gene10169 GeneID:105283771 LOC105283771 Gene LOC105283771 protein_coding NA
character(0) character(0) NW_011925323.1 2111421 2111426 6 * 6 1000
NW_011925351.1 11934 15244 3311 - Gnomon gene NA NA gene10387 GeneID:105284041 LOC105284041 Gene LOC105284041 protein_coding NA character(0)
character(0) NW_011925351.1 14519 14524 6 * 6 1000
NW_011925353.1 26250 32260 6011 + Gnomon gene NA NA gene10402 GeneID:105284053 LOC105284053 Gene LOC105284053 protein_coding NA character(0)
character(0) NW_011925353.1 27469 27474 6 * 6 1000
NW_011925353.1 40476 43146 2671 - Gnomon gene NA NA gene10405 GeneID:105284057 LOC105284057 Gene LOC105284057 protein_coding NA character(0)
character(0) NW_011925353.1 41948 41953 6 * 6 1000
NW_011925355.1 581482 584005 2524 + Gnomon gene NA NA gene10482 GeneID:105284137 LOC105284137 Gene LOC105284137 protein_coding NA
character(0) character(0) NW_011925355.1 582446 582451 6 * 6 1000
NW_011925356.1 704771 710120 5350 - Gnomon gene NA NA gene10539 GeneID:105284201 LOC105284201 Gene LOC105284201 protein_coding NA
character(0) character(0) NW_011925356.1 708176 708181 6 * 6 1000
NW_011925356.1 1573184 1579703 6520 - Gnomon gene NA NA gene10596 GeneID:105284262 LOC105284262 Gene LOC105284262 protein_coding NA
character(0) character(0) NW_011925356.1 1573650 1573655 6 * 6 1000
NW_011925367.1 1119922 1121887 1966 + Gnomon gene NA NA gene10714 GeneID:105284396 LOC105284396 Gene LOC105284396 protein_coding NA
character(0) character(0) NW_011925367.1 1120960 1120965 6 * 6 1000
NW_011925371.1 1203185 1204659 1475 - Gnomon gene NA NA gene10886 GeneID:105284582 LOC105284582 Gene LOC105284582 protein_coding NA
character(0) character(0) NW_011925371.1 1203884 1203889 6 * 6 1000
NW_011925387.1 39150 46759 7610 + Gnomon gene NA NA gene10928 GeneID:105284644 LOC105284644 Gene LOC105284644 protein_coding NA character(0)
character(0) NW_011925387.1 40039 40044 6 * 6 1000
NW_011925387.1 271672 273220 1549 + Gnomon gene NA NA gene10957 GeneID:105284676 LOC105284676 Gene LOC105284676 protein_coding NA

character(0) character(0) NW_011925387.1 272491 272496 6 * 6 1000
NW_011925401.1 10153 19053 8901 - Gnomon gene NA NA gene11099 GeneID:105284845 LOC105284845 Gene LOC105284845 protein_coding NA character(0)
character(0) NW_011925401.1 17311 17316 6 * 6 1000
NW_011925438.1 122592 124529 1938 + Gnomon gene NA NA gene11397 GeneID:105285205 LOC105285205 Gene LOC105285205 protein_coding NA
character(0) character(0) NW_011925438.1 123277 123282 6 * 6 1000
NW_011925442.1 70936 77041 6106 + Gnomon gene NA NA gene11466 GeneID:105285277 LOC105285277 Gene LOC105285277 protein_coding NA character(0)
character(0) NW_011925442.1 71492 71497 6 * 6 1000
NW_011925442.1 70999 75078 4080 - Gnomon gene NA NA gene11467 GeneID:105285278 LOC105285278 Gene LOC105285278 protein_coding NA character(0)
character(0) NW_011925442.1 71492 71497 6 * 6 1000
NW_011925459.1 186244 187554 1311 - Gnomon gene NA NA gene11569 GeneID:105285394 LOC105285394 Gene LOC105285394 protein_coding NA
character(0) character(0) NW_011925459.1 186501 186506 6 * 6 1000
NW_011925470.1 613911 617551 3641 - Gnomon gene NA NA gene11628 GeneID:105285465 LOC105285465 Gene LOC105285465 protein_coding NA
character(0) character(0) NW_011925470.1 616616 616621 6 * 6 1000
NW_011925490.1 137354 140755 3402 + Gnomon gene NA NA gene11689 GeneID:105285534 LOC105285534 Gene LOC105285534 protein_coding NA
character(0) character(0) NW_011925490.1 138283 138288 6 * 6 1000
NW_011925516.1 389602 391122 1521 - Gnomon gene NA NA gene11817 GeneID:105285670 LOC105285670 Gene LOC105285670 protein_coding NA
character(0) character(0) NW_011925516.1 390371 390376 6 * 6 1000
NW_011925595.1 1 4548 4548 - Gnomon gene NA NA gene12102 GeneID:105286003 LOC105286003 Gene LOC105286003 protein_coding NA character(0)
character(0) NW_011925595.1 3717 3722 6 * 6 1000
NW_011925653.1 98728 102889 4162 - Gnomon gene NA NA gene12259 GeneID:105286160 LOC105286160 Gene LOC105286160 protein_coding NA
character(0) character(0) NW_011925653.1 101522 101527 6 * 6 1000
NW_011925653.1 98728 102889 4162 - Gnomon gene NA NA gene12259 GeneID:105286160 LOC105286160 Gene LOC105286160 protein_coding NA
character(0) character(0) NW_011925653.1 101541 101546 6 * 6 1000
NW_011925664.1 38296 39300 1005 + Gnomon gene NA NA gene12311 GeneID:105286212 LOC105286212 Gene LOC105286212 protein_coding NA character(0)
character(0) NW_011925664.1 39054 39061 8 * 8 1000
NW_011925664.1 277932 283313 5382 + Gnomon gene NA NA gene12359 GeneID:105286364 LOC105286364 Gene LOC105286364 protein_coding NA
character(0) character(0) NW_011925664.1 279108 279113 6 * 6 1000
NW_011925664.1 580837 582496 1660 - Gnomon gene NA NA gene12411 GeneID:105286320 LOC105286320 Gene LOC105286320 protein_coding NA
character(0) character(0) NW_011925664.1 581347 581352 6 * 6 1000
NW_011925672.1 39885 43167 3283 - Gnomon gene NA NA gene12467 GeneID:105286395 LOC105286395 Gene LOC105286395 protein_coding NA character(0)
character(0) NW_011925672.1 42017 42022 6 * 6 1000
NW_011925715.1 952620 956416 3797 - Gnomon gene NA NA gene12567 GeneID:105286509 LOC105286509 Gene LOC105286509 protein_coding NA
character(0) character(0) NW_011925715.1 955653 955658 6 * 6 1000
NW_011925810.1 23502 27182 3681 + Gnomon gene NA NA gene12829 GeneID:105286828 LOC105286828 Gene LOC105286828 protein_coding true character(0)
c("27182", ".") NW_011925810.1 24132 24137 6 * 6 1000
NW_011925984.1 48360 54191 5832 + Gnomon gene NA NA gene12992 GeneID:105287013 LOC105287013 Gene LOC105287013 protein_coding NA character(0)
character(0) NW_011925984.1 49836 49841 6 * 6 1000
NW_011926135.1 70608 74554 3947 + Gnomon gene NA NA gene13076 GeneID:105287104 LOC105287104 Gene LOC105287104 protein_coding NA character(0)
character(0) NW_011926135.1 71338 71343 6 * 6 1000
NW_011929339.1 7 1883 1877 + Gnomon gene NA NA gene13384 GeneID:105287447 LOC105287447 Gene LOC105287447 protein_coding true character(0)
c("1883", ".") NW_011929339.1 1129 1134 6 * 6 1000
NW_011929339.1 7 1883 1877 + Gnomon gene NA NA gene13384 GeneID:105287447 LOC105287447 Gene LOC105287447 protein_coding true character(0)
c("1883", ".") NW_011929339.1 1180 1185 6 * 6 1000

Directory: MRPR File: gwr-Cb_LI_Rphase.txt

| seqnames | start | end | width | strand | source | type | score | phase | ID | Dbxref | Name | gbkey | gene | gene_biotype | partial | start_range | end_range | seqnames | start | end | width | strand | NbrSites | Sdnsty |
|---|---|---|---|---|---|---|---|---|---|---|---|---|---|---|---|---|---|---|---|---|---|---|---|---|
| NW_011924887.1 | 1546715 | 1553169 | 6455 | + | Gnomon | gene | NA | NA | gene209 | GeneID:105274475 | LOC105274475 | Gene | LOC105274475 | protein_coding | NA | character(0) | character(0) | NW_011924887.1 | 1550201 | 1550206 | 6 | * | 6 | 1000 |
| NW_011924887.1 | 1817915 | 1820166 | 2252 | - | Gnomon | gene | NA | NA | gene234 | GeneID:105274775 | LOC105274775 | Gene | LOC105274775 | protein_coding | NA | character(0) | character(0) | NW_011924887.1 | 1819270 | 1819275 | 6 | * | 6 | 1000 |
| NW_011924888.1 | 66076 | 68170 | 2095 | + | Gnomon | gene | NA | NA | gene272 | GeneID:105275200 | LOC105275200 | Gene | LOC105275200 | protein_coding | NA | character(0) | character(0) | NW_011924888.1 | 67012 | 67017 | 6 | * | 6 | 1000 |
| NW_011924888.1 | 77334 | 78632 | 1299 | - | Gnomon | gene | NA | NA | gene276 | GeneID:105275214 | LOC105275214 | Gene | LOC105275214 | protein_coding | NA | character(0) | character(0) | NW_011924888.1 | 77560 | 77565 | 6 | * | 6 | 1000 |
| NW_011924888.1 | 368049 | 370372 | 2324 | - | Gnomon | gene | NA | NA | gene315 | GeneID:105275731 | LOC105275731 | Gene | LOC105275731 | protein_coding | NA | character(0) | character(0) | NW_011924888.1 | 369704 | 369709 | 6 | * | 6 | 1000 |
| NW_011924888.1 | 389413 | 407214 | 17802 | - | Gnomon | gene | NA | NA | gene320 | GeneID:105275770 | LOC105275770 | Gene | LOC105275770 | protein_coding | NA | character(0) | character(0) | NW_011924888.1 | 405832 | 405837 | 6 | * | 6 | 1000 |
| NW_011924888.1 | 1237213 | 1240061 | 2849 | + | Gnomon | gene | NA | NA | gene363 | GeneID:105276248 | LOC105276248 | Gene | LOC105276248 | protein_coding | NA | character(0) | character(0) | NW_011924888.1 | 1237975 | 1237981 | 7 | * | 7 | 1000 |
| NW_011924889.1 | 138414 | 140469 | 2056 | - | Gnomon | gene | NA | NA | gene455 | GeneID:105277237 | LOC105277237 | Gene | LOC105277237 | protein_coding | NA | character(0) | character(0) | NW_011924889.1 | 138461 | 138466 | 6 | * | 6 | 1000 |
| NW_011924889.1 | 237492 | 240474 | 2983 | + | Gnomon | gene | NA | NA | gene478 | GeneID:105277523 | LOC105277523 | Gene | LOC105277523 | protein_coding | NA | character(0) | character(0) | NW_011924889.1 | 238756 | 238761 | 6 | * | 6 | 1000 |
| NW_011924892.1 | 1124810 | 1127611 | 2802 | - | Gnomon | gene | NA | NA | gene564 | GeneID:105278478 | LOC105278478 | Gene | LOC105278478 | protein_coding | NA | character(0) | character(0) | NW_011924892.1 | 1125399 | 1125404 | 6 | * | 6 | 1000 |
| NW_011924892.1 | 1158089 | 1160303 | 2215 | - | Gnomon | gene | NA | NA | gene569 | GeneID:105278527 | LOC105278527 | Gene | LOC105278527 | protein_coding | NA | character(0) | character(0) | NW_011924892.1 | 1159381 | 1159386 | 6 | * | 6 | 1000 |
| NW_011924904.1 | 855071 | 858056 | 2986 | - | Gnomon | gene | NA | NA | gene755 | GeneID:105280619 | LOC105280619 | Gene | LOC105280619 | protein_coding | NA | character(0) | character(0) | NW_011924904.1 | 856949 | 856954 | 6 | * | 6 | 1000 |
| NW_011924905.1 | 521528 | 526485 | 4958 | - | Gnomon | gene | NA | NA | gene796 | GeneID:105281111 | LOC105281111 | Gene | LOC105281111 | protein_coding | NA | character(0) | character(0) | NW_011924905.1 | 525144 | 525149 | 6 | * | 6 | 1000 |
| NW_011924906.1 | 8168 | 13592 | 5425 | - | Gnomon | gene | NA | NA | gene801 | GeneID:105281166 | LOC105281166 | Gene | LOC105281166 | protein_coding | NA | character(0) | character(0) | NW_011924906.1 | 12384 | 12389 | 6 | * | 6 | 1000 |
| NW_011924907.1 | 997121 | 998515 | 1395 | - | Gnomon | gene | NA | NA | gene870 | GeneID:105281848 | LOC105281848 | Gene | LOC105281848 | protein_coding | NA | character(0) | character(0) | NW_011924907.1 | 997828 | 997833 | 6 | * | 6 | 1000 |
| NW_011924914.1 | 259948 | 262322 | 2375 | - | Gnomon | gene | NA | NA | gene952 | GeneID:105283016 | LOC105283016 | Gene | LOC105283016 | protein_coding | NA | character(0) | character(0) | NW_011924914.1 | 261398 | 261403 | 6 | * | 6 | 1000 |
| NW_011924931.1 | 214221 | 216344 | 2124 | + | Gnomon | gene | NA | NA | gene1375 | GeneID:105287333 | LOC105287333 | Gene | LOC105287333 | protein_coding | NA | character(0) | character(0) | NW_011924931.1 | 215617 | 215622 | 6 | * | 6 | 1000 |
| NW_011924931.1 | 2973218 | 2974707 | 1490 | + | Gnomon | gene | NA | NA | gene1499 | GeneID:105287627 | LOC105287627 | Gene | LOC105287627 | protein_coding | NA | character(0) | character(0) | NW_011924931.1 | 2974343 | 2974348 | 6 | * | 6 | 1000 |
| NW_011924931.1 | 3318371 | 3321702 | 3332 | - | Gnomon | gene | NA | NA | gene1504 | GeneID:105287632 | LOC105287632 | Gene | LOC105287632 | protein_coding | NA | character(0) | character(0) | NW_011924931.1 | 3320449 | 3320454 | 6 | * | 6 | 1000 |
| NW_011924931.1 | 3689781 | 3693711 | 3931 | + | Gnomon | gene | NA | NA | gene1514 | GeneID:105287646 | LOC105287646 | Gene | LOC105287646 | protein_coding | NA | character(0) | character(0) | NW_011924931.1 | 3690689 | 3690694 | 6 | * | 6 | 1000 |
| NW_011924931.1 | 3778005 | 3781577 | 3573 | - | Gnomon | gene | NA | NA | gene1536 | GeneID:105287663 | LOC105287663 | Gene | LOC105287663 | protein_coding | NA | character(0) | character(0) | NW_011924931.1 | 3780672 | 3780677 | 6 | * | 6 | 1000 |
| NW_011924931.1 | 3888281 | 3891433 | 3153 | + | Gnomon | gene | NA | NA | gene1547 | GeneID:105287677 | LOC105287677 | Gene | LOC105287677 | protein_coding | NA | character(0) | character(0) | NW_011924931.1 | 3889175 | 3889180 | 6 | * | 6 | 1000 |
| NW_011924931.1 | 3952621 | 3973270 | 20650 | + | Gnomon | gene | NA | NA | gene1560 | GeneID:105287694 | LOC105287694 | Gene | LOC105287694 | protein_coding | NA | character(0) | character(0) | NW_011924931.1 | 3966620 | 3966625 | 6 | * | 6 | 1000 |
| NW_011924931.1 | 3966137 | 3967627 | 1491 | - | Gnomon | gene | NA | NA | gene1562 | GeneID:105287692 | LOC105287692 | Gene | LOC105287692 | protein_coding | NA | character(0) | character(0) | NW_011924931.1 | 3966620 | 3966625 | 6 | * | 6 | 1000 |
| NW_011924931.1 | 4065100 | 4067558 | 2459 | - | Gnomon | gene | NA | NA | gene1584 | GeneID:105287715 | LOC105287715 | Gene | LOC105287715 | protein_coding | NA | character(0) | character(0) | NW_011924931.1 | 4066201 | 4066206 | 6 | * | 6 | 1000 |
| NW_011924931.1 | 4488185 | 4490767 | 2583 | + | Gnomon | gene | NA | NA | gene1620 | GeneID:105287754 | LOC105287754 | Gene | LOC105287754 | protein_coding | NA | character(0) | character(0) | NW_011924931.1 | 4489292 | 4489297 | 6 | * | 6 | 1000 |
| NW_011924932.1 | 543289 | 544100 | 812 | + | Gnomon | gene | NA | NA | gene1656 | GeneID:105287814 | LOC105287814 | Gene | LOC105287814 | protein_coding | NA | character(0) | character(0) | NW_011924932.1 | 543633 | 543638 | 6 | * | 6 | 1000 |
| NW_011924940.1 | 334573 | 349050 | 14478 | - | Gnomon | gene | NA | NA | gene1863 | GeneID:105288044 | LOC105288044 | Gene | LOC105288044 | protein_coding | NA | character(0) | character(0) | NW_011924940.1 | 347385 | 347390 | 6 | * | 6 | 1000 |
| NW_011924945.1 | 16609 | 22779 | 6171 | - | Gnomon | gene | NA | NA | gene1878 | GeneID:105288065 | LOC105288065 | Gene | LOC105288065 | protein_coding | NA | character(0) | character(0) | NW_011924945.1 | 20640 | 20645 | 6 | * | 6 | 1000 |
| NW_011924946.1 | 775753 | 777201 | 1449 | - | Gnomon | gene | NA | NA | gene1993 | GeneID:105288193 | LOC105288193 | Gene | LOC105288193 | protein_coding | NA | | | | | | | | | |

character(0) character(0) NW_011924946.1 776368 776373 6 * 6 1000
NW_011924946.1 786370 788739 2370 - Gnomon gene NA NA gene1995 GeneID:105288196 LOC105288196 Gene LOC105288196 protein_coding NA
character(0) character(0) NW_011924946.1 787444 787449 6 * 6 1000
NW_011924946.1 1411689 1414880 3192 - Gnomon gene NA NA gene2041 GeneID:105288248 LOC105288248 Gene LOC105288248 protein_coding NA
character(0) character(0) NW_011924946.1 1413935 1413940 6 * 6 1000
NW_011924946.1 1502005 1507040 5036 - Gnomon gene NA NA gene2047 GeneID:105274446 LOC105274446 Gene LOC105274446 protein_coding NA
character(0) character(0) NW_011924946.1 1502183 1502188 6 * 6 1000
NW_011924946.1 2832086 2841975 9890 - Gnomon gene NA NA gene2162 GeneID:105274578 LOC105274578 Gene LOC105274578 protein_coding NA
character(0) character(0) NW_011924946.1 2839967 2839972 6 * 6 1000
NW_011924946.1 3044550 3047207 2658 + Gnomon gene NA NA gene2194 GeneID:105274613 LOC105274613 Gene LOC105274613 protein_coding NA
character(0) character(0) NW_011924946.1 3045669 3045674 6 * 6 1000
NW_011924946.1 3306018 3309376 3359 - Gnomon gene NA NA gene2251 GeneID:105274673 LOC105274673 Gene LOC105274673 protein_coding NA
character(0) character(0) NW_011924946.1 3307400 3307405 6 * 6 1000
NW_011924951.1 64945 71389 6445 - Gnomon gene NA NA gene2323 GeneID:105274768 LOC105274768 Gene LOC105274768 protein_coding NA character(0)
character(0) NW_011924951.1 69822 69827 6 * 6 1000
NW_011924955.1 604490 614233 9744 - Gnomon gene NA NA gene2480 GeneID:105275005 LOC105275005 Gene LOC105275005 protein_coding NA
character(0) character(0) NW_011924955.1 613098 613103 6 * 6 1000
NW_011924956.1 717757 722032 4276 + Gnomon gene NA NA gene2570 GeneID:105275036 LOC105275036 Gene LOC105275036 protein_coding NA
character(0) character(0) NW_011924956.1 719335 719344 10 * 10 1000
NW_011924956.1 789246 793927 4682 + Gnomon gene NA NA gene2578 GeneID:105275047 LOC105275047 Gene LOC105275047 protein_coding NA
character(0) character(0) NW_011924956.1 790656 790661 6 * 6 1000
NW_011924973.1 236871 239530 2660 - Gnomon gene NA NA gene2751 GeneID:105275264 LOC105275264 Gene LOC105275264 protein_coding NA
character(0) character(0) NW_011924973.1 238411 238416 6 * 6 1000
NW_011924973.1 244133 245537 1405 + Gnomon gene NA NA gene2754 GeneID:105275270 LOC105275270 Gene LOC105275270 protein_coding NA
character(0) character(0) NW_011924973.1 244951 244956 6 * 6 1000
NW_011924973.1 828831 835120 6290 + Gnomon gene NA NA gene2799 GeneID:105275316 LOC105275316 Gene LOC105275316 protein_coding NA
character(0) character(0) NW_011924973.1 829674 829679 6 * 6 1000
NW_011924973.1 935440 945867 10428 + Gnomon gene NA NA gene2820 GeneID:105275342 LOC105275342 Gene LOC105275342 protein_coding NA
character(0) character(0) NW_011924973.1 936776 936781 6 * 6 1000
NW_011924973.1 983252 986808 3557 - Gnomon gene NA NA gene2832 GeneID:105275350 LOC105275350 Gene LOC105275350 protein_coding NA
character(0) character(0) NW_011924973.1 985890 985895 6 * 6 1000
NW_011924973.1 1001098 1003969 2872 + Gnomon gene NA NA gene2837 GeneID:105275355 LOC105275355 Gene LOC105275355 protein_coding NA
character(0) character(0) NW_011924973.1 1002105 1002110 6 * 6 1000
NW_011924977.1 31430 34619 3190 + Gnomon gene NA NA gene2916 GeneID:109610741 LOC109610741 Gene LOC109610741 protein_coding NA character(0)
character(0) NW_011924977.1 32395 32400 6 * 6 1000
NW_011924977.1 516991 520258 3268 + Gnomon gene NA NA gene2981 GeneID:105275526 LOC105275526 Gene LOC105275526 protein_coding NA
character(0) character(0) NW_011924977.1 519073 519078 6 * 6 1000
NW_011924977.1 662427 665182 2756 + Gnomon gene NA NA gene3008 GeneID:105275550 LOC105275550 Gene LOC105275550 protein_coding NA
character(0) character(0) NW_011924977.1 663534 663539 6 * 6 1000
NW_011924977.1 1463641 1466501 2861 + Gnomon gene NA NA gene3074 GeneID:105275617 LOC105275617 Gene LOC105275617 protein_coding NA
character(0) character(0) NW_011924977.1 1465037 1465042 6 * 6 1000
NW_011924977.1 3723831 3727852 4022 - Gnomon gene NA NA gene3212 GeneID:105275760 LOC105275760 Gene LOC105275760 protein_coding NA
character(0) character(0) NW_011924977.1 3727104 3727109 6 * 6 1000
NW_011924977.1 3728770 3731404 2635 + Gnomon gene NA NA gene3213 GeneID:105275761 LOC105275761 Gene LOC105275761 protein_coding NA
character(0) character(0) NW_011924977.1 3729892 3729897 6 * 6 1000
NW_011924979.1 16258 20038 3781 + Gnomon gene NA NA gene3218 GeneID:105275781 LOC105275781 Gene LOC105275781 protein_coding NA character(0)
character(0) NW_011924979.1 17774 17779 6 * 6 1000
NW_011924979.1 1319139 1322882 3744 - Gnomon gene NA NA gene3286 GeneID:105275848 LOC105275848 Gene LOC105275848 protein_coding NA
character(0) character(0) NW_011924979.1 1321281 1321286 6 * 6 1000
NW_011924979.1 1593634 1596428 2795 + Gnomon gene NA NA gene3297 GeneID:105275861 LOC105275861 Gene LOC105275861 protein_coding NA
character(0) character(0) NW_011924979.1 1594463 1594468 6 * 6 1000
NW_011924979.1 1988584 1990897 2314 + Gnomon gene NA NA gene3314 GeneID:105275878 LOC105275878 Gene LOC105275878 protein_coding NA
character(0) character(0) NW_011924979.1 1989444 1989449 6 * 6 1000
NW_011924979.1 2744383 2746071 1689 + Gnomon gene NA NA gene3334 GeneID:105275897 LOC105275897 Gene LOC105275897 protein_coding NA
character(0) character(0) NW_011924979.1 2745453 2745458 6 * 6 1000
NW_011924996.1 75277 77259 1983 + Gnomon gene NA NA gene3743 GeneID:105276382 LOC105276382 Gene LOC105276382 protein_coding NA character(0)
character(0) NW_011924996.1 76074 76079 6 * 6 1000
NW_011924996.1 1822758 1830685 7928 - Gnomon gene NA NA gene3885 GeneID:105276527 LOC105276527 Gene LOC105276527 protein_coding NA
character(0) character(0) NW_011924996.1 1829379 1829384 6 * 6 1000
NW_011924999.1 2422 6583 4162 + Gnomon gene NA NA gene3908 GeneID:105276559 LOC105276559 Gene LOC105276559 protein_coding NA character(0)
character(0) NW_011924999.1 3525 3530 6 * 6 1000
NW_011925007.1 926256 929694 3439 + Gnomon gene NA NA gene4194 GeneID:105276881 LOC105276881 Gene LOC105276881 protein_coding NA
character(0) character(0) NW_011925007.1 927466 927471 6 * 6 1000
NW_011925007.1 1370227 1386638 16412 - Gnomon gene NA NA gene4246 GeneID:105276941 LOC105276941 Gene LOC105276941 protein_coding NA

character(0) character(0) NW_011925007.1 1385156 1385161 6 * 6 1000
NW_011925007.1 1454403 1457778 3376 + Gnomon gene NA NA gene4253 GeneID:105276950 LOC105276950 Gene LOC105276950 protein_coding NA
character(0) character(0) NW_011925007.1 1455553 1455558 6 * 6 1000
NW_011925007.1 3769589 3777100 7512 + Gnomon gene NA NA gene4385 GeneID:105277094 LOC105277094 Gene LOC105277094 protein_coding NA
character(0) character(0) NW_011925007.1 3770514 3770519 6 * 6 1000
NW_011925012.1 1356105 1358935 2831 + Gnomon gene NA NA gene4498 GeneID:105277239 LOC105277239 Gene LOC105277239 protein_coding NA
character(0) character(0) NW_011925012.1 1357276 1357281 6 * 6 1000
NW_011925016.1 800369 803160 2792 - Gnomon gene NA NA gene4610 GeneID:105277357 LOC105277357 Gene LOC105277357 protein_coding NA
character(0) character(0) NW_011925016.1 801483 801490 8 * 8 1000
NW_011925019.1 1195331 1199707 4377 - Gnomon gene NA NA gene4693 GeneID:105277452 LOC105277452 Gene LOC105277452 protein_coding NA
character(0) character(0) NW_011925019.1 1198913 1198918 6 * 6 1000
NW_011925020.1 469190 474352 5163 + Gnomon gene NA NA gene4780 GeneID:105277552 LOC105277552 Gene LOC105277552 protein_coding NA
character(0) character(0) NW_011925020.1 470416 470421 6 * 6 1000
NW_011925020.1 3366359 3368752 2394 + Gnomon gene NA NA gene4926 GeneID:105277697 LOC105277697 Gene LOC105277697 protein_coding NA
character(0) character(0) NW_011925020.1 3367088 3367093 6 * 6 1000
NW_011925022.1 386851 387976 1126 - Gnomon gene NA NA gene5000 GeneID:105277804 LOC105277804 Gene LOC105277804 protein_coding NA
character(0) character(0) NW_011925022.1 387077 387082 6 * 6 1000
NW_011925035.1 818580 820886 2307 - Gnomon gene NA NA gene5379 GeneID:105278234 LOC105278234 Gene LOC105278234 protein_coding NA
character(0) character(0) NW_011925035.1 820260 820265 6 * 6 1000
NW_011925050.1 266014 271068 5055 + Gnomon gene NA NA gene5758 GeneID:105278681 LOC105278681 Gene LOC105278681 protein_coding NA
character(0) character(0) NW_011925050.1 266840 266845 6 * 6 1000
NW_011925050.1 692537 695061 2525 - Gnomon gene NA NA gene5769 GeneID:105278683 LOC105278683 Gene LOC105278683 protein_coding NA
character(0) character(0) NW_011925050.1 693967 693972 6 * 6 1000
NW_011925051.1 754067 756870 2804 - Gnomon gene NA NA gene5860 GeneID:105278789 LOC105278789 Gene LOC105278789 protein_coding NA
character(0) character(0) NW_011925051.1 755643 755648 6 * 6 1000
NW_011925053.1 636493 648405 11913 - Gnomon gene NA NA gene5881 GeneID:105278811 LOC105278811 Gene LOC105278811 protein_coding NA
character(0) character(0) NW_011925053.1 647315 647320 6 * 6 1000
NW_011925053.1 649052 656133 7082 - Gnomon gene NA NA gene5882 GeneID:105278812 LOC105278812 Gene LOC105278812 protein_coding NA
character(0) character(0) NW_011925053.1 655139 655144 6 * 6 1000
NW_011925053.1 1481875 1483457 1583 - Gnomon gene NA NA gene5932 GeneID:105278862 LOC105278862 Gene LOC105278862 protein_coding NA
character(0) character(0) NW_011925053.1 1482901 1482906 6 * 6 1000
NW_011925067.1 201362 204847 3486 - Gnomon gene NA NA gene6034 GeneID:105279004 LOC105279004 Gene LOC105279004 protein_coding NA
character(0) character(0) NW_011925067.1 203691 203696 6 * 6 1000
NW_011925075.1 793827 800147 6321 + Gnomon gene NA NA gene6165 GeneID:105279155 LOC105279155 Gene LOC105279155 protein_coding NA
character(0) character(0) NW_011925075.1 794882 794887 6 * 6 1000
NW_011925086.1 128430 135139 6710 + Gnomon gene NA NA gene6311 GeneID:105279333 LOC105279333 Gene LOC105279333 protein_coding NA
character(0) character(0) NW_011925086.1 129637 129644 8 * 8 1000
NW_011925086.1 604599 609266 4668 + Gnomon gene NA NA gene6342 GeneID:105279367 LOC105279367 Gene LOC105279367 protein_coding NA
character(0) character(0) NW_011925086.1 606157 606162 6 * 6 1000
NW_011925099.1 12321 14914 2594 + Gnomon gene NA NA gene6401 GeneID:105279437 LOC105279437 Gene LOC105279437 protein_coding NA character(0)
character(0) NW_011925099.1 14885 14890 6 * 6 1000
NW_011925106.1 487858 493874 6017 + Gnomon gene NA NA gene6649 GeneID:105279717 LOC105279717 Gene LOC105279717 protein_coding NA
character(0) character(0) NW_011925106.1 489460 489465 6 * 6 1000
NW_011925106.1 1997159 1999737 2579 - Gnomon gene NA NA gene6753 GeneID:105279838 LOC105279838 Gene LOC105279838 protein_coding NA
character(0) character(0) NW_011925106.1 1998173 1998178 6 * 6 1000
NW_011925106.1 2086959 2096661 9703 + Gnomon gene NA NA gene6771 GeneID:105279854 LOC105279854 Gene LOC105279854 protein_coding NA
character(0) character(0) NW_011925106.1 2088114 2088121 8 * 8 1000
NW_011925106.1 2086959 2096661 9703 + Gnomon gene NA NA gene6771 GeneID:105279854 LOC105279854 Gene LOC105279854 protein_coding NA
character(0) character(0) NW_011925106.1 2088216 2088221 6 * 6 1000
NW_011925106.1 3275310 3277500 2191 - Gnomon gene NA NA gene6816 GeneID:105279900 LOC105279900 Gene LOC105279900 protein_coding NA
character(0) character(0) NW_011925106.1 3276041 3276046 6 * 6 1000
NW_011925106.1 3331206 3338136 6931 - Gnomon gene NA NA gene6822 GeneID:105279903 LOC105279903 Gene LOC105279903 protein_coding NA
character(0) character(0) NW_011925106.1 3336364 3336369 6 * 6 1000
NW_011925117.1 94064 101904 7841 - Gnomon gene NA NA gene6900 GeneID:105280035 LOC105280035 Gene LOC105280035 protein_coding NA
character(0) character(0) NW_011925117.1 100585 100590 6 * 6 1000
NW_011925118.1 771350 783967 12618 - Gnomon gene NA NA gene6991 GeneID:105280128 LOC105280128 Gene LOC105280128 protein_coding NA
character(0) character(0) NW_011925118.1 783338 783343 6 * 6 1000
NW_011925128.1 1875336 1880845 5510 + Gnomon gene NA NA gene7225 GeneID:105280395 LOC105280395 Gene LOC105280395 protein_coding NA
character(0) character(0) NW_011925128.1 1876634 1876639 6 * 6 1000
NW_011925128.1 2152768 2155243 2476 - Gnomon gene NA NA gene7252 GeneID:105280424 LOC105280424 Gene LOC105280424 protein_coding NA
character(0) character(0) NW_011925128.1 2153538 2153543 6 * 6 1000
NW_011925142.1 743409 745933 2525 - Gnomon gene NA NA gene7568 GeneID:105280786 LOC105280786 Gene LOC105280786 protein_coding NA
character(0) character(0) NW_011925142.1 744366 744371 6 * 6 1000
NW_011925142.1 777887 780161 2275 - Gnomon gene NA NA gene7574 GeneID:105280791 LOC105280791 Gene LOC105280791 protein_coding NA

character(0) character(0) NW_011925142.1 779002 779007 6 * 6 1000
NW_011925150.1 76140 79045 2906 - Gnomon gene NA NA gene7626 GeneID:105280856 LOC105280856 Gene LOC105280856 protein_coding NA character(0)
character(0) NW_011925150.1 78377 78382 6 * 6 1000
NW_011925151.1 38146 39177 1032 - Gnomon gene NA NA gene7635 GeneID:105280860 LOC105280860 Gene LOC105280860 protein_coding NA character(0)
character(0) NW_011925151.1 38257 38262 6 * 6 1000
NW_011925161.1 128427 131377 2951 + Gnomon gene NA NA gene7706 GeneID:105280950 LOC105280950 Gene LOC105280950 protein_coding NA
character(0) character(0) NW_011925161.1 129875 129880 6 * 6 1000
NW_011925161.1 145622 148482 2861 - Gnomon gene NA NA gene7710 GeneID:105280955 LOC105280955 Gene LOC105280955 protein_coding NA
character(0) character(0) NW_011925161.1 147171 147176 6 * 6 1000
NW_011925161.1 1702331 1707999 5669 - Gnomon gene NA NA gene7821 GeneID:105281067 LOC105281067 Gene LOC105281067 protein_coding NA
character(0) character(0) NW_011925161.1 1707165 1707170 6 * 6 1000
NW_011925161.1 1705906 1731780 25875 + Gnomon gene NA NA gene7822 GeneID:105281066 LOC105281066 Gene LOC105281066 protein_coding NA
character(0) character(0) NW_011925161.1 1707165 1707170 6 * 6 1000
NW_011925161.1 1984768 1994445 9678 + Gnomon gene NA NA gene7841 GeneID:105281080 LOC105281080 Gene LOC105281080 protein_coding NA
character(0) character(0) NW_011925161.1 1986883 1986888 6 * 6 1000
NW_011925161.1 2209807 2215262 5456 - Gnomon gene NA NA gene7866 GeneID:105281115 LOC105281115 Gene LOC105281115 protein_coding NA
character(0) character(0) NW_011925161.1 2212367 2212372 6 * 6 1000
NW_011925163.1 470647 473680 3034 + Gnomon gene NA NA gene8040 GeneID:105281341 LOC105281341 Gene LOC105281341 protein_coding NA
character(0) character(0) NW_011925163.1 471213 471218 6 * 6 1000
NW_011925182.1 57229 59555 2327 - Gnomon gene NA NA gene8209 GeneID:105281549 LOC105281549 Gene LOC105281549 protein_coding NA character(0)
character(0) NW_011925182.1 58922 58927 6 * 6 1000
NW_011925182.1 162285 170775 8491 + Gnomon gene NA NA gene8218 GeneID:105281562 LOC105281562 Gene LOC105281562 protein_coding NA
character(0) character(0) NW_011925182.1 163572 163577 6 * 6 1000
NW_011925183.1 39511 42630 3120 + Gnomon gene NA NA gene8236 GeneID:105281577 LOC105281577 Gene LOC105281577 protein_coding NA character(0)
character(0) NW_011925183.1 40051 40056 6 * 6 1000
NW_011925188.1 199014 202632 3619 - Gnomon gene NA NA gene8257 GeneID:105281603 LOC105281603 Gene LOC105281603 protein_coding NA
character(0) character(0) NW_011925188.1 201626 201631 6 * 6 1000
NW_011925189.1 165659 175663 10005 - Gnomon gene NA NA gene8294 GeneID:105281645 LOC105281645 Gene LOC105281645 protein_coding NA
character(0) character(0) NW_011925189.1 174276 174281 6 * 6 1000
NW_011925189.1 575609 577709 2101 - Gnomon gene NA NA gene8332 GeneID:105281690 LOC105281690 Gene LOC105281690 protein_coding NA
character(0) character(0) NW_011925189.1 577082 577087 6 * 6 1000
NW_011925189.1 633407 636090 2684 + Gnomon gene NA NA gene8339 GeneID:105281698 LOC105281698 Gene LOC105281698 protein_coding NA
character(0) character(0) NW_011925189.1 634540 634545 6 * 6 1000
NW_011925203.1 63038 97839 34802 + Gnomon gene NA NA gene8521 GeneID:105281906 LOC105281906 Gene LOC105281906 protein_coding NA character(0)
character(0) NW_011925203.1 64603 64608 6 * 6 1000
NW_011925216.1 136610 140476 3867 - Gnomon gene NA NA gene8735 GeneID:105282144 LOC105282144 Gene LOC105282144 protein_coding NA
character(0) character(0) NW_011925216.1 137522 137527 6 * 6 1000
NW_011925216.1 136795 145548 8754 + Gnomon gene NA NA gene8736 GeneID:105282142 LOC105282142 Gene LOC105282142 protein_coding NA
character(0) character(0) NW_011925216.1 137522 137527 6 * 6 1000
NW_011925216.1 156691 162086 5396 + Gnomon gene NA NA gene8741 GeneID:105282150 LOC105282150 Gene LOC105282150 protein_coding NA
character(0) character(0) NW_011925216.1 157541 157546 6 * 6 1000
NW_011925232.1 152153 159881 7729 + Gnomon gene NA NA gene8919 GeneID:105282356 LOC105282356 Gene LOC105282356 protein_coding NA
character(0) character(0) NW_011925232.1 153164 153169 6 * 6 1000
NW_011925239.1 3979 7808 3830 + Gnomon gene NA NA gene9001 GeneID:105282461 LOC105282461 Gene LOC105282461 protein_coding NA character(0)
character(0) NW_011925239.1 4877 4882 6 * 6 1000
NW_011925240.1 757237 760222 2986 - Gnomon gene NA NA gene9059 GeneID:105282525 LOC105282525 Gene LOC105282525 protein_coding NA
character(0) character(0) NW_011925240.1 758844 758849 6 * 6 1000
NW_011925240.1 2490948 2493602 2655 - Gnomon gene NA NA gene9134 GeneID:105282589 LOC105282589 Gene LOC105282589 protein_coding NA
character(0) character(0) NW_011925240.1 2492742 2492747 6 * 6 1000
NW_011925241.1 428008 431512 3505 + Gnomon gene NA NA gene9196 GeneID:105282662 LOC105282662 Gene LOC105282662 protein_coding NA
character(0) character(0) NW_011925241.1 428600 428605 6 * 6 1000
NW_011925258.1 22829 32812 9984 + Gnomon gene NA NA gene9308 GeneID:105282804 LOC105282804 Gene LOC105282804 protein_coding NA character(0)
character(0) NW_011925258.1 28153 28158 6 * 6 1000
NW_011925258.1 404301 408788 4488 - Gnomon gene NA NA gene9347 GeneID:105282847 LOC105282847 Gene LOC105282847 protein_coding NA
character(0) character(0) NW_011925258.1 407771 407776 6 * 6 1000
NW_011925304.1 378599 380691 2093 - Gnomon gene NA NA gene9814 GeneID:105283373 LOC105283373 Gene LOC105283373 protein_coding NA
character(0) character(0) NW_011925304.1 379611 379616 6 * 6 1000
NW_011925304.1 468966 476624 7659 - Gnomon gene NA NA gene9824 GeneID:105283383 LOC105283383 Gene LOC105283383 protein_coding NA
character(0) character(0) NW_011925304.1 475523 475528 6 * 6 1000
NW_011925321.1 975185 978704 3520 - Gnomon gene NA NA gene10031 GeneID:105283620 LOC105283620 Gene LOC105283620 protein_coding NA
character(0) character(0) NW_011925321.1 977422 977427 6 * 6 1000
NW_011925321.1 1107715 1109901 2187 - Gnomon gene NA NA gene10043 GeneID:105283634 LOC105283634 Gene LOC105283634 protein_coding NA
character(0) character(0) NW_011925321.1 1108690 1108695 6 * 6 1000
NW_011925323.1 2107588 2112140 4553 - Gnomon gene NA NA gene10169 GeneID:105283771 LOC105283771 Gene LOC105283771 protein_coding NA

character(0) character(0) NW_011925323.1 2111421 2111426 6 * 6 1000
NW_011925351.1 11934 15244 3311 - Gnomon gene NA NA gene10387 GeneID:105284041 LOC105284041 Gene LOC105284041 protein_coding NA character(0)
character(0) NW_011925351.1 14519 14524 6 * 6 1000
NW_011925353.1 26250 32260 6011 + Gnomon gene NA NA gene10402 GeneID:105284053 LOC105284053 Gene LOC105284053 protein_coding NA character(0)
character(0) NW_011925353.1 27469 27474 6 * 6 1000
NW_011925353.1 40476 43146 2671 - Gnomon gene NA NA gene10405 GeneID:105284057 LOC105284057 Gene LOC105284057 protein_coding NA character(0)
character(0) NW_011925353.1 41948 41953 6 * 6 1000
NW_011925355.1 581482 584005 2524 + Gnomon gene NA NA gene10482 GeneID:105284137 LOC105284137 Gene LOC105284137 protein_coding NA
character(0) character(0) NW_011925355.1 582446 582451 6 * 6 1000
NW_011925356.1 704771 710120 5350 - Gnomon gene NA NA gene10539 GeneID:105284201 LOC105284201 Gene LOC105284201 protein_coding NA
character(0) character(0) NW_011925356.1 708176 708181 6 * 6 1000
NW_011925356.1 1573184 1579703 6520 - Gnomon gene NA NA gene10596 GeneID:105284262 LOC105284262 Gene LOC105284262 protein_coding NA
character(0) character(0) NW_011925356.1 1573650 1573655 6 * 6 1000
NW_011925367.1 1119922 1121887 1966 + Gnomon gene NA NA gene10714 GeneID:105284396 LOC105284396 Gene LOC105284396 protein_coding NA
character(0) character(0) NW_011925367.1 1120960 1120965 6 * 6 1000
NW_011925371.1 357596 361737 4142 + Gnomon gene NA NA gene10820 GeneID:105284595 LOC105284595 Gene LOC105284595 protein_coding NA
character(0) character(0) NW_011925371.1 359444 359449 6 * 6 1000
NW_011925371.1 1203185 1204659 1475 - Gnomon gene NA NA gene10886 GeneID:105284582 LOC105284582 Gene LOC105284582 protein_coding NA
character(0) character(0) NW_011925371.1 1203884 1203889 6 * 6 1000
NW_011925387.1 39150 46759 7610 + Gnomon gene NA NA gene10928 GeneID:105284644 LOC105284644 Gene LOC105284644 protein_coding NA character(0)
character(0) NW_011925387.1 40039 40044 6 * 6 1000
NW_011925387.1 271672 273220 1549 + Gnomon gene NA NA gene10957 GeneID:105284676 LOC105284676 Gene LOC105284676 protein_coding NA
character(0) character(0) NW_011925387.1 272491 272496 6 * 6 1000
NW_011925401.1 10153 19053 8901 - Gnomon gene NA NA gene11099 GeneID:105284845 LOC105284845 Gene LOC105284845 protein_coding NA character(0)
character(0) NW_011925401.1 17311 17316 6 * 6 1000
NW_011925438.1 122592 124529 1938 + Gnomon gene NA NA gene11397 GeneID:105285205 LOC105285205 Gene LOC105285205 protein_coding NA
character(0) character(0) NW_011925438.1 123277 123282 6 * 6 1000
NW_011925442.1 70936 77041 6106 + Gnomon gene NA NA gene11466 GeneID:105285277 LOC105285277 Gene LOC105285277 protein_coding NA character(0)
character(0) NW_011925442.1 71492 71497 6 * 6 1000
NW_011925442.1 70999 75078 4080 - Gnomon gene NA NA gene11467 GeneID:105285278 LOC105285278 Gene LOC105285278 protein_coding NA character(0)
character(0) NW_011925442.1 71492 71497 6 * 6 1000
NW_011925459.1 186244 187554 1311 - Gnomon gene NA NA gene11569 GeneID:105285394 LOC105285394 Gene LOC105285394 protein_coding NA
character(0) character(0) NW_011925459.1 186501 186506 6 * 6 1000
NW_011925470.1 613911 617551 3641 - Gnomon gene NA NA gene11628 GeneID:105285465 LOC105285465 Gene LOC105285465 protein_coding NA
character(0) character(0) NW_011925470.1 616616 616621 6 * 6 1000
NW_011925490.1 137354 140755 3402 + Gnomon gene NA NA gene11689 GeneID:105285534 LOC105285534 Gene LOC105285534 protein_coding NA
character(0) character(0) NW_011925490.1 138283 138288 6 * 6 1000
NW_011925516.1 389602 391122 1521 - Gnomon gene NA NA gene11817 GeneID:105285670 LOC105285670 Gene LOC105285670 protein_coding NA
character(0) character(0) NW_011925516.1 390371 390376 6 * 6 1000
NW_011925595.1 1 4548 4548 - Gnomon gene NA NA gene12102 GeneID:105286003 LOC105286003 Gene LOC105286003 protein_coding NA character(0)
character(0) NW_011925595.1 3717 3722 6 * 6 1000
NW_011925653.1 98728 102889 4162 - Gnomon gene NA NA gene12259 GeneID:105286160 LOC105286160 Gene LOC105286160 protein_coding NA
character(0) character(0) NW_011925653.1 101522 101527 6 * 6 1000
NW_011925653.1 98728 102889 4162 - Gnomon gene NA NA gene12259 GeneID:105286160 LOC105286160 Gene LOC105286160 protein_coding NA
character(0) character(0) NW_011925653.1 101541 101546 6 * 6 1000
NW_011925664.1 38296 39300 1005 + Gnomon gene NA NA gene12311 GeneID:105286212 LOC105286212 Gene LOC105286212 protein_coding NA character(0)
character(0) NW_011925664.1 39054 39061 8 * 8 1000
NW_011925664.1 277932 283313 5382 + Gnomon gene NA NA gene12359 GeneID:105286364 LOC105286364 Gene LOC105286364 protein_coding NA
character(0) character(0) NW_011925664.1 279108 279113 6 * 6 1000
NW_011925664.1 580837 582496 1660 - Gnomon gene NA NA gene12411 GeneID:105286320 LOC105286320 Gene LOC105286320 protein_coding NA
character(0) character(0) NW_011925664.1 581347 581352 6 * 6 1000
NW_011925715.1 952620 956416 3797 - Gnomon gene NA NA gene12567 GeneID:105286509 LOC105286509 Gene LOC105286509 protein_coding NA
character(0) character(0) NW_011925715.1 955653 955658 6 * 6 1000
NW_011925810.1 23502 27182 3681 + Gnomon gene NA NA gene12829 GeneID:105286828 LOC105286828 Gene LOC105286828 protein_coding true character(0)
c("27182", ".") NW_011925810.1 24132 24137 6 * 6 1000
NW_011925919.1 45093 46724 1632 + Gnomon gene NA NA gene12928 GeneID:105286948 LOC105286948 Gene LOC105286948 protein_coding NA character(0)
character(0) NW_011925919.1 45973 45978 6 * 6 1000
NW_011925984.1 48360 54191 5832 + Gnomon gene NA NA gene12992 GeneID:105287013 LOC105287013 Gene LOC105287013 protein_coding NA character(0)
character(0) NW_011925984.1 49836 49841 6 * 6 1000
NW_011926135.1 70608 74554 3947 + Gnomon gene NA NA gene13076 GeneID:105287104 LOC105287104 Gene LOC105287104 protein_coding NA character(0)
character(0) NW_011926135.1 71338 71343 6 * 6 1000
NW_011929339.1 7 1883 1877 + Gnomon gene NA NA gene13384 GeneID:105287447 LOC105287447 Gene LOC105287447 protein_coding true character(0)
c("1883", ".") NW_011929339.1 1129 1134 6 * 6 1000
NW_011929339.1 7 1883 1877 + Gnomon gene NA NA gene13384 GeneID:105287447 LOC105287447 Gene LOC105287447 protein_coding true character(0)

| | | | | | | | |
|---|---|---|---|---|---|---|---|
| c("1883", ".") | NW_011929339.1 | 1180 | 1185 | 6 | * | 6 | 1000 |

```
Directory: MRPR File: rmp-Cb_LI_BCphase.txt

Overlap of methylation-rich regions with genome features for Cb sample Cb_LI_BCphase

Total size of the methylation-rich regions of Cb_LI_BCphase:                                       890 bp (  0.0% of genome)
Size of overlap of methylation-rich regions of Cb Cb_LI_BCphase with genic regions:                   866 bp ( 97.3% of total size)   ( 1.36 O/E)
Size of overlap of methylation-rich regions of Cb Cb_LI_BCphase with   exon regions:                    760 bp ( 85.4% of total size) ( 0.64 O/E)
Size of overlap of methylation-rich regions of Cb Cb_LI_BCphase with   intron regions:                  106 bp ( 11.9% of total size) ( 0.22 O/E)
Size of overlap of methylation-rich regions of Cb Cb_LI_BCphase with intergenic regions:               24 bp (  2.7% of total size)   ( 3.44 O/E)
Size of overlap of methylation-rich regions of Cb Cb_LI_BCphase with   promoter regions:                  0 bp (  0.0% of total size) ( 0.00 O/E)
Size of overlap of methylation-rich regions of Cb Cb_LI_BCphase with   other intergenic regions:         24 bp (  2.7% of total size) ( 0.11 O/E)

Overlap of methylation-poor regions with genome features for Cb sample Cb_LI_BCphase

Total size of the methylation-poor regions of Cb_LI_BCphase:                                   4616352 bp (  2.2% of genome)
Size of overlap of methylation-poor regions of Cb Cb_LI_BCphase with genic regions:               2907433 bp ( 63.0% of total size)   ( 0.88 O/E)
Size of overlap of methylation-poor regions of Cb Cb_LI_BCphase with   exon regions:                 635232 bp ( 13.8% of total size) ( 2.63 O/E)
Size of overlap of methylation-poor regions of Cb Cb_LI_BCphase with   intron regions:              2272201 bp ( 49.2% of total size) ( 0.93 O/E)
Size of overlap of methylation-poor regions of Cb Cb_LI_BCphase with intergenic regions:          1708919 bp ( 37.0% of total size)   ( 2.23 O/E)
Size of overlap of methylation-poor regions of Cb Cb_LI_BCphase with   promoter regions:              72740 bp (  1.6% of total size) ( 0.51 O/E)
Size of overlap of methylation-poor regions of Cb Cb_LI_BCphase with   other intergenic regions:    1636179 bp ( 35.4% of total size) ( 1.41 O/E)
```

```
Directory: MRPR File: rmp-Cb_LI_Rphase.txt

Overlap of methylation-rich regions with genome features for Cb sample Cb_LI_Rphase

Total size of the methylation-rich regions of Cb_LI_Rphase:                                       955 bp (  0.0% of genome)
Size of overlap of methylation-rich regions of Cb Cb_LI_Rphase with genic regions:                   937 bp ( 98.1% of total size)   ( 1.37 O/E)
Size of overlap of methylation-rich regions of Cb Cb_LI_Rphase with   exon regions:                    819 bp ( 85.8% of total size) ( 0.66 O/E)
Size of overlap of methylation-rich regions of Cb Cb_LI_Rphase with   intron regions:                  118 bp ( 12.4% of total size) ( 0.23 O/E)
Size of overlap of methylation-rich regions of Cb Cb_LI_Rphase with intergenic regions:               18 bp (  1.9% of total size)   ( 3.47 O/E)
Size of overlap of methylation-rich regions of Cb Cb_LI_Rphase with   promoter regions:                  0 bp (  0.0% of total size) ( 0.00 O/E)
Size of overlap of methylation-rich regions of Cb Cb_LI_Rphase with   other intergenic regions:         18 bp (  1.9% of total size) ( 0.07 O/E)

Overlap of methylation-poor regions with genome features for Cb sample Cb_LI_Rphase

Total size of the methylation-poor regions of Cb_LI_Rphase:                                   3699834 bp (  1.7% of genome)
Size of overlap of methylation-poor regions of Cb Cb_LI_Rphase with genic regions:               2452008 bp ( 66.3% of total size)   ( 0.92 O/E)
Size of overlap of methylation-poor regions of Cb Cb_LI_Rphase with   exon regions:                 405382 bp ( 11.0% of total size) ( 2.96 O/E)
Size of overlap of methylation-poor regions of Cb Cb_LI_Rphase with   intron regions:              2046626 bp ( 55.3% of total size) ( 1.04 O/E)
Size of overlap of methylation-poor regions of Cb Cb_LI_Rphase with intergenic regions:          1247826 bp ( 33.7% of total size)   ( 2.35 O/E)
Size of overlap of methylation-poor regions of Cb Cb_LI_Rphase with   promoter regions:               65741 bp (  1.8% of total size) ( 0.58 O/E)
Size of overlap of methylation-poor regions of Cb Cb_LI_Rphase with   other intergenic regions:     1182085 bp ( 31.9% of total size) ( 1.27 O/E)
```

Cb_LI_BCphase
1−distance distributions

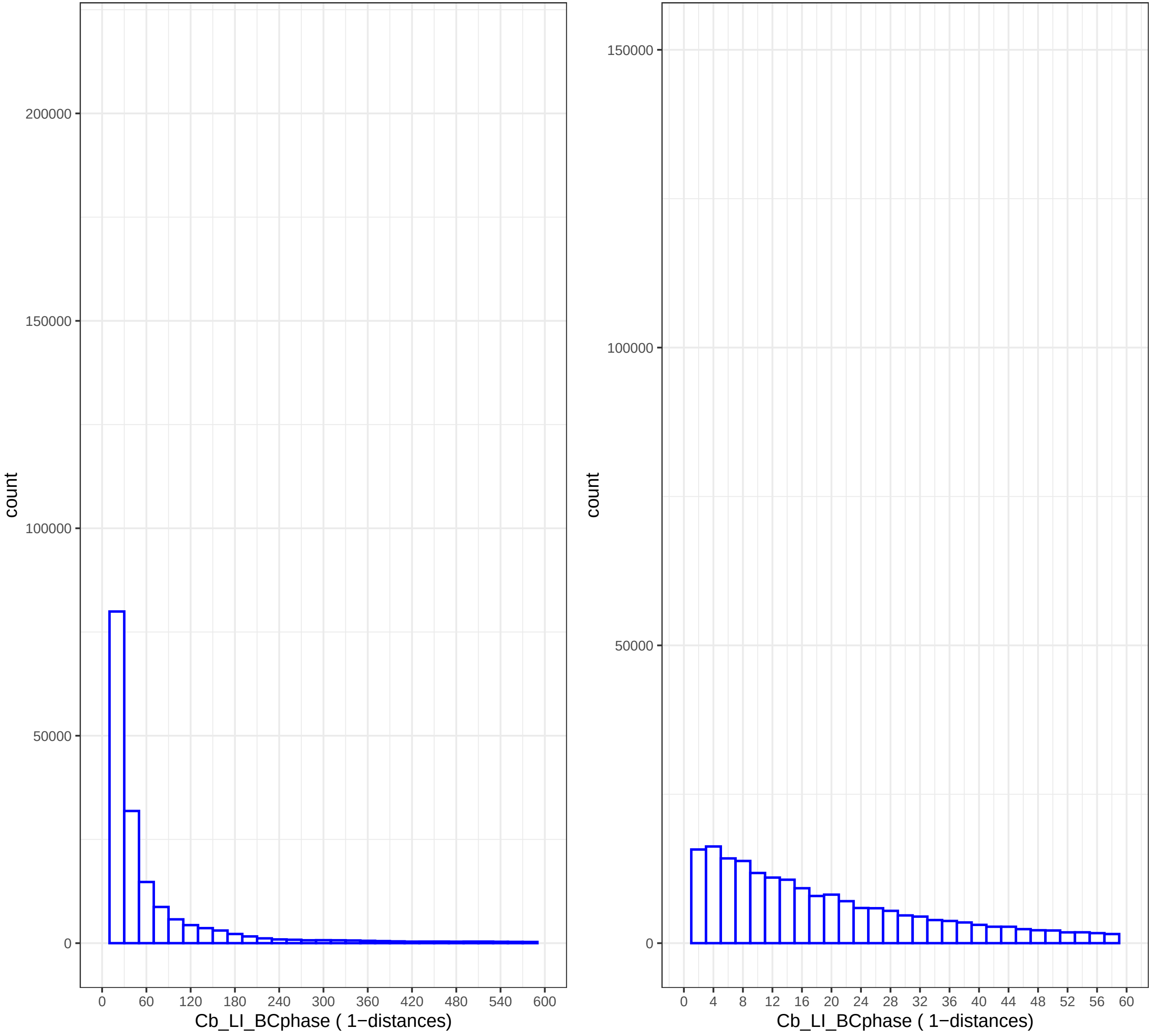

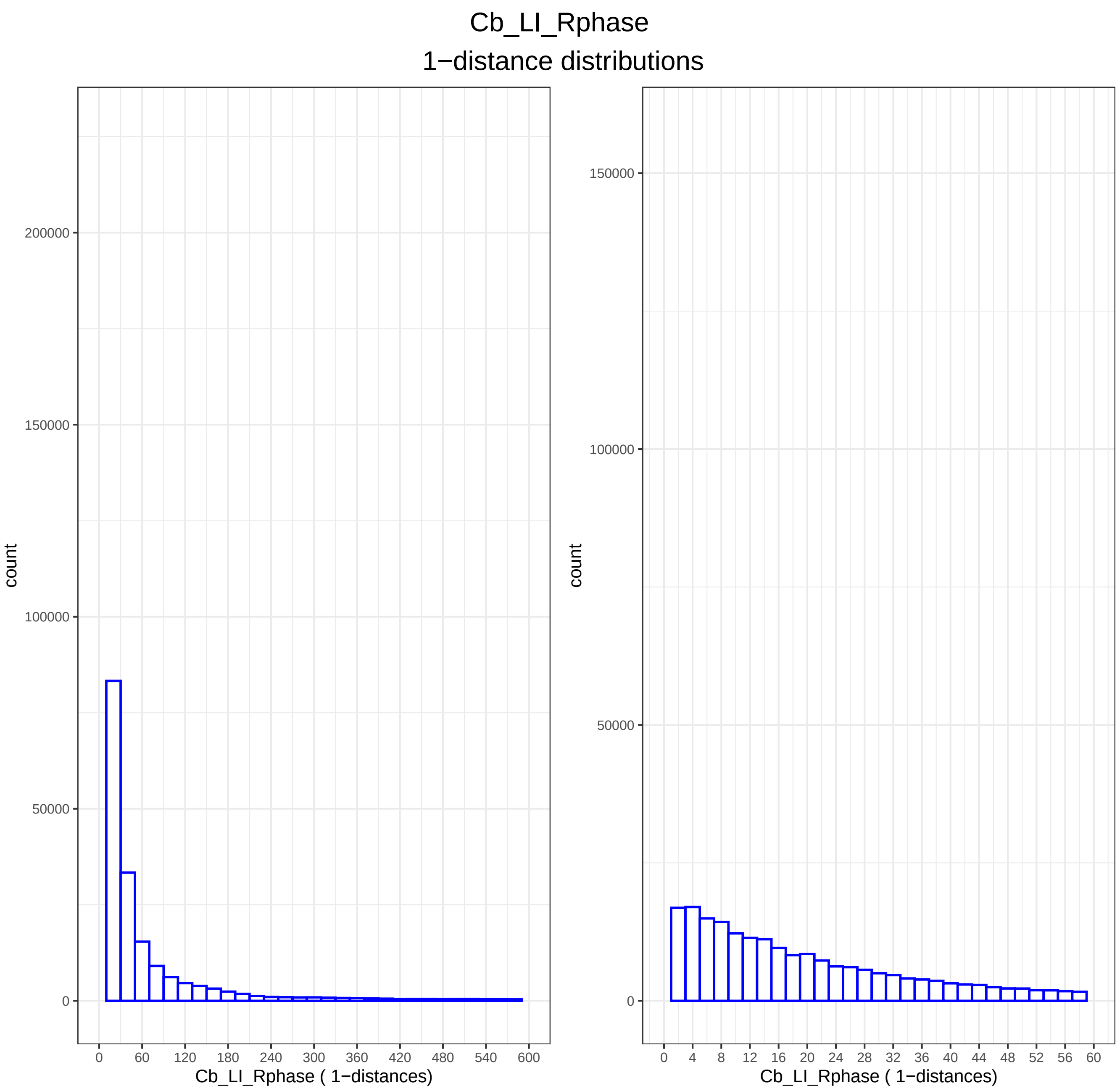

Cb_LI_Rphase
1−distance distributions
count
Cb_LI_Rphase ( 1−distances)
0
50000
100000
150000
200000
0
60
120
180
240
300
360
420
480
540
600
count
Cb_LI_Rphase ( 1−distances)
0
50000
100000
150000
0
4
8
12
16
20
24
28
32
36
40
44
48
52
56
60

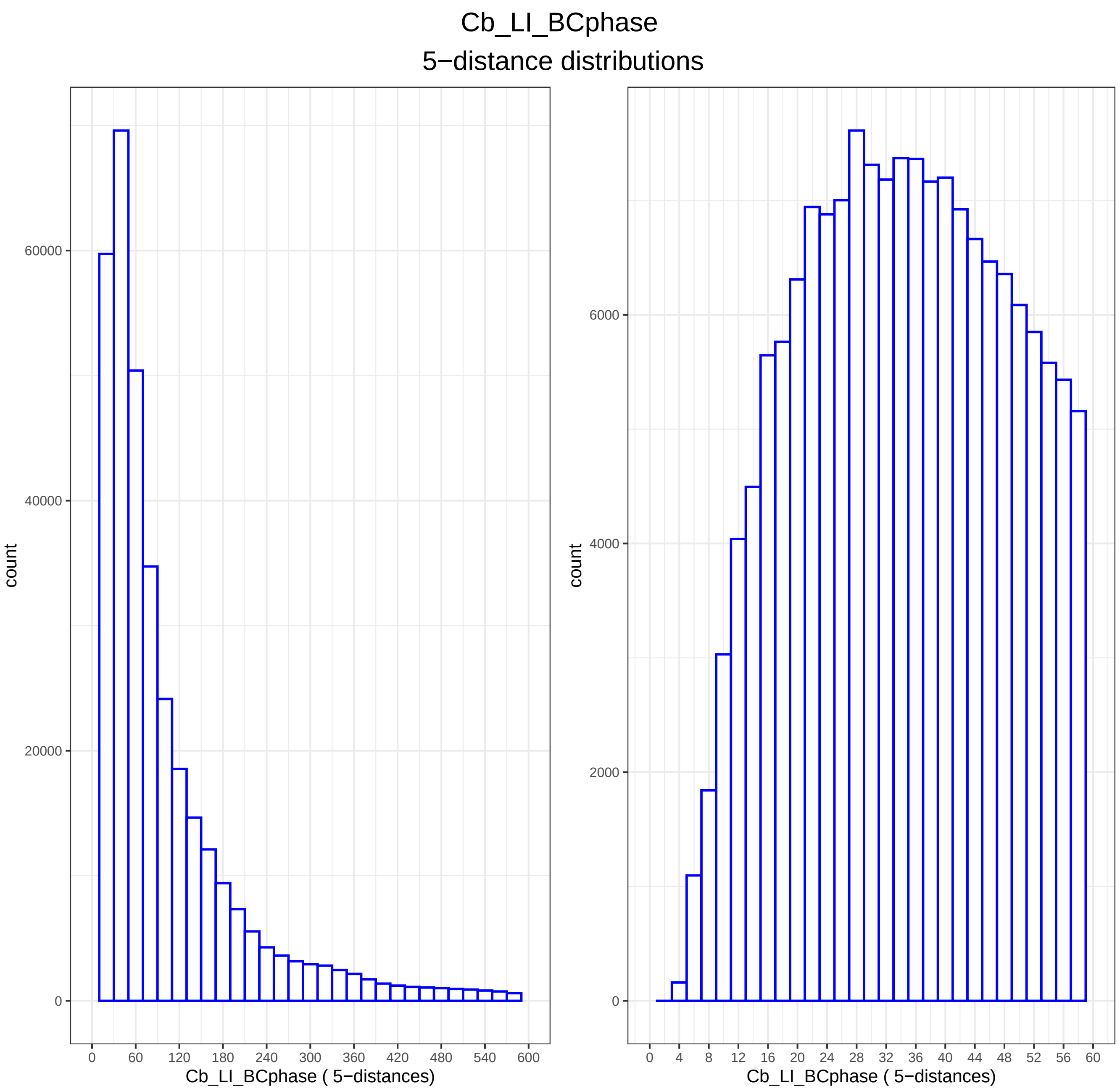

Cb_LI_BCphase
5−distance distributions
count
60000
40000
20000
0
0 60 120 180 240 300 360 420 480 540 600
Cb_LI_BCphase ( 5−distances)
count
6000
4000
2000
0
0 4 8 12 16 20 24 28 32 36 40 44 48 52 56 60
Cb_LI_BCphase ( 5−distances)

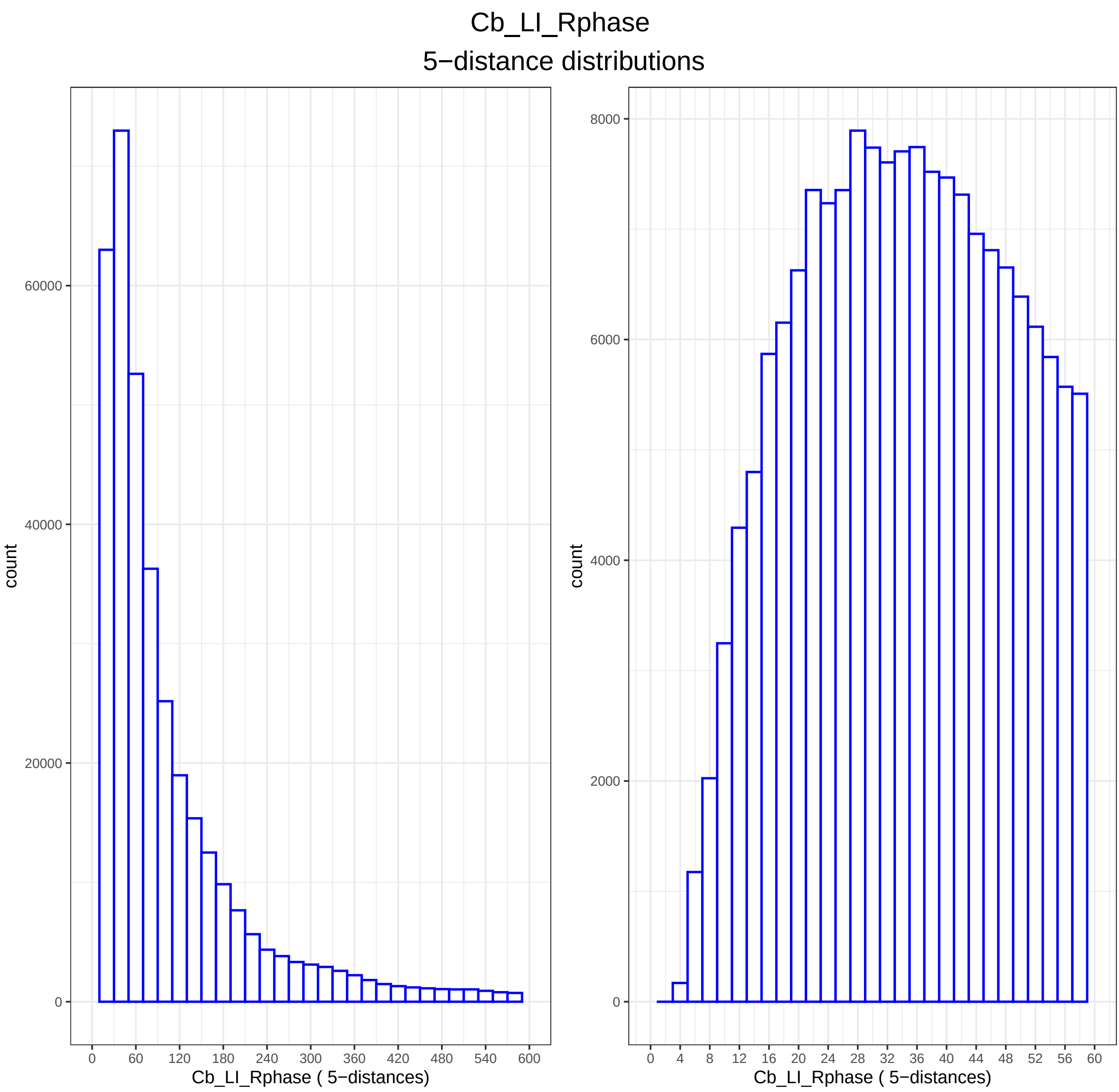

Cb_LI_Rphase
5−distance distributions
count
60000
40000
20000
0
0 60 120 180 240 300 360 420 480 540 600
Cb_LI_Rphase ( 5−distances)
count
8000
6000
4000
2000
0
0 4 8 12 16 20 24 28 32 36 40 44 48 52 56 60
Cb_LI_Rphase ( 5−distances)

```
Directory: DMT File: 0READMEdmt

DMT - Differentially methylated tiles.

Input: studymc wsize stepsize threshold qvalue
Output: files dmt-*.txt dmg-*.txt

Notes: Output files dmt-*.txt and dmg-*.txt are generated by BWASPR::det_dmt() and show the
        differentially methylated tiles and genes as determined by methylKit::getMethylDiff with
        parameters difference=threshold and qvalue, as provided in the Cb_LIp.conf
        configuration file.  Tiles refer to sliding windows along the genome within which
        methylation calls are cumulated by methylKit::tileMethylCounts.
        The dmg-*.txt files show all genes with at least one differentially methylated tile.

        A positive meth.diff value in comparison A.vs.B means that the B methylation percentage
        in that tile is higher than the A methylation percentage.
```

Directory: DMT File: dmg-Cb_LIp.txt

| seqnames | start | end | width | strand | source | type | score | phase | ID | Dbxref | Name | gbkey | gene | gene_biotype | partial | start_range | end_range | comparison |
|---|---|---|---|---|---|---|---|---|---|---|---|---|---|---|---|---|---|---|
| NW_011925163.1 | 518428 | 530854 | 12427 | - | Gnomon | gene | NA | NA | gene8045 | GeneID:105281345 | LOC105281345 | Gene | LOC105281345 | protein_coding | NA | character(0) | character(0) | BCphase.vs.Rphase |
| NW_011925362.1 | 60444 | 65208 | 4765 | + | Gnomon | gene | NA | NA | gene10649 | GeneID:105284325 | LOC105284325 | Gene | LOC105284325 | protein_coding | NA | character(0) | character(0) | BCphase.vs.Rphase |

Directory: DMT File: dmt-Cb_LIp.txt

| seqnames | start | end | width | strand | pvalue | qvalue | meth.diff | comparison |
|---|---|---|---|---|---|---|---|---|
| NW_011925163.1 | 530001 | 530500 | 500 | * | 3.09541760182134e-07 | 0.006 | -58.82 | BCphase.vs.Rphase |
| NW_011925362.1 | 64501 | 65000 | 500 | * | 7.05004578641165e-07 | 0.011 | -39.88 | BCphase.vs.Rphase |

```
Directory: DMSG File: 0READMEdmsg

DMSG - Differentially methylated sites and genes.

Input: studyhc threshold qvalue
       The studyhc methylKit raw object contains all CpGscd sites with coverage at least
        20 reads.
Output: files dms-*.txt dmg-*.txt  dmg-*details.txt dmg-*heatmaps.pdf

Notes: Output files dms-*.txt and dmg-*.txt are generated by BWASPR::det_dmsg() and show the
        differentially methylated sites and genes as determined by methylKit::getMethylDiff with
        parameters difference=threshold and qvalue, as provided in the Cb_LIp.conf
        configuration file.  The table of differentially methylated genes contains all genes
        with at least one differentially methylated site.

       The files dmg-*details.txt, generated by BWASPR::show_dmsg(), show all CpGscd sites in
        the differentially methylated genes, with coverage numbers and methylation percentages.

       The files dmg-*heatmaps.pdf, generated by BWASPR::show_dmsg(), give heatmap displays for
        genes meeting the following criteria: 1) there are between minNsites and maxNsites common
        CpGscd sites; 2) at least minPdmsites % of these sites are differentially methylated sites.
        The parameters are set in the Cb_LIp.conf configuration file.
```

Directory: DMSG File: dmg-Cb_LIp.txt

| seqnames | start | end | width | strand | source | type | score | phase | ID | Dbxref | Name | gbkey | gene | gene_biotype | partial | start_range | end_range | comparison |
|---|---|---|---|---|---|---|---|---|---|---|---|---|---|---|---|---|---|---|
| NW_011924881.1 | 39621 | 40962 | 1342 | - | Gnomon | gene | NA | NA | gene17 | GeneID:105286636 | LOC105286636 | Gene | LOC105286636 | protein_coding | NA | character(0) | character(0) | BCphase.vs.Rphase |
| NW_011924881.1 | 501948 | 588270 | 86323 | - | Gnomon | gene | NA | NA | gene29 | GeneID:105274746 | LOC105274746 | Gene | LOC105274746 | lncRNA | NA | character(0) | character(0) | BCphase.vs.Rphase |
| NW_011924881.1 | 751669 | 759027 | 7359 | - | Gnomon | gene | NA | NA | gene44 | GeneID:105275709 | LOC105275709 | Gene | LOC105275709 | protein_coding | NA | character(0) | character(0) | BCphase.vs.Rphase |
| NW_011924881.1 | 766631 | 768231 | 1601 | - | Gnomon | gene | NA | NA | gene48 | GeneID:105276896 | LOC105276896 | Gene | LOC105276896 | protein_coding | NA | character(0) | character(0) | BCphase.vs.Rphase |
| NW_011924881.1 | 989753 | 994960 | 5208 | + | Gnomon | gene | NA | NA | gene61 | GeneID:105277580 | LOC105277580 | Gene | LOC105277580 | protein_coding | NA | character(0) | character(0) | BCphase.vs.Rphase |
| NW_011924883.1 | 23752 | 46738 | 22987 | + | Gnomon | gene | NA | NA | gene70 | GeneID:105279190 | LOC105279190 | Gene | LOC105279190 | protein_coding | NA | character(0) | character(0) | BCphase.vs.Rphase |
| NW_011924887.1 | 111874 | 119948 | 8075 | - | Gnomon | gene | NA | NA | gene111 | GeneID:105284718 | LOC105284718 | Gene | LOC105284718 | protein_coding | NA | character(0) | character(0) | BCphase.vs.Rphase |
| NW_011924887.1 | 116318 | 118918 | 2601 | + | Gnomon | gene | NA | NA | gene112 | GeneID:105284841 | LOC105284841 | Gene | LOC105284841 | protein_coding | NA | character(0) | character(0) | BCphase.vs.Rphase |
| NW_011924887.1 | 120279 | 132526 | 12248 | + | Gnomon | gene | NA | NA | gene113 | GeneID:105285339 | LOC105285339 | Gene | LOC105285339 | protein_coding | NA | character(0) | character(0) | BCphase.vs.Rphase |
| NW_011924887.1 | 142565 | 143792 | 1228 | + | Gnomon | gene | NA | NA | gene117 | GeneID:105285795 | LOC105285795 | Gene | LOC105285795 | protein_coding | NA | character(0) | character(0) | BCphase.vs.Rphase |
| NW_011924887.1 | 143670 | 150430 | 6761 | - | Gnomon | gene | NA | NA | gene118 | GeneID:105285614 | LOC105285614 | Gene | LOC105285614 | protein_coding | NA | character(0) | character(0) | BCphase.vs.Rphase |
| NW_011924887.1 | 683254 | 689485 | 6232 | - | Gnomon | gene | NA | NA | gene155 | GeneID:105287785 | LOC105287785 | Gene | LOC105287785 | protein_coding | NA | character(0) | character(0) | BCphase.vs.Rphase |
| NW_011924887.1 | 705774 | 710286 | 4513 | + | Gnomon | gene | NA | NA | gene159 | GeneID:105287796 | LOC105287796 | Gene | LOC105287796 | protein_coding | NA | character(0) | character(0) | BCphase.vs.Rphase |
| NW_011924887.1 | 724203 | 726021 | 1819 | - | Gnomon | gene | NA | NA | gene163 | GeneID:105287838 | LOC105287838 | Gene | LOC105287838 | protein_coding | NA | character(0) | character(0) | BCphase.vs.Rphase |
| NW_011924887.1 | 949487 | 958414 | 8928 | + | Gnomon | gene | NA | NA | gene176 | GeneID:105288014 | LOC105288014 | Gene | LOC105288014 | protein_coding | NA | character(0) | character(0) | BCphase.vs.Rphase |
| NW_011924887.1 | 958098 | 992012 | 33915 | - | Gnomon | gene | NA | NA | gene177 | GeneID:105288005 | LOC105288005 | Gene | LOC105288005 | protein_coding | NA | character(0) | character(0) | BCphase.vs.Rphase |
| NW_011924887.1 | 1134654 | 1138168 | 3515 | - | Gnomon | gene | NA | NA | gene183 | GeneID:105288058 | LOC105288058 | Gene | LOC105288058 | protein_coding | NA | character(0) | character(0) | BCphase.vs.Rphase |
| NW_011924887.1 | 1274363 | 1356124 | 81762 | + | Gnomon | gene | NA | NA | gene194 | GeneID:105288172 | LOC105288172 | Gene | LOC105288172 | protein_coding | NA | character(0) | character(0) | BCphase.vs.Rphase |
| NW_011924887.1 | 1509692 | 1510490 | 799 | - | Gnomon | gene | NA | NA | gene200 | GeneID:105288225 | LOC105288225 | Gene | LOC105288225 | protein_coding | NA | character(0) | character(0) | BCphase.vs.Rphase |
| NW_011924887.1 | 1546715 | 1553169 | 6455 | + | Gnomon | gene | NA | NA | gene209 | GeneID:105274475 | LOC105274475 | Gene | LOC105274475 | protein_coding | NA | character(0) | character(0) | BCphase.vs.Rphase |
| NW_011924887.1 | 1594363 | 1599359 | 4997 | - | Gnomon | gene | NA | NA | gene218 | GeneID:105274600 | LOC105274600 | Gene | LOC105274600 | protein_coding | NA | character(0) | character(0) | BCphase.vs.Rphase |
| NW_011924887.1 | 1817915 | 1820166 | 2252 | - | Gnomon | gene | NA | NA | gene234 | GeneID:105274775 | LOC105274775 | Gene | LOC105274775 | protein_coding | NA | character(0) | character(0) | BCphase.vs.Rphase |
| NW_011924887.1 | 1845614 | 1848814 | 3201 | - | Gnomon | gene | NA | NA | gene239 | GeneID:105274807 | LOC105274807 | Gene | LOC105274807 | protein_coding | NA | character(0) | character(0) | BCphase.vs.Rphase |
| NW_011924888.1 | 96821 | 100703 | 3883 | - | Gnomon | gene | NA | NA | gene278 | GeneID:105275240 | LOC105275240 | Gene | LOC105275240 | protein_coding | NA | character(0) | character(0) | BCphase.vs.Rphase |

...

... only first 25 lines shown ...

Directory: DMSG File: dmg-Cb_LIp_BCphase.vs.Rphase_details.txt

| seqnames | start | end | width | strand | coverage1 | numCs1 | numTs1 | coverage2 | numCs2 | numTs2 | BCphase | Rphase | gene_seqnames | gene_start | gene_end | gene_width | gene_strand | gene_source | gene_type | gene_ID | gene_Dbxref | gene_Name | gene_gbkey | gene_gene | gene_gene_biotype | gene_partial | gene_start_range | gene_end_range | gene_comparison | is.dm |
|---|---|---|---|---|---|---|---|---|---|---|---|---|---|---|---|---|---|---|---|---|---|---|---|---|---|---|---|---|---|---|
| NW_011924881.1 | 39625 | 39625 | 1 | + | 77 | 4 | 73 | 96 | 2 | 94 | 5.19 | 2.08 | NW_011924881.1 | 39621 | 40962 | 1342 | - | Gnomon | gene | gene17 | GeneID:105286636 | LOC105286636 | Gene | LOC105286636 | protein_coding | NA | character(0) | character(0) | BCphase.vs.Rphase | FALSE |
| NW_011924881.1 | 39633 | 39633 | 1 | + | 79 | 7 | 72 | 92 | 6 | 86 | 8.86 | 6.52 | NW_011924881.1 | 39621 | 40962 | 1342 | - | Gnomon | gene | gene17 | GeneID:105286636 | LOC105286636 | Gene | LOC105286636 | protein_coding | NA | character(0) | character(0) | BCphase.vs.Rphase | FALSE |
| NW_011924881.1 | 39706 | 39706 | 1 | + | 49 | 5 | 44 | 52 | 1 | 51 | 10.2 | 1.92 | NW_011924881.1 | 39621 | 40962 | 1342 | - | Gnomon | gene | gene17 | GeneID:105286636 | LOC105286636 | Gene | LOC105286636 | protein_coding | NA | character(0) | character(0) | BCphase.vs.Rphase | FALSE |
| NW_011924881.1 | 39711 | 39711 | 1 | + | 48 | 3 | 45 | 47 | 1 | 46 | 6.25 | 2.13 | NW_011924881.1 | 39621 | 40962 | 1342 | - | Gnomon | gene | gene17 | GeneID:105286636 | LOC105286636 | Gene | LOC105286636 | protein_coding | NA | character(0) | character(0) | BCphase.vs.Rphase | FALSE |
| NW_011924881.1 | 39723 | 39723 | 1 | + | 51 | 1 | 50 | 48 | 1 | 47 | 1.96 | 2.08 | NW_011924881.1 | 39621 | 40962 | 1342 | - | Gnomon | gene | gene17 | GeneID:105286636 | LOC105286636 | Gene | LOC105286636 | protein_coding | NA | character(0) | character(0) | BCphase.vs.Rphase | FALSE |
| NW_011924881.1 | 39743 | 39743 | 1 | + | 51 | 1 | 50 | 51 | 4 | 47 | 1.96 | 7.84 | NW_011924881.1 | 39621 | 40962 | 1342 | - | Gnomon | gene | gene17 | GeneID:105286636 | LOC105286636 | Gene | LOC105286636 | protein_coding | NA | character(0) | character(0) | BCphase.vs.Rphase | FALSE |
| NW_011924881.1 | 39767 | 39767 | 1 | + | 49 | 8 | 41 | 48 | 6 | 42 | 16.33 | 12.5 | NW_011924881.1 | 39621 | 40962 | 1342 | - | Gnomon | gene | gene17 | GeneID:105286636 | LOC105286636 | Gene | LOC105286636 | protein_coding | NA | character(0) | character(0) | BCphase.vs.Rphase | FALSE |
| NW_011924881.1 | 39782 | 39782 | 1 | + | 27 | 4 | 23 | 22 | 4 | 18 | 14.81 | 18.18 | NW_011924881.1 | 39621 | 40962 | 1342 | - | Gnomon | gene | gene17 | GeneID:105286636 | LOC105286636 | Gene | LOC105286636 | protein_coding | NA | character(0) | character(0) | BCphase.vs.Rphase | FALSE |
| NW_011924881.1 | 39790 | 39790 | 1 | + | 29 | 2 | 27 | 25 | 0 | 25 | 6.9 | 0 | NW_011924881.1 | 39621 | 40962 | 1342 | - | Gnomon | gene | gene17 | GeneID:105286636 | LOC105286636 | Gene | LOC105286636 | protein_coding | NA | character(0) | character(0) | BCphase.vs.Rphase | FALSE |
| NW_011924881.1 | 39798 | 39798 | 1 | + | 51 | 1 | 50 | 47 | 1 | 46 | 1.96 | 2.13 | NW_011924881.1 | 39621 | 40962 | 1342 | - | Gnomon | gene | gene17 | GeneID:105286636 | LOC105286636 | Gene | LOC105286636 | protein_coding | NA | character(0) | character(0) | BCphase.vs.Rphase | FALSE |
| NW_011924881.1 | 39806 | 39806 | 1 | + | 52 | 0 | 52 | 45 | 0 | 45 | 0 | 0 | NW_011924881.1 | 39621 | 40962 | 1342 | - | Gnomon | gene | gene17 | GeneID:105286636 | LOC105286636 | Gene | LOC105286636 | protein_coding | NA | character(0) | character(0) | BCphase.vs.Rphase | FALSE |
| NW_011924881.1 | 39844 | 39844 | 1 | + | 44 | 7 | 37 | 35 | 11 | 24 | 15.91 | 31.43 | NW_011924881.1 | 39621 | 40962 | 1342 | - | Gnomon | gene | gene17 | GeneID:105286636 | LOC105286636 | Gene | LOC105286636 | protein_coding | NA | character(0) | character(0) | BCphase.vs.Rphase | FALSE |
| NW_011924881.1 | 39857 | 39857 | 1 | + | 61 | 2 | 59 | 31 | 4 | 27 | 3.28 | 12.9 | NW_011924881.1 | 39621 | 40962 | 1342 | - | Gnomon | gene | gene17 | GeneID:105286636 | LOC105286636 | Gene | LOC105286636 | protein_coding | NA | character(0) | character(0) | BCphase.vs.Rphase | FALSE |
| NW_011924881.1 | 39863 | 39863 | 1 | + | 57 | 1 | 56 | 30 | 1 | 29 | 1.75 | 3.33 | NW_011924881.1 | 39621 | 40962 | 1342 | - | Gnomon | gene | gene17 | GeneID:105286636 | LOC105286636 | Gene | LOC105286636 | protein_coding | NA | character(0) | character(0) | BCphase.vs.Rphase | FALSE |
| NW_011924881.1 | 39967 | 39967 | 1 | + | 48 | 38 | 10 | 40 | 28 | 12 | 79.17 | 70 | NW_011924881.1 | 39621 | 40962 | 1342 | - | Gnomon | gene | gene17 | GeneID:105286636 | LOC105286636 | Gene | LOC105286636 | protein_coding | NA | character(0) | character(0) | BCphase.vs.Rphase | FALSE |
| NW_011924881.1 | 39987 | 39987 | 1 | + | 58 | 30 | 28 | 55 | 31 | 24 | 51.72 | 56.36 | NW_011924881.1 | 39621 | 40962 | 1342 | - | Gnomon | gene | gene17 | GeneID:105286636 | LOC105286636 | Gene | LOC105286636 | protein_coding | NA | character(0) | character(0) | BCphase.vs.Rphase | FALSE |
| NW_011924881.1 | 40005 | 40005 | 1 | + | 71 | 5 | 66 | 63 | 6 | 57 | 7.04 | 9.52 | NW_011924881.1 | 39621 | 40962 | 1342 | - | Gnomon | gene | gene17 | GeneID:105286636 | LOC105286636 | Gene | LOC105286636 | protein_coding | NA | character(0) | character(0) | BCphase.vs.Rphase | FALSE |
| NW_011924881.1 | 40021 | 40021 | 1 | + | 70 | 9 | 61 | 65 | 7 | 58 | 12.86 | 10.77 | NW_011924881.1 | 39621 | 40962 | 1342 | - | Gnomon | gene | gene17 | GeneID:105286636 | LOC105286636 | Gene | LOC105286636 | protein_coding | NA | character(0) | character(0) | BCphase.vs.Rphase | FALSE |
| NW_011924881.1 | 40035 | 40035 | 1 | + | 82 | 15 | 67 | 78 | 13 | 65 | 18.29 | 16.67 | NW_011924881.1 | 39621 | 40962 | 1342 | - | Gnomon | gene | gene17 | GeneID:105286636 | LOC105286636 | Gene | LOC105286636 | protein_coding | NA | character(0) | character(0) | BCphase.vs.Rphase | FALSE |
| NW_011924881.1 | 40061 | 40061 | 1 | + | 49 | 34 | 15 | 49 | 45 | 4 | 69.39 | 91.84 | NW_011924881.1 | 39621 | 40962 | 1342 | - | Gnomon | gene | gene17 | GeneID:105286636 | LOC105286636 | Gene | LOC105286636 | protein_coding | NA | character(0) | character(0) | BCphase.vs.Rphase | TRUE |
| NW_011924881.1 | 40072 | 40072 | 1 | + | 45 | 42 | 3 | 51 | 50 | 1 | 93.33 | 98.04 | NW_011924881.1 | 39621 | 40962 | 1342 | - | Gnomon | gene | gene17 | GeneID:105286636 | LOC105286636 | Gene | LOC105286636 | protein_coding | NA | character(0) | character(0) | BCphase.vs.Rphase | FALSE |
| NW_011924881.1 | 40095 | 40095 | 1 | + | 60 | 30 | 30 | 71 | 39 | 32 | 50 | 54.93 | NW_011924881.1 | 39621 | 40962 | 1342 | - | Gnomon | gene | gene17 | GeneID:105286636 | LOC105286636 | Gene | LOC105286636 | protein_coding | NA | character(0) | character(0) | BCphase.vs.Rphase | FALSE |
| NW_011924881.1 | 40120 | 40120 | 1 | + | 58 | 44 | 14 | 99 | 64 | 35 | 75.86 | 64.65 | NW_011924881.1 | 39621 | 40962 | 1342 | - | Gnomon | gene | gene17 | GeneID:105286636 | LOC105286636 | Gene | LOC105286636 | protein_coding | NA | character(0) | character(0) | BCphase.vs.Rphase | FALSE |
| NW_011924881.1 | 40159 | 40159 | 1 | + | 48 | 27 | 21 | 71 | 46 | 25 | 56.25 | 64.79 | NW_011924881.1 | 39621 | 40962 | 1342 | - | Gnomon | gene | gene17 | GeneID:105286636 | LOC105286636 | Gene | LOC105286636 | protein_coding | NA | character(0) | character(0) | BCphase.vs.Rphase | FALSE |

...
... only first 25 lines shown ...

Directory: DMSG File: dms-Cb_LIp.txt

| seqnames | start | end | width | strand | pvalue | qvalue | meth.diff | comparison |
|---|---|---|---|---|---|---|---|---|
| NW_011924881.1 | 40061 | 40061 | 1 | + | 0.00932174524770422 | 0.032 | 22.45 | BCphase.vs.Rphase |
| NW_011924881.1 | 322701 | 322701 | 1 | + | 0.022289766970618 | 0.032 | -20.83 | BCphase.vs.Rphase |
| NW_011924881.1 | 502438 | 502438 | 1 | + | 0.00130822983503945 | 0.032 | -21.6 | BCphase.vs.Rphase |
| NW_011924881.1 | 758111 | 758111 | 1 | + | 0.0649554512710637 | 0.032 | 21.84 | BCphase.vs.Rphase |
| NW_011924881.1 | 766996 | 766996 | 1 | + | 0.0258381002114838 | 0.032 | 20.97 | BCphase.vs.Rphase |
| NW_011924881.1 | 968662 | 968662 | 1 | + | 0.0533104697272171 | 0.032 | -21 | BCphase.vs.Rphase |
| NW_011924881.1 | 990307 | 990307 | 1 | + | 0.0507485560143629 | 0.032 | -26.67 | BCphase.vs.Rphase |
| NW_011924881.1 | 991519 | 991519 | 1 | + | 0.026288595737396 | 0.032 | 29.87 | BCphase.vs.Rphase |
| NW_011924883.1 | 25514 | 25514 | 1 | + | 0.109143837804694 | 0.032 | -21.17 | BCphase.vs.Rphase |
| NW_011924885.1 | 202111 | 202111 | 1 | + | 0.00630025908783233 | 0.032 | -20.51 | BCphase.vs.Rphase |
| NW_011924887.1 | 118860 | 118860 | 1 | + | 0.154104550362431 | 0.032 | -21.57 | BCphase.vs.Rphase |
| NW_011924887.1 | 121567 | 121567 | 1 | + | 0.0515162767433556 | 0.032 | -26.51 | BCphase.vs.Rphase |
| NW_011924887.1 | 143757 | 143757 | 1 | + | 0.0500728618830326 | 0.032 | 20.91 | BCphase.vs.Rphase |
| NW_011924887.1 | 260029 | 260029 | 1 | + | 0.0309244977521026 | 0.032 | -25.81 | BCphase.vs.Rphase |
| NW_011924887.1 | 687007 | 687007 | 1 | + | 0.00466548997965979 | 0.032 | -22.56 | BCphase.vs.Rphase |
| NW_011924887.1 | 706978 | 706978 | 1 | + | 0.0139403462580346 | 0.032 | -40.22 | BCphase.vs.Rphase |
| NW_011924887.1 | 724642 | 724642 | 1 | + | 0.121870575690829 | 0.032 | -24.54 | BCphase.vs.Rphase |
| NW_011924887.1 | 796793 | 796793 | 1 | + | 0.00454940267680876 | 0.032 | -22.68 | BCphase.vs.Rphase |
| NW_011924887.1 | 950614 | 950614 | 1 | + | 0.0689935471511017 | 0.032 | 30.22 | BCphase.vs.Rphase |
| NW_011924887.1 | 985609 | 985609 | 1 | + | 0.00589044728009387 | 0.032 | 21.36 | BCphase.vs.Rphase |
| NW_011924887.1 | 1136535 | 1136535 | 1 | + | 0.0102244593048954 | 0.032 | -20.05 | BCphase.vs.Rphase |
| NW_011924887.1 | 1321288 | 1321288 | 1 | + | 0.00324154819160368 | 0.032 | -26.92 | BCphase.vs.Rphase |
| NW_011924887.1 | 1509753 | 1509753 | 1 | + | 0.00609873540936131 | 0.032 | 29.09 | BCphase.vs.Rphase |
| NW_011924887.1 | 1528900 | 1528900 | 1 | + | 0.0125226035492142 | 0.032 | 20.15 | BCphase.vs.Rphase |

...
... only first 25 lines shown ...

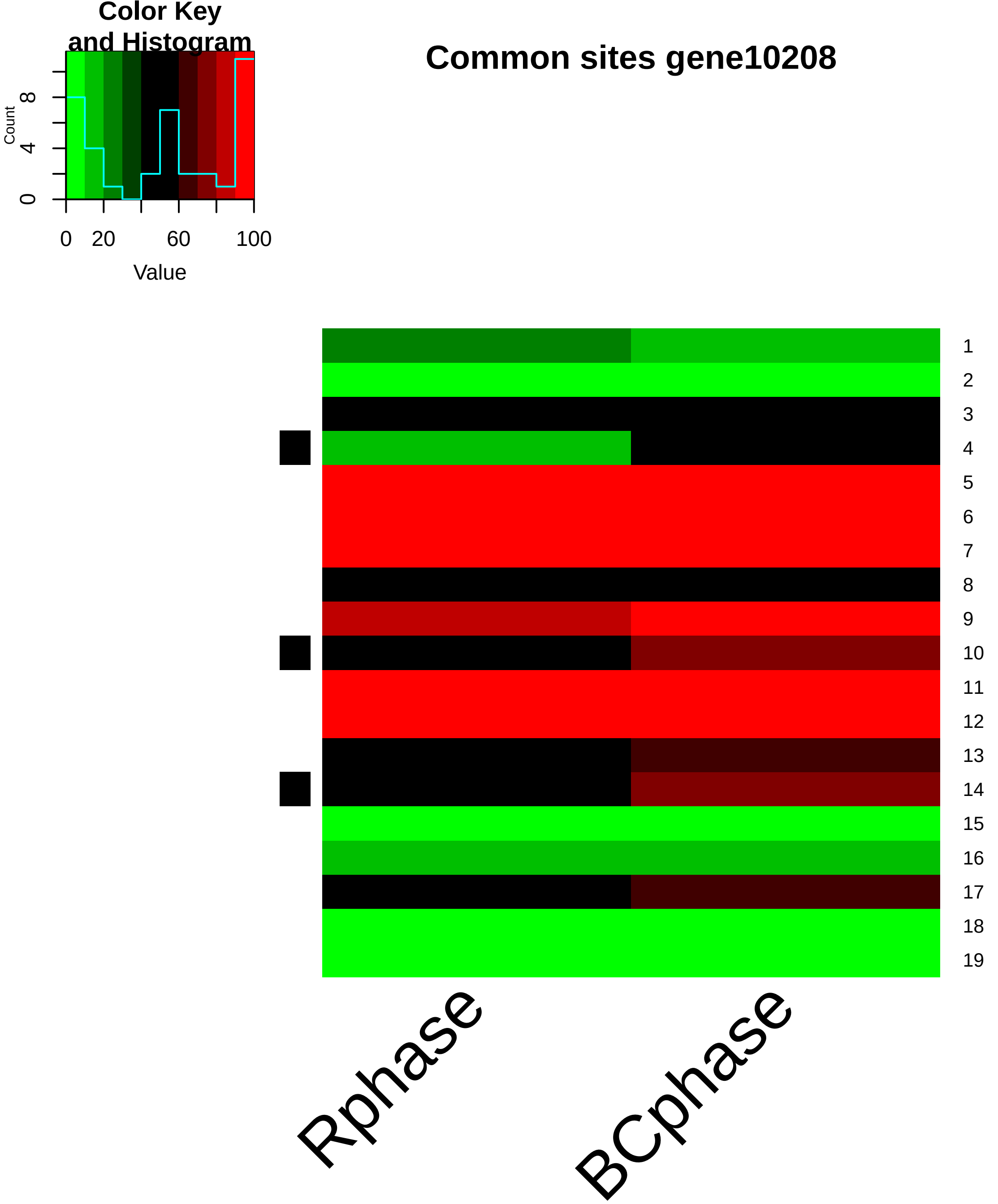
Color Key
and Histogram
Count
0
4
8
0
20
60
100
Value
Common sites gene10208
Rphase
BCphase

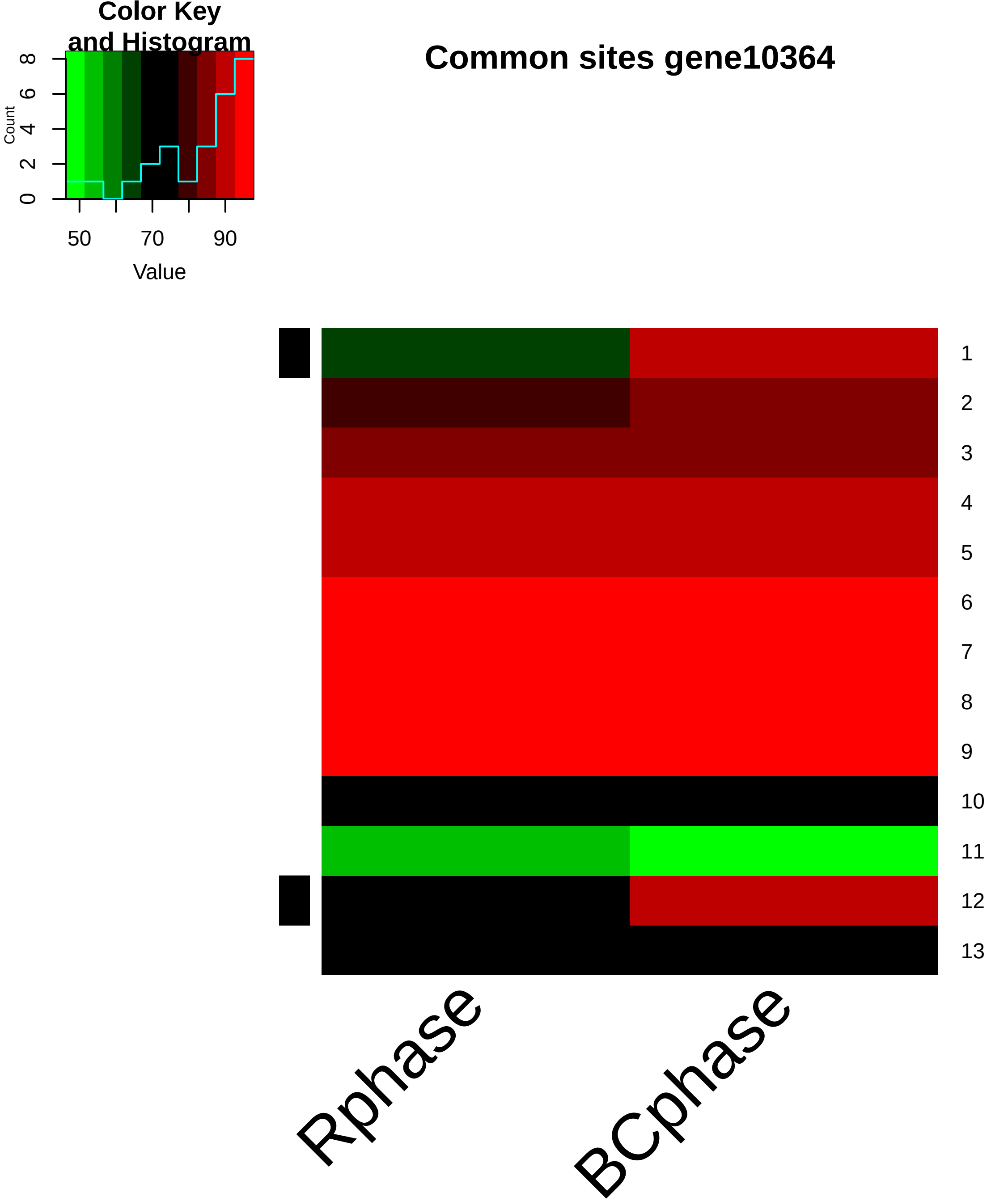
Color Key
and Histogram
Count
0
2
4
6
8
50
70
90
Value
Common sites gene10364
1
2
3
4
5
6
7
8
9
10
11
12
13
Rphase
BCphase

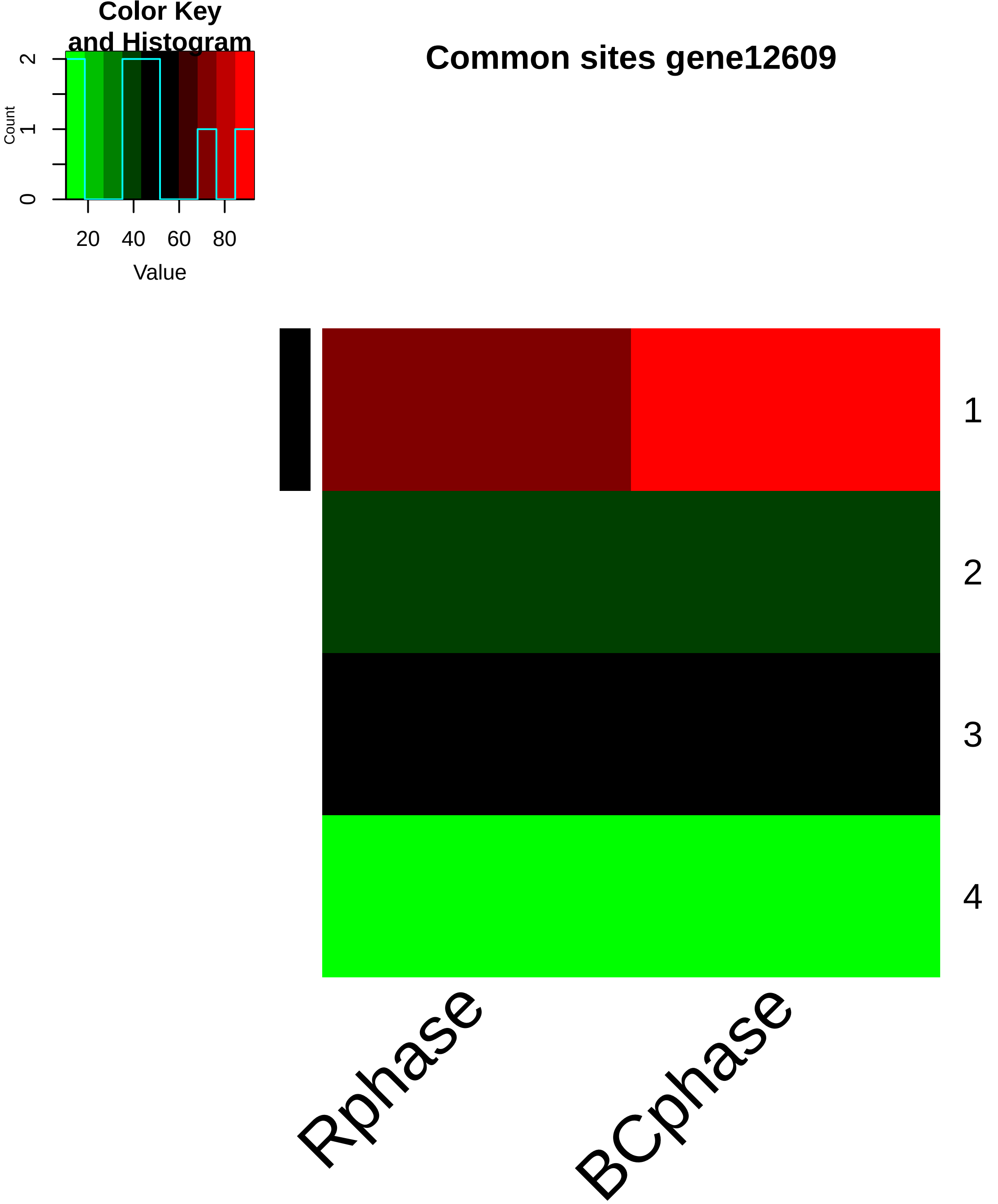
Color Key
and Histogram
Count
0
1
2
20
40
60
80
Value
Common sites gene12609
1
2
3
4
Rphase
BCphase

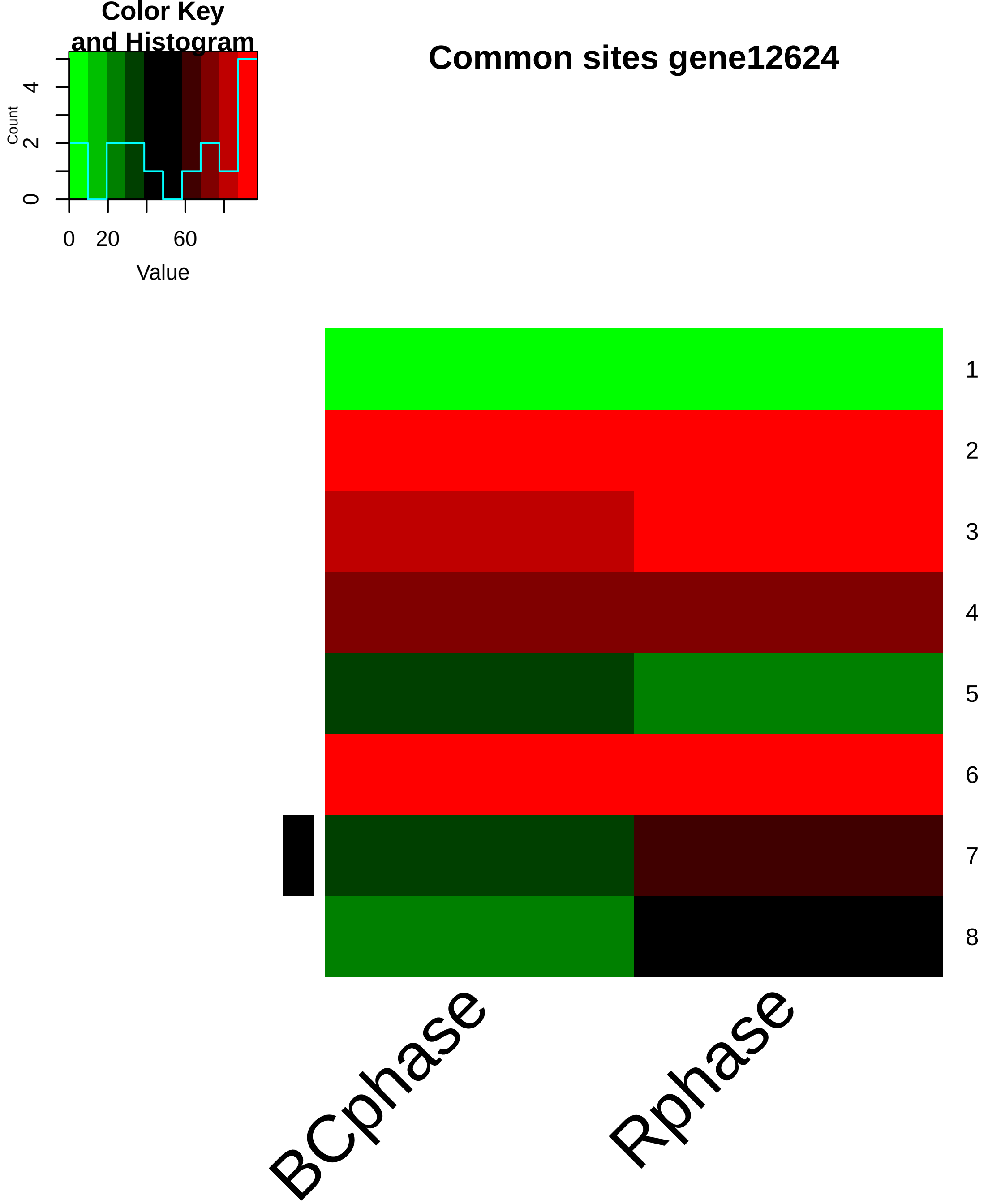

Color Key
and Histogram
Count
0
2
4
0
20
60
Value
Common sites gene12624
1
2
3
4
5
6
7
8
BCphase
Rphase

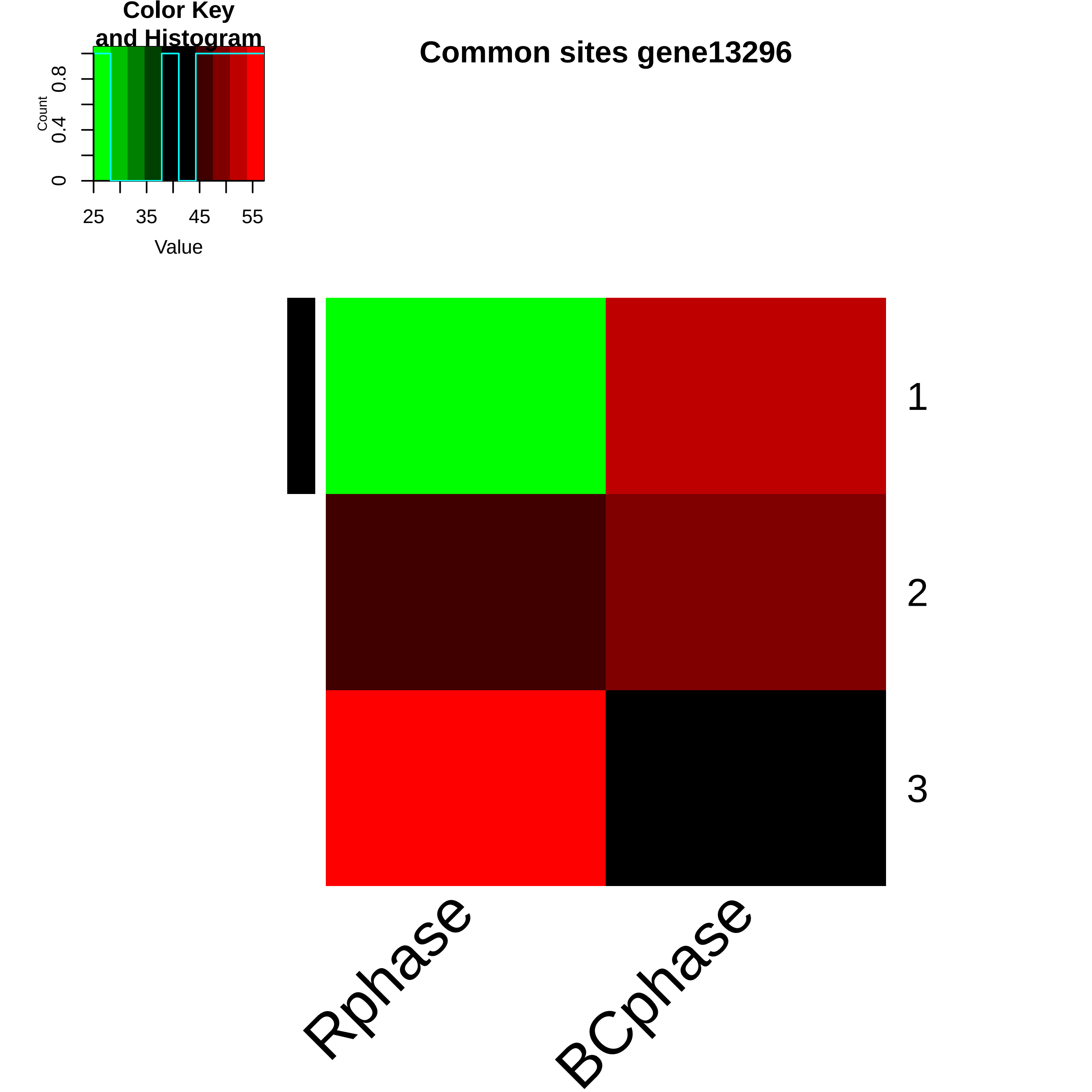
Color Key
and Histogram
Count
0
0.4
0.8
25
35
45
55
Value
Common sites gene13296
1
2
3
Rphase
BCphase

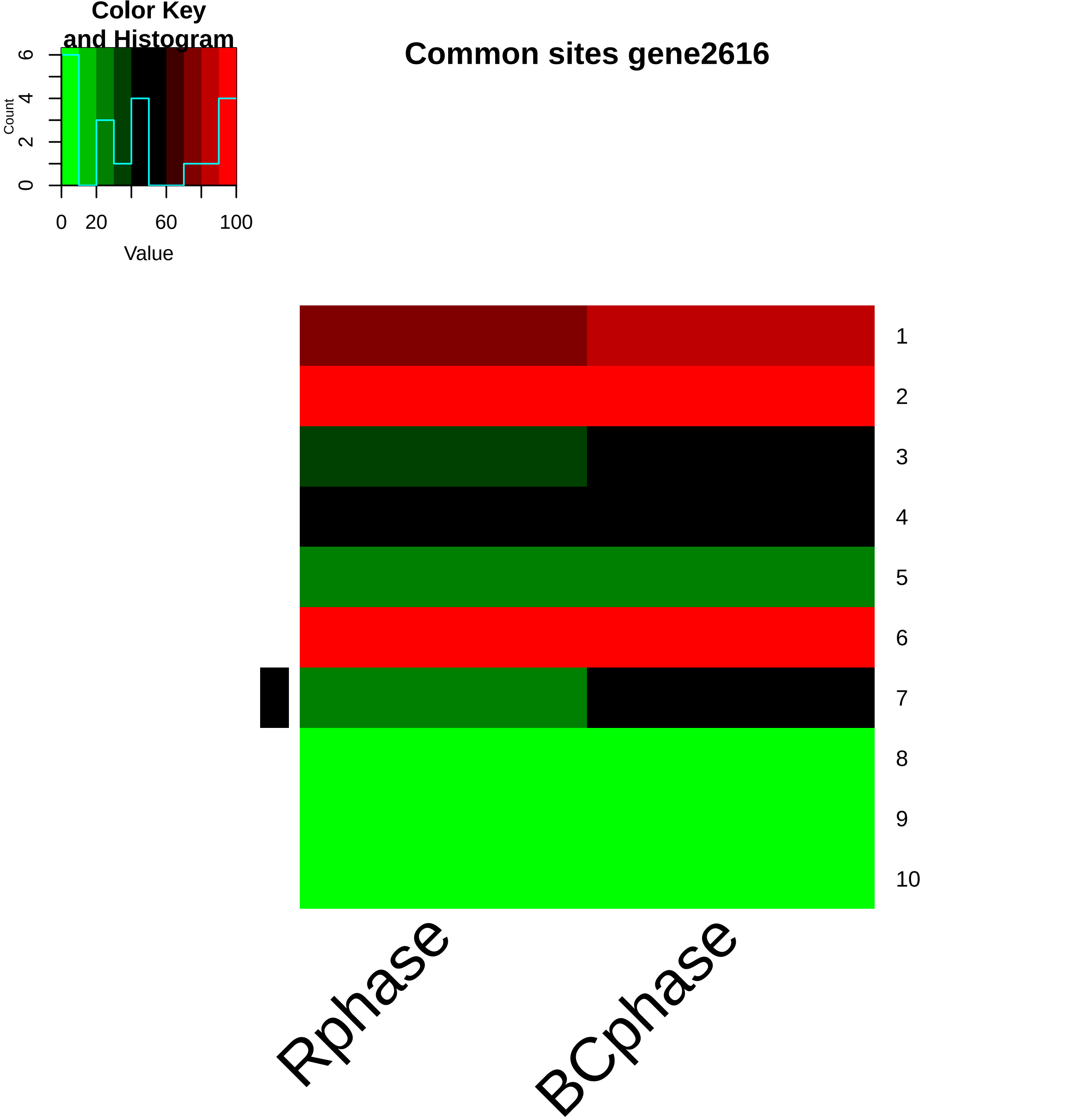
Color Key
and Histogram
Count
0
2
4
6
0
20
60
100
Value
Common sites gene2616
1
2
3
4
5
6
7
8
9
10
Rphase
BCphase

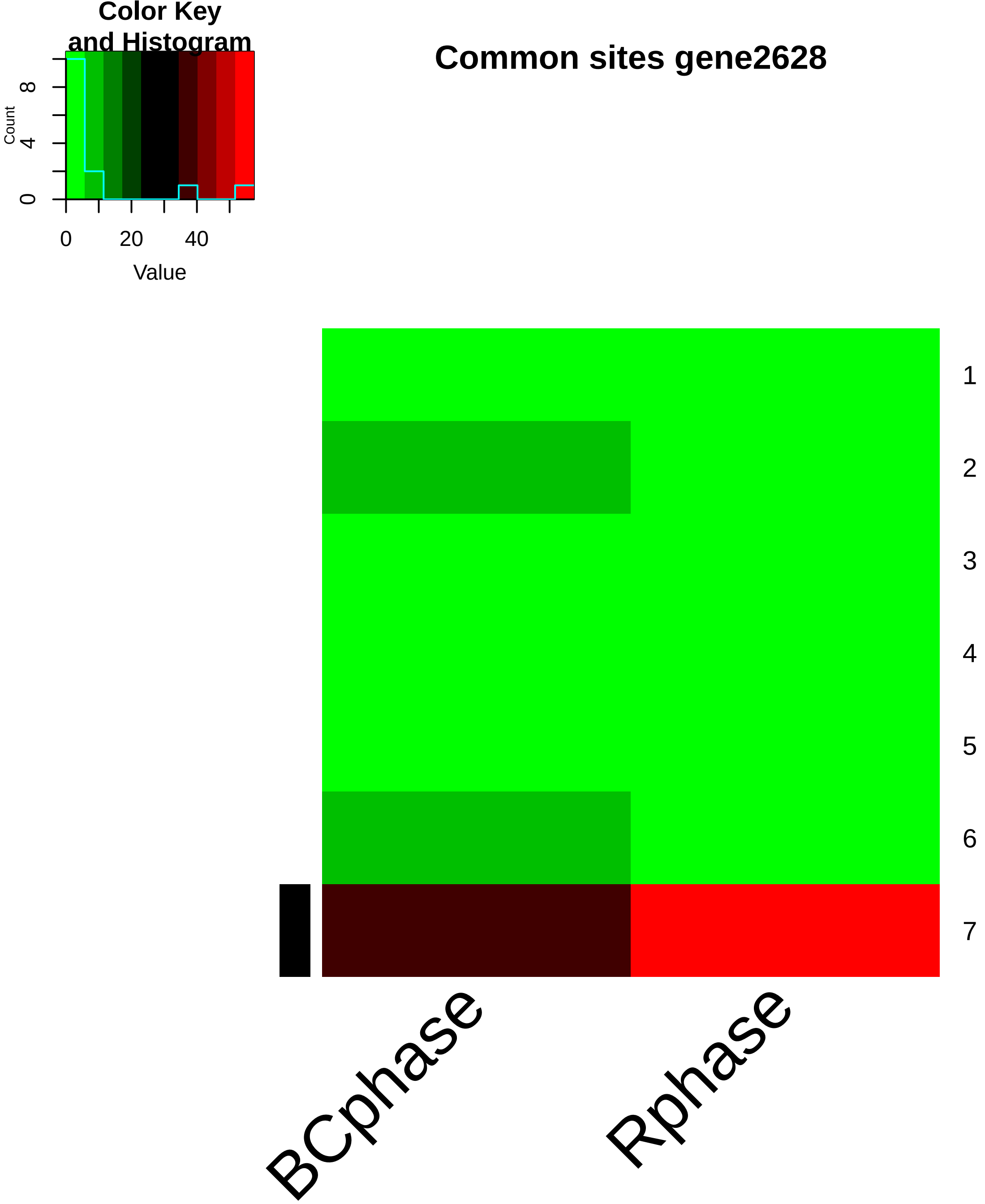

Color Key
and Histogram
Count
0
4
8
0
20
40
Value
Common sites gene2628
1
2
3
4
5
6
7
BCphase
Rphase

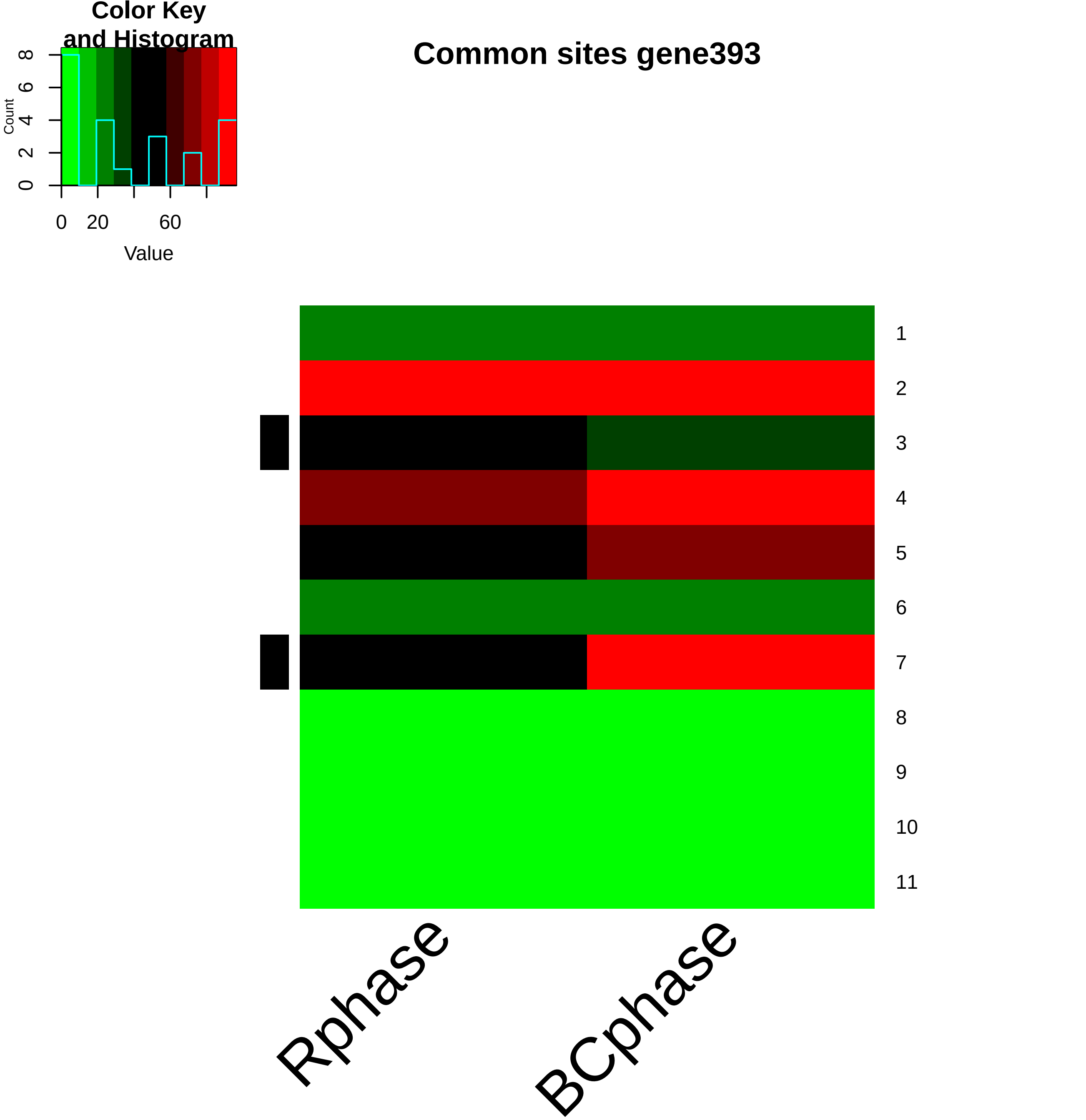
Color Key
and Histogram
Count
0
2
4
6
8
0
20
60
Value
Common sites gene393
1
2
3
4
5
6
7
8
9
10
11
Rphase
BCphase

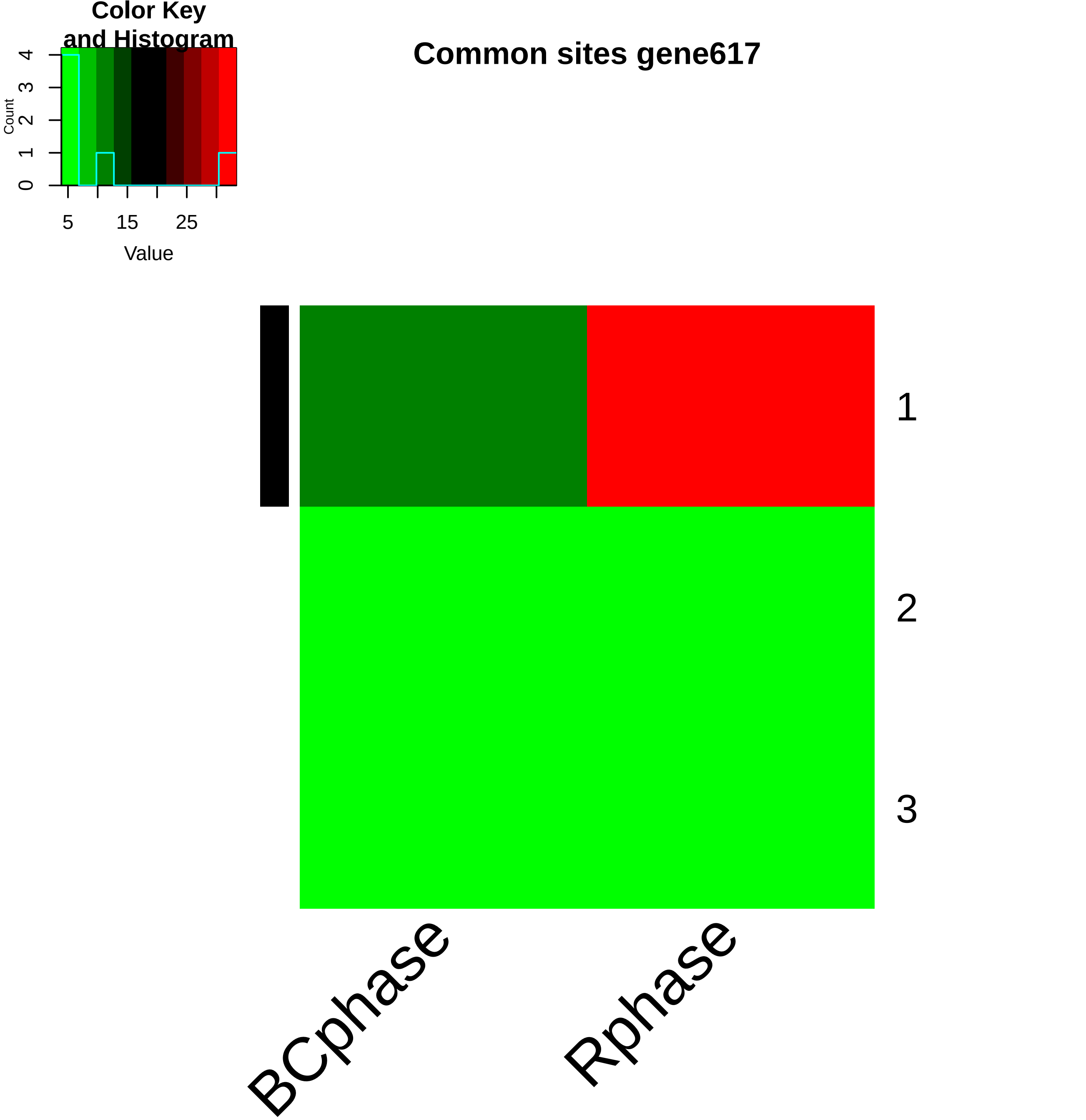
Color Key
and Histogram
Count
0
1
2
3
4
5
15
25
Value
Common sites gene617
1
2
3
BCphase
Rphase

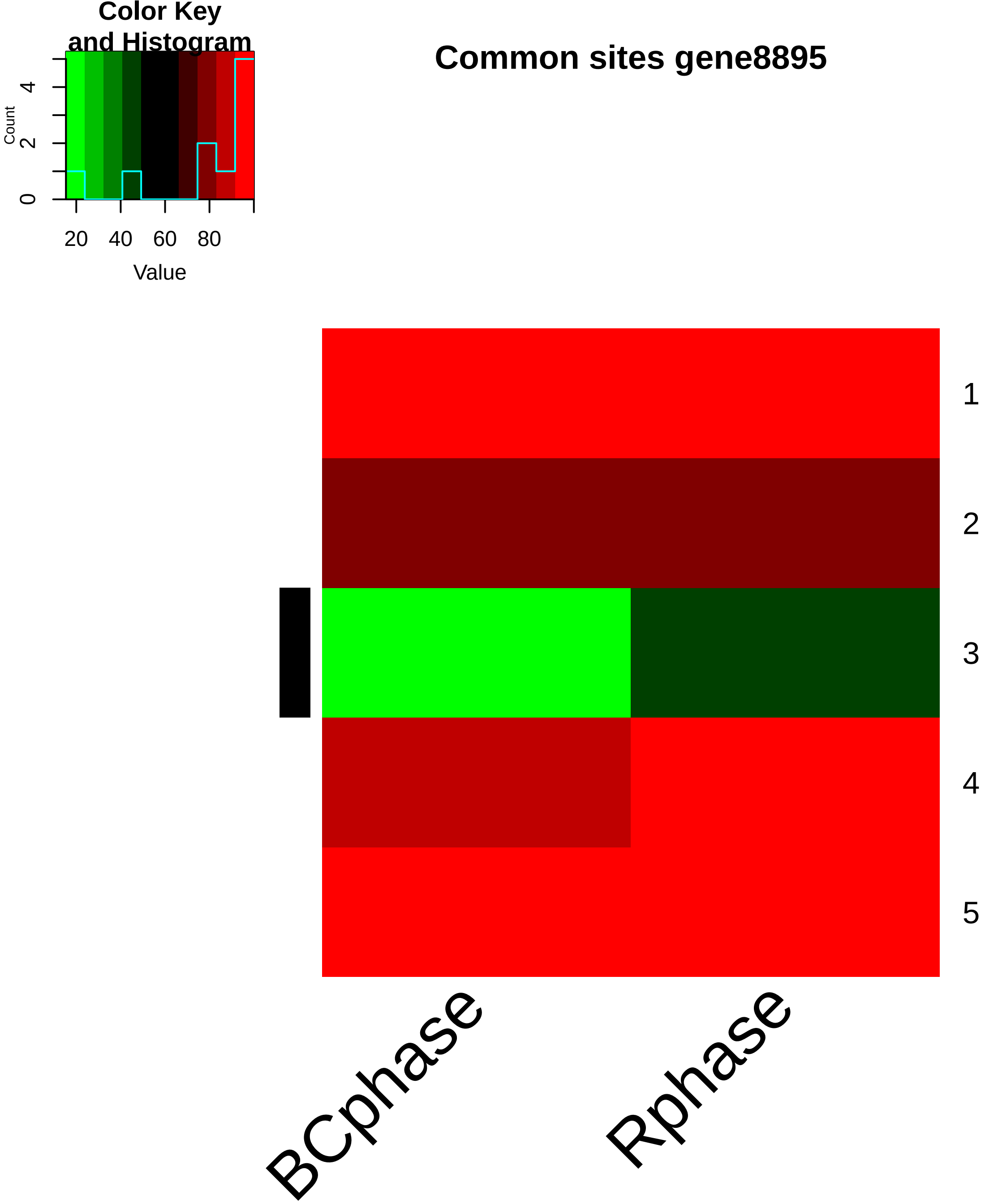
Color Key
and Histogram
Count
Value
20
40
60
80
0
2
4
Common sites gene8895
1
2
3
4
5
BCphase
Rphase

```
Directory: OGL File: 0READMEogl

OGL - Ordered gene lists.

Input: studyhc annotation   maxgwidth minnbrdmsites
       dmgprp (= output of BWASPR:show_dmsg())

Output: files ogl-*.txt rnk-dmg-*.txt
         ogl-<study>_<sample>.txt ogl-<study>_<sample1>.vs.<sample2>.txt
         rnk-dmg--<study>_<sample1>.vs.<sample2>.txt rnk-dmg--<study>_<sample1>.vs.<sample2>.pdf
         wrt-<study>.txt

Notes: Output files ogl-<study>_<sample>.txt give tables for each sample with columns

        gene_ID gwidth #Sites #per10Kb %perSite %pNucl

       ordered by %pNucl.  If available, a link to an NCBI entry of gene_ID is inserted as second column.
       Abbreviations used: %perSite, percent methylation per site
                           %perNucl, percent methylation per nucleotide of the gene

       Output files ogl-<study>_<sample1>.vs.<sample2>.txt give tables for each comparison with columns

        gene_ID gwidth #Sites #per10Kb #dmSites #dmsp10kb %dmSites %pSite1 %pSite2 DMpSite ADMpSite DMpNucl ADMpNucl

       ordered by DMpSite.  If available, a link to an NCBI entry of gene_ID is inserted as second column.
       Abbreviations used: %dmSites, percent sites that are differentially methylated
                           %pSite1, average per site % methylation for sample1
                           %pSite2, average per site % methylation for sample2
                           DMpSite, average per site difference in % methylation between sample1 and sample2
                           ADMpSite, absolute value of DMpSite
                           DMpNucl, average per nucleotide difference in % methylation between sample1 and sample2
                           ADMpNucl, absolute value of DMpNucl

       Output files rnk-dmg-<study>_<sample1>.vs.<sample2>.txt are equivalent to files
        ogl-<study>_<sample1>.vs.<sample2>.txt but ordered by ADMpNucl.
       Output files rnk-dmg-<study>_<sample1>.vs.<sample2>.pdf provide visualization of the distribution
        of ADMpNucl values.

       Output file wrt-<study>.txt gives results of the Wilcoxon signed rank test comparing
        the %pSite1 and %pSite2 vectors

       In output directory OGL, tables are restricted to genes with gwidth <= maxgwidth and, for
        pairwise comparisons, #dmSites >= minnbrdmsites

       If either maxgwidth or minnbrdmsites is set, then output directory OGLall will show the full tables
        with all genes (for reference).
```

Directory: OGL File: ogl-Cb_LIp_BCphase.txt

| gene_ID | gene_link | gwidth | #Sites | #per10Kb | %perSite | %pNucl |
|---|---|---|---|---|---|---|
| gene11569 | https://www.ncbi.nlm.nih.gov/gene/?term=LOC105285394 | 1311 | 149 | 1136.54 | 49.95 | 5.68 |
| gene10219 | https://www.ncbi.nlm.nih.gov/gene/?term=LOC105283830 | 235 | 32 | 1361.7 | 36.97 | 5.03 |
| gene4878 | https://www.ncbi.nlm.nih.gov/gene/?term=LOC105277648 | 675 | 53 | 785.19 | 62.39 | 4.9 |
| gene13384 | https://www.ncbi.nlm.nih.gov/gene/?term=LOC105287447 | 1877 | 356 | 1896.64 | 25.55 | 4.85 |
| gene5917 | https://www.ncbi.nlm.nih.gov/gene/?term=LOC105278849 | 823 | 163 | 1980.56 | 24.16 | 4.79 |
| gene12602 | https://www.ncbi.nlm.nih.gov/gene/?term=LOC105286562 | 1399 | 128 | 914.94 | 51.55 | 4.72 |
| gene5714 | https://www.ncbi.nlm.nih.gov/gene/?term=LOC105278729 | 710 | 54 | 760.56 | 61.16 | 4.65 |
| gene3719 | https://www.ncbi.nlm.nih.gov/gene/?term=LOC105276360 | 1471 | 239 | 1624.75 | 28.35 | 4.61 |
| gene12538 | https://www.ncbi.nlm.nih.gov/gene/?term=LOC105286531 | 372 | 46 | 1236.56 | 36.97 | 4.57 |
| gene1894 | https://www.ncbi.nlm.nih.gov/gene/?term=LOC105288083 | 445 | 36 | 808.99 | 56.38 | 4.56 |
| gene12364 | https://www.ncbi.nlm.nih.gov/gene/?term=LOC105286269 | 1403 | 156 | 1111.9 | 40.37 | 4.49 |
| gene10943 | https://www.ncbi.nlm.nih.gov/gene/?term=LOC105284659 | 1759 | 196 | 1114.27 | 40.19 | 4.48 |
| gene12403 | https://www.ncbi.nlm.nih.gov/gene/?term=LOC105286311 | 1413 | 129 | 912.95 | 46.44 | 4.24 |
| gene1171 | https://www.ncbi.nlm.nih.gov/gene/?term=LOC105285174 | 844 | 95 | 1125.59 | 37.4 | 4.21 |
| gene2797 | https://www.ncbi.nlm.nih.gov/gene/?term=LOC105275314 | 1489 | 189 | 1269.31 | 33.05 | 4.2 |
| gene5309 | https://www.ncbi.nlm.nih.gov/gene/?term=LOC105278160 | 1704 | 201 | 1179.58 | 35.46 | 4.18 |
| gene6195 | https://www.ncbi.nlm.nih.gov/gene/?term=LOC105279187 | 2221 | 235 | 1058.08 | 39.44 | 4.17 |
| gene211 | https://www.ncbi.nlm.nih.gov/gene/?term=LOC105274534 | 1280 | 140 | 1093.75 | 37.99 | 4.16 |
| gene12330 | https://www.ncbi.nlm.nih.gov/gene/?term=LOC105286235 | 1501 | 149 | 992.67 | 41.83 | 4.15 |
| gene12652 | https://www.ncbi.nlm.nih.gov/gene/?term=LOC109610640 | 1122 | 93 | 828.88 | 50.01 | 4.14 |
| gene4635 | https://www.ncbi.nlm.nih.gov/gene/?term=LOC105277390 | 2067 | 232 | 1122.4 | 36.6 | 4.11 |
| gene1654 | https://www.ncbi.nlm.nih.gov/gene/?term=LOC105287811 | 2285 | 259 | 1133.48 | 36.08 | 4.09 |
| gene5412 | https://www.ncbi.nlm.nih.gov/gene/?term=LOC105278266 | 1868 | 259 | 1386.51 | 29.43 | 4.08 |
| gene11566 | https://www.ncbi.nlm.nih.gov/gene/?term=LOC105285392 | 1603 | 230 | 1434.81 | 28.17 | 4.04 |

...

... only first 25 lines shown ...

Directory: OGL File: ogl-Cb_LIp_BCphase.vs.Rphase.txt

| gene_ID | gene_link | gwidth | #Sites | #per10Kb | #dmSites | #dmsp10kb | %dmSites | %pSite1 | %pSite2 | DMpSite | ADMpSite | DMpNucl | ADMpNucl |
|---|---|---|---|---|---|---|---|---|---|---|---|---|---|
| gene8087 | https://www.ncbi.nlm.nih.gov/gene/?term=LOC105281386 | 3923 | 20 | 50.98 | 1 | 2.55 | 5 | 60.39 | 54.32 | 6.08 | 6.08 | 0.03 | 0.03 |
| gene13296 | https://www.ncbi.nlm.nih.gov/gene/?term=LOC109611531 | 578 | 3 | 51.9 | 1 | 17.3 | 33.33 | 47.46 | 42.32 | 5.14 | 5.14 | 0.03 | 0.03 |
| gene12609 | https://www.ncbi.nlm.nih.gov/gene/?term=LOC105286567 | 1498 | 4 | 26.7 | 1 | 6.68 | 25 | 46.34 | 41.34 | 5.01 | 5.01 | 0.01 | 0.01 |
| gene13317 | https://www.ncbi.nlm.nih.gov/gene/?term=LOC105287372 | 874 | 23 | 263.16 | 1 | 11.44 | 4.35 | 18.83 | 14.08 | 4.75 | 4.75 | 0.12 | 0.12 |
| gene393 | https://www.ncbi.nlm.nih.gov/gene/?term=LOC105276566 | 1123 | 11 | 97.95 | 2 | 17.81 | 18.18 | 39.24 | 34.93 | 4.31 | 4.31 | 0.04 | 0.04 |
| gene10208 | https://www.ncbi.nlm.nih.gov/gene/?term=LOC105283817 | 1335 | 19 | 142.32 | 3 | 22.47 | 15.79 | 54.74 | 50.44 | 4.29 | 4.29 | 0.06 | 0.06 |
| gene8172 | https://www.ncbi.nlm.nih.gov/gene/?term=LOC105281507 | 946 | 25 | 264.27 | 2 | 21.14 | 8 | 23.6 | 19.87 | 3.73 | 3.73 | 0.1 | 0.1 |
| gene2616 | https://www.ncbi.nlm.nih.gov/gene/?term=LOC105275097 | 1073 | 10 | 93.2 | 1 | 9.32 | 10 | 44.18 | 41.09 | 3.09 | 3.09 | 0.03 | 0.03 |
| gene13119 | https://www.ncbi.nlm.nih.gov/gene/?term=LOC105287152 | 1525 | 18 | 118.03 | 1 | 6.56 | 5.56 | 36.95 | 33.99 | 2.96 | 2.96 | 0.03 | 0.03 |
| gene10189 | https://www.ncbi.nlm.nih.gov/gene/?term=LOC105283800 | 871 | 25 | 287.03 | 1 | 11.48 | 4 | 23.42 | 20.63 | 2.79 | 2.79 | 0.08 | 0.08 |
| gene1025 | https://www.ncbi.nlm.nih.gov/gene/?term=LOC105283760 | 1140 | 19 | 166.67 | 1 | 8.77 | 5.26 | 35.33 | 32.72 | 2.61 | 2.61 | 0.04 | 0.04 |
| gene10364 | https://www.ncbi.nlm.nih.gov/gene/?term=LOC105284008 | 560 | 13 | 232.14 | 2 | 35.71 | 15.38 | 84.57 | 82 | 2.56 | 2.56 | 0.06 | 0.06 |
| gene12876 | https://www.ncbi.nlm.nih.gov/gene/?term=LOC105286884 | 944 | 23 | 243.64 | 1 | 10.59 | 4.35 | 6.57 | 4.54 | 2.02 | 2.02 | 0.05 | 0.05 |
| gene4112 | https://www.ncbi.nlm.nih.gov/gene/?term=LOC105276797 | 4145 | 48 | 115.8 | 1 | 2.41 | 2.08 | 7.91 | 5.93 | 1.99 | 1.99 | 0.02 | 0.02 |
| gene6457 | https://www.ncbi.nlm.nih.gov/gene/?term=LOC105279501 | 2130 | 40 | 187.79 | 1 | 4.69 | 2.5 | 51.82 | 49.97 | 1.84 | 1.84 | 0.03 | 0.03 |
| gene2855 | https://www.ncbi.nlm.nih.gov/gene/?term=LOC105275378 | 1189 | 35 | 294.37 | 1 | 8.41 | 2.86 | 45.71 | 43.88 | 1.83 | 1.83 | 0.05 | 0.05 |
| gene3562 | https://www.ncbi.nlm.nih.gov/gene/?term=LOC105276157 | 1591 | 44 | 276.56 | 1 | 6.29 | 2.27 | 27.02 | 25.31 | 1.71 | 1.71 | 0.05 | 0.05 |
| gene8909 | https://www.ncbi.nlm.nih.gov/gene/?term=LOC105282348 | 2254 | 39 | 173.03 | 1 | 4.44 | 2.56 | 7.4 | 5.7 | 1.7 | 1.7 | 0.03 | 0.03 |
| gene11598 | https://www.ncbi.nlm.nih.gov/gene/?term=LOC105285429 | 1324 | 36 | 271.9 | 1 | 7.55 | 2.78 | 40.59 | 38.95 | 1.64 | 1.64 | 0.04 | 0.04 |
| gene1359 | https://www.ncbi.nlm.nih.gov/gene/?term=LOC105287127 | 2231 | 70 | 313.76 | 1 | 4.48 | 1.43 | 6.15 | 4.55 | 1.6 | 1.6 | 0.05 | 0.05 |
| gene12258 | https://www.ncbi.nlm.nih.gov/gene/?term=LOC105286157 | 1983 | 72 | 363.09 | 2 | 10.09 | 2.78 | 24.73 | 23.14 | 1.59 | 1.59 | 0.06 | 0.06 |
| gene7909 | https://www.ncbi.nlm.nih.gov/gene/?term=LOC105281162 | 2177 | 27 | 124.02 | 1 | 4.59 | 3.7 | 32.45 | 30.88 | 1.57 | 1.57 | 0.02 | 0.02 |
| gene11850 | https://www.ncbi.nlm.nih.gov/gene/?term=LOC105285707 | 2053 | 70 | 340.96 | 1 | 4.87 | 1.43 | 32.1 | 30.54 | 1.55 | 1.55 | 0.05 | 0.05 |
| gene7212 | https://www.ncbi.nlm.nih.gov/gene/?term=LOC105280380 | 1416 | 38 | 268.36 | 1 | 7.06 | 2.63 | 6.59 | 5.06 | 1.54 | 1.54 | 0.04 | 0.04 |

...
... only first 25 lines shown ...

Directory: OGL File: ogl-Cb_LIp_Rphase.txt

| gene_ID | gene_link | gwidth | #Sites | #per10Kb | %perSite | %pNucl |
|---|---|---|---|---|---|---|
| gene11569 | https://www.ncbi.nlm.nih.gov/gene/?term=LOC105285394 | 1311 | 150 | 1144.16 | 51.14 | 5.85 |
| gene10219 | https://www.ncbi.nlm.nih.gov/gene/?term=LOC105283830 | 235 | 32 | 1361.7 | 37.49 | 5.1 |
| gene13384 | https://www.ncbi.nlm.nih.gov/gene/?term=LOC105287447 | 1877 | 357 | 1901.97 | 25.33 | 4.82 |
| gene5917 | https://www.ncbi.nlm.nih.gov/gene/?term=LOC105278849 | 823 | 163 | 1980.56 | 24.04 | 4.76 |
| gene4878 | https://www.ncbi.nlm.nih.gov/gene/?term=LOC105277648 | 675 | 52 | 770.37 | 61.25 | 4.72 |
| gene5714 | https://www.ncbi.nlm.nih.gov/gene/?term=LOC105278729 | 710 | 54 | 760.56 | 61.99 | 4.71 |
| gene12602 | https://www.ncbi.nlm.nih.gov/gene/?term=LOC105286562 | 1399 | 134 | 957.83 | 48.86 | 4.68 |
| gene3719 | https://www.ncbi.nlm.nih.gov/gene/?term=LOC105276360 | 1471 | 239 | 1624.75 | 28.29 | 4.6 |
| gene10943 | https://www.ncbi.nlm.nih.gov/gene/?term=LOC105284659 | 1759 | 197 | 1119.95 | 40.8 | 4.57 |
| gene12364 | https://www.ncbi.nlm.nih.gov/gene/?term=LOC105286269 | 1403 | 155 | 1104.78 | 40.99 | 4.53 |
| gene12538 | https://www.ncbi.nlm.nih.gov/gene/?term=LOC105286531 | 372 | 46 | 1236.56 | 36.36 | 4.5 |
| gene211 | https://www.ncbi.nlm.nih.gov/gene/?term=LOC105274534 | 1280 | 144 | 1125 | 38.51 | 4.33 |
| gene2797 | https://www.ncbi.nlm.nih.gov/gene/?term=LOC105275314 | 1489 | 193 | 1296.17 | 33.09 | 4.29 |
| gene12403 | https://www.ncbi.nlm.nih.gov/gene/?term=LOC105286311 | 1413 | 128 | 905.87 | 47.27 | 4.28 |
| gene5309 | https://www.ncbi.nlm.nih.gov/gene/?term=LOC105278160 | 1704 | 206 | 1208.92 | 35.12 | 4.25 |
| gene5412 | https://www.ncbi.nlm.nih.gov/gene/?term=LOC105278266 | 1868 | 277 | 1482.87 | 28.68 | 4.25 |
| gene1894 | https://www.ncbi.nlm.nih.gov/gene/?term=LOC105288083 | 445 | 35 | 786.52 | 52.99 | 4.17 |
| gene1171 | https://www.ncbi.nlm.nih.gov/gene/?term=LOC105285174 | 844 | 94 | 1113.74 | 37.32 | 4.16 |
| gene4635 | https://www.ncbi.nlm.nih.gov/gene/?term=LOC105277390 | 2067 | 234 | 1132.08 | 36.45 | 4.13 |
| gene1654 | https://www.ncbi.nlm.nih.gov/gene/?term=LOC105287811 | 2285 | 260 | 1137.86 | 36.15 | 4.11 |
| gene12652 | https://www.ncbi.nlm.nih.gov/gene/?term=LOC109610640 | 1122 | 90 | 802.14 | 50.98 | 4.09 |
| gene12330 | https://www.ncbi.nlm.nih.gov/gene/?term=LOC105286235 | 1501 | 148 | 986.01 | 41.13 | 4.06 |
| gene6195 | https://www.ncbi.nlm.nih.gov/gene/?term=LOC105279187 | 2221 | 231 | 1040.07 | 38.82 | 4.04 |
| gene5911 | https://www.ncbi.nlm.nih.gov/gene/?term=LOC105278844 | 943 | 135 | 1431.6 | 28.12 | 4.03 |

...
... only first 25 lines shown ...

Directory: OGL File: rnk-dmg-Cb_LIp_BCphase.vs.Rphase.txt

| gene_ID | gene_link | gwidth | #Sites | #per10Kb | #dmSites | #dmsp10kb | %dmSites | %pSite1 | %pSite2 | DMpSite | ADMpSite | DMpNucl | ADMpNucl |
|---|---|---|---|---|---|---|---|---|---|---|---|---|---|
| gene13317 | https://www.ncbi.nlm.nih.gov/gene/?term=LOC105287372 | 874 | 23 | 263.16 | 1 | 11.44 | 4.35 | 18.83 | 14.08 | 4.75 | 4.75 | 0.12 | 0.12 |
| gene8172 | https://www.ncbi.nlm.nih.gov/gene/?term=LOC105281507 | 946 | 25 | 264.27 | 2 | 21.14 | 8 | 23.6 | 19.87 | 3.73 | 3.73 | 0.1 | 0.1 |
| gene6221 | https://www.ncbi.nlm.nih.gov/gene/?term=LOC105279226 | 784 | 27 | 344.39 | 1 | 12.76 | 3.7 | 17.88 | 20.88 | -3 | 3 | -0.1 | 0.1 |
| gene10784 | https://www.ncbi.nlm.nih.gov/gene/?term=LOC105284474 | 481 | 13 | 270.27 | 1 | 20.79 | 7.69 | 21.47 | 24.75 | -3.28 | 3.28 | -0.09 | 0.09 |
| gene10189 | https://www.ncbi.nlm.nih.gov/gene/?term=LOC105283800 | 871 | 25 | 287.03 | 1 | 11.48 | 4 | 23.42 | 20.63 | 2.79 | 2.79 | 0.08 | 0.08 |
| gene5859 | https://www.ncbi.nlm.nih.gov/gene/?term=LOC105278790 | 1169 | 85 | 727.12 | 1 | 8.55 | 1.18 | 23.37 | 22.28 | 1.09 | 1.09 | 0.08 | 0.08 |
| gene6658 | https://www.ncbi.nlm.nih.gov/gene/?term=LOC105279725 | 1518 | 93 | 612.65 | 2 | 13.18 | 2.15 | 21.31 | 20.15 | 1.16 | 1.16 | 0.07 | 0.07 |
| gene10156 | https://www.ncbi.nlm.nih.gov/gene/?term=LOC105283758 | 1649 | 97 | 588.24 | 1 | 6.06 | 1.03 | 22.8 | 21.67 | 1.12 | 1.12 | 0.07 | 0.07 |
| gene5932 | https://www.ncbi.nlm.nih.gov/gene/?term=LOC105278862 | 1583 | 99 | 625.39 | 1 | 6.32 | 1.01 | 28.64 | 27.53 | 1.11 | 1.11 | 0.07 | 0.07 |
| gene11940 | https://www.ncbi.nlm.nih.gov/gene/?term=LOC105285817 | 1432 | 69 | 481.84 | 1 | 6.98 | 1.45 | 16.71 | 18.17 | -1.46 | 1.46 | -0.07 | 0.07 |
| gene13020 | https://www.ncbi.nlm.nih.gov/gene/?term=LOC105287040 | 2622 | 121 | 461.48 | 3 | 11.44 | 2.48 | 5.32 | 6.8 | -1.49 | 1.49 | -0.07 | 0.07 |
| gene4574 | https://www.ncbi.nlm.nih.gov/gene/?term=LOC105277322 | 1768 | 67 | 378.96 | 1 | 5.66 | 1.49 | 36.26 | 38.02 | -1.76 | 1.76 | -0.07 | 0.07 |
| gene2709 | https://www.ncbi.nlm.nih.gov/gene/?term=LOC105275219 | 999 | 36 | 360.36 | 1 | 10.01 | 2.78 | 41.82 | 43.85 | -2.02 | 2.02 | -0.07 | 0.07 |
| gene13311 | https://www.ncbi.nlm.nih.gov/gene/?term=LOC105287364 | 812 | 22 | 270.94 | 1 | 12.32 | 4.55 | 17.25 | 19.87 | -2.62 | 2.62 | -0.07 | 0.07 |
| gene12733 | https://www.ncbi.nlm.nih.gov/gene/?term=LOC109611472 | 1351 | 30 | 222.06 | 1 | 7.4 | 3.33 | 36.52 | 39.58 | -3.05 | 3.05 | -0.07 | 0.07 |
| gene8259 | https://www.ncbi.nlm.nih.gov/gene/?term=LOC105281606 | 1606 | 23 | 143.21 | 2 | 12.45 | 8.7 | 35.16 | 39.91 | -4.74 | 4.74 | -0.07 | 0.07 |
| gene10208 | https://www.ncbi.nlm.nih.gov/gene/?term=LOC105283817 | 1335 | 19 | 142.32 | 3 | 22.47 | 15.79 | 54.74 | 50.44 | 4.29 | 4.29 | 0.06 | 0.06 |
| gene10364 | https://www.ncbi.nlm.nih.gov/gene/?term=LOC105284008 | 560 | 13 | 232.14 | 2 | 35.71 | 15.38 | 84.57 | 82 | 2.56 | 2.56 | 0.06 | 0.06 |
| gene12258 | https://www.ncbi.nlm.nih.gov/gene/?term=LOC105286157 | 1983 | 72 | 363.09 | 2 | 10.09 | 2.78 | 24.73 | 23.14 | 1.59 | 1.59 | 0.06 | 0.06 |
| gene8256 | https://www.ncbi.nlm.nih.gov/gene/?term=LOC105281604 | 2613 | 133 | 508.99 | 2 | 7.65 | 1.5 | 14.92 | 13.78 | 1.14 | 1.14 | 0.06 | 0.06 |
| gene5809 | https://www.ncbi.nlm.nih.gov/gene/?term=LOC105278737 | 1159 | 93 | 802.42 | 1 | 8.63 | 1.08 | 16.5 | 17.22 | -0.71 | 0.71 | -0.06 | 0.06 |
| gene12660 | https://www.ncbi.nlm.nih.gov/gene/?term=LOC105286633 | 1293 | 91 | 703.79 | 1 | 7.73 | 1.1 | 16.75 | 17.57 | -0.82 | 0.82 | -0.06 | 0.06 |
| gene9444 | https://www.ncbi.nlm.nih.gov/gene/?term=LOC105282952 | 1653 | 108 | 653.36 | 2 | 12.1 | 1.85 | 15.9 | 16.75 | -0.85 | 0.85 | -0.06 | 0.06 |
| gene2541 | https://www.ncbi.nlm.nih.gov/gene/?term=LOC105274993 | 2623 | 170 | 648.11 | 1 | 3.81 | 0.59 | 12.06 | 12.92 | -0.86 | 0.86 | -0.06 | 0.06 |

...

... only first 25 lines shown ...

```
Directory: OGL File: wrt-Cb_LIp.txt

... comparing  BCphase.vs.Rphase ...

Number of genes <=  20000  and with #dmsites >=  1 :  1284
sample1     sum of percent methylation: 20177.02     average:    15.71
sample2     sum of percent methylation: 20306.86     average:    15.82

sum difference:  -129.97      average difference:  -0.1012

      Wilcoxon signed rank test with continuity correction

data:  pw_summary$`%pSite1` and pw_summary$`%pSite2`
V = 3e+05, p-value = 3e-13
alternative hypothesis: true location shift is not equal to 0
```

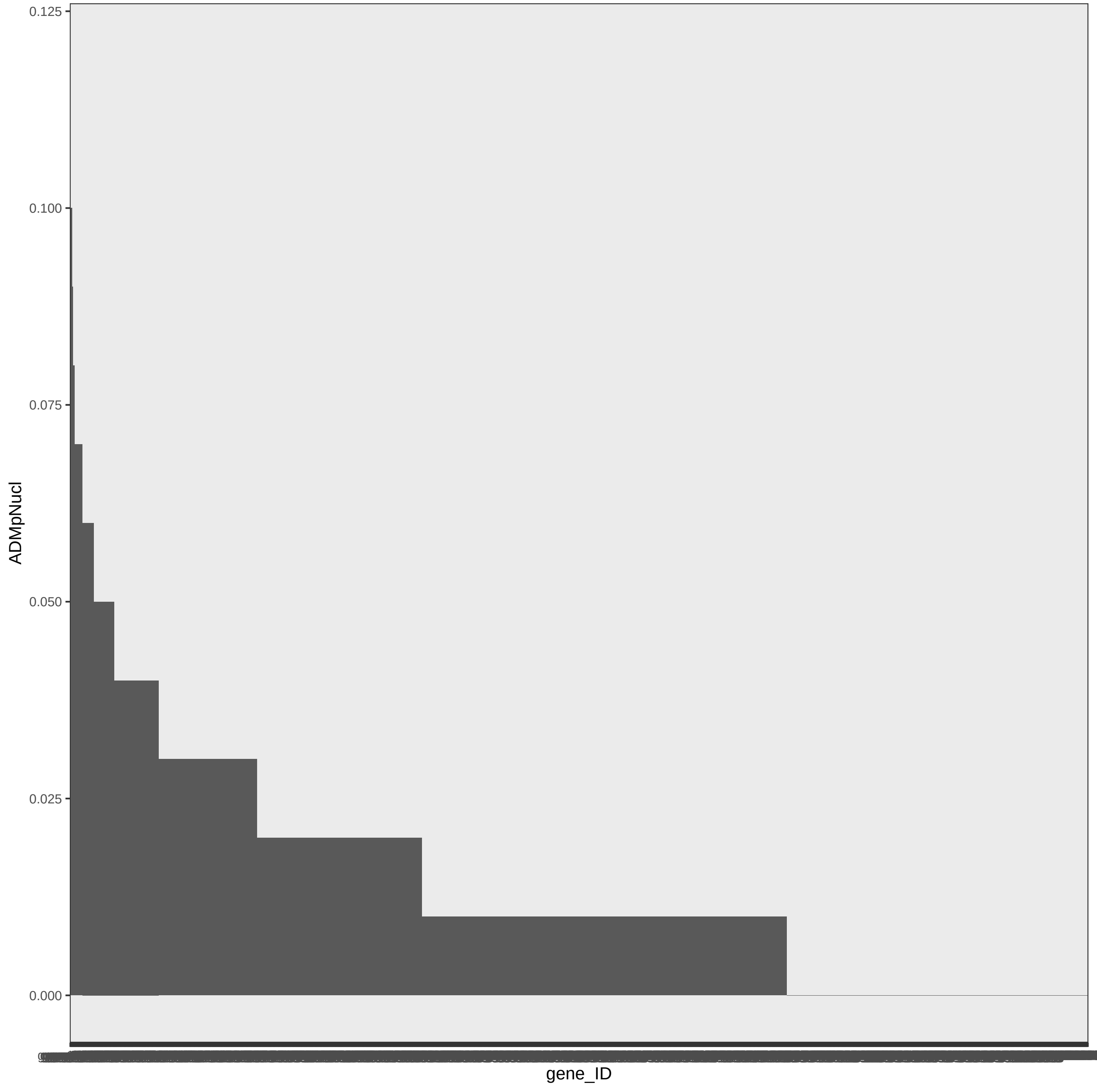

0.125
0.100
0.075
0.050
0.025
0.000
ADMpNucl
gene_ID

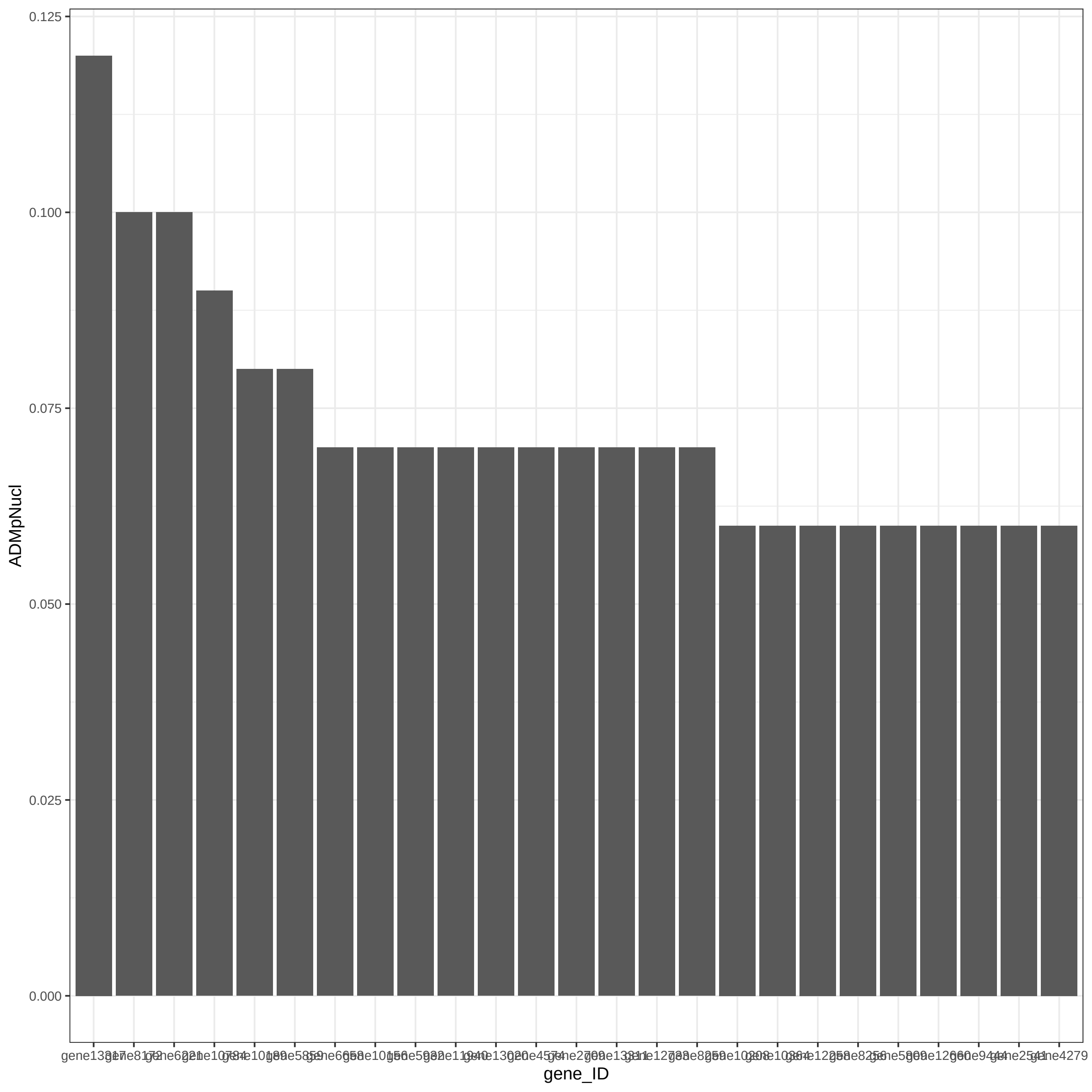

0.125
0.100
0.075
0.050
0.025
0.000
ADMpNucl
gene4279
gene_ID

```
Directory: OGLall File: 0READMEogl

OGL - Ordered gene lists.

Input: studyhc annotation   maxgwidth minnbrdmsites
       dmgprp (= output of BWASPR:show_dmsg())

Output: files ogl-*.txt rnk-dmg-*.txt
         ogl-<study>_<sample>.txt ogl-<study>_<sample1>.vs.<sample2>.txt
         rnk-dmg--<study>_<sample1>.vs.<sample2>.txt rnk-dmg--<study>_<sample1>.vs.<sample2>.pdf
         wrt-<study>.txt

Notes: Output files ogl-<study>_<sample>.txt give tables for each sample with columns

        gene_ID gwidth #Sites #per10Kb %perSite %pNucl

       ordered by %pNucl.  If available, a link to an NCBI entry of gene_ID is inserted as second column.
       Abbreviations used: %perSite, percent methylation per site
                           %perNucl, percent methylation per nucleotide of the gene

       Output files ogl-<study>_<sample1>.vs.<sample2>.txt give tables for each comparison with columns

        gene_ID gwidth #Sites #per10Kb #dmSites #dmsp10kb %dmSites %pSite1 %pSite2 DMpSite ADMpSite DMpNucl ADMpNucl

       ordered by DMpSite.  If available, a link to an NCBI entry of gene_ID is inserted as second column.
       Abbreviations used: %dmSites, percent sites that are differentially methylated
                           %pSite1, average per site % methylation for sample1
                           %pSite2, average per site % methylation for sample2
                           DMpSite, average per site difference in % methylation between sample1 and sample2
                           ADMpSite, absolute value of DMpSite
                           DMpNucl, average per nucleotide difference in % methylation between sample1 and sample2
                           ADMpNucl, absolute value of DMpNucl

       Output files rnk-dmg-<study>_<sample1>.vs.<sample2>.txt are equivalent to files
        ogl-<study>_<sample1>.vs.<sample2>.txt but ordered by ADMpNucl.
       Output files rnk-dmg-<study>_<sample1>.vs.<sample2>.pdf provide visualization of the distribution
        of ADMpNucl values.

       Output file wrt-<study>.txt gives results of the Wilcoxon signed rank test comparing
        the %pSite1 and %pSite2 vectors

       In output directory OGL, tables are restricted to genes with gwidth <= maxgwidth and, for
        pairwise comparisons, #dmSites >= minnbrdmsites

       If either maxgwidth or minnbrdmsites is set, then output directory OGLall will show the full tables
        with all genes (for reference).
```

Directory: OGLall File: ogl-Cb_LIp_BCphase.txt

| gene_ID | gene_link | gwidth | #Sites | #per10Kb | %perSite | %pNucl |
|---|---|---|---|---|---|---|
| gene11569 | https://www.ncbi.nlm.nih.gov/gene/?term=LOC105285394 | 1311 | 149 | 1136.54 | 49.95 | 5.68 |
| gene10219 | https://www.ncbi.nlm.nih.gov/gene/?term=LOC105283830 | 235 | 32 | 1361.7 | 36.97 | 5.03 |
| gene4878 | https://www.ncbi.nlm.nih.gov/gene/?term=LOC105277648 | 675 | 53 | 785.19 | 62.39 | 4.9 |
| gene13384 | https://www.ncbi.nlm.nih.gov/gene/?term=LOC105287447 | 1877 | 356 | 1896.64 | 25.55 | 4.85 |
| gene5917 | https://www.ncbi.nlm.nih.gov/gene/?term=LOC105278849 | 823 | 163 | 1980.56 | 24.16 | 4.79 |
| gene12602 | https://www.ncbi.nlm.nih.gov/gene/?term=LOC105286562 | 1399 | 128 | 914.94 | 51.55 | 4.72 |
| gene5714 | https://www.ncbi.nlm.nih.gov/gene/?term=LOC105278729 | 710 | 54 | 760.56 | 61.16 | 4.65 |
| gene3719 | https://www.ncbi.nlm.nih.gov/gene/?term=LOC105276360 | 1471 | 239 | 1624.75 | 28.35 | 4.61 |
| gene12538 | https://www.ncbi.nlm.nih.gov/gene/?term=LOC105286531 | 372 | 46 | 1236.56 | 36.97 | 4.57 |
| gene1894 | https://www.ncbi.nlm.nih.gov/gene/?term=LOC105288083 | 445 | 36 | 808.99 | 56.38 | 4.56 |
| gene12364 | https://www.ncbi.nlm.nih.gov/gene/?term=LOC105286269 | 1403 | 156 | 1111.9 | 40.37 | 4.49 |
| gene10943 | https://www.ncbi.nlm.nih.gov/gene/?term=LOC105284659 | 1759 | 196 | 1114.27 | 40.19 | 4.48 |
| gene12403 | https://www.ncbi.nlm.nih.gov/gene/?term=LOC105286311 | 1413 | 129 | 912.95 | 46.44 | 4.24 |
| gene1171 | https://www.ncbi.nlm.nih.gov/gene/?term=LOC105285174 | 844 | 95 | 1125.59 | 37.4 | 4.21 |
| gene2797 | https://www.ncbi.nlm.nih.gov/gene/?term=LOC105275314 | 1489 | 189 | 1269.31 | 33.05 | 4.2 |
| gene5309 | https://www.ncbi.nlm.nih.gov/gene/?term=LOC105278160 | 1704 | 201 | 1179.58 | 35.46 | 4.18 |
| gene6195 | https://www.ncbi.nlm.nih.gov/gene/?term=LOC105279187 | 2221 | 235 | 1058.08 | 39.44 | 4.17 |
| gene211 | https://www.ncbi.nlm.nih.gov/gene/?term=LOC105274534 | 1280 | 140 | 1093.75 | 37.99 | 4.16 |
| gene12330 | https://www.ncbi.nlm.nih.gov/gene/?term=LOC105286235 | 1501 | 149 | 992.67 | 41.83 | 4.15 |
| gene12652 | https://www.ncbi.nlm.nih.gov/gene/?term=LOC109610640 | 1122 | 93 | 828.88 | 50.01 | 4.14 |
| gene4635 | https://www.ncbi.nlm.nih.gov/gene/?term=LOC105277390 | 2067 | 232 | 1122.4 | 36.6 | 4.11 |
| gene1654 | https://www.ncbi.nlm.nih.gov/gene/?term=LOC105287811 | 2285 | 259 | 1133.48 | 36.08 | 4.09 |
| gene5412 | https://www.ncbi.nlm.nih.gov/gene/?term=LOC105278266 | 1868 | 259 | 1386.51 | 29.43 | 4.08 |
| gene11566 | https://www.ncbi.nlm.nih.gov/gene/?term=LOC105285392 | 1603 | 230 | 1434.81 | 28.17 | 4.04 |

...

... only first 25 lines shown ...

Directory: OGLall File: ogl-Cb_LIp_BCphase.vs.Rphase.txt

| gene_ID | gene_link | gwidth | #Sites | #per10Kb | #dmSites | #dmsp10kb | %dmSites | %pSite1 | %pSite2 | DMpSite | ADMpSite | DMpNucl | ADMpNucl |
|---|---|---|---|---|---|---|---|---|---|---|---|---|---|
| gene8087 | https://www.ncbi.nlm.nih.gov/gene/?term=LOC105281386 | 3923 | 20 | 50.98 | 1 | 2.55 | 5 | 60.39 | 54.32 | 6.08 | 6.08 | 0.03 | 0.03 |
| gene13296 | https://www.ncbi.nlm.nih.gov/gene/?term=LOC109611531 | 578 | 3 | 51.9 | 1 | 17.3 | 33.33 | 47.46 | 42.32 | 5.14 | 5.14 | 0.03 | 0.03 |
| gene12609 | https://www.ncbi.nlm.nih.gov/gene/?term=LOC105286567 | 1498 | 4 | 26.7 | 1 | 6.68 | 25 | 46.34 | 41.34 | 5.01 | 5.01 | 0.01 | 0.01 |
| gene13317 | https://www.ncbi.nlm.nih.gov/gene/?term=LOC105287372 | 874 | 23 | 263.16 | 1 | 11.44 | 4.35 | 18.83 | 14.08 | 4.75 | 4.75 | 0.12 | 0.12 |
| gene393 | https://www.ncbi.nlm.nih.gov/gene/?term=LOC105276566 | 1123 | 11 | 97.95 | 2 | 17.81 | 18.18 | 39.24 | 34.93 | 4.31 | 4.31 | 0.04 | 0.04 |
| gene10208 | https://www.ncbi.nlm.nih.gov/gene/?term=LOC105283817 | 1335 | 19 | 142.32 | 3 | 22.47 | 15.79 | 54.74 | 50.44 | 4.29 | 4.29 | 0.06 | 0.06 |
| gene8172 | https://www.ncbi.nlm.nih.gov/gene/?term=LOC105281507 | 946 | 25 | 264.27 | 2 | 21.14 | 8 | 23.6 | 19.87 | 3.73 | 3.73 | 0.1 | 0.1 |
| gene2616 | https://www.ncbi.nlm.nih.gov/gene/?term=LOC105275097 | 1073 | 10 | 93.2 | 1 | 9.32 | 10 | 44.18 | 41.09 | 3.09 | 3.09 | 0.03 | 0.03 |
| gene13119 | https://www.ncbi.nlm.nih.gov/gene/?term=LOC105287152 | 1525 | 18 | 118.03 | 1 | 6.56 | 5.56 | 36.95 | 33.99 | 2.96 | 2.96 | 0.03 | 0.03 |
| gene10189 | https://www.ncbi.nlm.nih.gov/gene/?term=LOC105283800 | 871 | 25 | 287.03 | 1 | 11.48 | 4 | 23.42 | 20.63 | 2.79 | 2.79 | 0.08 | 0.08 |
| gene1025 | https://www.ncbi.nlm.nih.gov/gene/?term=LOC105283760 | 1140 | 19 | 166.67 | 1 | 8.77 | 5.26 | 35.33 | 32.72 | 2.61 | 2.61 | 0.04 | 0.04 |
| gene10364 | https://www.ncbi.nlm.nih.gov/gene/?term=LOC105284008 | 560 | 13 | 232.14 | 2 | 35.71 | 15.38 | 84.57 | 82 | 2.56 | 2.56 | 0.06 | 0.06 |
| gene12876 | https://www.ncbi.nlm.nih.gov/gene/?term=LOC105286884 | 944 | 23 | 243.64 | 1 | 10.59 | 4.35 | 6.57 | 4.54 | 2.02 | 2.02 | 0.05 | 0.05 |
| gene4112 | https://www.ncbi.nlm.nih.gov/gene/?term=LOC105276797 | 4145 | 48 | 115.8 | 1 | 2.41 | 2.08 | 7.91 | 5.93 | 1.99 | 1.99 | 0.02 | 0.02 |
| gene6457 | https://www.ncbi.nlm.nih.gov/gene/?term=LOC105279501 | 2130 | 40 | 187.79 | 1 | 4.69 | 2.5 | 51.82 | 49.97 | 1.84 | 1.84 | 0.03 | 0.03 |
| gene2855 | https://www.ncbi.nlm.nih.gov/gene/?term=LOC105275378 | 1189 | 35 | 294.37 | 1 | 8.41 | 2.86 | 45.71 | 43.88 | 1.83 | 1.83 | 0.05 | 0.05 |
| gene3562 | https://www.ncbi.nlm.nih.gov/gene/?term=LOC105276157 | 1591 | 44 | 276.56 | 1 | 6.29 | 2.27 | 27.02 | 25.31 | 1.71 | 1.71 | 0.05 | 0.05 |
| gene8909 | https://www.ncbi.nlm.nih.gov/gene/?term=LOC105282348 | 2254 | 39 | 173.03 | 1 | 4.44 | 2.56 | 7.4 | 5.7 | 1.7 | 1.7 | 0.03 | 0.03 |
| gene11598 | https://www.ncbi.nlm.nih.gov/gene/?term=LOC105285429 | 1324 | 36 | 271.9 | 1 | 7.55 | 2.78 | 40.59 | 38.95 | 1.64 | 1.64 | 0.04 | 0.04 |
| gene1359 | https://www.ncbi.nlm.nih.gov/gene/?term=LOC105287127 | 2231 | 70 | 313.76 | 1 | 4.48 | 1.43 | 6.15 | 4.55 | 1.6 | 1.6 | 0.05 | 0.05 |
| gene12258 | https://www.ncbi.nlm.nih.gov/gene/?term=LOC105286157 | 1983 | 72 | 363.09 | 2 | 10.09 | 2.78 | 24.73 | 23.14 | 1.59 | 1.59 | 0.06 | 0.06 |
| gene7909 | https://www.ncbi.nlm.nih.gov/gene/?term=LOC105281162 | 2177 | 27 | 124.02 | 1 | 4.59 | 3.7 | 32.45 | 30.88 | 1.57 | 1.57 | 0.02 | 0.02 |
| gene11850 | https://www.ncbi.nlm.nih.gov/gene/?term=LOC105285707 | 2053 | 70 | 340.96 | 1 | 4.87 | 1.43 | 32.1 | 30.54 | 1.55 | 1.55 | 0.05 | 0.05 |
| gene7212 | https://www.ncbi.nlm.nih.gov/gene/?term=LOC105280380 | 1416 | 38 | 268.36 | 1 | 7.06 | 2.63 | 6.59 | 5.06 | 1.54 | 1.54 | 0.04 | 0.04 |

...
... only first 25 lines shown ...

Directory: OGLall File: ogl-Cb_LIp_Rphase.txt

| gene_ID | gene_link | gwidth | #Sites | #per10Kb | %perSite | %pNucl |
|---|---|---|---|---|---|---|
| gene11569 | https://www.ncbi.nlm.nih.gov/gene/?term=LOC105285394 | 1311 | 150 | 1144.16 | 51.14 | 5.85 |
| gene10219 | https://www.ncbi.nlm.nih.gov/gene/?term=LOC105283830 | 235 | 32 | 1361.7 | 37.49 | 5.1 |
| gene13384 | https://www.ncbi.nlm.nih.gov/gene/?term=LOC105287447 | 1877 | 357 | 1901.97 | 25.33 | 4.82 |
| gene5917 | https://www.ncbi.nlm.nih.gov/gene/?term=LOC105278849 | 823 | 163 | 1980.56 | 24.04 | 4.76 |
| gene4878 | https://www.ncbi.nlm.nih.gov/gene/?term=LOC105277648 | 675 | 52 | 770.37 | 61.25 | 4.72 |
| gene5714 | https://www.ncbi.nlm.nih.gov/gene/?term=LOC105278729 | 710 | 54 | 760.56 | 61.99 | 4.71 |
| gene12602 | https://www.ncbi.nlm.nih.gov/gene/?term=LOC105286562 | 1399 | 134 | 957.83 | 48.86 | 4.68 |
| gene3719 | https://www.ncbi.nlm.nih.gov/gene/?term=LOC105276360 | 1471 | 239 | 1624.75 | 28.29 | 4.6 |
| gene10943 | https://www.ncbi.nlm.nih.gov/gene/?term=LOC105284659 | 1759 | 197 | 1119.95 | 40.8 | 4.57 |
| gene12364 | https://www.ncbi.nlm.nih.gov/gene/?term=LOC105286269 | 1403 | 155 | 1104.78 | 40.99 | 4.53 |
| gene12538 | https://www.ncbi.nlm.nih.gov/gene/?term=LOC105286531 | 372 | 46 | 1236.56 | 36.36 | 4.5 |
| gene211 | https://www.ncbi.nlm.nih.gov/gene/?term=LOC105274534 | 1280 | 144 | 1125 38.51 | 4.33 | |
| gene2797 | https://www.ncbi.nlm.nih.gov/gene/?term=LOC105275314 | 1489 | 193 | 1296.17 | 33.09 | 4.29 |
| gene12403 | https://www.ncbi.nlm.nih.gov/gene/?term=LOC105286311 | 1413 | 128 | 905.87 | 47.27 | 4.28 |
| gene5309 | https://www.ncbi.nlm.nih.gov/gene/?term=LOC105278160 | 1704 | 206 | 1208.92 | 35.12 | 4.25 |
| gene5412 | https://www.ncbi.nlm.nih.gov/gene/?term=LOC105278266 | 1868 | 277 | 1482.87 | 28.68 | 4.25 |
| gene1894 | https://www.ncbi.nlm.nih.gov/gene/?term=LOC105288083 | 445 | 35 | 786.52 | 52.99 | 4.17 |
| gene1171 | https://www.ncbi.nlm.nih.gov/gene/?term=LOC105285174 | 844 | 94 | 1113.74 | 37.32 | 4.16 |
| gene4635 | https://www.ncbi.nlm.nih.gov/gene/?term=LOC105277390 | 2067 | 234 | 1132.08 | 36.45 | 4.13 |
| gene1654 | https://www.ncbi.nlm.nih.gov/gene/?term=LOC105287811 | 2285 | 260 | 1137.86 | 36.15 | 4.11 |
| gene12652 | https://www.ncbi.nlm.nih.gov/gene/?term=LOC109610640 | 1122 | 90 | 802.14 | 50.98 | 4.09 |
| gene12330 | https://www.ncbi.nlm.nih.gov/gene/?term=LOC105286235 | 1501 | 148 | 986.01 | 41.13 | 4.06 |
| gene6195 | https://www.ncbi.nlm.nih.gov/gene/?term=LOC105279187 | 2221 | 231 | 1040.07 | 38.82 | 4.04 |
| gene5911 | https://www.ncbi.nlm.nih.gov/gene/?term=LOC105278844 | 943 | 135 | 1431.6 | 28.12 | 4.03 |

...
... only first 25 lines shown ...

Directory: OGLall File: rnk-dmg-Cb_LIp_BCphase.vs.Rphase.txt

| gene_ID | gene_link | gwidth | #Sites | #per10Kb | #dmSites | #dmsp10kb | %dmSites | %pSite1 | %pSite2 | DMpSite | ADMpSite | DMpNucl | ADMpNucl |
|---|---|---|---|---|---|---|---|---|---|---|---|---|---|
| gene13317 | https://www.ncbi.nlm.nih.gov/gene/?term=LOC105287372 | 874 | 23 | 263.16 | 1 | 11.44 | 4.35 | 18.83 | 14.08 | 4.75 | 4.75 | 0.12 | 0.12 |
| gene8172 | https://www.ncbi.nlm.nih.gov/gene/?term=LOC105281507 | 946 | 25 | 264.27 | 2 | 21.14 | 8 | 23.6 | 19.87 | 3.73 | 3.73 | 0.1 | 0.1 |
| gene6221 | https://www.ncbi.nlm.nih.gov/gene/?term=LOC105279226 | 784 | 27 | 344.39 | 1 | 12.76 | 3.7 | 17.88 | 20.88 | -3 | 3 | -0.1 | 0.1 |
| gene10784 | https://www.ncbi.nlm.nih.gov/gene/?term=LOC105284474 | 481 | 13 | 270.27 | 1 | 20.79 | 7.69 | 21.47 | 24.75 | -3.28 | 3.28 | -0.09 | 0.09 |
| gene10189 | https://www.ncbi.nlm.nih.gov/gene/?term=LOC105283800 | 871 | 25 | 287.03 | 1 | 11.48 | 4 | 23.42 | 20.63 | 2.79 | 2.79 | 0.08 | 0.08 |
| gene5859 | https://www.ncbi.nlm.nih.gov/gene/?term=LOC105278790 | 1169 | 85 | 727.12 | 1 | 8.55 | 1.18 | 23.37 | 22.28 | 1.09 | 1.09 | 0.08 | 0.08 |
| gene6658 | https://www.ncbi.nlm.nih.gov/gene/?term=LOC105279725 | 1518 | 93 | 612.65 | 2 | 13.18 | 2.15 | 21.31 | 20.15 | 1.16 | 1.16 | 0.07 | 0.07 |
| gene10156 | https://www.ncbi.nlm.nih.gov/gene/?term=LOC105283758 | 1649 | 97 | 588.24 | 1 | 6.06 | 1.03 | 22.8 | 21.67 | 1.12 | 1.12 | 0.07 | 0.07 |
| gene5932 | https://www.ncbi.nlm.nih.gov/gene/?term=LOC105278862 | 1583 | 99 | 625.39 | 1 | 6.32 | 1.01 | 28.64 | 27.53 | 1.11 | 1.11 | 0.07 | 0.07 |
| gene11940 | https://www.ncbi.nlm.nih.gov/gene/?term=LOC105285817 | 1432 | 69 | 481.84 | 1 | 6.98 | 1.45 | 16.71 | 18.17 | -1.46 | 1.46 | -0.07 | 0.07 |
| gene13020 | https://www.ncbi.nlm.nih.gov/gene/?term=LOC105287040 | 2622 | 121 | 461.48 | 3 | 11.44 | 2.48 | 5.32 | 6.8 | -1.49 | 1.49 | -0.07 | 0.07 |
| gene4574 | https://www.ncbi.nlm.nih.gov/gene/?term=LOC105277322 | 1768 | 67 | 378.96 | 1 | 5.66 | 1.49 | 36.26 | 38.02 | -1.76 | 1.76 | -0.07 | 0.07 |
| gene2709 | https://www.ncbi.nlm.nih.gov/gene/?term=LOC105275219 | 999 | 36 | 360.36 | 1 | 10.01 | 2.78 | 41.82 | 43.85 | -2.02 | 2.02 | -0.07 | 0.07 |
| gene13311 | https://www.ncbi.nlm.nih.gov/gene/?term=LOC105287364 | 812 | 22 | 270.94 | 1 | 12.32 | 4.55 | 17.25 | 19.87 | -2.62 | 2.62 | -0.07 | 0.07 |
| gene12733 | https://www.ncbi.nlm.nih.gov/gene/?term=LOC109611472 | 1351 | 30 | 222.06 | 1 | 7.4 | 3.33 | 36.52 | 39.58 | -3.05 | 3.05 | -0.07 | 0.07 |
| gene8259 | https://www.ncbi.nlm.nih.gov/gene/?term=LOC105281606 | 1606 | 23 | 143.21 | 2 | 12.45 | 8.7 | 35.16 | 39.91 | -4.74 | 4.74 | -0.07 | 0.07 |
| gene10208 | https://www.ncbi.nlm.nih.gov/gene/?term=LOC105283817 | 1335 | 19 | 142.32 | 3 | 22.47 | 15.79 | 54.74 | 50.44 | 4.29 | 4.29 | 0.06 | 0.06 |
| gene10364 | https://www.ncbi.nlm.nih.gov/gene/?term=LOC105284008 | 560 | 13 | 232.14 | 2 | 35.71 | 15.38 | 84.57 | 82 | 2.56 | 2.56 | 0.06 | 0.06 |
| gene12258 | https://www.ncbi.nlm.nih.gov/gene/?term=LOC105286157 | 1983 | 72 | 363.09 | 2 | 10.09 | 2.78 | 24.73 | 23.14 | 1.59 | 1.59 | 0.06 | 0.06 |
| gene8256 | https://www.ncbi.nlm.nih.gov/gene/?term=LOC105281604 | 2613 | 133 | 508.99 | 2 | 7.65 | 1.5 | 14.92 | 13.78 | 1.14 | 1.14 | 0.06 | 0.06 |
| gene5809 | https://www.ncbi.nlm.nih.gov/gene/?term=LOC105278737 | 1159 | 93 | 802.42 | 1 | 8.63 | 1.08 | 16.5 | 17.22 | -0.71 | 0.71 | -0.06 | 0.06 |
| gene12660 | https://www.ncbi.nlm.nih.gov/gene/?term=LOC105286633 | 1293 | 91 | 703.79 | 1 | 7.73 | 1.1 | 16.75 | 17.57 | -0.82 | 0.82 | -0.06 | 0.06 |
| gene9444 | https://www.ncbi.nlm.nih.gov/gene/?term=LOC105282952 | 1653 | 108 | 653.36 | 2 | 12.1 | 1.85 | 15.9 | 16.75 | -0.85 | 0.85 | -0.06 | 0.06 |
| gene2541 | https://www.ncbi.nlm.nih.gov/gene/?term=LOC105274993 | 2623 | 170 | 648.11 | 1 | 3.81 | 0.59 | 12.06 | 12.92 | -0.86 | 0.86 | -0.06 | 0.06 |

...

... only first 25 lines shown ...

```
Directory: OGLall File: wrt-Cb_LIp.txt

... comparing  BCphase.vs.Rphase ...

Number of genes with #dmsites >=  1 :  1533

sample1     sum of percent methylation: 20408.72      average:    13.31
sample2     sum of percent methylation: 20542.75      average:    13.40

sum difference:  -134.18      average difference:  -0.0875

      Wilcoxon signed rank test with continuity correction

data:  pw_summary$`%pSite1` and pw_summary$`%pSite2`
V = 4e+05, p-value = 6e-16
alternative hypothesis: true location shift is not equal to 0
```

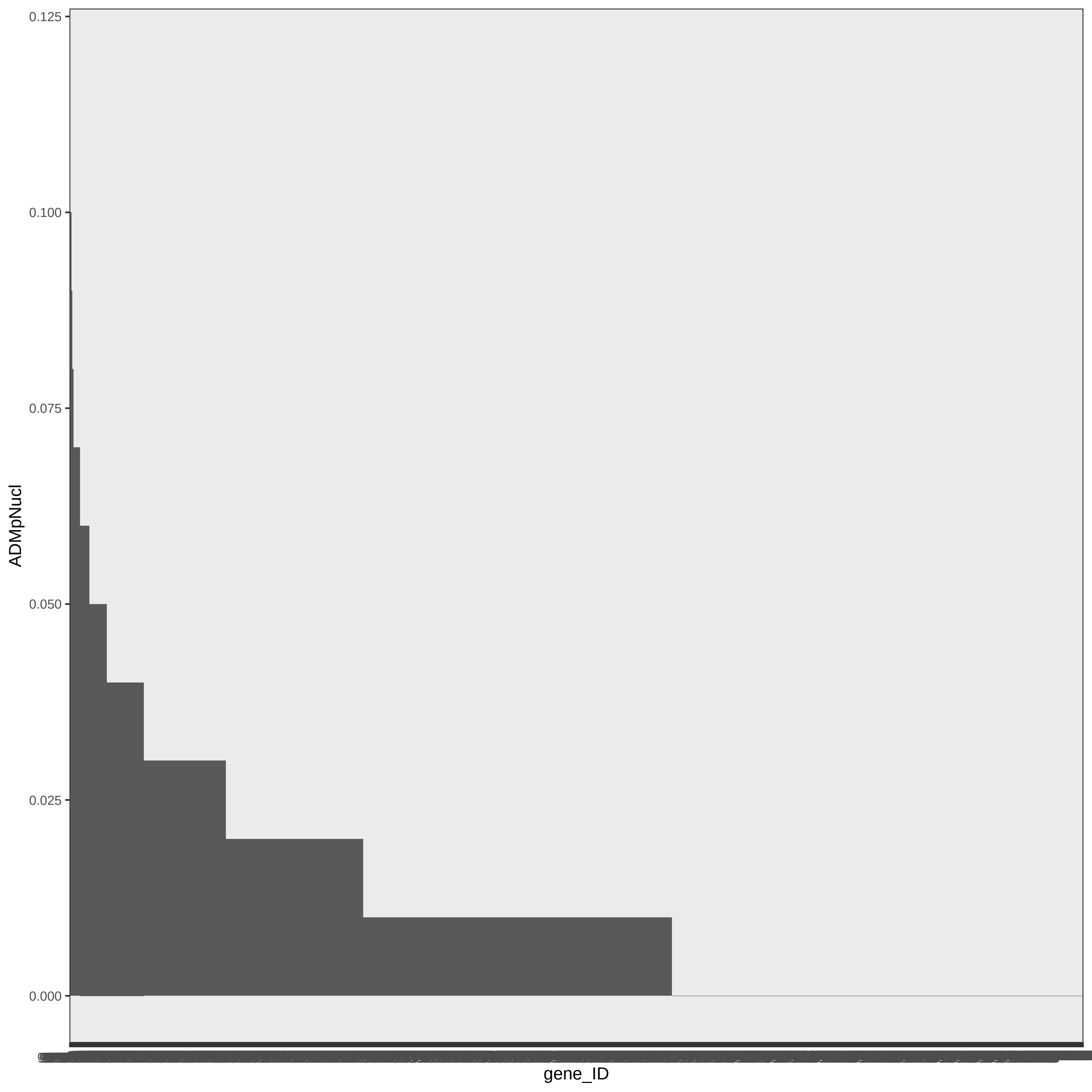

0.125
0.100
0.075
0.050
0.025
0.000
ADMpNucl
gene_ID

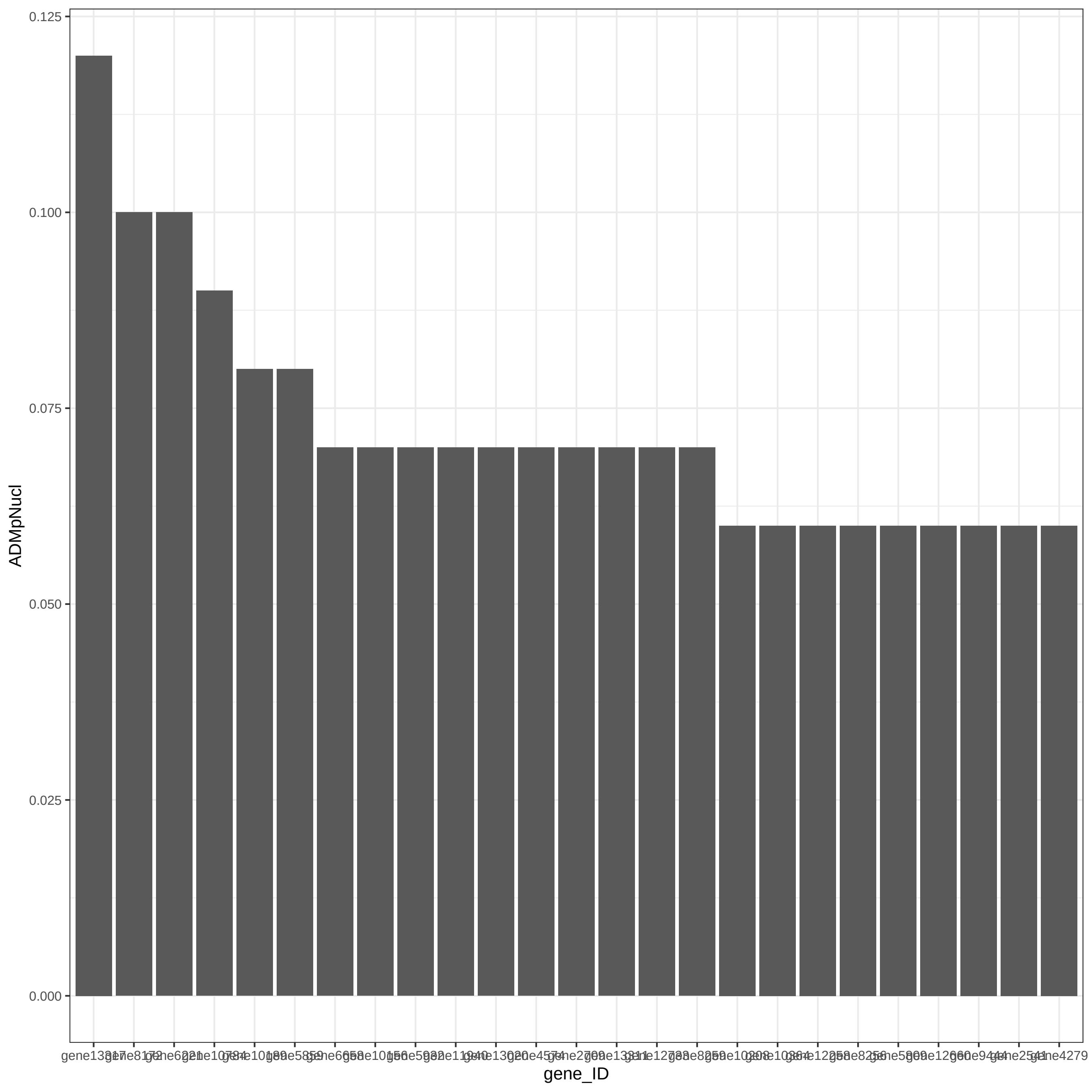
0.125
0.100
0.075
0.050
0.025
0.000
ADMpNucl
gene4279
gene_ID
