## Supplementary material for "Tools and applications for integrative analysis of DNA methylation in social insects": Dataset S18

Cb_LI.conf

```
#Customize variables here:#####################################################
###############################################################################

#Load the files:
#
infiles       <- setup_BWASPR(datafile="./Cb.dat",
                              parfile="./Cb.par")

#Set the study and samples in the study:
#
species       <- "Cb"
study         <- "LI"
samplelist    <- list("BCphaseA1", "BCphaseA2", "BCphaseB1", "BCphaseB2", "RphaseA1", "RphaseA2", "RphaseB1", "RphaseB2")
## The following two variables are used for output file labeling:
studyLabel    <- "LI"
sampleLabels  <- list("Cb_LI_BCphaseA1", "Cb_LI_BCphaseA2", "Cb_LI_BCphaseB1", "Cb_LI_BCphaseB2", "Cb_LI_RphaseA1", "Cb_LI_RphaseA2", "Cb_LI_RphaseB1", "Cb_LI_RphaseB2")

hasreplicates <- FALSE
type          <- "CpG"
destrand      <- TRUE
covlist       <- c(6,20,50)
locount       <- 100
hicount       <- 1000
repcovlist    <- c(6,10,15)
replocount    <- 10
rephicount    <- 100
hheight       <- 0.035
nbrpnts       <- 5000

## The following four variables are sent to rank_rbm() and determine what data
#  points get plotted:
minnbrmsprm   <- 5      # minimum number of methylation sites per promoter
mingenewidth  <- 500    # minimum gene width
maxgenewidth  <- 5000   # maximum gene width
minnbrmsgene  <- 5      # minimum number of methylation sites per gene

## Other parameters (see Rscript.BWASPR for usage notes):
#
highcoverage  <- 20     # high read coverage threshold for studyhc methylRawList object
threshold     <- 20.0   # "difference" threshold for getMethylDiff(), called by det_dmsg()
qvalue        <- 0.05   # "qvalue" setting for getMethylDiff(), called by det_dmsg()
wsize         <- 500    # "win.size" parameter for tileMethylCounts() in det_dmt()
stepsize      <- 500    # "step.size" parameter for tileMethylCounts() in det_dmt()

minNsites     <-  1     # minimum number of hc sites in a gene to be heatmapped in show_dmsg()
maxNsites     <- 60     # maximum number of hc sites in a gene to be heatmapped in show_dmsg()
minPdmsites   <- 10     # minimum ratio of dm/hc (in %) in a gene to be heatmapped in show_dmsg()

maxgwidth     <- 20000  # maximal gene width for a gene to be considered by explore_dmsg()
minnbrdmsites <- 1      # minimum number of differentially methylated sites for a gene to be
                        #  considered in sample comparisons by explore_dmsg()
glink         <- "NCBIgene"   # URL to show in explore_dmsg(); options: "" or "NCBIgene"

#Set the number of processors to use:
#
numprc <- 6

#Determine what analyses to run:
#
RUNload    <- FALSE
```

```
RUNcms     <- TRUE
RUNpwc     <- TRUE
RUNcrl     <- TRUE

RUNrepcms  <- FALSE
RUNrepcrl  <- FALSE

RUNmmp     <- TRUE
RUNacs     <- TRUE
RUNrnk     <- TRUE
RUNmrpr    <- TRUE

RUNdmt     <- TRUE
RUNdmsg    <- TRUE
RUNdmgdtls <- TRUE
RUNogl     <- TRUE

RUNsave    <- TRUE

mymessage <- sprintf("\nAnalyzing %s study %s for type %s\n\n",species,studyLabel,type)
message(mymessage)

################################################################################
#End of typical customization.##################################################
```

Cb.par

```
SPECIESNAME Cerapachys biroi
TOTALNBRPMSITES   26684764
ASSEMBLYVERSION   CerBir1.0
GENOMESIZE  212825769
SPECIESGFF3DIR    ./MCALLS/Cbir/genome/GFF3DIR
GENELISTGFF3      Cbir.gene.gff3
EXONLISTGFF3      Cbir.exon.gff3
PCGEXNLISTGFF3    Cbir.pcg-exon.gff3
PROMOTRLISTGFF3   Cbir.promoter.gff3
CDSLISTGFF3 Cbir.pcg-CDS.gff3
UTRFLAGSET  1
5UTRLISTGFF3      Cbir.pcg-5pUTR.gff3
3UTRLISTGFF3      Cbir.pcg-3pUTR.gff3
```

Cb.dat

```
# Samples from Libbrecht et al. (2016) Current Biology:
#

### 1. Comparisons between all 8 data sets (no replicates) ###
#

Cb    LI    BCphaseA1   0     CpGhsm      ./MCALLS/Cbir/Libbrecht2016/BCphaseA/replicate1/BCphaseA1.CpGhsm.mcalls
Cb    LI    BCphaseA1   0     CpGscd      ./MCALLS/Cbir/Libbrecht2016/BCphaseA/replicate1/BCphaseA1.CpGscd.mcalls
Cb    LI    BCphaseA2   0     CpGhsm      ./MCALLS/Cbir/Libbrecht2016/BCphaseA/replicate2/BCphaseA2.CpGhsm.mcalls
Cb    LI    BCphaseA2   0     CpGscd      ./MCALLS/Cbir/Libbrecht2016/BCphaseA/replicate2/BCphaseA2.CpGscd.mcalls

Cb    LI    BCphaseB1   0     CpGhsm      ./MCALLS/Cbir/Libbrecht2016/BCphaseB/replicate1/BCphaseB1.CpGhsm.mcalls
Cb    LI    BCphaseB1   0     CpGscd      ./MCALLS/Cbir/Libbrecht2016/BCphaseB/replicate1/BCphaseB1.CpGscd.mcalls
Cb    LI    BCphaseB2   0     CpGhsm      ./MCALLS/Cbir/Libbrecht2016/BCphaseB/replicate2/BCphaseB2.CpGhsm.mcalls
Cb    LI    BCphaseB2   0     CpGscd      ./MCALLS/Cbir/Libbrecht2016/BCphaseB/replicate2/BCphaseB2.CpGscd.mcalls

Cb    LI    RphaseA1    0     CpGhsm      ./MCALLS/Cbir/Libbrecht2016/RphaseA/replicate1/RphaseA1.CpGhsm.mcalls
Cb    LI    RphaseA1    0     CpGscd      ./MCALLS/Cbir/Libbrecht2016/RphaseA/replicate1/RphaseA1.CpGscd.mcalls
Cb    LI    RphaseA2    0     CpGhsm      ./MCALLS/Cbir/Libbrecht2016/RphaseA/replicate2/RphaseA2.CpGhsm.mcalls
Cb    LI    RphaseA2    0     CpGscd      ./MCALLS/Cbir/Libbrecht2016/RphaseA/replicate2/RphaseA2.CpGscd.mcalls

Cb    LI    RphaseB1    0     CpGhsm      ./MCALLS/Cbir/Libbrecht2016/RphaseB/replicate1/RphaseB1.CpGhsm.mcalls
Cb    LI    RphaseB1    0     CpGscd      ./MCALLS/Cbir/Libbrecht2016/RphaseB/replicate1/RphaseB1.CpGscd.mcalls
Cb    LI    RphaseB2    0     CpGhsm      ./MCALLS/Cbir/Libbrecht2016/RphaseB/replicate2/RphaseB2.CpGhsm.mcalls
Cb    LI    RphaseB2    0     CpGscd      ./MCALLS/Cbir/Libbrecht2016/RphaseB/replicate2/RphaseB2.CpGscd.mcalls

### 2. Pairwise omparisons between data sets split accorinding to phase, or line, or batch (2 samples, each with 4 replicates) ###
### 2.1 Comparing BCphase versus Rphase ###
#
# BCphase
Cb    LI    BCphase     0     CpGhsm      ./MCALLS/Cbir/Libbrecht2016/PhaseBCvsR/PhaseBC/PhaseBC.CpGhsm.mcalls
Cb    LI    BCphase     0     CpGscd      ./MCALLS/Cbir/Libbrecht2016/PhaseBCvsR/PhaseBC/PhaseBC.CpGscd.mcalls

# ...with replicates
Cb    LI    BCphase     1     CpGhsm      ./MCALLS/Cbir/Libbrecht2016/BCphaseA/replicate1/BCphaseA1.CpGhsm.mcalls
Cb    LI    BCphase     1     CpGscd      ./MCALLS/Cbir/Libbrecht2016/BCphaseA/replicate1/BCphaseA1.CpGscd.mcalls
Cb    LI    BCphase     2     CpGhsm      ./MCALLS/Cbir/Libbrecht2016/BCphaseA/replicate2/BCphaseA2.CpGhsm.mcalls
Cb    LI    BCphase     2     CpGscd      ./MCALLS/Cbir/Libbrecht2016/BCphaseA/replicate2/BCphaseA2.CpGscd.mcalls
Cb    LI    BCphase     3     CpGhsm      ./MCALLS/Cbir/Libbrecht2016/BCphaseB/replicate1/BCphaseB1.CpGhsm.mcalls
Cb    LI    BCphase     3     CpGscd      ./MCALLS/Cbir/Libbrecht2016/BCphaseB/replicate1/BCphaseB1.CpGscd.mcalls
Cb    LI    BCphase     4     CpGhsm      ./MCALLS/Cbir/Libbrecht2016/BCphaseB/replicate2/BCphaseB2.CpGhsm.mcalls
Cb    LI    BCphase     4     CpGscd      ./MCALLS/Cbir/Libbrecht2016/BCphaseB/replicate2/BCphaseB2.CpGscd.mcalls

# Rphase
Cb    LI    Rphase      0     CpGhsm      ./MCALLS/Cbir/Libbrecht2016/PhaseBCvsR/PhaseR/PhaseR.CpGhsm.mcalls
Cb    LI    Rphase      0     CpGscd      ./MCALLS/Cbir/Libbrecht2016/PhaseBCvsR/PhaseR/PhaseR.CpGscd.mcalls

# ...with replicates
Cb    LI    Rphase      1     CpGhsm      ./MCALLS/Cbir/Libbrecht2016/RphaseA/replicate1/RphaseA1.CpGhsm.mcalls
Cb    LI    Rphase      1     CpGscd      ./MCALLS/Cbir/Libbrecht2016/RphaseA/replicate1/RphaseA1.CpGscd.mcalls
Cb    LI    Rphase      2     CpGhsm      ./MCALLS/Cbir/Libbrecht2016/RphaseA/replicate2/RphaseA2.CpGhsm.mcalls
Cb    LI    Rphase      2     CpGscd      ./MCALLS/Cbir/Libbrecht2016/RphaseA/replicate2/RphaseA2.CpGscd.mcalls
Cb    LI    Rphase      3     CpGhsm      ./MCALLS/Cbir/Libbrecht2016/RphaseB/replicate1/RphaseB1.CpGhsm.mcalls
Cb    LI    Rphase      3     CpGscd      ./MCALLS/Cbir/Libbrecht2016/RphaseB/replicate1/RphaseB1.CpGscd.mcalls
Cb    LI    Rphase      4     CpGhsm      ./MCALLS/Cbir/Libbrecht2016/RphaseB/replicate2/RphaseB2.CpGhsm.mcalls
Cb    LI    Rphase      4     CpGscd      ./MCALLS/Cbir/Libbrecht2016/RphaseB/replicate2/RphaseB2.CpGscd.mcalls

### 2.2 Comparing LineA versus LineB ###
```

```
#
# LineA
Cb    LI    LineA 0     CpGhsm      ./MCALLS/Cbir/Libbrecht2016/LineAvsB/LineA/LineA.CpGhsm.mcalls
Cb    LI    LineA 0     CpGscd      ./MCALLS/Cbir/Libbrecht2016/LineAvsB/LineA/LineA.CpGscd.mcalls

# ...with replicates
Cb    LI    LineA 1     CpGhsm      ./MCALLS/Cbir/Libbrecht2016/BCphaseA/replicate1/BCphaseA1.CpGhsm.mcalls
Cb    LI    LineA 1     CpGscd      ./MCALLS/Cbir/Libbrecht2016/BCphaseA/replicate1/BCphaseA1.CpGscd.mcalls
Cb    LI    LineA 2     CpGhsm      ./MCALLS/Cbir/Libbrecht2016/BCphaseA/replicate2/BCphaseA2.CpGhsm.mcalls
Cb    LI    LineA 2     CpGscd      ./MCALLS/Cbir/Libbrecht2016/BCphaseA/replicate2/BCphaseA2.CpGscd.mcalls
Cb    LI    LineA 3     CpGhsm      ./MCALLS/Cbir/Libbrecht2016/RphaseA/replicate1/RphaseA1.CpGhsm.mcalls
Cb    LI    LineA 3     CpGscd      ./MCALLS/Cbir/Libbrecht2016/RphaseA/replicate1/RphaseA1.CpGscd.mcalls
Cb    LI    LineA 4     CpGhsm      ./MCALLS/Cbir/Libbrecht2016/RphaseA/replicate2/RphaseA2.CpGhsm.mcalls
Cb    LI    LineA 4     CpGscd      ./MCALLS/Cbir/Libbrecht2016/RphaseA/replicate2/RphaseA2.CpGscd.mcalls

# LineB
Cb    LI    LineB 0     CpGhsm      ./MCALLS/Cbir/Libbrecht2016/LineAvsB/LineB/LineB.CpGhsm.mcalls
Cb    LI    LineB 0     CpGscd      ./MCALLS/Cbir/Libbrecht2016/LineAvsB/LineB/LineB.CpGscd.mcalls

# ...with replicates
Cb    LI    LineB 1     CpGhsm      ./MCALLS/Cbir/Libbrecht2016/BCphaseB/replicate1/BCphaseB1.CpGhsm.mcalls
Cb    LI    LineB 1     CpGscd      ./MCALLS/Cbir/Libbrecht2016/BCphaseB/replicate1/BCphaseB1.CpGscd.mcalls
Cb    LI    LineB 2     CpGhsm      ./MCALLS/Cbir/Libbrecht2016/BCphaseB/replicate2/BCphaseB2.CpGhsm.mcalls
Cb    LI    LineB 2     CpGscd      ./MCALLS/Cbir/Libbrecht2016/BCphaseB/replicate2/BCphaseB2.CpGscd.mcalls
Cb    LI    LineB 3     CpGhsm      ./MCALLS/Cbir/Libbrecht2016/RphaseB/replicate1/RphaseB1.CpGhsm.mcalls
Cb    LI    LineB 3     CpGscd      ./MCALLS/Cbir/Libbrecht2016/RphaseB/replicate1/RphaseB1.CpGscd.mcalls
Cb    LI    LineB 4     CpGhsm      ./MCALLS/Cbir/Libbrecht2016/RphaseB/replicate2/RphaseB2.CpGhsm.mcalls
Cb    LI    LineB 4     CpGscd      ./MCALLS/Cbir/Libbrecht2016/RphaseB/replicate2/RphaseB2.CpGscd.mcalls

### 2.2 2.2 Comparing Batch1 versus Batch2 ###
#
# Batch1
Cb    LI    Batch1      0     CpGhsm      ./MCALLS/Cbir/Libbrecht2016/Batch1vs2/Batch1/Batch1.CpGhsm.mcalls
Cb    LI    Batch1      0     CpGscd      ./MCALLS/Cbir/Libbrecht2016/Batch1vs2/Batch1/Batch1.CpGscd.mcalls

# ...with replicates
Cb    LI    Batch1      1     CpGhsm      ./MCALLS/Cbir/Libbrecht2016/BCphaseA/replicate1/BCphaseA1.CpGhsm.mcalls
Cb    LI    Batch1      1     CpGscd      ./MCALLS/Cbir/Libbrecht2016/BCphaseA/replicate1/BCphaseA1.CpGscd.mcalls
Cb    LI    Batch1      2     CpGhsm      ./MCALLS/Cbir/Libbrecht2016/BCphaseB/replicate1/BCphaseB1.CpGhsm.mcalls
Cb    LI    Batch1      2     CpGscd      ./MCALLS/Cbir/Libbrecht2016/BCphaseB/replicate1/BCphaseB1.CpGscd.mcalls
Cb    LI    Batch1      3     CpGhsm      ./MCALLS/Cbir/Libbrecht2016/RphaseA/replicate1/RphaseA1.CpGhsm.mcalls
Cb    LI    Batch1      3     CpGscd      ./MCALLS/Cbir/Libbrecht2016/RphaseA/replicate1/RphaseA1.CpGscd.mcalls
Cb    LI    Batch1      4     CpGhsm      ./MCALLS/Cbir/Libbrecht2016/RphaseB/replicate1/RphaseB1.CpGhsm.mcalls
Cb    LI    Batch1      4     CpGscd      ./MCALLS/Cbir/Libbrecht2016/RphaseB/replicate1/RphaseB1.CpGscd.mcalls

# Batch2
Cb    LI    Batch2      0     CpGhsm      ./MCALLS/Cbir/Libbrecht2016/Batch1vs2/Batch2/Batch2.CpGhsm.mcalls
Cb    LI    Batch2      0     CpGscd      ./MCALLS/Cbir/Libbrecht2016/Batch1vs2/Batch2/Batch2.CpGscd.mcalls

# ...with replicates
Cb    LI    Batch2      1     CpGhsm      ./MCALLS/Cbir/Libbrecht2016/BCphaseA/replicate2/BCphaseA2.CpGhsm.mcalls
Cb    LI    Batch2      1     CpGscd      ./MCALLS/Cbir/Libbrecht2016/BCphaseA/replicate2/BCphaseA2.CpGscd.mcalls
Cb    LI    Batch2      2     CpGhsm      ./MCALLS/Cbir/Libbrecht2016/BCphaseB/replicate2/BCphaseB2.CpGhsm.mcalls
Cb    LI    Batch2      2     CpGscd      ./MCALLS/Cbir/Libbrecht2016/BCphaseB/replicate2/BCphaseB2.CpGscd.mcalls
Cb    LI    Batch2      3     CpGhsm      ./MCALLS/Cbir/Libbrecht2016/RphaseA/replicate2/RphaseA2.CpGhsm.mcalls
Cb    LI    Batch2      3     CpGscd      ./MCALLS/Cbir/Libbrecht2016/RphaseA/replicate2/RphaseA2.CpGscd.mcalls
Cb    LI    Batch2      4     CpGhsm      ./MCALLS/Cbir/Libbrecht2016/RphaseB/replicate2/RphaseB2.CpGhsm.mcalls
Cb    LI    Batch2      4     CpGscd      ./MCALLS/Cbir/Libbrecht2016/RphaseB/replicate2/RphaseB2.CpGscd.mcalls

### 3. Pairwise comparisons between BCphaseA, BCphaseB, RphaseA, and RphaseB (2 replicates each, corresponding to batches 1 and 2) ###
#
```

```
# BCphaseA
Cb    LI    BCphaseA    0     CpGhsm      ./MCALLS/Cbir/Libbrecht2016/BCphaseA/BCphaseA.CpGhsm.mcalls
Cb    LI    BCphaseA    0     CpGscd      ./MCALLS/Cbir/Libbrecht2016/BCphaseA/BCphaseA.CpGscd.mcalls

# ... with replicates:
Cb    LI    BCphaseA    1     CpGhsm      ./MCALLS/Cbir/Libbrecht2016/BCphaseA/replicate1/BCphaseA1.CpGhsm.mcalls
Cb    LI    BCphaseA    1     CpGscd      ./MCALLS/Cbir/Libbrecht2016/BCphaseA/replicate1/BCphaseA1.CpGscd.mcalls
Cb    LI    BCphaseA    2     CpGhsm      ./MCALLS/Cbir/Libbrecht2016/BCphaseA/replicate2/BCphaseA2.CpGhsm.mcalls
Cb    LI    BCphaseA    2     CpGscd      ./MCALLS/Cbir/Libbrecht2016/BCphaseA/replicate2/BCphaseA2.CpGscd.mcalls

# BCphaseB
Cb    LI    BCphaseB    0     CpGhsm      ./MCALLS/Cbir/Libbrecht2016/BCphaseB/BCphaseB.CpGhsm.mcalls
Cb    LI    BCphaseB    0     CpGscd      ./MCALLS/Cbir/Libbrecht2016/BCphaseB/BCphaseB.CpGscd.mcalls

# ... with replicates:
Cb    LI    BCphaseB    1     CpGhsm      ./MCALLS/Cbir/Libbrecht2016/BCphaseB/replicate1/BCphaseB1.CpGhsm.mcalls
Cb    LI    BCphaseB    1     CpGscd      ./MCALLS/Cbir/Libbrecht2016/BCphaseB/replicate1/BCphaseB1.CpGscd.mcalls
Cb    LI    BCphaseB    2     CpGhsm      ./MCALLS/Cbir/Libbrecht2016/BCphaseB/replicate2/BCphaseB2.CpGhsm.mcalls
Cb    LI    BCphaseB    2     CpGscd      ./MCALLS/Cbir/Libbrecht2016/BCphaseB/replicate2/BCphaseB2.CpGscd.mcalls

# RphaseA
Cb    LI    RphaseA     0     CpGhsm      ./MCALLS/Cbir/Libbrecht2016/RphaseA/RphaseA.CpGhsm.mcalls
Cb    LI    RphaseA     0     CpGscd      ./MCALLS/Cbir/Libbrecht2016/RphaseA/RphaseA.CpGscd.mcalls

# ... with replicates:
Cb    LI    RphaseA     1     CpGhsm      ./MCALLS/Cbir/Libbrecht2016/RphaseA/replicate1/RphaseA1.CpGhsm.mcalls
Cb    LI    RphaseA     1     CpGscd      ./MCALLS/Cbir/Libbrecht2016/RphaseA/replicate1/RphaseA1.CpGscd.mcalls
Cb    LI    RphaseA     2     CpGhsm      ./MCALLS/Cbir/Libbrecht2016/RphaseA/replicate2/RphaseA2.CpGhsm.mcalls
Cb    LI    RphaseA     2     CpGscd      ./MCALLS/Cbir/Libbrecht2016/RphaseA/replicate2/RphaseA2.CpGscd.mcalls

# RphaseB
Cb    LI    RphaseB     0     CpGhsm      ./MCALLS/Cbir/Libbrecht2016/RphaseB/RphaseB.CpGhsm.mcalls
Cb    LI    RphaseB     0     CpGscd      ./MCALLS/Cbir/Libbrecht2016/RphaseB/RphaseB.CpGscd.mcalls

# ... with replicates:
Cb    LI    RphaseB     1     CpGhsm      ./MCALLS/Cbir/Libbrecht2016/RphaseB/replicate1/RphaseB1.CpGhsm.mcalls
Cb    LI    RphaseB     1     CpGscd      ./MCALLS/Cbir/Libbrecht2016/RphaseB/replicate1/RphaseB1.CpGscd.mcalls
Cb    LI    RphaseB     2     CpGhsm      ./MCALLS/Cbir/Libbrecht2016/RphaseB/replicate2/RphaseB2.CpGhsm.mcalls
Cb    LI    RphaseB     2     CpGscd      ./MCALLS/Cbir/Libbrecht2016/RphaseB/replicate2/RphaseB2.CpGscd.mcalls
```

```
0README

Rscript.BWASPR output for .conf file Cb_LI.conf.

Data input are the specified *.mcalls files and parameters set in Cb_LI.conf.
The *.mcalls input data are saved in the following data structures:

studymk - a methylKit methylRaw(List) object storing the CpGhsm site data for all samples
studymc - a methylKit methylRaw(List) object storing the CpGscd site data for all samples
studyhc - a methylKit methylRaw(List) object storing the CpGscd site data for all samples,
           restricted to sites with high coverage (here set to 20)
mkrd    - a methylKit methylRawList object storing the CpGhsm site data for all
           replicates of a given sample

Output of the analysis is stored in the following subdirectories (if the corresponding
 function calls were specified in Cb_LI.conf; please consult the 0README
 files in the subdirectories for details on the output files):

CMS - Coverage and methylation statistics for aggregate samples
PWC - Pairwise comparisons between all samples
CRL - Correlations between aggregate samples
REPCMS - Coverage and methylation statistics for replicate samples
REPCRL - Correlations between replicates
MMP - Mapping of methylation sites on genome annotation
ACS - Annotation of conserved methylation sites
RNK - Genomic feature regions ranked by CpGhsm statistics
MRPR - Methylation-rich and -poor regions
DMSG - Differentially methylateds sites and genes
OGL - Ordered gene lists
```

Directory: CMS File: 0READMEcms

CMS - Coverage and methylation statistics for aggregate samples.

Input: studymk, covlist, locount, hicount

Output: files cms-*.txt and cms-*.pdf

Notes: The statistics are given for CpGhsm sites restricted to minimum or higher level coverage. At the minimum level, the number of sites will be the number of lines of the corresponding *CpGhsm.mcalls file minus one (consistency check). The distribution of methylation levels is really only interesting for high coverage because at low coverage there will be a strong bias towards high methylation levels (by definition of hsm sites). The same bias at high coverage would suggest that the sample includes a preponderance of consistently methylated genomic sites.

locount and hicount set bounds on the coverage to exclude sites with too few or too many covering reads to provide statistics on a typical range.

```
Directory: CMS File: cms-Cb_LI_BCphaseA1.txt

Number of "BCphaseA1" CpGhsm-sites with minimal and higher level coverage:

  number of "BCphaseA1" CpGhsm-sites with coverage >=  5: 198408
  number of "BCphaseA1" CpGhsm-sites with coverage >=  6: 195355
  number of "BCphaseA1" CpGhsm-sites with coverage >= 20: 113393
  number of "BCphaseA1" CpGhsm-sites with coverage >= 50:  11021

Coverage and methylation statistics for "BCphaseA1" CpGhsm-sites at different levels of minimum coverage:

methylKit::getCoverageStats output for "BCphaseA1" CpGhsm-sites (#: 198408) at minimum coverage 5    - read coverage statistics per base
summary:
   Min. 1st Qu.  Median    Mean 3rd Qu.    Max. 
      5      14      22      25      33    1266 
percentiles:
   0%   10%   20%   30%   40%   50%   60%   70%   80%   90%   95%   99% 99.5% 99.9%  100% 
    5     9    12    15    19    22    26    31    36    44    50    64    69    84  1266 

methylKit::getMethylationStats output for "BCphaseA1" CpGhsm-sites (#: 198408) at minimum coverage 5 - methylation statistics per base
summary:
   Min. 1st Qu.  Median    Mean 3rd Qu.    Max. 
    2.2    46.7    71.4    67.8    90.5   100.0 
percentiles:
    0%    10%    20%    30%    40%    50%    60%    70%    80%    90%    95%    99%  99.5%  99.9%   100% 
  2.21  31.71  41.40  52.00  61.54  71.43  80.00  87.50  93.55 100.00 100.00 100.00 100.00 100.00 100.00 

methylKit::getCoverageStats output for "BCphaseA1" CpGhsm-sites (#: 195355) at minimum coverage 6    - read coverage statistics per base
summary:
   Min. 1st Qu.  Median    Mean 3rd Qu.    Max. 
      6      14      23      25      33    1266 
percentiles:
   0%   10%   20%   30%   40%   50%   60%   70%   80%   90%   95%   99% 99.5% 99.9%  100% 
    6     9    12    16    19    23    27    31    36    44    51    64    69    84  1266 

methylKit::getMethylationStats output for "BCphaseA1" CpGhsm-sites (#: 195355) at minimum coverage 6 - methylation statistics per base
summary:
   Min. 1st Qu.  Median    Mean 3rd Qu.    Max. 
    2.2    46.2    70.4    67.3    90.0   100.0 
percentiles:
    0%    10%    20%    30%    40%    50%    60%    70%    80%    90%    95%    99%  99.5%  99.9%   100% 
  2.21  31.58  41.18  51.22  60.96  70.37  79.31  86.67  92.86 100.00 100.00 100.00 100.00 100.00 100.00 

methylKit::getCoverageStats output for "BCphaseA1" CpGhsm-sites (#: 113393) at minimum coverage 20   - read coverage statistics per base
summary:
   Min. 1st Qu.  Median    Mean 3rd Qu.    Max. 
     20      25      31      34      40    1266 
percentiles:
   0%   10%   20%   30%   40%   50%   60%   70%   80%   90%   95%   99% 99.5% 99.9%  100% 
   20    22    24    26    29    31    34    38    43    49    55    68    73    92  1266 

methylKit::getMethylationStats output for "BCphaseA1" CpGhsm-sites (#: 113393) at minimum coverage 20      - methylation statistics per base
summary:
   Min. 1st Qu.  Median    Mean 3rd Qu.    Max. 
    2.2    37.2    60.0    61.0    85.7   100.0 
percentiles:
    0%    10%    20%    30%    40%    50%    60%    70%    80%    90%    95%    99%  99.5%  99.9%   100% 
```

```
  2.21  26.67  33.33  41.03  50.00  60.00  70.97  81.25  90.00  95.83  98.41 100.00 100.00 100.00 100.00

methylKit::getCoverageStats output for "BCphaseA1" CpGhsm-sites (#: 11021) at minimum coverage 50   - read coverage statistics per base
summary:
   Min. 1st Qu.  Median    Mean 3rd Qu.    Max.
     50      52      55      61      61    1266
percentiles:
    0%    10%    20%    30%    40%    50%    60%    70%    80%    90%    95%    99%  99.5%  99.9%   100%
  50.0   50.0   51.0   53.0   54.0   55.0   57.0   60.0   63.0   68.0   73.0   92.8  186.5 1118.3 1266.0

methylKit::getMethylationStats output for "BCphaseA1" CpGhsm-sites (#: 11021) at minimum coverage 50 - methylation statistics per base
summary:
   Min. 1st Qu.  Median    Mean 3rd Qu.    Max.
    2.2    33.3    61.4    59.5    86.7   100.0
percentiles:
    0%    10%    20%    30%    40%    50%    60%    70%    80%    90%    95%    99%  99.5%  99.9%   100%
  2.21  20.00  28.33  38.18  49.09  61.36  72.31  82.46  90.38  96.00  98.08 100.00 100.00 100.00 100.00

methylKit::getCoverageStats output for "BCphaseA1" CpGhsm-sites (#: 77) in coverage range [100-1000] - read coverage statistics per base
summary:
   Min. 1st Qu.  Median    Mean 3rd Qu.    Max.
    100     130     194     286     320     974
percentiles:
   0%   10%   20%   30%   40%   50%   60%   70%   80%   90%   95%   99% 99.5% 99.9%  100%
  100   109   120   137   154   194   256   280   407   665   816   971   972   974   974

methylKit::getMethylationStats output for "BCphaseA1" CpGhsm-sites (#: 77) in coverage range [100-1000]   - methylation statistics per base
summary:
   Min. 1st Qu.  Median    Mean 3rd Qu.    Max.
    2.8     5.6     8.5    20.3    18.5    99.0
percentiles:
   0%   10%   20%   30%   40%   50%   60%   70%   80%   90%   95%   99% 99.5% 99.9%  100%
 2.83  3.96  5.06  6.34  7.85  8.55 10.24 15.72 36.53 63.73 77.02 89.40 94.21 98.06 99.02
```

Directory: CMS File: cms-Cb_LI_BCphaseA2.txt

Number of "BCphaseA2" CpGhsm-sites with minimal and higher level coverage:

```
  number of "BCphaseA2" CpGhsm-sites with coverage >=  5: 213448
  number of "BCphaseA2" CpGhsm-sites with coverage >=  6: 210593
  number of "BCphaseA2" CpGhsm-sites with coverage >= 20: 137831
  number of "BCphaseA2" CpGhsm-sites with coverage >= 50:  39262
```

Coverage and methylation statistics for "BCphaseA2" CpGhsm-sites at different levels of minimum coverage:

```
methylKit::getCoverageStats output for "BCphaseA2" CpGhsm-sites (#: 213448) at minimum coverage 5    - read coverage statistics per base
summary:
   Min. 1st Qu.  Median    Mean 3rd Qu.    Max.
      5      15      27      32      43    1272
percentiles:
   0%   10%   20%   30%   40%   50%   60%   70%   80%   90%   95%   99% 99.5% 99.9%  100%
    5     9    13    17    22    27    32    39    48    61    72    96   106   131  1272

methylKit::getMethylationStats output for "BCphaseA2" CpGhsm-sites (#: 213448) at minimum coverage 5 - methylation statistics per base
summary:
   Min. 1st Qu.  Median    Mean 3rd Qu.    Max.
    2.5    42.9    68.8    65.6    90.0   100.0
percentiles:
    0%    10%    20%    30%    40%    50%    60%    70%    80%    90%    95%    99%  99.5%  99.9%   100%
  2.52  27.59  37.50  47.83  58.33  68.75  78.26  86.49  92.86 100.00 100.00 100.00 100.00 100.00 100.00

methylKit::getCoverageStats output for "BCphaseA2" CpGhsm-sites (#: 210593) at minimum coverage 6    - read coverage statistics per base
summary:
   Min. 1st Qu.  Median    Mean 3rd Qu.    Max.
      6      16      27      32      43    1272
percentiles:
   0%   10%   20%   30%   40%   50%   60%   70%   80%   90%   95%   99% 99.5% 99.9%  100%
    6    10    14    18    22    27    33    39    48    61    72    96   106   131  1272

methylKit::getMethylationStats output for "BCphaseA2" CpGhsm-sites (#: 210593) at minimum coverage 6 - methylation statistics per base
summary:
   Min. 1st Qu.  Median    Mean 3rd Qu.    Max.
    2.5    42.4    67.9    65.1    89.2   100.0
percentiles:
    0%    10%    20%    30%    40%    50%    60%    70%    80%    90%    95%    99%  99.5%  99.9%   100%
  2.52  27.50  37.50  47.37  57.89  67.92  77.78  85.71  92.31 100.00 100.00 100.00 100.00 100.00 100.00

methylKit::getCoverageStats output for "BCphaseA2" CpGhsm-sites (#: 137831) at minimum coverage 20   - read coverage statistics per base
summary:
   Min. 1st Qu.  Median    Mean 3rd Qu.    Max.
     20      27      37      42      52    1272
percentiles:
   0%   10%   20%   30%   40%   50%   60%   70%   80%   90%   95%   99% 99.5% 99.9%  100%
   20    23    26    29    33    37    42    48    56    68    79   102   113   138  1272

methylKit::getMethylationStats output for "BCphaseA2" CpGhsm-sites (#: 137831) at minimum coverage 20     - methylation statistics per base
summary:
   Min. 1st Qu.  Median    Mean 3rd Qu.    Max.
    2.5    34.5    57.7    59.0    85.1   100.0
percentiles:
    0%    10%    20%    30%    40%    50%    60%    70%    80%    90%    95%    99%  99.5%  99.9%   100%
```

```
  2.52  23.53  30.77  38.24  47.46  57.69  69.23  80.00  89.47  95.65  97.92 100.00 100.00 100.00 100.00

methylKit::getCoverageStats output for "BCphaseA2" CpGhsm-sites (#: 39262) at minimum coverage 50   - read coverage statistics per base
summary:
   Min. 1st Qu.  Median    Mean 3rd Qu.    Max.
     50      55      62      67      74    1272
percentiles:
   0%   10%   20%   30%   40%   50%   60%   70%   80%   90%   95%   99% 99.5% 99.9%  100%
   50    52    54    56    59    62    66    71    77    87    97   121   132   186  1272

methylKit::getMethylationStats output for "BCphaseA2" CpGhsm-sites (#: 39262) at minimum coverage 50 - methylation statistics per base
summary:
   Min. 1st Qu.  Median    Mean 3rd Qu.    Max.
    2.5    27.3    52.3    54.5    82.5   100.0
percentiles:
    0%    10%    20%    30%    40%    50%    60%    70%    80%    90%    95%    99%  99.5%  99.9%   100%
  2.52  17.54  23.64  31.43  41.07  52.27  64.29  76.86  87.30  94.64  97.06 100.00 100.00 100.00 100.00

methylKit::getCoverageStats output for "BCphaseA2" CpGhsm-sites (#: 1678) in coverage range [100-1000]    - read coverage statistics per base
summary:
   Min. 1st Qu.  Median    Mean 3rd Qu.    Max.
    100     103     110     121     120     969
percentiles:
   0%   10%   20%   30%   40%   50%   60%   70%   80%   90%   95%   99% 99.5% 99.9%  100%
  100   101   103   105   107   110   113   117   123   134   148   335   708   927   969

methylKit::getMethylationStats output for "BCphaseA2" CpGhsm-sites (#: 1678) in coverage range [100-1000] - methylation statistics per base
summary:
   Min. 1st Qu.  Median    Mean 3rd Qu.    Max.
    3.0    28.3    56.0    55.4    84.0   100.0
percentiles:
    0%    10%    20%    30%    40%    50%    60%    70%    80%    90%    95%    99%  99.5%  99.9%   100%
  3.01  14.56  24.22  33.36  43.78  56.00  66.98  78.83  88.43  94.59  97.04  99.06  99.25 100.00 100.00
```

```
Directory: CMS File: cms-Cb_LI_BCphaseB1.txt

Number of "BCphaseB1" CpGhsm-sites with minimal and higher level coverage:

  number of "BCphaseB1" CpGhsm-sites with coverage >=  5: 226815
  number of "BCphaseB1" CpGhsm-sites with coverage >=  6: 224272
  number of "BCphaseB1" CpGhsm-sites with coverage >= 20: 150096
  number of "BCphaseB1" CpGhsm-sites with coverage >= 50:  29629

Coverage and methylation statistics for "BCphaseB1" CpGhsm-sites at different levels of minimum coverage:

methylKit::getCoverageStats output for "BCphaseB1" CpGhsm-sites (#: 226815) at minimum coverage 5    - read coverage statistics per base
summary:
   Min. 1st Qu.  Median    Mean 3rd Qu.    Max.
      5      16      27      30      40    1524
percentiles:
   0%   10%   20%   30%   40%   50%   60%   70%   80%   90%   95%   99% 99.5% 99.9%  100%
    5    10    14    18    22    27    32    37    44    53    61    77    83   102  1524

methylKit::getMethylationStats output for "BCphaseB1" CpGhsm-sites (#: 226815) at minimum coverage 5 - methylation statistics per base
summary:
   Min. 1st Qu.  Median    Mean 3rd Qu.    Max.
    2.4    41.9    66.7    65.0    89.3   100.0
percentiles:
    0%    10%    20%    30%    40%    50%    60%    70%    80%    90%    95%    99%  99.5%  99.9%   100%
  2.37  27.87  37.18  46.67  57.14  66.67  77.19  85.71  92.50 100.00 100.00 100.00 100.00 100.00 100.00

methylKit::getCoverageStats output for "BCphaseB1" CpGhsm-sites (#: 224272) at minimum coverage 6    - read coverage statistics per base
summary:
   Min. 1st Qu.  Median    Mean 3rd Qu.    Max.
      6      16      27      30      40    1524
percentiles:
   0%   10%   20%   30%   40%   50%   60%   70%   80%   90%   95%   99% 99.5% 99.9%  100%
    6    10    14    18    23    27    32    37    44    53    61    77    83   102  1524

methylKit::getMethylationStats output for "BCphaseB1" CpGhsm-sites (#: 224272) at minimum coverage 6 - methylation statistics per base
summary:
   Min. 1st Qu.  Median    Mean 3rd Qu.    Max.
    2.4    41.7    66.7    64.6    88.9   100.0
percentiles:
    0%    10%    20%    30%    40%    50%    60%    70%    80%    90%    95%    99%  99.5%  99.9%   100%
  2.37  27.78  36.96  46.67  56.41  66.67  76.47  85.37  91.94 100.00 100.00 100.00 100.00 100.00 100.00

methylKit::getCoverageStats output for "BCphaseB1" CpGhsm-sites (#: 150096) at minimum coverage 20   - read coverage statistics per base
summary:
   Min. 1st Qu.  Median    Mean 3rd Qu.    Max.
     20      27      35      38      46    1524
percentiles:
   0%   10%   20%   30%   40%   50%   60%   70%   80%   90%   95%   99% 99.5% 99.9%  100%
   20    22    26    29    32    35    39    44    49    57    65    81    88   109  1524

methylKit::getMethylationStats output for "BCphaseB1" CpGhsm-sites (#: 150096) at minimum coverage 20      - methylation statistics per base
summary:
   Min. 1st Qu.  Median    Mean 3rd Qu.    Max.
    2.4    34.6    57.1    59.0    85.0   100.0
percentiles:
    0%    10%    20%    30%    40%    50%    60%    70%    80%    90%    95%    99%  99.5%  99.9%   100%
```

```
  2.37  24.24  31.03  38.24  47.06  57.14  68.75  80.00  89.29  95.65  97.96 100.00 100.00 100.00 100.00

methylKit::getCoverageStats output for "BCphaseB1" CpGhsm-sites (#: 29629) at minimum coverage 50   - read coverage statistics per base
summary:
   Min. 1st Qu.  Median    Mean 3rd Qu.    Max. 
     50      53      58      62      65    1524 
percentiles:
   0%   10%   20%   30%   40%   50%   60%   70%   80%   90%   95%   99% 99.5% 99.9%  100% 
   50    51    52    54    56    58    60    63    67    74    81    98   109   678  1524 

methylKit::getMethylationStats output for "BCphaseB1" CpGhsm-sites (#: 29629) at minimum coverage 50 - methylation statistics per base
summary:
   Min. 1st Qu.  Median    Mean 3rd Qu.    Max. 
    2.4    29.2    54.8    56.1    84.1   100.0 
percentiles:
    0%    10%    20%    30%    40%    50%    60%    70%    80%    90%    95%    99%  99.5%  99.9%   100% 
  2.37  18.57  25.00  33.33  43.33  54.79  66.67  78.85  88.46  94.83  97.30 100.00 100.00 100.00 100.00

methylKit::getCoverageStats output for "BCphaseB1" CpGhsm-sites (#: 250) in coverage range [100-1000]     - read coverage statistics per base
summary:
   Min. 1st Qu.  Median    Mean 3rd Qu.    Max. 
    100     103     109     177     149     906 
percentiles:
   0%   10%   20%   30%   40%   50%   60%   70%   80%   90%   95%   99% 99.5% 99.9%  100% 
  100   101   103   104   106   109   120   132   177   348   638   864   873   898   906 

methylKit::getMethylationStats output for "BCphaseB1" CpGhsm-sites (#: 250) in coverage range [100-1000]  - methylation statistics per base
summary:
   Min. 1st Qu.  Median    Mean 3rd Qu.    Max. 
    2.8    10.4    34.9    42.1    73.6    99.0 
percentiles:
   0%   10%   20%   30%   40%   50%   60%   70%   80%   90%   95%   99% 99.5% 99.9%  100% 
 2.76  5.61  8.51 11.73 20.68 34.88 53.77 65.70 79.00 91.04 94.03 98.06 98.14 98.80 99.01 
```

```
Directory: CMS File: cms-Cb_LI_BCphaseB2.txt

Number of "BCphaseB2" CpGhsm-sites with minimal and higher level coverage:

  number of "BCphaseB2" CpGhsm-sites with coverage >=  5: 212569
  number of "BCphaseB2" CpGhsm-sites with coverage >=  6: 209742
  number of "BCphaseB2" CpGhsm-sites with coverage >= 20: 136735
  number of "BCphaseB2" CpGhsm-sites with coverage >= 50:  32573

Coverage and methylation statistics for "BCphaseB2" CpGhsm-sites at different levels of minimum coverage:

methylKit::getCoverageStats output for "BCphaseB2" CpGhsm-sites (#: 212569) at minimum coverage 5    - read coverage statistics per base
summary:
   Min. 1st Qu.  Median    Mean 3rd Qu.    Max.
      5      15      26      30      41    1287
percentiles:
   0%   10%   20%   30%   40%   50%   60%   70%   80%   90%   95%   99% 99.5% 99.9%  100%
    5     9    13    17    21    26    31    37    45    56    66    86    94   115  1287

methylKit::getMethylationStats output for "BCphaseB2" CpGhsm-sites (#: 212569) at minimum coverage 5 - methylation statistics per base
summary:
   Min. 1st Qu.  Median    Mean 3rd Qu.    Max.
    2.4    42.9    68.4    65.6    89.7   100.0
percentiles:
    0%    10%    20%    30%    40%    50%    60%    70%    80%    90%    95%    99%  99.5%  99.9%   100%
  2.37  28.21  37.93  48.00  58.33  68.42  77.78  86.11  92.86 100.00 100.00 100.00 100.00 100.00 100.00

methylKit::getCoverageStats output for "BCphaseB2" CpGhsm-sites (#: 209742) at minimum coverage 6    - read coverage statistics per base
summary:
   Min. 1st Qu.  Median    Mean 3rd Qu.    Max.
      6      16      27      30      41    1287
percentiles:
   0%   10%   20%   30%   40%   50%   60%   70%   80%   90%   95%   99% 99.5% 99.9%  100%
    6    10    14    18    22    27    32    38    45    56    66    86    94   115  1287

methylKit::getMethylationStats output for "BCphaseB2" CpGhsm-sites (#: 209742) at minimum coverage 6 - methylation statistics per base
summary:
   Min. 1st Qu.  Median    Mean 3rd Qu.    Max.
    2.4    42.9    67.6    65.2    88.9   100.0
percentiles:
    0%    10%    20%    30%    40%    50%    60%    70%    80%    90%    95%    99%  99.5%  99.9%   100%
  2.37  28.07  37.50  47.62  57.89  67.65  77.27  85.71  92.11 100.00 100.00 100.00 100.00 100.00 100.00

methylKit::getCoverageStats output for "BCphaseB2" CpGhsm-sites (#: 136735) at minimum coverage 20   - read coverage statistics per base
summary:
   Min. 1st Qu.  Median    Mean 3rd Qu.    Max.
     20      27      36      40      49    1287
percentiles:
   0%   10%   20%   30%   40%   50%   60%   70%   80%   90%   95%   99% 99.5% 99.9%  100%
   20    22    25    29    32    36    40    46    52    63    72    91    99   121  1287

methylKit::getMethylationStats output for "BCphaseB2" CpGhsm-sites (#: 136735) at minimum coverage 20     - methylation statistics per base
summary:
   Min. 1st Qu.  Median    Mean 3rd Qu.    Max.
    2.4    34.9    58.1    59.3    85.0   100.0
percentiles:
    0%    10%    20%    30%    40%    50%    60%    70%    80%    90%    95%    99%  99.5%  99.9%   100%
```

```
  2.37  24.32  31.37  38.71  47.83  58.14  69.23  80.00  89.29  95.56  97.96 100.00 100.00 100.00 100.00

methylKit::getCoverageStats output for "BCphaseB2" CpGhsm-sites (#: 32573) at minimum coverage 50   - read coverage statistics per base
summary:
   Min. 1st Qu.  Median    Mean 3rd Qu.    Max. 
     50      54      60      64      70    1287 
percentiles:
   0%   10%   20%   30%   40%   50%   60%   70%   80%   90%   95%   99% 99.5% 99.9%  100% 
   50    51    53    55    57    60    63    67    72    81    89   109   118   259  1287 

methylKit::getMethylationStats output for "BCphaseB2" CpGhsm-sites (#: 32573) at minimum coverage 50 - methylation statistics per base
summary:
   Min. 1st Qu.  Median    Mean 3rd Qu.    Max. 
    2.4    28.8    54.2    55.8    83.7   100.0 
percentiles:
    0%    10%    20%    30%    40%    50%    60%    70%    80%    90%    95%    99%  99.5%  99.9%   100% 
  2.37  18.30  24.69  33.33  43.10  54.17  66.18  78.43  88.06  94.83  97.26 100.00 100.00 100.00 100.00

methylKit::getCoverageStats output for "BCphaseB2" CpGhsm-sites (#: 669) in coverage range [100-1000]     - read coverage statistics per base
summary:
   Min. 1st Qu.  Median    Mean 3rd Qu.    Max. 
    100     103     108     126     117     922 
percentiles:
   0%   10%   20%   30%   40%   50%   60%   70%   80%   90%   95%   99% 99.5% 99.9%  100% 
  100   101   102   103   105   108   111   115   120   139   199   628   748   905   922 

methylKit::getMethylationStats output for "BCphaseB2" CpGhsm-sites (#: 669) in coverage range [100-1000]  - methylation statistics per base
summary:
   Min. 1st Qu.  Median    Mean 3rd Qu.    Max. 
    3.0    27.3    55.6    55.3    85.4   100.0 
percentiles:
    0%    10%    20%    30%    40%    50%    60%    70%    80%    90%    95%    99%  99.5%  99.9%   100% 
  3.01  13.84  22.00  34.77  44.08  55.56  68.44  79.94  88.38  95.11  97.03  99.05  99.12 100.00 100.00 
```

```
Directory: CMS File: cms-Cb_LI_RphaseA1.txt

Number of "RphaseA1" CpGhsm-sites with minimal and higher level coverage:

  number of "RphaseA1" CpGhsm-sites with coverage >=  5: 213610
  number of "RphaseA1" CpGhsm-sites with coverage >=  6: 210877
  number of "RphaseA1" CpGhsm-sites with coverage >= 20: 133847
  number of "RphaseA1" CpGhsm-sites with coverage >= 50:  18249

Coverage and methylation statistics for "RphaseA1" CpGhsm-sites at different levels of minimum coverage:

methylKit::getCoverageStats output for "RphaseA1" CpGhsm-sites (#: 213610) at minimum coverage 5    - read coverage statistics per base
summary:
   Min. 1st Qu.  Median    Mean 3rd Qu.    Max. 
      5      15      25      27      37    1360 
percentiles:
   0%   10%   20%   30%   40%   50%   60%   70%   80%   90%   95%   99% 99.5% 99.9%  100% 
    5     9    13    17    21    25    29    34    40    48    54    69    74    91  1360 

methylKit::getMethylationStats output for "RphaseA1" CpGhsm-sites (#: 213610) at minimum coverage 5  - methylation statistics per base
summary:
   Min. 1st Qu.  Median    Mean 3rd Qu.    Max. 
    2.5    44.4    69.6    66.5    90.0   100.0 
percentiles:
    0%    10%    20%    30%    40%    50%    60%    70%    80%    90%    95%    99%  99.5%  99.9%   100% 
  2.48  29.73  39.39  50.00  59.52  69.57  78.79  86.84  93.10 100.00 100.00 100.00 100.00 100.00 100.00 

methylKit::getCoverageStats output for "RphaseA1" CpGhsm-sites (#: 210877) at minimum coverage 6    - read coverage statistics per base
summary:
   Min. 1st Qu.  Median    Mean 3rd Qu.    Max. 
      6      15      25      27      37    1360 
percentiles:
   0%   10%   20%   30%   40%   50%   60%   70%   80%   90%   95%   99% 99.5% 99.9%  100% 
    6    10    13    17    21    25    30    34    40    48    55    69    74    91  1360 

methylKit::getMethylationStats output for "RphaseA1" CpGhsm-sites (#: 210877) at minimum coverage 6  - methylation statistics per base
summary:
   Min. 1st Qu.  Median    Mean 3rd Qu.    Max. 
    2.5    43.8    68.8    66.1    89.5   100.0 
percentiles:
    0%    10%    20%    30%    40%    50%    60%    70%    80%    90%    95%    99%  99.5%  99.9%   100% 
  2.48  29.63  39.13  50.00  58.82  68.75  77.78  86.05  92.59 100.00 100.00 100.00 100.00 100.00 100.00 

methylKit::getCoverageStats output for "RphaseA1" CpGhsm-sites (#: 133847) at minimum coverage 20   - read coverage statistics per base
summary:
   Min. 1st Qu.  Median    Mean 3rd Qu.    Max. 
     20      26      33      36      43    1360 
percentiles:
   0%   10%   20%   30%   40%   50%   60%   70%   80%   90%   95%   99% 99.5% 99.9%  100% 
   20    22    25    28    30    33    37    41    45    52    59    73    78    96  1360 

methylKit::getMethylationStats output for "RphaseA1" CpGhsm-sites (#: 133847) at minimum coverage 20 - methylation statistics per base
summary:
   Min. 1st Qu.  Median    Mean 3rd Qu.    Max. 
    2.5    36.0    59.1    60.2    85.7   100.0 
percentiles:
    0%    10%    20%    30%    40%    50%    60%    70%    80%    90%    95%    99%  99.5%  99.9%   100% 
```

```
  2.48  25.64  32.43  40.00  48.72  59.09  70.00  80.95  90.00  95.83  98.11 100.00 100.00 100.00 100.00

methylKit::getCoverageStats output for "RphaseA1" CpGhsm-sites (#: 18249) at minimum coverage 50    - read coverage statistics per base
summary:
   Min. 1st Qu.  Median    Mean 3rd Qu.    Max.
     50      52      56      60      62    1360
percentiles:
   0%   10%   20%   30%   40%   50%   60%   70%   80%   90%   95%   99% 99.5% 99.9%  100%
   50    51    52    53    54    56    58    60    64    70    76    92   106   654  1360

methylKit::getMethylationStats output for "RphaseA1" CpGhsm-sites (#: 18249) at minimum coverage 50  - methylation statistics per base
summary:
   Min. 1st Qu.  Median    Mean 3rd Qu.    Max.
    2.5    31.9    58.0    58.0    85.4   100.0
percentiles:
    0%    10%    20%    30%    40%    50%    60%    70%    80%    90%    95%    99%  99.5%  99.9%   100%
  2.48  20.00  27.45  36.23  47.06  58.00  69.64  80.60  89.29  95.45  98.04 100.00 100.00 100.00 100.00

methylKit::getCoverageStats output for "RphaseA1" CpGhsm-sites (#: 100) in coverage range [100-1000]- read coverage statistics per base
summary:
   Min. 1st Qu.  Median    Mean 3rd Qu.    Max.
    100     108     124     252     272     972
percentiles:
   0%   10%   20%   30%   40%   50%   60%   70%   80%   90%   95%   99% 99.5% 99.9%  100%
  100   102   106   110   115   124   154   221   360   633   875   938   955   969   972

methylKit::getMethylationStats output for "RphaseA1" CpGhsm-sites (#: 100) in coverage range [100-1000]   - methylation statistics per base
summary:
   Min. 1st Qu.  Median    Mean 3rd Qu.    Max.
    2.8     6.5    16.2    31.9    58.5    98.1
percentiles:
   0%   10%   20%   30%   40%   50%   60%   70%   80%   90%   95%   99% 99.5% 99.9%  100%
 2.78  4.41  5.68  7.10 10.32 16.20 27.33 50.90 61.46 82.27 94.79 98.00 98.06 98.10 98.11
```

```
Directory: CMS File: cms-Cb_LI_RphaseA2.txt

Number of "RphaseA2" CpGhsm-sites with minimal and higher level coverage:

  number of "RphaseA2" CpGhsm-sites with coverage >=  5: 265952
  number of "RphaseA2" CpGhsm-sites with coverage >=  6: 263685
  number of "RphaseA2" CpGhsm-sites with coverage >= 20: 201127
  number of "RphaseA2" CpGhsm-sites with coverage >= 50:  88662

Coverage and methylation statistics for "RphaseA2" CpGhsm-sites at different levels of minimum coverage:

methylKit::getCoverageStats output for "RphaseA2" CpGhsm-sites (#: 265952) at minimum coverage 5     - read coverage statistics per base
summary:
   Min. 1st Qu.  Median    Mean 3rd Qu.    Max.
      5      20      36      42      58    1345
percentiles:
   0%   10%   20%   30%   40%   50%   60%   70%   80%   90%   95%   99% 99.5% 99.9%  100%
    5    11    17    23    29    36    44    53    65    83    98   130   143   175  1345

methylKit::getMethylationStats output for "RphaseA2" CpGhsm-sites (#: 265952) at minimum coverage 5  - methylation statistics per base
summary:
   Min. 1st Qu.  Median    Mean 3rd Qu.    Max.
    2.7    35.4    62.1    60.9    87.5   100.0
percentiles:
    0%    10%    20%    30%    40%    50%    60%    70%    80%    90%    95%    99%  99.5%  99.9%   100%
  2.68  22.45  31.25  40.00  50.67  62.07  73.08  83.33  91.23  97.37 100.00 100.00 100.00 100.00 100.00

methylKit::getCoverageStats output for "RphaseA2" CpGhsm-sites (#: 263685) at minimum coverage 6     - read coverage statistics per base
summary:
   Min. 1st Qu.  Median    Mean 3rd Qu.    Max.
      6      20      36      43      59    1345
percentiles:
   0%   10%   20%   30%   40%   50%   60%   70%   80%   90%   95%   99% 99.5% 99.9%  100%
    6    12    18    23    29    36    44    53    65    83    99   130   143   175  1345

methylKit::getMethylationStats output for "RphaseA2" CpGhsm-sites (#: 263685) at minimum coverage 6  - methylation statistics per base
summary:
   Min. 1st Qu.  Median    Mean 3rd Qu.    Max.
    2.7    35.3    61.5    60.6    87.2   100.0
percentiles:
    0%    10%    20%    30%    40%    50%    60%    70%    80%    90%    95%    99%  99.5%  99.9%   100%
  2.68  22.35  31.03  40.00  50.00  61.54  72.55  83.33  90.91  96.88 100.00 100.00 100.00 100.00 100.00

methylKit::getCoverageStats output for "RphaseA2" CpGhsm-sites (#: 201127) at minimum coverage 20    - read coverage statistics per base
summary:
   Min. 1st Qu.  Median    Mean 3rd Qu.    Max.
     20      31      45      52      66    1345
percentiles:
   0%   10%   20%   30%   40%   50%   60%   70%   80%   90%   95%   99% 99.5% 99.9%  100%
   20    24    29    34    39    45    53    61    73    89   104   135   148   183  1345

methylKit::getMethylationStats output for "RphaseA2" CpGhsm-sites (#: 201127) at minimum coverage 20 - methylation statistics per base
summary:
   Min. 1st Qu.  Median    Mean 3rd Qu.    Max.
    2.7    30.2    53.0    55.9    83.3   100.0
percentiles:
    0%    10%    20%    30%    40%    50%    60%    70%    80%    90%    95%    99%  99.5%  99.9%   100%
```

```
  2.68  20.45  26.76  33.33  42.42  53.03  65.12  77.42  88.00  95.24  97.50 100.00 100.00 100.00 100.00
```

```
methylKit::getCoverageStats output for "RphaseA2" CpGhsm-sites (#: 88662) at minimum coverage 50    - read coverage statistics per base
summary:
   Min. 1st Qu.  Median    Mean 3rd Qu.    Max. 
     50      58      70      76      87    1345 
percentiles:
   0%   10%   20%   30%   40%   50%   60%   70%   80%   90%   95%   99% 99.5% 99.9%  100% 
   50    53    56    60    65    70    76    83    92   107   120   151   164   218  1345 

methylKit::getMethylationStats output for "RphaseA2" CpGhsm-sites (#: 88662) at minimum coverage 50  - methylation statistics per base
summary:
   Min. 1st Qu.  Median    Mean 3rd Qu.    Max. 
    2.7    24.3    48.2    52.2    80.6   100.0 
percentiles:
    0%    10%    20%    30%    40%    50%    60%    70%    80%    90%    95%    99%  99.5%  99.9%   100% 
  2.68  16.00  21.21  28.00  37.10  48.21  61.11  74.32  86.27  94.35  96.94 100.00 100.00 100.00 100.00
```

```
methylKit::getCoverageStats output for "RphaseA2" CpGhsm-sites (#: 12653) in coverage range [100-1000]    - read coverage statistics per base
summary:
   Min. 1st Qu.  Median    Mean 3rd Qu.    Max. 
    100     105     113     120     127     984 
percentiles:
   0%   10%   20%   30%   40%   50%   60%   70%   80%   90%   95%   99% 99.5% 99.9%  100% 
  100   102   104   107   110   113   117   123   131   144   157   196   230   695   984 

methylKit::getMethylationStats output for "RphaseA2" CpGhsm-sites (#: 12653) in coverage range [100-1000] - methylation statistics per base
summary:
   Min. 1st Qu.  Median    Mean 3rd Qu.    Max. 
    2.9    25.0    52.2    53.5    83.0   100.0 
percentiles:
    0%    10%    20%    30%    40%    50%    60%    70%    80%    90%    95%    99%  99.5%  99.9%   100% 
  2.85  13.76  20.77  29.52  40.28  52.17  64.76  77.57  87.62  94.61  97.03  99.04  99.21 100.00 100.00
```

```
Directory: CMS File: cms-Cb_LI_RphaseB1.txt

Number of "RphaseB1" CpGhsm-sites with minimal and higher level coverage:

  number of "RphaseB1" CpGhsm-sites with coverage >=  5: 218593
  number of "RphaseB1" CpGhsm-sites with coverage >=  6: 215760
  number of "RphaseB1" CpGhsm-sites with coverage >= 20: 139277
  number of "RphaseB1" CpGhsm-sites with coverage >= 50:  23245

Coverage and methylation statistics for "RphaseB1" CpGhsm-sites at different levels of minimum coverage:

methylKit::getCoverageStats output for "RphaseB1" CpGhsm-sites (#: 218593) at minimum coverage 5     - read coverage statistics per base
summary:
   Min. 1st Qu.  Median    Mean 3rd Qu.    Max.
      5      15      26      28      38    1983
percentiles:
   0%   10%   20%   30%   40%   50%   60%   70%   80%   90%   95%   99% 99.5% 99.9%  100%
    5     9    13    17    21    26    30    35    41    50    58    73    80    98  1983

methylKit::getMethylationStats output for "RphaseB1" CpGhsm-sites (#: 218593) at minimum coverage 5  - methylation statistics per base
summary:
   Min. 1st Qu.  Median    Mean 3rd Qu.    Max.
    2.0    43.3    68.4    65.9    89.7   100.0
percentiles:
   0%   10%   20%   30%   40%   50%   60%   70%   80%   90%   95%   99% 99.5% 99.9%  100%
  2.0  29.0  38.5  48.4  58.3  68.4  77.8  86.2  92.9 100.0 100.0 100.0 100.0 100.0 100.0

methylKit::getCoverageStats output for "RphaseB1" CpGhsm-sites (#: 215760) at minimum coverage 6     - read coverage statistics per base
summary:
   Min. 1st Qu.  Median    Mean 3rd Qu.    Max.
      6      16      26      29      38    1983
percentiles:
   0%   10%   20%   30%   40%   50%   60%   70%   80%   90%   95%   99% 99.5% 99.9%  100%
    6    10    14    17    21    26    30    35    42    50    58    73    80    98  1983

methylKit::getMethylationStats output for "RphaseB1" CpGhsm-sites (#: 215760) at minimum coverage 6  - methylation statistics per base
summary:
   Min. 1st Qu.  Median    Mean 3rd Qu.    Max.
    2.0    42.9    67.6    65.4    88.9   100.0
percentiles:
   0%   10%   20%   30%   40%   50%   60%   70%   80%   90%   95%   99% 99.5% 99.9%  100%
  2.0  28.9  38.1  47.8  57.9  67.6  77.3  85.7  92.3 100.0 100.0 100.0 100.0 100.0 100.0

methylKit::getCoverageStats output for "RphaseB1" CpGhsm-sites (#: 139277) at minimum coverage 20    - read coverage statistics per base
summary:
   Min. 1st Qu.  Median    Mean 3rd Qu.    Max.
     20      27      34      37      45    1983
percentiles:
   0%   10%   20%   30%   40%   50%   60%   70%   80%   90%   95%   99% 99.5% 99.9%  100%
   20    22    25    28    31    34    38    42    47    55    62    78    84   108  1983

methylKit::getMethylationStats output for "RphaseB1" CpGhsm-sites (#: 139277) at minimum coverage 20 - methylation statistics per base
summary:
   Min. 1st Qu.  Median    Mean 3rd Qu.    Max.
    2.0    35.3    58.3    59.7    85.2   100.0
percentiles:
   0%   10%   20%   30%   40%   50%   60%   70%   80%   90%   95%   99% 99.5% 99.9%  100%
```

```
  2.0  25.0  32.0  39.3  48.1  58.3  69.6  80.0  89.5  95.7  98.1 100.0 100.0 100.0 100.0

methylKit::getCoverageStats output for "RphaseB1" CpGhsm-sites (#: 23245) at minimum coverage 50    - read coverage statistics per base
summary:
   Min. 1st Qu.  Median    Mean 3rd Qu.    Max.
     50      53      57      62      64    1983
percentiles:
   0%   10%   20%   30%   40%   50%   60%   70%   80%   90%   95%   99% 99.5% 99.9%  100%
   50    51    52    54    55    57    59    62    66    73    79    97   116  1274  1983

methylKit::getMethylationStats output for "RphaseB1" CpGhsm-sites (#: 23245) at minimum coverage 50  - methylation statistics per base
summary:
   Min. 1st Qu.  Median    Mean 3rd Qu.    Max.
    2.0    30.6    56.9    57.3    84.6   100.0
percentiles:
   0%   10%   20%   30%   40%   50%   60%   70%   80%   90%   95%   99% 99.5% 99.9%  100%
  2.0  19.3  26.4  35.3  45.6  56.9  68.9  79.7  88.7  95.2  98.0 100.0 100.0 100.0 100.0

methylKit::getCoverageStats output for "RphaseB1" CpGhsm-sites (#: 156) in coverage range [100-1000]- read coverage statistics per base
summary:
   Min. 1st Qu.  Median    Mean 3rd Qu.    Max.
    100     106     120     216     200     918
percentiles:
   0%   10%   20%   30%   40%   50%   60%   70%   80%   90%   95%   99% 99.5% 99.9%  100%
  100   102   104   107   112   120   140   171   267   562   760   885   902   915   918

methylKit::getMethylationStats output for "RphaseB1" CpGhsm-sites (#: 156) in coverage range [100-1000]   - methylation statistics per base
summary:
   Min. 1st Qu.  Median    Mean 3rd Qu.    Max.
    2.6     8.3    34.1    39.9    71.5    98.4
percentiles:
   0%   10%   20%   30%   40%   50%   60%   70%   80%   90%   95%   99% 99.5% 99.9%  100%
 2.63  4.55  7.45 11.02 17.43 34.10 48.55 61.70 75.93 89.81 96.13 97.62 98.18 98.36 98.40
```

```
Directory: CMS File: cms-Cb_LI_RphaseB2.txt

Number of "RphaseB2" CpGhsm-sites with minimal and higher level coverage:

  number of "RphaseB2" CpGhsm-sites with coverage >=  5: 200707
  number of "RphaseB2" CpGhsm-sites with coverage >=  6: 197722
  number of "RphaseB2" CpGhsm-sites with coverage >= 20: 121851
  number of "RphaseB2" CpGhsm-sites with coverage >= 50:  25387

Coverage and methylation statistics for "RphaseB2" CpGhsm-sites at different levels of minimum coverage:

methylKit::getCoverageStats output for "RphaseB2" CpGhsm-sites (#: 200707) at minimum coverage 5     - read coverage statistics per base
summary:
   Min. 1st Qu.  Median    Mean 3rd Qu.    Max.
      5      14      24      28      38    1405
percentiles:
   0%   10%   20%   30%   40%   50%   60%   70%   80%   90%   95%   99% 99.5% 99.9%  100%
    5     9    12    16    20    24    29    35    42    53    63    82    90   110  1405

methylKit::getMethylationStats output for "RphaseB2" CpGhsm-sites (#: 200707) at minimum coverage 5  - methylation statistics per base
summary:
   Min. 1st Qu.  Median    Mean 3rd Qu.    Max.
    2.1    44.9    70.0    66.8    90.0   100.0
percentiles:
    0%    10%    20%    30%    40%    50%    60%    70%    80%    90%    95%    99%  99.5%  99.9%   100%
  2.06  29.79  40.00  50.00  60.00  70.00  79.31  87.10  93.33 100.00 100.00 100.00 100.00 100.00 100.00

methylKit::getCoverageStats output for "RphaseB2" CpGhsm-sites (#: 197722) at minimum coverage 6     - read coverage statistics per base
summary:
   Min. 1st Qu.  Median    Mean 3rd Qu.    Max.
      6      15      25      29      38    1405
percentiles:
   0%   10%   20%   30%   40%   50%   60%   70%   80%   90%   95%   99% 99.5% 99.9%  100%
    6     9    13    16    20    25    29    35    42    53    63    82    90   110  1405

methylKit::getMethylationStats output for "RphaseB2" CpGhsm-sites (#: 197722) at minimum coverage 6  - methylation statistics per base
summary:
   Min. 1st Qu.  Median    Mean 3rd Qu.    Max.
    2.1    44.4    69.4    66.3    89.5   100.0
percentiles:
    0%    10%    20%    30%    40%    50%    60%    70%    80%    90%    95%    99%  99.5%  99.9%   100%
  2.06  29.63  39.58  50.00  60.00  69.44  78.43  86.11  92.45 100.00 100.00 100.00 100.00 100.00 100.00

methylKit::getCoverageStats output for "RphaseB2" CpGhsm-sites (#: 121851) at minimum coverage 20    - read coverage statistics per base
summary:
   Min. 1st Qu.  Median    Mean 3rd Qu.    Max.
     20      26      34      39      47    1405
percentiles:
   0%   10%   20%   30%   40%   50%   60%   70%   80%   90%   95%   99% 99.5% 99.9%  100%
   20    22    25    28    31    34    39    44    50    60    69    88    96   119  1405

methylKit::getMethylationStats output for "RphaseB2" CpGhsm-sites (#: 121851) at minimum coverage 20 - methylation statistics per base
summary:
   Min. 1st Qu.  Median    Mean 3rd Qu.    Max.
    2.1    35.7    59.3    60.0    85.7   100.0
percentiles:
    0%    10%    20%    30%    40%    50%    60%    70%    80%    90%    95%    99%  99.5%  99.9%   100%
```

```
  2.06  25.00  32.26  40.00  48.84  59.26  70.00  80.77  89.66  95.65  98.04 100.00 100.00 100.00 100.00

methylKit::getCoverageStats output for "RphaseB2" CpGhsm-sites (#: 25387) at minimum coverage 50     - read coverage statistics per base
summary:
   Min. 1st Qu.  Median    Mean 3rd Qu.    Max.
     50      54      59      64      68    1405
percentiles:
   0%   10%   20%   30%   40%   50%   60%   70%   80%   90%   95%   99% 99.5% 99.9%  100%
   50    51    53    55    57    59    62    66    71    79    87   106   118   356  1405

methylKit::getMethylationStats output for "RphaseB2" CpGhsm-sites (#: 25387) at minimum coverage 50  - methylation statistics per base
summary:
   Min. 1st Qu.  Median    Mean 3rd Qu.    Max.
    2.1    29.8    55.4    56.4    83.9   100.0
percentiles:
    0%    10%    20%    30%    40%    50%    60%    70%    80%    90%    95%    99%  99.5%  99.9%   100%
  2.06  18.64  25.76  34.29  44.29  55.45  67.24  78.67  88.33  94.81  97.33 100.00 100.00 100.00 100.00

methylKit::getCoverageStats output for "RphaseB2" CpGhsm-sites (#: 436) in coverage range [100-1000]- read coverage statistics per base
summary:
   Min. 1st Qu.  Median    Mean 3rd Qu.    Max.
    100     103     107     140     119     994
percentiles:
   0%   10%   20%   30%   40%   50%   60%   70%   80%   90%   95%   99% 99.5% 99.9%  100%
  100   101   102   103   105   107   111   116   124   154   280   808   865   946   994

methylKit::getMethylationStats output for "RphaseB2" CpGhsm-sites (#: 436) in coverage range [100-1000]    - methylation statistics per base
summary:
   Min. 1st Qu.  Median    Mean 3rd Qu.    Max.
    3.1    28.5    62.5    57.2    87.8   100.0
percentiles:
    0%    10%    20%    30%    40%    50%    60%    70%    80%    90%    95%    99%  99.5%  99.9%   100%
  3.05  10.84  22.88  36.02  48.08  62.46  73.39  82.33  91.09  95.41  97.18  99.09  99.25 100.00 100.00
```

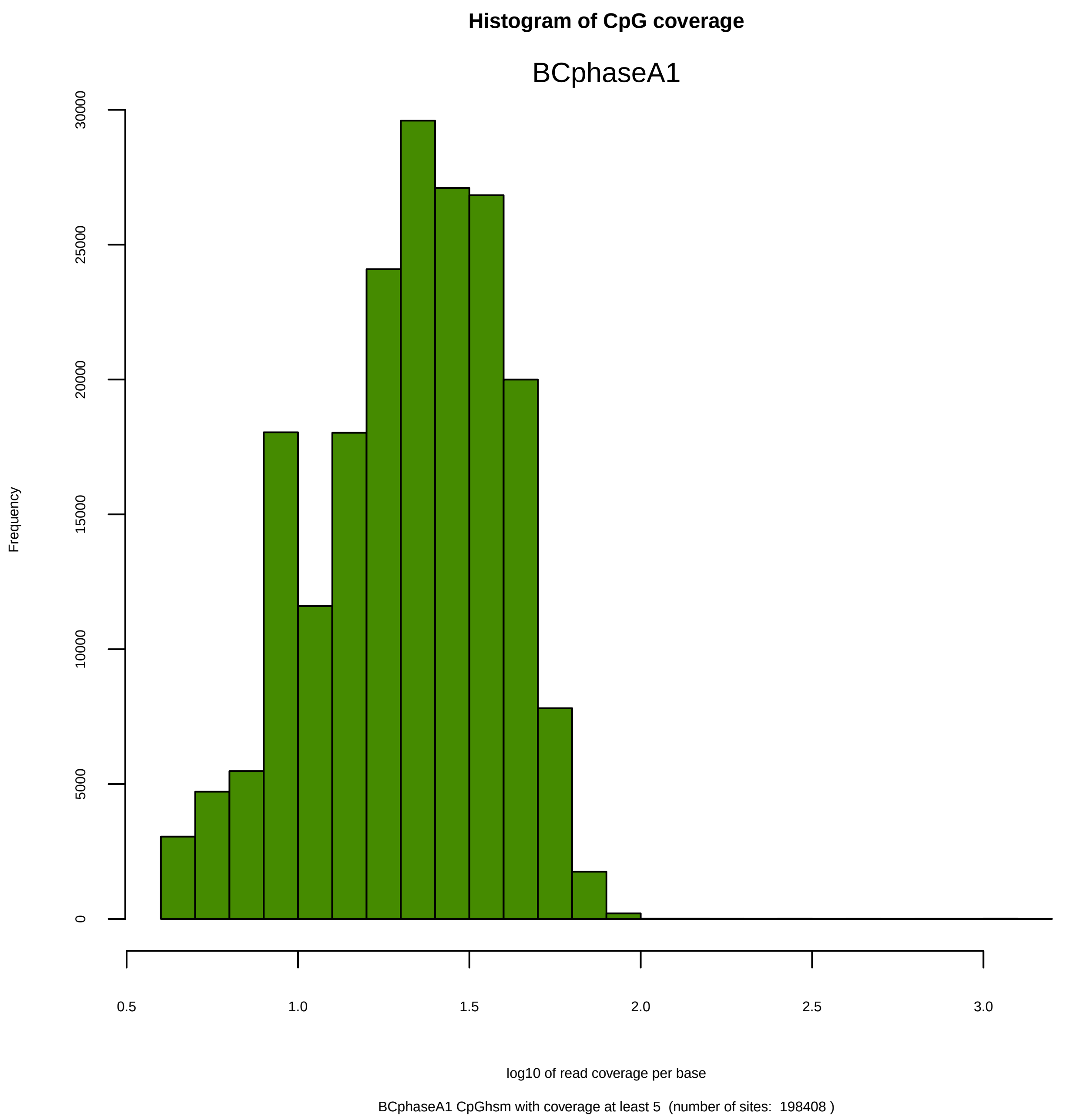

Histogram of CpG coverage
BCphaseA1
Frequency
0
5000
10000
15000
20000
25000
30000
0.5
1.0
1.5
2.0
2.5
3.0
log10 of read coverage per base
BCphaseA1 CpGhsm with coverage at least 5 (number of sites: 198408 )

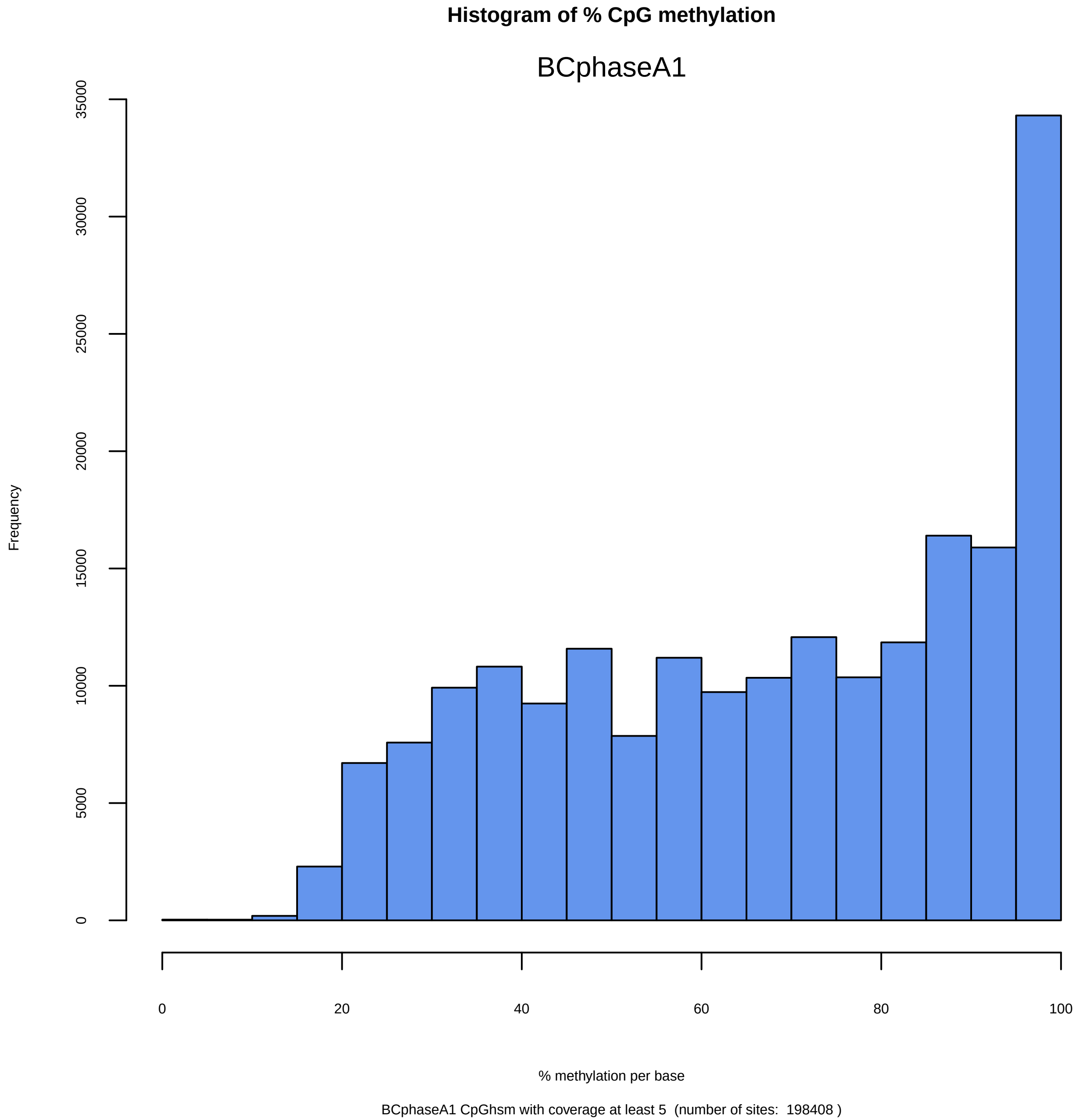

Histogram of % CpG methylation
BCphaseA1
Frequency
0
5000
10000
15000
20000
25000
30000
35000
0
20
40
60
80
100
% methylation per base
BCphaseA1 CpGhsm with coverage at least 5 (number of sites: 198408 )

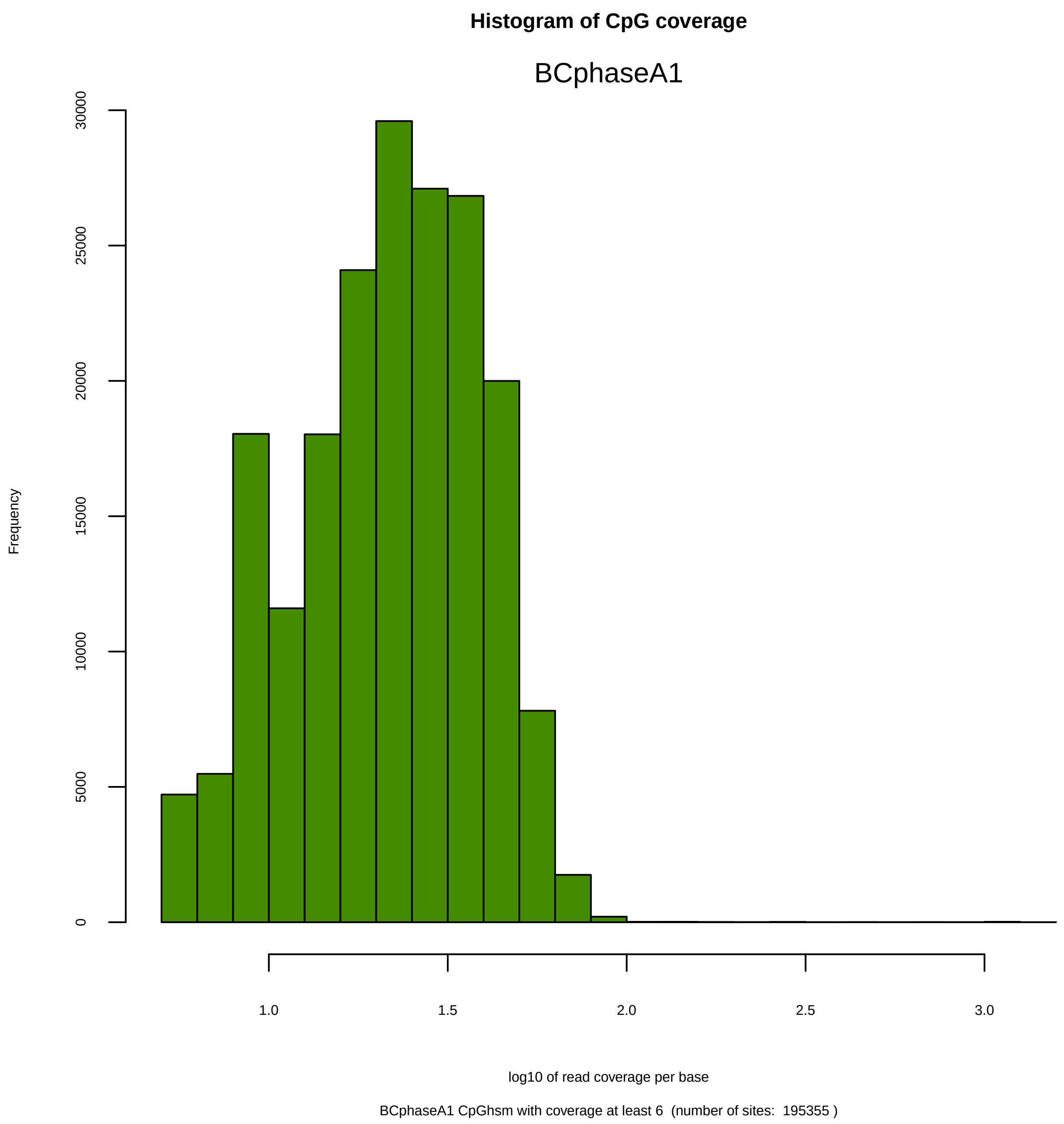

Histogram of CpG coverage
BCphaseA1
Frequency
0
5000
10000
15000
20000
25000
30000
1.0
1.5
2.0
2.5
3.0
log10 of read coverage per base
BCphaseA1 CpGhsm with coverage at least 6 (number of sites: 195355 )

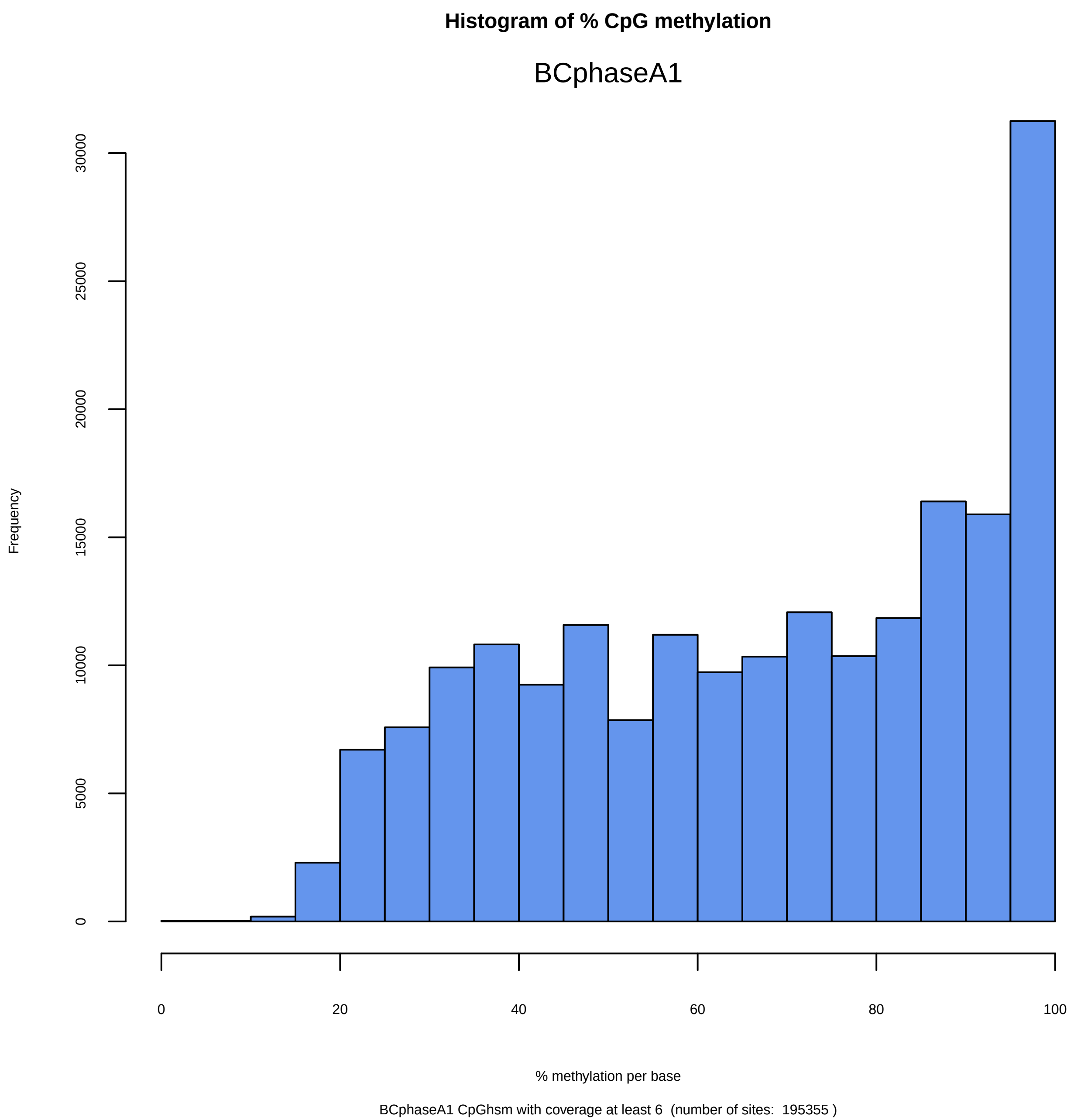

Histogram of % CpG methylation
BCphaseA1
Frequency
0
5000
10000
15000
20000
25000
30000
0
20
40
60
80
100
% methylation per base
BCphaseA1 CpGhsm with coverage at least 6  (number of sites:  195355 )

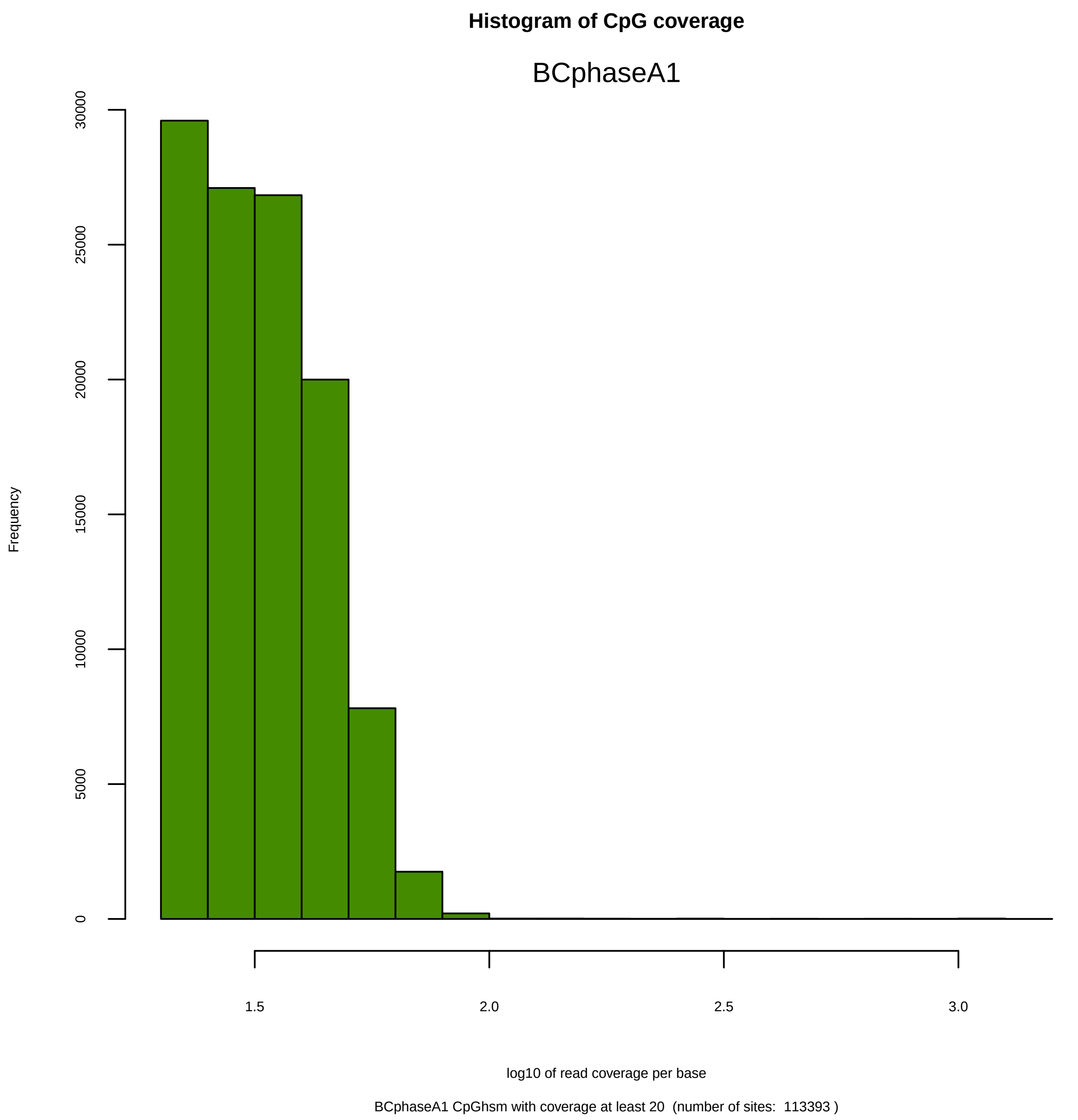
Histogram of CpG coverage
BCphaseA1
Frequency
0
5000
10000
15000
20000
25000
30000
1.5
2.0
2.5
3.0
log10 of read coverage per base
BCphaseA1 CpGhsm with coverage at least 20  (number of sites:  113393 )

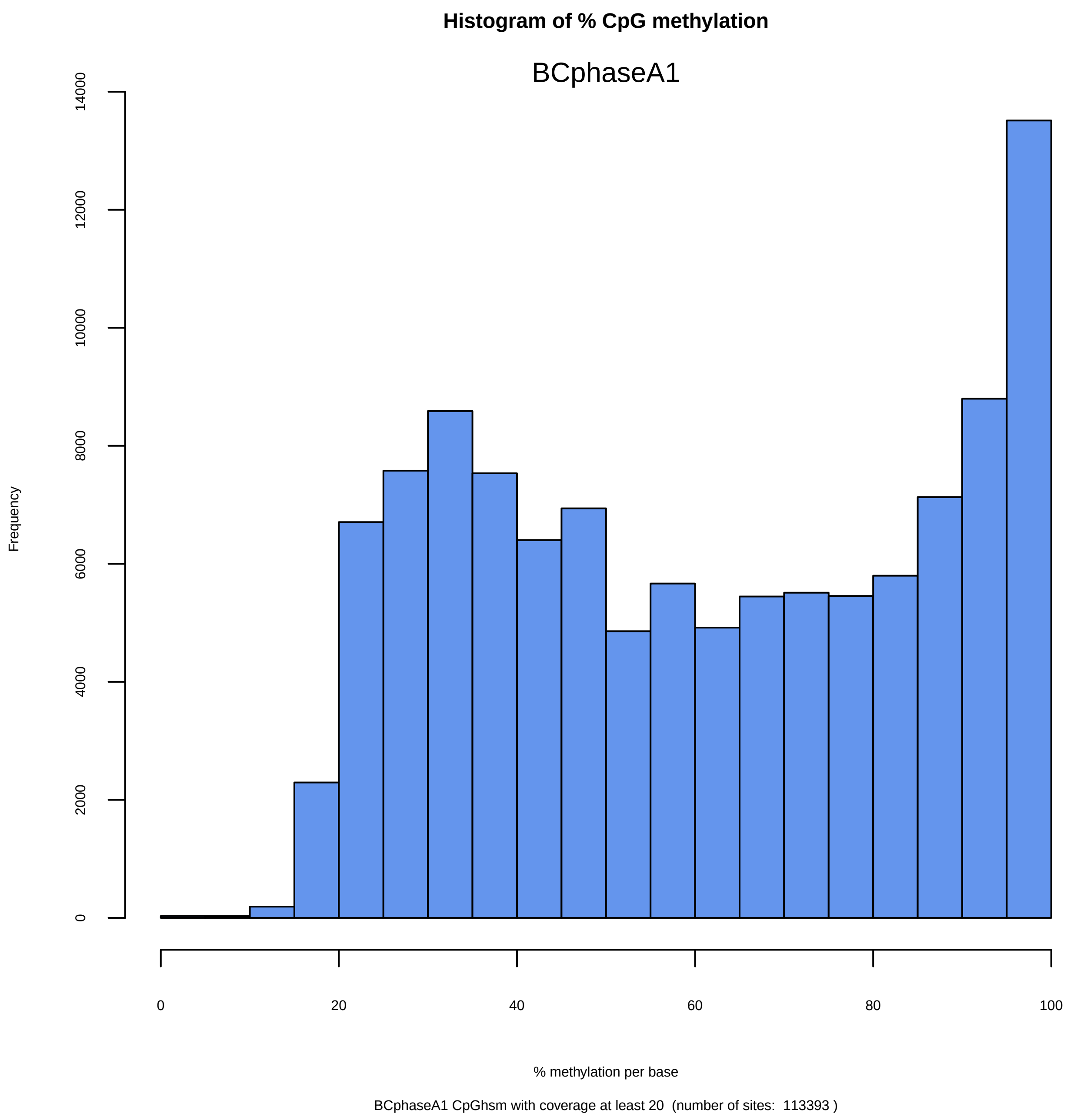

Histogram of % CpG methylation
BCphaseA1
Frequency
0
2000
4000
6000
8000
10000
12000
14000
0
20
40
60
80
100
% methylation per base
BCphaseA1 CpGhsm with coverage at least 20 (number of sites: 113393 )

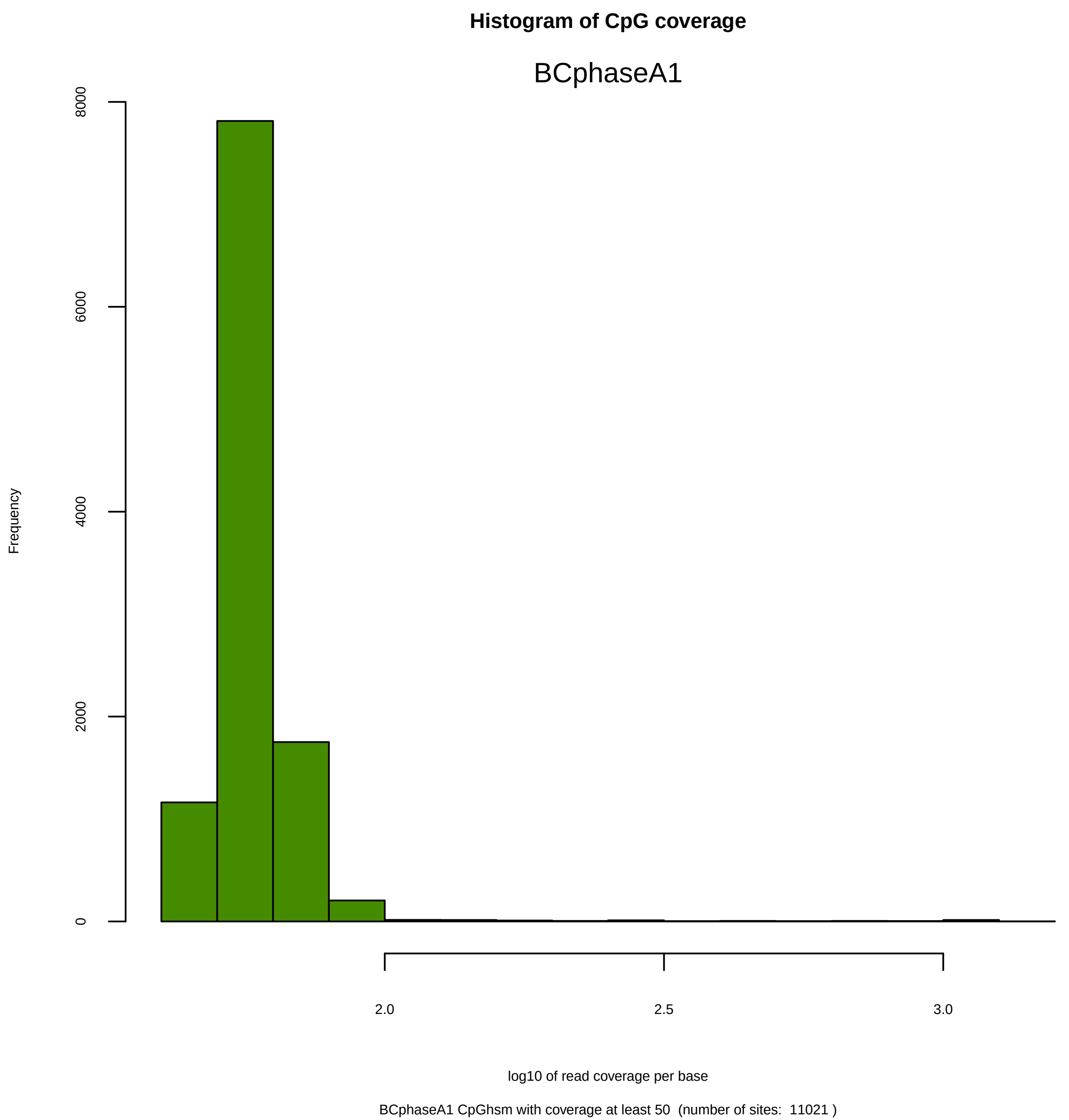

Histogram of CpG coverage
BCphaseA1
Frequency
0
2000
4000
6000
8000
2.0
2.5
3.0
log10 of read coverage per base
BCphaseA1 CpGhsm with coverage at least 50 (number of sites: 11021 )

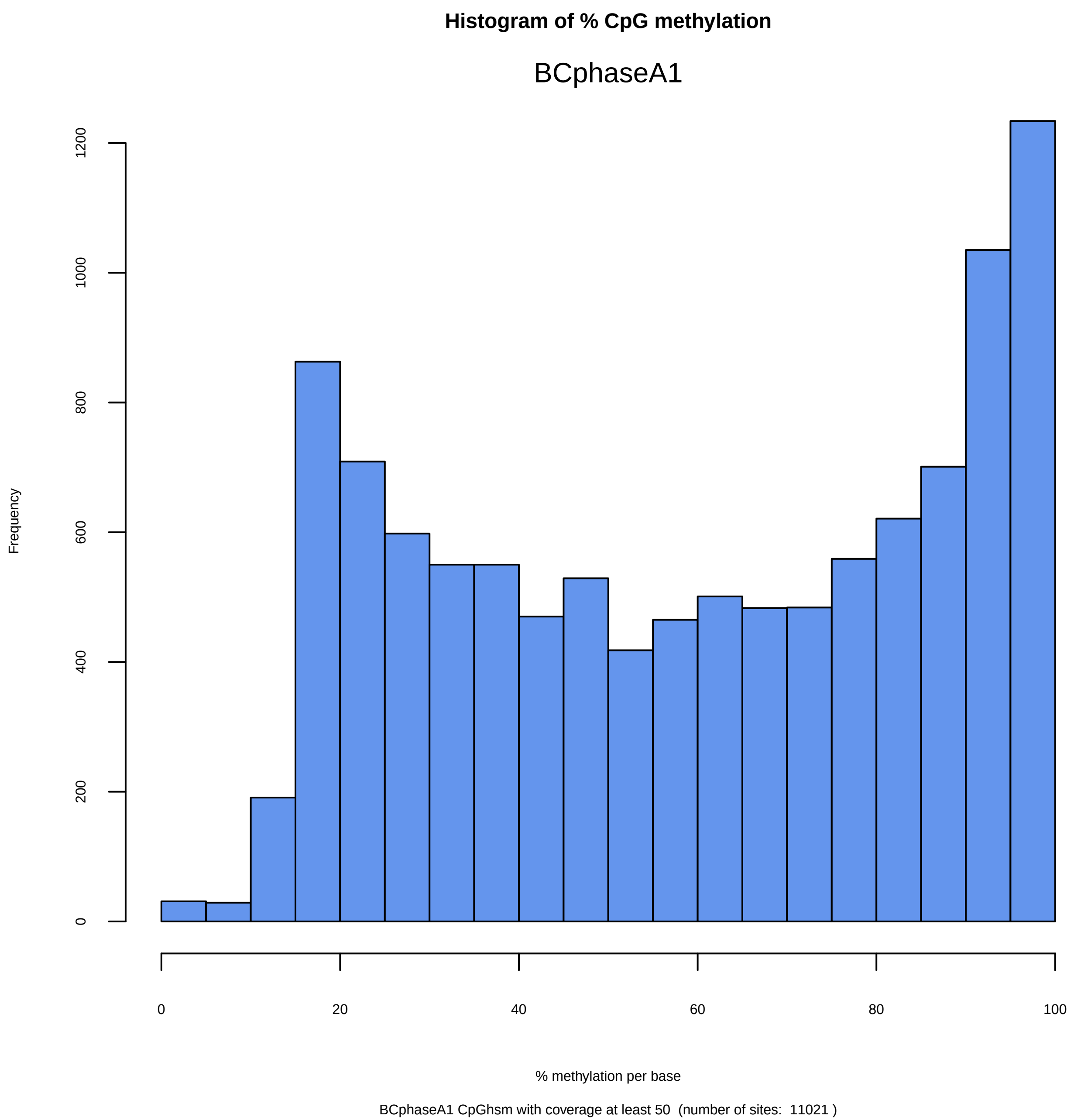

Histogram of % CpG methylation
BCphaseA1
Frequency
0
200
400
600
800
1000
1200
0
20
40
60
80
100
% methylation per base
BCphaseA1 CpGhsm with coverage at least 50 (number of sites: 11021 )

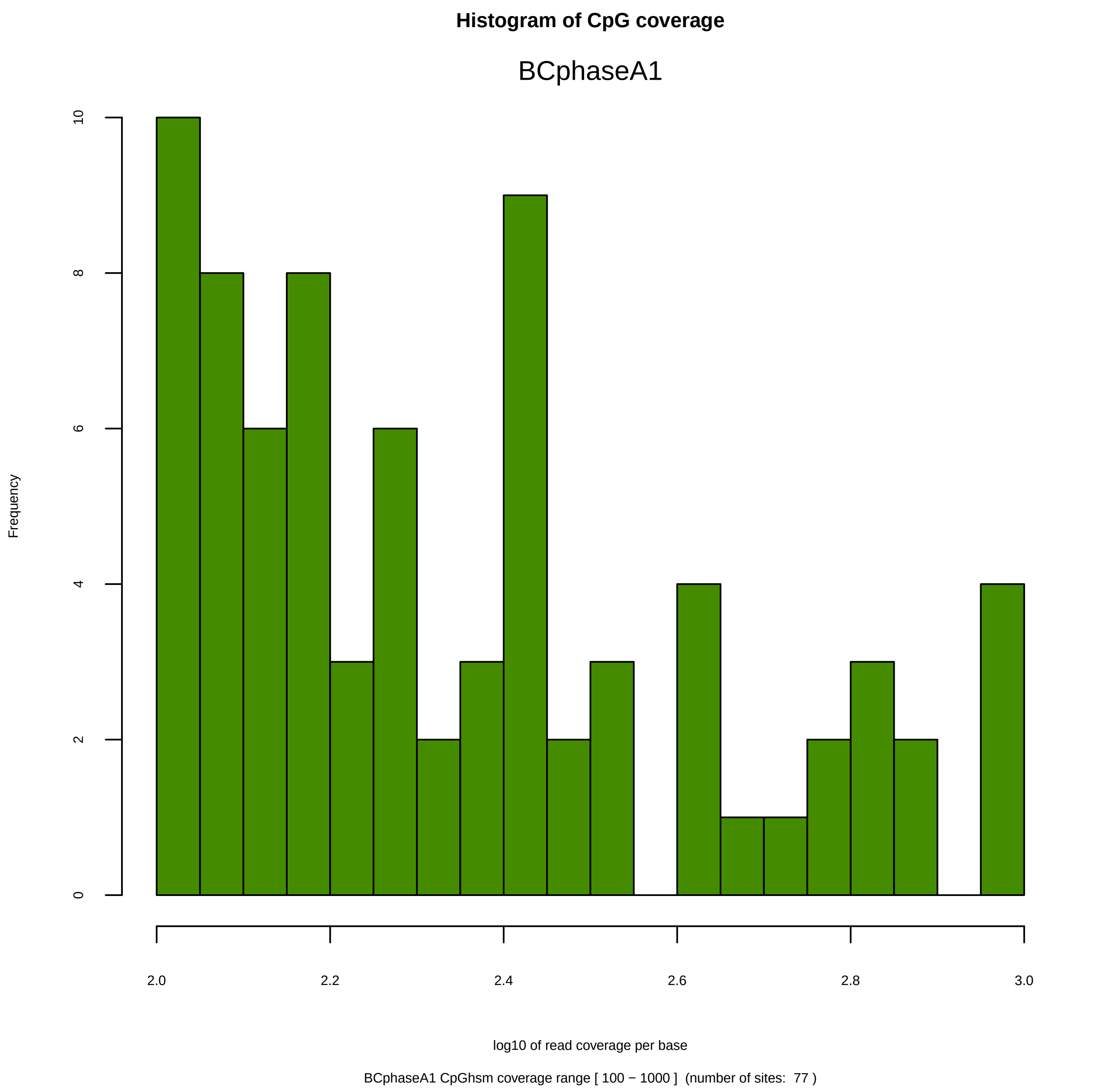
Histogram of CpG coverage
BCphaseA1
Frequency
0
2
4
6
8
10
2.0
2.2
2.4
2.6
2.8
3.0
log10 of read coverage per base
BCphaseA1 CpGhsm coverage range [ 100 − 1000 ] (number of sites: 77 )

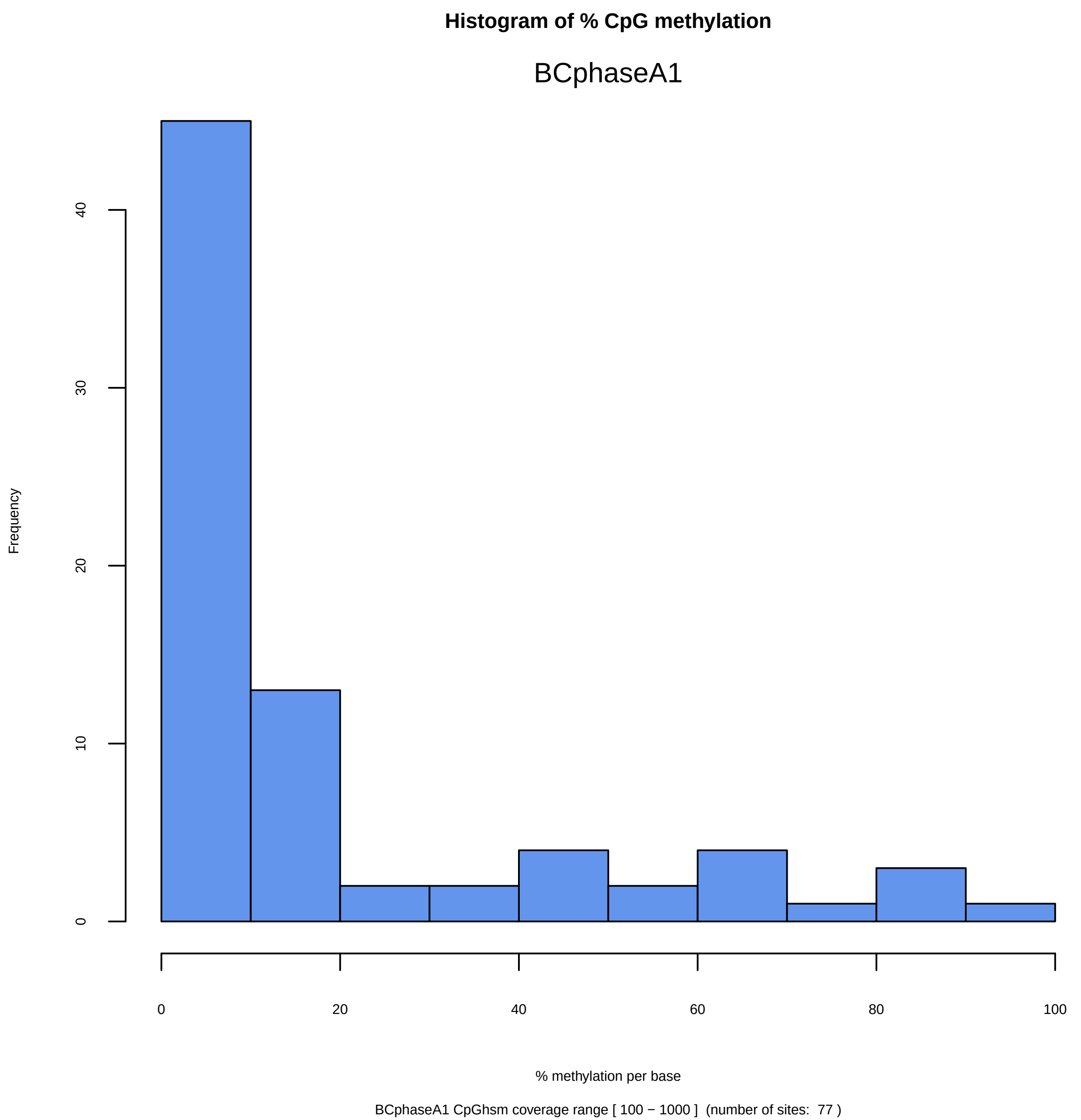

Histogram of % CpG methylation
BCphaseA1
Frequency
0
10
20
30
40
0
20
40
60
80
100
% methylation per base
BCphaseA1 CpGhsm coverage range [ 100 − 1000 ]  (number of sites:  77 )

#### Histogram of CpG coverage

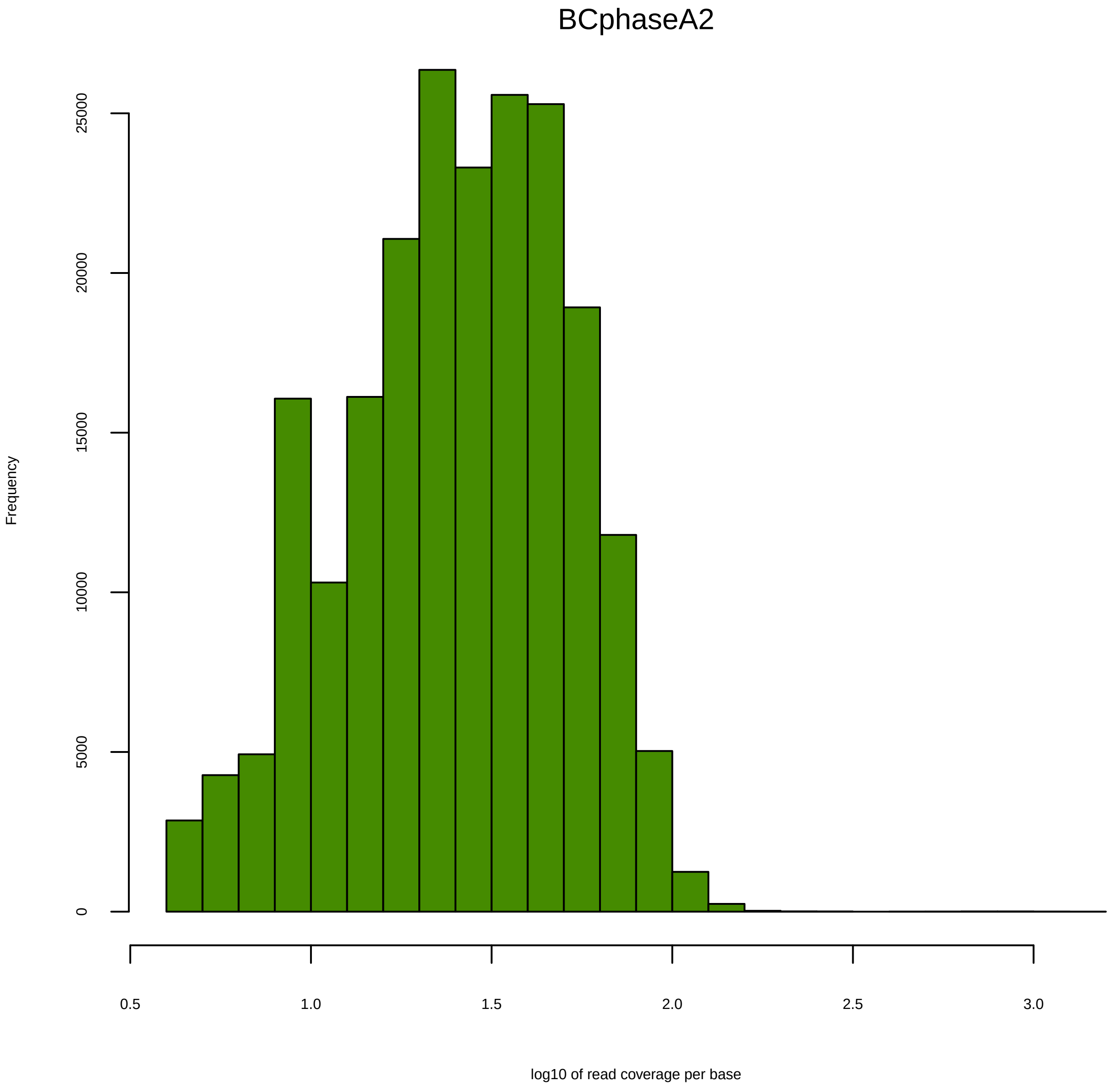

BCphaseA2 CpGhsm with coverage at least 5 (number of sites: 213448 )

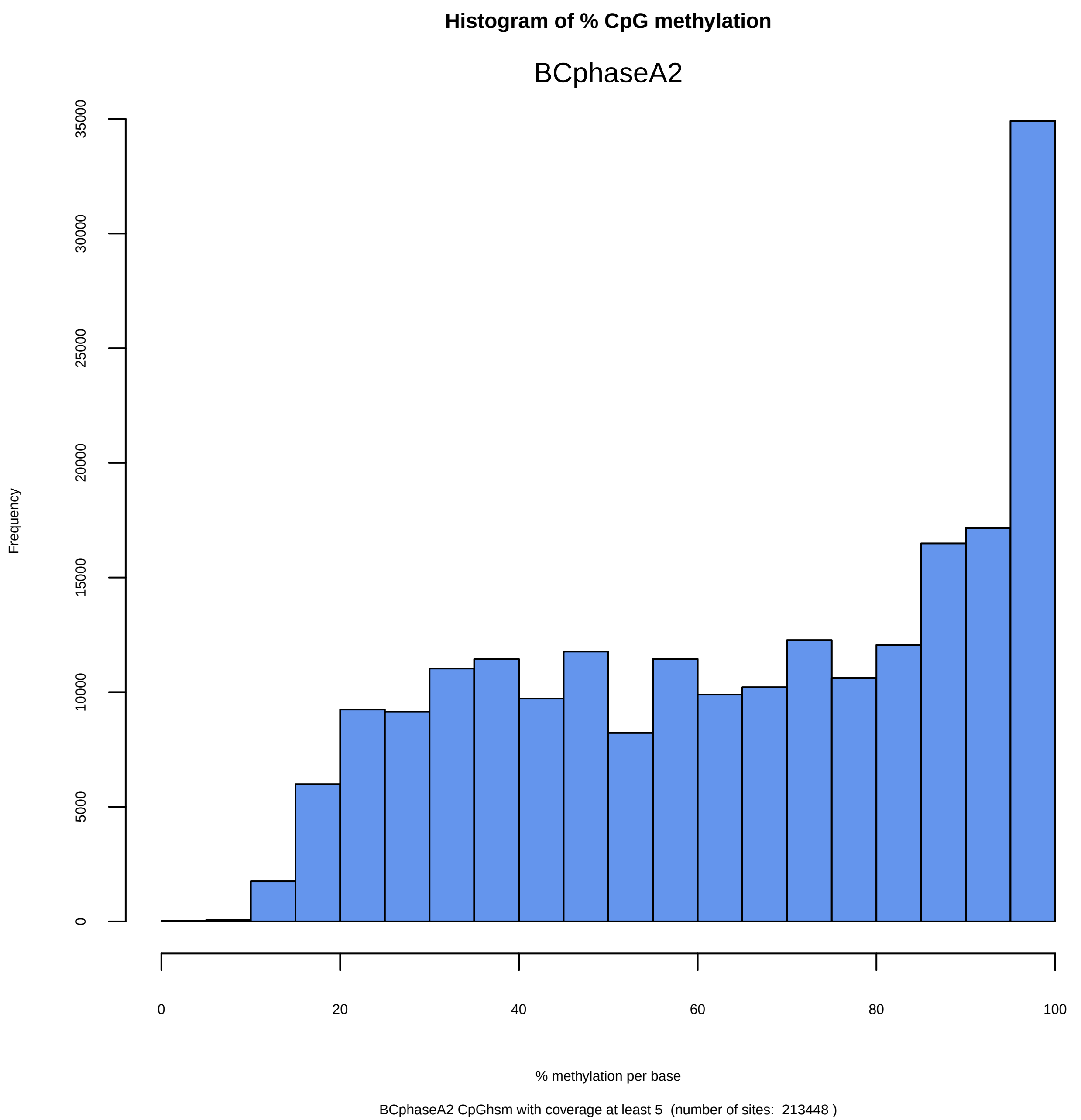

Histogram of % CpG methylation
BCphaseA2
Frequency
0
5000
10000
15000
20000
25000
30000
35000
0
20
40
60
80
100
% methylation per base
BCphaseA2 CpGhsm with coverage at least 5 (number of sites: 213448 )

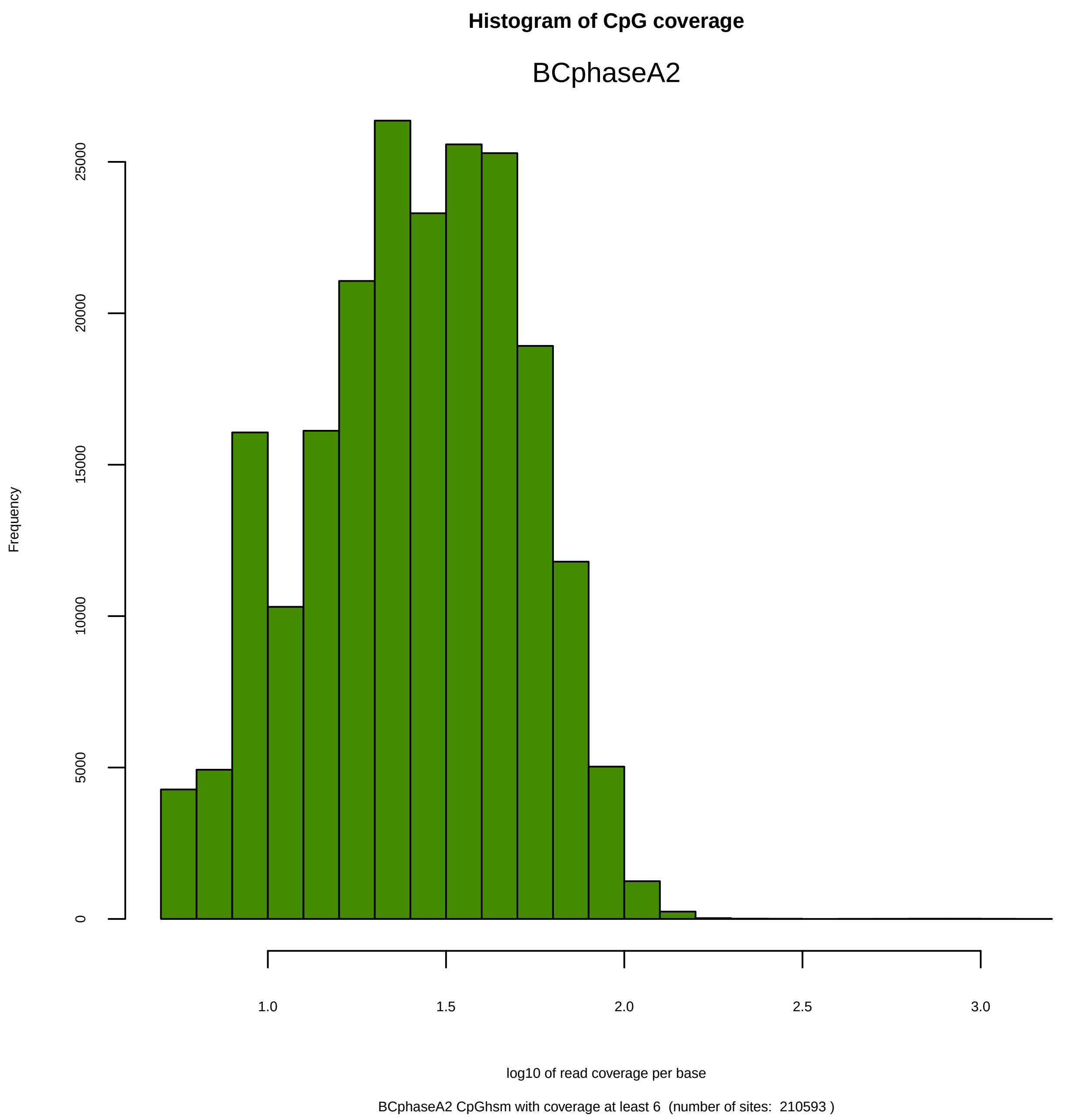

Histogram of CpG coverage
BCphaseA2
Frequency
0
5000
10000
15000
20000
25000
1.0
1.5
2.0
2.5
3.0
log10 of read coverage per base
BCphaseA2 CpGhsm with coverage at least 6 (number of sites: 210593 )

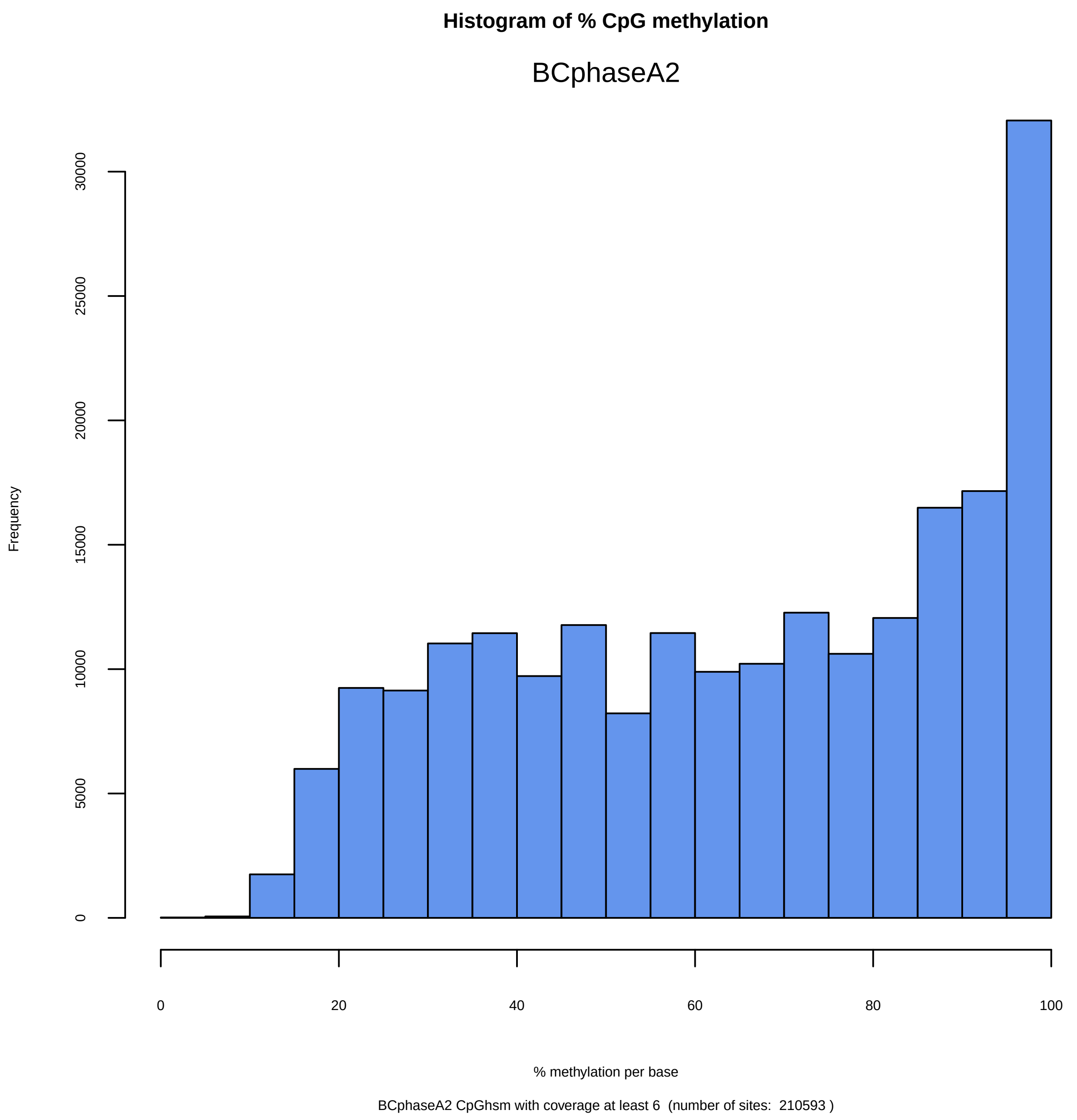

Histogram of % CpG methylation
BCphaseA2
Frequency
0
5000
10000
15000
20000
25000
30000
0
20
40
60
80
100
% methylation per base
BCphaseA2 CpGhsm with coverage at least 6 (number of sites: 210593 )

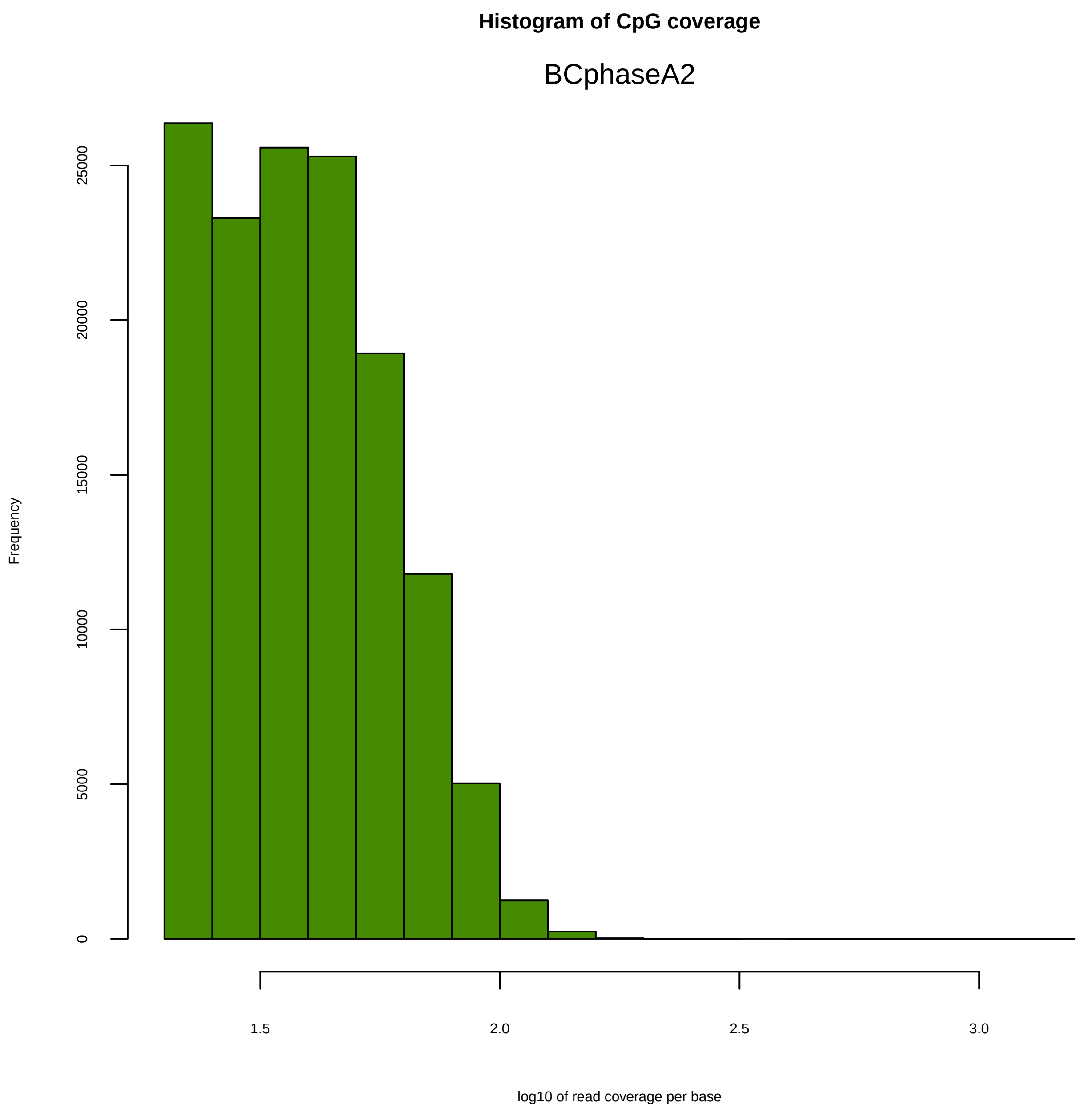

BCphaseA2 CpGhsm with coverage at least 20  (number of sites:  137831 )

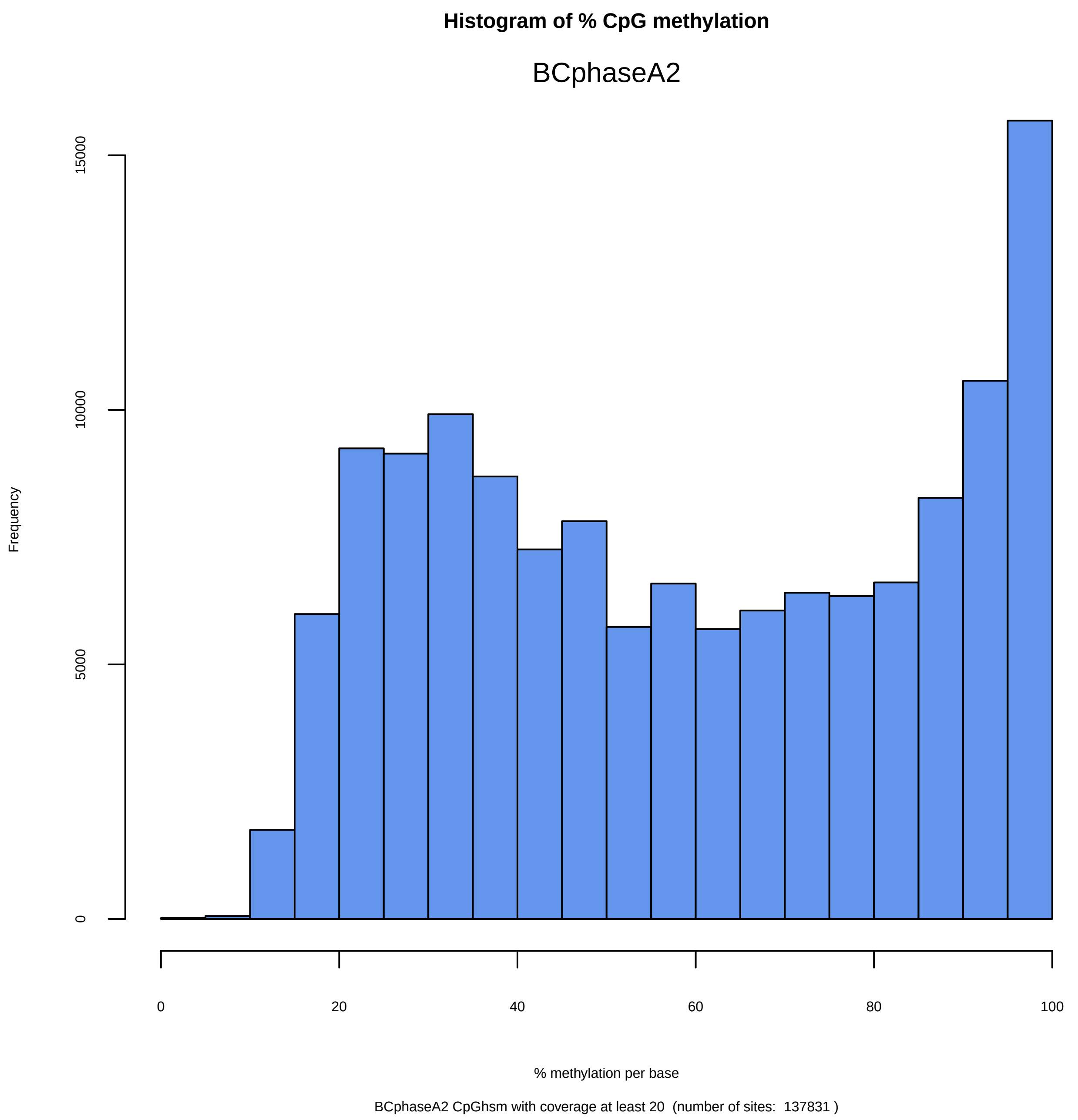
Histogram of % CpG methylation
BCphaseA2
Frequency
0
5000
10000
15000
0
20
40
60
80
100
% methylation per base
BCphaseA2 CpGhsm with coverage at least 20 (number of sites: 137831 )

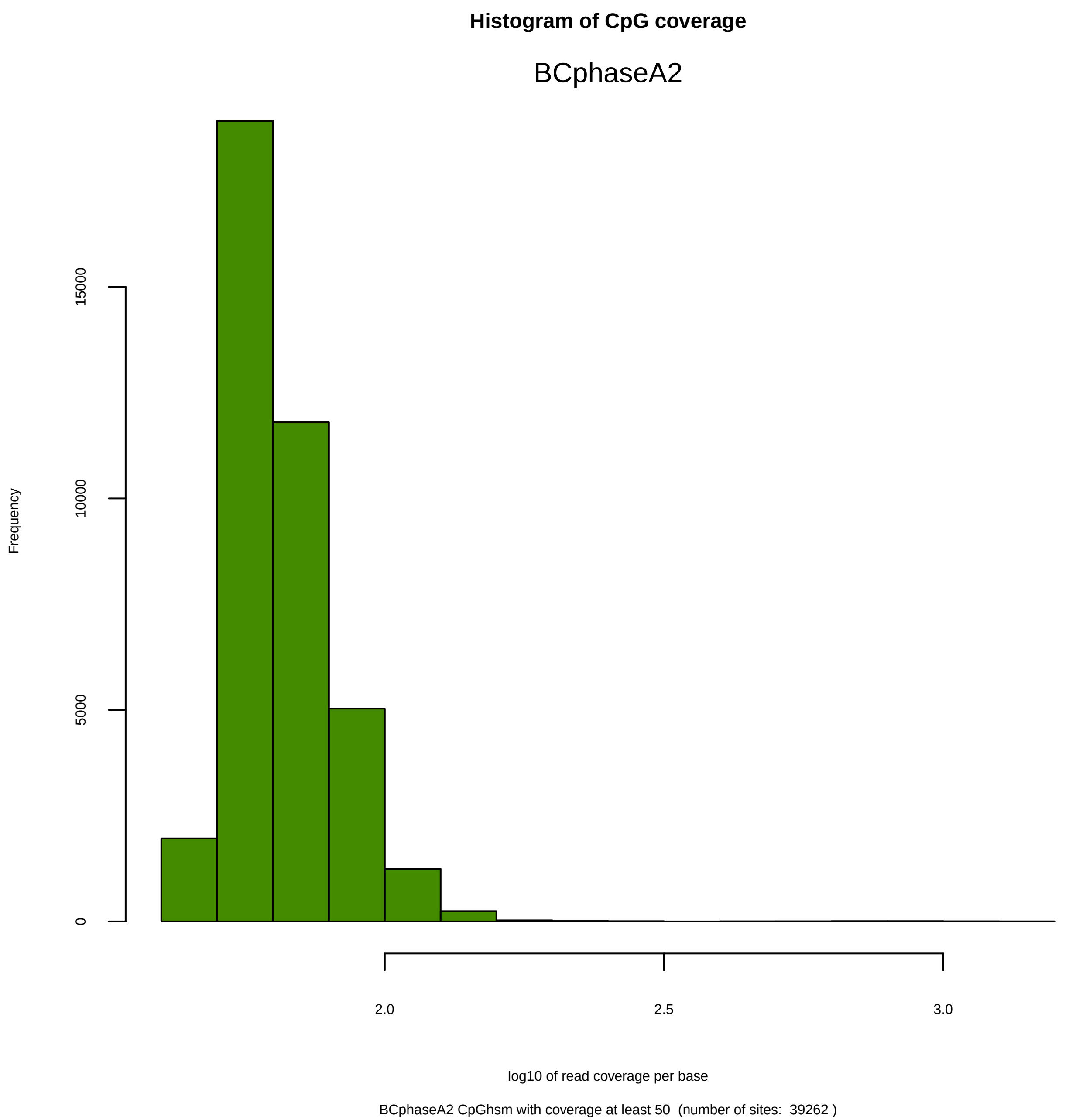

Histogram of CpG coverage
BCphaseA2
Frequency
0
5000
10000
15000
2.0
2.5
3.0
log10 of read coverage per base
BCphaseA2 CpGhsm with coverage at least 50 (number of sites: 39262 )

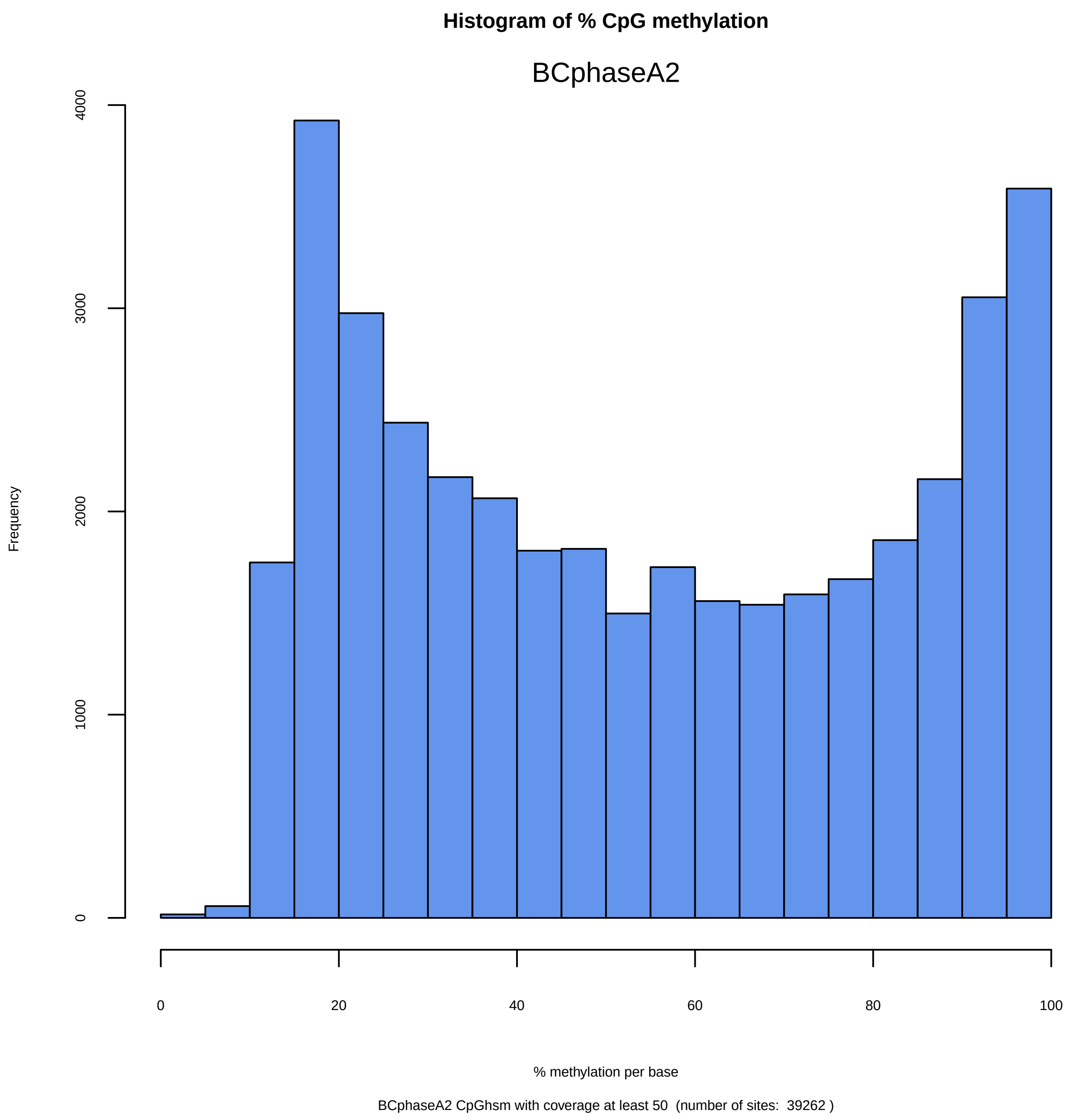
Histogram of % CpG methylation
BCphaseA2
Frequency
0
1000
2000
3000
4000
0
20
40
60
80
100
% methylation per base
BCphaseA2 CpGhsm with coverage at least 50 (number of sites: 39262 )

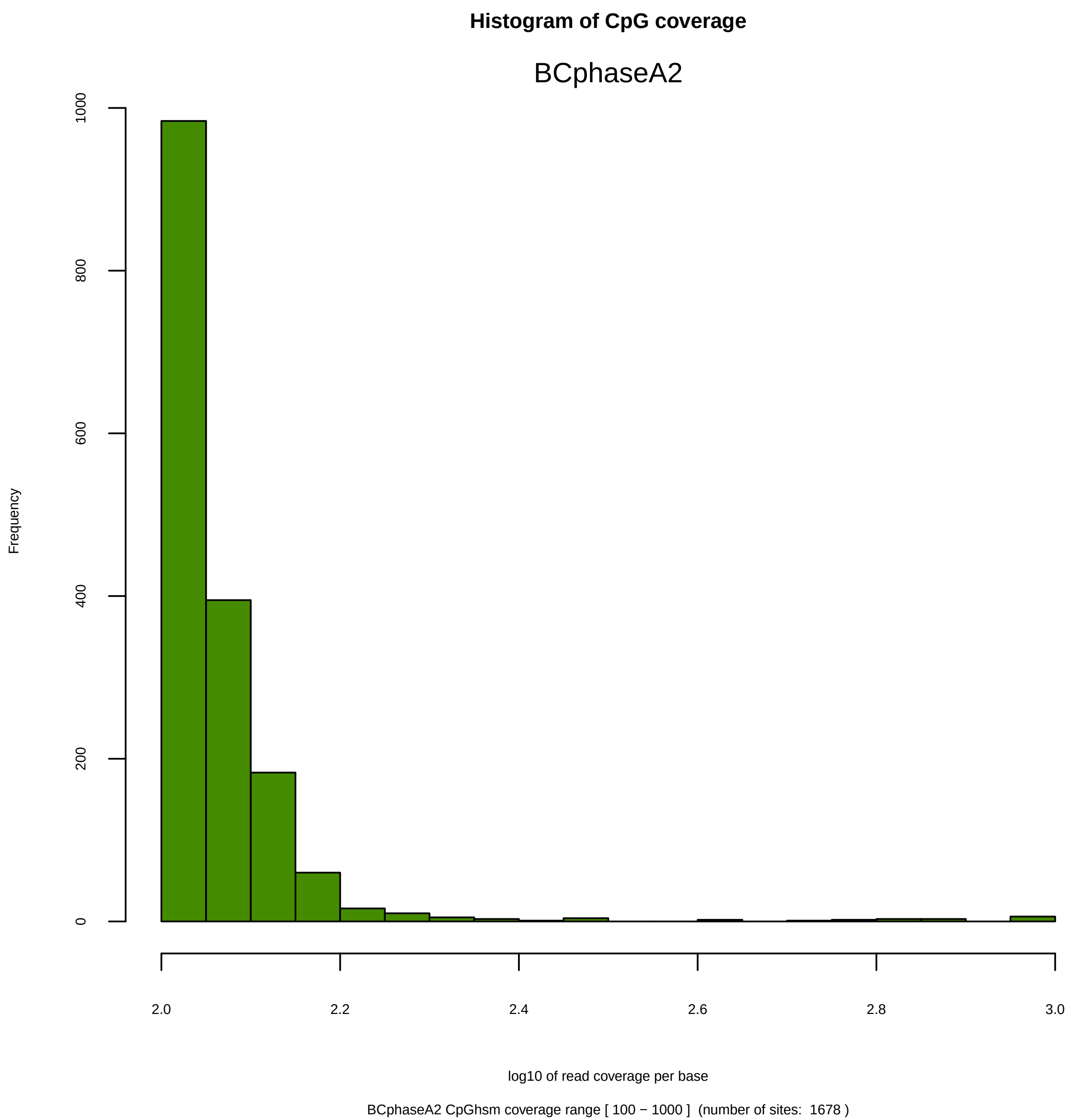
Histogram of CpG coverage
BCphaseA2
Frequency
0
200
400
600
800
1000
2.0
2.2
2.4
2.6
2.8
3.0
log10 of read coverage per base
BCphaseA2 CpGhsm coverage range [ 100 − 1000 ]  (number of sites:  1678 )

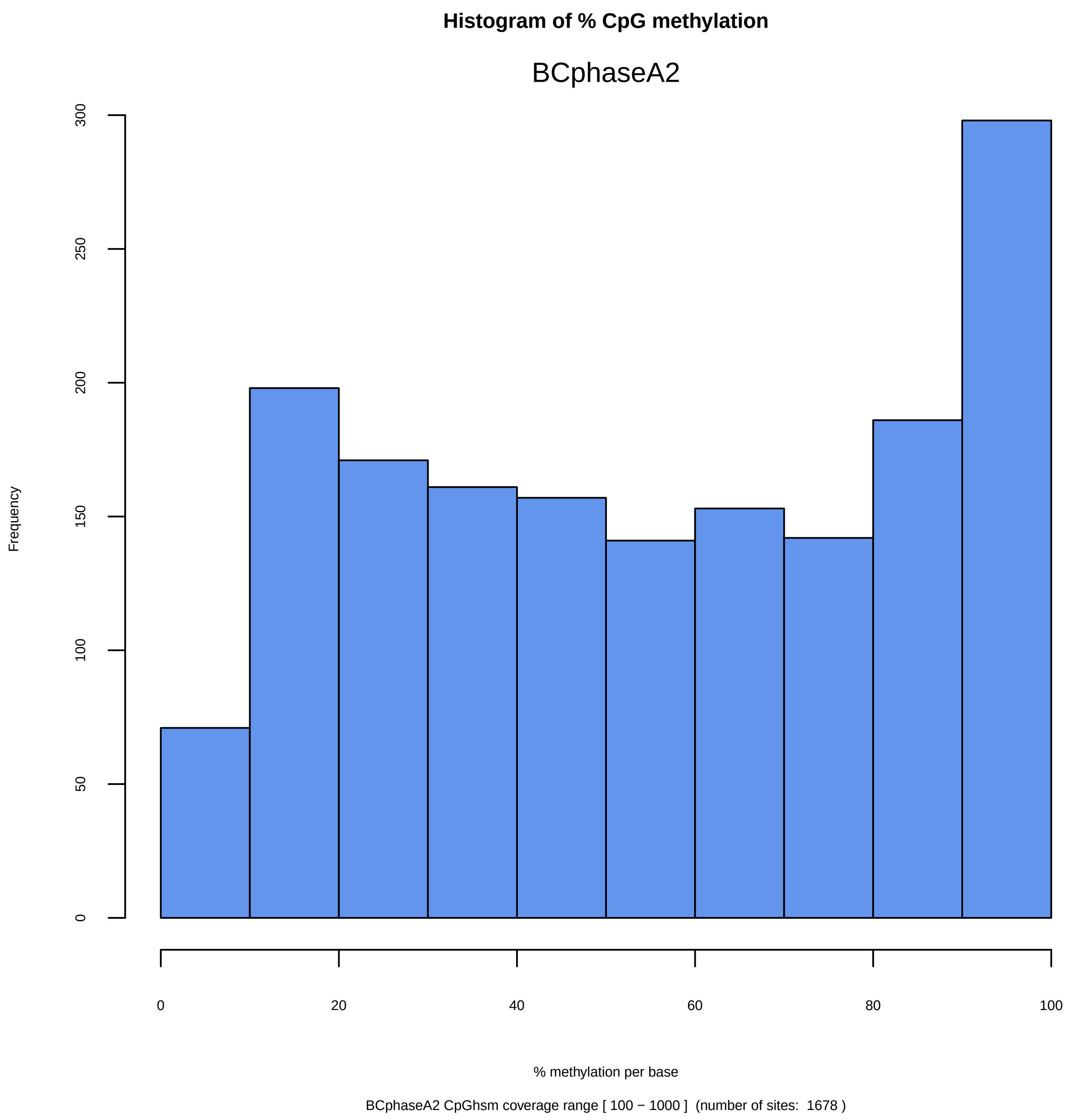

Histogram of % CpG methylation
BCphaseA2
Frequency
0
50
100
150
200
250
300
0
20
40
60
80
100
% methylation per base
BCphaseA2 CpGhsm coverage range [ 100 − 1000 ] (number of sites: 1678 )

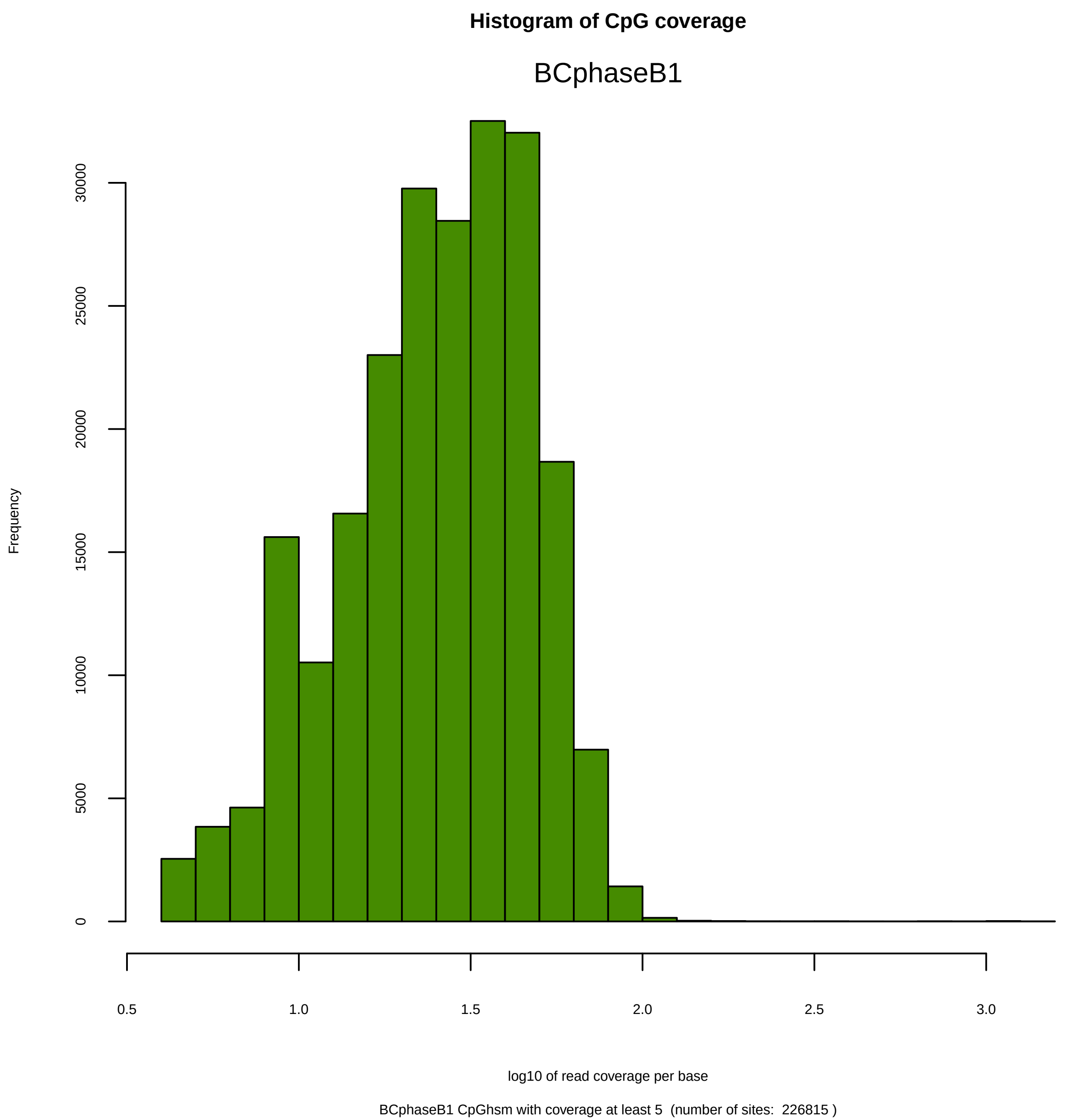
Histogram of CpG coverage
BCphaseB1
Frequency
0
5000
10000
15000
20000
25000
30000
0.5
1.0
1.5
2.0
2.5
3.0
log10 of read coverage per base
BCphaseB1 CpGhsm with coverage at least 5  (number of sites:  226815 )

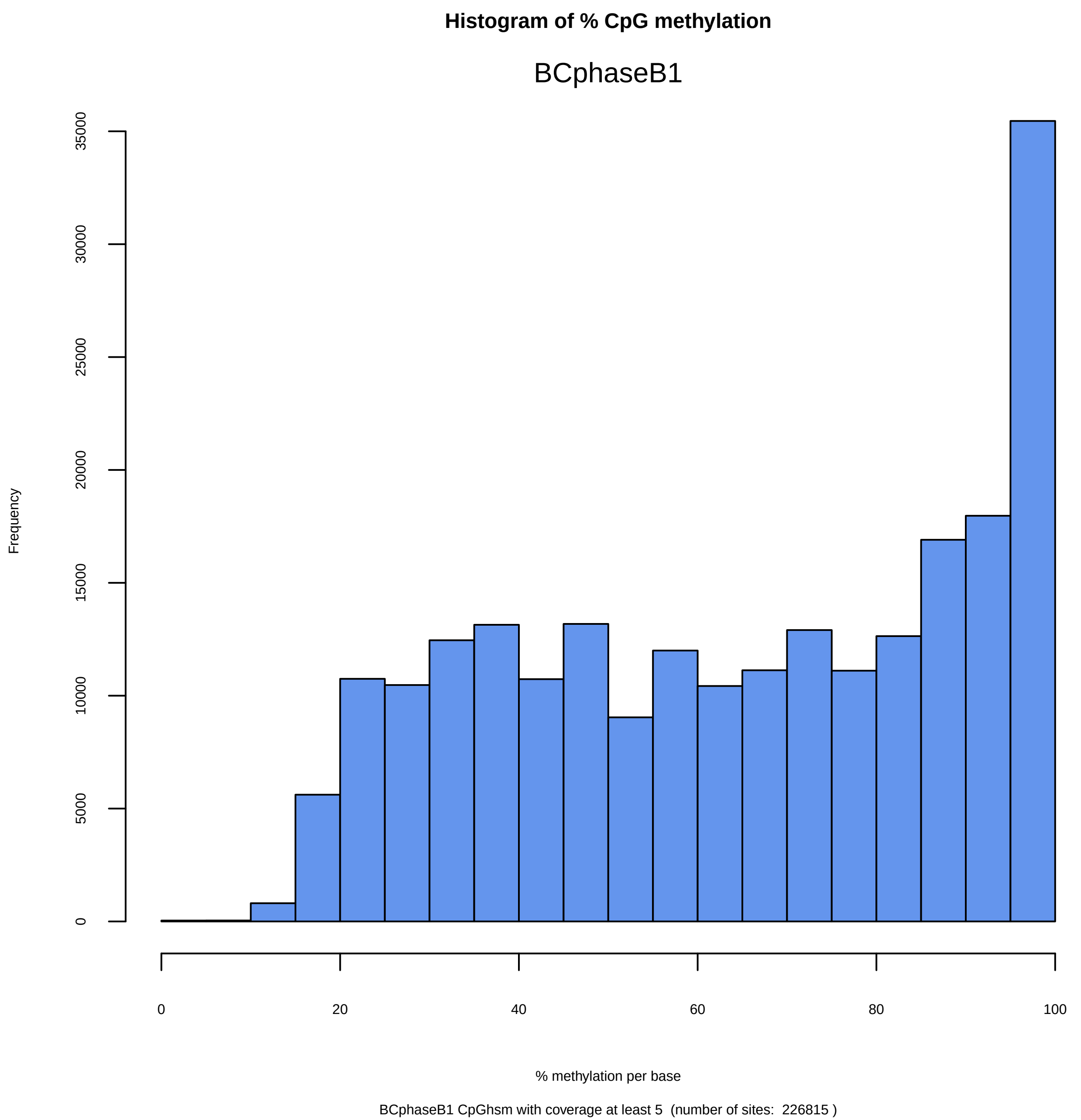

Histogram of % CpG methylation
BCphaseB1
Frequency
0
5000
10000
15000
20000
25000
30000
35000
0
20
40
60
80
100
% methylation per base
BCphaseB1 CpGhsm with coverage at least 5  (number of sites:  226815 )

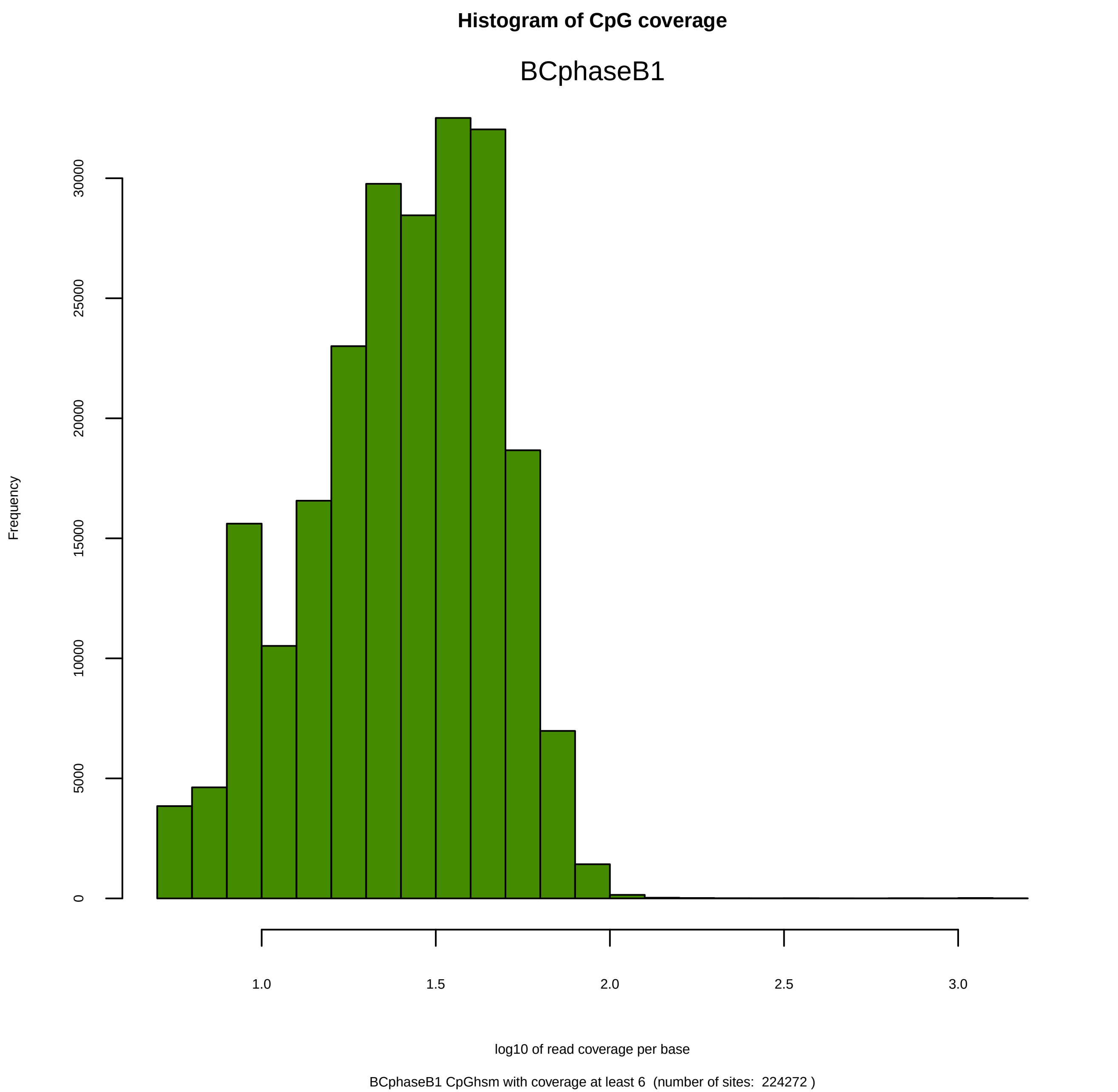

Histogram of CpG coverage
BCphaseB1
Frequency
0
5000
10000
15000
20000
25000
30000
1.0
1.5
2.0
2.5
3.0
log10 of read coverage per base
BCphaseB1 CpGhsm with coverage at least 6 (number of sites: 224272 )

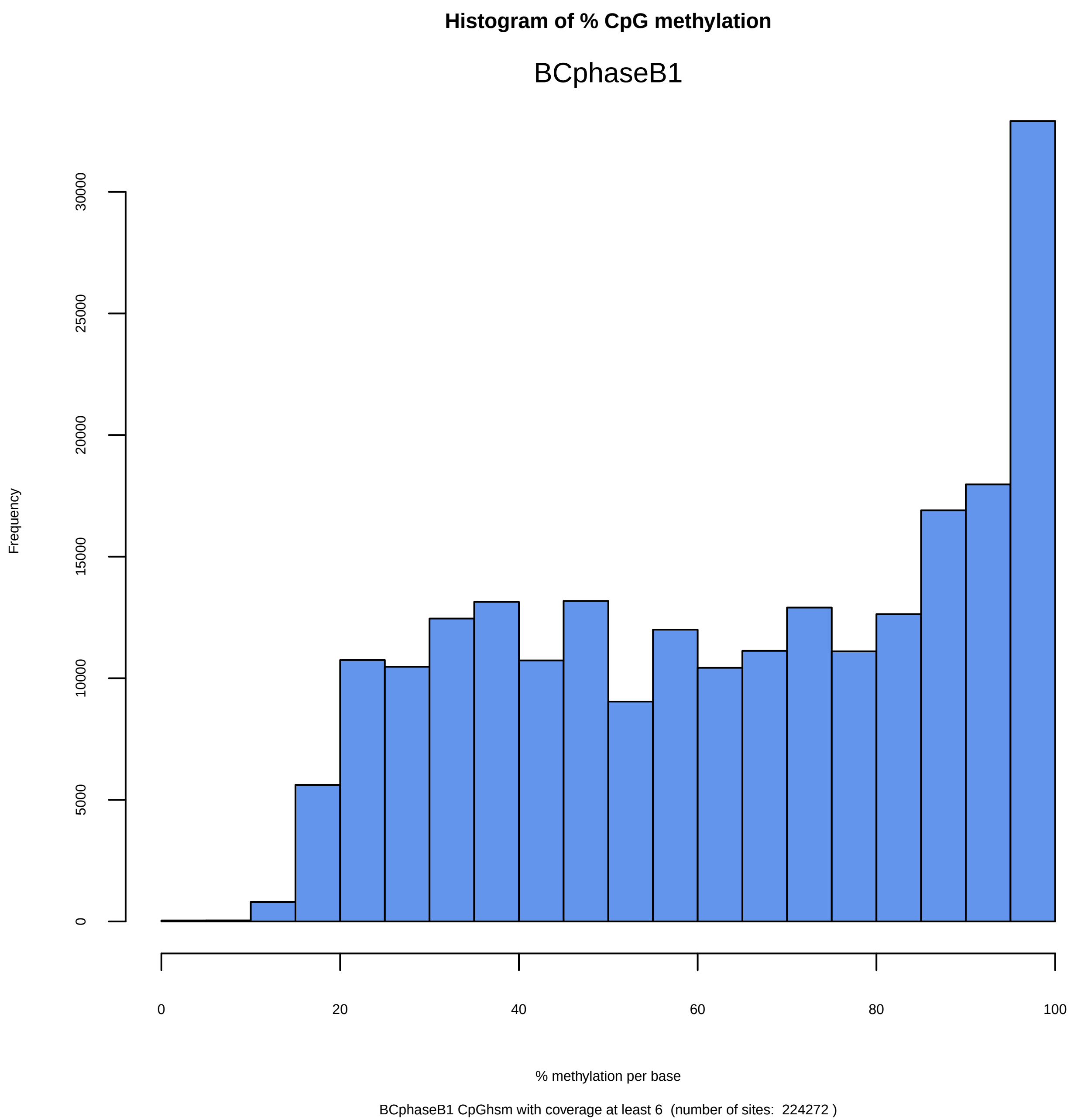

Histogram of % CpG methylation
BCphaseB1
Frequency
0
5000
10000
15000
20000
25000
30000
0
20
40
60
80
100
% methylation per base
BCphaseB1 CpGhsm with coverage at least 6 (number of sites: 224272 )

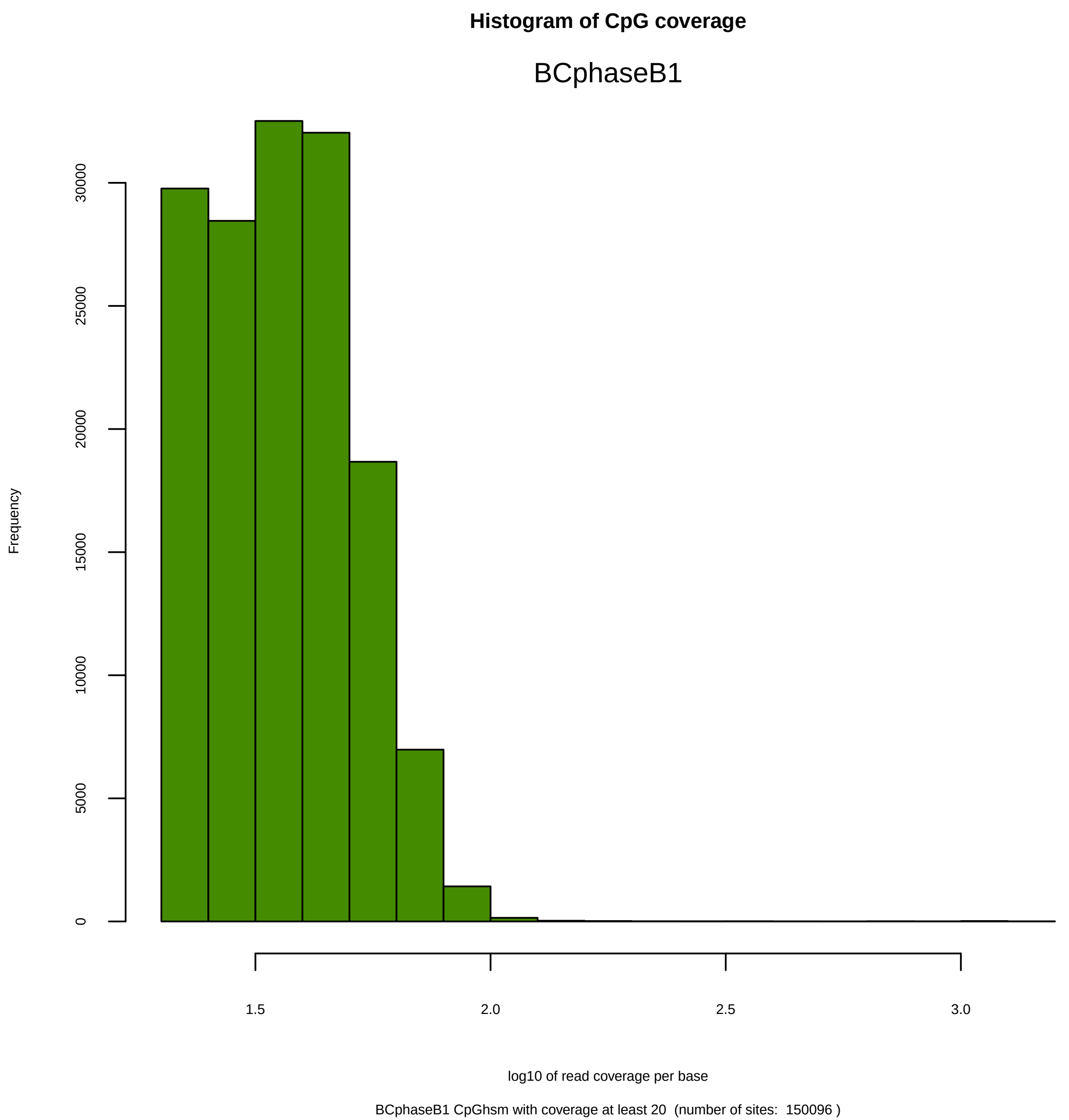

Histogram of CpG coverage
BCphaseB1
Frequency
0
5000
10000
15000
20000
25000
30000
1.5
2.0
2.5
3.0
log10 of read coverage per base
BCphaseB1 CpGhsm with coverage at least 20 (number of sites: 150096 )

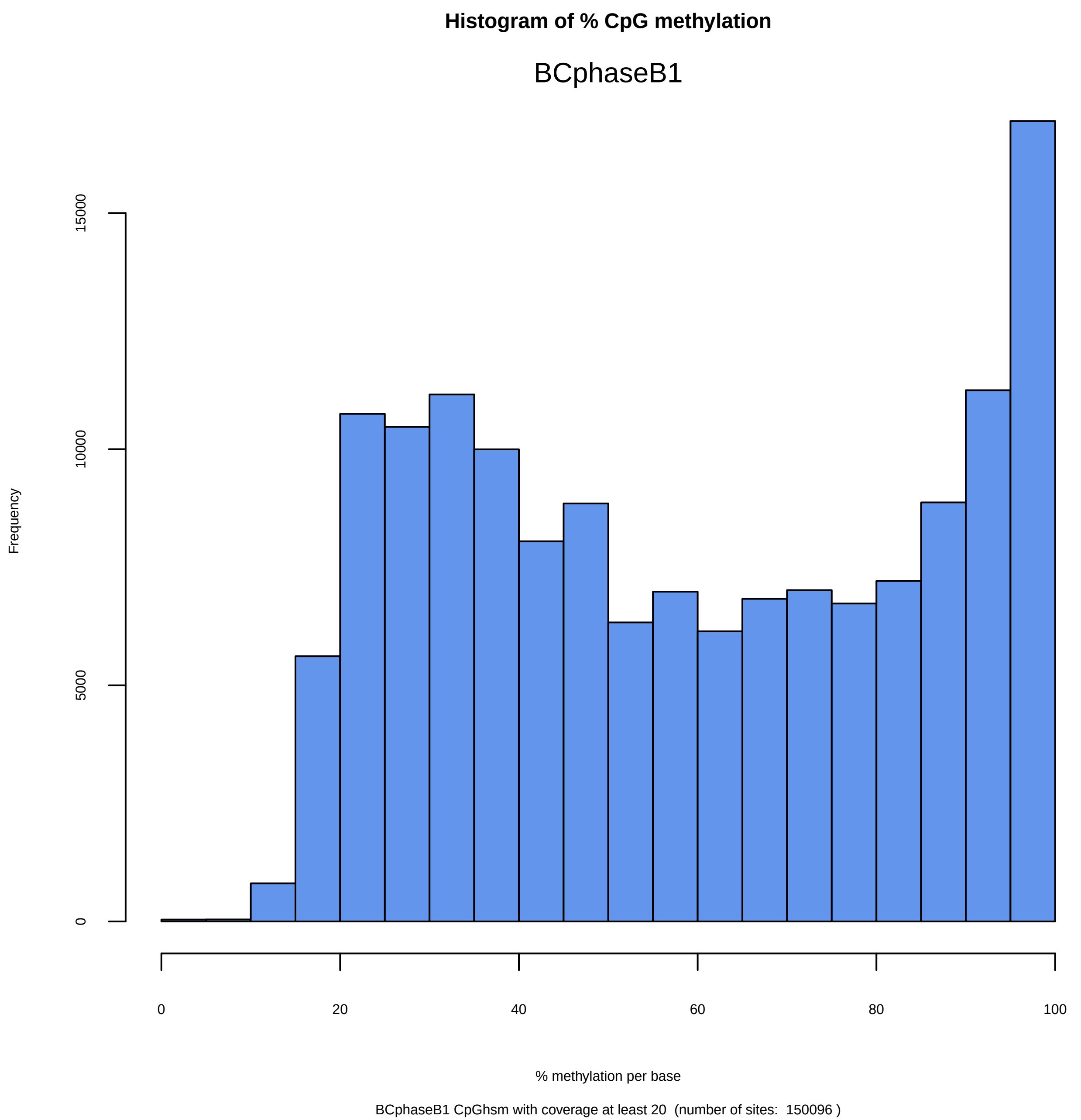
Histogram of % CpG methylation
BCphaseB1
Frequency
0
5000
10000
15000
0
20
40
60
80
100
% methylation per base
BCphaseB1 CpGhsm with coverage at least 20  (number of sites:  150096 )

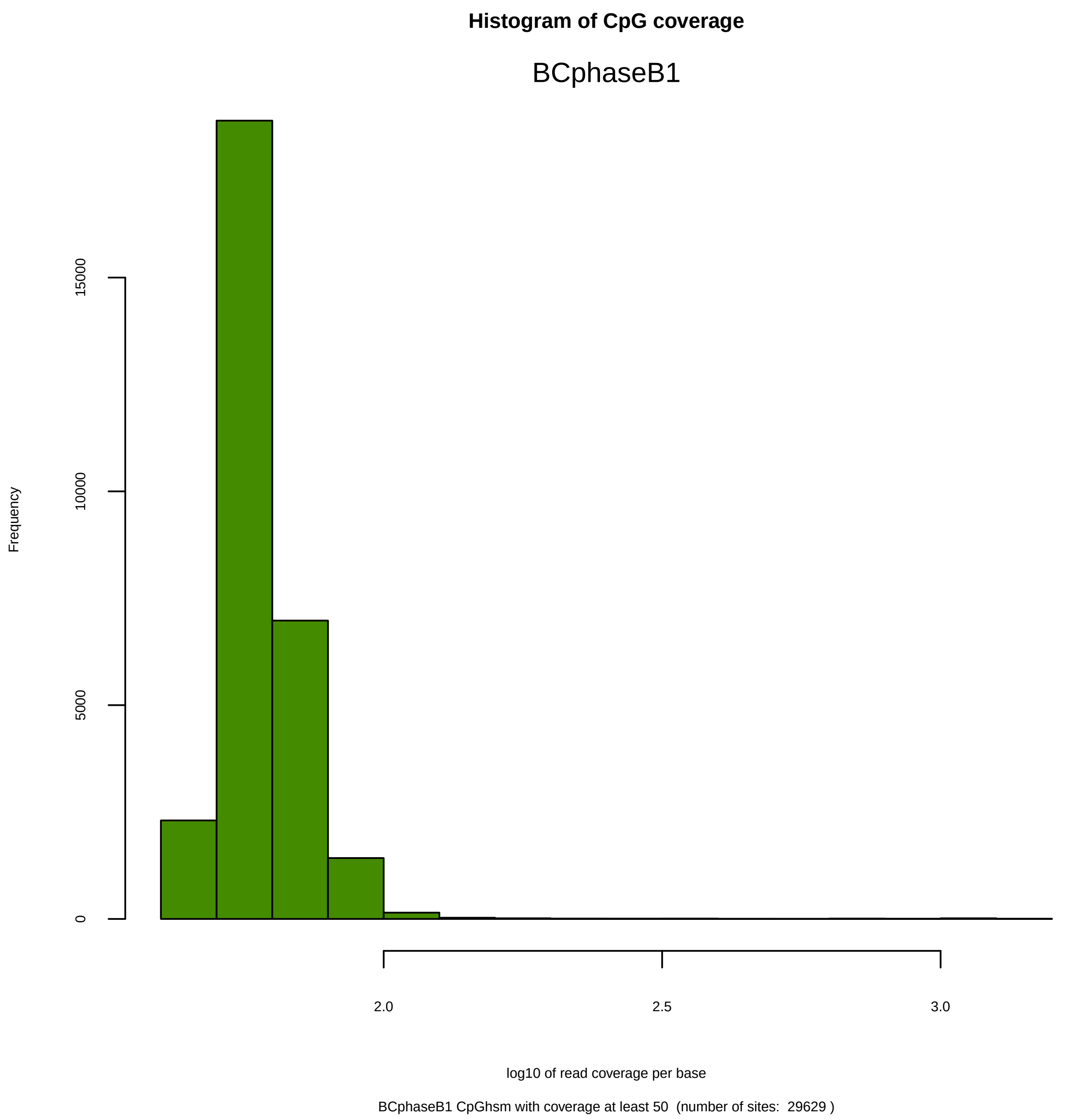
Histogram of CpG coverage
BCphaseB1
Frequency
0
5000
10000
15000
2.0
2.5
3.0
log10 of read coverage per base
BCphaseB1 CpGhsm with coverage at least 50 (number of sites: 29629 )

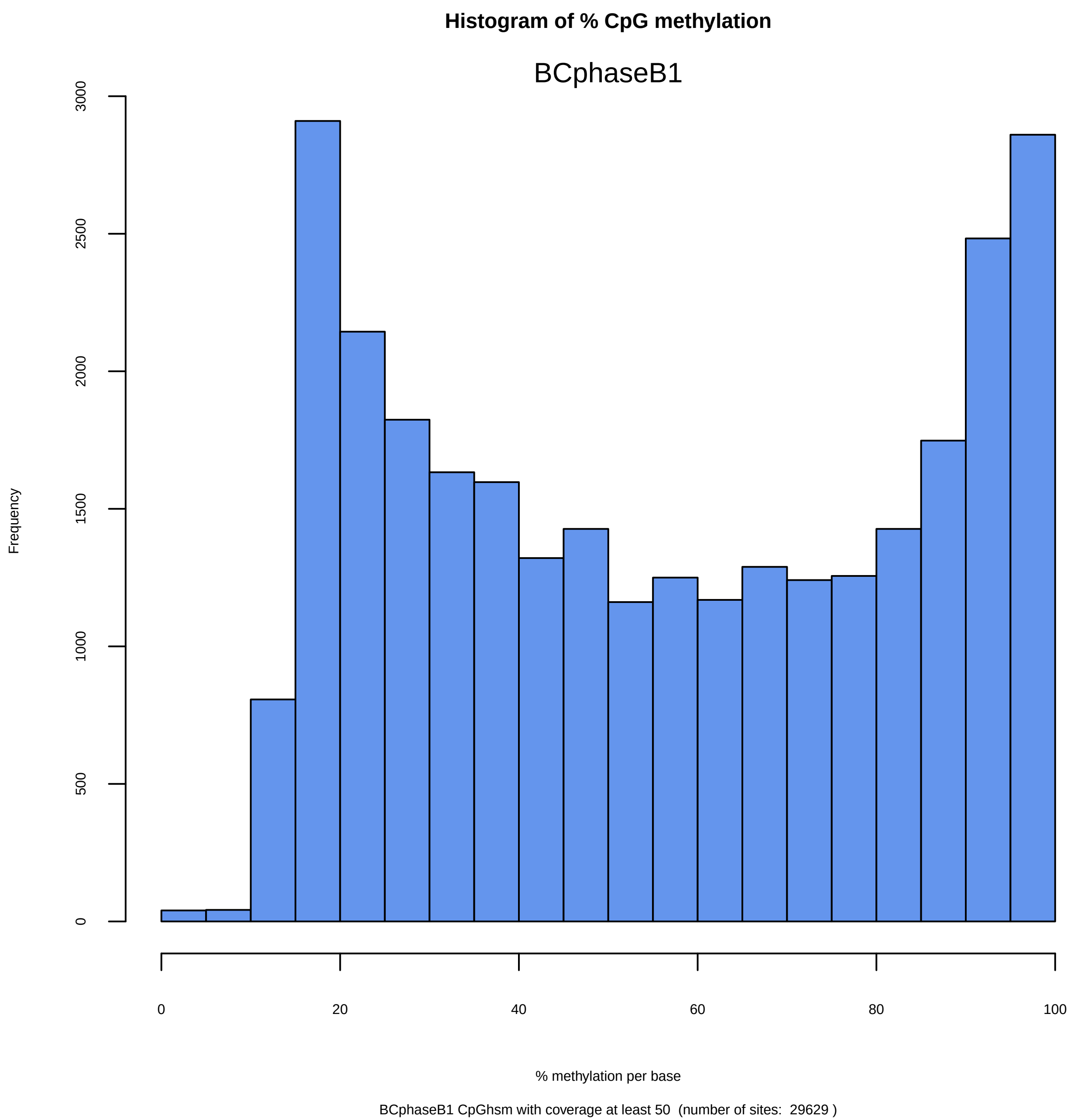

Histogram of % CpG methylation
BCphaseB1
Frequency
0
500
1000
1500
2000
2500
3000
0
20
40
60
80
100
% methylation per base
BCphaseB1 CpGhsm with coverage at least 50  (number of sites:  29629 )

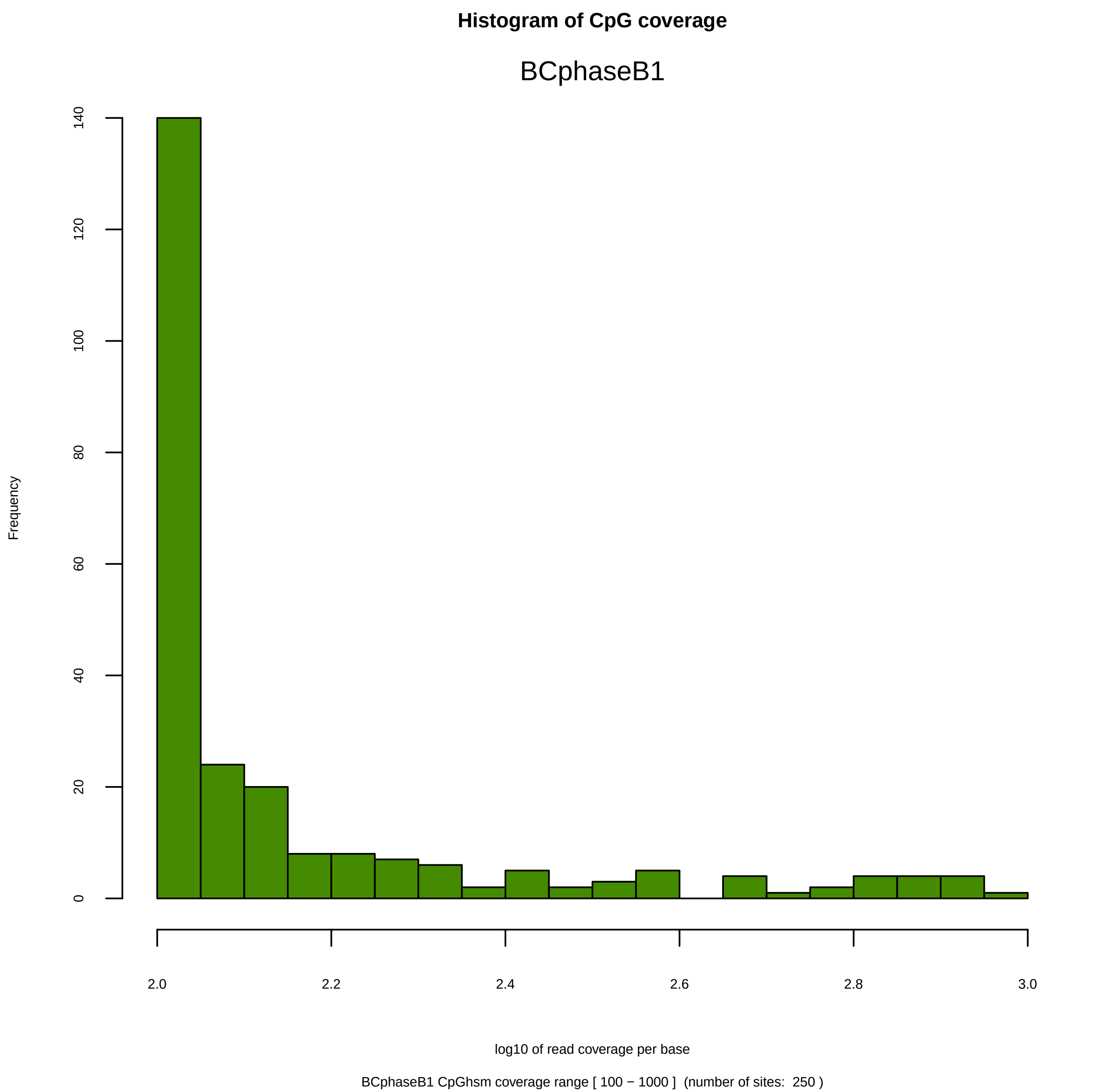
Histogram of CpG coverage
BCphaseB1
Frequency
0
20
40
60
80
100
120
140
2.0
2.2
2.4
2.6
2.8
3.0
log10 of read coverage per base
BCphaseB1 CpGhsm coverage range [ 100 − 1000 ]  (number of sites:  250 )

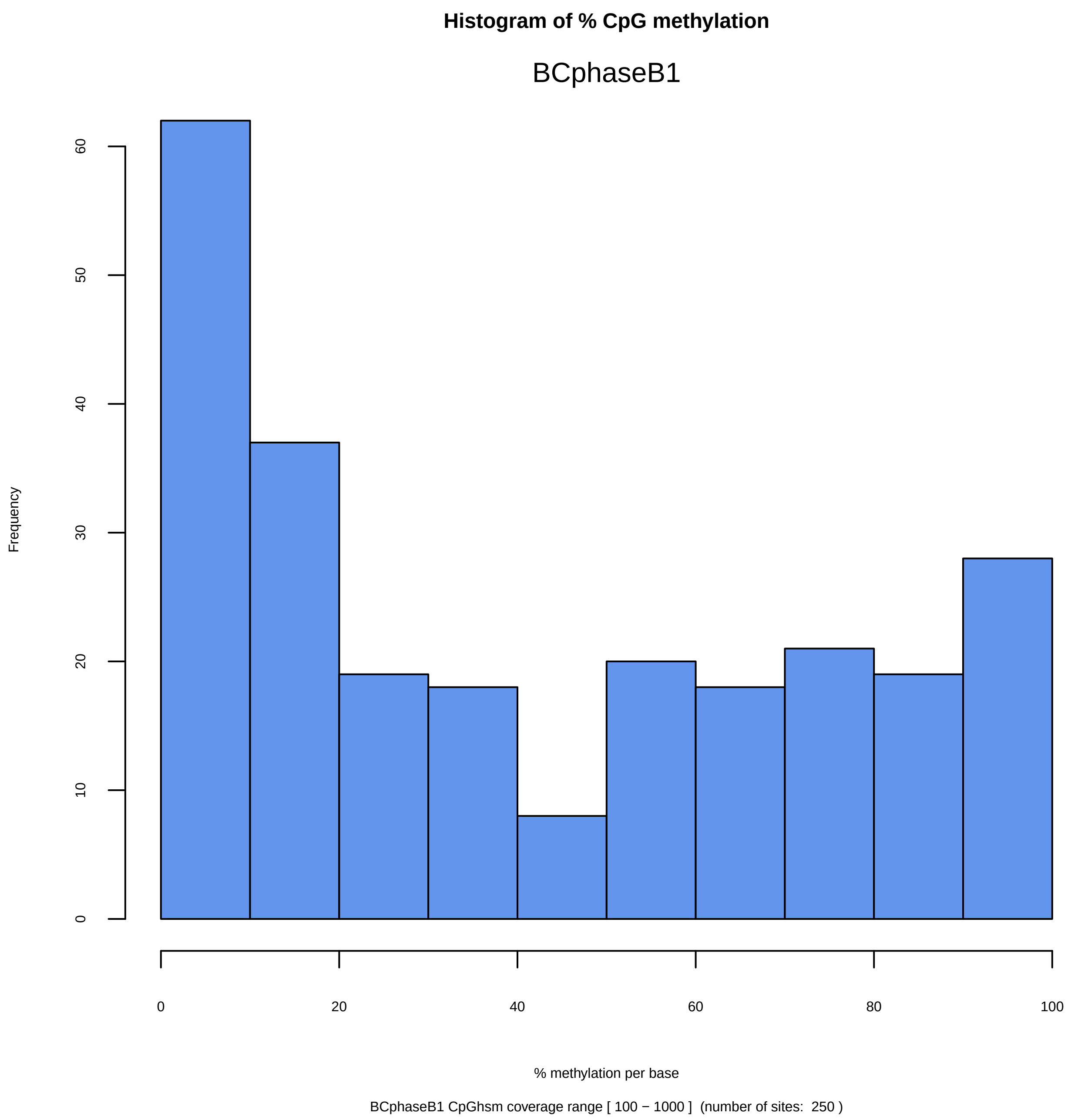

Histogram of % CpG methylation
BCphaseB1
Frequency
0
10
20
30
40
50
60
0
20
40
60
80
100
% methylation per base
BCphaseB1 CpGhsm coverage range [ 100 − 1000 ]  (number of sites:  250 )

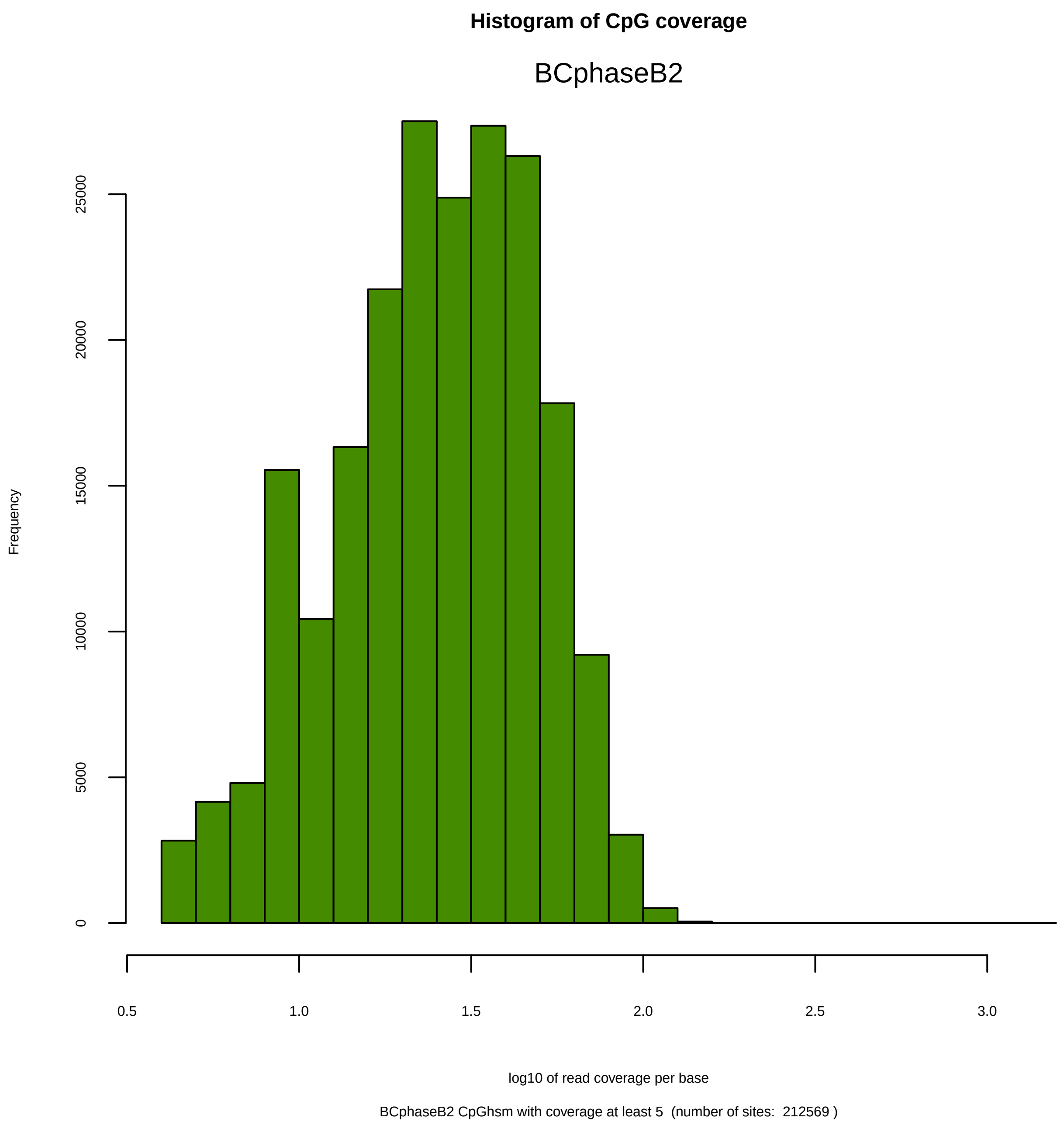
Histogram of CpG coverage
BCphaseB2
Frequency
0
5000
10000
15000
20000
25000
0.5
1.0
1.5
2.0
2.5
3.0
log10 of read coverage per base
BCphaseB2 CpGhsm with coverage at least 5  (number of sites:  212569 )

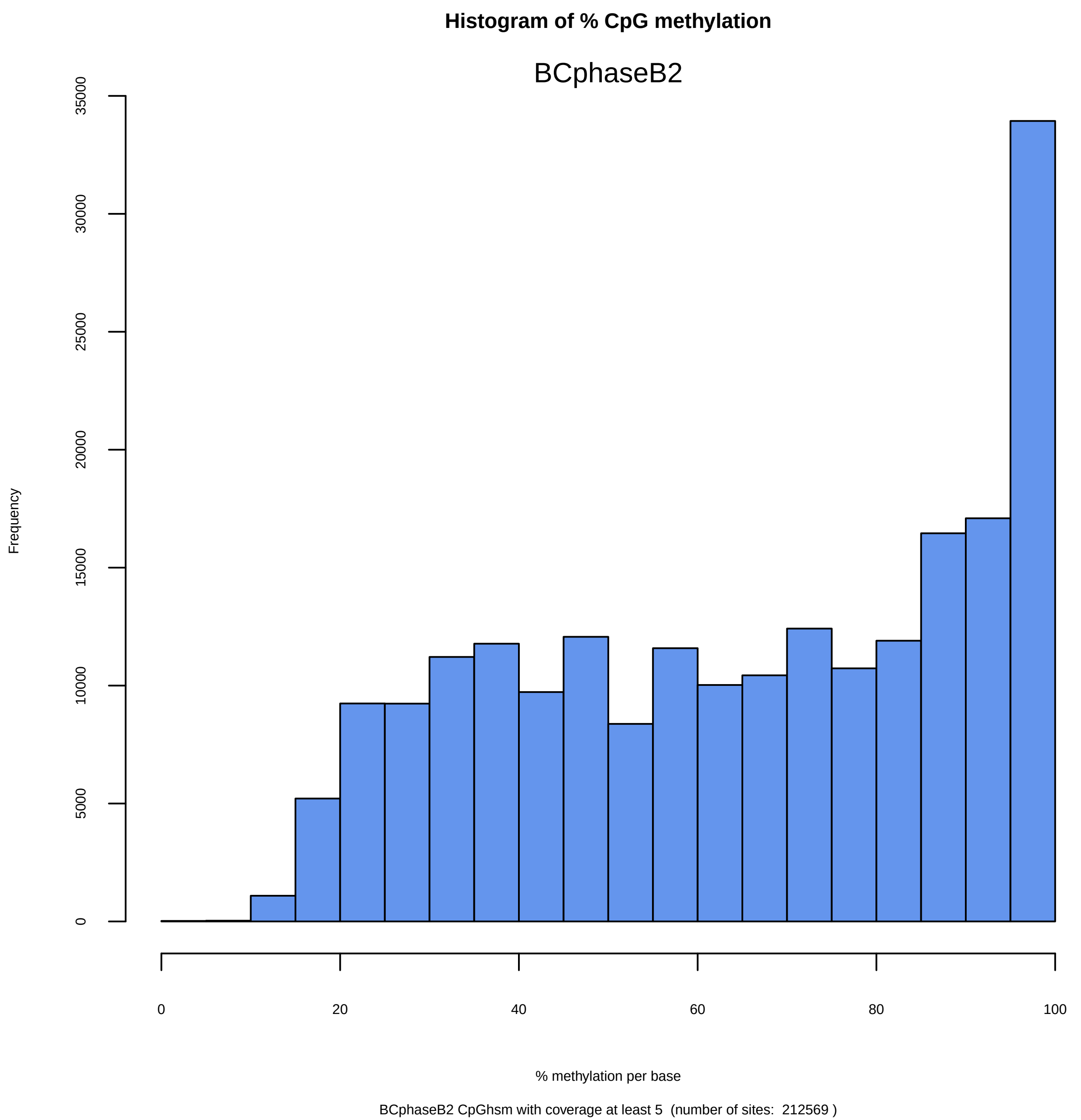
Histogram of % CpG methylation
BCphaseB2
Frequency
0
5000
10000
15000
20000
25000
30000
35000
0
20
40
60
80
100
% methylation per base
BCphaseB2 CpGhsm with coverage at least 5  (number of sites:  212569 )

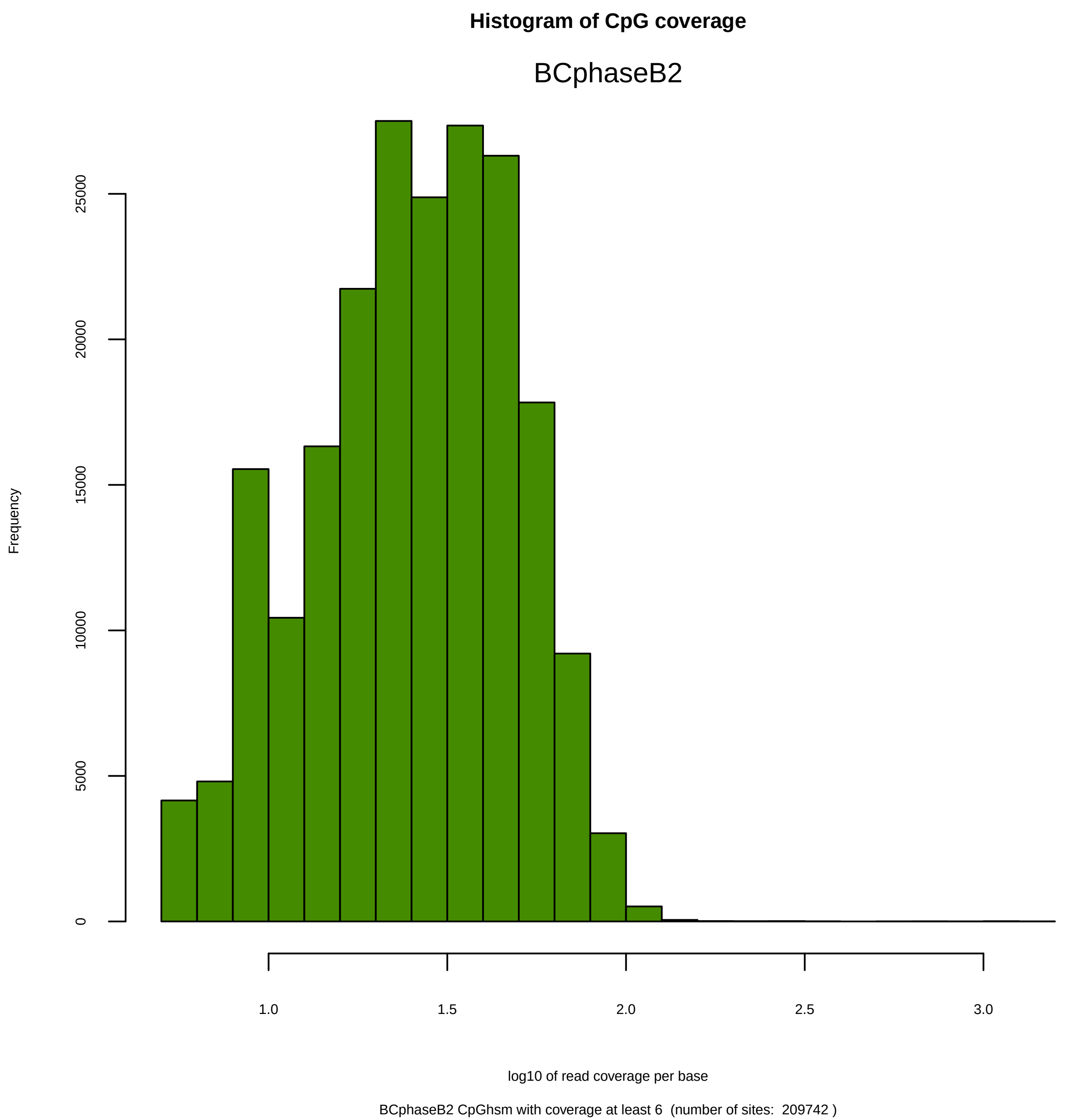
Histogram of CpG coverage
BCphaseB2
Frequency
0
5000
10000
15000
20000
25000
1.0
1.5
2.0
2.5
3.0
log10 of read coverage per base
BCphaseB2 CpGhsm with coverage at least 6  (number of sites:  209742 )

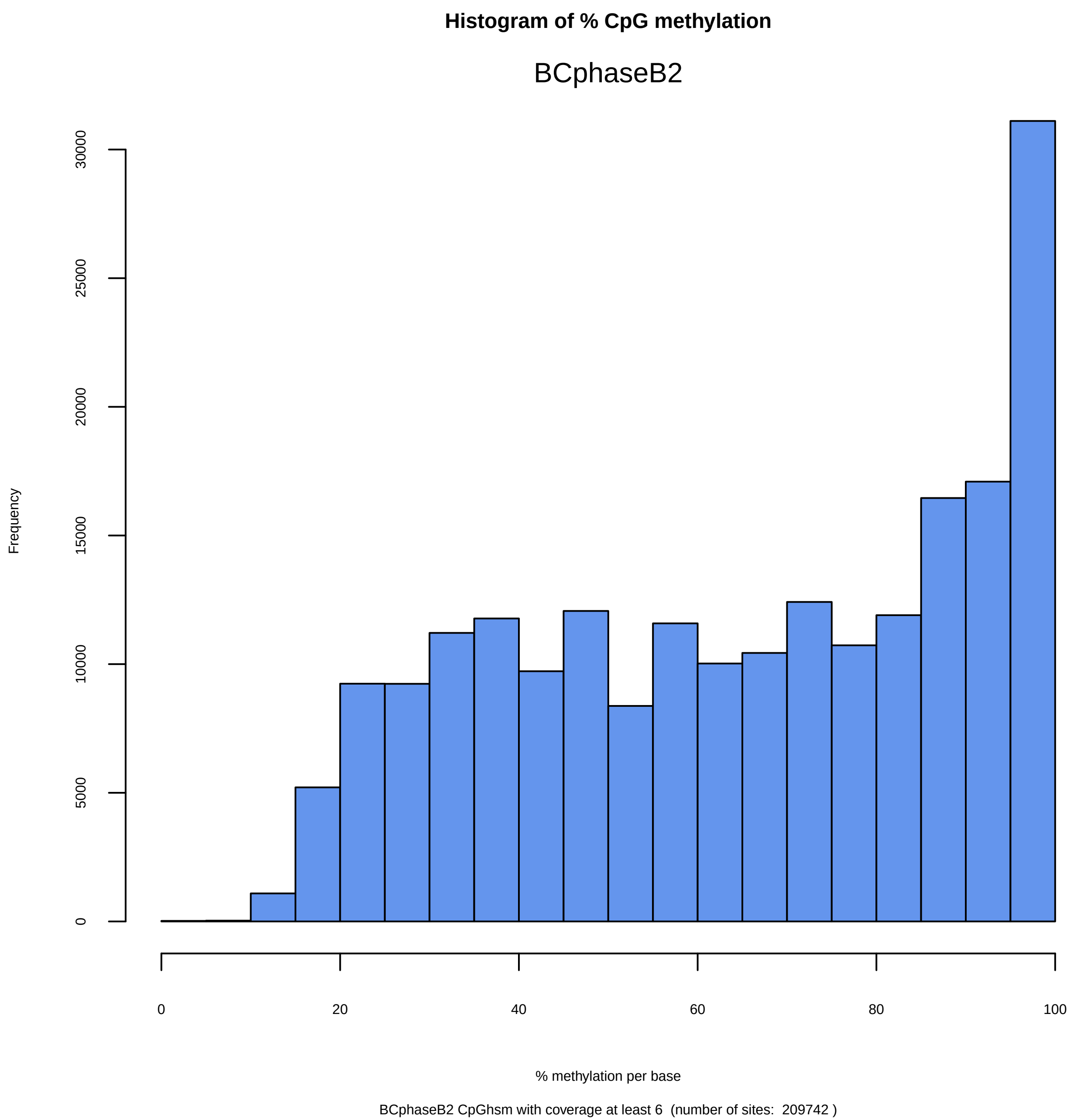

Histogram of % CpG methylation
BCphaseB2
Frequency
0
5000
10000
15000
20000
25000
30000
0
20
40
60
80
100
% methylation per base
BCphaseB2 CpGhsm with coverage at least 6  (number of sites:  209742 )

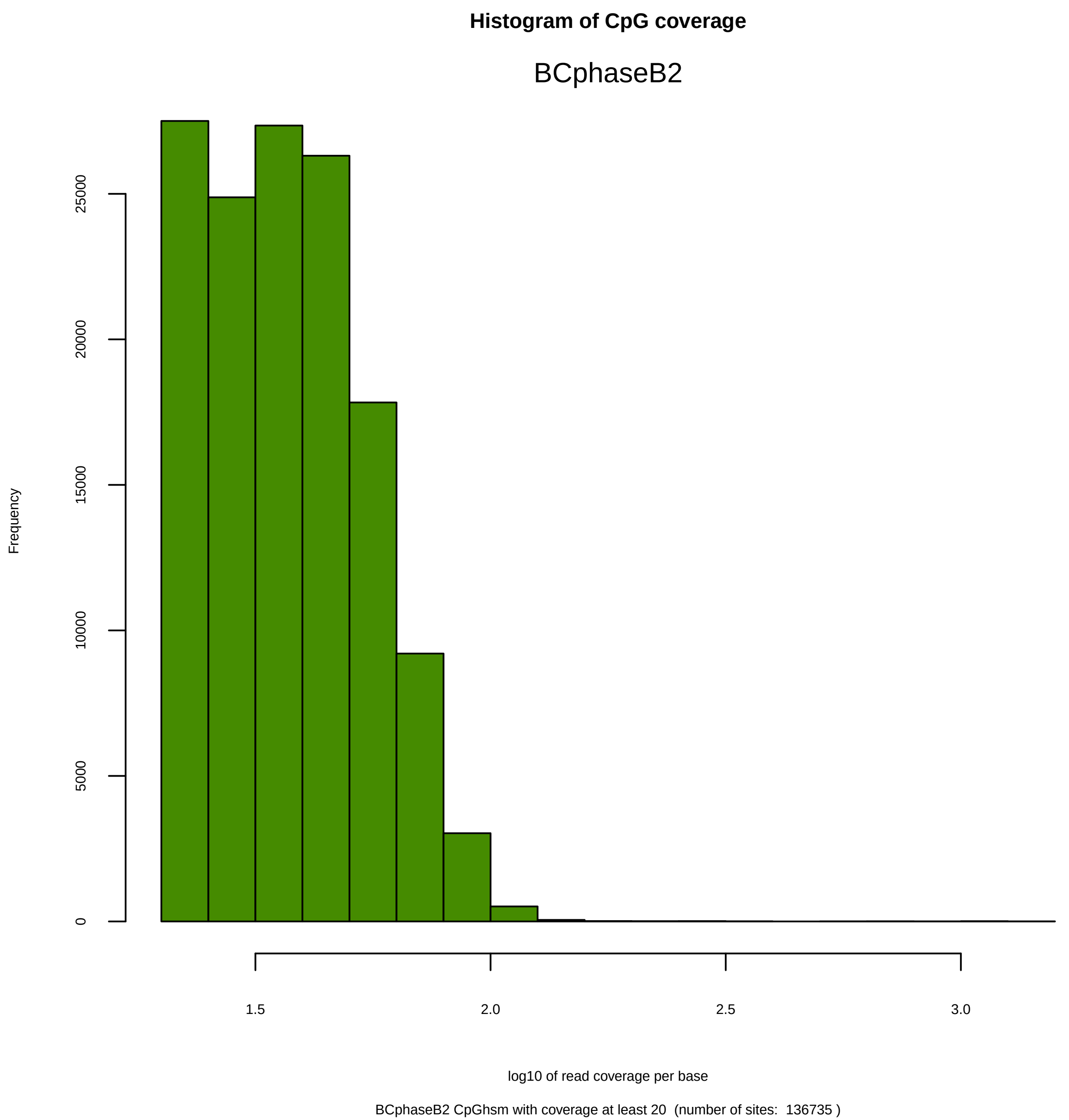

Histogram of CpG coverage
BCphaseB2
Frequency
0
5000
10000
15000
20000
25000
1.5
2.0
2.5
3.0
log10 of read coverage per base
BCphaseB2 CpGhsm with coverage at least 20  (number of sites:  136735 )

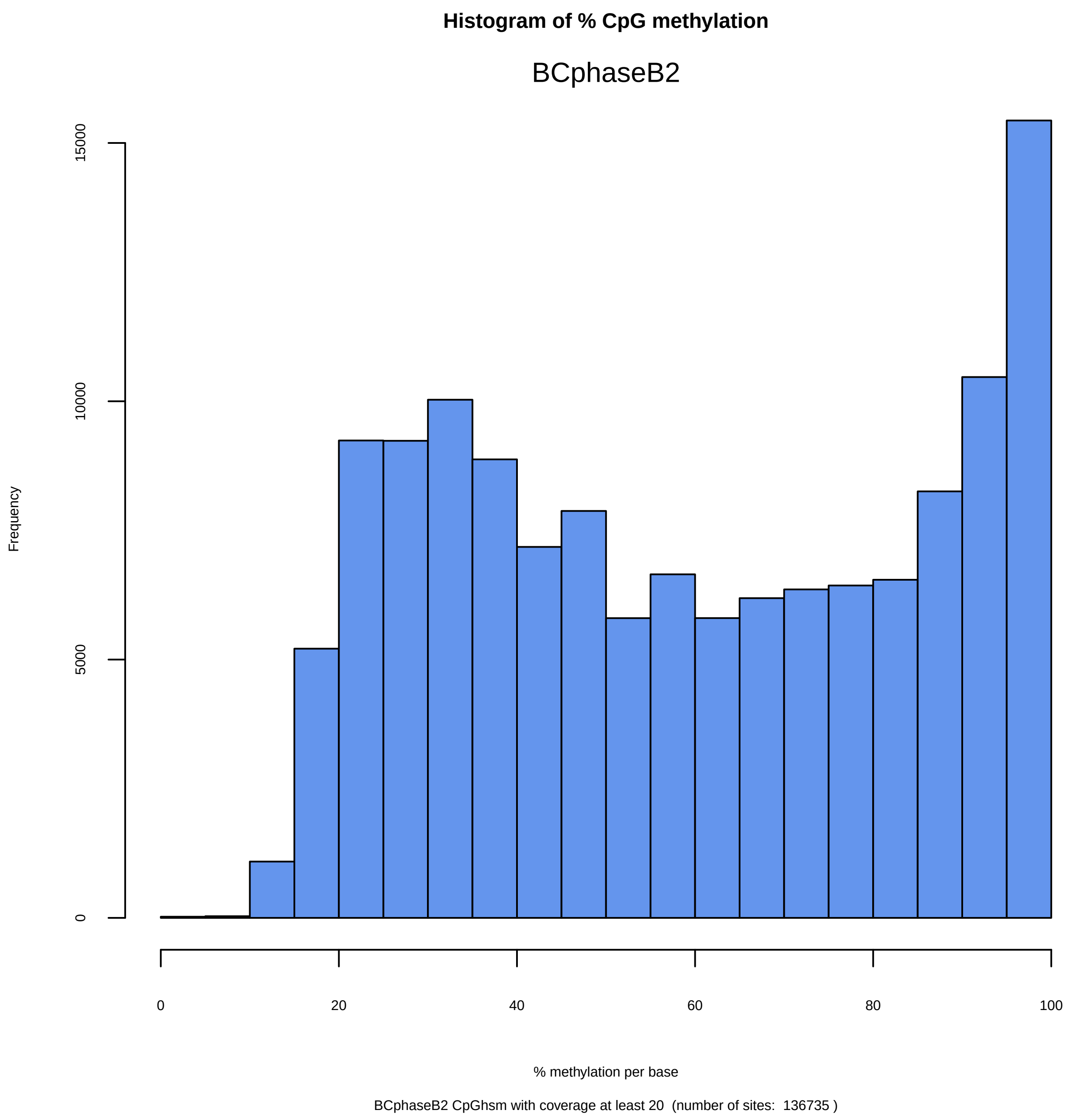
Histogram of % CpG methylation
BCphaseB2
Frequency
0
5000
10000
15000
0
20
40
60
80
100
% methylation per base
BCphaseB2 CpGhsm with coverage at least 20 (number of sites: 136735 )

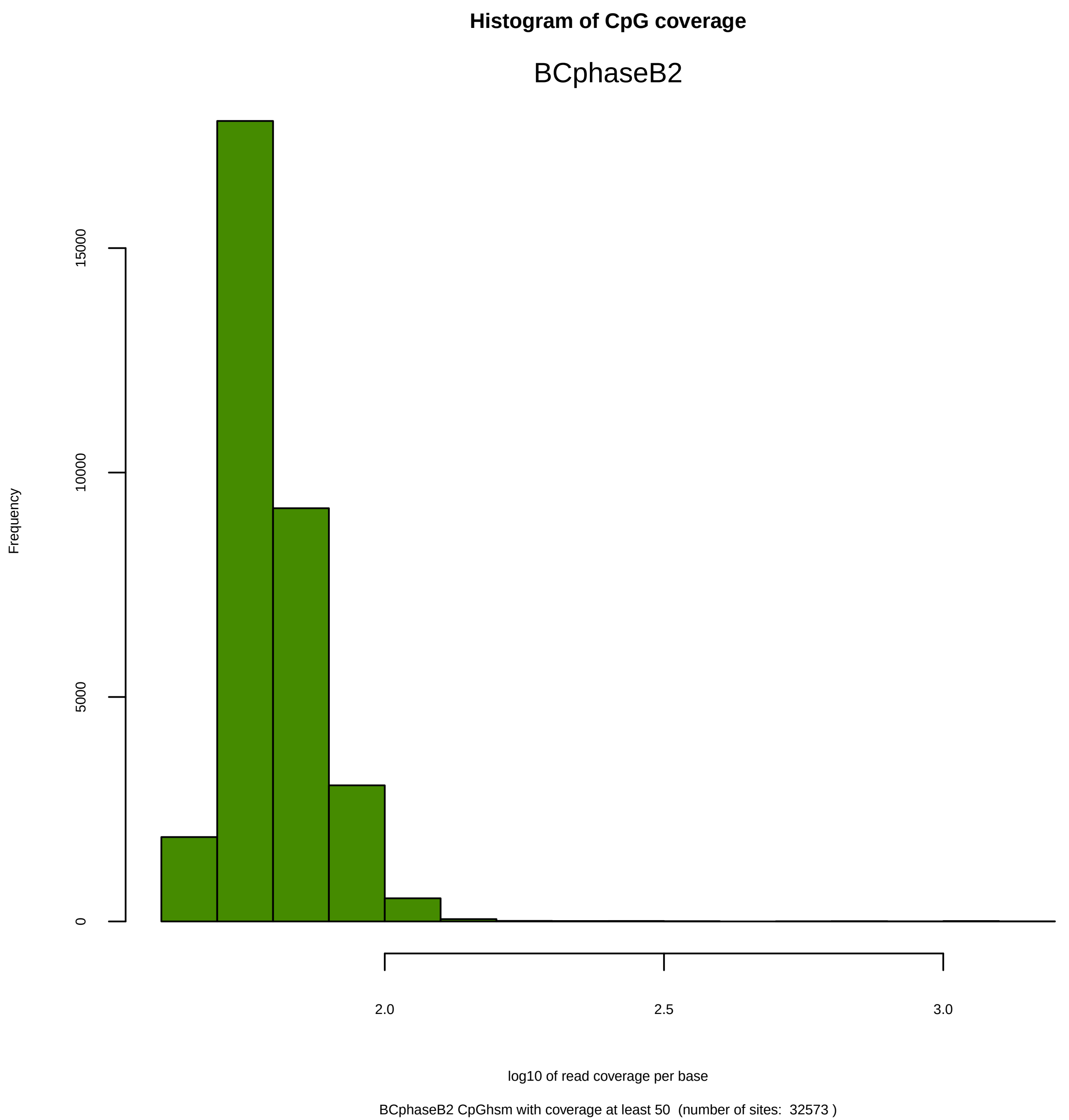
Histogram of CpG coverage
BCphaseB2
Frequency
0
5000
10000
15000
2.0
2.5
3.0
log10 of read coverage per base
BCphaseB2 CpGhsm with coverage at least 50  (number of sites:  32573 )

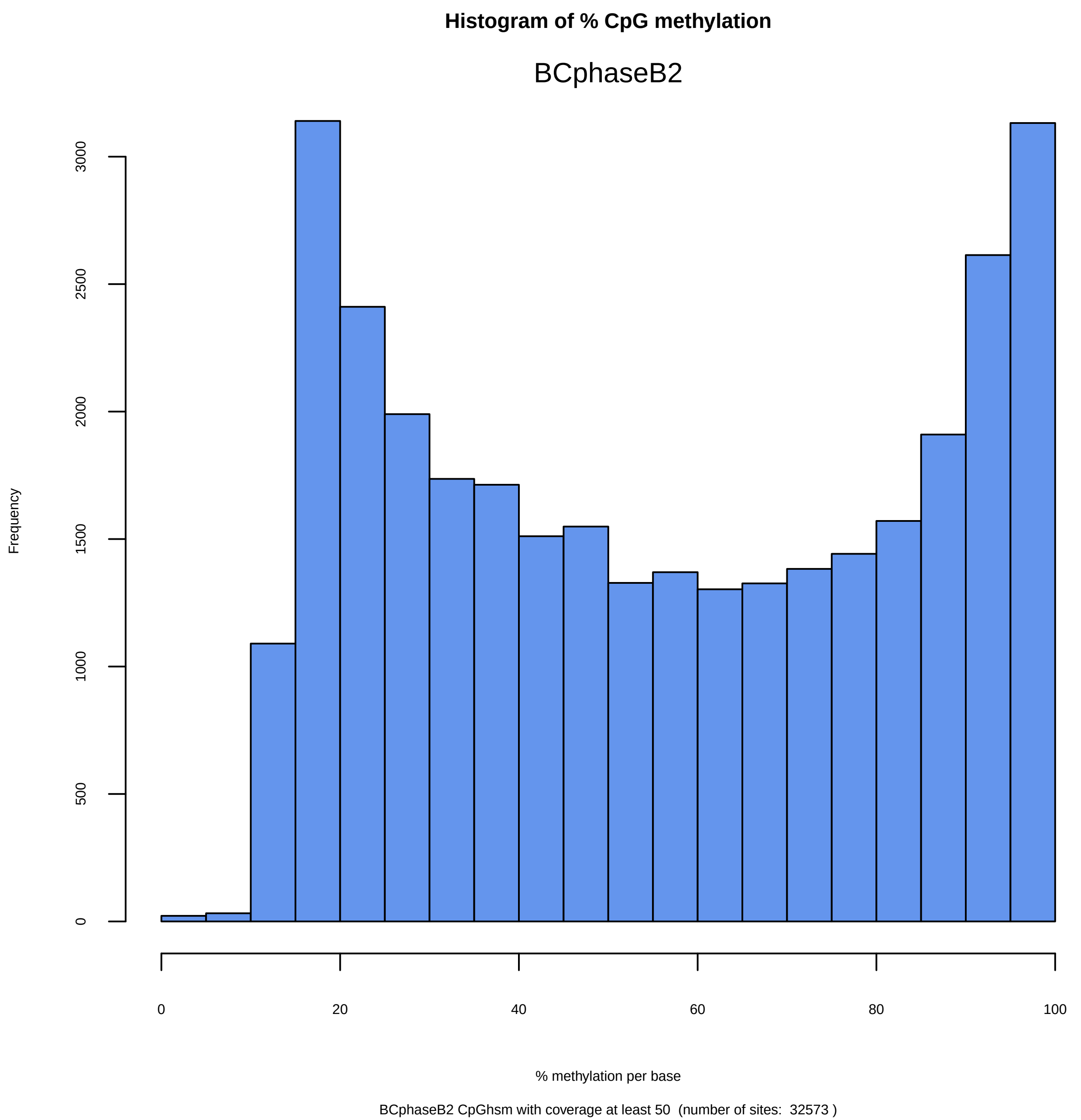
Histogram of % CpG methylation
BCphaseB2
Frequency
0
500
1000
1500
2000
2500
3000
0
20
40
60
80
100
% methylation per base
BCphaseB2 CpGhsm with coverage at least 50  (number of sites:  32573 )

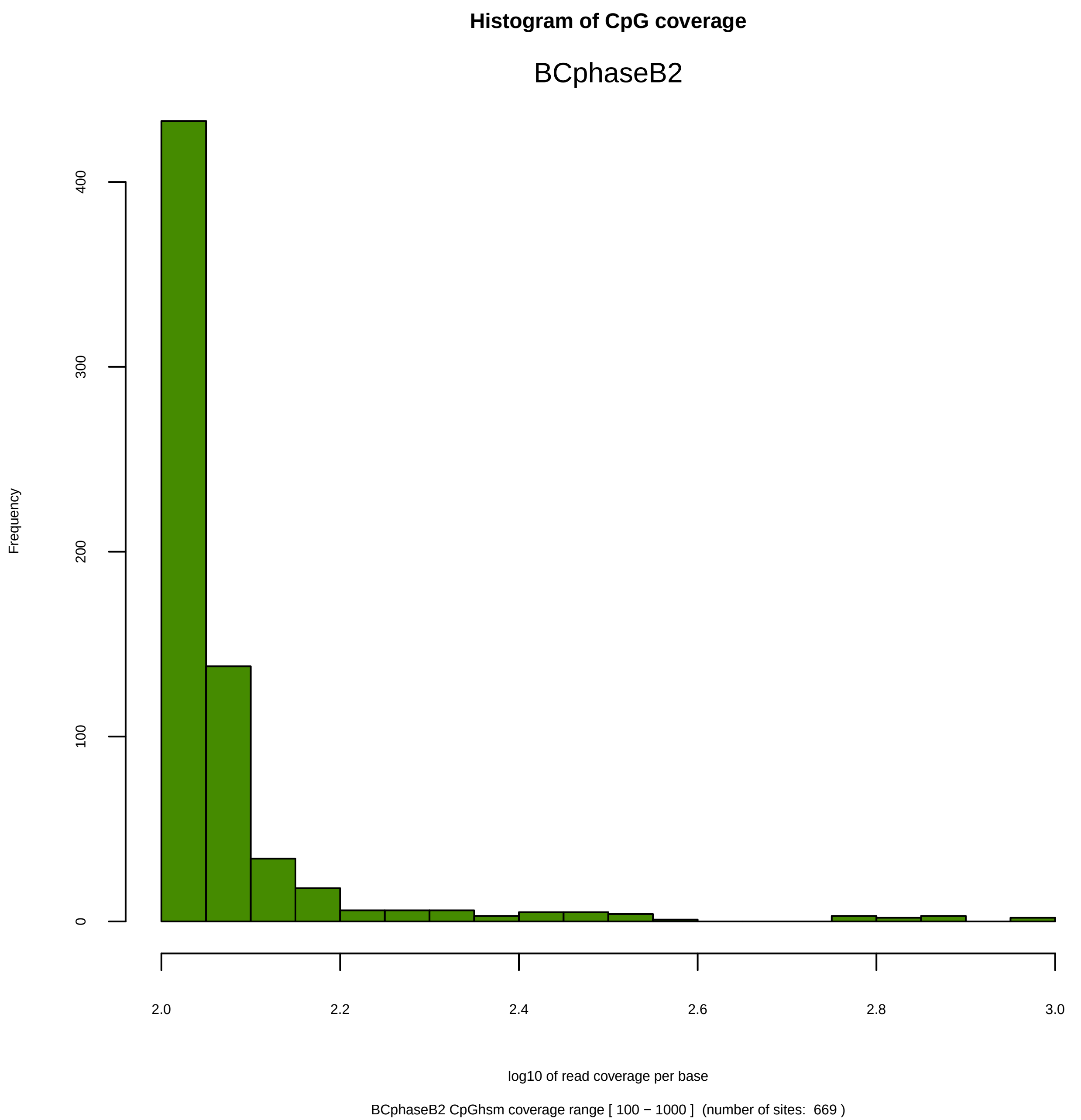

Histogram of CpG coverage
BCphaseB2
Frequency
0
100
200
300
400
2.0
2.2
2.4
2.6
2.8
3.0
log10 of read coverage per base
BCphaseB2 CpGhsm coverage range [ 100 − 1000 ] (number of sites: 669 )

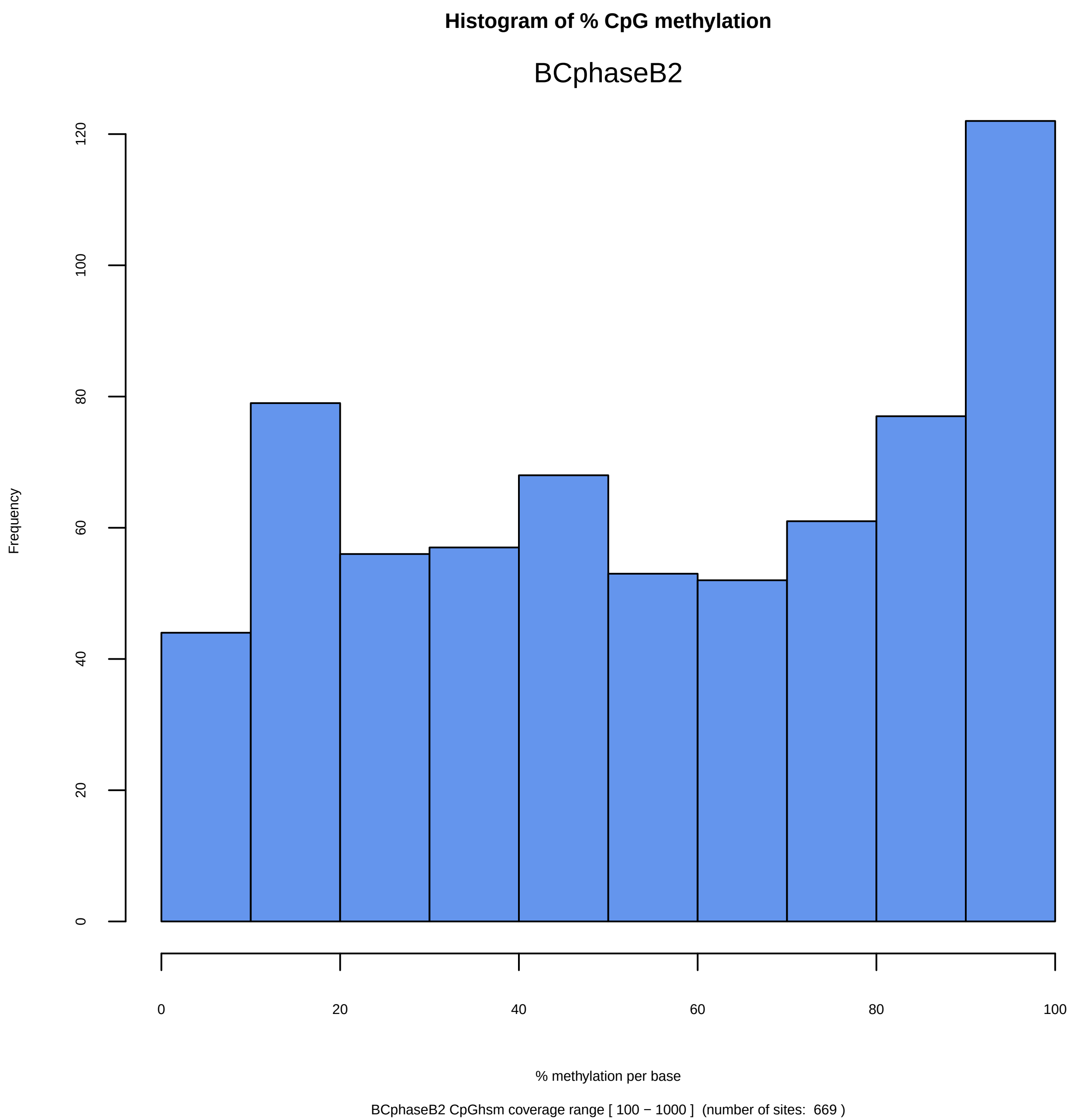
Histogram of % CpG methylation
BCphaseB2
Frequency
0
20
40
60
80
100
120
0
20
40
60
80
100
% methylation per base
BCphaseB2 CpGhsm coverage range [ 100 − 1000 ]  (number of sites:  669 )

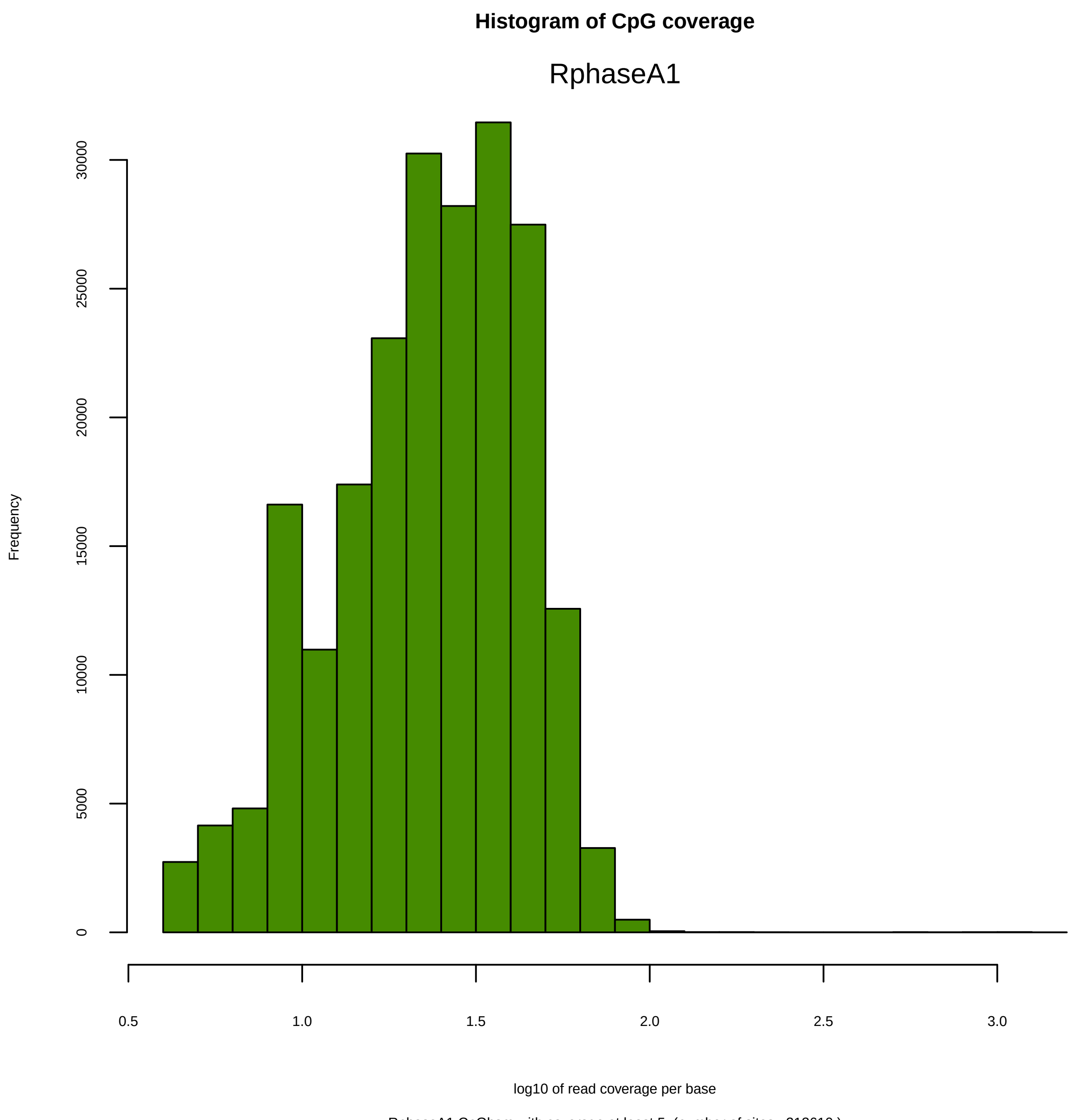
Histogram of CpG coverage
RphaseA1
Frequency
0
5000
10000
15000
20000
25000
30000
0.5
1.0
1.5
2.0
2.5
3.0
log10 of read coverage per base
RphaseA1 CpGhsm with coverage at least 5  (number of sites:  213610 )

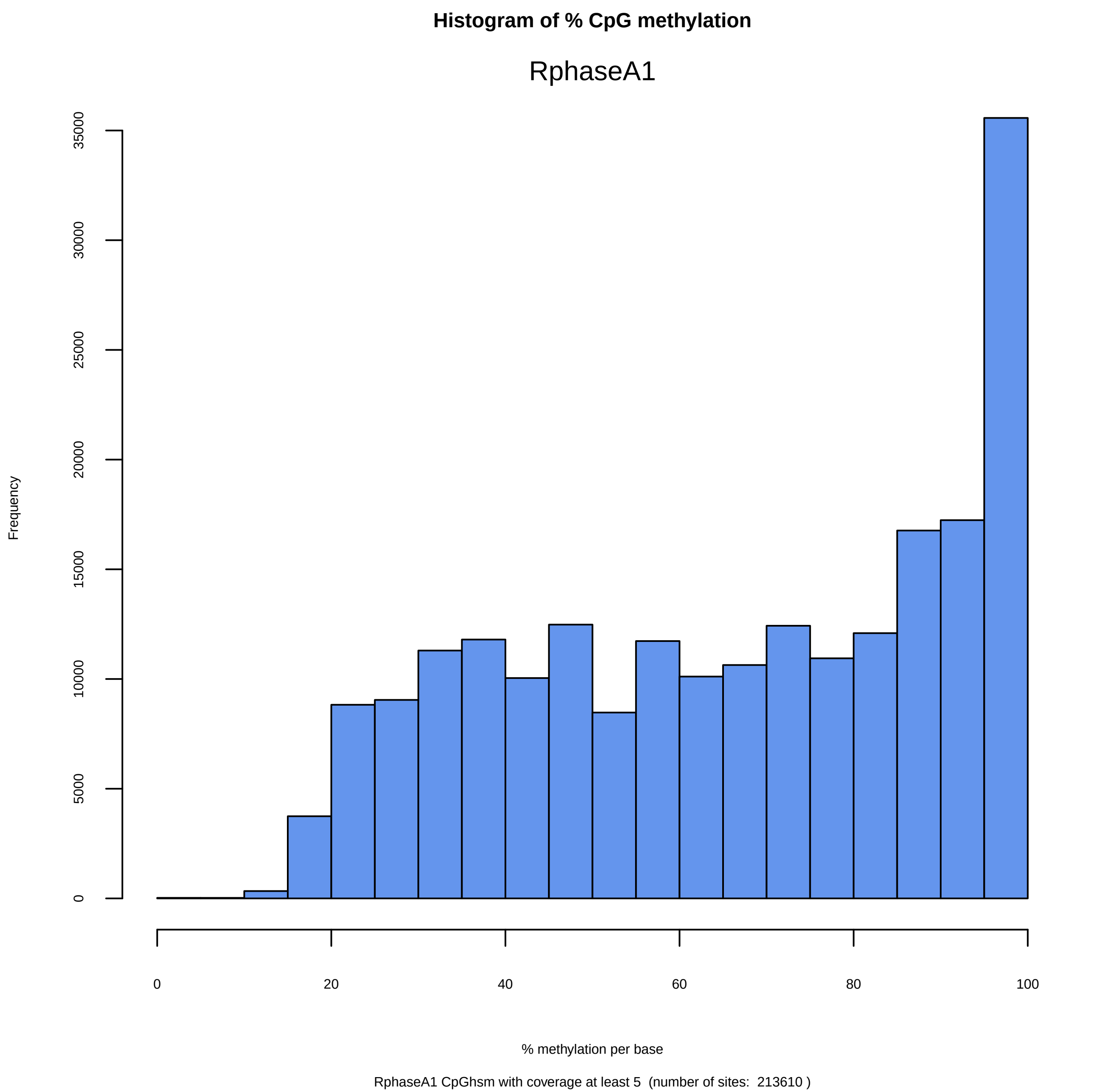
Histogram of % CpG methylation
RphaseA1
Frequency
0
5000
10000
15000
20000
25000
30000
35000
0
20
40
60
80
100
% methylation per base
RphaseA1 CpGhsm with coverage at least 5  (number of sites:  213610 )

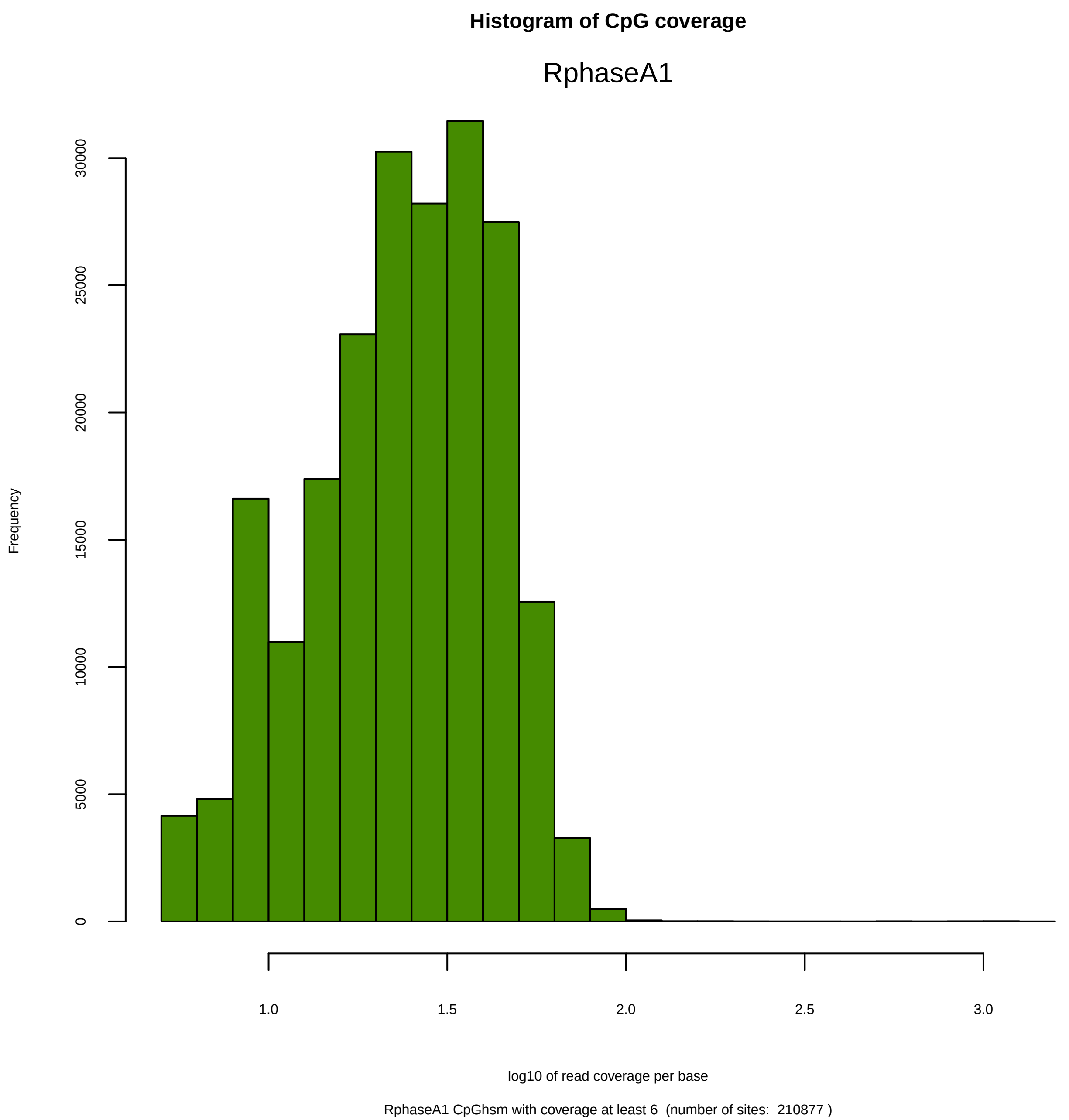
Histogram of CpG coverage
RphaseA1
Frequency
0
5000
10000
15000
20000
25000
30000
1.0
1.5
2.0
2.5
3.0
log10 of read coverage per base
RphaseA1 CpGhsm with coverage at least 6 (number of sites: 210877 )

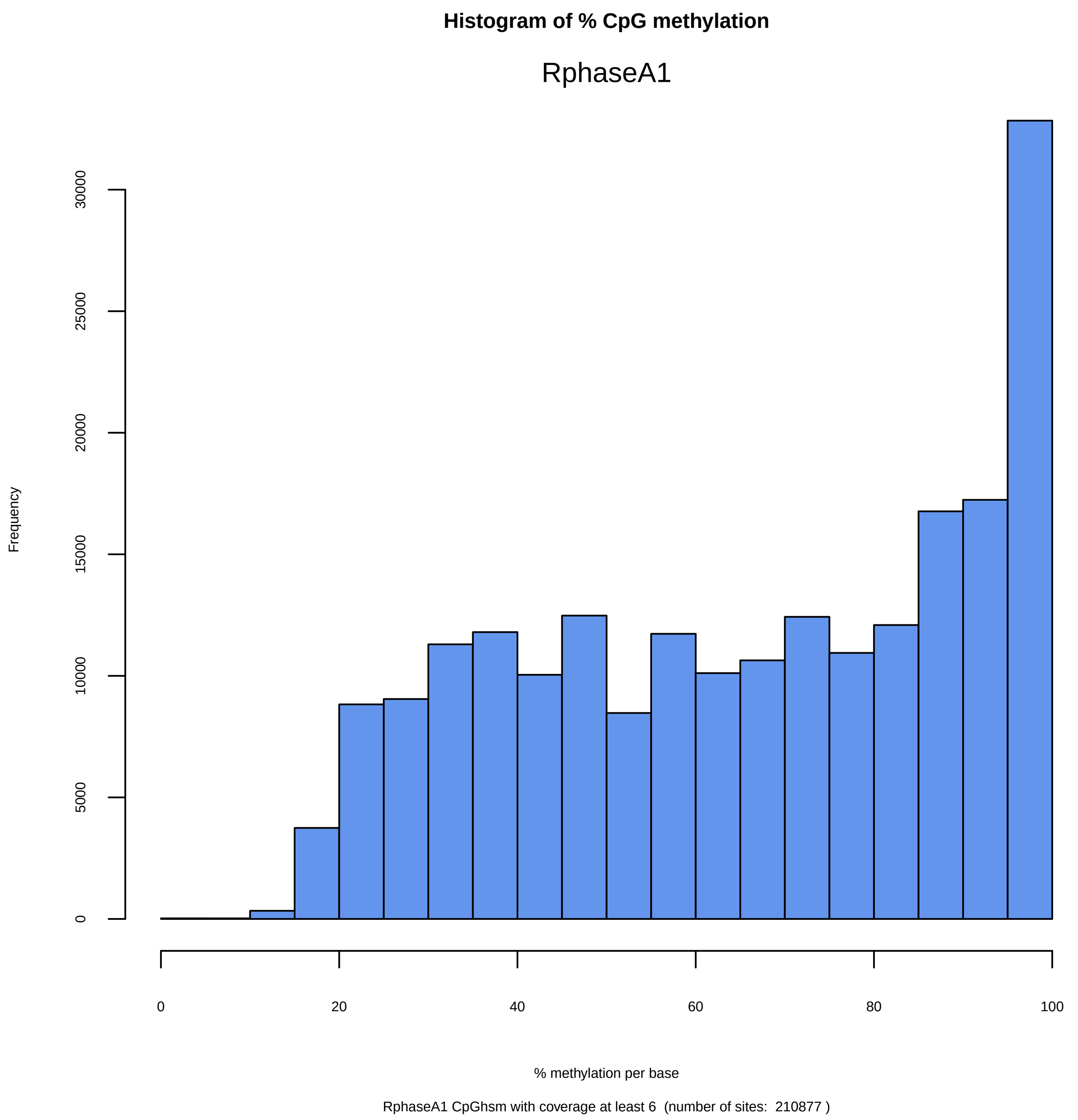

Histogram of % CpG methylation
RphaseA1
Frequency
0
5000
10000
15000
20000
25000
30000
0
20
40
60
80
100
% methylation per base
RphaseA1 CpGhsm with coverage at least 6 (number of sites: 210877 )

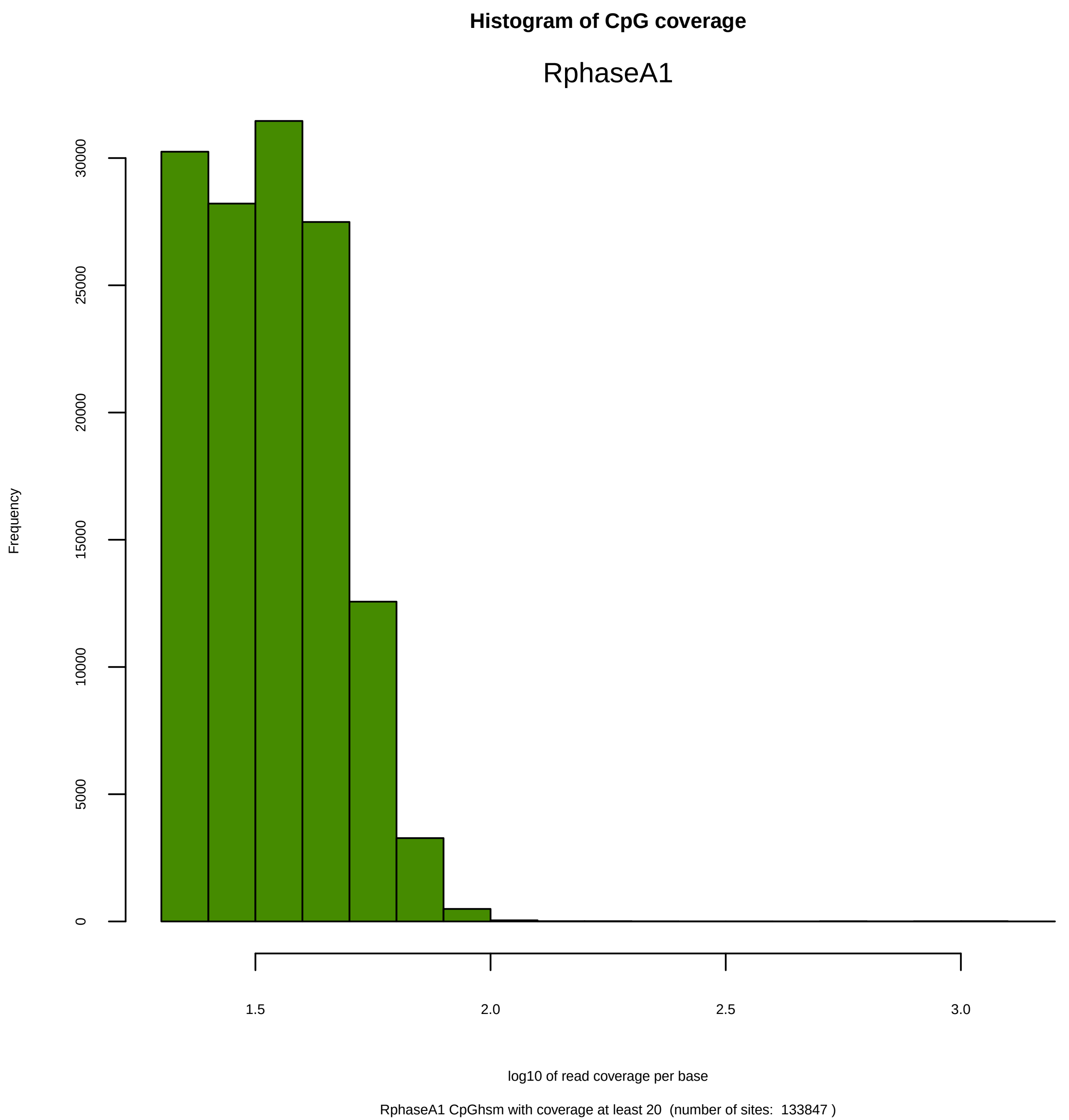
Histogram of CpG coverage
RphaseA1
Frequency
0
5000
10000
15000
20000
25000
30000
1.5
2.0
2.5
3.0
log10 of read coverage per base
RphaseA1 CpGhsm with coverage at least 20 (number of sites: 133847 )

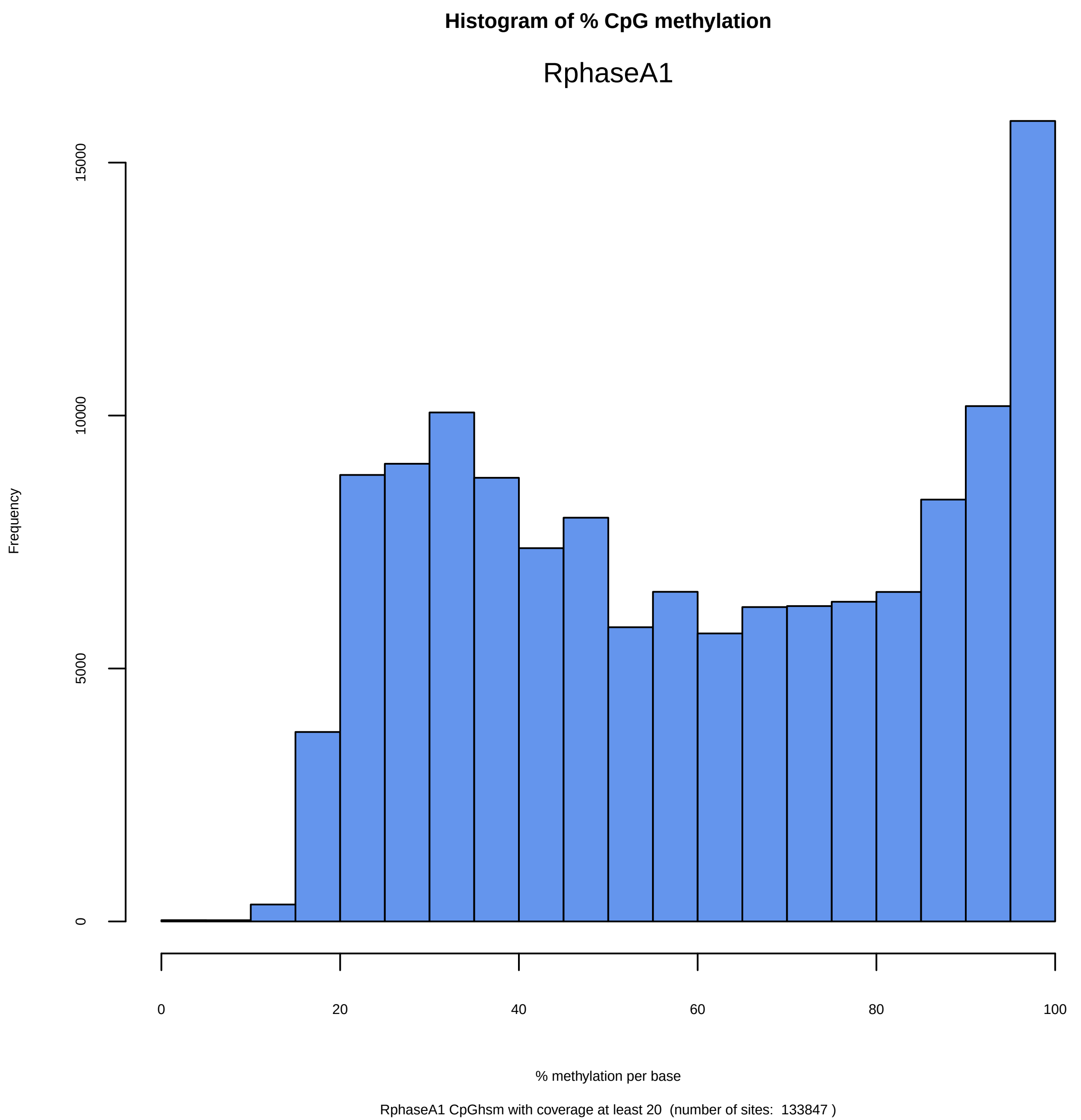

Histogram of % CpG methylation
RphaseA1
Frequency
0
5000
10000
15000
0
20
40
60
80
100
% methylation per base
RphaseA1 CpGhsm with coverage at least 20 (number of sites: 133847 )

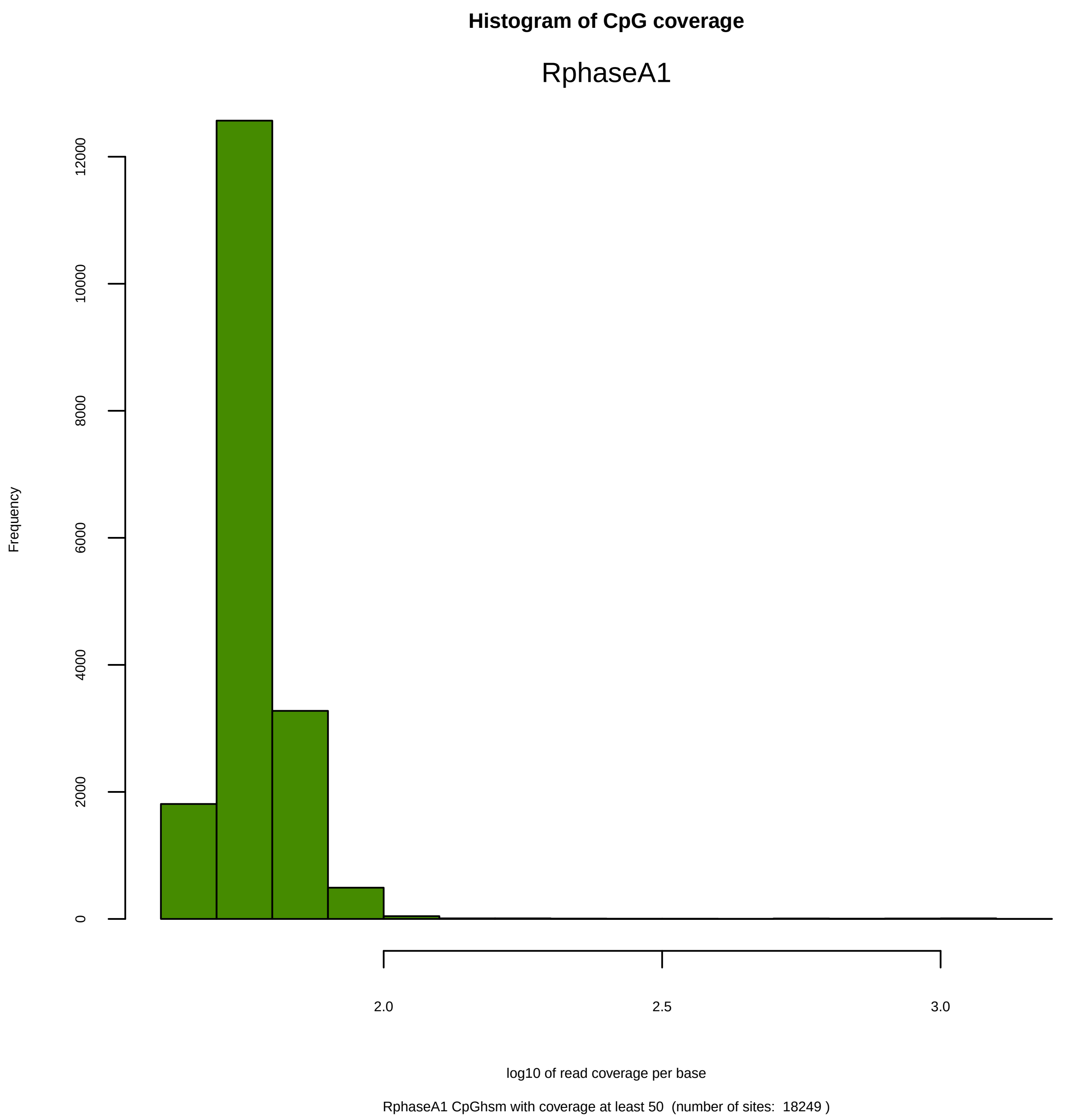

Histogram of CpG coverage
RphaseA1
Frequency
0
2000
4000
6000
8000
10000
12000
2.0
2.5
3.0
log10 of read coverage per base
RphaseA1 CpGhsm with coverage at least 50 (number of sites: 18249 )

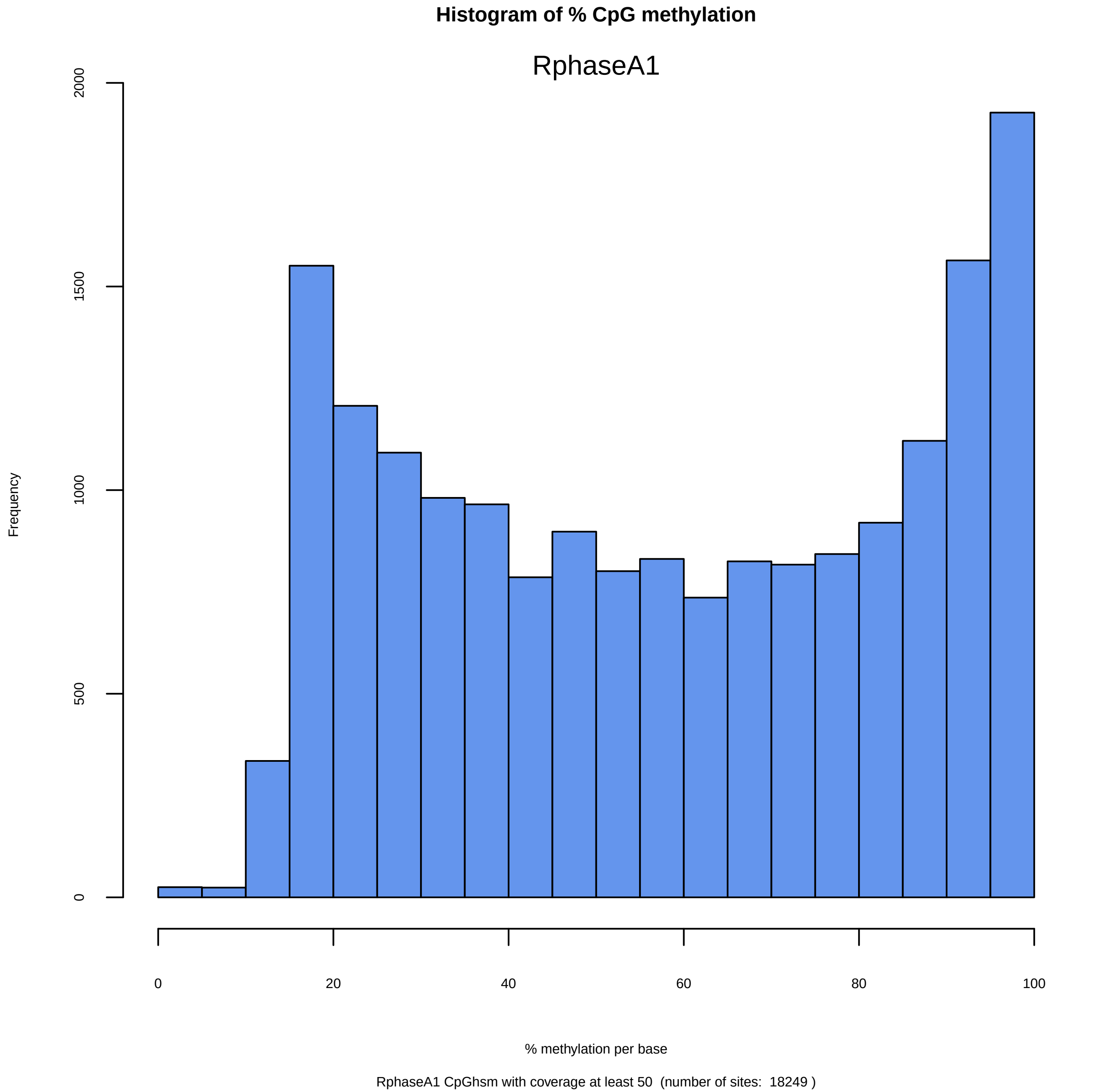

Histogram of % CpG methylation
RphaseA1
Frequency
0
500
1000
1500
2000
0
20
40
60
80
100
% methylation per base
RphaseA1 CpGhsm with coverage at least 50  (number of sites:  18249 )

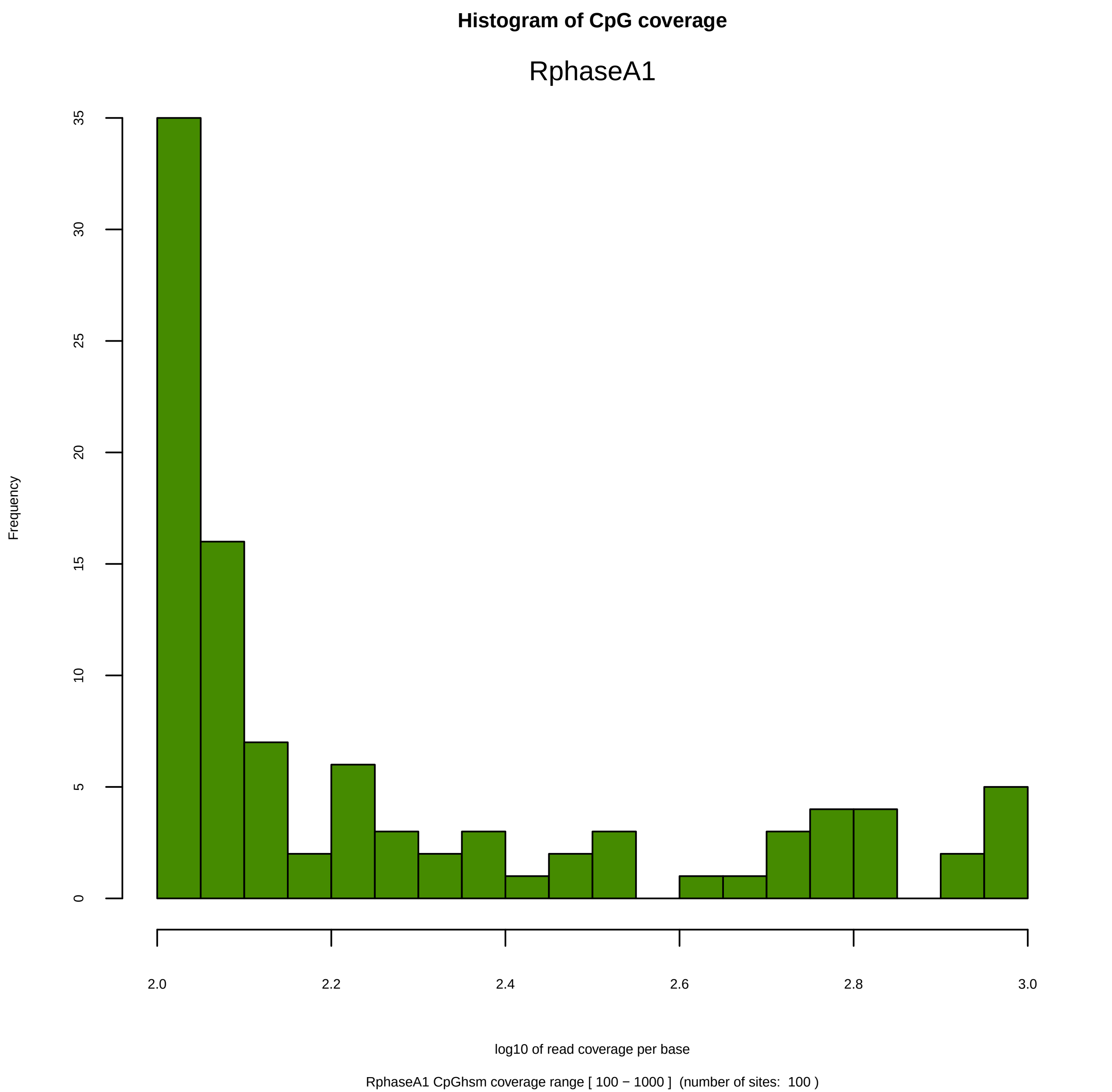

Histogram of CpG coverage
RphaseA1
Frequency
0
5
10
15
20
25
30
35
2.0
2.2
2.4
2.6
2.8
3.0
log10 of read coverage per base
RphaseA1 CpGhsm coverage range [ 100 − 1000 ]  (number of sites:  100 )

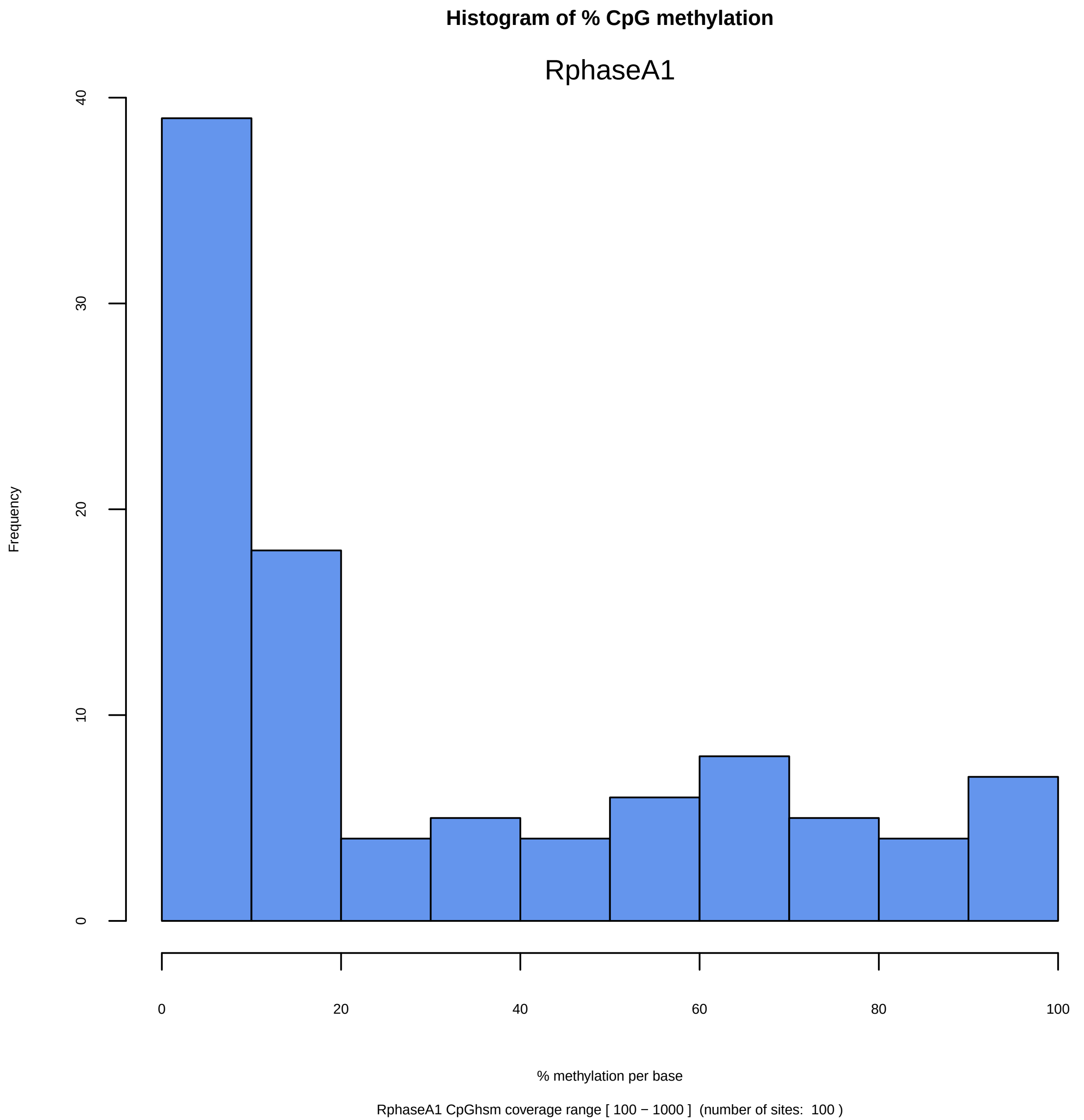

Histogram of % CpG methylation
RphaseA1
Frequency
0
10
20
30
40
0
20
40
60
80
100
% methylation per base
RphaseA1 CpGhsm coverage range [ 100 − 1000 ]  (number of sites:  100 )

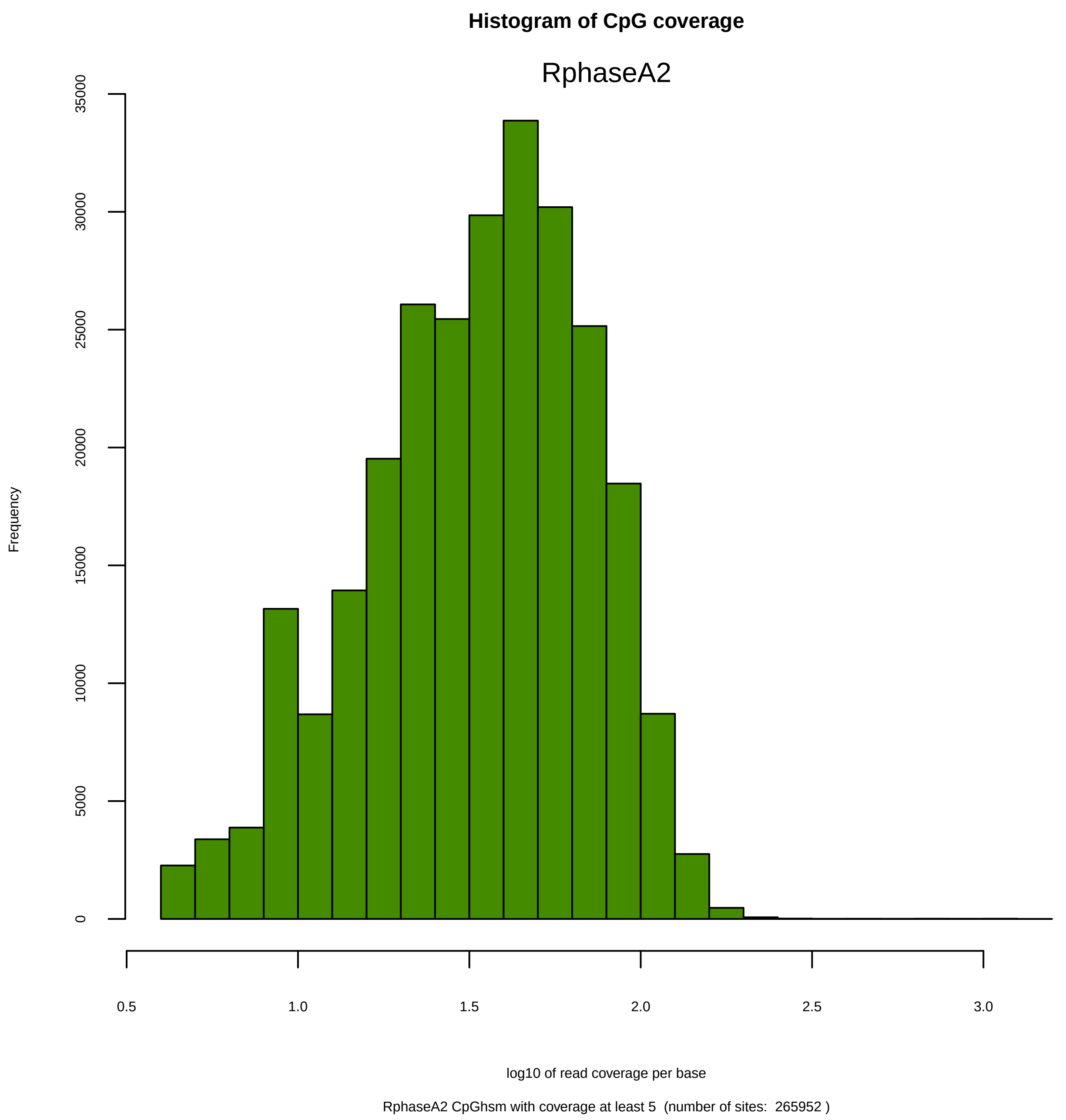

Histogram of CpG coverage
RphaseA2
Frequency
0
5000
10000
15000
20000
25000
30000
35000
0.5
1.0
1.5
2.0
2.5
3.0
log10 of read coverage per base
RphaseA2 CpGhsm with coverage at least 5 (number of sites: 265952 )

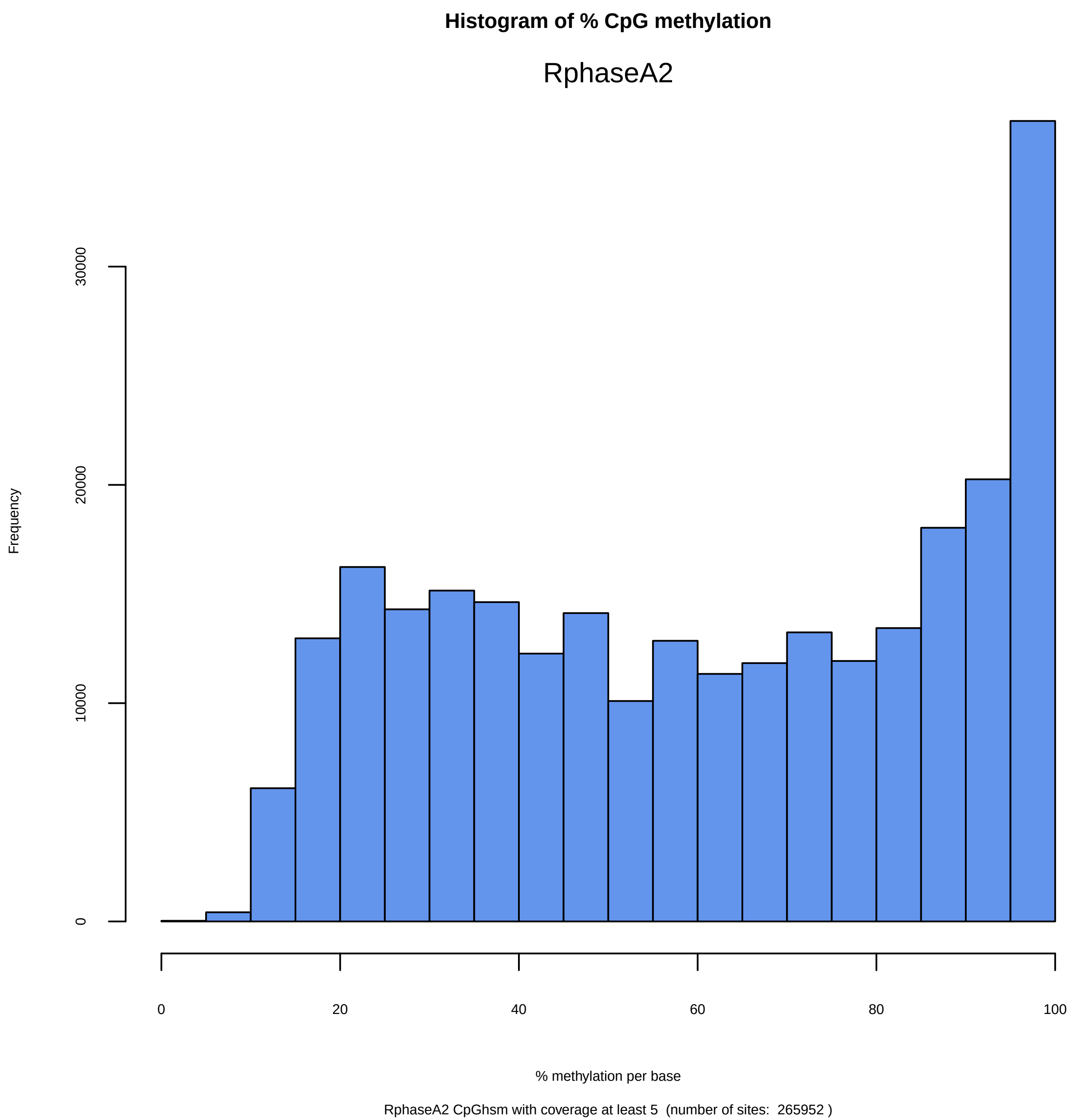

Histogram of % CpG methylation
RphaseA2
Frequency
0
10000
20000
30000
0
20
40
60
80
100
% methylation per base
RphaseA2 CpGhsm with coverage at least 5  (number of sites:  265952 )

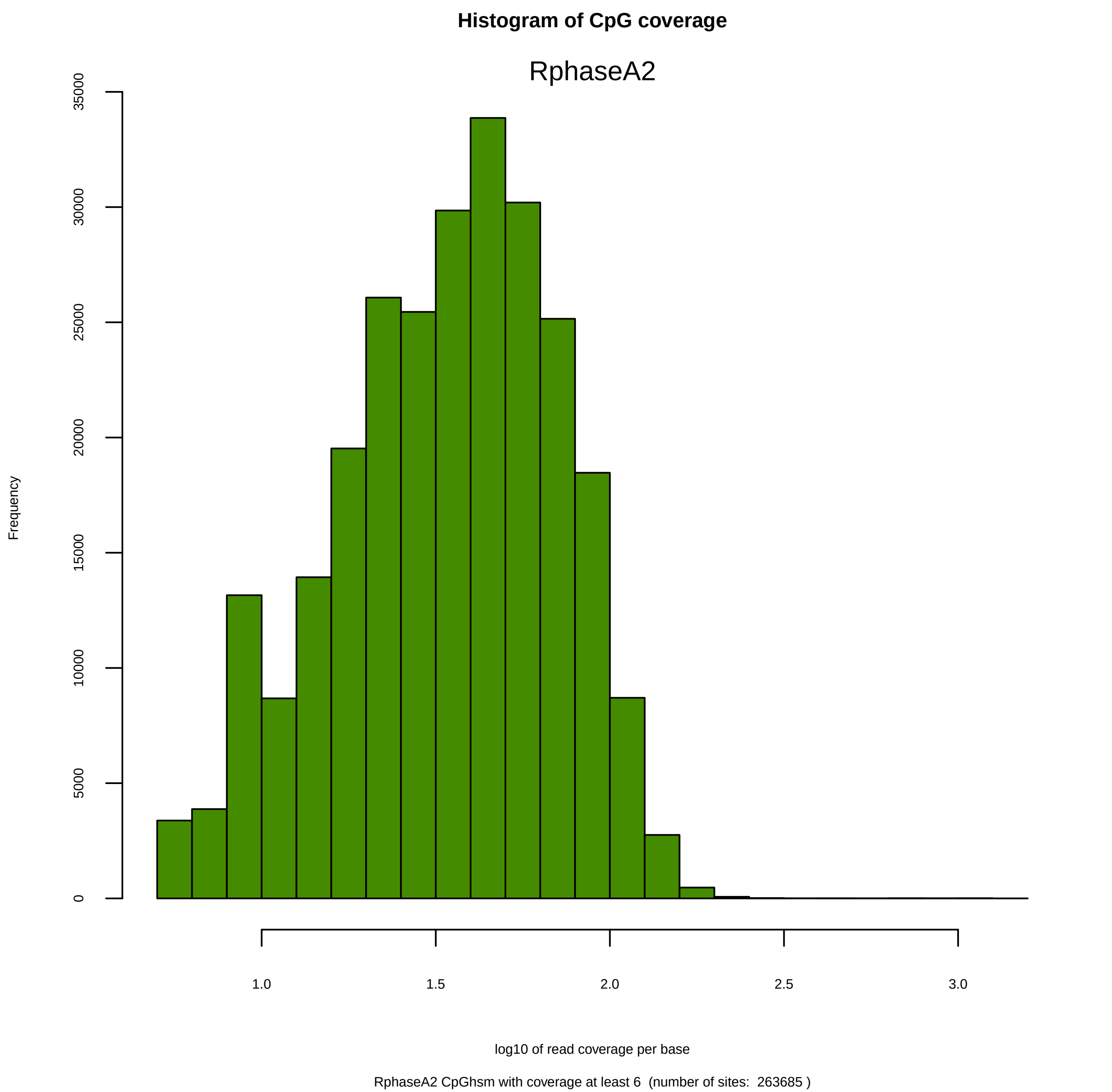
Histogram of CpG coverage
RphaseA2
Frequency
0
5000
10000
15000
20000
25000
30000
35000
1.0
1.5
2.0
2.5
3.0
log10 of read coverage per base
RphaseA2 CpGhsm with coverage at least 6 (number of sites: 263685 )

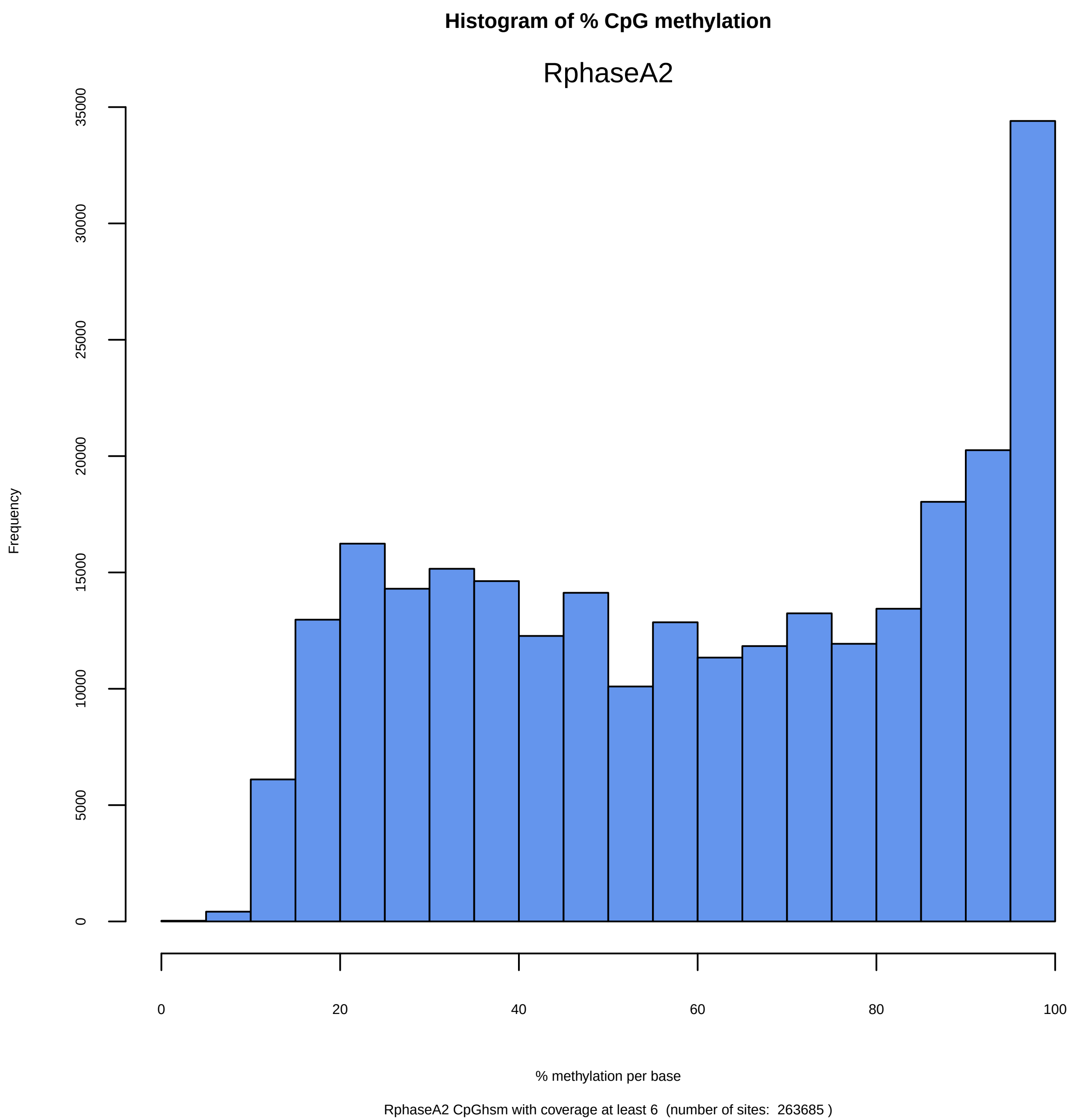
Histogram of % CpG methylation
RphaseA2
Frequency
0
5000
10000
15000
20000
25000
30000
35000
0
20
40
60
80
100
% methylation per base
RphaseA2 CpGhsm with coverage at least 6  (number of sites:  263685 )

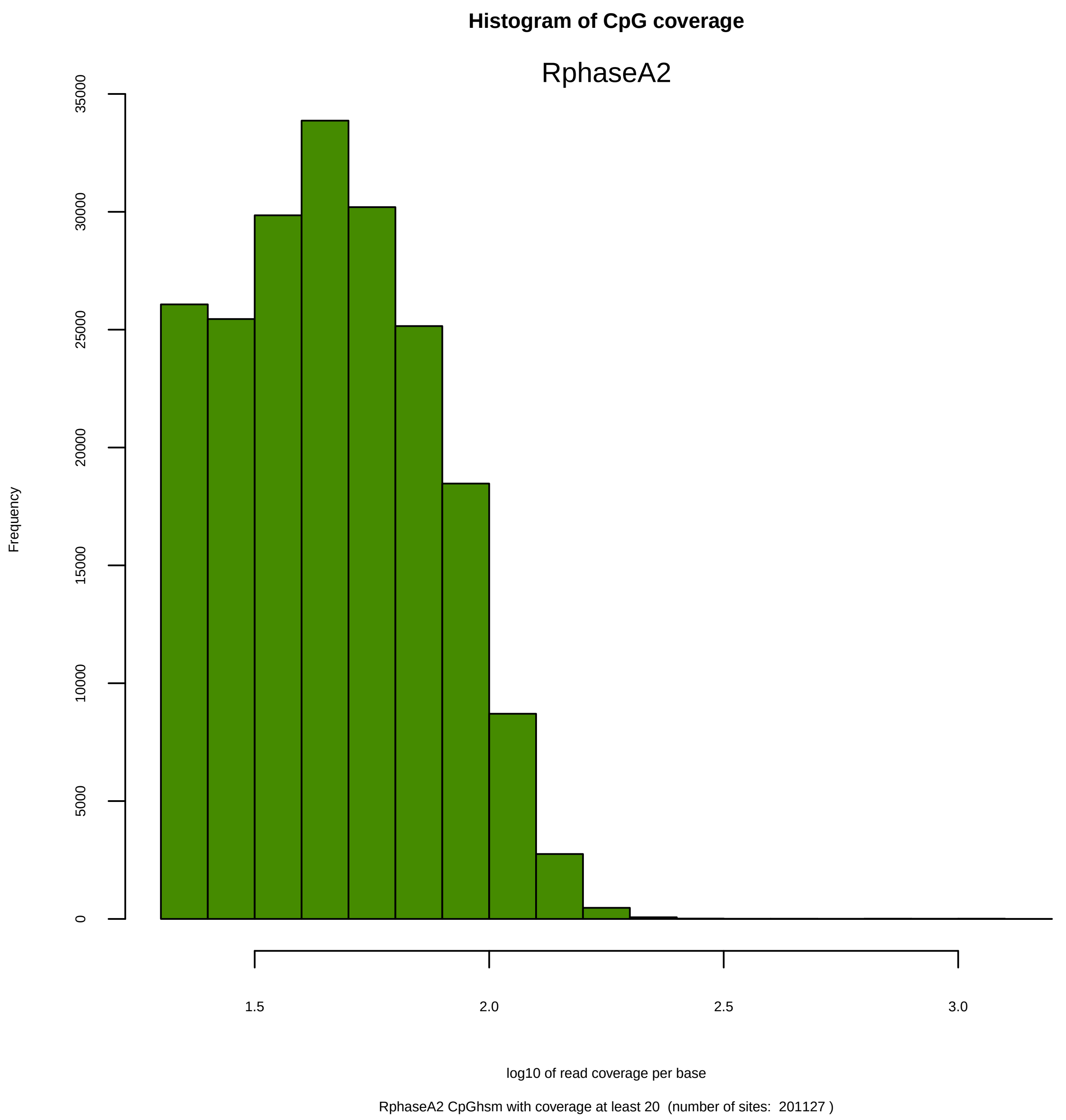
Histogram of CpG coverage
RphaseA2
Frequency
0
5000
10000
15000
20000
25000
30000
35000
1.5
2.0
2.5
3.0
log10 of read coverage per base
RphaseA2 CpGhsm with coverage at least 20 (number of sites: 201127 )

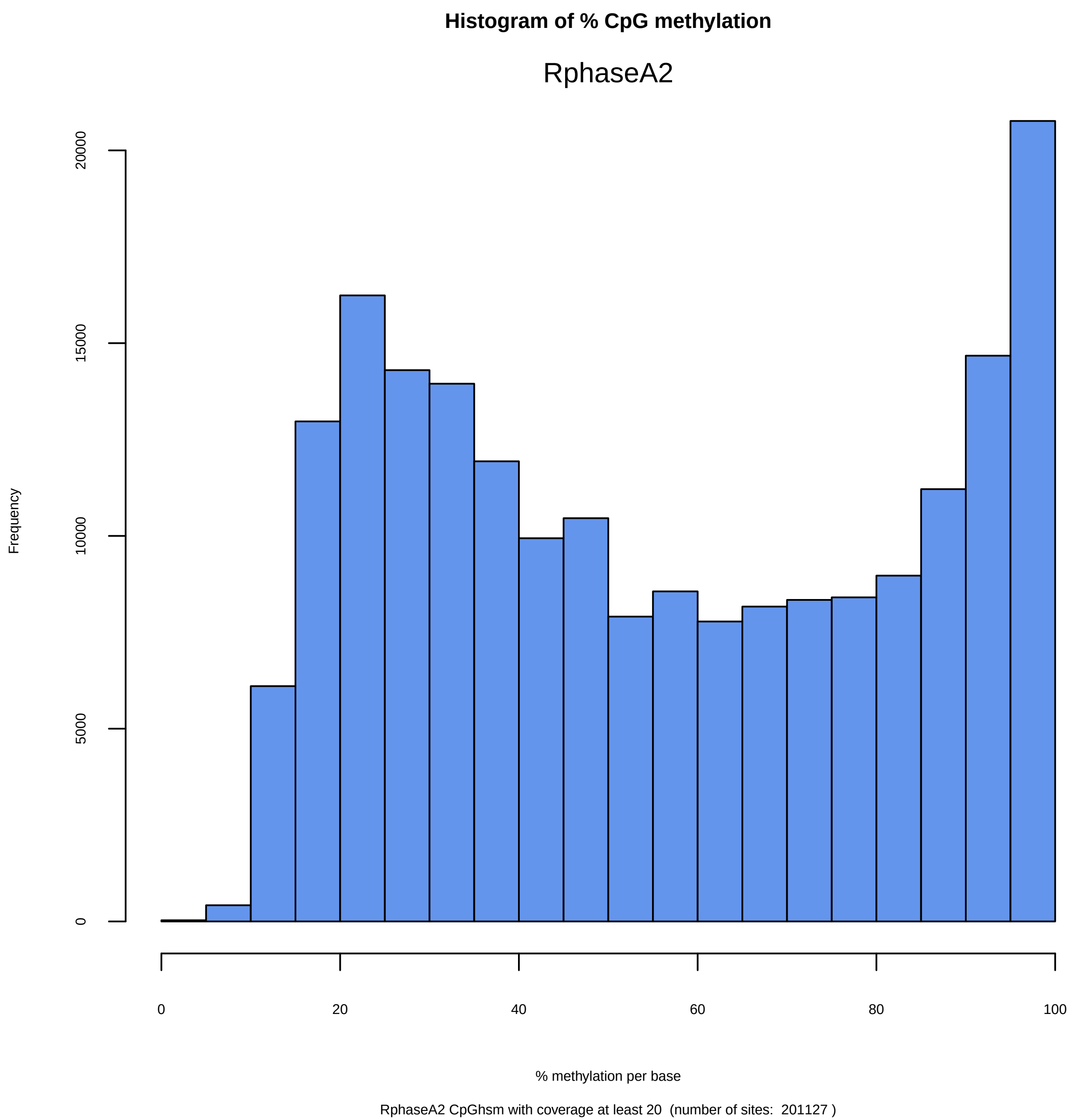
Histogram of % CpG methylation
RphaseA2
Frequency
0
5000
10000
15000
20000
0
20
40
60
80
100
% methylation per base
RphaseA2 CpGhsm with coverage at least 20 (number of sites: 201127 )

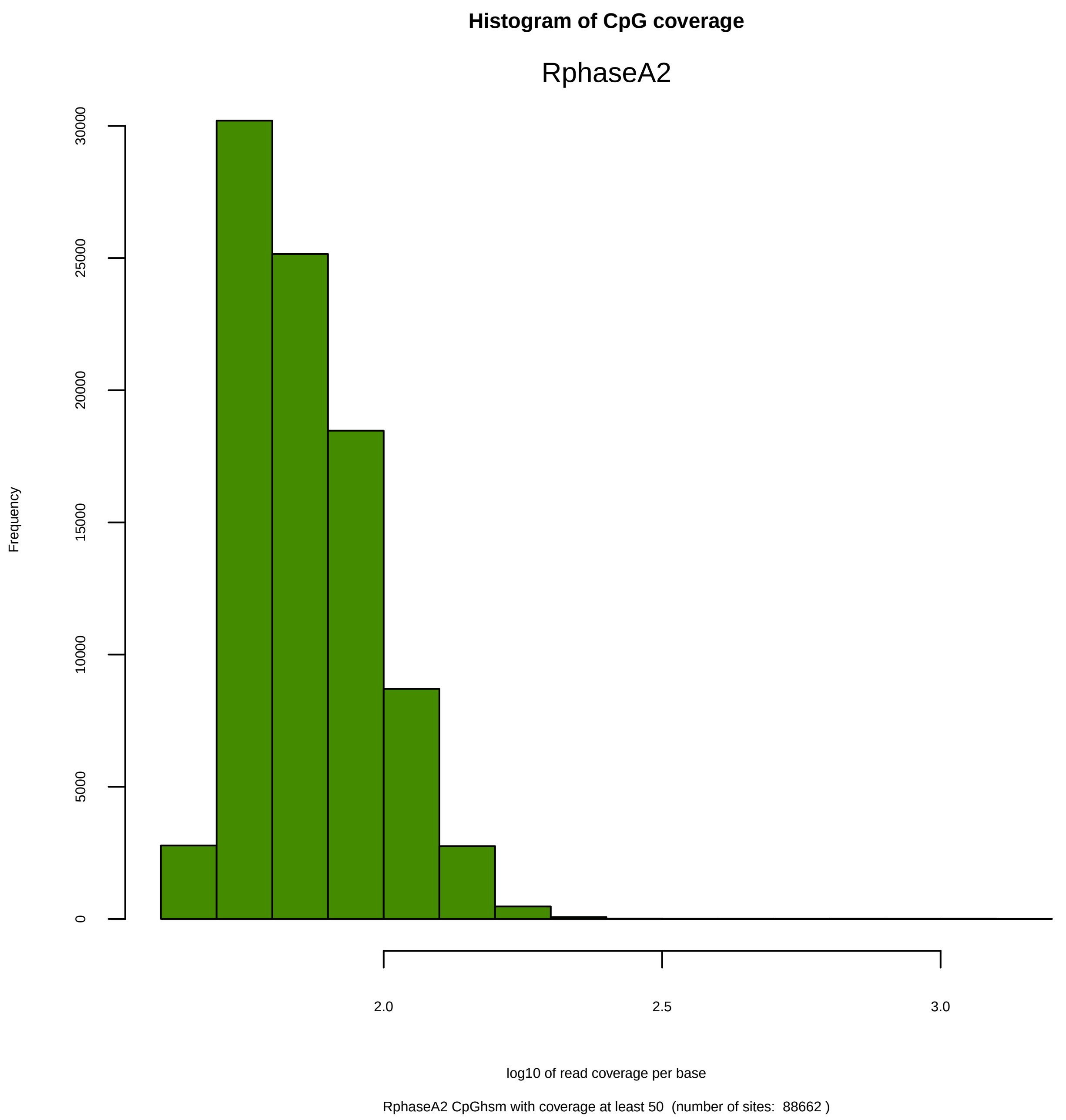
Histogram of CpG coverage
RphaseA2
Frequency
0
5000
10000
15000
20000
25000
30000
2.0
2.5
3.0
log10 of read coverage per base
RphaseA2 CpGhsm with coverage at least 50  (number of sites:  88662 )

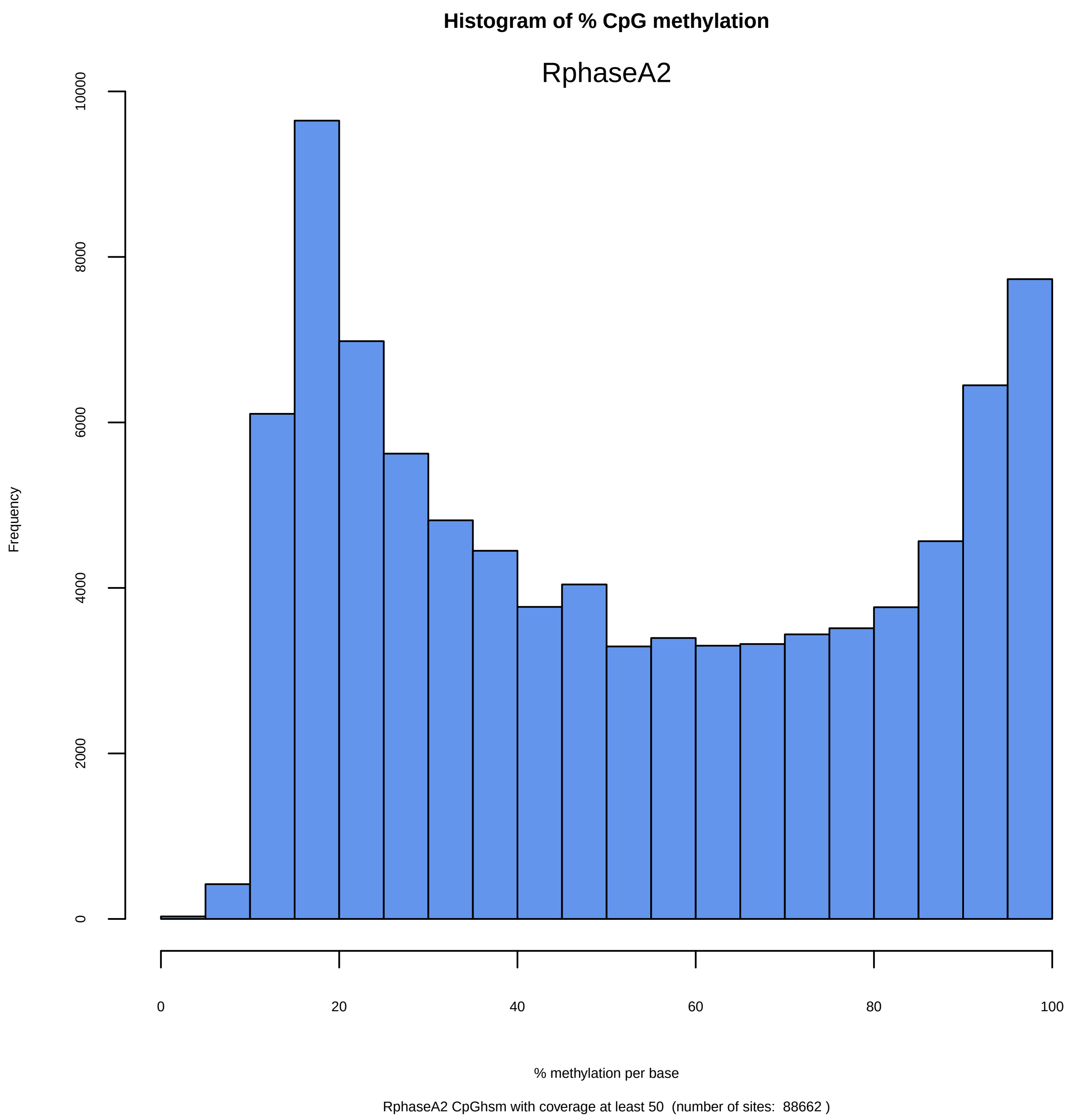
Histogram of % CpG methylation
RphaseA2
Frequency
0
2000
4000
6000
8000
10000
0
20
40
60
80
100
% methylation per base
RphaseA2 CpGhsm with coverage at least 50  (number of sites:  88662 )

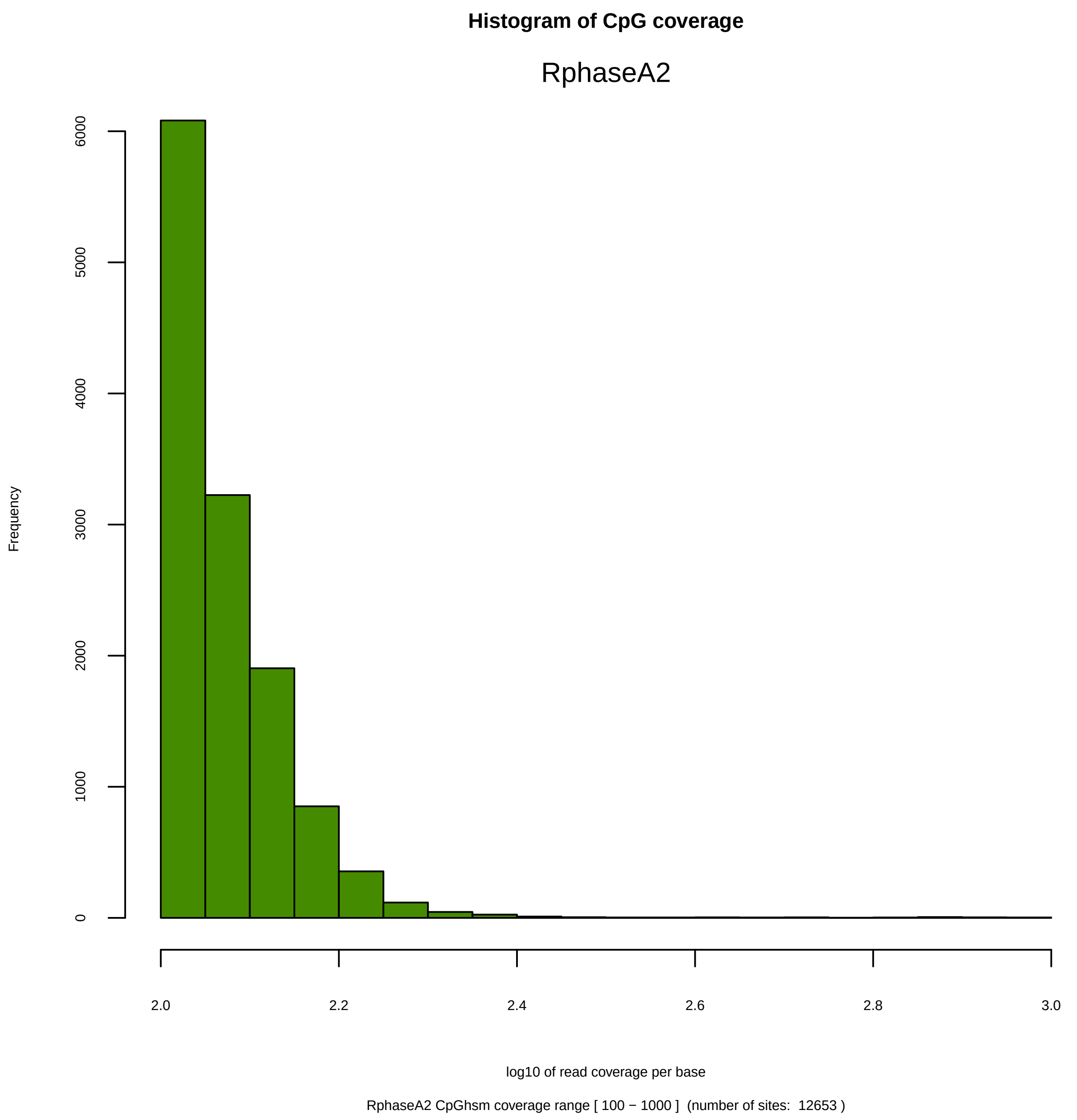

Histogram of CpG coverage
RphaseA2
Frequency
0
1000
2000
3000
4000
5000
6000
2.0
2.2
2.4
2.6
2.8
3.0
log10 of read coverage per base
RphaseA2 CpGhsm coverage range [ 100 − 1000 ] (number of sites: 12653 )

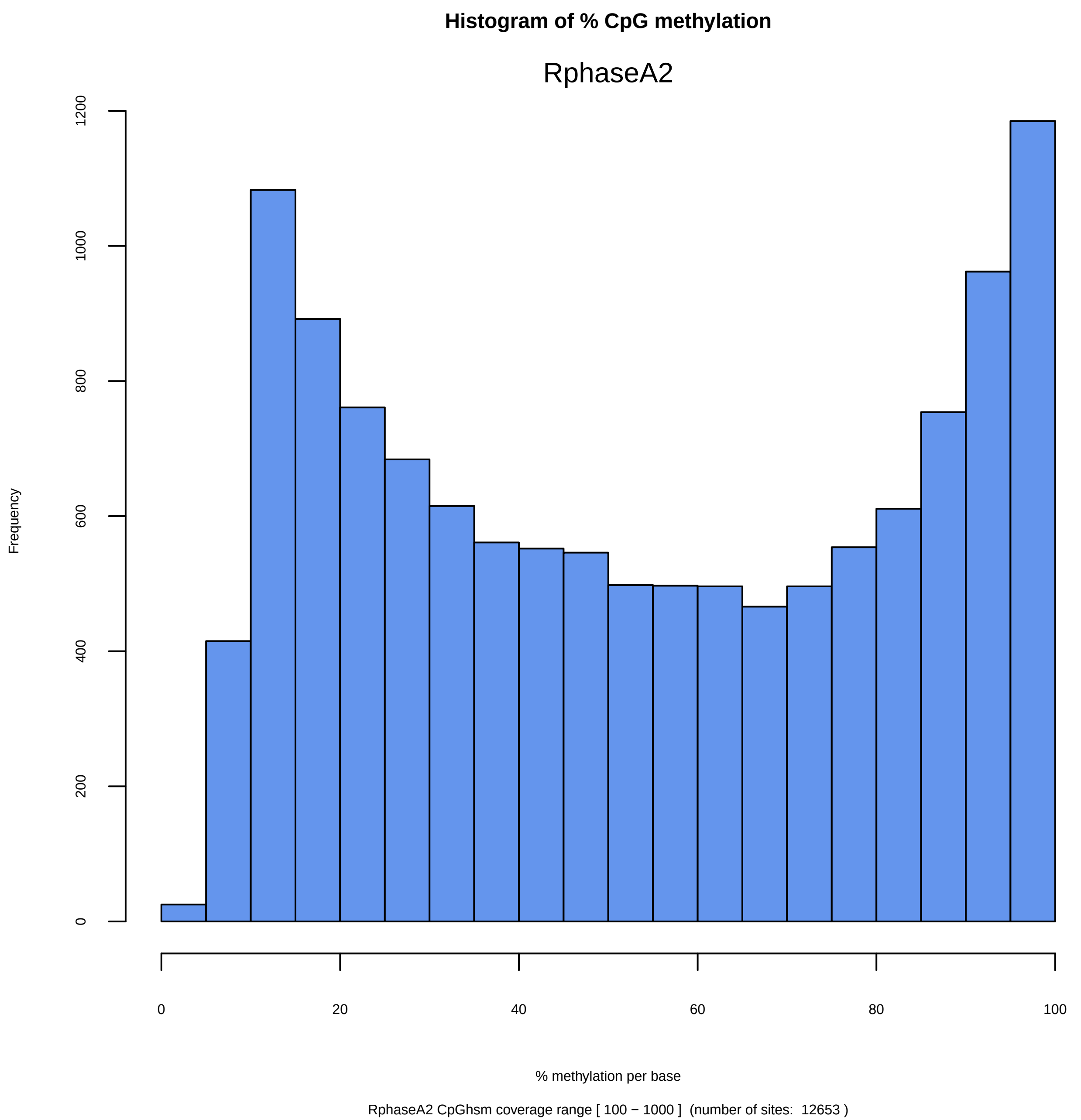
Histogram of % CpG methylation
RphaseA2
Frequency
0
200
400
600
800
1000
1200
0
20
40
60
80
100
% methylation per base
RphaseA2 CpGhsm coverage range [ 100 − 1000 ]  (number of sites:  12653 )

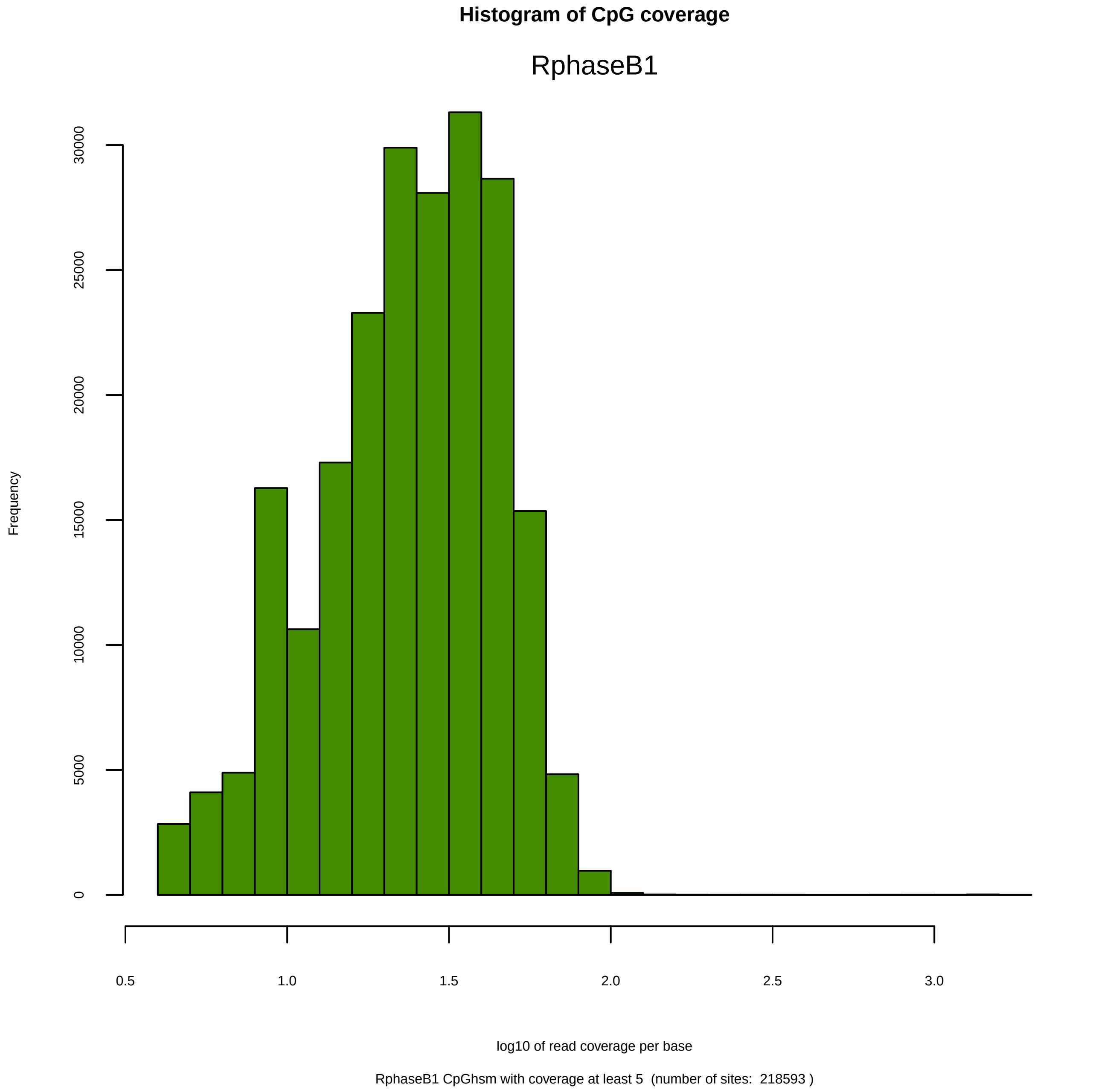
Histogram of CpG coverage
RphaseB1
Frequency
0
5000
10000
15000
20000
25000
30000
0.5
1.0
1.5
2.0
2.5
3.0
log10 of read coverage per base
RphaseB1 CpGhsm with coverage at least 5 (number of sites: 218593 )

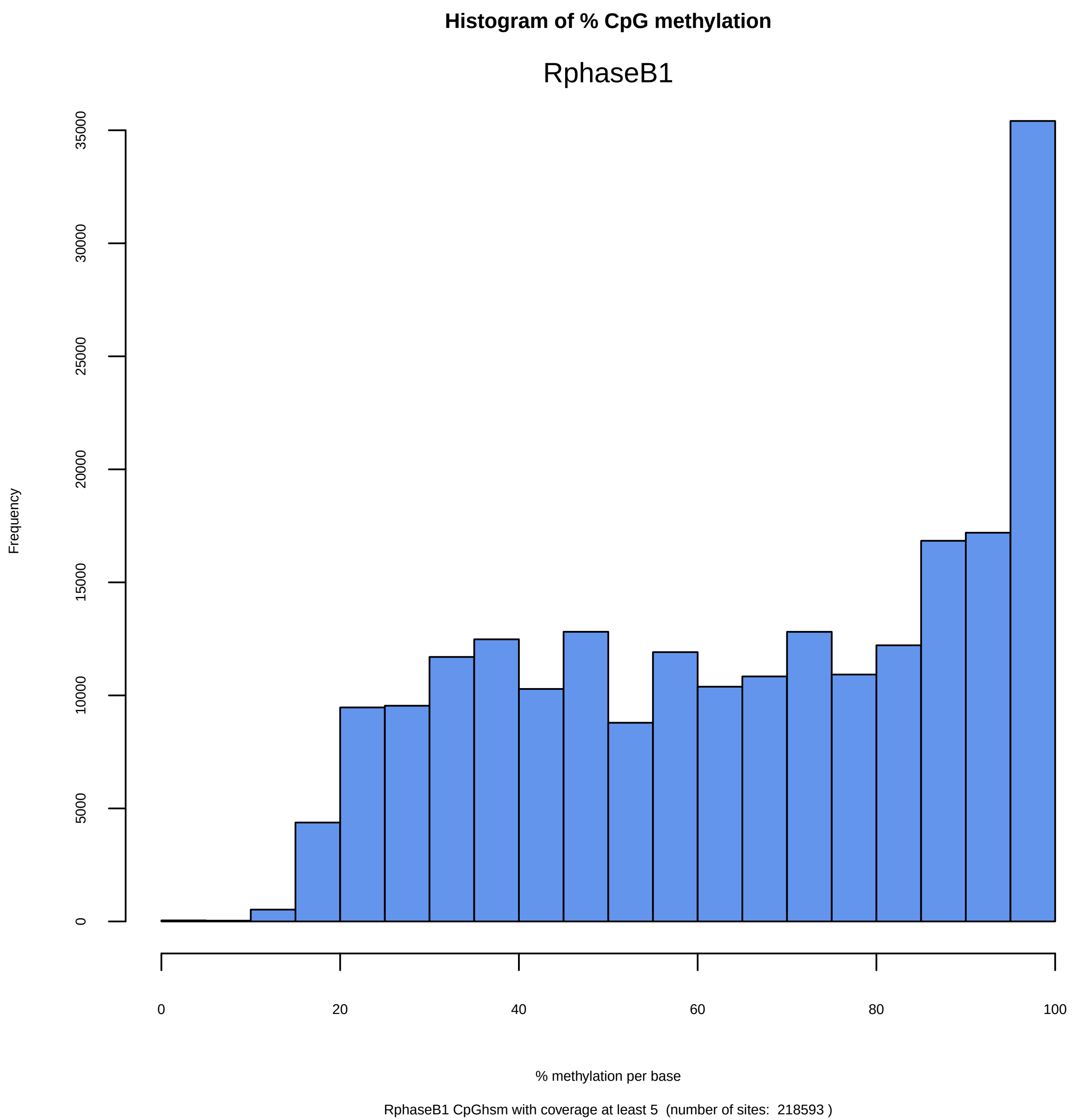

Histogram of % CpG methylation
RphaseB1
Frequency
0
5000
10000
15000
20000
25000
30000
35000
0
20
40
60
80
100
% methylation per base
RphaseB1 CpGhsm with coverage at least 5  (number of sites:  218593 )

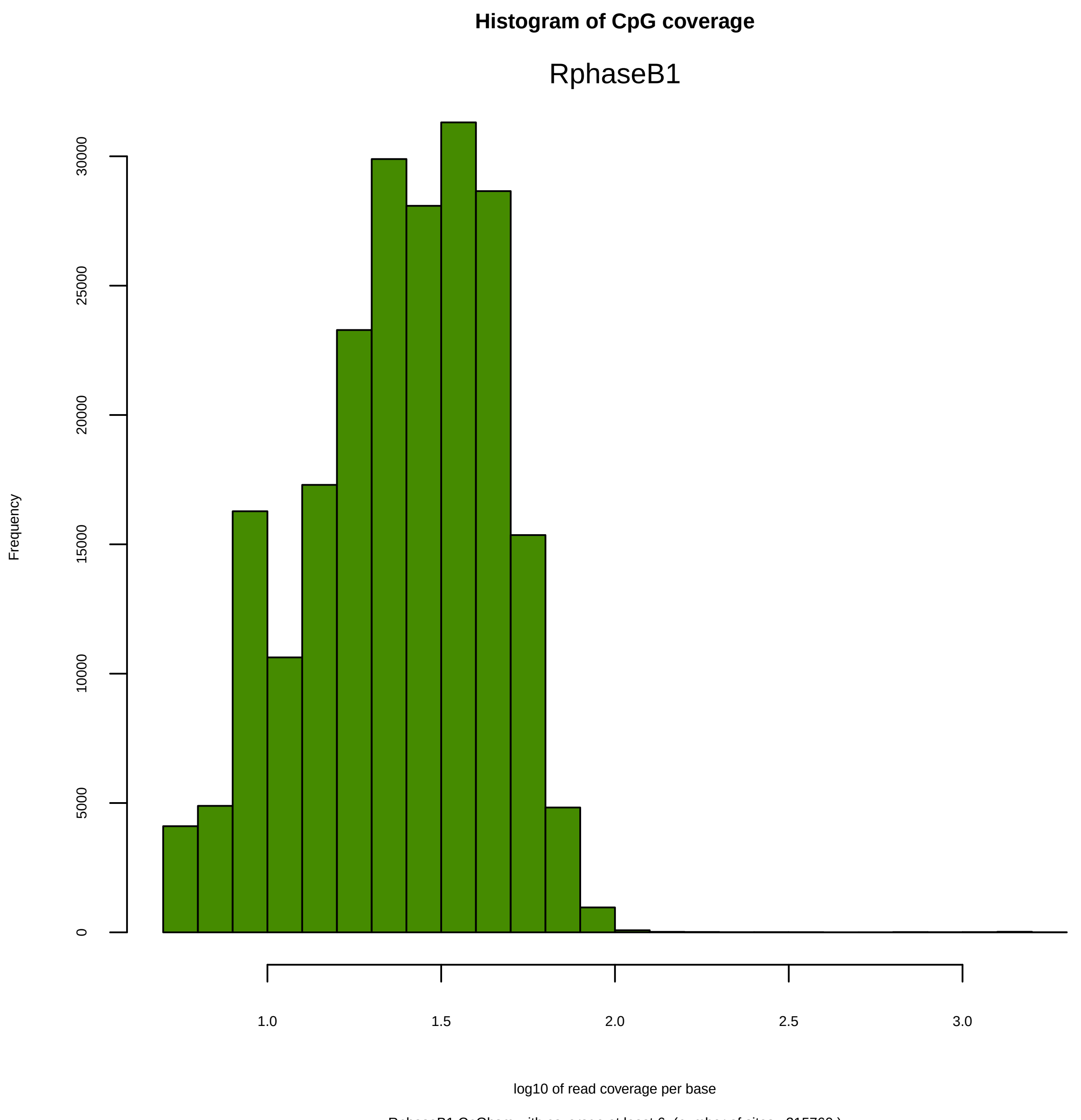
Histogram of CpG coverage
RphaseB1
Frequency
0
5000
10000
15000
20000
25000
30000
1.0
1.5
2.0
2.5
3.0
log10 of read coverage per base
RphaseB1 CpGhsm with coverage at least 6 (number of sites: 215760 )

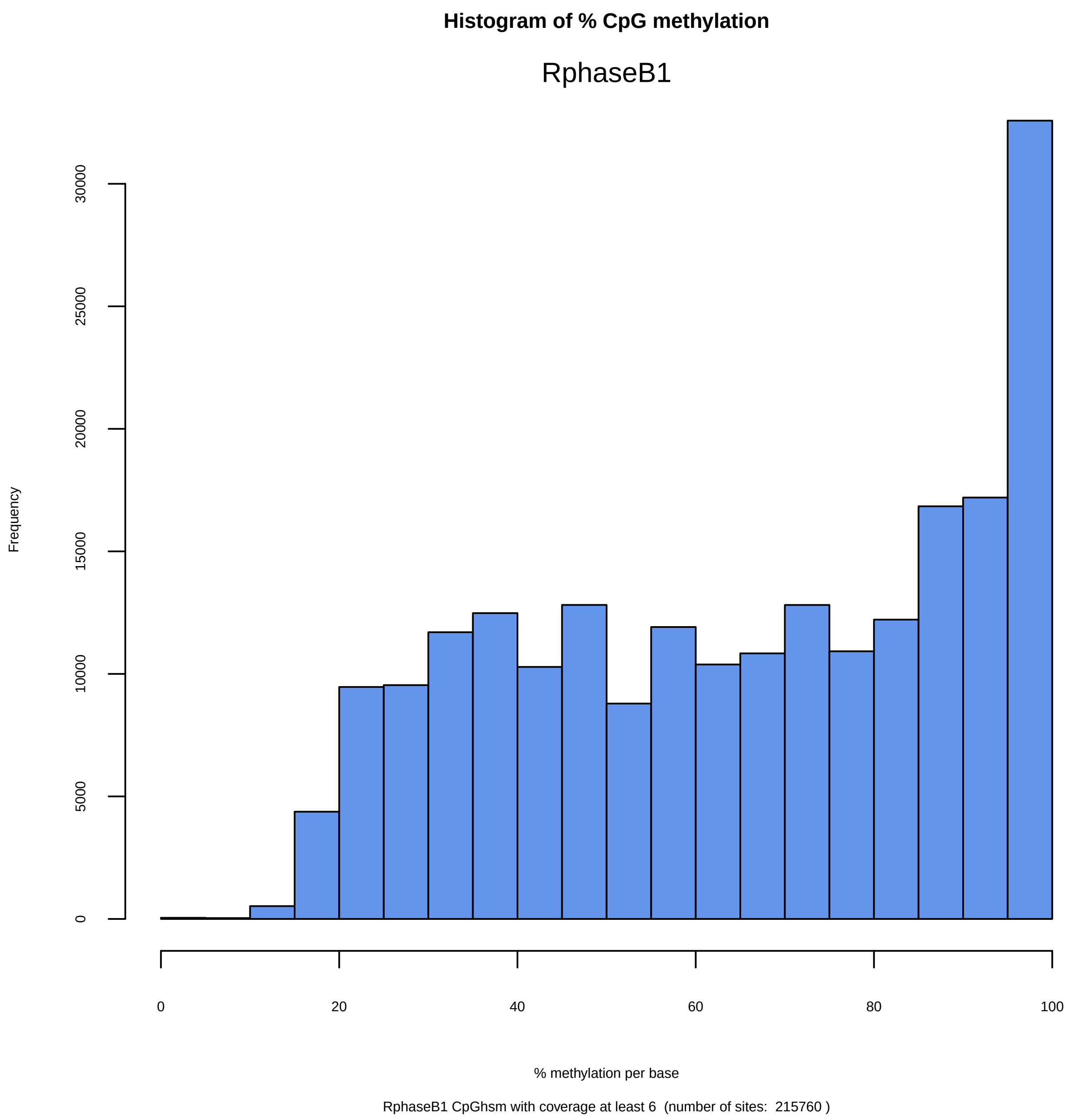

Histogram of % CpG methylation
RphaseB1
Frequency
0
5000
10000
15000
20000
25000
30000
0
20
40
60
80
100
% methylation per base
RphaseB1 CpGhsm with coverage at least 6 (number of sites: 215760 )

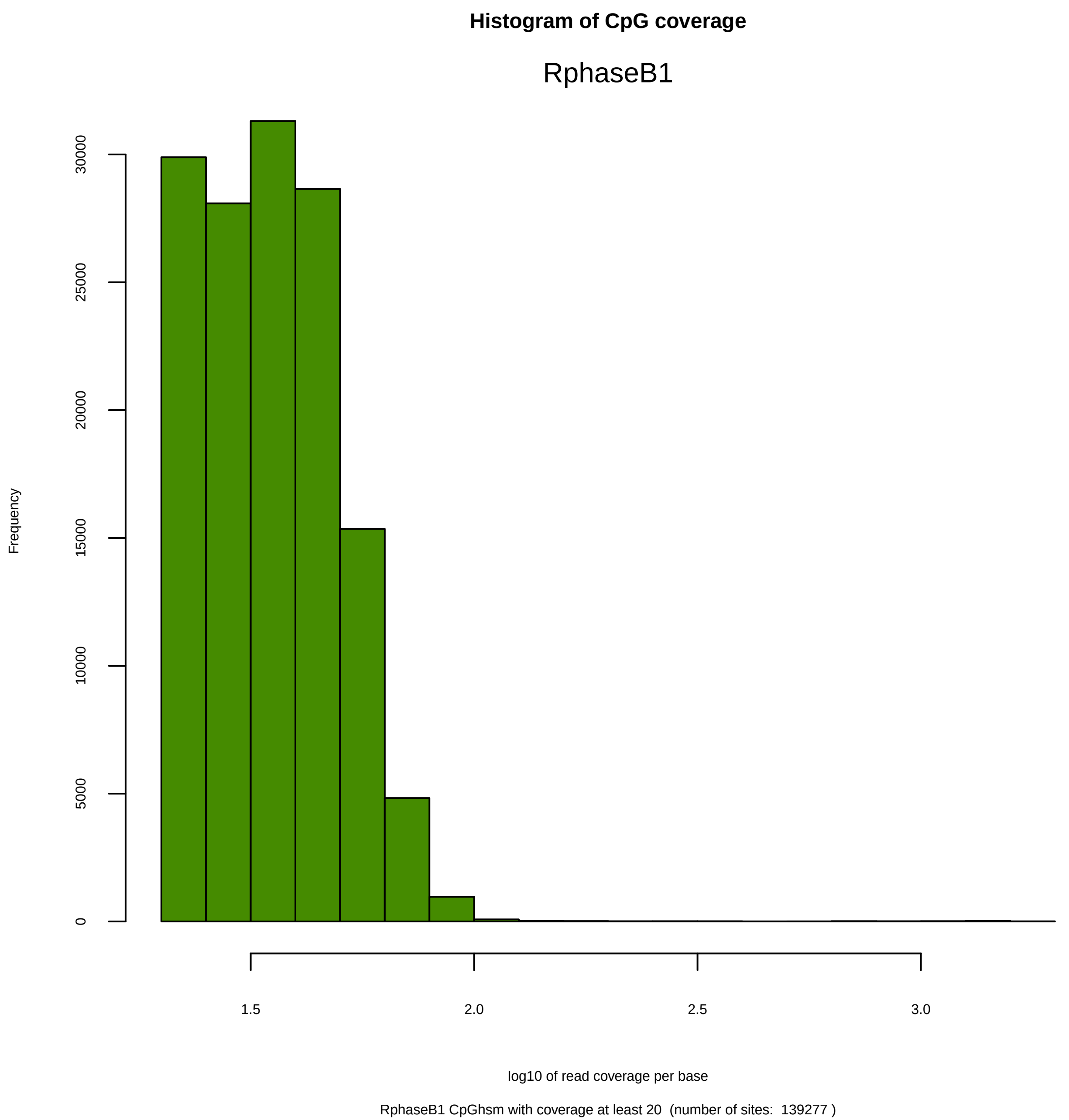
Histogram of CpG coverage
RphaseB1
Frequency
0
5000
10000
15000
20000
25000
30000
1.5
2.0
2.5
3.0
log10 of read coverage per base
RphaseB1 CpGhsm with coverage at least 20 (number of sites: 139277 )

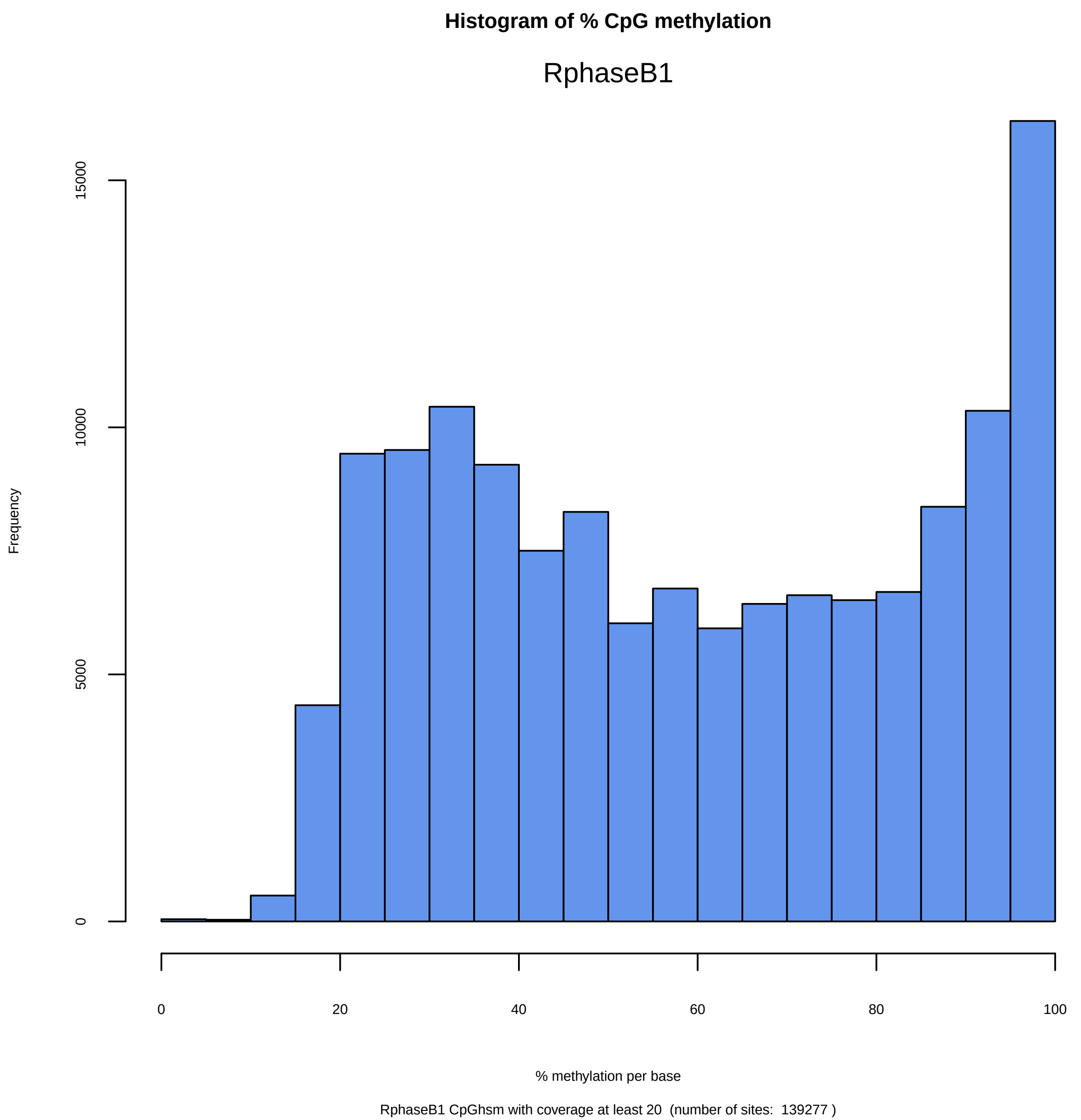
Histogram of % CpG methylation
RphaseB1
Frequency
0
5000
10000
15000
0
20
40
60
80
100
% methylation per base
RphaseB1 CpGhsm with coverage at least 20 (number of sites: 139277 )

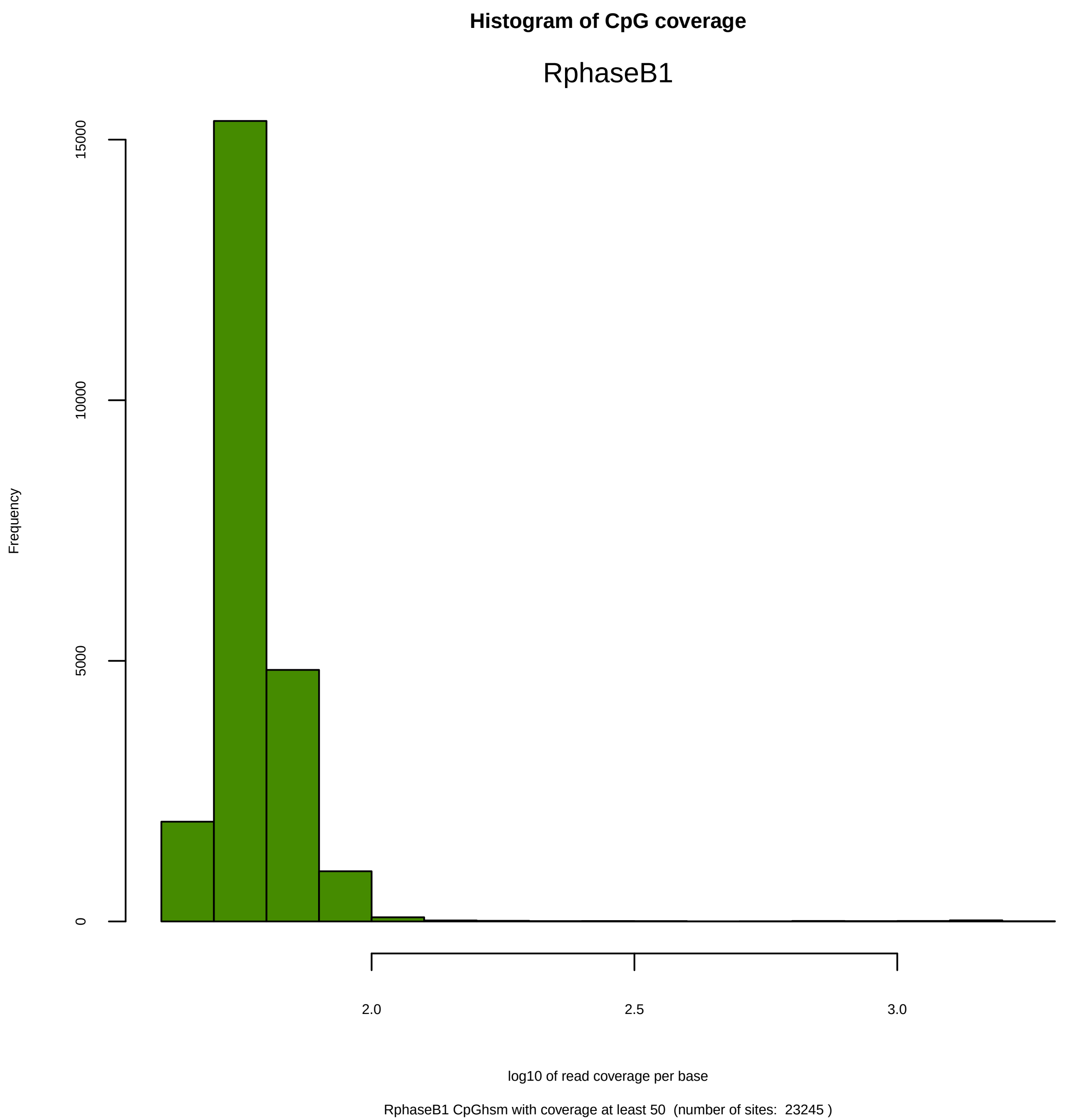

Histogram of CpG coverage
RphaseB1
Frequency
0
5000
10000
15000
2.0
2.5
3.0
log10 of read coverage per base
RphaseB1 CpGhsm with coverage at least 50  (number of sites:  23245 )

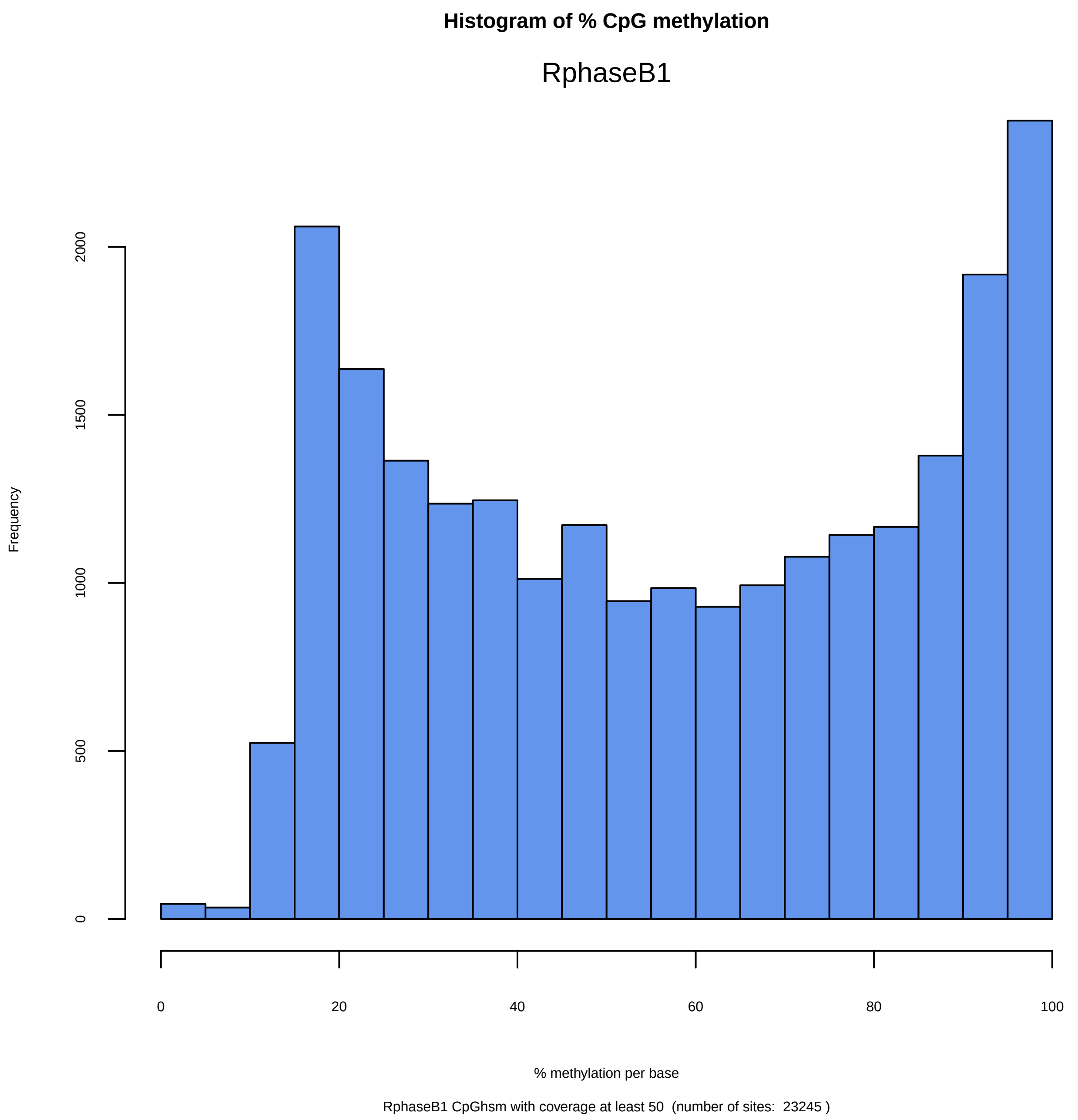
Histogram of % CpG methylation
RphaseB1
Frequency
0
500
1000
1500
2000
0
20
40
60
80
100
% methylation per base
RphaseB1 CpGhsm with coverage at least 50  (number of sites:  23245 )

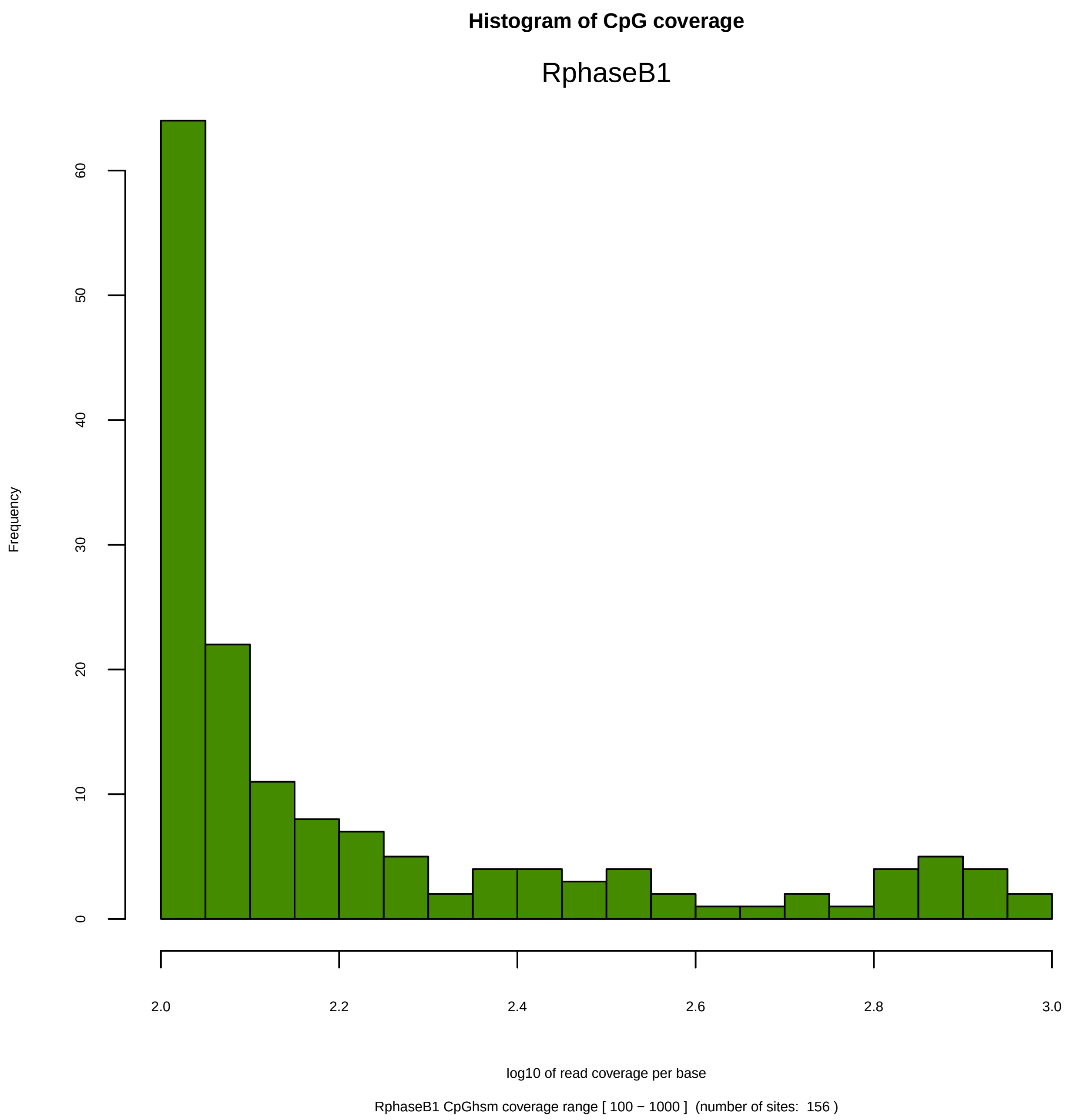

Histogram of CpG coverage
RphaseB1
Frequency
0
10
20
30
40
50
60
2.0
2.2
2.4
2.6
2.8
3.0
log10 of read coverage per base
RphaseB1 CpGhsm coverage range [ 100 − 1000 ] (number of sites: 156 )

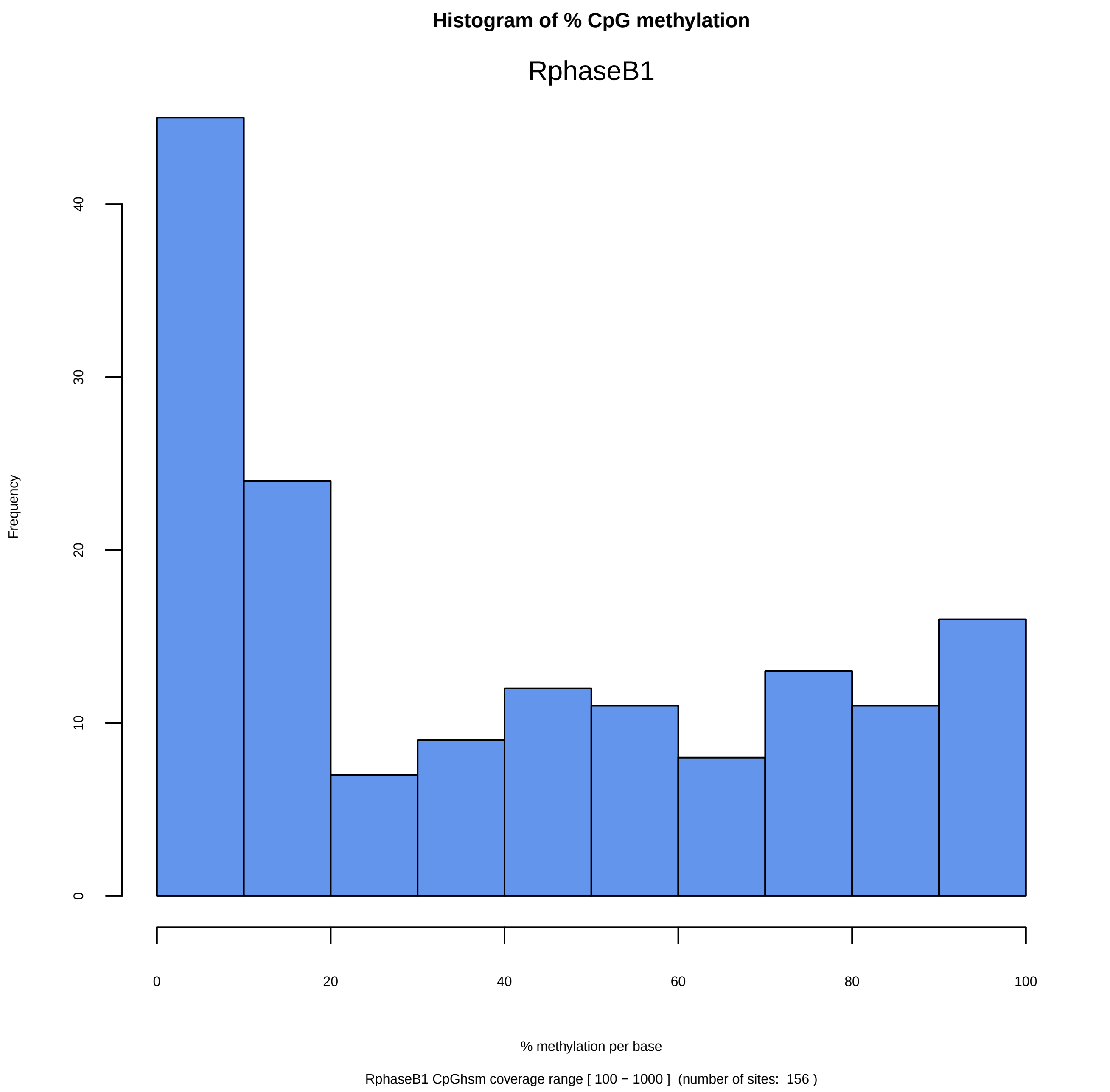
Histogram of % CpG methylation
RphaseB1
Frequency
0
10
20
30
40
0
20
40
60
80
100
% methylation per base
RphaseB1 CpGhsm coverage range [ 100 − 1000 ] (number of sites: 156 )

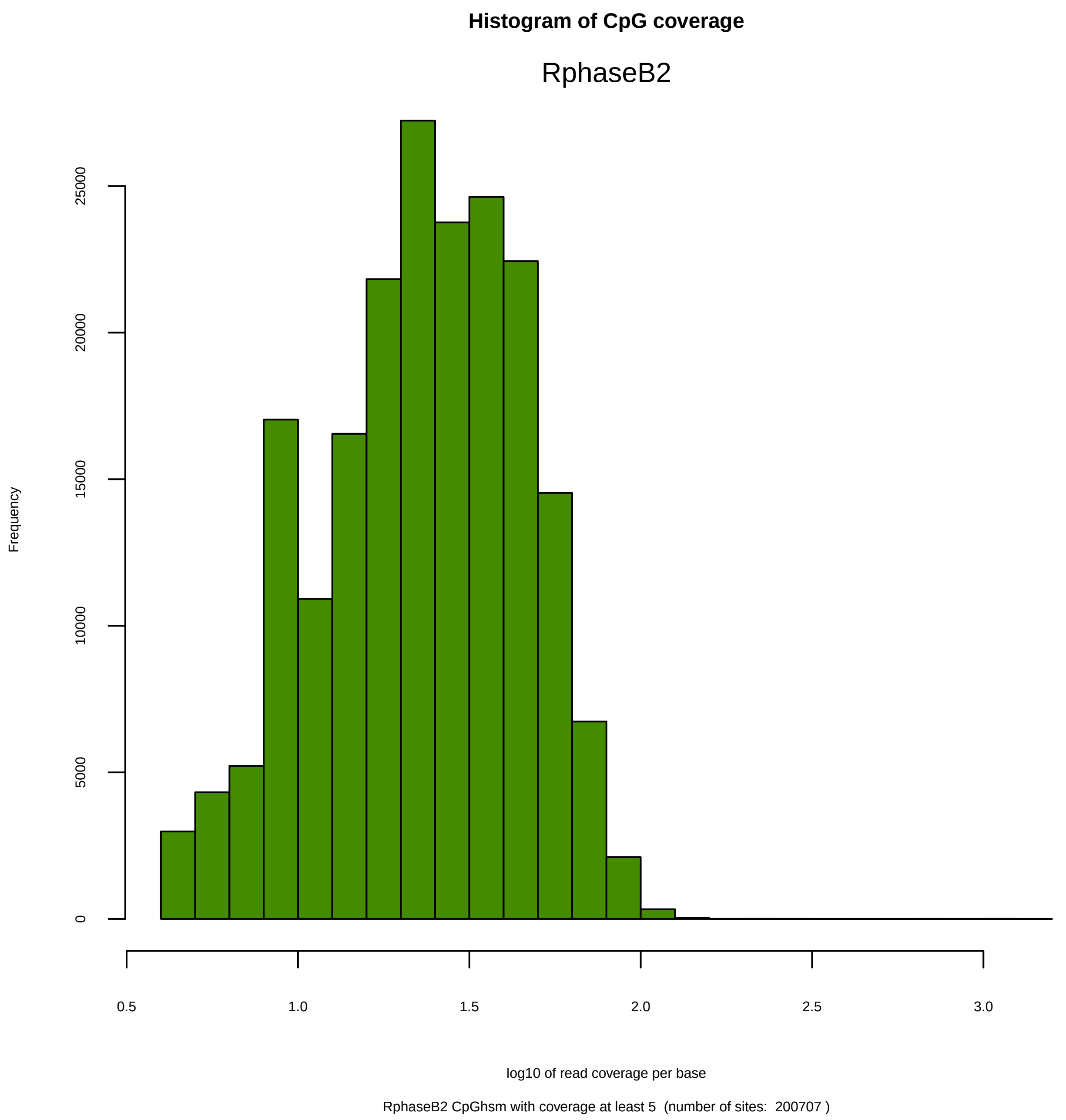
Histogram of CpG coverage
RphaseB2
Frequency
0
5000
10000
15000
20000
25000
0.5
1.0
1.5
2.0
2.5
3.0
log10 of read coverage per base
RphaseB2 CpGhsm with coverage at least 5 (number of sites: 200707 )

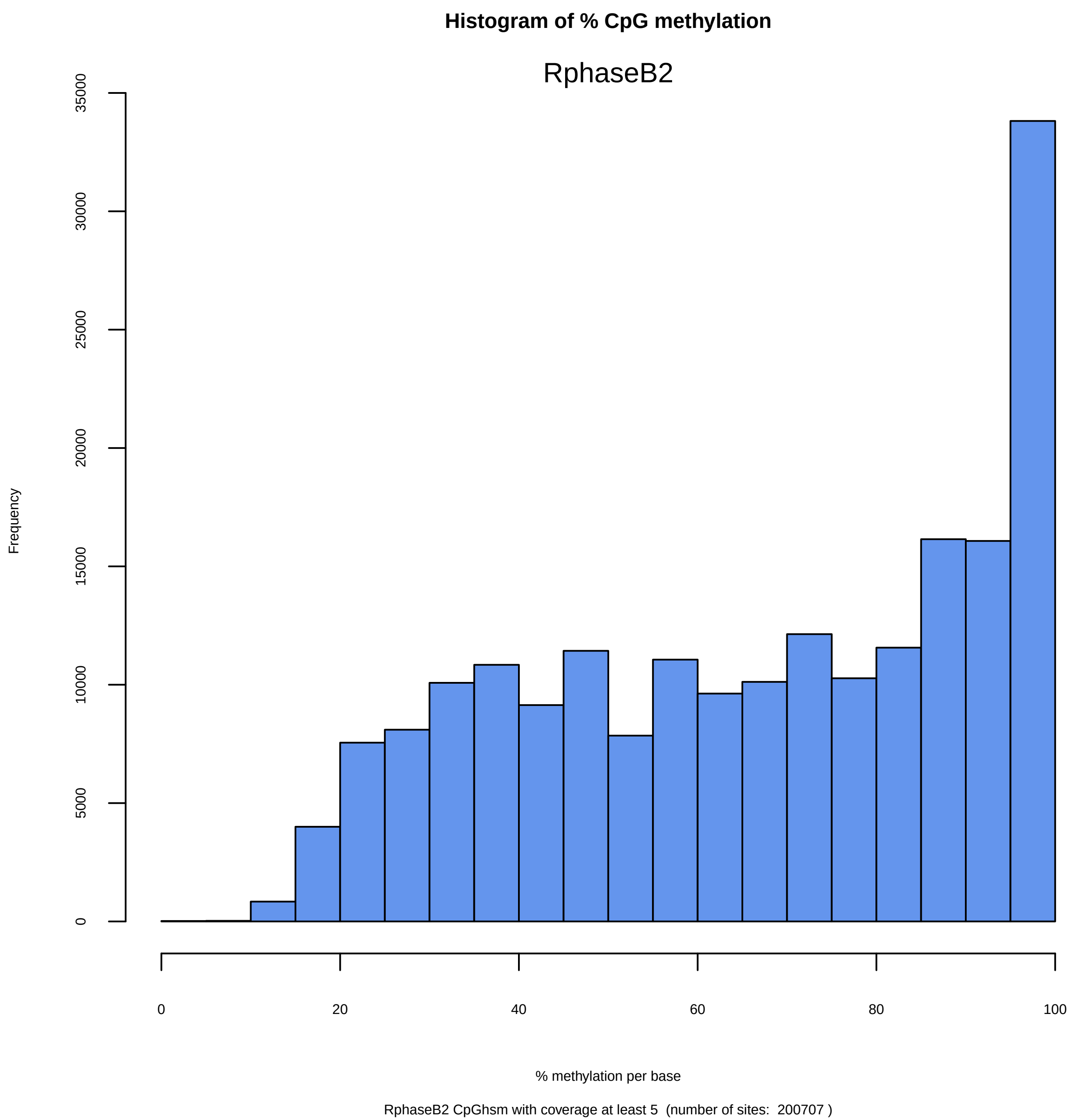

Histogram of % CpG methylation
RphaseB2
Frequency
0
5000
10000
15000
20000
25000
30000
35000
0
20
40
60
80
100
% methylation per base
RphaseB2 CpGhsm with coverage at least 5 (number of sites: 200707 )

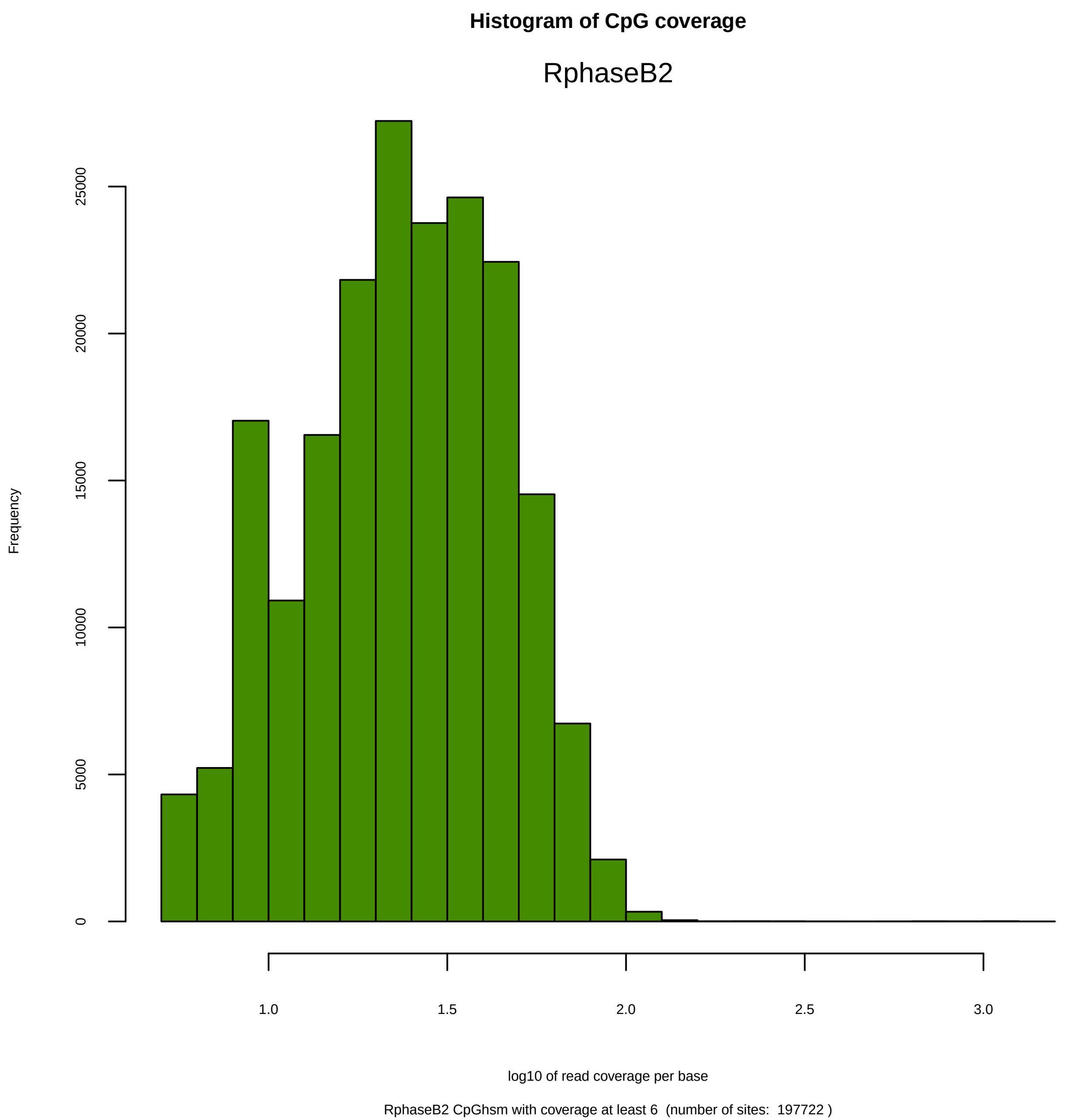
Histogram of CpG coverage
RphaseB2
Frequency
0
5000
10000
15000
20000
25000
1.0
1.5
2.0
2.5
3.0
log10 of read coverage per base
RphaseB2 CpGhsm with coverage at least 6 (number of sites: 197722 )

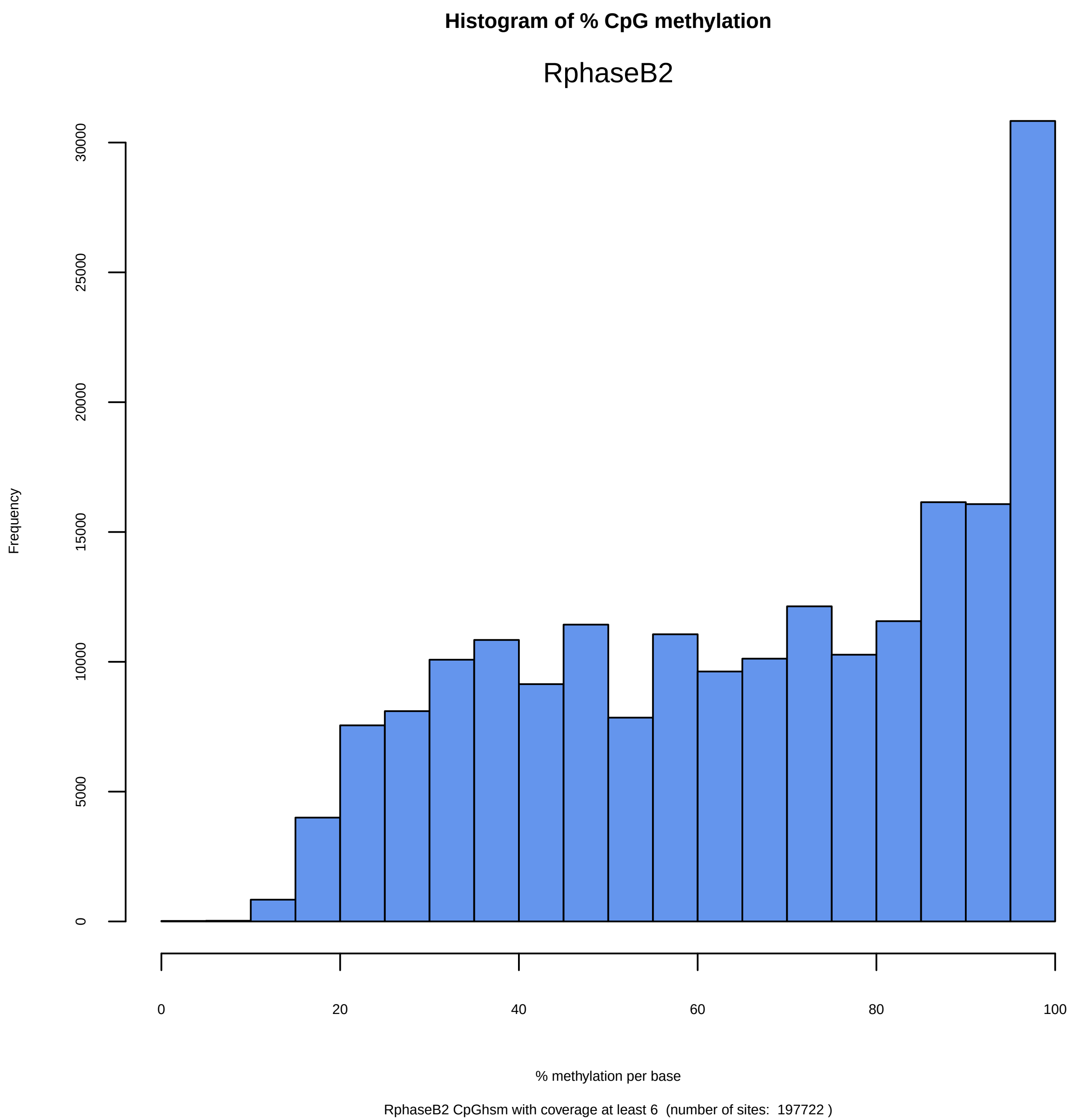
Histogram of % CpG methylation
RphaseB2
Frequency
0
5000
10000
15000
20000
25000
30000
0
20
40
60
80
100
% methylation per base
RphaseB2 CpGhsm with coverage at least 6  (number of sites:  197722 )

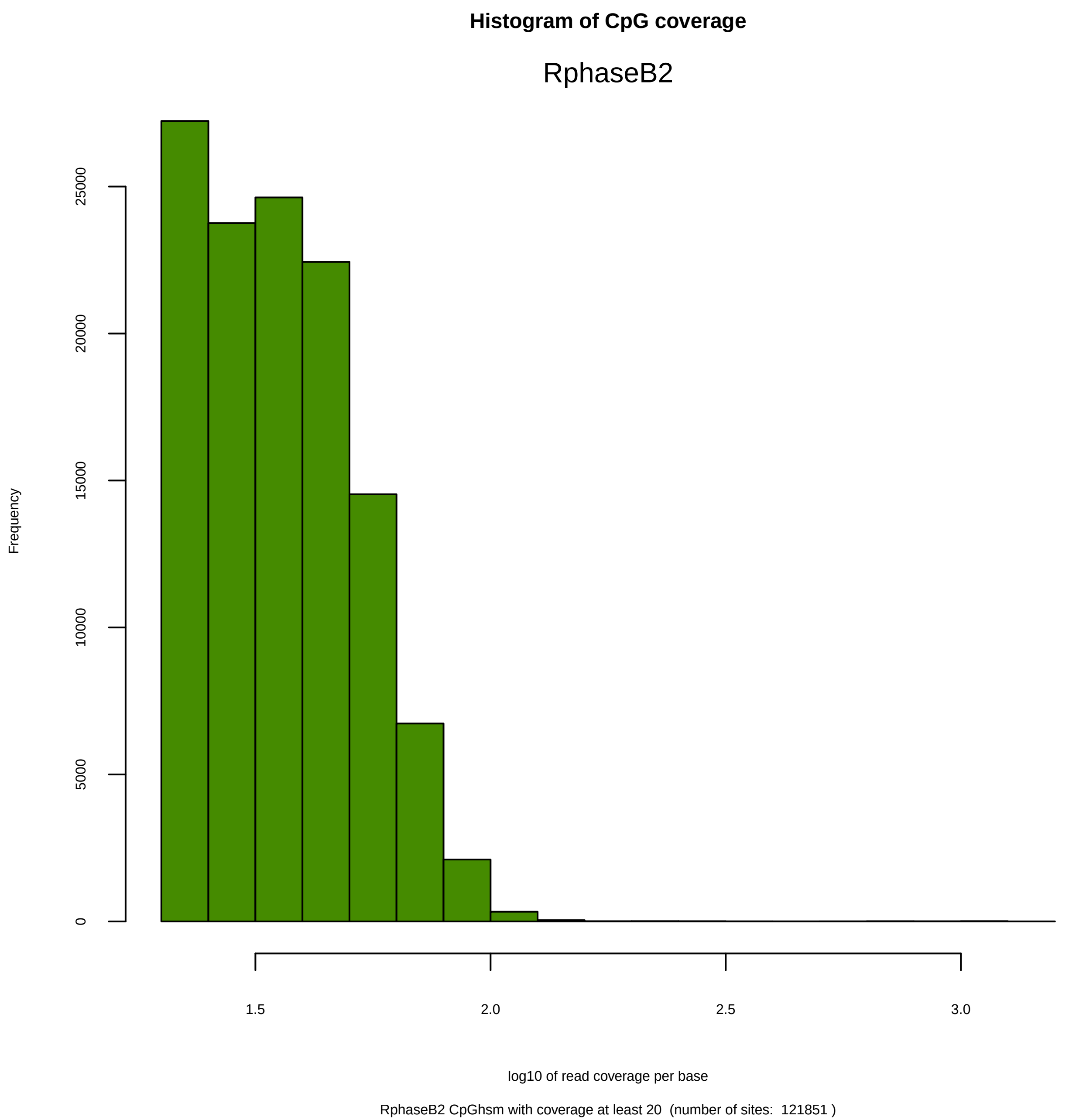
Histogram of CpG coverage
RphaseB2
Frequency
0
5000
10000
15000
20000
25000
1.5
2.0
2.5
3.0
log10 of read coverage per base
RphaseB2 CpGhsm with coverage at least 20  (number of sites:  121851 )

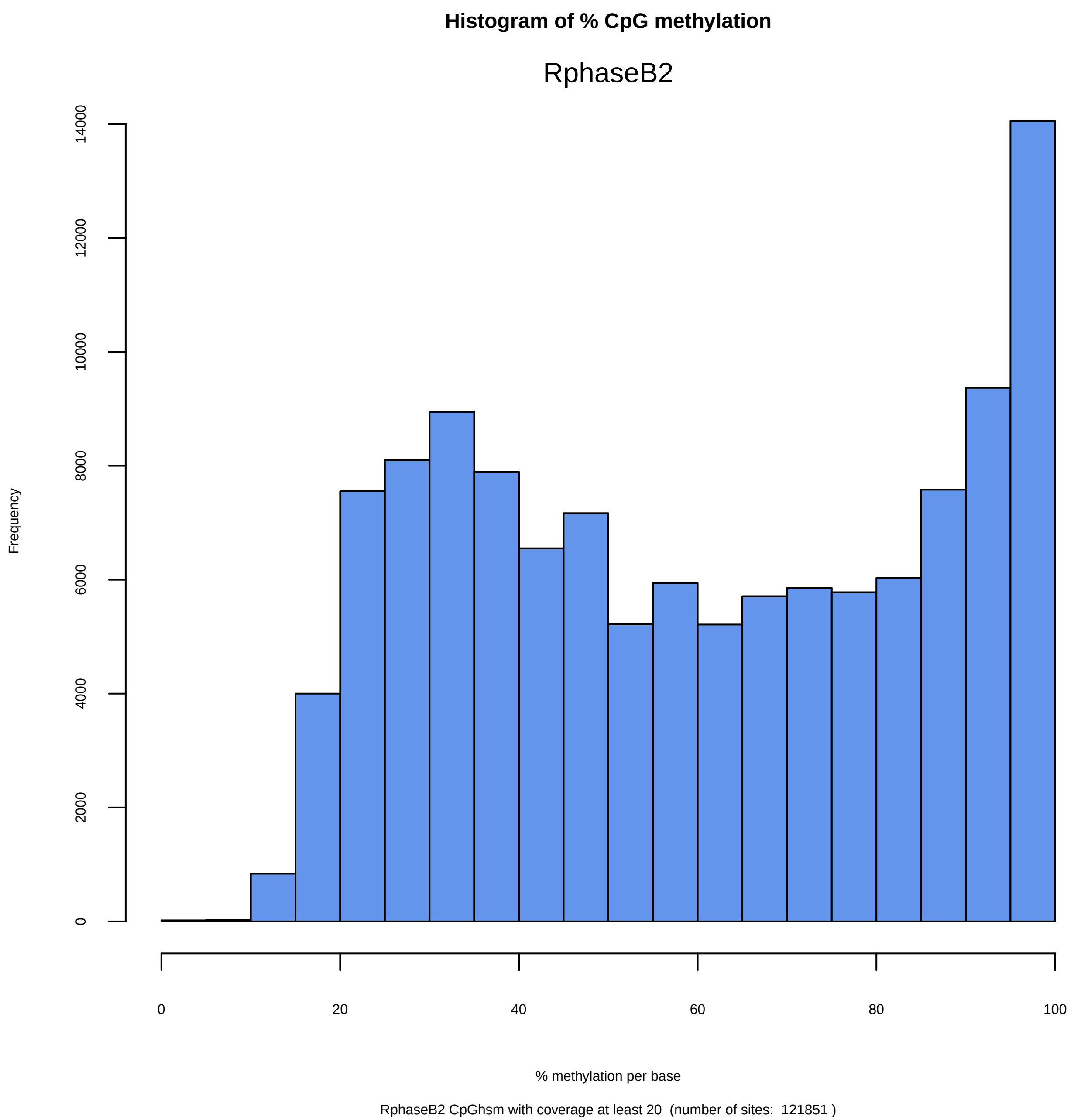

Histogram of % CpG methylation
RphaseB2
Frequency
0
2000
4000
6000
8000
10000
12000
14000
0
20
40
60
80
100
% methylation per base
RphaseB2 CpGhsm with coverage at least 20  (number of sites:  121851 )

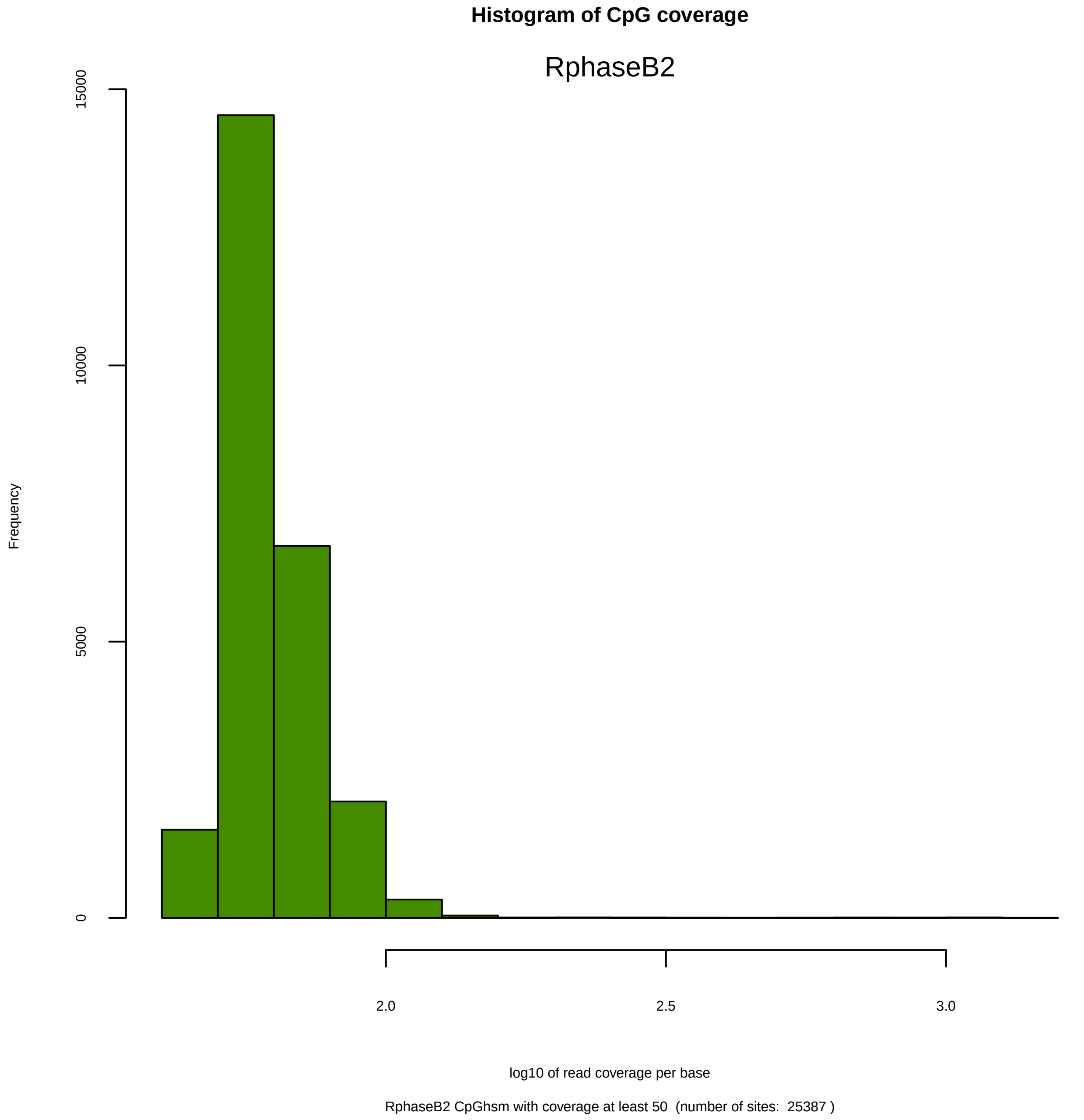

Histogram of CpG coverage
RphaseB2
Frequency
15000
10000
5000
0
2.0
2.5
3.0
log10 of read coverage per base
RphaseB2 CpGhsm with coverage at least 50  (number of sites:  25387 )

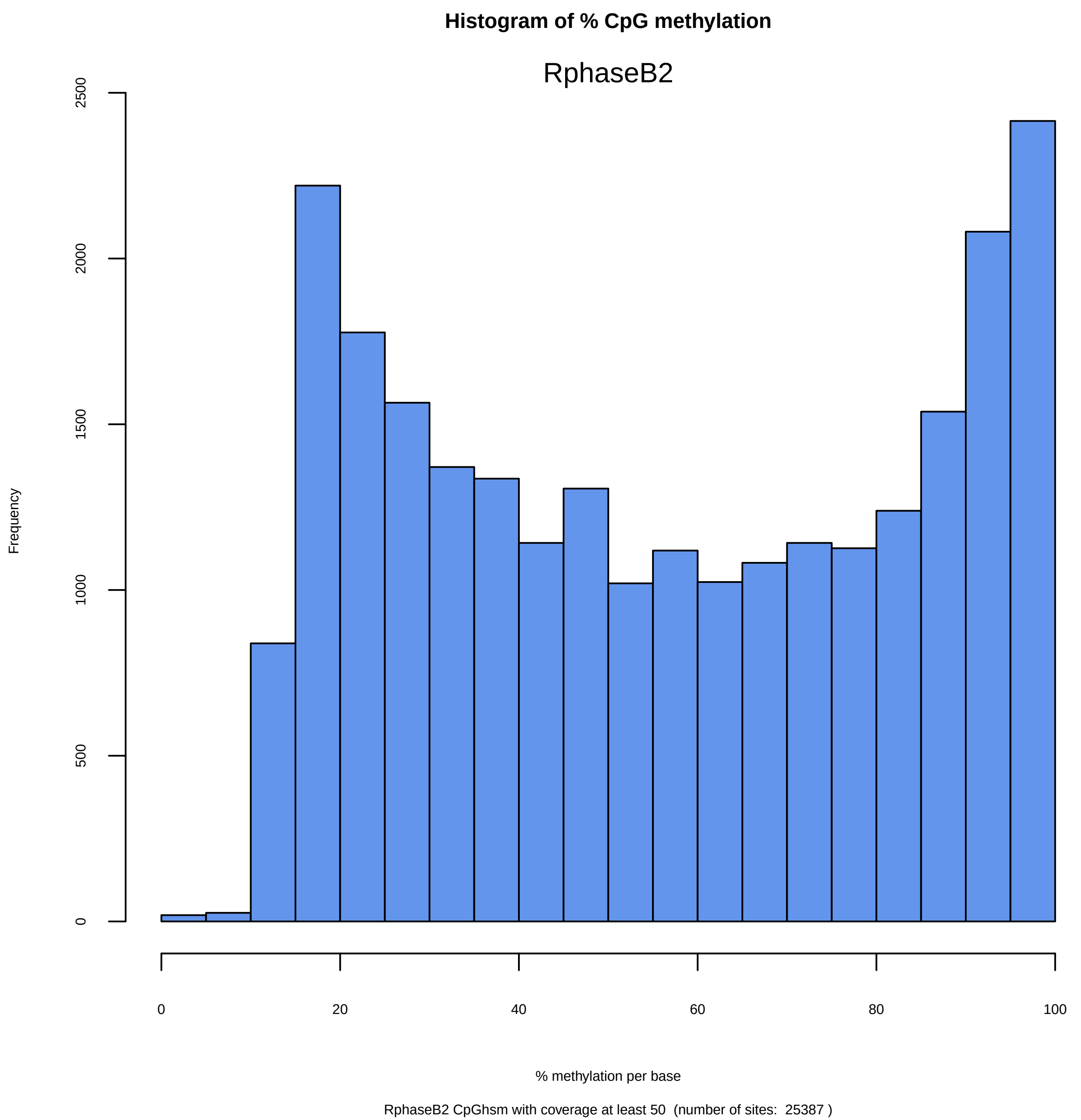

Histogram of % CpG methylation
RphaseB2
Frequency
0
500
1000
1500
2000
2500
0
20
40
60
80
100
% methylation per base
RphaseB2 CpGhsm with coverage at least 50  (number of sites:  25387 )

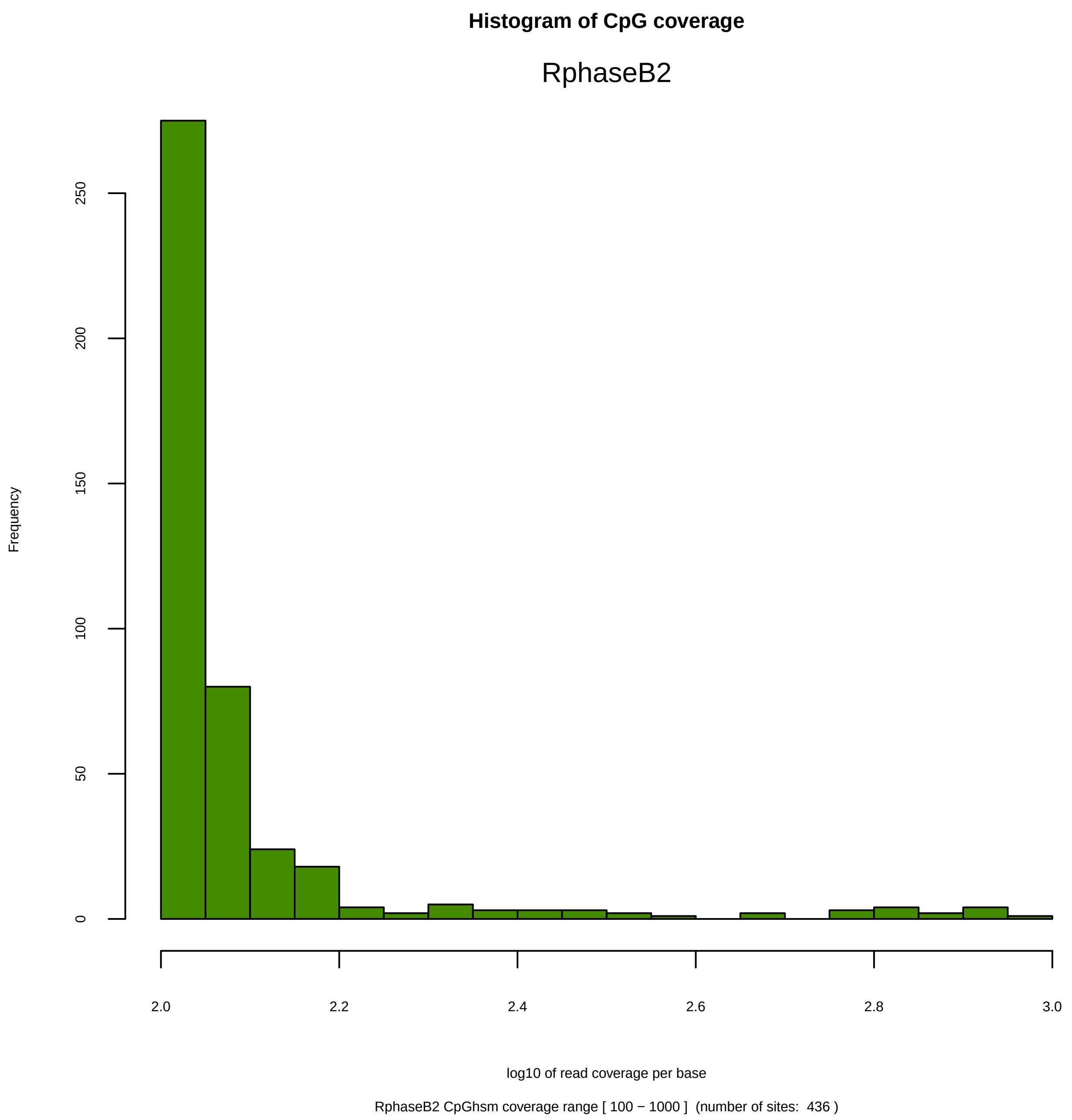

Histogram of CpG coverage
RphaseB2
Frequency
0
50
100
150
200
250
2.0
2.2
2.4
2.6
2.8
3.0
log10 of read coverage per base
RphaseB2 CpGhsm coverage range [ 100 − 1000 ]  (number of sites:  436 )

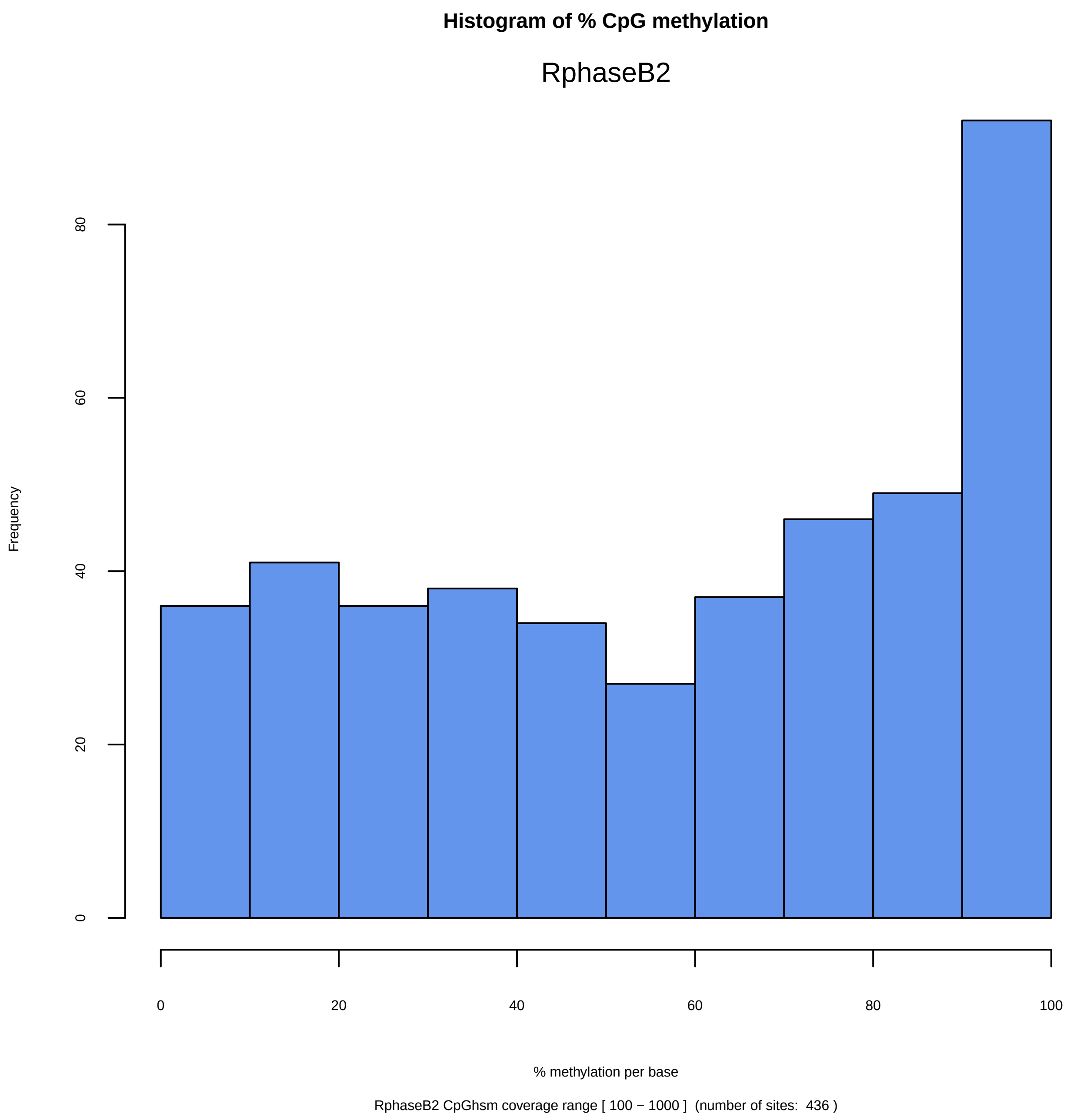

Histogram of % CpG methylation
RphaseB2
Frequency
0
20
40
60
80
0
20
40
60
80
100
% methylation per base
RphaseB2 CpGhsm coverage range [ 100 − 1000 ] (number of sites: 436 )

```
Directory: PWC File: 0READMEpwc

PWC - Pairwise comparisons between all samples.

Input: studymk, studymc, nbrpms, hheight, nbrpnts

Output: files pwc-*.txt pwc-.*pdf

Notes: The output is generated by BWASPR::cmpSites(), which determines for each pairwise
        comparison the numbers of common and distinct sites.  The function also calculates
        the overlap index between the samples and estimates the size of the common pool of
        potential methylation sites.

       The CpGscd data input (studymc) is necessary to determine sites that are detectable
        in both samples, and nbrpms (listed in the *.par file specified in the Cb_LI.conf
        configuration file as argument to TOTALNBRPMSITES) provides the total number of
        potential methylation sites (typically, all CpG sites), a necessary parameter for
        the estimations.
       hheight (default: 0.10) specifies the y-axis limit in the histogram plots.
       nbrpnts (default: 5000) specifies the number of common sites to be plotted in the
        methylation levels scatter plot.
```

```
Directory: PWC File: pwc-Cb_LI_BCphaseA1.vs.Cb_LI_BCphaseA2.txt

Numbers of common and distinct sites comparing Cb_LI_BCphaseA1 versus Cb_LI_BCphaseA2
================================================================================

  total number of potential sites:   26684764

  number of "Cb_LI_BCphaseA1_hsm"         sites:     198408
  number of "Cb_LI_BCphaseA2_hsm"         sites:     213448
    number of "Cb_LI_BCphaseA1_hsm"-unique sites:     28570
    number of common sites:        169838
    number of "Cb_LI_BCphaseA2_hsm"-unique sites:     43610
  total number of "Cb_LI_BCphaseA1_hsm+Cb_LI_BCphaseA2_hsm"-sites observed:  242018

  number of "Cb_LI_BCphaseA1_scd"        sites: 18501067 ( 69.33% of total)
  number of "Cb_LI_BCphaseA2_scd"        sites: 18589454 ( 69.66% of total)
    number of "Cb_LI_BCphaseA1_scd"-unique sites:      1613553
    number of sites in common:      16887514   (Expected:  12888431; O/E:       1.3)
    number of "Cb_LI_BCphaseA2_scd"-unique sites:      1701940
  total number of "Cb_LI_BCphaseA1_scd+Cb_LI_BCphaseA2_scd"-sites observed:   20203007

  number of sites in "Cb_LI_BCphaseA1_hsm" that are not  detectable in "Cb_LI_BCphaseA2_hsm":    6824
  number of sites in "Cb_LI_BCphaseA1_hsm" that are also detectable in "Cb_LI_BCphaseA2_hsm":  191584
    number of sites unique to "Cb_LI_BCphaseA1_hsm" although detectable in "Cb_LI_BCphaseA2_hsm":          21746
    number of "Cb_LI_BCphaseA1_hsm" / "Cb_LI_BCphaseA2_hsm" common sites                        :         169838      (Expected:   2341; O/E:      72.5)

  number of sites in "Cb_LI_BCphaseA2_hsm" that are not  detectable in "Cb_LI_BCphaseA1_hsm":    7102
  number of sites in "Cb_LI_BCphaseA2_hsm" that are also detectable in "Cb_LI_BCphaseA1_hsm":  206346
    number of sites unique to "Cb_LI_BCphaseA2_hsm" although detectable in "Cb_LI_BCphaseA1_hsm":          36508
    number of "Cb_LI_BCphaseA2_hsm" / "Cb_LI_BCphaseA1_hsm" common sites                        :         169838      (Expected:   2341; O/E:      72.5)

  Overlap index of "Cb_LI_BCphaseA1_hsm" with "Cb_LI_BCphaseA2_hsm":         0.802
  Estimated number of "Cb_LI_BCphaseA1_hsm" = "Cb_LI_BCphaseA2_hsm" sites (assuming sampling from one population):             232766
  Adjusted population size of "Cb_LI_BCphaseA1_hsm" = "Cb_LI_BCphaseA2_hsm" sites (assuming all sites detectable):             367805 (  1.52x of observed)
```

```
Directory: PWC File: pwc-Cb_LI_BCphaseA1.vs.Cb_LI_BCphaseB1.txt

Numbers of common and distinct sites comparing Cb_LI_BCphaseA1 versus Cb_LI_BCphaseB1
================================================================================

  total number of potential sites:   26684764

  number of "Cb_LI_BCphaseA1_hsm"         sites:     198408
  number of "Cb_LI_BCphaseB1_hsm"         sites:     226815
    number of "Cb_LI_BCphaseA1_hsm"-unique sites:     22669
    number of common sites:        175739
    number of "Cb_LI_BCphaseB1_hsm"-unique sites:     51076
  total number of "Cb_LI_BCphaseA1_hsm+Cb_LI_BCphaseB1_hsm"-sites observed:  249484

  number of "Cb_LI_BCphaseA1_scd"        sites: 18501067 ( 69.33% of total)
  number of "Cb_LI_BCphaseB1_scd"        sites: 20321376 ( 76.15% of total)
    number of "Cb_LI_BCphaseA1_scd"-unique sites:       713441
    number of sites in common:       17787626   (Expected:  14089206; O/E:       1.3)
    number of "Cb_LI_BCphaseB1_scd"-unique sites:      2533750
  total number of "Cb_LI_BCphaseA1_scd+Cb_LI_BCphaseB1_scd"-sites observed:   21034817

  number of sites in "Cb_LI_BCphaseA1_hsm" that are not  detectable in "Cb_LI_BCphaseB1_hsm":    3258
  number of sites in "Cb_LI_BCphaseA1_hsm" that are also detectable in "Cb_LI_BCphaseB1_hsm":  195150
    number of sites unique to "Cb_LI_BCphaseA1_hsm" although detectable in "Cb_LI_BCphaseB1_hsm":          19411
    number of "Cb_LI_BCphaseA1_hsm" / "Cb_LI_BCphaseB1_hsm" common sites                        :         175739      (Expected:   2376; O/E:      74.0)

  number of sites in "Cb_LI_BCphaseB1_hsm" that are not  detectable in "Cb_LI_BCphaseA1_hsm":   10223
  number of sites in "Cb_LI_BCphaseB1_hsm" that are also detectable in "Cb_LI_BCphaseA1_hsm":  216592
    number of sites unique to "Cb_LI_BCphaseB1_hsm" although detectable in "Cb_LI_BCphaseA1_hsm":          40853
    number of "Cb_LI_BCphaseB1_hsm" / "Cb_LI_BCphaseA1_hsm" common sites                        :         175739      (Expected:   2376; O/E:      74.0)

  Overlap index of "Cb_LI_BCphaseA1_hsm" with "Cb_LI_BCphaseB1_hsm":         0.826
  Estimated number of "Cb_LI_BCphaseA1_hsm" = "Cb_LI_BCphaseB1_hsm" sites (assuming sampling from one population):             240515
  Adjusted population size of "Cb_LI_BCphaseA1_hsm" = "Cb_LI_BCphaseB1_hsm" sites (assuming all sites detectable):             360818 (  1.45x of observed)
```

```
Directory: PWC File: pwc-Cb_LI_BCphaseA1.vs.Cb_LI_BCphaseB2.txt

Numbers of common and distinct sites comparing Cb_LI_BCphaseA1 versus Cb_LI_BCphaseB2
================================================================================

  total number of potential sites:  26684764

  number of "Cb_LI_BCphaseA1_hsm"         sites:     198408
  number of "Cb_LI_BCphaseB2_hsm"         sites:     212569
    number of "Cb_LI_BCphaseA1_hsm"-unique sites:     28088
    number of common sites:        170320
    number of "Cb_LI_BCphaseB2_hsm"-unique sites:     42249
  total number of "Cb_LI_BCphaseA1_hsm+Cb_LI_BCphaseB2_hsm"-sites observed:  240657

  number of "Cb_LI_BCphaseA1_scd"        sites: 18501067 ( 69.33% of total)
  number of "Cb_LI_BCphaseB2_scd"        sites: 19192933 ( 71.92% of total)
    number of "Cb_LI_BCphaseA1_scd"-unique sites:      1273021
    number of sites in common:       17228046   (Expected:  13306835; O/E:       1.3)
    number of "Cb_LI_BCphaseB2_scd"-unique sites:      1964887
  total number of "Cb_LI_BCphaseA1_scd+Cb_LI_BCphaseB2_scd"-sites observed:   20465954

  number of sites in "Cb_LI_BCphaseA1_hsm" that are not  detectable in "Cb_LI_BCphaseB2_hsm":    5496
  number of sites in "Cb_LI_BCphaseA1_hsm" that are also detectable in "Cb_LI_BCphaseB2_hsm":  192912
    number of sites unique to "Cb_LI_BCphaseA1_hsm" although detectable in "Cb_LI_BCphaseB2_hsm":          22592
    number of "Cb_LI_BCphaseA1_hsm" / "Cb_LI_BCphaseB2_hsm" common sites                        :         170320      (Expected:   2292; O/E:      74.3)

  number of sites in "Cb_LI_BCphaseB2_hsm" that are not  detectable in "Cb_LI_BCphaseA1_hsm":    7912
  number of sites in "Cb_LI_BCphaseB2_hsm" that are also detectable in "Cb_LI_BCphaseA1_hsm":  204657
    number of sites unique to "Cb_LI_BCphaseB2_hsm" although detectable in "Cb_LI_BCphaseA1_hsm":          34337
    number of "Cb_LI_BCphaseB2_hsm" / "Cb_LI_BCphaseA1_hsm" common sites                        :         170320      (Expected:   2292; O/E:      74.3)

  Overlap index of "Cb_LI_BCphaseA1_hsm" with "Cb_LI_BCphaseB2_hsm":         0.795
  Estimated number of "Cb_LI_BCphaseA1_hsm" = "Cb_LI_BCphaseB2_hsm" sites (assuming sampling from one population):             231804
  Adjusted population size of "Cb_LI_BCphaseA1_hsm" = "Cb_LI_BCphaseB2_hsm" sites (assuming all sites detectable):             359044 (  1.49x of observed)
```

Directory: PWC File: pwc-Cb_LI_BCphaseA1.vs.Cb_LI_RphaseA1.txt

```
Numbers of common and distinct sites comparing Cb_LI_BCphaseA1 versus Cb_LI_RphaseA1
================================================================================

  total number of potential sites:  26684764

  number of "Cb_LI_BCphaseA1_hsm"         sites:     198408
  number of "Cb_LI_RphaseA1_hsm"         sites:213610
    number of "Cb_LI_BCphaseA1_hsm"-unique sites:     27451
    number of common sites:        170957
    number of "Cb_LI_RphaseA1_hsm"-unique sites:      42653
  total number of "Cb_LI_BCphaseA1_hsm+Cb_LI_RphaseA1_hsm"-sites observed:   241061

  number of "Cb_LI_BCphaseA1_scd"        sites: 18501067 ( 69.33% of total)
  number of "Cb_LI_RphaseA1_scd"        sites:  19659217 ( 73.67% of total)
    number of "Cb_LI_BCphaseA1_scd"-unique sites:      1006096
    number of sites in common:      17494971   (Expected:  13630118; O/E:       1.3)
    number of "Cb_LI_RphaseA1_scd"-unique sites:       2164246
  total number of "Cb_LI_BCphaseA1_scd+Cb_LI_RphaseA1_scd"-sites observed:    20665313

  number of sites in "Cb_LI_BCphaseA1_hsm" that are not  detectable in "Cb_LI_RphaseA1_hsm":     4321
  number of sites in "Cb_LI_BCphaseA1_hsm" that are also detectable in "Cb_LI_RphaseA1_hsm":   194087
    number of sites unique to "Cb_LI_BCphaseA1_hsm" although detectable in "Cb_LI_RphaseA1_hsm":           23130
    number of "Cb_LI_BCphaseA1_hsm" / "Cb_LI_RphaseA1_hsm" common sites                        :          170957      (Expected:   2272; O/E:      75.2)

  number of sites in "Cb_LI_RphaseA1_hsm" that are not  detectable in "Cb_LI_BCphaseA1_hsm":     8799
  number of sites in "Cb_LI_RphaseA1_hsm" that are also detectable in "Cb_LI_BCphaseA1_hsm":   204811
    number of sites unique to "Cb_LI_RphaseA1_hsm" although detectable in "Cb_LI_BCphaseA1_hsm":           33854
    number of "Cb_LI_RphaseA1_hsm" / "Cb_LI_BCphaseA1_hsm" common sites                        :          170957      (Expected:   2272; O/E:      75.2)

  Overlap index of "Cb_LI_BCphaseA1_hsm" with "Cb_LI_RphaseA1_hsm":          0.791
  Estimated number of "Cb_LI_BCphaseA1_hsm" = "Cb_LI_RphaseA1_hsm" sites (assuming sampling from one population):              232521
  Adjusted population size of "Cb_LI_BCphaseA1_hsm" = "Cb_LI_RphaseA1_hsm" sites (assuming all sites detectable):              354661 (  1.47x of observed)
```

```
Directory: PWC File: pwc-Cb_LI_BCphaseA1.vs.Cb_LI_RphaseA2.txt

Numbers of common and distinct sites comparing Cb_LI_BCphaseA1 versus Cb_LI_RphaseA2
================================================================================

  total number of potential sites:  26684764

  number of "Cb_LI_BCphaseA1_hsm"         sites:     198408
  number of "Cb_LI_RphaseA2_hsm"         sites:265952
    number of "Cb_LI_BCphaseA1_hsm"-unique sites:     13716
    number of common sites:        184692
    number of "Cb_LI_RphaseA2_hsm"-unique sites:      81260
  total number of "Cb_LI_BCphaseA1_hsm+Cb_LI_RphaseA2_hsm"-sites observed:   279668

  number of "Cb_LI_BCphaseA1_scd"        sites: 18501067 ( 69.33% of total)
  number of "Cb_LI_RphaseA2_scd"        sites:  20435277 ( 76.58% of total)
    number of "Cb_LI_BCphaseA1_scd"-unique sites:       710114
    number of sites in common:      17790953   (Expected:  14168176; O/E:       1.3)
    number of "Cb_LI_RphaseA2_scd"-unique sites:       2644324
  total number of "Cb_LI_BCphaseA1_scd+Cb_LI_RphaseA2_scd"-sites observed:    21145391

  number of sites in "Cb_LI_BCphaseA1_hsm" that are not  detectable in "Cb_LI_RphaseA2_hsm":     2190
  number of sites in "Cb_LI_BCphaseA1_hsm" that are also detectable in "Cb_LI_RphaseA2_hsm":   196218
    number of sites unique to "Cb_LI_BCphaseA1_hsm" although detectable in "Cb_LI_RphaseA2_hsm":           11526
    number of "Cb_LI_BCphaseA1_hsm" / "Cb_LI_RphaseA2_hsm" common sites                        :          184692      (Expected:   2778; O/E:      66.5)

  number of sites in "Cb_LI_RphaseA2_hsm" that are not  detectable in "Cb_LI_BCphaseA1_hsm":    14073
  number of sites in "Cb_LI_RphaseA2_hsm" that are also detectable in "Cb_LI_BCphaseA1_hsm":   251879
    number of sites unique to "Cb_LI_RphaseA2_hsm" although detectable in "Cb_LI_BCphaseA1_hsm":           67187
    number of "Cb_LI_RphaseA2_hsm" / "Cb_LI_BCphaseA1_hsm" common sites                        :          184692      (Expected:   2778; O/E:      66.5)

  Overlap index of "Cb_LI_BCphaseA1_hsm" with "Cb_LI_RphaseA2_hsm":          0.900
  Estimated number of "Cb_LI_BCphaseA1_hsm" = "Cb_LI_RphaseA2_hsm" sites (assuming sampling from one population):              267598
  Adjusted population size of "Cb_LI_BCphaseA1_hsm" = "Cb_LI_RphaseA2_hsm" sites (assuming all sites detectable):              401372 (  1.44x of observed)
```

```
Directory: PWC File: pwc-Cb_LI_BCphaseA1.vs.Cb_LI_RphaseB1.txt

Numbers of common and distinct sites comparing Cb_LI_BCphaseA1 versus Cb_LI_RphaseB1
================================================================================

  total number of potential sites:  26684764

  number of "Cb_LI_BCphaseA1_hsm"         sites:     198408
  number of "Cb_LI_RphaseB1_hsm"         sites: 218593
    number of "Cb_LI_BCphaseA1_hsm"-unique sites:     25173
    number of common sites:        173235
    number of "Cb_LI_RphaseB1_hsm"-unique sites:      45358
  total number of "Cb_LI_BCphaseA1_hsm+Cb_LI_RphaseB1_hsm"-sites observed:   243766

  number of "Cb_LI_BCphaseA1_scd"        sites: 18501067 ( 69.33% of total)
  number of "Cb_LI_RphaseB1_scd"        sites:  19576474 ( 73.36% of total)
    number of "Cb_LI_BCphaseA1_scd"-unique sites:      1014280
    number of sites in common:      17486787   (Expected:  13572751; O/E:       1.3)
    number of "Cb_LI_RphaseB1_scd"-unique sites:       2089687
  total number of "Cb_LI_BCphaseA1_scd+Cb_LI_RphaseB1_scd"-sites observed:    20590754

  number of sites in "Cb_LI_BCphaseA1_hsm" that are not  detectable in "Cb_LI_RphaseB1_hsm":     3900
  number of sites in "Cb_LI_BCphaseA1_hsm" that are also detectable in "Cb_LI_RphaseB1_hsm":   194508
    number of sites unique to "Cb_LI_BCphaseA1_hsm" although detectable in "Cb_LI_RphaseB1_hsm":           21273
    number of "Cb_LI_BCphaseA1_hsm" / "Cb_LI_RphaseB1_hsm" common sites                        :          173235      (Expected:   2332; O/E:      74.3)

  number of sites in "Cb_LI_RphaseB1_hsm" that are not  detectable in "Cb_LI_BCphaseA1_hsm":     8903
  number of sites in "Cb_LI_RphaseB1_hsm" that are also detectable in "Cb_LI_BCphaseA1_hsm":   209690
    number of sites unique to "Cb_LI_RphaseB1_hsm" although detectable in "Cb_LI_BCphaseA1_hsm":           36455
    number of "Cb_LI_RphaseB1_hsm" / "Cb_LI_BCphaseA1_hsm" common sites                        :          173235      (Expected:   2332; O/E:      74.3)

  Overlap index of "Cb_LI_BCphaseA1_hsm" with "Cb_LI_RphaseB1_hsm":          0.809
  Estimated number of "Cb_LI_BCphaseA1_hsm" = "Cb_LI_RphaseB1_hsm" sites (assuming sampling from one population):              235440
  Adjusted population size of "Cb_LI_BCphaseA1_hsm" = "Cb_LI_RphaseB1_hsm" sites (assuming all sites detectable):              359280 (  1.47x of observed)
```

```
Directory: PWC File: pwc-Cb_LI_BCphaseA1.vs.Cb_LI_RphaseB2.txt

Numbers of common and distinct sites comparing Cb_LI_BCphaseA1 versus Cb_LI_RphaseB2
================================================================================

  total number of potential sites:   26684764

  number of "Cb_LI_BCphaseA1_hsm"         sites:     198408
  number of "Cb_LI_RphaseB2_hsm"         sites:200707
    number of "Cb_LI_BCphaseA1_hsm"-unique sites:     33263
    number of common sites:        165145
    number of "Cb_LI_RphaseB2_hsm"-unique sites:      35562
  total number of "Cb_LI_BCphaseA1_hsm+Cb_LI_RphaseB2_hsm"-sites observed:   233970

  number of "Cb_LI_BCphaseA1_scd"        sites: 18501067 ( 69.33% of total)
  number of "Cb_LI_RphaseB2_scd"        sites:  18335682 ( 68.71% of total)
    number of "Cb_LI_BCphaseA1_scd"-unique sites:      1760590
    number of sites in common:      16740477   (Expected:  12712486; O/E:       1.3)
    number of "Cb_LI_RphaseB2_scd"-unique sites:       1595205
  total number of "Cb_LI_BCphaseA1_scd+Cb_LI_RphaseB2_scd"-sites observed:    20096272

  number of sites in "Cb_LI_BCphaseA1_hsm" that are not  detectable in "Cb_LI_RphaseB2_hsm":     7233
  number of sites in "Cb_LI_BCphaseA1_hsm" that are also detectable in "Cb_LI_RphaseB2_hsm":   191175
    number of sites unique to "Cb_LI_BCphaseA1_hsm" although detectable in "Cb_LI_RphaseB2_hsm":           26030
    number of "Cb_LI_BCphaseA1_hsm" / "Cb_LI_RphaseB2_hsm" common sites                        :          165145      (Expected:   2216; O/E:      74.5)

  number of sites in "Cb_LI_RphaseB2_hsm" that are not  detectable in "Cb_LI_BCphaseA1_hsm":     6654
  number of sites in "Cb_LI_RphaseB2_hsm" that are also detectable in "Cb_LI_BCphaseA1_hsm":   194053
    number of sites unique to "Cb_LI_RphaseB2_hsm" although detectable in "Cb_LI_BCphaseA1_hsm":           28908
    number of "Cb_LI_RphaseB2_hsm" / "Cb_LI_BCphaseA1_hsm" common sites                        :          165145      (Expected:   2216; O/E:      74.5)

  Overlap index of "Cb_LI_BCphaseA1_hsm" with "Cb_LI_RphaseB2_hsm":          0.762
  Estimated number of "Cb_LI_BCphaseA1_hsm" = "Cb_LI_RphaseB2_hsm" sites (assuming sampling from one population):              224639
  Adjusted population size of "Cb_LI_BCphaseA1_hsm" = "Cb_LI_RphaseB2_hsm" sites (assuming all sites detectable):              358081 (  1.53x of observed)
```

```
Directory: PWC File: pwc-Cb_LI_BCphaseA2.vs.Cb_LI_BCphaseB1.txt

Numbers of common and distinct sites comparing Cb_LI_BCphaseA2 versus Cb_LI_BCphaseB1
================================================================================

  total number of potential sites:  26684764

  number of "Cb_LI_BCphaseA2_hsm"         sites:     213448
  number of "Cb_LI_BCphaseB1_hsm"         sites:     226815
    number of "Cb_LI_BCphaseA2_hsm"-unique sites:     29469
    number of common sites:        183979
    number of "Cb_LI_BCphaseB1_hsm"-unique sites:     42836
  total number of "Cb_LI_BCphaseA2_hsm+Cb_LI_BCphaseB1_hsm"-sites observed:  256284

  number of "Cb_LI_BCphaseA2_scd"        sites: 18589454 ( 69.66% of total)
  number of "Cb_LI_BCphaseB1_scd"        sites: 20321376 ( 76.15% of total)
    number of "Cb_LI_BCphaseA2_scd"-unique sites:       765929
    number of sites in common:      17823525   (Expected:  14156516; O/E:       1.3)
    number of "Cb_LI_BCphaseB1_scd"-unique sites:      2497851
  total number of "Cb_LI_BCphaseA2_scd+Cb_LI_BCphaseB1_scd"-sites observed:   21087305

  number of sites in "Cb_LI_BCphaseA2_hsm" that are not  detectable in "Cb_LI_BCphaseB1_hsm":    3451
  number of sites in "Cb_LI_BCphaseA2_hsm" that are also detectable in "Cb_LI_BCphaseB1_hsm":  209997
    number of sites unique to "Cb_LI_BCphaseA2_hsm" although detectable in "Cb_LI_BCphaseB1_hsm":          26018
    number of "Cb_LI_BCphaseA2_hsm" / "Cb_LI_BCphaseB1_hsm" common sites                        :         183979      (Expected:   2550; O/E:      72.1)

  number of sites in "Cb_LI_BCphaseB1_hsm" that are not  detectable in "Cb_LI_BCphaseA2_hsm":   10377
  number of sites in "Cb_LI_BCphaseB1_hsm" that are also detectable in "Cb_LI_BCphaseA2_hsm":  216438
    number of sites unique to "Cb_LI_BCphaseB1_hsm" although detectable in "Cb_LI_BCphaseA2_hsm":          32459
    number of "Cb_LI_BCphaseB1_hsm" / "Cb_LI_BCphaseA2_hsm" common sites                        :         183979      (Expected:   2550; O/E:      72.1)

  Overlap index of "Cb_LI_BCphaseA2_hsm" with "Cb_LI_BCphaseB1_hsm":         0.782
  Estimated number of "Cb_LI_BCphaseA2_hsm" = "Cb_LI_BCphaseB1_hsm" sites (assuming sampling from one population):             247046
  Adjusted population size of "Cb_LI_BCphaseA2_hsm" = "Cb_LI_BCphaseB1_hsm" sites (assuming all sites detectable):             369869 (  1.44x of observed)
```

```
Directory: PWC File: pwc-Cb_LI_BCphaseA2.vs.Cb_LI_BCphaseB2.txt

Numbers of common and distinct sites comparing Cb_LI_BCphaseA2 versus Cb_LI_BCphaseB2
================================================================================

  total number of potential sites:  26684764

  number of "Cb_LI_BCphaseA2_hsm"         sites:     213448
  number of "Cb_LI_BCphaseB2_hsm"         sites:     212569
    number of "Cb_LI_BCphaseA2_hsm"-unique sites:     34388
    number of common sites:        179060
    number of "Cb_LI_BCphaseB2_hsm"-unique sites:     33509
  total number of "Cb_LI_BCphaseA2_hsm+Cb_LI_BCphaseB2_hsm"-sites observed:  246957

  number of "Cb_LI_BCphaseA2_scd"        sites: 18589454 ( 69.66% of total)
  number of "Cb_LI_BCphaseB2_scd"        sites: 19192933 ( 71.92% of total)
    number of "Cb_LI_BCphaseA2_scd"-unique sites:      1209274
    number of sites in common:      17380180   (Expected:  13370407; O/E:       1.3)
    number of "Cb_LI_BCphaseB2_scd"-unique sites:      1812753
  total number of "Cb_LI_BCphaseA2_scd+Cb_LI_BCphaseB2_scd"-sites observed:   20402207

  number of sites in "Cb_LI_BCphaseA2_hsm" that are not  detectable in "Cb_LI_BCphaseB2_hsm":    5044
  number of sites in "Cb_LI_BCphaseA2_hsm" that are also detectable in "Cb_LI_BCphaseB2_hsm":  208404
    number of sites unique to "Cb_LI_BCphaseA2_hsm" although detectable in "Cb_LI_BCphaseB2_hsm":          29344
    number of "Cb_LI_BCphaseA2_hsm" / "Cb_LI_BCphaseB2_hsm" common sites                        :         179060      (Expected:   2462; O/E:      72.7)

  number of sites in "Cb_LI_BCphaseB2_hsm" that are not  detectable in "Cb_LI_BCphaseA2_hsm":    7206
  number of sites in "Cb_LI_BCphaseB2_hsm" that are also detectable in "Cb_LI_BCphaseA2_hsm":  205363
    number of sites unique to "Cb_LI_BCphaseB2_hsm" although detectable in "Cb_LI_BCphaseA2_hsm":          26303
    number of "Cb_LI_BCphaseB2_hsm" / "Cb_LI_BCphaseA2_hsm" common sites                        :         179060      (Expected:   2462; O/E:      72.7)

  Overlap index of "Cb_LI_BCphaseA2_hsm" with "Cb_LI_BCphaseB2_hsm":         0.774
  Estimated number of "Cb_LI_BCphaseA2_hsm" = "Cb_LI_BCphaseB2_hsm" sites (assuming sampling from one population):             239017
  Adjusted population size of "Cb_LI_BCphaseA2_hsm" = "Cb_LI_BCphaseB2_hsm" sites (assuming all sites detectable):             366977 (  1.49x of observed)
```

```
Directory: PWC File: pwc-Cb_LI_BCphaseA2.vs.Cb_LI_RphaseA1.txt

Numbers of common and distinct sites comparing Cb_LI_BCphaseA2 versus Cb_LI_RphaseA1
================================================================================

  total number of potential sites:   26684764

  number of "Cb_LI_BCphaseA2_hsm"         sites:     213448
  number of "Cb_LI_RphaseA1_hsm"         sites:213610
    number of "Cb_LI_BCphaseA2_hsm"-unique sites:     35191
    number of common sites:         178257
    number of "Cb_LI_RphaseA1_hsm"-unique sites:      35353
  total number of "Cb_LI_BCphaseA2_hsm+Cb_LI_RphaseA1_hsm"-sites observed:   248801

  number of "Cb_LI_BCphaseA2_scd"        sites: 18589454 ( 69.66% of total)
  number of "Cb_LI_RphaseA1_scd"        sites:  19659217 ( 73.67% of total)
    number of "Cb_LI_BCphaseA2_scd"-unique sites:      1071008
    number of sites in common:      17518446   (Expected:  13695235; O/E:       1.3)
    number of "Cb_LI_RphaseA1_scd"-unique sites:       2140771
  total number of "Cb_LI_BCphaseA2_scd+Cb_LI_RphaseA1_scd"-sites observed:    20730225

  number of sites in "Cb_LI_BCphaseA2_hsm" that are not  detectable in "Cb_LI_RphaseA1_hsm":     4666
  number of sites in "Cb_LI_BCphaseA2_hsm" that are also detectable in "Cb_LI_RphaseA1_hsm":   208782
    number of sites unique to "Cb_LI_BCphaseA2_hsm" although detectable in "Cb_LI_RphaseA1_hsm":           30525
    number of "Cb_LI_BCphaseA2_hsm" / "Cb_LI_RphaseA1_hsm" common sites                        :          178257      (Expected:   2439; O/E:      73.1)

  number of sites in "Cb_LI_RphaseA1_hsm" that are not  detectable in "Cb_LI_BCphaseA2_hsm":     8926
  number of sites in "Cb_LI_RphaseA1_hsm" that are also detectable in "Cb_LI_BCphaseA2_hsm":   204684
    number of sites unique to "Cb_LI_RphaseA1_hsm" although detectable in "Cb_LI_BCphaseA2_hsm":           26427
    number of "Cb_LI_RphaseA1_hsm" / "Cb_LI_BCphaseA2_hsm" common sites                        :          178257      (Expected:   2439; O/E:      73.1)

  Overlap index of "Cb_LI_BCphaseA2_hsm" with "Cb_LI_RphaseA1_hsm":          0.773
  Estimated number of "Cb_LI_BCphaseA2_hsm" = "Cb_LI_RphaseA1_hsm" sites (assuming sampling from one population):              239734
  Adjusted population size of "Cb_LI_BCphaseA2_hsm" = "Cb_LI_RphaseA1_hsm" sites (assuming all sites detectable):              365173 (  1.47x of observed)
```

Directory: PWC File: pwc-Cb_LI_BCphaseA2.vs.Cb_LI_RphaseA2.txt

```
Numbers of common and distinct sites comparing Cb_LI_BCphaseA2 versus Cb_LI_RphaseA2
================================================================================

  total number of potential sites:  26684764

  number of "Cb_LI_BCphaseA2_hsm"         sites:     213448
  number of "Cb_LI_RphaseA2_hsm"         sites:265952
    number of "Cb_LI_BCphaseA2_hsm"-unique sites:     15126
    number of common sites:        198322
    number of "Cb_LI_RphaseA2_hsm"-unique sites:      67630
  total number of "Cb_LI_BCphaseA2_hsm+Cb_LI_RphaseA2_hsm"-sites observed:   281078

  number of "Cb_LI_BCphaseA2_scd"        sites: 18589454 ( 69.66% of total)
  number of "Cb_LI_RphaseA2_scd"        sites:  20435277 ( 76.58% of total)
    number of "Cb_LI_BCphaseA2_scd"-unique sites:       627181
    number of sites in common:      17962273   (Expected:  14235863; O/E:       1.3)
    number of "Cb_LI_RphaseA2_scd"-unique sites:       2473004
  total number of "Cb_LI_BCphaseA2_scd+Cb_LI_RphaseA2_scd"-sites observed:    21062458

  number of sites in "Cb_LI_BCphaseA2_hsm" that are not  detectable in "Cb_LI_RphaseA2_hsm":     1786
  number of sites in "Cb_LI_BCphaseA2_hsm" that are also detectable in "Cb_LI_RphaseA2_hsm":   211662
    number of sites unique to "Cb_LI_BCphaseA2_hsm" although detectable in "Cb_LI_RphaseA2_hsm":           13340
    number of "Cb_LI_BCphaseA2_hsm" / "Cb_LI_RphaseA2_hsm" common sites                        :          198322      (Expected:   2980; O/E:      66.6)

  number of sites in "Cb_LI_RphaseA2_hsm" that are not  detectable in "Cb_LI_BCphaseA2_hsm":    13053
  number of sites in "Cb_LI_RphaseA2_hsm" that are also detectable in "Cb_LI_BCphaseA2_hsm":   252899
    number of sites unique to "Cb_LI_RphaseA2_hsm" although detectable in "Cb_LI_BCphaseA2_hsm":           54577
    number of "Cb_LI_RphaseA2_hsm" / "Cb_LI_BCphaseA2_hsm" common sites                        :          198322      (Expected:   2980; O/E:      66.6)

  Overlap index of "Cb_LI_BCphaseA2_hsm" with "Cb_LI_RphaseA2_hsm":          0.890
  Estimated number of "Cb_LI_BCphaseA2_hsm" = "Cb_LI_RphaseA2_hsm" sites (assuming sampling from one population):              269910
  Adjusted population size of "Cb_LI_BCphaseA2_hsm" = "Cb_LI_RphaseA2_hsm" sites (assuming all sites detectable):              400979 (  1.43x of observed)
```

```
Directory: PWC File: pwc-Cb_LI_BCphaseA2.vs.Cb_LI_RphaseB1.txt

Numbers of common and distinct sites comparing Cb_LI_BCphaseA2 versus Cb_LI_RphaseB1
================================================================================

  total number of potential sites:   26684764

  number of "Cb_LI_BCphaseA2_hsm"         sites:     213448
  number of "Cb_LI_RphaseB1_hsm"         sites:218593
    number of "Cb_LI_BCphaseA2_hsm"-unique sites:     33848
    number of common sites:        179600
    number of "Cb_LI_RphaseB1_hsm"-unique sites:      38993
  total number of "Cb_LI_BCphaseA2_hsm+Cb_LI_RphaseB1_hsm"-sites observed:   252441

  number of "Cb_LI_BCphaseA2_scd"        sites: 18589454 ( 69.66% of total)
  number of "Cb_LI_RphaseB1_scd"        sites:  19576474 ( 73.36% of total)
    number of "Cb_LI_BCphaseA2_scd"-unique sites:      1107737
    number of sites in common:      17481717   (Expected:  13637593; O/E:       1.3)
    number of "Cb_LI_RphaseB1_scd"-unique sites:       2094757
  total number of "Cb_LI_BCphaseA2_scd+Cb_LI_RphaseB1_scd"-sites observed:    20684211

  number of sites in "Cb_LI_BCphaseA2_hsm" that are not  detectable in "Cb_LI_RphaseB1_hsm":     4515
  number of sites in "Cb_LI_BCphaseA2_hsm" that are also detectable in "Cb_LI_RphaseB1_hsm":   208933
    number of sites unique to "Cb_LI_BCphaseA2_hsm" although detectable in "Cb_LI_RphaseB1_hsm":           29333
    number of "Cb_LI_BCphaseA2_hsm" / "Cb_LI_RphaseB1_hsm" common sites                        :          179600      (Expected:   2502; O/E:      71.8)

  number of sites in "Cb_LI_RphaseB1_hsm" that are not  detectable in "Cb_LI_BCphaseA2_hsm":     9264
  number of sites in "Cb_LI_RphaseB1_hsm" that are also detectable in "Cb_LI_BCphaseA2_hsm":   209329
    number of sites unique to "Cb_LI_RphaseB1_hsm" although detectable in "Cb_LI_BCphaseA2_hsm":           29729
    number of "Cb_LI_RphaseB1_hsm" / "Cb_LI_BCphaseA2_hsm" common sites                        :          179600      (Expected:   2502; O/E:      71.8)

  Overlap index of "Cb_LI_BCphaseA2_hsm" with "Cb_LI_RphaseB1_hsm":          0.754
  Estimated number of "Cb_LI_BCphaseA2_hsm" = "Cb_LI_RphaseB1_hsm" sites (assuming sampling from one population):              243517
  Adjusted population size of "Cb_LI_BCphaseA2_hsm" = "Cb_LI_RphaseB1_hsm" sites (assuming all sites detectable):              371714 (  1.47x of observed)
```

```
Directory: PWC File: pwc-Cb_LI_BCphaseA2.vs.Cb_LI_RphaseB2.txt

Numbers of common and distinct sites comparing Cb_LI_BCphaseA2 versus Cb_LI_RphaseB2
================================================================================

  total number of potential sites:   26684764

  number of "Cb_LI_BCphaseA2_hsm"         sites:     213448
  number of "Cb_LI_RphaseB2_hsm"         sites:200707
    number of "Cb_LI_BCphaseA2_hsm"-unique sites:     39989
    number of common sites:        173459
    number of "Cb_LI_RphaseB2_hsm"-unique sites:      27248
  total number of "Cb_LI_BCphaseA2_hsm+Cb_LI_RphaseB2_hsm"-sites observed:   240696

  number of "Cb_LI_BCphaseA2_scd"        sites: 18589454 ( 69.66% of total)
  number of "Cb_LI_RphaseB2_scd"        sites:  18335682 ( 68.71% of total)
    number of "Cb_LI_BCphaseA2_scd"-unique sites:      1689835
    number of sites in common:      16899619   (Expected:  12773218; O/E:       1.3)
    number of "Cb_LI_RphaseB2_scd"-unique sites:       1436063
  total number of "Cb_LI_BCphaseA2_scd+Cb_LI_RphaseB2_scd"-sites observed:    20025517

  number of sites in "Cb_LI_BCphaseA2_hsm" that are not  detectable in "Cb_LI_RphaseB2_hsm":     6726
  number of sites in "Cb_LI_BCphaseA2_hsm" that are also detectable in "Cb_LI_RphaseB2_hsm":   206722
    number of sites unique to "Cb_LI_BCphaseA2_hsm" although detectable in "Cb_LI_RphaseB2_hsm":           33263
    number of "Cb_LI_BCphaseA2_hsm" / "Cb_LI_RphaseB2_hsm" common sites                        :          173459      (Expected:   2385; O/E:      72.7)

  number of sites in "Cb_LI_RphaseB2_hsm" that are not  detectable in "Cb_LI_BCphaseA2_hsm":     5769
  number of sites in "Cb_LI_RphaseB2_hsm" that are also detectable in "Cb_LI_BCphaseA2_hsm":   194938
    number of sites unique to "Cb_LI_RphaseB2_hsm" although detectable in "Cb_LI_BCphaseA2_hsm":           21479
    number of "Cb_LI_RphaseB2_hsm" / "Cb_LI_BCphaseA2_hsm" common sites                        :          173459      (Expected:   2385; O/E:      72.7)

  Overlap index of "Cb_LI_BCphaseA2_hsm" with "Cb_LI_RphaseB2_hsm":          0.806
  Estimated number of "Cb_LI_BCphaseA2_hsm" = "Cb_LI_RphaseB2_hsm" sites (assuming sampling from one population):              232320
  Adjusted population size of "Cb_LI_BCphaseA2_hsm" = "Cb_LI_RphaseB2_hsm" sites (assuming all sites detectable):              366837 (  1.52x of observed)
```

Directory: PWC File: pwc-Cb_LI_BCphaseB1.vs.Cb_LI_BCphaseB2.txt

```
Numbers of common and distinct sites comparing Cb_LI_BCphaseB1 versus Cb_LI_BCphaseB2
================================================================================

  total number of potential sites:   26684764

  number of "Cb_LI_BCphaseB1_hsm"         sites:     226815
  number of "Cb_LI_BCphaseB2_hsm"         sites:     212569
    number of "Cb_LI_BCphaseB1_hsm"-unique sites:     43589
    number of common sites:        183226
    number of "Cb_LI_BCphaseB2_hsm"-unique sites:     29343
  total number of "Cb_LI_BCphaseB1_hsm+Cb_LI_BCphaseB2_hsm"-sites observed:  256158

  number of "Cb_LI_BCphaseB1_scd"        sites: 20321376 ( 76.15% of total)
  number of "Cb_LI_BCphaseB2_scd"        sites: 19192933 ( 71.92% of total)
    number of "Cb_LI_BCphaseB1_scd"-unique sites:      2070149
    number of sites in common:      18251227   (Expected:  14616086; O/E:       1.2)
    number of "Cb_LI_BCphaseB2_scd"-unique sites:       941706
  total number of "Cb_LI_BCphaseB1_scd+Cb_LI_BCphaseB2_scd"-sites observed:   21263082

  number of sites in "Cb_LI_BCphaseB1_hsm" that are not  detectable in "Cb_LI_BCphaseB2_hsm":    8809
  number of sites in "Cb_LI_BCphaseB1_hsm" that are also detectable in "Cb_LI_BCphaseB2_hsm":  218006
    number of sites unique to "Cb_LI_BCphaseB1_hsm" although detectable in "Cb_LI_BCphaseB2_hsm":          34780
    number of "Cb_LI_BCphaseB1_hsm" / "Cb_LI_BCphaseB2_hsm" common sites                        :         183226      (Expected:   2489; O/E:      73.6)

  number of sites in "Cb_LI_BCphaseB2_hsm" that are not  detectable in "Cb_LI_BCphaseB1_hsm":    4196
  number of sites in "Cb_LI_BCphaseB2_hsm" that are also detectable in "Cb_LI_BCphaseB1_hsm":  208373
    number of sites unique to "Cb_LI_BCphaseB2_hsm" although detectable in "Cb_LI_BCphaseB1_hsm":          25147
    number of "Cb_LI_BCphaseB2_hsm" / "Cb_LI_BCphaseB1_hsm" common sites                        :         183226      (Expected:   2489; O/E:      73.6)

  Overlap index of "Cb_LI_BCphaseB1_hsm" with "Cb_LI_BCphaseB2_hsm":         0.788
  Estimated number of "Cb_LI_BCphaseB1_hsm" = "Cb_LI_BCphaseB2_hsm" sites (assuming sampling from one population):             247926
  Adjusted population size of "Cb_LI_BCphaseB1_hsm" = "Cb_LI_BCphaseB2_hsm" sites (assuming all sites detectable):             362488 (  1.42x of observed)
```

```
Directory: PWC File: pwc-Cb_LI_BCphaseB1.vs.Cb_LI_RphaseA1.txt

Numbers of common and distinct sites comparing Cb_LI_BCphaseB1 versus Cb_LI_RphaseA1
================================================================================

  total number of potential sites:   26684764

  number of "Cb_LI_BCphaseB1_hsm"         sites:     226815
  number of "Cb_LI_RphaseA1_hsm"         sites: 213610
    number of "Cb_LI_BCphaseB1_hsm"-unique sites:     41481
    number of common sites:        185334
    number of "Cb_LI_RphaseA1_hsm"-unique sites:      28276
  total number of "Cb_LI_BCphaseB1_hsm+Cb_LI_RphaseA1_hsm"-sites observed:   255091

  number of "Cb_LI_BCphaseB1_scd"        sites: 20321376 ( 76.15% of total)
  number of "Cb_LI_RphaseA1_scd"        sites:  19659217 ( 73.67% of total)
    number of "Cb_LI_BCphaseB1_scd"-unique sites:      1684770
    number of sites in common:      18636606   (Expected:  14971178; O/E:       1.2)
    number of "Cb_LI_RphaseA1_scd"-unique sites:       1022611
  total number of "Cb_LI_BCphaseB1_scd+Cb_LI_RphaseA1_scd"-sites observed:    21343987

  number of sites in "Cb_LI_BCphaseB1_hsm" that are not  detectable in "Cb_LI_RphaseA1_hsm":     6978
  number of sites in "Cb_LI_BCphaseB1_hsm" that are also detectable in "Cb_LI_RphaseA1_hsm":   219837
    number of sites unique to "Cb_LI_BCphaseB1_hsm" although detectable in "Cb_LI_RphaseA1_hsm":           34503
    number of "Cb_LI_BCphaseB1_hsm" / "Cb_LI_RphaseA1_hsm" common sites                        :          185334      (Expected:   2465; O/E:      75.2)

  number of sites in "Cb_LI_RphaseA1_hsm" that are not  detectable in "Cb_LI_BCphaseB1_hsm":     4604
  number of sites in "Cb_LI_RphaseA1_hsm" that are also detectable in "Cb_LI_BCphaseB1_hsm":   209006
    number of sites unique to "Cb_LI_RphaseA1_hsm" although detectable in "Cb_LI_BCphaseB1_hsm":           23672
    number of "Cb_LI_RphaseA1_hsm" / "Cb_LI_BCphaseB1_hsm" common sites                        :          185334      (Expected:   2465; O/E:      75.2)

  Overlap index of "Cb_LI_BCphaseB1_hsm" with "Cb_LI_RphaseA1_hsm":          0.801
  Estimated number of "Cb_LI_BCphaseB1_hsm" = "Cb_LI_RphaseA1_hsm" sites (assuming sampling from one population):              247916
  Adjusted population size of "Cb_LI_BCphaseB1_hsm" = "Cb_LI_RphaseA1_hsm" sites (assuming all sites detectable):              354978 (  1.39x of observed)
```

```
Directory: PWC File: pwc-Cb_LI_BCphaseB1.vs.Cb_LI_RphaseA2.txt

Numbers of common and distinct sites comparing Cb_LI_BCphaseB1 versus Cb_LI_RphaseA2
================================================================================

  total number of potential sites:  26684764

  number of "Cb_LI_BCphaseB1_hsm"         sites:     226815
  number of "Cb_LI_RphaseA2_hsm"         sites:265952
    number of "Cb_LI_BCphaseB1_hsm"-unique sites:     22274
    number of common sites:        204541
    number of "Cb_LI_RphaseA2_hsm"-unique sites:      61411
  total number of "Cb_LI_BCphaseB1_hsm+Cb_LI_RphaseA2_hsm"-sites observed:   288226

  number of "Cb_LI_BCphaseB1_scd"        sites: 20321376 ( 76.15% of total)
  number of "Cb_LI_RphaseA2_scd"        sites:  20435277 ( 76.58% of total)
    number of "Cb_LI_BCphaseB1_scd"-unique sites:      1247854
    number of sites in common:      19073522   (Expected:  15562174; O/E:       1.2)
    number of "Cb_LI_RphaseA2_scd"-unique sites:       1361755
  total number of "Cb_LI_BCphaseB1_scd+Cb_LI_RphaseA2_scd"-sites observed:    21683131

  number of sites in "Cb_LI_BCphaseB1_hsm" that are not  detectable in "Cb_LI_RphaseA2_hsm":     3797
  number of sites in "Cb_LI_BCphaseB1_hsm" that are also detectable in "Cb_LI_RphaseA2_hsm":   223018
    number of sites unique to "Cb_LI_BCphaseB1_hsm" although detectable in "Cb_LI_RphaseA2_hsm":           18477
    number of "Cb_LI_BCphaseB1_hsm" / "Cb_LI_RphaseA2_hsm" common sites                        :          204541      (Expected:   3016; O/E:      67.8)

  number of sites in "Cb_LI_RphaseA2_hsm" that are not  detectable in "Cb_LI_BCphaseB1_hsm":     7997
  number of sites in "Cb_LI_RphaseA2_hsm" that are also detectable in "Cb_LI_BCphaseB1_hsm":   257955
    number of sites unique to "Cb_LI_RphaseA2_hsm" although detectable in "Cb_LI_BCphaseB1_hsm":           53414
    number of "Cb_LI_RphaseA2_hsm" / "Cb_LI_BCphaseB1_hsm" common sites                        :          204541      (Expected:   3016; O/E:      67.8)

  Overlap index of "Cb_LI_BCphaseB1_hsm" with "Cb_LI_RphaseA2_hsm":          0.856
  Estimated number of "Cb_LI_BCphaseB1_hsm" = "Cb_LI_RphaseA2_hsm" sites (assuming sampling from one population):              281257
  Adjusted population size of "Cb_LI_BCphaseB1_hsm" = "Cb_LI_RphaseA2_hsm" sites (assuming all sites detectable):              393492 (  1.37x of observed)
```

```
Directory: PWC File: pwc-Cb_LI_BCphaseB1.vs.Cb_LI_RphaseB1.txt

Numbers of common and distinct sites comparing Cb_LI_BCphaseB1 versus Cb_LI_RphaseB1
================================================================================

  total number of potential sites:  26684764

  number of "Cb_LI_BCphaseB1_hsm"         sites:     226815
  number of "Cb_LI_RphaseB1_hsm"         sites:218593
    number of "Cb_LI_BCphaseB1_hsm"-unique sites:     39842
    number of common sites:        186973
    number of "Cb_LI_RphaseB1_hsm"-unique sites:      31620
  total number of "Cb_LI_BCphaseB1_hsm+Cb_LI_RphaseB1_hsm"-sites observed:   258435

  number of "Cb_LI_BCphaseB1_scd"        sites: 20321376 ( 76.15% of total)
  number of "Cb_LI_RphaseB1_scd"        sites:  19576474 ( 73.36% of total)
    number of "Cb_LI_BCphaseB1_scd"-unique sites:      1745022
    number of sites in common:      18576354   (Expected:  14908166; O/E:       1.2)
    number of "Cb_LI_RphaseB1_scd"-unique sites:       1000120
  total number of "Cb_LI_BCphaseB1_scd+Cb_LI_RphaseB1_scd"-sites observed:    21321496

  number of sites in "Cb_LI_BCphaseB1_hsm" that are not  detectable in "Cb_LI_RphaseB1_hsm":     6736
  number of sites in "Cb_LI_BCphaseB1_hsm" that are also detectable in "Cb_LI_RphaseB1_hsm":   220079
    number of sites unique to "Cb_LI_BCphaseB1_hsm" although detectable in "Cb_LI_RphaseB1_hsm":           33106
    number of "Cb_LI_BCphaseB1_hsm" / "Cb_LI_RphaseB1_hsm" common sites                        :          186973      (Expected:   2534; O/E:      73.8)

  number of sites in "Cb_LI_RphaseB1_hsm" that are not  detectable in "Cb_LI_BCphaseB1_hsm":     4711
  number of sites in "Cb_LI_RphaseB1_hsm" that are also detectable in "Cb_LI_BCphaseB1_hsm":   213882
    number of sites unique to "Cb_LI_RphaseB1_hsm" although detectable in "Cb_LI_BCphaseB1_hsm":           26909
    number of "Cb_LI_RphaseB1_hsm" / "Cb_LI_BCphaseB1_hsm" common sites                        :          186973      (Expected:   2534; O/E:      73.8)

  Overlap index of "Cb_LI_BCphaseB1_hsm" with "Cb_LI_RphaseB1_hsm":          0.779
  Estimated number of "Cb_LI_BCphaseB1_hsm" = "Cb_LI_RphaseB1_hsm" sites (assuming sampling from one population):              251753
  Adjusted population size of "Cb_LI_BCphaseB1_hsm" = "Cb_LI_RphaseB1_hsm" sites (assuming all sites detectable):              361640 (  1.40x of observed)
```

```
Directory: PWC File: pwc-Cb_LI_BCphaseB1.vs.Cb_LI_RphaseB2.txt

Numbers of common and distinct sites comparing Cb_LI_BCphaseB1 versus Cb_LI_RphaseB2
================================================================================

  total number of potential sites:   26684764

  number of "Cb_LI_BCphaseB1_hsm"         sites:     226815
  number of "Cb_LI_RphaseB2_hsm"         sites:200707
    number of "Cb_LI_BCphaseB1_hsm"-unique sites:     50149
    number of common sites:        176666
    number of "Cb_LI_RphaseB2_hsm"-unique sites:      24041
  total number of "Cb_LI_BCphaseB1_hsm+Cb_LI_RphaseB2_hsm"-sites observed:   250856

  number of "Cb_LI_BCphaseB1_scd"        sites: 20321376 ( 76.15% of total)
  number of "Cb_LI_RphaseB2_scd"        sites:  18335682 ( 68.71% of total)
    number of "Cb_LI_BCphaseB1_scd"-unique sites:      2719354
    number of sites in common:      17602022   (Expected:  13963260; O/E:       1.3)
    number of "Cb_LI_RphaseB2_scd"-unique sites:        733660
  total number of "Cb_LI_BCphaseB1_scd+Cb_LI_RphaseB2_scd"-sites observed:    21055036

  number of sites in "Cb_LI_BCphaseB1_hsm" that are not  detectable in "Cb_LI_RphaseB2_hsm":    11250
  number of sites in "Cb_LI_BCphaseB1_hsm" that are also detectable in "Cb_LI_RphaseB2_hsm":   215565
    number of sites unique to "Cb_LI_BCphaseB1_hsm" although detectable in "Cb_LI_RphaseB2_hsm":           38899
    number of "Cb_LI_BCphaseB1_hsm" / "Cb_LI_RphaseB2_hsm" common sites                        :          176666      (Expected:   2416; O/E:      73.1)

  number of sites in "Cb_LI_RphaseB2_hsm" that are not  detectable in "Cb_LI_BCphaseB1_hsm":     3429
  number of sites in "Cb_LI_RphaseB2_hsm" that are also detectable in "Cb_LI_BCphaseB1_hsm":   197278
    number of sites unique to "Cb_LI_RphaseB2_hsm" although detectable in "Cb_LI_BCphaseB1_hsm":           20612
    number of "Cb_LI_RphaseB2_hsm" / "Cb_LI_BCphaseB1_hsm" common sites                        :          176666      (Expected:   2416; O/E:      73.1)

  Overlap index of "Cb_LI_BCphaseB1_hsm" with "Cb_LI_RphaseB2_hsm":          0.817
  Estimated number of "Cb_LI_BCphaseB1_hsm" = "Cb_LI_RphaseB2_hsm" sites (assuming sampling from one population):              240715
  Adjusted population size of "Cb_LI_BCphaseB1_hsm" = "Cb_LI_RphaseB2_hsm" sites (assuming all sites detectable):              364926 (  1.45x of observed)
```

```
Directory: PWC File: pwc-Cb_LI_BCphaseB2.vs.Cb_LI_RphaseA1.txt

Numbers of common and distinct sites comparing Cb_LI_BCphaseB2 versus Cb_LI_RphaseA1
================================================================================

  total number of potential sites:  26684764

  number of "Cb_LI_BCphaseB2_hsm"         sites:     212569
  number of "Cb_LI_RphaseA1_hsm"         sites:213610
    number of "Cb_LI_BCphaseB2_hsm"-unique sites:     34924
    number of common sites:        177645
    number of "Cb_LI_RphaseA1_hsm"-unique sites:      35965
  total number of "Cb_LI_BCphaseB2_hsm+Cb_LI_RphaseA1_hsm"-sites observed:   248534

  number of "Cb_LI_BCphaseB2_scd"        sites: 19192933 ( 71.92% of total)
  number of "Cb_LI_RphaseA1_scd"        sites:  19659217 ( 73.67% of total)
    number of "Cb_LI_BCphaseB2_scd"-unique sites:      1286227
    number of sites in common:      17906706   (Expected:  14139830; O/E:       1.3)
    number of "Cb_LI_RphaseA1_scd"-unique sites:       1752511
  total number of "Cb_LI_BCphaseB2_scd+Cb_LI_RphaseA1_scd"-sites observed:    20945444

  number of sites in "Cb_LI_BCphaseB2_hsm" that are not  detectable in "Cb_LI_RphaseA1_hsm":     5500
  number of sites in "Cb_LI_BCphaseB2_hsm" that are also detectable in "Cb_LI_RphaseA1_hsm":   207069
    number of sites unique to "Cb_LI_BCphaseB2_hsm" although detectable in "Cb_LI_RphaseA1_hsm":           29424
    number of "Cb_LI_BCphaseB2_hsm" / "Cb_LI_RphaseA1_hsm" common sites                        :          177645      (Expected:   2384; O/E:      74.5)

  number of sites in "Cb_LI_RphaseA1_hsm" that are not  detectable in "Cb_LI_BCphaseB2_hsm":     7490
  number of sites in "Cb_LI_RphaseA1_hsm" that are also detectable in "Cb_LI_BCphaseB2_hsm":   206120
    number of sites unique to "Cb_LI_RphaseA1_hsm" although detectable in "Cb_LI_BCphaseB2_hsm":           28475
    number of "Cb_LI_RphaseA1_hsm" / "Cb_LI_BCphaseB2_hsm" common sites                        :          177645      (Expected:   2384; O/E:      74.5)

  Overlap index of "Cb_LI_BCphaseB2_hsm" with "Cb_LI_RphaseA1_hsm":          0.758
  Estimated number of "Cb_LI_BCphaseB2_hsm" = "Cb_LI_RphaseA1_hsm" sites (assuming sampling from one population):              240260
  Adjusted population size of "Cb_LI_BCphaseB2_hsm" = "Cb_LI_RphaseA1_hsm" sites (assuming all sites detectable):              358039 (  1.44x of observed)
```

```
Directory: PWC File: pwc-Cb_LI_BCphaseB2.vs.Cb_LI_RphaseA2.txt

Numbers of common and distinct sites comparing Cb_LI_BCphaseB2 versus Cb_LI_RphaseA2
================================================================================

  total number of potential sites:  26684764

  number of "Cb_LI_BCphaseB2_hsm"         sites:     212569
  number of "Cb_LI_RphaseA2_hsm"         sites:265952
    number of "Cb_LI_BCphaseB2_hsm"-unique sites:     15763
    number of common sites:        196806
    number of "Cb_LI_RphaseA2_hsm"-unique sites:      69146
  total number of "Cb_LI_BCphaseB2_hsm+Cb_LI_RphaseA2_hsm"-sites observed:   281715

  number of "Cb_LI_BCphaseB2_scd"        sites: 19192933 ( 71.92% of total)
  number of "Cb_LI_RphaseA2_scd"        sites:  20435277 ( 76.58% of total)
    number of "Cb_LI_BCphaseB2_scd"-unique sites:       797027
    number of sites in common:      18395906   (Expected:  14698009; O/E:       1.3)
    number of "Cb_LI_RphaseA2_scd"-unique sites:       2039371
  total number of "Cb_LI_BCphaseB2_scd+Cb_LI_RphaseA2_scd"-sites observed:    21232304

  number of sites in "Cb_LI_BCphaseB2_hsm" that are not  detectable in "Cb_LI_RphaseA2_hsm":     2154
  number of sites in "Cb_LI_BCphaseB2_hsm" that are also detectable in "Cb_LI_RphaseA2_hsm":   210415
    number of sites unique to "Cb_LI_BCphaseB2_hsm" although detectable in "Cb_LI_RphaseA2_hsm":           13609
    number of "Cb_LI_BCphaseB2_hsm" / "Cb_LI_RphaseA2_hsm" common sites                        :          196806      (Expected:   2917; O/E:      67.5)

  number of sites in "Cb_LI_RphaseA2_hsm" that are not  detectable in "Cb_LI_BCphaseB2_hsm":    10896
  number of sites in "Cb_LI_RphaseA2_hsm" that are also detectable in "Cb_LI_BCphaseB2_hsm":   255056
    number of sites unique to "Cb_LI_RphaseA2_hsm" although detectable in "Cb_LI_BCphaseB2_hsm":           58250
    number of "Cb_LI_RphaseA2_hsm" / "Cb_LI_BCphaseB2_hsm" common sites                        :          196806      (Expected:   2917; O/E:      67.5)

  Overlap index of "Cb_LI_BCphaseB2_hsm" with "Cb_LI_RphaseA2_hsm":          0.888
  Estimated number of "Cb_LI_BCphaseB2_hsm" = "Cb_LI_RphaseA2_hsm" sites (assuming sampling from one population):              272693
  Adjusted population size of "Cb_LI_BCphaseB2_hsm" = "Cb_LI_RphaseA2_hsm" sites (assuming all sites detectable):              395563 (  1.40x of observed)
```

```
Directory: PWC File: pwc-Cb_LI_BCphaseB2.vs.Cb_LI_RphaseB1.txt

Numbers of common and distinct sites comparing Cb_LI_BCphaseB2 versus Cb_LI_RphaseB1
================================================================================

  total number of potential sites:   26684764

  number of "Cb_LI_BCphaseB2_hsm"         sites:     212569
  number of "Cb_LI_RphaseB1_hsm"         sites: 218593
    number of "Cb_LI_BCphaseB2_hsm"-unique sites:     32301
    number of common sites:        180268
    number of "Cb_LI_RphaseB1_hsm"-unique sites:      38325
  total number of "Cb_LI_BCphaseB2_hsm+Cb_LI_RphaseB1_hsm"-sites observed:   250894

  number of "Cb_LI_BCphaseB2_scd"        sites: 19192933 ( 71.92% of total)
  number of "Cb_LI_RphaseB1_scd"        sites:  19576474 ( 73.36% of total)
    number of "Cb_LI_BCphaseB2_scd"-unique sites:      1305678
    number of sites in common:      17887255   (Expected:  14080318; O/E:       1.3)
    number of "Cb_LI_RphaseB1_scd"-unique sites:       1689219
  total number of "Cb_LI_BCphaseB2_scd+Cb_LI_RphaseB1_scd"-sites observed:    20882152

  number of sites in "Cb_LI_BCphaseB2_hsm" that are not  detectable in "Cb_LI_RphaseB1_hsm":     4970
  number of sites in "Cb_LI_BCphaseB2_hsm" that are also detectable in "Cb_LI_RphaseB1_hsm":   207599
    number of sites unique to "Cb_LI_BCphaseB2_hsm" although detectable in "Cb_LI_RphaseB1_hsm":           27331
    number of "Cb_LI_BCphaseB2_hsm" / "Cb_LI_RphaseB1_hsm" common sites                        :          180268      (Expected:   2451; O/E:      73.5)

  number of sites in "Cb_LI_RphaseB1_hsm" that are not  detectable in "Cb_LI_BCphaseB2_hsm":     7443
  number of sites in "Cb_LI_RphaseB1_hsm" that are also detectable in "Cb_LI_BCphaseB2_hsm":   211150
    number of sites unique to "Cb_LI_RphaseB1_hsm" although detectable in "Cb_LI_BCphaseB2_hsm":           30882
    number of "Cb_LI_RphaseB1_hsm" / "Cb_LI_BCphaseB2_hsm" common sites                        :          180268      (Expected:   2451; O/E:      73.5)

  Overlap index of "Cb_LI_BCphaseB2_hsm" with "Cb_LI_RphaseB1_hsm":          0.769
  Estimated number of "Cb_LI_BCphaseB2_hsm" = "Cb_LI_RphaseB1_hsm" sites (assuming sampling from one population):              243163
  Adjusted population size of "Cb_LI_BCphaseB2_hsm" = "Cb_LI_RphaseB1_hsm" sites (assuming all sites detectable):              362758 (  1.45x of observed)
```

```
Directory: PWC File: pwc-Cb_LI_BCphaseB2.vs.Cb_LI_RphaseB2.txt

Numbers of common and distinct sites comparing Cb_LI_BCphaseB2 versus Cb_LI_RphaseB2
================================================================================

  total number of potential sites:   26684764

  number of "Cb_LI_BCphaseB2_hsm"         sites:     212569
  number of "Cb_LI_RphaseB2_hsm"         sites:200707
    number of "Cb_LI_BCphaseB2_hsm"-unique sites:     38811
    number of common sites:        173758
    number of "Cb_LI_RphaseB2_hsm"-unique sites:      26949
  total number of "Cb_LI_BCphaseB2_hsm+Cb_LI_RphaseB2_hsm"-sites observed:   239518

  number of "Cb_LI_BCphaseB2_scd"        sites: 19192933 ( 71.92% of total)
  number of "Cb_LI_RphaseB2_scd"        sites:  18335682 ( 68.71% of total)
    number of "Cb_LI_BCphaseB2_scd"-unique sites:      1982156
    number of sites in common:      17210777   (Expected:  13187882; O/E:       1.3)
    number of "Cb_LI_RphaseB2_scd"-unique sites:       1124905
  total number of "Cb_LI_BCphaseB2_scd+Cb_LI_RphaseB2_scd"-sites observed:    20317838

  number of sites in "Cb_LI_BCphaseB2_hsm" that are not  detectable in "Cb_LI_RphaseB2_hsm":     7532
  number of sites in "Cb_LI_BCphaseB2_hsm" that are also detectable in "Cb_LI_RphaseB2_hsm":   205037
    number of sites unique to "Cb_LI_BCphaseB2_hsm" although detectable in "Cb_LI_RphaseB2_hsm":           31279
    number of "Cb_LI_BCphaseB2_hsm" / "Cb_LI_RphaseB2_hsm" common sites                        :          173758      (Expected:   2337; O/E:      74.4)

  number of sites in "Cb_LI_RphaseB2_hsm" that are not  detectable in "Cb_LI_BCphaseB2_hsm":     4505
  number of sites in "Cb_LI_RphaseB2_hsm" that are also detectable in "Cb_LI_BCphaseB2_hsm":   196202
    number of sites unique to "Cb_LI_RphaseB2_hsm" although detectable in "Cb_LI_BCphaseB2_hsm":           22444
    number of "Cb_LI_RphaseB2_hsm" / "Cb_LI_BCphaseB2_hsm" common sites                        :          173758      (Expected:   2337; O/E:      74.4)

  Overlap index of "Cb_LI_BCphaseB2_hsm" with "Cb_LI_RphaseB2_hsm":          0.798
  Estimated number of "Cb_LI_BCphaseB2_hsm" = "Cb_LI_RphaseB2_hsm" sites (assuming sampling from one population):              231521
  Adjusted population size of "Cb_LI_BCphaseB2_hsm" = "Cb_LI_RphaseB2_hsm" sites (assuming all sites detectable):              358966 (  1.50x of observed)
```

```
Directory: PWC File: pwc-Cb_LI_RphaseA1.vs.Cb_LI_RphaseA2.txt

Numbers of common and distinct sites comparing Cb_LI_RphaseA1 versus Cb_LI_RphaseA2
================================================================================

  total number of potential sites:  26684764

  number of "Cb_LI_RphaseA1_hsm"         sites:213610
  number of "Cb_LI_RphaseA2_hsm"         sites:265952
    number of "Cb_LI_RphaseA1_hsm"-unique sites:      17654
    number of common sites:        195956
    number of "Cb_LI_RphaseA2_hsm"-unique sites:      69996
  total number of "Cb_LI_RphaseA1_hsm+Cb_LI_RphaseA2_hsm"-sites observed:    283606

  number of "Cb_LI_RphaseA1_scd"        sites:  19659217 ( 73.67% of total)
  number of "Cb_LI_RphaseA2_scd"        sites:  20435277 ( 76.58% of total)
    number of "Cb_LI_RphaseA1_scd"-unique sites:       1024788
    number of sites in common:      18634429   (Expected:  15055091; O/E:       1.2)
    number of "Cb_LI_RphaseA2_scd"-unique sites:       1800848
  total number of "Cb_LI_RphaseA1_scd+Cb_LI_RphaseA2_scd"-sites observed:     21460065

  number of sites in "Cb_LI_RphaseA1_hsm" that are not  detectable in "Cb_LI_RphaseA2_hsm":      3267
  number of sites in "Cb_LI_RphaseA1_hsm" that are also detectable in "Cb_LI_RphaseA2_hsm":    210343
    number of sites unique to "Cb_LI_RphaseA1_hsm" although detectable in "Cb_LI_RphaseA2_hsm":            14387
    number of "Cb_LI_RphaseA1_hsm" / "Cb_LI_RphaseA2_hsm" common sites                        :           195956      (Expected:   2886; O/E:      67.9)

  number of sites in "Cb_LI_RphaseA2_hsm" that are not  detectable in "Cb_LI_RphaseA1_hsm":     10240
  number of sites in "Cb_LI_RphaseA2_hsm" that are also detectable in "Cb_LI_RphaseA1_hsm":    255712
    number of sites unique to "Cb_LI_RphaseA2_hsm" although detectable in "Cb_LI_RphaseA1_hsm":            59756
    number of "Cb_LI_RphaseA2_hsm" / "Cb_LI_RphaseA1_hsm" common sites                        :           195956      (Expected:   2886; O/E:      67.9)

  Overlap index of "Cb_LI_RphaseA1_hsm" with "Cb_LI_RphaseA2_hsm":           0.882
  Estimated number of "Cb_LI_RphaseA1_hsm" = "Cb_LI_RphaseA2_hsm" sites (assuming sampling from one population):         274486
  Adjusted population size of "Cb_LI_RphaseA1_hsm" = "Cb_LI_RphaseA2_hsm" sites (assuming all sites detectable):         393068 (  1.39x of observed)
```

```
Directory: PWC File: pwc-Cb_LI_RphaseA1.vs.Cb_LI_RphaseB1.txt

Numbers of common and distinct sites comparing Cb_LI_RphaseA1 versus Cb_LI_RphaseB1
================================================================================

  total number of potential sites:  26684764

  number of "Cb_LI_RphaseA1_hsm"         sites:213610
  number of "Cb_LI_RphaseB1_hsm"         sites:218593
    number of "Cb_LI_RphaseA1_hsm"-unique sites:      32439
    number of common sites:        181171
    number of "Cb_LI_RphaseB1_hsm"-unique sites:      37422
  total number of "Cb_LI_RphaseA1_hsm+Cb_LI_RphaseB1_hsm"-sites observed:    251032

  number of "Cb_LI_RphaseA1_scd"        sites:  19659217 ( 73.67% of total)
  number of "Cb_LI_RphaseB1_scd"        sites:  19576474 ( 73.36% of total)
    number of "Cb_LI_RphaseA1_scd"-unique sites:       1453473
    number of sites in common:      18205744   (Expected:  14422393; O/E:       1.3)
    number of "Cb_LI_RphaseB1_scd"-unique sites:       1370730
  total number of "Cb_LI_RphaseA1_scd+Cb_LI_RphaseB1_scd"-sites observed:     21029947

  number of sites in "Cb_LI_RphaseA1_hsm" that are not  detectable in "Cb_LI_RphaseB1_hsm":      5672
  number of sites in "Cb_LI_RphaseA1_hsm" that are also detectable in "Cb_LI_RphaseB1_hsm":    207938
    number of sites unique to "Cb_LI_RphaseA1_hsm" although detectable in "Cb_LI_RphaseB1_hsm":            26767
    number of "Cb_LI_RphaseA1_hsm" / "Cb_LI_RphaseB1_hsm" common sites                        :           181171      (Expected:   2427; O/E:      74.6)

  number of sites in "Cb_LI_RphaseB1_hsm" that are not  detectable in "Cb_LI_RphaseA1_hsm":      6134
  number of sites in "Cb_LI_RphaseB1_hsm" that are also detectable in "Cb_LI_RphaseA1_hsm":    212459
    number of sites unique to "Cb_LI_RphaseB1_hsm" although detectable in "Cb_LI_RphaseA1_hsm":            31288
    number of "Cb_LI_RphaseB1_hsm" / "Cb_LI_RphaseA1_hsm" common sites                        :           181171      (Expected:   2427; O/E:      74.6)

  Overlap index of "Cb_LI_RphaseA1_hsm" with "Cb_LI_RphaseB1_hsm":           0.774
  Estimated number of "Cb_LI_RphaseA1_hsm" = "Cb_LI_RphaseB1_hsm" sites (assuming sampling from one population):         243849
  Adjusted population size of "Cb_LI_RphaseA1_hsm" = "Cb_LI_RphaseB1_hsm" sites (assuming all sites detectable):         357417 (  1.42x of observed)
```

```
Directory: PWC File: pwc-Cb_LI_RphaseA1.vs.Cb_LI_RphaseB2.txt

Numbers of common and distinct sites comparing Cb_LI_RphaseA1 versus Cb_LI_RphaseB2
================================================================================

  total number of potential sites:   26684764

  number of "Cb_LI_RphaseA1_hsm"         sites: 213610
  number of "Cb_LI_RphaseB2_hsm"         sites: 200707
    number of "Cb_LI_RphaseA1_hsm"-unique sites:      42005
    number of common sites:         171605
    number of "Cb_LI_RphaseB2_hsm"-unique sites:      29102
  total number of "Cb_LI_RphaseA1_hsm+Cb_LI_RphaseB2_hsm"-sites observed:    242712

  number of "Cb_LI_RphaseA1_scd"        sites:  19659217 ( 73.67% of total)
  number of "Cb_LI_RphaseB2_scd"        sites:  18335682 ( 68.71% of total)
    number of "Cb_LI_RphaseA1_scd"-unique sites:       2346831
    number of sites in common:       17312386   (Expected:  13508276; O/E:       1.3)
    number of "Cb_LI_RphaseB2_scd"-unique sites:       1023296
  total number of "Cb_LI_RphaseA1_scd+Cb_LI_RphaseB2_scd"-sites observed:     20682513

  number of sites in "Cb_LI_RphaseA1_hsm" that are not  detectable in "Cb_LI_RphaseB2_hsm":      9760
  number of sites in "Cb_LI_RphaseA1_hsm" that are also detectable in "Cb_LI_RphaseB2_hsm":    203850
    number of sites unique to "Cb_LI_RphaseA1_hsm" although detectable in "Cb_LI_RphaseB2_hsm":            32245
    number of "Cb_LI_RphaseA1_hsm" / "Cb_LI_RphaseB2_hsm" common sites                        :           171605      (Expected:   2310; O/E:      74.3)

  number of sites in "Cb_LI_RphaseB2_hsm" that are not  detectable in "Cb_LI_RphaseA1_hsm":      4495
  number of sites in "Cb_LI_RphaseB2_hsm" that are also detectable in "Cb_LI_RphaseA1_hsm":    196212
    number of sites unique to "Cb_LI_RphaseB2_hsm" although detectable in "Cb_LI_RphaseA1_hsm":            24607
    number of "Cb_LI_RphaseB2_hsm" / "Cb_LI_RphaseA1_hsm" common sites                        :           171605      (Expected:   2310; O/E:      74.3)

  Overlap index of "Cb_LI_RphaseA1_hsm" with "Cb_LI_RphaseB2_hsm":           0.780
  Estimated number of "Cb_LI_RphaseA1_hsm" = "Cb_LI_RphaseB2_hsm" sites (assuming sampling from one population):         233081
  Adjusted population size of "Cb_LI_RphaseA1_hsm" = "Cb_LI_RphaseB2_hsm" sites (assuming all sites detectable):         359263 (  1.48x of observed)
```

```
Directory: PWC File: pwc-Cb_LI_RphaseA2.vs.Cb_LI_RphaseB1.txt

Numbers of common and distinct sites comparing Cb_LI_RphaseA2 versus Cb_LI_RphaseB1
================================================================================

  total number of potential sites:   26684764

  number of "Cb_LI_RphaseA2_hsm"         sites: 265952
  number of "Cb_LI_RphaseB1_hsm"         sites: 218593
    number of "Cb_LI_RphaseA2_hsm"-unique sites:      67311
    number of common sites:        198641
    number of "Cb_LI_RphaseB1_hsm"-unique sites:      19952
  total number of "Cb_LI_RphaseA2_hsm+Cb_LI_RphaseB1_hsm"-sites observed:    285904

  number of "Cb_LI_RphaseA2_scd"        sites:  20435277 ( 76.58% of total)
  number of "Cb_LI_RphaseB1_scd"        sites:  19576474 ( 73.36% of total)
    number of "Cb_LI_RphaseA2_scd"-unique sites:       1848137
    number of sites in common:       18587140   (Expected:  14991726; O/E:       1.2)
    number of "Cb_LI_RphaseB1_scd"-unique sites:        989334
  total number of "Cb_LI_RphaseA2_scd+Cb_LI_RphaseB1_scd"-sites observed:     21424611

  number of sites in "Cb_LI_RphaseA2_hsm" that are not  detectable in "Cb_LI_RphaseB1_hsm":      9626
  number of sites in "Cb_LI_RphaseA2_hsm" that are also detectable in "Cb_LI_RphaseB1_hsm":    256326
    number of sites unique to "Cb_LI_RphaseA2_hsm" although detectable in "Cb_LI_RphaseB1_hsm":            57685
    number of "Cb_LI_RphaseA2_hsm" / "Cb_LI_RphaseB1_hsm" common sites                        :           198641      (Expected:   2969; O/E:      66.9)

  number of sites in "Cb_LI_RphaseB1_hsm" that are not  detectable in "Cb_LI_RphaseA2_hsm":      3267
  number of sites in "Cb_LI_RphaseB1_hsm" that are also detectable in "Cb_LI_RphaseA2_hsm":    215326
    number of sites unique to "Cb_LI_RphaseB1_hsm" although detectable in "Cb_LI_RphaseA2_hsm":            16685
    number of "Cb_LI_RphaseB1_hsm" / "Cb_LI_RphaseA2_hsm" common sites                        :           198641      (Expected:   2969; O/E:      66.9)

  Overlap index of "Cb_LI_RphaseA2_hsm" with "Cb_LI_RphaseB1_hsm":           0.866
  Estimated number of "Cb_LI_RphaseA2_hsm" = "Cb_LI_RphaseB1_hsm" sites (assuming sampling from one population):         277856
  Adjusted population size of "Cb_LI_RphaseA2_hsm" = "Cb_LI_RphaseB1_hsm" sites (assuming all sites detectable):         398906 (  1.40x of observed)
```

```
Directory: PWC File: pwc-Cb_LI_RphaseA2.vs.Cb_LI_RphaseB2.txt

Numbers of common and distinct sites comparing Cb_LI_RphaseA2 versus Cb_LI_RphaseB2
================================================================================

  total number of potential sites:  26684764

  number of "Cb_LI_RphaseA2_hsm"         sites:265952
  number of "Cb_LI_RphaseB2_hsm"         sites:200707
    number of "Cb_LI_RphaseA2_hsm"-unique sites:      77593
    number of common sites:        188359
    number of "Cb_LI_RphaseB2_hsm"-unique sites:      12348
  total number of "Cb_LI_RphaseA2_hsm+Cb_LI_RphaseB2_hsm"-sites observed:    278300

  number of "Cb_LI_RphaseA2_scd"        sites:  20435277 ( 76.58% of total)
  number of "Cb_LI_RphaseB2_scd"        sites:  18335682 ( 68.71% of total)
    number of "Cb_LI_RphaseA2_scd"-unique sites:       2695681
    number of sites in common:      17739596   (Expected:  14041523; O/E:       1.3)
    number of "Cb_LI_RphaseB2_scd"-unique sites:        596086
  total number of "Cb_LI_RphaseA2_scd+Cb_LI_RphaseB2_scd"-sites observed:     21031363

  number of sites in "Cb_LI_RphaseA2_hsm" that are not  detectable in "Cb_LI_RphaseB2_hsm":     13896
  number of sites in "Cb_LI_RphaseA2_hsm" that are also detectable in "Cb_LI_RphaseB2_hsm":    252056
    number of sites unique to "Cb_LI_RphaseA2_hsm" although detectable in "Cb_LI_RphaseB2_hsm":            63697
    number of "Cb_LI_RphaseA2_hsm" / "Cb_LI_RphaseB2_hsm" common sites                        :           188359      (Expected:   2827; O/E:      66.6)

  number of sites in "Cb_LI_RphaseB2_hsm" that are not  detectable in "Cb_LI_RphaseA2_hsm":      1709
  number of sites in "Cb_LI_RphaseB2_hsm" that are also detectable in "Cb_LI_RphaseA2_hsm":    198998
    number of sites unique to "Cb_LI_RphaseB2_hsm" although detectable in "Cb_LI_RphaseA2_hsm":            10639
    number of "Cb_LI_RphaseB2_hsm" / "Cb_LI_RphaseA2_hsm" common sites                        :           188359      (Expected:   2827; O/E:      66.6)

  Overlap index of "Cb_LI_RphaseA2_hsm" with "Cb_LI_RphaseB2_hsm":           0.908
  Estimated number of "Cb_LI_RphaseA2_hsm" = "Cb_LI_RphaseB2_hsm" sites (assuming sampling from one population):         266293
  Adjusted population size of "Cb_LI_RphaseA2_hsm" = "Cb_LI_RphaseB2_hsm" sites (assuming all sites detectable):         400571 (  1.44x of observed)
```

```
Directory: PWC File: pwc-Cb_LI_RphaseB1.vs.Cb_LI_RphaseB2.txt

Numbers of common and distinct sites comparing Cb_LI_RphaseB1 versus Cb_LI_RphaseB2
================================================================================

  total number of potential sites:  26684764

  number of "Cb_LI_RphaseB1_hsm"         sites:218593
  number of "Cb_LI_RphaseB2_hsm"         sites:200707
    number of "Cb_LI_RphaseB1_hsm"-unique sites:      44553
    number of common sites:        174040
    number of "Cb_LI_RphaseB2_hsm"-unique sites:      26667
  total number of "Cb_LI_RphaseB1_hsm+Cb_LI_RphaseB2_hsm"-sites observed:    245260

  number of "Cb_LI_RphaseB1_scd"        sites:  19576474 ( 73.36% of total)
  number of "Cb_LI_RphaseB2_scd"        sites:  18335682 ( 68.71% of total)
    number of "Cb_LI_RphaseB1_scd"-unique sites:       2272926
    number of sites in common:      17303548   (Expected:  13451421; O/E:       1.3)
    number of "Cb_LI_RphaseB2_scd"-unique sites:       1032134
  total number of "Cb_LI_RphaseB1_scd+Cb_LI_RphaseB2_scd"-sites observed:     20608608

  number of sites in "Cb_LI_RphaseB1_hsm" that are not  detectable in "Cb_LI_RphaseB2_hsm":      9698
  number of sites in "Cb_LI_RphaseB1_hsm" that are also detectable in "Cb_LI_RphaseB2_hsm":    208895
    number of sites unique to "Cb_LI_RphaseB1_hsm" although detectable in "Cb_LI_RphaseB2_hsm":            34855
    number of "Cb_LI_RphaseB1_hsm" / "Cb_LI_RphaseB2_hsm" common sites                        :           174040      (Expected:   2374; O/E:      73.3)

  number of sites in "Cb_LI_RphaseB2_hsm" that are not  detectable in "Cb_LI_RphaseB1_hsm":      4076
  number of sites in "Cb_LI_RphaseB2_hsm" that are also detectable in "Cb_LI_RphaseB1_hsm":    196631
    number of sites unique to "Cb_LI_RphaseB2_hsm" although detectable in "Cb_LI_RphaseB1_hsm":            22591
    number of "Cb_LI_RphaseB2_hsm" / "Cb_LI_RphaseB1_hsm" common sites                        :           174040      (Expected:   2374; O/E:      73.3)

  Overlap index of "Cb_LI_RphaseB1_hsm" with "Cb_LI_RphaseB2_hsm":           0.799
  Estimated number of "Cb_LI_RphaseB1_hsm" = "Cb_LI_RphaseB2_hsm" sites (assuming sampling from one population):         236010
  Adjusted population size of "Cb_LI_RphaseB1_hsm" = "Cb_LI_RphaseB2_hsm" sites (assuming all sites detectable):         363965 (  1.48x of observed)
```

Directory: CRL File: 0READMEcrl

CRL - Correlations between aggregate samples.

Input: studymk

Output: files crl-*.txt crl-*.pdf

```
Notes: The output is generated by BWASPR::cmpSamples(), which uses methylKit::unite()
        to determine the sites common to all samples and applies methylKit::getCorrelation()
        and methylKit::PCASamples().  The *.txt files show the correlations in a table, and
        the *.pdf files show graphics.

       Beware that the number of common sites (shown at the bottom of the *.txt files) may be
        small, which may make the correlations less informative.
```

Directory: CRL File: crl-Cb_LI.txt

| | BCphaseA1 | BCphaseA2 | BCphaseB1 | BCphaseB2 | RphaseA1 | RphaseA2 | RphaseB1 | RphaseB2 |
|---|---|---|---|---|---|---|---|---|
| BCphaseA1 | 1.000 | 0.905 | 0.908 | 0.913 | 0.905 | 0.915 | 0.914 | 0.909 |
| BCphaseA2 | 0.905 | 1.000 | 0.922 | 0.913 | 0.918 | 0.934 | 0.912 | 0.910 |
| BCphaseB1 | 0.908 | 0.922 | 1.000 | 0.914 | 0.923 | 0.933 | 0.915 | 0.911 |
| BCphaseB2 | 0.913 | 0.913 | 0.914 | 1.000 | 0.910 | 0.923 | 0.919 | 0.919 |
| RphaseA1 | 0.905 | 0.918 | 0.923 | 0.910 | 1.000 | 0.929 | 0.912 | 0.908 |
| RphaseA2 | 0.915 | 0.934 | 0.933 | 0.923 | 0.929 | 1.000 | 0.922 | 0.920 |
| RphaseB1 | 0.914 | 0.912 | 0.915 | 0.919 | 0.912 | 0.922 | 1.000 | 0.916 |
| RphaseB2 | 0.909 | 0.910 | 0.911 | 0.919 | 0.908 | 0.920 | 0.916 | 1.000 |

The number of conserved sites is 104770.

CpG base pearson cor.

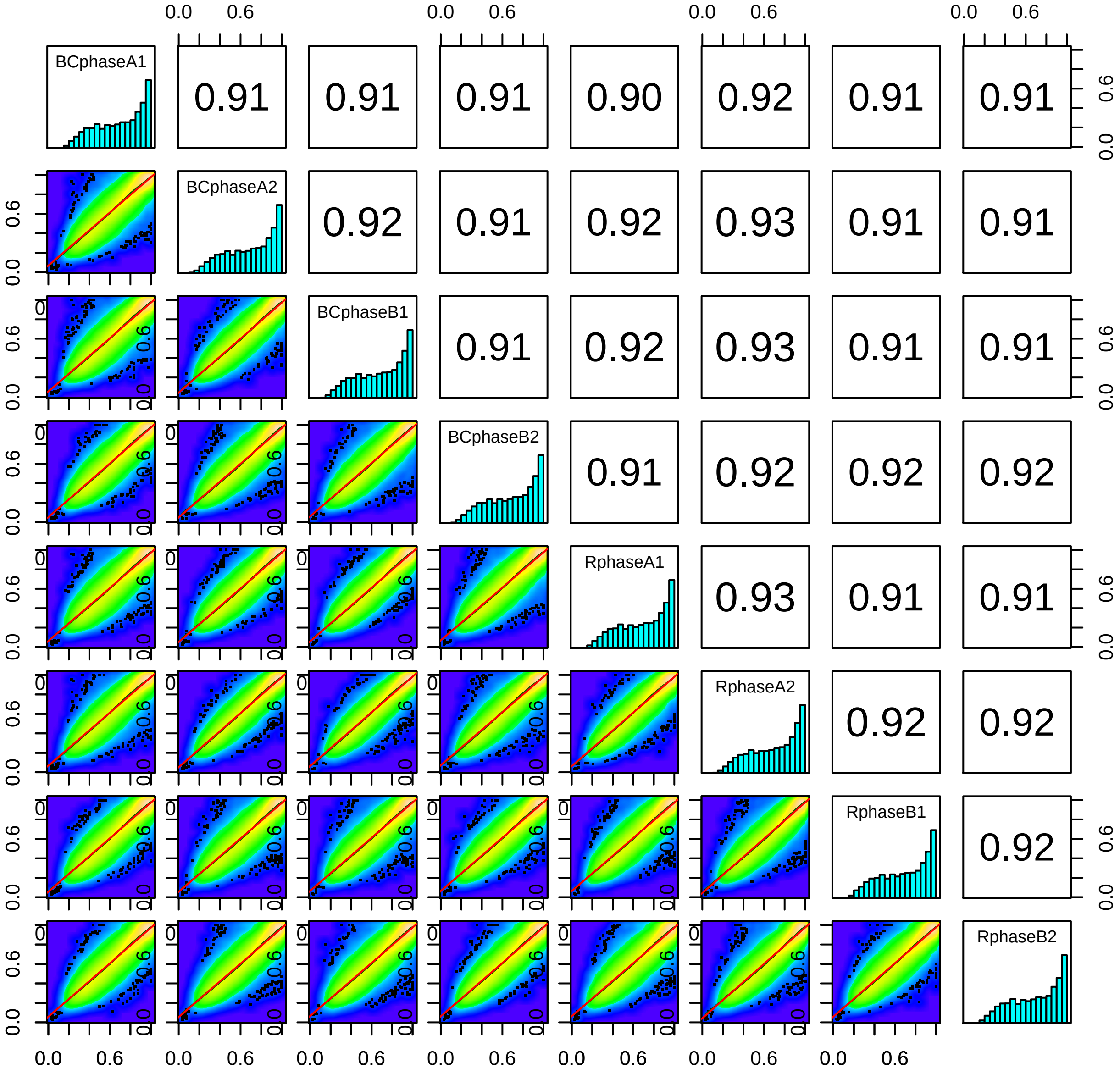

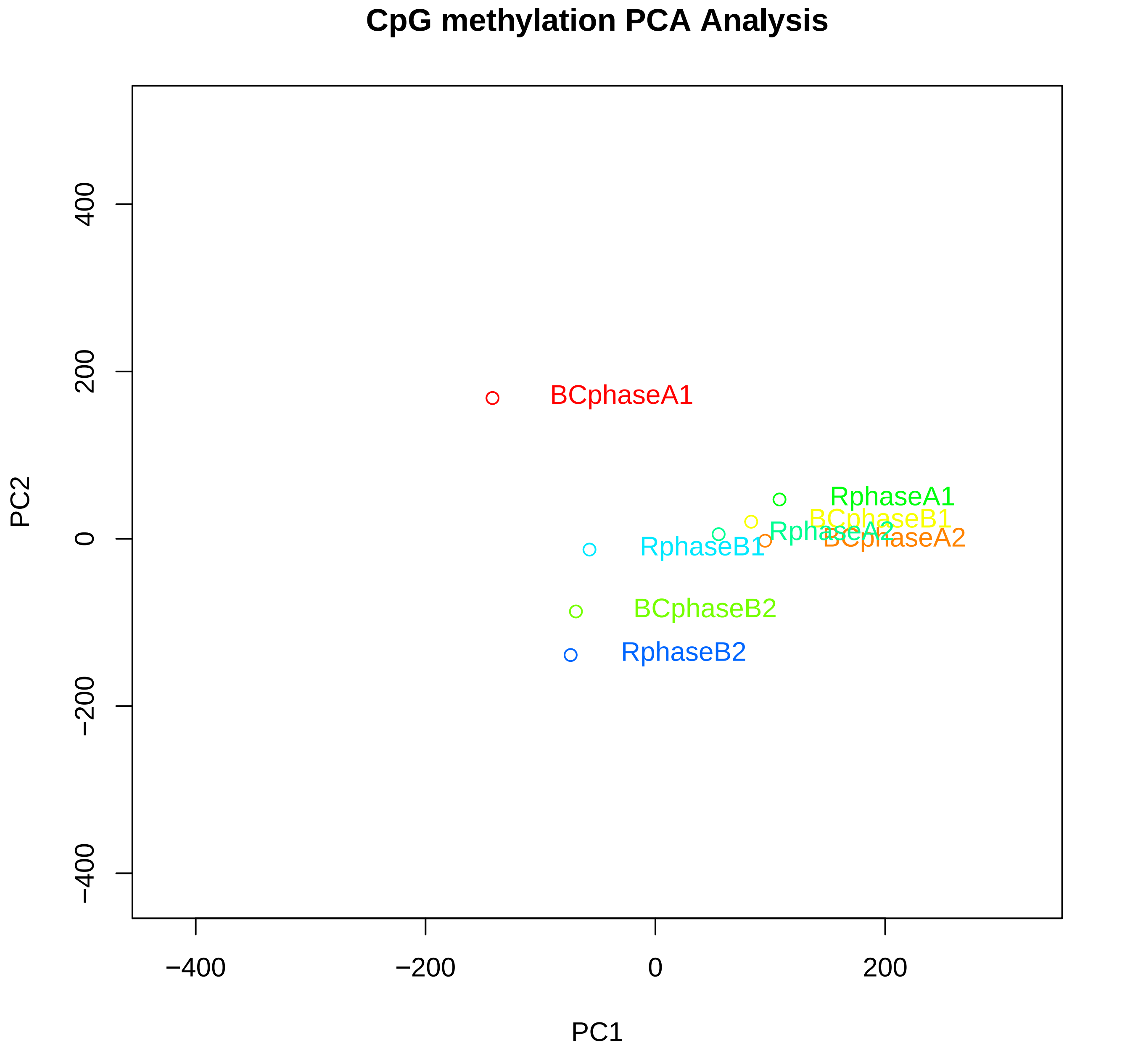
CpG methylation PCA Analysis
PC1
PC2
−400
−200
0
200
400
BCphaseA1
RphaseA1
BCphaseB1
RphaseA2
BCphaseA2
RphaseB1
BCphaseB2
RphaseB2

```
Directory: MMP File: 0READMEmmp

MMP - Mapping of methylation sites on genome annotation.

Input: studymk, studymc, genome_ann (a list of GRanges objects providing annotated region labels
        and boundaries; output of BWASPR::get_genome_annotation())

Output: files mmp-*.txt

Notes: The output is generated by BWASPR::map_methylome() and gives an accounting for every sample
        as to where the CpGhsm and CpGscd (control) sites reside relative to the genome annotation.
       Abbreviations used: O/E, observed over expected
                           (expected percentages are based on the respective feature proportions
                            in the genome, as annotated).
```

```
Directory: MMP File: mmp-Cb_LI_BCphaseA1.txt

================================================================================
Genomic composition in terms of feature regions
================================================================================

Cb genome size:                         212825769 bp
  Cb genic region size:                   152693430 bp ( 71.7%)
    Cb exon region size:                     39755998 bp ( 18.7%)
      Cb CDS region size:                      19364035 bp ( 48.7%)
      Cb five-prime UTR region size:            5122247 bp ( 12.9%)
      Cb three-prime UTR region size:           8663897 bp ( 21.8%)
      Cb other exon region size:                6605819 bp ( 16.6%)
    Cb intron region size:                 112937432 bp ( 53.1%)
  Cb intergenic region size:               60132339 bp ( 28.3%)
    Cb promoter region size:                  6533556 bp (  3.1%)
    Cb other intergenic region size:         53598783 bp ( 25.2%)

================================================================================
Methylation sites in genomic feature regions
================================================================================

Number and density (per 10kb) of sites in Cb_LI_BCphaseA1_scd

Number of sites identified in Cb Cb_LI_BCphaseA1_scd:                             18501067     (  869.31)
Number of sites identified in Cb Cb_LI_BCphaseA1_scd genic regions:               12914469     (  845.78)                       69.80%     ( 0.97 O/E)
Number of sites identified in Cb Cb_LI_BCphaseA1_scd   exon regions:                 3634575   (  914.22)                         19.65%   ( 1.05 O/E)
Number of sites identified in Cb Cb_LI_BCphaseA1_scd     CDS regions:                  2146608 ( 1108.55)                           59.06% ( 1.21 O/E)
Number of sites identified in Cb Cb_LI_BCphaseA1_scd     five-prime UTR regions:        592623 ( 1156.96)                           16.31% ( 1.27 O/E)
Number of sites identified in Cb Cb_LI_BCphaseA1_scd     three-prime UTR regions:       552901 (  638.17)                           15.21% ( 0.70 O/E)
Number of sites identified in Cb Cb_LI_BCphaseA1_scd     other exon regions:            342443 (  518.40)                            9.42% ( 0.57 O/E)
Number of sites identified in Cb Cb_LI_BCphaseA1_scd   intron regions:               9279894   (  821.68)                         50.16%   ( 0.95 O/E)
Number of sites identified in Cb Cb_LI_BCphaseA1_scd intergenic regions:           5586598     (  929.05)                       30.20%     ( 1.07 O/E)
Number of sites identified in Cb Cb_LI_BCphaseA1_scd   promoter regions:              297619   (  455.52)                          1.61%   ( 0.52 O/E)
Number of sites identified in Cb Cb_LI_BCphaseA1_scd   other intergenic regions:     5288979   (  986.77)                         28.59%   ( 1.14 O/E)

Number and density (per 10kb) of sites in Cb_LI_BCphaseA1_hsm

Number of sites identified in Cb Cb_LI_BCphaseA1_hsm:                               198408     (    9.32) (  1.07% of control)
Number of sites identified in Cb Cb_LI_BCphaseA1_hsm genic regions:                 195381     (   12.80) (  1.51% of control)  98.47%     ( 1.37 O/E)
Number of sites identified in Cb Cb_LI_BCphaseA1_hsm   exon regions:                  160476   (   40.37) (  4.42% of control)    80.88%   ( 4.33 O/E)
Number of sites identified in Cb Cb_LI_BCphaseA1_hsm     CDS regions:                   153138 (   79.08) (  7.13% of control)      95.43% ( 1.96 O/E)
Number of sites identified in Cb Cb_LI_BCphaseA1_hsm     five-prime UTR regions:          2594 (    5.06) (  0.44% of control)       1.62% ( 0.13 O/E)
Number of sites identified in Cb Cb_LI_BCphaseA1_hsm     three-prime UTR regions:         3786 (    4.37) (  0.68% of control)       2.36% ( 0.11 O/E)
Number of sites identified in Cb Cb_LI_BCphaseA1_hsm     other exon regions:               958 (    1.45) (  0.28% of control)       0.60% ( 0.04 O/E)
Number of sites identified in Cb Cb_LI_BCphaseA1_hsm   intron regions:                 34905   (    3.09) (  0.38% of control)    17.59%   ( 0.33 O/E)
Number of sites identified in Cb Cb_LI_BCphaseA1_hsm intergenic regions:              3027     (    0.50) (  0.05% of control)   1.53%     ( 0.05 O/E)
Number of sites identified in Cb Cb_LI_BCphaseA1_hsm   promoter regions:                 509   (    0.78) (  0.17% of control)     0.26%   ( 0.08 O/E)
Number of sites identified in Cb Cb_LI_BCphaseA1_hsm   other intergenic regions:        2518   (    0.47) (  0.05% of control)     1.27%   ( 0.05 O/E)
```

```
Directory: MMP File: mmp-Cb_LI_BCphaseA2.txt

================================================================================
Genomic composition in terms of feature regions
================================================================================

Cb genome size:                        212825769 bp
  Cb genic region size:                  152693430 bp ( 71.7%)
    Cb exon region size:                    39755998 bp ( 18.7%)
      Cb CDS region size:                     19364035 bp ( 48.7%)
      Cb five-prime UTR region size:           5122247 bp ( 12.9%)
      Cb three-prime UTR region size:          8663897 bp ( 21.8%)
      Cb other exon region size:               6605819 bp ( 16.6%)
    Cb intron region size:                 112937432 bp ( 53.1%)
  Cb intergenic region size:              60132339 bp ( 28.3%)
    Cb promoter region size:                 6533556 bp (  3.1%)
    Cb other intergenic region size:        53598783 bp ( 25.2%)

================================================================================
Methylation sites in genomic feature regions
================================================================================

Number and density (per 10kb) of sites in Cb_LI_BCphaseA2_scd

Number of sites identified in Cb Cb_LI_BCphaseA2_scd:                             18589454     (  873.46)                         
Number of sites identified in Cb Cb_LI_BCphaseA2_scd genic regions:               12989006     (  850.66)                           69.87%     ( 0.97 O/E)
Number of sites identified in Cb Cb_LI_BCphaseA2_scd   exon regions:                 3635773   (  914.52)                             19.56%   ( 1.05 O/E)
Number of sites identified in Cb Cb_LI_BCphaseA2_scd     CDS regions:                  2145780 ( 1108.13)                               59.02% ( 1.21 O/E)
Number of sites identified in Cb Cb_LI_BCphaseA2_scd     five-prime UTR regions:        597306 ( 1166.10)                               16.43% ( 1.28 O/E)
Number of sites identified in Cb Cb_LI_BCphaseA2_scd     three-prime UTR regions:       548814 (  633.45)                               15.09% ( 0.69 O/E)
Number of sites identified in Cb Cb_LI_BCphaseA2_scd     other exon regions:            343873 (  520.56)                                9.46% ( 0.57 O/E)
Number of sites identified in Cb Cb_LI_BCphaseA2_scd   intron regions:               9353233   (  828.18)                             50.31%   ( 0.95 O/E)
Number of sites identified in Cb Cb_LI_BCphaseA2_scd intergenic regions:           5600448     (  931.35)                           30.13%     ( 1.07 O/E)
Number of sites identified in Cb Cb_LI_BCphaseA2_scd   promoter regions:              300711   (  460.26)                              1.62%   ( 0.53 O/E)
Number of sites identified in Cb Cb_LI_BCphaseA2_scd   other intergenic regions:     5299737   (  988.78)                             28.51%   ( 1.13 O/E)

Number and density (per 10kb) of sites in Cb_LI_BCphaseA2_hsm

Number of sites identified in Cb Cb_LI_BCphaseA2_hsm:                               213448     (   10.03) (  1.15% of control)
Number of sites identified in Cb Cb_LI_BCphaseA2_hsm genic regions:                 210355     (   13.78) (  1.62% of control)  98.55%     ( 1.37 O/E)
Number of sites identified in Cb Cb_LI_BCphaseA2_hsm   exon regions:                  173143   (   43.55) (  4.76% of control)    81.12%   ( 4.34 O/E)
Number of sites identified in Cb Cb_LI_BCphaseA2_hsm     CDS regions:                   165297 (   85.36) (  7.70% of control)      95.47% ( 1.96 O/E)
Number of sites identified in Cb Cb_LI_BCphaseA2_hsm     five-prime UTR regions:          2821 (    5.51) (  0.47% of control)       1.63% ( 0.13 O/E)
Number of sites identified in Cb Cb_LI_BCphaseA2_hsm     three-prime UTR regions:         4071 (    4.70) (  0.74% of control)       2.35% ( 0.11 O/E)
Number of sites identified in Cb Cb_LI_BCphaseA2_hsm     other exon regions:               954 (    1.44) (  0.28% of control)       0.55% ( 0.03 O/E)
Number of sites identified in Cb Cb_LI_BCphaseA2_hsm   intron regions:                 37212   (    3.29) (  0.40% of control)    17.43%   ( 0.33 O/E)
Number of sites identified in Cb Cb_LI_BCphaseA2_hsm intergenic regions:              3093     (    0.51) (  0.06% of control)   1.45%     ( 0.05 O/E)
Number of sites identified in Cb Cb_LI_BCphaseA2_hsm   promoter regions:                 518   (    0.79) (  0.17% of control)     0.24%   ( 0.08 O/E)
Number of sites identified in Cb Cb_LI_BCphaseA2_hsm   other intergenic regions:        2575   (    0.48) (  0.05% of control)     1.21%   ( 0.05 O/E)
```

```
Directory: MMP File: mmp-Cb_LI_BCphaseB1.txt

================================================================================
Genomic composition in terms of feature regions
================================================================================

Cb genome size:                         212825769 bp
  Cb genic region size:                   152693430 bp ( 71.7%)
    Cb exon region size:                    39755998 bp ( 18.7%)
      Cb CDS region size:                     19364035 bp ( 48.7%)
      Cb five-prime UTR region size:           5122247 bp ( 12.9%)
      Cb three-prime UTR region size:          8663897 bp ( 21.8%)
      Cb other exon region size:               6605819 bp ( 16.6%)
    Cb intron region size:                 112937432 bp ( 53.1%)
  Cb intergenic region size:               60132339 bp ( 28.3%)
    Cb promoter region size:                 6533556 bp (  3.1%)
    Cb other intergenic region size:        53598783 bp ( 25.2%)

================================================================================
Methylation sites in genomic feature regions
================================================================================

Number and density (per 10kb) of sites in Cb_LI_BCphaseB1_scd

Number of sites identified in Cb Cb_LI_BCphaseB1_scd:                             20321376     (  954.84)                       69.77%     ( 0.97 O/E)
Number of sites identified in Cb Cb_LI_BCphaseB1_scd genic regions:               14177457     (  928.49)                       69.77%     ( 0.97 O/E)
Number of sites identified in Cb Cb_LI_BCphaseB1_scd   exon regions:                3925477   (  987.39)                         19.32%   ( 1.03 O/E)
Number of sites identified in Cb Cb_LI_BCphaseB1_scd     CDS regions:                 2288236 ( 1181.69)                           58.29% ( 1.20 O/E)
Number of sites identified in Cb Cb_LI_BCphaseB1_scd     five-prime UTR regions:       639088 ( 1247.67)                           16.28% ( 1.26 O/E)
Number of sites identified in Cb Cb_LI_BCphaseB1_scd     three-prime UTR regions:      619174 (  714.66)                           15.77% ( 0.72 O/E)
Number of sites identified in Cb Cb_LI_BCphaseB1_scd     other exon regions:           378979 (  573.70)                            9.65% ( 0.58 O/E)
Number of sites identified in Cb Cb_LI_BCphaseB1_scd   intron regions:             10251980   (  907.76)                         50.45%   ( 0.95 O/E)
Number of sites identified in Cb Cb_LI_BCphaseB1_scd intergenic regions:           6143919     ( 1021.73)                       30.23%     ( 1.07 O/E)
Number of sites identified in Cb Cb_LI_BCphaseB1_scd   promoter regions:             325804   (  498.66)                          1.60%   ( 0.52 O/E)
Number of sites identified in Cb Cb_LI_BCphaseB1_scd   other intergenic regions:    5818115   ( 1085.49)                         28.63%   ( 1.14 O/E)

Number and density (per 10kb) of sites in Cb_LI_BCphaseB1_hsm

Number of sites identified in Cb Cb_LI_BCphaseB1_hsm:                               226815     (   10.66) (  1.12% of control)
Number of sites identified in Cb Cb_LI_BCphaseB1_hsm genic regions:                 222990     (   14.60) (  1.57% of control)  98.31%     ( 1.37 O/E)
Number of sites identified in Cb Cb_LI_BCphaseB1_hsm   exon regions:                  182513   (   45.91) (  4.65% of control)   80.47%   ( 4.31 O/E)
Number of sites identified in Cb Cb_LI_BCphaseB1_hsm     CDS regions:                   173686 (   89.70) (  7.59% of control)     95.16% ( 1.95 O/E)
Number of sites identified in Cb Cb_LI_BCphaseB1_hsm     five-prime UTR regions:          2987 (    5.83) (  0.47% of control)      1.64% ( 0.13 O/E)
Number of sites identified in Cb Cb_LI_BCphaseB1_hsm     three-prime UTR regions:         4719 (    5.45) (  0.76% of control)      2.59% ( 0.12 O/E)
Number of sites identified in Cb Cb_LI_BCphaseB1_hsm     other exon regions:              1121 (    1.70) (  0.30% of control)      0.61% ( 0.04 O/E)
Number of sites identified in Cb Cb_LI_BCphaseB1_hsm   intron regions:                 40477   (    3.58) (  0.39% of control)   17.85%   ( 0.34 O/E)
Number of sites identified in Cb Cb_LI_BCphaseB1_hsm intergenic regions:              3825     (    0.64) (  0.06% of control)  1.69%     ( 0.06 O/E)
Number of sites identified in Cb Cb_LI_BCphaseB1_hsm   promoter regions:                590   (    0.90) (  0.18% of control)    0.26%   ( 0.08 O/E)
Number of sites identified in Cb Cb_LI_BCphaseB1_hsm   other intergenic regions:       3235   (    0.60) (  0.06% of control)    1.43%   ( 0.06 O/E)
```

Directory: MMP File: mmp-Cb_LI_BCphaseB2.txt

================================================================================

Genomic composition in terms of feature regions

================================================================================

| | | |
|---|---|---|
| Cb genome size: | 212825769 bp | |
| Cb genic region size: | 152693430 bp | ( 71.7%) |
| Cb exon region size: | 39755998 bp | ( 18.7%) |
| Cb CDS region size: | 19364035 bp | ( 48.7%) |
| Cb five-prime UTR region size: | 5122247 bp | ( 12.9%) |
| Cb three-prime UTR region size: | 8663897 bp | ( 21.8%) |
| Cb other exon region size: | 6605819 bp | ( 16.6%) |
| Cb intron region size: | 112937432 bp | ( 53.1%) |
| Cb intergenic region size: | 60132339 bp | ( 28.3%) |
| Cb promoter region size: | 6533556 bp | ( 3.1%) |
| Cb other intergenic region size: | 53598783 bp | ( 25.2%) |

================================================================================

Methylation sites in genomic feature regions

================================================================================

Number and density (per 10kb) of sites in Cb_LI_BCphaseB2_scd

| | | | | |
|---|---|---|---|---|
| Number of sites identified in Cb Cb_LI_BCphaseB2_scd: | 19192933 | ( 901.81) | | |
| Number of sites identified in Cb Cb_LI_BCphaseB2_scd genic regions: | 13390750 | ( 876.97) | 69.77% | ( 0.97 O/E) |
| Number of sites identified in Cb Cb_LI_BCphaseB2_scd exon regions: | 3722251 | ( 936.27) | 19.39% | ( 1.04 O/E) |
| Number of sites identified in Cb Cb_LI_BCphaseB2_scd CDS regions: | 2185149 | ( 1128.46) | 58.71% | ( 1.21 O/E) |
| Number of sites identified in Cb Cb_LI_BCphaseB2_scd five-prime UTR regions: | 610145 | ( 1191.17) | 16.39% | ( 1.27 O/E) |
| Number of sites identified in Cb Cb_LI_BCphaseB2_scd three-prime UTR regions: | 572080 | ( 660.30) | 15.37% | ( 0.71 O/E) |
| Number of sites identified in Cb Cb_LI_BCphaseB2_scd other exon regions: | 354877 | ( 537.22) | 9.53% | ( 0.57 O/E) |
| Number of sites identified in Cb Cb_LI_BCphaseB2_scd intron regions: | 9668499 | ( 856.09) | 50.38% | ( 0.95 O/E) |
| Number of sites identified in Cb Cb_LI_BCphaseB2_scd intergenic regions: | 5802183 | ( 964.90) | 30.23% | ( 1.07 O/E) |
| Number of sites identified in Cb Cb_LI_BCphaseB2_scd promoter regions: | 308616 | ( 472.36) | 1.61% | ( 0.52 O/E) |
| Number of sites identified in Cb Cb_LI_BCphaseB2_scd other intergenic regions: | 5493567 | ( 1024.94) | 28.62% | ( 1.14 O/E) |

Number and density (per 10kb) of sites in Cb_LI_BCphaseB2_hsm

| | | | | | |
|---|---|---|---|---|---|
| Number of sites identified in Cb Cb_LI_BCphaseB2_hsm: | 212569 | ( 9.99) | ( 1.11% of control) | | |
| Number of sites identified in Cb Cb_LI_BCphaseB2_hsm genic regions: | 209300 | ( 13.71) | ( 1.56% of control) | 98.46% | ( 1.37 O/E) |
| Number of sites identified in Cb Cb_LI_BCphaseB2_hsm exon regions: | 171901 | ( 43.24) | ( 4.62% of control) | 80.87% | ( 4.33 O/E) |
| Number of sites identified in Cb Cb_LI_BCphaseB2_hsm CDS regions: | 163912 | ( 84.65) | ( 7.50% of control) | 95.35% | ( 1.96 O/E) |
| Number of sites identified in Cb Cb_LI_BCphaseB2_hsm five-prime UTR regions: | 2720 | ( 5.31) | ( 0.45% of control) | 1.58% | ( 0.12 O/E) |
| Number of sites identified in Cb Cb_LI_BCphaseB2_hsm three-prime UTR regions: | 4237 | ( 4.89) | ( 0.74% of control) | 2.46% | ( 0.11 O/E) |
| Number of sites identified in Cb Cb_LI_BCphaseB2_hsm other exon regions: | 1032 | ( 1.56) | ( 0.29% of control) | 0.60% | ( 0.04 O/E) |
| Number of sites identified in Cb Cb_LI_BCphaseB2_hsm intron regions: | 37399 | ( 3.31) | ( 0.39% of control) | 17.59% | ( 0.33 O/E) |
| Number of sites identified in Cb Cb_LI_BCphaseB2_hsm intergenic regions: | 3269 | ( 0.54) | ( 0.06% of control) | 1.54% | ( 0.05 O/E) |
| Number of sites identified in Cb Cb_LI_BCphaseB2_hsm promoter regions: | 514 | ( 0.79) | ( 0.17% of control) | 0.24% | ( 0.08 O/E) |
| Number of sites identified in Cb Cb_LI_BCphaseB2_hsm other intergenic regions: | 2755 | ( 0.51) | ( 0.05% of control) | 1.30% | ( 0.05 O/E) |

```
Directory: MMP File: mmp-Cb_LI_RphaseA1.txt

================================================================================
Genomic composition in terms of feature regions
================================================================================

Cb genome size:                        212825769 bp
  Cb genic region size:                  152693430 bp ( 71.7%)
    Cb exon region size:                    39755998 bp ( 18.7%)
      Cb CDS region size:                     19364035 bp ( 48.7%)
      Cb five-prime UTR region size:           5122247 bp ( 12.9%)
      Cb three-prime UTR region size:          8663897 bp ( 21.8%)
      Cb other exon region size:               6605819 bp ( 16.6%)
    Cb intron region size:                 112937432 bp ( 53.1%)
  Cb intergenic region size:              60132339 bp ( 28.3%)
    Cb promoter region size:                 6533556 bp (  3.1%)
    Cb other intergenic region size:        53598783 bp ( 25.2%)

================================================================================
Methylation sites in genomic feature regions
================================================================================

Number and density (per 10kb) of sites in Cb_LI_RphaseA1_scd

Number of sites identified in Cb Cb_LI_RphaseA1_scd:                      19659217     (  923.72)                          
Number of sites identified in Cb Cb_LI_RphaseA1_scd genic regions:        13724777     (  898.85)                        69.81%     ( 0.97 O/E)
Number of sites identified in Cb Cb_LI_RphaseA1_scd   exon regions:          3830012   (  963.38)                          19.48%   ( 1.04 O/E)
Number of sites identified in Cb Cb_LI_RphaseA1_scd     CDS regions:           2244781 ( 1159.25)                            58.61% ( 1.20 O/E)
Number of sites identified in Cb Cb_LI_RphaseA1_scd     five-prime UTR regions:   623455 ( 1217.15)                          16.28% ( 1.26 O/E)
Number of sites identified in Cb Cb_LI_RphaseA1_scd     three-prime UTR regions:  595755 (  687.63)                          15.55% ( 0.71 O/E)
Number of sites identified in Cb Cb_LI_RphaseA1_scd     other exon regions:       366021 (  554.09)                           9.56% ( 0.58 O/E)
Number of sites identified in Cb Cb_LI_RphaseA1_scd   intron regions:        9894765   (  876.13)                          50.33%   ( 0.95 O/E)
Number of sites identified in Cb Cb_LI_RphaseA1_scd intergenic regions:    5934440     (  986.90)                        30.19%     ( 1.07 O/E)
Number of sites identified in Cb Cb_LI_RphaseA1_scd   promoter regions:       315515   (  482.91)                           1.60%   ( 0.52 O/E)
Number of sites identified in Cb Cb_LI_RphaseA1_scd   other intergenic regions:  5618925   ( 1048.33)                      28.58%   ( 1.13 O/E)

Number and density (per 10kb) of sites in Cb_LI_RphaseA1_hsm

Number of sites identified in Cb Cb_LI_RphaseA1_hsm:                        213610     (   10.04) (  1.09% of control)
Number of sites identified in Cb Cb_LI_RphaseA1_hsm genic regions:          210359     (   13.78) (  1.53% of control)  98.48%     ( 1.37 O/E)
Number of sites identified in Cb Cb_LI_RphaseA1_hsm   exon regions:            172454   (   43.38) (  4.50% of control)   80.73%   ( 4.32 O/E)
Number of sites identified in Cb Cb_LI_RphaseA1_hsm     CDS regions:             164286 (   84.84) (  7.32% of control)     95.26% ( 1.96 O/E)
Number of sites identified in Cb Cb_LI_RphaseA1_hsm     five-prime UTR regions:     2896 (    5.65) (  0.46% of control)      1.68% ( 0.13 O/E)
Number of sites identified in Cb Cb_LI_RphaseA1_hsm     three-prime UTR regions:    4235 (    4.89) (  0.71% of control)      2.46% ( 0.11 O/E)
Number of sites identified in Cb Cb_LI_RphaseA1_hsm     other exon regions:         1037 (    1.57) (  0.28% of control)      0.60% ( 0.04 O/E)
Number of sites identified in Cb Cb_LI_RphaseA1_hsm   intron regions:           37905   (    3.36) (  0.38% of control)   17.74%   ( 0.33 O/E)
Number of sites identified in Cb Cb_LI_RphaseA1_hsm intergenic regions:        3251     (    0.54) (  0.05% of control)  1.52%      ( 0.05 O/E)
Number of sites identified in Cb Cb_LI_RphaseA1_hsm   promoter regions:           538   (    0.82) (  0.17% of control)    0.25%   ( 0.08 O/E)
Number of sites identified in Cb Cb_LI_RphaseA1_hsm   other intergenic regions:      2713   (    0.51) (  0.05% of control)    1.27%   ( 0.05 O/E)
```

```
Directory: MMP File: mmp-Cb_LI_RphaseA2.txt

================================================================================
Genomic composition in terms of feature regions
================================================================================

Cb genome size:                        212825769 bp
  Cb genic region size:                  152693430 bp ( 71.7%)
    Cb exon region size:                    39755998 bp ( 18.7%)
      Cb CDS region size:                     19364035 bp ( 48.7%)
      Cb five-prime UTR region size:           5122247 bp ( 12.9%)
      Cb three-prime UTR region size:          8663897 bp ( 21.8%)
      Cb other exon region size:               6605819 bp ( 16.6%)
    Cb intron region size:                 112937432 bp ( 53.1%)
  Cb intergenic region size:              60132339 bp ( 28.3%)
    Cb promoter region size:                 6533556 bp (  3.1%)
    Cb other intergenic region size:        53598783 bp ( 25.2%)

================================================================================
Methylation sites in genomic feature regions
================================================================================

Number and density (per 10kb) of sites in Cb_LI_RphaseA2_scd

Number of sites identified in Cb Cb_LI_RphaseA2_scd:                            20435277     (  960.19)
Number of sites identified in Cb Cb_LI_RphaseA2_scd genic regions:              14291364     (  935.95)                        69.93%     ( 0.97 O/E)
Number of sites identified in Cb Cb_LI_RphaseA2_scd   exon regions:                3958110   (  995.60)                          19.37%   ( 1.04 O/E)
Number of sites identified in Cb Cb_LI_RphaseA2_scd     CDS regions:                 2300288 ( 1187.92)                            58.12% ( 1.19 O/E)
Number of sites identified in Cb Cb_LI_RphaseA2_scd     five-prime UTR regions:       646001 ( 1261.17)                            16.32% ( 1.27 O/E)
Number of sites identified in Cb Cb_LI_RphaseA2_scd     three-prime UTR regions:      629789 (  726.91)                            15.91% ( 0.73 O/E)
Number of sites identified in Cb Cb_LI_RphaseA2_scd     other exon regions:           382032 (  578.33)                             9.65% ( 0.58 O/E)
Number of sites identified in Cb Cb_LI_RphaseA2_scd   intron regions:             10333254   (  914.95)                          50.57%   ( 0.95 O/E)
Number of sites identified in Cb Cb_LI_RphaseA2_scd intergenic regions:          6143913     ( 1021.73)                        30.07%     ( 1.06 O/E)
Number of sites identified in Cb Cb_LI_RphaseA2_scd   promoter regions:             330350   (  505.62)                           1.62%   ( 0.53 O/E)
Number of sites identified in Cb Cb_LI_RphaseA2_scd   other intergenic regions:    5813563   ( 1084.64)                          28.45%   ( 1.13 O/E)

Number and density (per 10kb) of sites in Cb_LI_RphaseA2_hsm

Number of sites identified in Cb Cb_LI_RphaseA2_hsm:                              265952     (   12.50) (  1.30% of control)
Number of sites identified in Cb Cb_LI_RphaseA2_hsm genic regions:                260429     (   17.06) (  1.82% of control)  97.92%     ( 1.36 O/E)
Number of sites identified in Cb Cb_LI_RphaseA2_hsm   exon regions:                 211437   (   53.18) (  5.34% of control)    79.50%   ( 4.26 O/E)
Number of sites identified in Cb Cb_LI_RphaseA2_hsm     CDS regions:                  200311 (  103.44) (  8.71% of control)      94.74% ( 1.95 O/E)
Number of sites identified in Cb Cb_LI_RphaseA2_hsm     five-prime UTR regions:         3574 (    6.98) (  0.55% of control)       1.69% ( 0.13 O/E)
Number of sites identified in Cb Cb_LI_RphaseA2_hsm     three-prime UTR regions:        6177 (    7.13) (  0.98% of control)       2.92% ( 0.13 O/E)
Number of sites identified in Cb Cb_LI_RphaseA2_hsm     other exon regions:             1375 (    2.08) (  0.36% of control)       0.65% ( 0.04 O/E)
Number of sites identified in Cb Cb_LI_RphaseA2_hsm   intron regions:               48992   (    4.34) (  0.47% of control)    18.42%   ( 0.35 O/E)
Number of sites identified in Cb Cb_LI_RphaseA2_hsm intergenic regions:            5523     (    0.92) (  0.09% of control)   2.08%     ( 0.07 O/E)
Number of sites identified in Cb Cb_LI_RphaseA2_hsm   promoter regions:               811   (    1.24) (  0.25% of control)     0.30%   ( 0.10 O/E)
Number of sites identified in Cb Cb_LI_RphaseA2_hsm   other intergenic regions:      4712   (    0.88) (  0.08% of control)     1.77%   ( 0.07 O/E)
```

```
Directory: MMP File: mmp-Cb_LI_RphaseB1.txt

================================================================================
Genomic composition in terms of feature regions
================================================================================

Cb genome size:                        212825769 bp
  Cb genic region size:                  152693430 bp ( 71.7%)
    Cb exon region size:                    39755998 bp ( 18.7%)
      Cb CDS region size:                     19364035 bp ( 48.7%)
      Cb five-prime UTR region size:           5122247 bp ( 12.9%)
      Cb three-prime UTR region size:          8663897 bp ( 21.8%)
      Cb other exon region size:               6605819 bp ( 16.6%)
    Cb intron region size:                 112937432 bp ( 53.1%)
  Cb intergenic region size:              60132339 bp ( 28.3%)
    Cb promoter region size:                 6533556 bp (  3.1%)
    Cb other intergenic region size:        53598783 bp ( 25.2%)

================================================================================
Methylation sites in genomic feature regions
================================================================================

Number and density (per 10kb) of sites in Cb_LI_RphaseB1_scd

Number of sites identified in Cb Cb_LI_RphaseB1_scd:                            19576474     (  919.84)                         
Number of sites identified in Cb Cb_LI_RphaseB1_scd genic regions:              13665978     (  894.99)                       69.81%     ( 0.97 O/E)
Number of sites identified in Cb Cb_LI_RphaseB1_scd   exon regions:               3815488   (  959.73)                         19.49%   ( 1.04 O/E)
Number of sites identified in Cb Cb_LI_RphaseB1_scd     CDS regions:                2238558 ( 1156.04)                           58.67% ( 1.20 O/E)
Number of sites identified in Cb Cb_LI_RphaseB1_scd     five-prime UTR regions:      621655 ( 1213.64)                           16.29% ( 1.26 O/E)
Number of sites identified in Cb Cb_LI_RphaseB1_scd     three-prime UTR regions:     592005 (  683.30)                           15.52% ( 0.71 O/E)
Number of sites identified in Cb Cb_LI_RphaseB1_scd     other exon regions:          363270 (  549.92)                            9.52% ( 0.57 O/E)
Number of sites identified in Cb Cb_LI_RphaseB1_scd   intron regions:             9850490   (  872.21)                         50.32%   ( 0.95 O/E)
Number of sites identified in Cb Cb_LI_RphaseB1_scd intergenic regions:          5910496     (  982.91)                       30.19%     ( 1.07 O/E)
Number of sites identified in Cb Cb_LI_RphaseB1_scd   promoter regions:            315507   (  482.90)                          1.61%   ( 0.52 O/E)
Number of sites identified in Cb Cb_LI_RphaseB1_scd   other intergenic regions:   5594989   ( 1043.86)                         28.58%   ( 1.13 O/E)

Number and density (per 10kb) of sites in Cb_LI_RphaseB1_hsm

Number of sites identified in Cb Cb_LI_RphaseB1_hsm:                              218593     (   10.27) (  1.12% of control)
Number of sites identified in Cb Cb_LI_RphaseB1_hsm genic regions:                215039     (   14.08) (  1.57% of control)  98.37%     ( 1.37 O/E)
Number of sites identified in Cb Cb_LI_RphaseB1_hsm   exon regions:                 175978   (   44.26) (  4.61% of control)    80.50%   ( 4.31 O/E)
Number of sites identified in Cb Cb_LI_RphaseB1_hsm     CDS regions:                  167612 (   86.56) (  7.49% of control)      95.25% ( 1.96 O/E)
Number of sites identified in Cb Cb_LI_RphaseB1_hsm     five-prime UTR regions:         2853 (    5.57) (  0.46% of control)       1.62% ( 0.13 O/E)
Number of sites identified in Cb Cb_LI_RphaseB1_hsm     three-prime UTR regions:        4436 (    5.12) (  0.75% of control)       2.52% ( 0.12 O/E)
Number of sites identified in Cb Cb_LI_RphaseB1_hsm     other exon regions:             1077 (    1.63) (  0.30% of control)       0.61% ( 0.04 O/E)
Number of sites identified in Cb Cb_LI_RphaseB1_hsm   intron regions:                39061   (    3.46) (  0.40% of control)    17.87%   ( 0.34 O/E)
Number of sites identified in Cb Cb_LI_RphaseB1_hsm intergenic regions:            3554     (    0.59) (  0.06% of control)   1.63%     ( 0.06 O/E)
Number of sites identified in Cb Cb_LI_RphaseB1_hsm   promoter regions:               566   (    0.87) (  0.18% of control)     0.26%   ( 0.08 O/E)
Number of sites identified in Cb Cb_LI_RphaseB1_hsm   other intergenic regions:      2988   (    0.56) (  0.05% of control)     1.37%   ( 0.05 O/E)
```

```
Directory: MMP File: mmp-Cb_LI_RphaseB2.txt

================================================================================
Genomic composition in terms of feature regions
================================================================================

Cb genome size:                        212825769 bp
  Cb genic region size:                  152693430 bp ( 71.7%)
    Cb exon region size:                    39755998 bp ( 18.7%)
      Cb CDS region size:                     19364035 bp ( 48.7%)
      Cb five-prime UTR region size:           5122247 bp ( 12.9%)
      Cb three-prime UTR region size:          8663897 bp ( 21.8%)
      Cb other exon region size:               6605819 bp ( 16.6%)
    Cb intron region size:                 112937432 bp ( 53.1%)
  Cb intergenic region size:              60132339 bp ( 28.3%)
    Cb promoter region size:                 6533556 bp (  3.1%)
    Cb other intergenic region size:        53598783 bp ( 25.2%)

================================================================================
Methylation sites in genomic feature regions
================================================================================

Number and density (per 10kb) of sites in Cb_LI_RphaseB2_scd

Number of sites identified in Cb Cb_LI_RphaseB2_scd:                        18335682     (  861.53)                          
Number of sites identified in Cb Cb_LI_RphaseB2_scd genic regions:          12803181     (  838.49)                          69.83%     ( 0.97 O/E)
Number of sites identified in Cb Cb_LI_RphaseB2_scd   exon regions:            3587938   (  902.49)                            19.57%   ( 1.05 O/E)
Number of sites identified in Cb Cb_LI_RphaseB2_scd     CDS regions:             2117142 ( 1093.34)                              59.01% ( 1.21 O/E)
Number of sites identified in Cb Cb_LI_RphaseB2_scd     five-prime UTR regions:   588411 ( 1148.74)                              16.40% ( 1.27 O/E)
Number of sites identified in Cb Cb_LI_RphaseB2_scd     three-prime UTR regions:  543642 (  627.48)                              15.15% ( 0.70 O/E)
Number of sites identified in Cb Cb_LI_RphaseB2_scd     other exon regions:       338743 (  512.79)                               9.44% ( 0.57 O/E)
Number of sites identified in Cb Cb_LI_RphaseB2_scd   intron regions:          9215243   (  815.96)                            50.26%   ( 0.95 O/E)
Number of sites identified in Cb Cb_LI_RphaseB2_scd intergenic regions:      5532501     (  920.05)                          30.17%     ( 1.07 O/E)
Number of sites identified in Cb Cb_LI_RphaseB2_scd   promoter regions:         296653   (  454.05)                             1.62%   ( 0.53 O/E)
Number of sites identified in Cb Cb_LI_RphaseB2_scd   other intergenic regions: 5235848  (  976.86)                            28.56%   ( 1.13 O/E)

Number and density (per 10kb) of sites in Cb_LI_RphaseB2_hsm

Number of sites identified in Cb Cb_LI_RphaseB2_hsm:                          200707     (    9.43) (  1.09% of control)
Number of sites identified in Cb Cb_LI_RphaseB2_hsm genic regions:            197784     (   12.95) (  1.54% of control)  98.54%     ( 1.37 O/E)
Number of sites identified in Cb Cb_LI_RphaseB2_hsm   exon regions:             162470   (   40.87) (  4.53% of control)    80.95%   ( 4.33 O/E)
Number of sites identified in Cb Cb_LI_RphaseB2_hsm     CDS regions:              155084 (   80.09) (  7.33% of control)      95.45% ( 1.96 O/E)
Number of sites identified in Cb Cb_LI_RphaseB2_hsm     five-prime UTR regions:     2563 (    5.00) (  0.44% of control)       1.58% ( 0.12 O/E)
Number of sites identified in Cb Cb_LI_RphaseB2_hsm     three-prime UTR regions:    3869 (    4.47) (  0.71% of control)       2.38% ( 0.11 O/E)
Number of sites identified in Cb Cb_LI_RphaseB2_hsm     other exon regions:          954 (    1.44) (  0.28% of control)       0.59% ( 0.04 O/E)
Number of sites identified in Cb Cb_LI_RphaseB2_hsm   intron regions:            35314   (    3.13) (  0.38% of control)    17.59%   ( 0.33 O/E)
Number of sites identified in Cb Cb_LI_RphaseB2_hsm intergenic regions:         2923     (    0.49) (  0.05% of control)   1.46%     ( 0.05 O/E)
Number of sites identified in Cb Cb_LI_RphaseB2_hsm   promoter regions:            470   (    0.72) (  0.16% of control)     0.23%   ( 0.08 O/E)
Number of sites identified in Cb Cb_LI_RphaseB2_hsm   other intergenic regions:   2453   (    0.46) (  0.05% of control)     1.22%   ( 0.05 O/E)
```

```
Directory: ACS File: 0READMEacs

ACS - Annotation of conserved methylation sites.

Input: studymk, genome_ann (a list of GRanges objects providing annotated region labels and bounds;
                            output of BWASPR::get_genome_annotation())

Output: file acs-*.txt

Notes: The output is generated by BWASPR::annotate_methylome(), which uses methylKit::unite()
        to determine the sites common to all experiments and applies genomation::annotateWithFeature().
        The acs-*.txt files show the common sites with methylation levels in all samples
        and columns with booleans indicating whether the site fits a specific annotation feature.

       Abbreviations used: pc, protein coding; nc, non-coding; fp, five prime; tp, three prime.
                           UTR, untranslated region; unique, not overlapping with other categories.
```

Directory: ACS File: acs-Cb_LI.txt

| BCphaseA1 | BCphaseA2 | BCphaseB1 | BCphaseB2 | RphaseA1 | RphaseA2 | RphaseB1 | RphaseB2 | chr | start | end | strand | gene | exon | pcexon | promoter | CDS | fpUTR | tpUTR | fpUTRnotCDS | tpUTRnotCDS | tpUTRunique | ncexon |
|---|---|---|---|---|---|---|---|---|---|---|---|---|---|---|---|---|---|---|---|---|---|---|
| 61.54 | 88.89 | 61.54 | 71.43 | 88.89 | 93.33 | 92.86 | 90.91 | NW_011924881.1 | 40061 | 40061 | + | 1 | 1 | 1 | 0 | 0 | 0 | 1 | 0 | 1 | 1 | 0 |
| 81.82 | 100 | 100 | 92.31 | 100 | 100 | 100 | 92.31 | NW_011924881.1 | 40072 | 40072 | + | 1 | 1 | 1 | 0 | 0 | 0 | 1 | 0 | 1 | 1 | 0 |
| 63.64 | 90 | 80 | 70.59 | 70.97 | 62.07 | 68 | 50 | NW_011924881.1 | 40120 | 40120 | + | 1 | 1 | 1 | 0 | 0 | 0 | 1 | 0 | 1 | 1 | 0 |
| 78.95 | 92.31 | 73.91 | 83.33 | 81.25 | 89.74 | 78.26 | 100 | NW_011924881.1 | 40256 | 40256 | + | 1 | 1 | 1 | 0 | 1 | 0 | 0 | 0 | 0 | 0 | 0 |
| 89.47 | 96.43 | 91.3 | 83.33 | 100 | 90.32 | 90.48 | 100 | NW_011924881.1 | 40259 | 40259 | + | 1 | 1 | 1 | 0 | 1 | 0 | 0 | 0 | 0 | 0 | 0 |
| 76.47 | 86.67 | 86.96 | 91.3 | 83.33 | 88.57 | 90.48 | 100 | NW_011924881.1 | 40270 | 40270 | + | 1 | 1 | 1 | 0 | 1 | 0 | 0 | 0 | 0 | 0 | 0 |
| 82.35 | 95 | 95.65 | 84 | 90 | 93.33 | 87.5 | 94.12 | NW_011924881.1 | 40359 | 40359 | + | 1 | 1 | 1 | 0 | 1 | 0 | 0 | 0 | 0 | 0 | 0 |
| 93.75 | 88.89 | 95.83 | 91.67 | 100 | 96.55 | 93.33 | 100 | NW_011924881.1 | 40371 | 40371 | + | 1 | 1 | 1 | 0 | 1 | 0 | 0 | 0 | 0 | 0 | 0 |
| 37.31 | 35.37 | 31.33 | 37.61 | 45.45 | 42.96 | 48.57 | 42.22 | NW_011924881.1 | 447836 | 447836 | + | 1 | 1 | 1 | 0 | 1 | 0 | 0 | 0 | 0 | 0 | 0 |
| 87.1 | 84.62 | 78.12 | 82.76 | 89.47 | 86.96 | 91.67 | 93.1 | NW_011924881.1 | 447945 | 447945 | + | 1 | 0 | 0 | 0 | 0 | 0 | 0 | 0 | 0 | 0 | 0 |
| 74.07 | 93.75 | 80 | 61.54 | 88.89 | 97.22 | 85.71 | 72.73 | NW_011924881.1 | 447952 | 447952 | + | 1 | 0 | 0 | 0 | 0 | 0 | 0 | 0 | 0 | 0 | 0 |
| 92.31 | 84 | 87.5 | 86.21 | 100 | 92.68 | 96.55 | 87.5 | NW_011924881.1 | 447958 | 447958 | + | 1 | 0 | 0 | 0 | 0 | 0 | 0 | 0 | 0 | 0 | 0 |
| 85.71 | 72.73 | 82.86 | 80.56 | 82.35 | 75.51 | 79.49 | 78.95 | NW_011924881.1 | 448007 | 448007 | + | 1 | 0 | 0 | 0 | 0 | 0 | 0 | 0 | 0 | 0 | 0 |
| 66.67 | 47.83 | 61.54 | 61.36 | 62.07 | 64.81 | 68.18 | 54.84 | NW_011924881.1 | 448050 | 448050 | + | 1 | 1 | 1 | 0 | 1 | 0 | 0 | 0 | 0 | 0 | 0 |
| 78.26 | 71.7 | 70.27 | 72.92 | 71.43 | 79.41 | 75.51 | 48.72 | NW_011924881.1 | 448056 | 448056 | + | 1 | 1 | 1 | 0 | 1 | 0 | 0 | 0 | 0 | 0 | 0 |
| 94 | 100 | 92.31 | 89.58 | 97.96 | 98.15 | 95.52 | 88.37 | NW_011924881.1 | 448083 | 448083 | + | 1 | 1 | 1 | 0 | 1 | 0 | 0 | 0 | 0 | 0 | 0 |
| 94.87 | 96.55 | 95.56 | 97.73 | 100 | 94.83 | 94.83 | 89.58 | NW_011924881.1 | 448109 | 448109 | + | 1 | 1 | 1 | 0 | 1 | 0 | 0 | 0 | 0 | 0 | 0 |
| 94.74 | 92.86 | 95.35 | 97.56 | 100 | 100 | 94.34 | 94.12 | NW_011924881.1 | 448125 | 448125 | + | 1 | 1 | 1 | 0 | 1 | 0 | 0 | 0 | 0 | 0 | 0 |
| 100 | 98.21 | 97.37 | 97.96 | 100 | 97.01 | 98.08 | 100 | NW_011924881.1 | 448136 | 448136 | + | 1 | 1 | 1 | 0 | 1 | 0 | 0 | 0 | 0 | 0 | 0 |
| 82.35 | 73.47 | 89.47 | 95.92 | 85.71 | 87.5 | 92.73 | 86.54 | NW_011924881.1 | 448140 | 448140 | + | 1 | 1 | 1 | 0 | 1 | 0 | 0 | 0 | 0 | 0 | 0 |
| 82.14 | 88.57 | 90 | 93.1 | 91.67 | 95.35 | 87.5 | 89.74 | NW_011924881.1 | 448152 | 448152 | + | 1 | 1 | 1 | 0 | 1 | 0 | 0 | 0 | 0 | 0 | 0 |
| 90.91 | 97.56 | 100 | 100 | 89.66 | 100 | 94.44 | 100 | NW_011924881.1 | 448167 | 448167 | + | 1 | 1 | 1 | 0 | 1 | 0 | 0 | 0 | 0 | 0 | 0 |
| 100 | 96.15 | 89.29 | 90.48 | 80 | 97.56 | 90 | 80 | NW_011924881.1 | 448200 | 448200 | + | 1 | 1 | 1 | 0 | 1 | 0 | 0 | 0 | 0 | 0 | 0 |
| 71.43 | 53.85 | 74.07 | 55.56 | 70.37 | 73.81 | 52.94 | 56 | NW_011924881.1 | 448247 | 448247 | + | 1 | 1 | 1 | 0 | 0 | 0 | 1 | 0 | 1 | 1 | 0 |

...
... only first 25 lines shown ...

```
Directory: RNK File: 0READMErnk

RNK - Genomic feature regions ranked by CpG methylation and hsm statistics.

Input: studymc, studymk, region.gr (GRanges object providing annotated region labels and bounds)

Output: files ranked-genes-*.txt sites-in-genes-*.txt
              ranked-promoters-*.txt sites-in-promoters-*.txt
              plot*.pdf

Notes: The output is generated by BWASPR::rank_rbm().
       The files ranked-genes-*.txt provide ranked lists of genes based on
        overall methylation percentage or the occurrence of CpGhms sites
        within the annotated gene bounds.  The output columns include
          region_ID    (= region/gene name)
          rwidth       (= region/gene width)
          nbrsites     (= number of CpGhsm sites in the region/gene)
          nbrper10kb   (= number of CpGhsm sites in the region/gene normalized to 10kb width)
          pmrpersite   (= average % methylation per CpGhsm site in the region/gene)
          pmrpernucl   (= average % CpGhsm methylation per nucleotide in the region/gene)
          prcntM       (= overall % CpGhsm methylation in the region/gene)

       The tables are sorted by prcntM (*byPrcntM* files) or by nbper10kb
         (*bySiteDensity* files) in descending order.

       The ranked-promoters-*.txt files provide analogous tables for promoter regions.
        If the promoter annotation was derived with BWASP, then these regions are simply
        defined here as 500 nucleotides upstream of the 5'-end of the gene (shorter if the
        scaffold ends before).

       The files sites-*.txt provide details for the CpGhsm sites in the respective
        regions.
       The files plot-*.pdf explore concordance of the two methylation measures.
       The optional parameters minrwidth, maxrwidth, and minnbrsites specify
       the length range and minimal number of methylation sites for regions
       to be included in the plot.
```

Directory: RNK File: ranked-genes-byPrcntM-Cb_LI_BCphaseA1.txt

| region_ID | seqnames | start | end | width | strand | coverage | numCs | numTs | prcntM |
|---|---|---|---|---|---|---|---|---|---|
| gene13336 | NW_011929072.1 | 30 | 517 | 488 | + | 71 | 69 | 2 | 97.18 |
| gene8895 | NW_011925229.1 | 67030 | 70140 | 3111 | - | 42 | 38 | 4 | 90.48 |
| gene13247 | NW_011926712.1 | 6 | 783 | 778 | + | 10 | 9 | 1 | 90 |
| gene1643 | NW_011924932.1 | 342040 | 343422 | 1383 | - | 19 | 17 | 2 | 89.47 |
| gene10364 | NW_011925342.1 | 111577 | 112136 | 560 | + | 372 | 332 | 40 | 89.25 |
| gene13380 | NW_011929316.1 | 135 | 1735 | 1601 | - | 69 | 60 | 9 | 86.96 |
| gene903 | NW_011924914.1 | 17973 | 18954 | 982 | - | 125 | 106 | 19 | 84.8 |
| gene5000 | NW_011925022.1 | 386851 | 387976 | 1126 | - | 416 | 345 | 71 | 82.93 |
| gene10653 | NW_011925362.1 | 80044 | 81082 | 1039 | - | 145 | 119 | 26 | 82.07 |
| gene12557 | NW_011925715.1 | 916490 | 917391 | 902 | - | 26 | 21 | 5 | 80.77 |
| gene2956 | NW_011924977.1 | 323781 | 325133 | 1353 | + | 484 | 372 | 112 | 76.86 |
| gene13327 | NW_011929009.1 | 6 | 1008 | 1003 | + | 56 | 43 | 13 | 76.79 |
| gene6356 | NW_011925092.1 | 13079 | 14878 | 1800 | + | 152 | 116 | 36 | 76.32 |
| gene405 | NW_011924888.1 | 1707554 | 1708995 | 1442 | - | 961 | 730 | 231 | 75.96 |
| gene8458 | NW_011925192.1 | 234647 | 235699 | 1053 | - | 512 | 385 | 127 | 75.2 |
| gene3015 | NW_011924977.1 | 732939 | 733749 | 811 | - | 614 | 460 | 154 | 74.92 |
| gene4579 | NW_011925016.1 | 246765 | 248208 | 1444 | - | 616 | 458 | 158 | 74.35 |
| gene9952 | NW_011925321.1 | 87267 | 88468 | 1202 | + | 537 | 398 | 139 | 74.12 |
| gene10196 | NW_011925327.1 | 165617 | 166924 | 1308 | + | 246 | 182 | 64 | 73.98 |
| gene8315 | NW_011925189.1 | 490017 | 490870 | 854 | + | 482 | 354 | 128 | 73.44 |
| gene10163 | NW_011925323.1 | 2060801 | 2062147 | 1347 | - | 679 | 490 | 189 | 72.16 |
| gene9525 | NW_011925273.1 | 781582 | 783179 | 1598 | + | 843 | 608 | 235 | 72.12 |
| gene3362 | NW_011924987.1 | 68555 | 70011 | 1457 | - | 342 | 246 | 96 | 71.93 |
| gene2249 | NW_011924946.1 | 3301258 | 3302569 | 1312 | - | 404 | 290 | 114 | 71.78 |

...

... only first 25 lines shown ...

Directory: RNK File: ranked-genes-byPrcntM-Cb_LI_BCphaseA2.txt

| region_ID | seqnames | start | end | width | strand | coverage | numCs | numTs | prcntM |
|---|---|---|---|---|---|---|---|---|---|
| gene8895 | NW_011925229.1 | 67030 | 70140 | 3111 | - | 34 | 34 | 0 | 100 |
| gene13336 | NW_011929072.1 | 30 | 517 | 488 | + | 150 | 140 | 10 | 93.33 |
| gene10364 | NW_011925342.1 | 111577 | 112136 | 560 | + | 285 | 254 | 31 | 89.12 |
| gene13213 | NW_011926558.1 | 4 | 1118 | 1115 | - | 44 | 39 | 5 | 88.64 |
| gene2219 | NW_011924946.1 | 3167076 | 3167848 | 773 | - | 30 | 26 | 4 | 86.67 |
| gene10653 | NW_011925362.1 | 80044 | 81082 | 1039 | - | 120 | 104 | 16 | 86.67 |
| gene1643 | NW_011924932.1 | 342040 | 343422 | 1383 | - | 27 | 23 | 4 | 85.19 |
| gene5000 | NW_011925022.1 | 386851 | 387976 | 1126 | - | 388 | 314 | 74 | 80.93 |
| gene12557 | NW_011925715.1 | 916490 | 917391 | 902 | - | 36 | 29 | 7 | 80.56 |
| gene8203 | NW_011925182.1 | 19245 | 20253 | 1009 | + | 457 | 364 | 93 | 79.65 |
| gene2956 | NW_011924977.1 | 323781 | 325133 | 1353 | + | 471 | 374 | 97 | 79.41 |
| gene1677 | NW_011924932.1 | 641367 | 642280 | 914 | - | 126 | 99 | 27 | 78.57 |
| gene3362 | NW_011924987.1 | 68555 | 70011 | 1457 | - | 242 | 188 | 54 | 77.69 |
| gene10196 | NW_011925327.1 | 165617 | 166924 | 1308 | + | 322 | 249 | 73 | 77.33 |
| gene266 | NW_011924888.1 | 34060 | 35504 | 1445 | + | 160 | 123 | 37 | 76.88 |
| gene8330 | NW_011925189.1 | 572782 | 573960 | 1179 | + | 233 | 179 | 54 | 76.82 |
| gene6173 | NW_011925075.1 | 824766 | 825641 | 876 | + | 207 | 157 | 50 | 75.85 |
| gene3015 | NW_011924977.1 | 732939 | 733749 | 811 | - | 610 | 457 | 153 | 74.92 |
| gene9312 | NW_011925258.1 | 34384 | 35413 | 1030 | + | 211 | 158 | 53 | 74.88 |
| gene4579 | NW_011925016.1 | 246765 | 248208 | 1444 | - | 665 | 495 | 170 | 74.44 |
| gene405 | NW_011924888.1 | 1707554 | 1708995 | 1442 | - | 1165 | 857 | 308 | 73.56 |
| gene13380 | NW_011929316.1 | 135 | 1735 | 1601 | - | 34 | 25 | 9 | 73.53 |
| gene6356 | NW_011925092.1 | 13079 | 14878 | 1800 | + | 133 | 96 | 37 | 72.18 |
| gene8315 | NW_011925189.1 | 490017 | 490870 | 854 | + | 671 | 481 | 190 | 71.68 |

...
... only first 25 lines shown ...

Directory: RNK File: ranked-genes-byPrcntM-Cb_LI_BCphaseB1.txt

| region_ID | seqnames | start | end | width | strand | coverage | numCs | numTs | prcntM |
|---|---|---|---|---|---|---|---|---|---|
| gene13213 | NW_011926558.1 | 4 | 1118 | 1115 | - | 18 | 18 | 0 | 100 |
| gene13336 | NW_011929072.1 | 30 | 517 | 488 | + | 95 | 90 | 5 | 94.74 |
| gene13380 | NW_011929316.1 | 135 | 1735 | 1601 | - | 41 | 37 | 4 | 90.24 |
| gene1643 | NW_011924932.1 | 342040 | 343422 | 1383 | - | 18 | 16 | 2 | 88.89 |
| gene10364 | NW_011925342.1 | 111577 | 112136 | 560 | + | 378 | 325 | 53 | 85.98 |
| gene8895 | NW_011925229.1 | 67030 | 70140 | 3111 | - | 72 | 61 | 11 | 84.72 |
| gene5000 | NW_011925022.1 | 386851 | 387976 | 1126 | - | 460 | 374 | 86 | 81.3 |
| gene10653 | NW_011925362.1 | 80044 | 81082 | 1039 | - | 173 | 136 | 37 | 78.61 |
| gene10196 | NW_011925327.1 | 165617 | 166924 | 1308 | + | 370 | 281 | 89 | 75.95 |
| gene2956 | NW_011924977.1 | 323781 | 325133 | 1353 | + | 697 | 524 | 173 | 75.18 |
| gene2219 | NW_011924946.1 | 3167076 | 3167848 | 773 | - | 16 | 12 | 4 | 75 |
| gene4579 | NW_011925016.1 | 246765 | 248208 | 1444 | - | 691 | 508 | 183 | 73.52 |
| gene8203 | NW_011925182.1 | 19245 | 20253 | 1009 | + | 452 | 331 | 121 | 73.23 |
| gene8315 | NW_011925189.1 | 490017 | 490870 | 854 | + | 651 | 476 | 175 | 73.12 |
| gene3015 | NW_011924977.1 | 732939 | 733749 | 811 | - | 864 | 629 | 235 | 72.8 |
| gene405 | NW_011924888.1 | 1707554 | 1708995 | 1442 | - | 1451 | 1051 | 400 | 72.43 |
| gene2249 | NW_011924946.1 | 3301258 | 3302569 | 1312 | - | 387 | 280 | 107 | 72.35 |
| gene9952 | NW_011925321.1 | 87267 | 88468 | 1202 | + | 697 | 501 | 196 | 71.88 |
| gene903 | NW_011924914.1 | 17973 | 18954 | 982 | - | 132 | 94 | 38 | 71.21 |
| gene9412 | NW_011925267.1 | 159703 | 161346 | 1644 | - | 699 | 489 | 210 | 69.96 |
| gene4880 | NW_011925020.1 | 2602046 | 2603046 | 1001 | + | 757 | 516 | 241 | 68.16 |
| gene12557 | NW_011925715.1 | 916490 | 917391 | 902 | - | 56 | 38 | 18 | 67.86 |
| gene3362 | NW_011924987.1 | 68555 | 70011 | 1457 | - | 267 | 180 | 87 | 67.42 |
| gene9525 | NW_011925273.1 | 781582 | 783179 | 1598 | + | 1035 | 697 | 338 | 67.34 |

...

... only first 25 lines shown ...

Directory: RNK File: ranked-genes-byPrcntM-Cb_LI_BCphaseB2.txt

| region_ID | seqnames | start | end | width | strand | coverage | numCs | numTs | prcntM |
|---|---|---|---|---|---|---|---|---|---|
| gene13336 | NW_011929072.1 | 30 | 517 | 488 | + | 97 | 97 | 0 | 100 |
| gene13213 | NW_011926558.1 | 4 | 1118 | 1115 | - | 30 | 29 | 1 | 96.67 |
| gene13247 | NW_011926712.1 | 6 | 783 | 778 | + | 16 | 15 | 1 | 93.75 |
| gene10364 | NW_011925342.1 | 111577 | 112136 | 560 | + | 342 | 297 | 45 | 86.84 |
| gene13380 | NW_011929316.1 | 135 | 1735 | 1601 | - | 30 | 25 | 5 | 83.33 |
| gene10653 | NW_011925362.1 | 80044 | 81082 | 1039 | - | 158 | 128 | 30 | 81.01 |
| gene10196 | NW_011925327.1 | 165617 | 166924 | 1308 | + | 280 | 226 | 54 | 80.71 |
| gene2956 | NW_011924977.1 | 323781 | 325133 | 1353 | + | 587 | 467 | 120 | 79.56 |
| gene8203 | NW_011925182.1 | 19245 | 20253 | 1009 | + | 453 | 360 | 93 | 79.47 |
| gene8895 | NW_011925229.1 | 67030 | 70140 | 3111 | - | 85 | 67 | 18 | 78.82 |
| gene5000 | NW_011925022.1 | 386851 | 387976 | 1126 | - | 428 | 337 | 91 | 78.74 |
| gene3015 | NW_011924977.1 | 732939 | 733749 | 811 | - | 697 | 529 | 168 | 75.9 |
| gene9525 | NW_011925273.1 | 781582 | 783179 | 1598 | + | 910 | 684 | 226 | 75.16 |
| gene8055 | NW_011925163.1 | 580633 | 581209 | 577 | - | 19 | 14 | 5 | 73.68 |
| gene2249 | NW_011924946.1 | 3301258 | 3302569 | 1312 | - | 525 | 383 | 142 | 72.95 |
| gene4579 | NW_011925016.1 | 246765 | 248208 | 1444 | - | 639 | 465 | 174 | 72.77 |
| gene13327 | NW_011929009.1 | 6 | 1008 | 1003 | + | 77 | 56 | 21 | 72.73 |
| gene903 | NW_011924914.1 | 17973 | 18954 | 982 | - | 105 | 76 | 29 | 72.38 |
| gene8458 | NW_011925192.1 | 234647 | 235699 | 1053 | - | 616 | 439 | 177 | 71.27 |
| gene405 | NW_011924888.1 | 1707554 | 1708995 | 1442 | - | 1205 | 847 | 358 | 70.29 |
| gene1780 | NW_011924937.1 | 525027 | 527068 | 2042 | - | 999 | 693 | 306 | 69.37 |
| gene6356 | NW_011925092.1 | 13079 | 14878 | 1800 | + | 153 | 106 | 47 | 69.28 |
| gene8946 | NW_011925232.1 | 235912 | 237534 | 1623 | - | 1955 | 1352 | 603 | 69.16 |
| gene4906 | NW_011925020.1 | 2960255 | 2961622 | 1368 | - | 1133 | 782 | 351 | 69.02 |

...

... only first 25 lines shown ...

Directory: RNK File: ranked-genes-byPrcntM-Cb_LI_RphaseA1.txt

| region_ID | seqnames | start | end | width | strand | coverage | numCs | numTs | prcntM |
|---|---|---|---|---|---|---|---|---|---|
| gene13213 | NW_011926558.1 | 4 | 1118 | 1115 | - | 25 | 24 | 1 | 96 |
| gene1643 | NW_011924932.1 | 342040 | 343422 | 1383 | - | 38 | 36 | 2 | 94.74 |
| gene13336 | NW_011929072.1 | 30 | 517 | 488 | + | 132 | 123 | 9 | 93.18 |
| gene8895 | NW_011925229.1 | 67030 | 70140 | 3111 | - | 26 | 22 | 4 | 84.62 |
| gene13247 | NW_011926712.1 | 6 | 783 | 778 | + | 13 | 11 | 2 | 84.62 |
| gene13380 | NW_011929316.1 | 135 | 1735 | 1601 | - | 51 | 43 | 8 | 84.31 |
| gene10364 | NW_011925342.1 | 111577 | 112136 | 560 | + | 383 | 316 | 67 | 82.51 |
| gene10653 | NW_011925362.1 | 80044 | 81082 | 1039 | - | 112 | 88 | 24 | 78.57 |
| gene3362 | NW_011924987.1 | 68555 | 70011 | 1457 | - | 273 | 213 | 60 | 78.02 |
| gene903 | NW_011924914.1 | 17973 | 18954 | 982 | - | 146 | 113 | 33 | 77.4 |
| gene5000 | NW_011925022.1 | 386851 | 387976 | 1126 | - | 347 | 267 | 80 | 76.95 |
| gene2956 | NW_011924977.1 | 323781 | 325133 | 1353 | + | 508 | 382 | 126 | 75.2 |
| gene4579 | NW_011925016.1 | 246765 | 248208 | 1444 | - | 602 | 448 | 154 | 74.42 |
| gene4880 | NW_011925020.1 | 2602046 | 2603046 | 1001 | + | 621 | 459 | 162 | 73.91 |
| gene10196 | NW_011925327.1 | 165617 | 166924 | 1308 | + | 328 | 239 | 89 | 72.87 |
| gene4906 | NW_011925020.1 | 2960255 | 2961622 | 1368 | - | 777 | 566 | 211 | 72.84 |
| gene3015 | NW_011924977.1 | 732939 | 733749 | 811 | - | 661 | 481 | 180 | 72.77 |
| gene8315 | NW_011925189.1 | 490017 | 490870 | 854 | + | 691 | 498 | 193 | 72.07 |
| gene2249 | NW_011924946.1 | 3301258 | 3302569 | 1312 | - | 331 | 230 | 101 | 69.49 |
| gene4879 | NW_011925020.1 | 2601782 | 2603159 | 1378 | - | 795 | 550 | 245 | 69.18 |
| gene5374 | NW_011925035.1 | 691928 | 694456 | 2529 | - | 327 | 225 | 102 | 68.81 |
| gene405 | NW_011924888.1 | 1707554 | 1708995 | 1442 | - | 1298 | 893 | 405 | 68.8 |
| gene8203 | NW_011925182.1 | 19245 | 20253 | 1009 | + | 456 | 313 | 143 | 68.64 |
| gene6356 | NW_011925092.1 | 13079 | 14878 | 1800 | + | 227 | 155 | 72 | 68.28 |

...

... only first 25 lines shown ...

Directory: RNK File: ranked-genes-byPrcntM-Cb_LI_RphaseA2.txt

| region_ID | seqnames | start | end | width | strand | coverage | numCs | numTs | prcntM |
|---|---|---|---|---|---|---|---|---|---|
| gene13336 | NW_011929072.1 | 30 | 517 | 488 | + | 176 | 168 | 8 | 95.45 |
| gene10364 | NW_011925342.1 | 111577 | 112136 | 560 | + | 561 | 500 | 61 | 89.13 |
| gene13213 | NW_011926558.1 | 4 | 1118 | 1115 | - | 59 | 52 | 7 | 88.14 |
| gene10653 | NW_011925362.1 | 80044 | 81082 | 1039 | - | 234 | 198 | 36 | 84.62 |
| gene1643 | NW_011924932.1 | 342040 | 343422 | 1383 | - | 36 | 30 | 6 | 83.33 |
| gene8895 | NW_011925229.1 | 67030 | 70140 | 3111 | - | 46 | 38 | 8 | 82.61 |
| gene2956 | NW_011924977.1 | 323781 | 325133 | 1353 | + | 854 | 679 | 175 | 79.51 |
| gene5000 | NW_011925022.1 | 386851 | 387976 | 1126 | - | 660 | 523 | 137 | 79.24 |
| gene13380 | NW_011929316.1 | 135 | 1735 | 1601 | - | 89 | 70 | 19 | 78.65 |
| gene10196 | NW_011925327.1 | 165617 | 166924 | 1308 | + | 510 | 398 | 112 | 78.04 |
| gene3015 | NW_011924977.1 | 732939 | 733749 | 811 | - | 870 | 662 | 208 | 76.09 |
| gene405 | NW_011924888.1 | 1707554 | 1708995 | 1442 | - | 2009 | 1512 | 497 | 75.26 |
| gene903 | NW_011924914.1 | 17973 | 18954 | 982 | - | 317 | 238 | 79 | 75.08 |
| gene8203 | NW_011925182.1 | 19245 | 20253 | 1009 | + | 664 | 498 | 166 | 75 |
| gene13247 | NW_011926712.1 | 6 | 783 | 778 | + | 32 | 24 | 8 | 75 |
| gene12557 | NW_011925715.1 | 916490 | 917391 | 902 | - | 61 | 45 | 16 | 73.77 |
| gene2249 | NW_011924946.1 | 3301258 | 3302569 | 1312 | - | 583 | 428 | 155 | 73.41 |
| gene8315 | NW_011925189.1 | 490017 | 490870 | 854 | + | 900 | 660 | 240 | 73.33 |
| gene4579 | NW_011925016.1 | 246765 | 248208 | 1444 | - | 1150 | 838 | 312 | 72.87 |
| gene4880 | NW_011925020.1 | 2602046 | 2603046 | 1001 | + | 821 | 594 | 227 | 72.35 |
| gene9525 | NW_011925273.1 | 781582 | 783179 | 1598 | + | 1285 | 923 | 362 | 71.83 |
| gene13327 | NW_011929009.1 | 6 | 1008 | 1003 | + | 206 | 146 | 60 | 70.87 |
| gene3362 | NW_011924987.1 | 68555 | 70011 | 1457 | - | 384 | 272 | 112 | 70.83 |
| gene6356 | NW_011925092.1 | 13079 | 14878 | 1800 | + | 284 | 200 | 84 | 70.42 |

...

... only first 25 lines shown ...

Directory: RNK File: ranked-genes-byPrcntM-Cb_LI_RphaseB1.txt

| region_ID | seqnames | start | end | width | strand | coverage | numCs | numTs | prcntM |
|---|---|---|---|---|---|---|---|---|---|
| gene13336 | NW_011929072.1 | 30 | 517 | 488 | + | 101 | 96 | 5 | 95.05 |
| gene13213 | NW_011926558.1 | 4 | 1118 | 1115 | - | 30 | 28 | 2 | 93.33 |
| gene8895 | NW_011925229.1 | 67030 | 70140 | 3111 | - | 26 | 24 | 2 | 92.31 |
| gene10364 | NW_011925342.1 | 111577 | 112136 | 560 | + | 420 | 362 | 58 | 86.19 |
| gene10653 | NW_011925362.1 | 80044 | 81082 | 1039 | - | 157 | 135 | 22 | 85.99 |
| gene903 | NW_011924914.1 | 17973 | 18954 | 982 | - | 50 | 42 | 8 | 84 |
| gene1643 | NW_011924932.1 | 342040 | 343422 | 1383 | - | 30 | 25 | 5 | 83.33 |
| gene8055 | NW_011925163.1 | 580633 | 581209 | 577 | - | 24 | 20 | 4 | 83.33 |
| gene5000 | NW_011925022.1 | 386851 | 387976 | 1126 | - | 386 | 306 | 80 | 79.27 |
| gene8203 | NW_011925182.1 | 19245 | 20253 | 1009 | + | 443 | 338 | 105 | 76.3 |
| gene2956 | NW_011924977.1 | 323781 | 325133 | 1353 | + | 616 | 469 | 147 | 76.14 |
| gene9312 | NW_011925258.1 | 34384 | 35413 | 1030 | + | 292 | 217 | 75 | 74.32 |
| gene10196 | NW_011925327.1 | 165617 | 166924 | 1308 | + | 317 | 235 | 82 | 74.13 |
| gene9952 | NW_011925321.1 | 87267 | 88468 | 1202 | + | 593 | 439 | 154 | 74.03 |
| gene3015 | NW_011924977.1 | 732939 | 733749 | 811 | - | 732 | 536 | 196 | 73.22 |
| gene8330 | NW_011925189.1 | 572782 | 573960 | 1179 | + | 265 | 194 | 71 | 73.21 |
| gene4579 | NW_011925016.1 | 246765 | 248208 | 1444 | - | 712 | 514 | 198 | 72.19 |
| gene5374 | NW_011925035.1 | 691928 | 694456 | 2529 | - | 298 | 215 | 83 | 72.15 |
| gene12557 | NW_011925715.1 | 916490 | 917391 | 902 | - | 32 | 23 | 9 | 71.88 |
| gene9525 | NW_011925273.1 | 781582 | 783179 | 1598 | + | 816 | 575 | 241 | 70.47 |
| gene405 | NW_011924888.1 | 1707554 | 1708995 | 1442 | - | 1244 | 873 | 371 | 70.18 |
| gene266 | NW_011924888.1 | 34060 | 35504 | 1445 | + | 156 | 109 | 47 | 69.87 |
| gene6356 | NW_011925092.1 | 13079 | 14878 | 1800 | + | 246 | 170 | 76 | 69.11 |
| gene8315 | NW_011925189.1 | 490017 | 490870 | 854 | + | 630 | 434 | 196 | 68.89 |

...
... only first 25 lines shown ...

Directory: RNK File: ranked-genes-byPrcntM-Cb_LI_RphaseB2.txt

| region_ID | seqnames | start | end | width | strand | coverage | numCs | numTs | prcntM |
|---|---|---|---|---|---|---|---|---|---|
| gene1643 | NW_011924932.1 | 342040 | 343422 | 1383 | - | 17 | 17 | 0 | 100 |
| gene13213 | NW_011926558.1 | 4 | 1118 | 1115 | - | 21 | 20 | 1 | 95.24 |
| gene8895 | NW_011925229.1 | 67030 | 70140 | 3111 | - | 33 | 31 | 2 | 93.94 |
| gene13336 | NW_011929072.1 | 30 | 517 | 488 | + | 116 | 108 | 8 | 93.1 |
| gene12557 | NW_011925715.1 | 916490 | 917391 | 902 | - | 21 | 19 | 2 | 90.48 |
| gene10653 | NW_011925362.1 | 80044 | 81082 | 1039 | - | 139 | 120 | 19 | 86.33 |
| gene13380 | NW_011929316.1 | 135 | 1735 | 1601 | - | 20 | 17 | 3 | 85 |
| gene10364 | NW_011925342.1 | 111577 | 112136 | 560 | + | 305 | 259 | 46 | 84.92 |
| gene5000 | NW_011925022.1 | 386851 | 387976 | 1126 | - | 361 | 306 | 55 | 84.76 |
| gene13247 | NW_011926712.1 | 6 | 783 | 778 | + | 28 | 23 | 5 | 82.14 |
| gene2956 | NW_011924977.1 | 323781 | 325133 | 1353 | + | 484 | 394 | 90 | 81.4 |
| gene10196 | NW_011925327.1 | 165617 | 166924 | 1308 | + | 270 | 219 | 51 | 81.11 |
| gene8203 | NW_011925182.1 | 19245 | 20253 | 1009 | + | 457 | 369 | 88 | 80.74 |
| gene9312 | NW_011925258.1 | 34384 | 35413 | 1030 | + | 211 | 167 | 44 | 79.15 |
| gene2249 | NW_011924946.1 | 3301258 | 3302569 | 1312 | - | 366 | 285 | 81 | 77.87 |
| gene9525 | NW_011925273.1 | 781582 | 783179 | 1598 | + | 782 | 601 | 181 | 76.85 |
| gene3015 | NW_011924977.1 | 732939 | 733749 | 811 | - | 656 | 487 | 169 | 74.24 |
| gene4880 | NW_011925020.1 | 2602046 | 2603046 | 1001 | + | 556 | 405 | 151 | 72.84 |
| gene9952 | NW_011925321.1 | 87267 | 88468 | 1202 | + | 468 | 333 | 135 | 71.15 |
| gene5374 | NW_011925035.1 | 691928 | 694456 | 2529 | - | 286 | 202 | 84 | 70.63 |
| gene405 | NW_011924888.1 | 1707554 | 1708995 | 1442 | - | 1030 | 723 | 307 | 70.19 |
| gene6356 | NW_011925092.1 | 13079 | 14878 | 1800 | + | 201 | 141 | 60 | 70.15 |
| gene2274 | NW_011924947.1 | 109214 | 110680 | 1467 | - | 2064 | 1447 | 617 | 70.11 |
| gene7935 | NW_011925161.1 | 3201027 | 3202301 | 1275 | - | 833 | 584 | 249 | 70.11 |

...
... only first 25 lines shown ...

Directory: RNK File: ranked-genes-bySiteDensity-Cb_LI_BCphaseA1.txt

| region_ID | rwidth | nbrsites | nbrper10kb | pmpersite | pmpernucl | pglink | seqnames | start | end | strand | coverage | numCs | numTs | prcntM |
|---|---|---|---|---|---|---|---|---|---|---|---|---|---|---|
| gene11569 | 1311 | 92 | 701.75 | 74.9 | 5.26 | https://www.ncbi.nlm.nih.gov/gene/?term=LOC105285394 | NW_011925459.1 | 186244 | 187554 | - | 2833 | 1687 | 1146 | 59.55 |
| gene10219 | 235 | 16 | 680.85 | 65.39 | 4.45 | https://www.ncbi.nlm.nih.gov/gene/?term=LOC105283830 | NW_011925327.1 | 293521 | 293755 | + | 695 | 278 | 417 | 40 |
| gene13384 | 1877 | 124 | 660.63 | 53.27 | 3.52 | https://www.ncbi.nlm.nih.gov/gene/?term=LOC105287447 | NW_011929339.1 | 7 | 1883 | + | 8410 | 2426 | 5984 | 28.85 |
| gene5917 | 823 | 53 | 643.99 | 42.31 | 2.72 | https://www.ncbi.nlm.nih.gov/gene/?term=LOC105278849 | NW_011925053.1 | 1371807 | 1372629 | + | 4237 | 1013 | 3224 | 23.91 |
| gene5309 | 1704 | 107 | 627.93 | 53.57 | 3.36 | https://www.ncbi.nlm.nih.gov/gene/?term=LOC105278160 | NW_011925031.1 | 150501 | 152204 | - | 5674 | 2146 | 3528 | 37.82 |
| gene211 | 1280 | 77 | 601.56 | 63.58 | 3.82 | https://www.ncbi.nlm.nih.gov/gene/?term=LOC105274534 | NW_011924887.1 | 1565391 | 1566670 | + | 3697 | 1566 | 2131 | 42.36 |
| gene5911 | 943 | 56 | 593.85 | 63.63 | 3.78 | https://www.ncbi.nlm.nih.gov/gene/?term=LOC105278844 | NW_011925053.1 | 1331236 | 1332178 | - | 3938 | 1371 | 2567 | 34.81 |
| gene12364 | 1403 | 83 | 591.59 | 67.3 | 3.98 | https://www.ncbi.nlm.nih.gov/gene/?term=LOC105286269 | NW_011925664.1 | 291441 | 292843 | - | 3486 | 1682 | 1804 | 48.25 |
| gene4878 | 675 | 39 | 577.78 | 77.75 | 4.49 | https://www.ncbi.nlm.nih.gov/gene/?term=LOC105277648 | NW_011925020.1 | 2601147 | 2601821 | - | 1222 | 737 | 485 | 60.31 |
| gene1562 | 1491 | 85 | 570.09 | 58.01 | 3.31 | https://www.ncbi.nlm.nih.gov/gene/?term=LOC105287692 | NW_011924931.1 | 3966137 | 3967627 | - | 5013 | 1543 | 3470 | 30.78 |
| gene3719 | 1471 | 82 | 557.44 | 69.7 | 3.89 | https://www.ncbi.nlm.nih.gov/gene/?term=LOC105276360 | NW_011924994.1 | 94735 | 96205 | - | 5289 | 1702 | 3587 | 32.18 |
| gene3731 | 865 | 48 | 554.91 | 59.87 | 3.32 | https://www.ncbi.nlm.nih.gov/gene/?term=LOC105276371 | NW_011924994.1 | 459736 | 460600 | - | 2341 | 990 | 1351 | 42.29 |
| gene2797 | 1489 | 82 | 550.71 | 65.25 | 3.59 | https://www.ncbi.nlm.nih.gov/gene/?term=LOC105275314 | NW_011924973.1 | 823242 | 824730 | + | 3826 | 1545 | 2281 | 40.38 |
| gene570 | 1673 | 91 | 543.93 | 63.03 | 3.43 | https://www.ncbi.nlm.nih.gov/gene/?term=LOC105278549 | NW_011924892.1 | 1160484 | 1162156 | + | 4357 | 1716 | 2641 | 39.38 |
| gene11566 | 1603 | 87 | 542.73 | 58.21 | 3.16 | https://www.ncbi.nlm.nih.gov/gene/?term=LOC105285392 | NW_011925459.1 | 162939 | 164541 | + | 4181 | 1491 | 2690 | 35.66 |
| gene1894 | 445 | 24 | 539.33 | 69.07 | 3.73 | https://www.ncbi.nlm.nih.gov/gene/?term=LOC105288083 | NW_011924945.1 | 117665 | 118109 | + | 633 | 351 | 282 | 55.45 |
| gene12538 | 372 | 20 | 537.63 | 67.08 | 3.61 | https://www.ncbi.nlm.nih.gov/gene/?term=LOC105286531 | NW_011925715.1 | 412527 | 412898 | - | 1217 | 424 | 793 | 34.84 |
| gene12652 | 1122 | 60 | 534.76 | 68.25 | 3.65 | https://www.ncbi.nlm.nih.gov/gene/?term=LOC109610640 | NW_011925762.1 | 557576 | 558697 | + | 2312 | 1182 | 1130 | 51.12 |
| gene1223 | 1048 | 56 | 534.35 | 64.89 | 3.47 | https://www.ncbi.nlm.nih.gov/gene/?term=LOC105285713 | NW_011924922.1 | 2517529 | 2518576 | - | 2099 | 986 | 1113 | 46.97 |
| gene1654 | 2285 | 121 | 529.54 | 65.7 | 3.48 | https://www.ncbi.nlm.nih.gov/gene/?term=LOC105287811 | NW_011924932.1 | 539418 | 541702 | - | 6306 | 2727 | 3579 | 43.24 |
| gene12602 | 1399 | 73 | 521.8 | 76.82 | 4.01 | https://www.ncbi.nlm.nih.gov/gene/?term=LOC105286562 | NW_011925739.1 | 19908 | 21306 | + | 1963 | 1007 | 956 | 51.3 |
| gene5714 | 710 | 37 | 521.13 | 87.71 | 4.57 | https://www.ncbi.nlm.nih.gov/gene/?term=LOC105278729 | NW_011925050.1 | 7095 | 7804 | - | 833 | 509 | 324 | 61.1 |
| gene12330 | 1501 | 78 | 519.65 | 65.45 | 3.4 | https://www.ncbi.nlm.nih.gov/gene/?term=LOC105286235 | NW_011925664.1 | 138328 | 139828 | - | 3911 | 1654 | 2257 | 42.29 |
| gene10943 | 1759 | 91 | 517.34 | 68.07 | 3.52 | https://www.ncbi.nlm.nih.gov/gene/?term=LOC105284659 | NW_011925387.1 | 118203 | 119961 | - | 3593 | 1606 | 1987 | 44.7 |

...
... only first 25 lines shown ...

Directory: RNK File: ranked-genes-bySiteDensity-Cb_LI_BCphaseA2.txt

| region_ID | rwidth | nbrsites | nbrper10kb | pmpersite | pmpernucl | pglink | seqnames | start | end | strand | coverage | numCs | numTs | prcntM |
|---|---|---|---|---|---|---|---|---|---|---|---|---|---|---|
| gene5917 | 823 | 81 | 984.2 | 39.76 | 3.91 | https://www.ncbi.nlm.nih.gov/gene/?term=LOC105278849 | NW_011925053.1 | 1371807 | 1372629 | + | 7068 | 1725 | 5343 | 24.41 |
| gene13384 | 1877 | 164 | 873.73 | 49.49 | 4.32 | https://www.ncbi.nlm.nih.gov/gene/?term=LOC105287447 | NW_011929339.1 | 7 | 1883 | + | 12571 | 4014 | 8557 | 31.93 |
| gene5309 | 1704 | 126 | 739.44 | 52.83 | 3.91 | https://www.ncbi.nlm.nih.gov/gene/?term=LOC105278160 | NW_011925031.1 | 150501 | 152204 | - | 8379 | 3329 | 5050 | 39.73 |
| gene11569 | 1311 | 92 | 701.75 | 77.64 | 5.45 | https://www.ncbi.nlm.nih.gov/gene/?term=LOC105285394 | NW_011925459.1 | 186244 | 187554 | - | 3733 | 2333 | 1400 | 62.5 |
| gene12538 | 372 | 25 | 672.04 | 61.42 | 4.13 | https://www.ncbi.nlm.nih.gov/gene/?term=LOC105286531 | NW_011925715.1 | 412527 | 412898 | - | 1839 | 685 | 1154 | 37.25 |
| gene11566 | 1603 | 105 | 655.02 | 55.03 | 3.6 | https://www.ncbi.nlm.nih.gov/gene/?term=LOC105285392 | NW_011925459.1 | 162939 | 164541 | + | 5989 | 2272 | 3717 | 37.94 |
| gene12364 | 1403 | 91 | 648.61 | 66.37 | 4.3 | https://www.ncbi.nlm.nih.gov/gene/?term=LOC105286269 | NW_011925664.1 | 291441 | 292843 | - | 4276 | 2147 | 2129 | 50.21 |
| gene211 | 1280 | 81 | 632.81 | 61.78 | 3.91 | https://www.ncbi.nlm.nih.gov/gene/?term=LOC105274534 | NW_011924887.1 | 1565391 | 1566670 | + | 4336 | 1936 | 2400 | 44.65 |
| gene1894 | 445 | 27 | 606.74 | 63.65 | 3.86 | https://www.ncbi.nlm.nih.gov/gene/?term=LOC105288083 | NW_011924945.1 | 117665 | 118109 | + | 681 | 367 | 314 | 53.89 |
| gene5911 | 943 | 57 | 604.45 | 62 | 3.75 | https://www.ncbi.nlm.nih.gov/gene/?term=LOC105278844 | NW_011925053.1 | 1331236 | 1332178 | - | 5790 | 2198 | 3592 | 37.96 |
| gene570 | 1673 | 101 | 603.71 | 60.08 | 3.63 | https://www.ncbi.nlm.nih.gov/gene/?term=LOC105278549 | NW_011924892.1 | 1160484 | 1162156 | + | 5642 | 2230 | 3412 | 39.52 |
| gene12330 | 1501 | 90 | 599.6 | 62.31 | 3.74 | https://www.ncbi.nlm.nih.gov/gene/?term=LOC105286235 | NW_011925664.1 | 138328 | 139828 | - | 5153 | 2321 | 2832 | 45.04 |
| gene2797 | 1489 | 89 | 597.72 | 64.83 | 3.87 | https://www.ncbi.nlm.nih.gov/gene/?term=LOC105275314 | NW_011924973.1 | 823242 | 824730 | + | 5033 | 2280 | 2753 | 45.3 |
| gene4635 | 2067 | 123 | 595.07 | 63.09 | 3.75 | https://www.ncbi.nlm.nih.gov/gene/?term=LOC105277390 | NW_011925019.1 | 5988 | 8054 | - | 6715 | 3009 | 3706 | 44.81 |
| gene4878 | 675 | 40 | 592.59 | 80.1 | 4.75 | https://www.ncbi.nlm.nih.gov/gene/?term=LOC105277648 | NW_011925020.1 | 2601147 | 2601821 | - | 1475 | 883 | 592 | 59.86 |
| gene5301 | 1881 | 111 | 590.11 | 51.46 | 3.04 | https://www.ncbi.nlm.nih.gov/gene/?term=LOC105278148 | NW_011925031.1 | 114108 | 115988 | + | 9100 | 2470 | 6630 | 27.14 |
| gene10742 | 776 | 45 | 579.9 | 63.05 | 3.66 | https://www.ncbi.nlm.nih.gov/gene/?term=LOC109611309 | NW_011925370.1 | 1617 | 2392 | + | 2269 | 931 | 1338 | 41.03 |
| gene10943 | 1759 | 102 | 579.87 | 65.14 | 3.78 | https://www.ncbi.nlm.nih.gov/gene/?term=LOC105284659 | NW_011925387.1 | 118203 | 119961 | - | 5599 | 2629 | 2970 | 46.95 |
| gene3731 | 865 | 50 | 578.03 | 59.38 | 3.43 | https://www.ncbi.nlm.nih.gov/gene/?term=LOC105276371 | NW_011924994.1 | 459736 | 460600 | - | 2407 | 1050 | 1357 | 43.62 |
| gene3719 | 1471 | 84 | 571.04 | 70.56 | 4.03 | https://www.ncbi.nlm.nih.gov/gene/?term=LOC105276360 | NW_011924994.1 | 94735 | 96205 | - | 6779 | 2253 | 4526 | 33.23 |
| gene1562 | 1491 | 85 | 570.09 | 56.72 | 3.23 | https://www.ncbi.nlm.nih.gov/gene/?term=LOC105287692 | NW_011924931.1 | 3966137 | 3967627 | - | 6285 | 2051 | 4234 | 32.63 |
| gene1654 | 2285 | 130 | 568.93 | 66.29 | 3.77 | https://www.ncbi.nlm.nih.gov/gene/?term=LOC105287811 | NW_011924932.1 | 539418 | 541702 | - | 8300 | 3696 | 4604 | 44.53 |
| gene10232 | 1591 | 90 | 565.68 | 48.51 | 2.74 | https://www.ncbi.nlm.nih.gov/gene/?term=LOC105283847 | NW_011925327.1 | 546966 | 548556 | + | 6353 | 2176 | 4177 | 34.25 |
| gene12602 | 1399 | 79 | 564.69 | 75.22 | 4.25 | https://www.ncbi.nlm.nih.gov/gene/?term=LOC105286562 | NW_011925739.1 | 19908 | 21306 | + | 2465 | 1340 | 1125 | 54.36 |

...
... only first 25 lines shown ...

Directory: RNK File: ranked-genes-bySiteDensity-Cb_LI_BCphaseB1.txt

| region_ID | rwidth | nbrsites | nbrper10kb | pmpersite | pmpernucl | pglink | seqnames | start | end | strand | coverage | numCs | numTs | prcntM |
|---|---|---|---|---|---|---|---|---|---|---|---|---|---|---|
| gene5917 | 823 | 79 | 959.9 | 42.51 | 4.08 | https://www.ncbi.nlm.nih.gov/gene/?term=LOC105278849 | NW_011925053.1 | 1371807 | 1372629 | + | 5427 | 1351 | 4076 | 24.89 |
| gene13384 | 1877 | 153 | 815.13 | 52.38 | 4.27 | https://www.ncbi.nlm.nih.gov/gene/?term=LOC105287447 | NW_011929339.1 | 7 | 1883 | + | 10903 | 3243 | 7660 | 29.74 |
| gene5309 | 1704 | 127 | 745.31 | 54.07 | 4.03 | https://www.ncbi.nlm.nih.gov/gene/?term=LOC105278160 | NW_011925031.1 | 150501 | 152204 | - | 7369 | 2990 | 4379 | 40.58 |
| gene11569 | 1311 | 91 | 694.13 | 76.92 | 5.34 | https://www.ncbi.nlm.nih.gov/gene/?term=LOC105285394 | NW_011925459.1 | 186244 | 187554 | - | 3258 | 1960 | 1298 | 60.16 |
| gene12538 | 372 | 25 | 672.04 | 64.86 | 4.36 | https://www.ncbi.nlm.nih.gov/gene/?term=LOC105286531 | NW_011925715.1 | 412527 | 412898 | - | 1770 | 643 | 1127 | 36.33 |
| gene211 | 1280 | 83 | 648.44 | 59.17 | 3.84 | https://www.ncbi.nlm.nih.gov/gene/?term=LOC105274534 | NW_011924887.1 | 1565391 | 1566670 | + | 4583 | 1829 | 2754 | 39.91 |
| gene1894 | 445 | 28 | 629.21 | 62.94 | 3.96 | https://www.ncbi.nlm.nih.gov/gene/?term=LOC105288083 | NW_011924945.1 | 117665 | 118109 | + | 858 | 470 | 388 | 54.78 |
| gene12364 | 1403 | 88 | 627.23 | 67.09 | 4.21 | https://www.ncbi.nlm.nih.gov/gene/?term=LOC105286269 | NW_011925664.1 | 291441 | 292843 | - | 4073 | 1970 | 2103 | 48.37 |
| gene4878 | 675 | 42 | 622.22 | 79.8 | 4.97 | https://www.ncbi.nlm.nih.gov/gene/?term=LOC105277648 | NW_011925020.1 | 2601147 | 2601821 | - | 1744 | 1042 | 702 | 59.75 |
| gene12330 | 1501 | 93 | 619.59 | 62.46 | 3.87 | https://www.ncbi.nlm.nih.gov/gene/?term=LOC105286235 | NW_011925664.1 | 138328 | 139828 | - | 5529 | 2329 | 3200 | 42.12 |
| gene10232 | 1591 | 98 | 615.96 | 49.05 | 3.02 | https://www.ncbi.nlm.nih.gov/gene/?term=LOC105283847 | NW_011925327.1 | 546966 | 548556 | + | 6029 | 2031 | 3998 | 33.69 |
| gene12602 | 1399 | 86 | 614.72 | 71.97 | 4.42 | https://www.ncbi.nlm.nih.gov/gene/?term=LOC105286562 | NW_011925739.1 | 19908 | 21306 | + | 2947 | 1532 | 1415 | 51.99 |
| gene3719 | 1471 | 90 | 611.83 | 66.77 | 4.09 | https://www.ncbi.nlm.nih.gov/gene/?term=LOC105276360 | NW_011924994.1 | 94735 | 96205 | - | 7059 | 2146 | 4913 | 30.4 |
| gene5911 | 943 | 57 | 604.45 | 62.5 | 3.78 | https://www.ncbi.nlm.nih.gov/gene/?term=LOC105278844 | NW_011925053.1 | 1331236 | 1332178 | - | 4781 | 1733 | 3048 | 36.25 |
| gene570 | 1673 | 101 | 603.71 | 60.72 | 3.67 | https://www.ncbi.nlm.nih.gov/gene/?term=LOC105278549 | NW_011924892.1 | 1160484 | 1162156 | + | 5685 | 2277 | 3408 | 40.05 |
| gene10943 | 1759 | 105 | 596.93 | 65.29 | 3.9 | https://www.ncbi.nlm.nih.gov/gene/?term=LOC105284659 | NW_011925387.1 | 118203 | 119961 | - | 4822 | 2170 | 2652 | 45 |
| gene11566 | 1603 | 95 | 592.64 | 58.11 | 3.44 | https://www.ncbi.nlm.nih.gov/gene/?term=LOC105285392 | NW_011925459.1 | 162939 | 164541 | + | 6006 | 2095 | 3911 | 34.88 |
| gene1562 | 1491 | 88 | 590.21 | 55.24 | 3.26 | https://www.ncbi.nlm.nih.gov/gene/?term=LOC105287692 | NW_011924931.1 | 3966137 | 3967627 | - | 5819 | 1882 | 3937 | 32.34 |
| gene5301 | 1881 | 108 | 574.16 | 51.89 | 2.98 | https://www.ncbi.nlm.nih.gov/gene/?term=LOC105278148 | NW_011925031.1 | 114108 | 115988 | + | 8326 | 2323 | 6003 | 27.9 |
| gene2797 | 1489 | 84 | 564.14 | 64.48 | 3.64 | https://www.ncbi.nlm.nih.gov/gene/?term=LOC105275314 | NW_011924973.1 | 823242 | 824730 | + | 4912 | 1965 | 2947 | 40 |
| gene5412 | 1868 | 104 | 556.75 | 56.68 | 3.16 | https://www.ncbi.nlm.nih.gov/gene/?term=LOC105278266 | NW_011925035.1 | 1477936 | 1479803 | + | 6612 | 2104 | 4508 | 31.82 |
| gene2279 | 2037 | 113 | 554.74 | 63.44 | 3.52 | https://www.ncbi.nlm.nih.gov/gene/?term=LOC105274711 | NW_011924947.1 | 141424 | 143460 | + | 5737 | 2574 | 3163 | 44.87 |
| gene12652 | 1122 | 62 | 552.58 | 66.57 | 3.68 | https://www.ncbi.nlm.nih.gov/gene/?term=LOC109610640 | NW_011925762.1 | 557576 | 558697 | + | 2911 | 1423 | 1488 | 48.88 |
| gene4635 | 2067 | 114 | 551.52 | 64.07 | 3.53 | https://www.ncbi.nlm.nih.gov/gene/?term=LOC105277390 | NW_011925019.1 | 5988 | 8054 | - | 6098 | 2458 | 3640 | 40.31 |

...

... only first 25 lines shown ...

Directory: RNK File: ranked-genes-bySiteDensity-Cb_LI_BCphaseB2.txt

| region_ID | rwidth | nbrsites | nbrper10kb | pmpersite | pmpernucl | pglink | seqnames | start | end | strand | coverage | numCs | numTs | prcntM |
|---|---|---|---|---|---|---|---|---|---|---|---|---|---|---|
| gene5917 | 823 | 72 | 874.85 | 37.91 | 3.32 | https://www.ncbi.nlm.nih.gov/gene/?term=LOC105278849 | NW_011925053.1 | 1371807 | 1372629 | + | 6189 | 1391 | 4798 | 22.48 |
| gene13384 | 1877 | 153 | 815.13 | 50.8 | 4.14 | https://www.ncbi.nlm.nih.gov/gene/?term=LOC105287447 | NW_011929339.1 | 7 | 1883 | + | 11689 | 3556 | 8133 | 30.42 |
| gene5309 | 1704 | 118 | 692.49 | 51.39 | 3.56 | https://www.ncbi.nlm.nih.gov/gene/?term=LOC105278160 | NW_011925031.1 | 150501 | 152204 | - | 7261 | 2804 | 4457 | 38.62 |
| gene11569 | 1311 | 89 | 678.87 | 74.63 | 5.07 | https://www.ncbi.nlm.nih.gov/gene/?term=LOC105285394 | NW_011925459.1 | 186244 | 187554 | - | 3427 | 2074 | 1353 | 60.52 |
| gene1562 | 1491 | 95 | 637.16 | 55.48 | 3.53 | https://www.ncbi.nlm.nih.gov/gene/?term=LOC105287692 | NW_011924931.1 | 3966137 | 3967627 | - | 5852 | 2025 | 3827 | 34.6 |
| gene1894 | 445 | 28 | 629.21 | 64.12 | 4.03 | https://www.ncbi.nlm.nih.gov/gene/?term=LOC105288083 | NW_011924945.1 | 117665 | 118109 | + | 771 | 443 | 328 | 57.46 |
| gene5911 | 943 | 59 | 625.66 | 61.03 | 3.82 | https://www.ncbi.nlm.nih.gov/gene/?term=LOC105278844 | NW_011925053.1 | 1331236 | 1332178 | - | 4983 | 1977 | 3006 | 39.67 |
| gene12538 | 372 | 23 | 618.28 | 63.84 | 3.95 | https://www.ncbi.nlm.nih.gov/gene/?term=LOC105286531 | NW_011925715.1 | 412527 | 412898 | - | 1801 | 643 | 1158 | 35.7 |
| gene211 | 1280 | 79 | 617.19 | 64.56 | 3.98 | https://www.ncbi.nlm.nih.gov/gene/?term=LOC105274534 | NW_011924887.1 | 1565391 | 1566670 | + | 4636 | 2015 | 2621 | 43.46 |
| gene3719 | 1471 | 90 | 611.83 | 68.46 | 4.19 | https://www.ncbi.nlm.nih.gov/gene/?term=LOC105276360 | NW_011924994.1 | 94735 | 96205 | - | 7176 | 2253 | 4923 | 31.4 |
| gene12364 | 1403 | 85 | 605.84 | 67.3 | 4.08 | https://www.ncbi.nlm.nih.gov/gene/?term=LOC105286269 | NW_011925664.1 | 291441 | 292843 | - | 4065 | 1948 | 2117 | 47.92 |
| gene2797 | 1489 | 89 | 597.72 | 64.87 | 3.88 | https://www.ncbi.nlm.nih.gov/gene/?term=LOC105275314 | NW_011924973.1 | 823242 | 824730 | + | 4905 | 2009 | 2896 | 40.96 |
| gene12602 | 1399 | 82 | 586.13 | 73.4 | 4.3 | https://www.ncbi.nlm.nih.gov/gene/?term=LOC105286562 | NW_011925739.1 | 19908 | 21306 | + | 2810 | 1462 | 1348 | 52.03 |
| gene4878 | 675 | 39 | 577.78 | 75.33 | 4.35 | https://www.ncbi.nlm.nih.gov/gene/?term=LOC105277648 | NW_011925020.1 | 2601147 | 2601821 | - | 1384 | 787 | 597 | 56.86 |
| gene11566 | 1603 | 92 | 573.92 | 57.49 | 3.3 | https://www.ncbi.nlm.nih.gov/gene/?term=LOC105285392 | NW_011925459.1 | 162939 | 164541 | + | 5843 | 2054 | 3789 | 35.15 |
| gene4635 | 2067 | 118 | 570.88 | 61.88 | 3.53 | https://www.ncbi.nlm.nih.gov/gene/?term=LOC105277390 | NW_011925019.1 | 5988 | 8054 | - | 6821 | 2822 | 3999 | 41.37 |
| gene10943 | 1759 | 100 | 568.5 | 66 | 3.75 | https://www.ncbi.nlm.nih.gov/gene/?term=LOC105284659 | NW_011925387.1 | 118203 | 119961 | - | 4388 | 2074 | 2314 | 47.27 |
| gene3731 | 865 | 49 | 566.47 | 57.7 | 3.27 | https://www.ncbi.nlm.nih.gov/gene/?term=LOC105276371 | NW_011924994.1 | 459736 | 460600 | - | 2458 | 954 | 1504 | 38.81 |
| gene1654 | 2285 | 128 | 560.18 | 65.44 | 3.67 | https://www.ncbi.nlm.nih.gov/gene/?term=LOC105287811 | NW_011924932.1 | 539418 | 541702 | - | 7900 | 3372 | 4528 | 42.68 |
| gene12330 | 1501 | 84 | 559.63 | 63.25 | 3.54 | https://www.ncbi.nlm.nih.gov/gene/?term=LOC105286235 | NW_011925664.1 | 138328 | 139828 | - | 4719 | 2011 | 2708 | 42.61 |
| gene5301 | 1881 | 105 | 558.21 | 51.35 | 2.87 | https://www.ncbi.nlm.nih.gov/gene/?term=LOC105278148 | NW_011925031.1 | 114108 | 115988 | + | 8331 | 2248 | 6083 | 26.98 |
| gene5412 | 1868 | 104 | 556.75 | 54.88 | 3.06 | https://www.ncbi.nlm.nih.gov/gene/?term=LOC105278266 | NW_011925035.1 | 1477936 | 1479803 | + | 7288 | 2211 | 5077 | 30.34 |
| gene570 | 1673 | 93 | 555.89 | 62.37 | 3.47 | https://www.ncbi.nlm.nih.gov/gene/?term=LOC105278549 | NW_011924892.1 | 1160484 | 1162156 | + | 5135 | 2068 | 3067 | 40.27 |
| gene9471 | 1605 | 86 | 535.83 | 58.99 | 3.16 | https://www.ncbi.nlm.nih.gov/gene/?term=LOC105282986 | NW_011925273.1 | 229004 | 230608 | + | 6567 | 2027 | 4540 | 30.87 |

...
... only first 25 lines shown ...

Directory: RNK File: ranked-genes-bySiteDensity-Cb_LI_RphaseA1.txt

| region_ID | rwidth | nbrsites | nbrper10kb | pmpersite | pmpernucl | pglink | seqnames | start | end | strand | coverage | numCs | numTs | prcntM |
|---|---|---|---|---|---|---|---|---|---|---|---|---|---|---|
| gene13384 | 1877 | 152 | 809.8 | 51.2 | 4.15 | https://www.ncbi.nlm.nih.gov/gene/?term=LOC105287447 | NW_011929339.1 | 7 | 1883 | + | 10357 | 3038 | 7319 | 29.33 |
| gene5917 | 823 | 60 | 729.04 | 44.01 | 3.21 | https://www.ncbi.nlm.nih.gov/gene/?term=LOC105278849 | NW_011925053.1 | 1371807 | 1372629 | + | 4476 | 1068 | 3408 | 23.86 |
| gene5309 | 1704 | 115 | 674.88 | 55.23 | 3.73 | https://www.ncbi.nlm.nih.gov/gene/?term=LOC105278160 | NW_011925031.1 | 150501 | 152204 | - | 6813 | 2568 | 4245 | 37.69 |
| gene11569 | 1311 | 84 | 640.73 | 77.72 | 4.98 | https://www.ncbi.nlm.nih.gov/gene/?term=LOC105285394 | NW_011925459.1 | 186244 | 187554 | - | 3335 | 2004 | 1331 | 60.09 |
| gene12364 | 1403 | 88 | 627.23 | 66.88 | 4.19 | https://www.ncbi.nlm.nih.gov/gene/?term=LOC105286269 | NW_011925664.1 | 291441 | 292843 | - | 4029 | 1958 | 2071 | 48.6 |
| gene211 | 1280 | 80 | 625 | 63.97 | 4 | https://www.ncbi.nlm.nih.gov/gene/?term=LOC105274534 | NW_011924887.1 | 1565391 | 1566670 | + | 4214 | 1808 | 2406 | 42.9 |
| gene12538 | 372 | 23 | 618.28 | 64.99 | 4.02 | https://www.ncbi.nlm.nih.gov/gene/?term=LOC105286531 | NW_011925715.1 | 412527 | 412898 | - | 1418 | 504 | 914 | 35.54 |
| gene5911 | 943 | 56 | 593.85 | 64.01 | 3.8 | https://www.ncbi.nlm.nih.gov/gene/?term=LOC105278844 | NW_011925053.1 | 1331236 | 1332178 | - | 4009 | 1373 | 2636 | 34.25 |
| gene11566 | 1603 | 95 | 592.64 | 54.33 | 3.22 | https://www.ncbi.nlm.nih.gov/gene/?term=LOC105285392 | NW_011925459.1 | 162939 | 164541 | + | 5744 | 1933 | 3811 | 33.65 |
| gene10943 | 1759 | 103 | 585.56 | 64.13 | 3.76 | https://www.ncbi.nlm.nih.gov/gene/?term=LOC105284659 | NW_011925387.1 | 118203 | 119961 | - | 4357 | 1913 | 2444 | 43.91 |
| gene4878 | 675 | 39 | 577.78 | 80.24 | 4.64 | https://www.ncbi.nlm.nih.gov/gene/?term=LOC105277648 | NW_011925020.1 | 2601147 | 2601821 | - | 1536 | 958 | 578 | 62.37 |
| gene12330 | 1501 | 86 | 572.95 | 64.27 | 3.68 | https://www.ncbi.nlm.nih.gov/gene/?term=LOC105286235 | NW_011925664.1 | 138328 | 139828 | - | 4624 | 1945 | 2679 | 42.06 |
| gene1993 | 1449 | 82 | 565.91 | 52.71 | 2.98 | https://www.ncbi.nlm.nih.gov/gene/?term=LOC105288193 | NW_011924946.1 | 775753 | 777201 | - | 5039 | 1409 | 3630 | 27.96 |
| gene12602 | 1399 | 79 | 564.69 | 72.06 | 4.07 | https://www.ncbi.nlm.nih.gov/gene/?term=LOC105286562 | NW_011925739.1 | 19908 | 21306 | + | 2472 | 1223 | 1249 | 49.47 |
| gene570 | 1673 | 94 | 561.86 | 62.56 | 3.51 | https://www.ncbi.nlm.nih.gov/gene/?term=LOC105278549 | NW_011924892.1 | 1160484 | 1162156 | + | 4947 | 1896 | 3051 | 38.33 |
| gene1894 | 445 | 25 | 561.8 | 63.33 | 3.56 | https://www.ncbi.nlm.nih.gov/gene/?term=LOC105288083 | NW_011924945.1 | 117665 | 118109 | + | 688 | 359 | 329 | 52.18 |
| gene3719 | 1471 | 82 | 557.44 | 68.3 | 3.81 | https://www.ncbi.nlm.nih.gov/gene/?term=LOC105276360 | NW_011924994.1 | 94735 | 96205 | - | 6342 | 1907 | 4435 | 30.07 |
| gene2797 | 1489 | 83 | 557.42 | 65.53 | 3.65 | https://www.ncbi.nlm.nih.gov/gene/?term=LOC105275314 | NW_011924973.1 | 823242 | 824730 | + | 4368 | 1859 | 2509 | 42.56 |
| gene10219 | 235 | 13 | 553.19 | 70.96 | 3.93 | https://www.ncbi.nlm.nih.gov/gene/?term=LOC105283830 | NW_011925327.1 | 293521 | 293755 | + | 670 | 238 | 432 | 35.52 |
| gene1654 | 2285 | 126 | 551.42 | 65.05 | 3.59 | https://www.ncbi.nlm.nih.gov/gene/?term=LOC105287811 | NW_011924932.1 | 539418 | 541702 | - | 7363 | 3210 | 4153 | 43.6 |
| gene5412 | 1868 | 102 | 546.04 | 59.25 | 3.24 | https://www.ncbi.nlm.nih.gov/gene/?term=LOC105278266 | NW_011925035.1 | 1477936 | 1479803 | + | 6566 | 1976 | 4590 | 30.09 |
| gene12652 | 1122 | 61 | 543.67 | 66.1 | 3.59 | https://www.ncbi.nlm.nih.gov/gene/?term=LOC109610640 | NW_011925762.1 | 557576 | 558697 | + | 2703 | 1396 | 1307 | 51.65 |
| gene10232 | 1591 | 86 | 540.54 | 47.45 | 2.56 | https://www.ncbi.nlm.nih.gov/gene/?term=LOC105283847 | NW_011925327.1 | 546966 | 548556 | + | 5122 | 1632 | 3490 | 31.86 |
| gene1562 | 1491 | 80 | 536.55 | 59.8 | 3.21 | https://www.ncbi.nlm.nih.gov/gene/?term=LOC105287692 | NW_011924931.1 | 3966137 | 3967627 | - | 5082 | 1672 | 3410 | 32.9 |

...

... only first 25 lines shown ...

Directory: RNK File: ranked-genes-bySiteDensity-Cb_LI_RphaseA2.txt

| region_ID | rwidth | nbrsites | nbrper10kb | pmpersite | pmpernucl | pglink | seqnames | start | end | strand | coverage | numCs | numTs | prcntM |
|---|---|---|---|---|---|---|---|---|---|---|---|---|---|---|
| gene5917 | 823 | 102 | 1239.37 | 33.57 | 4.16 | https://www.ncbi.nlm.nih.gov/gene/?term=LOC105278849 | NW_011925053.1 | 1371807 | 1372629 | + | 10685 | 2555 | 8130 | 23.91 |
| gene13384 | 1877 | 175 | 932.34 | 46.29 | 4.32 | https://www.ncbi.nlm.nih.gov/gene/?term=LOC105287447 | NW_011929339.1 | 7 | 1883 | + | 17714 | 5351 | 12363 | 30.21 |
| gene5309 | 1704 | 135 | 792.25 | 50.44 | 4 | https://www.ncbi.nlm.nih.gov/gene/?term=LOC105278160 | NW_011925031.1 | 150501 | 152204 | - | 11574 | 4537 | 7037 | 39.2 |
| gene11569 | 1311 | 102 | 778.03 | 75.91 | 5.91 | https://www.ncbi.nlm.nih.gov/gene/?term=LOC105285394 | NW_011925459.1 | 186244 | 187554 | - | 5164 | 3333 | 1831 | 64.54 |
| gene10219 | 235 | 18 | 765.96 | 59.08 | 4.53 | https://www.ncbi.nlm.nih.gov/gene/?term=LOC105283830 | NW_011925327.1 | 293521 | 293755 | + | 1023 | 386 | 637 | 37.73 |
| gene211 | 1280 | 91 | 710.94 | 59.28 | 4.21 | https://www.ncbi.nlm.nih.gov/gene/?term=LOC105274534 | NW_011924887.1 | 1565391 | 1566670 | + | 7060 | 3129 | 3931 | 44.32 |
| gene5937 | 1902 | 134 | 704.52 | 40.07 | 2.82 | https://www.ncbi.nlm.nih.gov/gene/?term=LOC105278873 | NW_011925053.1 | 1492486 | 1494387 | - | 16447 | 3556 | 12891 | 21.62 |
| gene1993 | 1449 | 102 | 703.93 | 46.1 | 3.25 | https://www.ncbi.nlm.nih.gov/gene/?term=LOC105288193 | NW_011924946.1 | 775753 | 777201 | - | 8656 | 2593 | 6063 | 29.96 |
| gene3719 | 1471 | 103 | 700.2 | 64.53 | 4.52 | https://www.ncbi.nlm.nih.gov/gene/?term=LOC105276360 | NW_011924994.1 | 94735 | 96205 | - | 10626 | 3521 | 7105 | 33.14 |
| gene12364 | 1403 | 98 | 698.5 | 63.68 | 4.45 | https://www.ncbi.nlm.nih.gov/gene/?term=LOC105286269 | NW_011925664.1 | 291441 | 292843 | - | 6095 | 3089 | 3006 | 50.68 |
| gene10943 | 1759 | 122 | 693.58 | 63.08 | 4.38 | https://www.ncbi.nlm.nih.gov/gene/?term=LOC105284659 | NW_011925387.1 | 118203 | 119961 | - | 7759 | 3843 | 3916 | 49.53 |
| gene1562 | 1491 | 103 | 690.81 | 53.02 | 3.66 | https://www.ncbi.nlm.nih.gov/gene/?term=LOC105287692 | NW_011924931.1 | 3966137 | 3967627 | - | 9836 | 3460 | 6376 | 35.18 |
| gene5412 | 1868 | 127 | 679.87 | 51.95 | 3.53 | https://www.ncbi.nlm.nih.gov/gene/?term=LOC105278266 | NW_011925035.1 | 1477936 | 1479803 | + | 10650 | 3326 | 7324 | 31.23 |
| gene1894 | 445 | 30 | 674.16 | 57.72 | 3.89 | https://www.ncbi.nlm.nih.gov/gene/?term=LOC105288083 | NW_011924945.1 | 117665 | 118109 | + | 1144 | 606 | 538 | 52.97 |
| gene11566 | 1603 | 108 | 673.74 | 52.56 | 3.54 | https://www.ncbi.nlm.nih.gov/gene/?term=LOC105285392 | NW_011925459.1 | 162939 | 164541 | + | 8693 | 3278 | 5415 | 37.71 |
| gene10232 | 1591 | 107 | 672.53 | 46.19 | 3.11 | https://www.ncbi.nlm.nih.gov/gene/?term=LOC105283847 | NW_011925327.1 | 546966 | 548556 | + | 8501 | 3096 | 5405 | 36.42 |
| gene12538 | 372 | 25 | 672.04 | 61.6 | 4.14 | https://www.ncbi.nlm.nih.gov/gene/?term=LOC105286531 | NW_011925715.1 | 412527 | 412898 | - | 2425 | 869 | 1556 | 35.84 |
| gene5301 | 1881 | 126 | 669.86 | 47.19 | 3.16 | https://www.ncbi.nlm.nih.gov/gene/?term=LOC105278148 | NW_011925031.1 | 114108 | 115988 | + | 12972 | 3637 | 9335 | 28.04 |
| gene12602 | 1399 | 93 | 664.76 | 72.35 | 4.81 | https://www.ncbi.nlm.nih.gov/gene/?term=LOC105286562 | NW_011925739.1 | 19908 | 21306 | + | 4090 | 2262 | 1828 | 55.31 |
| gene5911 | 943 | 61 | 646.87 | 62.79 | 4.06 | https://www.ncbi.nlm.nih.gov/gene/?term=LOC105278844 | NW_011925053.1 | 1331236 | 1332178 | - | 7490 | 2978 | 4512 | 39.76 |
| gene2797 | 1489 | 96 | 644.73 | 62.31 | 4.02 | https://www.ncbi.nlm.nih.gov/gene/?term=LOC105275314 | NW_011924973.1 | 823242 | 824730 | + | 7455 | 3164 | 4291 | 42.44 |
| gene1654 | 2285 | 147 | 643.33 | 60.4 | 3.89 | https://www.ncbi.nlm.nih.gov/gene/?term=LOC105287811 | NW_011924932.1 | 539418 | 541702 | - | 12199 | 5318 | 6881 | 43.59 |
| gene570 | 1673 | 107 | 639.57 | 59.59 | 3.81 | https://www.ncbi.nlm.nih.gov/gene/?term=LOC105278549 | NW_011924892.1 | 1160484 | 1162156 | + | 8287 | 3324 | 4963 | 40.11 |
| gene4635 | 2067 | 132 | 638.61 | 59.89 | 3.82 | https://www.ncbi.nlm.nih.gov/gene/?term=LOC105277390 | NW_011925019.1 | 5988 | 8054 | - | 9946 | 4242 | 5704 | 42.65 |

...

... only first 25 lines shown ...

Directory: RNK File: ranked-genes-bySiteDensity-Cb_LI_RphaseB1.txt

| region_ID | rwidth | nbrsites | nbrper10kb | pmpersite | pmpernucl | pglink | seqnames | start | end | strand | coverage | numCs | numTs | prcntM |
|---|---|---|---|---|---|---|---|---|---|---|---|---|---|---|
| gene5917 | 823 | 64 | 777.64 | 38.44 | 2.99 | https://www.ncbi.nlm.nih.gov/gene/?term=LOC105278849 | NW_011925053.1 | 1371807 | 1372629 | + | 4901 | 1050 | 3851 | 21.42 |
| gene13384 | 1877 | 141 | 751.2 | 53.46 | 4.02 | https://www.ncbi.nlm.nih.gov/gene/?term=LOC105287447 | NW_011929339.1 | 7 | 1883 | + | 10660 | 3206 | 7454 | 30.08 |
| gene11569 | 1311 | 89 | 678.87 | 75.99 | 5.16 | https://www.ncbi.nlm.nih.gov/gene/?term=LOC105285394 | NW_011925459.1 | 186244 | 187554 | - | 3154 | 1890 | 1264 | 59.92 |
| gene12364 | 1403 | 95 | 677.12 | 63.53 | 4.3 | https://www.ncbi.nlm.nih.gov/gene/?term=LOC105286269 | NW_011925664.1 | 291441 | 292843 | - | 4205 | 2105 | 2100 | 50.06 |
| gene5309 | 1704 | 114 | 669.01 | 51.01 | 3.41 | https://www.ncbi.nlm.nih.gov/gene/?term=LOC105278160 | NW_011925031.1 | 150501 | 152204 | - | 7036 | 2519 | 4517 | 35.8 |
| gene1894 | 445 | 29 | 651.69 | 60.17 | 3.92 | https://www.ncbi.nlm.nih.gov/gene/?term=LOC105288083 | NW_011924945.1 | 117665 | 118109 | + | 762 | 387 | 375 | 50.79 |
| gene211 | 1280 | 82 | 640.62 | 62.9 | 4.03 | https://www.ncbi.nlm.nih.gov/gene/?term=LOC105274534 | NW_011924887.1 | 1565391 | 1566670 | + | 4423 | 1924 | 2499 | 43.5 |
| gene12538 | 372 | 23 | 618.28 | 62.34 | 3.85 | https://www.ncbi.nlm.nih.gov/gene/?term=LOC105286531 | NW_011925715.1 | 412527 | 412898 | - | 1632 | 551 | 1081 | 33.76 |
| gene570 | 1673 | 101 | 603.71 | 60.31 | 3.64 | https://www.ncbi.nlm.nih.gov/gene/?term=LOC105278549 | NW_011924892.1 | 1160484 | 1162156 | + | 5477 | 2245 | 3232 | 40.99 |
| gene12602 | 1399 | 84 | 600.43 | 73.12 | 4.39 | https://www.ncbi.nlm.nih.gov/gene/?term=LOC105286562 | NW_011925739.1 | 19908 | 21306 | + | 2547 | 1276 | 1271 | 50.1 |
| gene3719 | 1471 | 88 | 598.23 | 68.28 | 4.08 | https://www.ncbi.nlm.nih.gov/gene/?term=LOC105276360 | NW_011924994.1 | 94735 | 96205 | - | 6631 | 2093 | 4538 | 31.56 |
| gene10219 | 235 | 14 | 595.74 | 72.78 | 4.34 | https://www.ncbi.nlm.nih.gov/gene/?term=LOC105283830 | NW_011925327.1 | 293521 | 293755 | + | 655 | 257 | 398 | 39.24 |
| gene1562 | 1491 | 87 | 583.5 | 56.45 | 3.29 | https://www.ncbi.nlm.nih.gov/gene/?term=LOC105287692 | NW_011924931.1 | 3966137 | 3967627 | - | 5966 | 1907 | 4059 | 31.96 |
| gene5911 | 943 | 55 | 583.24 | 64.42 | 3.76 | https://www.ncbi.nlm.nih.gov/gene/?term=LOC105278844 | NW_011925053.1 | 1331236 | 1332178 | - | 4514 | 1504 | 3010 | 33.32 |
| gene4878 | 675 | 39 | 577.78 | 75.44 | 4.36 | https://www.ncbi.nlm.nih.gov/gene/?term=LOC105277648 | NW_011925020.1 | 2601147 | 2601821 | - | 1482 | 882 | 600 | 59.51 |
| gene12330 | 1501 | 86 | 572.95 | 63.93 | 3.66 | https://www.ncbi.nlm.nih.gov/gene/?term=LOC105286235 | NW_011925664.1 | 138328 | 139828 | - | 4577 | 2008 | 2569 | 43.87 |
| gene2797 | 1489 | 85 | 570.85 | 64.88 | 3.7 | https://www.ncbi.nlm.nih.gov/gene/?term=LOC105275314 | NW_011924973.1 | 823242 | 824730 | + | 4719 | 1967 | 2752 | 41.68 |
| gene3731 | 865 | 47 | 543.35 | 58.92 | 3.2 | https://www.ncbi.nlm.nih.gov/gene/?term=LOC105276371 | NW_011924994.1 | 459736 | 460600 | - | 2474 | 993 | 1481 | 40.14 |
| gene11566 | 1603 | 87 | 542.73 | 58.13 | 3.16 | https://www.ncbi.nlm.nih.gov/gene/?term=LOC105285392 | NW_011925459.1 | 162939 | 164541 | + | 5456 | 2005 | 3451 | 36.75 |
| gene1654 | 2285 | 124 | 542.67 | 65.33 | 3.55 | https://www.ncbi.nlm.nih.gov/gene/?term=LOC105287811 | NW_011924932.1 | 539418 | 541702 | - | 7155 | 3267 | 3888 | 45.66 |
| gene4635 | 2067 | 112 | 541.85 | 66 | 3.58 | https://www.ncbi.nlm.nih.gov/gene/?term=LOC105277390 | NW_011925019.1 | 5988 | 8054 | - | 5603 | 2376 | 3227 | 42.41 |
| gene12652 | 1122 | 60 | 534.76 | 68.09 | 3.64 | https://www.ncbi.nlm.nih.gov/gene/?term=LOC109610640 | NW_011925762.1 | 557576 | 558697 | + | 2589 | 1361 | 1228 | 52.57 |
| gene1223 | 1048 | 56 | 534.35 | 64.54 | 3.45 | https://www.ncbi.nlm.nih.gov/gene/?term=LOC105285713 | NW_011924922.1 | 2517529 | 2518576 | - | 2528 | 1163 | 1365 | 46 |
| gene3806 | 2385 | 127 | 532.49 | 60.6 | 3.23 | https://www.ncbi.nlm.nih.gov/gene/?term=LOC105276439 | NW_011924996.1 | 675985 | 678369 | + | 7115 | 2711 | 4404 | 38.1 |

...
... only first 25 lines shown ...

Directory: RNK File: ranked-genes-bySiteDensity-Cb_LI_RphaseB2.txt

| region_ID | rwidth | nbrsites | nbrper10kb | pmpersite | pmpernucl | pglink | seqnames | start | end | strand | coverage | numCs | numTs | prcntM |
|---|---|---|---|---|---|---|---|---|---|---|---|---|---|---|
| gene5917 | 823 | 67 | 814.09 | 39.8 | 3.24 | https://www.ncbi.nlm.nih.gov/gene/?term=LOC105278849 | NW_011925053.1 | 1371807 | 1372629 | + | 5249 | 1222 | 4027 | 23.28 |
| gene13384 | 1877 | 143 | 761.85 | 52.16 | 3.97 | https://www.ncbi.nlm.nih.gov/gene/?term=LOC105287447 | NW_011929339.1 | 7 | 1883 | + | 10632 | 3145 | 7487 | 29.58 |
| gene5309 | 1704 | 118 | 692.49 | 53.46 | 3.7 | https://www.ncbi.nlm.nih.gov/gene/?term=LOC105278160 | NW_011925031.1 | 150501 | 152204 | - | 7398 | 2908 | 4490 | 39.31 |
| gene11569 | 1311 | 86 | 655.99 | 75.08 | 4.92 | https://www.ncbi.nlm.nih.gov/gene/?term=LOC105285394 | NW_011925459.1 | 186244 | 187554 | - | 3138 | 1985 | 1153 | 63.26 |
| gene211 | 1280 | 81 | 632.81 | 62.78 | 3.97 | https://www.ncbi.nlm.nih.gov/gene/?term=LOC105274534 | NW_011924887.1 | 1565391 | 1566670 | + | 4363 | 1950 | 2413 | 44.69 |
| gene12538 | 372 | 23 | 618.28 | 63.53 | 3.93 | https://www.ncbi.nlm.nih.gov/gene/?term=LOC105286531 | NW_011925715.1 | 412527 | 412898 | - | 1559 | 543 | 1016 | 34.83 |
| gene5911 | 943 | 57 | 604.45 | 61.57 | 3.72 | https://www.ncbi.nlm.nih.gov/gene/?term=LOC105278844 | NW_011925053.1 | 1331236 | 1332178 | - | 4499 | 1718 | 2781 | 38.19 |
| gene2797 | 1489 | 89 | 597.72 | 63.22 | 3.78 | https://www.ncbi.nlm.nih.gov/gene/?term=LOC105275314 | NW_011924973.1 | 823242 | 824730 | + | 4716 | 2098 | 2618 | 44.49 |
| gene10219 | 235 | 14 | 595.74 | 70.11 | 4.18 | https://www.ncbi.nlm.nih.gov/gene/?term=LOC105283830 | NW_011925327.1 | 293521 | 293755 | + | 441 | 168 | 273 | 38.1 |
| gene3719 | 1471 | 85 | 577.84 | 69.22 | 4 | https://www.ncbi.nlm.nih.gov/gene/?term=LOC105276360 | NW_011924994.1 | 94735 | 96205 | - | 6046 | 1998 | 4048 | 33.05 |
| gene1562 | 1491 | 85 | 570.09 | 56.03 | 3.19 | https://www.ncbi.nlm.nih.gov/gene/?term=LOC105287692 | NW_011924931.1 | 3966137 | 3967627 | - | 5892 | 2023 | 3869 | 34.33 |
| gene1654 | 2285 | 130 | 568.93 | 64.4 | 3.66 | https://www.ncbi.nlm.nih.gov/gene/?term=LOC105287811 | NW_011924932.1 | 539418 | 541702 | - | 7293 | 3219 | 4074 | 44.14 |
| gene11566 | 1603 | 91 | 567.69 | 56.06 | 3.18 | https://www.ncbi.nlm.nih.gov/gene/?term=LOC105285392 | NW_011925459.1 | 162939 | 164541 | + | 5465 | 2011 | 3454 | 36.8 |
| gene5301 | 1881 | 105 | 558.21 | 51.96 | 2.9 | https://www.ncbi.nlm.nih.gov/gene/?term=LOC105278148 | NW_011925031.1 | 114108 | 115988 | + | 7619 | 2018 | 5601 | 26.49 |
| gene3731 | 865 | 48 | 554.91 | 58.57 | 3.25 | https://www.ncbi.nlm.nih.gov/gene/?term=LOC105276371 | NW_011924994.1 | 459736 | 460600 | - | 2436 | 1019 | 1417 | 41.83 |
| gene4635 | 2067 | 114 | 551.52 | 63.65 | 3.51 | https://www.ncbi.nlm.nih.gov/gene/?term=LOC105277390 | NW_011925019.1 | 5988 | 8054 | - | 5972 | 2523 | 3449 | 42.25 |
| gene10943 | 1759 | 97 | 551.45 | 66.13 | 3.65 | https://www.ncbi.nlm.nih.gov/gene/?term=LOC105284659 | NW_011925387.1 | 118203 | 119961 | - | 4322 | 1987 | 2335 | 45.97 |
| gene12364 | 1403 | 77 | 548.82 | 71.2 | 3.91 | https://www.ncbi.nlm.nih.gov/gene/?term=LOC105286269 | NW_011925664.1 | 291441 | 292843 | - | 3388 | 1685 | 1703 | 49.73 |
| gene10232 | 1591 | 87 | 546.83 | 47.24 | 2.58 | https://www.ncbi.nlm.nih.gov/gene/?term=LOC105283847 | NW_011925327.1 | 546966 | 548556 | + | 4811 | 1620 | 3191 | 33.67 |
| gene570 | 1673 | 91 | 543.93 | 61.67 | 3.35 | https://www.ncbi.nlm.nih.gov/gene/?term=LOC105278549 | NW_011924892.1 | 1160484 | 1162156 | + | 4509 | 1788 | 2721 | 39.65 |
| gene12330 | 1501 | 81 | 539.64 | 64.21 | 3.46 | https://www.ncbi.nlm.nih.gov/gene/?term=LOC105286235 | NW_011925664.1 | 138328 | 139828 | - | 4303 | 1915 | 2388 | 44.5 |
| gene12602 | 1399 | 75 | 536.1 | 73.75 | 3.95 | https://www.ncbi.nlm.nih.gov/gene/?term=LOC105286562 | NW_011925739.1 | 19908 | 21306 | + | 2234 | 1219 | 1015 | 54.57 |
| gene5412 | 1868 | 100 | 535.33 | 56.79 | 3.04 | https://www.ncbi.nlm.nih.gov/gene/?term=LOC105278266 | NW_011925035.1 | 1477936 | 1479803 | + | 6499 | 2067 | 4432 | 31.8 |
| gene4878 | 675 | 36 | 533.33 | 75.02 | 4 | https://www.ncbi.nlm.nih.gov/gene/?term=LOC105277648 | NW_011925020.1 | 2601147 | 2601821 | - | 1265 | 730 | 535 | 57.71 |

...
... only first 25 lines shown ...

Directory: RNK File: ranked-promoters-byPrcntM-Cb_LI_BCphaseA1.txt

| region_ID | seqnames | start | end | width | strand | coverage | numCs | numTs | prcntM |
|---|---|---|---|---|---|---|---|---|---|
| gene12609 | NW_011925739.1 | 39939 | 40438 | 500 | + | 5 | 5 | 0 | 100 |
| gene8259 | NW_011925188.1 | 207457 | 207956 | 500 | - | 33 | 32 | 1 | 96.97 |
| gene9311 | NW_011925258.1 | 34560 | 35059 | 500 | - | 63 | 60 | 3 | 95.24 |
| gene5713 | NW_011925050.1 | 7072 | 7571 | 500 | - | 330 | 303 | 27 | 91.82 |
| gene13190 | NW_011926458.1 | 36701 | 37200 | 500 | - | 33 | 30 | 3 | 90.91 |
| gene4016 | NW_011925003.1 | 66624 | 67123 | 500 | + | 754 | 676 | 78 | 89.66 |
| gene5601 | NW_011925036.1 | 147325 | 147824 | 500 | - | 335 | 300 | 35 | 89.55 |
| gene3546 | NW_011924987.1 | 1804967 | 1805466 | 500 | - | 64 | 57 | 7 | 89.06 |
| gene4040 | NW_011925003.1 | 277741 | 278240 | 500 | + | 18 | 16 | 2 | 88.89 |
| gene6765 | NW_011925106.1 | 2070681 | 2071180 | 500 | - | 92 | 81 | 11 | 88.04 |
| gene11598 | NW_011925467.1 | 116917 | 117416 | 500 | + | 307 | 268 | 39 | 87.3 |
| gene9313 | NW_011925258.1 | 34883 | 35382 | 500 | + | 178 | 153 | 25 | 85.96 |
| gene8331 | NW_011925189.1 | 573460 | 573959 | 500 | + | 117 | 99 | 18 | 84.62 |
| gene772 | NW_011924904.1 | 1310349 | 1310848 | 500 | + | 608 | 500 | 108 | 82.24 |
| gene3373 | NW_011924987.1 | 206378 | 206877 | 500 | - | 132 | 107 | 25 | 81.06 |
| gene2337 | NW_011924951.1 | 247713 | 248212 | 500 | + | 108 | 87 | 21 | 80.56 |
| gene9622 | NW_011925287.1 | 414625 | 415124 | 500 | - | 784 | 631 | 153 | 80.48 |
| gene617 | NW_011924894.1 | 393331 | 393830 | 500 | - | 90 | 72 | 18 | 80 |
| gene2749 | NW_011924973.1 | 232730 | 233229 | 500 | + | 10 | 8 | 2 | 80 |
| gene3478 | NW_011924987.1 | 1323515 | 1324014 | 500 | - | 200 | 158 | 42 | 79 |
| gene10656 | NW_011925362.1 | 86544 | 87043 | 500 | + | 237 | 187 | 50 | 78.9 |
| gene2675 | NW_011924970.1 | 296515 | 297014 | 500 | - | 257 | 201 | 56 | 78.21 |
| gene12926 | NW_011925919.1 | 43401 | 43900 | 500 | - | 679 | 529 | 150 | 77.91 |
| gene4521 | NW_011925013.1 | 44114 | 44613 | 500 | + | 217 | 169 | 48 | 77.88 |

...
... only first 25 lines shown ...

Directory: RNK File: ranked-promoters-byPrcntM-Cb_LI_BCphaseA2.txt

| region_ID | seqnames | start | end | width | strand | coverage | numCs | numTs | prcntM |
|---|---|---|---|---|---|---|---|---|---|
| gene7105 | NW_011925128.1 | 181746 | 182245 | 500 | + | 6 | 6 | 0 | 100 |
| gene3546 | NW_011924987.1 | 1804967 | 1805466 | 500 | - | 49 | 48 | 1 | 97.96 |
| gene4040 | NW_011925003.1 | 277741 | 278240 | 500 | + | 48 | 47 | 1 | 97.92 |
| gene4016 | NW_011925003.1 | 66624 | 67123 | 500 | + | 926 | 858 | 68 | 92.66 |
| gene5713 | NW_011925050.1 | 7072 | 7571 | 500 | - | 449 | 410 | 39 | 91.31 |
| gene2337 | NW_011924951.1 | 247713 | 248212 | 500 | + | 97 | 87 | 10 | 89.69 |
| gene11598 | NW_011925467.1 | 116917 | 117416 | 500 | + | 271 | 243 | 28 | 89.67 |
| gene8331 | NW_011925189.1 | 573460 | 573959 | 500 | + | 204 | 179 | 25 | 87.75 |
| gene3373 | NW_011924987.1 | 206378 | 206877 | 500 | - | 94 | 82 | 12 | 87.23 |
| gene5601 | NW_011925036.1 | 147325 | 147824 | 500 | - | 310 | 266 | 44 | 85.81 |
| gene779 | NW_011924904.1 | 1346112 | 1346611 | 500 | - | 48 | 41 | 7 | 85.42 |
| gene12926 | NW_011925919.1 | 43401 | 43900 | 500 | - | 839 | 710 | 129 | 84.62 |
| gene9313 | NW_011925258.1 | 34883 | 35382 | 500 | + | 189 | 158 | 31 | 83.6 |
| gene11827 | NW_011925516.1 | 416246 | 416745 | 500 | + | 6 | 5 | 1 | 83.33 |
| gene6595 | NW_011925106.1 | 124931 | 125430 | 500 | + | 183 | 152 | 31 | 83.06 |
| gene9311 | NW_011925258.1 | 34560 | 35059 | 500 | - | 72 | 59 | 13 | 81.94 |
| gene8077 | NW_011925163.1 | 1005320 | 1005819 | 500 | - | 11 | 9 | 2 | 81.82 |
| gene617 | NW_011924894.1 | 393331 | 393830 | 500 | - | 74 | 60 | 14 | 81.08 |
| gene2124 | NW_011924946.1 | 2553236 | 2553735 | 500 | + | 73 | 59 | 14 | 80.82 |
| gene772 | NW_011924904.1 | 1310349 | 1310848 | 500 | + | 641 | 518 | 123 | 80.81 |
| gene8259 | NW_011925188.1 | 207457 | 207956 | 500 | - | 26 | 21 | 5 | 80.77 |
| gene9485 | NW_011925273.1 | 340332 | 340831 | 500 | - | 31 | 25 | 6 | 80.65 |
| gene9622 | NW_011925287.1 | 414625 | 415124 | 500 | - | 788 | 634 | 154 | 80.46 |
| gene3298 | NW_011924979.1 | 1594523 | 1595022 | 500 | + | 121 | 97 | 24 | 80.17 |

...
... only first 25 lines shown ...

Directory: RNK File: ranked-promoters-byPrcntM-Cb_LI_BCphaseB1.txt

| region_ID | seqnames | start | end | width | strand | coverage | numCs | numTs | prcntM |
|---|---|---|---|---|---|---|---|---|---|
| gene4040 | NW_011925003.1 | 277741 | 278240 | 500 | + | 30 | 28 | 2 | 93.33 |
| gene3546 | NW_011924987.1 | 1804967 | 1805466 | 500 | - | 65 | 60 | 5 | 92.31 |
| gene5713 | NW_011925050.1 | 7072 | 7571 | 500 | - | 494 | 456 | 38 | 92.31 |
| gene4016 | NW_011925003.1 | 66624 | 67123 | 500 | + | 913 | 824 | 89 | 90.25 |
| gene11598 | NW_011925467.1 | 116917 | 117416 | 500 | + | 385 | 344 | 41 | 89.35 |
| gene2337 | NW_011924951.1 | 247713 | 248212 | 500 | + | 117 | 104 | 13 | 88.89 |
| gene5601 | NW_011925036.1 | 147325 | 147824 | 500 | - | 517 | 446 | 71 | 86.27 |
| gene779 | NW_011924904.1 | 1346112 | 1346611 | 500 | - | 84 | 71 | 13 | 84.52 |
| gene6765 | NW_011925106.1 | 2070681 | 2071180 | 500 | - | 128 | 108 | 20 | 84.38 |
| gene12609 | NW_011925739.1 | 39939 | 40438 | 500 | + | 6 | 5 | 1 | 83.33 |
| gene3478 | NW_011924987.1 | 1323515 | 1324014 | 500 | - | 277 | 229 | 48 | 82.67 |
| gene617 | NW_011924894.1 | 393331 | 393830 | 500 | - | 83 | 68 | 15 | 81.93 |
| gene12926 | NW_011925919.1 | 43401 | 43900 | 500 | - | 612 | 501 | 111 | 81.86 |
| gene772 | NW_011924904.1 | 1310349 | 1310848 | 500 | + | 822 | 670 | 152 | 81.51 |
| gene9622 | NW_011925287.1 | 414625 | 415124 | 500 | - | 746 | 606 | 140 | 81.23 |
| gene4521 | NW_011925013.1 | 44114 | 44613 | 500 | + | 235 | 190 | 45 | 80.85 |
| gene3373 | NW_011924987.1 | 206378 | 206877 | 500 | - | 142 | 114 | 28 | 80.28 |
| gene11827 | NW_011925516.1 | 416246 | 416745 | 500 | + | 5 | 4 | 1 | 80 |
| gene3238 | NW_011924979.1 | 133011 | 133510 | 500 | + | 69 | 54 | 15 | 78.26 |
| gene8331 | NW_011925189.1 | 573460 | 573959 | 500 | + | 225 | 175 | 50 | 77.78 |
| gene1675 | NW_011924932.1 | 638769 | 639268 | 500 | - | 324 | 250 | 74 | 77.16 |
| gene9313 | NW_011925258.1 | 34883 | 35382 | 500 | + | 239 | 184 | 55 | 76.99 |
| gene5463 | NW_011925035.1 | 1933485 | 1933984 | 500 | - | 377 | 288 | 89 | 76.39 |
| gene9311 | NW_011925258.1 | 34560 | 35059 | 500 | - | 70 | 53 | 17 | 75.71 |

...

... only first 25 lines shown ...

Directory: RNK File: ranked-promoters-byPrcntM-Cb_LI_BCphaseB2.txt

```
region_ID   seqnames    start end   width strand      coverage    numCs numTs prcntM
gene4040    NW_011925003.1    277741      278240      500   +     30    30    0     100
gene8748    NW_011925216.1    183604      184103      500   -     5     5     0     100
gene4016    NW_011925003.1    66624 67123 500   +     852   778   74    91.31
gene9311    NW_011925258.1    34560 35059 500   -     56    51    5     91.07
gene3546    NW_011924987.1    1804967     1805466     500   -     65    59    6     90.77
gene11598   NW_011925467.1    116917      117416      500   +     231   208   23    90.04
gene5601    NW_011925036.1    147325      147824      500   -     347   312   35    89.91
gene779     NW_011924904.1    1346112     1346611     500   -     34    30    4     88.24
gene13190   NW_011926458.1    36701 37200 500   -     32    28    4     87.5
gene5713    NW_011925050.1    7072  7571  500   -     442   382   60    86.43
gene12609   NW_011925739.1    39939 40438 500   +     21    18    3     85.71
gene8331    NW_011925189.1    573460      573959      500   +     181   152   29    83.98
gene3478    NW_011924987.1    1323515     1324014     500   -     282   232   50    82.27
gene617     NW_011924894.1    393331      393830      500   -     87    71    16    81.61
gene2337    NW_011924951.1    247713      248212      500   +     100   81    19    81
gene9313    NW_011925258.1    34883 35382 500   +     192   155   37    80.73
gene12926   NW_011925919.1    43401 43900 500   -     838   676   162   80.67
gene9622    NW_011925287.1    414625      415124      500   -     776   622   154   80.15
gene8259    NW_011925188.1    207457      207956      500   -     25    20    5     80
gene9485    NW_011925273.1    340332      340831      500   -     40    32    8     80
gene2032    NW_011924946.1    1344986     1345485     500   -     24    19    5     79.17
gene772     NW_011924904.1    1310349     1310848     500   +     691   537   154   77.71
gene3373    NW_011924987.1    206378      206877      500   -     137   106   31    77.37
gene2675    NW_011924970.1    296515      297014      500   -     297   226   71    76.09
...
... only first 25 lines shown ...
```

Directory: RNK File: ranked-promoters-byPrcntM-Cb_LI_RphaseA1.txt

| region_ID | seqnames | start | end | width | strand | coverage | numCs | numTs | prcntM |
|---|---|---|---|---|---|---|---|---|---|
| gene536 | NW_011924892.1 | 67669 | 68168 | 500 | - | 5 | 5 | 0 | 100 |
| gene8259 | NW_011925188.1 | 207457 | 207956 | 500 | - | 29 | 29 | 0 | 100 |
| gene8748 | NW_011925216.1 | 183604 | 184103 | 500 | - | 9 | 9 | 0 | 100 |
| gene4040 | NW_011925003.1 | 277741 | 278240 | 500 | + | 42 | 41 | 1 | 97.62 |
| gene3546 | NW_011924987.1 | 1804967 | 1805466 | 500 | - | 75 | 72 | 3 | 96 |
| gene13190 | NW_011926458.1 | 36701 | 37200 | 500 | - | 23 | 21 | 2 | 91.3 |
| gene4016 | NW_011925003.1 | 66624 | 67123 | 500 | + | 880 | 798 | 82 | 90.68 |
| gene5713 | NW_011925050.1 | 7072 | 7571 | 500 | - | 394 | 357 | 37 | 90.61 |
| gene11598 | NW_011925467.1 | 116917 | 117416 | 500 | + | 295 | 265 | 30 | 89.83 |
| gene2337 | NW_011924951.1 | 247713 | 248212 | 500 | + | 132 | 117 | 15 | 88.64 |
| gene5601 | NW_011925036.1 | 147325 | 147824 | 500 | - | 448 | 390 | 58 | 87.05 |
| gene12926 | NW_011925919.1 | 43401 | 43900 | 500 | - | 634 | 529 | 105 | 83.44 |
| gene11827 | NW_011925516.1 | 416246 | 416745 | 500 | + | 6 | 5 | 1 | 83.33 |
| gene3373 | NW_011924987.1 | 206378 | 206877 | 500 | - | 128 | 105 | 23 | 82.03 |
| gene3478 | NW_011924987.1 | 1323515 | 1324014 | 500 | - | 280 | 223 | 57 | 79.64 |
| gene9311 | NW_011925258.1 | 34560 | 35059 | 500 | - | 68 | 54 | 14 | 79.41 |
| gene772 | NW_011924904.1 | 1310349 | 1310848 | 500 | + | 674 | 533 | 141 | 79.08 |
| gene8331 | NW_011925189.1 | 573460 | 573959 | 500 | + | 193 | 151 | 42 | 78.24 |
| gene3929 | NW_011924999.1 | 140417 | 140916 | 500 | - | 640 | 497 | 143 | 77.66 |
| gene4521 | NW_011925013.1 | 44114 | 44613 | 500 | + | 241 | 187 | 54 | 77.59 |
| gene9622 | NW_011925287.1 | 414625 | 415124 | 500 | - | 790 | 612 | 178 | 77.47 |
| gene9313 | NW_011925258.1 | 34883 | 35382 | 500 | + | 204 | 158 | 46 | 77.45 |
| gene6595 | NW_011925106.1 | 124931 | 125430 | 500 | + | 154 | 119 | 35 | 77.27 |
| gene7677 | NW_011925160.1 | 14541 | 15040 | 500 | - | 171 | 132 | 39 | 77.19 |

...

... only first 25 lines shown ...

Directory: RNK File: ranked-promoters-byPrcntM-Cb_LI_RphaseA2.txt

| region_ID | seqnames | start | end | width | strand | coverage | numCs | numTs | prcntM |
|---|---|---|---|---|---|---|---|---|---|
| gene13190 | NW_011926458.1 | 36701 | 37200 | 500 | - | 22 | 21 | 1 | 95.45 |
| gene3546 | NW_011924987.1 | 1804967 | 1805466 | 500 | - | 61 | 57 | 4 | 93.44 |
| gene4040 | NW_011925003.1 | 277741 | 278240 | 500 | + | 60 | 56 | 4 | 93.33 |
| gene4016 | NW_011925003.1 | 66624 | 67123 | 500 | + | 1484 | 1383 | 101 | 93.19 |
| gene5713 | NW_011925050.1 | 7072 | 7571 | 500 | - | 678 | 620 | 58 | 91.45 |
| gene617 | NW_011924894.1 | 393331 | 393830 | 500 | - | 146 | 131 | 15 | 89.73 |
| gene11598 | NW_011925467.1 | 116917 | 117416 | 500 | + | 416 | 372 | 44 | 89.42 |
| gene9311 | NW_011925258.1 | 34560 | 35059 | 500 | - | 111 | 98 | 13 | 88.29 |
| gene5601 | NW_011925036.1 | 147325 | 147824 | 500 | - | 584 | 509 | 75 | 87.16 |
| gene2337 | NW_011924951.1 | 247713 | 248212 | 500 | + | 122 | 105 | 17 | 86.07 |
| gene6595 | NW_011925106.1 | 124931 | 125430 | 500 | + | 202 | 172 | 30 | 85.15 |
| gene9313 | NW_011925258.1 | 34883 | 35382 | 500 | + | 297 | 248 | 49 | 83.5 |
| gene536 | NW_011924892.1 | 67669 | 68168 | 500 | - | 6 | 5 | 1 | 83.33 |
| gene12926 | NW_011925919.1 | 43401 | 43900 | 500 | - | 1168 | 970 | 198 | 83.05 |
| gene2032 | NW_011924946.1 | 1344986 | 1345485 | 500 | - | 63 | 52 | 11 | 82.54 |
| gene3478 | NW_011924987.1 | 1323515 | 1324014 | 500 | - | 347 | 281 | 66 | 80.98 |
| gene3238 | NW_011924979.1 | 133011 | 133510 | 500 | + | 72 | 58 | 14 | 80.56 |
| gene3373 | NW_011924987.1 | 206378 | 206877 | 500 | - | 170 | 136 | 34 | 80 |
| gene7331 | NW_011925129.1 | 188998 | 189497 | 500 | - | 5 | 4 | 1 | 80 |
| gene10311 | NW_011925335.1 | 493532 | 494031 | 500 | + | 5 | 4 | 1 | 80 |
| gene12609 | NW_011925739.1 | 39939 | 40438 | 500 | + | 15 | 12 | 3 | 80 |
| gene772 | NW_011924904.1 | 1310349 | 1310848 | 500 | + | 1028 | 820 | 208 | 79.77 |
| gene8331 | NW_011925189.1 | 573460 | 573959 | 500 | + | 303 | 240 | 63 | 79.21 |
| gene9622 | NW_011925287.1 | 414625 | 415124 | 500 | - | 1009 | 799 | 210 | 79.19 |

...
... only first 25 lines shown ...

Directory: RNK File: ranked-promoters-byPrcntM-Cb_LI_RphaseB1.txt

| region_ID | seqnames | start | end | width | strand | coverage | numCs | numTs | prcntM |
|---|---|---|---|---|---|---|---|---|---|
| gene4040 | NW_011925003.1 | 277741 | 278240 | 500 | + | 28 | 27 | 1 | 96.43 |
| gene13190 | NW_011926458.1 | 36701 | 37200 | 500 | - | 18 | 17 | 1 | 94.44 |
| gene4016 | NW_011925003.1 | 66624 | 67123 | 500 | + | 900 | 828 | 72 | 92 |
| gene3546 | NW_011924987.1 | 1804967 | 1805466 | 500 | - | 51 | 45 | 6 | 88.24 |
| gene5713 | NW_011925050.1 | 7072 | 7571 | 500 | - | 357 | 315 | 42 | 88.24 |
| gene11598 | NW_011925467.1 | 116917 | 117416 | 500 | + | 379 | 334 | 45 | 88.13 |
| gene12609 | NW_011925739.1 | 39939 | 40438 | 500 | + | 16 | 14 | 2 | 87.5 |
| gene9311 | NW_011925258.1 | 34560 | 35059 | 500 | - | 83 | 71 | 12 | 85.54 |
| gene617 | NW_011924894.1 | 393331 | 393830 | 500 | - | 111 | 94 | 17 | 84.68 |
| gene3373 | NW_011924987.1 | 206378 | 206877 | 500 | - | 141 | 119 | 22 | 84.4 |
| gene8331 | NW_011925189.1 | 573460 | 573959 | 500 | + | 230 | 194 | 36 | 84.35 |
| gene3238 | NW_011924979.1 | 133011 | 133510 | 500 | + | 70 | 59 | 11 | 84.29 |
| gene5601 | NW_011925036.1 | 147325 | 147824 | 500 | - | 402 | 338 | 64 | 84.08 |
| gene8259 | NW_011925188.1 | 207457 | 207956 | 500 | - | 35 | 29 | 6 | 82.86 |
| gene3478 | NW_011924987.1 | 1323515 | 1324014 | 500 | - | 285 | 234 | 51 | 82.11 |
| gene2337 | NW_011924951.1 | 247713 | 248212 | 500 | + | 112 | 89 | 23 | 79.46 |
| gene779 | NW_011924904.1 | 1346112 | 1346611 | 500 | - | 63 | 50 | 13 | 79.37 |
| gene9781 | NW_011925300.1 | 457658 | 458157 | 500 | + | 218 | 173 | 45 | 79.36 |
| gene12926 | NW_011925919.1 | 43401 | 43900 | 500 | - | 627 | 496 | 131 | 79.11 |
| gene6595 | NW_011925106.1 | 124931 | 125430 | 500 | + | 148 | 117 | 31 | 79.05 |
| gene6765 | NW_011925106.1 | 2070681 | 2071180 | 500 | - | 100 | 79 | 21 | 79 |
| gene9865 | NW_011925315.1 | 89622 | 90121 | 500 | + | 28 | 22 | 6 | 78.57 |
| gene5463 | NW_011925035.1 | 1933485 | 1933984 | 500 | - | 390 | 304 | 86 | 77.95 |
| gene772 | NW_011924904.1 | 1310349 | 1310848 | 500 | + | 772 | 601 | 171 | 77.85 |

...
... only first 25 lines shown ...

Directory: RNK File: ranked-promoters-byPrcntM-Cb_LI_RphaseB2.txt

| region_ID | seqnames | start | end | width | strand | coverage | numCs | numTs | prcntM |
|---|---|---|---|---|---|---|---|---|---|
| gene8748 | NW_011925216.1 | 183604 | 184103 | 500 | - | 5 | 5 | 0 | 100 |
| gene11598 | NW_011925467.1 | 116917 | 117416 | 500 | + | 260 | 243 | 17 | 93.46 |
| gene3546 | NW_011924987.1 | 1804967 | 1805466 | 500 | - | 43 | 40 | 3 | 93.02 |
| gene4016 | NW_011925003.1 | 66624 | 67123 | 500 | + | 795 | 725 | 70 | 91.19 |
| gene5713 | NW_011925050.1 | 7072 | 7571 | 500 | - | 371 | 327 | 44 | 88.14 |
| gene5601 | NW_011925036.1 | 147325 | 147824 | 500 | - | 355 | 312 | 43 | 87.89 |
| gene4040 | NW_011925003.1 | 277741 | 278240 | 500 | + | 40 | 35 | 5 | 87.5 |
| gene9313 | NW_011925258.1 | 34883 | 35382 | 500 | + | 197 | 167 | 30 | 84.77 |
| gene779 | NW_011924904.1 | 1346112 | 1346611 | 500 | - | 51 | 43 | 8 | 84.31 |
| gene617 | NW_011924894.1 | 393331 | 393830 | 500 | - | 77 | 64 | 13 | 83.12 |
| gene3373 | NW_011924987.1 | 206378 | 206877 | 500 | - | 78 | 64 | 14 | 82.05 |
| gene6595 | NW_011925106.1 | 124931 | 125430 | 500 | + | 168 | 137 | 31 | 81.55 |
| gene13190 | NW_011926458.1 | 36701 | 37200 | 500 | - | 16 | 13 | 3 | 81.25 |
| gene2532 | NW_011924955.1 | 1998697 | 1999196 | 500 | - | 200 | 162 | 38 | 81 |
| gene12926 | NW_011925919.1 | 43401 | 43900 | 500 | - | 881 | 710 | 171 | 80.59 |
| gene9311 | NW_011925258.1 | 34560 | 35059 | 500 | - | 70 | 56 | 14 | 80 |
| gene3478 | NW_011924987.1 | 1323515 | 1324014 | 500 | - | 203 | 161 | 42 | 79.31 |
| gene8331 | NW_011925189.1 | 573460 | 573959 | 500 | + | 178 | 141 | 37 | 79.21 |
| gene10656 | NW_011925362.1 | 86544 | 87043 | 500 | + | 367 | 290 | 77 | 79.02 |
| gene9485 | NW_011925273.1 | 340332 | 340831 | 500 | - | 44 | 34 | 10 | 77.27 |
| gene2337 | NW_011924951.1 | 247713 | 248212 | 500 | + | 118 | 91 | 27 | 77.12 |
| gene772 | NW_011924904.1 | 1310349 | 1310848 | 500 | + | 593 | 450 | 143 | 75.89 |
| gene8259 | NW_011925188.1 | 207457 | 207956 | 500 | - | 12 | 9 | 3 | 75 |
| gene9622 | NW_011925287.1 | 414625 | 415124 | 500 | - | 589 | 441 | 148 | 74.87 |

...

... only first 25 lines shown ...

Directory: RNK File: ranked-promoters-bySiteDensity-Cb_LI_BCphaseA1.txt

| region_ID | rwidth | nbrsites | nbrper10kb | pmpersite | pmpernucl | pglink | seqnames | start | end | strand | coverage | numCs | numTs | prcntM |
|---|---|---|---|---|---|---|---|---|---|---|---|---|---|---|
| gene4515 | 500 | 37 | 740 | 75.66 | 5.6 | https://www.ncbi.nlm.nih.gov/gene/?term=LOC109610839 | NW_011925013.1 | 19534 | 20033 | + | 1160 | 766 | 394 | 66.03 |
| gene12926 | 500 | 36 | 720 | 84.69 | 6.1 | https://www.ncbi.nlm.nih.gov/gene/?term=LOC105286944 | NW_011925919.1 | 43401 | 43900 | - | 679 | 529 | 150 | 77.91 |
| gene8964 | 500 | 35 | 700 | 60.17 | 4.21 | https://www.ncbi.nlm.nih.gov/gene/?term=LOC105282404 | NW_011925232.1 | 562782 | 563281 | - | 1239 | 554 | 685 | 44.71 |
| gene505 | 500 | 34 | 680 | 59.33 | 4.03 | https://www.ncbi.nlm.nih.gov/gene/?term=LOC105277808 | NW_011924889.1 | 477908 | 478407 | + | 1897 | 567 | 1330 | 29.89 |
| gene5753 | 500 | 32 | 640 | 74.85 | 4.79 | https://www.ncbi.nlm.nih.gov/gene/?term=LOC105278673 | NW_011925050.1 | 215094 | 215593 | + | 1046 | 695 | 351 | 66.44 |
| gene7634 | 500 | 32 | 640 | 73.61 | 4.71 | https://www.ncbi.nlm.nih.gov/gene/?term=LOC105280861 | NW_011925151.1 | 38041 | 38540 | - | 1280 | 658 | 622 | 51.41 |
| gene10220 | 500 | 31 | 620 | 66.72 | 4.14 | https://www.ncbi.nlm.nih.gov/gene/?term=LOC105283833 | NW_011925327.1 | 293294 | 293793 | + | 1277 | 528 | 749 | 41.35 |
| gene465 | 500 | 31 | 620 | 70.5 | 4.37 | https://www.ncbi.nlm.nih.gov/gene/?term=LOC105277304 | NW_011924889.1 | 171845 | 172344 | + | 1150 | 613 | 537 | 53.3 |
| gene772 | 500 | 30 | 600 | 82.9 | 4.97 | https://www.ncbi.nlm.nih.gov/gene/?term=LOC105280935 | NW_011924904.1 | 1310349 | 1310848 | + | 608 | 500 | 108 | 82.24 |
| gene10147 | 500 | 29 | 580 | 84.36 | 4.89 | https://www.ncbi.nlm.nih.gov/gene/?term=LOC105283749 | NW_011925323.1 | 1722554 | 1723053 | + | 1003 | 599 | 404 | 59.72 |
| gene12363 | 500 | 28 | 560 | 69.55 | 3.89 | https://www.ncbi.nlm.nih.gov/gene/?term=LOC105286266 | NW_011925664.1 | 291330 | 291829 | - | 847 | 387 | 460 | 45.69 |
| gene11980 | 500 | 27 | 540 | 75.19 | 4.06 | https://www.ncbi.nlm.nih.gov/gene/?term=LOC109611395 | NW_011925546.1 | 60031 | 60530 | + | 765 | 533 | 232 | 69.67 |
| gene128 | 500 | 27 | 540 | 56.69 | 3.06 | https://www.ncbi.nlm.nih.gov/gene/?term=LOC105286681 | NW_011924887.1 | 225822 | 226321 | + | 1015 | 450 | 565 | 44.33 |
| gene9622 | 500 | 27 | 540 | 84.94 | 4.59 | https://www.ncbi.nlm.nih.gov/gene/?term=LOC105283159 | NW_011925287.1 | 414625 | 415124 | - | 784 | 631 | 153 | 80.48 |
| gene12925 | 500 | 26 | 520 | 53.8 | 2.8 | https://www.ncbi.nlm.nih.gov/gene/?term=LOC109611487 | NW_011925919.1 | 41730 | 42229 | - | 1706 | 436 | 1270 | 25.56 |
| gene4016 | 500 | 26 | 520 | 89.4 | 4.65 | https://www.ncbi.nlm.nih.gov/gene/?term=LOC105276685 | NW_011925003.1 | 66624 | 67123 | + | 754 | 676 | 78 | 89.66 |
| gene12539 | 500 | 25 | 500 | 69.76 | 3.49 | https://www.ncbi.nlm.nih.gov/gene/?term=LOC105286478 | NW_011925715.1 | 412371 | 412870 | + | 1210 | 480 | 730 | 39.67 |
| gene3929 | 500 | 23 | 460 | 80.01 | 3.68 | https://www.ncbi.nlm.nih.gov/gene/?term=LOC105276585 | NW_011924999.1 | 140417 | 140916 | - | 496 | 357 | 139 | 71.98 |
| gene525 | 500 | 23 | 460 | 57.73 | 2.66 | https://www.ncbi.nlm.nih.gov/gene/?term=LOC105278116 | NW_011924889.1 | 635990 | 636489 | + | 1207 | 360 | 847 | 29.83 |
| gene9243 | 500 | 23 | 460 | 73.25 | 3.37 | https://www.ncbi.nlm.nih.gov/gene/?term=LOC105282715 | NW_011925242.1 | 392772 | 393271 | - | 1244 | 477 | 767 | 38.34 |
| gene1025 | 500 | 22 | 440 | 74.88 | 3.29 | https://www.ncbi.nlm.nih.gov/gene/?term=LOC105283760 | NW_011924920.1 | 14737 | 15236 | + | 430 | 302 | 128 | 70.23 |
| gene1895 | 500 | 22 | 440 | 69.68 | 3.07 | https://www.ncbi.nlm.nih.gov/gene/?term=LOC105288085 | NW_011924945.1 | 117559 | 118058 | + | 605 | 350 | 255 | 57.85 |
| gene2223 | 500 | 22 | 440 | 87.84 | 3.87 | https://www.ncbi.nlm.nih.gov/gene/?term=LOC109610674 | NW_011924946.1 | 3172465 | 3172964 | - | 690 | 456 | 234 | 66.09 |
| gene4878 | 500 | 22 | 440 | 78.98 | 3.48 | https://www.ncbi.nlm.nih.gov/gene/?term=LOC105277648 | NW_011925020.1 | 2601822 | 2602321 | - | 451 | 309 | 142 | 68.51 |

...

... only first 25 lines shown ...

Directory: RNK File: ranked-promoters-bySiteDensity-Cb_LI_BCphaseA2.txt

| region_ID | rwidth | nbrsites | nbrper10kb | pmpersite | pmpernucl | pglink | seqnames | start | end | strand | coverage | numCs | numTs | prcntM |
|---|---|---|---|---|---|---|---|---|---|---|---|---|---|---|
| gene505 | 500 | 39 | 780 | 57.58 | 4.49 | https://www.ncbi.nlm.nih.gov/gene/?term=LOC105277808 | NW_011924889.1 | 477908 | 478407 | + | 3124 | 995 | 2129 | 31.85 |
| gene4515 | 500 | 37 | 740 | 72.16 | 5.34 | https://www.ncbi.nlm.nih.gov/gene/?term=LOC109610839 | NW_011925013.1 | 19534 | 20033 | + | 1698 | 1093 | 605 | 64.37 |
| gene1399 | 500 | 35 | 700 | 38.24 | 2.68 | https://www.ncbi.nlm.nih.gov/gene/?term=LOC105287518 | NW_011924931.1 | 442908 | 443407 | - | 2483 | 681 | 1802 | 27.43 |
| gene8964 | 500 | 34 | 680 | 59.31 | 4.03 | https://www.ncbi.nlm.nih.gov/gene/?term=LOC105282404 | NW_011925232.1 | 562782 | 563281 | - | 1591 | 700 | 891 | 44 |
| gene12926 | 500 | 32 | 640 | 88.97 | 5.69 | https://www.ncbi.nlm.nih.gov/gene/?term=LOC105286944 | NW_011925919.1 | 43401 | 43900 | - | 839 | 710 | 129 | 84.62 |
| gene7634 | 500 | 32 | 640 | 74.76 | 4.78 | https://www.ncbi.nlm.nih.gov/gene/?term=LOC105280861 | NW_011925151.1 | 38041 | 38540 | - | 1316 | 691 | 625 | 52.51 |
| gene12331 | 500 | 31 | 620 | 52.05 | 3.23 | https://www.ncbi.nlm.nih.gov/gene/?term=LOC105286234 | NW_011925664.1 | 139161 | 139660 | + | 1980 | 686 | 1294 | 34.65 |
| gene12925 | 500 | 31 | 620 | 52.42 | 3.25 | https://www.ncbi.nlm.nih.gov/gene/?term=LOC109611487 | NW_011925919.1 | 41730 | 42229 | - | 2612 | 686 | 1926 | 26.26 |
| gene465 | 500 | 31 | 620 | 72.55 | 4.5 | https://www.ncbi.nlm.nih.gov/gene/?term=LOC105277304 | NW_011924889.1 | 171845 | 172344 | + | 1418 | 670 | 748 | 47.25 |
| gene5753 | 500 | 30 | 600 | 78.39 | 4.7 | https://www.ncbi.nlm.nih.gov/gene/?term=LOC105278673 | NW_011925050.1 | 215094 | 215593 | + | 1104 | 797 | 307 | 72.19 |
| gene772 | 500 | 30 | 600 | 86.1 | 5.17 | https://www.ncbi.nlm.nih.gov/gene/?term=LOC105280935 | NW_011924904.1 | 1310349 | 1310848 | + | 641 | 518 | 123 | 80.81 |
| gene10147 | 500 | 29 | 580 | 85.6 | 4.96 | https://www.ncbi.nlm.nih.gov/gene/?term=LOC105283749 | NW_011925323.1 | 1722554 | 1723053 | + | 1280 | 734 | 546 | 57.34 |
| gene10220 | 500 | 29 | 580 | 67.1 | 3.89 | https://www.ncbi.nlm.nih.gov/gene/?term=LOC105283833 | NW_011925327.1 | 293294 | 293793 | + | 1224 | 494 | 730 | 40.36 |
| gene10441 | 500 | 28 | 560 | 39.48 | 2.21 | https://www.ncbi.nlm.nih.gov/gene/?term=LOC105284094 | NW_011925353.1 | 595996 | 596495 | - | 2602 | 532 | 2070 | 20.45 |
| gene12363 | 500 | 28 | 560 | 68.81 | 3.85 | https://www.ncbi.nlm.nih.gov/gene/?term=LOC105286266 | NW_011925664.1 | 291330 | 291829 | - | 1115 | 509 | 606 | 45.65 |
| gene12539 | 500 | 28 | 560 | 62.86 | 3.52 | https://www.ncbi.nlm.nih.gov/gene/?term=LOC105286478 | NW_011925715.1 | 412371 | 412870 | + | 1757 | 719 | 1038 | 40.92 |
| gene128 | 500 | 27 | 540 | 57.61 | 3.11 | https://www.ncbi.nlm.nih.gov/gene/?term=LOC105286681 | NW_011924887.1 | 225822 | 226321 | + | 1120 | 482 | 638 | 43.04 |
| gene9622 | 500 | 27 | 540 | 87.74 | 4.74 | https://www.ncbi.nlm.nih.gov/gene/?term=LOC105283159 | NW_011925287.1 | 414625 | 415124 | - | 788 | 634 | 154 | 80.46 |
| gene11980 | 500 | 26 | 520 | 75.79 | 3.94 | https://www.ncbi.nlm.nih.gov/gene/?term=LOC109611395 | NW_011925546.1 | 60031 | 60530 | + | 982 | 716 | 266 | 72.91 |
| gene3929 | 500 | 26 | 520 | 86.03 | 4.47 | https://www.ncbi.nlm.nih.gov/gene/?term=LOC105276585 | NW_011924999.1 | 140417 | 140916 | - | 705 | 506 | 199 | 71.77 |
| gene4016 | 500 | 26 | 520 | 91.87 | 4.78 | https://www.ncbi.nlm.nih.gov/gene/?term=LOC105276685 | NW_011925003.1 | 66624 | 67123 | + | 926 | 858 | 68 | 92.66 |
| gene12537 | 500 | 25 | 500 | 61.42 | 3.07 | https://www.ncbi.nlm.nih.gov/gene/?term=LOC105286479 | NW_011925715.1 | 412531 | 413030 | - | 2365 | 695 | 1670 | 29.39 |
| gene1855 | 500 | 25 | 500 | 64.17 | 3.21 | https://www.ncbi.nlm.nih.gov/gene/?term=LOC105288036 | NW_011924940.1 | 293742 | 294241 | - | 2989 | 626 | 2363 | 20.94 |
| gene723 | 500 | 25 | 500 | 53.87 | 2.69 | https://www.ncbi.nlm.nih.gov/gene/?term=LOC105280278 | NW_011924902.1 | 275704 | 276203 | + | 1050 | 436 | 614 | 41.52 |

...

... only first 25 lines shown ...

Directory: RNK File: ranked-promoters-bySiteDensity-Cb_LI_BCphaseB1.txt

| region_ID | rwidth | nbrsites | nbrper10kb | pmpersite | pmpernucl | pglink | seqnames | start | end | strand | coverage | numCs | numTs | prcntM |
|---|---|---|---|---|---|---|---|---|---|---|---|---|---|---|
| gene12925 | 500 | 43 | 860 | 43.33 | 3.73 | https://www.ncbi.nlm.nih.gov/gene/?term=LOC109611487 | NW_011925919.1 | 41730 | 42229 | - | 3109 | 762 | 2347 | 24.51 |
| gene505 | 500 | 39 | 780 | 58.34 | 4.55 | https://www.ncbi.nlm.nih.gov/gene/?term=LOC105277808 | NW_011924889.1 | 477908 | 478407 | + | 2575 | 843 | 1732 | 32.74 |
| gene4515 | 500 | 37 | 740 | 73.61 | 5.45 | https://www.ncbi.nlm.nih.gov/gene/?term=LOC109610839 | NW_011925013.1 | 19534 | 20033 | + | 1454 | 932 | 522 | 64.1 |
| gene8964 | 500 | 37 | 740 | 50.57 | 3.74 | https://www.ncbi.nlm.nih.gov/gene/?term=LOC105282404 | NW_011925232.1 | 562782 | 563281 | - | 1698 | 651 | 1047 | 38.34 |
| gene465 | 500 | 36 | 720 | 72.24 | 5.2 | https://www.ncbi.nlm.nih.gov/gene/?term=LOC105277304 | NW_011924889.1 | 171845 | 172344 | + | 1385 | 671 | 714 | 48.45 |
| gene7634 | 500 | 36 | 720 | 72.51 | 5.22 | https://www.ncbi.nlm.nih.gov/gene/?term=LOC105280861 | NW_011925151.1 | 38041 | 38540 | - | 1554 | 728 | 826 | 46.85 |
| gene12926 | 500 | 33 | 660 | 87.9 | 5.8 | https://www.ncbi.nlm.nih.gov/gene/?term=LOC105286944 | NW_011925919.1 | 43401 | 43900 | - | 612 | 501 | 111 | 81.86 |
| gene12331 | 500 | 32 | 640 | 51.75 | 3.31 | https://www.ncbi.nlm.nih.gov/gene/?term=LOC105286234 | NW_011925664.1 | 139161 | 139660 | + | 2175 | 685 | 1490 | 31.49 |
| gene5753 | 500 | 32 | 640 | 75.72 | 4.85 | https://www.ncbi.nlm.nih.gov/gene/?term=LOC105278673 | NW_011925050.1 | 215094 | 215593 | + | 1076 | 723 | 353 | 67.19 |
| gene12539 | 500 | 31 | 620 | 67.58 | 4.19 | https://www.ncbi.nlm.nih.gov/gene/?term=LOC105286478 | NW_011925715.1 | 412371 | 412870 | + | 1723 | 707 | 1016 | 41.03 |
| gene2406 | 500 | 31 | 620 | 40.37 | 2.5 | https://www.ncbi.nlm.nih.gov/gene/?term=LOC105274855 | NW_011924953.1 | 426755 | 427254 | - | 1872 | 438 | 1434 | 23.4 |
| gene10147 | 500 | 29 | 580 | 84.77 | 4.92 | https://www.ncbi.nlm.nih.gov/gene/?term=LOC105283749 | NW_011925323.1 | 1722554 | 1723053 | + | 1335 | 797 | 538 | 59.7 |
| gene1025 | 500 | 29 | 580 | 70.32 | 4.08 | https://www.ncbi.nlm.nih.gov/gene/?term=LOC105283760 | NW_011924920.1 | 14737 | 15236 | + | 692 | 448 | 244 | 64.74 |
| gene772 | 500 | 29 | 580 | 84.73 | 4.91 | https://www.ncbi.nlm.nih.gov/gene/?term=LOC105280935 | NW_011924904.1 | 1310349 | 1310848 | + | 822 | 670 | 152 | 81.51 |
| gene10220 | 500 | 27 | 540 | 70.81 | 3.82 | https://www.ncbi.nlm.nih.gov/gene/?term=LOC105283833 | NW_011925327.1 | 293294 | 293793 | + | 1409 | 559 | 850 | 39.67 |
| gene11980 | 500 | 27 | 540 | 72.77 | 3.93 | https://www.ncbi.nlm.nih.gov/gene/?term=LOC109611395 | NW_011925546.1 | 60031 | 60530 | + | 756 | 526 | 230 | 69.58 |
| gene12363 | 500 | 27 | 540 | 73.44 | 3.97 | https://www.ncbi.nlm.nih.gov/gene/?term=LOC105286266 | NW_011925664.1 | 291330 | 291829 | - | 917 | 437 | 480 | 47.66 |
| gene3929 | 500 | 27 | 540 | 77.17 | 4.17 | https://www.ncbi.nlm.nih.gov/gene/?term=LOC105276585 | NW_011924999.1 | 140417 | 140916 | - | 753 | 505 | 248 | 67.07 |
| gene4878 | 500 | 27 | 540 | 78.62 | 4.25 | https://www.ncbi.nlm.nih.gov/gene/?term=LOC105277648 | NW_011925020.1 | 2601822 | 2602321 | - | 652 | 443 | 209 | 67.94 |
| gene12537 | 500 | 26 | 520 | 64.28 | 3.34 | https://www.ncbi.nlm.nih.gov/gene/?term=LOC105286479 | NW_011925715.1 | 412531 | 413030 | - | 2380 | 671 | 1709 | 28.19 |
| gene128 | 500 | 26 | 520 | 59.8 | 3.11 | https://www.ncbi.nlm.nih.gov/gene/?term=LOC105286681 | NW_011924887.1 | 225822 | 226321 | + | 1074 | 458 | 616 | 42.64 |
| gene1399 | 500 | 26 | 520 | 39.58 | 2.06 | https://www.ncbi.nlm.nih.gov/gene/?term=LOC105287518 | NW_011924931.1 | 442908 | 443407 | - | 1927 | 533 | 1394 | 27.66 |
| gene4016 | 500 | 26 | 520 | 90.67 | 4.71 | https://www.ncbi.nlm.nih.gov/gene/?term=LOC105276685 | NW_011925003.1 | 66624 | 67123 | + | 913 | 824 | 89 | 90.25 |
| gene525 | 500 | 26 | 520 | 56.36 | 2.93 | https://www.ncbi.nlm.nih.gov/gene/?term=LOC105278116 | NW_011924889.1 | 635990 | 636489 | + | 1274 | 407 | 867 | 31.95 |

...
... only first 25 lines shown ...

Directory: RNK File: ranked-promoters-bySiteDensity-Cb_LI_BCphaseB2.txt

| region_ID | rwidth | nbrsites | nbrper10kb | pmpersite | pmpernucl | pglink | seqnames | start | end | strand | coverage | numCs | numTs | prcntM |
|---|---|---|---|---|---|---|---|---|---|---|---|---|---|---|
| gene7634 | 500 | 39 | 780 | 68.87 | 5.37 | https://www.ncbi.nlm.nih.gov/gene/?term=LOC105280861 | NW_011925151.1 | 38041 | 38540 | - | 1602 | 799 | 803 | 49.88 |
| gene12925 | 500 | 37 | 740 | 46.07 | 3.41 | https://www.ncbi.nlm.nih.gov/gene/?term=LOC109611487 | NW_011925919.1 | 41730 | 42229 | - | 2560 | 659 | 1901 | 25.74 |
| gene505 | 500 | 37 | 740 | 55.98 | 4.14 | https://www.ncbi.nlm.nih.gov/gene/?term=LOC105277808 | NW_011924889.1 | 477908 | 478407 | + | 3183 | 958 | 2225 | 30.1 |
| gene5753 | 500 | 37 | 740 | 77.53 | 5.74 | https://www.ncbi.nlm.nih.gov/gene/?term=LOC105278673 | NW_011925050.1 | 215094 | 215593 | + | 1045 | 731 | 314 | 69.95 |
| gene128 | 500 | 34 | 680 | 60.57 | 4.12 | https://www.ncbi.nlm.nih.gov/gene/?term=LOC105286681 | NW_011924887.1 | 225822 | 226321 | + | 1124 | 514 | 610 | 45.73 |
| gene4515 | 500 | 34 | 680 | 71.67 | 4.87 | https://www.ncbi.nlm.nih.gov/gene/?term=LOC109610839 | NW_011925013.1 | 19534 | 20033 | + | 1580 | 1025 | 555 | 64.87 |
| gene8964 | 500 | 33 | 660 | 56.19 | 3.71 | https://www.ncbi.nlm.nih.gov/gene/?term=LOC105282404 | NW_011925232.1 | 562782 | 563281 | - | 1710 | 717 | 993 | 41.93 |
| gene12926 | 500 | 32 | 640 | 86.9 | 5.56 | https://www.ncbi.nlm.nih.gov/gene/?term=LOC105286944 | NW_011925919.1 | 43401 | 43900 | - | 838 | 676 | 162 | 80.67 |
| gene465 | 500 | 31 | 620 | 71.76 | 4.45 | https://www.ncbi.nlm.nih.gov/gene/?term=LOC105277304 | NW_011924889.1 | 171845 | 172344 | + | 1287 | 618 | 669 | 48.02 |
| gene10147 | 500 | 30 | 600 | 81.9 | 4.91 | https://www.ncbi.nlm.nih.gov/gene/?term=LOC105283749 | NW_011925323.1 | 1722554 | 1723053 | + | 1245 | 728 | 517 | 58.47 |
| gene12363 | 500 | 30 | 600 | 66.4 | 3.98 | https://www.ncbi.nlm.nih.gov/gene/?term=LOC105286266 | NW_011925664.1 | 291330 | 291829 | - | 1089 | 531 | 558 | 48.76 |
| gene1399 | 500 | 30 | 600 | 39.97 | 2.4 | https://www.ncbi.nlm.nih.gov/gene/?term=LOC105287518 | NW_011924931.1 | 442908 | 443407 | - | 2470 | 603 | 1867 | 24.41 |
| gene3929 | 500 | 30 | 600 | 81.21 | 4.87 | https://www.ncbi.nlm.nih.gov/gene/?term=LOC105276585 | NW_011924999.1 | 140417 | 140916 | - | 692 | 522 | 170 | 75.43 |
| gene12331 | 500 | 29 | 580 | 50.62 | 2.94 | https://www.ncbi.nlm.nih.gov/gene/?term=LOC105286234 | NW_011925664.1 | 139161 | 139660 | + | 1685 | 497 | 1188 | 29.5 |
| gene772 | 500 | 29 | 580 | 78 | 4.52 | https://www.ncbi.nlm.nih.gov/gene/?term=LOC105280935 | NW_011924904.1 | 1310349 | 1310848 | + | 691 | 537 | 154 | 77.71 |
| gene2406 | 500 | 27 | 540 | 40.65 | 2.19 | https://www.ncbi.nlm.nih.gov/gene/?term=LOC105274855 | NW_011924953.1 | 426755 | 427254 | - | 1831 | 444 | 1387 | 24.25 |
| gene4016 | 500 | 27 | 540 | 90.76 | 4.9 | https://www.ncbi.nlm.nih.gov/gene/?term=LOC105276685 | NW_011925003.1 | 66624 | 67123 | + | 852 | 778 | 74 | 91.31 |
| gene9622 | 500 | 27 | 540 | 86.65 | 4.68 | https://www.ncbi.nlm.nih.gov/gene/?term=LOC105283159 | NW_011925287.1 | 414625 | 415124 | - | 776 | 622 | 154 | 80.15 |
| gene1829 | 500 | 26 | 520 | 60.29 | 3.14 | https://www.ncbi.nlm.nih.gov/gene/?term=LOC105288013 | NW_011924940.1 | 83218 | 83717 | + | 667 | 283 | 384 | 42.43 |
| gene1895 | 500 | 26 | 520 | 66.08 | 3.44 | https://www.ncbi.nlm.nih.gov/gene/?term=LOC105288085 | NW_011924945.1 | 117559 | 118058 | + | 774 | 468 | 306 | 60.47 |
| gene10220 | 500 | 25 | 500 | 65.76 | 3.29 | https://www.ncbi.nlm.nih.gov/gene/?term=LOC105283833 | NW_011925327.1 | 293294 | 293793 | + | 1310 | 477 | 833 | 36.41 |
| gene10441 | 500 | 25 | 500 | 36.76 | 1.84 | https://www.ncbi.nlm.nih.gov/gene/?term=LOC105284094 | NW_011925353.1 | 595996 | 596495 | - | 2232 | 423 | 1809 | 18.95 |
| gene11980 | 500 | 25 | 500 | 71.64 | 3.58 | https://www.ncbi.nlm.nih.gov/gene/?term=LOC109611395 | NW_011925546.1 | 60031 | 60530 | + | 759 | 505 | 254 | 66.53 |
| gene12539 | 500 | 25 | 500 | 65.7 | 3.28 | https://www.ncbi.nlm.nih.gov/gene/?term=LOC105286478 | NW_011925715.1 | 412371 | 412870 | + | 1716 | 668 | 1048 | 38.93 |

...
... only first 25 lines shown ...

Directory: RNK File: ranked-promoters-bySiteDensity-Cb_LI_RphaseA1.txt

| region_ID | rwidth | nbrsites | nbrper10kb | pmpersite | pmpernucl | pglink | seqnames | start | end | strand | coverage | numCs | numTs | prcntM |
|---|---|---|---|---|---|---|---|---|---|---|---|---|---|---|
| gene4515 | 500 | 41 | 820 | 76.8 | 6.3 | https://www.ncbi.nlm.nih.gov/gene/?term=LOC109610839 | NW_011925013.1 | 19534 | 20033 | + | 1440 | 957 | 483 | 66.46 |
| gene12925 | 500 | 37 | 740 | 50.61 | 3.75 | https://www.ncbi.nlm.nih.gov/gene/?term=LOC109611487 | NW_011925919.1 | 41730 | 42229 | - | 2395 | 674 | 1721 | 28.14 |
| gene505 | 500 | 37 | 740 | 60.43 | 4.47 | https://www.ncbi.nlm.nih.gov/gene/?term=LOC105277808 | NW_011924889.1 | 477908 | 478407 | + | 2685 | 872 | 1813 | 32.48 |
| gene7634 | 500 | 35 | 700 | 73.59 | 5.15 | https://www.ncbi.nlm.nih.gov/gene/?term=LOC105280861 | NW_011925151.1 | 38041 | 38540 | - | 1301 | 583 | 718 | 44.81 |
| gene12926 | 500 | 34 | 680 | 86.5 | 5.88 | https://www.ncbi.nlm.nih.gov/gene/?term=LOC105286944 | NW_011925919.1 | 43401 | 43900 | - | 634 | 529 | 105 | 83.44 |
| gene5753 | 500 | 33 | 660 | 79.01 | 5.21 | https://www.ncbi.nlm.nih.gov/gene/?term=LOC105278673 | NW_011925050.1 | 215094 | 215593 | + | 1026 | 710 | 316 | 69.2 |
| gene8964 | 500 | 33 | 660 | 51.32 | 3.39 | https://www.ncbi.nlm.nih.gov/gene/?term=LOC105282404 | NW_011925232.1 | 562782 | 563281 | - | 1211 | 433 | 778 | 35.76 |
| gene10220 | 500 | 31 | 620 | 66.28 | 4.11 | https://www.ncbi.nlm.nih.gov/gene/?term=LOC105283833 | NW_011925327.1 | 293294 | 293793 | + | 1314 | 547 | 767 | 41.63 |
| gene3929 | 500 | 31 | 620 | 81.46 | 5.05 | https://www.ncbi.nlm.nih.gov/gene/?term=LOC105276585 | NW_011924999.1 | 140417 | 140916 | - | 640 | 497 | 143 | 77.66 |
| gene10147 | 500 | 29 | 580 | 82.41 | 4.78 | https://www.ncbi.nlm.nih.gov/gene/?term=LOC105283749 | NW_011925323.1 | 1722554 | 1723053 | + | 1219 | 756 | 463 | 62.02 |
| gene12331 | 500 | 29 | 580 | 52.3 | 3.03 | https://www.ncbi.nlm.nih.gov/gene/?term=LOC105286234 | NW_011925664.1 | 139161 | 139660 | + | 1759 | 525 | 1234 | 29.85 |
| gene12539 | 500 | 29 | 580 | 67.78 | 3.93 | https://www.ncbi.nlm.nih.gov/gene/?term=LOC105286478 | NW_011925715.1 | 412371 | 412870 | + | 1398 | 560 | 838 | 40.06 |
| gene772 | 500 | 29 | 580 | 81.49 | 4.73 | https://www.ncbi.nlm.nih.gov/gene/?term=LOC105280935 | NW_011924904.1 | 1310349 | 1310848 | + | 674 | 533 | 141 | 79.08 |
| gene128 | 500 | 28 | 560 | 58.11 | 3.25 | https://www.ncbi.nlm.nih.gov/gene/?term=LOC105286681 | NW_011924887.1 | 225822 | 226321 | + | 1016 | 430 | 586 | 42.32 |
| gene1010 | 500 | 27 | 540 | 75.93 | 4.1 | https://www.ncbi.nlm.nih.gov/gene/?term=LOC105283498 | NW_011924914.1 | 831409 | 831908 | - | 1198 | 657 | 541 | 54.84 |
| gene12363 | 500 | 27 | 540 | 73.62 | 3.98 | https://www.ncbi.nlm.nih.gov/gene/?term=LOC105286266 | NW_011925664.1 | 291330 | 291829 | - | 1098 | 524 | 574 | 47.72 |
| gene1399 | 500 | 27 | 540 | 41.24 | 2.23 | https://www.ncbi.nlm.nih.gov/gene/?term=LOC105287518 | NW_011924931.1 | 442908 | 443407 | - | 1748 | 449 | 1299 | 25.69 |
| gene525 | 500 | 27 | 540 | 55.25 | 2.98 | https://www.ncbi.nlm.nih.gov/gene/?term=LOC105278116 | NW_011924889.1 | 635990 | 636489 | + | 1464 | 488 | 976 | 33.33 |
| gene1025 | 500 | 26 | 520 | 65.97 | 3.43 | https://www.ncbi.nlm.nih.gov/gene/?term=LOC105283760 | NW_011924920.1 | 14737 | 15236 | + | 657 | 417 | 240 | 63.47 |
| gene4016 | 500 | 26 | 520 | 91.14 | 4.74 | https://www.ncbi.nlm.nih.gov/gene/?term=LOC105276685 | NW_011925003.1 | 66624 | 67123 | + | 880 | 798 | 82 | 90.68 |
| gene9622 | 500 | 26 | 520 | 83.51 | 4.34 | https://www.ncbi.nlm.nih.gov/gene/?term=LOC105283159 | NW_011925287.1 | 414625 | 415124 | - | 790 | 612 | 178 | 77.47 |
| gene465 | 500 | 25 | 500 | 72.12 | 3.61 | https://www.ncbi.nlm.nih.gov/gene/?term=LOC105277304 | NW_011924889.1 | 171845 | 172344 | + | 1235 | 586 | 649 | 47.45 |
| gene10219 | 500 | 24 | 480 | 65.69 | 3.15 | https://www.ncbi.nlm.nih.gov/gene/?term=LOC105283830 | NW_011925327.1 | 293021 | 293520 | + | 955 | 410 | 545 | 42.93 |
| gene2406 | 500 | 24 | 480 | 43.71 | 2.1 | https://www.ncbi.nlm.nih.gov/gene/?term=LOC105274855 | NW_011924953.1 | 426755 | 427254 | - | 1844 | 451 | 1393 | 24.46 |

...

... only first 25 lines shown ...

Directory: RNK File: ranked-promoters-bySiteDensity-Cb_LI_RphaseA2.txt

| region_ID | rwidth | nbrsites | nbrper10kb | pmpersite | pmpernucl | pglink | seqnames | start | end | strand | coverage | numCs | numTs | prcntM |
|---|---|---|---|---|---|---|---|---|---|---|---|---|---|---|
| gene12925 | 500 | 48 | 960 | 45.53 | 4.37 | https://www.ncbi.nlm.nih.gov/gene/?term=LOC109611487 | NW_011925919.1 | 41730 | 42229 | - | 4216 | 1167 | 3049 | 27.68 |
| gene505 | 500 | 46 | 920 | 55.88 | 5.14 | https://www.ncbi.nlm.nih.gov/gene/?term=LOC105277808 | NW_011924889.1 | 477908 | 478407 | + | 4302 | 1503 | 2799 | 34.94 |
| gene1399 | 500 | 45 | 900 | 35.61 | 3.2 | https://www.ncbi.nlm.nih.gov/gene/?term=LOC105287518 | NW_011924931.1 | 442908 | 443407 | - | 3536 | 1027 | 2509 | 29.04 |
| gene8964 | 500 | 43 | 860 | 50.12 | 4.31 | https://www.ncbi.nlm.nih.gov/gene/?term=LOC105282404 | NW_011925232.1 | 562782 | 563281 | - | 2413 | 1025 | 1388 | 42.48 |
| gene7634 | 500 | 42 | 840 | 69.2 | 5.81 | https://www.ncbi.nlm.nih.gov/gene/?term=LOC105280861 | NW_011925151.1 | 38041 | 38540 | - | 2205 | 1108 | 1097 | 50.25 |
| gene2406 | 500 | 41 | 820 | 38.81 | 3.18 | https://www.ncbi.nlm.nih.gov/gene/?term=LOC105274855 | NW_011924953.1 | 426755 | 427254 | - | 3082 | 828 | 2254 | 26.87 |
| gene5753 | 500 | 40 | 800 | 75.83 | 6.07 | https://www.ncbi.nlm.nih.gov/gene/?term=LOC105278673 | NW_011925050.1 | 215094 | 215593 | + | 1561 | 1050 | 511 | 67.26 |
| gene12926 | 500 | 39 | 780 | 87.36 | 6.81 | https://www.ncbi.nlm.nih.gov/gene/?term=LOC105286944 | NW_011925919.1 | 43401 | 43900 | - | 1168 | 970 | 198 | 83.05 |
| gene4515 | 500 | 38 | 760 | 72.59 | 5.52 | https://www.ncbi.nlm.nih.gov/gene/?term=LOC109610839 | NW_011925013.1 | 19534 | 20033 | + | 2269 | 1495 | 774 | 65.89 |
| gene465 | 500 | 37 | 740 | 68.01 | 5.03 | https://www.ncbi.nlm.nih.gov/gene/?term=LOC105277304 | NW_011924889.1 | 171845 | 172344 | + | 2027 | 1026 | 1001 | 50.62 |
| gene128 | 500 | 36 | 720 | 58.87 | 4.24 | https://www.ncbi.nlm.nih.gov/gene/?term=LOC105286681 | NW_011924887.1 | 225822 | 226321 | + | 1514 | 677 | 837 | 44.72 |
| gene3929 | 500 | 34 | 680 | 80.62 | 5.48 | https://www.ncbi.nlm.nih.gov/gene/?term=LOC105276585 | NW_011924999.1 | 140417 | 140916 | - | 1118 | 848 | 270 | 75.85 |
| gene10220 | 500 | 33 | 660 | 63.8 | 4.21 | https://www.ncbi.nlm.nih.gov/gene/?term=LOC105283833 | NW_011925327.1 | 293294 | 293793 | + | 1891 | 788 | 1103 | 41.67 |
| gene772 | 500 | 33 | 660 | 81.04 | 5.35 | https://www.ncbi.nlm.nih.gov/gene/?term=LOC105280935 | NW_011924904.1 | 1310349 | 1310848 | + | 1028 | 820 | 208 | 79.77 |
| gene1010 | 500 | 32 | 640 | 66.26 | 4.24 | https://www.ncbi.nlm.nih.gov/gene/?term=LOC105283498 | NW_011924914.1 | 831409 | 831908 | - | 2637 | 1440 | 1197 | 54.61 |
| gene12331 | 500 | 32 | 640 | 48.96 | 3.13 | https://www.ncbi.nlm.nih.gov/gene/?term=LOC105286234 | NW_011925664.1 | 139161 | 139660 | + | 2728 | 895 | 1833 | 32.81 |
| gene10441 | 500 | 31 | 620 | 34.83 | 2.16 | https://www.ncbi.nlm.nih.gov/gene/?term=LOC105284094 | NW_011925353.1 | 595996 | 596495 | - | 3571 | 711 | 2860 | 19.91 |
| gene12363 | 500 | 31 | 620 | 67.1 | 4.16 | https://www.ncbi.nlm.nih.gov/gene/?term=LOC105286266 | NW_011925664.1 | 291330 | 291829 | - | 1621 | 774 | 847 | 47.75 |
| gene12539 | 500 | 31 | 620 | 64.17 | 3.98 | https://www.ncbi.nlm.nih.gov/gene/?term=LOC105286478 | NW_011925715.1 | 412371 | 412870 | + | 2381 | 941 | 1440 | 39.52 |
| gene1179 | 500 | 30 | 600 | 56.51 | 3.39 | https://www.ncbi.nlm.nih.gov/gene/?term=LOC105285207 | NW_011924922.1 | 2259192 | 2259691 | - | 1899 | 584 | 1315 | 30.75 |
| gene763 | 500 | 30 | 600 | 43.49 | 2.61 | https://www.ncbi.nlm.nih.gov/gene/?term=LOC105280696 | NW_011924904.1 | 1268134 | 1268633 | + | 3285 | 786 | 2499 | 23.93 |
| gene10147 | 500 | 29 | 580 | 84.49 | 4.9 | https://www.ncbi.nlm.nih.gov/gene/?term=LOC105283749 | NW_011925323.1 | 1722554 | 1723053 | + | 1583 | 949 | 634 | 59.95 |
| gene10397 | 500 | 29 | 580 | 58.33 | 3.38 | https://www.ncbi.nlm.nih.gov/gene/?term=LOC105284046 | NW_011925353.1 | 2217 | 2716 | + | 1481 | 539 | 942 | 36.39 |
| gene4878 | 500 | 29 | 580 | 77.18 | 4.48 | https://www.ncbi.nlm.nih.gov/gene/?term=LOC105277648 | NW_011925020.1 | 2601822 | 2602321 | - | 736 | 498 | 238 | 67.66 |

...

... only first 25 lines shown ...

Directory: RNK File: ranked-promoters-bySiteDensity-Cb_LI_RphaseB1.txt

| region_ID | rwidth | nbrsites | nbrper10kb | pmpersite | pmpernucl | pglink | seqnames | start | end | strand | coverage | numCs | numTs | prcntM |
|---|---|---|---|---|---|---|---|---|---|---|---|---|---|---|
| gene4515 | 500 | 40 | 800 | 73.92 | 5.91 | https://www.ncbi.nlm.nih.gov/gene/?term=LOC109610839 | NW_011925013.1 | 19534 | 20033 | + | 1555 | 1031 | 524 | 66.3 |
| gene505 | 500 | 39 | 780 | 56.58 | 4.41 | https://www.ncbi.nlm.nih.gov/gene/?term=LOC105277808 | NW_011924889.1 | 477908 | 478407 | + | 2814 | 869 | 1945 | 30.88 |
| gene465 | 500 | 38 | 760 | 68.31 | 5.19 | https://www.ncbi.nlm.nih.gov/gene/?term=LOC105277304 | NW_011924889.1 | 171845 | 172344 | + | 1531 | 753 | 778 | 49.18 |
| gene8964 | 500 | 35 | 700 | 55.5 | 3.89 | https://www.ncbi.nlm.nih.gov/gene/?term=LOC105282404 | NW_011925232.1 | 562782 | 563281 | - | 1626 | 632 | 994 | 38.87 |
| gene12925 | 500 | 34 | 680 | 46.84 | 3.18 | https://www.ncbi.nlm.nih.gov/gene/?term=LOC109611487 | NW_011925919.1 | 41730 | 42229 | - | 2639 | 673 | 1966 | 25.5 |
| gene12926 | 500 | 33 | 660 | 86.77 | 5.73 | https://www.ncbi.nlm.nih.gov/gene/?term=LOC105286944 | NW_011925919.1 | 43401 | 43900 | - | 627 | 496 | 131 | 79.11 |
| gene1399 | 500 | 31 | 620 | 42.39 | 2.63 | https://www.ncbi.nlm.nih.gov/gene/?term=LOC105287518 | NW_011924931.1 | 442908 | 443407 | - | 1775 | 494 | 1281 | 27.83 |
| gene128 | 500 | 30 | 600 | 58.02 | 3.48 | https://www.ncbi.nlm.nih.gov/gene/?term=LOC105286681 | NW_011924887.1 | 225822 | 226321 | + | 1223 | 509 | 714 | 41.62 |
| gene772 | 500 | 30 | 600 | 78.25 | 4.7 | https://www.ncbi.nlm.nih.gov/gene/?term=LOC105280935 | NW_011924904.1 | 1310349 | 1310848 | + | 772 | 601 | 171 | 77.85 |
| gene10220 | 500 | 29 | 580 | 69.4 | 4.03 | https://www.ncbi.nlm.nih.gov/gene/?term=LOC105283833 | NW_011925327.1 | 293294 | 293793 | + | 1233 | 524 | 709 | 42.5 |
| gene3929 | 500 | 29 | 580 | 82.1 | 4.76 | https://www.ncbi.nlm.nih.gov/gene/?term=LOC105276585 | NW_011924999.1 | 140417 | 140916 | - | 655 | 502 | 153 | 76.64 |
| gene5753 | 500 | 29 | 580 | 71.59 | 4.15 | https://www.ncbi.nlm.nih.gov/gene/?term=LOC105278673 | NW_011925050.1 | 215094 | 215593 | + | 956 | 641 | 315 | 67.05 |
| gene10147 | 500 | 28 | 560 | 85.36 | 4.78 | https://www.ncbi.nlm.nih.gov/gene/?term=LOC105283749 | NW_011925323.1 | 1722554 | 1723053 | + | 1269 | 736 | 533 | 58 |
| gene12363 | 500 | 28 | 560 | 71.24 | 3.99 | https://www.ncbi.nlm.nih.gov/gene/?term=LOC105286266 | NW_011925664.1 | 291330 | 291829 | - | 985 | 493 | 492 | 50.05 |
| gene12539 | 500 | 28 | 560 | 66 | 3.7 | https://www.ncbi.nlm.nih.gov/gene/?term=LOC105286478 | NW_011925715.1 | 412371 | 412870 | + | 1595 | 612 | 983 | 38.37 |
| gene7634 | 500 | 28 | 560 | 74.2 | 4.16 | https://www.ncbi.nlm.nih.gov/gene/?term=LOC105280861 | NW_011925151.1 | 38041 | 38540 | - | 1273 | 561 | 712 | 44.07 |
| gene11980 | 500 | 27 | 540 | 75.78 | 4.09 | https://www.ncbi.nlm.nih.gov/gene/?term=LOC109611395 | NW_011925546.1 | 60031 | 60530 | + | 824 | 592 | 232 | 71.84 |
| gene4016 | 500 | 27 | 540 | 91.87 | 4.96 | https://www.ncbi.nlm.nih.gov/gene/?term=LOC105276685 | NW_011925003.1 | 66624 | 67123 | + | 900 | 828 | 72 | 92 |
| gene9622 | 500 | 27 | 540 | 85.16 | 4.6 | https://www.ncbi.nlm.nih.gov/gene/?term=LOC105283159 | NW_011925287.1 | 414625 | 415124 | - | 788 | 611 | 177 | 77.54 |
| gene1010 | 500 | 26 | 520 | 65.7 | 3.42 | https://www.ncbi.nlm.nih.gov/gene/?term=LOC105283498 | NW_011924914.1 | 831409 | 831908 | - | 1273 | 550 | 723 | 43.21 |
| gene12331 | 500 | 26 | 520 | 50.19 | 2.61 | https://www.ncbi.nlm.nih.gov/gene/?term=LOC105286234 | NW_011925664.1 | 139161 | 139660 | + | 1644 | 471 | 1173 | 28.65 |
| gene1895 | 500 | 25 | 500 | 63.9 | 3.19 | https://www.ncbi.nlm.nih.gov/gene/?term=LOC105288085 | NW_011924945.1 | 117559 | 118058 | + | 678 | 375 | 303 | 55.31 |
| gene9243 | 500 | 24 | 480 | 74.47 | 3.57 | https://www.ncbi.nlm.nih.gov/gene/?term=LOC105282715 | NW_011925242.1 | 392772 | 393271 | - | 1351 | 573 | 778 | 42.41 |
| gene1025 | 500 | 23 | 460 | 74.18 | 3.41 | https://www.ncbi.nlm.nih.gov/gene/?term=LOC105283760 | NW_011924920.1 | 14737 | 15236 | + | 533 | 376 | 157 | 70.54 |

...

... only first 25 lines shown ...

Directory: RNK File: ranked-promoters-bySiteDensity-Cb_LI_RphaseB2.txt

| region_ID | rwidth | nbrsites | nbrper10kb | pmpersite | pmpernucl | pglink | seqnames | start | end | strand | coverage | numCs | numTs | prcntM |
|---|---|---|---|---|---|---|---|---|---|---|---|---|---|---|
| gene505 | 500 | 36 | 720 | 57.91 | 4.17 | https://www.ncbi.nlm.nih.gov/gene/?term=LOC105277808 | NW_011924889.1 | 477908 | 478407 | + | 2540 | 775 | 1765 | 30.51 |
| gene12925 | 500 | 35 | 700 | 47.48 | 3.32 | https://www.ncbi.nlm.nih.gov/gene/?term=LOC109611487 | NW_011925919.1 | 41730 | 42229 | - | 2545 | 655 | 1890 | 25.74 |
| gene12926 | 500 | 34 | 680 | 88.15 | 5.99 | https://www.ncbi.nlm.nih.gov/gene/?term=LOC105286944 | NW_011925919.1 | 43401 | 43900 | - | 881 | 710 | 171 | 80.59 |
| gene4515 | 500 | 34 | 680 | 70.16 | 4.77 | https://www.ncbi.nlm.nih.gov/gene/?term=LOC109610839 | NW_011925013.1 | 19534 | 20033 | + | 1424 | 890 | 534 | 62.5 |
| gene7634 | 500 | 31 | 620 | 75.89 | 4.7 | https://www.ncbi.nlm.nih.gov/gene/?term=LOC105280861 | NW_011925151.1 | 38041 | 38540 | - | 1365 | 644 | 721 | 47.18 |
| gene8964 | 500 | 30 | 600 | 55.48 | 3.33 | https://www.ncbi.nlm.nih.gov/gene/?term=LOC105282404 | NW_011925232.1 | 562782 | 563281 | - | 1367 | 558 | 809 | 40.82 |
| gene465 | 500 | 29 | 580 | 79.04 | 4.58 | https://www.ncbi.nlm.nih.gov/gene/?term=LOC105277304 | NW_011924889.1 | 171845 | 172344 | + | 1216 | 612 | 604 | 50.33 |
| gene10220 | 500 | 28 | 560 | 69.61 | 3.9 | https://www.ncbi.nlm.nih.gov/gene/?term=LOC105283833 | NW_011925327.1 | 293294 | 293793 | + | 970 | 410 | 560 | 42.27 |
| gene1010 | 500 | 27 | 540 | 67.03 | 3.62 | https://www.ncbi.nlm.nih.gov/gene/?term=LOC105283498 | NW_011924914.1 | 831409 | 831908 | - | 1459 | 670 | 789 | 45.92 |
| gene10147 | 500 | 27 | 540 | 83.57 | 4.51 | https://www.ncbi.nlm.nih.gov/gene/?term=LOC105283749 | NW_011925323.1 | 1722554 | 1723053 | + | 1049 | 635 | 414 | 60.53 |
| gene4016 | 500 | 27 | 540 | 88.49 | 4.78 | https://www.ncbi.nlm.nih.gov/gene/?term=LOC105276685 | NW_011925003.1 | 66624 | 67123 | + | 795 | 725 | 70 | 91.19 |
| gene5753 | 500 | 27 | 540 | 78.9 | 4.26 | https://www.ncbi.nlm.nih.gov/gene/?term=LOC105278673 | NW_011925050.1 | 215094 | 215593 | + | 949 | 625 | 324 | 65.86 |
| gene772 | 500 | 27 | 540 | 81.38 | 4.39 | https://www.ncbi.nlm.nih.gov/gene/?term=LOC105280935 | NW_011924904.1 | 1310349 | 1310848 | + | 593 | 450 | 143 | 75.89 |
| gene11980 | 500 | 26 | 520 | 69.15 | 3.6 | https://www.ncbi.nlm.nih.gov/gene/?term=LOC109611395 | NW_011925546.1 | 60031 | 60530 | + | 822 | 541 | 281 | 65.82 |
| gene12539 | 500 | 26 | 520 | 65.62 | 3.41 | https://www.ncbi.nlm.nih.gov/gene/?term=LOC105286478 | NW_011925715.1 | 412371 | 412870 | + | 1513 | 573 | 940 | 37.87 |
| gene128 | 500 | 26 | 520 | 57.92 | 3.01 | https://www.ncbi.nlm.nih.gov/gene/?term=LOC105286681 | NW_011924887.1 | 225822 | 226321 | + | 998 | 415 | 583 | 41.58 |
| gene12331 | 500 | 25 | 500 | 51.66 | 2.58 | https://www.ncbi.nlm.nih.gov/gene/?term=LOC105286234 | NW_011925664.1 | 139161 | 139660 | + | 1489 | 493 | 996 | 33.11 |
| gene3929 | 500 | 25 | 500 | 79.29 | 3.96 | https://www.ncbi.nlm.nih.gov/gene/?term=LOC105276585 | NW_011924999.1 | 140417 | 140916 | - | 661 | 483 | 178 | 73.07 |
| gene1399 | 500 | 24 | 480 | 42.71 | 2.05 | https://www.ncbi.nlm.nih.gov/gene/?term=LOC105287518 | NW_011924931.1 | 442908 | 443407 | - | 1654 | 439 | 1215 | 26.54 |
| gene2223 | 500 | 24 | 480 | 83.36 | 4 | https://www.ncbi.nlm.nih.gov/gene/?term=LOC109610674 | NW_011924946.1 | 3172465 | 3172964 | - | 714 | 470 | 244 | 65.83 |
| gene9622 | 500 | 24 | 480 | 82.96 | 3.98 | https://www.ncbi.nlm.nih.gov/gene/?term=LOC105283159 | NW_011925287.1 | 414625 | 415124 | - | 589 | 441 | 148 | 74.87 |
| gene12363 | 500 | 23 | 460 | 74.36 | 3.42 | https://www.ncbi.nlm.nih.gov/gene/?term=LOC105286266 | NW_011925664.1 | 291330 | 291829 | - | 885 | 428 | 457 | 48.36 |
| gene12537 | 500 | 23 | 460 | 63.53 | 2.92 | https://www.ncbi.nlm.nih.gov/gene/?term=LOC105286479 | NW_011925715.1 | 412531 | 413030 | - | 1923 | 558 | 1365 | 29.02 |
| gene1895 | 500 | 23 | 460 | 67.12 | 3.09 | https://www.ncbi.nlm.nih.gov/gene/?term=LOC105288085 | NW_011924945.1 | 117559 | 118058 | + | 649 | 372 | 277 | 57.32 |

...
... only first 25 lines shown ...

Directory: RNK File: sites-in-genes-Cb_LI_BCphaseA1.txt

```
seqnames	start	end	width	strand	coverage	numCs	numTs	perc_meth	region_seqnames	region_start	region_end	region_width	region_strand	region_source
	region_type	region_score	region_phase	region_ID	region_Dbxref	region_Name	region_gbkey	region_gene	region_gene_biotype	region_partial
	region_start_range	region_end_range
NW_011924881.1	39967	39967	1	+	9	8	1	88.8888888888889	NW_011924881.1	39621	40962	1342	-	Gnomon	gene	NA	NA	gene17	GeneID:105286636
	LOC105286636	Gene	LOC105286636	protein_coding	NA	character(0)	character(0)
NW_011924881.1	39987	39987	1	+	11	6	5	54.5454545454545	NW_011924881.1	39621	40962	1342	-	Gnomon	gene	NA	NA	gene17	GeneID:105286636
	LOC105286636	Gene	LOC105286636	protein_coding	NA	character(0)	character(0)
NW_011924881.1	40061	40061	1	+	13	8	5	61.5384615384615	NW_011924881.1	39621	40962	1342	-	Gnomon	gene	NA	NA	gene17	GeneID:105286636
	LOC105286636	Gene	LOC105286636	protein_coding	NA	character(0)	character(0)
NW_011924881.1	40072	40072	1	+	11	9	2	81.8181818181818	NW_011924881.1	39621	40962	1342	-	Gnomon	gene	NA	NA	gene17	GeneID:105286636
	LOC105286636	Gene	LOC105286636	protein_coding	NA	character(0)	character(0)
NW_011924881.1	40095	40095	1	+	13	6	7	46.1538461538462	NW_011924881.1	39621	40962	1342	-	Gnomon	gene	NA	NA	gene17	GeneID:105286636
	LOC105286636	Gene	LOC105286636	protein_coding	NA	character(0)	character(0)
NW_011924881.1	40120	40120	1	+	11	7	4	63.6363636363636	NW_011924881.1	39621	40962	1342	-	Gnomon	gene	NA	NA	gene17	GeneID:105286636
	LOC105286636	Gene	LOC105286636	protein_coding	NA	character(0)	character(0)
NW_011924881.1	40257	40257	1	-	19	15	4	78.9473684210526	NW_011924881.1	39621	40962	1342	-	Gnomon	gene	NA	NA	gene17	GeneID:105286636
	LOC105286636	Gene	LOC105286636	protein_coding	NA	character(0)	character(0)
NW_011924881.1	40260	40260	1	-	19	17	2	89.4736842105263	NW_011924881.1	39621	40962	1342	-	Gnomon	gene	NA	NA	gene17	GeneID:105286636
	LOC105286636	Gene	LOC105286636	protein_coding	NA	character(0)	character(0)
NW_011924881.1	40271	40271	1	-	17	13	4	76.4705882352941	NW_011924881.1	39621	40962	1342	-	Gnomon	gene	NA	NA	gene17	GeneID:105286636
	LOC105286636	Gene	LOC105286636	protein_coding	NA	character(0)	character(0)
NW_011924881.1	40335	40335	1	-	12	7	5	58.3333333333333	NW_011924881.1	39621	40962	1342	-	Gnomon	gene	NA	NA	gene17	GeneID:105286636
	LOC105286636	Gene	LOC105286636	protein_coding	NA	character(0)	character(0)
NW_011924881.1	40350	40350	1	-	10	6	4	60	NW_011924881.1	39621	40962	1342	-	Gnomon	gene	NA	NA	gene17	GeneID:105286636	LOC105286636
	Gene	LOC105286636	protein_coding	NA	character(0)	character(0)
NW_011924881.1	40360	40360	1	-	17	14	3	82.3529411764706	NW_011924881.1	39621	40962	1342	-	Gnomon	gene	NA	NA	gene17	GeneID:105286636
	LOC105286636	Gene	LOC105286636	protein_coding	NA	character(0)	character(0)
NW_011924881.1	40372	40372	1	-	16	15	1	93.75	NW_011924881.1	39621	40962	1342	-	Gnomon	gene	NA	NA	gene17	GeneID:105286636	LOC105286636
	Gene	LOC105286636	protein_coding	NA	character(0)	character(0)
NW_011924881.1	40385	40385	1	-	13	13	0	100	NW_011924881.1	39621	40962	1342	-	Gnomon	gene	NA	NA	gene17	GeneID:105286636	LOC105286636
	Gene	LOC105286636	protein_coding	NA	character(0)	character(0)
NW_011924881.1	40390	40390	1	-	13	13	0	100	NW_011924881.1	39621	40962	1342	-	Gnomon	gene	NA	NA	gene17	GeneID:105286636	LOC105286636
	Gene	LOC105286636	protein_coding	NA	character(0)	character(0)
NW_011924881.1	40404	40404	1	-	10	8	2	80	NW_011924881.1	39621	40962	1342	-	Gnomon	gene	NA	NA	gene17	GeneID:105286636	LOC105286636
	Gene	LOC105286636	protein_coding	NA	character(0)	character(0)
NW_011924881.1	40406	40406	1	-	10	7	3	70	NW_011924881.1	39621	40962	1342	-	Gnomon	gene	NA	NA	gene17	GeneID:105286636	LOC105286636
	Gene	LOC105286636	protein_coding	NA	character(0)	character(0)
NW_011924881.1	247875	247875	1	+	28	8	20	28.5714285714286	NW_011924881.1	195683	286417	90735	-	Gnomon	gene	NA	NA	gene22
	GeneID:105288120	LOC105288120	Gene	LOC105288120	protein_coding	NA	character(0)	character(0)
NW_011924881.1	401787	401787	1	+	26	7	19	26.9230769230769	NW_011924881.1	369231	418861	49631	-	Gnomon	gene	NA	NA	gene24
	GeneID:109611552	LOC109611552	Gene	LOC109611552	protein_coding	NA	character(0)	character(0)
NW_011924881.1	447772	447772	1	+	35	7	28	20	NW_011924881.1	447189	448306	1118	+	Gnomon	gene	NA	NA	gene25
	GeneID:105274524	LOC105274524	Gene	LOC105274524	protein_coding	NA	character(0)	character(0)
NW_011924881.1	447836	447836	1	+	44	16	28	36.3636363636364	NW_011924881.1	447189	448306	1118	+	Gnomon	gene	NA	NA	gene25
	GeneID:105274524	LOC105274524	Gene	LOC105274524	protein_coding	NA	character(0)	character(0)
NW_011924881.1	447837	447837	1	-	23	9	14	39.1304347826087	NW_011924881.1	447189	448306	1118	+	Gnomon	gene	NA	NA	gene25
	GeneID:105274524	LOC105274524	Gene	LOC105274524	protein_coding	NA	character(0)	character(0)
NW_011924881.1	447927	447927	1	+	25	7	18	28	NW_011924881.1	447189	448306	1118	+	Gnomon	gene	NA	NA	gene25
	GeneID:105274524	LOC105274524	Gene	LOC105274524	protein_coding	NA	character(0)	character(0)
NW_011924881.1	447945	447945	1	+	22	19	3	86.3636363636364	NW_011924881.1	447189	448306	1118	+	Gnomon	gene	NA	NA	gene25
	GeneID:105274524	LOC105274524	Gene	LOC105274524	protein_coding	NA	character(0)	character(0)
...
... only first 25 lines shown ...
```

Directory: RNK File: sites-in-genes-Cb_LI_BCphaseA2.txt

```
seqnames	start	end	width	strand	coverage	numCs	numTs	perc_meth	region_seqnames	region_start	region_end	region_width	region_strand	region_source
	region_type	region_score	region_phase	region_ID	region_Dbxref	region_Name	region_gbkey	region_gene	region_gene_biotype	region_partial
	region_start_range	region_end_range
NW_011924879.1	2162	2162	1	+	31	10	21	32.258064516129	NW_011924879.1	7	3079	3073	+	Gnomon	gene	NA	NA	gene5	GeneID:105275124
	LOC105275124	Gene	LOC105275124	protein_coding	NA	character(0)	character(0)
NW_011924881.1	39987	39987	1	+	9	6	3	66.6666666666667	NW_011924881.1	39621	40962	1342	-	Gnomon	gene	NA	NA	gene17	GeneID:105286636
	LOC105286636	Gene	LOC105286636	protein_coding	NA	character(0)	character(0)
NW_011924881.1	40061	40061	1	+	9	8	1	88.8888888888889	NW_011924881.1	39621	40962	1342	-	Gnomon	gene	NA	NA	gene17	GeneID:105286636
	LOC105286636	Gene	LOC105286636	protein_coding	NA	character(0)	character(0)
NW_011924881.1	40072	40072	1	+	10	10	0	100	NW_011924881.1	39621	40962	1342	-	Gnomon	gene	NA	NA	gene17	GeneID:105286636	LOC105286636
	Gene	LOC105286636	protein_coding	NA	character(0)	character(0)
NW_011924881.1	40120	40120	1	+	10	9	1	90	NW_011924881.1	39621	40962	1342	-	Gnomon	gene	NA	NA	gene17	GeneID:105286636	LOC105286636
	Gene	LOC105286636	protein_coding	NA	character(0)	character(0)
NW_011924881.1	40257	40257	1	-	26	24	2	92.3076923076923	NW_011924881.1	39621	40962	1342	-	Gnomon	gene	NA	NA	gene17	GeneID:105286636
	LOC105286636	Gene	LOC105286636	protein_coding	NA	character(0)	character(0)
NW_011924881.1	40259	40259	1	+	5	5	0	100	NW_011924881.1	39621	40962	1342	-	Gnomon	gene	NA	NA	gene17	GeneID:105286636	LOC105286636
	Gene	LOC105286636	protein_coding	NA	character(0)	character(0)
NW_011924881.1	40260	40260	1	-	23	22	1	95.6521739130435	NW_011924881.1	39621	40962	1342	-	Gnomon	gene	NA	NA	gene17	GeneID:105286636
	LOC105286636	Gene	LOC105286636	protein_coding	NA	character(0)	character(0)
NW_011924881.1	40270	40270	1	+	6	5	1	83.3333333333333	NW_011924881.1	39621	40962	1342	-	Gnomon	gene	NA	NA	gene17	GeneID:105286636
	LOC105286636	Gene	LOC105286636	protein_coding	NA	character(0)	character(0)
NW_011924881.1	40271	40271	1	-	24	21	3	87.5	NW_011924881.1	39621	40962	1342	-	Gnomon	gene	NA	NA	gene17	GeneID:105286636	LOC105286636
	Gene	LOC105286636	protein_coding	NA	character(0)	character(0)
NW_011924881.1	40335	40335	1	-	14	8	6	57.1428571428571	NW_011924881.1	39621	40962	1342	-	Gnomon	gene	NA	NA	gene17	GeneID:105286636
	LOC105286636	Gene	LOC105286636	protein_coding	NA	character(0)	character(0)
NW_011924881.1	40350	40350	1	-	14	13	1	92.8571428571429	NW_011924881.1	39621	40962	1342	-	Gnomon	gene	NA	NA	gene17	GeneID:105286636
	LOC105286636	Gene	LOC105286636	protein_coding	NA	character(0)	character(0)
NW_011924881.1	40360	40360	1	-	20	19	1	95	NW_011924881.1	39621	40962	1342	-	Gnomon	gene	NA	NA	gene17	GeneID:105286636	LOC105286636
	Gene	LOC105286636	protein_coding	NA	character(0)	character(0)
NW_011924881.1	40372	40372	1	-	18	16	2	88.8888888888889	NW_011924881.1	39621	40962	1342	-	Gnomon	gene	NA	NA	gene17	GeneID:105286636
	LOC105286636	Gene	LOC105286636	protein_coding	NA	character(0)	character(0)
NW_011924881.1	40385	40385	1	-	14	14	0	100	NW_011924881.1	39621	40962	1342	-	Gnomon	gene	NA	NA	gene17	GeneID:105286636	LOC105286636
	Gene	LOC105286636	protein_coding	NA	character(0)	character(0)
NW_011924881.1	40390	40390	1	-	14	14	0	100	NW_011924881.1	39621	40962	1342	-	Gnomon	gene	NA	NA	gene17	GeneID:105286636	LOC105286636
	Gene	LOC105286636	protein_coding	NA	character(0)	character(0)
NW_011924881.1	145546	145546	1	-	19	6	13	31.5789473684211	NW_011924881.1	140800	154070	13271	-	Gnomon	gene	NA	NA	gene19
	GeneID:105287639	LOC105287639	Gene	LOC105287639	protein_coding	NA	character(0)	character(0)
NW_011924881.1	447836	447836	1	+	36	14	22	38.8888888888889	NW_011924881.1	447189	448306	1118	+	Gnomon	gene	NA	NA	gene25
	GeneID:105274524	LOC105274524	Gene	LOC105274524	protein_coding	NA	character(0)	character(0)
NW_011924881.1	447837	447837	1	-	46	15	31	32.6086956521739	NW_011924881.1	447189	448306	1118	+	Gnomon	gene	NA	NA	gene25
	GeneID:105274524	LOC105274524	Gene	LOC105274524	protein_coding	NA	character(0)	character(0)
NW_011924881.1	447927	447927	1	+	22	9	13	40.9090909090909	NW_011924881.1	447189	448306	1118	+	Gnomon	gene	NA	NA	gene25
	GeneID:105274524	LOC105274524	Gene	LOC105274524	protein_coding	NA	character(0)	character(0)
NW_011924881.1	447928	447928	1	-	15	7	8	46.6666666666667	NW_011924881.1	447189	448306	1118	+	Gnomon	gene	NA	NA	gene25
	GeneID:105274524	LOC105274524	Gene	LOC105274524	protein_coding	NA	character(0)	character(0)
NW_011924881.1	447945	447945	1	+	18	16	2	88.8888888888889	NW_011924881.1	447189	448306	1118	+	Gnomon	gene	NA	NA	gene25
	GeneID:105274524	LOC105274524	Gene	LOC105274524	protein_coding	NA	character(0)	character(0)
NW_011924881.1	447946	447946	1	-	8	6	2	75	NW_011924881.1	447189	448306	1118	+	Gnomon	gene	NA	NA	gene25
	GeneID:105274524	LOC105274524	Gene	LOC105274524	protein_coding	NA	character(0)	character(0)
NW_011924881.1	447952	447952	1	+	16	15	1	93.75	NW_011924881.1	447189	448306	1118	+	Gnomon	gene	NA	NA	gene25
	GeneID:105274524	LOC105274524	Gene	LOC105274524	protein_coding	NA	character(0)	character(0)
...
... only first 25 lines shown ...
```

Directory: RNK File: sites-in-genes-Cb_LI_BCphaseB1.txt

```
seqnames	start	end	width	strand	coverage	numCs	numTs	perc_meth	region_seqnames	region_start	region_end	region_width	region_strand	region_source
	region_type	region_score	region_phase	region_ID	region_Dbxref	region_Name	region_gbkey	region_gene	region_gene_biotype	region_partial
	region_start_range	region_end_range
NW_011924881.1	39967	39967	1	+	16	14	2	87.5	NW_011924881.1	39621	40962	1342	-	Gnomon	gene	NA	NA	gene17	GeneID:105286636	LOC105286636
	Gene	LOC105286636	protein_coding	NA	character(0)	character(0)
NW_011924881.1	39987	39987	1	+	23	12	11	52.1739130434783	NW_011924881.1	39621	40962	1342	-	Gnomon	gene	NA	NA	gene17	GeneID:105286636
	LOC105286636	Gene	LOC105286636	protein_coding	NA	character(0)	character(0)
NW_011924881.1	40061	40061	1	+	13	8	5	61.5384615384615	NW_011924881.1	39621	40962	1342	-	Gnomon	gene	NA	NA	gene17	GeneID:105286636
	LOC105286636	Gene	LOC105286636	protein_coding	NA	character(0)	character(0)
NW_011924881.1	40072	40072	1	+	11	11	0	100	NW_011924881.1	39621	40962	1342	-	Gnomon	gene	NA	NA	gene17	GeneID:105286636	LOC105286636
	Gene	LOC105286636	protein_coding	NA	character(0)	character(0)
NW_011924881.1	40095	40095	1	+	16	10	6	62.5	NW_011924881.1	39621	40962	1342	-	Gnomon	gene	NA	NA	gene17	GeneID:105286636	LOC105286636
	Gene	LOC105286636	protein_coding	NA	character(0)	character(0)
NW_011924881.1	40120	40120	1	+	20	16	4	80	NW_011924881.1	39621	40962	1342	-	Gnomon	gene	NA	NA	gene17	GeneID:105286636	LOC105286636
	Gene	LOC105286636	protein_coding	NA	character(0)	character(0)
NW_011924881.1	40159	40159	1	+	20	11	9	55	NW_011924881.1	39621	40962	1342	-	Gnomon	gene	NA	NA	gene17	GeneID:105286636	LOC105286636
	Gene	LOC105286636	protein_coding	NA	character(0)	character(0)
NW_011924881.1	40257	40257	1	-	23	17	6	73.9130434782609	NW_011924881.1	39621	40962	1342	-	Gnomon	gene	NA	NA	gene17	GeneID:105286636
	LOC105286636	Gene	LOC105286636	protein_coding	NA	character(0)	character(0)
NW_011924881.1	40260	40260	1	-	23	21	2	91.304347826087	NW_011924881.1	39621	40962	1342	-	Gnomon	gene	NA	NA	gene17	GeneID:105286636
	LOC105286636	Gene	LOC105286636	protein_coding	NA	character(0)	character(0)
NW_011924881.1	40271	40271	1	-	23	20	3	86.9565217391304	NW_011924881.1	39621	40962	1342	-	Gnomon	gene	NA	NA	gene17	GeneID:105286636
	LOC105286636	Gene	LOC105286636	protein_coding	NA	character(0)	character(0)
NW_011924881.1	40335	40335	1	-	13	8	5	61.5384615384615	NW_011924881.1	39621	40962	1342	-	Gnomon	gene	NA	NA	gene17	GeneID:105286636
	LOC105286636	Gene	LOC105286636	protein_coding	NA	character(0)	character(0)
NW_011924881.1	40350	40350	1	-	15	11	4	73.3333333333333	NW_011924881.1	39621	40962	1342	-	Gnomon	gene	NA	NA	gene17	GeneID:105286636
	LOC105286636	Gene	LOC105286636	protein_coding	NA	character(0)	character(0)
NW_011924881.1	40360	40360	1	-	23	22	1	95.6521739130435	NW_011924881.1	39621	40962	1342	-	Gnomon	gene	NA	NA	gene17	GeneID:105286636
	LOC105286636	Gene	LOC105286636	protein_coding	NA	character(0)	character(0)
NW_011924881.1	40372	40372	1	-	24	23	1	95.8333333333333	NW_011924881.1	39621	40962	1342	-	Gnomon	gene	NA	NA	gene17	GeneID:105286636
	LOC105286636	Gene	LOC105286636	protein_coding	NA	character(0)	character(0)
NW_011924881.1	40385	40385	1	-	18	18	0	100	NW_011924881.1	39621	40962	1342	-	Gnomon	gene	NA	NA	gene17	GeneID:105286636	LOC105286636
	Gene	LOC105286636	protein_coding	NA	character(0)	character(0)
NW_011924881.1	40390	40390	1	-	16	15	1	93.75	NW_011924881.1	39621	40962	1342	-	Gnomon	gene	NA	NA	gene17	GeneID:105286636	LOC105286636
	Gene	LOC105286636	protein_coding	NA	character(0)	character(0)
NW_011924881.1	40406	40406	1	-	13	6	7	46.1538461538462	NW_011924881.1	39621	40962	1342	-	Gnomon	gene	NA	NA	gene17	GeneID:105286636
	LOC105286636	Gene	LOC105286636	protein_coding	NA	character(0)	character(0)
NW_011924881.1	45873	45873	1	-	14	7	7	50	NW_011924881.1	42715	135016	92302	-	Gnomon	gene	NA	NA	gene18	GeneID:105287290
	LOC105287290	Gene	LOC105287290	protein_coding	NA	character(0)	character(0)
NW_011924881.1	62383	62383	1	+	17	6	11	35.2941176470588	NW_011924881.1	42715	135016	92302	-	Gnomon	gene	NA	NA	gene18
	GeneID:105287290	LOC105287290	Gene	LOC105287290	protein_coding	NA	character(0)	character(0)
NW_011924881.1	125434	125434	1	-	13	6	7	46.1538461538462	NW_011924881.1	42715	135016	92302	-	Gnomon	gene	NA	NA	gene18
	GeneID:105287290	LOC105287290	Gene	LOC105287290	protein_coding	NA	character(0)	character(0)
NW_011924881.1	145080	145080	1	-	29	8	21	27.5862068965517	NW_011924881.1	140800	154070	13271	-	Gnomon	gene	NA	NA	gene19
	GeneID:105287639	LOC105287639	Gene	LOC105287639	protein_coding	NA	character(0)	character(0)
NW_011924881.1	227838	227838	1	-	14	7	7	50	NW_011924881.1	195683	286417	90735	-	Gnomon	gene	NA	NA	gene22
	GeneID:105288120	LOC105288120	Gene	LOC105288120	protein_coding	NA	character(0)	character(0)
NW_011924881.1	447836	447836	1	+	50	17	33	34	NW_011924881.1	447189	448306	1118	+	Gnomon	gene	NA	NA	gene25
	GeneID:105274524	LOC105274524	Gene	LOC105274524	protein_coding	NA	character(0)	character(0)
NW_011924881.1	447837	447837	1	-	33	9	24	27.2727272727273	NW_011924881.1	447189	448306	1118	+	Gnomon	gene	NA	NA	gene25
	GeneID:105274524	LOC105274524	Gene	LOC105274524	protein_coding	NA	character(0)	character(0)
...
... only first 25 lines shown ...
```

Directory: RNK File: sites-in-genes-Cb_LI_BCphaseB2.txt

| seqnames | start | end | width | strand | coverage | numCs | numTs | perc_meth | region_seqnames | region_start | region_end | region_width | region_strand | region_source | region_type | region_score | region_phase | region_ID | region_Dbxref | region_Name | region_gbkey | region_gene | region_gene_biotype | region_partial | region_start_range | region_end_range |
|---|---|---|---|---|---|---|---|---|---|---|---|---|---|---|---|---|---|---|---|---|---|---|---|---|---|---|
| NW_011924881.1 | 39967 | 39967 | 1 | + | 15 | 12 | 3 | 80 | NW_011924881.1 | 39621 | 40962 | 1342 | - | Gnomon | gene | NA | NA | gene17 | GeneID:105286636 | LOC105286636 | Gene | LOC105286636 | protein_coding | NA | character(0) | character(0) |
| NW_011924881.1 | 39987 | 39987 | 1 | + | 15 | 6 | 9 | 40 | NW_011924881.1 | 39621 | 40962 | 1342 | - | Gnomon | gene | NA | NA | gene17 | GeneID:105286636 | LOC105286636 | Gene | LOC105286636 | protein_coding | NA | character(0) | character(0) |
| NW_011924881.1 | 40061 | 40061 | 1 | + | 14 | 10 | 4 | 71.4285714285714 | NW_011924881.1 | 39621 | 40962 | 1342 | - | Gnomon | gene | NA | NA | gene17 | GeneID:105286636 | LOC105286636 | Gene | LOC105286636 | protein_coding | NA | character(0) | character(0) |
| NW_011924881.1 | 40072 | 40072 | 1 | + | 13 | 12 | 1 | 92.3076923076923 | NW_011924881.1 | 39621 | 40962 | 1342 | - | Gnomon | gene | NA | NA | gene17 | GeneID:105286636 | LOC105286636 | Gene | LOC105286636 | protein_coding | NA | character(0) | character(0) |
| NW_011924881.1 | 40095 | 40095 | 1 | + | 17 | 9 | 8 | 52.9411764705882 | NW_011924881.1 | 39621 | 40962 | 1342 | - | Gnomon | gene | NA | NA | gene17 | GeneID:105286636 | LOC105286636 | Gene | LOC105286636 | protein_coding | NA | character(0) | character(0) |
| NW_011924881.1 | 40120 | 40120 | 1 | + | 17 | 12 | 5 | 70.5882352941177 | NW_011924881.1 | 39621 | 40962 | 1342 | - | Gnomon | gene | NA | NA | gene17 | GeneID:105286636 | LOC105286636 | Gene | LOC105286636 | protein_coding | NA | character(0) | character(0) |
| NW_011924881.1 | 40159 | 40159 | 1 | + | 13 | 9 | 4 | 69.2307692307692 | NW_011924881.1 | 39621 | 40962 | 1342 | - | Gnomon | gene | NA | NA | gene17 | GeneID:105286636 | LOC105286636 | Gene | LOC105286636 | protein_coding | NA | character(0) | character(0) |
| NW_011924881.1 | 40257 | 40257 | 1 | - | 24 | 20 | 4 | 83.3333333333333 | NW_011924881.1 | 39621 | 40962 | 1342 | - | Gnomon | gene | NA | NA | gene17 | GeneID:105286636 | LOC105286636 | Gene | LOC105286636 | protein_coding | NA | character(0) | character(0) |
| NW_011924881.1 | 40260 | 40260 | 1 | - | 24 | 20 | 4 | 83.3333333333333 | NW_011924881.1 | 39621 | 40962 | 1342 | - | Gnomon | gene | NA | NA | gene17 | GeneID:105286636 | LOC105286636 | Gene | LOC105286636 | protein_coding | NA | character(0) | character(0) |
| NW_011924881.1 | 40271 | 40271 | 1 | - | 23 | 21 | 2 | 91.304347826087 | NW_011924881.1 | 39621 | 40962 | 1342 | - | Gnomon | gene | NA | NA | gene17 | GeneID:105286636 | LOC105286636 | Gene | LOC105286636 | protein_coding | NA | character(0) | character(0) |
| NW_011924881.1 | 40335 | 40335 | 1 | - | 13 | 6 | 7 | 46.1538461538462 | NW_011924881.1 | 39621 | 40962 | 1342 | - | Gnomon | gene | NA | NA | gene17 | GeneID:105286636 | LOC105286636 | Gene | LOC105286636 | protein_coding | NA | character(0) | character(0) |
| NW_011924881.1 | 40350 | 40350 | 1 | - | 19 | 16 | 3 | 84.2105263157895 | NW_011924881.1 | 39621 | 40962 | 1342 | - | Gnomon | gene | NA | NA | gene17 | GeneID:105286636 | LOC105286636 | Gene | LOC105286636 | protein_coding | NA | character(0) | character(0) |
| NW_011924881.1 | 40360 | 40360 | 1 | - | 25 | 21 | 4 | 84 | NW_011924881.1 | 39621 | 40962 | 1342 | - | Gnomon | gene | NA | NA | gene17 | GeneID:105286636 | LOC105286636 | Gene | LOC105286636 | protein_coding | NA | character(0) | character(0) |
| NW_011924881.1 | 40372 | 40372 | 1 | - | 24 | 22 | 2 | 91.6666666666667 | NW_011924881.1 | 39621 | 40962 | 1342 | - | Gnomon | gene | NA | NA | gene17 | GeneID:105286636 | LOC105286636 | Gene | LOC105286636 | protein_coding | NA | character(0) | character(0) |
| NW_011924881.1 | 40385 | 40385 | 1 | - | 20 | 20 | 0 | 100 | NW_011924881.1 | 39621 | 40962 | 1342 | - | Gnomon | gene | NA | NA | gene17 | GeneID:105286636 | LOC105286636 | Gene | LOC105286636 | protein_coding | NA | character(0) | character(0) |
| NW_011924881.1 | 40390 | 40390 | 1 | - | 18 | 18 | 0 | 100 | NW_011924881.1 | 39621 | 40962 | 1342 | - | Gnomon | gene | NA | NA | gene17 | GeneID:105286636 | LOC105286636 | Gene | LOC105286636 | protein_coding | NA | character(0) | character(0) |
| NW_011924881.1 | 40403 | 40403 | 1 | + | 5 | 5 | 0 | 100 | NW_011924881.1 | 39621 | 40962 | 1342 | - | Gnomon | gene | NA | NA | gene17 | GeneID:105286636 | LOC105286636 | Gene | LOC105286636 | protein_coding | NA | character(0) | character(0) |
| NW_011924881.1 | 40404 | 40404 | 1 | - | 17 | 8 | 9 | 47.0588235294118 | NW_011924881.1 | 39621 | 40962 | 1342 | - | Gnomon | gene | NA | NA | gene17 | GeneID:105286636 | LOC105286636 | Gene | LOC105286636 | protein_coding | NA | character(0) | character(0) |
| NW_011924881.1 | 40406 | 40406 | 1 | - | 17 | 10 | 7 | 58.8235294117647 | NW_011924881.1 | 39621 | 40962 | 1342 | - | Gnomon | gene | NA | NA | gene17 | GeneID:105286636 | LOC105286636 | Gene | LOC105286636 | protein_coding | NA | character(0) | character(0) |
| NW_011924881.1 | 152633 | 152633 | 1 | - | 16 | 6 | 10 | 37.5 | NW_011924881.1 | 140800 | 154070 | 13271 | - | Gnomon | gene | NA | NA | gene19 | GeneID:105287639 | LOC105287639 | Gene | LOC105287639 | protein_coding | NA | character(0) | character(0) |
| NW_011924881.1 | 447772 | 447772 | 1 | + | 35 | 11 | 24 | 31.4285714285714 | NW_011924881.1 | 447189 | 448306 | 1118 | + | Gnomon | gene | NA | NA | gene25 | GeneID:105274524 | LOC105274524 | Gene | LOC105274524 | protein_coding | NA | character(0) | character(0) |
| NW_011924881.1 | 447836 | 447836 | 1 | + | 54 | 17 | 37 | 31.4814814814815 | NW_011924881.1 | 447189 | 448306 | 1118 | + | Gnomon | gene | NA | NA | gene25 | GeneID:105274524 | LOC105274524 | Gene | LOC105274524 | protein_coding | NA | character(0) | character(0) |
| NW_011924881.1 | 447837 | 447837 | 1 | - | 55 | 24 | 31 | 43.6363636363636 | NW_011924881.1 | 447189 | 448306 | 1118 | + | Gnomon | gene | NA | NA | gene25 | GeneID:105274524 | LOC105274524 | Gene | LOC105274524 | protein_coding | NA | character(0) | character(0) |
| NW_011924881.1 | 447945 | 447945 | 1 | + | 13 | 9 | 4 | 69.2307692307692 | NW_011924881.1 | 447189 | 448306 | 1118 | + | Gnomon | gene | NA | NA | gene25 | GeneID:105274524 | LOC105274524 | Gene | LOC105274524 | protein_coding | NA | character(0) | character(0) |

...
... only first 25 lines shown ...

```
Directory: RNK File: sites-in-genes-Cb_LI_RphaseA1.txt

seqnames	start	end	width	strand	coverage	numCs	numTs	perc_meth	region_seqnames	region_start	region_end	region_width	region_strand	region_source
	region_type	region_score	region_phase	region_ID	region_Dbxref	region_Name	region_gbkey	region_gene	region_gene_biotype	region_partial
	region_start_range	region_end_range
NW_011924881.1	39967	39967	1	+	8	7	1	87.5	NW_011924881.1	39621	40962	1342	-	Gnomon	gene	NA	NA	gene17	GeneID:105286636	LOC105286636
	Gene	LOC105286636	protein_coding	NA	character(0)	character(0)
NW_011924881.1	39987	39987	1	+	8	5	3	62.5	NW_011924881.1	39621	40962	1342	-	Gnomon	gene	NA	NA	gene17	GeneID:105286636	LOC105286636
	Gene	LOC105286636	protein_coding	NA	character(0)	character(0)
NW_011924881.1	40061	40061	1	+	9	8	1	88.8888888888889	NW_011924881.1	39621	40962	1342	-	Gnomon	gene	NA	NA	gene17	GeneID:105286636
	LOC105286636	Gene	LOC105286636	protein_coding	NA	character(0)	character(0)
NW_011924881.1	40072	40072	1	+	15	15	0	100	NW_011924881.1	39621	40962	1342	-	Gnomon	gene	NA	NA	gene17	GeneID:105286636	LOC105286636
	Gene	LOC105286636	protein_coding	NA	character(0)	character(0)
NW_011924881.1	40095	40095	1	+	20	12	8	60	NW_011924881.1	39621	40962	1342	-	Gnomon	gene	NA	NA	gene17	GeneID:105286636	LOC105286636
	Gene	LOC105286636	protein_coding	NA	character(0)	character(0)
NW_011924881.1	40120	40120	1	+	22	15	7	68.1818181818182	NW_011924881.1	39621	40962	1342	-	Gnomon	gene	NA	NA	gene17	GeneID:105286636
	LOC105286636	Gene	LOC105286636	protein_coding	NA	character(0)	character(0)
NW_011924881.1	40121	40121	1	-	9	7	2	77.7777777777778	NW_011924881.1	39621	40962	1342	-	Gnomon	gene	NA	NA	gene17	GeneID:105286636
	LOC105286636	Gene	LOC105286636	protein_coding	NA	character(0)	character(0)
NW_011924881.1	40159	40159	1	+	16	11	5	68.75	NW_011924881.1	39621	40962	1342	-	Gnomon	gene	NA	NA	gene17	GeneID:105286636	LOC105286636
	Gene	LOC105286636	protein_coding	NA	character(0)	character(0)
NW_011924881.1	40160	40160	1	-	10	6	4	60	NW_011924881.1	39621	40962	1342	-	Gnomon	gene	NA	NA	gene17	GeneID:105286636	LOC105286636
	Gene	LOC105286636	protein_coding	NA	character(0)	character(0)
NW_011924881.1	40257	40257	1	-	16	13	3	81.25	NW_011924881.1	39621	40962	1342	-	Gnomon	gene	NA	NA	gene17	GeneID:105286636	LOC105286636
	Gene	LOC105286636	protein_coding	NA	character(0)	character(0)
NW_011924881.1	40260	40260	1	-	13	13	0	100	NW_011924881.1	39621	40962	1342	-	Gnomon	gene	NA	NA	gene17	GeneID:105286636	LOC105286636
	Gene	LOC105286636	protein_coding	NA	character(0)	character(0)
NW_011924881.1	40270	40270	1	+	5	5	0	100	NW_011924881.1	39621	40962	1342	-	Gnomon	gene	NA	NA	gene17	GeneID:105286636	LOC105286636
	Gene	LOC105286636	protein_coding	NA	character(0)	character(0)
NW_011924881.1	40271	40271	1	-	13	10	3	76.9230769230769	NW_011924881.1	39621	40962	1342	-	Gnomon	gene	NA	NA	gene17	GeneID:105286636
	LOC105286636	Gene	LOC105286636	protein_coding	NA	character(0)	character(0)
NW_011924881.1	40335	40335	1	-	13	7	6	53.8461538461538	NW_011924881.1	39621	40962	1342	-	Gnomon	gene	NA	NA	gene17	GeneID:105286636
	LOC105286636	Gene	LOC105286636	protein_coding	NA	character(0)	character(0)
NW_011924881.1	40360	40360	1	-	10	9	1	90	NW_011924881.1	39621	40962	1342	-	Gnomon	gene	NA	NA	gene17	GeneID:105286636	LOC105286636
	Gene	LOC105286636	protein_coding	NA	character(0)	character(0)
NW_011924881.1	40372	40372	1	-	10	10	0	100	NW_011924881.1	39621	40962	1342	-	Gnomon	gene	NA	NA	gene17	GeneID:105286636	LOC105286636
	Gene	LOC105286636	protein_coding	NA	character(0)	character(0)
NW_011924881.1	447836	447836	1	+	34	15	19	44.1176470588235	NW_011924881.1	447189	448306	1118	+	Gnomon	gene	NA	NA	gene25
	GeneID:105274524	LOC105274524	Gene	LOC105274524	protein_coding	NA	character(0)	character(0)
NW_011924881.1	447837	447837	1	-	21	10	11	47.6190476190476	NW_011924881.1	447189	448306	1118	+	Gnomon	gene	NA	NA	gene25
	GeneID:105274524	LOC105274524	Gene	LOC105274524	protein_coding	NA	character(0)	character(0)
NW_011924881.1	447945	447945	1	+	10	10	0	100	NW_011924881.1	447189	448306	1118	+	Gnomon	gene	NA	NA	gene25
	GeneID:105274524	LOC105274524	Gene	LOC105274524	protein_coding	NA	character(0)	character(0)
NW_011924881.1	447946	447946	1	-	9	7	2	77.7777777777778	NW_011924881.1	447189	448306	1118	+	Gnomon	gene	NA	NA	gene25
	GeneID:105274524	LOC105274524	Gene	LOC105274524	protein_coding	NA	character(0)	character(0)
NW_011924881.1	447952	447952	1	+	10	10	0	100	NW_011924881.1	447189	448306	1118	+	Gnomon	gene	NA	NA	gene25
	GeneID:105274524	LOC105274524	Gene	LOC105274524	protein_coding	NA	character(0)	character(0)
NW_011924881.1	447953	447953	1	-	8	6	2	75	NW_011924881.1	447189	448306	1118	+	Gnomon	gene	NA	NA	gene25
	GeneID:105274524	LOC105274524	Gene	LOC105274524	protein_coding	NA	character(0)	character(0)
NW_011924881.1	447958	447958	1	+	10	10	0	100	NW_011924881.1	447189	448306	1118	+	Gnomon	gene	NA	NA	gene25
	GeneID:105274524	LOC105274524	Gene	LOC105274524	protein_coding	NA	character(0)	character(0)
NW_011924881.1	447959	447959	1	-	7	7	0	100	NW_011924881.1	447189	448306	1118	+	Gnomon	gene	NA	NA	gene25
	GeneID:105274524	LOC105274524	Gene	LOC105274524	protein_coding	NA	character(0)	character(0)
...
... only first 25 lines shown ...
```

Directory: RNK File: sites-in-genes-Cb_LI_RphaseA2.txt

| seqnames | start | end | width | strand | coverage | numCs | numTs | perc_meth | region_seqnames | region_start | region_end | region_width | region_strand | region_source | region_type | region_score | region_phase | region_ID | region_Dbxref | region_Name | region_gbkey | region_gene | region_gene_biotype | region_partial | region_start_range | region_end_range |
|---|---|---|---|---|---|---|---|---|---|---|---|---|---|---|---|---|---|---|---|---|---|---|---|---|---|---|
| NW_011924879.1 | 2162 | 2162 | 1 | + | 38 | 11 | 27 | 28.9473684210526 | NW_011924879.1 | 7 | 3079 | 3073 | + | Gnomon | gene | NA | NA | gene5 | GeneID:105275124 | LOC105275124 | Gene | LOC105275124 | protein_coding | NA | character(0) | character(0) |
| NW_011924881.1 | 39967 | 39967 | 1 | + | 17 | 10 | 7 | 58.8235294117647 | NW_011924881.1 | 39621 | 40962 | 1342 | - | Gnomon | gene | NA | NA | gene17 | GeneID:105286636 | LOC105286636 | Gene | LOC105286636 | protein_coding | NA | character(0) | character(0) |
| NW_011924881.1 | 39987 | 39987 | 1 | + | 21 | 14 | 7 | 66.6666666666667 | NW_011924881.1 | 39621 | 40962 | 1342 | - | Gnomon | gene | NA | NA | gene17 | GeneID:105286636 | LOC105286636 | Gene | LOC105286636 | protein_coding | NA | character(0) | character(0) |
| NW_011924881.1 | 40061 | 40061 | 1 | + | 15 | 14 | 1 | 93.3333333333333 | NW_011924881.1 | 39621 | 40962 | 1342 | - | Gnomon | gene | NA | NA | gene17 | GeneID:105286636 | LOC105286636 | Gene | LOC105286636 | protein_coding | NA | character(0) | character(0) |
| NW_011924881.1 | 40072 | 40072 | 1 | + | 14 | 14 | 0 | 100 | NW_011924881.1 | 39621 | 40962 | 1342 | - | Gnomon | gene | NA | NA | gene17 | GeneID:105286636 | LOC105286636 | Gene | LOC105286636 | protein_coding | NA | character(0) | character(0) |
| NW_011924881.1 | 40095 | 40095 | 1 | + | 22 | 13 | 9 | 59.0909090909091 | NW_011924881.1 | 39621 | 40962 | 1342 | - | Gnomon | gene | NA | NA | gene17 | GeneID:105286636 | LOC105286636 | Gene | LOC105286636 | protein_coding | NA | character(0) | character(0) |
| NW_011924881.1 | 40120 | 40120 | 1 | + | 22 | 13 | 9 | 59.0909090909091 | NW_011924881.1 | 39621 | 40962 | 1342 | - | Gnomon | gene | NA | NA | gene17 | GeneID:105286636 | LOC105286636 | Gene | LOC105286636 | protein_coding | NA | character(0) | character(0) |
| NW_011924881.1 | 40121 | 40121 | 1 | - | 7 | 5 | 2 | 71.4285714285714 | NW_011924881.1 | 39621 | 40962 | 1342 | - | Gnomon | gene | NA | NA | gene17 | GeneID:105286636 | LOC105286636 | Gene | LOC105286636 | protein_coding | NA | character(0) | character(0) |
| NW_011924881.1 | 40160 | 40160 | 1 | - | 7 | 6 | 1 | 85.7142857142857 | NW_011924881.1 | 39621 | 40962 | 1342 | - | Gnomon | gene | NA | NA | gene17 | GeneID:105286636 | LOC105286636 | Gene | LOC105286636 | protein_coding | NA | character(0) | character(0) |
| NW_011924881.1 | 40256 | 40256 | 1 | + | 5 | 5 | 0 | 100 | NW_011924881.1 | 39621 | 40962 | 1342 | - | Gnomon | gene | NA | NA | gene17 | GeneID:105286636 | LOC105286636 | Gene | LOC105286636 | protein_coding | NA | character(0) | character(0) |
| NW_011924881.1 | 40257 | 40257 | 1 | - | 34 | 30 | 4 | 88.2352941176471 | NW_011924881.1 | 39621 | 40962 | 1342 | - | Gnomon | gene | NA | NA | gene17 | GeneID:105286636 | LOC105286636 | Gene | LOC105286636 | protein_coding | NA | character(0) | character(0) |
| NW_011924881.1 | 40260 | 40260 | 1 | - | 31 | 28 | 3 | 90.3225806451613 | NW_011924881.1 | 39621 | 40962 | 1342 | - | Gnomon | gene | NA | NA | gene17 | GeneID:105286636 | LOC105286636 | Gene | LOC105286636 | protein_coding | NA | character(0) | character(0) |
| NW_011924881.1 | 40270 | 40270 | 1 | + | 6 | 5 | 1 | 83.3333333333333 | NW_011924881.1 | 39621 | 40962 | 1342 | - | Gnomon | gene | NA | NA | gene17 | GeneID:105286636 | LOC105286636 | Gene | LOC105286636 | protein_coding | NA | character(0) | character(0) |
| NW_011924881.1 | 40271 | 40271 | 1 | - | 29 | 26 | 3 | 89.6551724137931 | NW_011924881.1 | 39621 | 40962 | 1342 | - | Gnomon | gene | NA | NA | gene17 | GeneID:105286636 | LOC105286636 | Gene | LOC105286636 | protein_coding | NA | character(0) | character(0) |
| NW_011924881.1 | 40305 | 40305 | 1 | - | 23 | 7 | 16 | 30.4347826086957 | NW_011924881.1 | 39621 | 40962 | 1342 | - | Gnomon | gene | NA | NA | gene17 | GeneID:105286636 | LOC105286636 | Gene | LOC105286636 | protein_coding | NA | character(0) | character(0) |
| NW_011924881.1 | 40350 | 40350 | 1 | - | 18 | 11 | 7 | 61.1111111111111 | NW_011924881.1 | 39621 | 40962 | 1342 | - | Gnomon | gene | NA | NA | gene17 | GeneID:105286636 | LOC105286636 | Gene | LOC105286636 | protein_coding | NA | character(0) | character(0) |
| NW_011924881.1 | 40360 | 40360 | 1 | - | 30 | 28 | 2 | 93.3333333333333 | NW_011924881.1 | 39621 | 40962 | 1342 | - | Gnomon | gene | NA | NA | gene17 | GeneID:105286636 | LOC105286636 | Gene | LOC105286636 | protein_coding | NA | character(0) | character(0) |
| NW_011924881.1 | 40372 | 40372 | 1 | - | 29 | 28 | 1 | 96.551724137931 | NW_011924881.1 | 39621 | 40962 | 1342 | - | Gnomon | gene | NA | NA | gene17 | GeneID:105286636 | LOC105286636 | Gene | LOC105286636 | protein_coding | NA | character(0) | character(0) |
| NW_011924881.1 | 40385 | 40385 | 1 | - | 23 | 23 | 0 | 100 | NW_011924881.1 | 39621 | 40962 | 1342 | - | Gnomon | gene | NA | NA | gene17 | GeneID:105286636 | LOC105286636 | Gene | LOC105286636 | protein_coding | NA | character(0) | character(0) |
| NW_011924881.1 | 40390 | 40390 | 1 | - | 23 | 23 | 0 | 100 | NW_011924881.1 | 39621 | 40962 | 1342 | - | Gnomon | gene | NA | NA | gene17 | GeneID:105286636 | LOC105286636 | Gene | LOC105286636 | protein_coding | NA | character(0) | character(0) |
| NW_011924881.1 | 40404 | 40404 | 1 | - | 18 | 11 | 7 | 61.1111111111111 | NW_011924881.1 | 39621 | 40962 | 1342 | - | Gnomon | gene | NA | NA | gene17 | GeneID:105286636 | LOC105286636 | Gene | LOC105286636 | protein_coding | NA | character(0) | character(0) |
| NW_011924881.1 | 40406 | 40406 | 1 | - | 17 | 13 | 4 | 76.4705882352941 | NW_011924881.1 | 39621 | 40962 | 1342 | - | Gnomon | gene | NA | NA | gene17 | GeneID:105286636 | LOC105286636 | Gene | LOC105286636 | protein_coding | NA | character(0) | character(0) |
| NW_011924881.1 | 40417 | 40417 | 1 | - | 20 | 8 | 12 | 40 | NW_011924881.1 | 39621 | 40962 | 1342 | - | Gnomon | gene | NA | NA | gene17 | GeneID:105286636 | LOC105286636 | Gene | LOC105286636 | protein_coding | NA | character(0) | character(0) |
| NW_011924881.1 | 152633 | 152633 | 1 | - | 42 | 13 | 29 | 30.952380952381 | NW_011924881.1 | 140800 | 154070 | 13271 | - | Gnomon | gene | NA | NA | gene19 | GeneID:105287639 | LOC105287639 | Gene | LOC105287639 | protein_coding | NA | character(0) | character(0) |

...
... only first 25 lines shown ...

Directory: RNK File: sites-in-genes-Cb_LI_RphaseB1.txt

```
seqnames	start	end	width	strand	coverage	numCs	numTs	perc_meth	region_seqnames	region_start	region_end	region_width	region_strand	region_source
	region_type	region_score	region_phase	region_ID	region_Dbxref	region_Name	region_gbkey	region_gene	region_gene_biotype	region_partial
	region_start_range	region_end_range
NW_011924881.1	39967	39967	1	+	9	6	3	66.6666666666667	NW_011924881.1	39621	40962	1342	-	Gnomon	gene	NA	NA	gene17	GeneID:105286636
	LOC105286636	Gene	LOC105286636	protein_coding	NA	character(0)	character(0)
NW_011924881.1	39968	39968	1	-	5	5	0	100	NW_011924881.1	39621	40962	1342	-	Gnomon	gene	NA	NA	gene17	GeneID:105286636	LOC105286636
	Gene	LOC105286636	protein_coding	NA	character(0)	character(0)
NW_011924881.1	39987	39987	1	+	15	7	8	46.6666666666667	NW_011924881.1	39621	40962	1342	-	Gnomon	gene	NA	NA	gene17	GeneID:105286636
	LOC105286636	Gene	LOC105286636	protein_coding	NA	character(0)	character(0)
NW_011924881.1	40061	40061	1	+	14	13	1	92.8571428571429	NW_011924881.1	39621	40962	1342	-	Gnomon	gene	NA	NA	gene17	GeneID:105286636
	LOC105286636	Gene	LOC105286636	protein_coding	NA	character(0)	character(0)
NW_011924881.1	40072	40072	1	+	9	9	0	100	NW_011924881.1	39621	40962	1342	-	Gnomon	gene	NA	NA	gene17	GeneID:105286636	LOC105286636
	Gene	LOC105286636	protein_coding	NA	character(0)	character(0)
NW_011924881.1	40095	40095	1	+	14	7	7	50	NW_011924881.1	39621	40962	1342	-	Gnomon	gene	NA	NA	gene17	GeneID:105286636	LOC105286636
	Gene	LOC105286636	protein_coding	NA	character(0)	character(0)
NW_011924881.1	40120	40120	1	+	17	11	6	64.7058823529412	NW_011924881.1	39621	40962	1342	-	Gnomon	gene	NA	NA	gene17	GeneID:105286636
	LOC105286636	Gene	LOC105286636	protein_coding	NA	character(0)	character(0)
NW_011924881.1	40121	40121	1	-	8	6	2	75	NW_011924881.1	39621	40962	1342	-	Gnomon	gene	NA	NA	gene17	GeneID:105286636	LOC105286636
	Gene	LOC105286636	protein_coding	NA	character(0)	character(0)
NW_011924881.1	40159	40159	1	+	14	9	5	64.2857142857143	NW_011924881.1	39621	40962	1342	-	Gnomon	gene	NA	NA	gene17	GeneID:105286636
	LOC105286636	Gene	LOC105286636	protein_coding	NA	character(0)	character(0)
NW_011924881.1	40160	40160	1	-	7	5	2	71.4285714285714	NW_011924881.1	39621	40962	1342	-	Gnomon	gene	NA	NA	gene17	GeneID:105286636
	LOC105286636	Gene	LOC105286636	protein_coding	NA	character(0)	character(0)
NW_011924881.1	40257	40257	1	-	23	18	5	78.2608695652174	NW_011924881.1	39621	40962	1342	-	Gnomon	gene	NA	NA	gene17	GeneID:105286636
	LOC105286636	Gene	LOC105286636	protein_coding	NA	character(0)	character(0)
NW_011924881.1	40260	40260	1	-	21	19	2	90.4761904761905	NW_011924881.1	39621	40962	1342	-	Gnomon	gene	NA	NA	gene17	GeneID:105286636
	LOC105286636	Gene	LOC105286636	protein_coding	NA	character(0)	character(0)
NW_011924881.1	40271	40271	1	-	21	19	2	90.4761904761905	NW_011924881.1	39621	40962	1342	-	Gnomon	gene	NA	NA	gene17	GeneID:105286636
	LOC105286636	Gene	LOC105286636	protein_coding	NA	character(0)	character(0)
NW_011924881.1	40335	40335	1	-	16	10	6	62.5	NW_011924881.1	39621	40962	1342	-	Gnomon	gene	NA	NA	gene17	GeneID:105286636	LOC105286636
	Gene	LOC105286636	protein_coding	NA	character(0)	character(0)
NW_011924881.1	40350	40350	1	-	15	12	3	80	NW_011924881.1	39621	40962	1342	-	Gnomon	gene	NA	NA	gene17	GeneID:105286636	LOC105286636
	Gene	LOC105286636	protein_coding	NA	character(0)	character(0)
NW_011924881.1	40360	40360	1	-	16	14	2	87.5	NW_011924881.1	39621	40962	1342	-	Gnomon	gene	NA	NA	gene17	GeneID:105286636	LOC105286636
	Gene	LOC105286636	protein_coding	NA	character(0)	character(0)
NW_011924881.1	40372	40372	1	-	15	14	1	93.3333333333333	NW_011924881.1	39621	40962	1342	-	Gnomon	gene	NA	NA	gene17	GeneID:105286636
	LOC105286636	Gene	LOC105286636	protein_coding	NA	character(0)	character(0)
NW_011924881.1	40385	40385	1	-	11	11	0	100	NW_011924881.1	39621	40962	1342	-	Gnomon	gene	NA	NA	gene17	GeneID:105286636	LOC105286636
	Gene	LOC105286636	protein_coding	NA	character(0)	character(0)
NW_011924881.1	40390	40390	1	-	11	10	1	90.9090909090909	NW_011924881.1	39621	40962	1342	-	Gnomon	gene	NA	NA	gene17	GeneID:105286636
	LOC105286636	Gene	LOC105286636	protein_coding	NA	character(0)	character(0)
NW_011924881.1	40404	40404	1	-	10	6	4	60	NW_011924881.1	39621	40962	1342	-	Gnomon	gene	NA	NA	gene17	GeneID:105286636	LOC105286636
	Gene	LOC105286636	protein_coding	NA	character(0)	character(0)
NW_011924881.1	40406	40406	1	-	9	5	4	55.5555555555556	NW_011924881.1	39621	40962	1342	-	Gnomon	gene	NA	NA	gene17	GeneID:105286636
	LOC105286636	Gene	LOC105286636	protein_coding	NA	character(0)	character(0)
NW_011924881.1	53486	53486	1	+	11	6	5	54.5454545454545	NW_011924881.1	42715	135016	92302	-	Gnomon	gene	NA	NA	gene18
	GeneID:105287290	LOC105287290	Gene	LOC105287290	protein_coding	NA	character(0)	character(0)
NW_011924881.1	106949	106949	1	+	14	7	7	50	NW_011924881.1	42715	135016	92302	-	Gnomon	gene	NA	NA	gene18
	GeneID:105287290	LOC105287290	Gene	LOC105287290	protein_coding	NA	character(0)	character(0)
NW_011924881.1	247875	247875	1	+	25	8	17	32	NW_011924881.1	195683	286417	90735	-	Gnomon	gene	NA	NA	gene22
	GeneID:105288120	LOC105288120	Gene	LOC105288120	protein_coding	NA	character(0)	character(0)
...
... only first 25 lines shown ...
```

Directory: RNK File: sites-in-genes-Cb_LI_RphaseB2.txt

seqnames start end width strand coverage numCs numTs perc_meth region_seqnames region_start region_end region_width region_strand region_source
region_type region_score region_phase region_ID region_Dbxref region_Name region_gbkey region_gene region_gene_biotype region_partial
region_start_range region_end_range
NW_011924879.1 2162 2162 1 + 34 9 25 26.4705882352941 NW_011924879.1 7 3079 3073 + Gnomon gene NA NA gene5 GeneID:105275124
LOC105275124 Gene LOC105275124 protein_coding NA character(0) character(0)
NW_011924881.1 39967 39967 1 + 6 5 1 83.3333333333333 NW_011924881.1 39621 40962 1342 - Gnomon gene NA NA gene17 GeneID:105286636
LOC105286636 Gene LOC105286636 protein_coding NA character(0) character(0)
NW_011924881.1 40061 40061 1 + 11 10 1 90.9090909090909 NW_011924881.1 39621 40962 1342 - Gnomon gene NA NA gene17 GeneID:105286636
LOC105286636 Gene LOC105286636 protein_coding NA character(0) character(0)
NW_011924881.1 40072 40072 1 + 13 12 1 92.3076923076923 NW_011924881.1 39621 40962 1342 - Gnomon gene NA NA gene17 GeneID:105286636
LOC105286636 Gene LOC105286636 protein_coding NA character(0) character(0)
NW_011924881.1 40095 40095 1 + 15 7 8 46.6666666666667 NW_011924881.1 39621 40962 1342 - Gnomon gene NA NA gene17 GeneID:105286636
LOC105286636 Gene LOC105286636 protein_coding NA character(0) character(0)
NW_011924881.1 40120 40120 1 + 14 7 7 50 NW_011924881.1 39621 40962 1342 - Gnomon gene NA NA gene17 GeneID:105286636 LOC105286636
Gene LOC105286636 protein_coding NA character(0) character(0)
NW_011924881.1 40257 40257 1 - 13 13 0 100 NW_011924881.1 39621 40962 1342 - Gnomon gene NA NA gene17 GeneID:105286636 LOC105286636
Gene LOC105286636 protein_coding NA character(0) character(0)
NW_011924881.1 40260 40260 1 - 11 11 0 100 NW_011924881.1 39621 40962 1342 - Gnomon gene NA NA gene17 GeneID:105286636 LOC105286636
Gene LOC105286636 protein_coding NA character(0) character(0)
NW_011924881.1 40271 40271 1 - 11 11 0 100 NW_011924881.1 39621 40962 1342 - Gnomon gene NA NA gene17 GeneID:105286636 LOC105286636
Gene LOC105286636 protein_coding NA character(0) character(0)
NW_011924881.1 40350 40350 1 - 9 7 2 77.7777777777778 NW_011924881.1 39621 40962 1342 - Gnomon gene NA NA gene17 GeneID:105286636
LOC105286636 Gene LOC105286636 protein_coding NA character(0) character(0)
NW_011924881.1 40360 40360 1 - 17 16 1 94.1176470588235 NW_011924881.1 39621 40962 1342 - Gnomon gene NA NA gene17 GeneID:105286636
LOC105286636 Gene LOC105286636 protein_coding NA character(0) character(0)
NW_011924881.1 40372 40372 1 - 16 16 0 100 NW_011924881.1 39621 40962 1342 - Gnomon gene NA NA gene17 GeneID:105286636 LOC105286636
Gene LOC105286636 protein_coding NA character(0) character(0)
NW_011924881.1 40385 40385 1 - 15 15 0 100 NW_011924881.1 39621 40962 1342 - Gnomon gene NA NA gene17 GeneID:105286636 LOC105286636
Gene LOC105286636 protein_coding NA character(0) character(0)
NW_011924881.1 40390 40390 1 - 15 13 2 86.6666666666667 NW_011924881.1 39621 40962 1342 - Gnomon gene NA NA gene17 GeneID:105286636
LOC105286636 Gene LOC105286636 protein_coding NA character(0) character(0)
NW_011924881.1 40406 40406 1 - 16 6 10 37.5 NW_011924881.1 39621 40962 1342 - Gnomon gene NA NA gene17 GeneID:105286636 LOC105286636
Gene LOC105286636 protein_coding NA character(0) character(0)
NW_011924881.1 379413 379413 1 + 16 6 10 37.5 NW_011924881.1 369231 418861 49631 - Gnomon gene NA NA gene24
GeneID:109611552 LOC109611552 Gene LOC109611552 protein_coding NA character(0) character(0)
NW_011924881.1 447836 447836 1 + 36 17 19 47.2222222222222 NW_011924881.1 447189 448306 1118 + Gnomon gene NA NA gene25
GeneID:105274524 LOC105274524 Gene LOC105274524 protein_coding NA character(0) character(0)
NW_011924881.1 447837 447837 1 - 54 21 33 38.8888888888889 NW_011924881.1 447189 448306 1118 + Gnomon gene NA NA gene25
GeneID:105274524 LOC105274524 Gene LOC105274524 protein_coding NA character(0) character(0)
NW_011924881.1 447945 447945 1 + 15 14 1 93.3333333333333 NW_011924881.1 447189 448306 1118 + Gnomon gene NA NA gene25
GeneID:105274524 LOC105274524 Gene LOC105274524 protein_coding NA character(0) character(0)
NW_011924881.1 447946 447946 1 - 14 13 1 92.8571428571429 NW_011924881.1 447189 448306 1118 + Gnomon gene NA NA gene25
GeneID:105274524 LOC105274524 Gene LOC105274524 protein_coding NA character(0) character(0)
NW_011924881.1 447952 447952 1 + 12 9 3 75 NW_011924881.1 447189 448306 1118 + Gnomon gene NA NA gene25
GeneID:105274524 LOC105274524 Gene LOC105274524 protein_coding NA character(0) character(0)
NW_011924881.1 447953 447953 1 - 10 7 3 70 NW_011924881.1 447189 448306 1118 + Gnomon gene NA NA gene25
GeneID:105274524 LOC105274524 Gene LOC105274524 protein_coding NA character(0) character(0)
NW_011924881.1 447958 447958 1 + 14 13 1 92.8571428571429 NW_011924881.1 447189 448306 1118 + Gnomon gene NA NA gene25
GeneID:105274524 LOC105274524 Gene LOC105274524 protein_coding NA character(0) character(0)
NW_011924881.1 447959 447959 1 - 10 8 2 80 NW_011924881.1 447189 448306 1118 + Gnomon gene NA NA gene25
GeneID:105274524 LOC105274524 Gene LOC105274524 protein_coding NA character(0) character(0)
...
... only first 25 lines shown ...

Directory: RNK File: sites-in-promoters-Cb_LI_BCphaseA1.txt

| seqnames | start | end | width | strand | coverage | numCs | numTs | perc_meth | region_seqnames | region_start | region_end | region_width | region_strand | region_source | region_type | region_score | region_phase | region_ID | region_Dbxref | region_Name | region_gbkey | region_gene | region_gene_biotype | region_partial | region_start_range | region_end_range |
|---|---|---|---|---|---|---|---|---|---|---|---|---|---|---|---|---|---|---|---|---|---|---|---|---|---|---|
| NW_011924881.1 | 745918 | 745918 | 1 | + | 45 | 34 | 11 | 75.5555555555556 | NW_011924881.1 | 745918 | 746417 | 500 | + | Gnomon | promoter | NA | NA | gene42 | GeneID:105275579 | LOC105275579 | Gene | LOC105275579 | protein_coding | NA | character(0) | character(0) |
| NW_011924887.1 | 221098 | 221098 | 1 | - | 25 | 15 | 10 | 60 | NW_011924887.1 | 220963 | 221462 | 500 | + | Gnomon | promoter | NA | NA | gene126 | GeneID:105286831 | LOC105286831 | Gene | LOC105286831 | protein_coding | NA | character(0) | character(0) |
| NW_011924887.1 | 221214 | 221214 | 1 | - | 14 | 6 | 8 | 42.8571428571429 | NW_011924887.1 | 220963 | 221462 | 500 | + | Gnomon | promoter | NA | NA | gene126 | GeneID:105286831 | LOC105286831 | Gene | LOC105286831 | protein_coding | NA | character(0) | character(0) |
| NW_011924887.1 | 225897 | 225897 | 1 | - | 16 | 6 | 10 | 37.5 | NW_011924887.1 | 225822 | 226321 | 500 | + | Gnomon | promoter | NA | NA | gene128 | GeneID:105286681 | LOC105286681 | Gene | LOC105286681 | lncRNA | NA | character(0) | character(0) |
| NW_011924887.1 | 225945 | 225945 | 1 | + | 19 | 14 | 5 | 73.6842105263158 | NW_011924887.1 | 225822 | 226321 | 500 | + | Gnomon | promoter | NA | NA | gene128 | GeneID:105286681 | LOC105286681 | Gene | LOC105286681 | lncRNA | NA | character(0) | character(0) |
| NW_011924887.1 | 225946 | 225946 | 1 | - | 14 | 13 | 1 | 92.8571428571429 | NW_011924887.1 | 225822 | 226321 | 500 | + | Gnomon | promoter | NA | NA | gene128 | GeneID:105286681 | LOC105286681 | Gene | LOC105286681 | lncRNA | NA | character(0) | character(0) |
| NW_011924887.1 | 225950 | 225950 | 1 | + | 21 | 17 | 4 | 80.9523809523809 | NW_011924887.1 | 225822 | 226321 | 500 | + | Gnomon | promoter | NA | NA | gene128 | GeneID:105286681 | LOC105286681 | Gene | LOC105286681 | lncRNA | NA | character(0) | character(0) |
| NW_011924887.1 | 225951 | 225951 | 1 | - | 12 | 9 | 3 | 75 | NW_011924887.1 | 225822 | 226321 | 500 | + | Gnomon | promoter | NA | NA | gene128 | GeneID:105286681 | LOC105286681 | Gene | LOC105286681 | lncRNA | NA | character(0) | character(0) |
| NW_011924887.1 | 225973 | 225973 | 1 | + | 26 | 18 | 8 | 69.2307692307692 | NW_011924887.1 | 225822 | 226321 | 500 | + | Gnomon | promoter | NA | NA | gene128 | GeneID:105286681 | LOC105286681 | Gene | LOC105286681 | lncRNA | NA | character(0) | character(0) |
| NW_011924887.1 | 225974 | 225974 | 1 | - | 9 | 8 | 1 | 88.8888888888889 | NW_011924887.1 | 225822 | 226321 | 500 | + | Gnomon | promoter | NA | NA | gene128 | GeneID:105286681 | LOC105286681 | Gene | LOC105286681 | lncRNA | NA | character(0) | character(0) |
| NW_011924887.1 | 225981 | 225981 | 1 | + | 28 | 20 | 8 | 71.4285714285714 | NW_011924887.1 | 225822 | 226321 | 500 | + | Gnomon | promoter | NA | NA | gene128 | GeneID:105286681 | LOC105286681 | Gene | LOC105286681 | lncRNA | NA | character(0) | character(0) |
| NW_011924887.1 | 226007 | 226007 | 1 | + | 29 | 12 | 17 | 41.3793103448276 | NW_011924887.1 | 225822 | 226321 | 500 | + | Gnomon | promoter | NA | NA | gene128 | GeneID:105286681 | LOC105286681 | Gene | LOC105286681 | lncRNA | NA | character(0) | character(0) |
| NW_011924887.1 | 226008 | 226008 | 1 | - | 24 | 7 | 17 | 29.1666666666667 | NW_011924887.1 | 225822 | 226321 | 500 | + | Gnomon | promoter | NA | NA | gene128 | GeneID:105286681 | LOC105286681 | Gene | LOC105286681 | lncRNA | NA | character(0) | character(0) |
| NW_011924887.1 | 226055 | 226055 | 1 | + | 23 | 10 | 13 | 43.4782608695652 | NW_011924887.1 | 225822 | 226321 | 500 | + | Gnomon | promoter | NA | NA | gene128 | GeneID:105286681 | LOC105286681 | Gene | LOC105286681 | lncRNA | NA | character(0) | character(0) |
| NW_011924887.1 | 226056 | 226056 | 1 | - | 51 | 18 | 33 | 35.2941176470588 | NW_011924887.1 | 225822 | 226321 | 500 | + | Gnomon | promoter | NA | NA | gene128 | GeneID:105286681 | LOC105286681 | Gene | LOC105286681 | lncRNA | NA | character(0) | character(0) |
| NW_011924887.1 | 226086 | 226086 | 1 | + | 15 | 7 | 8 | 46.6666666666667 | NW_011924887.1 | 225822 | 226321 | 500 | + | Gnomon | promoter | NA | NA | gene128 | GeneID:105286681 | LOC105286681 | Gene | LOC105286681 | lncRNA | NA | character(0) | character(0) |
| NW_011924887.1 | 226087 | 226087 | 1 | - | 73 | 30 | 43 | 41.0958904109589 | NW_011924887.1 | 225822 | 226321 | 500 | + | Gnomon | promoter | NA | NA | gene128 | GeneID:105286681 | LOC105286681 | Gene | LOC105286681 | lncRNA | NA | character(0) | character(0) |
| NW_011924887.1 | 226089 | 226089 | 1 | - | 71 | 16 | 55 | 22.5352112676056 | NW_011924887.1 | 225822 | 226321 | 500 | + | Gnomon | promoter | NA | NA | gene128 | GeneID:105286681 | LOC105286681 | Gene | LOC105286681 | lncRNA | NA | character(0) | character(0) |
| NW_011924887.1 | 226099 | 226099 | 1 | - | 65 | 11 | 54 | 16.9230769230769 | NW_011924887.1 | 225822 | 226321 | 500 | + | Gnomon | promoter | NA | NA | gene128 | GeneID:105286681 | LOC105286681 | Gene | LOC105286681 | lncRNA | NA | character(0) | character(0) |
| NW_011924887.1 | 226100 | 226100 | 1 | + | 8 | 5 | 3 | 62.5 | NW_011924887.1 | 225822 | 226321 | 500 | + | Gnomon | promoter | NA | NA | gene128 | GeneID:105286681 | LOC105286681 | Gene | LOC105286681 | lncRNA | NA | character(0) | character(0) |
| NW_011924887.1 | 226101 | 226101 | 1 | - | 58 | 40 | 18 | 68.9655172413793 | NW_011924887.1 | 225822 | 226321 | 500 | + | Gnomon | promoter | NA | NA | gene128 | GeneID:105286681 | LOC105286681 | Gene | LOC105286681 | lncRNA | NA | character(0) | character(0) |
| NW_011924887.1 | 226117 | 226117 | 1 | - | 49 | 20 | 29 | 40.8163265306122 | NW_011924887.1 | 225822 | 226321 | 500 | + | Gnomon | promoter | NA | NA | gene128 | GeneID:105286681 | LOC105286681 | Gene | LOC105286681 | lncRNA | NA | character(0) | character(0) |
| NW_011924887.1 | 226128 | 226128 | 1 | - | 45 | 32 | 13 | 71.1111111111111 | NW_011924887.1 | 225822 | 226321 | 500 | + | Gnomon | promoter | NA | NA | gene128 | GeneID:105286681 | LOC105286681 | Gene | LOC105286681 | lncRNA | NA | character(0) | character(0) |
| NW_011924887.1 | 226149 | 226149 | 1 | - | 41 | 24 | 17 | 58.5365853658537 | NW_011924887.1 | 225822 | 226321 | 500 | + | Gnomon | promoter | NA | NA | gene128 | GeneID:105286681 | LOC105286681 | Gene | LOC105286681 | lncRNA | NA | character(0) | character(0) |

...
... only first 25 lines shown ...

Directory: RNK File: sites-in-promoters-Cb_LI_BCphaseA2.txt

| seqnames | start | end | width | strand | coverage | numCs | numTs | perc_meth | region_seqnames | region_start | region_end | region_width | region_strand | region_source | region_type | region_score | region_phase | region_ID | region_Dbxref | region_Name | region_gbkey | region_gene | region_gene_biotype | region_partial | region_start_range | region_end_range |
|---|---|---|---|---|---|---|---|---|---|---|---|---|---|---|---|---|---|---|---|---|---|---|---|---|---|---|
| NW_011924881.1 | 745918 | 745918 | 1 | + | 25 | 22 | 3 | 88 | NW_011924881.1 | 745918 | 746417 | 500 | + | Gnomon | promoter | NA | NA | gene42 | GeneID:105275579 | LOC105275579 | Gene | LOC105275579 | protein_coding | NA | character(0) | character(0) |
| NW_011924881.1 | 989262 | 989262 | 1 | - | 6 | 6 | 0 | 100 | NW_011924881.1 | 989253 | 989752 | 500 | + | Gnomon | promoter | NA | NA | gene61 | GeneID:105277580 | LOC105277580 | Gene | LOC105277580 | protein_coding | NA | character(0) | character(0) |
| NW_011924887.1 | 221082 | 221082 | 1 | - | 31 | 11 | 20 | 35.4838709677419 | NW_011924887.1 | 220963 | 221462 | 500 | + | Gnomon | promoter | NA | NA | gene126 | GeneID:105286831 | LOC105286831 | Gene | LOC105286831 | protein_coding | NA | character(0) | character(0) |
| NW_011924887.1 | 221097 | 221097 | 1 | + | 5 | 5 | 0 | 100 | NW_011924887.1 | 220963 | 221462 | 500 | + | Gnomon | promoter | NA | NA | gene126 | GeneID:105286831 | LOC105286831 | Gene | LOC105286831 | protein_coding | NA | character(0) | character(0) |
| NW_011924887.1 | 221098 | 221098 | 1 | - | 28 | 18 | 10 | 64.2857142857143 | NW_011924887.1 | 220963 | 221462 | 500 | + | Gnomon | promoter | NA | NA | gene126 | GeneID:105286831 | LOC105286831 | Gene | LOC105286831 | protein_coding | NA | character(0) | character(0) |
| NW_011924887.1 | 221105 | 221105 | 1 | - | 29 | 8 | 21 | 27.5862068965517 | NW_011924887.1 | 220963 | 221462 | 500 | + | Gnomon | promoter | NA | NA | gene126 | GeneID:105286831 | LOC105286831 | Gene | LOC105286831 | protein_coding | NA | character(0) | character(0) |
| NW_011924887.1 | 221253 | 221253 | 1 | - | 14 | 7 | 7 | 50 | NW_011924887.1 | 220963 | 221462 | 500 | + | Gnomon | promoter | NA | NA | gene126 | GeneID:105286831 | LOC105286831 | Gene | LOC105286831 | protein_coding | NA | character(0) | character(0) |
| NW_011924887.1 | 225945 | 225945 | 1 | + | 20 | 15 | 5 | 75 | NW_011924887.1 | 225822 | 226321 | 500 | + | Gnomon | promoter | NA | NA | gene128 | GeneID:105286681 | LOC105286681 | Gene | LOC105286681 | lncRNA | NA | character(0) | character(0) |
| NW_011924887.1 | 225946 | 225946 | 1 | - | 14 | 11 | 3 | 78.5714285714286 | NW_011924887.1 | 225822 | 226321 | 500 | + | Gnomon | promoter | NA | NA | gene128 | GeneID:105286681 | LOC105286681 | Gene | LOC105286681 | lncRNA | NA | character(0) | character(0) |
| NW_011924887.1 | 225950 | 225950 | 1 | + | 20 | 12 | 8 | 60 | NW_011924887.1 | 225822 | 226321 | 500 | + | Gnomon | promoter | NA | NA | gene128 | GeneID:105286681 | LOC105286681 | Gene | LOC105286681 | lncRNA | NA | character(0) | character(0) |
| NW_011924887.1 | 225951 | 225951 | 1 | - | 14 | 6 | 8 | 42.8571428571429 | NW_011924887.1 | 225822 | 226321 | 500 | + | Gnomon | promoter | NA | NA | gene128 | GeneID:105286681 | LOC105286681 | Gene | LOC105286681 | lncRNA | NA | character(0) | character(0) |
| NW_011924887.1 | 225973 | 225973 | 1 | + | 30 | 25 | 5 | 83.3333333333333 | NW_011924887.1 | 225822 | 226321 | 500 | + | Gnomon | promoter | NA | NA | gene128 | GeneID:105286681 | LOC105286681 | Gene | LOC105286681 | lncRNA | NA | character(0) | character(0) |
| NW_011924887.1 | 225974 | 225974 | 1 | - | 12 | 10 | 2 | 83.3333333333333 | NW_011924887.1 | 225822 | 226321 | 500 | + | Gnomon | promoter | NA | NA | gene128 | GeneID:105286681 | LOC105286681 | Gene | LOC105286681 | lncRNA | NA | character(0) | character(0) |
| NW_011924887.1 | 225981 | 225981 | 1 | + | 31 | 23 | 8 | 74.1935483870968 | NW_011924887.1 | 225822 | 226321 | 500 | + | Gnomon | promoter | NA | NA | gene128 | GeneID:105286681 | LOC105286681 | Gene | LOC105286681 | lncRNA | NA | character(0) | character(0) |
| NW_011924887.1 | 225982 | 225982 | 1 | - | 13 | 6 | 7 | 46.1538461538462 | NW_011924887.1 | 225822 | 226321 | 500 | + | Gnomon | promoter | NA | NA | gene128 | GeneID:105286681 | LOC105286681 | Gene | LOC105286681 | lncRNA | NA | character(0) | character(0) |
| NW_011924887.1 | 226007 | 226007 | 1 | + | 43 | 21 | 22 | 48.8372093023256 | NW_011924887.1 | 225822 | 226321 | 500 | + | Gnomon | promoter | NA | NA | gene128 | GeneID:105286681 | LOC105286681 | Gene | LOC105286681 | lncRNA | NA | character(0) | character(0) |
| NW_011924887.1 | 226055 | 226055 | 1 | + | 27 | 9 | 18 | 33.3333333333333 | NW_011924887.1 | 225822 | 226321 | 500 | + | Gnomon | promoter | NA | NA | gene128 | GeneID:105286681 | LOC105286681 | Gene | LOC105286681 | lncRNA | NA | character(0) | character(0) |
| NW_011924887.1 | 226056 | 226056 | 1 | - | 46 | 20 | 26 | 43.4782608695652 | NW_011924887.1 | 225822 | 226321 | 500 | + | Gnomon | promoter | NA | NA | gene128 | GeneID:105286681 | LOC105286681 | Gene | LOC105286681 | lncRNA | NA | character(0) | character(0) |
| NW_011924887.1 | 226087 | 226087 | 1 | - | 77 | 22 | 55 | 28.5714285714286 | NW_011924887.1 | 225822 | 226321 | 500 | + | Gnomon | promoter | NA | NA | gene128 | GeneID:105286681 | LOC105286681 | Gene | LOC105286681 | lncRNA | NA | character(0) | character(0) |
| NW_011924887.1 | 226088 | 226088 | 1 | + | 13 | 6 | 7 | 46.1538461538462 | NW_011924887.1 | 225822 | 226321 | 500 | + | Gnomon | promoter | NA | NA | gene128 | GeneID:105286681 | LOC105286681 | Gene | LOC105286681 | lncRNA | NA | character(0) | character(0) |
| NW_011924887.1 | 226089 | 226089 | 1 | - | 76 | 14 | 62 | 18.4210526315789 | NW_011924887.1 | 225822 | 226321 | 500 | + | Gnomon | promoter | NA | NA | gene128 | GeneID:105286681 | LOC105286681 | Gene | LOC105286681 | lncRNA | NA | character(0) | character(0) |
| NW_011924887.1 | 226100 | 226100 | 1 | + | 13 | 6 | 7 | 46.1538461538462 | NW_011924887.1 | 225822 | 226321 | 500 | + | Gnomon | promoter | NA | NA | gene128 | GeneID:105286681 | LOC105286681 | Gene | LOC105286681 | lncRNA | NA | character(0) | character(0) |
| NW_011924887.1 | 226101 | 226101 | 1 | - | 63 | 41 | 22 | 65.0793650793651 | NW_011924887.1 | 225822 | 226321 | 500 | + | Gnomon | promoter | NA | NA | gene128 | GeneID:105286681 | LOC105286681 | Gene | LOC105286681 | lncRNA | NA | character(0) | character(0) |
| NW_011924887.1 | 226117 | 226117 | 1 | - | 54 | 26 | 28 | 48.1481481481481 | NW_011924887.1 | 225822 | 226321 | 500 | + | Gnomon | promoter | NA | NA | gene128 | GeneID:105286681 | LOC105286681 | Gene | LOC105286681 | lncRNA | NA | character(0) | character(0) |

...
... only first 25 lines shown ...

Directory: RNK File: sites-in-promoters-Cb_LI_BCphaseB1.txt

| seqnames | start | end | width | strand | coverage | numCs | numTs | perc_meth | region_seqnames | region_start | region_end | region_width | region_strand | region_source | region_type | region_score | region_phase | region_ID | region_Dbxref | region_Name | region_gbkey | region_gene | region_gene_biotype | region_partial | region_start_range | region_end_range |
|---|---|---|---|---|---|---|---|---|---|---|---|---|---|---|---|---|---|---|---|---|---|---|---|---|---|---|
| NW_011924881.1 | 745918 | 745918 | 1 | + | 47 | 35 | 12 | 74.468085106383 | NW_011924881.1 | 745918 | 746417 | 500 | + | Gnomon | promoter | NA | NA | gene42 | GeneID:105275579 | LOC105275579 | Gene | LOC105275579 | protein_coding | NA | character(0) | character(0) |
| NW_011924881.1 | 989262 | 989262 | 1 | - | 6 | 5 | 1 | 83.3333333333333 | NW_011924881.1 | 989253 | 989752 | 500 | + | Gnomon | promoter | NA | NA | gene61 | GeneID:105277580 | LOC105277580 | Gene | LOC105277580 | protein_coding | NA | character(0) | character(0) |
| NW_011924887.1 | 221082 | 221082 | 1 | - | 16 | 6 | 10 | 37.5 | NW_011924887.1 | 220963 | 221462 | 500 | + | Gnomon | promoter | NA | NA | gene126 | GeneID:105286831 | LOC105286831 | Gene | LOC105286831 | protein_coding | NA | character(0) | character(0) |
| NW_011924887.1 | 221097 | 221097 | 1 | + | 9 | 7 | 2 | 77.7777777777778 | NW_011924887.1 | 220963 | 221462 | 500 | + | Gnomon | promoter | NA | NA | gene126 | GeneID:105286831 | LOC105286831 | Gene | LOC105286831 | protein_coding | NA | character(0) | character(0) |
| NW_011924887.1 | 221098 | 221098 | 1 | - | 34 | 18 | 16 | 52.9411764705882 | NW_011924887.1 | 220963 | 221462 | 500 | + | Gnomon | promoter | NA | NA | gene126 | GeneID:105286831 | LOC105286831 | Gene | LOC105286831 | protein_coding | NA | character(0) | character(0) |
| NW_011924887.1 | 221105 | 221105 | 1 | - | 34 | 7 | 27 | 20.5882352941176 | NW_011924887.1 | 220963 | 221462 | 500 | + | Gnomon | promoter | NA | NA | gene126 | GeneID:105286831 | LOC105286831 | Gene | LOC105286831 | protein_coding | NA | character(0) | character(0) |
| NW_011924887.1 | 221292 | 221292 | 1 | - | 23 | 7 | 16 | 30.4347826086957 | NW_011924887.1 | 220963 | 221462 | 500 | + | Gnomon | promoter | NA | NA | gene126 | GeneID:105286831 | LOC105286831 | Gene | LOC105286831 | protein_coding | NA | character(0) | character(0) |
| NW_011924887.1 | 225896 | 225896 | 1 | + | 25 | 11 | 14 | 44 | NW_011924887.1 | 225822 | 226321 | 500 | + | Gnomon | promoter | NA | NA | gene128 | GeneID:105286681 | LOC105286681 | Gene | LOC105286681 | lncRNA | NA | character(0) | character(0) |
| NW_011924887.1 | 225945 | 225945 | 1 | + | 29 | 23 | 6 | 79.3103448275862 | NW_011924887.1 | 225822 | 226321 | 500 | + | Gnomon | promoter | NA | NA | gene128 | GeneID:105286681 | LOC105286681 | Gene | LOC105286681 | lncRNA | NA | character(0) | character(0) |
| NW_011924887.1 | 225946 | 225946 | 1 | - | 19 | 15 | 4 | 78.9473684210526 | NW_011924887.1 | 225822 | 226321 | 500 | + | Gnomon | promoter | NA | NA | gene128 | GeneID:105286681 | LOC105286681 | Gene | LOC105286681 | lncRNA | NA | character(0) | character(0) |
| NW_011924887.1 | 225950 | 225950 | 1 | + | 32 | 17 | 15 | 53.125 | NW_011924887.1 | 225822 | 226321 | 500 | + | Gnomon | promoter | NA | NA | gene128 | GeneID:105286681 | LOC105286681 | Gene | LOC105286681 | lncRNA | NA | character(0) | character(0) |
| NW_011924887.1 | 225951 | 225951 | 1 | - | 14 | 9 | 5 | 64.2857142857143 | NW_011924887.1 | 225822 | 226321 | 500 | + | Gnomon | promoter | NA | NA | gene128 | GeneID:105286681 | LOC105286681 | Gene | LOC105286681 | lncRNA | NA | character(0) | character(0) |
| NW_011924887.1 | 225973 | 225973 | 1 | + | 36 | 31 | 5 | 86.1111111111111 | NW_011924887.1 | 225822 | 226321 | 500 | + | Gnomon | promoter | NA | NA | gene128 | GeneID:105286681 | LOC105286681 | Gene | LOC105286681 | lncRNA | NA | character(0) | character(0) |
| NW_011924887.1 | 225974 | 225974 | 1 | - | 11 | 8 | 3 | 72.7272727272727 | NW_011924887.1 | 225822 | 226321 | 500 | + | Gnomon | promoter | NA | NA | gene128 | GeneID:105286681 | LOC105286681 | Gene | LOC105286681 | lncRNA | NA | character(0) | character(0) |
| NW_011924887.1 | 225981 | 225981 | 1 | + | 37 | 28 | 9 | 75.6756756756757 | NW_011924887.1 | 225822 | 226321 | 500 | + | Gnomon | promoter | NA | NA | gene128 | GeneID:105286681 | LOC105286681 | Gene | LOC105286681 | lncRNA | NA | character(0) | character(0) |
| NW_011924887.1 | 225982 | 225982 | 1 | - | 10 | 9 | 1 | 90 | NW_011924887.1 | 225822 | 226321 | 500 | + | Gnomon | promoter | NA | NA | gene128 | GeneID:105286681 | LOC105286681 | Gene | LOC105286681 | lncRNA | NA | character(0) | character(0) |
| NW_011924887.1 | 226007 | 226007 | 1 | + | 37 | 9 | 28 | 24.3243243243243 | NW_011924887.1 | 225822 | 226321 | 500 | + | Gnomon | promoter | NA | NA | gene128 | GeneID:105286681 | LOC105286681 | Gene | LOC105286681 | lncRNA | NA | character(0) | character(0) |
| NW_011924887.1 | 226008 | 226008 | 1 | - | 21 | 11 | 10 | 52.3809523809524 | NW_011924887.1 | 225822 | 226321 | 500 | + | Gnomon | promoter | NA | NA | gene128 | GeneID:105286681 | LOC105286681 | Gene | LOC105286681 | lncRNA | NA | character(0) | character(0) |
| NW_011924887.1 | 226055 | 226055 | 1 | + | 30 | 15 | 15 | 50 | NW_011924887.1 | 225822 | 226321 | 500 | + | Gnomon | promoter | NA | NA | gene128 | GeneID:105286681 | LOC105286681 | Gene | LOC105286681 | lncRNA | NA | character(0) | character(0) |
| NW_011924887.1 | 226056 | 226056 | 1 | - | 46 | 18 | 28 | 39.1304347826087 | NW_011924887.1 | 225822 | 226321 | 500 | + | Gnomon | promoter | NA | NA | gene128 | GeneID:105286681 | LOC105286681 | Gene | LOC105286681 | lncRNA | NA | character(0) | character(0) |
| NW_011924887.1 | 226086 | 226086 | 1 | + | 16 | 6 | 10 | 37.5 | NW_011924887.1 | 225822 | 226321 | 500 | + | Gnomon | promoter | NA | NA | gene128 | GeneID:105286681 | LOC105286681 | Gene | LOC105286681 | lncRNA | NA | character(0) | character(0) |
| NW_011924887.1 | 226087 | 226087 | 1 | - | 63 | 17 | 46 | 26.984126984127 | NW_011924887.1 | 225822 | 226321 | 500 | + | Gnomon | promoter | NA | NA | gene128 | GeneID:105286681 | LOC105286681 | Gene | LOC105286681 | lncRNA | NA | character(0) | character(0) |
| NW_011924887.1 | 226101 | 226101 | 1 | - | 49 | 31 | 18 | 63.265306122449 | NW_011924887.1 | 225822 | 226321 | 500 | + | Gnomon | promoter | NA | NA | gene128 | GeneID:105286681 | LOC105286681 | Gene | LOC105286681 | lncRNA | NA | character(0) | character(0) |
| NW_011924887.1 | 226117 | 226117 | 1 | - | 50 | 20 | 30 | 40 | NW_011924887.1 | 225822 | 226321 | 500 | + | Gnomon | promoter | NA | NA | gene128 | GeneID:105286681 | LOC105286681 | Gene | LOC105286681 | lncRNA | NA | character(0) | character(0) |

...

... only first 25 lines shown ...

Directory: RNK File: sites-in-promoters-Cb_LI_BCphaseB2.txt

```
seqnames	start	end	width	strand	coverage	numCs	numTs	perc_meth	region_seqnames	region_start	region_end	region_width	region_strand	region_source	region_type	region_score	region_phase	region_ID	region_Dbxref	region_Name	region_gbkey	region_gene	region_gene_biotype	region_partial	region_start_range	region_end_range
NW_011924881.1	745918	745918	1	+	19	15	4	78.9473684210526	NW_011924881.1	745918	746417	500	+	Gnomon	promoter	NA	NA	gene42	GeneID:105275579	LOC105275579	Gene	LOC105275579	protein_coding	NA	character(0)	character(0)
NW_011924881.1	745919	745919	1	-	5	5	0	100	NW_011924881.1	745918	746417	500	+	Gnomon	promoter	NA	NA	gene42	GeneID:105275579	LOC105275579	Gene	LOC105275579	protein_coding	NA	character(0)	character(0)
NW_011924881.1	746036	746036	1	-	33	9	24	27.2727272727273	NW_011924881.1	745918	746417	500	+	Gnomon	promoter	NA	NA	gene42	GeneID:105275579	LOC105275579	Gene	LOC105275579	protein_coding	NA	character(0)	character(0)
NW_011924881.1	1012572	1012572	1	+	25	7	18	28	NW_011924881.1	1012159	1012658	500	+	Gnomon	promoter	NA	NA	gene65	GeneID:105277639	LOC105277639	Gene	LOC105277639	protein_coding	NA	character(0)	character(0)
NW_011924887.1	221082	221082	1	-	34	7	27	20.5882352941176	NW_011924887.1	220963	221462	500	+	Gnomon	promoter	NA	NA	gene126	GeneID:105286831	LOC105286831	Gene	LOC105286831	protein_coding	NA	character(0)	character(0)
NW_011924887.1	221097	221097	1	+	6	5	1	83.3333333333333	NW_011924887.1	220963	221462	500	+	Gnomon	promoter	NA	NA	gene126	GeneID:105286831	LOC105286831	Gene	LOC105286831	protein_coding	NA	character(0)	character(0)
NW_011924887.1	221098	221098	1	-	32	20	12	62.5	NW_011924887.1	220963	221462	500	+	Gnomon	promoter	NA	NA	gene126	GeneID:105286831	LOC105286831	Gene	LOC105286831	protein_coding	NA	character(0)	character(0)
NW_011924887.1	221292	221292	1	-	16	8	8	50	NW_011924887.1	220963	221462	500	+	Gnomon	promoter	NA	NA	gene126	GeneID:105286831	LOC105286831	Gene	LOC105286831	protein_coding	NA	character(0)	character(0)
NW_011924887.1	225896	225896	1	+	24	9	15	37.5	NW_011924887.1	225822	226321	500	+	Gnomon	promoter	NA	NA	gene128	GeneID:105286681	LOC105286681	Gene	LOC105286681	lncRNA	NA	character(0)	character(0)
NW_011924887.1	225897	225897	1	-	19	8	11	42.1052631578947	NW_011924887.1	225822	226321	500	+	Gnomon	promoter	NA	NA	gene128	GeneID:105286681	LOC105286681	Gene	LOC105286681	lncRNA	NA	character(0)	character(0)
NW_011924887.1	225945	225945	1	+	27	19	8	70.3703703703704	NW_011924887.1	225822	226321	500	+	Gnomon	promoter	NA	NA	gene128	GeneID:105286681	LOC105286681	Gene	LOC105286681	lncRNA	NA	character(0)	character(0)
NW_011924887.1	225946	225946	1	-	14	12	2	85.7142857142857	NW_011924887.1	225822	226321	500	+	Gnomon	promoter	NA	NA	gene128	GeneID:105286681	LOC105286681	Gene	LOC105286681	lncRNA	NA	character(0)	character(0)
NW_011924887.1	225950	225950	1	+	30	18	12	60	NW_011924887.1	225822	226321	500	+	Gnomon	promoter	NA	NA	gene128	GeneID:105286681	LOC105286681	Gene	LOC105286681	lncRNA	NA	character(0)	character(0)
NW_011924887.1	225951	225951	1	-	12	10	2	83.3333333333333	NW_011924887.1	225822	226321	500	+	Gnomon	promoter	NA	NA	gene128	GeneID:105286681	LOC105286681	Gene	LOC105286681	lncRNA	NA	character(0)	character(0)
NW_011924887.1	225973	225973	1	+	29	25	4	86.2068965517241	NW_011924887.1	225822	226321	500	+	Gnomon	promoter	NA	NA	gene128	GeneID:105286681	LOC105286681	Gene	LOC105286681	lncRNA	NA	character(0)	character(0)
NW_011924887.1	225974	225974	1	-	7	7	0	100	NW_011924887.1	225822	226321	500	+	Gnomon	promoter	NA	NA	gene128	GeneID:105286681	LOC105286681	Gene	LOC105286681	lncRNA	NA	character(0)	character(0)
NW_011924887.1	225981	225981	1	+	31	20	11	64.5161290322581	NW_011924887.1	225822	226321	500	+	Gnomon	promoter	NA	NA	gene128	GeneID:105286681	LOC105286681	Gene	LOC105286681	lncRNA	NA	character(0)	character(0)
NW_011924887.1	225982	225982	1	-	9	6	3	66.6666666666667	NW_011924887.1	225822	226321	500	+	Gnomon	promoter	NA	NA	gene128	GeneID:105286681	LOC105286681	Gene	LOC105286681	lncRNA	NA	character(0)	character(0)
NW_011924887.1	226007	226007	1	+	36	12	24	33.3333333333333	NW_011924887.1	225822	226321	500	+	Gnomon	promoter	NA	NA	gene128	GeneID:105286681	LOC105286681	Gene	LOC105286681	lncRNA	NA	character(0)	character(0)
NW_011924887.1	226008	226008	1	-	28	7	21	25	NW_011924887.1	225822	226321	500	+	Gnomon	promoter	NA	NA	gene128	GeneID:105286681	LOC105286681	Gene	LOC105286681	lncRNA	NA	character(0)	character(0)
NW_011924887.1	226055	226055	1	+	24	11	13	45.8333333333333	NW_011924887.1	225822	226321	500	+	Gnomon	promoter	NA	NA	gene128	GeneID:105286681	LOC105286681	Gene	LOC105286681	lncRNA	NA	character(0)	character(0)
NW_011924887.1	226056	226056	1	-	54	24	30	44.4444444444444	NW_011924887.1	225822	226321	500	+	Gnomon	promoter	NA	NA	gene128	GeneID:105286681	LOC105286681	Gene	LOC105286681	lncRNA	NA	character(0)	character(0)
NW_011924887.1	226087	226087	1	-	71	24	47	33.8028169014084	NW_011924887.1	225822	226321	500	+	Gnomon	promoter	NA	NA	gene128	GeneID:105286681	LOC105286681	Gene	LOC105286681	lncRNA	NA	character(0)	character(0)
NW_011924887.1	226088	226088	1	+	18	7	11	38.8888888888889	NW_011924887.1	225822	226321	500	+	Gnomon	promoter	NA	NA	gene128	GeneID:105286681	LOC105286681	Gene	LOC105286681	lncRNA	NA	character(0)	character(0)
...
... only first 25 lines shown ...
```

Directory: RNK File: sites-in-promoters-Cb_LI_RphaseA1.txt

| seqnames | start | end | width | strand | coverage | numCs | numTs | perc_meth | region_seqnames | region_start | region_end | region_width | region_strand | region_source | region_type | region_score | region_phase | region_ID | region_Dbxref | region_Name | region_gbkey | region_gene | region_gene_biotype | region_partial | region_start_range | region_end_range |
|---|---|---|---|---|---|---|---|---|---|---|---|---|---|---|---|---|---|---|---|---|---|---|---|---|---|---|
| NW_011924881.1 | 744707 | 744707 | 1 | + | 19 | 6 | 13 | 31.5789473684211 | NW_011924881.1 | 744598 | 745097 | 500 | - | Gnomon | promoter | NA | NA | gene40 | GeneID:105275462 | LOC105275462 | Gene | LOC105275462 | protein_coding | NA | character(0) | character(0) |
| NW_011924881.1 | 745918 | 745918 | 1 | + | 42 | 38 | 4 | 90.4761904761905 | NW_011924881.1 | 745918 | 746417 | 500 | + | Gnomon | promoter | NA | NA | gene42 | GeneID:105275579 | LOC105275579 | Gene | LOC105275579 | protein_coding | NA | character(0) | character(0) |
| NW_011924881.1 | 746036 | 746036 | 1 | - | 26 | 7 | 19 | 26.9230769230769 | NW_011924881.1 | 745918 | 746417 | 500 | + | Gnomon | promoter | NA | NA | gene42 | GeneID:105275579 | LOC105275579 | Gene | LOC105275579 | protein_coding | NA | character(0) | character(0) |
| NW_011924881.1 | 807838 | 807838 | 1 | + | 14 | 6 | 8 | 42.8571428571429 | NW_011924881.1 | 807523 | 808022 | 500 | - | Gnomon | promoter | NA | NA | gene52 | GeneID:105276184 | LOC105276184 | Gene | LOC105276184 | protein_coding | NA | character(0) | character(0) |
| NW_011924881.1 | 989262 | 989262 | 1 | - | 6 | 5 | 1 | 83.3333333333333 | NW_011924881.1 | 989253 | 989752 | 500 | + | Gnomon | promoter | NA | NA | gene61 | GeneID:105277580 | LOC105277580 | Gene | LOC105277580 | protein_coding | NA | character(0) | character(0) |
| NW_011924887.1 | 221097 | 221097 | 1 | + | 10 | 8 | 2 | 80 | NW_011924887.1 | 220963 | 221462 | 500 | + | Gnomon | promoter | NA | NA | gene126 | GeneID:105286831 | LOC105286831 | Gene | LOC105286831 | protein_coding | NA | character(0) | character(0) |
| NW_011924887.1 | 221098 | 221098 | 1 | - | 28 | 14 | 14 | 50 | NW_011924887.1 | 220963 | 221462 | 500 | + | Gnomon | promoter | NA | NA | gene126 | GeneID:105286831 | LOC105286831 | Gene | LOC105286831 | protein_coding | NA | character(0) | character(0) |
| NW_011924887.1 | 225896 | 225896 | 1 | + | 29 | 11 | 18 | 37.9310344827586 | NW_011924887.1 | 225822 | 226321 | 500 | + | Gnomon | promoter | NA | NA | gene128 | GeneID:105286681 | LOC105286681 | Gene | LOC105286681 | lncRNA | NA | character(0) | character(0) |
| NW_011924887.1 | 225897 | 225897 | 1 | - | 19 | 6 | 13 | 31.5789473684211 | NW_011924887.1 | 225822 | 226321 | 500 | + | Gnomon | promoter | NA | NA | gene128 | GeneID:105286681 | LOC105286681 | Gene | LOC105286681 | lncRNA | NA | character(0) | character(0) |
| NW_011924887.1 | 225945 | 225945 | 1 | + | 20 | 12 | 8 | 60 | NW_011924887.1 | 225822 | 226321 | 500 | + | Gnomon | promoter | NA | NA | gene128 | GeneID:105286681 | LOC105286681 | Gene | LOC105286681 | lncRNA | NA | character(0) | character(0) |
| NW_011924887.1 | 225946 | 225946 | 1 | - | 19 | 15 | 4 | 78.9473684210526 | NW_011924887.1 | 225822 | 226321 | 500 | + | Gnomon | promoter | NA | NA | gene128 | GeneID:105286681 | LOC105286681 | Gene | LOC105286681 | lncRNA | NA | character(0) | character(0) |
| NW_011924887.1 | 225950 | 225950 | 1 | + | 23 | 13 | 10 | 56.5217391304348 | NW_011924887.1 | 225822 | 226321 | 500 | + | Gnomon | promoter | NA | NA | gene128 | GeneID:105286681 | LOC105286681 | Gene | LOC105286681 | lncRNA | NA | character(0) | character(0) |
| NW_011924887.1 | 225951 | 225951 | 1 | - | 14 | 9 | 5 | 64.2857142857143 | NW_011924887.1 | 225822 | 226321 | 500 | + | Gnomon | promoter | NA | NA | gene128 | GeneID:105286681 | LOC105286681 | Gene | LOC105286681 | lncRNA | NA | character(0) | character(0) |
| NW_011924887.1 | 225973 | 225973 | 1 | + | 30 | 24 | 6 | 80 | NW_011924887.1 | 225822 | 226321 | 500 | + | Gnomon | promoter | NA | NA | gene128 | GeneID:105286681 | LOC105286681 | Gene | LOC105286681 | lncRNA | NA | character(0) | character(0) |
| NW_011924887.1 | 225974 | 225974 | 1 | - | 12 | 12 | 0 | 100 | NW_011924887.1 | 225822 | 226321 | 500 | + | Gnomon | promoter | NA | NA | gene128 | GeneID:105286681 | LOC105286681 | Gene | LOC105286681 | lncRNA | NA | character(0) | character(0) |
| NW_011924887.1 | 225981 | 225981 | 1 | + | 36 | 20 | 16 | 55.5555555555556 | NW_011924887.1 | 225822 | 226321 | 500 | + | Gnomon | promoter | NA | NA | gene128 | GeneID:105286681 | LOC105286681 | Gene | LOC105286681 | lncRNA | NA | character(0) | character(0) |
| NW_011924887.1 | 225982 | 225982 | 1 | - | 9 | 6 | 3 | 66.6666666666667 | NW_011924887.1 | 225822 | 226321 | 500 | + | Gnomon | promoter | NA | NA | gene128 | GeneID:105286681 | LOC105286681 | Gene | LOC105286681 | lncRNA | NA | character(0) | character(0) |
| NW_011924887.1 | 226007 | 226007 | 1 | + | 45 | 14 | 31 | 31.1111111111111 | NW_011924887.1 | 225822 | 226321 | 500 | + | Gnomon | promoter | NA | NA | gene128 | GeneID:105286681 | LOC105286681 | Gene | LOC105286681 | lncRNA | NA | character(0) | character(0) |
| NW_011924887.1 | 226008 | 226008 | 1 | - | 20 | 8 | 12 | 40 | NW_011924887.1 | 225822 | 226321 | 500 | + | Gnomon | promoter | NA | NA | gene128 | GeneID:105286681 | LOC105286681 | Gene | LOC105286681 | lncRNA | NA | character(0) | character(0) |
| NW_011924887.1 | 226055 | 226055 | 1 | + | 29 | 14 | 15 | 48.2758620689655 | NW_011924887.1 | 225822 | 226321 | 500 | + | Gnomon | promoter | NA | NA | gene128 | GeneID:105286681 | LOC105286681 | Gene | LOC105286681 | lncRNA | NA | character(0) | character(0) |
| NW_011924887.1 | 226056 | 226056 | 1 | - | 43 | 22 | 21 | 51.1627906976744 | NW_011924887.1 | 225822 | 226321 | 500 | + | Gnomon | promoter | NA | NA | gene128 | GeneID:105286681 | LOC105286681 | Gene | LOC105286681 | lncRNA | NA | character(0) | character(0) |
| NW_011924887.1 | 226087 | 226087 | 1 | - | 54 | 20 | 34 | 37.037037037037 | NW_011924887.1 | 225822 | 226321 | 500 | + | Gnomon | promoter | NA | NA | gene128 | GeneID:105286681 | LOC105286681 | Gene | LOC105286681 | lncRNA | NA | character(0) | character(0) |
| NW_011924887.1 | 226089 | 226089 | 1 | - | 56 | 15 | 41 | 26.7857142857143 | NW_011924887.1 | 225822 | 226321 | 500 | + | Gnomon | promoter | NA | NA | gene128 | GeneID:105286681 | LOC105286681 | Gene | LOC105286681 | lncRNA | NA | character(0) | character(0) |
| NW_011924887.1 | 226100 | 226100 | 1 | + | 9 | 9 | 0 | 100 | NW_011924887.1 | 225822 | 226321 | 500 | + | Gnomon | promoter | NA | NA | gene128 | GeneID:105286681 | LOC105286681 | Gene | LOC105286681 | lncRNA | NA | character(0) | character(0) |

...
... only first 25 lines shown ...

Directory: RNK File: sites-in-promoters-Cb_LI_RphaseA2.txt

| seqnames | start | end | width | strand | coverage | numCs | numTs | perc_meth | region_seqnames | region_start | region_end | region_width | region_strand | region_source | region_type | region_score | region_phase | region_ID | region_Dbxref | region_Name | region_gbkey | region_gene | region_gene_biotype | region_partial | region_start_range | region_end_range |
|---|---|---|---|---|---|---|---|---|---|---|---|---|---|---|---|---|---|---|---|---|---|---|---|---|---|---|
| NW_011924881.1 | 745918 | 745918 | 1 | + | 41 | 33 | 8 | 80.4878048780488 | NW_011924881.1 | 745918 | 746417 | 500 | + | Gnomon | promoter | NA | NA | gene42 | GeneID:105275579 | LOC105275579 | Gene | LOC105275579 | protein_coding | NA | character(0) | character(0) |
| NW_011924881.1 | 746036 | 746036 | 1 | - | 41 | 15 | 26 | 36.5853658536585 | NW_011924881.1 | 745918 | 746417 | 500 | + | Gnomon | promoter | NA | NA | gene42 | GeneID:105275579 | LOC105275579 | Gene | LOC105275579 | protein_coding | NA | character(0) | character(0) |
| NW_011924881.1 | 746261 | 746261 | 1 | - | 46 | 8 | 38 | 17.3913043478261 | NW_011924881.1 | 745918 | 746417 | 500 | + | Gnomon | promoter | NA | NA | gene42 | GeneID:105275579 | LOC105275579 | Gene | LOC105275579 | protein_coding | NA | character(0) | character(0) |
| NW_011924881.1 | 989262 | 989262 | 1 | - | 6 | 6 | 0 | 100 | NW_011924881.1 | 989253 | 989752 | 500 | + | Gnomon | promoter | NA | NA | gene61 | GeneID:105277580 | LOC105277580 | Gene | LOC105277580 | protein_coding | NA | character(0) | character(0) |
| NW_011924881.1 | 1012572 | 1012572 | 1 | + | 52 | 8 | 44 | 15.3846153846154 | NW_011924881.1 | 1012159 | 1012658 | 500 | + | Gnomon | promoter | NA | NA | gene65 | GeneID:105277639 | LOC105277639 | Gene | LOC105277639 | protein_coding | NA | character(0) | character(0) |
| NW_011924887.1 | 221082 | 221082 | 1 | - | 39 | 9 | 30 | 23.0769230769231 | NW_011924887.1 | 220963 | 221462 | 500 | + | Gnomon | promoter | NA | NA | gene126 | GeneID:105286831 | LOC105286831 | Gene | LOC105286831 | protein_coding | NA | character(0) | character(0) |
| NW_011924887.1 | 221097 | 221097 | 1 | + | 10 | 8 | 2 | 80 | NW_011924887.1 | 220963 | 221462 | 500 | + | Gnomon | promoter | NA | NA | gene126 | GeneID:105286831 | LOC105286831 | Gene | LOC105286831 | protein_coding | NA | character(0) | character(0) |
| NW_011924887.1 | 221098 | 221098 | 1 | - | 35 | 17 | 18 | 48.5714285714286 | NW_011924887.1 | 220963 | 221462 | 500 | + | Gnomon | promoter | NA | NA | gene126 | GeneID:105286831 | LOC105286831 | Gene | LOC105286831 | protein_coding | NA | character(0) | character(0) |
| NW_011924887.1 | 221105 | 221105 | 1 | - | 33 | 7 | 26 | 21.2121212121212 | NW_011924887.1 | 220963 | 221462 | 500 | + | Gnomon | promoter | NA | NA | gene126 | GeneID:105286831 | LOC105286831 | Gene | LOC105286831 | protein_coding | NA | character(0) | character(0) |
| NW_011924887.1 | 221253 | 221253 | 1 | - | 26 | 8 | 18 | 30.7692307692308 | NW_011924887.1 | 220963 | 221462 | 500 | + | Gnomon | promoter | NA | NA | gene126 | GeneID:105286831 | LOC105286831 | Gene | LOC105286831 | protein_coding | NA | character(0) | character(0) |
| NW_011924887.1 | 225896 | 225896 | 1 | + | 20 | 8 | 12 | 40 | NW_011924887.1 | 225822 | 226321 | 500 | + | Gnomon | promoter | NA | NA | gene128 | GeneID:105286681 | LOC105286681 | Gene | LOC105286681 | lncRNA | NA | character(0) | character(0) |
| NW_011924887.1 | 225897 | 225897 | 1 | - | 22 | 7 | 15 | 31.8181818181818 | NW_011924887.1 | 225822 | 226321 | 500 | + | Gnomon | promoter | NA | NA | gene128 | GeneID:105286681 | LOC105286681 | Gene | LOC105286681 | lncRNA | NA | character(0) | character(0) |
| NW_011924887.1 | 225945 | 225945 | 1 | + | 33 | 26 | 7 | 78.7878787878788 | NW_011924887.1 | 225822 | 226321 | 500 | + | Gnomon | promoter | NA | NA | gene128 | GeneID:105286681 | LOC105286681 | Gene | LOC105286681 | lncRNA | NA | character(0) | character(0) |
| NW_011924887.1 | 225946 | 225946 | 1 | - | 13 | 12 | 1 | 92.3076923076923 | NW_011924887.1 | 225822 | 226321 | 500 | + | Gnomon | promoter | NA | NA | gene128 | GeneID:105286681 | LOC105286681 | Gene | LOC105286681 | lncRNA | NA | character(0) | character(0) |
| NW_011924887.1 | 225950 | 225950 | 1 | + | 35 | 25 | 10 | 71.4285714285714 | NW_011924887.1 | 225822 | 226321 | 500 | + | Gnomon | promoter | NA | NA | gene128 | GeneID:105286681 | LOC105286681 | Gene | LOC105286681 | lncRNA | NA | character(0) | character(0) |
| NW_011924887.1 | 225951 | 225951 | 1 | - | 14 | 7 | 7 | 50 | NW_011924887.1 | 225822 | 226321 | 500 | + | Gnomon | promoter | NA | NA | gene128 | GeneID:105286681 | LOC105286681 | Gene | LOC105286681 | lncRNA | NA | character(0) | character(0) |
| NW_011924887.1 | 225973 | 225973 | 1 | + | 41 | 35 | 6 | 85.3658536585366 | NW_011924887.1 | 225822 | 226321 | 500 | + | Gnomon | promoter | NA | NA | gene128 | GeneID:105286681 | LOC105286681 | Gene | LOC105286681 | lncRNA | NA | character(0) | character(0) |
| NW_011924887.1 | 225974 | 225974 | 1 | - | 13 | 10 | 3 | 76.9230769230769 | NW_011924887.1 | 225822 | 226321 | 500 | + | Gnomon | promoter | NA | NA | gene128 | GeneID:105286681 | LOC105286681 | Gene | LOC105286681 | lncRNA | NA | character(0) | character(0) |
| NW_011924887.1 | 225981 | 225981 | 1 | + | 42 | 31 | 11 | 73.8095238095238 | NW_011924887.1 | 225822 | 226321 | 500 | + | Gnomon | promoter | NA | NA | gene128 | GeneID:105286681 | LOC105286681 | Gene | LOC105286681 | lncRNA | NA | character(0) | character(0) |
| NW_011924887.1 | 225982 | 225982 | 1 | - | 14 | 12 | 2 | 85.7142857142857 | NW_011924887.1 | 225822 | 226321 | 500 | + | Gnomon | promoter | NA | NA | gene128 | GeneID:105286681 | LOC105286681 | Gene | LOC105286681 | lncRNA | NA | character(0) | character(0) |
| NW_011924887.1 | 226007 | 226007 | 1 | + | 60 | 28 | 32 | 46.6666666666667 | NW_011924887.1 | 225822 | 226321 | 500 | + | Gnomon | promoter | NA | NA | gene128 | GeneID:105286681 | LOC105286681 | Gene | LOC105286681 | lncRNA | NA | character(0) | character(0) |
| NW_011924887.1 | 226008 | 226008 | 1 | - | 43 | 17 | 26 | 39.5348837209302 | NW_011924887.1 | 225822 | 226321 | 500 | + | Gnomon | promoter | NA | NA | gene128 | GeneID:105286681 | LOC105286681 | Gene | LOC105286681 | lncRNA | NA | character(0) | character(0) |
| NW_011924887.1 | 226014 | 226014 | 1 | + | 62 | 10 | 52 | 16.1290322580645 | NW_011924887.1 | 225822 | 226321 | 500 | + | Gnomon | promoter | NA | NA | gene128 | GeneID:105286681 | LOC105286681 | Gene | LOC105286681 | lncRNA | NA | character(0) | character(0) |
| NW_011924887.1 | 226055 | 226055 | 1 | + | 45 | 19 | 26 | 42.2222222222222 | NW_011924887.1 | 225822 | 226321 | 500 | + | Gnomon | promoter | NA | NA | gene128 | GeneID:105286681 | LOC105286681 | Gene | LOC105286681 | lncRNA | NA | character(0) | character(0) |

...
... only first 25 lines shown ...

Directory: RNK File: sites-in-promoters-Cb_LI_RphaseB1.txt

| seqnames | start | end | width | strand | coverage | numCs | numTs | perc_meth | region_seqnames | region_start | region_end | region_width | region_strand | region_source | region_type | region_score | region_phase | region_ID | region_Dbxref | region_Name | region_gbkey | region_gene | region_gene_biotype | region_partial | region_start_range | region_end_range |
|---|---|---|---|---|---|---|---|---|---|---|---|---|---|---|---|---|---|---|---|---|---|---|---|---|---|---|
| NW_011924881.1 | 745918 | 745918 | 1 | + | 37 | 24 | 13 | 64.8648648648649 | NW_011924881.1 | 745918 | 746417 | 500 | + | Gnomon | promoter | NA | NA | gene42 | GeneID:105275579 | LOC105275579 | Gene | LOC105275579 | protein_coding | NA | character(0) | character(0) |
| NW_011924881.1 | 746036 | 746036 | 1 | - | 19 | 7 | 12 | 36.8421052631579 | NW_011924881.1 | 745918 | 746417 | 500 | + | Gnomon | promoter | NA | NA | gene42 | GeneID:105275579 | LOC105275579 | Gene | LOC105275579 | protein_coding | NA | character(0) | character(0) |
| NW_011924881.1 | 989262 | 989262 | 1 | - | 9 | 8 | 1 | 88.8888888888889 | NW_011924881.1 | 989253 | 989752 | 500 | + | Gnomon | promoter | NA | NA | gene61 | GeneID:105277580 | LOC105277580 | Gene | LOC105277580 | protein_coding | NA | character(0) | character(0) |
| NW_011924887.1 | 221098 | 221098 | 1 | - | 35 | 16 | 19 | 45.7142857142857 | NW_011924887.1 | 220963 | 221462 | 500 | + | Gnomon | promoter | NA | NA | gene126 | GeneID:105286831 | LOC105286831 | Gene | LOC105286831 | protein_coding | NA | character(0) | character(0) |
| NW_011924887.1 | 221105 | 221105 | 1 | - | 37 | 11 | 26 | 29.7297297297297 | NW_011924887.1 | 220963 | 221462 | 500 | + | Gnomon | promoter | NA | NA | gene126 | GeneID:105286831 | LOC105286831 | Gene | LOC105286831 | protein_coding | NA | character(0) | character(0) |
| NW_011924887.1 | 221253 | 221253 | 1 | - | 36 | 8 | 28 | 22.2222222222222 | NW_011924887.1 | 220963 | 221462 | 500 | + | Gnomon | promoter | NA | NA | gene126 | GeneID:105286831 | LOC105286831 | Gene | LOC105286831 | protein_coding | NA | character(0) | character(0) |
| NW_011924887.1 | 221292 | 221292 | 1 | - | 28 | 9 | 19 | 32.1428571428571 | NW_011924887.1 | 220963 | 221462 | 500 | + | Gnomon | promoter | NA | NA | gene126 | GeneID:105286831 | LOC105286831 | Gene | LOC105286831 | protein_coding | NA | character(0) | character(0) |
| NW_011924887.1 | 225896 | 225896 | 1 | + | 20 | 7 | 13 | 35 | NW_011924887.1 | 225822 | 226321 | 500 | + | Gnomon | promoter | NA | NA | gene128 | GeneID:105286681 | LOC105286681 | Gene | LOC105286681 | lncRNA | NA | character(0) | character(0) |
| NW_011924887.1 | 225945 | 225945 | 1 | + | 30 | 19 | 11 | 63.3333333333333 | NW_011924887.1 | 225822 | 226321 | 500 | + | Gnomon | promoter | NA | NA | gene128 | GeneID:105286681 | LOC105286681 | Gene | LOC105286681 | lncRNA | NA | character(0) | character(0) |
| NW_011924887.1 | 225946 | 225946 | 1 | - | 20 | 16 | 4 | 80 | NW_011924887.1 | 225822 | 226321 | 500 | + | Gnomon | promoter | NA | NA | gene128 | GeneID:105286681 | LOC105286681 | Gene | LOC105286681 | lncRNA | NA | character(0) | character(0) |
| NW_011924887.1 | 225950 | 225950 | 1 | + | 36 | 22 | 14 | 61.1111111111111 | NW_011924887.1 | 225822 | 226321 | 500 | + | Gnomon | promoter | NA | NA | gene128 | GeneID:105286681 | LOC105286681 | Gene | LOC105286681 | lncRNA | NA | character(0) | character(0) |
| NW_011924887.1 | 225951 | 225951 | 1 | - | 12 | 6 | 6 | 50 | NW_011924887.1 | 225822 | 226321 | 500 | + | Gnomon | promoter | NA | NA | gene128 | GeneID:105286681 | LOC105286681 | Gene | LOC105286681 | lncRNA | NA | character(0) | character(0) |
| NW_011924887.1 | 225973 | 225973 | 1 | + | 38 | 32 | 6 | 84.2105263157895 | NW_011924887.1 | 225822 | 226321 | 500 | + | Gnomon | promoter | NA | NA | gene128 | GeneID:105286681 | LOC105286681 | Gene | LOC105286681 | lncRNA | NA | character(0) | character(0) |
| NW_011924887.1 | 225974 | 225974 | 1 | - | 10 | 8 | 2 | 80 | NW_011924887.1 | 225822 | 226321 | 500 | + | Gnomon | promoter | NA | NA | gene128 | GeneID:105286681 | LOC105286681 | Gene | LOC105286681 | lncRNA | NA | character(0) | character(0) |
| NW_011924887.1 | 225981 | 225981 | 1 | + | 44 | 34 | 10 | 77.2727272727273 | NW_011924887.1 | 225822 | 226321 | 500 | + | Gnomon | promoter | NA | NA | gene128 | GeneID:105286681 | LOC105286681 | Gene | LOC105286681 | lncRNA | NA | character(0) | character(0) |
| NW_011924887.1 | 225982 | 225982 | 1 | - | 10 | 7 | 3 | 70 | NW_011924887.1 | 225822 | 226321 | 500 | + | Gnomon | promoter | NA | NA | gene128 | GeneID:105286681 | LOC105286681 | Gene | LOC105286681 | lncRNA | NA | character(0) | character(0) |
| NW_011924887.1 | 226007 | 226007 | 1 | + | 50 | 18 | 32 | 36 | NW_011924887.1 | 225822 | 226321 | 500 | + | Gnomon | promoter | NA | NA | gene128 | GeneID:105286681 | LOC105286681 | Gene | LOC105286681 | lncRNA | NA | character(0) | character(0) |
| NW_011924887.1 | 226008 | 226008 | 1 | - | 35 | 9 | 26 | 25.7142857142857 | NW_011924887.1 | 225822 | 226321 | 500 | + | Gnomon | promoter | NA | NA | gene128 | GeneID:105286681 | LOC105286681 | Gene | LOC105286681 | lncRNA | NA | character(0) | character(0) |
| NW_011924887.1 | 226055 | 226055 | 1 | + | 37 | 9 | 28 | 24.3243243243243 | NW_011924887.1 | 225822 | 226321 | 500 | + | Gnomon | promoter | NA | NA | gene128 | GeneID:105286681 | LOC105286681 | Gene | LOC105286681 | lncRNA | NA | character(0) | character(0) |
| NW_011924887.1 | 226056 | 226056 | 1 | - | 47 | 18 | 29 | 38.2978723404255 | NW_011924887.1 | 225822 | 226321 | 500 | + | Gnomon | promoter | NA | NA | gene128 | GeneID:105286681 | LOC105286681 | Gene | LOC105286681 | lncRNA | NA | character(0) | character(0) |
| NW_011924887.1 | 226087 | 226087 | 1 | - | 65 | 16 | 49 | 24.6153846153846 | NW_011924887.1 | 225822 | 226321 | 500 | + | Gnomon | promoter | NA | NA | gene128 | GeneID:105286681 | LOC105286681 | Gene | LOC105286681 | lncRNA | NA | character(0) | character(0) |
| NW_011924887.1 | 226089 | 226089 | 1 | - | 65 | 10 | 55 | 15.3846153846154 | NW_011924887.1 | 225822 | 226321 | 500 | + | Gnomon | promoter | NA | NA | gene128 | GeneID:105286681 | LOC105286681 | Gene | LOC105286681 | lncRNA | NA | character(0) | character(0) |
| NW_011924887.1 | 226100 | 226100 | 1 | + | 9 | 5 | 4 | 55.5555555555556 | NW_011924887.1 | 225822 | 226321 | 500 | + | Gnomon | promoter | NA | NA | gene128 | GeneID:105286681 | LOC105286681 | Gene | LOC105286681 | lncRNA | NA | character(0) | character(0) |
| NW_011924887.1 | 226101 | 226101 | 1 | - | 55 | 42 | 13 | 76.3636363636364 | NW_011924887.1 | 225822 | 226321 | 500 | + | Gnomon | promoter | NA | NA | gene128 | GeneID:105286681 | LOC105286681 | Gene | LOC105286681 | lncRNA | NA | character(0) | character(0) |

...
... only first 25 lines shown ...

Directory: RNK File: sites-in-promoters-Cb_LI_RphaseB2.txt

| seqnames | start | end | width | strand | coverage | numCs | numTs | perc_meth | region_seqnames | region_start | region_end | region_width | region_strand | region_source | region_type | region_score | region_phase | region_ID | region_Dbxref | region_Name | region_gbkey | region_gene | region_gene_biotype | region_partial | region_start_range | region_end_range |
|---|---|---|---|---|---|---|---|---|---|---|---|---|---|---|---|---|---|---|---|---|---|---|---|---|---|---|
| NW_011924881.1 | 745918 | 745918 | 1 | + | 34 | 27 | 7 | 79.4117647058823 | NW_011924881.1 | 745918 | 746417 | 500 | + | Gnomon | promoter | NA | NA | gene42 | GeneID:105275579 | LOC105275579 | Gene | LOC105275579 | protein_coding | NA | character(0) | character(0) |
| NW_011924881.1 | 746261 | 746261 | 1 | - | 42 | 12 | 30 | 28.5714285714286 | NW_011924881.1 | 745918 | 746417 | 500 | + | Gnomon | promoter | NA | NA | gene42 | GeneID:105275579 | LOC105275579 | Gene | LOC105275579 | protein_coding | NA | character(0) | character(0) |
| NW_011924887.1 | 221082 | 221082 | 1 | - | 29 | 7 | 22 | 24.1379310344828 | NW_011924887.1 | 220963 | 221462 | 500 | + | Gnomon | promoter | NA | NA | gene126 | GeneID:105286831 | LOC105286831 | Gene | LOC105286831 | protein_coding | NA | character(0) | character(0) |
| NW_011924887.1 | 221097 | 221097 | 1 | + | 7 | 5 | 2 | 71.4285714285714 | NW_011924887.1 | 220963 | 221462 | 500 | + | Gnomon | promoter | NA | NA | gene126 | GeneID:105286831 | LOC105286831 | Gene | LOC105286831 | protein_coding | NA | character(0) | character(0) |
| NW_011924887.1 | 221098 | 221098 | 1 | - | 27 | 19 | 8 | 70.3703703703704 | NW_011924887.1 | 220963 | 221462 | 500 | + | Gnomon | promoter | NA | NA | gene126 | GeneID:105286831 | LOC105286831 | Gene | LOC105286831 | protein_coding | NA | character(0) | character(0) |
| NW_011924887.1 | 221105 | 221105 | 1 | - | 27 | 7 | 20 | 25.9259259259259 | NW_011924887.1 | 220963 | 221462 | 500 | + | Gnomon | promoter | NA | NA | gene126 | GeneID:105286831 | LOC105286831 | Gene | LOC105286831 | protein_coding | NA | character(0) | character(0) |
| NW_011924887.1 | 225945 | 225945 | 1 | + | 21 | 12 | 9 | 57.1428571428571 | NW_011924887.1 | 225822 | 226321 | 500 | + | Gnomon | promoter | NA | NA | gene128 | GeneID:105286681 | LOC105286681 | Gene | LOC105286681 | lncRNA | NA | character(0) | character(0) |
| NW_011924887.1 | 225950 | 225950 | 1 | + | 21 | 14 | 7 | 66.6666666666667 | NW_011924887.1 | 225822 | 226321 | 500 | + | Gnomon | promoter | NA | NA | gene128 | GeneID:105286681 | LOC105286681 | Gene | LOC105286681 | lncRNA | NA | character(0) | character(0) |
| NW_011924887.1 | 225973 | 225973 | 1 | + | 18 | 11 | 7 | 61.1111111111111 | NW_011924887.1 | 225822 | 226321 | 500 | + | Gnomon | promoter | NA | NA | gene128 | GeneID:105286681 | LOC105286681 | Gene | LOC105286681 | lncRNA | NA | character(0) | character(0) |
| NW_011924887.1 | 225974 | 225974 | 1 | - | 7 | 6 | 1 | 85.7142857142857 | NW_011924887.1 | 225822 | 226321 | 500 | + | Gnomon | promoter | NA | NA | gene128 | GeneID:105286681 | LOC105286681 | Gene | LOC105286681 | lncRNA | NA | character(0) | character(0) |
| NW_011924887.1 | 225981 | 225981 | 1 | + | 18 | 10 | 8 | 55.5555555555556 | NW_011924887.1 | 225822 | 226321 | 500 | + | Gnomon | promoter | NA | NA | gene128 | GeneID:105286681 | LOC105286681 | Gene | LOC105286681 | lncRNA | NA | character(0) | character(0) |
| NW_011924887.1 | 225982 | 225982 | 1 | - | 7 | 6 | 1 | 85.7142857142857 | NW_011924887.1 | 225822 | 226321 | 500 | + | Gnomon | promoter | NA | NA | gene128 | GeneID:105286681 | LOC105286681 | Gene | LOC105286681 | lncRNA | NA | character(0) | character(0) |
| NW_011924887.1 | 226007 | 226007 | 1 | + | 27 | 10 | 17 | 37.037037037037 | NW_011924887.1 | 225822 | 226321 | 500 | + | Gnomon | promoter | NA | NA | gene128 | GeneID:105286681 | LOC105286681 | Gene | LOC105286681 | lncRNA | NA | character(0) | character(0) |
| NW_011924887.1 | 226008 | 226008 | 1 | - | 24 | 9 | 15 | 37.5 | NW_011924887.1 | 225822 | 226321 | 500 | + | Gnomon | promoter | NA | NA | gene128 | GeneID:105286681 | LOC105286681 | Gene | LOC105286681 | lncRNA | NA | character(0) | character(0) |
| NW_011924887.1 | 226055 | 226055 | 1 | + | 27 | 11 | 16 | 40.7407407407407 | NW_011924887.1 | 225822 | 226321 | 500 | + | Gnomon | promoter | NA | NA | gene128 | GeneID:105286681 | LOC105286681 | Gene | LOC105286681 | lncRNA | NA | character(0) | character(0) |
| NW_011924887.1 | 226056 | 226056 | 1 | - | 39 | 17 | 22 | 43.5897435897436 | NW_011924887.1 | 225822 | 226321 | 500 | + | Gnomon | promoter | NA | NA | gene128 | GeneID:105286681 | LOC105286681 | Gene | LOC105286681 | lncRNA | NA | character(0) | character(0) |
| NW_011924887.1 | 226087 | 226087 | 1 | - | 65 | 26 | 39 | 40 | NW_011924887.1 | 225822 | 226321 | 500 | + | Gnomon | promoter | NA | NA | gene128 | GeneID:105286681 | LOC105286681 | Gene | LOC105286681 | lncRNA | NA | character(0) | character(0) |
| NW_011924887.1 | 226089 | 226089 | 1 | - | 62 | 19 | 43 | 30.6451612903226 | NW_011924887.1 | 225822 | 226321 | 500 | + | Gnomon | promoter | NA | NA | gene128 | GeneID:105286681 | LOC105286681 | Gene | LOC105286681 | lncRNA | NA | character(0) | character(0) |
| NW_011924887.1 | 226100 | 226100 | 1 | + | 19 | 11 | 8 | 57.8947368421053 | NW_011924887.1 | 225822 | 226321 | 500 | + | Gnomon | promoter | NA | NA | gene128 | GeneID:105286681 | LOC105286681 | Gene | LOC105286681 | lncRNA | NA | character(0) | character(0) |
| NW_011924887.1 | 226101 | 226101 | 1 | - | 52 | 37 | 15 | 71.1538461538462 | NW_011924887.1 | 225822 | 226321 | 500 | + | Gnomon | promoter | NA | NA | gene128 | GeneID:105286681 | LOC105286681 | Gene | LOC105286681 | lncRNA | NA | character(0) | character(0) |
| NW_011924887.1 | 226117 | 226117 | 1 | - | 46 | 18 | 28 | 39.1304347826087 | NW_011924887.1 | 225822 | 226321 | 500 | + | Gnomon | promoter | NA | NA | gene128 | GeneID:105286681 | LOC105286681 | Gene | LOC105286681 | lncRNA | NA | character(0) | character(0) |
| NW_011924887.1 | 226127 | 226127 | 1 | + | 12 | 7 | 5 | 58.3333333333333 | NW_011924887.1 | 225822 | 226321 | 500 | + | Gnomon | promoter | NA | NA | gene128 | GeneID:105286681 | LOC105286681 | Gene | LOC105286681 | lncRNA | NA | character(0) | character(0) |
| NW_011924887.1 | 226128 | 226128 | 1 | - | 48 | 28 | 20 | 58.3333333333333 | NW_011924887.1 | 225822 | 226321 | 500 | + | Gnomon | promoter | NA | NA | gene128 | GeneID:105286681 | LOC105286681 | Gene | LOC105286681 | lncRNA | NA | character(0) | character(0) |
| NW_011924887.1 | 226148 | 226148 | 1 | + | 9 | 5 | 4 | 55.5555555555556 | NW_011924887.1 | 225822 | 226321 | 500 | + | Gnomon | promoter | NA | NA | gene128 | GeneID:105286681 | LOC105286681 | Gene | LOC105286681 | lncRNA | NA | character(0) | character(0) |

...
... only first 25 lines shown ...

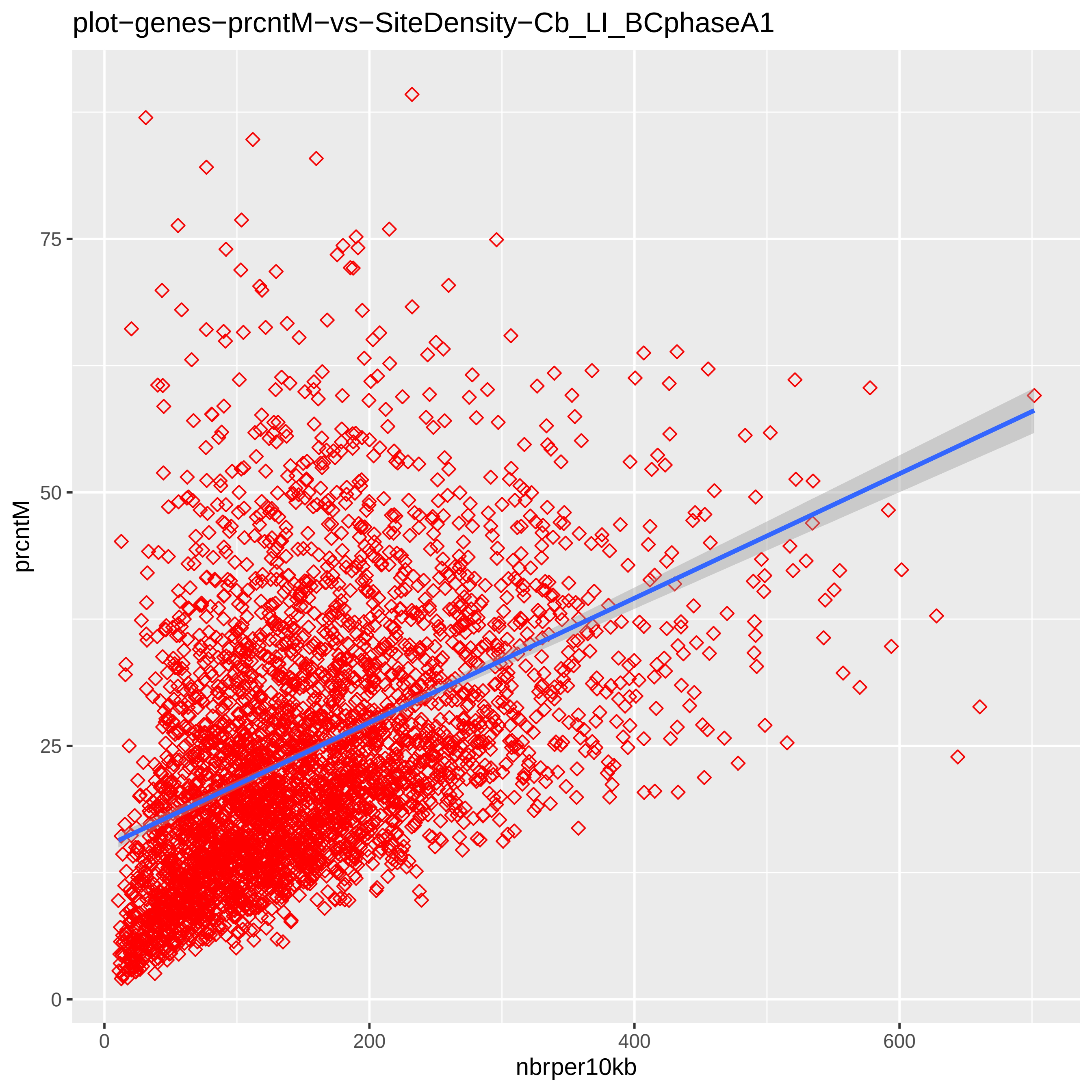
plot−genes−prcntM−vs−SiteDensity−Cb_LI_BCphaseA1
prcntM
nbrper10kb
0
25
50
75
0
200
400
600

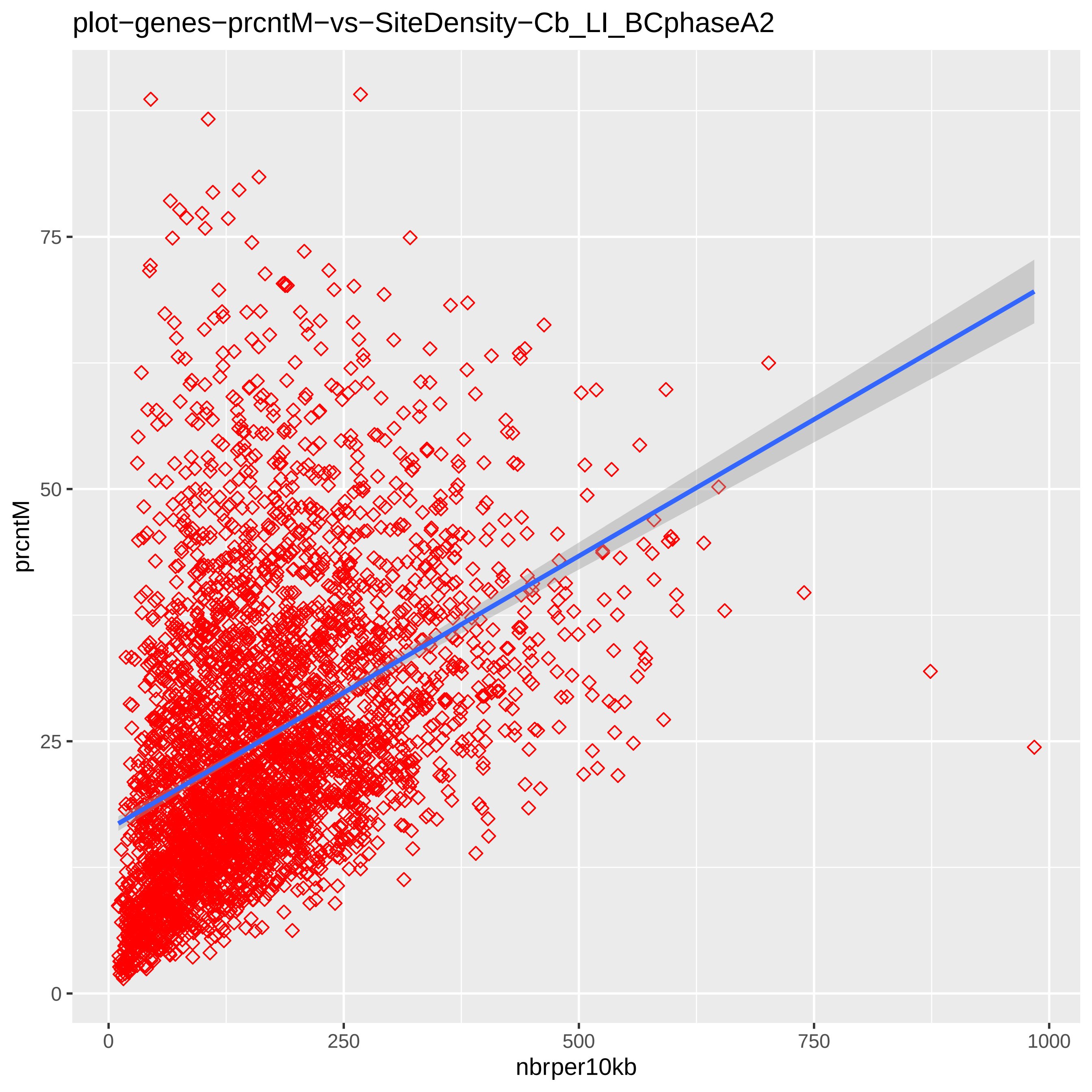
plot−genes−prcntM−vs−SiteDensity−Cb_LI_BCphaseA2
prcntM
nbrper10kb
0
25
50
75
0
250
500
750
1000

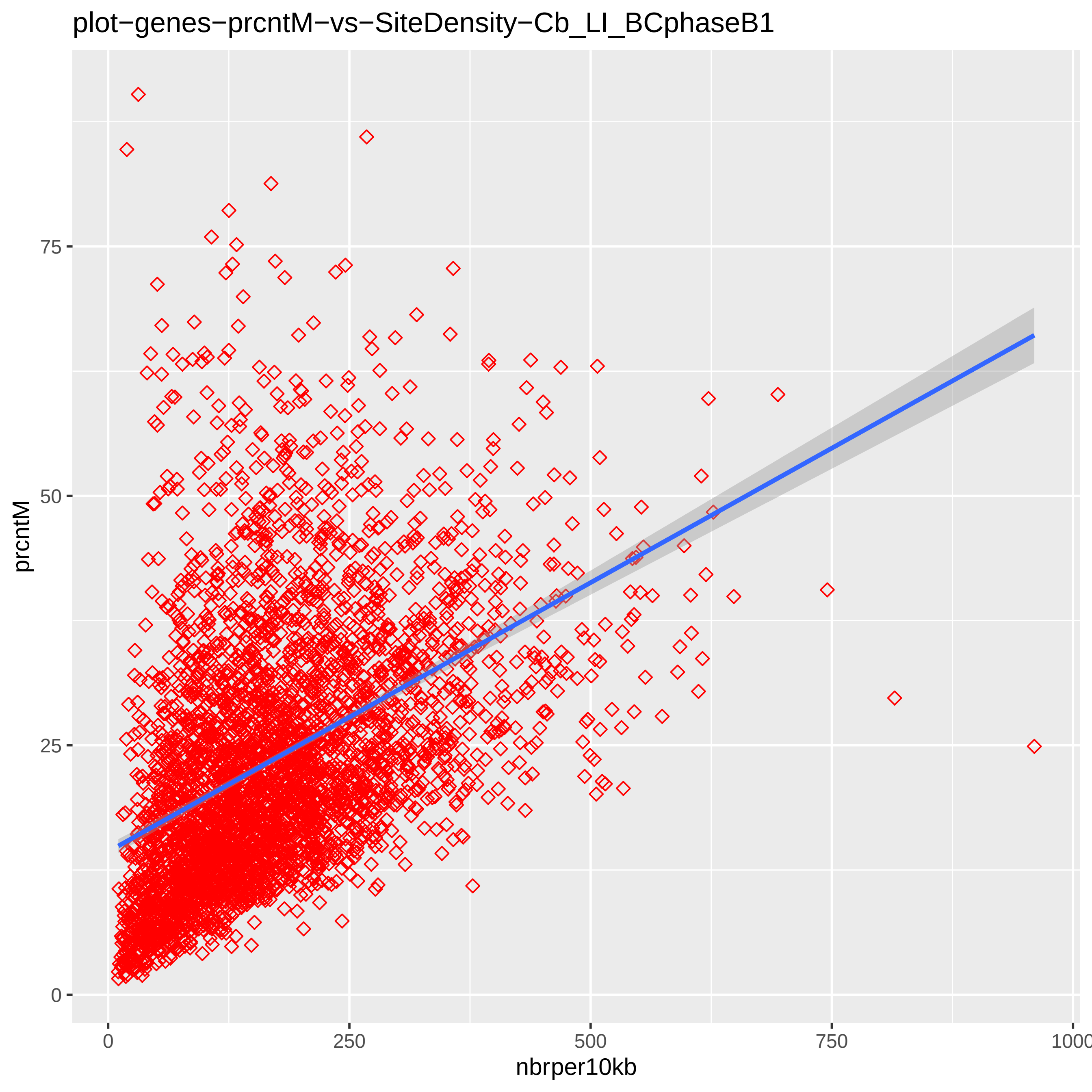

plot−genes−prcntM−vs−SiteDensity−Cb_LI_BCphaseB1
prcntM
nbrper10kb
0
25
50
75
0
250
500
750
1000

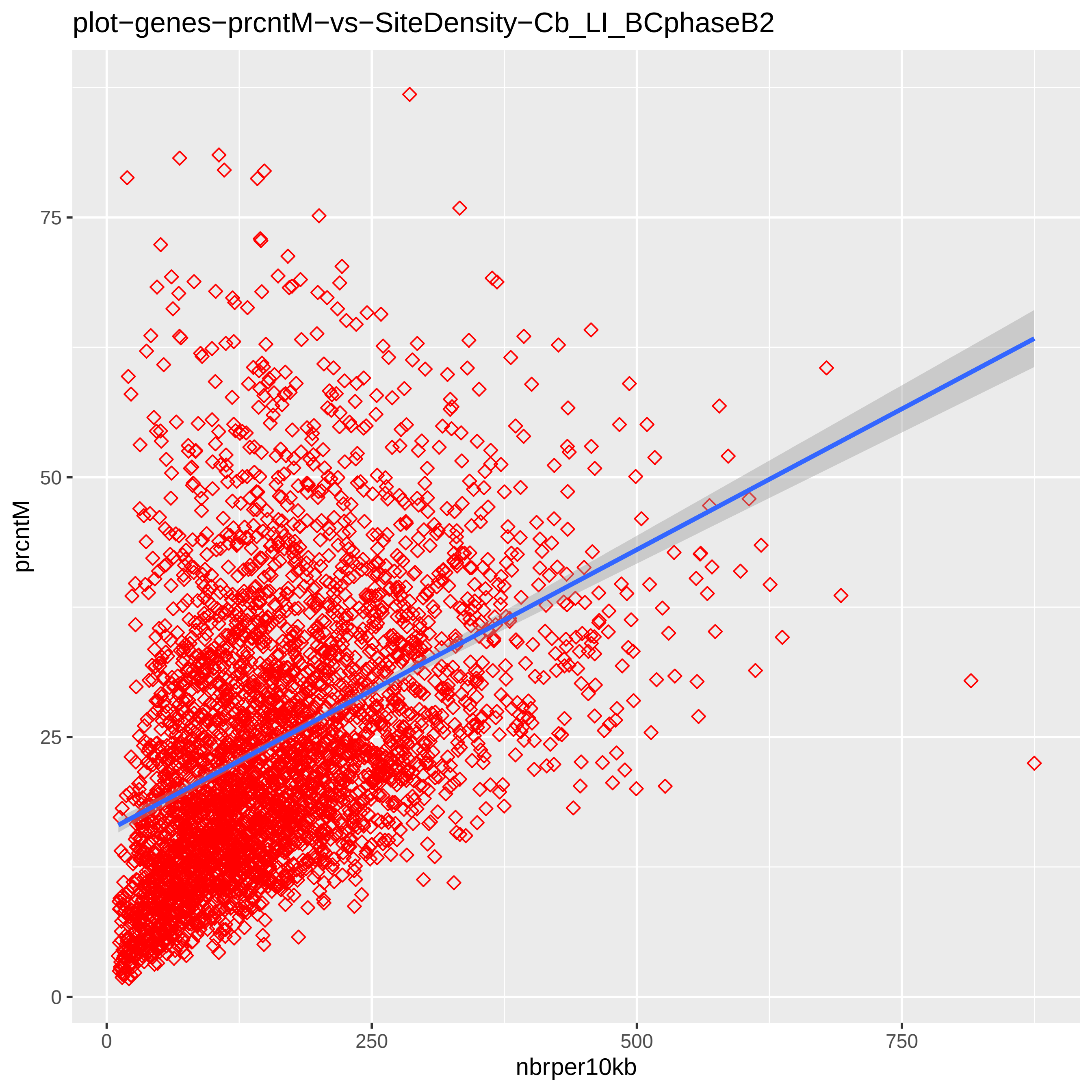

plot−genes−prcntM−vs−SiteDensity−Cb_LI_BCphaseB2
prcntM
nbrper10kb
0
25
50
75
0
250
500
750

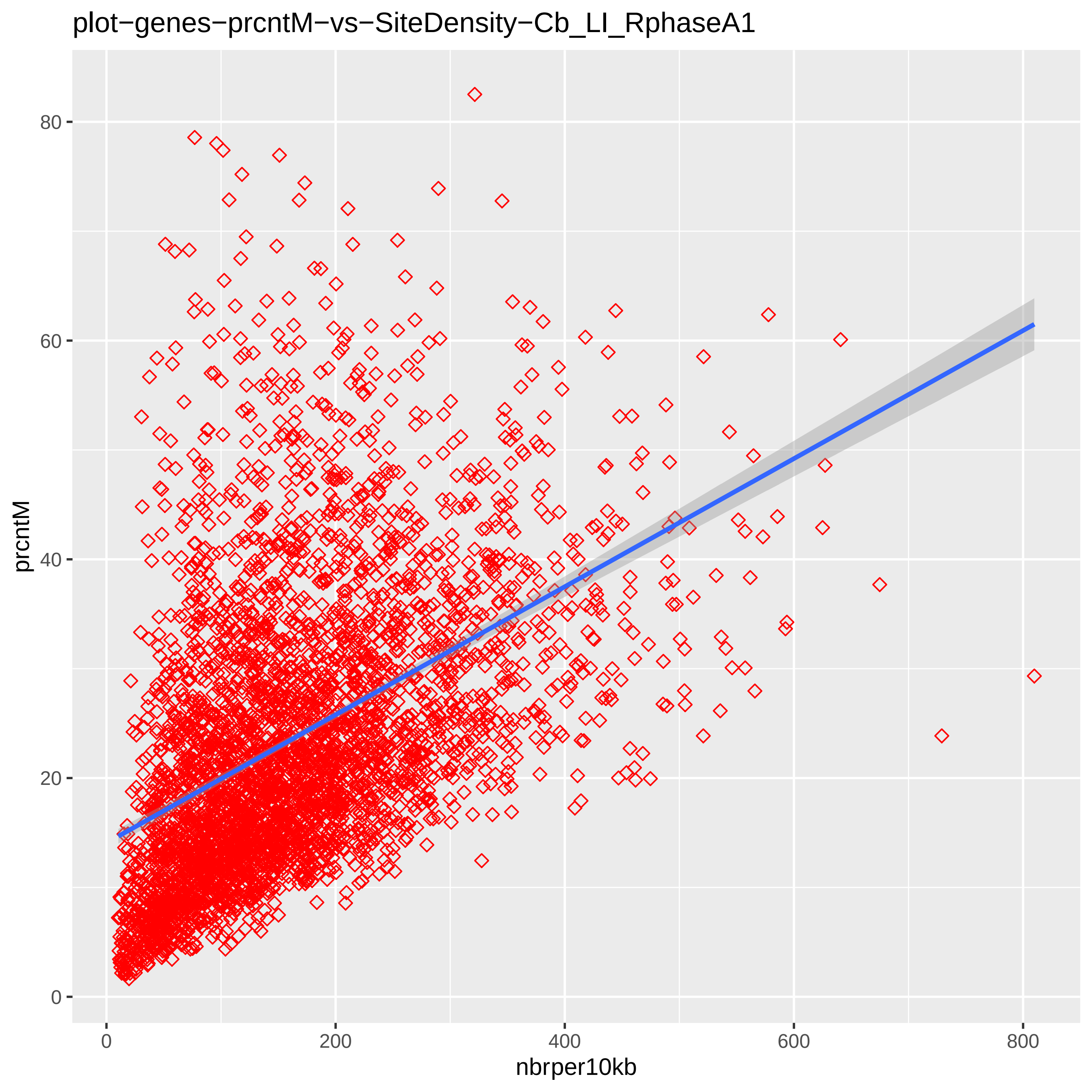

plot−genes−prcntM−vs−SiteDensity−Cb_LI_RphaseA1
prcntM
nbrper10kb
0
20
40
60
80
0
200
400
600
800

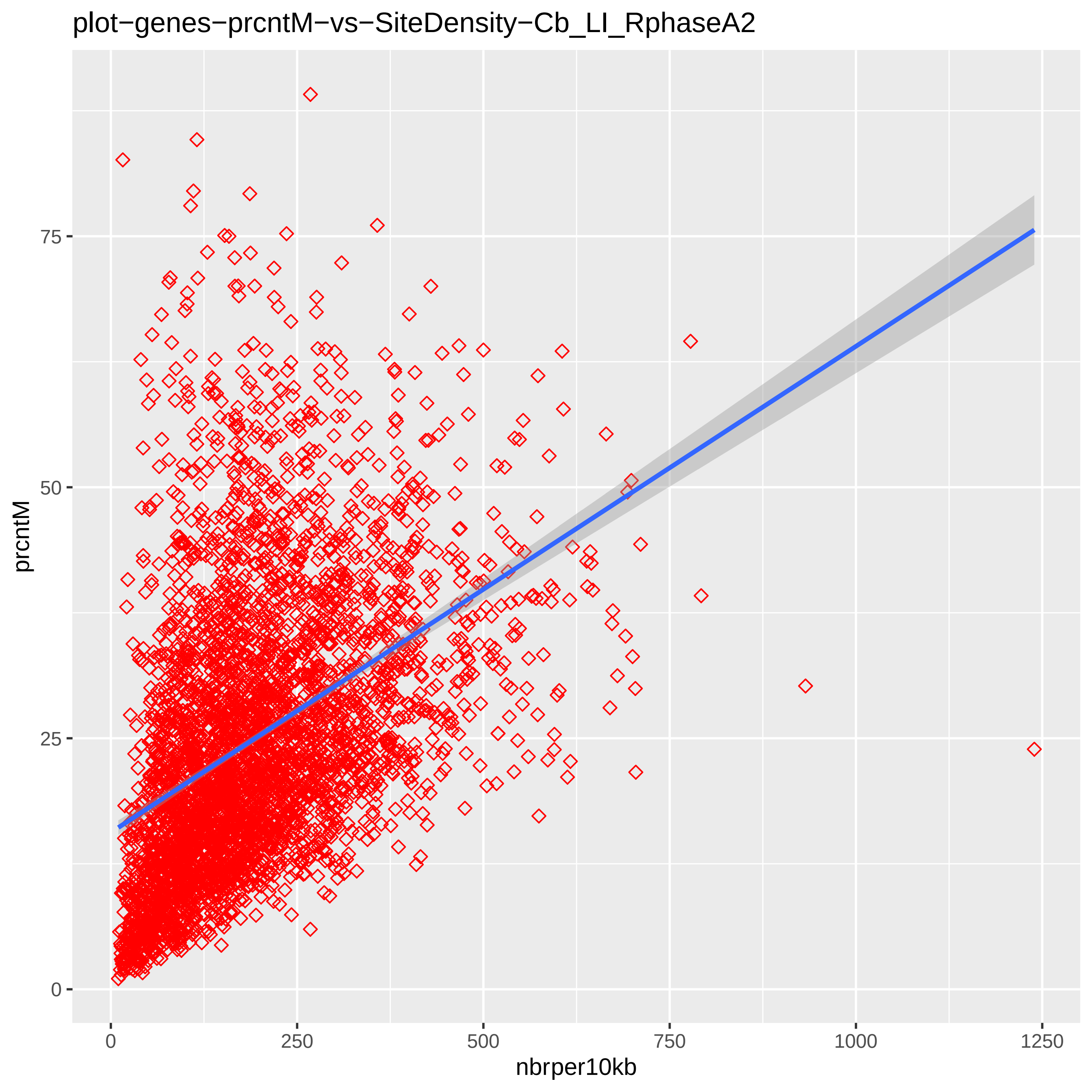

plot−genes−prcntM−vs−SiteDensity−Cb_LI_RphaseA2
prcntM
75
50
25
0
nbrper10kb
0
250
500
750
1000
1250

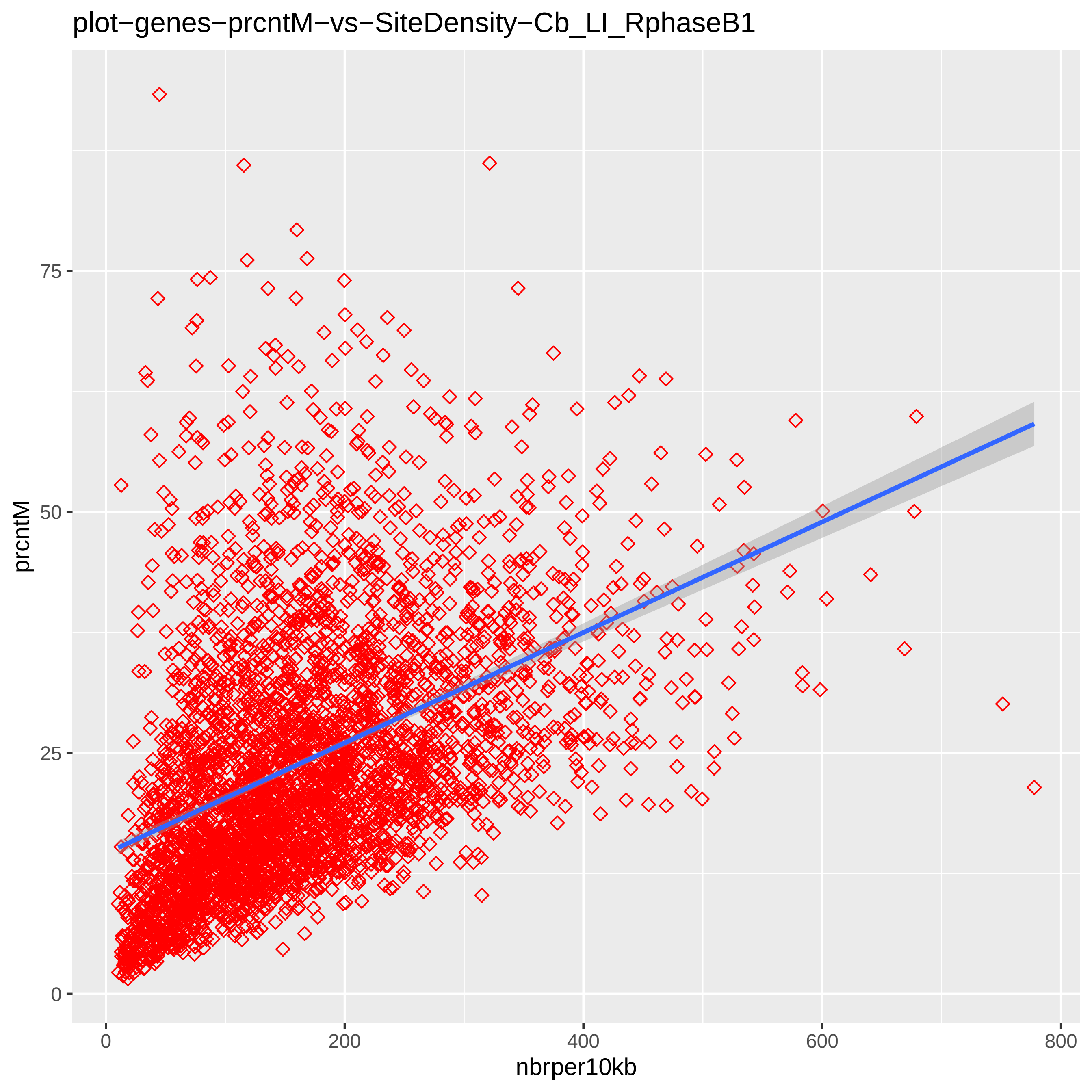
plot−genes−prcntM−vs−SiteDensity−Cb_LI_RphaseB1
prcntM
0
25
50
75
0
200
400
600
800
nbrper10kb

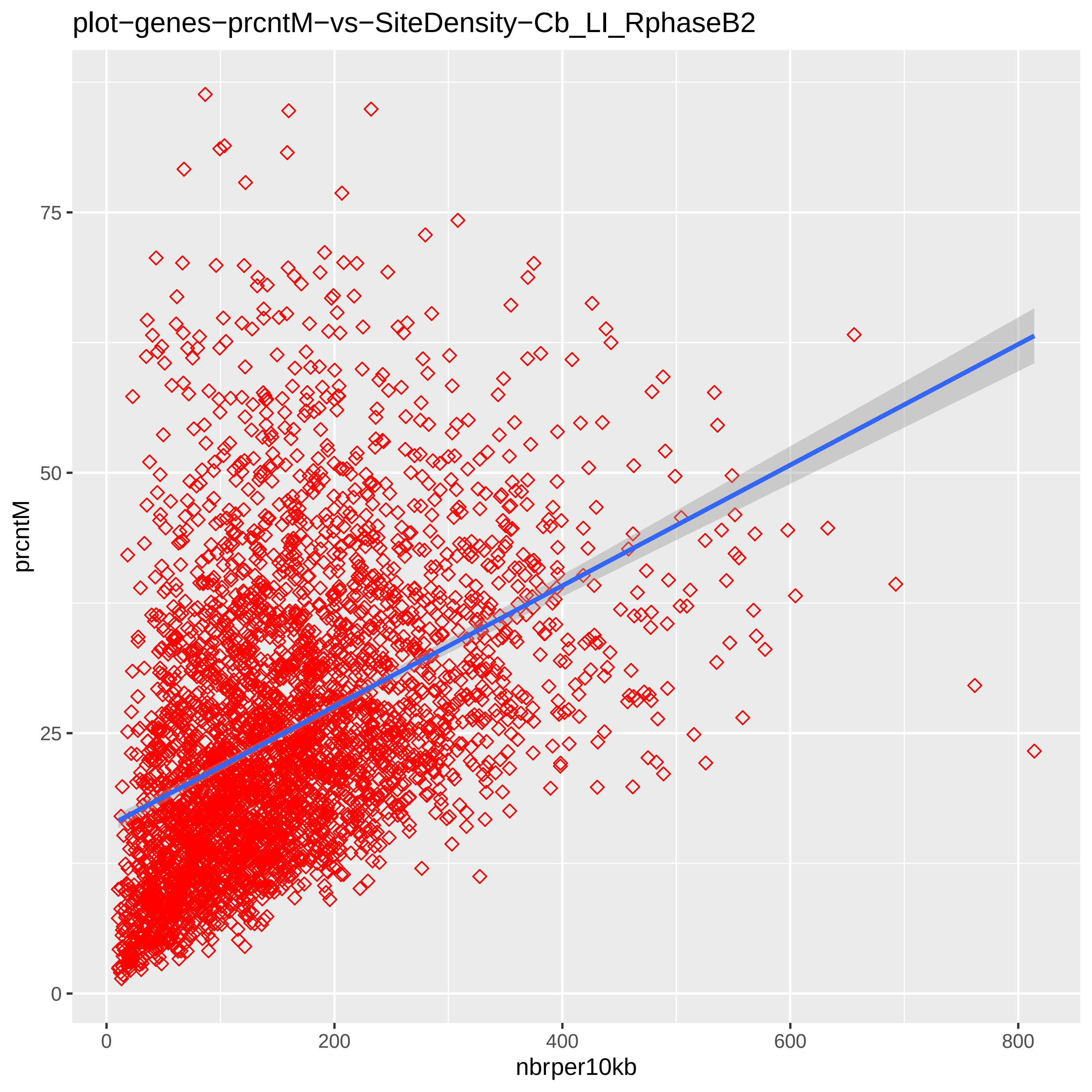
plot−genes−prcntM−vs−SiteDensity−Cb_LI_RphaseB2
prcntM
0
25
50
75
nbrper10kb
0
200
400
600
800

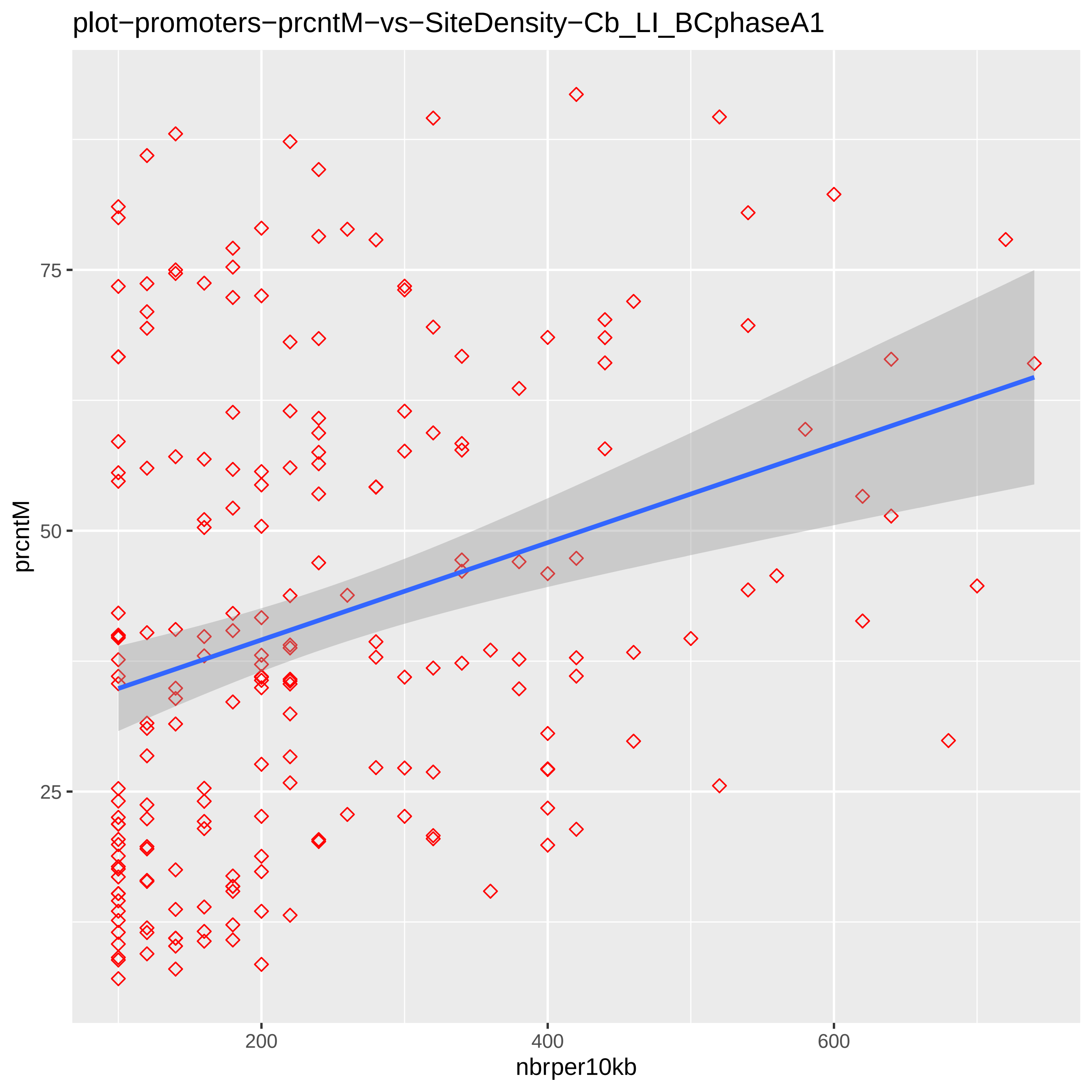
plot−promoters−prcntM−vs−SiteDensity−Cb_LI_BCphaseA1
prcntM
75
50
25
200
400
600
nbrper10kb

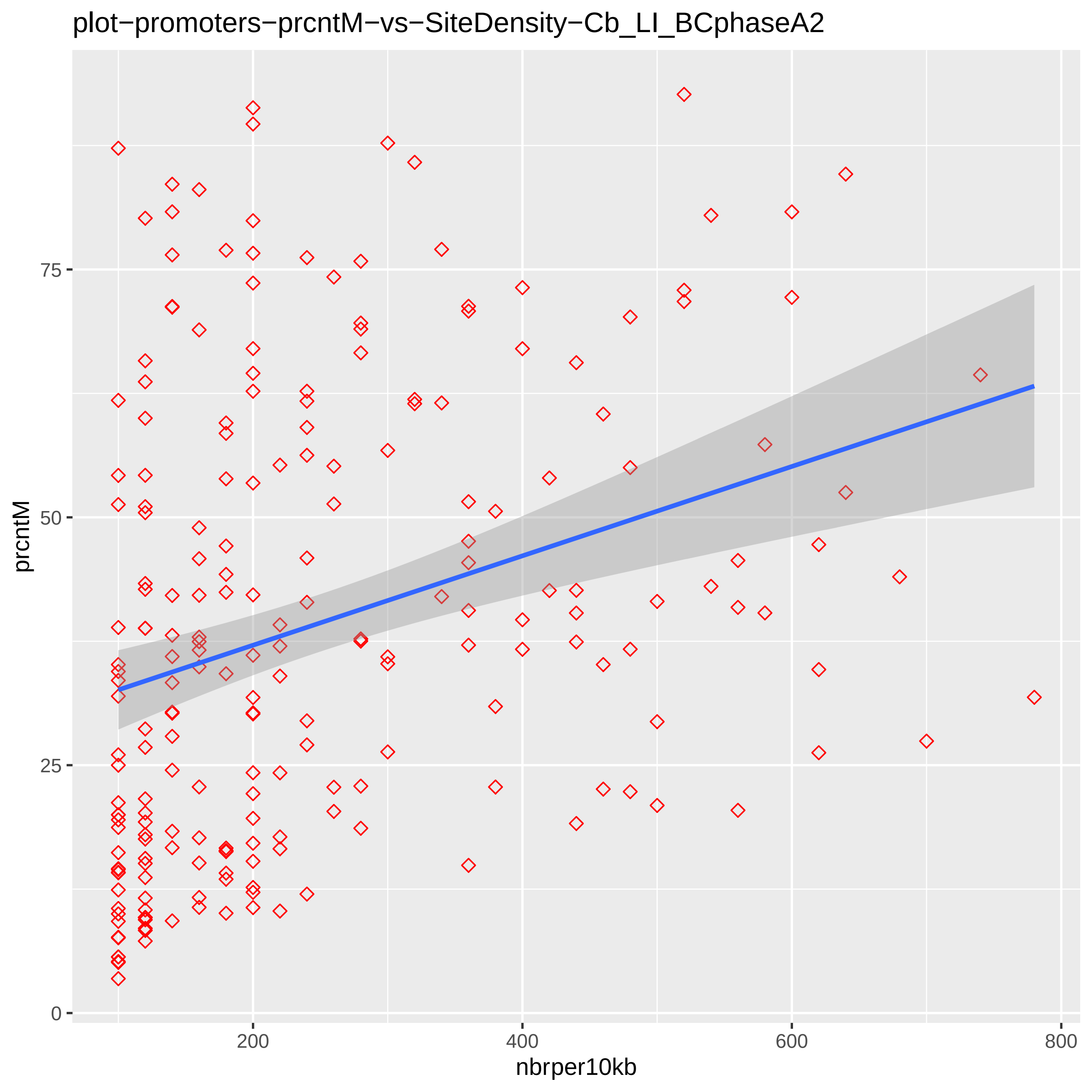

plot−promoters−prcntM−vs−SiteDensity−Cb_LI_BCphaseA2
prcntM
nbrper10kb
0
25
50
75
200
400
600
800

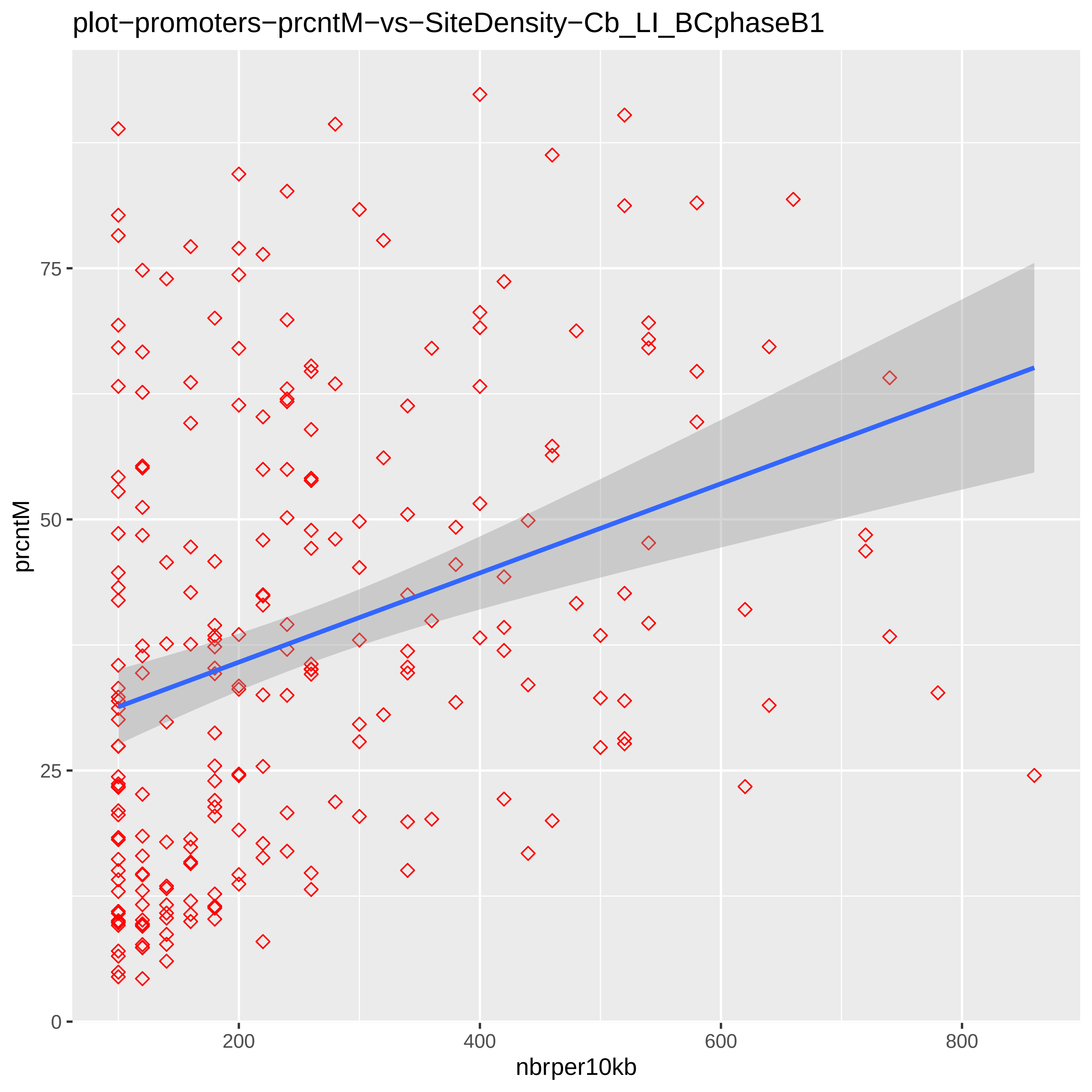
plot−promoters−prcntM−vs−SiteDensity−Cb_LI_BCphaseB1
prcntM
75
50
25
0
200
400
600
800
nbrper10kb

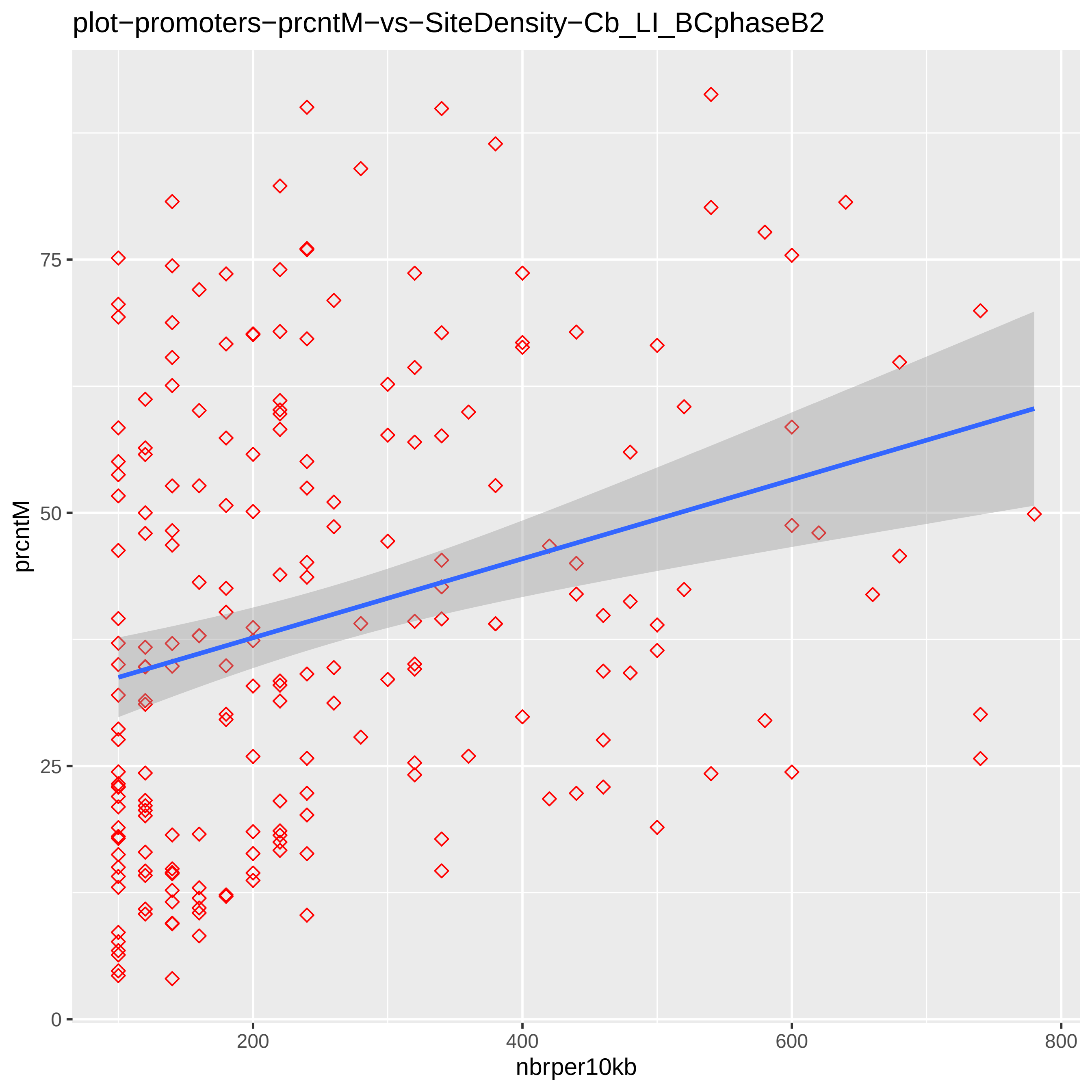
plot−promoters−prcntM−vs−SiteDensity−Cb_LI_BCphaseB2
prcntM
0
25
50
75
200
400
600
800
nbrper10kb

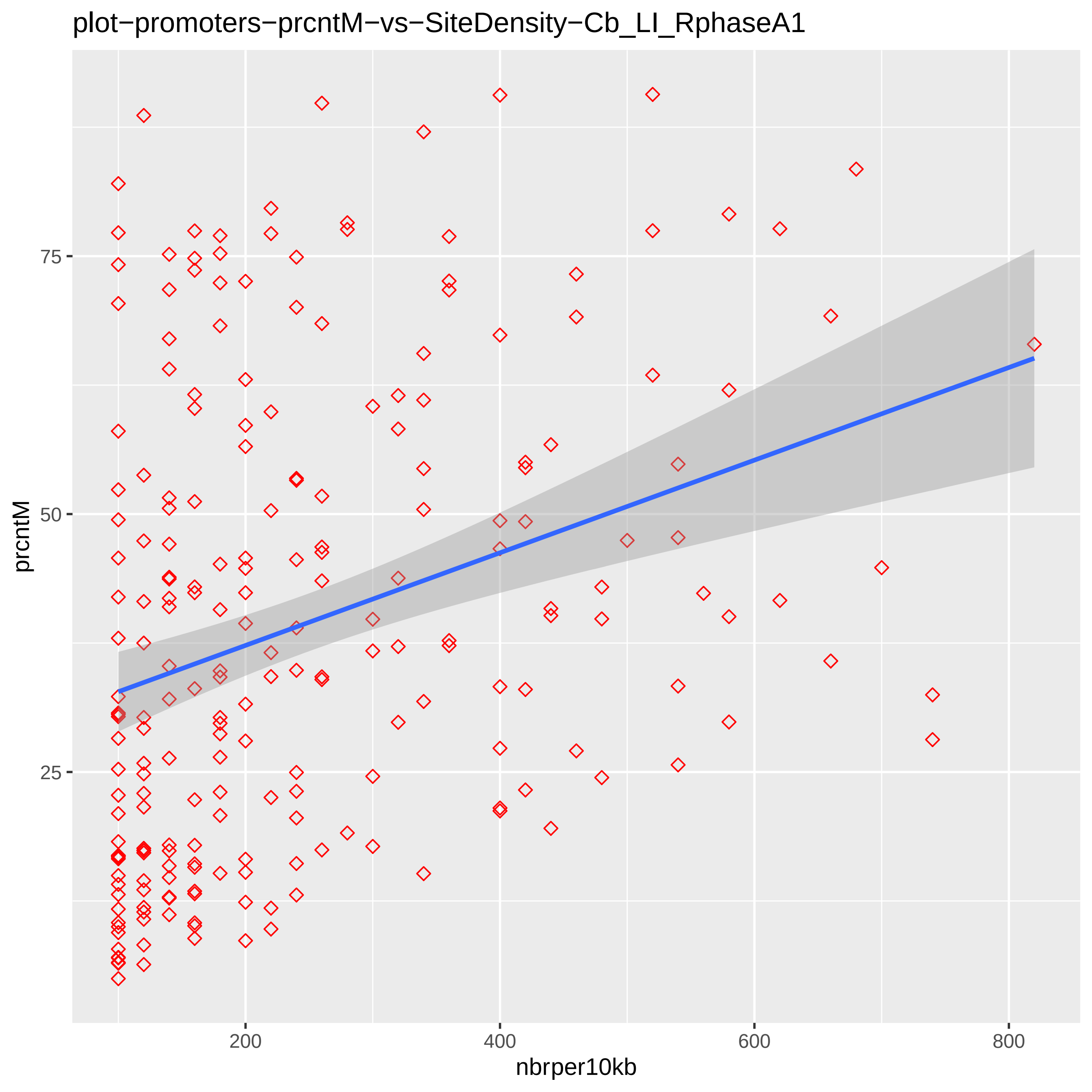
plot−promoters−prcntM−vs−SiteDensity−Cb_LI_RphaseA1
prcntM
75
50
25
200
400
600
800
nbrper10kb

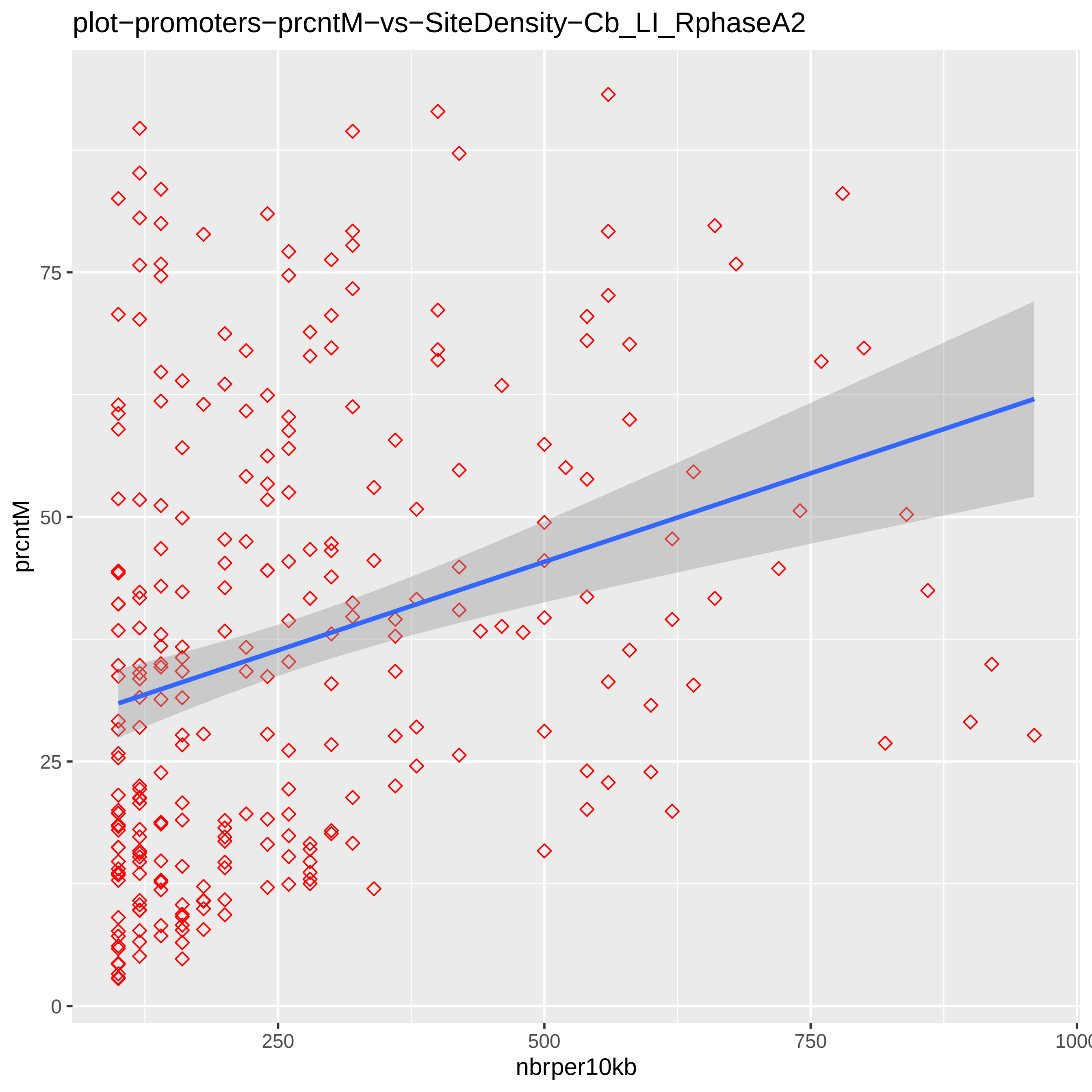
plot−promoters−prcntM−vs−SiteDensity−Cb_LI_RphaseA2
prcntM
nbrper10kb
0
25
50
75
250
500
750
1000

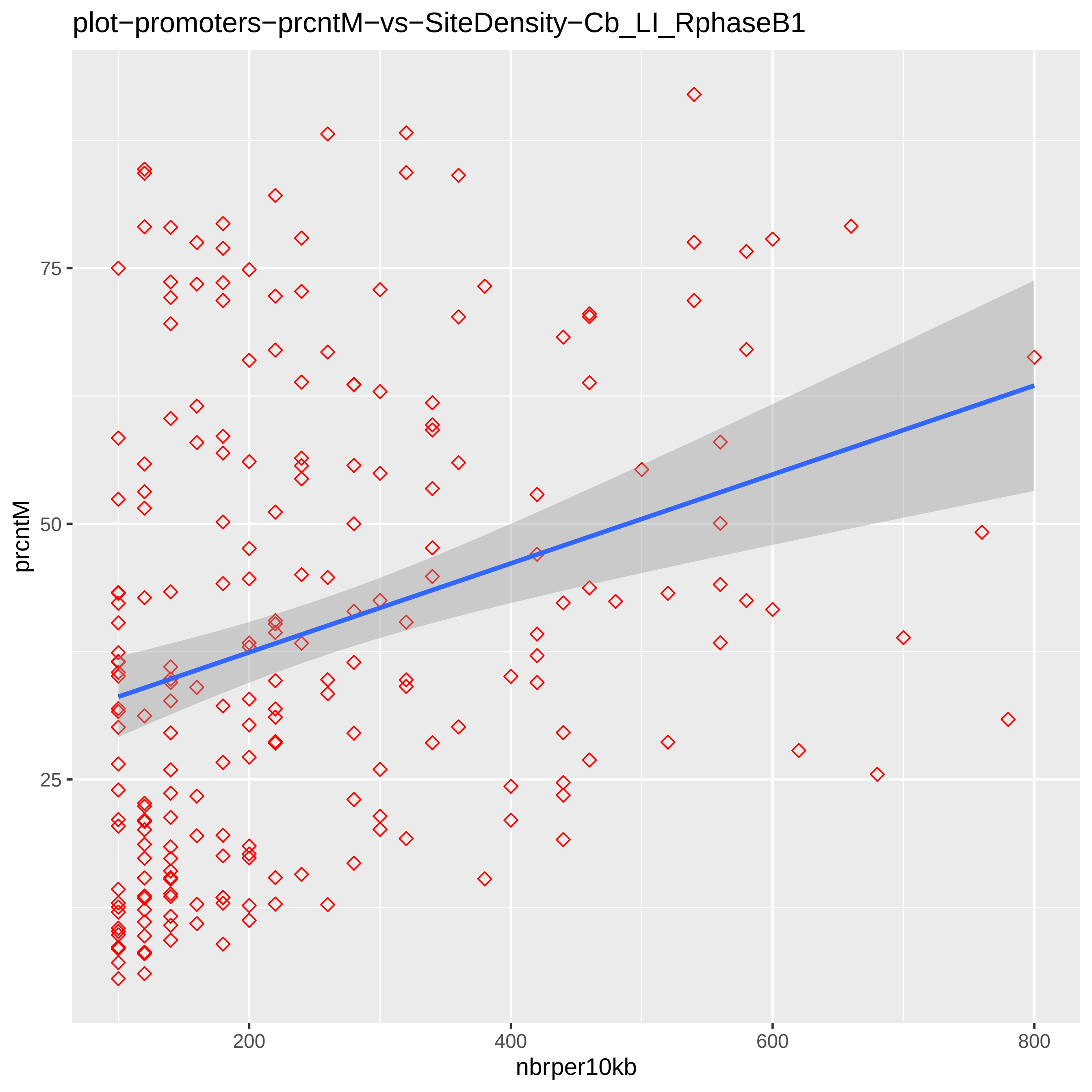

plot−promoters−prcntM−vs−SiteDensity−Cb_LI_RphaseB1
prcntM
75
50
25
200
400
600
800
nbrper10kb

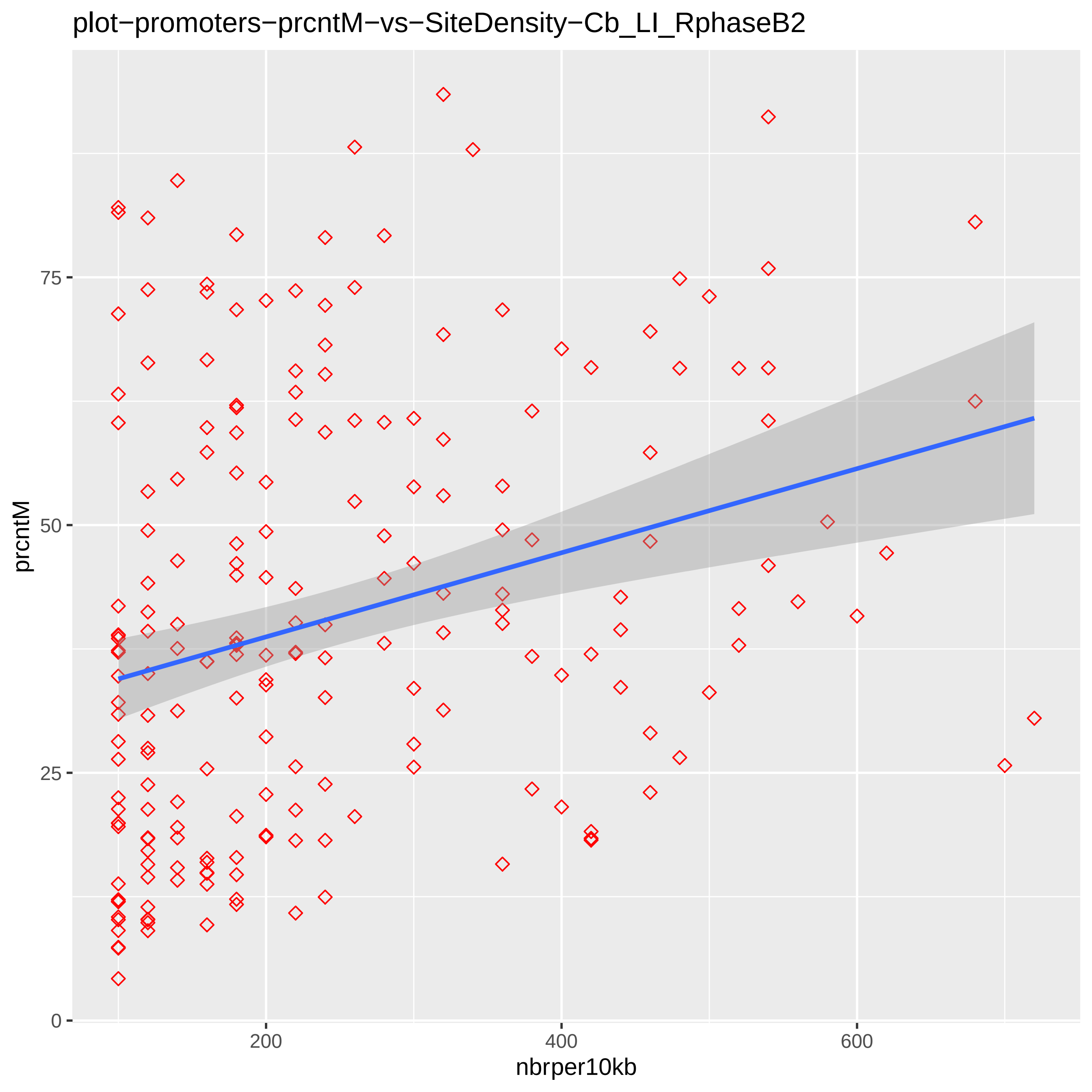
plot−promoters−prcntM−vs−SiteDensity−Cb_LI_RphaseB2
75
50
25
0
prcntM
200
400
600
nbrper10kb

```
Directory: MRPR File: 0READMEmrpr

MRPR - Methylation-rich and -poor regions.

Input: studymk, genome_ann (a list of GRanges objects providing annotated region labels and bounds;
                           output of BWASPR::get_genome_annotation())
       nbrxtrms (specifies the number of richest and poorest regions to be displayed; default: 100)

Output: files dst-*.txt 1ds-*.pdf 5ds-*.pdf    mdr-*.bed   rmp-* gwp-* gwr-*

Notes: The output is generated by BWASPR functions det_mrpr() and map_mrpr().
       Methylation-rich and -poor regions are determined based on the spacing between
        neighboring CpGhsm sites.  If sites occur at positions a, b, c, d, e, and f
        (and not in between), then b-a, c-b, ... are 1-distances, and f-a is a 5-distance.
        The file dst-*.txt shows the empirical distribution of d-distances for the sample.
        Methylation-poor regions are determined as long 1-distances (in the top <nbrxtrms>; merged
        if adjacent 1-distances are both in the top <nbrxtrms>).  Methylation-rich regions are determined
        as short 5-distances (in the low <nbrxtrms>; merged if adjacent 5-distances are both in the low
        <nbrxtrms>).  Parts of the distribution are plotted in the 1ds-*.pdf and 5ds-*.pdf files.

       The methylation-rich and -poor regions are listed in files dst-*.txt and mdr-*.bed,
        the latter in BED format for potential display in genome browsers.

       The overlap of methylation regions with genome features is summarized in files rmp-*.txt.
        Files gwr-*.txt show genes overlapping with methylation-rich regions, ordered by site
        density in the methylation-rich region.
        Files gwp-*.txt show genes overlapping with methylation-poor regions, ordered by site
        density in the methylation-poor region.
```

Directory: MRPR File: dst-Cb_LI_BCphaseA1.txt

Analysis of 1-distances for sample "Cb_LI_BCphaseA1" (198408 sites):

Total number of 1-distances: 197942
Median: 11.0 Mean: 828.3 Std: 10304.0
Quantiles:

| 0% | 5% | 10% | 15% | 20% | 25% | 30% | 35% | 40% | 45% | 50% | 55% | 60% | 65% | 70% | 75% | 80% | 85% | 90% | 95% | 100% |
|---|---|---|---|---|---|---|---|---|---|---|---|---|---|---|---|---|---|---|---|---|
| 1 | 1 | 1 | 1 | 1 | 1 | 1 | 3 | 6 | 8 | 11 | 14 | 18 | 22 | 28 | 35 | 46 | 67 | 113 | 305 | 554497 |

Low density region distance cutoff (lowest 100): 228717
High density region distance cutoff (highest 100): 1

Ordered lists of methylation-poor regions:

Top 100 (out of 96) methylation-poor regions in "Cb_LI_BCphaseA1":
(Sdnsty = sites per 1kb)

| | Rtype | Sample | SeqID | From | To | Rlgth | NbrSites | Sdnsty |
|---|---|---|---|---|---|---|---|---|
| 1 | Poor | BCphaseA1 | NW_011924956.1 | 107369 | 661866 | 554498 | 2 | 0.00361 |
| 2 | Poor | BCphaseA1 | NW_011925216.1 | 184346 | 723298 | 538953 | 2 | 0.00371 |
| 3 | Poor | BCphaseA1 | NW_011924931.1 | 535781 | 1038333 | 502553 | 2 | 0.00398 |
| 4 | Poor | BCphaseA1 | NW_011925438.1 | 816165 | 1308645 | 492481 | 2 | 0.00406 |
| 5 | Poor | BCphaseA1 | NW_011925020.1 | 1500346 | 1984956 | 484611 | 2 | 0.00413 |
| 6 | Poor | BCphaseA1 | NW_011925012.1 | 710605 | 1186542 | 475938 | 2 | 0.00420 |

...
... only first 25 lines shown ...

Directory: MRPR File: dst-Cb_LI_BCphaseA2.txt

Analysis of 1-distances for sample "Cb_LI_BCphaseA2" (213448 sites):

Total number of 1-distances: 213040
Median: 10.0 Mean: 736.4 Std: 11292.2
Quantiles:

| 0% | 5% | 10% | 15% | 20% | 25% | 30% | 35% | 40% | 45% | 50% | 55% | 60% | 65% | 70% | 75% | 80% | 85% | 90% | 95% | 100% |
|---|---|---|---|---|---|---|---|---|---|---|---|---|---|---|---|---|---|---|---|---|
| 1 | 1 | 1 | 1 | 1 | 1 | 1 | 3 | 5 | 8 | 10 | 13 | 17 | 21 | 26 | 33 | 42 | 60 | 100 | 246 | 899037 |

Low density region distance cutoff (lowest 100): 258414
High density region distance cutoff (highest 100): 1

Ordered lists of methylation-poor regions:

Top 100 (out of 97) methylation-poor regions in "Cb_LI_BCphaseA2":
(Sdnsty = sites per 1kb)

| | Rtype | Sample | SeqID | From | To | Rlgth | NbrSites | Sdnsty |
|---|---|---|---|---|---|---|---|---|
| 1 | Poor | BCphaseA2 | NW_011924892.1 | 1201246 | 2100283 | 899038 | 2 | 0.00222 |
| 2 | Poor | BCphaseA2 | NW_011924892.1 | 284899 | 1097469 | 812571 | 2 | 0.00246 |
| 3 | Poor | BCphaseA2 | NW_011925216.1 | 389005 | 1092286 | 703282 | 2 | 0.00284 |
| 4 | Poor | BCphaseA2 | NW_011925036.1 | 463239 | 1155855 | 692617 | 2 | 0.00289 |
| 5 | Poor | BCphaseA2 | NW_011925438.1 | 651775 | 1308432 | 656658 | 2 | 0.00305 |
| 6 | Poor | BCphaseA2 | NW_011925387.1 | 685943 | 1328713 | 642771 | 2 | 0.00311 |

...
... only first 25 lines shown ...

Directory: MRPR File: dst-Cb_LI_BCphaseB1.txt

Analysis of 1-distances for sample "Cb_LI_BCphaseB1" (226815 sites):

Total number of 1-distances: 226290
Median: 10.0 Mean: 742.8 Std: 8906.9
Quantiles:

| 0% | 5% | 10% | 15% | 20% | 25% | 30% | 35% | 40% | 45% | 50% | 55% | 60% | 65% | 70% | 75% | 80% | 85% | 90% | 95% | 100% |
|---|---|---|---|---|---|---|---|---|---|---|---|---|---|---|---|---|---|---|---|---|
| 1 | 1 | 1 | 1 | 1 | 1 | 1 | 2 | 5 | 7 | 10 | 13 | 17 | 21 | 26 | 34 | 44 | 64 | 108 | 287 | 515472 |

Low density region distance cutoff (lowest 100): 199442
High density region distance cutoff (highest 100): 1

Ordered lists of methylation-poor regions:

Top 100 (out of 98) methylation-poor regions in "Cb_LI_BCphaseB1":
(Sdnsty = sites per 1kb)

| | Rtype | Sample | SeqID | From | To | Rlgth | NbrSites | Sdnsty |
|---|---|---|---|---|---|---|---|---|
| 1 | Poor | BCphaseB1 | NW_011924931.1 | 535781 | 1051253 | 515473 | 2 | 0.00388 |
| 2 | Poor | BCphaseB1 | NW_011924888.1 | 408597 | 897056 | 488460 | 2 | 0.00409 |
| 3 | Poor | BCphaseB1 | NW_011925053.1 | 681984 | 1153435 | 471452 | 2 | 0.00424 |
| 4 | Poor | BCphaseB1 | NW_011925443.1 | 308926 | 765268 | 456343 | 2 | 0.00438 |
| 5 | Poor | BCphaseB1 | NW_011925036.1 | 590572 | 1041090 | 450519 | 2 | 0.00444 |
| 6 | Poor | BCphaseB1 | NW_011925106.1 | 2601151 | 3047459 | 446309 | 2 | 0.00448 |

...
... only first 25 lines shown ...

```
Directory: MRPR File: dst-Cb_LI_BCphaseB2.txt

Analysis of  1-distances for sample "Cb_LI_BCphaseB2" (212569 sites):

  Total number of  1-distances:     212126
  Median:    10.0 Mean:   774.5 Std:  9483.3
  Quantiles:
    0%     5%    10%    15%    20%    25%    30%    35%    40%    45%    50%    55%    60%    65%    70%    75%    80%    85%    90%    95%   100%
     1      1      1      1      1      1      1      3      5      8     10     13     17     21     26     33     44     63    106    282 645758

  Low  density region distance cutoff (lowest  100):  204232
  High density region distance cutoff (highest 100):       1

Ordered lists of methylation-poor regions:

 Top 100 (out of    97) methylation-poor regions in "Cb_LI_BCphaseB2":
 (Sdnsty = sites per 1kb)

   Rtype    Sample          SeqID    From      To  Rlgth NbrSites  Sdnsty
1   Poor BCphaseB2 NW_011924892.1  284893  930651 645759        2 0.00310
2   Poor BCphaseB2 NW_011925082.1   53097  666977 613881        2 0.00326
3   Poor BCphaseB2 NW_011924937.1  528587 1038579 509993        2 0.00392
4   Poor BCphaseB2 NW_011925438.1  713751 1203971 490221        2 0.00408
5   Poor BCphaseB2 NW_011924899.1  885423 1375424 490002        2 0.00408
6   Poor BCphaseB2 NW_011925020.1 1690022 2150743 460722        2 0.00434
...
... only first 25 lines shown ...
```

Directory: MRPR File: dst-Cb_LI_RphaseA1.txt

Analysis of 1-distances for sample "Cb_LI_RphaseA1" (213610 sites):

Total number of 1-distances: 213171
Median: 10.0 Mean: 734.1 Std: 10922.5
Quantiles:

| 0% | 5% | 10% | 15% | 20% | 25% | 30% | 35% | 40% | 45% | 50% | 55% | 60% | 65% | 70% | 75% | 80% | 85% | 90% | 95% | 100% |
|---|---|---|---|---|---|---|---|---|---|---|---|---|---|---|---|---|---|---|---|---|
| 1 | 1 | 1 | 1 | 1 | 1 | 1 | 2 | 5 | 7 | 10 | 13 | 17 | 21 | 26 | 34 | 44 | 63 | 106 | 259 | 756940 |

Low density region distance cutoff (lowest 100): 253535
High density region distance cutoff (highest 100): 1

Ordered lists of methylation-poor regions:

Top 100 (out of 97) methylation-poor regions in "Cb_LI_RphaseA1":
(Sdnsty = sites per 1kb)

| | Rtype | Sample | SeqID | From | To | Rlgth | NbrSites | Sdnsty |
|---|---|---|---|---|---|---|---|---|
| 1 | Poor | RphaseA1 | NW_011925315.1 | 741815 | 1498755 | 756941 | 2 | 0.00264 |
| 2 | Poor | RphaseA1 | NW_011925036.1 | 463225 | 1155855 | 692631 | 2 | 0.00289 |
| 3 | Poor | RphaseA1 | NW_011924987.1 | 3659357 | 4326160 | 666804 | 2 | 0.00300 |
| 4 | Poor | RphaseA1 | NW_011924945.1 | 324927 | 958341 | 633415 | 2 | 0.00316 |
| 5 | Poor | RphaseA1 | NW_011924956.1 | 36360 | 661866 | 625507 | 2 | 0.00320 |
| 6 | Poor | RphaseA1 | NW_011925387.1 | 685943 | 1275933 | 589991 | 2 | 0.00339 |

...
... only first 25 lines shown ...

Directory: MRPR File: dst-Cb_LI_RphaseA2.txt

Analysis of 1-distances for sample "Cb_LI_RphaseA2" (265952 sites):

Total number of 1-distances: 265356
Median: 9.0 Mean: 671.5 Std: 5821.6
Quantiles:

| 0% | 5% | 10% | 15% | 20% | 25% | 30% | 35% | 40% | 45% | 50% | 55% | 60% | 65% | 70% | 75% | 80% | 85% | 90% | 95% | 100% |
|---|---|---|---|---|---|---|---|---|---|---|---|---|---|---|---|---|---|---|---|---|
| 1 | 1 | 1 | 1 | 1 | 1 | 1 | 2 | 4 | 7 | 9 | 13 | 16 | 20 | 26 | 33 | 43 | 63 | 113 | 388 | 320202 |

Low density region distance cutoff (lowest 100): 122021
High density region distance cutoff (highest 100): 1

Ordered lists of methylation-poor regions:

Top 100 (out of 99) methylation-poor regions in "Cb_LI_RphaseA2":
(Sdnsty = sites per 1kb)

| | Rtype | Sample | SeqID | From | To | Rlgth | NbrSites | Sdnsty |
|---|---|---|---|---|---|---|---|---|
| 1 | Poor | RphaseA2 | NW_011924996.1 | 1277657 | 1597859 | 320203 | 2 | 0.00625 |
| 2 | Poor | RphaseA2 | NW_011925007.1 | 96864 | 381604 | 284741 | 2 | 0.00702 |
| 3 | Poor | RphaseA2 | NW_011925300.1 | 134597 | 401637 | 267041 | 2 | 0.00749 |
| 4 | Poor | RphaseA2 | NW_011925099.1 | 1006421 | 1228934 | 222514 | 2 | 0.00899 |
| 5 | Poor | RphaseA2 | NW_011925027.1 | 689188 | 897305 | 208118 | 2 | 0.00961 |
| 6 | Poor | RphaseA2 | NW_011925059.1 | 361653 | 563982 | 202330 | 2 | 0.00988 |

...
... only first 25 lines shown ...

Directory: MRPR File: dst-Cb_LI_RphaseB1.txt

Analysis of 1-distances for sample "Cb_LI_RphaseB1" (218593 sites):

Total number of 1-distances: 218089
Median: 10.0 Mean: 765.0 Std: 8675.1
Quantiles:

| 0% | 5% | 10% | 15% | 20% | 25% | 30% | 35% | 40% | 45% | 50% | 55% | 60% | 65% | 70% | 75% | 80% | 85% | 90% | 95% | 100% |
|---|---|---|---|---|---|---|---|---|---|---|---|---|---|---|---|---|---|---|---|---|
| 1 | 1 | 1 | 1 | 1 | 1 | 1 | 3 | 5 | 8 | 10 | 13 | 17 | 21 | 27 | 34 | 45 | 65 | 111 | 300 | 393633 |

Low density region distance cutoff (lowest 100): 189238
High density region distance cutoff (highest 100): 1

Ordered lists of methylation-poor regions:

Top 100 (out of 97) methylation-poor regions in "Cb_LI_RphaseB1":
(Sdnsty = sites per 1kb)

| | Rtype | Sample | SeqID | From | To | Rlgth | NbrSites | Sdnsty |
|---|---|---|---|---|---|---|---|---|
| 1 | Poor | RphaseB1 | NW_011925387.1 | 685943 | 1328666 | 642724 | 3 | 0.00467 |
| 2 | Poor | RphaseB1 | NW_011924987.1 | 3688298 | 4326160 | 637863 | 3 | 0.00470 |
| 3 | Poor | RphaseB1 | NW_011925367.1 | 152381 | 546014 | 393634 | 2 | 0.00508 |
| 4 | Poor | RphaseB1 | NW_011924888.1 | 408597 | 992521 | 583925 | 3 | 0.00514 |
| 5 | Poor | RphaseB1 | NW_011925241.1 | 534433 | 910312 | 375880 | 2 | 0.00532 |
| 6 | Poor | RphaseB1 | NW_011925053.1 | 658977 | 1034593 | 375617 | 2 | 0.00532 |

...
... only first 25 lines shown ...

Directory: MRPR File: dst-Cb_LI_RphaseB2.txt

Analysis of 1-distances for sample "Cb_LI_RphaseB2" (200707 sites):

Total number of 1-distances: 200274
Median: 11.0 Mean: 798.6 Std: 10646.0
Quantiles:

| 0% | 5% | 10% | 15% | 20% | 25% | 30% | 35% | 40% | 45% | 50% | 55% | 60% | 65% | 70% | 75% | 80% | 85% | 90% | 95% | 100% |
|---|---|---|---|---|---|---|---|---|---|---|---|---|---|---|---|---|---|---|---|---|
| 1 | 1 | 1 | 1 | 1 | 1 | 1 | 3 | 6 | 8 | 11 | 14 | 17 | 22 | 27 | 34 | 45 | 64 | 107 | 274 | 876231 |

Low density region distance cutoff (lowest 100): 227715
High density region distance cutoff (highest 100): 1

Ordered lists of methylation-poor regions:

Top 100 (out of 97) methylation-poor regions in "Cb_LI_RphaseB2":
(Sdnsty = sites per 1kb)

| | Rtype | Sample | SeqID | From | To | Rlgth | NbrSites | Sdnsty |
|---|---|---|---|---|---|---|---|---|
| 1 | Poor | RphaseB2 | NW_011925216.1 | 216055 | 1092286 | 876232 | 2 | 0.00228 |
| 2 | Poor | RphaseB2 | NW_011925036.1 | 566786 | 1156139 | 589354 | 2 | 0.00339 |
| 3 | Poor | RphaseB2 | NW_011925020.1 | 1500346 | 2352286 | 851941 | 3 | 0.00352 |
| 4 | Poor | RphaseB2 | NW_011924975.1 | 94387 | 654441 | 560055 | 2 | 0.00357 |
| 5 | Poor | RphaseB2 | NW_011924931.1 | 2329425 | 2877401 | 547977 | 2 | 0.00365 |
| 6 | Poor | RphaseB2 | NW_011925082.1 | 53097 | 600439 | 547343 | 2 | 0.00365 |

...
... only first 25 lines shown ...

Directory: MRPR File: gwp-Cb_LI_BCphaseA1.txt

```
seqnames	start	end	width	strand	source	type	score	phase	ID	Dbxref	Name	gbkey	gene	gene_biotype	partial	start_range	end_range	seqnames	start
	end	width	strand	NbrSites	Sdnsty
NW_011924956.1	107962	344153	236192	-	Gnomon	gene	NA	NA	gene2555	GeneID:105275019	LOC105275019	Gene	LOC105275019	protein_coding	NA
	character(0)	character(0)	NW_011924956.1	107369	661866	554498	*	2	0.00360686603017504
NW_011924956.1	140571	150861	10291	+	Gnomon	gene	NA	NA	gene2556	GeneID:105275018	LOC105275018	Gene	LOC105275018	lncRNA	NA
	character(0)	character(0)	NW_011924956.1	107369	661866	554498	*	2	0.00360686603017504
NW_011924956.1	415268	419160	3893	+	Gnomon	gene	NA	NA	gene2557	GeneID:105275021	LOC105275021	Gene	LOC105275021	protein_coding	NA
	character(0)	character(0)	NW_011924956.1	107369	661866	554498	*	2	0.00360686603017504
NW_011924956.1	428078	657058	228981	-	Gnomon	gene	NA	NA	gene2558	GeneID:105275023	LOC105275023	Gene	LOC105275023	protein_coding	NA
	character(0)	character(0)	NW_011924956.1	107369	661866	554498	*	2	0.00360686603017504
NW_011924956.1	661360	663256	1897	+	Gnomon	gene	NA	NA	gene2559	GeneID:105275030	LOC105275030	Gene	LOC105275030	protein_coding	NA
	character(0)	character(0)	NW_011924956.1	107369	661866	554498	*	2	0.00360686603017504
NW_011925216.1	184882	185741	860	-	Gnomon	gene	NA	NA	gene8749	GeneID:105282158	LOC105282158	Gene	LOC105282158	lncRNA	NA
	character(0)	character(0)	NW_011925216.1	184346	723298	538953	*	2	0.00371089872400747
NW_011925216.1	190952	205333	14382	-	Gnomon	gene	NA	NA	gene8750	GeneID:105282160	LOC105282160	Gene	LOC105282160	protein_coding	NA
	character(0)	character(0)	NW_011925216.1	184346	723298	538953	*	2	0.00371089872400747
NW_011925216.1	219999	246618	26620	-	Gnomon	gene	NA	NA	gene8751	GeneID:105282161	LOC105282161	Gene	LOC105282161	protein_coding	NA
	character(0)	character(0)	NW_011925216.1	184346	723298	538953	*	2	0.00371089872400747
NW_011925216.1	277106	281176	4071	-	Gnomon	gene	NA	NA	gene8752	GeneID:105282162	LOC105282162	Gene	LOC105282162	protein_coding	NA
	character(0)	character(0)	NW_011925216.1	184346	723298	538953	*	2	0.00371089872400747
NW_011925216.1	290094	291816	1723	+	Gnomon	gene	NA	NA	gene8753	GeneID:105282163	LOC105282163	Gene	LOC105282163	lncRNA	NA
	character(0)	character(0)	NW_011925216.1	184346	723298	538953	*	2	0.00371089872400747
NW_011925216.1	313949	321221	7273	-	Gnomon	gene	NA	NA	gene8754	GeneID:105282165	LOC105282165	Gene	LOC105282165	protein_coding	NA
	character(0)	character(0)	NW_011925216.1	184346	723298	538953	*	2	0.00371089872400747
NW_011925216.1	337863	347845	9983	-	Gnomon	gene	NA	NA	gene8755	GeneID:105282166	LOC105282166	Gene	LOC105282166	lncRNA	NA
	character(0)	character(0)	NW_011925216.1	184346	723298	538953	*	2	0.00371089872400747
NW_011925216.1	367045	368882	1838	-	Gnomon	gene	NA	NA	gene8756	GeneID:105282192	LOC105282192	Gene	LOC105282192	protein_coding	NA
	character(0)	character(0)	NW_011925216.1	184346	723298	538953	*	2	0.00371089872400747
NW_011925216.1	382786	404605	21820	-	Gnomon	gene	NA	NA	gene8757	GeneID:105282169	LOC105282169	Gene	LOC105282169	protein_coding	NA
	character(0)	character(0)	NW_011925216.1	184346	723298	538953	*	2	0.00371089872400747
NW_011925216.1	406528	409862	3335	-	Gnomon	gene	NA	NA	gene8758	GeneID:105282168	LOC105282168	Gene	LOC105282168	protein_coding	NA
	character(0)	character(0)	NW_011925216.1	184346	723298	538953	*	2	0.00371089872400747
NW_011925216.1	411413	433190	21778	+	Gnomon	gene	NA	NA	gene8759	GeneID:105282167	LOC105282167	Gene	LOC105282167	lncRNA	NA
	character(0)	character(0)	NW_011925216.1	184346	723298	538953	*	2	0.00371089872400747
NW_011925216.1	446916	480871	33956	-	Gnomon	gene	NA	NA	gene8760	GeneID:109611151	LOC109611151	Gene	LOC109611151	lncRNA	NA
	character(0)	character(0)	NW_011925216.1	184346	723298	538953	*	2	0.00371089872400747
NW_011925216.1	496030	584409	88380	-	Gnomon	gene	NA	NA	gene8761	GeneID:105282171	LOC105282171	Gene	LOC105282171	protein_coding	NA
	character(0)	character(0)	NW_011925216.1	184346	723298	538953	*	2	0.00371089872400747
NW_011925216.1	611697	703889	92193	-	Gnomon	gene	NA	NA	gene8762	GeneID:105282173	LOC105282173	Gene	LOC105282173	protein_coding	NA
	character(0)	character(0)	NW_011925216.1	184346	723298	538953	*	2	0.00371089872400747
NW_011924931.1	534549	536012	1464	+	Gnomon	gene	NA	NA	gene1415	GeneID:105287536	LOC105287536	Gene	LOC105287536	protein_coding	NA
	character(0)	character(0)	NW_011924931.1	535781	1038333	502553	*	2	0.00397967975517 01
NW_011924931.1	536217	855035	318819	-	Gnomon	gene	NA	NA	gene1416	GeneID:105287539	LOC105287539	Gene	LOC105287539	protein_coding	NA
	character(0)	character(0)	NW_011924931.1	535781	1038333	502553	*	2	0.0039796797551701
NW_011924931.1	1014634	1019043	4410	-	Gnomon	gene	NA	NA	gene1417	GeneID:105287538	LOC105287538	Gene	LOC105287538	protein_coding	NA
	character(0)	character(0)	NW_011924931.1	535781	1038333	502553	*	2	0.0039796797551701
NW_011925438.1	752792	844175	91384	-	Gnomon	gene	NA	NA	gene11416	GeneID:105285218	LOC105285218	Gene	LOC105285218	protein_coding	NA
	character(0)	character(0)	NW_011925438.1	816165	1308645	492481	*	2	0.00406107037631909
NW_011925438.1	864395	874202	9808	+	Gnomon	gene	NA	NA	gene11417	GeneID:105285217	LOC105285217	Gene	LOC105285217	protein_coding	NA
	character(0)	character(0)	NW_011925438.1	816165	1308645	492481	*	2	0.00406107037631909
...
... only first 25 lines shown ...
```

Directory: MRPR File: gwp-Cb_LI_BCphaseA2.txt

| seqnames | start | end | width | strand | source | type | score | phase | ID | Dbxref | Name | gbkey | gene | gene_biotype | partial | start_range | end_range | seqnames | start | end | width | strand | NbrSites | Sdnsty |
|---|---|---|---|---|---|---|---|---|---|---|---|---|---|---|---|---|---|---|---|---|---|---|---|---|
| NW_011924892.1 | 1206093 | 1223652 | 17560 | - | Gnomon | gene | NA | NA | gene574 | GeneID:105278577 | LOC105278577 | Gene | LOC105278577 | protein_coding | NA | character(0) | character(0) | NW_011924892.1 | 1201246 | 2100283 | 899038 | * | 2 | 0.00222460007252196 |
| NW_011924892.1 | 1369781 | 1567211 | 197431 | + | Gnomon | gene | NA | NA | gene575 | GeneID:105278587 | LOC105278587 | Gene | LOC105278587 | protein_coding | NA | character(0) | character(0) | NW_011924892.1 | 1201246 | 2100283 | 899038 | * | 2 | 0.00222460007252196 |
| NW_011924892.1 | 1679442 | 1682114 | 2673 | - | Gnomon | gene | NA | NA | gene576 | GeneID:105278916 | LOC105278916 | Gene | LOC105278916 | protein_coding | NA | character(0) | character(0) | NW_011924892.1 | 1201246 | 2100283 | 899038 | * | 2 | 0.00222460007252196 |
| NW_011924892.1 | 1695292 | 1776980 | 81689 | + | Gnomon | gene | NA | NA | gene577 | GeneID:105278599 | LOC105278599 | Gene | LOC105278599 | protein_coding | NA | character(0) | character(0) | NW_011924892.1 | 1201246 | 2100283 | 899038 | * | 2 | 0.00222460007252196 |
| NW_011924892.1 | 1706057 | 1732187 | 26131 | - | Gnomon | gene | NA | NA | gene578 | GeneID:105278631 | LOC105278631 | Gene | LOC105278631 | protein_coding | NA | character(0) | character(0) | NW_011924892.1 | 1201246 | 2100283 | 899038 | * | 2 | 0.00222460007252196 |
| NW_011924892.1 | 1733989 | 1736080 | 2092 | + | Gnomon | gene | NA | NA | gene579 | GeneID:105278624 | LOC105278624 | Gene | LOC105278624 | protein_coding | NA | character(0) | character(0) | NW_011924892.1 | 1201246 | 2100283 | 899038 | * | 2 | 0.00222460007252196 |
| NW_011924892.1 | 1736532 | 1783302 | 46771 | - | Gnomon | gene | NA | NA | gene580 | GeneID:105278615 | LOC105278615 | Gene | LOC105278615 | protein_coding | NA | character(0) | character(0) | NW_011924892.1 | 1201246 | 2100283 | 899038 | * | 2 | 0.00222460007252196 |
| NW_011924892.1 | 1791449 | 1820341 | 28893 | + | Gnomon | gene | NA | NA | gene581 | GeneID:105278660 | LOC105278660 | Gene | LOC105278660 | protein_coding | NA | character(0) | character(0) | NW_011924892.1 | 1201246 | 2100283 | 899038 | * | 2 | 0.00222460007252196 |
| NW_011924892.1 | 1808558 | 1809484 | 927 | - | Gnomon | gene | NA | NA | gene582 | GeneID:105278679 | LOC105278679 | Gene | LOC105278679 | lncRNA | NA | character(0) | character(0) | NW_011924892.1 | 1201246 | 2100283 | 899038 | * | 2 | 0.00222460007252196 |
| NW_011924892.1 | 1819808 | 1868151 | 48344 | - | Gnomon | gene | NA | NA | gene583 | GeneID:105278653 | LOC105278653 | Gene | LOC105278653 | protein_coding | NA | character(0) | character(0) | NW_011924892.1 | 1201246 | 2100283 | 899038 | * | 2 | 0.00222460007252196 |
| NW_011924892.1 | 1933978 | 1947699 | 13722 | - | Gnomon | gene | NA | NA | gene584 | GeneID:105278693 | LOC105278693 | Gene | LOC105278693 | protein_coding | NA | character(0) | character(0) | NW_011924892.1 | 1201246 | 2100283 | 899038 | * | 2 | 0.00222460007252196 |
| NW_011924892.1 | 1949247 | 1951469 | 2223 | - | Gnomon | gene | NA | NA | gene585 | GeneID:105278705 | LOC105278705 | Gene | LOC105278705 | protein_coding | NA | character(0) | character(0) | NW_011924892.1 | 1201246 | 2100283 | 899038 | * | 2 | 0.00222460007252196 |
| NW_011924892.1 | 1951575 | 1954903 | 3329 | - | Gnomon | gene | NA | NA | gene586 | GeneID:105278699 | LOC105278699 | Gene | LOC105278699 | protein_coding | NA | character(0) | character(0) | NW_011924892.1 | 1201246 | 2100283 | 899038 | * | 2 | 0.00222460007252196 |
| NW_011924892.1 | 1955005 | 2048428 | 93424 | - | Gnomon | gene | NA | NA | gene587 | GeneID:105278716 | LOC105278716 | Gene | LOC105278716 | protein_coding | NA | character(0) | character(0) | NW_011924892.1 | 1201246 | 2100283 | 899038 | * | 2 | 0.00222460007252196 |
| NW_011924892.1 | 2067467 | 2077511 | 10045 | - | Gnomon | gene | NA | NA | gene588 | GeneID:105278925 | LOC105278925 | Gene | LOC105278925 | protein_coding | NA | character(0) | character(0) | NW_011924892.1 | 1201246 | 2100283 | 899038 | * | 2 | 0.00222460007252196 |
| NW_011924892.1 | 2074929 | 2079592 | 4664 | - | Gnomon | gene | NA | NA | gene589 | GeneID:105278751 | LOC105278751 | Gene | LOC105278751 | protein_coding | NA | character(0) | character(0) | NW_011924892.1 | 1201246 | 2100283 | 899038 | * | 2 | 0.00222460007252196 |
| NW_011924892.1 | 2081041 | 2099417 | 18377 | - | Gnomon | gene | NA | NA | gene590 | GeneID:105278759 | LOC105278759 | Gene | LOC105278759 | protein_coding | NA | character(0) | character(0) | NW_011924892.1 | 1201246 | 2100283 | 899038 | * | 2 | 0.00222460007252196 |
| NW_011924892.1 | 2099866 | 2105958 | 6093 | + | Gnomon | gene | NA | NA | gene591 | GeneID:105278782 | LOC105278782 | Gene | LOC105278782 | protein_coding | NA | character(0) | character(0) | NW_011924892.1 | 1201246 | 2100283 | 899038 | * | 2 | 0.00222460007252196 |
| NW_011924892.1 | 274786 | 285690 | 10905 | - | Gnomon | gene | NA | NA | gene544 | GeneID:105278288 | LOC105278288 | Gene | LOC105278288 | protein_coding | NA | character(0) | character(0) | NW_011924892.1 | 284899 | 1097469 | 812571 | * | 2 | 0.00246132337974159 |
| NW_011924892.1 | 302199 | 519970 | 217772 | + | Gnomon | gene | NA | NA | gene545 | GeneID:105278329 | LOC105278329 | Gene | LOC105278329 | protein_coding | NA | character(0) | character(0) | NW_011924892.1 | 284899 | 1097469 | 812571 | * | 2 | 0.00246132337974159 |
| NW_011924892.1 | 378175 | 382124 | 3950 | + | Gnomon | gene | NA | NA | gene546 | GeneID:105278325 | LOC105278325 | Gene | LOC105278325 | lncRNA | NA | character(0) | character(0) | NW_011924892.1 | 284899 | 1097469 | 812571 | * | 2 | 0.00246132337974159 |
| NW_011924892.1 | 446522 | 450804 | 4283 | - | Gnomon | gene | NA | NA | gene547 | GeneID:105278319 | LOC105278319 | Gene | LOC105278319 | protein_coding | NA | character(0) | character(0) | NW_011924892.1 | 284899 | 1097469 | 812571 | * | 2 | 0.00246132337974159 |
| NW_011924892.1 | 458698 | 471223 | 12526 | - | Gnomon | gene | NA | NA | gene548 | GeneID:105278305 | LOC105278305 | Gene | LOC105278305 | protein_coding | NA | character(0) | character(0) | NW_011924892.1 | 284899 | 1097469 | 812571 | * | 2 | 0.00246132337974159 |
| NW_011924892.1 | 604295 | 664845 | 60551 | + | Gnomon | gene | NA | NA | gene549 | GeneID:105278340 | LOC105278340 | Gene | LOC105278340 | protein_coding | NA | character(0) | character(0) | NW_011924892.1 | 284899 | 1097469 | 812571 | * | 2 | 0.00246132337974159 |

...

... only first 25 lines shown ...

Directory: MRPR File: gwp-Cb_LI_BCphaseB1.txt

| seqnames | start | end | width | strand | source | type | score | phase | ID | Dbxref | Name | gbkey | gene | gene_biotype | partial | start_range | end_range | seqnames | start | end | width | strand | NbrSites | Sdnsty |
|---|---|---|---|---|---|---|---|---|---|---|---|---|---|---|---|---|---|---|---|---|---|---|---|---|
| NW_011924931.1 | 534549 | 536012 | 1464 | + | Gnomon | gene | NA | NA | gene1415 | GeneID:105287536 | LOC105287536 | Gene | LOC105287536 | protein_coding | NA | character(0) | character(0) | NW_011924931.1 | 535781 | 1051253 | 515473 | * | 2 | 0.00387993163560458 |
| NW_011924931.1 | 536217 | 855035 | 318819 | - | Gnomon | gene | NA | NA | gene1416 | GeneID:105287539 | LOC105287539 | Gene | LOC105287539 | protein_coding | NA | character(0) | character(0) | NW_011924931.1 | 535781 | 1051253 | 515473 | * | 2 | 0.00387993163560458 |
| NW_011924931.1 | 1014634 | 1019043 | 4410 | - | Gnomon | gene | NA | NA | gene1417 | GeneID:105287538 | LOC105287538 | Gene | LOC105287538 | protein_coding | NA | character(0) | character(0) | NW_011924931.1 | 535781 | 1051253 | 515473 | * | 2 | 0.00387993163560458 |
| NW_011924888.1 | 407815 | 409420 | 1606 | - | Gnomon | gene | NA | NA | gene322 | GeneID:105275777 | LOC105275777 | Gene | LOC105275777 | protein_coding | NA | character(0) | character(0) | NW_011924888.1 | 408597 | 897056 | 488460 | * | 2 | 0.00409450108504279 |
| NW_011924888.1 | 413553 | 425756 | 12204 | + | Gnomon | gene | NA | NA | gene323 | GeneID:105277093 | LOC105277093 | Gene | LOC105277093 | protein_coding | NA | character(0) | character(0) | NW_011924888.1 | 408597 | 897056 | 488460 | * | 2 | 0.00409450108504279 |
| NW_011924888.1 | 422756 | 424085 | 1330 | - | Gnomon | gene | NA | NA | gene324 | GeneID:105275786 | LOC105275786 | Gene | LOC105275786 | protein_coding | NA | character(0) | character(0) | NW_011924888.1 | 408597 | 897056 | 488460 | * | 2 | 0.00409450108504279 |
| NW_011924888.1 | 425900 | 425972 | 73 | + | tRNAscan-SE | gene | NA | NA | gene325 | GeneID:109610824 | Trnav-uac | Gene | Trnav-uac | tRNA | NA | character(0) | character(0) | NW_011924888.1 | 408597 | 897056 | 488460 | * | 2 | 0.00409450108504279 |
| NW_011924888.1 | 428309 | 444138 | 15830 | + | Gnomon | gene | NA | NA | gene326 | GeneID:105275823 | LOC105275823 | Gene | LOC105275823 | protein_coding | NA | character(0) | character(0) | NW_011924888.1 | 408597 | 897056 | 488460 | * | 2 | 0.00409450108504279 |
| NW_011924888.1 | 452858 | 539134 | 86277 | - | Gnomon | gene | NA | NA | gene327 | GeneID:105275815 | LOC105275815 | Gene | LOC105275815 | protein_coding | NA | character(0) | character(0) | NW_011924888.1 | 408597 | 897056 | 488460 | * | 2 | 0.00409450108504279 |
| NW_011924888.1 | 558196 | 568778 | 10583 | + | Gnomon | gene | NA | NA | gene328 | GeneID:105275807 | LOC105275807 | Gene | LOC105275807 | protein_coding | NA | character(0) | character(0) | NW_011924888.1 | 408597 | 897056 | 488460 | * | 2 | 0.00409450108504279 |
| NW_011924888.1 | 569356 | 570956 | 1601 | + | Gnomon | gene | NA | NA | gene329 | GeneID:105275794 | LOC105275794 | Gene | LOC105275794 | protein_coding | NA | character(0) | character(0) | NW_011924888.1 | 408597 | 897056 | 488460 | * | 2 | 0.00409450108504279 |
| NW_011924888.1 | 571293 | 679878 | 108586 | + | Gnomon | gene | NA | NA | gene330 | GeneID:105275842 | LOC105275842 | Gene | LOC105275842 | protein_coding | NA | character(0) | character(0) | NW_011924888.1 | 408597 | 897056 | 488460 | * | 2 | 0.00409450108504279 |
| NW_011924888.1 | 700961 | 713940 | 12980 | + | Gnomon | gene | NA | NA | gene331 | GeneID:105275867 | LOC105275867 | Gene | LOC105275867 | protein_coding | NA | character(0) | character(0) | NW_011924888.1 | 408597 | 897056 | 488460 | * | 2 | 0.00409450108504279 |
| NW_011924888.1 | 715828 | 720753 | 4926 | + | Gnomon | gene | NA | NA | gene332 | GeneID:109610823 | LOC109610823 | Gene | LOC109610823 | lncRNA | NA | character(0) | character(0) | NW_011924888.1 | 408597 | 897056 | 488460 | * | 2 | 0.00409450108504279 |
| NW_011924888.1 | 788065 | 793095 | 5031 | - | Gnomon | gene | NA | NA | gene333 | GeneID:105275873 | LOC105275873 | Gene | LOC105275873 | protein_coding | NA | character(0) | character(0) | NW_011924888.1 | 408597 | 897056 | 488460 | * | 2 | 0.00409450108504279 |
| NW_011924888.1 | 848004 | 855282 | 7279 | + | Gnomon | gene | NA | NA | gene334 | GeneID:105275890 | LOC105275890 | Gene | LOC105275890 | protein_coding | NA | character(0) | character(0) | NW_011924888.1 | 408597 | 897056 | 488460 | * | 2 | 0.00409450108504279 |
| NW_011924888.1 | 855465 | 858826 | 3362 | + | Gnomon | gene | NA | NA | gene335 | GeneID:109610755 | LOC109610755 | Gene | LOC109610755 | protein_coding | NA | character(0) | character(0) | NW_011924888.1 | 408597 | 897056 | 488460 | * | 2 | 0.00409450108504279 |
| NW_011924888.1 | 858153 | 985759 | 127607 | - | Gnomon | gene | NA | NA | gene336 | GeneID:105275903 | LOC105275903 | Gene | LOC105275903 | protein_coding | NA | character(0) | character(0) | NW_011924888.1 | 408597 | 897056 | 488460 | * | 2 | 0.00409450108504279 |
| NW_011925053.1 | 1014381 | 1042267 | 27887 | - | Gnomon | gene | NA | NA | gene5886 | GeneID:105278816 | LOC105278816 | Gene | LOC105278816 | protein_coding | NA | character(0) | character(0) | NW_011925053.1 | 681984 | 1153435 | 471452 | * | 2 | 0.0042422134172726 |
| NW_011925053.1 | 1060400 | 1064769 | 4370 | - | Gnomon | gene | NA | NA | gene5887 | GeneID:105278817 | LOC105278817 | Gene | LOC105278817 | protein_coding | NA | character(0) | character(0) | NW_011925053.1 | 681984 | 1153435 | 471452 | * | 2 | 0.0042422134172726 |
| NW_011925053.1 | 1103568 | 1106298 | 2731 | - | Gnomon | gene | NA | NA | gene5888 | GeneID:105278887 | LOC105278887 | Gene | LOC105278887 | protein_coding | NA | character(0) | character(0) | NW_011925053.1 | 681984 | 1153435 | 471452 | * | 2 | 0.0042422134172726 |
| NW_011925053.1 | 1106355 | 1107818 | 1464 | - | Gnomon | gene | NA | NA | gene5889 | GeneID:105278823 | LOC105278823 | Gene | LOC105278823 | protein_coding | NA | character(0) | character(0) | NW_011925053.1 | 681984 | 1153435 | 471452 | * | 2 | 0.0042422134172726 |
| NW_011925053.1 | 1110412 | 1112281 | 1870 | + | Gnomon | gene | NA | NA | gene5890 | GeneID:109610933 | LOC109610933 | Gene | LOC109610933 | protein_coding | NA | character(0) | character(0) | NW_011925053.1 | 681984 | 1153435 | 471452 | * | 2 | 0.0042422134172726 |
| NW_011925053.1 | 1119431 | 1126904 | 7474 | + | Gnomon | gene | NA | NA | gene5891 | GeneID:105278821 | LOC105278821 | Gene | LOC105278821 | protein_coding | NA | character(0) | character(0) | NW_011925053.1 | 681984 | 1153435 | 471452 | * | 2 | 0.0042422134172726 |

...
... only first 25 lines shown ...

Directory: MRPR File: gwp-Cb_LI_BCphaseB2.txt

seqnames start end width strand source type score phase ID Dbxref Name gbkey gene gene_biotype partial start_range end_range seqnames start
end width strand NbrSites Sdnsty
NW_011924892.1 274786 285690 10905 - Gnomon gene NA NA gene544 GeneID:105278288 LOC105278288 Gene LOC105278288 protein_coding NA
character(0) character(0) NW_011924892.1 284893 930651 645759 * 2 0.00309713066329699
NW_011924892.1 302199 519970 217772 + Gnomon gene NA NA gene545 GeneID:105278329 LOC105278329 Gene LOC105278329 protein_coding NA
character(0) character(0) NW_011924892.1 284893 930651 645759 * 2 0.00309713066329699
NW_011924892.1 378175 382124 3950 + Gnomon gene NA NA gene546 GeneID:105278325 LOC105278325 Gene LOC105278325 lncRNA NA
character(0) character(0) NW_011924892.1 284893 930651 645759 * 2 0.00309713066329699
NW_011924892.1 446522 450804 4283 - Gnomon gene NA NA gene547 GeneID:105278319 LOC105278319 Gene LOC105278319 protein_coding NA
character(0) character(0) NW_011924892.1 284893 930651 645759 * 2 0.00309713066329699
NW_011924892.1 458698 471223 12526 - Gnomon gene NA NA gene548 GeneID:105278305 LOC105278305 Gene LOC105278305 protein_coding NA
character(0) character(0) NW_011924892.1 284893 930651 645759 * 2 0.00309713066329699
NW_011924892.1 604295 664845 60551 + Gnomon gene NA NA gene549 GeneID:105278340 LOC105278340 Gene LOC105278340 protein_coding NA
character(0) character(0) NW_011924892.1 284893 930651 645759 * 2 0.00309713066329699
NW_011924892.1 720318 1032506 312189 + Gnomon gene NA NA gene550 GeneID:105278394 LOC105278394 Gene LOC105278394 protein_coding NA
character(0) character(0) NW_011924892.1 284893 930651 645759 * 2 0.00309713066329699
NW_011924892.1 859826 860748 923 + Gnomon gene NA NA gene551 GeneID:105278378 LOC105278378 Gene LOC105278378 protein_coding NA
character(0) character(0) NW_011924892.1 284893 930651 645759 * 2 0.00309713066329699
NW_011924892.1 895456 911391 15936 - Gnomon gene NA NA gene552 GeneID:105278371 LOC105278371 Gene LOC105278371 lncRNA NA
character(0) character(0) NW_011924892.1 284893 930651 645759 * 2 0.00309713066329699
NW_011925082.1 50833 53693 2861 - Gnomon gene NA NA gene6286 GeneID:105279298 LOC105279298 Gene LOC105279298 protein_coding NA character(0)
character(0) NW_011925082.1 53097 666977 613881 * 2 0.00325796041903887
NW_011925082.1 54202 179969 125768 + Gnomon gene NA NA gene6287 GeneID:105279302 LOC105279302 Gene LOC105279302 protein_coding NA
character(0) character(0) NW_011925082.1 53097 666977 613881 * 2 0.00325796041903887
NW_011925082.1 157598 334193 176596 + Gnomon gene NA NA gene6288 GeneID:105279301 LOC105279301 Gene LOC105279301 protein_coding NA
character(0) character(0) NW_011925082.1 53097 666977 613881 * 2 0.00325796041903887
NW_011925082.1 304108 306517 2410 - Gnomon gene NA NA gene6289 GeneID:105279300 LOC105279300 Gene LOC105279300 protein_coding NA
character(0) character(0) NW_011925082.1 53097 666977 613881 * 2 0.00325796041903887
NW_011925082.1 308223 310906 2684 - Gnomon gene NA NA gene6290 GeneID:105279299 LOC105279299 Gene LOC105279299 protein_coding NA
character(0) character(0) NW_011925082.1 53097 666977 613881 * 2 0.00325796041903887
NW_011925082.1 338811 368308 29498 - Gnomon gene NA NA gene6291 GeneID:105279304 LOC105279304 Gene LOC105279304 protein_coding NA
character(0) character(0) NW_011925082.1 53097 666977 613881 * 2 0.00325796041903887
NW_011925082.1 372157 401370 29214 - Gnomon gene NA NA gene6292 GeneID:105279306 LOC105279306 Gene LOC105279306 protein_coding NA
character(0) character(0) NW_011925082.1 53097 666977 613881 * 2 0.00325796041903887
NW_011925082.1 402393 414452 12060 - Gnomon gene NA NA gene6293 GeneID:105279308 LOC105279308 Gene LOC105279308 protein_coding NA
character(0) character(0) NW_011925082.1 53097 666977 613881 * 2 0.00325796041903887
NW_011925082.1 403950 406160 2211 + Gnomon gene NA NA gene6294 GeneID:105279307 LOC105279307 Gene LOC105279307 protein_coding NA
character(0) character(0) NW_011925082.1 53097 666977 613881 * 2 0.00325796041903887
NW_011925082.1 425182 450421 25240 + Gnomon gene NA NA gene6295 GeneID:105279309 LOC105279309 Gene LOC105279309 protein_coding NA
character(0) character(0) NW_011925082.1 53097 666977 613881 * 2 0.00325796041903887
NW_011925082.1 457116 650005 192890 + Gnomon gene NA NA gene6296 GeneID:105279311 LOC105279311 Gene LOC105279311 protein_coding NA
character(0) character(0) NW_011925082.1 53097 666977 613881 * 2 0.00325796041903887
NW_011925082.1 650138 655780 5643 - Gnomon gene NA NA gene6297 GeneID:105279310 LOC105279310 Gene LOC105279310 protein_coding NA
character(0) character(0) NW_011925082.1 53097 666977 613881 * 2 0.00325796041903887
NW_011925082.1 656023 664801 8779 - Gnomon gene NA NA gene6298 GeneID:105279312 LOC105279312 Gene LOC105279312 protein_coding NA
character(0) character(0) NW_011925082.1 53097 666977 613881 * 2 0.00325796041903887
NW_011925082.1 664899 668532 3634 - Gnomon gene NA NA gene6299 GeneID:105279313 LOC105279313 Gene LOC105279313 protein_coding NA
character(0) character(0) NW_011925082.1 53097 666977 613881 * 2 0.00325796041903887
NW_011924937.1 527309 528853 1545 + Gnomon gene NA NA gene1781 GeneID:105287956 LOC105287956 Gene LOC105287956 protein_coding NA
character(0) character(0) NW_011924937.1 528587 1038579 509993 * 2 0.00392162245364152
...
... only first 25 lines shown ...

Directory: MRPR File: gwp-Cb_LI_RphaseA1.txt

| seqnames | start | end | width | strand | source | type | score | phase | ID | Dbxref | Name | gbkey | gene | gene_biotype | partial | start_range | end_range | seqnames | start | end | width | strand | NbrSites | Sdnsty |
|---|---|---|---|---|---|---|---|---|---|---|---|---|---|---|---|---|---|---|---|---|---|---|---|---|
| NW_011925315.1 | 753750 | 761692 | 7943 | - | Gnomon | gene | NA | NA | gene9929 | GeneID:105283502 | LOC105283502 | Gene | LOC105283502 | lncRNA | NA | character(0) | character(0) | NW_011925315.1 | 741815 | 1498755 | 756941 | * | 2 | 0.00264221385814746 |
| NW_011925315.1 | 889570 | 1485837 | 596268 | + | Gnomon | gene | NA | NA | gene9930 | GeneID:105283503 | LOC105283503 | Gene | LOC105283503 | protein_coding | NA | character(0) | character(0) | NW_011925315.1 | 741815 | 1498755 | 756941 | * | 2 | 0.00264221385814746 |
| NW_011925315.1 | 993535 | 999713 | 6179 | - | Gnomon | gene | NA | NA | gene9931 | GeneID:105283505 | LOC105283505 | Gene | LOC105283505 | protein_coding | NA | character(0) | character(0) | NW_011925315.1 | 741815 | 1498755 | 756941 | * | 2 | 0.00264221385814746 |
| NW_011925315.1 | 1003778 | 1006949 | 3172 | + | Gnomon | gene | NA | NA | gene9932 | GeneID:105283504 | LOC105283504 | Gene | LOC105283504 | protein_coding | NA | character(0) | character(0) | NW_011925315.1 | 741815 | 1498755 | 756941 | * | 2 | 0.00264221385814746 |
| NW_011925315.1 | 1493432 | 1494004 | 573 | + | Gnomon | gene | NA | NA | gene9933 | GeneID:105283508 | LOC105283508 | Gene | LOC105283508 | protein_coding | NA | character(0) | character(0) | NW_011925315.1 | 741815 | 1498755 | 756941 | * | 2 | 0.00264221385814746 |
| NW_011925315.1 | 1493819 | 1497141 | 3323 | - | Gnomon | gene | NA | NA | gene9934 | GeneID:105283506 | LOC105283506 | Gene | LOC105283506 | lncRNA | NA | character(0) | character(0) | NW_011925315.1 | 741815 | 1498755 | 756941 | * | 2 | 0.00264221385814746 |
| NW_011925315.1 | 1497840 | 1501619 | 3780 | + | Gnomon | gene | NA | NA | gene9935 | GeneID:105283509 | LOC105283509 | Gene | LOC105283509 | protein_coding | NA | character(0) | character(0) | NW_011925315.1 | 741815 | 1498755 | 756941 | * | 2 | 0.00264221385814746 |
| NW_011925036.1 | 461196 | 469719 | 8524 | + | Gnomon | gene | NA | NA | gene5613 | GeneID:105278504 | LOC105278504 | Gene | LOC105278504 | protein_coding | NA | character(0) | character(0) | NW_011925036.1 | 463225 | 1155855 | 692631 | * | 2 | 0.00288754040751858 |
| NW_011925036.1 | 474043 | 476463 | 2421 | - | Gnomon | gene | NA | NA | gene5614 | GeneID:105278505 | LOC105278505 | Gene | LOC105278505 | protein_coding | NA | character(0) | character(0) | NW_011925036.1 | 463225 | 1155855 | 692631 | * | 2 | 0.00288754040751858 |
| NW_011925036.1 | 477004 | 482029 | 5026 | + | Gnomon | gene | NA | NA | gene5615 | GeneID:105278506 | LOC105278506 | Gene | LOC105278506 | protein_coding | NA | character(0) | character(0) | NW_011925036.1 | 463225 | 1155855 | 692631 | * | 2 | 0.00288754040751858 |
| NW_011925036.1 | 489591 | 492420 | 2830 | - | Gnomon | gene | NA | NA | gene5616 | GeneID:105278560 | LOC105278560 | Gene | LOC105278560 | protein_coding | NA | character(0) | character(0) | NW_011925036.1 | 463225 | 1155855 | 692631 | * | 2 | 0.00288754040751858 |
| NW_011925036.1 | 524857 | 527309 | 2453 | - | Gnomon | gene | NA | NA | gene5617 | GeneID:105278507 | LOC105278507 | Gene | LOC105278507 | lncRNA | NA | character(0) | character(0) | NW_011925036.1 | 463225 | 1155855 | 692631 | * | 2 | 0.00288754040751858 |
| NW_011925036.1 | 544456 | 608309 | 63854 | + | Gnomon | gene | NA | NA | gene5618 | GeneID:109610912 | LOC109610912 | Gene | LOC109610912 | lncRNA | NA | character(0) | character(0) | NW_011925036.1 | 463225 | 1155855 | 692631 | * | 2 | 0.00288754040751858 |
| NW_011925036.1 | 608861 | 873355 | 264495 | + | Gnomon | gene | NA | NA | gene5619 | GeneID:105278508 | LOC105278508 | Gene | LOC105278508 | protein_coding | NA | character(0) | character(0) | NW_011925036.1 | 463225 | 1155855 | 692631 | * | 2 | 0.00288754040751858 |
| NW_011925036.1 | 927483 | 928885 | 1403 | + | Gnomon | gene | NA | NA | gene5620 | GeneID:105278511 | LOC105278511 | Gene | LOC105278511 | protein_coding | NA | character(0) | character(0) | NW_011925036.1 | 463225 | 1155855 | 692631 | * | 2 | 0.00288754040751858 |
| NW_011925036.1 | 929651 | 932446 | 2796 | + | Gnomon | gene | NA | NA | gene5621 | GeneID:105278510 | LOC105278510 | Gene | LOC105278510 | protein_coding | NA | character(0) | character(0) | NW_011925036.1 | 463225 | 1155855 | 692631 | * | 2 | 0.00288754040751858 |
| NW_011925036.1 | 1006642 | 1011165 | 4524 | + | Gnomon | gene | NA | NA | gene5622 | GeneID:105278519 | LOC105278519 | Gene | LOC105278519 | protein_coding | NA | character(0) | character(0) | NW_011925036.1 | 463225 | 1155855 | 692631 | * | 2 | 0.00288754040751858 |
| NW_011925036.1 | 1011369 | 1143497 | 132129 | - | Gnomon | gene | NA | NA | gene5623 | GeneID:105278518 | LOC105278518 | Gene | LOC105278518 | protein_coding | NA | character(0) | character(0) | NW_011925036.1 | 463225 | 1155855 | 692631 | * | 2 | 0.00288754040751858 |
| NW_011925036.1 | 1093799 | 1100136 | 6338 | + | Gnomon | gene | NA | NA | gene5624 | GeneID:105278516 | LOC105278516 | Gene | LOC105278516 | protein_coding | NA | character(0) | character(0) | NW_011925036.1 | 463225 | 1155855 | 692631 | * | 2 | 0.00288754040751858 |
| NW_011925036.1 | 1114815 | 1119640 | 4826 | + | Gnomon | gene | NA | NA | gene5625 | GeneID:105278515 | LOC105278515 | Gene | LOC105278515 | protein_coding | NA | character(0) | character(0) | NW_011925036.1 | 463225 | 1155855 | 692631 | * | 2 | 0.00288754040751858 |
| NW_011925036.1 | 1120601 | 1124464 | 3864 | - | Gnomon | gene | NA | NA | gene5626 | GeneID:105278514 | LOC105278514 | Gene | LOC105278514 | lncRNA | NA | character(0) | character(0) | NW_011925036.1 | 463225 | 1155855 | 692631 | * | 2 | 0.00288754040751858 |
| NW_011925036.1 | 1124968 | 1126812 | 1845 | + | Gnomon | gene | NA | NA | gene5627 | GeneID:105278513 | LOC105278513 | Gene | LOC105278513 | protein_coding | NA | character(0) | character(0) | NW_011925036.1 | 463225 | 1155855 | 692631 | * | 2 | 0.00288754040751858 |
| NW_011925036.1 | 1131653 | 1134176 | 2524 | + | Gnomon | gene | NA | NA | gene5628 | GeneID:105278512 | LOC105278512 | Gene | LOC105278512 | protein_coding | NA | character(0) | character(0) | NW_011925036.1 | 463225 | 1155855 | 692631 | * | 2 | 0.00288754040751858 |
| NW_011925036.1 | 1143942 | 1157567 | 13626 | - | Gnomon | gene | NA | NA | gene5629 | GeneID:105278520 | LOC105278520 | Gene | LOC105278520 | protein_coding | NA | character(0) | character(0) | NW_011925036.1 | 463225 | 1155855 | 692631 | * | 2 | 0.00288754040751858 |

...
... only first 25 lines shown ...

Directory: MRPR File: gwp-Cb_LI_RphaseA2.txt

```
seqnames    start end   width strand      source      type  score phase ID    Dbxref      Name  gbkey gene  gene_biotype      partial     start_range end_range   seqnames    start
      end   width strand      NbrSites    Sdnsty
NW_011924996.1    1257761     1305220     47460 +     Gnomon      gene  NA    NA    gene3840    GeneID:105276485  LOC105276485      Gene  LOC105276485      protein_coding    NA
      character(0)      character(0)      NW_011924996.1    1277657     1597859     320203      *     2     0.00624603766985319
NW_011924996.1    1306100     1429542     123443      -     Gnomon      gene  NA    NA    gene3841    GeneID:105276484  LOC105276484      Gene  LOC105276484      protein_coding    NA
      character(0)      character(0)      NW_011924996.1    1277657     1597859     320203      *     2     0.00624603766985319
NW_011924996.1    1355655     1356219     565   -     Gnomon      gene  NA    NA    gene3842    GeneID:105276483  LOC105276483      Gene  LOC105276483      lncRNA      NA
      character(0)      character(0)      NW_011924996.1    1277657     1597859     320203      *     2     0.00624603766985319
NW_011924996.1    1450370     1584562     134193      -     Gnomon      gene  NA    NA    gene3843    GeneID:105276481  LOC105276481      Gene  LOC105276481      protein_coding    NA
      character(0)      character(0)      NW_011924996.1    1277657     1597859     320203      *     2     0.00624603766985319
NW_011924996.1    1488645     1492750     4106  -     Gnomon      gene  NA    NA    gene3844    GeneID:105276480  LOC105276480      Gene  LOC105276480      protein_coding    NA
      character(0)      character(0)      NW_011924996.1    1277657     1597859     320203      *     2     0.00624603766985319
NW_011924996.1    1493046     1497958     4913  -     Gnomon      gene  NA    NA    gene3845    GeneID:105276478  LOC105276478      Gene  LOC105276478      protein_coding    NA
      character(0)      character(0)      NW_011924996.1    1277657     1597859     320203      *     2     0.00624603766985319
NW_011924996.1    1584788     1593914     9127  +     Gnomon      gene  NA    NA    gene3846    GeneID:105276477  LOC105276477      Gene  LOC105276477      protein_coding    NA
      character(0)      character(0)      NW_011924996.1    1277657     1597859     320203      *     2     0.00624603766985319
NW_011924996.1    1593872     1600328     6457  -     Gnomon      gene  NA    NA    gene3847    GeneID:105276487  LOC105276487      Gene  LOC105276487      protein_coding    NA
      character(0)      character(0)      NW_011924996.1    1277657     1597859     320203      *     2     0.00624603766985319
NW_011925007.1    86240 97994 11755 -     Gnomon      gene  NA    NA    gene4157    GeneID:105276841  LOC105276841      Gene  LOC105276841      protein_coding    NA    character(0)
      character(0)      NW_011925007.1    96864 381604      284741      *     2     0.00702392700735054
NW_011925007.1    117788      336278      218491      -     Gnomon      gene  NA    NA    gene4158    GeneID:105276843  LOC105276843      Gene  LOC105276843      protein_coding    NA
      character(0)      character(0)      NW_011925007.1    96864 381604      284741      *     2     0.00702392700735054
NW_011925007.1    191348      196652      5305  +     Gnomon      gene  NA    NA    gene4159    GeneID:105276842  LOC105276842      Gene  LOC105276842      lncRNA      NA
      character(0)      character(0)      NW_011925007.1    96864 381604      284741      *     2     0.00702392700735054
NW_011925300.1    106737      174288      67552 +     Gnomon      gene  NA    NA    gene9770    GeneID:105283320  LOC105283320      Gene  LOC105283320      protein_coding    NA
      character(0)      character(0)      NW_011925300.1    134597      401637      267041      *     2     0.00748948663313873
NW_011925300.1    177556      235020      57465 -     Gnomon      gene  NA    NA    gene9771    GeneID:105283321  LOC105283321      Gene  LOC105283321      protein_coding    NA
      character(0)      character(0)      NW_011925300.1    134597      401637      267041      *     2     0.00748948663313873
NW_011925300.1    236613      251599      14987 -     Gnomon      gene  NA    NA    gene9772    GeneID:105283347  LOC105283347      Gene  LOC105283347      protein_coding    NA
      character(0)      character(0)      NW_011925300.1    134597      401637      267041      *     2     0.00748948663313873
NW_011925300.1    241190      245222      4033  +     Gnomon      gene  NA    NA    gene9773    GeneID:105283322  LOC105283322      Gene  LOC105283322      protein_coding    NA
      character(0)      character(0)      NW_011925300.1    134597      401637      267041      *     2     0.00748948663313873
NW_011925300.1    252785      259998      7214  -     Gnomon      gene  NA    NA    gene9774    GeneID:105283348  LOC105283348      Gene  LOC105283348      protein_coding    NA
      character(0)      character(0)      NW_011925300.1    134597      401637      267041      *     2     0.00748948663313873
NW_011925300.1    277125      425165      148041      -     Gnomon      gene  NA    NA    gene9775    GeneID:105283323  LOC105283323      Gene  LOC105283323      protein_coding    NA
      character(0)      character(0)      NW_011925300.1    134597      401637      267041      *     2     0.00748948663313873
NW_011925300.1    356733      369726      12994 +     Gnomon      gene  NA    NA    gene9776    GeneID:105283325  LOC105283325      Gene  LOC105283325      protein_coding    NA
      character(0)      character(0)      NW_011925300.1    134597      401637      267041      *     2     0.00748948663313873
NW_011925300.1    378676      396625      17950 -     Gnomon      gene  NA    NA    gene9777    GeneID:105283349  LOC105283349      Gene  LOC105283349      protein_coding    NA
      character(0)      character(0)      NW_011925300.1    134597      401637      267041      *     2     0.00748948663313873
NW_011925099.1    958705      1058161     99457 +     Gnomon      gene  NA    NA    gene6467    GeneID:105279513  LOC105279513      Gene  LOC105279513      protein_coding    NA
      character(0)      character(0)      NW_011925099.1    1006421     1228934     222514      *     2     0.00898819849537557
NW_011925099.1    1028256     1031094     2839  +     Gnomon      gene  NA    NA    gene6469    GeneID:109610978  LOC109610978      Gene  LOC109610978      lncRNA      NA
      character(0)      character(0)      NW_011925099.1    1006421     1228934     222514      *     2     0.00898819849537557
NW_011925099.1    1061474     1084552     23079 +     Gnomon      gene  NA    NA    gene6470    GeneID:105279514  LOC105279514      Gene  LOC105279514      protein_coding    NA
      character(0)      character(0)      NW_011925099.1    1006421     1228934     222514      *     2     0.00898819849537557
NW_011925099.1    1100140     1106592     6453  +     Gnomon      gene  NA    NA    gene6471    GeneID:105279515  LOC105279515      Gene  LOC105279515      lncRNA      NA
      character(0)      character(0)      NW_011925099.1    1006421     1228934     222514      *     2     0.00898819849537557
NW_011925027.1    687169      695552      8384  -     Gnomon      gene  NA    NA    gene5153    GeneID:105277971  LOC105277971      Gene  LOC105277971      protein_coding    NA
      character(0)      character(0)      NW_011925027.1    689188      897305      208118      *     2     0.00960993282656954
...
... only first 25 lines shown ...
```

Directory: MRPR File: gwp-Cb_LI_RphaseB1.txt

```
seqnames	start	end	width	strand	source	type	score	phase	ID	Dbxref	Name	gbkey	gene	gene_biotype	partial	start_range	end_range	seqnames	start
	end	width	strand	NbrSites	Sdnsty
NW_011925387.1	683388	700925	17538	+	Gnomon	gene	NA	NA	gene10985	GeneID:105284708	LOC105284708	Gene	LOC105284708	protein_coding	NA
	character(0)	character(0)	NW_011925387.1	685943	1328666	642724	*	3	0.00466763338540338
NW_011925387.1	686855	697617	10763	-	Gnomon	gene	NA	NA	gene10986	GeneID:105284707	LOC105284707	Gene	LOC105284707	protein_coding	NA
	character(0)	character(0)	NW_011925387.1	685943	1328666	642724	*	3	0.00466763338540338
NW_011925387.1	719208	725289	6082	+	Gnomon	gene	NA	NA	gene10987	GeneID:105284731	LOC105284731	Gene	LOC105284731	protein_coding	NA
	character(0)	character(0)	NW_011925387.1	685943	1328666	642724	*	3	0.00466763338540338
NW_011925387.1	745822	766017	20196	-	Gnomon	gene	NA	NA	gene10988	GeneID:105284709	LOC105284709	Gene	LOC105284709	protein_coding	NA
	character(0)	character(0)	NW_011925387.1	685943	1328666	642724	*	3	0.00466763338540338
NW_011925387.1	830435	979579	149145	-	Gnomon	gene	NA	NA	gene10989	GeneID:105284711	LOC105284711	Gene	LOC105284711	protein_coding	NA
	character(0)	character(0)	NW_011925387.1	685943	1328666	642724	*	3	0.00466763338540338
NW_011925387.1	1173433	1275017	101585	-	Gnomon	gene	NA	NA	gene10990	GeneID:105284714	LOC105284714	Gene	LOC105284714	protein_coding	NA
	character(0)	character(0)	NW_011925387.1	685943	1328666	642724	*	3	0.00466763338540338
NW_011925387.1	1325075	1330400	5326	-	Gnomon	gene	NA	NA	gene10991	GeneID:105284713	LOC105284713	Gene	LOC105284713	protein_coding	NA
	character(0)	character(0)	NW_011925387.1	685943	1328666	642724	*	3	0.00466763338540338
NW_011924987.1	3614126	3750507	136382	-	Gnomon	gene	NA	NA	gene3684	GeneID:105276288	LOC105276288	Gene	LOC105276288	protein_coding	NA
	character(0)	character(0)	NW_011924987.1	3688298	4326160	637863	*	3	0.00470320429308488
NW_011924987.1	3944593	4206024	261432	-	Gnomon	gene	NA	NA	gene3685	GeneID:105276294	LOC105276294	Gene	LOC105276294	protein_coding	NA
	character(0)	character(0)	NW_011924987.1	3688298	4326160	637863	*	3	0.00470320429308488
NW_011924987.1	4104169	4104978	810	+	Gnomon	gene	NA	NA	gene3686	GeneID:105276293	LOC105276293	Gene	LOC105276293	lncRNA	NA
	character(0)	character(0)	NW_011924987.1	3688298	4326160	637863	*	3	0.00470320429308488
NW_011924987.1	4108870	4111526	2657	-	Gnomon	gene	NA	NA	gene3687	GeneID:105276292	LOC105276292	Gene	LOC105276292	protein_coding	NA
	character(0)	character(0)	NW_011924987.1	3688298	4326160	637863	*	3	0.00470320429308488
NW_011924987.1	4193029	4199116	6088	+	Gnomon	gene	NA	NA	gene3688	GeneID:105276291	LOC105276291	Gene	LOC105276291	lncRNA	NA
	character(0)	character(0)	NW_011924987.1	3688298	4326160	637863	*	3	0.00470320429308488
NW_011924987.1	4306305	4325191	18887	-	Gnomon	gene	NA	NA	gene3689	GeneID:109610771	LOC109610771	Gene	LOC109610771	lncRNA	NA
	character(0)	character(0)	NW_011924987.1	3688298	4326160	637863	*	3	0.00470320429308488
NW_011924987.1	4324236	4324309	74	+	tRNAscan-SE	gene	NA	NA	gene3690	GeneID:109610778	Trnan-guu	Gene	Trnan-guu	tRNA	NA	character(0)
	character(0)	NW_011924987.1	3688298	4326160	637863	*	3	0.00470320429308488
NW_011924987.1	4324725	4324798	74	+	tRNAscan-SE	gene	NA	NA	gene3691	GeneID:109610779	Trnat-agu	Gene	Trnat-agu	tRNA	NA	character(0)
	character(0)	NW_011924987.1	3688298	4326160	637863	*	3	0.00470320429308488
NW_011924987.1	4325126	4385544	60419	+	Gnomon	gene	NA	NA	gene3692	GeneID:105276290	LOC105276290	Gene	LOC105276290	protein_coding	NA
	character(0)	character(0)	NW_011924987.1	3688298	4326160	637863	*	3	0.00470320429308488
NW_011925367.1	143267	668644	525378	-	Gnomon	gene	NA	NA	gene10696	GeneID:105284382	LOC105284382	Gene	LOC105284382	protein_coding	NA
	character(0)	character(0)	NW_011925367.1	152381	546014	393634	*	2	0.00508086191741567
NW_011925367.1	531411	536651	5241	-	Gnomon	gene	NA	NA	gene10697	GeneID:105284381	LOC105284381	Gene	LOC105284381	protein_coding	NA
	character(0)	character(0)	NW_011925367.1	152381	546014	393634	*	2	0.00508086191741567
NW_011924888.1	407815	409420	1606	-	Gnomon	gene	NA	NA	gene322	GeneID:105275777	LOC105275777	Gene	LOC105275777	protein_coding	NA
	character(0)	character(0)	NW_011924888.1	408597	992521	583925	*	3	0.00513764610181102
NW_011924888.1	413553	425756	12204	+	Gnomon	gene	NA	NA	gene323	GeneID:105277093	LOC105277093	Gene	LOC105277093	protein_coding	NA
	character(0)	character(0)	NW_011924888.1	408597	992521	583925	*	3	0.00513764610181102
NW_011924888.1	422756	424085	1330	-	Gnomon	gene	NA	NA	gene324	GeneID:105275786	LOC105275786	Gene	LOC105275786	protein_coding	NA
	character(0)	character(0)	NW_011924888.1	408597	992521	583925	*	3	0.00513764610181102
NW_011924888.1	425900	425972	73	+	tRNAscan-SE	gene	NA	NA	gene325	GeneID:109610824	Trnav-uac	Gene	Trnav-uac	tRNA	NA	character(0)
	character(0)	NW_011924888.1	408597	992521	583925	*	3	0.00513764610181102
NW_011924888.1	428309	444138	15830	+	Gnomon	gene	NA	NA	gene326	GeneID:105275823	LOC105275823	Gene	LOC105275823	protein_coding	NA
	character(0)	character(0)	NW_011924888.1	408597	992521	583925	*	3	0.00513764610181102
NW_011924888.1	452858	539134	86277	-	Gnomon	gene	NA	NA	gene327	GeneID:105275815	LOC105275815	Gene	LOC105275815	protein_coding	NA
	character(0)	character(0)	NW_011924888.1	408597	992521	583925	*	3	0.00513764610181102
...
... only first 25 lines shown ...
```

Directory: MRPR File: gwp-Cb_LI_RphaseB2.txt

| seqnames | start | end | width | strand | source | type | score | phase | ID | Dbxref | Name | gbkey | gene | gene_biotype | partial | start_range | end_range | seqnames | start | end | width | strand | NbrSites | Sdnsty |
|---|---|---|---|---|---|---|---|---|---|---|---|---|---|---|---|---|---|---|---|---|---|---|---|---|
| NW_011925216.1 | 219999 | 246618 | 26620 | - | Gnomon | gene | NA | NA | gene8751 | GeneID:105282161 | LOC105282161 | Gene | LOC105282161 | protein_coding | NA | character(0) | character(0) | NW_011925216.1 | 216055 | 1092286 | 876232 | * | 2 | 0.00228250052497512 |
| NW_011925216.1 | 277106 | 281176 | 4071 | - | Gnomon | gene | NA | NA | gene8752 | GeneID:105282162 | LOC105282162 | Gene | LOC105282162 | protein_coding | NA | character(0) | character(0) | NW_011925216.1 | 216055 | 1092286 | 876232 | * | 2 | 0.00228250052497512 |
| NW_011925216.1 | 290094 | 291816 | 1723 | + | Gnomon | gene | NA | NA | gene8753 | GeneID:105282163 | LOC105282163 | Gene | LOC105282163 | lncRNA | NA | character(0) | character(0) | NW_011925216.1 | 216055 | 1092286 | 876232 | * | 2 | 0.00228250052497512 |
| NW_011925216.1 | 313949 | 321221 | 7273 | - | Gnomon | gene | NA | NA | gene8754 | GeneID:105282165 | LOC105282165 | Gene | LOC105282165 | protein_coding | NA | character(0) | character(0) | NW_011925216.1 | 216055 | 1092286 | 876232 | * | 2 | 0.00228250052497512 |
| NW_011925216.1 | 337863 | 347845 | 9983 | - | Gnomon | gene | NA | NA | gene8755 | GeneID:105282166 | LOC105282166 | Gene | LOC105282166 | lncRNA | NA | character(0) | character(0) | NW_011925216.1 | 216055 | 1092286 | 876232 | * | 2 | 0.00228250052497512 |
| NW_011925216.1 | 367045 | 368882 | 1838 | - | Gnomon | gene | NA | NA | gene8756 | GeneID:105282192 | LOC105282192 | Gene | LOC105282192 | protein_coding | NA | character(0) | character(0) | NW_011925216.1 | 216055 | 1092286 | 876232 | * | 2 | 0.00228250052497512 |
| NW_011925216.1 | 382786 | 404605 | 21820 | - | Gnomon | gene | NA | NA | gene8757 | GeneID:105282169 | LOC105282169 | Gene | LOC105282169 | protein_coding | NA | character(0) | character(0) | NW_011925216.1 | 216055 | 1092286 | 876232 | * | 2 | 0.00228250052497512 |
| NW_011925216.1 | 406528 | 409862 | 3335 | - | Gnomon | gene | NA | NA | gene8758 | GeneID:105282168 | LOC105282168 | Gene | LOC105282168 | protein_coding | NA | character(0) | character(0) | NW_011925216.1 | 216055 | 1092286 | 876232 | * | 2 | 0.00228250052497512 |
| NW_011925216.1 | 411413 | 433190 | 21778 | + | Gnomon | gene | NA | NA | gene8759 | GeneID:105282167 | LOC105282167 | Gene | LOC105282167 | lncRNA | NA | character(0) | character(0) | NW_011925216.1 | 216055 | 1092286 | 876232 | * | 2 | 0.00228250052497512 |
| NW_011925216.1 | 446916 | 480871 | 33956 | - | Gnomon | gene | NA | NA | gene8760 | GeneID:109611151 | LOC109611151 | Gene | LOC109611151 | lncRNA | NA | character(0) | character(0) | NW_011925216.1 | 216055 | 1092286 | 876232 | * | 2 | 0.00228250052497512 |
| NW_011925216.1 | 496030 | 584409 | 88380 | - | Gnomon | gene | NA | NA | gene8761 | GeneID:105282171 | LOC105282171 | Gene | LOC105282171 | protein_coding | NA | character(0) | character(0) | NW_011925216.1 | 216055 | 1092286 | 876232 | * | 2 | 0.00228250052497512 |
| NW_011925216.1 | 611697 | 703889 | 92193 | - | Gnomon | gene | NA | NA | gene8762 | GeneID:105282173 | LOC105282173 | Gene | LOC105282173 | protein_coding | NA | character(0) | character(0) | NW_011925216.1 | 216055 | 1092286 | 876232 | * | 2 | 0.00228250052497512 |
| NW_011925216.1 | 796968 | 820657 | 23690 | - | Gnomon | gene | NA | NA | gene8763 | GeneID:105282174 | LOC105282174 | Gene | LOC105282174 | protein_coding | NA | character(0) | character(0) | NW_011925216.1 | 216055 | 1092286 | 876232 | * | 2 | 0.00228250052497512 |
| NW_011925216.1 | 821894 | 915458 | 93565 | - | Gnomon | gene | NA | NA | gene8764 | GeneID:105282175 | LOC105282175 | Gene | LOC105282175 | lncRNA | NA | character(0) | character(0) | NW_011925216.1 | 216055 | 1092286 | 876232 | * | 2 | 0.00228250052497512 |
| NW_011925216.1 | 958187 | 960788 | 2602 | + | Gnomon | gene | NA | NA | gene8765 | GeneID:105282176 | LOC105282176 | Gene | LOC105282176 | lncRNA | NA | character(0) | character(0) | NW_011925216.1 | 216055 | 1092286 | 876232 | * | 2 | 0.00228250052497512 |
| NW_011925216.1 | 984296 | 991766 | 7471 | - | Gnomon | gene | NA | NA | gene8766 | GeneID:105282177 | LOC105282177 | Gene | LOC105282177 | protein_coding | NA | character(0) | character(0) | NW_011925216.1 | 216055 | 1092286 | 876232 | * | 2 | 0.00228250052497512 |
| NW_011925216.1 | 1091748 | 1092721 | 974 | + | Gnomon | gene | NA | NA | gene8767 | GeneID:105282179 | LOC105282179 | Gene | LOC105282179 | protein_coding | NA | character(0) | character(0) | NW_011925216.1 | 216055 | 1092286 | 876232 | * | 2 | 0.00228250052497512 |
| NW_011925036.1 | 544456 | 608309 | 63854 | + | Gnomon | gene | NA | NA | gene5618 | GeneID:109610912 | LOC109610912 | Gene | LOC109610912 | lncRNA | NA | character(0) | character(0) | NW_011925036.1 | 566786 | 1156139 | 589354 | * | 2 | 0.00339354615392447 |
| NW_011925036.1 | 608861 | 873355 | 264495 | + | Gnomon | gene | NA | NA | gene5619 | GeneID:105278508 | LOC105278508 | Gene | LOC105278508 | protein_coding | NA | character(0) | character(0) | NW_011925036.1 | 566786 | 1156139 | 589354 | * | 2 | 0.00339354615392447 |
| NW_011925036.1 | 927483 | 928885 | 1403 | + | Gnomon | gene | NA | NA | gene5620 | GeneID:105278511 | LOC105278511 | Gene | LOC105278511 | protein_coding | NA | character(0) | character(0) | NW_011925036.1 | 566786 | 1156139 | 589354 | * | 2 | 0.00339354615392447 |
| NW_011925036.1 | 929651 | 932446 | 2796 | + | Gnomon | gene | NA | NA | gene5621 | GeneID:105278510 | LOC105278510 | Gene | LOC105278510 | protein_coding | NA | character(0) | character(0) | NW_011925036.1 | 566786 | 1156139 | 589354 | * | 2 | 0.00339354615392447 |
| NW_011925036.1 | 1006642 | 1011165 | 4524 | + | Gnomon | gene | NA | NA | gene5622 | GeneID:105278519 | LOC105278519 | Gene | LOC105278519 | protein_coding | NA | character(0) | character(0) | NW_011925036.1 | 566786 | 1156139 | 589354 | * | 2 | 0.00339354615392447 |
| NW_011925036.1 | 1011369 | 1143497 | 132129 | - | Gnomon | gene | NA | NA | gene5623 | GeneID:105278518 | LOC105278518 | Gene | LOC105278518 | protein_coding | NA | character(0) | character(0) | NW_011925036.1 | 566786 | 1156139 | 589354 | * | 2 | 0.00339354615392447 |
| NW_011925036.1 | 1093799 | 1100136 | 6338 | + | Gnomon | gene | NA | NA | gene5624 | GeneID:105278516 | LOC105278516 | Gene | LOC105278516 | protein_coding | NA | character(0) | character(0) | NW_011925036.1 | 566786 | 1156139 | 589354 | * | 2 | 0.00339354615392447 |

...
... only first 25 lines shown ...

Directory: MRPR File: gwr-Cb_LI_BCphaseA1.txt

| seqnames | start | end | width | strand | source | type | score | phase | ID | Dbxref | Name | gbkey | gene | gene_biotype | partial | start_range | end_range | seqnames | start | end | width | strand | NbrSites | Sdnsty |
|---|---|---|---|---|---|---|---|---|---|---|---|---|---|---|---|---|---|---|---|---|---|---|---|---|
| NW_011924887.1 | 1817915 | 1820166 | 2252 | - | Gnomon | gene | NA | NA | gene234 | GeneID:105274775 | LOC105274775 | Gene | LOC105274775 | protein_coding | NA | character(0) | character(0) | NW_011924887.1 | 1819270 | 1819275 | 6 | * | 6 | 1000 |
| NW_011924888.1 | 389413 | 407214 | 17802 | - | Gnomon | gene | NA | NA | gene320 | GeneID:105275770 | LOC105275770 | Gene | LOC105275770 | protein_coding | NA | character(0) | character(0) | NW_011924888.1 | 405832 | 405837 | 6 | * | 6 | 1000 |
| NW_011924892.1 | 1158089 | 1160303 | 2215 | - | Gnomon | gene | NA | NA | gene569 | GeneID:105278527 | LOC105278527 | Gene | LOC105278527 | protein_coding | NA | character(0) | character(0) | NW_011924892.1 | 1159381 | 1159386 | 6 | * | 6 | 1000 |
| NW_011924973.1 | 828831 | 835120 | 6290 | + | Gnomon | gene | NA | NA | gene2799 | GeneID:105275316 | LOC105275316 | Gene | LOC105275316 | protein_coding | NA | character(0) | character(0) | NW_011924973.1 | 829674 | 829679 | 6 | * | 6 | 1000 |
| NW_011924973.1 | 1001098 | 1003969 | 2872 | + | Gnomon | gene | NA | NA | gene2837 | GeneID:105275355 | LOC105275355 | Gene | LOC105275355 | protein_coding | NA | character(0) | character(0) | NW_011924973.1 | 1002105 | 1002110 | 6 | * | 6 | 1000 |
| NW_011924977.1 | 662427 | 665182 | 2756 | + | Gnomon | gene | NA | NA | gene3008 | GeneID:105275550 | LOC105275550 | Gene | LOC105275550 | protein_coding | NA | character(0) | character(0) | NW_011924977.1 | 663534 | 663539 | 6 | * | 6 | 1000 |
| NW_011924996.1 | 75277 | 77259 | 1983 | + | Gnomon | gene | NA | NA | gene3743 | GeneID:105276382 | LOC105276382 | Gene | LOC105276382 | protein_coding | NA | character(0) | character(0) | NW_011924996.1 | 76074 | 76079 | 6 | * | 6 | 1000 |
| NW_011924996.1 | 1822758 | 1830685 | 7928 | - | Gnomon | gene | NA | NA | gene3885 | GeneID:105276527 | LOC105276527 | Gene | LOC105276527 | protein_coding | NA | character(0) | character(0) | NW_011924996.1 | 1829379 | 1829384 | 6 | * | 6 | 1000 |
| NW_011925007.1 | 926256 | 929694 | 3439 | + | Gnomon | gene | NA | NA | gene4194 | GeneID:105276881 | LOC105276881 | Gene | LOC105276881 | protein_coding | NA | character(0) | character(0) | NW_011925007.1 | 927466 | 927471 | 6 | * | 6 | 1000 |
| NW_011925022.1 | 386851 | 387976 | 1126 | - | Gnomon | gene | NA | NA | gene5000 | GeneID:105277804 | LOC105277804 | Gene | LOC105277804 | protein_coding | NA | character(0) | character(0) | NW_011925022.1 | 387077 | 387082 | 6 | * | 6 | 1000 |
| NW_011925053.1 | 649052 | 656133 | 7082 | - | Gnomon | gene | NA | NA | gene5882 | GeneID:105278812 | LOC105278812 | Gene | LOC105278812 | protein_coding | NA | character(0) | character(0) | NW_011925053.1 | 655139 | 655144 | 6 | * | 6 | 1000 |
| NW_011925142.1 | 777887 | 780161 | 2275 | - | Gnomon | gene | NA | NA | gene7574 | GeneID:105280791 | LOC105280791 | Gene | LOC105280791 | protein_coding | NA | character(0) | character(0) | NW_011925142.1 | 779002 | 779007 | 6 | * | 6 | 1000 |
| NW_011925163.1 | 470647 | 473680 | 3034 | + | Gnomon | gene | NA | NA | gene8040 | GeneID:105281341 | LOC105281341 | Gene | LOC105281341 | protein_coding | NA | character(0) | character(0) | NW_011925163.1 | 471213 | 471218 | 6 | * | 6 | 1000 |
| NW_011925183.1 | 39511 | 42630 | 3120 | + | Gnomon | gene | NA | NA | gene8236 | GeneID:105281577 | LOC105281577 | Gene | LOC105281577 | protein_coding | NA | character(0) | character(0) | NW_011925183.1 | 40051 | 40056 | 6 | * | 6 | 1000 |
| NW_011925188.1 | 199014 | 202632 | 3619 | - | Gnomon | gene | NA | NA | gene8257 | GeneID:105281603 | LOC105281603 | Gene | LOC105281603 | protein_coding | NA | character(0) | character(0) | NW_011925188.1 | 201626 | 201631 | 6 | * | 6 | 1000 |
| NW_011925203.1 | 63038 | 97839 | 34802 | + | Gnomon | gene | NA | NA | gene8521 | GeneID:105281906 | LOC105281906 | Gene | LOC105281906 | protein_coding | NA | character(0) | character(0) | NW_011925203.1 | 64603 | 64608 | 6 | * | 6 | 1000 |
| NW_011925216.1 | 156691 | 162086 | 5396 | + | Gnomon | gene | NA | NA | gene8741 | GeneID:105282150 | LOC105282150 | Gene | LOC105282150 | protein_coding | NA | character(0) | character(0) | NW_011925216.1 | 157541 | 157546 | 6 | * | 6 | 1000 |
| NW_011925258.1 | 404301 | 408788 | 4488 | - | Gnomon | gene | NA | NA | gene9347 | GeneID:105282847 | LOC105282847 | Gene | LOC105282847 | protein_coding | NA | character(0) | character(0) | NW_011925258.1 | 407771 | 407776 | 6 | * | 6 | 1000 |
| NW_011925304.1 | 378599 | 380691 | 2093 | - | Gnomon | gene | NA | NA | gene9814 | GeneID:105283373 | LOC105283373 | Gene | LOC105283373 | protein_coding | NA | character(0) | character(0) | NW_011925304.1 | 379611 | 379616 | 6 | * | 6 | 1000 |
| NW_011925321.1 | 975185 | 978704 | 3520 | - | Gnomon | gene | NA | NA | gene10031 | GeneID:105283620 | LOC105283620 | Gene | LOC105283620 | protein_coding | NA | character(0) | character(0) | NW_011925321.1 | 977422 | 977427 | 6 | * | 6 | 1000 |
| NW_011925321.1 | 1107715 | 1109901 | 2187 | - | Gnomon | gene | NA | NA | gene10043 | GeneID:105283634 | LOC105283634 | Gene | LOC105283634 | protein_coding | NA | character(0) | character(0) | NW_011925321.1 | 1108690 | 1108695 | 6 | * | 6 | 1000 |
| NW_011925353.1 | 40476 | 43146 | 2671 | - | Gnomon | gene | NA | NA | gene10405 | GeneID:105284057 | LOC105284057 | Gene | LOC105284057 | protein_coding | NA | character(0) | character(0) | NW_011925353.1 | 41948 | 41953 | 6 | * | 6 | 1000 |
| NW_011925356.1 | 1573184 | 1579703 | 6520 | - | Gnomon | gene | NA | NA | gene10596 | GeneID:105284262 | LOC105284262 | Gene | LOC105284262 | protein_coding | NA | character(0) | character(0) | NW_011925356.1 | 1573650 | 1573655 | 6 | * | 6 | 1000 |
| NW_011925367.1 | 1119922 | 1121887 | 1966 | + | Gnomon | gene | NA | NA | gene10714 | GeneID:105284396 | LOC105284396 | Gene | LOC105284396 | protein_coding | NA | character(0) | character(0) | NW_011925367.1 | 1120960 | 1120965 | 6 | * | 6 | 1000 |
| NW_011925387.1 | 39150 | 46759 | 7610 | + | Gnomon | gene | NA | NA | gene10928 | GeneID:105284644 | LOC105284644 | Gene | LOC105284644 | protein_coding | NA | character(0) | character(0) | NW_011925387.1 | 40039 | 40044 | 6 | * | 6 | 1000 |
| NW_011925470.1 | 613911 | 617551 | 3641 | - | Gnomon | gene | NA | NA | gene11628 | GeneID:105285465 | LOC105285465 | Gene | LOC105285465 | protein_coding | NA | character(0) | character(0) | NW_011925470.1 | 616616 | 616621 | 6 | * | 6 | 1000 |
| NW_011925490.1 | 137354 | 140755 | 3402 | + | Gnomon | gene | NA | NA | gene11689 | GeneID:105285534 | LOC105285534 | Gene | LOC105285534 | protein_coding | NA | character(0) | character(0) | NW_011925490.1 | 138283 | 138288 | 6 | * | 6 | 1000 |
| NW_011925595.1 | 1 | 4548 | 4548 | - | Gnomon | gene | NA | NA | gene12102 | GeneID:105286003 | LOC105286003 | Gene | LOC105286003 | protein_coding | NA | character(0) | character(0) | NW_011925595.1 | 3717 | 3722 | 6 | * | 6 | 1000 |
| NW_011925086.1 | 128430 | 135139 | 6710 | + | Gnomon | gene | NA | NA | gene6311 | GeneID:105279333 | LOC105279333 | Gene | LOC105279333 | protein_coding | NA | character(0) | character(0) | NW_011925086.1 | 129637 | 129647 | 11 | * | 10 | 909.090909090909 |
| NW_011924956.1 | 717757 | 722032 | 4276 | + | Gnomon | gene | NA | NA | gene2570 | GeneID:105275036 | LOC105275036 | Gene | LOC105275036 | protein_coding | NA | | | | | | | | | |

character(0) character(0) NW_011924956.1 719335 719344 10 * 9 900
NW_011924914.1 259948 262322 2375 - Gnomon gene NA NA gene952 GeneID:105283016 LOC105283016 Gene LOC105283016 protein_coding NA
character(0) character(0) NW_011924914.1 261398 261406 9 * 8 888.888888888889
NW_011924967.1 91700 93046 1347 - Gnomon gene NA NA gene2644 GeneID:105275137 LOC105275137 Gene LOC105275137 protein_coding NA character(0)
character(0) NW_011924967.1 92497 92505 9 * 8 888.888888888889
NW_011924977.1 3728770 3731404 2635 + Gnomon gene NA NA gene3213 GeneID:105275761 LOC105275761 Gene LOC105275761 protein_coding NA
character(0) character(0) NW_011924977.1 3729890 3729897 8 * 7 875
NW_011925664.1 38296 39300 1005 + Gnomon gene NA NA gene12311 GeneID:105286212 LOC105286212 Gene LOC105286212 protein_coding NA character(0)
character(0) NW_011925664.1 39054 39061 8 * 7 875
NW_011924887.1 547855 552902 5048 + Gnomon gene NA NA gene149 GeneID:105287687 LOC105287687 Gene LOC105287687 protein_coding NA
character(0) character(0) NW_011924887.1 549006 549012 7 * 6 857.142857142857
NW_011924888.1 30738 31847 1110 + Gnomon gene NA NA gene264 GeneID:105275153 LOC105275153 Gene LOC105275153 protein_coding NA character(0)
character(0) NW_011924888.1 31453 31459 7 * 6 857.142857142857
NW_011924889.1 476236 478868 2633 - Gnomon gene NA NA gene504 GeneID:105277817 LOC105277817 Gene LOC105277817 protein_coding NA
character(0) character(0) NW_011924889.1 477980 477986 7 * 6 857.142857142857
NW_011924889.1 606387 610322 3936 - Gnomon gene NA NA gene519 GeneID:105278022 LOC105278022 Gene LOC105278022 protein_coding NA
character(0) character(0) NW_011924889.1 609130 609136 7 * 6 857.142857142857
NW_011924889.1 606387 610322 3936 - Gnomon gene NA NA gene519 GeneID:105278022 LOC105278022 Gene LOC105278022 protein_coding NA
character(0) character(0) NW_011924889.1 609259 609265 7 * 6 857.142857142857
NW_011924892.1 274786 285690 10905 - Gnomon gene NA NA gene544 GeneID:105278288 LOC105278288 Gene LOC105278288 protein_coding NA
character(0) character(0) NW_011924892.1 284836 284842 7 * 6 857.142857142857
NW_011924907.1 889683 896841 7159 - Gnomon gene NA NA gene861 GeneID:105281750 LOC105281750 Gene LOC105281750 protein_coding NA
character(0) character(0) NW_011924907.1 896035 896041 7 * 6 857.142857142857
NW_011924914.1 87641 92698 5058 - Gnomon gene NA NA gene915 GeneID:105282459 LOC105282459 Gene LOC105282459 protein_coding NA character(0)
character(0) NW_011924914.1 90894 90900 7 * 6 857.142857142857
NW_011924914.1 827084 831408 4325 - Gnomon gene NA NA gene1010 GeneID:105283498 LOC105283498 Gene LOC105283498 protein_coding NA
character(0) character(0) NW_011924914.1 831099 831105 7 * 6 857.142857142857
NW_011924922.1 2390151 2392888 2738 - Gnomon gene NA NA gene1198 GeneID:105285507 LOC105285507 Gene LOC105285507 protein_coding NA
character(0) character(0) NW_011924922.1 2392009 2392015 7 * 6 857.142857142857
NW_011924931.1 201740 204575 2836 - Gnomon gene NA NA gene1371 GeneID:105287279 LOC105287279 Gene LOC105287279 protein_coding NA
character(0) character(0) NW_011924931.1 203497 203503 7 * 6 857.142857142857
NW_011924931.1 242950 250477 7528 - Gnomon gene NA NA gene1378 GeneID:105287371 LOC105287371 Gene LOC105287371 protein_coding NA
character(0) character(0) NW_011924931.1 249191 249197 7 * 6 857.142857142857
NW_011924931.1 347408 353524 6117 + Gnomon gene NA NA gene1391 GeneID:105287511 LOC105287511 Gene LOC105287511 protein_coding NA
character(0) character(0) NW_011924931.1 348868 348874 7 * 6 857.142857142857
NW_011924931.1 1565109 1566883 1775 + Gnomon gene NA NA gene1444 GeneID:105287569 LOC105287569 Gene LOC105287569 protein_coding NA
character(0) character(0) NW_011924931.1 1565961 1565967 7 * 6 857.142857142857
NW_011924931.1 2028839 2031812 2974 + Gnomon gene NA NA gene1470 GeneID:105287597 LOC105287597 Gene LOC105287597 protein_coding NA
character(0) character(0) NW_011924931.1 2030239 2030245 7 * 6 857.142857142857
NW_011924931.1 3979390 3986328 6939 + Gnomon gene NA NA gene1566 GeneID:105287698 LOC105287698 Gene LOC105287698 protein_coding NA
character(0) character(0) NW_011924931.1 3983680 3983686 7 * 6 857.142857142857
NW_011924931.1 4058265 4059684 1420 - Gnomon gene NA NA gene1581 GeneID:105287710 LOC105287710 Gene LOC105287710 protein_coding NA
character(0) character(0) NW_011924931.1 4058767 4058773 7 * 6 857.142857142857
NW_011924932.1 569126 573789 4664 + Gnomon gene NA NA gene1661 GeneID:105287820 LOC105287820 Gene LOC105287820 protein_coding NA
character(0) character(0) NW_011924932.1 570776 570782 7 * 6 857.142857142857
NW_011924938.1 72334 83258 10925 + Gnomon gene NA NA gene1805 GeneID:105287985 LOC105287985 Gene LOC105287985 protein_coding NA character(0)
character(0) NW_011924938.1 72914 72920 7 * 6 857.142857142857
NW_011924946.1 775753 777201 1449 - Gnomon gene NA NA gene1993 GeneID:105288193 LOC105288193 Gene LOC105288193 protein_coding NA
character(0) character(0) NW_011924946.1 776237 776243 7 * 6 857.142857142857
NW_011924946.1 2914862 2919248 4387 - Gnomon gene NA NA gene2173 GeneID:105274590 LOC105274590 Gene LOC105274590 protein_coding NA
character(0) character(0) NW_011924946.1 2918055 2918061 7 * 6 857.142857142857
NW_011924946.1 3060317 3062791 2475 - Gnomon gene NA NA gene2198 GeneID:105274618 LOC105274618 Gene LOC105274618 protein_coding NA
character(0) character(0) NW_011924946.1 3060957 3060963 7 * 6 857.142857142857
NW_011924946.1 3145253 3148790 3538 - Gnomon gene NA NA gene2215 GeneID:105274638 LOC105274638 Gene LOC105274638 protein_coding NA
character(0) character(0) NW_011924946.1 3147722 3147728 7 * 6 857.142857142857
NW_011924947.1 141424 143460 2037 + Gnomon gene NA NA gene2279 GeneID:105274711 LOC105274711 Gene LOC105274711 protein_coding NA
character(0) character(0) NW_011924947.1 142119 142125 7 * 6 857.142857142857
NW_011924953.1 401032 402582 1551 - Gnomon gene NA NA gene2400 GeneID:105274850 LOC105274850 Gene LOC105274850 protein_coding NA
character(0) character(0) NW_011924953.1 401754 401760 7 * 6 857.142857142857
NW_011924956.1 701839 703915 2077 - Gnomon gene NA NA gene2566 GeneID:105275031 LOC105275031 Gene LOC105275031 protein_coding NA
character(0) character(0) NW_011924956.1 702713 702719 7 * 6 857.142857142857
NW_011924967.1 11923 12765 843 - Gnomon gene NA NA gene2630 GeneID:105275119 LOC105275119 Gene LOC105275119 protein_coding NA character(0)
character(0) NW_011924967.1 12053 12059 7 * 6 857.142857142857
NW_011924967.1 91700 93046 1347 - Gnomon gene NA NA gene2644 GeneID:105275137 LOC105275137 Gene LOC105275137 protein_coding NA character(0)

character(0) NW_011924967.1 92468 92474 7 * 6 857.142857142857
NW_011924973.1 774415 775843 1429 - Gnomon gene NA NA gene2790 GeneID:105275307 LOC105275307 Gene LOC105275307 protein_coding NA
character(0) character(0) NW_011924973.1 775075 775081 7 * 6 857.142857142857
NW_011924974.1 483919 487626 3708 + Gnomon gene NA NA gene2873 GeneID:105275401 LOC105275401 Gene LOC105275401 protein_coding NA
character(0) character(0) NW_011924974.1 484702 484708 7 * 6 857.142857142857
NW_011924977.1 1460154 1463690 3537 - Gnomon gene NA NA gene3072 GeneID:105275615 LOC105275615 Gene LOC105275615 protein_coding NA
character(0) character(0) NW_011924977.1 1461887 1461893 7 * 6 857.142857142857
NW_011924977.1 1460154 1463690 3537 - Gnomon gene NA NA gene3072 GeneID:105275615 LOC105275615 Gene LOC105275615 protein_coding NA
character(0) character(0) NW_011924977.1 1462188 1462194 7 * 6 857.142857142857
NW_011924977.1 2573287 2581453 8167 - Gnomon gene NA NA gene3147 GeneID:105275689 LOC105275689 Gene LOC105275689 protein_coding NA
character(0) character(0) NW_011924977.1 2580506 2580512 7 * 6 857.142857142857
NW_011924977.1 3678529 3682684 4156 - Gnomon gene NA NA gene3205 GeneID:105275753 LOC105275753 Gene LOC105275753 protein_coding NA
character(0) character(0) NW_011924977.1 3681359 3681365 7 * 6 857.142857142857
NW_011924977.1 3678529 3682684 4156 - Gnomon gene NA NA gene3205 GeneID:105275753 LOC105275753 Gene LOC105275753 protein_coding NA
character(0) character(0) NW_011924977.1 3681568 3681574 7 * 6 857.142857142857
NW_011924977.1 3682584 3686057 3474 - Gnomon gene NA NA gene3206 GeneID:105275752 LOC105275752 Gene LOC105275752 protein_coding NA
character(0) character(0) NW_011924977.1 3684978 3684984 7 * 6 857.142857142857
NW_011924979.1 2744383 2746071 1689 + Gnomon gene NA NA gene3334 GeneID:105275897 LOC105275897 Gene LOC105275897 protein_coding NA
character(0) character(0) NW_011924979.1 2745351 2745357 7 * 6 857.142857142857
NW_011924987.1 2638171 2648657 10487 - Gnomon gene NA NA gene3636 GeneID:105276233 LOC105276233 Gene LOC105276233 protein_coding NA
character(0) character(0) NW_011924987.1 2647465 2647471 7 * 6 857.142857142857
NW_011924996.1 292526 303470 10945 + Gnomon gene NA NA gene3775 GeneID:105276410 LOC105276410 Gene LOC105276410 protein_coding NA
character(0) character(0) NW_011924996.1 293762 293768 7 * 6 857.142857142857
NW_011924996.1 292526 303470 10945 + Gnomon gene NA NA gene3775 GeneID:105276410 LOC105276410 Gene LOC105276410 protein_coding NA
character(0) character(0) NW_011924996.1 293819 293825 7 * 6 857.142857142857
NW_011924996.1 641332 646889 5558 - Gnomon gene NA NA gene3801 GeneID:105276437 LOC105276437 Gene LOC105276437 protein_coding NA
character(0) character(0) NW_011924996.1 645416 645422 7 * 6 857.142857142857
NW_011925007.1 1417723 1421435 3713 + Gnomon gene NA NA gene4248 GeneID:105276948 LOC105276948 Gene LOC105276948 protein_coding NA
character(0) character(0) NW_011925007.1 1419353 1419359 7 * 6 857.142857142857
NW_011925007.1 1452162 1454301 2140 + Gnomon gene NA NA gene4252 GeneID:105276949 LOC105276949 Gene LOC105276949 protein_coding NA
character(0) character(0) NW_011925007.1 1453109 1453115 7 * 6 857.142857142857
NW_011925007.1 3760218 3764466 4249 - Gnomon gene NA NA gene4383 GeneID:105277128 LOC105277128 Gene LOC105277128 protein_coding NA
character(0) character(0) NW_011925007.1 3760305 3760311 7 * 6 857.142857142857
NW_011925012.1 1316386 1318477 2092 + Gnomon gene NA NA gene4492 GeneID:105277232 LOC105277232 Gene LOC105277232 protein_coding NA
character(0) character(0) NW_011925012.1 1317469 1317475 7 * 6 857.142857142857
NW_011925013.1 5041 7730 2690 - Gnomon gene NA NA gene4511 GeneID:105277256 LOC105277256 Gene LOC105277256 protein_coding NA character(0)
character(0) NW_011925013.1 6764 6770 7 * 6 857.142857142857
NW_011925019.1 475 2900 2426 + Gnomon gene NA NA gene4633 GeneID:105277389 LOC105277389 Gene LOC105277389 protein_coding NA character(0)
character(0) NW_011925019.1 1670 1676 7 * 6 857.142857142857
NW_011925019.1 1427431 1429968 2538 - Gnomon gene NA NA gene4709 GeneID:109610850 LOC109610850 Gene LOC109610850 protein_coding NA
character(0) character(0) NW_011925019.1 1429240 1429246 7 * 6 857.142857142857
NW_011925022.1 1610155 1615436 5282 + Gnomon gene NA NA gene5106 GeneID:105277909 LOC105277909 Gene LOC105277909 protein_coding NA
character(0) character(0) NW_011925022.1 1611777 1611783 7 * 6 857.142857142857
NW_011925029.1 39174 43410 4237 + Gnomon gene NA NA gene5243 GeneID:105278106 LOC105278106 Gene LOC105278106 protein_coding NA character(0)
character(0) NW_011925029.1 40261 40267 7 * 6 857.142857142857
NW_011925035.1 3102848 3104698 1851 + Gnomon gene NA NA gene5557 GeneID:105278431 LOC105278431 Gene LOC105278431 protein_coding NA
character(0) character(0) NW_011925035.1 3103902 3103908 7 * 6 857.142857142857
NW_011925036.1 1418708 1424550 5843 + Gnomon gene NA NA gene5653 GeneID:105278544 LOC105278544 Gene LOC105278544 protein_coding NA
character(0) character(0) NW_011925036.1 1420178 1420184 7 * 6 857.142857142857
NW_011925050.1 126141 128598 2458 - Gnomon gene NA NA gene5737 GeneID:105278656 LOC105278656 Gene LOC105278656 protein_coding NA
character(0) character(0) NW_011925050.1 127417 127423 7 * 6 857.142857142857
NW_011925067.1 5971 15150 9180 + Gnomon gene NA NA gene6024 GeneID:105278996 LOC105278996 Gene LOC105278996 protein_coding NA character(0)
character(0) NW_011925067.1 7054 7060 7 * 6 857.142857142857
NW_011925082.1 664899 668532 3634 - Gnomon gene NA NA gene6299 GeneID:105279313 LOC105279313 Gene LOC105279313 protein_coding NA
character(0) character(0) NW_011925082.1 667658 667664 7 * 6 857.142857142857
NW_011925086.1 250638 255287 4650 - Gnomon gene NA NA gene6324 GeneID:105279346 LOC105279346 Gene LOC105279346 protein_coding NA
character(0) character(0) NW_011925086.1 254146 254152 7 * 6 857.142857142857
NW_011925106.1 154197 158224 4028 + Gnomon gene NA NA gene6601 GeneID:105279664 LOC105279664 Gene LOC105279664 protein_coding NA
character(0) character(0) NW_011925106.1 155492 155498 7 * 6 857.142857142857
NW_011925106.1 431654 436050 4397 + Gnomon gene NA NA gene6641 GeneID:105279709 LOC105279709 Gene LOC105279709 protein_coding NA
character(0) character(0) NW_011925106.1 432897 432903 7 * 6 857.142857142857
NW_011925106.1 512988 515818 2831 - Gnomon gene NA NA gene6652 GeneID:105279719 LOC105279719 Gene LOC105279719 protein_coding NA
character(0) character(0) NW_011925106.1 514935 514941 7 * 6 857.142857142857
NW_011925106.1 629782 633106 3325 - Gnomon gene NA NA gene6668 GeneID:105279740 LOC105279740 Gene LOC105279740 protein_coding NA

character(0) character(0) NW_011925106.1 631881 631887 7 * 6 857.142857142857
NW_011925106.1 779647 785206 5560 + Gnomon gene NA NA gene6679 GeneID:105279748 LOC105279748 Gene LOC105279748 protein_coding NA
character(0) character(0) NW_011925106.1 780733 780739 7 * 6 857.142857142857
NW_011925106.1 2086959 2096661 9703 + Gnomon gene NA NA gene6771 GeneID:105279854 LOC105279854 Gene LOC105279854 protein_coding NA
character(0) character(0) NW_011925106.1 2087848 2087854 7 * 6 857.142857142857
NW_011925118.1 457689 459947 2259 + Gnomon gene NA NA gene6953 GeneID:105280092 LOC105280092 Gene LOC105280092 protein_coding NA
character(0) character(0) NW_011925118.1 458682 458688 7 * 6 857.142857142857
NW_011925118.1 481415 484075 2661 + Gnomon gene NA NA gene6956 GeneID:105280102 LOC105280102 Gene LOC105280102 protein_coding NA
character(0) character(0) NW_011925118.1 482695 482701 7 * 6 857.142857142857
NW_011925128.1 7781 9472 1692 + Gnomon gene NA NA gene7085 GeneID:105280236 LOC105280236 Gene LOC105280236 protein_coding NA character(0)
character(0) NW_011925128.1 8930 8936 7 * 6 857.142857142857
NW_011925128.1 800543 802328 1786 - Gnomon gene NA NA gene7160 GeneID:105280319 LOC105280319 Gene LOC105280319 protein_coding NA
character(0) character(0) NW_011925128.1 801605 801611 7 * 6 857.142857142857
NW_011925142.1 423375 426060 2686 - Gnomon gene NA NA gene7527 GeneID:105280739 LOC105280739 Gene LOC105280739 protein_coding NA
character(0) character(0) NW_011925142.1 424101 424107 7 * 6 857.142857142857
NW_011925142.1 423375 426060 2686 - Gnomon gene NA NA gene7527 GeneID:105280739 LOC105280739 Gene LOC105280739 protein_coding NA
character(0) character(0) NW_011925142.1 424227 424233 7 * 6 857.142857142857
NW_011925161.1 210886 215244 4359 + Gnomon gene NA NA gene7720 GeneID:105280964 LOC105280964 Gene LOC105280964 protein_coding NA
character(0) character(0) NW_011925161.1 213136 213142 7 * 6 857.142857142857
NW_011925161.1 3740915 3743520 2606 + Gnomon gene NA NA gene7963 GeneID:105281227 LOC105281227 Gene LOC105281227 protein_coding NA
character(0) character(0) NW_011925161.1 3741983 3741989 7 * 6 857.142857142857
NW_011925163.1 477881 479636 1756 - Gnomon gene NA NA gene8042 GeneID:105281339 LOC105281339 Gene LOC105281339 protein_coding NA
character(0) character(0) NW_011925163.1 478478 478484 7 * 6 857.142857142857
NW_011925183.1 39511 42630 3120 + Gnomon gene NA NA gene8236 GeneID:105281577 LOC105281577 Gene LOC105281577 protein_coding NA character(0)
character(0) NW_011925183.1 40353 40359 7 * 6 857.142857142857
NW_011925188.1 199014 202632 3619 - Gnomon gene NA NA gene8257 GeneID:105281603 LOC105281603 Gene LOC105281603 protein_coding NA
character(0) character(0) NW_011925188.1 201441 201447 7 * 6 857.142857142857
NW_011925189.1 165659 175663 10005 - Gnomon gene NA NA gene8294 GeneID:105281645 LOC105281645 Gene LOC105281645 protein_coding NA
character(0) character(0) NW_011925189.1 174278 174284 7 * 6 857.142857142857
NW_011925203.1 23205 24951 1747 - Gnomon gene NA NA gene8516 GeneID:105281896 LOC105281896 Gene LOC105281896 protein_coding NA character(0)
character(0) NW_011925203.1 24180 24186 7 * 6 857.142857142857
NW_011925209.1 85655 90207 4553 - Gnomon gene NA NA gene8551 GeneID:105281943 LOC105281943 Gene LOC105281943 protein_coding NA character(0)
character(0) NW_011925209.1 86875 86881 7 * 6 857.142857142857
NW_011925209.1 169877 173749 3873 - Gnomon gene NA NA gene8560 GeneID:105281949 LOC105281949 Gene LOC105281949 protein_coding NA
character(0) character(0) NW_011925209.1 172697 172703 7 * 6 857.142857142857
NW_011925221.1 474109 476408 2300 + Gnomon gene NA NA gene8798 GeneID:105282220 LOC105282220 Gene LOC105282220 protein_coding NA
character(0) character(0) NW_011925221.1 474855 474861 7 * 6 857.142857142857
NW_011925224.1 208794 211425 2632 + Gnomon gene NA NA gene8852 GeneID:105282278 LOC105282278 Gene LOC105282278 protein_coding NA
character(0) character(0) NW_011925224.1 209730 209736 7 * 6 857.142857142857
NW_011925224.1 219079 219935 857 - Gnomon gene NA NA gene8854 GeneID:105282280 LOC105282280 Gene LOC105282280 protein_coding NA
character(0) character(0) NW_011925224.1 219257 219263 7 * 6 857.142857142857
NW_011925224.1 327268 339256 11989 - Gnomon gene NA NA gene8872 GeneID:105282298 LOC105282298 Gene LOC105282298 protein_coding NA
character(0) character(0) NW_011925224.1 338306 338312 7 * 6 857.142857142857
NW_011925240.1 1526960 1531289 4330 + Gnomon gene NA NA gene9107 GeneID:105282559 LOC105282559 Gene LOC105282559 protein_coding NA
character(0) character(0) NW_011925240.1 1527897 1527903 7 * 6 857.142857142857
NW_011925241.1 397336 409261 11926 - Gnomon gene NA NA gene9191 GeneID:105282655 LOC105282655 Gene LOC105282655 protein_coding NA
character(0) character(0) NW_011925241.1 408196 408202 7 * 6 857.142857142857
NW_011925257.1 52519 55399 2881 - Gnomon gene NA NA gene9303 GeneID:105282793 LOC105282793 Gene LOC105282793 protein_coding NA character(0)
character(0) NW_011925257.1 54841 54847 7 * 6 857.142857142857
NW_011925267.1 224087 232805 8719 + Gnomon gene NA NA gene9418 GeneID:105282921 LOC105282921 Gene LOC105282921 protein_coding NA
character(0) character(0) NW_011925267.1 225570 225576 7 * 6 857.142857142857
NW_011925315.1 538327 543302 4976 + Gnomon gene NA NA gene9909 GeneID:105283484 LOC105283484 Gene LOC105283484 protein_coding NA
character(0) character(0) NW_011925315.1 539621 539627 7 * 6 857.142857142857
NW_011925323.1 1444062 1448038 3977 - Gnomon gene NA NA gene10138 GeneID:105283741 LOC105283741 Gene LOC105283741 protein_coding NA
character(0) character(0) NW_011925323.1 1445991 1445997 7 * 6 857.142857142857
NW_011925327.1 214150 224738 10589 + Gnomon gene NA NA gene10207 GeneID:105283818 LOC105283818 Gene LOC105283818 protein_coding NA
character(0) character(0) NW_011925327.1 216086 216092 7 * 6 857.142857142857
NW_011925336.1 92449 95536 3088 + Gnomon gene NA NA gene10324 GeneID:105283961 LOC105283961 Gene LOC105283961 protein_coding NA character(0)
character(0) NW_011925336.1 93305 93311 7 * 6 857.142857142857
NW_011925336.1 100271 103076 2806 - Gnomon gene NA NA gene10327 GeneID:105283964 LOC105283964 Gene LOC105283964 protein_coding NA
character(0) character(0) NW_011925336.1 102423 102429 7 * 6 857.142857142857
NW_011925353.1 16223 20332 4110 + Gnomon gene NA NA gene10401 GeneID:105284052 LOC105284052 Gene LOC105284052 protein_coding NA character(0)
character(0) NW_011925353.1 19139 19145 7 * 6 857.142857142857
NW_011925356.1 701727 704945 3219 + Gnomon gene NA NA gene10538 GeneID:105284202 LOC105284202 Gene LOC105284202 protein_coding NA

character(0) character(0) NW_011925356.1 703128 703134 7 * 6 857.142857142857
NW_011925367.1 1119922 1121887 1966 + Gnomon gene NA NA gene10714 GeneID:105284396 LOC105284396 Gene LOC105284396 protein_coding NA
character(0) character(0) NW_011925367.1 1120846 1120852 7 * 6 857.142857142857
NW_011925371.1 899674 901634 1961 - Gnomon gene NA NA gene10880 GeneID:105284576 LOC105284576 Gene LOC105284576 protein_coding NA
character(0) character(0) NW_011925371.1 900958 900964 7 * 6 857.142857142857
NW_011925387.1 118203 119961 1759 - Gnomon gene NA NA gene10943 GeneID:105284659 LOC105284659 Gene LOC105284659 protein_coding NA
character(0) character(0) NW_011925387.1 119303 119309 7 * 6 857.142857142857
NW_011925434.1 120720 130419 9700 + Gnomon gene NA NA gene11344 GeneID:105285135 LOC105285135 Gene LOC105285135 protein_coding NA
character(0) character(0) NW_011925434.1 123131 123137 7 * 6 857.142857142857
NW_011925446.1 246855 250131 3277 + Gnomon gene NA NA gene11531 GeneID:105285345 LOC105285345 Gene LOC105285345 protein_coding NA
character(0) character(0) NW_011925446.1 248422 248428 7 * 6 857.142857142857
NW_011925459.1 186244 187554 1311 - Gnomon gene NA NA gene11569 GeneID:105285394 LOC105285394 Gene LOC105285394 protein_coding NA
character(0) character(0) NW_011925459.1 186700 186706 7 * 6 857.142857142857
NW_011925465.1 71398 78050 6653 - Gnomon gene NA NA gene11580 GeneID:105285410 LOC105285410 Gene LOC105285410 protein_coding NA character(0)
character(0) NW_011925465.1 77157 77163 7 * 6 857.142857142857
NW_011925471.1 1427 3141 1715 - Gnomon gene NA NA gene11632 GeneID:109611363 LOC109611363 Gene LOC109611363 lncRNA NA character(0)
character(0) NW_011925471.1 2780 2786 7 * 6 857.142857142857
NW_011925516.1 278917 286323 7407 - Gnomon gene NA NA gene11797 GeneID:105285648 LOC105285648 Gene LOC105285648 protein_coding NA
character(0) character(0) NW_011925516.1 285124 285130 7 * 6 857.142857142857
NW_011925546.1 59043 60354 1312 + Gnomon gene NA NA gene11979 GeneID:105285864 LOC105285864 Gene LOC105285864 protein_coding NA character(0)
character(0) NW_011925546.1 59986 59992 7 * 6 857.142857142857
NW_011925549.1 267852 270480 2629 - Gnomon gene NA NA gene12010 GeneID:105285889 LOC105285889 Gene LOC105285889 protein_coding NA
character(0) character(0) NW_011925549.1 269441 269447 7 * 6 857.142857142857
NW_011925664.1 495388 526731 31344 + Gnomon gene NA NA gene12399 GeneID:105286310 LOC105286310 Gene LOC105286310 lncRNA NA
character(0) character(0) NW_011925664.1 514139 514145 7 * 6 857.142857142857
NW_011925664.1 513347 514759 1413 - Gnomon gene NA NA gene12403 GeneID:105286311 LOC105286311 Gene LOC105286311 protein_coding NA
character(0) character(0) NW_011925664.1 514139 514145 7 * 6 857.142857142857
NW_011925664.1 845127 848378 3252 - Gnomon gene NA NA gene12441 GeneID:105286355 LOC105286355 Gene LOC105286355 protein_coding NA
character(0) character(0) NW_011925664.1 847281 847287 7 * 6 857.142857142857
NW_011925715.1 251551 257367 5817 - Gnomon gene NA NA gene12526 GeneID:105286461 LOC105286461 Gene LOC105286461 protein_coding NA
character(0) character(0) NW_011925715.1 256038 256044 7 * 6 857.142857142857
NW_011926194.1 11132 18740 7609 + Gnomon gene NA NA gene13114 GeneID:105287147 LOC105287147 Gene LOC105287147 protein_coding NA character(0)
character(0) NW_011926194.1 12247 12253 7 * 6 857.142857142857
NW_011924899.1 718895 720452 1558 + Gnomon gene NA NA gene664 GeneID:105279718 LOC105279718 Gene LOC105279718 protein_coding NA
character(0) character(0) NW_011924899.1 719598 719607 10 * 8 800
NW_011924951.1 73901 76560 2660 + Gnomon gene NA NA gene2325 GeneID:105274767 LOC105274767 Gene LOC105274767 protein_coding NA character(0)
character(0) NW_011924951.1 74986 74995 10 * 8 800
NW_011925007.1 3759369 3760273 905 + Gnomon gene NA NA gene4382 GeneID:105277091 LOC105277091 Gene LOC105277091 protein_coding NA
character(0) character(0) NW_011925007.1 3760113 3760122 10 * 8 800

Directory: MRPR File: gwr-Cb_LI_BCphaseA2.txt

```
seqnames    start end   width strand      source      type  score phase ID    Dbxref      Name  gbkey gene  gene_biotype      partial     start_range end_range   seqnames    start
      end   width strand      NbrSites    Sdnsty
NW_011924887.1    1546715     1553169     6455  +     Gnomon      gene  NA    NA    gene209     GeneID:105274475  LOC105274475      Gene  LOC105274475      protein_coding    NA
      character(0)      character(0)      NW_011924887.1    1550201     1550206     6     *     6     1000
NW_011924887.1    1817915     1820166     2252  -     Gnomon      gene  NA    NA    gene234     GeneID:105274775  LOC105274775      Gene  LOC105274775      protein_coding    NA
      character(0)      character(0)      NW_011924887.1    1819270     1819275     6     *     6     1000
NW_011924888.1    389413      407214      17802 -     Gnomon      gene  NA    NA    gene320     GeneID:105275770  LOC105275770      Gene  LOC105275770      protein_coding    NA
      character(0)      character(0)      NW_011924888.1    405832      405837      6     *     6     1000
NW_011924892.1    1158089     1160303     2215  -     Gnomon      gene  NA    NA    gene569     GeneID:105278527  LOC105278527      Gene  LOC105278527      protein_coding    NA
      character(0)      character(0)      NW_011924892.1    1159381     1159386     6     *     6     1000
NW_011924931.1    3778005     3781577     3573  -     Gnomon      gene  NA    NA    gene1536    GeneID:105287663  LOC105287663      Gene  LOC105287663      protein_coding    NA
      character(0)      character(0)      NW_011924931.1    3780672     3780677     6     *     6     1000
NW_011924931.1    4488185     4490767     2583  +     Gnomon      gene  NA    NA    gene1620    GeneID:105287754  LOC105287754      Gene  LOC105287754      protein_coding    NA
      character(0)      character(0)      NW_011924931.1    4489292     4489297     6     *     6     1000
NW_011924940.1    334573      349050      14478 -     Gnomon      gene  NA    NA    gene1863    GeneID:105288044  LOC105288044      Gene  LOC105288044      protein_coding    NA
      character(0)      character(0)      NW_011924940.1    347385      347390      6     *     6     1000
NW_011924946.1    1502005     1507040     5036  -     Gnomon      gene  NA    NA    gene2047    GeneID:105274446  LOC105274446      Gene  LOC105274446      protein_coding    NA
      character(0)      character(0)      NW_011924946.1    1502183     1502188     6     *     6     1000
NW_011924956.1    717757      722032      4276  +     Gnomon      gene  NA    NA    gene2570    GeneID:105275036  LOC105275036      Gene  LOC105275036      protein_coding    NA
      character(0)      character(0)      NW_011924956.1    719335      719344      10    *     10    1000
NW_011924973.1    828831      835120      6290  +     Gnomon      gene  NA    NA    gene2799    GeneID:105275316  LOC105275316      Gene  LOC105275316      protein_coding    NA
      character(0)      character(0)      NW_011924973.1    829674      829679      6     *     6     1000
NW_011924973.1    1001098     1003969     2872  +     Gnomon      gene  NA    NA    gene2837    GeneID:105275355  LOC105275355      Gene  LOC105275355      protein_coding    NA
      character(0)      character(0)      NW_011924973.1    1002105     1002110     6     *     6     1000
NW_011924977.1    31430 34619 3190  +     Gnomon      gene  NA    NA    gene2916    GeneID:109610741  LOC109610741      Gene  LOC109610741      protein_coding    NA    character(0)
      character(0)      NW_011924977.1    32395 32400 6     *     6     1000
NW_011924977.1    662427      665182      2756  +     Gnomon      gene  NA    NA    gene3008    GeneID:105275550  LOC105275550      Gene  LOC105275550      protein_coding    NA
      character(0)      character(0)      NW_011924977.1    663534      663539      6     *     6     1000
NW_011924979.1    2744383     2746071     1689  +     Gnomon      gene  NA    NA    gene3334    GeneID:105275897  LOC105275897      Gene  LOC105275897      protein_coding    NA
      character(0)      character(0)      NW_011924979.1    2745453     2745458     6     *     6     1000
NW_011924996.1    75277 77259 1983  +     Gnomon      gene  NA    NA    gene3743    GeneID:105276382  LOC105276382      Gene  LOC105276382      protein_coding    NA    character(0)
      character(0)      NW_011924996.1    76074 76079 6     *     6     1000
NW_011925007.1    926256      929694      3439  +     Gnomon      gene  NA    NA    gene4194    GeneID:105276881  LOC105276881      Gene  LOC105276881      protein_coding    NA
      character(0)      character(0)      NW_011925007.1    927466      927471      6     *     6     1000
NW_011925007.1    1370227     1386638     16412 -     Gnomon      gene  NA    NA    gene4246    GeneID:105276941  LOC105276941      Gene  LOC105276941      protein_coding    NA
      character(0)      character(0)      NW_011925007.1    1385156     1385161     6     *     6     1000
NW_011925022.1    386851      387976      1126  -     Gnomon      gene  NA    NA    gene5000    GeneID:105277804  LOC105277804      Gene  LOC105277804      protein_coding    NA
      character(0)      character(0)      NW_011925022.1    387077      387082      6     *     6     1000
NW_011925053.1    649052      656133      7082  -     Gnomon      gene  NA    NA    gene5882    GeneID:105278812  LOC105278812      Gene  LOC105278812      protein_coding    NA
      character(0)      character(0)      NW_011925053.1    655139      655144      6     *     6     1000
NW_011925053.1    1481875     1483457     1583  -     Gnomon      gene  NA    NA    gene5932    GeneID:105278862  LOC105278862      Gene  LOC105278862      protein_coding    NA
      character(0)      character(0)      NW_011925053.1    1482901     1482906     6     *     6     1000
NW_011925128.1    1875336     1880845     5510  +     Gnomon      gene  NA    NA    gene7225    GeneID:105280395  LOC105280395      Gene  LOC105280395      protein_coding    NA
      character(0)      character(0)      NW_011925128.1    1876634     1876639     6     *     6     1000
NW_011925142.1    777887      780161      2275  -     Gnomon      gene  NA    NA    gene7574    GeneID:105280791  LOC105280791      Gene  LOC105280791      protein_coding    NA
      character(0)      character(0)      NW_011925142.1    779002      779007      6     *     6     1000
NW_011925183.1    39511 42630 3120  +     Gnomon      gene  NA    NA    gene8236    GeneID:105281577  LOC105281577      Gene  LOC105281577      protein_coding    NA    character(0)
      character(0)      NW_011925183.1    40051 40056 6     *     6     1000
NW_011925188.1    199014      202632      3619  -     Gnomon      gene  NA    NA    gene8257    GeneID:105281603  LOC105281603      Gene  LOC105281603      protein_coding    NA
      character(0)      character(0)      NW_011925188.1    201626      201631      6     *     6     1000
...
... only first 25 lines shown ...
```

Directory: MRPR File: gwr-Cb_LI_BCphaseB1.txt

```
seqnames	start	end	width	strand	source	type	score	phase	ID	Dbxref	Name	gbkey	gene	gene_biotype	partial	start_range	end_range	seqnames	start
	end	width	strand	NbrSites	Sdnsty
NW_011924887.1	1817915	1820166	2252	-	Gnomon	gene	NA	NA	gene234	GeneID:105274775	LOC105274775	Gene	LOC105274775	protein_coding	NA
	character(0)	character(0)	NW_011924887.1	1819270	1819275	6	*	6	1000
NW_011924888.1	389413	407214	17802	-	Gnomon	gene	NA	NA	gene320	GeneID:105275770	LOC105275770	Gene	LOC105275770	protein_coding	NA
	character(0)	character(0)	NW_011924888.1	405832	405837	6	*	6	1000
NW_011924892.1	1158089	1160303	2215	-	Gnomon	gene	NA	NA	gene569	GeneID:105278527	LOC105278527	Gene	LOC105278527	protein_coding	NA
	character(0)	character(0)	NW_011924892.1	1159381	1159386	6	*	6	1000
NW_011924931.1	4488185	4490767	2583	+	Gnomon	gene	NA	NA	gene1620	GeneID:105287754	LOC105287754	Gene	LOC105287754	protein_coding	NA
	character(0)	character(0)	NW_011924931.1	4489292	4489297	6	*	6	1000
NW_011924946.1	1411689	1414880	3192	-	Gnomon	gene	NA	NA	gene2041	GeneID:105288248	LOC105288248	Gene	LOC105288248	protein_coding	NA
	character(0)	character(0)	NW_011924946.1	1413935	1413940	6	*	6	1000
NW_011924956.1	717757	722032	4276	+	Gnomon	gene	NA	NA	gene2570	GeneID:105275036	LOC105275036	Gene	LOC105275036	protein_coding	NA
	character(0)	character(0)	NW_011924956.1	719335	719344	10	*	10	1000
NW_011924973.1	244133	245537	1405	+	Gnomon	gene	NA	NA	gene2754	GeneID:105275270	LOC105275270	Gene	LOC105275270	protein_coding	NA
	character(0)	character(0)	NW_011924973.1	244951	244956	6	*	6	1000
NW_011924973.1	828831	835120	6290	+	Gnomon	gene	NA	NA	gene2799	GeneID:105275316	LOC105275316	Gene	LOC105275316	protein_coding	NA
	character(0)	character(0)	NW_011924973.1	829674	829679	6	*	6	1000
NW_011924973.1	1001098	1003969	2872	+	Gnomon	gene	NA	NA	gene2837	GeneID:105275355	LOC105275355	Gene	LOC105275355	protein_coding	NA
	character(0)	character(0)	NW_011924973.1	1002105	1002110	6	*	6	1000
NW_011924977.1	31430	34619	3190	+	Gnomon	gene	NA	NA	gene2916	GeneID:109610741	LOC109610741	Gene	LOC109610741	protein_coding	NA	character(0)
	character(0)	NW_011924977.1	32395	32400	6	*	6	1000
NW_011924977.1	662427	665182	2756	+	Gnomon	gene	NA	NA	gene3008	GeneID:105275550	LOC105275550	Gene	LOC105275550	protein_coding	NA
	character(0)	character(0)	NW_011924977.1	663534	663539	6	*	6	1000
NW_011924996.1	75277	77259	1983	+	Gnomon	gene	NA	NA	gene3743	GeneID:105276382	LOC105276382	Gene	LOC105276382	protein_coding	NA	character(0)
	character(0)	NW_011924996.1	76074	76079	6	*	6	1000
NW_011924996.1	1822758	1830685	7928	-	Gnomon	gene	NA	NA	gene3885	GeneID:105276527	LOC105276527	Gene	LOC105276527	protein_coding	NA
	character(0)	character(0)	NW_011924996.1	1829379	1829384	6	*	6	1000
NW_011925007.1	926256	929694	3439	+	Gnomon	gene	NA	NA	gene4194	GeneID:105276881	LOC105276881	Gene	LOC105276881	protein_coding	NA
	character(0)	character(0)	NW_011925007.1	927466	927471	6	*	6	1000
NW_011925022.1	386851	387976	1126	-	Gnomon	gene	NA	NA	gene5000	GeneID:105277804	LOC105277804	Gene	LOC105277804	protein_coding	NA
	character(0)	character(0)	NW_011925022.1	387077	387082	6	*	6	1000
NW_011925053.1	649052	656133	7082	-	Gnomon	gene	NA	NA	gene5882	GeneID:105278812	LOC105278812	Gene	LOC105278812	protein_coding	NA
	character(0)	character(0)	NW_011925053.1	655139	655144	6	*	6	1000
NW_011925053.1	1481875	1483457	1583	-	Gnomon	gene	NA	NA	gene5932	GeneID:105278862	LOC105278862	Gene	LOC105278862	protein_coding	NA
	character(0)	character(0)	NW_011925053.1	1482901	1482906	6	*	6	1000
NW_011925128.1	1875336	1880845	5510	+	Gnomon	gene	NA	NA	gene7225	GeneID:105280395	LOC105280395	Gene	LOC105280395	protein_coding	NA
	character(0)	character(0)	NW_011925128.1	1876634	1876639	6	*	6	1000
NW_011925142.1	777887	780161	2275	-	Gnomon	gene	NA	NA	gene7574	GeneID:105280791	LOC105280791	Gene	LOC105280791	protein_coding	NA
	character(0)	character(0)	NW_011925142.1	779002	779007	6	*	6	1000
NW_011925150.1	76140	79045	2906	-	Gnomon	gene	NA	NA	gene7626	GeneID:105280856	LOC105280856	Gene	LOC105280856	protein_coding	NA	character(0)
	character(0)	NW_011925150.1	78377	78382	6	*	6	1000
NW_011925151.1	38146	39177	1032	-	Gnomon	gene	NA	NA	gene7635	GeneID:105280860	LOC105280860	Gene	LOC105280860	protein_coding	NA	character(0)
	character(0)	NW_011925151.1	38257	38262	6	*	6	1000
NW_011925163.1	470647	473680	3034	+	Gnomon	gene	NA	NA	gene8040	GeneID:105281341	LOC105281341	Gene	LOC105281341	protein_coding	NA
	character(0)	character(0)	NW_011925163.1	471213	471218	6	*	6	1000
NW_011925183.1	39511	42630	3120	+	Gnomon	gene	NA	NA	gene8236	GeneID:105281577	LOC105281577	Gene	LOC105281577	protein_coding	NA	character(0)
	character(0)	NW_011925183.1	40051	40056	6	*	6	1000
NW_011925188.1	199014	202632	3619	-	Gnomon	gene	NA	NA	gene8257	GeneID:105281603	LOC105281603	Gene	LOC105281603	protein_coding	NA
	character(0)	character(0)	NW_011925188.1	201626	201631	6	*	6	1000
...
... only first 25 lines shown ...
```

Directory: MRPR File: gwr-Cb_LI_BCphaseB2.txt

seqnames start end width strand source type score phase ID Dbxref Name gbkey gene gene_biotype partial start_range end_range seqnames start
end width strand NbrSites Sdnsty
NW_011924887.1 1817915 1820166 2252 - Gnomon gene NA NA gene234 GeneID:105274775 LOC105274775 Gene LOC105274775 protein_coding NA
character(0) character(0) NW_011924887.1 1819270 1819275 6 * 6 1000
NW_011924888.1 389413 407214 17802 - Gnomon gene NA NA gene320 GeneID:105275770 LOC105275770 Gene LOC105275770 protein_coding NA
character(0) character(0) NW_011924888.1 405832 405837 6 * 6 1000
NW_011924892.1 1158089 1160303 2215 - Gnomon gene NA NA gene569 GeneID:105278527 LOC105278527 Gene LOC105278527 protein_coding NA
character(0) character(0) NW_011924892.1 1159381 1159386 6 * 6 1000
NW_011924931.1 4488185 4490767 2583 + Gnomon gene NA NA gene1620 GeneID:105287754 LOC105287754 Gene LOC105287754 protein_coding NA
character(0) character(0) NW_011924931.1 4489292 4489297 6 * 6 1000
NW_011924956.1 717757 722032 4276 + Gnomon gene NA NA gene2570 GeneID:105275036 LOC105275036 Gene LOC105275036 protein_coding NA
character(0) character(0) NW_011924956.1 719335 719344 10 * 10 1000
NW_011924973.1 828831 835120 6290 + Gnomon gene NA NA gene2799 GeneID:105275316 LOC105275316 Gene LOC105275316 protein_coding NA
character(0) character(0) NW_011924973.1 829674 829679 6 * 6 1000
NW_011924973.1 1001098 1003969 2872 + Gnomon gene NA NA gene2837 GeneID:105275355 LOC105275355 Gene LOC105275355 protein_coding NA
character(0) character(0) NW_011924973.1 1002105 1002110 6 * 6 1000
NW_011924977.1 662427 665182 2756 + Gnomon gene NA NA gene3008 GeneID:105275550 LOC105275550 Gene LOC105275550 protein_coding NA
character(0) character(0) NW_011924977.1 663534 663539 6 * 6 1000
NW_011924996.1 75277 77259 1983 + Gnomon gene NA NA gene3743 GeneID:105276382 LOC105276382 Gene LOC105276382 protein_coding NA character(0)
character(0) NW_011924996.1 76074 76079 6 * 6 1000
NW_011924996.1 1822758 1830685 7928 - Gnomon gene NA NA gene3885 GeneID:105276527 LOC105276527 Gene LOC105276527 protein_coding NA
character(0) character(0) NW_011924996.1 1829379 1829384 6 * 6 1000
NW_011925007.1 926256 929694 3439 + Gnomon gene NA NA gene4194 GeneID:105276881 LOC105276881 Gene LOC105276881 protein_coding NA
character(0) character(0) NW_011925007.1 927466 927471 6 * 6 1000
NW_011925007.1 1370227 1386638 16412 - Gnomon gene NA NA gene4246 GeneID:105276941 LOC105276941 Gene LOC105276941 protein_coding NA
character(0) character(0) NW_011925007.1 1385156 1385161 6 * 6 1000
NW_011925022.1 386851 387976 1126 - Gnomon gene NA NA gene5000 GeneID:105277804 LOC105277804 Gene LOC105277804 protein_coding NA
character(0) character(0) NW_011925022.1 387077 387082 6 * 6 1000
NW_011925053.1 649052 656133 7082 - Gnomon gene NA NA gene5882 GeneID:105278812 LOC105278812 Gene LOC105278812 protein_coding NA
character(0) character(0) NW_011925053.1 655139 655144 6 * 6 1000
NW_011925086.1 604599 609266 4668 + Gnomon gene NA NA gene6342 GeneID:105279367 LOC105279367 Gene LOC105279367 protein_coding NA
character(0) character(0) NW_011925086.1 606157 606162 6 * 6 1000
NW_011925128.1 1875336 1880845 5510 + Gnomon gene NA NA gene7225 GeneID:105280395 LOC105280395 Gene LOC105280395 protein_coding NA
character(0) character(0) NW_011925128.1 1876634 1876639 6 * 6 1000
NW_011925142.1 777887 780161 2275 - Gnomon gene NA NA gene7574 GeneID:105280791 LOC105280791 Gene LOC105280791 protein_coding NA
character(0) character(0) NW_011925142.1 779002 779007 6 * 6 1000
NW_011925150.1 76140 79045 2906 - Gnomon gene NA NA gene7626 GeneID:105280856 LOC105280856 Gene LOC105280856 protein_coding NA character(0)
character(0) NW_011925150.1 78377 78382 6 * 6 1000
NW_011925151.1 38146 39177 1032 - Gnomon gene NA NA gene7635 GeneID:105280860 LOC105280860 Gene LOC105280860 protein_coding NA character(0)
character(0) NW_011925151.1 38257 38262 6 * 6 1000
NW_011925163.1 470647 473680 3034 + Gnomon gene NA NA gene8040 GeneID:105281341 LOC105281341 Gene LOC105281341 protein_coding NA
character(0) character(0) NW_011925163.1 471213 471218 6 * 6 1000
NW_011925183.1 39511 42630 3120 + Gnomon gene NA NA gene8236 GeneID:105281577 LOC105281577 Gene LOC105281577 protein_coding NA character(0)
character(0) NW_011925183.1 40051 40056 6 * 6 1000
NW_011925188.1 199014 202632 3619 - Gnomon gene NA NA gene8257 GeneID:105281603 LOC105281603 Gene LOC105281603 protein_coding NA
character(0) character(0) NW_011925188.1 201626 201631 6 * 6 1000
NW_011925203.1 63038 97839 34802 + Gnomon gene NA NA gene8521 GeneID:105281906 LOC105281906 Gene LOC105281906 protein_coding NA character(0)
character(0) NW_011925203.1 64603 64608 6 * 6 1000
NW_011925216.1 156691 162086 5396 + Gnomon gene NA NA gene8741 GeneID:105282150 LOC105282150 Gene LOC105282150 protein_coding NA
character(0) character(0) NW_011925216.1 157541 157546 6 * 6 1000
NW_011925240.1 2490948 2493602 2655 - Gnomon gene NA NA gene9134 GeneID:105282589 LOC105282589 Gene LOC105282589 protein_coding NA
character(0) character(0) NW_011925240.1 2492742 2492747 6 * 6 1000
NW_011925304.1 378599 380691 2093 - Gnomon gene NA NA gene9814 GeneID:105283373 LOC105283373 Gene LOC105283373 protein_coding NA
character(0) character(0) NW_011925304.1 379611 379616 6 * 6 1000
NW_011925304.1 468966 476624 7659 - Gnomon gene NA NA gene9824 GeneID:105283383 LOC105283383 Gene LOC105283383 protein_coding NA
character(0) character(0) NW_011925304.1 475523 475528 6 * 6 1000
NW_011925321.1 975185 978704 3520 - Gnomon gene NA NA gene10031 GeneID:105283620 LOC105283620 Gene LOC105283620 protein_coding NA
character(0) character(0) NW_011925321.1 977422 977427 6 * 6 1000
NW_011925321.1 1107715 1109901 2187 - Gnomon gene NA NA gene10043 GeneID:105283634 LOC105283634 Gene LOC105283634 protein_coding NA
character(0) character(0) NW_011925321.1 1108690 1108695 6 * 6 1000
NW_011925353.1 40476 43146 2671 - Gnomon gene NA NA gene10405 GeneID:105284057 LOC105284057 Gene LOC105284057 protein_coding NA character(0)

character(0) NW_011925353.1 41948 41953 6 * 6 1000
NW_011925356.1 1573184 1579703 6520 - Gnomon gene NA NA gene10596 GeneID:105284262 LOC105284262 Gene LOC105284262 protein_coding NA
character(0) character(0) NW_011925356.1 1573650 1573655 6 * 6 1000
NW_011925367.1 1119922 1121887 1966 + Gnomon gene NA NA gene10714 GeneID:105284396 LOC105284396 Gene LOC105284396 protein_coding NA
character(0) character(0) NW_011925367.1 1120960 1120965 6 * 6 1000
NW_011925387.1 39150 46759 7610 + Gnomon gene NA NA gene10928 GeneID:105284644 LOC105284644 Gene LOC105284644 protein_coding NA character(0)
character(0) NW_011925387.1 40039 40044 6 * 6 1000
NW_011925401.1 10153 19053 8901 - Gnomon gene NA NA gene11099 GeneID:105284845 LOC105284845 Gene LOC105284845 protein_coding NA character(0)
character(0) NW_011925401.1 17311 17316 6 * 6 1000
NW_011925470.1 613911 617551 3641 - Gnomon gene NA NA gene11628 GeneID:105285465 LOC105285465 Gene LOC105285465 protein_coding NA
character(0) character(0) NW_011925470.1 616616 616621 6 * 6 1000
NW_011925490.1 137354 140755 3402 + Gnomon gene NA NA gene11689 GeneID:105285534 LOC105285534 Gene LOC105285534 protein_coding NA
character(0) character(0) NW_011925490.1 138283 138288 6 * 6 1000
NW_011925595.1 1 4548 4548 - Gnomon gene NA NA gene12102 GeneID:105286003 LOC105286003 Gene LOC105286003 protein_coding NA character(0)
character(0) NW_011925595.1 3717 3722 6 * 6 1000
NW_011925664.1 277932 283313 5382 + Gnomon gene NA NA gene12359 GeneID:105286364 LOC105286364 Gene LOC105286364 protein_coding NA
character(0) character(0) NW_011925664.1 279108 279113 6 * 6 1000
NW_011925715.1 952620 956416 3797 - Gnomon gene NA NA gene12567 GeneID:105286509 LOC105286509 Gene LOC105286509 protein_coding NA
character(0) character(0) NW_011925715.1 955653 955658 6 * 6 1000
NW_011925086.1 128430 135139 6710 + Gnomon gene NA NA gene6311 GeneID:105279333 LOC105279333 Gene LOC105279333 protein_coding NA
character(0) character(0) NW_011925086.1 129637 129647 11 * 10 909.090909090909
NW_011924914.1 259948 262322 2375 - Gnomon gene NA NA gene952 GeneID:105283016 LOC105283016 Gene LOC105283016 protein_coding NA
character(0) character(0) NW_011924914.1 261398 261406 9 * 8 888.888888888889
NW_011924967.1 91700 93046 1347 - Gnomon gene NA NA gene2644 GeneID:105275137 LOC105275137 Gene LOC105275137 protein_coding NA character(0)
character(0) NW_011924967.1 92497 92505 9 * 8 888.888888888889
NW_011924979.1 1988584 1990897 2314 + Gnomon gene NA NA gene3314 GeneID:105275878 LOC105275878 Gene LOC105275878 protein_coding NA
character(0) character(0) NW_011924979.1 1989441 1989449 9 * 8 888.888888888889
NW_011925239.1 3979 7808 3830 + Gnomon gene NA NA gene9001 GeneID:105282461 LOC105282461 Gene LOC105282461 protein_coding NA character(0)
character(0) NW_011925239.1 4874 4882 9 * 8 888.888888888889
NW_011924977.1 3728770 3731404 2635 + Gnomon gene NA NA gene3213 GeneID:105275761 LOC105275761 Gene LOC105275761 protein_coding NA
character(0) character(0) NW_011924977.1 3729890 3729897 8 * 7 875
NW_011924888.1 30738 31847 1110 + Gnomon gene NA NA gene264 GeneID:105275153 LOC105275153 Gene LOC105275153 protein_coding NA character(0)
character(0) NW_011924888.1 31453 31459 7 * 6 857.142857142857
NW_011924889.1 476236 478868 2633 - Gnomon gene NA NA gene504 GeneID:105277817 LOC105277817 Gene LOC105277817 protein_coding NA
character(0) character(0) NW_011924889.1 477980 477986 7 * 6 857.142857142857
NW_011924889.1 493449 499545 6097 + Gnomon gene NA NA gene508 GeneID:105277915 LOC105277915 Gene LOC105277915 protein_coding NA
character(0) character(0) NW_011924889.1 494195 494201 7 * 6 857.142857142857
NW_011924892.1 274786 285690 10905 - Gnomon gene NA NA gene544 GeneID:105278288 LOC105278288 Gene LOC105278288 protein_coding NA
character(0) character(0) NW_011924892.1 284836 284842 7 * 6 857.142857142857
NW_011924899.1 718895 720452 1558 + Gnomon gene NA NA gene664 GeneID:105279718 LOC105279718 Gene LOC105279718 protein_coding NA
character(0) character(0) NW_011924899.1 719601 719607 7 * 6 857.142857142857
NW_011924902.1 81345 99446 18102 - Gnomon gene NA NA gene702 GeneID:105280058 LOC105280058 Gene LOC105280058 protein_coding NA character(0)
character(0) NW_011924902.1 98752 98758 7 * 6 857.142857142857
NW_011924905.1 26269 321616 295348 - Gnomon gene NA NA gene787 GeneID:105280980 LOC105280980 Gene LOC105280980 protein_coding NA
character(0) character(0) NW_011924905.1 319741 319747 7 * 6 857.142857142857
NW_011924907.1 889683 896841 7159 - Gnomon gene NA NA gene861 GeneID:105281750 LOC105281750 Gene LOC105281750 protein_coding NA
character(0) character(0) NW_011924907.1 896035 896041 7 * 6 857.142857142857
NW_011924914.1 827084 831408 4325 - Gnomon gene NA NA gene1010 GeneID:105283498 LOC105283498 Gene LOC105283498 protein_coding NA
character(0) character(0) NW_011924914.1 831099 831105 7 * 6 857.142857142857
NW_011924922.1 2390151 2392888 2738 - Gnomon gene NA NA gene1198 GeneID:105285507 LOC105285507 Gene LOC105285507 protein_coding NA
character(0) character(0) NW_011924922.1 2392009 2392015 7 * 6 857.142857142857
NW_011924929.1 931843 943599 11757 + Gnomon gene NA NA gene1329 GeneID:105286844 LOC105286844 Gene LOC105286844 protein_coding NA
character(0) character(0) NW_011924929.1 933320 933326 7 * 6 857.142857142857
NW_011924931.1 122002 124933 2932 + Gnomon gene NA NA gene1364 GeneID:105287208 LOC105287208 Gene LOC105287208 protein_coding NA
character(0) character(0) NW_011924931.1 123017 123023 7 * 6 857.142857142857
NW_011924931.1 242950 250477 7528 - Gnomon gene NA NA gene1378 GeneID:105287371 LOC105287371 Gene LOC105287371 protein_coding NA
character(0) character(0) NW_011924931.1 249191 249197 7 * 6 857.142857142857
NW_011924931.1 290674 293586 2913 + Gnomon gene NA NA gene1388 GeneID:105287473 LOC105287473 Gene LOC105287473 protein_coding NA
character(0) character(0) NW_011924931.1 291628 291634 7 * 6 857.142857142857
NW_011924931.1 1565109 1566883 1775 + Gnomon gene NA NA gene1444 GeneID:105287569 LOC105287569 Gene LOC105287569 protein_coding NA
character(0) character(0) NW_011924931.1 1565961 1565967 7 * 6 857.142857142857
NW_011924931.1 2028839 2031812 2974 + Gnomon gene NA NA gene1470 GeneID:105287597 LOC105287597 Gene LOC105287597 protein_coding NA
character(0) character(0) NW_011924931.1 2030239 2030245 7 * 6 857.142857142857
NW_011924931.1 2871689 2878941 7253 - Gnomon gene NA NA gene1492 GeneID:105287620 LOC105287620 Gene LOC105287620 protein_coding NA

character(0) character(0) NW_011924931.1 2878145 2878151 7 * 6 857.142857142857
NW_011924931.1 3929932 3932866 2935 + Gnomon gene NA NA gene1555 GeneID:105287684 LOC105287684 Gene LOC105287684 protein_coding NA
character(0) character(0) NW_011924931.1 3930880 3930886 7 * 6 857.142857142857
NW_011924931.1 3979390 3986328 6939 + Gnomon gene NA NA gene1566 GeneID:105287698 LOC105287698 Gene LOC105287698 protein_coding NA
character(0) character(0) NW_011924931.1 3983680 3983686 7 * 6 857.142857142857
NW_011924931.1 3996443 4002577 6135 + Gnomon gene NA NA gene1569 GeneID:105287702 LOC105287702 Gene LOC105287702 protein_coding NA
character(0) character(0) NW_011924931.1 3997573 3997579 7 * 6 857.142857142857
NW_011924931.1 4058265 4059684 1420 - Gnomon gene NA NA gene1581 GeneID:105287710 LOC105287710 Gene LOC105287710 protein_coding NA
character(0) character(0) NW_011924931.1 4058767 4058773 7 * 6 857.142857142857
NW_011924932.1 1156872 1160705 3834 + Gnomon gene NA NA gene1695 GeneID:105287855 LOC105287855 Gene LOC105287855 protein_coding NA
character(0) character(0) NW_011924932.1 1157735 1157741 7 * 6 857.142857142857
NW_011924938.1 72334 83258 10925 + Gnomon gene NA NA gene1805 GeneID:105287985 LOC105287985 Gene LOC105287985 protein_coding NA character(0)
character(0) NW_011924938.1 72914 72920 7 * 6 857.142857142857
NW_011924946.1 1634653 1637620 2968 - Gnomon gene NA NA gene2065 GeneID:105274464 LOC105274464 Gene LOC105274464 protein_coding NA
character(0) character(0) NW_011924946.1 1636892 1636898 7 * 6 857.142857142857
NW_011924946.1 2342632 2344089 1458 - Gnomon gene NA NA gene2088 GeneID:105274497 LOC105274497 Gene LOC105274497 protein_coding NA
character(0) character(0) NW_011924946.1 2343055 2343061 7 * 6 857.142857142857
NW_011924946.1 2914862 2919248 4387 - Gnomon gene NA NA gene2173 GeneID:105274590 LOC105274590 Gene LOC105274590 protein_coding NA
character(0) character(0) NW_011924946.1 2918055 2918061 7 * 6 857.142857142857
NW_011924946.1 3145253 3148790 3538 - Gnomon gene NA NA gene2215 GeneID:105274638 LOC105274638 Gene LOC105274638 protein_coding NA
character(0) character(0) NW_011924946.1 3147722 3147728 7 * 6 857.142857142857
NW_011924947.1 141424 143460 2037 + Gnomon gene NA NA gene2279 GeneID:105274711 LOC105274711 Gene LOC105274711 protein_coding NA
character(0) character(0) NW_011924947.1 142119 142125 7 * 6 857.142857142857
NW_011924953.1 401032 402582 1551 - Gnomon gene NA NA gene2400 GeneID:105274850 LOC105274850 Gene LOC105274850 protein_coding NA
character(0) character(0) NW_011924953.1 401754 401760 7 * 6 857.142857142857
NW_011924955.1 473199 477158 3960 - Gnomon gene NA NA gene2462 GeneID:105274912 LOC105274912 Gene LOC105274912 protein_coding NA
character(0) character(0) NW_011924955.1 475912 475918 7 * 6 857.142857142857
NW_011924956.1 701839 703915 2077 - Gnomon gene NA NA gene2566 GeneID:105275031 LOC105275031 Gene LOC105275031 protein_coding NA
character(0) character(0) NW_011924956.1 702713 702719 7 * 6 857.142857142857
NW_011924967.1 11923 12765 843 - Gnomon gene NA NA gene2630 GeneID:105275119 LOC105275119 Gene LOC105275119 protein_coding NA character(0)
character(0) NW_011924967.1 12053 12059 7 * 6 857.142857142857
NW_011924967.1 91700 93046 1347 - Gnomon gene NA NA gene2644 GeneID:105275137 LOC105275137 Gene LOC105275137 protein_coding NA character(0)
character(0) NW_011924967.1 92468 92474 7 * 6 857.142857142857
NW_011924973.1 774415 775843 1429 - Gnomon gene NA NA gene2790 GeneID:105275307 LOC105275307 Gene LOC105275307 protein_coding NA
character(0) character(0) NW_011924973.1 775075 775081 7 * 6 857.142857142857
NW_011924974.1 483919 487626 3708 + Gnomon gene NA NA gene2873 GeneID:105275401 LOC105275401 Gene LOC105275401 protein_coding NA
character(0) character(0) NW_011924974.1 484702 484708 7 * 6 857.142857142857
NW_011924977.1 1460154 1463690 3537 - Gnomon gene NA NA gene3072 GeneID:105275615 LOC105275615 Gene LOC105275615 protein_coding NA
character(0) character(0) NW_011924977.1 1462188 1462194 7 * 6 857.142857142857
NW_011924977.1 1835450 1838251 2802 + Gnomon gene NA NA gene3114 GeneID:105275656 LOC105275656 Gene LOC105275656 protein_coding NA
character(0) character(0) NW_011924977.1 1836976 1836982 7 * 6 857.142857142857
NW_011924977.1 2573287 2581453 8167 - Gnomon gene NA NA gene3147 GeneID:105275689 LOC105275689 Gene LOC105275689 protein_coding NA
character(0) character(0) NW_011924977.1 2580506 2580512 7 * 6 857.142857142857
NW_011924977.1 3678529 3682684 4156 - Gnomon gene NA NA gene3205 GeneID:105275753 LOC105275753 Gene LOC105275753 protein_coding NA
character(0) character(0) NW_011924977.1 3681359 3681365 7 * 6 857.142857142857
NW_011924977.1 3678529 3682684 4156 - Gnomon gene NA NA gene3205 GeneID:105275753 LOC105275753 Gene LOC105275753 protein_coding NA
character(0) character(0) NW_011924977.1 3681568 3681574 7 * 6 857.142857142857
NW_011924977.1 3682584 3686057 3474 - Gnomon gene NA NA gene3206 GeneID:105275752 LOC105275752 Gene LOC105275752 protein_coding NA
character(0) character(0) NW_011924977.1 3684978 3684984 7 * 6 857.142857142857
NW_011924979.1 211193 212598 1406 - Gnomon gene NA NA gene3249 GeneID:105275908 LOC105275908 Gene LOC105275908 protein_coding NA
character(0) character(0) NW_011924979.1 212057 212063 7 * 6 857.142857142857
NW_011924979.1 2744383 2746071 1689 + Gnomon gene NA NA gene3334 GeneID:105275897 LOC105275897 Gene LOC105275897 protein_coding NA
character(0) character(0) NW_011924979.1 2745351 2745357 7 * 6 857.142857142857
NW_011924987.1 611928 613454 1527 - Gnomon gene NA NA gene3407 GeneID:105275991 LOC105275991 Gene LOC105275991 protein_coding NA
character(0) character(0) NW_011924987.1 612398 612404 7 * 6 857.142857142857
NW_011924987.1 1323431 1335995 12565 + Gnomon gene NA NA gene3479 GeneID:105276073 LOC105276073 Gene LOC105276073 protein_coding NA
character(0) character(0) NW_011924987.1 1329036 1329042 7 * 6 857.142857142857
NW_011924987.1 1327762 1332816 5055 + Gnomon gene NA NA gene3482 GeneID:105276070 LOC105276070 Gene LOC105276070 protein_coding NA
character(0) character(0) NW_011924987.1 1329036 1329042 7 * 6 857.142857142857
NW_011924987.1 2224897 2228137 3241 + Gnomon gene NA NA gene3607 GeneID:105276205 LOC105276205 Gene LOC105276205 protein_coding NA
character(0) character(0) NW_011924987.1 2225943 2225949 7 * 6 857.142857142857
NW_011924987.1 2638171 2648657 10487 - Gnomon gene NA NA gene3636 GeneID:105276233 LOC105276233 Gene LOC105276233 protein_coding NA
character(0) character(0) NW_011924987.1 2647465 2647471 7 * 6 857.142857142857
NW_011924996.1 292526 303470 10945 + Gnomon gene NA NA gene3775 GeneID:105276410 LOC105276410 Gene LOC105276410 protein_coding NA

character(0) character(0) NW_011924996.1 293762 293768 7 * 6 857.142857142857
NW_011924996.1 292526 303470 10945 + Gnomon gene NA NA gene3775 GeneID:105276410 LOC105276410 Gene LOC105276410 protein_coding NA
character(0) character(0) NW_011924996.1 293819 293825 7 * 6 857.142857142857
NW_011924996.1 448616 452114 3499 - Gnomon gene NA NA gene3788 GeneID:105276422 LOC105276422 Gene LOC105276422 protein_coding NA
character(0) character(0) NW_011924996.1 450468 450474 7 * 6 857.142857142857
NW_011924996.1 641332 646889 5558 - Gnomon gene NA NA gene3801 GeneID:105276437 LOC105276437 Gene LOC105276437 protein_coding NA
character(0) character(0) NW_011924996.1 645416 645422 7 * 6 857.142857142857
NW_011924996.1 1790083 1793499 3417 - Gnomon gene NA NA gene3877 GeneID:105276521 LOC105276521 Gene LOC105276521 protein_coding NA
character(0) character(0) NW_011924996.1 1792422 1792428 7 * 6 857.142857142857
NW_011925007.1 1417723 1421435 3713 + Gnomon gene NA NA gene4248 GeneID:105276948 LOC105276948 Gene LOC105276948 protein_coding NA
character(0) character(0) NW_011925007.1 1419353 1419359 7 * 6 857.142857142857
NW_011925007.1 1452162 1454301 2140 + Gnomon gene NA NA gene4252 GeneID:105276949 LOC105276949 Gene LOC105276949 protein_coding NA
character(0) character(0) NW_011925007.1 1453109 1453115 7 * 6 857.142857142857
NW_011925007.1 3760218 3764466 4249 - Gnomon gene NA NA gene4383 GeneID:105277128 LOC105277128 Gene LOC105277128 protein_coding NA
character(0) character(0) NW_011925007.1 3760305 3760311 7 * 6 857.142857142857
NW_011925012.1 1316386 1318477 2092 + Gnomon gene NA NA gene4492 GeneID:105277232 LOC105277232 Gene LOC105277232 protein_coding NA
character(0) character(0) NW_011925012.1 1317469 1317475 7 * 6 857.142857142857
NW_011925012.1 1360007 1362714 2708 - Gnomon gene NA NA gene4500 GeneID:105277238 LOC105277238 Gene LOC105277238 protein_coding NA
character(0) character(0) NW_011925012.1 1361666 1361672 7 * 6 857.142857142857
NW_011925013.1 5041 7730 2690 - Gnomon gene NA NA gene4511 GeneID:105277256 LOC105277256 Gene LOC105277256 protein_coding NA character(0)
character(0) NW_011925013.1 6764 6770 7 * 6 857.142857142857
NW_011925016.1 935483 939614 4132 + Gnomon gene NA NA gene4624 GeneID:105277376 LOC105277376 Gene LOC105277376 protein_coding NA
character(0) character(0) NW_011925016.1 936674 936680 7 * 6 857.142857142857
NW_011925019.1 475 2900 2426 + Gnomon gene NA NA gene4633 GeneID:105277389 LOC105277389 Gene LOC105277389 protein_coding NA character(0)
character(0) NW_011925019.1 1670 1676 7 * 6 857.142857142857
NW_011925019.1 2803 5896 3094 - Gnomon gene NA NA gene4634 GeneID:105277388 LOC105277388 Gene LOC105277388 protein_coding NA character(0)
character(0) NW_011925019.1 4685 4691 7 * 6 857.142857142857
NW_011925020.1 469190 474352 5163 + Gnomon gene NA NA gene4780 GeneID:105277552 LOC105277552 Gene LOC105277552 protein_coding NA
character(0) character(0) NW_011925020.1 470270 470276 7 * 6 857.142857142857
NW_011925027.1 1297549 1301844 4296 + Gnomon gene NA NA gene5179 GeneID:105278005 LOC105278005 Gene LOC105278005 protein_coding NA
character(0) character(0) NW_011925027.1 1298524 1298530 7 * 6 857.142857142857
NW_011925029.1 39174 43410 4237 + Gnomon gene NA NA gene5243 GeneID:105278106 LOC105278106 Gene LOC105278106 protein_coding NA character(0)
character(0) NW_011925029.1 40261 40267 7 * 6 857.142857142857
NW_011925029.1 78846 81944 3099 - Gnomon gene NA NA gene5249 GeneID:105278098 LOC105278098 Gene LOC105278098 protein_coding NA character(0)
character(0) NW_011925029.1 80103 80109 7 * 6 857.142857142857
NW_011925035.1 1492483 1496530 4048 - Gnomon gene NA NA gene5417 GeneID:105278277 LOC105278277 Gene LOC105278277 protein_coding NA
character(0) character(0) NW_011925035.1 1495407 1495413 7 * 6 857.142857142857
NW_011925035.1 1494546 1496299 1754 - Gnomon gene NA NA gene5418 GeneID:105278275 LOC105278275 Gene LOC105278275 protein_coding NA
character(0) character(0) NW_011925035.1 1495407 1495413 7 * 6 857.142857142857
NW_011925035.1 1812544 1818171 5628 - Gnomon gene NA NA gene5451 GeneID:105278309 LOC105278309 Gene LOC105278309 protein_coding NA
character(0) character(0) NW_011925035.1 1817088 1817094 7 * 6 857.142857142857
NW_011925035.1 3102848 3104698 1851 + Gnomon gene NA NA gene5557 GeneID:105278431 LOC105278431 Gene LOC105278431 protein_coding NA
character(0) character(0) NW_011925035.1 3103902 3103908 7 * 6 857.142857142857
NW_011925036.1 1418708 1424550 5843 + Gnomon gene NA NA gene5653 GeneID:105278544 LOC105278544 Gene LOC105278544 protein_coding NA
character(0) character(0) NW_011925036.1 1420178 1420184 7 * 6 857.142857142857
NW_011925067.1 5971 15150 9180 + Gnomon gene NA NA gene6024 GeneID:105278996 LOC105278996 Gene LOC105278996 protein_coding NA character(0)
character(0) NW_011925067.1 7054 7060 7 * 6 857.142857142857
NW_011925067.1 5971 15150 9180 + Gnomon gene NA NA gene6024 GeneID:105278996 LOC105278996 Gene LOC105278996 protein_coding NA character(0)
character(0) NW_011925067.1 7213 7219 7 * 6 857.142857142857
NW_011925068.1 10281 12328 2048 + Gnomon gene NA NA gene6111 GeneID:105279088 LOC105279088 Gene LOC105279088 protein_coding NA character(0)
character(0) NW_011925068.1 11234 11240 7 * 6 857.142857142857
NW_011925086.1 250638 255287 4650 - Gnomon gene NA NA gene6324 GeneID:105279346 LOC105279346 Gene LOC105279346 protein_coding NA
character(0) character(0) NW_011925086.1 254146 254152 7 * 6 857.142857142857
NW_011925106.1 154197 158224 4028 + Gnomon gene NA NA gene6601 GeneID:105279664 LOC105279664 Gene LOC105279664 protein_coding NA
character(0) character(0) NW_011925106.1 155492 155498 7 * 6 857.142857142857
NW_011925106.1 431654 436050 4397 + Gnomon gene NA NA gene6641 GeneID:105279709 LOC105279709 Gene LOC105279709 protein_coding NA
character(0) character(0) NW_011925106.1 432897 432903 7 * 6 857.142857142857
NW_011925106.1 512988 515818 2831 - Gnomon gene NA NA gene6652 GeneID:105279719 LOC105279719 Gene LOC105279719 protein_coding NA
character(0) character(0) NW_011925106.1 514935 514941 7 * 6 857.142857142857
NW_011925106.1 629782 633106 3325 - Gnomon gene NA NA gene6668 GeneID:105279740 LOC105279740 Gene LOC105279740 protein_coding NA
character(0) character(0) NW_011925106.1 631881 631887 7 * 6 857.142857142857
NW_011925106.1 779647 785206 5560 + Gnomon gene NA NA gene6679 GeneID:105279748 LOC105279748 Gene LOC105279748 protein_coding NA
character(0) character(0) NW_011925106.1 780733 780739 7 * 6 857.142857142857
NW_011925106.1 2086959 2096661 9703 + Gnomon gene NA NA gene6771 GeneID:105279854 LOC105279854 Gene LOC105279854 protein_coding NA

character(0) character(0) NW_011925106.1 2087848 2087854 7 * 6 857.142857142857
NW_011925118.1 457689 459947 2259 + Gnomon gene NA NA gene6953 GeneID:105280092 LOC105280092 Gene LOC105280092 protein_coding NA
character(0) character(0) NW_011925118.1 458682 458688 7 * 6 857.142857142857
NW_011925118.1 481415 484075 2661 + Gnomon gene NA NA gene6956 GeneID:105280102 LOC105280102 Gene LOC105280102 protein_coding NA
character(0) character(0) NW_011925118.1 482695 482701 7 * 6 857.142857142857
NW_011925128.1 7781 9472 1692 + Gnomon gene NA NA gene7085 GeneID:105280236 LOC105280236 Gene LOC105280236 protein_coding NA character(0)
character(0) NW_011925128.1 8930 8936 7 * 6 857.142857142857
NW_011925128.1 251790 262534 10745 - Gnomon gene NA NA gene7113 GeneID:105280269 LOC105280269 Gene LOC105280269 protein_coding NA
character(0) character(0) NW_011925128.1 259852 259858 7 * 6 857.142857142857
NW_011925128.1 800543 802328 1786 - Gnomon gene NA NA gene7160 GeneID:105280319 LOC105280319 Gene LOC105280319 protein_coding NA
character(0) character(0) NW_011925128.1 801605 801611 7 * 6 857.142857142857
NW_011925128.1 2453513 2456143 2631 + Gnomon gene NA NA gene7299 GeneID:105280469 LOC105280469 Gene LOC105280469 protein_coding NA
character(0) character(0) NW_011925128.1 2454945 2454951 7 * 6 857.142857142857
NW_011925142.1 423375 426060 2686 - Gnomon gene NA NA gene7527 GeneID:105280739 LOC105280739 Gene LOC105280739 protein_coding NA
character(0) character(0) NW_011925142.1 424227 424233 7 * 6 857.142857142857
NW_011925161.1 210886 215244 4359 + Gnomon gene NA NA gene7720 GeneID:105280964 LOC105280964 Gene LOC105280964 protein_coding NA
character(0) character(0) NW_011925161.1 213136 213142 7 * 6 857.142857142857
NW_011925161.1 3740915 3743520 2606 + Gnomon gene NA NA gene7963 GeneID:105281227 LOC105281227 Gene LOC105281227 protein_coding NA
character(0) character(0) NW_011925161.1 3741983 3741989 7 * 6 857.142857142857
NW_011925183.1 39511 42630 3120 + Gnomon gene NA NA gene8236 GeneID:105281577 LOC105281577 Gene LOC105281577 protein_coding NA character(0)
character(0) NW_011925183.1 40353 40359 7 * 6 857.142857142857
NW_011925188.1 199014 202632 3619 - Gnomon gene NA NA gene8257 GeneID:105281603 LOC105281603 Gene LOC105281603 protein_coding NA
character(0) character(0) NW_011925188.1 201441 201447 7 * 6 857.142857142857
NW_011925209.1 85655 90207 4553 - Gnomon gene NA NA gene8551 GeneID:105281943 LOC105281943 Gene LOC105281943 protein_coding NA character(0)
character(0) NW_011925209.1 86875 86881 7 * 6 857.142857142857
NW_011925209.1 169877 173749 3873 - Gnomon gene NA NA gene8560 GeneID:105281949 LOC105281949 Gene LOC105281949 protein_coding NA
character(0) character(0) NW_011925209.1 172697 172703 7 * 6 857.142857142857
NW_011925209.1 175864 177807 1944 + Gnomon gene NA NA gene8561 GeneID:105281950 LOC105281950 Gene LOC105281950 protein_coding NA
character(0) character(0) NW_011925209.1 176967 176973 7 * 6 857.142857142857
NW_011925216.1 130033 131250 1218 + Gnomon gene NA NA gene8733 GeneID:105282147 LOC105282147 Gene LOC105282147 protein_coding NA
character(0) character(0) NW_011925216.1 130799 130805 7 * 6 857.142857142857
NW_011925221.1 474109 476408 2300 + Gnomon gene NA NA gene8798 GeneID:105282220 LOC105282220 Gene LOC105282220 protein_coding NA
character(0) character(0) NW_011925221.1 474855 474861 7 * 6 857.142857142857
NW_011925224.1 208794 211425 2632 + Gnomon gene NA NA gene8852 GeneID:105282278 LOC105282278 Gene LOC105282278 protein_coding NA
character(0) character(0) NW_011925224.1 209730 209736 7 * 6 857.142857142857
NW_011925224.1 219079 219935 857 - Gnomon gene NA NA gene8854 GeneID:105282280 LOC105282280 Gene LOC105282280 protein_coding NA
character(0) character(0) NW_011925224.1 219257 219263 7 * 6 857.142857142857
NW_011925224.1 327268 339256 11989 - Gnomon gene NA NA gene8872 GeneID:105282298 LOC105282298 Gene LOC105282298 protein_coding NA
character(0) character(0) NW_011925224.1 338306 338312 7 * 6 857.142857142857
NW_011925241.1 397336 409261 11926 - Gnomon gene NA NA gene9191 GeneID:105282655 LOC105282655 Gene LOC105282655 protein_coding NA
character(0) character(0) NW_011925241.1 408196 408202 7 * 6 857.142857142857
NW_011925257.1 52519 55399 2881 - Gnomon gene NA NA gene9303 GeneID:105282793 LOC105282793 Gene LOC105282793 protein_coding NA character(0)
character(0) NW_011925257.1 54841 54847 7 * 6 857.142857142857
NW_011925258.1 360040 361894 1855 - Gnomon gene NA NA gene9341 GeneID:105282841 LOC105282841 Gene LOC105282841 protein_coding NA
character(0) character(0) NW_011925258.1 361089 361095 7 * 6 857.142857142857
NW_011925267.1 224087 232805 8719 + Gnomon gene NA NA gene9418 GeneID:105282921 LOC105282921 Gene LOC105282921 protein_coding NA
character(0) character(0) NW_011925267.1 225570 225576 7 * 6 857.142857142857
NW_011925304.1 443636 447978 4343 + Gnomon gene NA NA gene9820 GeneID:105283378 LOC105283378 Gene LOC105283378 protein_coding NA
character(0) character(0) NW_011925304.1 445224 445230 7 * 6 857.142857142857
NW_011925323.1 390910 395897 4988 - Gnomon gene NA NA gene10091 GeneID:105283697 LOC105283697 Gene LOC105283697 protein_coding NA
character(0) character(0) NW_011925323.1 394351 394357 7 * 6 857.142857142857
NW_011925323.1 1444062 1448038 3977 - Gnomon gene NA NA gene10138 GeneID:105283741 LOC105283741 Gene LOC105283741 protein_coding NA
character(0) character(0) NW_011925323.1 1445991 1445997 7 * 6 857.142857142857
NW_011925327.1 214150 224738 10589 + Gnomon gene NA NA gene10207 GeneID:105283818 LOC105283818 Gene LOC105283818 protein_coding NA
character(0) character(0) NW_011925327.1 216086 216092 7 * 6 857.142857142857
NW_011925330.1 58606 82143 23538 + Gnomon gene NA NA gene10255 GeneID:105283878 LOC105283878 Gene LOC105283878 protein_coding NA character(0)
character(0) NW_011925330.1 59800 59806 7 * 6 857.142857142857
NW_011925336.1 92449 95536 3088 + Gnomon gene NA NA gene10324 GeneID:105283961 LOC105283961 Gene LOC105283961 protein_coding NA character(0)
character(0) NW_011925336.1 93305 93311 7 * 6 857.142857142857
NW_011925336.1 100271 103076 2806 - Gnomon gene NA NA gene10327 GeneID:105283964 LOC105283964 Gene LOC105283964 protein_coding NA
character(0) character(0) NW_011925336.1 102423 102429 7 * 6 857.142857142857
NW_011925353.1 16223 20332 4110 + Gnomon gene NA NA gene10401 GeneID:105284052 LOC105284052 Gene LOC105284052 protein_coding NA character(0)
character(0) NW_011925353.1 19139 19145 7 * 6 857.142857142857
NW_011925356.1 701727 704945 3219 + Gnomon gene NA NA gene10538 GeneID:105284202 LOC105284202 Gene LOC105284202 protein_coding NA

character(0) character(0) NW_011925356.1 703128 703134 7 * 6 857.142857142857
NW_011925367.1 1119922 1121887 1966 + Gnomon gene NA NA gene10714 GeneID:105284396 LOC105284396 Gene LOC105284396 protein_coding NA
character(0) character(0) NW_011925367.1 1120846 1120852 7 * 6 857.142857142857
NW_011925371.1 899674 901634 1961 - Gnomon gene NA NA gene10880 GeneID:105284576 LOC105284576 Gene LOC105284576 protein_coding NA
character(0) character(0) NW_011925371.1 900958 900964 7 * 6 857.142857142857
NW_011925371.1 904647 906111 1465 - Gnomon gene NA NA gene10882 GeneID:105284579 LOC105284579 Gene LOC105284579 protein_coding NA
character(0) character(0) NW_011925371.1 905523 905529 7 * 6 857.142857142857
NW_011925387.1 118203 119961 1759 - Gnomon gene NA NA gene10943 GeneID:105284659 LOC105284659 Gene LOC105284659 protein_coding NA
character(0) character(0) NW_011925387.1 119303 119309 7 * 6 857.142857142857
NW_011925434.1 120720 130419 9700 + Gnomon gene NA NA gene11344 GeneID:105285135 LOC105285135 Gene LOC105285135 protein_coding NA
character(0) character(0) NW_011925434.1 123131 123137 7 * 6 857.142857142857
NW_011925446.1 246855 250131 3277 + Gnomon gene NA NA gene11531 GeneID:105285345 LOC105285345 Gene LOC105285345 protein_coding NA
character(0) character(0) NW_011925446.1 248422 248428 7 * 6 857.142857142857
NW_011925459.1 162939 164541 1603 + Gnomon gene NA NA gene11566 GeneID:105285392 LOC105285392 Gene LOC105285392 protein_coding NA
character(0) character(0) NW_011925459.1 163954 163960 7 * 6 857.142857142857
NW_011925459.1 186244 187554 1311 - Gnomon gene NA NA gene11569 GeneID:105285394 LOC105285394 Gene LOC105285394 protein_coding NA
character(0) character(0) NW_011925459.1 186700 186706 7 * 6 857.142857142857
NW_011925465.1 71398 78050 6653 - Gnomon gene NA NA gene11580 GeneID:105285410 LOC105285410 Gene LOC105285410 protein_coding NA character(0)
character(0) NW_011925465.1 77157 77163 7 * 6 857.142857142857
NW_011925467.1 95772 98981 3210 - Gnomon gene NA NA gene11594 GeneID:105285418 LOC105285418 Gene LOC105285418 protein_coding NA character(0)
character(0) NW_011925467.1 97582 97588 7 * 6 857.142857142857
NW_011925471.1 1427 3141 1715 - Gnomon gene NA NA gene11632 GeneID:109611363 LOC109611363 Gene LOC109611363 lncRNA NA character(0)
character(0) NW_011925471.1 2780 2786 7 * 6 857.142857142857
NW_011925490.1 207564 212389 4826 + Gnomon gene NA NA gene11699 GeneID:105285546 LOC105285546 Gene LOC105285546 protein_coding NA
character(0) character(0) NW_011925490.1 208609 208615 7 * 6 857.142857142857
NW_011925516.1 278917 286323 7407 - Gnomon gene NA NA gene11797 GeneID:105285648 LOC105285648 Gene LOC105285648 protein_coding NA
character(0) character(0) NW_011925516.1 285124 285130 7 * 6 857.142857142857
NW_011925538.1 63312 65618 2307 - Gnomon gene NA NA gene11931 GeneID:105285808 LOC105285808 Gene LOC105285808 protein_coding NA character(0)
character(0) NW_011925538.1 64230 64236 7 * 6 857.142857142857
NW_011925546.1 59043 60354 1312 + Gnomon gene NA NA gene11979 GeneID:105285864 LOC105285864 Gene LOC105285864 protein_coding NA character(0)
character(0) NW_011925546.1 59986 59992 7 * 6 857.142857142857
NW_011925546.1 78656 85452 6797 - Gnomon gene NA NA gene11986 GeneID:105285867 LOC105285867 Gene LOC105285867 protein_coding NA character(0)
character(0) NW_011925546.1 82761 82767 7 * 6 857.142857142857
NW_011925549.1 267852 270480 2629 - Gnomon gene NA NA gene12010 GeneID:105285889 LOC105285889 Gene LOC105285889 protein_coding NA
character(0) character(0) NW_011925549.1 269441 269447 7 * 6 857.142857142857
NW_011925664.1 188474 193249 4776 + Gnomon gene NA NA gene12339 GeneID:105286242 LOC105286242 Gene LOC105286242 protein_coding NA
character(0) character(0) NW_011925664.1 189517 189523 7 * 6 857.142857142857
NW_011925715.1 251551 257367 5817 - Gnomon gene NA NA gene12526 GeneID:105286461 LOC105286461 Gene LOC105286461 protein_coding NA
character(0) character(0) NW_011925715.1 256038 256044 7 * 6 857.142857142857
NW_011925715.1 412871 419485 6615 + Gnomon gene NA NA gene12539 GeneID:105286478 LOC105286478 Gene LOC105286478 protein_coding NA
character(0) character(0) NW_011925715.1 414243 414249 7 * 6 857.142857142857
NW_011926135.1 70608 74554 3947 + Gnomon gene NA NA gene13076 GeneID:105287104 LOC105287104 Gene LOC105287104 protein_coding NA character(0)
character(0) NW_011926135.1 71339 71345 7 * 6 857.142857142857
NW_011926194.1 11132 18740 7609 + Gnomon gene NA NA gene13114 GeneID:105287147 LOC105287147 Gene LOC105287147 protein_coding NA character(0)
character(0) NW_011926194.1 12247 12253 7 * 6 857.142857142857
NW_011924914.1 87641 92698 5058 - Gnomon gene NA NA gene915 GeneID:105282459 LOC105282459 Gene LOC105282459 protein_coding NA character(0)
character(0) NW_011924914.1 90891 90900 10 * 8 800
NW_011924951.1 73901 76560 2660 + Gnomon gene NA NA gene2325 GeneID:105274767 LOC105274767 Gene LOC105274767 protein_coding NA character(0)
character(0) NW_011924951.1 74986 74995 10 * 8 800
NW_011925007.1 3759369 3760273 905 + Gnomon gene NA NA gene4382 GeneID:105277091 LOC105277091 Gene LOC105277091 protein_coding NA
character(0) character(0) NW_011925007.1 3760113 3760122 10 * 8 800

Directory: MRPR File: gwr-Cb_LI_RphaseA1.txt

| seqnames | start | end | width | strand | source | type | score | phase | ID | Dbxref | Name | gbkey | gene | gene_biotype | partial | start_range | end_range | seqnames | start | end | width | strand | NbrSites | Sdnsty |
|---|---|---|---|---|---|---|---|---|---|---|---|---|---|---|---|---|---|---|---|---|---|---|---|---|
| NW_011924887.1 | 1817915 | 1820166 | 2252 | - | Gnomon | gene | NA | NA | gene234 | GeneID:105274775 | LOC105274775 | Gene | LOC105274775 | protein_coding | NA | character(0) | character(0) | NW_011924887.1 | 1819270 | 1819275 | 6 | * | 6 | 1000 |
| NW_011924888.1 | 77334 | 78632 | 1299 | - | Gnomon | gene | NA | NA | gene276 | GeneID:105275214 | LOC105275214 | Gene | LOC105275214 | protein_coding | NA | character(0) | character(0) | NW_011924888.1 | 77560 | 77565 | 6 | * | 6 | 1000 |
| NW_011924888.1 | 389413 | 407214 | 17802 | - | Gnomon | gene | NA | NA | gene320 | GeneID:105275770 | LOC105275770 | Gene | LOC105275770 | protein_coding | NA | character(0) | character(0) | NW_011924888.1 | 405832 | 405837 | 6 | * | 6 | 1000 |
| NW_011924892.1 | 1158089 | 1160303 | 2215 | - | Gnomon | gene | NA | NA | gene569 | GeneID:105278527 | LOC105278527 | Gene | LOC105278527 | protein_coding | NA | character(0) | character(0) | NW_011924892.1 | 1159381 | 1159386 | 6 | * | 6 | 1000 |
| NW_011924931.1 | 4488185 | 4490767 | 2583 | + | Gnomon | gene | NA | NA | gene1620 | GeneID:105287754 | LOC105287754 | Gene | LOC105287754 | protein_coding | NA | character(0) | character(0) | NW_011924931.1 | 4489292 | 4489297 | 6 | * | 6 | 1000 |
| NW_011924946.1 | 786370 | 788739 | 2370 | - | Gnomon | gene | NA | NA | gene1995 | GeneID:105288196 | LOC105288196 | Gene | LOC105288196 | protein_coding | NA | character(0) | character(0) | NW_011924946.1 | 787444 | 787449 | 6 | * | 6 | 1000 |
| NW_011924946.1 | 1411689 | 1414880 | 3192 | - | Gnomon | gene | NA | NA | gene2041 | GeneID:105288248 | LOC105288248 | Gene | LOC105288248 | protein_coding | NA | character(0) | character(0) | NW_011924946.1 | 1413935 | 1413940 | 6 | * | 6 | 1000 |
| NW_011924956.1 | 717757 | 722032 | 4276 | + | Gnomon | gene | NA | NA | gene2570 | GeneID:105275036 | LOC105275036 | Gene | LOC105275036 | protein_coding | NA | character(0) | character(0) | NW_011924956.1 | 719335 | 719344 | 10 | * | 10 | 1000 |
| NW_011924973.1 | 828831 | 835120 | 6290 | + | Gnomon | gene | NA | NA | gene2799 | GeneID:105275316 | LOC105275316 | Gene | LOC105275316 | protein_coding | NA | character(0) | character(0) | NW_011924973.1 | 829674 | 829679 | 6 | * | 6 | 1000 |
| NW_011924973.1 | 1001098 | 1003969 | 2872 | + | Gnomon | gene | NA | NA | gene2837 | GeneID:105275355 | LOC105275355 | Gene | LOC105275355 | protein_coding | NA | character(0) | character(0) | NW_011924973.1 | 1002105 | 1002110 | 6 | * | 6 | 1000 |
| NW_011924977.1 | 31430 | 34619 | 3190 | + | Gnomon | gene | NA | NA | gene2916 | GeneID:109610741 | LOC109610741 | Gene | LOC109610741 | protein_coding | NA | character(0) | character(0) | NW_011924977.1 | 32395 | 32400 | 6 | * | 6 | 1000 |
| NW_011924977.1 | 662427 | 665182 | 2756 | + | Gnomon | gene | NA | NA | gene3008 | GeneID:105275550 | LOC105275550 | Gene | LOC105275550 | protein_coding | NA | character(0) | character(0) | NW_011924977.1 | 663534 | 663539 | 6 | * | 6 | 1000 |
| NW_011924979.1 | 16258 | 20038 | 3781 | + | Gnomon | gene | NA | NA | gene3218 | GeneID:105275781 | LOC105275781 | Gene | LOC105275781 | protein_coding | NA | character(0) | character(0) | NW_011924979.1 | 17774 | 17779 | 6 | * | 6 | 1000 |
| NW_011924996.1 | 75277 | 77259 | 1983 | + | Gnomon | gene | NA | NA | gene3743 | GeneID:105276382 | LOC105276382 | Gene | LOC105276382 | protein_coding | NA | character(0) | character(0) | NW_011924996.1 | 76074 | 76079 | 6 | * | 6 | 1000 |
| NW_011925007.1 | 926256 | 929694 | 3439 | + | Gnomon | gene | NA | NA | gene4194 | GeneID:105276881 | LOC105276881 | Gene | LOC105276881 | protein_coding | NA | character(0) | character(0) | NW_011925007.1 | 927466 | 927471 | 6 | * | 6 | 1000 |
| NW_011925022.1 | 386851 | 387976 | 1126 | - | Gnomon | gene | NA | NA | gene5000 | GeneID:105277804 | LOC105277804 | Gene | LOC105277804 | protein_coding | NA | character(0) | character(0) | NW_011925022.1 | 387077 | 387082 | 6 | * | 6 | 1000 |
| NW_011925053.1 | 649052 | 656133 | 7082 | - | Gnomon | gene | NA | NA | gene5882 | GeneID:105278812 | LOC105278812 | Gene | LOC105278812 | protein_coding | NA | character(0) | character(0) | NW_011925053.1 | 655139 | 655144 | 6 | * | 6 | 1000 |
| NW_011925128.1 | 1875336 | 1880845 | 5510 | + | Gnomon | gene | NA | NA | gene7225 | GeneID:105280395 | LOC105280395 | Gene | LOC105280395 | protein_coding | NA | character(0) | character(0) | NW_011925128.1 | 1876634 | 1876639 | 6 | * | 6 | 1000 |
| NW_011925142.1 | 777887 | 780161 | 2275 | - | Gnomon | gene | NA | NA | gene7574 | GeneID:105280791 | LOC105280791 | Gene | LOC105280791 | protein_coding | NA | character(0) | character(0) | NW_011925142.1 | 779002 | 779007 | 6 | * | 6 | 1000 |
| NW_011925150.1 | 76140 | 79045 | 2906 | - | Gnomon | gene | NA | NA | gene7626 | GeneID:105280856 | LOC105280856 | Gene | LOC105280856 | protein_coding | NA | character(0) | character(0) | NW_011925150.1 | 78377 | 78382 | 6 | * | 6 | 1000 |
| NW_011925151.1 | 38146 | 39177 | 1032 | - | Gnomon | gene | NA | NA | gene7635 | GeneID:105280860 | LOC105280860 | Gene | LOC105280860 | protein_coding | NA | character(0) | character(0) | NW_011925151.1 | 38257 | 38262 | 6 | * | 6 | 1000 |
| NW_011925183.1 | 39511 | 42630 | 3120 | + | Gnomon | gene | NA | NA | gene8236 | GeneID:105281577 | LOC105281577 | Gene | LOC105281577 | protein_coding | NA | character(0) | character(0) | NW_011925183.1 | 40051 | 40056 | 6 | * | 6 | 1000 |
| NW_011925188.1 | 199014 | 202632 | 3619 | - | Gnomon | gene | NA | NA | gene8257 | GeneID:105281603 | LOC105281603 | Gene | LOC105281603 | protein_coding | NA | character(0) | character(0) | NW_011925188.1 | 201626 | 201631 | 6 | * | 6 | 1000 |
| NW_011925203.1 | 63038 | 97839 | 34802 | + | Gnomon | gene | NA | NA | gene8521 | GeneID:105281906 | LOC105281906 | Gene | LOC105281906 | protein_coding | NA | character(0) | character(0) | NW_011925203.1 | 64603 | 64608 | 6 | * | 6 | 1000 |

...
... only first 25 lines shown ...

Directory: MRPR File: gwr-Cb_LI_RphaseA2.txt

```
seqnames    start end   width strand      source      type  score phase ID    Dbxref      Name  gbkey gene  gene_biotype      partial     start_range end_range   seqnames    start
      end   width strand      NbrSites    Sdnsty
NW_011924887.1    1546715     1553169     6455  +     Gnomon      gene  NA    NA    gene209     GeneID:105274475  LOC105274475      Gene  LOC105274475      protein_coding    NA
      character(0)      character(0)      NW_011924887.1    1550201     1550206     6     *     6     1000
NW_011924887.1    1817915     1820166     2252  -     Gnomon      gene  NA    NA    gene234     GeneID:105274775  LOC105274775      Gene  LOC105274775      protein_coding    NA
      character(0)      character(0)      NW_011924887.1    1819270     1819275     6     *     6     1000
NW_011924888.1    77334 78632 1299  -     Gnomon      gene  NA    NA    gene276     GeneID:105275214  LOC105275214      Gene  LOC105275214      protein_coding    NA    character(0)
      character(0)      NW_011924888.1    77560 77565 6     *     6     1000
NW_011924888.1    389413      407214      17802 -     Gnomon      gene  NA    NA    gene320     GeneID:105275770  LOC105275770      Gene  LOC105275770      protein_coding    NA
      character(0)      character(0)      NW_011924888.1    405832      405837      6     *     6     1000
NW_011924892.1    1158089     1160303     2215  -     Gnomon      gene  NA    NA    gene569     GeneID:105278527  LOC105278527      Gene  LOC105278527      protein_coding    NA
      character(0)      character(0)      NW_011924892.1    1159381     1159386     6     *     6     1000
NW_011924931.1    214221      216344      2124  +     Gnomon      gene  NA    NA    gene1375    GeneID:105287333  LOC105287333      Gene  LOC105287333      protein_coding    NA
      character(0)      character(0)      NW_011924931.1    215617      215622      6     *     6     1000
NW_011924931.1    3952621     3973270     20650 +     Gnomon      gene  NA    NA    gene1560    GeneID:105287694  LOC105287694      Gene  LOC105287694      protein_coding    NA
      character(0)      character(0)      NW_011924931.1    3966620     3966625     6     *     6     1000
NW_011924931.1    3966137     3967627     1491  -     Gnomon      gene  NA    NA    gene1562    GeneID:105287692  LOC105287692      Gene  LOC105287692      protein_coding    NA
      character(0)      character(0)      NW_011924931.1    3966620     3966625     6     *     6     1000
NW_011924931.1    4488185     4490767     2583  +     Gnomon      gene  NA    NA    gene1620    GeneID:105287754  LOC105287754      Gene  LOC105287754      protein_coding    NA
      character(0)      character(0)      NW_011924931.1    4489292     4489297     6     *     6     1000
NW_011924940.1    334573      349050      14478 -     Gnomon      gene  NA    NA    gene1863    GeneID:105288044  LOC105288044      Gene  LOC105288044      protein_coding    NA
      character(0)      character(0)      NW_011924940.1    347385      347390      6     *     6     1000
NW_011924946.1    775753      777201      1449  -     Gnomon      gene  NA    NA    gene1993    GeneID:105288193  LOC105288193      Gene  LOC105288193      protein_coding    NA
      character(0)      character(0)      NW_011924946.1    776368      776373      6     *     6     1000
NW_011924946.1    1411689     1414880     3192  -     Gnomon      gene  NA    NA    gene2041    GeneID:105288248  LOC105288248      Gene  LOC105288248      protein_coding    NA
      character(0)      character(0)      NW_011924946.1    1413935     1413940     6     *     6     1000
NW_011924956.1    717757      722032      4276  +     Gnomon      gene  NA    NA    gene2570    GeneID:105275036  LOC105275036      Gene  LOC105275036      protein_coding    NA
      character(0)      character(0)      NW_011924956.1    719335      719344      10    *     10    1000
NW_011924973.1    244133      245537      1405  +     Gnomon      gene  NA    NA    gene2754    GeneID:105275270  LOC105275270      Gene  LOC105275270      protein_coding    NA
      character(0)      character(0)      NW_011924973.1    244951      244956      6     *     6     1000
NW_011924973.1    828831      835120      6290  +     Gnomon      gene  NA    NA    gene2799    GeneID:105275316  LOC105275316      Gene  LOC105275316      protein_coding    NA
      character(0)      character(0)      NW_011924973.1    829674      829679      6     *     6     1000
NW_011924973.1    935440      945867      10428 +     Gnomon      gene  NA    NA    gene2820    GeneID:105275342  LOC105275342      Gene  LOC105275342      protein_coding    NA
      character(0)      character(0)      NW_011924973.1    936776      936781      6     *     6     1000
NW_011924973.1    1001098     1003969     2872  +     Gnomon      gene  NA    NA    gene2837    GeneID:105275355  LOC105275355      Gene  LOC105275355      protein_coding    NA
      character(0)      character(0)      NW_011924973.1    1002105     1002110     6     *     6     1000
NW_011924977.1    31430 34619 3190  +     Gnomon      gene  NA    NA    gene2916    GeneID:109610741  LOC109610741      Gene  LOC109610741      protein_coding    NA    character(0)
      character(0)      NW_011924977.1    32395 32400 6     *     6     1000
NW_011924977.1    662427      665182      2756  +     Gnomon      gene  NA    NA    gene3008    GeneID:105275550  LOC105275550      Gene  LOC105275550      protein_coding    NA
      character(0)      character(0)      NW_011924977.1    663534      663539      6     *     6     1000
NW_011924979.1    16258 20038 3781  +     Gnomon      gene  NA    NA    gene3218    GeneID:105275781  LOC105275781      Gene  LOC105275781      protein_coding    NA    character(0)
      character(0)      NW_011924979.1    17774 17779 6     *     6     1000
NW_011924979.1    2744383     2746071     1689  +     Gnomon      gene  NA    NA    gene3334    GeneID:105275897  LOC105275897      Gene  LOC105275897      protein_coding    NA
      character(0)      character(0)      NW_011924979.1    2745453     2745458     6     *     6     1000
NW_011924996.1    75277 77259 1983  +     Gnomon      gene  NA    NA    gene3743    GeneID:105276382  LOC105276382      Gene  LOC105276382      protein_coding    NA    character(0)
      character(0)      NW_011924996.1    76074 76079 6     *     6     1000
NW_011924996.1    1822758     1830685     7928  -     Gnomon      gene  NA    NA    gene3885    GeneID:105276527  LOC105276527      Gene  LOC105276527      protein_coding    NA
      character(0)      character(0)      NW_011924996.1    1829379     1829384     6     *     6     1000
NW_011925007.1    926256      929694      3439  +     Gnomon      gene  NA    NA    gene4194    GeneID:105276881  LOC105276881      Gene  LOC105276881      protein_coding    NA
      character(0)      character(0)      NW_011925007.1    927466      927471      6     *     6     1000
...
... only first 25 lines shown ...
```

Directory: MRPR File: gwr-Cb_LI_RphaseB1.txt

| seqnames | start | end | width | strand | source | type | score | phase | ID | Dbxref | Name | gbkey | gene | gene_biotype | partial | start_range | end_range | seqnames | start | end | width | strand | NbrSites | Sdnsty |
|---|---|---|---|---|---|---|---|---|---|---|---|---|---|---|---|---|---|---|---|---|---|---|---|---|
| NW_011924887.1 | 1546715 | 1553169 | 6455 | + | Gnomon | gene | NA | NA | gene209 | GeneID:105274475 | LOC105274475 | Gene | LOC105274475 | protein_coding | NA | character(0) | character(0) | NW_011924887.1 | 1550201 | 1550206 | 6 | * | 6 | 1000 |
| NW_011924887.1 | 1817915 | 1820166 | 2252 | - | Gnomon | gene | NA | NA | gene234 | GeneID:105274775 | LOC105274775 | Gene | LOC105274775 | protein_coding | NA | character(0) | character(0) | NW_011924887.1 | 1819270 | 1819275 | 6 | * | 6 | 1000 |
| NW_011924888.1 | 77334 | 78632 | 1299 | - | Gnomon | gene | NA | NA | gene276 | GeneID:105275214 | LOC105275214 | Gene | LOC105275214 | protein_coding | NA | character(0) | character(0) | NW_011924888.1 | 77560 | 77565 | 6 | * | 6 | 1000 |
| NW_011924888.1 | 389413 | 407214 | 17802 | - | Gnomon | gene | NA | NA | gene320 | GeneID:105275770 | LOC105275770 | Gene | LOC105275770 | protein_coding | NA | character(0) | character(0) | NW_011924888.1 | 405832 | 405837 | 6 | * | 6 | 1000 |
| NW_011924892.1 | 1158089 | 1160303 | 2215 | - | Gnomon | gene | NA | NA | gene569 | GeneID:105278527 | LOC105278527 | Gene | LOC105278527 | protein_coding | NA | character(0) | character(0) | NW_011924892.1 | 1159381 | 1159386 | 6 | * | 6 | 1000 |
| NW_011924931.1 | 3778005 | 3781577 | 3573 | - | Gnomon | gene | NA | NA | gene1536 | GeneID:105287663 | LOC105287663 | Gene | LOC105287663 | protein_coding | NA | character(0) | character(0) | NW_011924931.1 | 3780672 | 3780677 | 6 | * | 6 | 1000 |
| NW_011924931.1 | 4488185 | 4490767 | 2583 | + | Gnomon | gene | NA | NA | gene1620 | GeneID:105287754 | LOC105287754 | Gene | LOC105287754 | protein_coding | NA | character(0) | character(0) | NW_011924931.1 | 4489292 | 4489297 | 6 | * | 6 | 1000 |
| NW_011924973.1 | 244133 | 245537 | 1405 | + | Gnomon | gene | NA | NA | gene2754 | GeneID:105275270 | LOC105275270 | Gene | LOC105275270 | protein_coding | NA | character(0) | character(0) | NW_011924973.1 | 244951 | 244956 | 6 | * | 6 | 1000 |
| NW_011924973.1 | 828831 | 835120 | 6290 | + | Gnomon | gene | NA | NA | gene2799 | GeneID:105275316 | LOC105275316 | Gene | LOC105275316 | protein_coding | NA | character(0) | character(0) | NW_011924973.1 | 829674 | 829679 | 6 | * | 6 | 1000 |
| NW_011924973.1 | 935440 | 945867 | 10428 | + | Gnomon | gene | NA | NA | gene2820 | GeneID:105275342 | LOC105275342 | Gene | LOC105275342 | protein_coding | NA | character(0) | character(0) | NW_011924973.1 | 936776 | 936781 | 6 | * | 6 | 1000 |
| NW_011924973.1 | 1001098 | 1003969 | 2872 | + | Gnomon | gene | NA | NA | gene2837 | GeneID:105275355 | LOC105275355 | Gene | LOC105275355 | protein_coding | NA | character(0) | character(0) | NW_011924973.1 | 1002105 | 1002110 | 6 | * | 6 | 1000 |
| NW_011924977.1 | 31430 | 34619 | 3190 | + | Gnomon | gene | NA | NA | gene2916 | GeneID:109610741 | LOC109610741 | Gene | LOC109610741 | protein_coding | NA | character(0) | character(0) | NW_011924977.1 | 32395 | 32400 | 6 | * | 6 | 1000 |
| NW_011924977.1 | 662427 | 665182 | 2756 | + | Gnomon | gene | NA | NA | gene3008 | GeneID:105275550 | LOC105275550 | Gene | LOC105275550 | protein_coding | NA | character(0) | character(0) | NW_011924977.1 | 663534 | 663539 | 6 | * | 6 | 1000 |
| NW_011924979.1 | 16258 | 20038 | 3781 | + | Gnomon | gene | NA | NA | gene3218 | GeneID:105275781 | LOC105275781 | Gene | LOC105275781 | protein_coding | NA | character(0) | character(0) | NW_011924979.1 | 17774 | 17779 | 6 | * | 6 | 1000 |
| NW_011924996.1 | 75277 | 77259 | 1983 | + | Gnomon | gene | NA | NA | gene3743 | GeneID:105276382 | LOC105276382 | Gene | LOC105276382 | protein_coding | NA | character(0) | character(0) | NW_011924996.1 | 76074 | 76079 | 6 | * | 6 | 1000 |
| NW_011924996.1 | 1822758 | 1830685 | 7928 | - | Gnomon | gene | NA | NA | gene3885 | GeneID:105276527 | LOC105276527 | Gene | LOC105276527 | protein_coding | NA | character(0) | character(0) | NW_011924996.1 | 1829379 | 1829384 | 6 | * | 6 | 1000 |
| NW_011925022.1 | 386851 | 387976 | 1126 | - | Gnomon | gene | NA | NA | gene5000 | GeneID:105277804 | LOC105277804 | Gene | LOC105277804 | protein_coding | NA | character(0) | character(0) | NW_011925022.1 | 387077 | 387082 | 6 | * | 6 | 1000 |
| NW_011925053.1 | 649052 | 656133 | 7082 | - | Gnomon | gene | NA | NA | gene5882 | GeneID:105278812 | LOC105278812 | Gene | LOC105278812 | protein_coding | NA | character(0) | character(0) | NW_011925053.1 | 655139 | 655144 | 6 | * | 6 | 1000 |
| NW_011925053.1 | 1481875 | 1483457 | 1583 | - | Gnomon | gene | NA | NA | gene5932 | GeneID:105278862 | LOC105278862 | Gene | LOC105278862 | protein_coding | NA | character(0) | character(0) | NW_011925053.1 | 1482901 | 1482906 | 6 | * | 6 | 1000 |
| NW_011925086.1 | 604599 | 609266 | 4668 | + | Gnomon | gene | NA | NA | gene6342 | GeneID:105279367 | LOC105279367 | Gene | LOC105279367 | protein_coding | NA | character(0) | character(0) | NW_011925086.1 | 606157 | 606162 | 6 | * | 6 | 1000 |
| NW_011925128.1 | 1875336 | 1880845 | 5510 | + | Gnomon | gene | NA | NA | gene7225 | GeneID:105280395 | LOC105280395 | Gene | LOC105280395 | protein_coding | NA | character(0) | character(0) | NW_011925128.1 | 1876634 | 1876639 | 6 | * | 6 | 1000 |
| NW_011925142.1 | 777887 | 780161 | 2275 | - | Gnomon | gene | NA | NA | gene7574 | GeneID:105280791 | LOC105280791 | Gene | LOC105280791 | protein_coding | NA | character(0) | character(0) | NW_011925142.1 | 779002 | 779007 | 6 | * | 6 | 1000 |
| NW_011925150.1 | 76140 | 79045 | 2906 | - | Gnomon | gene | NA | NA | gene7626 | GeneID:105280856 | LOC105280856 | Gene | LOC105280856 | protein_coding | NA | character(0) | character(0) | NW_011925150.1 | 78377 | 78382 | 6 | * | 6 | 1000 |
| NW_011925163.1 | 470647 | 473680 | 3034 | + | Gnomon | gene | NA | NA | gene8040 | GeneID:105281341 | LOC105281341 | Gene | LOC105281341 | protein_coding | NA | character(0) | character(0) | NW_011925163.1 | 471213 | 471218 | 6 | * | 6 | 1000 |
| NW_011925182.1 | 57229 | 59555 | 2327 | - | Gnomon | gene | NA | NA | gene8209 | GeneID:105281549 | LOC105281549 | Gene | LOC105281549 | protein_coding | NA | character(0) | character(0) | NW_011925182.1 | 58922 | 58927 | 6 | * | 6 | 1000 |
| NW_011925183.1 | 39511 | 42630 | 3120 | + | Gnomon | gene | NA | NA | gene8236 | GeneID:105281577 | LOC105281577 | Gene | LOC105281577 | protein_coding | NA | character(0) | character(0) | NW_011925183.1 | 40051 | 40056 | 6 | * | 6 | 1000 |
| NW_011925203.1 | 63038 | 97839 | 34802 | + | Gnomon | gene | NA | NA | gene8521 | GeneID:105281906 | LOC105281906 | Gene | LOC105281906 | protein_coding | NA | character(0) | character(0) | NW_011925203.1 | 64603 | 64608 | 6 | * | 6 | 1000 |
| NW_011925216.1 | 136610 | 140476 | 3867 | - | Gnomon | gene | NA | NA | gene8735 | GeneID:105282144 | LOC105282144 | Gene | LOC105282144 | protein_coding | NA | character(0) | character(0) | NW_011925216.1 | 137522 | 137527 | 6 | * | 6 | 1000 |
| NW_011925216.1 | 136795 | 145548 | 8754 | + | Gnomon | gene | NA | NA | gene8736 | GeneID:105282142 | LOC105282142 | Gene | LOC105282142 | protein_coding | NA | character(0) | character(0) | NW_011925216.1 | 137522 | 137527 | 6 | * | 6 | 1000 |
| NW_011925216.1 | 156691 | 162086 | 5396 | + | Gnomon | gene | NA | NA | gene8741 | GeneID:105282150 | LOC105282150 | Gene | LOC105282150 | protein_coding | NA | | | | | | | | | |

character(0) character(0) NW_011925216.1 157541 157546 6 * 6 1000
NW_011925240.1 2490948 2493602 2655 - Gnomon gene NA NA gene9134 GeneID:105282589 LOC105282589 Gene LOC105282589 protein_coding NA
character(0) character(0) NW_011925240.1 2492742 2492747 6 * 6 1000
NW_011925258.1 404301 408788 4488 - Gnomon gene NA NA gene9347 GeneID:105282847 LOC105282847 Gene LOC105282847 protein_coding NA
character(0) character(0) NW_011925258.1 407771 407776 6 * 6 1000
NW_011925304.1 378599 380691 2093 - Gnomon gene NA NA gene9814 GeneID:105283373 LOC105283373 Gene LOC105283373 protein_coding NA
character(0) character(0) NW_011925304.1 379611 379616 6 * 6 1000
NW_011925304.1 468966 476624 7659 - Gnomon gene NA NA gene9824 GeneID:105283383 LOC105283383 Gene LOC105283383 protein_coding NA
character(0) character(0) NW_011925304.1 475523 475528 6 * 6 1000
NW_011925321.1 1107715 1109901 2187 - Gnomon gene NA NA gene10043 GeneID:105283634 LOC105283634 Gene LOC105283634 protein_coding NA
character(0) character(0) NW_011925321.1 1108690 1108695 6 * 6 1000
NW_011925353.1 40476 43146 2671 - Gnomon gene NA NA gene10405 GeneID:105284057 LOC105284057 Gene LOC105284057 protein_coding NA character(0)
character(0) NW_011925353.1 41948 41953 6 * 6 1000
NW_011925356.1 1573184 1579703 6520 - Gnomon gene NA NA gene10596 GeneID:105284262 LOC105284262 Gene LOC105284262 protein_coding NA
character(0) character(0) NW_011925356.1 1573650 1573655 6 * 6 1000
NW_011925387.1 39150 46759 7610 + Gnomon gene NA NA gene10928 GeneID:105284644 LOC105284644 Gene LOC105284644 protein_coding NA character(0)
character(0) NW_011925387.1 40039 40044 6 * 6 1000
NW_011925401.1 10153 19053 8901 - Gnomon gene NA NA gene11099 GeneID:105284845 LOC105284845 Gene LOC105284845 protein_coding NA character(0)
character(0) NW_011925401.1 17311 17316 6 * 6 1000
NW_011925470.1 613911 617551 3641 - Gnomon gene NA NA gene11628 GeneID:105285465 LOC105285465 Gene LOC105285465 protein_coding NA
character(0) character(0) NW_011925470.1 616616 616621 6 * 6 1000
NW_011925595.1 1 4548 4548 - Gnomon gene NA NA gene12102 GeneID:105286003 LOC105286003 Gene LOC105286003 protein_coding NA character(0)
character(0) NW_011925595.1 3717 3722 6 * 6 1000
NW_011925664.1 277932 283313 5382 + Gnomon gene NA NA gene12359 GeneID:105286364 LOC105286364 Gene LOC105286364 protein_coding NA
character(0) character(0) NW_011925664.1 279108 279113 6 * 6 1000
NW_011925715.1 952620 956416 3797 - Gnomon gene NA NA gene12567 GeneID:105286509 LOC105286509 Gene LOC105286509 protein_coding NA
character(0) character(0) NW_011925715.1 955653 955658 6 * 6 1000
NW_011929339.1 7 1883 1877 + Gnomon gene NA NA gene13384 GeneID:105287447 LOC105287447 Gene LOC105287447 protein_coding true character(0)
c("1883", ".") NW_011929339.1 1180 1185 6 * 6 1000
NW_011925086.1 128430 135139 6710 + Gnomon gene NA NA gene6311 GeneID:105279333 LOC105279333 Gene LOC105279333 protein_coding NA
character(0) character(0) NW_011925086.1 129637 129647 11 * 10 909.090909090909
NW_011924956.1 717757 722032 4276 + Gnomon gene NA NA gene2570 GeneID:105275036 LOC105275036 Gene LOC105275036 protein_coding NA
character(0) character(0) NW_011924956.1 719335 719344 10 * 9 900
NW_011924914.1 259948 262322 2375 - Gnomon gene NA NA gene952 GeneID:105283016 LOC105283016 Gene LOC105283016 protein_coding NA
character(0) character(0) NW_011924914.1 261398 261406 9 * 8 888.888888888889
NW_011924967.1 91700 93046 1347 - Gnomon gene NA NA gene2644 GeneID:105275137 LOC105275137 Gene LOC105275137 protein_coding NA character(0)
character(0) NW_011924967.1 92497 92505 9 * 8 888.888888888889
NW_011924977.1 3728770 3731404 2635 + Gnomon gene NA NA gene3213 GeneID:105275761 LOC105275761 Gene LOC105275761 protein_coding NA
character(0) character(0) NW_011924977.1 3729889 3729897 9 * 8 888.888888888889
NW_011925189.1 165659 175663 10005 - Gnomon gene NA NA gene8294 GeneID:105281645 LOC105281645 Gene LOC105281645 protein_coding NA
character(0) character(0) NW_011925189.1 174276 174284 9 * 8 888.888888888889
NW_011925239.1 3979 7808 3830 + Gnomon gene NA NA gene9001 GeneID:105282461 LOC105282461 Gene LOC105282461 protein_coding NA character(0)
character(0) NW_011925239.1 4874 4882 9 * 8 888.888888888889
NW_011925106.1 2086959 2096661 9703 + Gnomon gene NA NA gene6771 GeneID:105279854 LOC105279854 Gene LOC105279854 protein_coding NA
character(0) character(0) NW_011925106.1 2088114 2088121 8 * 7 875
NW_011925664.1 38296 39300 1005 + Gnomon gene NA NA gene12311 GeneID:105286212 LOC105286212 Gene LOC105286212 protein_coding NA character(0)
character(0) NW_011925664.1 39054 39061 8 * 7 875
NW_011926135.1 70608 74554 3947 + Gnomon gene NA NA gene13076 GeneID:105287104 LOC105287104 Gene LOC105287104 protein_coding NA character(0)
character(0) NW_011926135.1 71339 71346 8 * 7 875
NW_011924887.1 547855 552902 5048 + Gnomon gene NA NA gene149 GeneID:105287687 LOC105287687 Gene LOC105287687 protein_coding NA
character(0) character(0) NW_011924887.1 549006 549012 7 * 6 857.142857142857
NW_011924888.1 30738 31847 1110 + Gnomon gene NA NA gene264 GeneID:105275153 LOC105275153 Gene LOC105275153 protein_coding NA character(0)
character(0) NW_011924888.1 31453 31459 7 * 6 857.142857142857
NW_011924889.1 476236 478868 2633 - Gnomon gene NA NA gene504 GeneID:105277817 LOC105277817 Gene LOC105277817 protein_coding NA
character(0) character(0) NW_011924889.1 477980 477986 7 * 6 857.142857142857
NW_011924889.1 493449 499545 6097 + Gnomon gene NA NA gene508 GeneID:105277915 LOC105277915 Gene LOC105277915 protein_coding NA
character(0) character(0) NW_011924889.1 494195 494201 7 * 6 857.142857142857
NW_011924889.1 606387 610322 3936 - Gnomon gene NA NA gene519 GeneID:105278022 LOC105278022 Gene LOC105278022 protein_coding NA
character(0) character(0) NW_011924889.1 609088 609094 7 * 6 857.142857142857
NW_011924889.1 606387 610322 3936 - Gnomon gene NA NA gene519 GeneID:105278022 LOC105278022 Gene LOC105278022 protein_coding NA
character(0) character(0) NW_011924889.1 609259 609265 7 * 6 857.142857142857
NW_011924892.1 274786 285690 10905 - Gnomon gene NA NA gene544 GeneID:105278288 LOC105278288 Gene LOC105278288 protein_coding NA
character(0) character(0) NW_011924892.1 284836 284842 7 * 6 857.142857142857
NW_011924907.1 889683 896841 7159 - Gnomon gene NA NA gene861 GeneID:105281750 LOC105281750 Gene LOC105281750 protein_coding NA

character(0) character(0) NW_011924907.1 896035 896041 7 * 6 857.142857142857
NW_011924914.1 827084 831408 4325 - Gnomon gene NA NA gene1010 GeneID:105283498 LOC105283498 Gene LOC105283498 protein_coding NA
character(0) character(0) NW_011924914.1 831099 831105 7 * 6 857.142857142857
NW_011924929.1 931843 943599 11757 + Gnomon gene NA NA gene1329 GeneID:105286844 LOC105286844 Gene LOC105286844 protein_coding NA
character(0) character(0) NW_011924929.1 933320 933326 7 * 6 857.142857142857
NW_011924931.1 116678 120078 3401 + Gnomon gene NA NA gene1363 GeneID:105287185 LOC105287185 Gene LOC105287185 protein_coding NA
character(0) character(0) NW_011924931.1 118254 118260 7 * 6 857.142857142857
NW_011924931.1 201740 204575 2836 - Gnomon gene NA NA gene1371 GeneID:105287279 LOC105287279 Gene LOC105287279 protein_coding NA
character(0) character(0) NW_011924931.1 203497 203503 7 * 6 857.142857142857
NW_011924931.1 242950 250477 7528 - Gnomon gene NA NA gene1378 GeneID:105287371 LOC105287371 Gene LOC105287371 protein_coding NA
character(0) character(0) NW_011924931.1 249191 249197 7 * 6 857.142857142857
NW_011924931.1 290674 293586 2913 + Gnomon gene NA NA gene1388 GeneID:105287473 LOC105287473 Gene LOC105287473 protein_coding NA
character(0) character(0) NW_011924931.1 291628 291634 7 * 6 857.142857142857
NW_011924931.1 347408 353524 6117 + Gnomon gene NA NA gene1391 GeneID:105287511 LOC105287511 Gene LOC105287511 protein_coding NA
character(0) character(0) NW_011924931.1 348868 348874 7 * 6 857.142857142857
NW_011924931.1 1565109 1566883 1775 + Gnomon gene NA NA gene1444 GeneID:105287569 LOC105287569 Gene LOC105287569 protein_coding NA
character(0) character(0) NW_011924931.1 1565961 1565967 7 * 6 857.142857142857
NW_011924931.1 2028839 2031812 2974 + Gnomon gene NA NA gene1470 GeneID:105287597 LOC105287597 Gene LOC105287597 protein_coding NA
character(0) character(0) NW_011924931.1 2030239 2030245 7 * 6 857.142857142857
NW_011924931.1 2871689 2878941 7253 - Gnomon gene NA NA gene1492 GeneID:105287620 LOC105287620 Gene LOC105287620 protein_coding NA
character(0) character(0) NW_011924931.1 2878145 2878151 7 * 6 857.142857142857
NW_011924931.1 3929932 3932866 2935 + Gnomon gene NA NA gene1555 GeneID:105287684 LOC105287684 Gene LOC105287684 protein_coding NA
character(0) character(0) NW_011924931.1 3930880 3930886 7 * 6 857.142857142857
NW_011924931.1 3979390 3986328 6939 + Gnomon gene NA NA gene1566 GeneID:105287698 LOC105287698 Gene LOC105287698 protein_coding NA
character(0) character(0) NW_011924931.1 3983680 3983686 7 * 6 857.142857142857
NW_011924931.1 4058265 4059684 1420 - Gnomon gene NA NA gene1581 GeneID:105287710 LOC105287710 Gene LOC105287710 protein_coding NA
character(0) character(0) NW_011924931.1 4058767 4058773 7 * 6 857.142857142857
NW_011924932.1 1156872 1160705 3834 + Gnomon gene NA NA gene1695 GeneID:105287855 LOC105287855 Gene LOC105287855 protein_coding NA
character(0) character(0) NW_011924932.1 1157735 1157741 7 * 6 857.142857142857
NW_011924938.1 72334 83258 10925 + Gnomon gene NA NA gene1805 GeneID:105287985 LOC105287985 Gene LOC105287985 protein_coding NA character(0)
character(0) NW_011924938.1 72914 72920 7 * 6 857.142857142857
NW_011924946.1 2342632 2344089 1458 - Gnomon gene NA NA gene2088 GeneID:105274497 LOC105274497 Gene LOC105274497 protein_coding NA
character(0) character(0) NW_011924946.1 2343055 2343061 7 * 6 857.142857142857
NW_011924946.1 2914862 2919248 4387 - Gnomon gene NA NA gene2173 GeneID:105274590 LOC105274590 Gene LOC105274590 protein_coding NA
character(0) character(0) NW_011924946.1 2917571 2917577 7 * 6 857.142857142857
NW_011924946.1 2914862 2919248 4387 - Gnomon gene NA NA gene2173 GeneID:105274590 LOC105274590 Gene LOC105274590 protein_coding NA
character(0) character(0) NW_011924946.1 2918055 2918061 7 * 6 857.142857142857
NW_011924946.1 3145253 3148790 3538 - Gnomon gene NA NA gene2215 GeneID:105274638 LOC105274638 Gene LOC105274638 protein_coding NA
character(0) character(0) NW_011924946.1 3147722 3147728 7 * 6 857.142857142857
NW_011924947.1 86691 90381 3691 - Gnomon gene NA NA gene2272 GeneID:105274705 LOC105274705 Gene LOC105274705 protein_coding NA character(0)
character(0) NW_011924947.1 89757 89763 7 * 6 857.142857142857
NW_011924947.1 141424 143460 2037 + Gnomon gene NA NA gene2279 GeneID:105274711 LOC105274711 Gene LOC105274711 protein_coding NA
character(0) character(0) NW_011924947.1 142119 142125 7 * 6 857.142857142857
NW_011924951.1 73901 76560 2660 + Gnomon gene NA NA gene2325 GeneID:105274767 LOC105274767 Gene LOC105274767 protein_coding NA character(0)
character(0) NW_011924951.1 74989 74995 7 * 6 857.142857142857
NW_011924951.1 223619 227563 3945 + Gnomon gene NA NA gene2335 GeneID:105274778 LOC105274778 Gene LOC105274778 protein_coding NA
character(0) character(0) NW_011924951.1 224683 224689 7 * 6 857.142857142857
NW_011924956.1 701839 703915 2077 - Gnomon gene NA NA gene2566 GeneID:105275031 LOC105275031 Gene LOC105275031 protein_coding NA
character(0) character(0) NW_011924956.1 702713 702719 7 * 6 857.142857142857
NW_011924967.1 11923 12765 843 - Gnomon gene NA NA gene2630 GeneID:105275119 LOC105275119 Gene LOC105275119 protein_coding NA character(0)
character(0) NW_011924967.1 12053 12059 7 * 6 857.142857142857
NW_011924973.1 774415 775843 1429 - Gnomon gene NA NA gene2790 GeneID:105275307 LOC105275307 Gene LOC105275307 protein_coding NA
character(0) character(0) NW_011924973.1 775075 775081 7 * 6 857.142857142857
NW_011924974.1 483919 487626 3708 + Gnomon gene NA NA gene2873 GeneID:105275401 LOC105275401 Gene LOC105275401 protein_coding NA
character(0) character(0) NW_011924974.1 484702 484708 7 * 6 857.142857142857
NW_011924977.1 1460154 1463690 3537 - Gnomon gene NA NA gene3072 GeneID:105275615 LOC105275615 Gene LOC105275615 protein_coding NA
character(0) character(0) NW_011924977.1 1461887 1461893 7 * 6 857.142857142857
NW_011924977.1 1460154 1463690 3537 - Gnomon gene NA NA gene3072 GeneID:105275615 LOC105275615 Gene LOC105275615 protein_coding NA
character(0) character(0) NW_011924977.1 1462188 1462194 7 * 6 857.142857142857
NW_011924977.1 2573287 2581453 8167 - Gnomon gene NA NA gene3147 GeneID:105275689 LOC105275689 Gene LOC105275689 protein_coding NA
character(0) character(0) NW_011924977.1 2580506 2580512 7 * 6 857.142857142857
NW_011924977.1 3678529 3682684 4156 - Gnomon gene NA NA gene3205 GeneID:105275753 LOC105275753 Gene LOC105275753 protein_coding NA
character(0) character(0) NW_011924977.1 3681359 3681365 7 * 6 857.142857142857
NW_011924977.1 3678529 3682684 4156 - Gnomon gene NA NA gene3205 GeneID:105275753 LOC105275753 Gene LOC105275753 protein_coding NA

character(0) character(0) NW_011924977.1 3681568 3681574 7 * 6 857.142857142857
NW_011924977.1 3682584 3686057 3474 - Gnomon gene NA NA gene3206 GeneID:105275752 LOC105275752 Gene LOC105275752 protein_coding NA
character(0) character(0) NW_011924977.1 3684978 3684984 7 * 6 857.142857142857
NW_011924979.1 211193 212598 1406 - Gnomon gene NA NA gene3249 GeneID:105275908 LOC105275908 Gene LOC105275908 protein_coding NA
character(0) character(0) NW_011924979.1 212057 212063 7 * 6 857.142857142857
NW_011924979.1 2744383 2746071 1689 + Gnomon gene NA NA gene3334 GeneID:105275897 LOC105275897 Gene LOC105275897 protein_coding NA
character(0) character(0) NW_011924979.1 2745351 2745357 7 * 6 857.142857142857
NW_011924987.1 611928 613454 1527 - Gnomon gene NA NA gene3407 GeneID:105275991 LOC105275991 Gene LOC105275991 protein_coding NA
character(0) character(0) NW_011924987.1 612398 612404 7 * 6 857.142857142857
NW_011924987.1 2638171 2648657 10487 - Gnomon gene NA NA gene3636 GeneID:105276233 LOC105276233 Gene LOC105276233 protein_coding NA
character(0) character(0) NW_011924987.1 2647465 2647471 7 * 6 857.142857142857
NW_011924996.1 292526 303470 10945 + Gnomon gene NA NA gene3775 GeneID:105276410 LOC105276410 Gene LOC105276410 protein_coding NA
character(0) character(0) NW_011924996.1 293762 293768 7 * 6 857.142857142857
NW_011924996.1 292526 303470 10945 + Gnomon gene NA NA gene3775 GeneID:105276410 LOC105276410 Gene LOC105276410 protein_coding NA
character(0) character(0) NW_011924996.1 293819 293825 7 * 6 857.142857142857
NW_011924996.1 439124 441322 2199 - Gnomon gene NA NA gene3784 GeneID:105276420 LOC105276420 Gene LOC105276420 protein_coding NA
character(0) character(0) NW_011924996.1 440297 440303 7 * 6 857.142857142857
NW_011924996.1 641332 646889 5558 - Gnomon gene NA NA gene3801 GeneID:105276437 LOC105276437 Gene LOC105276437 protein_coding NA
character(0) character(0) NW_011924996.1 645416 645422 7 * 6 857.142857142857
NW_011924996.1 1790083 1793499 3417 - Gnomon gene NA NA gene3877 GeneID:105276521 LOC105276521 Gene LOC105276521 protein_coding NA
character(0) character(0) NW_011924996.1 1792422 1792428 7 * 6 857.142857142857
NW_011925007.1 1417723 1421435 3713 + Gnomon gene NA NA gene4248 GeneID:105276948 LOC105276948 Gene LOC105276948 protein_coding NA
character(0) character(0) NW_011925007.1 1419353 1419359 7 * 6 857.142857142857
NW_011925007.1 1452162 1454301 2140 + Gnomon gene NA NA gene4252 GeneID:105276949 LOC105276949 Gene LOC105276949 protein_coding NA
character(0) character(0) NW_011925007.1 1453109 1453115 7 * 6 857.142857142857
NW_011925012.1 1316386 1318477 2092 + Gnomon gene NA NA gene4492 GeneID:105277232 LOC105277232 Gene LOC105277232 protein_coding NA
character(0) character(0) NW_011925012.1 1317469 1317475 7 * 6 857.142857142857
NW_011925019.1 475 2900 2426 + Gnomon gene NA NA gene4633 GeneID:105277389 LOC105277389 Gene LOC105277389 protein_coding NA character(0)
character(0) NW_011925019.1 1670 1676 7 * 6 857.142857142857
NW_011925019.1 2803 5896 3094 - Gnomon gene NA NA gene4634 GeneID:105277388 LOC105277388 Gene LOC105277388 protein_coding NA character(0)
character(0) NW_011925019.1 4685 4691 7 * 6 857.142857142857
NW_011925019.1 1195331 1199707 4377 - Gnomon gene NA NA gene4693 GeneID:105277452 LOC105277452 Gene LOC105277452 protein_coding NA
character(0) character(0) NW_011925019.1 1198454 1198460 7 * 6 857.142857142857
NW_011925019.1 1427431 1429968 2538 - Gnomon gene NA NA gene4709 GeneID:109610850 LOC109610850 Gene LOC109610850 protein_coding NA
character(0) character(0) NW_011925019.1 1429240 1429246 7 * 6 857.142857142857
NW_011925020.1 469190 474352 5163 + Gnomon gene NA NA gene4780 GeneID:105277552 LOC105277552 Gene LOC105277552 protein_coding NA
character(0) character(0) NW_011925020.1 470270 470276 7 * 6 857.142857142857
NW_011925027.1 1297549 1301844 4296 + Gnomon gene NA NA gene5179 GeneID:105278005 LOC105278005 Gene LOC105278005 protein_coding NA
character(0) character(0) NW_011925027.1 1298524 1298530 7 * 6 857.142857142857
NW_011925029.1 39174 43410 4237 + Gnomon gene NA NA gene5243 GeneID:105278106 LOC105278106 Gene LOC105278106 protein_coding NA character(0)
character(0) NW_011925029.1 40261 40267 7 * 6 857.142857142857
NW_011925031.1 146315 150217 3903 + Gnomon gene NA NA gene5308 GeneID:105278157 LOC105278157 Gene LOC105278157 protein_coding NA
character(0) character(0) NW_011925031.1 148598 148604 7 * 6 857.142857142857
NW_011925035.1 1812544 1818171 5628 - Gnomon gene NA NA gene5451 GeneID:105278309 LOC105278309 Gene LOC105278309 protein_coding NA
character(0) character(0) NW_011925035.1 1817088 1817094 7 * 6 857.142857142857
NW_011925035.1 3102848 3104698 1851 + Gnomon gene NA NA gene5557 GeneID:105278431 LOC105278431 Gene LOC105278431 protein_coding NA
character(0) character(0) NW_011925035.1 3103902 3103908 7 * 6 857.142857142857
NW_011925036.1 1418708 1424550 5843 + Gnomon gene NA NA gene5653 GeneID:105278544 LOC105278544 Gene LOC105278544 protein_coding NA
character(0) character(0) NW_011925036.1 1420178 1420184 7 * 6 857.142857142857
NW_011925067.1 5971 15150 9180 + Gnomon gene NA NA gene6024 GeneID:105278996 LOC105278996 Gene LOC105278996 protein_coding NA character(0)
character(0) NW_011925067.1 7054 7060 7 * 6 857.142857142857
NW_011925067.1 5971 15150 9180 + Gnomon gene NA NA gene6024 GeneID:105278996 LOC105278996 Gene LOC105278996 protein_coding NA character(0)
character(0) NW_011925067.1 7213 7219 7 * 6 857.142857142857
NW_011925067.1 111367 117057 5691 + Gnomon gene NA NA gene6031 GeneID:105279000 LOC105279000 Gene LOC105279000 protein_coding NA
character(0) character(0) NW_011925067.1 112547 112553 7 * 6 857.142857142857
NW_011925068.1 10281 12328 2048 + Gnomon gene NA NA gene6111 GeneID:105279088 LOC105279088 Gene LOC105279088 protein_coding NA character(0)
character(0) NW_011925068.1 11234 11240 7 * 6 857.142857142857
NW_011925082.1 664899 668532 3634 - Gnomon gene NA NA gene6299 GeneID:105279313 LOC105279313 Gene LOC105279313 protein_coding NA
character(0) character(0) NW_011925082.1 667658 667664 7 * 6 857.142857142857
NW_011925086.1 250638 255287 4650 - Gnomon gene NA NA gene6324 GeneID:105279346 LOC105279346 Gene LOC105279346 protein_coding NA
character(0) character(0) NW_011925086.1 254146 254152 7 * 6 857.142857142857
NW_011925106.1 154197 158224 4028 + Gnomon gene NA NA gene6601 GeneID:105279664 LOC105279664 Gene LOC105279664 protein_coding NA
character(0) character(0) NW_011925106.1 155492 155498 7 * 6 857.142857142857
NW_011925106.1 431654 436050 4397 + Gnomon gene NA NA gene6641 GeneID:105279709 LOC105279709 Gene LOC105279709 protein_coding NA

character(0) character(0) NW_011925106.1 432897 432903 7 * 6 857.142857142857
NW_011925106.1 512988 515818 2831 - Gnomon gene NA NA gene6652 GeneID:105279719 LOC105279719 Gene LOC105279719 protein_coding NA
character(0) character(0) NW_011925106.1 514935 514941 7 * 6 857.142857142857
NW_011925106.1 629782 633106 3325 - Gnomon gene NA NA gene6668 GeneID:105279740 LOC105279740 Gene LOC105279740 protein_coding NA
character(0) character(0) NW_011925106.1 631881 631887 7 * 6 857.142857142857
NW_011925106.1 2086959 2096661 9703 + Gnomon gene NA NA gene6771 GeneID:105279854 LOC105279854 Gene LOC105279854 protein_coding NA
character(0) character(0) NW_011925106.1 2087848 2087854 7 * 6 857.142857142857
NW_011925118.1 457689 459947 2259 + Gnomon gene NA NA gene6953 GeneID:105280092 LOC105280092 Gene LOC105280092 protein_coding NA
character(0) character(0) NW_011925118.1 458682 458688 7 * 6 857.142857142857
NW_011925118.1 481415 484075 2661 + Gnomon gene NA NA gene6956 GeneID:105280102 LOC105280102 Gene LOC105280102 protein_coding NA
character(0) character(0) NW_011925118.1 482695 482701 7 * 6 857.142857142857
NW_011925128.1 7781 9472 1692 + Gnomon gene NA NA gene7085 GeneID:105280236 LOC105280236 Gene LOC105280236 protein_coding NA character(0)
character(0) NW_011925128.1 8930 8936 7 * 6 857.142857142857
NW_011925128.1 709798 711703 1906 - Gnomon gene NA NA gene7149 GeneID:105280306 LOC105280306 Gene LOC105280306 protein_coding NA
character(0) character(0) NW_011925128.1 711187 711193 7 * 6 857.142857142857
NW_011925128.1 800543 802328 1786 - Gnomon gene NA NA gene7160 GeneID:105280319 LOC105280319 Gene LOC105280319 protein_coding NA
character(0) character(0) NW_011925128.1 801605 801611 7 * 6 857.142857142857
NW_011925142.1 423375 426060 2686 - Gnomon gene NA NA gene7527 GeneID:105280739 LOC105280739 Gene LOC105280739 protein_coding NA
character(0) character(0) NW_011925142.1 424101 424107 7 * 6 857.142857142857
NW_011925142.1 423375 426060 2686 - Gnomon gene NA NA gene7527 GeneID:105280739 LOC105280739 Gene LOC105280739 protein_coding NA
character(0) character(0) NW_011925142.1 424227 424233 7 * 6 857.142857142857
NW_011925142.1 781512 784836 3325 + Gnomon gene NA NA gene7575 GeneID:105280793 LOC105280793 Gene LOC105280793 protein_coding NA
character(0) character(0) NW_011925142.1 782345 782351 7 * 6 857.142857142857
NW_011925161.1 210886 215244 4359 + Gnomon gene NA NA gene7720 GeneID:105280964 LOC105280964 Gene LOC105280964 protein_coding NA
character(0) character(0) NW_011925161.1 213136 213142 7 * 6 857.142857142857
NW_011925161.1 3740915 3743520 2606 + Gnomon gene NA NA gene7963 GeneID:105281227 LOC105281227 Gene LOC105281227 protein_coding NA
character(0) character(0) NW_011925161.1 3741983 3741989 7 * 6 857.142857142857
NW_011925182.1 26861 30576 3716 - Gnomon gene NA NA gene8205 GeneID:105281545 LOC105281545 Gene LOC105281545 protein_coding NA character(0)
character(0) NW_011925182.1 29339 29345 7 * 6 857.142857142857
NW_011925183.1 39511 42630 3120 + Gnomon gene NA NA gene8236 GeneID:105281577 LOC105281577 Gene LOC105281577 protein_coding NA character(0)
character(0) NW_011925183.1 40353 40359 7 * 6 857.142857142857
NW_011925188.1 199014 202632 3619 - Gnomon gene NA NA gene8257 GeneID:105281603 LOC105281603 Gene LOC105281603 protein_coding NA
character(0) character(0) NW_011925188.1 201441 201447 7 * 6 857.142857142857
NW_011925203.1 23205 24951 1747 - Gnomon gene NA NA gene8516 GeneID:105281896 LOC105281896 Gene LOC105281896 protein_coding NA character(0)
character(0) NW_011925203.1 24180 24186 7 * 6 857.142857142857
NW_011925209.1 85655 90207 4553 - Gnomon gene NA NA gene8551 GeneID:105281943 LOC105281943 Gene LOC105281943 protein_coding NA character(0)
character(0) NW_011925209.1 86875 86881 7 * 6 857.142857142857
NW_011925209.1 169877 173749 3873 - Gnomon gene NA NA gene8560 GeneID:105281949 LOC105281949 Gene LOC105281949 protein_coding NA
character(0) character(0) NW_011925209.1 172697 172703 7 * 6 857.142857142857
NW_011925216.1 130033 131250 1218 + Gnomon gene NA NA gene8733 GeneID:105282147 LOC105282147 Gene LOC105282147 protein_coding NA
character(0) character(0) NW_011925216.1 130799 130805 7 * 6 857.142857142857
NW_011925224.1 208794 211425 2632 + Gnomon gene NA NA gene8852 GeneID:105282278 LOC105282278 Gene LOC105282278 protein_coding NA
character(0) character(0) NW_011925224.1 209730 209736 7 * 6 857.142857142857
NW_011925224.1 219079 219935 857 - Gnomon gene NA NA gene8854 GeneID:105282280 LOC105282280 Gene LOC105282280 protein_coding NA
character(0) character(0) NW_011925224.1 219257 219263 7 * 6 857.142857142857
NW_011925240.1 1526960 1531289 4330 + Gnomon gene NA NA gene9107 GeneID:105282559 LOC105282559 Gene LOC105282559 protein_coding NA
character(0) character(0) NW_011925240.1 1527897 1527903 7 * 6 857.142857142857
NW_011925240.1 1575290 1580897 5608 - Gnomon gene NA NA gene9116 GeneID:105282568 LOC105282568 Gene LOC105282568 protein_coding NA
character(0) character(0) NW_011925240.1 1579255 1579261 7 * 6 857.142857142857
NW_011925241.1 397336 409261 11926 - Gnomon gene NA NA gene9191 GeneID:105282655 LOC105282655 Gene LOC105282655 protein_coding NA
character(0) character(0) NW_011925241.1 408196 408202 7 * 6 857.142857142857
NW_011925257.1 52519 55399 2881 - Gnomon gene NA NA gene9303 GeneID:105282793 LOC105282793 Gene LOC105282793 protein_coding NA character(0)
character(0) NW_011925257.1 54841 54847 7 * 6 857.142857142857
NW_011925258.1 360040 361894 1855 - Gnomon gene NA NA gene9341 GeneID:105282841 LOC105282841 Gene LOC105282841 protein_coding NA
character(0) character(0) NW_011925258.1 361089 361095 7 * 6 857.142857142857
NW_011925267.1 224087 232805 8719 + Gnomon gene NA NA gene9418 GeneID:105282921 LOC105282921 Gene LOC105282921 protein_coding NA
character(0) character(0) NW_011925267.1 225570 225576 7 * 6 857.142857142857
NW_011925315.1 538327 543302 4976 + Gnomon gene NA NA gene9909 GeneID:105283484 LOC105283484 Gene LOC105283484 protein_coding NA
character(0) character(0) NW_011925315.1 539621 539627 7 * 6 857.142857142857
NW_011925323.1 390910 395897 4988 - Gnomon gene NA NA gene10091 GeneID:105283697 LOC105283697 Gene LOC105283697 protein_coding NA
character(0) character(0) NW_011925323.1 394351 394357 7 * 6 857.142857142857
NW_011925323.1 1444062 1448038 3977 - Gnomon gene NA NA gene10138 GeneID:105283741 LOC105283741 Gene LOC105283741 protein_coding NA
character(0) character(0) NW_011925323.1 1445991 1445997 7 * 6 857.142857142857
NW_011925330.1 58606 82143 23538 + Gnomon gene NA NA gene10255 GeneID:105283878 LOC105283878 Gene LOC105283878 protein_coding NA character(0)

character(0) NW_011925330.1 59800 59806 7 * 6 857.142857142857
NW_011925336.1 89625 92404 2780 - Gnomon gene NA NA gene10323 GeneID:105283960 LOC105283960 Gene LOC105283960 protein_coding NA character(0)
character(0) NW_011925336.1 91240 91246 7 * 6 857.142857142857
NW_011925336.1 92449 95536 3088 + Gnomon gene NA NA gene10324 GeneID:105283961 LOC105283961 Gene LOC105283961 protein_coding NA character(0)
character(0) NW_011925336.1 93305 93311 7 * 6 857.142857142857
NW_011925353.1 16223 20332 4110 + Gnomon gene NA NA gene10401 GeneID:105284052 LOC105284052 Gene LOC105284052 protein_coding NA character(0)
character(0) NW_011925353.1 19139 19145 7 * 6 857.142857142857
NW_011925353.1 584566 593506 8941 - Gnomon gene NA NA gene10439 GeneID:105284096 LOC105284096 Gene LOC105284096 protein_coding NA
character(0) character(0) NW_011925353.1 592218 592224 7 * 6 857.142857142857
NW_011925356.1 701727 704945 3219 + Gnomon gene NA NA gene10538 GeneID:105284202 LOC105284202 Gene LOC105284202 protein_coding NA
character(0) character(0) NW_011925356.1 703128 703134 7 * 6 857.142857142857
NW_011925367.1 1119922 1121887 1966 + Gnomon gene NA NA gene10714 GeneID:105284396 LOC105284396 Gene LOC105284396 protein_coding NA
character(0) character(0) NW_011925367.1 1120846 1120852 7 * 6 857.142857142857
NW_011925371.1 341773 343746 1974 - Gnomon gene NA NA gene10816 GeneID:105284502 LOC105284502 Gene LOC105284502 protein_coding NA
character(0) character(0) NW_011925371.1 342828 342834 7 * 6 857.142857142857
NW_011925371.1 899674 901634 1961 - Gnomon gene NA NA gene10880 GeneID:105284576 LOC105284576 Gene LOC105284576 protein_coding NA
character(0) character(0) NW_011925371.1 900958 900964 7 * 6 857.142857142857
NW_011925387.1 118203 119961 1759 - Gnomon gene NA NA gene10943 GeneID:105284659 LOC105284659 Gene LOC105284659 protein_coding NA
character(0) character(0) NW_011925387.1 119303 119309 7 * 6 857.142857142857
NW_011925390.1 59041 77340 18300 + Gnomon gene NA NA gene11018 GeneID:105284754 LOC105284754 Gene LOC105284754 protein_coding NA character(0)
character(0) NW_011925390.1 60352 60358 7 * 6 857.142857142857
NW_011925434.1 120720 130419 9700 + Gnomon gene NA NA gene11344 GeneID:105285135 LOC105285135 Gene LOC105285135 protein_coding NA
character(0) character(0) NW_011925434.1 123131 123137 7 * 6 857.142857142857
NW_011925459.1 162939 164541 1603 + Gnomon gene NA NA gene11566 GeneID:105285392 LOC105285392 Gene LOC105285392 protein_coding NA
character(0) character(0) NW_011925459.1 163954 163960 7 * 6 857.142857142857
NW_011925459.1 186244 187554 1311 - Gnomon gene NA NA gene11569 GeneID:105285394 LOC105285394 Gene LOC105285394 protein_coding NA
character(0) character(0) NW_011925459.1 186700 186706 7 * 6 857.142857142857
NW_011925465.1 71398 78050 6653 - Gnomon gene NA NA gene11580 GeneID:105285410 LOC105285410 Gene LOC105285410 protein_coding NA character(0)
character(0) NW_011925465.1 77157 77163 7 * 6 857.142857142857
NW_011925471.1 1427 3141 1715 - Gnomon gene NA NA gene11632 GeneID:109611363 LOC109611363 Gene LOC109611363 lncRNA NA character(0)
character(0) NW_011925471.1 2780 2786 7 * 6 857.142857142857
NW_011925490.1 207564 212389 4826 + Gnomon gene NA NA gene11699 GeneID:105285546 LOC105285546 Gene LOC105285546 protein_coding NA
character(0) character(0) NW_011925490.1 208609 208615 7 * 6 857.142857142857
NW_011925516.1 278917 286323 7407 - Gnomon gene NA NA gene11797 GeneID:105285648 LOC105285648 Gene LOC105285648 protein_coding NA
character(0) character(0) NW_011925516.1 285124 285130 7 * 6 857.142857142857
NW_011925520.1 152066 157750 5685 + Gnomon gene NA NA gene11859 GeneID:105285719 LOC105285719 Gene LOC105285719 protein_coding NA
character(0) character(0) NW_011925520.1 152944 152950 7 * 6 857.142857142857
NW_011925538.1 63312 65618 2307 - Gnomon gene NA NA gene11931 GeneID:105285808 LOC105285808 Gene LOC105285808 protein_coding NA character(0)
character(0) NW_011925538.1 64230 64236 7 * 6 857.142857142857
NW_011925546.1 59043 60354 1312 + Gnomon gene NA NA gene11979 GeneID:105285864 LOC105285864 Gene LOC105285864 protein_coding NA character(0)
character(0) NW_011925546.1 59986 59992 7 * 6 857.142857142857
NW_011925549.1 267852 270480 2629 - Gnomon gene NA NA gene12010 GeneID:105285889 LOC105285889 Gene LOC105285889 protein_coding NA
character(0) character(0) NW_011925549.1 269441 269447 7 * 6 857.142857142857
NW_011925664.1 188474 193249 4776 + Gnomon gene NA NA gene12339 GeneID:105286242 LOC105286242 Gene LOC105286242 protein_coding NA
character(0) character(0) NW_011925664.1 189517 189523 7 * 6 857.142857142857
NW_011925664.1 495388 526731 31344 + Gnomon gene NA NA gene12399 GeneID:105286310 LOC105286310 Gene LOC105286310 lncRNA NA
character(0) character(0) NW_011925664.1 514139 514145 7 * 6 857.142857142857
NW_011925664.1 513347 514759 1413 - Gnomon gene NA NA gene12403 GeneID:105286311 LOC105286311 Gene LOC105286311 protein_coding NA
character(0) character(0) NW_011925664.1 514139 514145 7 * 6 857.142857142857
NW_011925715.1 251551 257367 5817 - Gnomon gene NA NA gene12526 GeneID:105286461 LOC105286461 Gene LOC105286461 protein_coding NA
character(0) character(0) NW_011925715.1 256038 256044 7 * 6 857.142857142857
NW_011925715.1 412871 419485 6615 + Gnomon gene NA NA gene12539 GeneID:105286478 LOC105286478 Gene LOC105286478 protein_coding NA
character(0) character(0) NW_011925715.1 414243 414249 7 * 6 857.142857142857
NW_011925715.1 1168275 1170855 2581 - Gnomon gene NA NA gene12582 GeneID:105286526 LOC105286526 Gene LOC105286526 protein_coding NA
character(0) character(0) NW_011925715.1 1169658 1169664 7 * 6 857.142857142857
NW_011926194.1 11132 18740 7609 + Gnomon gene NA NA gene13114 GeneID:105287147 LOC105287147 Gene LOC105287147 protein_coding NA character(0)
character(0) NW_011926194.1 12247 12253 7 * 6 857.142857142857
NW_011929444.1 7 5430 5424 + Gnomon gene NA NA gene13428 GeneID:105287499 LOC105287499 Gene LOC105287499 protein_coding NA character(0)
character(0) NW_011929444.1 1627 1633 7 * 6 857.142857142857
NW_011924899.1 718895 720452 1558 + Gnomon gene NA NA gene664 GeneID:105279718 LOC105279718 Gene LOC105279718 protein_coding NA
character(0) character(0) NW_011924899.1 719598 719607 10 * 8 800
NW_011925007.1 3759369 3760273 905 + Gnomon gene NA NA gene4382 GeneID:105277091 LOC105277091 Gene LOC105277091 protein_coding NA
character(0) character(0) NW_011925007.1 3760113 3760122 10 * 8 800
NW_011925477.1 19131 23385 4255 + Gnomon gene NA NA gene11657 GeneID:105285496 LOC105285496 Gene LOC105285496 protein_coding NA character(0)

| | | | | | | | |
|---|---|---|---|---|---|---|---|
| character(0) | NW_011925477.1 | 19843 | 19852 | 10 | * | 8 | 800 |

Directory: MRPR File: gwr-Cb_LI_RphaseB2.txt

| seqnames | start | end | width | strand | source | type | score | phase | ID | Dbxref | Name | gbkey | gene | gene_biotype | partial | start_range | end_range | seqnames | start | end | width | strand | NbrSites | Sdnsty |
|---|---|---|---|---|---|---|---|---|---|---|---|---|---|---|---|---|---|---|---|---|---|---|---|---|
| NW_011924887.1 | 1817915 | 1820166 | 2252 | - | Gnomon | gene | NA | NA | gene234 | GeneID:105274775 | LOC105274775 | Gene | LOC105274775 | protein_coding | NA | character(0) | character(0) | NW_011924887.1 | 1819270 | 1819275 | 6 | * | 6 | 1000 |
| NW_011924888.1 | 77334 | 78632 | 1299 | - | Gnomon | gene | NA | NA | gene276 | GeneID:105275214 | LOC105275214 | Gene | LOC105275214 | protein_coding | NA | character(0) | character(0) | NW_011924888.1 | 77560 | 77565 | 6 | * | 6 | 1000 |
| NW_011924888.1 | 389413 | 407214 | 17802 | - | Gnomon | gene | NA | NA | gene320 | GeneID:105275770 | LOC105275770 | Gene | LOC105275770 | protein_coding | NA | character(0) | character(0) | NW_011924888.1 | 405832 | 405837 | 6 | * | 6 | 1000 |
| NW_011924892.1 | 1158089 | 1160303 | 2215 | - | Gnomon | gene | NA | NA | gene569 | GeneID:105278527 | LOC105278527 | Gene | LOC105278527 | protein_coding | NA | character(0) | character(0) | NW_011924892.1 | 1159381 | 1159386 | 6 | * | 6 | 1000 |
| NW_011924931.1 | 4488185 | 4490767 | 2583 | + | Gnomon | gene | NA | NA | gene1620 | GeneID:105287754 | LOC105287754 | Gene | LOC105287754 | protein_coding | NA | character(0) | character(0) | NW_011924931.1 | 4489292 | 4489297 | 6 | * | 6 | 1000 |
| NW_011924946.1 | 1411689 | 1414880 | 3192 | - | Gnomon | gene | NA | NA | gene2041 | GeneID:105288248 | LOC105288248 | Gene | LOC105288248 | protein_coding | NA | character(0) | character(0) | NW_011924946.1 | 1413935 | 1413940 | 6 | * | 6 | 1000 |
| NW_011924946.1 | 2832086 | 2841975 | 9890 | - | Gnomon | gene | NA | NA | gene2162 | GeneID:105274578 | LOC105274578 | Gene | LOC105274578 | protein_coding | NA | character(0) | character(0) | NW_011924946.1 | 2839967 | 2839972 | 6 | * | 6 | 1000 |
| NW_011924973.1 | 1001098 | 1003969 | 2872 | + | Gnomon | gene | NA | NA | gene2837 | GeneID:105275355 | LOC105275355 | Gene | LOC105275355 | protein_coding | NA | character(0) | character(0) | NW_011924973.1 | 1002105 | 1002110 | 6 | * | 6 | 1000 |
| NW_011924977.1 | 662427 | 665182 | 2756 | + | Gnomon | gene | NA | NA | gene3008 | GeneID:105275550 | LOC105275550 | Gene | LOC105275550 | protein_coding | NA | character(0) | character(0) | NW_011924977.1 | 663534 | 663539 | 6 | * | 6 | 1000 |
| NW_011924996.1 | 75277 | 77259 | 1983 | + | Gnomon | gene | NA | NA | gene3743 | GeneID:105276382 | LOC105276382 | Gene | LOC105276382 | protein_coding | NA | character(0) | character(0) | NW_011924996.1 | 76074 | 76079 | 6 | * | 6 | 1000 |
| NW_011924996.1 | 1822758 | 1830685 | 7928 | - | Gnomon | gene | NA | NA | gene3885 | GeneID:105276527 | LOC105276527 | Gene | LOC105276527 | protein_coding | NA | character(0) | character(0) | NW_011924996.1 | 1829379 | 1829384 | 6 | * | 6 | 1000 |
| NW_011925022.1 | 386851 | 387976 | 1126 | - | Gnomon | gene | NA | NA | gene5000 | GeneID:105277804 | LOC105277804 | Gene | LOC105277804 | protein_coding | NA | character(0) | character(0) | NW_011925022.1 | 387077 | 387082 | 6 | * | 6 | 1000 |
| NW_011925053.1 | 649052 | 656133 | 7082 | - | Gnomon | gene | NA | NA | gene5882 | GeneID:105278812 | LOC105278812 | Gene | LOC105278812 | protein_coding | NA | character(0) | character(0) | NW_011925053.1 | 655139 | 655144 | 6 | * | 6 | 1000 |
| NW_011925142.1 | 777887 | 780161 | 2275 | - | Gnomon | gene | NA | NA | gene7574 | GeneID:105280791 | LOC105280791 | Gene | LOC105280791 | protein_coding | NA | character(0) | character(0) | NW_011925142.1 | 779002 | 779007 | 6 | * | 6 | 1000 |
| NW_011925150.1 | 76140 | 79045 | 2906 | - | Gnomon | gene | NA | NA | gene7626 | GeneID:105280856 | LOC105280856 | Gene | LOC105280856 | protein_coding | NA | character(0) | character(0) | NW_011925150.1 | 78377 | 78382 | 6 | * | 6 | 1000 |
| NW_011925151.1 | 38146 | 39177 | 1032 | - | Gnomon | gene | NA | NA | gene7635 | GeneID:105280860 | LOC105280860 | Gene | LOC105280860 | protein_coding | NA | character(0) | character(0) | NW_011925151.1 | 38257 | 38262 | 6 | * | 6 | 1000 |
| NW_011925163.1 | 470647 | 473680 | 3034 | + | Gnomon | gene | NA | NA | gene8040 | GeneID:105281341 | LOC105281341 | Gene | LOC105281341 | protein_coding | NA | character(0) | character(0) | NW_011925163.1 | 471213 | 471218 | 6 | * | 6 | 1000 |
| NW_011925183.1 | 39511 | 42630 | 3120 | + | Gnomon | gene | NA | NA | gene8236 | GeneID:105281577 | LOC105281577 | Gene | LOC105281577 | protein_coding | NA | character(0) | character(0) | NW_011925183.1 | 40051 | 40056 | 6 | * | 6 | 1000 |
| NW_011925189.1 | 575609 | 577709 | 2101 | - | Gnomon | gene | NA | NA | gene8332 | GeneID:105281690 | LOC105281690 | Gene | LOC105281690 | protein_coding | NA | character(0) | character(0) | NW_011925189.1 | 577082 | 577087 | 6 | * | 6 | 1000 |
| NW_011925203.1 | 63038 | 97839 | 34802 | + | Gnomon | gene | NA | NA | gene8521 | GeneID:105281906 | LOC105281906 | Gene | LOC105281906 | protein_coding | NA | character(0) | character(0) | NW_011925203.1 | 64603 | 64608 | 6 | * | 6 | 1000 |
| NW_011925216.1 | 156691 | 162086 | 5396 | + | Gnomon | gene | NA | NA | gene8741 | GeneID:105282150 | LOC105282150 | Gene | LOC105282150 | protein_coding | NA | character(0) | character(0) | NW_011925216.1 | 157541 | 157546 | 6 | * | 6 | 1000 |
| NW_011925304.1 | 378599 | 380691 | 2093 | - | Gnomon | gene | NA | NA | gene9814 | GeneID:105283373 | LOC105283373 | Gene | LOC105283373 | protein_coding | NA | character(0) | character(0) | NW_011925304.1 | 379611 | 379616 | 6 | * | 6 | 1000 |
| NW_011925304.1 | 468966 | 476624 | 7659 | - | Gnomon | gene | NA | NA | gene9824 | GeneID:105283383 | LOC105283383 | Gene | LOC105283383 | protein_coding | NA | character(0) | character(0) | NW_011925304.1 | 475523 | 475528 | 6 | * | 6 | 1000 |
| NW_011925321.1 | 1107715 | 1109901 | 2187 | - | Gnomon | gene | NA | NA | gene10043 | GeneID:105283634 | LOC105283634 | Gene | LOC105283634 | protein_coding | NA | character(0) | character(0) | NW_011925321.1 | 1108690 | 1108695 | 6 | * | 6 | 1000 |
| NW_011925356.1 | 704771 | 710120 | 5350 | - | Gnomon | gene | NA | NA | gene10539 | GeneID:105284201 | LOC105284201 | Gene | LOC105284201 | protein_coding | NA | character(0) | character(0) | NW_011925356.1 | 708176 | 708181 | 6 | * | 6 | 1000 |
| NW_011925356.1 | 1573184 | 1579703 | 6520 | - | Gnomon | gene | NA | NA | gene10596 | GeneID:105284262 | LOC105284262 | Gene | LOC105284262 | protein_coding | NA | character(0) | character(0) | NW_011925356.1 | 1573650 | 1573655 | 6 | * | 6 | 1000 |
| NW_011925367.1 | 1119922 | 1121887 | 1966 | + | Gnomon | gene | NA | NA | gene10714 | GeneID:105284396 | LOC105284396 | Gene | LOC105284396 | protein_coding | NA | character(0) | character(0) | NW_011925367.1 | 1120960 | 1120965 | 6 | * | 6 | 1000 |
| NW_011925387.1 | 39150 | 46759 | 7610 | + | Gnomon | gene | NA | NA | gene10928 | GeneID:105284644 | LOC105284644 | Gene | LOC105284644 | protein_coding | NA | character(0) | character(0) | NW_011925387.1 | 40039 | 40044 | 6 | * | 6 | 1000 |
| NW_011925401.1 | 10153 | 19053 | 8901 | - | Gnomon | gene | NA | NA | gene11099 | GeneID:105284845 | LOC105284845 | Gene | LOC105284845 | protein_coding | NA | character(0) | character(0) | NW_011925401.1 | 17311 | 17316 | 6 | * | 6 | 1000 |
| NW_011925470.1 | 613911 | 617551 | 3641 | - | Gnomon | gene | NA | NA | gene11628 | GeneID:105285465 | LOC105285465 | Gene | LOC105285465 | protein_coding | NA | | | | | | | | | |

character(0) character(0) NW_011925470.1 616616 616621 6 * 6 1000
NW_011925490.1 137354 140755 3402 + Gnomon gene NA NA gene11689 GeneID:105285534 LOC105285534 Gene LOC105285534 protein_coding NA
character(0) character(0) NW_011925490.1 138283 138288 6 * 6 1000
NW_011925595.1 1 4548 4548 - Gnomon gene NA NA gene12102 GeneID:105286003 LOC105286003 Gene LOC105286003 protein_coding NA character(0)
character(0) NW_011925595.1 3717 3722 6 * 6 1000
NW_011925664.1 38296 39300 1005 + Gnomon gene NA NA gene12311 GeneID:105286212 LOC105286212 Gene LOC105286212 protein_coding NA character(0)
character(0) NW_011925664.1 39054 39061 8 * 8 1000
NW_011925664.1 277932 283313 5382 + Gnomon gene NA NA gene12359 GeneID:105286364 LOC105286364 Gene LOC105286364 protein_coding NA
character(0) character(0) NW_011925664.1 279108 279113 6 * 6 1000
NW_011925664.1 580837 582496 1660 - Gnomon gene NA NA gene12411 GeneID:105286320 LOC105286320 Gene LOC105286320 protein_coding NA
character(0) character(0) NW_011925664.1 581347 581352 6 * 6 1000
NW_011925715.1 952620 956416 3797 - Gnomon gene NA NA gene12567 GeneID:105286509 LOC105286509 Gene LOC105286509 protein_coding NA
character(0) character(0) NW_011925715.1 955653 955658 6 * 6 1000
NW_011925086.1 128430 135139 6710 + Gnomon gene NA NA gene6311 GeneID:105279333 LOC105279333 Gene LOC105279333 protein_coding NA
character(0) character(0) NW_011925086.1 129637 129647 11 * 10 909.090909090909
NW_011924956.1 717757 722032 4276 + Gnomon gene NA NA gene2570 GeneID:105275036 LOC105275036 Gene LOC105275036 protein_coding NA
character(0) character(0) NW_011924956.1 719335 719344 10 * 9 900
NW_011924914.1 259948 262322 2375 - Gnomon gene NA NA gene952 GeneID:105283016 LOC105283016 Gene LOC105283016 protein_coding NA
character(0) character(0) NW_011924914.1 261398 261406 9 * 8 888.888888888889
NW_011924967.1 91700 93046 1347 - Gnomon gene NA NA gene2644 GeneID:105275137 LOC105275137 Gene LOC105275137 protein_coding NA character(0)
character(0) NW_011924967.1 92497 92505 9 * 8 888.888888888889
NW_011924979.1 1988584 1990897 2314 + Gnomon gene NA NA gene3314 GeneID:105275878 LOC105275878 Gene LOC105275878 protein_coding NA
character(0) character(0) NW_011924979.1 1989441 1989449 9 * 8 888.888888888889
NW_011925239.1 3979 7808 3830 + Gnomon gene NA NA gene9001 GeneID:105282461 LOC105282461 Gene LOC105282461 protein_coding NA character(0)
character(0) NW_011925239.1 4874 4882 9 * 8 888.888888888889
NW_011924887.1 1205421 1208016 2596 + Gnomon gene NA NA gene191 GeneID:105288114 LOC105288114 Gene LOC105288114 protein_coding NA
character(0) character(0) NW_011924887.1 1206843 1206850 8 * 7 875
NW_011924977.1 3728770 3731404 2635 + Gnomon gene NA NA gene3213 GeneID:105275761 LOC105275761 Gene LOC105275761 protein_coding NA
character(0) character(0) NW_011924977.1 3729890 3729897 8 * 7 875
NW_011924887.1 748542 750844 2303 - Gnomon gene NA NA gene168 GeneID:105287895 LOC105287895 Gene LOC105287895 protein_coding NA
character(0) character(0) NW_011924887.1 749895 749901 7 * 6 857.142857142857
NW_011924888.1 30738 31847 1110 + Gnomon gene NA NA gene264 GeneID:105275153 LOC105275153 Gene LOC105275153 protein_coding NA character(0)
character(0) NW_011924888.1 31453 31459 7 * 6 857.142857142857
NW_011924889.1 476236 478868 2633 - Gnomon gene NA NA gene504 GeneID:105277817 LOC105277817 Gene LOC105277817 protein_coding NA
character(0) character(0) NW_011924889.1 477980 477986 7 * 6 857.142857142857
NW_011924889.1 493449 499545 6097 + Gnomon gene NA NA gene508 GeneID:105277915 LOC105277915 Gene LOC105277915 protein_coding NA
character(0) character(0) NW_011924889.1 494195 494201 7 * 6 857.142857142857
NW_011924889.1 606387 610322 3936 - Gnomon gene NA NA gene519 GeneID:105278022 LOC105278022 Gene LOC105278022 protein_coding NA
character(0) character(0) NW_011924889.1 609130 609136 7 * 6 857.142857142857
NW_011924892.1 274786 285690 10905 - Gnomon gene NA NA gene544 GeneID:105278288 LOC105278288 Gene LOC105278288 protein_coding NA
character(0) character(0) NW_011924892.1 284836 284842 7 * 6 857.142857142857
NW_011924902.1 81345 99446 18102 - Gnomon gene NA NA gene702 GeneID:105280058 LOC105280058 Gene LOC105280058 protein_coding NA character(0)
character(0) NW_011924902.1 98752 98758 7 * 6 857.142857142857
NW_011924905.1 26269 321616 295348 - Gnomon gene NA NA gene787 GeneID:105280980 LOC105280980 Gene LOC105280980 protein_coding NA
character(0) character(0) NW_011924905.1 319741 319747 7 * 6 857.142857142857
NW_011924907.1 889683 896841 7159 - Gnomon gene NA NA gene861 GeneID:105281750 LOC105281750 Gene LOC105281750 protein_coding NA
character(0) character(0) NW_011924907.1 896035 896041 7 * 6 857.142857142857
NW_011924914.1 827084 831408 4325 - Gnomon gene NA NA gene1010 GeneID:105283498 LOC105283498 Gene LOC105283498 protein_coding NA
character(0) character(0) NW_011924914.1 831099 831105 7 * 6 857.142857142857
NW_011924922.1 1096844 1105846 9003 + Gnomon gene NA NA gene1085 GeneID:105284317 LOC105284317 Gene LOC105284317 protein_coding NA
character(0) character(0) NW_011924922.1 1098044 1098050 7 * 6 857.142857142857
NW_011924922.1 2390151 2392888 2738 - Gnomon gene NA NA gene1198 GeneID:105285507 LOC105285507 Gene LOC105285507 protein_coding NA
character(0) character(0) NW_011924922.1 2392009 2392015 7 * 6 857.142857142857
NW_011924931.1 122002 124933 2932 + Gnomon gene NA NA gene1364 GeneID:105287208 LOC105287208 Gene LOC105287208 protein_coding NA
character(0) character(0) NW_011924931.1 123017 123023 7 * 6 857.142857142857
NW_011924931.1 201740 204575 2836 - Gnomon gene NA NA gene1371 GeneID:105287279 LOC105287279 Gene LOC105287279 protein_coding NA
character(0) character(0) NW_011924931.1 203497 203503 7 * 6 857.142857142857
NW_011924931.1 242950 250477 7528 - Gnomon gene NA NA gene1378 GeneID:105287371 LOC105287371 Gene LOC105287371 protein_coding NA
character(0) character(0) NW_011924931.1 249191 249197 7 * 6 857.142857142857
NW_011924931.1 290674 293586 2913 + Gnomon gene NA NA gene1388 GeneID:105287473 LOC105287473 Gene LOC105287473 protein_coding NA
character(0) character(0) NW_011924931.1 291628 291634 7 * 6 857.142857142857
NW_011924931.1 1565109 1566883 1775 + Gnomon gene NA NA gene1444 GeneID:105287569 LOC105287569 Gene LOC105287569 protein_coding NA
character(0) character(0) NW_011924931.1 1565961 1565967 7 * 6 857.142857142857
NW_011924931.1 2028839 2031812 2974 + Gnomon gene NA NA gene1470 GeneID:105287597 LOC105287597 Gene LOC105287597 protein_coding NA

character(0)	character(0)	NW_011924931.1	2030239	2030245	7	*	6	857.142857142857
NW_011924931.1	2871689	2878941	7253	-	Gnomon	gene	NA	NA	gene1492	GeneID:105287620	LOC105287620	Gene	LOC105287620	protein_coding	NA
character(0)	character(0)	NW_011924931.1	2878145	2878151	7	*	6	857.142857142857
NW_011924931.1	3929932	3932866	2935	+	Gnomon	gene	NA	NA	gene1555	GeneID:105287684	LOC105287684	Gene	LOC105287684	protein_coding	NA
character(0)	character(0)	NW_011924931.1	3930880	3930886	7	*	6	857.142857142857
NW_011924931.1	3979390	3986328	6939	+	Gnomon	gene	NA	NA	gene1566	GeneID:105287698	LOC105287698	Gene	LOC105287698	protein_coding	NA
character(0)	character(0)	NW_011924931.1	3983680	3983686	7	*	6	857.142857142857
NW_011924931.1	4058265	4059684	1420	-	Gnomon	gene	NA	NA	gene1581	GeneID:105287710	LOC105287710	Gene	LOC105287710	protein_coding	NA
character(0)	character(0)	NW_011924931.1	4058767	4058773	7	*	6	857.142857142857
NW_011924932.1	539418	541702	2285	-	Gnomon	gene	NA	NA	gene1654	GeneID:105287811	LOC105287811	Gene	LOC105287811	protein_coding	NA
character(0)	character(0)	NW_011924932.1	541032	541038	7	*	6	857.142857142857
NW_011924938.1	72334	83258	10925	+	Gnomon	gene	NA	NA	gene1805	GeneID:105287985	LOC105287985	Gene	LOC105287985	protein_coding	NA	character(0)
character(0)	NW_011924938.1	72914	72920	7	*	6	857.142857142857
NW_011924946.1	786370	788739	2370	-	Gnomon	gene	NA	NA	gene1995	GeneID:105288196	LOC105288196	Gene	LOC105288196	protein_coding	NA
character(0)	character(0)	NW_011924946.1	787628	787634	7	*	6	857.142857142857
NW_011924946.1	1634653	1637620	2968	-	Gnomon	gene	NA	NA	gene2065	GeneID:105274464	LOC105274464	Gene	LOC105274464	protein_coding	NA
character(0)	character(0)	NW_011924946.1	1636892	1636898	7	*	6	857.142857142857
NW_011924946.1	2342632	2344089	1458	-	Gnomon	gene	NA	NA	gene2088	GeneID:105274497	LOC105274497	Gene	LOC105274497	protein_coding	NA
character(0)	character(0)	NW_011924946.1	2343055	2343061	7	*	6	857.142857142857
NW_011924946.1	2914862	2919248	4387	-	Gnomon	gene	NA	NA	gene2173	GeneID:105274590	LOC105274590	Gene	LOC105274590	protein_coding	NA
character(0)	character(0)	NW_011924946.1	2917571	2917577	7	*	6	857.142857142857
NW_011924946.1	2914862	2919248	4387	-	Gnomon	gene	NA	NA	gene2173	GeneID:105274590	LOC105274590	Gene	LOC105274590	protein_coding	NA
character(0)	character(0)	NW_011924946.1	2918055	2918061	7	*	6	857.142857142857
NW_011924946.1	3145253	3148790	3538	-	Gnomon	gene	NA	NA	gene2215	GeneID:105274638	LOC105274638	Gene	LOC105274638	protein_coding	NA
character(0)	character(0)	NW_011924946.1	3147722	3147728	7	*	6	857.142857142857
NW_011924947.1	141424	143460	2037	+	Gnomon	gene	NA	NA	gene2279	GeneID:105274711	LOC105274711	Gene	LOC105274711	protein_coding	NA
character(0)	character(0)	NW_011924947.1	142119	142125	7	*	6	857.142857142857
NW_011924953.1	401032	402582	1551	-	Gnomon	gene	NA	NA	gene2400	GeneID:105274850	LOC105274850	Gene	LOC105274850	protein_coding	NA
character(0)	character(0)	NW_011924953.1	401754	401760	7	*	6	857.142857142857
NW_011924955.1	473199	477158	3960	-	Gnomon	gene	NA	NA	gene2462	GeneID:105274912	LOC105274912	Gene	LOC105274912	protein_coding	NA
character(0)	character(0)	NW_011924955.1	475912	475918	7	*	6	857.142857142857
NW_011924956.1	667547	676817	9271	+	Gnomon	gene	NA	NA	gene2561	GeneID:105275064	LOC105275064	Gene	LOC105275064	protein_coding	NA
character(0)	character(0)	NW_011924956.1	668424	668430	7	*	6	857.142857142857
NW_011924956.1	701839	703915	2077	-	Gnomon	gene	NA	NA	gene2566	GeneID:105275031	LOC105275031	Gene	LOC105275031	protein_coding	NA
character(0)	character(0)	NW_011924956.1	702713	702719	7	*	6	857.142857142857
NW_011924967.1	11923	12765	843	-	Gnomon	gene	NA	NA	gene2630	GeneID:105275119	LOC105275119	Gene	LOC105275119	protein_coding	NA	character(0)
character(0)	NW_011924967.1	12053	12059	7	*	6	857.142857142857
NW_011924967.1	91700	93046	1347	-	Gnomon	gene	NA	NA	gene2644	GeneID:105275137	LOC105275137	Gene	LOC105275137	protein_coding	NA	character(0)
character(0)	NW_011924967.1	92468	92474	7	*	6	857.142857142857
NW_011924973.1	774415	775843	1429	-	Gnomon	gene	NA	NA	gene2790	GeneID:105275307	LOC105275307	Gene	LOC105275307	protein_coding	NA
character(0)	character(0)	NW_011924973.1	775075	775081	7	*	6	857.142857142857
NW_011924974.1	483919	487626	3708	+	Gnomon	gene	NA	NA	gene2873	GeneID:105275401	LOC105275401	Gene	LOC105275401	protein_coding	NA
character(0)	character(0)	NW_011924974.1	484702	484708	7	*	6	857.142857142857
NW_011924977.1	1460154	1463690	3537	-	Gnomon	gene	NA	NA	gene3072	GeneID:105275615	LOC105275615	Gene	LOC105275615	protein_coding	NA
character(0)	character(0)	NW_011924977.1	1461887	1461893	7	*	6	857.142857142857
NW_011924977.1	1460154	1463690	3537	-	Gnomon	gene	NA	NA	gene3072	GeneID:105275615	LOC105275615	Gene	LOC105275615	protein_coding	NA
character(0)	character(0)	NW_011924977.1	1462188	1462194	7	*	6	857.142857142857
NW_011924977.1	2573287	2581453	8167	-	Gnomon	gene	NA	NA	gene3147	GeneID:105275689	LOC105275689	Gene	LOC105275689	protein_coding	NA
character(0)	character(0)	NW_011924977.1	2580506	2580512	7	*	6	857.142857142857
NW_011924977.1	3678529	3682684	4156	-	Gnomon	gene	NA	NA	gene3205	GeneID:105275753	LOC105275753	Gene	LOC105275753	protein_coding	NA
character(0)	character(0)	NW_011924977.1	3681359	3681365	7	*	6	857.142857142857
NW_011924977.1	3678529	3682684	4156	-	Gnomon	gene	NA	NA	gene3205	GeneID:105275753	LOC105275753	Gene	LOC105275753	protein_coding	NA
character(0)	character(0)	NW_011924977.1	3681568	3681574	7	*	6	857.142857142857
NW_011924977.1	3682584	3686057	3474	-	Gnomon	gene	NA	NA	gene3206	GeneID:105275752	LOC105275752	Gene	LOC105275752	protein_coding	NA
character(0)	character(0)	NW_011924977.1	3684978	3684984	7	*	6	857.142857142857
NW_011924979.1	211193	212598	1406	-	Gnomon	gene	NA	NA	gene3249	GeneID:105275908	LOC105275908	Gene	LOC105275908	protein_coding	NA
character(0)	character(0)	NW_011924979.1	212057	212063	7	*	6	857.142857142857
NW_011924996.1	292526	303470	10945	+	Gnomon	gene	NA	NA	gene3775	GeneID:105276410	LOC105276410	Gene	LOC105276410	protein_coding	NA
character(0)	character(0)	NW_011924996.1	293762	293768	7	*	6	857.142857142857
NW_011924996.1	292526	303470	10945	+	Gnomon	gene	NA	NA	gene3775	GeneID:105276410	LOC105276410	Gene	LOC105276410	protein_coding	NA
character(0)	character(0)	NW_011924996.1	293819	293825	7	*	6	857.142857142857
NW_011924996.1	448616	452114	3499	-	Gnomon	gene	NA	NA	gene3788	GeneID:105276422	LOC105276422	Gene	LOC105276422	protein_coding	NA
character(0)	character(0)	NW_011924996.1	450468	450474	7	*	6	857.142857142857
NW_011924996.1	641332	646889	5558	-	Gnomon	gene	NA	NA	gene3801	GeneID:105276437	LOC105276437	Gene	LOC105276437	protein_coding	NA

character(0) character(0) NW_011924996.1 645416 645422 7 * 6 857.142857142857
NW_011924996.1 1790083 1793499 3417 - Gnomon gene NA NA gene3877 GeneID:105276521 LOC105276521 Gene LOC105276521 protein_coding NA
character(0) character(0) NW_011924996.1 1792422 1792428 7 * 6 857.142857142857
NW_011925007.1 1417723 1421435 3713 + Gnomon gene NA NA gene4248 GeneID:105276948 LOC105276948 Gene LOC105276948 protein_coding NA
character(0) character(0) NW_011925007.1 1419353 1419359 7 * 6 857.142857142857
NW_011925007.1 1452162 1454301 2140 + Gnomon gene NA NA gene4252 GeneID:105276949 LOC105276949 Gene LOC105276949 protein_coding NA
character(0) character(0) NW_011925007.1 1453109 1453115 7 * 6 857.142857142857
NW_011925007.1 3760218 3764466 4249 - Gnomon gene NA NA gene4383 GeneID:105277128 LOC105277128 Gene LOC105277128 protein_coding NA
character(0) character(0) NW_011925007.1 3760305 3760311 7 * 6 857.142857142857
NW_011925012.1 1316386 1318477 2092 + Gnomon gene NA NA gene4492 GeneID:105277232 LOC105277232 Gene LOC105277232 protein_coding NA
character(0) character(0) NW_011925012.1 1317469 1317475 7 * 6 857.142857142857
NW_011925012.1 1360007 1362714 2708 - Gnomon gene NA NA gene4500 GeneID:105277238 LOC105277238 Gene LOC105277238 protein_coding NA
character(0) character(0) NW_011925012.1 1361666 1361672 7 * 6 857.142857142857
NW_011925013.1 5041 7730 2690 - Gnomon gene NA NA gene4511 GeneID:105277256 LOC105277256 Gene LOC105277256 protein_coding NA character(0)
character(0) NW_011925013.1 6764 6770 7 * 6 857.142857142857
NW_011925019.1 475 2900 2426 + Gnomon gene NA NA gene4633 GeneID:105277389 LOC105277389 Gene LOC105277389 protein_coding NA character(0)
character(0) NW_011925019.1 1670 1676 7 * 6 857.142857142857
NW_011925019.1 2803 5896 3094 - Gnomon gene NA NA gene4634 GeneID:105277388 LOC105277388 Gene LOC105277388 protein_coding NA character(0)
character(0) NW_011925019.1 4685 4691 7 * 6 857.142857142857
NW_011925019.1 1195331 1199707 4377 - Gnomon gene NA NA gene4693 GeneID:105277452 LOC105277452 Gene LOC105277452 protein_coding NA
character(0) character(0) NW_011925019.1 1198454 1198460 7 * 6 857.142857142857
NW_011925020.1 469190 474352 5163 + Gnomon gene NA NA gene4780 GeneID:105277552 LOC105277552 Gene LOC105277552 protein_coding NA
character(0) character(0) NW_011925020.1 470270 470276 7 * 6 857.142857142857
NW_011925027.1 1297549 1301844 4296 + Gnomon gene NA NA gene5179 GeneID:105278005 LOC105278005 Gene LOC105278005 protein_coding NA
character(0) character(0) NW_011925027.1 1298524 1298530 7 * 6 857.142857142857
NW_011925029.1 39174 43410 4237 + Gnomon gene NA NA gene5243 GeneID:105278106 LOC105278106 Gene LOC105278106 protein_coding NA character(0)
character(0) NW_011925029.1 40261 40267 7 * 6 857.142857142857
NW_011925035.1 1492483 1496530 4048 - Gnomon gene NA NA gene5417 GeneID:105278277 LOC105278277 Gene LOC105278277 protein_coding NA
character(0) character(0) NW_011925035.1 1495407 1495413 7 * 6 857.142857142857
NW_011925035.1 1494546 1496299 1754 - Gnomon gene NA NA gene5418 GeneID:105278275 LOC105278275 Gene LOC105278275 protein_coding NA
character(0) character(0) NW_011925035.1 1495407 1495413 7 * 6 857.142857142857
NW_011925035.1 3102848 3104698 1851 + Gnomon gene NA NA gene5557 GeneID:105278431 LOC105278431 Gene LOC105278431 protein_coding NA
character(0) character(0) NW_011925035.1 3103902 3103908 7 * 6 857.142857142857
NW_011925036.1 1418708 1424550 5843 + Gnomon gene NA NA gene5653 GeneID:105278544 LOC105278544 Gene LOC105278544 protein_coding NA
character(0) character(0) NW_011925036.1 1420178 1420184 7 * 6 857.142857142857
NW_011925050.1 126141 128598 2458 - Gnomon gene NA NA gene5737 GeneID:105278656 LOC105278656 Gene LOC105278656 protein_coding NA
character(0) character(0) NW_011925050.1 127417 127423 7 * 6 857.142857142857
NW_011925053.1 1491435 1517044 25610 - Gnomon gene NA NA gene5936 GeneID:105278874 LOC105278874 Gene LOC105278874 protein_coding NA
character(0) character(0) NW_011925053.1 1493590 1493596 7 * 6 857.142857142857
NW_011925053.1 1492486 1494387 1902 - Gnomon gene NA NA gene5937 GeneID:105278873 LOC105278873 Gene LOC105278873 protein_coding NA
character(0) character(0) NW_011925053.1 1493590 1493596 7 * 6 857.142857142857
NW_011925067.1 5971 15150 9180 + Gnomon gene NA NA gene6024 GeneID:105278996 LOC105278996 Gene LOC105278996 protein_coding NA character(0)
character(0) NW_011925067.1 7054 7060 7 * 6 857.142857142857
NW_011925067.1 111367 117057 5691 + Gnomon gene NA NA gene6031 GeneID:105279000 LOC105279000 Gene LOC105279000 protein_coding NA
character(0) character(0) NW_011925067.1 112547 112553 7 * 6 857.142857142857
NW_011925068.1 10281 12328 2048 + Gnomon gene NA NA gene6111 GeneID:105279088 LOC105279088 Gene LOC105279088 protein_coding NA character(0)
character(0) NW_011925068.1 11234 11240 7 * 6 857.142857142857
NW_011925086.1 250638 255287 4650 - Gnomon gene NA NA gene6324 GeneID:105279346 LOC105279346 Gene LOC105279346 protein_coding NA
character(0) character(0) NW_011925086.1 254146 254152 7 * 6 857.142857142857
NW_011925103.1 430898 432277 1380 + Gnomon gene NA NA gene6564 GeneID:105279626 LOC105279626 Gene LOC105279626 protein_coding NA
character(0) character(0) NW_011925103.1 431700 431706 7 * 6 857.142857142857
NW_011925106.1 154197 158224 4028 + Gnomon gene NA NA gene6601 GeneID:105279664 LOC105279664 Gene LOC105279664 protein_coding NA
character(0) character(0) NW_011925106.1 155492 155498 7 * 6 857.142857142857
NW_011925106.1 431654 436050 4397 + Gnomon gene NA NA gene6641 GeneID:105279709 LOC105279709 Gene LOC105279709 protein_coding NA
character(0) character(0) NW_011925106.1 432897 432903 7 * 6 857.142857142857
NW_011925106.1 512988 515818 2831 - Gnomon gene NA NA gene6652 GeneID:105279719 LOC105279719 Gene LOC105279719 protein_coding NA
character(0) character(0) NW_011925106.1 514935 514941 7 * 6 857.142857142857
NW_011925106.1 629782 633106 3325 - Gnomon gene NA NA gene6668 GeneID:105279740 LOC105279740 Gene LOC105279740 protein_coding NA
character(0) character(0) NW_011925106.1 631881 631887 7 * 6 857.142857142857
NW_011925106.1 779647 785206 5560 + Gnomon gene NA NA gene6679 GeneID:105279748 LOC105279748 Gene LOC105279748 protein_coding NA
character(0) character(0) NW_011925106.1 780733 780739 7 * 6 857.142857142857
NW_011925106.1 2086959 2096661 9703 + Gnomon gene NA NA gene6771 GeneID:105279854 LOC105279854 Gene LOC105279854 protein_coding NA
character(0) character(0) NW_011925106.1 2087848 2087854 7 * 6 857.142857142857
NW_011925106.1 2086959 2096661 9703 + Gnomon gene NA NA gene6771 GeneID:105279854 LOC105279854 Gene LOC105279854 protein_coding NA

character(0) character(0) NW_011925106.1 2088114 2088120 7 * 6 857.142857142857
NW_011925118.1 457689 459947 2259 + Gnomon gene NA NA gene6953 GeneID:105280092 LOC105280092 Gene LOC105280092 protein_coding NA
character(0) character(0) NW_011925118.1 458682 458688 7 * 6 857.142857142857
NW_011925118.1 481415 484075 2661 + Gnomon gene NA NA gene6956 GeneID:105280102 LOC105280102 Gene LOC105280102 protein_coding NA
character(0) character(0) NW_011925118.1 482695 482701 7 * 6 857.142857142857
NW_011925128.1 7781 9472 1692 + Gnomon gene NA NA gene7085 GeneID:105280236 LOC105280236 Gene LOC105280236 protein_coding NA character(0)
character(0) NW_011925128.1 8930 8936 7 * 6 857.142857142857
NW_011925128.1 800543 802328 1786 - Gnomon gene NA NA gene7160 GeneID:105280319 LOC105280319 Gene LOC105280319 protein_coding NA
character(0) character(0) NW_011925128.1 801605 801611 7 * 6 857.142857142857
NW_011925142.1 423375 426060 2686 - Gnomon gene NA NA gene7527 GeneID:105280739 LOC105280739 Gene LOC105280739 protein_coding NA
character(0) character(0) NW_011925142.1 424101 424107 7 * 6 857.142857142857
NW_011925142.1 423375 426060 2686 - Gnomon gene NA NA gene7527 GeneID:105280739 LOC105280739 Gene LOC105280739 protein_coding NA
character(0) character(0) NW_011925142.1 424227 424233 7 * 6 857.142857142857
NW_011925161.1 210886 215244 4359 + Gnomon gene NA NA gene7720 GeneID:105280964 LOC105280964 Gene LOC105280964 protein_coding NA
character(0) character(0) NW_011925161.1 213136 213142 7 * 6 857.142857142857
NW_011925161.1 3740915 3743520 2606 + Gnomon gene NA NA gene7963 GeneID:105281227 LOC105281227 Gene LOC105281227 protein_coding NA
character(0) character(0) NW_011925161.1 3741983 3741989 7 * 6 857.142857142857
NW_011925183.1 39511 42630 3120 + Gnomon gene NA NA gene8236 GeneID:105281577 LOC105281577 Gene LOC105281577 protein_coding NA character(0)
character(0) NW_011925183.1 40353 40359 7 * 6 857.142857142857
NW_011925188.1 199014 202632 3619 - Gnomon gene NA NA gene8257 GeneID:105281603 LOC105281603 Gene LOC105281603 protein_coding NA
character(0) character(0) NW_011925188.1 201441 201447 7 * 6 857.142857142857
NW_011925203.1 23205 24951 1747 - Gnomon gene NA NA gene8516 GeneID:105281896 LOC105281896 Gene LOC105281896 protein_coding NA character(0)
character(0) NW_011925203.1 24180 24186 7 * 6 857.142857142857
NW_011925209.1 85655 90207 4553 - Gnomon gene NA NA gene8551 GeneID:105281943 LOC105281943 Gene LOC105281943 protein_coding NA character(0)
character(0) NW_011925209.1 86875 86881 7 * 6 857.142857142857
NW_011925216.1 130033 131250 1218 + Gnomon gene NA NA gene8733 GeneID:105282147 LOC105282147 Gene LOC105282147 protein_coding NA
character(0) character(0) NW_011925216.1 130799 130805 7 * 6 857.142857142857
NW_011925221.1 474109 476408 2300 + Gnomon gene NA NA gene8798 GeneID:105282220 LOC105282220 Gene LOC105282220 protein_coding NA
character(0) character(0) NW_011925221.1 474855 474861 7 * 6 857.142857142857
NW_011925224.1 219079 219935 857 - Gnomon gene NA NA gene8854 GeneID:105282280 LOC105282280 Gene LOC105282280 protein_coding NA
character(0) character(0) NW_011925224.1 219257 219263 7 * 6 857.142857142857
NW_011925224.1 327268 339256 11989 - Gnomon gene NA NA gene8872 GeneID:105282298 LOC105282298 Gene LOC105282298 protein_coding NA
character(0) character(0) NW_011925224.1 338306 338312 7 * 6 857.142857142857
NW_011925240.1 1526960 1531289 4330 + Gnomon gene NA NA gene9107 GeneID:105282559 LOC105282559 Gene LOC105282559 protein_coding NA
character(0) character(0) NW_011925240.1 1527897 1527903 7 * 6 857.142857142857
NW_011925240.1 1575290 1580897 5608 - Gnomon gene NA NA gene9116 GeneID:105282568 LOC105282568 Gene LOC105282568 protein_coding NA
character(0) character(0) NW_011925240.1 1579255 1579261 7 * 6 857.142857142857
NW_011925241.1 397336 409261 11926 - Gnomon gene NA NA gene9191 GeneID:105282655 LOC105282655 Gene LOC105282655 protein_coding NA
character(0) character(0) NW_011925241.1 408196 408202 7 * 6 857.142857142857
NW_011925257.1 52519 55399 2881 - Gnomon gene NA NA gene9303 GeneID:105282793 LOC105282793 Gene LOC105282793 protein_coding NA character(0)
character(0) NW_011925257.1 54841 54847 7 * 6 857.142857142857
NW_011925258.1 360040 361894 1855 - Gnomon gene NA NA gene9341 GeneID:105282841 LOC105282841 Gene LOC105282841 protein_coding NA
character(0) character(0) NW_011925258.1 361089 361095 7 * 6 857.142857142857
NW_011925267.1 224087 232805 8719 + Gnomon gene NA NA gene9418 GeneID:105282921 LOC105282921 Gene LOC105282921 protein_coding NA
character(0) character(0) NW_011925267.1 225570 225576 7 * 6 857.142857142857
NW_011925315.1 538327 543302 4976 + Gnomon gene NA NA gene9909 GeneID:105283484 LOC105283484 Gene LOC105283484 protein_coding NA
character(0) character(0) NW_011925315.1 539621 539627 7 * 6 857.142857142857
NW_011925323.1 390910 395897 4988 - Gnomon gene NA NA gene10091 GeneID:105283697 LOC105283697 Gene LOC105283697 protein_coding NA
character(0) character(0) NW_011925323.1 394351 394357 7 * 6 857.142857142857
NW_011925323.1 1254678 1256970 2293 + Gnomon gene NA NA gene10126 GeneID:105283727 LOC105283727 Gene LOC105283727 protein_coding NA
character(0) character(0) NW_011925323.1 1255392 1255398 7 * 6 857.142857142857
NW_011925330.1 58606 82143 23538 + Gnomon gene NA NA gene10255 GeneID:105283878 LOC105283878 Gene LOC105283878 protein_coding NA character(0)
character(0) NW_011925330.1 59800 59806 7 * 6 857.142857142857
NW_011925336.1 89625 92404 2780 - Gnomon gene NA NA gene10323 GeneID:105283960 LOC105283960 Gene LOC105283960 protein_coding NA character(0)
character(0) NW_011925336.1 91240 91246 7 * 6 857.142857142857
NW_011925336.1 92449 95536 3088 + Gnomon gene NA NA gene10324 GeneID:105283961 LOC105283961 Gene LOC105283961 protein_coding NA character(0)
character(0) NW_011925336.1 93305 93311 7 * 6 857.142857142857
NW_011925353.1 16223 20332 4110 + Gnomon gene NA NA gene10401 GeneID:105284052 LOC105284052 Gene LOC105284052 protein_coding NA character(0)
character(0) NW_011925353.1 19139 19145 7 * 6 857.142857142857
NW_011925356.1 701727 704945 3219 + Gnomon gene NA NA gene10538 GeneID:105284202 LOC105284202 Gene LOC105284202 protein_coding NA
character(0) character(0) NW_011925356.1 703128 703134 7 * 6 857.142857142857
NW_011925356.1 843920 846762 2843 + Gnomon gene NA NA gene10559 GeneID:105284223 LOC105284223 Gene LOC105284223 protein_coding NA
character(0) character(0) NW_011925356.1 845129 845135 7 * 6 857.142857142857
NW_011925367.1 1119922 1121887 1966 + Gnomon gene NA NA gene10714 GeneID:105284396 LOC105284396 Gene LOC105284396 protein_coding NA

character(0) character(0) NW_011925367.1 1120846 1120852 7 * 6 857.142857142857
NW_011925371.1 341773 343746 1974 - Gnomon gene NA NA gene10816 GeneID:105284502 LOC105284502 Gene LOC105284502 protein_coding NA
character(0) character(0) NW_011925371.1 342828 342834 7 * 6 857.142857142857
NW_011925371.1 899674 901634 1961 - Gnomon gene NA NA gene10880 GeneID:105284576 LOC105284576 Gene LOC105284576 protein_coding NA
character(0) character(0) NW_011925371.1 900958 900964 7 * 6 857.142857142857
NW_011925387.1 118203 119961 1759 - Gnomon gene NA NA gene10943 GeneID:105284659 LOC105284659 Gene LOC105284659 protein_coding NA
character(0) character(0) NW_011925387.1 119303 119309 7 * 6 857.142857142857
NW_011925434.1 120720 130419 9700 + Gnomon gene NA NA gene11344 GeneID:105285135 LOC105285135 Gene LOC105285135 protein_coding NA
character(0) character(0) NW_011925434.1 123131 123137 7 * 6 857.142857142857
NW_011925459.1 162939 164541 1603 + Gnomon gene NA NA gene11566 GeneID:105285392 LOC105285392 Gene LOC105285392 protein_coding NA
character(0) character(0) NW_011925459.1 163954 163960 7 * 6 857.142857142857
NW_011925459.1 186244 187554 1311 - Gnomon gene NA NA gene11569 GeneID:105285394 LOC105285394 Gene LOC105285394 protein_coding NA
character(0) character(0) NW_011925459.1 186700 186706 7 * 6 857.142857142857
NW_011925465.1 71398 78050 6653 - Gnomon gene NA NA gene11580 GeneID:105285410 LOC105285410 Gene LOC105285410 protein_coding NA character(0)
character(0) NW_011925465.1 77157 77163 7 * 6 857.142857142857
NW_011925471.1 1427 3141 1715 - Gnomon gene NA NA gene11632 GeneID:109611363 LOC109611363 Gene LOC109611363 lncRNA NA character(0)
character(0) NW_011925471.1 2780 2786 7 * 6 857.142857142857
NW_011925490.1 207564 212389 4826 + Gnomon gene NA NA gene11699 GeneID:105285546 LOC105285546 Gene LOC105285546 protein_coding NA
character(0) character(0) NW_011925490.1 208609 208615 7 * 6 857.142857142857
NW_011925516.1 278917 286323 7407 - Gnomon gene NA NA gene11797 GeneID:105285648 LOC105285648 Gene LOC105285648 protein_coding NA
character(0) character(0) NW_011925516.1 285124 285130 7 * 6 857.142857142857
NW_011925520.1 152066 157750 5685 + Gnomon gene NA NA gene11859 GeneID:105285719 LOC105285719 Gene LOC105285719 protein_coding NA
character(0) character(0) NW_011925520.1 152861 152867 7 * 6 857.142857142857
NW_011925546.1 78656 85452 6797 - Gnomon gene NA NA gene11986 GeneID:105285867 LOC105285867 Gene LOC105285867 protein_coding NA character(0)
character(0) NW_011925546.1 82761 82767 7 * 6 857.142857142857
NW_011925549.1 267852 270480 2629 - Gnomon gene NA NA gene12010 GeneID:105285889 LOC105285889 Gene LOC105285889 protein_coding NA
character(0) character(0) NW_011925549.1 269441 269447 7 * 6 857.142857142857
NW_011925715.1 251551 257367 5817 - Gnomon gene NA NA gene12526 GeneID:105286461 LOC105286461 Gene LOC105286461 protein_coding NA
character(0) character(0) NW_011925715.1 256038 256044 7 * 6 857.142857142857
NW_011925715.1 412871 419485 6615 + Gnomon gene NA NA gene12539 GeneID:105286478 LOC105286478 Gene LOC105286478 protein_coding NA
character(0) character(0) NW_011925715.1 414243 414249 7 * 6 857.142857142857
NW_011925715.1 1168275 1170855 2581 - Gnomon gene NA NA gene12582 GeneID:105286526 LOC105286526 Gene LOC105286526 protein_coding NA
character(0) character(0) NW_011925715.1 1169658 1169664 7 * 6 857.142857142857
NW_011926194.1 11132 18740 7609 + Gnomon gene NA NA gene13114 GeneID:105287147 LOC105287147 Gene LOC105287147 protein_coding NA character(0)
character(0) NW_011926194.1 12247 12253 7 * 6 857.142857142857
NW_011929339.1 7 1883 1877 + Gnomon gene NA NA gene13384 GeneID:105287447 LOC105287447 Gene LOC105287447 protein_coding true character(0)
c("1883", ".") NW_011929339.1 710 716 7 * 6 857.142857142857
NW_011924899.1 718895 720452 1558 + Gnomon gene NA NA gene664 GeneID:105279718 LOC105279718 Gene LOC105279718 protein_coding NA
character(0) character(0) NW_011924899.1 719598 719607 10 * 8 800
NW_011924951.1 73901 76560 2660 + Gnomon gene NA NA gene2325 GeneID:105274767 LOC105274767 Gene LOC105274767 protein_coding NA character(0)
character(0) NW_011924951.1 74986 74995 10 * 8 800
NW_011925477.1 19131 23385 4255 + Gnomon gene NA NA gene11657 GeneID:105285496 LOC105285496 Gene LOC105285496 protein_coding NA character(0)
character(0) NW_011925477.1 19843 19852 10 * 8 800

```
Directory: MRPR File: rmp-Cb_LI_BCphaseA1.txt

Overlap of methylation-rich regions with genome features for Cb sample Cb_LI_BCphaseA1

Total size of the methylation-rich regions of Cb_LI_BCphaseA1:                                       1009 bp (  0.0% of genome)
Size of overlap of methylation-rich regions of Cb Cb_LI_BCphaseA1 with genic regions:                   1002 bp ( 99.3% of total size)   ( 1.38 O/E)
Size of overlap of methylation-rich regions of Cb Cb_LI_BCphaseA1 with   exon regions:                     923 bp ( 91.5% of total size) ( 0.42 O/E)
Size of overlap of methylation-rich regions of Cb Cb_LI_BCphaseA1 with   intron regions:                    79 bp (  7.8% of total size) ( 0.15 O/E)
Size of overlap of methylation-rich regions of Cb Cb_LI_BCphaseA1 with intergenic regions:                 7 bp (  0.7% of total size)   ( 3.51 O/E)
Size of overlap of methylation-rich regions of Cb Cb_LI_BCphaseA1 with   promoter regions:                   0 bp (  0.0% of total size) ( 0.00 O/E)
Size of overlap of methylation-rich regions of Cb Cb_LI_BCphaseA1 with   other intergenic regions:           7 bp (  0.7% of total size) ( 0.03 O/E)

Overlap of methylation-poor regions with genome features for Cb sample Cb_LI_BCphaseA1

Total size of the methylation-poor regions of Cb_LI_BCphaseA1:                                   31153530 bp ( 14.6% of genome)
Size of overlap of methylation-poor regions of Cb Cb_LI_BCphaseA1 with genic regions:               22335867 bp ( 71.7% of total size)   ( 1.00 O/E)
Size of overlap of methylation-poor regions of Cb Cb_LI_BCphaseA1 with   exon regions:                 2291347 bp (  7.4% of total size) ( 3.44 O/E)
Size of overlap of methylation-poor regions of Cb Cb_LI_BCphaseA1 with   intron regions:              20044520 bp ( 64.3% of total size) ( 1.21 O/E)
Size of overlap of methylation-poor regions of Cb Cb_LI_BCphaseA1 with intergenic regions:           8817663 bp ( 28.3% of total size)   ( 2.54 O/E)
Size of overlap of methylation-poor regions of Cb Cb_LI_BCphaseA1 with   promoter regions:              208140 bp (  0.7% of total size) ( 0.22 O/E)
Size of overlap of methylation-poor regions of Cb Cb_LI_BCphaseA1 with   other intergenic regions:     8609523 bp ( 27.6% of total size) ( 1.10 O/E)
```

Directory: MRPR File: rmp-Cb_LI_BCphaseA2.txt

Overlap of methylation-rich regions with genome features for Cb sample Cb_LI_BCphaseA2

```
Total size of the methylation-rich regions of Cb_LI_BCphaseA2:                                       1370 bp (  0.0% of genome)
Size of overlap of methylation-rich regions of Cb Cb_LI_BCphaseA2 with genic regions:                   1363 bp ( 99.5% of total size)   ( 1.39 O/E)
Size of overlap of methylation-rich regions of Cb Cb_LI_BCphaseA2 with   exon regions:                    1282 bp ( 93.6% of total size) ( 0.32 O/E)
Size of overlap of methylation-rich regions of Cb Cb_LI_BCphaseA2 with   intron regions:                    81 bp (  5.9% of total size) ( 0.11 O/E)
Size of overlap of methylation-rich regions of Cb Cb_LI_BCphaseA2 with intergenic regions:                 7 bp (  0.5% of total size)   ( 3.52 O/E)
Size of overlap of methylation-rich regions of Cb Cb_LI_BCphaseA2 with   promoter regions:                   0 bp (  0.0% of total size) ( 0.00 O/E)
Size of overlap of methylation-rich regions of Cb Cb_LI_BCphaseA2 with   other intergenic regions:           7 bp (  0.5% of total size) ( 0.02 O/E)
```

Overlap of methylation-poor regions with genome features for Cb sample Cb_LI_BCphaseA2

```
Total size of the methylation-poor regions of Cb_LI_BCphaseA2:                                   38006225 bp ( 17.9% of genome)
Size of overlap of methylation-poor regions of Cb Cb_LI_BCphaseA2 with genic regions:               25581629 bp ( 67.3% of total size)   ( 0.94 O/E)
Size of overlap of methylation-poor regions of Cb Cb_LI_BCphaseA2 with   exon regions:                 2836067 bp (  7.5% of total size) ( 3.20 O/E)
Size of overlap of methylation-poor regions of Cb Cb_LI_BCphaseA2 with   intron regions:              22745562 bp ( 59.9% of total size) ( 1.13 O/E)
Size of overlap of methylation-poor regions of Cb Cb_LI_BCphaseA2 with intergenic regions:          12424596 bp ( 32.7% of total size)   ( 2.38 O/E)
Size of overlap of methylation-poor regions of Cb Cb_LI_BCphaseA2 with   promoter regions:              251083 bp (  0.7% of total size) ( 0.21 O/E)
Size of overlap of methylation-poor regions of Cb Cb_LI_BCphaseA2 with   other intergenic regions:    12173513 bp ( 32.0% of total size) ( 1.27 O/E)
```

```
Directory: MRPR File: rmp-Cb_LI_BCphaseB1.txt

Overlap of methylation-rich regions with genome features for Cb sample Cb_LI_BCphaseB1

Total size of the methylation-rich regions of Cb_LI_BCphaseB1:                                     1462 bp (  0.0% of genome)
Size of overlap of methylation-rich regions of Cb Cb_LI_BCphaseB1 with genic regions:                 1455 bp ( 99.5% of total size)   ( 1.39 O/E)
Size of overlap of methylation-rich regions of Cb Cb_LI_BCphaseB1 with   exon regions:                  1340 bp ( 91.7% of total size) ( 0.42 O/E)
Size of overlap of methylation-rich regions of Cb Cb_LI_BCphaseB1 with   intron regions:                 115 bp (  7.9% of total size) ( 0.15 O/E)
Size of overlap of methylation-rich regions of Cb Cb_LI_BCphaseB1 with intergenic regions:                7 bp (  0.5% of total size)   ( 3.52 O/E)
Size of overlap of methylation-rich regions of Cb Cb_LI_BCphaseB1 with   promoter regions:                  0 bp (  0.0% of total size) ( 0.00 O/E)
Size of overlap of methylation-rich regions of Cb Cb_LI_BCphaseB1 with   other intergenic regions:          7 bp (  0.5% of total size) ( 0.02 O/E)

Overlap of methylation-poor regions with genome features for Cb sample Cb_LI_BCphaseB1

Total size of the methylation-poor regions of Cb_LI_BCphaseB1:                                 27543451 bp ( 12.9% of genome)
Size of overlap of methylation-poor regions of Cb Cb_LI_BCphaseB1 with genic regions:             19112859 bp ( 69.4% of total size)   ( 0.97 O/E)
Size of overlap of methylation-poor regions of Cb Cb_LI_BCphaseB1 with   exon regions:               2060008 bp (  7.5% of total size) ( 3.31 O/E)
Size of overlap of methylation-poor regions of Cb Cb_LI_BCphaseB1 with   intron regions:            17052851 bp ( 61.9% of total size) ( 1.17 O/E)
Size of overlap of methylation-poor regions of Cb Cb_LI_BCphaseB1 with intergenic regions:          8430592 bp ( 30.6% of total size)   ( 2.46 O/E)
Size of overlap of methylation-poor regions of Cb Cb_LI_BCphaseB1 with   promoter regions:             177764 bp (  0.7% of total size) ( 0.21 O/E)
Size of overlap of methylation-poor regions of Cb Cb_LI_BCphaseB1 with   other intergenic regions:    8252828 bp ( 30.0% of total size) ( 1.19 O/E)
```

Directory: MRPR File: rmp-Cb_LI_BCphaseB2.txt

Overlap of methylation-rich regions with genome features for Cb sample Cb_LI_BCphaseB2

```
Total size of the methylation-rich regions of Cb_LI_BCphaseB2:                                       1261 bp (  0.0% of genome)
Size of overlap of methylation-rich regions of Cb Cb_LI_BCphaseB2 with genic regions:                   1254 bp ( 99.4% of total size)   ( 1.39 O/E)
Size of overlap of methylation-rich regions of Cb Cb_LI_BCphaseB2 with   exon regions:                    1168 bp ( 92.6% of total size) ( 0.37 O/E)
Size of overlap of methylation-rich regions of Cb Cb_LI_BCphaseB2 with   intron regions:                    86 bp (  6.8% of total size) ( 0.13 O/E)
Size of overlap of methylation-rich regions of Cb Cb_LI_BCphaseB2 with intergenic regions:                 7 bp (  0.6% of total size)   ( 3.52 O/E)
Size of overlap of methylation-rich regions of Cb Cb_LI_BCphaseB2 with   promoter regions:                   0 bp (  0.0% of total size) ( 0.00 O/E)
Size of overlap of methylation-rich regions of Cb Cb_LI_BCphaseB2 with   other intergenic regions:           7 bp (  0.6% of total size) ( 0.02 O/E)
```

Overlap of methylation-poor regions with genome features for Cb sample Cb_LI_BCphaseB2

```
Total size of the methylation-poor regions of Cb_LI_BCphaseB2:                                   28781387 bp ( 13.5% of genome)
Size of overlap of methylation-poor regions of Cb Cb_LI_BCphaseB2 with genic regions:               20218084 bp ( 70.2% of total size)   ( 0.98 O/E)
Size of overlap of methylation-poor regions of Cb Cb_LI_BCphaseB2 with   exon regions:                2325208 bp (  8.1% of total size) ( 3.33 O/E)
Size of overlap of methylation-poor regions of Cb Cb_LI_BCphaseB2 with   intron regions:             17892876 bp ( 62.2% of total size) ( 1.17 O/E)
Size of overlap of methylation-poor regions of Cb Cb_LI_BCphaseB2 with intergenic regions:           8563303 bp ( 29.8% of total size)   ( 2.49 O/E)
Size of overlap of methylation-poor regions of Cb Cb_LI_BCphaseB2 with   promoter regions:              206376 bp (  0.7% of total size) ( 0.23 O/E)
Size of overlap of methylation-poor regions of Cb Cb_LI_BCphaseB2 with   other intergenic regions:     8356927 bp ( 29.0% of total size) ( 1.15 O/E)
```

```
Directory: MRPR File: rmp-Cb_LI_RphaseA1.txt

Overlap of methylation-rich regions with genome features for Cb sample Cb_LI_RphaseA1

Total size of the methylation-rich regions of Cb_LI_RphaseA1:                                        1391 bp (  0.0% of genome)
Size of overlap of methylation-rich regions of Cb Cb_LI_RphaseA1 with genic regions:                    1384 bp ( 99.5% of total size)   ( 1.39 O/E)
Size of overlap of methylation-rich regions of Cb Cb_LI_RphaseA1 with   exon regions:                     1303 bp ( 93.7% of total size) ( 0.31 O/E)
Size of overlap of methylation-rich regions of Cb Cb_LI_RphaseA1 with   intron regions:                     81 bp (  5.8% of total size) ( 0.11 O/E)
Size of overlap of methylation-rich regions of Cb Cb_LI_RphaseA1 with intergenic regions:                  7 bp (  0.5% of total size)   ( 3.52 O/E)
Size of overlap of methylation-rich regions of Cb Cb_LI_RphaseA1 with   promoter regions:                    0 bp (  0.0% of total size) ( 0.00 O/E)
Size of overlap of methylation-rich regions of Cb Cb_LI_RphaseA1 with   other intergenic regions:           7 bp (  0.5% of total size) ( 0.02 O/E)

Overlap of methylation-poor regions with genome features for Cb sample Cb_LI_RphaseA1

Total size of the methylation-poor regions of Cb_LI_RphaseA1:                                    36779916 bp ( 17.3% of genome)
Size of overlap of methylation-poor regions of Cb Cb_LI_RphaseA1 with genic regions:                25141438 bp ( 68.4% of total size)   ( 0.95 O/E)
Size of overlap of methylation-poor regions of Cb Cb_LI_RphaseA1 with   exon regions:                  2850413 bp (  7.8% of total size) ( 3.24 O/E)
Size of overlap of methylation-poor regions of Cb Cb_LI_RphaseA1 with   intron regions:               22291025 bp ( 60.6% of total size) ( 1.14 O/E)
Size of overlap of methylation-poor regions of Cb Cb_LI_RphaseA1 with intergenic regions:          11638478 bp ( 31.6% of total size)   ( 2.42 O/E)
Size of overlap of methylation-poor regions of Cb Cb_LI_RphaseA1 with   promoter regions:               260337 bp (  0.7% of total size) ( 0.23 O/E)
Size of overlap of methylation-poor regions of Cb Cb_LI_RphaseA1 with   other intergenic regions:    11378141 bp ( 30.9% of total size) ( 1.23 O/E)
```

Directory: MRPR File: rmp-Cb_LI_RphaseA2.txt

Overlap of methylation-rich regions with genome features for Cb sample Cb_LI_RphaseA2

Total size of the methylation-rich regions of Cb_LI_RphaseA2: 2011 bp ( 0.0% of genome)
Size of overlap of methylation-rich regions of Cb Cb_LI_RphaseA2 with genic regions: 1998 bp ( 99.3% of total size) ( 1.38 O/E)
Size of overlap of methylation-rich regions of Cb Cb_LI_RphaseA2 with exon regions: 1857 bp ( 92.3% of total size) ( 0.38 O/E)
Size of overlap of methylation-rich regions of Cb Cb_LI_RphaseA2 with intron regions: 141 bp ( 7.0% of total size) ( 0.13 O/E)
Size of overlap of methylation-rich regions of Cb Cb_LI_RphaseA2 with intergenic regions: 13 bp ( 0.7% of total size) ( 3.52 O/E)
Size of overlap of methylation-rich regions of Cb Cb_LI_RphaseA2 with promoter regions: 0 bp ( 0.0% of total size) ( 0.00 O/E)
Size of overlap of methylation-rich regions of Cb Cb_LI_RphaseA2 with other intergenic regions: 13 bp ( 0.7% of total size) ( 0.03 O/E)

Overlap of methylation-poor regions with genome features for Cb sample Cb_LI_RphaseA2

Total size of the methylation-poor regions of Cb_LI_RphaseA2: 15143436 bp ( 7.1% of genome)
Size of overlap of methylation-poor regions of Cb Cb_LI_RphaseA2 with genic regions: 10457224 bp ( 69.0% of total size) ( 0.96 O/E)
Size of overlap of methylation-poor regions of Cb Cb_LI_RphaseA2 with exon regions: 1285103 bp ( 8.5% of total size) ( 3.24 O/E)
Size of overlap of methylation-poor regions of Cb Cb_LI_RphaseA2 with intron regions: 9172121 bp ( 60.6% of total size) ( 1.14 O/E)
Size of overlap of methylation-poor regions of Cb Cb_LI_RphaseA2 with intergenic regions: 4686212 bp ( 30.9% of total size) ( 2.44 O/E)
Size of overlap of methylation-poor regions of Cb Cb_LI_RphaseA2 with promoter regions: 134542 bp ( 0.9% of total size) ( 0.29 O/E)
Size of overlap of methylation-poor regions of Cb Cb_LI_RphaseA2 with other intergenic regions: 4551670 bp ( 30.1% of total size) ( 1.19 O/E)

Directory: MRPR File: rmp-Cb_LI_RphaseB1.txt

Overlap of methylation-rich regions with genome features for Cb sample Cb_LI_RphaseB1

```
Total size of the methylation-rich regions of Cb_LI_RphaseB1:                                       1309 bp (  0.0% of genome)
Size of overlap of methylation-rich regions of Cb Cb_LI_RphaseB1 with genic regions:                   1302 bp ( 99.5% of total size)   ( 1.39 O/E)
Size of overlap of methylation-rich regions of Cb Cb_LI_RphaseB1 with   exon regions:                    1215 bp ( 92.8% of total size) ( 0.36 O/E)
Size of overlap of methylation-rich regions of Cb Cb_LI_RphaseB1 with   intron regions:                    87 bp (  6.7% of total size) ( 0.13 O/E)
Size of overlap of methylation-rich regions of Cb Cb_LI_RphaseB1 with intergenic regions:                 7 bp (  0.5% of total size)   ( 3.52 O/E)
Size of overlap of methylation-rich regions of Cb Cb_LI_RphaseB1 with   promoter regions:                   0 bp (  0.0% of total size) ( 0.00 O/E)
Size of overlap of methylation-rich regions of Cb Cb_LI_RphaseB1 with   other intergenic regions:           7 bp (  0.5% of total size) ( 0.02 O/E)
```

Overlap of methylation-poor regions with genome features for Cb sample Cb_LI_RphaseB1

```
Total size of the methylation-poor regions of Cb_LI_RphaseB1:                                   25550773 bp ( 12.0% of genome)
Size of overlap of methylation-poor regions of Cb Cb_LI_RphaseB1 with genic regions:               17192910 bp ( 67.3% of total size)   ( 0.94 O/E)
Size of overlap of methylation-poor regions of Cb Cb_LI_RphaseB1 with   exon regions:                 1858321 bp (  7.3% of total size) ( 3.21 O/E)
Size of overlap of methylation-poor regions of Cb Cb_LI_RphaseB1 with   intron regions:              15334589 bp ( 60.0% of total size) ( 1.13 O/E)
Size of overlap of methylation-poor regions of Cb Cb_LI_RphaseB1 with intergenic regions:           8357863 bp ( 32.7% of total size)   ( 2.38 O/E)
Size of overlap of methylation-poor regions of Cb Cb_LI_RphaseB1 with   promoter regions:              181059 bp (  0.7% of total size) ( 0.23 O/E)
Size of overlap of methylation-poor regions of Cb Cb_LI_RphaseB1 with   other intergenic regions:     8176804 bp ( 32.0% of total size) ( 1.27 O/E)
```

Directory: MRPR File: rmp-Cb_LI_RphaseB2.txt

Overlap of methylation-rich regions with genome features for Cb sample Cb_LI_RphaseB2

```
Total size of the methylation-rich regions of Cb_LI_RphaseB2:                                       1238 bp (  0.0% of genome)
Size of overlap of methylation-rich regions of Cb Cb_LI_RphaseB2 with genic regions:                   1231 bp ( 99.4% of total size)   ( 1.39 O/E)
Size of overlap of methylation-rich regions of Cb Cb_LI_RphaseB2 with   exon regions:                    1152 bp ( 93.0% of total size) ( 0.34 O/E)
Size of overlap of methylation-rich regions of Cb Cb_LI_RphaseB2 with   intron regions:                    79 bp (  6.4% of total size) ( 0.12 O/E)
Size of overlap of methylation-rich regions of Cb Cb_LI_RphaseB2 with intergenic regions:                 7 bp (  0.6% of total size)   ( 3.52 O/E)
Size of overlap of methylation-rich regions of Cb Cb_LI_RphaseB2 with   promoter regions:                   0 bp (  0.0% of total size) ( 0.00 O/E)
Size of overlap of methylation-rich regions of Cb Cb_LI_RphaseB2 with   other intergenic regions:           7 bp (  0.6% of total size) ( 0.02 O/E)
```

Overlap of methylation-poor regions with genome features for Cb sample Cb_LI_RphaseB2

```
Total size of the methylation-poor regions of Cb_LI_RphaseB2:                                   33169521 bp ( 15.6% of genome)
Size of overlap of methylation-poor regions of Cb Cb_LI_RphaseB2 with genic regions:               22735557 bp ( 68.5% of total size)   ( 0.96 O/E)
Size of overlap of methylation-poor regions of Cb Cb_LI_RphaseB2 with   exon regions:                2706326 bp (  8.2% of total size) ( 3.23 O/E)
Size of overlap of methylation-poor regions of Cb Cb_LI_RphaseB2 with   intron regions:             20029231 bp ( 60.4% of total size) ( 1.14 O/E)
Size of overlap of methylation-poor regions of Cb Cb_LI_RphaseB2 with intergenic regions:          10433964 bp ( 31.5% of total size)   ( 2.43 O/E)
Size of overlap of methylation-poor regions of Cb Cb_LI_RphaseB2 with   promoter regions:             254922 bp (  0.8% of total size) ( 0.25 O/E)
Size of overlap of methylation-poor regions of Cb Cb_LI_RphaseB2 with   other intergenic regions:    10179042 bp ( 30.7% of total size) ( 1.22 O/E)
```

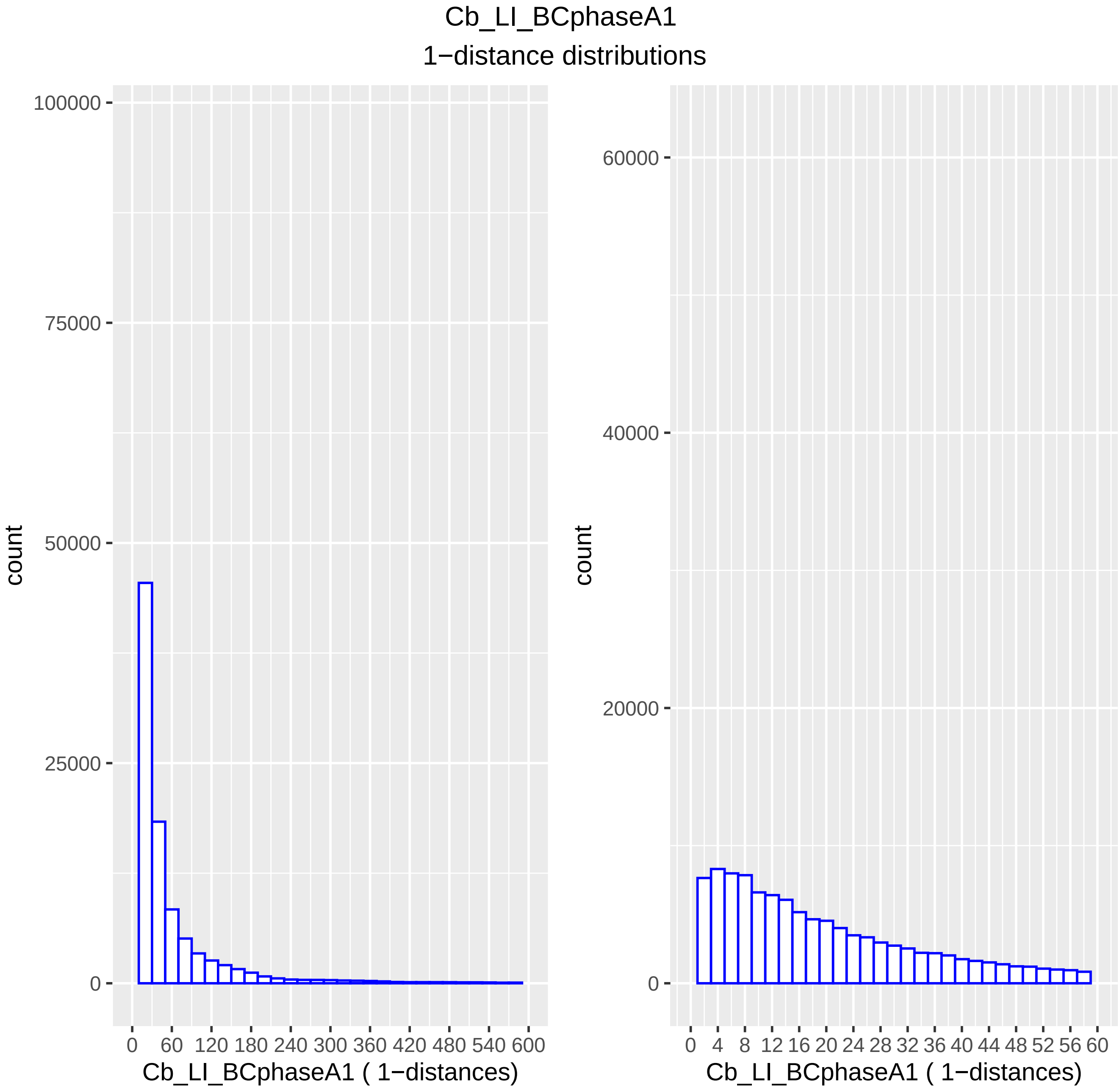
Cb_LI_BCphaseA1
1−distance distributions
count
100000
75000
50000
25000
0
0
60
120
180
240
300
360
420
480
540
600
Cb_LI_BCphaseA1 ( 1−distances)
count
60000
40000
20000
0
0
4
8
12
16
20
24
28
32
36
40
44
48
52
56
60
Cb_LI_BCphaseA1 ( 1−distances)

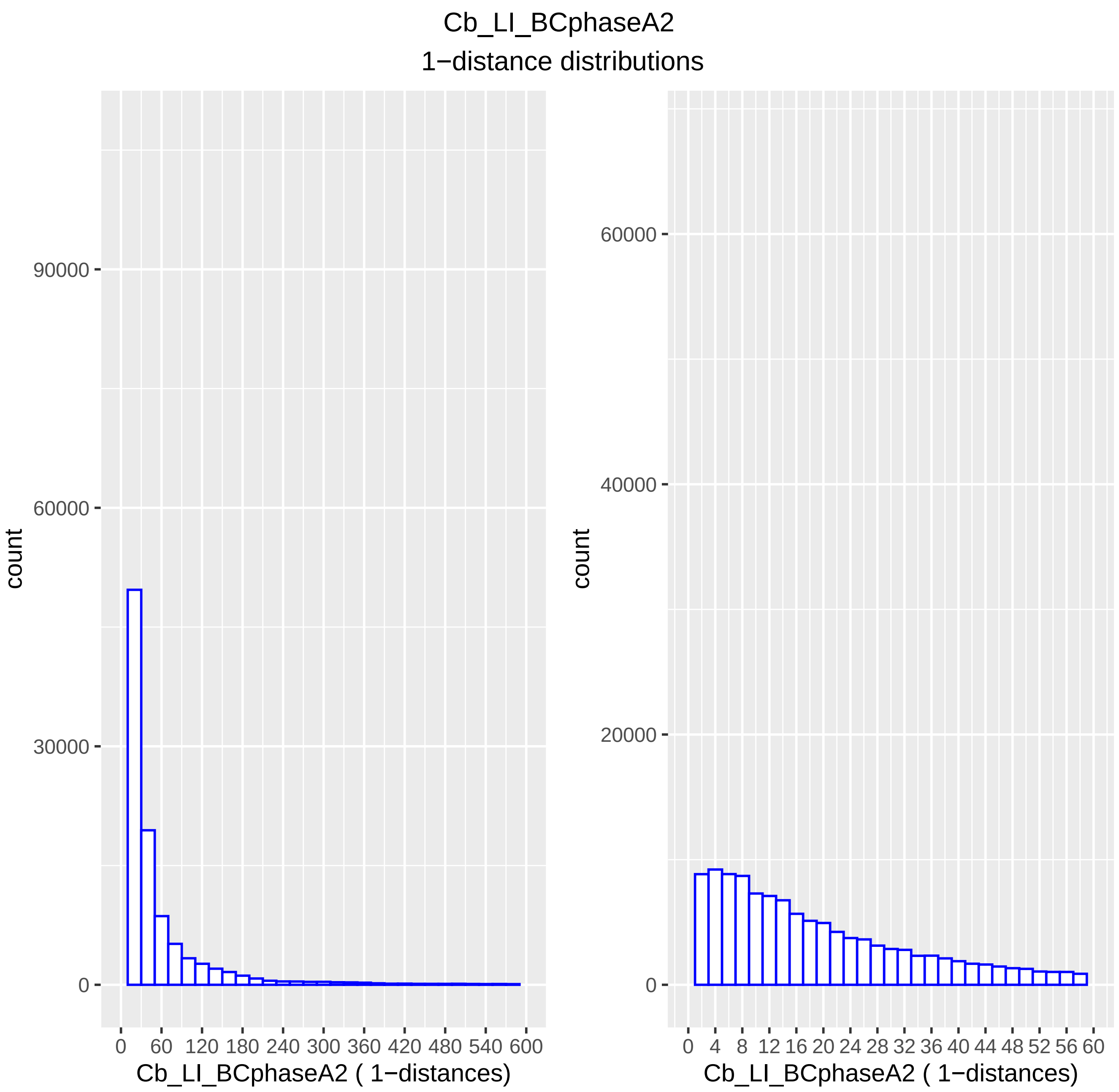

Cb_LI_BCphaseA2
1−distance distributions
count
90000
60000
30000
0
0 60 120 180 240 300 360 420 480 540 600
Cb_LI_BCphaseA2 ( 1−distances)
count
60000
40000
20000
0
0 4 8 12 16 20 24 28 32 36 40 44 48 52 56 60
Cb_LI_BCphaseA2 ( 1−distances)

Cb_LI_BCphaseB1
1−distance distributions

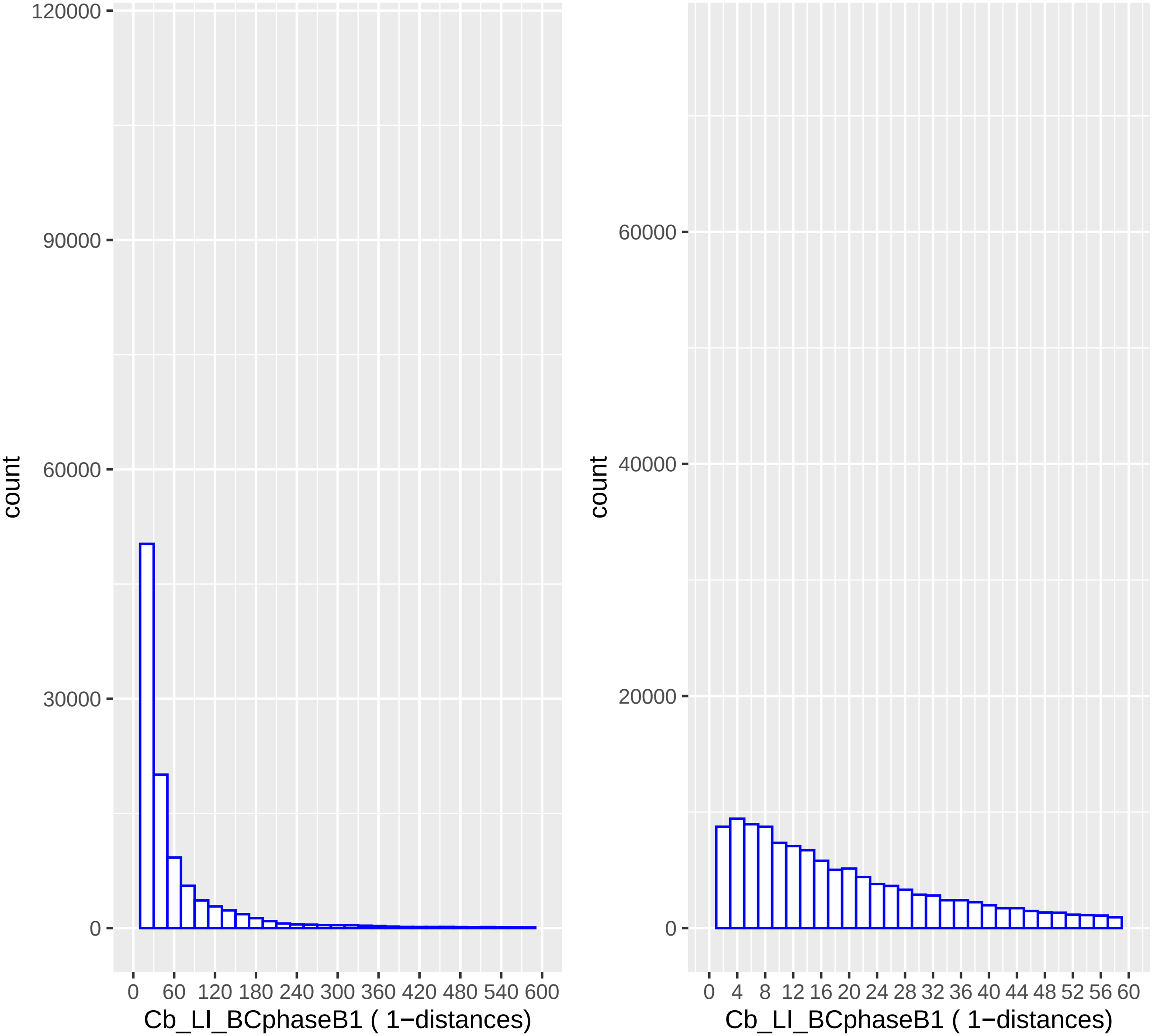

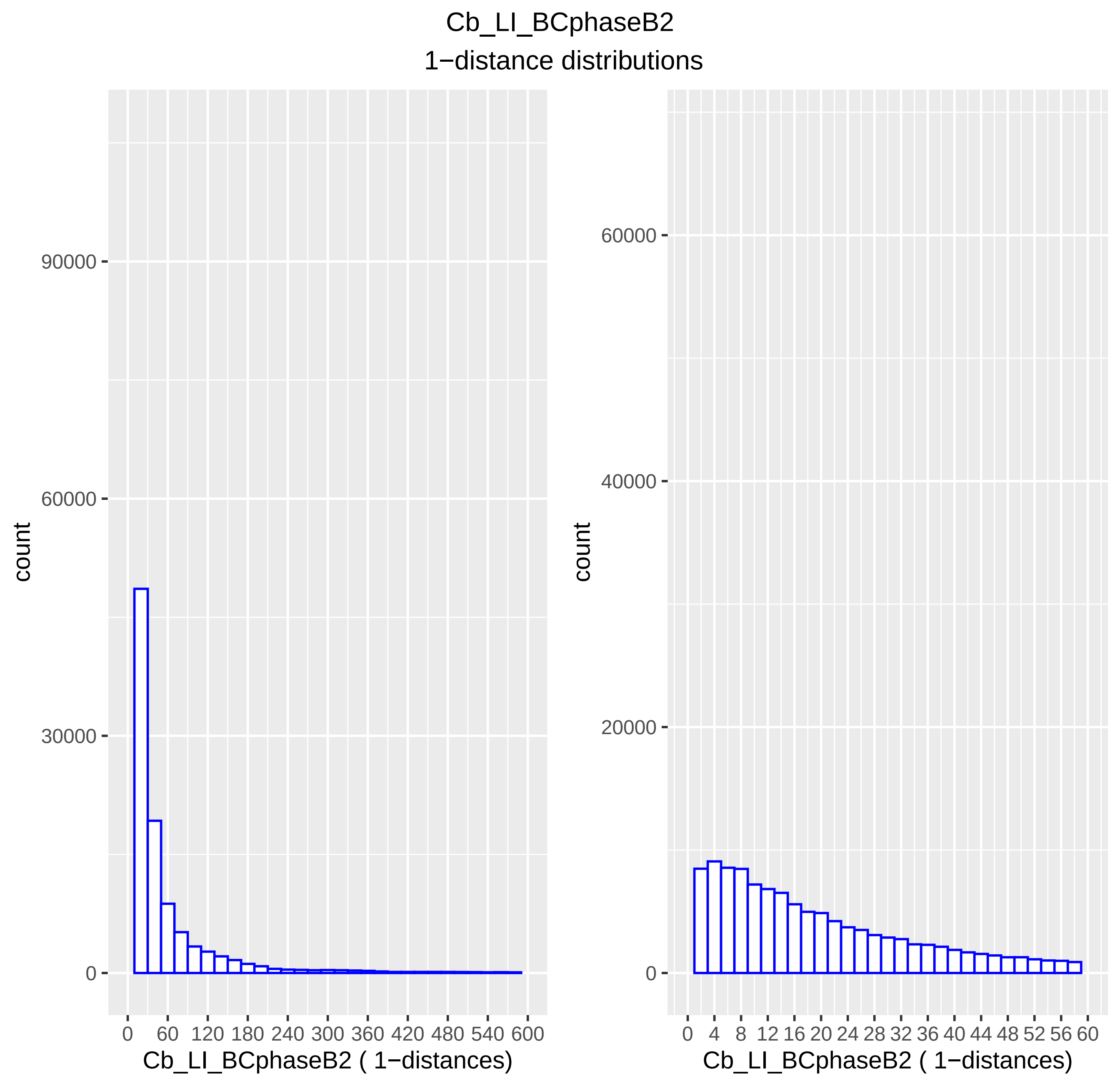
Cb_LI_BCphaseB2
1−distance distributions
count
90000
60000
30000
0
0 60 120 180 240 300 360 420 480 540 600
Cb_LI_BCphaseB2 ( 1−distances)
count
60000
40000
20000
0
0 4 8 12 16 20 24 28 32 36 40 44 48 52 56 60
Cb_LI_BCphaseB2 ( 1−distances)

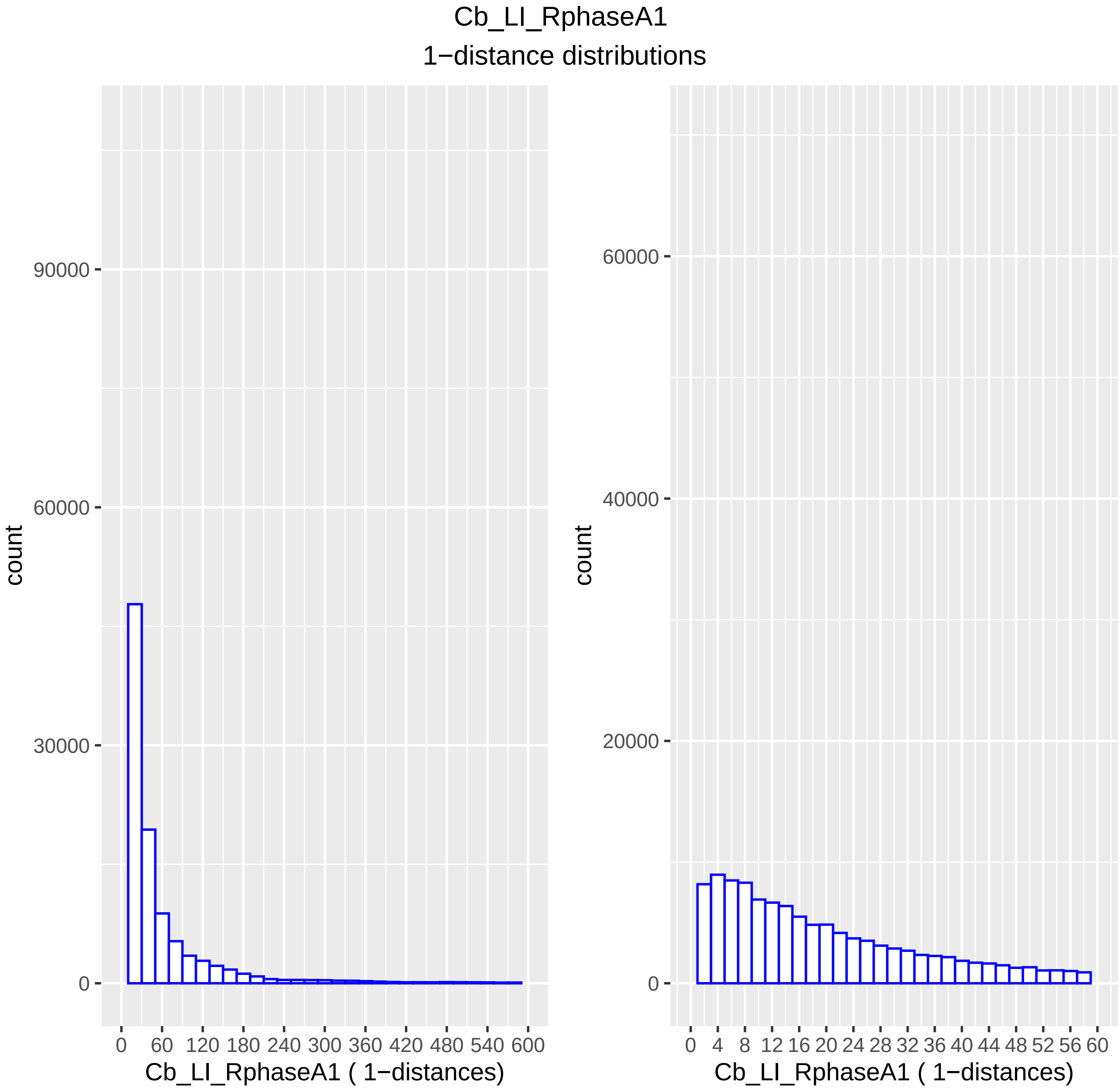
Cb_LI_RphaseA1
1−distance distributions
count
90000
60000
30000
0
0 60 120 180 240 300 360 420 480 540 600
Cb_LI_RphaseA1 ( 1−distances)
count
60000
40000
20000
0
0 4 8 12 16 20 24 28 32 36 40 44 48 52 56 60
Cb_LI_RphaseA1 ( 1−distances)

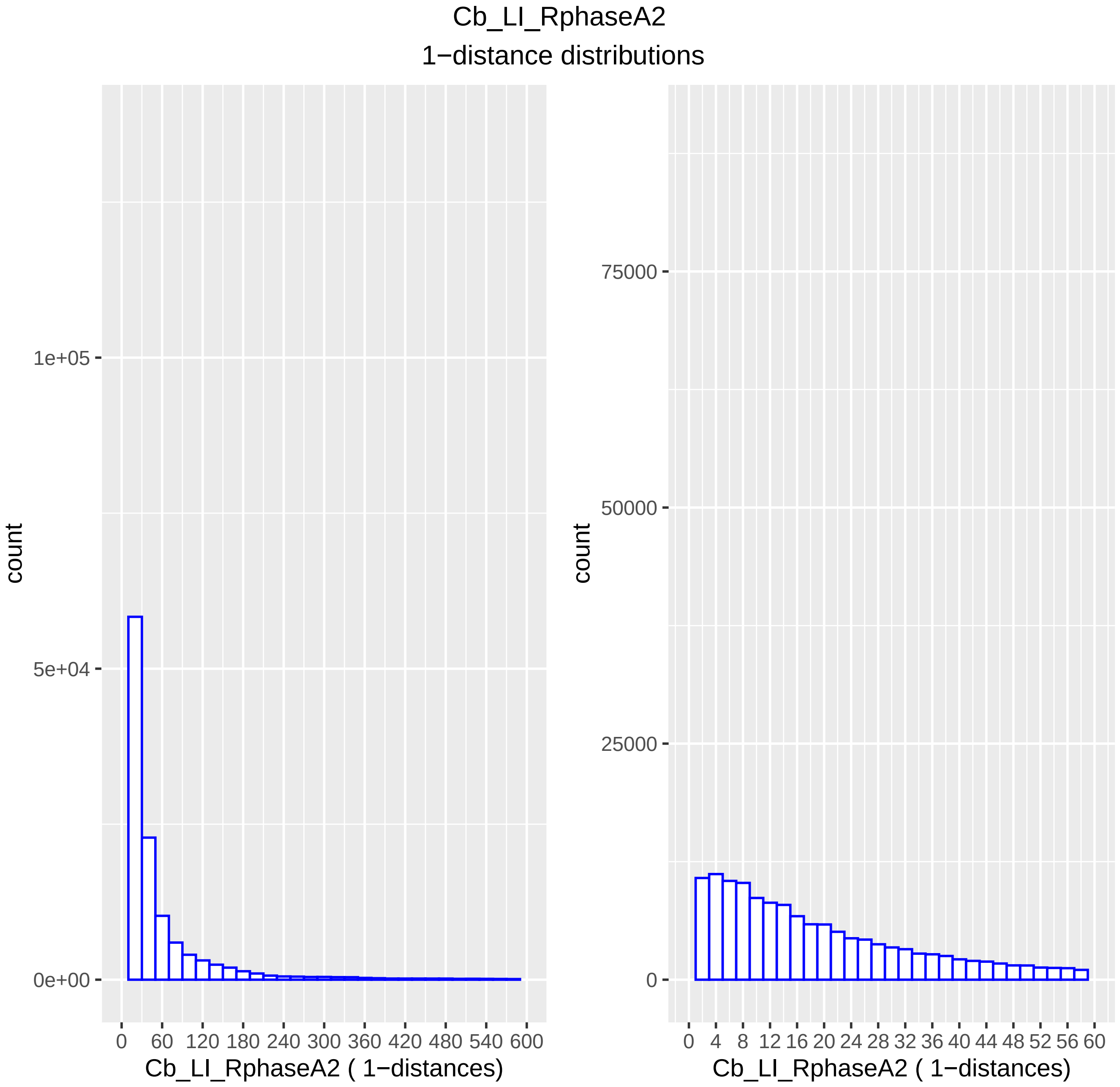

Cb_LI_RphaseA2
1−distance distributions
count
1e+05
5e+04
0e+00
0
60
120
180
240
300
360
420
480
540
600
Cb_LI_RphaseA2 ( 1−distances)
count
75000
50000
25000
0
0
4
8
12
16
20
24
28
32
36
40
44
48
52
56
60
Cb_LI_RphaseA2 ( 1−distances)

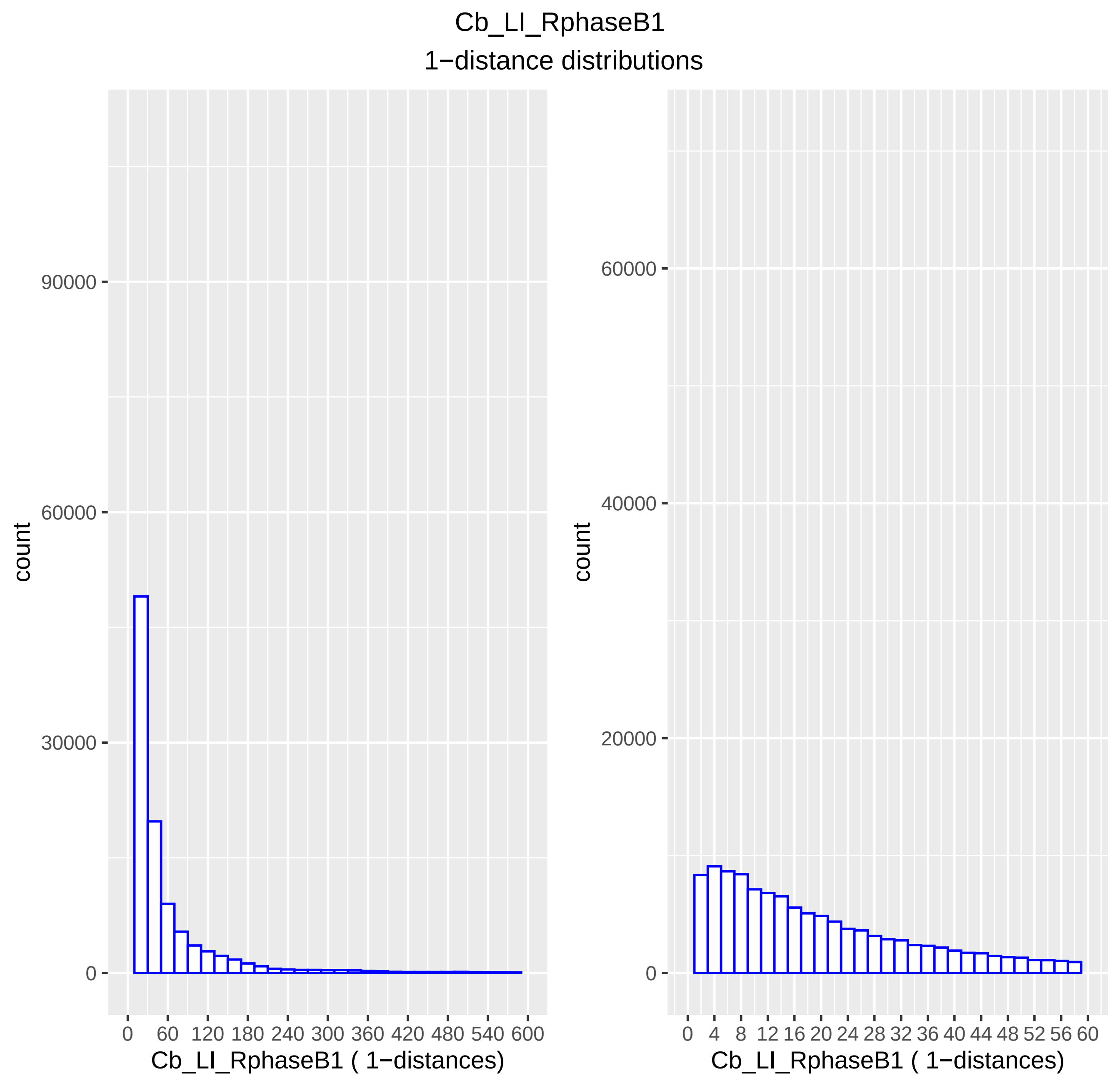

Cb_LI_RphaseB1
1−distance distributions
count
90000
60000
30000
0
0 60 120 180 240 300 360 420 480 540 600
Cb_LI_RphaseB1 ( 1−distances)
count
60000
40000
20000
0
0 4 8 12 16 20 24 28 32 36 40 44 48 52 56 60
Cb_LI_RphaseB1 ( 1−distances)

Cb_LI_RphaseB2
1−distance distributions

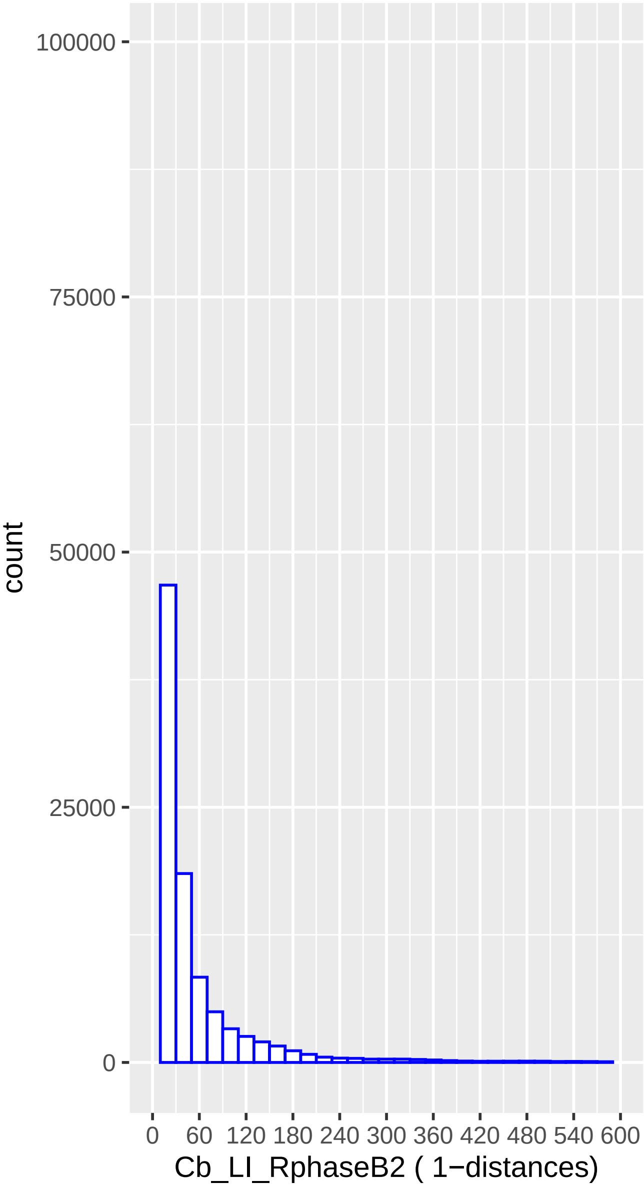

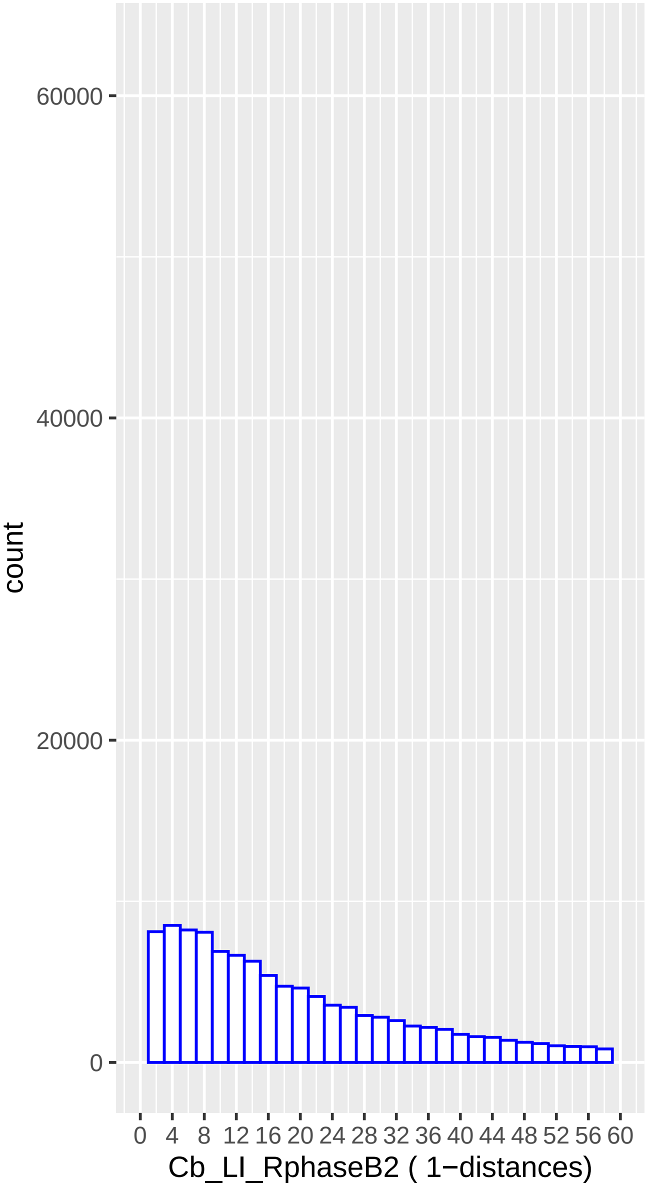

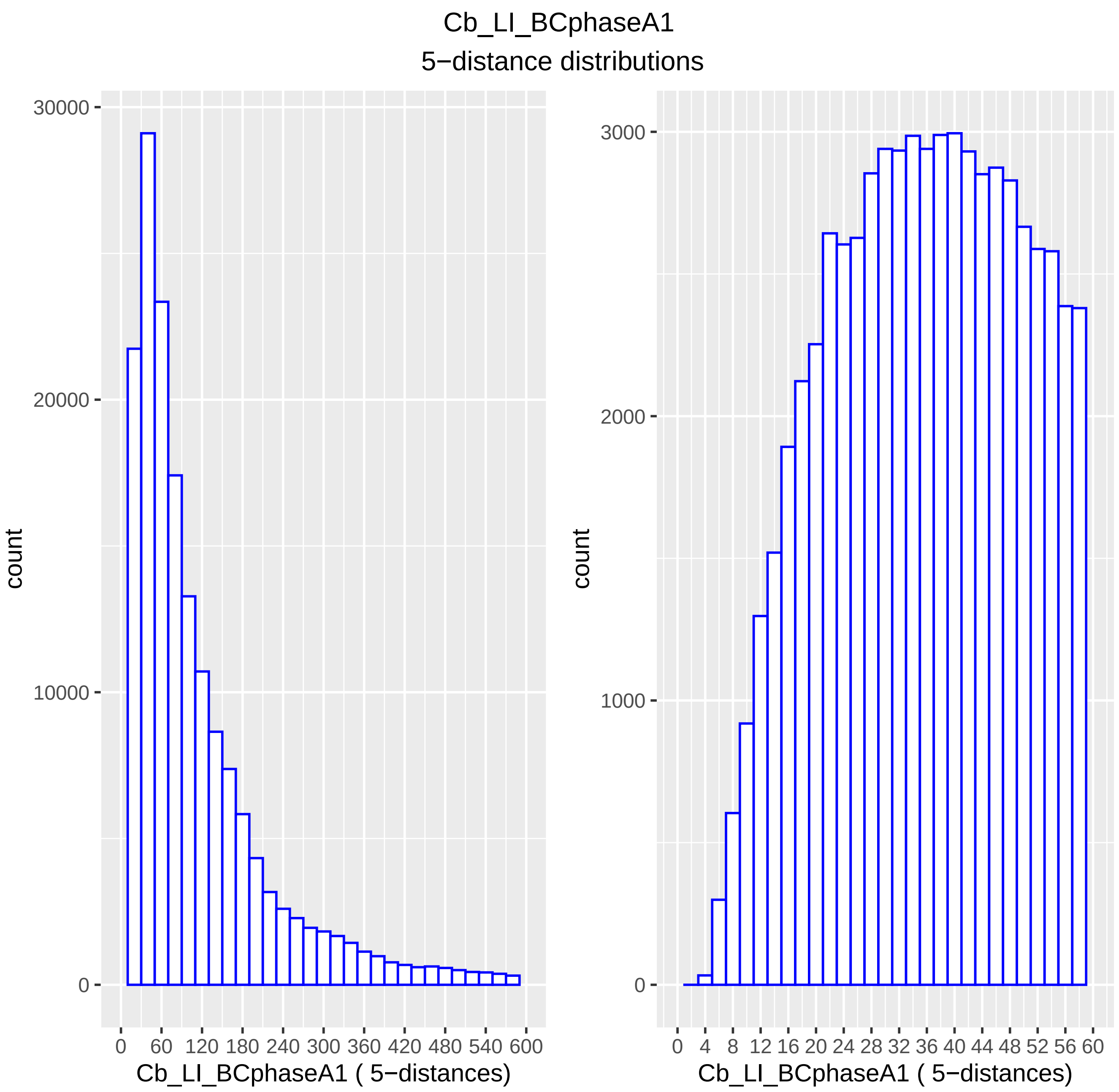
Cb_LI_BCphaseA1
5−distance distributions
count
30000
20000
10000
0
0 60 120 180 240 300 360 420 480 540 600
Cb_LI_BCphaseA1 ( 5−distances)
count
3000
2000
1000
0
0 4 8 12 16 20 24 28 32 36 40 44 48 52 56 60
Cb_LI_BCphaseA1 ( 5−distances)

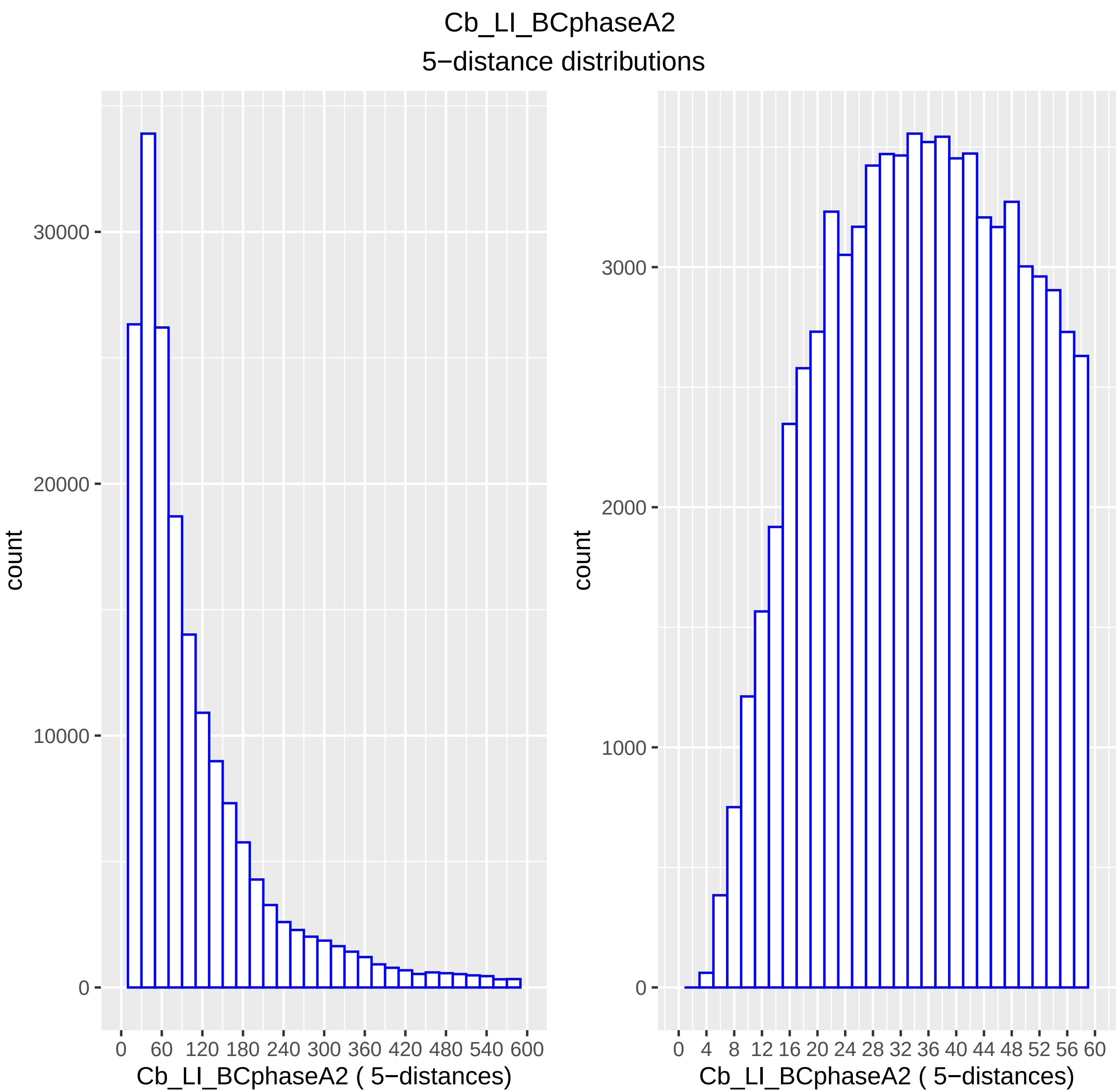
Cb_LI_BCphaseA2
5−distance distributions
count
30000
20000
10000
0
0 60 120 180 240 300 360 420 480 540 600
Cb_LI_BCphaseA2 ( 5−distances)
count
3000
2000
1000
0
0 4 8 12 16 20 24 28 32 36 40 44 48 52 56 60
Cb_LI_BCphaseA2 ( 5−distances)

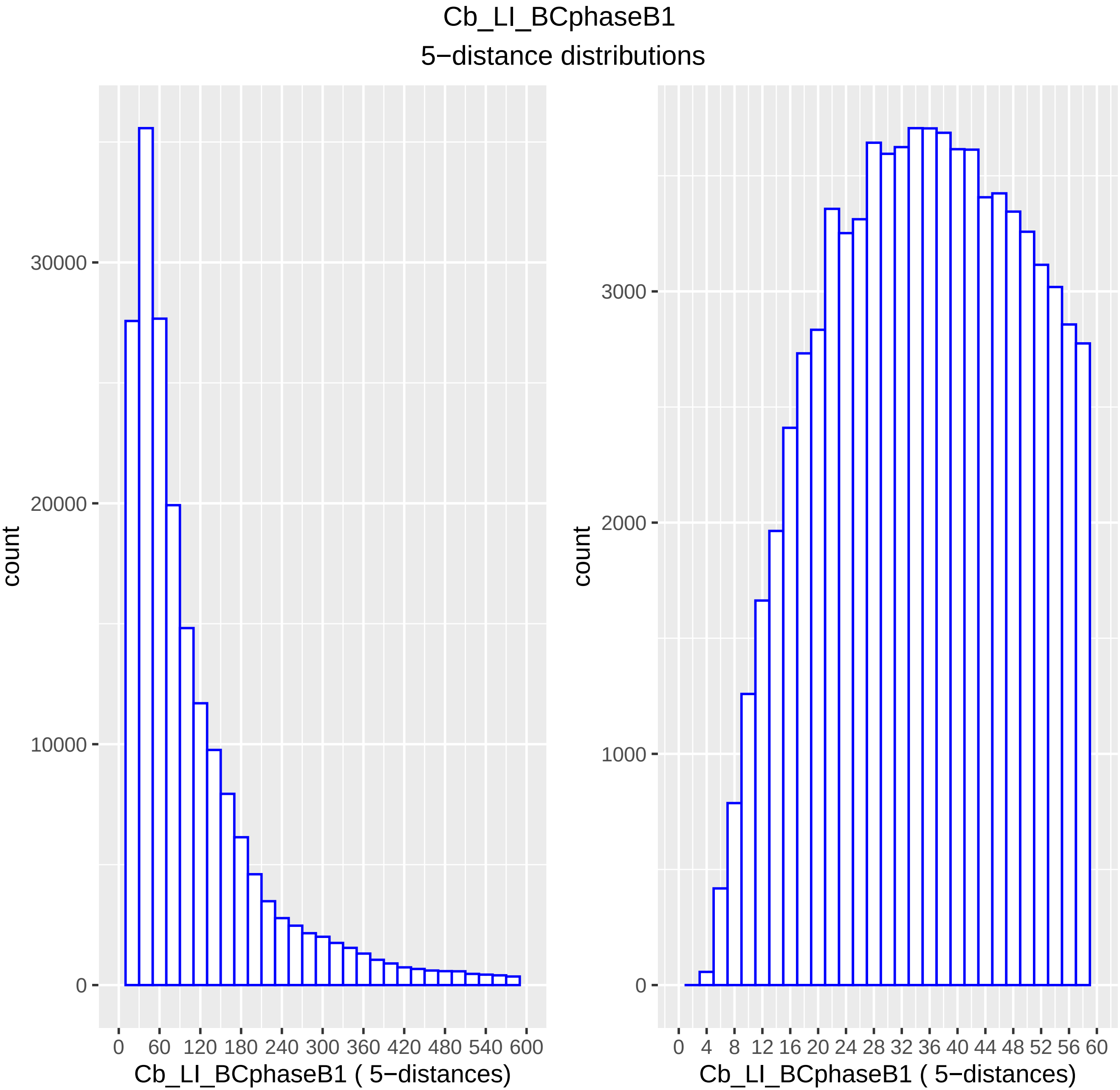

Cb_LI_BCphaseB1
5−distance distributions
count
30000
20000
10000
0
0 60 120 180 240 300 360 420 480 540 600
Cb_LI_BCphaseB1 ( 5−distances)
count
3000
2000
1000
0
0 4 8 12 16 20 24 28 32 36 40 44 48 52 56 60
Cb_LI_BCphaseB1 ( 5−distances)

Cb_LI_BCphaseB2
5−distance distributions

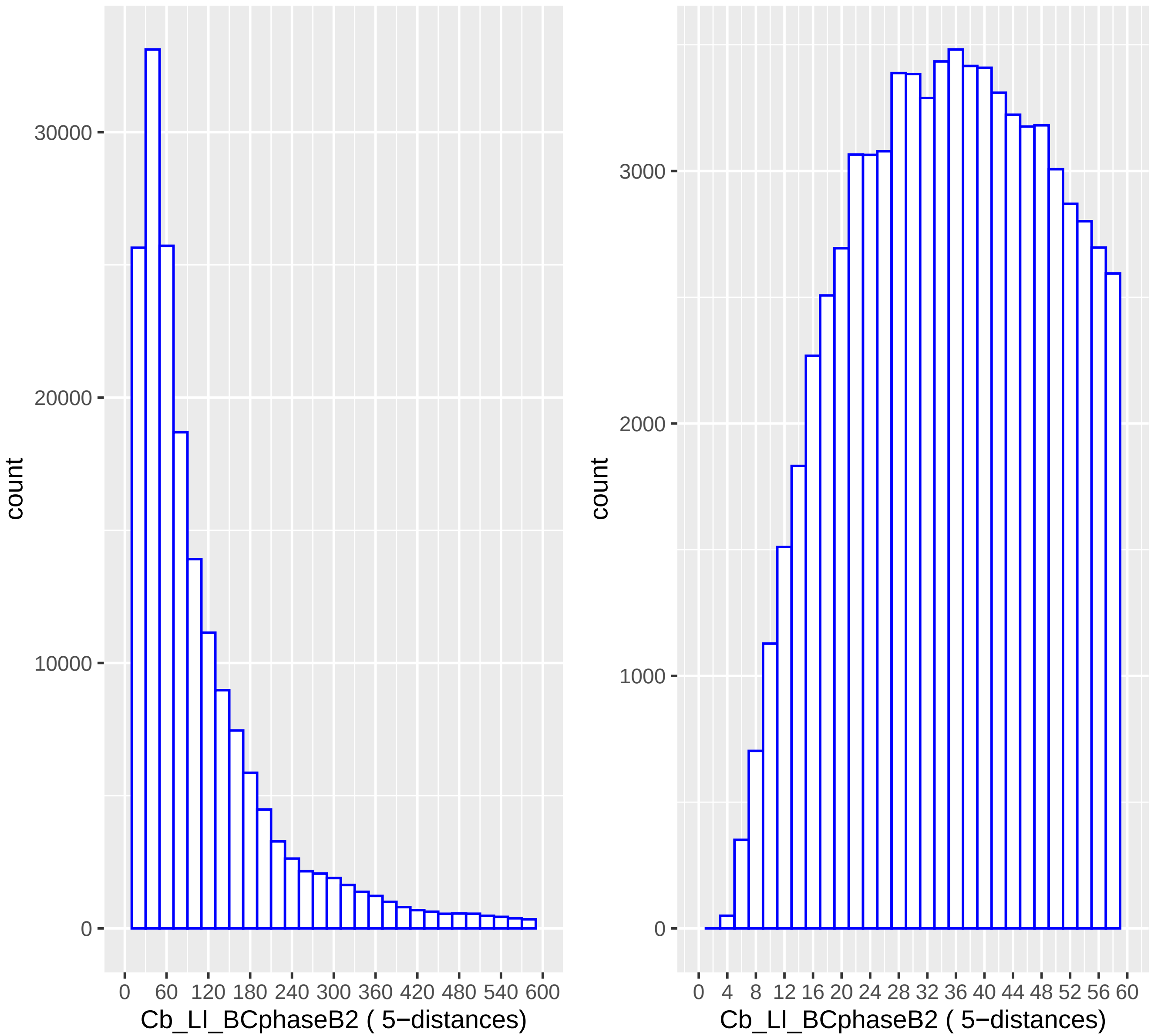

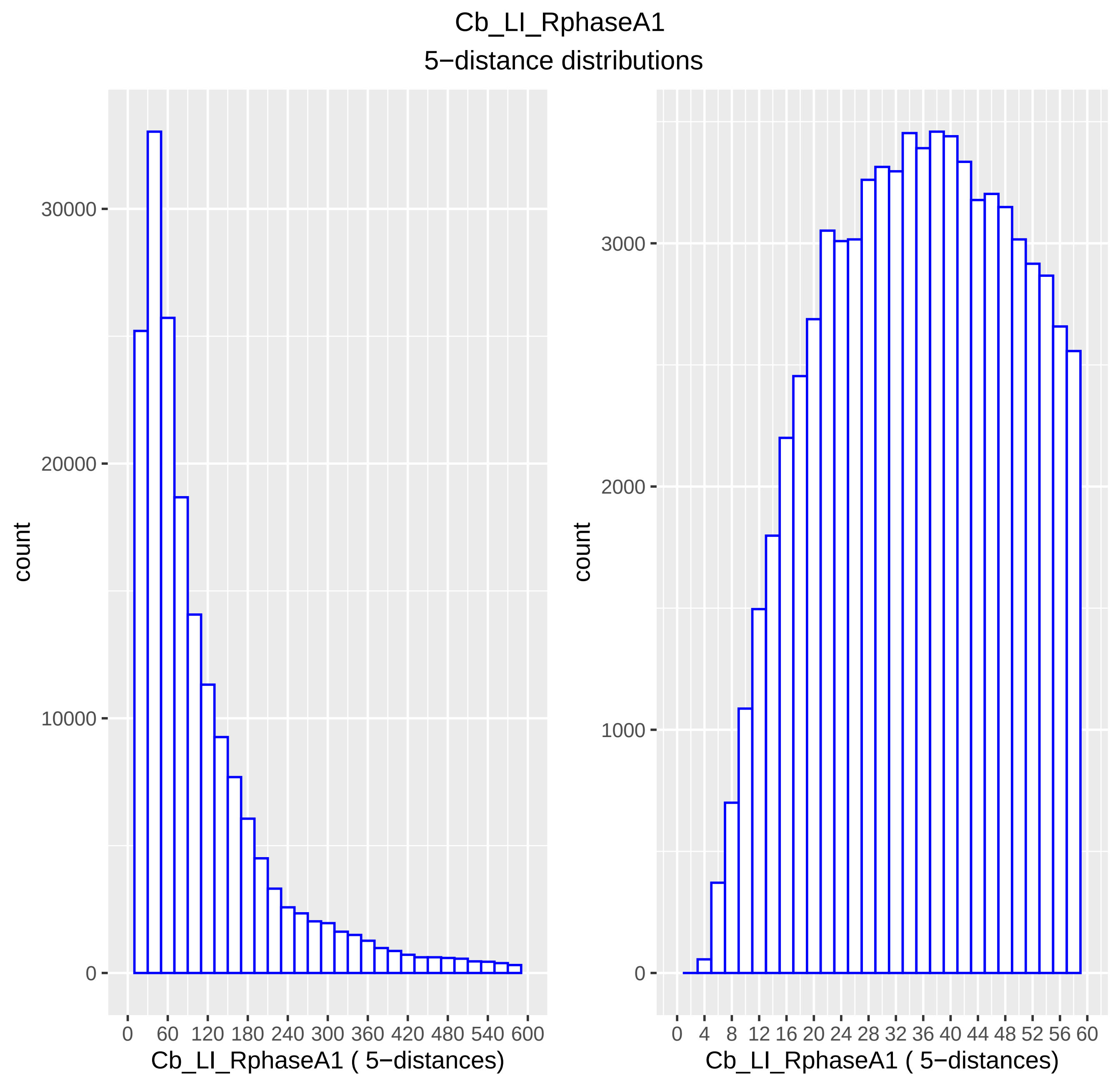
Cb_LI_RphaseA1
5−distance distributions
count
30000
20000
10000
0
0
60
120
180
240
300
360
420
480
540
600
Cb_LI_RphaseA1 ( 5−distances)
count
3000
2000
1000
0
0
4
8
12
16
20
24
28
32
36
40
44
48
52
56
60
Cb_LI_RphaseA1 ( 5−distances)

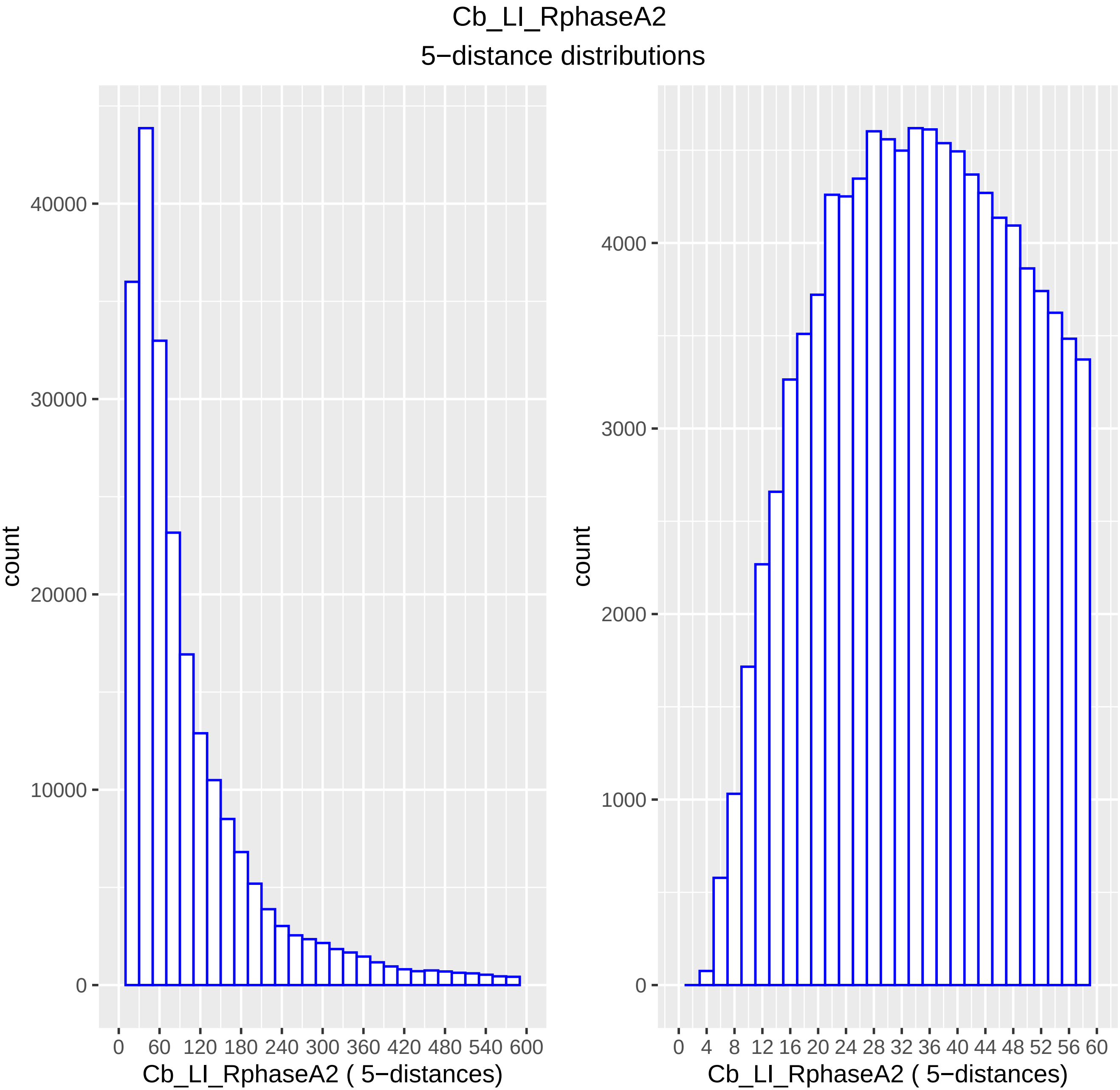

Cb_LI_RphaseA2
5−distance distributions
count
40000
30000
20000
10000
0
0
60
120
180
240
300
360
420
480
540
600
Cb_LI_RphaseA2 ( 5−distances)
count
4000
3000
2000
1000
0
0
4
8
12
16
20
24
28
32
36
40
44
48
52
56
60
Cb_LI_RphaseA2 ( 5−distances)

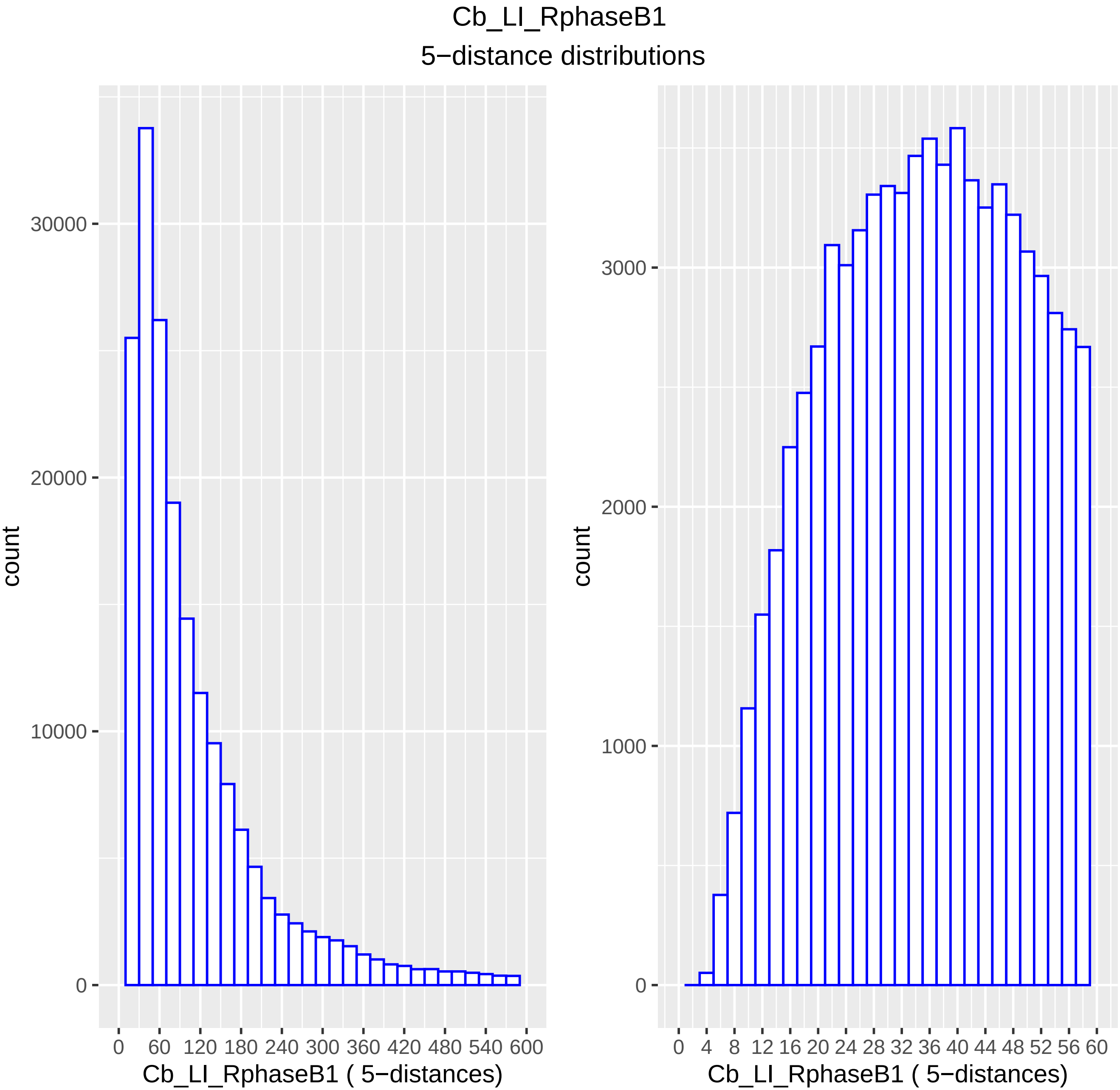

Cb_LI_RphaseB1
5−distance distributions
count
30000
20000
10000
0
0 60 120 180 240 300 360 420 480 540 600
Cb_LI_RphaseB1 ( 5−distances)
count
3000
2000
1000
0
0 4 8 12 16 20 24 28 32 36 40 44 48 52 56 60
Cb_LI_RphaseB1 ( 5−distances)

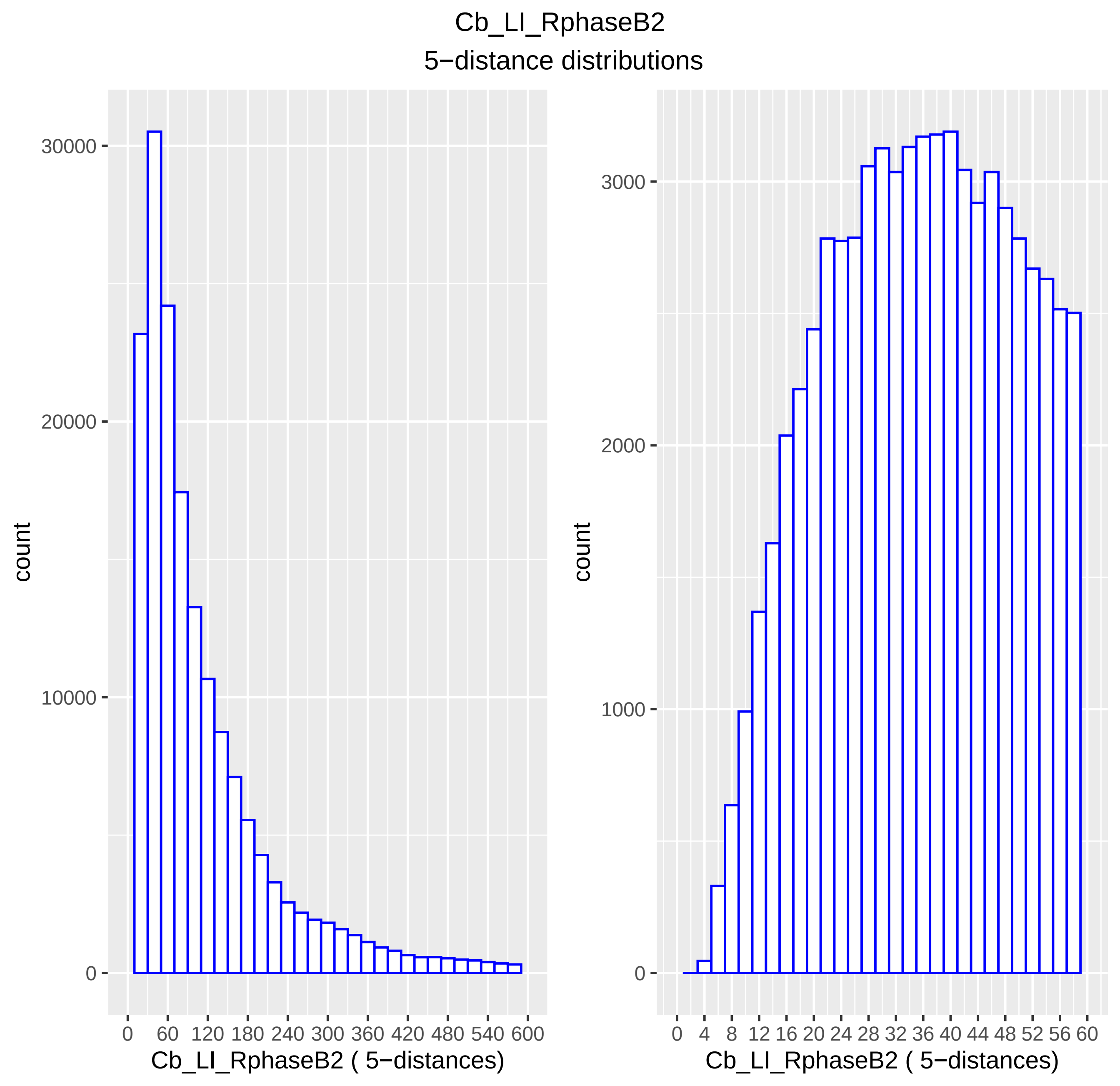

Cb_LI_RphaseB2
5−distance distributions
count
30000
20000
10000
0
0
60
120
180
240
300
360
420
480
540
600
Cb_LI_RphaseB2 ( 5−distances)
count
3000
2000
1000
0
0
4
8
12
16
20
24
28
32
36
40
44
48
52
56
60
Cb_LI_RphaseB2 ( 5−distances)

```
Directory: DMT File: 0READMEdmt

DMT - Differentially methylated tiles.

Input: studymc wsize stepsize threshold qvalue
Output: files dmt-*.txt dmg-*.txt

Notes: Output files dmt-*.txt and dmg-*.txt are generated by BWASPR::det_dmt() and show the
        differentially methylated tiles and genes as determined by methylKit::getMethylDiff with
        parameters difference=threshold and qvalue, as provided in the Cb_LI.conf
        configuration file.  Tiles refer to sliding windows along the genome within which
        methylation calls are cumulated by methylKit::tileMethylCounts.
        The dmg-*.txt files show all genes with at least one differentially methylated tile.

        A positive meth.diff value in comparison A.vs.B means that the B methylation percentage
        in that tile is higher than the A methylation percentage.
```

Directory: DMT File: dmg-Cb_LI.txt

| seqnames | start | end | width | strand | source | type | score | phase | ID | Dbxref | Name | gbkey | gene | gene_biotype | partial | start_range | end_range | comparison |
|---|---|---|---|---|---|---|---|---|---|---|---|---|---|---|---|---|---|---|
| NW_011924883.1 | 3349 | 9688 | 6340 | - | Gnomon | gene | NA | NA | gene67 | GeneID:105278822 | LOC105278822 | Gene | LOC105278822 | protein_coding | true | c(".", "3349") | character(0) | BCphaseA1.vs.BCphaseA2 |
| NW_011924922.1 | 2509149 | 2512601 | 3453 | + | Gnomon | gene | NA | NA | gene1220 | GeneID:105285742 | LOC105285742 | Gene | LOC105285742 | protein_coding | NA | character(0) | character(0) | BCphaseA1.vs.BCphaseA2 |
| NW_011924937.1 | 108248 | 110398 | 2151 | - | Gnomon | gene | NA | NA | gene1751 | GeneID:105287919 | LOC105287919 | Gene | LOC105287919 | protein_coding | NA | character(0) | character(0) | BCphaseA1.vs.BCphaseA2 |
| NW_011924946.1 | 2902307 | 2907088 | 4782 | - | Gnomon | gene | NA | NA | gene2170 | GeneID:105274589 | LOC105274589 | Gene | LOC105274589 | protein_coding | NA | character(0) | character(0) | BCphaseA1.vs.BCphaseA2 |
| NW_011924958.1 | 56714 | 58487 | 1774 | - | Gnomon | gene | NA | NA | gene2593 | GeneID:105275070 | LOC105275070 | Gene | LOC105275070 | protein_coding | NA | character(0) | character(0) | BCphaseA1.vs.BCphaseA2 |
| NW_011924977.1 | 3074052 | 3076425 | 2374 | - | Gnomon | gene | NA | NA | gene3165 | GeneID:105275706 | LOC105275706 | Gene | LOC105275706 | protein_coding | NA | character(0) | character(0) | BCphaseA1.vs.BCphaseA2 |
| NW_011924977.1 | 3614512 | 3642583 | 28072 | - | Gnomon | gene | NA | NA | gene3195 | GeneID:105275750 | LOC105275750 | Gene | LOC105275750 | protein_coding | NA | character(0) | character(0) | BCphaseA1.vs.BCphaseA2 |
| NW_011924977.1 | 3622479 | 3631189 | 8711 | - | Gnomon | gene | NA | NA | gene3196 | GeneID:105275748 | LOC105275748 | Gene | LOC105275748 | protein_coding | NA | character(0) | character(0) | BCphaseA1.vs.BCphaseA2 |
| NW_011924999.1 | 327071 | 336458 | 9388 | + | Gnomon | gene | NA | NA | gene3943 | GeneID:105276607 | LOC105276607 | Gene | LOC105276607 | protein_coding | NA | character(0) | character(0) | BCphaseA1.vs.BCphaseA2 |
| NW_011925007.1 | 3592727 | 3595751 | 3025 | + | Gnomon | gene | NA | NA | gene4366 | GeneID:105277073 | LOC105277073 | Gene | LOC105277073 | protein_coding | NA | character(0) | character(0) | BCphaseA1.vs.BCphaseA2 |
| NW_011925035.1 | 680867 | 687414 | 6548 | - | Gnomon | gene | NA | NA | gene5372 | GeneID:105278225 | LOC105278225 | Gene | LOC105278225 | protein_coding | NA | character(0) | character(0) | BCphaseA1.vs.BCphaseA2 |
| NW_011925061.1 | 83423 | 87193 | 3771 | + | Gnomon | gene | NA | NA | gene5985 | GeneID:105278934 | LOC105278934 | Gene | LOC105278934 | protein_coding | NA | character(0) | character(0) | BCphaseA1.vs.BCphaseA2 |
| NW_011925099.1 | 1724294 | 1725379 | 1086 | - | Gnomon | gene | NA | NA | gene6490 | GeneID:105279538 | LOC105279538 | Gene | LOC105279538 | protein_coding | NA | character(0) | character(0) | BCphaseA1.vs.BCphaseA2 |
| NW_011925106.1 | 3271050 | 3274726 | 3677 | + | Gnomon | gene | NA | NA | gene6815 | GeneID:105279899 | LOC105279899 | Gene | LOC105279899 | protein_coding | NA | character(0) | character(0) | BCphaseA1.vs.BCphaseA2 |
| NW_011925129.1 | 197807 | 211835 | 14029 | + | Gnomon | gene | NA | NA | gene7333 | GeneID:105280517 | LOC105280517 | Gene | LOC105280517 | protein_coding | NA | character(0) | character(0) | BCphaseA1.vs.BCphaseA2 |
| NW_011925162.1 | 17698 | 22041 | 4344 | - | Gnomon | gene | NA | NA | gene7987 | GeneID:105281280 | LOC105281280 | Gene | LOC105281280 | protein_coding | NA | character(0) | character(0) | BCphaseA1.vs.BCphaseA2 |
| NW_011925162.1 | 25062 | 28298 | 3237 | + | Gnomon | gene | NA | NA | gene7988 | GeneID:105281282 | LOC105281282 | Gene | LOC105281282 | protein_coding | NA | character(0) | character(0) | BCphaseA1.vs.BCphaseA2 |
| NW_011925163.1 | 1798313 | 1800209 | 1897 | + | Gnomon | gene | NA | NA | gene8111 | GeneID:105281418 | LOC105281418 | Gene | LOC105281418 | protein_coding | NA | character(0) | character(0) | BCphaseA1.vs.BCphaseA2 |
| NW_011925178.1 | 158465 | 173389 | 14925 | + | Gnomon | gene | NA | NA | gene8181 | GeneID:105281520 | LOC105281520 | Gene | LOC105281520 | protein_coding | NA | character(0) | character(0) | BCphaseA1.vs.BCphaseA2 |
| NW_011925232.1 | 185642 | 187490 | 1849 | - | Gnomon | gene | NA | NA | gene8929 | GeneID:105282371 | LOC105282371 | Gene | LOC105282371 | lncRNA | NA | character(0) | character(0) | BCphaseA1.vs.BCphaseA2 |
| NW_011925279.1 | 1541 | 25639 | 24099 | - | Gnomon | gene | NA | NA | gene9540 | GeneID:105283067 | LOC105283067 | Gene | LOC105283067 | protein_coding | NA | character(0) | character(0) | BCphaseA1.vs.BCphaseA2 |
| NW_011925279.1 | 14660 | 16170 | 1511 | + | Gnomon | gene | NA | NA | gene9542 | GeneID:105283065 | LOC105283065 | Gene | LOC105283065 | protein_coding | NA | character(0) | character(0) | BCphaseA1.vs.BCphaseA2 |
| NW_011925279.1 | 16317 | 25727 | 9411 | + | Gnomon | gene | NA | NA | gene9543 | GeneID:105283064 | LOC105283064 | Gene | LOC105283064 | protein_coding | NA | character(0) | character(0) | BCphaseA1.vs.BCphaseA2 |
| NW_011925284.1 | 24112 | 26626 | 2515 | - | Gnomon | gene | NA | NA | gene9559 | GeneID:105283091 | LOC105283091 | Gene | LOC105283091 | protein_coding | NA | character(0) | character(0) | BCphaseA1.vs.BCphaseA2 |

...
... only first 25 lines shown ...

Directory: DMT File: dmt-Cb_LI.txt

| seqnames | start | end | width | strand | pvalue | qvalue | meth.diff | comparison |
|---|---|---|---|---|---|---|---|---|
| NW_011924883.1 | 7001 | 7500 | 500 | * | 2.64917946375975e-06 | 0.007 | -63.83 | BCphaseA1.vs.BCphaseA2 |
| NW_011924914.1 | 216001 | 216500 | 500 | * | 1.49638979984964e-06 | 0.005 | -20.12 | BCphaseA1.vs.BCphaseA2 |
| NW_011924922.1 | 2510001 | 2510500 | 500 | * | 2.3284773946768e-06 | 0.006 | -23.91 | BCphaseA1.vs.BCphaseA2 |
| NW_011924937.1 | 109501 | 110000 | 500 | * | 3.39968732997585e-06 | 0.008 | -26.77 | BCphaseA1.vs.BCphaseA2 |
| NW_011924946.1 | 2906001 | 2906500 | 500 | * | 6.18182354927382e-07 | 0.002 | -23.37 | BCphaseA1.vs.BCphaseA2 |
| NW_011924958.1 | 57001 | 57500 | 500 | * | 1.45771221586136e-47 | 0 | 47.29 | BCphaseA1.vs.BCphaseA2 |
| NW_011924977.1 | 3074501 | 3075000 | 500 | * | 6.24914032067485e-32 | 0 | 31.78 | BCphaseA1.vs.BCphaseA2 |
| NW_011924977.1 | 3629501 | 3630000 | 500 | * | 8.90735365559623e-07 | 0.003 | 20.35 | BCphaseA1.vs.BCphaseA2 |
| NW_011924999.1 | 328001 | 328500 | 500 | * | 4.41028471943773e-05 | 0.05 | 21.79 | BCphaseA1.vs.BCphaseA2 |
| NW_011925007.1 | 3593501 | 3594000 | 500 | * | 1.27809653753215e-06 | 0.004 | -37.87 | BCphaseA1.vs.BCphaseA2 |
| NW_011925027.1 | 1407501 | 1408000 | 500 | * | 1.95048342571818e-05 | 0.029 | 21.81 | BCphaseA1.vs.BCphaseA2 |
| NW_011925035.1 | 686001 | 686500 | 500 | * | 2.60456775596222e-07 | 0.001 | -29.95 | BCphaseA1.vs.BCphaseA2 |
| NW_011925061.1 | 84001 | 84500 | 500 | * | 1.77340059543087e-07 | 0.001 | 23.8 | BCphaseA1.vs.BCphaseA2 |
| NW_011925099.1 | 1724001 | 1724500 | 500 | * | 4.27870698795725e-05 | 0.049 | 28.41 | BCphaseA1.vs.BCphaseA2 |
| NW_011925106.1 | 3271501 | 3272000 | 500 | * | 3.03811908469588e-05 | 0.039 | 45.06 | BCphaseA1.vs.BCphaseA2 |
| NW_011925129.1 | 199001 | 199500 | 500 | * | 1.88243830210409e-11 | 0 | 28.7 | BCphaseA1.vs.BCphaseA2 |
| NW_011925162.1 | 20501 | 21000 | 500 | * | 1.1145838772096e-06 | 0.004 | 25.31 | BCphaseA1.vs.BCphaseA2 |
| NW_011925162.1 | 27001 | 27500 | 500 | * | 5.16376754306592e-06 | 0.011 | 22.36 | BCphaseA1.vs.BCphaseA2 |
| NW_011925163.1 | 1798501 | 1799000 | 500 | * | 9.96175785983529e-09 | 0 | 29.99 | BCphaseA1.vs.BCphaseA2 |
| NW_011925178.1 | 161001 | 161500 | 500 | * | 2.67240731206555e-09 | 0 | 29.62 | BCphaseA1.vs.BCphaseA2 |
| NW_011925178.1 | 226501 | 227000 | 500 | * | 5.16071959899933e-18 | 0 | 22.67 | BCphaseA1.vs.BCphaseA2 |
| NW_011925215.1 | 700501 | 701000 | 500 | * | 4.66154773867494e-07 | 0.002 | -37.82 | BCphaseA1.vs.BCphaseA2 |
| NW_011925232.1 | 186001 | 186500 | 500 | * | 8.13054622215717e-09 | 0 | 31.79 | BCphaseA1.vs.BCphaseA2 |
| NW_011925279.1 | 16001 | 16500 | 500 | * | 1.27073794544147e-06 | 0.004 | 36.64 | BCphaseA1.vs.BCphaseA2 |

...
... only first 25 lines shown ...

```
Directory: DMSG File: 0READMEdmsg

DMSG - Differentially methylated sites and genes.

Input: studyhc threshold qvalue
       The studyhc methylKit raw object contains all CpGscd sites with coverage at least
        20 reads.
Output: files dms-*.txt dmg-*.txt  dmg-*details.txt dmg-*heatmaps.pdf

Notes: Output files dms-*.txt and dmg-*.txt are generated by BWASPR::det_dmsg() and show the
        differentially methylated sites and genes as determined by methylKit::getMethylDiff with
        parameters difference=threshold and qvalue, as provided in the Cb_LI.conf
        configuration file.  The table of differentially methylated genes contains all genes
        with at least one differentially methylated site.

       The files dmg-*details.txt, generated by BWASPR::show_dmsg(), show all CpGscd sites in
        the differentially methylated genes, with coverage numbers and methylation percentages.

       The files dmg-*heatmaps.pdf, generated by BWASPR::show_dmsg(), give heatmap displays for
        genes meeting the following criteria: 1) there are between minNsites and maxNsites common
        CpGscd sites; 2) at least minPdmsites % of these sites are differentially methylated sites.
        The parameters are set in the Cb_LI.conf configuration file.
```

Directory: DMSG File: dmg-Cb_LI.txt

| seqnames | start | end | width | strand | source | type | score | phase | ID | Dbxref | Name | gbkey | gene | gene_biotype | partial | start_range | end_range | comparison |
|---|---|---|---|---|---|---|---|---|---|---|---|---|---|---|---|---|---|---|
| NW_011924881.1 | 808343 | 827625 | 19283 | - | Gnomon | gene | NA | NA | gene53 | GeneID:105277148 | LOC105277148 | Gene | LOC105277148 | protein_coding | NA | character(0) | character(0) | BCphaseA1.vs.BCphaseA2 |
| NW_011924888.1 | 70798 | 74285 | 3488 | - | Gnomon | gene | NA | NA | gene274 | GeneID:105275190 | LOC105275190 | Gene | LOC105275190 | protein_coding | NA | character(0) | character(0) | BCphaseA1.vs.BCphaseA2 |
| NW_011924888.1 | 140602 | 143426 | 2825 | - | Gnomon | gene | NA | NA | gene292 | GeneID:105275398 | LOC105275398 | Gene | LOC105275398 | protein_coding | NA | character(0) | character(0) | BCphaseA1.vs.BCphaseA2 |
| NW_011924888.1 | 164054 | 166526 | 2473 | - | Gnomon | gene | NA | NA | gene298 | GeneID:105275460 | LOC105275460 | Gene | LOC105275460 | protein_coding | NA | character(0) | character(0) | BCphaseA1.vs.BCphaseA2 |
| NW_011924888.1 | 246411 | 249491 | 3081 | - | Gnomon | gene | NA | NA | gene303 | GeneID:105275584 | LOC105275584 | Gene | LOC105275584 | protein_coding | NA | character(0) | character(0) | BCphaseA1.vs.BCphaseA2 |
| NW_011924888.1 | 407815 | 409420 | 1606 | - | Gnomon | gene | NA | NA | gene322 | GeneID:105275777 | LOC105275777 | Gene | LOC105275777 | protein_coding | NA | character(0) | character(0) | BCphaseA1.vs.BCphaseA2 |
| NW_011924888.1 | 1582383 | 1583940 | 1558 | + | Gnomon | gene | NA | NA | gene388 | GeneID:105276538 | LOC105276538 | Gene | LOC105276538 | protein_coding | NA | character(0) | character(0) | BCphaseA1.vs.BCphaseA2 |
| NW_011924888.1 | 1598262 | 1599384 | 1123 | - | Gnomon | gene | NA | NA | gene393 | GeneID:105276566 | LOC105276566 | Gene | LOC105276566 | protein_coding | NA | character(0) | character(0) | BCphaseA1.vs.BCphaseA2 |
| NW_011924889.1 | 214123 | 217747 | 3625 | + | Gnomon | gene | NA | NA | gene474 | GeneID:105277435 | LOC105277435 | Gene | LOC105277435 | protein_coding | NA | character(0) | character(0) | BCphaseA1.vs.BCphaseA2 |
| NW_011924899.1 | 335137 | 341557 | 6421 | - | Gnomon | gene | NA | NA | gene657 | GeneID:105279667 | LOC105279667 | Gene | LOC105279667 | protein_coding | NA | character(0) | character(0) | BCphaseA1.vs.BCphaseA2 |
| NW_011924914.1 | 109715 | 110911 | 1197 | + | Gnomon | gene | NA | NA | gene920 | GeneID:105282518 | LOC105282518 | Gene | LOC105282518 | protein_coding | NA | character(0) | character(0) | BCphaseA1.vs.BCphaseA2 |
| NW_011924914.1 | 216572 | 220149 | 3578 | + | Gnomon | gene | NA | NA | gene940 | GeneID:105282794 | LOC105282794 | Gene | LOC105282794 | protein_coding | NA | character(0) | character(0) | BCphaseA1.vs.BCphaseA2 |
| NW_011924922.1 | 2078651 | 2082672 | 4022 | + | Gnomon | gene | NA | NA | gene1147 | GeneID:105284954 | LOC105284954 | Gene | LOC105284954 | protein_coding | NA | character(0) | character(0) | BCphaseA1.vs.BCphaseA2 |
| NW_011924922.1 | 2392503 | 2394722 | 2220 | - | Gnomon | gene | NA | NA | gene1199 | GeneID:105285498 | LOC105285498 | Gene | LOC105285498 | protein_coding | NA | character(0) | character(0) | BCphaseA1.vs.BCphaseA2 |
| NW_011924922.1 | 2509149 | 2512601 | 3453 | + | Gnomon | gene | NA | NA | gene1220 | GeneID:105285742 | LOC105285742 | Gene | LOC105285742 | protein_coding | NA | character(0) | character(0) | BCphaseA1.vs.BCphaseA2 |
| NW_011924929.1 | 732440 | 735021 | 2582 | + | Gnomon | gene | NA | NA | gene1310 | GeneID:105286661 | LOC105286661 | Gene | LOC105286661 | protein_coding | NA | character(0) | character(0) | BCphaseA1.vs.BCphaseA2 |
| NW_011924931.1 | 2028839 | 2031812 | 2974 | + | Gnomon | gene | NA | NA | gene1470 | GeneID:105287597 | LOC105287597 | Gene | LOC105287597 | protein_coding | NA | character(0) | character(0) | BCphaseA1.vs.BCphaseA2 |
| NW_011924945.1 | 67984 | 72152 | 4169 | + | Gnomon | gene | NA | NA | gene1884 | GeneID:105288072 | LOC105288072 | Gene | LOC105288072 | protein_coding | NA | character(0) | character(0) | BCphaseA1.vs.BCphaseA2 |
| NW_011924946.1 | 775753 | 777201 | 1449 | - | Gnomon | gene | NA | NA | gene1993 | GeneID:105288193 | LOC105288193 | Gene | LOC105288193 | protein_coding | NA | character(0) | character(0) | BCphaseA1.vs.BCphaseA2 |
| NW_011924946.1 | 1543495 | 1550078 | 6584 | - | Gnomon | gene | NA | NA | gene2056 | GeneID:105274452 | LOC105274452 | Gene | LOC105274452 | protein_coding | NA | character(0) | character(0) | BCphaseA1.vs.BCphaseA2 |
| NW_011924946.1 | 2914862 | 2919248 | 4387 | - | Gnomon | gene | NA | NA | gene2173 | GeneID:105274590 | LOC105274590 | Gene | LOC105274590 | protein_coding | NA | character(0) | character(0) | BCphaseA1.vs.BCphaseA2 |
| NW_011924947.1 | 85348 | 86795 | 1448 | + | Gnomon | gene | NA | NA | gene2271 | GeneID:105274706 | LOC105274706 | Gene | LOC105274706 | protein_coding | NA | character(0) | character(0) | BCphaseA1.vs.BCphaseA2 |
| NW_011924953.1 | 416151 | 417917 | 1767 | - | Gnomon | gene | NA | NA | gene2404 | GeneID:105274853 | LOC105274853 | Gene | LOC105274853 | protein_coding | NA | character(0) | character(0) | BCphaseA1.vs.BCphaseA2 |
| NW_011924958.1 | 56714 | 58487 | 1774 | - | Gnomon | gene | NA | NA | gene2593 | GeneID:105275070 | LOC105275070 | Gene | LOC105275070 | protein_coding | NA | character(0) | character(0) | BCphaseA1.vs.BCphaseA2 |

...
... only first 25 lines shown ...

Directory: DMSG File: dmg-Cb_LI_BCphaseA1.vs.BCphaseA2_details.txt

| seqnames | start | end | width | strand | coverage1 | numCs1 | numTs1 | coverage2 | numCs2 | numTs2 | BCphaseA1 | BCphaseA2 | gene_seqnames | gene_start | gene_end | gene_width | gene_strand | gene_source | gene_type | gene_ID | gene_Dbxref | gene_Name | gene_gbkey | gene_gene | gene_gene_biotype | gene_start_range | gene_end_range | gene_comparison | is.dm |
|---|---|---|---|---|---|---|---|---|---|---|---|---|---|---|---|---|---|---|---|---|---|---|---|---|---|---|---|---|---|
| NW_011924881.1 | 809481 | 809481 | 1 | + | 37 | 0 | 37 | 20 | 0 | 20 | 0 | 0 | NW_011924881.1 | 808343 | 827625 | 19283 | - | Gnomon | gene | gene53 | GeneID:105277148 | LOC105277148 | Gene | LOC105277148 | protein_coding | character(0) | character(0) | BCphaseA1.vs.BCphaseA2 | FALSE |
| NW_011924881.1 | 809504 | 809504 | 1 | + | 38 | 1 | 37 | 20 | 0 | 20 | 2.63 | 0 | NW_011924881.1 | 808343 | 827625 | 19283 | - | Gnomon | gene | gene53 | GeneID:105277148 | LOC105277148 | Gene | LOC105277148 | protein_coding | character(0) | character(0) | BCphaseA1.vs.BCphaseA2 | FALSE |
| NW_011924881.1 | 810155 | 810155 | 1 | + | 29 | 0 | 29 | 22 | 0 | 22 | 0 | 0 | NW_011924881.1 | 808343 | 827625 | 19283 | - | Gnomon | gene | gene53 | GeneID:105277148 | LOC105277148 | Gene | LOC105277148 | protein_coding | character(0) | character(0) | BCphaseA1.vs.BCphaseA2 | FALSE |
| NW_011924881.1 | 810182 | 810182 | 1 | + | 33 | 0 | 33 | 23 | 0 | 23 | 0 | 0 | NW_011924881.1 | 808343 | 827625 | 19283 | - | Gnomon | gene | gene53 | GeneID:105277148 | LOC105277148 | Gene | LOC105277148 | protein_coding | character(0) | character(0) | BCphaseA1.vs.BCphaseA2 | FALSE |
| NW_011924881.1 | 810335 | 810335 | 1 | + | 20 | 0 | 20 | 23 | 0 | 23 | 0 | 0 | NW_011924881.1 | 808343 | 827625 | 19283 | - | Gnomon | gene | gene53 | GeneID:105277148 | LOC105277148 | Gene | LOC105277148 | protein_coding | character(0) | character(0) | BCphaseA1.vs.BCphaseA2 | FALSE |
| NW_011924881.1 | 810337 | 810337 | 1 | + | 21 | 0 | 21 | 24 | 0 | 24 | 0 | 0 | NW_011924881.1 | 808343 | 827625 | 19283 | - | Gnomon | gene | gene53 | GeneID:105277148 | LOC105277148 | Gene | LOC105277148 | protein_coding | character(0) | character(0) | BCphaseA1.vs.BCphaseA2 | FALSE |
| NW_011924881.1 | 810348 | 810348 | 1 | + | 35 | 0 | 35 | 51 | 0 | 51 | 0 | 0 | NW_011924881.1 | 808343 | 827625 | 19283 | - | Gnomon | gene | gene53 | GeneID:105277148 | LOC105277148 | Gene | LOC105277148 | protein_coding | character(0) | character(0) | BCphaseA1.vs.BCphaseA2 | FALSE |
| NW_011924881.1 | 810391 | 810391 | 1 | + | 37 | 0 | 37 | 54 | 1 | 53 | 0 | 1.85 | NW_011924881.1 | 808343 | 827625 | 19283 | - | Gnomon | gene | gene53 | GeneID:105277148 | LOC105277148 | Gene | LOC105277148 | protein_coding | character(0) | character(0) | BCphaseA1.vs.BCphaseA2 | FALSE |
| NW_011924881.1 | 810409 | 810409 | 1 | + | 55 | 0 | 55 | 61 | 2 | 59 | 0 | 3.28 | NW_011924881.1 | 808343 | 827625 | 19283 | - | Gnomon | gene | gene53 | GeneID:105277148 | LOC105277148 | Gene | LOC105277148 | protein_coding | character(0) | character(0) | BCphaseA1.vs.BCphaseA2 | FALSE |
| NW_011924881.1 | 810412 | 810412 | 1 | + | 30 | 0 | 30 | 40 | 0 | 40 | 0 | 0 | NW_011924881.1 | 808343 | 827625 | 19283 | - | Gnomon | gene | gene53 | GeneID:105277148 | LOC105277148 | Gene | LOC105277148 | protein_coding | character(0) | character(0) | BCphaseA1.vs.BCphaseA2 | FALSE |
| NW_011924881.1 | 810417 | 810417 | 1 | + | 31 | 0 | 31 | 38 | 1 | 37 | 0 | 2.63 | NW_011924881.1 | 808343 | 827625 | 19283 | - | Gnomon | gene | gene53 | GeneID:105277148 | LOC105277148 | Gene | LOC105277148 | protein_coding | character(0) | character(0) | BCphaseA1.vs.BCphaseA2 | FALSE |
| NW_011924881.1 | 810424 | 810424 | 1 | + | 46 | 1 | 45 | 37 | 0 | 37 | 2.17 | 0 | NW_011924881.1 | 808343 | 827625 | 19283 | - | Gnomon | gene | gene53 | GeneID:105277148 | LOC105277148 | Gene | LOC105277148 | protein_coding | character(0) | character(0) | BCphaseA1.vs.BCphaseA2 | FALSE |
| NW_011924881.1 | 810430 | 810430 | 1 | + | 26 | 1 | 25 | 27 | 0 | 27 | 3.85 | 0 | NW_011924881.1 | 808343 | 827625 | 19283 | - | Gnomon | gene | gene53 | GeneID:105277148 | LOC105277148 | Gene | LOC105277148 | protein_coding | character(0) | character(0) | BCphaseA1.vs.BCphaseA2 | FALSE |
| NW_011924881.1 | 810436 | 810436 | 1 | + | 52 | 2 | 50 | 27 | 0 | 27 | 3.85 | 0 | NW_011924881.1 | 808343 | 827625 | 19283 | - | Gnomon | gene | gene53 | GeneID:105277148 | LOC105277148 | Gene | LOC105277148 | protein_coding | character(0) | character(0) | BCphaseA1.vs.BCphaseA2 | FALSE |
| NW_011924881.1 | 810505 | 810505 | 1 | + | 60 | 0 | 60 | 22 | 0 | 22 | 0 | 0 | NW_011924881.1 | 808343 | 827625 | 19283 | - | Gnomon | gene | gene53 | GeneID:105277148 | LOC105277148 | Gene | LOC105277148 | protein_coding | character(0) | character(0) | BCphaseA1.vs.BCphaseA2 | FALSE |
| NW_011924881.1 | 810511 | 810511 | 1 | + | 39 | 0 | 39 | 22 | 1 | 21 | 0 | 4.55 | NW_011924881.1 | 808343 | 827625 | 19283 | - | Gnomon | gene | gene53 | GeneID:105277148 | LOC105277148 | Gene | LOC105277148 | protein_coding | character(0) | character(0) | BCphaseA1.vs.BCphaseA2 | FALSE |
| NW_011924881.1 | 810516 | 810516 | 1 | + | 38 | 0 | 38 | 23 | 0 | 23 | 0 | 0 | NW_011924881.1 | 808343 | 827625 | 19283 | - | Gnomon | gene | gene53 | GeneID:105277148 | LOC105277148 | Gene | LOC105277148 | protein_coding | character(0) | character(0) | BCphaseA1.vs.BCphaseA2 | FALSE |
| NW_011924881.1 | 810520 | 810520 | 1 | + | 39 | 0 | 39 | 23 | 0 | 23 | 0 | 0 | NW_011924881.1 | 808343 | 827625 | 19283 | - | Gnomon | gene | gene53 | GeneID:105277148 | LOC105277148 | Gene | LOC105277148 | protein_coding | character(0) | character(0) | BCphaseA1.vs.BCphaseA2 | FALSE |
| NW_011924881.1 | 810633 | 810633 | 1 | + | 28 | 0 | 28 | 20 | 0 | 20 | 0 | 0 | NW_011924881.1 | 808343 | 827625 | 19283 | - | Gnomon | gene | gene53 | GeneID:105277148 | LOC105277148 | Gene | LOC105277148 | protein_coding | character(0) | character(0) | BCphaseA1.vs.BCphaseA2 | FALSE |
| NW_011924881.1 | 810645 | 810645 | 1 | + | 49 | 0 | 49 | 21 | 0 | 21 | 0 | 0 | NW_011924881.1 | 808343 | 827625 | 19283 | - | Gnomon | gene | gene53 | GeneID:105277148 | LOC105277148 | Gene | LOC105277148 | protein_coding | character(0) | character(0) | BCphaseA1.vs.BCphaseA2 | FALSE |
| NW_011924881.1 | 811042 | 811042 | 1 | + | 23 | 0 | 23 | 21 | 0 | 21 | 0 | 0 | NW_011924881.1 | 808343 | 827625 | 19283 | - | Gnomon | gene | gene53 | GeneID:105277148 | LOC105277148 | Gene | LOC105277148 | protein_coding | character(0) | character(0) | BCphaseA1.vs.BCphaseA2 | FALSE |
| NW_011924881.1 | 811083 | 811083 | 1 | + | 21 | 0 | 21 | 26 | 0 | 26 | 0 | 0 | NW_011924881.1 | 808343 | 827625 | 19283 | - | Gnomon | gene | gene53 | GeneID:105277148 | LOC105277148 | Gene | LOC105277148 | protein_coding | character(0) | character(0) | BCphaseA1.vs.BCphaseA2 | FALSE |
| NW_011924881.1 | 811089 | 811089 | 1 | + | 20 | 0 | 20 | 27 | 0 | 27 | 0 | 0 | NW_011924881.1 | 808343 | 827625 | 19283 | - | Gnomon | gene | gene53 | GeneID:105277148 | LOC105277148 | Gene | LOC105277148 | protein_coding | character(0) | character(0) | BCphaseA1.vs.BCphaseA2 | FALSE |
| NW_011924881.1 | 811424 | 811424 | 1 | + | 23 | 0 | 23 | 31 | 2 | 29 | 0 | 6.45 | NW_011924881.1 | 808343 | 827625 | 19283 | - | Gnomon | gene | gene53 | GeneID:105277148 | LOC105277148 | Gene | LOC105277148 | protein_coding | character(0) | character(0) | BCphaseA1.vs.BCphaseA2 | FALSE |

...
... only first 25 lines shown ...

Directory: DMSG File: dmg-Cb_LI_BCphaseA1.vs.BCphaseB1_details.txt

| seqnames | start | end | width | strand | coverage1 | numCs1 | numTs1 | coverage2 | numCs2 | numTs2 | BCphaseA1 | BCphaseB1 | gene_seqnames | gene_start | gene_end | gene_width | gene_strand | gene_source | gene_type | gene_ID | gene_Dbxref | gene_Name | gene_gbkey | gene_gene | gene_gene_biotype | gene_start_range | gene_end_range | gene_comparison | is.dm |
|---|---|---|---|---|---|---|---|---|---|---|---|---|---|---|---|---|---|---|---|---|---|---|---|---|---|---|---|---|---|
| NW_011924881.1 | 809447 | 809447 | 1 | + | 28 | 0 | 28 | 25 | 0 | 25 | 0 | 0 | NW_011924881.1 | 808343 | 827625 | 19283 | - | Gnomon | gene | gene53 | GeneID:105277148 | LOC105277148 | Gene | LOC105277148 | protein_coding | character(0) | character(0) | BCphaseA1.vs.BCphaseB1 | FALSE |
| NW_011924881.1 | 809481 | 809481 | 1 | + | 37 | 0 | 37 | 34 | 1 | 33 | 0 | 2.94 | NW_011924881.1 | 808343 | 827625 | 19283 | - | Gnomon | gene | gene53 | GeneID:105277148 | LOC105277148 | Gene | LOC105277148 | protein_coding | character(0) | character(0) | BCphaseA1.vs.BCphaseB1 | FALSE |
| NW_011924881.1 | 809504 | 809504 | 1 | + | 38 | 1 | 37 | 34 | 0 | 34 | 2.63 | 0 | NW_011924881.1 | 808343 | 827625 | 19283 | - | Gnomon | gene | gene53 | GeneID:105277148 | LOC105277148 | Gene | LOC105277148 | protein_coding | character(0) | character(0) | BCphaseA1.vs.BCphaseB1 | FALSE |
| NW_011924881.1 | 809520 | 809520 | 1 | + | 31 | 0 | 31 | 30 | 0 | 30 | 0 | 0 | NW_011924881.1 | 808343 | 827625 | 19283 | - | Gnomon | gene | gene53 | GeneID:105277148 | LOC105277148 | Gene | LOC105277148 | protein_coding | character(0) | character(0) | BCphaseA1.vs.BCphaseB1 | FALSE |
| NW_011924881.1 | 810123 | 810123 | 1 | + | 26 | 0 | 26 | 21 | 0 | 21 | 0 | 0 | NW_011924881.1 | 808343 | 827625 | 19283 | - | Gnomon | gene | gene53 | GeneID:105277148 | LOC105277148 | Gene | LOC105277148 | protein_coding | character(0) | character(0) | BCphaseA1.vs.BCphaseB1 | FALSE |
| NW_011924881.1 | 810155 | 810155 | 1 | + | 29 | 0 | 29 | 23 | 0 | 23 | 0 | 0 | NW_011924881.1 | 808343 | 827625 | 19283 | - | Gnomon | gene | gene53 | GeneID:105277148 | LOC105277148 | Gene | LOC105277148 | protein_coding | character(0) | character(0) | BCphaseA1.vs.BCphaseB1 | FALSE |
| NW_011924881.1 | 810182 | 810182 | 1 | + | 33 | 0 | 33 | 25 | 3 | 22 | 0 | 12 | NW_011924881.1 | 808343 | 827625 | 19283 | - | Gnomon | gene | gene53 | GeneID:105277148 | LOC105277148 | Gene | LOC105277148 | protein_coding | character(0) | character(0) | BCphaseA1.vs.BCphaseB1 | FALSE |
| NW_011924881.1 | 810337 | 810337 | 1 | + | 21 | 0 | 21 | 22 | 0 | 22 | 0 | 0 | NW_011924881.1 | 808343 | 827625 | 19283 | - | Gnomon | gene | gene53 | GeneID:105277148 | LOC105277148 | Gene | LOC105277148 | protein_coding | character(0) | character(0) | BCphaseA1.vs.BCphaseB1 | FALSE |
| NW_011924881.1 | 810348 | 810348 | 1 | + | 35 | 0 | 35 | 42 | 1 | 41 | 0 | 2.38 | NW_011924881.1 | 808343 | 827625 | 19283 | - | Gnomon | gene | gene53 | GeneID:105277148 | LOC105277148 | Gene | LOC105277148 | protein_coding | character(0) | character(0) | BCphaseA1.vs.BCphaseB1 | FALSE |
| NW_011924881.1 | 810391 | 810391 | 1 | + | 37 | 0 | 37 | 68 | 1 | 67 | 0 | 1.47 | NW_011924881.1 | 808343 | 827625 | 19283 | - | Gnomon | gene | gene53 | GeneID:105277148 | LOC105277148 | Gene | LOC105277148 | protein_coding | character(0) | character(0) | BCphaseA1.vs.BCphaseB1 | FALSE |
| NW_011924881.1 | 810409 | 810409 | 1 | + | 55 | 0 | 55 | 59 | 0 | 59 | 0 | 0 | NW_011924881.1 | 808343 | 827625 | 19283 | - | Gnomon | gene | gene53 | GeneID:105277148 | LOC105277148 | Gene | LOC105277148 | protein_coding | character(0) | character(0) | BCphaseA1.vs.BCphaseB1 | FALSE |
| NW_011924881.1 | 810412 | 810412 | 1 | + | 30 | 0 | 30 | 34 | 1 | 33 | 0 | 2.94 | NW_011924881.1 | 808343 | 827625 | 19283 | - | Gnomon | gene | gene53 | GeneID:105277148 | LOC105277148 | Gene | LOC105277148 | protein_coding | character(0) | character(0) | BCphaseA1.vs.BCphaseB1 | FALSE |
| NW_011924881.1 | 810417 | 810417 | 1 | + | 31 | 0 | 31 | 33 | 0 | 33 | 0 | 0 | NW_011924881.1 | 808343 | 827625 | 19283 | - | Gnomon | gene | gene53 | GeneID:105277148 | LOC105277148 | Gene | LOC105277148 | protein_coding | character(0) | character(0) | BCphaseA1.vs.BCphaseB1 | FALSE |
| NW_011924881.1 | 810424 | 810424 | 1 | + | 46 | 1 | 45 | 46 | 0 | 46 | 2.17 | 0 | NW_011924881.1 | 808343 | 827625 | 19283 | - | Gnomon | gene | gene53 | GeneID:105277148 | LOC105277148 | Gene | LOC105277148 | protein_coding | character(0) | character(0) | BCphaseA1.vs.BCphaseB1 | FALSE |
| NW_011924881.1 | 810430 | 810430 | 1 | + | 26 | 1 | 25 | 45 | 0 | 45 | 3.85 | 0 | NW_011924881.1 | 808343 | 827625 | 19283 | - | Gnomon | gene | gene53 | GeneID:105277148 | LOC105277148 | Gene | LOC105277148 | protein_coding | character(0) | character(0) | BCphaseA1.vs.BCphaseB1 | FALSE |
| NW_011924881.1 | 810436 | 810436 | 1 | + | 52 | 2 | 50 | 49 | 0 | 49 | 3.85 | 0 | NW_011924881.1 | 808343 | 827625 | 19283 | - | Gnomon | gene | gene53 | GeneID:105277148 | LOC105277148 | Gene | LOC105277148 | protein_coding | character(0) | character(0) | BCphaseA1.vs.BCphaseB1 | FALSE |
| NW_011924881.1 | 810482 | 810482 | 1 | + | 61 | 1 | 60 | 66 | 2 | 64 | 1.64 | 3.03 | NW_011924881.1 | 808343 | 827625 | 19283 | - | Gnomon | gene | gene53 | GeneID:105277148 | LOC105277148 | Gene | LOC105277148 | protein_coding | character(0) | character(0) | BCphaseA1.vs.BCphaseB1 | FALSE |
| NW_011924881.1 | 810505 | 810505 | 1 | + | 60 | 0 | 60 | 64 | 0 | 64 | 0 | 0 | NW_011924881.1 | 808343 | 827625 | 19283 | - | Gnomon | gene | gene53 | GeneID:105277148 | LOC105277148 | Gene | LOC105277148 | protein_coding | character(0) | character(0) | BCphaseA1.vs.BCphaseB1 | FALSE |
| NW_011924881.1 | 810511 | 810511 | 1 | + | 39 | 0 | 39 | 61 | 1 | 60 | 0 | 1.64 | NW_011924881.1 | 808343 | 827625 | 19283 | - | Gnomon | gene | gene53 | GeneID:105277148 | LOC105277148 | Gene | LOC105277148 | protein_coding | character(0) | character(0) | BCphaseA1.vs.BCphaseB1 | FALSE |
| NW_011924881.1 | 810516 | 810516 | 1 | + | 38 | 0 | 38 | 63 | 0 | 63 | 0 | 0 | NW_011924881.1 | 808343 | 827625 | 19283 | - | Gnomon | gene | gene53 | GeneID:105277148 | LOC105277148 | Gene | LOC105277148 | protein_coding | character(0) | character(0) | BCphaseA1.vs.BCphaseB1 | FALSE |
| NW_011924881.1 | 810520 | 810520 | 1 | + | 39 | 0 | 39 | 63 | 0 | 63 | 0 | 0 | NW_011924881.1 | 808343 | 827625 | 19283 | - | Gnomon | gene | gene53 | GeneID:105277148 | LOC105277148 | Gene | LOC105277148 | protein_coding | character(0) | character(0) | BCphaseA1.vs.BCphaseB1 | FALSE |
| NW_011924881.1 | 810577 | 810577 | 1 | + | 21 | 0 | 21 | 33 | 0 | 33 | 0 | 0 | NW_011924881.1 | 808343 | 827625 | 19283 | - | Gnomon | gene | gene53 | GeneID:105277148 | LOC105277148 | Gene | LOC105277148 | protein_coding | character(0) | character(0) | BCphaseA1.vs.BCphaseB1 | FALSE |
| NW_011924881.1 | 810581 | 810581 | 1 | + | 22 | 0 | 22 | 35 | 0 | 35 | 0 | 0 | NW_011924881.1 | 808343 | 827625 | 19283 | - | Gnomon | gene | gene53 | GeneID:105277148 | LOC105277148 | Gene | LOC105277148 | protein_coding | character(0) | character(0) | BCphaseA1.vs.BCphaseB1 | FALSE |
| NW_011924881.1 | 810583 | 810583 | 1 | + | 22 | 1 | 21 | 34 | 0 | 34 | 4.55 | 0 | NW_011924881.1 | 808343 | 827625 | 19283 | - | Gnomon | gene | gene53 | GeneID:105277148 | LOC105277148 | Gene | LOC105277148 | protein_coding | character(0) | character(0) | BCphaseA1.vs.BCphaseB1 | FALSE |

...
... only first 25 lines shown ...

Directory: DMSG File: dmg-Cb_LI_BCphaseA1.vs.BCphaseB2_details.txt

| seqnames | start | end | width | strand | coverage1 | numCs1 | numTs1 | coverage2 | numCs2 | numTs2 | BCphaseA1 | BCphaseB2 | gene_seqnames | gene_start | gene_end | gene_width | gene_strand | gene_source | gene_type | gene_ID | gene_Dbxref | gene_Name | gene_gbkey | gene_gene | gene_gene_biotype | gene_partial | gene_start_range | gene_end_range | gene_comparison | is.dm |
|---|---|---|---|---|---|---|---|---|---|---|---|---|---|---|---|---|---|---|---|---|---|---|---|---|---|---|---|---|---|---|
| NW_011924881.1 | 195703 | 195703 | 1 | + | 20 | 0 | 20 | 27 | 0 | 27 | 0 | 0 | NW_011924881.1 | 195683 | 286417 | 90735 | - | Gnomon | gene | gene22 | GeneID:105288120 | LOC105288120 | Gene | LOC105288120 | protein_coding | NA | character(0) | character(0) | BCphaseA1.vs.BCphaseB2 | FALSE |
| NW_011924881.1 | 195718 | 195718 | 1 | + | 24 | 0 | 24 | 36 | 0 | 36 | 0 | 0 | NW_011924881.1 | 195683 | 286417 | 90735 | - | Gnomon | gene | gene22 | GeneID:105288120 | LOC105288120 | Gene | LOC105288120 | protein_coding | NA | character(0) | character(0) | BCphaseA1.vs.BCphaseB2 | FALSE |
| NW_011924881.1 | 195729 | 195729 | 1 | + | 25 | 1 | 24 | 32 | 0 | 32 | 4 | 0 | NW_011924881.1 | 195683 | 286417 | 90735 | - | Gnomon | gene | gene22 | GeneID:105288120 | LOC105288120 | Gene | LOC105288120 | protein_coding | NA | character(0) | character(0) | BCphaseA1.vs.BCphaseB2 | FALSE |
| NW_011924881.1 | 195734 | 195734 | 1 | + | 28 | 1 | 27 | 33 | 0 | 33 | 3.57 | 0 | NW_011924881.1 | 195683 | 286417 | 90735 | - | Gnomon | gene | gene22 | GeneID:105288120 | LOC105288120 | Gene | LOC105288120 | protein_coding | NA | character(0) | character(0) | BCphaseA1.vs.BCphaseB2 | FALSE |
| NW_011924881.1 | 195747 | 195747 | 1 | + | 36 | 0 | 36 | 60 | 0 | 60 | 0 | 0 | NW_011924881.1 | 195683 | 286417 | 90735 | - | Gnomon | gene | gene22 | GeneID:105288120 | LOC105288120 | Gene | LOC105288120 | protein_coding | NA | character(0) | character(0) | BCphaseA1.vs.BCphaseB2 | FALSE |
| NW_011924881.1 | 195763 | 195763 | 1 | + | 58 | 0 | 58 | 102 | 0 | 102 | 0 | 0 | NW_011924881.1 | 195683 | 286417 | 90735 | - | Gnomon | gene | gene22 | GeneID:105288120 | LOC105288120 | Gene | LOC105288120 | protein_coding | NA | character(0) | character(0) | BCphaseA1.vs.BCphaseB2 | FALSE |
| NW_011924881.1 | 195773 | 195773 | 1 | + | 54 | 0 | 54 | 90 | 0 | 90 | 0 | 0 | NW_011924881.1 | 195683 | 286417 | 90735 | - | Gnomon | gene | gene22 | GeneID:105288120 | LOC105288120 | Gene | LOC105288120 | protein_coding | NA | character(0) | character(0) | BCphaseA1.vs.BCphaseB2 | FALSE |
| NW_011924881.1 | 195790 | 195790 | 1 | + | 52 | 0 | 52 | 91 | 0 | 91 | 0 | 0 | NW_011924881.1 | 195683 | 286417 | 90735 | - | Gnomon | gene | gene22 | GeneID:105288120 | LOC105288120 | Gene | LOC105288120 | protein_coding | NA | character(0) | character(0) | BCphaseA1.vs.BCphaseB2 | FALSE |
| NW_011924881.1 | 195812 | 195812 | 1 | + | 72 | 0 | 72 | 92 | 1 | 91 | 0 | 1.09 | NW_011924881.1 | 195683 | 286417 | 90735 | - | Gnomon | gene | gene22 | GeneID:105288120 | LOC105288120 | Gene | LOC105288120 | protein_coding | NA | character(0) | character(0) | BCphaseA1.vs.BCphaseB2 | FALSE |
| NW_011924881.1 | 195815 | 195815 | 1 | + | 72 | 0 | 72 | 92 | 0 | 92 | 0 | 0 | NW_011924881.1 | 195683 | 286417 | 90735 | - | Gnomon | gene | gene22 | GeneID:105288120 | LOC105288120 | Gene | LOC105288120 | protein_coding | NA | character(0) | character(0) | BCphaseA1.vs.BCphaseB2 | FALSE |
| NW_011924881.1 | 195830 | 195830 | 1 | + | 69 | 0 | 69 | 86 | 1 | 85 | 0 | 1.16 | NW_011924881.1 | 195683 | 286417 | 90735 | - | Gnomon | gene | gene22 | GeneID:105288120 | LOC105288120 | Gene | LOC105288120 | protein_coding | NA | character(0) | character(0) | BCphaseA1.vs.BCphaseB2 | FALSE |
| NW_011924881.1 | 195881 | 195881 | 1 | + | 20 | 0 | 20 | 29 | 1 | 28 | 0 | 3.45 | NW_011924881.1 | 195683 | 286417 | 90735 | - | Gnomon | gene | gene22 | GeneID:105288120 | LOC105288120 | Gene | LOC105288120 | protein_coding | NA | character(0) | character(0) | BCphaseA1.vs.BCphaseB2 | FALSE |
| NW_011924881.1 | 195905 | 195905 | 1 | + | 23 | 0 | 23 | 21 | 0 | 21 | 0 | 0 | NW_011924881.1 | 195683 | 286417 | 90735 | - | Gnomon | gene | gene22 | GeneID:105288120 | LOC105288120 | Gene | LOC105288120 | protein_coding | NA | character(0) | character(0) | BCphaseA1.vs.BCphaseB2 | FALSE |
| NW_011924881.1 | 195908 | 195908 | 1 | + | 24 | 0 | 24 | 31 | 0 | 31 | 0 | 0 | NW_011924881.1 | 195683 | 286417 | 90735 | - | Gnomon | gene | gene22 | GeneID:105288120 | LOC105288120 | Gene | LOC105288120 | protein_coding | NA | character(0) | character(0) | BCphaseA1.vs.BCphaseB2 | FALSE |
| NW_011924881.1 | 195914 | 195914 | 1 | + | 30 | 0 | 30 | 45 | 1 | 44 | 0 | 2.22 | NW_011924881.1 | 195683 | 286417 | 90735 | - | Gnomon | gene | gene22 | GeneID:105288120 | LOC105288120 | Gene | LOC105288120 | protein_coding | NA | character(0) | character(0) | BCphaseA1.vs.BCphaseB2 | FALSE |
| NW_011924881.1 | 195917 | 195917 | 1 | + | 32 | 0 | 32 | 46 | 0 | 46 | 0 | 0 | NW_011924881.1 | 195683 | 286417 | 90735 | - | Gnomon | gene | gene22 | GeneID:105288120 | LOC105288120 | Gene | LOC105288120 | protein_coding | NA | character(0) | character(0) | BCphaseA1.vs.BCphaseB2 | FALSE |
| NW_011924881.1 | 195923 | 195923 | 1 | + | 33 | 1 | 32 | 49 | 0 | 49 | 3.03 | 0 | NW_011924881.1 | 195683 | 286417 | 90735 | - | Gnomon | gene | gene22 | GeneID:105288120 | LOC105288120 | Gene | LOC105288120 | protein_coding | NA | character(0) | character(0) | BCphaseA1.vs.BCphaseB2 | FALSE |
| NW_011924881.1 | 195929 | 195929 | 1 | + | 34 | 0 | 34 | 49 | 0 | 49 | 0 | 0 | NW_011924881.1 | 195683 | 286417 | 90735 | - | Gnomon | gene | gene22 | GeneID:105288120 | LOC105288120 | Gene | LOC105288120 | protein_coding | NA | character(0) | character(0) | BCphaseA1.vs.BCphaseB2 | FALSE |
| NW_011924881.1 | 195935 | 195935 | 1 | + | 34 | 0 | 34 | 47 | 0 | 47 | 0 | 0 | NW_011924881.1 | 195683 | 286417 | 90735 | - | Gnomon | gene | gene22 | GeneID:105288120 | LOC105288120 | Gene | LOC105288120 | protein_coding | NA | character(0) | character(0) | BCphaseA1.vs.BCphaseB2 | FALSE |
| NW_011924881.1 | 195941 | 195941 | 1 | + | 35 | 0 | 35 | 47 | 0 | 47 | 0 | 0 | NW_011924881.1 | 195683 | 286417 | 90735 | - | Gnomon | gene | gene22 | GeneID:105288120 | LOC105288120 | Gene | LOC105288120 | protein_coding | NA | character(0) | character(0) | BCphaseA1.vs.BCphaseB2 | FALSE |
| NW_011924881.1 | 195946 | 195946 | 1 | + | 35 | 0 | 35 | 47 | 1 | 46 | 0 | 2.13 | NW_011924881.1 | 195683 | 286417 | 90735 | - | Gnomon | gene | gene22 | GeneID:105288120 | LOC105288120 | Gene | LOC105288120 | protein_coding | NA | character(0) | character(0) | BCphaseA1.vs.BCphaseB2 | FALSE |
| NW_011924881.1 | 195952 | 195952 | 1 | + | 34 | 0 | 34 | 47 | 0 | 47 | 0 | 0 | NW_011924881.1 | 195683 | 286417 | 90735 | - | Gnomon | gene | gene22 | GeneID:105288120 | LOC105288120 | Gene | LOC105288120 | protein_coding | NA | character(0) | character(0) | BCphaseA1.vs.BCphaseB2 | FALSE |
| NW_011924881.1 | 195961 | 195961 | 1 | + | 29 | 0 | 29 | 46 | 0 | 46 | 0 | 0 | NW_011924881.1 | 195683 | 286417 | 90735 | - | Gnomon | gene | gene22 | GeneID:105288120 | LOC105288120 | Gene | LOC105288120 | protein_coding | NA | character(0) | character(0) | BCphaseA1.vs.BCphaseB2 | FALSE |
| NW_011924881.1 | 195964 | 195964 | 1 | + | 29 | 0 | 29 | 45 | 0 | 45 | 0 | 0 | NW_011924881.1 | 195683 | 286417 | 90735 | - | Gnomon | gene | gene22 | GeneID:105288120 | LOC105288120 | Gene | LOC105288120 | protein_coding | NA | character(0) | character(0) | BCphaseA1.vs.BCphaseB2 | FALSE |

...
... only first 25 lines shown ...

Directory: DMSG File: dmg-Cb_LI_BCphaseA1.vs.RphaseA1_details.txt

| seqnames | start | end | width | strand | coverage1 | numCs1 | numTs1 | coverage2 | numCs2 | numTs2 | BCphaseA1 | RphaseA1 | gene_seqnames | gene_start | gene_end | gene_width | gene_strand | gene_source | gene_type | gene_ID | gene_Dbxref | gene_Name | gene_gbkey | gene_gene | gene_gene_biotype | gene_start_range | gene_end_range | gene_comparison | is.dm |
|---|---|---|---|---|---|---|---|---|---|---|---|---|---|---|---|---|---|---|---|---|---|---|---|---|---|---|---|---|---|
| NW_011924888.1 | 70881 | 70881 | 1 | + | 20 | 0 | 20 | 60 | 0 | 60 | 0 | 0 | NW_011924888.1 | 70798 | 74285 | 3488 | - | Gnomon | gene | gene274 | GeneID:105275190 | LOC105275190 | Gene | LOC105275190 | protein_coding | character(0) | character(0) | BCphaseA1.vs.RphaseA1 | FALSE |
| NW_011924888.1 | 70885 | 70885 | 1 | + | 20 | 0 | 20 | 60 | 1 | 59 | 0 | 1.67 | NW_011924888.1 | 70798 | 74285 | 3488 | - | Gnomon | gene | gene274 | GeneID:105275190 | LOC105275190 | Gene | LOC105275190 | protein_coding | character(0) | character(0) | BCphaseA1.vs.RphaseA1 | FALSE |
| NW_011924888.1 | 70922 | 70922 | 1 | + | 24 | 0 | 24 | 83 | 0 | 83 | 0 | 0 | NW_011924888.1 | 70798 | 74285 | 3488 | - | Gnomon | gene | gene274 | GeneID:105275190 | LOC105275190 | Gene | LOC105275190 | protein_coding | character(0) | character(0) | BCphaseA1.vs.RphaseA1 | FALSE |
| NW_011924888.1 | 70933 | 70933 | 1 | + | 47 | 1 | 46 | 93 | 0 | 93 | 2.13 | 0 | NW_011924888.1 | 70798 | 74285 | 3488 | - | Gnomon | gene | gene274 | GeneID:105275190 | LOC105275190 | Gene | LOC105275190 | protein_coding | character(0) | character(0) | BCphaseA1.vs.RphaseA1 | FALSE |
| NW_011924888.1 | 70949 | 70949 | 1 | + | 42 | 3 | 39 | 75 | 5 | 70 | 7.14 | 6.67 | NW_011924888.1 | 70798 | 74285 | 3488 | - | Gnomon | gene | gene274 | GeneID:105275190 | LOC105275190 | Gene | LOC105275190 | protein_coding | character(0) | character(0) | BCphaseA1.vs.RphaseA1 | FALSE |
| NW_011924888.1 | 70953 | 70953 | 1 | + | 23 | 0 | 23 | 76 | 0 | 76 | 0 | 0 | NW_011924888.1 | 70798 | 74285 | 3488 | - | Gnomon | gene | gene274 | GeneID:105275190 | LOC105275190 | Gene | LOC105275190 | protein_coding | character(0) | character(0) | BCphaseA1.vs.RphaseA1 | FALSE |
| NW_011924888.1 | 70955 | 70955 | 1 | + | 23 | 0 | 23 | 78 | 2 | 76 | 0 | 2.56 | NW_011924888.1 | 70798 | 74285 | 3488 | - | Gnomon | gene | gene274 | GeneID:105275190 | LOC105275190 | Gene | LOC105275190 | protein_coding | character(0) | character(0) | BCphaseA1.vs.RphaseA1 | FALSE |
| NW_011924888.1 | 70972 | 70972 | 1 | + | 21 | 0 | 21 | 67 | 1 | 66 | 0 | 1.49 | NW_011924888.1 | 70798 | 74285 | 3488 | - | Gnomon | gene | gene274 | GeneID:105275190 | LOC105275190 | Gene | LOC105275190 | protein_coding | character(0) | character(0) | BCphaseA1.vs.RphaseA1 | FALSE |
| NW_011924888.1 | 70981 | 70981 | 1 | + | 22 | 6 | 16 | 58 | 3 | 55 | 27.27 | 5.17 | NW_011924888.1 | 70798 | 74285 | 3488 | - | Gnomon | gene | gene274 | GeneID:105275190 | LOC105275190 | Gene | LOC105275190 | protein_coding | character(0) | character(0) | BCphaseA1.vs.RphaseA1 | FALSE |
| NW_011924888.1 | 71274 | 71274 | 1 | + | 24 | 0 | 24 | 24 | 1 | 23 | 0 | 4.17 | NW_011924888.1 | 70798 | 74285 | 3488 | - | Gnomon | gene | gene274 | GeneID:105275190 | LOC105275190 | Gene | LOC105275190 | protein_coding | character(0) | character(0) | BCphaseA1.vs.RphaseA1 | FALSE |
| NW_011924888.1 | 71283 | 71283 | 1 | + | 25 | 0 | 25 | 25 | 0 | 25 | 0 | 0 | NW_011924888.1 | 70798 | 74285 | 3488 | - | Gnomon | gene | gene274 | GeneID:105275190 | LOC105275190 | Gene | LOC105275190 | protein_coding | character(0) | character(0) | BCphaseA1.vs.RphaseA1 | FALSE |
| NW_011924888.1 | 71293 | 71293 | 1 | + | 20 | 0 | 20 | 22 | 0 | 22 | 0 | 0 | NW_011924888.1 | 70798 | 74285 | 3488 | - | Gnomon | gene | gene274 | GeneID:105275190 | LOC105275190 | Gene | LOC105275190 | protein_coding | character(0) | character(0) | BCphaseA1.vs.RphaseA1 | FALSE |
| NW_011924888.1 | 71323 | 71323 | 1 | + | 21 | 1 | 20 | 20 | 0 | 20 | 4.76 | 0 | NW_011924888.1 | 70798 | 74285 | 3488 | - | Gnomon | gene | gene274 | GeneID:105275190 | LOC105275190 | Gene | LOC105275190 | protein_coding | character(0) | character(0) | BCphaseA1.vs.RphaseA1 | FALSE |
| NW_011924888.1 | 71880 | 71880 | 1 | + | 22 | 0 | 22 | 24 | 0 | 24 | 0 | 0 | NW_011924888.1 | 70798 | 74285 | 3488 | - | Gnomon | gene | gene274 | GeneID:105275190 | LOC105275190 | Gene | LOC105275190 | protein_coding | character(0) | character(0) | BCphaseA1.vs.RphaseA1 | FALSE |
| NW_011924888.1 | 71914 | 71914 | 1 | + | 45 | 0 | 45 | 52 | 0 | 52 | 0 | 0 | NW_011924888.1 | 70798 | 74285 | 3488 | - | Gnomon | gene | gene274 | GeneID:105275190 | LOC105275190 | Gene | LOC105275190 | protein_coding | character(0) | character(0) | BCphaseA1.vs.RphaseA1 | FALSE |
| NW_011924888.1 | 72001 | 72001 | 1 | + | 22 | 1 | 21 | 32 | 1 | 31 | 4.55 | 3.12 | NW_011924888.1 | 70798 | 74285 | 3488 | - | Gnomon | gene | gene274 | GeneID:105275190 | LOC105275190 | Gene | LOC105275190 | protein_coding | character(0) | character(0) | BCphaseA1.vs.RphaseA1 | FALSE |
| NW_011924888.1 | 72012 | 72012 | 1 | + | 22 | 1 | 21 | 28 | 2 | 26 | 4.55 | 7.14 | NW_011924888.1 | 70798 | 74285 | 3488 | - | Gnomon | gene | gene274 | GeneID:105275190 | LOC105275190 | Gene | LOC105275190 | protein_coding | character(0) | character(0) | BCphaseA1.vs.RphaseA1 | FALSE |
| NW_011924888.1 | 72027 | 72027 | 1 | + | 24 | 1 | 23 | 32 | 0 | 32 | 4.17 | 0 | NW_011924888.1 | 70798 | 74285 | 3488 | - | Gnomon | gene | gene274 | GeneID:105275190 | LOC105275190 | Gene | LOC105275190 | protein_coding | character(0) | character(0) | BCphaseA1.vs.RphaseA1 | FALSE |
| NW_011924888.1 | 72336 | 72336 | 1 | + | 24 | 1 | 23 | 31 | 5 | 26 | 4.17 | 16.13 | NW_011924888.1 | 70798 | 74285 | 3488 | - | Gnomon | gene | gene274 | GeneID:105275190 | LOC105275190 | Gene | LOC105275190 | protein_coding | character(0) | character(0) | BCphaseA1.vs.RphaseA1 | FALSE |
| NW_011924888.1 | 72346 | 72346 | 1 | + | 27 | 2 | 25 | 31 | 1 | 30 | 7.41 | 3.23 | NW_011924888.1 | 70798 | 74285 | 3488 | - | Gnomon | gene | gene274 | GeneID:105275190 | LOC105275190 | Gene | LOC105275190 | protein_coding | character(0) | character(0) | BCphaseA1.vs.RphaseA1 | FALSE |
| NW_011924888.1 | 72368 | 72368 | 1 | + | 26 | 2 | 24 | 29 | 4 | 25 | 7.69 | 13.79 | NW_011924888.1 | 70798 | 74285 | 3488 | - | Gnomon | gene | gene274 | GeneID:105275190 | LOC105275190 | Gene | LOC105275190 | protein_coding | character(0) | character(0) | BCphaseA1.vs.RphaseA1 | FALSE |
| NW_011924888.1 | 72551 | 72551 | 1 | + | 20 | 1 | 19 | 20 | 2 | 18 | 5 | 10 | NW_011924888.1 | 70798 | 74285 | 3488 | - | Gnomon | gene | gene274 | GeneID:105275190 | LOC105275190 | Gene | LOC105275190 | protein_coding | character(0) | character(0) | BCphaseA1.vs.RphaseA1 | FALSE |
| NW_011924888.1 | 72555 | 72555 | 1 | + | 21 | 0 | 21 | 20 | 0 | 20 | 0 | 0 | NW_011924888.1 | 70798 | 74285 | 3488 | - | Gnomon | gene | gene274 | GeneID:105275190 | LOC105275190 | Gene | LOC105275190 | protein_coding | character(0) | character(0) | BCphaseA1.vs.RphaseA1 | FALSE |
| NW_011924888.1 | 72562 | 72562 | 1 | + | 20 | 1 | 19 | 20 | 2 | 18 | 5 | 10 | NW_011924888.1 | 70798 | 74285 | 3488 | - | Gnomon | gene | gene274 | GeneID:105275190 | LOC105275190 | Gene | LOC105275190 | protein_coding | character(0) | character(0) | BCphaseA1.vs.RphaseA1 | FALSE |

...
... only first 25 lines shown ...

Directory: DMSG File: dmg-Cb_LI_BCphaseA1.vs.RphaseA2_details.txt

| seqnames | start | end | width | strand | coverage1 | numCs1 | numTs1 | coverage2 | numCs2 | numTs2 | BCphaseA1 | RphaseA2 | gene_seqnames | gene_start | gene_end | gene_width | gene_strand | gene_source | gene_type | gene_ID | gene_Dbxref | gene_Name | gene_gbkey | gene_gene | gene_gene_biotype | gene_start_range | gene_end_range | gene_comparison | is.dm |
|---|---|---|---|---|---|---|---|---|---|---|---|---|---|---|---|---|---|---|---|---|---|---|---|---|---|---|---|---|---|
| NW_011924881.1 | 809447 | 809447 | 1 | + | 28 | 0 | 28 | 28 | 0 | 28 | 0 | 0 | NW_011924881.1 | 808343 | 827625 | 19283 | - | Gnomon | gene | gene53 | GeneID:105277148 | LOC105277148 | Gene | LOC105277148 | protein_coding | character(0) | character(0) | BCphaseA1.vs.RphaseA2 | FALSE |
| NW_011924881.1 | 809481 | 809481 | 1 | + | 37 | 0 | 37 | 38 | 1 | 37 | 0 | 2.63 | NW_011924881.1 | 808343 | 827625 | 19283 | - | Gnomon | gene | gene53 | GeneID:105277148 | LOC105277148 | Gene | LOC105277148 | protein_coding | character(0) | character(0) | BCphaseA1.vs.RphaseA2 | FALSE |
| NW_011924881.1 | 809504 | 809504 | 1 | + | 38 | 1 | 37 | 44 | 1 | 43 | 2.63 | 2.27 | NW_011924881.1 | 808343 | 827625 | 19283 | - | Gnomon | gene | gene53 | GeneID:105277148 | LOC105277148 | Gene | LOC105277148 | protein_coding | character(0) | character(0) | BCphaseA1.vs.RphaseA2 | FALSE |
| NW_011924881.1 | 809520 | 809520 | 1 | + | 31 | 0 | 31 | 37 | 1 | 36 | 0 | 2.7 | NW_011924881.1 | 808343 | 827625 | 19283 | - | Gnomon | gene | gene53 | GeneID:105277148 | LOC105277148 | Gene | LOC105277148 | protein_coding | character(0) | character(0) | BCphaseA1.vs.RphaseA2 | FALSE |
| NW_011924881.1 | 810123 | 810123 | 1 | + | 26 | 0 | 26 | 26 | 0 | 26 | 0 | 0 | NW_011924881.1 | 808343 | 827625 | 19283 | - | Gnomon | gene | gene53 | GeneID:105277148 | LOC105277148 | Gene | LOC105277148 | protein_coding | character(0) | character(0) | BCphaseA1.vs.RphaseA2 | FALSE |
| NW_011924881.1 | 810155 | 810155 | 1 | + | 29 | 0 | 29 | 36 | 0 | 36 | 0 | 0 | NW_011924881.1 | 808343 | 827625 | 19283 | - | Gnomon | gene | gene53 | GeneID:105277148 | LOC105277148 | Gene | LOC105277148 | protein_coding | character(0) | character(0) | BCphaseA1.vs.RphaseA2 | FALSE |
| NW_011924881.1 | 810182 | 810182 | 1 | + | 33 | 0 | 33 | 39 | 2 | 37 | 0 | 5.13 | NW_011924881.1 | 808343 | 827625 | 19283 | - | Gnomon | gene | gene53 | GeneID:105277148 | LOC105277148 | Gene | LOC105277148 | protein_coding | character(0) | character(0) | BCphaseA1.vs.RphaseA2 | FALSE |
| NW_011924881.1 | 810198 | 810198 | 1 | + | 20 | 0 | 20 | 23 | 0 | 23 | 0 | 0 | NW_011924881.1 | 808343 | 827625 | 19283 | - | Gnomon | gene | gene53 | GeneID:105277148 | LOC105277148 | Gene | LOC105277148 | protein_coding | character(0) | character(0) | BCphaseA1.vs.RphaseA2 | FALSE |
| NW_011924881.1 | 810335 | 810335 | 1 | + | 20 | 0 | 20 | 33 | 1 | 32 | 0 | 3.03 | NW_011924881.1 | 808343 | 827625 | 19283 | - | Gnomon | gene | gene53 | GeneID:105277148 | LOC105277148 | Gene | LOC105277148 | protein_coding | character(0) | character(0) | BCphaseA1.vs.RphaseA2 | FALSE |
| NW_011924881.1 | 810337 | 810337 | 1 | + | 21 | 0 | 21 | 32 | 0 | 32 | 0 | 0 | NW_011924881.1 | 808343 | 827625 | 19283 | - | Gnomon | gene | gene53 | GeneID:105277148 | LOC105277148 | Gene | LOC105277148 | protein_coding | character(0) | character(0) | BCphaseA1.vs.RphaseA2 | FALSE |
| NW_011924881.1 | 810348 | 810348 | 1 | + | 35 | 0 | 35 | 74 | 0 | 74 | 0 | 0 | NW_011924881.1 | 808343 | 827625 | 19283 | - | Gnomon | gene | gene53 | GeneID:105277148 | LOC105277148 | Gene | LOC105277148 | protein_coding | character(0) | character(0) | BCphaseA1.vs.RphaseA2 | FALSE |
| NW_011924881.1 | 810391 | 810391 | 1 | + | 37 | 0 | 37 | 93 | 0 | 93 | 0 | 0 | NW_011924881.1 | 808343 | 827625 | 19283 | - | Gnomon | gene | gene53 | GeneID:105277148 | LOC105277148 | Gene | LOC105277148 | protein_coding | character(0) | character(0) | BCphaseA1.vs.RphaseA2 | FALSE |
| NW_011924881.1 | 810409 | 810409 | 1 | + | 55 | 0 | 55 | 86 | 0 | 86 | 0 | 0 | NW_011924881.1 | 808343 | 827625 | 19283 | - | Gnomon | gene | gene53 | GeneID:105277148 | LOC105277148 | Gene | LOC105277148 | protein_coding | character(0) | character(0) | BCphaseA1.vs.RphaseA2 | FALSE |
| NW_011924881.1 | 810412 | 810412 | 1 | + | 30 | 0 | 30 | 84 | 0 | 84 | 0 | 0 | NW_011924881.1 | 808343 | 827625 | 19283 | - | Gnomon | gene | gene53 | GeneID:105277148 | LOC105277148 | Gene | LOC105277148 | protein_coding | character(0) | character(0) | BCphaseA1.vs.RphaseA2 | FALSE |
| NW_011924881.1 | 810417 | 810417 | 1 | + | 31 | 0 | 31 | 76 | 0 | 76 | 0 | 0 | NW_011924881.1 | 808343 | 827625 | 19283 | - | Gnomon | gene | gene53 | GeneID:105277148 | LOC105277148 | Gene | LOC105277148 | protein_coding | character(0) | character(0) | BCphaseA1.vs.RphaseA2 | FALSE |
| NW_011924881.1 | 810424 | 810424 | 1 | + | 46 | 1 | 45 | 74 | 1 | 73 | 2.17 | 1.35 | NW_011924881.1 | 808343 | 827625 | 19283 | - | Gnomon | gene | gene53 | GeneID:105277148 | LOC105277148 | Gene | LOC105277148 | protein_coding | character(0) | character(0) | BCphaseA1.vs.RphaseA2 | FALSE |
| NW_011924881.1 | 810430 | 810430 | 1 | + | 26 | 1 | 25 | 31 | 0 | 31 | 3.85 | 0 | NW_011924881.1 | 808343 | 827625 | 19283 | - | Gnomon | gene | gene53 | GeneID:105277148 | LOC105277148 | Gene | LOC105277148 | protein_coding | character(0) | character(0) | BCphaseA1.vs.RphaseA2 | FALSE |
| NW_011924881.1 | 810436 | 810436 | 1 | + | 52 | 2 | 50 | 32 | 0 | 32 | 3.85 | 0 | NW_011924881.1 | 808343 | 827625 | 19283 | - | Gnomon | gene | gene53 | GeneID:105277148 | LOC105277148 | Gene | LOC105277148 | protein_coding | character(0) | character(0) | BCphaseA1.vs.RphaseA2 | FALSE |
| NW_011924881.1 | 810482 | 810482 | 1 | + | 61 | 1 | 60 | 67 | 3 | 64 | 1.64 | 4.48 | NW_011924881.1 | 808343 | 827625 | 19283 | - | Gnomon | gene | gene53 | GeneID:105277148 | LOC105277148 | Gene | LOC105277148 | protein_coding | character(0) | character(0) | BCphaseA1.vs.RphaseA2 | FALSE |
| NW_011924881.1 | 810505 | 810505 | 1 | + | 60 | 0 | 60 | 71 | 0 | 71 | 0 | 0 | NW_011924881.1 | 808343 | 827625 | 19283 | - | Gnomon | gene | gene53 | GeneID:105277148 | LOC105277148 | Gene | LOC105277148 | protein_coding | character(0) | character(0) | BCphaseA1.vs.RphaseA2 | FALSE |
| NW_011924881.1 | 810511 | 810511 | 1 | + | 39 | 0 | 39 | 66 | 0 | 66 | 0 | 0 | NW_011924881.1 | 808343 | 827625 | 19283 | - | Gnomon | gene | gene53 | GeneID:105277148 | LOC105277148 | Gene | LOC105277148 | protein_coding | character(0) | character(0) | BCphaseA1.vs.RphaseA2 | FALSE |
| NW_011924881.1 | 810516 | 810516 | 1 | + | 38 | 0 | 38 | 64 | 0 | 64 | 0 | 0 | NW_011924881.1 | 808343 | 827625 | 19283 | - | Gnomon | gene | gene53 | GeneID:105277148 | LOC105277148 | Gene | LOC105277148 | protein_coding | character(0) | character(0) | BCphaseA1.vs.RphaseA2 | FALSE |
| NW_011924881.1 | 810520 | 810520 | 1 | + | 39 | 0 | 39 | 68 | 0 | 68 | 0 | 0 | NW_011924881.1 | 808343 | 827625 | 19283 | - | Gnomon | gene | gene53 | GeneID:105277148 | LOC105277148 | Gene | LOC105277148 | protein_coding | character(0) | character(0) | BCphaseA1.vs.RphaseA2 | FALSE |
| NW_011924881.1 | 810577 | 810577 | 1 | + | 21 | 0 | 21 | 25 | 0 | 25 | 0 | 0 | NW_011924881.1 | 808343 | 827625 | 19283 | - | Gnomon | gene | gene53 | GeneID:105277148 | LOC105277148 | Gene | LOC105277148 | protein_coding | character(0) | character(0) | BCphaseA1.vs.RphaseA2 | FALSE |

...

... only first 25 lines shown ...

Directory: DMSG File: dmg-Cb_LI_BCphaseA1.vs.RphaseB1_details.txt

| seqnames | start | end | width | strand | coverage1 | numCs1 | numTs1 | coverage2 | numCs2 | numTs2 | BCphaseA1 | RphaseB1 | gene_seqnames | gene_start | gene_end | gene_width | gene_strand | gene_source | gene_type | gene_ID | gene_Dbxref | gene_Name | gene_gbkey | gene_gene | gene_gene_biotype | gene_partial | gene_start_range | gene_end_range | gene_comparison | is.dm |
|---|---|---|---|---|---|---|---|---|---|---|---|---|---|---|---|---|---|---|---|---|---|---|---|---|---|---|---|---|---|---|
| NW_011924881.1 | 448501 | 448501 | 1 | + | 25 | 4 | 21 | 33 | 7 | 26 | 16 | 21.21 | NW_011924881.1 | 448307 | 454087 | 5781 | - | Gnomon | gene | gene26 | GeneID:105288189 | LOC105288189 | Gene | LOC105288189 | protein_coding | NA | character(0) | character(0) | BCphaseA1.vs.RphaseB1 | FALSE |
| NW_011924881.1 | 448524 | 448524 | 1 | + | 20 | 10 | 10 | 30 | 12 | 18 | 50 | 40 | NW_011924881.1 | 448307 | 454087 | 5781 | - | Gnomon | gene | gene26 | GeneID:105288189 | LOC105288189 | Gene | LOC105288189 | protein_coding | NA | character(0) | character(0) | BCphaseA1.vs.RphaseB1 | FALSE |
| NW_011924881.1 | 448684 | 448684 | 1 | + | 20 | 8 | 12 | 26 | 7 | 19 | 40 | 26.92 | NW_011924881.1 | 448307 | 454087 | 5781 | - | Gnomon | gene | gene26 | GeneID:105288189 | LOC105288189 | Gene | LOC105288189 | protein_coding | NA | character(0) | character(0) | BCphaseA1.vs.RphaseB1 | FALSE |
| NW_011924881.1 | 448698 | 448698 | 1 | + | 22 | 2 | 20 | 33 | 2 | 31 | 9.09 | 6.06 | NW_011924881.1 | 448307 | 454087 | 5781 | - | Gnomon | gene | gene26 | GeneID:105288189 | LOC105288189 | Gene | LOC105288189 | protein_coding | NA | character(0) | character(0) | BCphaseA1.vs.RphaseB1 | FALSE |
| NW_011924881.1 | 448707 | 448707 | 1 | + | 22 | 5 | 17 | 35 | 7 | 28 | 22.73 | 20 | NW_011924881.1 | 448307 | 454087 | 5781 | - | Gnomon | gene | gene26 | GeneID:105288189 | LOC105288189 | Gene | LOC105288189 | protein_coding | NA | character(0) | character(0) | BCphaseA1.vs.RphaseB1 | FALSE |
| NW_011924881.1 | 448725 | 448725 | 1 | + | 23 | 7 | 16 | 40 | 13 | 27 | 30.43 | 32.5 | NW_011924881.1 | 448307 | 454087 | 5781 | - | Gnomon | gene | gene26 | GeneID:105288189 | LOC105288189 | Gene | LOC105288189 | protein_coding | NA | character(0) | character(0) | BCphaseA1.vs.RphaseB1 | FALSE |
| NW_011924881.1 | 449119 | 449119 | 1 | + | 53 | 0 | 53 | 41 | 1 | 40 | 0 | 2.44 | NW_011924881.1 | 448307 | 454087 | 5781 | - | Gnomon | gene | gene26 | GeneID:105288189 | LOC105288189 | Gene | LOC105288189 | protein_coding | NA | character(0) | character(0) | BCphaseA1.vs.RphaseB1 | FALSE |
| NW_011924881.1 | 449147 | 449147 | 1 | + | 57 | 2 | 55 | 45 | 3 | 42 | 3.51 | 6.67 | NW_011924881.1 | 448307 | 454087 | 5781 | - | Gnomon | gene | gene26 | GeneID:105288189 | LOC105288189 | Gene | LOC105288189 | protein_coding | NA | character(0) | character(0) | BCphaseA1.vs.RphaseB1 | FALSE |
| NW_011924881.1 | 449151 | 449151 | 1 | + | 57 | 1 | 56 | 42 | 1 | 41 | 1.75 | 2.38 | NW_011924881.1 | 448307 | 454087 | 5781 | - | Gnomon | gene | gene26 | GeneID:105288189 | LOC105288189 | Gene | LOC105288189 | protein_coding | NA | character(0) | character(0) | BCphaseA1.vs.RphaseB1 | FALSE |
| NW_011924881.1 | 449161 | 449161 | 1 | + | 53 | 6 | 47 | 40 | 5 | 35 | 11.32 | 12.5 | NW_011924881.1 | 448307 | 454087 | 5781 | - | Gnomon | gene | gene26 | GeneID:105288189 | LOC105288189 | Gene | LOC105288189 | protein_coding | NA | character(0) | character(0) | BCphaseA1.vs.RphaseB1 | FALSE |
| NW_011924881.1 | 449168 | 449168 | 1 | + | 47 | 0 | 47 | 37 | 2 | 35 | 0 | 5.41 | NW_011924881.1 | 448307 | 454087 | 5781 | - | Gnomon | gene | gene26 | GeneID:105288189 | LOC105288189 | Gene | LOC105288189 | protein_coding | NA | character(0) | character(0) | BCphaseA1.vs.RphaseB1 | FALSE |
| NW_011924881.1 | 449174 | 449174 | 1 | + | 46 | 0 | 46 | 36 | 0 | 36 | 0 | 0 | NW_011924881.1 | 448307 | 454087 | 5781 | - | Gnomon | gene | gene26 | GeneID:105288189 | LOC105288189 | Gene | LOC105288189 | protein_coding | NA | character(0) | character(0) | BCphaseA1.vs.RphaseB1 | FALSE |
| NW_011924881.1 | 449192 | 449192 | 1 | + | 37 | 0 | 37 | 32 | 0 | 32 | 0 | 0 | NW_011924881.1 | 448307 | 454087 | 5781 | - | Gnomon | gene | gene26 | GeneID:105288189 | LOC105288189 | Gene | LOC105288189 | protein_coding | NA | character(0) | character(0) | BCphaseA1.vs.RphaseB1 | FALSE |
| NW_011924881.1 | 449197 | 449197 | 1 | + | 36 | 1 | 35 | 32 | 2 | 30 | 2.78 | 6.25 | NW_011924881.1 | 448307 | 454087 | 5781 | - | Gnomon | gene | gene26 | GeneID:105288189 | LOC105288189 | Gene | LOC105288189 | protein_coding | NA | character(0) | character(0) | BCphaseA1.vs.RphaseB1 | FALSE |
| NW_011924881.1 | 449201 | 449201 | 1 | + | 27 | 0 | 27 | 28 | 0 | 28 | 0 | 0 | NW_011924881.1 | 448307 | 454087 | 5781 | - | Gnomon | gene | gene26 | GeneID:105288189 | LOC105288189 | Gene | LOC105288189 | protein_coding | NA | character(0) | character(0) | BCphaseA1.vs.RphaseB1 | FALSE |
| NW_011924881.1 | 449208 | 449208 | 1 | + | 27 | 2 | 25 | 31 | 1 | 30 | 7.41 | 3.23 | NW_011924881.1 | 448307 | 454087 | 5781 | - | Gnomon | gene | gene26 | GeneID:105288189 | LOC105288189 | Gene | LOC105288189 | protein_coding | NA | character(0) | character(0) | BCphaseA1.vs.RphaseB1 | FALSE |
| NW_011924881.1 | 449220 | 449220 | 1 | + | 27 | 0 | 27 | 31 | 1 | 30 | 0 | 3.23 | NW_011924881.1 | 448307 | 454087 | 5781 | - | Gnomon | gene | gene26 | GeneID:105288189 | LOC105288189 | Gene | LOC105288189 | protein_coding | NA | character(0) | character(0) | BCphaseA1.vs.RphaseB1 | FALSE |
| NW_011924881.1 | 449229 | 449229 | 1 | + | 27 | 0 | 27 | 28 | 0 | 28 | 0 | 0 | NW_011924881.1 | 448307 | 454087 | 5781 | - | Gnomon | gene | gene26 | GeneID:105288189 | LOC105288189 | Gene | LOC105288189 | protein_coding | NA | character(0) | character(0) | BCphaseA1.vs.RphaseB1 | FALSE |
| NW_011924881.1 | 449250 | 449250 | 1 | + | 27 | 2 | 25 | 31 | 3 | 28 | 7.41 | 9.68 | NW_011924881.1 | 448307 | 454087 | 5781 | - | Gnomon | gene | gene26 | GeneID:105288189 | LOC105288189 | Gene | LOC105288189 | protein_coding | NA | character(0) | character(0) | BCphaseA1.vs.RphaseB1 | FALSE |
| NW_011924881.1 | 449293 | 449293 | 1 | + | 29 | 1 | 28 | 32 | 2 | 30 | 3.45 | 6.25 | NW_011924881.1 | 448307 | 454087 | 5781 | - | Gnomon | gene | gene26 | GeneID:105288189 | LOC105288189 | Gene | LOC105288189 | protein_coding | NA | character(0) | character(0) | BCphaseA1.vs.RphaseB1 | FALSE |
| NW_011924881.1 | 449296 | 449296 | 1 | + | 29 | 0 | 29 | 32 | 0 | 32 | 0 | 0 | NW_011924881.1 | 448307 | 454087 | 5781 | - | Gnomon | gene | gene26 | GeneID:105288189 | LOC105288189 | Gene | LOC105288189 | protein_coding | NA | character(0) | character(0) | BCphaseA1.vs.RphaseB1 | FALSE |
| NW_011924881.1 | 449314 | 449314 | 1 | + | 21 | 0 | 21 | 25 | 0 | 25 | 0 | 0 | NW_011924881.1 | 448307 | 454087 | 5781 | - | Gnomon | gene | gene26 | GeneID:105288189 | LOC105288189 | Gene | LOC105288189 | protein_coding | NA | character(0) | character(0) | BCphaseA1.vs.RphaseB1 | FALSE |
| NW_011924881.1 | 449318 | 449318 | 1 | + | 25 | 0 | 25 | 33 | 1 | 32 | 0 | 3.03 | NW_011924881.1 | 448307 | 454087 | 5781 | - | Gnomon | gene | gene26 | GeneID:105288189 | LOC105288189 | Gene | LOC105288189 | protein_coding | NA | character(0) | character(0) | BCphaseA1.vs.RphaseB1 | FALSE |
| NW_011924881.1 | 449321 | 449321 | 1 | + | 23 | 0 | 23 | 33 | 0 | 33 | 0 | 0 | NW_011924881.1 | 448307 | 454087 | 5781 | - | Gnomon | gene | gene26 | GeneID:105288189 | LOC105288189 | Gene | LOC105288189 | protein_coding | NA | character(0) | character(0) | BCphaseA1.vs.RphaseB1 | FALSE |

...
... only first 25 lines shown ...

Directory: DMSG File: dmg-Cb_LI_BCphaseA1.vs.RphaseB2_details.txt

```
seqnames	start	end	width	strand	coverage1	numCs1	numTs1	coverage2	numCs2	numTs2	BCphaseA1	RphaseB2	gene_seqnames	gene_start	gene_end
	gene_width	gene_strand	gene_source	gene_type	gene_ID	gene_Dbxref	gene_Name	gene_gbkey	gene_gene	gene_gene_biotype	gene_partial	gene_start_range	gene_end_range
	gene_comparison	is.dm
NW_011924879.1	1085	1085	1	+	23	0	23	32	0	32	0	0	NW_011924879.1	7	3079	3073	+	Gnomon	gene	gene5	GeneID:105275124
	LOC105275124	Gene	LOC105275124	protein_coding	NA	character(0)	character(0)	BCphaseA1.vs.RphaseB2	FALSE
NW_011924879.1	1110	1110	1	+	30	0	30	40	0	40	0	0	NW_011924879.1	7	3079	3073	+	Gnomon	gene	gene5	GeneID:105275124
	LOC105275124	Gene	LOC105275124	protein_coding	NA	character(0)	character(0)	BCphaseA1.vs.RphaseB2	FALSE
NW_011924879.1	1131	1131	1	+	32	0	32	45	0	45	0	0	NW_011924879.1	7	3079	3073	+	Gnomon	gene	gene5	GeneID:105275124
	LOC105275124	Gene	LOC105275124	protein_coding	NA	character(0)	character(0)	BCphaseA1.vs.RphaseB2	FALSE
NW_011924879.1	1148	1148	1	+	27	0	27	30	2	28	0	6.67	NW_011924879.1	7	3079	3073	+	Gnomon	gene	gene5	GeneID:105275124
	LOC105275124	Gene	LOC105275124	protein_coding	NA	character(0)	character(0)	BCphaseA1.vs.RphaseB2	FALSE
NW_011924879.1	1155	1155	1	+	27	0	27	24	0	24	0	0	NW_011924879.1	7	3079	3073	+	Gnomon	gene	gene5	GeneID:105275124
	LOC105275124	Gene	LOC105275124	protein_coding	NA	character(0)	character(0)	BCphaseA1.vs.RphaseB2	FALSE
NW_011924879.1	1177	1177	1	+	22	1	21	21	1	20	4.55	4.76	NW_011924879.1	7	3079	3073	+	Gnomon	gene	gene5	GeneID:105275124
	LOC105275124	Gene	LOC105275124	protein_coding	NA	character(0)	character(0)	BCphaseA1.vs.RphaseB2	FALSE
NW_011924879.1	1180	1180	1	+	25	1	24	20	2	18	4	10	NW_011924879.1	7	3079	3073	+	Gnomon	gene	gene5	GeneID:105275124
	LOC105275124	Gene	LOC105275124	protein_coding	NA	character(0)	character(0)	BCphaseA1.vs.RphaseB2	FALSE
NW_011924879.1	1309	1309	1	+	26	0	26	22	0	22	0	0	NW_011924879.1	7	3079	3073	+	Gnomon	gene	gene5	GeneID:105275124
	LOC105275124	Gene	LOC105275124	protein_coding	NA	character(0)	character(0)	BCphaseA1.vs.RphaseB2	FALSE
NW_011924879.1	1311	1311	1	+	26	0	26	23	0	23	0	0	NW_011924879.1	7	3079	3073	+	Gnomon	gene	gene5	GeneID:105275124
	LOC105275124	Gene	LOC105275124	protein_coding	NA	character(0)	character(0)	BCphaseA1.vs.RphaseB2	FALSE
NW_011924879.1	1313	1313	1	+	26	0	26	23	0	23	0	0	NW_011924879.1	7	3079	3073	+	Gnomon	gene	gene5	GeneID:105275124
	LOC105275124	Gene	LOC105275124	protein_coding	NA	character(0)	character(0)	BCphaseA1.vs.RphaseB2	FALSE
NW_011924879.1	1320	1320	1	+	27	0	27	25	0	25	0	0	NW_011924879.1	7	3079	3073	+	Gnomon	gene	gene5	GeneID:105275124
	LOC105275124	Gene	LOC105275124	protein_coding	NA	character(0)	character(0)	BCphaseA1.vs.RphaseB2	FALSE
NW_011924879.1	1332	1332	1	+	25	0	25	24	0	24	0	0	NW_011924879.1	7	3079	3073	+	Gnomon	gene	gene5	GeneID:105275124
	LOC105275124	Gene	LOC105275124	protein_coding	NA	character(0)	character(0)	BCphaseA1.vs.RphaseB2	FALSE
NW_011924879.1	1334	1334	1	+	24	0	24	24	0	24	0	0	NW_011924879.1	7	3079	3073	+	Gnomon	gene	gene5	GeneID:105275124
	LOC105275124	Gene	LOC105275124	protein_coding	NA	character(0)	character(0)	BCphaseA1.vs.RphaseB2	FALSE
NW_011924879.1	1337	1337	1	+	24	0	24	23	0	23	0	0	NW_011924879.1	7	3079	3073	+	Gnomon	gene	gene5	GeneID:105275124
	LOC105275124	Gene	LOC105275124	protein_coding	NA	character(0)	character(0)	BCphaseA1.vs.RphaseB2	FALSE
NW_011924879.1	1341	1341	1	+	24	0	24	22	0	22	0	0	NW_011924879.1	7	3079	3073	+	Gnomon	gene	gene5	GeneID:105275124
	LOC105275124	Gene	LOC105275124	protein_coding	NA	character(0)	character(0)	BCphaseA1.vs.RphaseB2	FALSE
NW_011924879.1	1344	1344	1	+	24	0	24	22	0	22	0	0	NW_011924879.1	7	3079	3073	+	Gnomon	gene	gene5	GeneID:105275124
	LOC105275124	Gene	LOC105275124	protein_coding	NA	character(0)	character(0)	BCphaseA1.vs.RphaseB2	FALSE
NW_011924879.1	2162	2162	1	+	22	1	21	34	9	25	4.55	26.47	NW_011924879.1	7	3079	3073	+	Gnomon	gene	gene5	GeneID:105275124
	LOC105275124	Gene	LOC105275124	protein_coding	NA	character(0)	character(0)	BCphaseA1.vs.RphaseB2	TRUE
NW_011924881.1	44105	44105	1	+	36	0	36	59	0	59	0	0	NW_011924881.1	42715	135016	92302	-	Gnomon	gene	gene18
	GeneID:105287290	LOC105287290	Gene	LOC105287290	protein_coding	NA	character(0)	character(0)	BCphaseA1.vs.RphaseB2	FALSE
NW_011924881.1	44112	44112	1	+	37	0	37	57	0	57	0	0	NW_011924881.1	42715	135016	92302	-	Gnomon	gene	gene18
	GeneID:105287290	LOC105287290	Gene	LOC105287290	protein_coding	NA	character(0)	character(0)	BCphaseA1.vs.RphaseB2	FALSE
NW_011924881.1	44121	44121	1	+	37	0	37	56	0	56	0	0	NW_011924881.1	42715	135016	92302	-	Gnomon	gene	gene18
	GeneID:105287290	LOC105287290	Gene	LOC105287290	protein_coding	NA	character(0)	character(0)	BCphaseA1.vs.RphaseB2	FALSE
NW_011924881.1	44123	44123	1	+	37	0	37	56	0	56	0	0	NW_011924881.1	42715	135016	92302	-	Gnomon	gene	gene18
	GeneID:105287290	LOC105287290	Gene	LOC105287290	protein_coding	NA	character(0)	character(0)	BCphaseA1.vs.RphaseB2	FALSE
NW_011924881.1	44125	44125	1	+	36	0	36	55	0	55	0	0	NW_011924881.1	42715	135016	92302	-	Gnomon	gene	gene18
	GeneID:105287290	LOC105287290	Gene	LOC105287290	protein_coding	NA	character(0)	character(0)	BCphaseA1.vs.RphaseB2	FALSE
NW_011924881.1	44129	44129	1	+	37	0	37	54	0	54	0	0	NW_011924881.1	42715	135016	92302	-	Gnomon	gene	gene18
	GeneID:105287290	LOC105287290	Gene	LOC105287290	protein_coding	NA	character(0)	character(0)	BCphaseA1.vs.RphaseB2	FALSE
NW_011924881.1	44138	44138	1	+	36	0	36	52	0	52	0	0	NW_011924881.1	42715	135016	92302	-	Gnomon	gene	gene18
	GeneID:105287290	LOC105287290	Gene	LOC105287290	protein_coding	NA	character(0)	character(0)	BCphaseA1.vs.RphaseB2	FALSE
...
... only first 25 lines shown ...
```

Directory: DMSG File: dmg-Cb_LI_BCphaseA2.vs.BCphaseB1_details.txt

| seqnames | start | end | width | strand | coverage1 | numCs1 | numTs1 | coverage2 | numCs2 | numTs2 | BCphaseA2 | BCphaseB1 | gene_seqnames | gene_start | gene_end | gene_width | gene_strand | gene_source | gene_type | gene_ID | gene_Dbxref | gene_Name | gene_gbkey | gene_gene | gene_gene_biotype | gene_start_range | gene_end_range | gene_comparison | is.dm |
|---|---|---|---|---|---|---|---|---|---|---|---|---|---|---|---|---|---|---|---|---|---|---|---|---|---|---|---|---|---|
| NW_011924892.1 | 1124938 | 1124938 | 1 | + | 36 | 17 | 19 | 37 | 16 | 21 | 47.22 | 43.24 | NW_011924892.1 | 1124810 | 1127611 | 2802 | - | Gnomon | gene | gene564 | GeneID:105278478 | LOC105278478 | Gene | LOC105278478 | protein_coding | character(0) | character(0) | BCphaseA2.vs.BCphaseB1 | FALSE |
| NW_011924892.1 | 1124976 | 1124976 | 1 | + | 54 | 5 | 49 | 58 | 1 | 57 | 9.26 | 1.72 | NW_011924892.1 | 1124810 | 1127611 | 2802 | - | Gnomon | gene | gene564 | GeneID:105278478 | LOC105278478 | Gene | LOC105278478 | protein_coding | character(0) | character(0) | BCphaseA2.vs.BCphaseB1 | FALSE |
| NW_011924892.1 | 1125023 | 1125023 | 1 | + | 36 | 12 | 24 | 39 | 20 | 19 | 33.33 | 51.28 | NW_011924892.1 | 1124810 | 1127611 | 2802 | - | Gnomon | gene | gene564 | GeneID:105278478 | LOC105278478 | Gene | LOC105278478 | protein_coding | character(0) | character(0) | BCphaseA2.vs.BCphaseB1 | FALSE |
| NW_011924892.1 | 1125031 | 1125031 | 1 | + | 32 | 11 | 21 | 35 | 17 | 18 | 34.38 | 48.57 | NW_011924892.1 | 1124810 | 1127611 | 2802 | - | Gnomon | gene | gene564 | GeneID:105278478 | LOC105278478 | Gene | LOC105278478 | protein_coding | character(0) | character(0) | BCphaseA2.vs.BCphaseB1 | FALSE |
| NW_011924892.1 | 1125137 | 1125137 | 1 | + | 61 | 24 | 37 | 62 | 1 | 61 | 39.34 | 1.61 | NW_011924892.1 | 1124810 | 1127611 | 2802 | - | Gnomon | gene | gene564 | GeneID:105278478 | LOC105278478 | Gene | LOC105278478 | protein_coding | character(0) | character(0) | BCphaseA2.vs.BCphaseB1 | TRUE |
| NW_011924892.1 | 1125159 | 1125159 | 1 | + | 53 | 2 | 51 | 58 | 1 | 57 | 3.77 | 1.72 | NW_011924892.1 | 1124810 | 1127611 | 2802 | - | Gnomon | gene | gene564 | GeneID:105278478 | LOC105278478 | Gene | LOC105278478 | protein_coding | character(0) | character(0) | BCphaseA2.vs.BCphaseB1 | FALSE |
| NW_011924892.1 | 1125173 | 1125173 | 1 | + | 52 | 3 | 49 | 48 | 4 | 44 | 5.77 | 8.33 | NW_011924892.1 | 1124810 | 1127611 | 2802 | - | Gnomon | gene | gene564 | GeneID:105278478 | LOC105278478 | Gene | LOC105278478 | protein_coding | character(0) | character(0) | BCphaseA2.vs.BCphaseB1 | FALSE |
| NW_011924892.1 | 1125176 | 1125176 | 1 | + | 51 | 6 | 45 | 48 | 9 | 39 | 11.76 | 18.75 | NW_011924892.1 | 1124810 | 1127611 | 2802 | - | Gnomon | gene | gene564 | GeneID:105278478 | LOC105278478 | Gene | LOC105278478 | protein_coding | character(0) | character(0) | BCphaseA2.vs.BCphaseB1 | FALSE |
| NW_011924892.1 | 1125182 | 1125182 | 1 | + | 48 | 6 | 42 | 46 | 5 | 41 | 12.5 | 10.87 | NW_011924892.1 | 1124810 | 1127611 | 2802 | - | Gnomon | gene | gene564 | GeneID:105278478 | LOC105278478 | Gene | LOC105278478 | protein_coding | character(0) | character(0) | BCphaseA2.vs.BCphaseB1 | FALSE |
| NW_011924892.1 | 1125188 | 1125188 | 1 | + | 42 | 5 | 37 | 43 | 6 | 37 | 11.9 | 13.95 | NW_011924892.1 | 1124810 | 1127611 | 2802 | - | Gnomon | gene | gene564 | GeneID:105278478 | LOC105278478 | Gene | LOC105278478 | protein_coding | character(0) | character(0) | BCphaseA2.vs.BCphaseB1 | FALSE |
| NW_011924892.1 | 1125200 | 1125200 | 1 | + | 43 | 19 | 24 | 43 | 15 | 28 | 44.19 | 34.88 | NW_011924892.1 | 1124810 | 1127611 | 2802 | - | Gnomon | gene | gene564 | GeneID:105278478 | LOC105278478 | Gene | LOC105278478 | protein_coding | character(0) | character(0) | BCphaseA2.vs.BCphaseB1 | FALSE |
| NW_011924892.1 | 1126219 | 1126219 | 1 | + | 23 | 0 | 23 | 31 | 0 | 31 | 0 | 0 | NW_011924892.1 | 1124810 | 1127611 | 2802 | - | Gnomon | gene | gene564 | GeneID:105278478 | LOC105278478 | Gene | LOC105278478 | protein_coding | character(0) | character(0) | BCphaseA2.vs.BCphaseB1 | FALSE |
| NW_011924892.1 | 1126228 | 1126228 | 1 | + | 25 | 0 | 25 | 31 | 0 | 31 | 0 | 0 | NW_011924892.1 | 1124810 | 1127611 | 2802 | - | Gnomon | gene | gene564 | GeneID:105278478 | LOC105278478 | Gene | LOC105278478 | protein_coding | character(0) | character(0) | BCphaseA2.vs.BCphaseB1 | FALSE |
| NW_011924892.1 | 1126242 | 1126242 | 1 | + | 26 | 0 | 26 | 27 | 1 | 26 | 0 | 3.7 | NW_011924892.1 | 1124810 | 1127611 | 2802 | - | Gnomon | gene | gene564 | GeneID:105278478 | LOC105278478 | Gene | LOC105278478 | protein_coding | character(0) | character(0) | BCphaseA2.vs.BCphaseB1 | FALSE |
| NW_011924892.1 | 1126244 | 1126244 | 1 | + | 25 | 0 | 25 | 27 | 0 | 27 | 0 | 0 | NW_011924892.1 | 1124810 | 1127611 | 2802 | - | Gnomon | gene | gene564 | GeneID:105278478 | LOC105278478 | Gene | LOC105278478 | protein_coding | character(0) | character(0) | BCphaseA2.vs.BCphaseB1 | FALSE |
| NW_011924892.1 | 1126248 | 1126248 | 1 | + | 24 | 0 | 24 | 27 | 0 | 27 | 0 | 0 | NW_011924892.1 | 1124810 | 1127611 | 2802 | - | Gnomon | gene | gene564 | GeneID:105278478 | LOC105278478 | Gene | LOC105278478 | protein_coding | character(0) | character(0) | BCphaseA2.vs.BCphaseB1 | FALSE |
| NW_011924892.1 | 1126254 | 1126254 | 1 | + | 24 | 0 | 24 | 27 | 0 | 27 | 0 | 0 | NW_011924892.1 | 1124810 | 1127611 | 2802 | - | Gnomon | gene | gene564 | GeneID:105278478 | LOC105278478 | Gene | LOC105278478 | protein_coding | character(0) | character(0) | BCphaseA2.vs.BCphaseB1 | FALSE |
| NW_011924892.1 | 1160882 | 1160882 | 1 | + | 37 | 1 | 36 | 22 | 0 | 22 | 2.7 | 0 | NW_011924892.1 | 1160484 | 1162156 | 1673 | + | Gnomon | gene | gene570 | GeneID:105278549 | LOC105278549 | Gene | LOC105278549 | protein_coding | character(0) | character(0) | BCphaseA2.vs.BCphaseB1 | FALSE |
| NW_011924892.1 | 1160884 | 1160884 | 1 | + | 36 | 2 | 34 | 23 | 2 | 21 | 5.56 | 8.7 | NW_011924892.1 | 1160484 | 1162156 | 1673 | + | Gnomon | gene | gene570 | GeneID:105278549 | LOC105278549 | Gene | LOC105278549 | protein_coding | character(0) | character(0) | BCphaseA2.vs.BCphaseB1 | FALSE |
| NW_011924892.1 | 1160898 | 1160898 | 1 | + | 36 | 0 | 36 | 23 | 1 | 22 | 0 | 4.35 | NW_011924892.1 | 1160484 | 1162156 | 1673 | + | Gnomon | gene | gene570 | GeneID:105278549 | LOC105278549 | Gene | LOC105278549 | protein_coding | character(0) | character(0) | BCphaseA2.vs.BCphaseB1 | FALSE |
| NW_011924892.1 | 1160905 | 1160905 | 1 | + | 62 | 0 | 62 | 49 | 1 | 48 | 0 | 2.04 | NW_011924892.1 | 1160484 | 1162156 | 1673 | + | Gnomon | gene | gene570 | GeneID:105278549 | LOC105278549 | Gene | LOC105278549 | protein_coding | character(0) | character(0) | BCphaseA2.vs.BCphaseB1 | FALSE |
| NW_011924892.1 | 1160910 | 1160910 | 1 | + | 60 | 2 | 58 | 58 | 2 | 56 | 3.33 | 3.45 | NW_011924892.1 | 1160484 | 1162156 | 1673 | + | Gnomon | gene | gene570 | GeneID:105278549 | LOC105278549 | Gene | LOC105278549 | protein_coding | character(0) | character(0) | BCphaseA2.vs.BCphaseB1 | FALSE |
| NW_011924892.1 | 1160928 | 1160928 | 1 | + | 63 | 0 | 63 | 60 | 0 | 60 | 0 | 0 | NW_011924892.1 | 1160484 | 1162156 | 1673 | + | Gnomon | gene | gene570 | GeneID:105278549 | LOC105278549 | Gene | LOC105278549 | protein_coding | character(0) | character(0) | BCphaseA2.vs.BCphaseB1 | FALSE |
| NW_011924892.1 | 1160953 | 1160953 | 1 | + | 96 | 5 | 91 | 63 | 2 | 61 | 5.21 | 3.17 | NW_011924892.1 | 1160484 | 1162156 | 1673 | + | Gnomon | gene | gene570 | GeneID:105278549 | LOC105278549 | Gene | LOC105278549 | protein_coding | character(0) | character(0) | BCphaseA2.vs.BCphaseB1 | FALSE |

...

... only first 25 lines shown ...

Directory: DMSG File: dmg-Cb_LI_BCphaseA2.vs.BCphaseB2_details.txt

| seqnames | start | end | width | strand | coverage1 | numCs1 | numTs1 | coverage2 | numCs2 | numTs2 | BCphaseA2 | BCphaseB2 | gene_seqnames | gene_start | gene_end | gene_width | gene_strand | gene_source | gene_type | gene_ID | gene_Dbxref | gene_Name | gene_gbkey | gene_gene | gene_gene_biotype | gene_start_range | gene_end_range | gene_comparison | is.dm |
|---|---|---|---|---|---|---|---|---|---|---|---|---|---|---|---|---|---|---|---|---|---|---|---|---|---|---|---|---|---|
| NW_011924881.1 | 809504 | 809504 | 1 | + | 20 | 0 | 20 | 23 | 0 | 23 | 0 | 0 | NW_011924881.1 | 808343 | 827625 | 19283 | - | Gnomon | gene | gene53 | GeneID:105277148 | LOC105277148 | Gene | LOC105277148 | protein_coding | character(0) | character(0) | BCphaseA2.vs.BCphaseB2 | FALSE |
| NW_011924881.1 | 810155 | 810155 | 1 | + | 22 | 0 | 22 | 25 | 0 | 25 | 0 | 0 | NW_011924881.1 | 808343 | 827625 | 19283 | - | Gnomon | gene | gene53 | GeneID:105277148 | LOC105277148 | Gene | LOC105277148 | protein_coding | character(0) | character(0) | BCphaseA2.vs.BCphaseB2 | FALSE |
| NW_011924881.1 | 810182 | 810182 | 1 | + | 23 | 0 | 23 | 28 | 2 | 26 | 0 | 7.14 | NW_011924881.1 | 808343 | 827625 | 19283 | - | Gnomon | gene | gene53 | GeneID:105277148 | LOC105277148 | Gene | LOC105277148 | protein_coding | character(0) | character(0) | BCphaseA2.vs.BCphaseB2 | FALSE |
| NW_011924881.1 | 810335 | 810335 | 1 | + | 23 | 0 | 23 | 20 | 0 | 20 | 0 | 0 | NW_011924881.1 | 808343 | 827625 | 19283 | - | Gnomon | gene | gene53 | GeneID:105277148 | LOC105277148 | Gene | LOC105277148 | protein_coding | character(0) | character(0) | BCphaseA2.vs.BCphaseB2 | FALSE |
| NW_011924881.1 | 810337 | 810337 | 1 | + | 24 | 0 | 24 | 20 | 0 | 20 | 0 | 0 | NW_011924881.1 | 808343 | 827625 | 19283 | - | Gnomon | gene | gene53 | GeneID:105277148 | LOC105277148 | Gene | LOC105277148 | protein_coding | character(0) | character(0) | BCphaseA2.vs.BCphaseB2 | FALSE |
| NW_011924881.1 | 810348 | 810348 | 1 | + | 51 | 0 | 51 | 43 | 0 | 43 | 0 | 0 | NW_011924881.1 | 808343 | 827625 | 19283 | - | Gnomon | gene | gene53 | GeneID:105277148 | LOC105277148 | Gene | LOC105277148 | protein_coding | character(0) | character(0) | BCphaseA2.vs.BCphaseB2 | FALSE |
| NW_011924881.1 | 810391 | 810391 | 1 | + | 54 | 1 | 53 | 38 | 0 | 38 | 1.85 | 0 | NW_011924881.1 | 808343 | 827625 | 19283 | - | Gnomon | gene | gene53 | GeneID:105277148 | LOC105277148 | Gene | LOC105277148 | protein_coding | character(0) | character(0) | BCphaseA2.vs.BCphaseB2 | FALSE |
| NW_011924881.1 | 810409 | 810409 | 1 | + | 61 | 2 | 59 | 32 | 0 | 32 | 3.28 | 0 | NW_011924881.1 | 808343 | 827625 | 19283 | - | Gnomon | gene | gene53 | GeneID:105277148 | LOC105277148 | Gene | LOC105277148 | protein_coding | character(0) | character(0) | BCphaseA2.vs.BCphaseB2 | FALSE |
| NW_011924881.1 | 810412 | 810412 | 1 | + | 40 | 0 | 40 | 32 | 1 | 31 | 0 | 3.12 | NW_011924881.1 | 808343 | 827625 | 19283 | - | Gnomon | gene | gene53 | GeneID:105277148 | LOC105277148 | Gene | LOC105277148 | protein_coding | character(0) | character(0) | BCphaseA2.vs.BCphaseB2 | FALSE |
| NW_011924881.1 | 810417 | 810417 | 1 | + | 38 | 1 | 37 | 35 | 0 | 35 | 2.63 | 0 | NW_011924881.1 | 808343 | 827625 | 19283 | - | Gnomon | gene | gene53 | GeneID:105277148 | LOC105277148 | Gene | LOC105277148 | protein_coding | character(0) | character(0) | BCphaseA2.vs.BCphaseB2 | FALSE |
| NW_011924881.1 | 810424 | 810424 | 1 | + | 37 | 0 | 37 | 36 | 0 | 36 | 0 | 0 | NW_011924881.1 | 808343 | 827625 | 19283 | - | Gnomon | gene | gene53 | GeneID:105277148 | LOC105277148 | Gene | LOC105277148 | protein_coding | character(0) | character(0) | BCphaseA2.vs.BCphaseB2 | FALSE |
| NW_011924881.1 | 810430 | 810430 | 1 | + | 27 | 0 | 27 | 31 | 0 | 31 | 0 | 0 | NW_011924881.1 | 808343 | 827625 | 19283 | - | Gnomon | gene | gene53 | GeneID:105277148 | LOC105277148 | Gene | LOC105277148 | protein_coding | character(0) | character(0) | BCphaseA2.vs.BCphaseB2 | FALSE |
| NW_011924881.1 | 810436 | 810436 | 1 | + | 27 | 0 | 27 | 32 | 0 | 32 | 0 | 0 | NW_011924881.1 | 808343 | 827625 | 19283 | - | Gnomon | gene | gene53 | GeneID:105277148 | LOC105277148 | Gene | LOC105277148 | protein_coding | character(0) | character(0) | BCphaseA2.vs.BCphaseB2 | FALSE |
| NW_011924881.1 | 810505 | 810505 | 1 | + | 22 | 0 | 22 | 55 | 0 | 55 | 0 | 0 | NW_011924881.1 | 808343 | 827625 | 19283 | - | Gnomon | gene | gene53 | GeneID:105277148 | LOC105277148 | Gene | LOC105277148 | protein_coding | character(0) | character(0) | BCphaseA2.vs.BCphaseB2 | FALSE |
| NW_011924881.1 | 810511 | 810511 | 1 | + | 22 | 1 | 21 | 54 | 0 | 54 | 4.55 | 0 | NW_011924881.1 | 808343 | 827625 | 19283 | - | Gnomon | gene | gene53 | GeneID:105277148 | LOC105277148 | Gene | LOC105277148 | protein_coding | character(0) | character(0) | BCphaseA2.vs.BCphaseB2 | FALSE |
| NW_011924881.1 | 810516 | 810516 | 1 | + | 23 | 0 | 23 | 53 | 0 | 53 | 0 | 0 | NW_011924881.1 | 808343 | 827625 | 19283 | - | Gnomon | gene | gene53 | GeneID:105277148 | LOC105277148 | Gene | LOC105277148 | protein_coding | character(0) | character(0) | BCphaseA2.vs.BCphaseB2 | FALSE |
| NW_011924881.1 | 810520 | 810520 | 1 | + | 23 | 0 | 23 | 51 | 1 | 50 | 0 | 1.96 | NW_011924881.1 | 808343 | 827625 | 19283 | - | Gnomon | gene | gene53 | GeneID:105277148 | LOC105277148 | Gene | LOC105277148 | protein_coding | character(0) | character(0) | BCphaseA2.vs.BCphaseB2 | FALSE |
| NW_011924881.1 | 810633 | 810633 | 1 | + | 20 | 0 | 20 | 37 | 0 | 37 | 0 | 0 | NW_011924881.1 | 808343 | 827625 | 19283 | - | Gnomon | gene | gene53 | GeneID:105277148 | LOC105277148 | Gene | LOC105277148 | protein_coding | character(0) | character(0) | BCphaseA2.vs.BCphaseB2 | FALSE |
| NW_011924881.1 | 810645 | 810645 | 1 | + | 21 | 0 | 21 | 37 | 0 | 37 | 0 | 0 | NW_011924881.1 | 808343 | 827625 | 19283 | - | Gnomon | gene | gene53 | GeneID:105277148 | LOC105277148 | Gene | LOC105277148 | protein_coding | character(0) | character(0) | BCphaseA2.vs.BCphaseB2 | FALSE |
| NW_011924881.1 | 811042 | 811042 | 1 | + | 21 | 0 | 21 | 21 | 0 | 21 | 0 | 0 | NW_011924881.1 | 808343 | 827625 | 19283 | - | Gnomon | gene | gene53 | GeneID:105277148 | LOC105277148 | Gene | LOC105277148 | protein_coding | character(0) | character(0) | BCphaseA2.vs.BCphaseB2 | FALSE |
| NW_011924881.1 | 811083 | 811083 | 1 | + | 26 | 0 | 26 | 29 | 0 | 29 | 0 | 0 | NW_011924881.1 | 808343 | 827625 | 19283 | - | Gnomon | gene | gene53 | GeneID:105277148 | LOC105277148 | Gene | LOC105277148 | protein_coding | character(0) | character(0) | BCphaseA2.vs.BCphaseB2 | FALSE |
| NW_011924881.1 | 811089 | 811089 | 1 | + | 27 | 0 | 27 | 29 | 0 | 29 | 0 | 0 | NW_011924881.1 | 808343 | 827625 | 19283 | - | Gnomon | gene | gene53 | GeneID:105277148 | LOC105277148 | Gene | LOC105277148 | protein_coding | character(0) | character(0) | BCphaseA2.vs.BCphaseB2 | FALSE |
| NW_011924881.1 | 811120 | 811120 | 1 | + | 23 | 0 | 23 | 24 | 0 | 24 | 0 | 0 | NW_011924881.1 | 808343 | 827625 | 19283 | - | Gnomon | gene | gene53 | GeneID:105277148 | LOC105277148 | Gene | LOC105277148 | protein_coding | character(0) | character(0) | BCphaseA2.vs.BCphaseB2 | FALSE |
| NW_011924881.1 | 811122 | 811122 | 1 | + | 23 | 0 | 23 | 25 | 0 | 25 | 0 | 0 | NW_011924881.1 | 808343 | 827625 | 19283 | - | Gnomon | gene | gene53 | GeneID:105277148 | LOC105277148 | Gene | LOC105277148 | protein_coding | character(0) | character(0) | BCphaseA2.vs.BCphaseB2 | FALSE |

...
... only first 25 lines shown ...

Directory: DMSG File: dmg-Cb_LI_BCphaseA2.vs.RphaseA1_details.txt

| seqnames | start | end | width | strand | coverage1 | numCs1 | numTs1 | coverage2 | numCs2 | numTs2 | BCphaseA2 | RphaseA1 | gene_seqnames | gene_start | gene_end | gene_width | gene_strand | gene_source | gene_type | gene_ID | gene_Dbxref | gene_Name | gene_gbkey | gene_gene | gene_gene_biotype | gene_partial | gene_start_range | gene_end_range | gene_comparison | is.dm |
|---|---|---|---|---|---|---|---|---|---|---|---|---|---|---|---|---|---|---|---|---|---|---|---|---|---|---|---|---|---|---|
| NW_011924881.1 | 448501 | 448501 | 1 | + | 26 | 9 | 17 | 28 | 6 | 22 | 34.62 | 21.43 | NW_011924881.1 | 448307 | 454087 | 5781 | - | Gnomon | gene | gene26 | GeneID:105288189 | LOC105288189 | Gene | LOC105288189 | protein_coding | NA | character(0) | character(0) | BCphaseA2.vs.RphaseA1 | FALSE |
| NW_011924881.1 | 448684 | 448684 | 1 | + | 21 | 9 | 12 | 25 | 6 | 19 | 42.86 | 24 | NW_011924881.1 | 448307 | 454087 | 5781 | - | Gnomon | gene | gene26 | GeneID:105288189 | LOC105288189 | Gene | LOC105288189 | protein_coding | NA | character(0) | character(0) | BCphaseA2.vs.RphaseA1 | FALSE |
| NW_011924881.1 | 448698 | 448698 | 1 | + | 26 | 2 | 24 | 30 | 4 | 26 | 7.69 | 13.33 | NW_011924881.1 | 448307 | 454087 | 5781 | - | Gnomon | gene | gene26 | GeneID:105288189 | LOC105288189 | Gene | LOC105288189 | protein_coding | NA | character(0) | character(0) | BCphaseA2.vs.RphaseA1 | FALSE |
| NW_011924881.1 | 448707 | 448707 | 1 | + | 26 | 2 | 24 | 31 | 6 | 25 | 7.69 | 19.35 | NW_011924881.1 | 448307 | 454087 | 5781 | - | Gnomon | gene | gene26 | GeneID:105288189 | LOC105288189 | Gene | LOC105288189 | protein_coding | NA | character(0) | character(0) | BCphaseA2.vs.RphaseA1 | FALSE |
| NW_011924881.1 | 448725 | 448725 | 1 | + | 29 | 9 | 20 | 32 | 6 | 26 | 31.03 | 18.75 | NW_011924881.1 | 448307 | 454087 | 5781 | - | Gnomon | gene | gene26 | GeneID:105288189 | LOC105288189 | Gene | LOC105288189 | protein_coding | NA | character(0) | character(0) | BCphaseA2.vs.RphaseA1 | FALSE |
| NW_011924881.1 | 448952 | 448952 | 1 | + | 21 | 0 | 21 | 22 | 0 | 22 | 0 | 0 | NW_011924881.1 | 448307 | 454087 | 5781 | - | Gnomon | gene | gene26 | GeneID:105288189 | LOC105288189 | Gene | LOC105288189 | protein_coding | NA | character(0) | character(0) | BCphaseA2.vs.RphaseA1 | FALSE |
| NW_011924881.1 | 449119 | 449119 | 1 | + | 47 | 1 | 46 | 36 | 1 | 35 | 2.13 | 2.78 | NW_011924881.1 | 448307 | 454087 | 5781 | - | Gnomon | gene | gene26 | GeneID:105288189 | LOC105288189 | Gene | LOC105288189 | protein_coding | NA | character(0) | character(0) | BCphaseA2.vs.RphaseA1 | FALSE |
| NW_011924881.1 | 449147 | 449147 | 1 | + | 45 | 1 | 44 | 37 | 1 | 36 | 2.22 | 2.7 | NW_011924881.1 | 448307 | 454087 | 5781 | - | Gnomon | gene | gene26 | GeneID:105288189 | LOC105288189 | Gene | LOC105288189 | protein_coding | NA | character(0) | character(0) | BCphaseA2.vs.RphaseA1 | FALSE |
| NW_011924881.1 | 449151 | 449151 | 1 | + | 44 | 1 | 43 | 36 | 0 | 36 | 2.27 | 0 | NW_011924881.1 | 448307 | 454087 | 5781 | - | Gnomon | gene | gene26 | GeneID:105288189 | LOC105288189 | Gene | LOC105288189 | protein_coding | NA | character(0) | character(0) | BCphaseA2.vs.RphaseA1 | FALSE |
| NW_011924881.1 | 449161 | 449161 | 1 | + | 43 | 1 | 42 | 37 | 6 | 31 | 2.33 | 16.22 | NW_011924881.1 | 448307 | 454087 | 5781 | - | Gnomon | gene | gene26 | GeneID:105288189 | LOC105288189 | Gene | LOC105288189 | protein_coding | NA | character(0) | character(0) | BCphaseA2.vs.RphaseA1 | FALSE |
| NW_011924881.1 | 449168 | 449168 | 1 | + | 43 | 0 | 43 | 36 | 0 | 36 | 0 | 0 | NW_011924881.1 | 448307 | 454087 | 5781 | - | Gnomon | gene | gene26 | GeneID:105288189 | LOC105288189 | Gene | LOC105288189 | protein_coding | NA | character(0) | character(0) | BCphaseA2.vs.RphaseA1 | FALSE |
| NW_011924881.1 | 449174 | 449174 | 1 | + | 41 | 0 | 41 | 36 | 0 | 36 | 0 | 0 | NW_011924881.1 | 448307 | 454087 | 5781 | - | Gnomon | gene | gene26 | GeneID:105288189 | LOC105288189 | Gene | LOC105288189 | protein_coding | NA | character(0) | character(0) | BCphaseA2.vs.RphaseA1 | FALSE |
| NW_011924881.1 | 449192 | 449192 | 1 | + | 27 | 0 | 27 | 30 | 0 | 30 | 0 | 0 | NW_011924881.1 | 448307 | 454087 | 5781 | - | Gnomon | gene | gene26 | GeneID:105288189 | LOC105288189 | Gene | LOC105288189 | protein_coding | NA | character(0) | character(0) | BCphaseA2.vs.RphaseA1 | FALSE |
| NW_011924881.1 | 449197 | 449197 | 1 | + | 27 | 2 | 25 | 22 | 2 | 20 | 7.41 | 9.09 | NW_011924881.1 | 448307 | 454087 | 5781 | - | Gnomon | gene | gene26 | GeneID:105288189 | LOC105288189 | Gene | LOC105288189 | protein_coding | NA | character(0) | character(0) | BCphaseA2.vs.RphaseA1 | FALSE |
| NW_011924881.1 | 449220 | 449220 | 1 | + | 30 | 0 | 30 | 20 | 0 | 20 | 0 | 0 | NW_011924881.1 | 448307 | 454087 | 5781 | - | Gnomon | gene | gene26 | GeneID:105288189 | LOC105288189 | Gene | LOC105288189 | protein_coding | NA | character(0) | character(0) | BCphaseA2.vs.RphaseA1 | FALSE |
| NW_011924881.1 | 449229 | 449229 | 1 | + | 32 | 0 | 32 | 23 | 1 | 22 | 0 | 4.35 | NW_011924881.1 | 448307 | 454087 | 5781 | - | Gnomon | gene | gene26 | GeneID:105288189 | LOC105288189 | Gene | LOC105288189 | protein_coding | NA | character(0) | character(0) | BCphaseA2.vs.RphaseA1 | FALSE |
| NW_011924881.1 | 449250 | 449250 | 1 | + | 30 | 0 | 30 | 30 | 1 | 29 | 0 | 3.33 | NW_011924881.1 | 448307 | 454087 | 5781 | - | Gnomon | gene | gene26 | GeneID:105288189 | LOC105288189 | Gene | LOC105288189 | protein_coding | NA | character(0) | character(0) | BCphaseA2.vs.RphaseA1 | FALSE |
| NW_011924881.1 | 449304 | 449304 | 1 | + | 33 | 1 | 32 | 34 | 0 | 34 | 3.03 | 0 | NW_011924881.1 | 448307 | 454087 | 5781 | - | Gnomon | gene | gene26 | GeneID:105288189 | LOC105288189 | Gene | LOC105288189 | protein_coding | NA | character(0) | character(0) | BCphaseA2.vs.RphaseA1 | FALSE |
| NW_011924881.1 | 449308 | 449308 | 1 | + | 32 | 0 | 32 | 32 | 0 | 32 | 0 | 0 | NW_011924881.1 | 448307 | 454087 | 5781 | - | Gnomon | gene | gene26 | GeneID:105288189 | LOC105288189 | Gene | LOC105288189 | protein_coding | NA | character(0) | character(0) | BCphaseA2.vs.RphaseA1 | FALSE |
| NW_011924881.1 | 449312 | 449312 | 1 | + | 52 | 1 | 51 | 30 | 0 | 30 | 1.92 | 0 | NW_011924881.1 | 448307 | 454087 | 5781 | - | Gnomon | gene | gene26 | GeneID:105288189 | LOC105288189 | Gene | LOC105288189 | protein_coding | NA | character(0) | character(0) | BCphaseA2.vs.RphaseA1 | FALSE |
| NW_011924881.1 | 449314 | 449314 | 1 | + | 64 | 0 | 64 | 34 | 1 | 33 | 0 | 2.94 | NW_011924881.1 | 448307 | 454087 | 5781 | - | Gnomon | gene | gene26 | GeneID:105288189 | LOC105288189 | Gene | LOC105288189 | protein_coding | NA | character(0) | character(0) | BCphaseA2.vs.RphaseA1 | FALSE |
| NW_011924881.1 | 449318 | 449318 | 1 | + | 64 | 0 | 64 | 41 | 1 | 40 | 0 | 2.44 | NW_011924881.1 | 448307 | 454087 | 5781 | - | Gnomon | gene | gene26 | GeneID:105288189 | LOC105288189 | Gene | LOC105288189 | protein_coding | NA | character(0) | character(0) | BCphaseA2.vs.RphaseA1 | FALSE |
| NW_011924881.1 | 449321 | 449321 | 1 | + | 64 | 1 | 63 | 41 | 1 | 40 | 1.56 | 2.44 | NW_011924881.1 | 448307 | 454087 | 5781 | - | Gnomon | gene | gene26 | GeneID:105288189 | LOC105288189 | Gene | LOC105288189 | protein_coding | NA | character(0) | character(0) | BCphaseA2.vs.RphaseA1 | FALSE |
| NW_011924881.1 | 449325 | 449325 | 1 | + | 68 | 2 | 66 | 43 | 0 | 43 | 2.94 | 0 | NW_011924881.1 | 448307 | 454087 | 5781 | - | Gnomon | gene | gene26 | GeneID:105288189 | LOC105288189 | Gene | LOC105288189 | protein_coding | NA | character(0) | character(0) | BCphaseA2.vs.RphaseA1 | FALSE |

...
... only first 25 lines shown ...

Directory: DMSG File: dmg-Cb_LI_BCphaseA2.vs.RphaseA2_details.txt

| seqnames | start | end | width | strand | coverage1 | numCs1 | numTs1 | coverage2 | numCs2 | numTs2 | BCphaseA2 | RphaseA2 | gene_seqnames | gene_start | gene_end | gene_width | gene_strand | gene_source | gene_type | gene_ID | gene_Dbxref | gene_Name | gene_gbkey | gene_gene | gene_gene_biotype | gene_partial | gene_start_range | gene_end_range | gene_comparison | is.dm |
|---|---|---|---|---|---|---|---|---|---|---|---|---|---|---|---|---|---|---|---|---|---|---|---|---|---|---|---|---|---|---|
| NW_011924881.1 | 43243 | 43243 | 1 | + | 24 | 0 | 24 | 33 | 0 | 33 | 0 | 0 | NW_011924881.1 | 42715 | 135016 | 92302 | - | Gnomon | gene | gene18 | GeneID:105287290 | LOC105287290 | Gene | LOC105287290 | protein_coding | NA | character(0) | character(0) | BCphaseA2.vs.RphaseA2 | FALSE |
| NW_011924881.1 | 44091 | 44091 | 1 | + | 20 | 0 | 20 | 58 | 2 | 56 | 0 | 3.45 | NW_011924881.1 | 42715 | 135016 | 92302 | - | Gnomon | gene | gene18 | GeneID:105287290 | LOC105287290 | Gene | LOC105287290 | protein_coding | NA | character(0) | character(0) | BCphaseA2.vs.RphaseA2 | FALSE |
| NW_011924881.1 | 44105 | 44105 | 1 | + | 65 | 1 | 64 | 87 | 1 | 86 | 1.54 | 1.15 | NW_011924881.1 | 42715 | 135016 | 92302 | - | Gnomon | gene | gene18 | GeneID:105287290 | LOC105287290 | Gene | LOC105287290 | protein_coding | NA | character(0) | character(0) | BCphaseA2.vs.RphaseA2 | FALSE |
| NW_011924881.1 | 44112 | 44112 | 1 | + | 65 | 0 | 65 | 84 | 1 | 83 | 0 | 1.19 | NW_011924881.1 | 42715 | 135016 | 92302 | - | Gnomon | gene | gene18 | GeneID:105287290 | LOC105287290 | Gene | LOC105287290 | protein_coding | NA | character(0) | character(0) | BCphaseA2.vs.RphaseA2 | FALSE |
| NW_011924881.1 | 44121 | 44121 | 1 | + | 65 | 0 | 65 | 84 | 1 | 83 | 0 | 1.19 | NW_011924881.1 | 42715 | 135016 | 92302 | - | Gnomon | gene | gene18 | GeneID:105287290 | LOC105287290 | Gene | LOC105287290 | protein_coding | NA | character(0) | character(0) | BCphaseA2.vs.RphaseA2 | FALSE |
| NW_011924881.1 | 44123 | 44123 | 1 | + | 65 | 1 | 64 | 83 | 0 | 83 | 1.54 | 0 | NW_011924881.1 | 42715 | 135016 | 92302 | - | Gnomon | gene | gene18 | GeneID:105287290 | LOC105287290 | Gene | LOC105287290 | protein_coding | NA | character(0) | character(0) | BCphaseA2.vs.RphaseA2 | FALSE |
| NW_011924881.1 | 44125 | 44125 | 1 | + | 62 | 0 | 62 | 83 | 2 | 81 | 0 | 2.41 | NW_011924881.1 | 42715 | 135016 | 92302 | - | Gnomon | gene | gene18 | GeneID:105287290 | LOC105287290 | Gene | LOC105287290 | protein_coding | NA | character(0) | character(0) | BCphaseA2.vs.RphaseA2 | FALSE |
| NW_011924881.1 | 44129 | 44129 | 1 | + | 59 | 0 | 59 | 76 | 0 | 76 | 0 | 0 | NW_011924881.1 | 42715 | 135016 | 92302 | - | Gnomon | gene | gene18 | GeneID:105287290 | LOC105287290 | Gene | LOC105287290 | protein_coding | NA | character(0) | character(0) | BCphaseA2.vs.RphaseA2 | FALSE |
| NW_011924881.1 | 44138 | 44138 | 1 | + | 57 | 0 | 57 | 65 | 0 | 65 | 0 | 0 | NW_011924881.1 | 42715 | 135016 | 92302 | - | Gnomon | gene | gene18 | GeneID:105287290 | LOC105287290 | Gene | LOC105287290 | protein_coding | NA | character(0) | character(0) | BCphaseA2.vs.RphaseA2 | FALSE |
| NW_011924881.1 | 44152 | 44152 | 1 | + | 66 | 0 | 66 | 75 | 1 | 74 | 0 | 1.33 | NW_011924881.1 | 42715 | 135016 | 92302 | - | Gnomon | gene | gene18 | GeneID:105287290 | LOC105287290 | Gene | LOC105287290 | protein_coding | NA | character(0) | character(0) | BCphaseA2.vs.RphaseA2 | FALSE |
| NW_011924881.1 | 44173 | 44173 | 1 | + | 75 | 0 | 75 | 79 | 0 | 79 | 0 | 0 | NW_011924881.1 | 42715 | 135016 | 92302 | - | Gnomon | gene | gene18 | GeneID:105287290 | LOC105287290 | Gene | LOC105287290 | protein_coding | NA | character(0) | character(0) | BCphaseA2.vs.RphaseA2 | FALSE |
| NW_011924881.1 | 44185 | 44185 | 1 | + | 61 | 1 | 60 | 66 | 0 | 66 | 1.64 | 0 | NW_011924881.1 | 42715 | 135016 | 92302 | - | Gnomon | gene | gene18 | GeneID:105287290 | LOC105287290 | Gene | LOC105287290 | protein_coding | NA | character(0) | character(0) | BCphaseA2.vs.RphaseA2 | FALSE |
| NW_011924881.1 | 44193 | 44193 | 1 | + | 56 | 0 | 56 | 60 | 2 | 58 | 0 | 3.33 | NW_011924881.1 | 42715 | 135016 | 92302 | - | Gnomon | gene | gene18 | GeneID:105287290 | LOC105287290 | Gene | LOC105287290 | protein_coding | NA | character(0) | character(0) | BCphaseA2.vs.RphaseA2 | FALSE |
| NW_011924881.1 | 44199 | 44199 | 1 | + | 57 | 0 | 57 | 60 | 1 | 59 | 0 | 1.67 | NW_011924881.1 | 42715 | 135016 | 92302 | - | Gnomon | gene | gene18 | GeneID:105287290 | LOC105287290 | Gene | LOC105287290 | protein_coding | NA | character(0) | character(0) | BCphaseA2.vs.RphaseA2 | FALSE |
| NW_011924881.1 | 44201 | 44201 | 1 | + | 57 | 0 | 57 | 61 | 0 | 61 | 0 | 0 | NW_011924881.1 | 42715 | 135016 | 92302 | - | Gnomon | gene | gene18 | GeneID:105287290 | LOC105287290 | Gene | LOC105287290 | protein_coding | NA | character(0) | character(0) | BCphaseA2.vs.RphaseA2 | FALSE |
| NW_011924881.1 | 44204 | 44204 | 1 | + | 56 | 0 | 56 | 63 | 2 | 61 | 0 | 3.17 | NW_011924881.1 | 42715 | 135016 | 92302 | - | Gnomon | gene | gene18 | GeneID:105287290 | LOC105287290 | Gene | LOC105287290 | protein_coding | NA | character(0) | character(0) | BCphaseA2.vs.RphaseA2 | FALSE |
| NW_011924881.1 | 44221 | 44221 | 1 | + | 49 | 1 | 48 | 55 | 0 | 55 | 2.04 | 0 | NW_011924881.1 | 42715 | 135016 | 92302 | - | Gnomon | gene | gene18 | GeneID:105287290 | LOC105287290 | Gene | LOC105287290 | protein_coding | NA | character(0) | character(0) | BCphaseA2.vs.RphaseA2 | FALSE |
| NW_011924881.1 | 44223 | 44223 | 1 | + | 47 | 0 | 47 | 50 | 0 | 50 | 0 | 0 | NW_011924881.1 | 42715 | 135016 | 92302 | - | Gnomon | gene | gene18 | GeneID:105287290 | LOC105287290 | Gene | LOC105287290 | protein_coding | NA | character(0) | character(0) | BCphaseA2.vs.RphaseA2 | FALSE |
| NW_011924881.1 | 44252 | 44252 | 1 | + | 32 | 0 | 32 | 44 | 0 | 44 | 0 | 0 | NW_011924881.1 | 42715 | 135016 | 92302 | - | Gnomon | gene | gene18 | GeneID:105287290 | LOC105287290 | Gene | LOC105287290 | protein_coding | NA | character(0) | character(0) | BCphaseA2.vs.RphaseA2 | FALSE |
| NW_011924881.1 | 44261 | 44261 | 1 | + | 30 | 0 | 30 | 41 | 1 | 40 | 0 | 2.44 | NW_011924881.1 | 42715 | 135016 | 92302 | - | Gnomon | gene | gene18 | GeneID:105287290 | LOC105287290 | Gene | LOC105287290 | protein_coding | NA | character(0) | character(0) | BCphaseA2.vs.RphaseA2 | FALSE |
| NW_011924881.1 | 44271 | 44271 | 1 | + | 30 | 0 | 30 | 44 | 0 | 44 | 0 | 0 | NW_011924881.1 | 42715 | 135016 | 92302 | - | Gnomon | gene | gene18 | GeneID:105287290 | LOC105287290 | Gene | LOC105287290 | protein_coding | NA | character(0) | character(0) | BCphaseA2.vs.RphaseA2 | FALSE |
| NW_011924881.1 | 44276 | 44276 | 1 | + | 33 | 0 | 33 | 53 | 0 | 53 | 0 | 0 | NW_011924881.1 | 42715 | 135016 | 92302 | - | Gnomon | gene | gene18 | GeneID:105287290 | LOC105287290 | Gene | LOC105287290 | protein_coding | NA | character(0) | character(0) | BCphaseA2.vs.RphaseA2 | FALSE |
| NW_011924881.1 | 44282 | 44282 | 1 | + | 29 | 0 | 29 | 47 | 0 | 47 | 0 | 0 | NW_011924881.1 | 42715 | 135016 | 92302 | - | Gnomon | gene | gene18 | GeneID:105287290 | LOC105287290 | Gene | LOC105287290 | protein_coding | NA | character(0) | character(0) | BCphaseA2.vs.RphaseA2 | FALSE |
| NW_011924881.1 | 45788 | 45788 | 1 | + | 20 | 0 | 20 | 34 | 0 | 34 | 0 | 0 | NW_011924881.1 | 42715 | 135016 | 92302 | - | Gnomon | gene | gene18 | GeneID:105287290 | LOC105287290 | Gene | LOC105287290 | protein_coding | NA | character(0) | character(0) | BCphaseA2.vs.RphaseA2 | FALSE |

...
... only first 25 lines shown ...

Directory: DMSG File: dmg-Cb_LI_BCphaseA2.vs.RphaseB1_details.txt

| seqnames | start | end | width | strand | coverage1 | numCs1 | numTs1 | coverage2 | numCs2 | numTs2 | BCphaseA2 | RphaseB1 | gene_seqnames | gene_start | gene_end | gene_width | gene_strand | gene_source | gene_type | gene_ID | gene_Dbxref | gene_Name | gene_gbkey | gene_gene | gene_gene_biotype | gene_start_range | gene_end_range | gene_comparison | is.dm |
|---|---|---|---|---|---|---|---|---|---|---|---|---|---|---|---|---|---|---|---|---|---|---|---|---|---|---|---|---|---|
| NW_011924881.1 | 809481 | 809481 | 1 | + | 20 | 0 | 20 | 30 | 0 | 30 | 0 | 0 | NW_011924881.1 | 808343 | 827625 | 19283 | - | Gnomon | gene | gene53 | GeneID:105277148 | LOC105277148 | Gene | LOC105277148 | protein_coding | character(0) | character(0) | BCphaseA2.vs.RphaseB1 | FALSE |
| NW_011924881.1 | 809504 | 809504 | 1 | + | 20 | 0 | 20 | 31 | 0 | 31 | 0 | 0 | NW_011924881.1 | 808343 | 827625 | 19283 | - | Gnomon | gene | gene53 | GeneID:105277148 | LOC105277148 | Gene | LOC105277148 | protein_coding | character(0) | character(0) | BCphaseA2.vs.RphaseB1 | FALSE |
| NW_011924881.1 | 810155 | 810155 | 1 | + | 22 | 0 | 22 | 25 | 0 | 25 | 0 | 0 | NW_011924881.1 | 808343 | 827625 | 19283 | - | Gnomon | gene | gene53 | GeneID:105277148 | LOC105277148 | Gene | LOC105277148 | protein_coding | character(0) | character(0) | BCphaseA2.vs.RphaseB1 | FALSE |
| NW_011924881.1 | 810182 | 810182 | 1 | + | 23 | 0 | 23 | 28 | 1 | 27 | 0 | 3.57 | NW_011924881.1 | 808343 | 827625 | 19283 | - | Gnomon | gene | gene53 | GeneID:105277148 | LOC105277148 | Gene | LOC105277148 | protein_coding | character(0) | character(0) | BCphaseA2.vs.RphaseB1 | FALSE |
| NW_011924881.1 | 810335 | 810335 | 1 | + | 23 | 0 | 23 | 28 | 0 | 28 | 0 | 0 | NW_011924881.1 | 808343 | 827625 | 19283 | - | Gnomon | gene | gene53 | GeneID:105277148 | LOC105277148 | Gene | LOC105277148 | protein_coding | character(0) | character(0) | BCphaseA2.vs.RphaseB1 | FALSE |
| NW_011924881.1 | 810337 | 810337 | 1 | + | 24 | 0 | 24 | 29 | 0 | 29 | 0 | 0 | NW_011924881.1 | 808343 | 827625 | 19283 | - | Gnomon | gene | gene53 | GeneID:105277148 | LOC105277148 | Gene | LOC105277148 | protein_coding | character(0) | character(0) | BCphaseA2.vs.RphaseB1 | FALSE |
| NW_011924881.1 | 810348 | 810348 | 1 | + | 51 | 0 | 51 | 41 | 1 | 40 | 0 | 2.44 | NW_011924881.1 | 808343 | 827625 | 19283 | - | Gnomon | gene | gene53 | GeneID:105277148 | LOC105277148 | Gene | LOC105277148 | protein_coding | character(0) | character(0) | BCphaseA2.vs.RphaseB1 | FALSE |
| NW_011924881.1 | 810391 | 810391 | 1 | + | 54 | 1 | 53 | 37 | 0 | 37 | 1.85 | 0 | NW_011924881.1 | 808343 | 827625 | 19283 | - | Gnomon | gene | gene53 | GeneID:105277148 | LOC105277148 | Gene | LOC105277148 | protein_coding | character(0) | character(0) | BCphaseA2.vs.RphaseB1 | FALSE |
| NW_011924881.1 | 810409 | 810409 | 1 | + | 61 | 2 | 59 | 32 | 1 | 31 | 3.28 | 3.12 | NW_011924881.1 | 808343 | 827625 | 19283 | - | Gnomon | gene | gene53 | GeneID:105277148 | LOC105277148 | Gene | LOC105277148 | protein_coding | character(0) | character(0) | BCphaseA2.vs.RphaseB1 | FALSE |
| NW_011924881.1 | 810412 | 810412 | 1 | + | 40 | 0 | 40 | 27 | 1 | 26 | 0 | 3.7 | NW_011924881.1 | 808343 | 827625 | 19283 | - | Gnomon | gene | gene53 | GeneID:105277148 | LOC105277148 | Gene | LOC105277148 | protein_coding | character(0) | character(0) | BCphaseA2.vs.RphaseB1 | FALSE |
| NW_011924881.1 | 810417 | 810417 | 1 | + | 38 | 1 | 37 | 24 | 1 | 23 | 2.63 | 4.17 | NW_011924881.1 | 808343 | 827625 | 19283 | - | Gnomon | gene | gene53 | GeneID:105277148 | LOC105277148 | Gene | LOC105277148 | protein_coding | character(0) | character(0) | BCphaseA2.vs.RphaseB1 | FALSE |
| NW_011924881.1 | 810424 | 810424 | 1 | + | 37 | 0 | 37 | 24 | 0 | 24 | 0 | 0 | NW_011924881.1 | 808343 | 827625 | 19283 | - | Gnomon | gene | gene53 | GeneID:105277148 | LOC105277148 | Gene | LOC105277148 | protein_coding | character(0) | character(0) | BCphaseA2.vs.RphaseB1 | FALSE |
| NW_011924881.1 | 810430 | 810430 | 1 | + | 27 | 0 | 27 | 26 | 0 | 26 | 0 | 0 | NW_011924881.1 | 808343 | 827625 | 19283 | - | Gnomon | gene | gene53 | GeneID:105277148 | LOC105277148 | Gene | LOC105277148 | protein_coding | character(0) | character(0) | BCphaseA2.vs.RphaseB1 | FALSE |
| NW_011924881.1 | 810436 | 810436 | 1 | + | 27 | 0 | 27 | 30 | 0 | 30 | 0 | 0 | NW_011924881.1 | 808343 | 827625 | 19283 | - | Gnomon | gene | gene53 | GeneID:105277148 | LOC105277148 | Gene | LOC105277148 | protein_coding | character(0) | character(0) | BCphaseA2.vs.RphaseB1 | FALSE |
| NW_011924881.1 | 810505 | 810505 | 1 | + | 22 | 0 | 22 | 50 | 0 | 50 | 0 | 0 | NW_011924881.1 | 808343 | 827625 | 19283 | - | Gnomon | gene | gene53 | GeneID:105277148 | LOC105277148 | Gene | LOC105277148 | protein_coding | character(0) | character(0) | BCphaseA2.vs.RphaseB1 | FALSE |
| NW_011924881.1 | 810511 | 810511 | 1 | + | 22 | 1 | 21 | 54 | 0 | 54 | 4.55 | 0 | NW_011924881.1 | 808343 | 827625 | 19283 | - | Gnomon | gene | gene53 | GeneID:105277148 | LOC105277148 | Gene | LOC105277148 | protein_coding | character(0) | character(0) | BCphaseA2.vs.RphaseB1 | FALSE |
| NW_011924881.1 | 810516 | 810516 | 1 | + | 23 | 0 | 23 | 55 | 1 | 54 | 0 | 1.82 | NW_011924881.1 | 808343 | 827625 | 19283 | - | Gnomon | gene | gene53 | GeneID:105277148 | LOC105277148 | Gene | LOC105277148 | protein_coding | character(0) | character(0) | BCphaseA2.vs.RphaseB1 | FALSE |
| NW_011924881.1 | 810520 | 810520 | 1 | + | 23 | 0 | 23 | 56 | 1 | 55 | 0 | 1.79 | NW_011924881.1 | 808343 | 827625 | 19283 | - | Gnomon | gene | gene53 | GeneID:105277148 | LOC105277148 | Gene | LOC105277148 | protein_coding | character(0) | character(0) | BCphaseA2.vs.RphaseB1 | FALSE |
| NW_011924881.1 | 810633 | 810633 | 1 | + | 20 | 0 | 20 | 67 | 0 | 67 | 0 | 0 | NW_011924881.1 | 808343 | 827625 | 19283 | - | Gnomon | gene | gene53 | GeneID:105277148 | LOC105277148 | Gene | LOC105277148 | protein_coding | character(0) | character(0) | BCphaseA2.vs.RphaseB1 | FALSE |
| NW_011924881.1 | 810645 | 810645 | 1 | + | 21 | 0 | 21 | 66 | 0 | 66 | 0 | 0 | NW_011924881.1 | 808343 | 827625 | 19283 | - | Gnomon | gene | gene53 | GeneID:105277148 | LOC105277148 | Gene | LOC105277148 | protein_coding | character(0) | character(0) | BCphaseA2.vs.RphaseB1 | FALSE |
| NW_011924881.1 | 811042 | 811042 | 1 | + | 21 | 0 | 21 | 31 | 1 | 30 | 0 | 3.23 | NW_011924881.1 | 808343 | 827625 | 19283 | - | Gnomon | gene | gene53 | GeneID:105277148 | LOC105277148 | Gene | LOC105277148 | protein_coding | character(0) | character(0) | BCphaseA2.vs.RphaseB1 | FALSE |
| NW_011924881.1 | 811083 | 811083 | 1 | + | 26 | 0 | 26 | 29 | 0 | 29 | 0 | 0 | NW_011924881.1 | 808343 | 827625 | 19283 | - | Gnomon | gene | gene53 | GeneID:105277148 | LOC105277148 | Gene | LOC105277148 | protein_coding | character(0) | character(0) | BCphaseA2.vs.RphaseB1 | FALSE |
| NW_011924881.1 | 811089 | 811089 | 1 | + | 27 | 0 | 27 | 28 | 0 | 28 | 0 | 0 | NW_011924881.1 | 808343 | 827625 | 19283 | - | Gnomon | gene | gene53 | GeneID:105277148 | LOC105277148 | Gene | LOC105277148 | protein_coding | character(0) | character(0) | BCphaseA2.vs.RphaseB1 | FALSE |
| NW_011924881.1 | 811120 | 811120 | 1 | + | 23 | 0 | 23 | 30 | 0 | 30 | 0 | 0 | NW_011924881.1 | 808343 | 827625 | 19283 | - | Gnomon | gene | gene53 | GeneID:105277148 | LOC105277148 | Gene | LOC105277148 | protein_coding | character(0) | character(0) | BCphaseA2.vs.RphaseB1 | FALSE |

...
... only first 25 lines shown ...

Directory: DMSG File: dmg-Cb_LI_BCphaseA2.vs.RphaseB2_details.txt

| seqnames | start | end | width | strand | coverage1 | numCs1 | numTs1 | coverage2 | numCs2 | numTs2 | BCphaseA2 | RphaseB2 | gene_seqnames | gene_start | gene_end | gene_width | gene_strand | gene_source | gene_type | gene_ID | gene_Dbxref | gene_Name | gene_gbkey | gene_gene | gene_gene_biotype | gene_start_range | gene_end_range | gene_comparison | is.dm |
|---|---|---|---|---|---|---|---|---|---|---|---|---|---|---|---|---|---|---|---|---|---|---|---|---|---|---|---|---|---|
| NW_011924881.1 | 724785 | 724785 | 1 | + | 34 | 0 | 34 | 28 | 0 | 28 | 0 | 0 | NW_011924881.1 | 724448 | 731567 | 7120 | - | Gnomon | gene | gene34 | GeneID:105278416 | LOC105278416 | Gene | LOC105278416 | protein_coding | character(0) | character(0) | BCphaseA2.vs.RphaseB2 | FALSE |
| NW_011924881.1 | 724788 | 724788 | 1 | + | 33 | 0 | 33 | 28 | 0 | 28 | 0 | 0 | NW_011924881.1 | 724448 | 731567 | 7120 | - | Gnomon | gene | gene34 | GeneID:105278416 | LOC105278416 | Gene | LOC105278416 | protein_coding | character(0) | character(0) | BCphaseA2.vs.RphaseB2 | FALSE |
| NW_011924881.1 | 724794 | 724794 | 1 | + | 33 | 0 | 33 | 28 | 0 | 28 | 0 | 0 | NW_011924881.1 | 724448 | 731567 | 7120 | - | Gnomon | gene | gene34 | GeneID:105278416 | LOC105278416 | Gene | LOC105278416 | protein_coding | character(0) | character(0) | BCphaseA2.vs.RphaseB2 | FALSE |
| NW_011924881.1 | 724802 | 724802 | 1 | + | 33 | 0 | 33 | 28 | 0 | 28 | 0 | 0 | NW_011924881.1 | 724448 | 731567 | 7120 | - | Gnomon | gene | gene34 | GeneID:105278416 | LOC105278416 | Gene | LOC105278416 | protein_coding | character(0) | character(0) | BCphaseA2.vs.RphaseB2 | FALSE |
| NW_011924881.1 | 724806 | 724806 | 1 | + | 33 | 0 | 33 | 29 | 0 | 29 | 0 | 0 | NW_011924881.1 | 724448 | 731567 | 7120 | - | Gnomon | gene | gene34 | GeneID:105278416 | LOC105278416 | Gene | LOC105278416 | protein_coding | character(0) | character(0) | BCphaseA2.vs.RphaseB2 | FALSE |
| NW_011924881.1 | 724815 | 724815 | 1 | + | 33 | 0 | 33 | 29 | 1 | 28 | 0 | 3.45 | NW_011924881.1 | 724448 | 731567 | 7120 | - | Gnomon | gene | gene34 | GeneID:105278416 | LOC105278416 | Gene | LOC105278416 | protein_coding | character(0) | character(0) | BCphaseA2.vs.RphaseB2 | FALSE |
| NW_011924881.1 | 724818 | 724818 | 1 | + | 32 | 0 | 32 | 29 | 0 | 29 | 0 | 0 | NW_011924881.1 | 724448 | 731567 | 7120 | - | Gnomon | gene | gene34 | GeneID:105278416 | LOC105278416 | Gene | LOC105278416 | protein_coding | character(0) | character(0) | BCphaseA2.vs.RphaseB2 | FALSE |
| NW_011924881.1 | 724839 | 724839 | 1 | + | 46 | 0 | 46 | 44 | 0 | 44 | 0 | 0 | NW_011924881.1 | 724448 | 731567 | 7120 | - | Gnomon | gene | gene34 | GeneID:105278416 | LOC105278416 | Gene | LOC105278416 | protein_coding | character(0) | character(0) | BCphaseA2.vs.RphaseB2 | FALSE |
| NW_011924881.1 | 724872 | 724872 | 1 | + | 36 | 0 | 36 | 36 | 0 | 36 | 0 | 0 | NW_011924881.1 | 724448 | 731567 | 7120 | - | Gnomon | gene | gene34 | GeneID:105278416 | LOC105278416 | Gene | LOC105278416 | protein_coding | character(0) | character(0) | BCphaseA2.vs.RphaseB2 | FALSE |
| NW_011924881.1 | 724889 | 724889 | 1 | + | 38 | 0 | 38 | 38 | 0 | 38 | 0 | 0 | NW_011924881.1 | 724448 | 731567 | 7120 | - | Gnomon | gene | gene34 | GeneID:105278416 | LOC105278416 | Gene | LOC105278416 | protein_coding | character(0) | character(0) | BCphaseA2.vs.RphaseB2 | FALSE |
| NW_011924881.1 | 724896 | 724896 | 1 | + | 38 | 0 | 38 | 36 | 0 | 36 | 0 | 0 | NW_011924881.1 | 724448 | 731567 | 7120 | - | Gnomon | gene | gene34 | GeneID:105278416 | LOC105278416 | Gene | LOC105278416 | protein_coding | character(0) | character(0) | BCphaseA2.vs.RphaseB2 | FALSE |
| NW_011924881.1 | 724898 | 724898 | 1 | + | 38 | 0 | 38 | 37 | 0 | 37 | 0 | 0 | NW_011924881.1 | 724448 | 731567 | 7120 | - | Gnomon | gene | gene34 | GeneID:105278416 | LOC105278416 | Gene | LOC105278416 | protein_coding | character(0) | character(0) | BCphaseA2.vs.RphaseB2 | FALSE |
| NW_011924881.1 | 724926 | 724926 | 1 | + | 50 | 0 | 50 | 58 | 1 | 57 | 0 | 1.72 | NW_011924881.1 | 724448 | 731567 | 7120 | - | Gnomon | gene | gene34 | GeneID:105278416 | LOC105278416 | Gene | LOC105278416 | protein_coding | character(0) | character(0) | BCphaseA2.vs.RphaseB2 | FALSE |
| NW_011924881.1 | 724934 | 724934 | 1 | + | 38 | 0 | 38 | 43 | 0 | 43 | 0 | 0 | NW_011924881.1 | 724448 | 731567 | 7120 | - | Gnomon | gene | gene34 | GeneID:105278416 | LOC105278416 | Gene | LOC105278416 | protein_coding | character(0) | character(0) | BCphaseA2.vs.RphaseB2 | FALSE |
| NW_011924881.1 | 724936 | 724936 | 1 | + | 38 | 0 | 38 | 44 | 0 | 44 | 0 | 0 | NW_011924881.1 | 724448 | 731567 | 7120 | - | Gnomon | gene | gene34 | GeneID:105278416 | LOC105278416 | Gene | LOC105278416 | protein_coding | character(0) | character(0) | BCphaseA2.vs.RphaseB2 | FALSE |
| NW_011924881.1 | 724943 | 724943 | 1 | + | 32 | 0 | 32 | 40 | 1 | 39 | 0 | 2.5 | NW_011924881.1 | 724448 | 731567 | 7120 | - | Gnomon | gene | gene34 | GeneID:105278416 | LOC105278416 | Gene | LOC105278416 | protein_coding | character(0) | character(0) | BCphaseA2.vs.RphaseB2 | FALSE |
| NW_011924881.1 | 724947 | 724947 | 1 | + | 33 | 0 | 33 | 42 | 0 | 42 | 0 | 0 | NW_011924881.1 | 724448 | 731567 | 7120 | - | Gnomon | gene | gene34 | GeneID:105278416 | LOC105278416 | Gene | LOC105278416 | protein_coding | character(0) | character(0) | BCphaseA2.vs.RphaseB2 | FALSE |
| NW_011924881.1 | 724961 | 724961 | 1 | + | 43 | 0 | 43 | 46 | 0 | 46 | 0 | 0 | NW_011924881.1 | 724448 | 731567 | 7120 | - | Gnomon | gene | gene34 | GeneID:105278416 | LOC105278416 | Gene | LOC105278416 | protein_coding | character(0) | character(0) | BCphaseA2.vs.RphaseB2 | FALSE |
| NW_011924881.1 | 724976 | 724976 | 1 | + | 53 | 0 | 53 | 54 | 1 | 53 | 0 | 1.85 | NW_011924881.1 | 724448 | 731567 | 7120 | - | Gnomon | gene | gene34 | GeneID:105278416 | LOC105278416 | Gene | LOC105278416 | protein_coding | character(0) | character(0) | BCphaseA2.vs.RphaseB2 | FALSE |
| NW_011924881.1 | 724982 | 724982 | 1 | + | 58 | 0 | 58 | 57 | 0 | 57 | 0 | 0 | NW_011924881.1 | 724448 | 731567 | 7120 | - | Gnomon | gene | gene34 | GeneID:105278416 | LOC105278416 | Gene | LOC105278416 | protein_coding | character(0) | character(0) | BCphaseA2.vs.RphaseB2 | FALSE |
| NW_011924881.1 | 724990 | 724990 | 1 | + | 60 | 0 | 60 | 58 | 0 | 58 | 0 | 0 | NW_011924881.1 | 724448 | 731567 | 7120 | - | Gnomon | gene | gene34 | GeneID:105278416 | LOC105278416 | Gene | LOC105278416 | protein_coding | character(0) | character(0) | BCphaseA2.vs.RphaseB2 | FALSE |
| NW_011924881.1 | 724994 | 724994 | 1 | + | 60 | 0 | 60 | 58 | 0 | 58 | 0 | 0 | NW_011924881.1 | 724448 | 731567 | 7120 | - | Gnomon | gene | gene34 | GeneID:105278416 | LOC105278416 | Gene | LOC105278416 | protein_coding | character(0) | character(0) | BCphaseA2.vs.RphaseB2 | FALSE |
| NW_011924881.1 | 725008 | 725008 | 1 | + | 41 | 1 | 40 | 42 | 0 | 42 | 2.44 | 0 | NW_011924881.1 | 724448 | 731567 | 7120 | - | Gnomon | gene | gene34 | GeneID:105278416 | LOC105278416 | Gene | LOC105278416 | protein_coding | character(0) | character(0) | BCphaseA2.vs.RphaseB2 | FALSE |
| NW_011924881.1 | 725014 | 725014 | 1 | + | 31 | 0 | 31 | 31 | 0 | 31 | 0 | 0 | NW_011924881.1 | 724448 | 731567 | 7120 | - | Gnomon | gene | gene34 | GeneID:105278416 | LOC105278416 | Gene | LOC105278416 | protein_coding | character(0) | character(0) | BCphaseA2.vs.RphaseB2 | FALSE |

...

... only first 25 lines shown ...

Directory: DMSG File: dmg-Cb_LI_BCphaseB1.vs.BCphaseB2_details.txt

| seqnames | start | end | width | strand | coverage1 | numCs1 | numTs1 | coverage2 | numCs2 | numTs2 | BCphaseB1 | BCphaseB2 | gene_seqnames | gene_start | gene_end | gene_width | gene_strand | gene_source | gene_type | gene_ID | gene_Dbxref | gene_Name | gene_gbkey | gene_gene | gene_gene_biotype | gene_start_range | gene_end_range | gene_comparison | is.dm |
|---|---|---|---|---|---|---|---|---|---|---|---|---|---|---|---|---|---|---|---|---|---|---|---|---|---|---|---|---|---|
| NW_011924881.1 | 724788 | 724788 | 1 | + | 37 | 0 | 37 | 31 | 0 | 31 | 0 | 0 | NW_011924881.1 | 724448 | 731567 | 7120 | - | Gnomon | gene | gene34 | GeneID:105278416 | LOC105278416 | Gene | LOC105278416 | protein_coding | character(0) | character(0) | BCphaseB1.vs.BCphaseB2 | FALSE |
| NW_011924881.1 | 724794 | 724794 | 1 | + | 35 | 0 | 35 | 31 | 0 | 31 | 0 | 0 | NW_011924881.1 | 724448 | 731567 | 7120 | - | Gnomon | gene | gene34 | GeneID:105278416 | LOC105278416 | Gene | LOC105278416 | protein_coding | character(0) | character(0) | BCphaseB1.vs.BCphaseB2 | FALSE |
| NW_011924881.1 | 724802 | 724802 | 1 | + | 33 | 0 | 33 | 30 | 0 | 30 | 0 | 0 | NW_011924881.1 | 724448 | 731567 | 7120 | - | Gnomon | gene | gene34 | GeneID:105278416 | LOC105278416 | Gene | LOC105278416 | protein_coding | character(0) | character(0) | BCphaseB1.vs.BCphaseB2 | FALSE |
| NW_011924881.1 | 724806 | 724806 | 1 | + | 33 | 1 | 32 | 30 | 0 | 30 | 3.03 | 0 | NW_011924881.1 | 724448 | 731567 | 7120 | - | Gnomon | gene | gene34 | GeneID:105278416 | LOC105278416 | Gene | LOC105278416 | protein_coding | character(0) | character(0) | BCphaseB1.vs.BCphaseB2 | FALSE |
| NW_011924881.1 | 724815 | 724815 | 1 | + | 33 | 0 | 33 | 30 | 0 | 30 | 0 | 0 | NW_011924881.1 | 724448 | 731567 | 7120 | - | Gnomon | gene | gene34 | GeneID:105278416 | LOC105278416 | Gene | LOC105278416 | protein_coding | character(0) | character(0) | BCphaseB1.vs.BCphaseB2 | FALSE |
| NW_011924881.1 | 724818 | 724818 | 1 | + | 33 | 0 | 33 | 30 | 0 | 30 | 0 | 0 | NW_011924881.1 | 724448 | 731567 | 7120 | - | Gnomon | gene | gene34 | GeneID:105278416 | LOC105278416 | Gene | LOC105278416 | protein_coding | character(0) | character(0) | BCphaseB1.vs.BCphaseB2 | FALSE |
| NW_011924881.1 | 724839 | 724839 | 1 | + | 33 | 0 | 33 | 23 | 0 | 23 | 0 | 0 | NW_011924881.1 | 724448 | 731567 | 7120 | - | Gnomon | gene | gene34 | GeneID:105278416 | LOC105278416 | Gene | LOC105278416 | protein_coding | character(0) | character(0) | BCphaseB1.vs.BCphaseB2 | FALSE |
| NW_011924881.1 | 724926 | 724926 | 1 | + | 34 | 0 | 34 | 37 | 0 | 37 | 0 | 0 | NW_011924881.1 | 724448 | 731567 | 7120 | - | Gnomon | gene | gene34 | GeneID:105278416 | LOC105278416 | Gene | LOC105278416 | protein_coding | character(0) | character(0) | BCphaseB1.vs.BCphaseB2 | FALSE |
| NW_011924881.1 | 724934 | 724934 | 1 | + | 36 | 0 | 36 | 34 | 0 | 34 | 0 | 0 | NW_011924881.1 | 724448 | 731567 | 7120 | - | Gnomon | gene | gene34 | GeneID:105278416 | LOC105278416 | Gene | LOC105278416 | protein_coding | character(0) | character(0) | BCphaseB1.vs.BCphaseB2 | FALSE |
| NW_011924881.1 | 724936 | 724936 | 1 | + | 36 | 0 | 36 | 35 | 0 | 35 | 0 | 0 | NW_011924881.1 | 724448 | 731567 | 7120 | - | Gnomon | gene | gene34 | GeneID:105278416 | LOC105278416 | Gene | LOC105278416 | protein_coding | character(0) | character(0) | BCphaseB1.vs.BCphaseB2 | FALSE |
| NW_011924881.1 | 724943 | 724943 | 1 | + | 33 | 0 | 33 | 33 | 0 | 33 | 0 | 0 | NW_011924881.1 | 724448 | 731567 | 7120 | - | Gnomon | gene | gene34 | GeneID:105278416 | LOC105278416 | Gene | LOC105278416 | protein_coding | character(0) | character(0) | BCphaseB1.vs.BCphaseB2 | FALSE |
| NW_011924881.1 | 724947 | 724947 | 1 | + | 31 | 0 | 31 | 36 | 0 | 36 | 0 | 0 | NW_011924881.1 | 724448 | 731567 | 7120 | - | Gnomon | gene | gene34 | GeneID:105278416 | LOC105278416 | Gene | LOC105278416 | protein_coding | character(0) | character(0) | BCphaseB1.vs.BCphaseB2 | FALSE |
| NW_011924881.1 | 724961 | 724961 | 1 | + | 36 | 1 | 35 | 43 | 0 | 43 | 2.78 | 0 | NW_011924881.1 | 724448 | 731567 | 7120 | - | Gnomon | gene | gene34 | GeneID:105278416 | LOC105278416 | Gene | LOC105278416 | protein_coding | character(0) | character(0) | BCphaseB1.vs.BCphaseB2 | FALSE |
| NW_011924881.1 | 724976 | 724976 | 1 | + | 43 | 0 | 43 | 52 | 0 | 52 | 0 | 0 | NW_011924881.1 | 724448 | 731567 | 7120 | - | Gnomon | gene | gene34 | GeneID:105278416 | LOC105278416 | Gene | LOC105278416 | protein_coding | character(0) | character(0) | BCphaseB1.vs.BCphaseB2 | FALSE |
| NW_011924881.1 | 724982 | 724982 | 1 | + | 44 | 0 | 44 | 54 | 0 | 54 | 0 | 0 | NW_011924881.1 | 724448 | 731567 | 7120 | - | Gnomon | gene | gene34 | GeneID:105278416 | LOC105278416 | Gene | LOC105278416 | protein_coding | character(0) | character(0) | BCphaseB1.vs.BCphaseB2 | FALSE |
| NW_011924881.1 | 724990 | 724990 | 1 | + | 44 | 0 | 44 | 54 | 0 | 54 | 0 | 0 | NW_011924881.1 | 724448 | 731567 | 7120 | - | Gnomon | gene | gene34 | GeneID:105278416 | LOC105278416 | Gene | LOC105278416 | protein_coding | character(0) | character(0) | BCphaseB1.vs.BCphaseB2 | FALSE |
| NW_011924881.1 | 724994 | 724994 | 1 | + | 45 | 0 | 45 | 53 | 0 | 53 | 0 | 0 | NW_011924881.1 | 724448 | 731567 | 7120 | - | Gnomon | gene | gene34 | GeneID:105278416 | LOC105278416 | Gene | LOC105278416 | protein_coding | character(0) | character(0) | BCphaseB1.vs.BCphaseB2 | FALSE |
| NW_011924881.1 | 725008 | 725008 | 1 | + | 31 | 1 | 30 | 43 | 0 | 43 | 3.23 | 0 | NW_011924881.1 | 724448 | 731567 | 7120 | - | Gnomon | gene | gene34 | GeneID:105278416 | LOC105278416 | Gene | LOC105278416 | protein_coding | character(0) | character(0) | BCphaseB1.vs.BCphaseB2 | FALSE |
| NW_011924881.1 | 725014 | 725014 | 1 | + | 58 | 2 | 56 | 35 | 0 | 35 | 3.45 | 0 | NW_011924881.1 | 724448 | 731567 | 7120 | - | Gnomon | gene | gene34 | GeneID:105278416 | LOC105278416 | Gene | LOC105278416 | protein_coding | character(0) | character(0) | BCphaseB1.vs.BCphaseB2 | FALSE |
| NW_011924881.1 | 725022 | 725022 | 1 | + | 36 | 3 | 33 | 24 | 0 | 24 | 8.33 | 0 | NW_011924881.1 | 724448 | 731567 | 7120 | - | Gnomon | gene | gene34 | GeneID:105278416 | LOC105278416 | Gene | LOC105278416 | protein_coding | character(0) | character(0) | BCphaseB1.vs.BCphaseB2 | FALSE |
| NW_011924881.1 | 725041 | 725041 | 1 | + | 50 | 0 | 50 | 33 | 0 | 33 | 0 | 0 | NW_011924881.1 | 724448 | 731567 | 7120 | - | Gnomon | gene | gene34 | GeneID:105278416 | LOC105278416 | Gene | LOC105278416 | protein_coding | character(0) | character(0) | BCphaseB1.vs.BCphaseB2 | FALSE |
| NW_011924881.1 | 725048 | 725048 | 1 | + | 54 | 0 | 54 | 38 | 0 | 38 | 0 | 0 | NW_011924881.1 | 724448 | 731567 | 7120 | - | Gnomon | gene | gene34 | GeneID:105278416 | LOC105278416 | Gene | LOC105278416 | protein_coding | character(0) | character(0) | BCphaseB1.vs.BCphaseB2 | FALSE |
| NW_011924881.1 | 725062 | 725062 | 1 | + | 81 | 0 | 81 | 68 | 0 | 68 | 0 | 0 | NW_011924881.1 | 724448 | 731567 | 7120 | - | Gnomon | gene | gene34 | GeneID:105278416 | LOC105278416 | Gene | LOC105278416 | protein_coding | character(0) | character(0) | BCphaseB1.vs.BCphaseB2 | FALSE |
| NW_011924881.1 | 725065 | 725065 | 1 | + | 82 | 0 | 82 | 69 | 0 | 69 | 0 | 0 | NW_011924881.1 | 724448 | 731567 | 7120 | - | Gnomon | gene | gene34 | GeneID:105278416 | LOC105278416 | Gene | LOC105278416 | protein_coding | character(0) | character(0) | BCphaseB1.vs.BCphaseB2 | FALSE |

...
... only first 25 lines shown ...

Directory: DMSG File: dmg-Cb_LI_BCphaseB1.vs.RphaseA1_details.txt

| seqnames | start | end | width | strand | coverage1 | numCs1 | numTs1 | coverage2 | numCs2 | numTs2 | BCphaseB1 | RphaseA1 | gene_seqnames | gene_start | gene_end | gene_width | gene_strand | gene_source | gene_type | gene_ID | gene_Dbxref | gene_Name | gene_gbkey | gene_gene | gene_gene_biotype | gene_partial | gene_start_range | gene_end_range | gene_comparison | is.dm |
|---|---|---|---|---|---|---|---|---|---|---|---|---|---|---|---|---|---|---|---|---|---|---|---|---|---|---|---|---|---|---|
| NW_011924881.1 | 447193 | 447193 | 1 | + | 52 | 0 | 52 | 20 | 0 | 20 | 0 | 0 | NW_011924881.1 | 447189 | 448306 | 1118 | + | Gnomon | gene | gene25 | GeneID:105274524 | LOC105274524 | Gene | LOC105274524 | protein_coding | NA | character(0) | character(0) | BCphaseB1.vs.RphaseA1 | FALSE |
| NW_011924881.1 | 447200 | 447200 | 1 | + | 51 | 0 | 51 | 41 | 1 | 40 | 0 | 2.44 | NW_011924881.1 | 447189 | 448306 | 1118 | + | Gnomon | gene | gene25 | GeneID:105274524 | LOC105274524 | Gene | LOC105274524 | protein_coding | NA | character(0) | character(0) | BCphaseB1.vs.RphaseA1 | FALSE |
| NW_011924881.1 | 447209 | 447209 | 1 | + | 51 | 1 | 50 | 24 | 0 | 24 | 1.96 | 0 | NW_011924881.1 | 447189 | 448306 | 1118 | + | Gnomon | gene | gene25 | GeneID:105274524 | LOC105274524 | Gene | LOC105274524 | protein_coding | NA | character(0) | character(0) | BCphaseB1.vs.RphaseA1 | FALSE |
| NW_011924881.1 | 447215 | 447215 | 1 | + | 32 | 0 | 32 | 23 | 0 | 23 | 0 | 0 | NW_011924881.1 | 447189 | 448306 | 1118 | + | Gnomon | gene | gene25 | GeneID:105274524 | LOC105274524 | Gene | LOC105274524 | protein_coding | NA | character(0) | character(0) | BCphaseB1.vs.RphaseA1 | FALSE |
| NW_011924881.1 | 447221 | 447221 | 1 | + | 35 | 0 | 35 | 23 | 0 | 23 | 0 | 0 | NW_011924881.1 | 447189 | 448306 | 1118 | + | Gnomon | gene | gene25 | GeneID:105274524 | LOC105274524 | Gene | LOC105274524 | protein_coding | NA | character(0) | character(0) | BCphaseB1.vs.RphaseA1 | FALSE |
| NW_011924881.1 | 447233 | 447233 | 1 | + | 37 | 0 | 37 | 24 | 0 | 24 | 0 | 0 | NW_011924881.1 | 447189 | 448306 | 1118 | + | Gnomon | gene | gene25 | GeneID:105274524 | LOC105274524 | Gene | LOC105274524 | protein_coding | NA | character(0) | character(0) | BCphaseB1.vs.RphaseA1 | FALSE |
| NW_011924881.1 | 447235 | 447235 | 1 | + | 37 | 0 | 37 | 24 | 0 | 24 | 0 | 0 | NW_011924881.1 | 447189 | 448306 | 1118 | + | Gnomon | gene | gene25 | GeneID:105274524 | LOC105274524 | Gene | LOC105274524 | protein_coding | NA | character(0) | character(0) | BCphaseB1.vs.RphaseA1 | FALSE |
| NW_011924881.1 | 447536 | 447536 | 1 | + | 22 | 0 | 22 | 23 | 0 | 23 | 0 | 0 | NW_011924881.1 | 447189 | 448306 | 1118 | + | Gnomon | gene | gene25 | GeneID:105274524 | LOC105274524 | Gene | LOC105274524 | protein_coding | NA | character(0) | character(0) | BCphaseB1.vs.RphaseA1 | FALSE |
| NW_011924881.1 | 447571 | 447571 | 1 | + | 23 | 0 | 23 | 23 | 0 | 23 | 0 | 0 | NW_011924881.1 | 447189 | 448306 | 1118 | + | Gnomon | gene | gene25 | GeneID:105274524 | LOC105274524 | Gene | LOC105274524 | protein_coding | NA | character(0) | character(0) | BCphaseB1.vs.RphaseA1 | FALSE |
| NW_011924881.1 | 447585 | 447585 | 1 | + | 22 | 0 | 22 | 23 | 0 | 23 | 0 | 0 | NW_011924881.1 | 447189 | 448306 | 1118 | + | Gnomon | gene | gene25 | GeneID:105274524 | LOC105274524 | Gene | LOC105274524 | protein_coding | NA | character(0) | character(0) | BCphaseB1.vs.RphaseA1 | FALSE |
| NW_011924881.1 | 447602 | 447602 | 1 | + | 26 | 0 | 26 | 26 | 0 | 26 | 0 | 0 | NW_011924881.1 | 447189 | 448306 | 1118 | + | Gnomon | gene | gene25 | GeneID:105274524 | LOC105274524 | Gene | LOC105274524 | protein_coding | NA | character(0) | character(0) | BCphaseB1.vs.RphaseA1 | FALSE |
| NW_011924881.1 | 447604 | 447604 | 1 | + | 27 | 0 | 27 | 26 | 0 | 26 | 0 | 0 | NW_011924881.1 | 447189 | 448306 | 1118 | + | Gnomon | gene | gene25 | GeneID:105274524 | LOC105274524 | Gene | LOC105274524 | protein_coding | NA | character(0) | character(0) | BCphaseB1.vs.RphaseA1 | FALSE |
| NW_011924881.1 | 447757 | 447757 | 1 | + | 20 | 1 | 19 | 23 | 1 | 22 | 5 | 4.35 | NW_011924881.1 | 447189 | 448306 | 1118 | + | Gnomon | gene | gene25 | GeneID:105274524 | LOC105274524 | Gene | LOC105274524 | protein_coding | NA | character(0) | character(0) | BCphaseB1.vs.RphaseA1 | FALSE |
| NW_011924881.1 | 447772 | 447772 | 1 | + | 36 | 6 | 30 | 31 | 6 | 25 | 16.67 | 19.35 | NW_011924881.1 | 447189 | 448306 | 1118 | + | Gnomon | gene | gene25 | GeneID:105274524 | LOC105274524 | Gene | LOC105274524 | protein_coding | NA | character(0) | character(0) | BCphaseB1.vs.RphaseA1 | FALSE |
| NW_011924881.1 | 447836 | 447836 | 1 | + | 83 | 26 | 57 | 55 | 25 | 30 | 31.33 | 45.45 | NW_011924881.1 | 447189 | 448306 | 1118 | + | Gnomon | gene | gene25 | GeneID:105274524 | LOC105274524 | Gene | LOC105274524 | protein_coding | NA | character(0) | character(0) | BCphaseB1.vs.RphaseA1 | FALSE |
| NW_011924881.1 | 447850 | 447850 | 1 | + | 73 | 6 | 67 | 36 | 2 | 34 | 8.22 | 5.56 | NW_011924881.1 | 447189 | 448306 | 1118 | + | Gnomon | gene | gene25 | GeneID:105274524 | LOC105274524 | Gene | LOC105274524 | protein_coding | NA | character(0) | character(0) | BCphaseB1.vs.RphaseA1 | FALSE |
| NW_011924881.1 | 447872 | 447872 | 1 | + | 65 | 7 | 58 | 35 | 5 | 30 | 10.77 | 14.29 | NW_011924881.1 | 447189 | 448306 | 1118 | + | Gnomon | gene | gene25 | GeneID:105274524 | LOC105274524 | Gene | LOC105274524 | protein_coding | NA | character(0) | character(0) | BCphaseB1.vs.RphaseA1 | FALSE |
| NW_011924881.1 | 447882 | 447882 | 1 | + | 62 | 9 | 53 | 31 | 4 | 27 | 14.52 | 12.9 | NW_011924881.1 | 447189 | 448306 | 1118 | + | Gnomon | gene | gene25 | GeneID:105274524 | LOC105274524 | Gene | LOC105274524 | protein_coding | NA | character(0) | character(0) | BCphaseB1.vs.RphaseA1 | FALSE |
| NW_011924881.1 | 448007 | 448007 | 1 | + | 24 | 20 | 4 | 22 | 18 | 4 | 83.33 | 81.82 | NW_011924881.1 | 447189 | 448306 | 1118 | + | Gnomon | gene | gene25 | GeneID:105274524 | LOC105274524 | Gene | LOC105274524 | protein_coding | NA | character(0) | character(0) | BCphaseB1.vs.RphaseA1 | FALSE |
| NW_011924881.1 | 448050 | 448050 | 1 | + | 21 | 6 | 15 | 21 | 12 | 9 | 28.57 | 57.14 | NW_011924881.1 | 447189 | 448306 | 1118 | + | Gnomon | gene | gene25 | GeneID:105274524 | LOC105274524 | Gene | LOC105274524 | protein_coding | NA | character(0) | character(0) | BCphaseB1.vs.RphaseA1 | TRUE |
| NW_011924881.1 | 448056 | 448056 | 1 | + | 28 | 18 | 10 | 20 | 13 | 7 | 64.29 | 65 | NW_011924881.1 | 447189 | 448306 | 1118 | + | Gnomon | gene | gene25 | GeneID:105274524 | LOC105274524 | Gene | LOC105274524 | protein_coding | NA | character(0) | character(0) | BCphaseB1.vs.RphaseA1 | FALSE |
| NW_011924881.1 | 448083 | 448083 | 1 | + | 39 | 36 | 3 | 41 | 40 | 1 | 92.31 | 97.56 | NW_011924881.1 | 447189 | 448306 | 1118 | + | Gnomon | gene | gene25 | GeneID:105274524 | LOC105274524 | Gene | LOC105274524 | protein_coding | NA | character(0) | character(0) | BCphaseB1.vs.RphaseA1 | FALSE |
| NW_011924881.1 | 448109 | 448109 | 1 | + | 39 | 37 | 2 | 35 | 35 | 0 | 94.87 | 100 | NW_011924881.1 | 447189 | 448306 | 1118 | + | Gnomon | gene | gene25 | GeneID:105274524 | LOC105274524 | Gene | LOC105274524 | protein_coding | NA | character(0) | character(0) | BCphaseB1.vs.RphaseA1 | FALSE |
| NW_011924881.1 | 448125 | 448125 | 1 | + | 36 | 34 | 2 | 34 | 34 | 0 | 94.44 | 100 | NW_011924881.1 | 447189 | 448306 | 1118 | + | Gnomon | gene | gene25 | GeneID:105274524 | LOC105274524 | Gene | LOC105274524 | protein_coding | NA | character(0) | character(0) | BCphaseB1.vs.RphaseA1 | FALSE |

...
... only first 25 lines shown ...

Directory: DMSG File: dmg-Cb_LI_BCphaseB1.vs.RphaseA2_details.txt

| seqnames | start | end | width | strand | coverage1 | numCs1 | numTs1 | coverage2 | numCs2 | numTs2 | BCphaseB1 | RphaseA2 | gene_seqnames | gene_start | gene_end | gene_width | gene_strand | gene_source | gene_type | gene_ID | gene_Dbxref | gene_Name | gene_gbkey | gene_gene | gene_gene_biotype | gene_partial | gene_start_range | gene_end_range | gene_comparison | is.dm |
|---|---|---|---|---|---|---|---|---|---|---|---|---|---|---|---|---|---|---|---|---|---|---|---|---|---|---|---|---|---|---|
| NW_011924881.1 | 39625 | 39625 | 1 | + | 22 | 2 | 20 | 26 | 1 | 25 | 9.09 | 3.85 | NW_011924881.1 | 39621 | 40962 | 1342 | - | Gnomon | gene | gene17 | GeneID:105286636 | LOC105286636 | Gene | LOC105286636 | protein_coding | NA | character(0) | character(0) | BCphaseB1.vs.RphaseA2 | FALSE |
| NW_011924881.1 | 39633 | 39633 | 1 | + | 25 | 2 | 23 | 25 | 1 | 24 | 8 | 4 | NW_011924881.1 | 39621 | 40962 | 1342 | - | Gnomon | gene | gene17 | GeneID:105286636 | LOC105286636 | Gene | LOC105286636 | protein_coding | NA | character(0) | character(0) | BCphaseB1.vs.RphaseA2 | FALSE |
| NW_011924881.1 | 39987 | 39987 | 1 | + | 23 | 12 | 11 | 21 | 14 | 7 | 52.17 | 66.67 | NW_011924881.1 | 39621 | 40962 | 1342 | - | Gnomon | gene | gene17 | GeneID:105286636 | LOC105286636 | Gene | LOC105286636 | protein_coding | NA | character(0) | character(0) | BCphaseB1.vs.RphaseA2 | FALSE |
| NW_011924881.1 | 40005 | 40005 | 1 | + | 24 | 1 | 23 | 23 | 4 | 19 | 4.17 | 17.39 | NW_011924881.1 | 39621 | 40962 | 1342 | - | Gnomon | gene | gene17 | GeneID:105286636 | LOC105286636 | Gene | LOC105286636 | protein_coding | NA | character(0) | character(0) | BCphaseB1.vs.RphaseA2 | FALSE |
| NW_011924881.1 | 40021 | 40021 | 1 | + | 24 | 2 | 22 | 24 | 4 | 20 | 8.33 | 16.67 | NW_011924881.1 | 39621 | 40962 | 1342 | - | Gnomon | gene | gene17 | GeneID:105286636 | LOC105286636 | Gene | LOC105286636 | protein_coding | NA | character(0) | character(0) | BCphaseB1.vs.RphaseA2 | FALSE |
| NW_011924881.1 | 40035 | 40035 | 1 | + | 25 | 6 | 19 | 30 | 5 | 25 | 24 | 16.67 | NW_011924881.1 | 39621 | 40962 | 1342 | - | Gnomon | gene | gene17 | GeneID:105286636 | LOC105286636 | Gene | LOC105286636 | protein_coding | NA | character(0) | character(0) | BCphaseB1.vs.RphaseA2 | FALSE |
| NW_011924881.1 | 40120 | 40120 | 1 | + | 20 | 16 | 4 | 22 | 13 | 9 | 80 | 59.09 | NW_011924881.1 | 39621 | 40962 | 1342 | - | Gnomon | gene | gene17 | GeneID:105286636 | LOC105286636 | Gene | LOC105286636 | protein_coding | NA | character(0) | character(0) | BCphaseB1.vs.RphaseA2 | TRUE |
| NW_011924881.1 | 40256 | 40256 | 1 | + | 23 | 17 | 6 | 34 | 30 | 4 | 73.91 | 88.24 | NW_011924881.1 | 39621 | 40962 | 1342 | - | Gnomon | gene | gene17 | GeneID:105286636 | LOC105286636 | Gene | LOC105286636 | protein_coding | NA | character(0) | character(0) | BCphaseB1.vs.RphaseA2 | FALSE |
| NW_011924881.1 | 40259 | 40259 | 1 | + | 23 | 21 | 2 | 31 | 28 | 3 | 91.3 | 90.32 | NW_011924881.1 | 39621 | 40962 | 1342 | - | Gnomon | gene | gene17 | GeneID:105286636 | LOC105286636 | Gene | LOC105286636 | protein_coding | NA | character(0) | character(0) | BCphaseB1.vs.RphaseA2 | FALSE |
| NW_011924881.1 | 40270 | 40270 | 1 | + | 23 | 20 | 3 | 29 | 26 | 3 | 86.96 | 89.66 | NW_011924881.1 | 39621 | 40962 | 1342 | - | Gnomon | gene | gene17 | GeneID:105286636 | LOC105286636 | Gene | LOC105286636 | protein_coding | NA | character(0) | character(0) | BCphaseB1.vs.RphaseA2 | FALSE |
| NW_011924881.1 | 40304 | 40304 | 1 | + | 20 | 3 | 17 | 23 | 7 | 16 | 15 | 30.43 | NW_011924881.1 | 39621 | 40962 | 1342 | - | Gnomon | gene | gene17 | GeneID:105286636 | LOC105286636 | Gene | LOC105286636 | protein_coding | NA | character(0) | character(0) | BCphaseB1.vs.RphaseA2 | FALSE |
| NW_011924881.1 | 40359 | 40359 | 1 | + | 23 | 22 | 1 | 30 | 28 | 2 | 95.65 | 93.33 | NW_011924881.1 | 39621 | 40962 | 1342 | - | Gnomon | gene | gene17 | GeneID:105286636 | LOC105286636 | Gene | LOC105286636 | protein_coding | NA | character(0) | character(0) | BCphaseB1.vs.RphaseA2 | FALSE |
| NW_011924881.1 | 40371 | 40371 | 1 | + | 24 | 23 | 1 | 29 | 28 | 1 | 95.83 | 96.55 | NW_011924881.1 | 39621 | 40962 | 1342 | - | Gnomon | gene | gene17 | GeneID:105286636 | LOC105286636 | Gene | LOC105286636 | protein_coding | NA | character(0) | character(0) | BCphaseB1.vs.RphaseA2 | FALSE |
| NW_011924881.1 | 40583 | 40583 | 1 | + | 23 | 0 | 23 | 26 | 1 | 25 | 0 | 3.85 | NW_011924881.1 | 39621 | 40962 | 1342 | - | Gnomon | gene | gene17 | GeneID:105286636 | LOC105286636 | Gene | LOC105286636 | protein_coding | NA | character(0) | character(0) | BCphaseB1.vs.RphaseA2 | FALSE |
| NW_011924881.1 | 40595 | 40595 | 1 | + | 25 | 0 | 25 | 27 | 1 | 26 | 0 | 3.7 | NW_011924881.1 | 39621 | 40962 | 1342 | - | Gnomon | gene | gene17 | GeneID:105286636 | LOC105286636 | Gene | LOC105286636 | protein_coding | NA | character(0) | character(0) | BCphaseB1.vs.RphaseA2 | FALSE |
| NW_011924881.1 | 40641 | 40641 | 1 | + | 26 | 0 | 26 | 23 | 0 | 23 | 0 | 0 | NW_011924881.1 | 39621 | 40962 | 1342 | - | Gnomon | gene | gene17 | GeneID:105286636 | LOC105286636 | Gene | LOC105286636 | protein_coding | NA | character(0) | character(0) | BCphaseB1.vs.RphaseA2 | FALSE |
| NW_011924881.1 | 40646 | 40646 | 1 | + | 26 | 0 | 26 | 21 | 0 | 21 | 0 | 0 | NW_011924881.1 | 39621 | 40962 | 1342 | - | Gnomon | gene | gene17 | GeneID:105286636 | LOC105286636 | Gene | LOC105286636 | protein_coding | NA | character(0) | character(0) | BCphaseB1.vs.RphaseA2 | FALSE |
| NW_011924881.1 | 40762 | 40762 | 1 | + | 22 | 0 | 22 | 20 | 0 | 20 | 0 | 0 | NW_011924881.1 | 39621 | 40962 | 1342 | - | Gnomon | gene | gene17 | GeneID:105286636 | LOC105286636 | Gene | LOC105286636 | protein_coding | NA | character(0) | character(0) | BCphaseB1.vs.RphaseA2 | FALSE |
| NW_011924881.1 | 40781 | 40781 | 1 | + | 33 | 0 | 33 | 23 | 0 | 23 | 0 | 0 | NW_011924881.1 | 39621 | 40962 | 1342 | - | Gnomon | gene | gene17 | GeneID:105286636 | LOC105286636 | Gene | LOC105286636 | protein_coding | NA | character(0) | character(0) | BCphaseB1.vs.RphaseA2 | FALSE |
| NW_011924881.1 | 40810 | 40810 | 1 | + | 30 | 0 | 30 | 26 | 0 | 26 | 0 | 0 | NW_011924881.1 | 39621 | 40962 | 1342 | - | Gnomon | gene | gene17 | GeneID:105286636 | LOC105286636 | Gene | LOC105286636 | protein_coding | NA | character(0) | character(0) | BCphaseB1.vs.RphaseA2 | FALSE |
| NW_011924881.1 | 40843 | 40843 | 1 | + | 25 | 0 | 25 | 24 | 0 | 24 | 0 | 0 | NW_011924881.1 | 39621 | 40962 | 1342 | - | Gnomon | gene | gene17 | GeneID:105286636 | LOC105286636 | Gene | LOC105286636 | protein_coding | NA | character(0) | character(0) | BCphaseB1.vs.RphaseA2 | FALSE |
| NW_011924881.1 | 40852 | 40852 | 1 | + | 28 | 0 | 28 | 32 | 0 | 32 | 0 | 0 | NW_011924881.1 | 39621 | 40962 | 1342 | - | Gnomon | gene | gene17 | GeneID:105286636 | LOC105286636 | Gene | LOC105286636 | protein_coding | NA | character(0) | character(0) | BCphaseB1.vs.RphaseA2 | FALSE |
| NW_011924881.1 | 40879 | 40879 | 1 | + | 44 | 0 | 44 | 28 | 0 | 28 | 0 | 0 | NW_011924881.1 | 39621 | 40962 | 1342 | - | Gnomon | gene | gene17 | GeneID:105286636 | LOC105286636 | Gene | LOC105286636 | protein_coding | NA | character(0) | character(0) | BCphaseB1.vs.RphaseA2 | FALSE |
| NW_011924881.1 | 40895 | 40895 | 1 | + | 28 | 0 | 28 | 48 | 0 | 48 | 0 | 0 | NW_011924881.1 | 39621 | 40962 | 1342 | - | Gnomon | gene | gene17 | GeneID:105286636 | LOC105286636 | Gene | LOC105286636 | protein_coding | NA | character(0) | character(0) | BCphaseB1.vs.RphaseA2 | FALSE |

...
... only first 25 lines shown ...

Directory: DMSG File: dmg-Cb_LI_BCphaseB1.vs.RphaseB1_details.txt

| seqnames | start | end | width | strand | coverage1 | numCs1 | numTs1 | coverage2 | numCs2 | numTs2 | BCphaseB1 | RphaseB1 | gene_seqnames | gene_start | gene_end | gene_width | gene_strand | gene_source | gene_type | gene_ID | gene_Dbxref | gene_Name | gene_gbkey | gene_gene | gene_gene_biotype | gene_start_range | gene_end_range | gene_comparison | is.dm |
|---|---|---|---|---|---|---|---|---|---|---|---|---|---|---|---|---|---|---|---|---|---|---|---|---|---|---|---|---|---|
| NW_011924881.1 | 809447 | 809447 | 1 | + | 25 | 0 | 25 | 23 | 0 | 23 | 0 | 0 | NW_011924881.1 | 808343 | 827625 | 19283 | - | Gnomon | gene | gene53 | GeneID:105277148 | LOC105277148 | Gene | LOC105277148 | protein_coding | character(0) | character(0) | BCphaseB1.vs.RphaseB1 | FALSE |
| NW_011924881.1 | 809481 | 809481 | 1 | + | 34 | 1 | 33 | 30 | 0 | 30 | 2.94 | 0 | NW_011924881.1 | 808343 | 827625 | 19283 | - | Gnomon | gene | gene53 | GeneID:105277148 | LOC105277148 | Gene | LOC105277148 | protein_coding | character(0) | character(0) | BCphaseB1.vs.RphaseB1 | FALSE |
| NW_011924881.1 | 809504 | 809504 | 1 | + | 34 | 0 | 34 | 31 | 0 | 31 | 0 | 0 | NW_011924881.1 | 808343 | 827625 | 19283 | - | Gnomon | gene | gene53 | GeneID:105277148 | LOC105277148 | Gene | LOC105277148 | protein_coding | character(0) | character(0) | BCphaseB1.vs.RphaseB1 | FALSE |
| NW_011924881.1 | 809520 | 809520 | 1 | + | 30 | 0 | 30 | 25 | 0 | 25 | 0 | 0 | NW_011924881.1 | 808343 | 827625 | 19283 | - | Gnomon | gene | gene53 | GeneID:105277148 | LOC105277148 | Gene | LOC105277148 | protein_coding | character(0) | character(0) | BCphaseB1.vs.RphaseB1 | FALSE |
| NW_011924881.1 | 810123 | 810123 | 1 | + | 21 | 0 | 21 | 25 | 0 | 25 | 0 | 0 | NW_011924881.1 | 808343 | 827625 | 19283 | - | Gnomon | gene | gene53 | GeneID:105277148 | LOC105277148 | Gene | LOC105277148 | protein_coding | character(0) | character(0) | BCphaseB1.vs.RphaseB1 | FALSE |
| NW_011924881.1 | 810155 | 810155 | 1 | + | 23 | 0 | 23 | 25 | 0 | 25 | 0 | 0 | NW_011924881.1 | 808343 | 827625 | 19283 | - | Gnomon | gene | gene53 | GeneID:105277148 | LOC105277148 | Gene | LOC105277148 | protein_coding | character(0) | character(0) | BCphaseB1.vs.RphaseB1 | FALSE |
| NW_011924881.1 | 810182 | 810182 | 1 | + | 25 | 3 | 22 | 28 | 1 | 27 | 12 | 3.57 | NW_011924881.1 | 808343 | 827625 | 19283 | - | Gnomon | gene | gene53 | GeneID:105277148 | LOC105277148 | Gene | LOC105277148 | protein_coding | character(0) | character(0) | BCphaseB1.vs.RphaseB1 | FALSE |
| NW_011924881.1 | 810337 | 810337 | 1 | + | 22 | 0 | 22 | 29 | 0 | 29 | 0 | 0 | NW_011924881.1 | 808343 | 827625 | 19283 | - | Gnomon | gene | gene53 | GeneID:105277148 | LOC105277148 | Gene | LOC105277148 | protein_coding | character(0) | character(0) | BCphaseB1.vs.RphaseB1 | FALSE |
| NW_011924881.1 | 810348 | 810348 | 1 | + | 42 | 1 | 41 | 41 | 1 | 40 | 2.38 | 2.44 | NW_011924881.1 | 808343 | 827625 | 19283 | - | Gnomon | gene | gene53 | GeneID:105277148 | LOC105277148 | Gene | LOC105277148 | protein_coding | character(0) | character(0) | BCphaseB1.vs.RphaseB1 | FALSE |
| NW_011924881.1 | 810391 | 810391 | 1 | + | 68 | 1 | 67 | 37 | 0 | 37 | 1.47 | 0 | NW_011924881.1 | 808343 | 827625 | 19283 | - | Gnomon | gene | gene53 | GeneID:105277148 | LOC105277148 | Gene | LOC105277148 | protein_coding | character(0) | character(0) | BCphaseB1.vs.RphaseB1 | FALSE |
| NW_011924881.1 | 810409 | 810409 | 1 | + | 59 | 0 | 59 | 32 | 1 | 31 | 0 | 3.12 | NW_011924881.1 | 808343 | 827625 | 19283 | - | Gnomon | gene | gene53 | GeneID:105277148 | LOC105277148 | Gene | LOC105277148 | protein_coding | character(0) | character(0) | BCphaseB1.vs.RphaseB1 | FALSE |
| NW_011924881.1 | 810412 | 810412 | 1 | + | 34 | 1 | 33 | 27 | 1 | 26 | 2.94 | 3.7 | NW_011924881.1 | 808343 | 827625 | 19283 | - | Gnomon | gene | gene53 | GeneID:105277148 | LOC105277148 | Gene | LOC105277148 | protein_coding | character(0) | character(0) | BCphaseB1.vs.RphaseB1 | FALSE |
| NW_011924881.1 | 810417 | 810417 | 1 | + | 33 | 0 | 33 | 24 | 1 | 23 | 0 | 4.17 | NW_011924881.1 | 808343 | 827625 | 19283 | - | Gnomon | gene | gene53 | GeneID:105277148 | LOC105277148 | Gene | LOC105277148 | protein_coding | character(0) | character(0) | BCphaseB1.vs.RphaseB1 | FALSE |
| NW_011924881.1 | 810424 | 810424 | 1 | + | 46 | 0 | 46 | 24 | 0 | 24 | 0 | 0 | NW_011924881.1 | 808343 | 827625 | 19283 | - | Gnomon | gene | gene53 | GeneID:105277148 | LOC105277148 | Gene | LOC105277148 | protein_coding | character(0) | character(0) | BCphaseB1.vs.RphaseB1 | FALSE |
| NW_011924881.1 | 810430 | 810430 | 1 | + | 45 | 0 | 45 | 26 | 0 | 26 | 0 | 0 | NW_011924881.1 | 808343 | 827625 | 19283 | - | Gnomon | gene | gene53 | GeneID:105277148 | LOC105277148 | Gene | LOC105277148 | protein_coding | character(0) | character(0) | BCphaseB1.vs.RphaseB1 | FALSE |
| NW_011924881.1 | 810436 | 810436 | 1 | + | 49 | 0 | 49 | 30 | 0 | 30 | 0 | 0 | NW_011924881.1 | 808343 | 827625 | 19283 | - | Gnomon | gene | gene53 | GeneID:105277148 | LOC105277148 | Gene | LOC105277148 | protein_coding | character(0) | character(0) | BCphaseB1.vs.RphaseB1 | FALSE |
| NW_011924881.1 | 810482 | 810482 | 1 | + | 66 | 2 | 64 | 47 | 1 | 46 | 3.03 | 2.13 | NW_011924881.1 | 808343 | 827625 | 19283 | - | Gnomon | gene | gene53 | GeneID:105277148 | LOC105277148 | Gene | LOC105277148 | protein_coding | character(0) | character(0) | BCphaseB1.vs.RphaseB1 | FALSE |
| NW_011924881.1 | 810505 | 810505 | 1 | + | 64 | 0 | 64 | 50 | 0 | 50 | 0 | 0 | NW_011924881.1 | 808343 | 827625 | 19283 | - | Gnomon | gene | gene53 | GeneID:105277148 | LOC105277148 | Gene | LOC105277148 | protein_coding | character(0) | character(0) | BCphaseB1.vs.RphaseB1 | FALSE |
| NW_011924881.1 | 810511 | 810511 | 1 | + | 61 | 1 | 60 | 54 | 0 | 54 | 1.64 | 0 | NW_011924881.1 | 808343 | 827625 | 19283 | - | Gnomon | gene | gene53 | GeneID:105277148 | LOC105277148 | Gene | LOC105277148 | protein_coding | character(0) | character(0) | BCphaseB1.vs.RphaseB1 | FALSE |
| NW_011924881.1 | 810516 | 810516 | 1 | + | 63 | 0 | 63 | 55 | 1 | 54 | 0 | 1.82 | NW_011924881.1 | 808343 | 827625 | 19283 | - | Gnomon | gene | gene53 | GeneID:105277148 | LOC105277148 | Gene | LOC105277148 | protein_coding | character(0) | character(0) | BCphaseB1.vs.RphaseB1 | FALSE |
| NW_011924881.1 | 810520 | 810520 | 1 | + | 63 | 0 | 63 | 56 | 1 | 55 | 0 | 1.79 | NW_011924881.1 | 808343 | 827625 | 19283 | - | Gnomon | gene | gene53 | GeneID:105277148 | LOC105277148 | Gene | LOC105277148 | protein_coding | character(0) | character(0) | BCphaseB1.vs.RphaseB1 | FALSE |
| NW_011924881.1 | 810532 | 810532 | 1 | + | 24 | 0 | 24 | 22 | 0 | 22 | 0 | 0 | NW_011924881.1 | 808343 | 827625 | 19283 | - | Gnomon | gene | gene53 | GeneID:105277148 | LOC105277148 | Gene | LOC105277148 | protein_coding | character(0) | character(0) | BCphaseB1.vs.RphaseB1 | FALSE |
| NW_011924881.1 | 810535 | 810535 | 1 | + | 24 | 1 | 23 | 23 | 0 | 23 | 4.17 | 0 | NW_011924881.1 | 808343 | 827625 | 19283 | - | Gnomon | gene | gene53 | GeneID:105277148 | LOC105277148 | Gene | LOC105277148 | protein_coding | character(0) | character(0) | BCphaseB1.vs.RphaseB1 | FALSE |
| NW_011924881.1 | 810577 | 810577 | 1 | + | 33 | 0 | 33 | 30 | 0 | 30 | 0 | 0 | NW_011924881.1 | 808343 | 827625 | 19283 | - | Gnomon | gene | gene53 | GeneID:105277148 | LOC105277148 | Gene | LOC105277148 | protein_coding | character(0) | character(0) | BCphaseB1.vs.RphaseB1 | FALSE |

...
... only first 25 lines shown ...

Directory: DMSG File: dmg-Cb_LI_BCphaseB1.vs.RphaseB2_details.txt

| seqnames | start | end | width | strand | coverage1 | numCs1 | numTs1 | coverage2 | numCs2 | numTs2 | BCphaseB1 | RphaseB2 | gene_seqnames | gene_start | gene_end | gene_width | gene_strand | gene_source | gene_type | gene_ID | gene_Dbxref | gene_Name | gene_gbkey | gene_gene | gene_gene_biotype | gene_start_range | gene_end_range | gene_comparison | is.dm |
|---|---|---|---|---|---|---|---|---|---|---|---|---|---|---|---|---|---|---|---|---|---|---|---|---|---|---|---|---|---|
| NW_011924881.1 | 724788 | 724788 | 1 | + | 37 | 0 | 37 | 28 | 0 | 28 | 0 | 0 | NW_011924881.1 | 724448 | 731567 | 7120 | - | Gnomon | gene | gene34 | GeneID:105278416 | LOC105278416 | Gene | LOC105278416 | protein_coding | character(0) | character(0) | BCphaseB1.vs.RphaseB2 | FALSE |
| NW_011924881.1 | 724794 | 724794 | 1 | + | 35 | 0 | 35 | 28 | 0 | 28 | 0 | 0 | NW_011924881.1 | 724448 | 731567 | 7120 | - | Gnomon | gene | gene34 | GeneID:105278416 | LOC105278416 | Gene | LOC105278416 | protein_coding | character(0) | character(0) | BCphaseB1.vs.RphaseB2 | FALSE |
| NW_011924881.1 | 724802 | 724802 | 1 | + | 33 | 0 | 33 | 28 | 0 | 28 | 0 | 0 | NW_011924881.1 | 724448 | 731567 | 7120 | - | Gnomon | gene | gene34 | GeneID:105278416 | LOC105278416 | Gene | LOC105278416 | protein_coding | character(0) | character(0) | BCphaseB1.vs.RphaseB2 | FALSE |
| NW_011924881.1 | 724806 | 724806 | 1 | + | 33 | 1 | 32 | 29 | 0 | 29 | 3.03 | 0 | NW_011924881.1 | 724448 | 731567 | 7120 | - | Gnomon | gene | gene34 | GeneID:105278416 | LOC105278416 | Gene | LOC105278416 | protein_coding | character(0) | character(0) | BCphaseB1.vs.RphaseB2 | FALSE |
| NW_011924881.1 | 724815 | 724815 | 1 | + | 33 | 0 | 33 | 29 | 1 | 28 | 0 | 3.45 | NW_011924881.1 | 724448 | 731567 | 7120 | - | Gnomon | gene | gene34 | GeneID:105278416 | LOC105278416 | Gene | LOC105278416 | protein_coding | character(0) | character(0) | BCphaseB1.vs.RphaseB2 | FALSE |
| NW_011924881.1 | 724818 | 724818 | 1 | + | 33 | 0 | 33 | 29 | 0 | 29 | 0 | 0 | NW_011924881.1 | 724448 | 731567 | 7120 | - | Gnomon | gene | gene34 | GeneID:105278416 | LOC105278416 | Gene | LOC105278416 | protein_coding | character(0) | character(0) | BCphaseB1.vs.RphaseB2 | FALSE |
| NW_011924881.1 | 724839 | 724839 | 1 | + | 33 | 0 | 33 | 44 | 0 | 44 | 0 | 0 | NW_011924881.1 | 724448 | 731567 | 7120 | - | Gnomon | gene | gene34 | GeneID:105278416 | LOC105278416 | Gene | LOC105278416 | protein_coding | character(0) | character(0) | BCphaseB1.vs.RphaseB2 | FALSE |
| NW_011924881.1 | 724872 | 724872 | 1 | + | 29 | 0 | 29 | 36 | 0 | 36 | 0 | 0 | NW_011924881.1 | 724448 | 731567 | 7120 | - | Gnomon | gene | gene34 | GeneID:105278416 | LOC105278416 | Gene | LOC105278416 | protein_coding | character(0) | character(0) | BCphaseB1.vs.RphaseB2 | FALSE |
| NW_011924881.1 | 724889 | 724889 | 1 | + | 28 | 0 | 28 | 38 | 0 | 38 | 0 | 0 | NW_011924881.1 | 724448 | 731567 | 7120 | - | Gnomon | gene | gene34 | GeneID:105278416 | LOC105278416 | Gene | LOC105278416 | protein_coding | character(0) | character(0) | BCphaseB1.vs.RphaseB2 | FALSE |
| NW_011924881.1 | 724896 | 724896 | 1 | + | 30 | 0 | 30 | 36 | 0 | 36 | 0 | 0 | NW_011924881.1 | 724448 | 731567 | 7120 | - | Gnomon | gene | gene34 | GeneID:105278416 | LOC105278416 | Gene | LOC105278416 | protein_coding | character(0) | character(0) | BCphaseB1.vs.RphaseB2 | FALSE |
| NW_011924881.1 | 724898 | 724898 | 1 | + | 29 | 0 | 29 | 37 | 0 | 37 | 0 | 0 | NW_011924881.1 | 724448 | 731567 | 7120 | - | Gnomon | gene | gene34 | GeneID:105278416 | LOC105278416 | Gene | LOC105278416 | protein_coding | character(0) | character(0) | BCphaseB1.vs.RphaseB2 | FALSE |
| NW_011924881.1 | 724926 | 724926 | 1 | + | 34 | 0 | 34 | 58 | 1 | 57 | 0 | 1.72 | NW_011924881.1 | 724448 | 731567 | 7120 | - | Gnomon | gene | gene34 | GeneID:105278416 | LOC105278416 | Gene | LOC105278416 | protein_coding | character(0) | character(0) | BCphaseB1.vs.RphaseB2 | FALSE |
| NW_011924881.1 | 724934 | 724934 | 1 | + | 36 | 0 | 36 | 43 | 0 | 43 | 0 | 0 | NW_011924881.1 | 724448 | 731567 | 7120 | - | Gnomon | gene | gene34 | GeneID:105278416 | LOC105278416 | Gene | LOC105278416 | protein_coding | character(0) | character(0) | BCphaseB1.vs.RphaseB2 | FALSE |
| NW_011924881.1 | 724936 | 724936 | 1 | + | 36 | 0 | 36 | 44 | 0 | 44 | 0 | 0 | NW_011924881.1 | 724448 | 731567 | 7120 | - | Gnomon | gene | gene34 | GeneID:105278416 | LOC105278416 | Gene | LOC105278416 | protein_coding | character(0) | character(0) | BCphaseB1.vs.RphaseB2 | FALSE |
| NW_011924881.1 | 724943 | 724943 | 1 | + | 33 | 0 | 33 | 40 | 1 | 39 | 0 | 2.5 | NW_011924881.1 | 724448 | 731567 | 7120 | - | Gnomon | gene | gene34 | GeneID:105278416 | LOC105278416 | Gene | LOC105278416 | protein_coding | character(0) | character(0) | BCphaseB1.vs.RphaseB2 | FALSE |
| NW_011924881.1 | 724947 | 724947 | 1 | + | 31 | 0 | 31 | 42 | 0 | 42 | 0 | 0 | NW_011924881.1 | 724448 | 731567 | 7120 | - | Gnomon | gene | gene34 | GeneID:105278416 | LOC105278416 | Gene | LOC105278416 | protein_coding | character(0) | character(0) | BCphaseB1.vs.RphaseB2 | FALSE |
| NW_011924881.1 | 724961 | 724961 | 1 | + | 36 | 1 | 35 | 46 | 0 | 46 | 2.78 | 0 | NW_011924881.1 | 724448 | 731567 | 7120 | - | Gnomon | gene | gene34 | GeneID:105278416 | LOC105278416 | Gene | LOC105278416 | protein_coding | character(0) | character(0) | BCphaseB1.vs.RphaseB2 | FALSE |
| NW_011924881.1 | 724976 | 724976 | 1 | + | 43 | 0 | 43 | 54 | 1 | 53 | 0 | 1.85 | NW_011924881.1 | 724448 | 731567 | 7120 | - | Gnomon | gene | gene34 | GeneID:105278416 | LOC105278416 | Gene | LOC105278416 | protein_coding | character(0) | character(0) | BCphaseB1.vs.RphaseB2 | FALSE |
| NW_011924881.1 | 724982 | 724982 | 1 | + | 44 | 0 | 44 | 57 | 0 | 57 | 0 | 0 | NW_011924881.1 | 724448 | 731567 | 7120 | - | Gnomon | gene | gene34 | GeneID:105278416 | LOC105278416 | Gene | LOC105278416 | protein_coding | character(0) | character(0) | BCphaseB1.vs.RphaseB2 | FALSE |
| NW_011924881.1 | 724990 | 724990 | 1 | + | 44 | 0 | 44 | 58 | 0 | 58 | 0 | 0 | NW_011924881.1 | 724448 | 731567 | 7120 | - | Gnomon | gene | gene34 | GeneID:105278416 | LOC105278416 | Gene | LOC105278416 | protein_coding | character(0) | character(0) | BCphaseB1.vs.RphaseB2 | FALSE |
| NW_011924881.1 | 724994 | 724994 | 1 | + | 45 | 0 | 45 | 58 | 0 | 58 | 0 | 0 | NW_011924881.1 | 724448 | 731567 | 7120 | - | Gnomon | gene | gene34 | GeneID:105278416 | LOC105278416 | Gene | LOC105278416 | protein_coding | character(0) | character(0) | BCphaseB1.vs.RphaseB2 | FALSE |
| NW_011924881.1 | 725008 | 725008 | 1 | + | 31 | 1 | 30 | 42 | 0 | 42 | 3.23 | 0 | NW_011924881.1 | 724448 | 731567 | 7120 | - | Gnomon | gene | gene34 | GeneID:105278416 | LOC105278416 | Gene | LOC105278416 | protein_coding | character(0) | character(0) | BCphaseB1.vs.RphaseB2 | FALSE |
| NW_011924881.1 | 725014 | 725014 | 1 | + | 58 | 2 | 56 | 31 | 0 | 31 | 3.45 | 0 | NW_011924881.1 | 724448 | 731567 | 7120 | - | Gnomon | gene | gene34 | GeneID:105278416 | LOC105278416 | Gene | LOC105278416 | protein_coding | character(0) | character(0) | BCphaseB1.vs.RphaseB2 | FALSE |
| NW_011924881.1 | 725022 | 725022 | 1 | + | 36 | 3 | 33 | 22 | 1 | 21 | 8.33 | 4.55 | NW_011924881.1 | 724448 | 731567 | 7120 | - | Gnomon | gene | gene34 | GeneID:105278416 | LOC105278416 | Gene | LOC105278416 | protein_coding | character(0) | character(0) | BCphaseB1.vs.RphaseB2 | FALSE |

...
... only first 25 lines shown ...

Directory: DMSG File: dmg-Cb_LI_BCphaseB2.vs.RphaseA1_details.txt

| seqnames | start | end | width | strand | coverage1 | numCs1 | numTs1 | coverage2 | numCs2 | numTs2 | BCphaseB2 | RphaseA1 | gene_seqnames | gene_start | gene_end | gene_width | gene_strand | gene_source | gene_type | gene_ID | gene_Dbxref | gene_Name | gene_gbkey | gene_gene | gene_gene_biotype | gene_start_range | gene_end_range | gene_comparison | is.dm |
|---|---|---|---|---|---|---|---|---|---|---|---|---|---|---|---|---|---|---|---|---|---|---|---|---|---|---|---|---|---|
| NW_011924881.1 | 724788 | 724788 | 1 | + | 31 | 0 | 31 | 21 | 0 | 21 | 0 | 0 | NW_011924881.1 | 724448 | 731567 | 7120 | - | Gnomon | gene | gene34 | GeneID:105278416 | LOC105278416 | Gene | LOC105278416 | protein_coding | character(0) | character(0) | BCphaseB2.vs.RphaseA1 | FALSE |
| NW_011924881.1 | 724926 | 724926 | 1 | + | 37 | 0 | 37 | 30 | 0 | 30 | 0 | 0 | NW_011924881.1 | 724448 | 731567 | 7120 | - | Gnomon | gene | gene34 | GeneID:105278416 | LOC105278416 | Gene | LOC105278416 | protein_coding | character(0) | character(0) | BCphaseB2.vs.RphaseA1 | FALSE |
| NW_011924881.1 | 724934 | 724934 | 1 | + | 34 | 0 | 34 | 26 | 0 | 26 | 0 | 0 | NW_011924881.1 | 724448 | 731567 | 7120 | - | Gnomon | gene | gene34 | GeneID:105278416 | LOC105278416 | Gene | LOC105278416 | protein_coding | character(0) | character(0) | BCphaseB2.vs.RphaseA1 | FALSE |
| NW_011924881.1 | 724936 | 724936 | 1 | + | 35 | 0 | 35 | 27 | 0 | 27 | 0 | 0 | NW_011924881.1 | 724448 | 731567 | 7120 | - | Gnomon | gene | gene34 | GeneID:105278416 | LOC105278416 | Gene | LOC105278416 | protein_coding | character(0) | character(0) | BCphaseB2.vs.RphaseA1 | FALSE |
| NW_011924881.1 | 724943 | 724943 | 1 | + | 33 | 0 | 33 | 24 | 0 | 24 | 0 | 0 | NW_011924881.1 | 724448 | 731567 | 7120 | - | Gnomon | gene | gene34 | GeneID:105278416 | LOC105278416 | Gene | LOC105278416 | protein_coding | character(0) | character(0) | BCphaseB2.vs.RphaseA1 | FALSE |
| NW_011924881.1 | 724947 | 724947 | 1 | + | 36 | 0 | 36 | 27 | 0 | 27 | 0 | 0 | NW_011924881.1 | 724448 | 731567 | 7120 | - | Gnomon | gene | gene34 | GeneID:105278416 | LOC105278416 | Gene | LOC105278416 | protein_coding | character(0) | character(0) | BCphaseB2.vs.RphaseA1 | FALSE |
| NW_011924881.1 | 724961 | 724961 | 1 | + | 43 | 0 | 43 | 34 | 0 | 34 | 0 | 0 | NW_011924881.1 | 724448 | 731567 | 7120 | - | Gnomon | gene | gene34 | GeneID:105278416 | LOC105278416 | Gene | LOC105278416 | protein_coding | character(0) | character(0) | BCphaseB2.vs.RphaseA1 | FALSE |
| NW_011924881.1 | 724976 | 724976 | 1 | + | 52 | 0 | 52 | 41 | 0 | 41 | 0 | 0 | NW_011924881.1 | 724448 | 731567 | 7120 | - | Gnomon | gene | gene34 | GeneID:105278416 | LOC105278416 | Gene | LOC105278416 | protein_coding | character(0) | character(0) | BCphaseB2.vs.RphaseA1 | FALSE |
| NW_011924881.1 | 724982 | 724982 | 1 | + | 54 | 0 | 54 | 41 | 2 | 39 | 0 | 4.88 | NW_011924881.1 | 724448 | 731567 | 7120 | - | Gnomon | gene | gene34 | GeneID:105278416 | LOC105278416 | Gene | LOC105278416 | protein_coding | character(0) | character(0) | BCphaseB2.vs.RphaseA1 | FALSE |
| NW_011924881.1 | 724990 | 724990 | 1 | + | 54 | 0 | 54 | 39 | 0 | 39 | 0 | 0 | NW_011924881.1 | 724448 | 731567 | 7120 | - | Gnomon | gene | gene34 | GeneID:105278416 | LOC105278416 | Gene | LOC105278416 | protein_coding | character(0) | character(0) | BCphaseB2.vs.RphaseA1 | FALSE |
| NW_011924881.1 | 724994 | 724994 | 1 | + | 53 | 0 | 53 | 39 | 0 | 39 | 0 | 0 | NW_011924881.1 | 724448 | 731567 | 7120 | - | Gnomon | gene | gene34 | GeneID:105278416 | LOC105278416 | Gene | LOC105278416 | protein_coding | character(0) | character(0) | BCphaseB2.vs.RphaseA1 | FALSE |
| NW_011924881.1 | 725008 | 725008 | 1 | + | 43 | 0 | 43 | 31 | 1 | 30 | 0 | 3.23 | NW_011924881.1 | 724448 | 731567 | 7120 | - | Gnomon | gene | gene34 | GeneID:105278416 | LOC105278416 | Gene | LOC105278416 | protein_coding | character(0) | character(0) | BCphaseB2.vs.RphaseA1 | FALSE |
| NW_011924881.1 | 725014 | 725014 | 1 | + | 35 | 0 | 35 | 33 | 1 | 32 | 0 | 3.03 | NW_011924881.1 | 724448 | 731567 | 7120 | - | Gnomon | gene | gene34 | GeneID:105278416 | LOC105278416 | Gene | LOC105278416 | protein_coding | character(0) | character(0) | BCphaseB2.vs.RphaseA1 | FALSE |
| NW_011924881.1 | 725022 | 725022 | 1 | + | 24 | 0 | 24 | 37 | 1 | 36 | 0 | 2.7 | NW_011924881.1 | 724448 | 731567 | 7120 | - | Gnomon | gene | gene34 | GeneID:105278416 | LOC105278416 | Gene | LOC105278416 | protein_coding | character(0) | character(0) | BCphaseB2.vs.RphaseA1 | FALSE |
| NW_011924881.1 | 725041 | 725041 | 1 | + | 33 | 0 | 33 | 41 | 0 | 41 | 0 | 0 | NW_011924881.1 | 724448 | 731567 | 7120 | - | Gnomon | gene | gene34 | GeneID:105278416 | LOC105278416 | Gene | LOC105278416 | protein_coding | character(0) | character(0) | BCphaseB2.vs.RphaseA1 | FALSE |
| NW_011924881.1 | 725048 | 725048 | 1 | + | 38 | 0 | 38 | 43 | 0 | 43 | 0 | 0 | NW_011924881.1 | 724448 | 731567 | 7120 | - | Gnomon | gene | gene34 | GeneID:105278416 | LOC105278416 | Gene | LOC105278416 | protein_coding | character(0) | character(0) | BCphaseB2.vs.RphaseA1 | FALSE |
| NW_011924881.1 | 725062 | 725062 | 1 | + | 68 | 0 | 68 | 44 | 0 | 44 | 0 | 0 | NW_011924881.1 | 724448 | 731567 | 7120 | - | Gnomon | gene | gene34 | GeneID:105278416 | LOC105278416 | Gene | LOC105278416 | protein_coding | character(0) | character(0) | BCphaseB2.vs.RphaseA1 | FALSE |
| NW_011924881.1 | 725065 | 725065 | 1 | + | 69 | 0 | 69 | 44 | 0 | 44 | 0 | 0 | NW_011924881.1 | 724448 | 731567 | 7120 | - | Gnomon | gene | gene34 | GeneID:105278416 | LOC105278416 | Gene | LOC105278416 | protein_coding | character(0) | character(0) | BCphaseB2.vs.RphaseA1 | FALSE |
| NW_011924881.1 | 725072 | 725072 | 1 | + | 69 | 0 | 69 | 45 | 0 | 45 | 0 | 0 | NW_011924881.1 | 724448 | 731567 | 7120 | - | Gnomon | gene | gene34 | GeneID:105278416 | LOC105278416 | Gene | LOC105278416 | protein_coding | character(0) | character(0) | BCphaseB2.vs.RphaseA1 | FALSE |
| NW_011924881.1 | 725074 | 725074 | 1 | + | 69 | 0 | 69 | 45 | 0 | 45 | 0 | 0 | NW_011924881.1 | 724448 | 731567 | 7120 | - | Gnomon | gene | gene34 | GeneID:105278416 | LOC105278416 | Gene | LOC105278416 | protein_coding | character(0) | character(0) | BCphaseB2.vs.RphaseA1 | FALSE |
| NW_011924881.1 | 725091 | 725091 | 1 | + | 54 | 0 | 54 | 45 | 0 | 45 | 0 | 0 | NW_011924881.1 | 724448 | 731567 | 7120 | - | Gnomon | gene | gene34 | GeneID:105278416 | LOC105278416 | Gene | LOC105278416 | protein_coding | character(0) | character(0) | BCphaseB2.vs.RphaseA1 | FALSE |
| NW_011924881.1 | 725099 | 725099 | 1 | + | 76 | 1 | 75 | 41 | 0 | 41 | 1.32 | 0 | NW_011924881.1 | 724448 | 731567 | 7120 | - | Gnomon | gene | gene34 | GeneID:105278416 | LOC105278416 | Gene | LOC105278416 | protein_coding | character(0) | character(0) | BCphaseB2.vs.RphaseA1 | FALSE |
| NW_011924881.1 | 725122 | 725122 | 1 | + | 77 | 0 | 77 | 39 | 0 | 39 | 0 | 0 | NW_011924881.1 | 724448 | 731567 | 7120 | - | Gnomon | gene | gene34 | GeneID:105278416 | LOC105278416 | Gene | LOC105278416 | protein_coding | character(0) | character(0) | BCphaseB2.vs.RphaseA1 | FALSE |
| NW_011924881.1 | 725126 | 725126 | 1 | + | 75 | 0 | 75 | 40 | 0 | 40 | 0 | 0 | NW_011924881.1 | 724448 | 731567 | 7120 | - | Gnomon | gene | gene34 | GeneID:105278416 | LOC105278416 | Gene | LOC105278416 | protein_coding | character(0) | character(0) | BCphaseB2.vs.RphaseA1 | FALSE |

...
... only first 25 lines shown ...

Directory: DMSG File: dmg-Cb_LI_BCphaseB2.vs.RphaseA2_details.txt

| seqnames | start | end | width | strand | coverage1 | numCs1 | numTs1 | coverage2 | numCs2 | numTs2 | BCphaseB2 | RphaseA2 | gene_seqnames | gene_start | gene_end | gene_width | gene_strand | gene_source | gene_type | gene_ID | gene_Dbxref | gene_Name | gene_gbkey | gene_gene | gene_gene_biotype | gene_start_range | gene_end_range | gene_comparison | is.dm |
|---|---|---|---|---|---|---|---|---|---|---|---|---|---|---|---|---|---|---|---|---|---|---|---|---|---|---|---|---|---|
| NW_011924881.1 | 724785 | 724785 | 1 | + | 34 | 0 | 34 | 65 | 0 | 65 | 0 | 0 | NW_011924881.1 | 724448 | 731567 | 7120 | - | Gnomon | gene | gene34 | GeneID:105278416 | LOC105278416 | Gene | LOC105278416 | protein_coding | character(0) | character(0) | BCphaseB2.vs.RphaseA2 | FALSE |
| NW_011924881.1 | 724788 | 724788 | 1 | + | 31 | 0 | 31 | 63 | 0 | 63 | 0 | 0 | NW_011924881.1 | 724448 | 731567 | 7120 | - | Gnomon | gene | gene34 | GeneID:105278416 | LOC105278416 | Gene | LOC105278416 | protein_coding | character(0) | character(0) | BCphaseB2.vs.RphaseA2 | FALSE |
| NW_011924881.1 | 724794 | 724794 | 1 | + | 31 | 0 | 31 | 63 | 0 | 63 | 0 | 0 | NW_011924881.1 | 724448 | 731567 | 7120 | - | Gnomon | gene | gene34 | GeneID:105278416 | LOC105278416 | Gene | LOC105278416 | protein_coding | character(0) | character(0) | BCphaseB2.vs.RphaseA2 | FALSE |
| NW_011924881.1 | 724802 | 724802 | 1 | + | 30 | 0 | 30 | 61 | 0 | 61 | 0 | 0 | NW_011924881.1 | 724448 | 731567 | 7120 | - | Gnomon | gene | gene34 | GeneID:105278416 | LOC105278416 | Gene | LOC105278416 | protein_coding | character(0) | character(0) | BCphaseB2.vs.RphaseA2 | FALSE |
| NW_011924881.1 | 724806 | 724806 | 1 | + | 30 | 0 | 30 | 61 | 1 | 60 | 0 | 1.64 | NW_011924881.1 | 724448 | 731567 | 7120 | - | Gnomon | gene | gene34 | GeneID:105278416 | LOC105278416 | Gene | LOC105278416 | protein_coding | character(0) | character(0) | BCphaseB2.vs.RphaseA2 | FALSE |
| NW_011924881.1 | 724815 | 724815 | 1 | + | 30 | 0 | 30 | 61 | 0 | 61 | 0 | 0 | NW_011924881.1 | 724448 | 731567 | 7120 | - | Gnomon | gene | gene34 | GeneID:105278416 | LOC105278416 | Gene | LOC105278416 | protein_coding | character(0) | character(0) | BCphaseB2.vs.RphaseA2 | FALSE |
| NW_011924881.1 | 724818 | 724818 | 1 | + | 30 | 0 | 30 | 62 | 0 | 62 | 0 | 0 | NW_011924881.1 | 724448 | 731567 | 7120 | - | Gnomon | gene | gene34 | GeneID:105278416 | LOC105278416 | Gene | LOC105278416 | protein_coding | character(0) | character(0) | BCphaseB2.vs.RphaseA2 | FALSE |
| NW_011924881.1 | 724839 | 724839 | 1 | + | 23 | 0 | 23 | 73 | 1 | 72 | 0 | 1.37 | NW_011924881.1 | 724448 | 731567 | 7120 | - | Gnomon | gene | gene34 | GeneID:105278416 | LOC105278416 | Gene | LOC105278416 | protein_coding | character(0) | character(0) | BCphaseB2.vs.RphaseA2 | FALSE |
| NW_011924881.1 | 724926 | 724926 | 1 | + | 37 | 0 | 37 | 62 | 0 | 62 | 0 | 0 | NW_011924881.1 | 724448 | 731567 | 7120 | - | Gnomon | gene | gene34 | GeneID:105278416 | LOC105278416 | Gene | LOC105278416 | protein_coding | character(0) | character(0) | BCphaseB2.vs.RphaseA2 | FALSE |
| NW_011924881.1 | 724934 | 724934 | 1 | + | 34 | 0 | 34 | 51 | 0 | 51 | 0 | 0 | NW_011924881.1 | 724448 | 731567 | 7120 | - | Gnomon | gene | gene34 | GeneID:105278416 | LOC105278416 | Gene | LOC105278416 | protein_coding | character(0) | character(0) | BCphaseB2.vs.RphaseA2 | FALSE |
| NW_011924881.1 | 724936 | 724936 | 1 | + | 35 | 0 | 35 | 51 | 0 | 51 | 0 | 0 | NW_011924881.1 | 724448 | 731567 | 7120 | - | Gnomon | gene | gene34 | GeneID:105278416 | LOC105278416 | Gene | LOC105278416 | protein_coding | character(0) | character(0) | BCphaseB2.vs.RphaseA2 | FALSE |
| NW_011924881.1 | 724943 | 724943 | 1 | + | 33 | 0 | 33 | 70 | 1 | 69 | 0 | 1.43 | NW_011924881.1 | 724448 | 731567 | 7120 | - | Gnomon | gene | gene34 | GeneID:105278416 | LOC105278416 | Gene | LOC105278416 | protein_coding | character(0) | character(0) | BCphaseB2.vs.RphaseA2 | FALSE |
| NW_011924881.1 | 724947 | 724947 | 1 | + | 36 | 0 | 36 | 72 | 0 | 72 | 0 | 0 | NW_011924881.1 | 724448 | 731567 | 7120 | - | Gnomon | gene | gene34 | GeneID:105278416 | LOC105278416 | Gene | LOC105278416 | protein_coding | character(0) | character(0) | BCphaseB2.vs.RphaseA2 | FALSE |
| NW_011924881.1 | 724961 | 724961 | 1 | + | 43 | 0 | 43 | 65 | 0 | 65 | 0 | 0 | NW_011924881.1 | 724448 | 731567 | 7120 | - | Gnomon | gene | gene34 | GeneID:105278416 | LOC105278416 | Gene | LOC105278416 | protein_coding | character(0) | character(0) | BCphaseB2.vs.RphaseA2 | FALSE |
| NW_011924881.1 | 724976 | 724976 | 1 | + | 52 | 0 | 52 | 75 | 0 | 75 | 0 | 0 | NW_011924881.1 | 724448 | 731567 | 7120 | - | Gnomon | gene | gene34 | GeneID:105278416 | LOC105278416 | Gene | LOC105278416 | protein_coding | character(0) | character(0) | BCphaseB2.vs.RphaseA2 | FALSE |
| NW_011924881.1 | 724982 | 724982 | 1 | + | 54 | 0 | 54 | 79 | 0 | 79 | 0 | 0 | NW_011924881.1 | 724448 | 731567 | 7120 | - | Gnomon | gene | gene34 | GeneID:105278416 | LOC105278416 | Gene | LOC105278416 | protein_coding | character(0) | character(0) | BCphaseB2.vs.RphaseA2 | FALSE |
| NW_011924881.1 | 724990 | 724990 | 1 | + | 54 | 0 | 54 | 82 | 0 | 82 | 0 | 0 | NW_011924881.1 | 724448 | 731567 | 7120 | - | Gnomon | gene | gene34 | GeneID:105278416 | LOC105278416 | Gene | LOC105278416 | protein_coding | character(0) | character(0) | BCphaseB2.vs.RphaseA2 | FALSE |
| NW_011924881.1 | 724994 | 724994 | 1 | + | 53 | 0 | 53 | 81 | 0 | 81 | 0 | 0 | NW_011924881.1 | 724448 | 731567 | 7120 | - | Gnomon | gene | gene34 | GeneID:105278416 | LOC105278416 | Gene | LOC105278416 | protein_coding | character(0) | character(0) | BCphaseB2.vs.RphaseA2 | FALSE |
| NW_011924881.1 | 725008 | 725008 | 1 | + | 43 | 0 | 43 | 53 | 0 | 53 | 0 | 0 | NW_011924881.1 | 724448 | 731567 | 7120 | - | Gnomon | gene | gene34 | GeneID:105278416 | LOC105278416 | Gene | LOC105278416 | protein_coding | character(0) | character(0) | BCphaseB2.vs.RphaseA2 | FALSE |
| NW_011924881.1 | 725014 | 725014 | 1 | + | 35 | 0 | 35 | 63 | 1 | 62 | 0 | 1.59 | NW_011924881.1 | 724448 | 731567 | 7120 | - | Gnomon | gene | gene34 | GeneID:105278416 | LOC105278416 | Gene | LOC105278416 | protein_coding | character(0) | character(0) | BCphaseB2.vs.RphaseA2 | FALSE |
| NW_011924881.1 | 725022 | 725022 | 1 | + | 24 | 0 | 24 | 48 | 2 | 46 | 0 | 4.17 | NW_011924881.1 | 724448 | 731567 | 7120 | - | Gnomon | gene | gene34 | GeneID:105278416 | LOC105278416 | Gene | LOC105278416 | protein_coding | character(0) | character(0) | BCphaseB2.vs.RphaseA2 | FALSE |
| NW_011924881.1 | 725041 | 725041 | 1 | + | 33 | 0 | 33 | 41 | 0 | 41 | 0 | 0 | NW_011924881.1 | 724448 | 731567 | 7120 | - | Gnomon | gene | gene34 | GeneID:105278416 | LOC105278416 | Gene | LOC105278416 | protein_coding | character(0) | character(0) | BCphaseB2.vs.RphaseA2 | FALSE |
| NW_011924881.1 | 725048 | 725048 | 1 | + | 38 | 0 | 38 | 51 | 0 | 51 | 0 | 0 | NW_011924881.1 | 724448 | 731567 | 7120 | - | Gnomon | gene | gene34 | GeneID:105278416 | LOC105278416 | Gene | LOC105278416 | protein_coding | character(0) | character(0) | BCphaseB2.vs.RphaseA2 | FALSE |
| NW_011924881.1 | 725062 | 725062 | 1 | + | 68 | 0 | 68 | 80 | 0 | 80 | 0 | 0 | NW_011924881.1 | 724448 | 731567 | 7120 | - | Gnomon | gene | gene34 | GeneID:105278416 | LOC105278416 | Gene | LOC105278416 | protein_coding | character(0) | character(0) | BCphaseB2.vs.RphaseA2 | FALSE |

...
... only first 25 lines shown ...

Directory: DMSG File: dmg-Cb_LI_BCphaseB2.vs.RphaseB1_details.txt

| seqnames | start | end | width | strand | coverage1 | numCs1 | numTs1 | coverage2 | numCs2 | numTs2 | BCphaseB2 | RphaseB1 | gene_seqnames | gene_start | gene_end | gene_width | gene_strand | gene_source | gene_type | gene_ID | gene_Dbxref | gene_Name | gene_gbkey | gene_gene | gene_gene_biotype | gene_start_range | gene_end_range | gene_comparison | is.dm |
|---|---|---|---|---|---|---|---|---|---|---|---|---|---|---|---|---|---|---|---|---|---|---|---|---|---|---|---|---|---|
| NW_011924931.1 | 2028841 | 2028841 | 1 | + | 24 | 0 | 24 | 21 | 0 | 21 | 0 | 0 | NW_011924931.1 | 2028839 | 2031812 | 2974 | + | Gnomon | gene | gene1470 | GeneID:105287597 | LOC105287597 | Gene | LOC105287597 | protein_coding | character(0) | character(0) | BCphaseB2.vs.RphaseB1 | FALSE |
| NW_011924931.1 | 2028882 | 2028882 | 1 | + | 20 | 0 | 20 | 20 | 0 | 20 | 0 | 0 | NW_011924931.1 | 2028839 | 2031812 | 2974 | + | Gnomon | gene | gene1470 | GeneID:105287597 | LOC105287597 | Gene | LOC105287597 | protein_coding | character(0) | character(0) | BCphaseB2.vs.RphaseB1 | FALSE |
| NW_011924931.1 | 2028885 | 2028885 | 1 | + | 21 | 0 | 21 | 21 | 0 | 21 | 0 | 0 | NW_011924931.1 | 2028839 | 2031812 | 2974 | + | Gnomon | gene | gene1470 | GeneID:105287597 | LOC105287597 | Gene | LOC105287597 | protein_coding | character(0) | character(0) | BCphaseB2.vs.RphaseB1 | FALSE |
| NW_011924931.1 | 2028887 | 2028887 | 1 | + | 20 | 0 | 20 | 21 | 0 | 21 | 0 | 0 | NW_011924931.1 | 2028839 | 2031812 | 2974 | + | Gnomon | gene | gene1470 | GeneID:105287597 | LOC105287597 | Gene | LOC105287597 | protein_coding | character(0) | character(0) | BCphaseB2.vs.RphaseB1 | FALSE |
| NW_011924931.1 | 2028897 | 2028897 | 1 | + | 21 | 0 | 21 | 22 | 0 | 22 | 0 | 0 | NW_011924931.1 | 2028839 | 2031812 | 2974 | + | Gnomon | gene | gene1470 | GeneID:105287597 | LOC105287597 | Gene | LOC105287597 | protein_coding | character(0) | character(0) | BCphaseB2.vs.RphaseB1 | FALSE |
| NW_011924931.1 | 2028908 | 2028908 | 1 | + | 20 | 1 | 19 | 24 | 1 | 23 | 5 | 4.17 | NW_011924931.1 | 2028839 | 2031812 | 2974 | + | Gnomon | gene | gene1470 | GeneID:105287597 | LOC105287597 | Gene | LOC105287597 | protein_coding | character(0) | character(0) | BCphaseB2.vs.RphaseB1 | FALSE |
| NW_011924931.1 | 2028911 | 2028911 | 1 | + | 20 | 0 | 20 | 23 | 0 | 23 | 0 | 0 | NW_011924931.1 | 2028839 | 2031812 | 2974 | + | Gnomon | gene | gene1470 | GeneID:105287597 | LOC105287597 | Gene | LOC105287597 | protein_coding | character(0) | character(0) | BCphaseB2.vs.RphaseB1 | FALSE |
| NW_011924931.1 | 2028927 | 2028927 | 1 | + | 33 | 0 | 33 | 28 | 0 | 28 | 0 | 0 | NW_011924931.1 | 2028839 | 2031812 | 2974 | + | Gnomon | gene | gene1470 | GeneID:105287597 | LOC105287597 | Gene | LOC105287597 | protein_coding | character(0) | character(0) | BCphaseB2.vs.RphaseB1 | FALSE |
| NW_011924931.1 | 2028930 | 2028930 | 1 | + | 51 | 0 | 51 | 28 | 0 | 28 | 0 | 0 | NW_011924931.1 | 2028839 | 2031812 | 2974 | + | Gnomon | gene | gene1470 | GeneID:105287597 | LOC105287597 | Gene | LOC105287597 | protein_coding | character(0) | character(0) | BCphaseB2.vs.RphaseB1 | FALSE |
| NW_011924931.1 | 2028932 | 2028932 | 1 | + | 32 | 0 | 32 | 28 | 0 | 28 | 0 | 0 | NW_011924931.1 | 2028839 | 2031812 | 2974 | + | Gnomon | gene | gene1470 | GeneID:105287597 | LOC105287597 | Gene | LOC105287597 | protein_coding | character(0) | character(0) | BCphaseB2.vs.RphaseB1 | FALSE |
| NW_011924931.1 | 2028934 | 2028934 | 1 | + | 32 | 0 | 32 | 28 | 0 | 28 | 0 | 0 | NW_011924931.1 | 2028839 | 2031812 | 2974 | + | Gnomon | gene | gene1470 | GeneID:105287597 | LOC105287597 | Gene | LOC105287597 | protein_coding | character(0) | character(0) | BCphaseB2.vs.RphaseB1 | FALSE |
| NW_011924931.1 | 2028940 | 2028940 | 1 | + | 32 | 0 | 32 | 28 | 0 | 28 | 0 | 0 | NW_011924931.1 | 2028839 | 2031812 | 2974 | + | Gnomon | gene | gene1470 | GeneID:105287597 | LOC105287597 | Gene | LOC105287597 | protein_coding | character(0) | character(0) | BCphaseB2.vs.RphaseB1 | FALSE |
| NW_011924931.1 | 2028946 | 2028946 | 1 | + | 30 | 0 | 30 | 27 | 0 | 27 | 0 | 0 | NW_011924931.1 | 2028839 | 2031812 | 2974 | + | Gnomon | gene | gene1470 | GeneID:105287597 | LOC105287597 | Gene | LOC105287597 | protein_coding | character(0) | character(0) | BCphaseB2.vs.RphaseB1 | FALSE |
| NW_011924931.1 | 2028951 | 2028951 | 1 | + | 30 | 0 | 30 | 27 | 0 | 27 | 0 | 0 | NW_011924931.1 | 2028839 | 2031812 | 2974 | + | Gnomon | gene | gene1470 | GeneID:105287597 | LOC105287597 | Gene | LOC105287597 | protein_coding | character(0) | character(0) | BCphaseB2.vs.RphaseB1 | FALSE |
| NW_011924931.1 | 2028957 | 2028957 | 1 | + | 30 | 0 | 30 | 29 | 0 | 29 | 0 | 0 | NW_011924931.1 | 2028839 | 2031812 | 2974 | + | Gnomon | gene | gene1470 | GeneID:105287597 | LOC105287597 | Gene | LOC105287597 | protein_coding | character(0) | character(0) | BCphaseB2.vs.RphaseB1 | FALSE |
| NW_011924931.1 | 2028964 | 2028964 | 1 | + | 28 | 0 | 28 | 51 | 1 | 50 | 0 | 1.96 | NW_011924931.1 | 2028839 | 2031812 | 2974 | + | Gnomon | gene | gene1470 | GeneID:105287597 | LOC105287597 | Gene | LOC105287597 | protein_coding | character(0) | character(0) | BCphaseB2.vs.RphaseB1 | FALSE |
| NW_011924931.1 | 2028969 | 2028969 | 1 | + | 25 | 0 | 25 | 49 | 0 | 49 | 0 | 0 | NW_011924931.1 | 2028839 | 2031812 | 2974 | + | Gnomon | gene | gene1470 | GeneID:105287597 | LOC105287597 | Gene | LOC105287597 | protein_coding | character(0) | character(0) | BCphaseB2.vs.RphaseB1 | FALSE |
| NW_011924931.1 | 2028973 | 2028973 | 1 | + | 28 | 0 | 28 | 47 | 0 | 47 | 0 | 0 | NW_011924931.1 | 2028839 | 2031812 | 2974 | + | Gnomon | gene | gene1470 | GeneID:105287597 | LOC105287597 | Gene | LOC105287597 | protein_coding | character(0) | character(0) | BCphaseB2.vs.RphaseB1 | FALSE |
| NW_011924931.1 | 2028977 | 2028977 | 1 | + | 28 | 0 | 28 | 22 | 0 | 22 | 0 | 0 | NW_011924931.1 | 2028839 | 2031812 | 2974 | + | Gnomon | gene | gene1470 | GeneID:105287597 | LOC105287597 | Gene | LOC105287597 | protein_coding | character(0) | character(0) | BCphaseB2.vs.RphaseB1 | FALSE |
| NW_011924931.1 | 2028979 | 2028979 | 1 | + | 29 | 0 | 29 | 22 | 0 | 22 | 0 | 0 | NW_011924931.1 | 2028839 | 2031812 | 2974 | + | Gnomon | gene | gene1470 | GeneID:105287597 | LOC105287597 | Gene | LOC105287597 | protein_coding | character(0) | character(0) | BCphaseB2.vs.RphaseB1 | FALSE |
| NW_011924931.1 | 2028982 | 2028982 | 1 | + | 37 | 0 | 37 | 23 | 0 | 23 | 0 | 0 | NW_011924931.1 | 2028839 | 2031812 | 2974 | + | Gnomon | gene | gene1470 | GeneID:105287597 | LOC105287597 | Gene | LOC105287597 | protein_coding | character(0) | character(0) | BCphaseB2.vs.RphaseB1 | FALSE |
| NW_011924931.1 | 2028984 | 2028984 | 1 | + | 39 | 0 | 39 | 23 | 1 | 22 | 0 | 4.35 | NW_011924931.1 | 2028839 | 2031812 | 2974 | + | Gnomon | gene | gene1470 | GeneID:105287597 | LOC105287597 | Gene | LOC105287597 | protein_coding | character(0) | character(0) | BCphaseB2.vs.RphaseB1 | FALSE |
| NW_011924931.1 | 2028987 | 2028987 | 1 | + | 58 | 0 | 58 | 23 | 0 | 23 | 0 | 0 | NW_011924931.1 | 2028839 | 2031812 | 2974 | + | Gnomon | gene | gene1470 | GeneID:105287597 | LOC105287597 | Gene | LOC105287597 | protein_coding | character(0) | character(0) | BCphaseB2.vs.RphaseB1 | FALSE |
| NW_011924931.1 | 2028989 | 2028989 | 1 | + | 63 | 0 | 63 | 24 | 0 | 24 | 0 | 0 | NW_011924931.1 | 2028839 | 2031812 | 2974 | + | Gnomon | gene | gene1470 | GeneID:105287597 | LOC105287597 | Gene | LOC105287597 | protein_coding | character(0) | character(0) | BCphaseB2.vs.RphaseB1 | FALSE |

...
... only first 25 lines shown ...

Directory: DMSG File: dmg-Cb_LI_BCphaseB2.vs.RphaseB2_details.txt

| seqnames | start | end | width | strand | coverage1 | numCs1 | numTs1 | coverage2 | numCs2 | numTs2 | BCphaseB2 | RphaseB2 | gene_seqnames | gene_start | gene_end | gene_width | gene_strand | gene_source | gene_type | gene_ID | gene_Dbxref | gene_Name | gene_gbkey | gene_gene | gene_gene_biotype | gene_partial | gene_start_range | gene_end_range | gene_comparison | is.dm |
|---|---|---|---|---|---|---|---|---|---|---|---|---|---|---|---|---|---|---|---|---|---|---|---|---|---|---|---|---|---|---|
| NW_011924881.1 | 43846 | 43846 | 1 | + | 25 | 0 | 25 | 24 | 0 | 24 | 0 | 0 | NW_011924881.1 | 42715 | 135016 | 92302 | - | Gnomon | gene | gene18 | GeneID:105287290 | LOC105287290 | Gene | LOC105287290 | protein_coding | NA | character(0) | character(0) | BCphaseB2.vs.RphaseB2 | FALSE |
| NW_011924881.1 | 43853 | 43853 | 1 | + | 24 | 0 | 24 | 22 | 0 | 22 | 0 | 0 | NW_011924881.1 | 42715 | 135016 | 92302 | - | Gnomon | gene | gene18 | GeneID:105287290 | LOC105287290 | Gene | LOC105287290 | protein_coding | NA | character(0) | character(0) | BCphaseB2.vs.RphaseB2 | FALSE |
| NW_011924881.1 | 43864 | 43864 | 1 | + | 22 | 0 | 22 | 22 | 0 | 22 | 0 | 0 | NW_011924881.1 | 42715 | 135016 | 92302 | - | Gnomon | gene | gene18 | GeneID:105287290 | LOC105287290 | Gene | LOC105287290 | protein_coding | NA | character(0) | character(0) | BCphaseB2.vs.RphaseB2 | FALSE |
| NW_011924881.1 | 43879 | 43879 | 1 | + | 22 | 0 | 22 | 21 | 0 | 21 | 0 | 0 | NW_011924881.1 | 42715 | 135016 | 92302 | - | Gnomon | gene | gene18 | GeneID:105287290 | LOC105287290 | Gene | LOC105287290 | protein_coding | NA | character(0) | character(0) | BCphaseB2.vs.RphaseB2 | FALSE |
| NW_011924881.1 | 44105 | 44105 | 1 | + | 45 | 0 | 45 | 59 | 0 | 59 | 0 | 0 | NW_011924881.1 | 42715 | 135016 | 92302 | - | Gnomon | gene | gene18 | GeneID:105287290 | LOC105287290 | Gene | LOC105287290 | protein_coding | NA | character(0) | character(0) | BCphaseB2.vs.RphaseB2 | FALSE |
| NW_011924881.1 | 44112 | 44112 | 1 | + | 42 | 0 | 42 | 57 | 0 | 57 | 0 | 0 | NW_011924881.1 | 42715 | 135016 | 92302 | - | Gnomon | gene | gene18 | GeneID:105287290 | LOC105287290 | Gene | LOC105287290 | protein_coding | NA | character(0) | character(0) | BCphaseB2.vs.RphaseB2 | FALSE |
| NW_011924881.1 | 44121 | 44121 | 1 | + | 41 | 0 | 41 | 56 | 0 | 56 | 0 | 0 | NW_011924881.1 | 42715 | 135016 | 92302 | - | Gnomon | gene | gene18 | GeneID:105287290 | LOC105287290 | Gene | LOC105287290 | protein_coding | NA | character(0) | character(0) | BCphaseB2.vs.RphaseB2 | FALSE |
| NW_011924881.1 | 44123 | 44123 | 1 | + | 40 | 0 | 40 | 56 | 0 | 56 | 0 | 0 | NW_011924881.1 | 42715 | 135016 | 92302 | - | Gnomon | gene | gene18 | GeneID:105287290 | LOC105287290 | Gene | LOC105287290 | protein_coding | NA | character(0) | character(0) | BCphaseB2.vs.RphaseB2 | FALSE |
| NW_011924881.1 | 44125 | 44125 | 1 | + | 41 | 0 | 41 | 55 | 0 | 55 | 0 | 0 | NW_011924881.1 | 42715 | 135016 | 92302 | - | Gnomon | gene | gene18 | GeneID:105287290 | LOC105287290 | Gene | LOC105287290 | protein_coding | NA | character(0) | character(0) | BCphaseB2.vs.RphaseB2 | FALSE |
| NW_011924881.1 | 44129 | 44129 | 1 | + | 38 | 0 | 38 | 54 | 0 | 54 | 0 | 0 | NW_011924881.1 | 42715 | 135016 | 92302 | - | Gnomon | gene | gene18 | GeneID:105287290 | LOC105287290 | Gene | LOC105287290 | protein_coding | NA | character(0) | character(0) | BCphaseB2.vs.RphaseB2 | FALSE |
| NW_011924881.1 | 44138 | 44138 | 1 | + | 35 | 0 | 35 | 52 | 0 | 52 | 0 | 0 | NW_011924881.1 | 42715 | 135016 | 92302 | - | Gnomon | gene | gene18 | GeneID:105287290 | LOC105287290 | Gene | LOC105287290 | protein_coding | NA | character(0) | character(0) | BCphaseB2.vs.RphaseB2 | FALSE |
| NW_011924881.1 | 44152 | 44152 | 1 | + | 36 | 0 | 36 | 52 | 0 | 52 | 0 | 0 | NW_011924881.1 | 42715 | 135016 | 92302 | - | Gnomon | gene | gene18 | GeneID:105287290 | LOC105287290 | Gene | LOC105287290 | protein_coding | NA | character(0) | character(0) | BCphaseB2.vs.RphaseB2 | FALSE |
| NW_011924881.1 | 44173 | 44173 | 1 | + | 42 | 0 | 42 | 51 | 1 | 50 | 0 | 1.96 | NW_011924881.1 | 42715 | 135016 | 92302 | - | Gnomon | gene | gene18 | GeneID:105287290 | LOC105287290 | Gene | LOC105287290 | protein_coding | NA | character(0) | character(0) | BCphaseB2.vs.RphaseB2 | FALSE |
| NW_011924881.1 | 44185 | 44185 | 1 | + | 34 | 0 | 34 | 37 | 0 | 37 | 0 | 0 | NW_011924881.1 | 42715 | 135016 | 92302 | - | Gnomon | gene | gene18 | GeneID:105287290 | LOC105287290 | Gene | LOC105287290 | protein_coding | NA | character(0) | character(0) | BCphaseB2.vs.RphaseB2 | FALSE |
| NW_011924881.1 | 44193 | 44193 | 1 | + | 33 | 0 | 33 | 29 | 0 | 29 | 0 | 0 | NW_011924881.1 | 42715 | 135016 | 92302 | - | Gnomon | gene | gene18 | GeneID:105287290 | LOC105287290 | Gene | LOC105287290 | protein_coding | NA | character(0) | character(0) | BCphaseB2.vs.RphaseB2 | FALSE |
| NW_011924881.1 | 44199 | 44199 | 1 | + | 35 | 0 | 35 | 31 | 0 | 31 | 0 | 0 | NW_011924881.1 | 42715 | 135016 | 92302 | - | Gnomon | gene | gene18 | GeneID:105287290 | LOC105287290 | Gene | LOC105287290 | protein_coding | NA | character(0) | character(0) | BCphaseB2.vs.RphaseB2 | FALSE |
| NW_011924881.1 | 44201 | 44201 | 1 | + | 35 | 0 | 35 | 31 | 0 | 31 | 0 | 0 | NW_011924881.1 | 42715 | 135016 | 92302 | - | Gnomon | gene | gene18 | GeneID:105287290 | LOC105287290 | Gene | LOC105287290 | protein_coding | NA | character(0) | character(0) | BCphaseB2.vs.RphaseB2 | FALSE |
| NW_011924881.1 | 44204 | 44204 | 1 | + | 36 | 1 | 35 | 30 | 0 | 30 | 2.78 | 0 | NW_011924881.1 | 42715 | 135016 | 92302 | - | Gnomon | gene | gene18 | GeneID:105287290 | LOC105287290 | Gene | LOC105287290 | protein_coding | NA | character(0) | character(0) | BCphaseB2.vs.RphaseB2 | FALSE |
| NW_011924881.1 | 44221 | 44221 | 1 | + | 34 | 0 | 34 | 26 | 0 | 26 | 0 | 0 | NW_011924881.1 | 42715 | 135016 | 92302 | - | Gnomon | gene | gene18 | GeneID:105287290 | LOC105287290 | Gene | LOC105287290 | protein_coding | NA | character(0) | character(0) | BCphaseB2.vs.RphaseB2 | FALSE |
| NW_011924881.1 | 44223 | 44223 | 1 | + | 32 | 0 | 32 | 24 | 0 | 24 | 0 | 0 | NW_011924881.1 | 42715 | 135016 | 92302 | - | Gnomon | gene | gene18 | GeneID:105287290 | LOC105287290 | Gene | LOC105287290 | protein_coding | NA | character(0) | character(0) | BCphaseB2.vs.RphaseB2 | FALSE |
| NW_011924881.1 | 44252 | 44252 | 1 | + | 24 | 0 | 24 | 26 | 0 | 26 | 0 | 0 | NW_011924881.1 | 42715 | 135016 | 92302 | - | Gnomon | gene | gene18 | GeneID:105287290 | LOC105287290 | Gene | LOC105287290 | protein_coding | NA | character(0) | character(0) | BCphaseB2.vs.RphaseB2 | FALSE |
| NW_011924881.1 | 44261 | 44261 | 1 | + | 20 | 0 | 20 | 24 | 1 | 23 | 0 | 4.17 | NW_011924881.1 | 42715 | 135016 | 92302 | - | Gnomon | gene | gene18 | GeneID:105287290 | LOC105287290 | Gene | LOC105287290 | protein_coding | NA | character(0) | character(0) | BCphaseB2.vs.RphaseB2 | FALSE |
| NW_011924881.1 | 44276 | 44276 | 1 | + | 27 | 0 | 27 | 29 | 0 | 29 | 0 | 0 | NW_011924881.1 | 42715 | 135016 | 92302 | - | Gnomon | gene | gene18 | GeneID:105287290 | LOC105287290 | Gene | LOC105287290 | protein_coding | NA | character(0) | character(0) | BCphaseB2.vs.RphaseB2 | FALSE |
| NW_011924881.1 | 44282 | 44282 | 1 | + | 25 | 0 | 25 | 26 | 0 | 26 | 0 | 0 | NW_011924881.1 | 42715 | 135016 | 92302 | - | Gnomon | gene | gene18 | GeneID:105287290 | LOC105287290 | Gene | LOC105287290 | protein_coding | NA | character(0) | character(0) | BCphaseB2.vs.RphaseB2 | FALSE |

...
... only first 25 lines shown ...

Directory: DMSG File: dmg-Cb_LI_RphaseA1.vs.RphaseA2_details.txt

| seqnames | start | end | width | strand | coverage1 | numCs1 | numTs1 | coverage2 | numCs2 | numTs2 | RphaseA1 | RphaseA2 | gene_seqnames | gene_start | gene_end | gene_width | gene_strand | gene_source | gene_type | gene_ID | gene_Dbxref | gene_Name | gene_gbkey | gene_gene | gene_gene_biotype | gene_partial | gene_start_range | gene_end_range | gene_comparison | is.dm |
|---|---|---|---|---|---|---|---|---|---|---|---|---|---|---|---|---|---|---|---|---|---|---|---|---|---|---|---|---|---|---|
| NW_011924881.1 | 724688 | 724688 | 1 | + | 23 | 0 | 23 | 23 | 0 | 23 | 0 | 0 | NW_011924881.1 | 724448 | 731567 | 7120 | - | Gnomon | gene | gene34 | GeneID:105278416 | LOC105278416 | Gene | LOC105278416 | protein_coding | NA | character(0) | character(0) | RphaseA1.vs.RphaseA2 | FALSE |
| NW_011924881.1 | 724692 | 724692 | 1 | + | 22 | 0 | 22 | 22 | 0 | 22 | 0 | 0 | NW_011924881.1 | 724448 | 731567 | 7120 | - | Gnomon | gene | gene34 | GeneID:105278416 | LOC105278416 | Gene | LOC105278416 | protein_coding | NA | character(0) | character(0) | RphaseA1.vs.RphaseA2 | FALSE |
| NW_011924881.1 | 724695 | 724695 | 1 | + | 22 | 0 | 22 | 22 | 0 | 22 | 0 | 0 | NW_011924881.1 | 724448 | 731567 | 7120 | - | Gnomon | gene | gene34 | GeneID:105278416 | LOC105278416 | Gene | LOC105278416 | protein_coding | NA | character(0) | character(0) | RphaseA1.vs.RphaseA2 | FALSE |
| NW_011924881.1 | 724788 | 724788 | 1 | + | 21 | 0 | 21 | 63 | 0 | 63 | 0 | 0 | NW_011924881.1 | 724448 | 731567 | 7120 | - | Gnomon | gene | gene34 | GeneID:105278416 | LOC105278416 | Gene | LOC105278416 | protein_coding | NA | character(0) | character(0) | RphaseA1.vs.RphaseA2 | FALSE |
| NW_011924881.1 | 724872 | 724872 | 1 | + | 32 | 0 | 32 | 62 | 0 | 62 | 0 | 0 | NW_011924881.1 | 724448 | 731567 | 7120 | - | Gnomon | gene | gene34 | GeneID:105278416 | LOC105278416 | Gene | LOC105278416 | protein_coding | NA | character(0) | character(0) | RphaseA1.vs.RphaseA2 | FALSE |
| NW_011924881.1 | 724889 | 724889 | 1 | + | 32 | 0 | 32 | 44 | 0 | 44 | 0 | 0 | NW_011924881.1 | 724448 | 731567 | 7120 | - | Gnomon | gene | gene34 | GeneID:105278416 | LOC105278416 | Gene | LOC105278416 | protein_coding | NA | character(0) | character(0) | RphaseA1.vs.RphaseA2 | FALSE |
| NW_011924881.1 | 724896 | 724896 | 1 | + | 32 | 0 | 32 | 42 | 0 | 42 | 0 | 0 | NW_011924881.1 | 724448 | 731567 | 7120 | - | Gnomon | gene | gene34 | GeneID:105278416 | LOC105278416 | Gene | LOC105278416 | protein_coding | NA | character(0) | character(0) | RphaseA1.vs.RphaseA2 | FALSE |
| NW_011924881.1 | 724898 | 724898 | 1 | + | 33 | 0 | 33 | 64 | 1 | 63 | 0 | 1.56 | NW_011924881.1 | 724448 | 731567 | 7120 | - | Gnomon | gene | gene34 | GeneID:105278416 | LOC105278416 | Gene | LOC105278416 | protein_coding | NA | character(0) | character(0) | RphaseA1.vs.RphaseA2 | FALSE |
| NW_011924881.1 | 724926 | 724926 | 1 | + | 30 | 0 | 30 | 62 | 0 | 62 | 0 | 0 | NW_011924881.1 | 724448 | 731567 | 7120 | - | Gnomon | gene | gene34 | GeneID:105278416 | LOC105278416 | Gene | LOC105278416 | protein_coding | NA | character(0) | character(0) | RphaseA1.vs.RphaseA2 | FALSE |
| NW_011924881.1 | 724934 | 724934 | 1 | + | 26 | 0 | 26 | 51 | 0 | 51 | 0 | 0 | NW_011924881.1 | 724448 | 731567 | 7120 | - | Gnomon | gene | gene34 | GeneID:105278416 | LOC105278416 | Gene | LOC105278416 | protein_coding | NA | character(0) | character(0) | RphaseA1.vs.RphaseA2 | FALSE |
| NW_011924881.1 | 724936 | 724936 | 1 | + | 27 | 0 | 27 | 51 | 0 | 51 | 0 | 0 | NW_011924881.1 | 724448 | 731567 | 7120 | - | Gnomon | gene | gene34 | GeneID:105278416 | LOC105278416 | Gene | LOC105278416 | protein_coding | NA | character(0) | character(0) | RphaseA1.vs.RphaseA2 | FALSE |
| NW_011924881.1 | 724943 | 724943 | 1 | + | 24 | 0 | 24 | 70 | 1 | 69 | 0 | 1.43 | NW_011924881.1 | 724448 | 731567 | 7120 | - | Gnomon | gene | gene34 | GeneID:105278416 | LOC105278416 | Gene | LOC105278416 | protein_coding | NA | character(0) | character(0) | RphaseA1.vs.RphaseA2 | FALSE |
| NW_011924881.1 | 724947 | 724947 | 1 | + | 27 | 0 | 27 | 72 | 0 | 72 | 0 | 0 | NW_011924881.1 | 724448 | 731567 | 7120 | - | Gnomon | gene | gene34 | GeneID:105278416 | LOC105278416 | Gene | LOC105278416 | protein_coding | NA | character(0) | character(0) | RphaseA1.vs.RphaseA2 | FALSE |
| NW_011924881.1 | 724961 | 724961 | 1 | + | 34 | 0 | 34 | 65 | 0 | 65 | 0 | 0 | NW_011924881.1 | 724448 | 731567 | 7120 | - | Gnomon | gene | gene34 | GeneID:105278416 | LOC105278416 | Gene | LOC105278416 | protein_coding | NA | character(0) | character(0) | RphaseA1.vs.RphaseA2 | FALSE |
| NW_011924881.1 | 724976 | 724976 | 1 | + | 41 | 0 | 41 | 75 | 0 | 75 | 0 | 0 | NW_011924881.1 | 724448 | 731567 | 7120 | - | Gnomon | gene | gene34 | GeneID:105278416 | LOC105278416 | Gene | LOC105278416 | protein_coding | NA | character(0) | character(0) | RphaseA1.vs.RphaseA2 | FALSE |
| NW_011924881.1 | 724982 | 724982 | 1 | + | 41 | 2 | 39 | 79 | 0 | 79 | 4.88 | 0 | NW_011924881.1 | 724448 | 731567 | 7120 | - | Gnomon | gene | gene34 | GeneID:105278416 | LOC105278416 | Gene | LOC105278416 | protein_coding | NA | character(0) | character(0) | RphaseA1.vs.RphaseA2 | FALSE |
| NW_011924881.1 | 724990 | 724990 | 1 | + | 39 | 0 | 39 | 82 | 0 | 82 | 0 | 0 | NW_011924881.1 | 724448 | 731567 | 7120 | - | Gnomon | gene | gene34 | GeneID:105278416 | LOC105278416 | Gene | LOC105278416 | protein_coding | NA | character(0) | character(0) | RphaseA1.vs.RphaseA2 | FALSE |
| NW_011924881.1 | 724994 | 724994 | 1 | + | 39 | 0 | 39 | 81 | 0 | 81 | 0 | 0 | NW_011924881.1 | 724448 | 731567 | 7120 | - | Gnomon | gene | gene34 | GeneID:105278416 | LOC105278416 | Gene | LOC105278416 | protein_coding | NA | character(0) | character(0) | RphaseA1.vs.RphaseA2 | FALSE |
| NW_011924881.1 | 725008 | 725008 | 1 | + | 31 | 1 | 30 | 53 | 0 | 53 | 3.23 | 0 | NW_011924881.1 | 724448 | 731567 | 7120 | - | Gnomon | gene | gene34 | GeneID:105278416 | LOC105278416 | Gene | LOC105278416 | protein_coding | NA | character(0) | character(0) | RphaseA1.vs.RphaseA2 | FALSE |
| NW_011924881.1 | 725014 | 725014 | 1 | + | 33 | 1 | 32 | 63 | 1 | 62 | 3.03 | 1.59 | NW_011924881.1 | 724448 | 731567 | 7120 | - | Gnomon | gene | gene34 | GeneID:105278416 | LOC105278416 | Gene | LOC105278416 | protein_coding | NA | character(0) | character(0) | RphaseA1.vs.RphaseA2 | FALSE |
| NW_011924881.1 | 725022 | 725022 | 1 | + | 37 | 1 | 36 | 48 | 2 | 46 | 2.7 | 4.17 | NW_011924881.1 | 724448 | 731567 | 7120 | - | Gnomon | gene | gene34 | GeneID:105278416 | LOC105278416 | Gene | LOC105278416 | protein_coding | NA | character(0) | character(0) | RphaseA1.vs.RphaseA2 | FALSE |
| NW_011924881.1 | 725041 | 725041 | 1 | + | 41 | 0 | 41 | 41 | 0 | 41 | 0 | 0 | NW_011924881.1 | 724448 | 731567 | 7120 | - | Gnomon | gene | gene34 | GeneID:105278416 | LOC105278416 | Gene | LOC105278416 | protein_coding | NA | character(0) | character(0) | RphaseA1.vs.RphaseA2 | FALSE |
| NW_011924881.1 | 725048 | 725048 | 1 | + | 43 | 0 | 43 | 51 | 0 | 51 | 0 | 0 | NW_011924881.1 | 724448 | 731567 | 7120 | - | Gnomon | gene | gene34 | GeneID:105278416 | LOC105278416 | Gene | LOC105278416 | protein_coding | NA | character(0) | character(0) | RphaseA1.vs.RphaseA2 | FALSE |
| NW_011924881.1 | 725062 | 725062 | 1 | + | 44 | 0 | 44 | 80 | 0 | 80 | 0 | 0 | NW_011924881.1 | 724448 | 731567 | 7120 | - | Gnomon | gene | gene34 | GeneID:105278416 | LOC105278416 | Gene | LOC105278416 | protein_coding | NA | character(0) | character(0) | RphaseA1.vs.RphaseA2 | FALSE |

...
... only first 25 lines shown ...

Directory: DMSG File: dmg-Cb_LI_RphaseA1.vs.RphaseB1_details.txt

| seqnames | start | end | width | strand | coverage1 | numCs1 | numTs1 | coverage2 | numCs2 | numTs2 | RphaseA1 | RphaseB1 | gene_seqnames | gene_start | gene_end | gene_width | gene_strand | gene_source | gene_type | gene_ID | gene_Dbxref | gene_Name | gene_gbkey | gene_gene | gene_gene_biotype | gene_start_range | gene_end_range | gene_comparison | is.dm |
|---|---|---|---|---|---|---|---|---|---|---|---|---|---|---|---|---|---|---|---|---|---|---|---|---|---|---|---|---|---|
| NW_011924881.1 | 809447 | 809447 | 1 | + | 22 | 0 | 22 | 23 | 0 | 23 | 0 | 0 | NW_011924881.1 | 808343 | 827625 | 19283 | - | Gnomon | gene | gene53 | GeneID:105277148 | LOC105277148 | Gene | LOC105277148 | protein_coding | character(0) | character(0) | RphaseA1.vs.RphaseB1 | FALSE |
| NW_011924881.1 | 809481 | 809481 | 1 | + | 35 | 1 | 34 | 30 | 0 | 30 | 2.86 | 0 | NW_011924881.1 | 808343 | 827625 | 19283 | - | Gnomon | gene | gene53 | GeneID:105277148 | LOC105277148 | Gene | LOC105277148 | protein_coding | character(0) | character(0) | RphaseA1.vs.RphaseB1 | FALSE |
| NW_011924881.1 | 809504 | 809504 | 1 | + | 34 | 0 | 34 | 31 | 0 | 31 | 0 | 0 | NW_011924881.1 | 808343 | 827625 | 19283 | - | Gnomon | gene | gene53 | GeneID:105277148 | LOC105277148 | Gene | LOC105277148 | protein_coding | character(0) | character(0) | RphaseA1.vs.RphaseB1 | FALSE |
| NW_011924881.1 | 809520 | 809520 | 1 | + | 30 | 1 | 29 | 25 | 0 | 25 | 3.33 | 0 | NW_011924881.1 | 808343 | 827625 | 19283 | - | Gnomon | gene | gene53 | GeneID:105277148 | LOC105277148 | Gene | LOC105277148 | protein_coding | character(0) | character(0) | RphaseA1.vs.RphaseB1 | FALSE |
| NW_011924881.1 | 810123 | 810123 | 1 | + | 28 | 0 | 28 | 25 | 0 | 25 | 0 | 0 | NW_011924881.1 | 808343 | 827625 | 19283 | - | Gnomon | gene | gene53 | GeneID:105277148 | LOC105277148 | Gene | LOC105277148 | protein_coding | character(0) | character(0) | RphaseA1.vs.RphaseB1 | FALSE |
| NW_011924881.1 | 810155 | 810155 | 1 | + | 29 | 0 | 29 | 25 | 0 | 25 | 0 | 0 | NW_011924881.1 | 808343 | 827625 | 19283 | - | Gnomon | gene | gene53 | GeneID:105277148 | LOC105277148 | Gene | LOC105277148 | protein_coding | character(0) | character(0) | RphaseA1.vs.RphaseB1 | FALSE |
| NW_011924881.1 | 810182 | 810182 | 1 | + | 28 | 2 | 26 | 28 | 1 | 27 | 7.14 | 3.57 | NW_011924881.1 | 808343 | 827625 | 19283 | - | Gnomon | gene | gene53 | GeneID:105277148 | LOC105277148 | Gene | LOC105277148 | protein_coding | character(0) | character(0) | RphaseA1.vs.RphaseB1 | FALSE |
| NW_011924881.1 | 810335 | 810335 | 1 | + | 21 | 0 | 21 | 28 | 0 | 28 | 0 | 0 | NW_011924881.1 | 808343 | 827625 | 19283 | - | Gnomon | gene | gene53 | GeneID:105277148 | LOC105277148 | Gene | LOC105277148 | protein_coding | character(0) | character(0) | RphaseA1.vs.RphaseB1 | FALSE |
| NW_011924881.1 | 810337 | 810337 | 1 | + | 24 | 1 | 23 | 29 | 0 | 29 | 4.17 | 0 | NW_011924881.1 | 808343 | 827625 | 19283 | - | Gnomon | gene | gene53 | GeneID:105277148 | LOC105277148 | Gene | LOC105277148 | protein_coding | character(0) | character(0) | RphaseA1.vs.RphaseB1 | FALSE |
| NW_011924881.1 | 810348 | 810348 | 1 | + | 59 | 0 | 59 | 41 | 1 | 40 | 0 | 2.44 | NW_011924881.1 | 808343 | 827625 | 19283 | - | Gnomon | gene | gene53 | GeneID:105277148 | LOC105277148 | Gene | LOC105277148 | protein_coding | character(0) | character(0) | RphaseA1.vs.RphaseB1 | FALSE |
| NW_011924881.1 | 810391 | 810391 | 1 | + | 66 | 0 | 66 | 37 | 0 | 37 | 0 | 0 | NW_011924881.1 | 808343 | 827625 | 19283 | - | Gnomon | gene | gene53 | GeneID:105277148 | LOC105277148 | Gene | LOC105277148 | protein_coding | character(0) | character(0) | RphaseA1.vs.RphaseB1 | FALSE |
| NW_011924881.1 | 810409 | 810409 | 1 | + | 57 | 0 | 57 | 32 | 1 | 31 | 0 | 3.12 | NW_011924881.1 | 808343 | 827625 | 19283 | - | Gnomon | gene | gene53 | GeneID:105277148 | LOC105277148 | Gene | LOC105277148 | protein_coding | character(0) | character(0) | RphaseA1.vs.RphaseB1 | FALSE |
| NW_011924881.1 | 810412 | 810412 | 1 | + | 26 | 0 | 26 | 27 | 1 | 26 | 0 | 3.7 | NW_011924881.1 | 808343 | 827625 | 19283 | - | Gnomon | gene | gene53 | GeneID:105277148 | LOC105277148 | Gene | LOC105277148 | protein_coding | character(0) | character(0) | RphaseA1.vs.RphaseB1 | FALSE |
| NW_011924881.1 | 810417 | 810417 | 1 | + | 29 | 1 | 28 | 24 | 1 | 23 | 3.45 | 4.17 | NW_011924881.1 | 808343 | 827625 | 19283 | - | Gnomon | gene | gene53 | GeneID:105277148 | LOC105277148 | Gene | LOC105277148 | protein_coding | character(0) | character(0) | RphaseA1.vs.RphaseB1 | FALSE |
| NW_011924881.1 | 810424 | 810424 | 1 | + | 40 | 0 | 40 | 24 | 0 | 24 | 0 | 0 | NW_011924881.1 | 808343 | 827625 | 19283 | - | Gnomon | gene | gene53 | GeneID:105277148 | LOC105277148 | Gene | LOC105277148 | protein_coding | character(0) | character(0) | RphaseA1.vs.RphaseB1 | FALSE |
| NW_011924881.1 | 810430 | 810430 | 1 | + | 21 | 0 | 21 | 26 | 0 | 26 | 0 | 0 | NW_011924881.1 | 808343 | 827625 | 19283 | - | Gnomon | gene | gene53 | GeneID:105277148 | LOC105277148 | Gene | LOC105277148 | protein_coding | character(0) | character(0) | RphaseA1.vs.RphaseB1 | FALSE |
| NW_011924881.1 | 810436 | 810436 | 1 | + | 46 | 1 | 45 | 30 | 0 | 30 | 2.17 | 0 | NW_011924881.1 | 808343 | 827625 | 19283 | - | Gnomon | gene | gene53 | GeneID:105277148 | LOC105277148 | Gene | LOC105277148 | protein_coding | character(0) | character(0) | RphaseA1.vs.RphaseB1 | FALSE |
| NW_011924881.1 | 810482 | 810482 | 1 | + | 38 | 0 | 38 | 47 | 1 | 46 | 0 | 2.13 | NW_011924881.1 | 808343 | 827625 | 19283 | - | Gnomon | gene | gene53 | GeneID:105277148 | LOC105277148 | Gene | LOC105277148 | protein_coding | character(0) | character(0) | RphaseA1.vs.RphaseB1 | FALSE |
| NW_011924881.1 | 810505 | 810505 | 1 | + | 63 | 0 | 63 | 50 | 0 | 50 | 0 | 0 | NW_011924881.1 | 808343 | 827625 | 19283 | - | Gnomon | gene | gene53 | GeneID:105277148 | LOC105277148 | Gene | LOC105277148 | protein_coding | character(0) | character(0) | RphaseA1.vs.RphaseB1 | FALSE |
| NW_011924881.1 | 810511 | 810511 | 1 | + | 61 | 0 | 61 | 54 | 0 | 54 | 0 | 0 | NW_011924881.1 | 808343 | 827625 | 19283 | - | Gnomon | gene | gene53 | GeneID:105277148 | LOC105277148 | Gene | LOC105277148 | protein_coding | character(0) | character(0) | RphaseA1.vs.RphaseB1 | FALSE |
| NW_011924881.1 | 810516 | 810516 | 1 | + | 41 | 0 | 41 | 55 | 1 | 54 | 0 | 1.82 | NW_011924881.1 | 808343 | 827625 | 19283 | - | Gnomon | gene | gene53 | GeneID:105277148 | LOC105277148 | Gene | LOC105277148 | protein_coding | character(0) | character(0) | RphaseA1.vs.RphaseB1 | FALSE |
| NW_011924881.1 | 810520 | 810520 | 1 | + | 38 | 0 | 38 | 56 | 1 | 55 | 0 | 1.79 | NW_011924881.1 | 808343 | 827625 | 19283 | - | Gnomon | gene | gene53 | GeneID:105277148 | LOC105277148 | Gene | LOC105277148 | protein_coding | character(0) | character(0) | RphaseA1.vs.RphaseB1 | FALSE |
| NW_011924881.1 | 810532 | 810532 | 1 | + | 20 | 0 | 20 | 22 | 0 | 22 | 0 | 0 | NW_011924881.1 | 808343 | 827625 | 19283 | - | Gnomon | gene | gene53 | GeneID:105277148 | LOC105277148 | Gene | LOC105277148 | protein_coding | character(0) | character(0) | RphaseA1.vs.RphaseB1 | FALSE |
| NW_011924881.1 | 810535 | 810535 | 1 | + | 20 | 0 | 20 | 23 | 0 | 23 | 0 | 0 | NW_011924881.1 | 808343 | 827625 | 19283 | - | Gnomon | gene | gene53 | GeneID:105277148 | LOC105277148 | Gene | LOC105277148 | protein_coding | character(0) | character(0) | RphaseA1.vs.RphaseB1 | FALSE |

...
... only first 25 lines shown ...

Directory: DMSG File: dmg-Cb_LI_RphaseA1.vs.RphaseB2_details.txt

| seqnames | start | end | width | strand | coverage1 | numCs1 | numTs1 | coverage2 | numCs2 | numTs2 | RphaseA1 | RphaseB2 | gene_seqnames | gene_start | gene_end | gene_width | gene_strand | gene_source | gene_type | gene_ID | gene_Dbxref | gene_Name | gene_gbkey | gene_gene | gene_gene_biotype | gene_start_range | gene_end_range | gene_comparison | is.dm |
|---|---|---|---|---|---|---|---|---|---|---|---|---|---|---|---|---|---|---|---|---|---|---|---|---|---|---|---|---|---|
| NW_011924881.1 | 724788 | 724788 | 1 | + | 21 | 0 | 21 | 28 | 0 | 28 | 0 | 0 | NW_011924881.1 | 724448 | 731567 | 7120 | - | Gnomon | gene | gene34 | GeneID:105278416 | LOC105278416 | Gene | LOC105278416 | protein_coding | character(0) | character(0) | RphaseA1.vs.RphaseB2 | FALSE |
| NW_011924881.1 | 724872 | 724872 | 1 | + | 32 | 0 | 32 | 36 | 0 | 36 | 0 | 0 | NW_011924881.1 | 724448 | 731567 | 7120 | - | Gnomon | gene | gene34 | GeneID:105278416 | LOC105278416 | Gene | LOC105278416 | protein_coding | character(0) | character(0) | RphaseA1.vs.RphaseB2 | FALSE |
| NW_011924881.1 | 724889 | 724889 | 1 | + | 32 | 0 | 32 | 38 | 0 | 38 | 0 | 0 | NW_011924881.1 | 724448 | 731567 | 7120 | - | Gnomon | gene | gene34 | GeneID:105278416 | LOC105278416 | Gene | LOC105278416 | protein_coding | character(0) | character(0) | RphaseA1.vs.RphaseB2 | FALSE |
| NW_011924881.1 | 724896 | 724896 | 1 | + | 32 | 0 | 32 | 36 | 0 | 36 | 0 | 0 | NW_011924881.1 | 724448 | 731567 | 7120 | - | Gnomon | gene | gene34 | GeneID:105278416 | LOC105278416 | Gene | LOC105278416 | protein_coding | character(0) | character(0) | RphaseA1.vs.RphaseB2 | FALSE |
| NW_011924881.1 | 724898 | 724898 | 1 | + | 33 | 0 | 33 | 37 | 0 | 37 | 0 | 0 | NW_011924881.1 | 724448 | 731567 | 7120 | - | Gnomon | gene | gene34 | GeneID:105278416 | LOC105278416 | Gene | LOC105278416 | protein_coding | character(0) | character(0) | RphaseA1.vs.RphaseB2 | FALSE |
| NW_011924881.1 | 724926 | 724926 | 1 | + | 30 | 0 | 30 | 58 | 1 | 57 | 0 | 1.72 | NW_011924881.1 | 724448 | 731567 | 7120 | - | Gnomon | gene | gene34 | GeneID:105278416 | LOC105278416 | Gene | LOC105278416 | protein_coding | character(0) | character(0) | RphaseA1.vs.RphaseB2 | FALSE |
| NW_011924881.1 | 724934 | 724934 | 1 | + | 26 | 0 | 26 | 43 | 0 | 43 | 0 | 0 | NW_011924881.1 | 724448 | 731567 | 7120 | - | Gnomon | gene | gene34 | GeneID:105278416 | LOC105278416 | Gene | LOC105278416 | protein_coding | character(0) | character(0) | RphaseA1.vs.RphaseB2 | FALSE |
| NW_011924881.1 | 724936 | 724936 | 1 | + | 27 | 0 | 27 | 44 | 0 | 44 | 0 | 0 | NW_011924881.1 | 724448 | 731567 | 7120 | - | Gnomon | gene | gene34 | GeneID:105278416 | LOC105278416 | Gene | LOC105278416 | protein_coding | character(0) | character(0) | RphaseA1.vs.RphaseB2 | FALSE |
| NW_011924881.1 | 724943 | 724943 | 1 | + | 24 | 0 | 24 | 40 | 1 | 39 | 0 | 2.5 | NW_011924881.1 | 724448 | 731567 | 7120 | - | Gnomon | gene | gene34 | GeneID:105278416 | LOC105278416 | Gene | LOC105278416 | protein_coding | character(0) | character(0) | RphaseA1.vs.RphaseB2 | FALSE |
| NW_011924881.1 | 724947 | 724947 | 1 | + | 27 | 0 | 27 | 42 | 0 | 42 | 0 | 0 | NW_011924881.1 | 724448 | 731567 | 7120 | - | Gnomon | gene | gene34 | GeneID:105278416 | LOC105278416 | Gene | LOC105278416 | protein_coding | character(0) | character(0) | RphaseA1.vs.RphaseB2 | FALSE |
| NW_011924881.1 | 724961 | 724961 | 1 | + | 34 | 0 | 34 | 46 | 0 | 46 | 0 | 0 | NW_011924881.1 | 724448 | 731567 | 7120 | - | Gnomon | gene | gene34 | GeneID:105278416 | LOC105278416 | Gene | LOC105278416 | protein_coding | character(0) | character(0) | RphaseA1.vs.RphaseB2 | FALSE |
| NW_011924881.1 | 724976 | 724976 | 1 | + | 41 | 0 | 41 | 54 | 1 | 53 | 0 | 1.85 | NW_011924881.1 | 724448 | 731567 | 7120 | - | Gnomon | gene | gene34 | GeneID:105278416 | LOC105278416 | Gene | LOC105278416 | protein_coding | character(0) | character(0) | RphaseA1.vs.RphaseB2 | FALSE |
| NW_011924881.1 | 724982 | 724982 | 1 | + | 41 | 2 | 39 | 57 | 0 | 57 | 4.88 | 0 | NW_011924881.1 | 724448 | 731567 | 7120 | - | Gnomon | gene | gene34 | GeneID:105278416 | LOC105278416 | Gene | LOC105278416 | protein_coding | character(0) | character(0) | RphaseA1.vs.RphaseB2 | FALSE |
| NW_011924881.1 | 724990 | 724990 | 1 | + | 39 | 0 | 39 | 58 | 0 | 58 | 0 | 0 | NW_011924881.1 | 724448 | 731567 | 7120 | - | Gnomon | gene | gene34 | GeneID:105278416 | LOC105278416 | Gene | LOC105278416 | protein_coding | character(0) | character(0) | RphaseA1.vs.RphaseB2 | FALSE |
| NW_011924881.1 | 724994 | 724994 | 1 | + | 39 | 0 | 39 | 58 | 0 | 58 | 0 | 0 | NW_011924881.1 | 724448 | 731567 | 7120 | - | Gnomon | gene | gene34 | GeneID:105278416 | LOC105278416 | Gene | LOC105278416 | protein_coding | character(0) | character(0) | RphaseA1.vs.RphaseB2 | FALSE |
| NW_011924881.1 | 725008 | 725008 | 1 | + | 31 | 1 | 30 | 42 | 0 | 42 | 3.23 | 0 | NW_011924881.1 | 724448 | 731567 | 7120 | - | Gnomon | gene | gene34 | GeneID:105278416 | LOC105278416 | Gene | LOC105278416 | protein_coding | character(0) | character(0) | RphaseA1.vs.RphaseB2 | FALSE |
| NW_011924881.1 | 725014 | 725014 | 1 | + | 33 | 1 | 32 | 31 | 0 | 31 | 3.03 | 0 | NW_011924881.1 | 724448 | 731567 | 7120 | - | Gnomon | gene | gene34 | GeneID:105278416 | LOC105278416 | Gene | LOC105278416 | protein_coding | character(0) | character(0) | RphaseA1.vs.RphaseB2 | FALSE |
| NW_011924881.1 | 725022 | 725022 | 1 | + | 37 | 1 | 36 | 22 | 1 | 21 | 2.7 | 4.55 | NW_011924881.1 | 724448 | 731567 | 7120 | - | Gnomon | gene | gene34 | GeneID:105278416 | LOC105278416 | Gene | LOC105278416 | protein_coding | character(0) | character(0) | RphaseA1.vs.RphaseB2 | FALSE |
| NW_011924881.1 | 725041 | 725041 | 1 | + | 41 | 0 | 41 | 32 | 0 | 32 | 0 | 0 | NW_011924881.1 | 724448 | 731567 | 7120 | - | Gnomon | gene | gene34 | GeneID:105278416 | LOC105278416 | Gene | LOC105278416 | protein_coding | character(0) | character(0) | RphaseA1.vs.RphaseB2 | FALSE |
| NW_011924881.1 | 725048 | 725048 | 1 | + | 43 | 0 | 43 | 37 | 1 | 36 | 0 | 2.7 | NW_011924881.1 | 724448 | 731567 | 7120 | - | Gnomon | gene | gene34 | GeneID:105278416 | LOC105278416 | Gene | LOC105278416 | protein_coding | character(0) | character(0) | RphaseA1.vs.RphaseB2 | FALSE |
| NW_011924881.1 | 725062 | 725062 | 1 | + | 44 | 0 | 44 | 43 | 0 | 43 | 0 | 0 | NW_011924881.1 | 724448 | 731567 | 7120 | - | Gnomon | gene | gene34 | GeneID:105278416 | LOC105278416 | Gene | LOC105278416 | protein_coding | character(0) | character(0) | RphaseA1.vs.RphaseB2 | FALSE |
| NW_011924881.1 | 725065 | 725065 | 1 | + | 44 | 0 | 44 | 44 | 0 | 44 | 0 | 0 | NW_011924881.1 | 724448 | 731567 | 7120 | - | Gnomon | gene | gene34 | GeneID:105278416 | LOC105278416 | Gene | LOC105278416 | protein_coding | character(0) | character(0) | RphaseA1.vs.RphaseB2 | FALSE |
| NW_011924881.1 | 725072 | 725072 | 1 | + | 45 | 0 | 45 | 45 | 0 | 45 | 0 | 0 | NW_011924881.1 | 724448 | 731567 | 7120 | - | Gnomon | gene | gene34 | GeneID:105278416 | LOC105278416 | Gene | LOC105278416 | protein_coding | character(0) | character(0) | RphaseA1.vs.RphaseB2 | FALSE |
| NW_011924881.1 | 725074 | 725074 | 1 | + | 45 | 0 | 45 | 45 | 0 | 45 | 0 | 0 | NW_011924881.1 | 724448 | 731567 | 7120 | - | Gnomon | gene | gene34 | GeneID:105278416 | LOC105278416 | Gene | LOC105278416 | protein_coding | character(0) | character(0) | RphaseA1.vs.RphaseB2 | FALSE |

...
... only first 25 lines shown ...

Directory: DMSG File: dmg-Cb_LI_RphaseA2.vs.RphaseB1_details.txt

| seqnames | start | end | width | strand | coverage1 | numCs1 | numTs1 | coverage2 | numCs2 | numTs2 | RphaseA2 | RphaseB1 | gene_seqnames | gene_start | gene_end | gene_width | gene_strand | gene_source | gene_type | gene_ID | gene_Dbxref | gene_Name | gene_gbkey | gene_gene | gene_gene_biotype | gene_start_range | gene_end_range | gene_comparison | is.dm |
|---|---|---|---|---|---|---|---|---|---|---|---|---|---|---|---|---|---|---|---|---|---|---|---|---|---|---|---|---|---|
| NW_011924881.1 | 809447 | 809447 | 1 | + | 28 | 0 | 28 | 23 | 0 | 23 | 0 | 0 | NW_011924881.1 | 808343 | 827625 | 19283 | - | Gnomon | gene | gene53 | GeneID:105277148 | LOC105277148 | Gene | LOC105277148 | protein_coding | character(0) | character(0) | RphaseA2.vs.RphaseB1 | FALSE |
| NW_011924881.1 | 809481 | 809481 | 1 | + | 38 | 1 | 37 | 30 | 0 | 30 | 2.63 | 0 | NW_011924881.1 | 808343 | 827625 | 19283 | - | Gnomon | gene | gene53 | GeneID:105277148 | LOC105277148 | Gene | LOC105277148 | protein_coding | character(0) | character(0) | RphaseA2.vs.RphaseB1 | FALSE |
| NW_011924881.1 | 809504 | 809504 | 1 | + | 44 | 1 | 43 | 31 | 0 | 31 | 2.27 | 0 | NW_011924881.1 | 808343 | 827625 | 19283 | - | Gnomon | gene | gene53 | GeneID:105277148 | LOC105277148 | Gene | LOC105277148 | protein_coding | character(0) | character(0) | RphaseA2.vs.RphaseB1 | FALSE |
| NW_011924881.1 | 809520 | 809520 | 1 | + | 37 | 1 | 36 | 25 | 0 | 25 | 2.7 | 0 | NW_011924881.1 | 808343 | 827625 | 19283 | - | Gnomon | gene | gene53 | GeneID:105277148 | LOC105277148 | Gene | LOC105277148 | protein_coding | character(0) | character(0) | RphaseA2.vs.RphaseB1 | FALSE |
| NW_011924881.1 | 810123 | 810123 | 1 | + | 26 | 0 | 26 | 25 | 0 | 25 | 0 | 0 | NW_011924881.1 | 808343 | 827625 | 19283 | - | Gnomon | gene | gene53 | GeneID:105277148 | LOC105277148 | Gene | LOC105277148 | protein_coding | character(0) | character(0) | RphaseA2.vs.RphaseB1 | FALSE |
| NW_011924881.1 | 810155 | 810155 | 1 | + | 36 | 0 | 36 | 25 | 0 | 25 | 0 | 0 | NW_011924881.1 | 808343 | 827625 | 19283 | - | Gnomon | gene | gene53 | GeneID:105277148 | LOC105277148 | Gene | LOC105277148 | protein_coding | character(0) | character(0) | RphaseA2.vs.RphaseB1 | FALSE |
| NW_011924881.1 | 810182 | 810182 | 1 | + | 39 | 2 | 37 | 28 | 1 | 27 | 5.13 | 3.57 | NW_011924881.1 | 808343 | 827625 | 19283 | - | Gnomon | gene | gene53 | GeneID:105277148 | LOC105277148 | Gene | LOC105277148 | protein_coding | character(0) | character(0) | RphaseA2.vs.RphaseB1 | FALSE |
| NW_011924881.1 | 810330 | 810330 | 1 | + | 32 | 2 | 30 | 22 | 3 | 19 | 6.25 | 13.64 | NW_011924881.1 | 808343 | 827625 | 19283 | - | Gnomon | gene | gene53 | GeneID:105277148 | LOC105277148 | Gene | LOC105277148 | protein_coding | character(0) | character(0) | RphaseA2.vs.RphaseB1 | FALSE |
| NW_011924881.1 | 810335 | 810335 | 1 | + | 33 | 1 | 32 | 28 | 0 | 28 | 3.03 | 0 | NW_011924881.1 | 808343 | 827625 | 19283 | - | Gnomon | gene | gene53 | GeneID:105277148 | LOC105277148 | Gene | LOC105277148 | protein_coding | character(0) | character(0) | RphaseA2.vs.RphaseB1 | FALSE |
| NW_011924881.1 | 810337 | 810337 | 1 | + | 32 | 0 | 32 | 29 | 0 | 29 | 0 | 0 | NW_011924881.1 | 808343 | 827625 | 19283 | - | Gnomon | gene | gene53 | GeneID:105277148 | LOC105277148 | Gene | LOC105277148 | protein_coding | character(0) | character(0) | RphaseA2.vs.RphaseB1 | FALSE |
| NW_011924881.1 | 810348 | 810348 | 1 | + | 74 | 0 | 74 | 41 | 1 | 40 | 0 | 2.44 | NW_011924881.1 | 808343 | 827625 | 19283 | - | Gnomon | gene | gene53 | GeneID:105277148 | LOC105277148 | Gene | LOC105277148 | protein_coding | character(0) | character(0) | RphaseA2.vs.RphaseB1 | FALSE |
| NW_011924881.1 | 810391 | 810391 | 1 | + | 93 | 0 | 93 | 37 | 0 | 37 | 0 | 0 | NW_011924881.1 | 808343 | 827625 | 19283 | - | Gnomon | gene | gene53 | GeneID:105277148 | LOC105277148 | Gene | LOC105277148 | protein_coding | character(0) | character(0) | RphaseA2.vs.RphaseB1 | FALSE |
| NW_011924881.1 | 810409 | 810409 | 1 | + | 86 | 0 | 86 | 32 | 1 | 31 | 0 | 3.12 | NW_011924881.1 | 808343 | 827625 | 19283 | - | Gnomon | gene | gene53 | GeneID:105277148 | LOC105277148 | Gene | LOC105277148 | protein_coding | character(0) | character(0) | RphaseA2.vs.RphaseB1 | FALSE |
| NW_011924881.1 | 810412 | 810412 | 1 | + | 84 | 0 | 84 | 27 | 1 | 26 | 0 | 3.7 | NW_011924881.1 | 808343 | 827625 | 19283 | - | Gnomon | gene | gene53 | GeneID:105277148 | LOC105277148 | Gene | LOC105277148 | protein_coding | character(0) | character(0) | RphaseA2.vs.RphaseB1 | FALSE |
| NW_011924881.1 | 810417 | 810417 | 1 | + | 76 | 0 | 76 | 24 | 1 | 23 | 0 | 4.17 | NW_011924881.1 | 808343 | 827625 | 19283 | - | Gnomon | gene | gene53 | GeneID:105277148 | LOC105277148 | Gene | LOC105277148 | protein_coding | character(0) | character(0) | RphaseA2.vs.RphaseB1 | FALSE |
| NW_011924881.1 | 810424 | 810424 | 1 | + | 74 | 1 | 73 | 24 | 0 | 24 | 1.35 | 0 | NW_011924881.1 | 808343 | 827625 | 19283 | - | Gnomon | gene | gene53 | GeneID:105277148 | LOC105277148 | Gene | LOC105277148 | protein_coding | character(0) | character(0) | RphaseA2.vs.RphaseB1 | FALSE |
| NW_011924881.1 | 810430 | 810430 | 1 | + | 31 | 0 | 31 | 26 | 0 | 26 | 0 | 0 | NW_011924881.1 | 808343 | 827625 | 19283 | - | Gnomon | gene | gene53 | GeneID:105277148 | LOC105277148 | Gene | LOC105277148 | protein_coding | character(0) | character(0) | RphaseA2.vs.RphaseB1 | FALSE |
| NW_011924881.1 | 810436 | 810436 | 1 | + | 32 | 0 | 32 | 30 | 0 | 30 | 0 | 0 | NW_011924881.1 | 808343 | 827625 | 19283 | - | Gnomon | gene | gene53 | GeneID:105277148 | LOC105277148 | Gene | LOC105277148 | protein_coding | character(0) | character(0) | RphaseA2.vs.RphaseB1 | FALSE |
| NW_011924881.1 | 810482 | 810482 | 1 | + | 67 | 3 | 64 | 47 | 1 | 46 | 4.48 | 2.13 | NW_011924881.1 | 808343 | 827625 | 19283 | - | Gnomon | gene | gene53 | GeneID:105277148 | LOC105277148 | Gene | LOC105277148 | protein_coding | character(0) | character(0) | RphaseA2.vs.RphaseB1 | FALSE |
| NW_011924881.1 | 810505 | 810505 | 1 | + | 71 | 0 | 71 | 50 | 0 | 50 | 0 | 0 | NW_011924881.1 | 808343 | 827625 | 19283 | - | Gnomon | gene | gene53 | GeneID:105277148 | LOC105277148 | Gene | LOC105277148 | protein_coding | character(0) | character(0) | RphaseA2.vs.RphaseB1 | FALSE |
| NW_011924881.1 | 810511 | 810511 | 1 | + | 66 | 0 | 66 | 54 | 0 | 54 | 0 | 0 | NW_011924881.1 | 808343 | 827625 | 19283 | - | Gnomon | gene | gene53 | GeneID:105277148 | LOC105277148 | Gene | LOC105277148 | protein_coding | character(0) | character(0) | RphaseA2.vs.RphaseB1 | FALSE |
| NW_011924881.1 | 810516 | 810516 | 1 | + | 64 | 0 | 64 | 55 | 1 | 54 | 0 | 1.82 | NW_011924881.1 | 808343 | 827625 | 19283 | - | Gnomon | gene | gene53 | GeneID:105277148 | LOC105277148 | Gene | LOC105277148 | protein_coding | character(0) | character(0) | RphaseA2.vs.RphaseB1 | FALSE |
| NW_011924881.1 | 810520 | 810520 | 1 | + | 68 | 0 | 68 | 56 | 1 | 55 | 0 | 1.79 | NW_011924881.1 | 808343 | 827625 | 19283 | - | Gnomon | gene | gene53 | GeneID:105277148 | LOC105277148 | Gene | LOC105277148 | protein_coding | character(0) | character(0) | RphaseA2.vs.RphaseB1 | FALSE |
| NW_011924881.1 | 810532 | 810532 | 1 | + | 27 | 0 | 27 | 22 | 0 | 22 | 0 | 0 | NW_011924881.1 | 808343 | 827625 | 19283 | - | Gnomon | gene | gene53 | GeneID:105277148 | LOC105277148 | Gene | LOC105277148 | protein_coding | character(0) | character(0) | RphaseA2.vs.RphaseB1 | FALSE |

...
... only first 25 lines shown ...

Directory: DMSG File: dmg-Cb_LI_RphaseA2.vs.RphaseB2_details.txt

| seqnames | start | end | width | strand | coverage1 | numCs1 | numTs1 | coverage2 | numCs2 | numTs2 | RphaseA2 | RphaseB2 | gene_seqnames | gene_start | gene_end | gene_width | gene_strand | gene_source | gene_type | gene_ID | gene_Dbxref | gene_Name | gene_gbkey | gene_gene | gene_gene_biotype | gene_start_range | gene_end_range | gene_comparison | is.dm |
|---|---|---|---|---|---|---|---|---|---|---|---|---|---|---|---|---|---|---|---|---|---|---|---|---|---|---|---|---|---|
| NW_011924881.1 | 724785 | 724785 | 1 | + | 65 | 0 | 65 | 28 | 0 | 28 | 0 | 0 | NW_011924881.1 | 724448 | 731567 | 7120 | - | Gnomon | gene | gene34 | GeneID:105278416 | LOC105278416 | Gene | LOC105278416 | protein_coding | character(0) | character(0) | RphaseA2.vs.RphaseB2 | FALSE |
| NW_011924881.1 | 724788 | 724788 | 1 | + | 63 | 0 | 63 | 28 | 0 | 28 | 0 | 0 | NW_011924881.1 | 724448 | 731567 | 7120 | - | Gnomon | gene | gene34 | GeneID:105278416 | LOC105278416 | Gene | LOC105278416 | protein_coding | character(0) | character(0) | RphaseA2.vs.RphaseB2 | FALSE |
| NW_011924881.1 | 724794 | 724794 | 1 | + | 63 | 0 | 63 | 28 | 0 | 28 | 0 | 0 | NW_011924881.1 | 724448 | 731567 | 7120 | - | Gnomon | gene | gene34 | GeneID:105278416 | LOC105278416 | Gene | LOC105278416 | protein_coding | character(0) | character(0) | RphaseA2.vs.RphaseB2 | FALSE |
| NW_011924881.1 | 724802 | 724802 | 1 | + | 61 | 0 | 61 | 28 | 0 | 28 | 0 | 0 | NW_011924881.1 | 724448 | 731567 | 7120 | - | Gnomon | gene | gene34 | GeneID:105278416 | LOC105278416 | Gene | LOC105278416 | protein_coding | character(0) | character(0) | RphaseA2.vs.RphaseB2 | FALSE |
| NW_011924881.1 | 724806 | 724806 | 1 | + | 61 | 1 | 60 | 29 | 0 | 29 | 1.64 | 0 | NW_011924881.1 | 724448 | 731567 | 7120 | - | Gnomon | gene | gene34 | GeneID:105278416 | LOC105278416 | Gene | LOC105278416 | protein_coding | character(0) | character(0) | RphaseA2.vs.RphaseB2 | FALSE |
| NW_011924881.1 | 724815 | 724815 | 1 | + | 61 | 0 | 61 | 29 | 1 | 28 | 0 | 3.45 | NW_011924881.1 | 724448 | 731567 | 7120 | - | Gnomon | gene | gene34 | GeneID:105278416 | LOC105278416 | Gene | LOC105278416 | protein_coding | character(0) | character(0) | RphaseA2.vs.RphaseB2 | FALSE |
| NW_011924881.1 | 724818 | 724818 | 1 | + | 62 | 0 | 62 | 29 | 0 | 29 | 0 | 0 | NW_011924881.1 | 724448 | 731567 | 7120 | - | Gnomon | gene | gene34 | GeneID:105278416 | LOC105278416 | Gene | LOC105278416 | protein_coding | character(0) | character(0) | RphaseA2.vs.RphaseB2 | FALSE |
| NW_011924881.1 | 724839 | 724839 | 1 | + | 73 | 1 | 72 | 44 | 0 | 44 | 1.37 | 0 | NW_011924881.1 | 724448 | 731567 | 7120 | - | Gnomon | gene | gene34 | GeneID:105278416 | LOC105278416 | Gene | LOC105278416 | protein_coding | character(0) | character(0) | RphaseA2.vs.RphaseB2 | FALSE |
| NW_011924881.1 | 724872 | 724872 | 1 | + | 62 | 0 | 62 | 36 | 0 | 36 | 0 | 0 | NW_011924881.1 | 724448 | 731567 | 7120 | - | Gnomon | gene | gene34 | GeneID:105278416 | LOC105278416 | Gene | LOC105278416 | protein_coding | character(0) | character(0) | RphaseA2.vs.RphaseB2 | FALSE |
| NW_011924881.1 | 724889 | 724889 | 1 | + | 44 | 0 | 44 | 38 | 0 | 38 | 0 | 0 | NW_011924881.1 | 724448 | 731567 | 7120 | - | Gnomon | gene | gene34 | GeneID:105278416 | LOC105278416 | Gene | LOC105278416 | protein_coding | character(0) | character(0) | RphaseA2.vs.RphaseB2 | FALSE |
| NW_011924881.1 | 724896 | 724896 | 1 | + | 42 | 0 | 42 | 36 | 0 | 36 | 0 | 0 | NW_011924881.1 | 724448 | 731567 | 7120 | - | Gnomon | gene | gene34 | GeneID:105278416 | LOC105278416 | Gene | LOC105278416 | protein_coding | character(0) | character(0) | RphaseA2.vs.RphaseB2 | FALSE |
| NW_011924881.1 | 724898 | 724898 | 1 | + | 64 | 1 | 63 | 37 | 0 | 37 | 1.56 | 0 | NW_011924881.1 | 724448 | 731567 | 7120 | - | Gnomon | gene | gene34 | GeneID:105278416 | LOC105278416 | Gene | LOC105278416 | protein_coding | character(0) | character(0) | RphaseA2.vs.RphaseB2 | FALSE |
| NW_011924881.1 | 724926 | 724926 | 1 | + | 62 | 0 | 62 | 58 | 1 | 57 | 0 | 1.72 | NW_011924881.1 | 724448 | 731567 | 7120 | - | Gnomon | gene | gene34 | GeneID:105278416 | LOC105278416 | Gene | LOC105278416 | protein_coding | character(0) | character(0) | RphaseA2.vs.RphaseB2 | FALSE |
| NW_011924881.1 | 724934 | 724934 | 1 | + | 51 | 0 | 51 | 43 | 0 | 43 | 0 | 0 | NW_011924881.1 | 724448 | 731567 | 7120 | - | Gnomon | gene | gene34 | GeneID:105278416 | LOC105278416 | Gene | LOC105278416 | protein_coding | character(0) | character(0) | RphaseA2.vs.RphaseB2 | FALSE |
| NW_011924881.1 | 724936 | 724936 | 1 | + | 51 | 0 | 51 | 44 | 0 | 44 | 0 | 0 | NW_011924881.1 | 724448 | 731567 | 7120 | - | Gnomon | gene | gene34 | GeneID:105278416 | LOC105278416 | Gene | LOC105278416 | protein_coding | character(0) | character(0) | RphaseA2.vs.RphaseB2 | FALSE |
| NW_011924881.1 | 724943 | 724943 | 1 | + | 70 | 1 | 69 | 40 | 1 | 39 | 1.43 | 2.5 | NW_011924881.1 | 724448 | 731567 | 7120 | - | Gnomon | gene | gene34 | GeneID:105278416 | LOC105278416 | Gene | LOC105278416 | protein_coding | character(0) | character(0) | RphaseA2.vs.RphaseB2 | FALSE |
| NW_011924881.1 | 724947 | 724947 | 1 | + | 72 | 0 | 72 | 42 | 0 | 42 | 0 | 0 | NW_011924881.1 | 724448 | 731567 | 7120 | - | Gnomon | gene | gene34 | GeneID:105278416 | LOC105278416 | Gene | LOC105278416 | protein_coding | character(0) | character(0) | RphaseA2.vs.RphaseB2 | FALSE |
| NW_011924881.1 | 724961 | 724961 | 1 | + | 65 | 0 | 65 | 46 | 0 | 46 | 0 | 0 | NW_011924881.1 | 724448 | 731567 | 7120 | - | Gnomon | gene | gene34 | GeneID:105278416 | LOC105278416 | Gene | LOC105278416 | protein_coding | character(0) | character(0) | RphaseA2.vs.RphaseB2 | FALSE |
| NW_011924881.1 | 724976 | 724976 | 1 | + | 75 | 0 | 75 | 54 | 1 | 53 | 0 | 1.85 | NW_011924881.1 | 724448 | 731567 | 7120 | - | Gnomon | gene | gene34 | GeneID:105278416 | LOC105278416 | Gene | LOC105278416 | protein_coding | character(0) | character(0) | RphaseA2.vs.RphaseB2 | FALSE |
| NW_011924881.1 | 724982 | 724982 | 1 | + | 79 | 0 | 79 | 57 | 0 | 57 | 0 | 0 | NW_011924881.1 | 724448 | 731567 | 7120 | - | Gnomon | gene | gene34 | GeneID:105278416 | LOC105278416 | Gene | LOC105278416 | protein_coding | character(0) | character(0) | RphaseA2.vs.RphaseB2 | FALSE |
| NW_011924881.1 | 724990 | 724990 | 1 | + | 82 | 0 | 82 | 58 | 0 | 58 | 0 | 0 | NW_011924881.1 | 724448 | 731567 | 7120 | - | Gnomon | gene | gene34 | GeneID:105278416 | LOC105278416 | Gene | LOC105278416 | protein_coding | character(0) | character(0) | RphaseA2.vs.RphaseB2 | FALSE |
| NW_011924881.1 | 724994 | 724994 | 1 | + | 81 | 0 | 81 | 58 | 0 | 58 | 0 | 0 | NW_011924881.1 | 724448 | 731567 | 7120 | - | Gnomon | gene | gene34 | GeneID:105278416 | LOC105278416 | Gene | LOC105278416 | protein_coding | character(0) | character(0) | RphaseA2.vs.RphaseB2 | FALSE |
| NW_011924881.1 | 725008 | 725008 | 1 | + | 53 | 0 | 53 | 42 | 0 | 42 | 0 | 0 | NW_011924881.1 | 724448 | 731567 | 7120 | - | Gnomon | gene | gene34 | GeneID:105278416 | LOC105278416 | Gene | LOC105278416 | protein_coding | character(0) | character(0) | RphaseA2.vs.RphaseB2 | FALSE |
| NW_011924881.1 | 725014 | 725014 | 1 | + | 63 | 1 | 62 | 31 | 0 | 31 | 1.59 | 0 | NW_011924881.1 | 724448 | 731567 | 7120 | - | Gnomon | gene | gene34 | GeneID:105278416 | LOC105278416 | Gene | LOC105278416 | protein_coding | character(0) | character(0) | RphaseA2.vs.RphaseB2 | FALSE |

...
... only first 25 lines shown ...

Directory: DMSG File: dmg-Cb_LI_RphaseB1.vs.RphaseB2_details.txt

| seqnames | start | end | width | strand | coverage1 | numCs1 | numTs1 | coverage2 | numCs2 | numTs2 | RphaseB1 | RphaseB2 | gene_seqnames | gene_start | gene_end | gene_width | gene_strand | gene_source | gene_type | gene_ID | gene_Dbxref | gene_Name | gene_gbkey | gene_gene | gene_gene_biotype | gene_partial | gene_start_range | gene_end_range | gene_comparison | is.dm |
|---|---|---|---|---|---|---|---|---|---|---|---|---|---|---|---|---|---|---|---|---|---|---|---|---|---|---|---|---|---|---|
| NW_011924881.1 | 43853 | 43853 | 1 | + | 30 | 0 | 30 | 22 | 0 | 22 | 0 | 0 | NW_011924881.1 | 42715 | 135016 | 92302 | - | Gnomon | gene | gene18 | GeneID:105287290 | LOC105287290 | Gene | LOC105287290 | protein_coding | NA | character(0) | character(0) | RphaseB1.vs.RphaseB2 | FALSE |
| NW_011924881.1 | 43864 | 43864 | 1 | + | 30 | 0 | 30 | 22 | 0 | 22 | 0 | 0 | NW_011924881.1 | 42715 | 135016 | 92302 | - | Gnomon | gene | gene18 | GeneID:105287290 | LOC105287290 | Gene | LOC105287290 | protein_coding | NA | character(0) | character(0) | RphaseB1.vs.RphaseB2 | FALSE |
| NW_011924881.1 | 43879 | 43879 | 1 | + | 28 | 0 | 28 | 21 | 0 | 21 | 0 | 0 | NW_011924881.1 | 42715 | 135016 | 92302 | - | Gnomon | gene | gene18 | GeneID:105287290 | LOC105287290 | Gene | LOC105287290 | protein_coding | NA | character(0) | character(0) | RphaseB1.vs.RphaseB2 | FALSE |
| NW_011924881.1 | 43901 | 43901 | 1 | + | 30 | 0 | 30 | 22 | 0 | 22 | 0 | 0 | NW_011924881.1 | 42715 | 135016 | 92302 | - | Gnomon | gene | gene18 | GeneID:105287290 | LOC105287290 | Gene | LOC105287290 | protein_coding | NA | character(0) | character(0) | RphaseB1.vs.RphaseB2 | FALSE |
| NW_011924881.1 | 43904 | 43904 | 1 | + | 27 | 0 | 27 | 21 | 0 | 21 | 0 | 0 | NW_011924881.1 | 42715 | 135016 | 92302 | - | Gnomon | gene | gene18 | GeneID:105287290 | LOC105287290 | Gene | LOC105287290 | protein_coding | NA | character(0) | character(0) | RphaseB1.vs.RphaseB2 | FALSE |
| NW_011924881.1 | 43907 | 43907 | 1 | + | 27 | 0 | 27 | 21 | 0 | 21 | 0 | 0 | NW_011924881.1 | 42715 | 135016 | 92302 | - | Gnomon | gene | gene18 | GeneID:105287290 | LOC105287290 | Gene | LOC105287290 | protein_coding | NA | character(0) | character(0) | RphaseB1.vs.RphaseB2 | FALSE |
| NW_011924881.1 | 43912 | 43912 | 1 | + | 24 | 0 | 24 | 20 | 0 | 20 | 0 | 0 | NW_011924881.1 | 42715 | 135016 | 92302 | - | Gnomon | gene | gene18 | GeneID:105287290 | LOC105287290 | Gene | LOC105287290 | protein_coding | NA | character(0) | character(0) | RphaseB1.vs.RphaseB2 | FALSE |
| NW_011924881.1 | 43914 | 43914 | 1 | + | 23 | 0 | 23 | 20 | 0 | 20 | 0 | 0 | NW_011924881.1 | 42715 | 135016 | 92302 | - | Gnomon | gene | gene18 | GeneID:105287290 | LOC105287290 | Gene | LOC105287290 | protein_coding | NA | character(0) | character(0) | RphaseB1.vs.RphaseB2 | FALSE |
| NW_011924881.1 | 44105 | 44105 | 1 | + | 32 | 0 | 32 | 59 | 0 | 59 | 0 | 0 | NW_011924881.1 | 42715 | 135016 | 92302 | - | Gnomon | gene | gene18 | GeneID:105287290 | LOC105287290 | Gene | LOC105287290 | protein_coding | NA | character(0) | character(0) | RphaseB1.vs.RphaseB2 | FALSE |
| NW_011924881.1 | 44112 | 44112 | 1 | + | 32 | 0 | 32 | 57 | 0 | 57 | 0 | 0 | NW_011924881.1 | 42715 | 135016 | 92302 | - | Gnomon | gene | gene18 | GeneID:105287290 | LOC105287290 | Gene | LOC105287290 | protein_coding | NA | character(0) | character(0) | RphaseB1.vs.RphaseB2 | FALSE |
| NW_011924881.1 | 44121 | 44121 | 1 | + | 30 | 0 | 30 | 56 | 0 | 56 | 0 | 0 | NW_011924881.1 | 42715 | 135016 | 92302 | - | Gnomon | gene | gene18 | GeneID:105287290 | LOC105287290 | Gene | LOC105287290 | protein_coding | NA | character(0) | character(0) | RphaseB1.vs.RphaseB2 | FALSE |
| NW_011924881.1 | 44123 | 44123 | 1 | + | 30 | 0 | 30 | 56 | 0 | 56 | 0 | 0 | NW_011924881.1 | 42715 | 135016 | 92302 | - | Gnomon | gene | gene18 | GeneID:105287290 | LOC105287290 | Gene | LOC105287290 | protein_coding | NA | character(0) | character(0) | RphaseB1.vs.RphaseB2 | FALSE |
| NW_011924881.1 | 44125 | 44125 | 1 | + | 30 | 0 | 30 | 55 | 0 | 55 | 0 | 0 | NW_011924881.1 | 42715 | 135016 | 92302 | - | Gnomon | gene | gene18 | GeneID:105287290 | LOC105287290 | Gene | LOC105287290 | protein_coding | NA | character(0) | character(0) | RphaseB1.vs.RphaseB2 | FALSE |
| NW_011924881.1 | 44129 | 44129 | 1 | + | 30 | 0 | 30 | 54 | 0 | 54 | 0 | 0 | NW_011924881.1 | 42715 | 135016 | 92302 | - | Gnomon | gene | gene18 | GeneID:105287290 | LOC105287290 | Gene | LOC105287290 | protein_coding | NA | character(0) | character(0) | RphaseB1.vs.RphaseB2 | FALSE |
| NW_011924881.1 | 44138 | 44138 | 1 | + | 29 | 0 | 29 | 52 | 0 | 52 | 0 | 0 | NW_011924881.1 | 42715 | 135016 | 92302 | - | Gnomon | gene | gene18 | GeneID:105287290 | LOC105287290 | Gene | LOC105287290 | protein_coding | NA | character(0) | character(0) | RphaseB1.vs.RphaseB2 | FALSE |
| NW_011924881.1 | 44152 | 44152 | 1 | + | 33 | 0 | 33 | 52 | 0 | 52 | 0 | 0 | NW_011924881.1 | 42715 | 135016 | 92302 | - | Gnomon | gene | gene18 | GeneID:105287290 | LOC105287290 | Gene | LOC105287290 | protein_coding | NA | character(0) | character(0) | RphaseB1.vs.RphaseB2 | FALSE |
| NW_011924881.1 | 44173 | 44173 | 1 | + | 37 | 0 | 37 | 51 | 1 | 50 | 0 | 1.96 | NW_011924881.1 | 42715 | 135016 | 92302 | - | Gnomon | gene | gene18 | GeneID:105287290 | LOC105287290 | Gene | LOC105287290 | protein_coding | NA | character(0) | character(0) | RphaseB1.vs.RphaseB2 | FALSE |
| NW_011924881.1 | 44185 | 44185 | 1 | + | 29 | 0 | 29 | 37 | 0 | 37 | 0 | 0 | NW_011924881.1 | 42715 | 135016 | 92302 | - | Gnomon | gene | gene18 | GeneID:105287290 | LOC105287290 | Gene | LOC105287290 | protein_coding | NA | character(0) | character(0) | RphaseB1.vs.RphaseB2 | FALSE |
| NW_011924881.1 | 44193 | 44193 | 1 | + | 27 | 0 | 27 | 29 | 0 | 29 | 0 | 0 | NW_011924881.1 | 42715 | 135016 | 92302 | - | Gnomon | gene | gene18 | GeneID:105287290 | LOC105287290 | Gene | LOC105287290 | protein_coding | NA | character(0) | character(0) | RphaseB1.vs.RphaseB2 | FALSE |
| NW_011924881.1 | 44199 | 44199 | 1 | + | 23 | 0 | 23 | 31 | 0 | 31 | 0 | 0 | NW_011924881.1 | 42715 | 135016 | 92302 | - | Gnomon | gene | gene18 | GeneID:105287290 | LOC105287290 | Gene | LOC105287290 | protein_coding | NA | character(0) | character(0) | RphaseB1.vs.RphaseB2 | FALSE |
| NW_011924881.1 | 44201 | 44201 | 1 | + | 26 | 0 | 26 | 31 | 0 | 31 | 0 | 0 | NW_011924881.1 | 42715 | 135016 | 92302 | - | Gnomon | gene | gene18 | GeneID:105287290 | LOC105287290 | Gene | LOC105287290 | protein_coding | NA | character(0) | character(0) | RphaseB1.vs.RphaseB2 | FALSE |
| NW_011924881.1 | 44204 | 44204 | 1 | + | 25 | 1 | 24 | 30 | 0 | 30 | 4 | 0 | NW_011924881.1 | 42715 | 135016 | 92302 | - | Gnomon | gene | gene18 | GeneID:105287290 | LOC105287290 | Gene | LOC105287290 | protein_coding | NA | character(0) | character(0) | RphaseB1.vs.RphaseB2 | FALSE |
| NW_011924881.1 | 44221 | 44221 | 1 | + | 25 | 1 | 24 | 26 | 0 | 26 | 4 | 0 | NW_011924881.1 | 42715 | 135016 | 92302 | - | Gnomon | gene | gene18 | GeneID:105287290 | LOC105287290 | Gene | LOC105287290 | protein_coding | NA | character(0) | character(0) | RphaseB1.vs.RphaseB2 | FALSE |
| NW_011924881.1 | 44223 | 44223 | 1 | + | 25 | 0 | 25 | 24 | 0 | 24 | 0 | 0 | NW_011924881.1 | 42715 | 135016 | 92302 | - | Gnomon | gene | gene18 | GeneID:105287290 | LOC105287290 | Gene | LOC105287290 | protein_coding | NA | character(0) | character(0) | RphaseB1.vs.RphaseB2 | FALSE |

...
... only first 25 lines shown ...

Directory: DMSG File: dms-Cb_LI.txt

| seqnames | start | end | width | strand | pvalue | qvalue | meth.diff | comparison |
|---|---|---|---|---|---|---|---|---|
| NW_011924881.1 | 821269 | 821269 | 1 | + | 2.65918081615226e-07 | 0.005 | -38.24 | BCphaseA1.vs.BCphaseA2 |
| NW_011924888.1 | 73339 | 73339 | 1 | + | 1.92295370159431e-06 | 0.027 | 53.66 | BCphaseA1.vs.BCphaseA2 |
| NW_011924888.1 | 142295 | 142295 | 1 | + | 2.72122700544346e-07 | 0.005 | -56.16 | BCphaseA1.vs.BCphaseA2 |
| NW_011924888.1 | 165750 | 165750 | 1 | + | 9.07639766429664e-12 | 0 | 46.99 | BCphaseA1.vs.BCphaseA2 |
| NW_011924888.1 | 248296 | 248296 | 1 | + | 8.22908913910749e-07 | 0.012 | 59.59 | BCphaseA1.vs.BCphaseA2 |
| NW_011924888.1 | 408443 | 408443 | 1 | + | 5.4059524894543e-09 | 0 | -54.84 | BCphaseA1.vs.BCphaseA2 |
| NW_011924888.1 | 1582933 | 1582933 | 1 | + | 3.34949007748681e-06 | 0.042 | -37.84 | BCphaseA1.vs.BCphaseA2 |
| NW_011924888.1 | 1583145 | 1583145 | 1 | + | 4.26359761773129e-14 | 0 | 49.61 | BCphaseA1.vs.BCphaseA2 |
| NW_011924888.1 | 1598633 | 1598633 | 1 | + | 3.25350819055164e-07 | 0.005 | -58.33 | BCphaseA1.vs.BCphaseA2 |
| NW_011924889.1 | 215297 | 215297 | 1 | + | 3.27593239984402e-07 | 0.005 | 62.94 | BCphaseA1.vs.BCphaseA2 |
| NW_011924899.1 | 340853 | 340853 | 1 | + | 8.35741651473022e-08 | 0.002 | -64.52 | BCphaseA1.vs.BCphaseA2 |
| NW_011924914.1 | 110806 | 110806 | 1 | + | 3.62977783354751e-17 | 0 | -93.1 | BCphaseA1.vs.BCphaseA2 |
| NW_011924914.1 | 216827 | 216827 | 1 | + | 3.62272750504221e-09 | 0 | 42.58 | BCphaseA1.vs.BCphaseA2 |
| NW_011924922.1 | 2079399 | 2079399 | 1 | + | 8.54485992285842e-12 | 0 | -51.98 | BCphaseA1.vs.BCphaseA2 |
| NW_011924922.1 | 2393776 | 2393776 | 1 | + | 1.09881883780235e-07 | 0.002 | -46.97 | BCphaseA1.vs.BCphaseA2 |
| NW_011924922.1 | 2510440 | 2510440 | 1 | + | 3.71569111196151e-12 | 0 | -100 | BCphaseA1.vs.BCphaseA2 |
| NW_011924929.1 | 733820 | 733820 | 1 | + | 4.09088981861159e-13 | 0 | -39.53 | BCphaseA1.vs.BCphaseA2 |
| NW_011924931.1 | 2030310 | 2030310 | 1 | + | 2.20866896389859e-29 | 0 | -95.35 | BCphaseA1.vs.BCphaseA2 |
| NW_011924945.1 | 68768 | 68768 | 1 | + | 1.57574067083735e-10 | 0 | 42.36 | BCphaseA1.vs.BCphaseA2 |
| NW_011924946.1 | 776599 | 776599 | 1 | + | 3.11981545968302e-07 | 0.005 | -57.84 | BCphaseA1.vs.BCphaseA2 |
| NW_011924946.1 | 1548965 | 1548965 | 1 | + | 5.04666914642911e-08 | 0.001 | -35 | BCphaseA1.vs.BCphaseA2 |
| NW_011924946.1 | 2918036 | 2918036 | 1 | + | 2.59941918016128e-08 | 0.001 | -51.28 | BCphaseA1.vs.BCphaseA2 |
| NW_011924947.1 | 86088 | 86088 | 1 | + | 6.01726773352562e-12 | 0 | 53.26 | BCphaseA1.vs.BCphaseA2 |
| NW_011924953.1 | 417309 | 417309 | 1 | + | 1.15993617050659e-16 | 0 | -90 | BCphaseA1.vs.BCphaseA2 |

...

... only first 25 lines shown ...

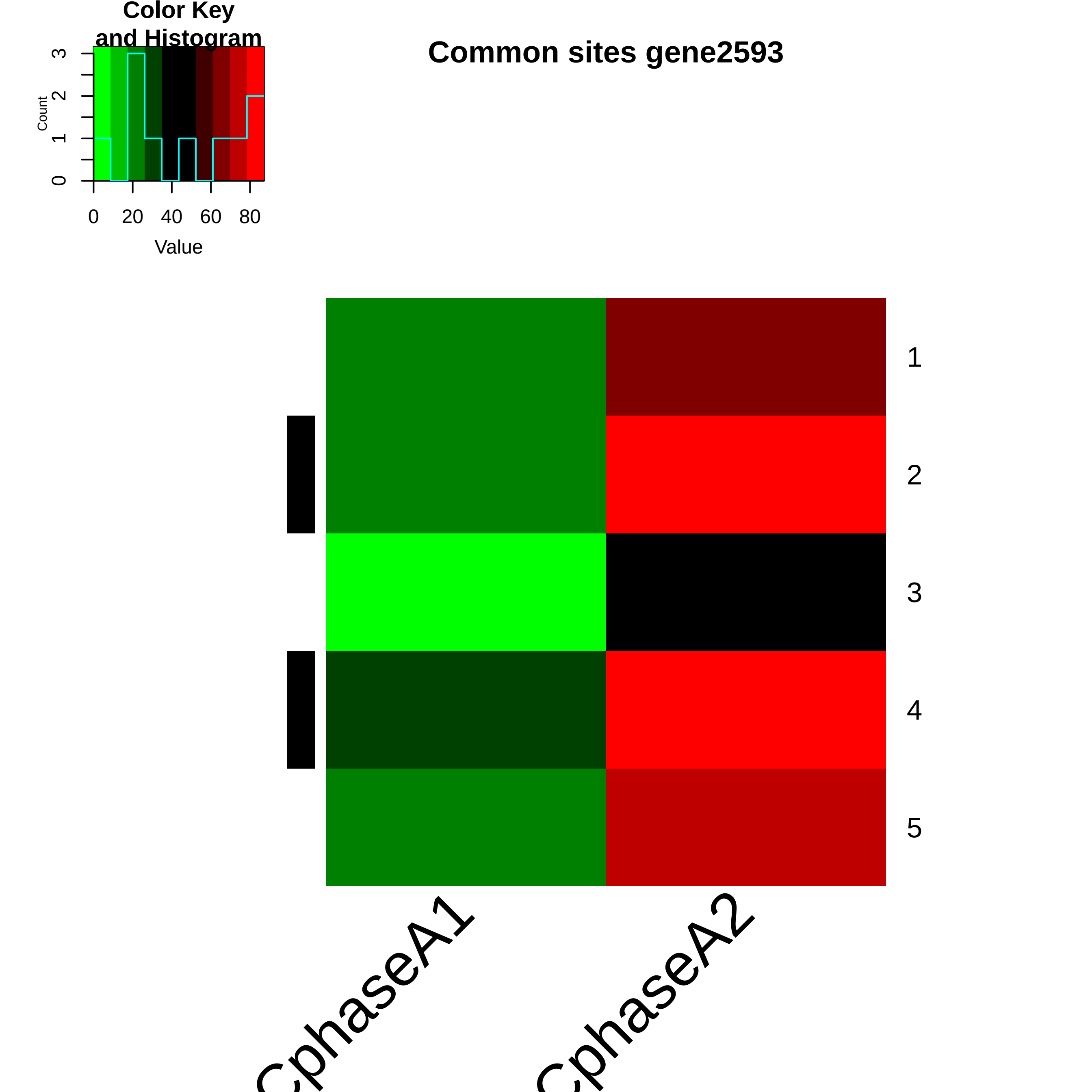

Color Key
and Histogram
Count
0
1
2
3
0
20
40
60
80
Value
Common sites gene2593
1
2
3
4
5
CphaseA1
CphaseA2

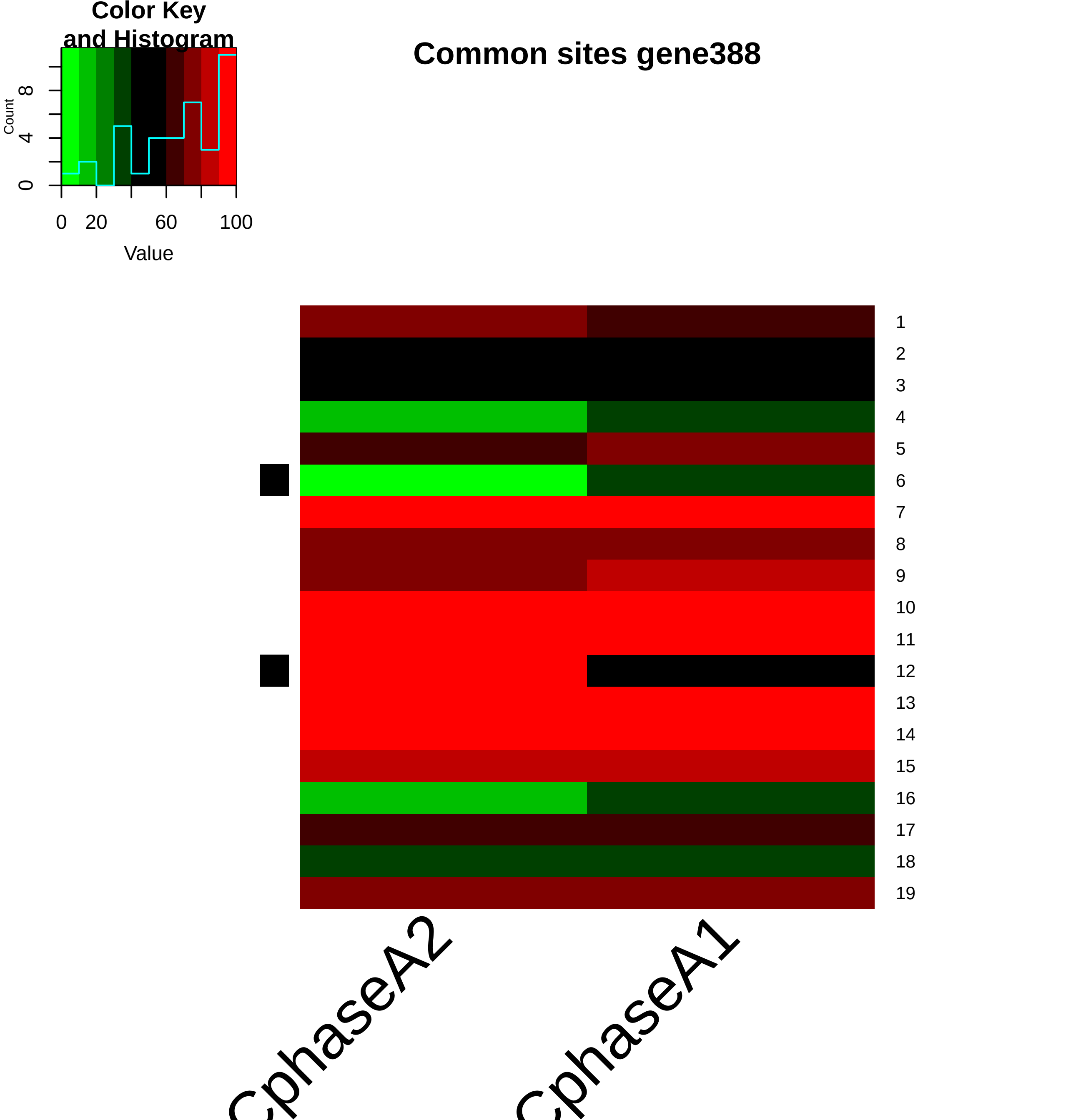

Color Key
and Histogram
Count
0
4
8
0
20
60
100
Value
Common sites gene388
CphaseA2
CphaseA1

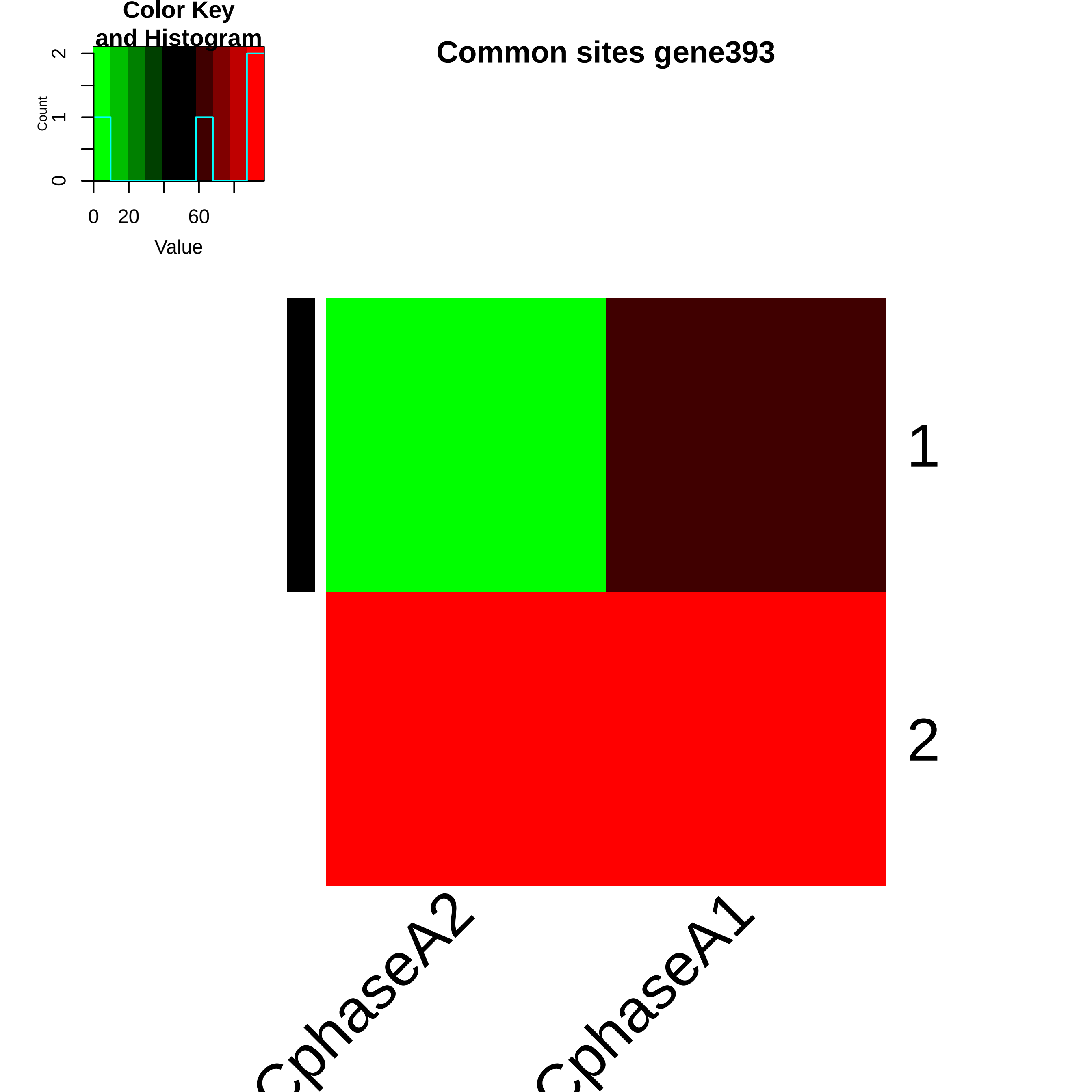

Color Key
and Histogram
Count
0
1
2
0
20
60
Value
Common sites gene393
1
2
CphaseA2
CphaseA1

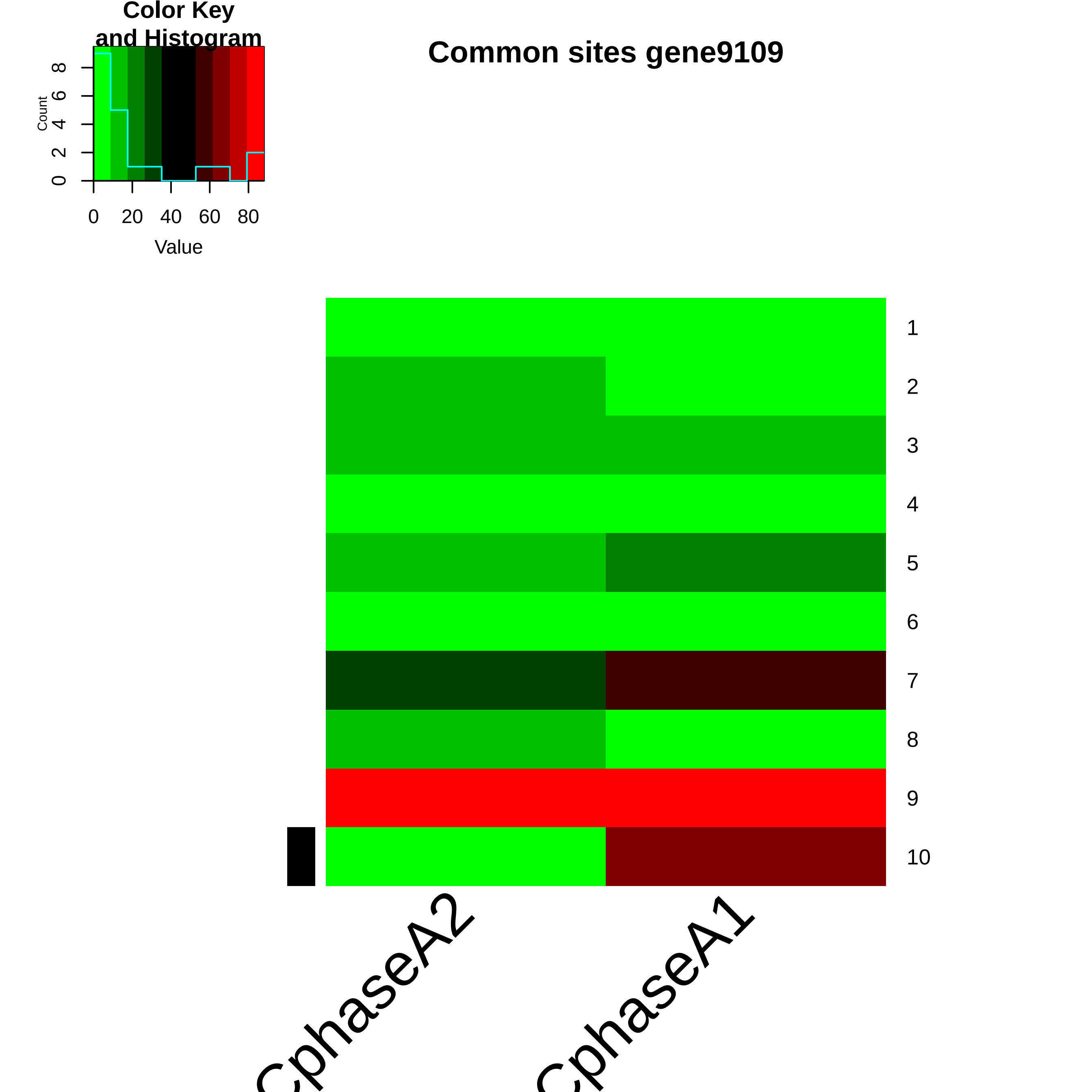
Color Key
and Histogram
Count
0
2
4
6
8
0
20
40
60
80
Value
Common sites gene9109
1
2
3
4
5
6
7
8
9
10
CphaseA2
CphaseA1

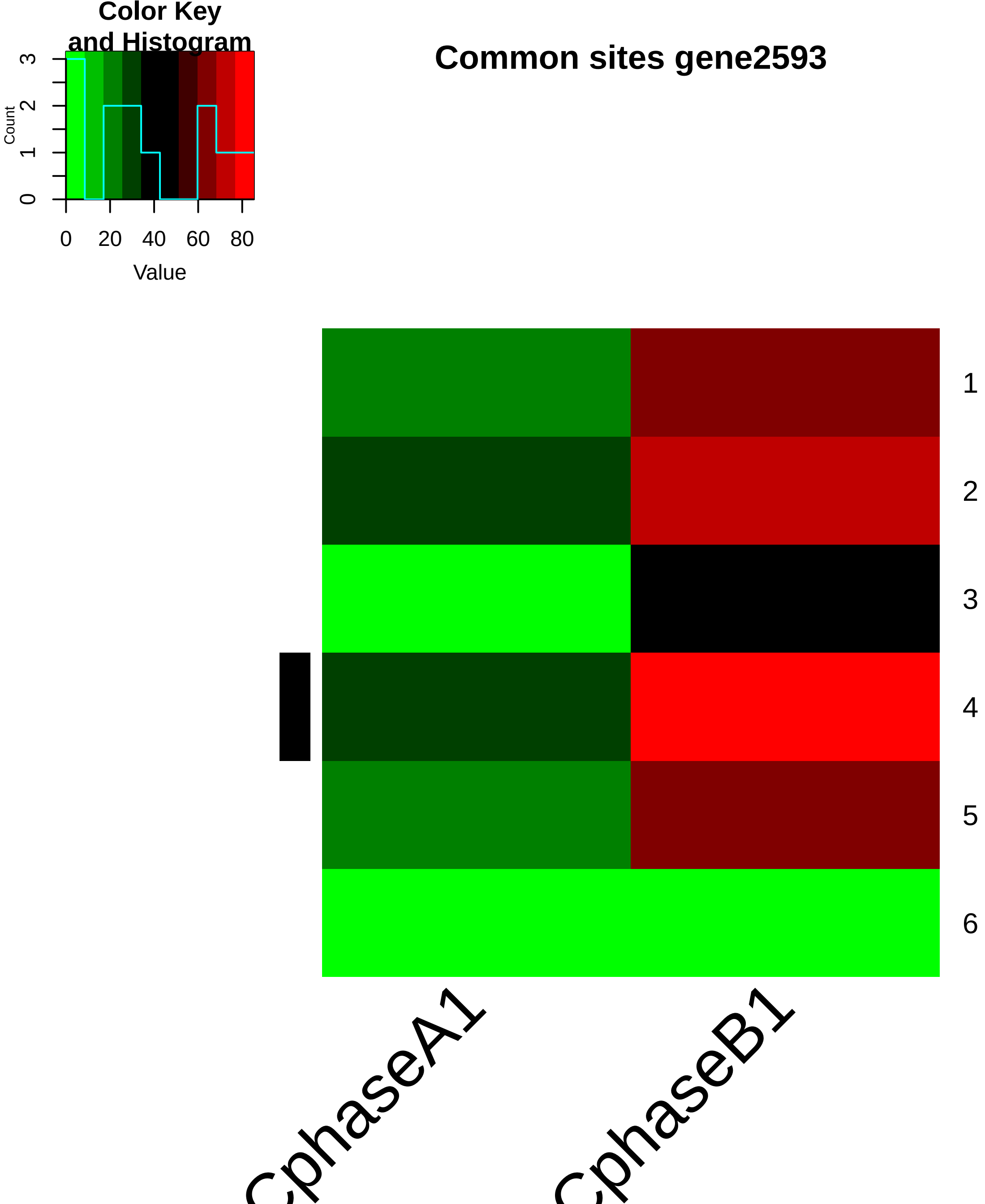

Color Key
and Histogram
Count
0
1
2
3
0
20
40
60
80
Value
Common sites gene2593
1
2
3
4
5
6
CphaseA1
CphaseB1

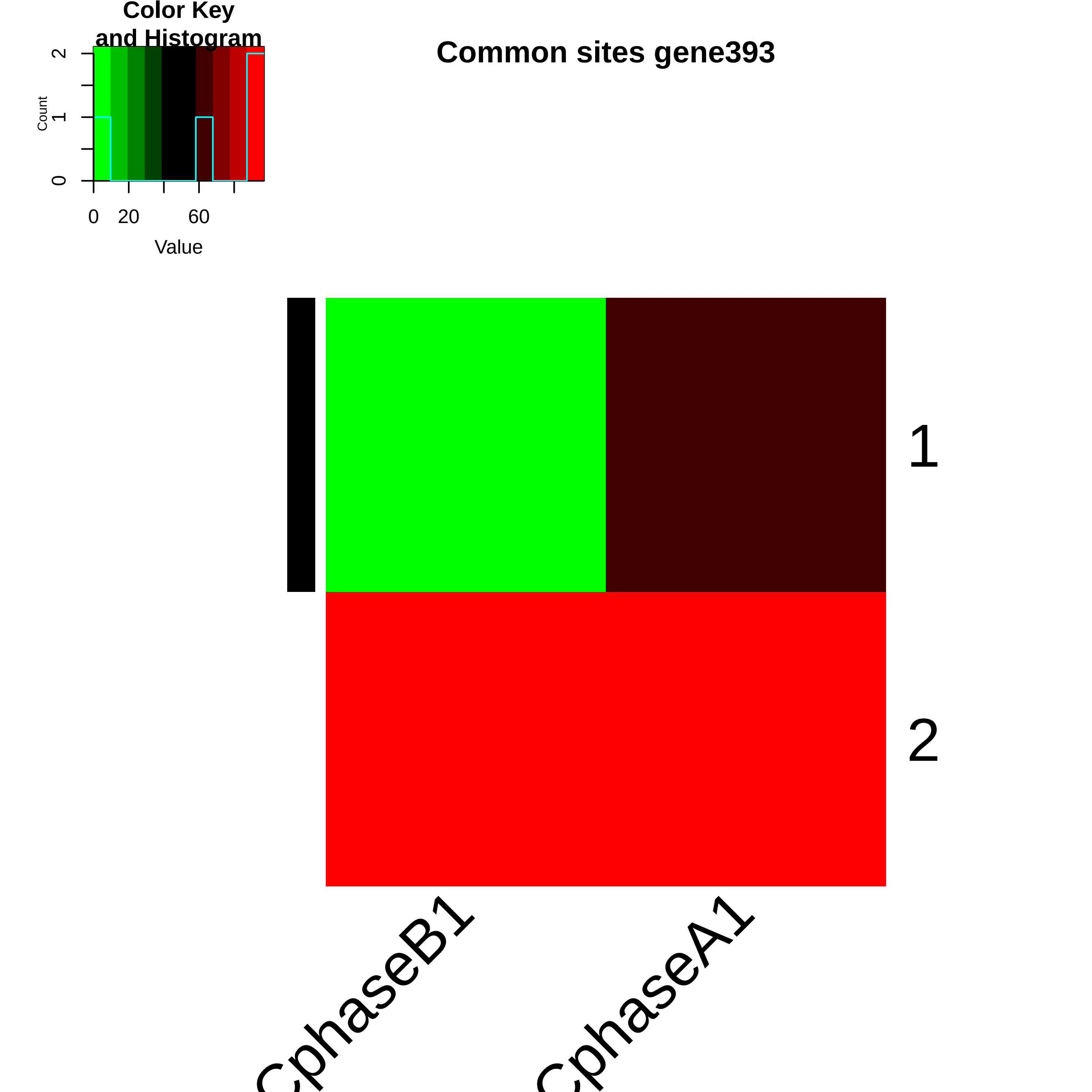

Color Key
and Histogram
Count
0
1
2
0
20
60
Value
Common sites gene393
1
2
CphaseB1
CphaseA1

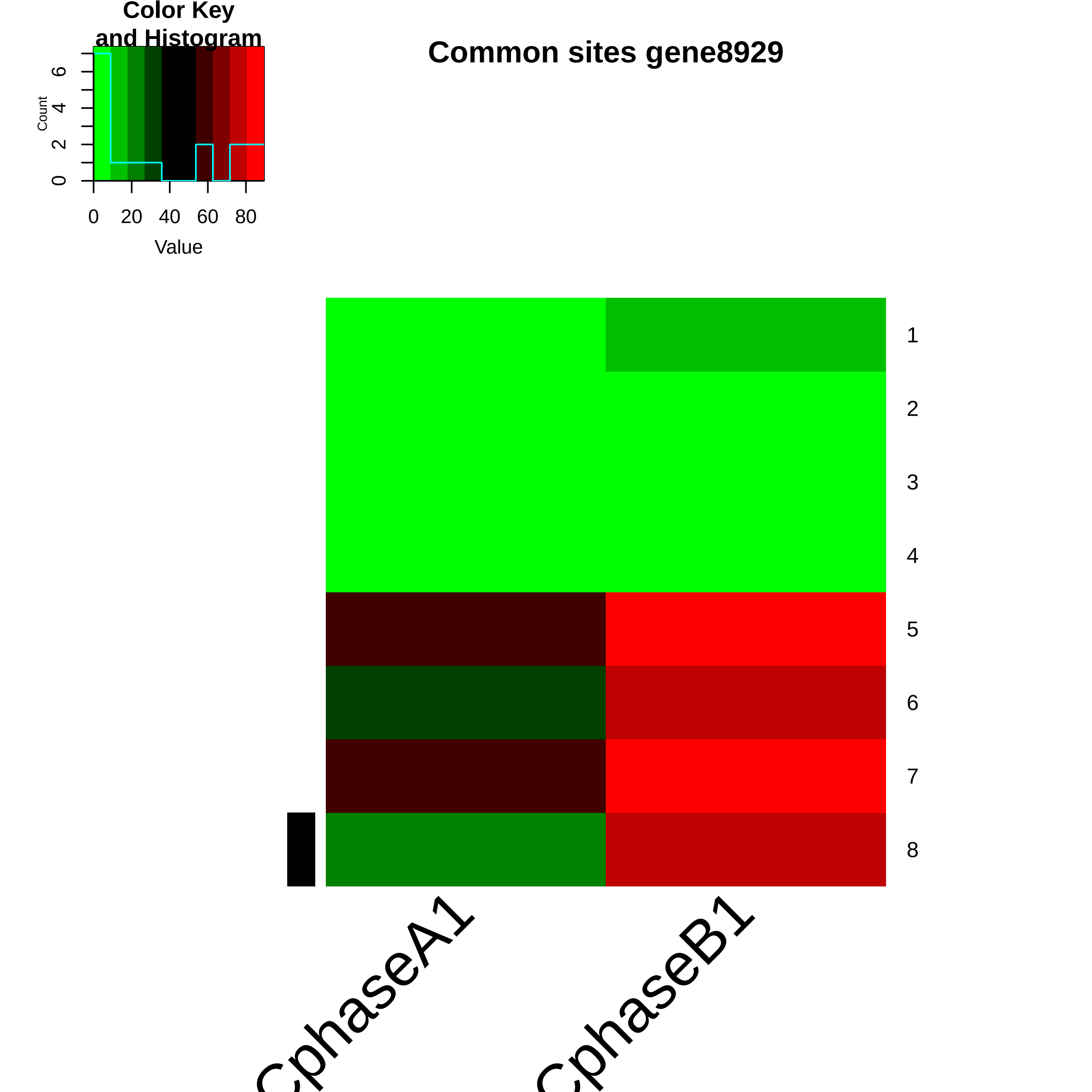

Color Key
and Histogram
Count
0
2
4
6
0
20
40
60
80
Value
Common sites gene8929
1
2
3
4
5
6
7
8
CphaseA1
CphaseB1

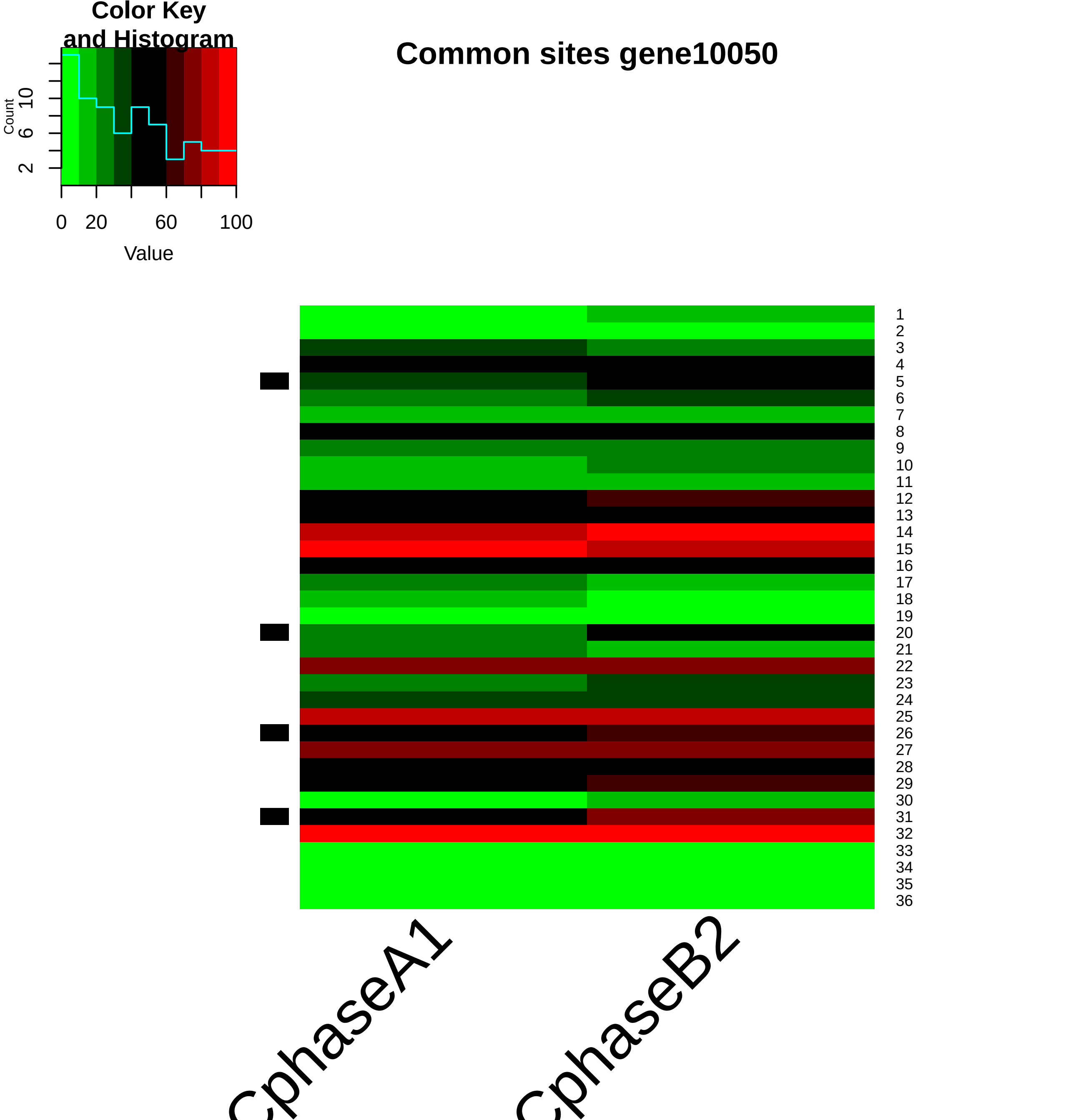

Color Key
and Histogram
Count
2
6
10
0
20
60
100
Value
Common sites gene10050
CphaseA1
CphaseB2

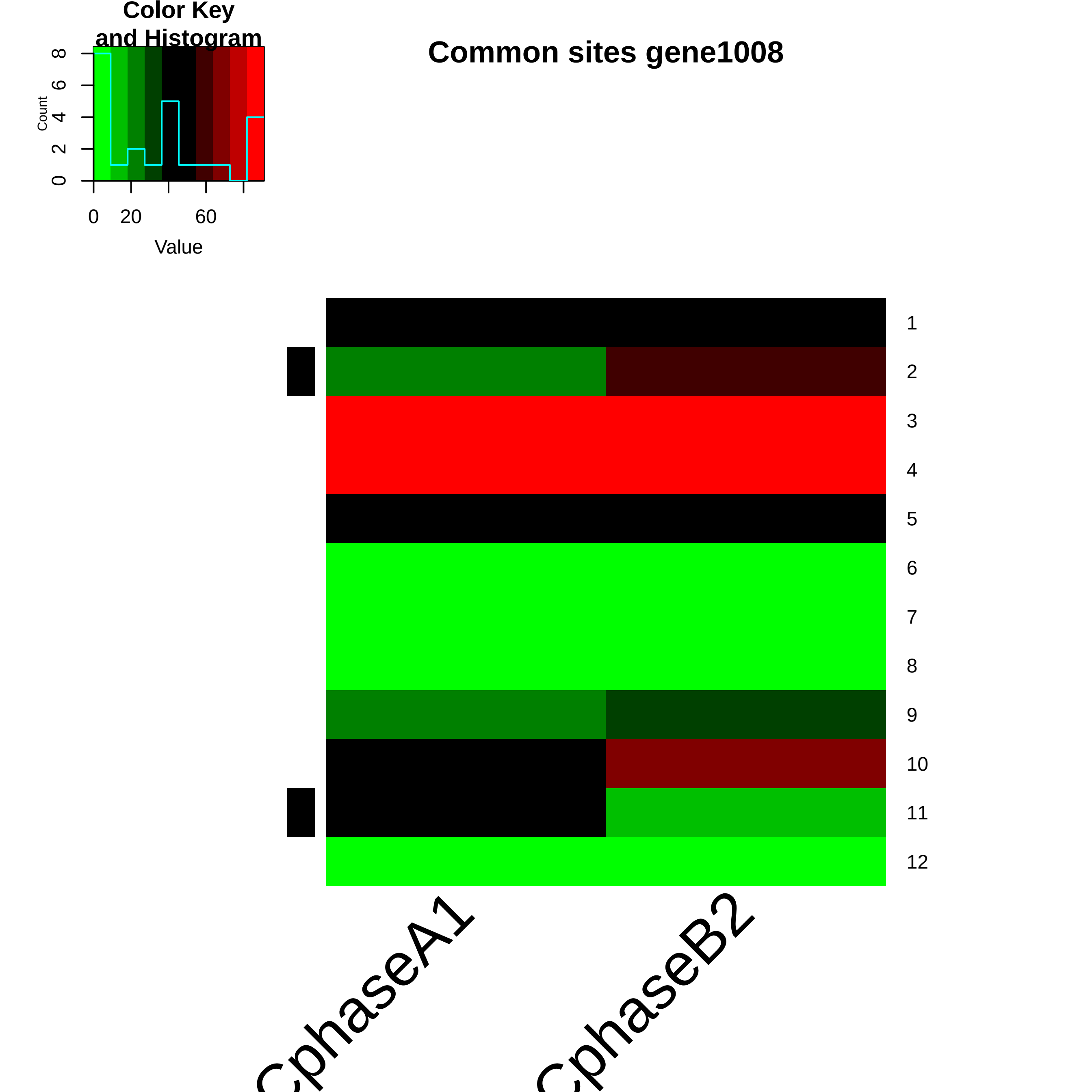
Color Key
and Histogram
Count
0
2
4
6
8
0
20
60
Value
Common sites gene1008
1
2
3
4
5
6
7
8
9
10
11
12
CphaseA1
CphaseB2

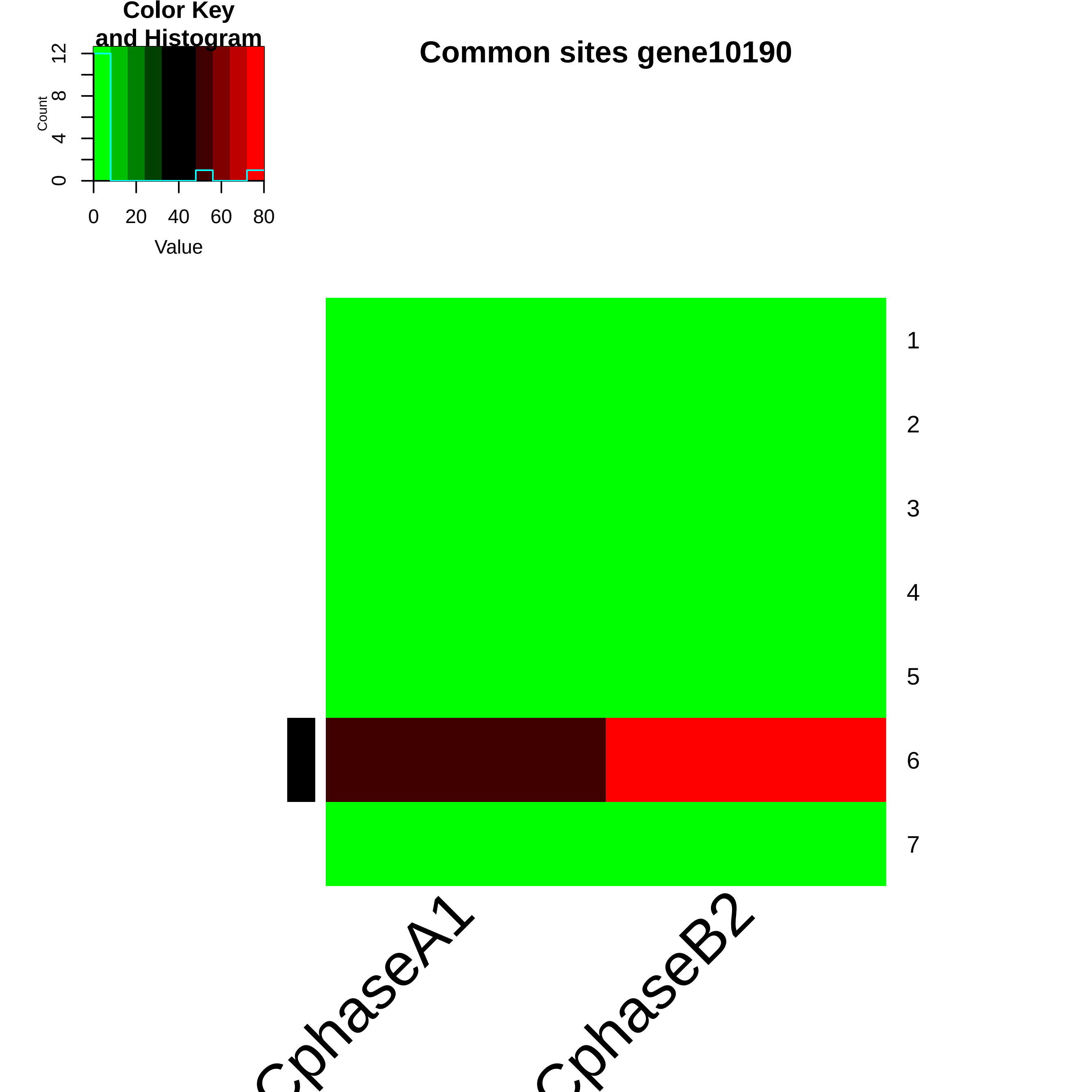
Color Key
and Histogram
Count
0
4
8
12
0
20
40
60
80
Value
Common sites gene10190
1
2
3
4
5
6
7
CphaseA1
CphaseB2

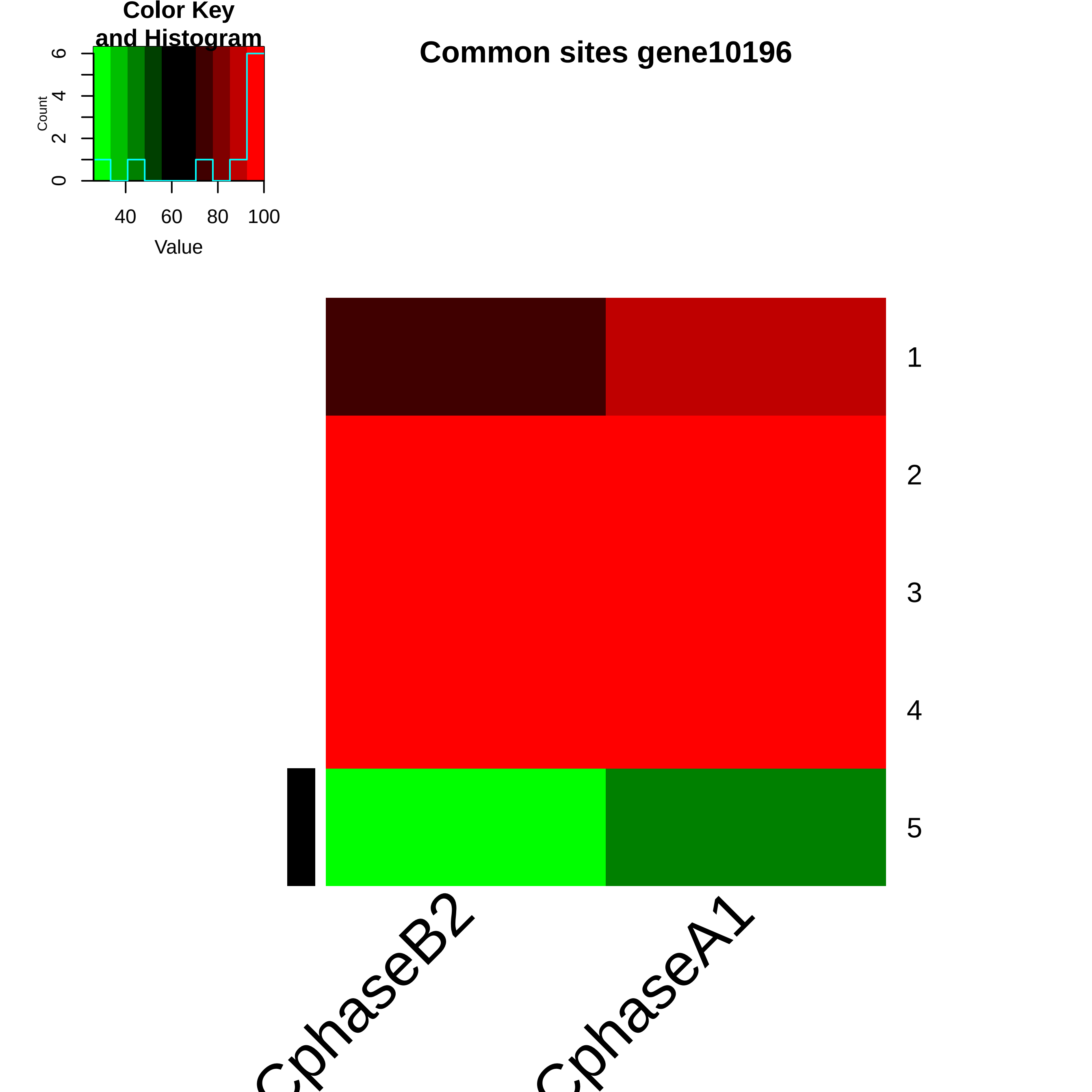

Color Key
and Histogram
Count
0
2
4
6
40
60
80
100
Value
Common sites gene10196
1
2
3
4
5
CphaseB2
CphaseA1

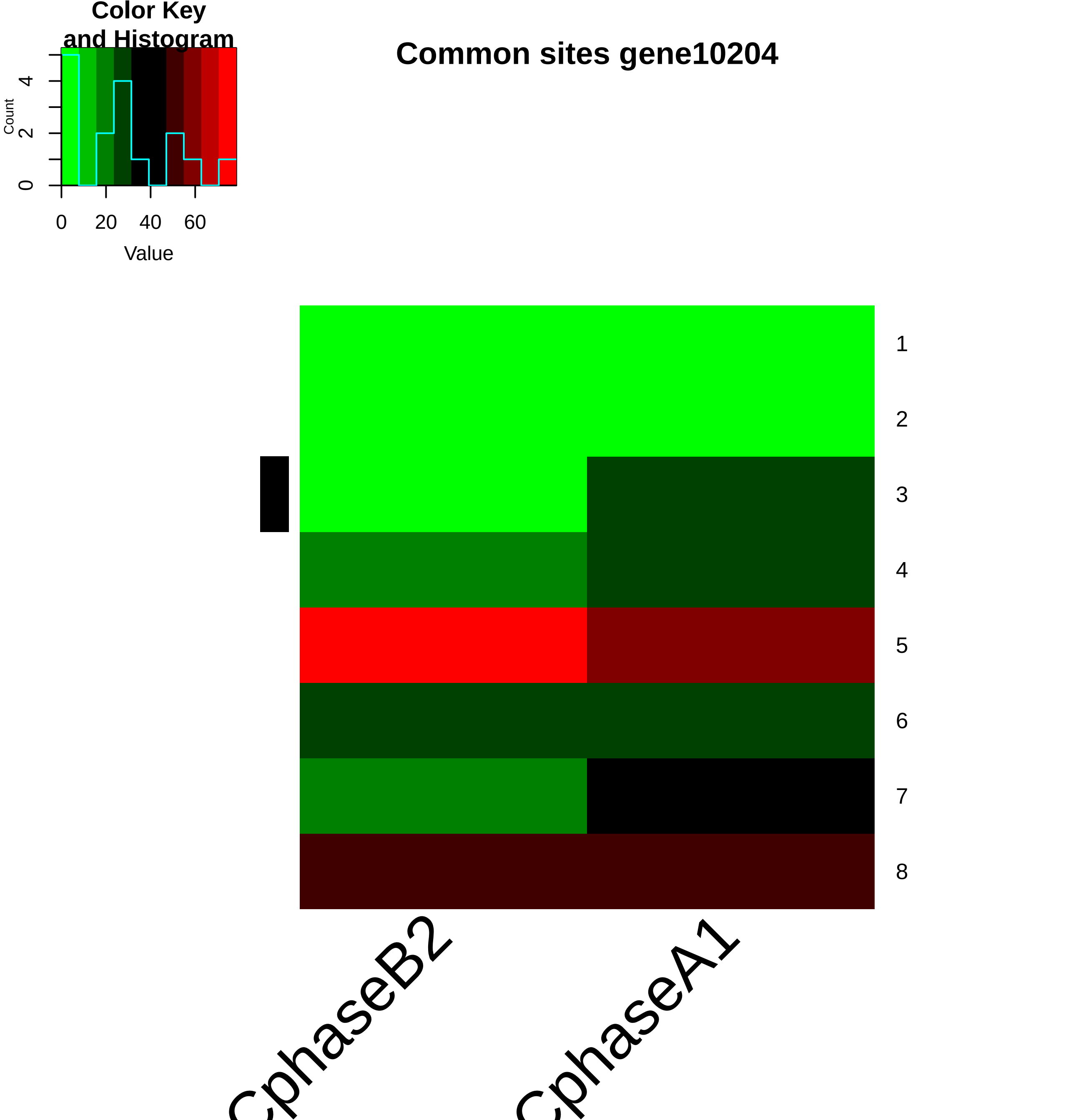
Color Key
and Histogram
Count
0
2
4
0
20
40
60
Value
Common sites gene10204
1
2
3
4
5
6
7
8
CphaseB2
CphaseA1

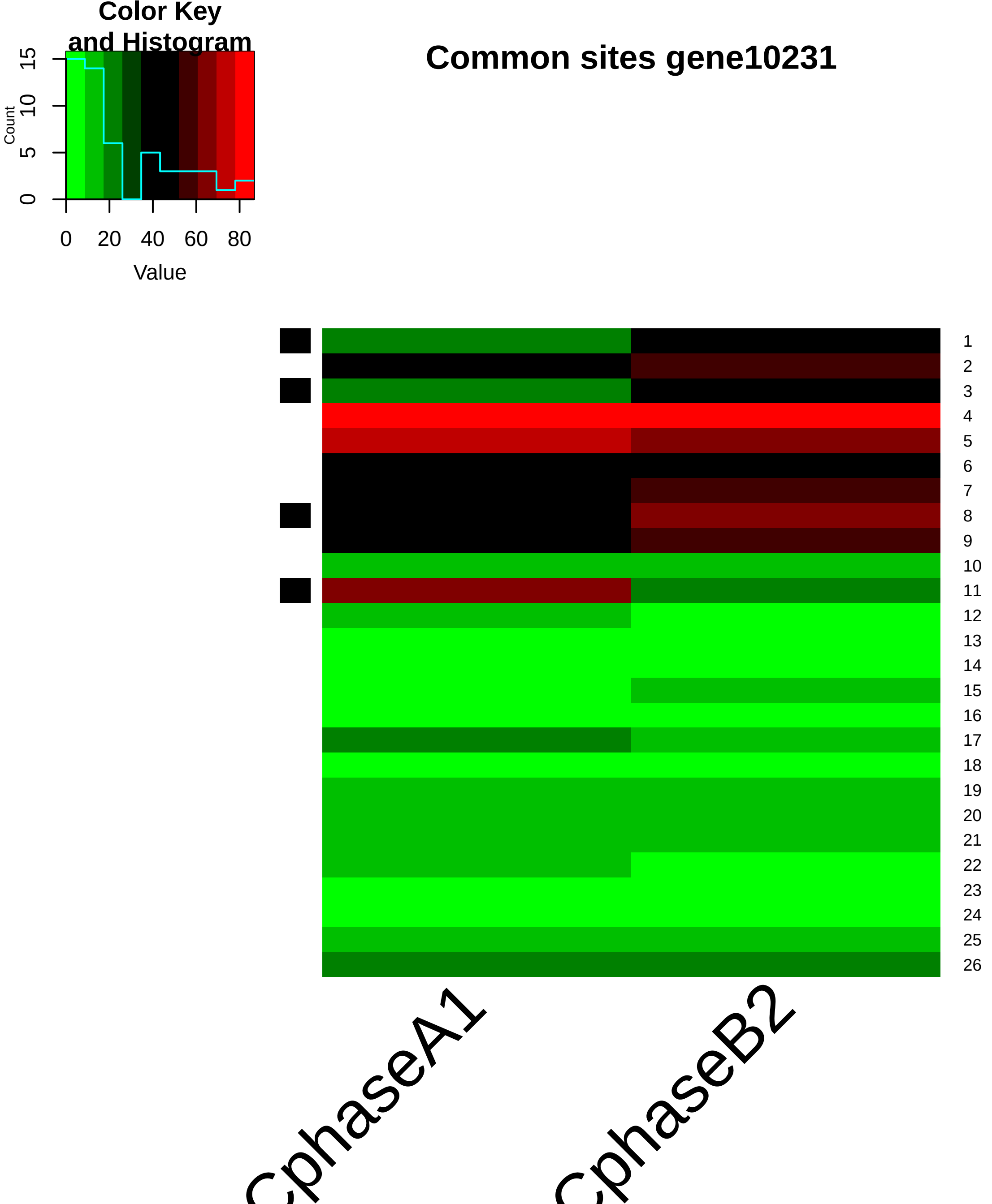

Color Key
and Histogram
Count
0
5
10
15
0
20
40
60
80
Value
Common sites gene10231
CphaseA1
CphaseB2

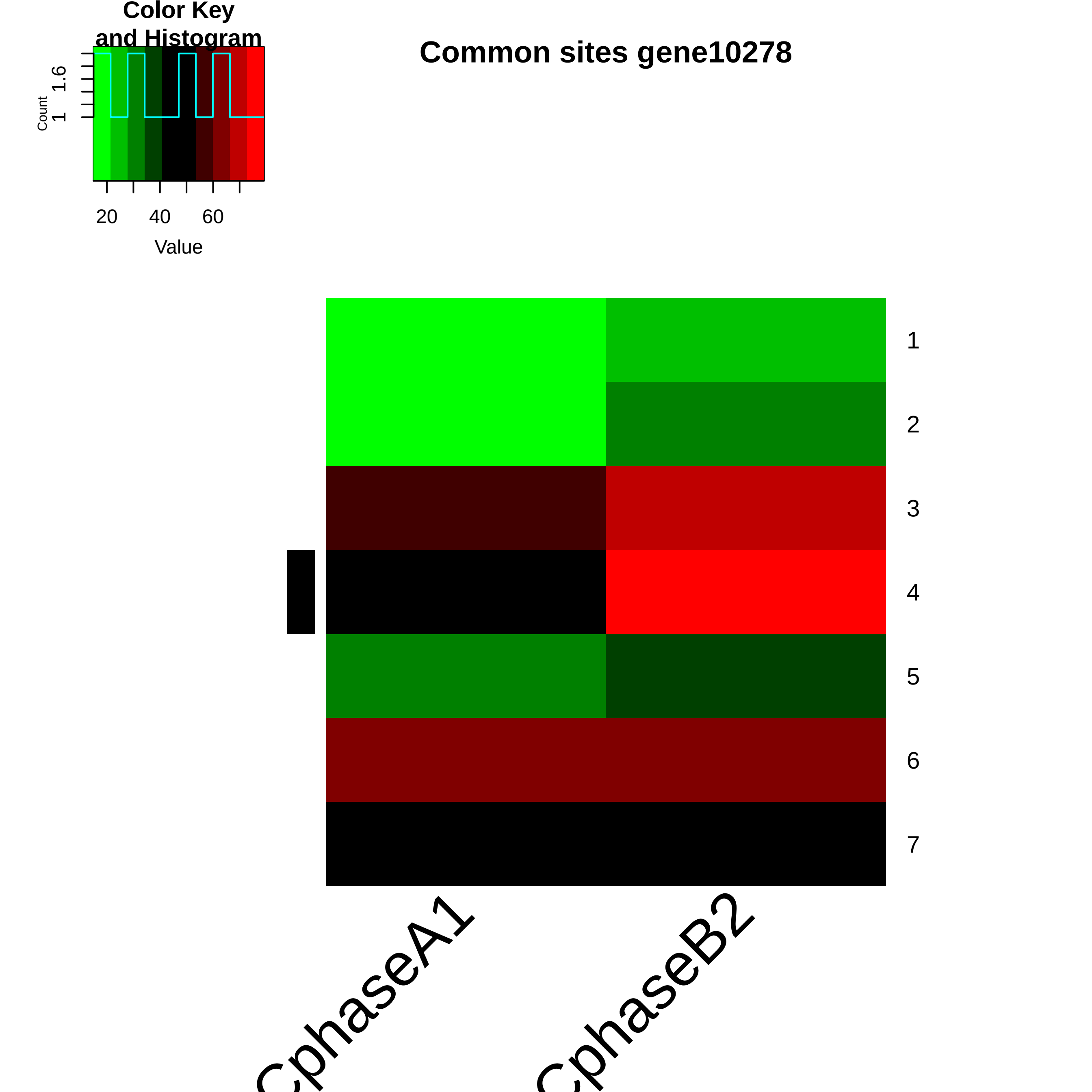

Color Key
and Histogram
Count
1
1.6
20
40
60
Value
Common sites gene10278
1
2
3
4
5
6
7
CphaseA1
CphaseB2

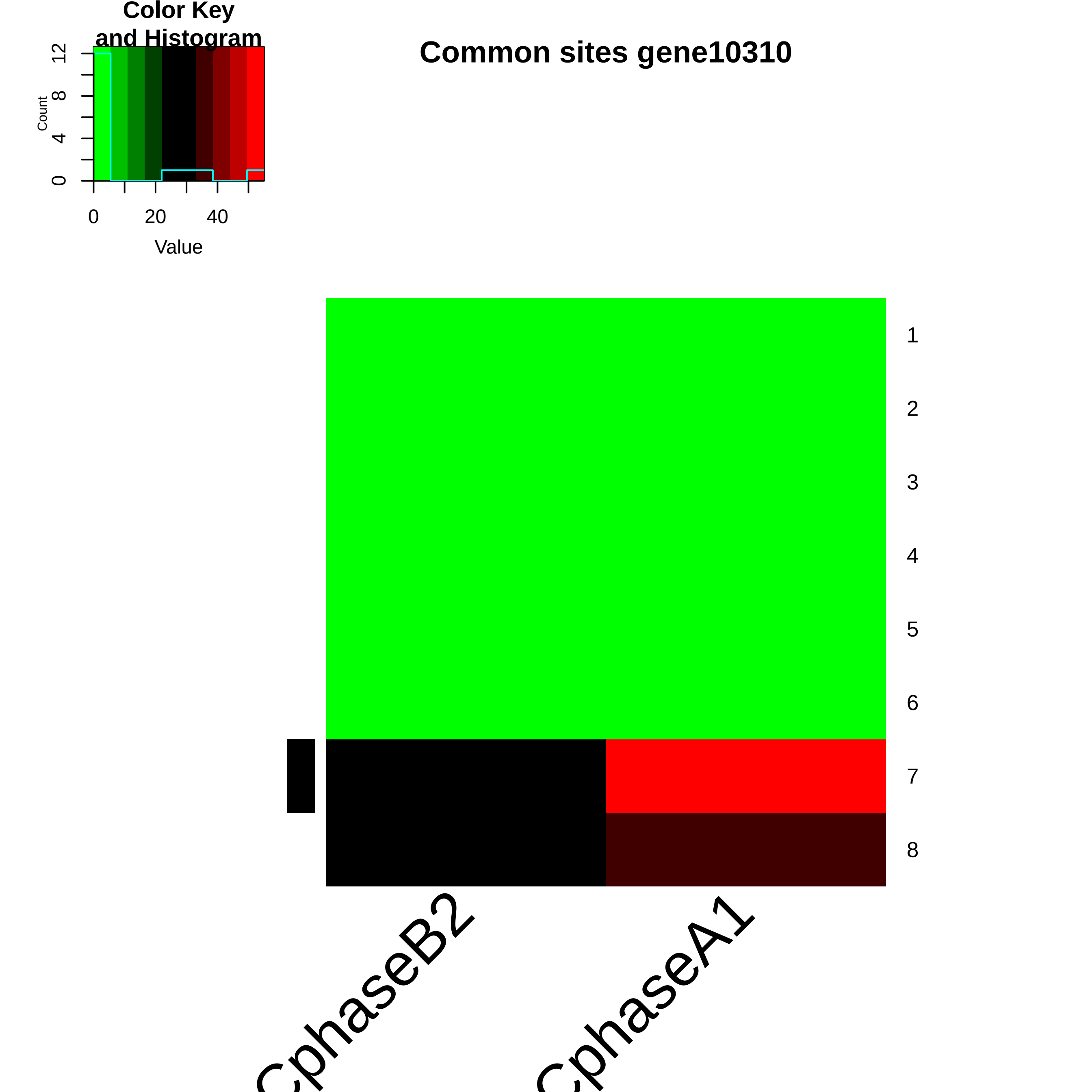
Color Key
and Histogram
Count
0
4
8
12
0
20
40
Value
Common sites gene10310
1
2
3
4
5
6
7
8
CphaseB2
CphaseA1

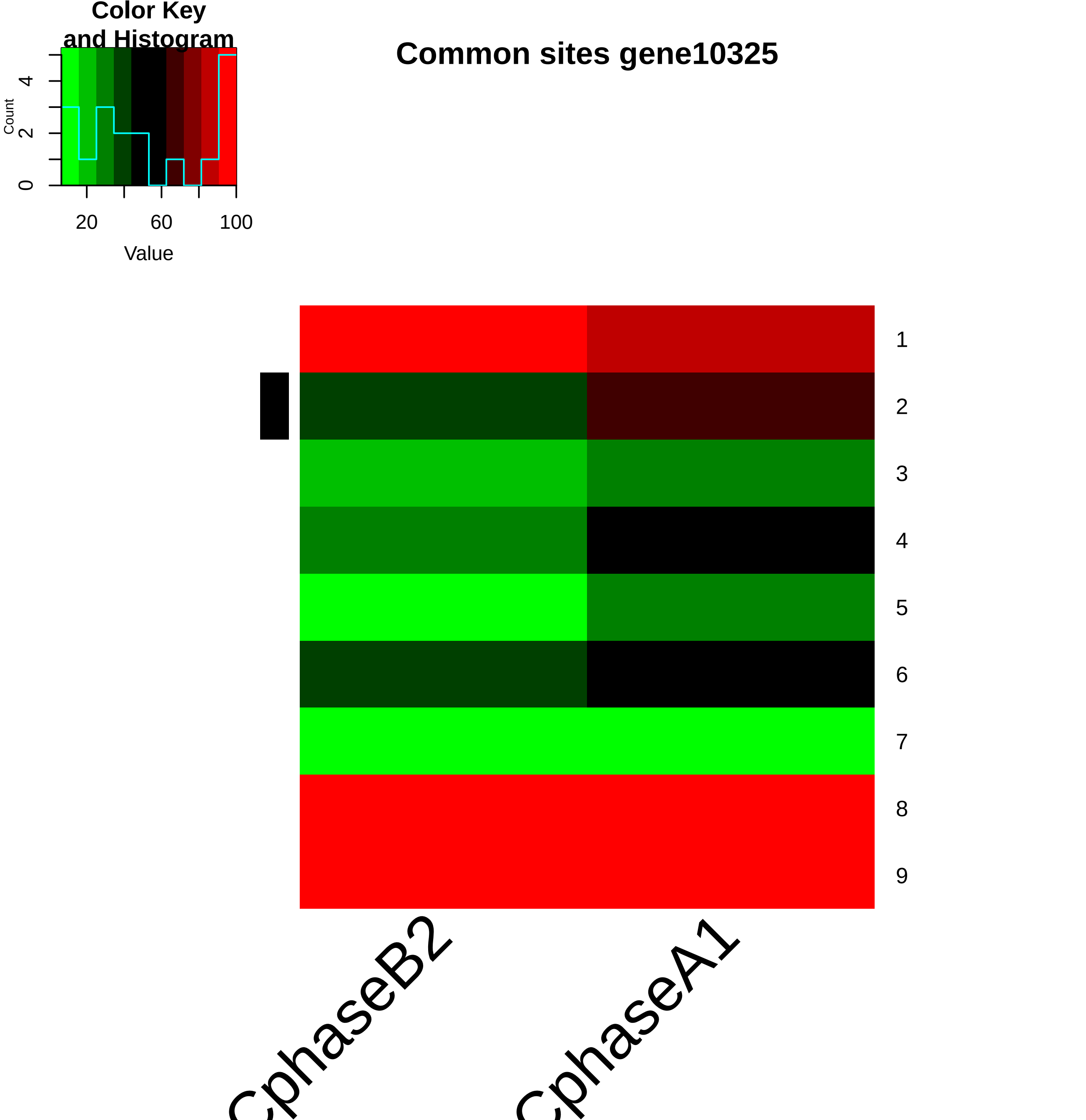
Color Key
and Histogram
Count
0
2
4
20
60
100
Value
Common sites gene10325
1
2
3
4
5
6
7
8
9
CphaseB2
CphaseA1

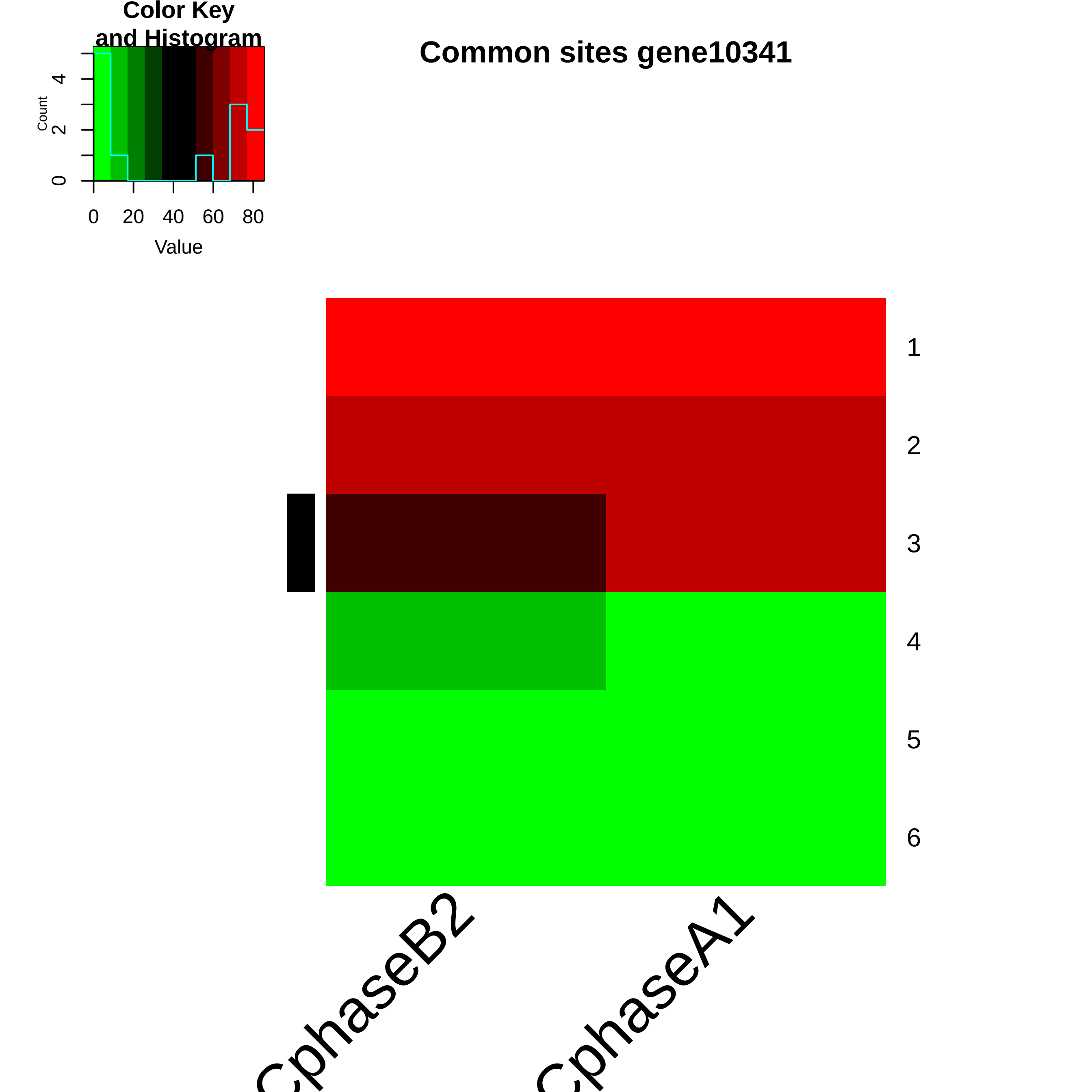

Color Key
and Histogram
Count
0
2
4
0
20
40
60
80
Value
Common sites gene10341
1
2
3
4
5
6
CphaseB2
CphaseA1

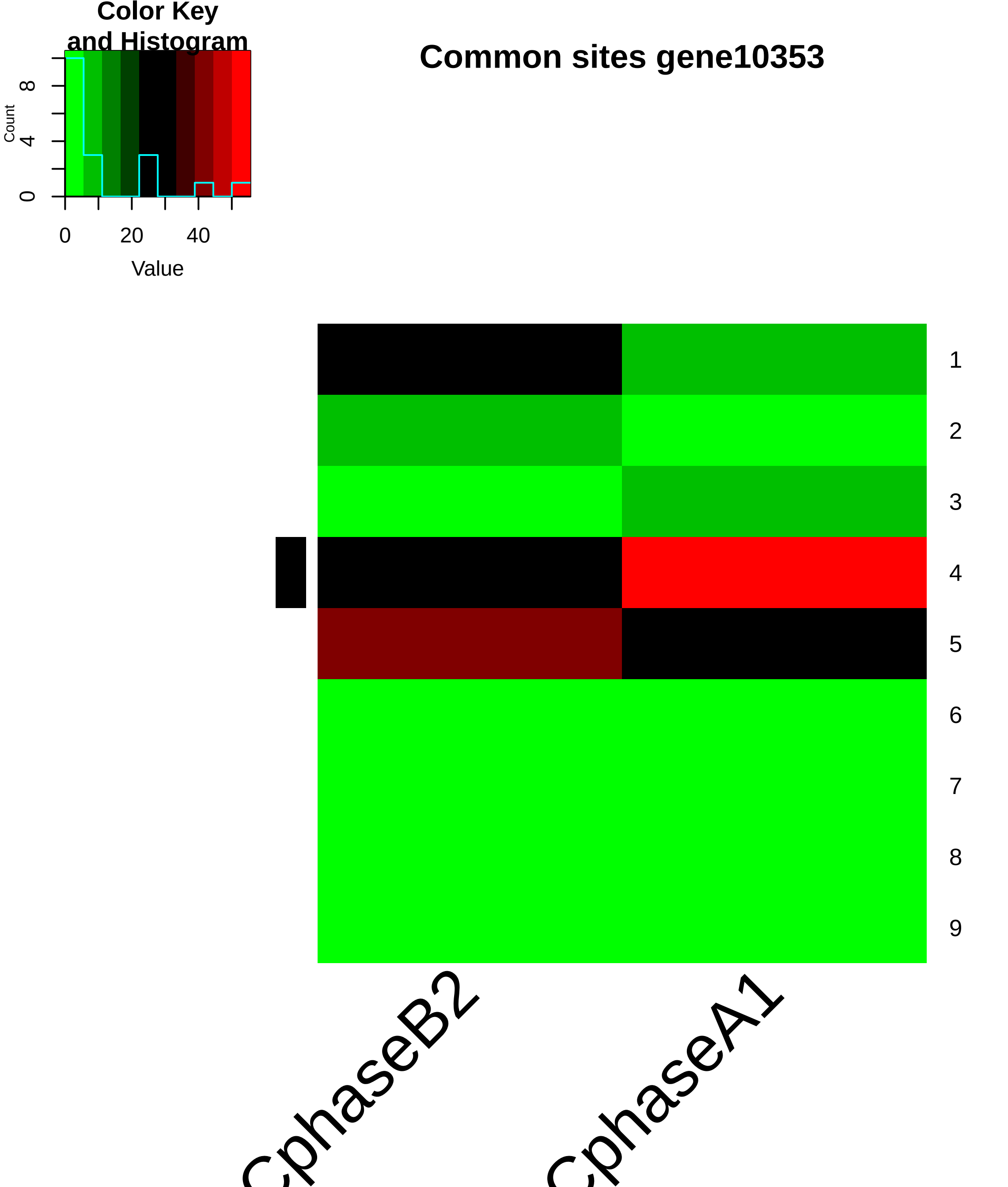
Color Key
and Histogram
Count
0
4
8
0
20
40
Value
Common sites gene10353
1
2
3
4
5
6
7
8
9
CphaseB2
CphaseA1

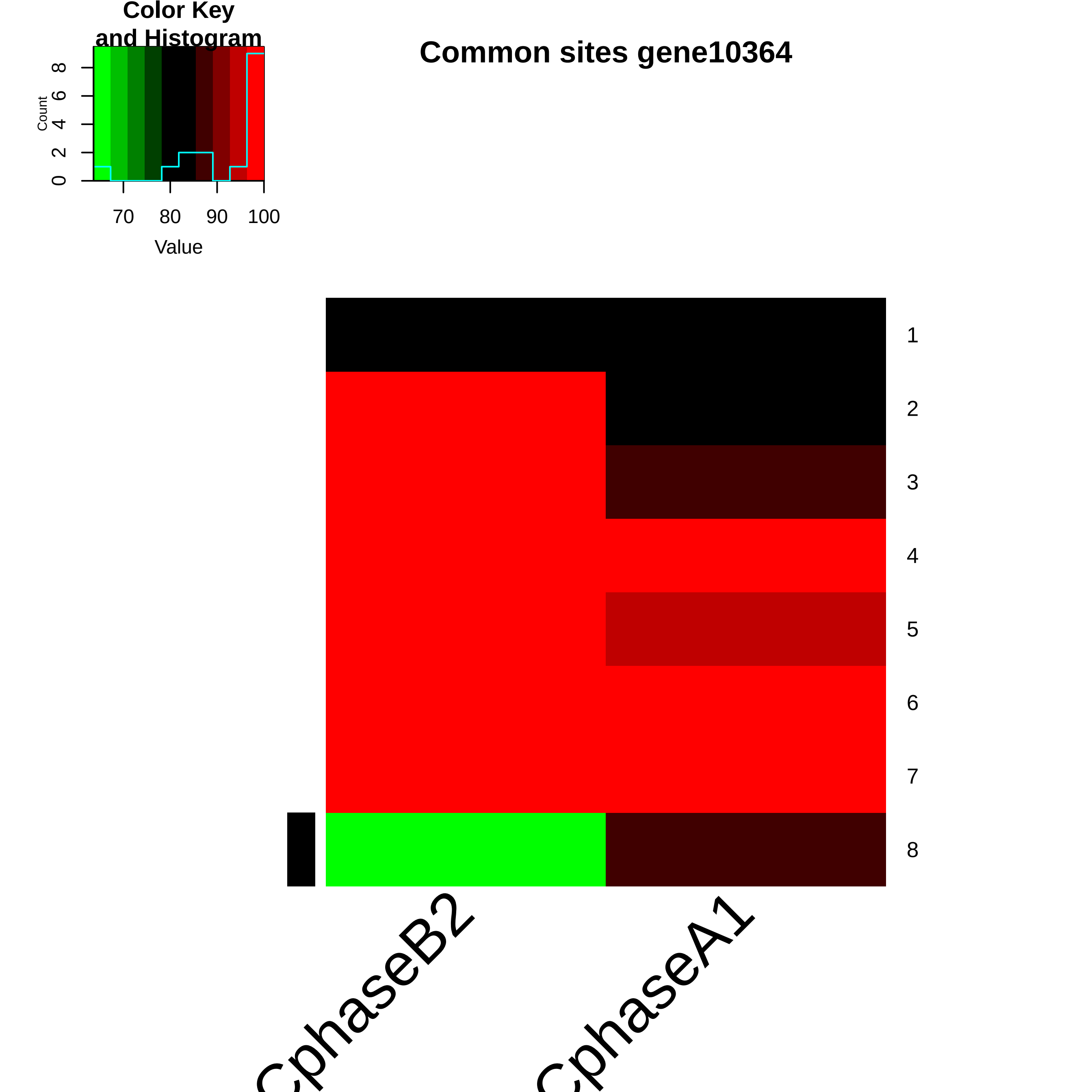
Color Key
and Histogram
Count
0 2 4 6 8
70 80 90 100
Value
Common sites gene10364
1
2
3
4
5
6
7
8
CphaseB2
CphaseA1

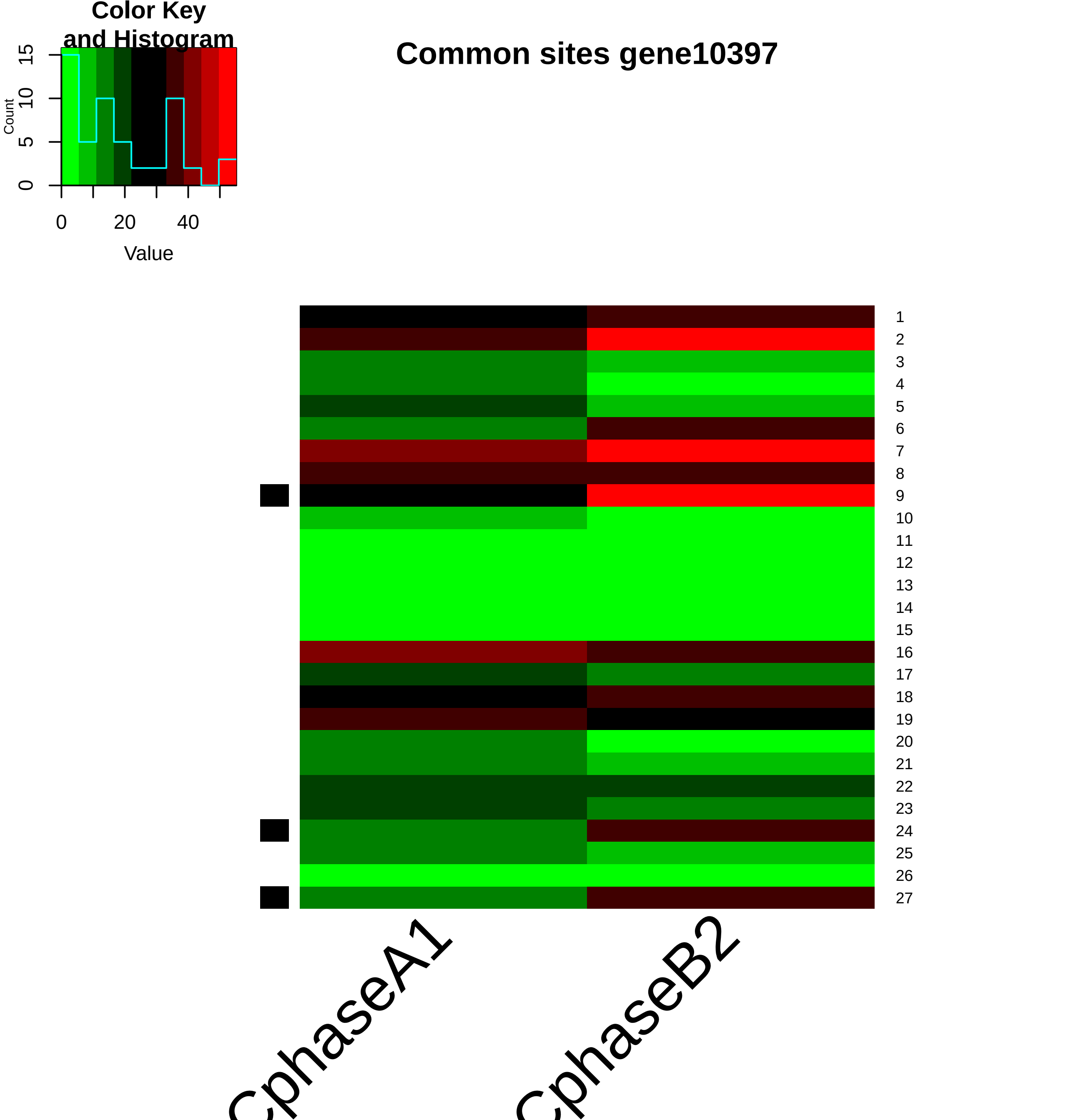
Color Key
and Histogram
Count
Value
Common sites gene10397
CphaseA1
CphaseB2

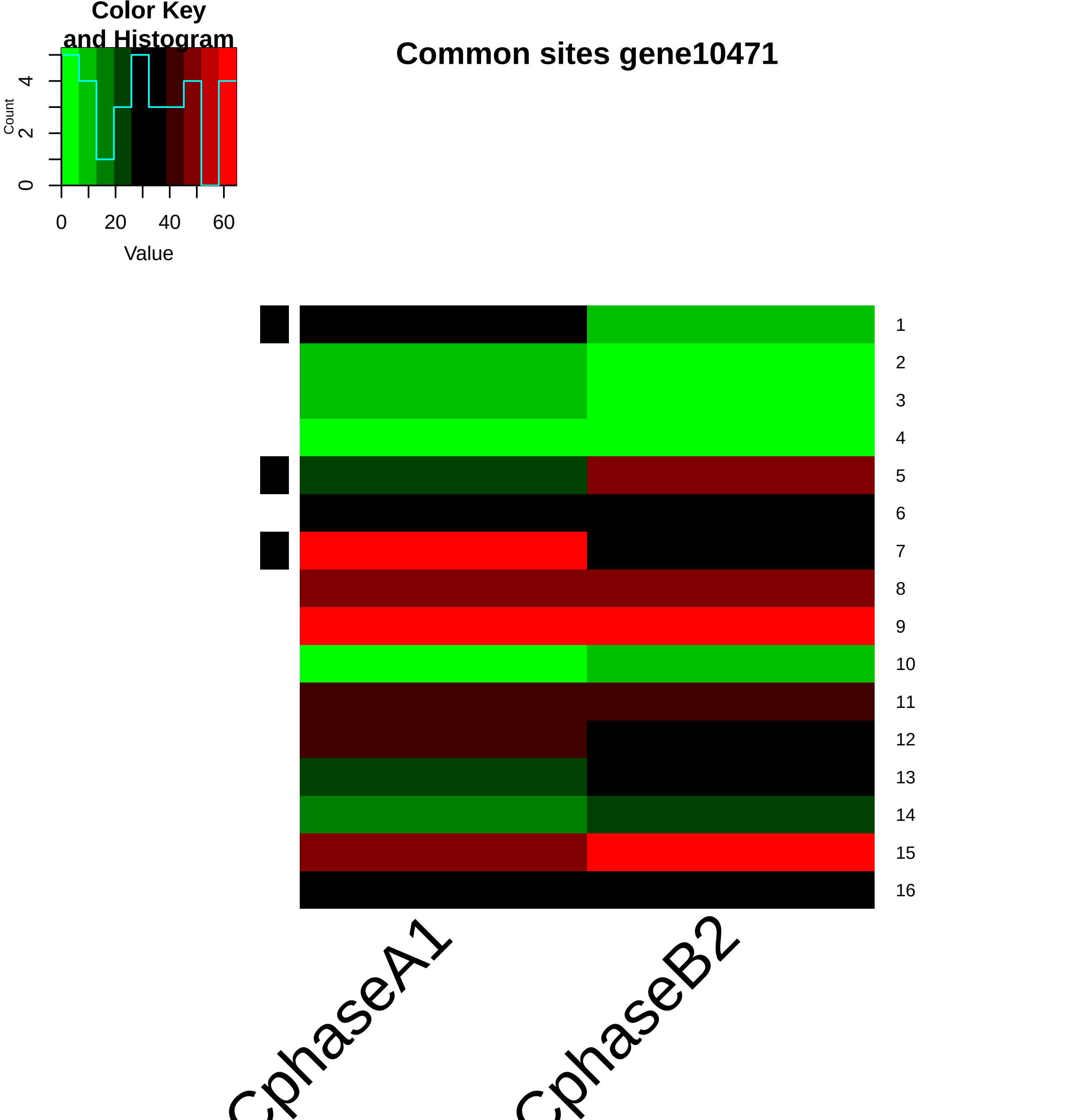
Color Key
and Histogram
Count
0
2
4
0
20
40
60
Value
Common sites gene10471
CphaseA1
CphaseB2

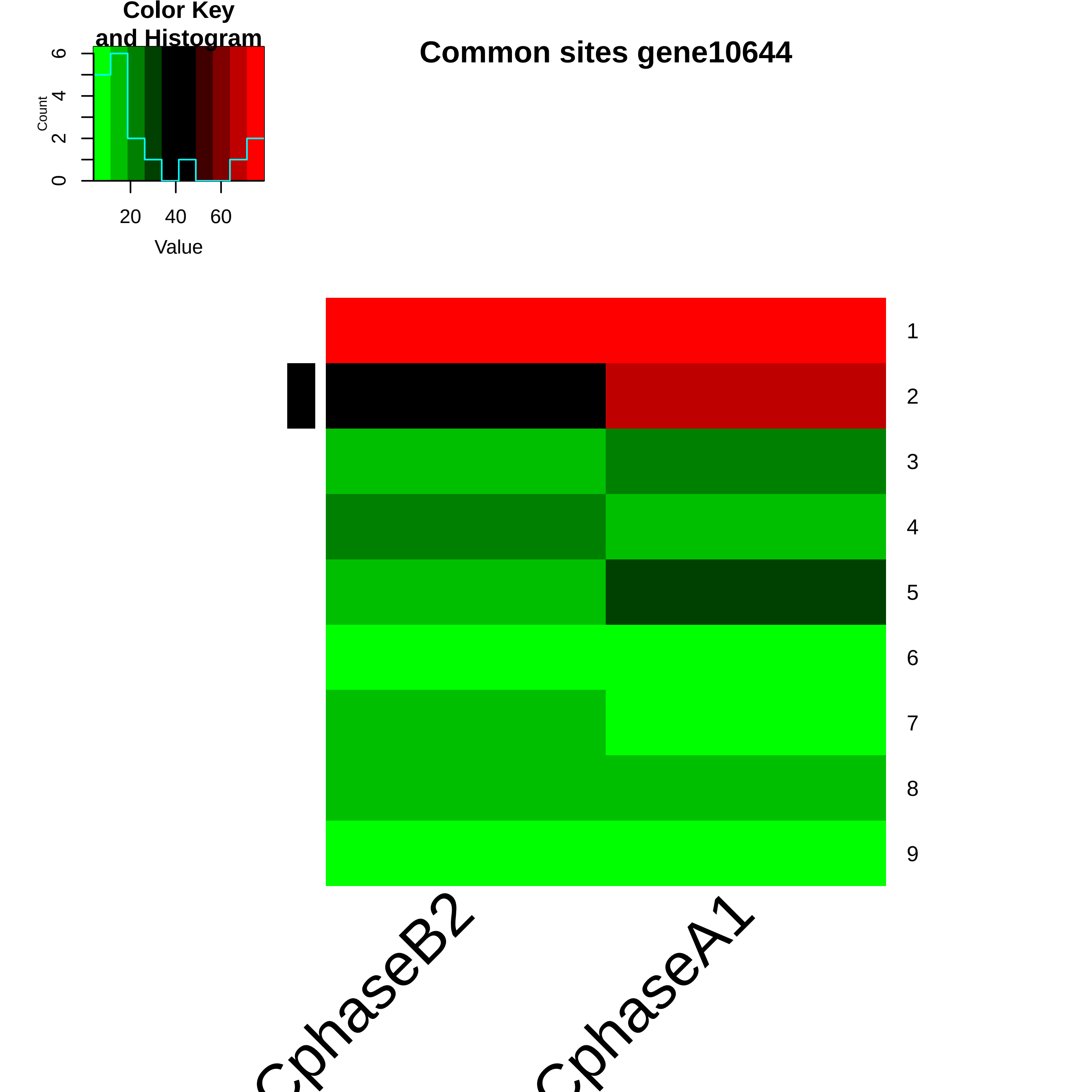
Color Key
and Histogram
Count
0
2
4
6
20
40
60
Value
Common sites gene10644
1
2
3
4
5
6
7
8
9
CphaseB2
CphaseA1

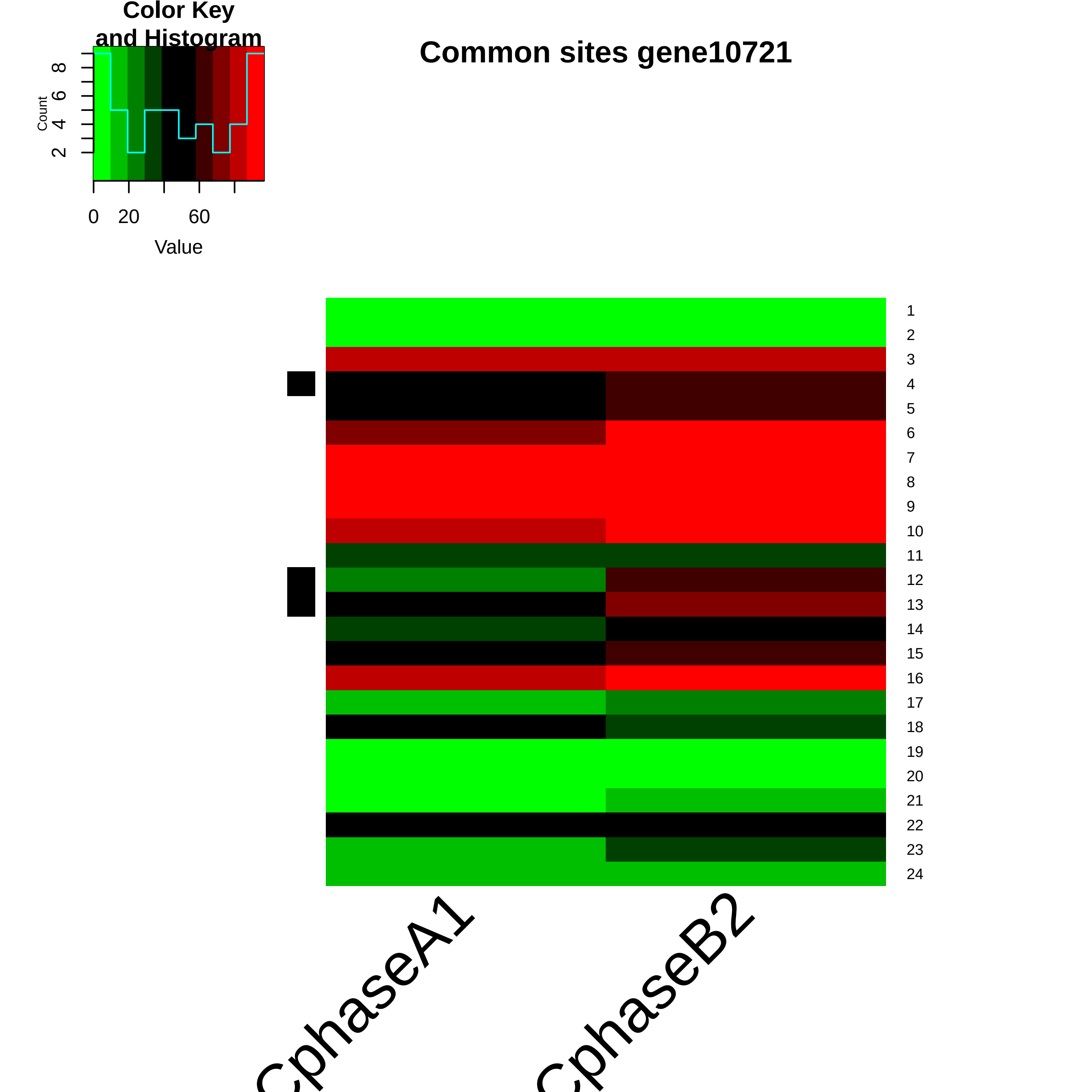

Color Key
and Histogram
Count
2
4
6
8
0
20
60
Value
Common sites gene10721
CphaseA1
CphaseB2

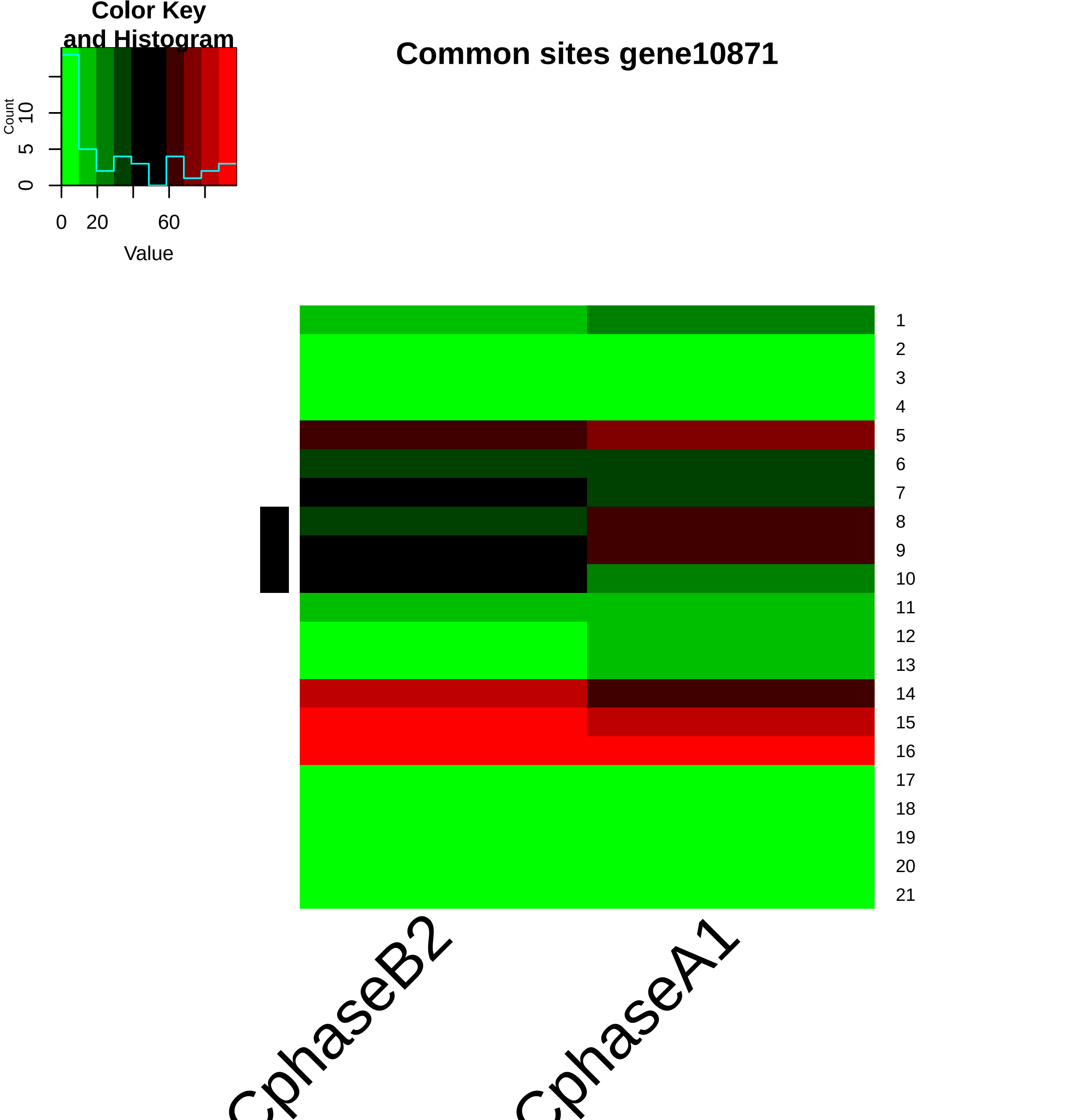

Color Key
and Histogram
Count
0
5
10
0
20
60
Value
Common sites gene10871
CphaseB2
CphaseA1

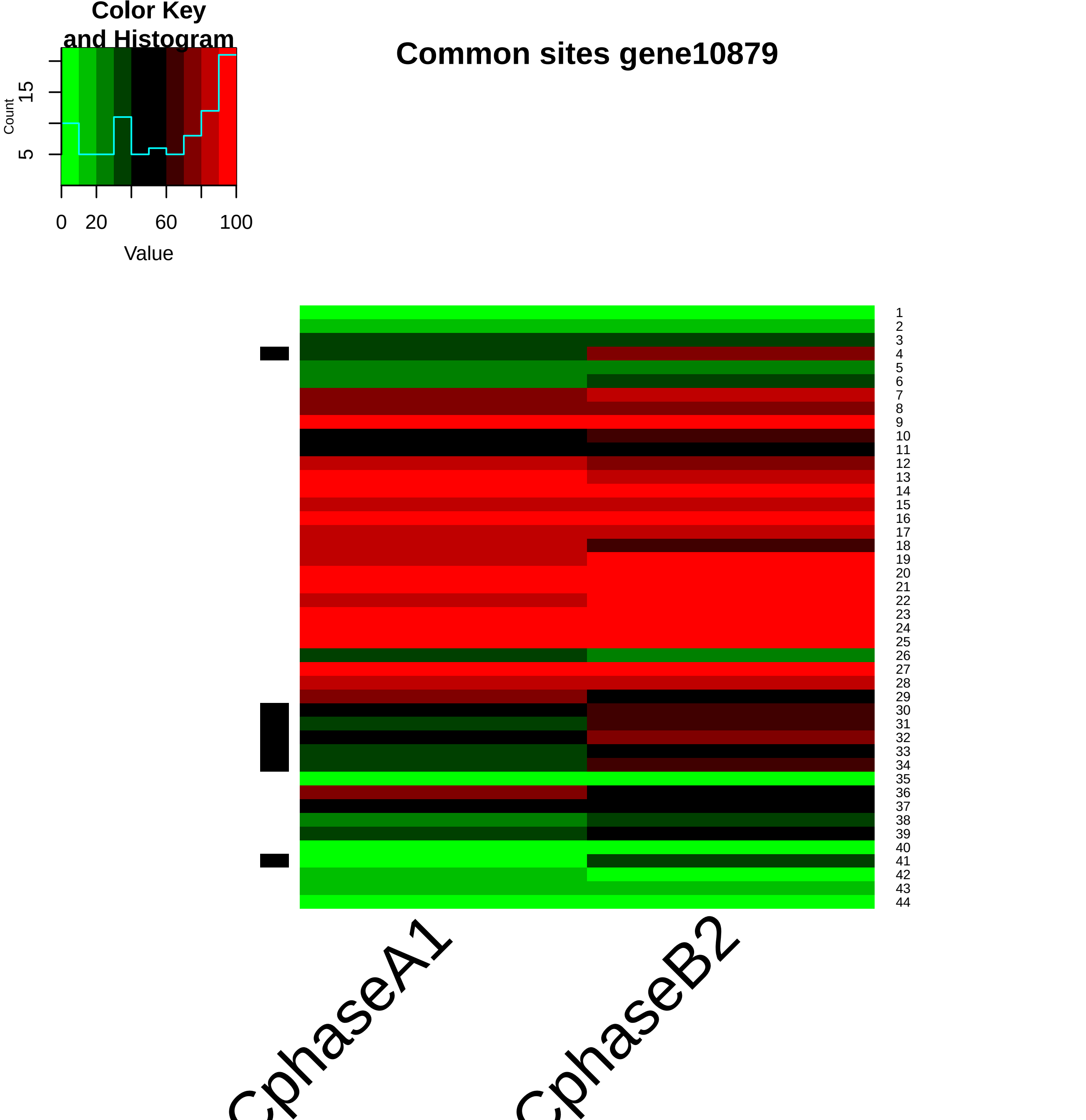
Color Key
and Histogram
Count
5
15
0
20
60
100
Value
Common sites gene10879
CphaseA1
CphaseB2

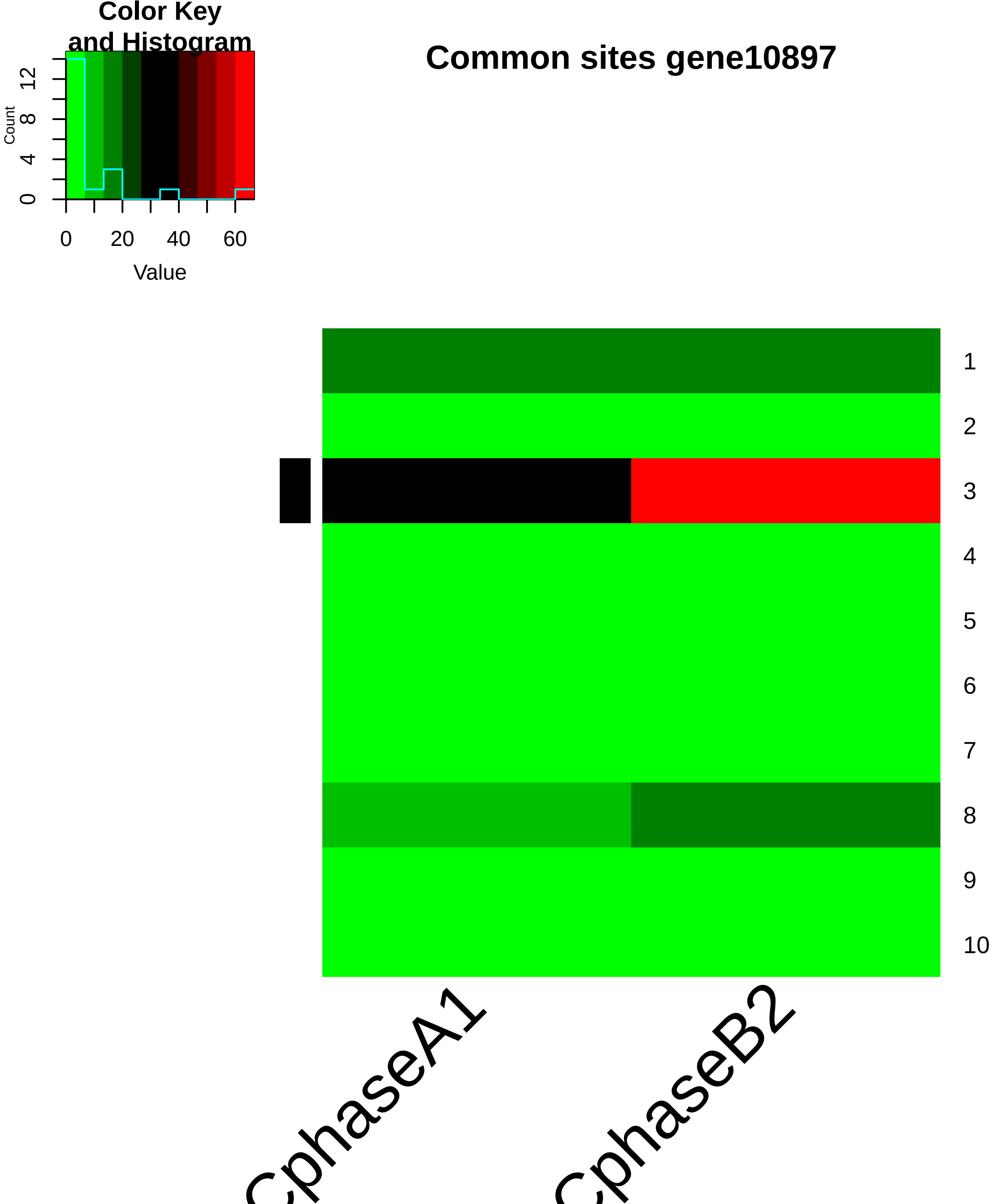
Color Key
and Histogram
Count
0
4
8
12
0
20
40
60
Value
Common sites gene10897
1
2
3
4
5
6
7
8
9
10
CphaseA1
CphaseB2

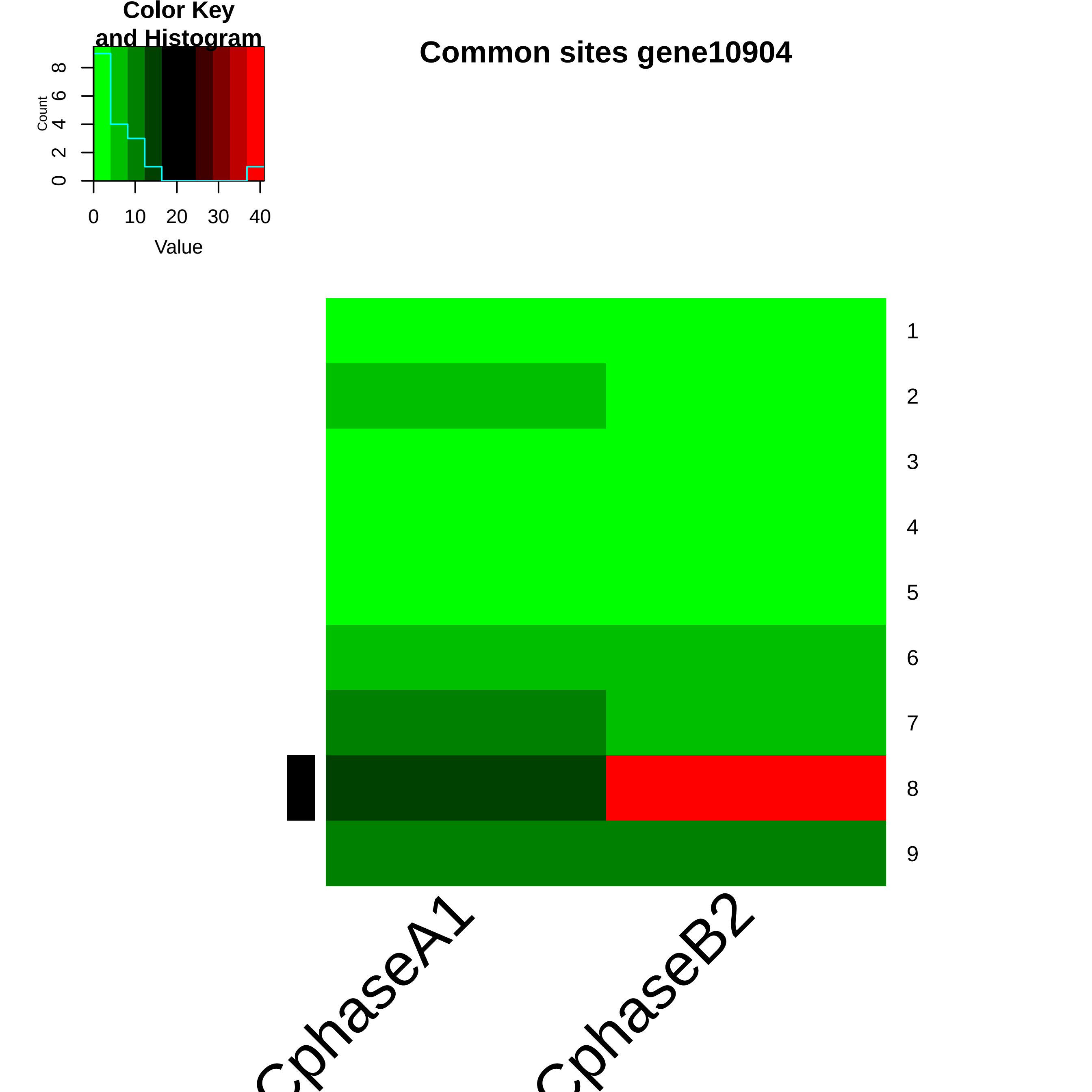
Color Key
and Histogram
Count
0
2
4
6
8
0
10
20
30
40
Value
Common sites gene10904
1
2
3
4
5
6
7
8
9
CphaseA1
CphaseB2

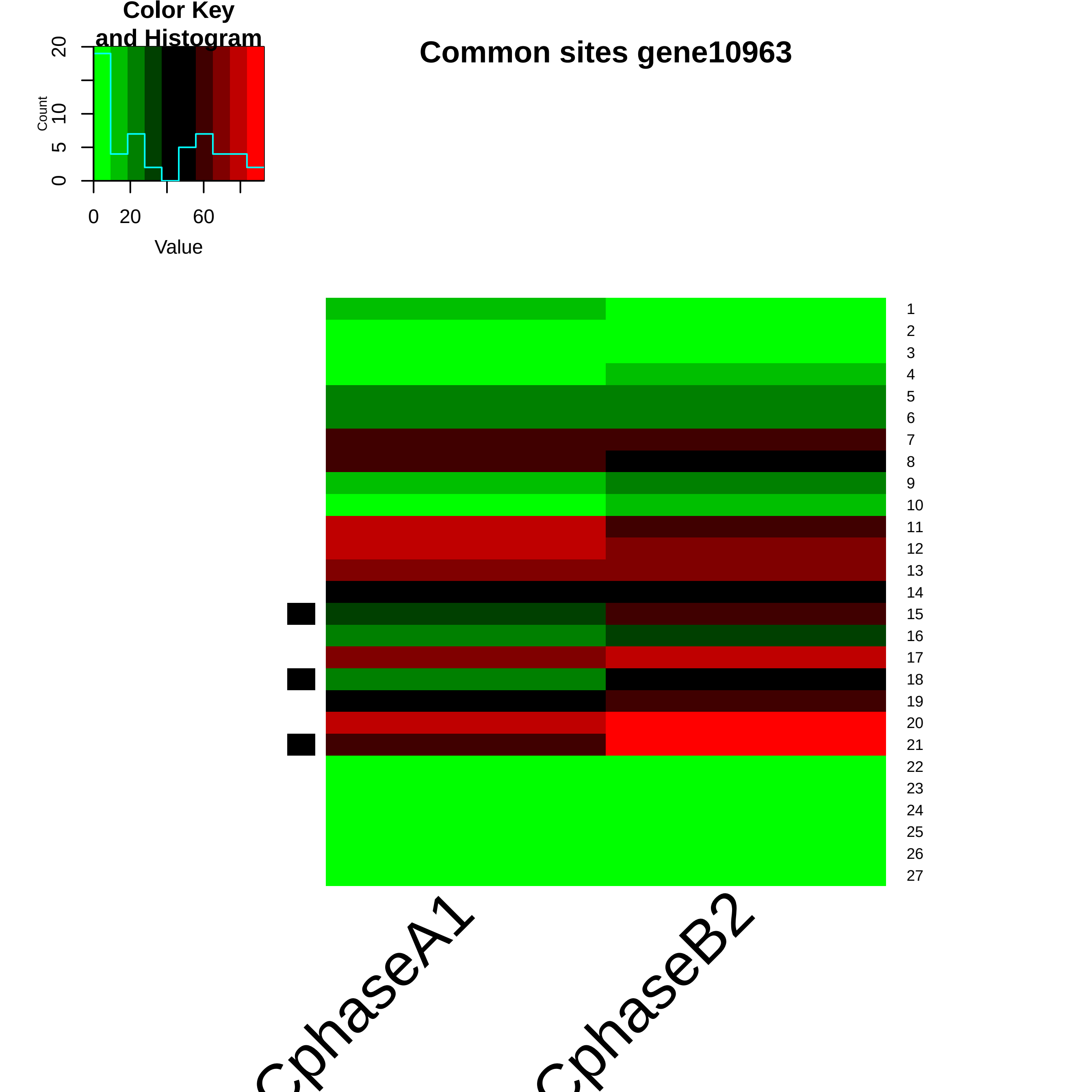
Common sites gene10963
Color Key
and Histogram
Count
Value
0
5
10
20
0
20
60
CphaseA1
CphaseB2

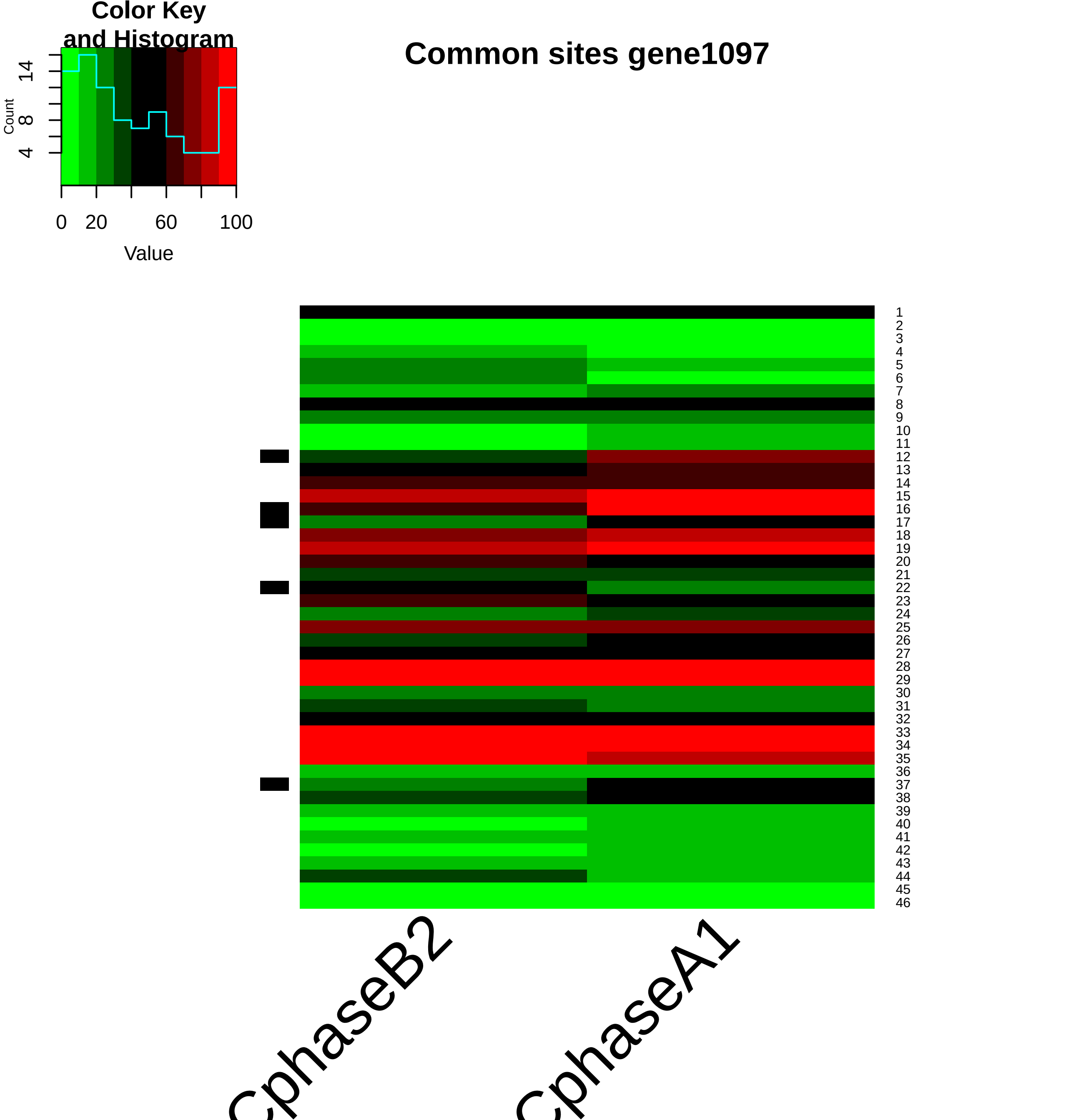

Color Key
and Histogram
Count
14
8
4
0
20
60
100
Value
Common sites gene1097
CphaseB2
CphaseA1

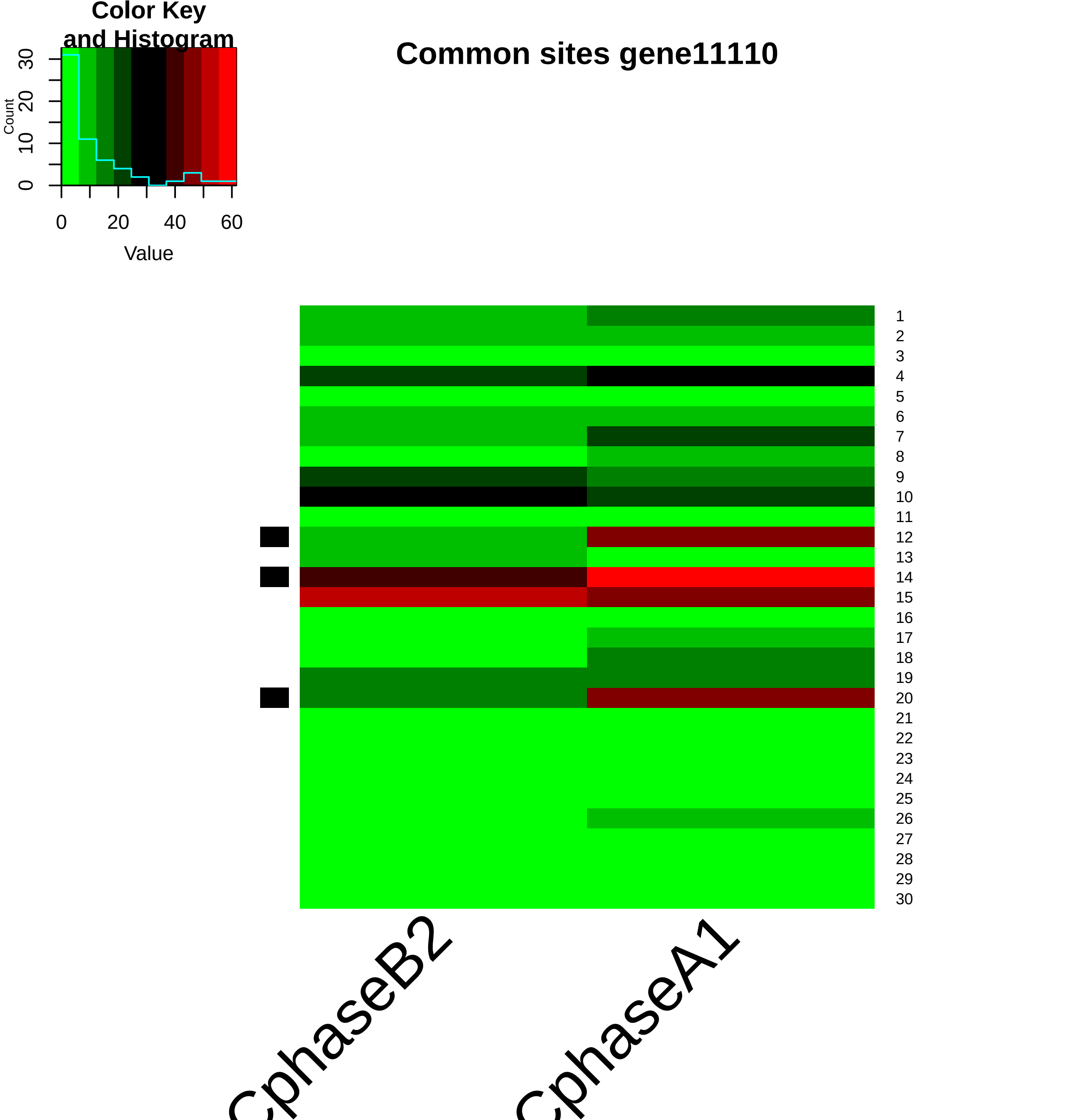

Color Key
and Histogram
Count
0
10
20
30
0
20
40
60
Value
Common sites gene11110
CphaseB2
CphaseA1

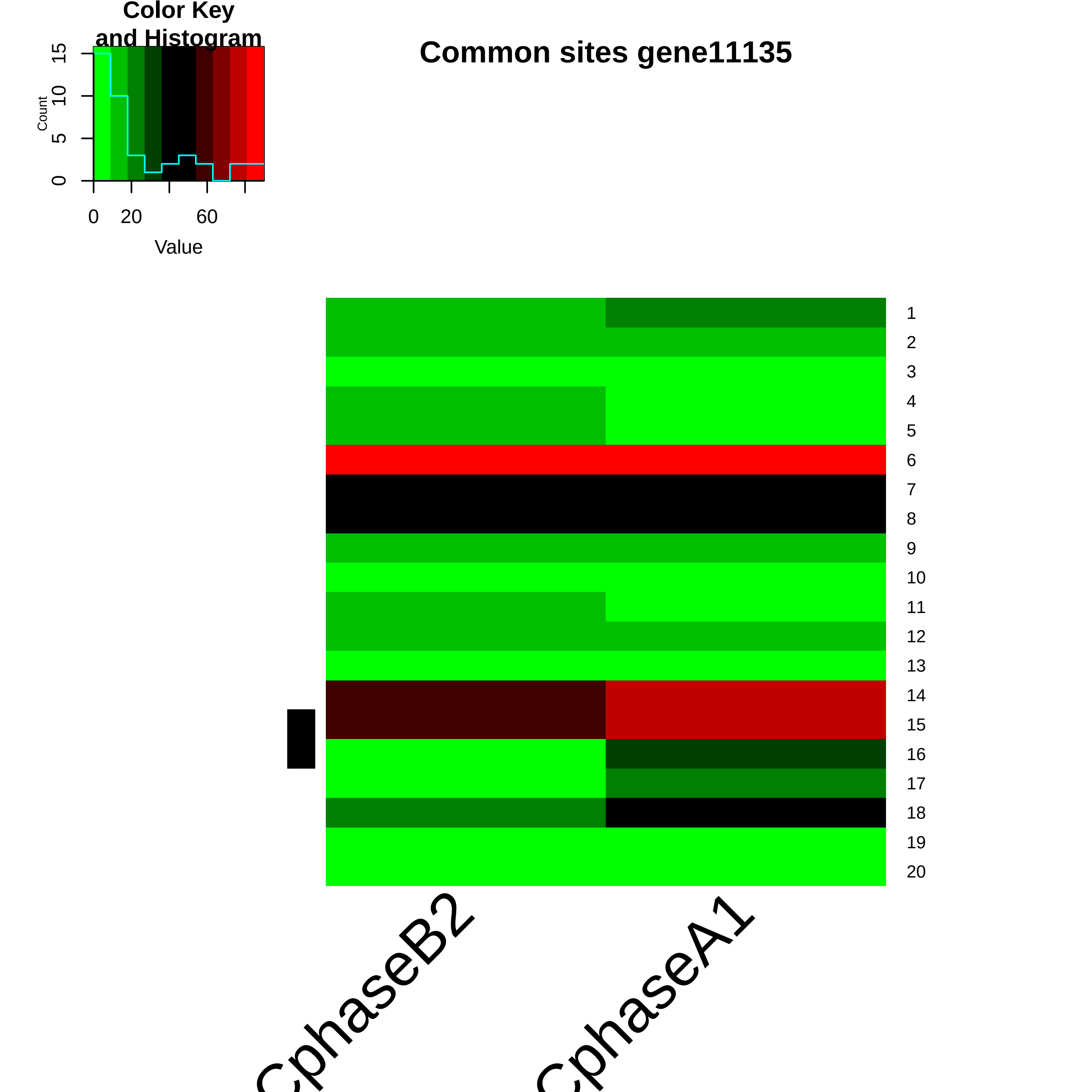

Color Key
and Histogram
Count
0
5
10
15
0
20
60
Value
Common sites gene11135
CphaseB2
CphaseA1

#### Common sites gene11235

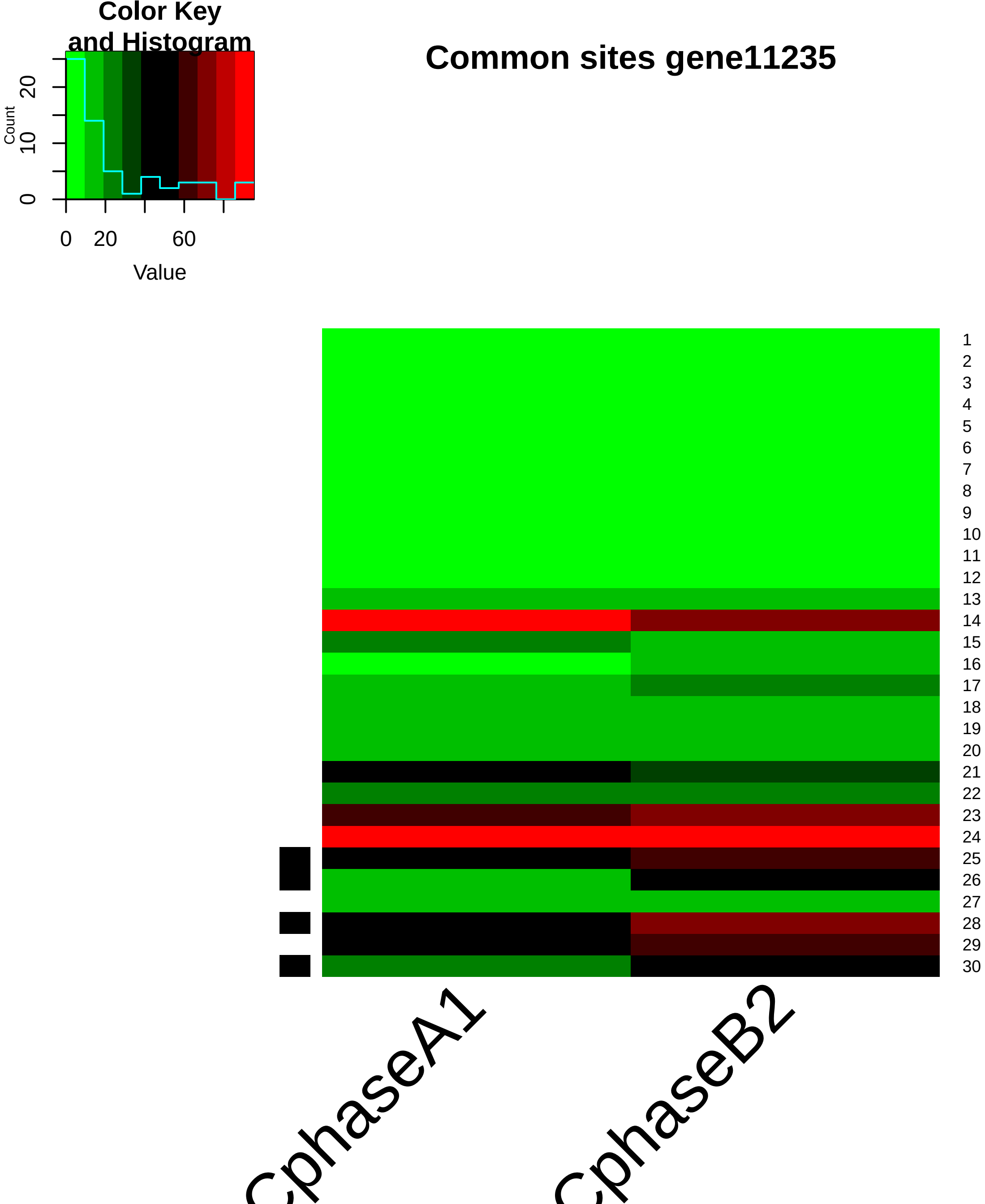

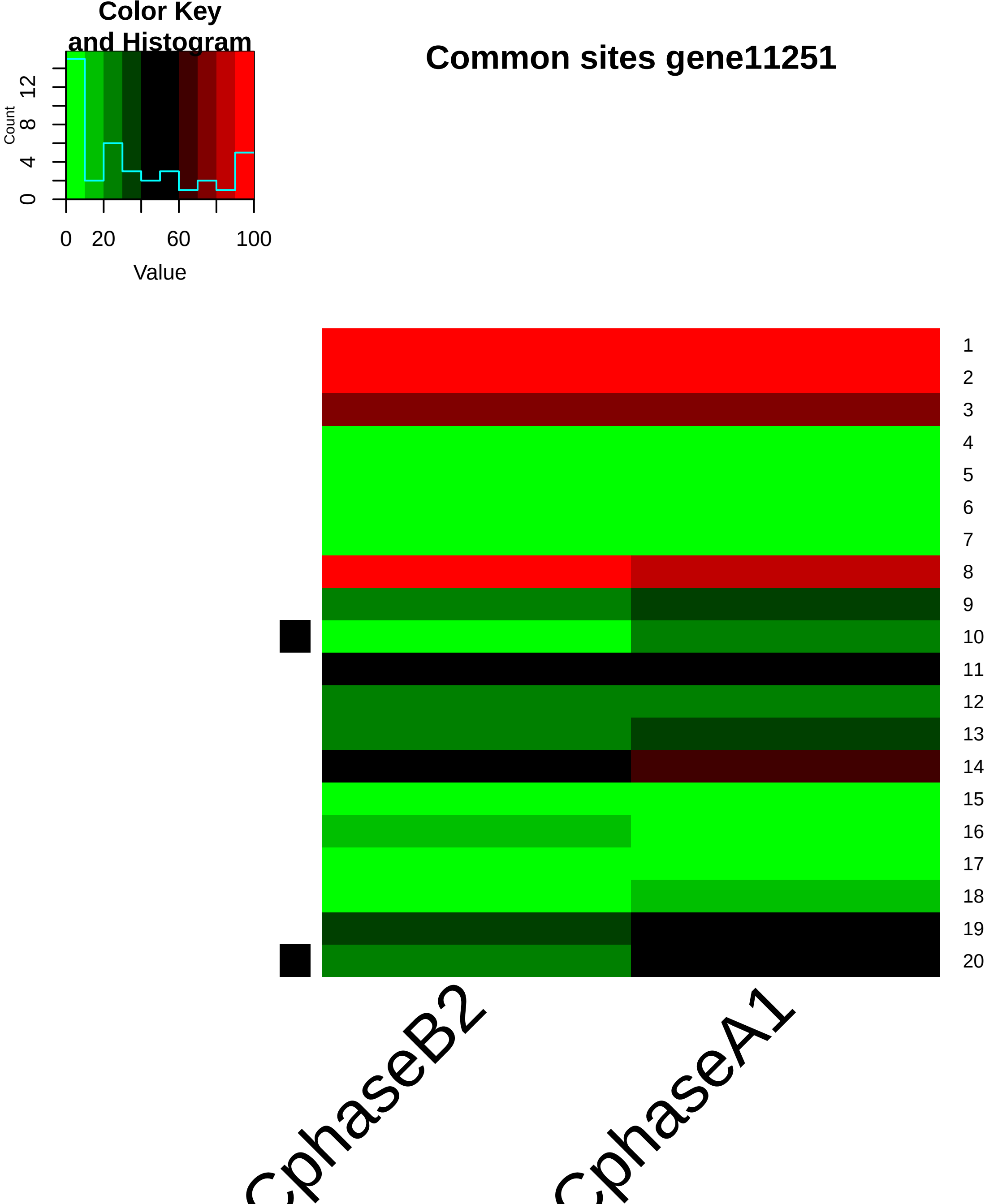
Color Key
and Histogram
Count
0
4
8
12
0
20
60
100
Value
Common sites gene11251
CphaseB2
CphaseA1

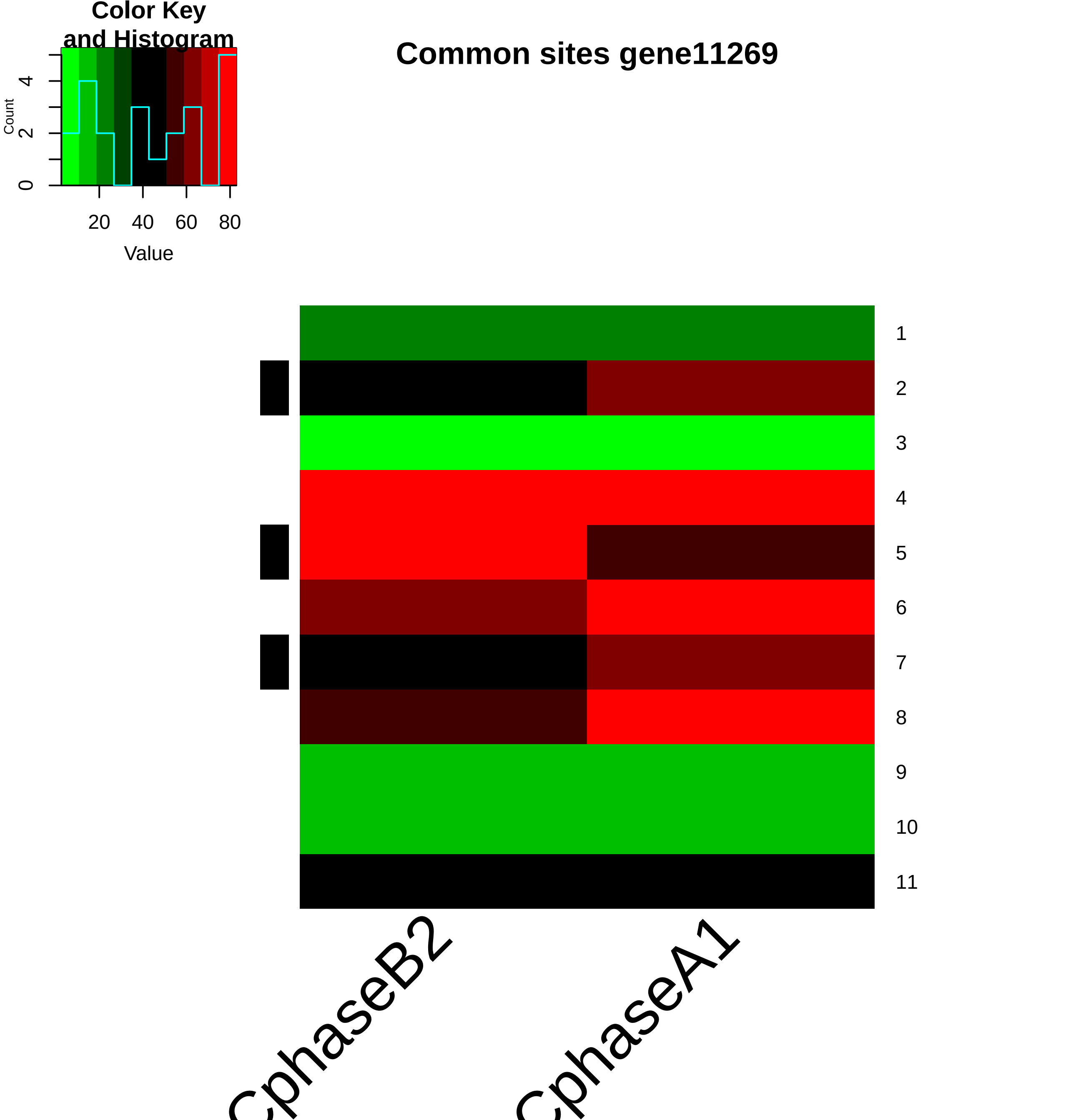

Color Key
and Histogram
Count
0
2
4
20
40
60
80
Value
Common sites gene11269
1
2
3
4
5
6
7
8
9
10
11
CphaseB2
CphaseA1

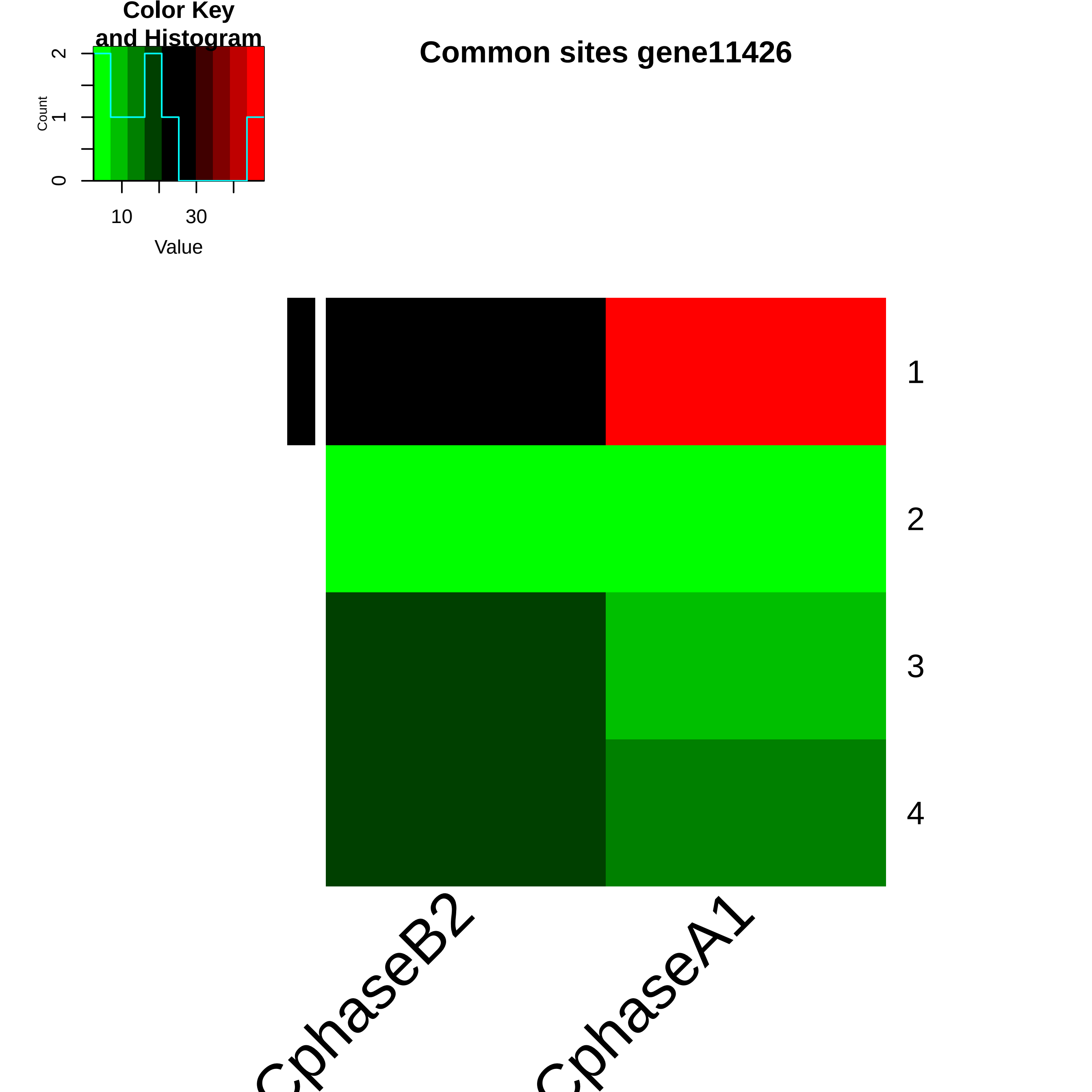
Color Key
and Histogram
Count
0
1
2
10
30
Value
Common sites gene11426
1
2
3
4
CphaseB2
CphaseA1

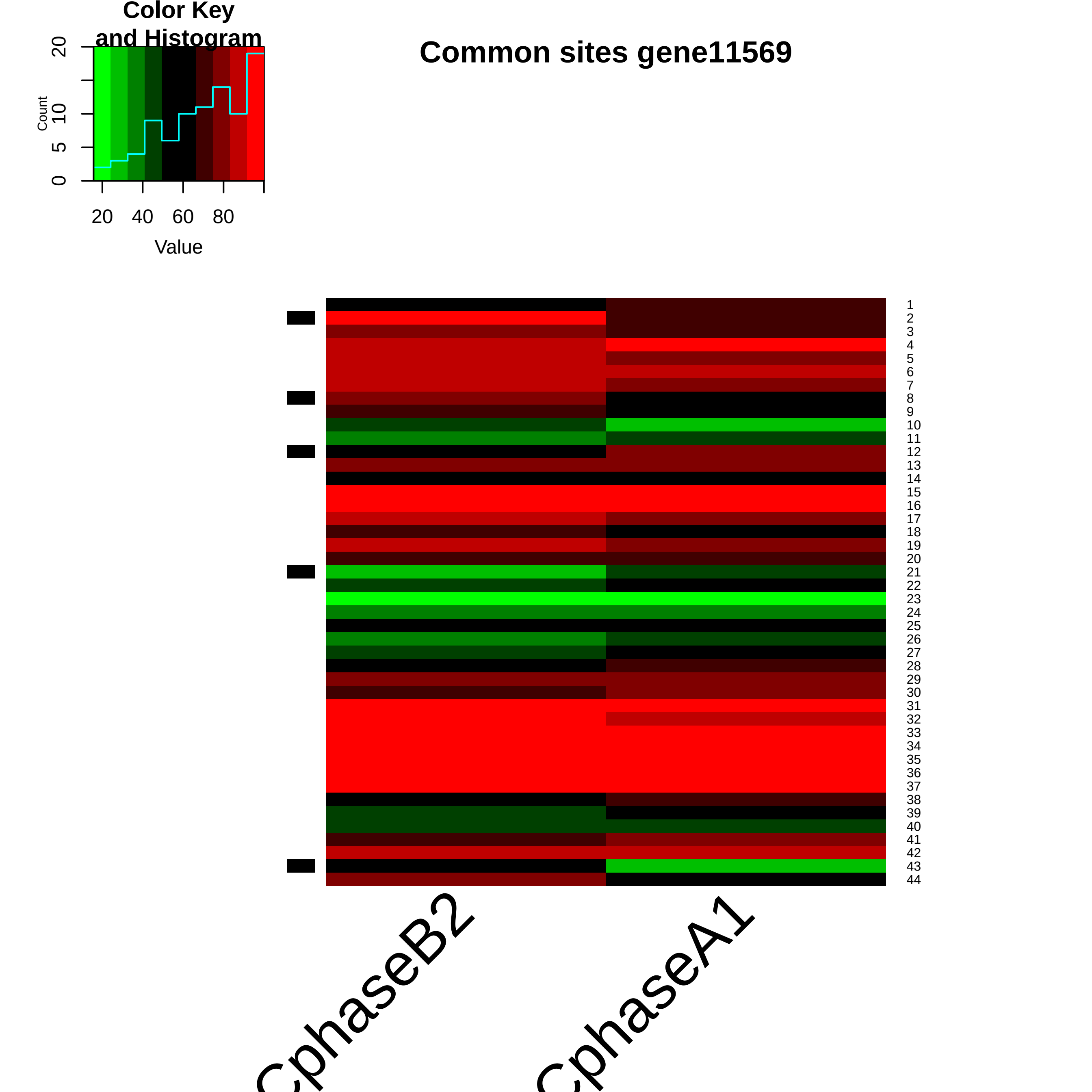
Color Key
and Histogram
Count
0
5
10
20
20
40
60
80
Value
Common sites gene11569
CphaseB2
CphaseA1

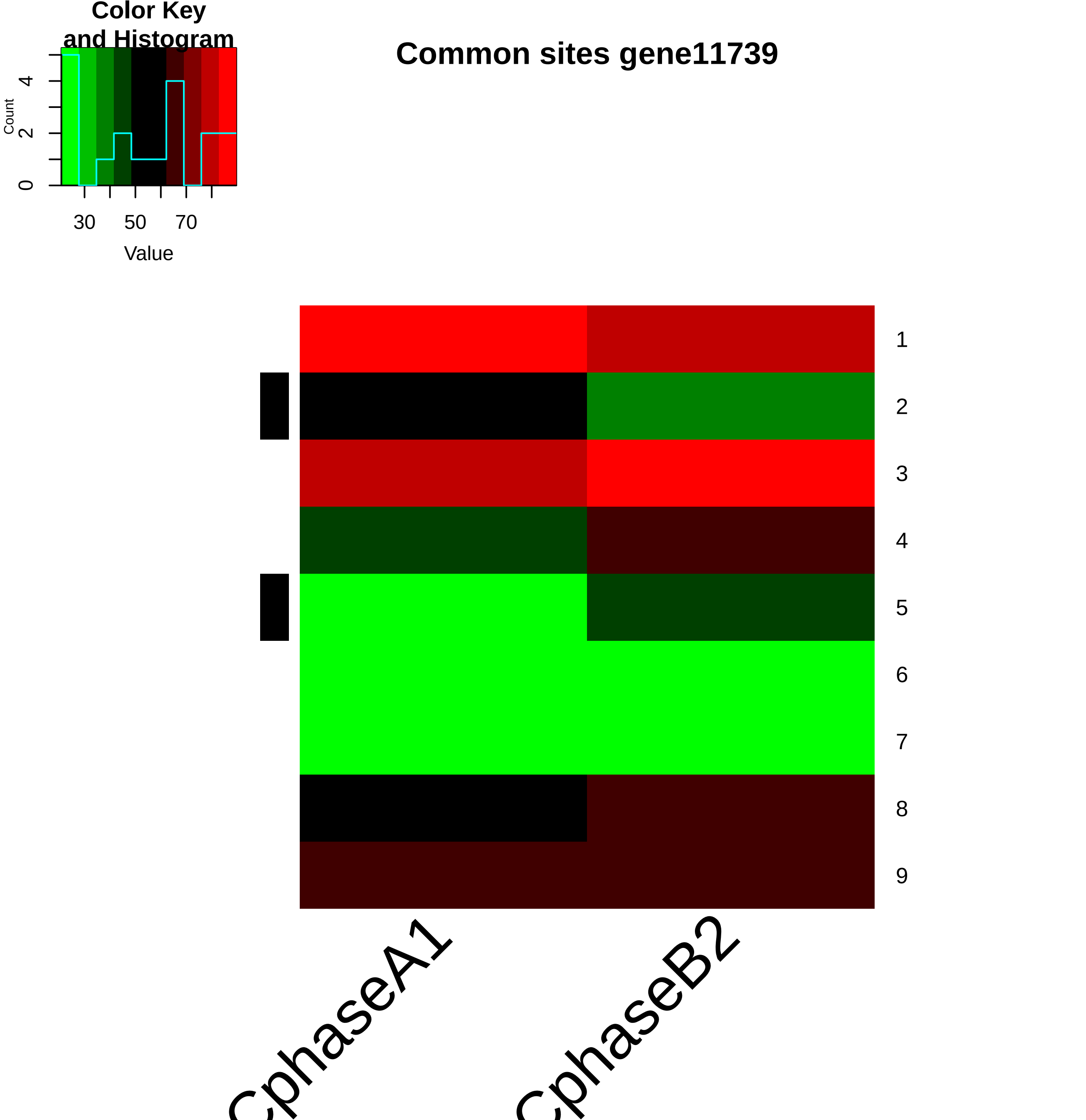
Color Key
and Histogram
Count
0
2
4
30
50
70
Value
Common sites gene11739
1
2
3
4
5
6
7
8
9
CphaseA1
CphaseB2

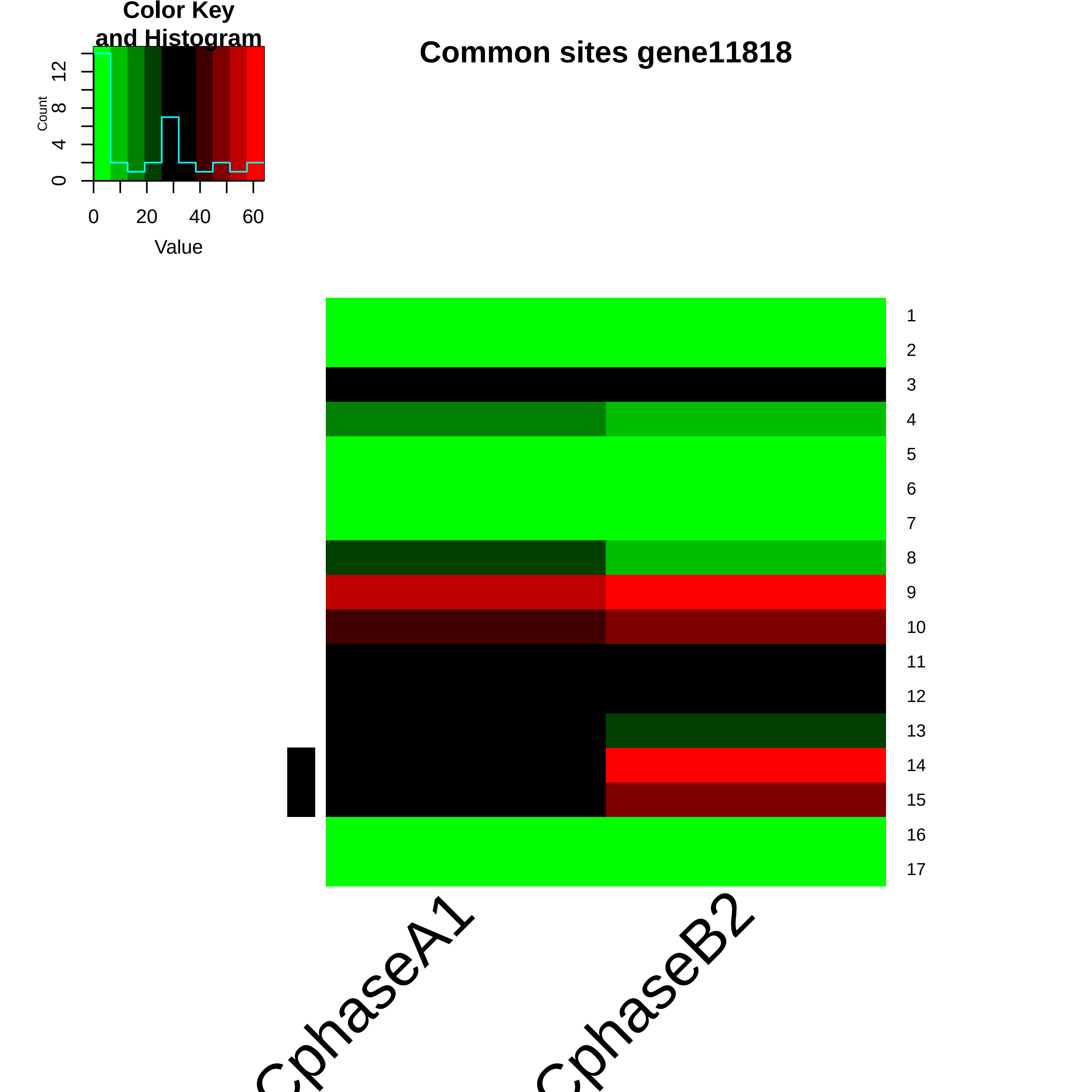
Color Key
and Histogram
Count
0
4
8
12
0
20
40
60
Value
Common sites gene11818
CphaseA1
CphaseB2

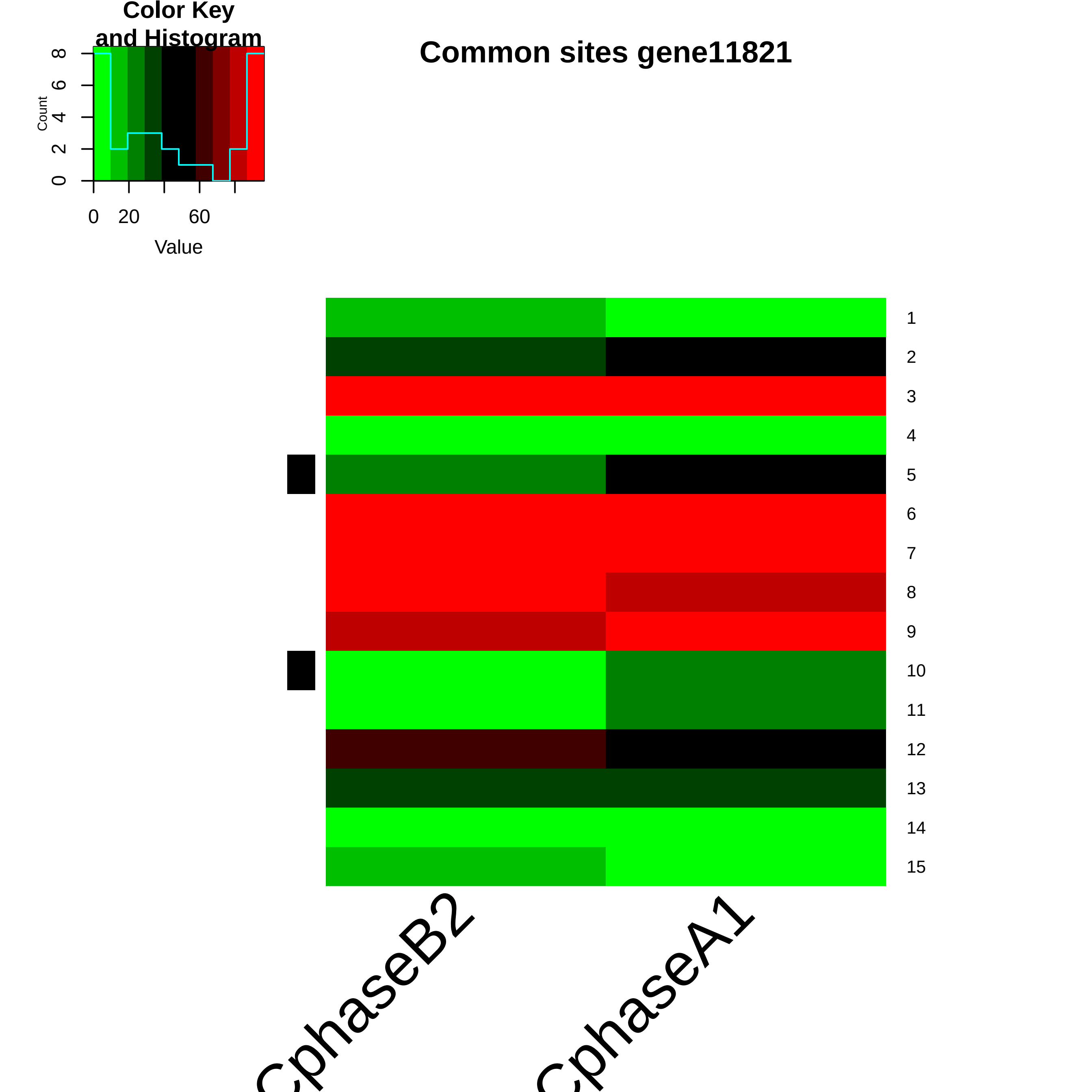
Color Key
and Histogram
Count
0
2
4
6
8
0
20
60
Value
Common sites gene11821
CphaseB2
CphaseA1

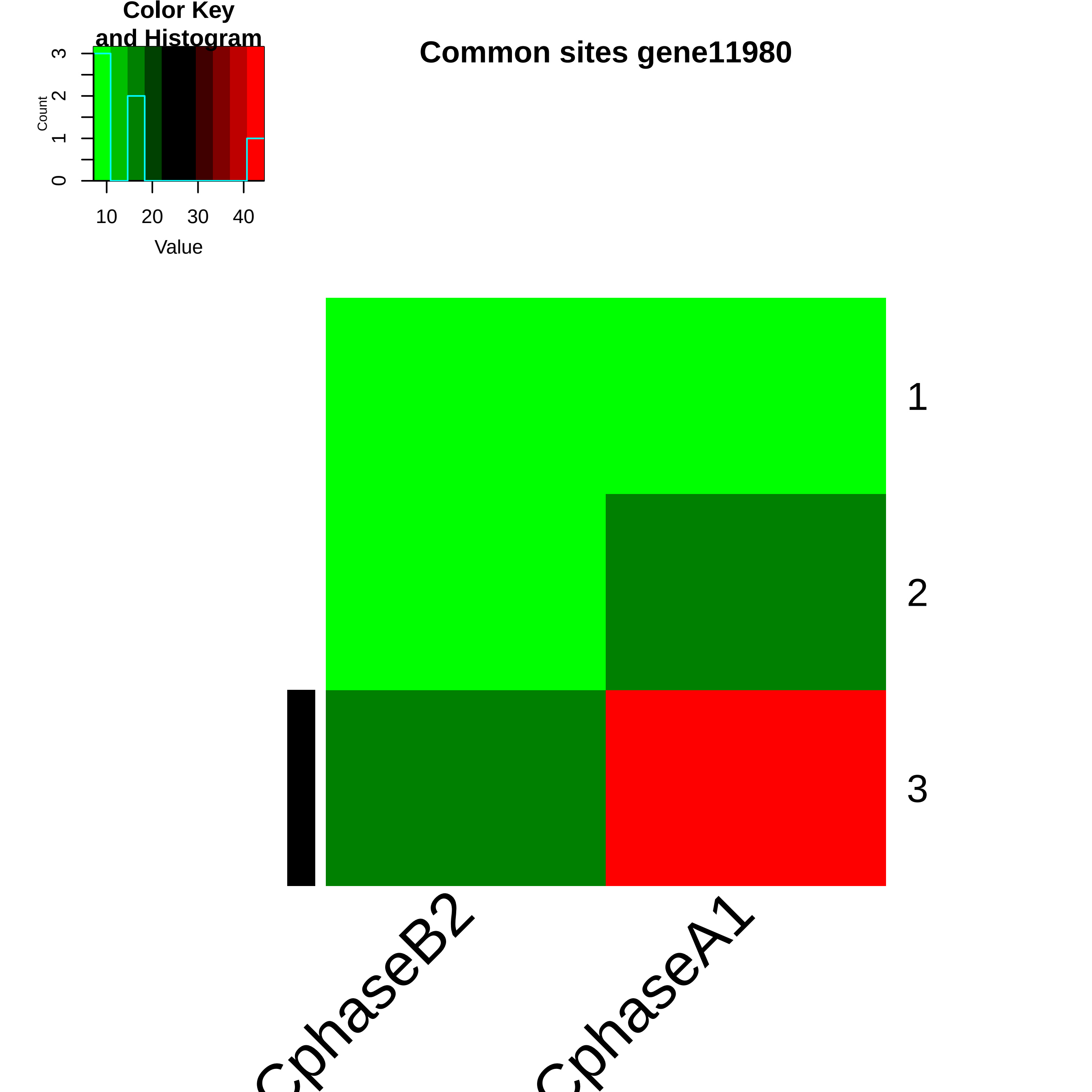
Color Key
and Histogram
Count
Value
10
20
30
40
0
1
2
3
Common sites gene11980
1
2
3
CphaseB2
CphaseA1

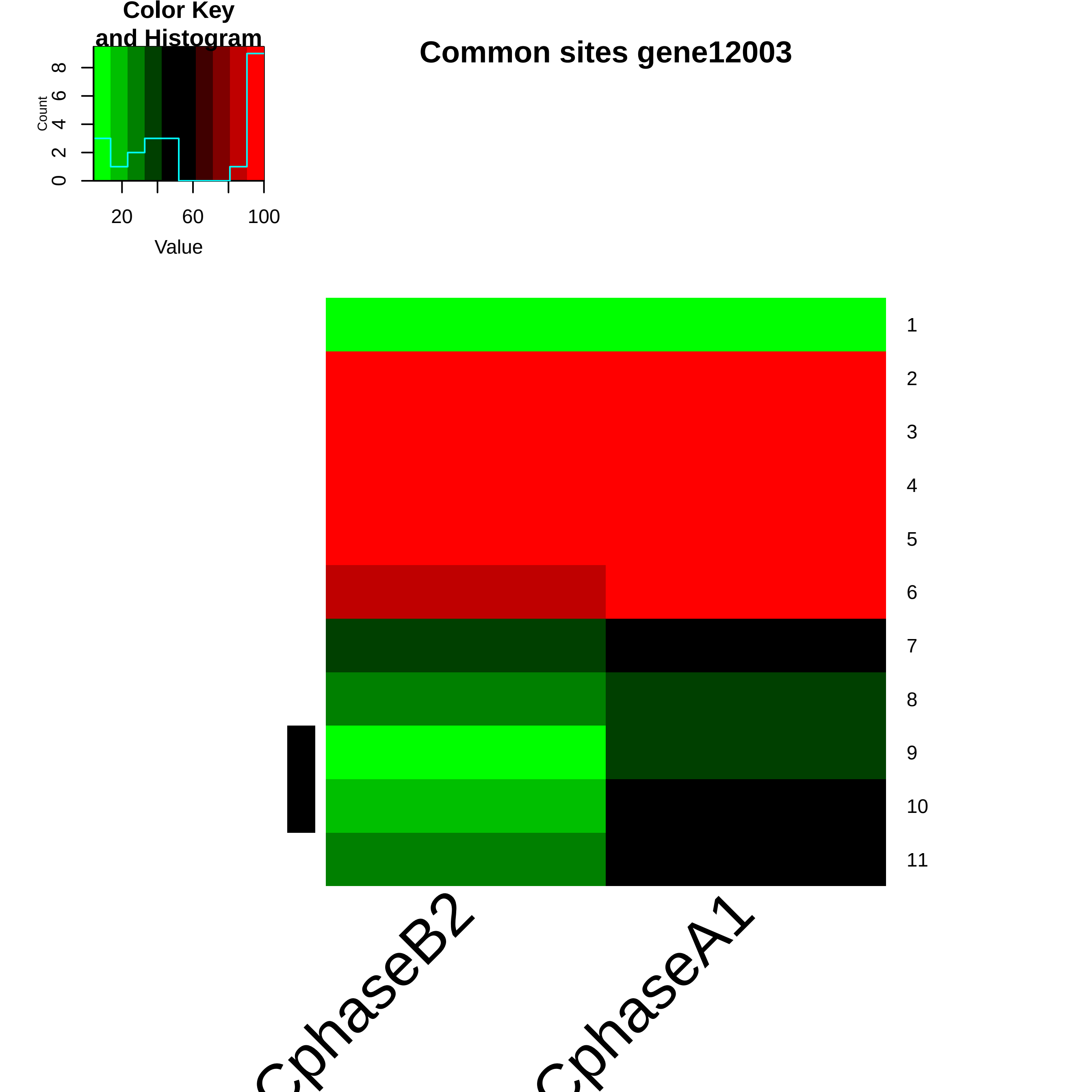

Color Key
and Histogram
Count
0
2
4
6
8
20
60
100
Value
Common sites gene12003
1
2
3
4
5
6
7
8
9
10
11
CphaseB2
CphaseA1

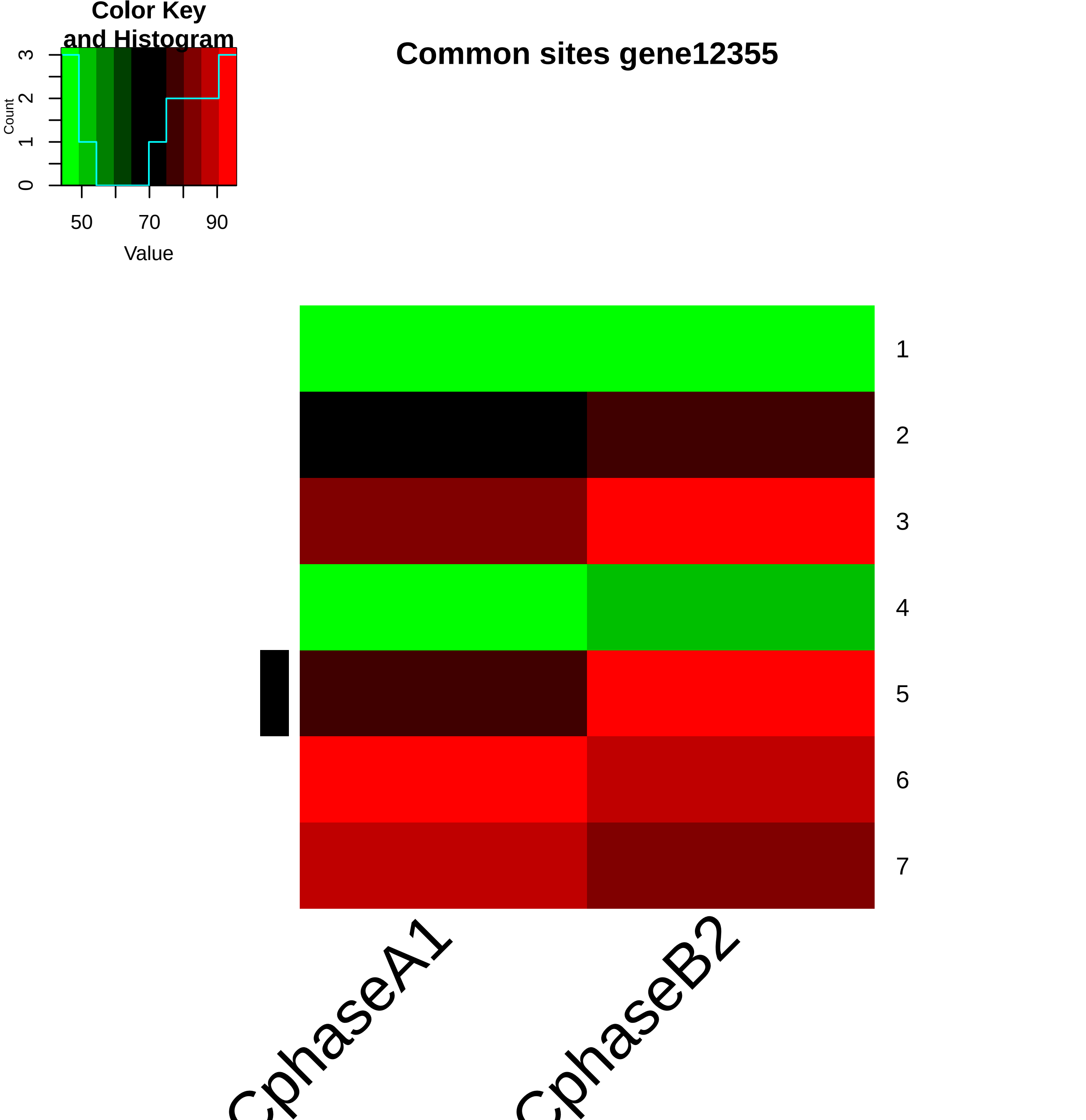
Color Key
and Histogram
Count
0
1
2
3
50
70
90
Value
Common sites gene12355
1
2
3
4
5
6
7
CphaseA1
CphaseB2

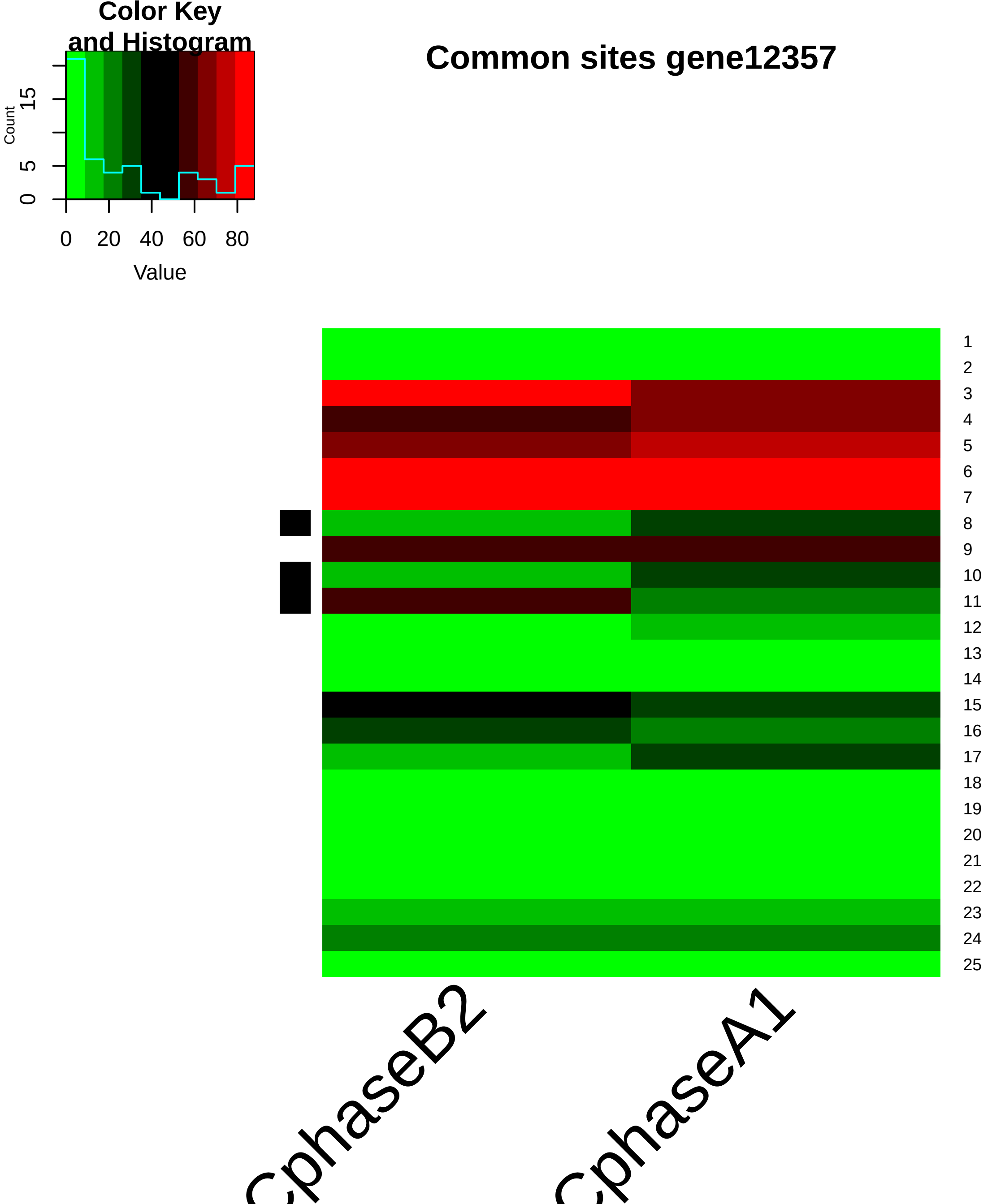
Color Key
and Histogram
Count
0
5
15
0
20
40
60
80
Value
Common sites gene12357
CphaseB2
CphaseA1

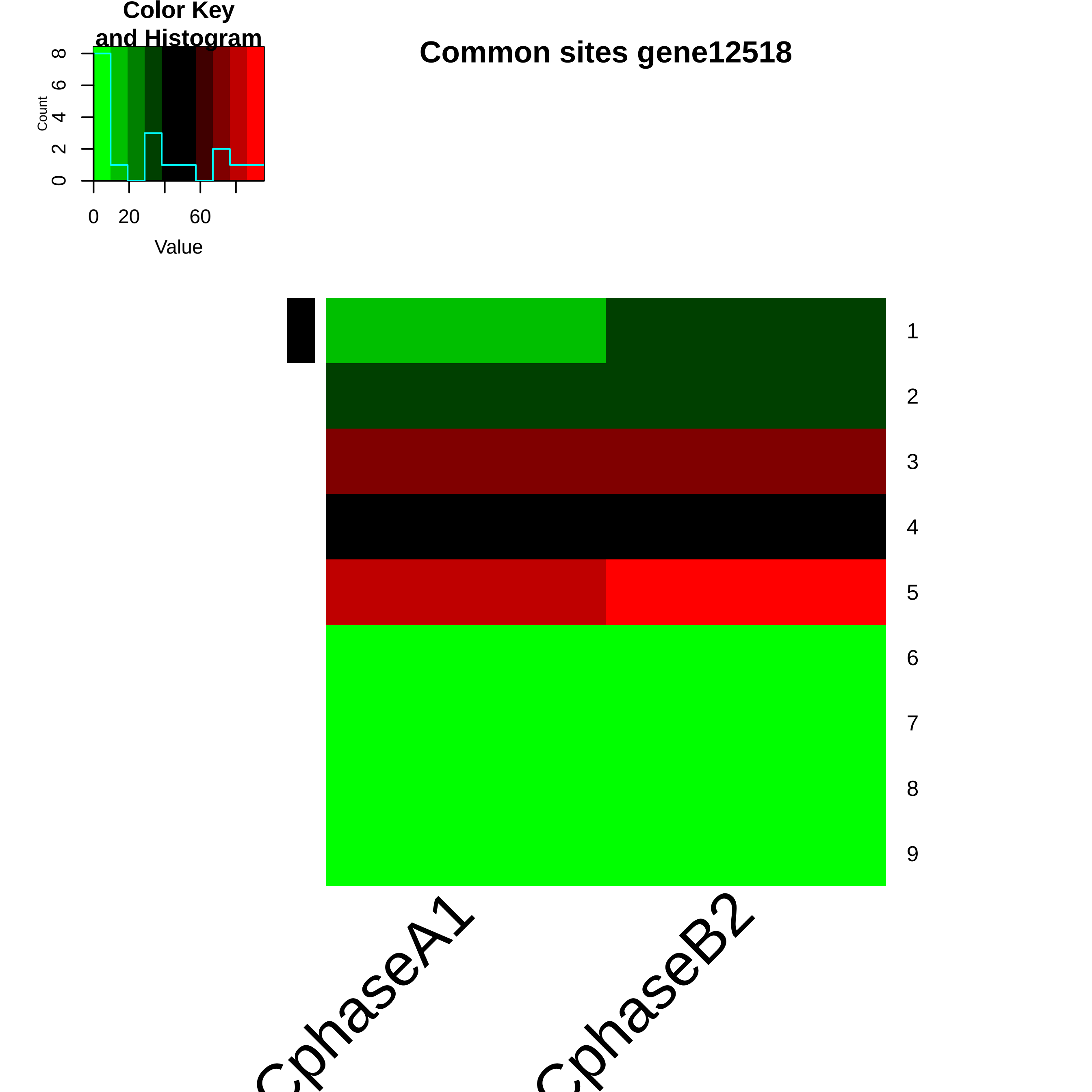

Color Key
and Histogram
Count
0
2
4
6
8
0
20
60
Value
Common sites gene12518
1
2
3
4
5
6
7
8
9
CphaseA1
CphaseB2

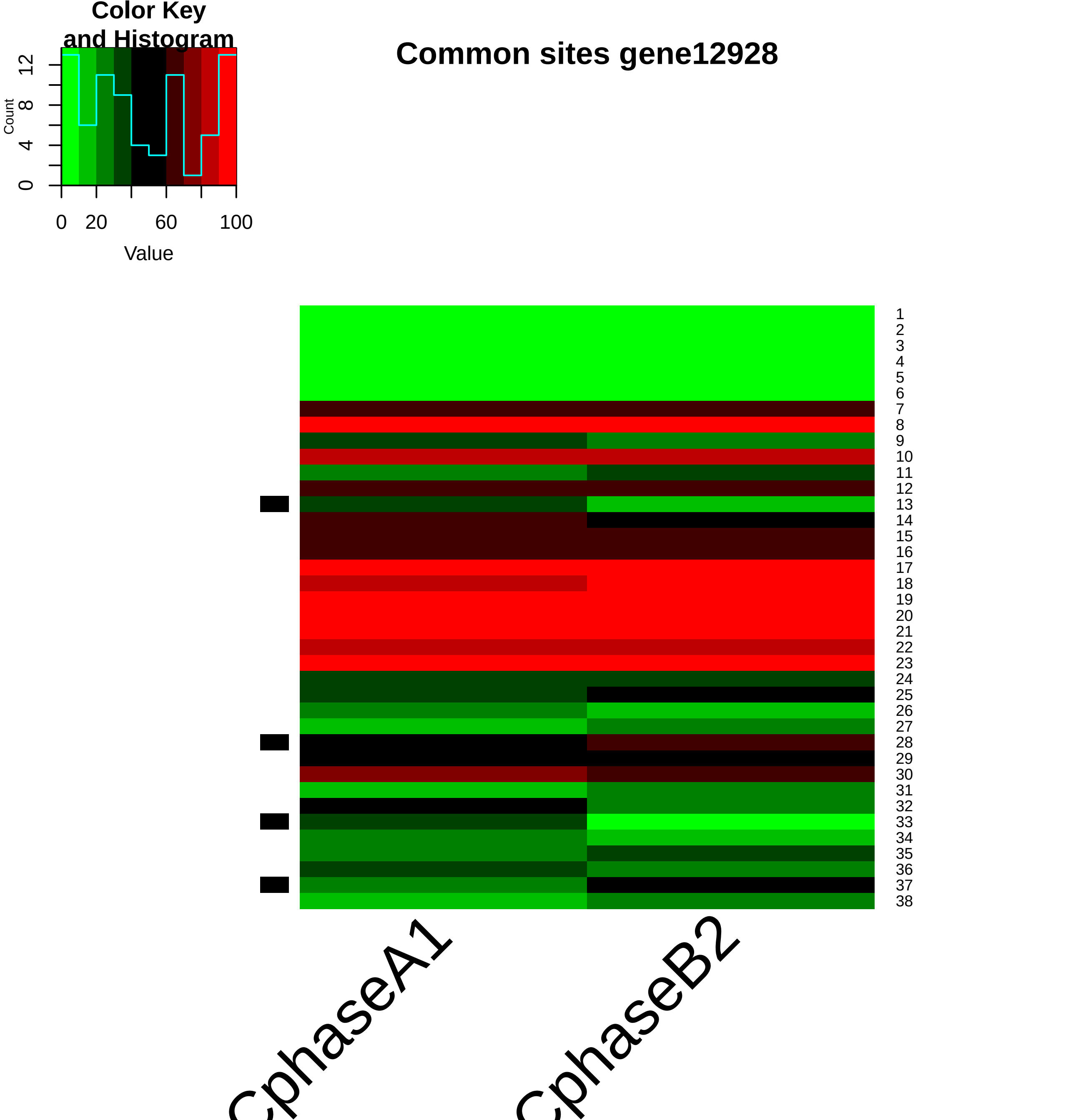

Color Key
and Histogram
Count
0
4
8
12
0
20
60
100
Value
Common sites gene12928
CphaseA1
CphaseB2

### Common sites gene13158

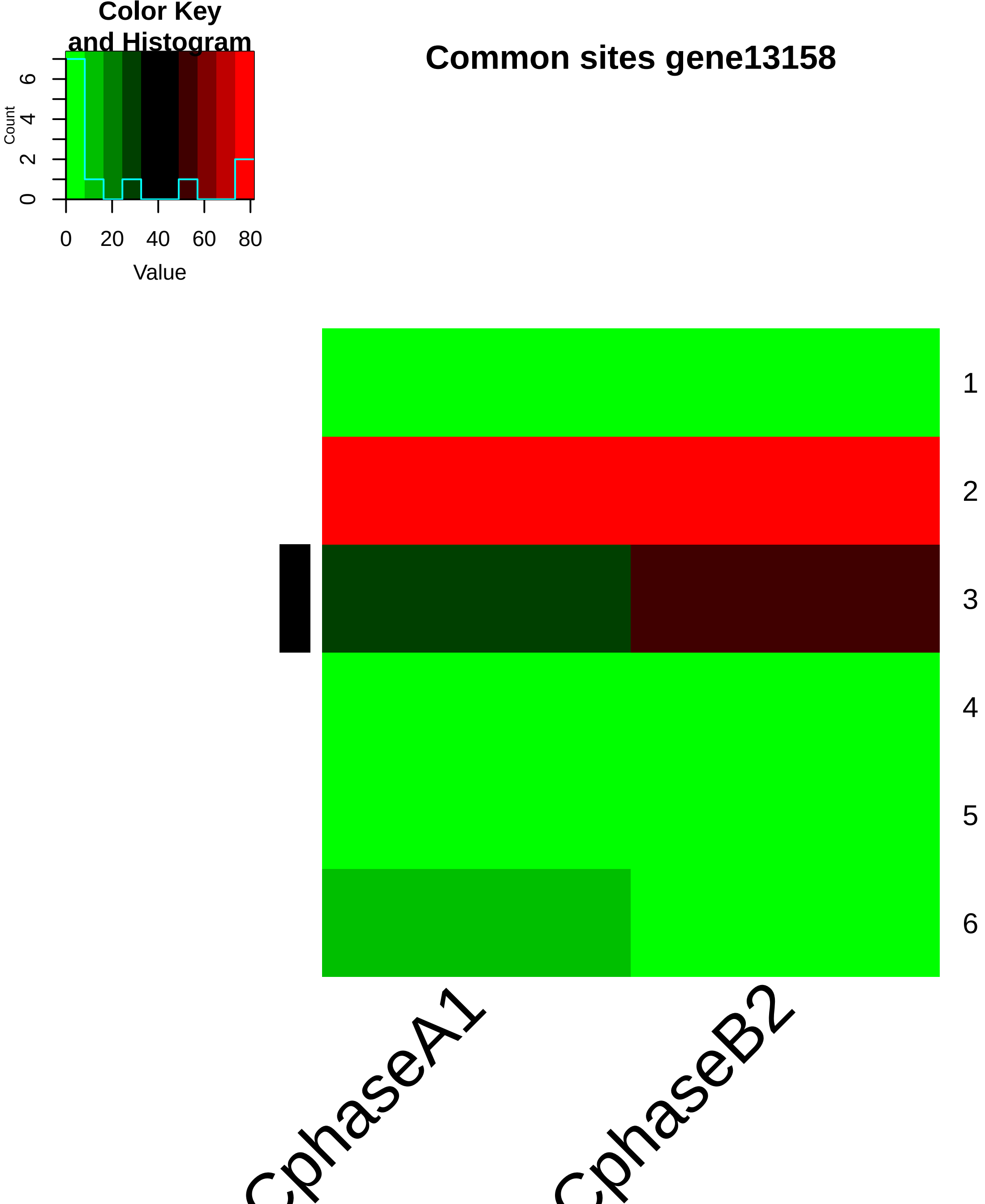

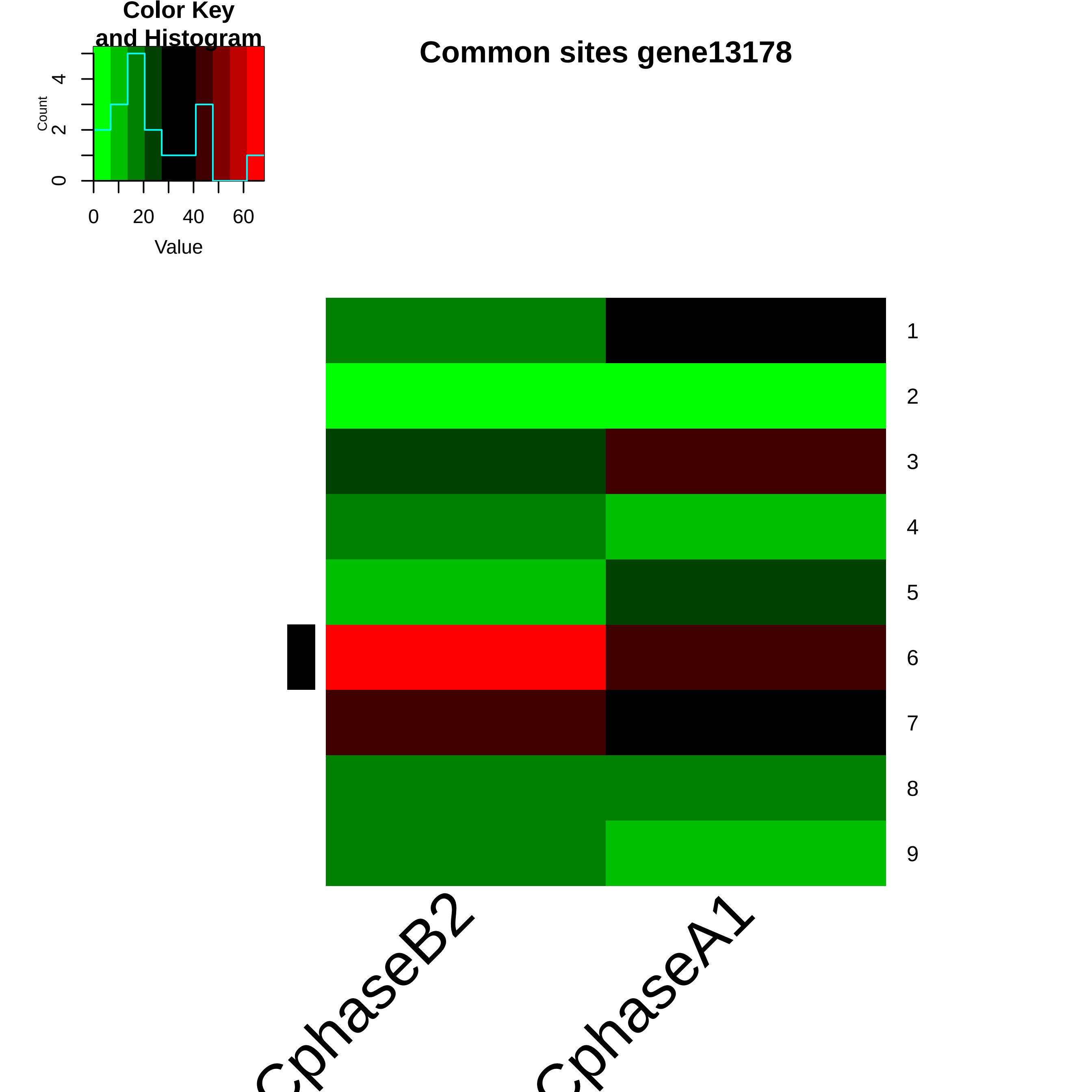
Color Key
and Histogram
Count
0
2
4
0
20
40
60
Value
Common sites gene13178
1
2
3
4
5
6
7
8
9
CphaseB2
CphaseA1

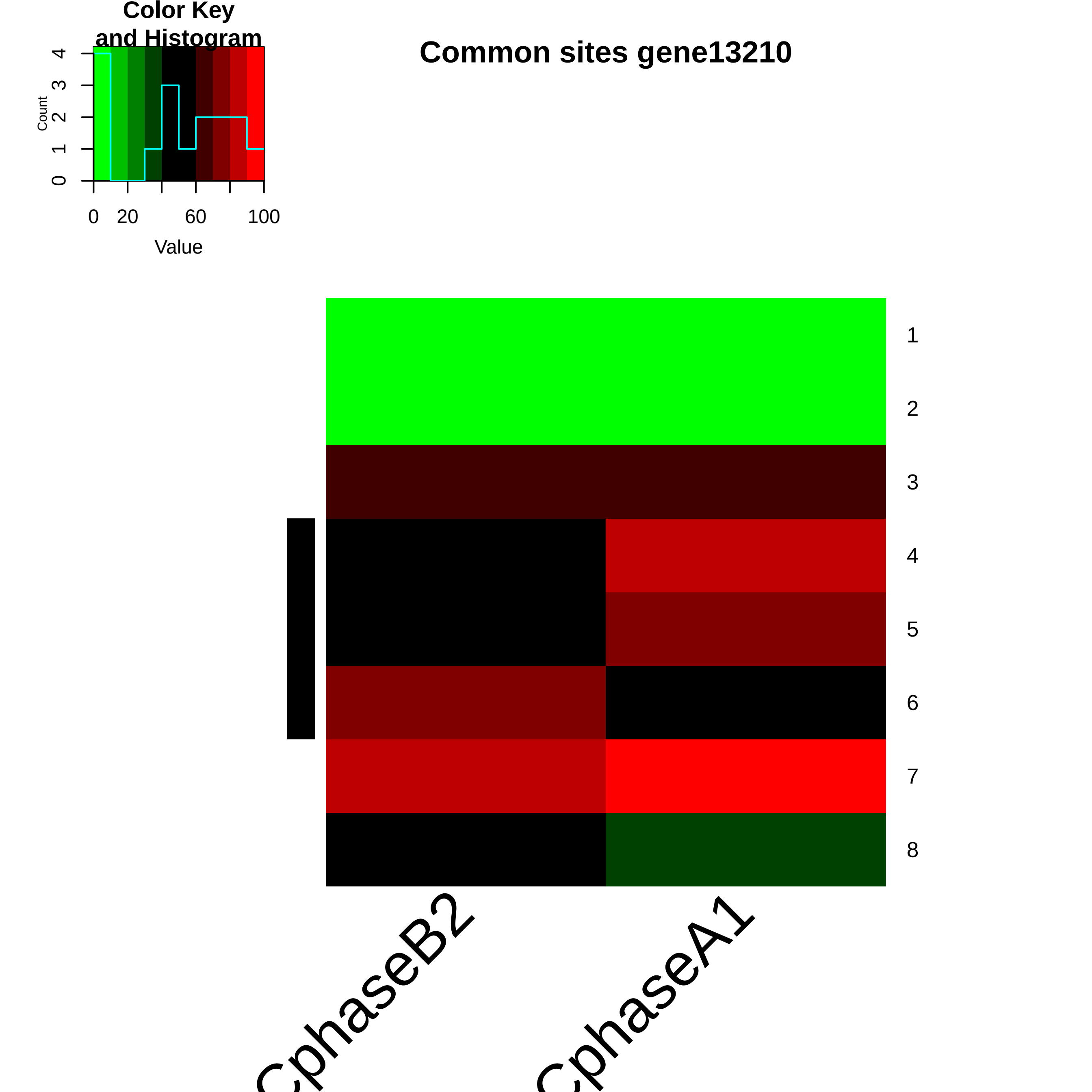

Color Key
and Histogram
Count
Value
0
20
60
100
0
1
2
3
4
Common sites gene13210
1
2
3
4
5
6
7
8
CphaseB2
CphaseA1

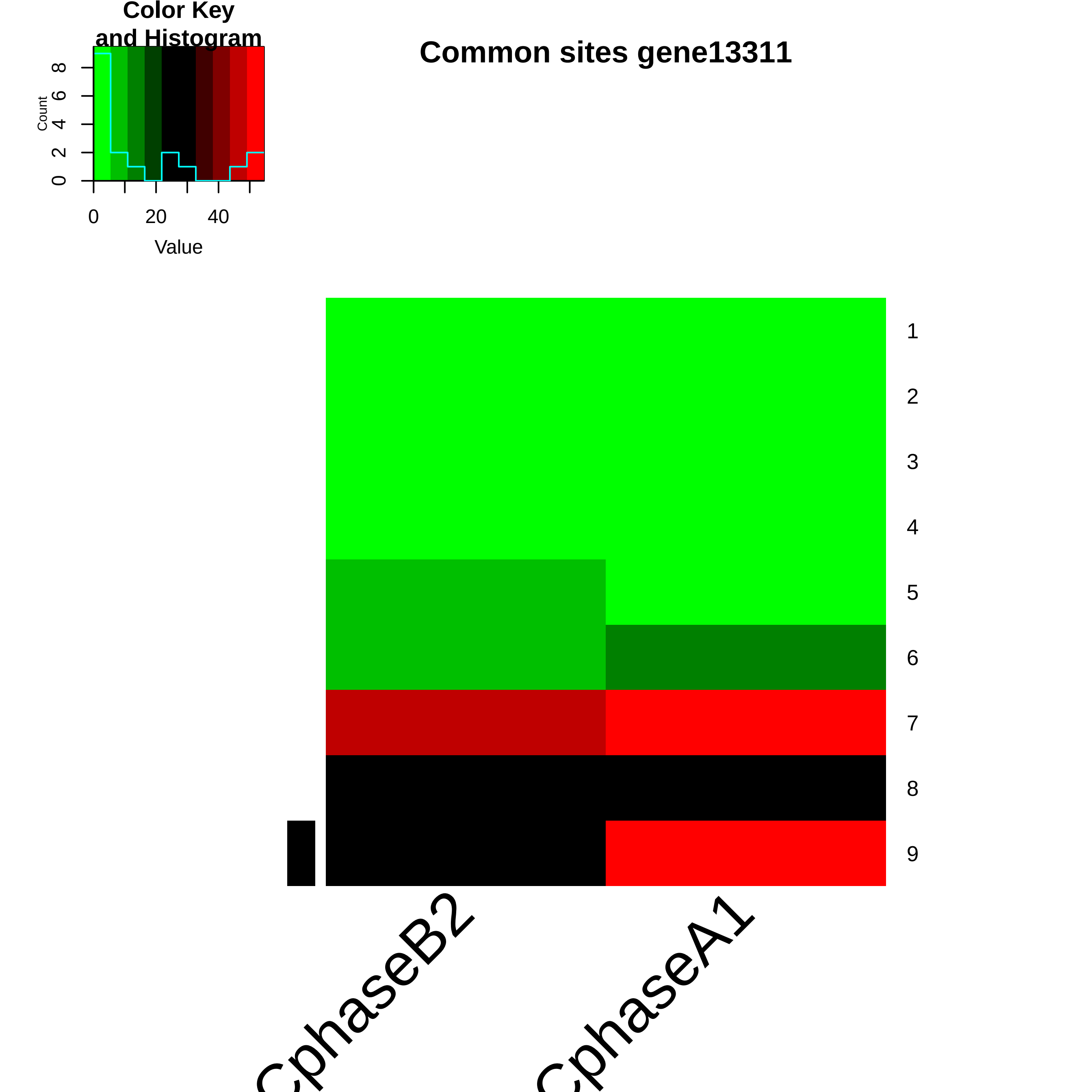
Color Key
and Histogram
Count
0
2
4
6
8
0
20
40
Value
Common sites gene13311
1
2
3
4
5
6
7
8
9
CphaseB2
CphaseA1

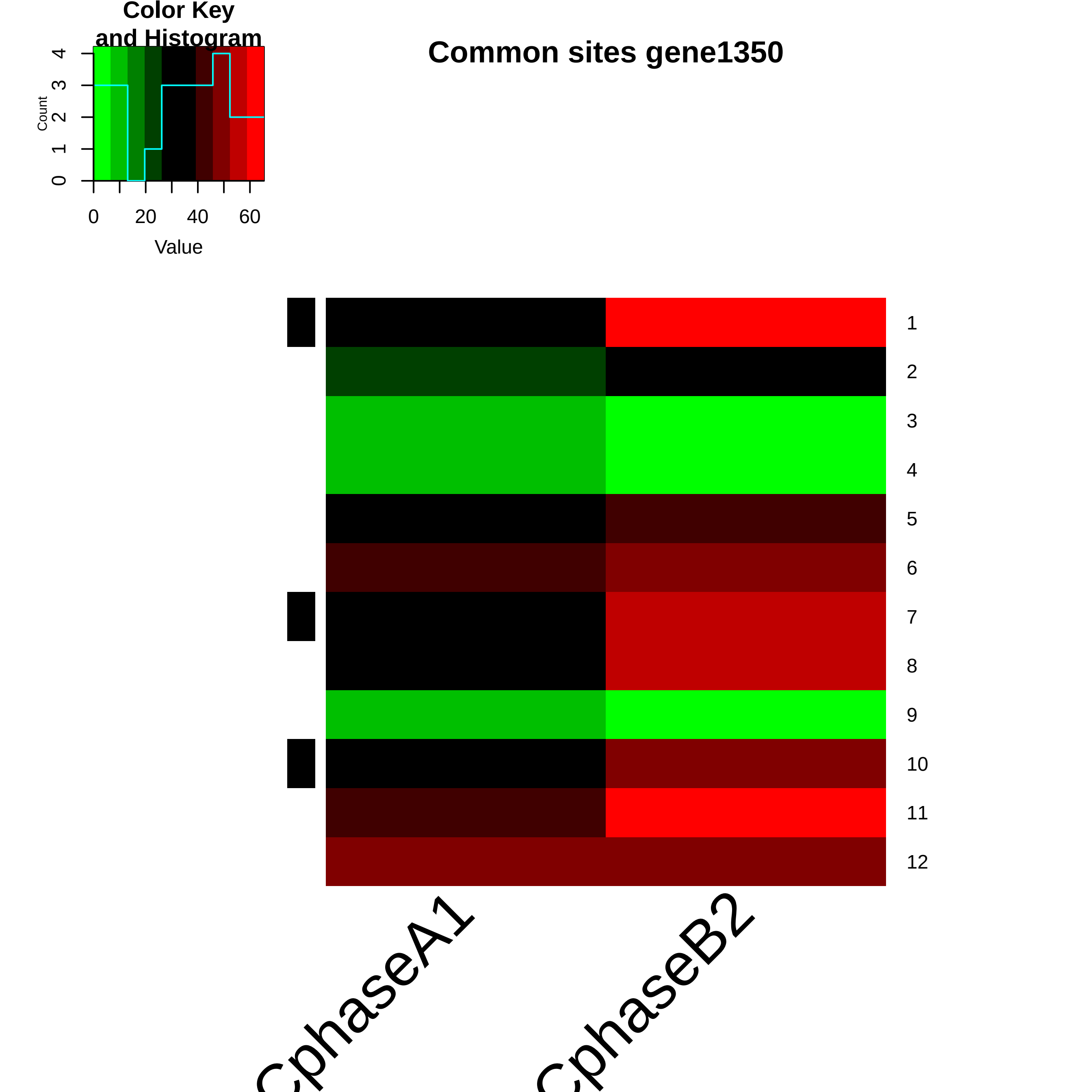
Color Key
and Histogram
Count
0
1
2
3
4
0
20
40
60
Value
Common sites gene1350
1
2
3
4
5
6
7
8
9
10
11
12
CphaseA1
CphaseB2

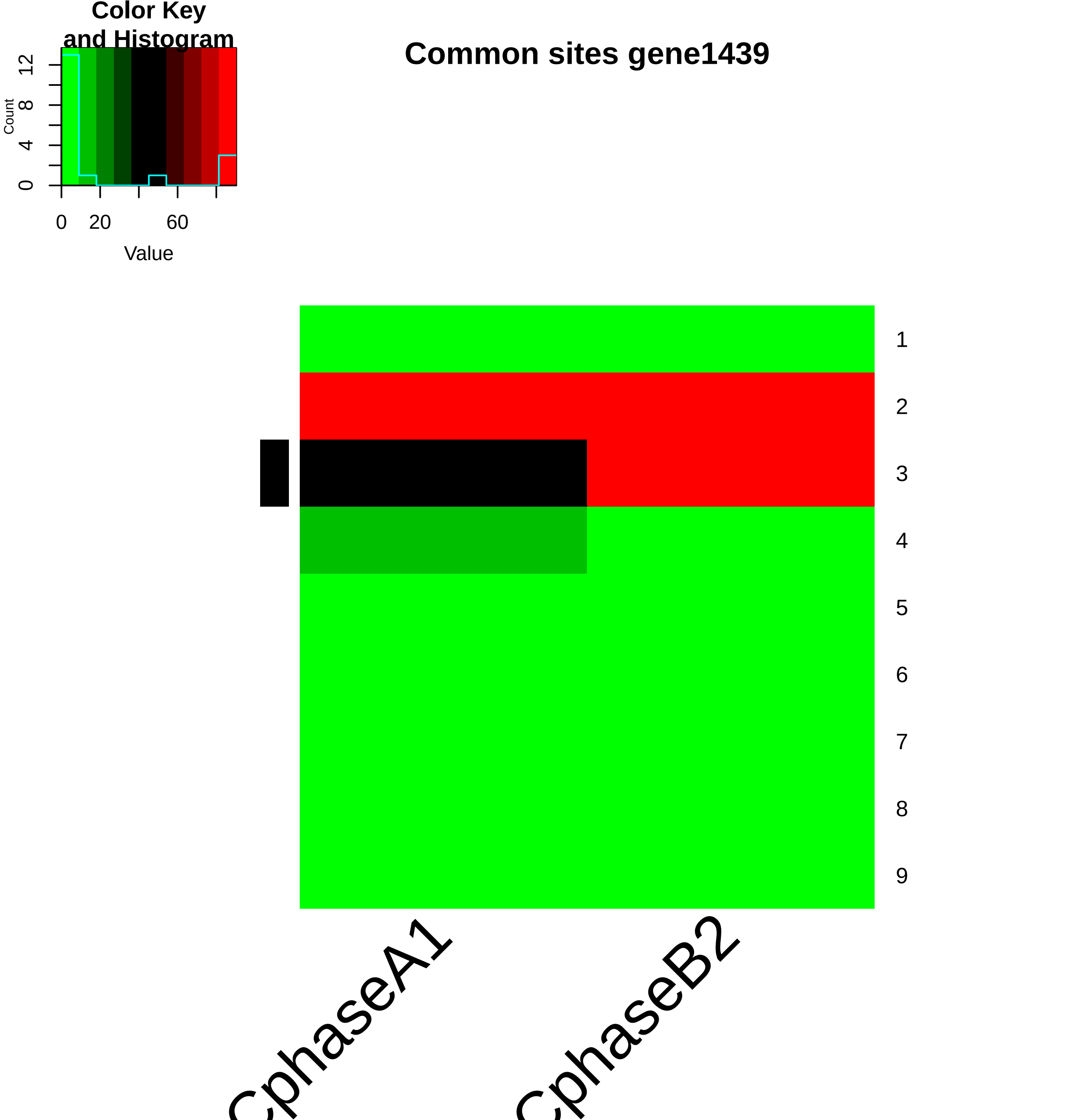
Color Key
and Histogram
Count
0
4
8
12
0
20
60
Value
Common sites gene1439
1
2
3
4
5
6
7
8
9
CphaseA1
CphaseB2

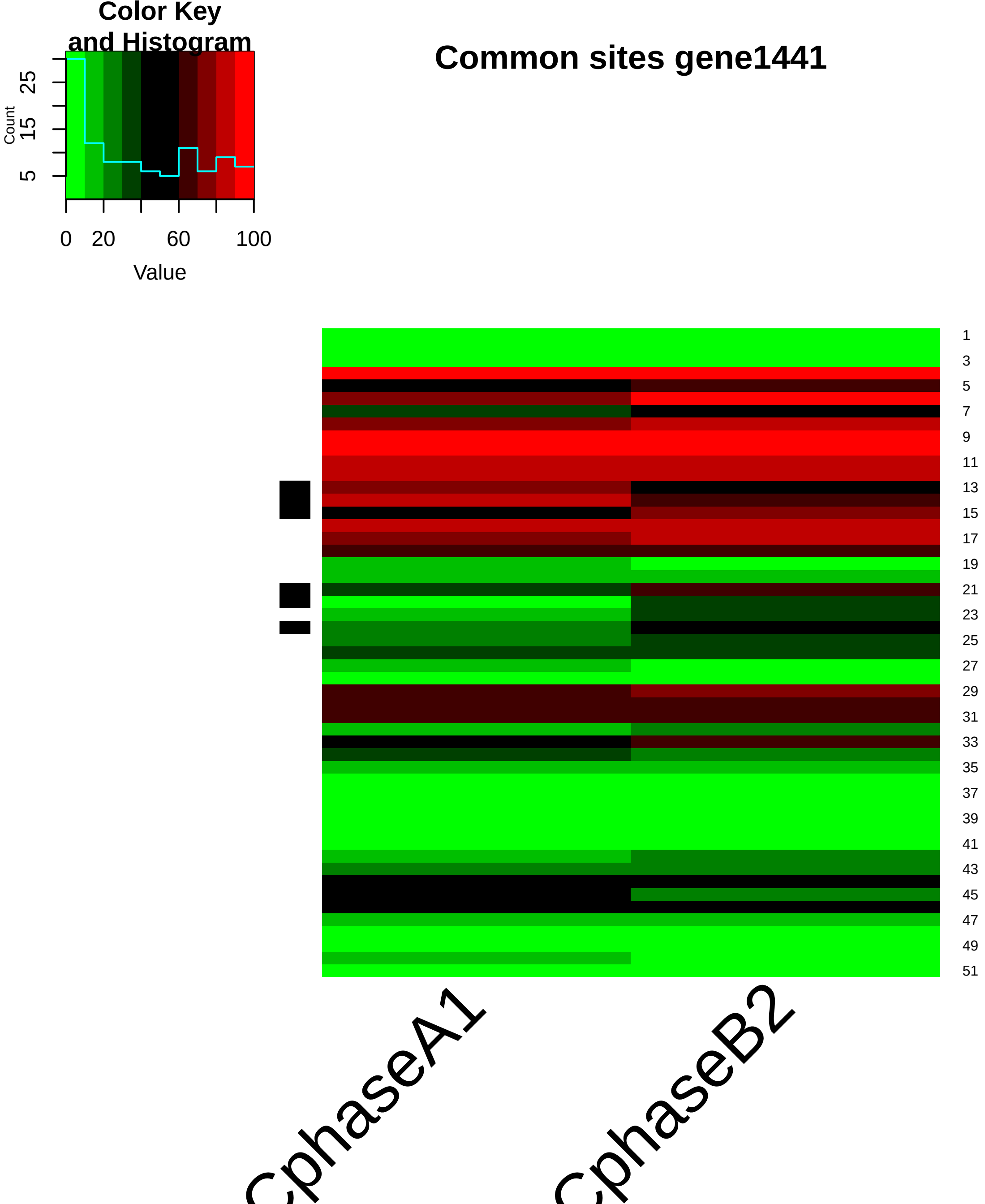

Color Key
and Histogram
Count
5
15
25
0
20
60
100
Value
Common sites gene1441
1
3
5
7
9
11
13
15
17
19
21
23
25
27
29
31
33
35
37
39
41
43
45
47
49
51
CphaseA1
CphaseB2

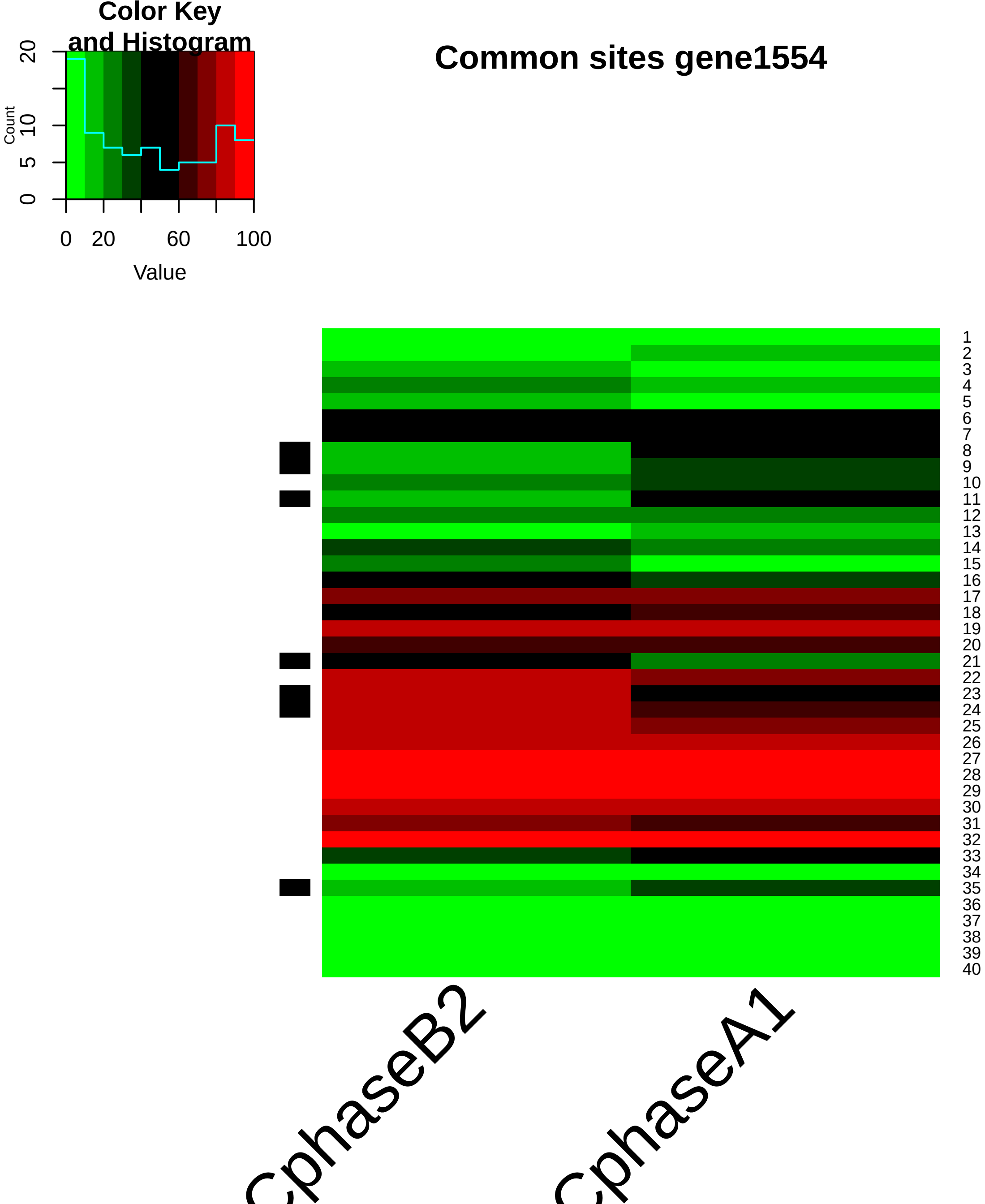
Color Key
and Histogram
Count
0
5
10
20
0
20
60
100
Value
Common sites gene1554
CphaseB2
CphaseA1

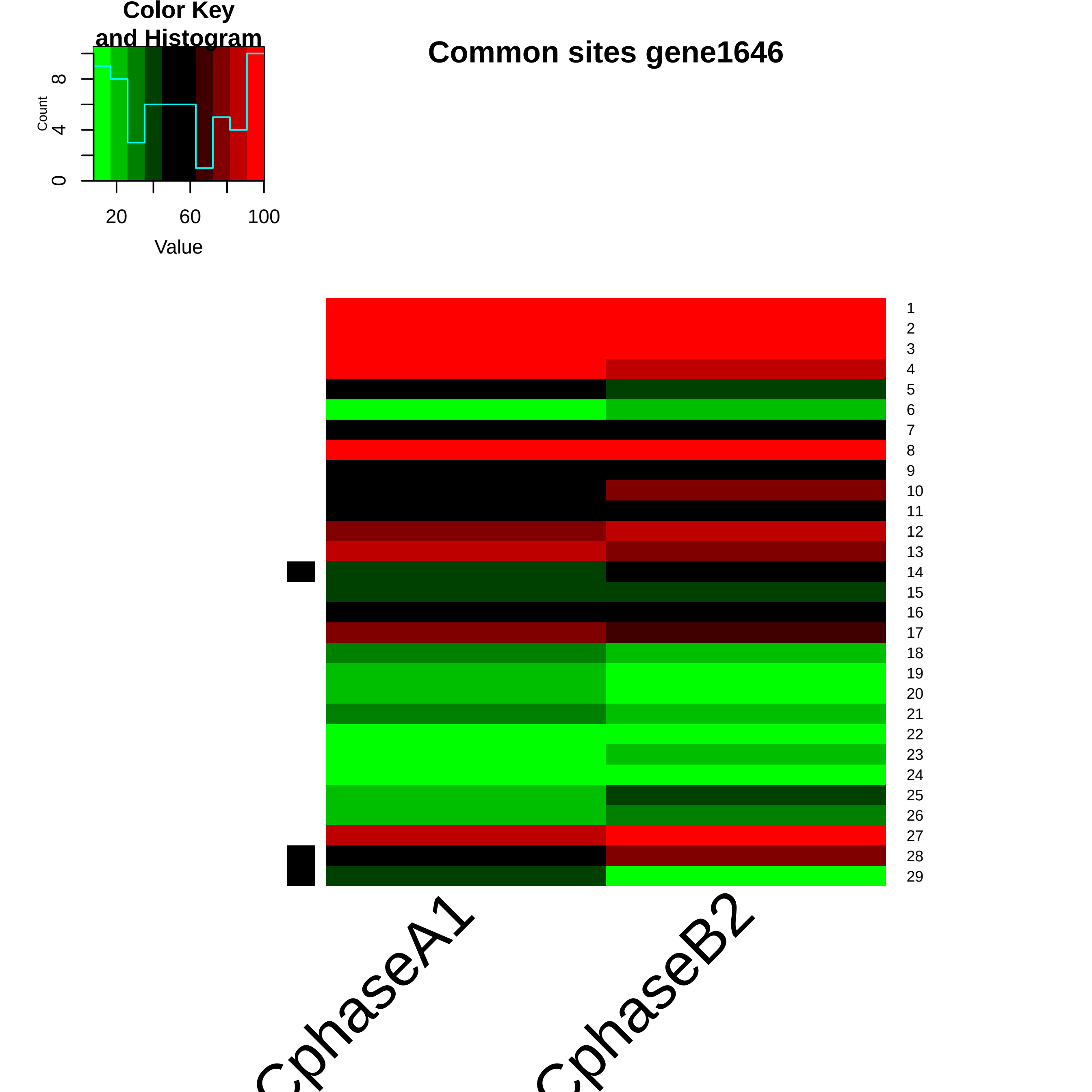

Color Key
and Histogram
Count
0
4
8
20
60
100
Value
Common sites gene1646
CphaseA1
CphaseB2

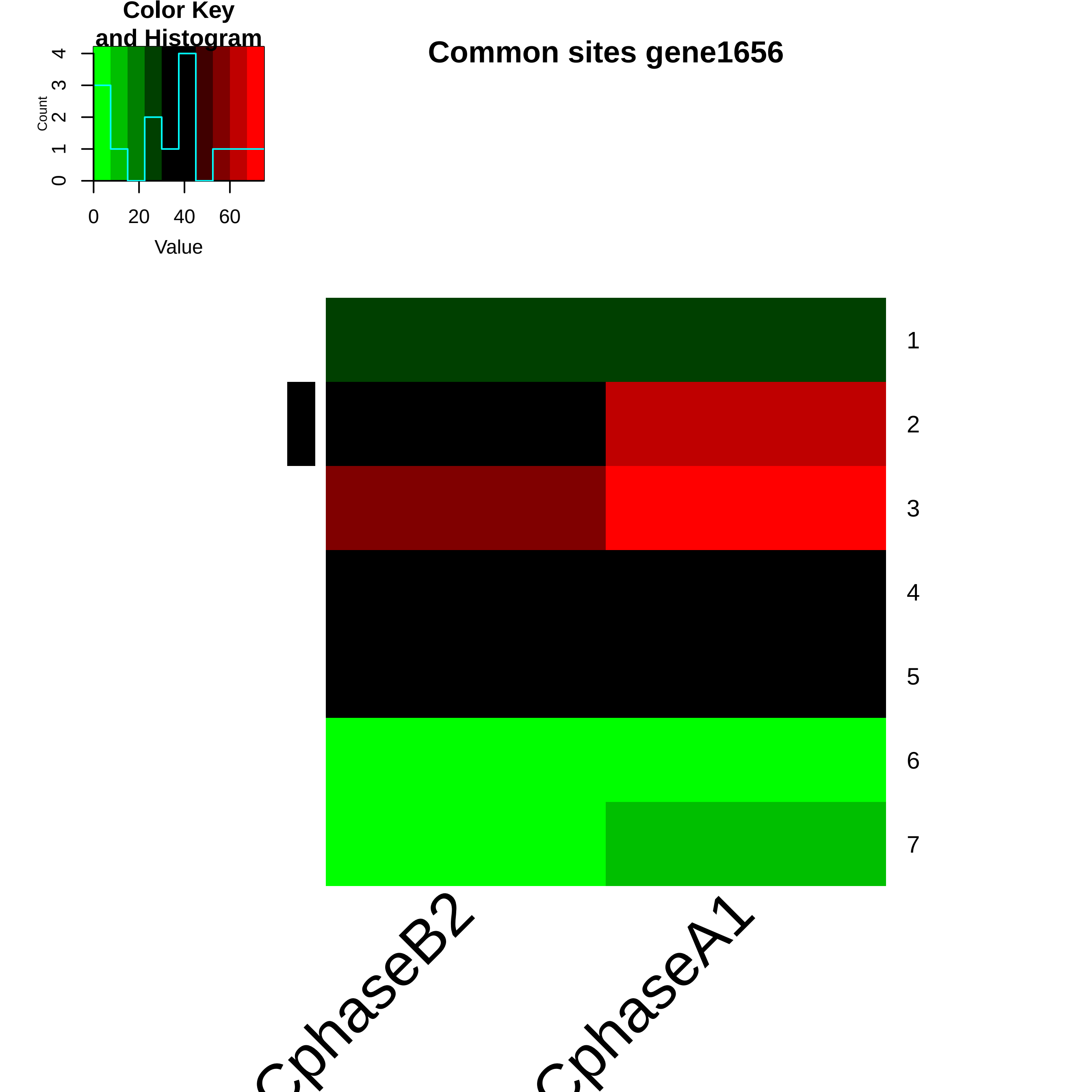

Color Key
and Histogram
Count
Value
0
20
40
60
0
1
2
3
4
Common sites gene1656
1
2
3
4
5
6
7
CphaseB2
CphaseA1

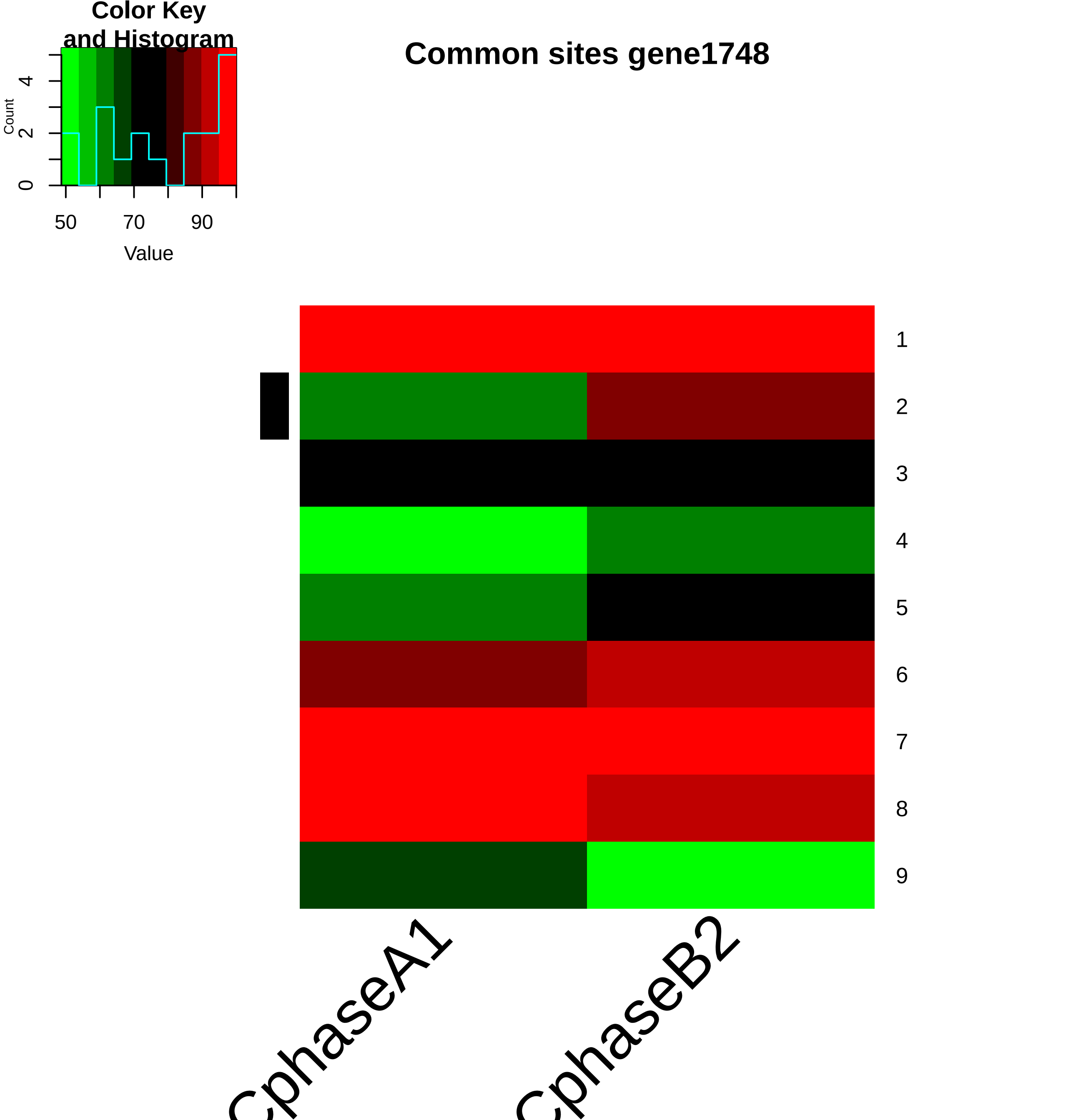
Color Key
and Histogram
Count
0
2
4
50
70
90
Value
Common sites gene1748
1
2
3
4
5
6
7
8
9
CphaseA1
CphaseB2

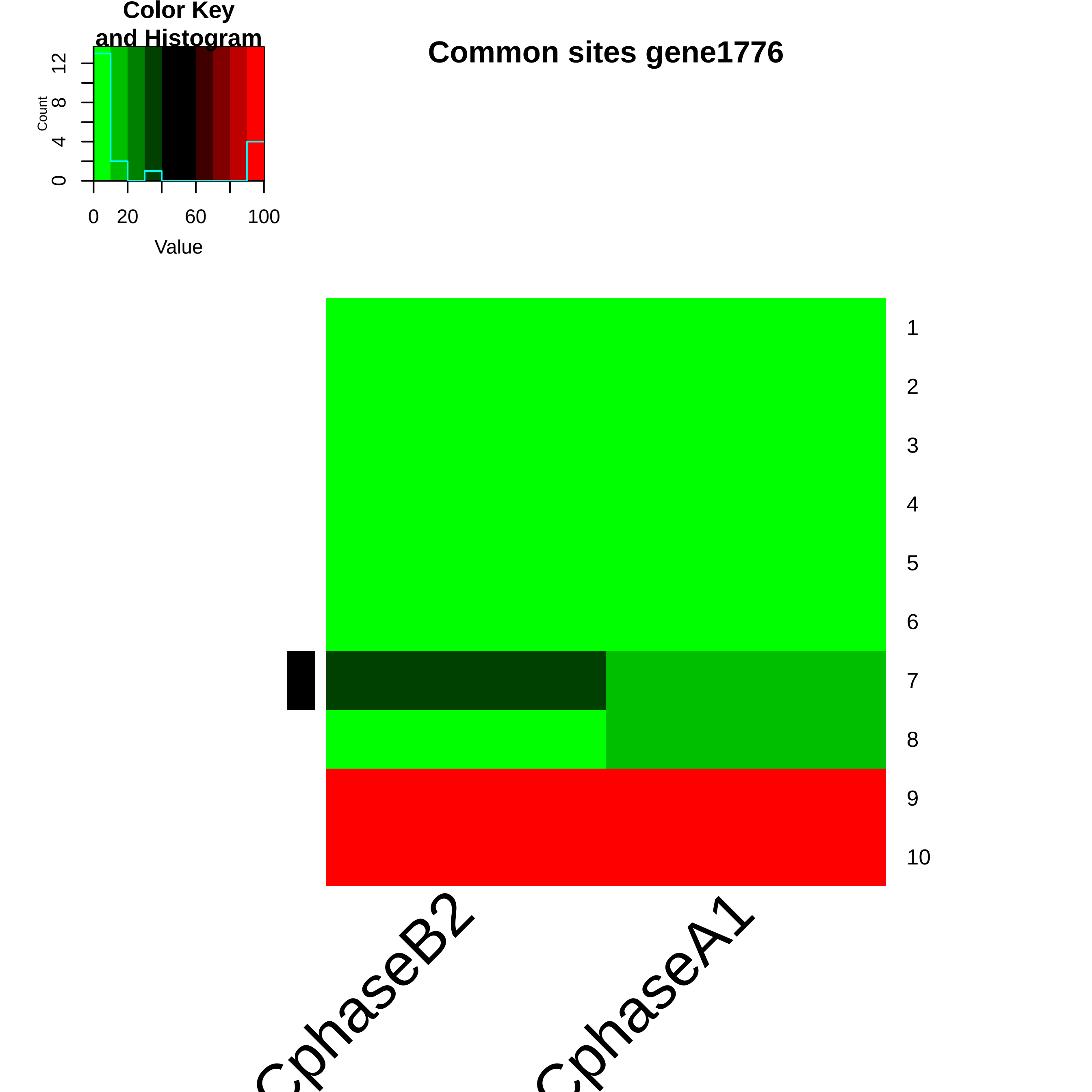

Color Key
and Histogram
Count
0
4
8
12
0
20
60
100
Value
Common sites gene1776
1
2
3
4
5
6
7
8
9
10
CphaseB2
CphaseA1

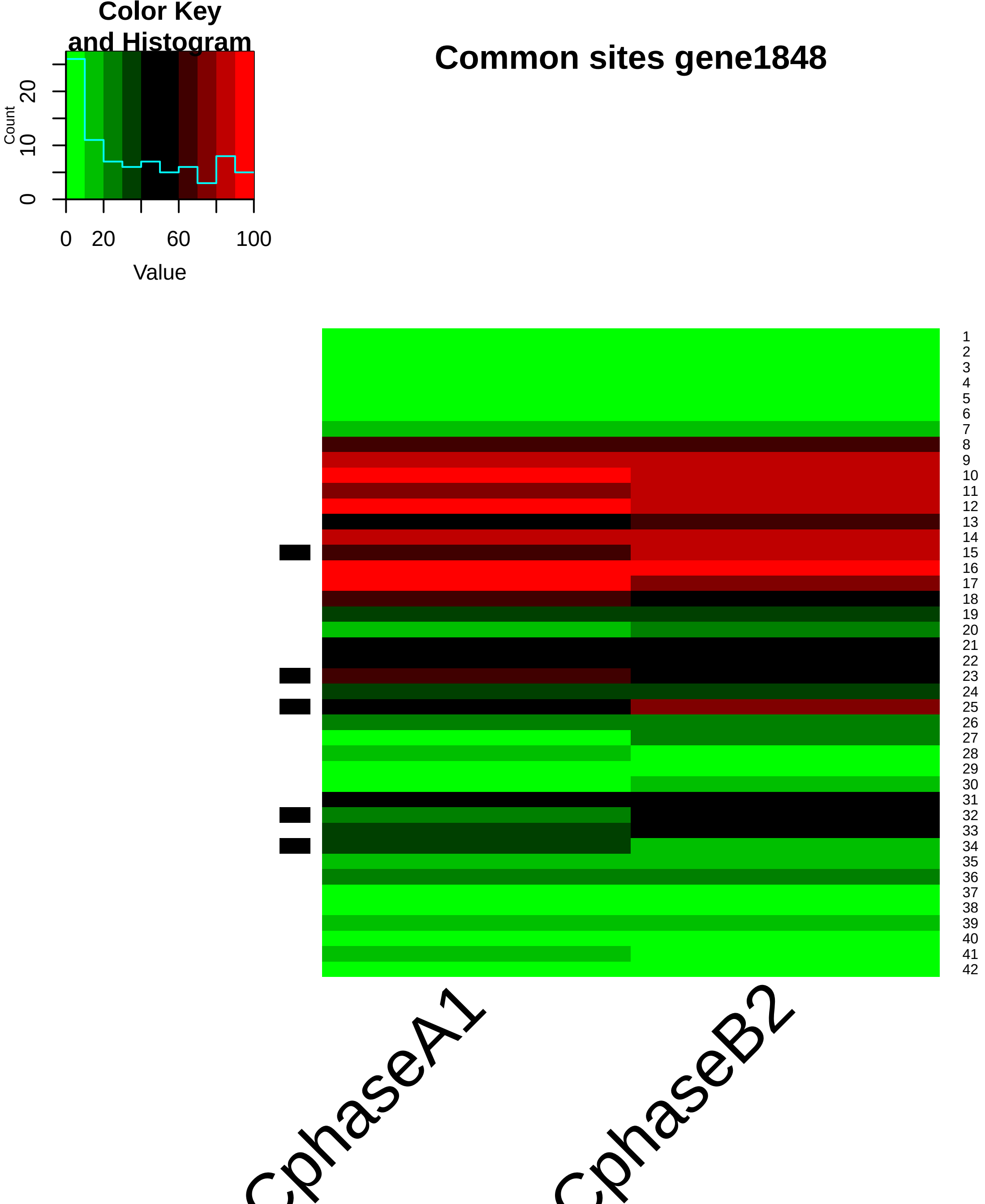
Color Key
and Histogram
Count
0
10
20
0
20
60
100
Value
Common sites gene1848
CphaseA1
CphaseB2

### Common sites gene1910

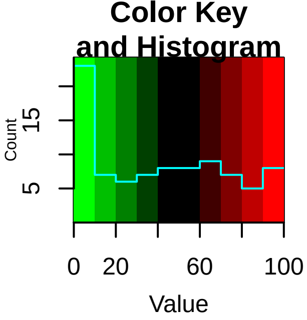

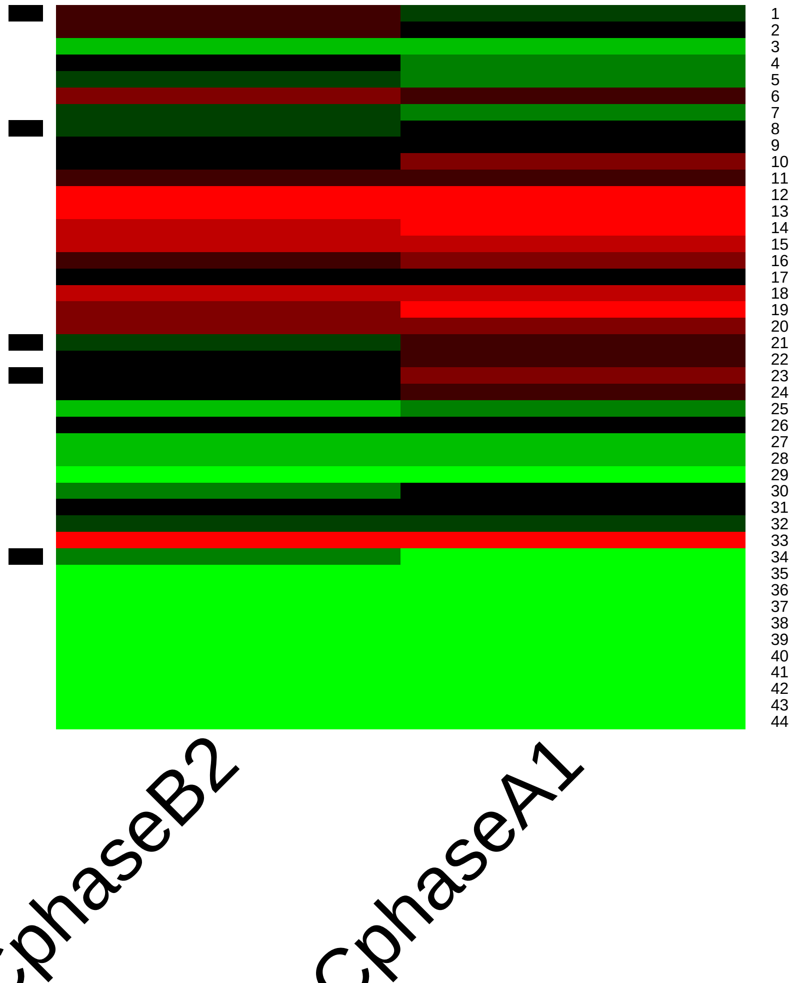

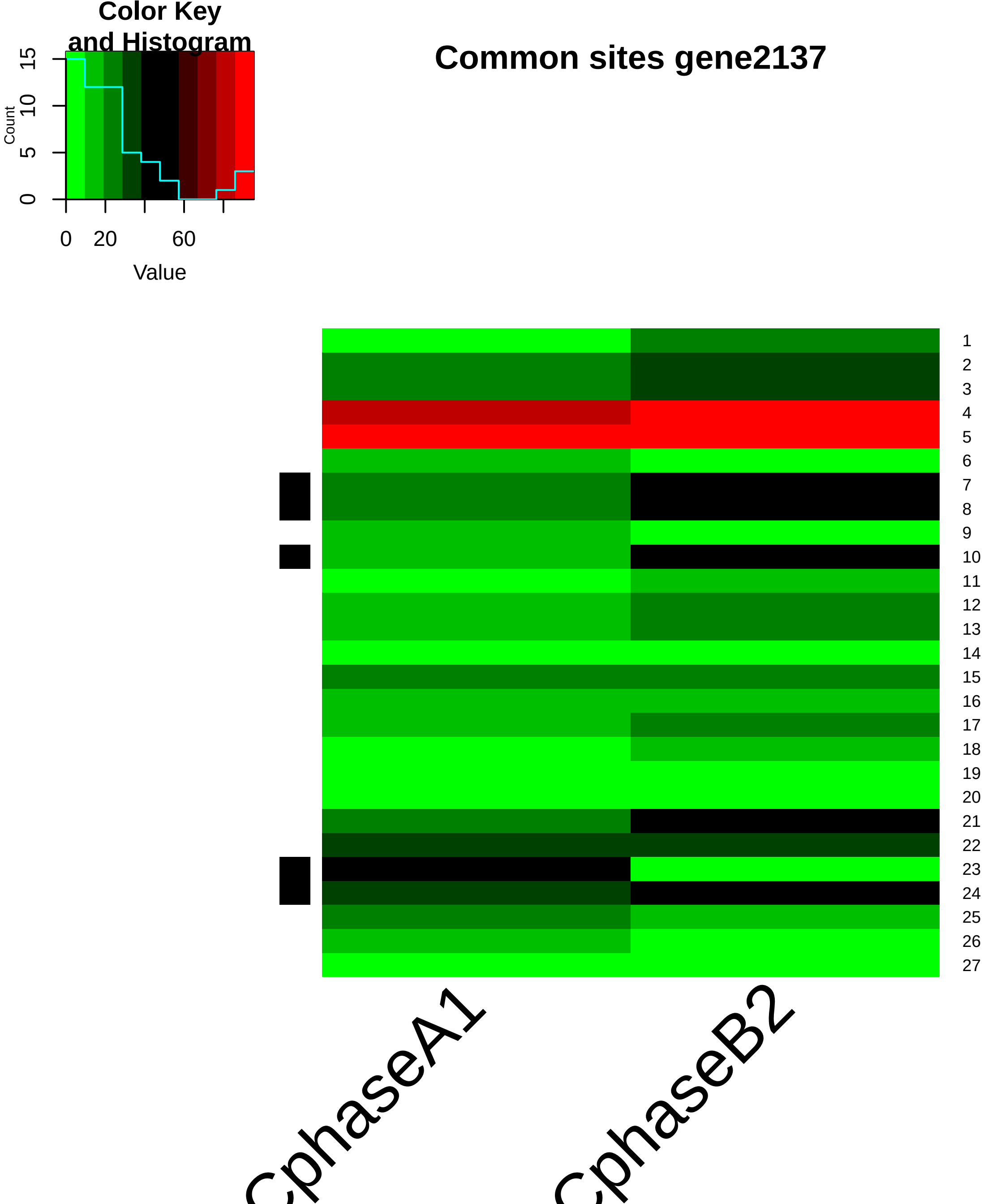
Color Key
and Histogram
Count
0
5
10
15
0
20
60
Value
Common sites gene2137
CphaseA1
CphaseB2

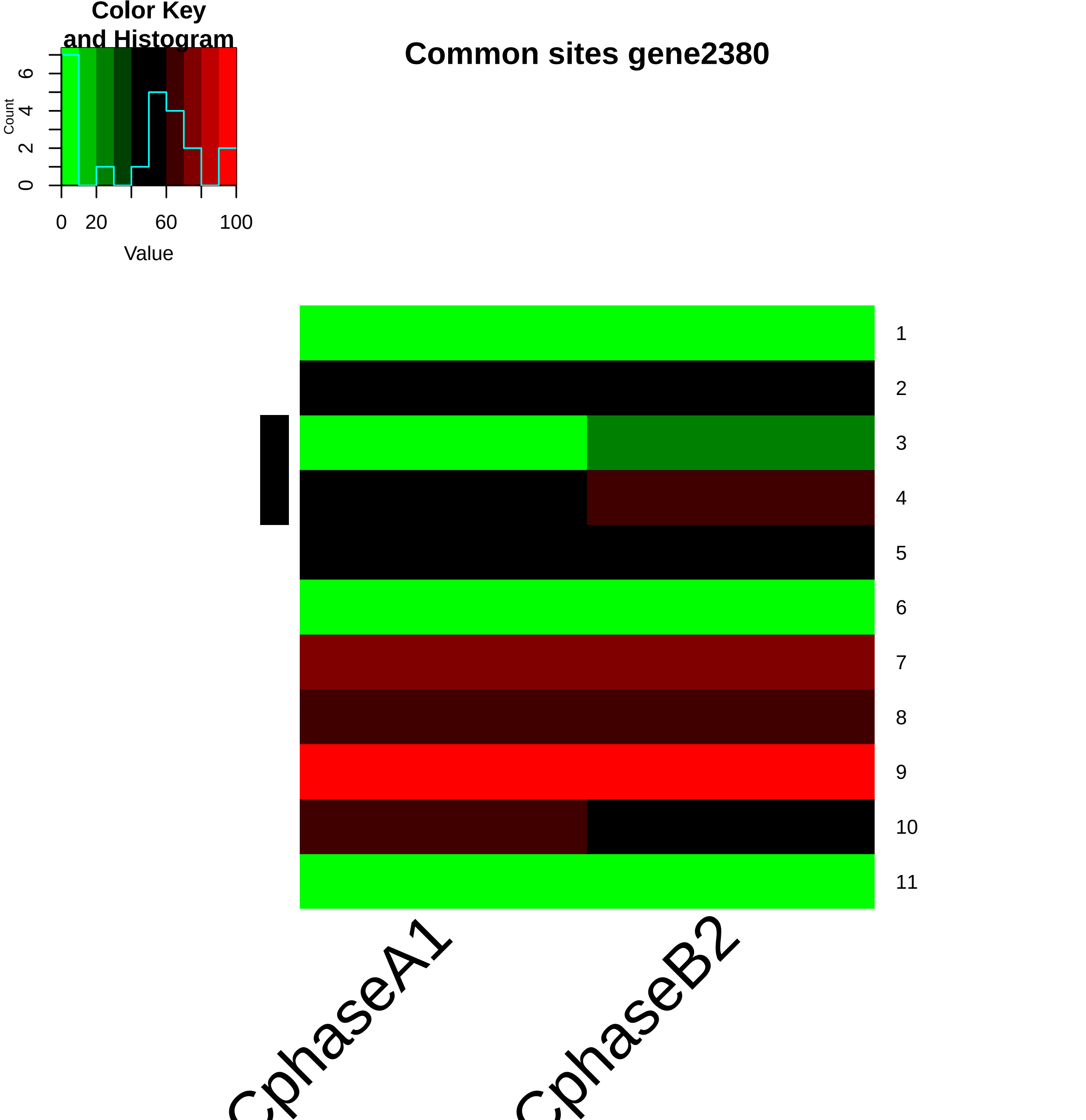
Color Key
and Histogram
Count
0
2
4
6
0
20
60
100
Value
Common sites gene2380
1
2
3
4
5
6
7
8
9
10
11
CphaseA1
CphaseB2

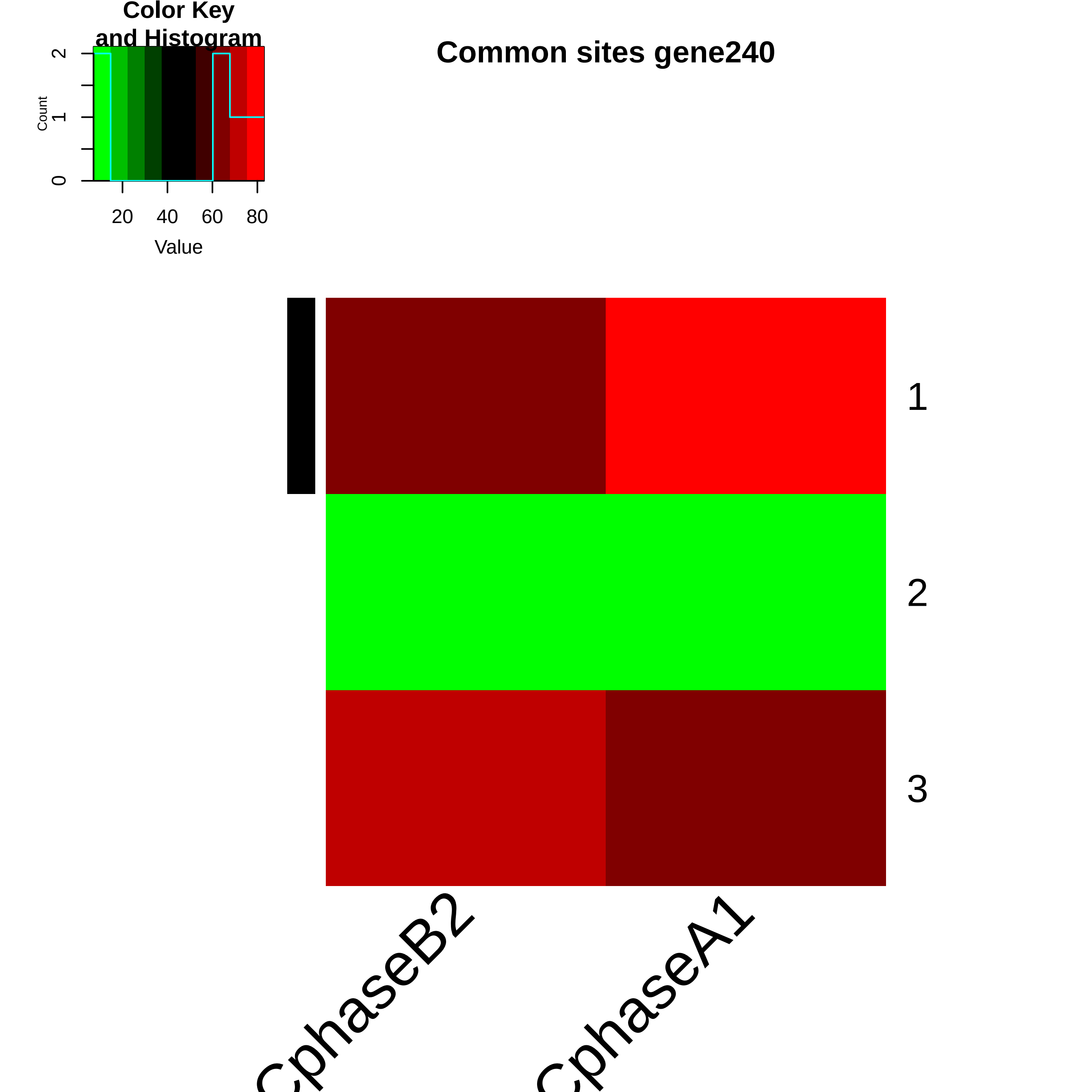
Color Key
and Histogram
Count
0
1
2
20
40
60
80
Value
Common sites gene240
1
2
3
CphaseB2
CphaseA1

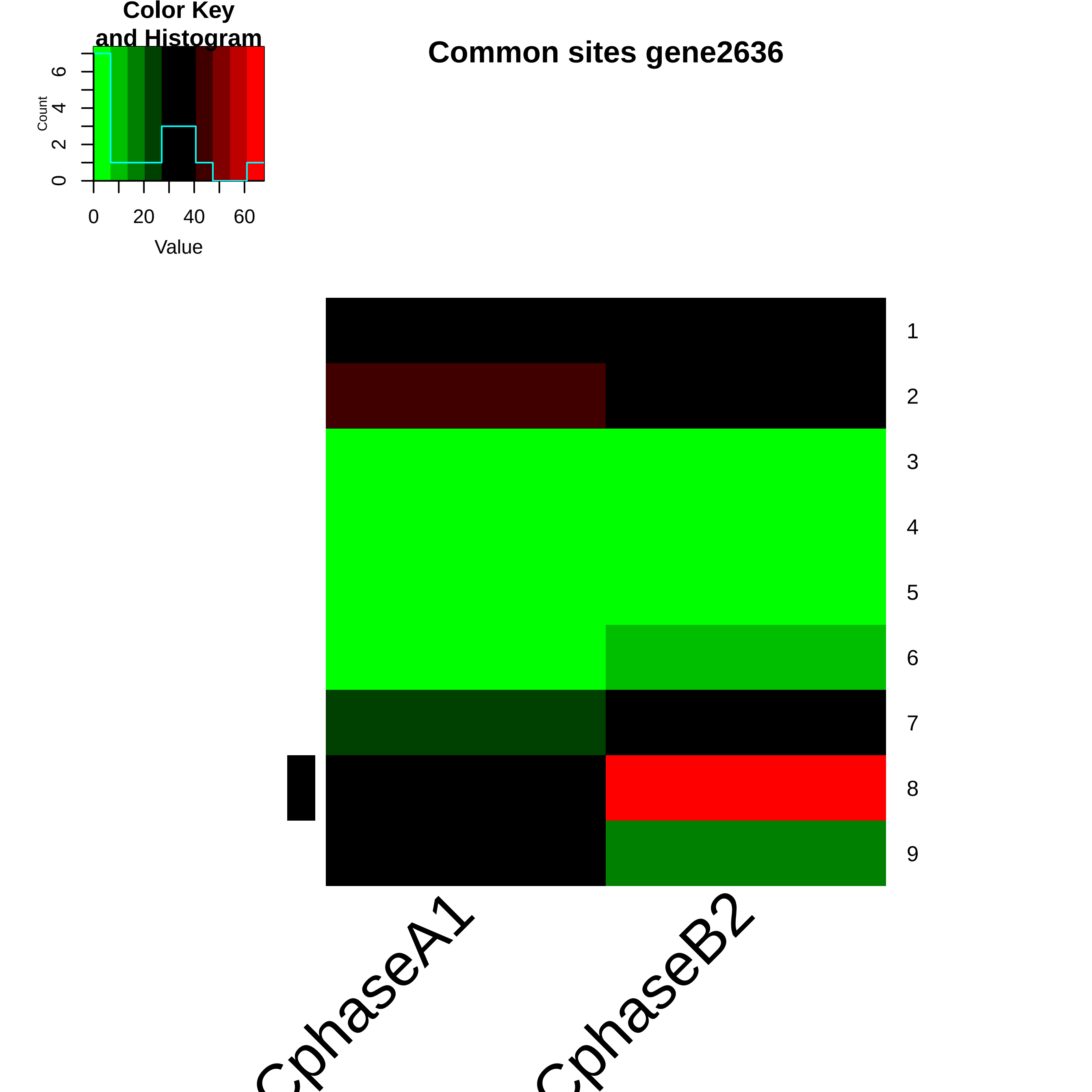
Color Key
and Histogram
Count
0
2
4
6
0
20
40
60
Value
Common sites gene2636
1
2
3
4
5
6
7
8
9
CphaseA1
CphaseB2

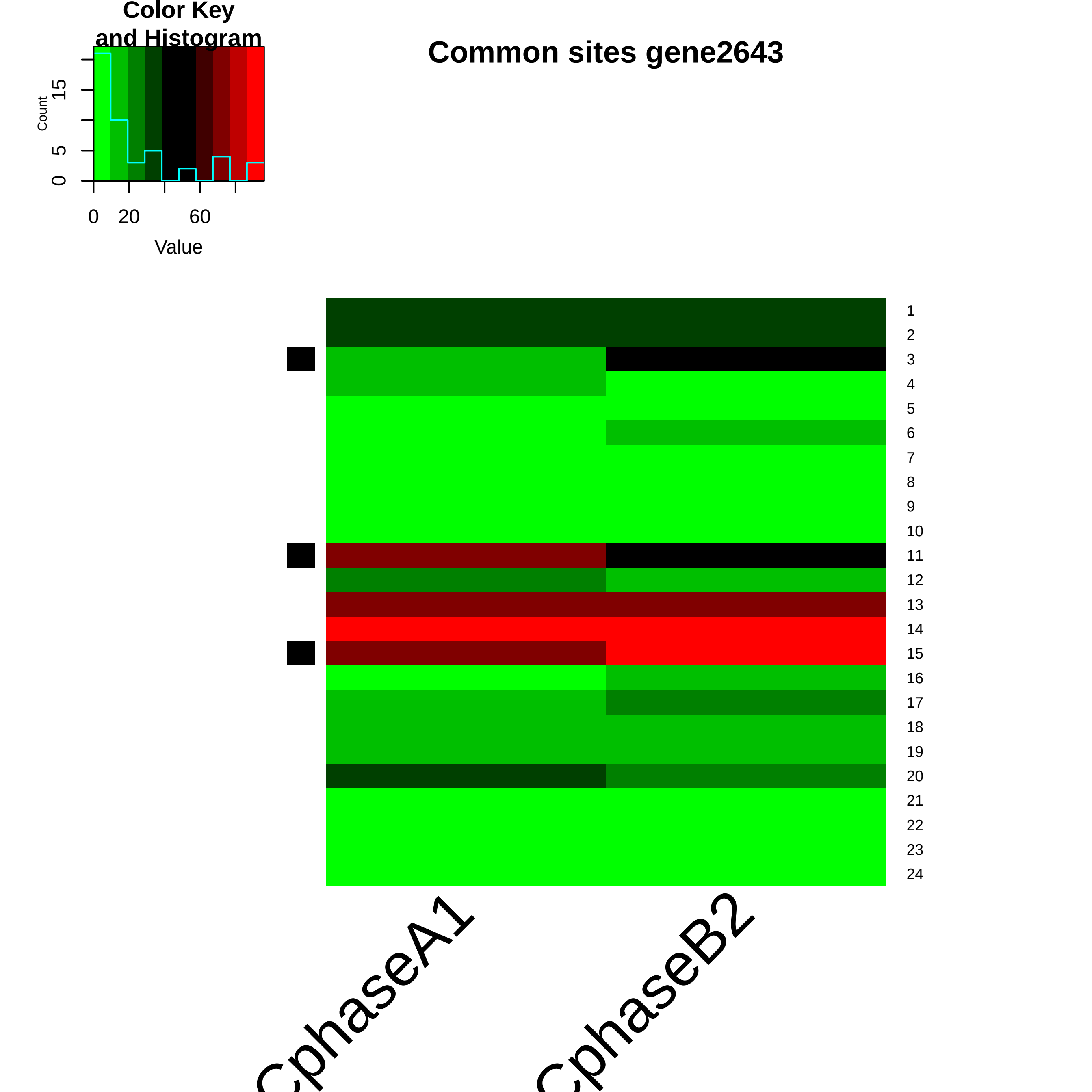
Color Key
and Histogram
Count
0
5
15
0
20
60
Value
Common sites gene2643
CphaseA1
CphaseB2

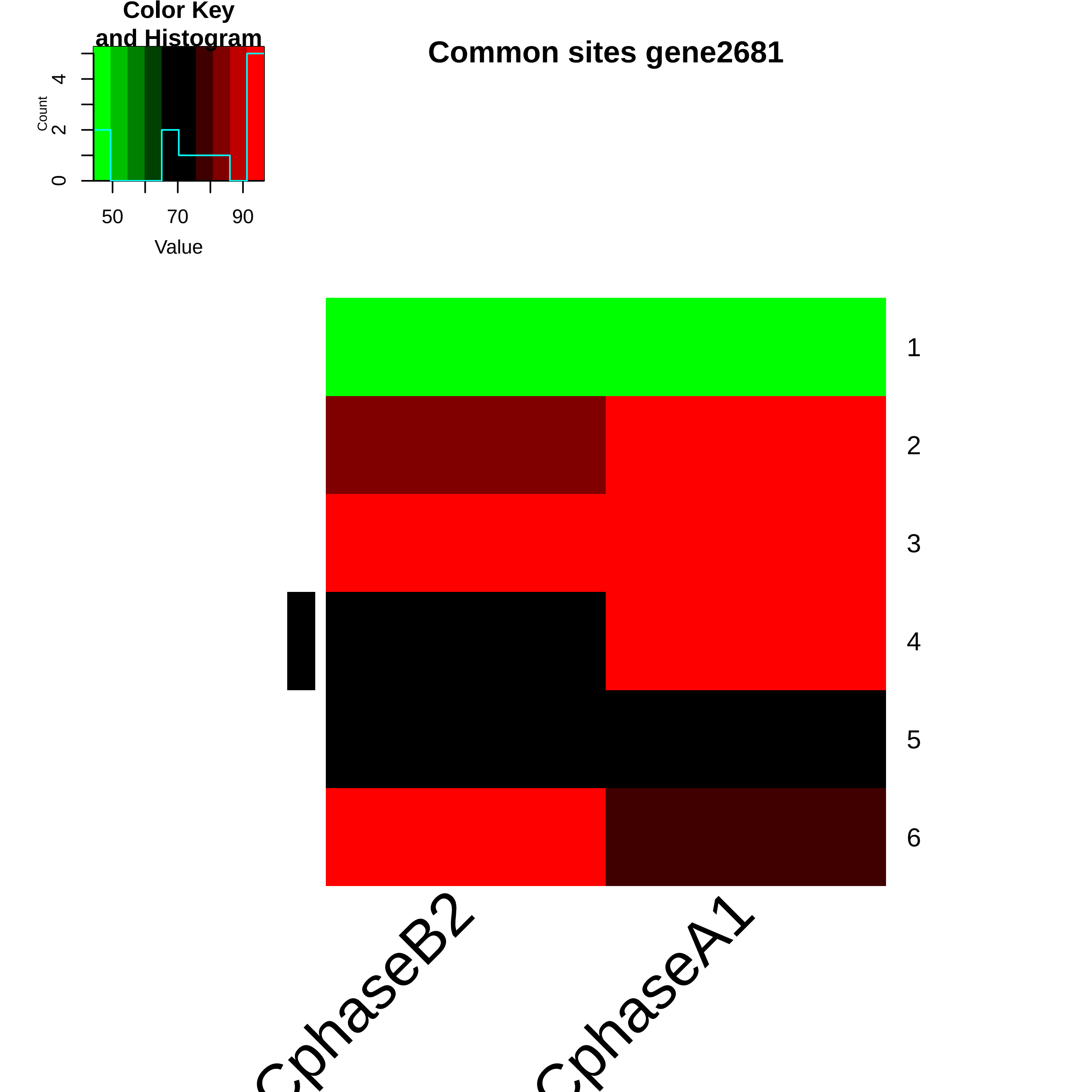

Color Key
and Histogram
Count
0
2
4
50
70
90
Value
Common sites gene2681
1
2
3
4
5
6
CphaseB2
CphaseA1

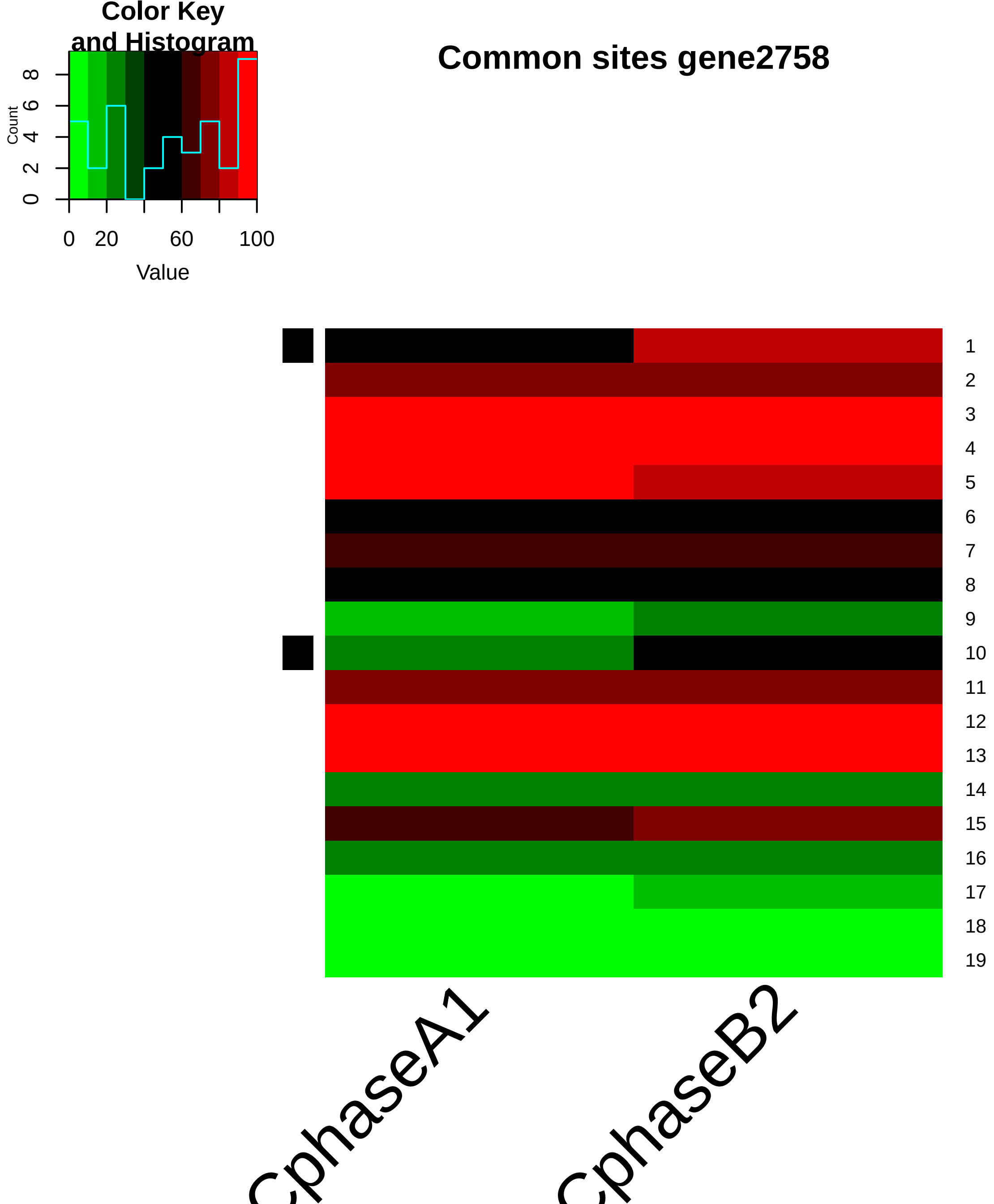

Color Key
and Histogram
Count
0 2 4 6 8
0 20 60 100
Value
Common sites gene2758
CphaseA1
CphaseB2

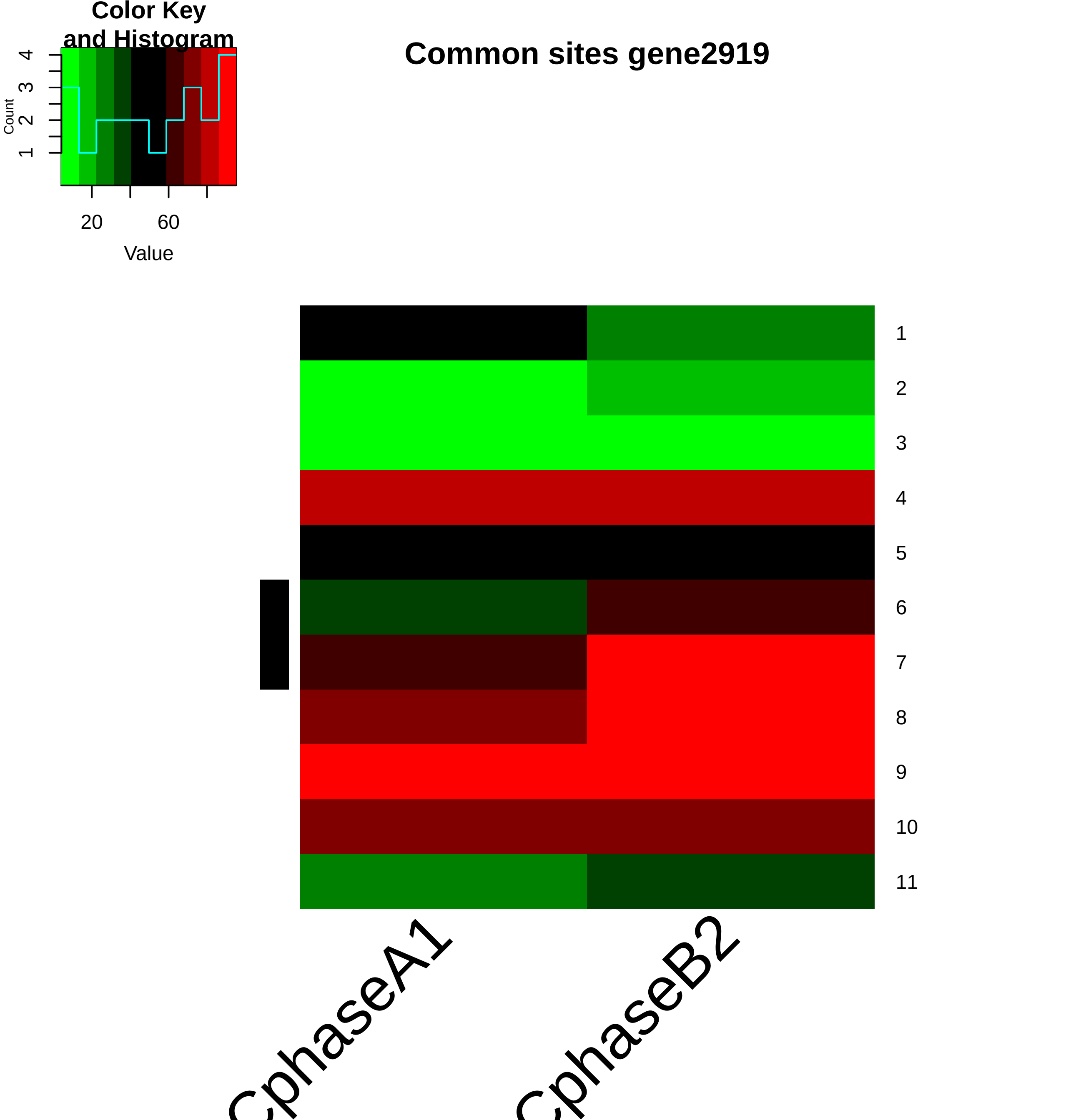
Color Key
and Histogram
Count
1
2
3
4
20
60
Value
Common sites gene2919
1
2
3
4
5
6
7
8
9
10
11
CphaseA1
CphaseB2

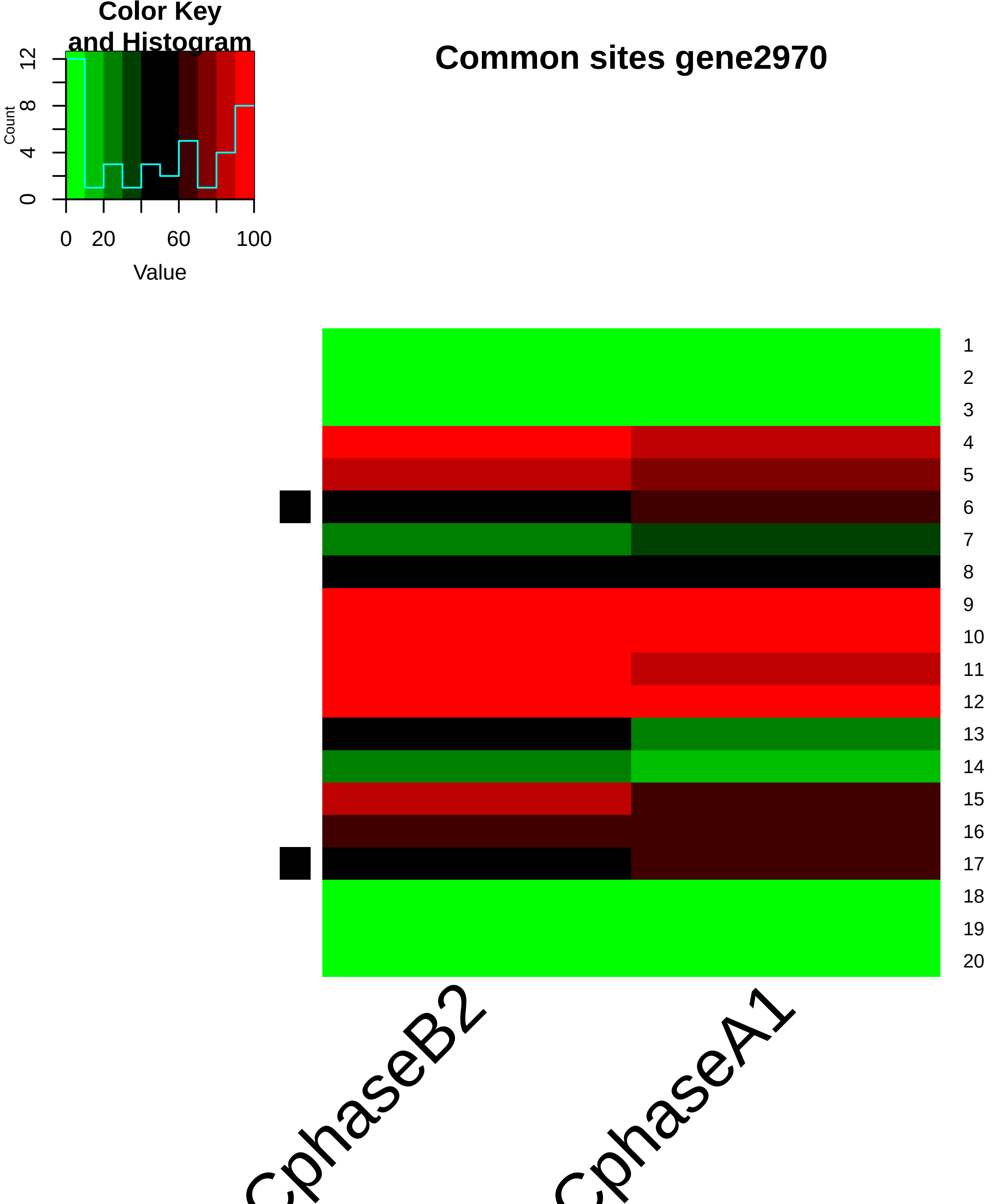
Color Key
and Histogram
Count
0
4
8
12
0
20
60
100
Value
Common sites gene2970
CphaseB2
CphaseA1

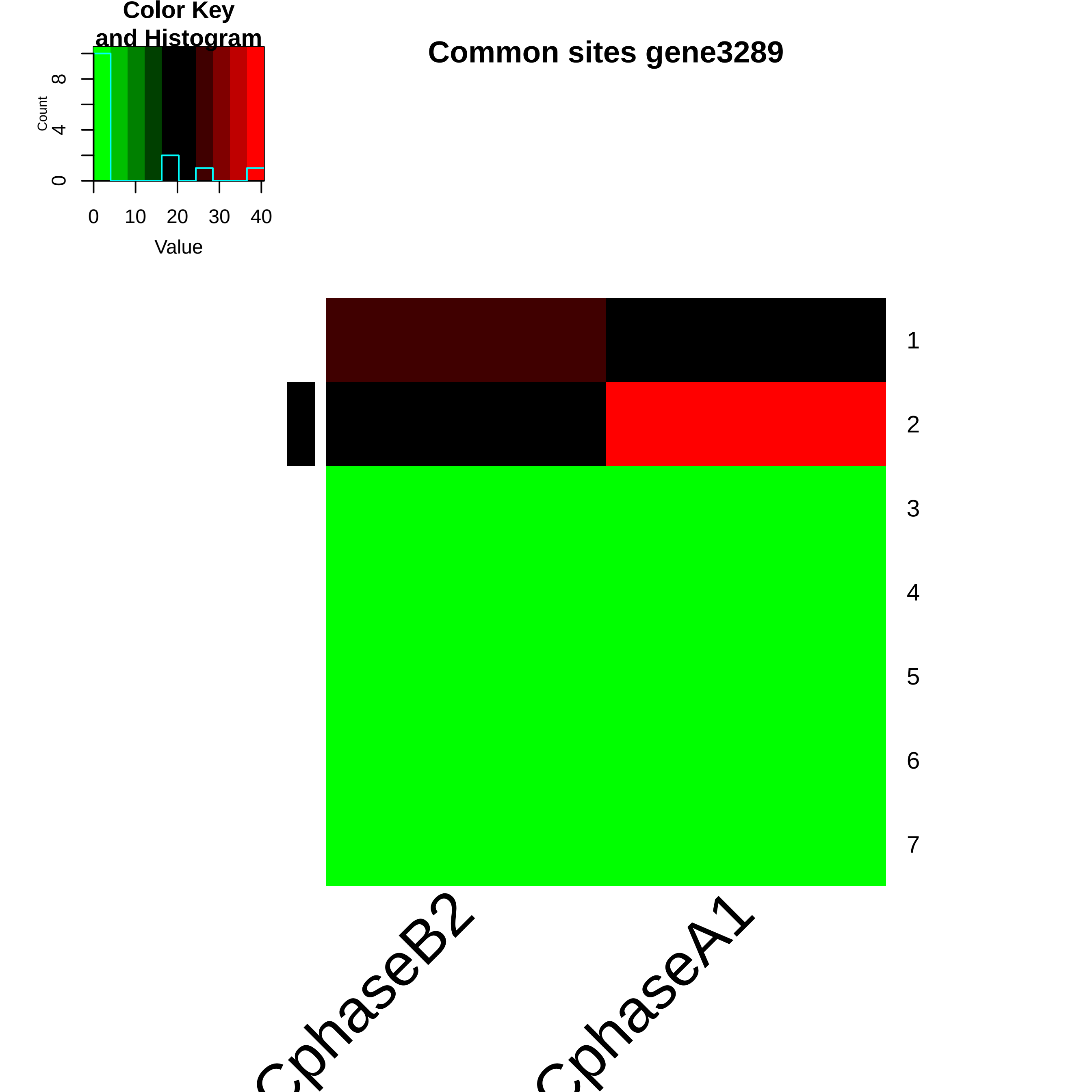
Color Key
and Histogram
Count
0
4
8
0
10
20
30
40
Value
Common sites gene3289
1
2
3
4
5
6
7
CphaseB2
CphaseA1

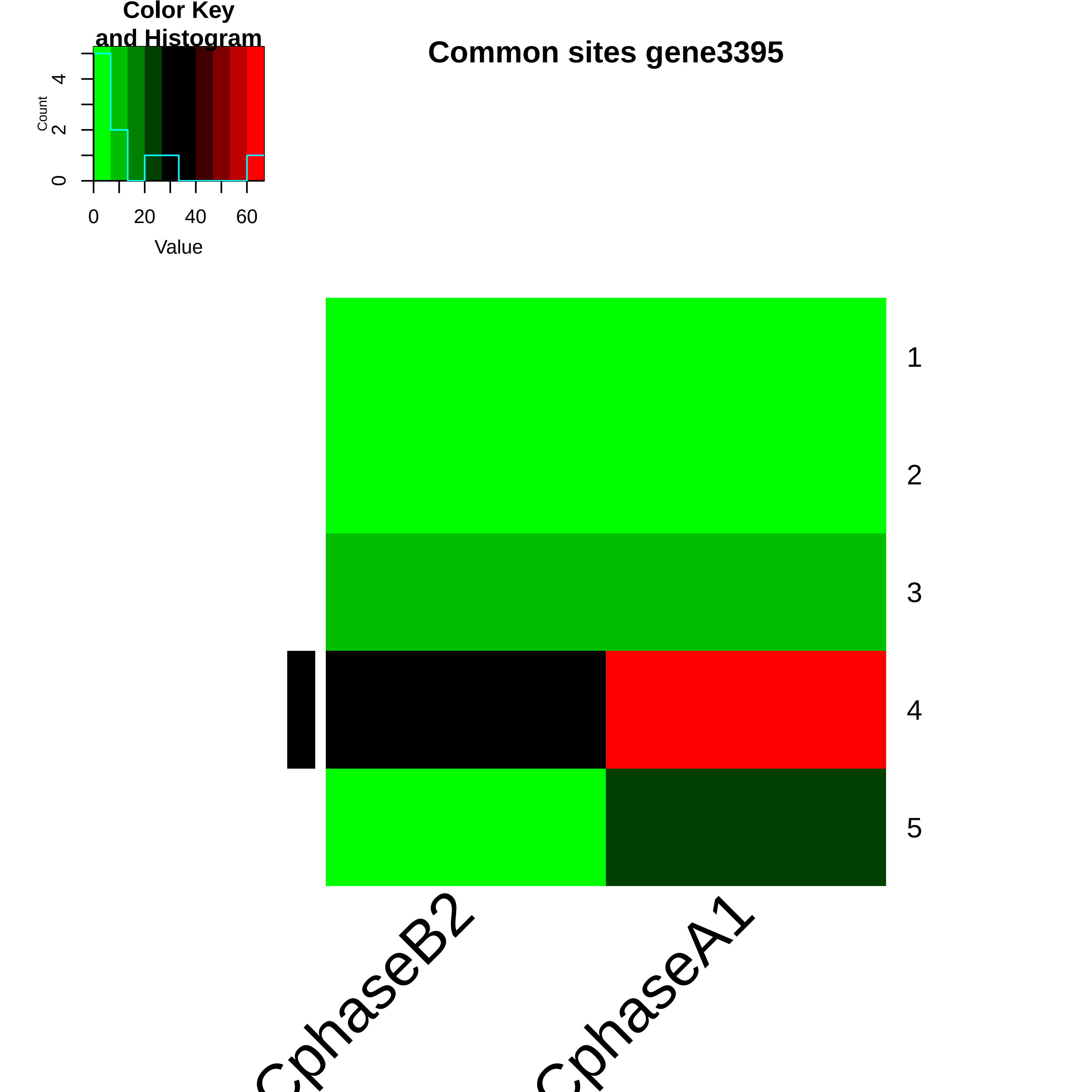
Color Key
and Histogram
Count
0
2
4
0
20
40
60
Value
Common sites gene3395
1
2
3
4
5
CphaseB2
CphaseA1

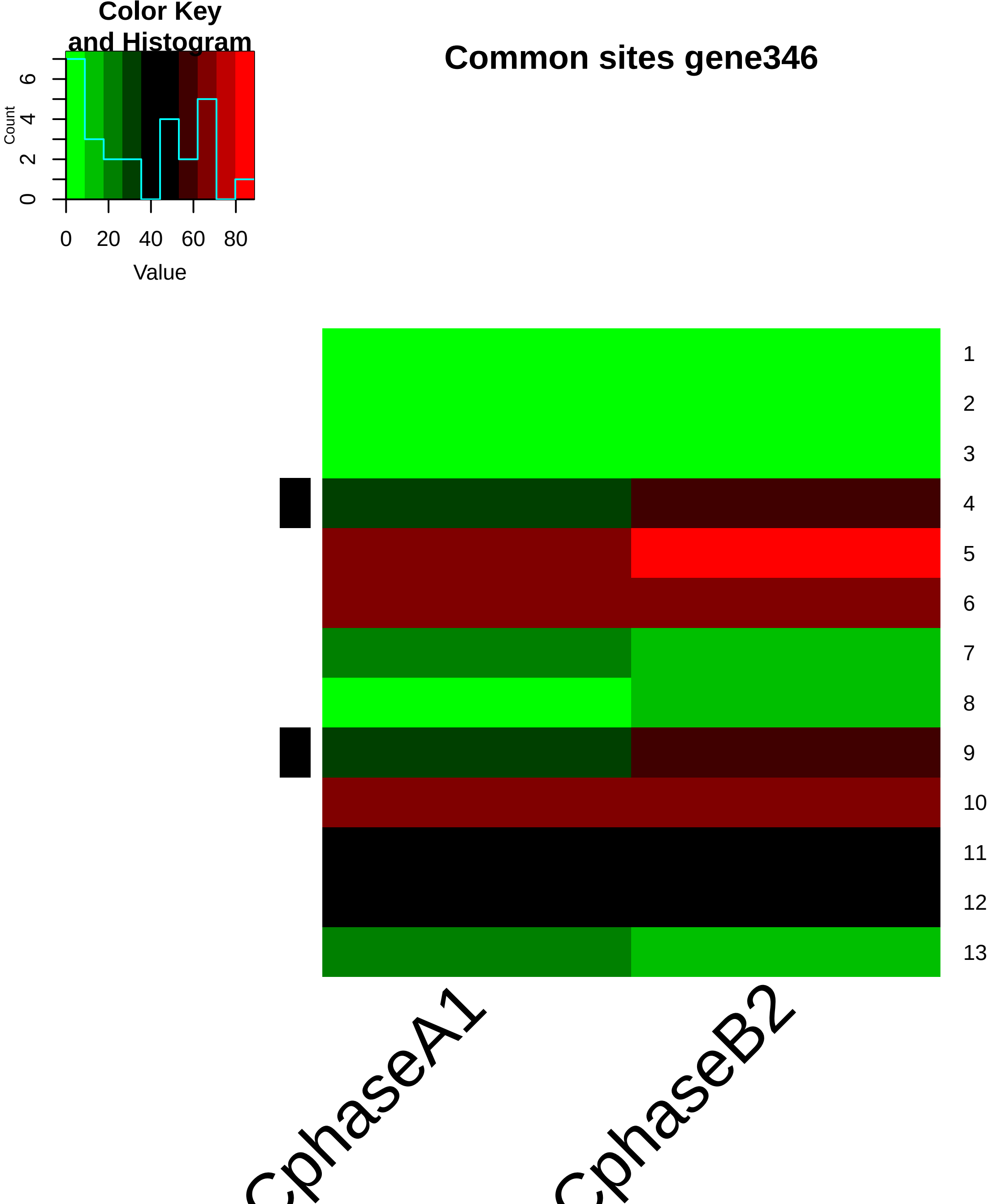
Color Key
and Histogram
Count
0
2
4
6
0
20
40
60
80
Value
Common sites gene346
1
2
3
4
5
6
7
8
9
10
11
12
13
CphaseA1
CphaseB2

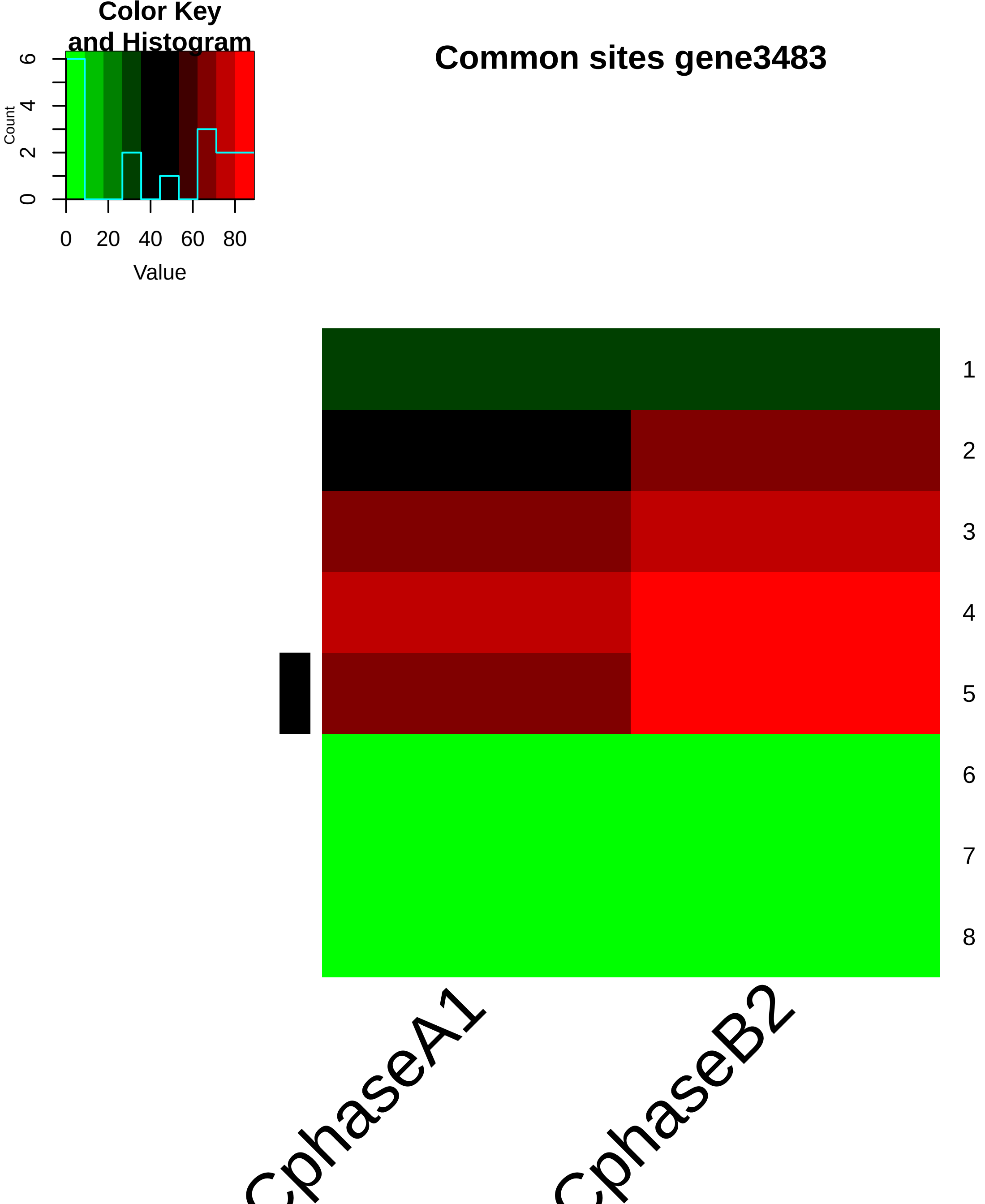
Color Key
and Histogram
Count
0
2
4
6
0
20
40
60
80
Value
Common sites gene3483
1
2
3
4
5
6
7
8
CphaseA1
CphaseB2

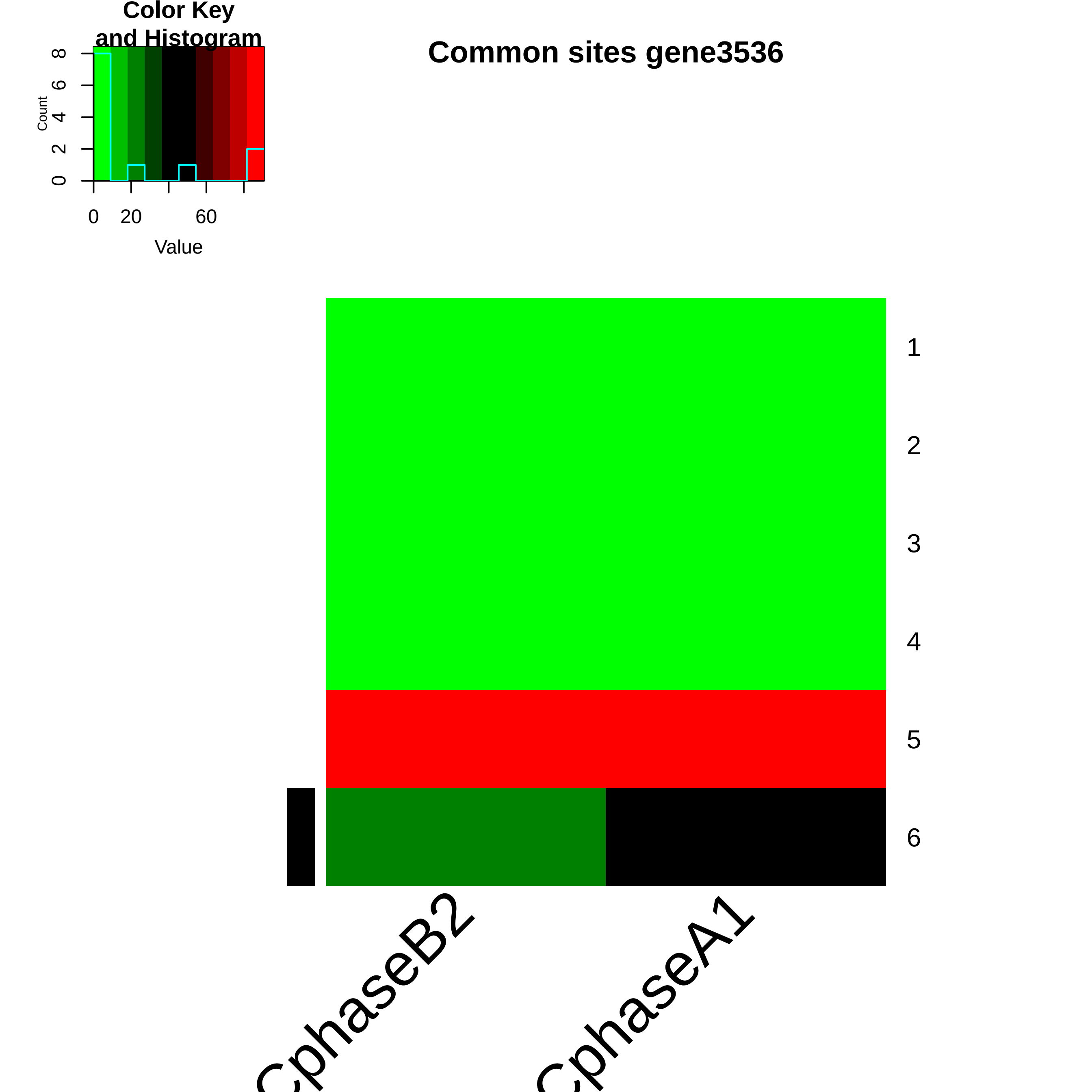

Color Key
and Histogram
Count
0
2
4
6
8
0
20
60
Value
Common sites gene3536
1
2
3
4
5
6
CphaseB2
CphaseA1

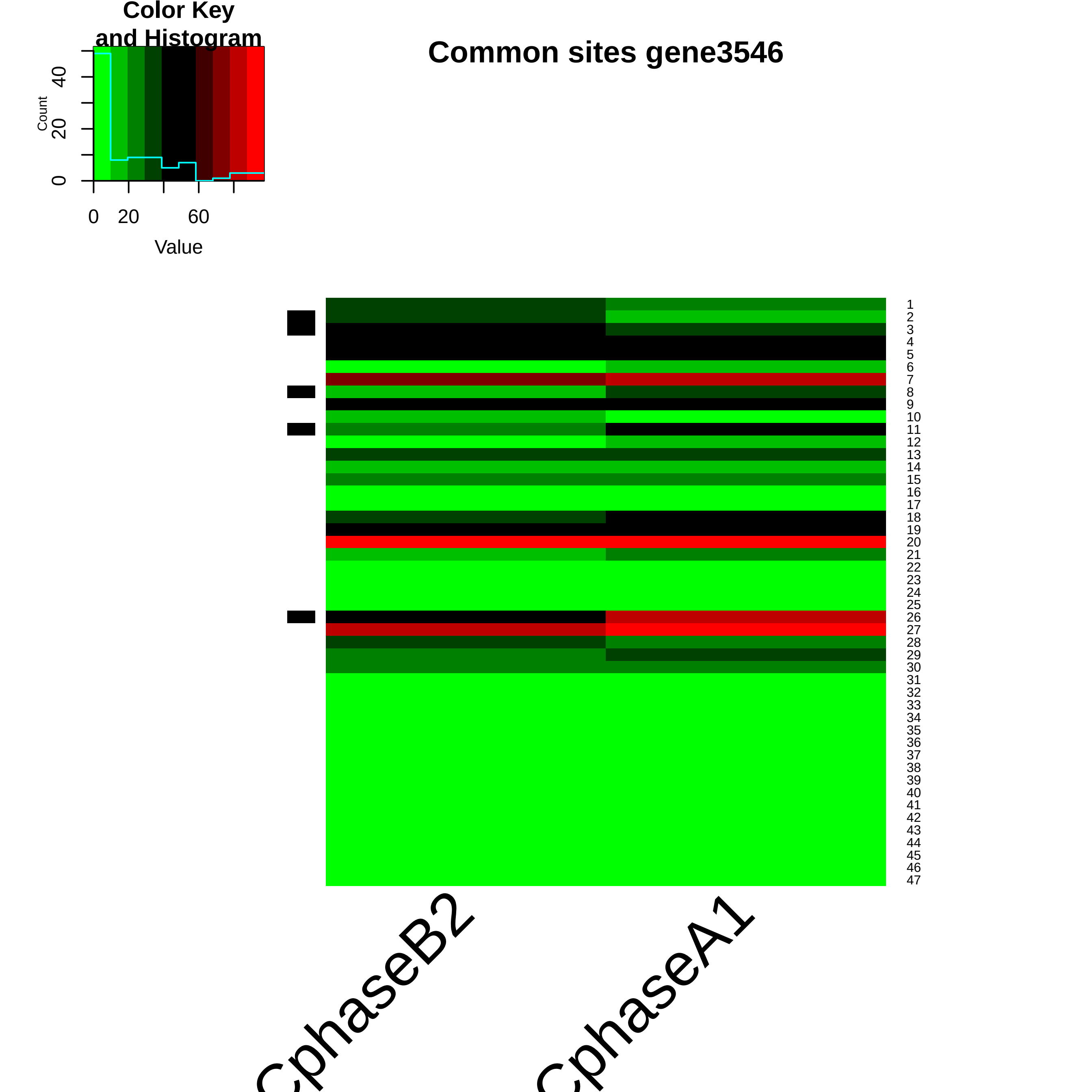
Color Key
and Histogram
Count
0
20
40
0
20
60
Value
Common sites gene3546
CphaseB2
CphaseA1

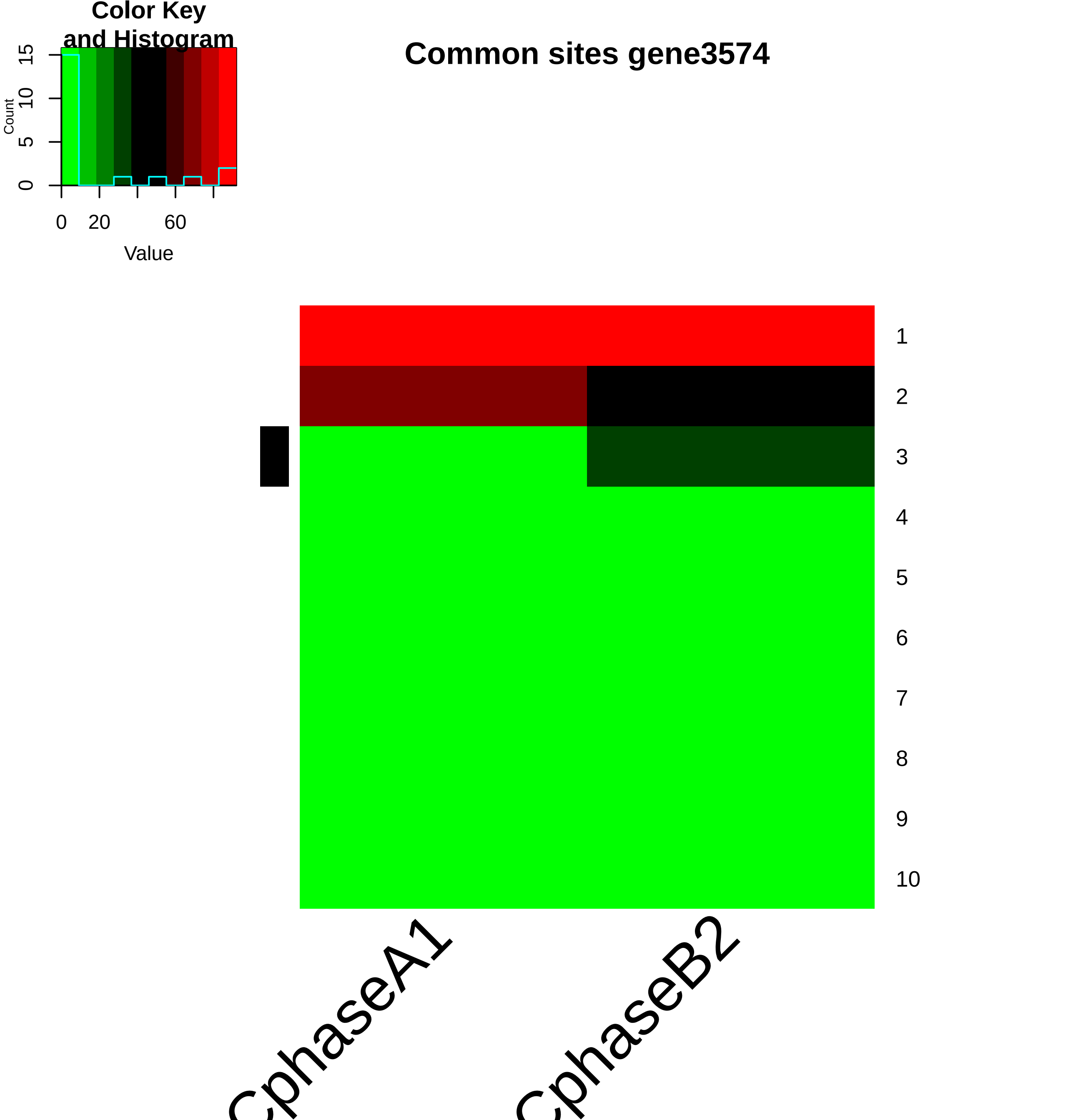

Color Key
and Histogram
Count
0
5
10
15
0
20
60
Value
Common sites gene3574
1
2
3
4
5
6
7
8
9
10
CphaseA1
CphaseB2

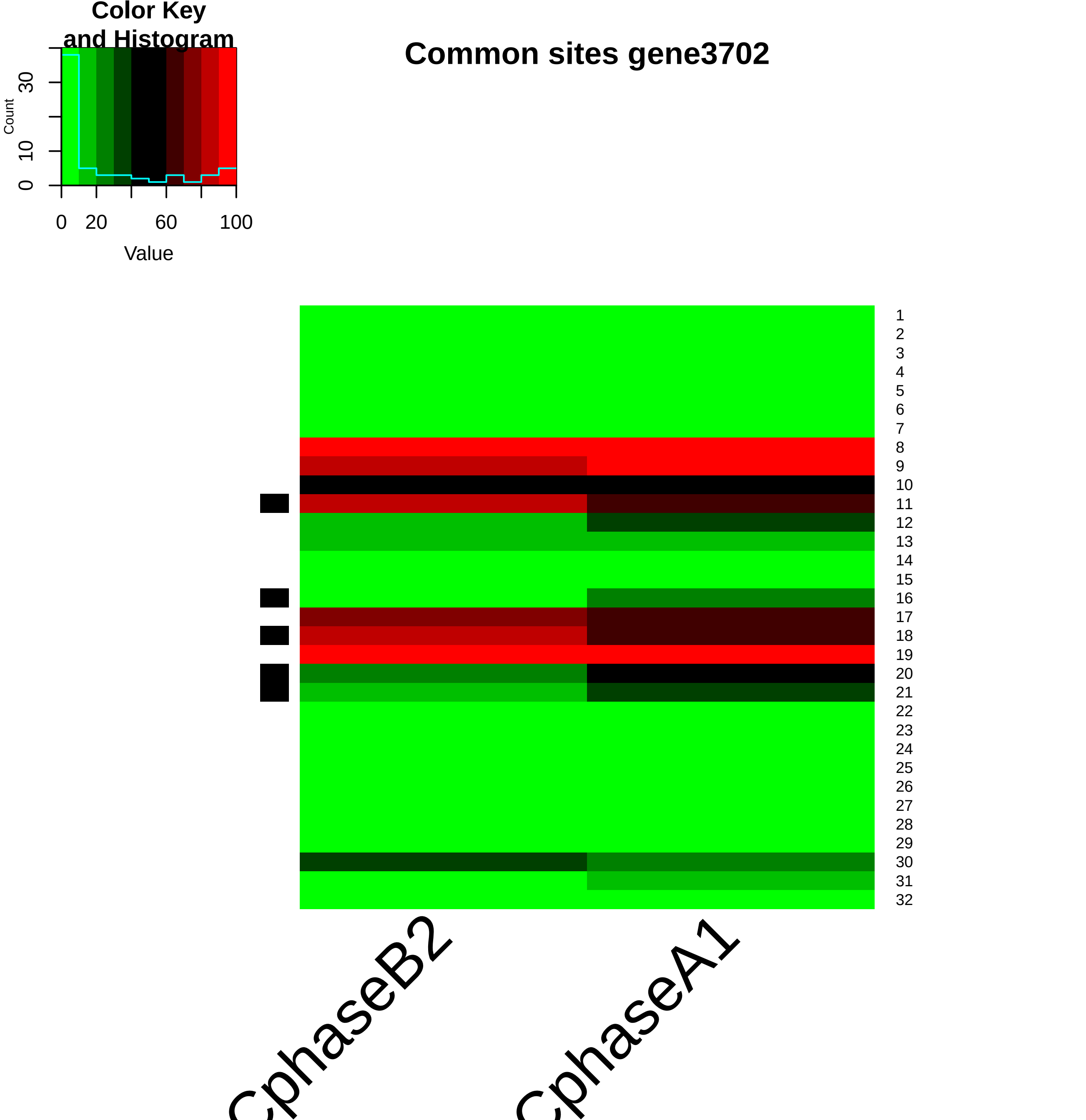

Color Key
and Histogram
Count
0
10
30
0
20
60
100
Value
Common sites gene3702
CphaseB2
CphaseA1

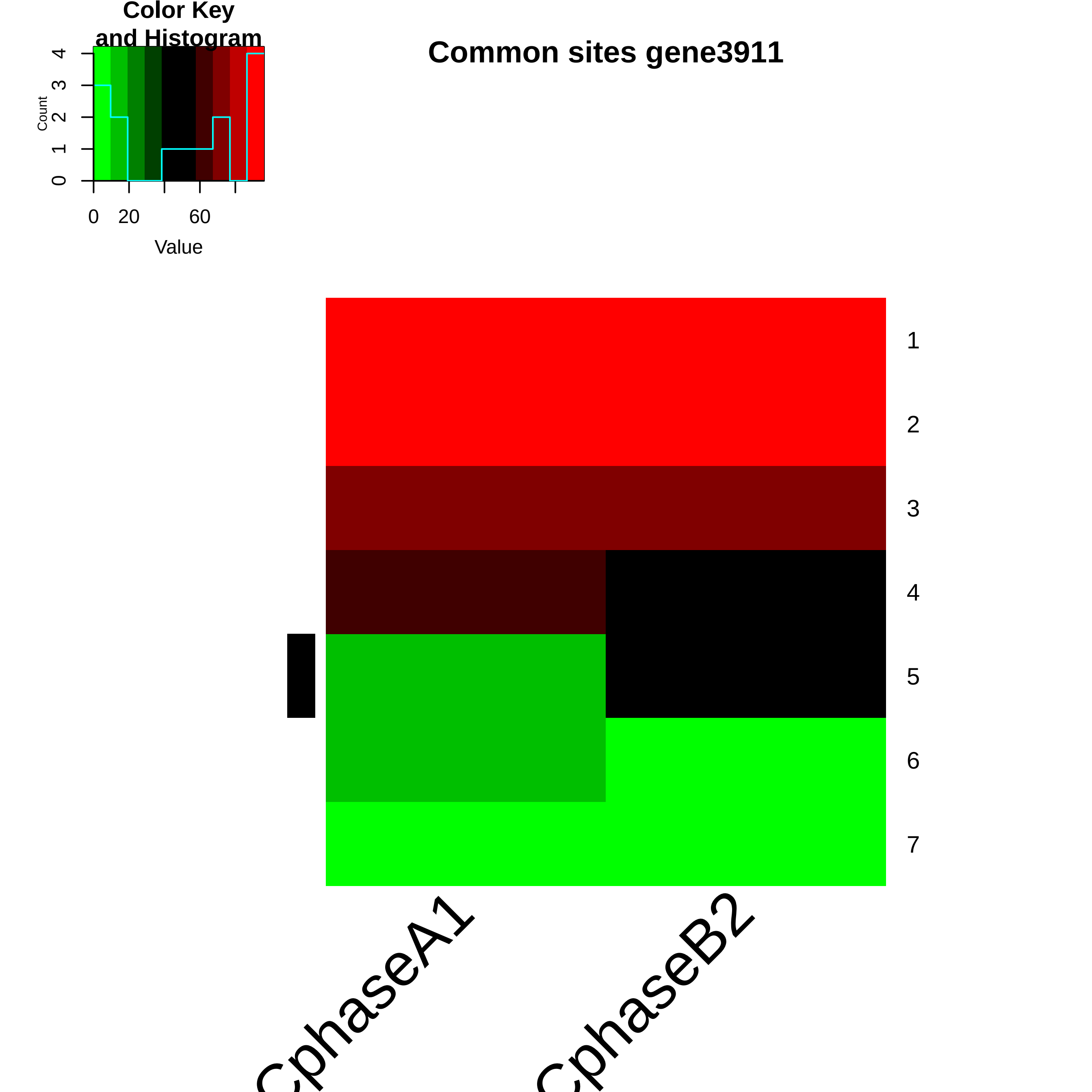

Color Key
and Histogram
Count
0
1
2
3
4
0
20
60
Value
Common sites gene3911
1
2
3
4
5
6
7
CphaseA1
CphaseB2

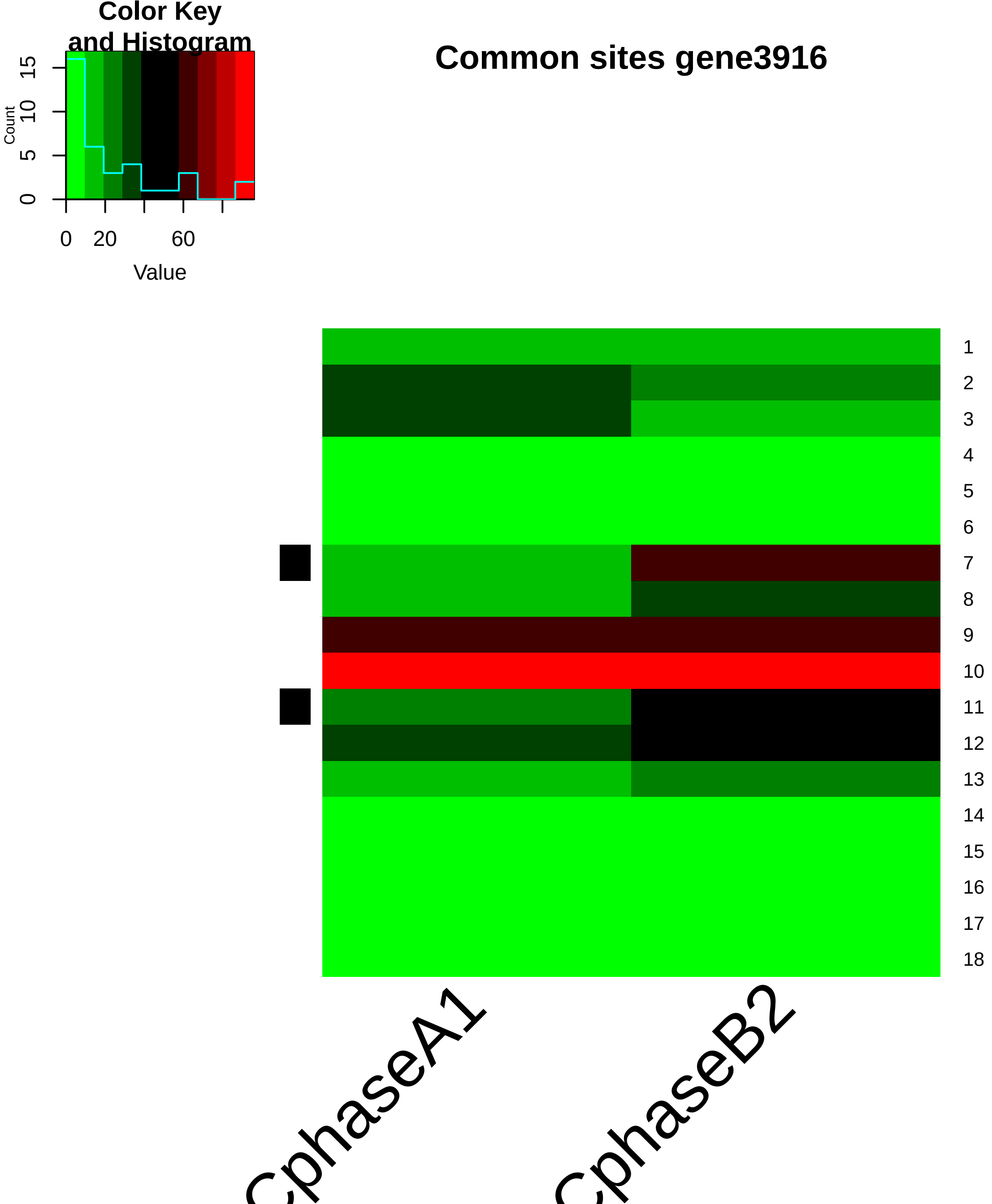
Color Key
and Histogram
Count
0
5
10
15
0
20
60
Value
Common sites gene3916
CphaseA1
CphaseB2

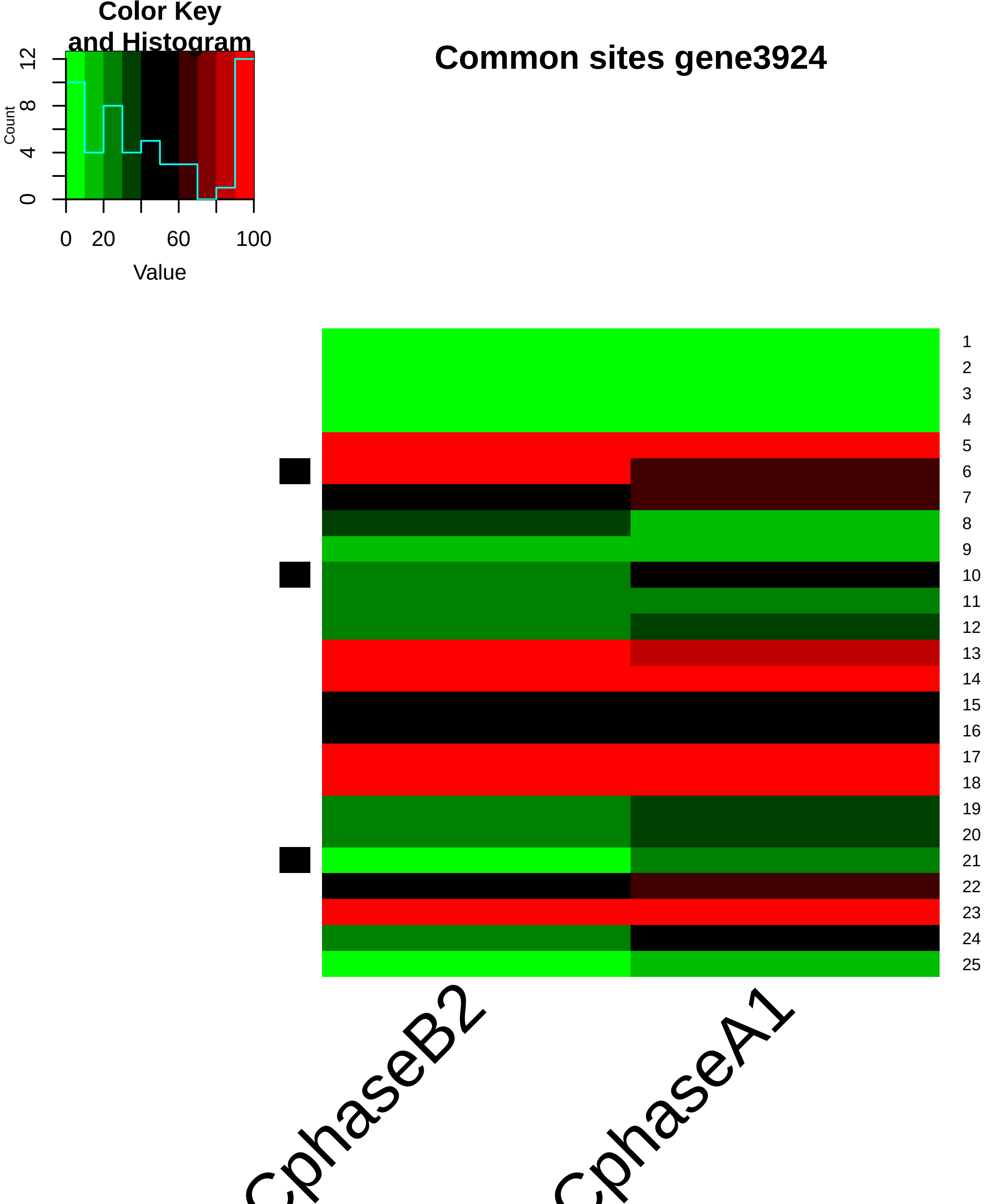
Color Key
and Histogram
Count
0
4
8
12
0
20
60
100
Value
Common sites gene3924
CphaseB2
CphaseA1

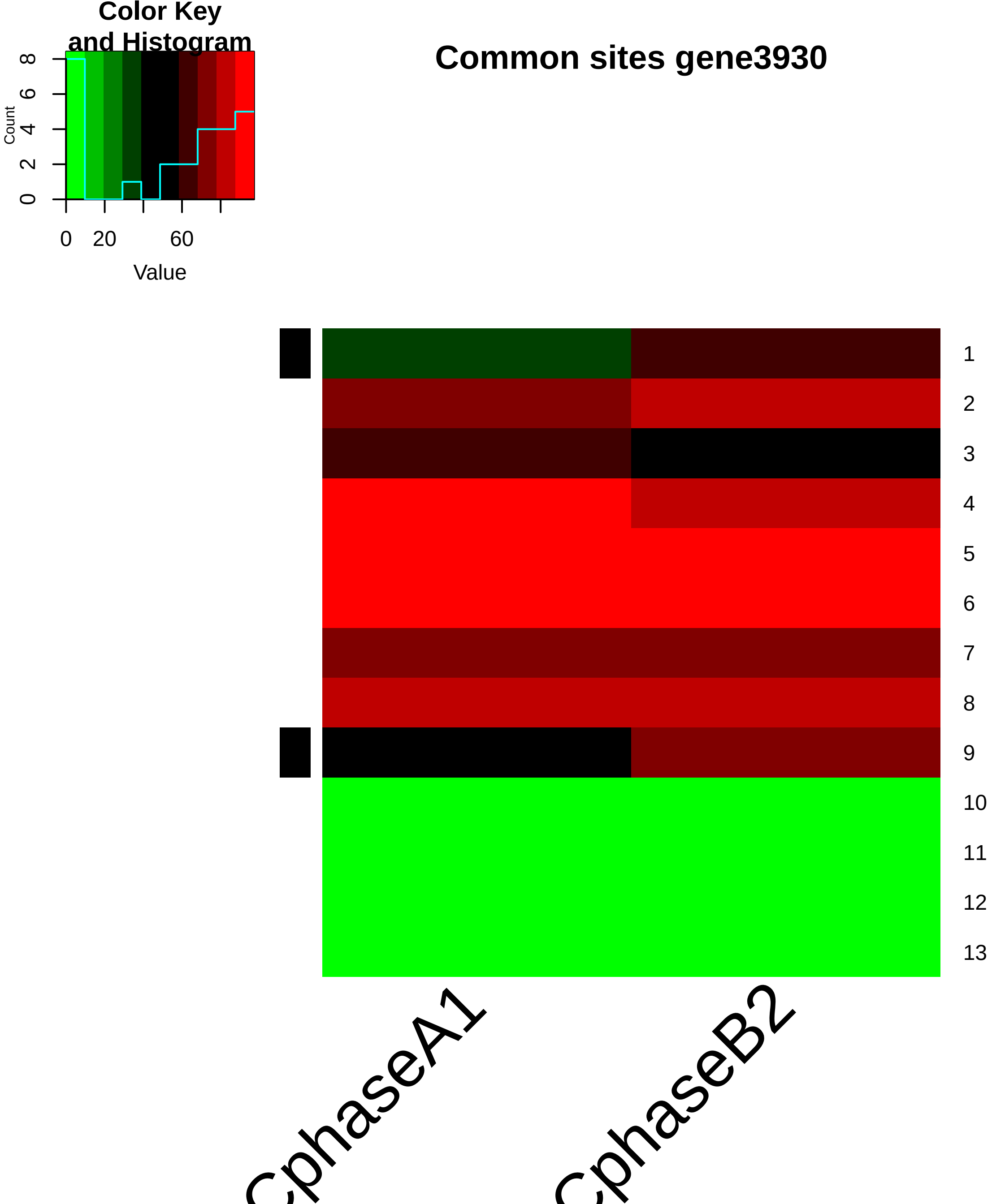

Color Key
and Histogram
Count
0
2
4
6
8
0
20
60
Value
Common sites gene3930
1
2
3
4
5
6
7
8
9
10
11
12
13
CphaseA1
CphaseB2

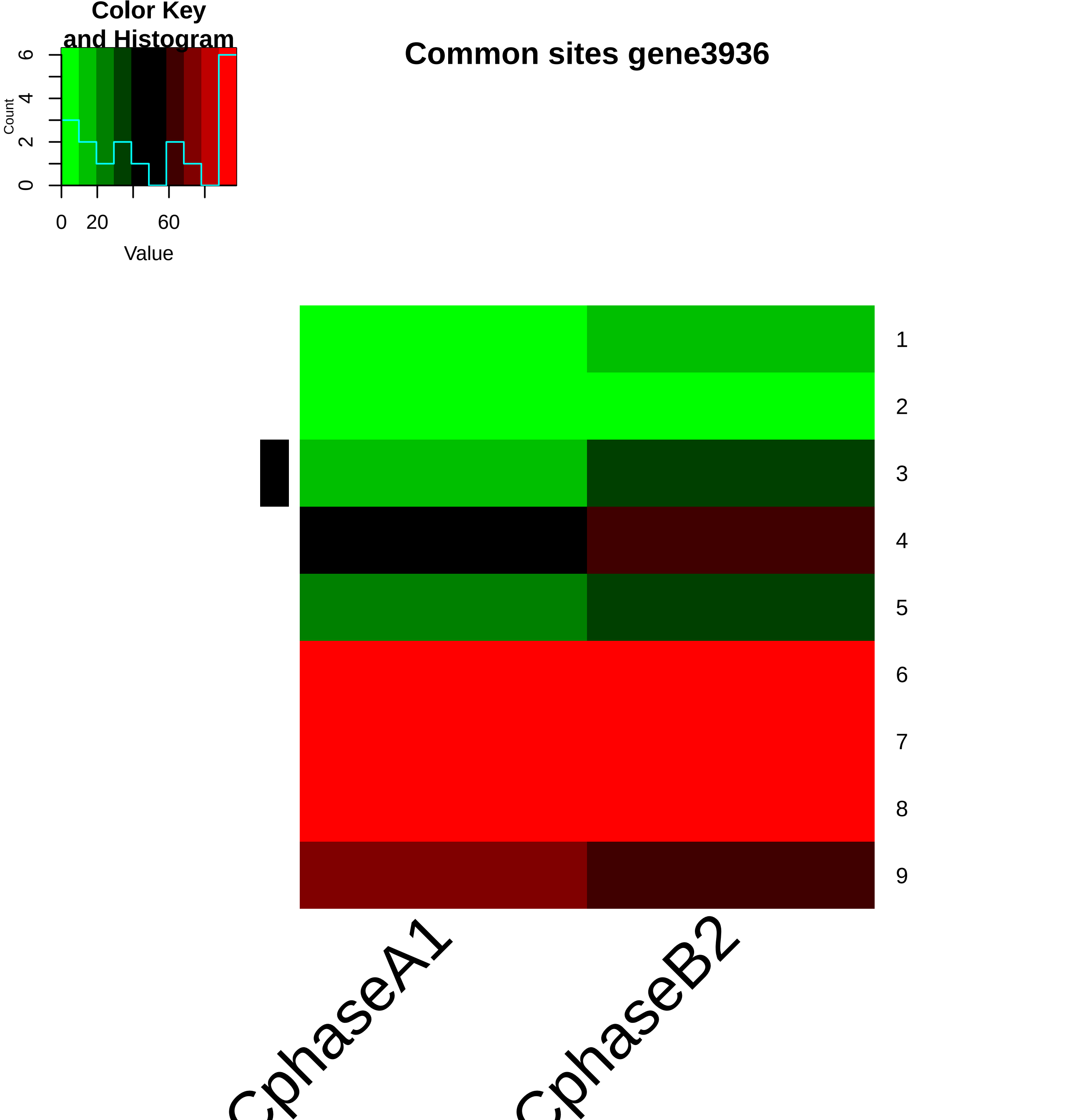

Color Key
and Histogram
Count
0
2
4
6
0
20
60
Value
Common sites gene3936
1
2
3
4
5
6
7
8
9
CphaseA1
CphaseB2

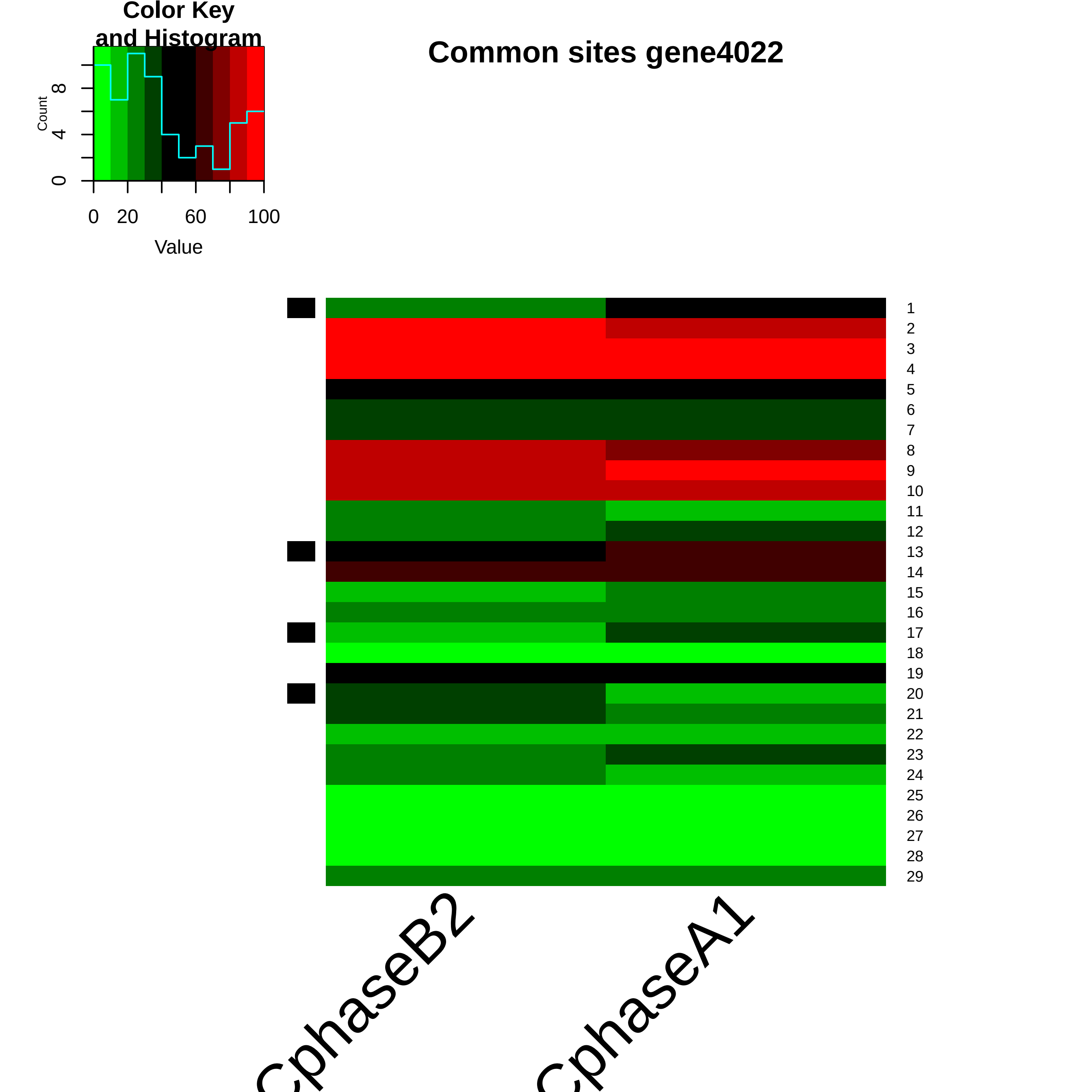
Color Key
and Histogram
Count
0
4
8
0
20
60
100
Value
Common sites gene4022
CphaseB2
CphaseA1

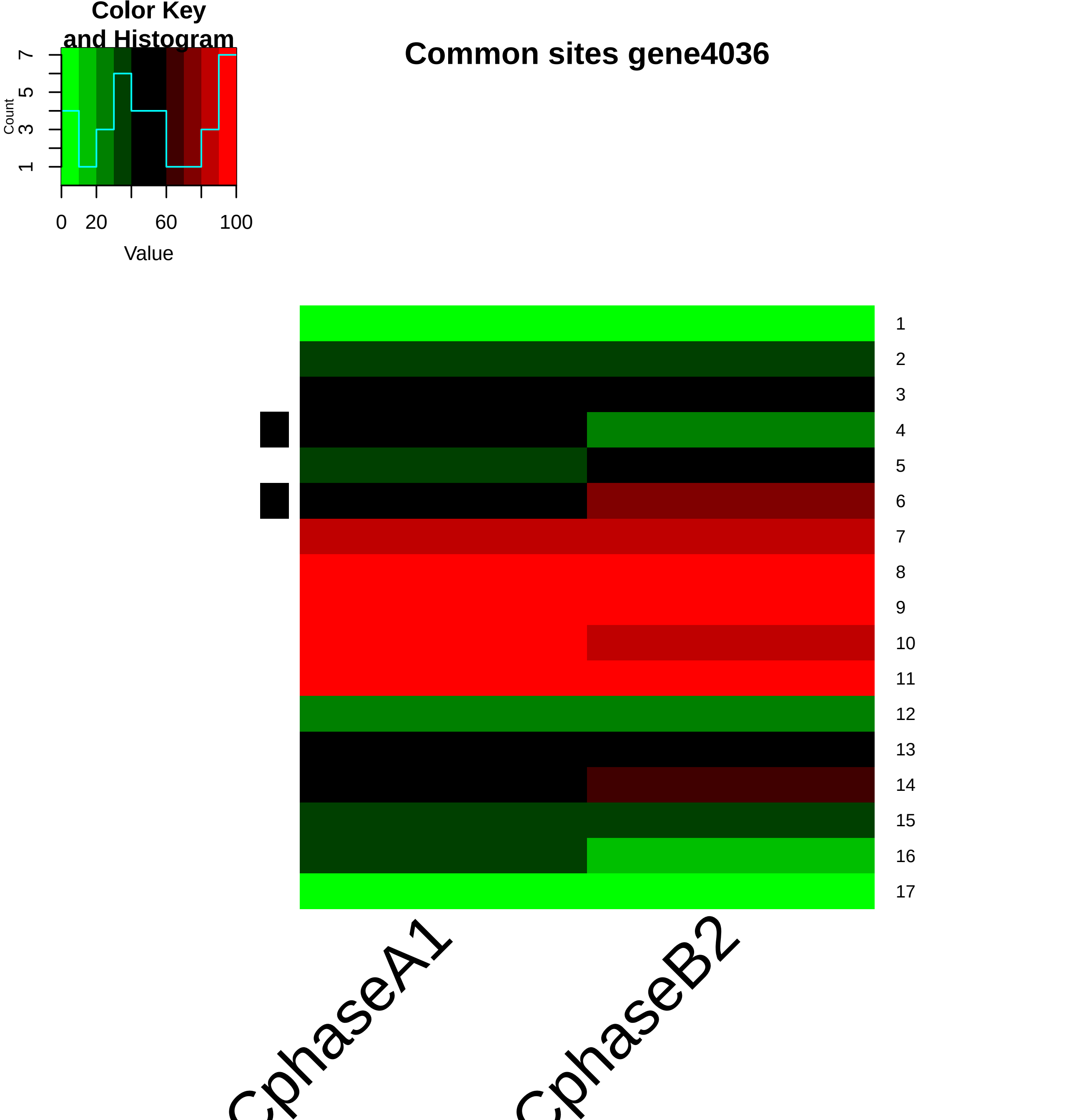

Color Key
and Histogram
Count
1
3
5
7
0
20
60
100
Value
Common sites gene4036
CphaseA1
CphaseB2

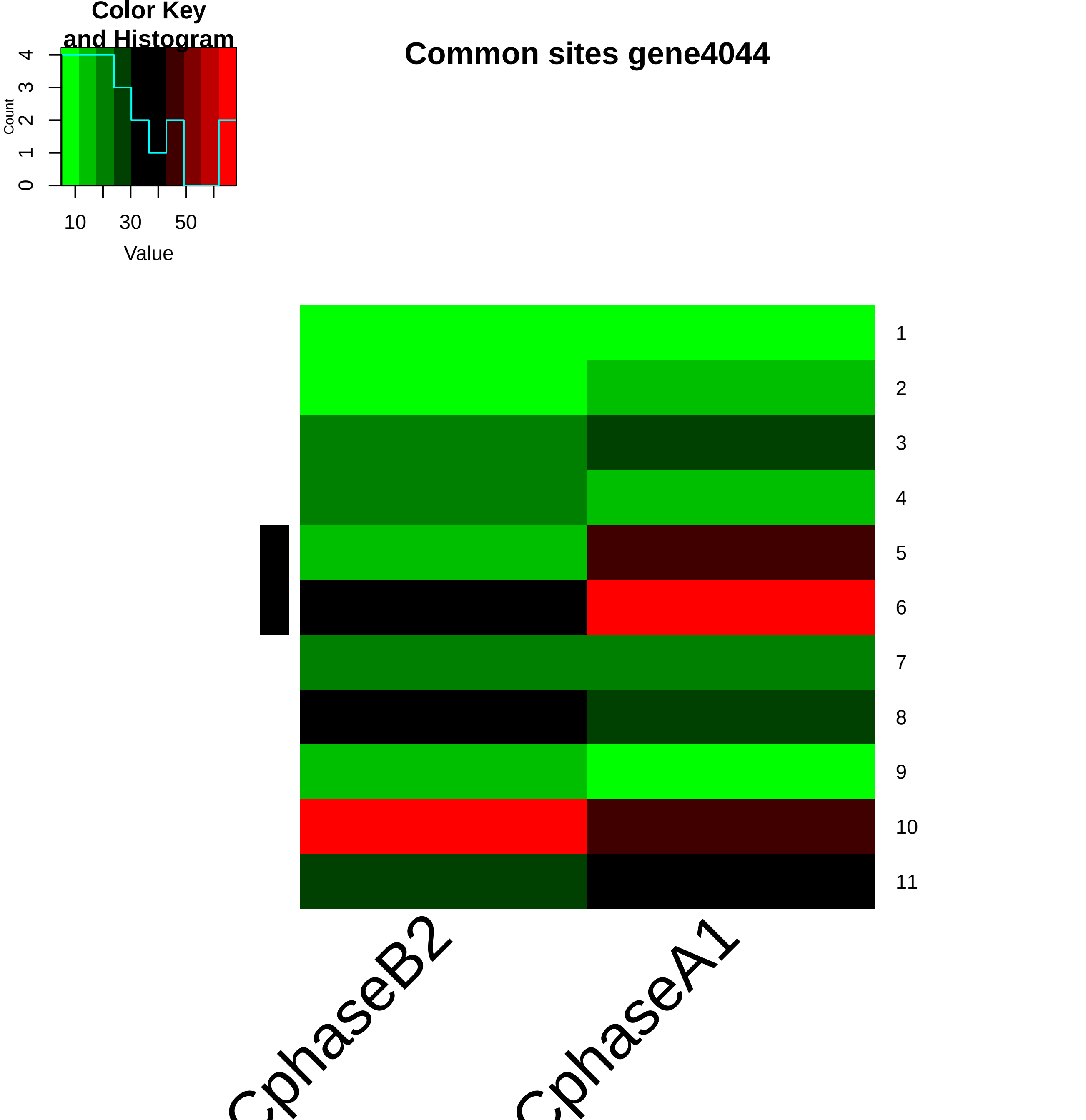

Color Key
and Histogram
Count
0
1
2
3
4
10
30
50
Value
Common sites gene4044
1
2
3
4
5
6
7
8
9
10
11
CphaseB2
CphaseA1

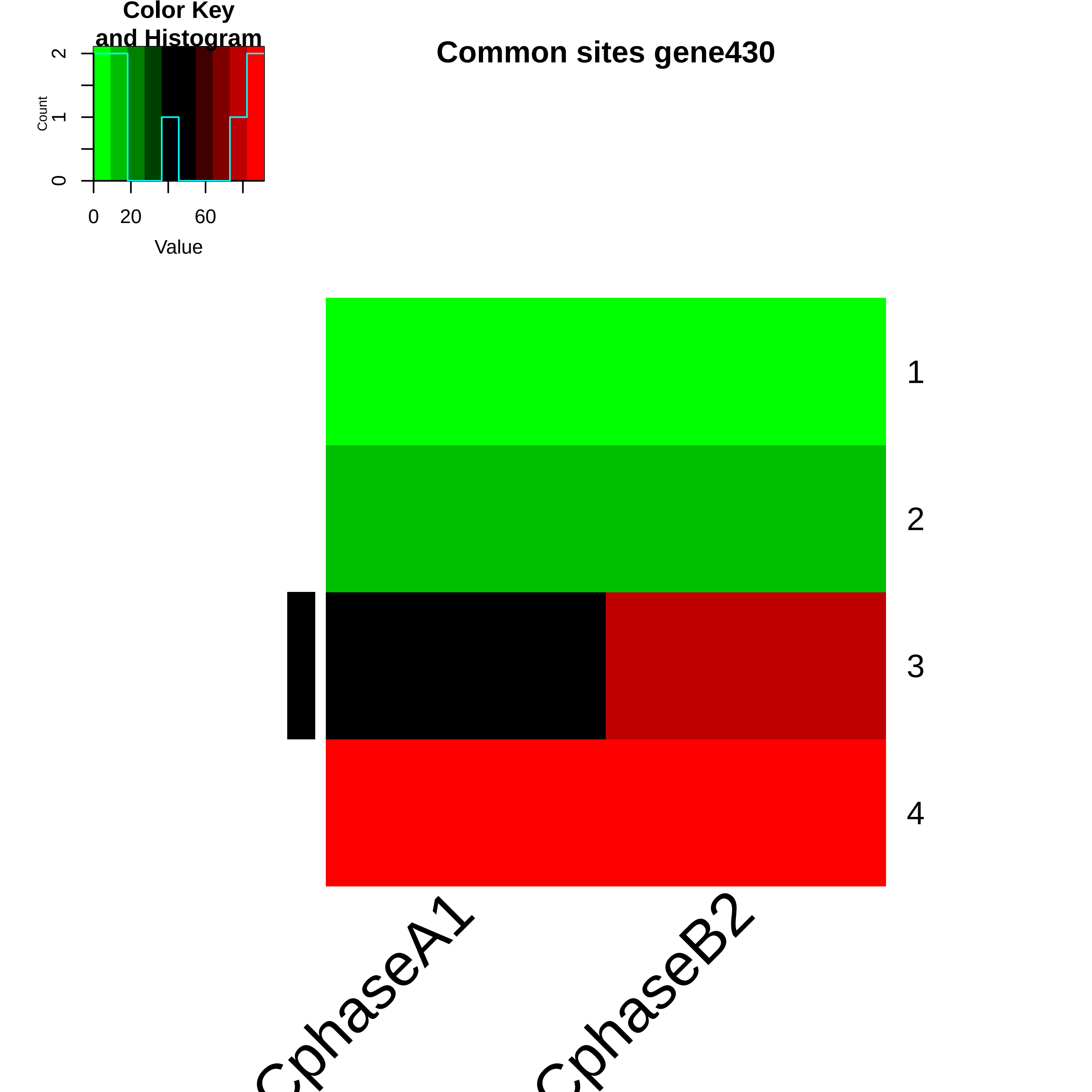
Color Key
and Histogram
Count
0
1
2
0
20
60
Value
Common sites gene430
1
2
3
4
CphaseA1
CphaseB2

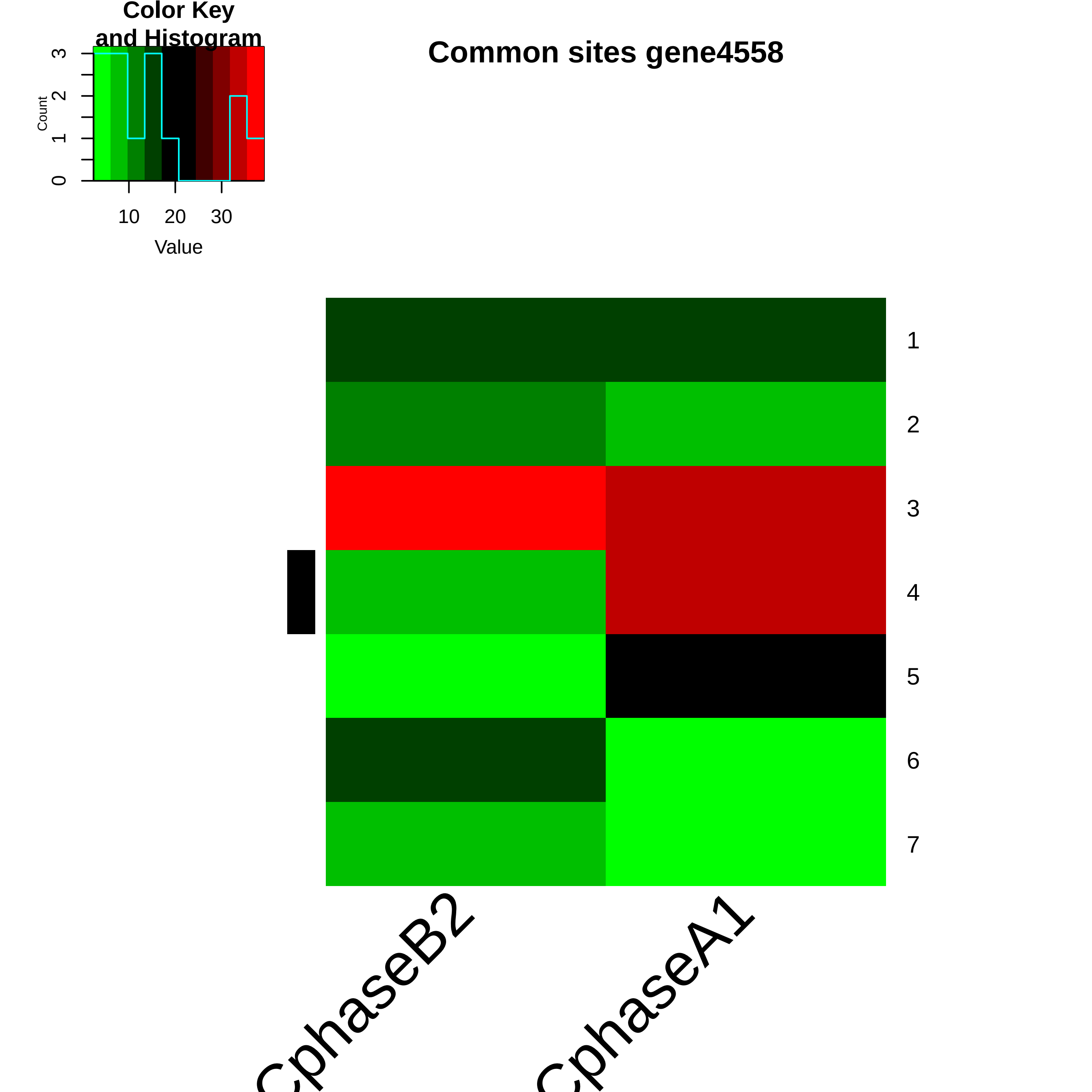

Color Key
and Histogram
Count
0
1
2
3
10
20
30
Value
Common sites gene4558
1
2
3
4
5
6
7
CphaseB2
CphaseA1

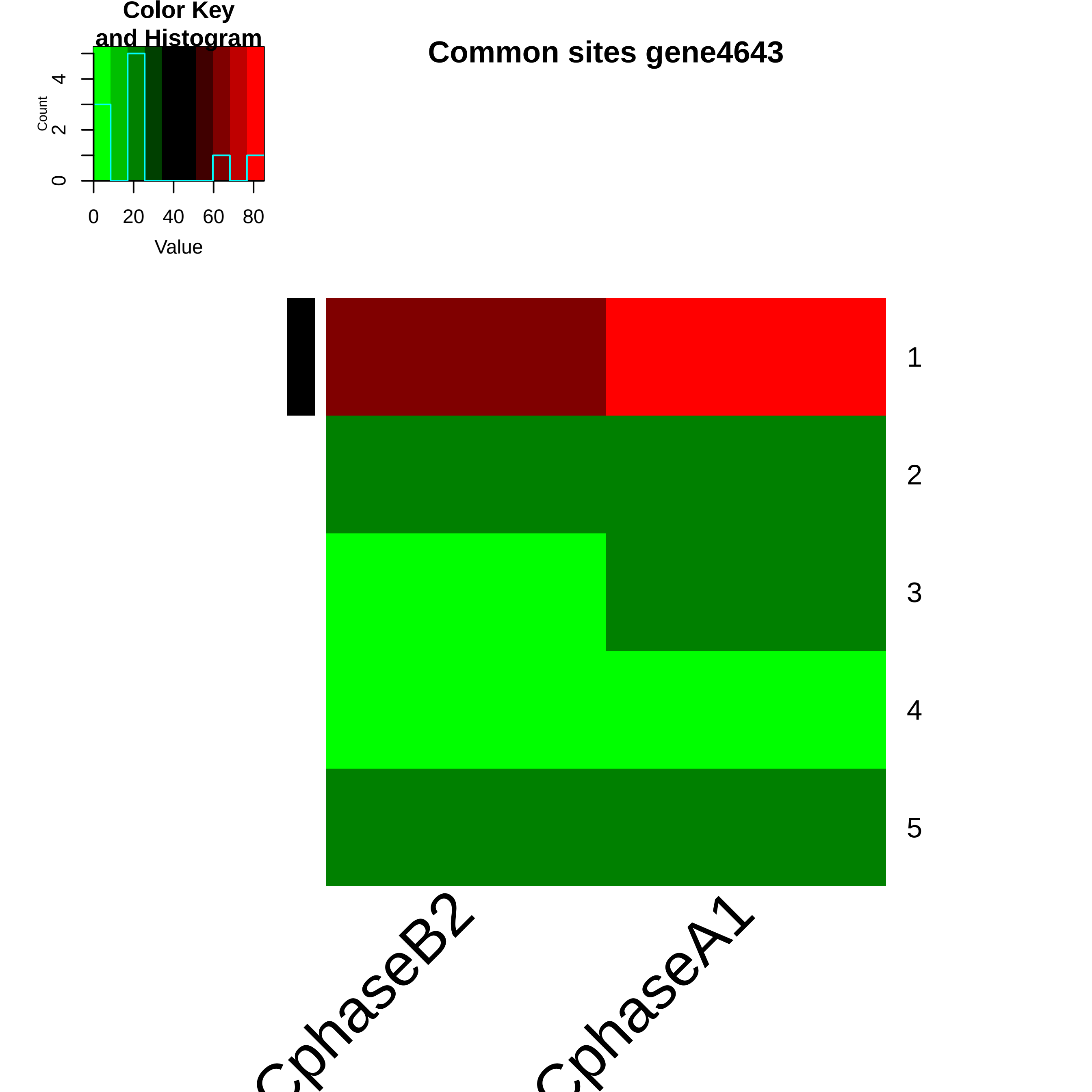

Color Key
and Histogram
Count
0
2
4
0
20
40
60
80
Value
Common sites gene4643
1
2
3
4
5
CphaseB2
CphaseA1

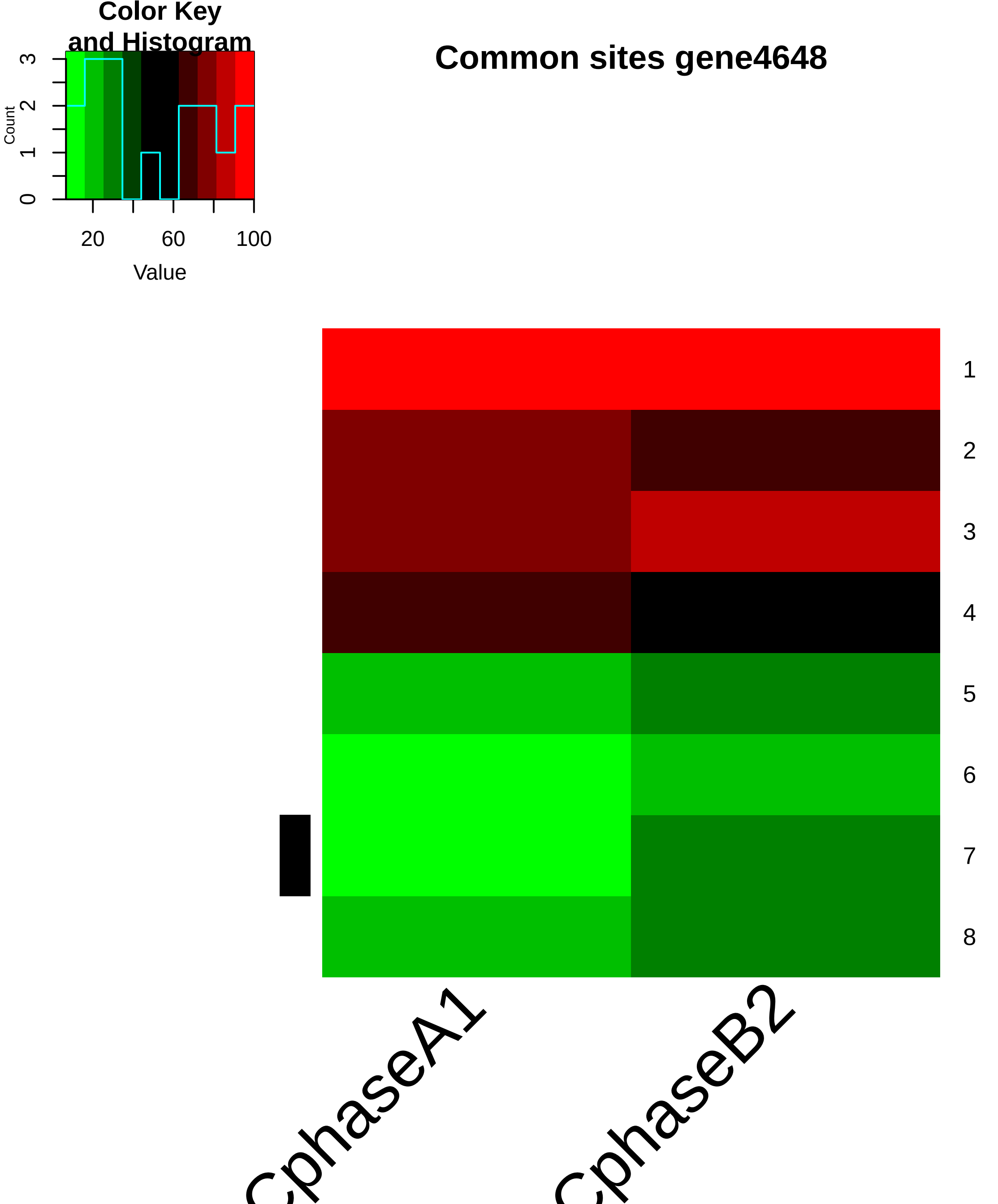

Color Key
and Histogram
Count
0
1
2
3
20
60
100
Value
Common sites gene4648
1
2
3
4
5
6
7
8
CphaseA1
CphaseB2

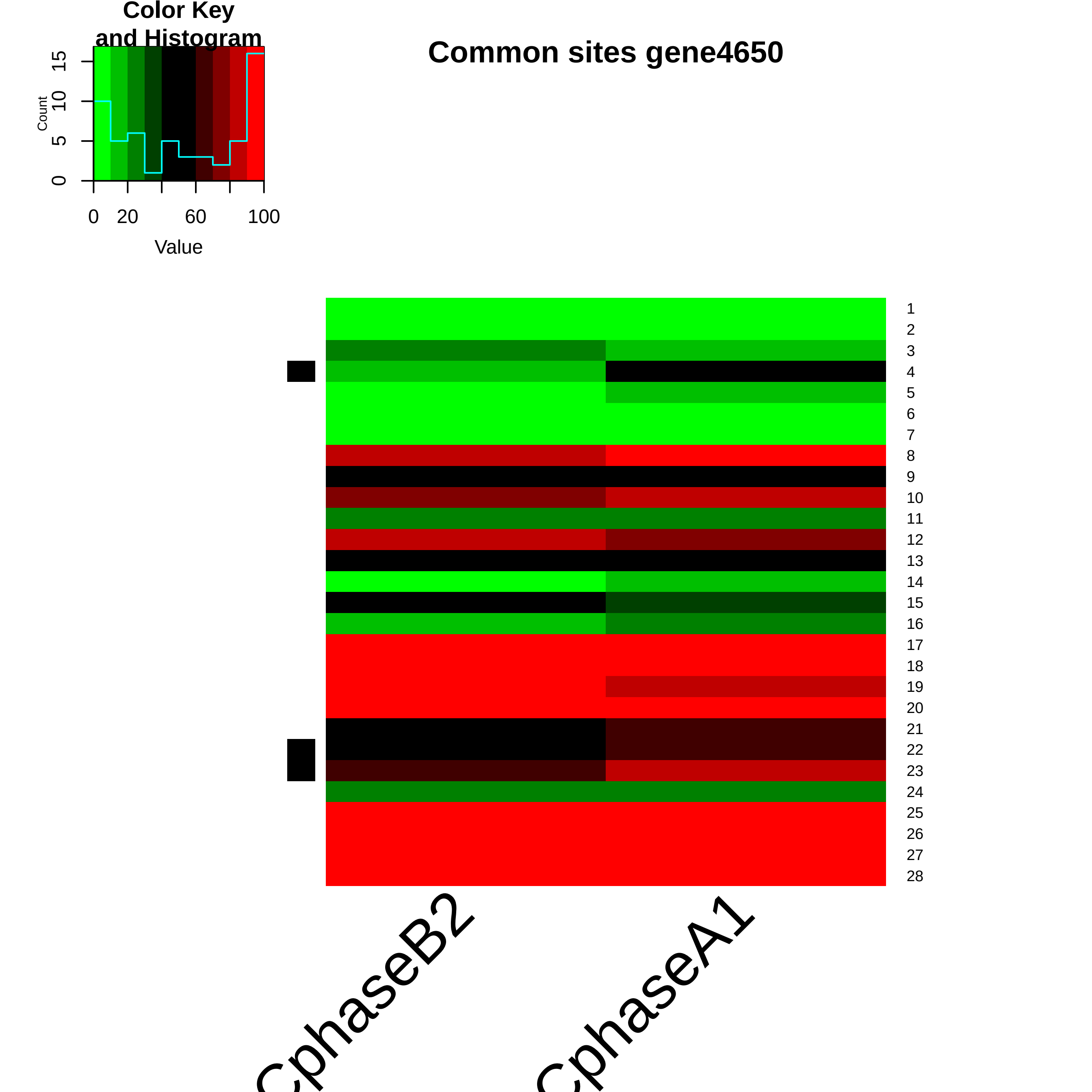
Color Key
and Histogram
Count
0
5
10
15
0
20
60
100
Value
Common sites gene4650
CphaseB2
CphaseA1

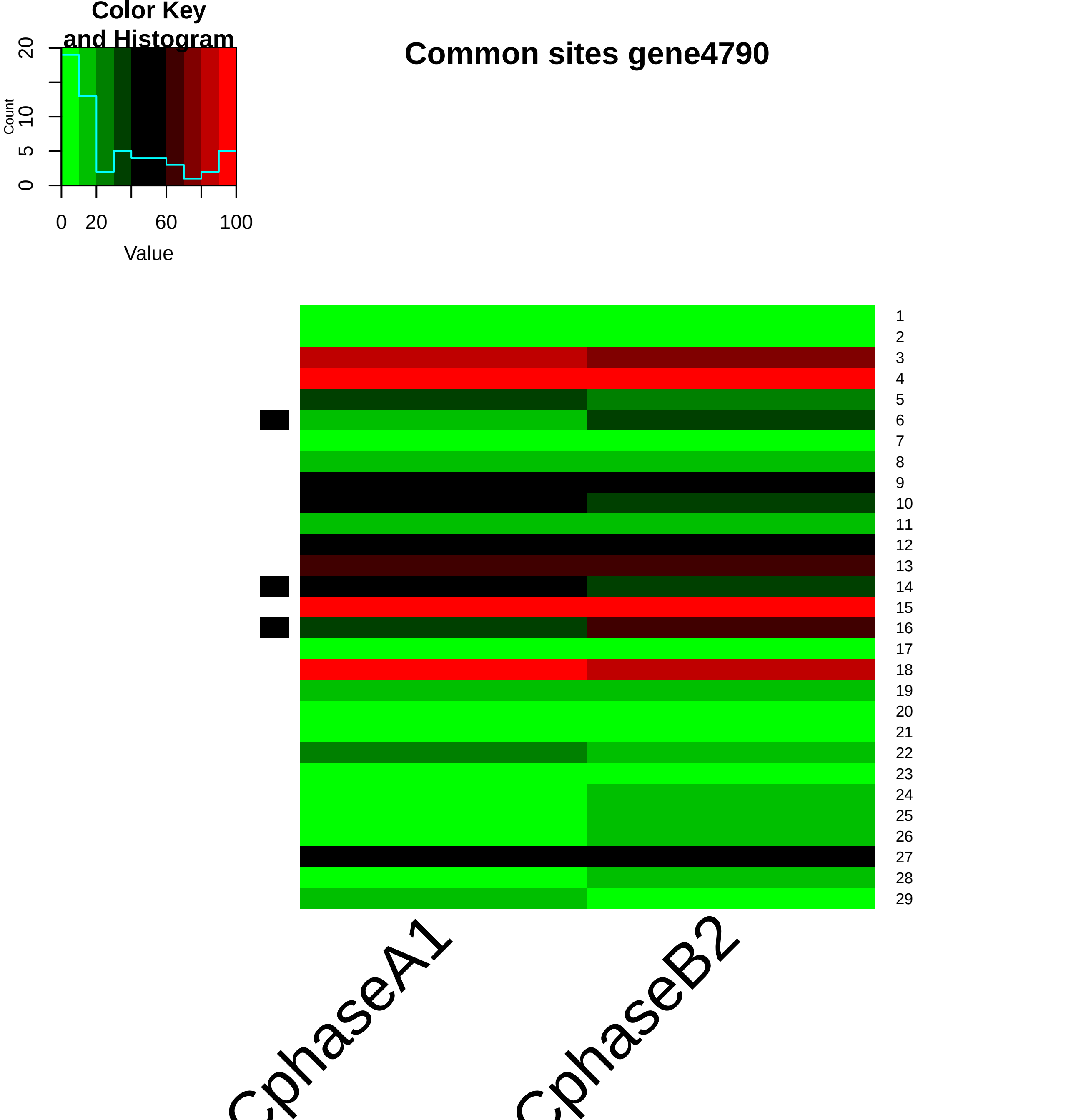
Color Key
and Histogram
Count
0
5
10
20
0
20
60
100
Value
Common sites gene4790
CphaseA1
CphaseB2

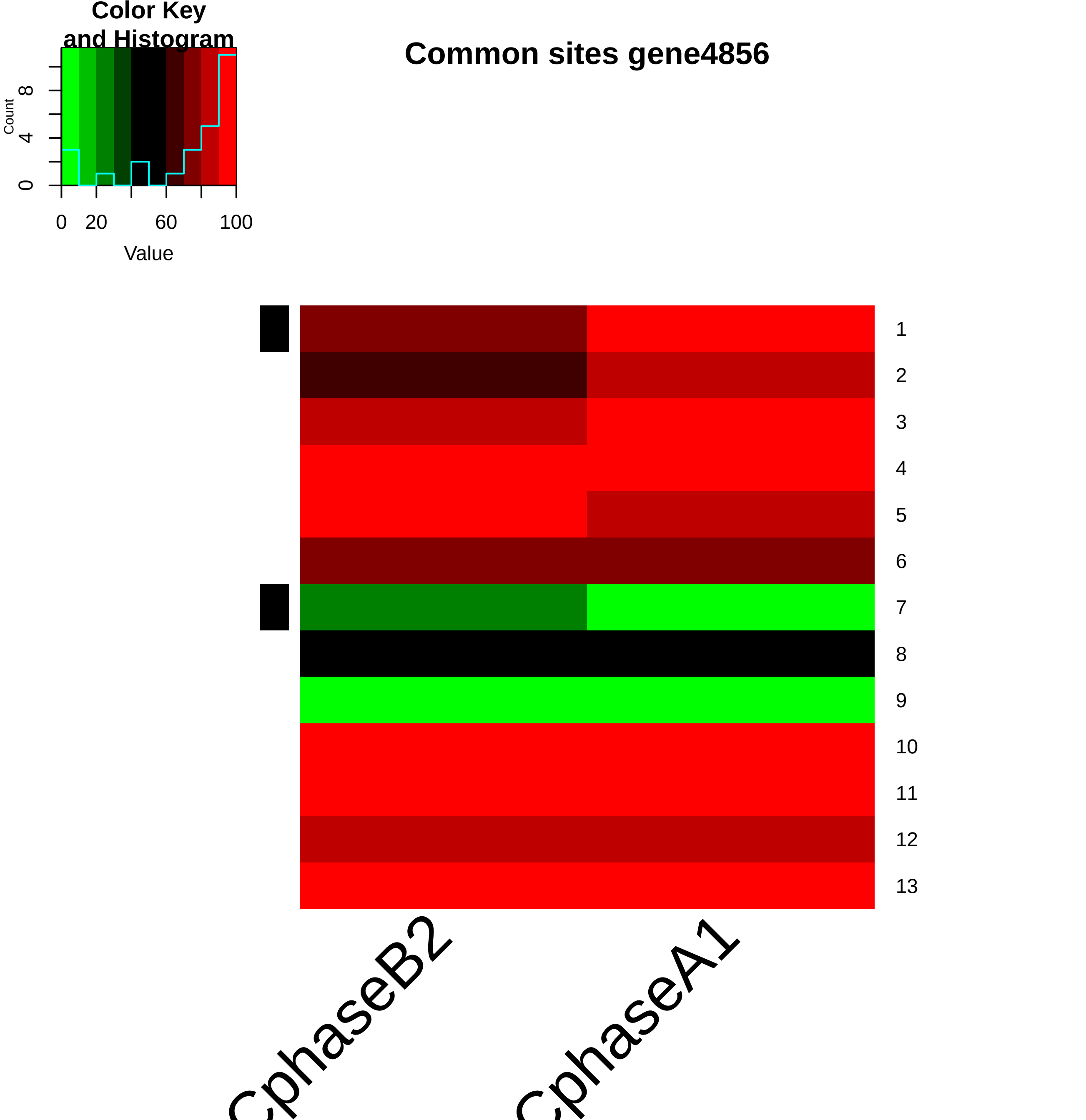
Color Key
and Histogram
Count
0
4
8
0
20
60
100
Value
Common sites gene4856
1
2
3
4
5
6
7
8
9
10
11
12
13
CphaseB2
CphaseA1

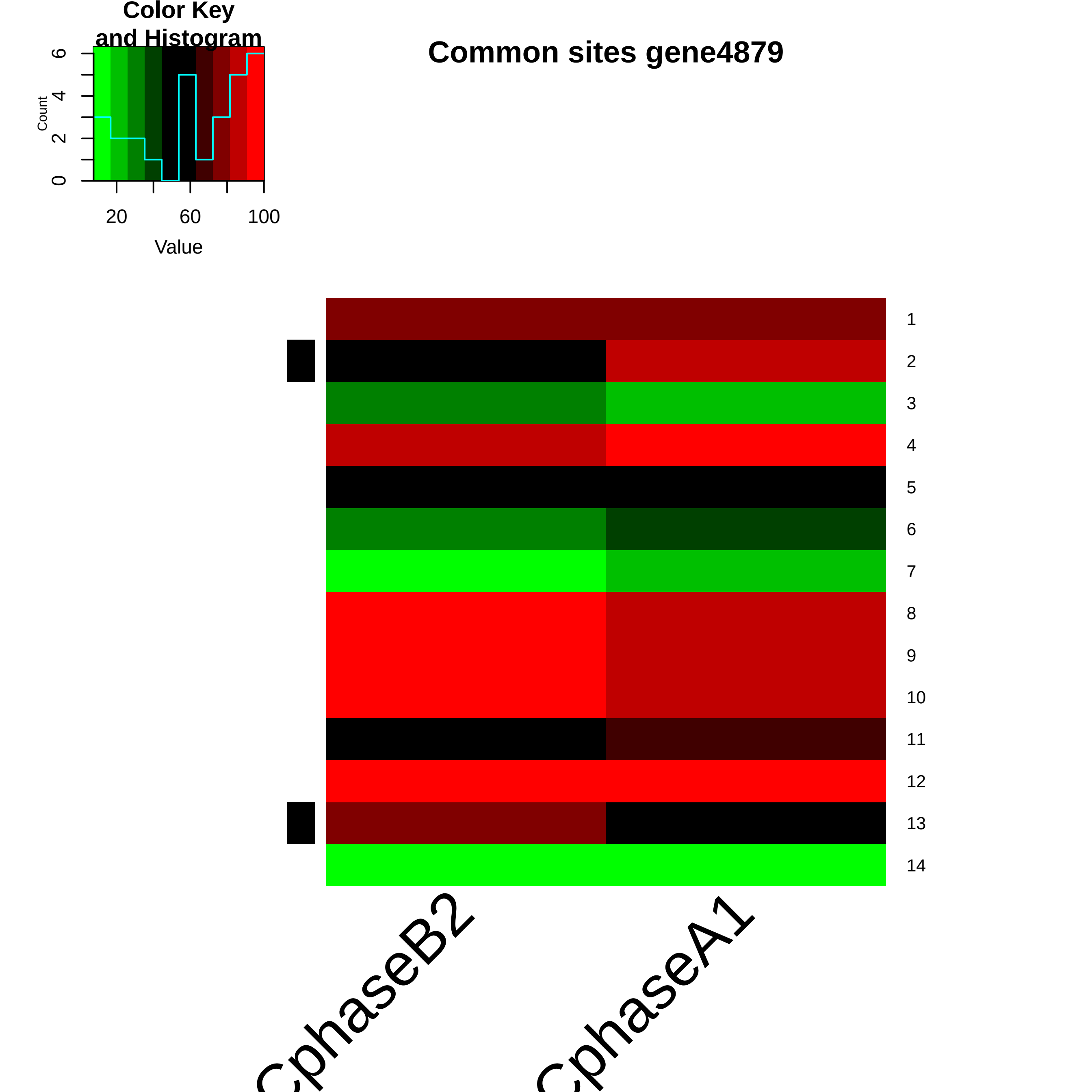

Color Key
and Histogram
Count
0
2
4
6
20
60
100
Value
Common sites gene4879
1
2
3
4
5
6
7
8
9
10
11
12
13
14
CphaseB2
CphaseA1

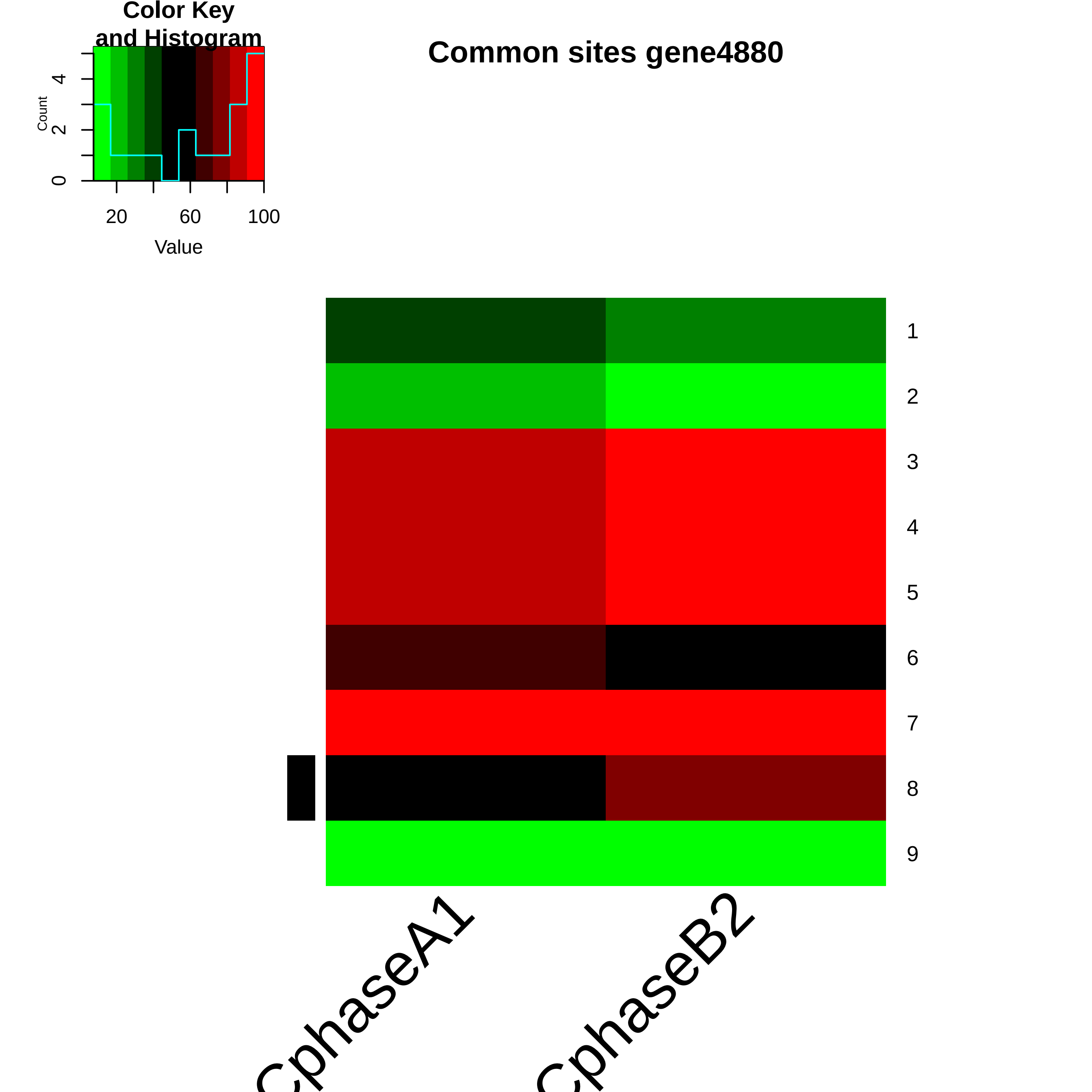
Color Key
and Histogram
Count
0
2
4
20
60
100
Value
Common sites gene4880
1
2
3
4
5
6
7
8
9
CphaseA1
CphaseB2

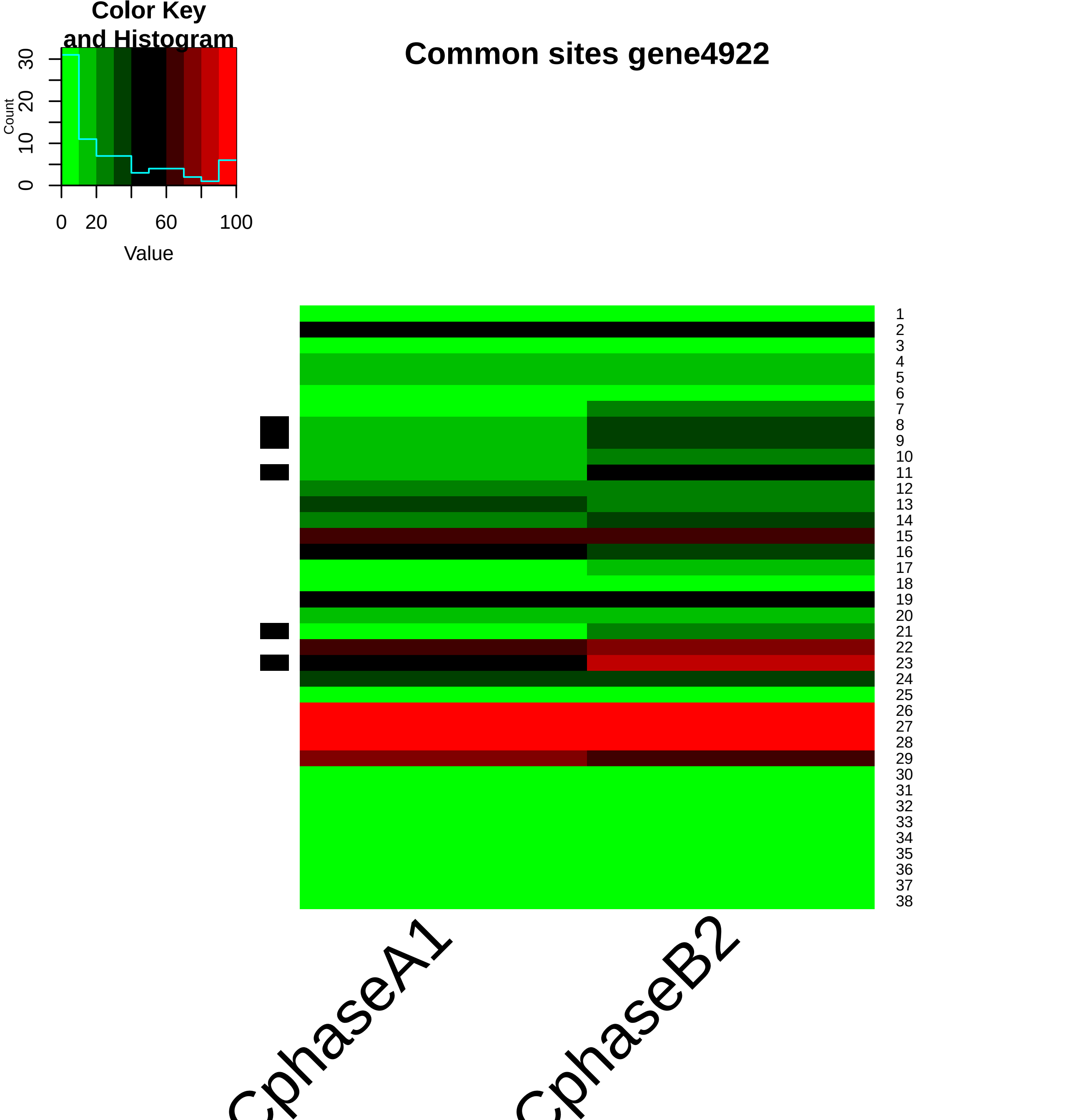
Color Key
and Histogram
Count
0
10
20
30
0
20
60
100
Value
Common sites gene4922
CphaseA1
CphaseB2

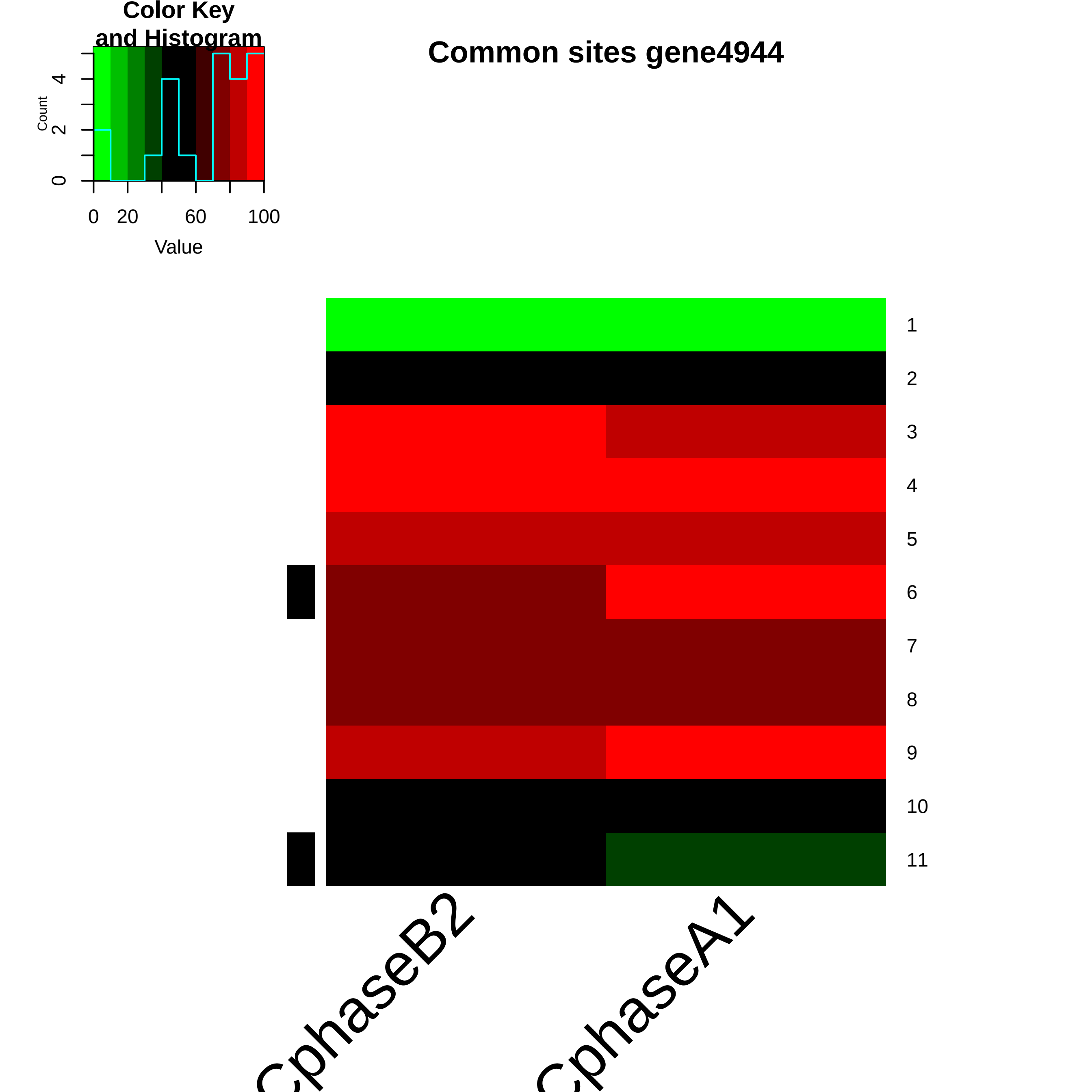
Color Key
and Histogram
Count
0
2
4
0
20
60
100
Value
Common sites gene4944
1
2
3
4
5
6
7
8
9
10
11
CphaseB2
CphaseA1

### Common sites gene4974

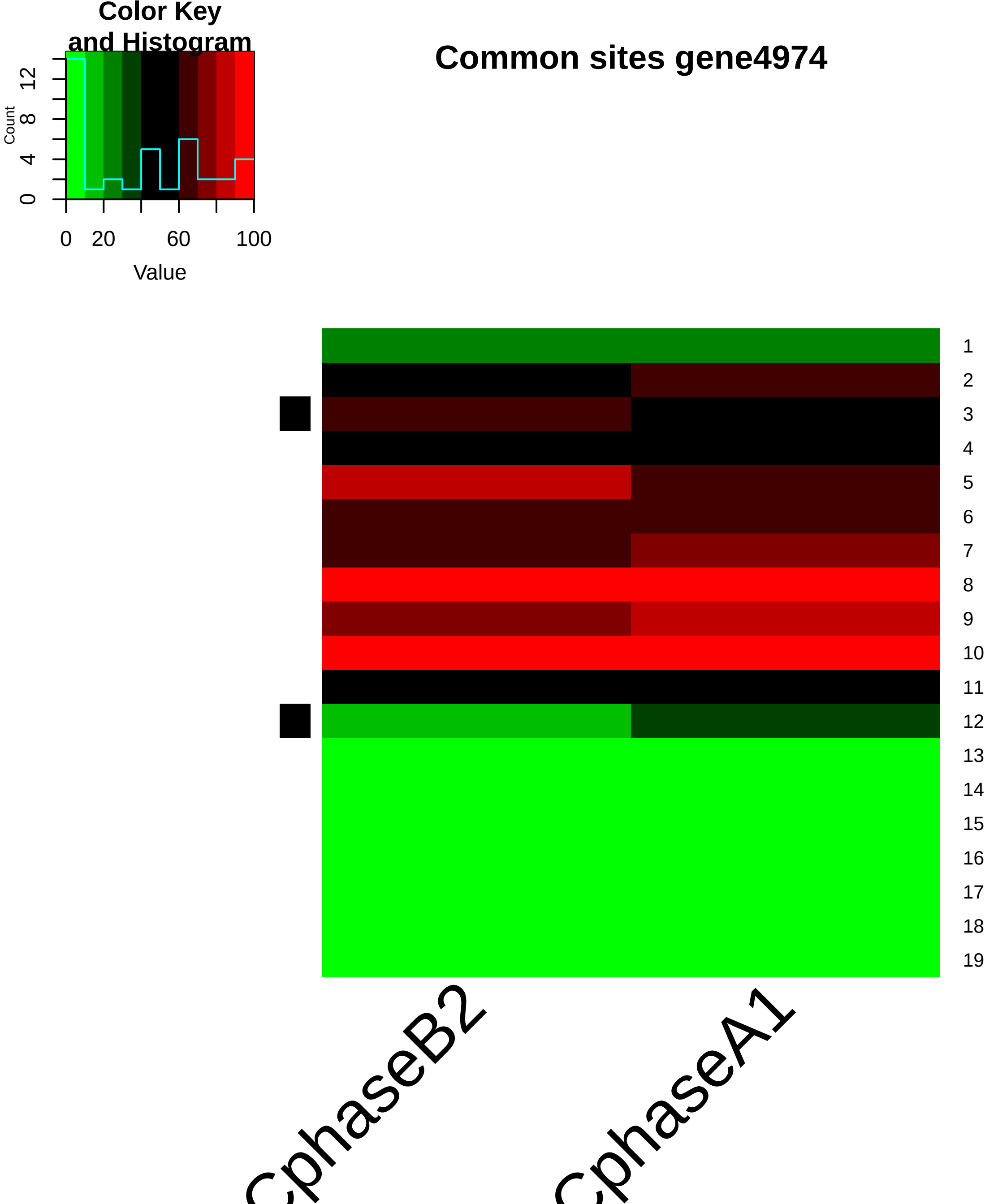

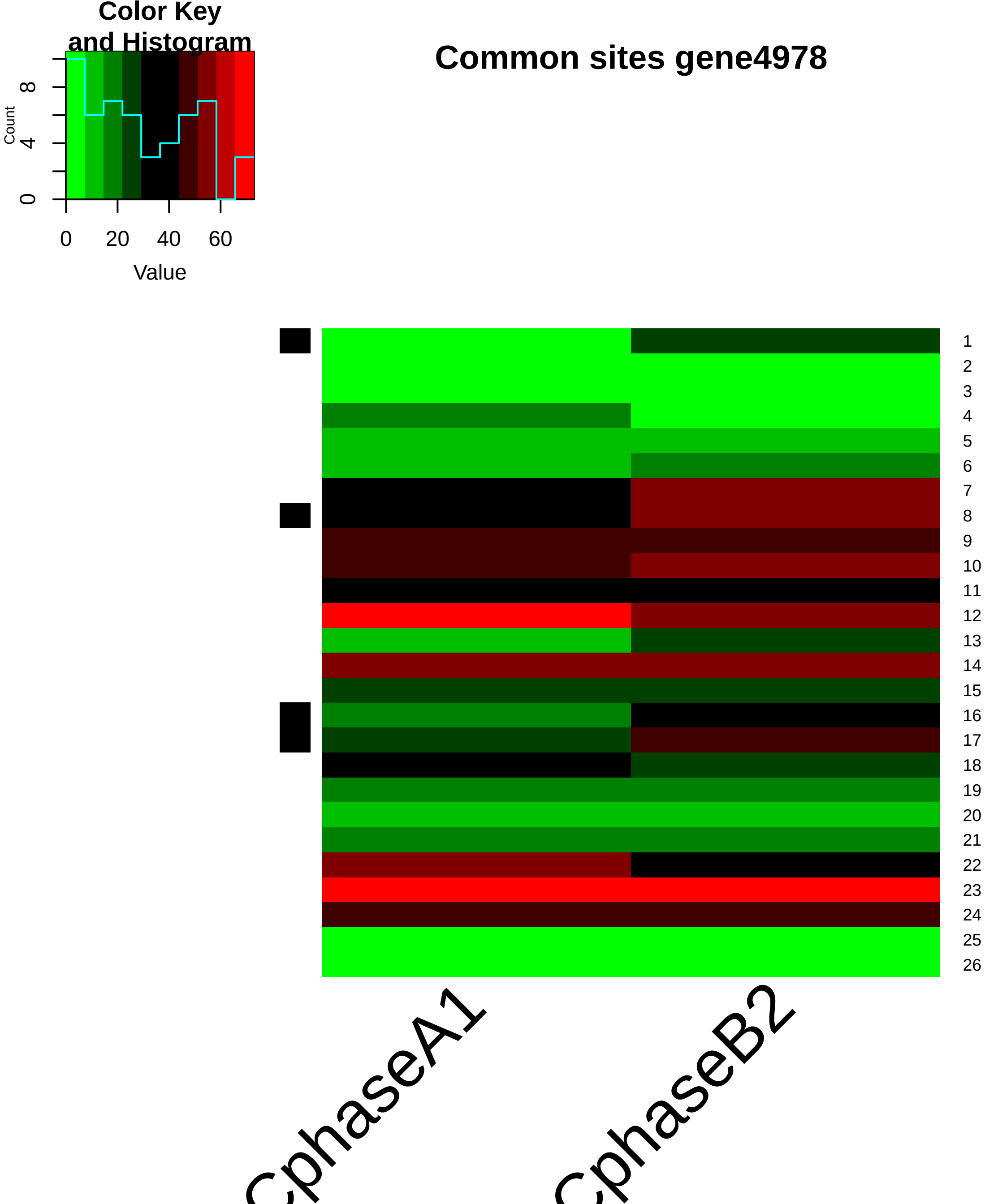
Color Key
and Histogram
Count
0
4
8
0
20
40
60
Value
Common sites gene4978
CphaseA1
CphaseB2

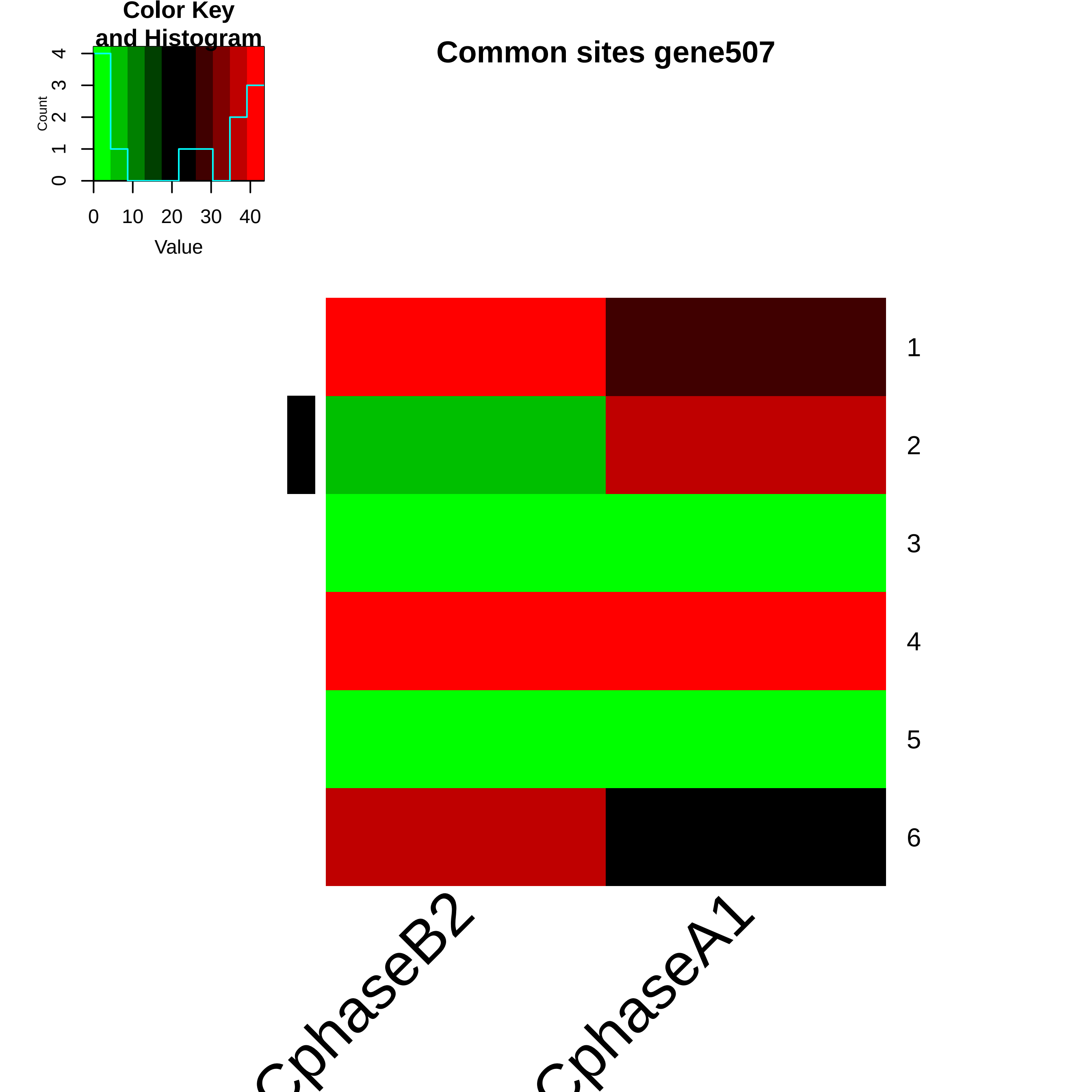
Color Key
and Histogram
Count
0
1
2
3
4
0
10
20
30
40
Value
Common sites gene507
1
2
3
4
5
6
CphaseB2
CphaseA1

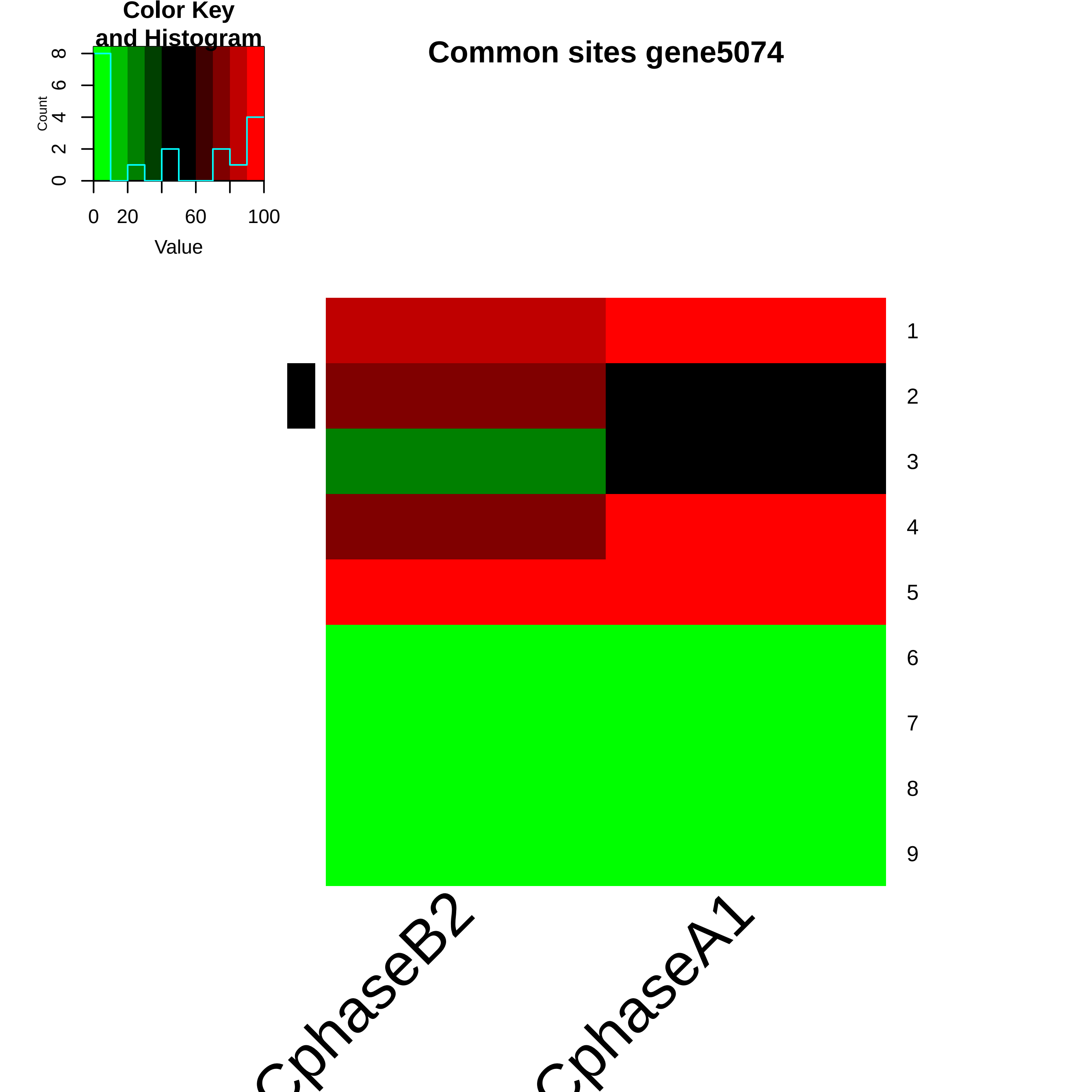

Color Key
and Histogram
Count
0
2
4
6
8
0
20
60
100
Value
Common sites gene5074
1
2
3
4
5
6
7
8
9
CphaseB2
CphaseA1

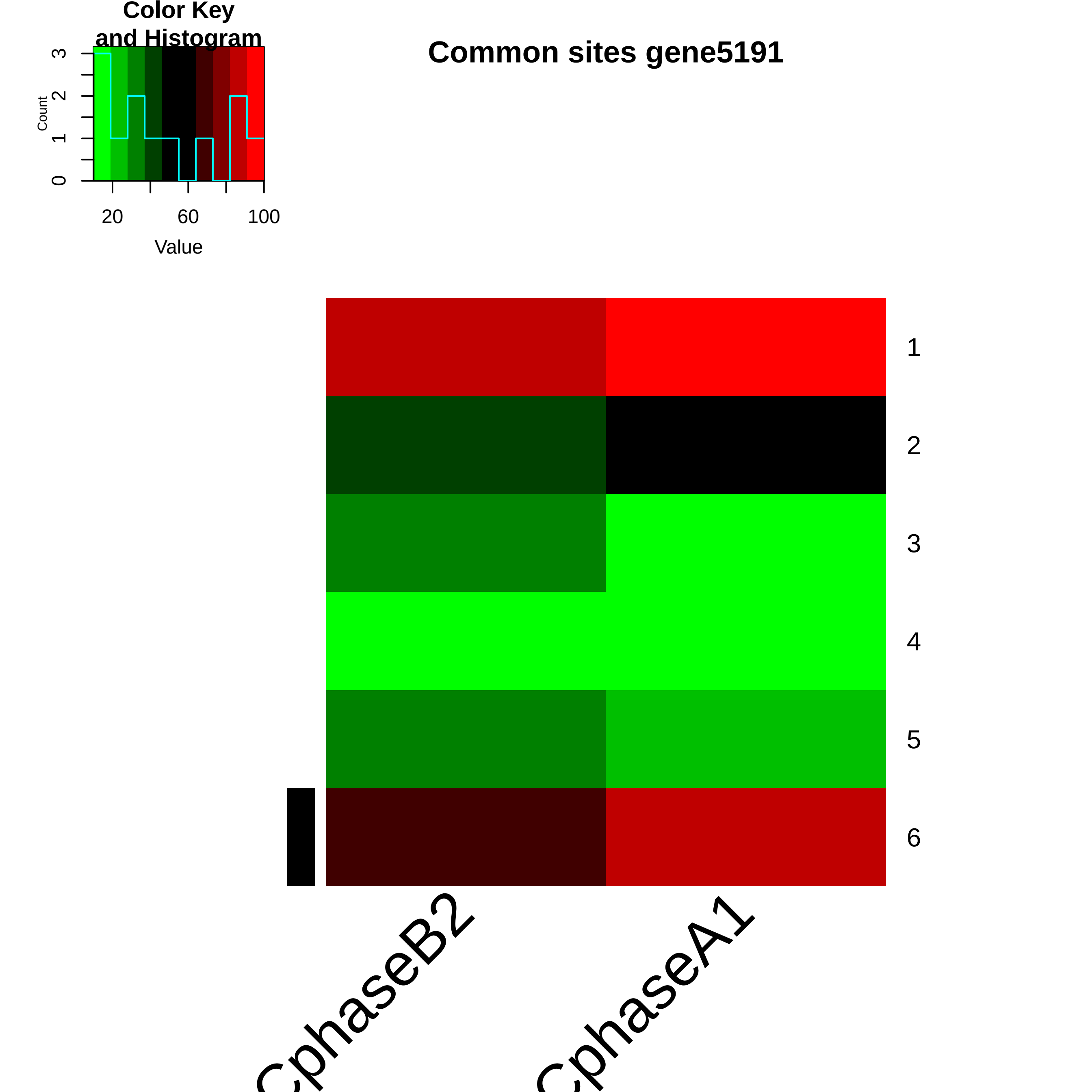
Color Key
and Histogram
Count
0
1
2
3
20
60
100
Value
Common sites gene5191
1
2
3
4
5
6
CphaseB2
CphaseA1

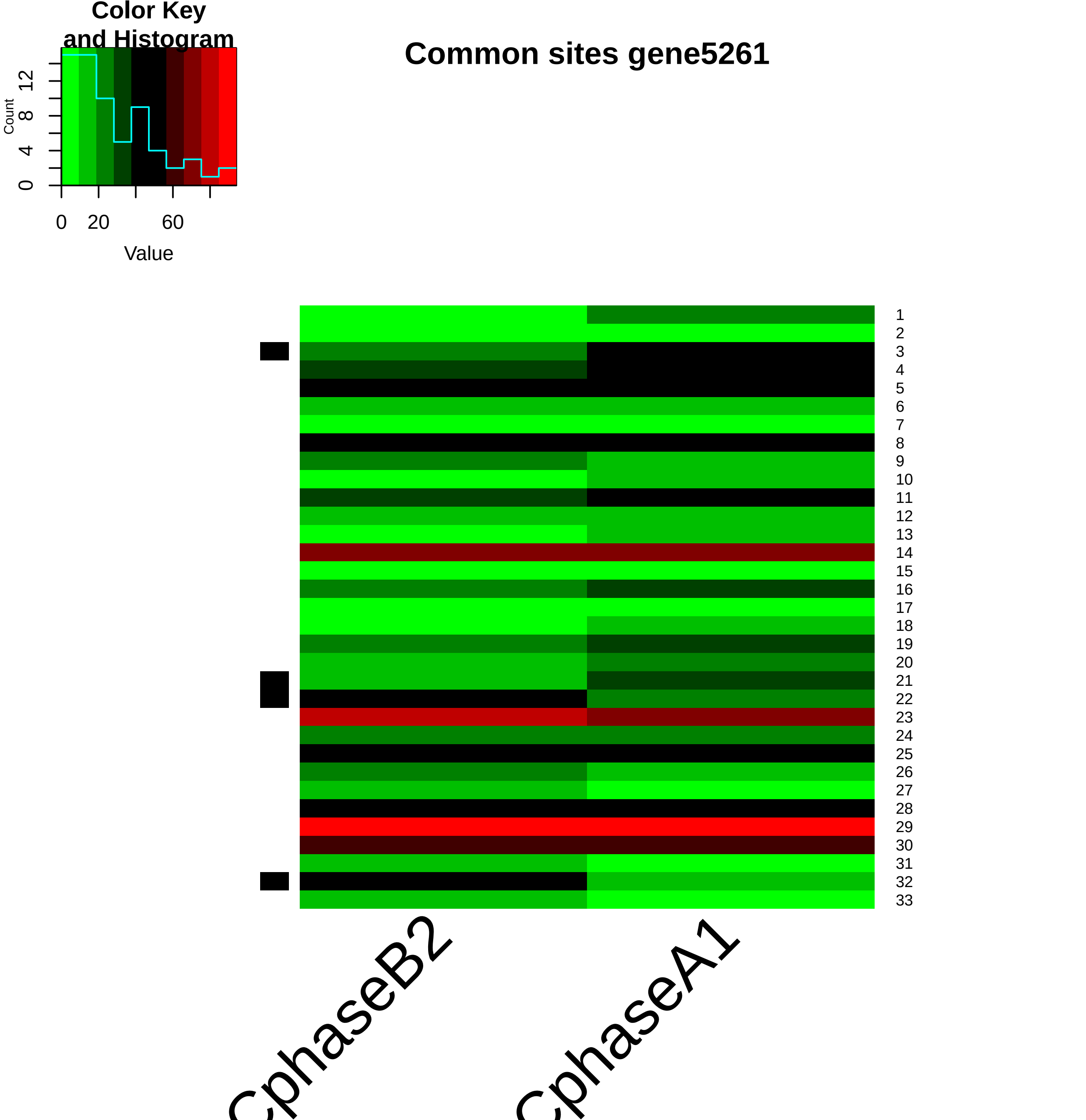
Color Key
and Histogram
Count
0
4
8
12
0
20
60
Value
Common sites gene5261
CphaseB2
CphaseA1

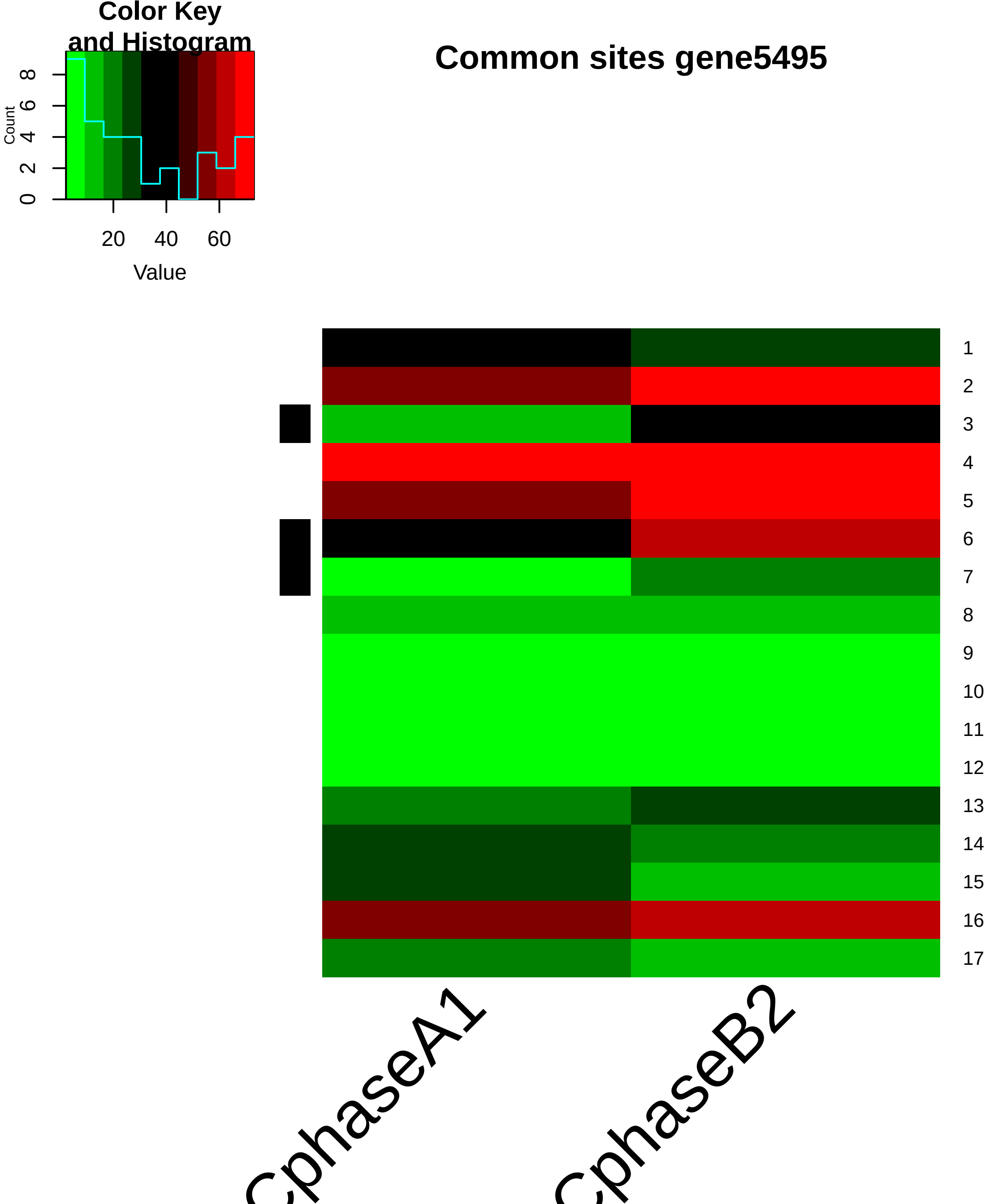
Color Key
and Histogram
Count
0
2
4
6
8
20
40
60
Value
Common sites gene5495
CphaseA1
CphaseB2

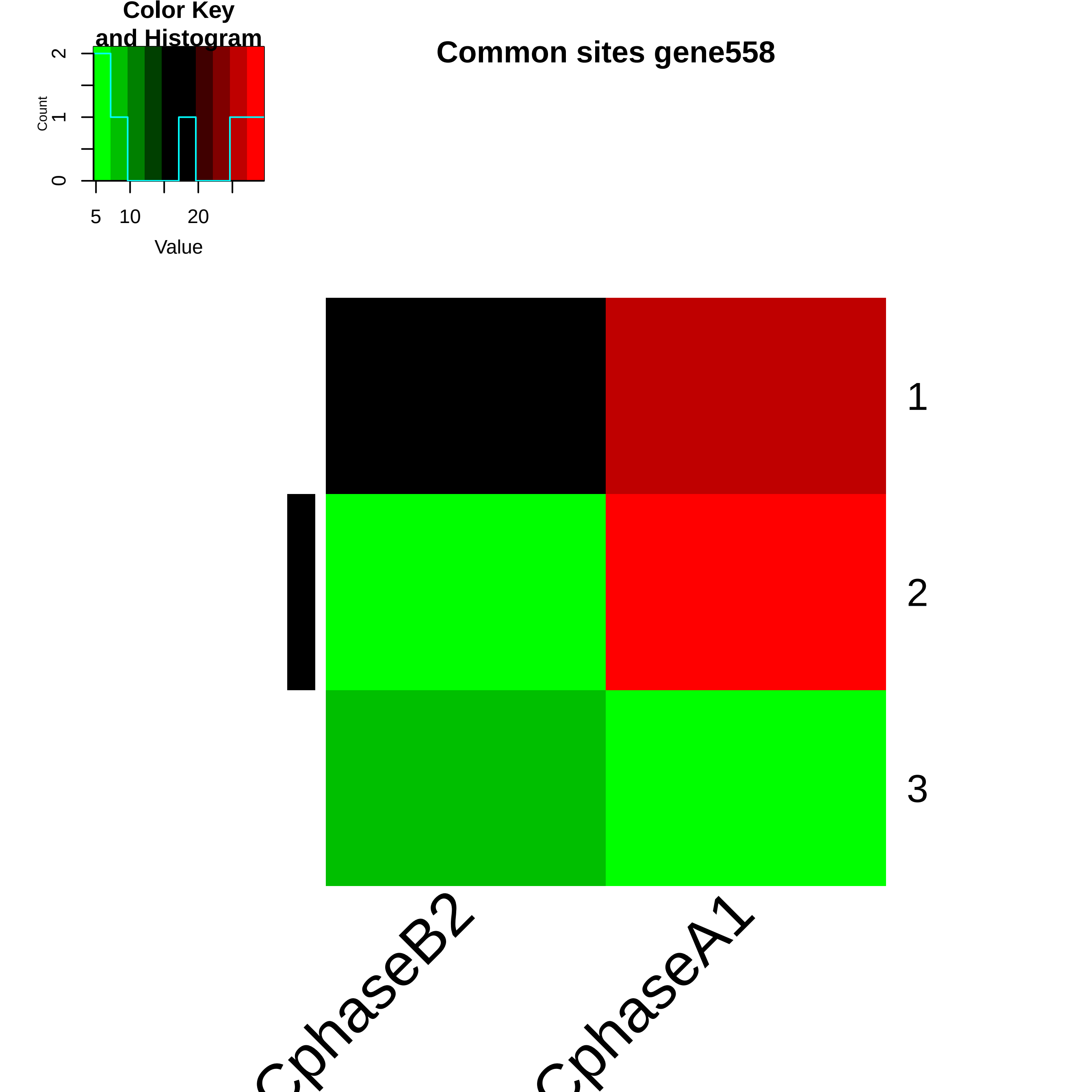
Color Key
and Histogram
Count
0
1
2
5
10
20
Value
Common sites gene558
1
2
3
CphaseB2
CphaseA1

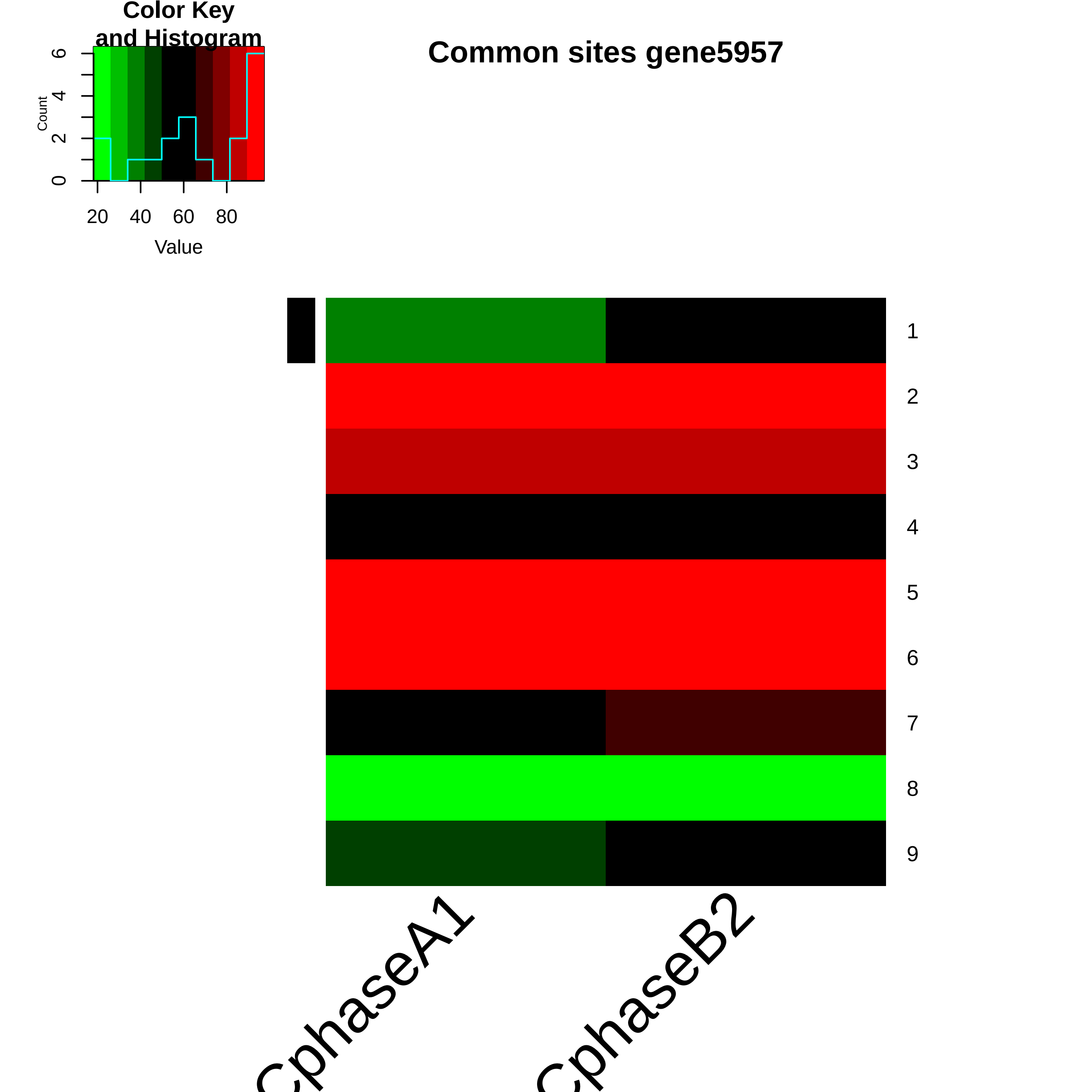
Color Key
and Histogram
Count
0
2
4
6
20
40
60
80
Value
Common sites gene5957
1
2
3
4
5
6
7
8
9
CphaseA1
CphaseB2

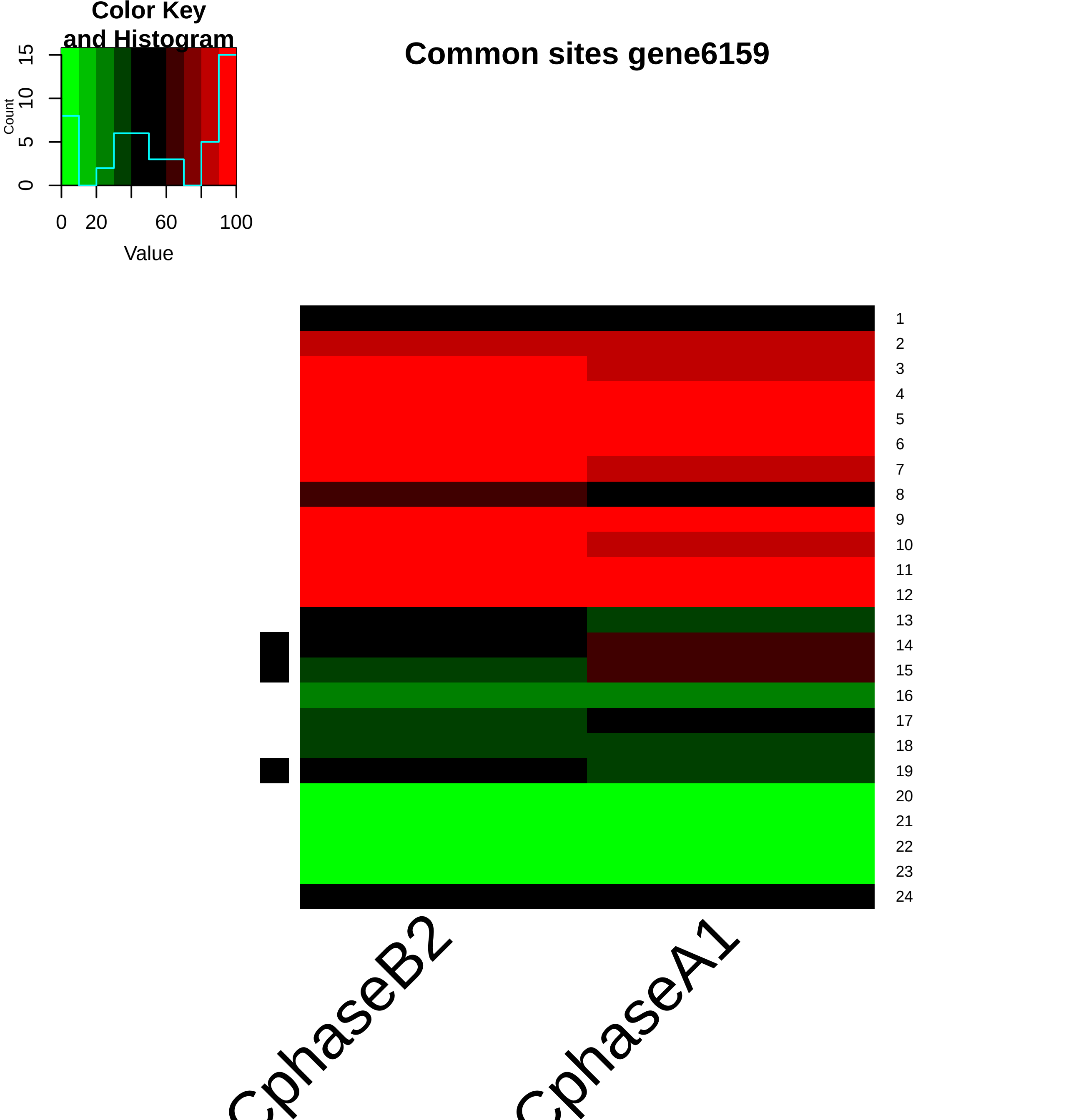

Color Key
and Histogram
Count
0
5
10
15
0
20
60
100
Value
Common sites gene6159
CphaseB2
CphaseA1

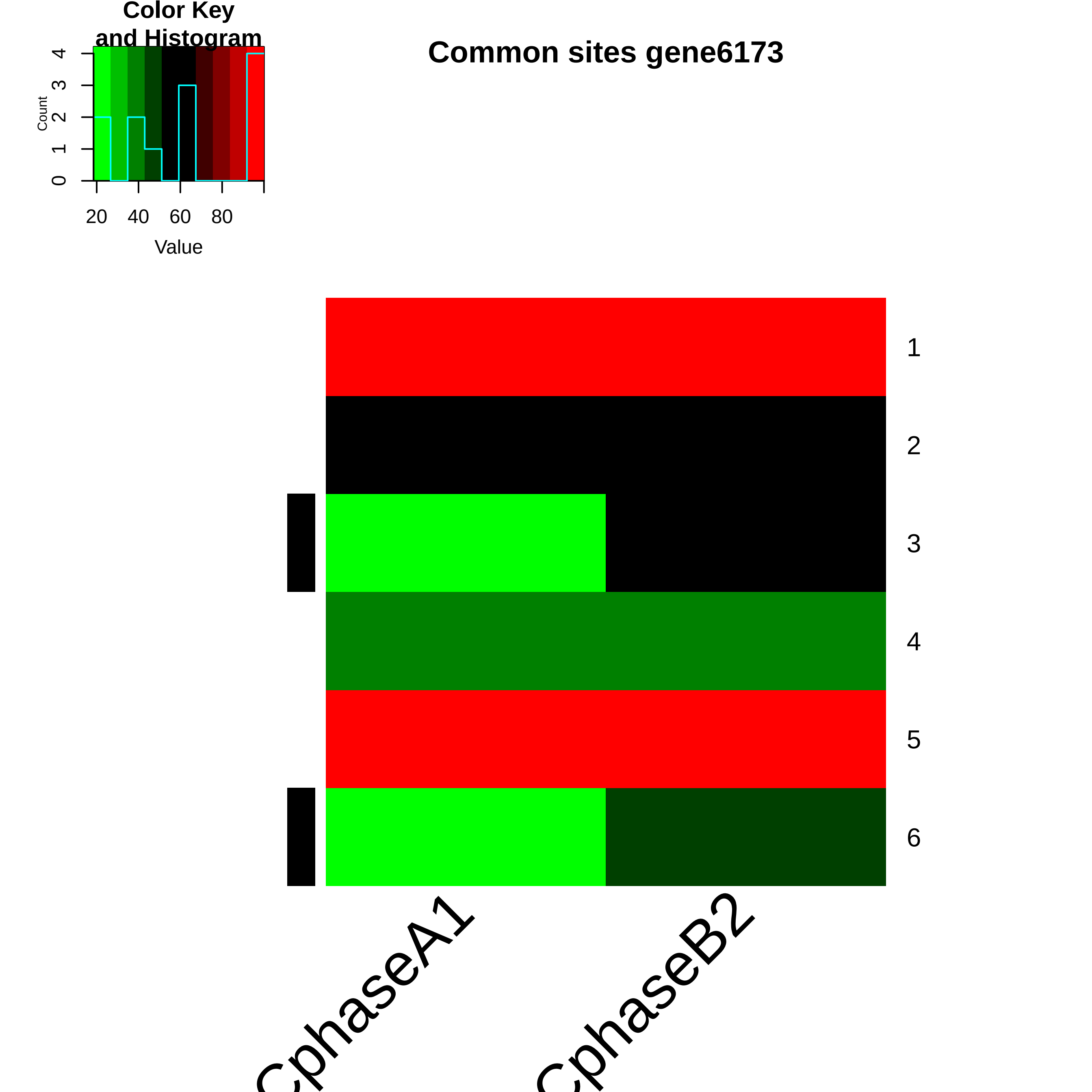
Color Key
and Histogram
Count
Value
20
40
60
80
0
1
2
3
4
Common sites gene6173
1
2
3
4
5
6
CphaseA1
CphaseB2

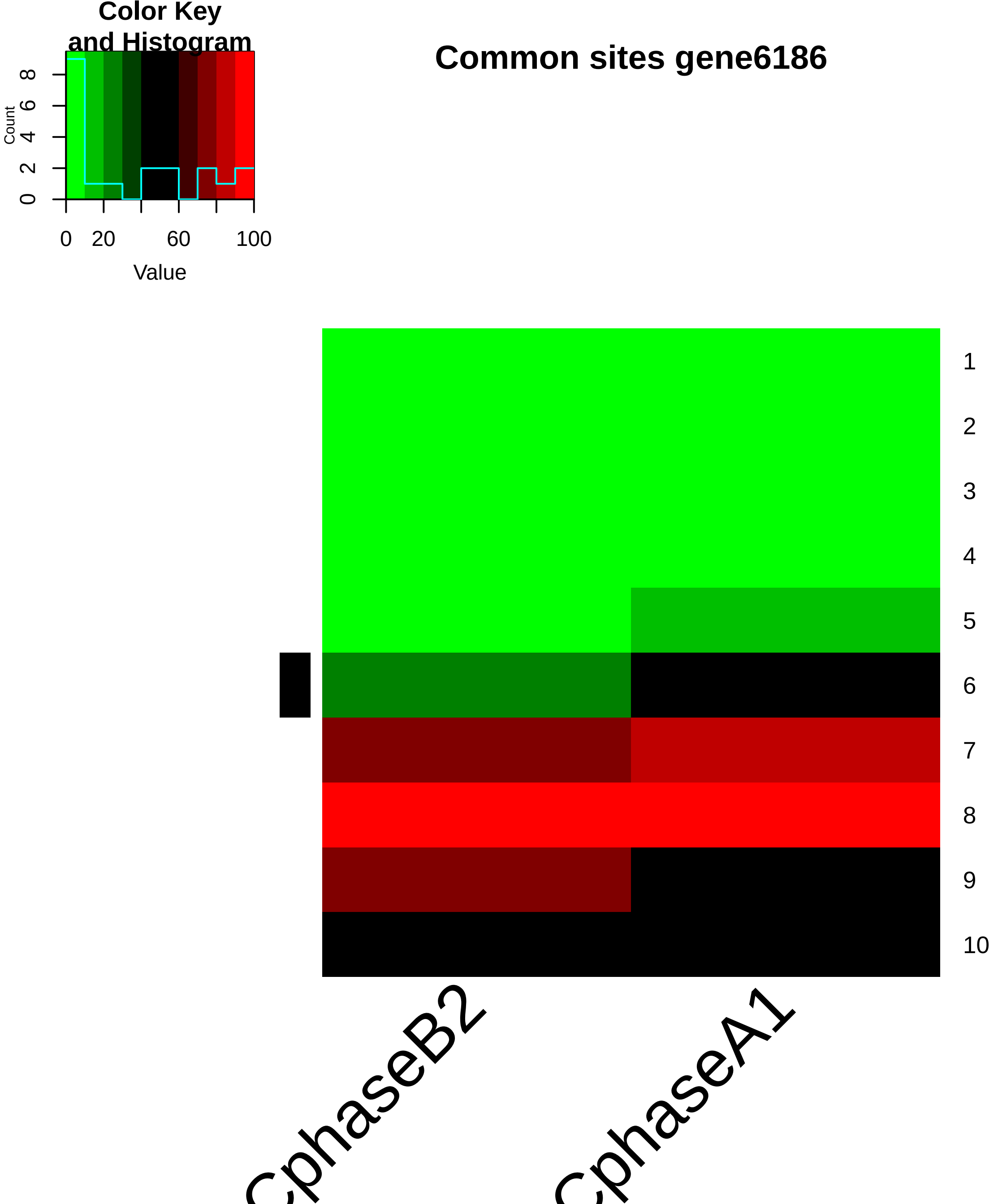

Color Key
and Histogram
Count
0
2
4
6
8
0
20
60
100
Value
Common sites gene6186
1
2
3
4
5
6
7
8
9
10
CphaseB2
CphaseA1

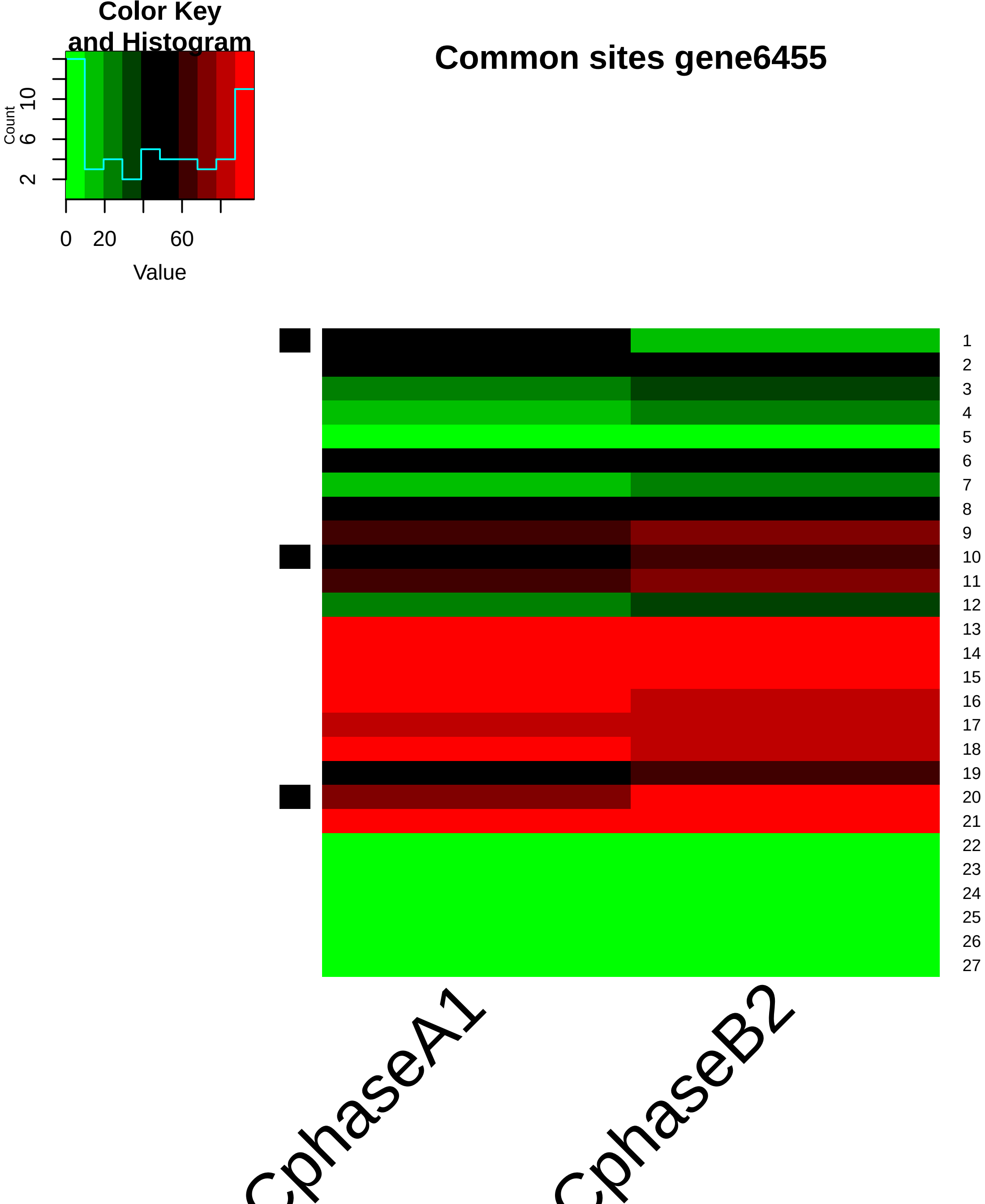

Color Key
and Histogram
Count
2
6
10
0
20
60
Value
Common sites gene6455
CphaseA1
CphaseB2

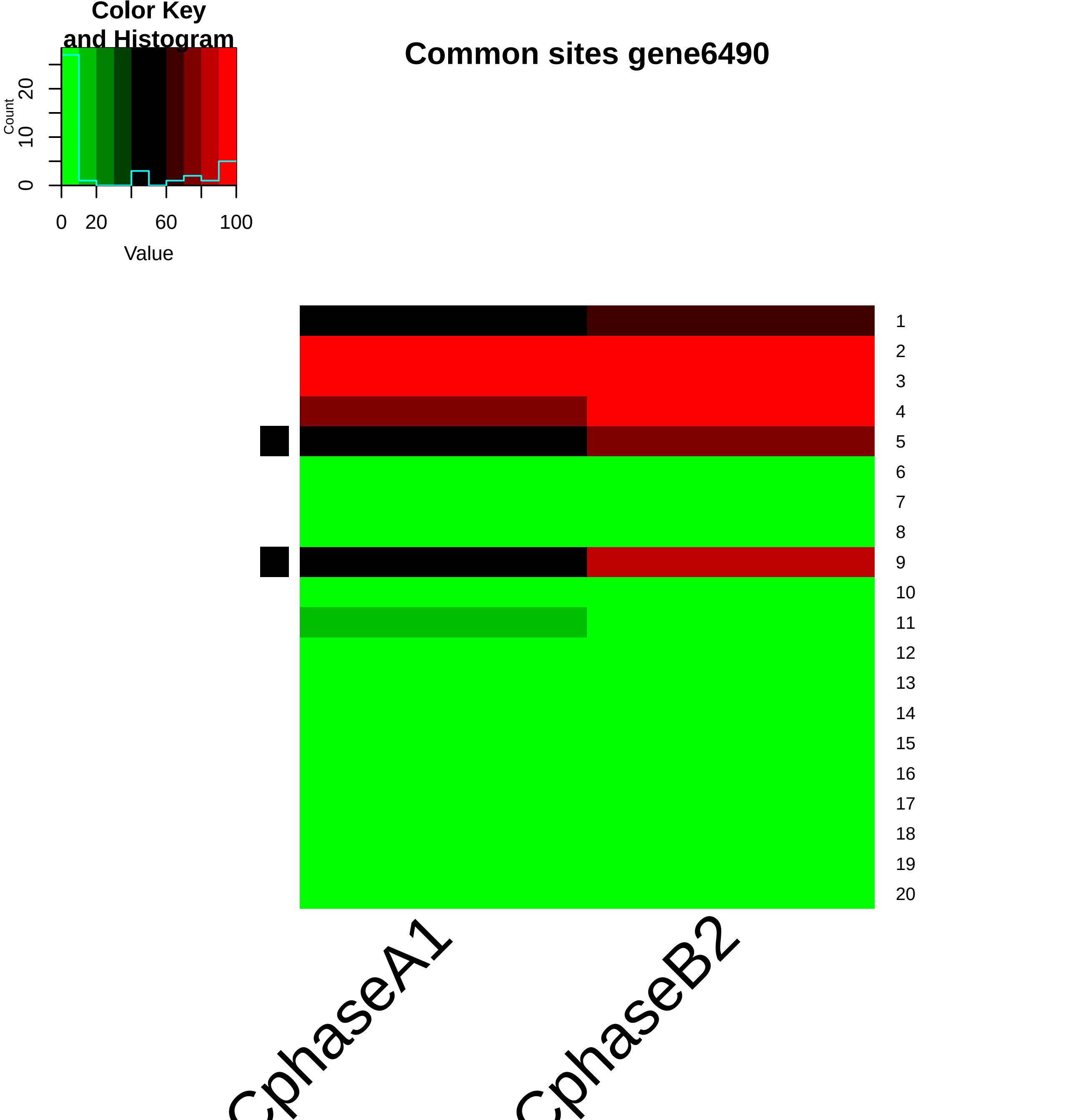
Color Key
and Histogram
Count
0
10
20
0
20
60
100
Value
Common sites gene6490
CphaseA1
CphaseB2

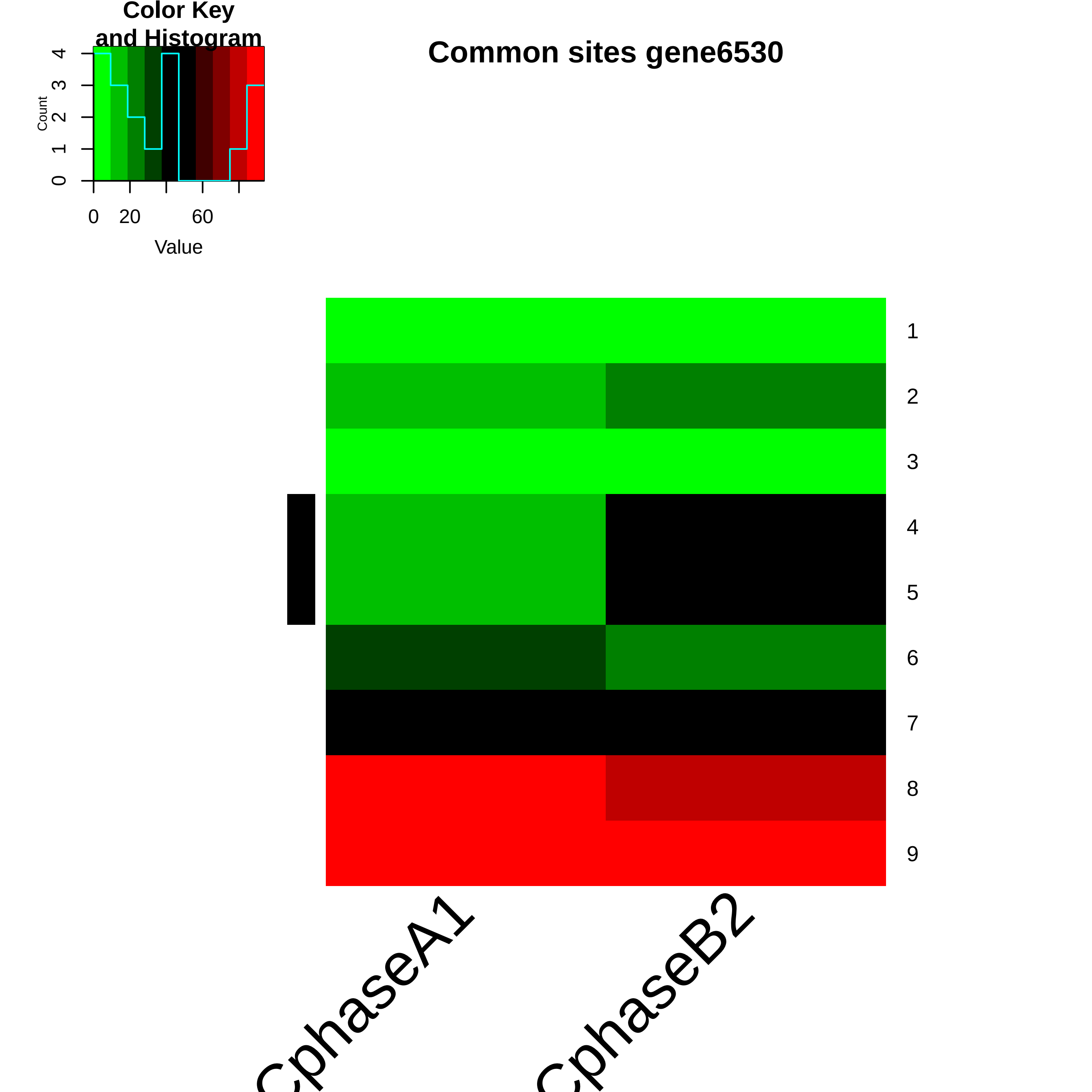
Color Key
and Histogram
Count
0
1
2
3
4
0
20
60
Value
Common sites gene6530
1
2
3
4
5
6
7
8
9
CphaseA1
CphaseB2

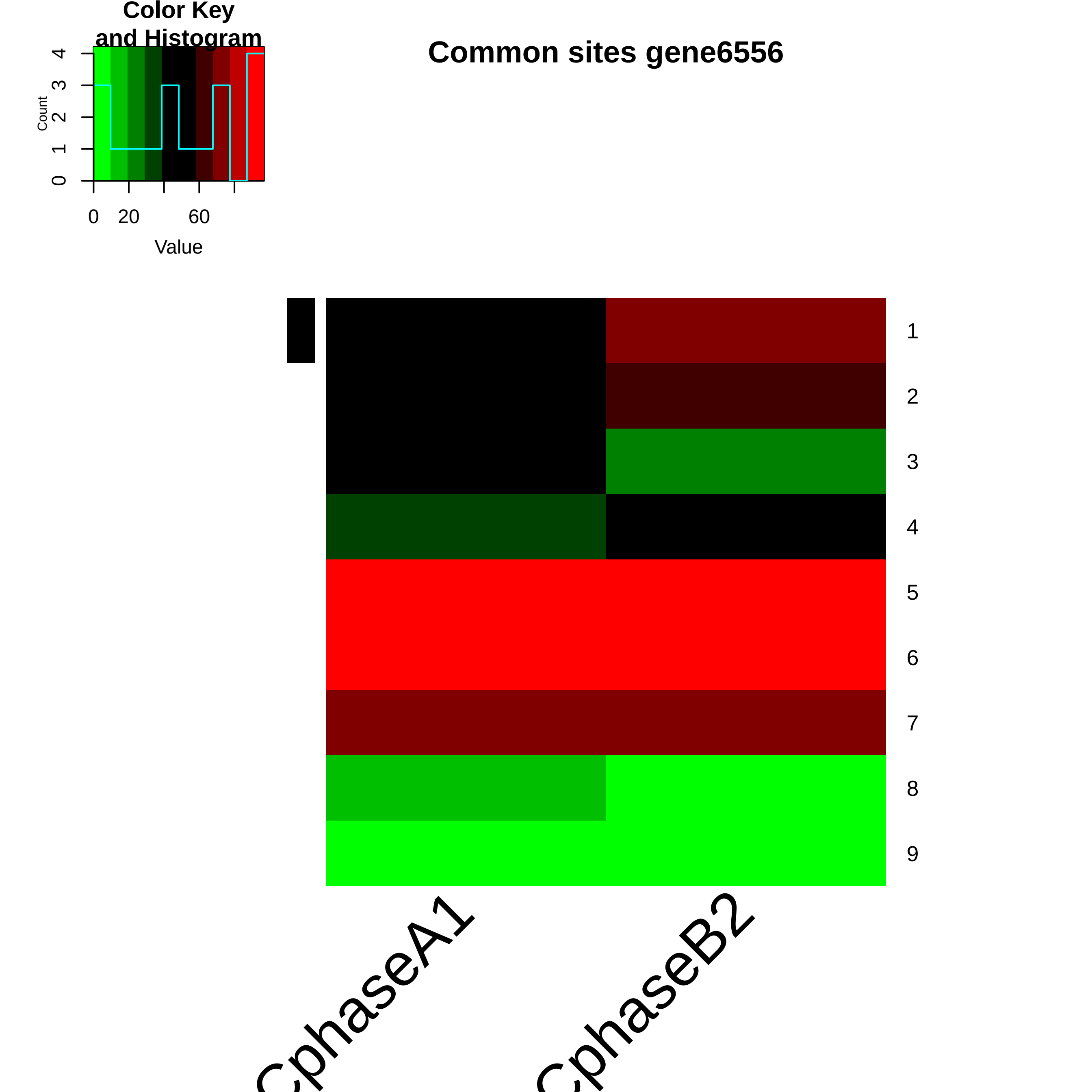
Color Key
and Histogram
Count
Value
0
20
60
0
1
2
3
4
Common sites gene6556
1
2
3
4
5
6
7
8
9
CphaseA1
CphaseB2

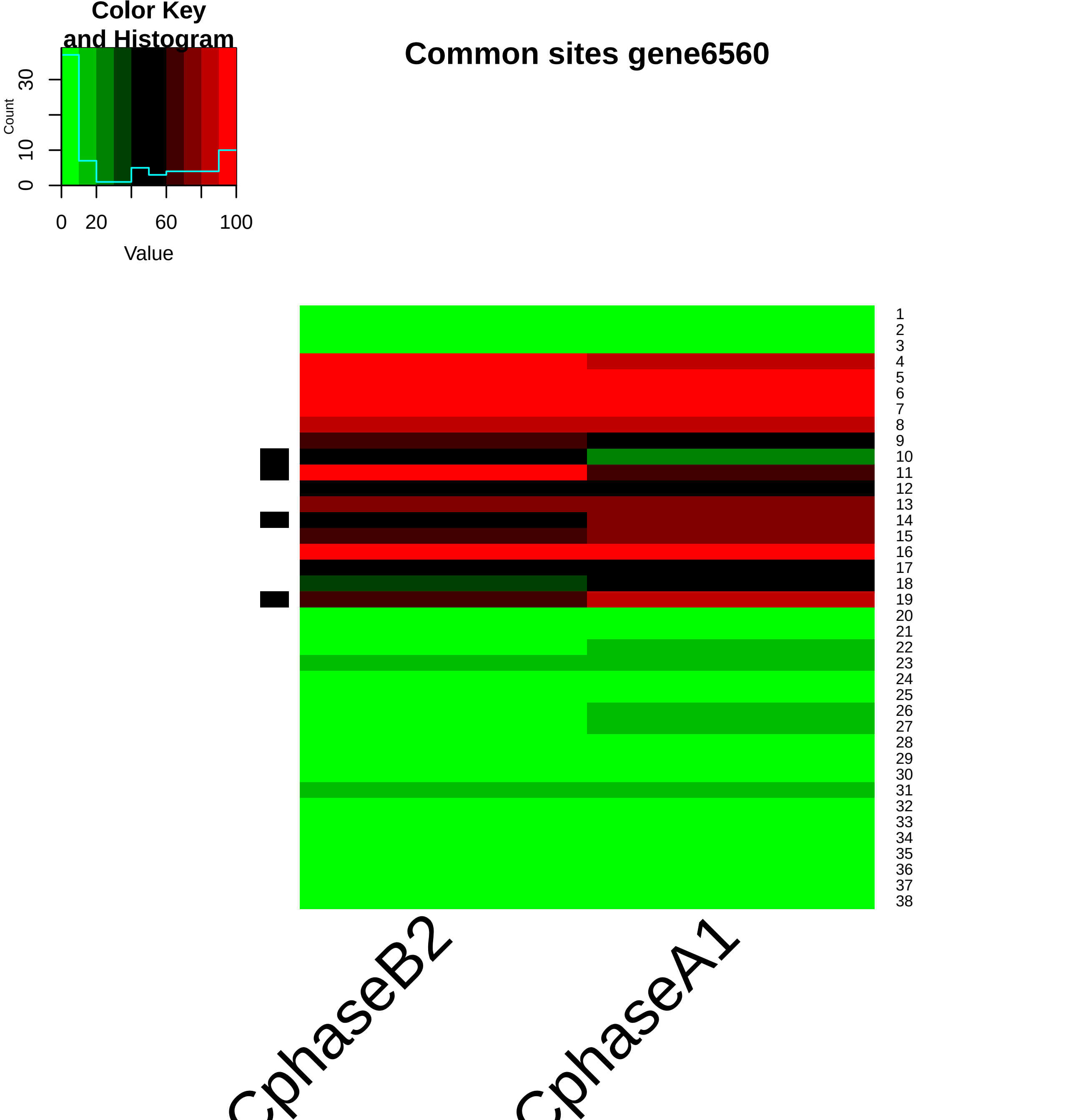

Color Key and Histogram
Count
0
10
30
0
20
60
100
Value
Common sites gene6560
CphaseB2
CphaseA1

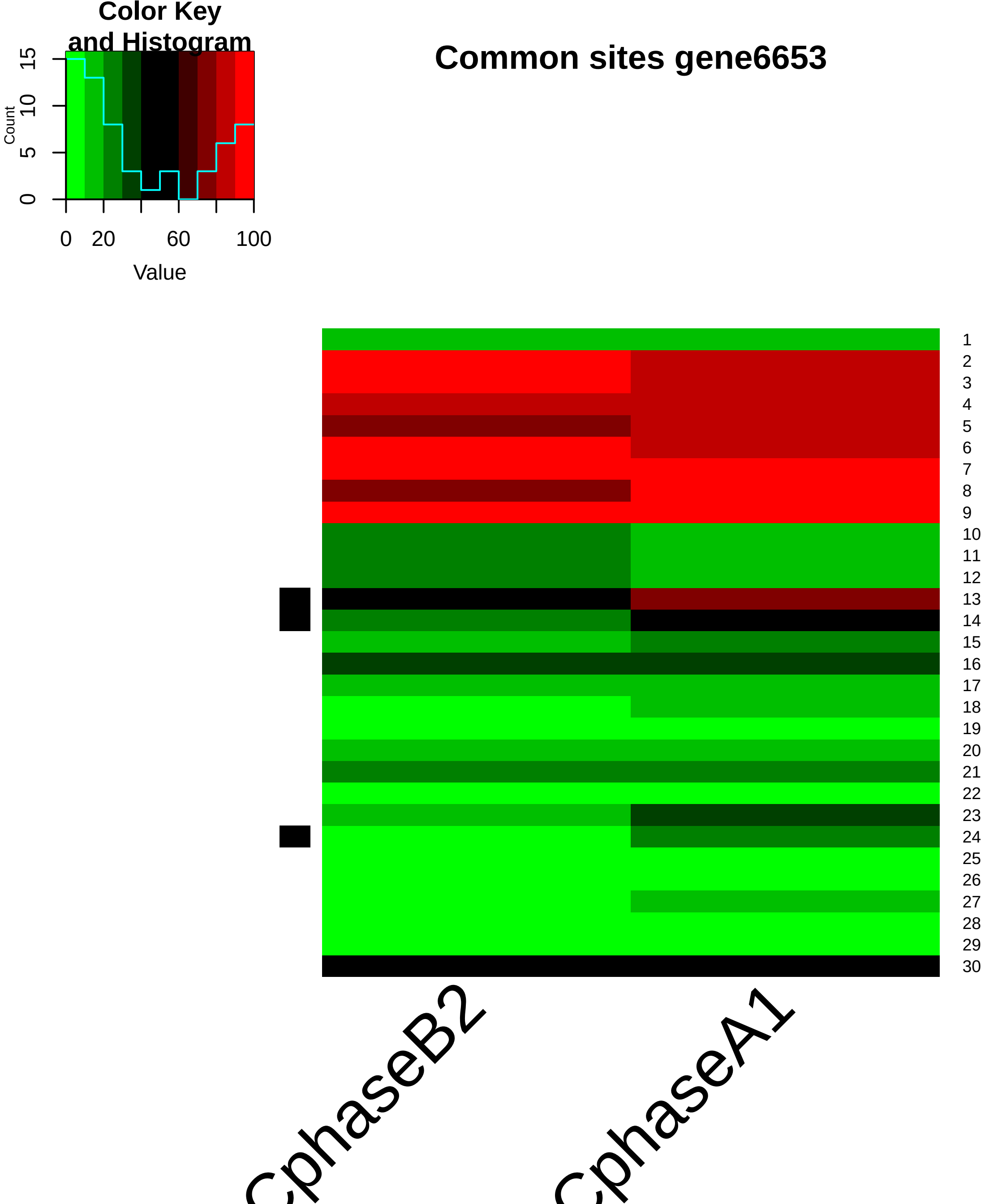
Color Key
and Histogram
Count
0
5
10
15
0
20
60
100
Value
Common sites gene6653
CphaseB2
CphaseA1

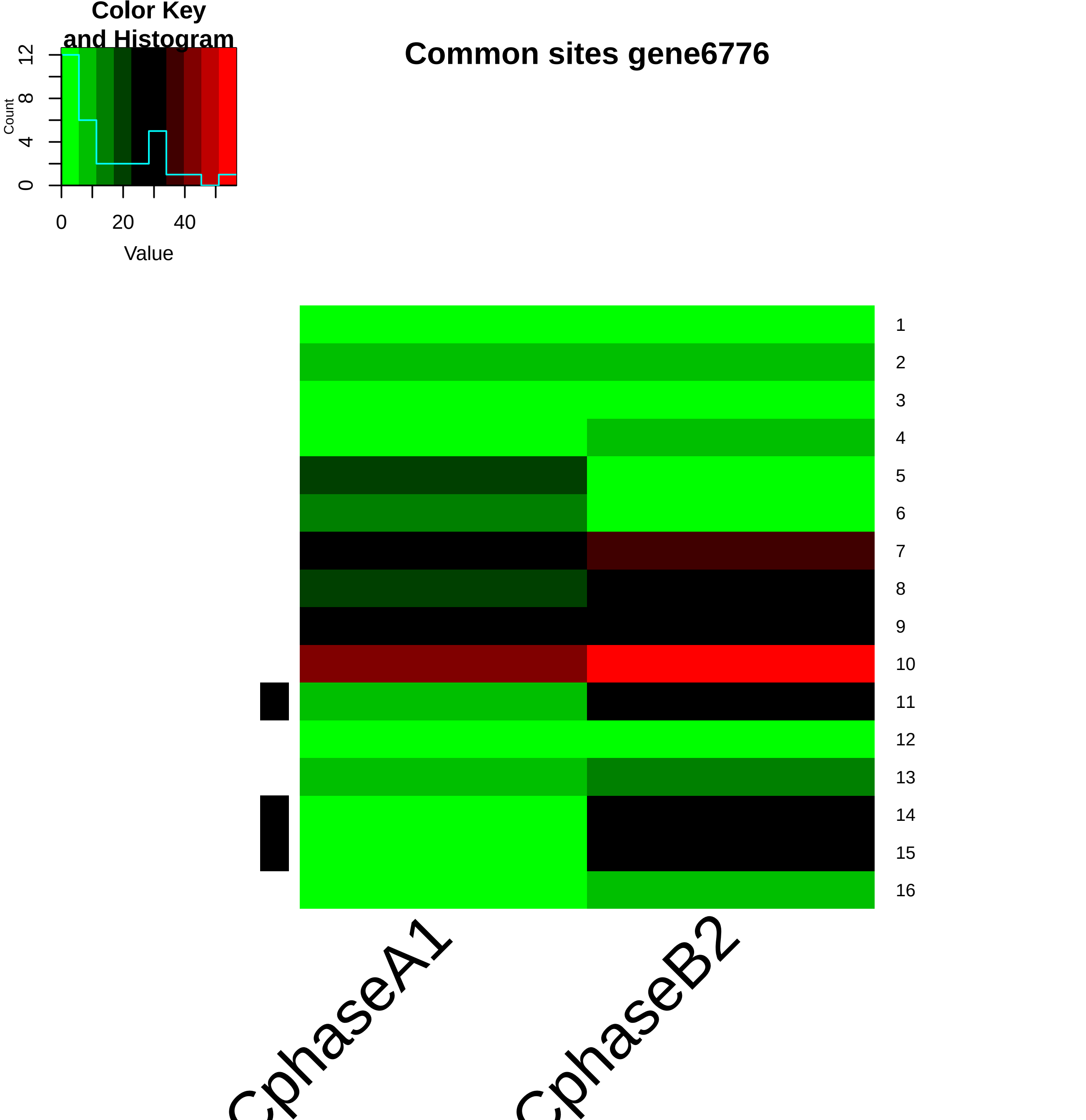

Color Key
and Histogram
Count
0
4
8
12
0
20
40
Value
Common sites gene6776
CphaseA1
CphaseB2

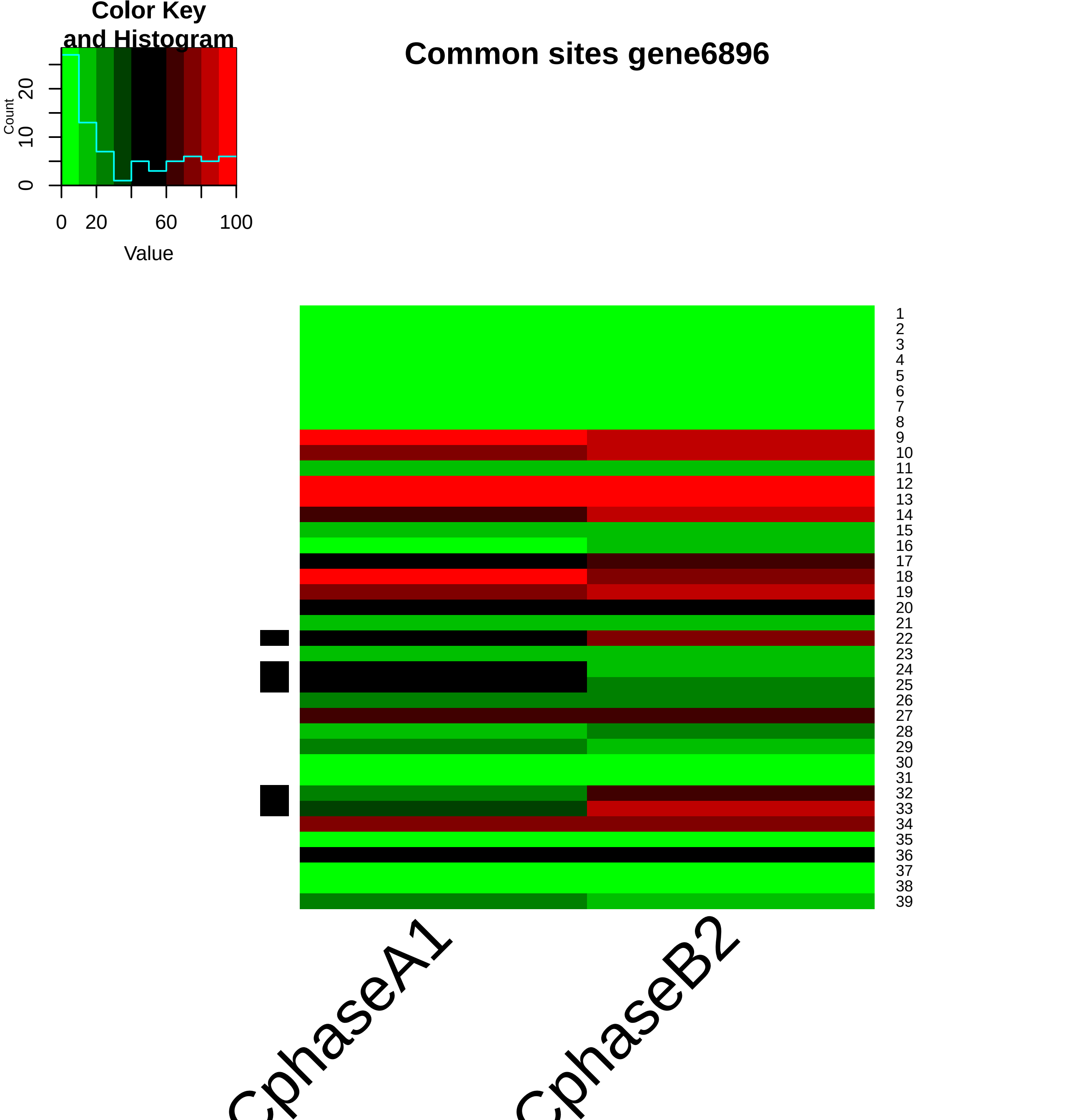

Color Key
and Histogram
Count
0
10
20
0
20
60
100
Value
Common sites gene6896
CphaseA1
CphaseB2

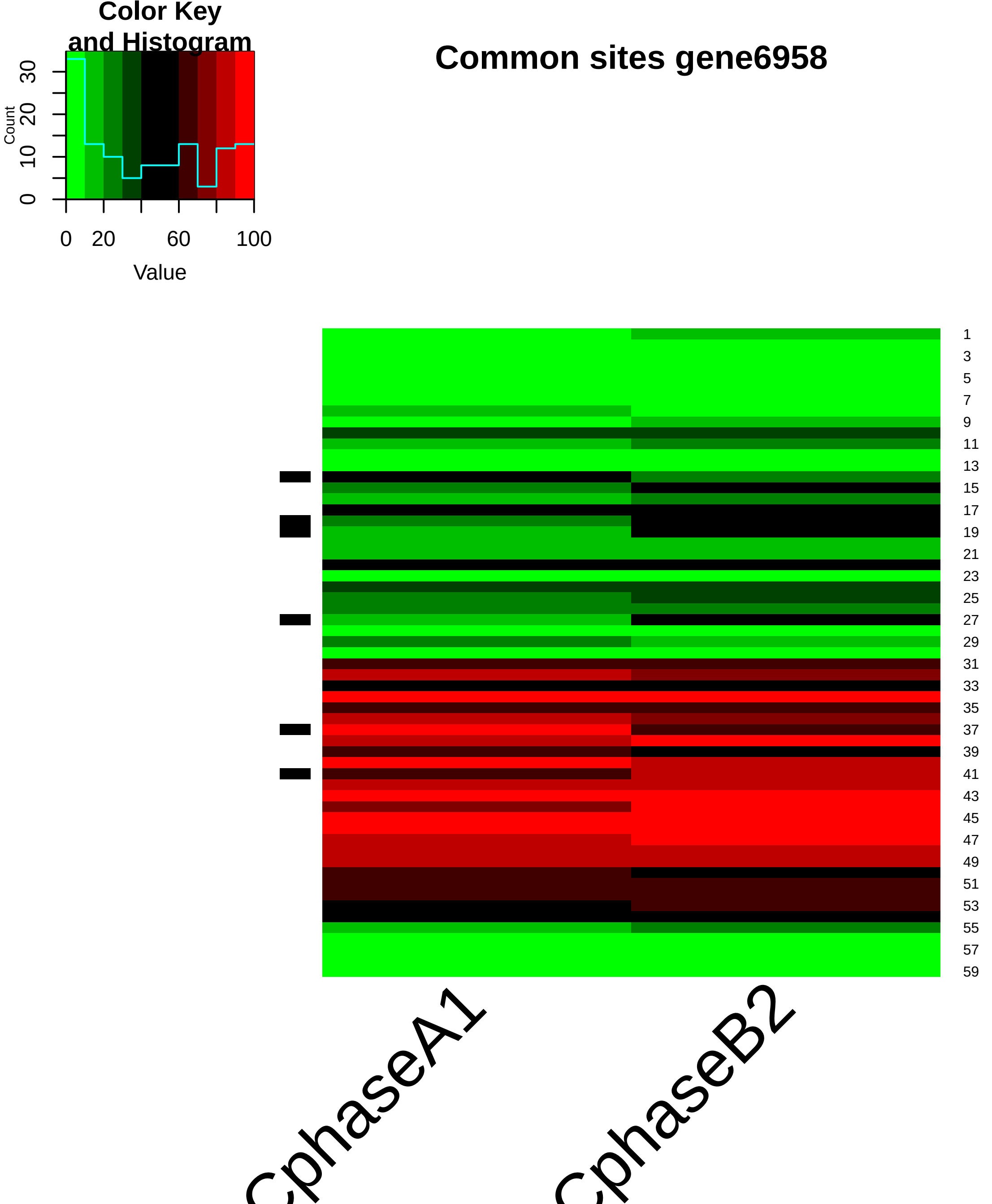

Color Key
and Histogram
Count
0
10
20
30
0
20
60
100
Value
Common sites gene6958
1
3
5
7
9
11
13
15
17
19
21
23
25
27
29
31
33
35
37
39
41
43
45
47
49
51
53
55
57
59
CphaseA1
CphaseB2

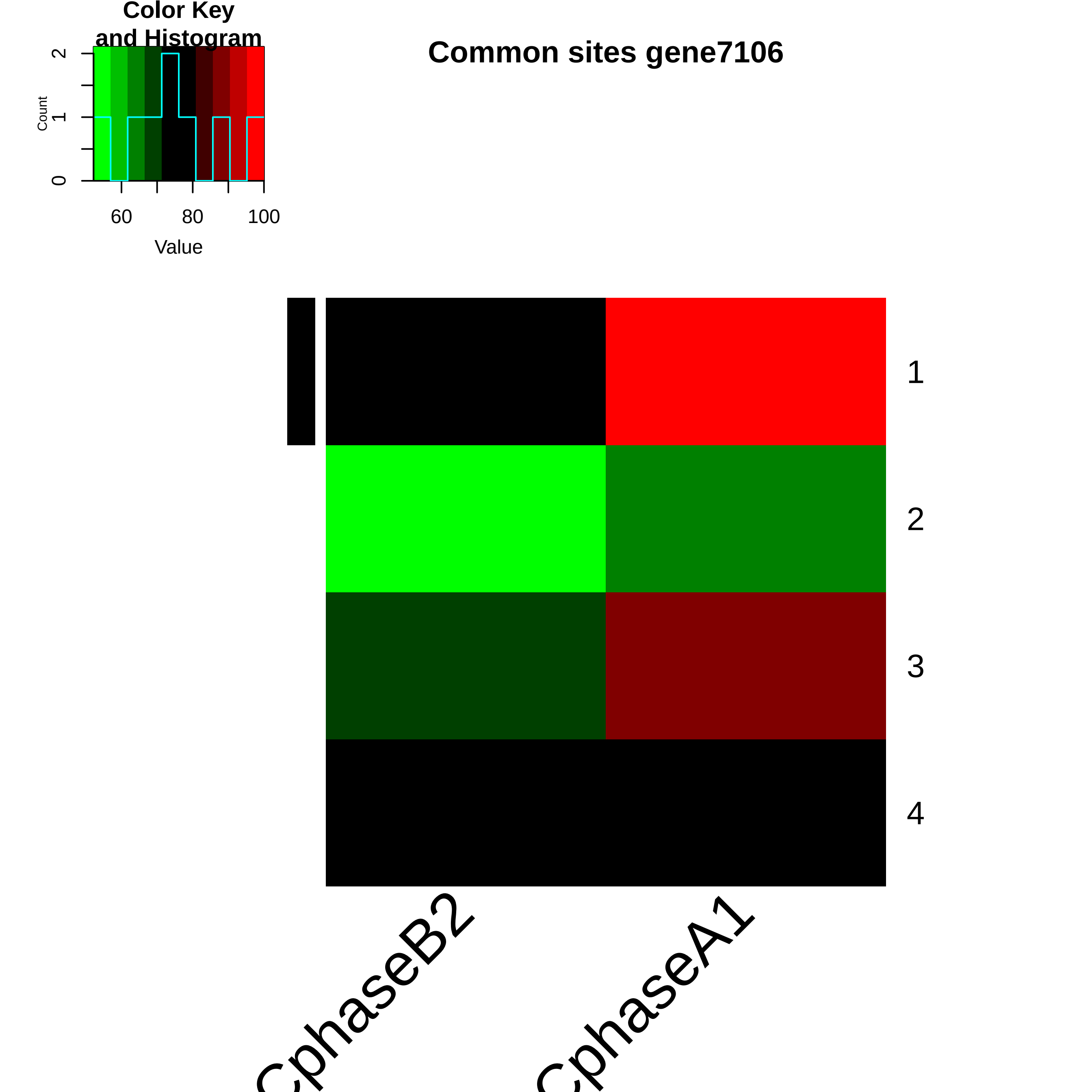

Color Key
and Histogram
Count
0
1
2
60
80
100
Value
Common sites gene7106
1
2
3
4
CphaseB2
CphaseA1

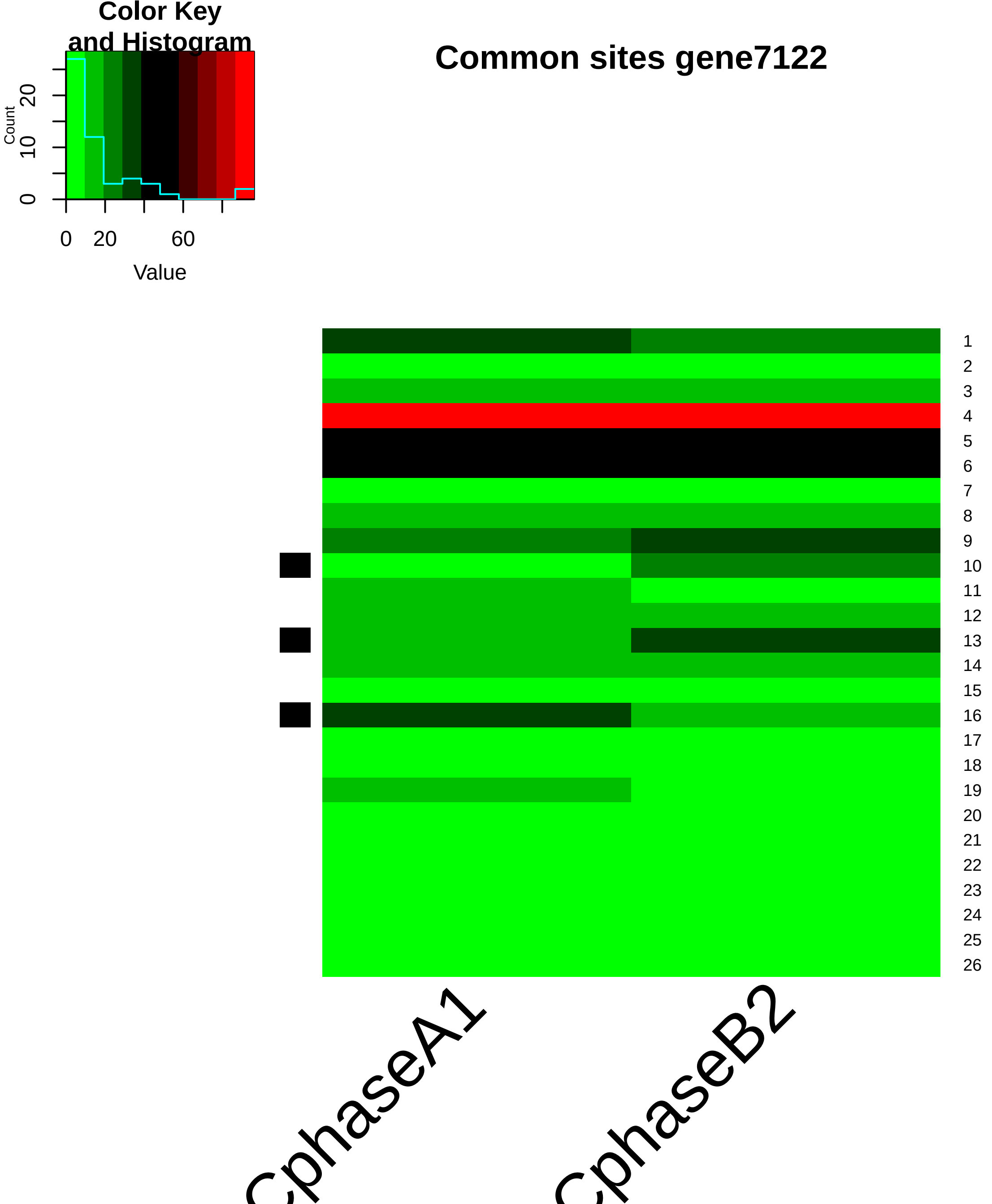
Color Key
and Histogram
Count
0
10
20
0
20
60
Value
Common sites gene7122
CphaseA1
CphaseB2

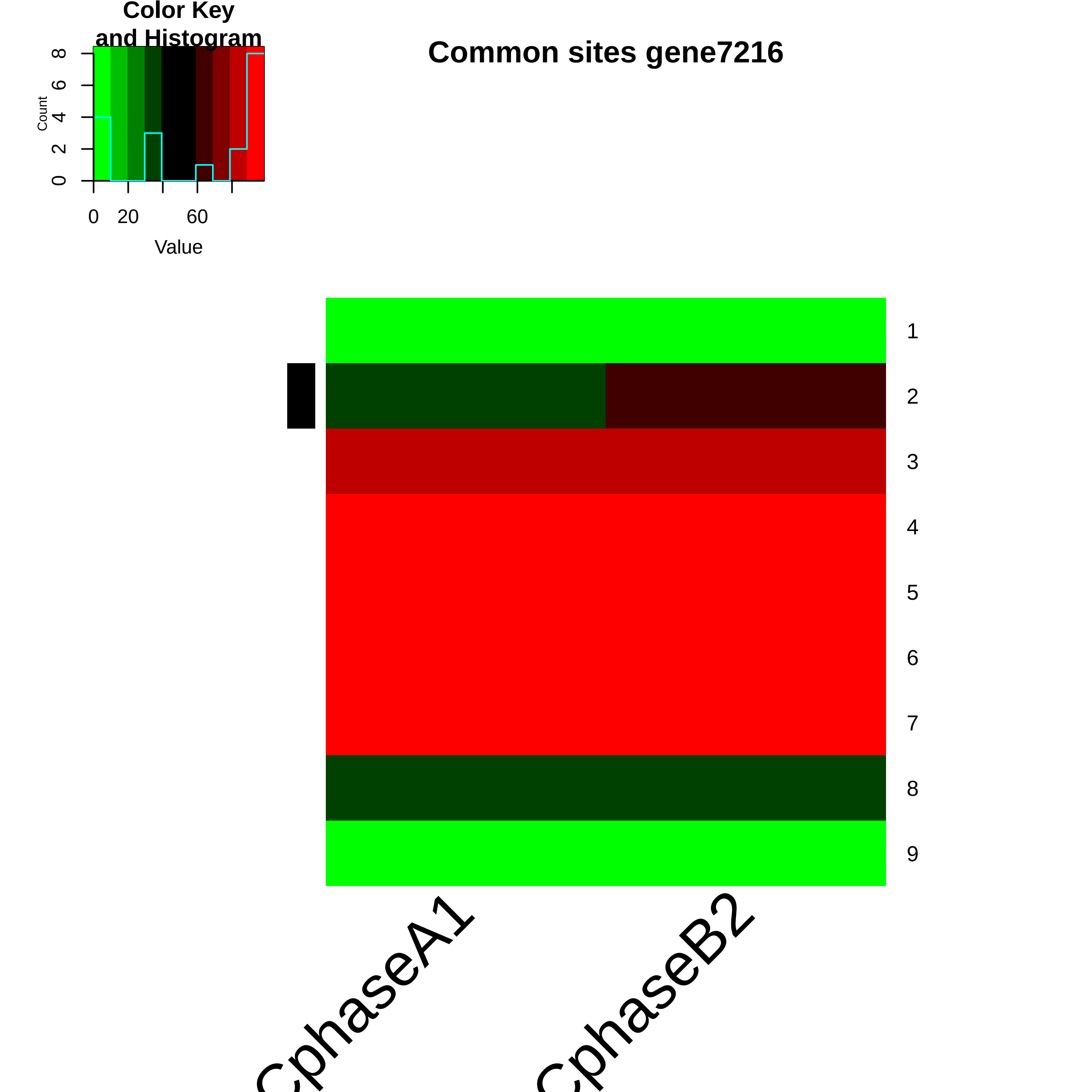

Color Key
and Histogram
Count
0
2
4
6
8
0
20
60
Value
Common sites gene7216
1
2
3
4
5
6
7
8
9
CphaseA1
CphaseB2

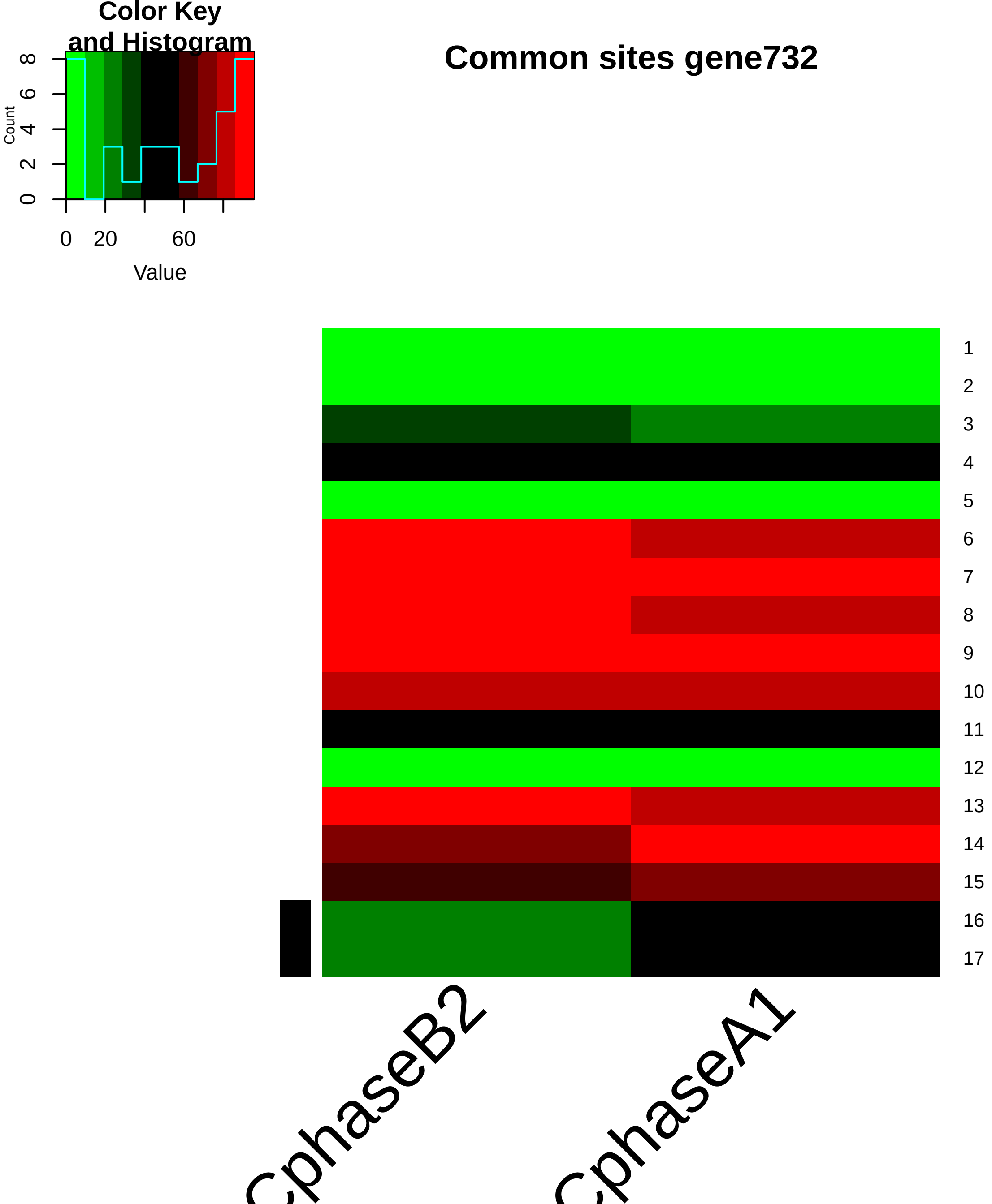
Color Key
and Histogram
Count
Value
0
20
60
0
2
4
6
8
Common sites gene732
CphaseB2
CphaseA1

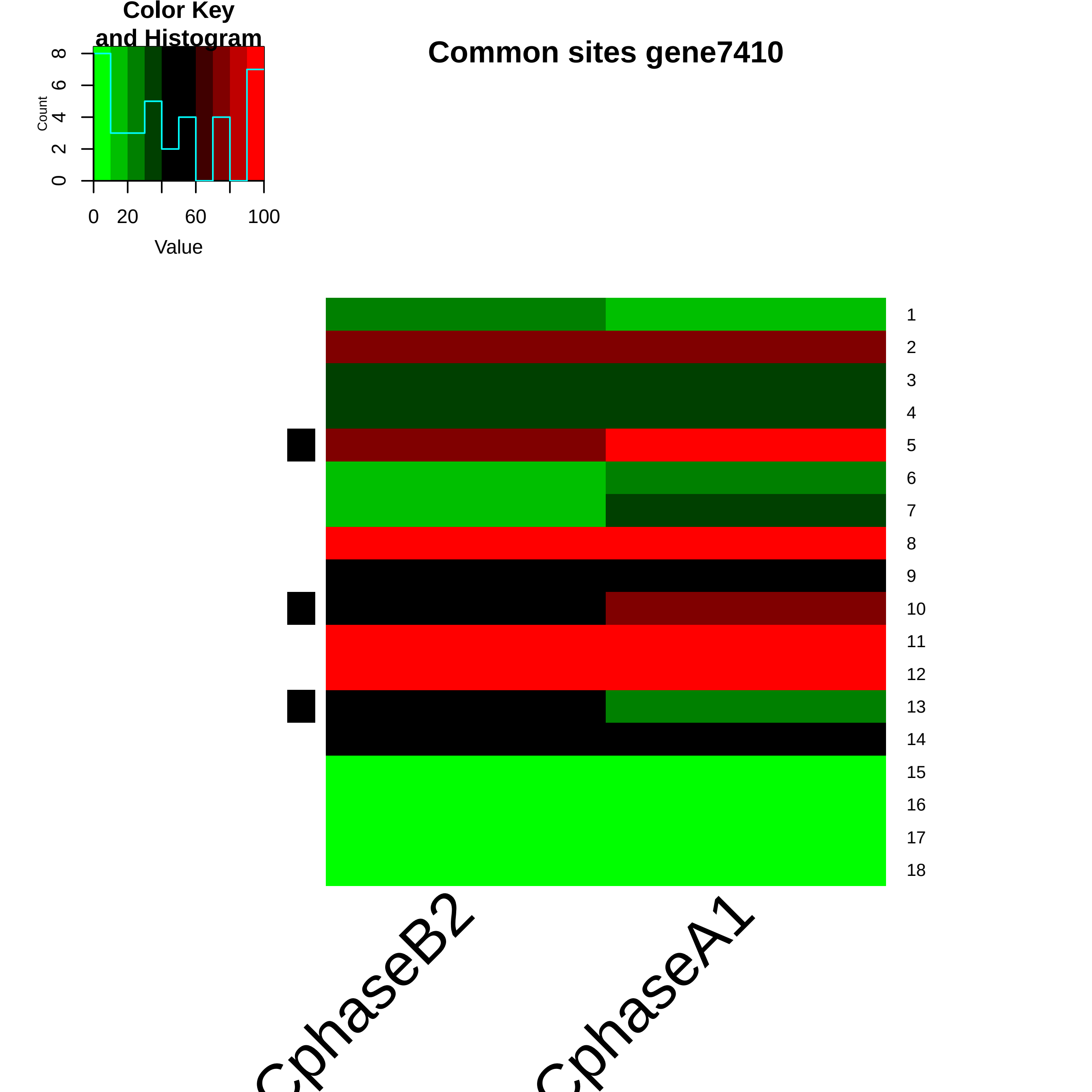

Color Key
and Histogram
Count
0
2
4
6
8
0
20
60
100
Value
Common sites gene7410
CphaseB2
CphaseA1

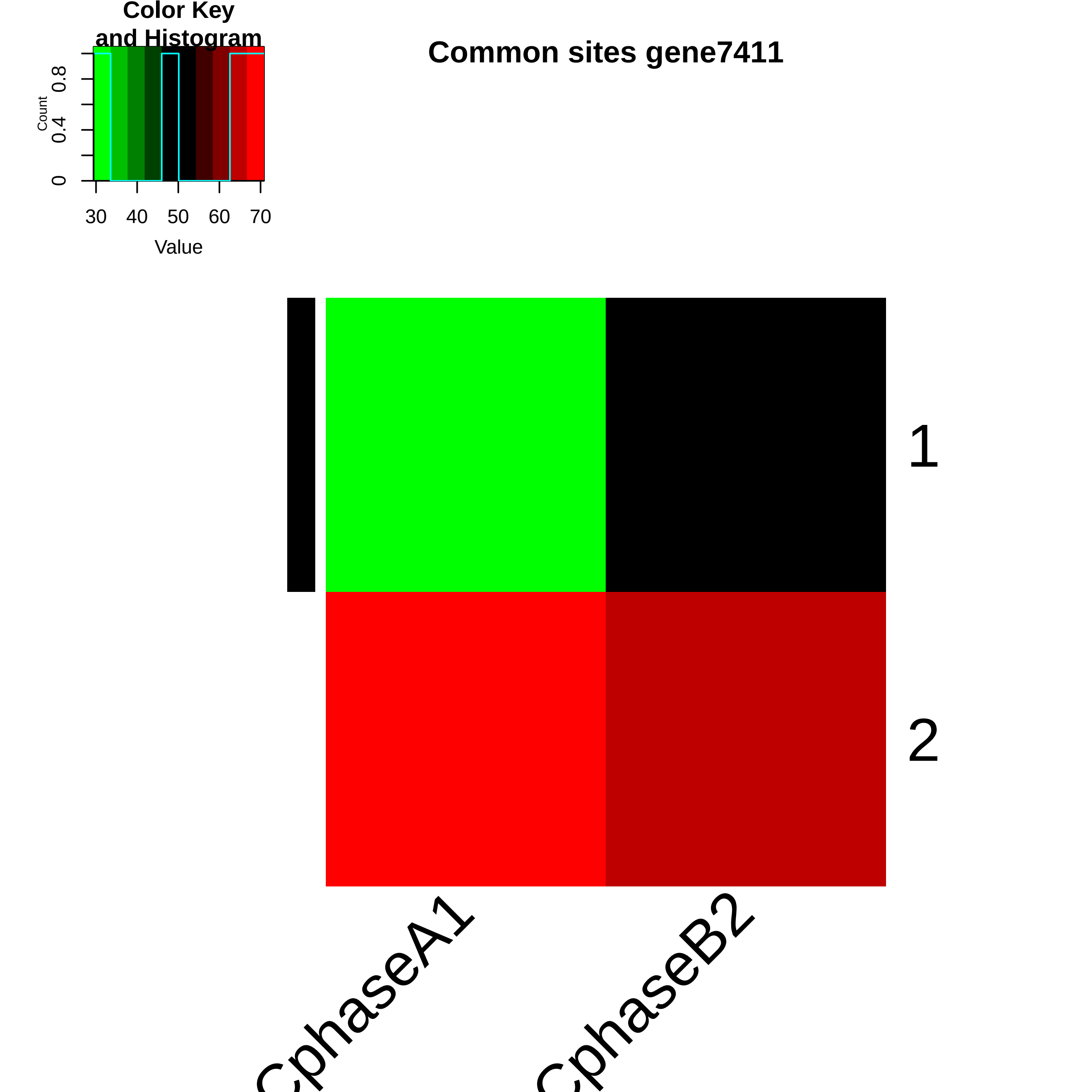
Color Key
and Histogram
Count
0
0.4
0.8
30
40
50
60
70
Value
Common sites gene7411
1
2
CphaseA1
CphaseB2

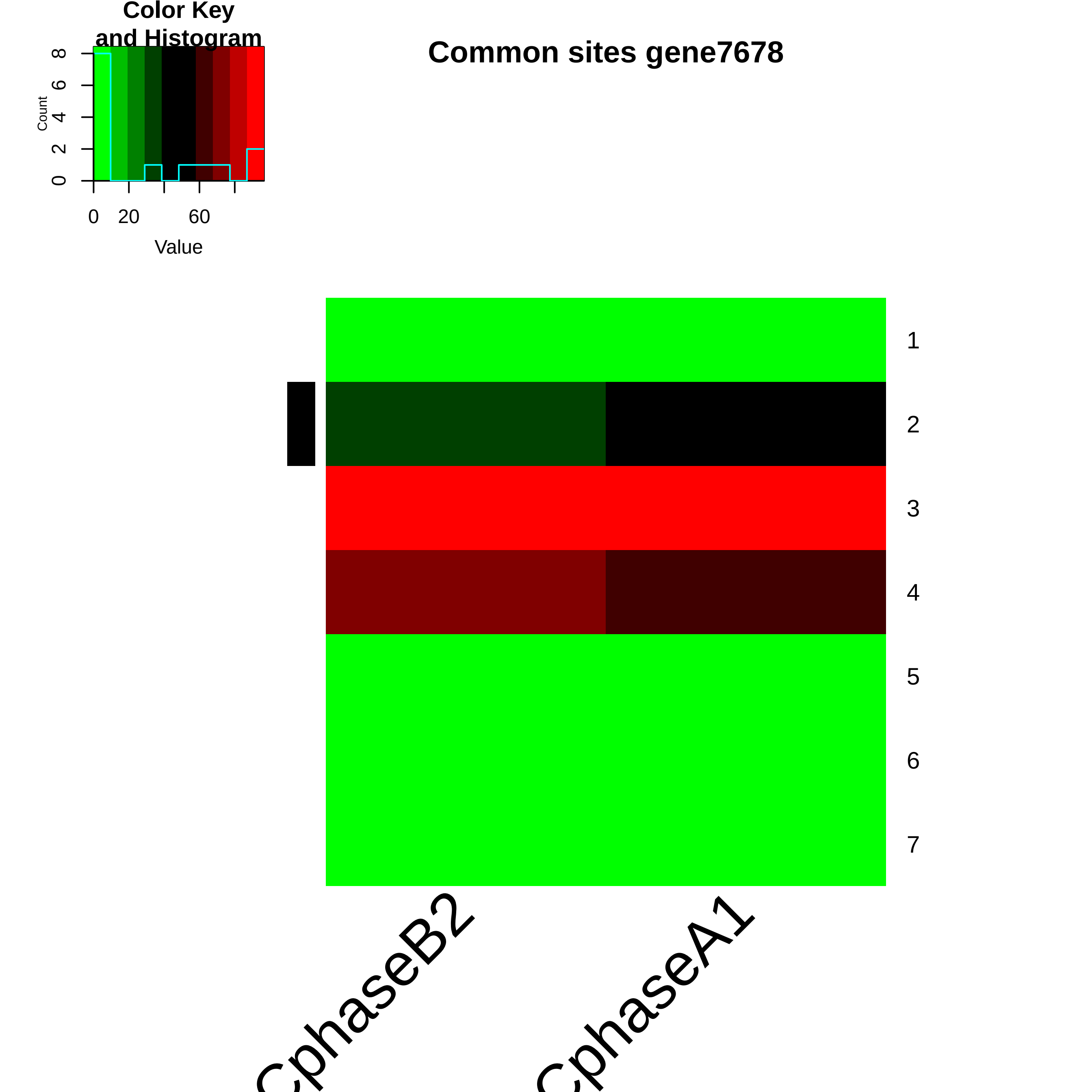

Color Key
and Histogram
Count
0
2
4
6
8
0
20
60
Value
Common sites gene7678
1
2
3
4
5
6
7
CphaseB2
CphaseA1

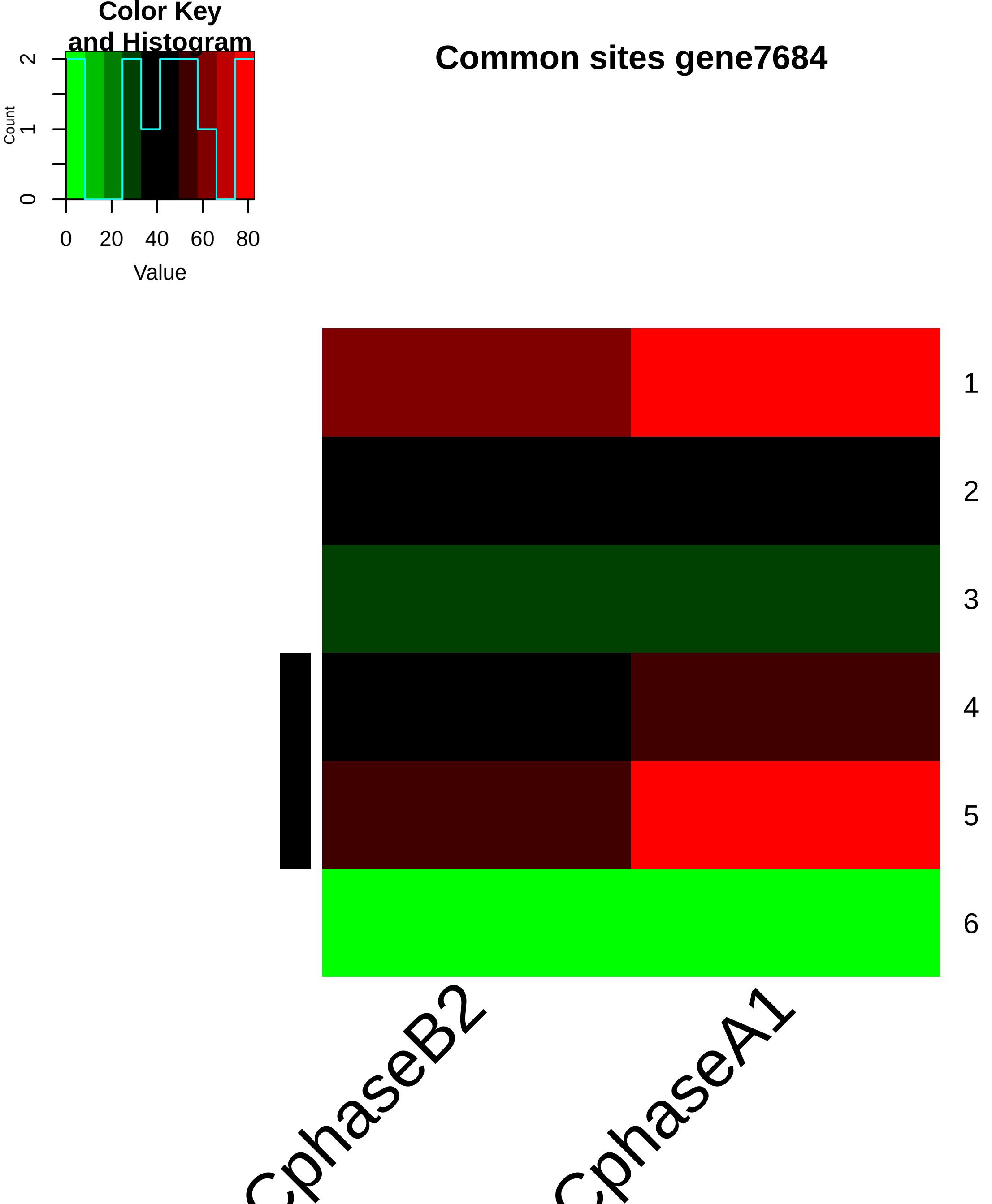
Color Key
and Histogram
Count
0
1
2
0
20
40
60
80
Value
Common sites gene7684
1
2
3
4
5
6
CphaseB2
CphaseA1

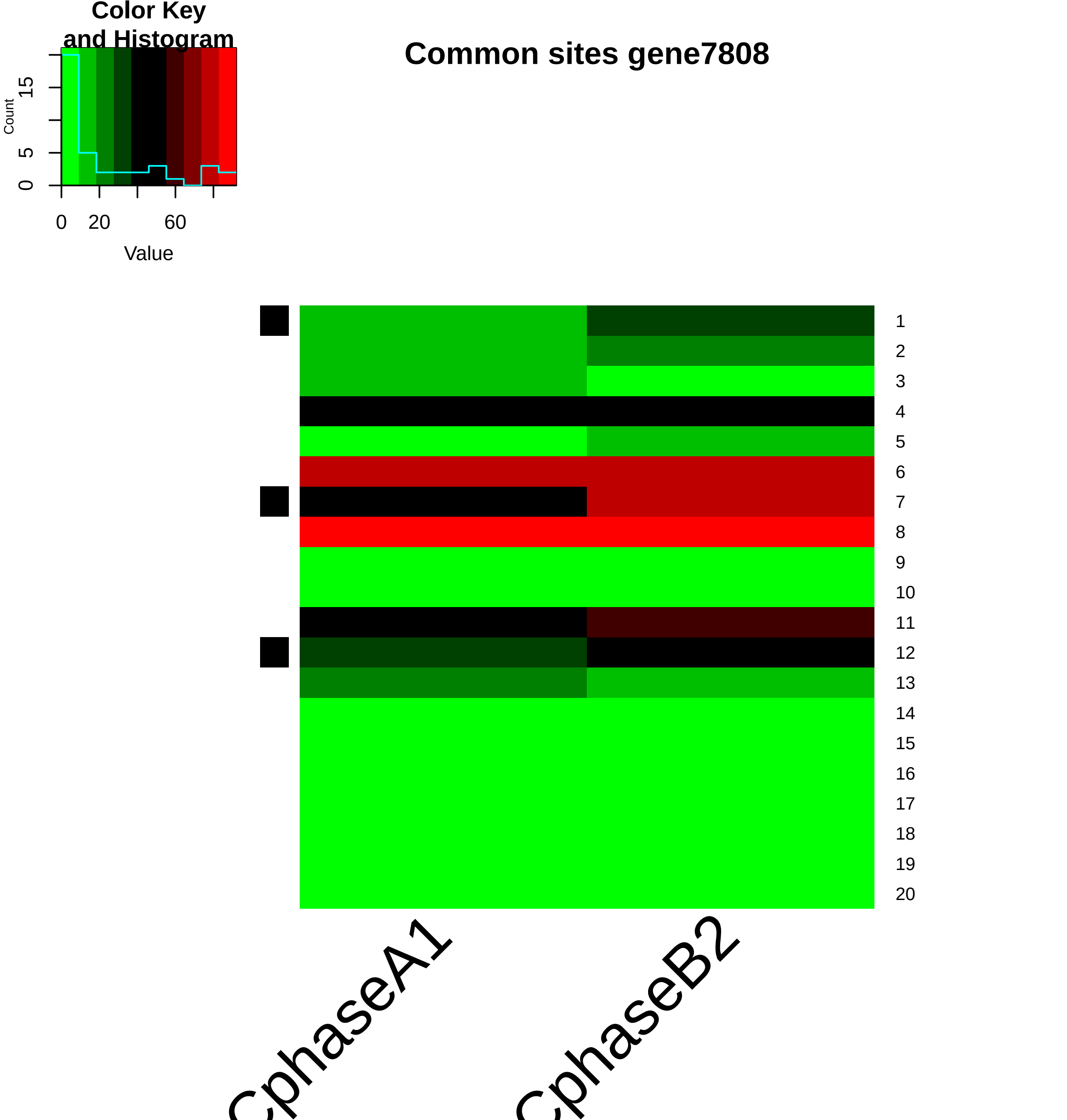

Color Key
and Histogram
Count
0
5
15
0
20
60
Value
Common sites gene7808
CphaseA1
CphaseB2

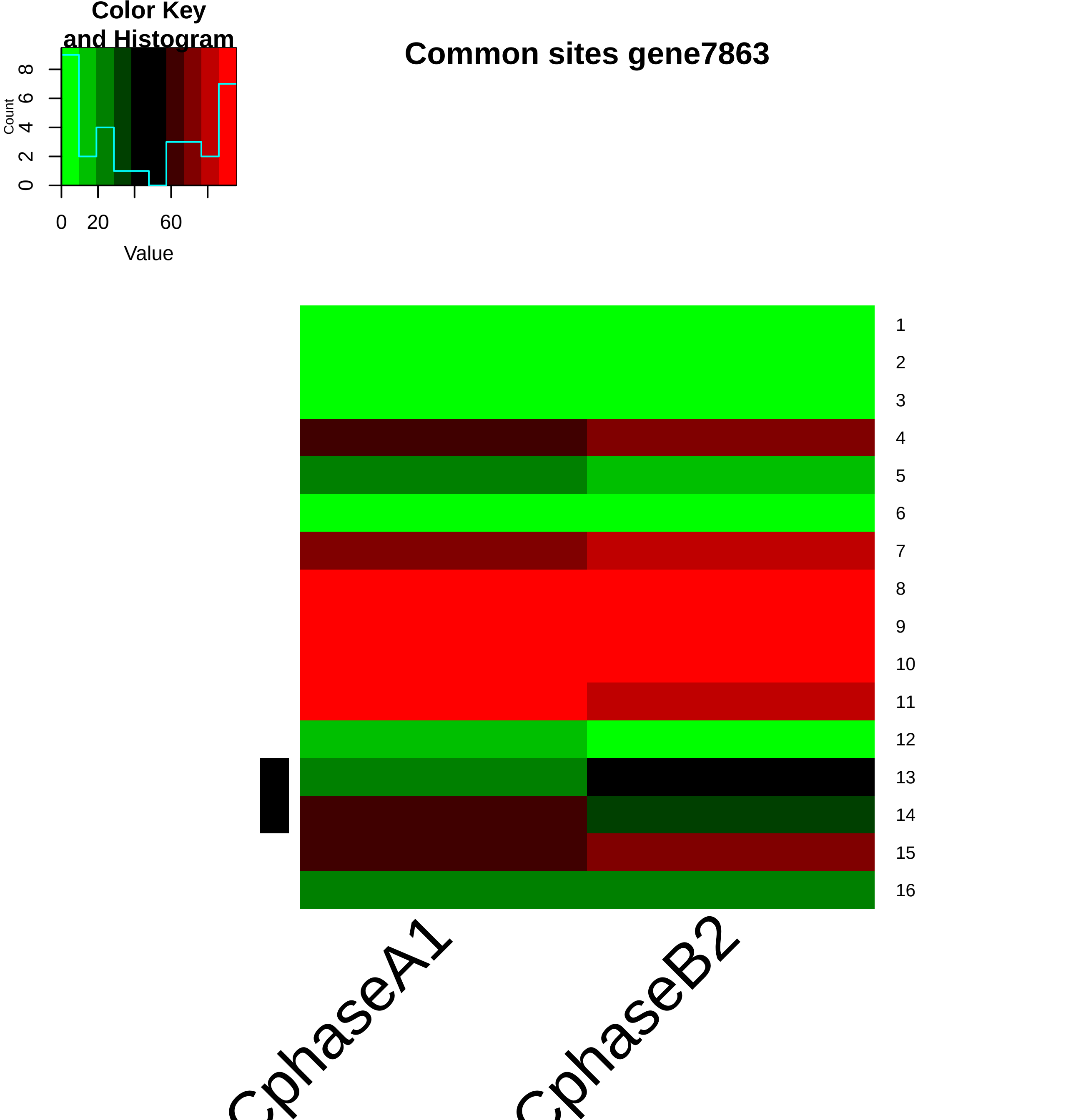

Color Key
and Histogram
Count
0
2
4
6
8
0
20
60
Value
Common sites gene7863
CphaseA1
CphaseB2

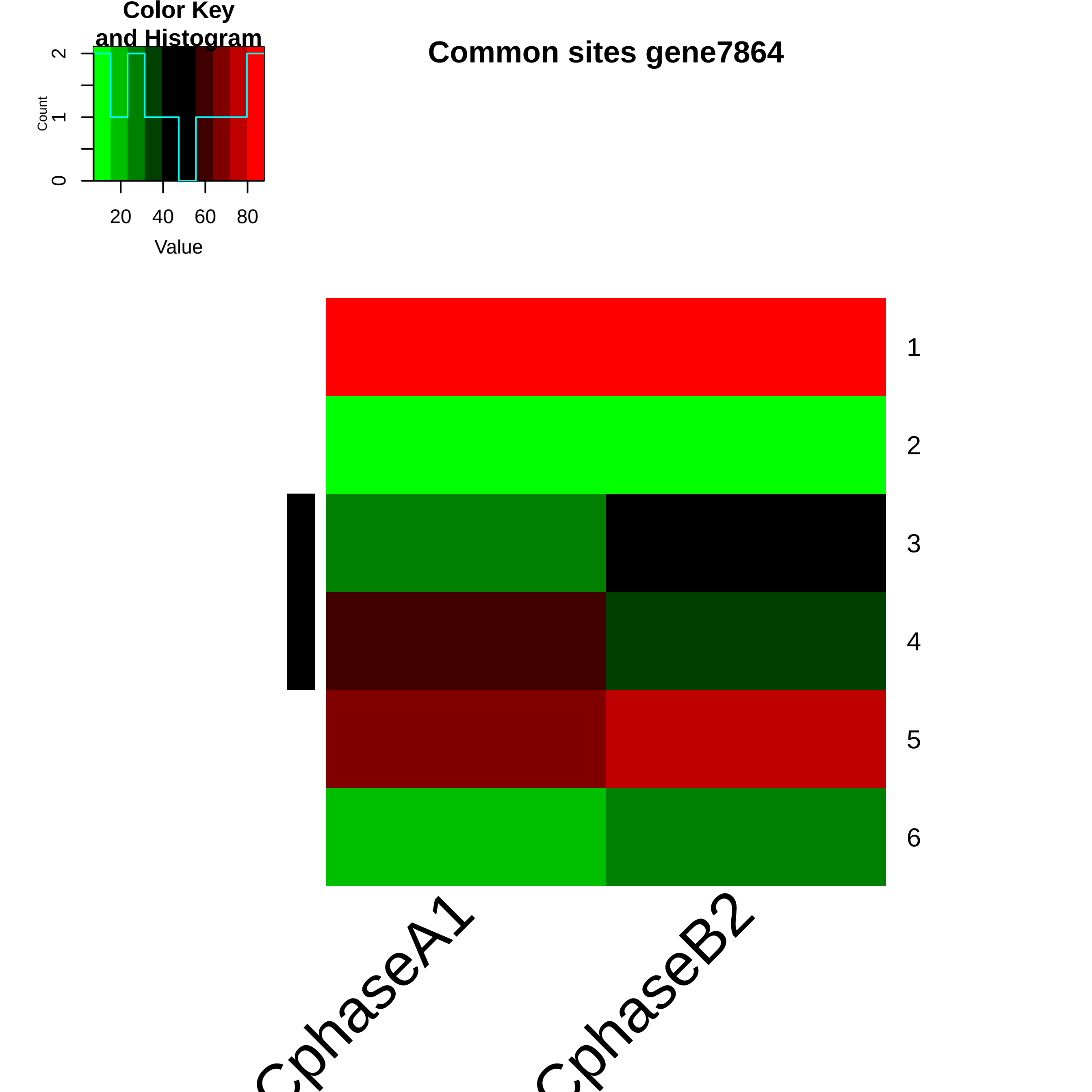

Color Key
and Histogram
Count
0
1
2
20
40
60
80
Value
Common sites gene7864
1
2
3
4
5
6
CphaseA1
CphaseB2

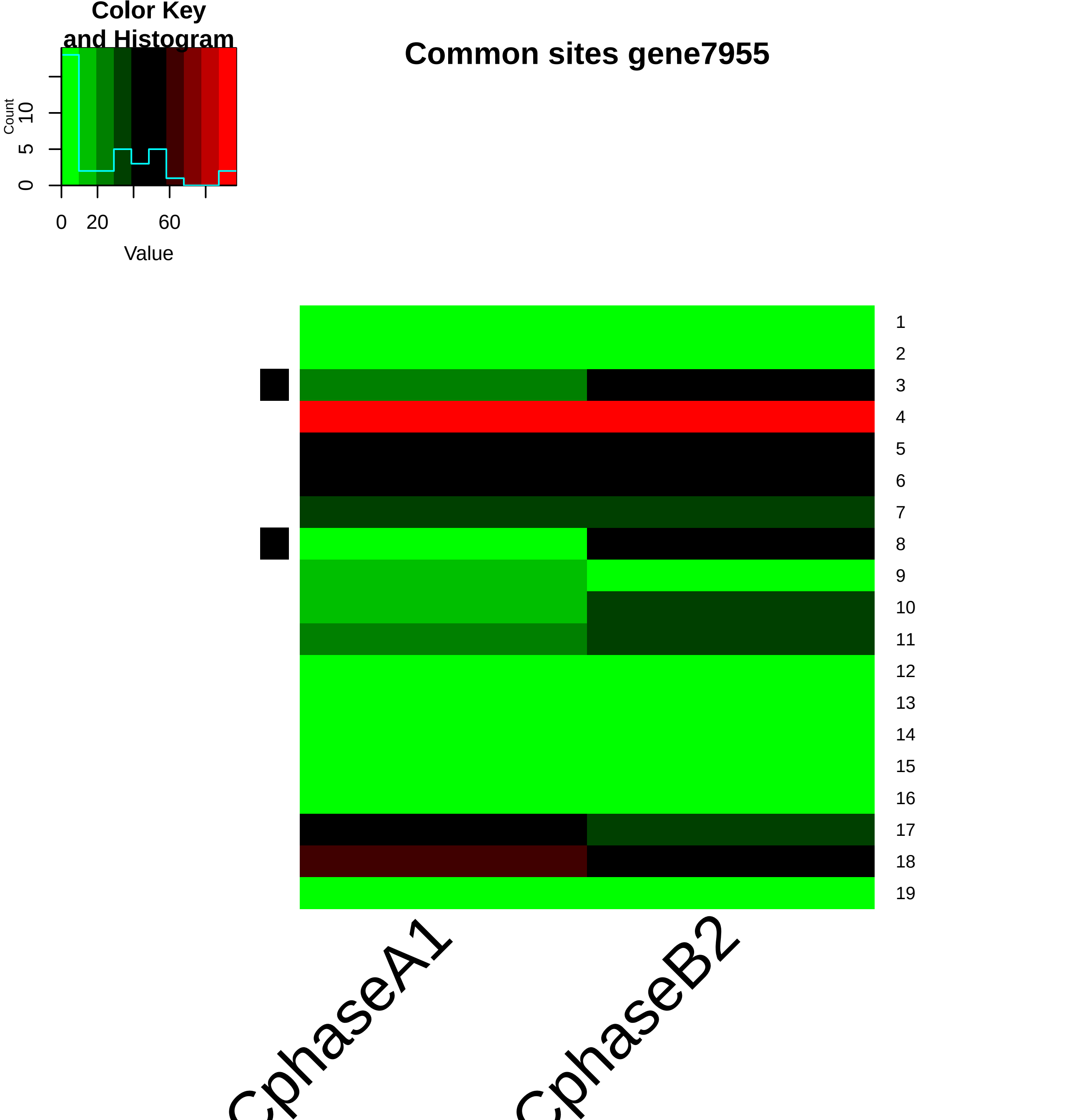
Color Key
and Histogram
Count
0
5
10
0
20
60
Value
Common sites gene7955
CphaseA1
CphaseB2

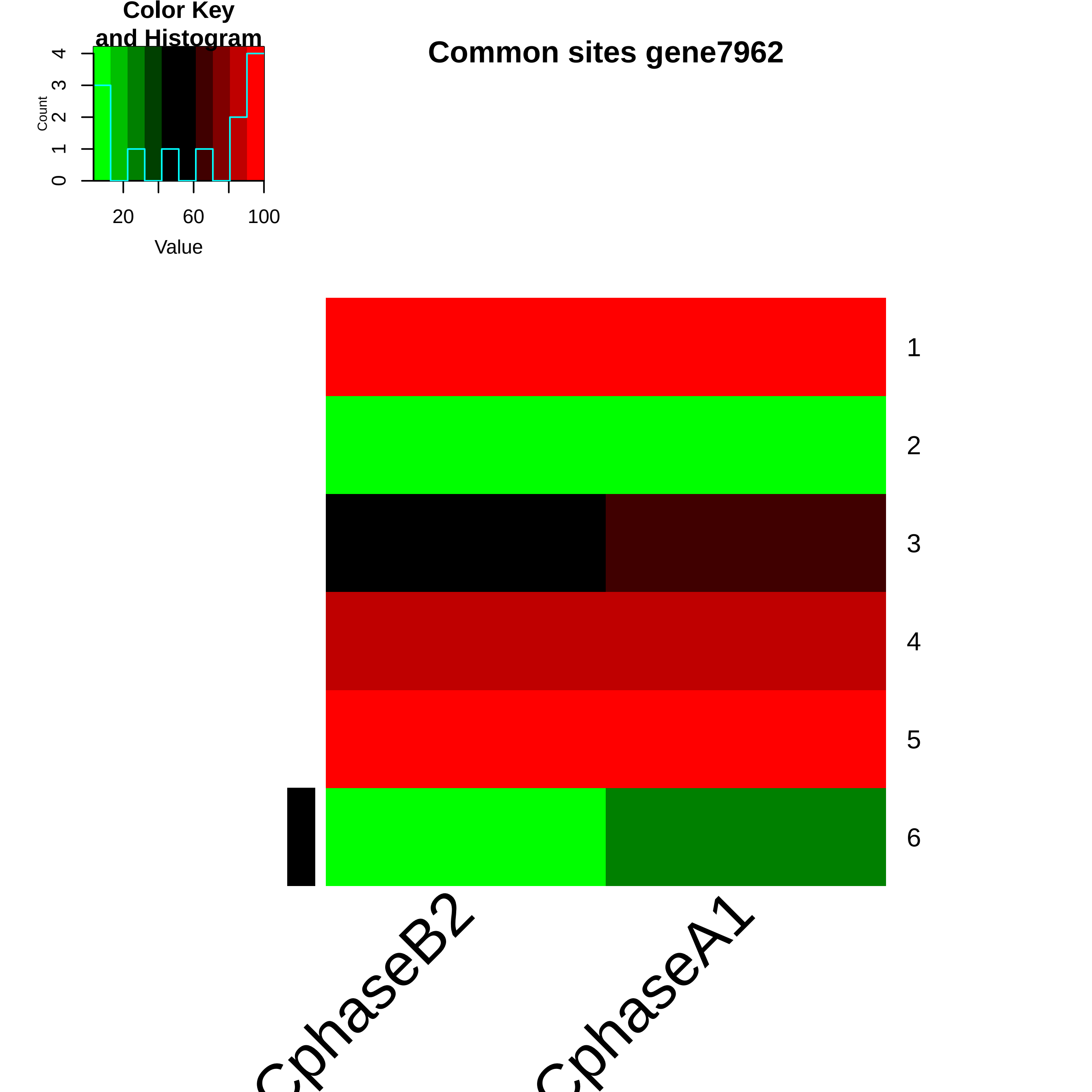
Color Key
and Histogram
Count
0
1
2
3
4
20
60
100
Value
Common sites gene7962
1
2
3
4
5
6
CphaseB2
CphaseA1

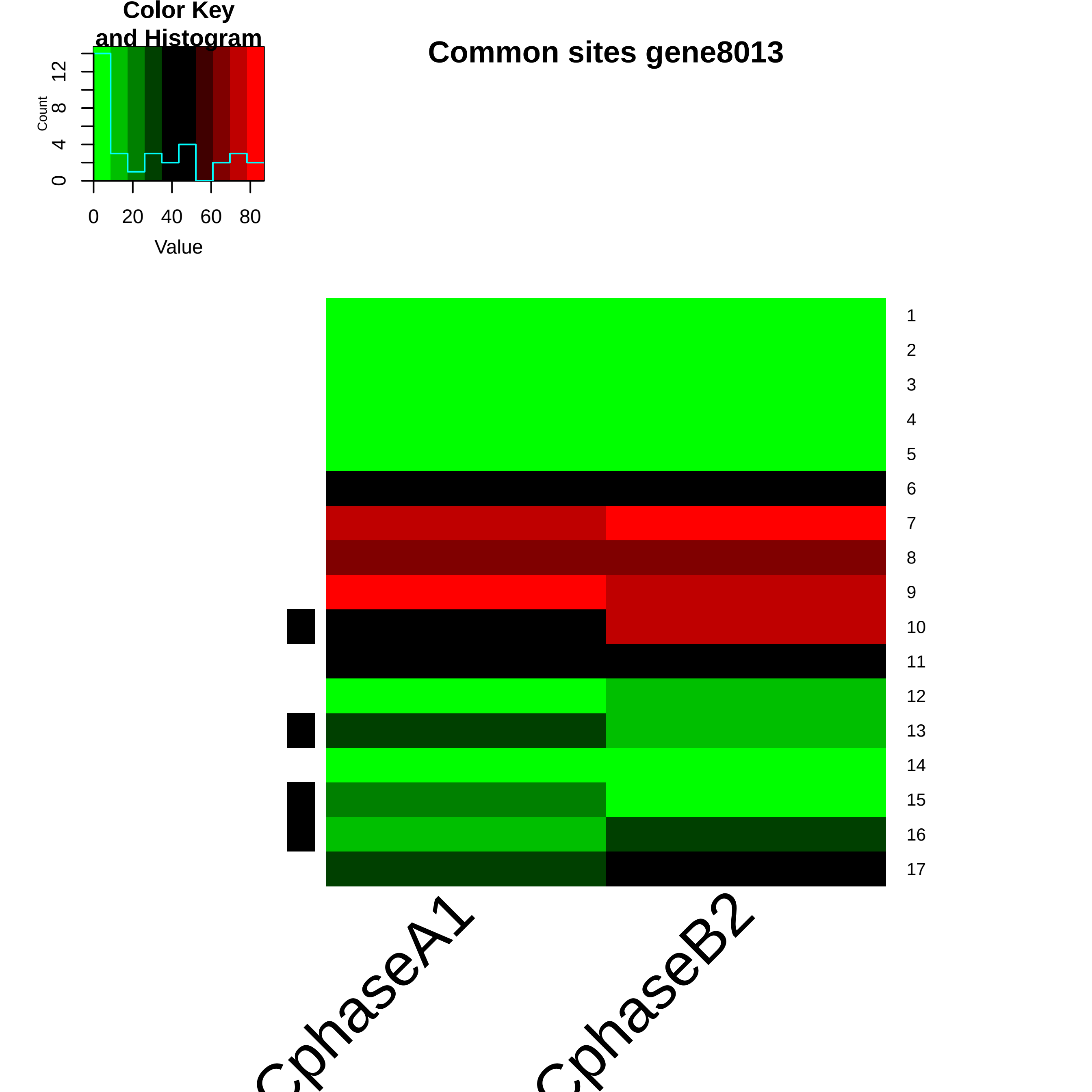
Color Key
and Histogram
Count
0
4
8
12
0
20
40
60
80
Value
Common sites gene8013
CphaseA1
CphaseB2

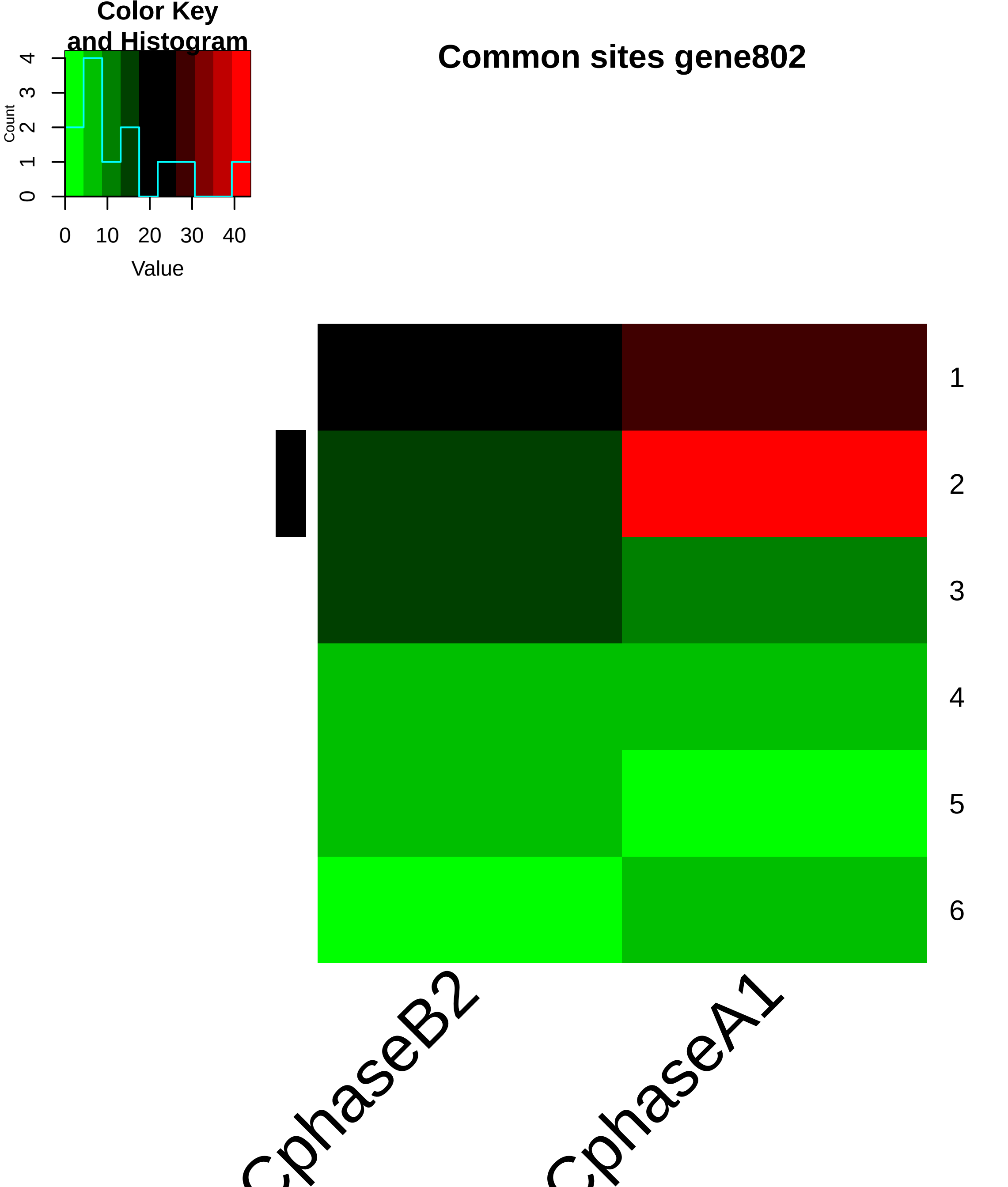
Color Key
and Histogram
Count
Value
0
10
20
30
40
0
1
2
3
4
Common sites gene802
1
2
3
4
5
6
CphaseB2
CphaseA1

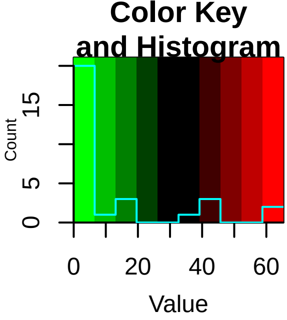

### Common sites gene8093

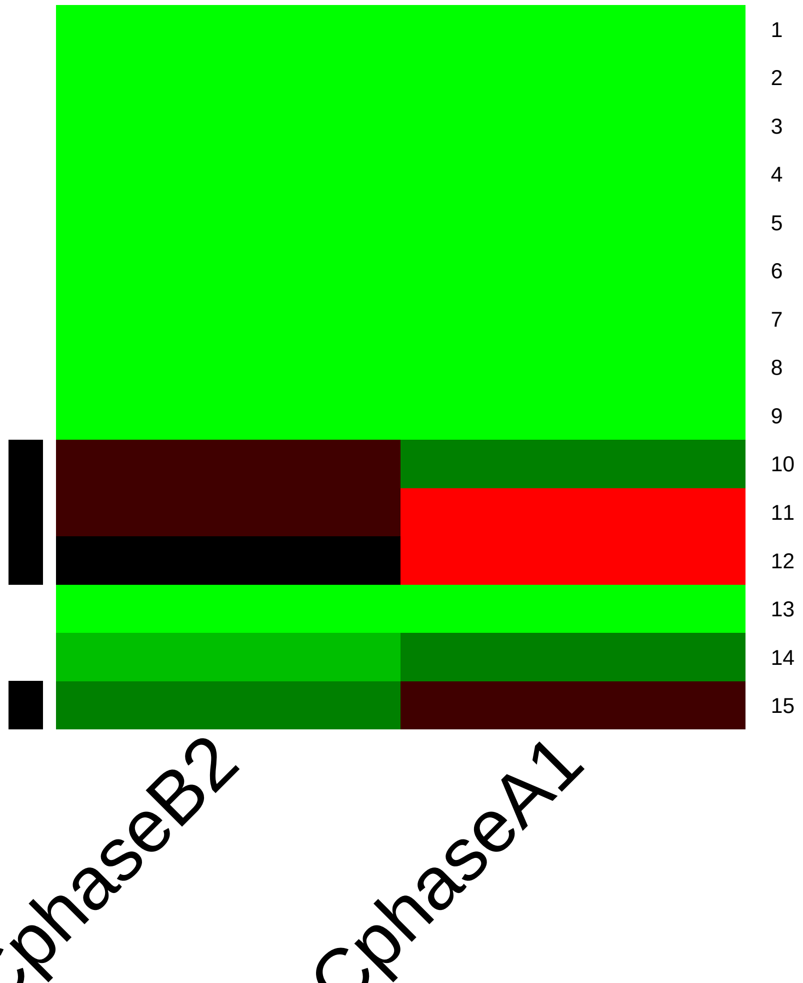

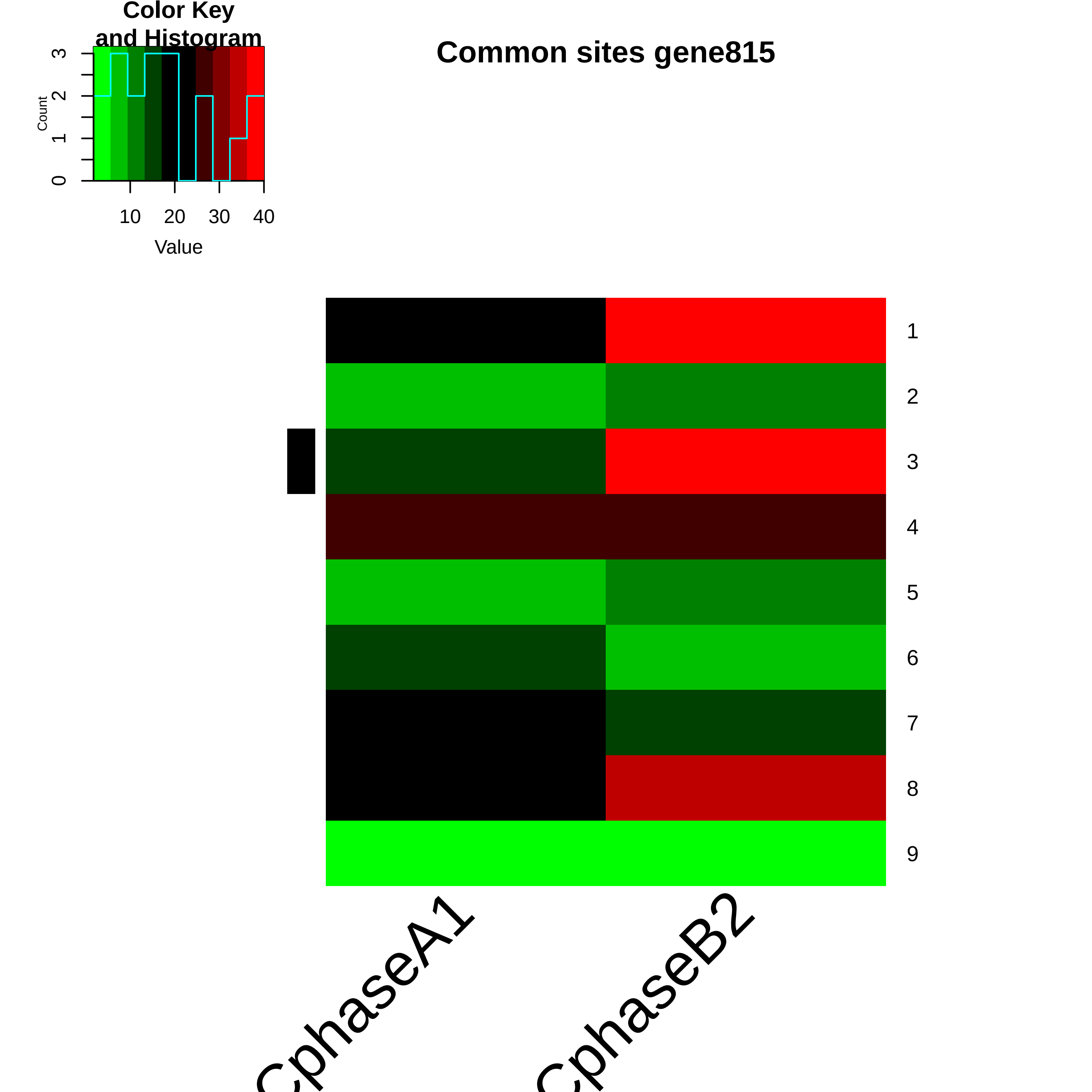

Color Key
and Histogram
Count
0
1
2
3
10
20
30
40
Value
Common sites gene815
1
2
3
4
5
6
7
8
9
CphaseA1
CphaseB2

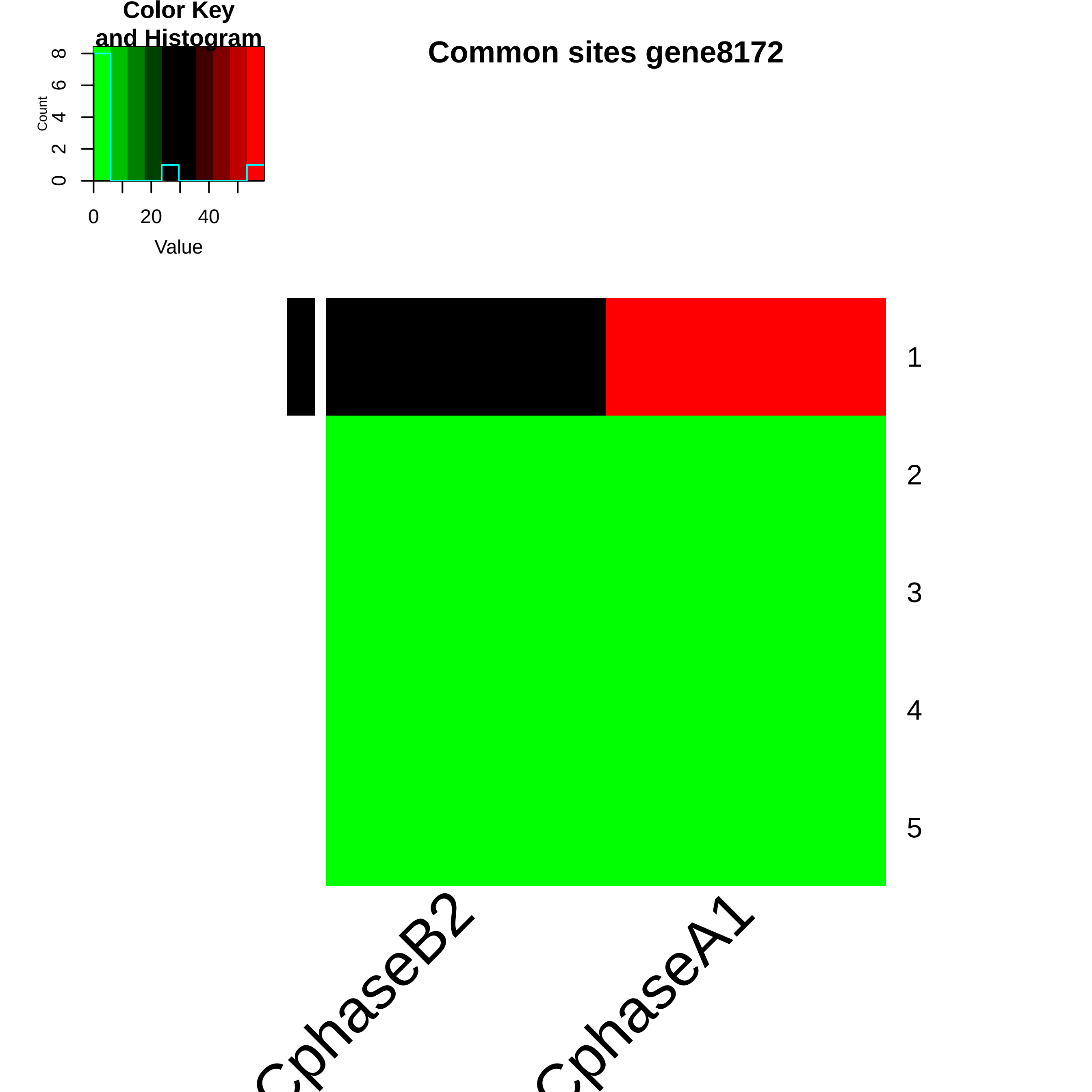

Color Key
and Histogram
Count
0
2
4
6
8
0
20
40
Value
Common sites gene8172
1
2
3
4
5
CphaseB2
CphaseA1

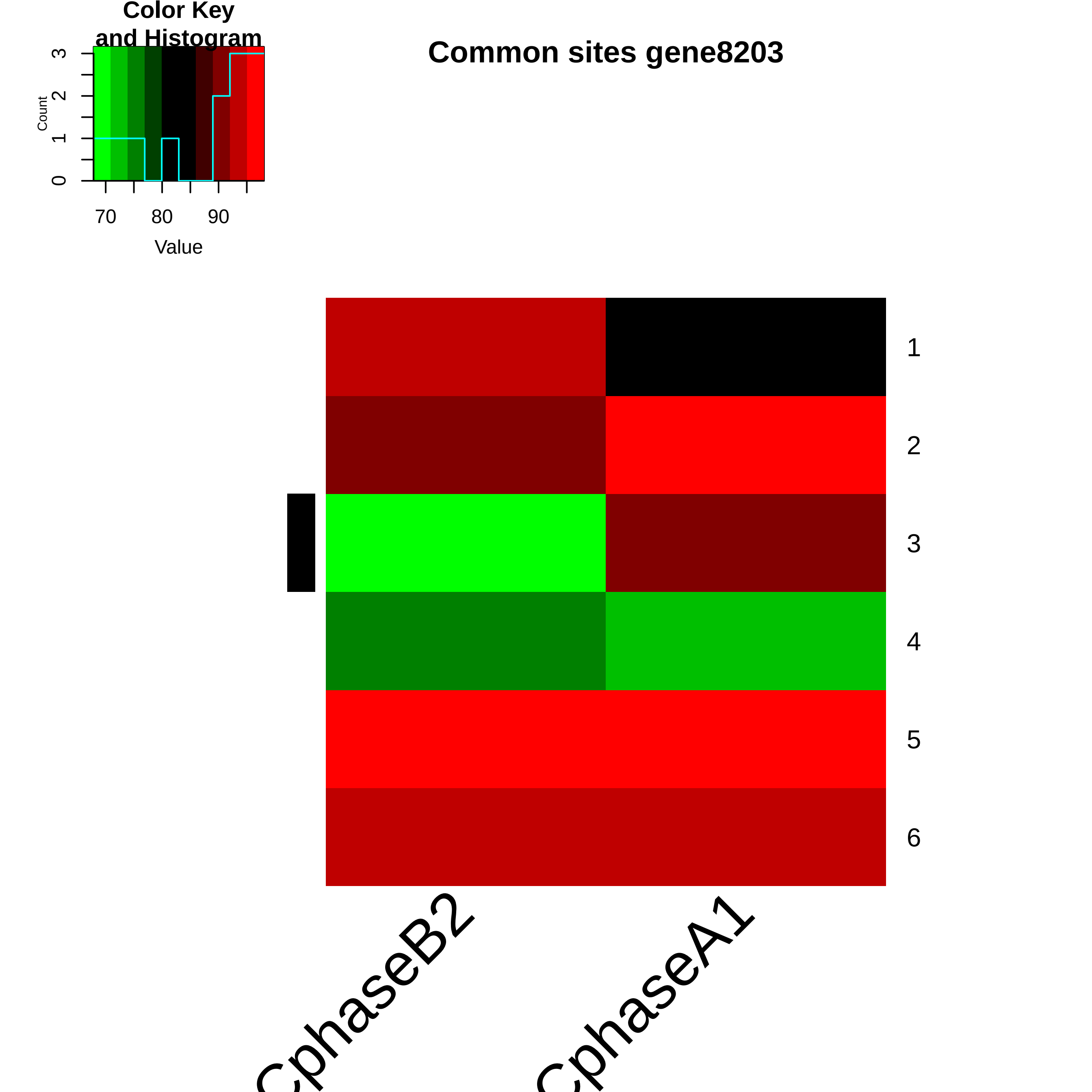
Color Key
and Histogram
Count
0
1
2
3
70
80
90
Value
Common sites gene8203
1
2
3
4
5
6
CphaseB2
CphaseA1

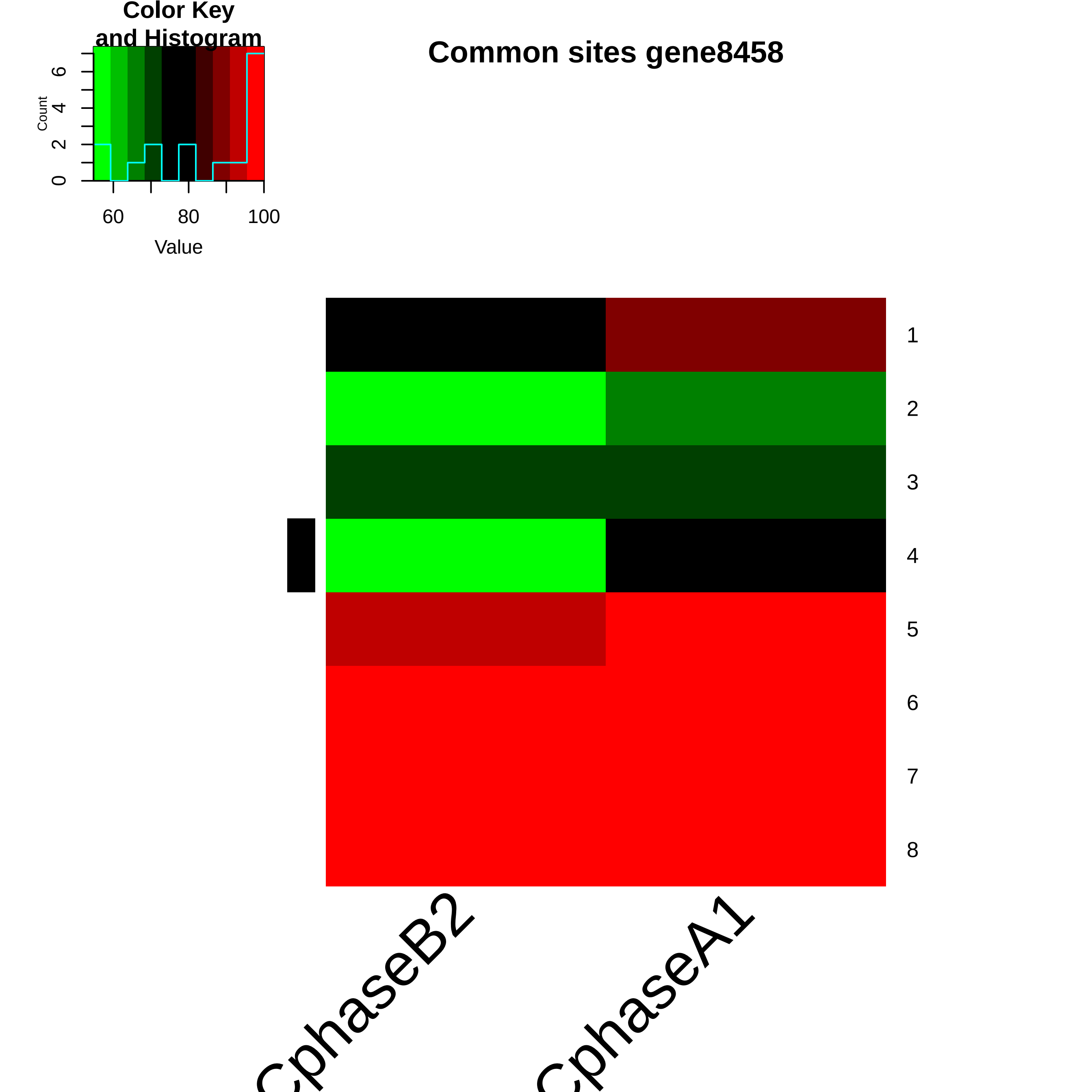

Color Key
and Histogram
Count
0
2
4
6
60
80
100
Value
Common sites gene8458
1
2
3
4
5
6
7
8
CphaseB2
CphaseA1

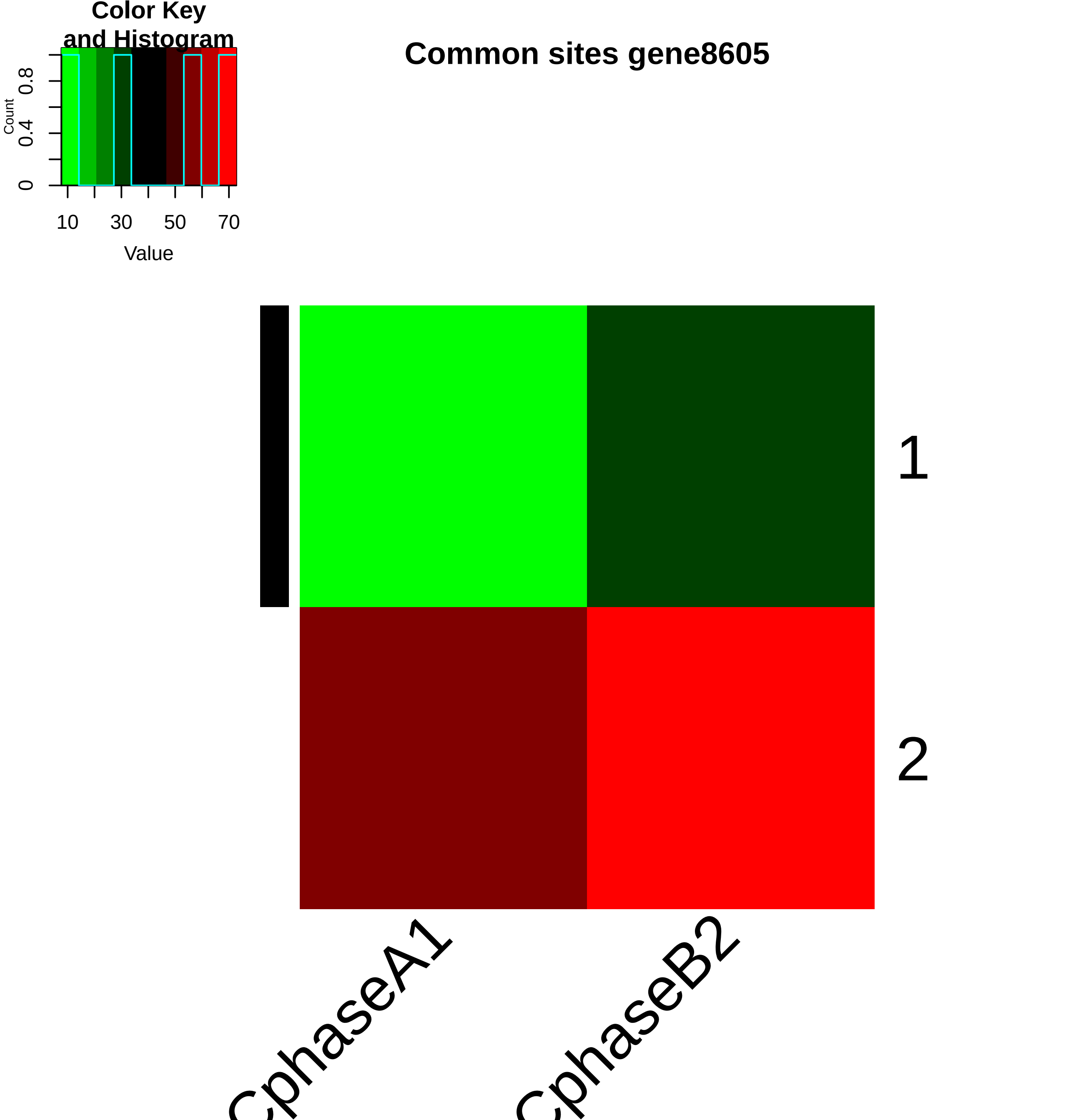

Color Key
and Histogram
Count
0
0.4
0.8
10
30
50
70
Value
Common sites gene8605
1
2
CphaseA1
CphaseB2

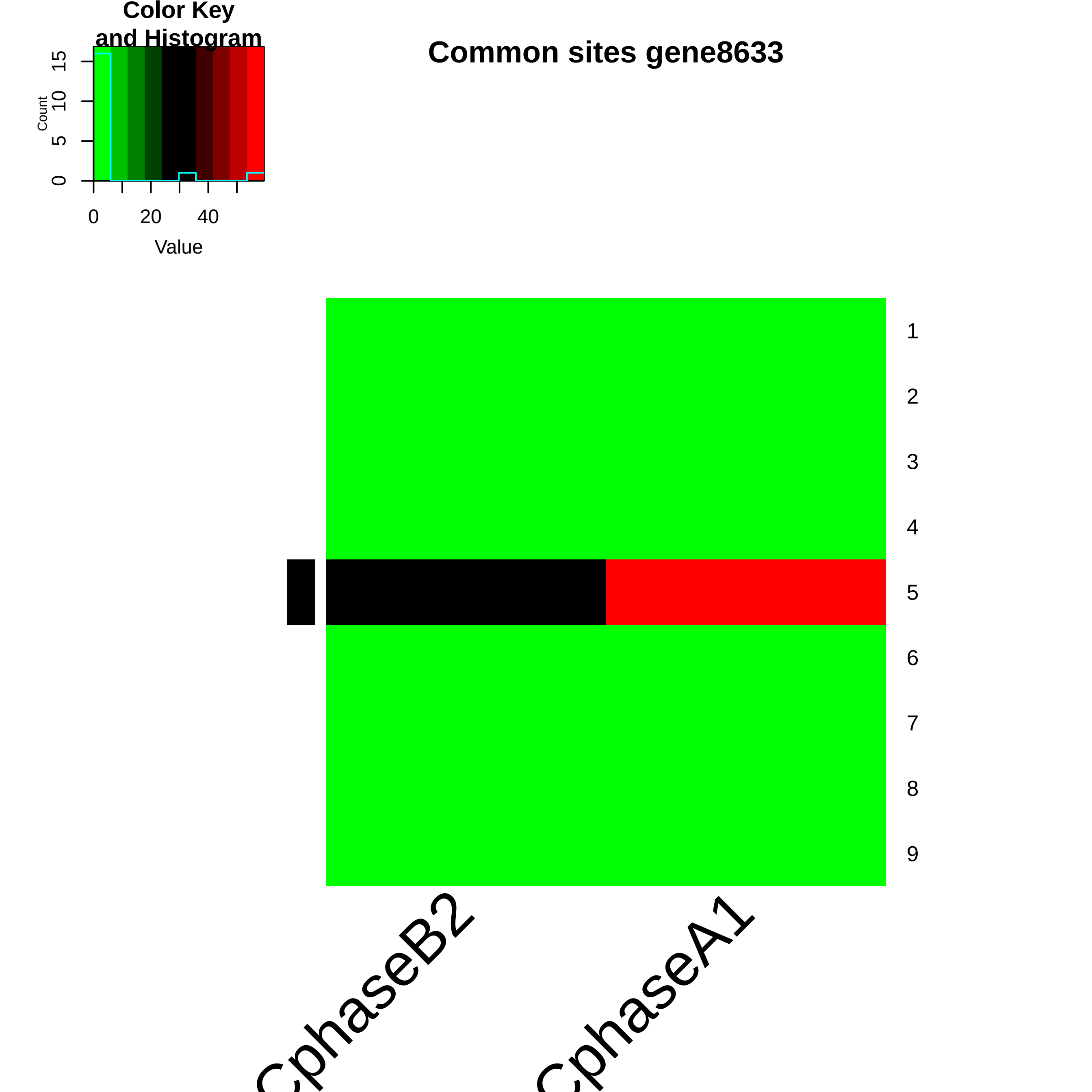
Color Key
and Histogram
Count
0
5
10
15
0
20
40
Value
Common sites gene8633
1
2
3
4
5
6
7
8
9
CphaseB2
CphaseA1

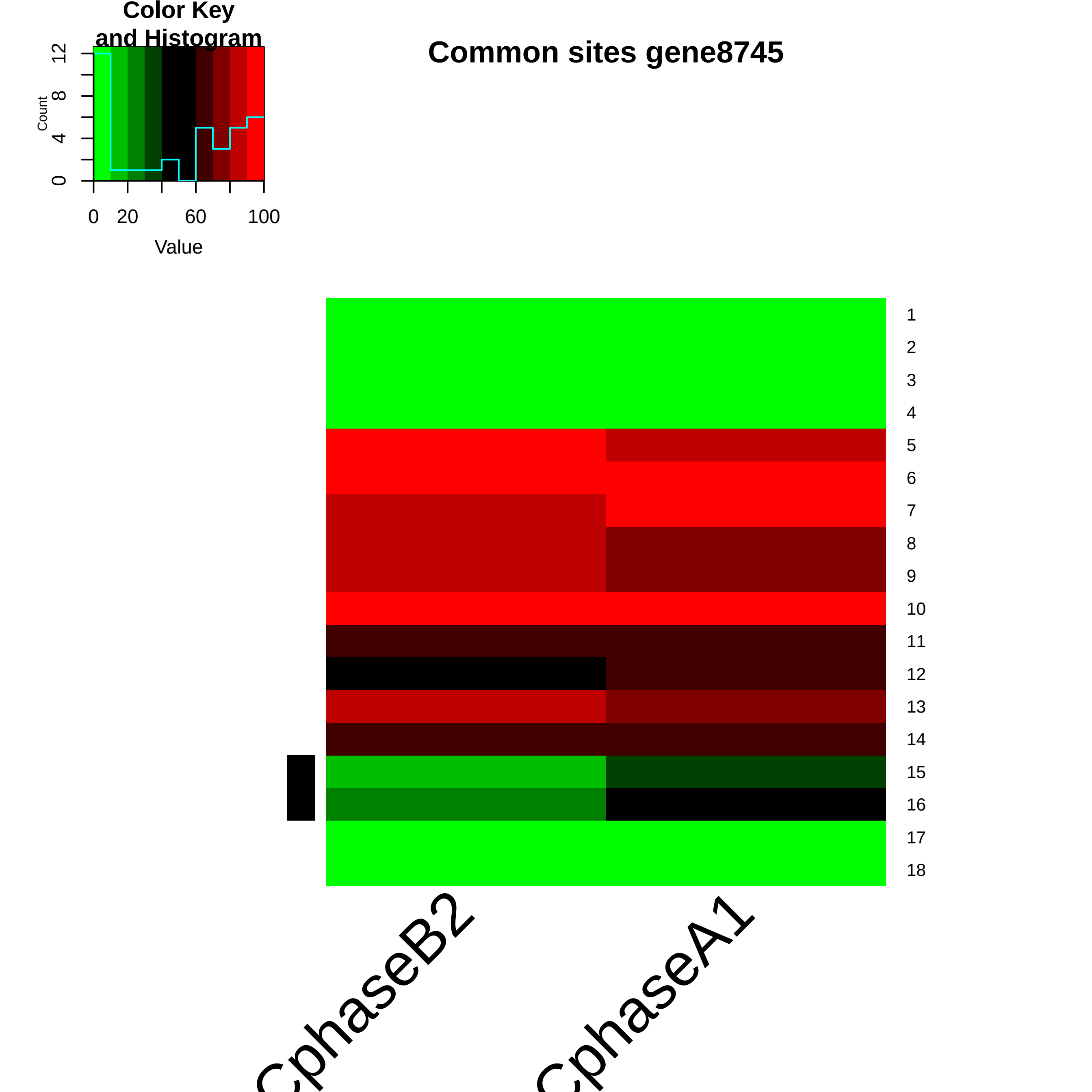
Color Key
and Histogram
Count
0
4
8
12
0
20
60
100
Value
Common sites gene8745
CphaseB2
CphaseA1

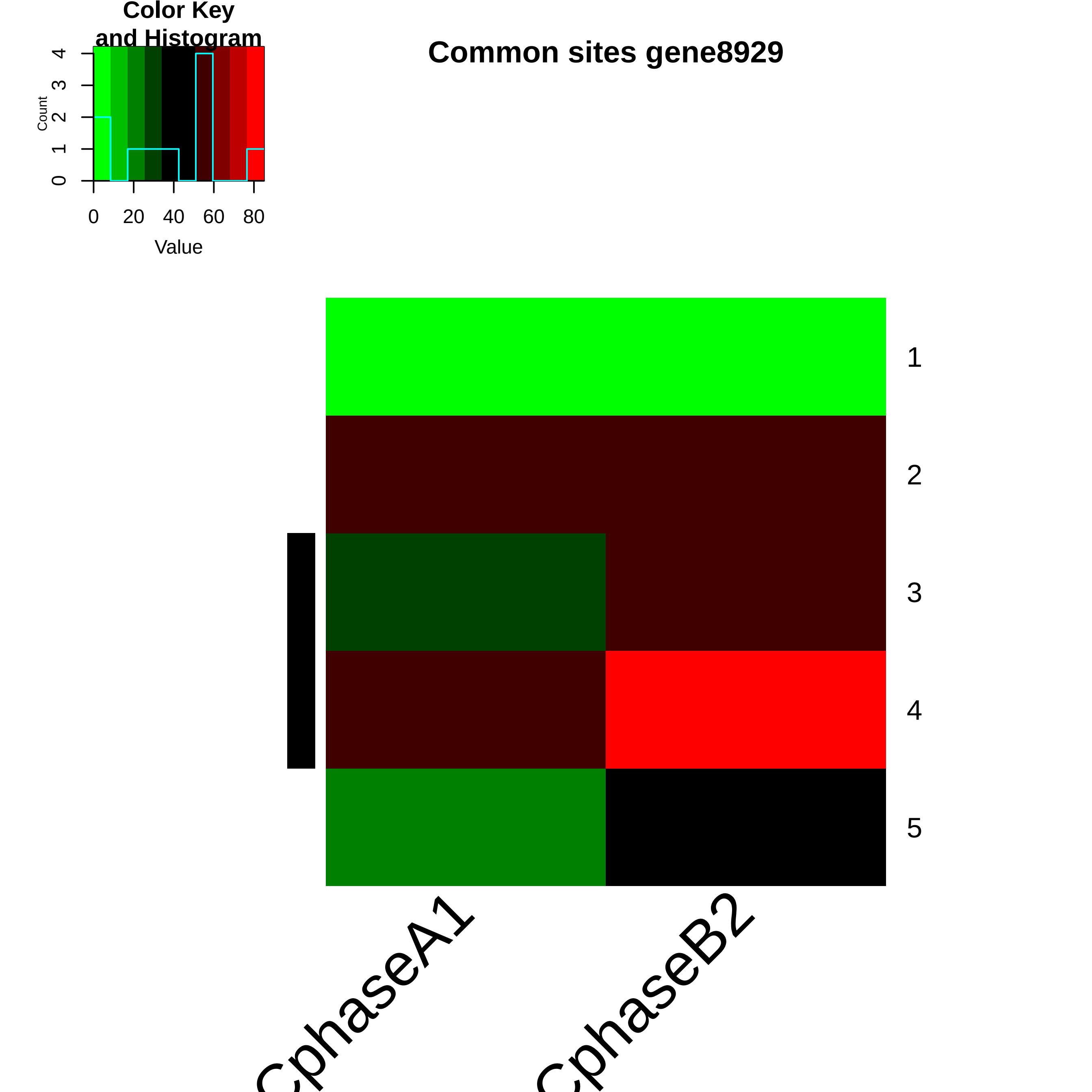

Color Key
and Histogram
Count
0
1
2
3
4
0
20
40
60
80
Value
Common sites gene8929
1
2
3
4
5
CphaseA1
CphaseB2

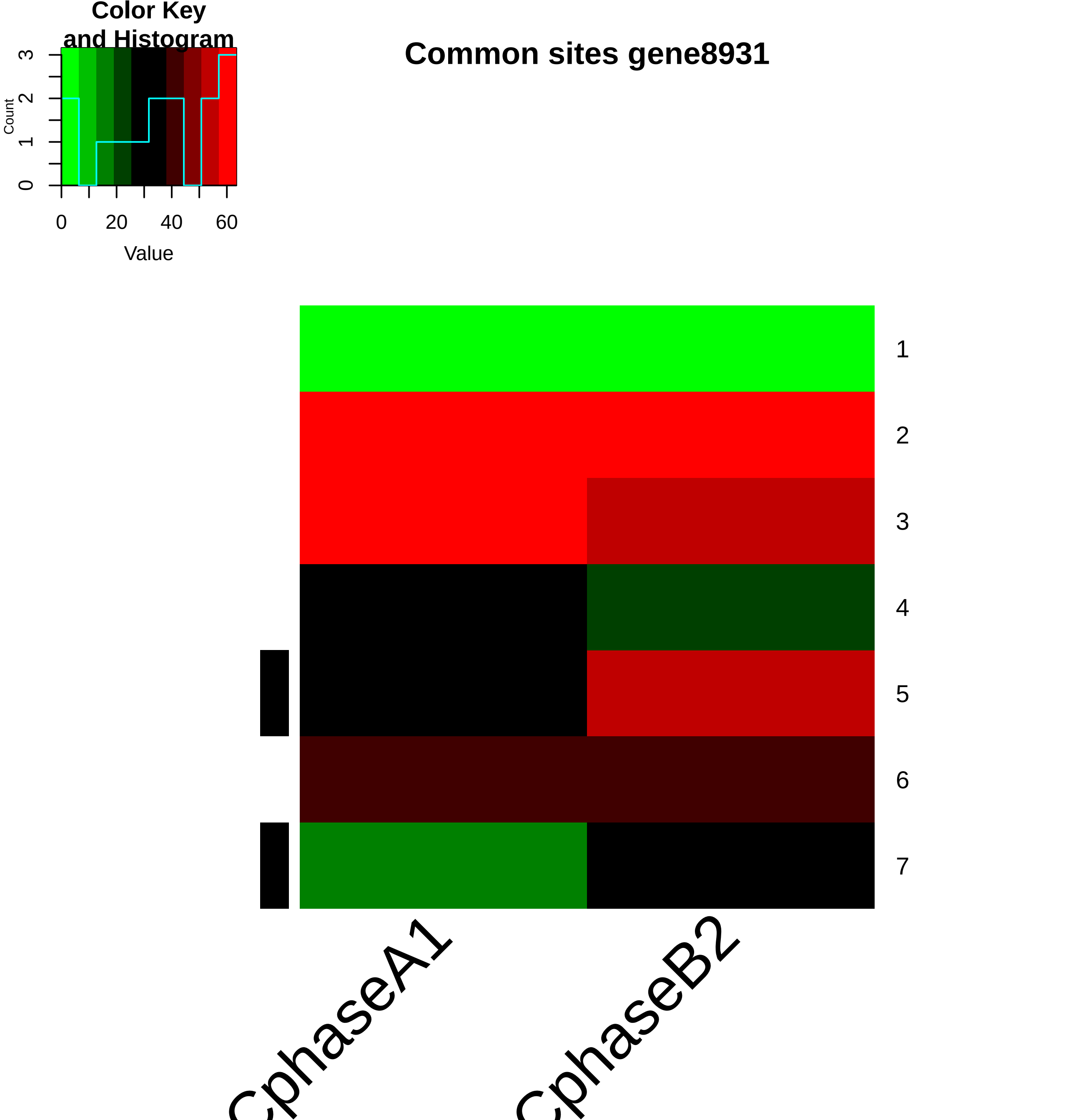

Color Key
and Histogram
Count
0
1
2
3
0
20
40
60
Value
Common sites gene8931
1
2
3
4
5
6
7
CphaseA1
CphaseB2

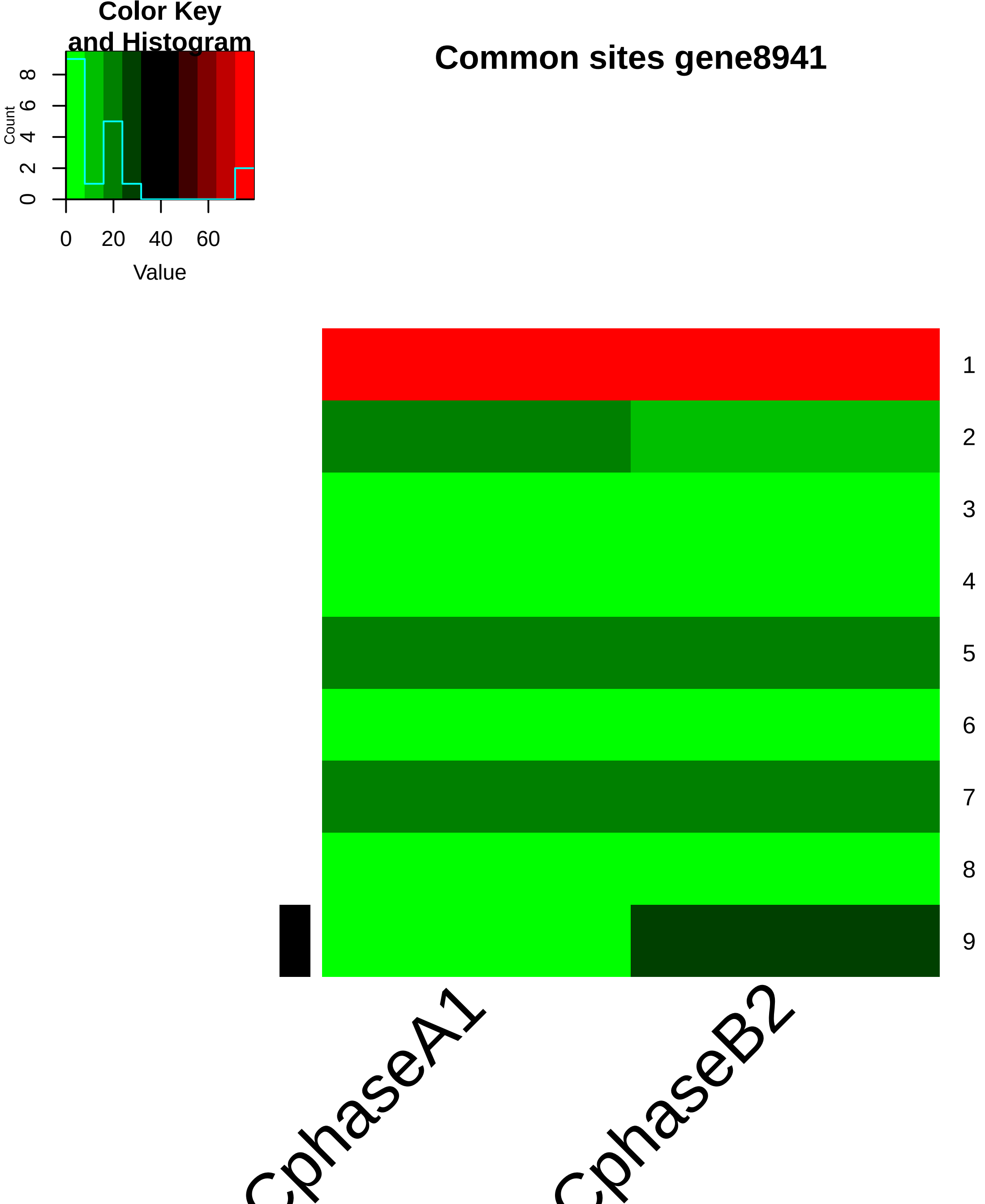

Color Key
and Histogram
Count
0 2 4 6 8
0 20 40 60
Value
Common sites gene8941
1
2
3
4
5
6
7
8
9
CphaseA1
CphaseB2

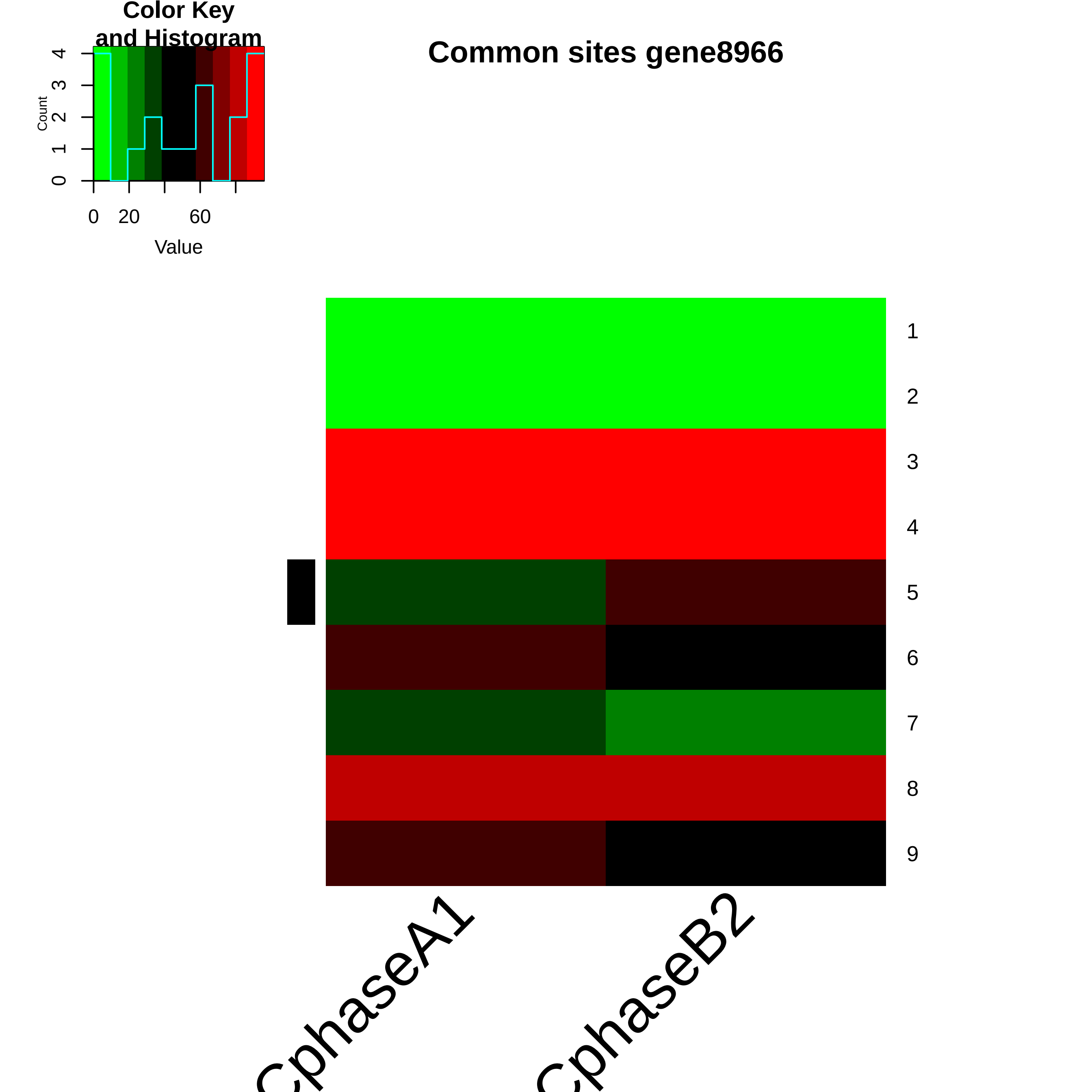

Color Key
and Histogram
Count
0
1
2
3
4
0
20
60
Value
Common sites gene8966
1
2
3
4
5
6
7
8
9
CphaseA1
CphaseB2

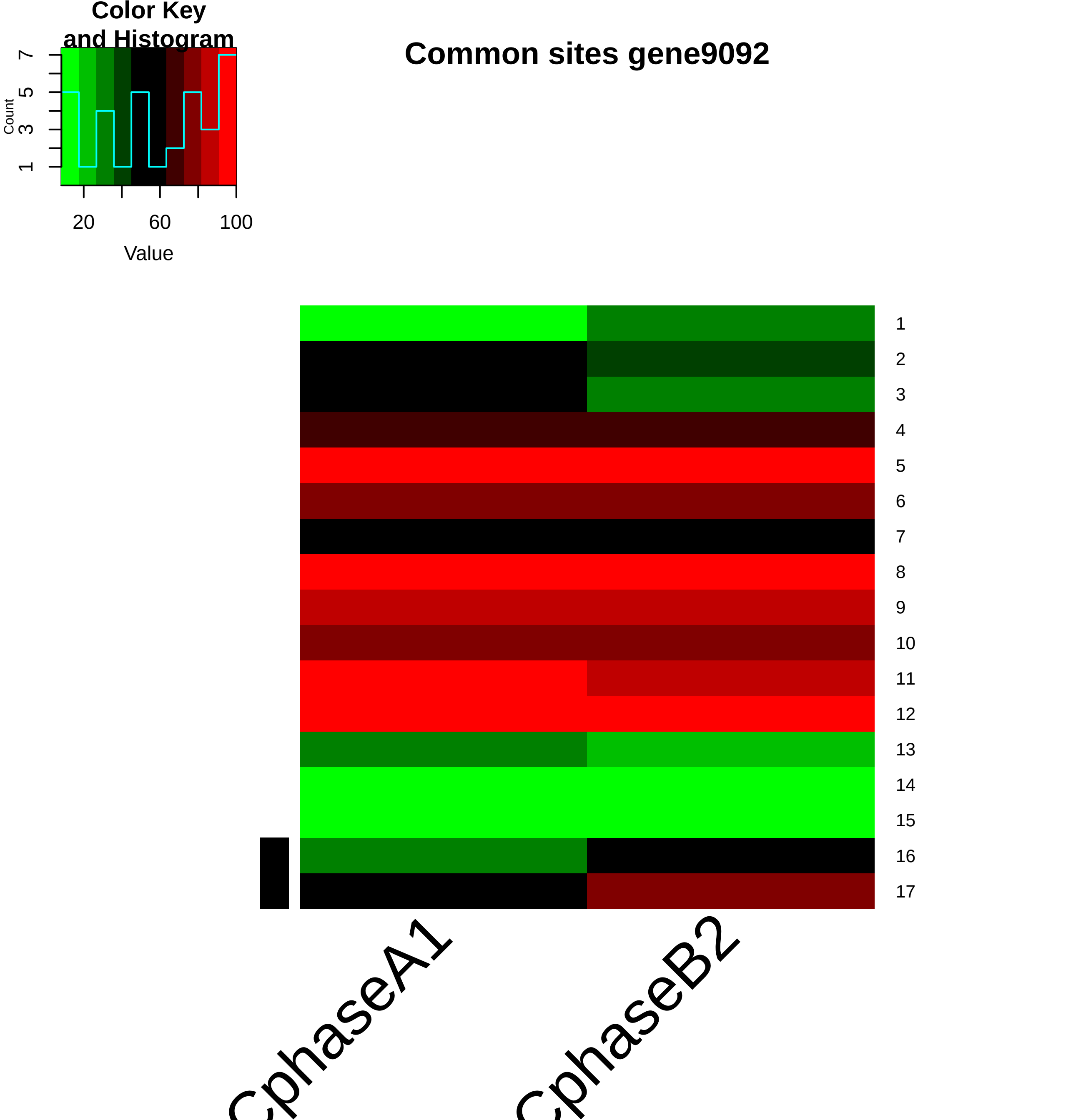
Color Key
and Histogram
Count
1
3
5
7
20
60
100
Value
Common sites gene9092
CphaseA1
CphaseB2

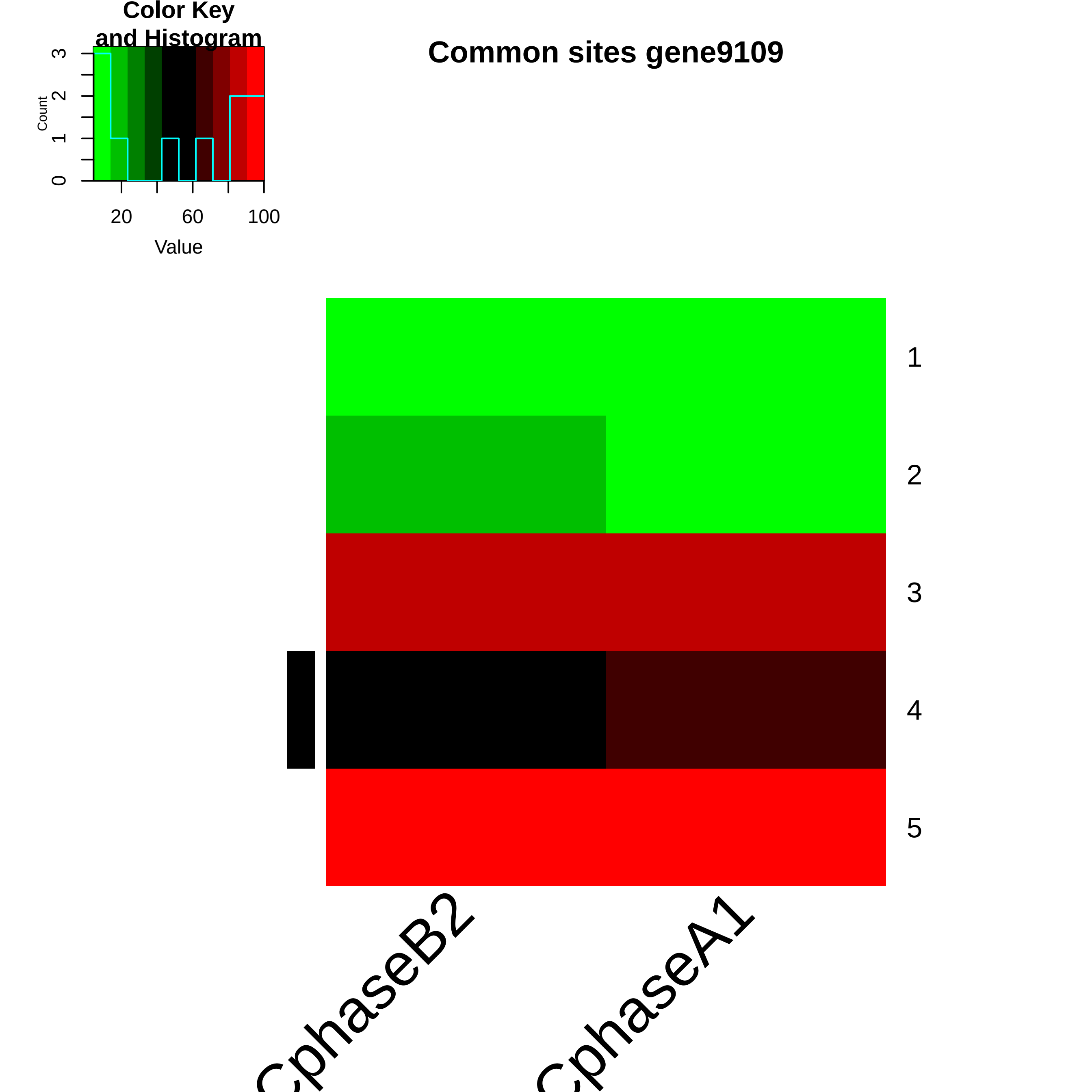

Color Key
and Histogram
Count
0
1
2
3
20
60
100
Value
Common sites gene9109
1
2
3
4
5
CphaseB2
CphaseA1

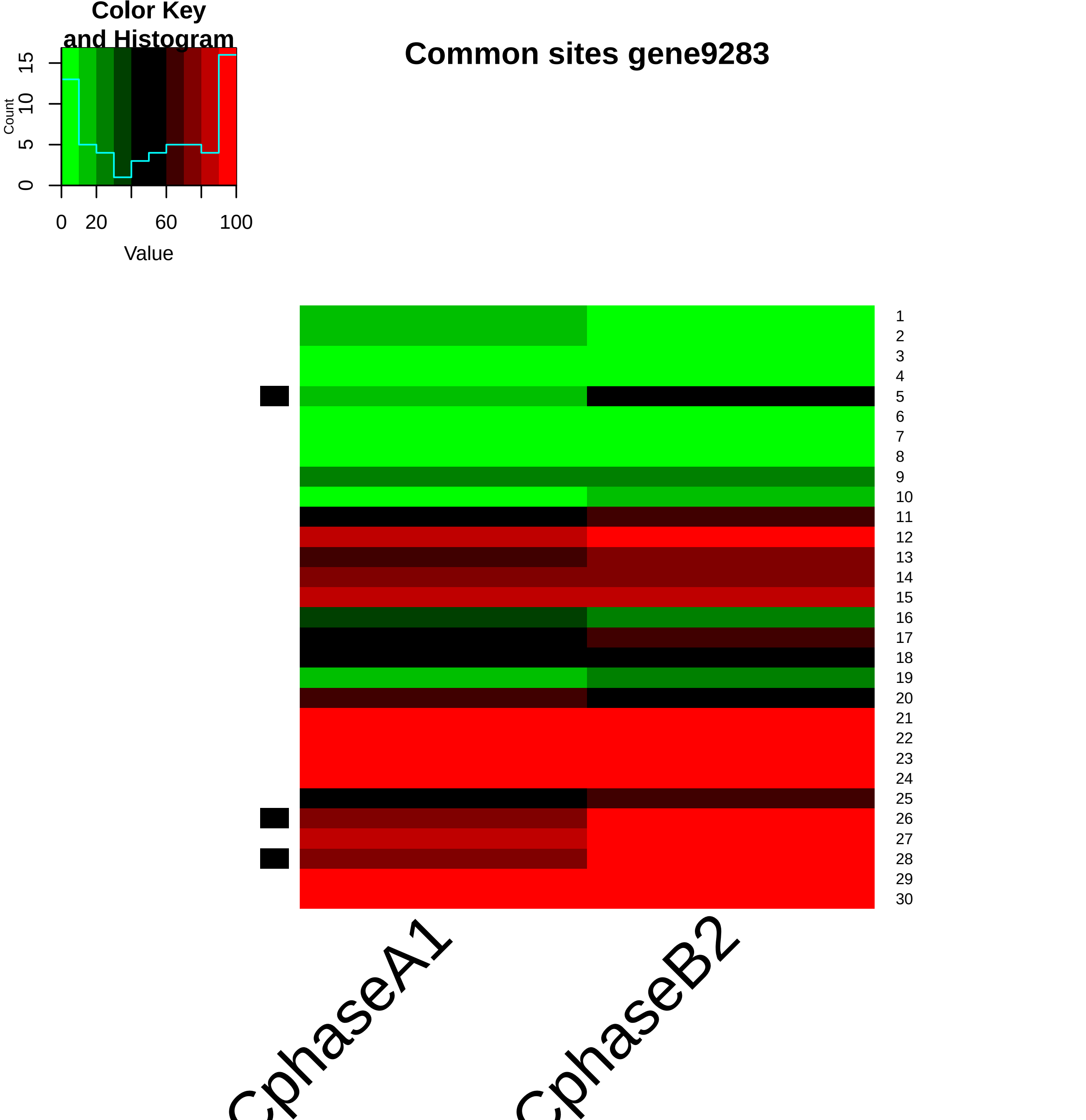

Color Key
and Histogram
Count
0
5
10
15
0
20
60
100
Value
Common sites gene9283
CphaseA1
CphaseB2

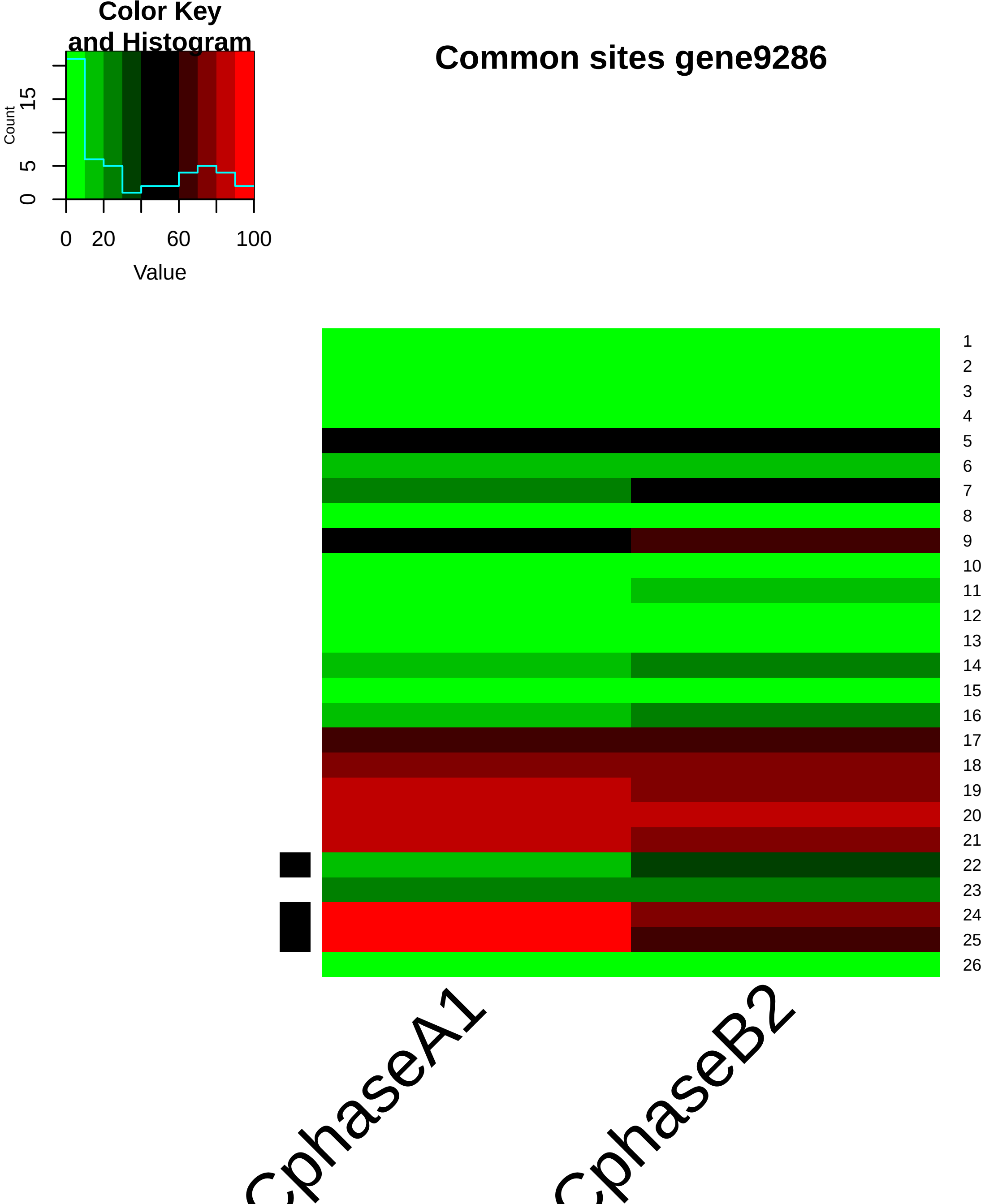

Color Key
and Histogram
Count
0
5
15
0
20
60
100
Value
Common sites gene9286
CphaseA1
CphaseB2

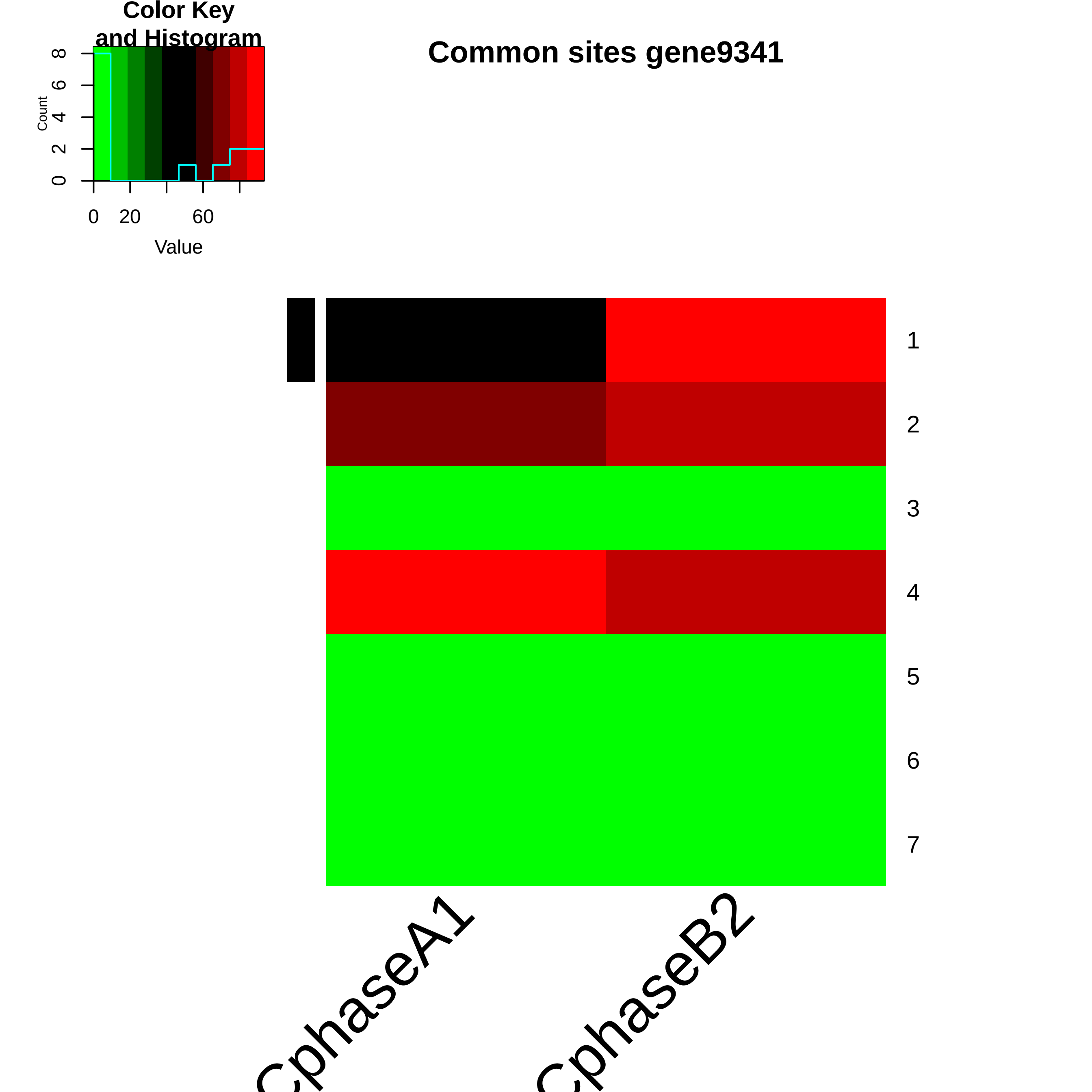
Color Key
and Histogram
Count
0
2
4
6
8
0
20
60
Value
Common sites gene9341
1
2
3
4
5
6
7
CphaseA1
CphaseB2

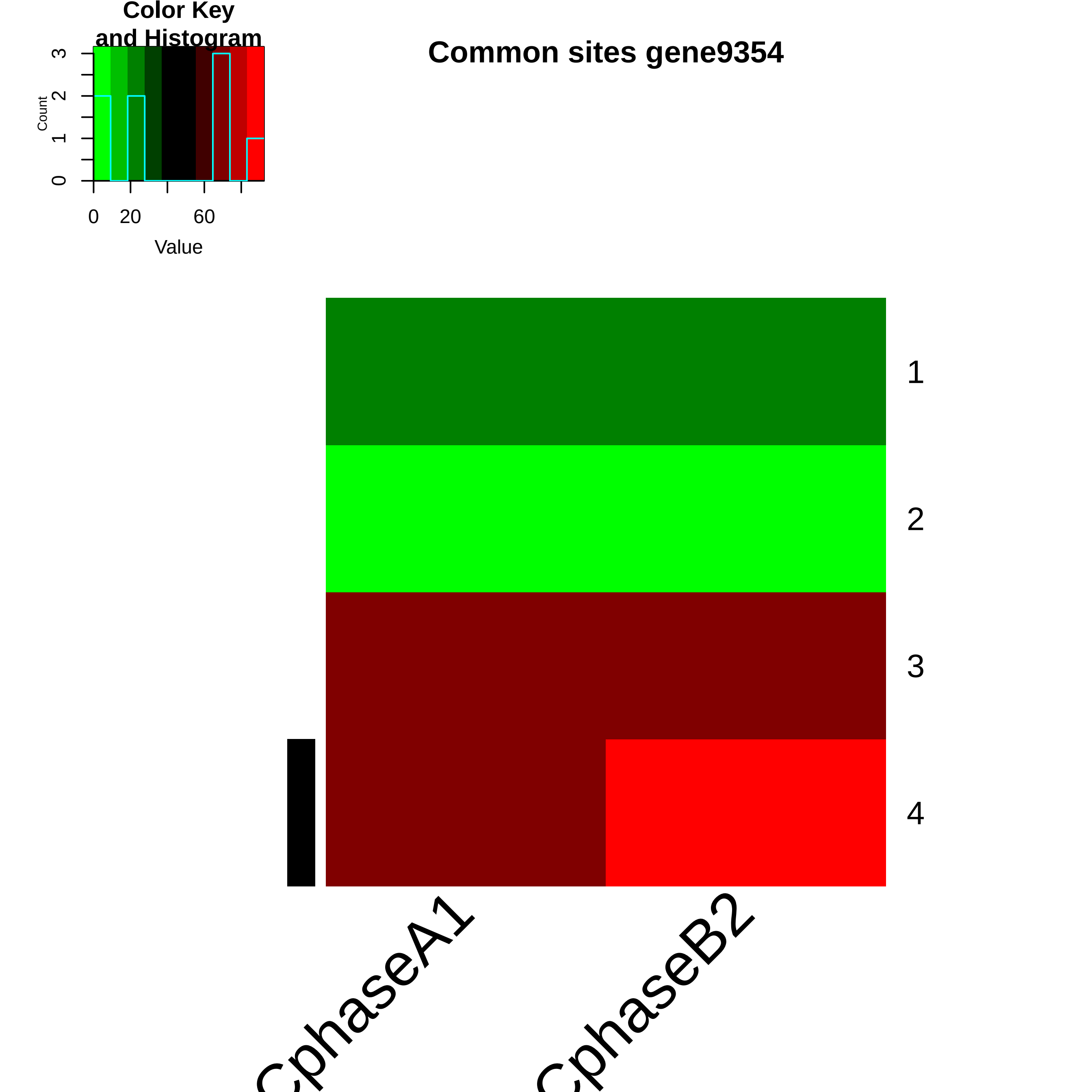

Color Key
and Histogram
Count
Value
Common sites gene9354
1
2
3
4
phaseA1
phaseB2

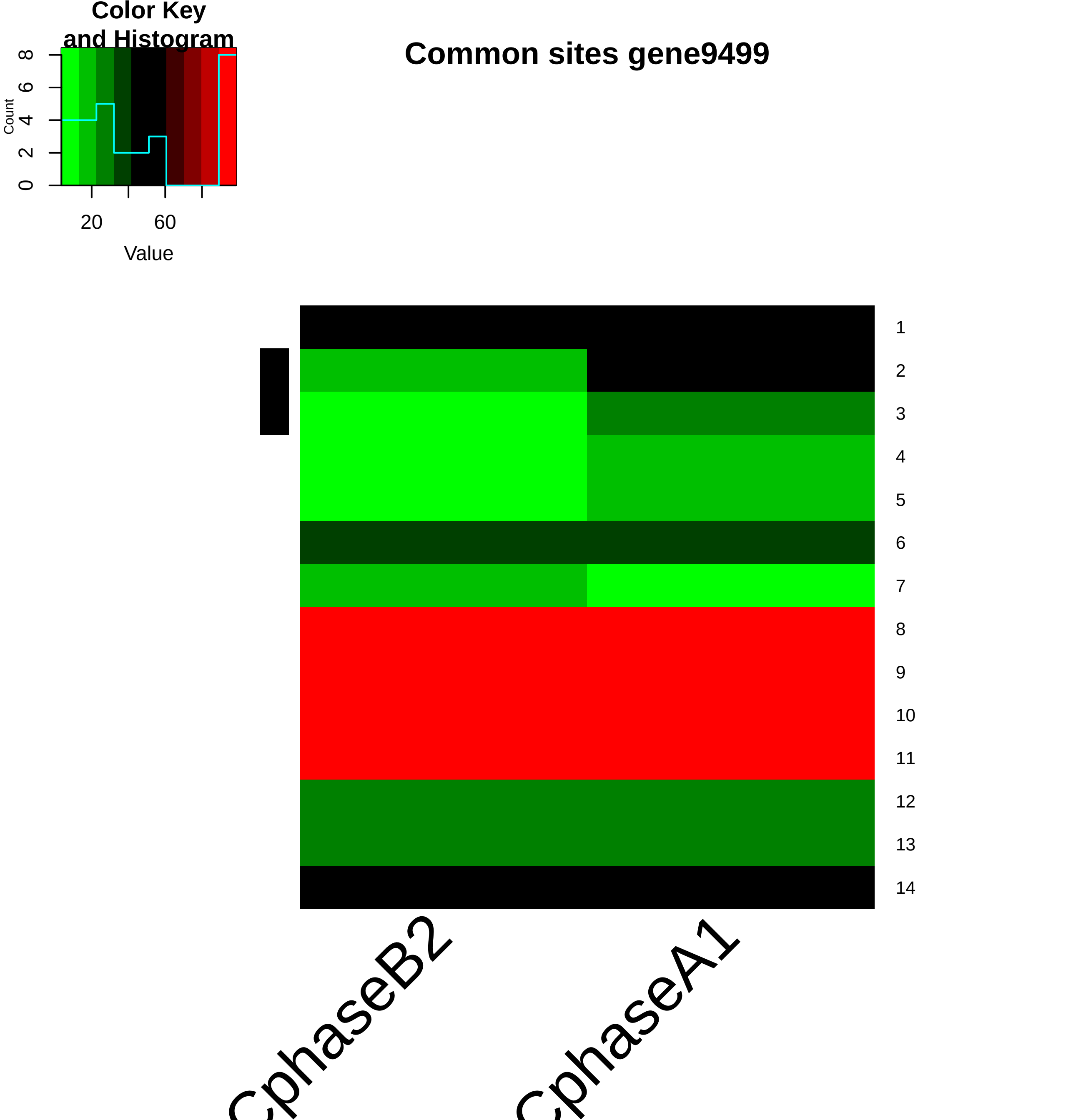
Color Key
and Histogram
Count
0
2
4
6
8
20
60
Value
Common sites gene9499
1
2
3
4
5
6
7
8
9
10
11
12
13
14
CphaseB2
CphaseA1

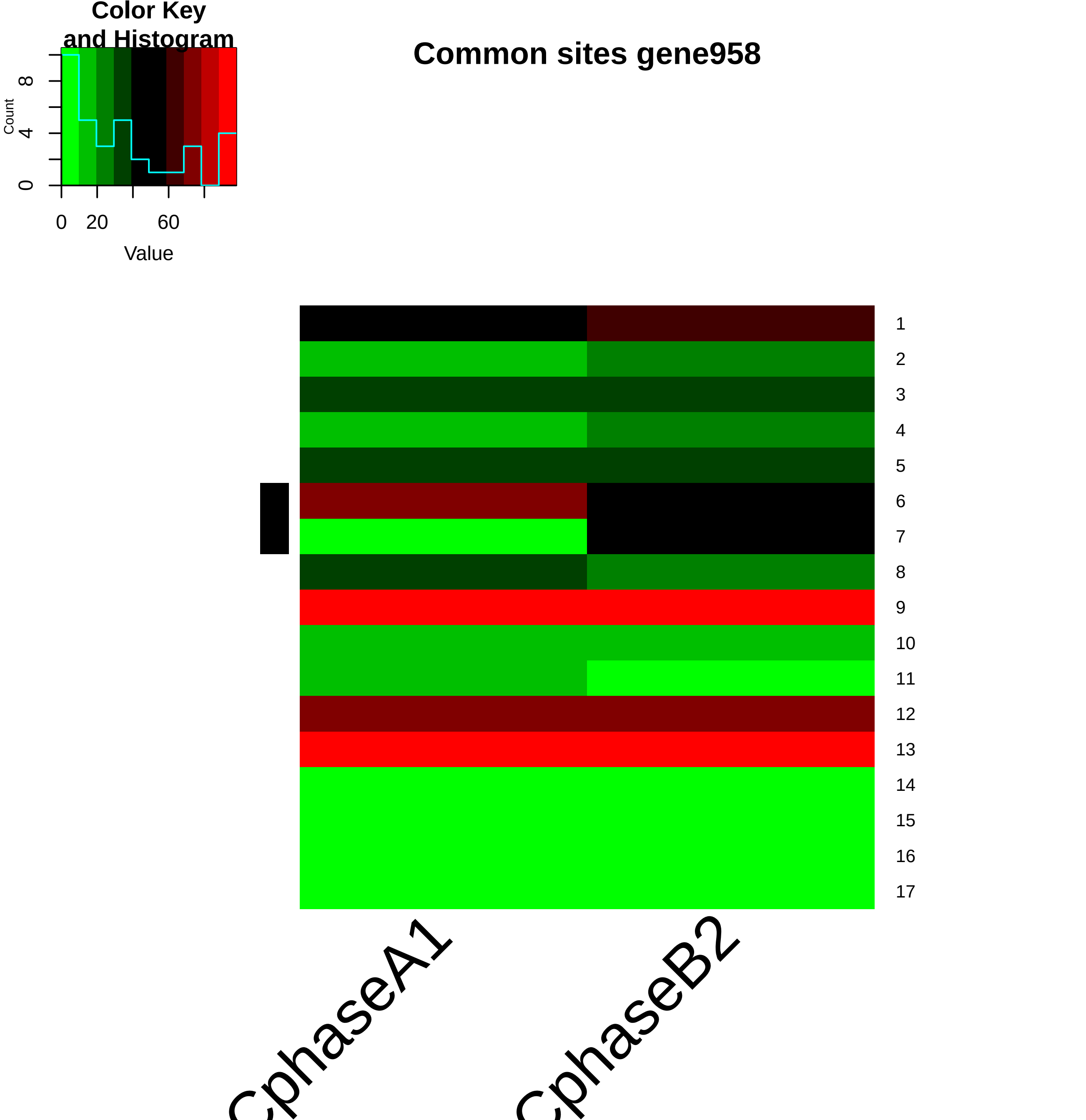

Color Key
and Histogram
Count
0
4
8
0
20
60
Value
Common sites gene958
CphaseA1
CphaseB2

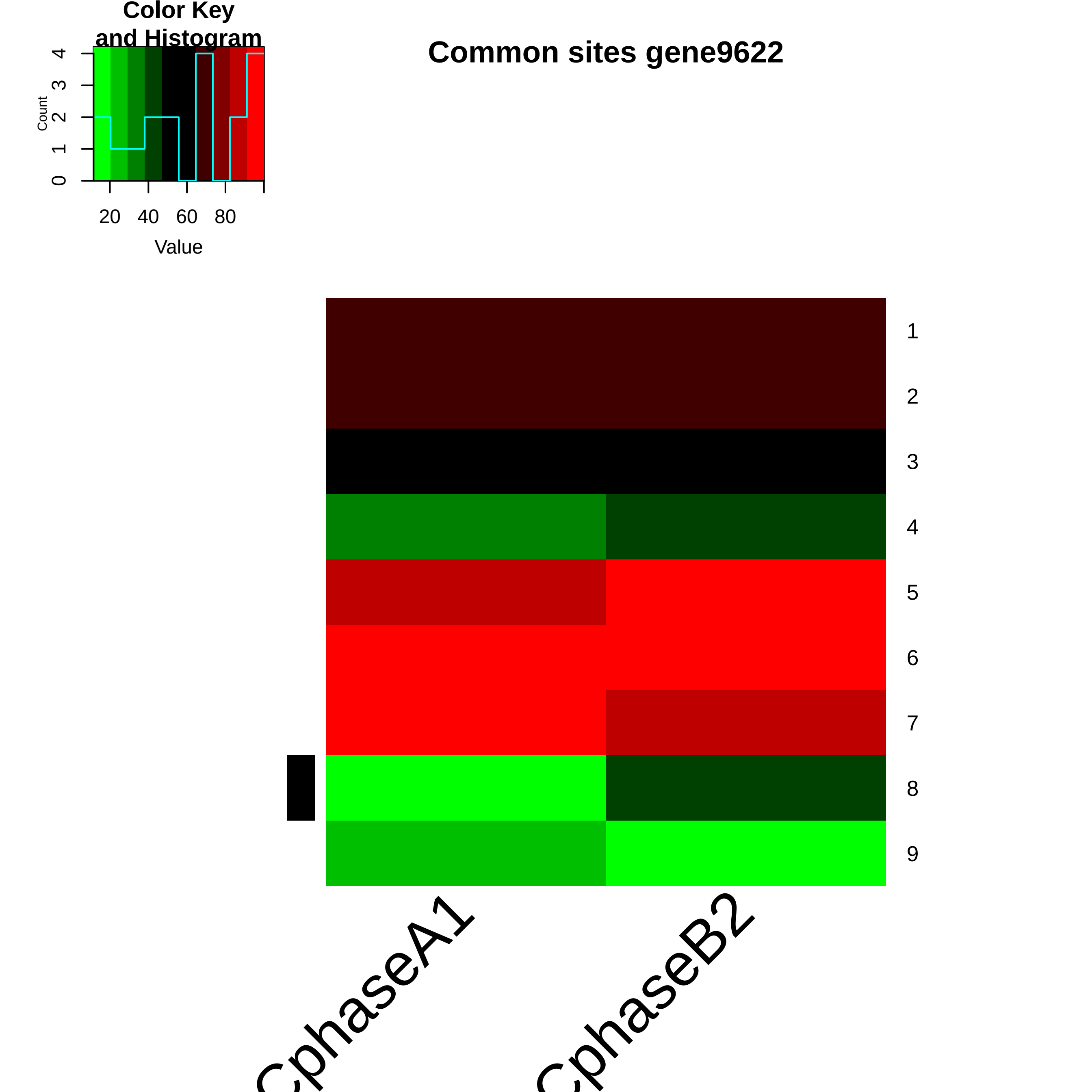
Color Key
and Histogram
Count
0
1
2
3
4
20
40
60
80
Value
Common sites gene9622
1
2
3
4
5
6
7
8
9
CphaseA1
CphaseB2

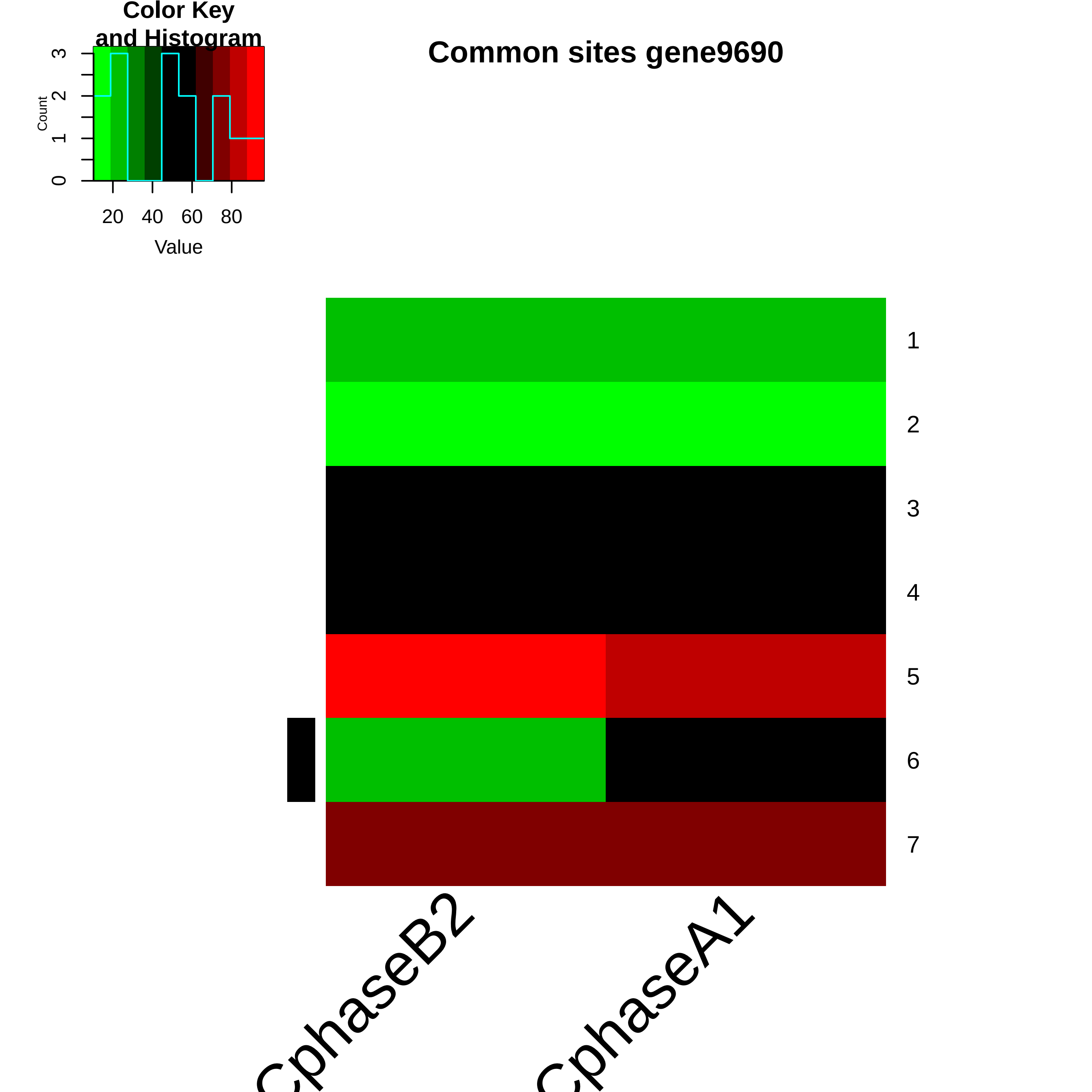
Color Key
and Histogram
Count
0
1
2
3
20
40
60
80
Value
Common sites gene9690
1
2
3
4
5
6
7
CphaseB2
CphaseA1

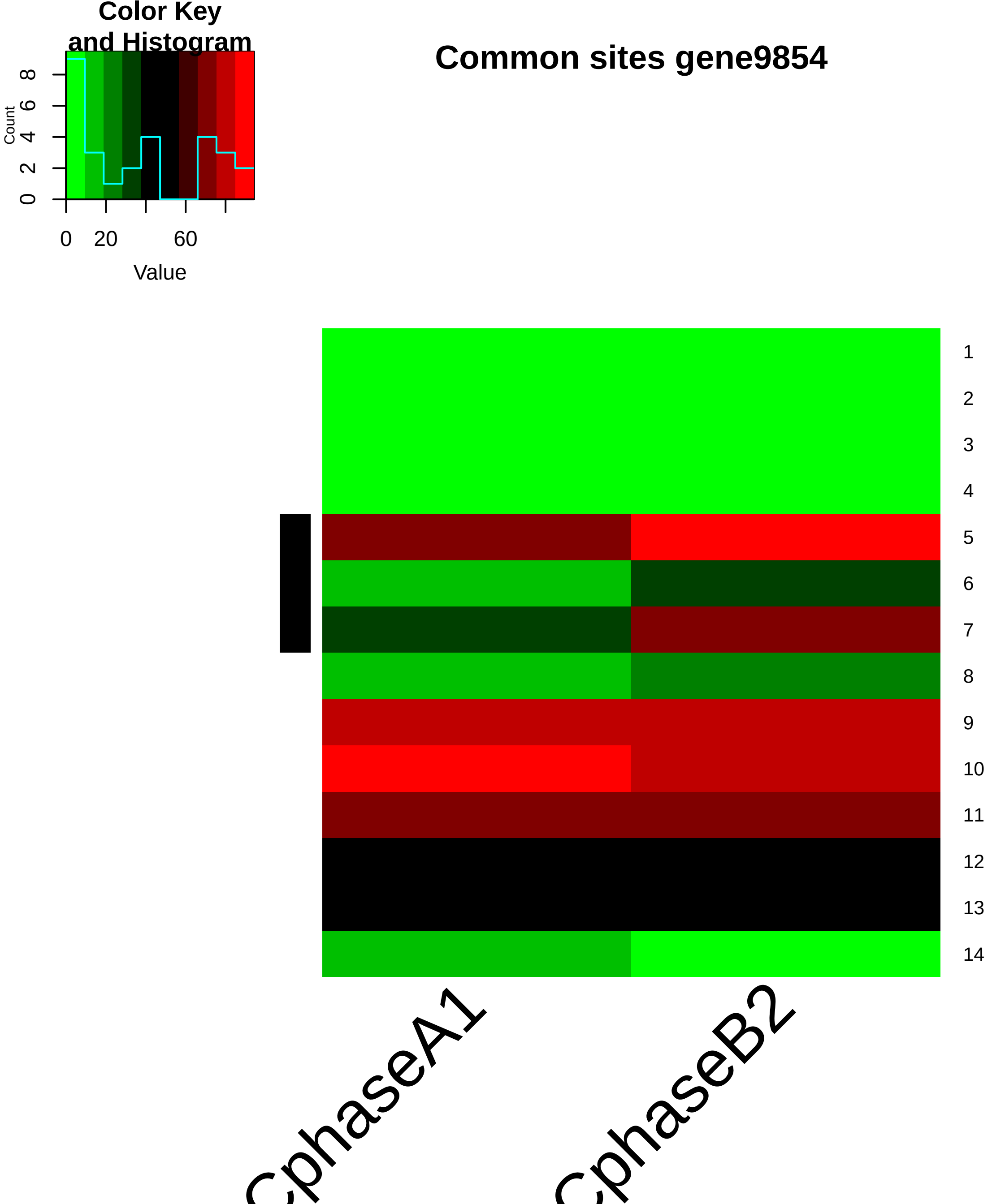

Color Key
and Histogram
Count
0
2
4
6
8
0
20
60
Value
Common sites gene9854
1
2
3
4
5
6
7
8
9
10
11
12
13
14
CphaseA1
CphaseB2

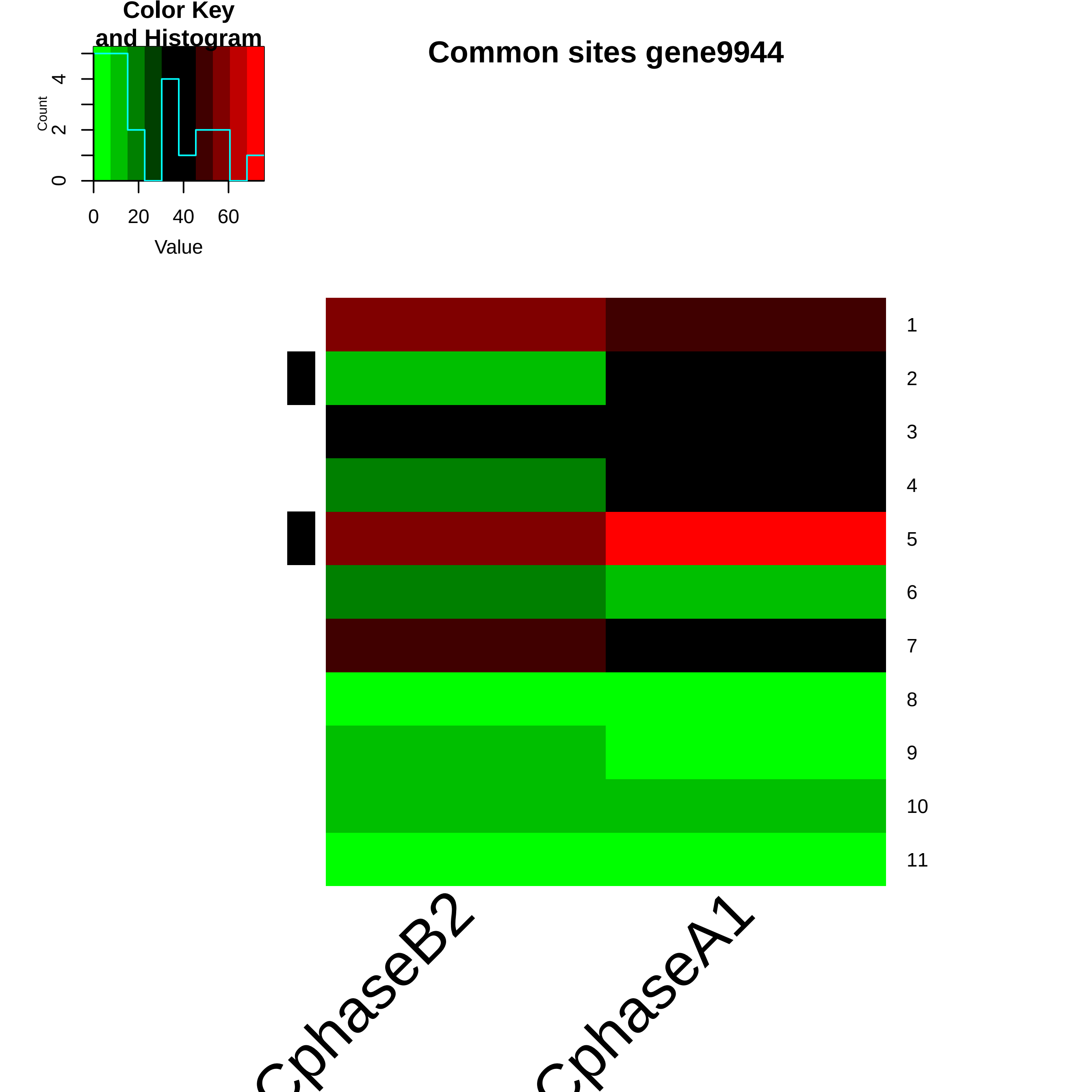
Color Key
and Histogram
Count
0
2
4
0
20
40
60
Value
Common sites gene9944
1
2
3
4
5
6
7
8
9
10
11
CphaseB2
CphaseA1

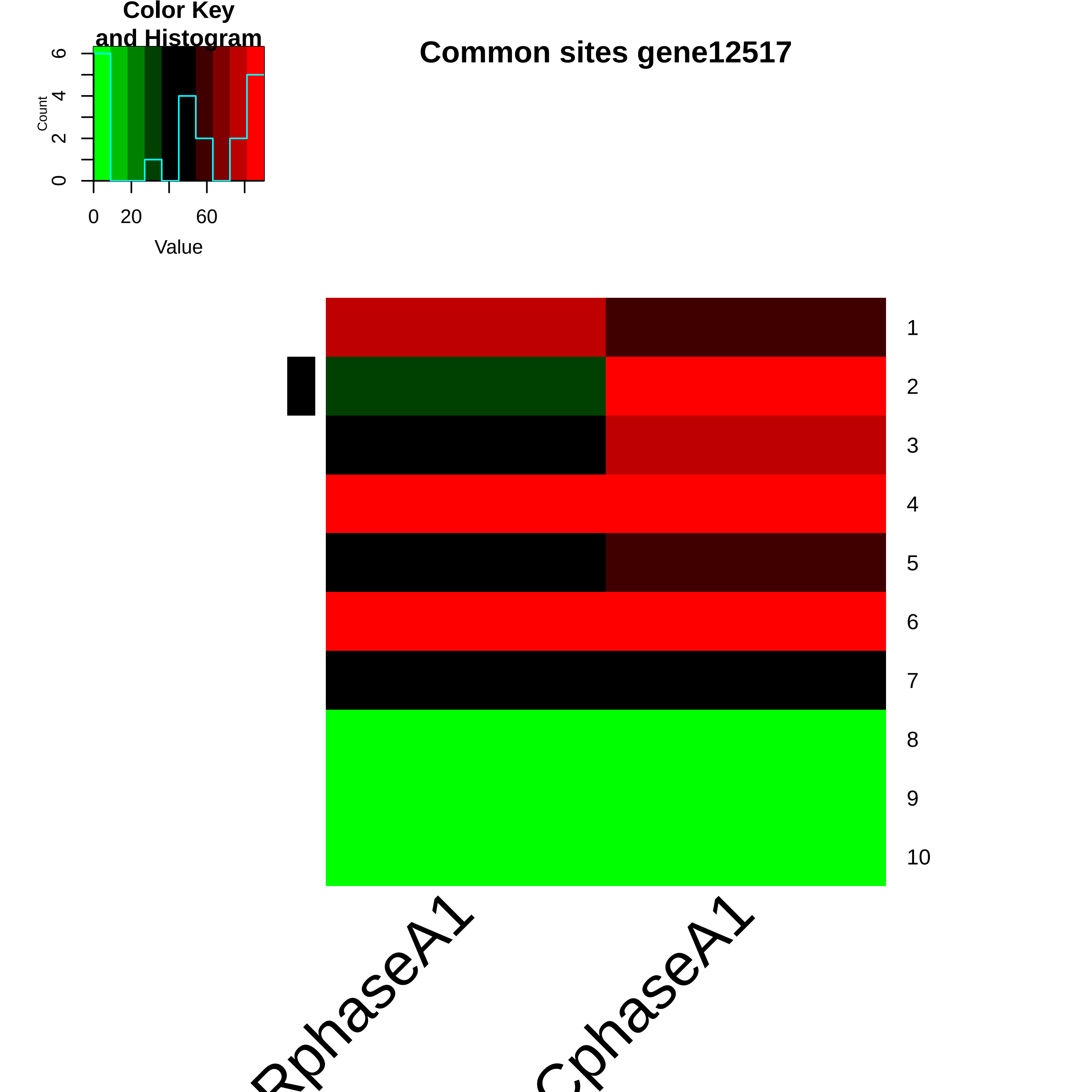
Color Key
and Histogram
Count
0
2
4
6
0
20
60
Value
Common sites gene12517
1
2
3
4
5
6
7
8
9
10
RphaseA1
CphaseA1

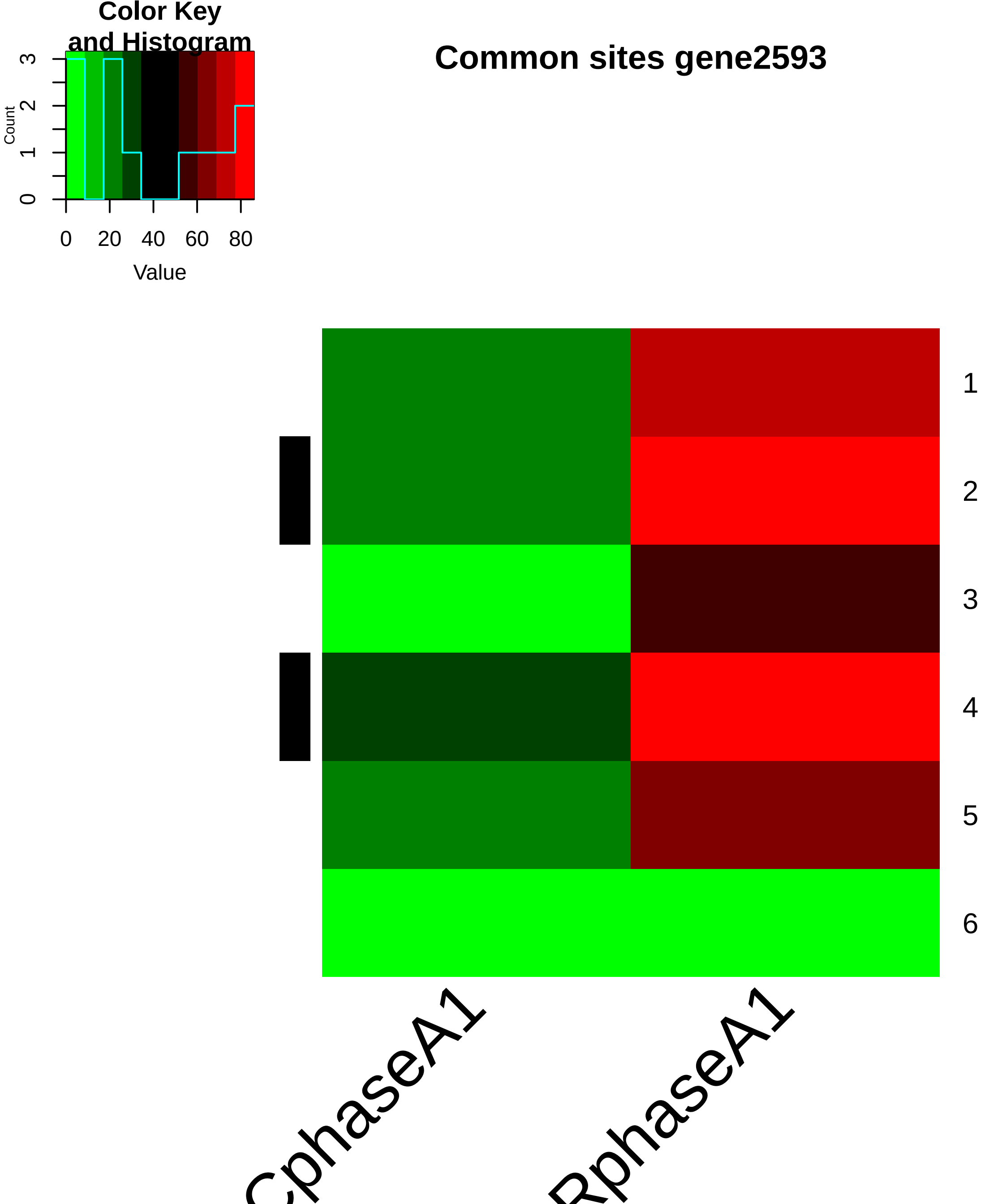
Color Key
and Histogram
Count
0
1
2
3
0
20
40
60
80
Value
Common sites gene2593
1
2
3
4
5
6
CphaseA1
RphaseA1

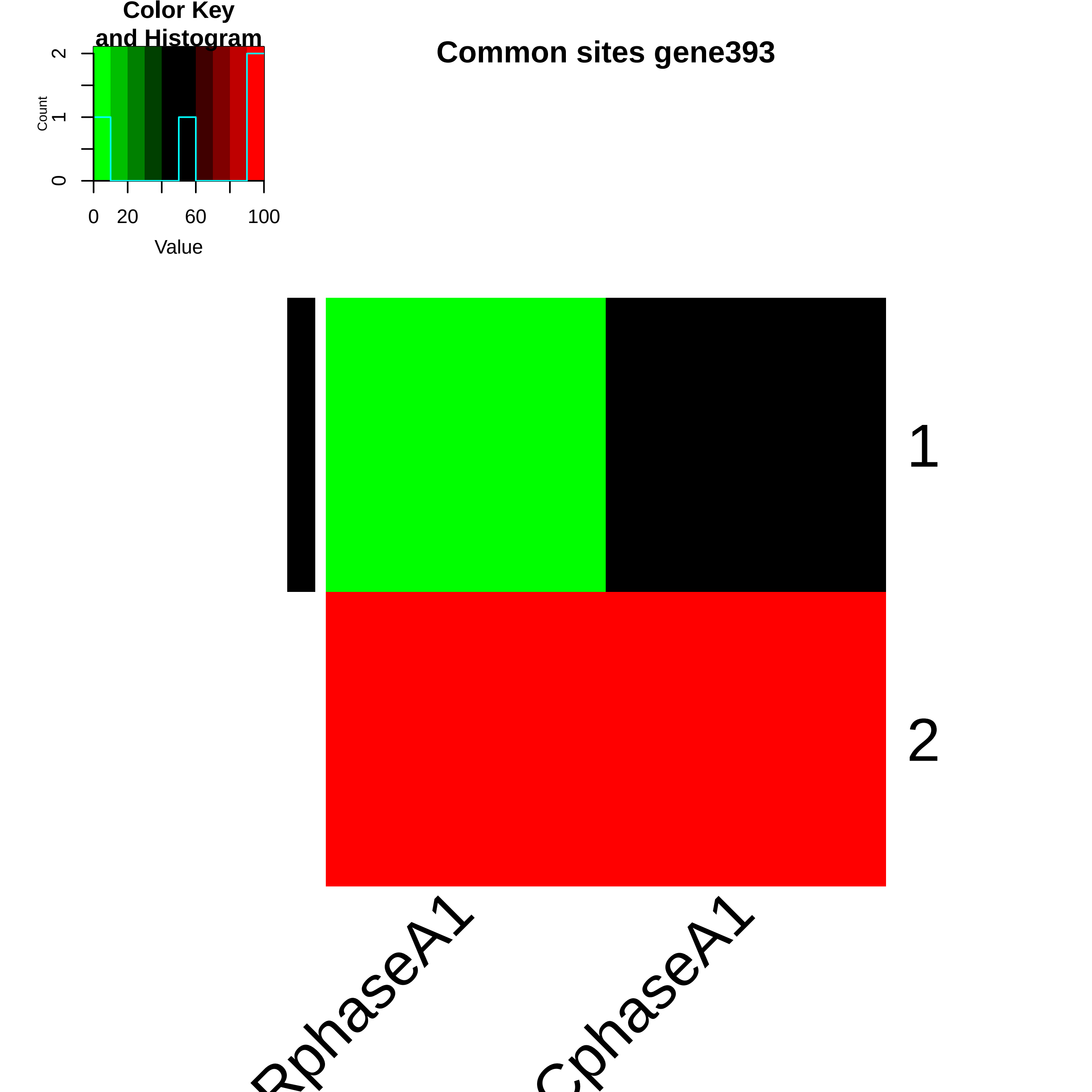

Color Key
and Histogram
Count
0
1
2
0
20
60
100
Value
Common sites gene393
1
2
RphaseA1
CphaseA1

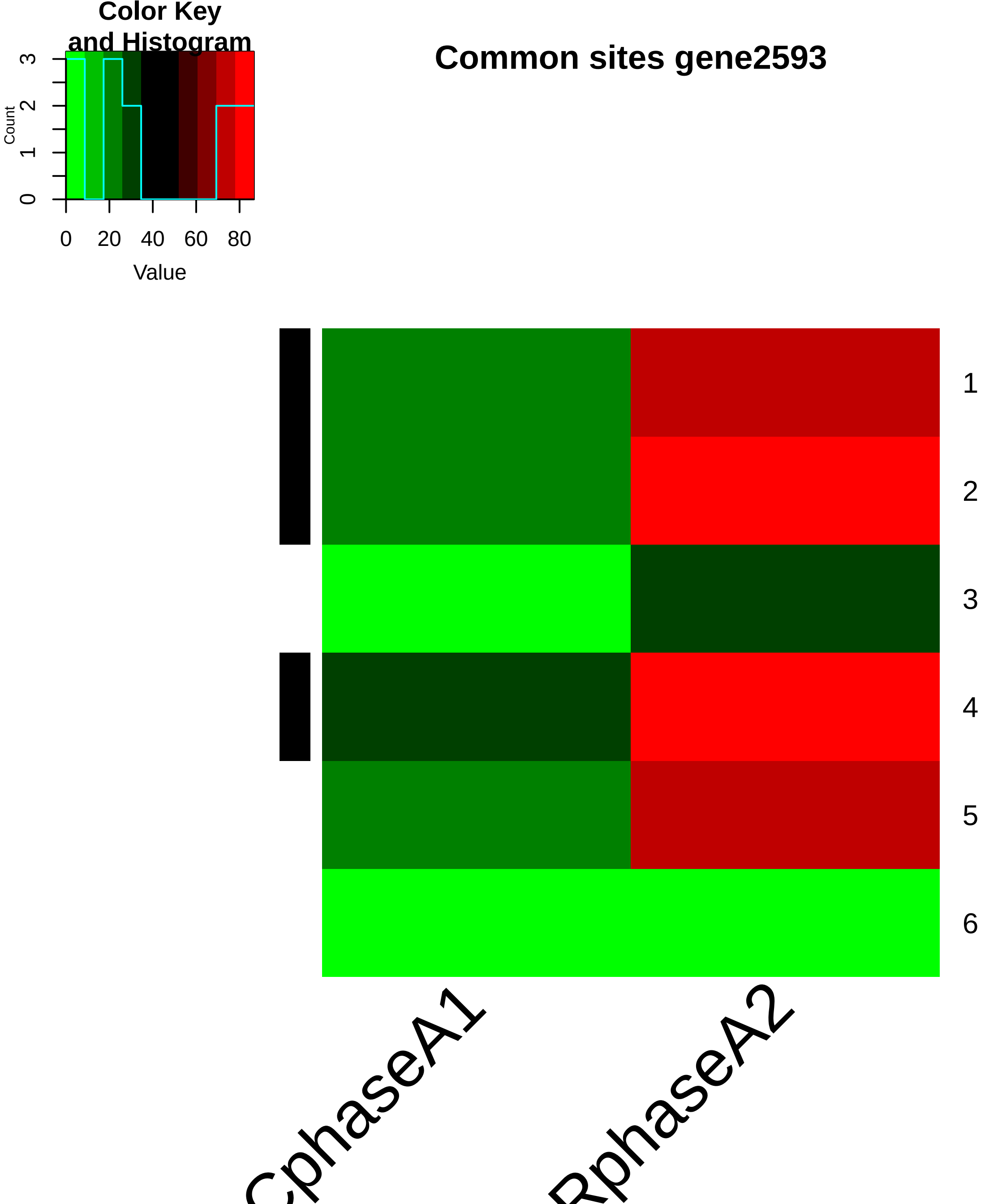
Color Key
and Histogram
Count
0
1
2
3
0
20
40
60
80
Value
Common sites gene2593
1
2
3
4
5
6
CphaseA1
RphaseA2

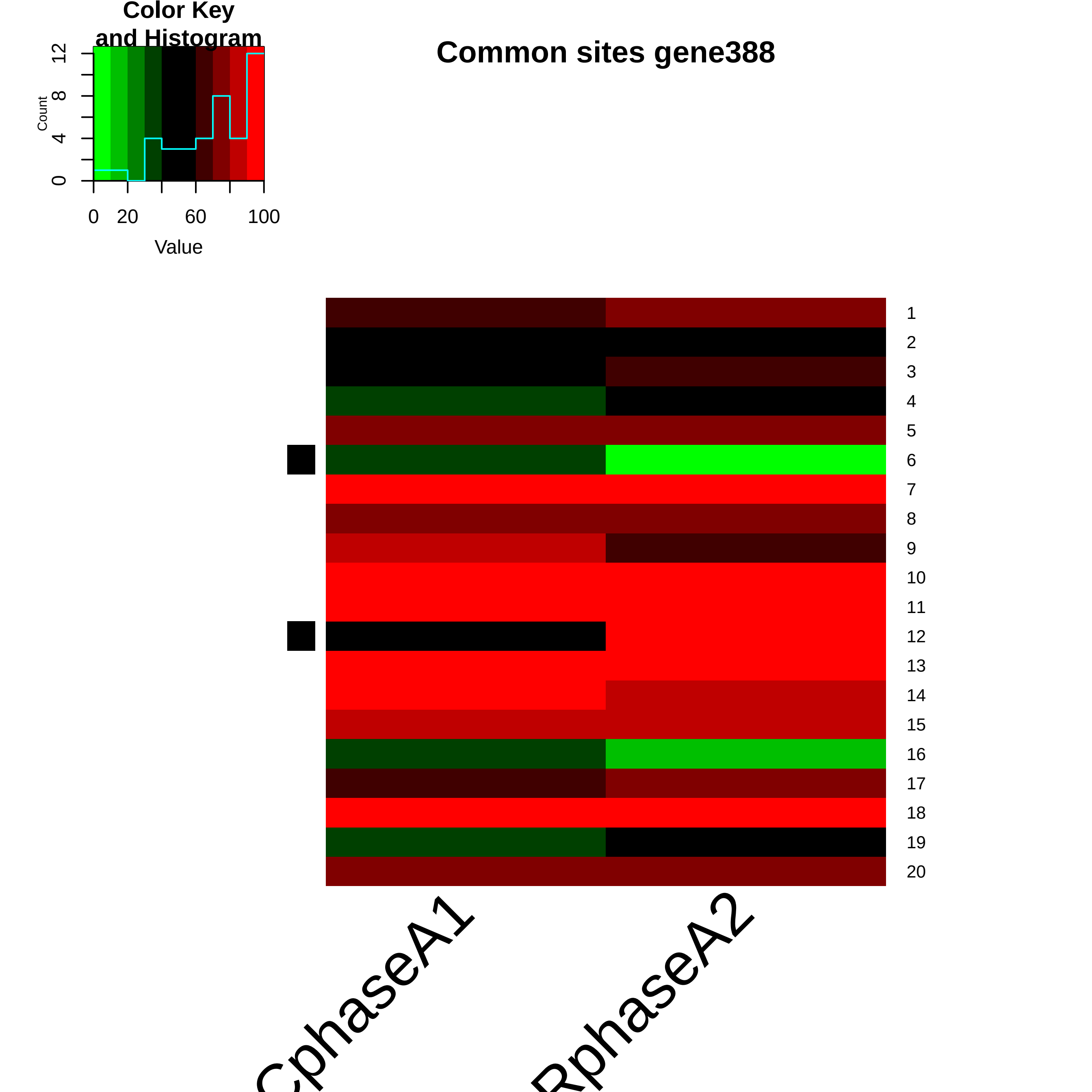
Color Key
and Histogram
Count
0
4
8
12
0
20
60
100
Value
Common sites gene388
CphaseA1
RphaseA2

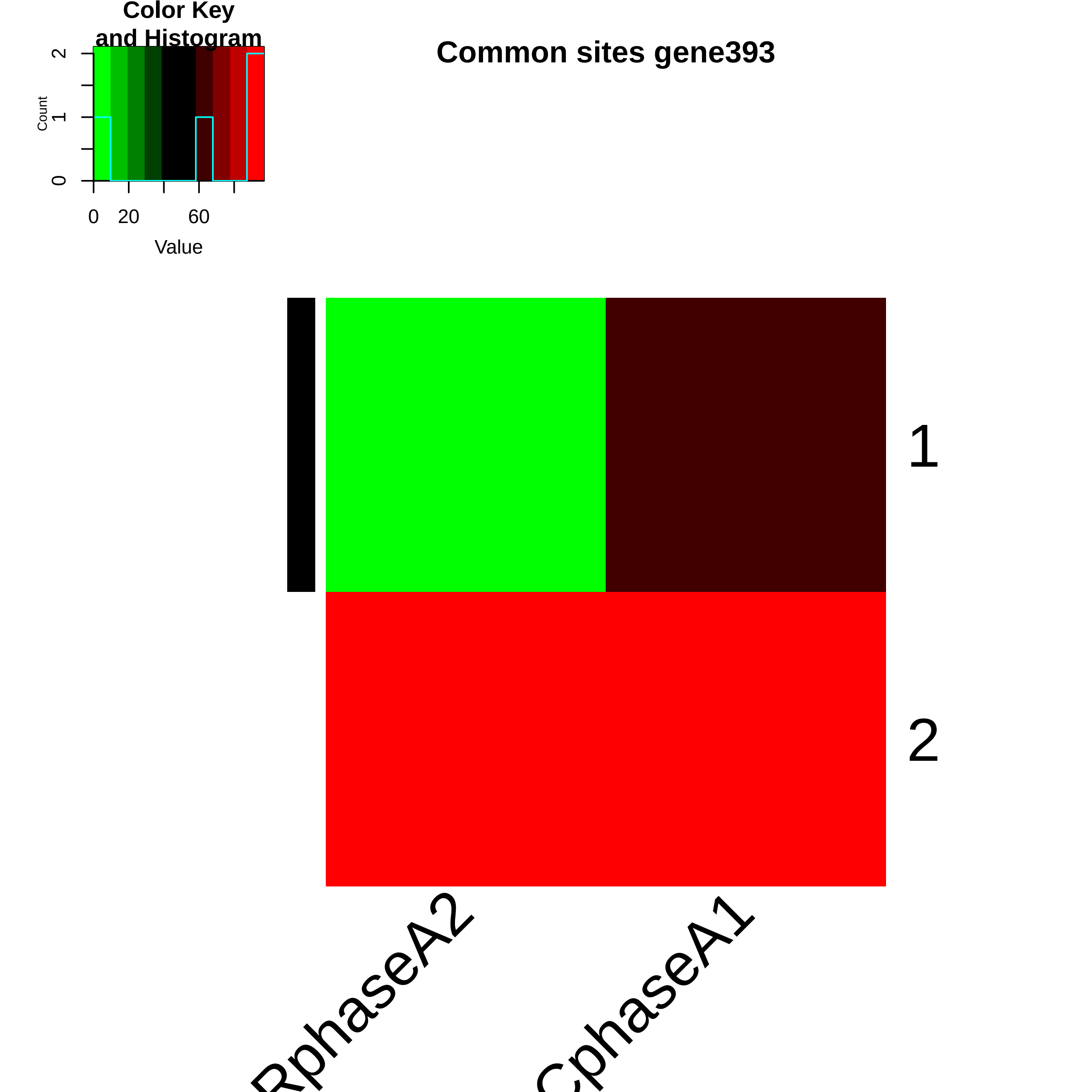

Color Key
and Histogram
Count
0
1
2
0
20
60
Value
Common sites gene393
1
2
RphaseA2
CphaseA1

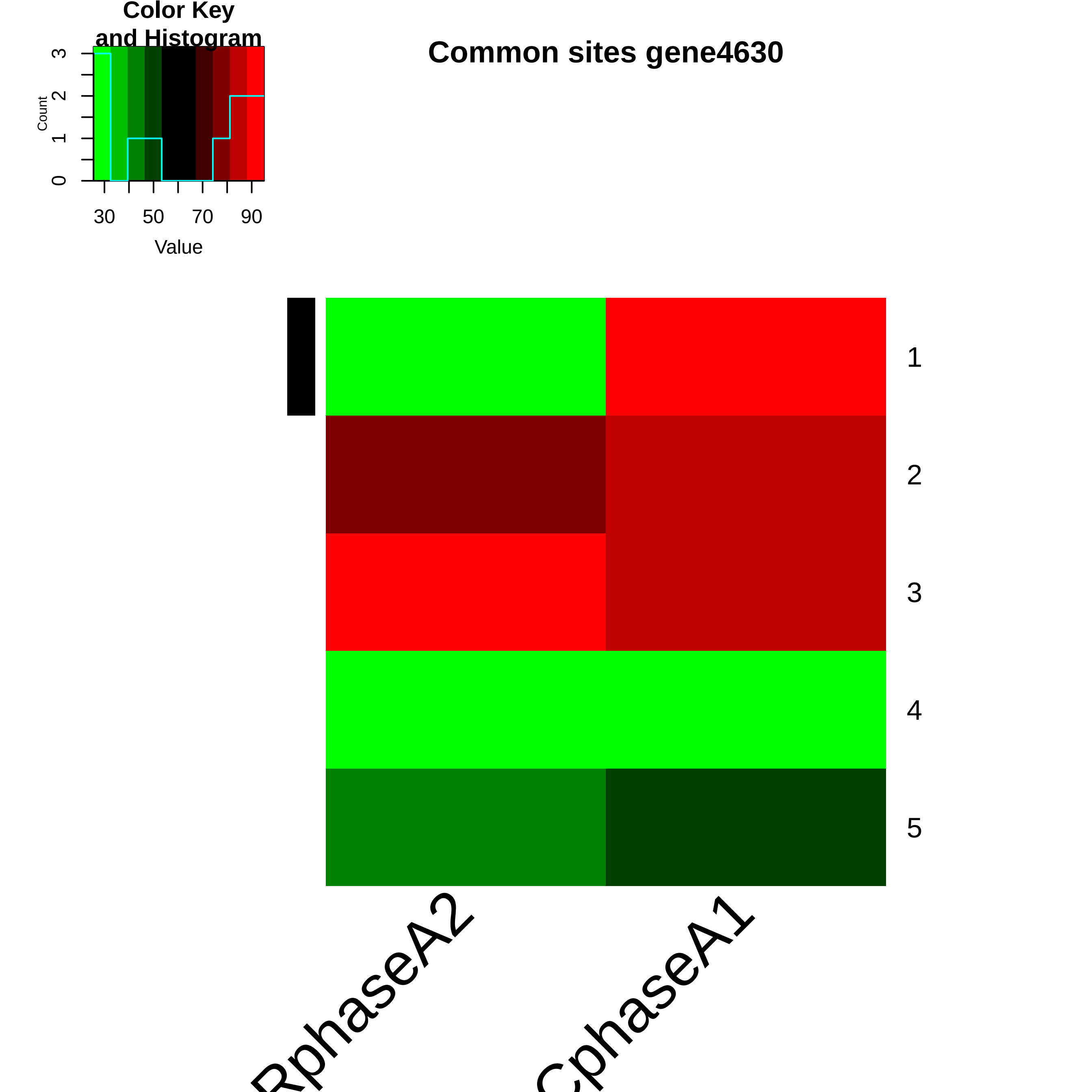

Color Key
and Histogram
Count
0
1
2
3
30
50
70
90
Value
Common sites gene4630
1
2
3
4
5
RphaseA2
CphaseA1

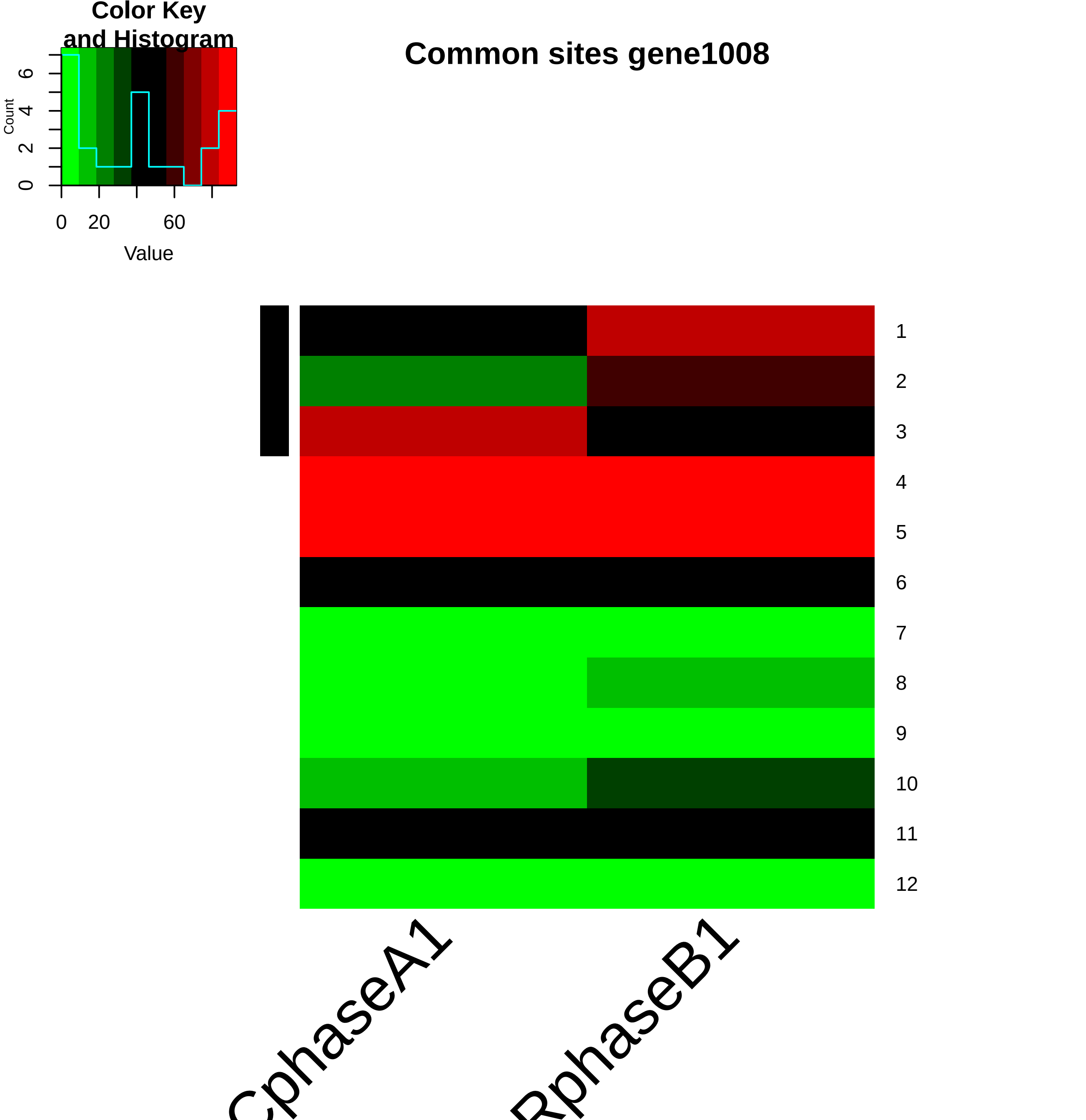
Color Key
and Histogram
Count
0
2
4
6
0
20
60
Value
Common sites gene1008
1
2
3
4
5
6
7
8
9
10
11
12
CphaseA1
RphaseB1

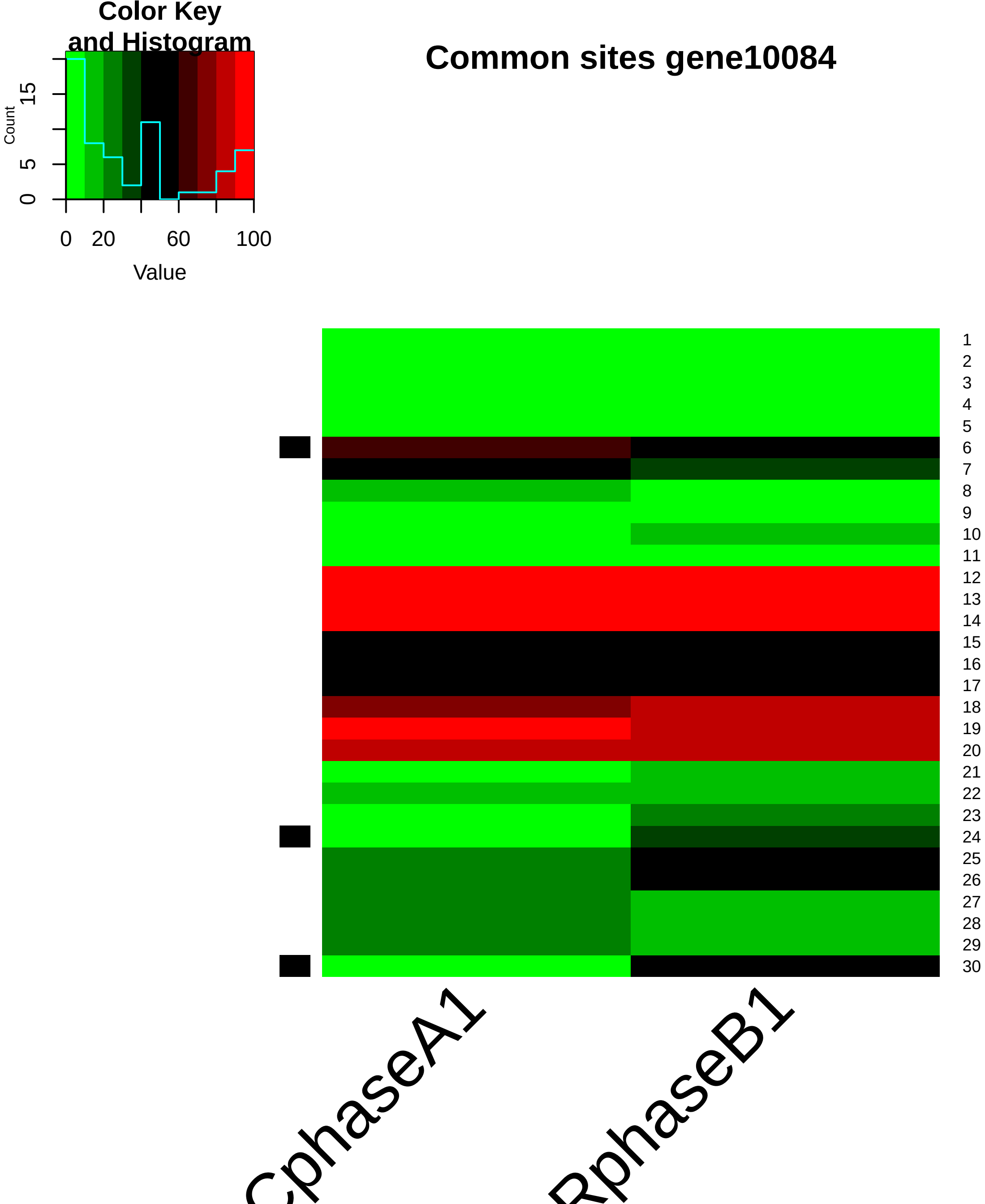
Color Key
and Histogram
Count
0
5
15
0
20
60
100
Value
Common sites gene10084
CphaseA1
RphaseB1

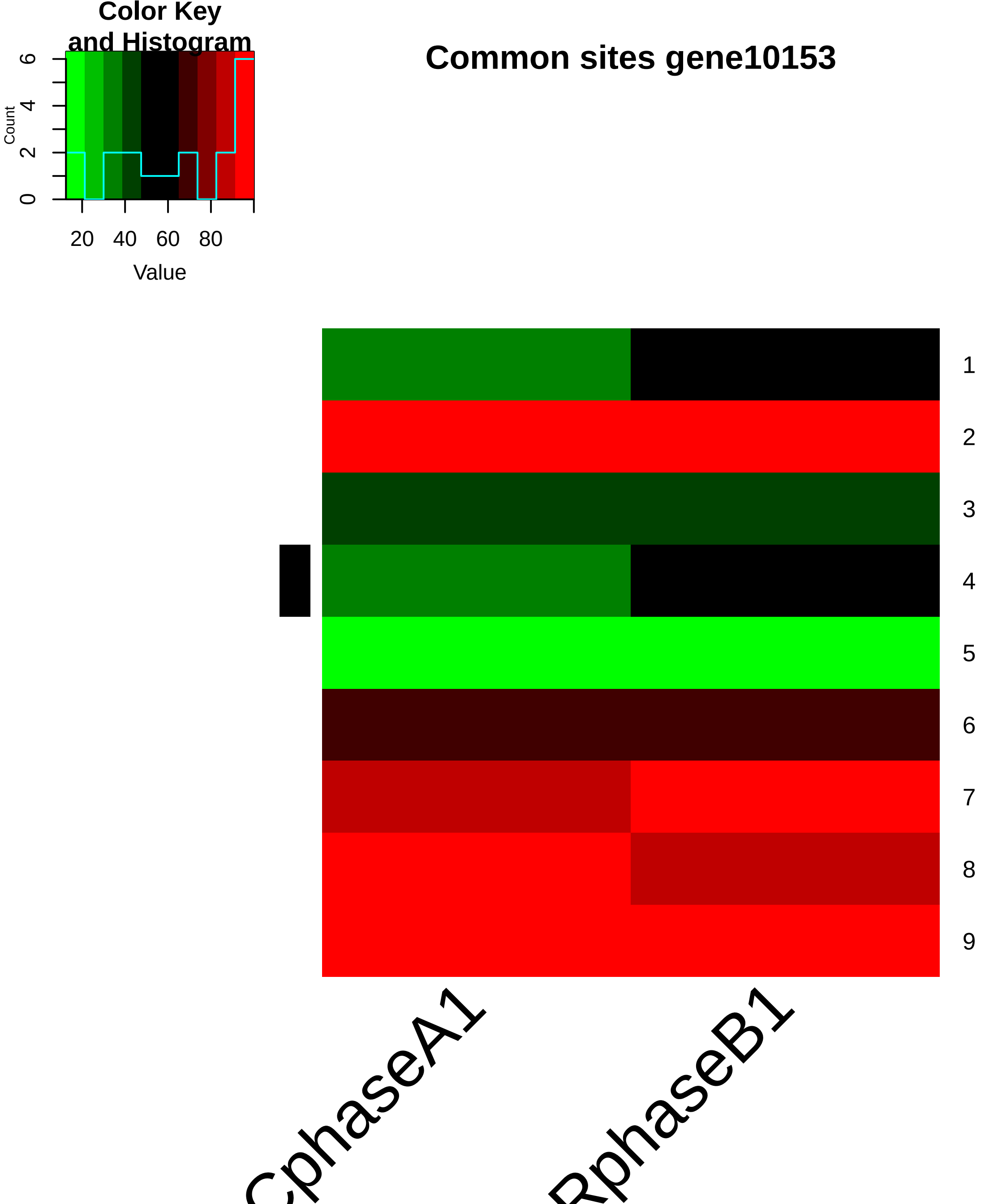
Color Key
and Histogram
Count
0
2
4
6
20
40
60
80
Value
Common sites gene10153
1
2
3
4
5
6
7
8
9
CphaseA1
RphaseB1

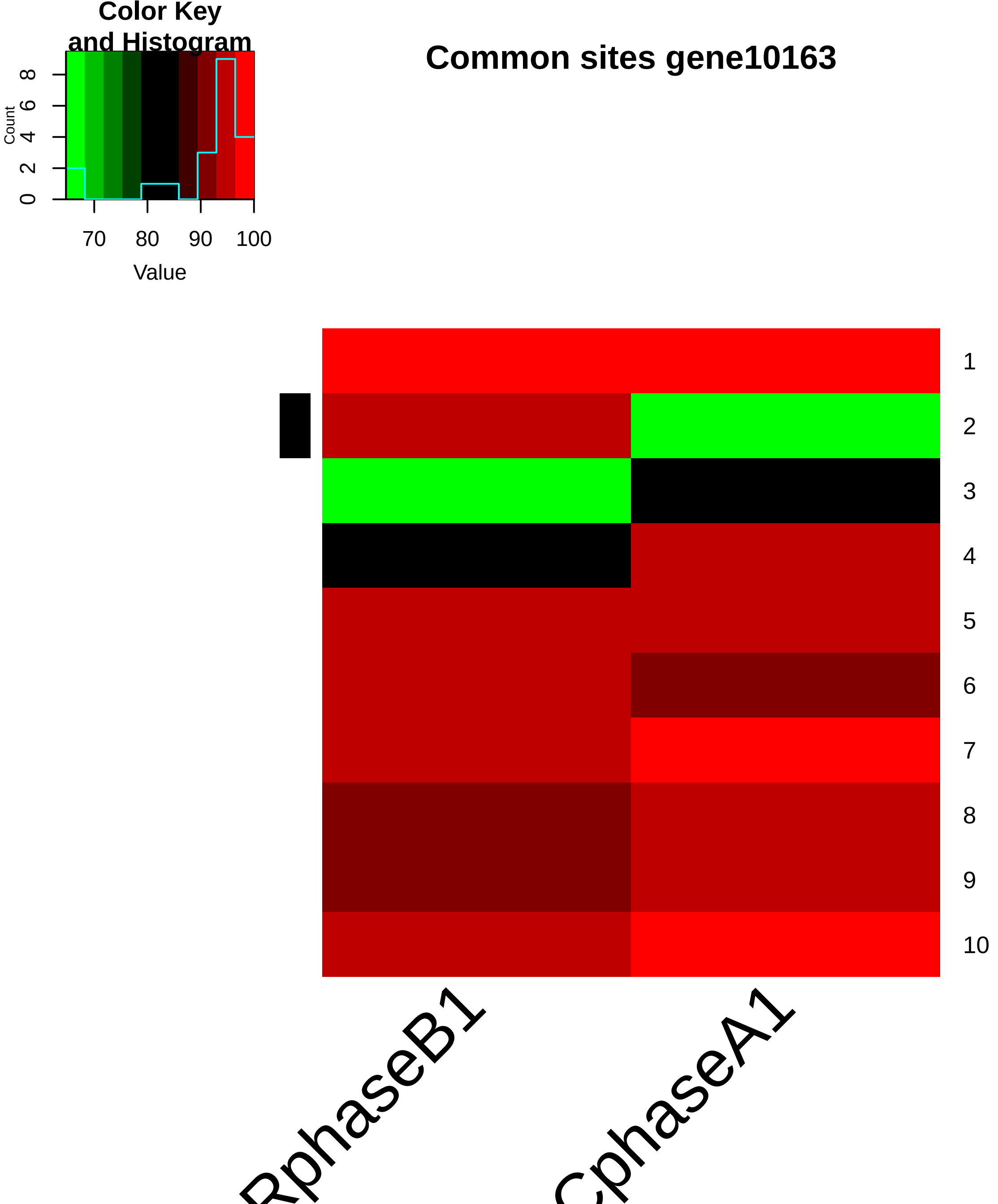
Color Key
and Histogram
Count
0
2
4
6
8
70
80
90
100
Value
Common sites gene10163
1
2
3
4
5
6
7
8
9
10
RphaseB1
CphaseA1

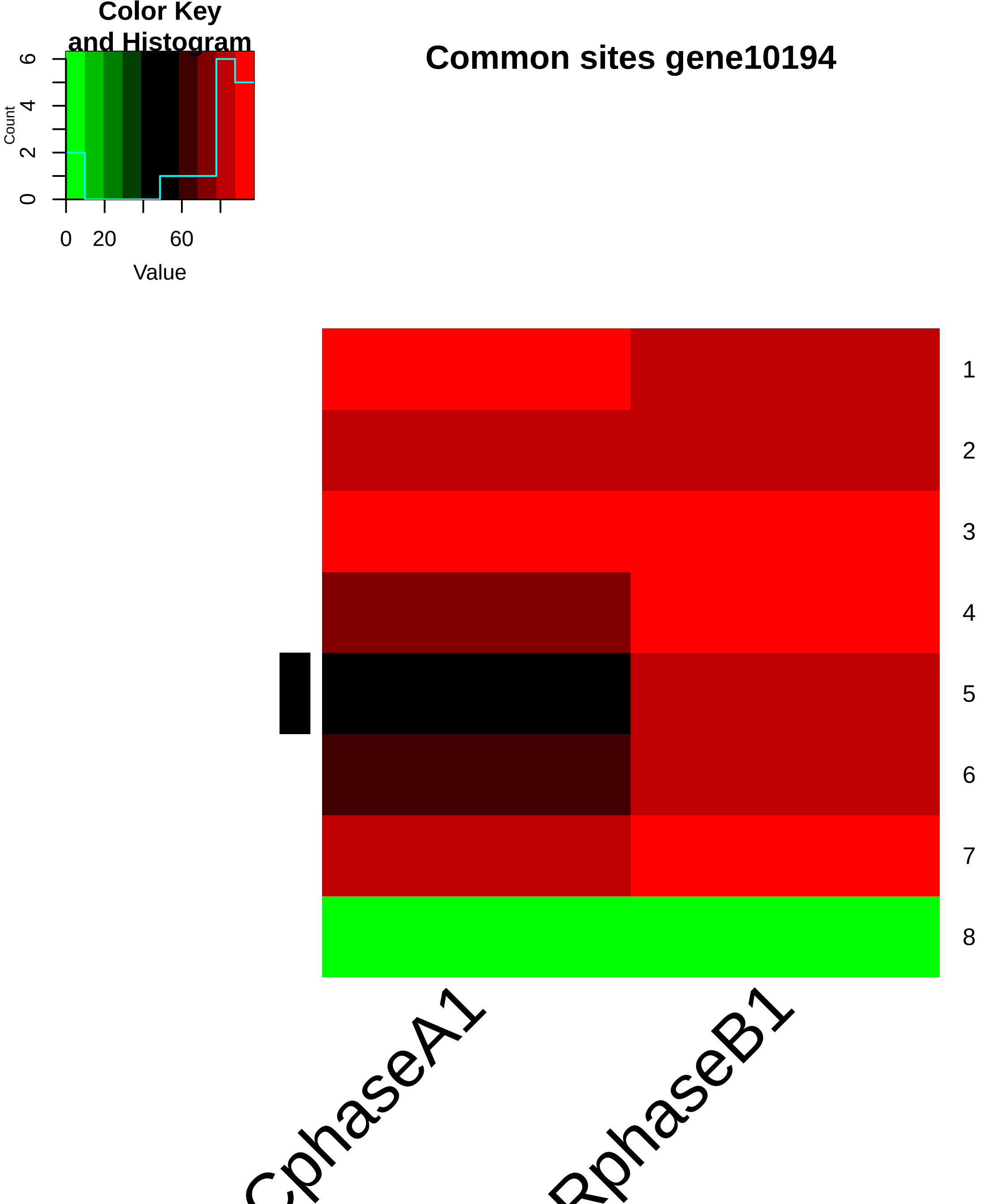
Color Key
and Histogram
Count
0
2
4
6
0
20
60
Value
Common sites gene10194
1
2
3
4
5
6
7
8
CphaseA1
RphaseB1

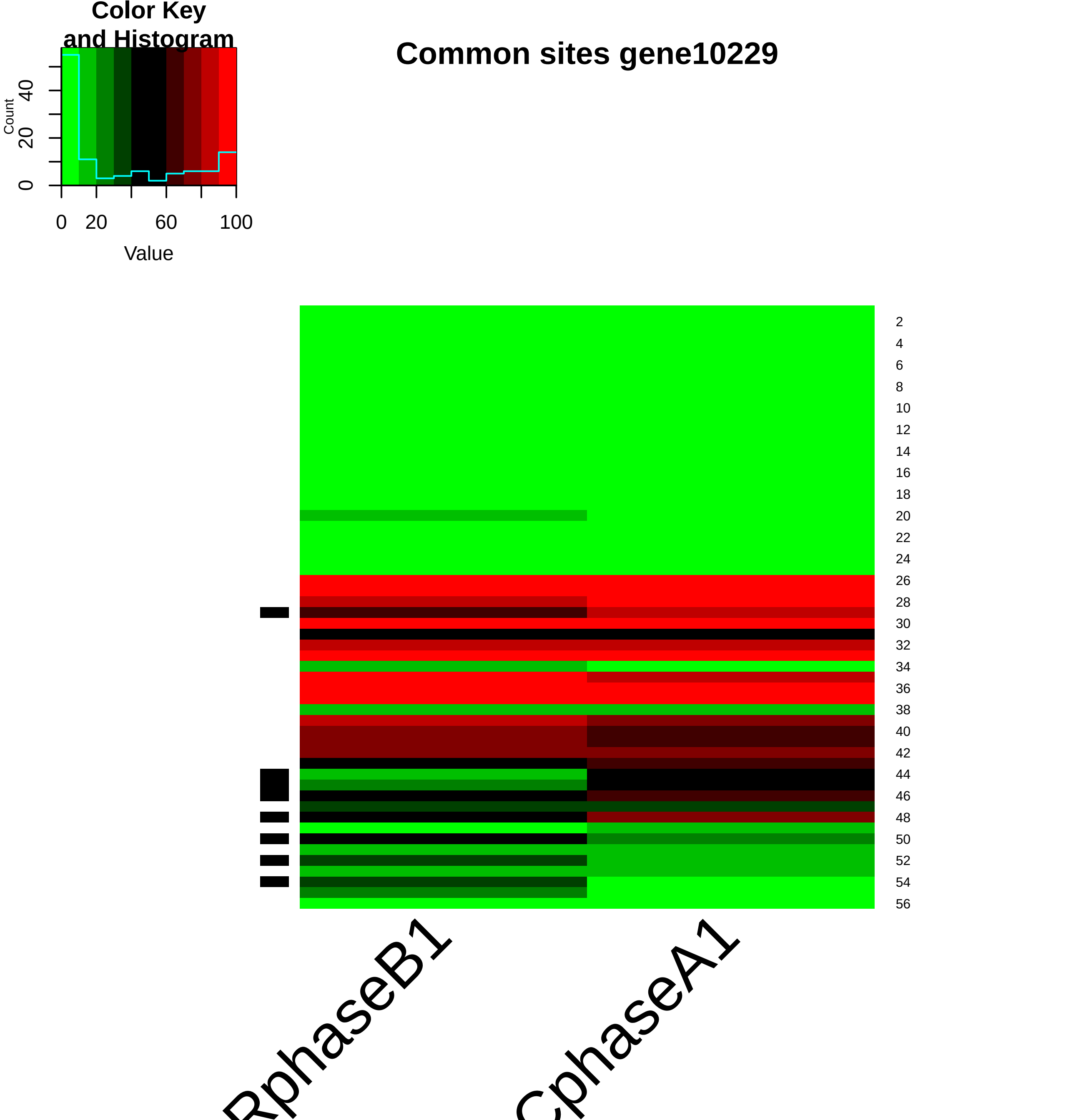

Color Key
and Histogram
Count
0
20
40
0
20
60
100
Value
Common sites gene10229
2
4
6
8
10
12
14
16
18
20
22
24
26
28
30
32
34
36
38
40
42
44
46
48
50
52
54
56
RphaseB1
CphaseA1

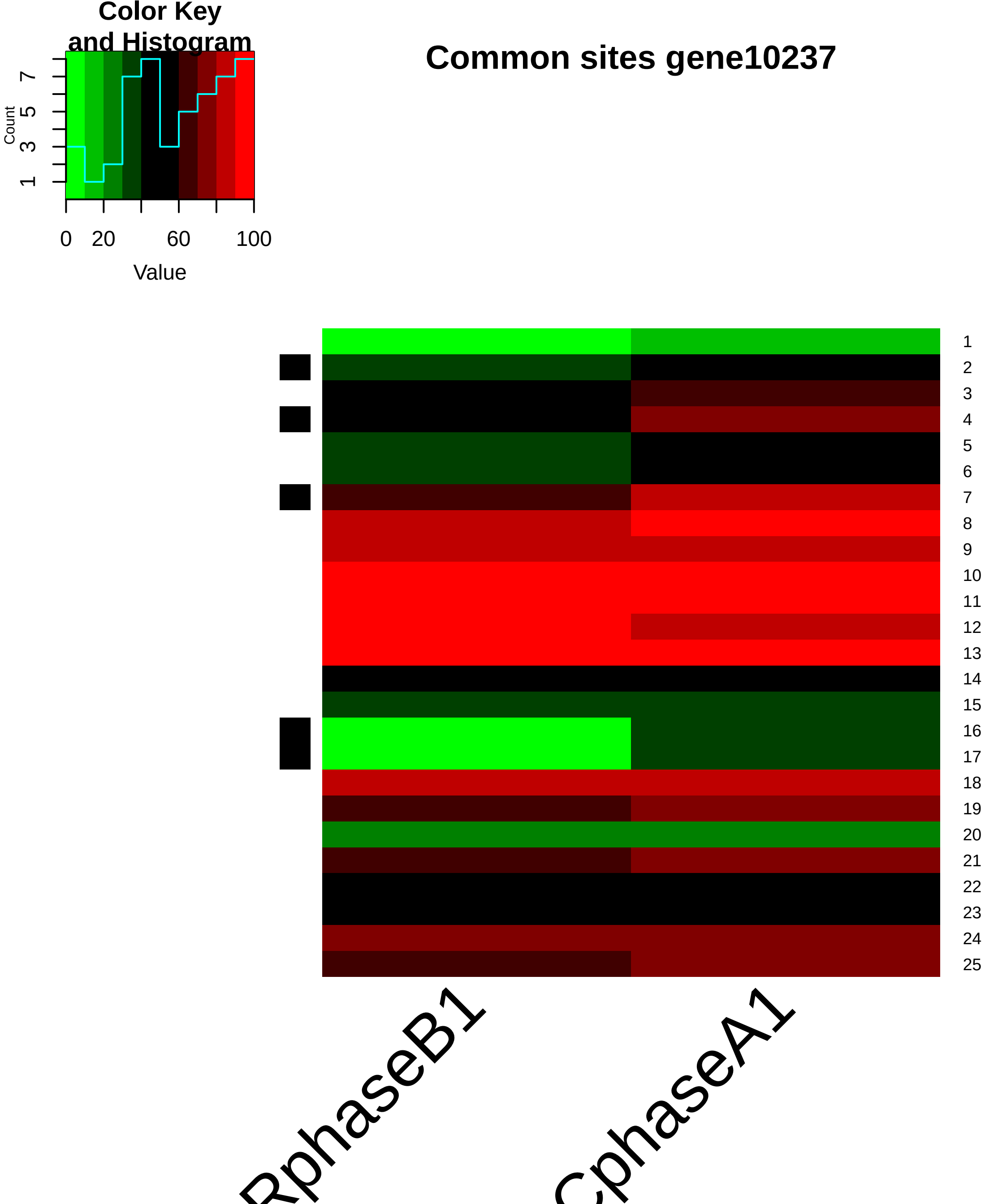

Color Key
and Histogram
Count
1
3
5
7
0
20
60
100
Value
Common sites gene10237
RphaseB1
CphaseA1

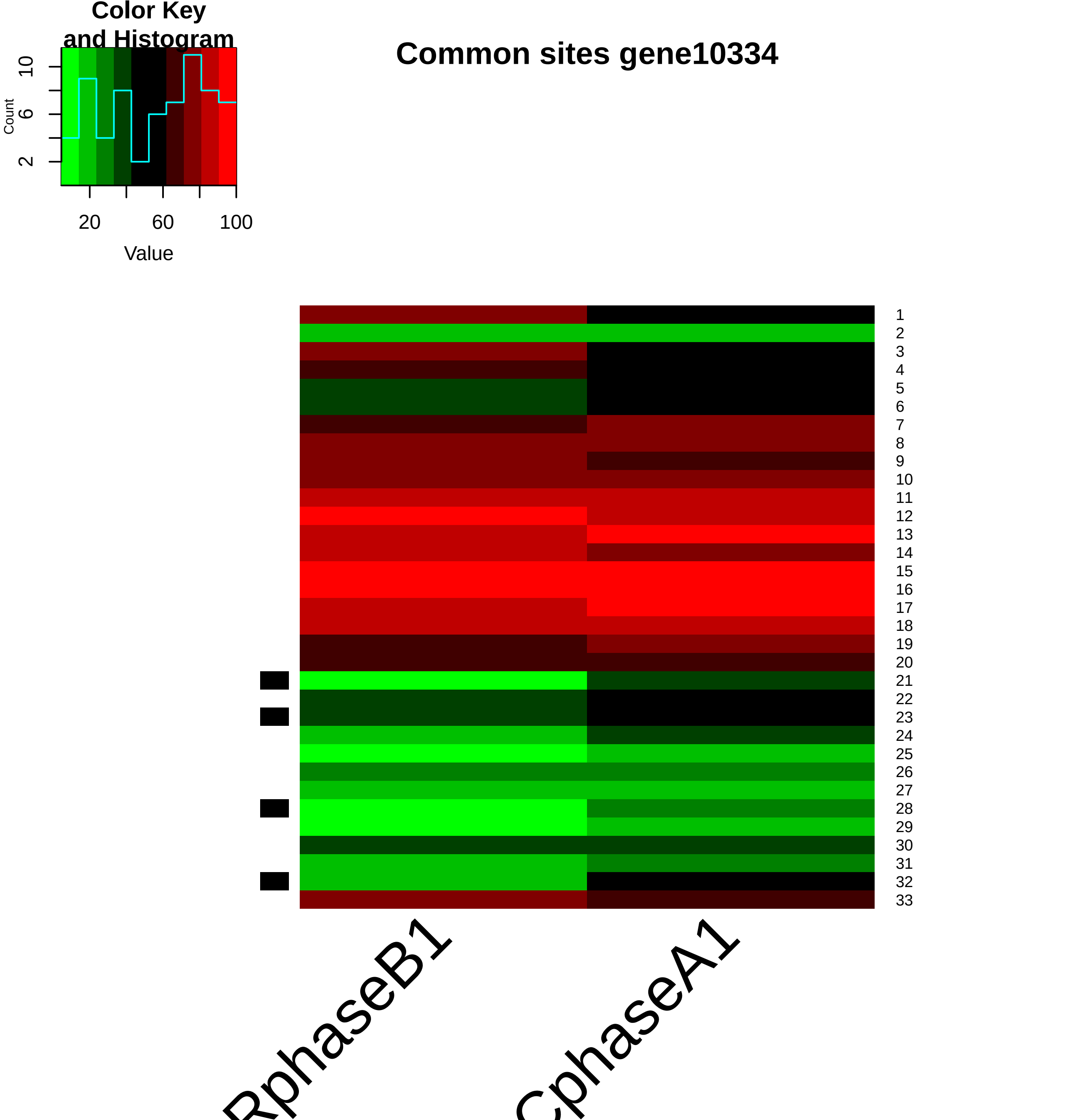

Color Key
and Histogram
Count
2
6
10
20
60
100
Value
Common sites gene10334
RphaseB1
CphaseA1

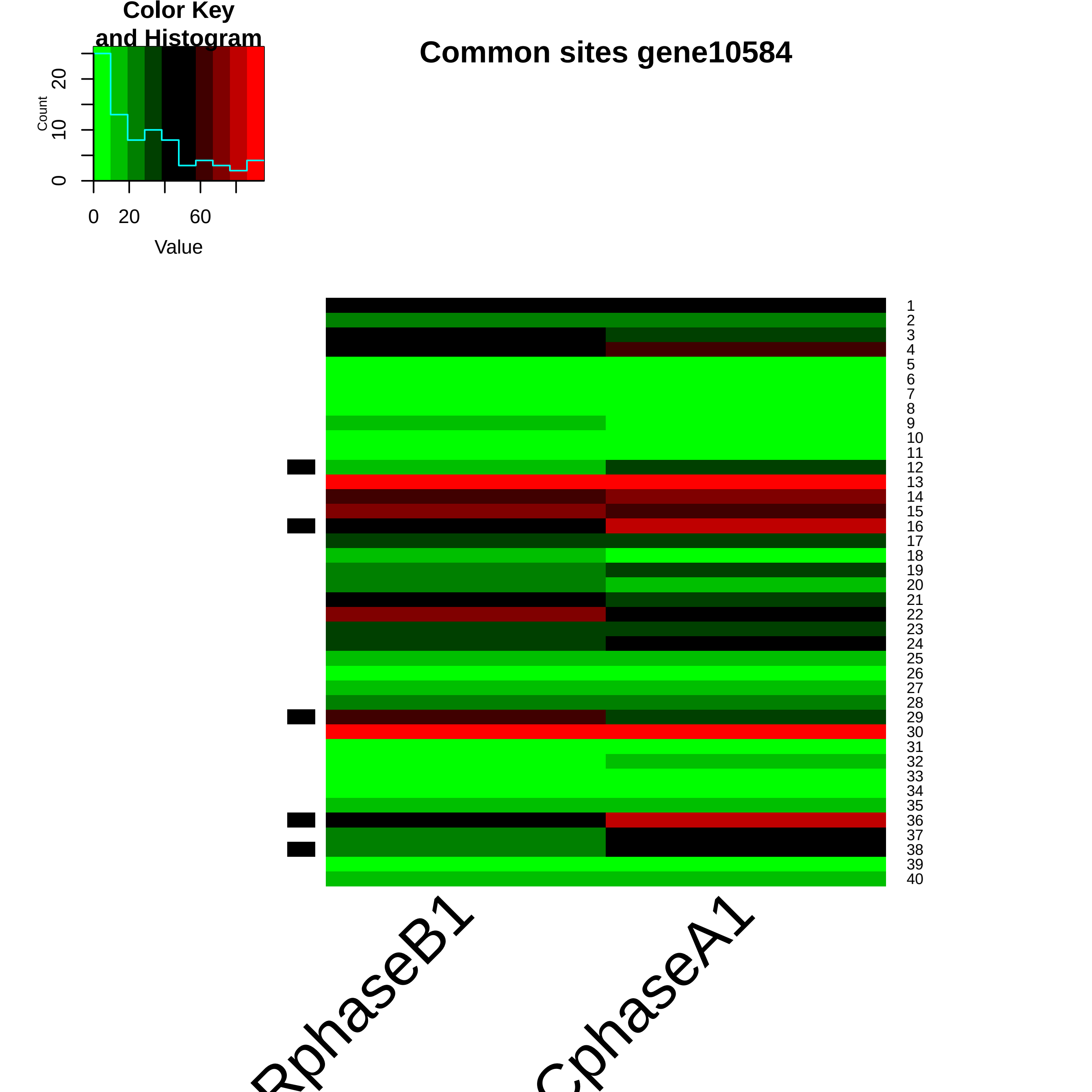

Color Key
and Histogram
Count
0
10
20
0
20
60
Value
Common sites gene10584
RphaseB1
CphaseA1

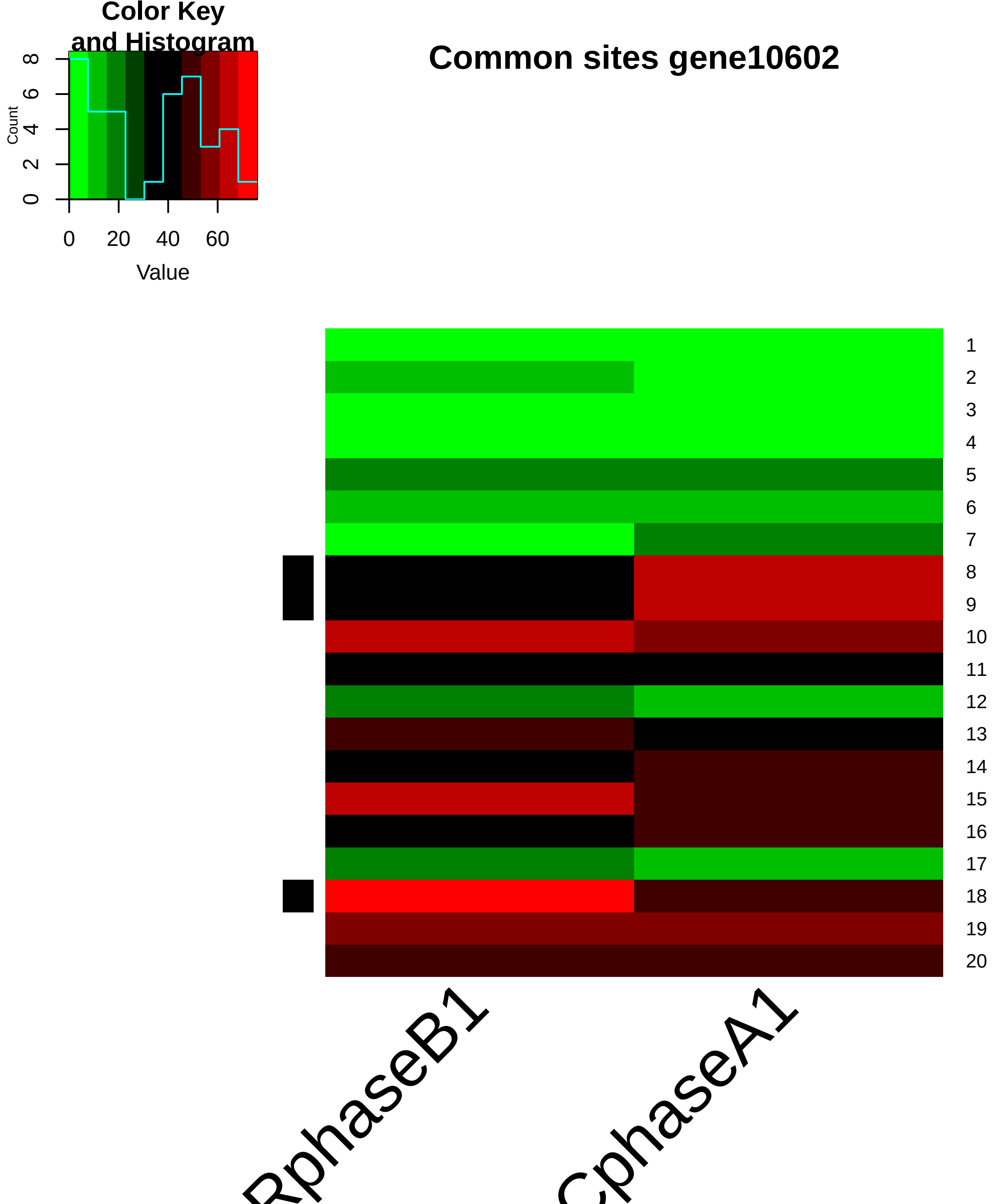

Color Key
and Histogram
Count
0
2
4
6
8
0
20
40
60
Value
Common sites gene10602
RphaseB1
CphaseA1

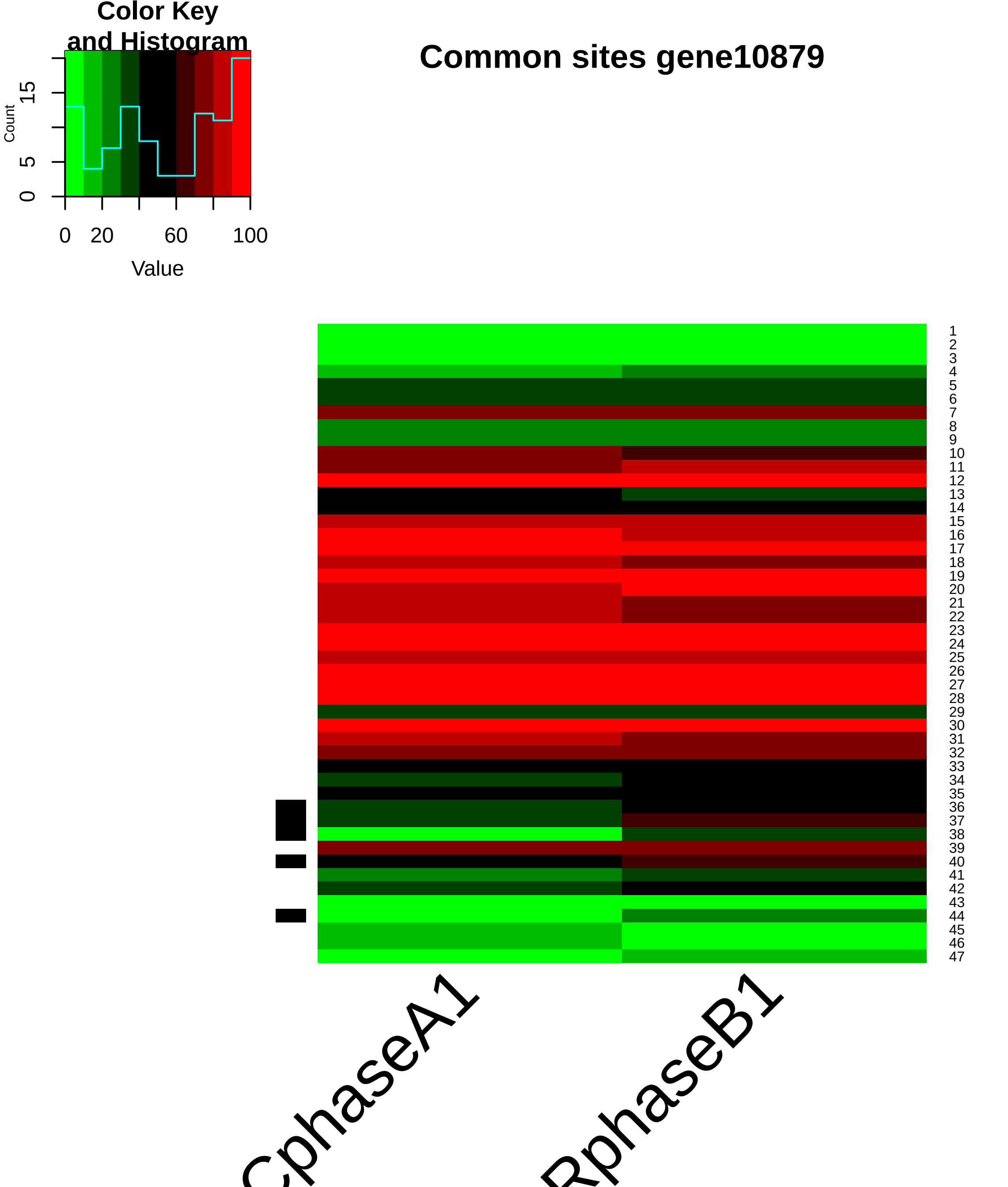
Color Key
and Histogram
Count
0
5
15
0
20
60
100
Value
Common sites gene10879
CphaseA1
RphaseB1

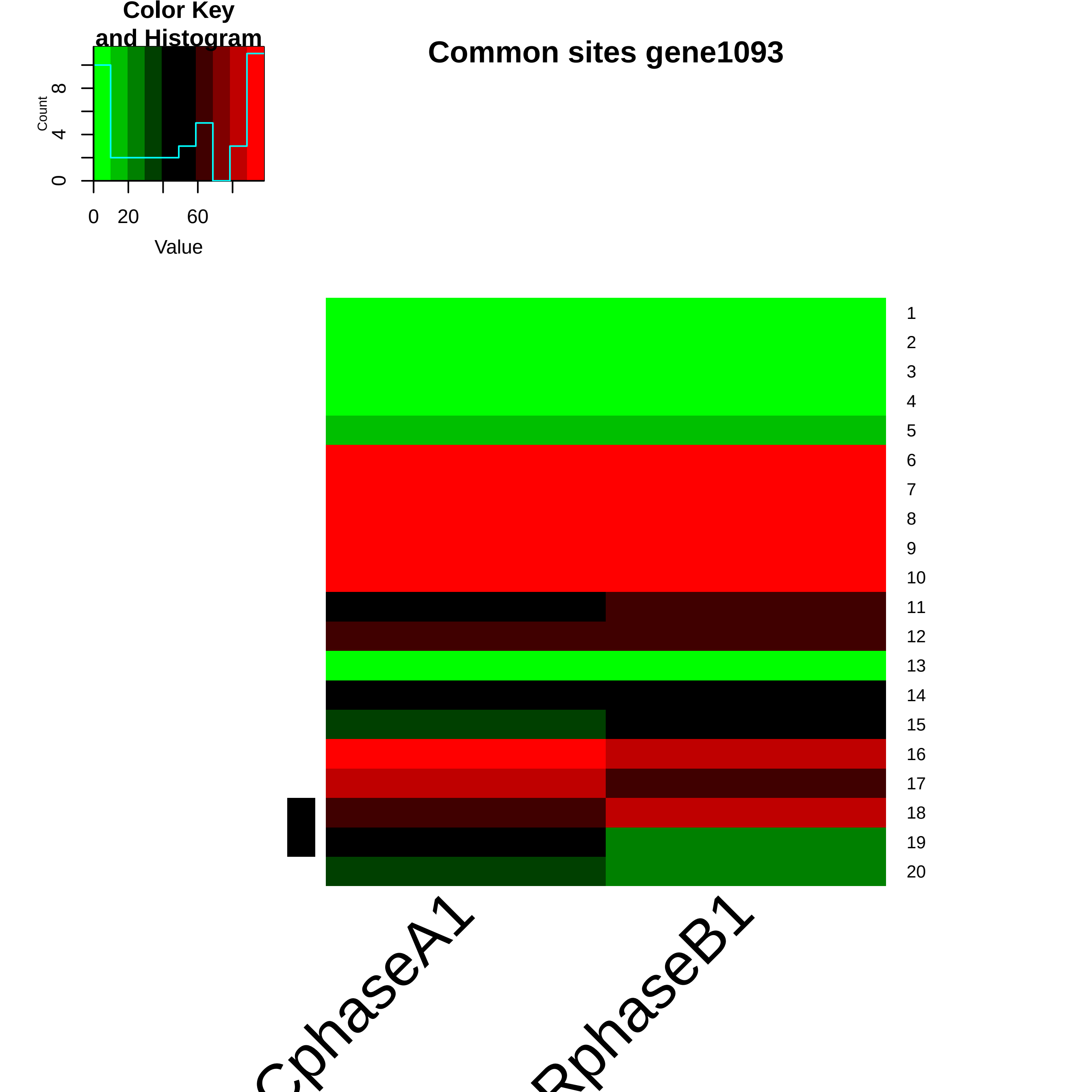
Color Key
and Histogram
Count
0
4
8
0
20
60
Value
Common sites gene1093
CphaseA1
RphaseB1

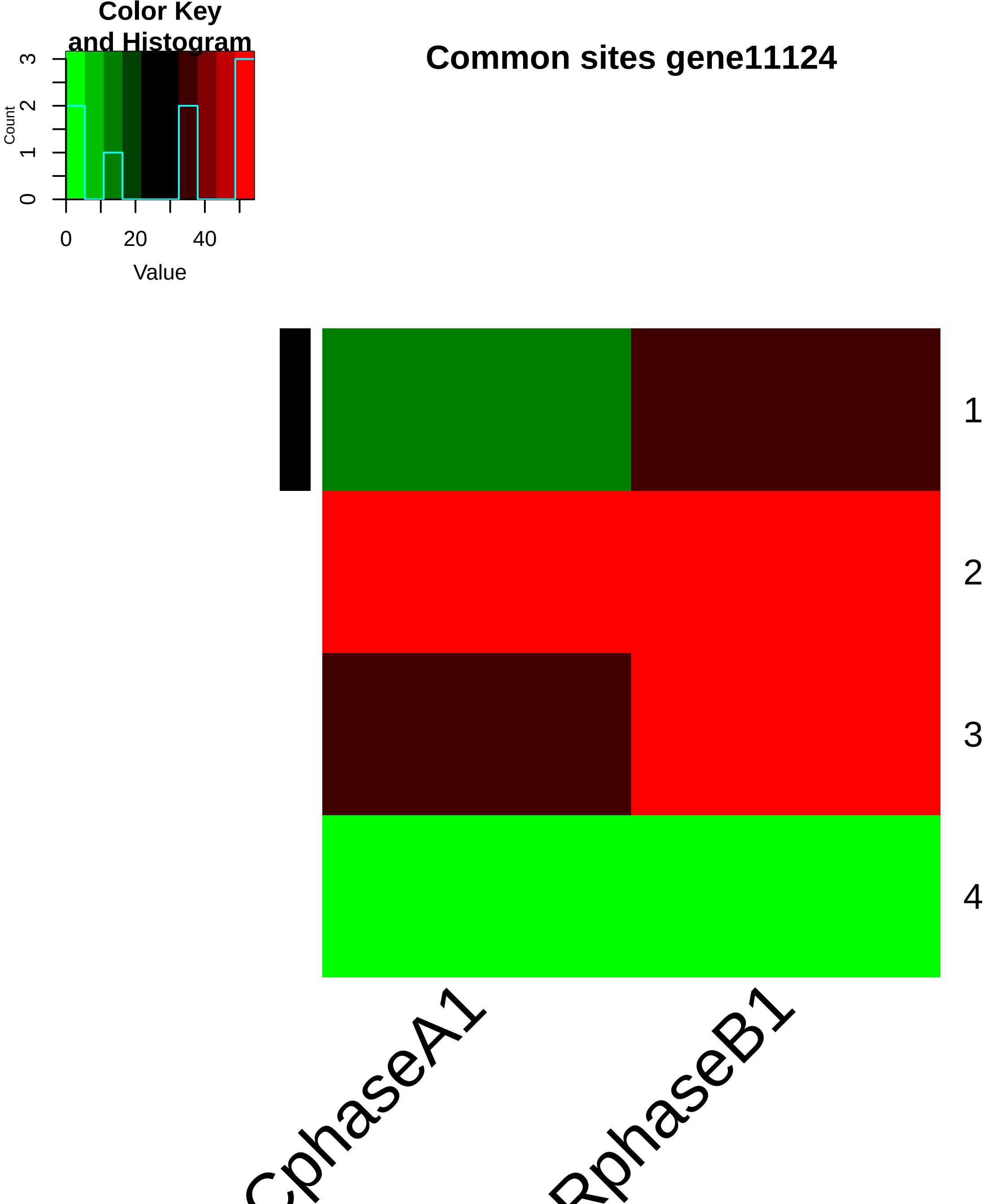

Color Key
and Histogram
Count
0
1
2
3
0
20
40
Value
Common sites gene11124
1
2
3
4
CphaseA1
RphaseB1

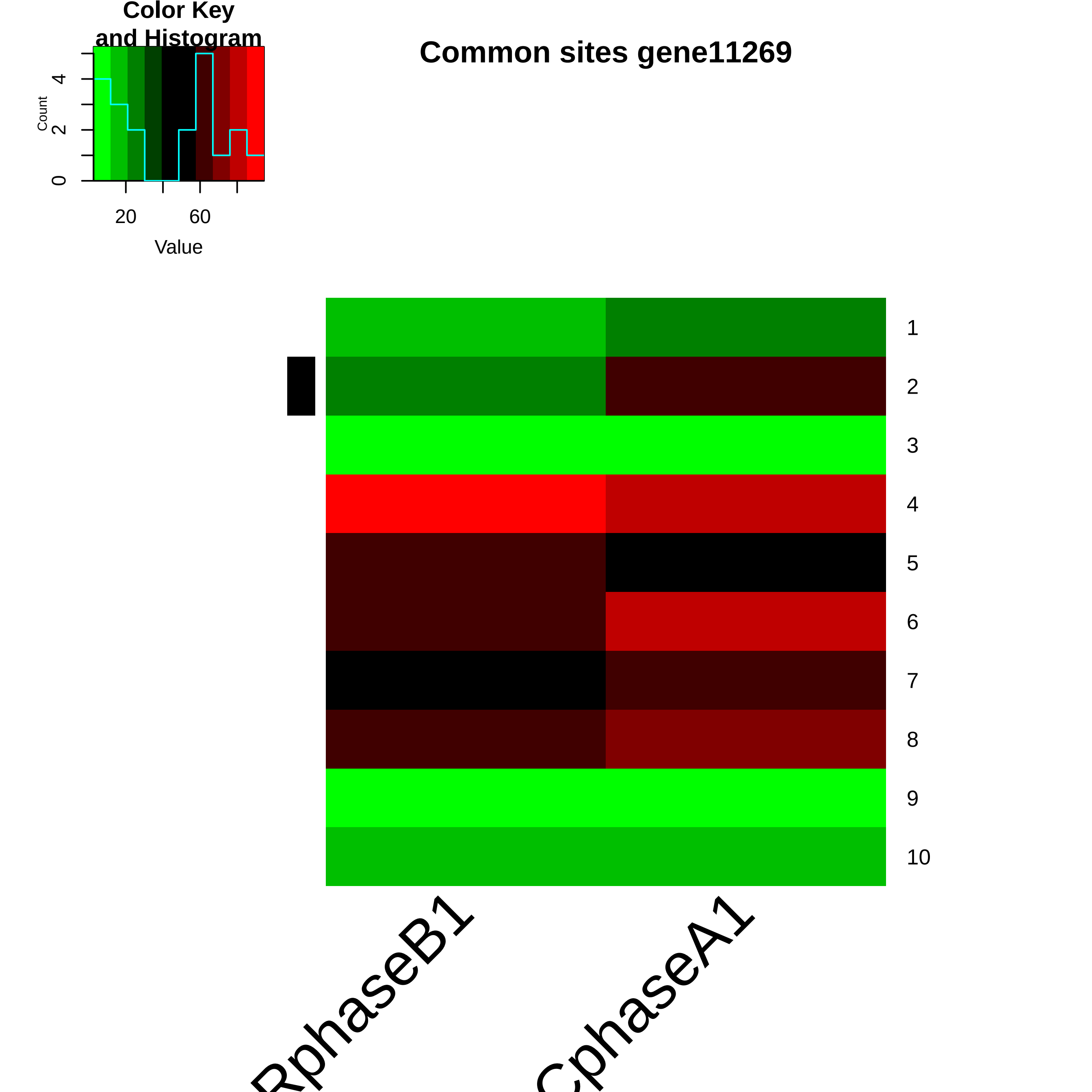
Color Key
and Histogram
Count
0
2
4
20
60
Value
Common sites gene11269
1
2
3
4
5
6
7
8
9
10
RphaseB1
CphaseA1

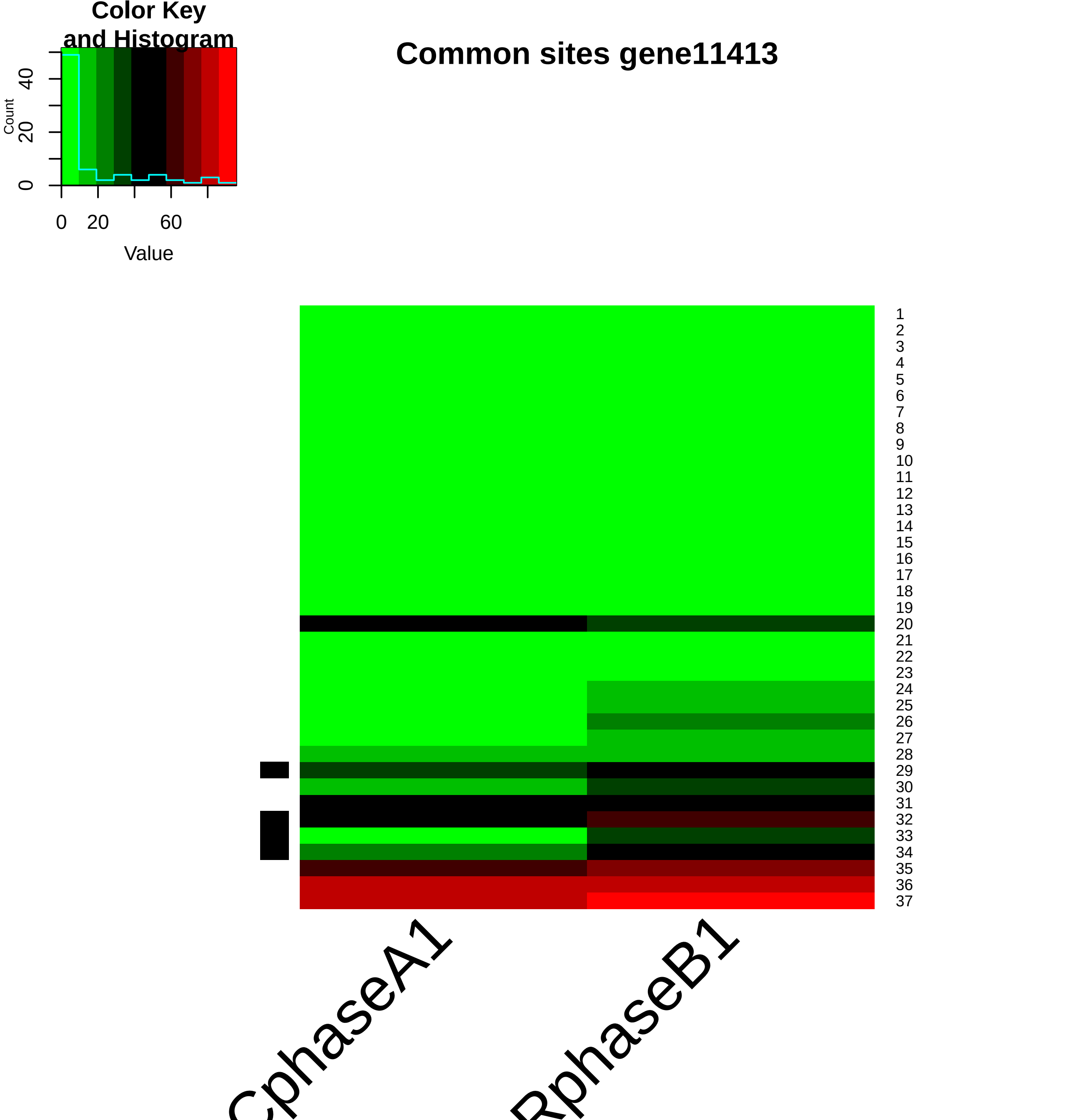
Color Key
and Histogram
Count
0
20
40
0
20
60
Value
Common sites gene11413
CphaseA1
RphaseB1

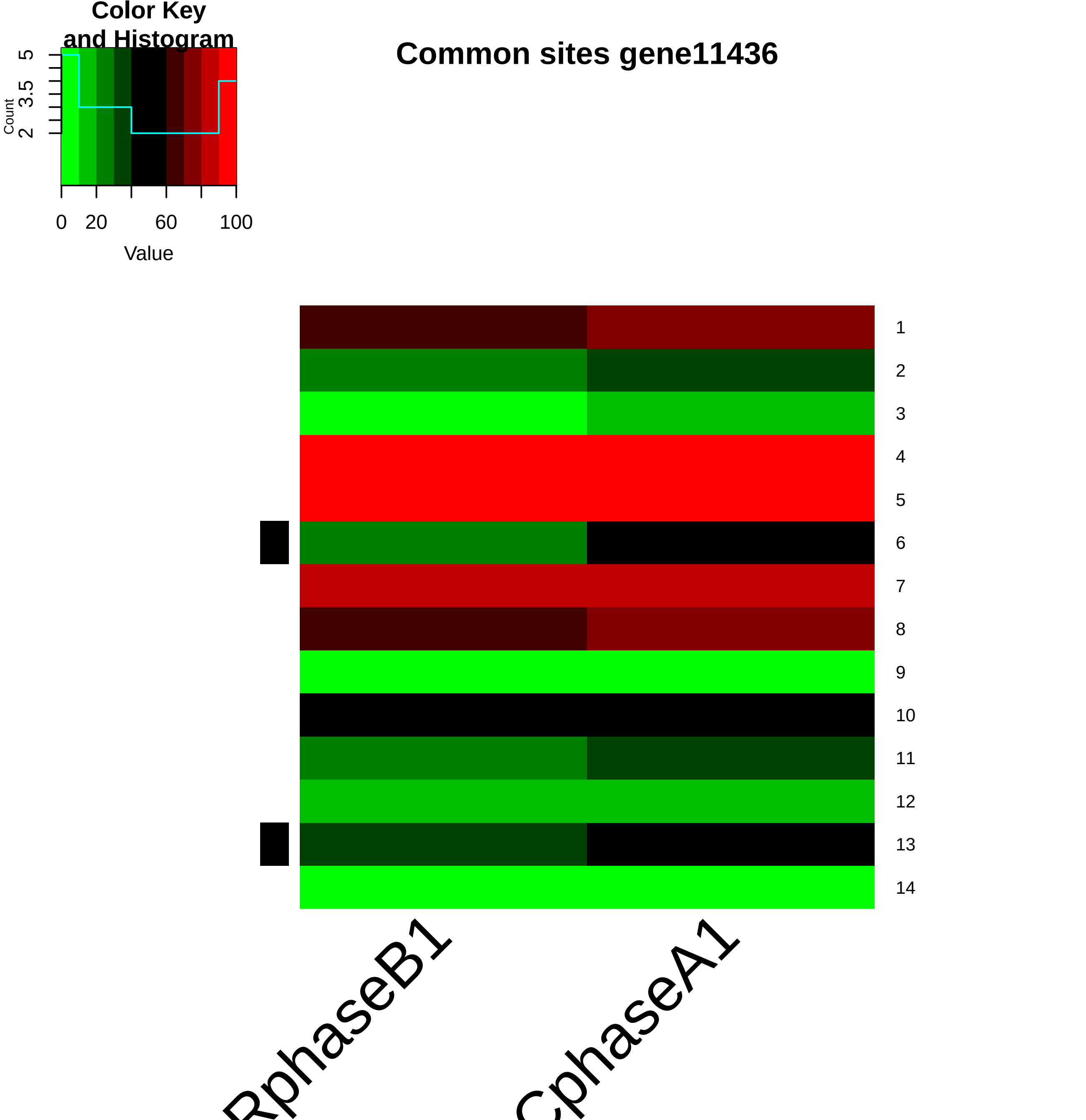

Color Key
and Histogram
Count
2
3.5
5
0
20
60
100
Value
Common sites gene11436
1
2
3
4
5
6
7
8
9
10
11
12
13
14
RphaseB1
CphaseA1

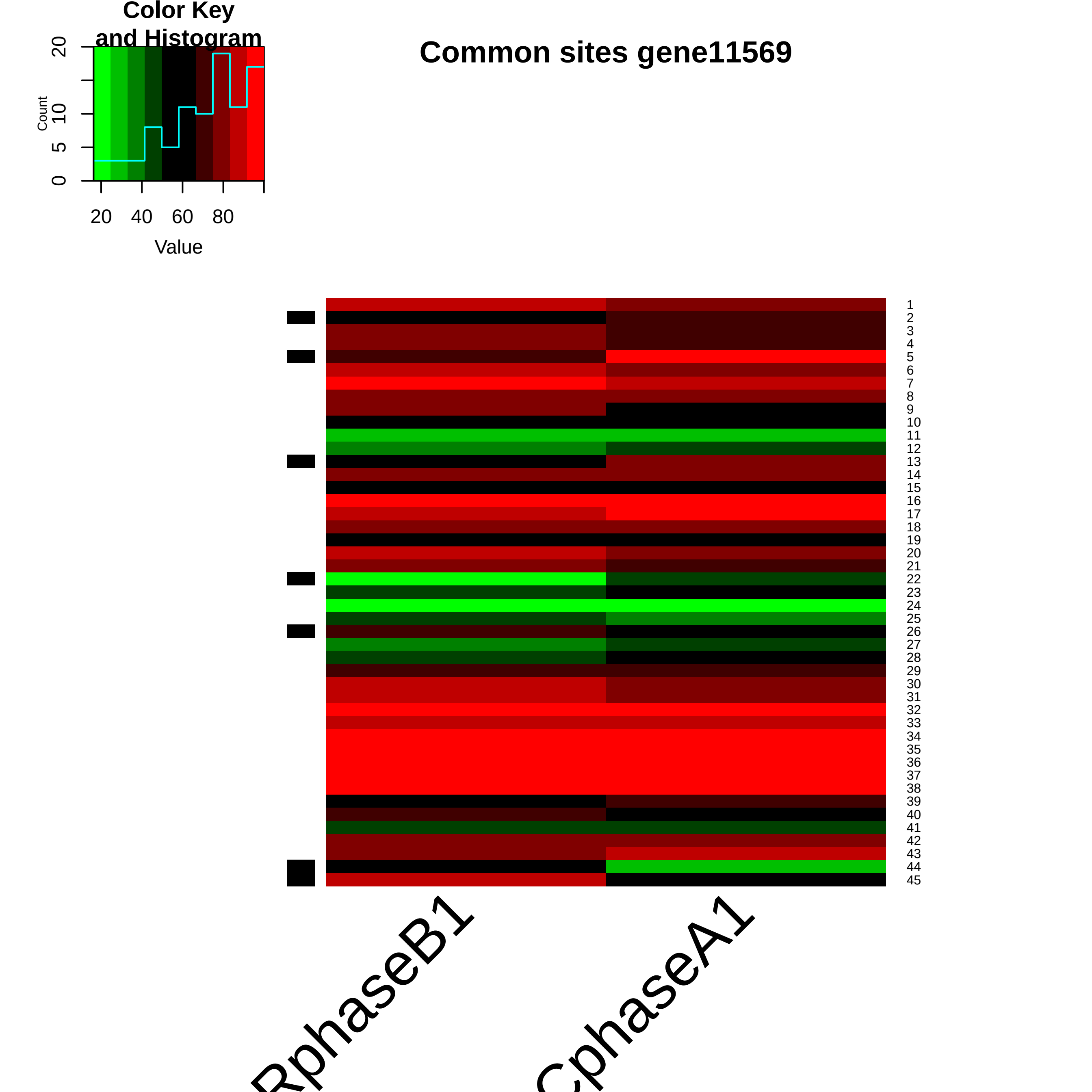

Color Key
and Histogram
Count
0
5
10
20
20
40
60
80
Value
Common sites gene11569
RphaseB1
CphaseA1

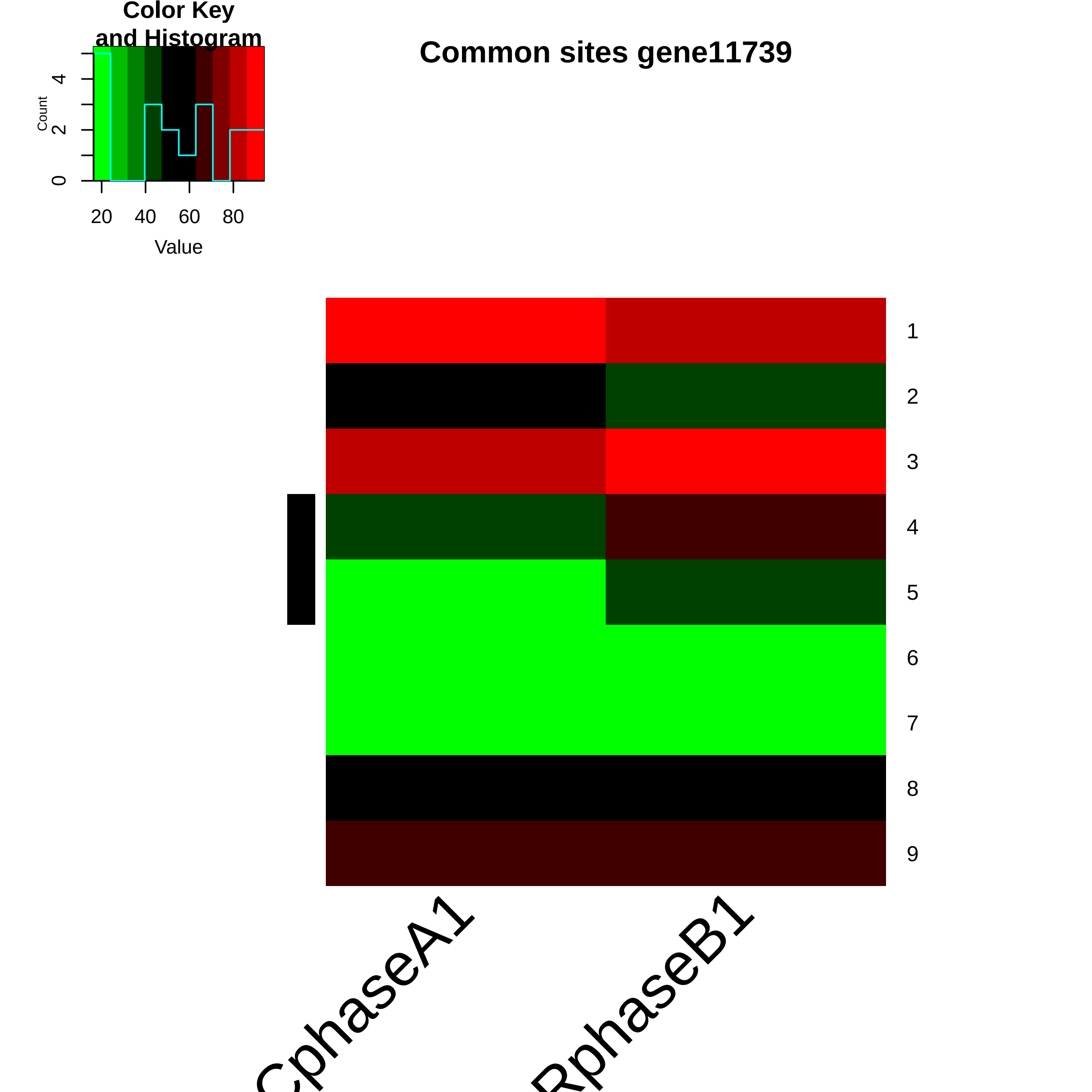

Color Key
and Histogram
Count
0
2
4
20
40
60
80
Value
Common sites gene11739
1
2
3
4
5
6
7
8
9

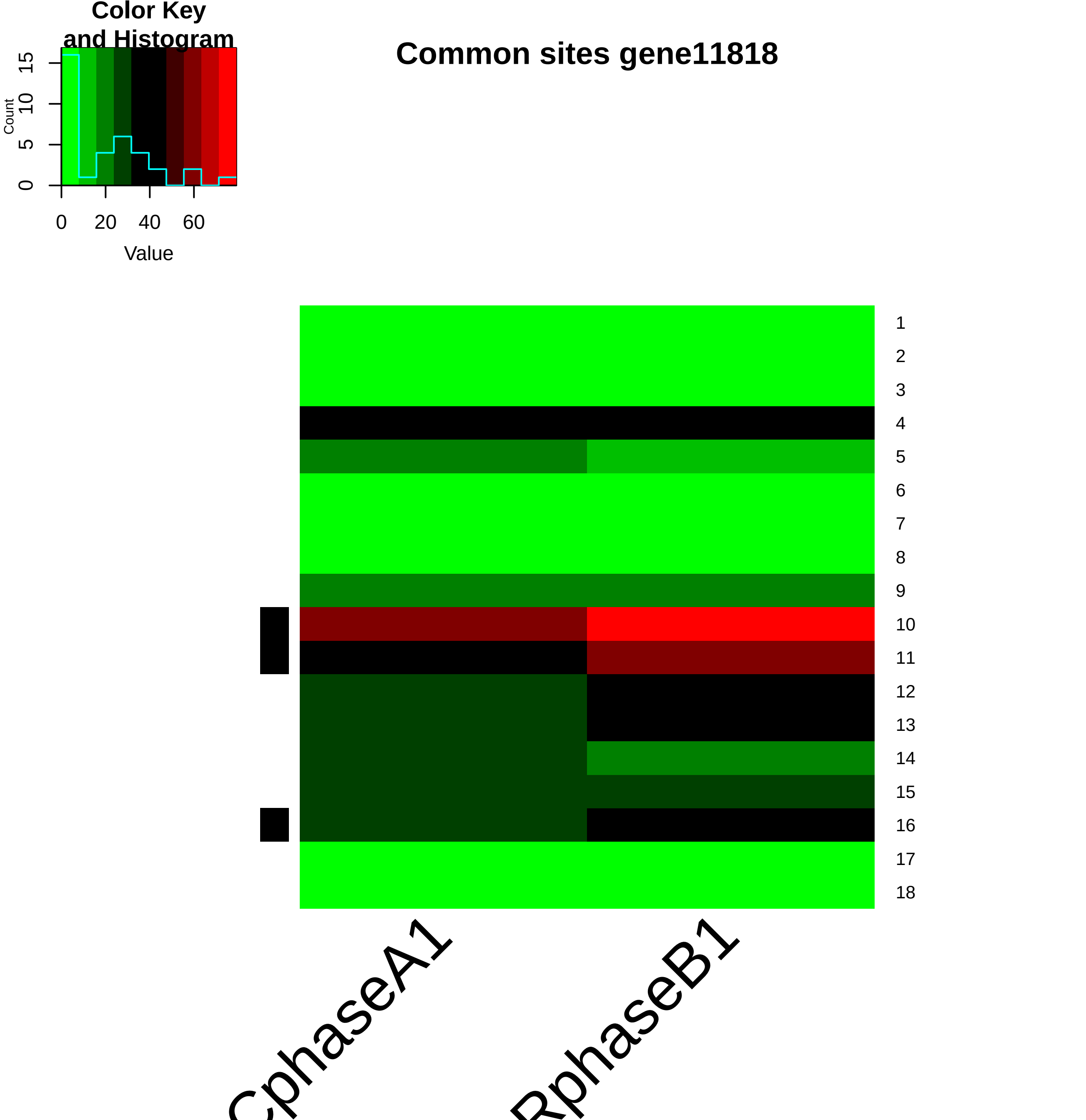

Color Key
and Histogram
Count
0
5
10
15
0
20
40
60
Value
Common sites gene11818
CphaseA1
RphaseB1

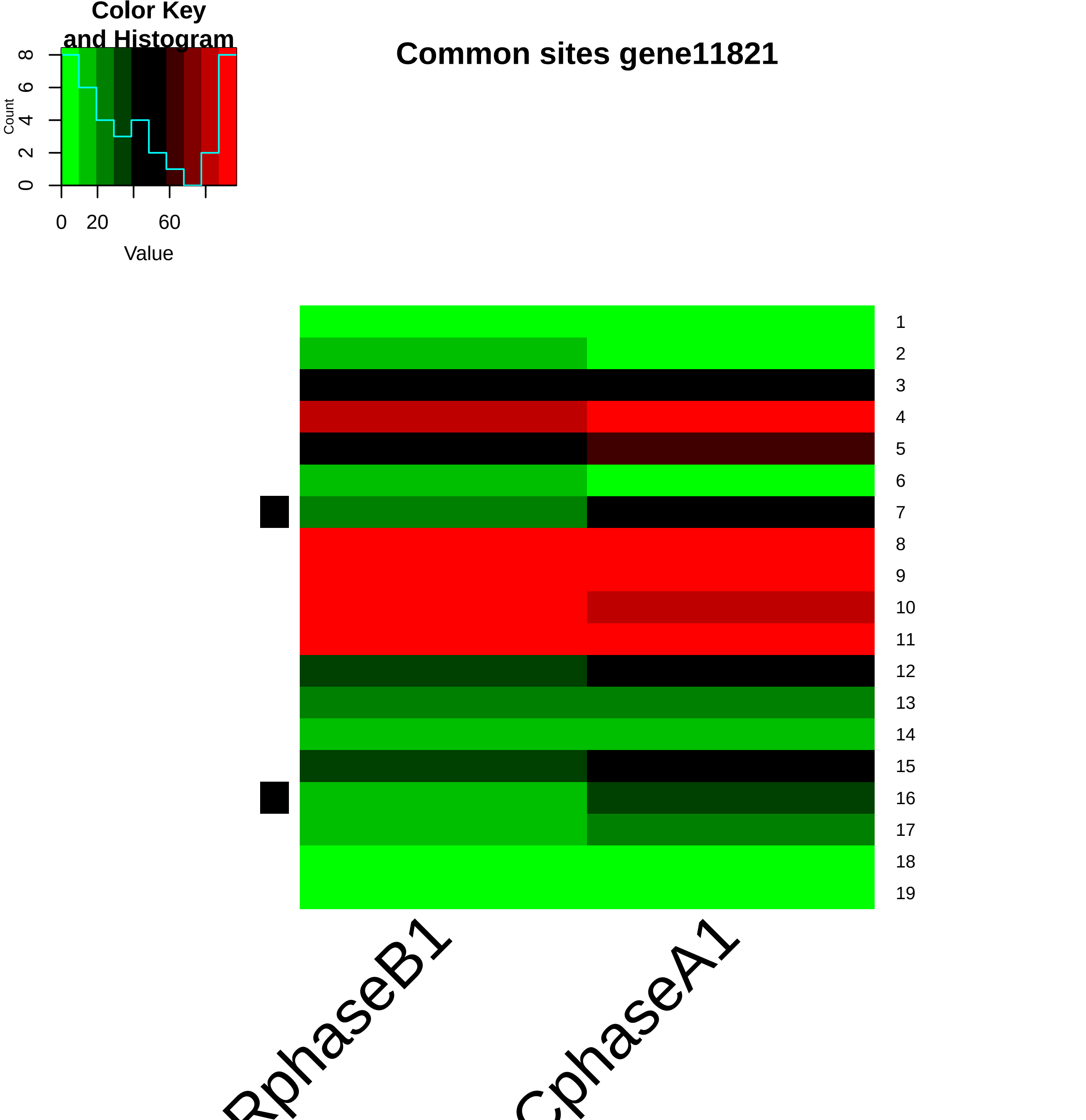
Color Key
and Histogram
Count
0
2
4
6
8
0
20
60
Value
Common sites gene11821
RphaseB1
CphaseA1

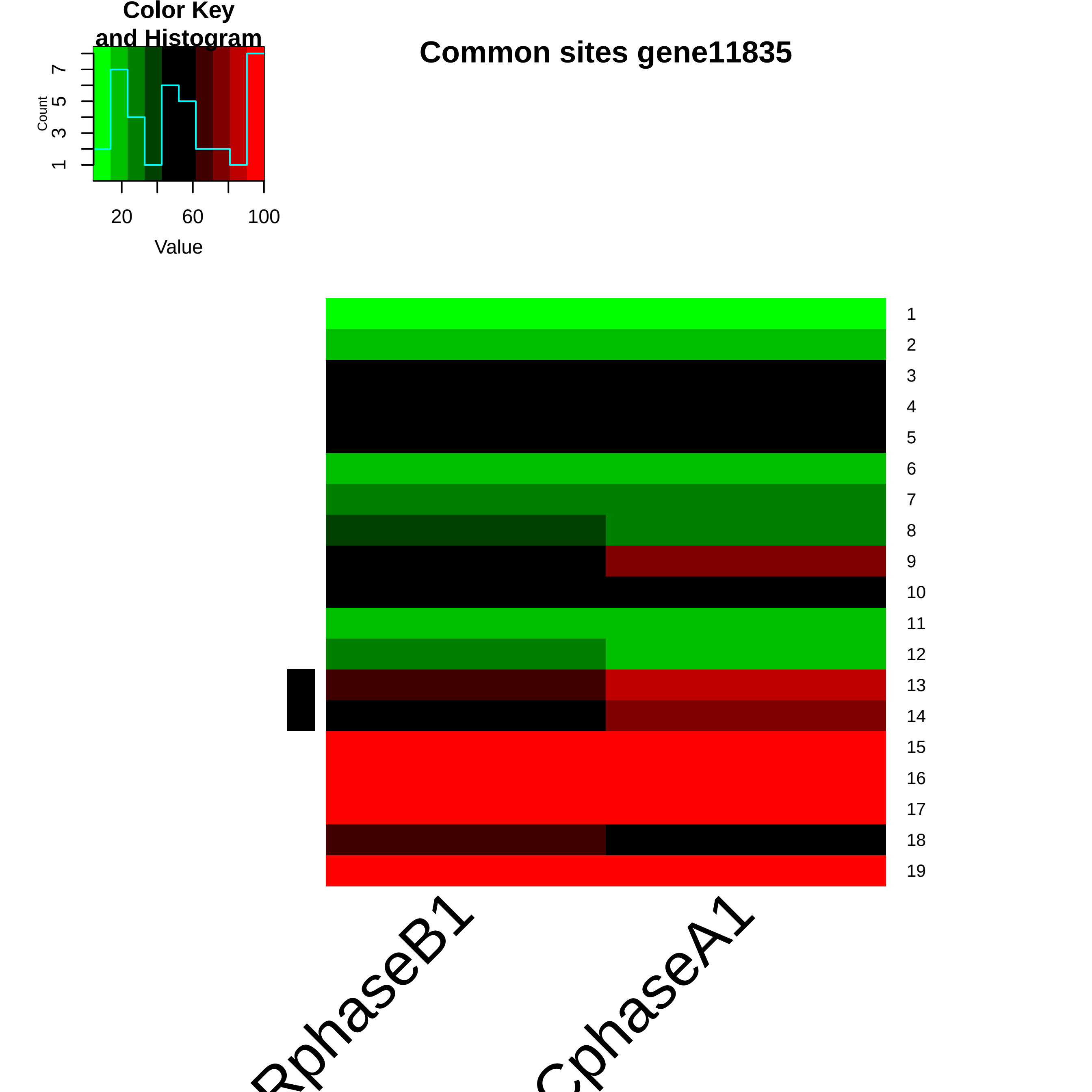

Color Key
and Histogram
Count
1
3
5
7
20
60
100
Value
Common sites gene11835
RphaseB1
CphaseA1

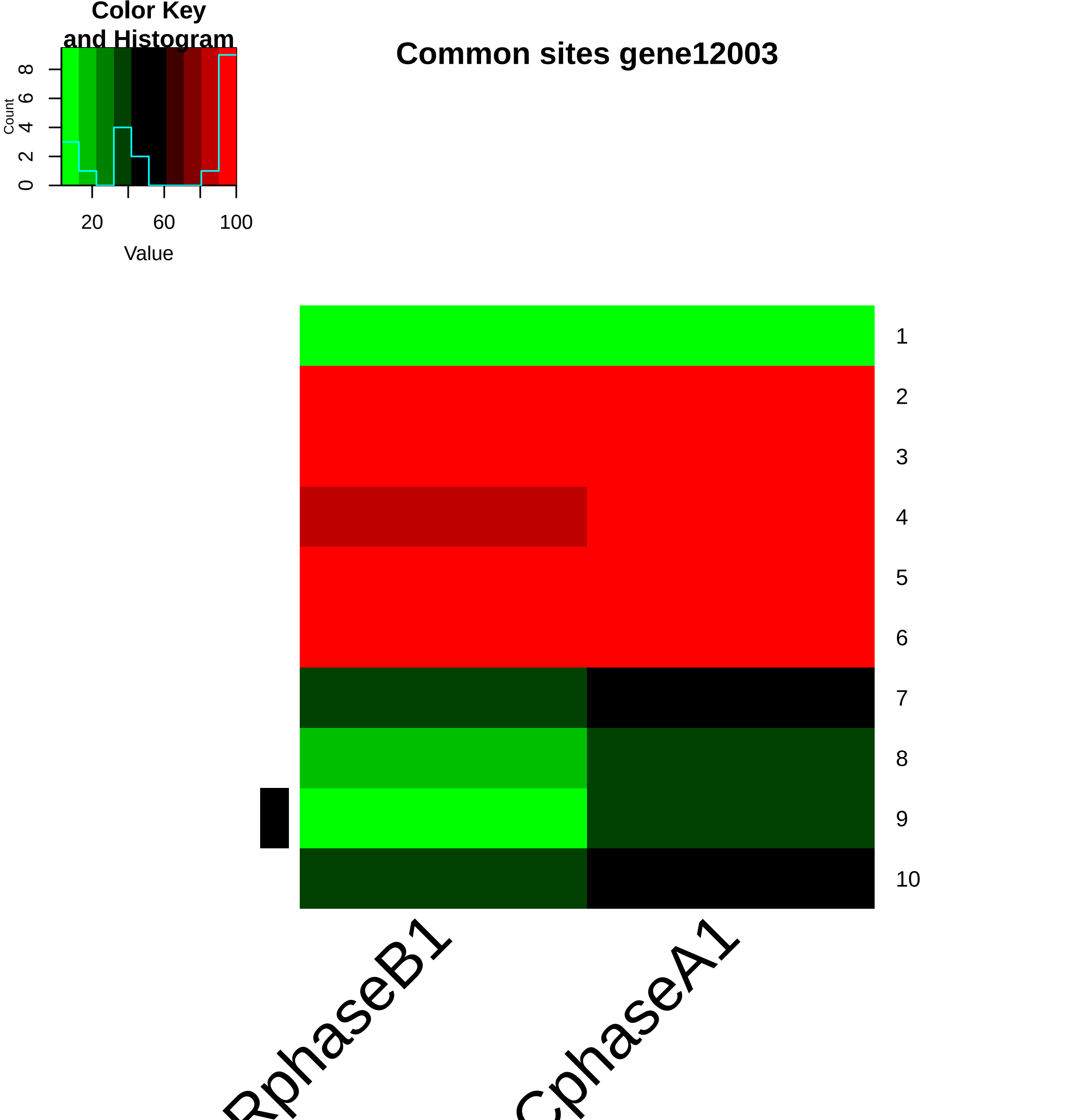
Color Key
and Histogram
Count
0
2
4
6
8
20
60
100
Value
Common sites gene12003
1
2
3
4
5
6
7
8
9
10
RphaseB1
CphaseA1

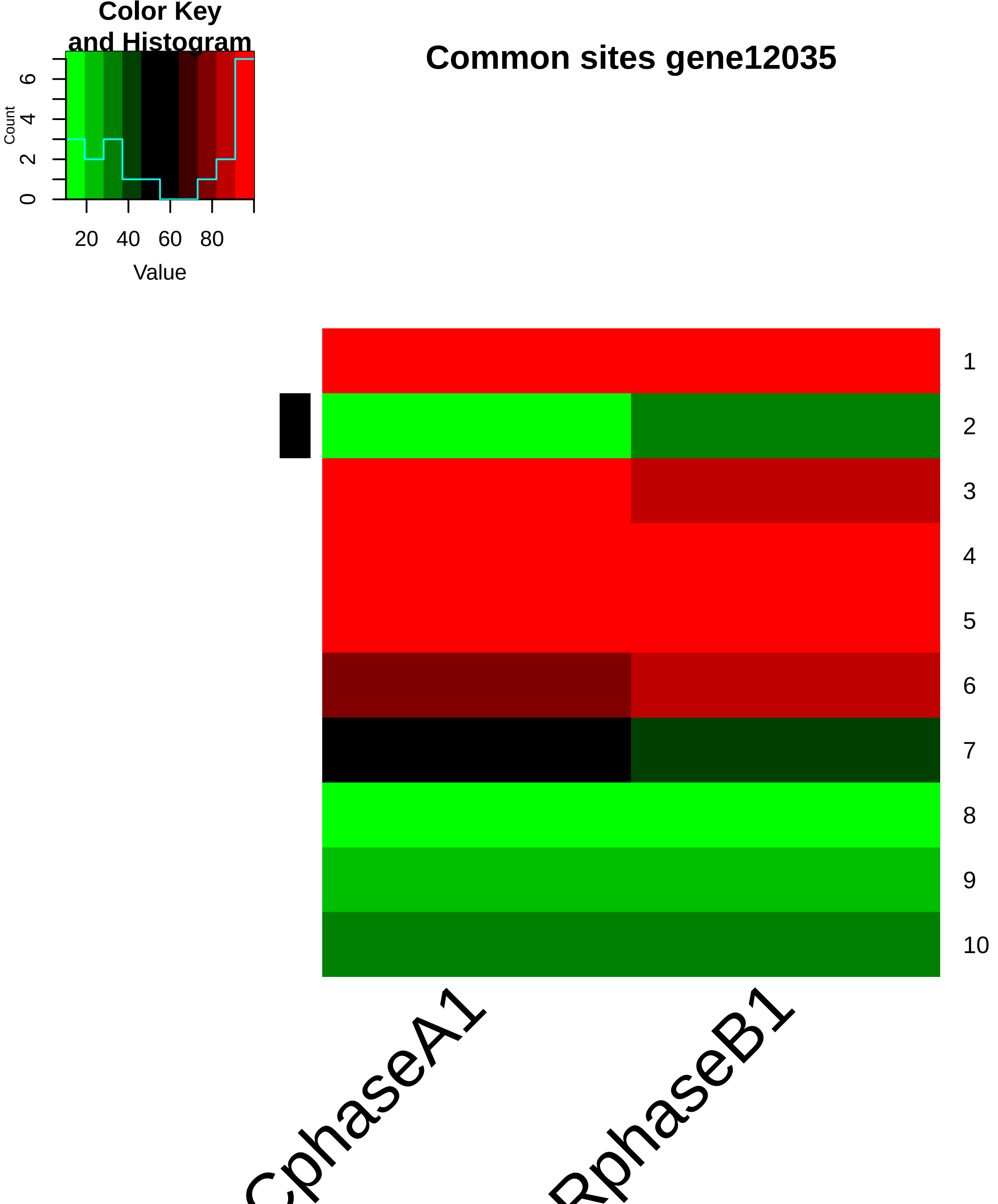
Color Key
and Histogram
Count
0
2
4
6
20
40
60
80
Value
Common sites gene12035
1
2
3
4
5
6
7
8
9
10
CphaseA1
RphaseB1

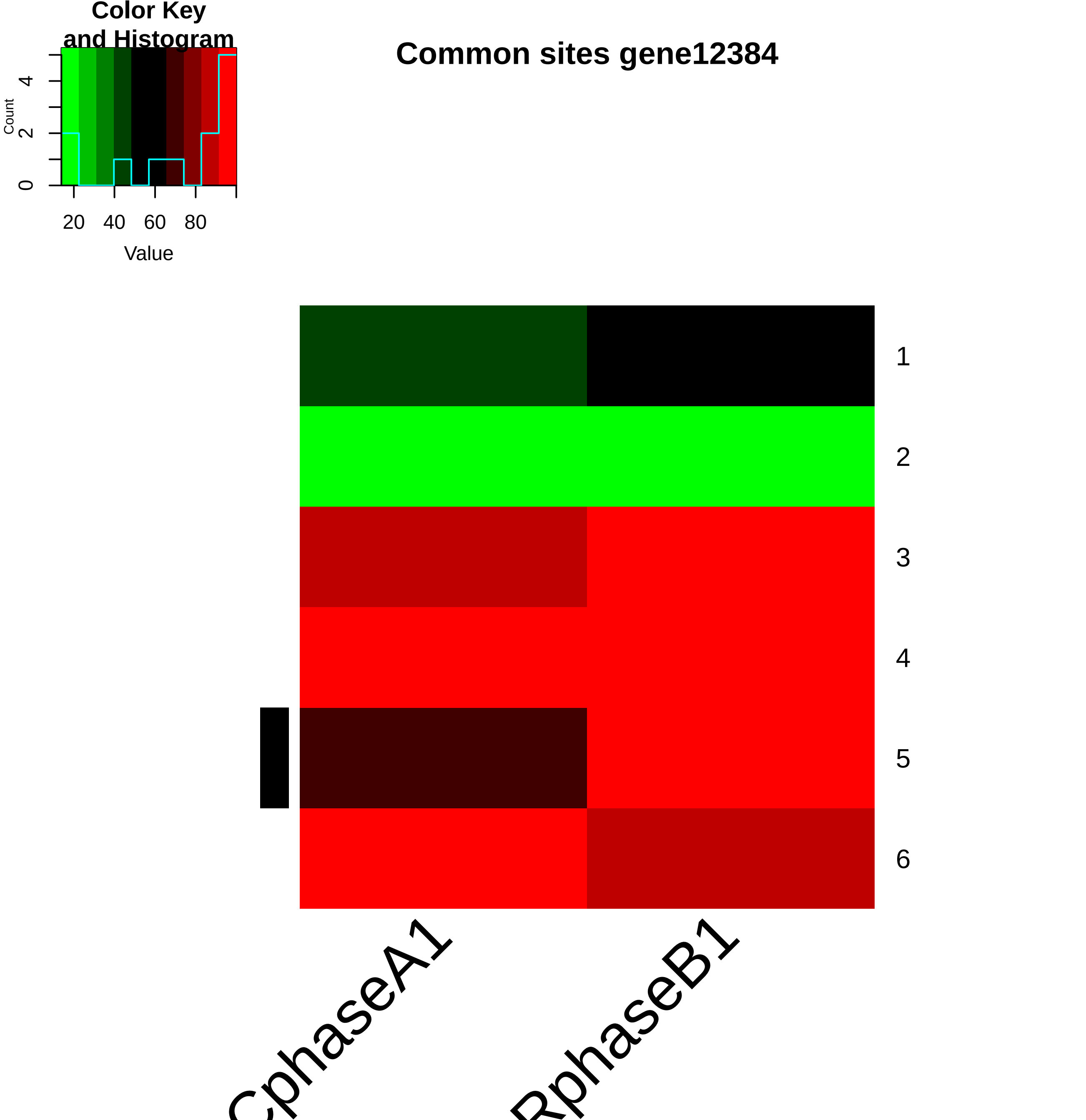
Color Key
and Histogram
Count
0
2
4
20
40
60
80
Value
Common sites gene12384
1
2
3
4
5
6
CphaseA1
RphaseB1

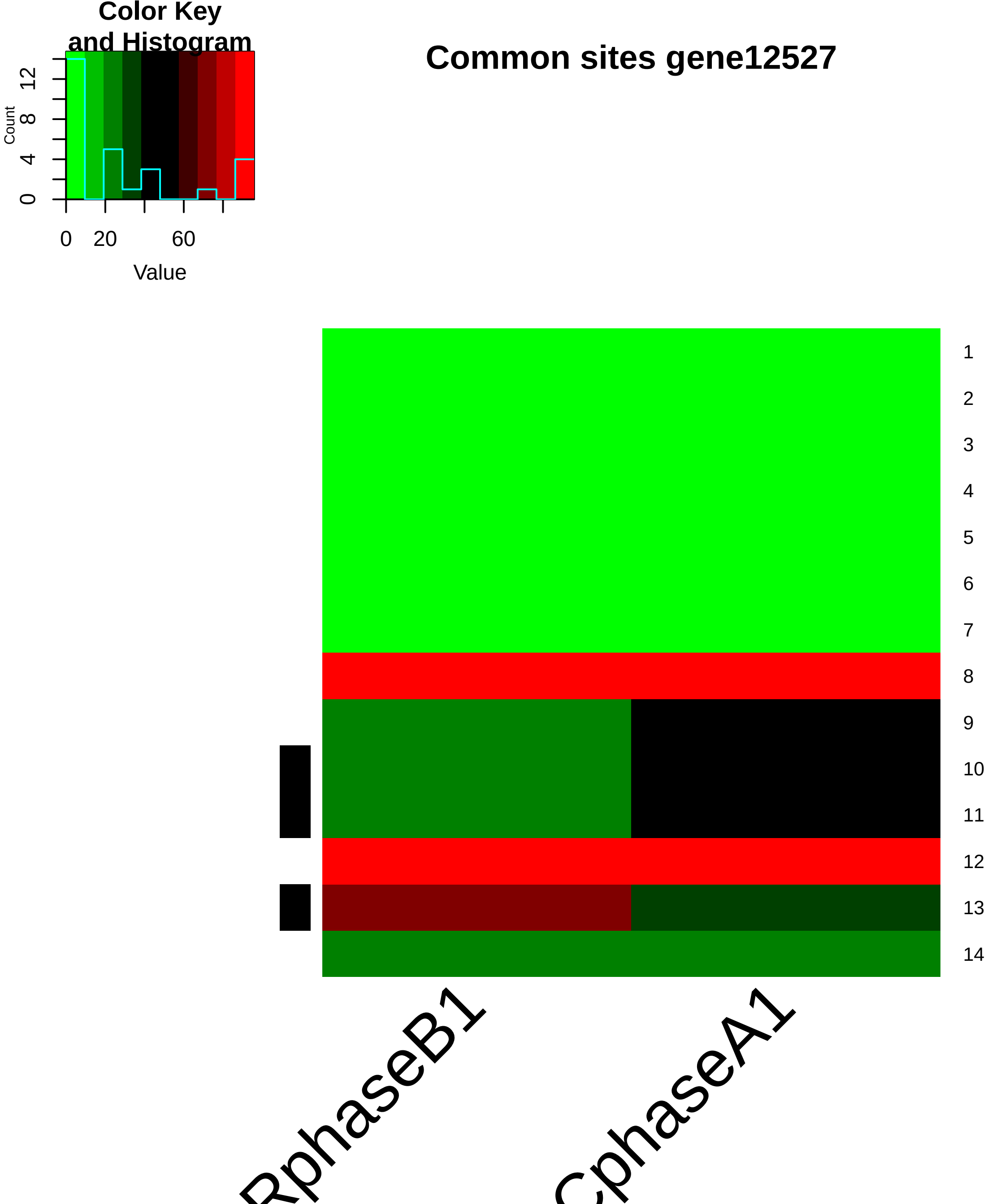

Color Key
and Histogram
Count
0
4
8
12
0
20
60
Value
Common sites gene12527
1
2
3
4
5
6
7
8
9
10
11
12
13
14
RphaseB1
CphaseA1

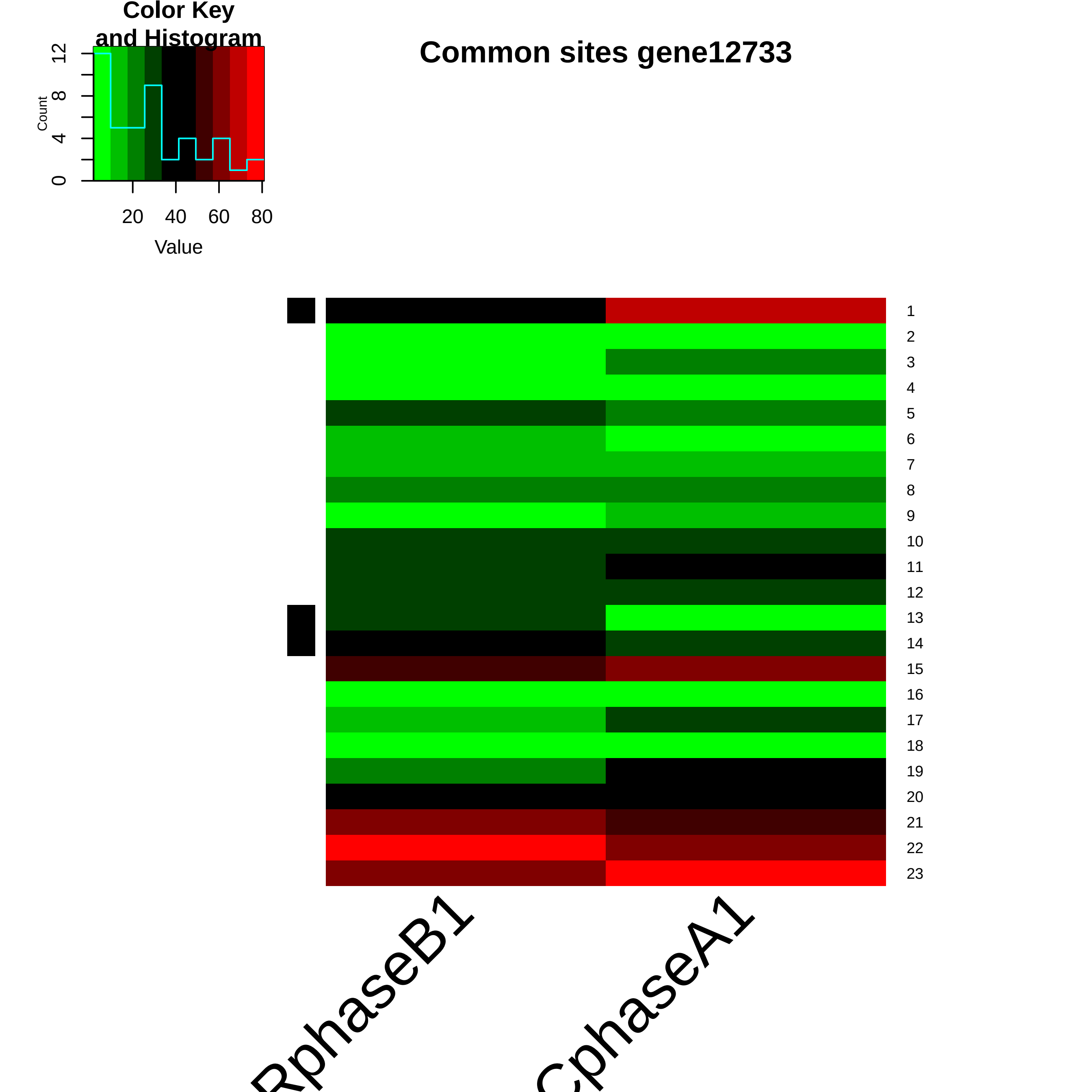
Color Key
and Histogram
Count
0
4
8
12
20
40
60
80
Value
Common sites gene12733
RphaseB1
CphaseA1

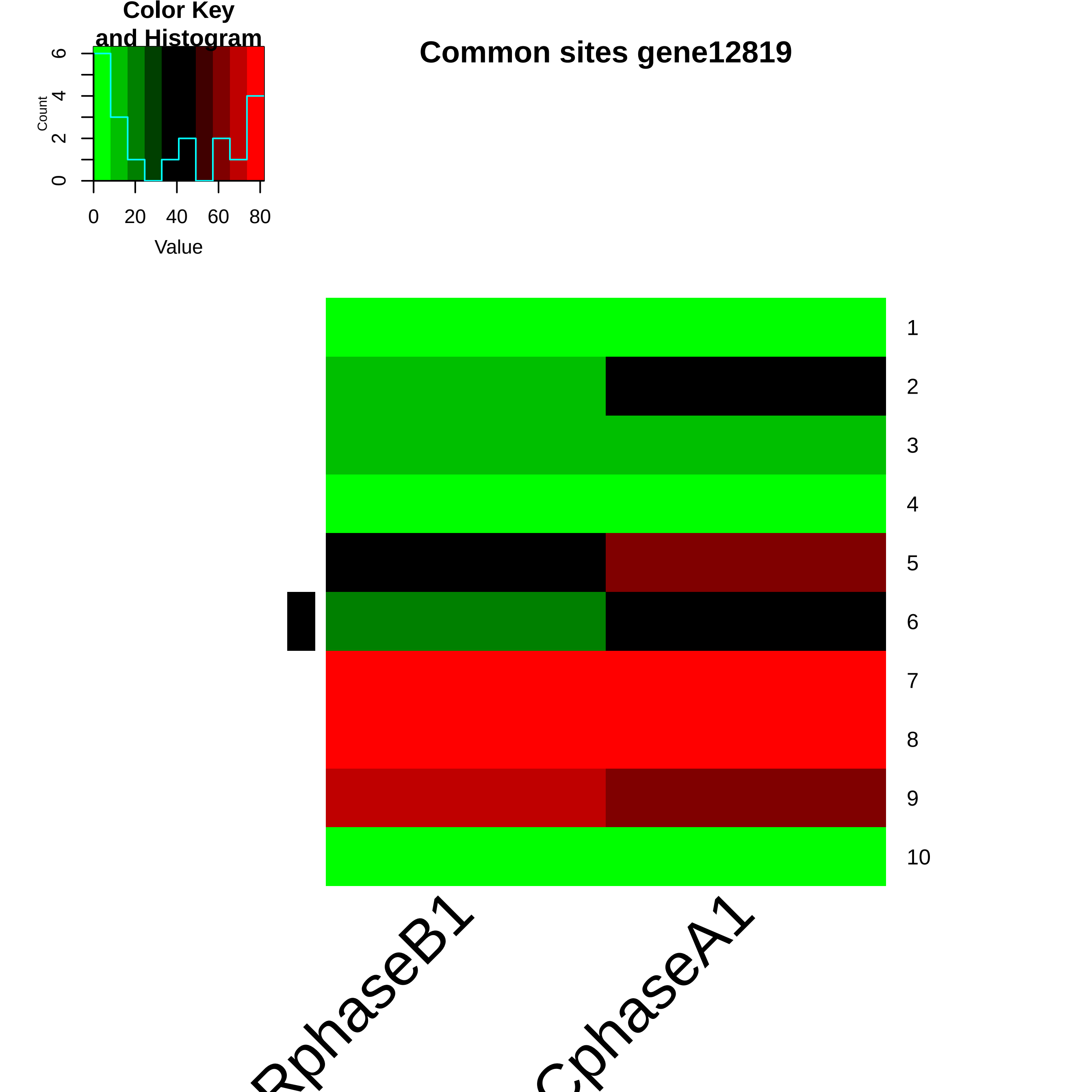
Color Key
and Histogram
Count
0
2
4
6
0
20
40
60
80
Value
Common sites gene12819
1
2
3
4
5
6
7
8
9
10
RphaseB1
CphaseA1

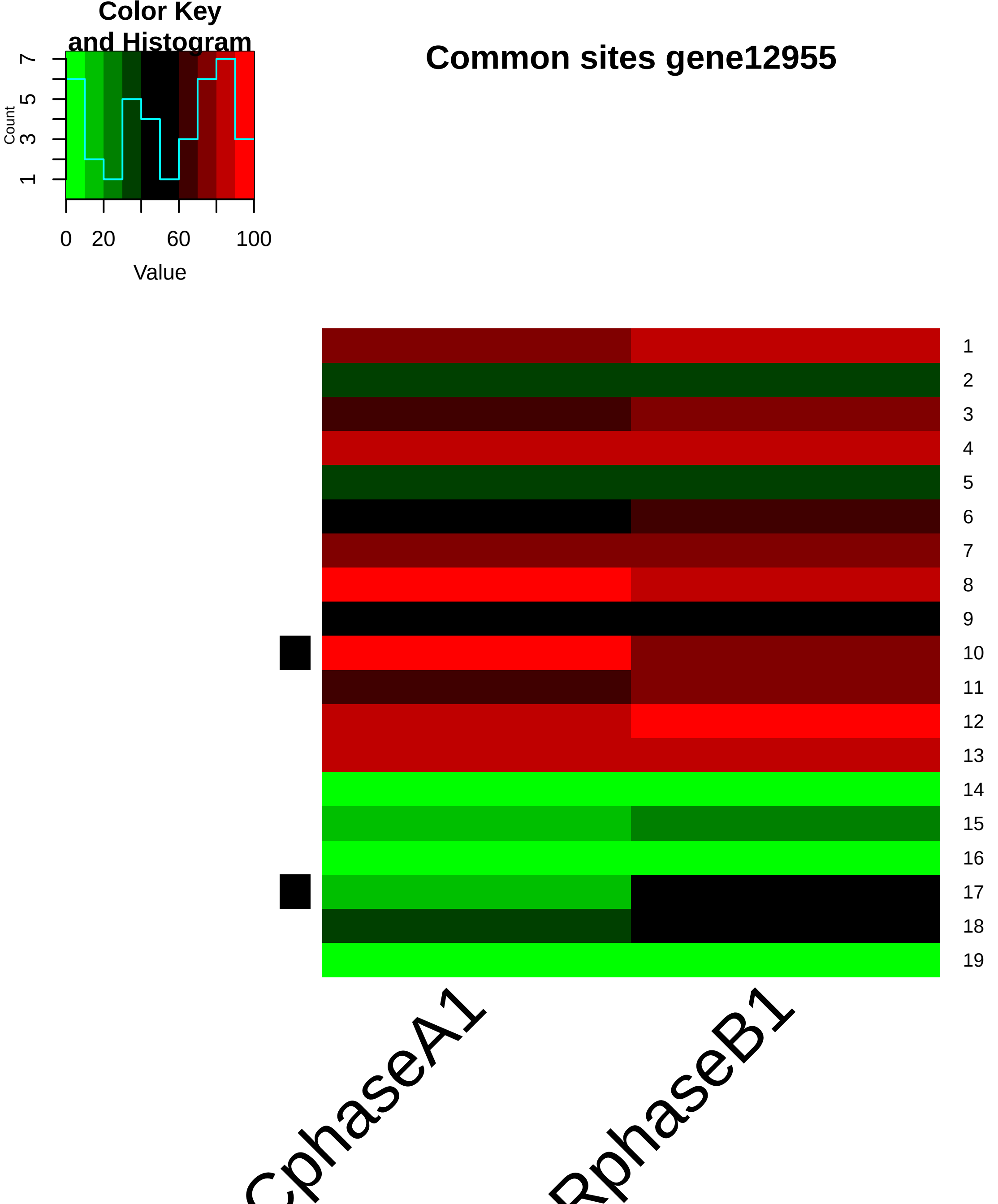

Color Key
and Histogram
Count
1
3
5
7
0
20
60
100
Value
Common sites gene12955
CphaseA1
RphaseB1

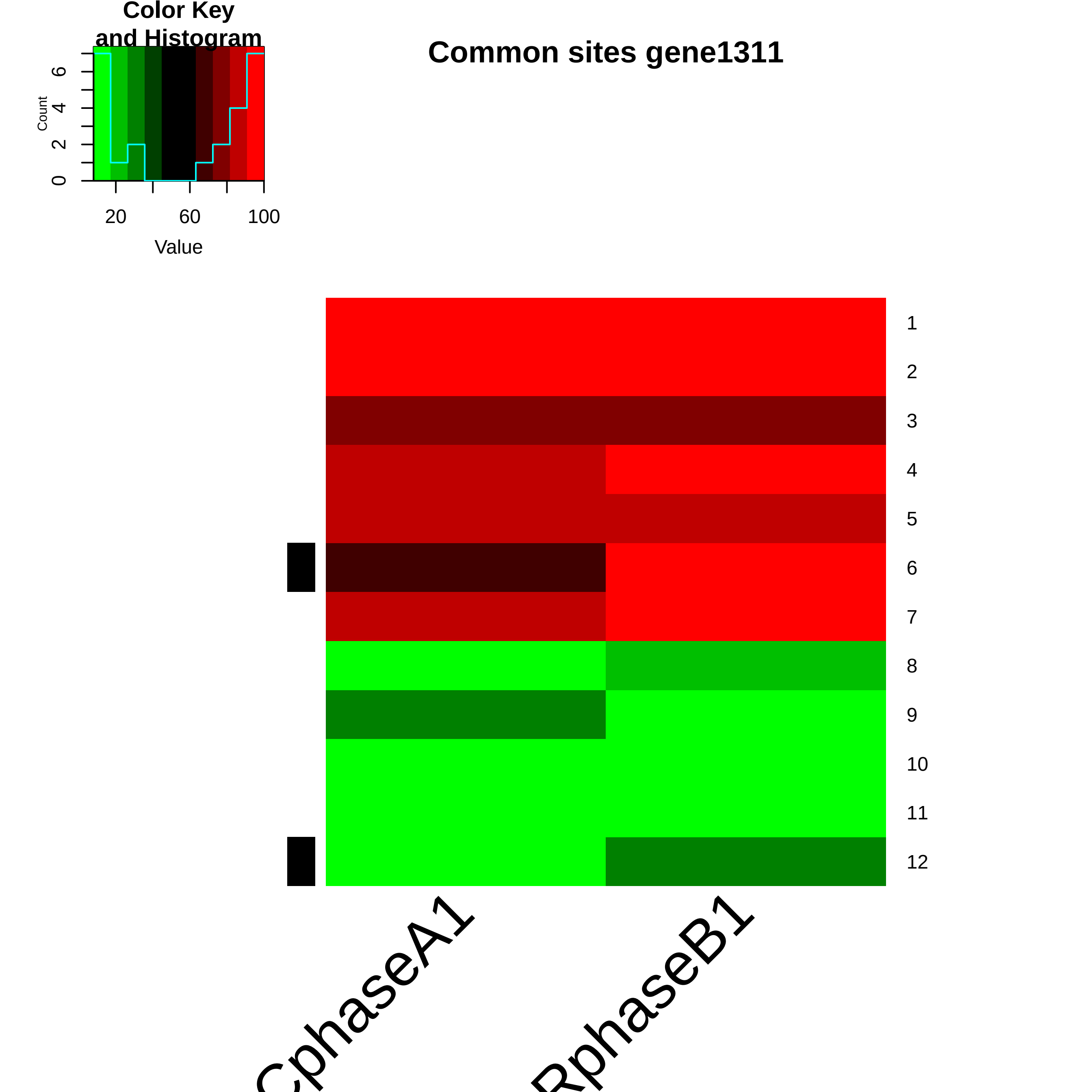

Color Key
and Histogram
Count
0
2
4
6
20
60
100
Value
Common sites gene1311
1
2
3
4
5
6
7
8
9
10
11
12
CphaseA1
RphaseB1

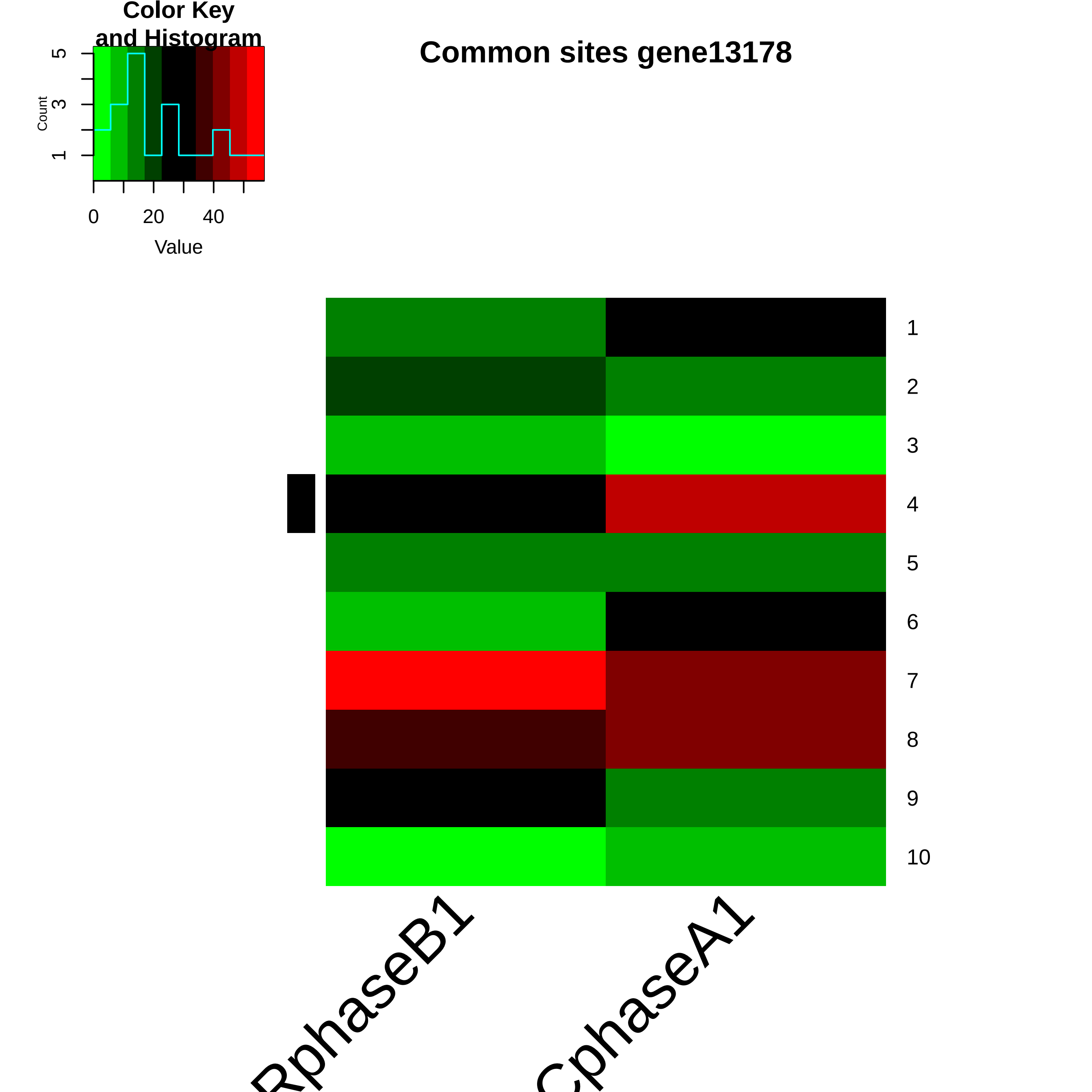
Color Key
and Histogram
Count
5
3
1
0
20
40
Value
Common sites gene13178
1
2
3
4
5
6
7
8
9
10
RphaseB1
CphaseA1

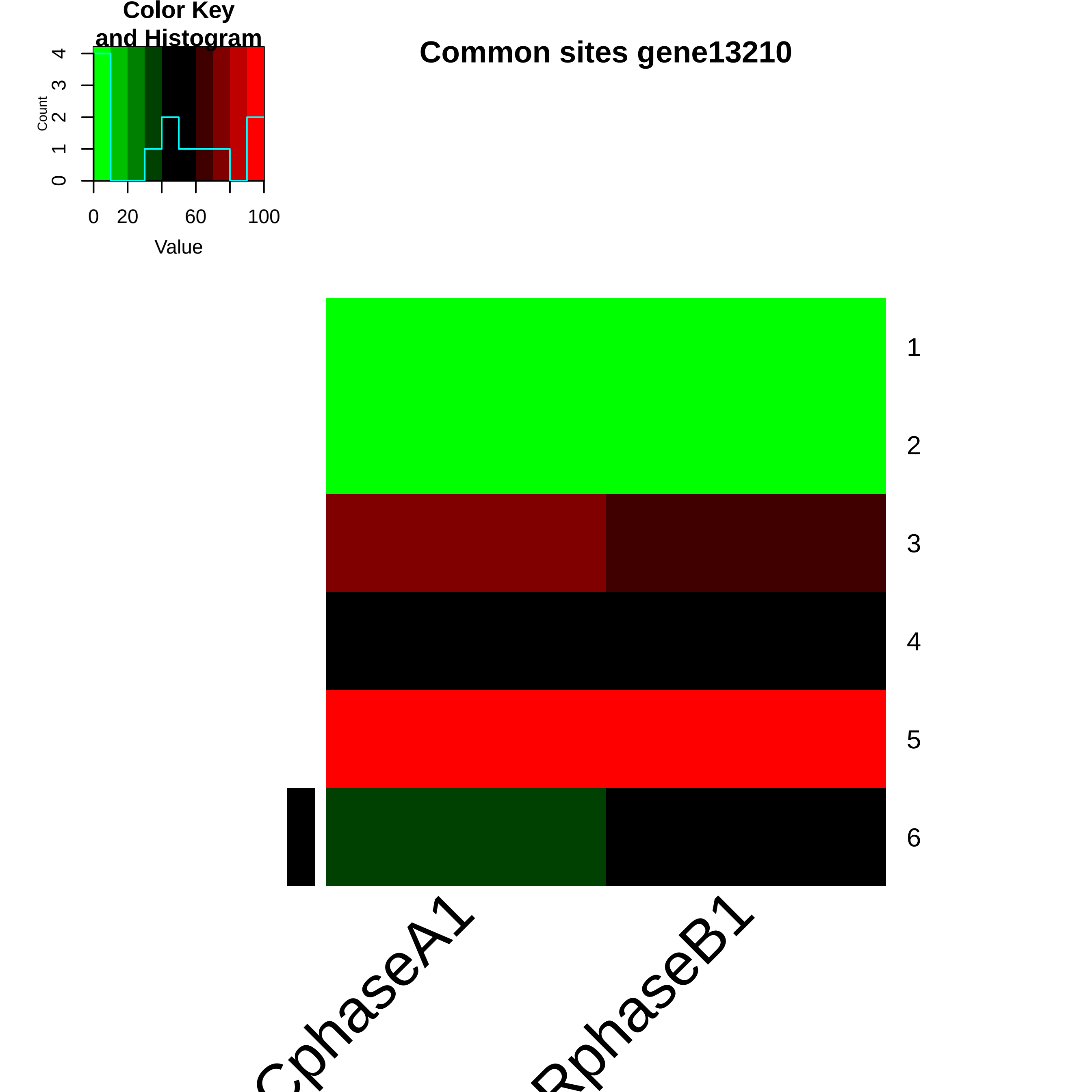

Color Key
and Histogram
Count
0
1
2
3
4
0
20
60
100
Value
Common sites gene13210
1
2
3
4
5
6
CphaseA1
RphaseB1

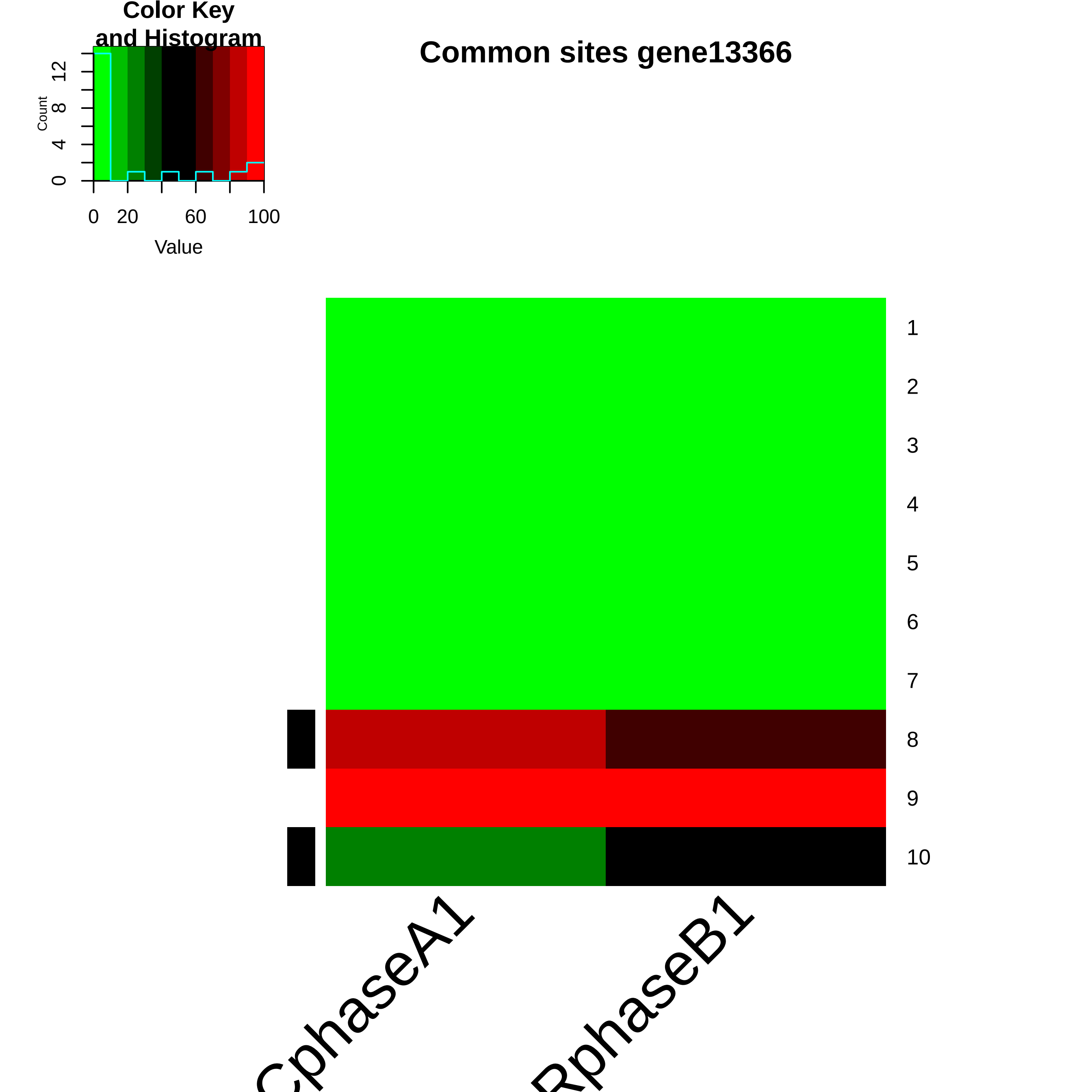

Color Key
and Histogram
Count
0
4
8
12
0
20
60
100
Value
Common sites gene13366
1
2
3
4
5
6
7
8
9
10
CphaseA1
RphaseB1

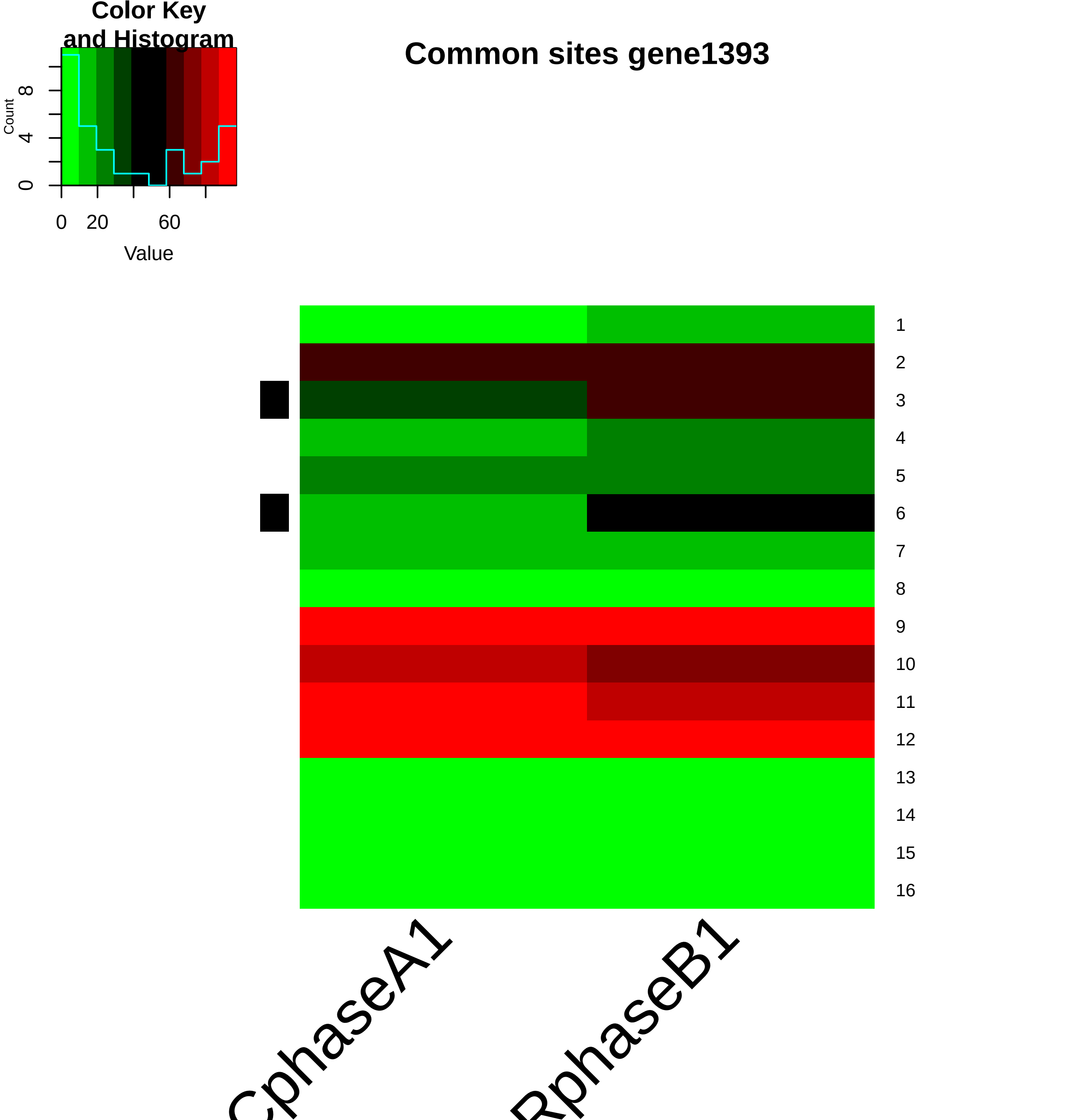
Color Key
and Histogram
Count
0
4
8
0
20
60
Value
Common sites gene1393
CphaseA1
RphaseB1

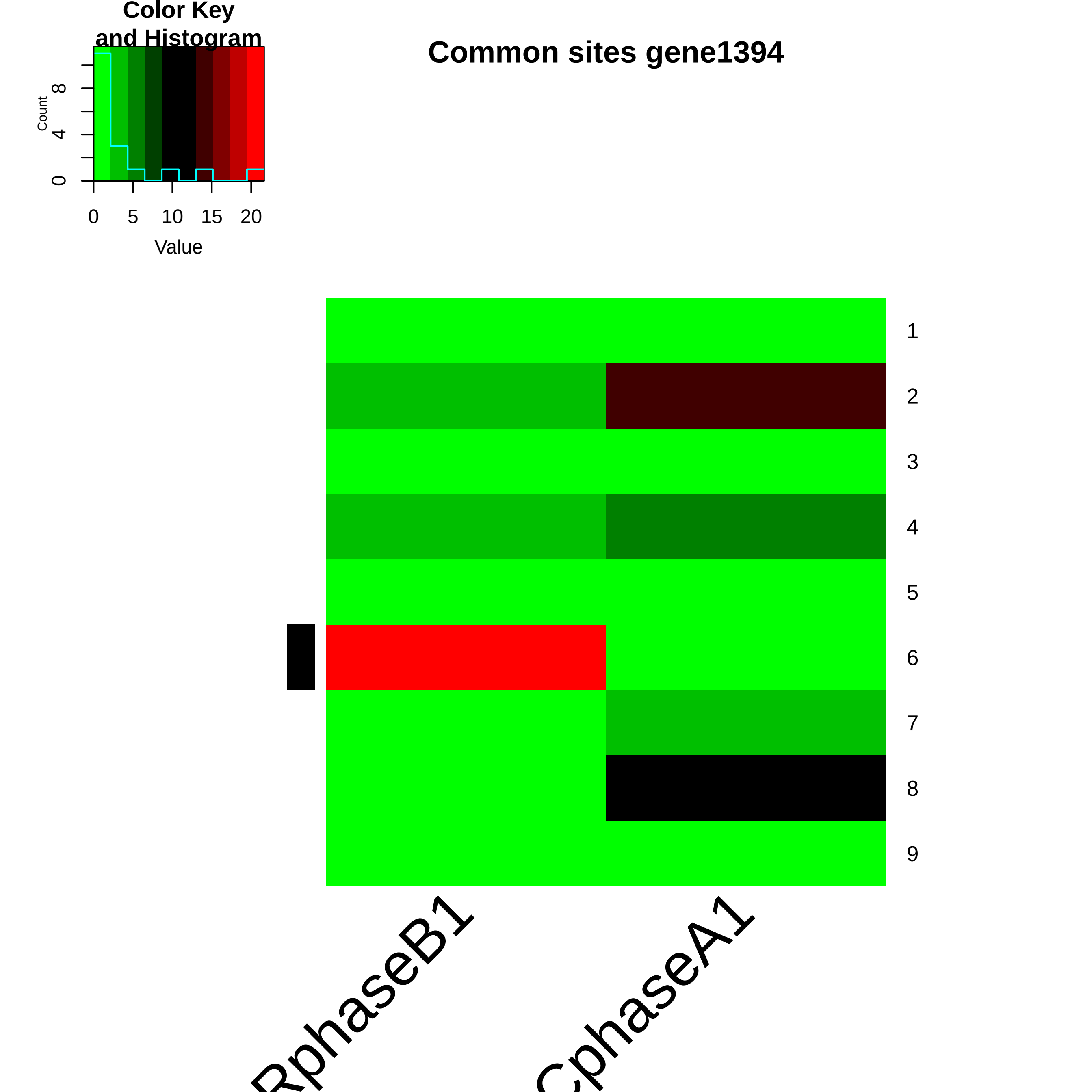
Color Key
and Histogram
Count
0
4
8
0
5
10
15
20
Value
Common sites gene1394
1
2
3
4
5
6
7
8
9
RphaseB1
CphaseA1

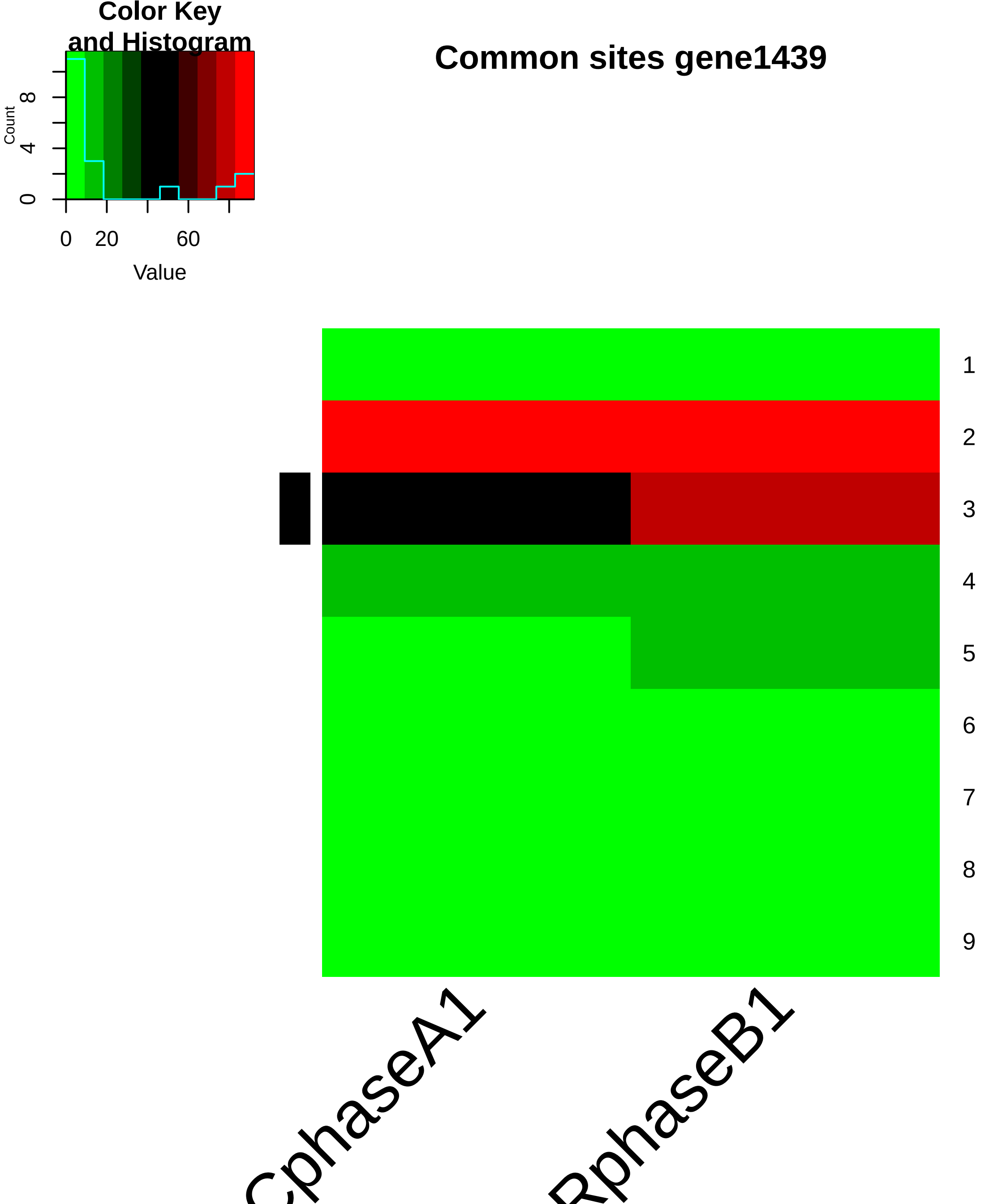
Color Key
and Histogram
Count
0
4
8
0
20
60
Value
Common sites gene1439
1
2
3
4
5
6
7
8
9
CphaseA1
RphaseB1

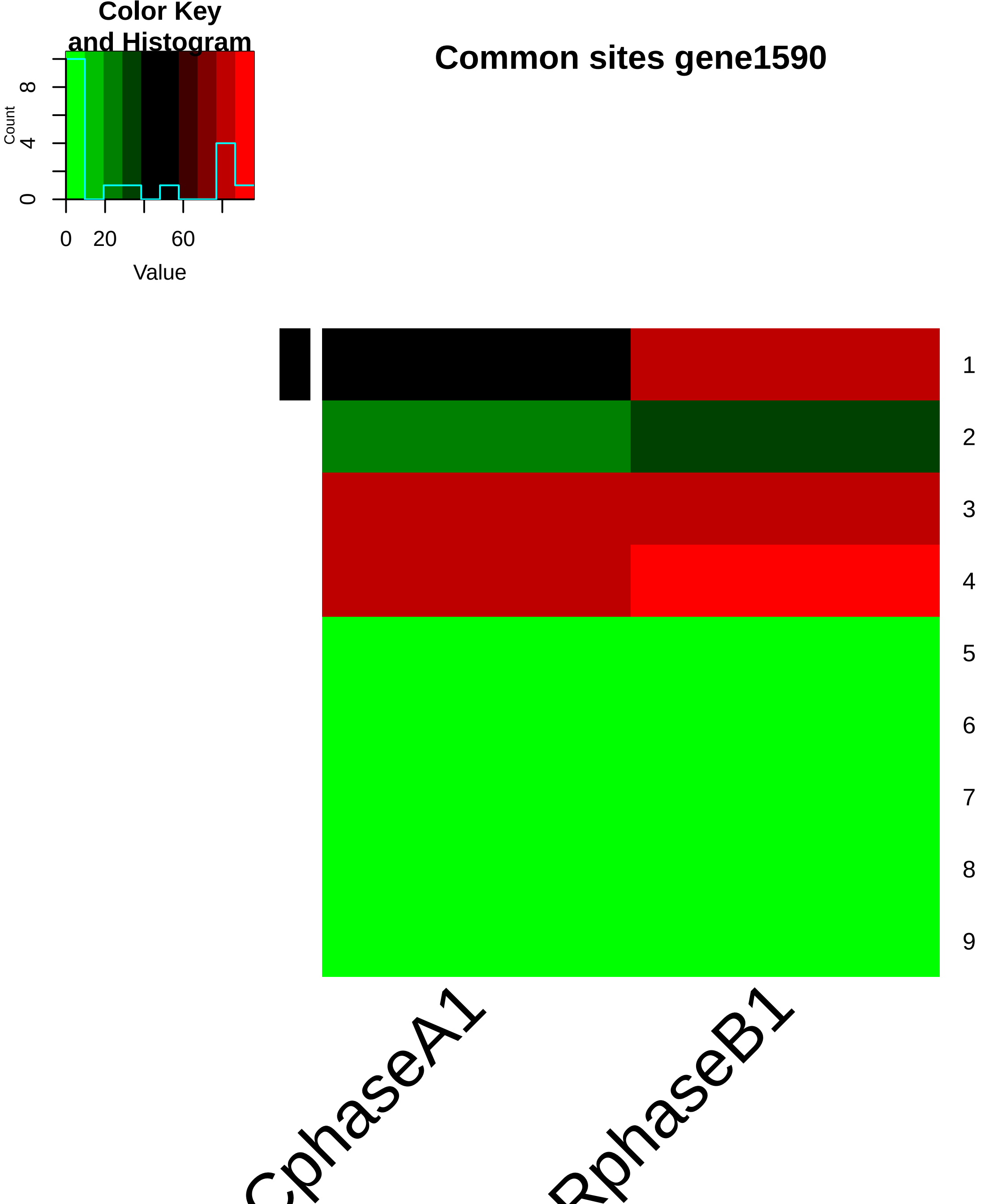

Color Key
and Histogram
Count
0
4
8
0
20
60
Value
Common sites gene1590
1
2
3
4
5
6
7
8
9
CphaseA1
RphaseB1

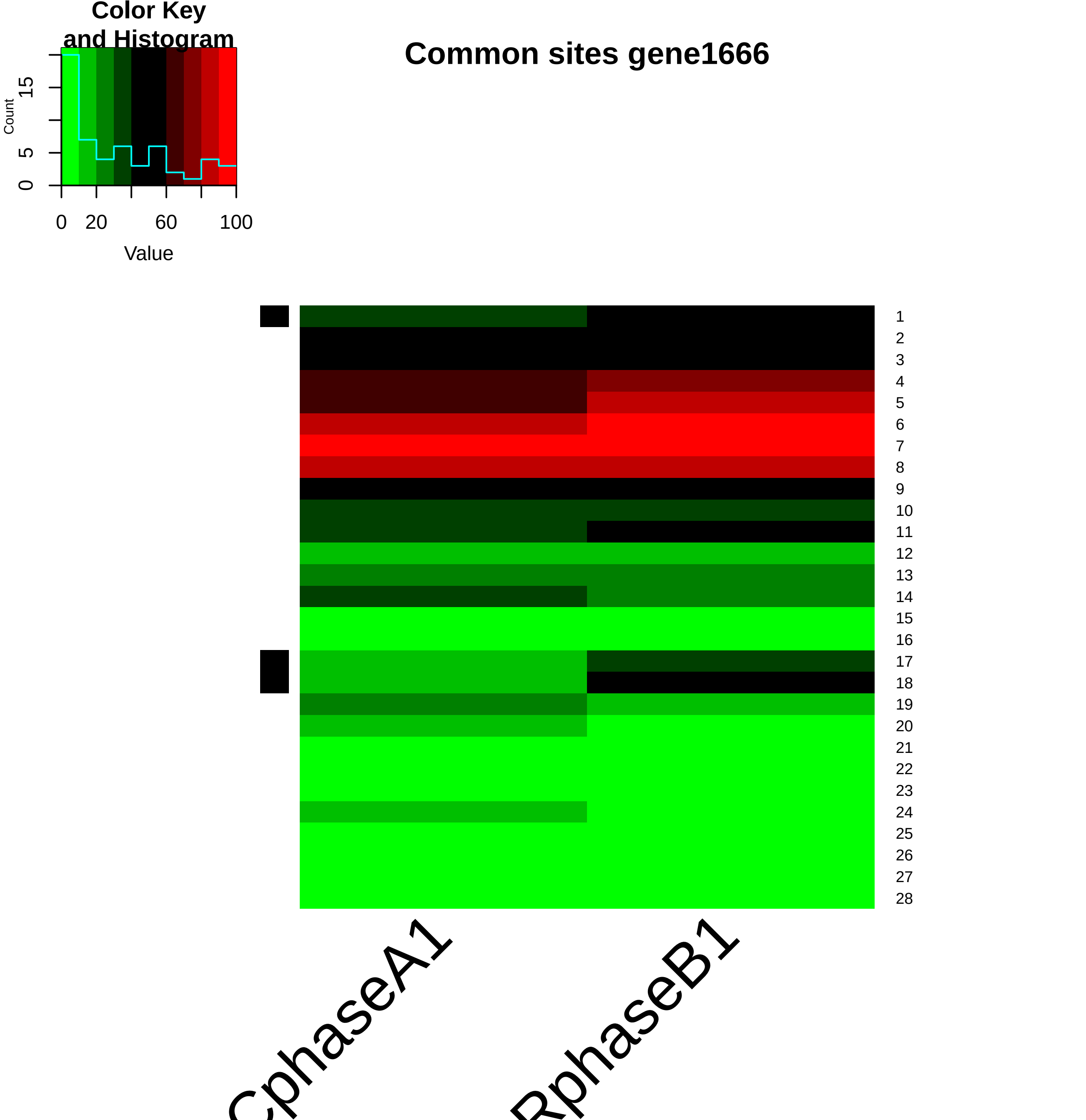

Color Key
and Histogram
Count
0
5
15
0
20
60
100
Value
Common sites gene1666
CphaseA1
RphaseB1

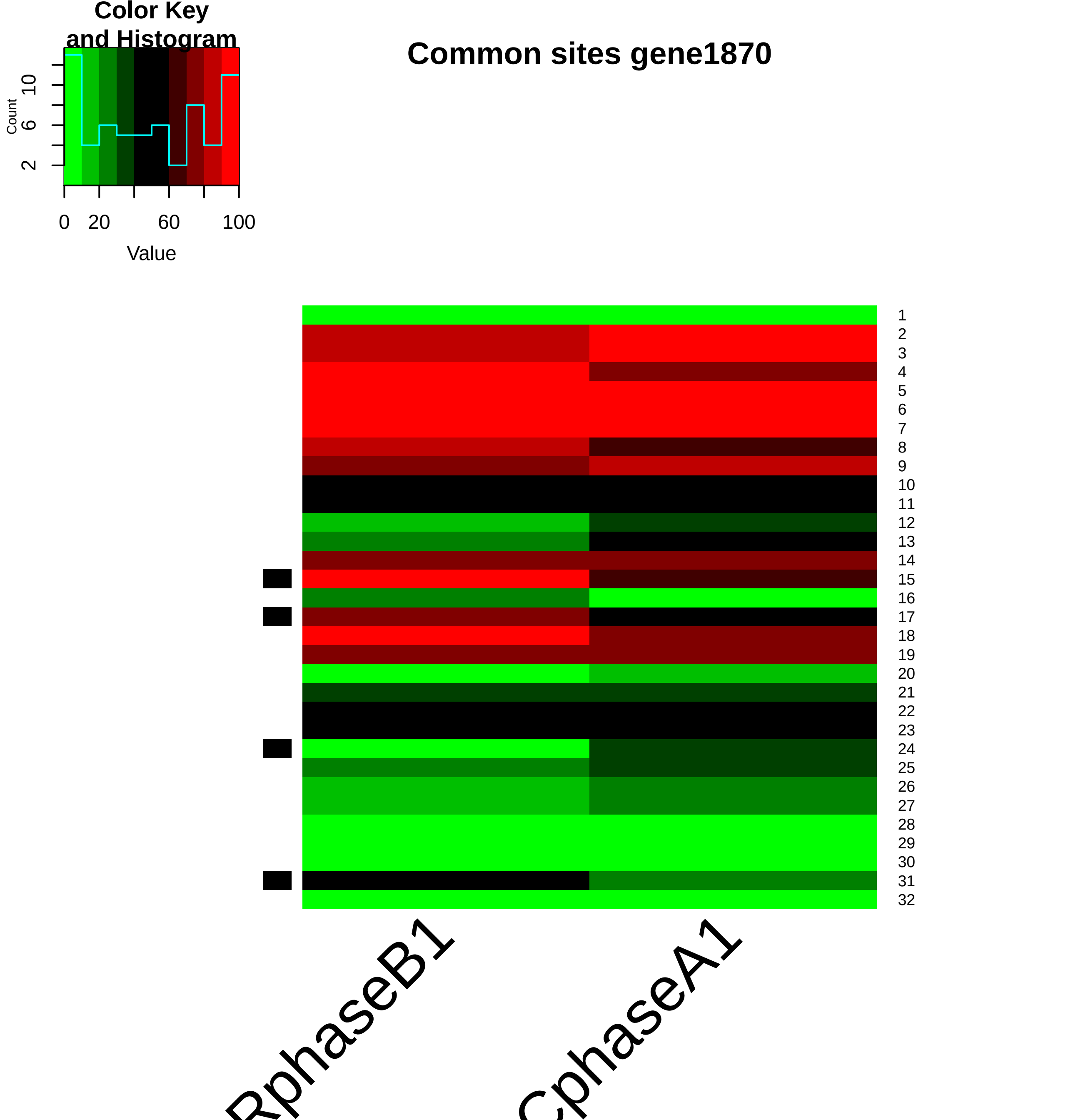
Color Key
and Histogram
Count
2
6
10
0
20
60
100
Value
Common sites gene1870
RphaseB1
CphaseA1

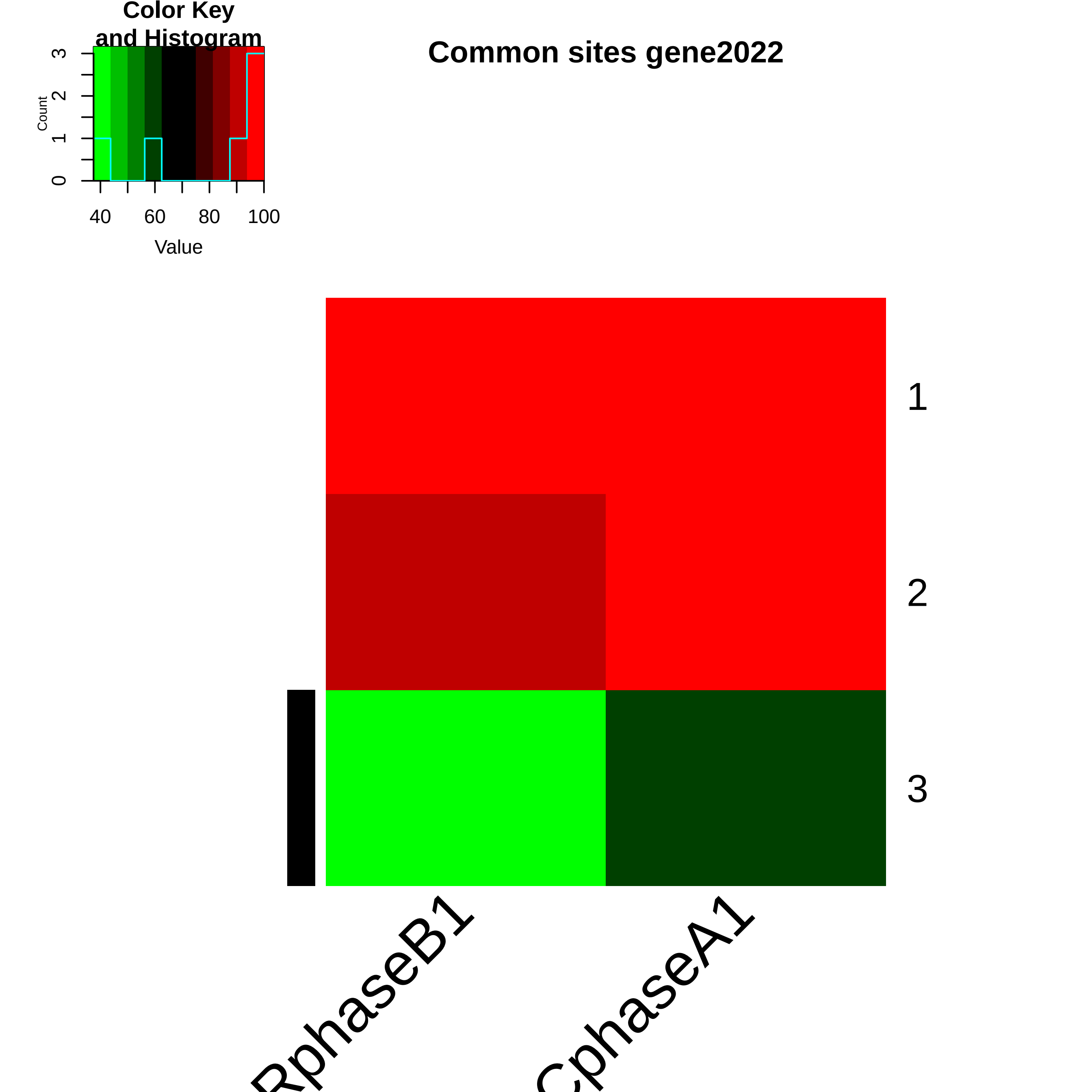
Color Key
and Histogram
Count
0
1
2
3
40
60
80
100
Value
Common sites gene2022
1
2
3
RphaseB1
CphaseA1

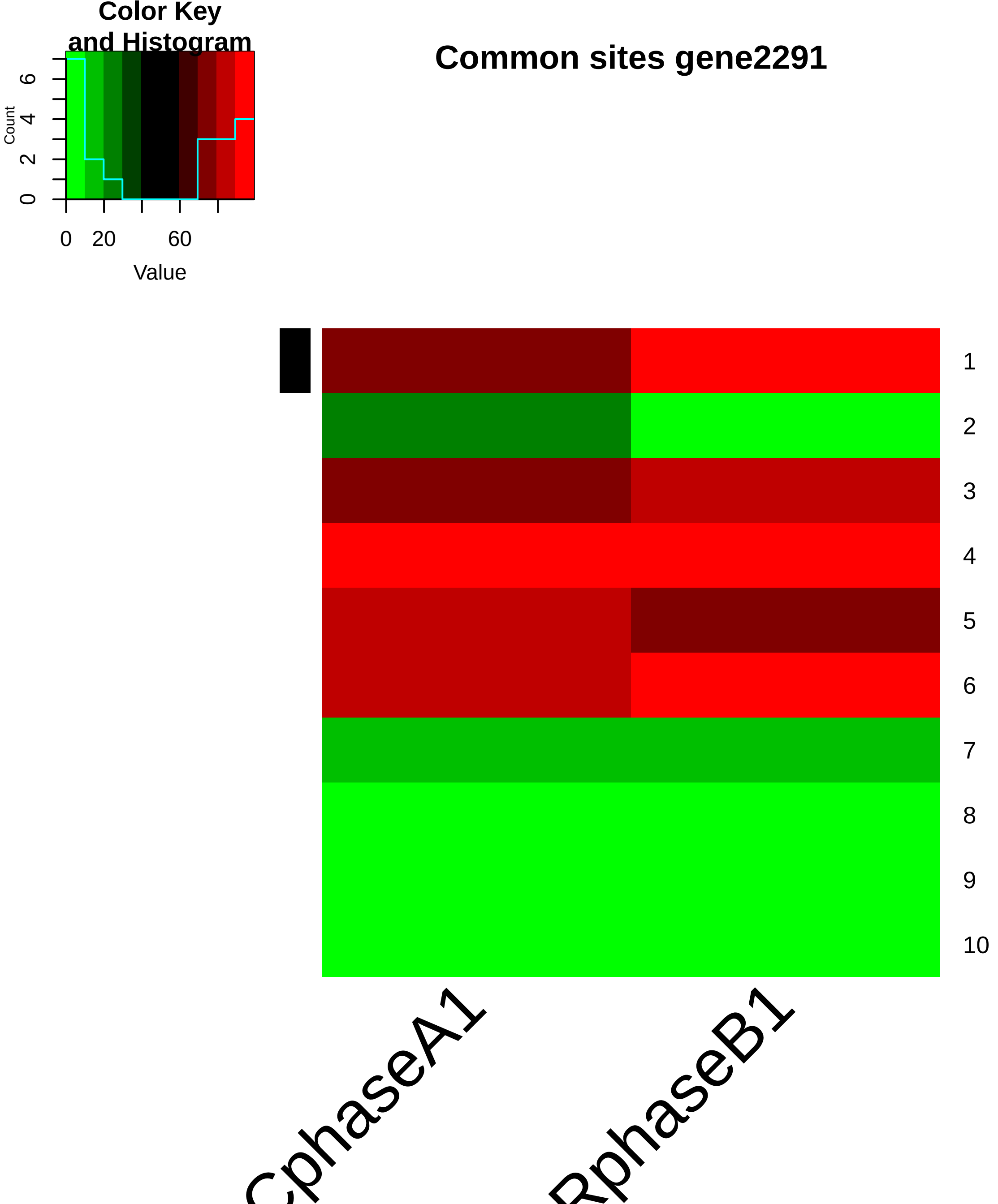
Color Key
and Histogram
Count
0
2
4
6
0
20
60
Value
Common sites gene2291
1
2
3
4
5
6
7
8
9
10
CphaseA1
RphaseB1

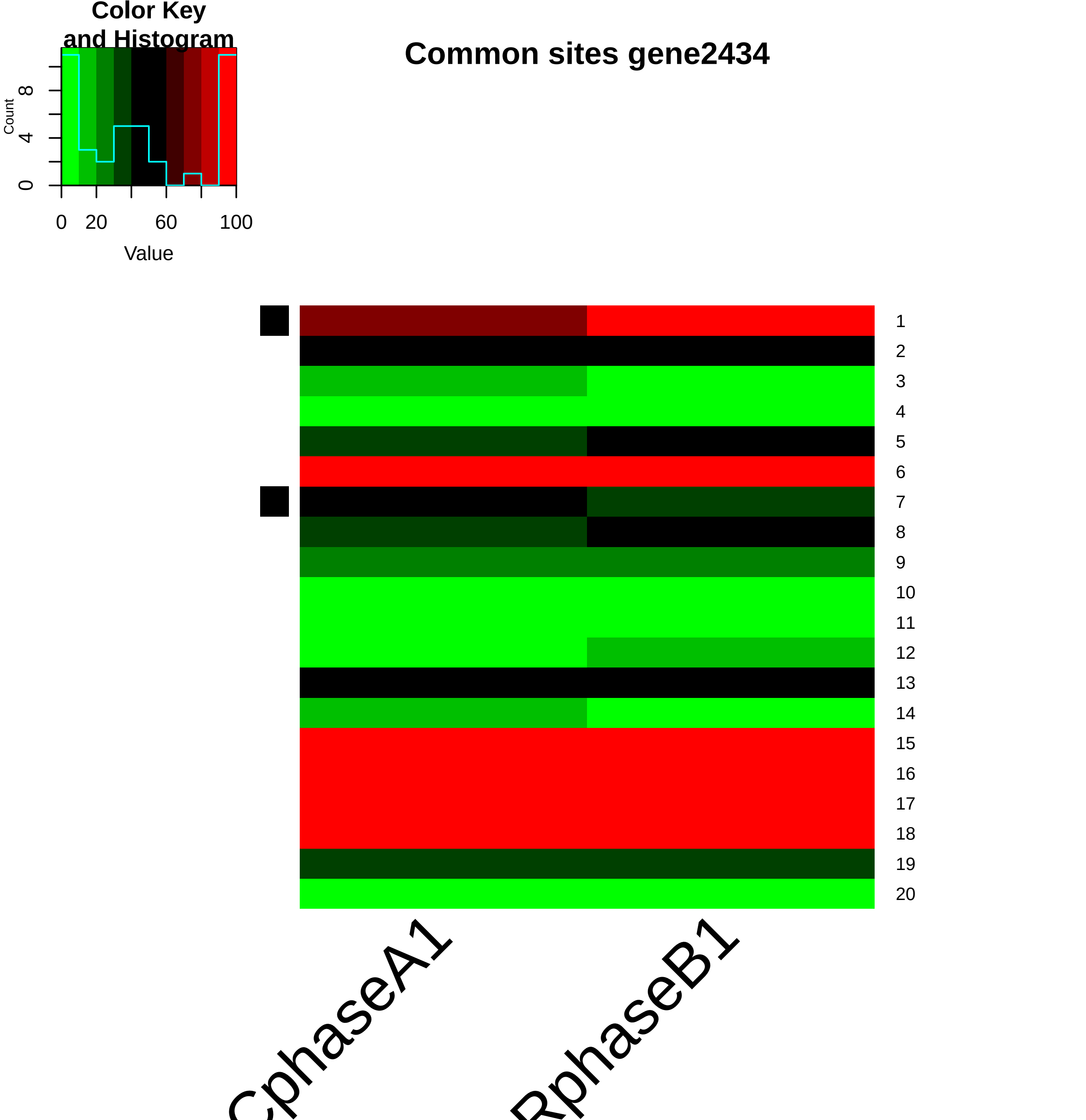

Color Key
and Histogram
Count
0
4
8
0
20
60
100
Value
Common sites gene2434
CphaseA1
RphaseB1

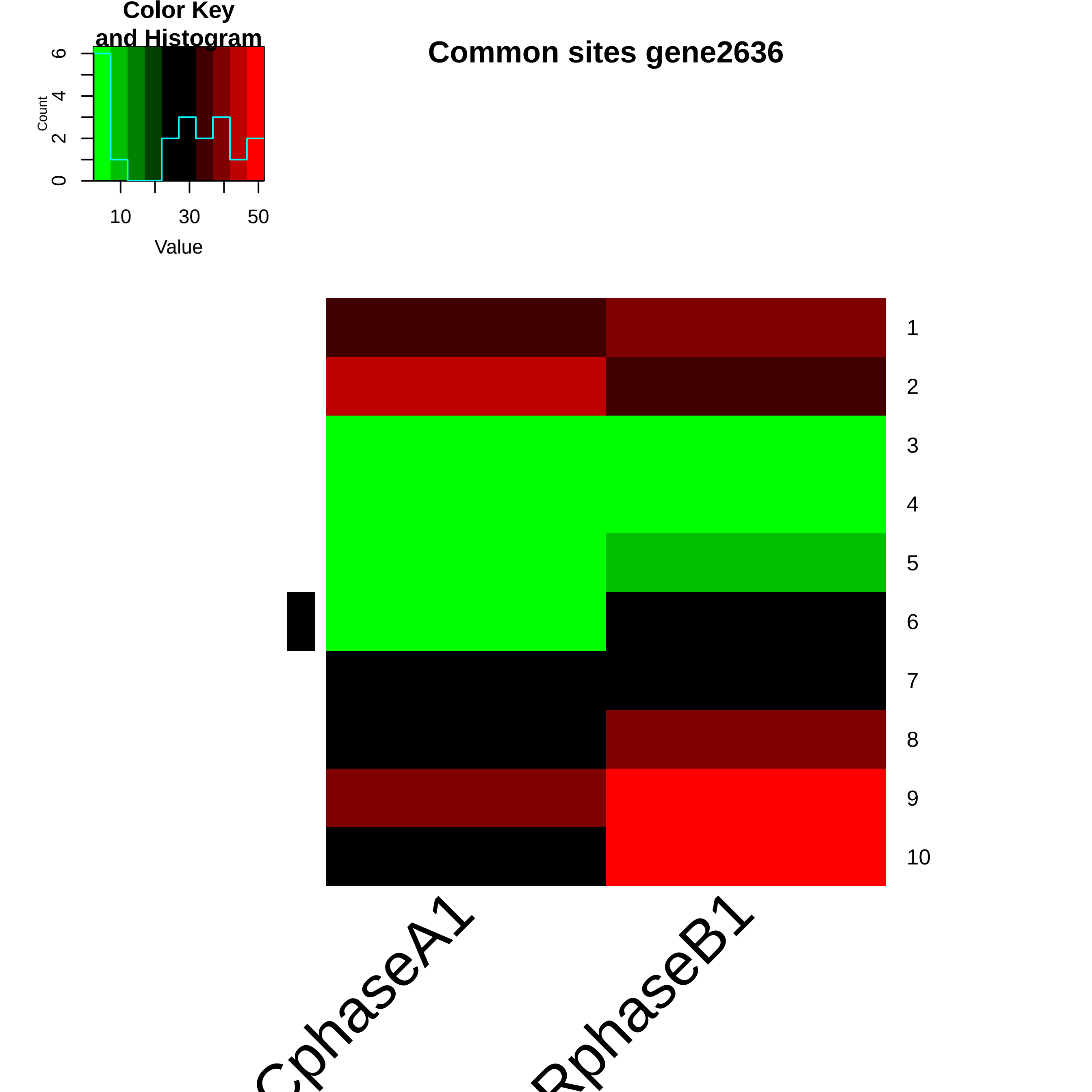
Color Key
and Histogram
Count
0
2
4
6
10
30
50
Value
Common sites gene2636
1
2
3
4
5
6
7
8
9
10
CphaseA1
RphaseB1

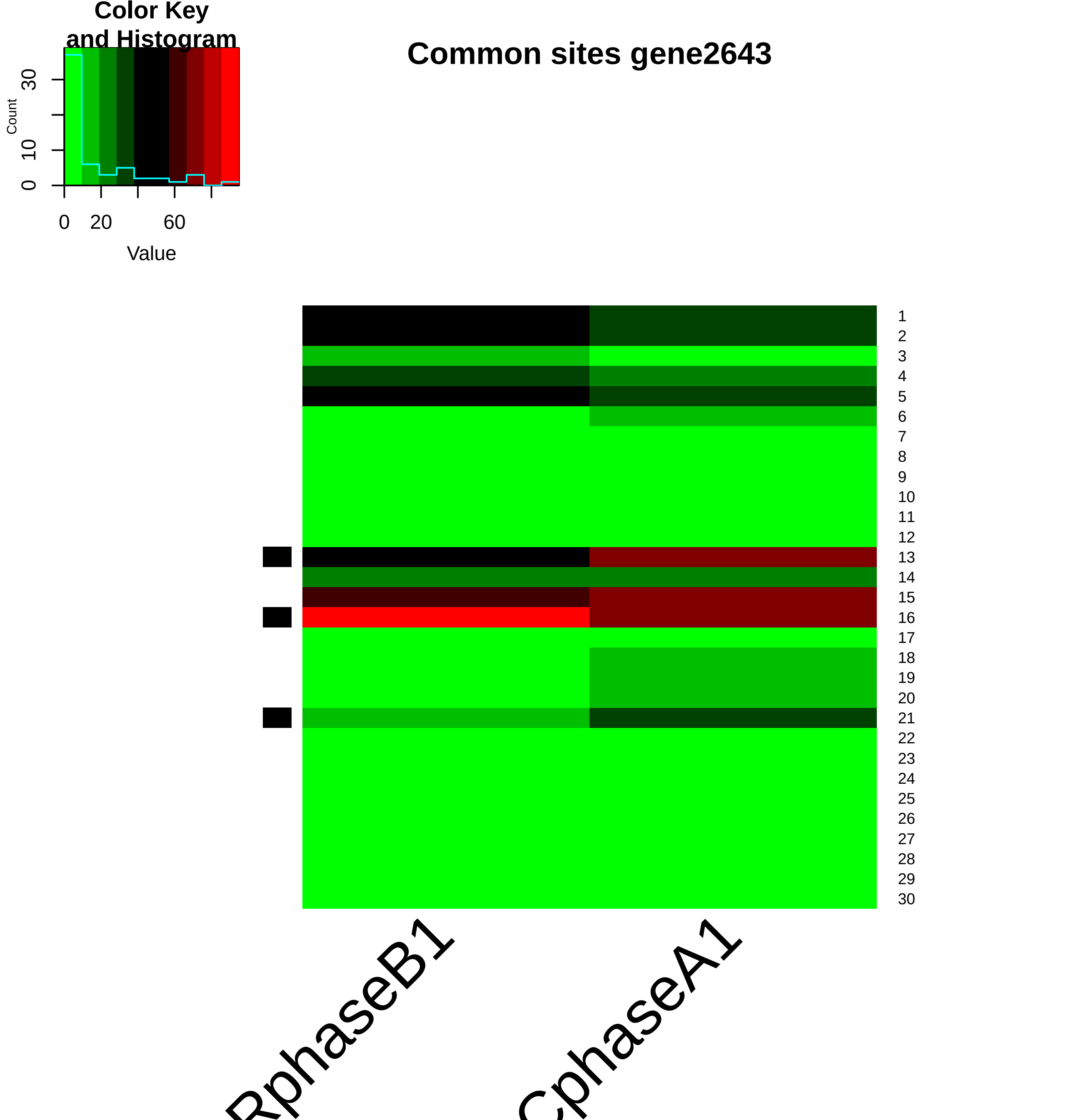
Color Key
and Histogram
Count
0
10
30
0
20
60
Value
Common sites gene2643
RphaseB1
CphaseA1

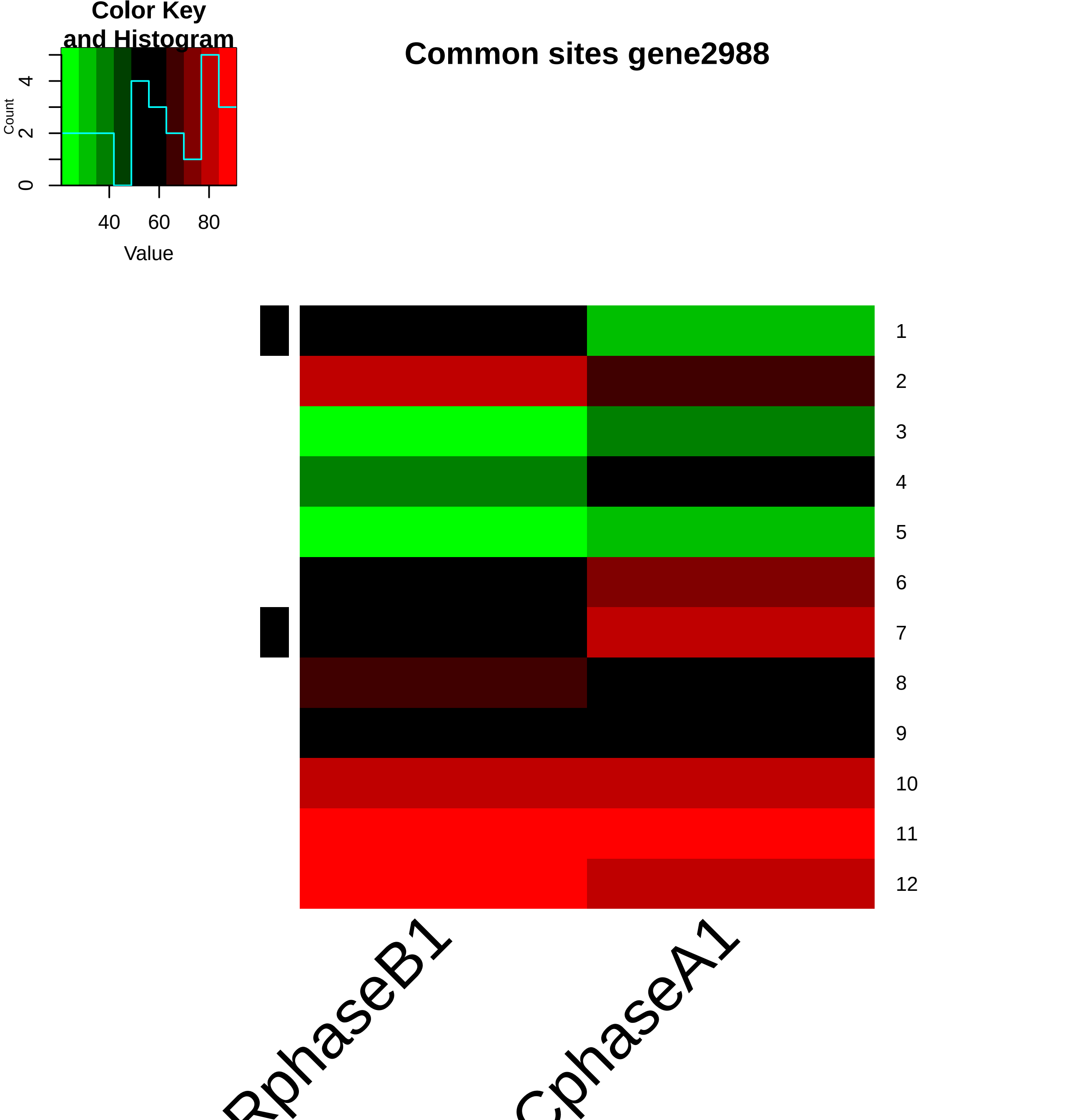
Color Key
and Histogram
Count
0
2
4
40
60
80
Value
Common sites gene2988
1
2
3
4
5
6
7
8
9
10
11
12
RphaseB1
CphaseA1

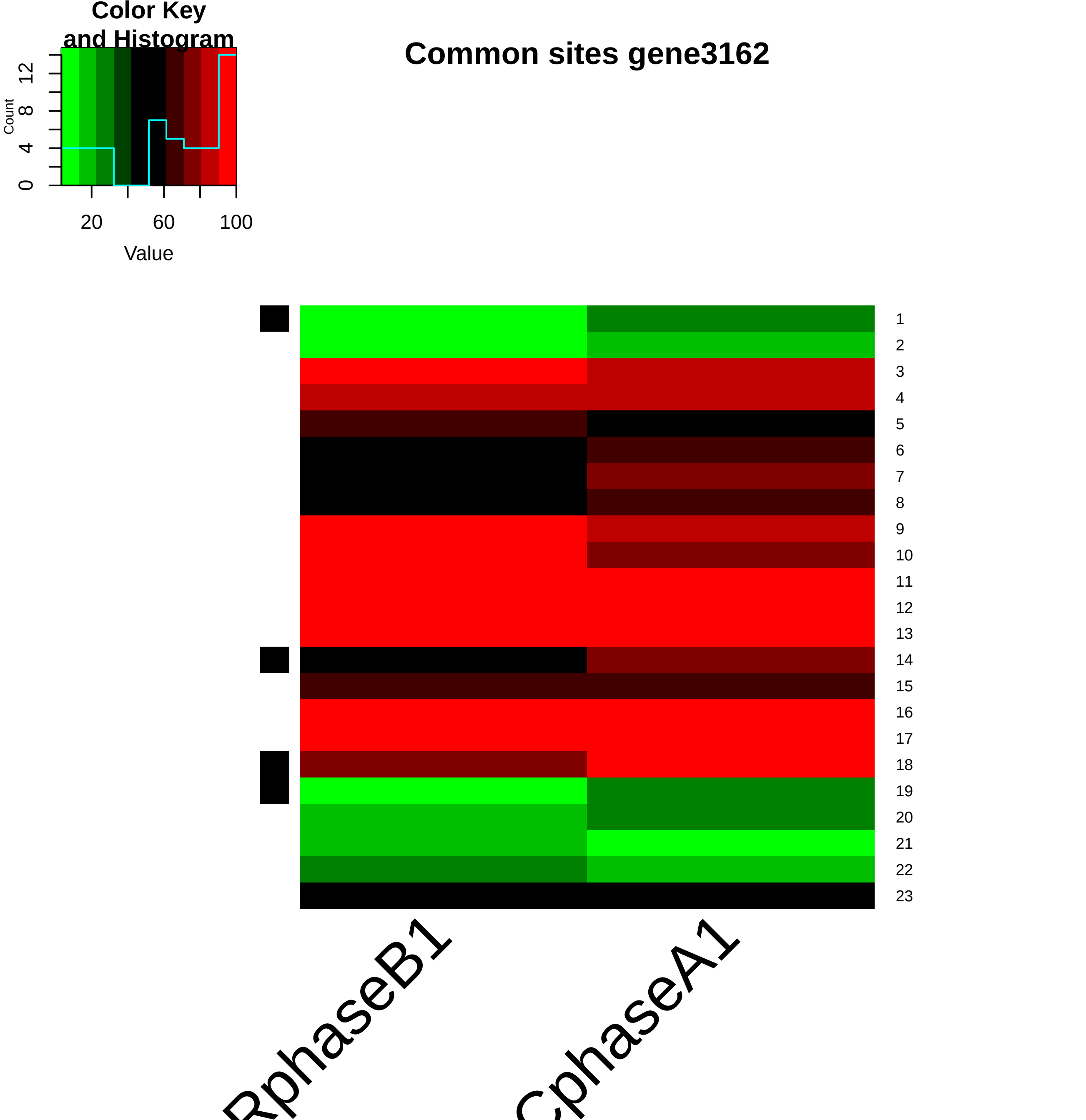

Color Key
and Histogram
Count
0
4
8
12
20
60
100
Value
Common sites gene3162
RphaseB1
CphaseA1

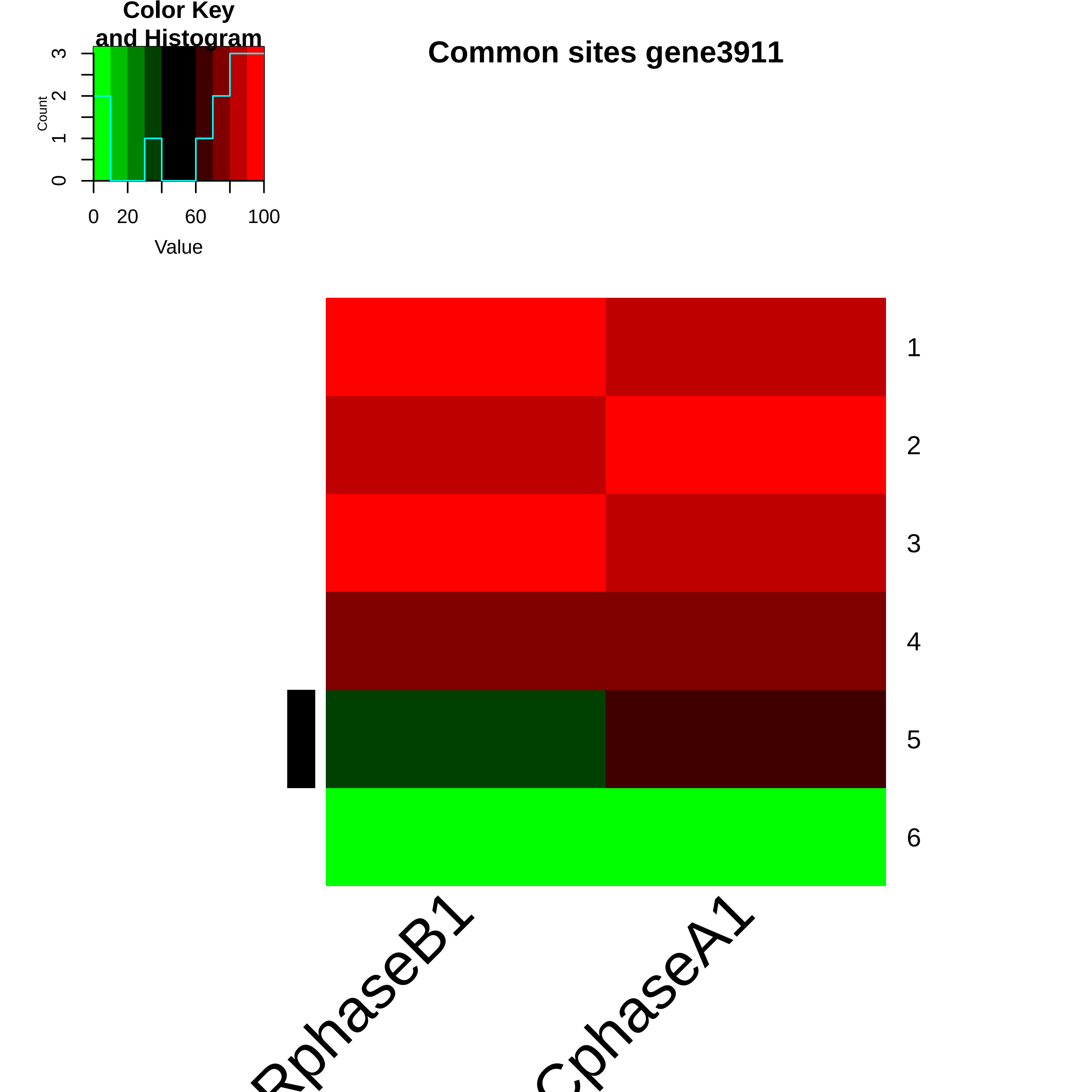
Color Key
and Histogram
Count
0
1
2
3
0
20
60
100
Value
Common sites gene3911
1
2
3
4
5
6
RphaseB1
CphaseA1

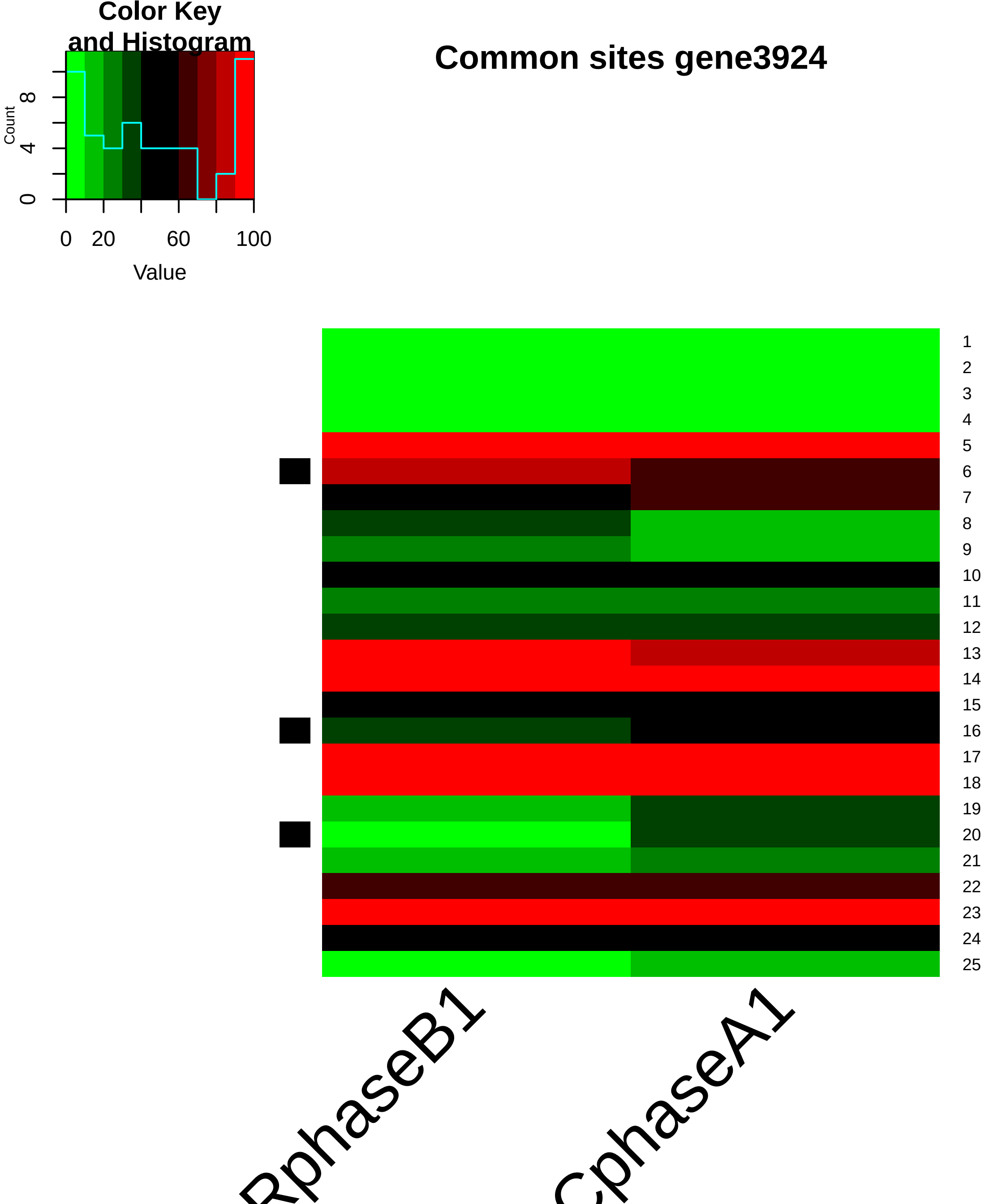
Color Key
and Histogram
Count
0
4
8
0
20
60
100
Value
Common sites gene3924
RphaseB1
CphaseA1

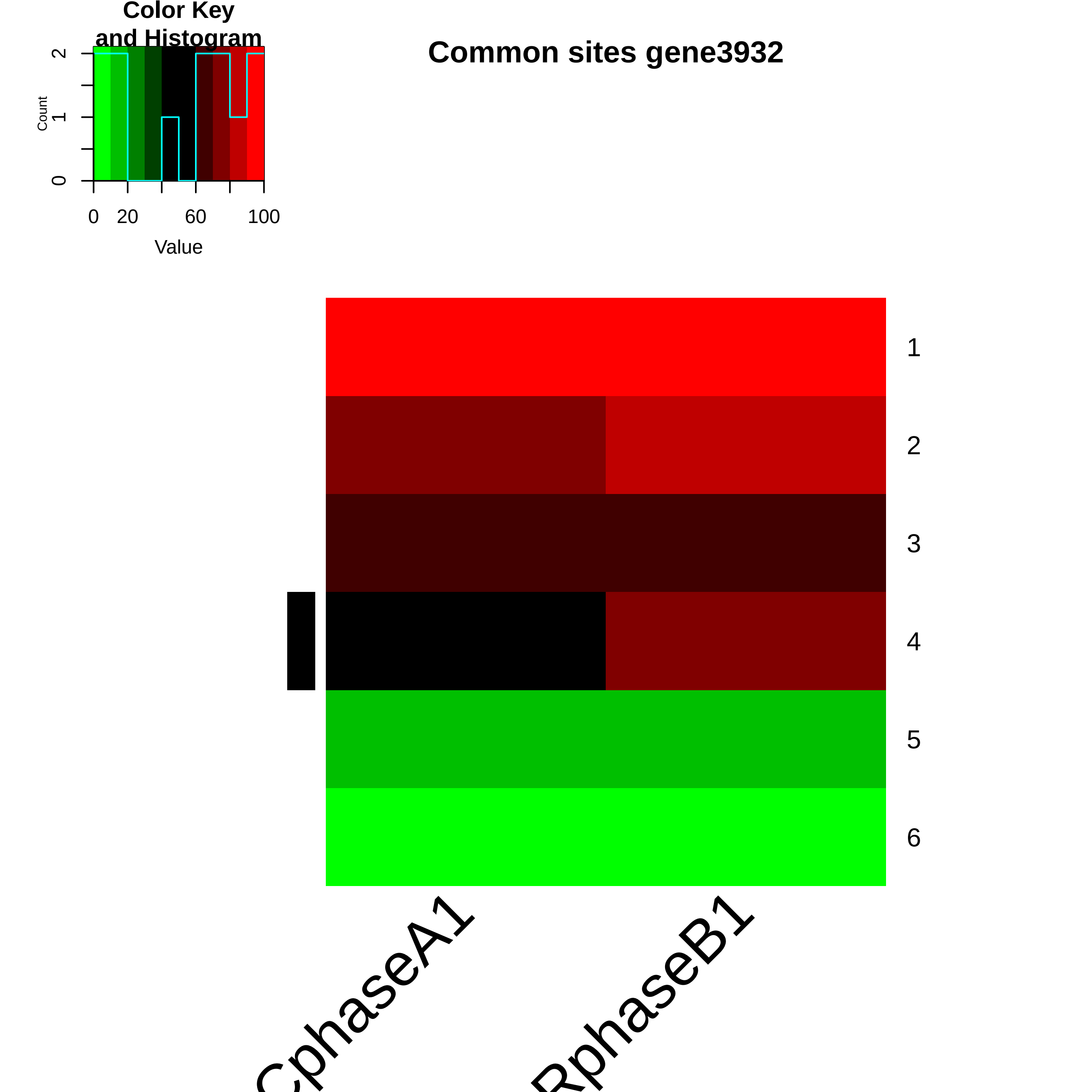

Color Key
and Histogram
Count
0
1
2
0
20
60
100
Value
Common sites gene3932
1
2
3
4
5
6
CphaseA1
RphaseB1

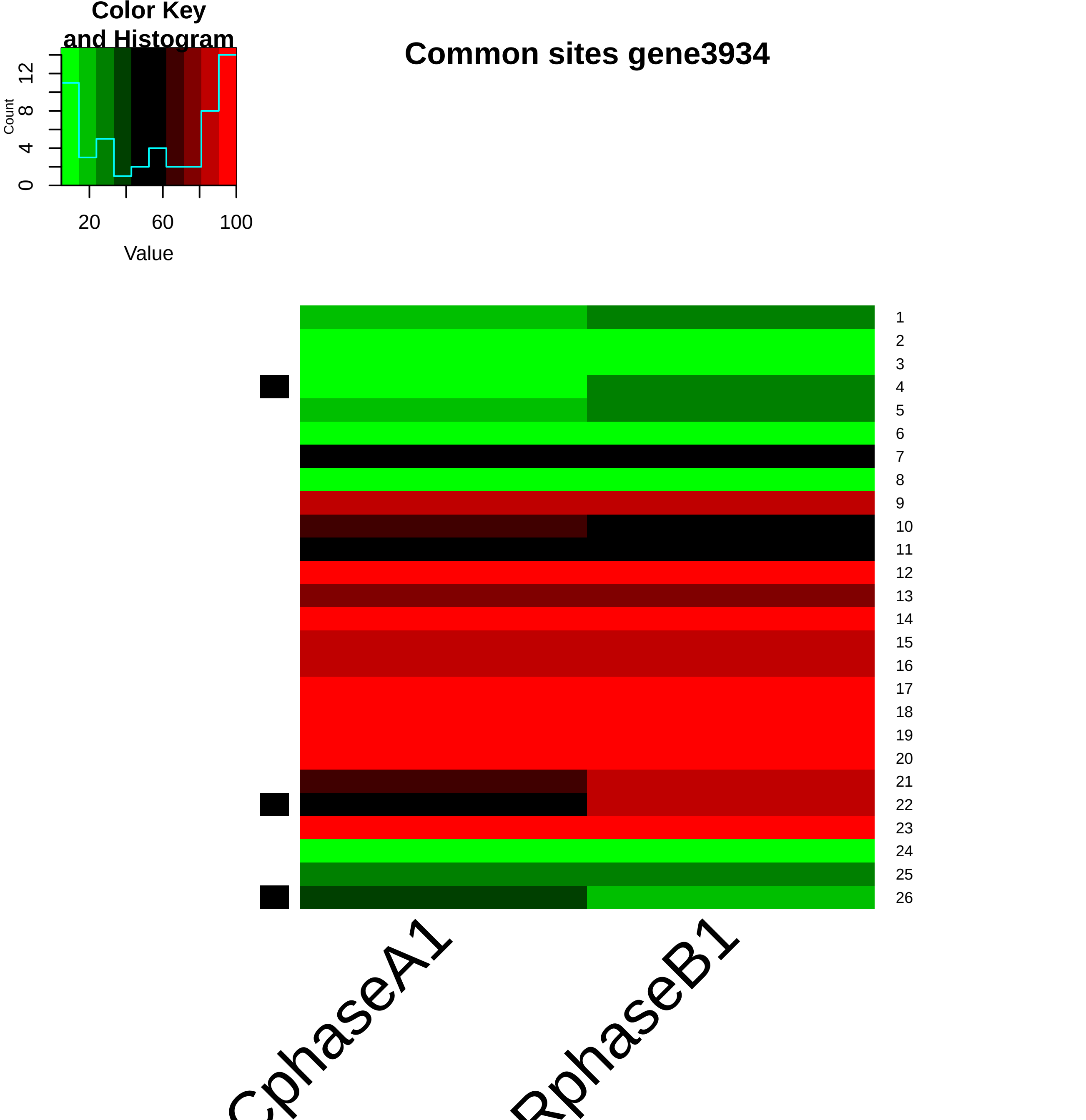
Color Key
and Histogram
Count
0
4
8
12
20
60
100
Value
Common sites gene3934
CphaseA1
RphaseB1

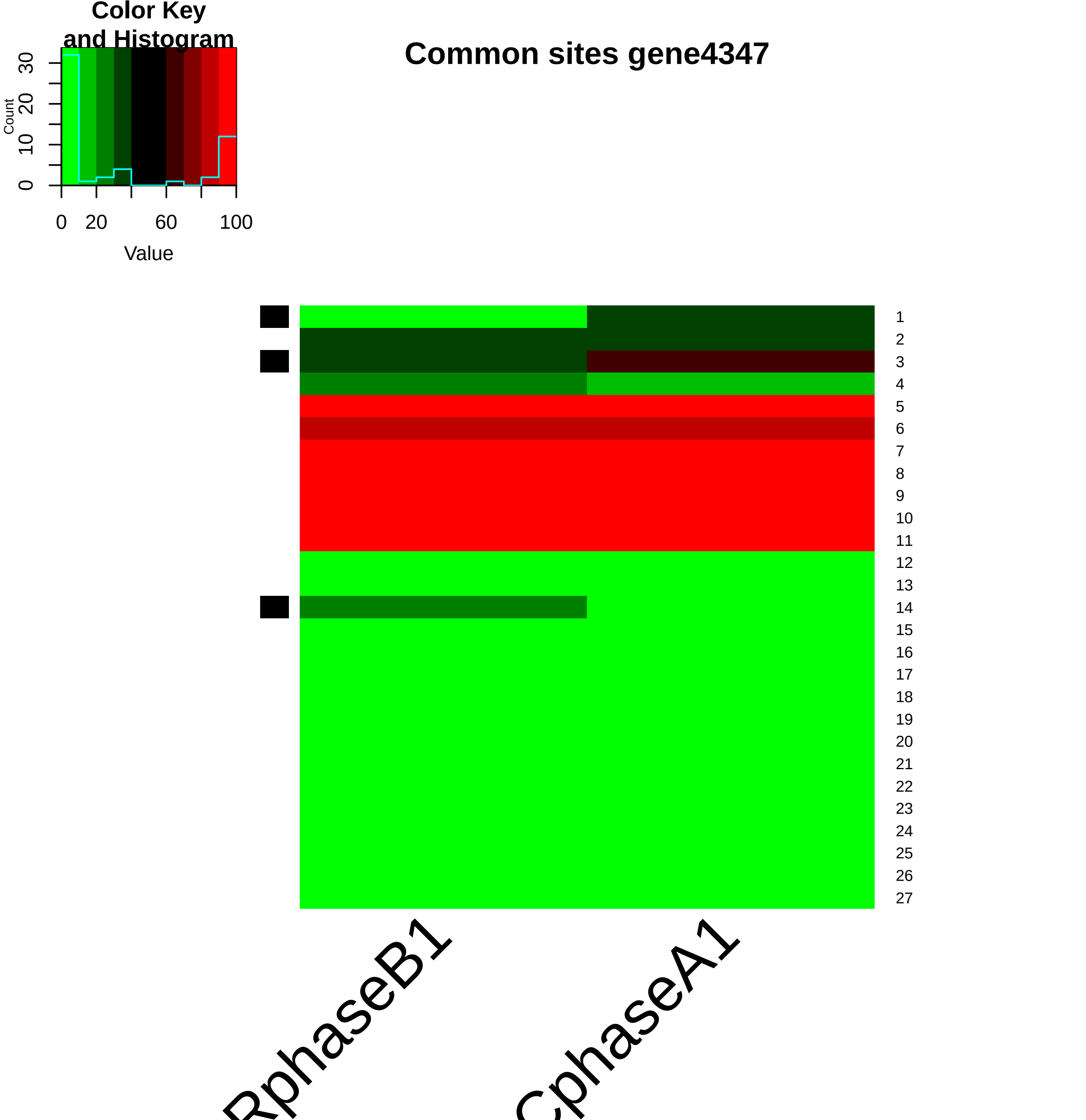
Color Key
and Histogram
Count
0
10
20
30
0
20
60
100
Value
Common sites gene4347
RphaseB1
CphaseA1

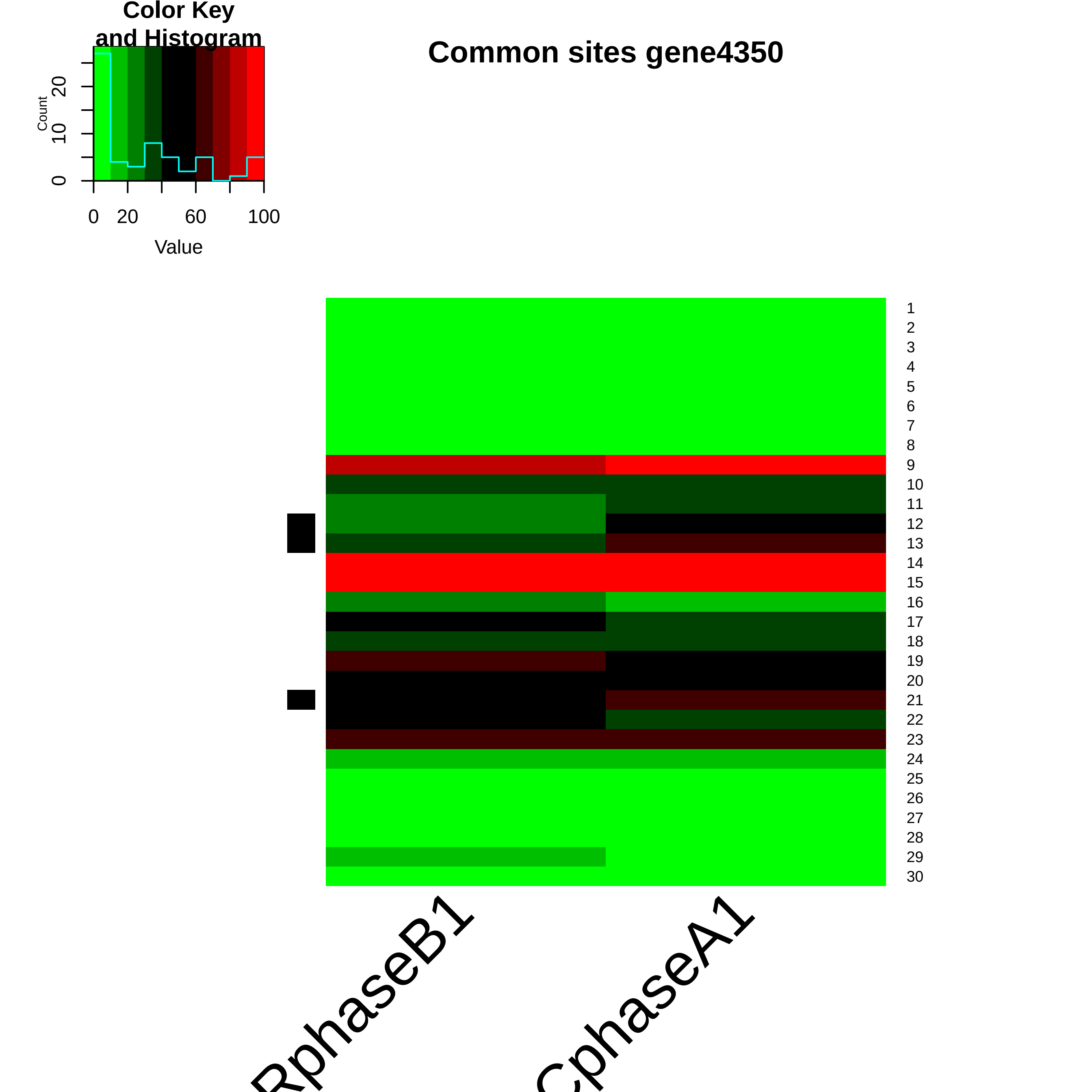
Color Key
and Histogram
Count
0
10
20
0
20
60
100
Value
Common sites gene4350
RphaseB1
CphaseA1

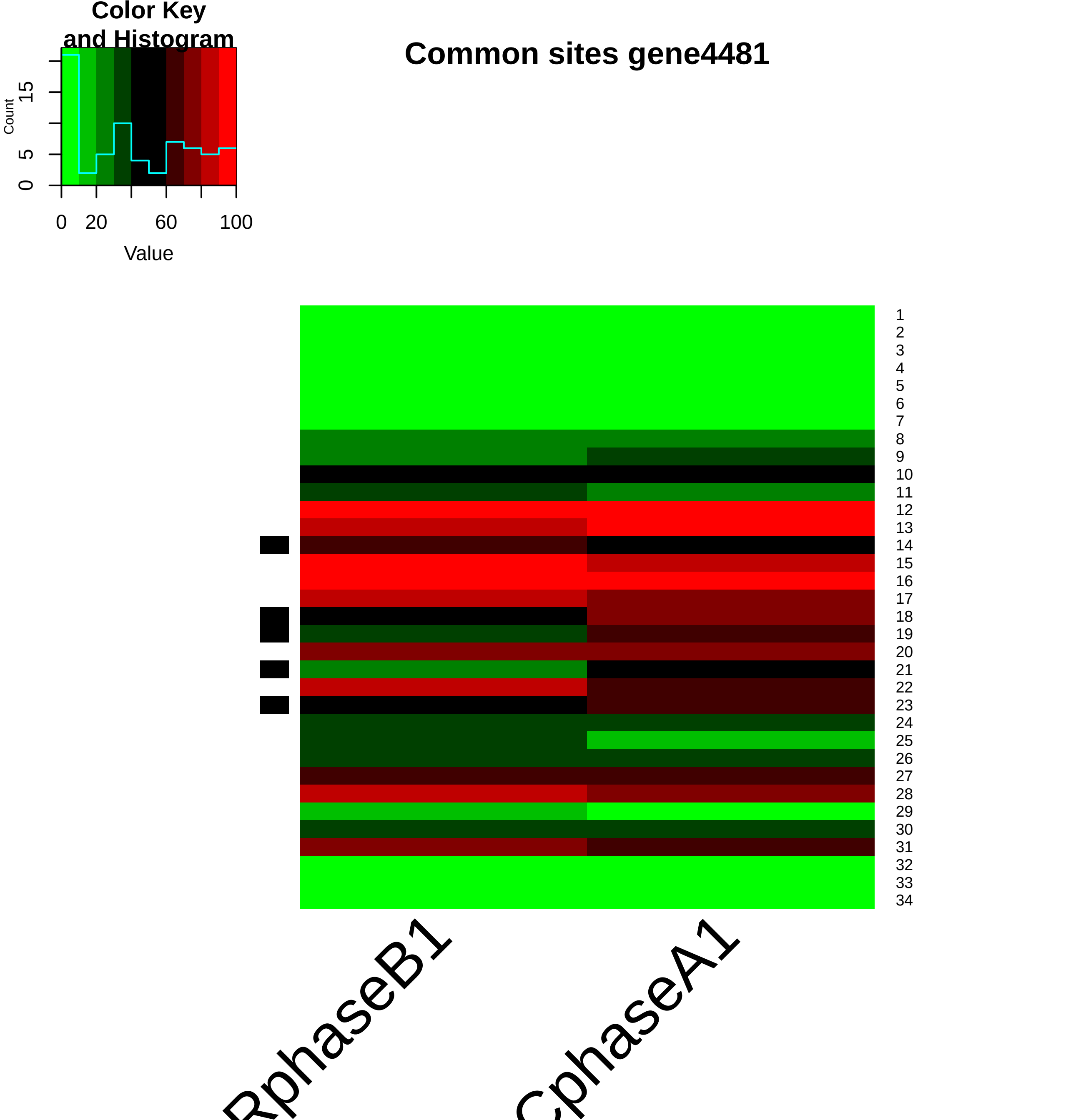

Color Key
and Histogram
Count
0
5
15
0
20
60
100
Value
Common sites gene4481
RphaseB1
CphaseA1

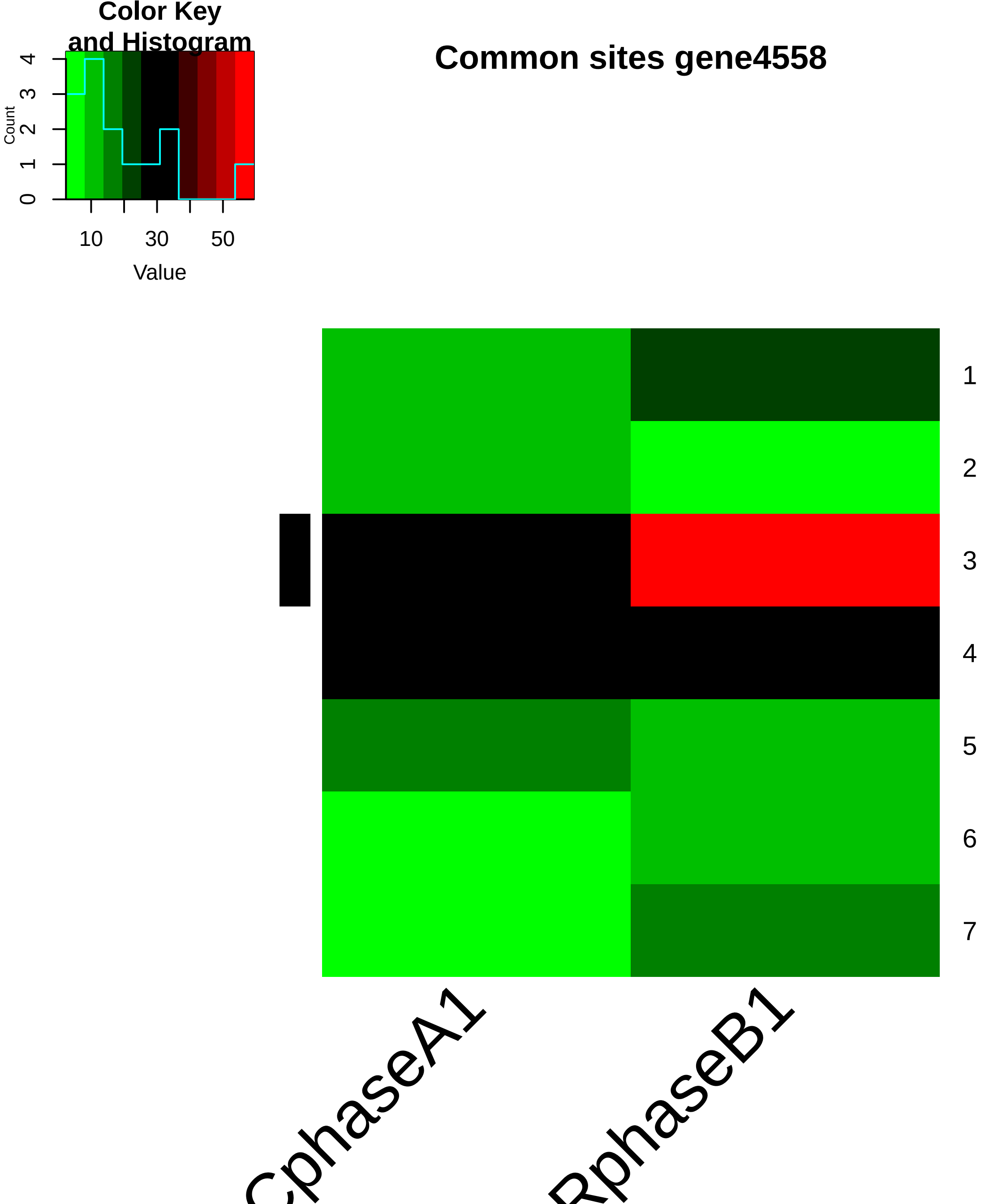
Color Key
and Histogram
Count
0
1
2
3
4
10
30
50
Value
Common sites gene4558
1
2
3
4
5
6
7
CphaseA1
RphaseB1

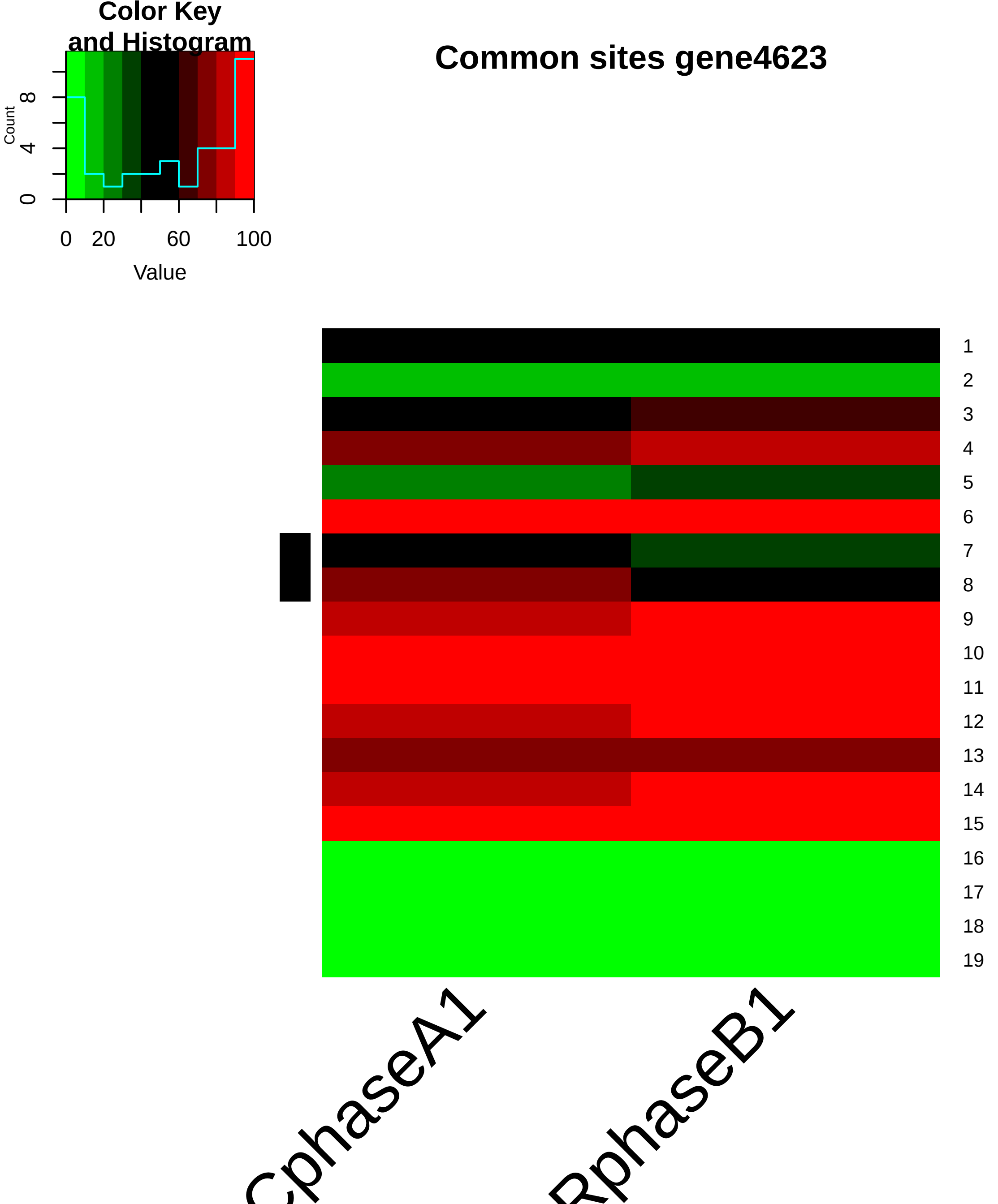

Color Key
and Histogram
Count
0
4
8
0
20
60
100
Value
Common sites gene4623
CphaseA1
RphaseB1

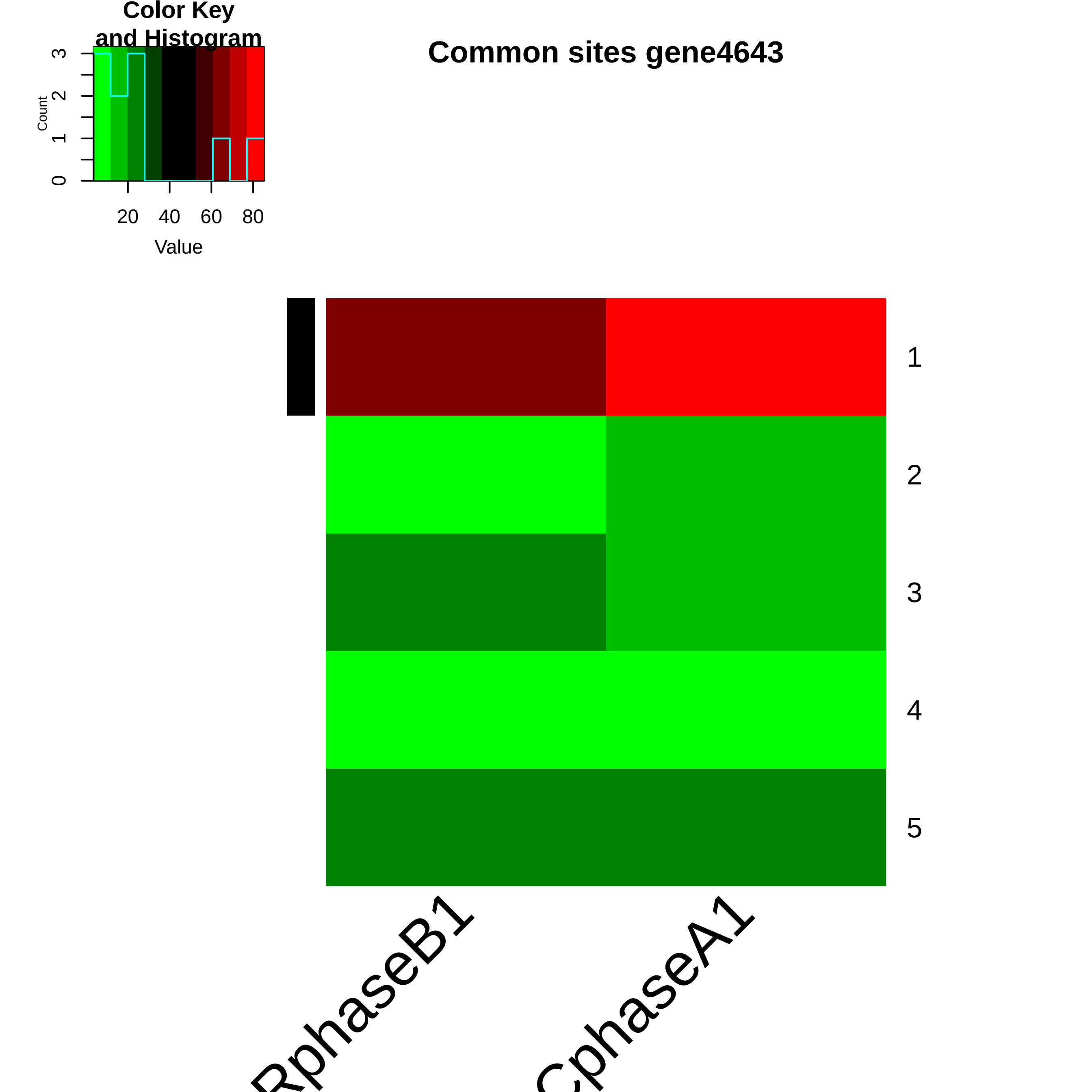
Color Key
and Histogram
Count
0
1
2
3
20
40
60
80
Value
Common sites gene4643
1
2
3
4
5

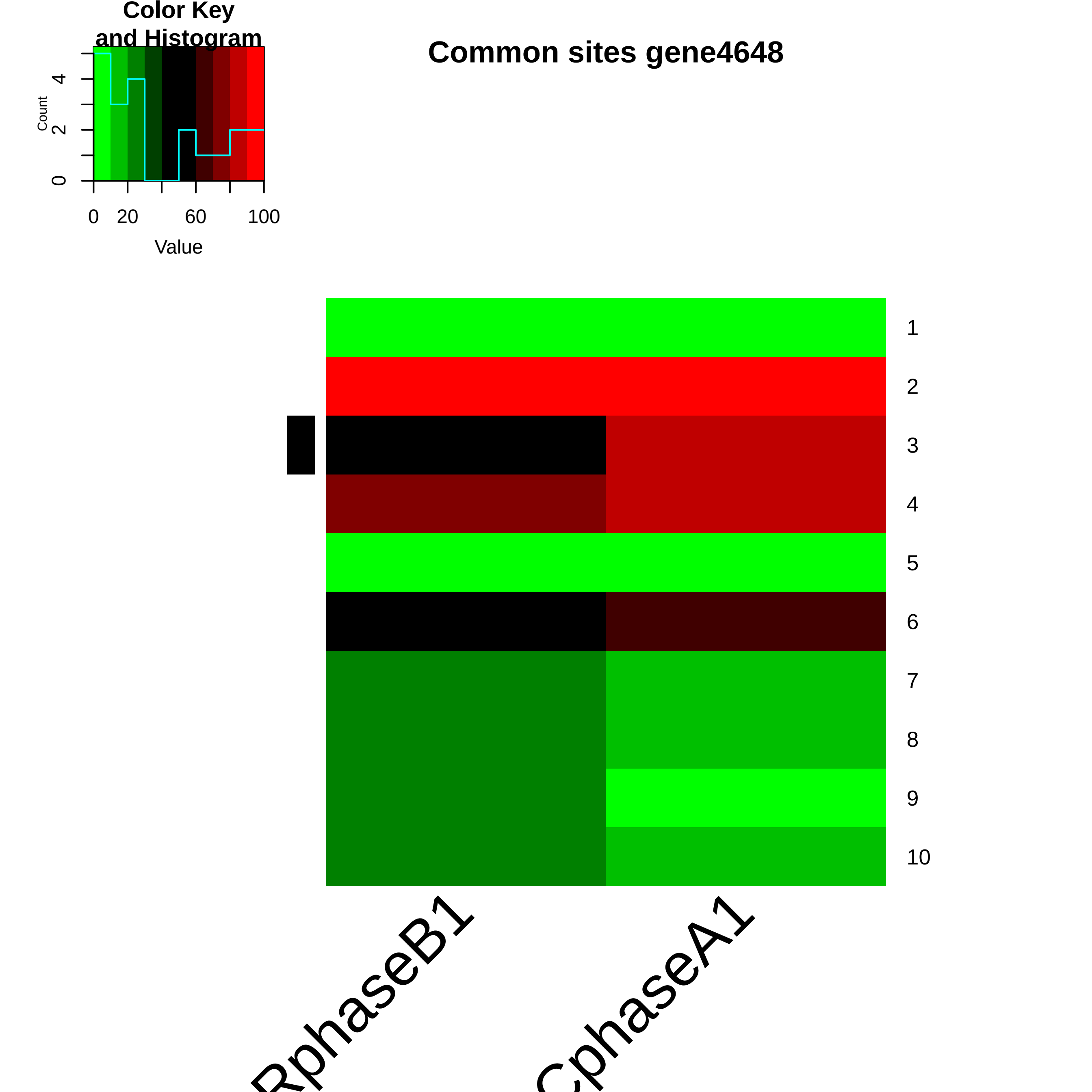
Color Key
and Histogram
Count
0
2
4
0
20
60
100
Value
Common sites gene4648
1
2
3
4
5
6
7
8
9
10
RphaseB1
CphaseA1

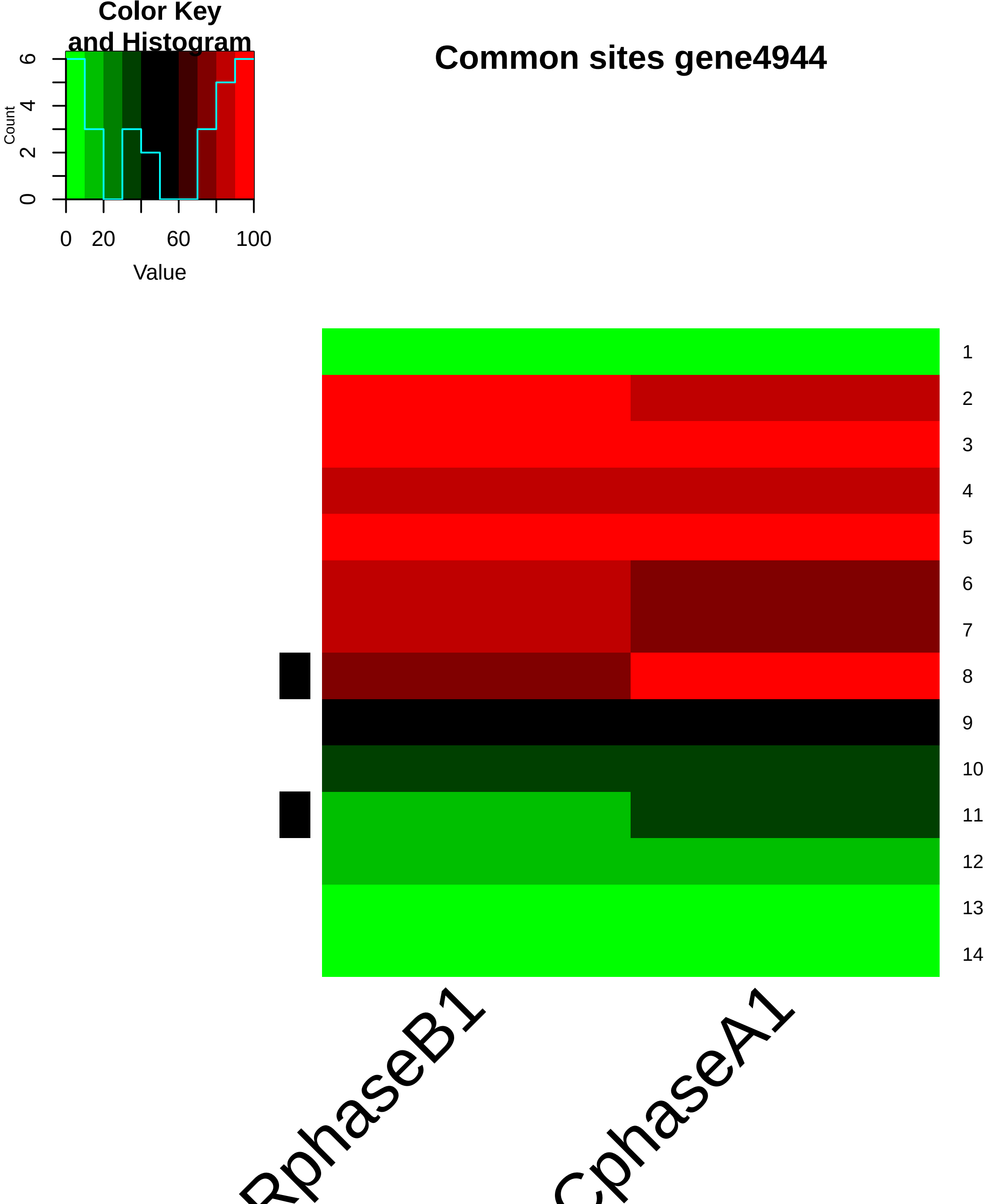
Color Key
and Histogram
Count
0
2
4
6
0
20
60
100
Value
Common sites gene4944
1
2
3
4
5
6
7
8
9
10
11
12
13
14
RphaseB1
CphaseA1

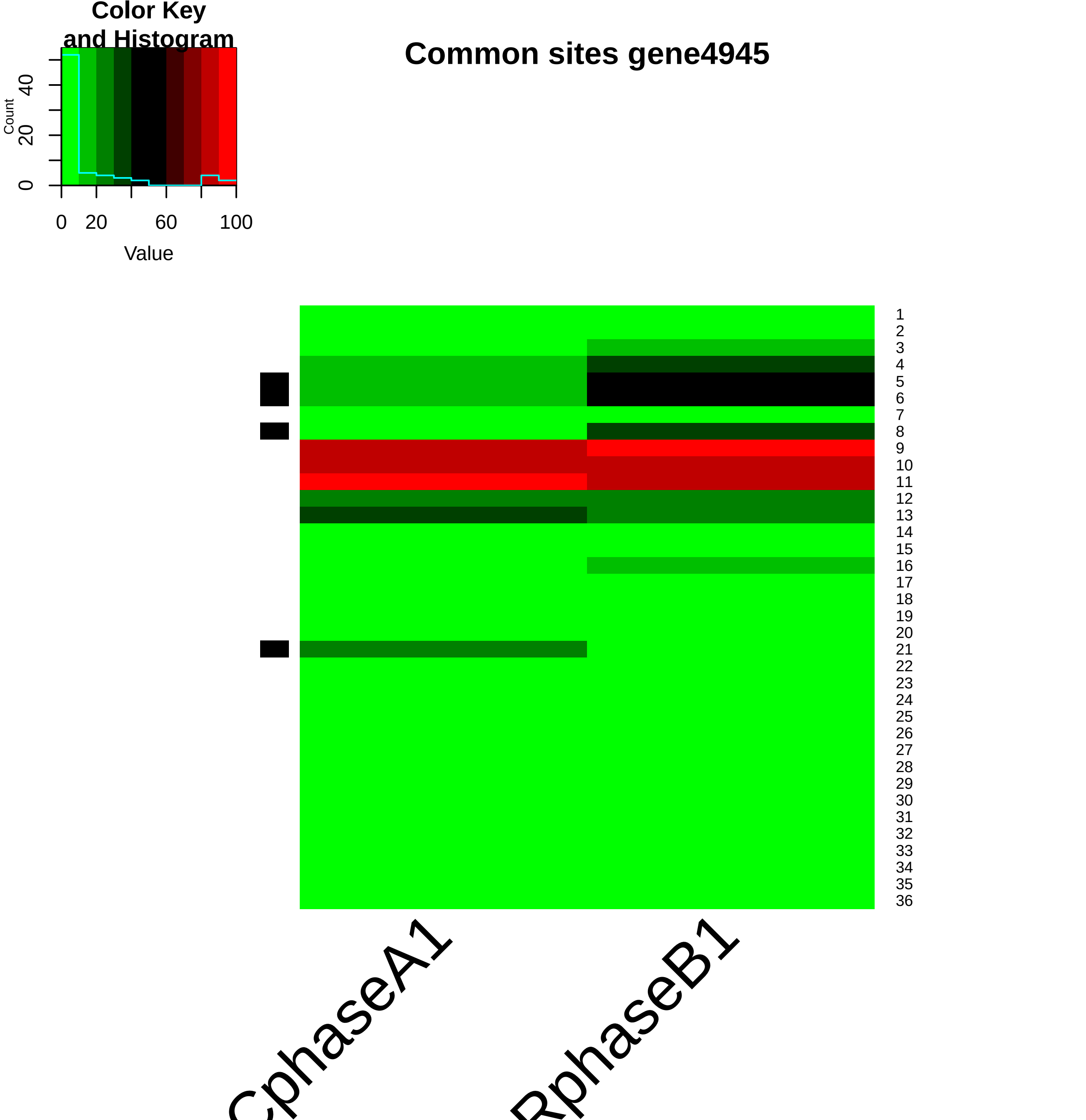
Color Key
and Histogram
Count
0
20
40
0
20
60
100
Value
Common sites gene4945
CphaseA1
RphaseB1

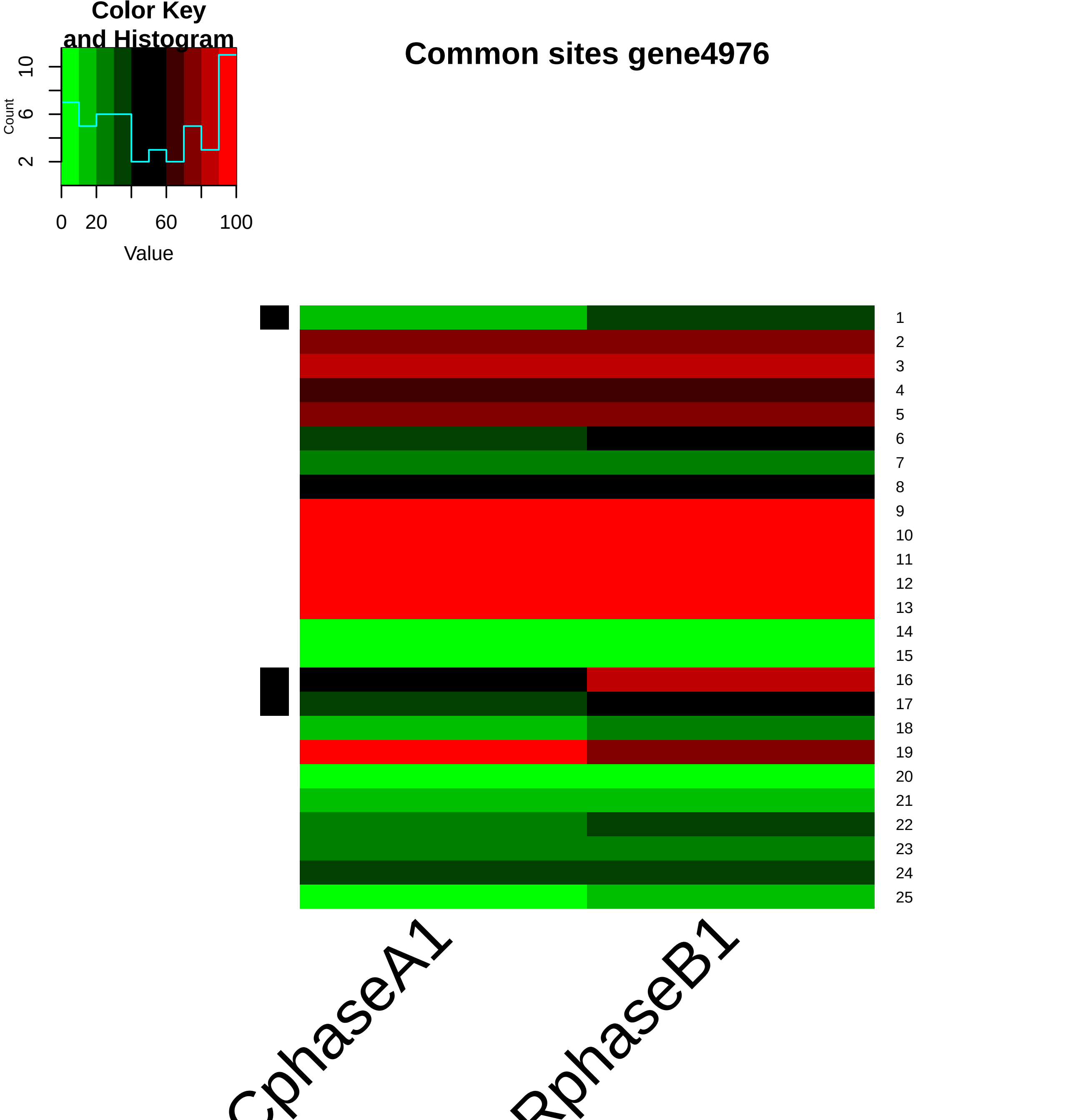

Color Key
and Histogram
Count
2
6
10
0
20
60
100
Value
Common sites gene4976
CphaseA1
RphaseB1

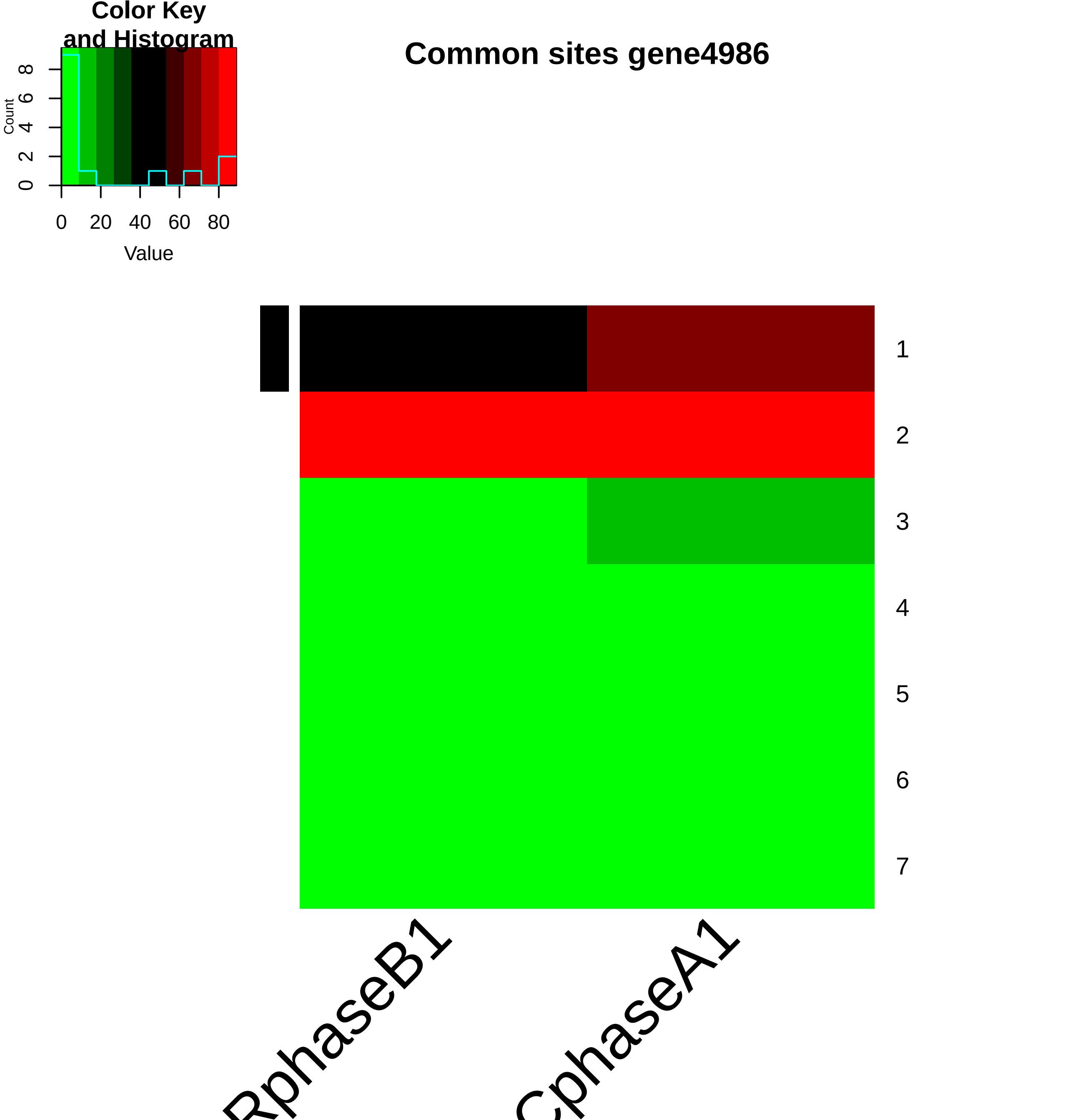

Color Key
and Histogram
Count
0
2
4
6
8
0
20
40
60
80
Value
Common sites gene4986
1
2
3
4
5
6
7
RphaseB1
CphaseA1

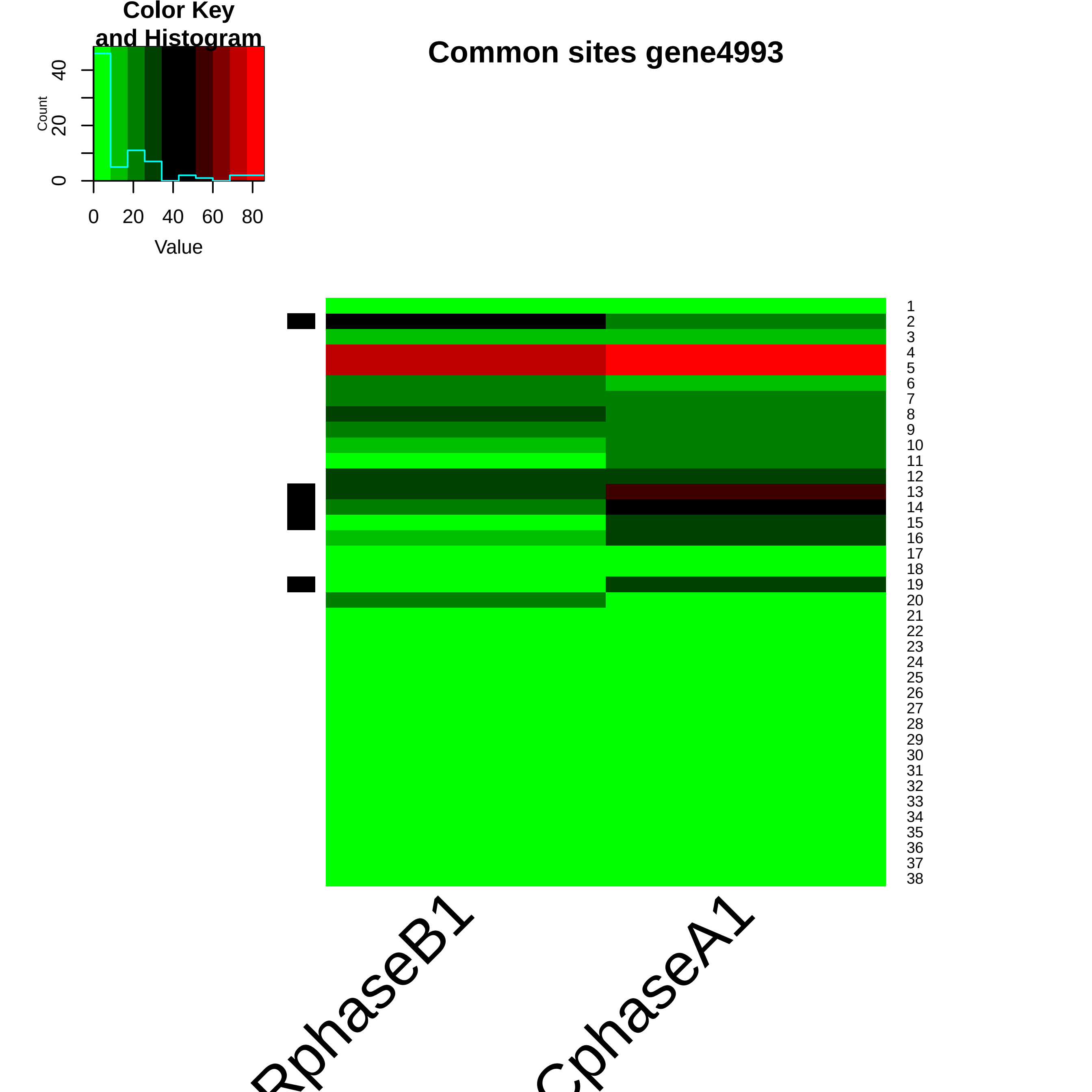

Color Key
and Histogram
Count
0
20
40
0
20
40
60
80
Value
Common sites gene4993
RphaseB1
CphaseA1

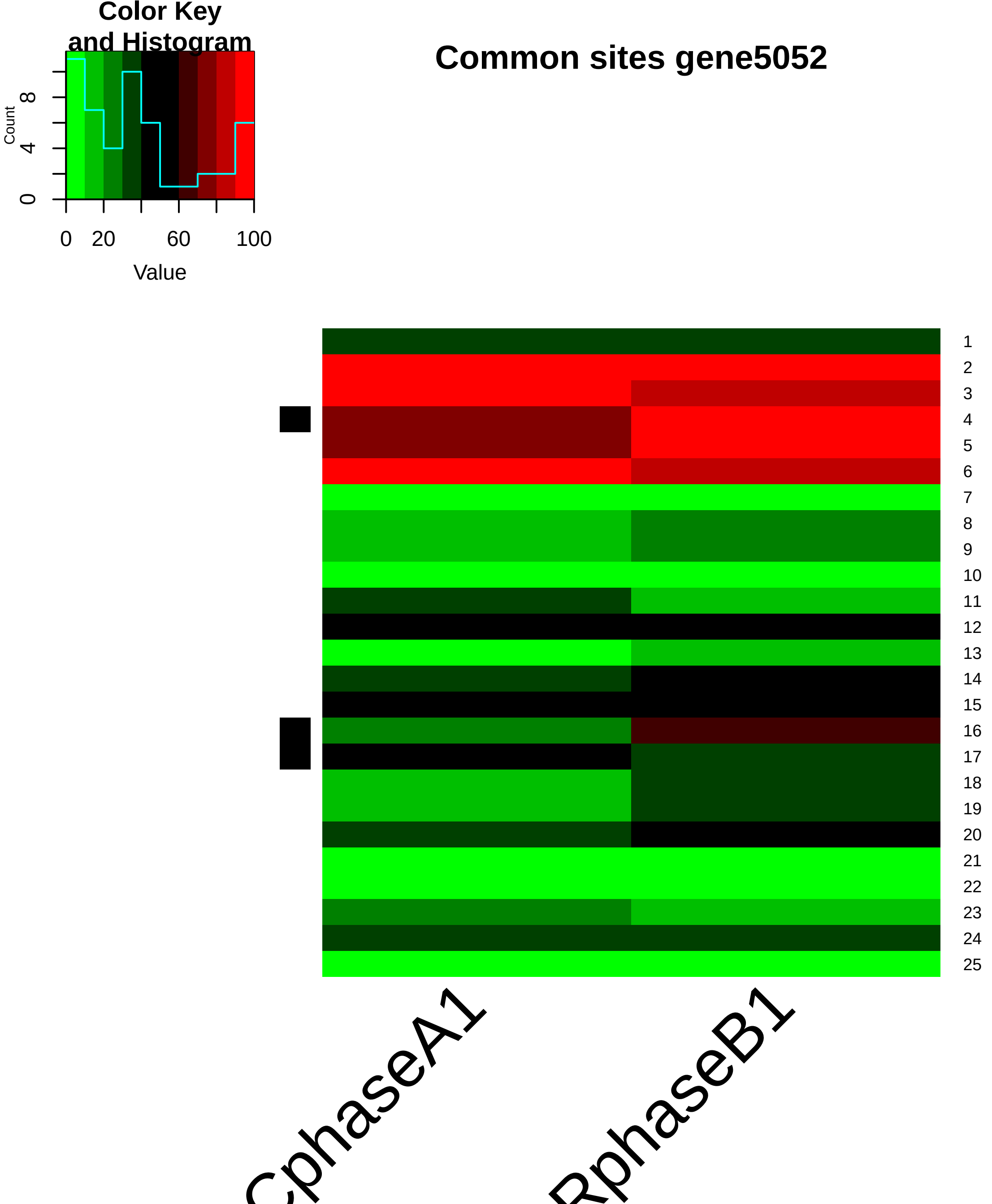
Color Key
and Histogram
Count
0
4
8
0
20
60
100
Value
Common sites gene5052
CphaseA1
RphaseB1

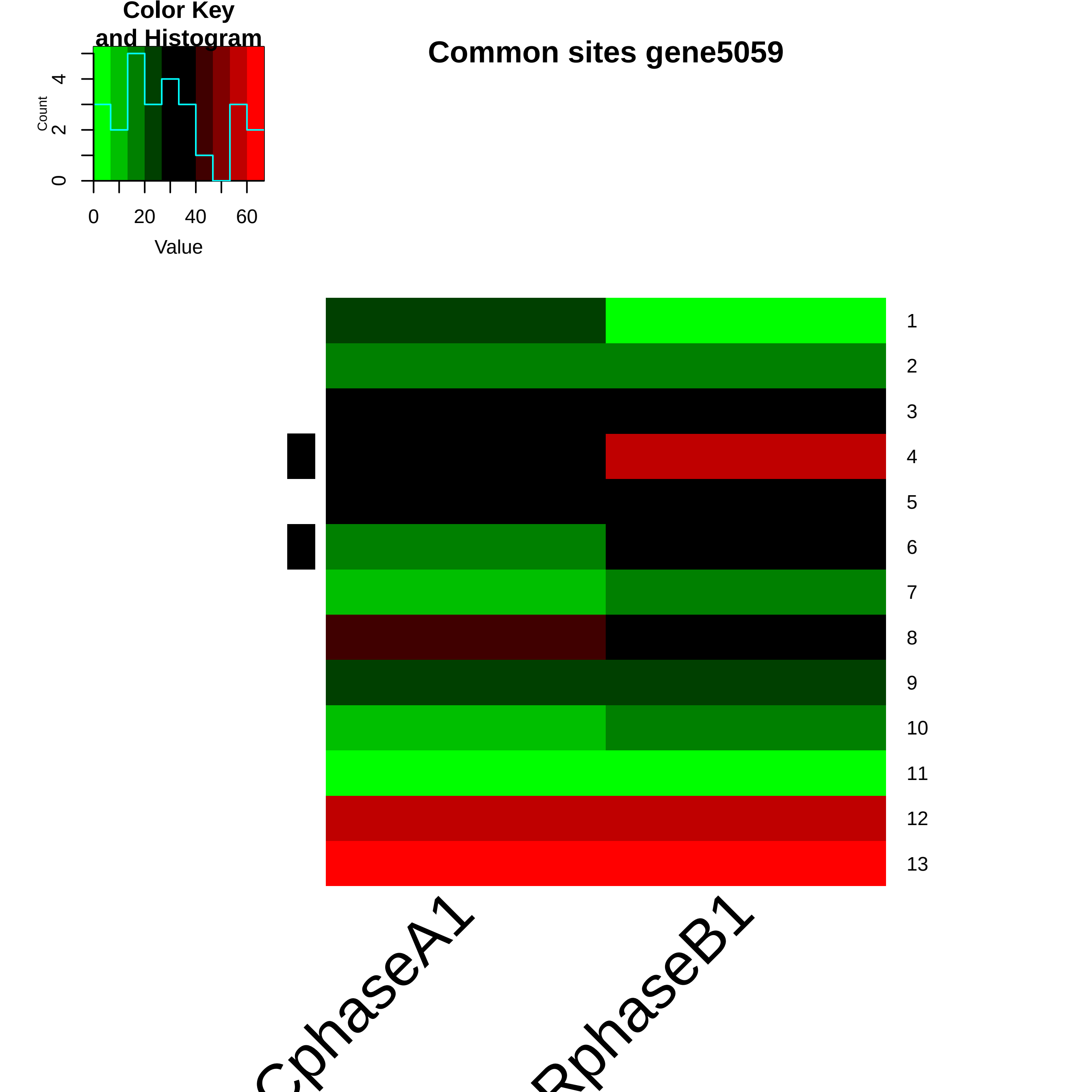
Color Key
and Histogram
Count
0
2
4
0
20
40
60
Value
Common sites gene5059
1
2
3
4
5
6
7
8
9
10
11
12
13
CphaseA1
RphaseB1

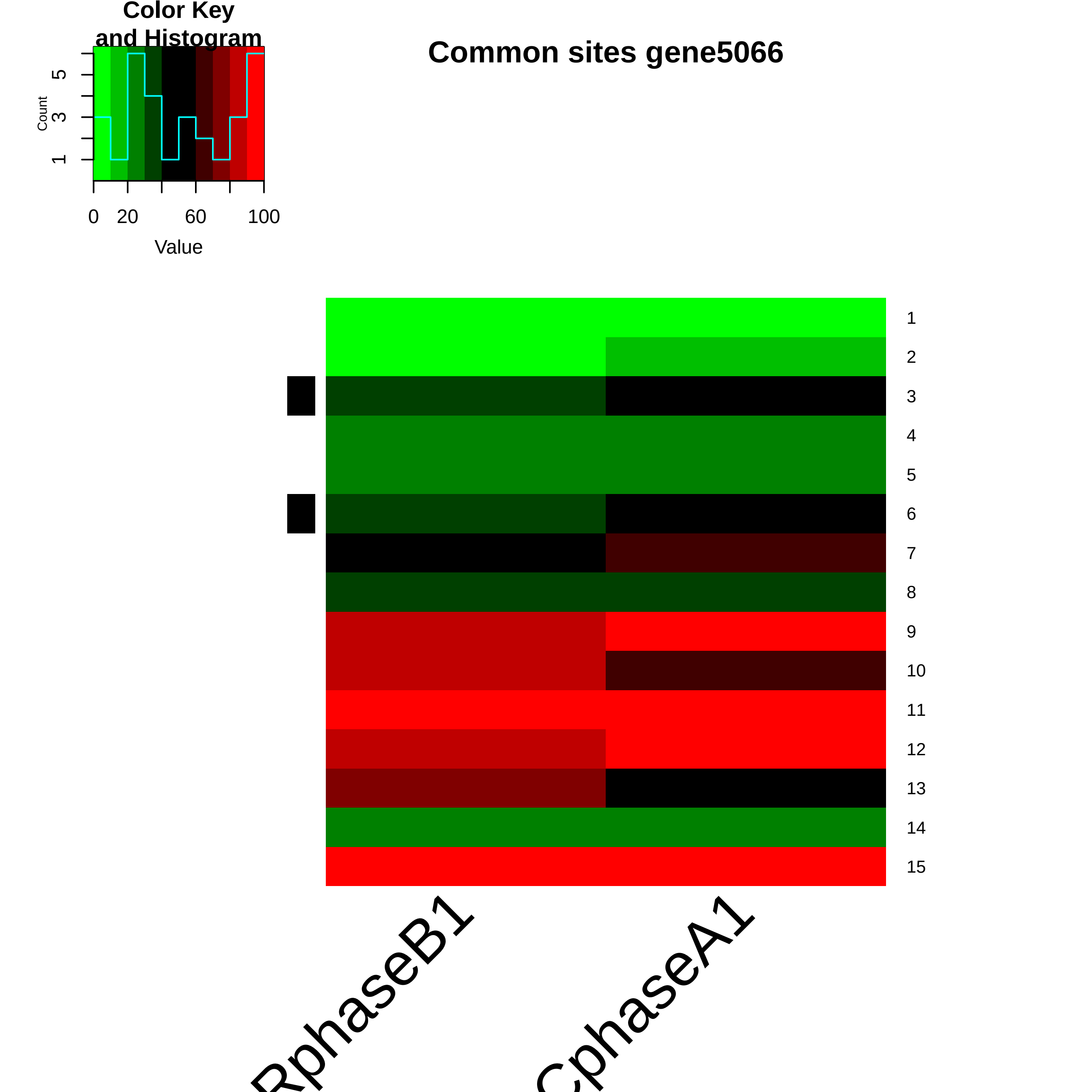
Color Key
and Histogram
Count
1
3
5
0
20
60
100
Value
Common sites gene5066
RphaseB1
CphaseA1

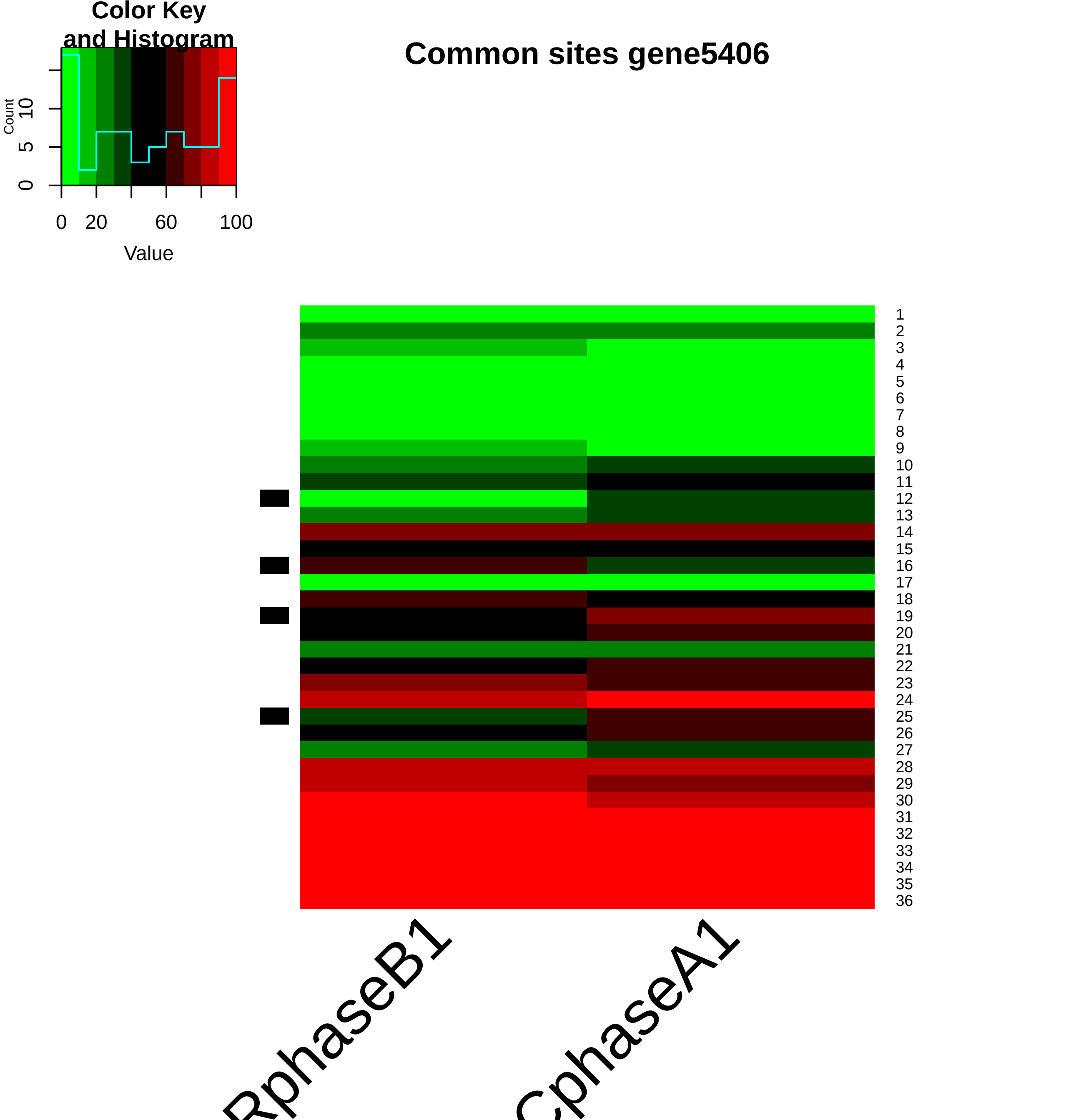
Color Key
and Histogram
Count
0
5
10
0
20
60
100
Value
Common sites gene5406
RphaseB1
CphaseA1

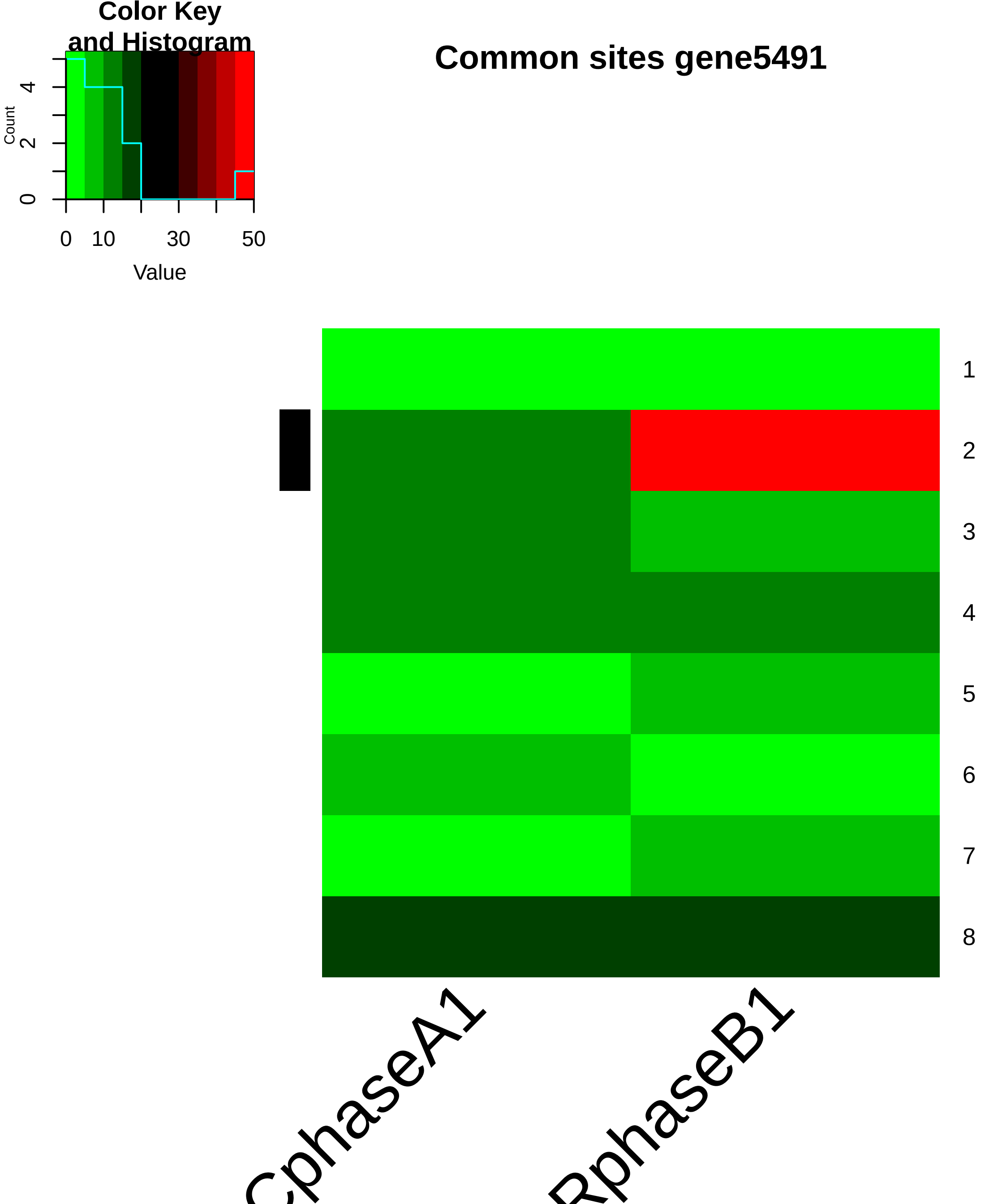
Color Key
and Histogram
Count
0
2
4
0
10
30
50
Value
Common sites gene5491
1
2
3
4
5
6
7
8

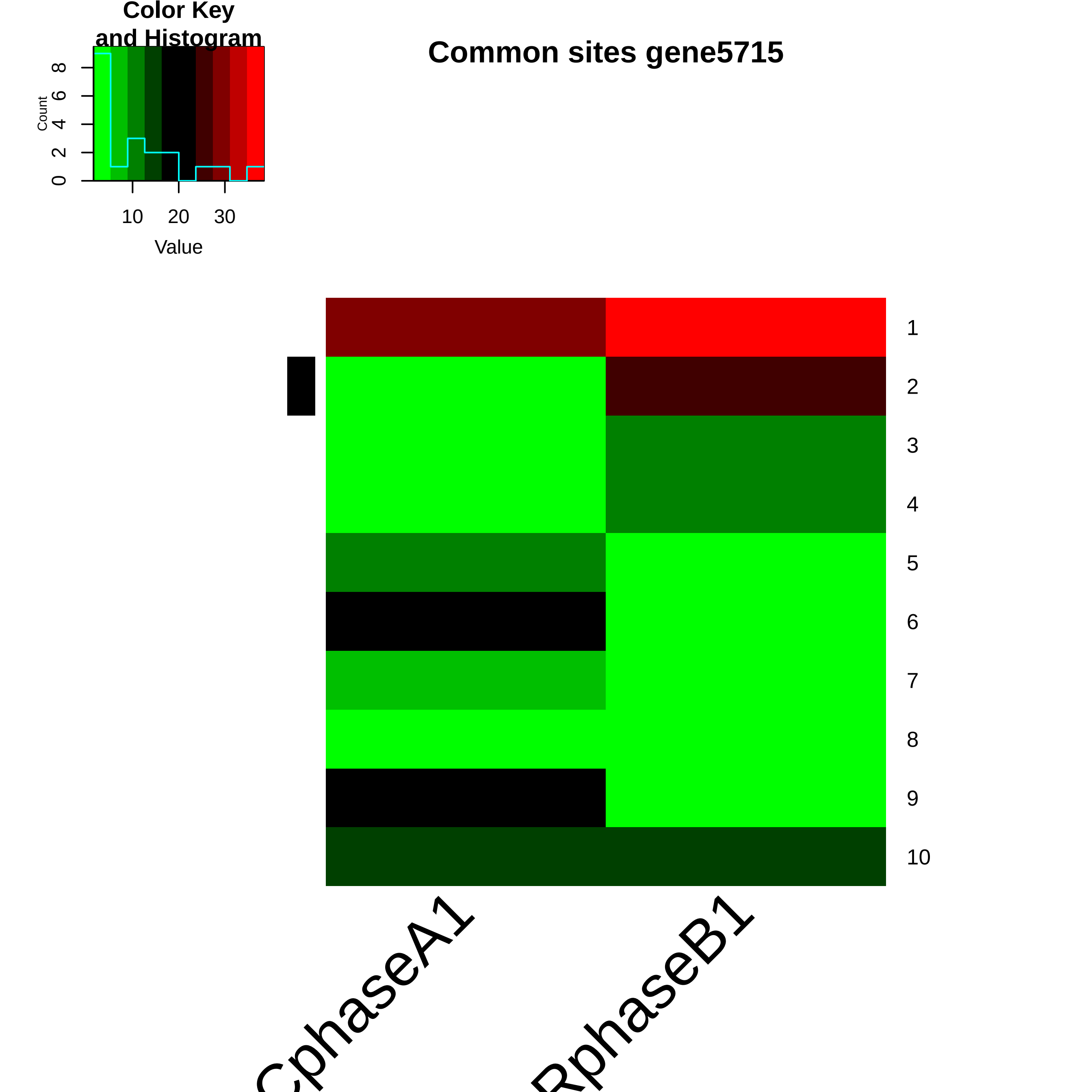
Color Key
and Histogram
Count
0
2
4
6
8
10
20
30
Value
Common sites gene5715
1
2
3
4
5
6
7
8
9
10
CphaseA1
RphaseB1

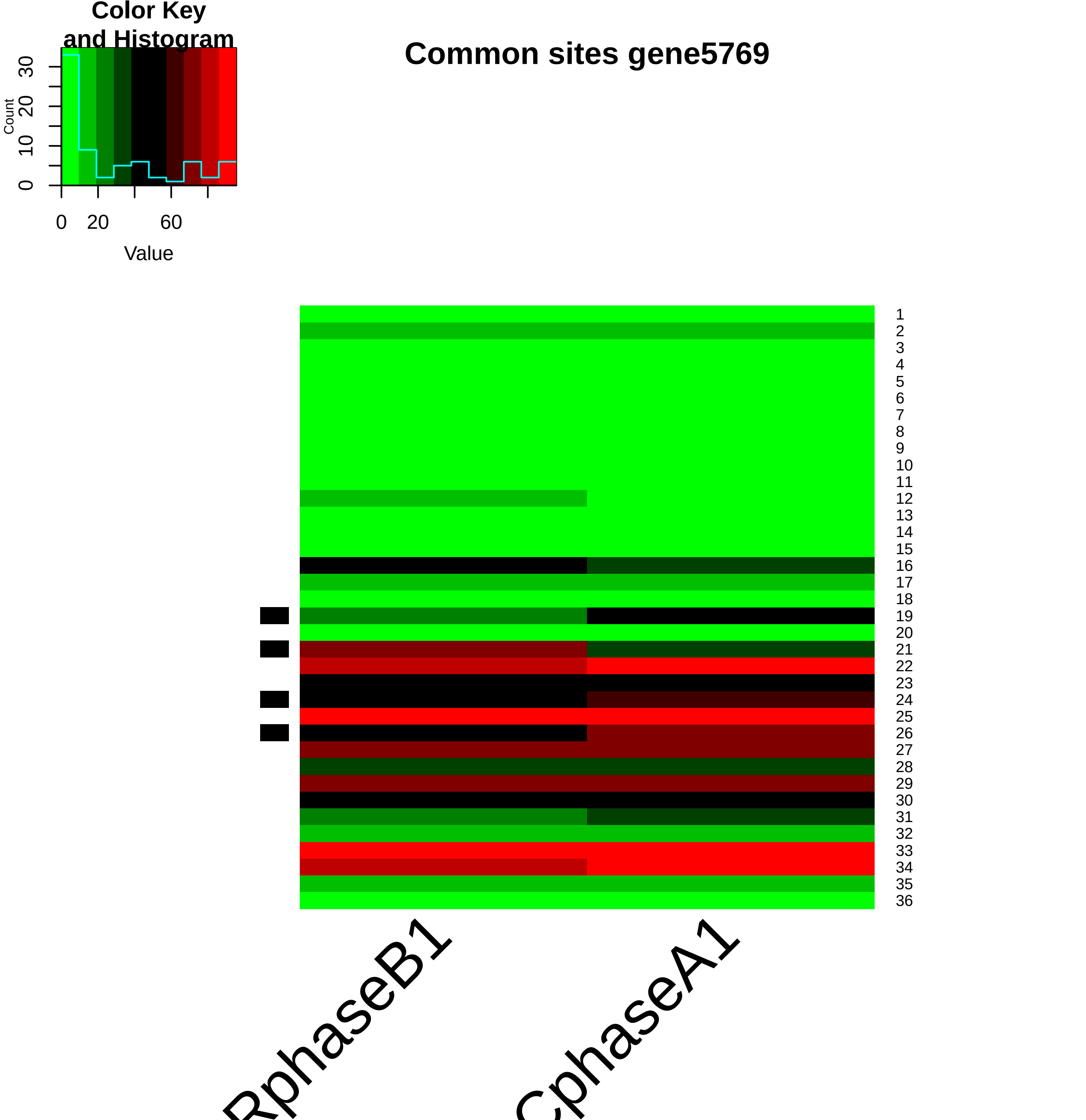
Color Key
and Histogram
Count
0
10
20
30
0
20
60
Value
Common sites gene5769
RphaseB1
CphaseA1

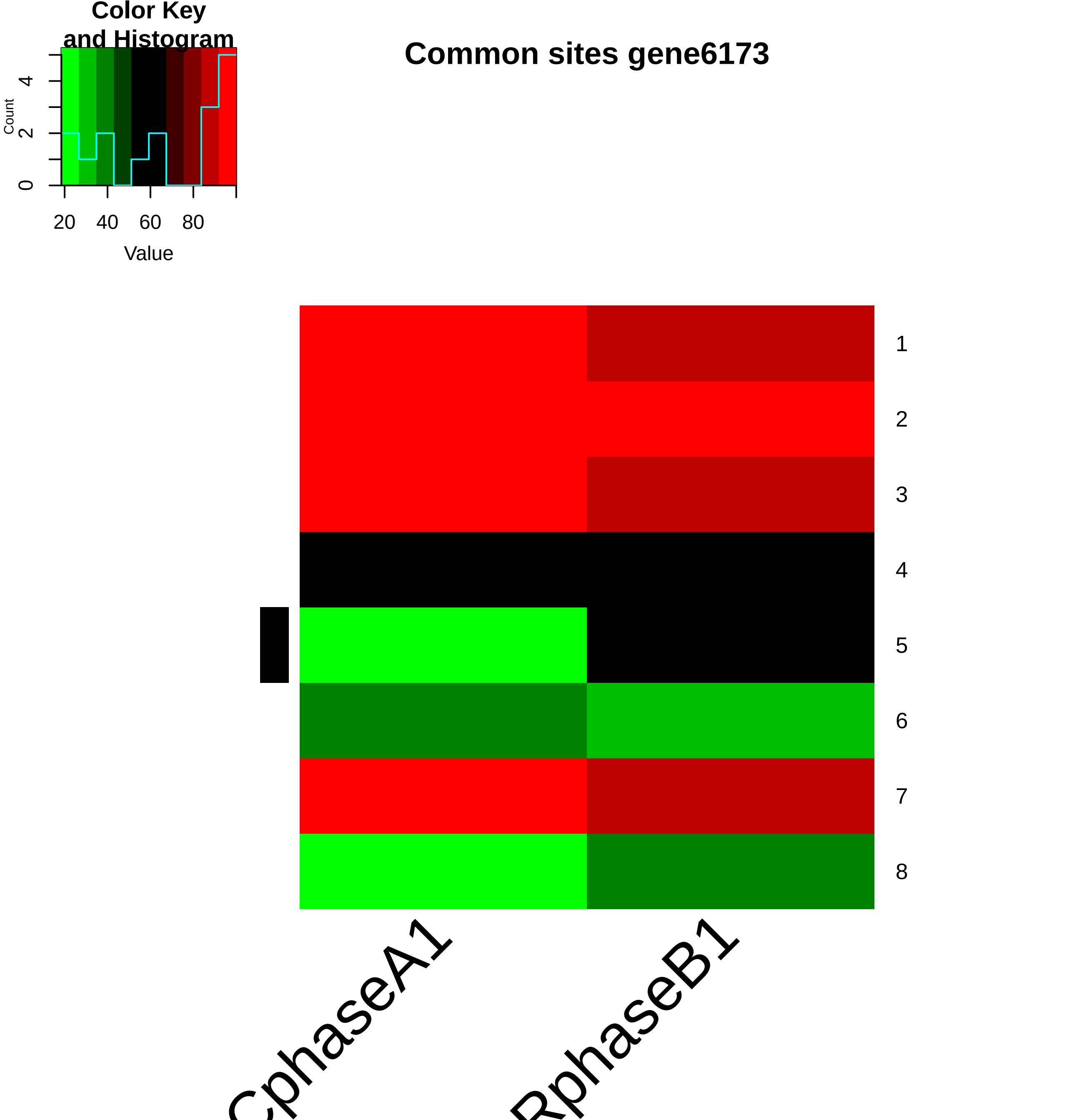
Color Key
and Histogram
Count
0
2
4
20
40
60
80
Value
Common sites gene6173
1
2
3
4
5
6
7
8
CphaseA1
RphaseB1

### Common sites gene6530

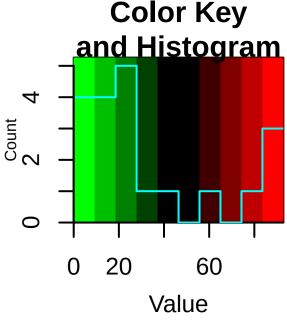

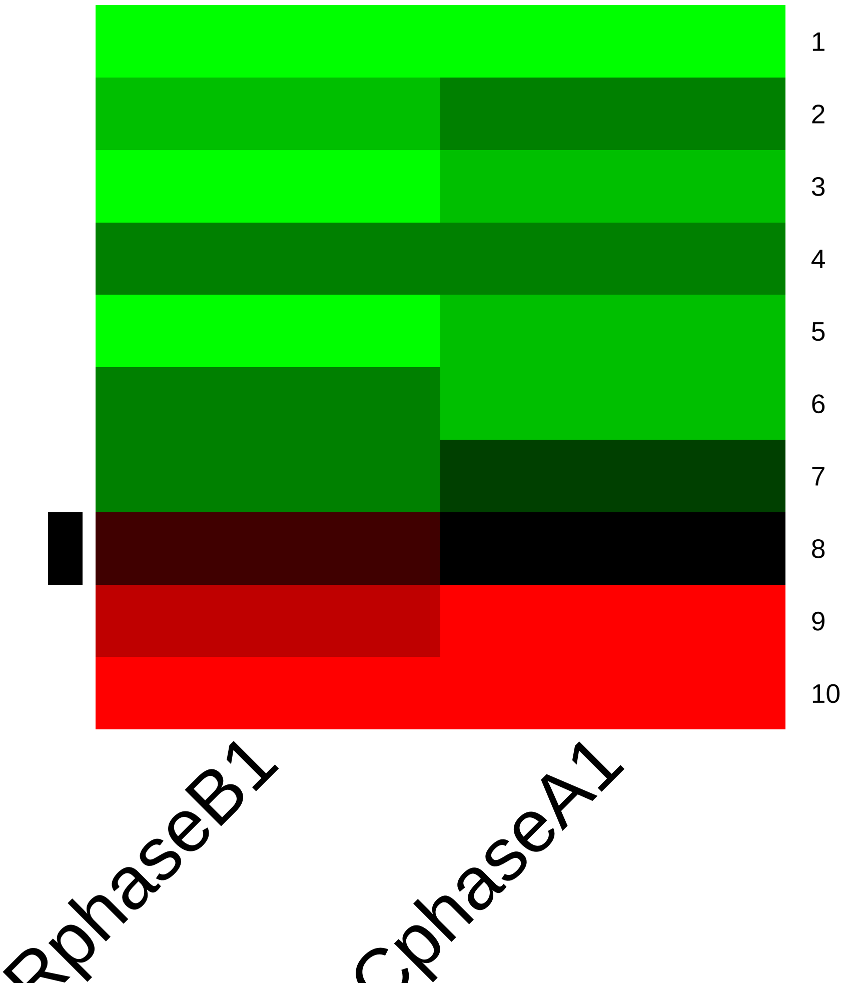

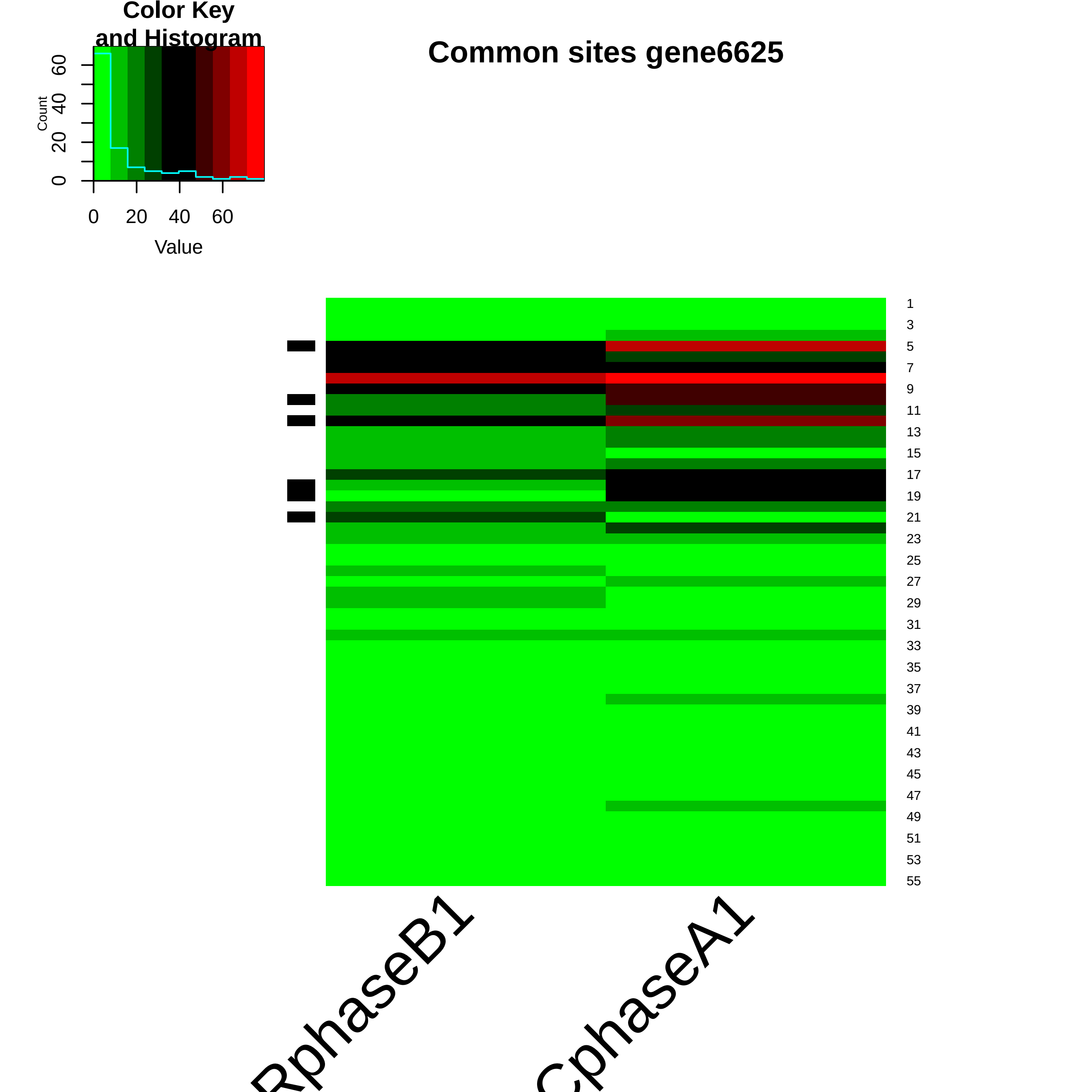
Color Key
and Histogram
Count
0
20
40
60
0
20
40
60
Value
Common sites gene6625
1
3
5
7
9
11
13
15
17
19
21
23
25
27
29
31
33
35
37
39
41
43
45
47
49
51
53
55
RphaseB1
CphaseA1

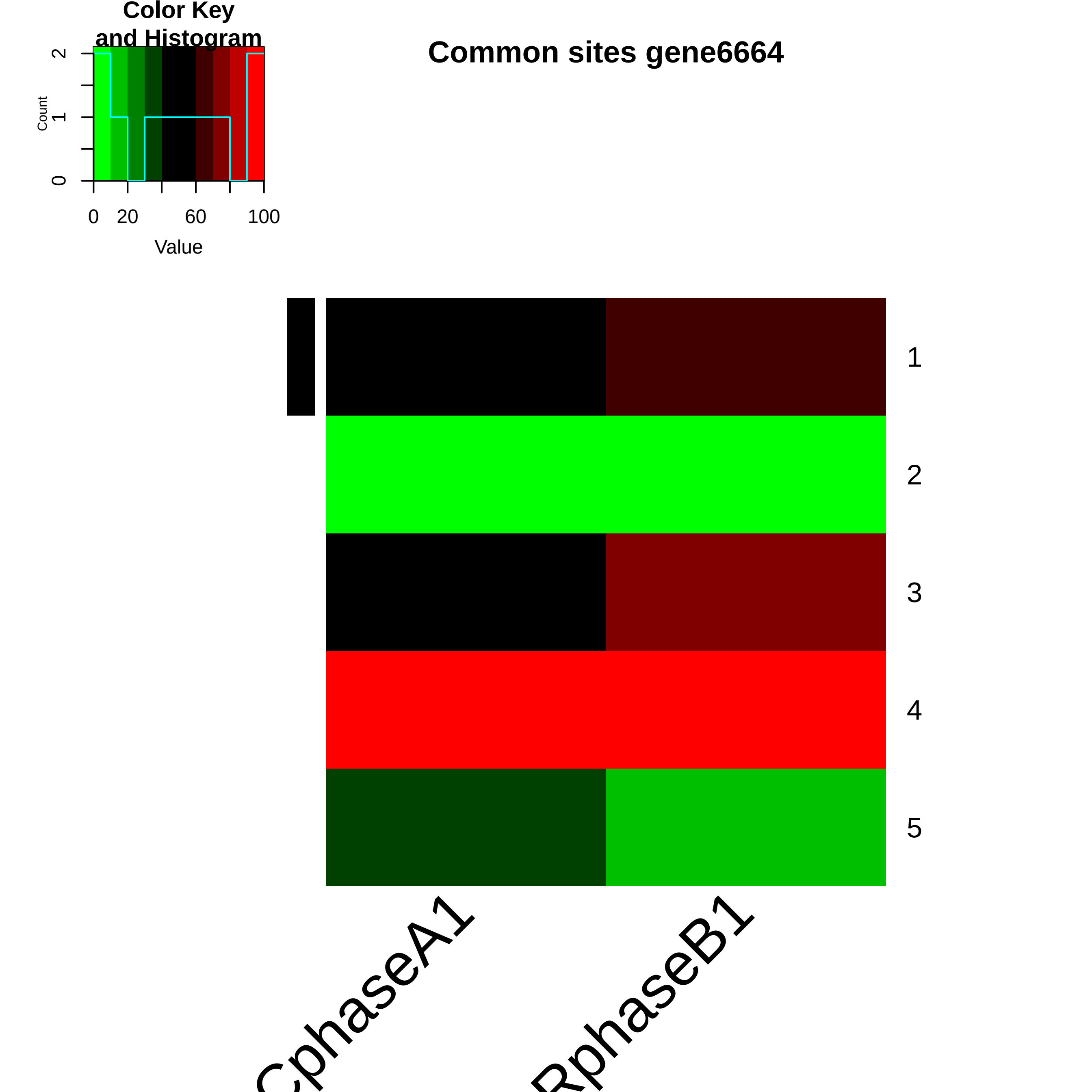

Color Key
and Histogram
Count
0
1
2
0
20
60
100
Value
Common sites gene6664
1
2
3
4
5
CphaseA1
RphaseB1

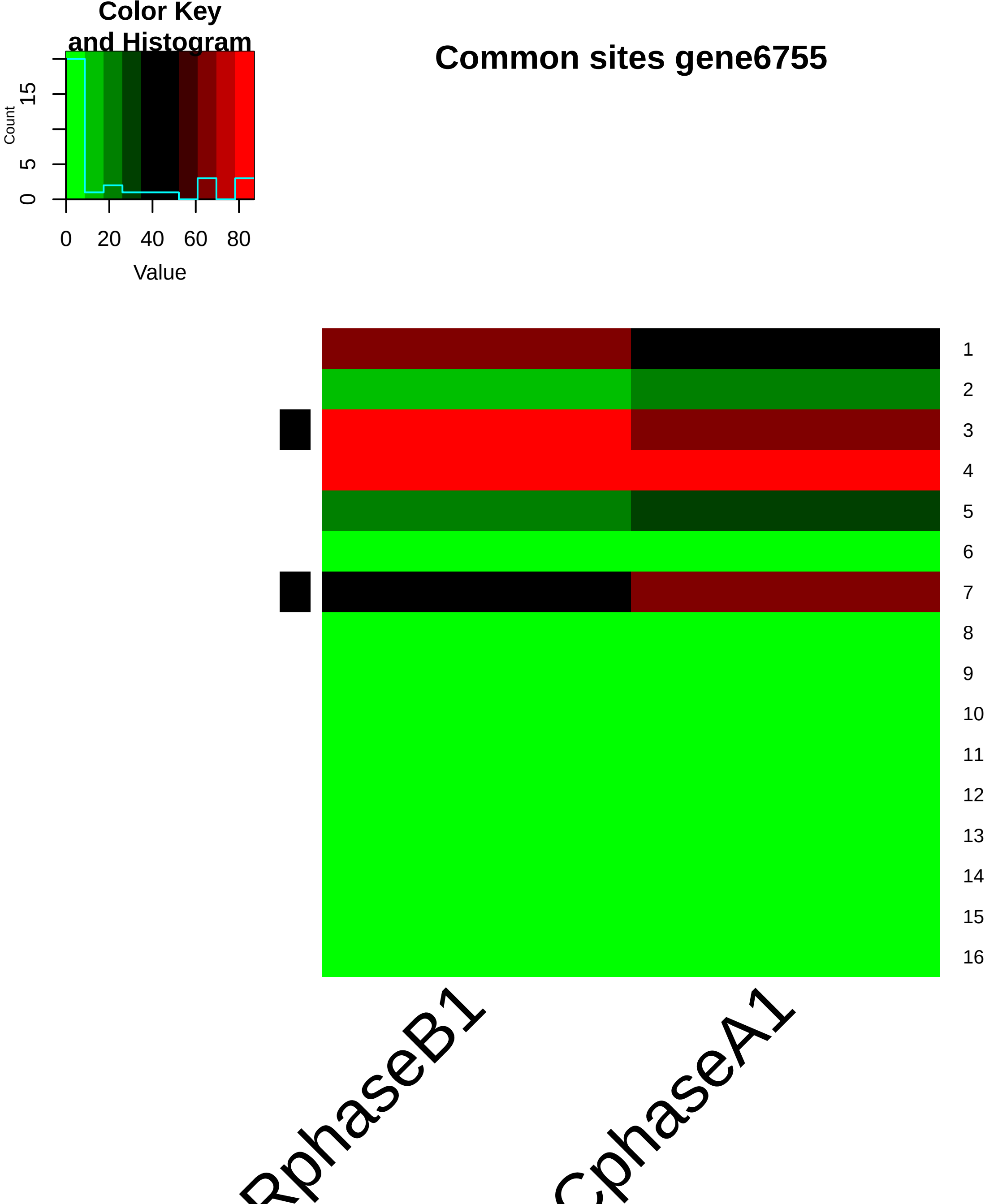

Color Key
and Histogram
Count
0
5
15
0
20
40
60
80
Value
Common sites gene6755
RphaseB1
CphaseA1

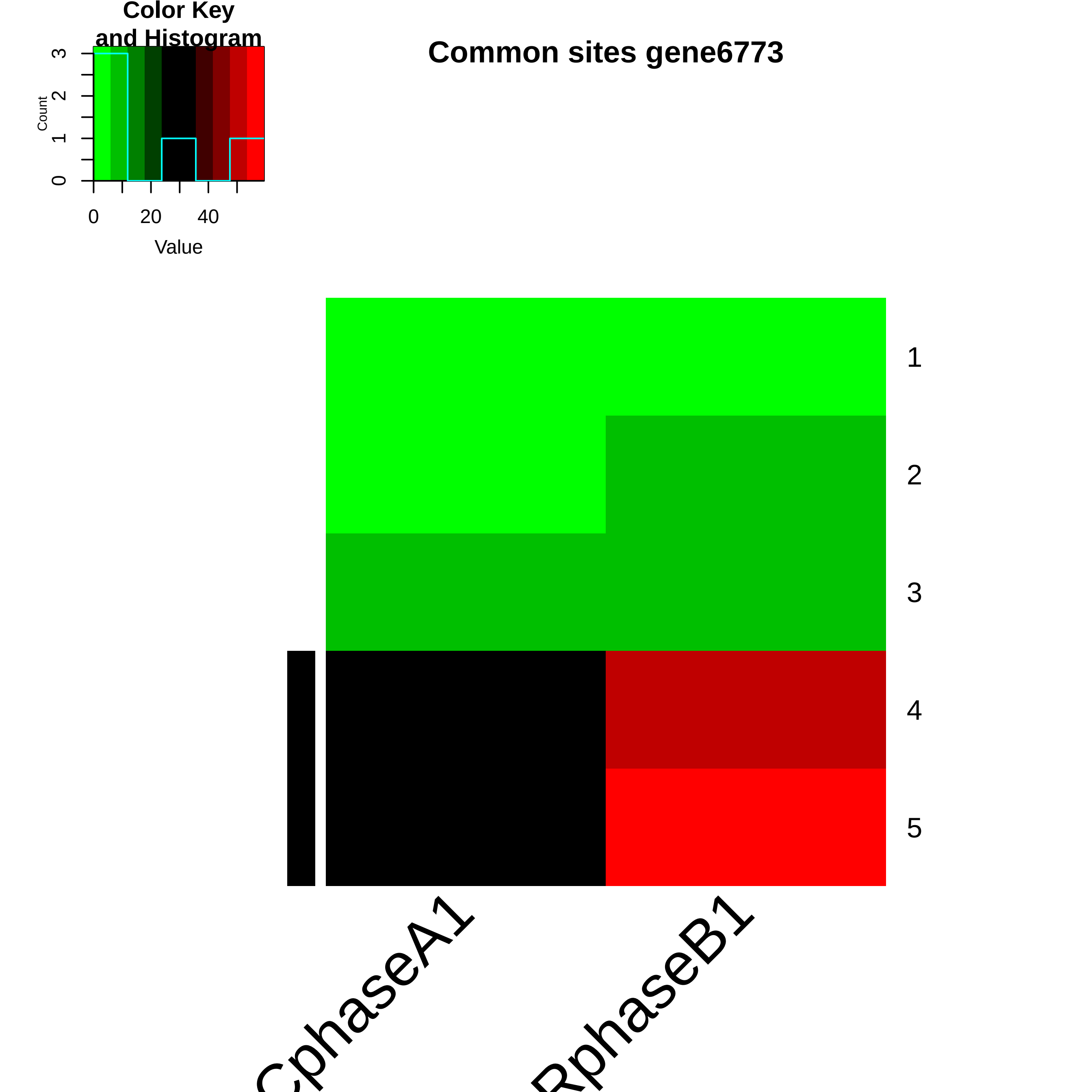

Color Key
and Histogram
Count
0
1
2
3
0
20
40
Value
Common sites gene6773
1
2
3
4
5
CphaseA1
RphaseB1

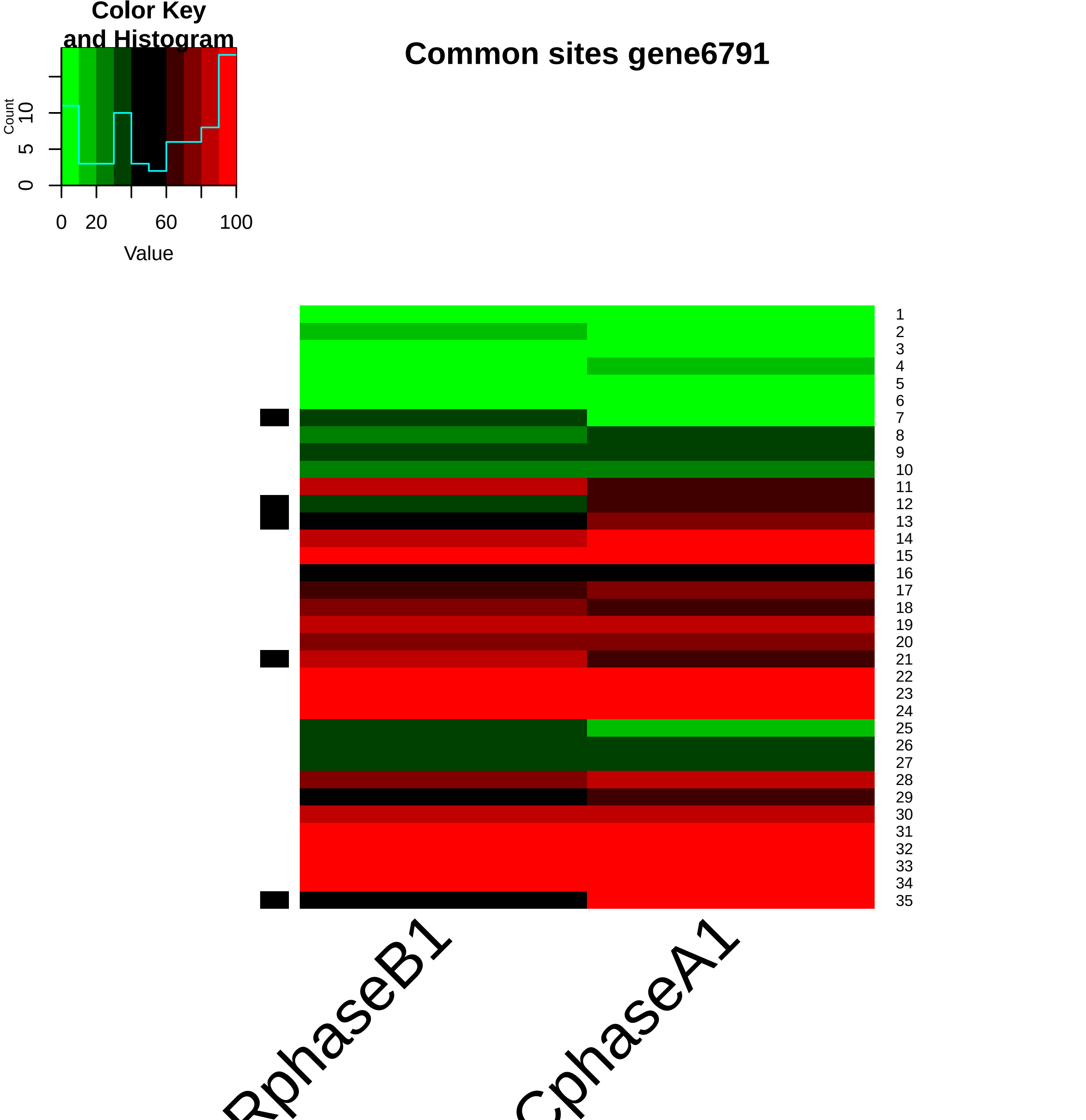

Color Key
and Histogram
Count
0
5
10
0
20
60
100
Value
Common sites gene6791
RphaseB1
CphaseA1

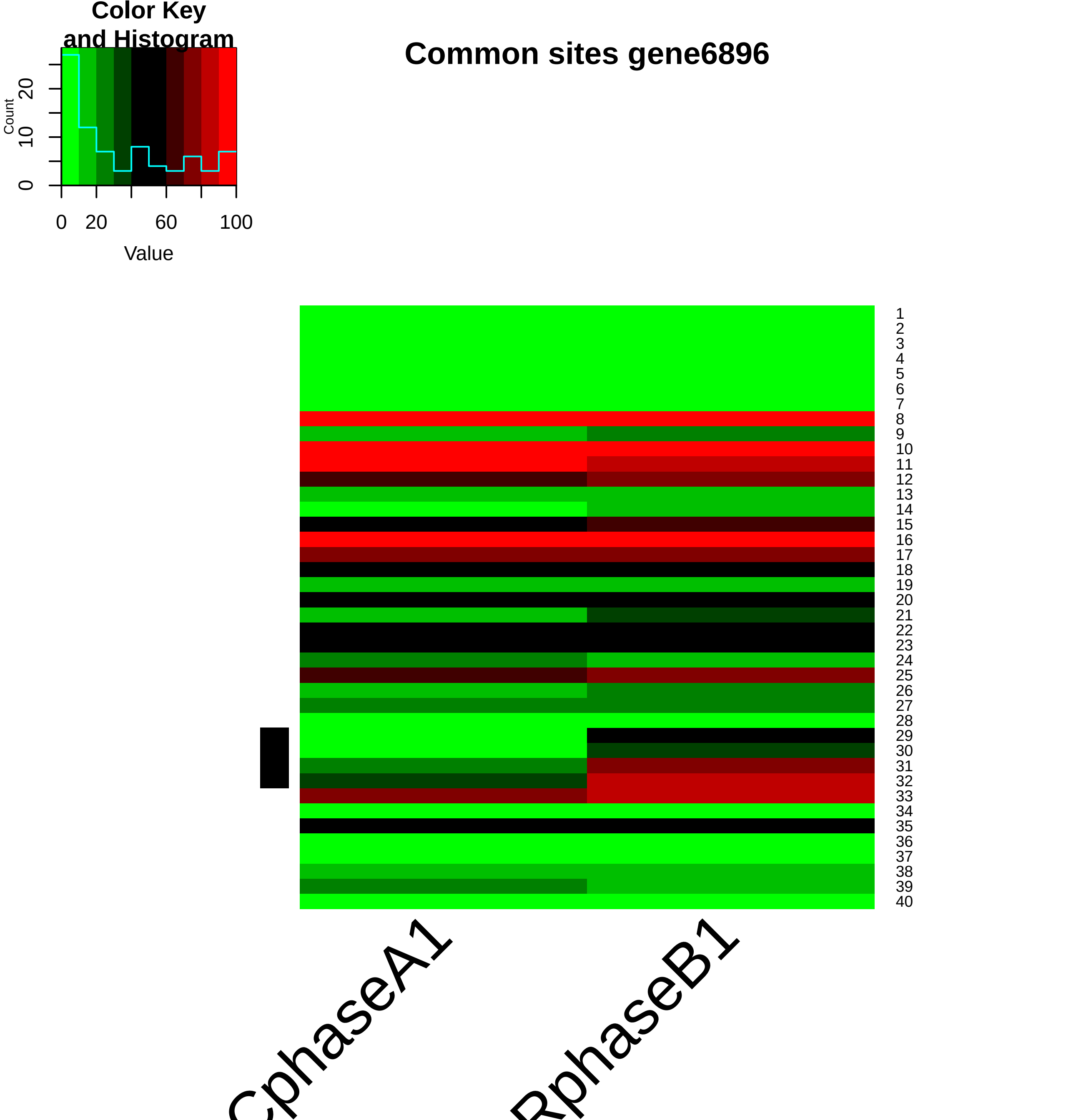

Color Key
and Histogram
Count
0
10
20
0
20
60
100
Value
Common sites gene6896
CphaseA1
RphaseB1

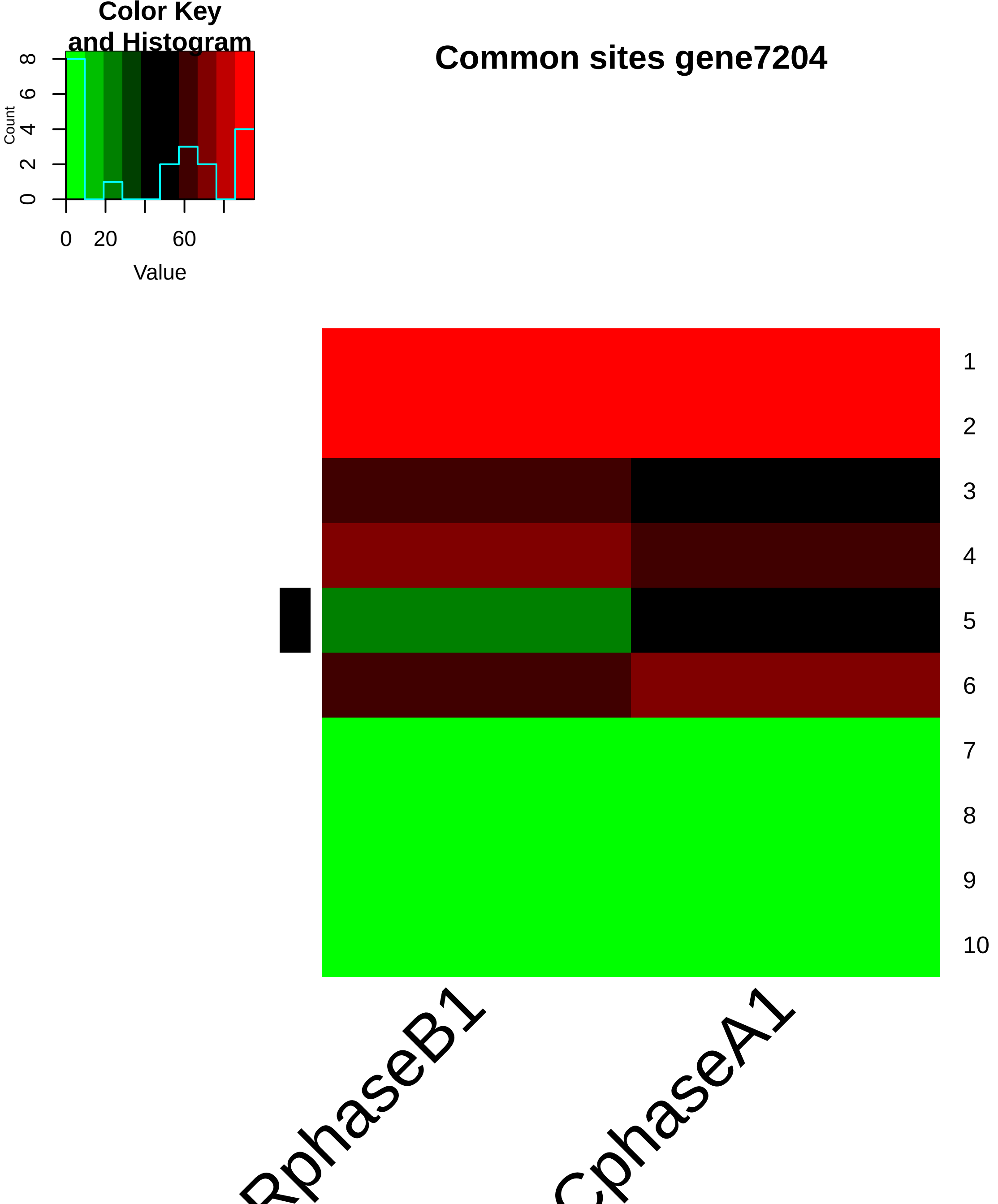
Color Key
and Histogram
Count
0
2
4
6
8
0
20
60
Value
Common sites gene7204
1
2
3
4
5
6
7
8
9
10
RphaseB1
CphaseA1

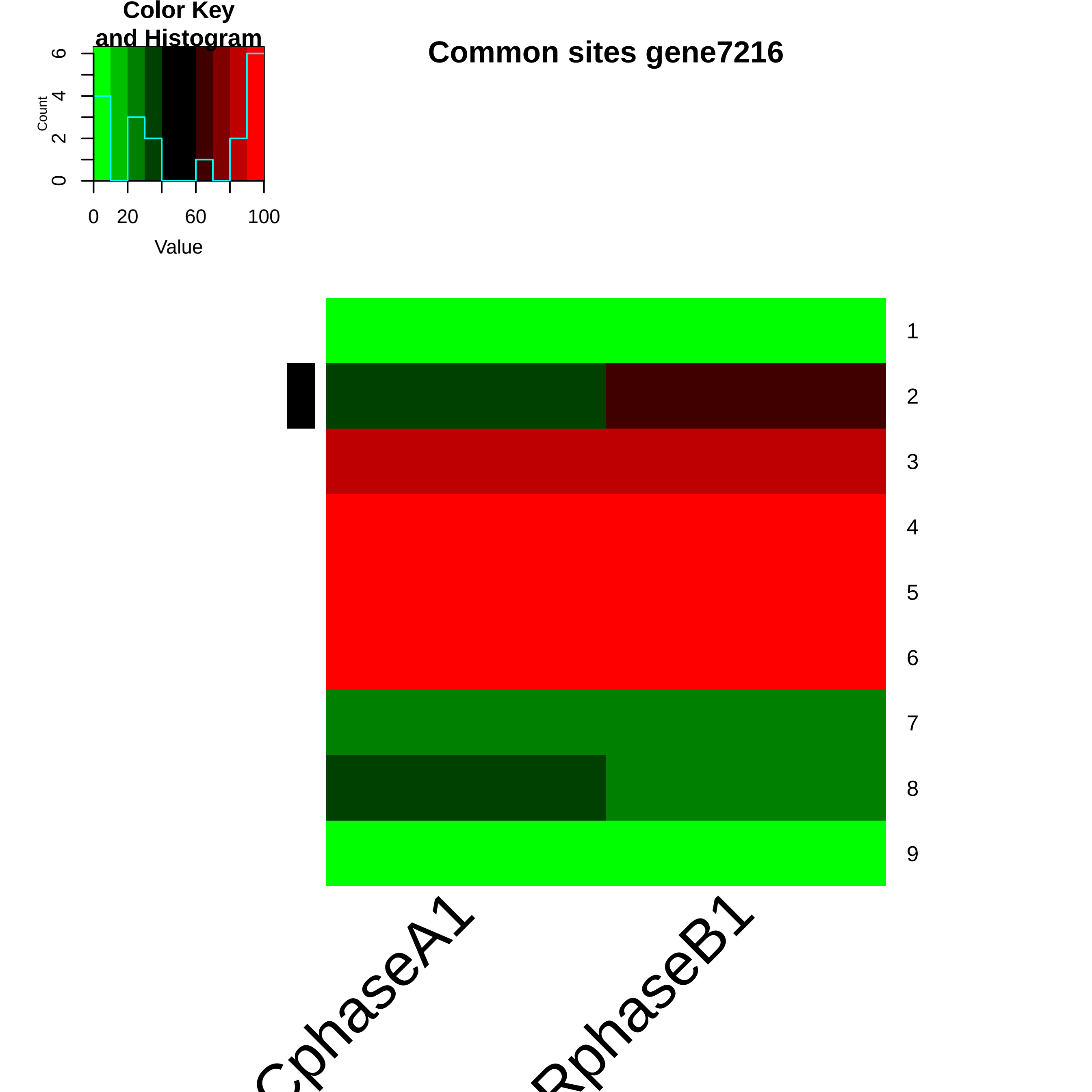
Color Key
and Histogram
Count
0
2
4
6
0
20
60
100
Value
Common sites gene7216
1
2
3
4
5
6
7
8
9

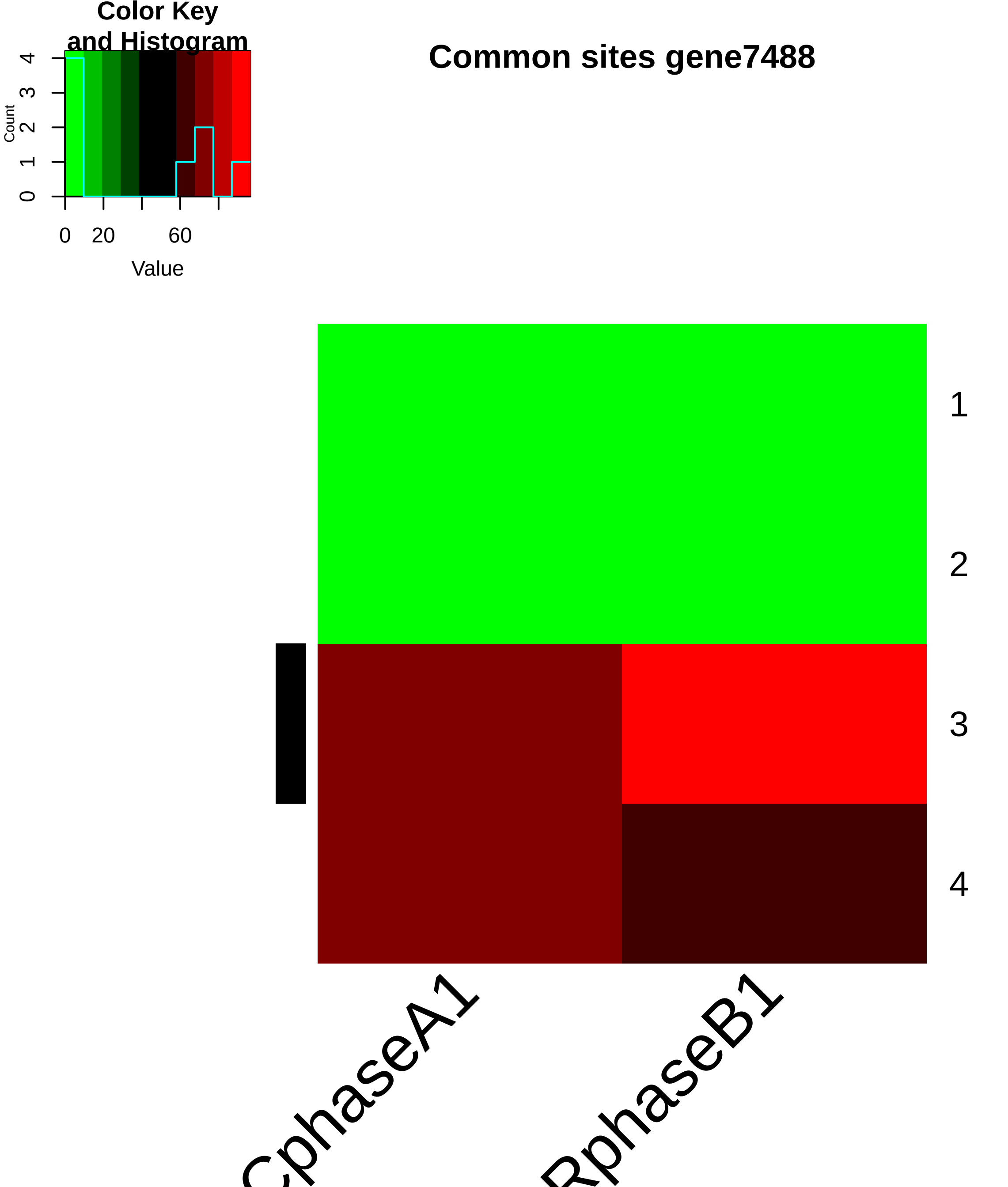
Color Key
and Histogram
Count
Value
0
20
60
0
1
2
3
4
Common sites gene7488
1
2
3
4
CphaseA1
RphaseB1

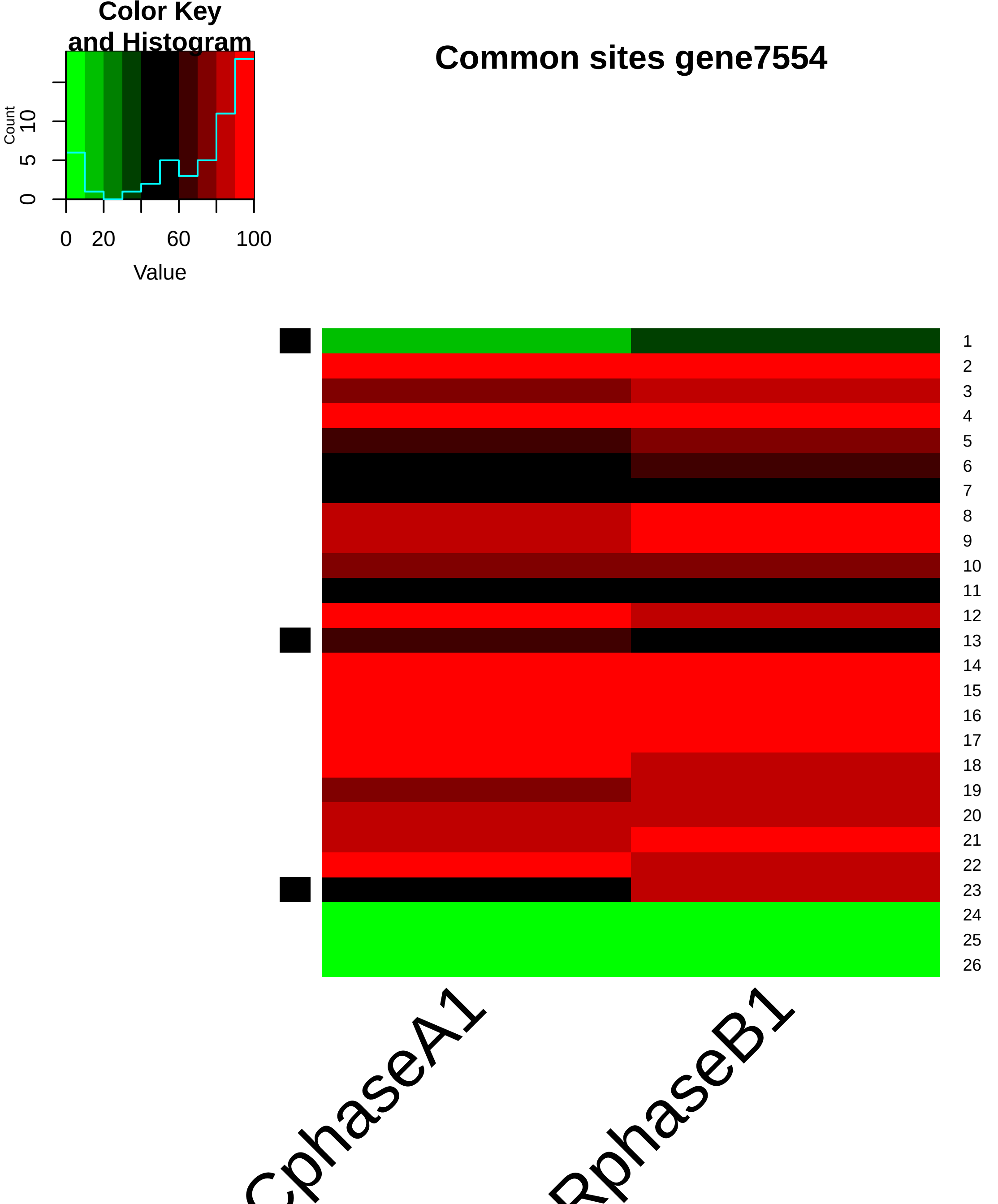
Color Key
and Histogram
Count
0
5
10
0
20
60
100
Value
Common sites gene7554
CphaseA1
RphaseB1

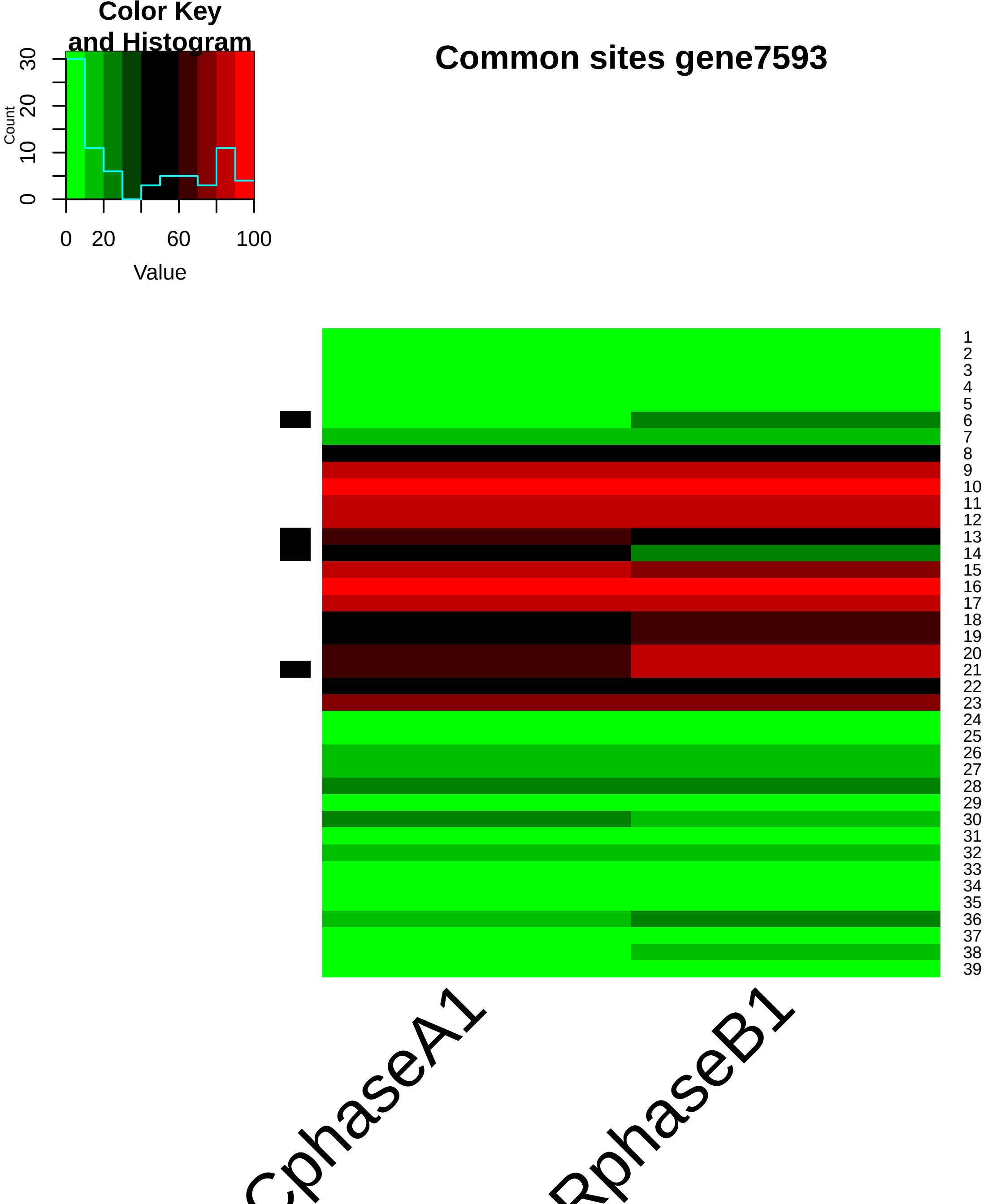

Color Key
and Histogram
Count
0
10
20
30
0
20
60
100
Value
Common sites gene7593
CphaseA1
RphaseB1

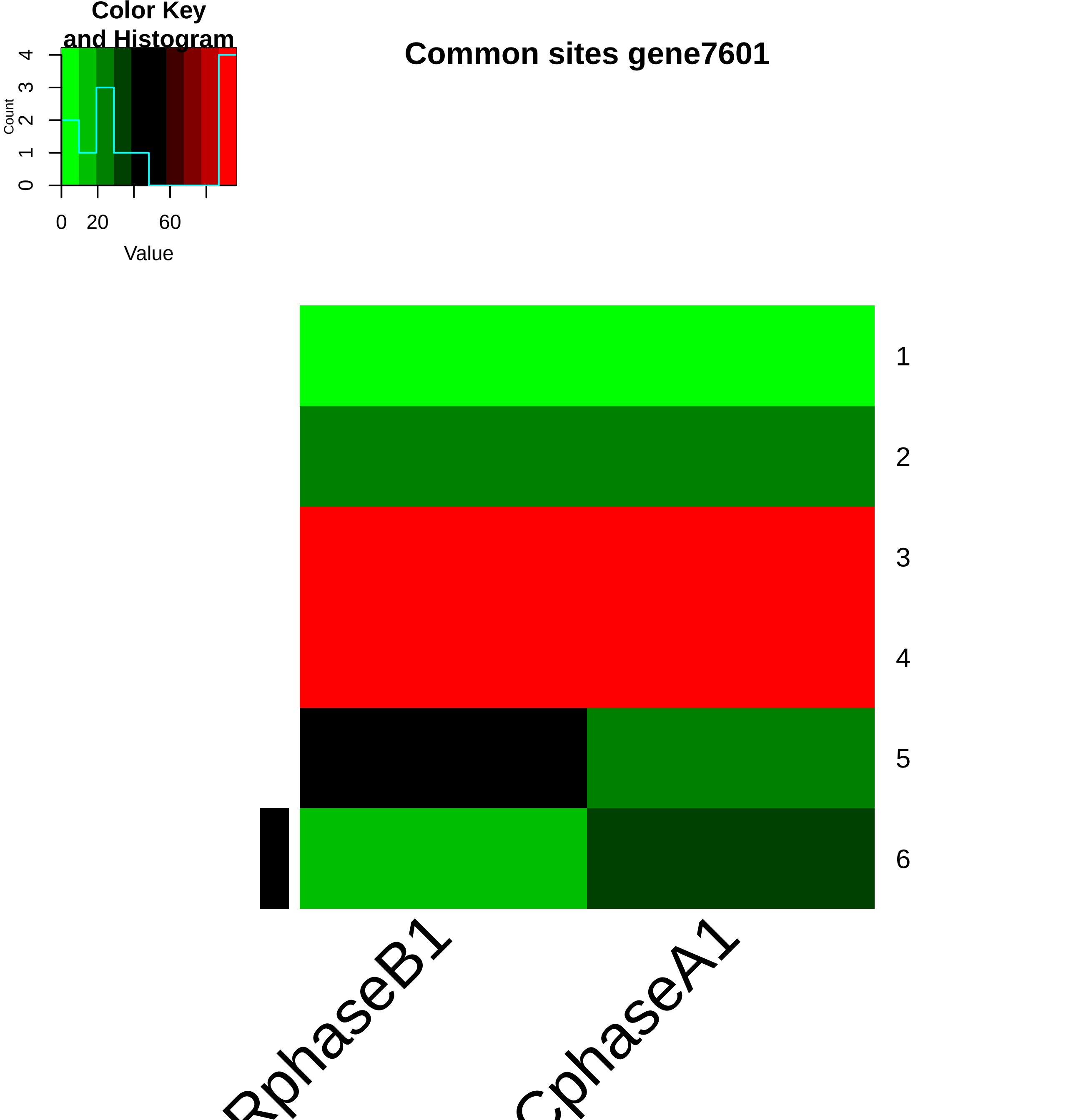
Color Key
and Histogram
Count
0
1
2
3
4
0
20
60
Value
Common sites gene7601
1
2
3
4
5
6
RphaseB1
CphaseA1

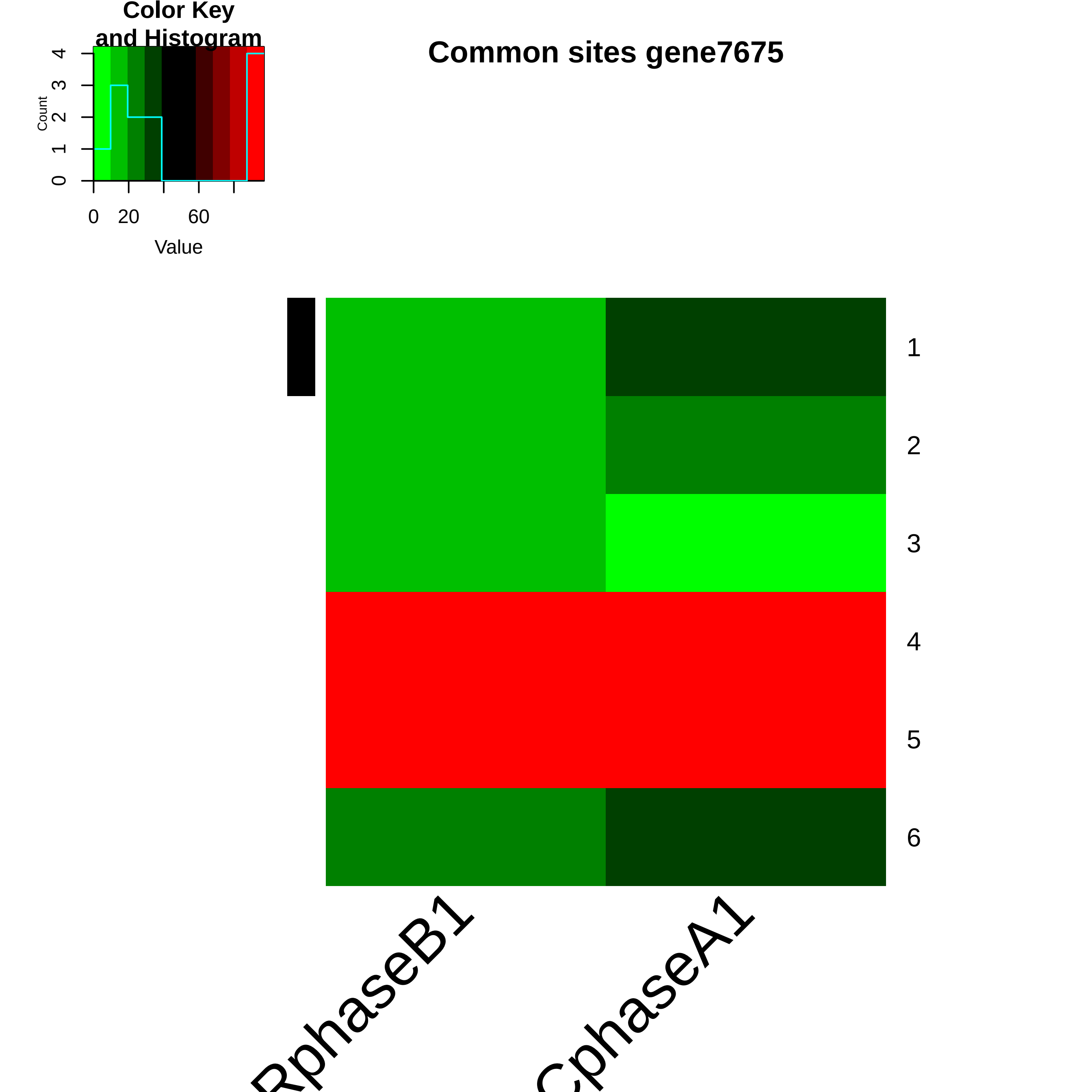
Color Key
and Histogram
Count
Value
0
20
60
0
1
2
3
4
Common sites gene7675
1
2
3
4
5
6
RphaseB1
CphaseA1

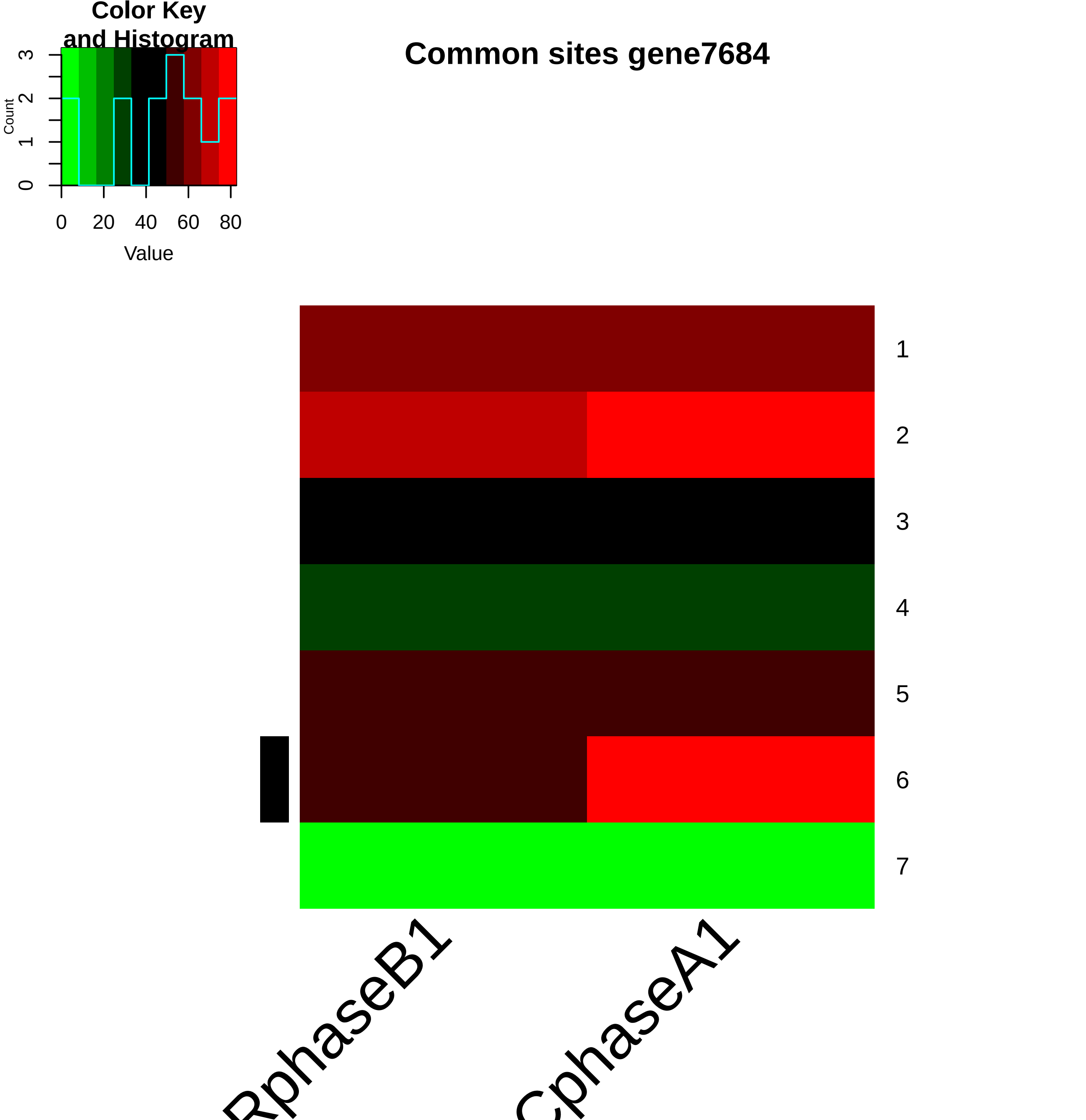
Color Key
and Histogram
Count
0
1
2
3
0
20
40
60
80
Value
Common sites gene7684
1
2
3
4
5
6
7
RphaseB1
CphaseA1

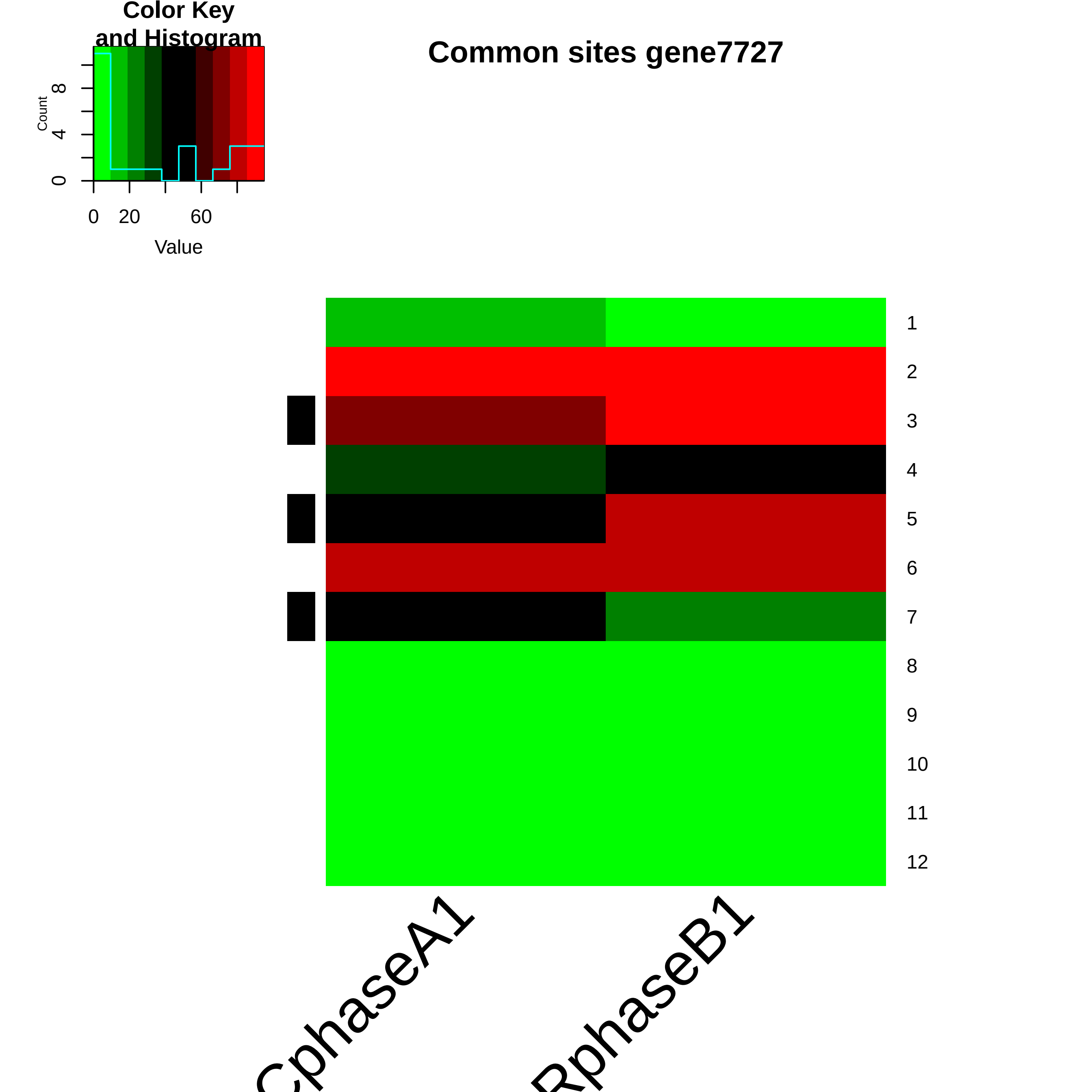
Color Key
and Histogram
Count
0
4
8
0
20
60
Value
Common sites gene7727
1
2
3
4
5
6
7
8
9
10
11
12
CphaseA1
RphaseB1

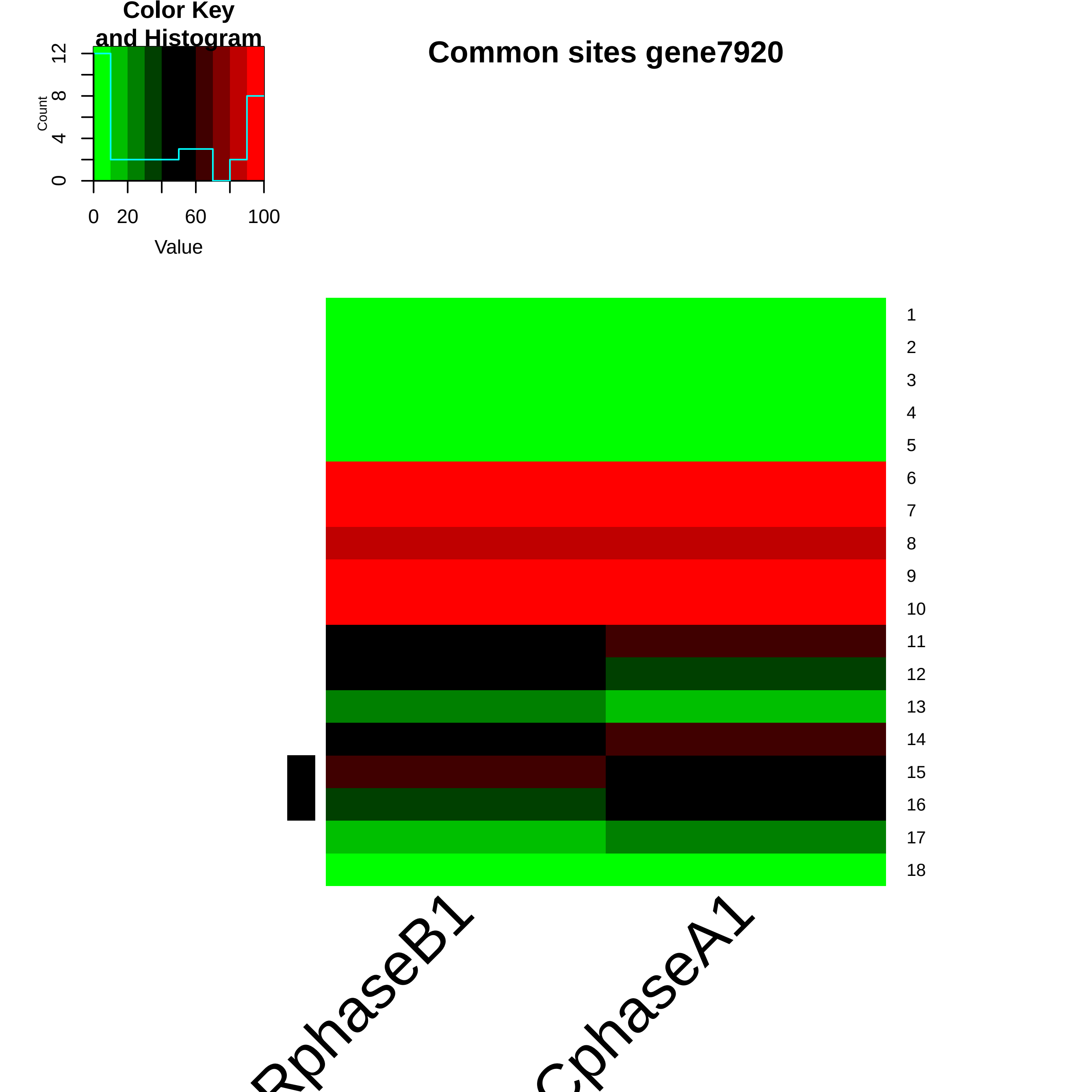

Color Key
and Histogram
Count
0
4
8
12
0
20
60
100
Value
Common sites gene7920
RphaseB1
CphaseA1

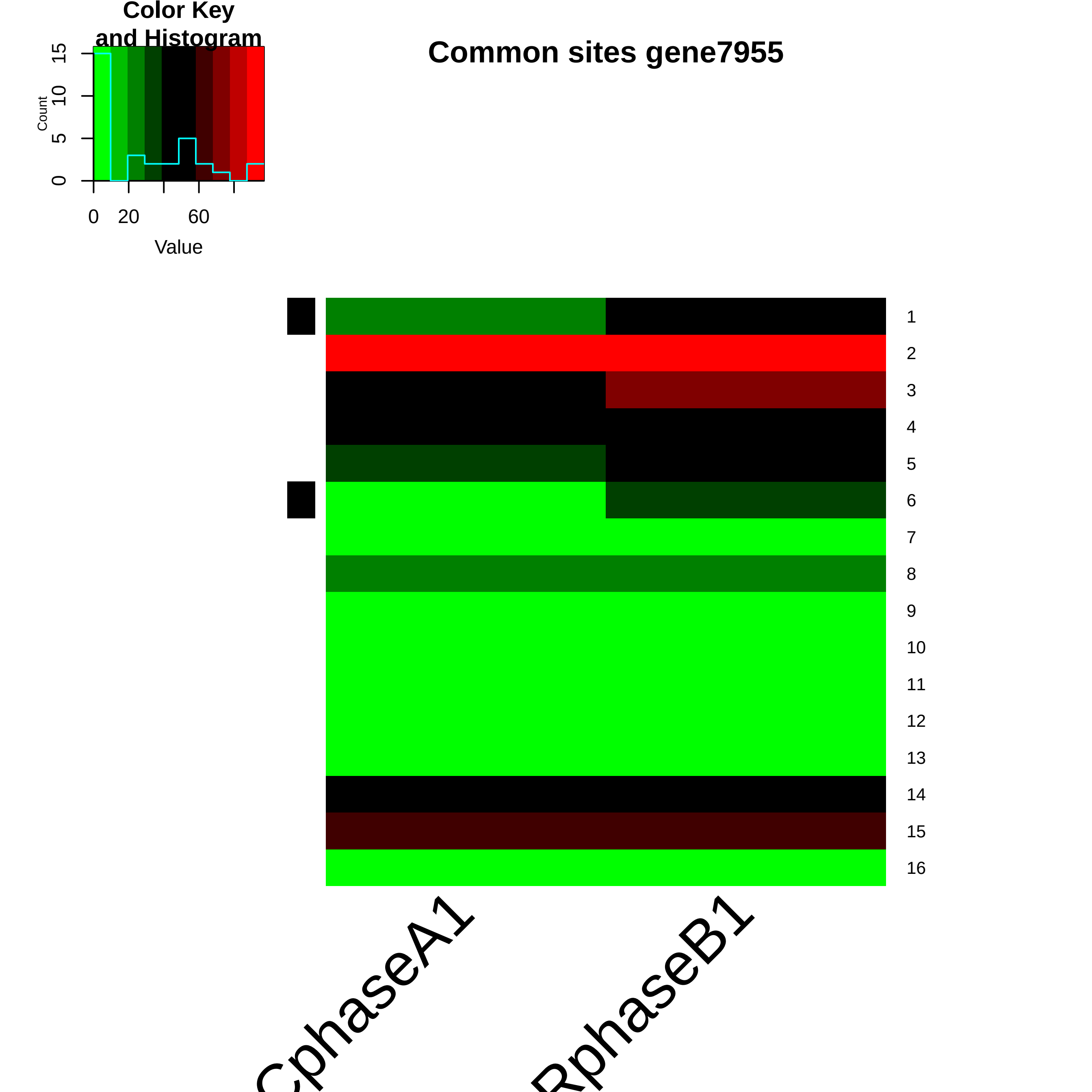

Color Key
and Histogram
Count
0
5
10
15
0
20
60
Value
Common sites gene7955
CphaseA1
RphaseB1

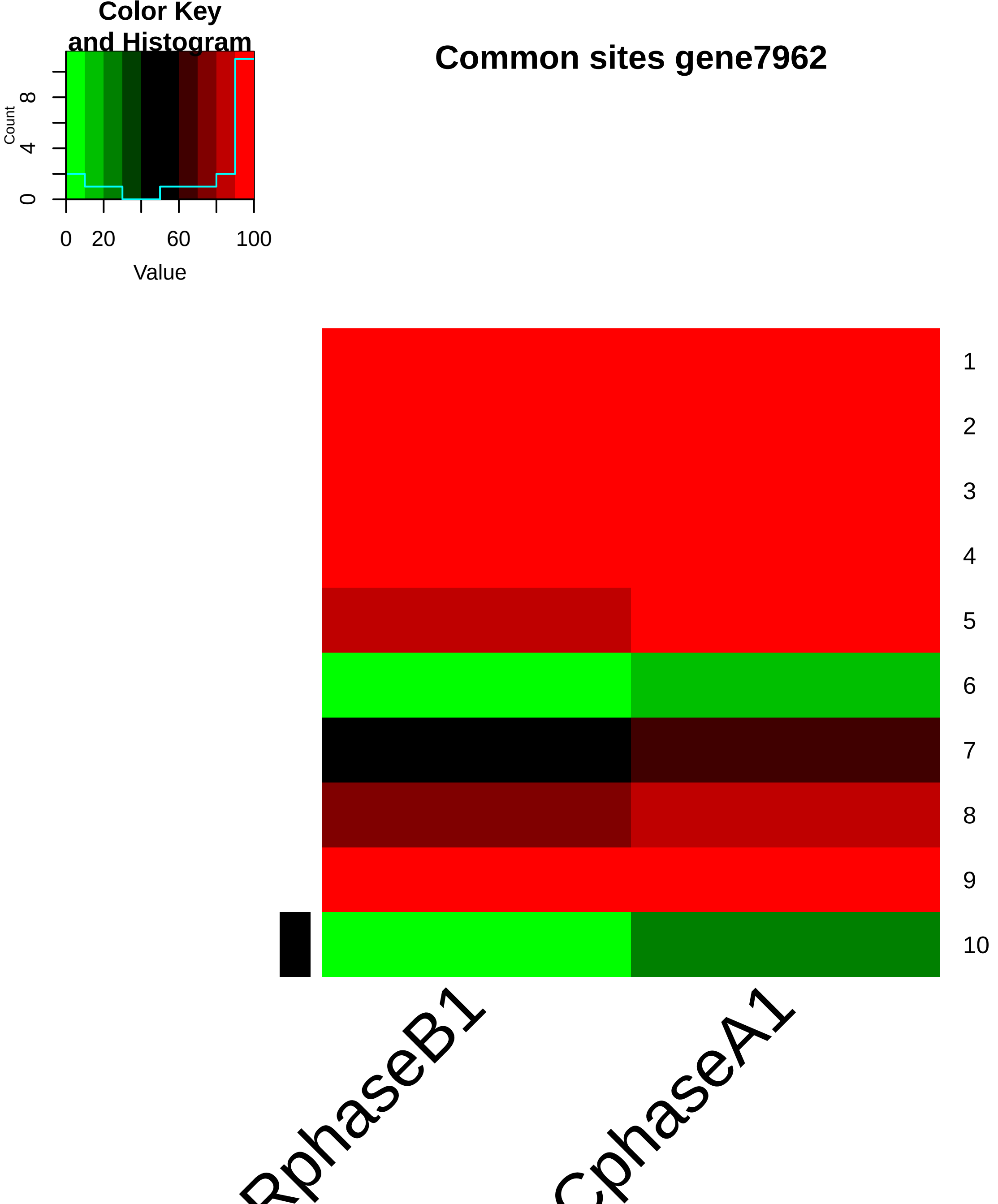

Color Key
and Histogram
Count
0
4
8
0
20
60
100
Value
Common sites gene7962
1
2
3
4
5
6
7
8
9
10
RphaseB1
CphaseA1

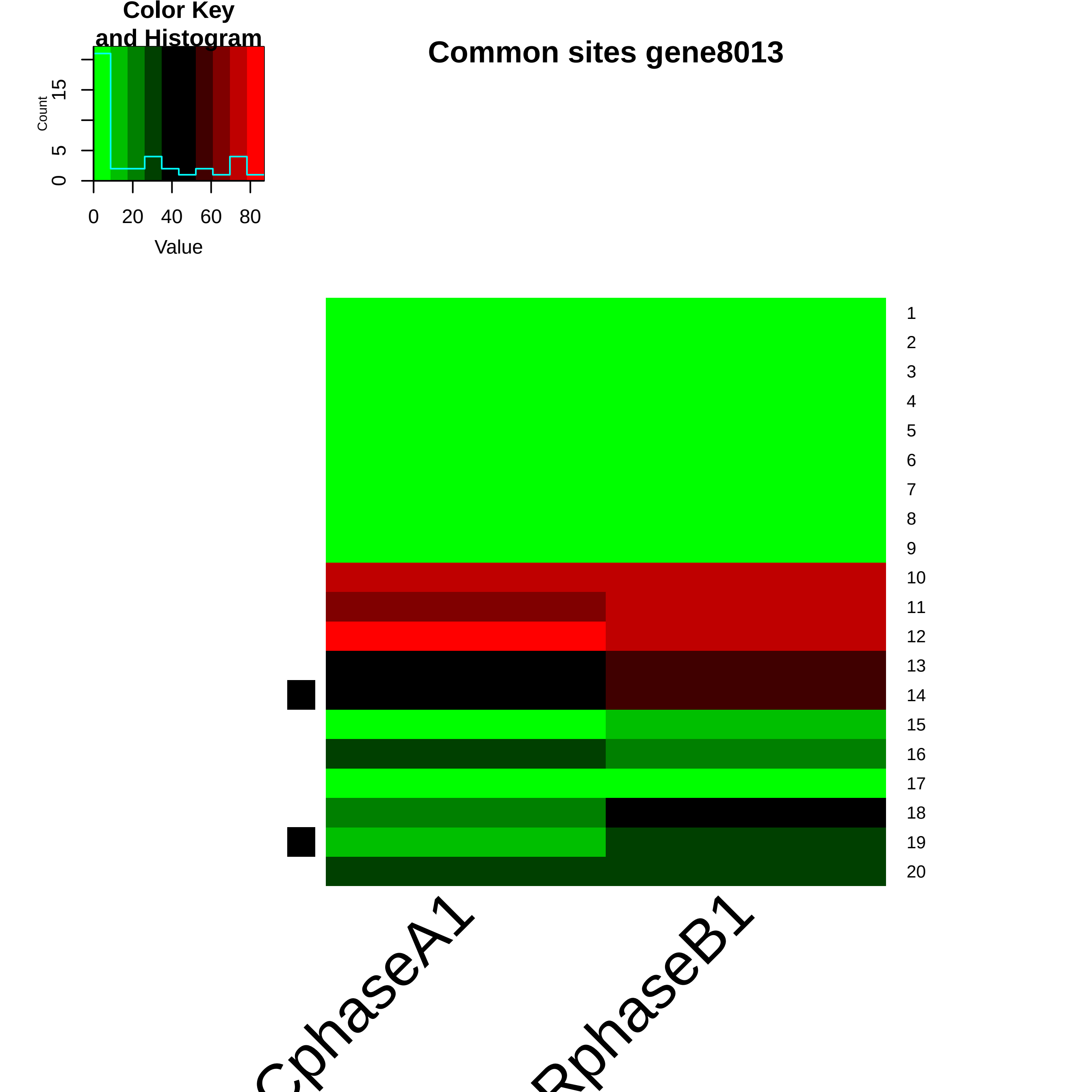
Color Key
and Histogram
Count
0
5
15
0
20
40
60
80
Value
Common sites gene8013
CphaseA1
RphaseB1

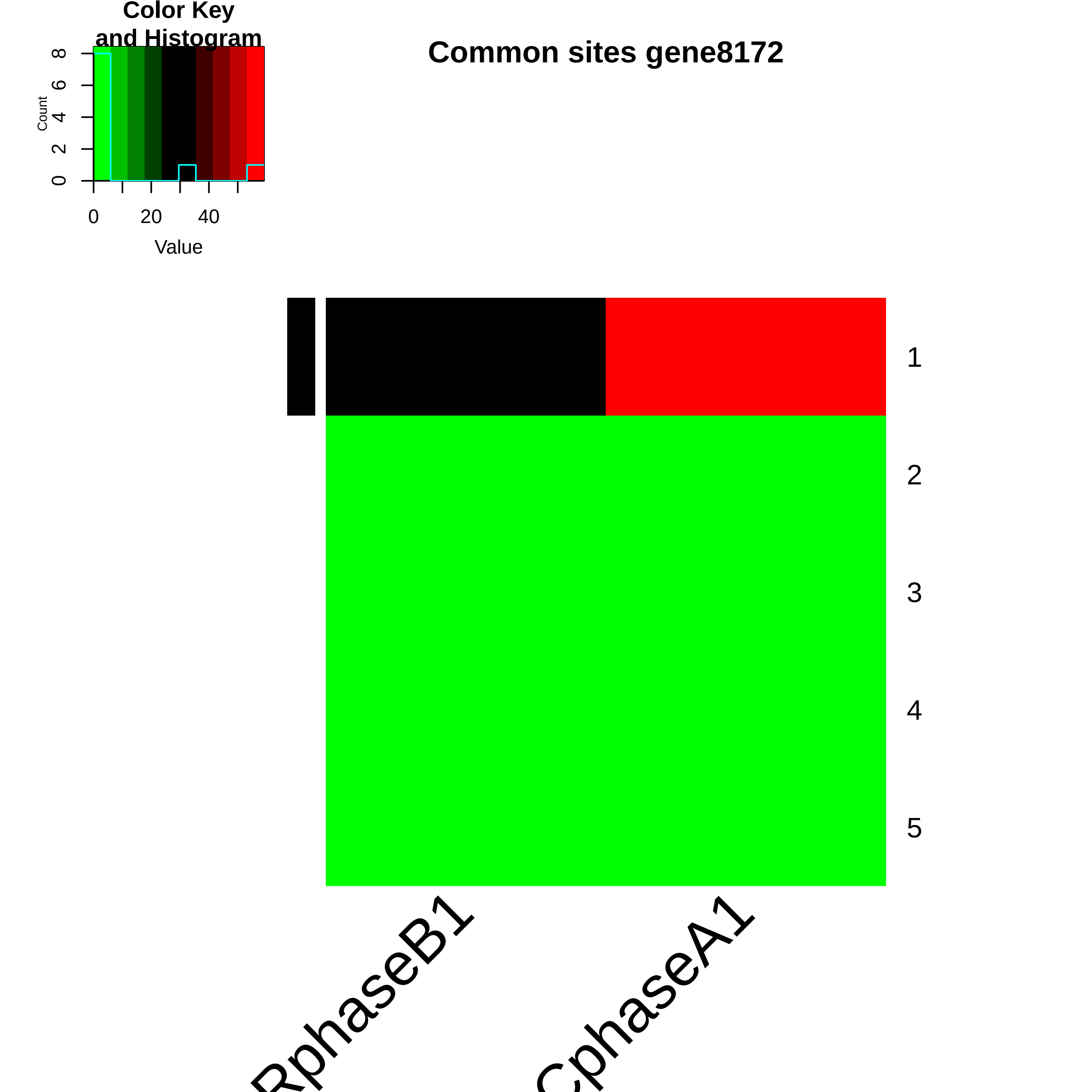
Color Key
and Histogram
Count
0
2
4
6
8
0
20
40
Value
Common sites gene8172
1
2
3
4
5
RphaseB1
CphaseA1

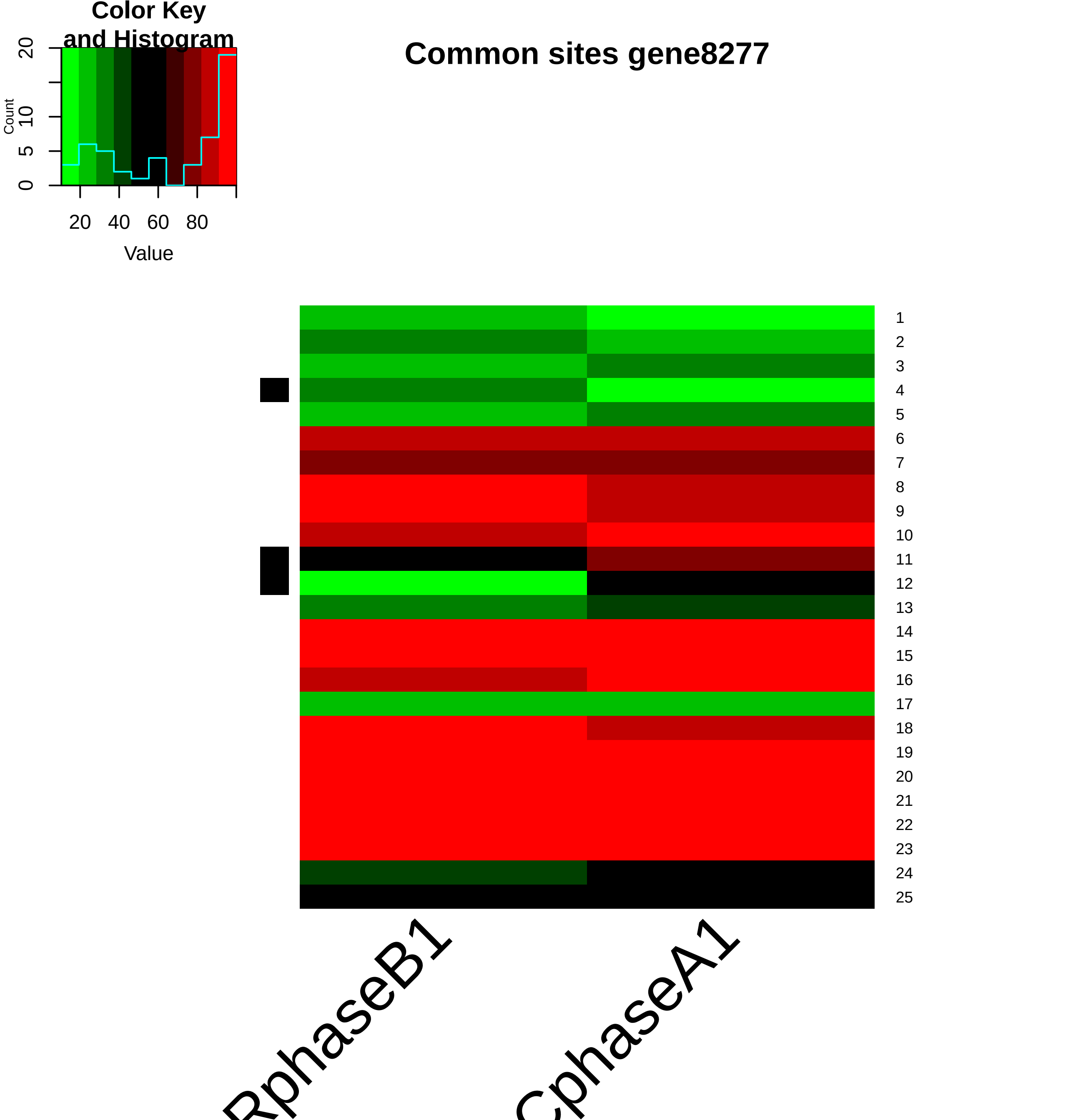

Color Key
and Histogram
Count
0
5
10
20
20
40
60
80
Value
Common sites gene8277
RphaseB1
CphaseA1

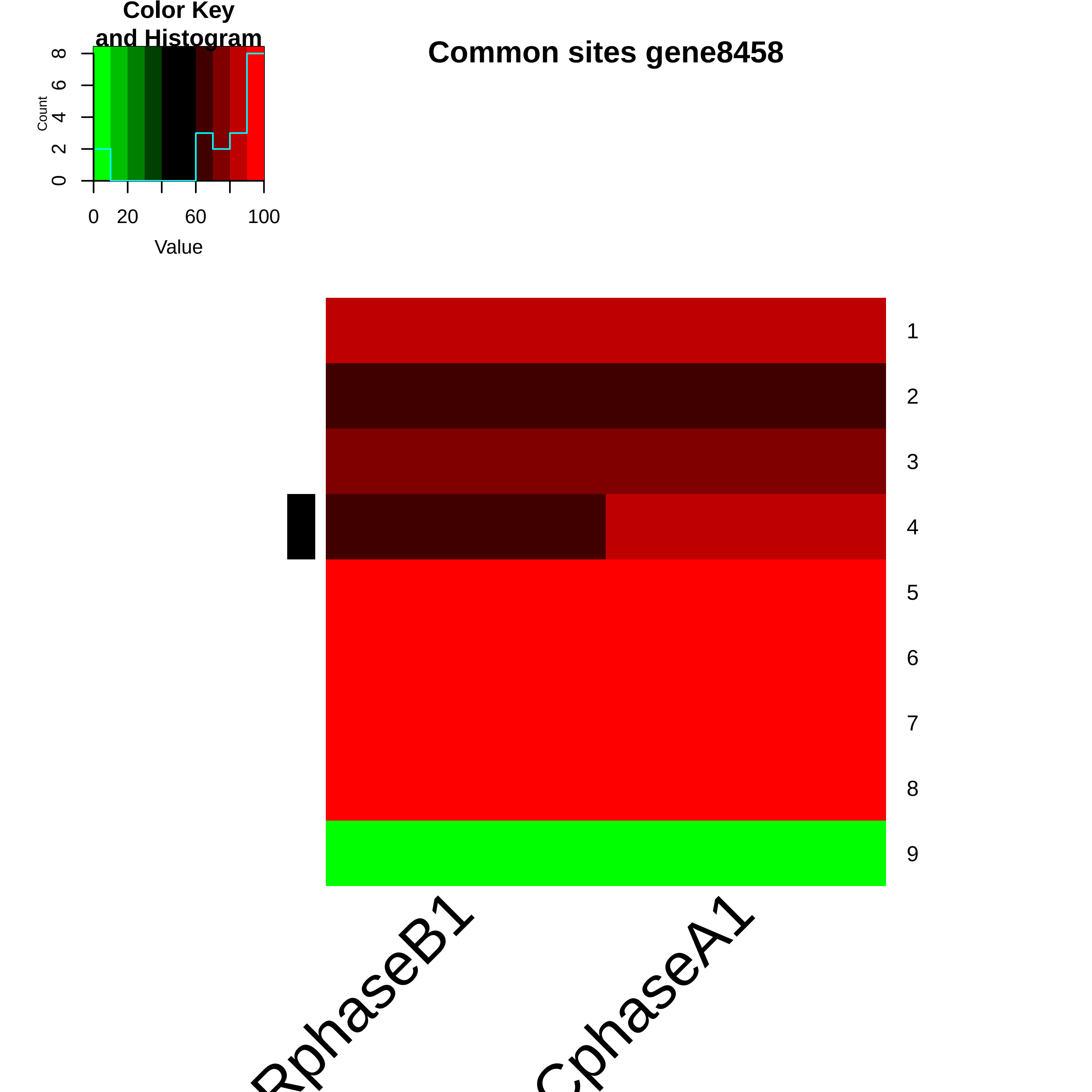

Color Key
and Histogram
Count
0
2
4
6
8
0
20
60
100
Value
Common sites gene8458
1
2
3
4
5
6
7
8
9
RphaseB1
CphaseA1

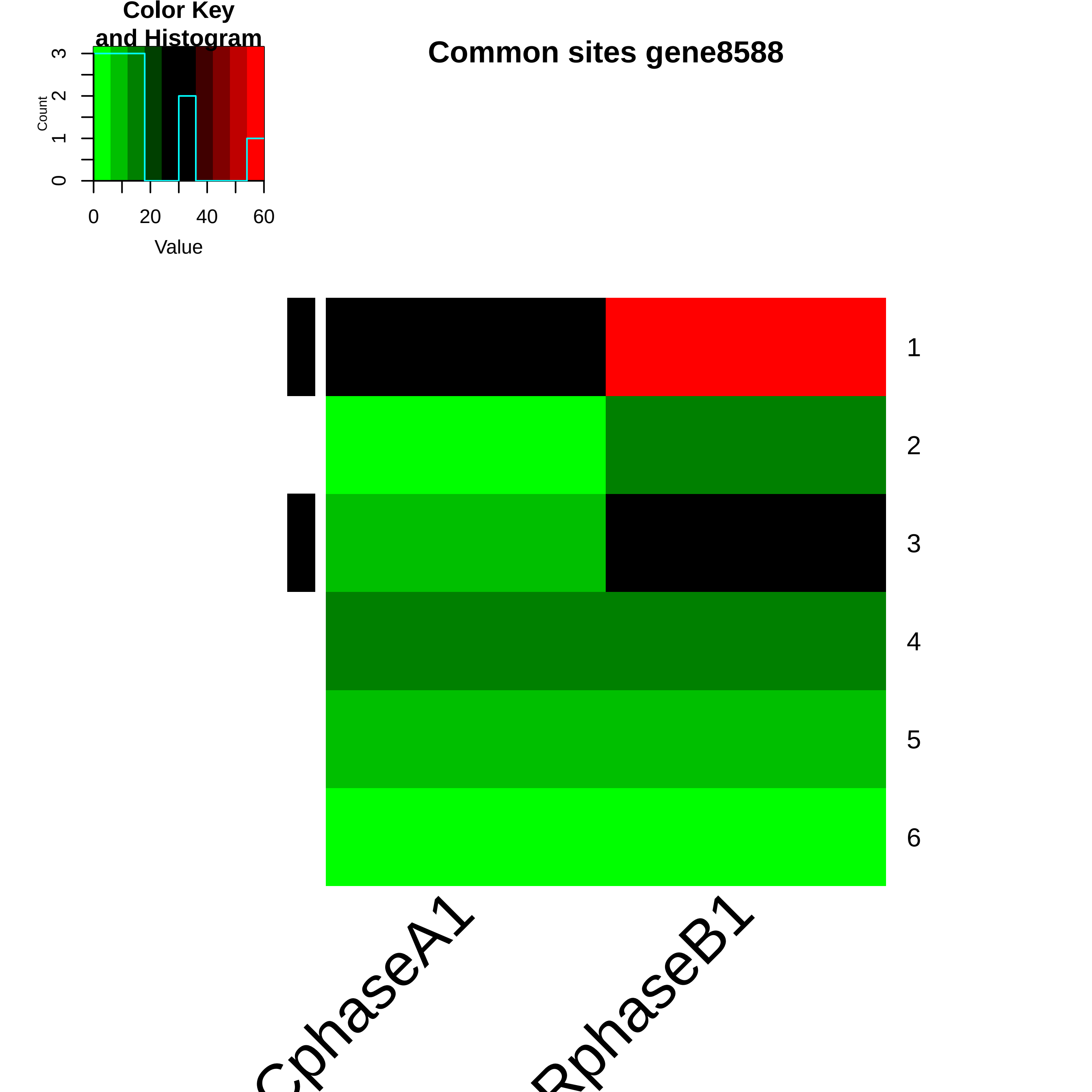

Color Key
and Histogram
Count
0
1
2
3
0
20
40
60
Value
Common sites gene8588
1
2
3
4
5
6
CphaseA1
RphaseB1

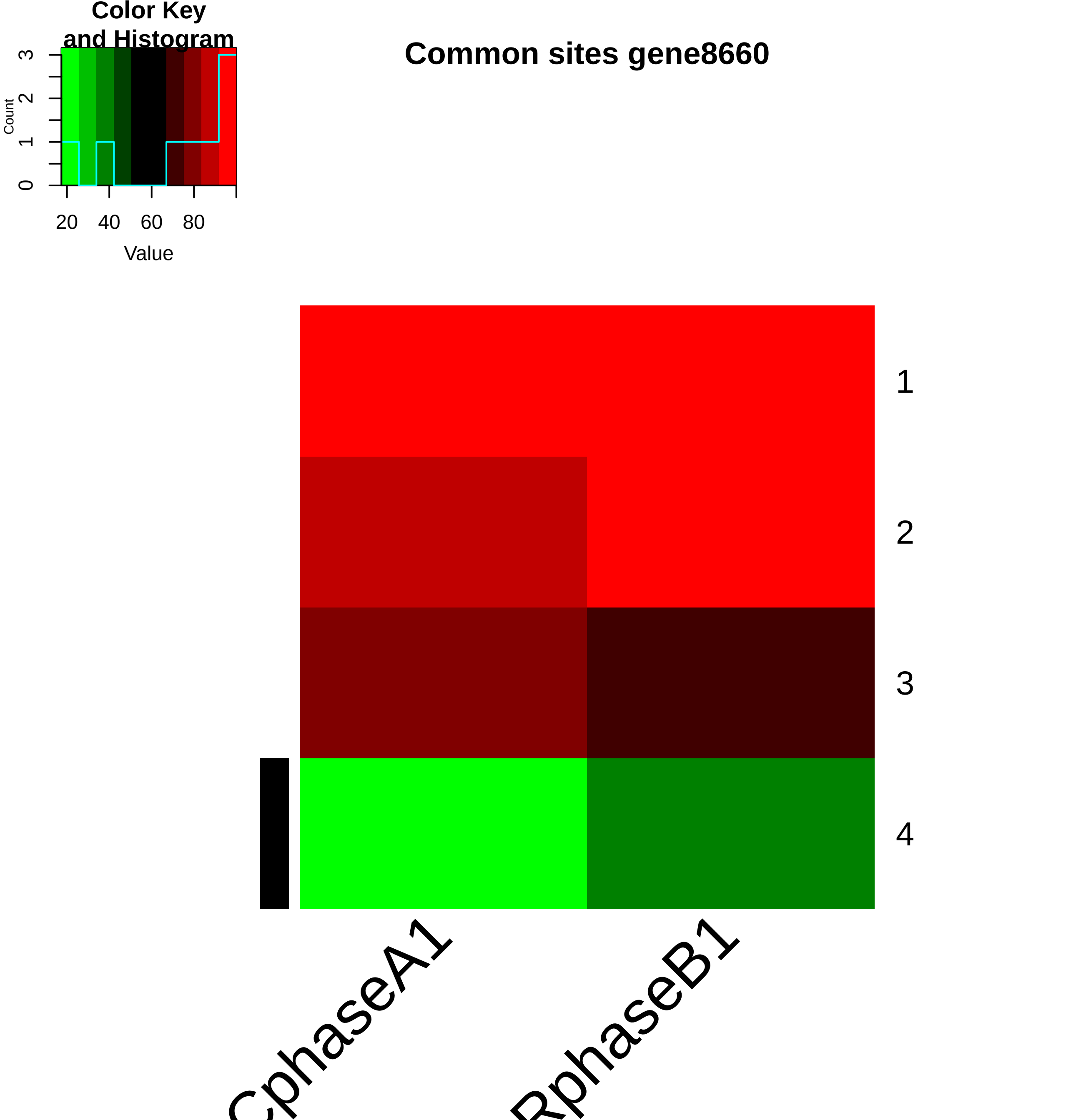

Color Key
and Histogram
Count
0
1
2
3
20
40
60
80
Value
Common sites gene8660
1
2
3
4
CphaseA1
RphaseB1

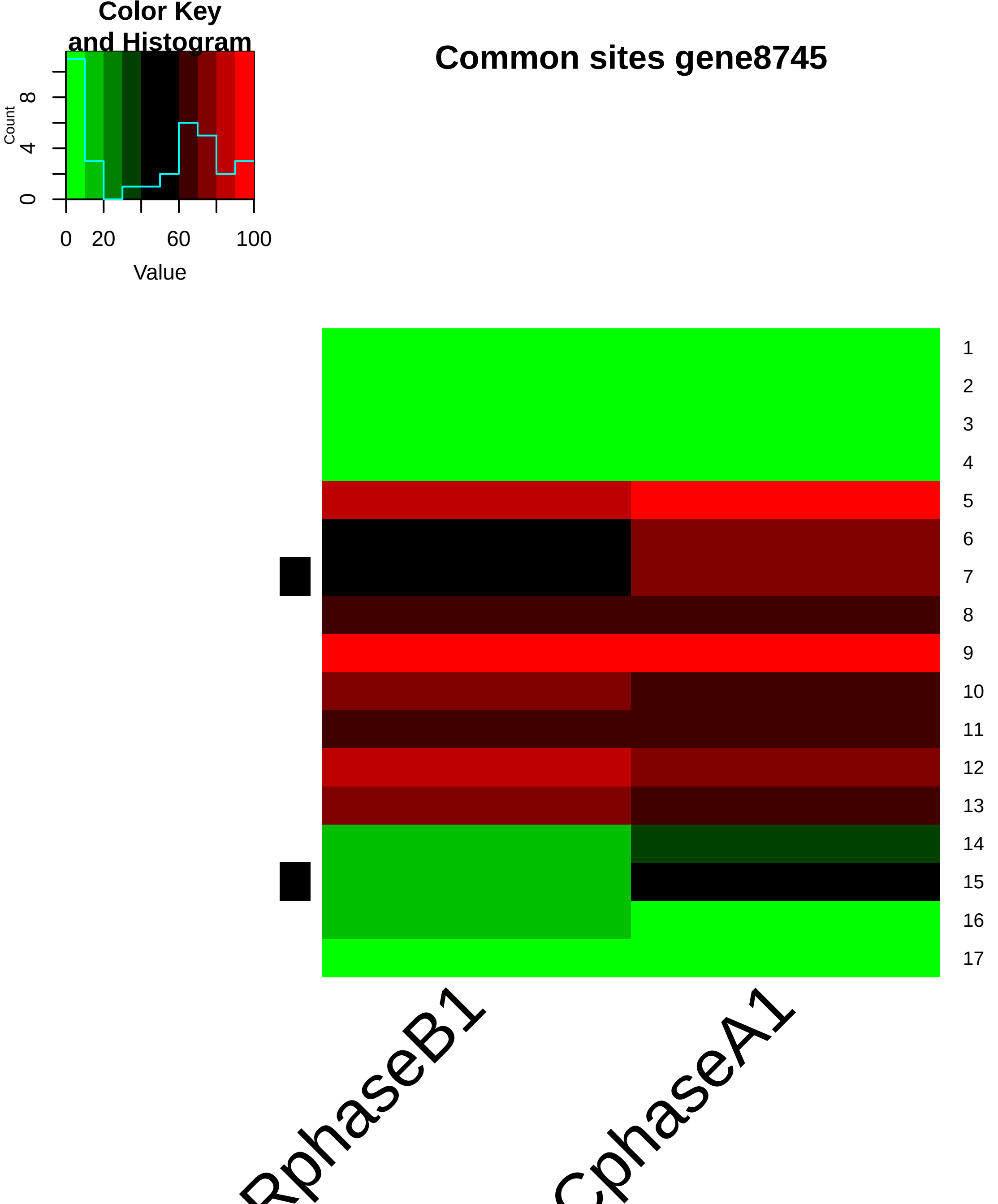

Color Key
and Histogram
Count
0
4
8
0
20
60
100
Value
Common sites gene8745
RphaseB1
CphaseA1

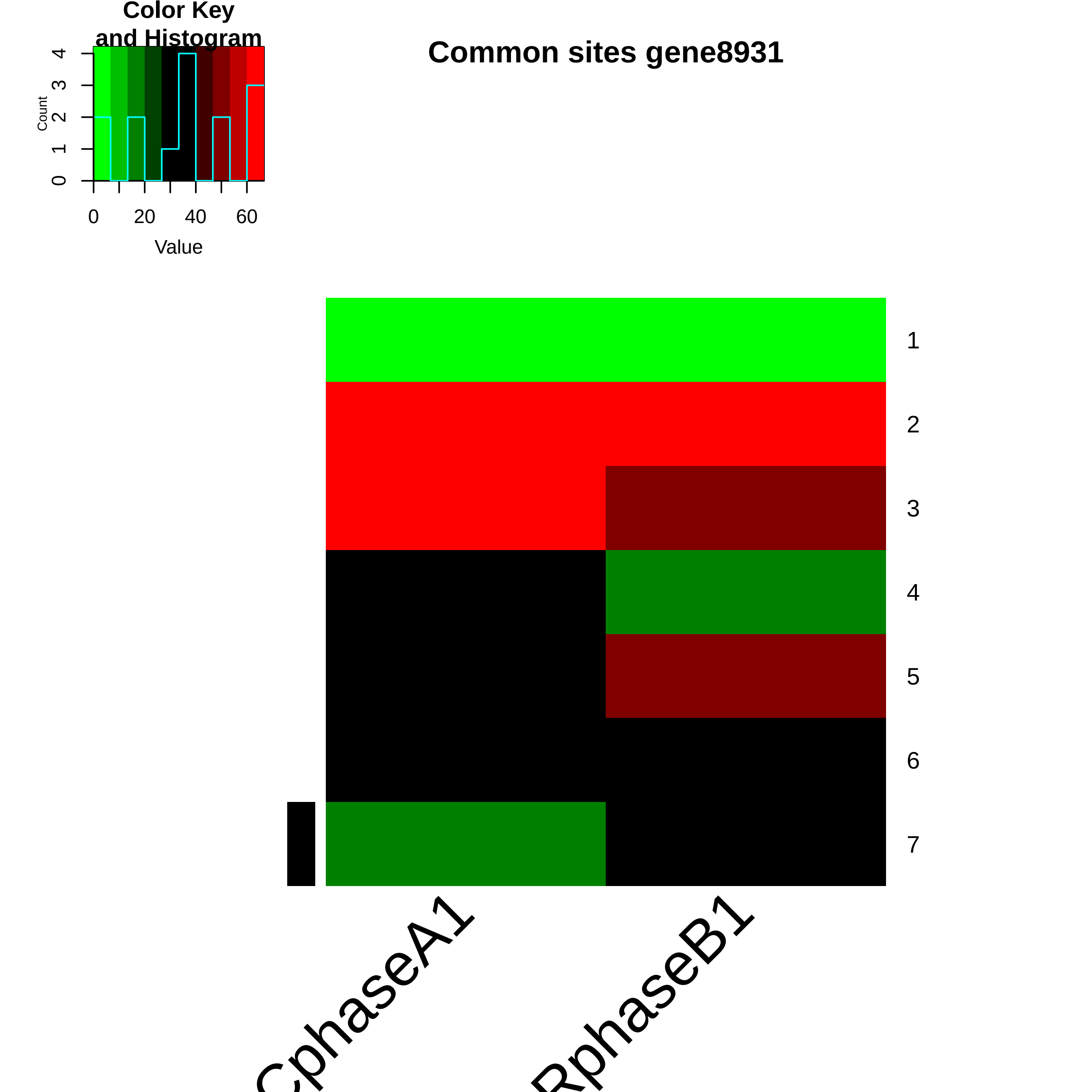
Color Key
and Histogram
Count
Value
0
20
40
60
0
1
2
3
4
Common sites gene8931
1
2
3
4
5
6
7
CphaseA1
RphaseB1

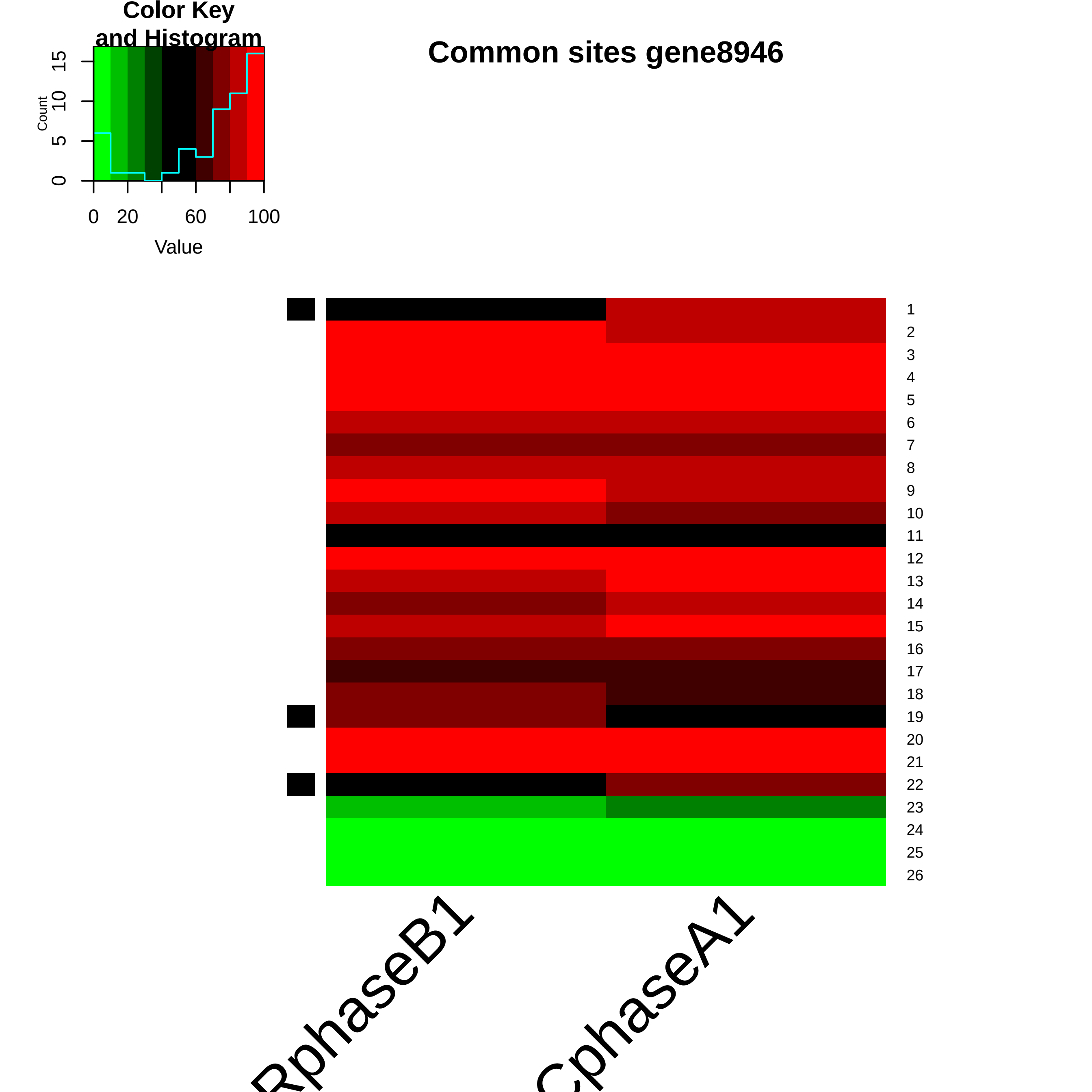

Color Key
and Histogram
Count
0
5
10
15
0
20
60
100
Value
Common sites gene8946
RphaseB1
CphaseA1

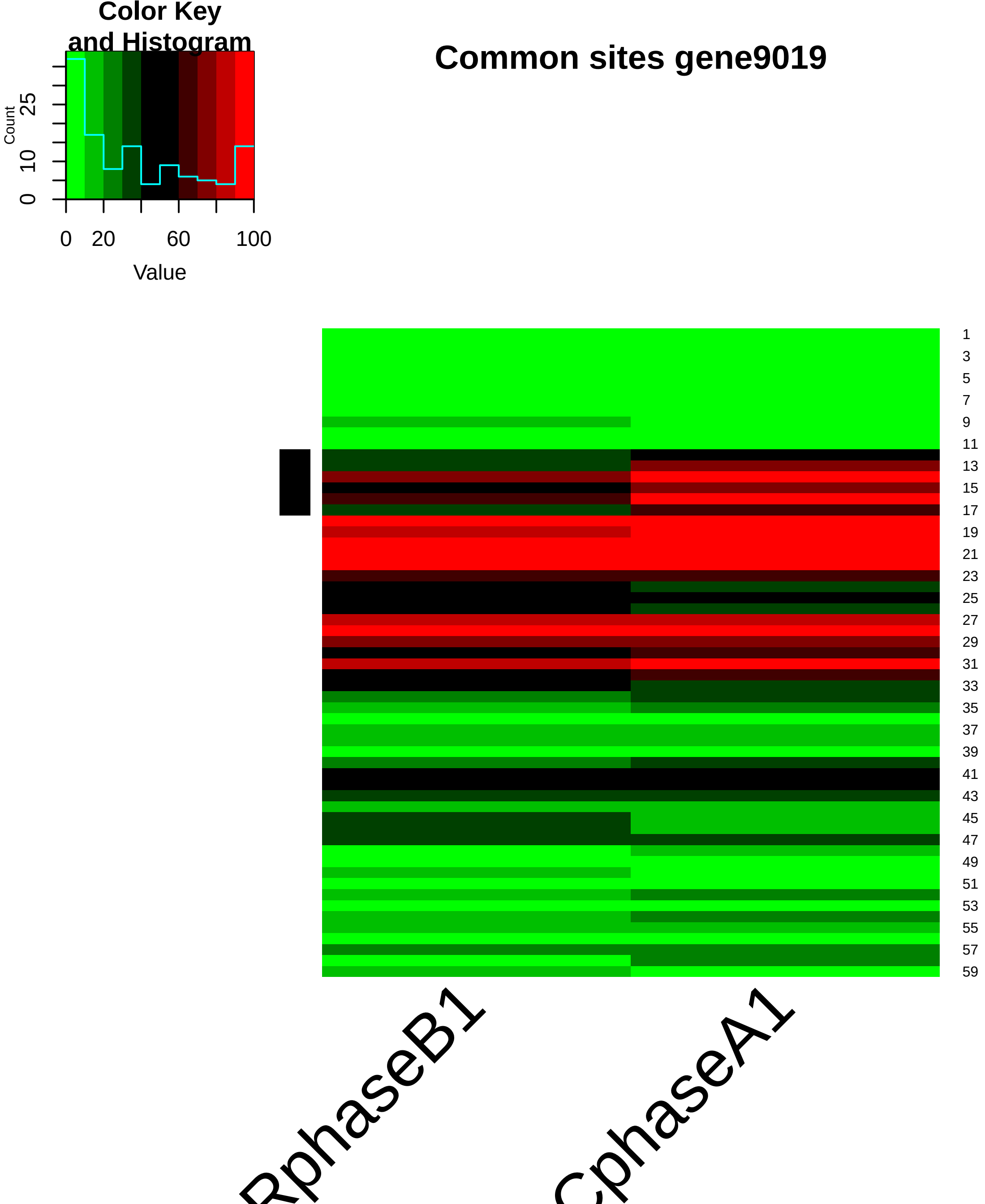
Color Key
and Histogram
Count
0
10
25
0
20
60
100
Value
Common sites gene9019
1
3
5
7
9
11
13
15
17
19
21
23
25
27
29
31
33
35
37
39
41
43
45
47
49
51
53
55
57
59
RphaseB1
CphaseA1

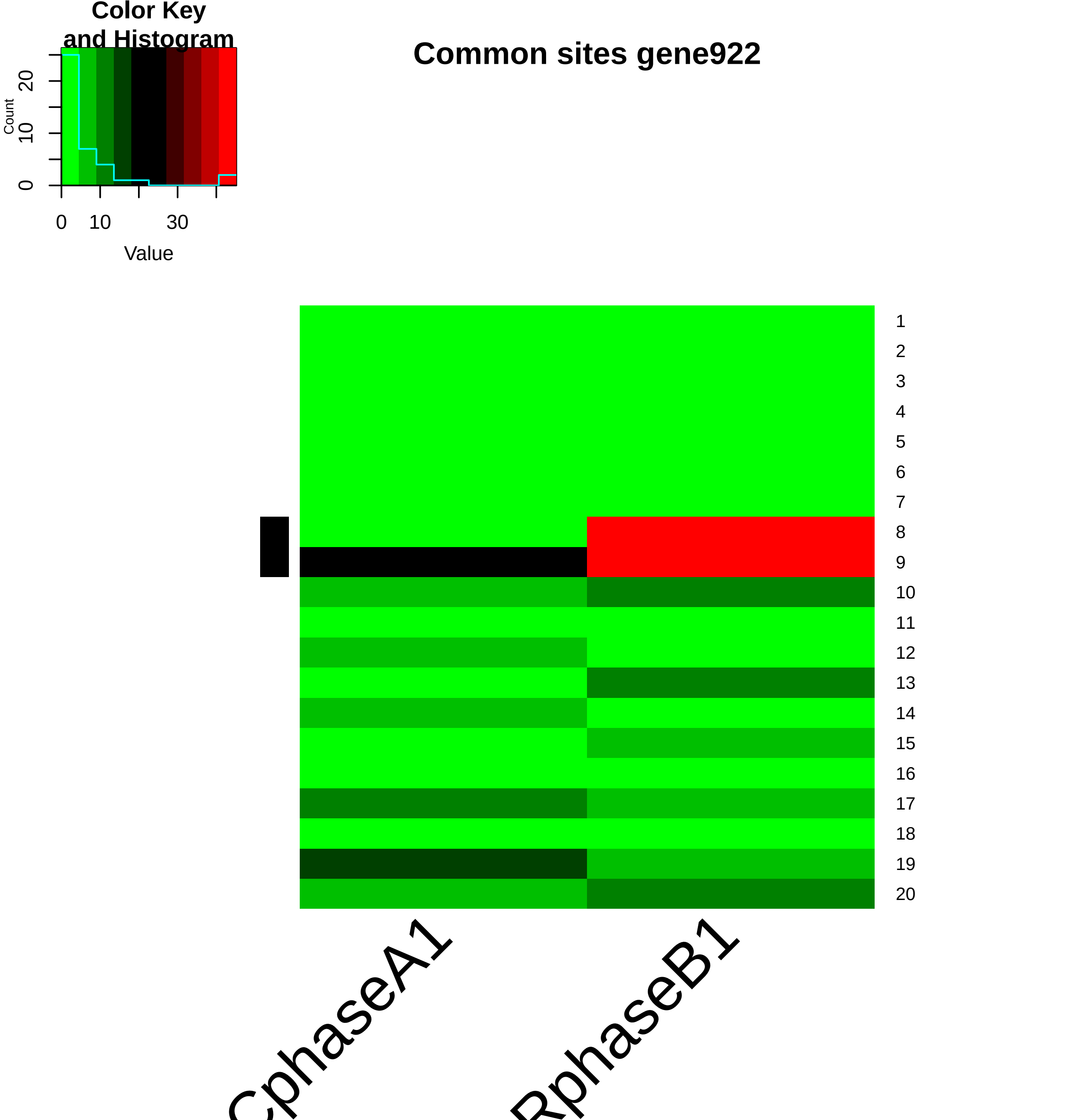
Color Key
and Histogram
Count
0
10
20
0
10
30
Value
Common sites gene922
CphaseA1
RphaseB1

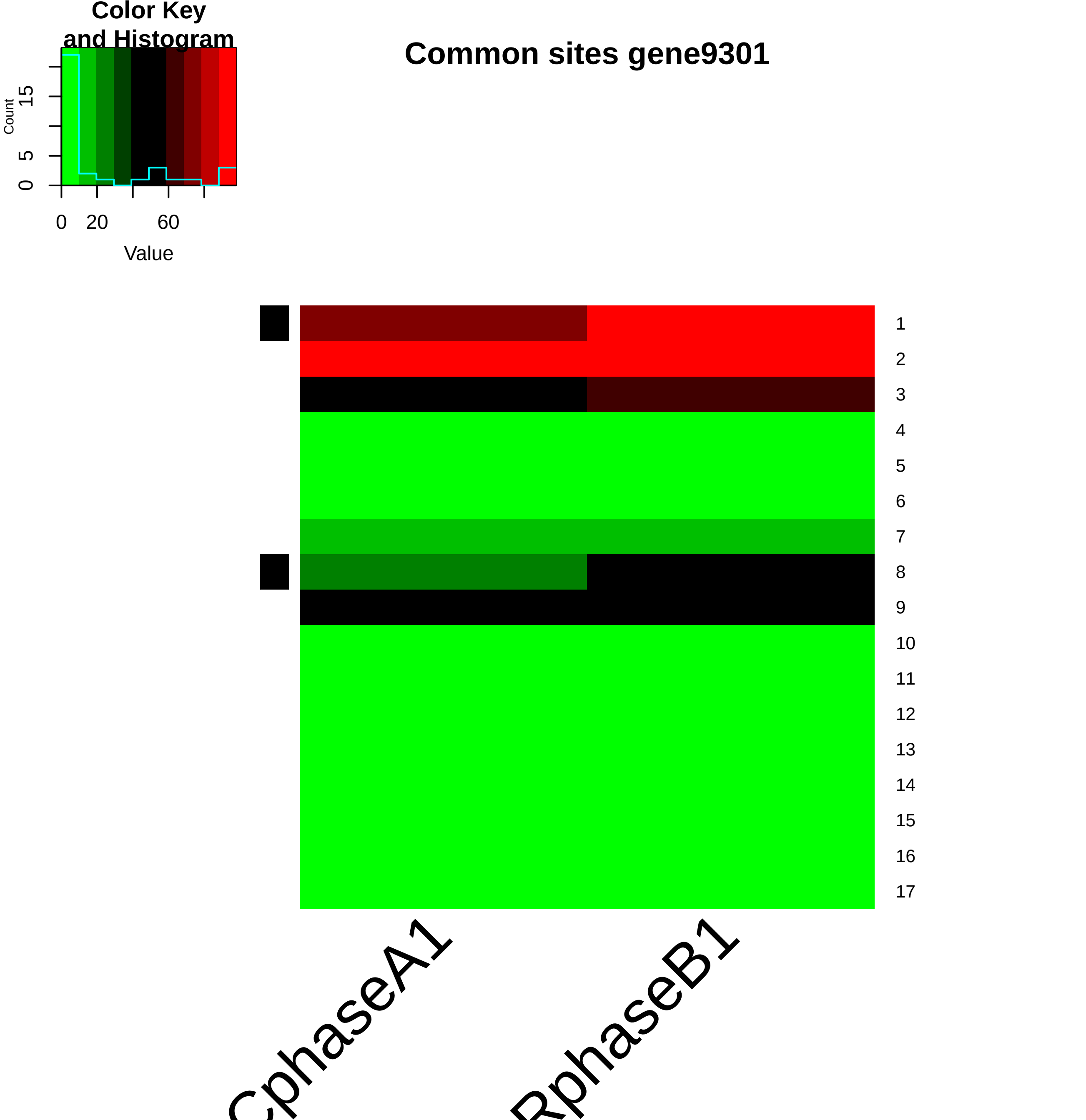

Color Key
and Histogram
Count
0
5
15
0
20
60
Value
Common sites gene9301
CphaseA1
RphaseB1

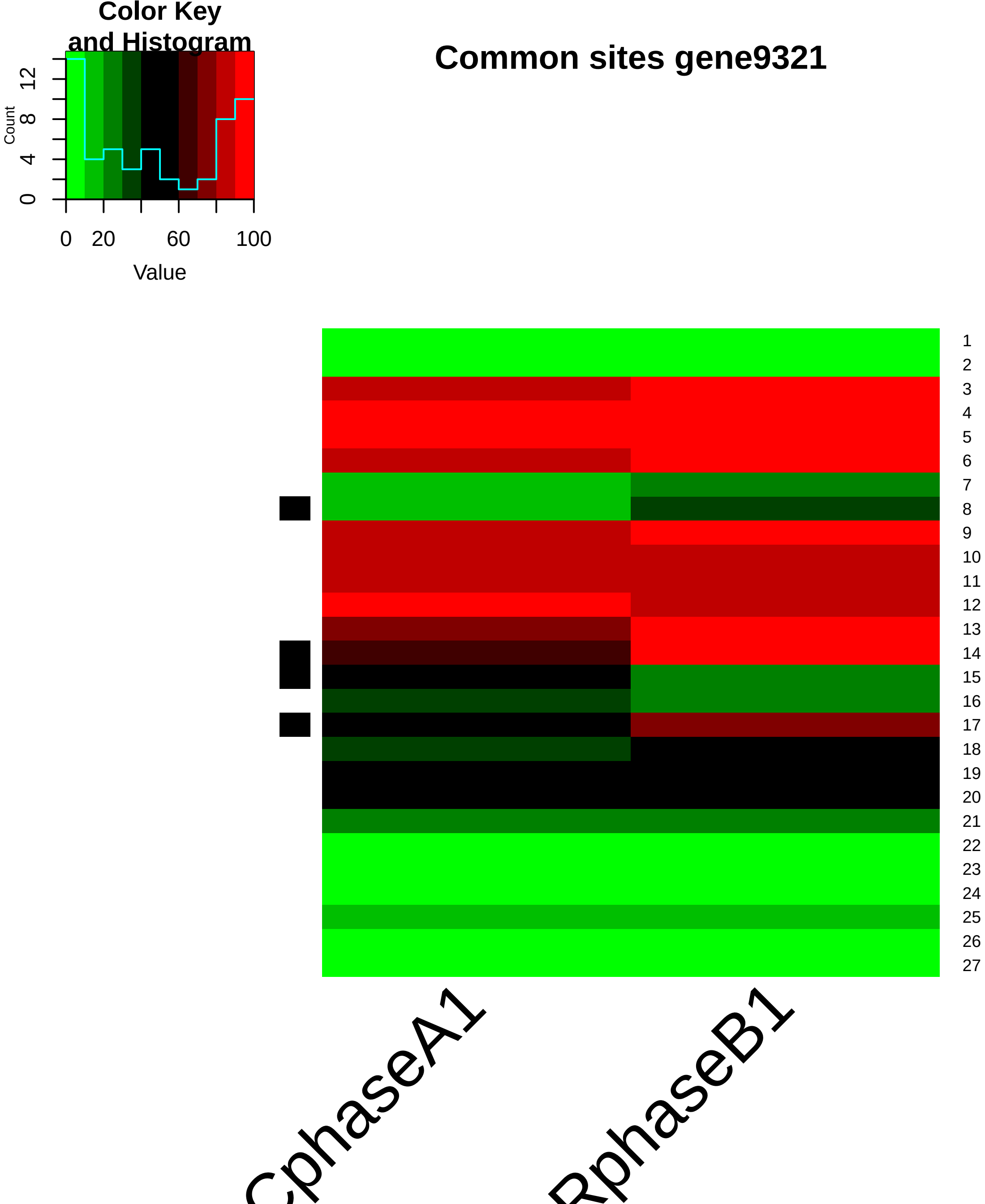
Color Key
and Histogram
Count
0
4
8
12
0
20
60
100
Value
Common sites gene9321
CphaseA1
RphaseB1

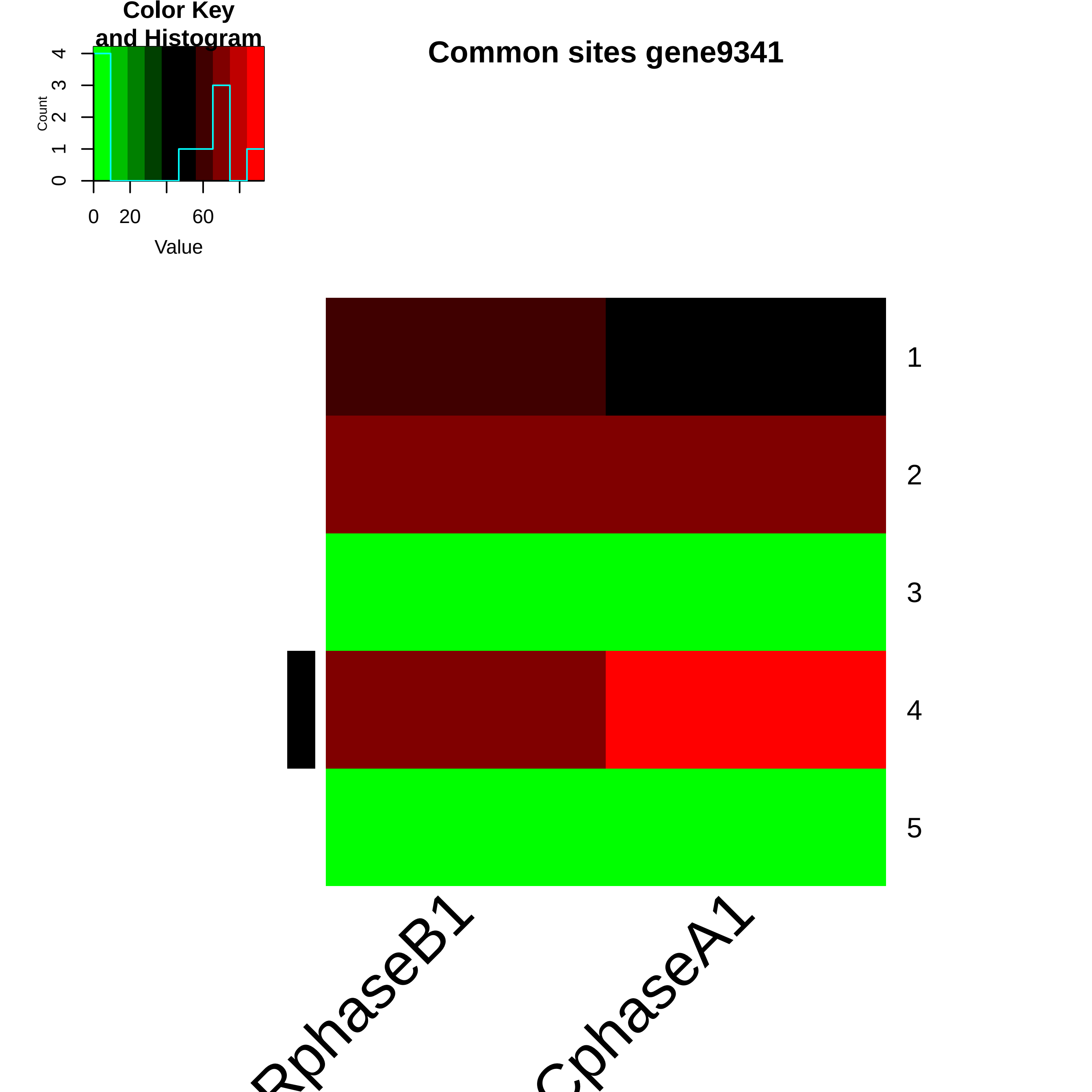

Color Key
and Histogram
Count
Value
0
20
60
0
1
2
3
4
Common sites gene9341
1
2
3
4
5
RphaseB1
CphaseA1

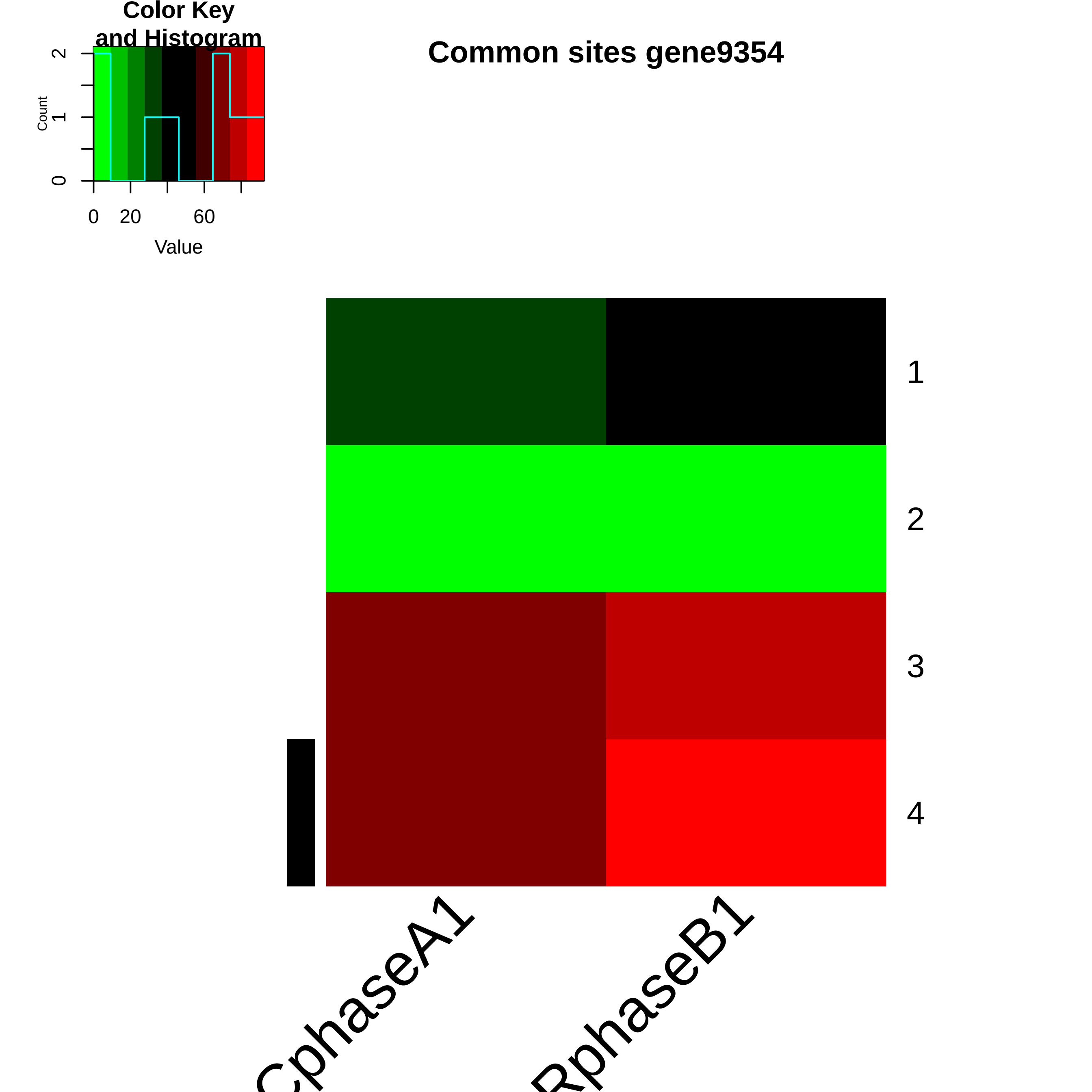

Color Key
and Histogram
Count
0
1
2
0
20
60
Value
Common sites gene9354
1
2
3
4
CphaseA1
RphaseB1

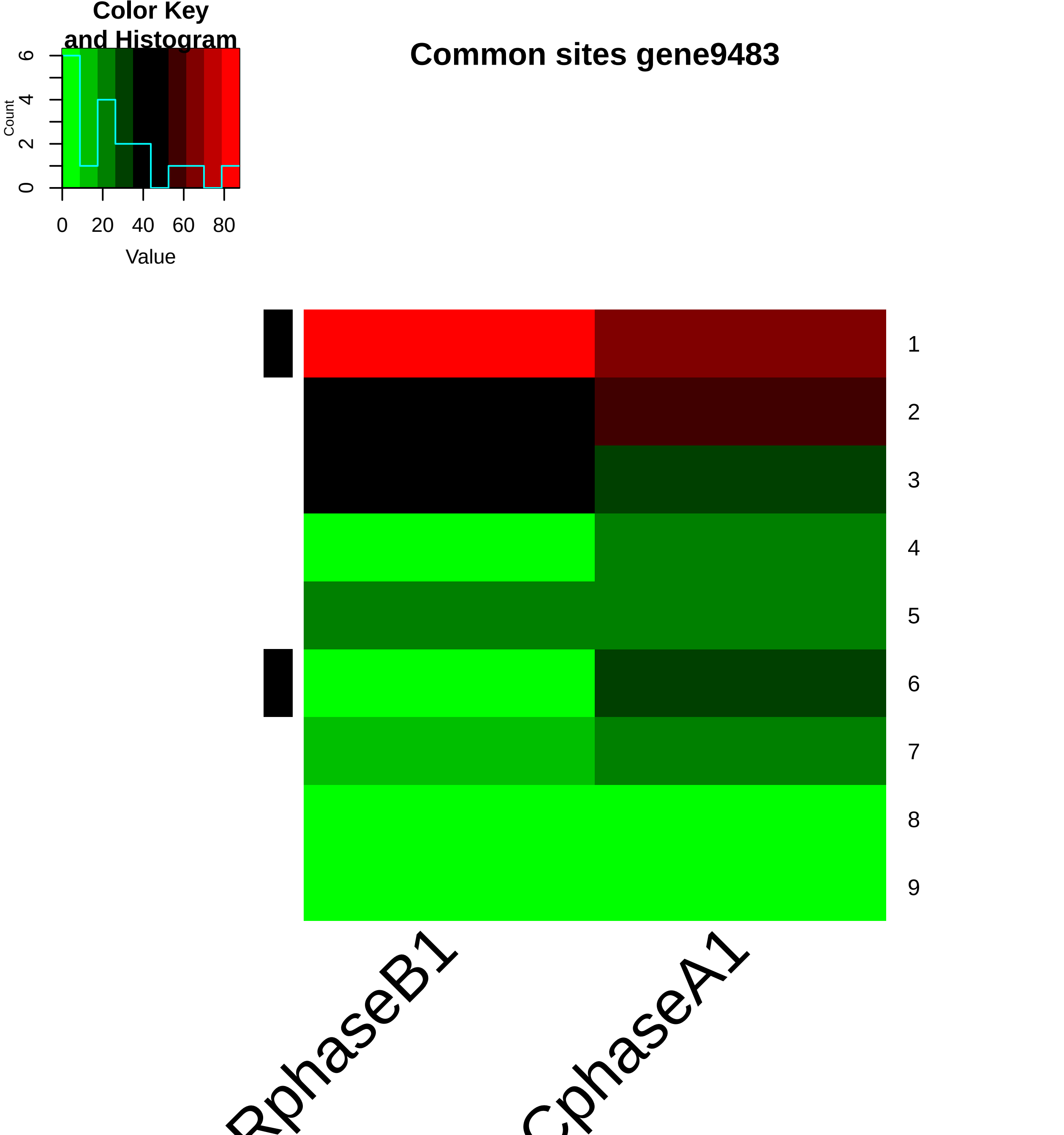
Color Key
and Histogram
Count
0
2
4
6
0
20
40
60
80
Value
Common sites gene9483
1
2
3
4
5
6
7
8
9
RphaseB1
CphaseA1

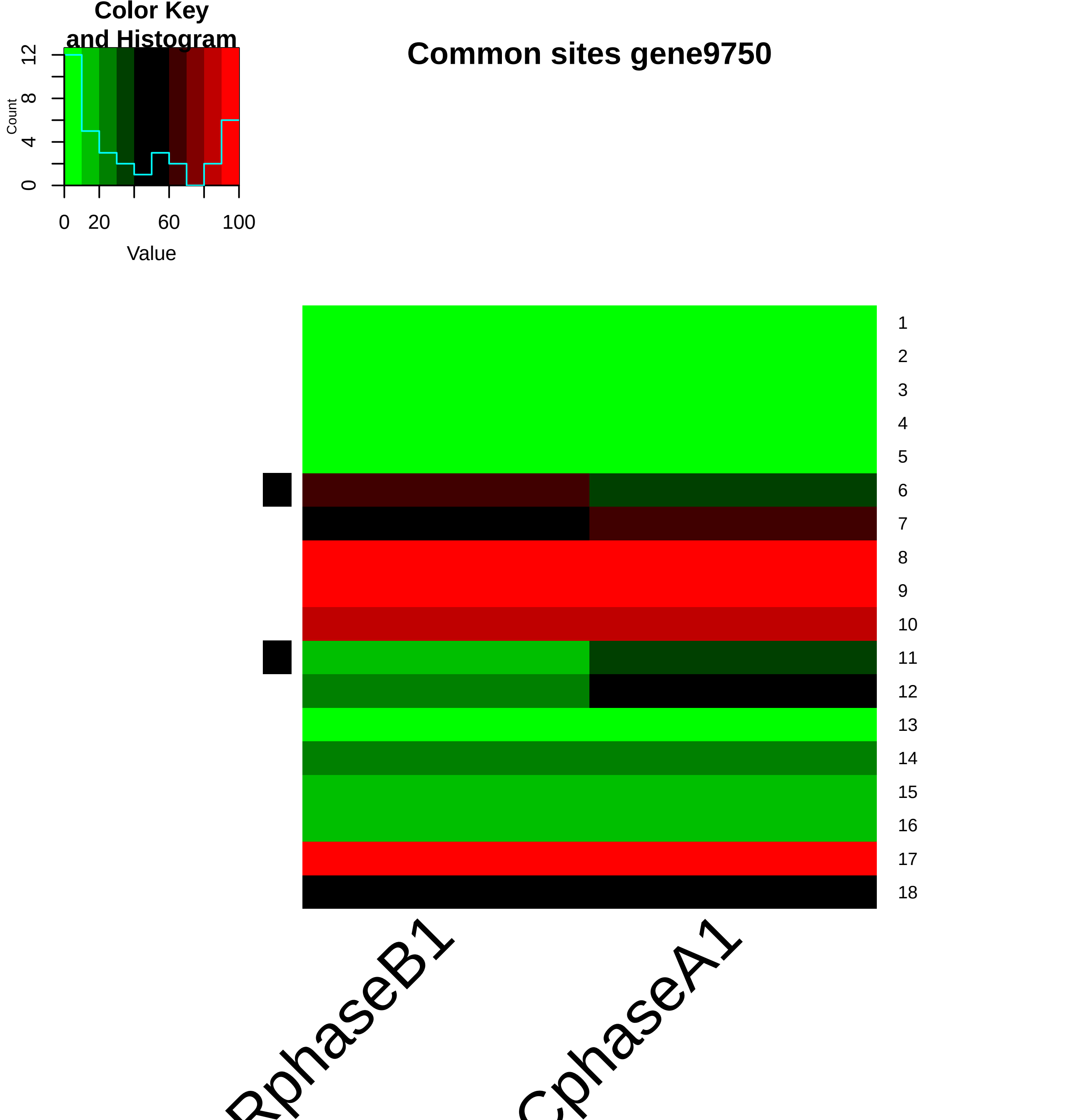

Color Key
and Histogram
Count
0
4
8
12
0
20
60
100
Value
Common sites gene9750
RphaseB1
CphaseA1

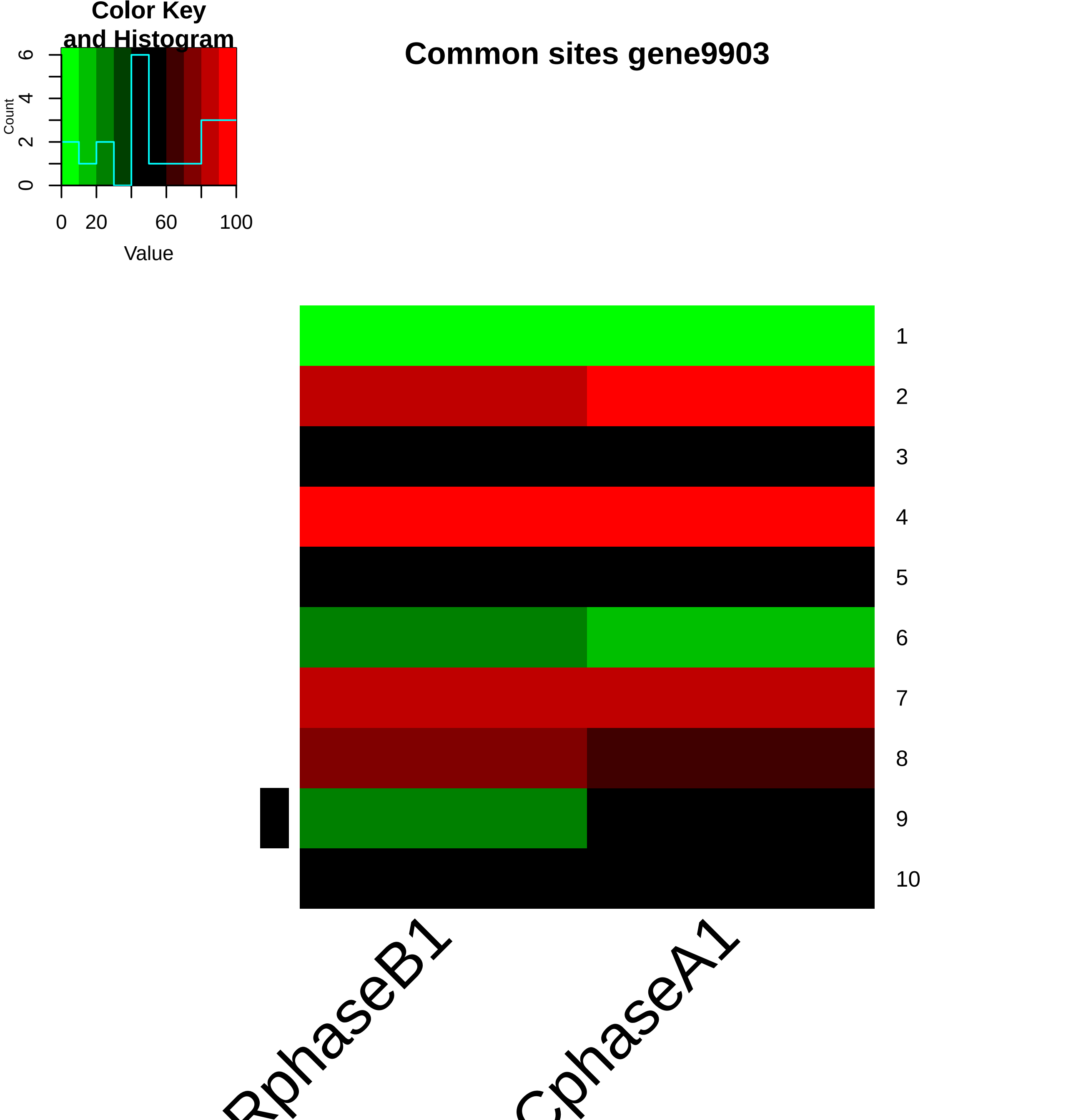
Color Key
and Histogram
Count
0
2
4
6
0
20
60
100
Value
Common sites gene9903
1
2
3
4
5
6
7
8
9
10
RphaseB1
CphaseA1

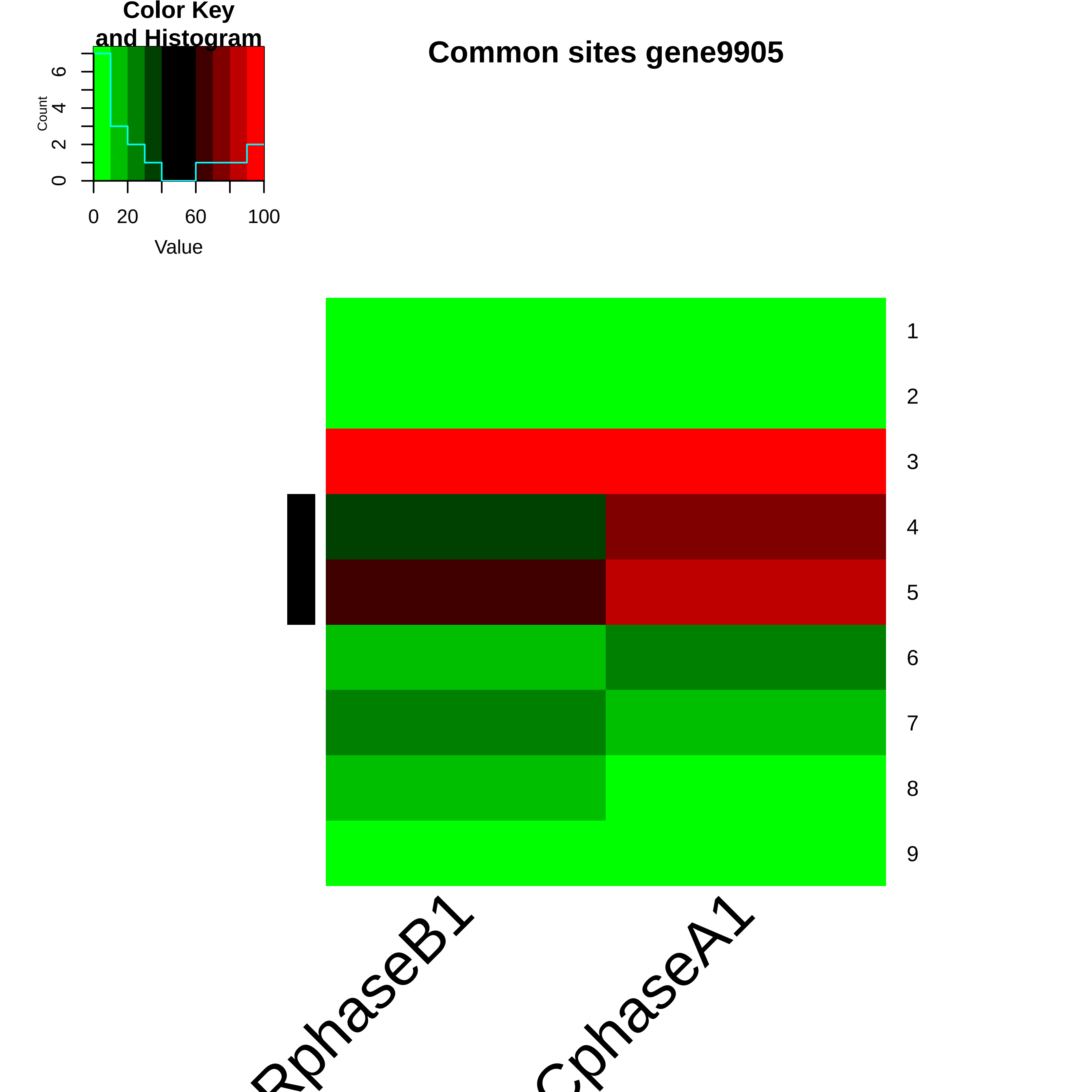
Color Key
and Histogram
Count
0
2
4
6
0
20
60
100
Value
Common sites gene9905
1
2
3
4
5
6
7
8
9
RphaseB1
CphaseA1

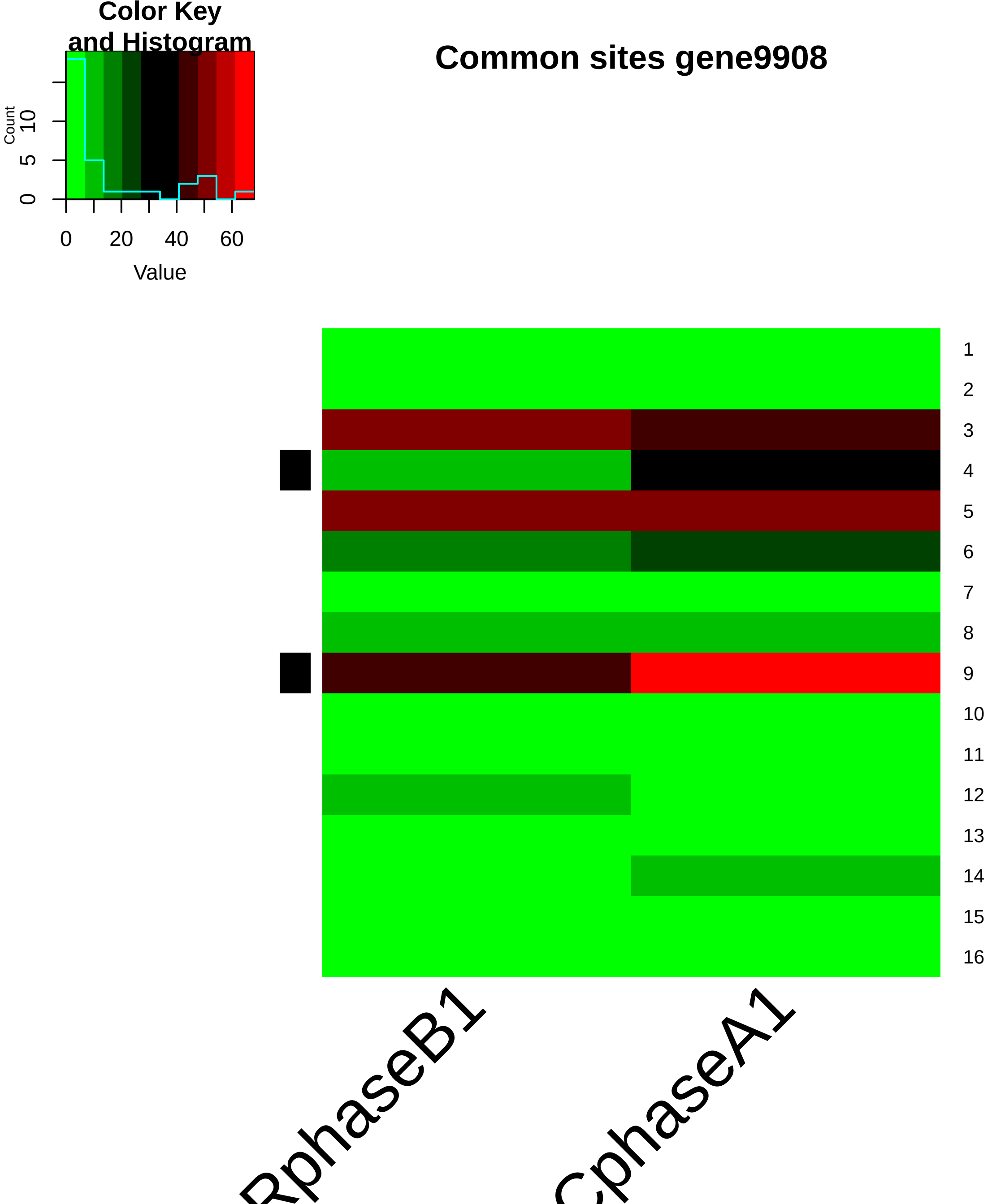

Color Key
and Histogram
Count
0
5
10
0
20
40
60
Value
Common sites gene9908
RphaseB1
CphaseA1

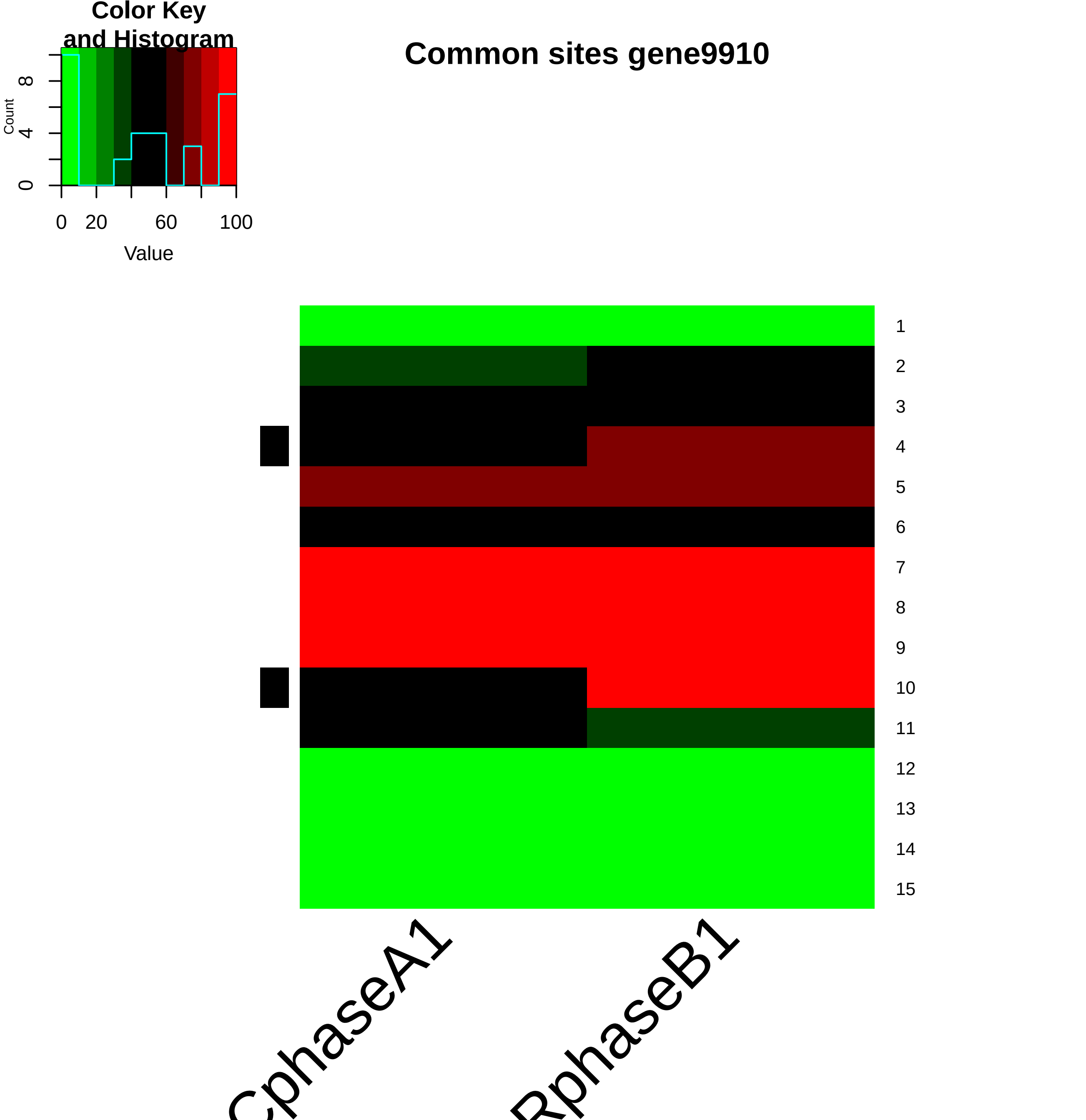
Color Key
and Histogram
Count
0
4
8
0
20
60
100
Value
Common sites gene9910
CphaseA1
RphaseB1

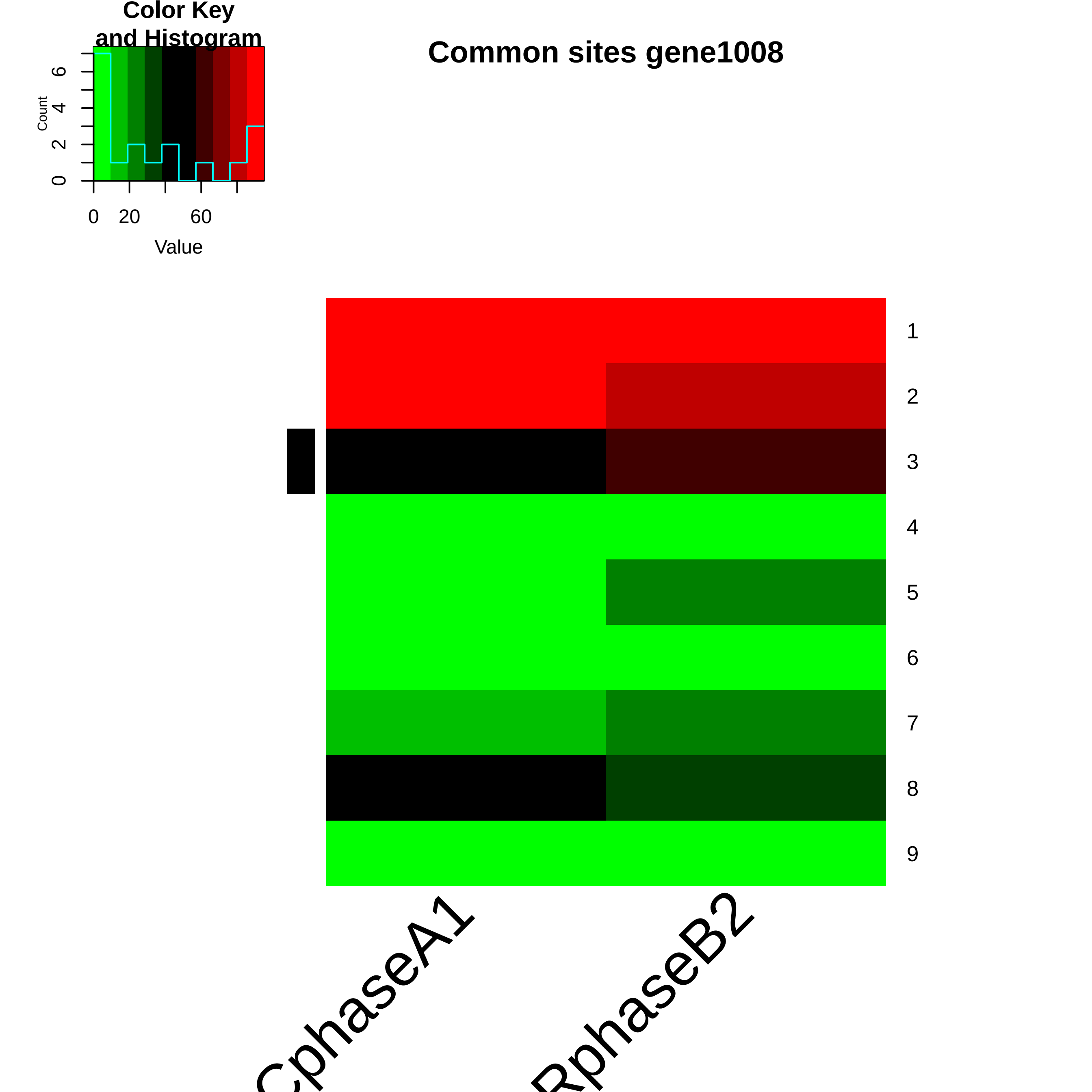

Color Key
and Histogram
Count
0
2
4
6
0
20
60
Value
Common sites gene1008
1
2
3
4
5
6
7
8
9
CphaseA1
RphaseB2

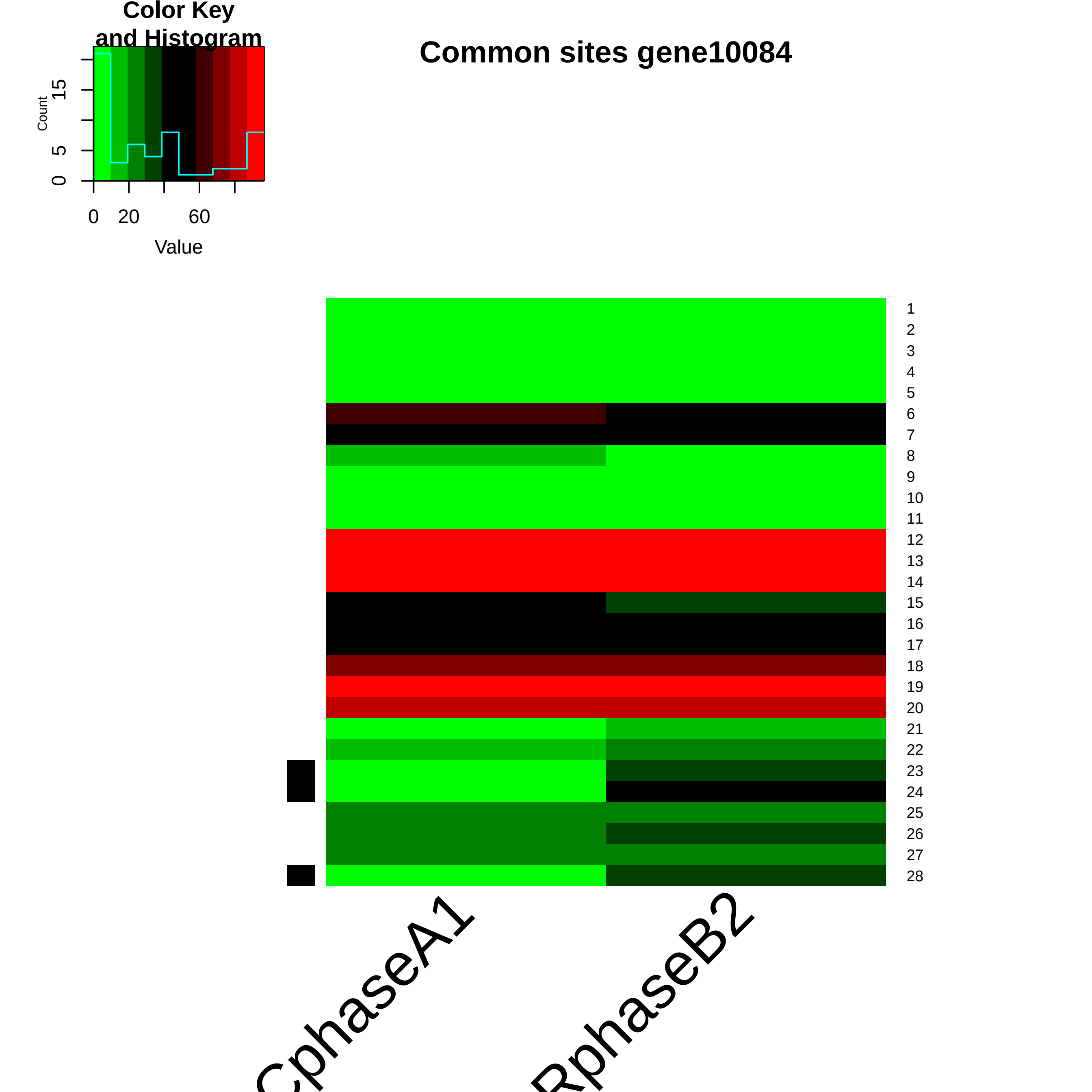
Color Key
and Histogram
Count
0
5
15
0
20
60
Value
Common sites gene10084
CphaseA1
RphaseB2

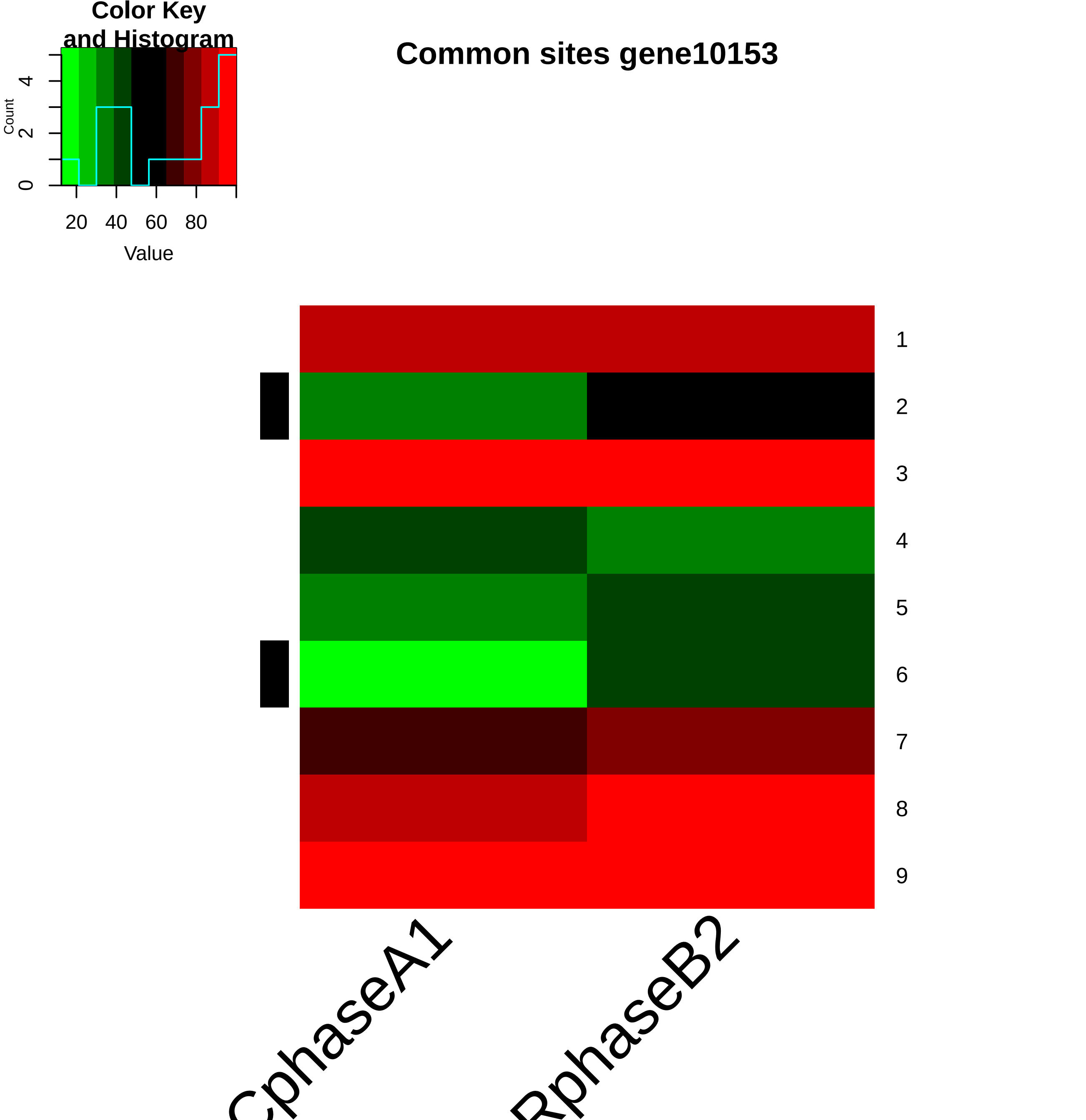
Color Key
and Histogram
Count
0
2
4
20
40
60
80
Value
Common sites gene10153
1
2
3
4
5
6
7
8
9

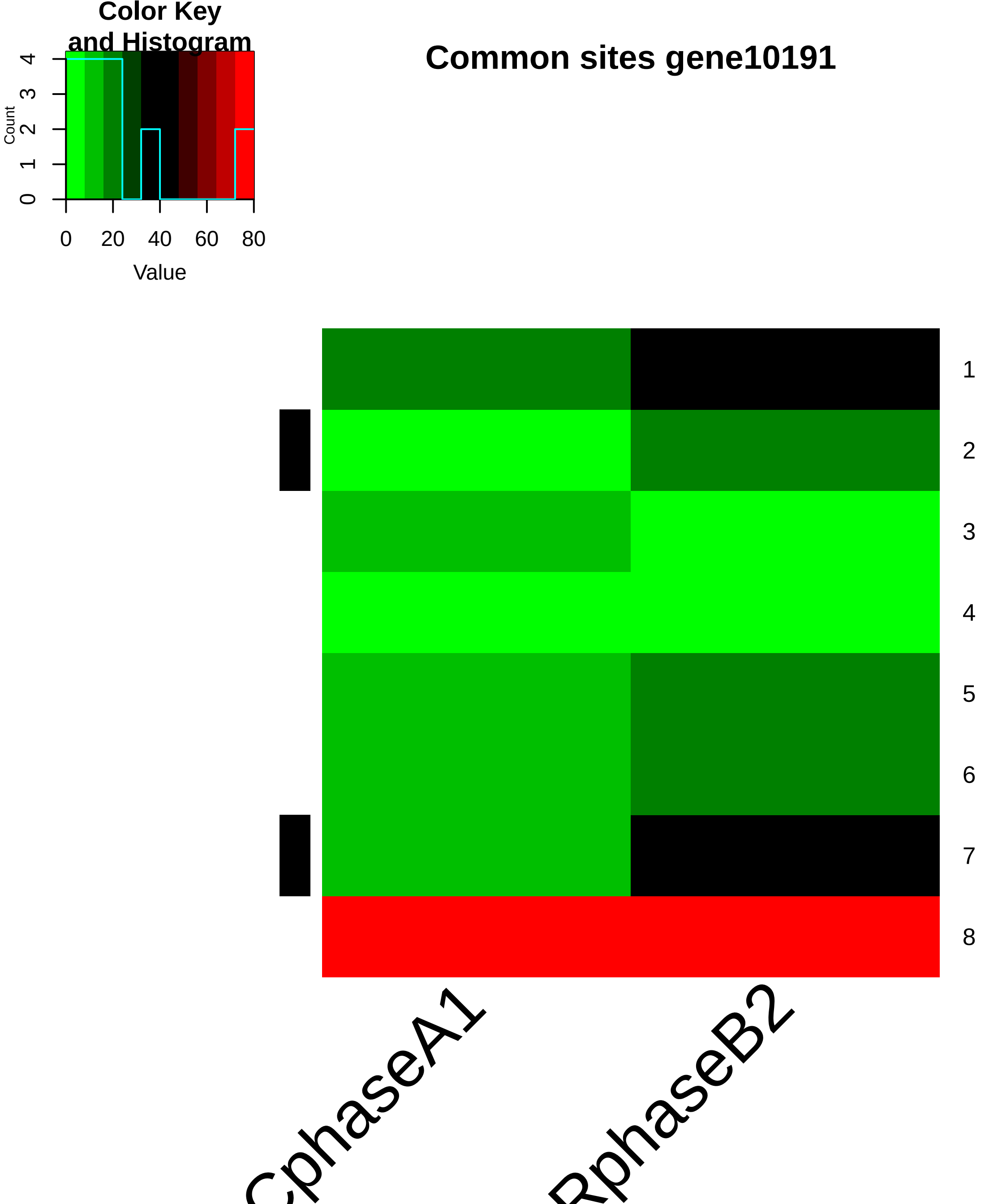
Color Key
and Histogram
Count
0
1
2
3
4
0
20
40
60
80
Value
Common sites gene10191
1
2
3
4
5
6
7
8
CphaseA1
RphaseB2

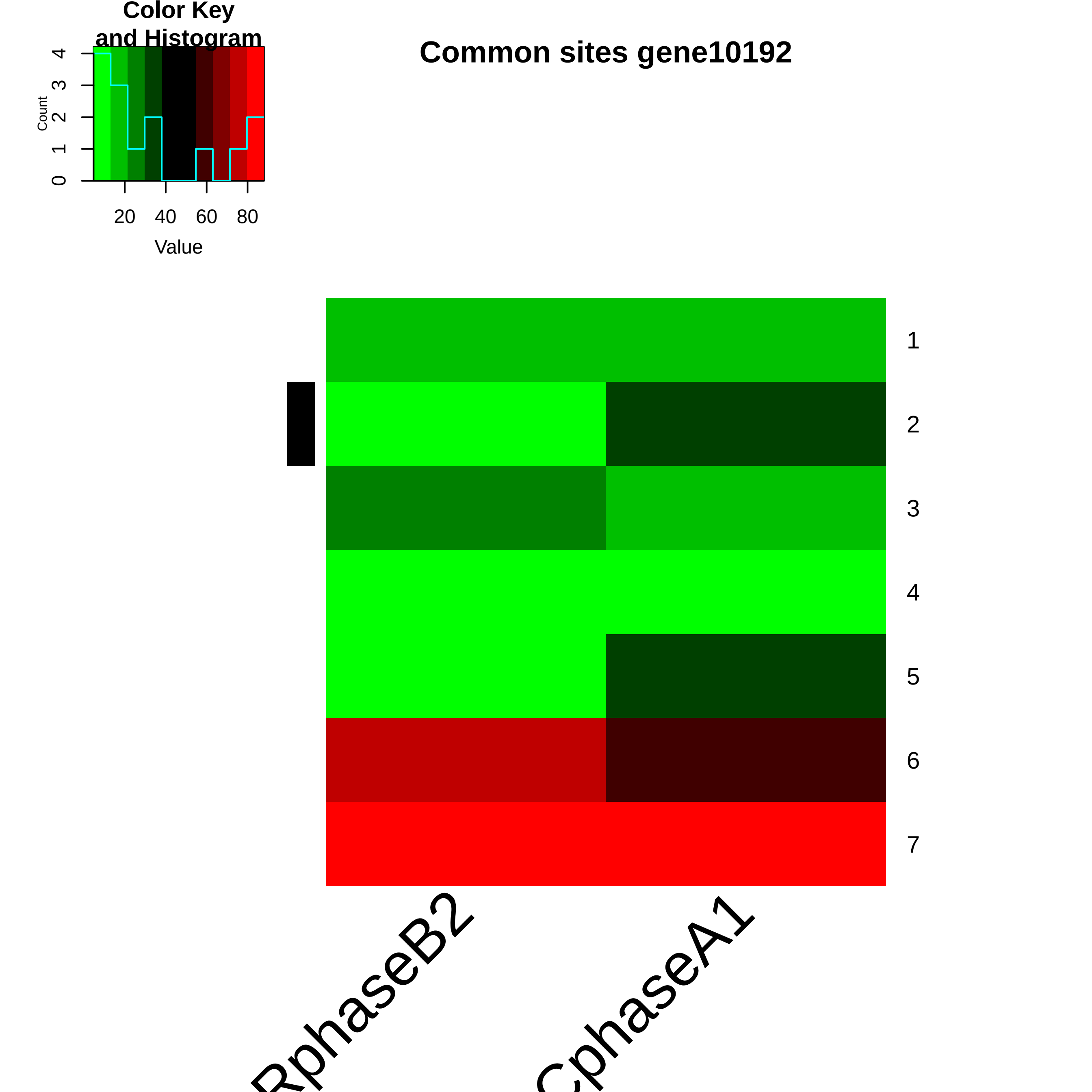

Color Key
and Histogram
Count
0
1
2
3
4
20
40
60
80
Value
Common sites gene10192
1
2
3
4
5
6
7
RphaseB2
CphaseA1

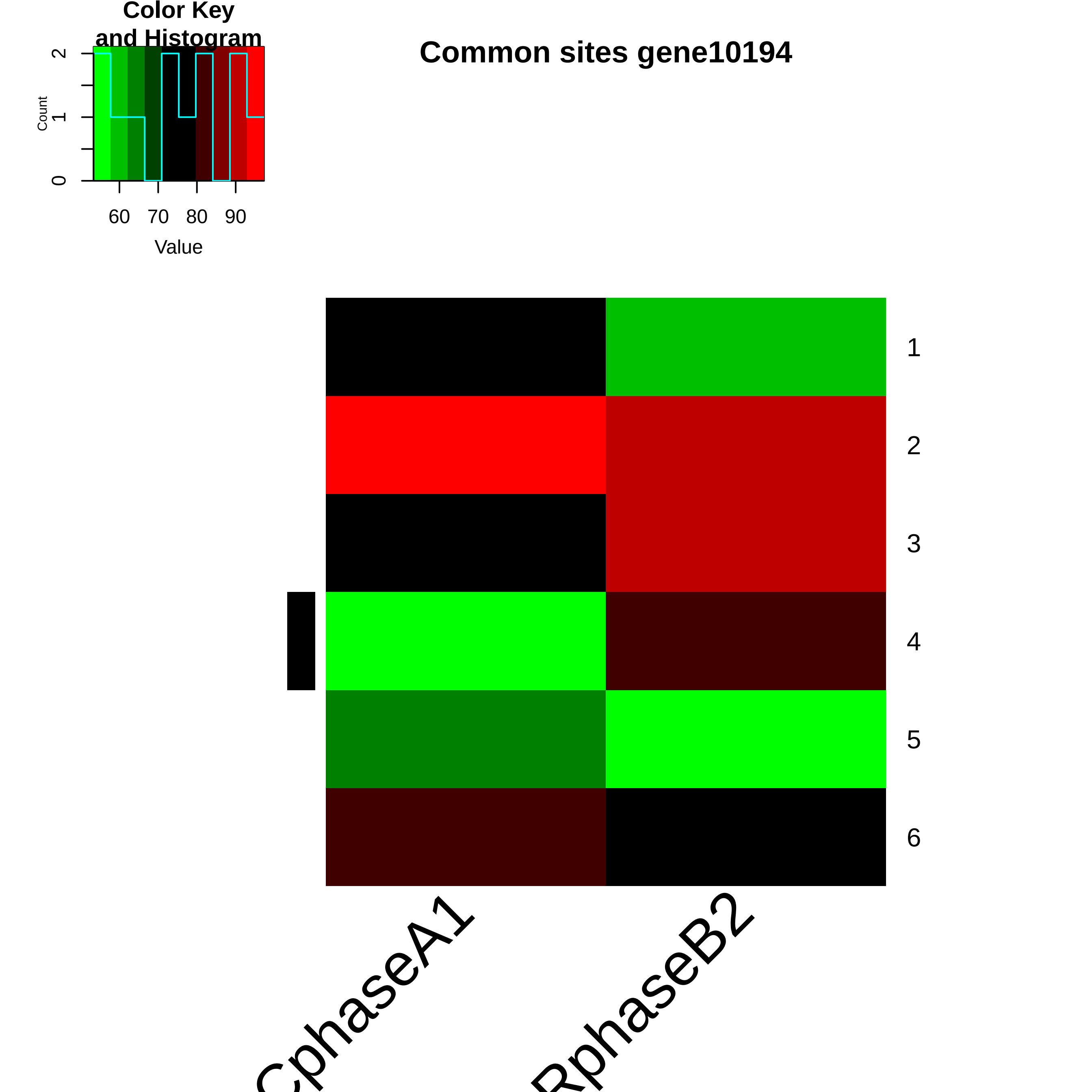

Color Key
and Histogram
Count
0
1
2
60
70
80
90
Value
Common sites gene10194
1
2
3
4
5
6
CphaseA1
RphaseB2

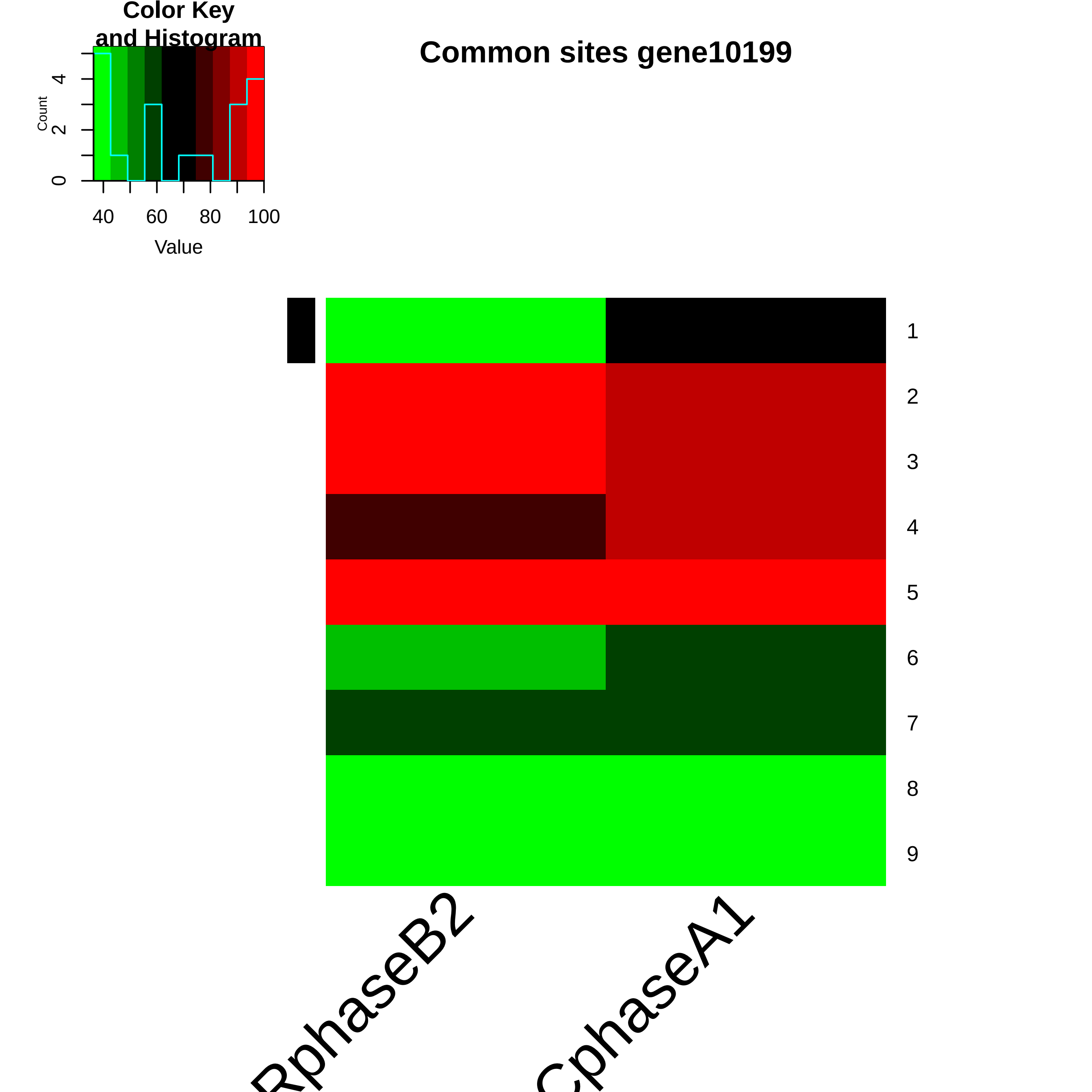
Color Key
and Histogram
Count
0
2
4
40
60
80
100
Value
Common sites gene10199
1
2
3
4
5
6
7
8
9
RphaseB2
CphaseA1

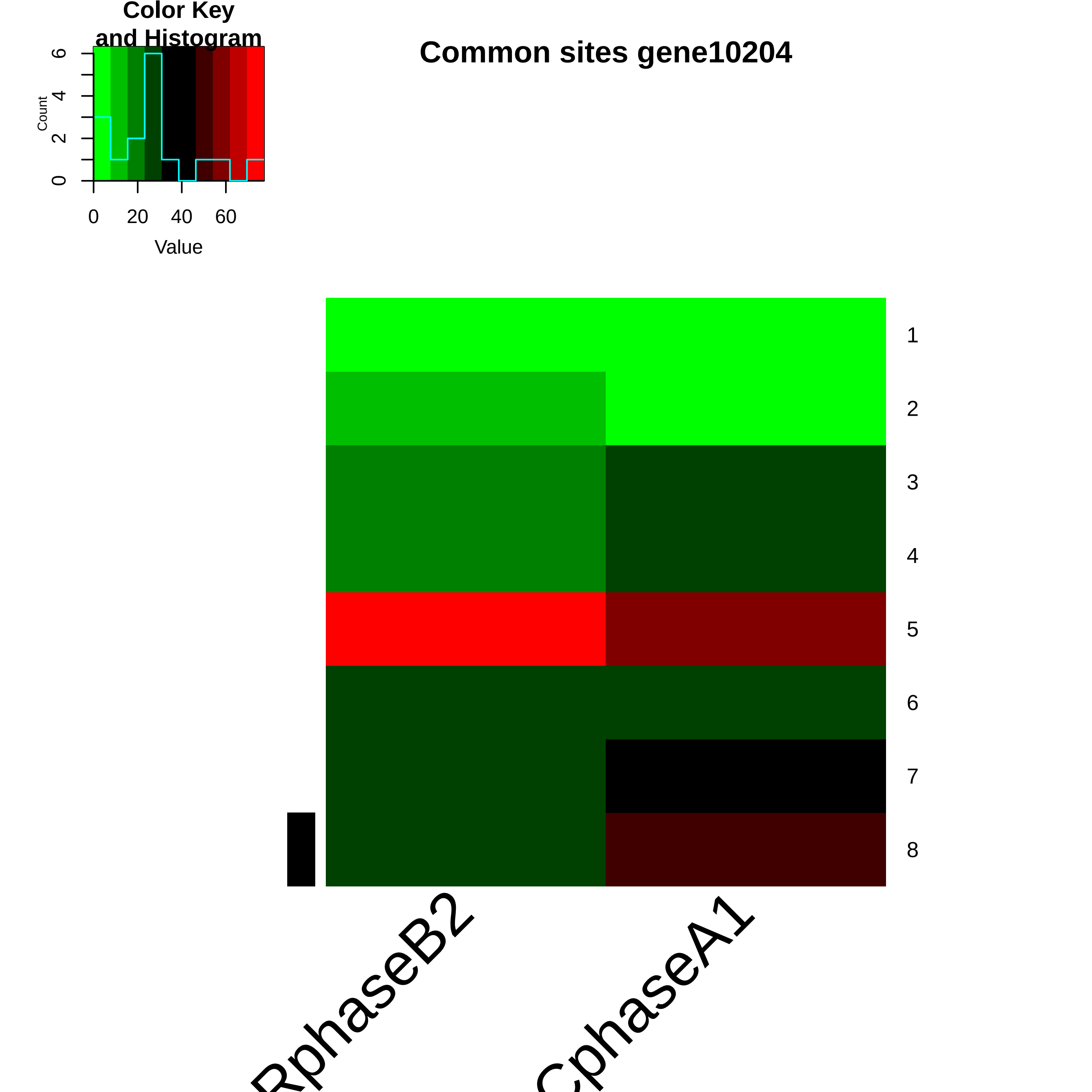
Color Key
and Histogram
Count
Value
0
20
40
60
0
2
4
6
Common sites gene10204
1
2
3
4
5
6
7
8
RphaseB2
CphaseA1

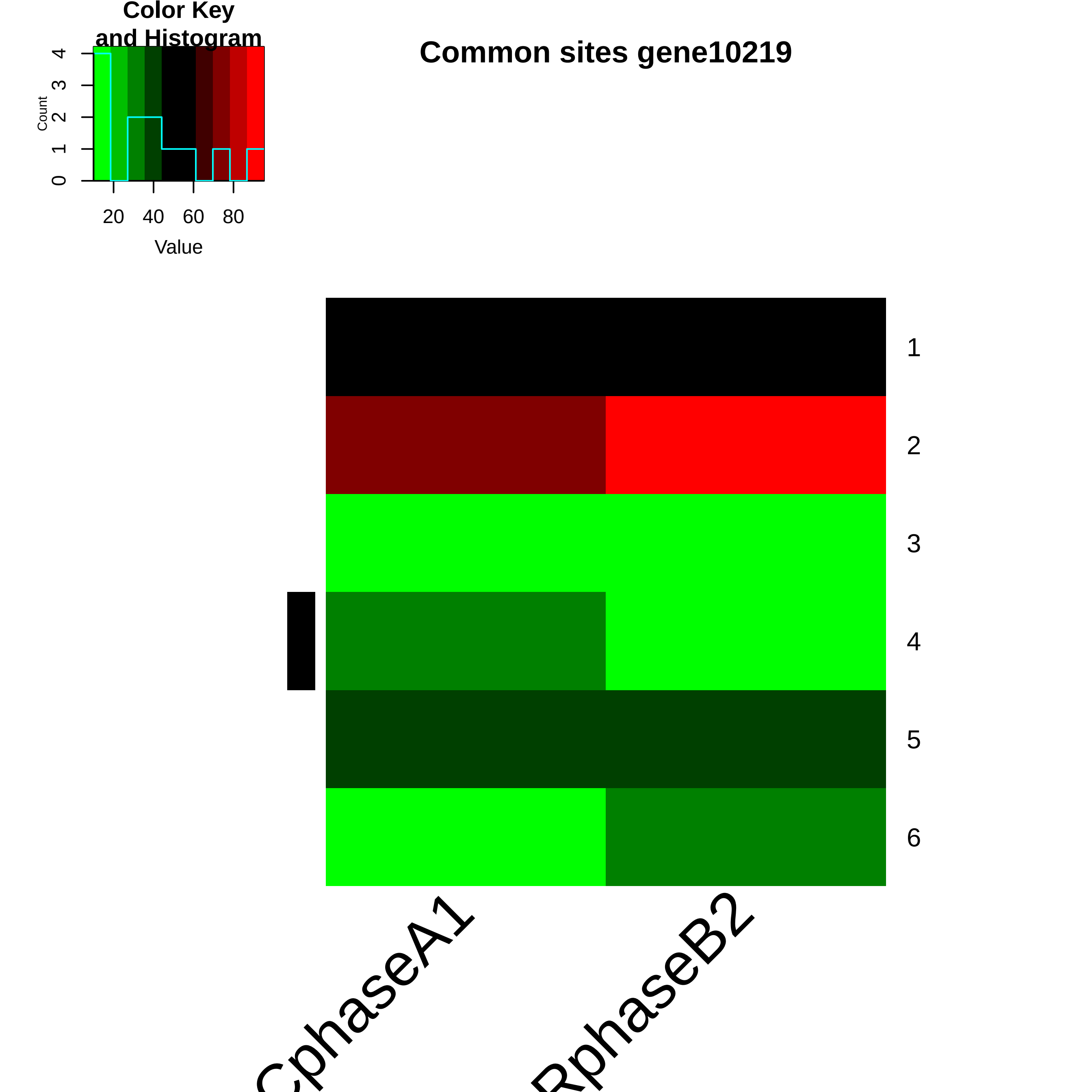

Color Key
and Histogram
Count
0
1
2
3
4
20
40
60
80
Value
Common sites gene10219
1
2
3
4
5
6
CphaseA1
RphaseB2

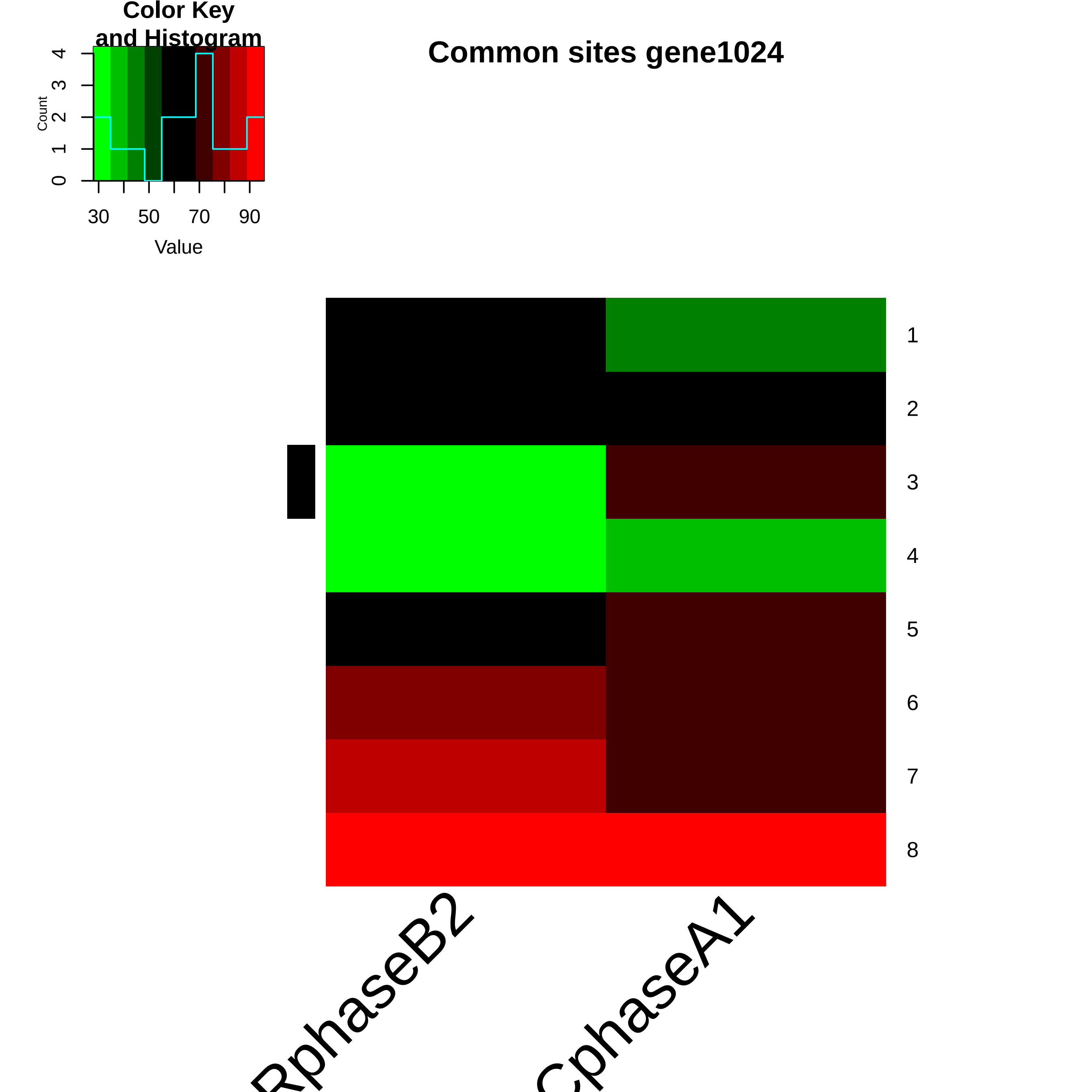
Color Key
and Histogram
Count
0
1
2
3
4
30
50
70
90
Value
Common sites gene1024
1
2
3
4
5
6
7
8
RphaseB2
CphaseA1

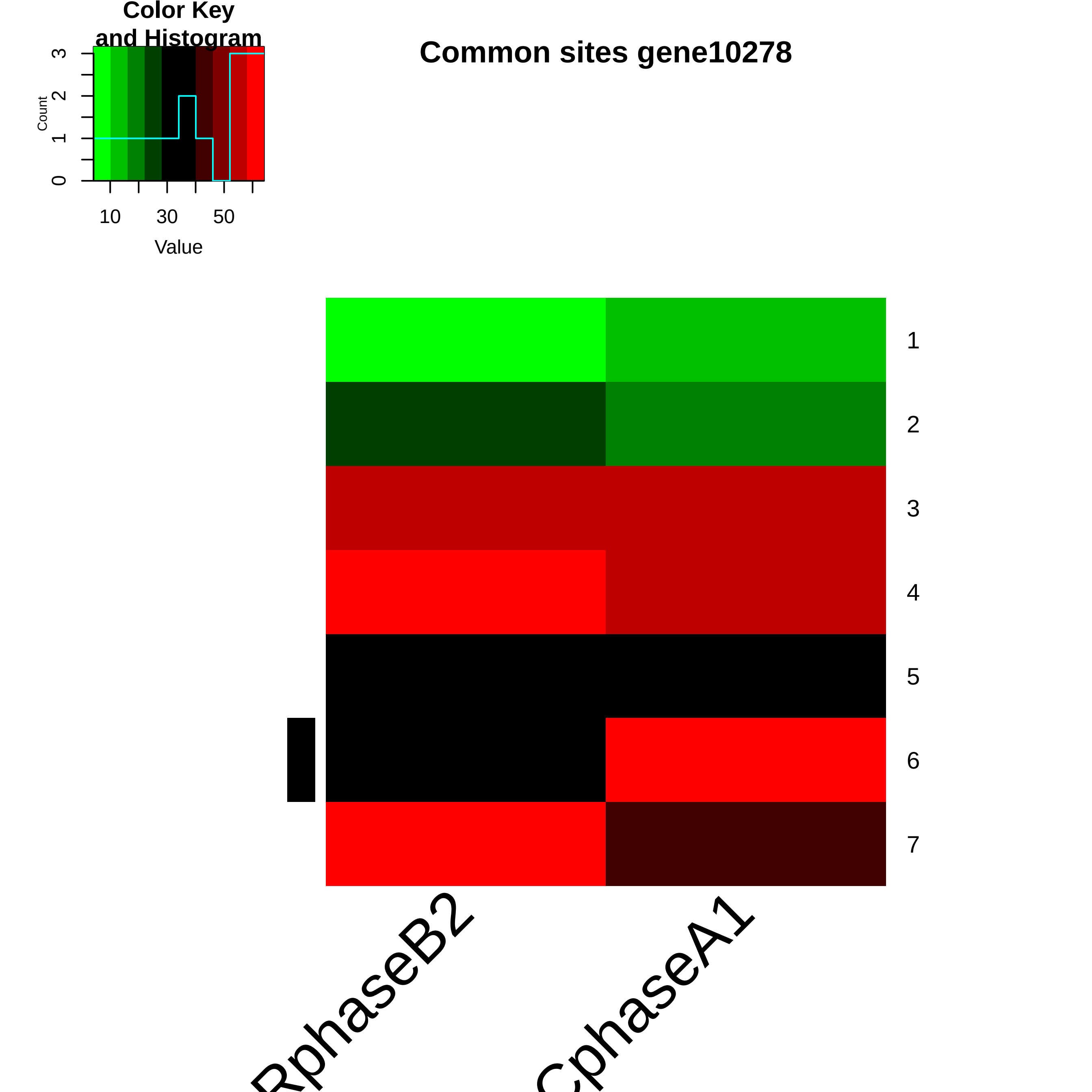
Color Key
and Histogram
Count
0
1
2
3
10
30
50
Value
Common sites gene10278
1
2
3
4
5
6
7
RphaseB2
CphaseA1

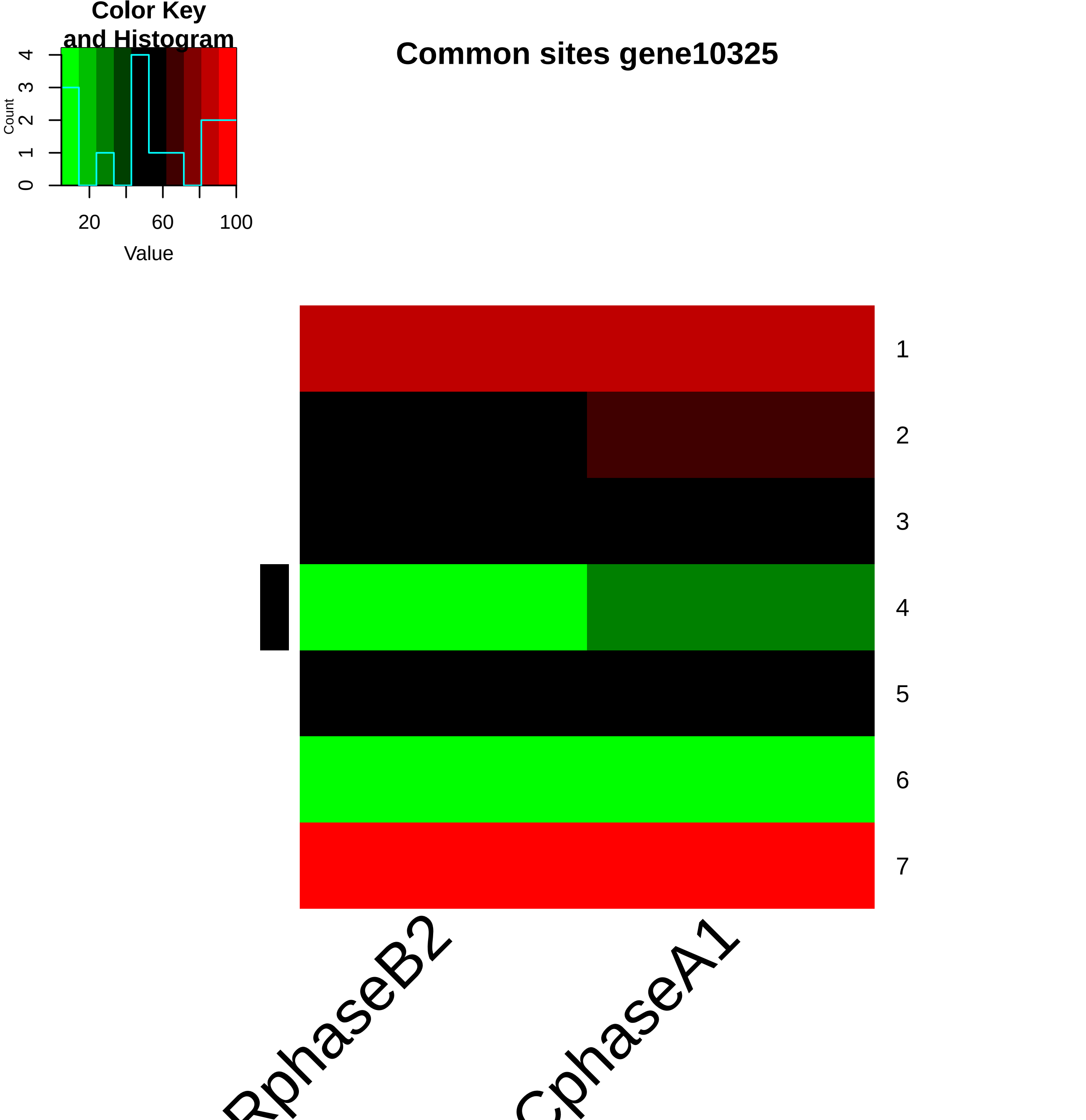
Color Key
and Histogram
Count
0
1
2
3
4
20
60
100
Value
Common sites gene10325
1
2
3
4
5
6
7
RphaseB2
CphaseA1

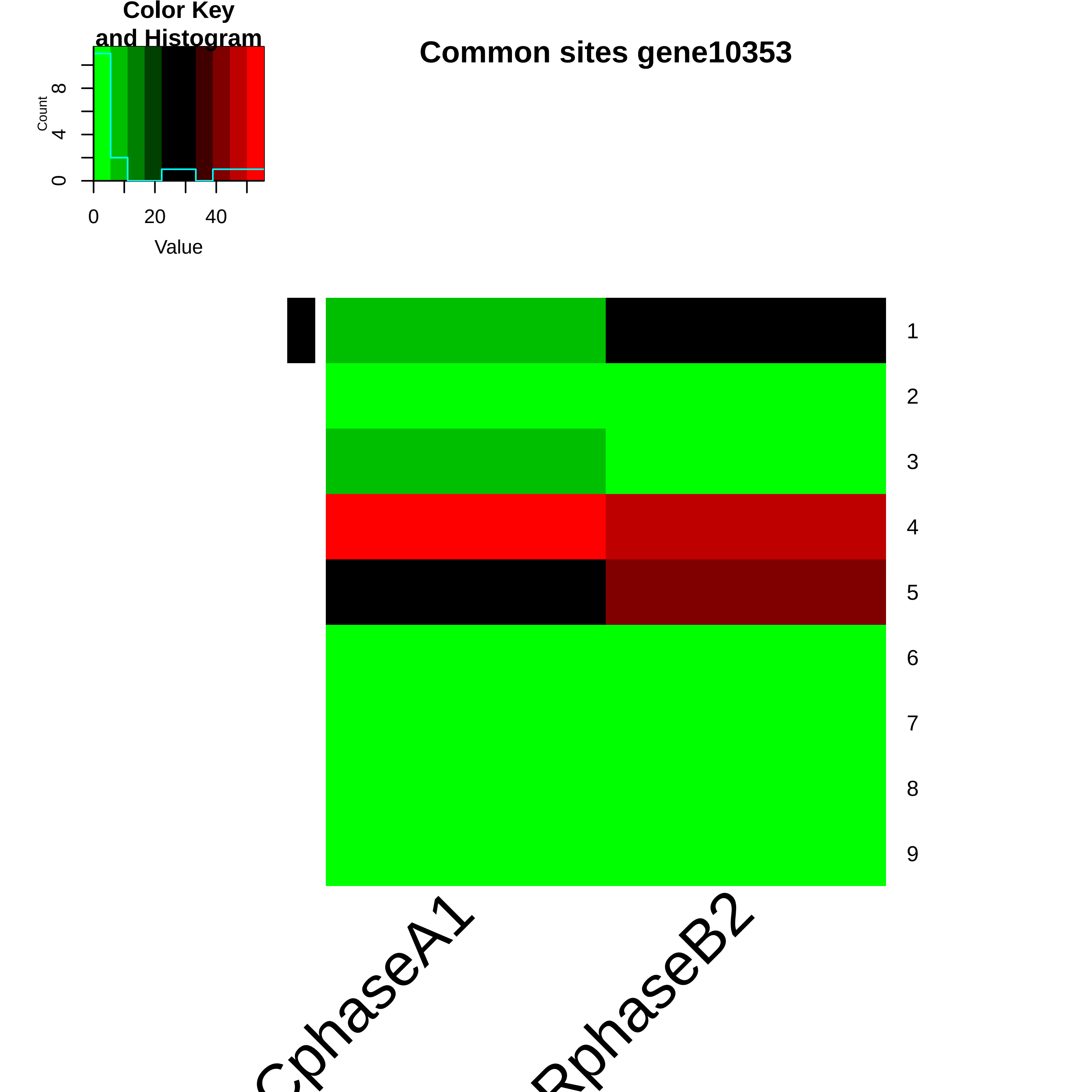
Color Key
and Histogram
Count
0
4
8
0
20
40
Value
Common sites gene10353
1
2
3
4
5
6
7
8
9
CphaseA1
RphaseB2

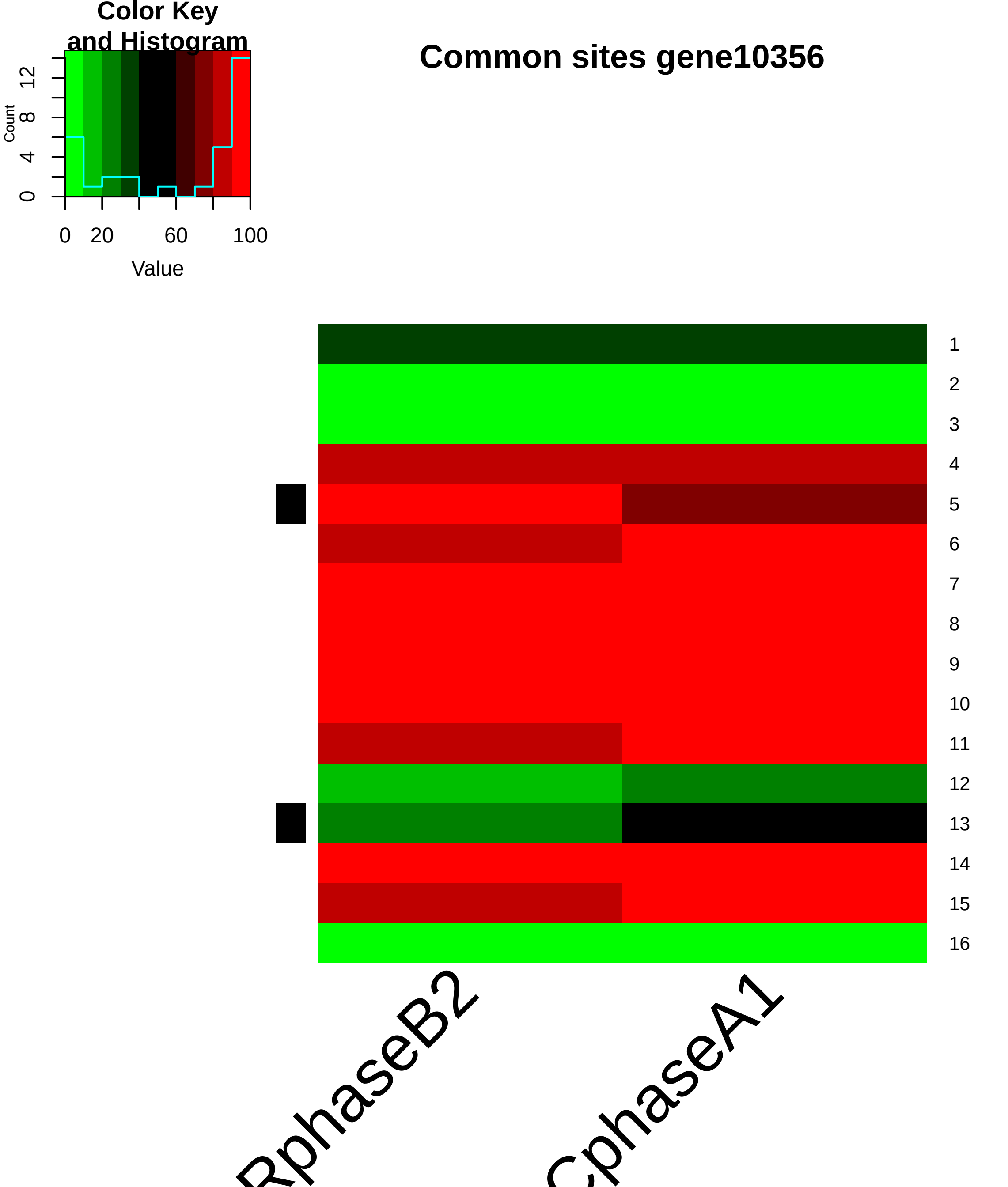

Color Key
and Histogram
Count
0
4
8
12
0
20
60
100
Value
Common sites gene10356
RphaseB2
CphaseA1

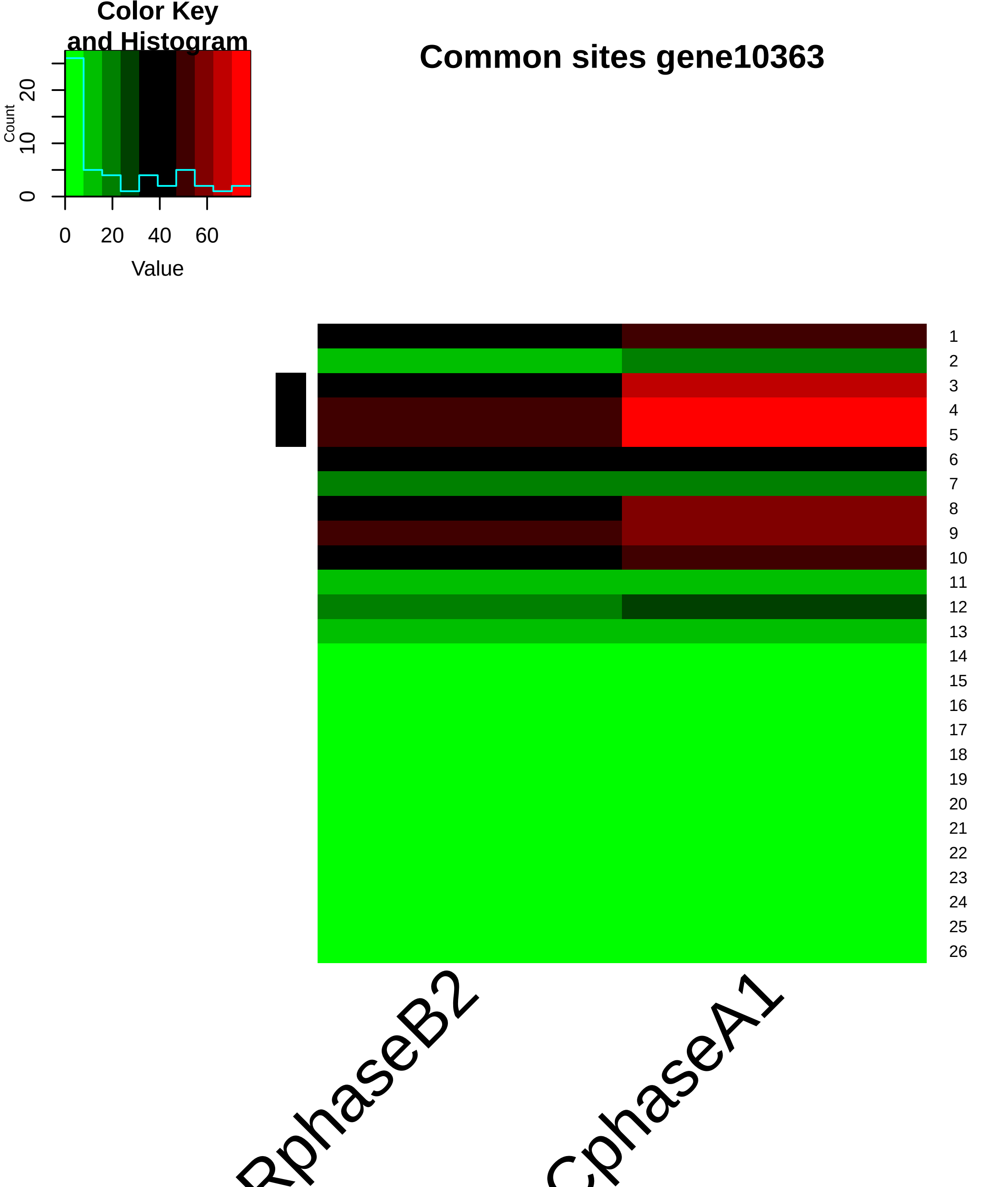
Color Key
and Histogram
Count
0
10
20
0
20
40
60
Value
Common sites gene10363
RphaseB2
CphaseA1

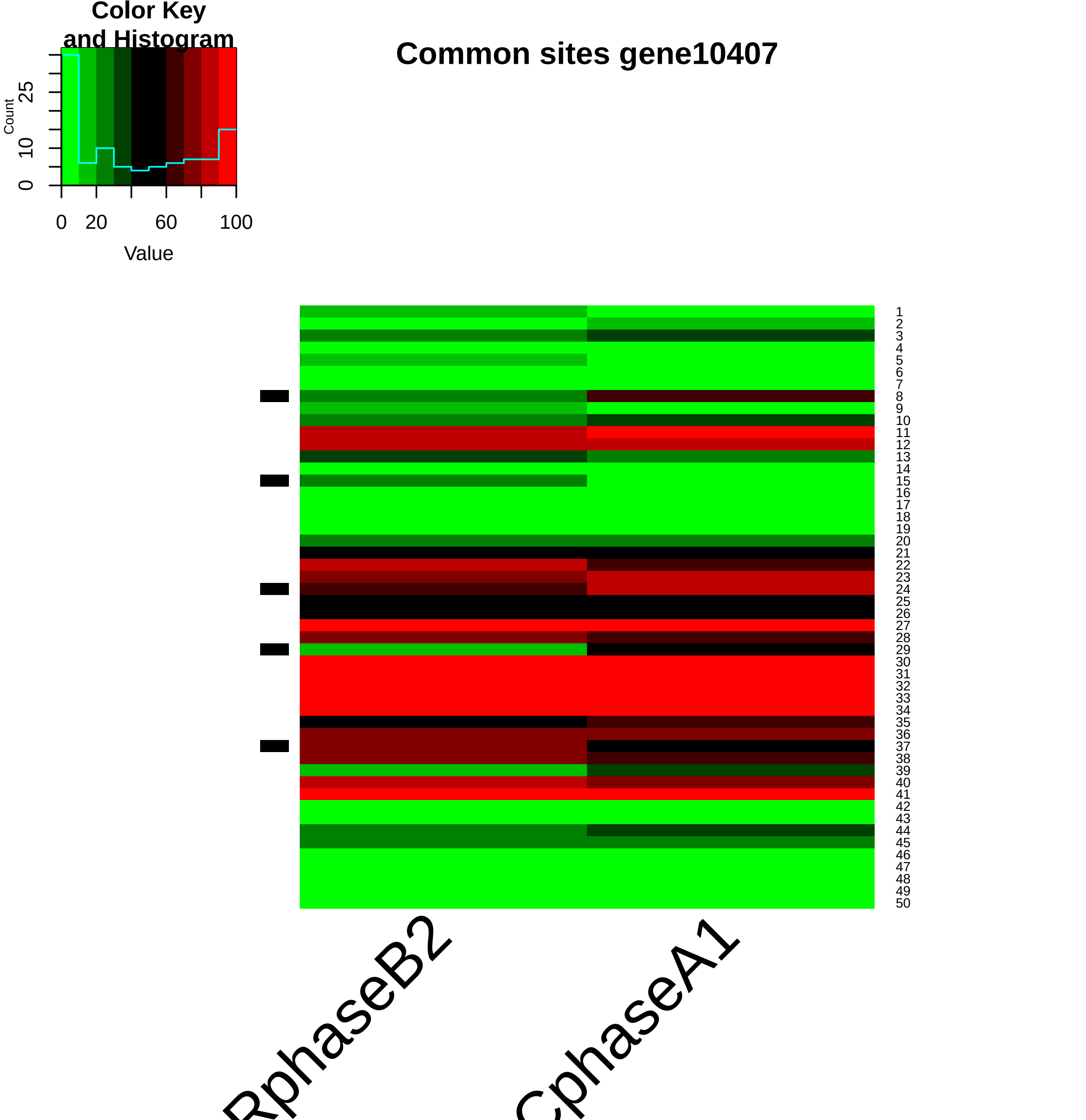
Color Key
and Histogram
Count
0
10
25
0
20
60
100
Value
Common sites gene10407
RphaseB2
CphaseA1

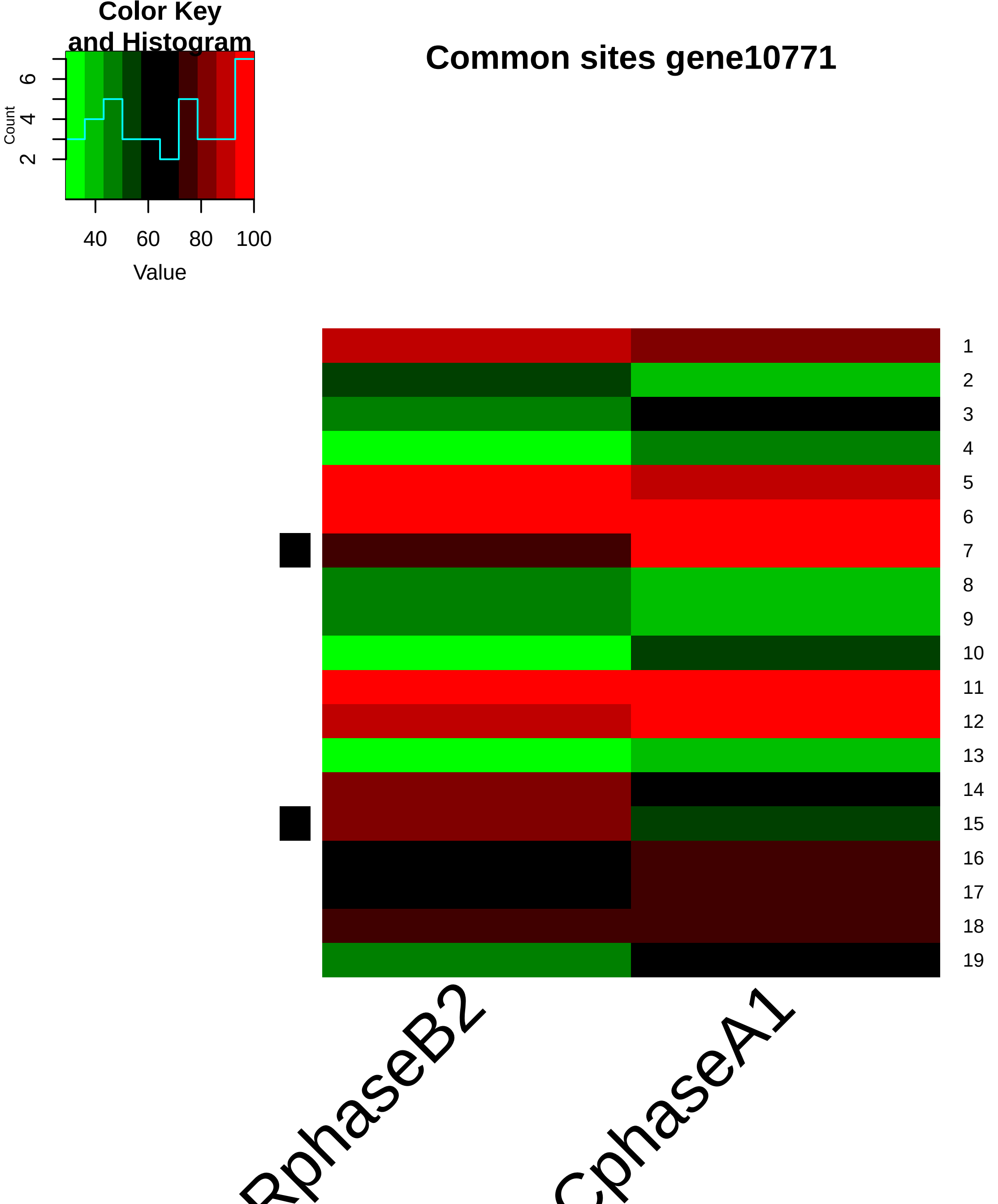
Color Key
and Histogram
Count
2
4
6
40
60
80
100
Value
Common sites gene10771
RphaseB2
CphaseA1

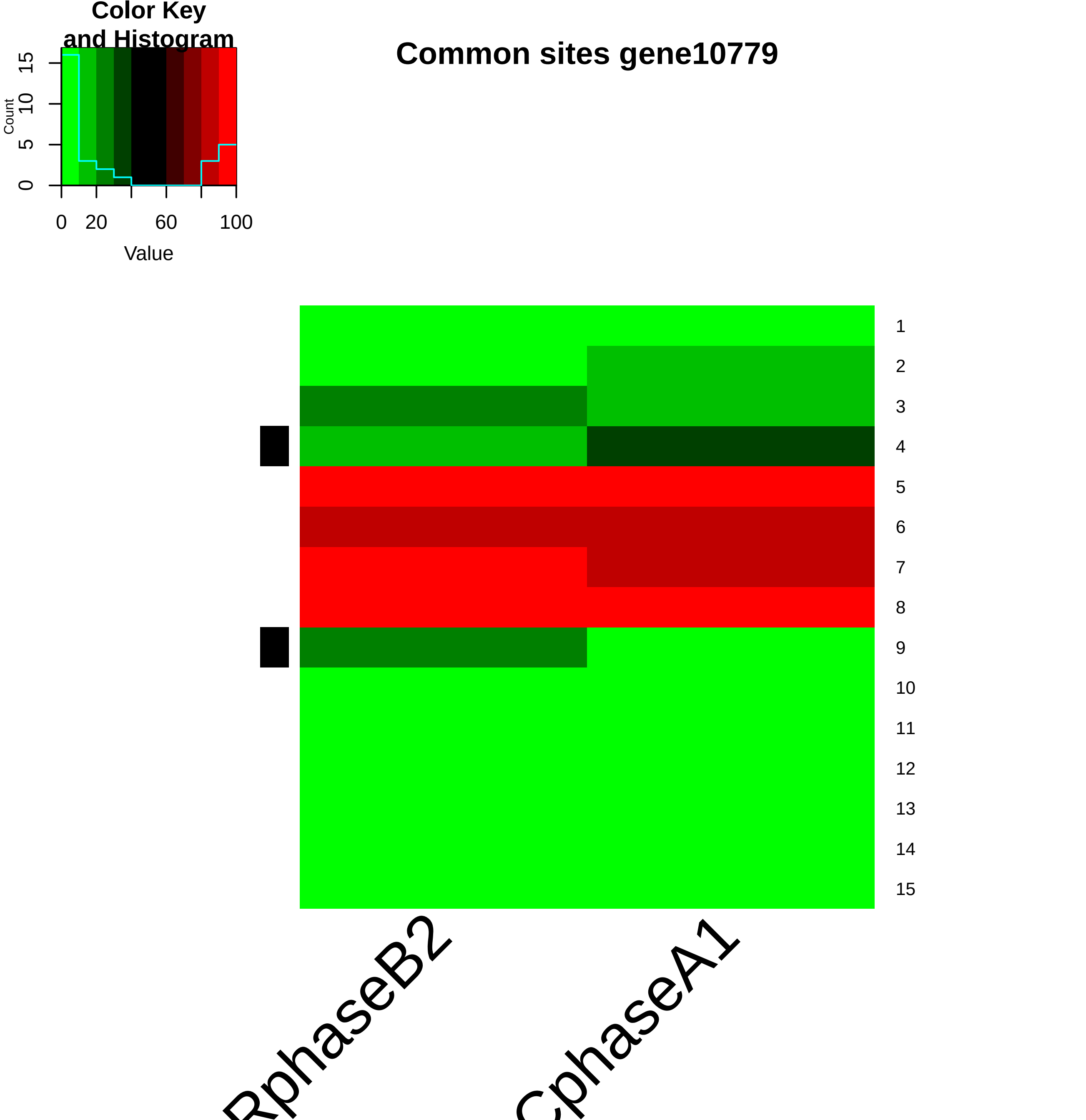
Color Key
and Histogram
Count
0
5
10
15
0
20
60
100
Value
Common sites gene10779
RphaseB2
CphaseA1

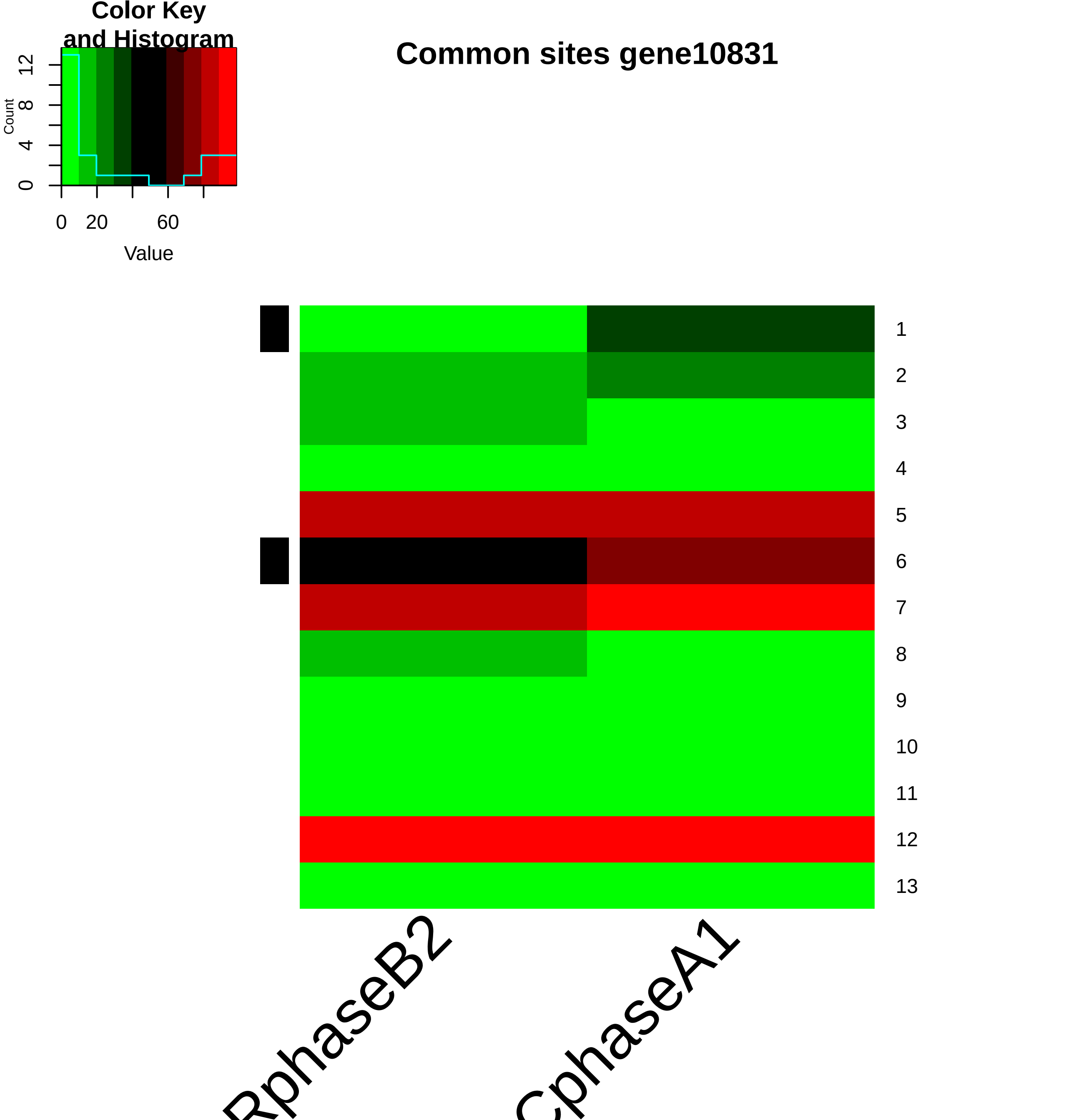

Color Key
and Histogram
Count
0
4
8
12
0
20
60
Value
Common sites gene10831
1
2
3
4
5
6
7
8
9
10
11
12
13
RphaseB2
CphaseA1

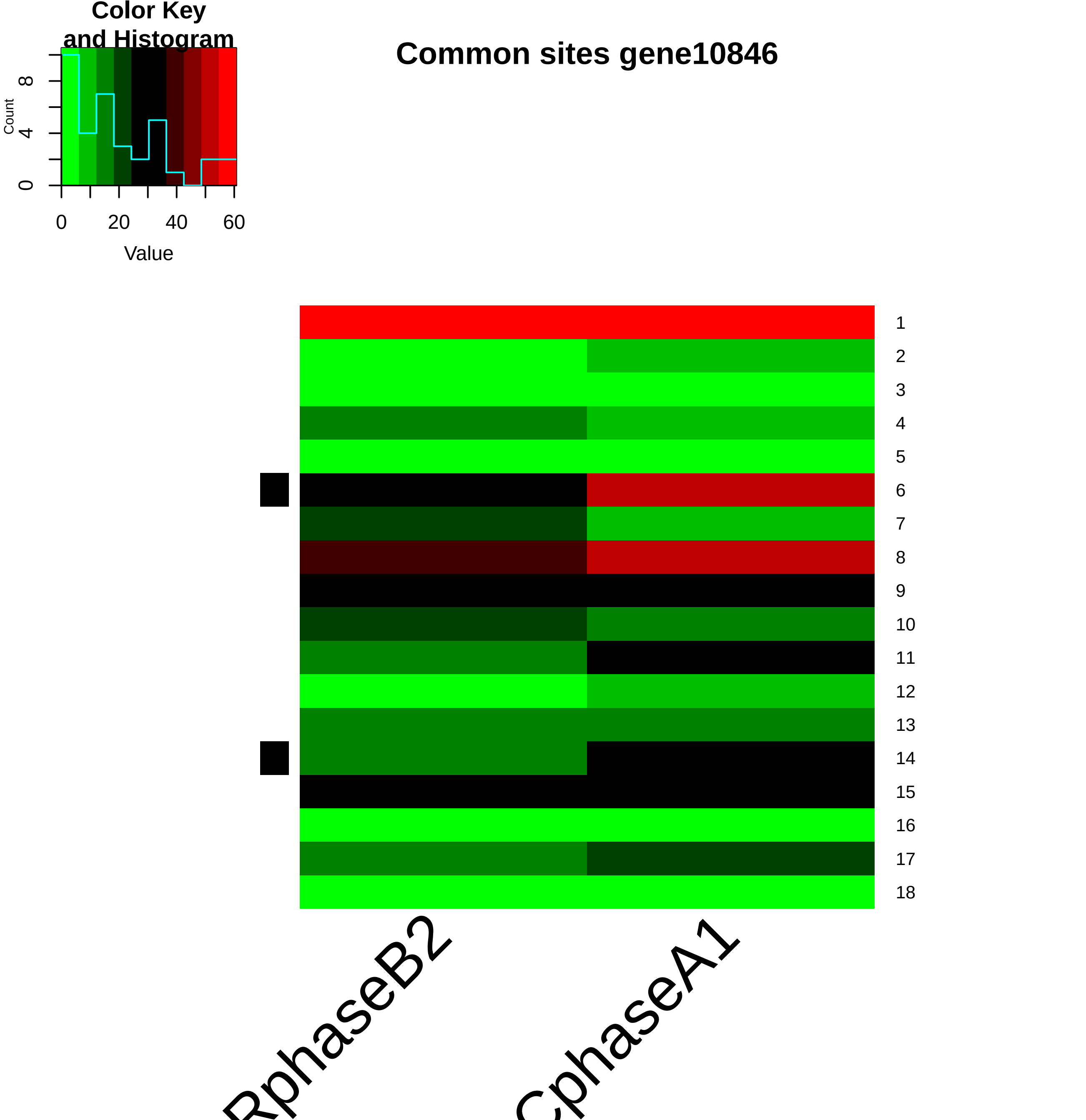
Color Key
and Histogram
Count
0
4
8
0
20
40
60
Value
Common sites gene10846
RphaseB2
CphaseA1

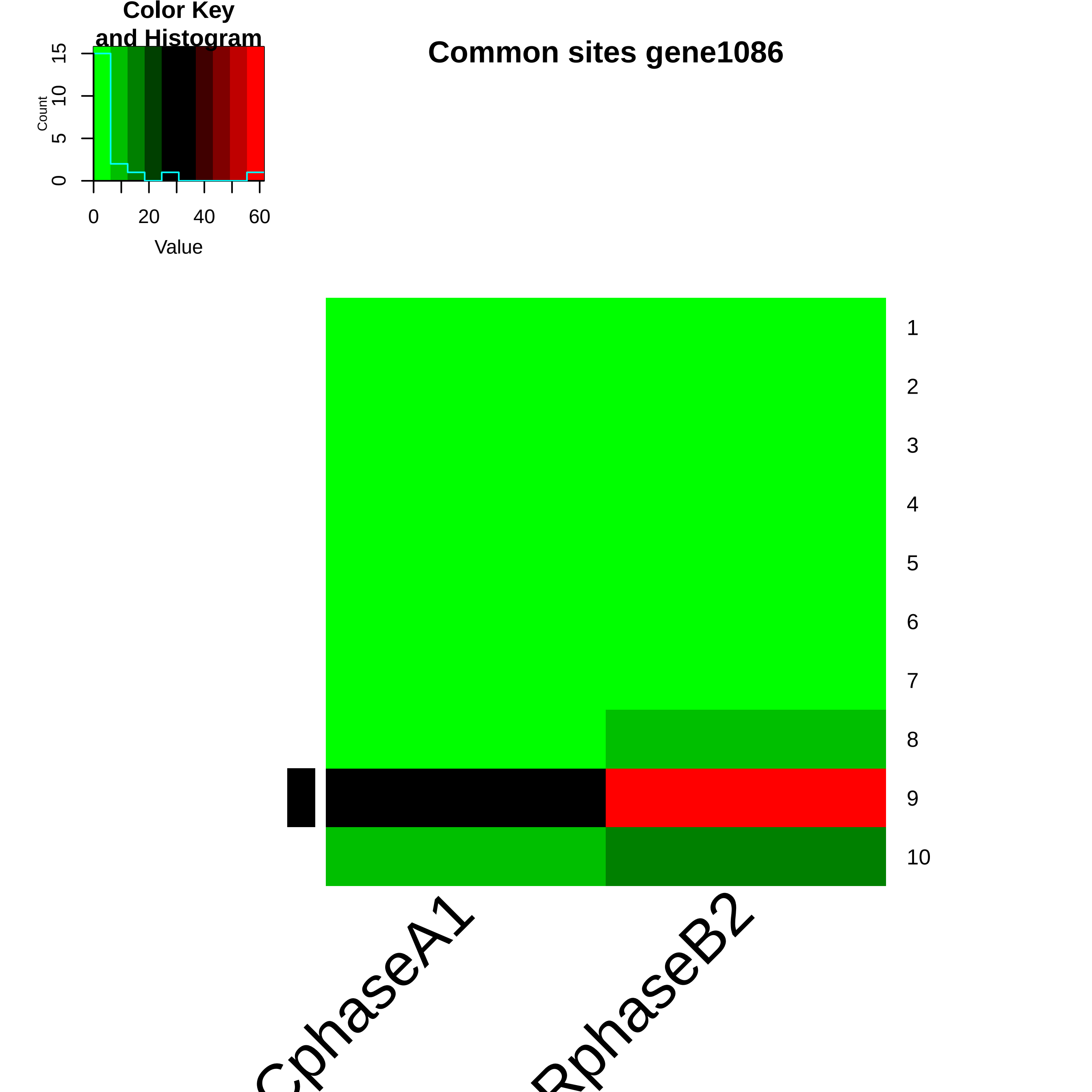

Color Key and Histogram
Count
0
5
10
15
0
20
40
60
Value
Common sites gene1086
1
2
3
4
5
6
7
8
9
10
CphaseA1
RphaseB2

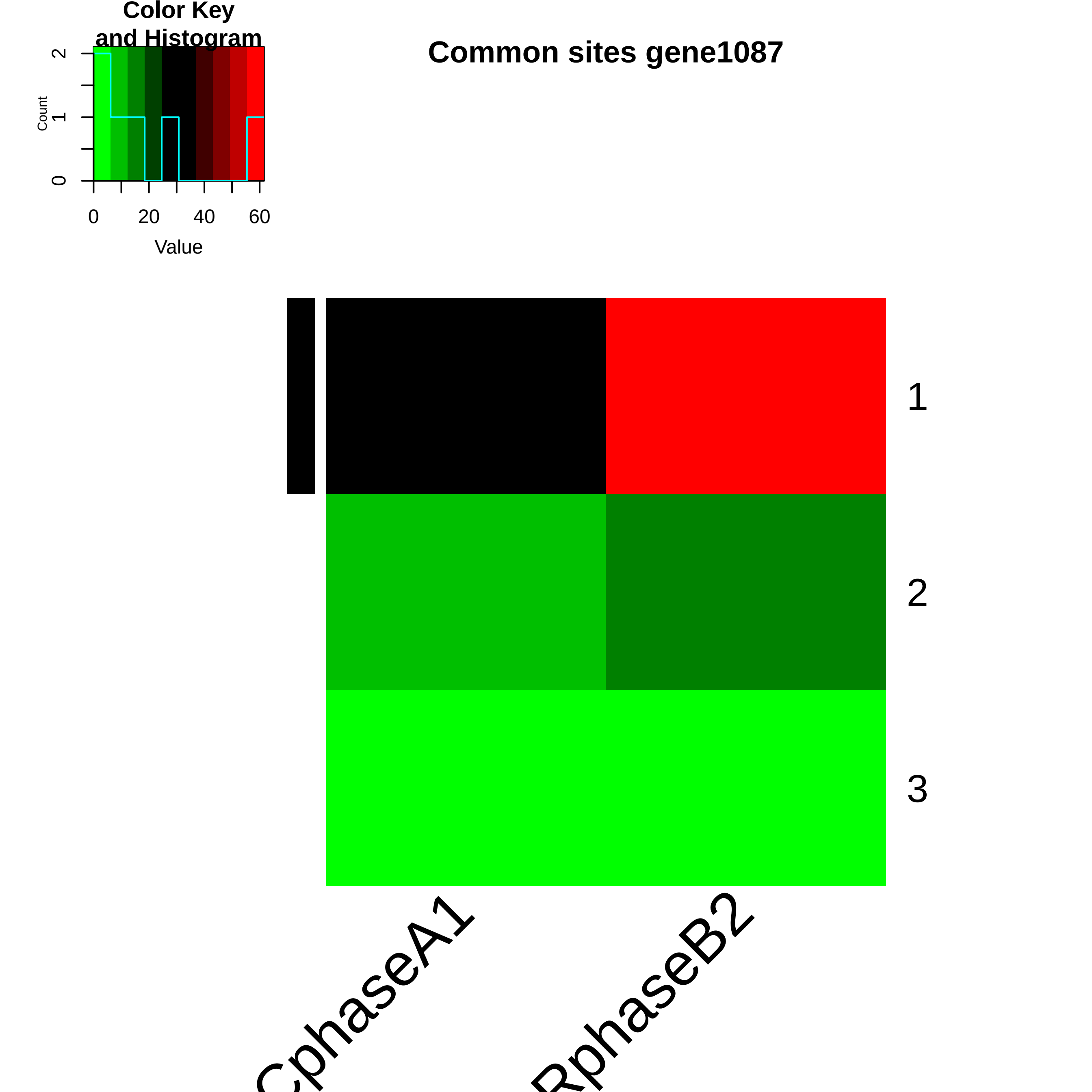

Color Key
and Histogram
Count
0
1
2
0
20
40
60
Value
Common sites gene1087
1
2
3
CphaseA1
RphaseB2

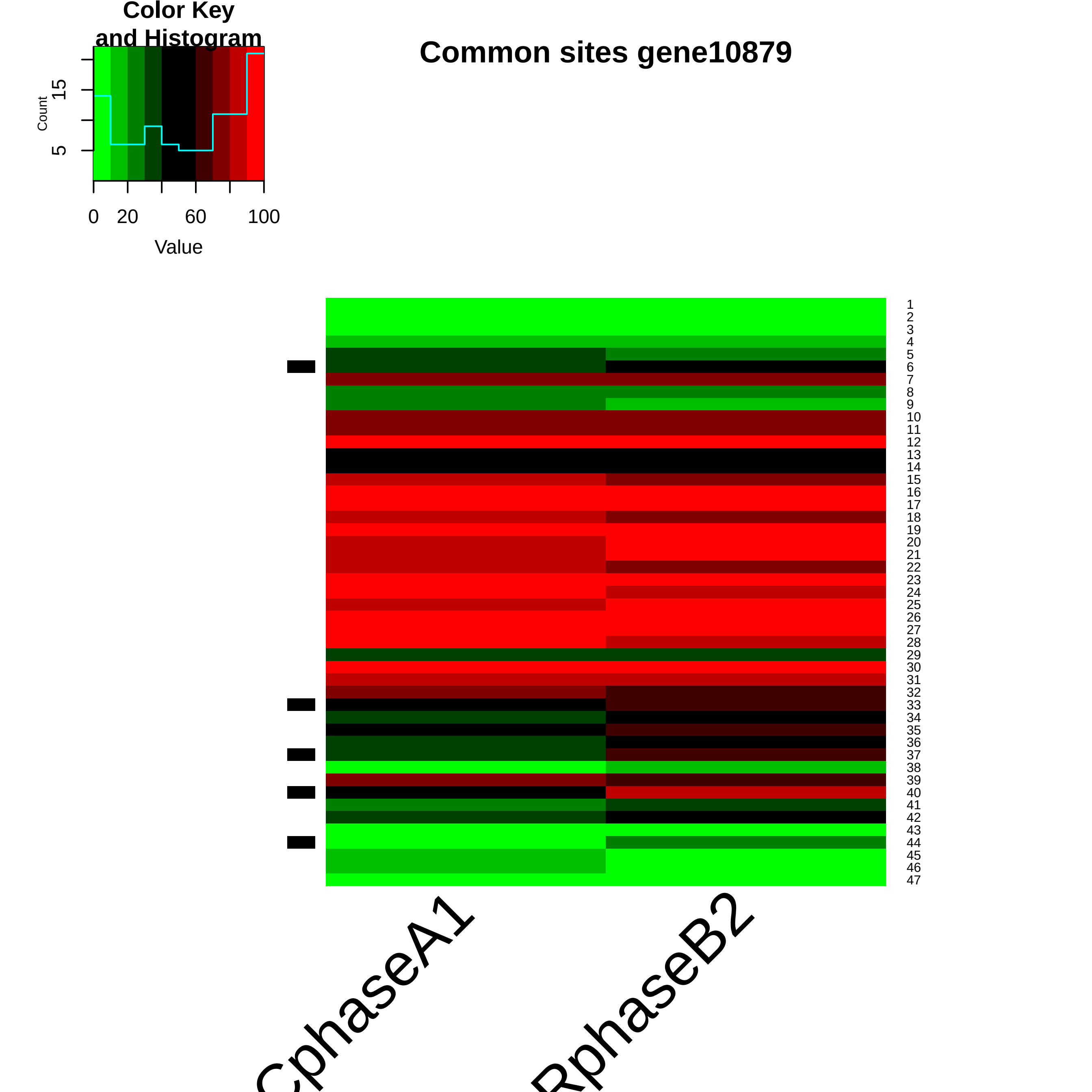
Color Key
and Histogram
Count
5
15
0
20
60
100
Value
Common sites gene10879
1 2 3 4 5 6 7 8 9 10 11 12 13 14 15 16 17 18 19 20 21 22 23 24 25 26 27 28 29 30 31 32 33 34 35 36 37 38 39 40 41 42 43 44 45 46 47
CphaseA1
RphaseB2

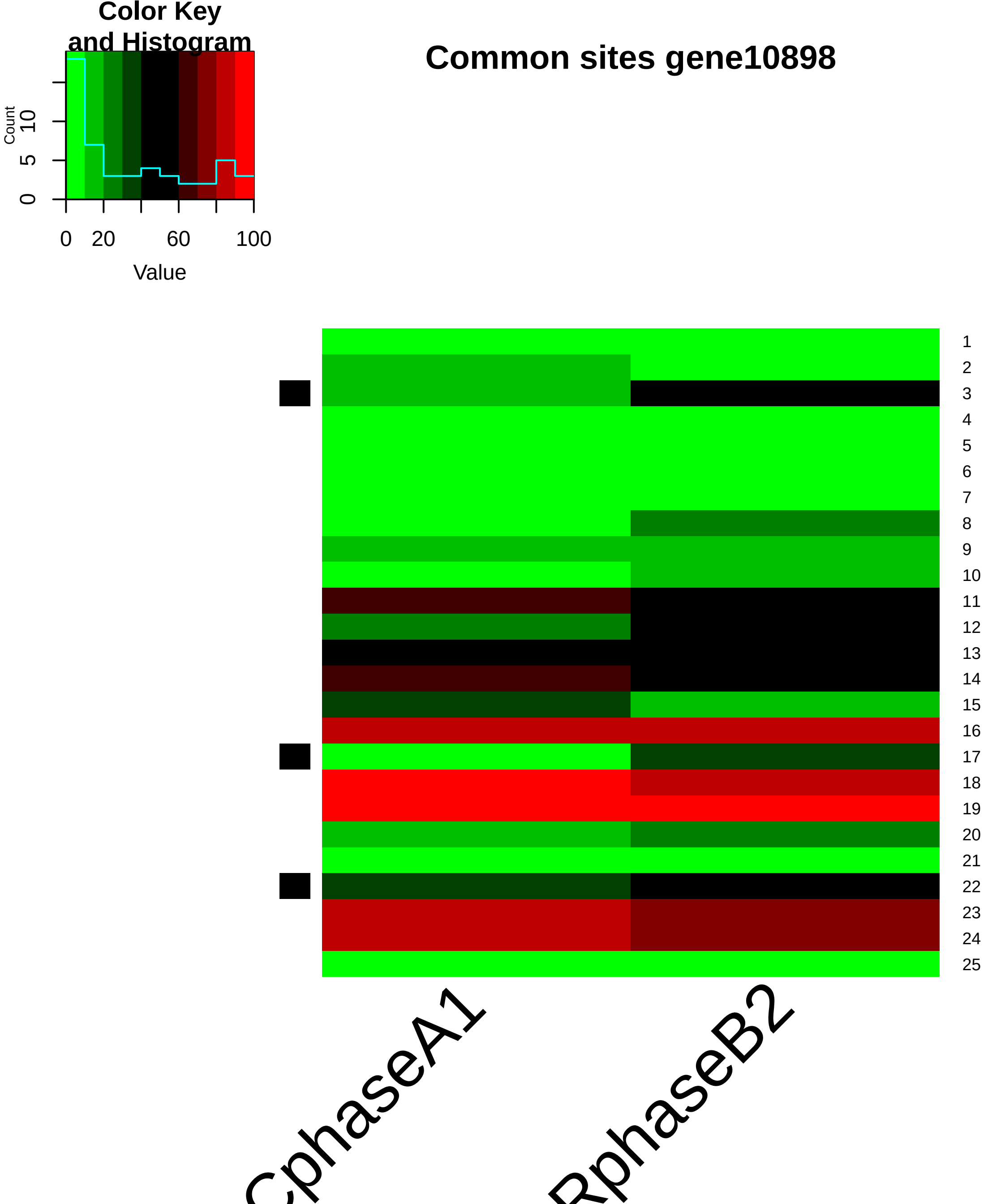
Color Key
and Histogram
Count
0
5
10
0
20
60
100
Value
Common sites gene10898
CphaseA1
RphaseB2

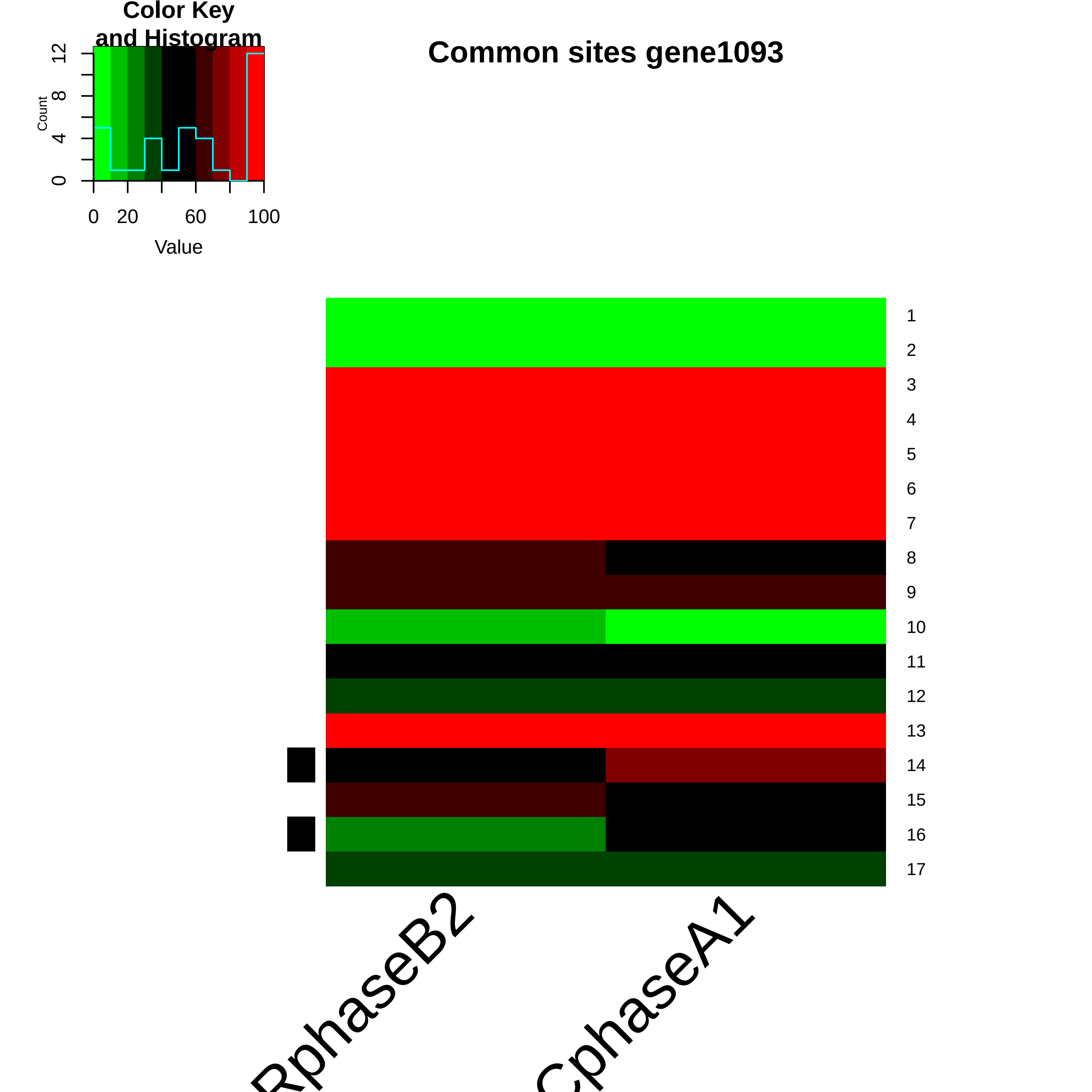
Color Key
and Histogram
Count
0
4
8
12
0
20
60
100
Value
Common sites gene1093
RphaseB2
CphaseA1

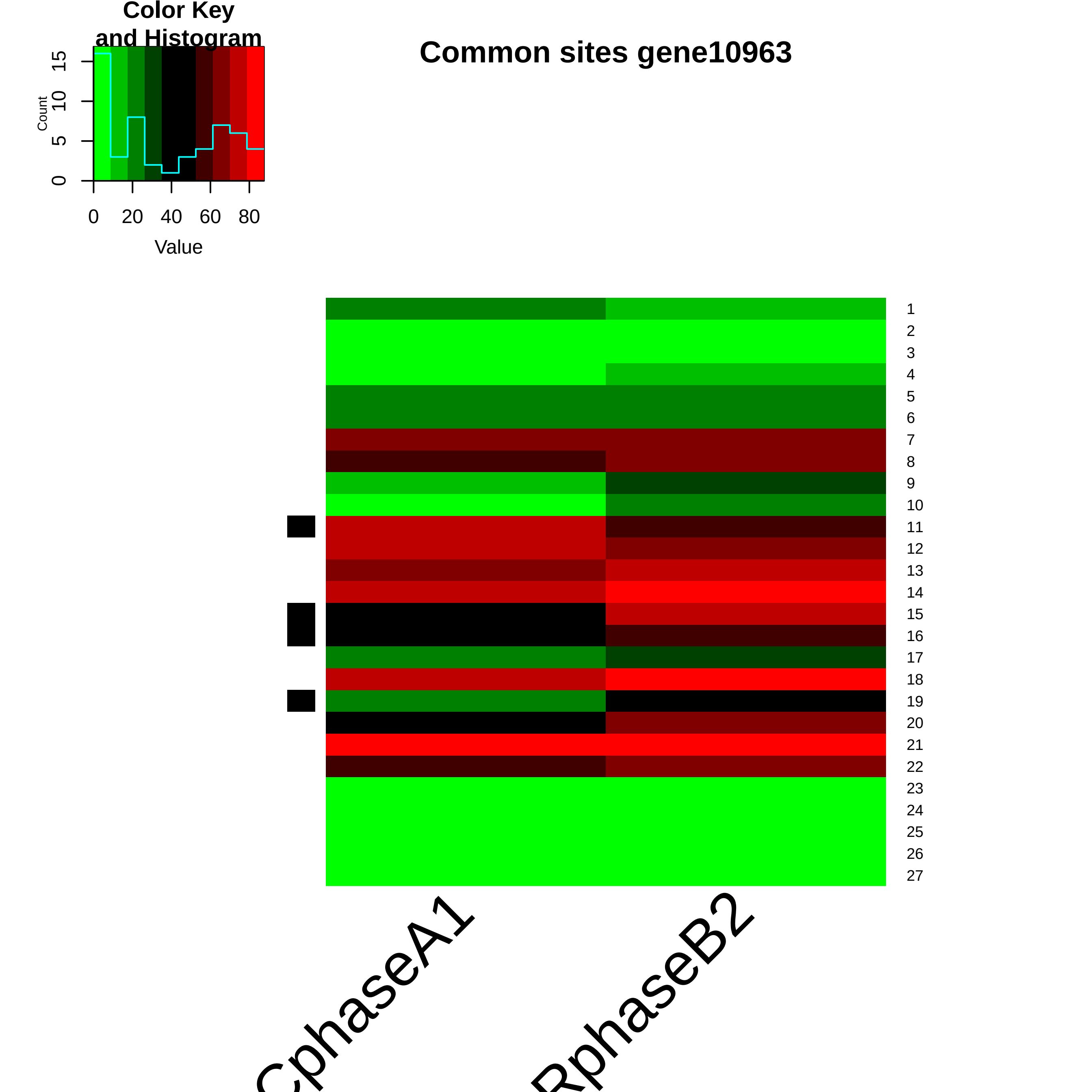
Color Key
and Histogram
Count
Value
0
20
40
60
80
0
5
10
15
Common sites gene10963
CphaseA1
RphaseB2

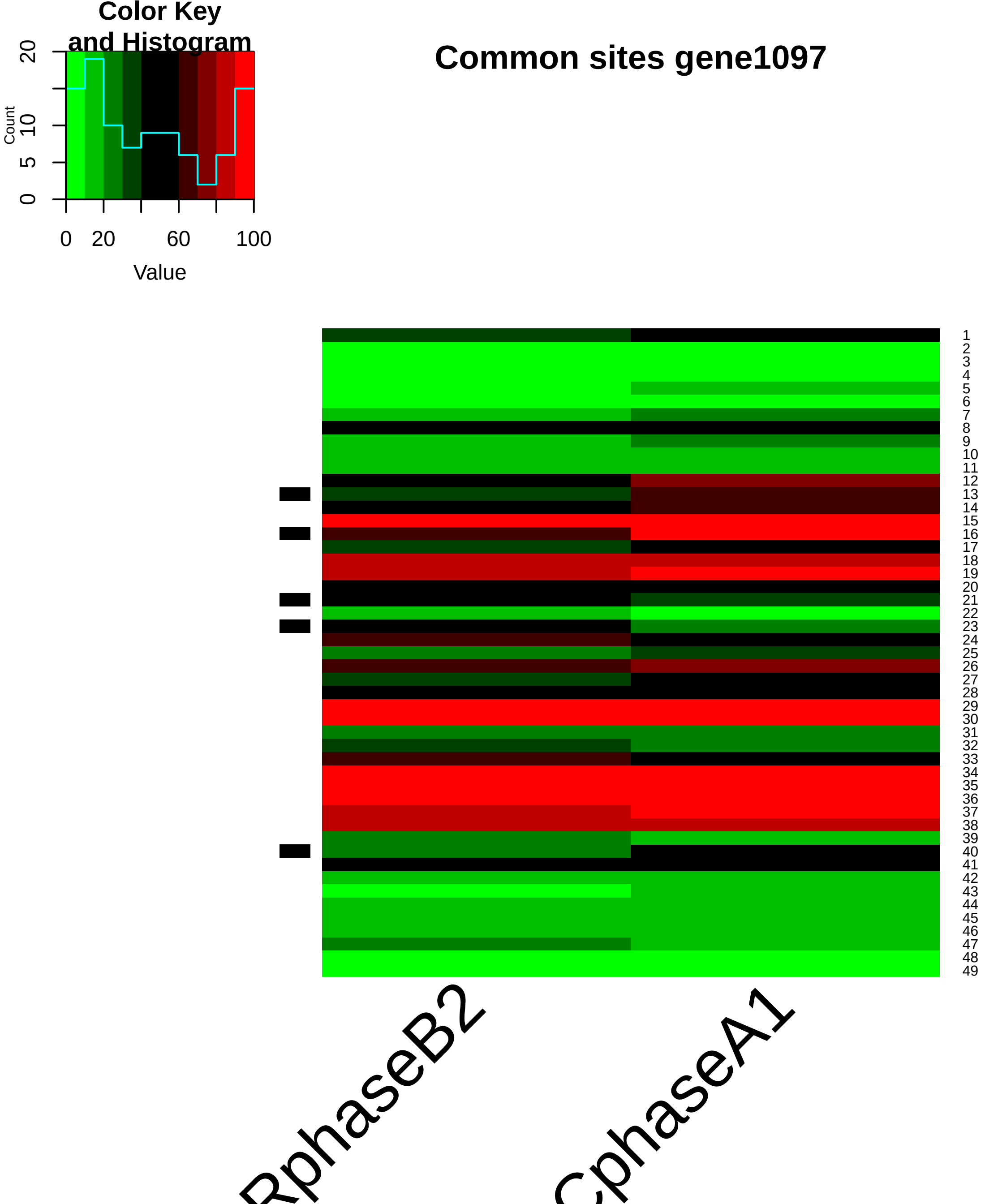

Color Key
and Histogram
Count
Value
0
20
60
100
0
5
10
20
Common sites gene1097
RphaseB2
CphaseA1

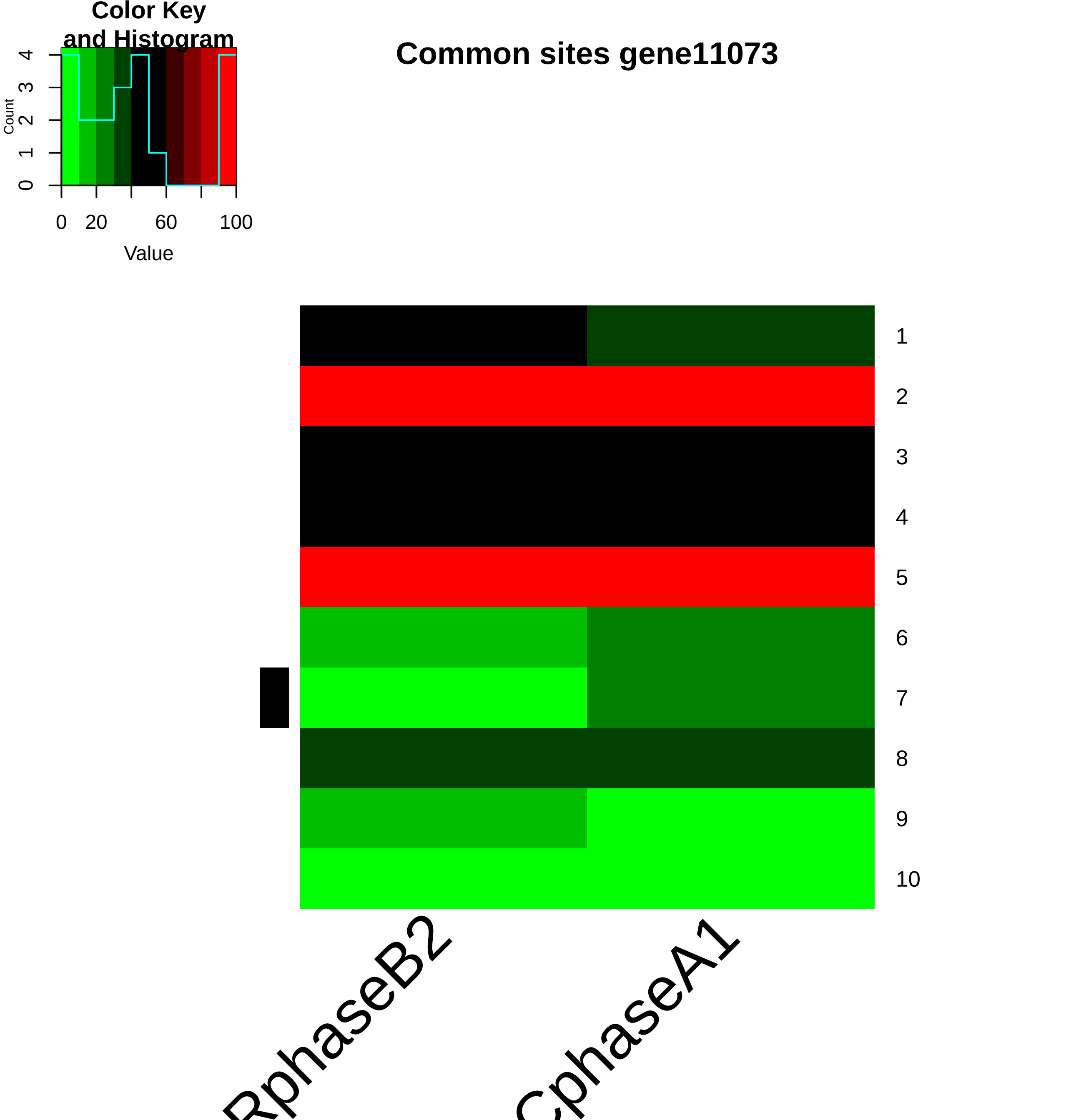
Color Key
and Histogram
Count
Value
0
20
60
100
0
1
2
3
4
Common sites gene11073
1
2
3
4
5
6
7
8
9
10
RphaseB2
CphaseA1

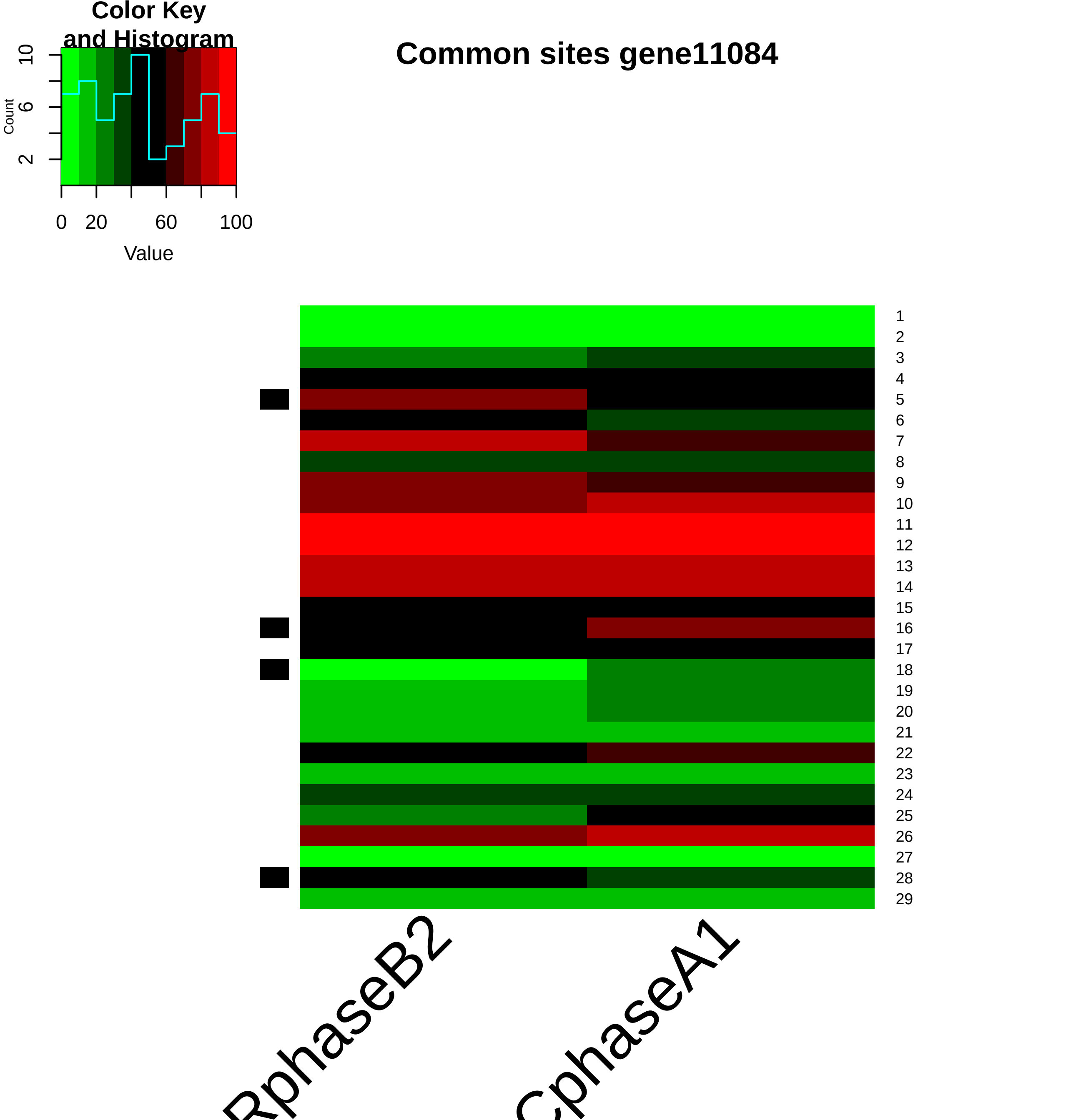
Color Key
and Histogram
Count
10
6
2
0
20
60
100
Value
Common sites gene11084
RphaseB2
CphaseA1

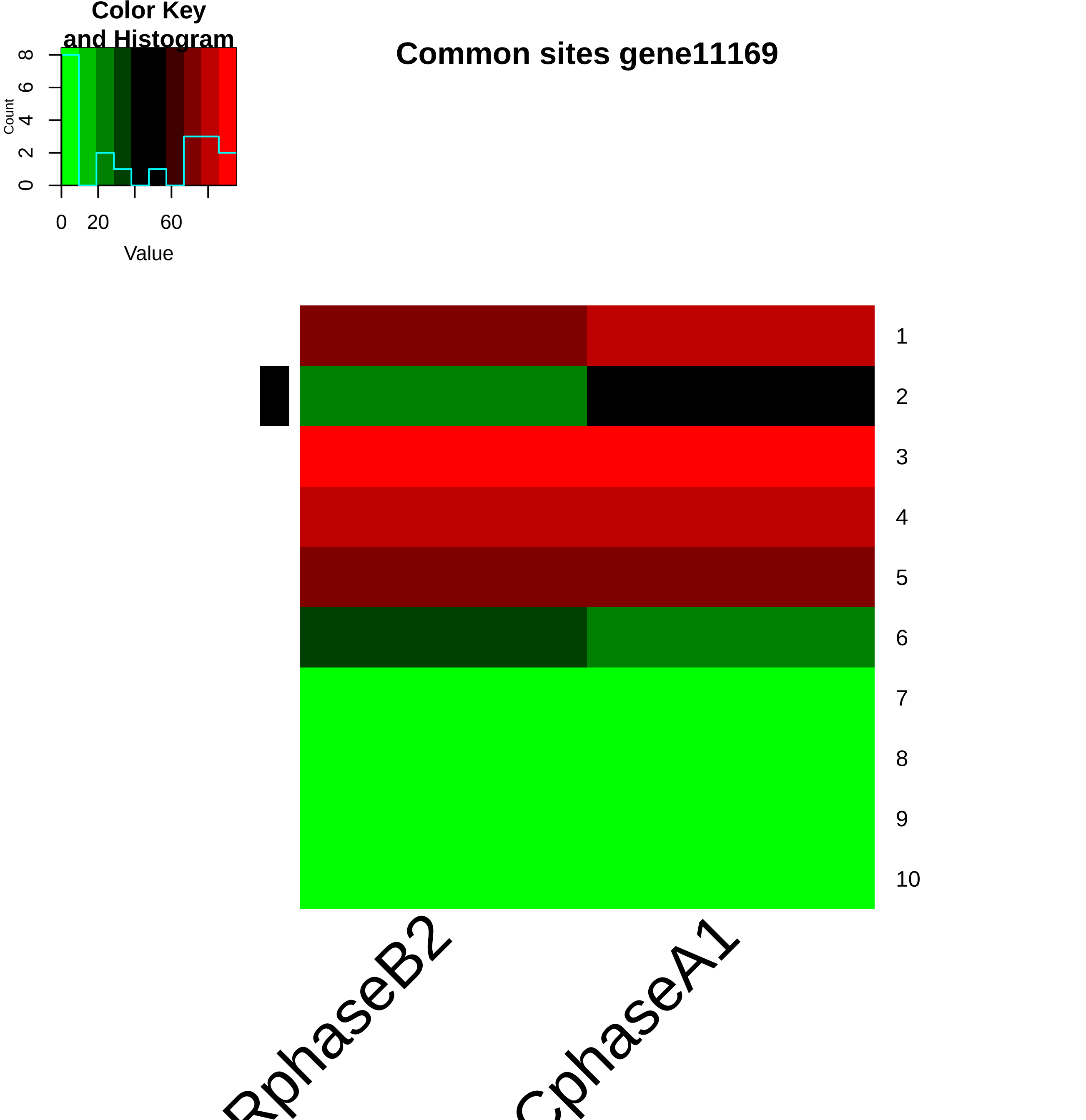

Color Key
and Histogram
Count
0
2
4
6
8
0
20
60
Value
Common sites gene11169
1
2
3
4
5
6
7
8
9
10
RphaseB2
CphaseA1

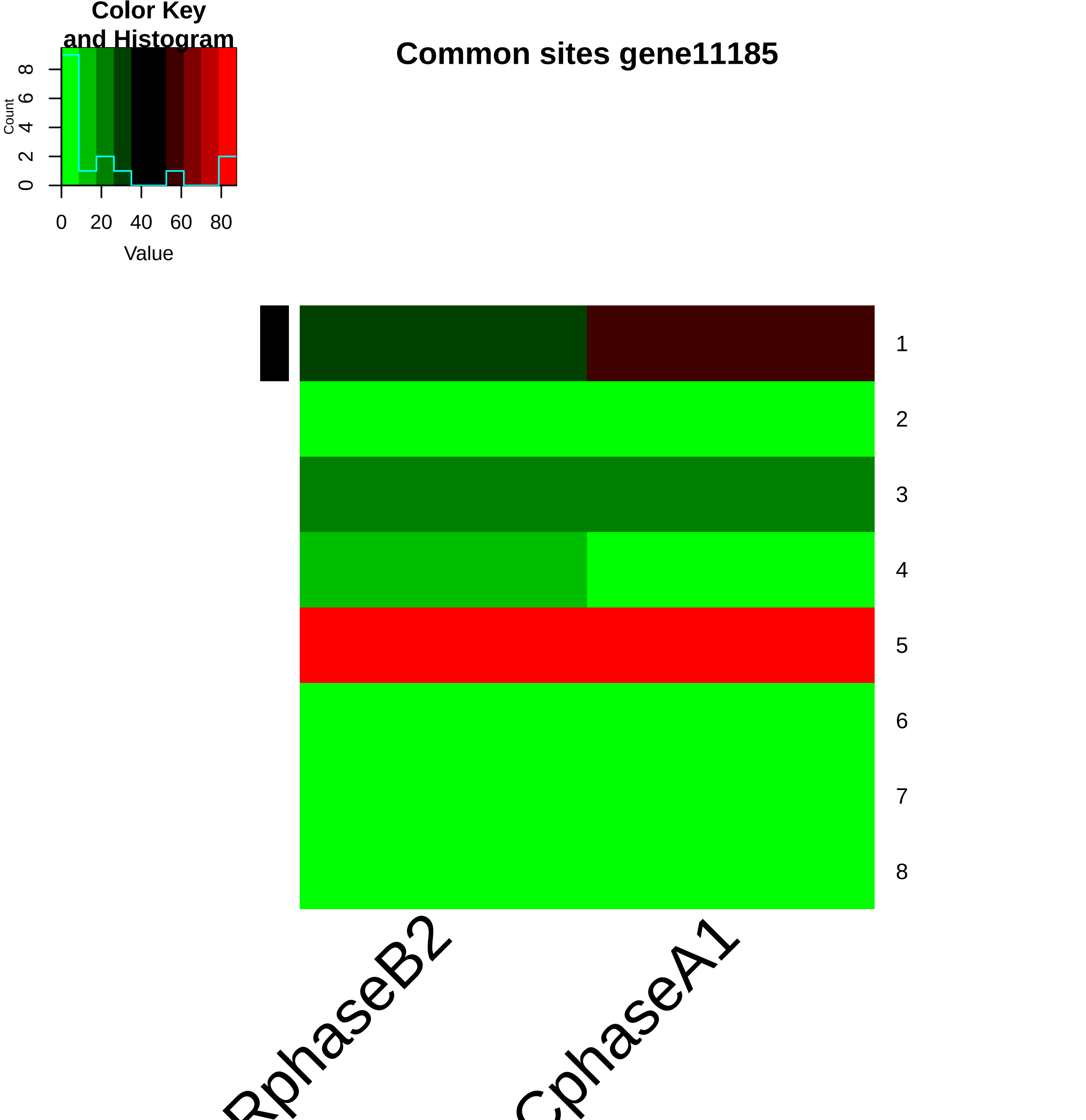
Color Key
and Histogram
Count
0
2
4
6
8
0
20
40
60
80
Value
Common sites gene11185
1
2
3
4
5
6
7
8
RphaseB2
CphaseA1

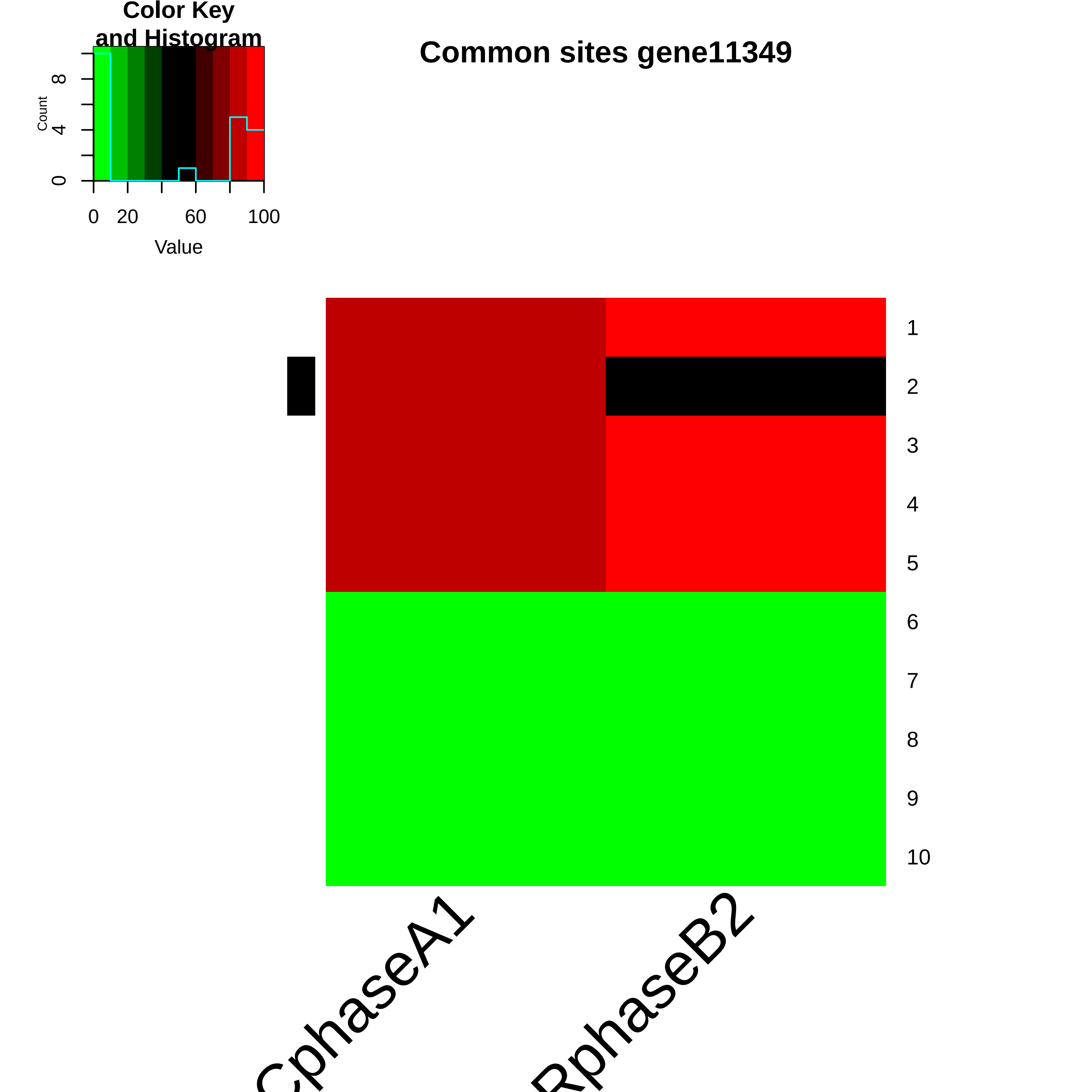

Color Key
and Histogram
Count
0
4
8
0
20
60
100
Value
Common sites gene11349
1
2
3
4
5
6
7
8
9
10
CphaseA1
RphaseB2

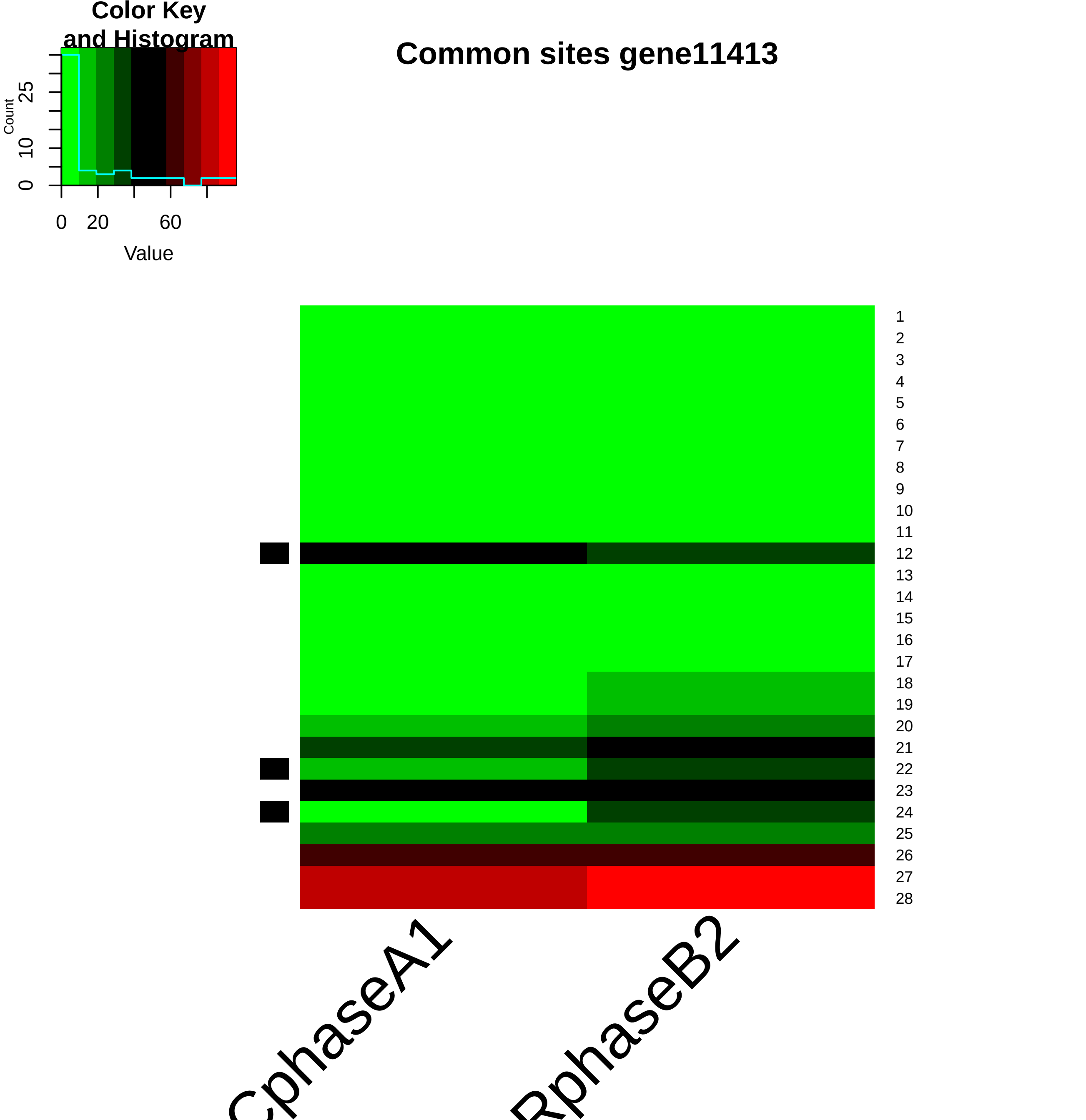
Color Key
and Histogram
Count
0
10
25
0
20
60
Value
Common sites gene11413
CphaseA1
RphaseB2

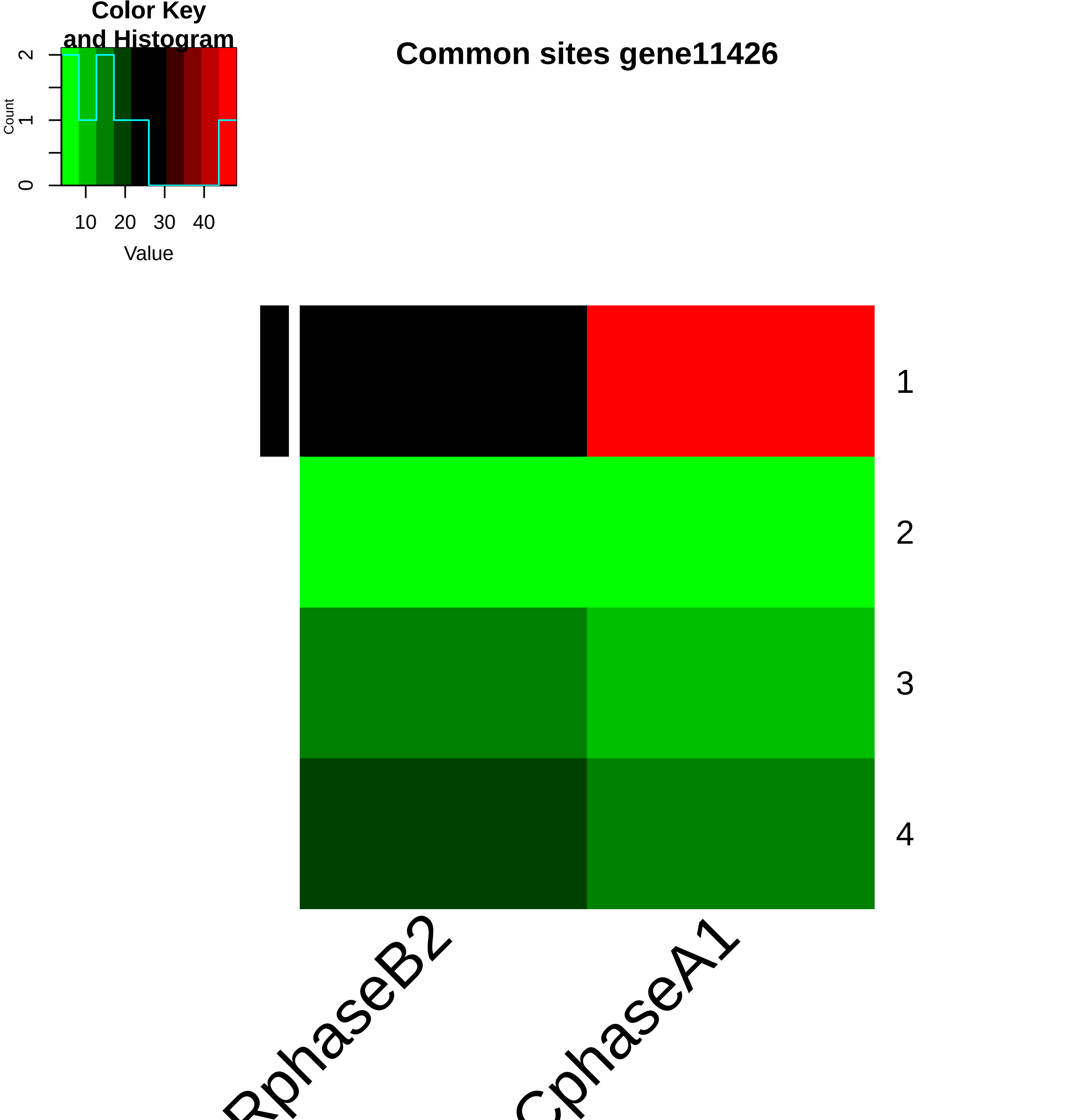
Color Key
and Histogram
Count
0
1
2
10
20
30
40
Value
Common sites gene11426
1
2
3
4
RphaseB2
CphaseA1

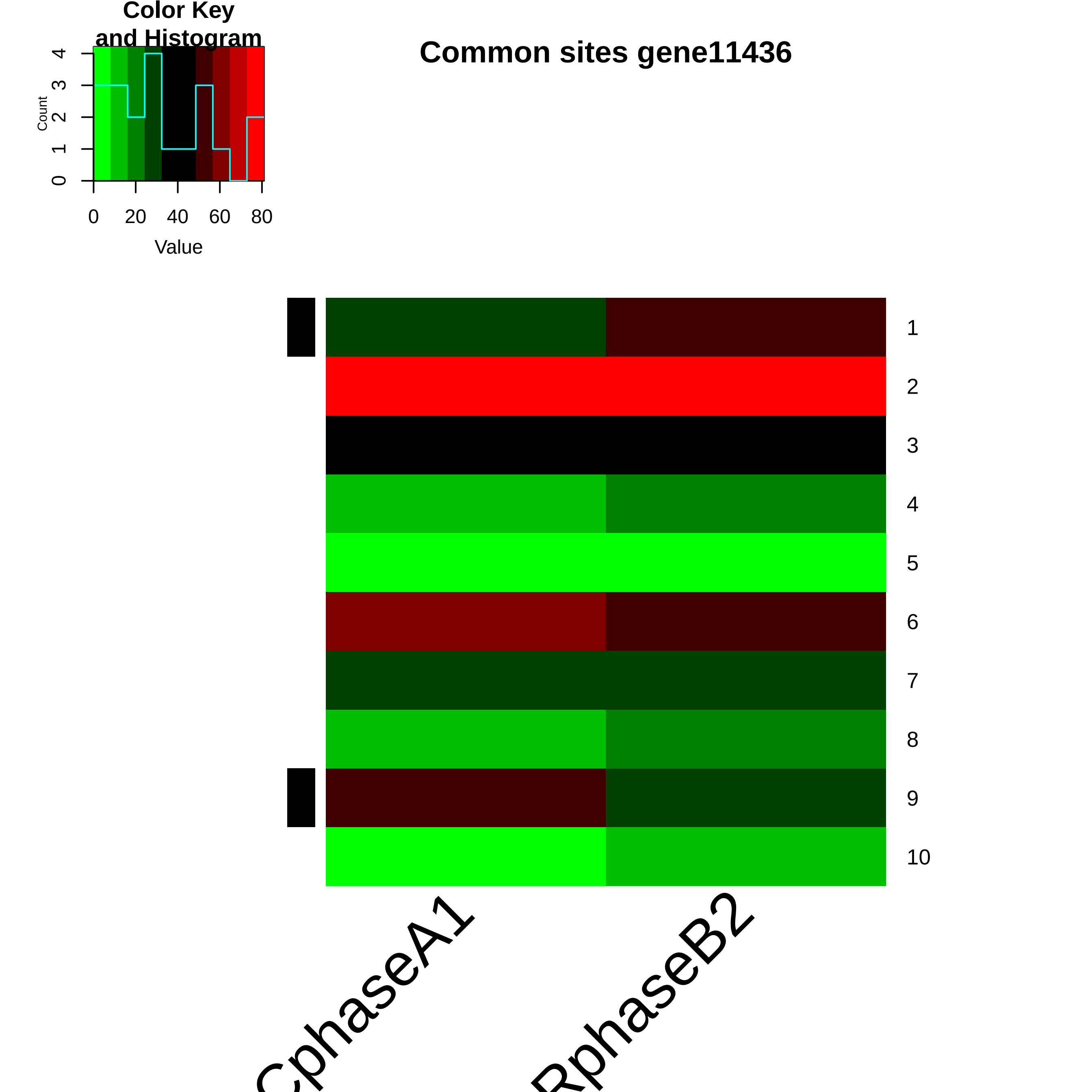
Color Key
and Histogram
Count
0
1
2
3
4
0
20
40
60
80
Value
Common sites gene11436
1
2
3
4
5
6
7
8
9
10
CphaseA1
RphaseB2

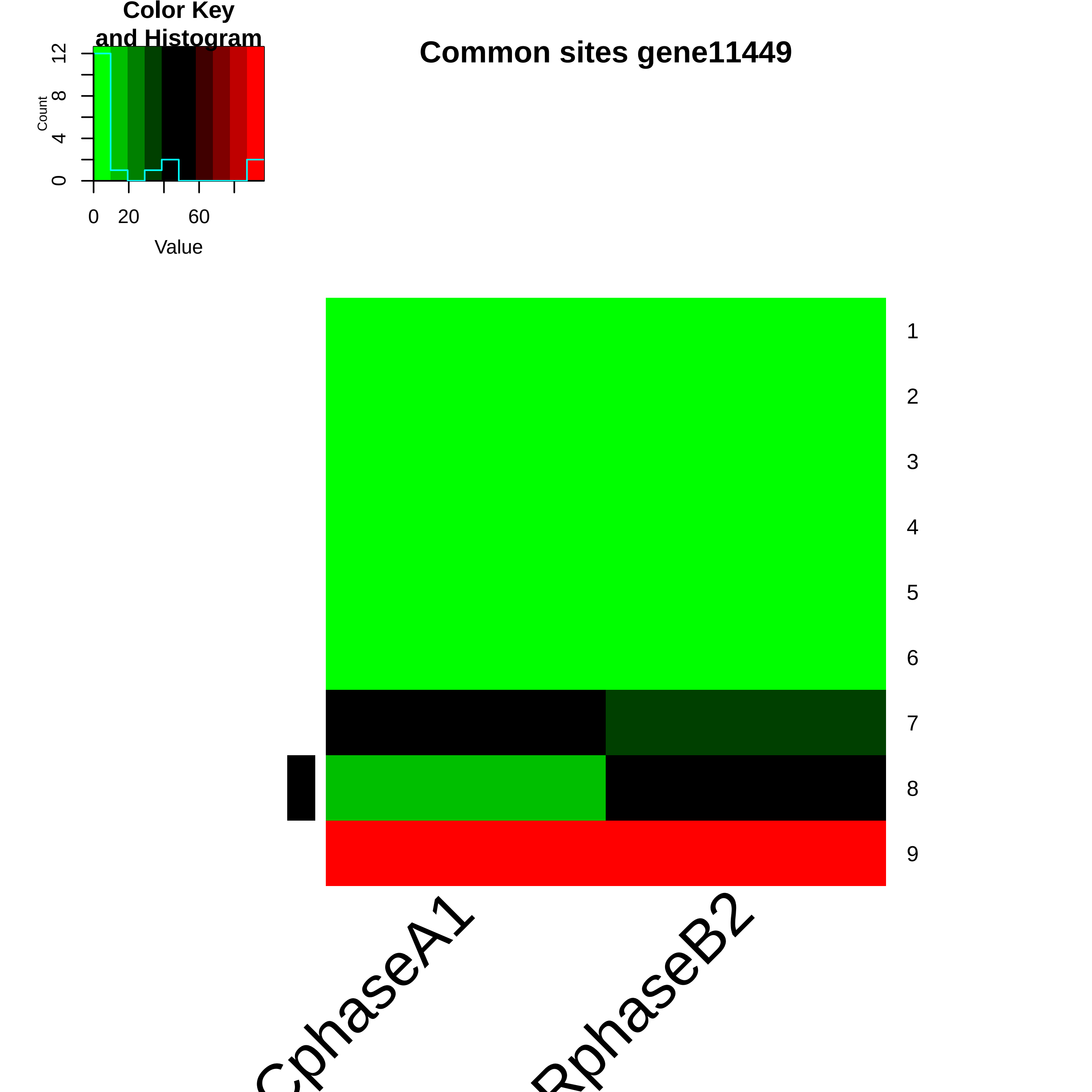
Color Key
and Histogram
Count
0
4
8
12
0
20
60
Value
Common sites gene11449
1
2
3
4
5
6
7
8
9
CphaseA1
RphaseB2

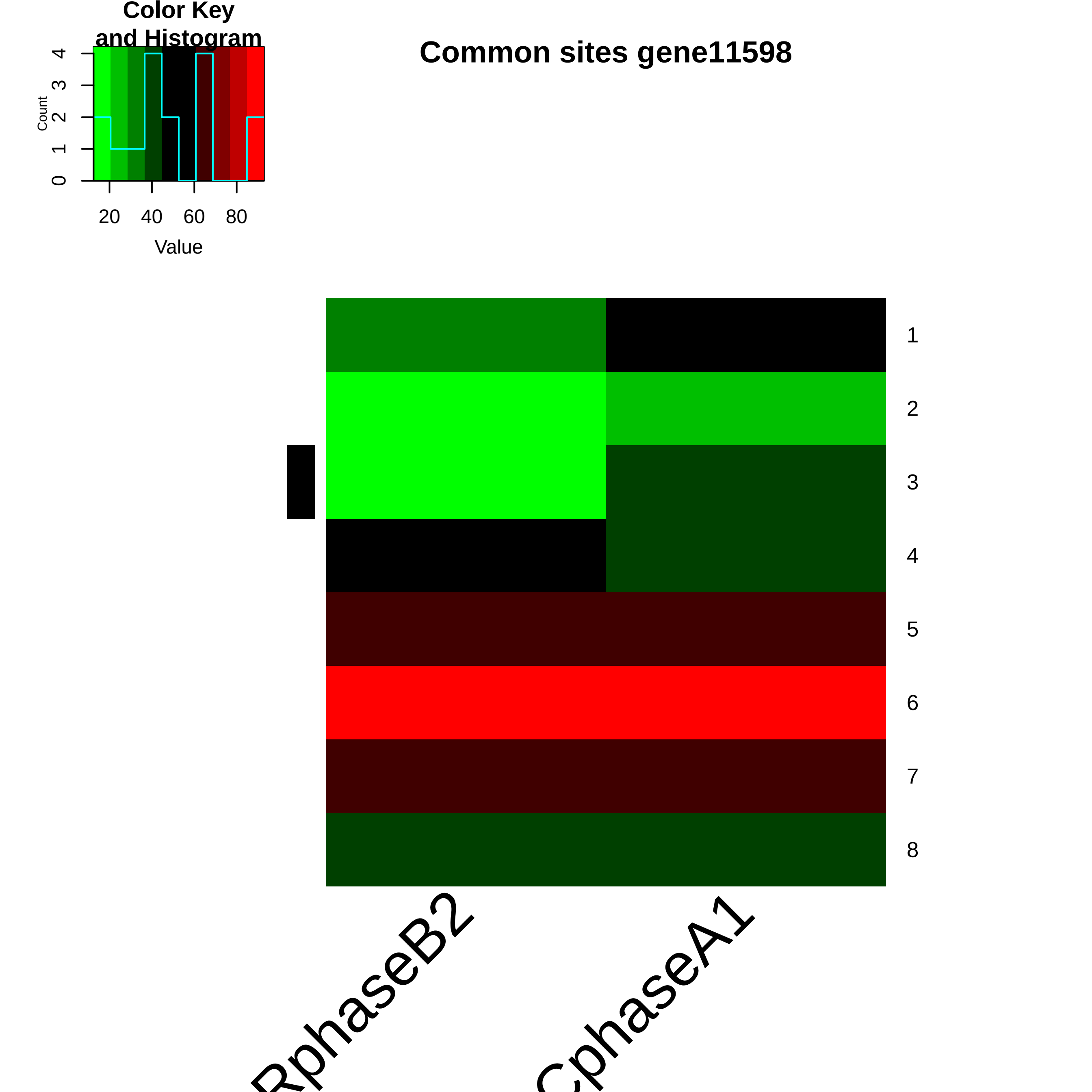
Color Key
and Histogram
Count
0
1
2
3
4
20
40
60
80
Value
Common sites gene11598
1
2
3
4
5
6
7
8
RphaseB2
CphaseA1

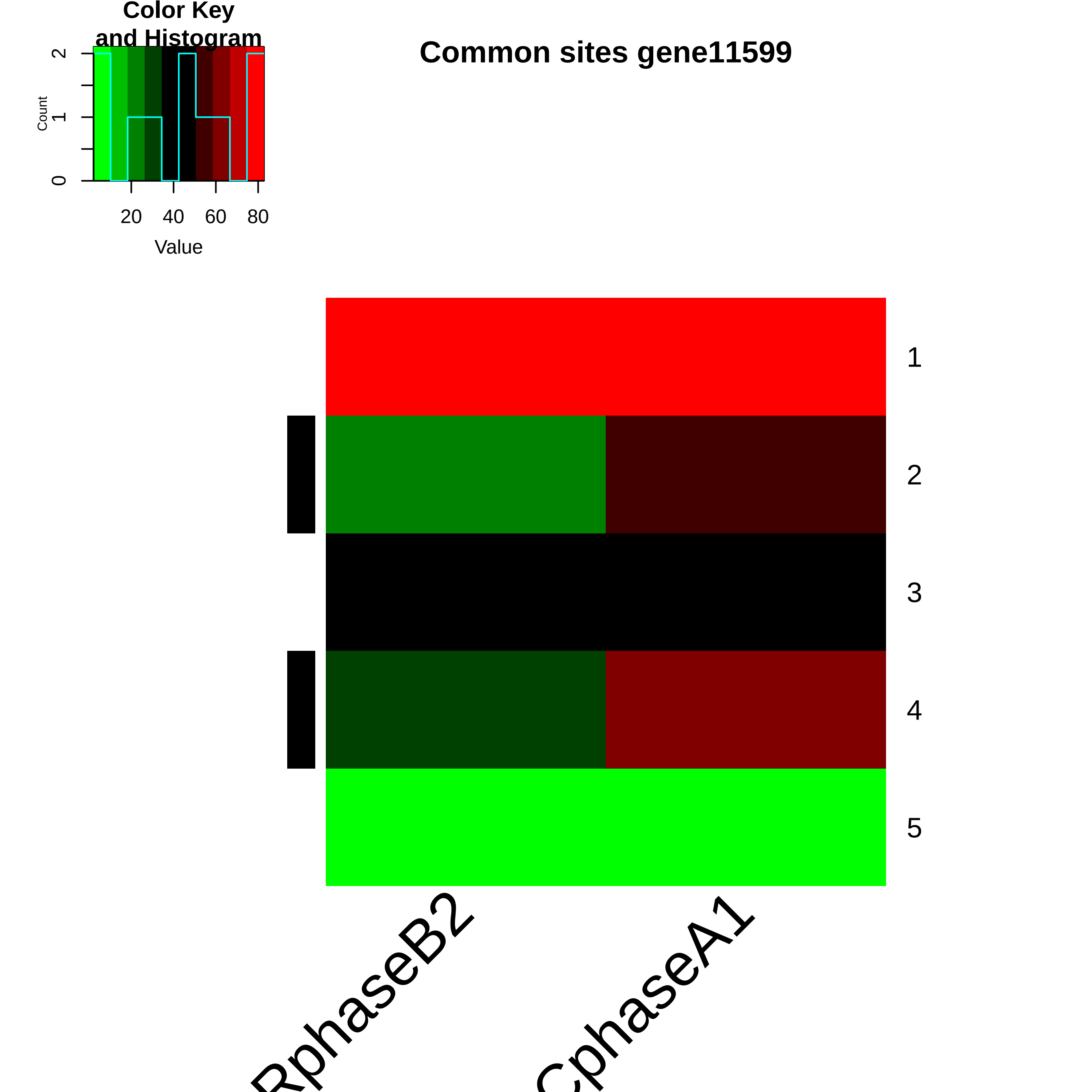

Color Key
and Histogram
Count
0
1
2
20
40
60
80
Value
Common sites gene11599
1
2
3
4
5
RphaseB2
CphaseA1

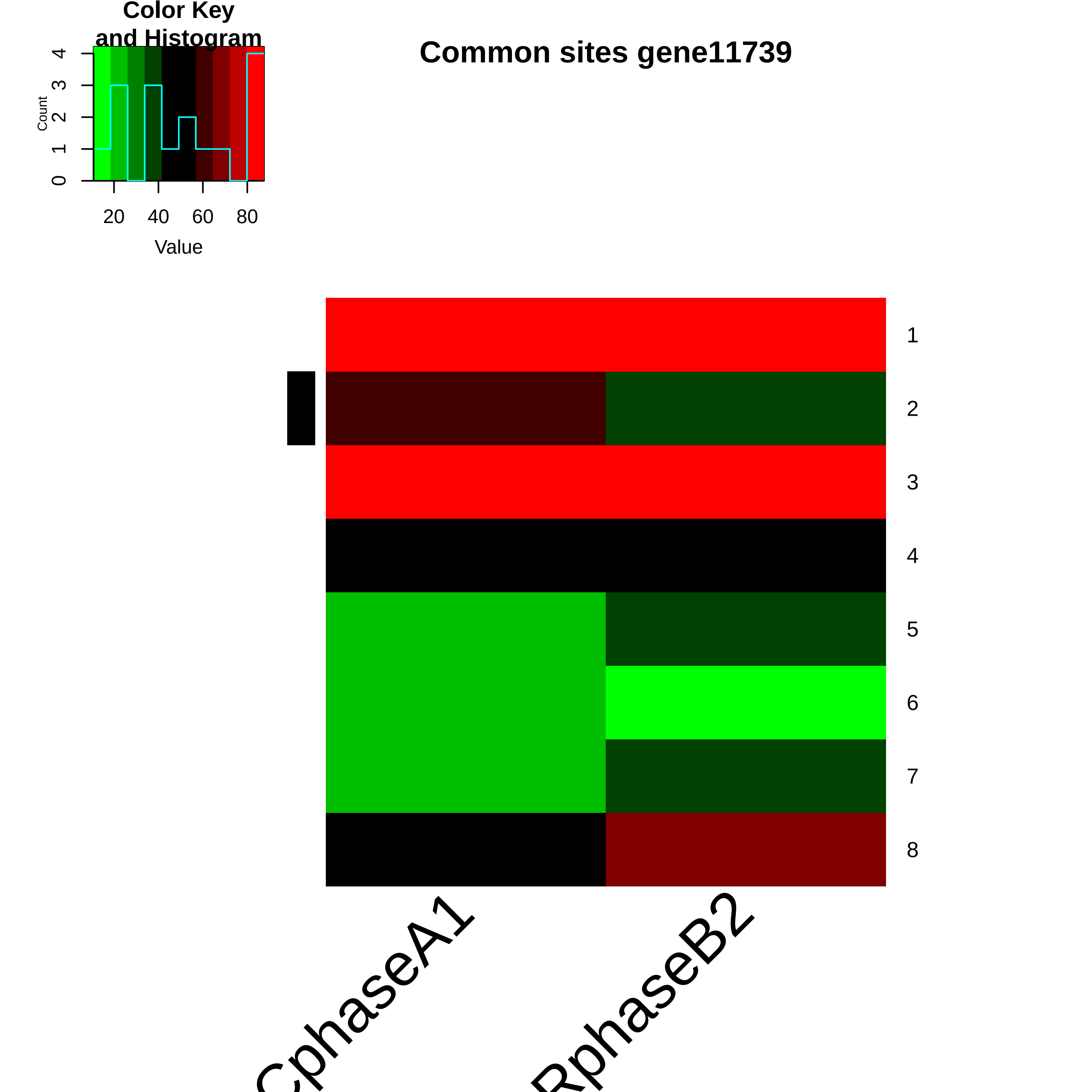

Color Key
and Histogram
Count
0
1
2
3
4
20
40
60
80
Value
Common sites gene11739
1
2
3
4
5
6
7
8
CphaseA1
RphaseB2

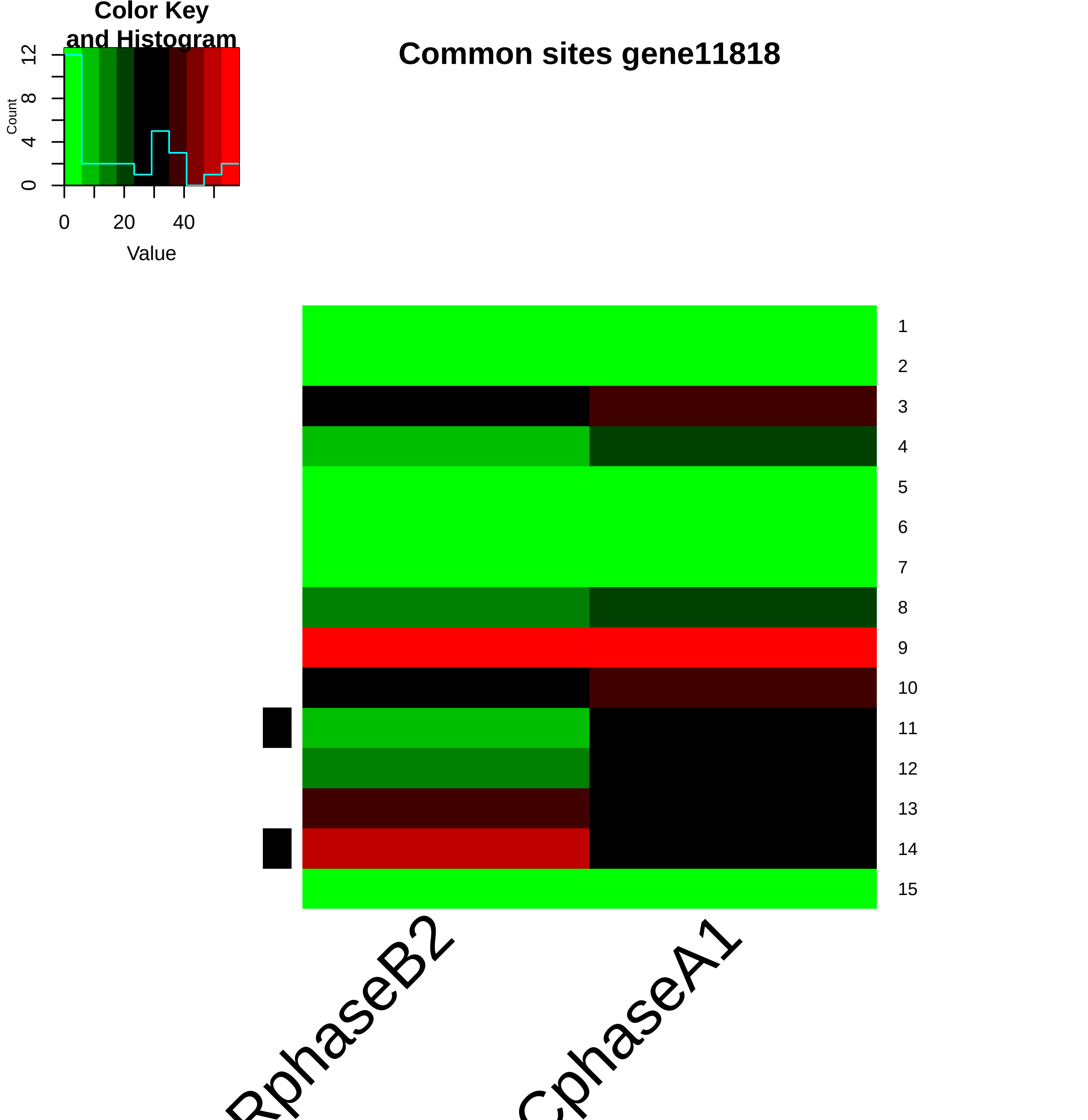
Color Key
and Histogram
Count
0
4
8
12
0
20
40
Value
Common sites gene11818
RphaseB2
CphaseA1

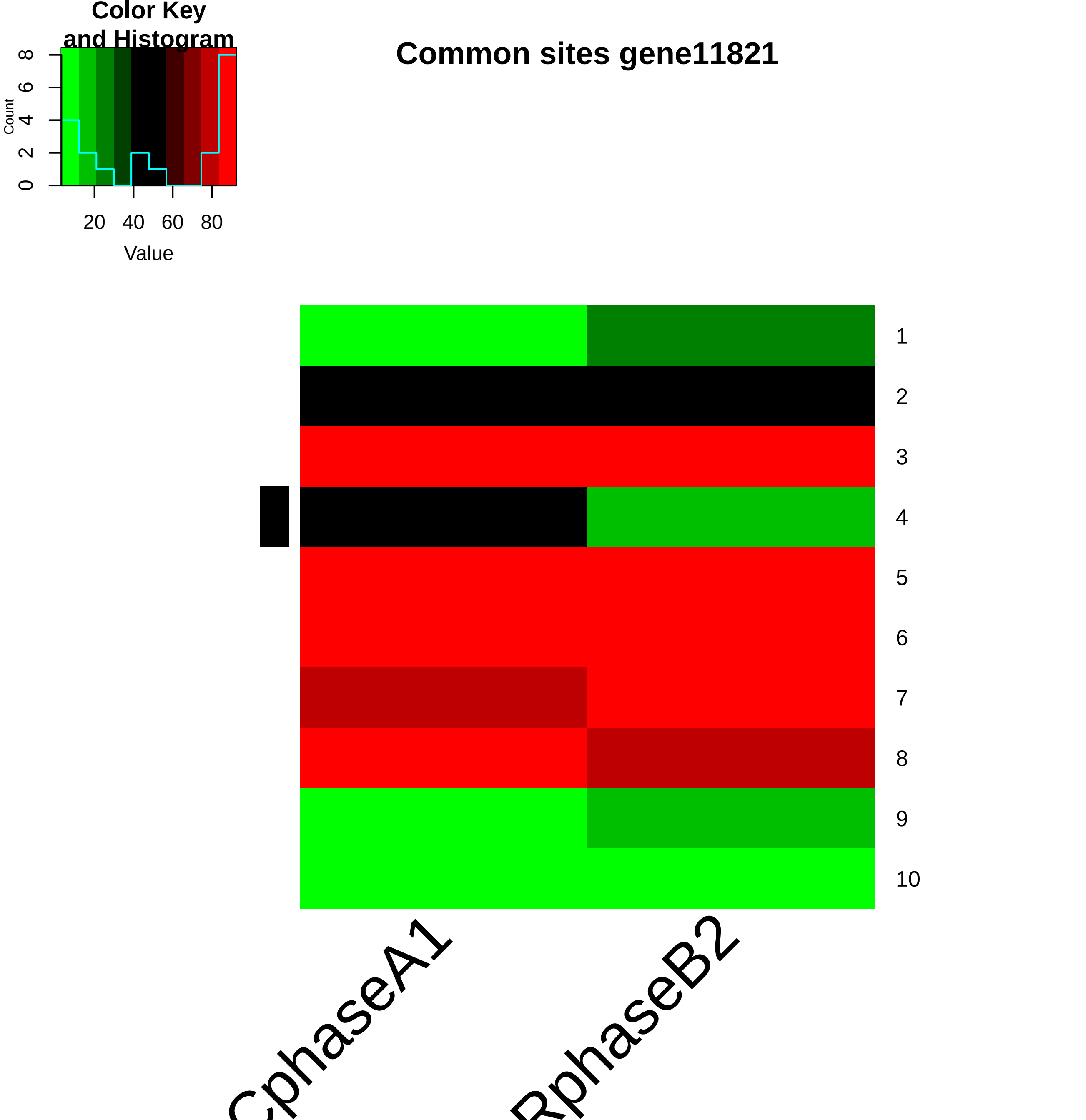
Color Key
and Histogram
Count
0 2 4 6 8
20 40 60 80
Value
Common sites gene11821
1
2
3
4
5
6
7
8
9
10
CphaseA1
RphaseB2

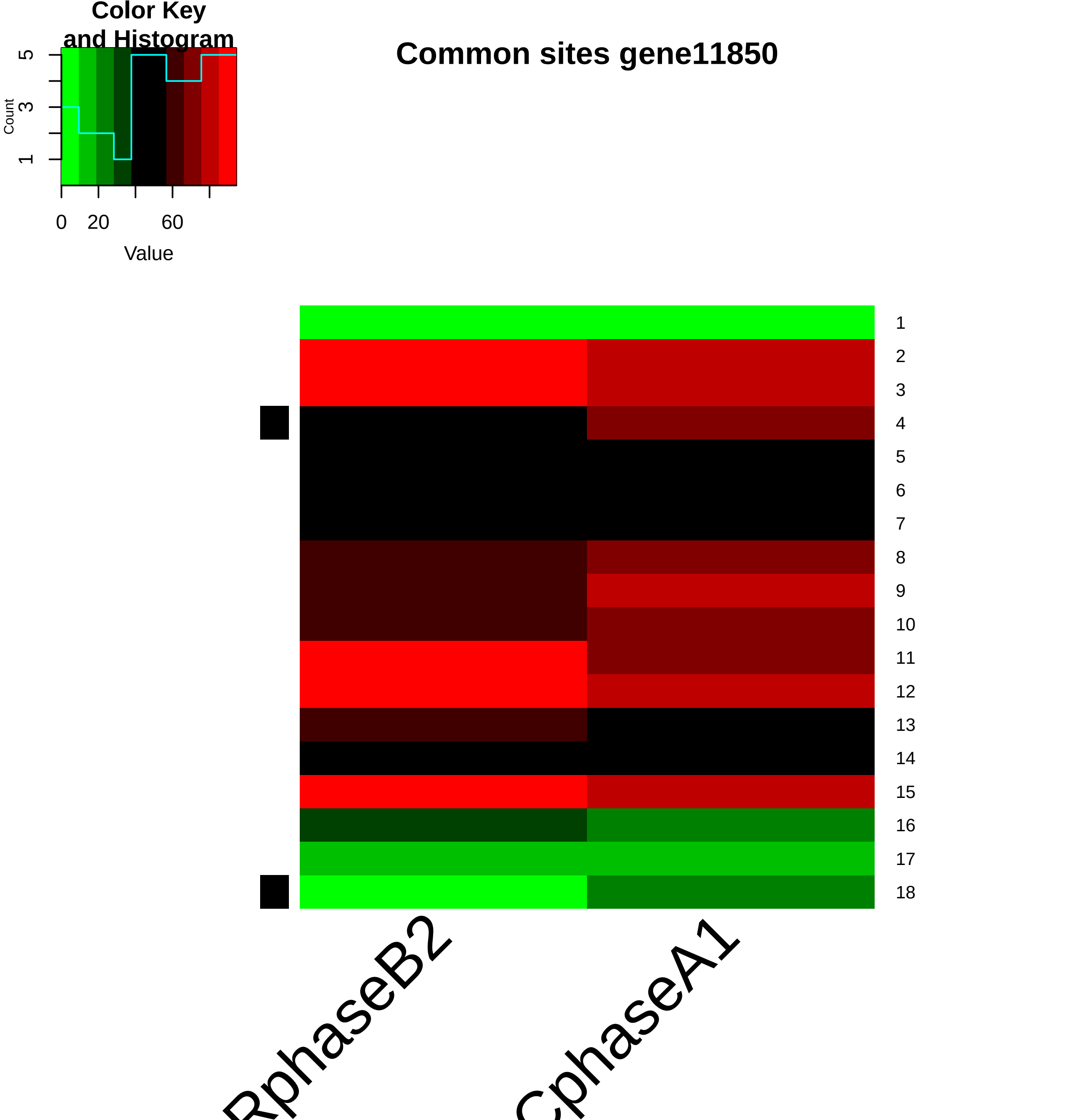
Color Key
and Histogram
Count
1
3
5
0
20
60
Value
Common sites gene11850
RphaseB2
CphaseA1

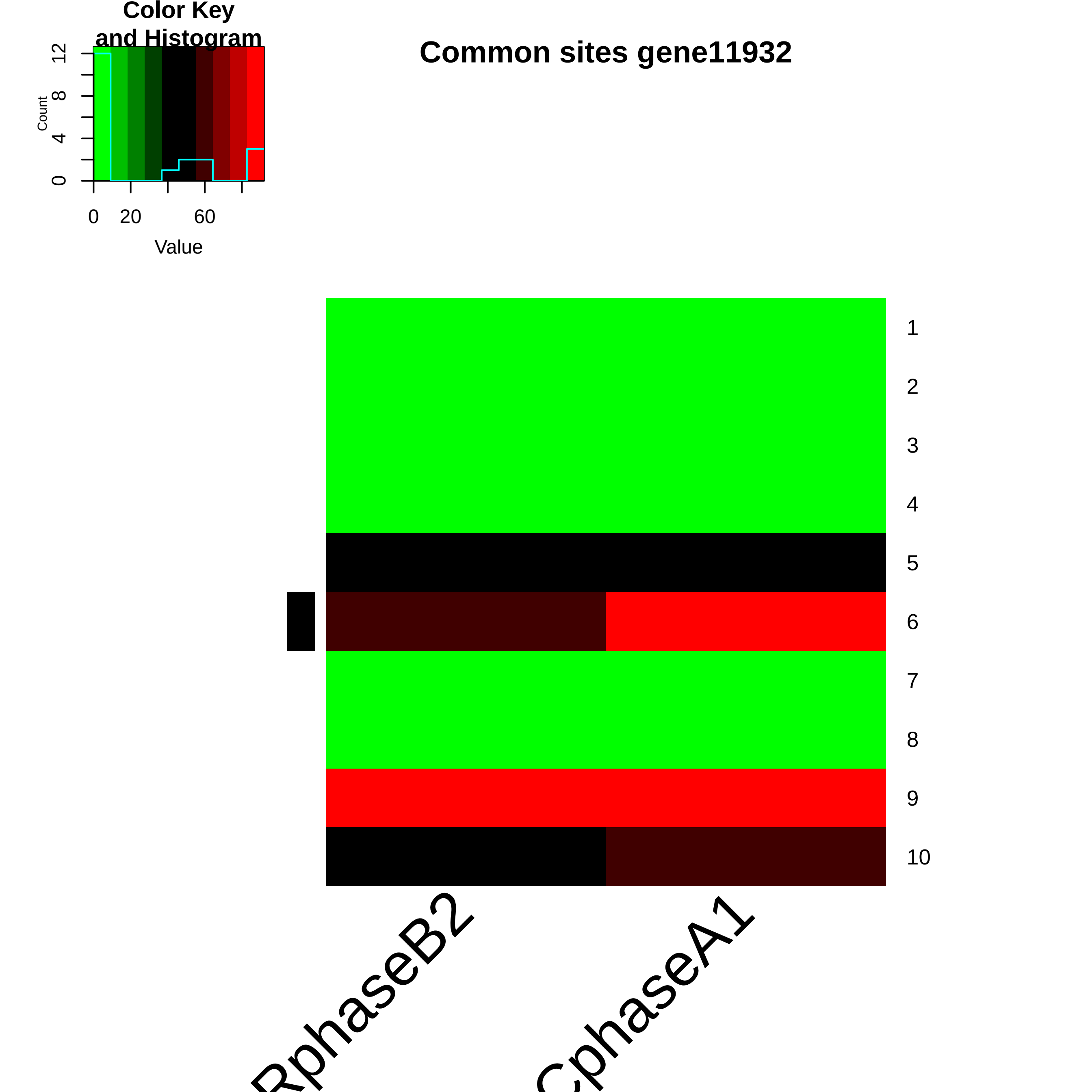
Color Key
and Histogram
Count
0
4
8
12
0
20
60
Value
Common sites gene11932
1
2
3
4
5
6
7
8
9
10
RphaseB2
CphaseA1

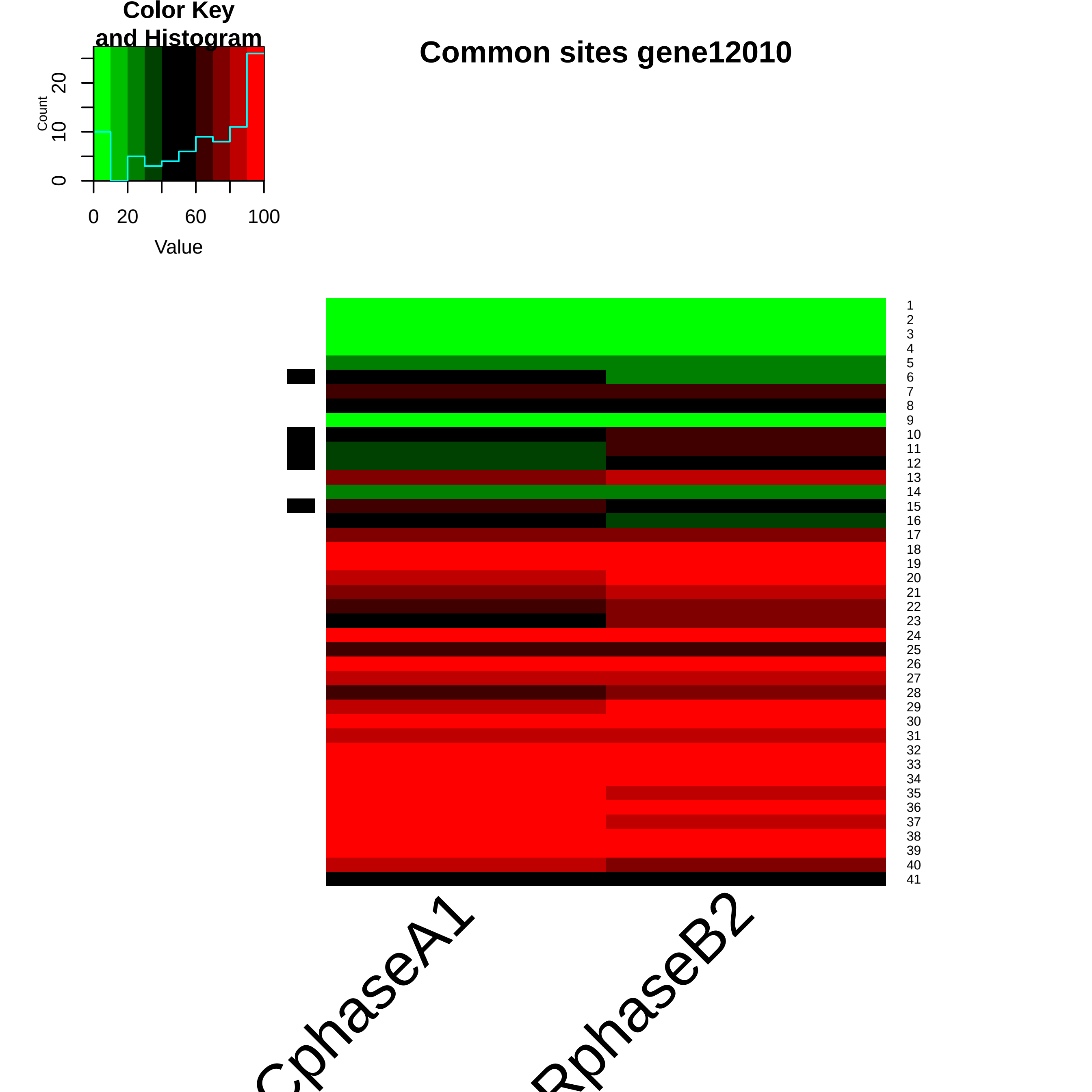
Color Key
and Histogram
Count
0
10
20
0
20
60
100
Value
Common sites gene12010
CphaseA1
RphaseB2

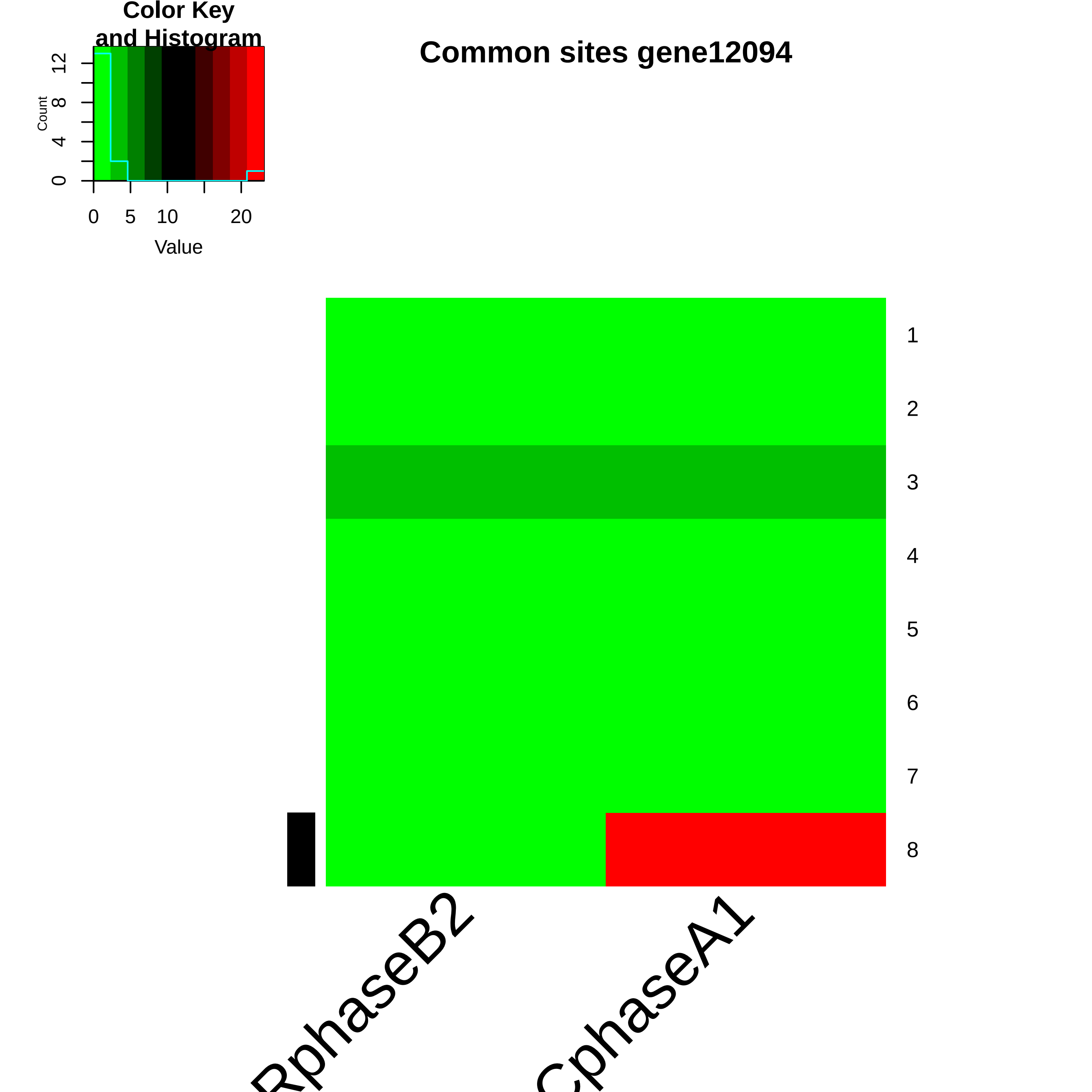

Color Key
and Histogram
Count
0
4
8
12
0
5
10
20
Value
Common sites gene12094
1
2
3
4
5
6
7
8
RphaseB2
CphaseA1

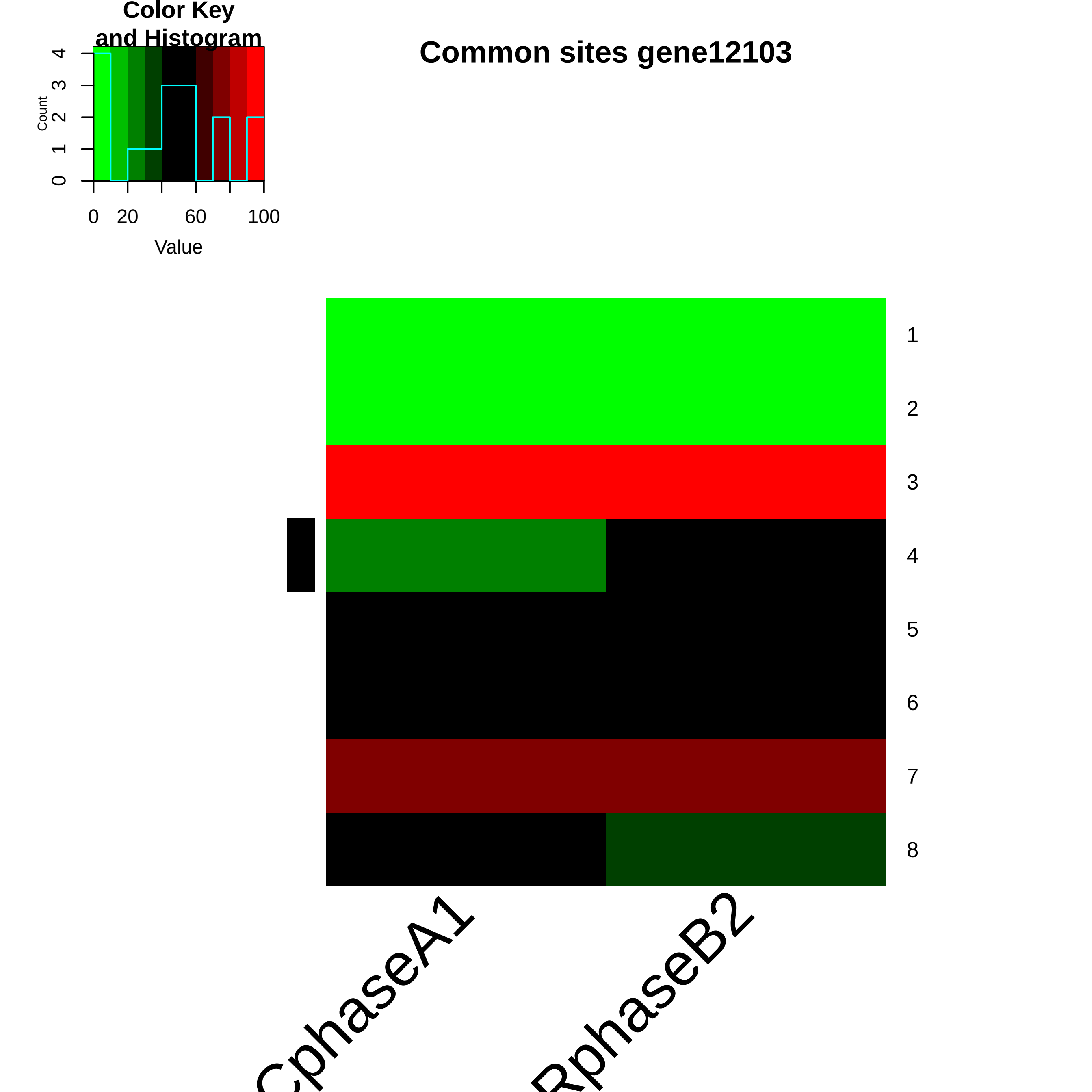

Color Key
and Histogram
Count
0
1
2
3
4
0
20
60
100
Value
Common sites gene12103
1
2
3
4
5
6
7
8
CphaseA1
RphaseB2

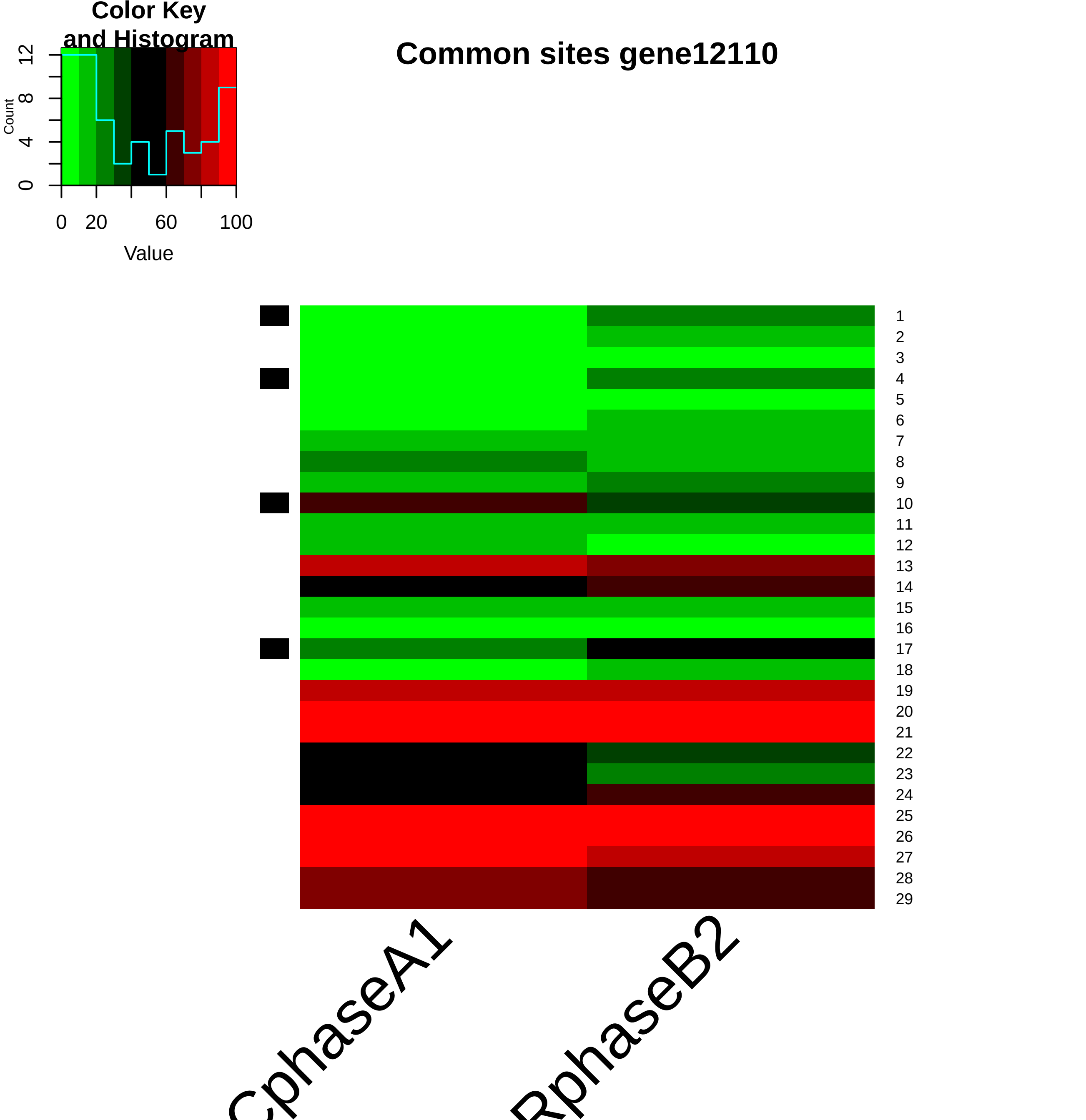
Color Key
and Histogram
Count
0
4
8
12
0
20
60
100
Value
Common sites gene12110
CphaseA1
RphaseB2

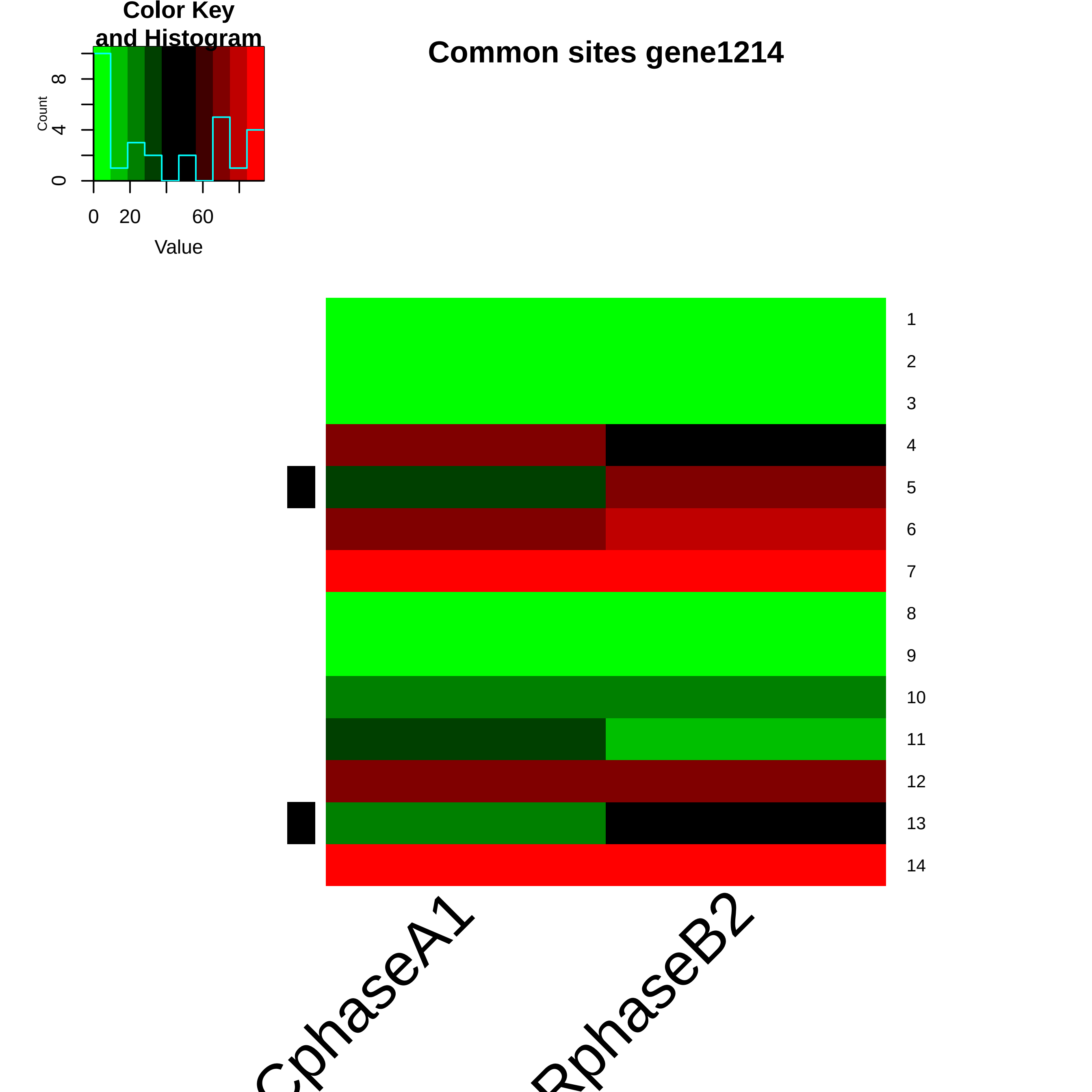
Color Key
and Histogram
Count
0
4
8
0
20
60
Value
Common sites gene1214
1
2
3
4
5
6
7
8
9
10
11
12
13
14
CphaseA1
RphaseB2

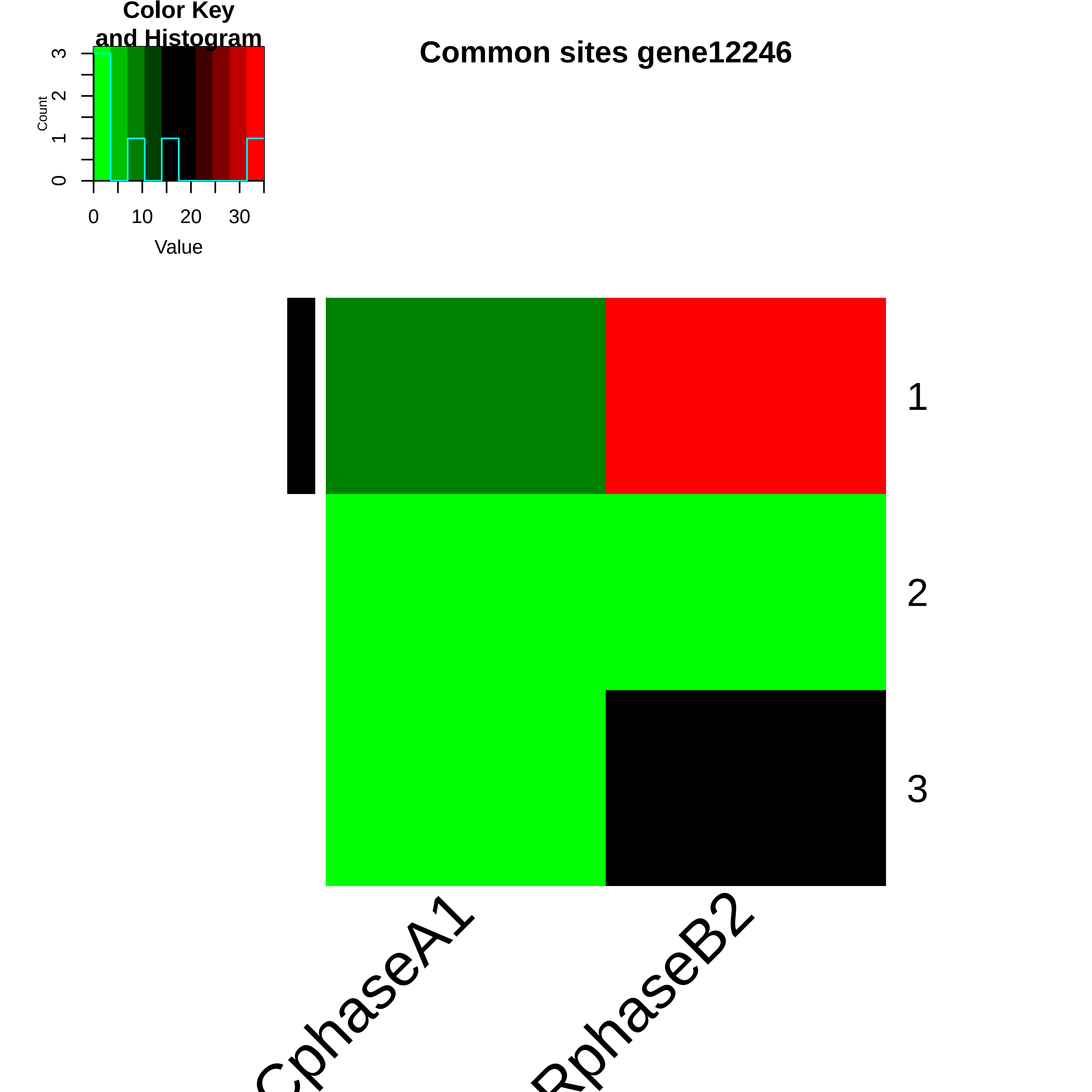
Color Key
and Histogram
Count
0
1
2
3
0
10
20
30
Value
Common sites gene12246
1
2
3
CphaseA1
RphaseB2

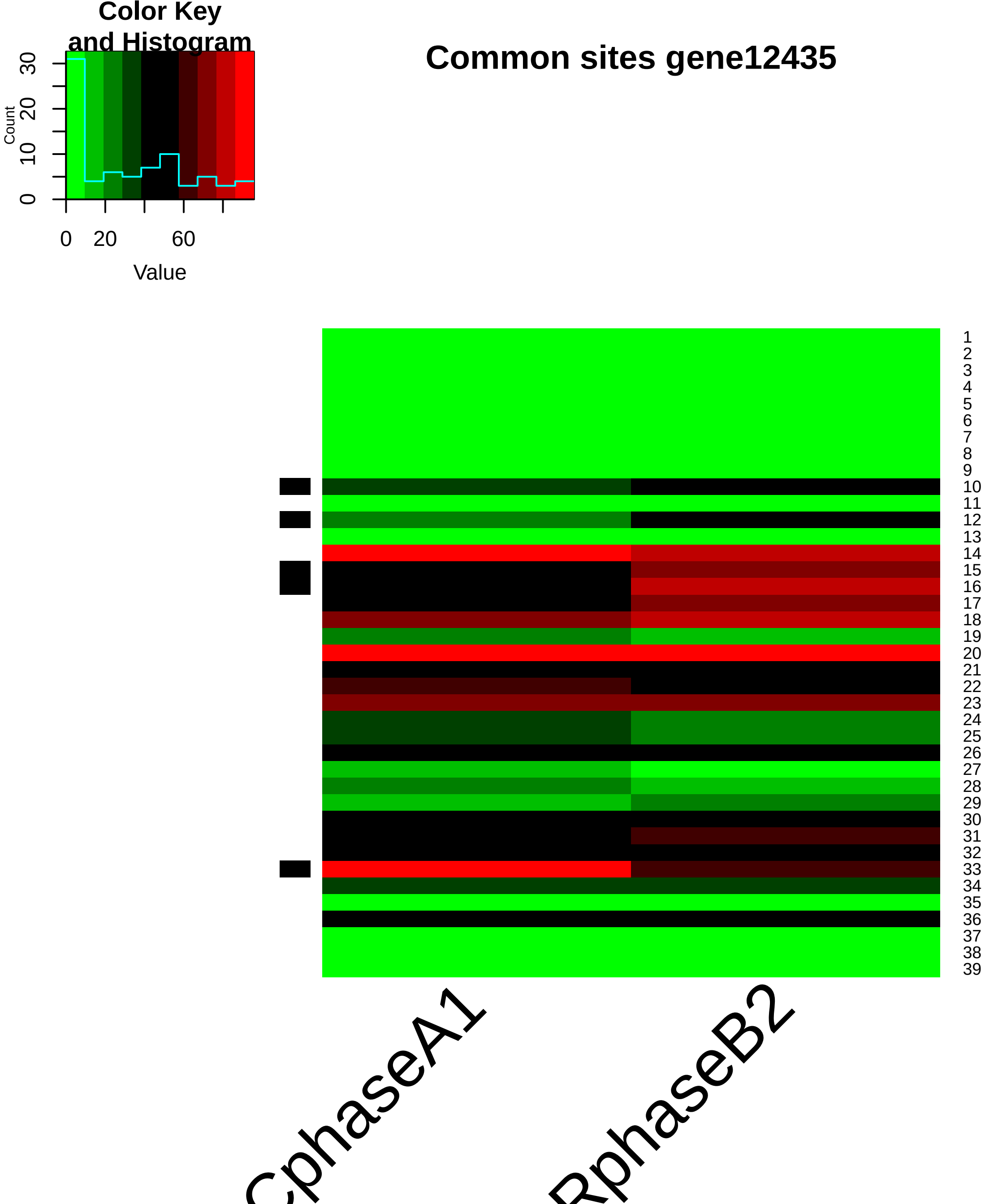
Color Key
and Histogram
Count
0
10
20
30
0
20
60
Value
Common sites gene12435
CphaseA1
RphaseB2

### Common sites gene12538

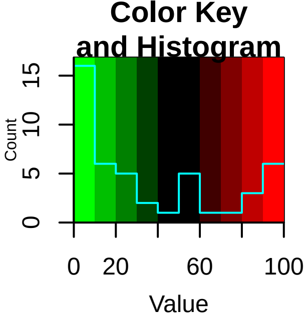

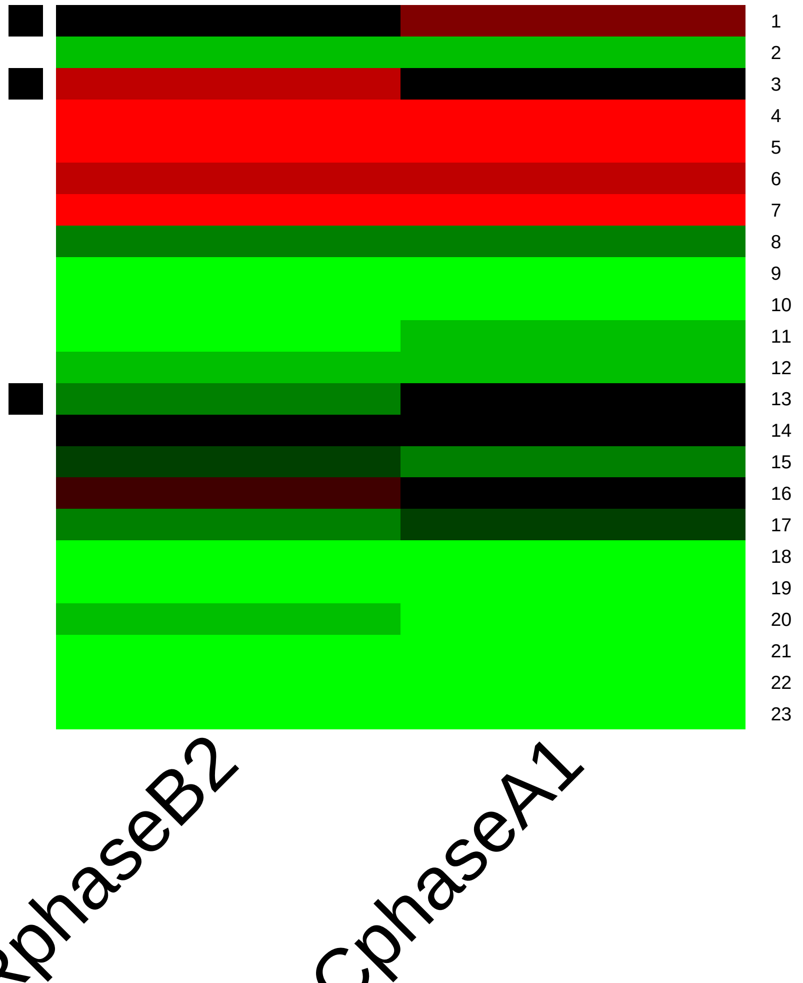

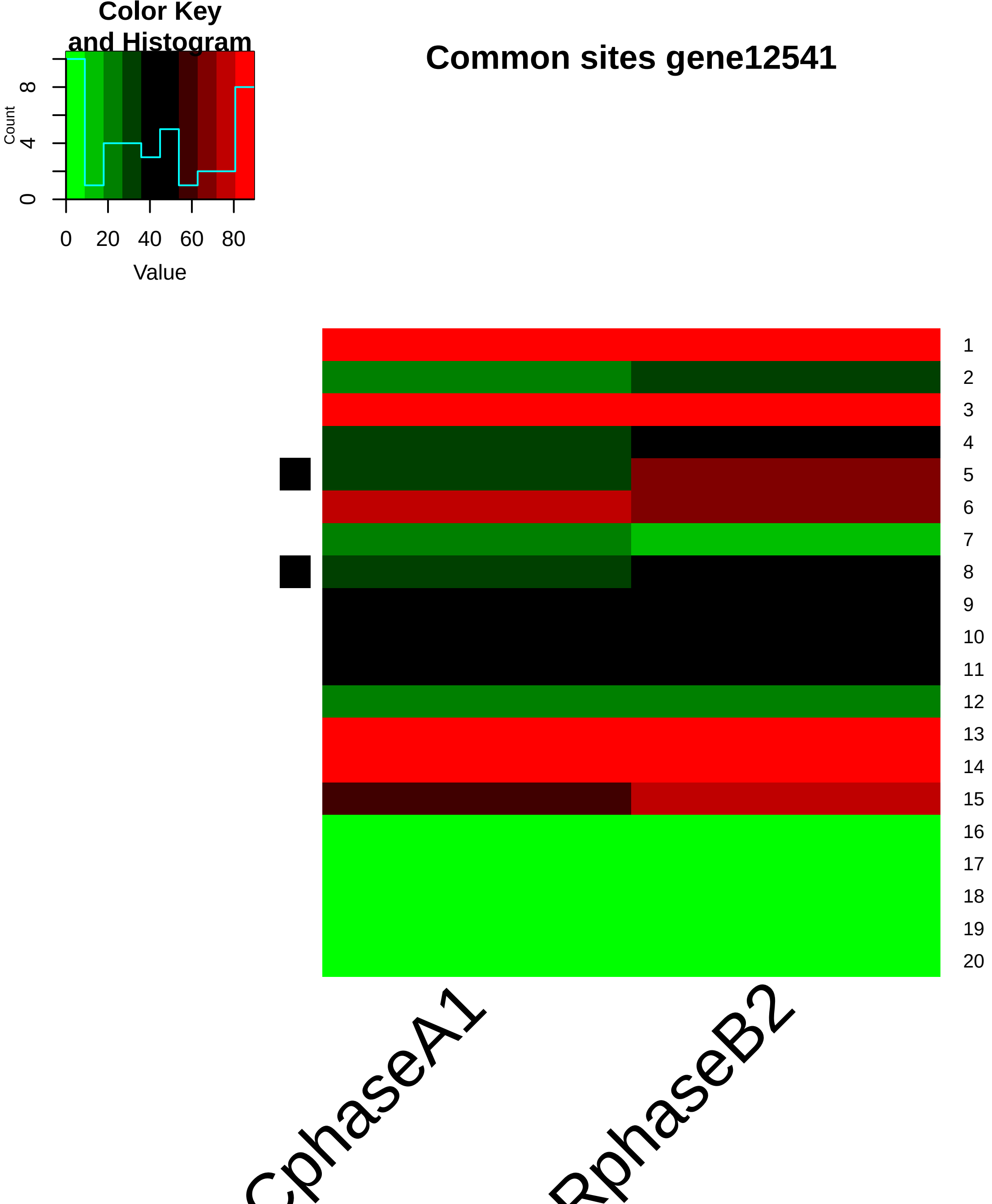
Color Key
and Histogram
Count
0
4
8
0
20
40
60
80
Value
Common sites gene12541
CphaseA1
RphaseB2

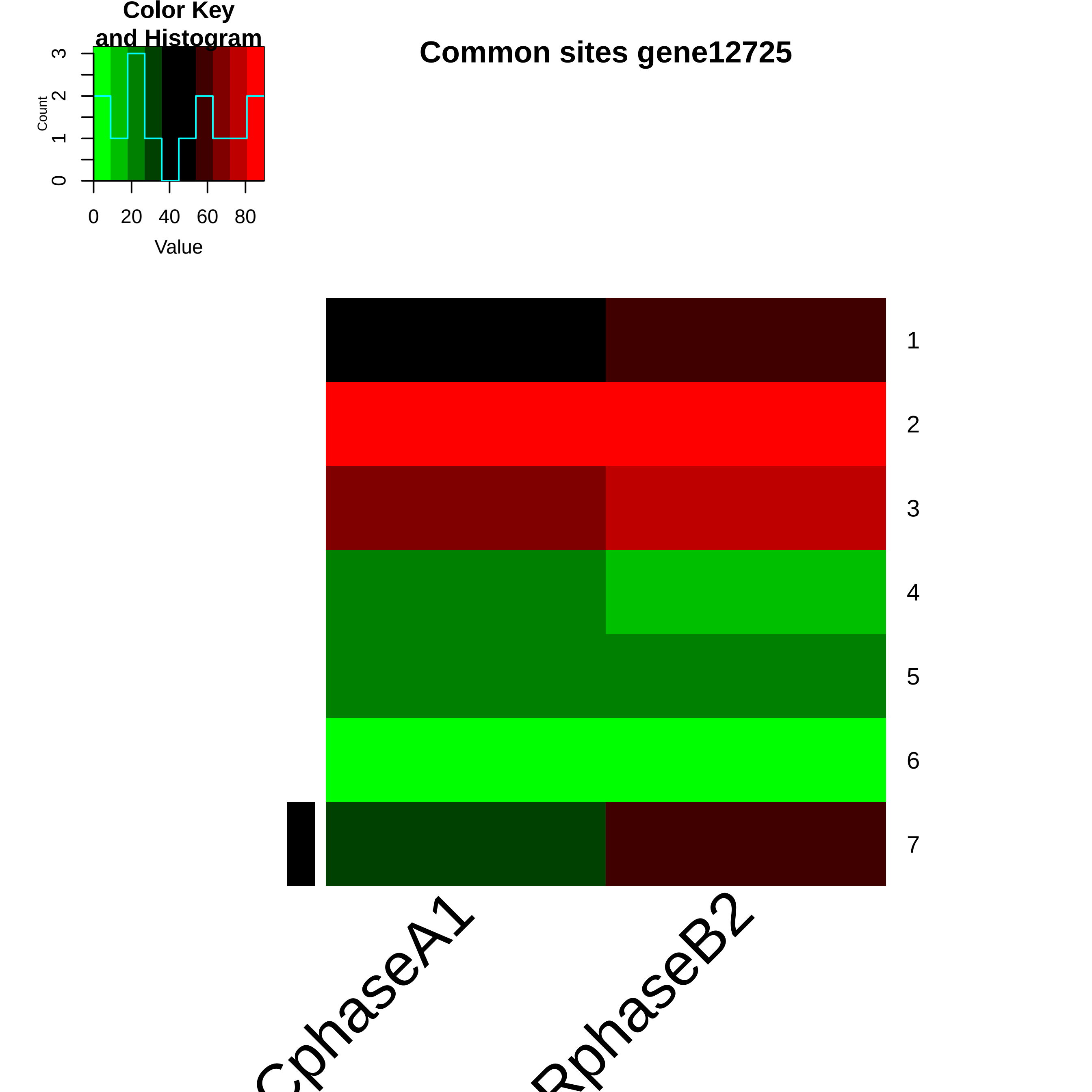
Color Key
and Histogram
Count
0
1
2
3
0
20
40
60
80
Value
Common sites gene12725
1
2
3
4
5
6
7
CphaseA1
RphaseB2

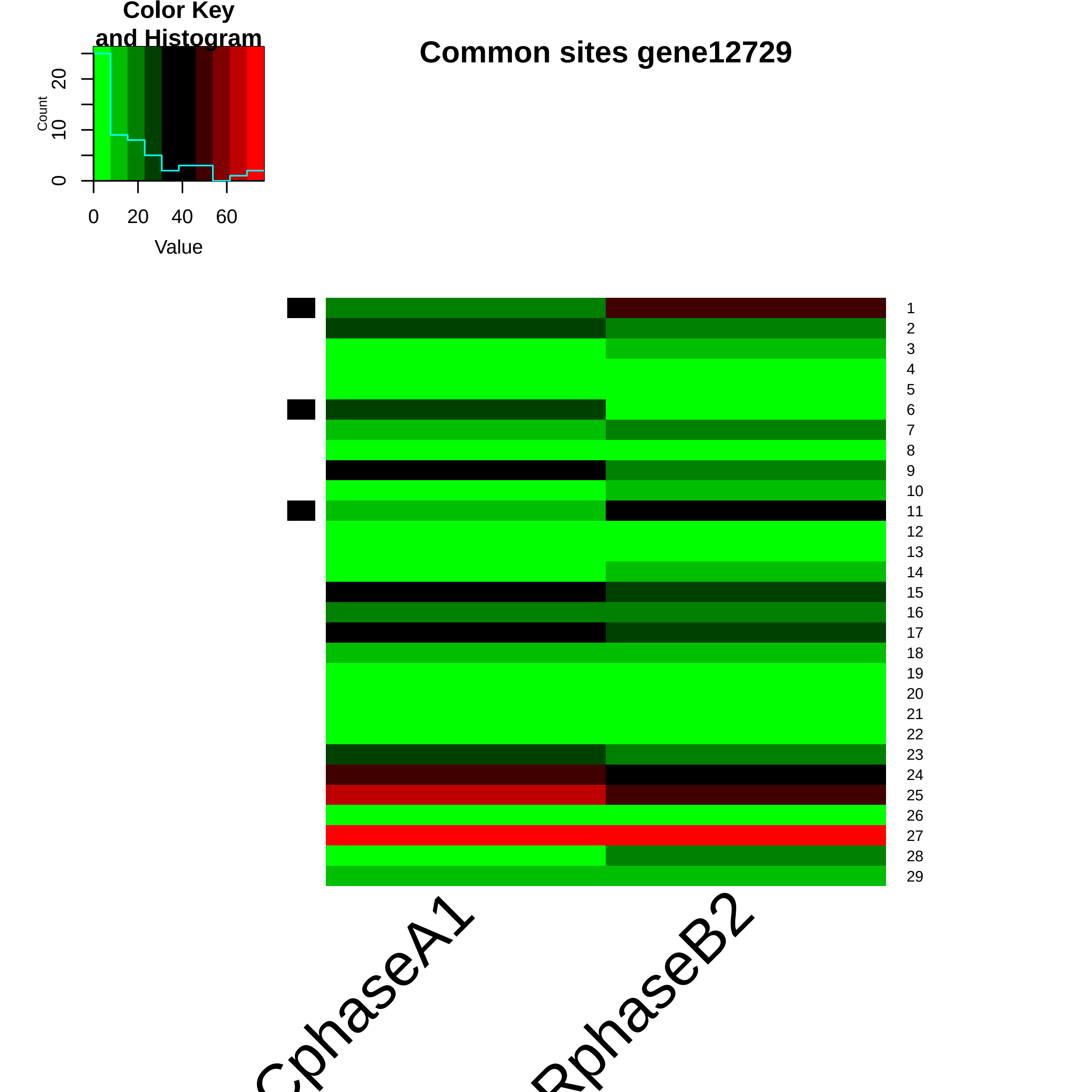

Color Key
and Histogram
Count
0
10
20
0
20
40
60
Value
Common sites gene12729
CphaseA1
RphaseB2

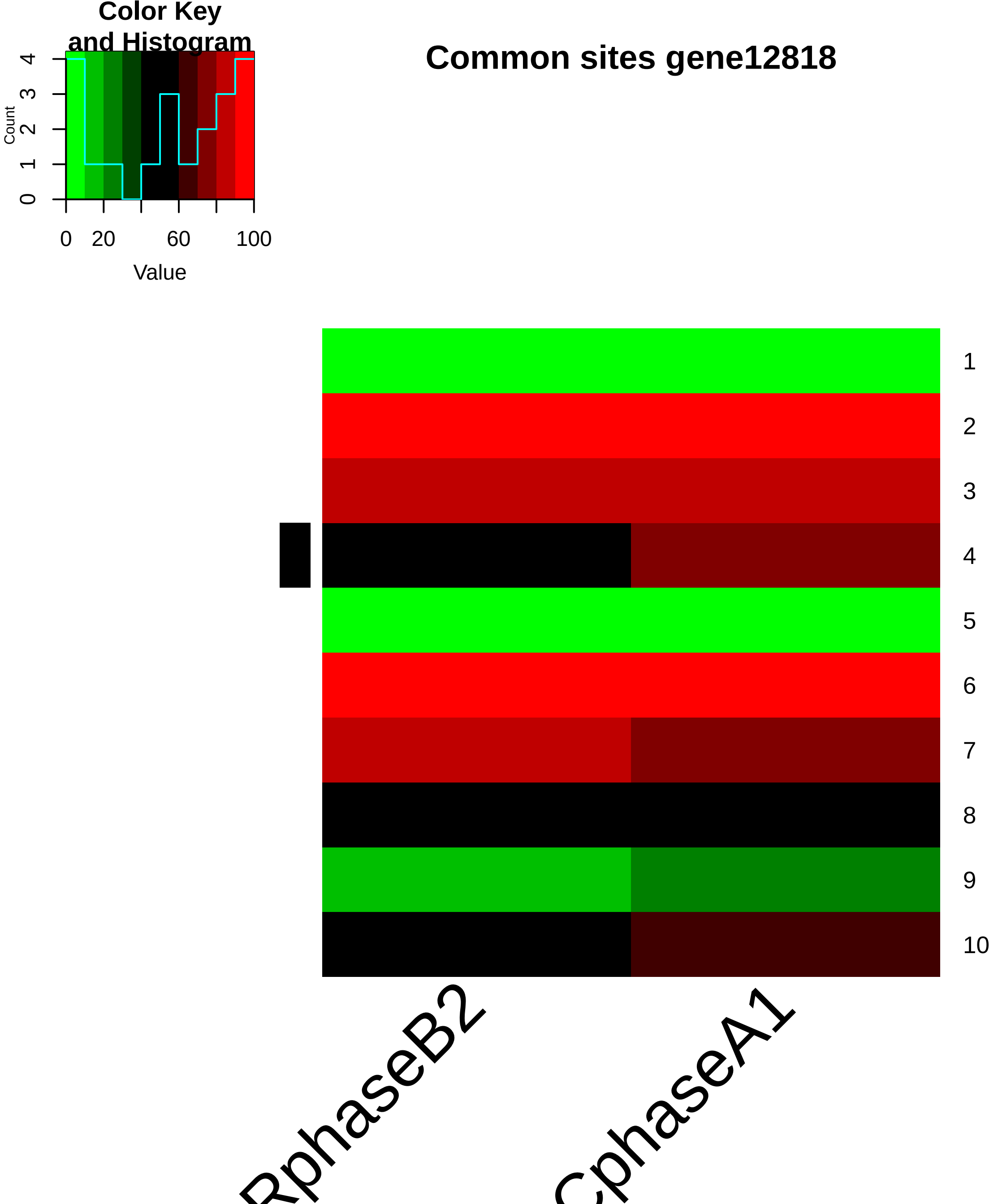
Color Key
and Histogram
Count
0
1
2
3
4
0
20
60
100
Value
Common sites gene12818
1
2
3
4
5
6
7
8
9
10
RphaseB2
CphaseA1

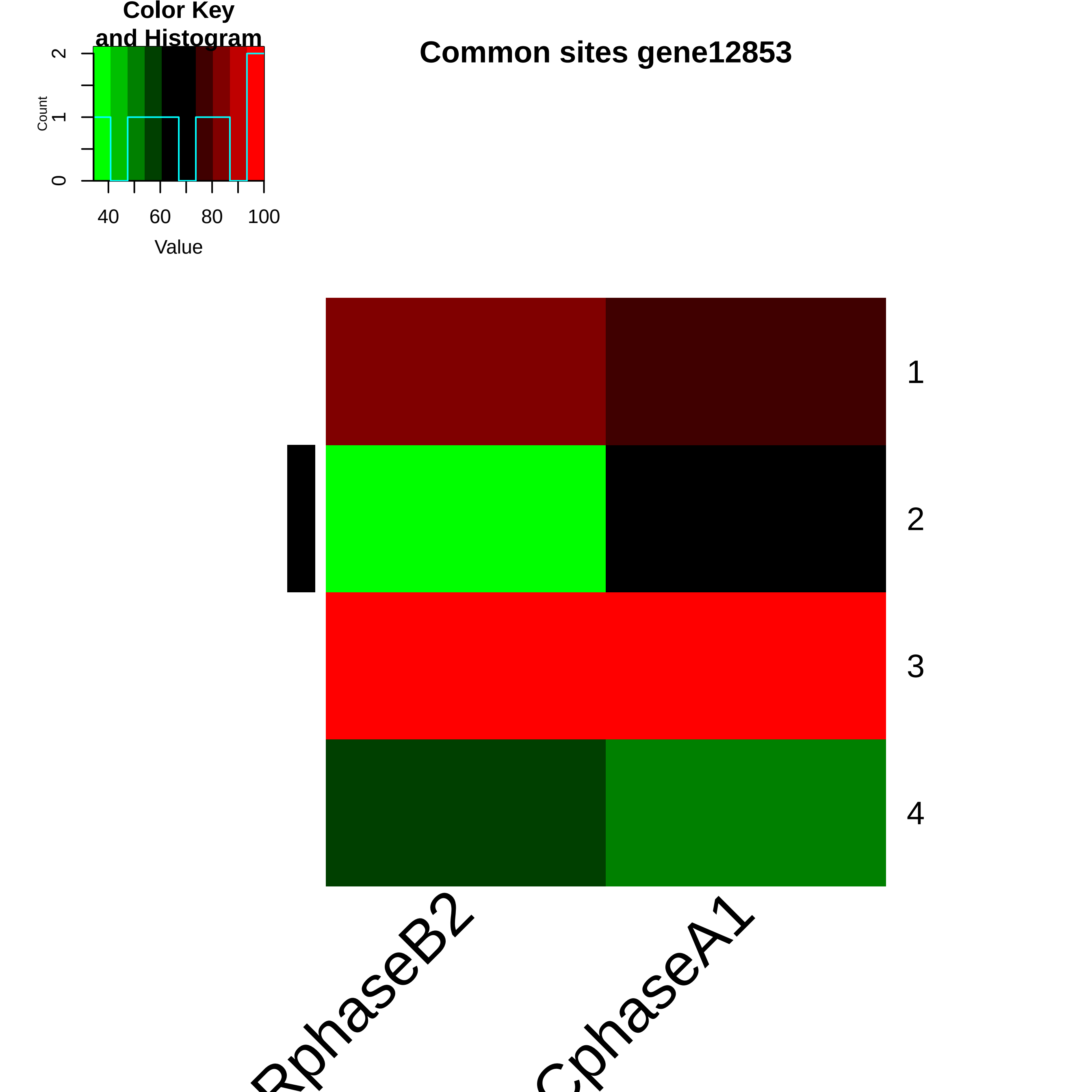
Color Key
and Histogram
Count
0
1
2
40
60
80
100
Value
Common sites gene12853
1
2
3
4
RphaseB2
CphaseA1

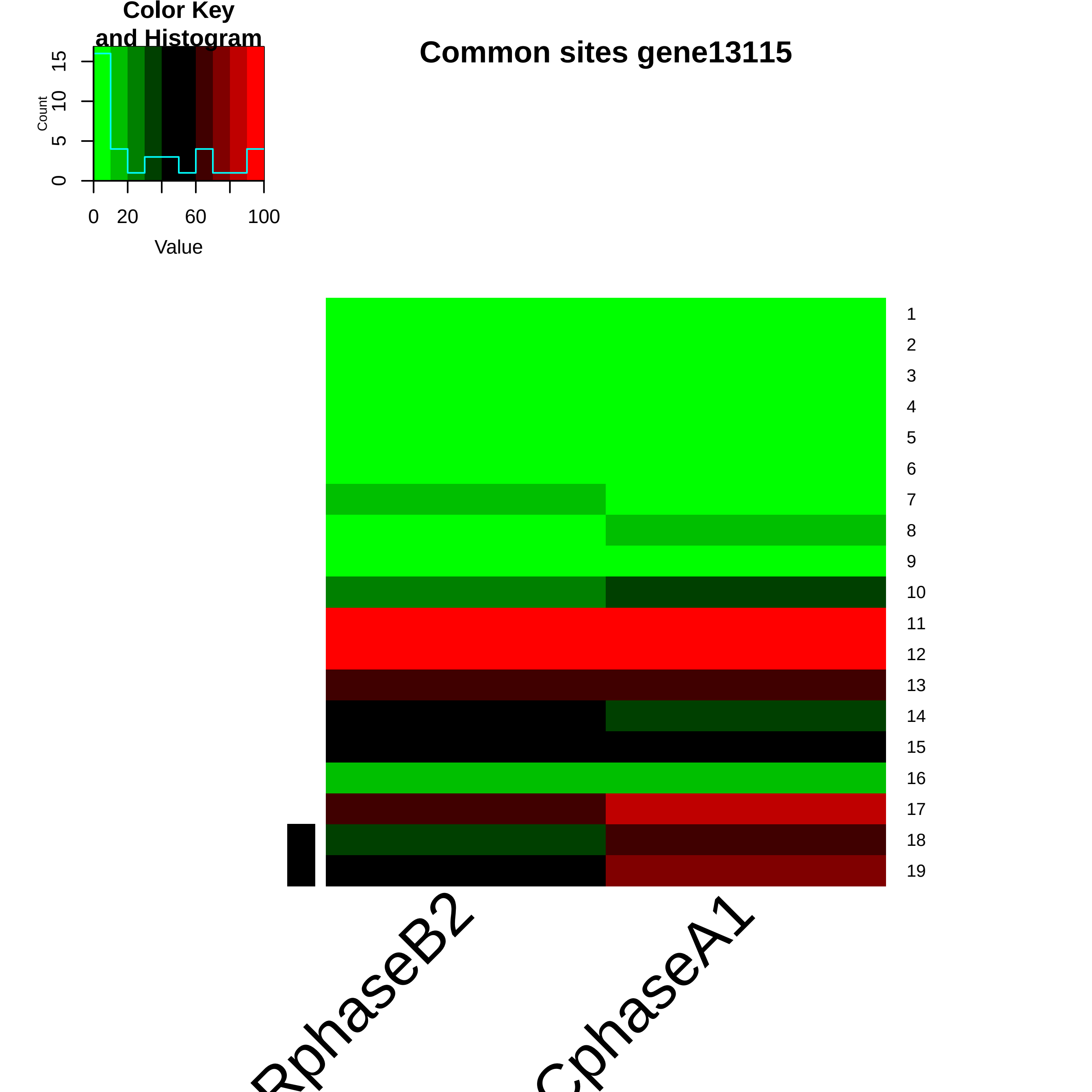
Color Key
and Histogram
Count
0
5
10
15
0
20
60
100
Value
Common sites gene13115
RphaseB2
CphaseA1

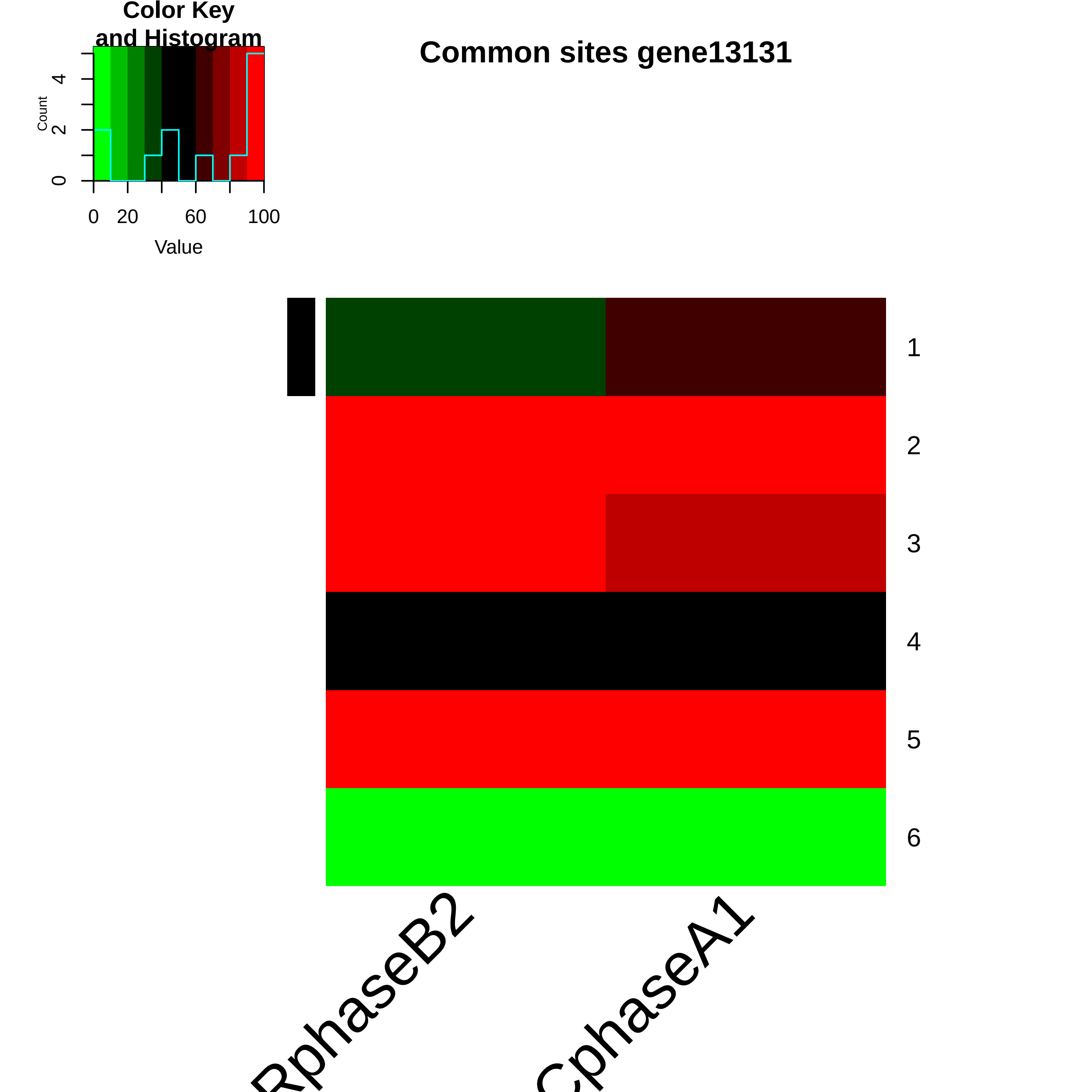

Color Key and Histogram
Count
0
2
4
0
20
60
100
Value
Common sites gene13131
1
2
3
4
5
6
RphaseB2
CphaseA1

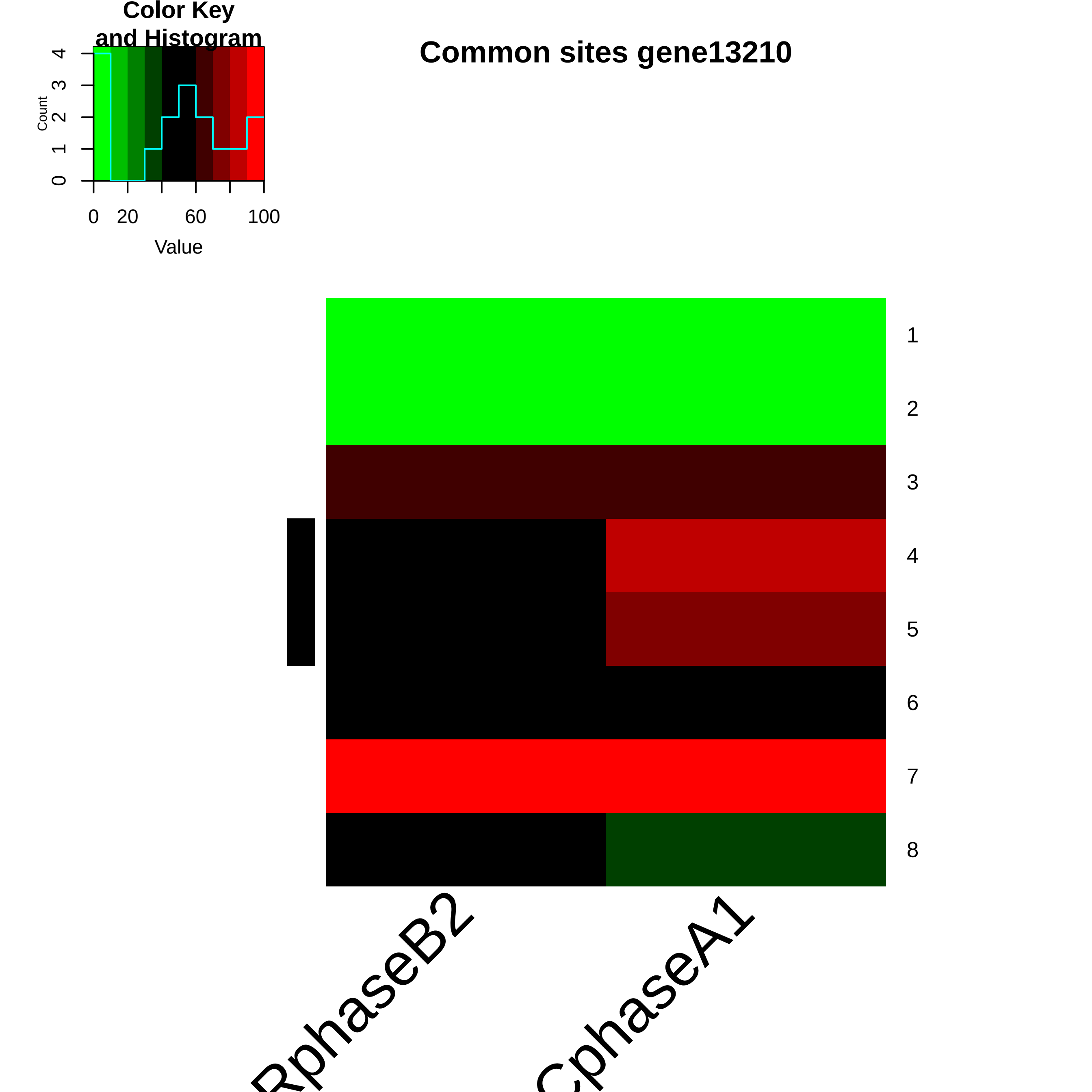

Color Key
and Histogram
Count
0
1
2
3
4
0
20
60
100
Value
Common sites gene13210
1
2
3
4
5
6
7
8
RphaseB2
CphaseA1

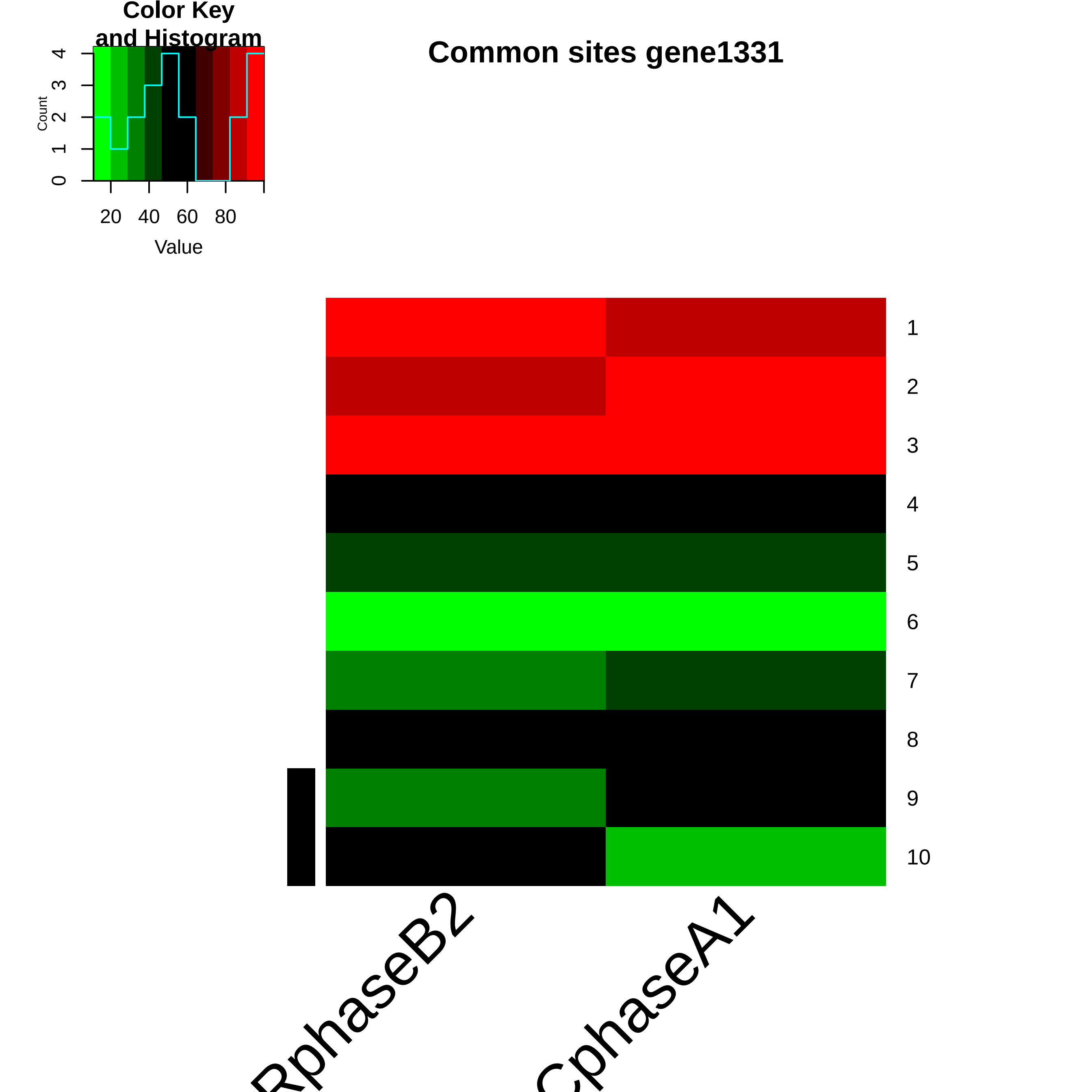

Color Key
and Histogram
Count
0
1
2
3
4
20
40
60
80
Value
Common sites gene1331
1
2
3
4
5
6
7
8
9
10
RphaseB2
CphaseA1

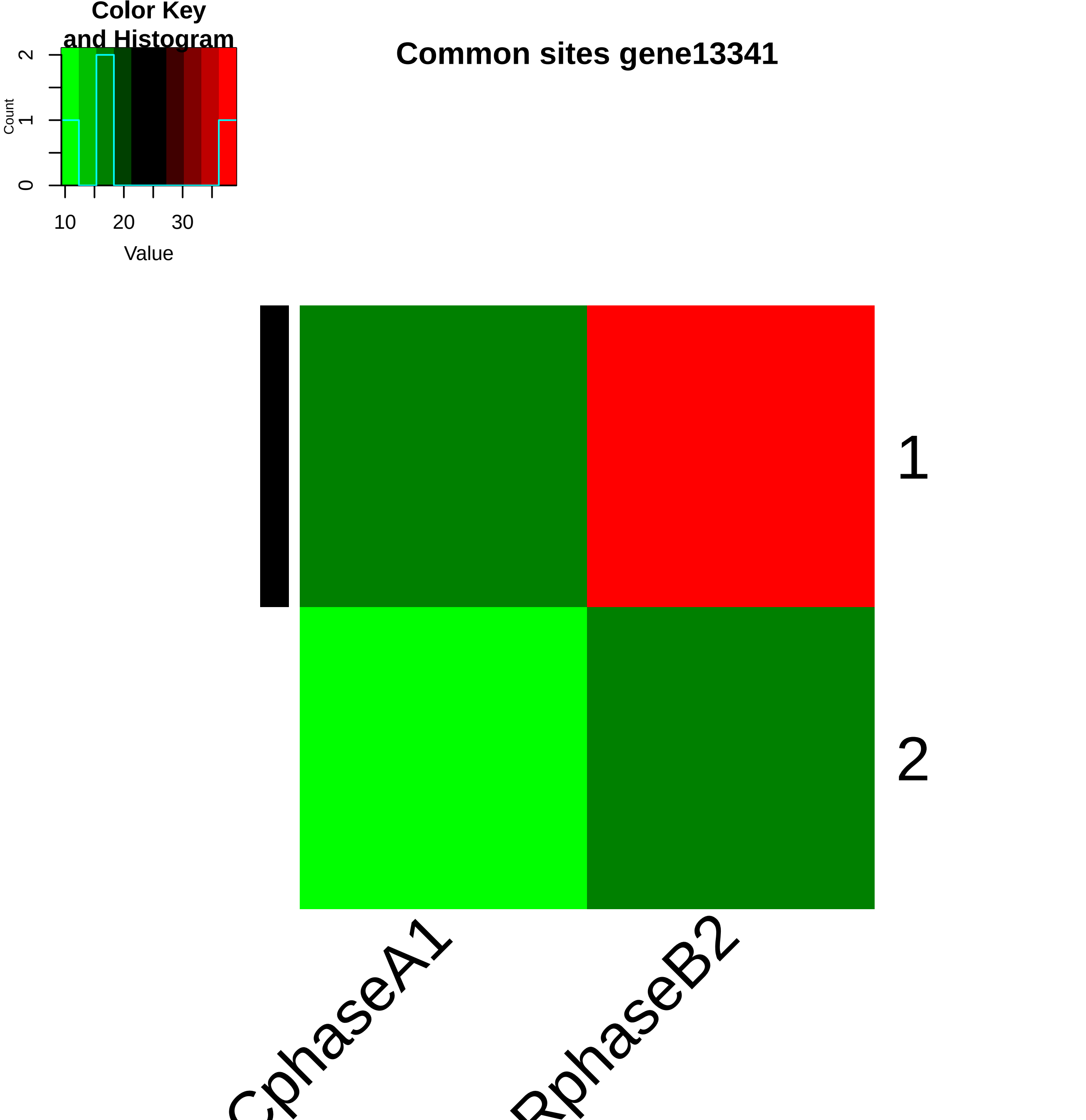
Color Key
and Histogram
Count
0
1
2
10
20
30
Value
Common sites gene13341
1
2
CphaseA1
RphaseB2

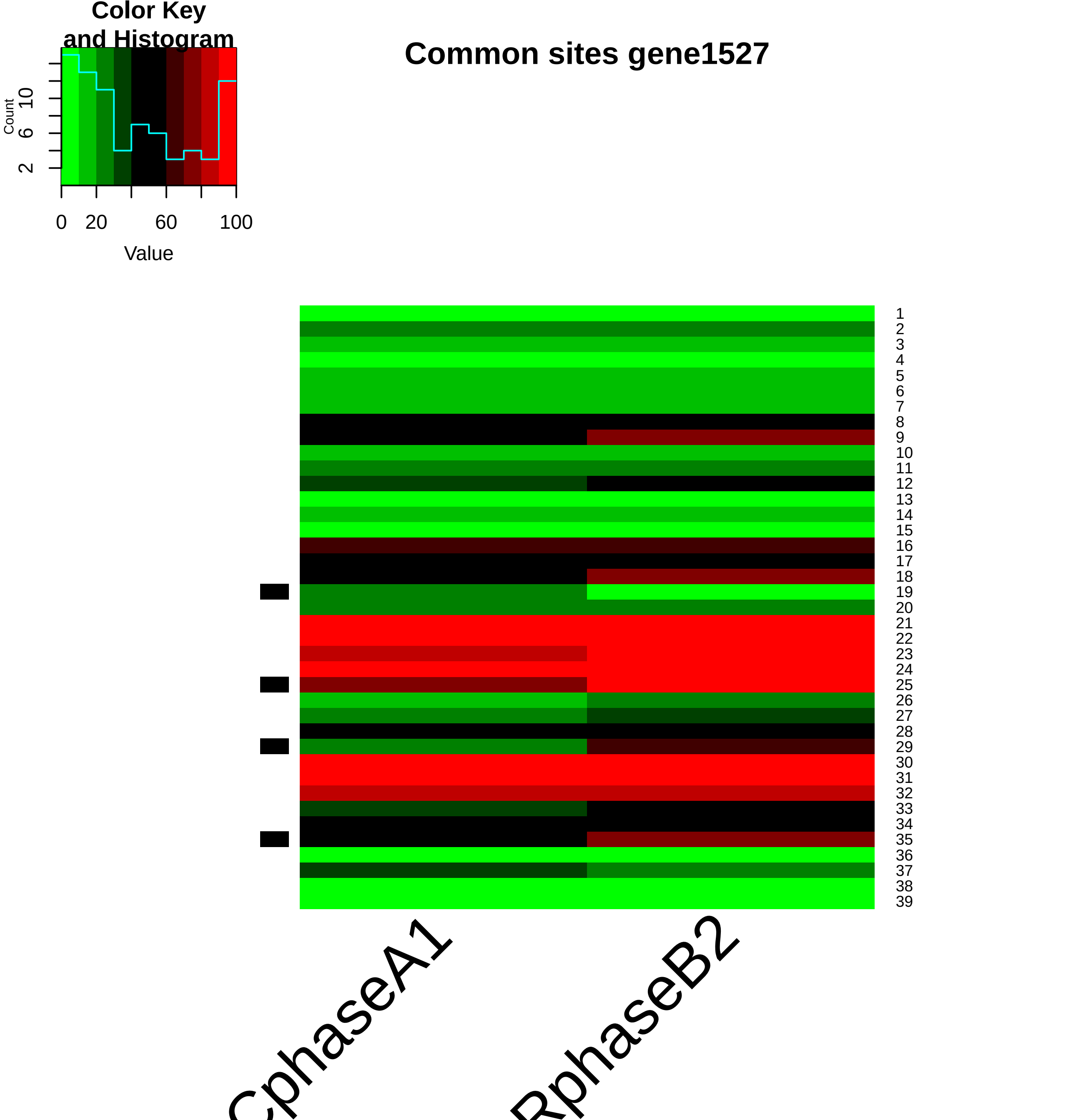
Color Key
and Histogram
Count
2
6
10
0
20
60
100
Value
Common sites gene1527
CphaseA1
RphaseB2

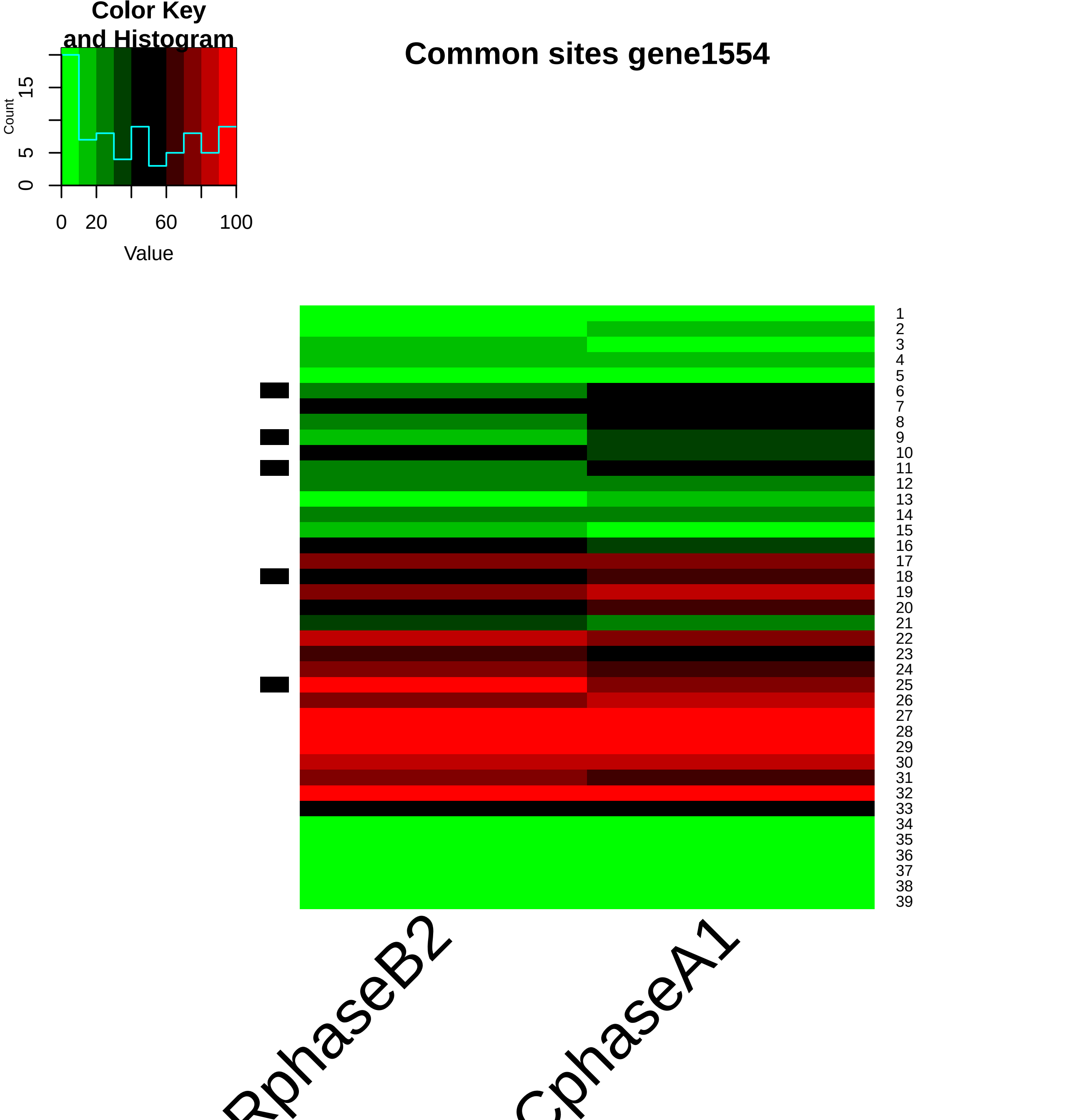
Color Key
and Histogram
Count
0
5
15
0
20
60
100
Value
Common sites gene1554
RphaseB2
CphaseA1

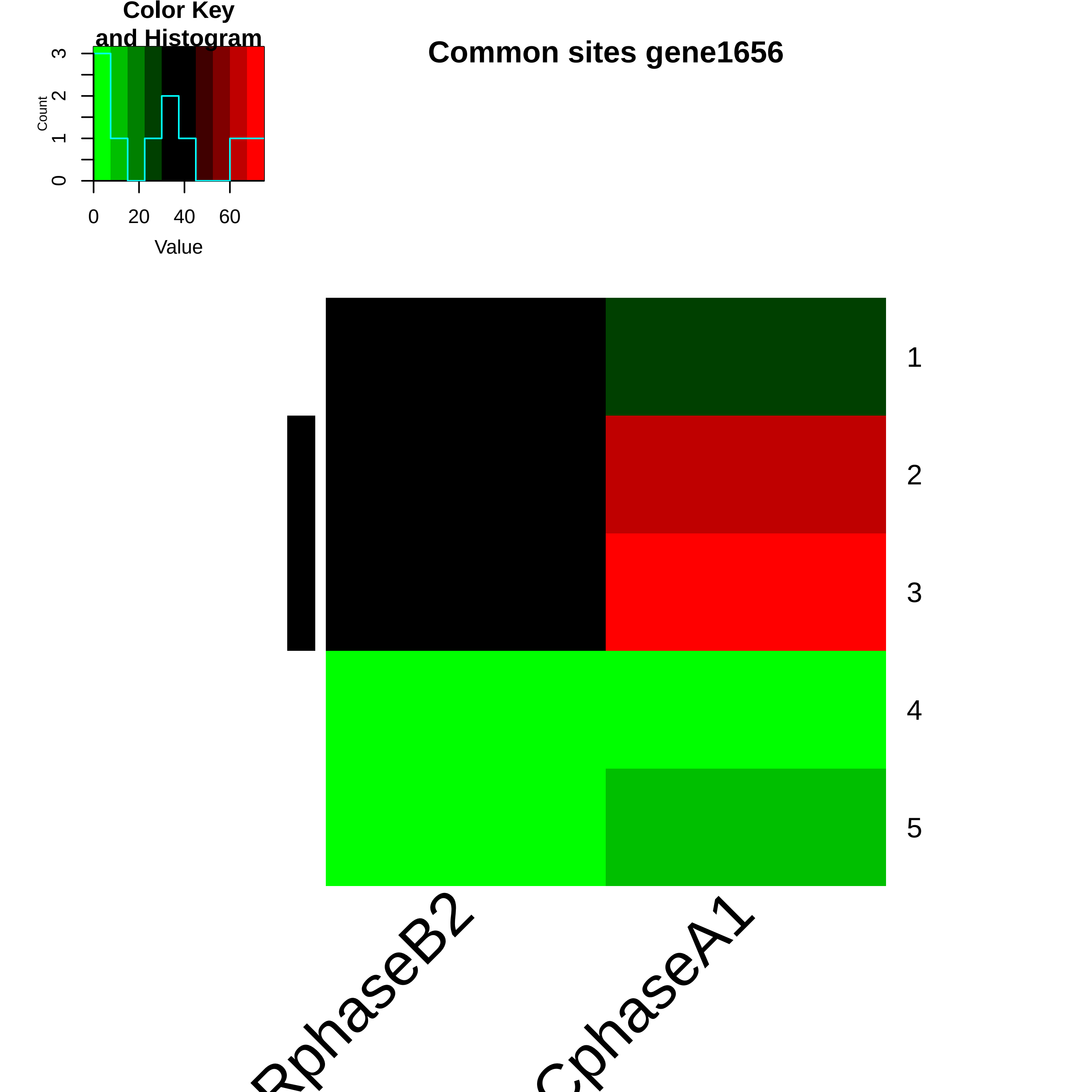

Color Key
and Histogram
Count
0
1
2
3
0
20
40
60
Value
Common sites gene1656
1
2
3
4
5
RphaseB2
CphaseA1

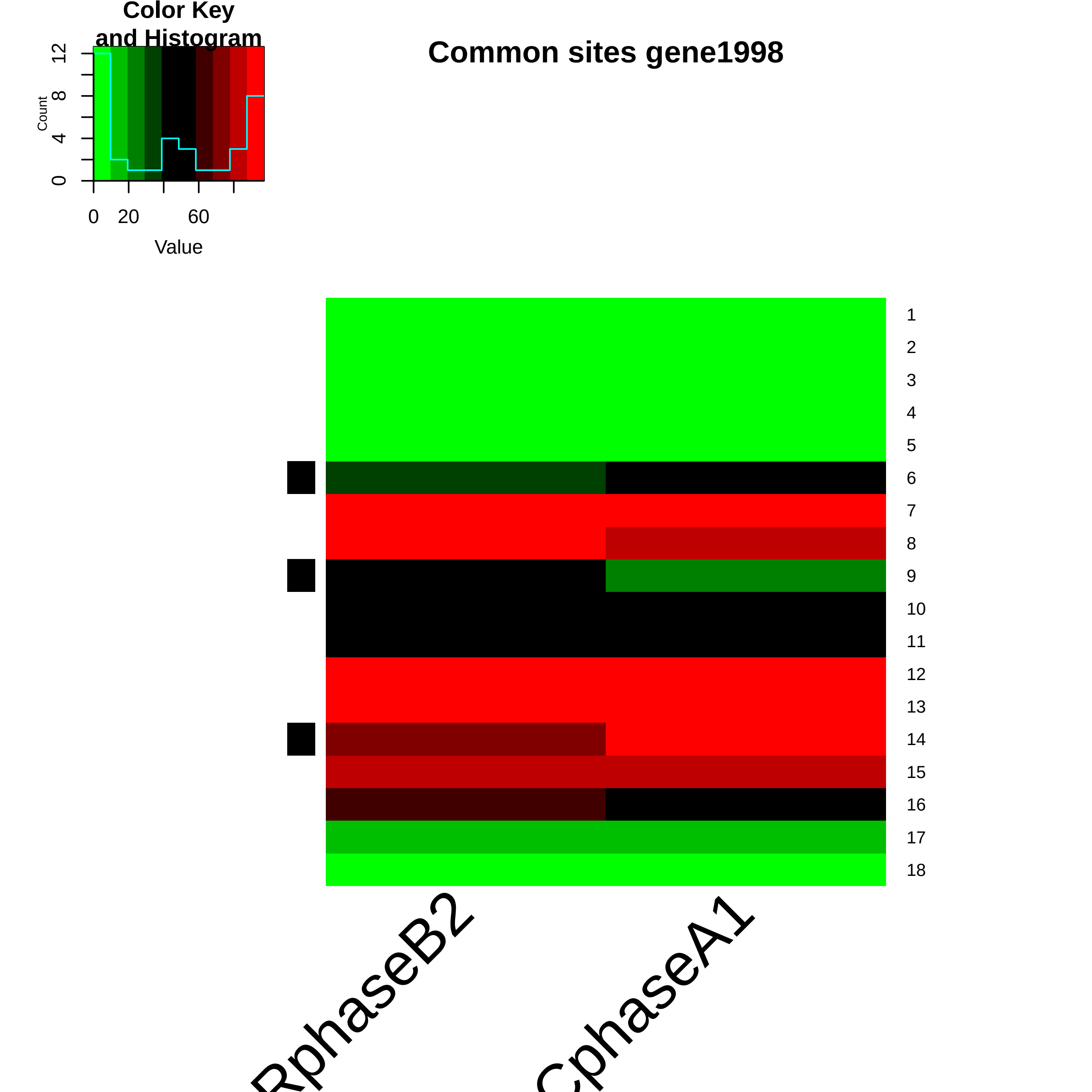
Color Key
and Histogram
Count
0
4
8
12
0
20
60
Value
Common sites gene1998
RphaseB2
CphaseA1

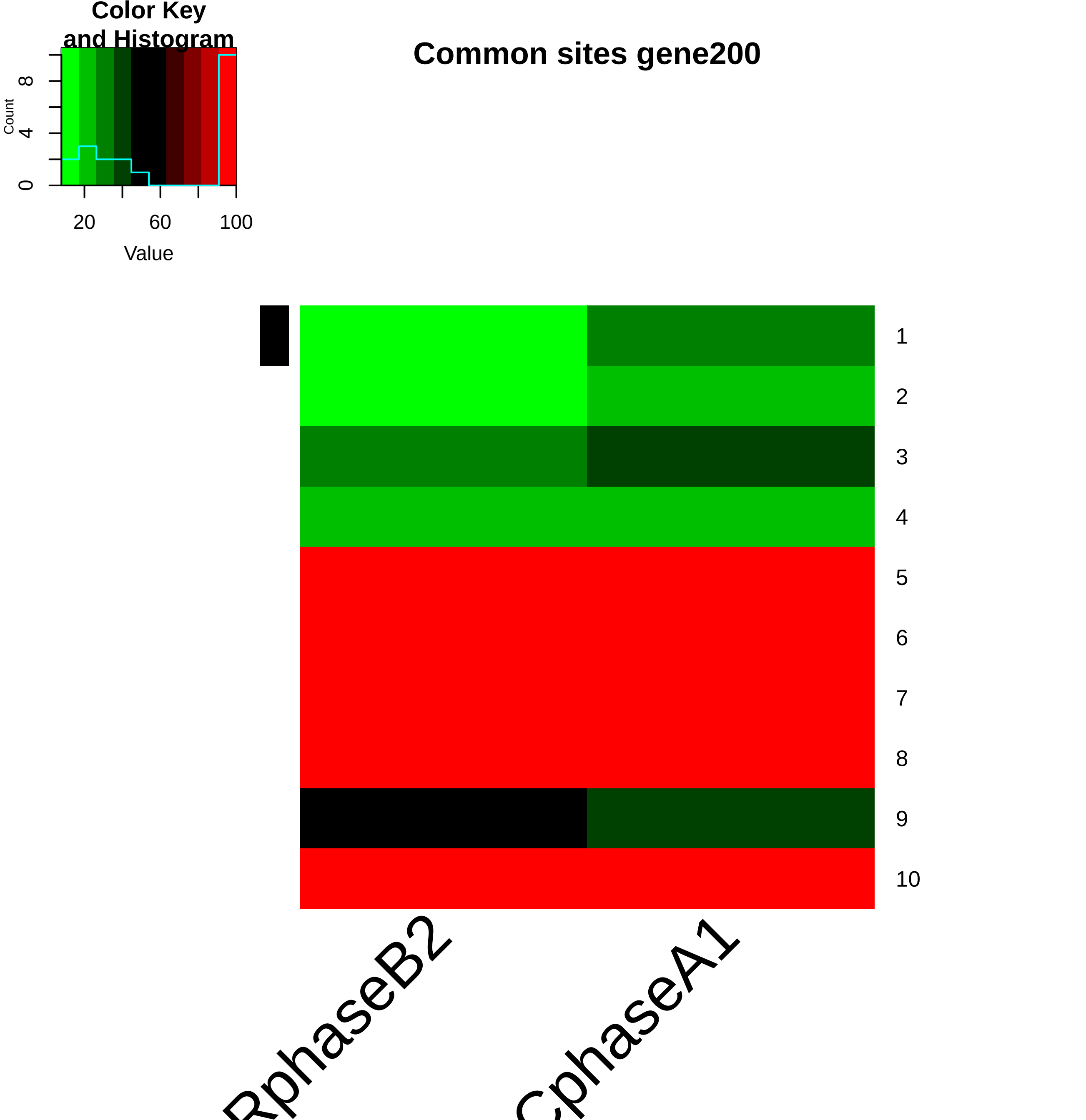
Color Key
and Histogram
Count
0
4
8
20
60
100
Value
Common sites gene200
1
2
3
4
5
6
7
8
9
10
RphaseB2
CphaseA1

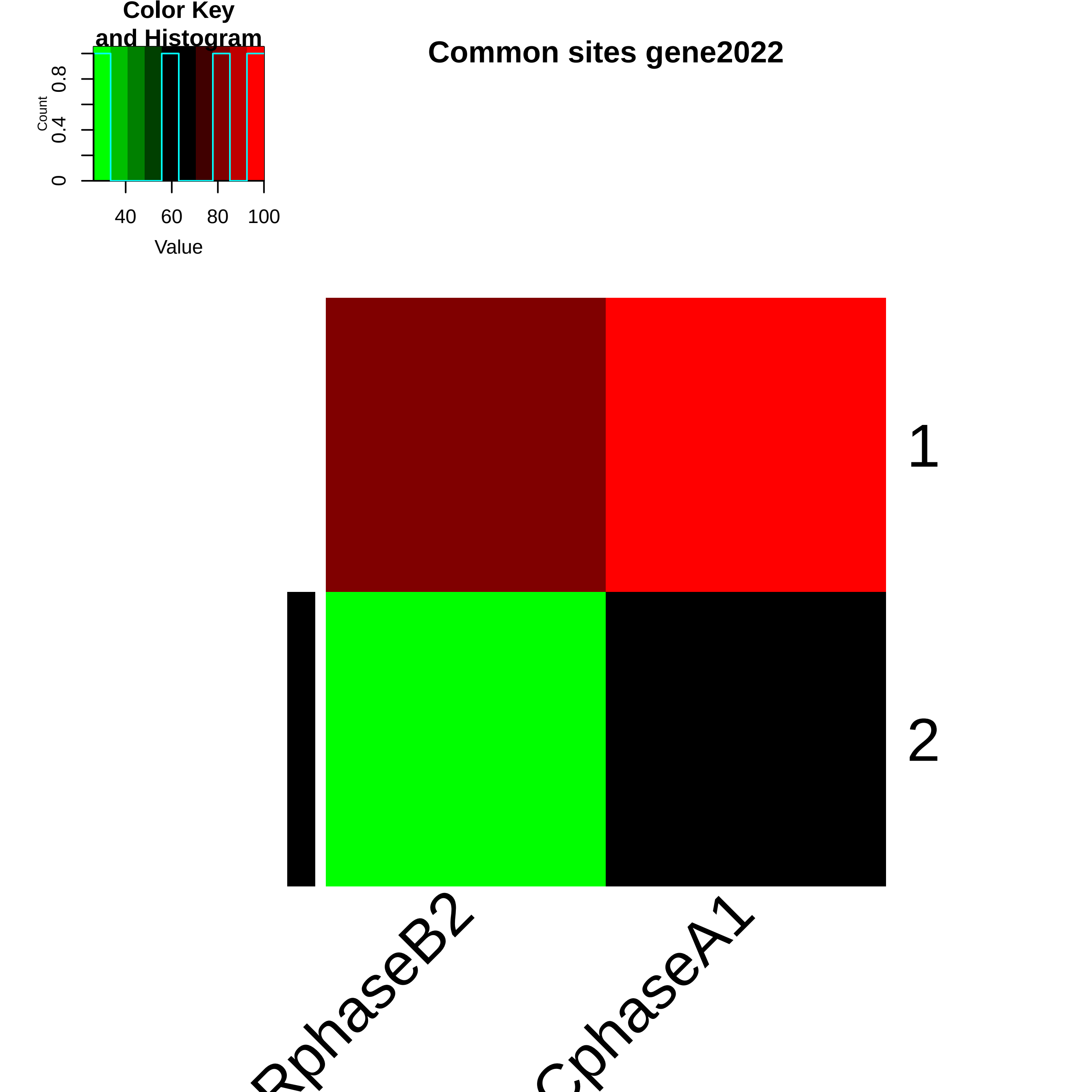

Color Key
and Histogram
Count
0
0.4
0.8
40
60
80
100
Value
Common sites gene2022
1
2
RphaseB2
CphaseA1

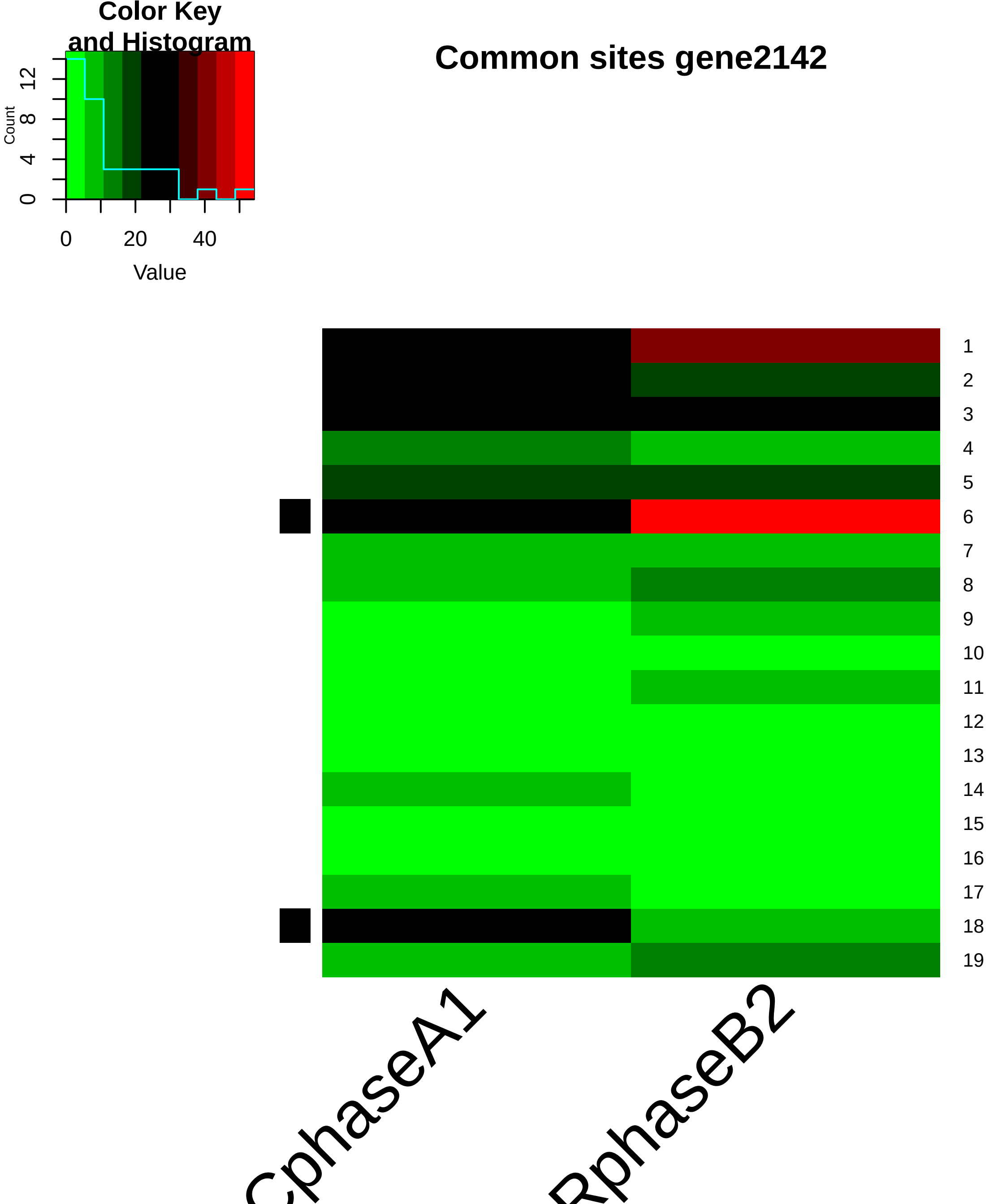

Color Key
and Histogram
Count
0
4
8
12
0
20
40
Value
Common sites gene2142
CphaseA1
RphaseB2

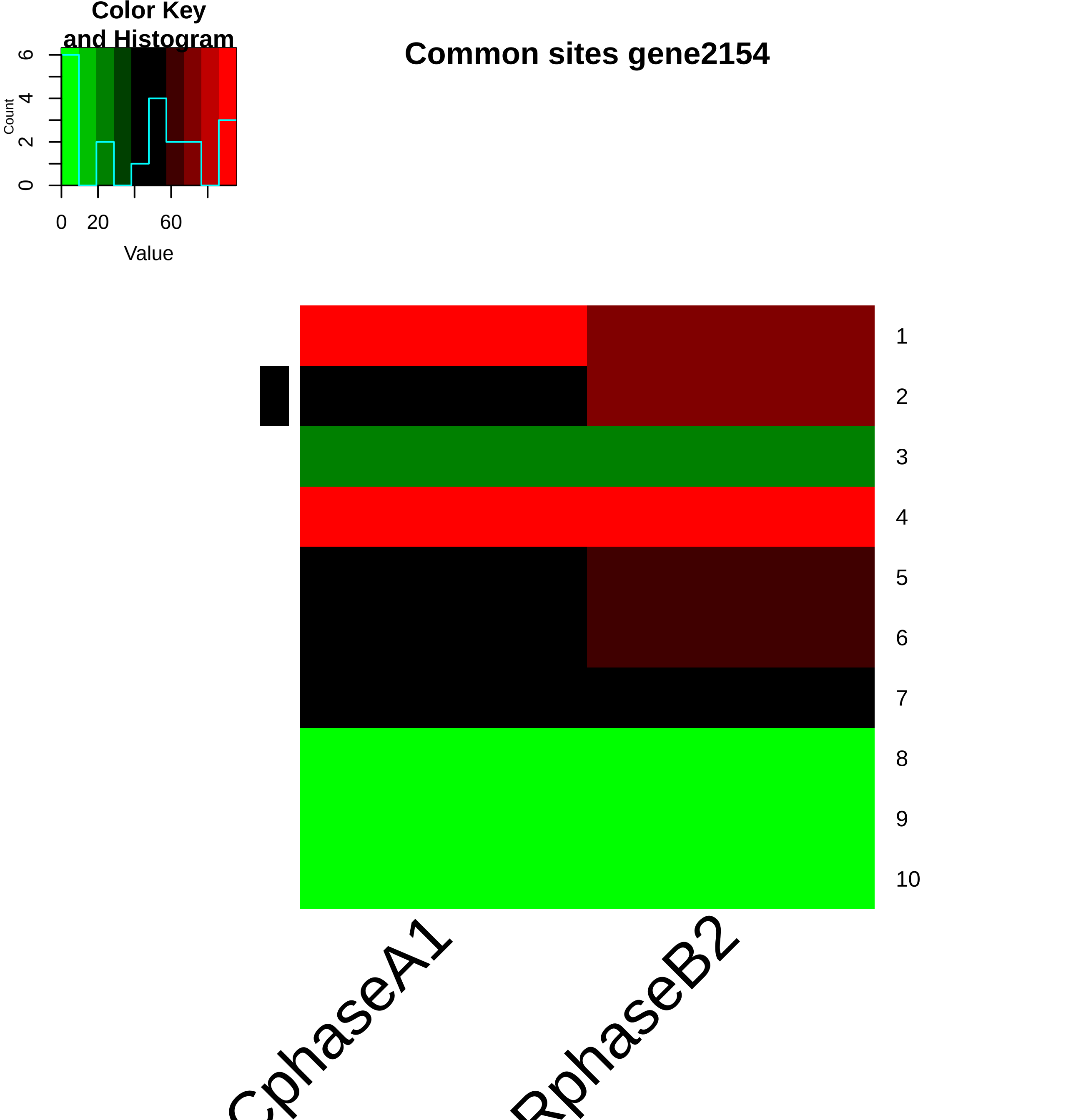

Color Key
and Histogram
Count
0
2
4
6
0
20
60
Value
Common sites gene2154
1
2
3
4
5
6
7
8
9
10
CphaseA1
RphaseB2

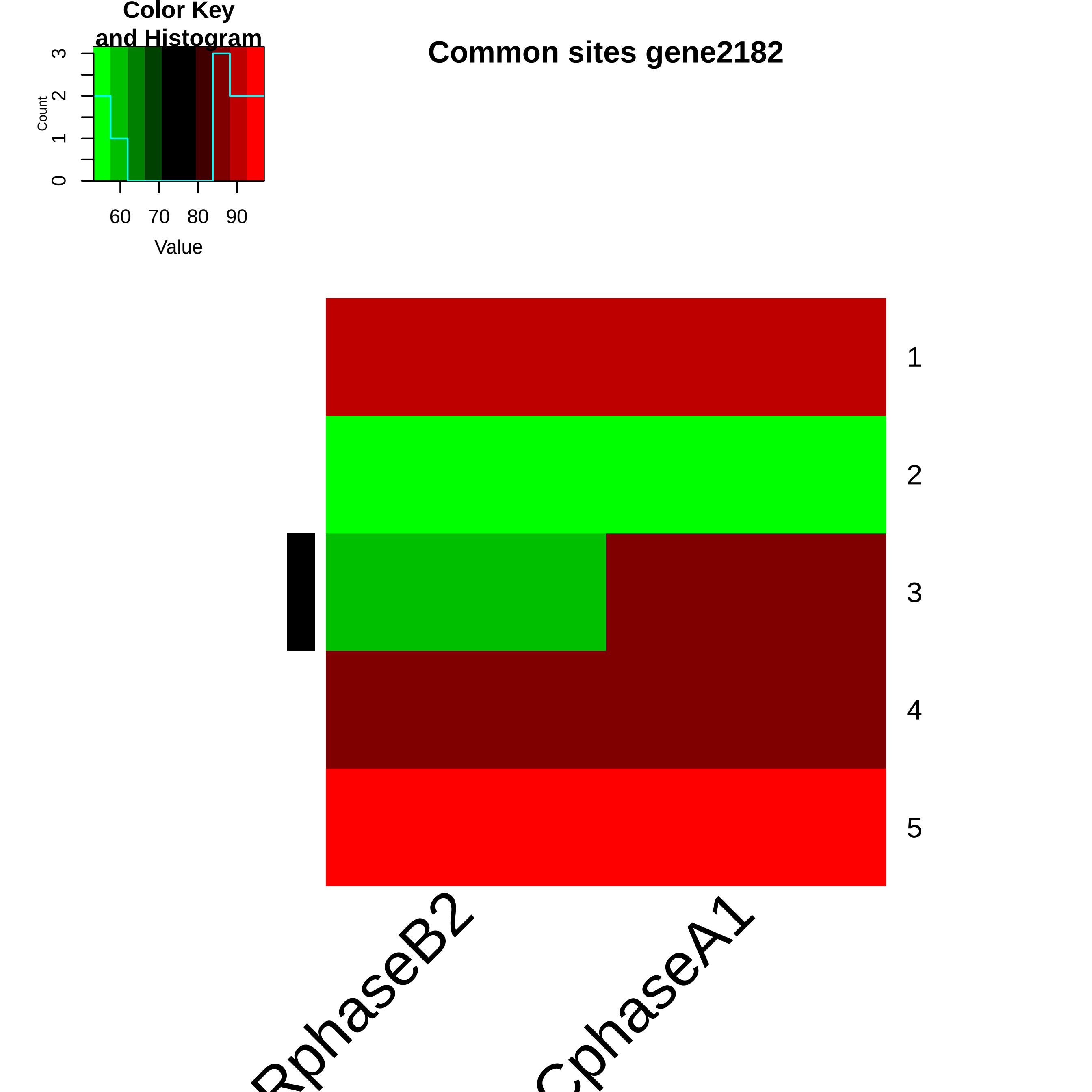

Color Key
and Histogram
Count
0
1
2
3
60
70
80
90
Value
Common sites gene2182
1
2
3
4
5
RphaseB2
CphaseA1

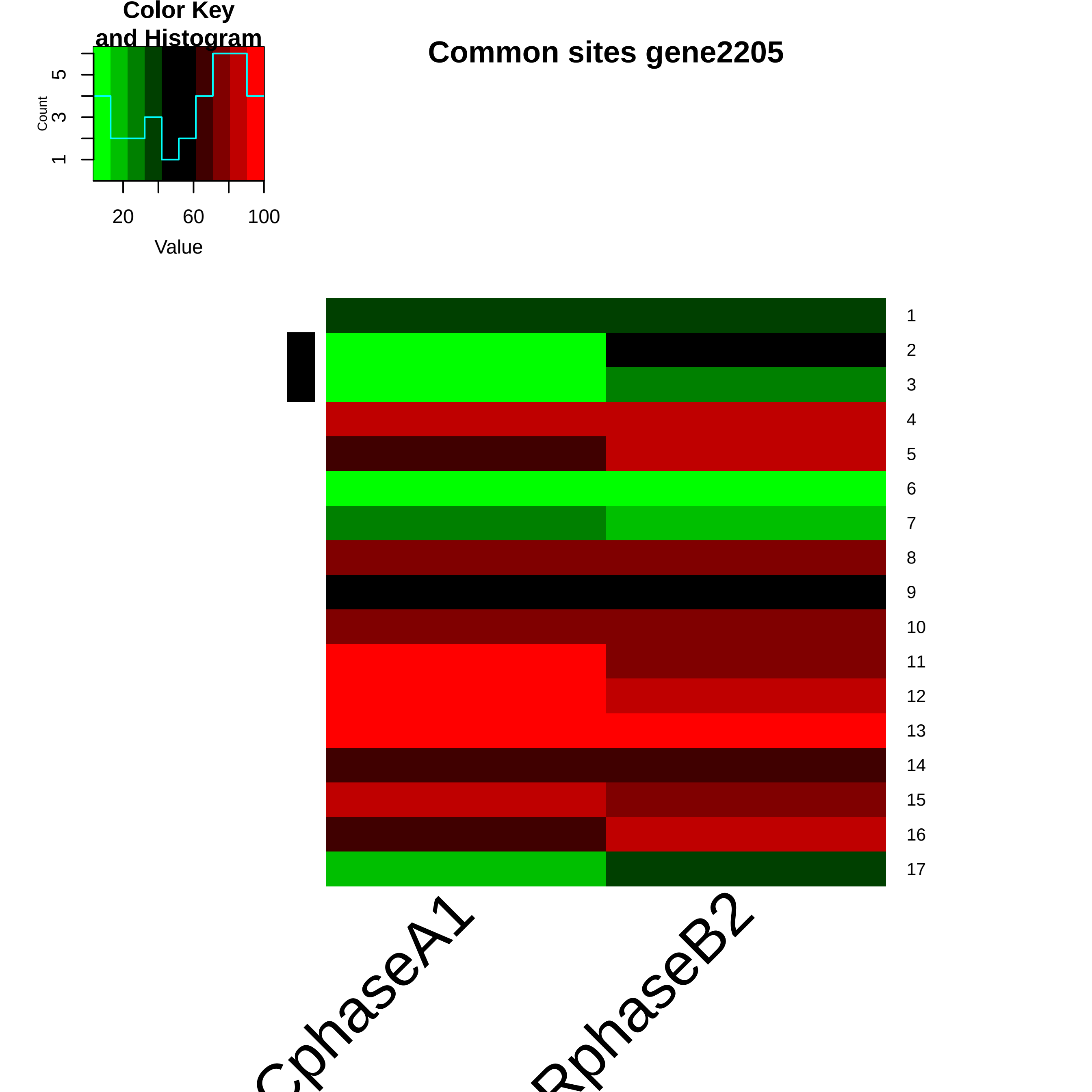

Color Key
and Histogram
Count
1
3
5
20
60
100
Value
Common sites gene2205
CphaseA1
RphaseB2

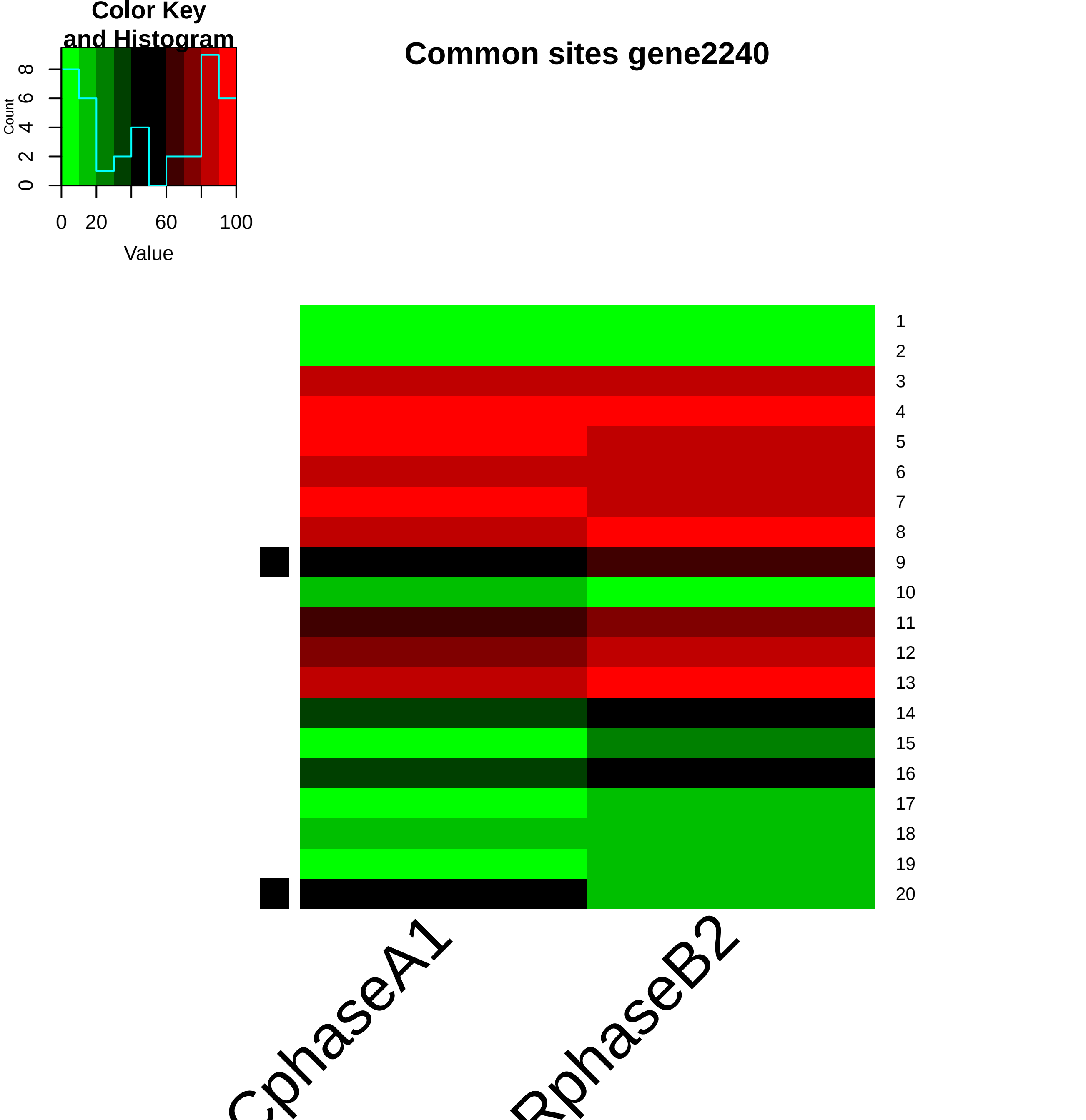

Color Key
and Histogram
Count
0
2
4
6
8
0
20
60
100
Value
Common sites gene2240
CphaseA1
RphaseB2

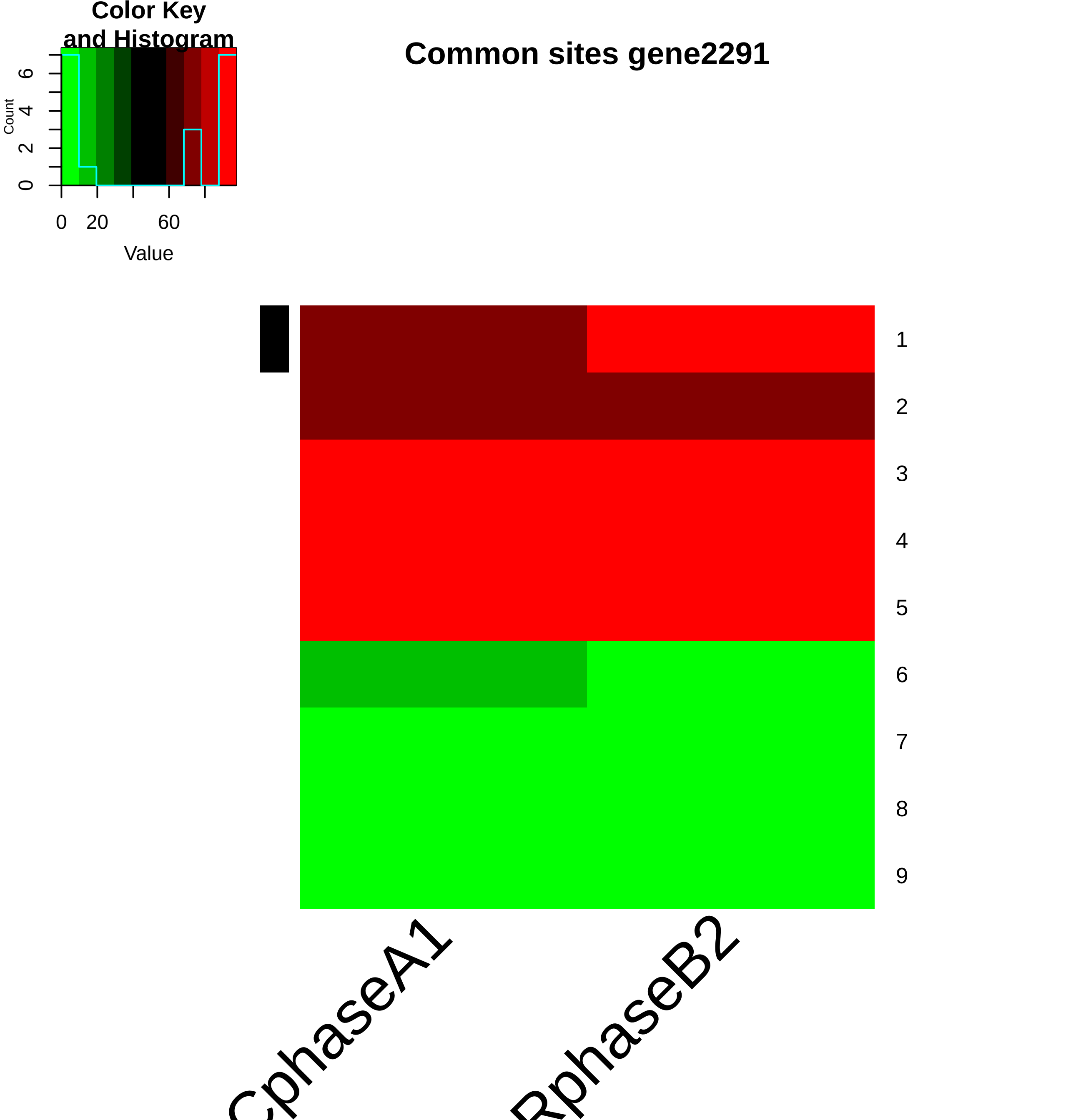
Color Key
and Histogram
Count
0
2
4
6
0
20
60
Value
Common sites gene2291
1
2
3
4
5
6
7
8
9
CphaseA1
RphaseB2

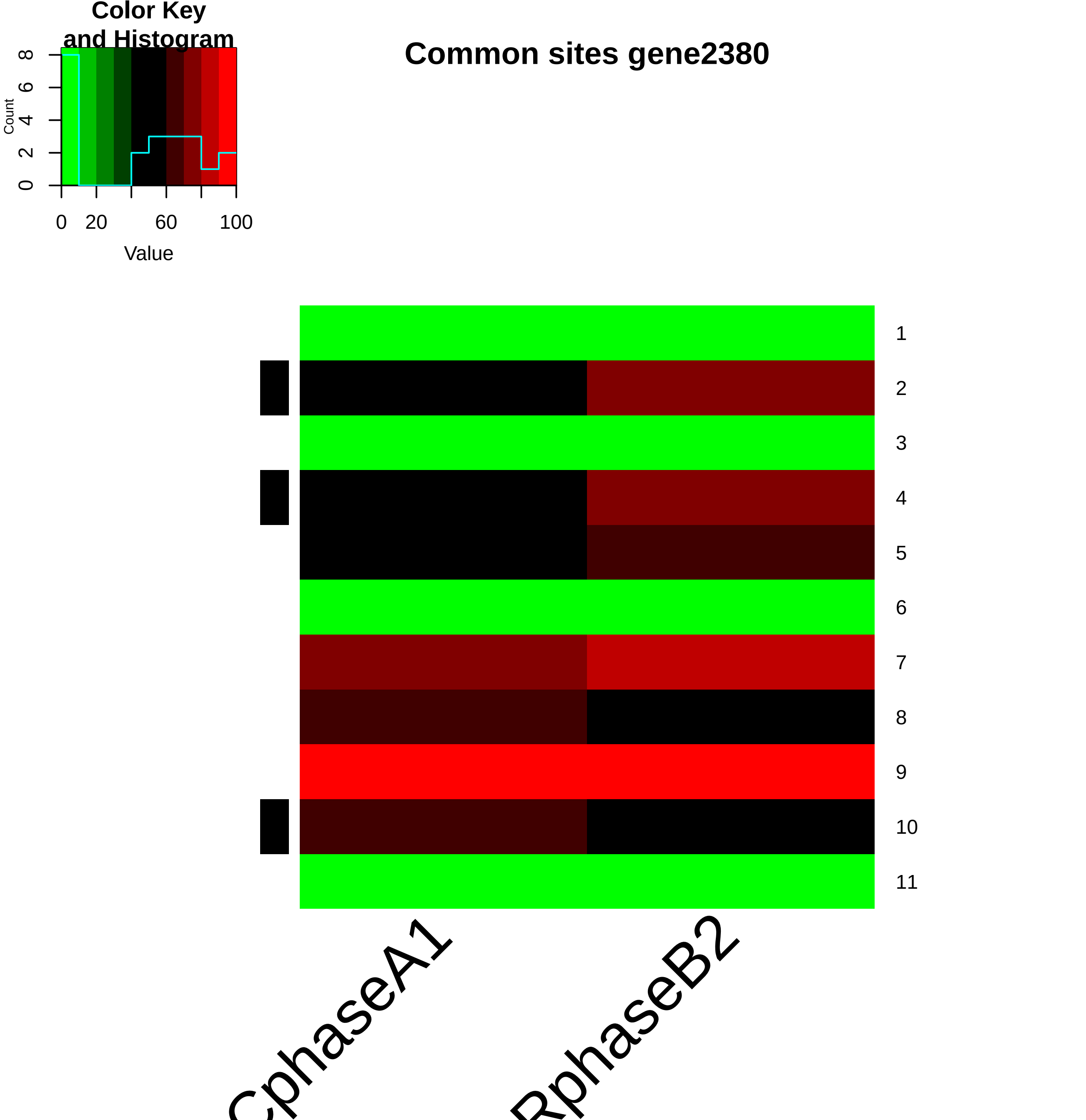
Color Key
and Histogram
Count
0
2
4
6
8
0
20
60
100
Value
Common sites gene2380
1
2
3
4
5
6
7
8
9
10
11
CphaseA1
RphaseB2

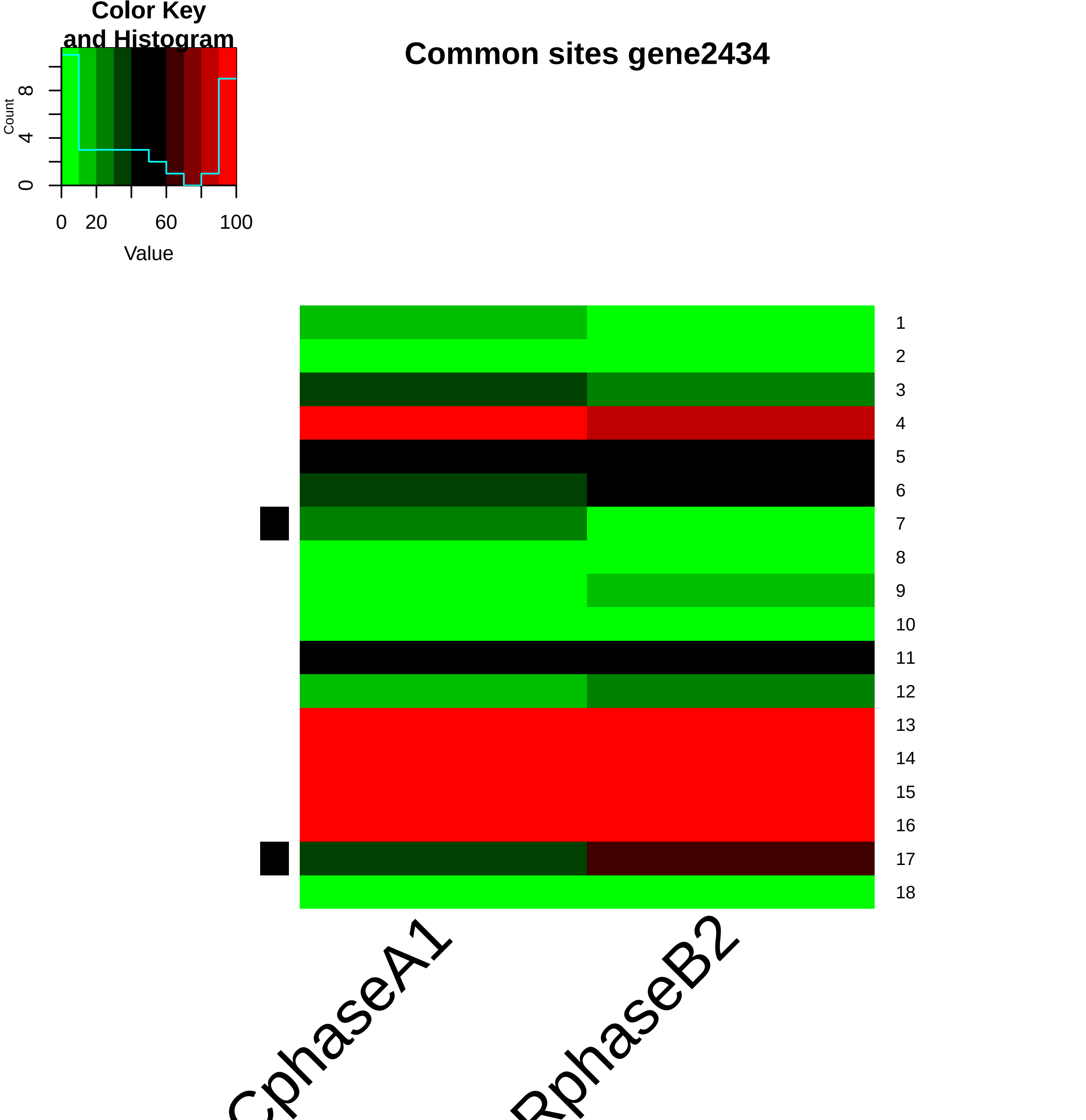
Color Key
and Histogram
Count
0
4
8
0
20
60
100
Value
Common sites gene2434
CphaseA1
RphaseB2

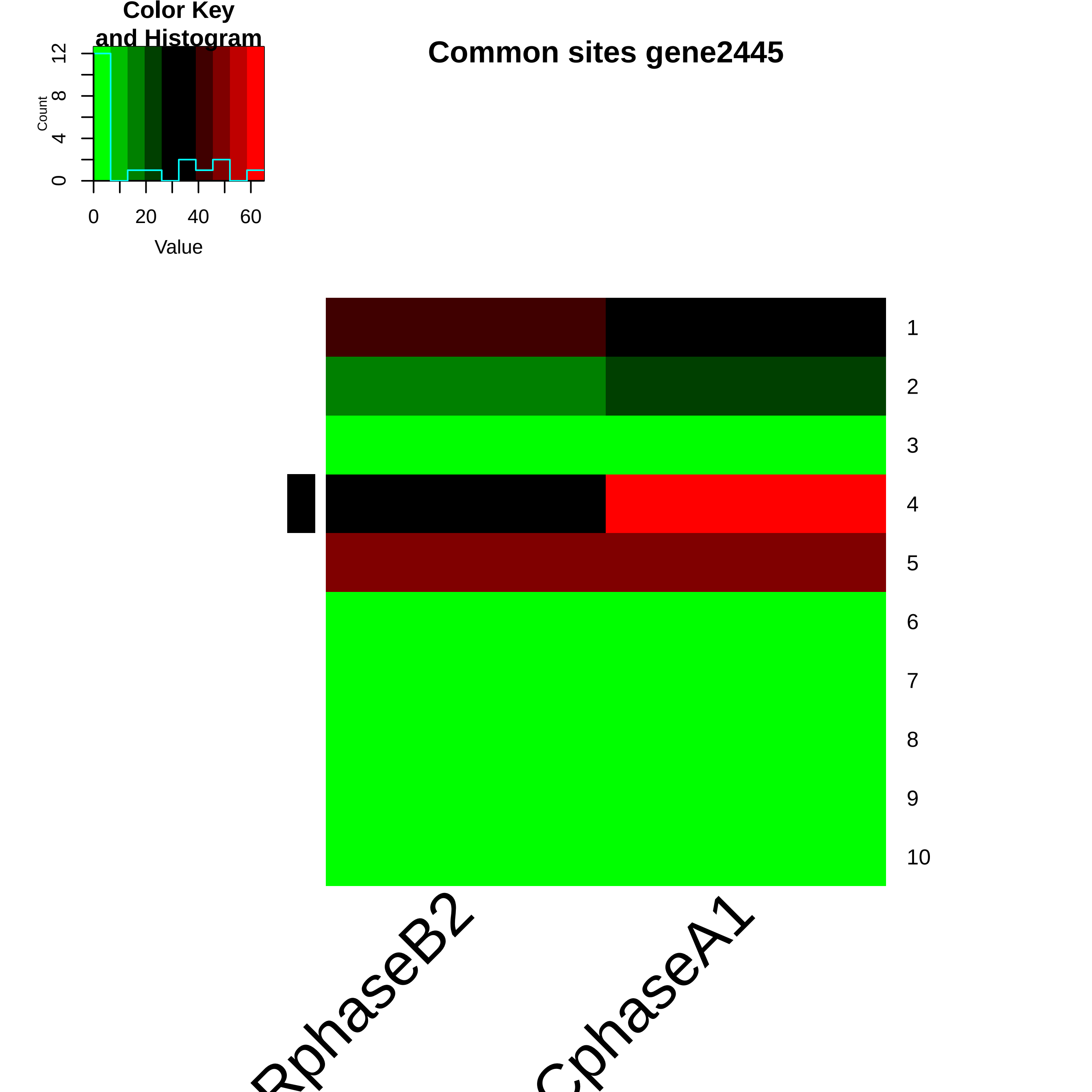

Color Key
and Histogram
Count
0
4
8
12
0
20
40
60
Value
Common sites gene2445
1
2
3
4
5
6
7
8
9
10
RphaseB2
CphaseA1

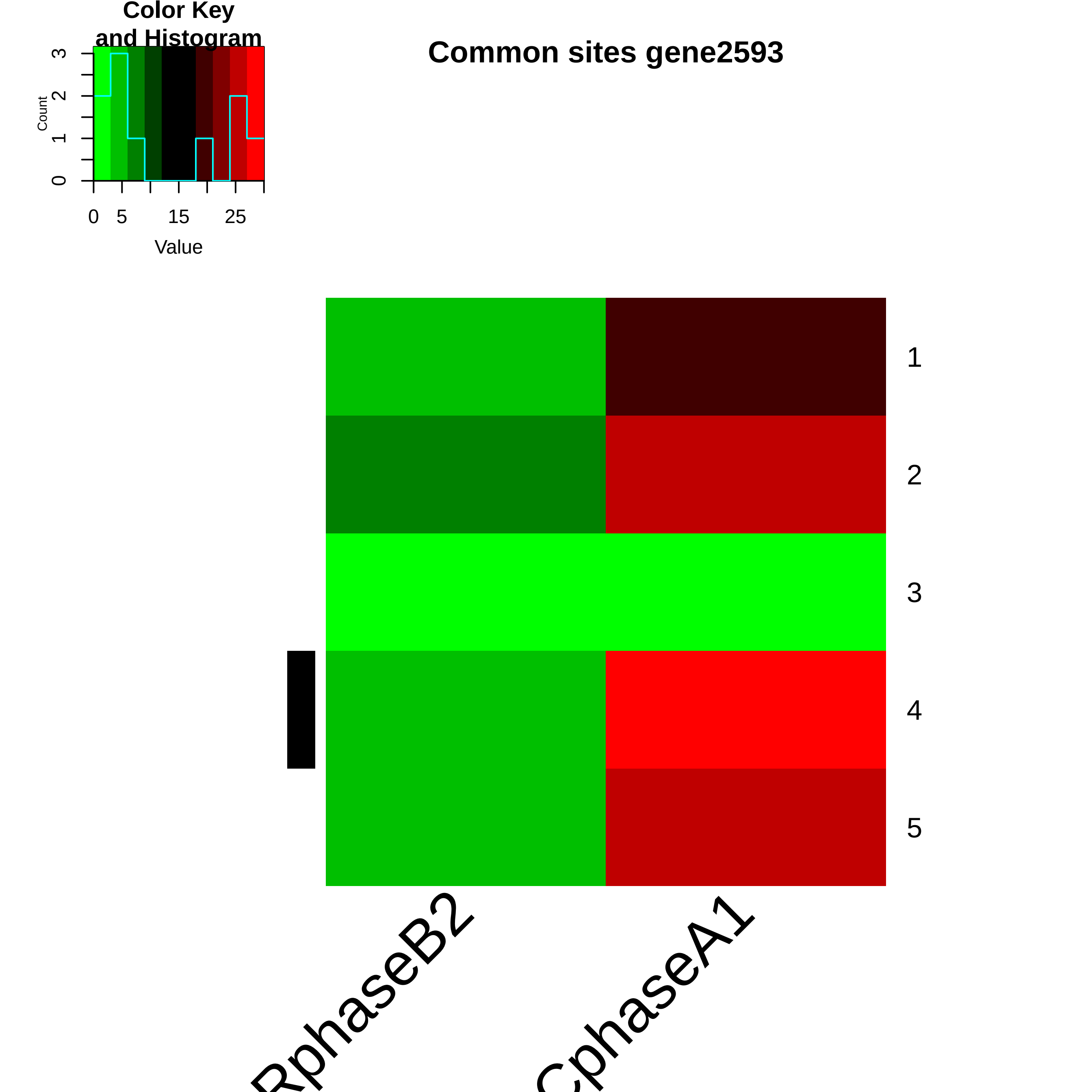

Color Key
and Histogram
Count
Value
0
5
15
25
0
1
2
3
Common sites gene2593
1
2
3
4
5
RphaseB2
CphaseA1

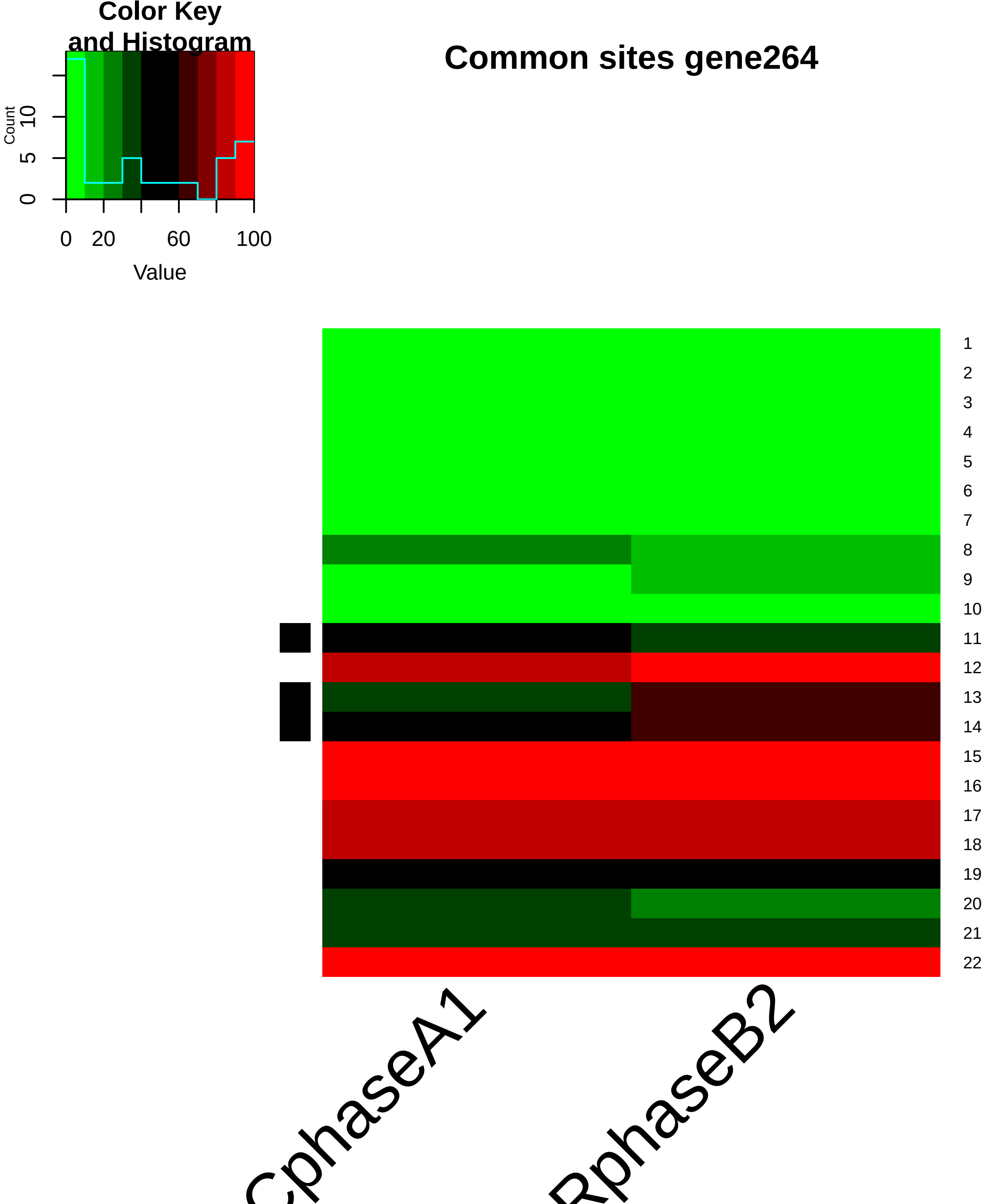
Color Key
and Histogram
Count
0
5
10
0
20
60
100
Value
Common sites gene264
CphaseA1
RphaseB2

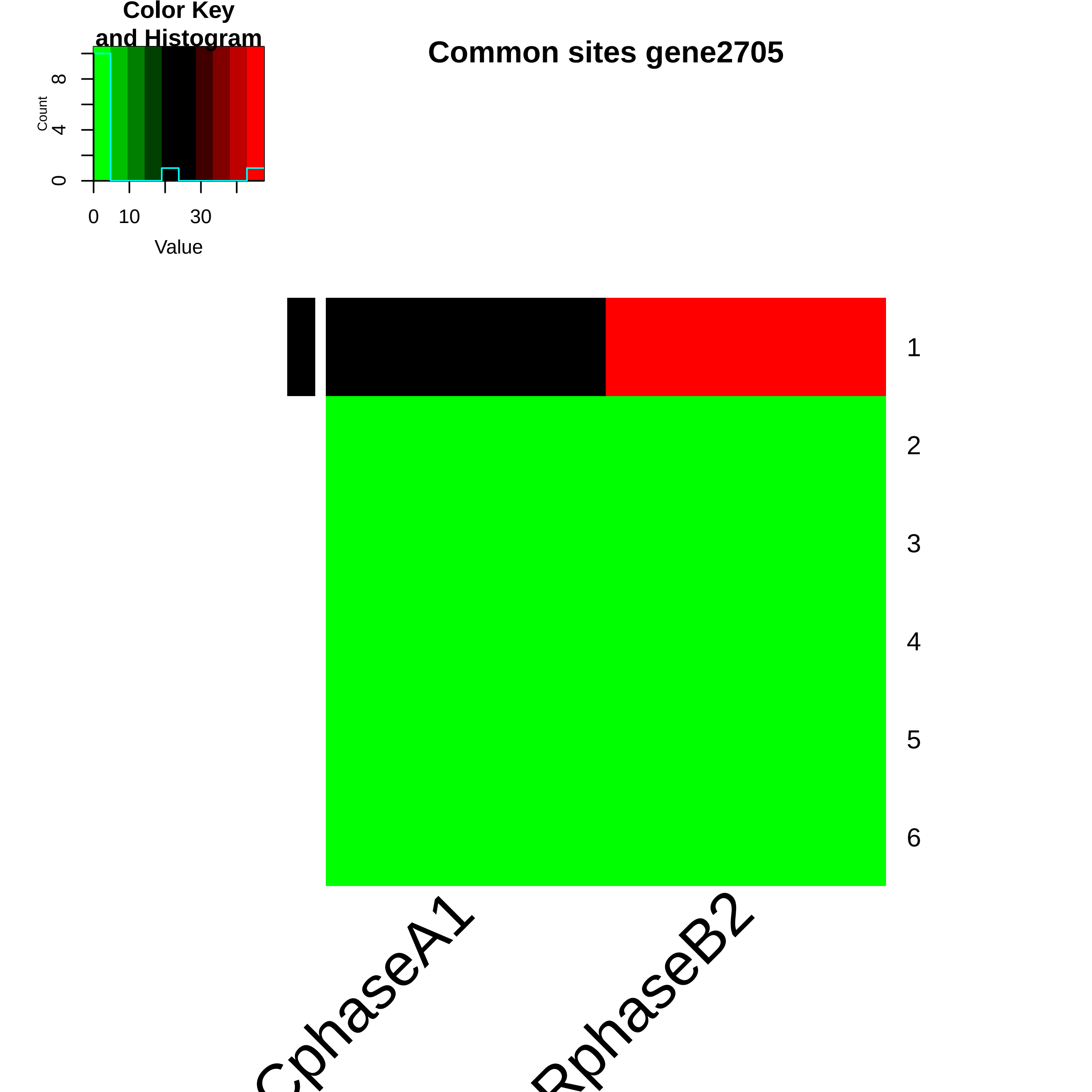
Color Key
and Histogram
Count
0
4
8
0
10
30
Value
Common sites gene2705
1
2
3
4
5
6

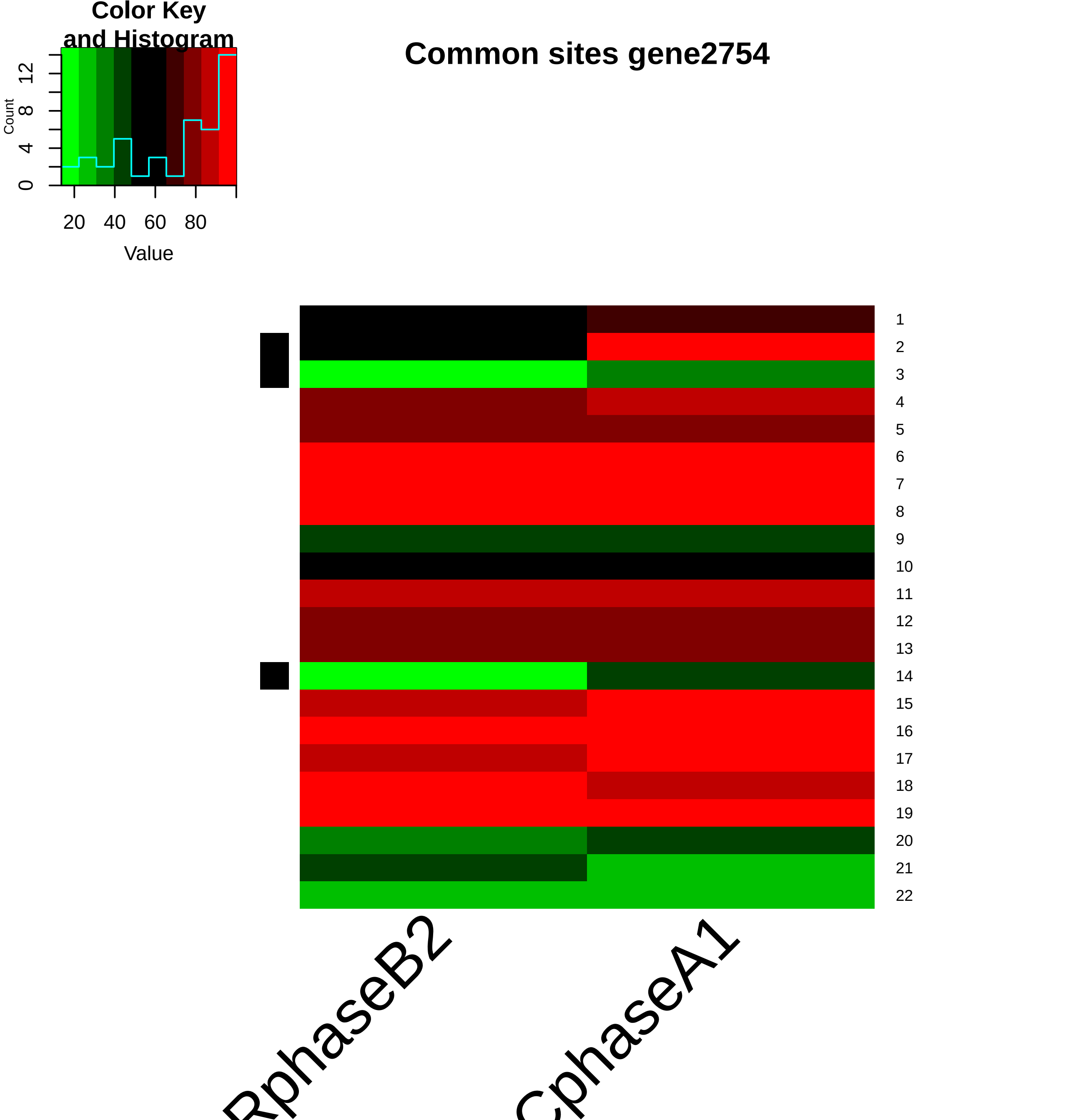
Color Key
and Histogram
Count
0
4
8
12
20
40
60
80
Value
Common sites gene2754
RphaseB2
CphaseA1

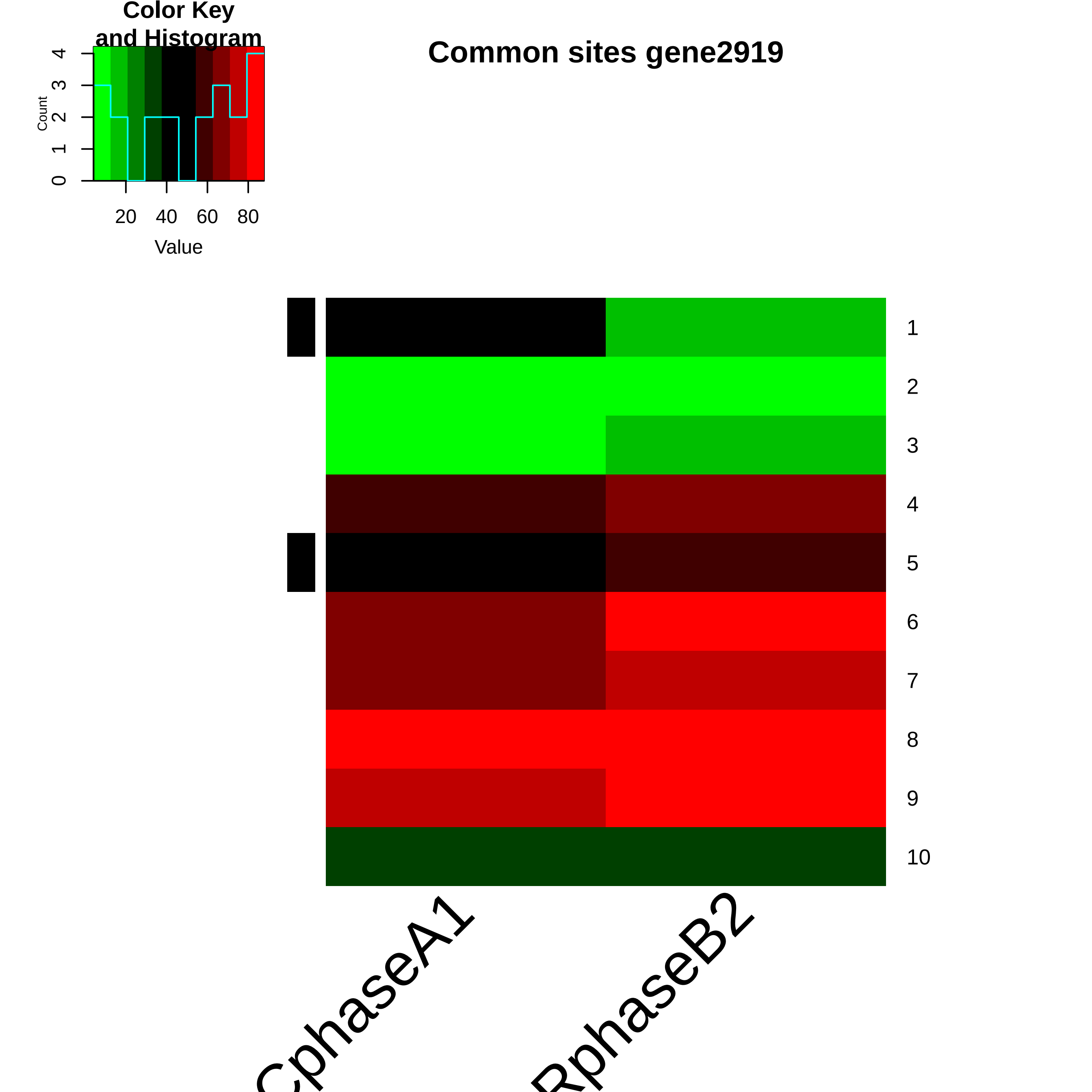

Color Key
and Histogram
Count
0
1
2
3
4
20
40
60
80
Value
Common sites gene2919
1
2
3
4
5
6
7
8
9
10
CphaseA1
RphaseB2

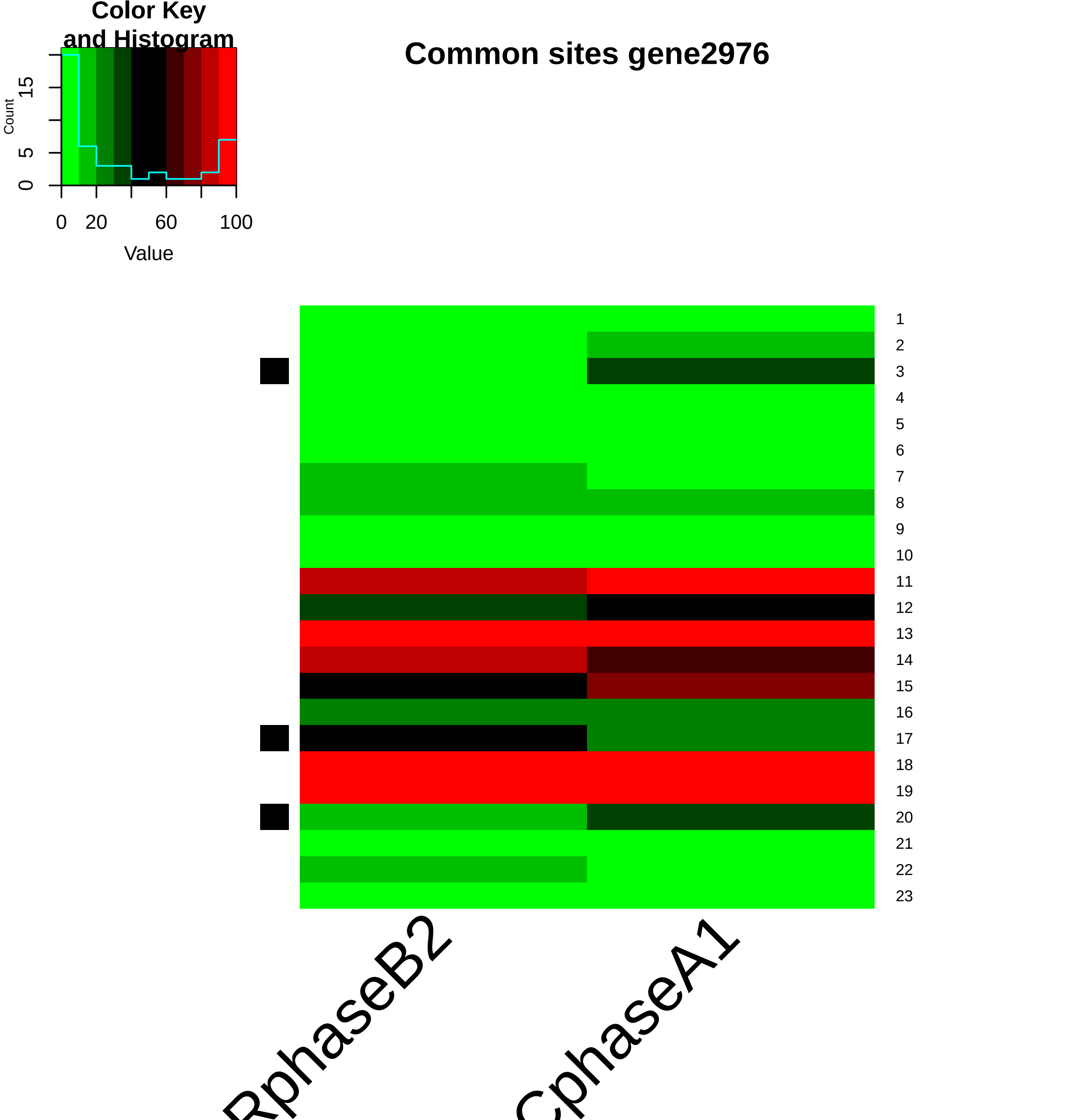

Color Key
and Histogram
Count
0
5
15
0
20
60
100
Value
Common sites gene2976
RphaseB2
CphaseA1

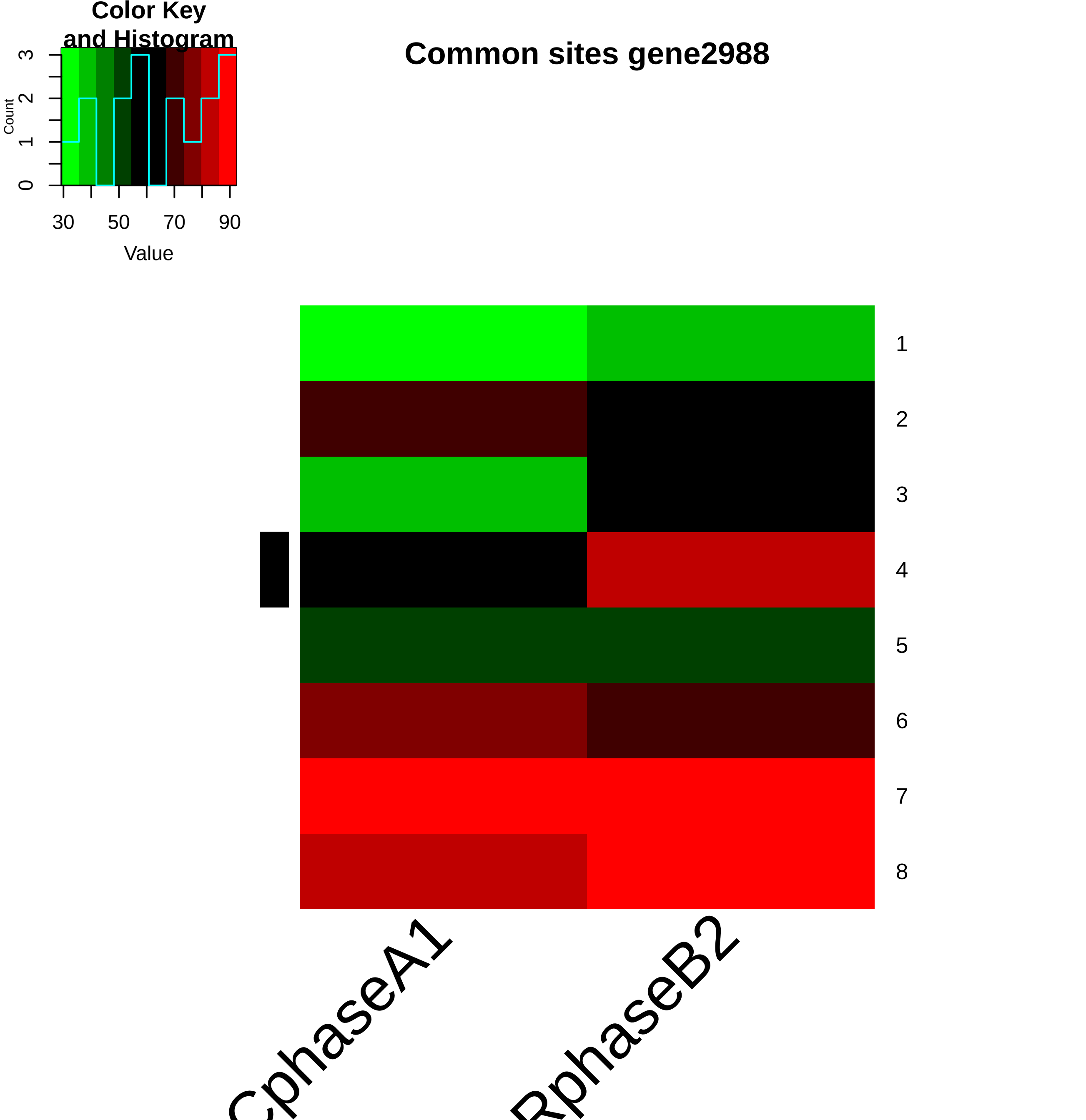
Color Key
and Histogram
Count
0
1
2
3
30
50
70
90
Value
Common sites gene2988
1
2
3
4
5
6
7
8
CphaseA1
RphaseB2

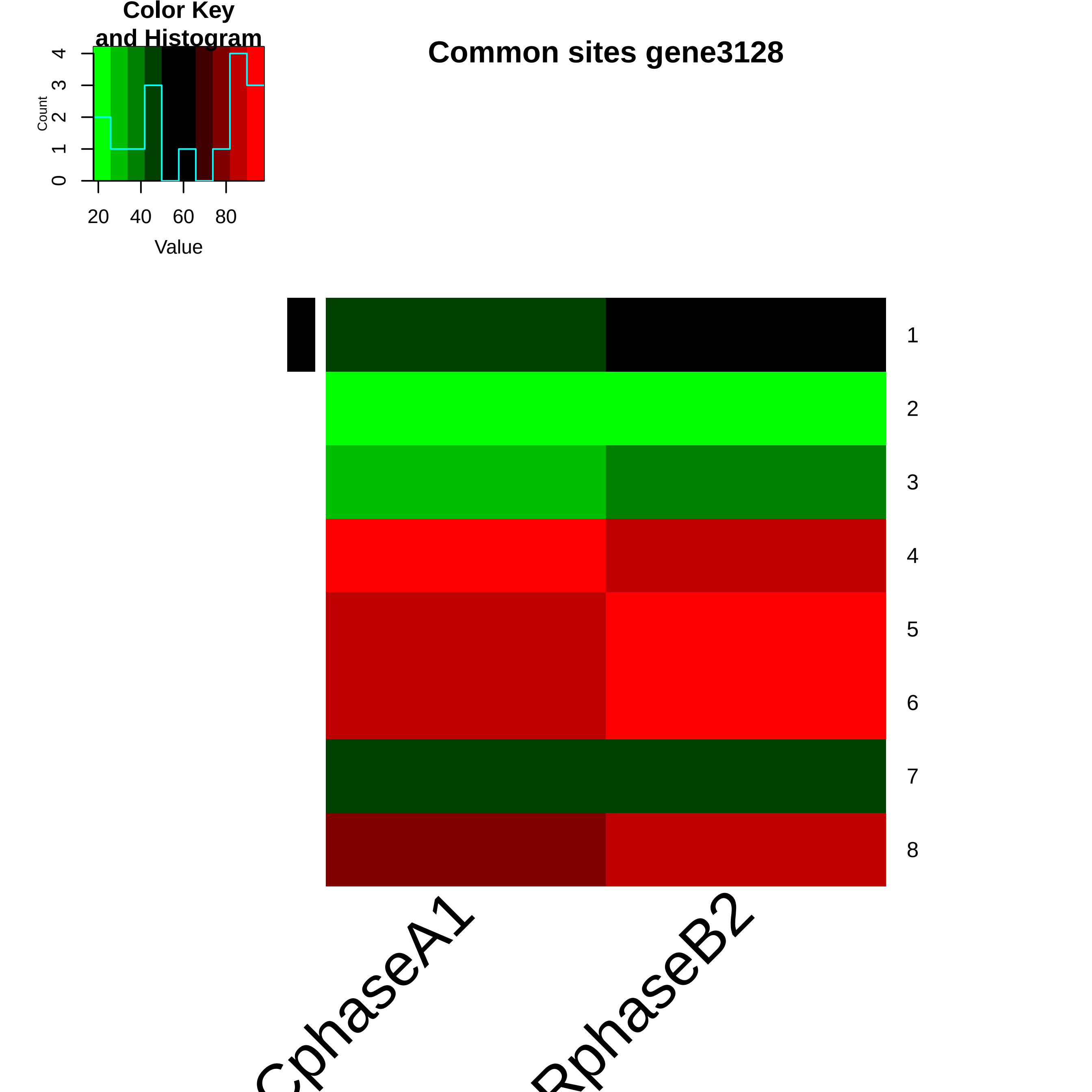
Color Key
and Histogram
Count
0
1
2
3
4
20
40
60
80
Value
Common sites gene3128
1
2
3
4
5
6
7
8
CphaseA1
RphaseB2

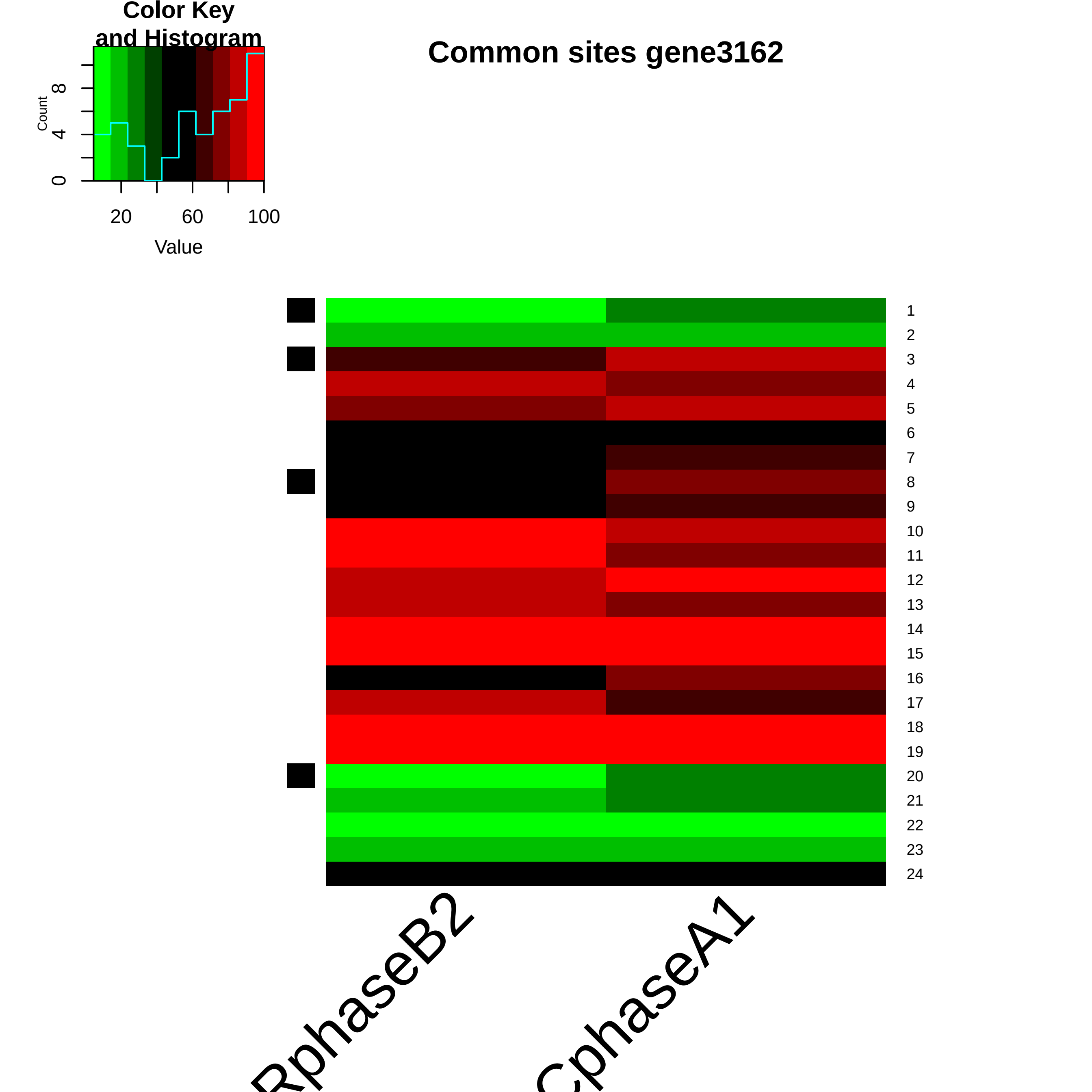

Color Key
and Histogram
Count
0
4
8
20
60
100
Value
Common sites gene3162
RphaseB2
CphaseA1

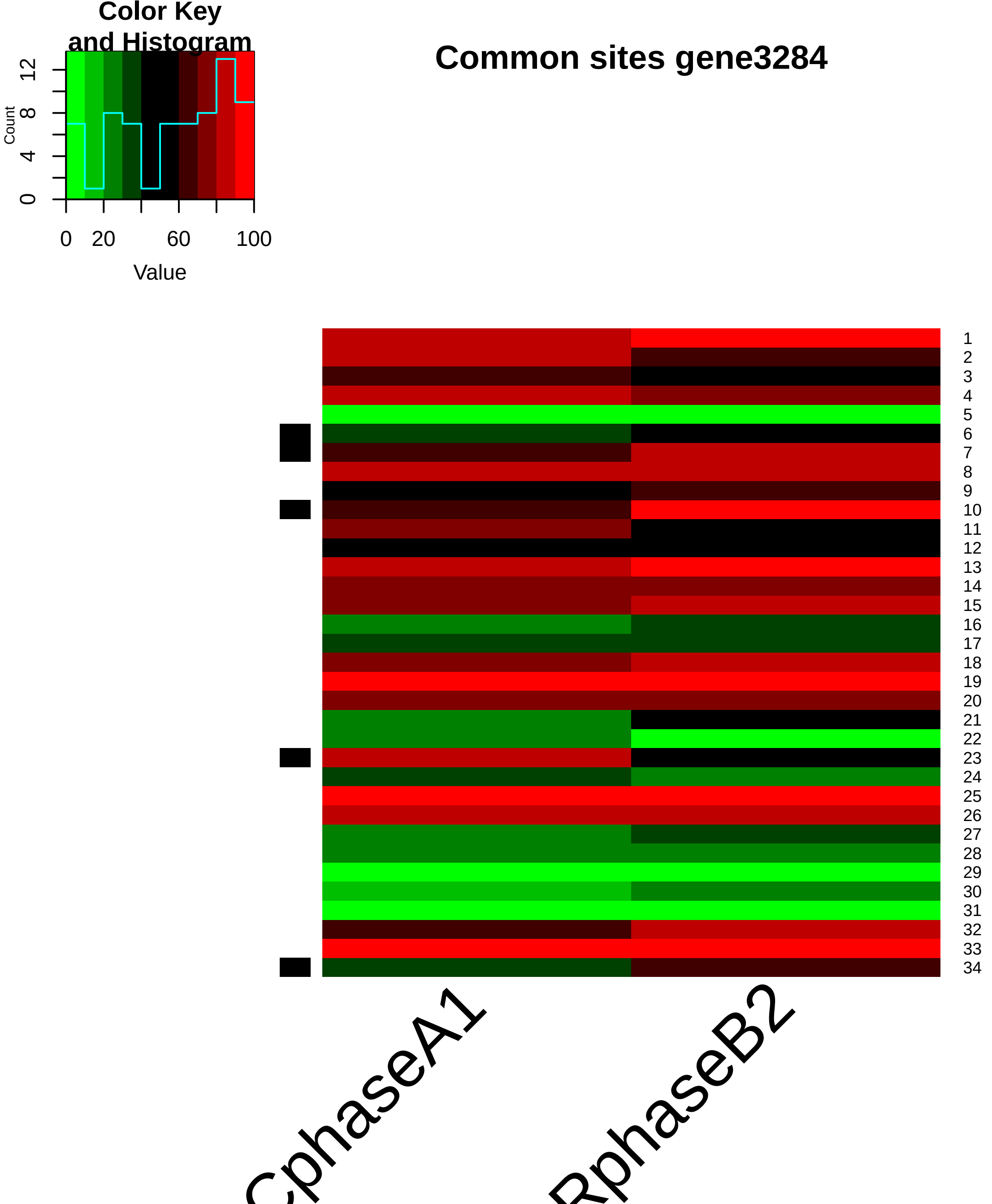
Color Key
and Histogram
Count
0
4
8
12
0
20
60
100
Value
Common sites gene3284
CphaseA1
RphaseB2

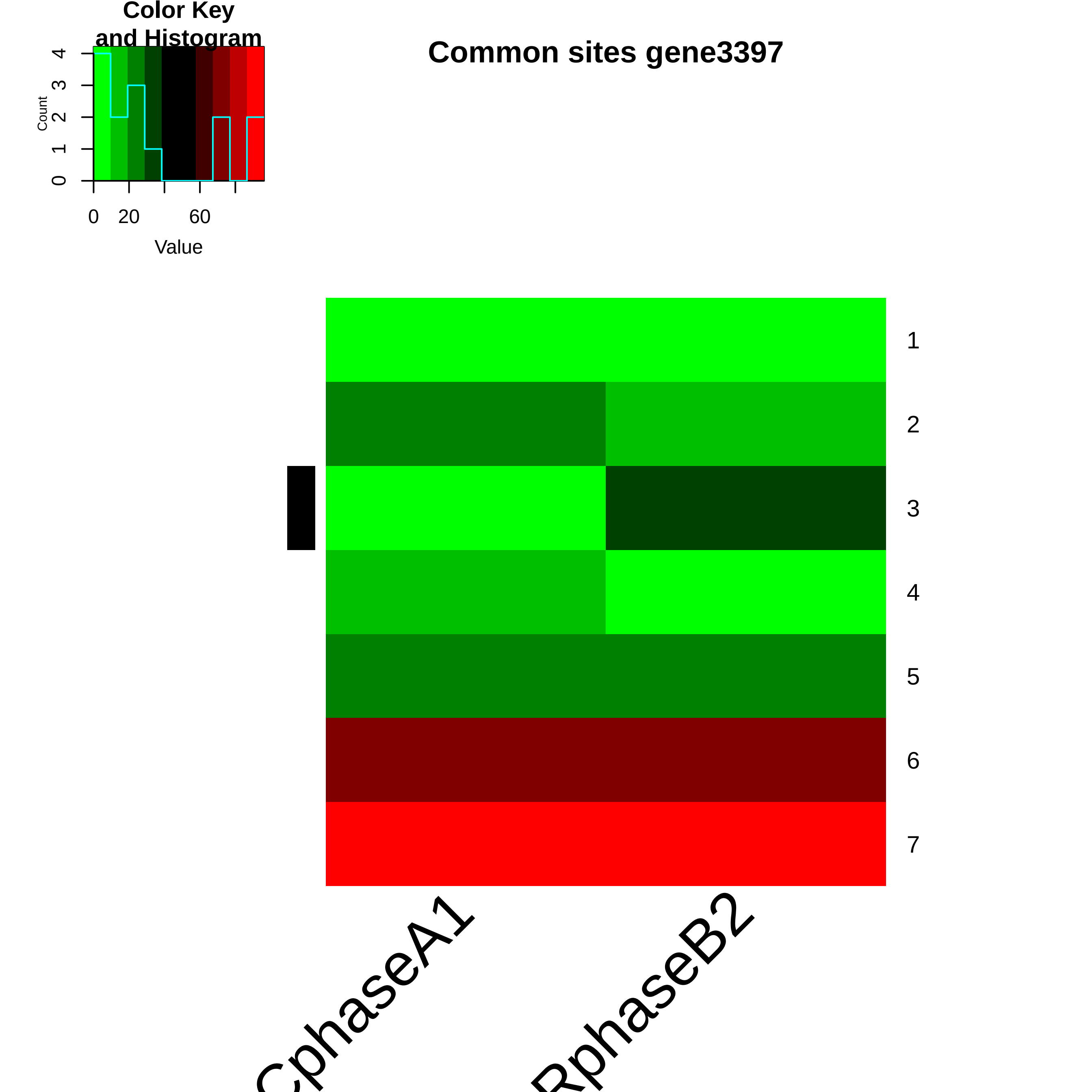

Color Key
and Histogram
Count
0
1
2
3
4
0
20
60
Value
Common sites gene3397
1
2
3
4
5
6
7
CphaseA1
RphaseB2

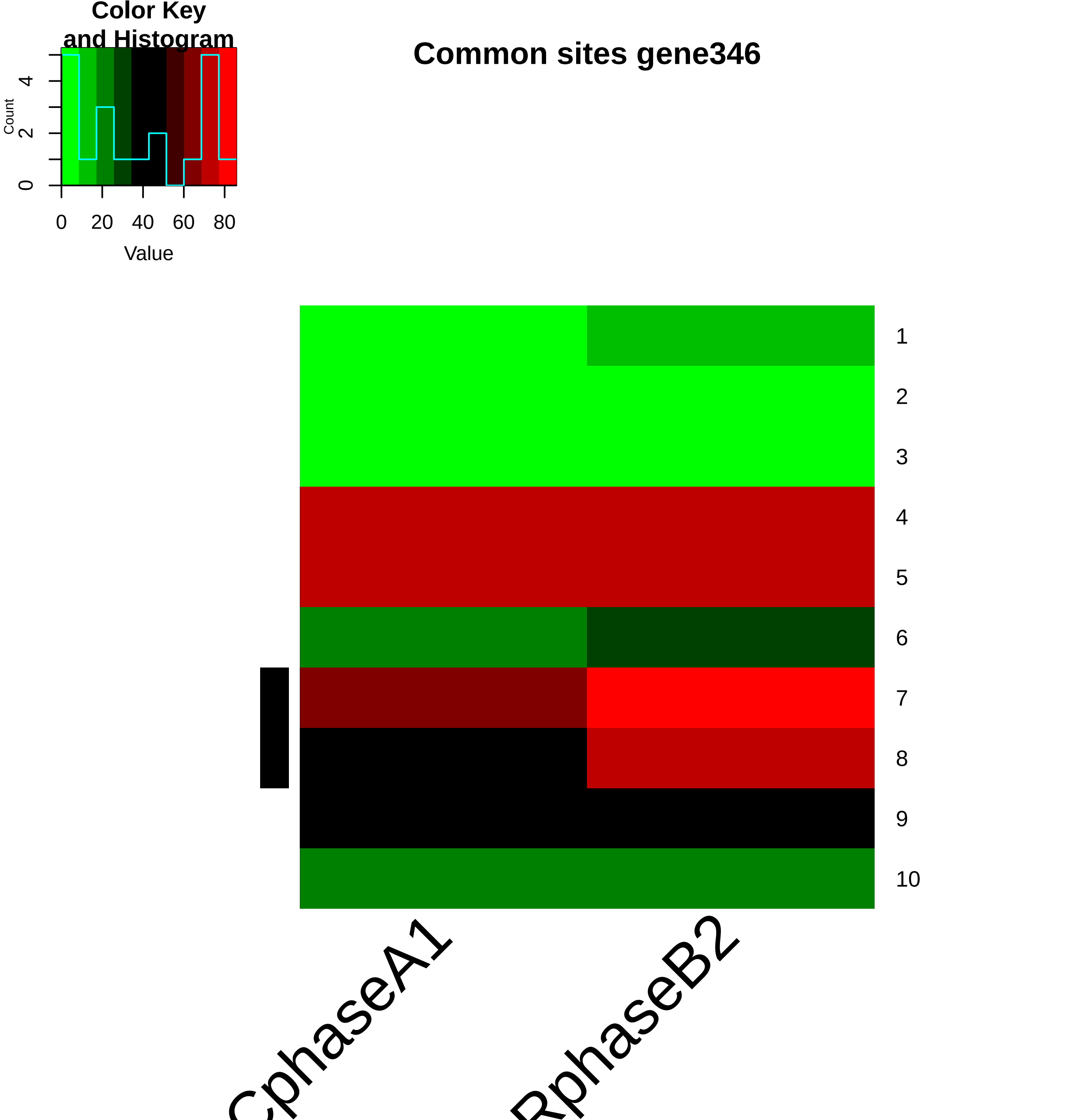
Color Key
and Histogram
Count
0
2
4
0
20
40
60
80
Value
Common sites gene346
1
2
3
4
5
6
7
8
9
10
CphaseA1
RphaseB2

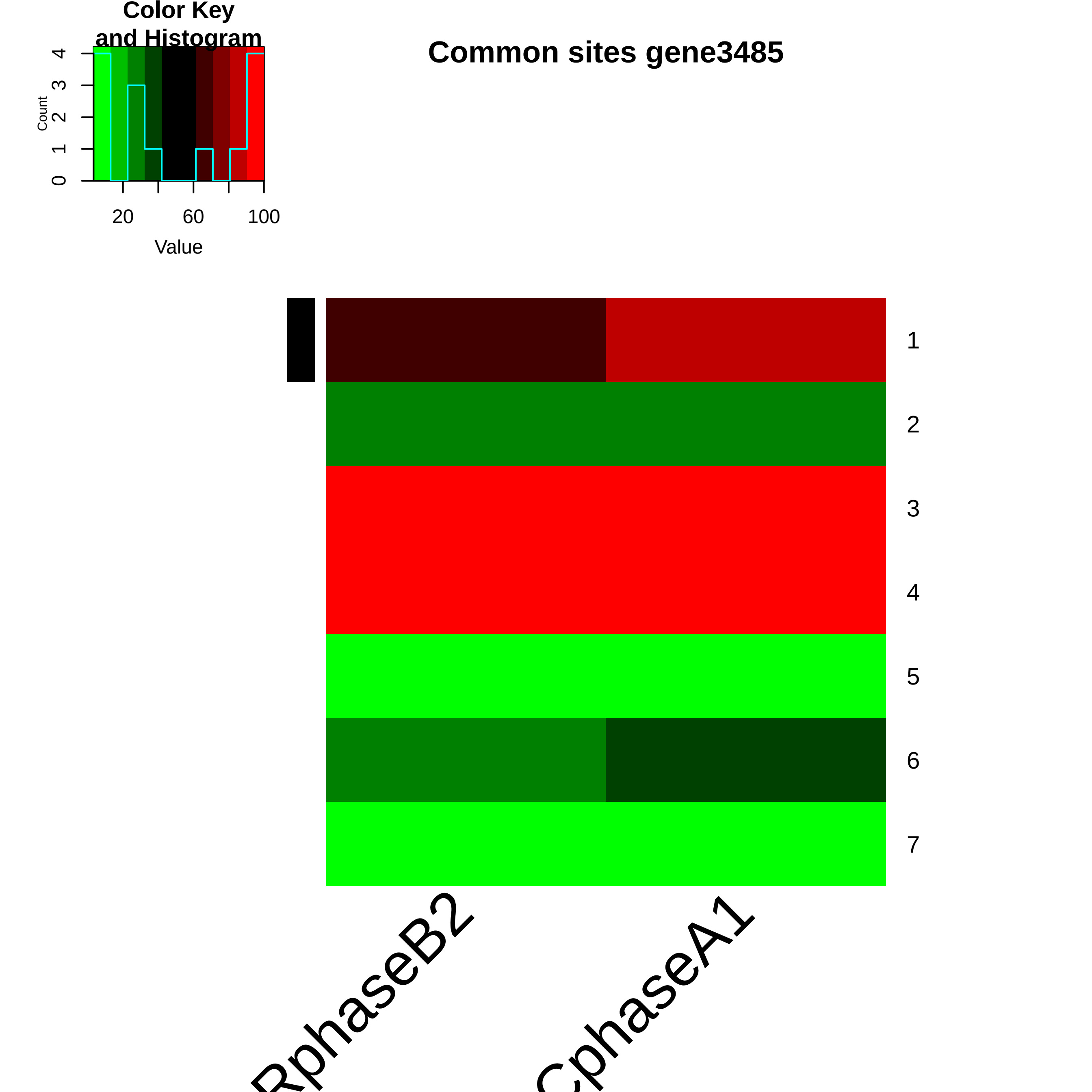
Color Key
and Histogram
Count
0
1
2
3
4
20
60
100
Value
Common sites gene3485
1
2
3
4
5
6
7
RphaseB2
CphaseA1

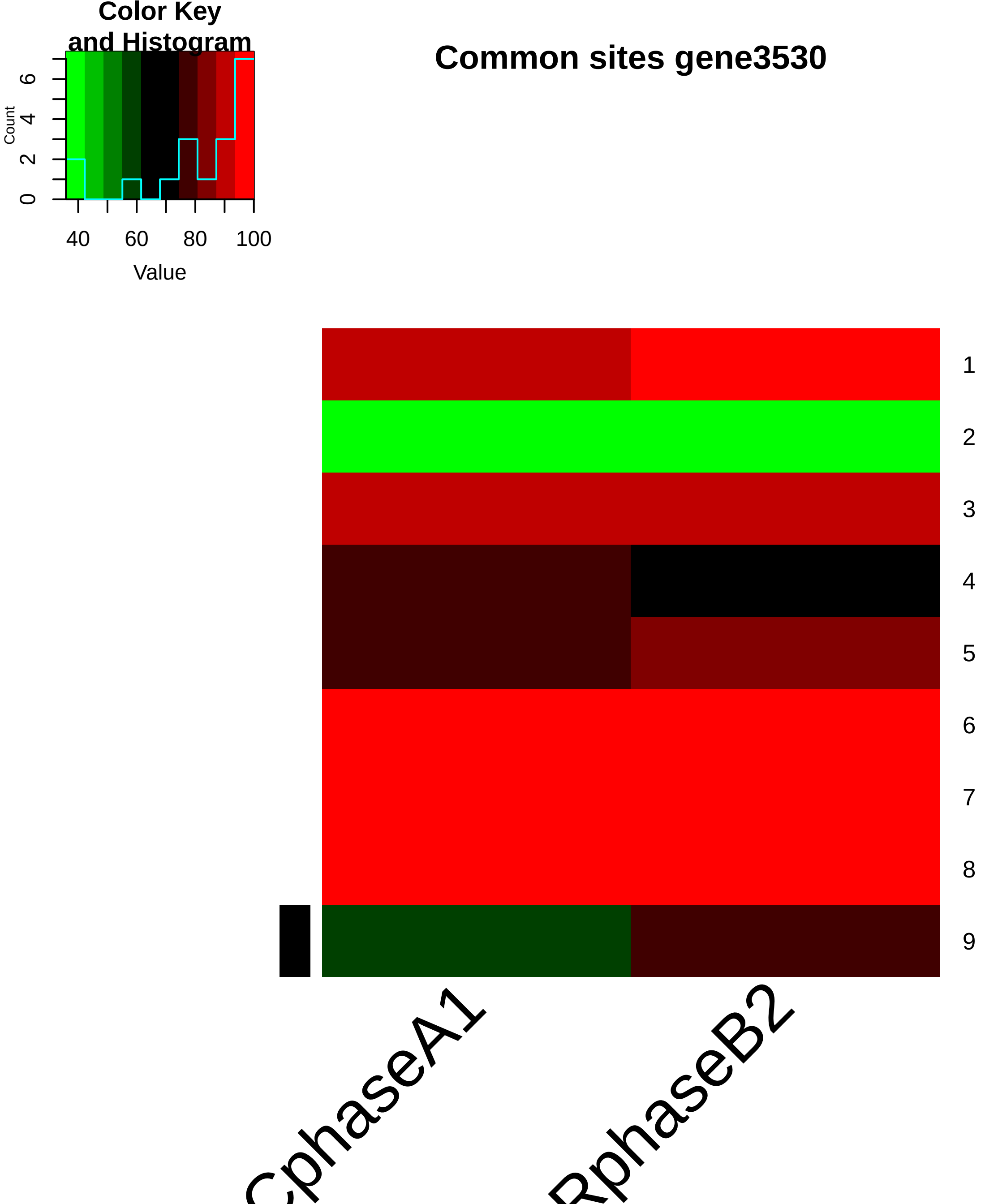
Color Key
and Histogram
Count
0
2
4
6
40
60
80
100
Value
Common sites gene3530
1
2
3
4
5
6
7
8
9
CphaseA1
RphaseB2

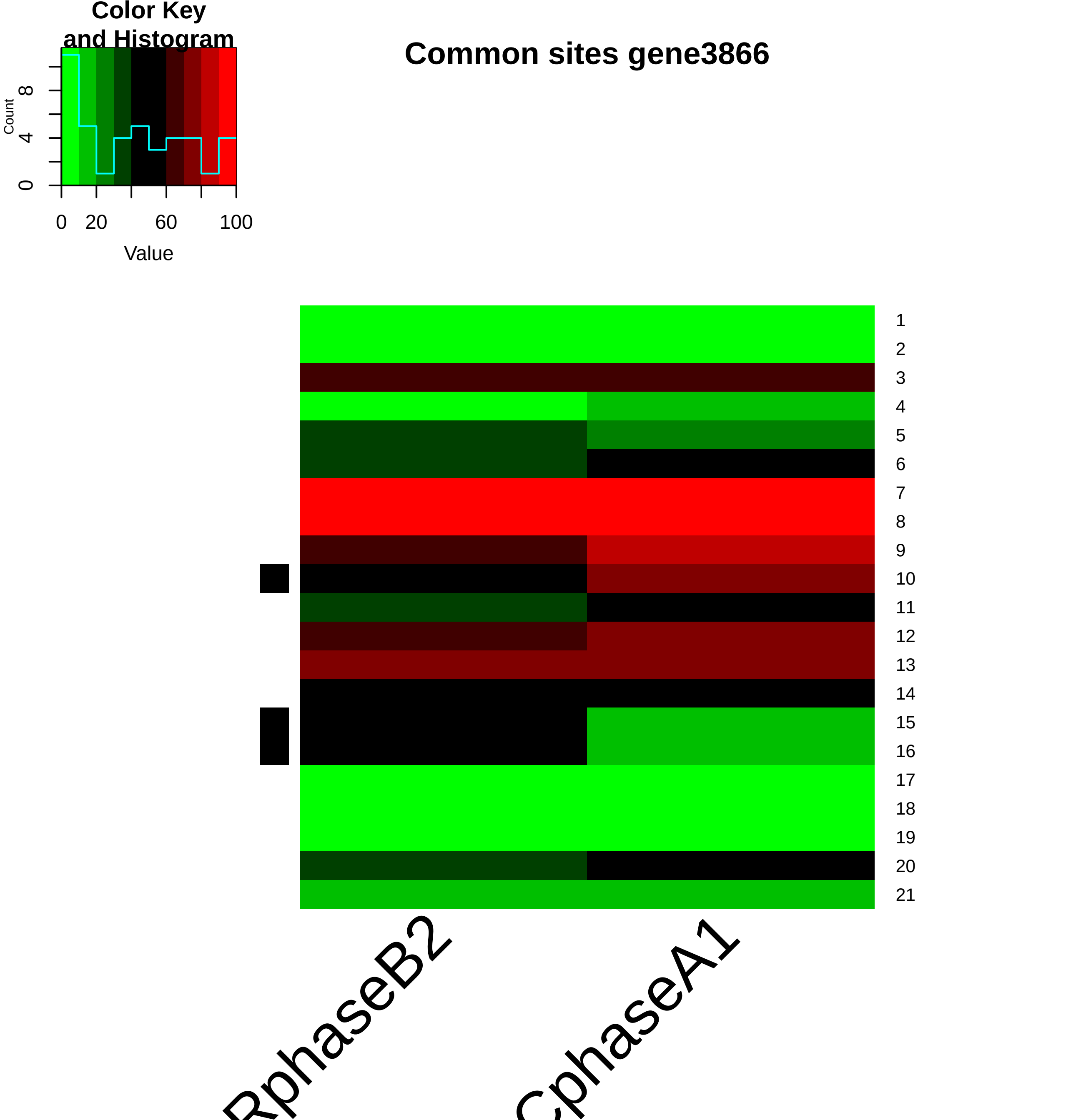

Color Key
and Histogram
Count
0
4
8
0
20
60
100
Value
Common sites gene3866
RphaseB2
CphaseA1

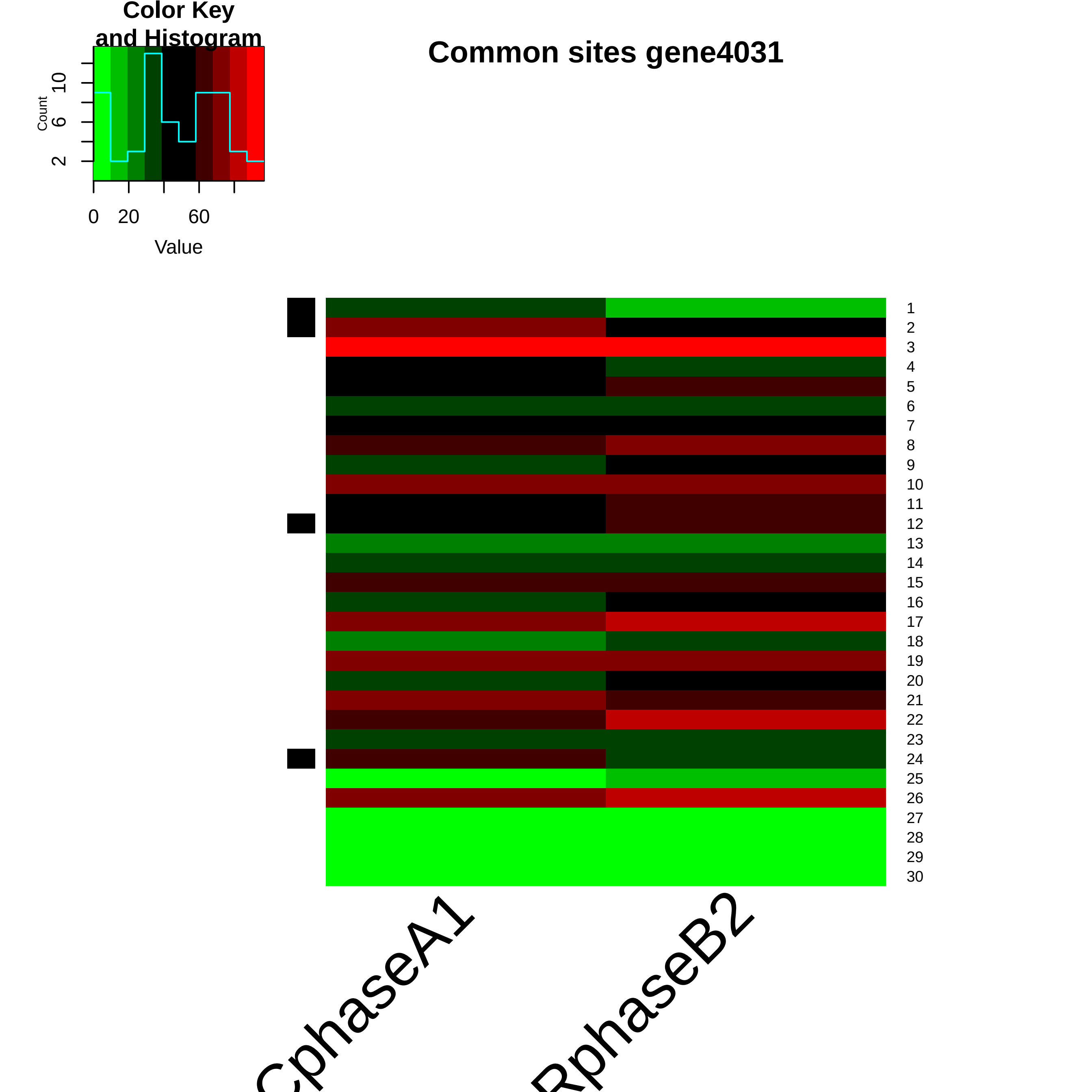

Color Key
and Histogram
Count
2
6
10
0
20
60
Value
Common sites gene4031
CphaseA1
RphaseB2

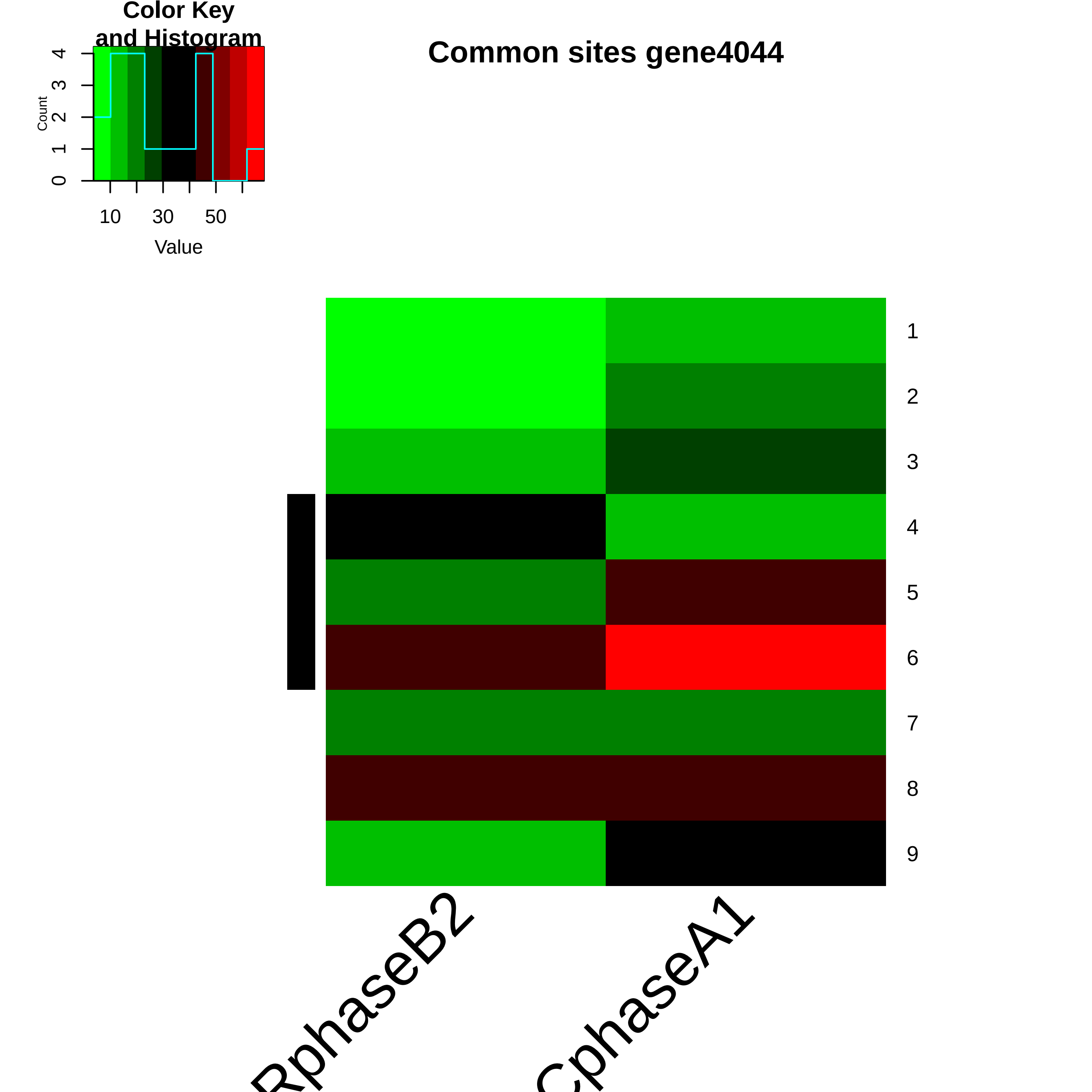
Color Key
and Histogram
Count
0
1
2
3
4
10
30
50
Value
Common sites gene4044
1
2
3
4
5
6
7
8
9
RphaseB2
CphaseA1

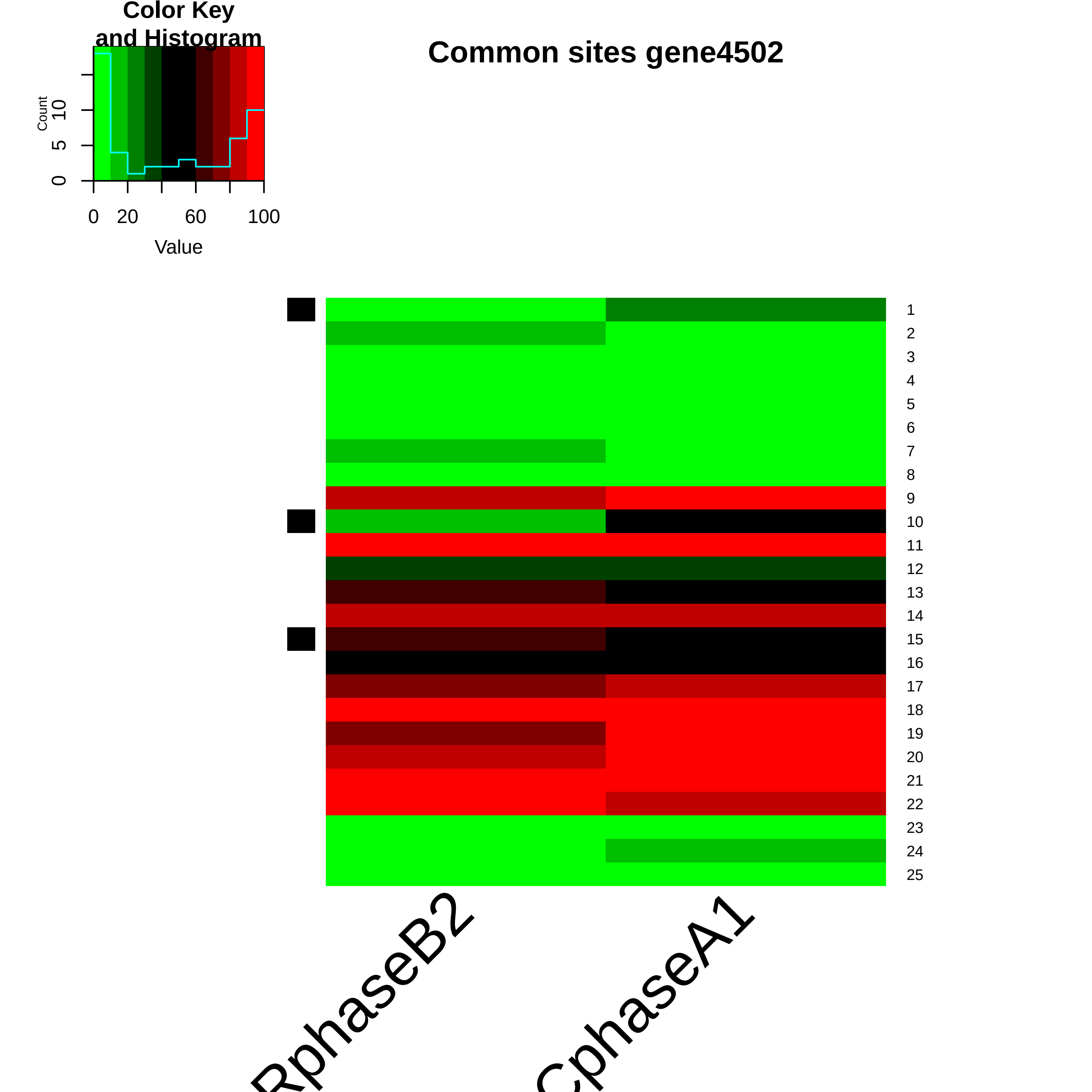

Color Key
and Histogram
Count
0
5
10
0
20
60
100
Value
Common sites gene4502
RphaseB2
CphaseA1

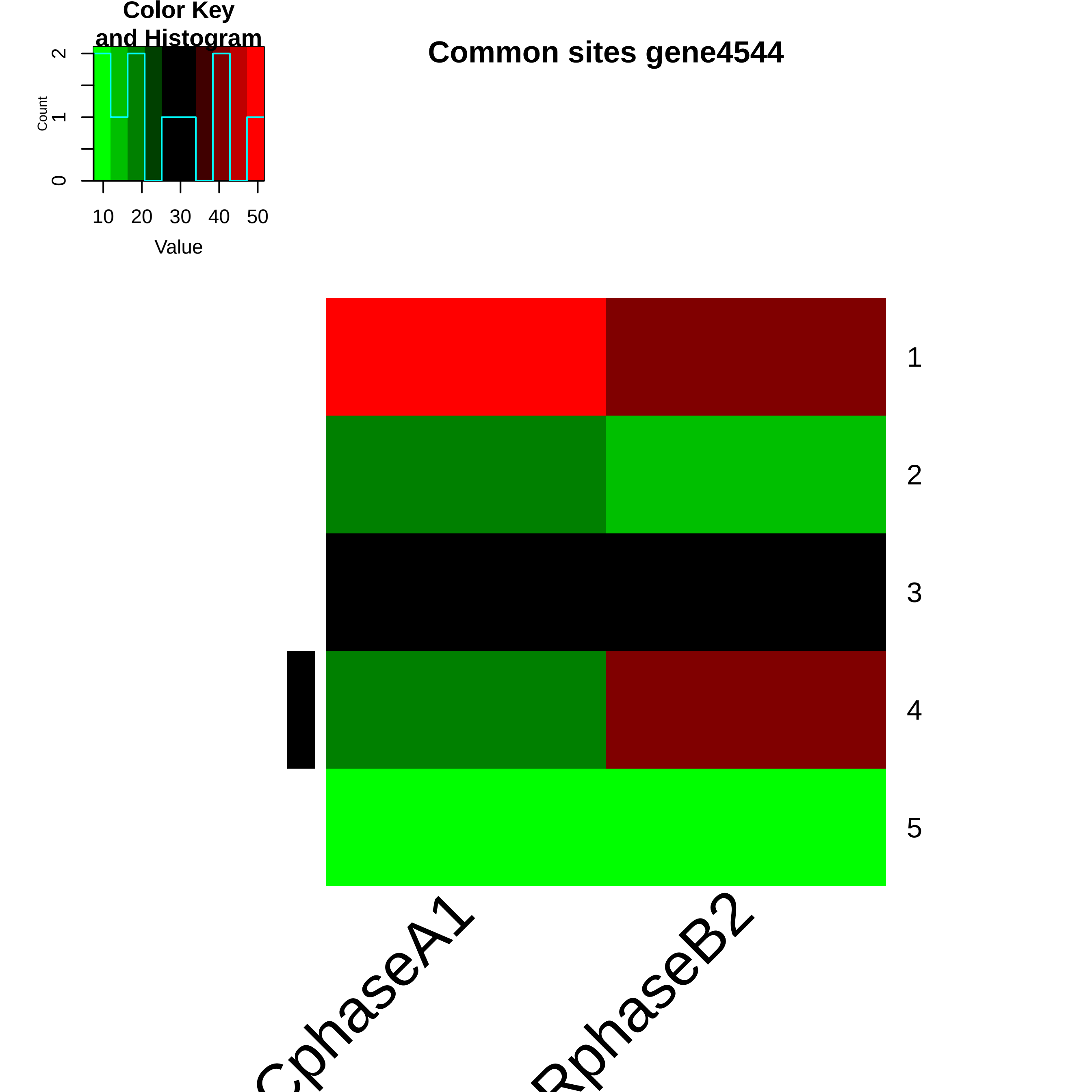

Color Key
and Histogram
Count
0
1
2
10
20
30
40
50
Value
Common sites gene4544
1
2
3
4
5
CphaseA1
RphaseB2

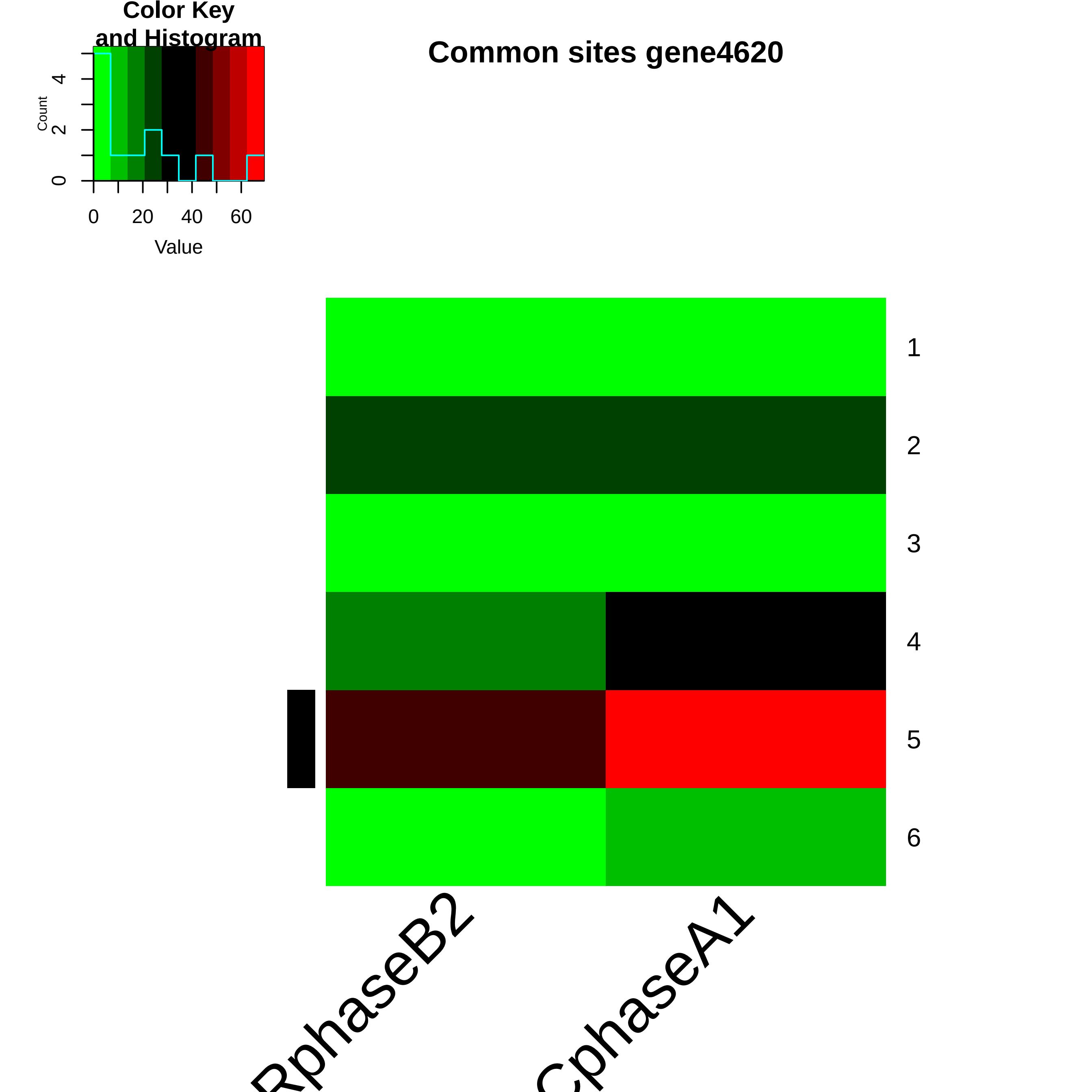
Color Key
and Histogram
Count
0
2
4
0
20
40
60
Value
Common sites gene4620
1
2
3
4
5
6

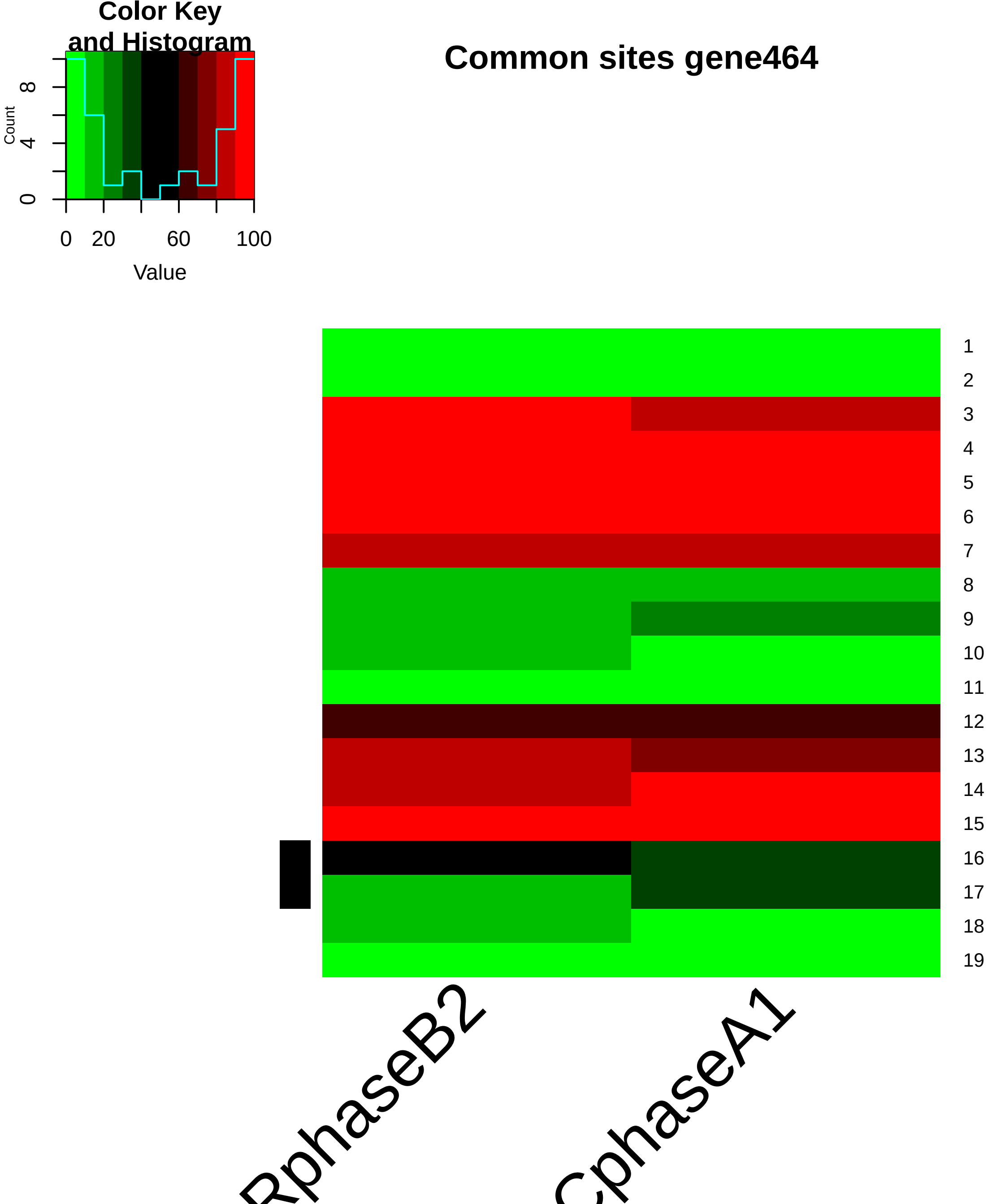

Color Key
and Histogram
Count
0
4
8
0
20
60
100
Value
Common sites gene464
RphaseB2
CphaseA1

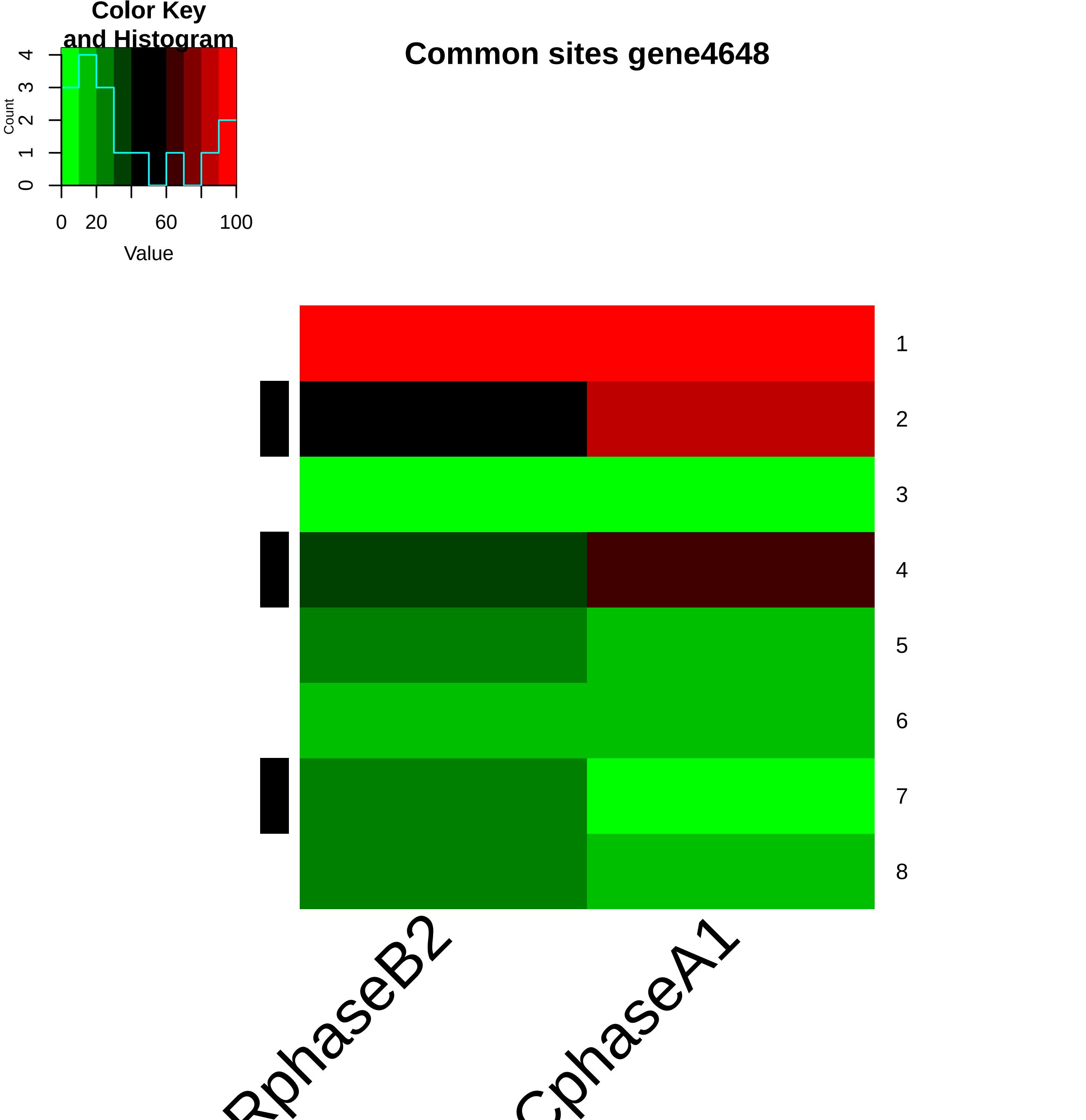
Color Key
and Histogram
Count
0
1
2
3
4
0
20
60
100
Value
Common sites gene4648
1
2
3
4
5
6
7
8
RphaseB2
CphaseA1

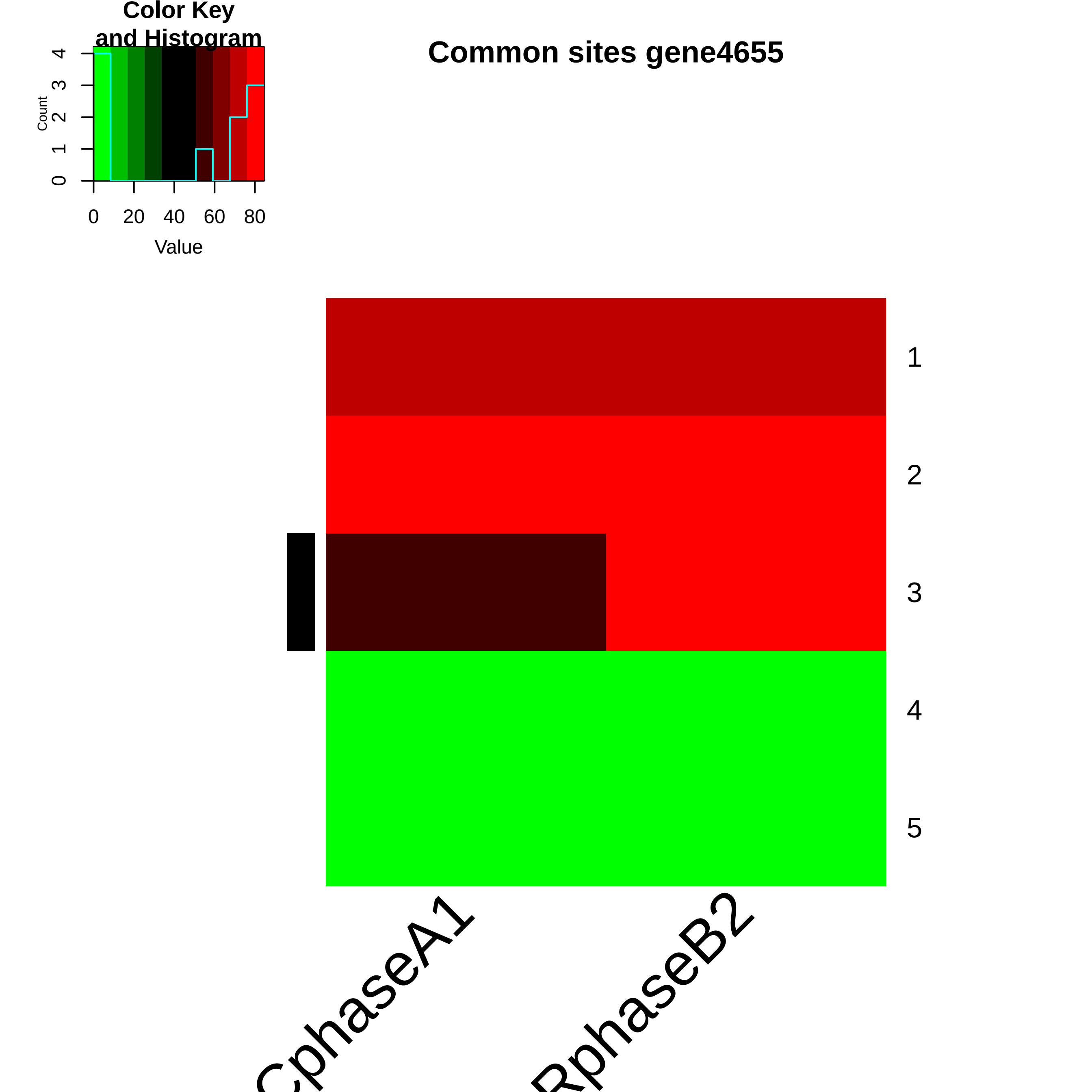

Color Key
and Histogram
Count
0
1
2
3
4
0
20
40
60
80
Value
Common sites gene4655
1
2
3
4
5
CphaseA1
RphaseB2

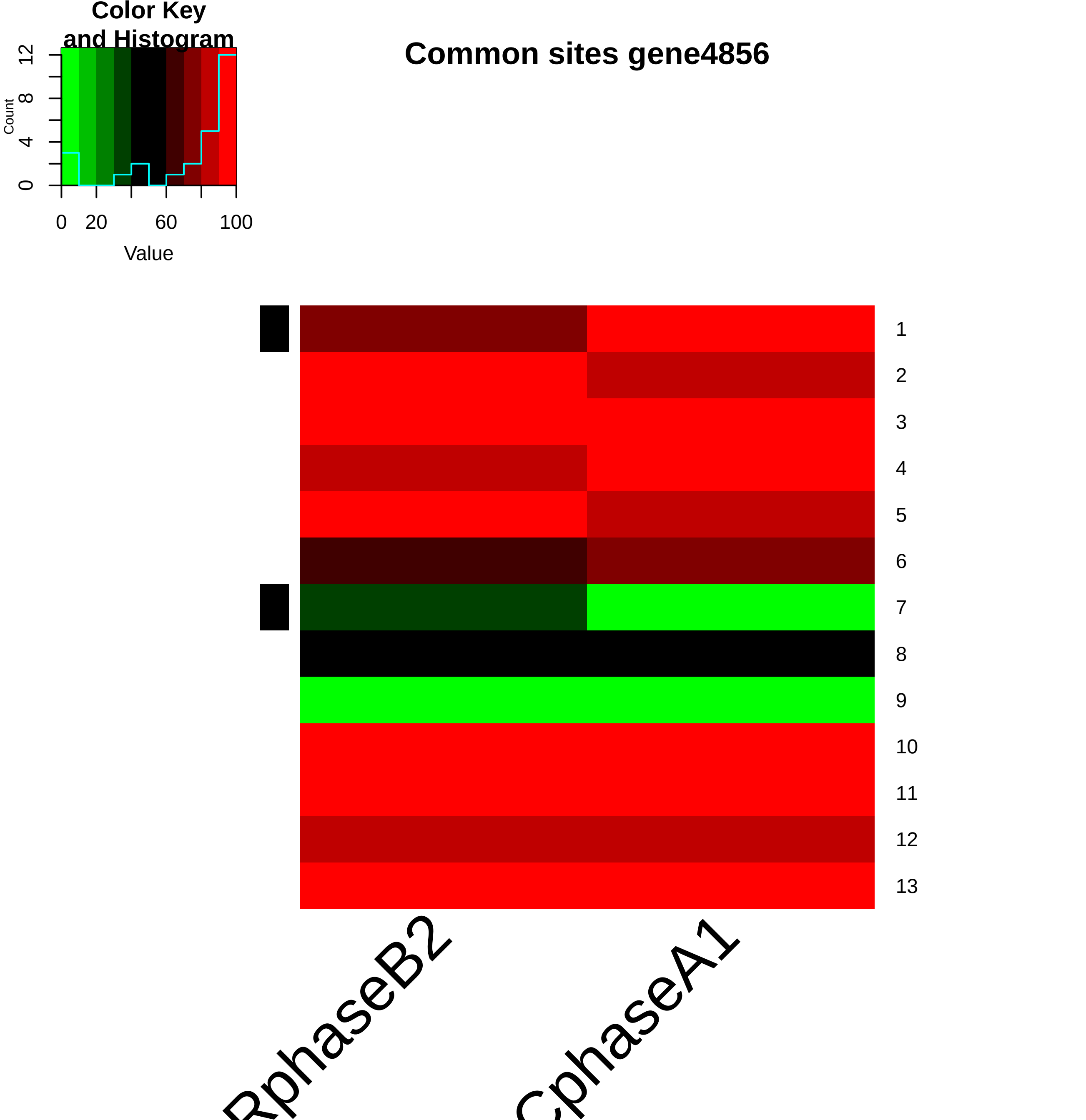
Color Key
and Histogram
Count
0
4
8
12
0
20
60
100
Value
Common sites gene4856
1
2
3
4
5
6
7
8
9
10
11
12
13
RphaseB2
CphaseA1

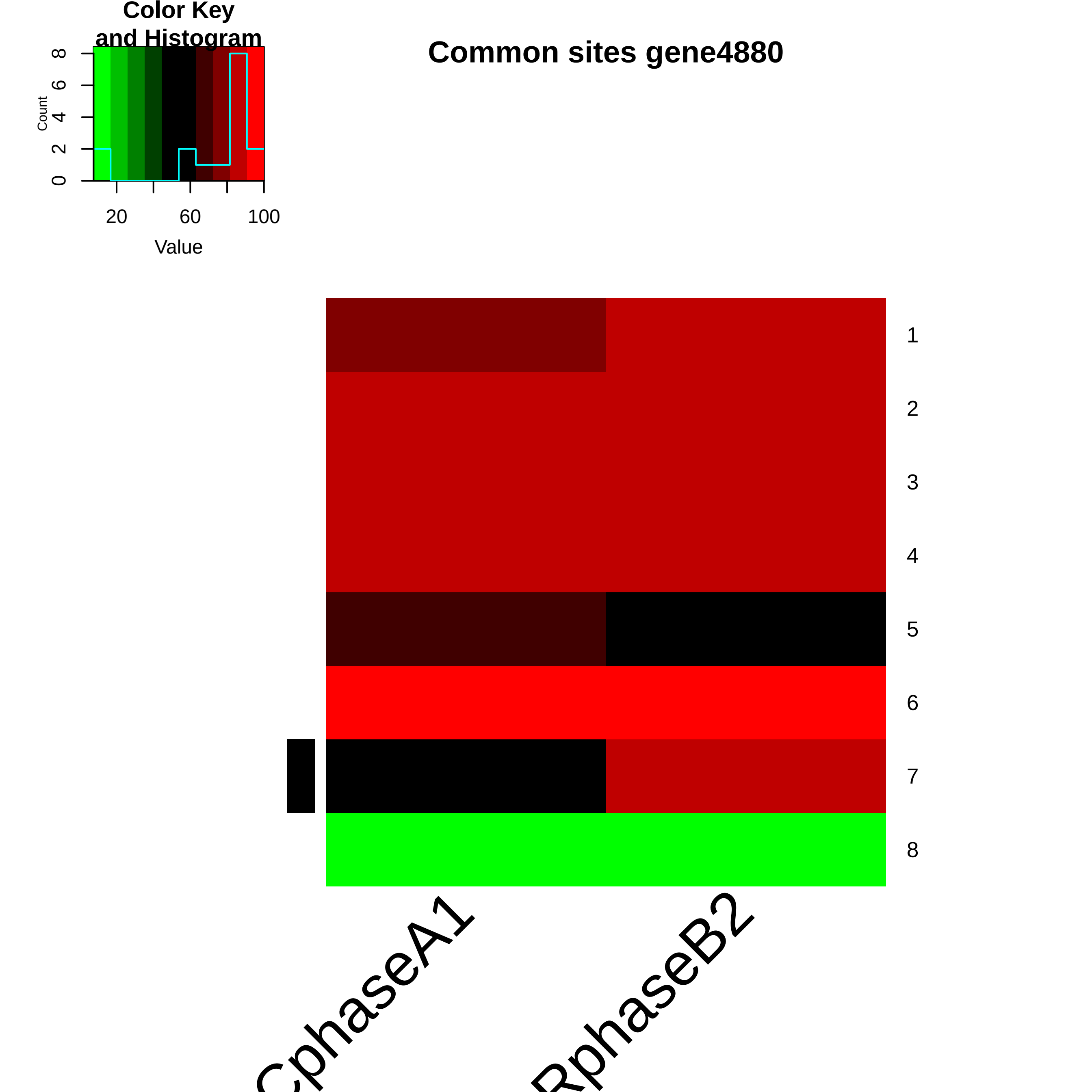
Color Key
and Histogram
Count
0
2
4
6
8
20
60
100
Value
Common sites gene4880
1
2
3
4
5
6
7
8
CphaseA1
RphaseB2

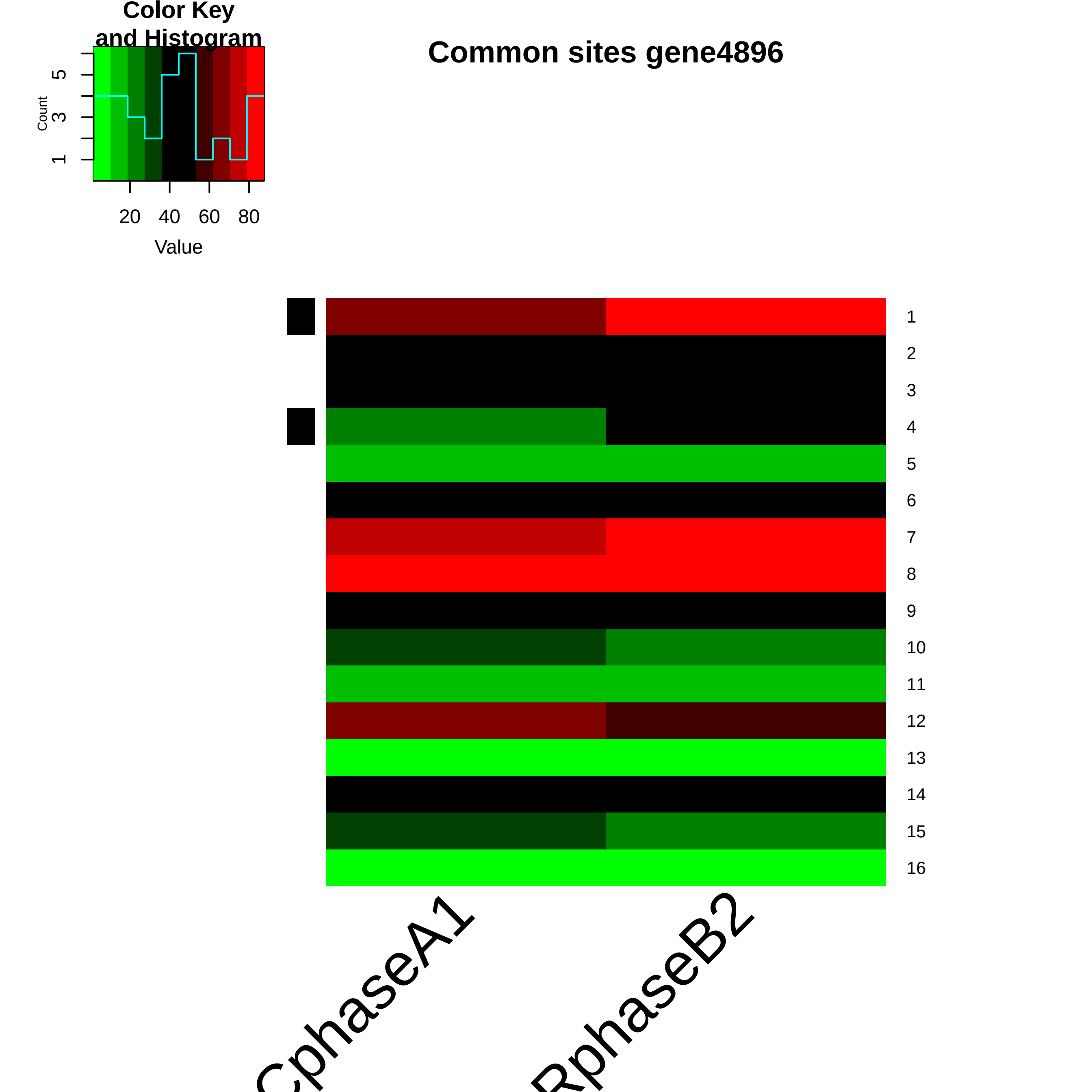

Color Key
and Histogram
Count
1
3
5
20
40
60
80
Value
Common sites gene4896
CphaseA1
RphaseB2

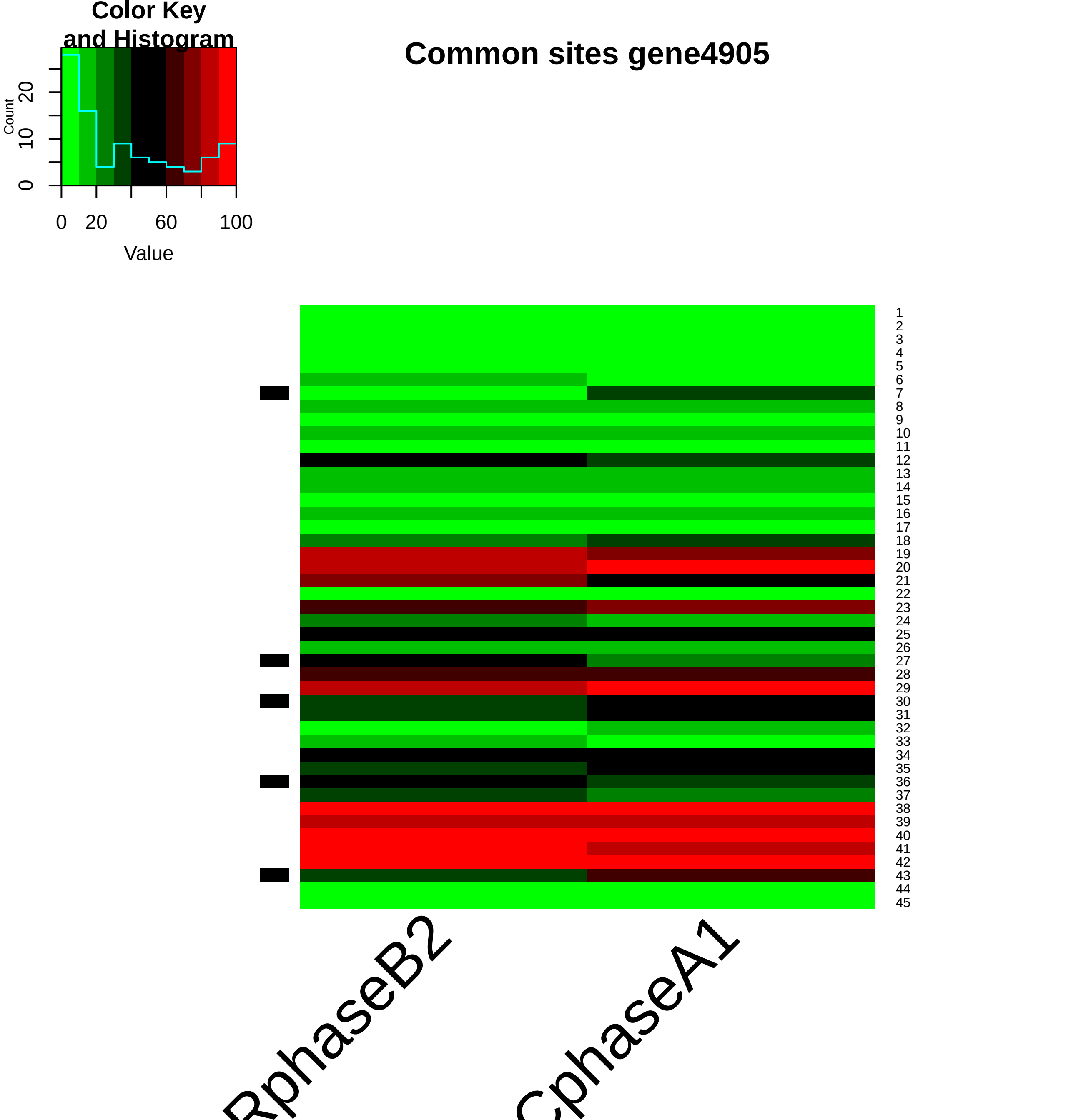

Color Key
and Histogram
Count
0
10
20
0
20
60
100
Value
Common sites gene4905
RphaseB2
CphaseA1

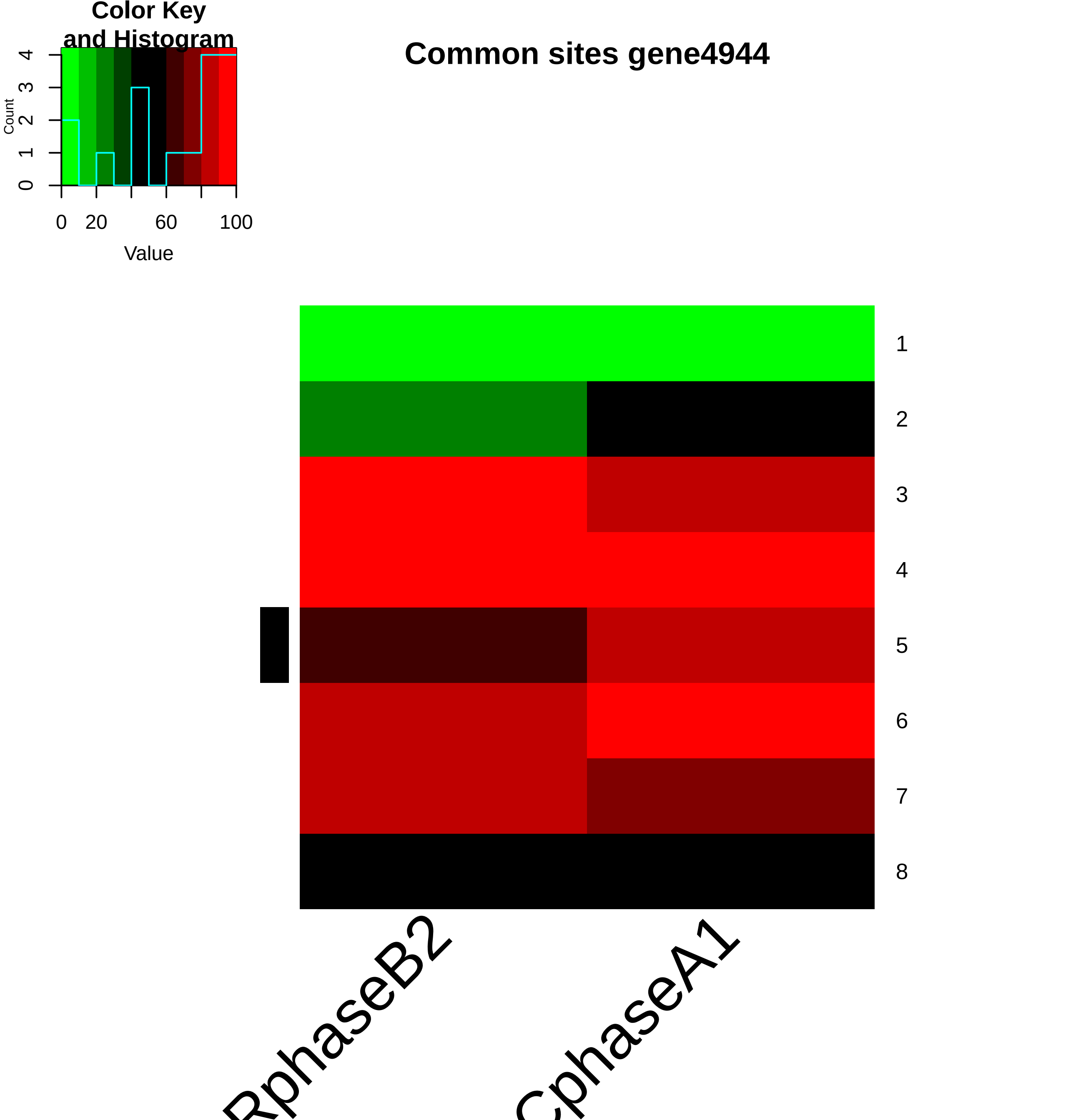
Color Key
and Histogram
Count
0
1
2
3
4
0
20
60
100
Value
Common sites gene4944
1
2
3
4
5
6
7
8
RphaseB2
CphaseA1

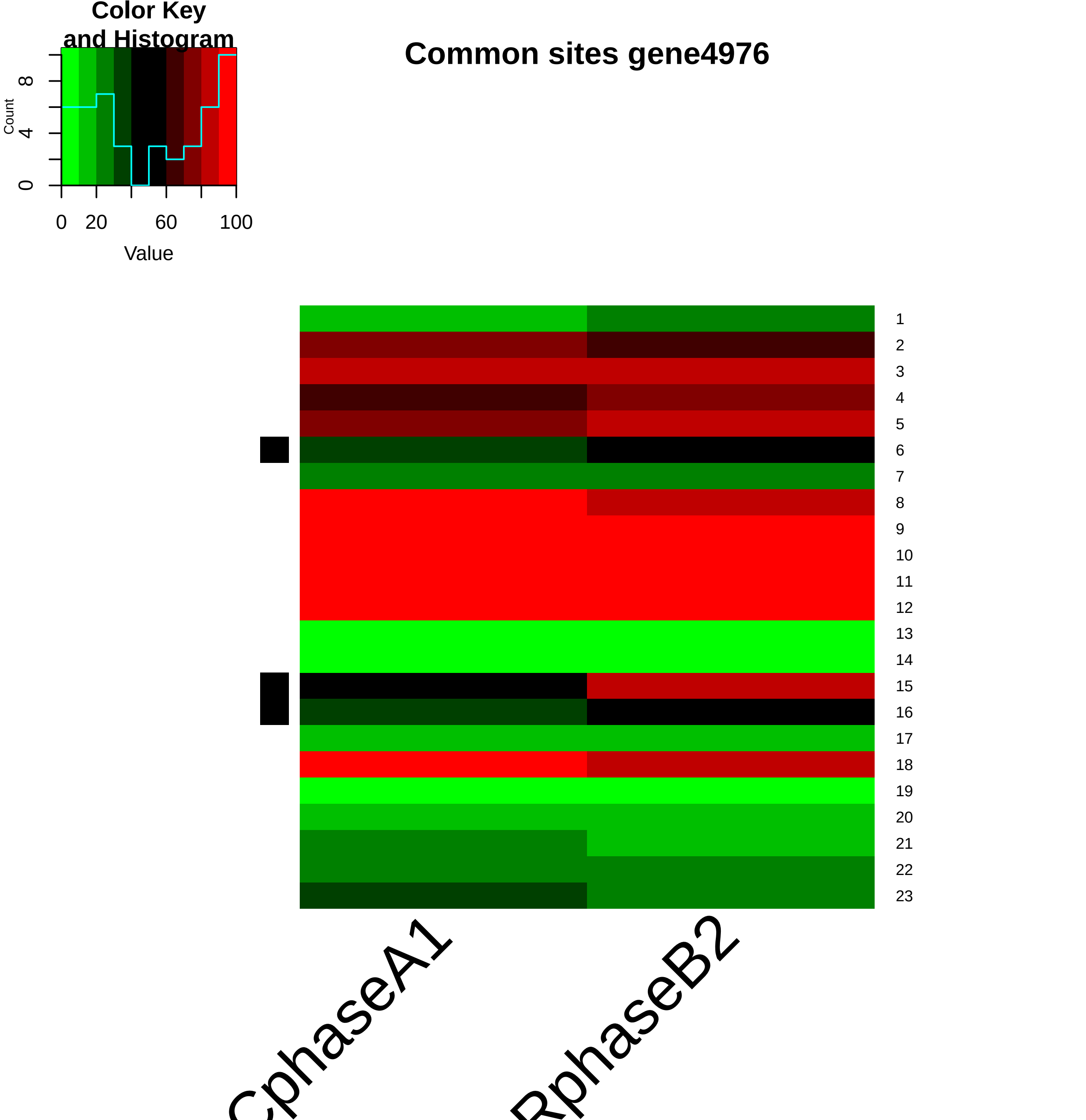

Color Key
and Histogram
Count
0
4
8
0
20
60
100
Value
Common sites gene4976
CphaseA1
RphaseB2

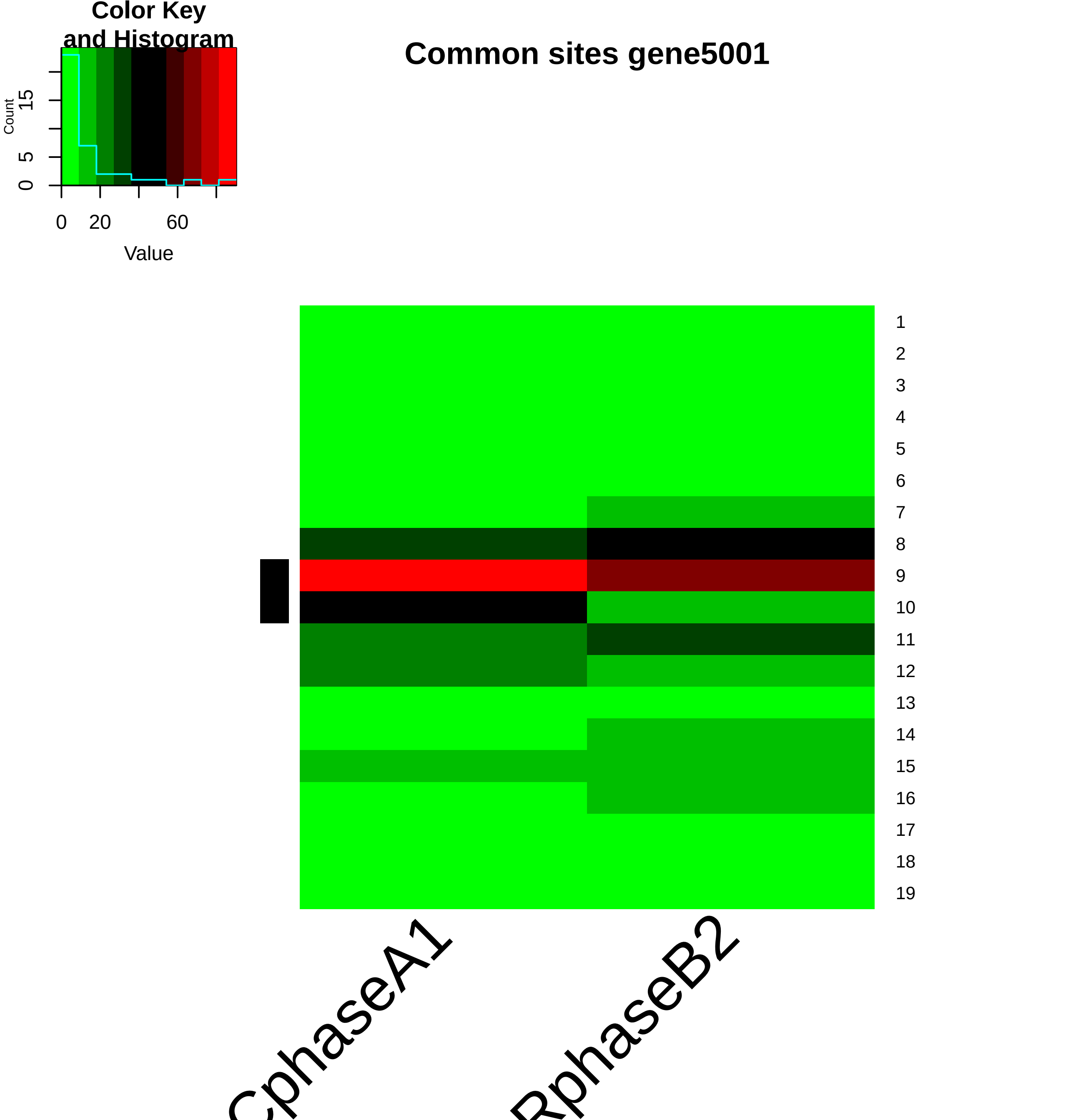
Color Key
and Histogram
Count
0
5
15
0
20
60
Value
Common sites gene5001
CphaseA1
RphaseB2

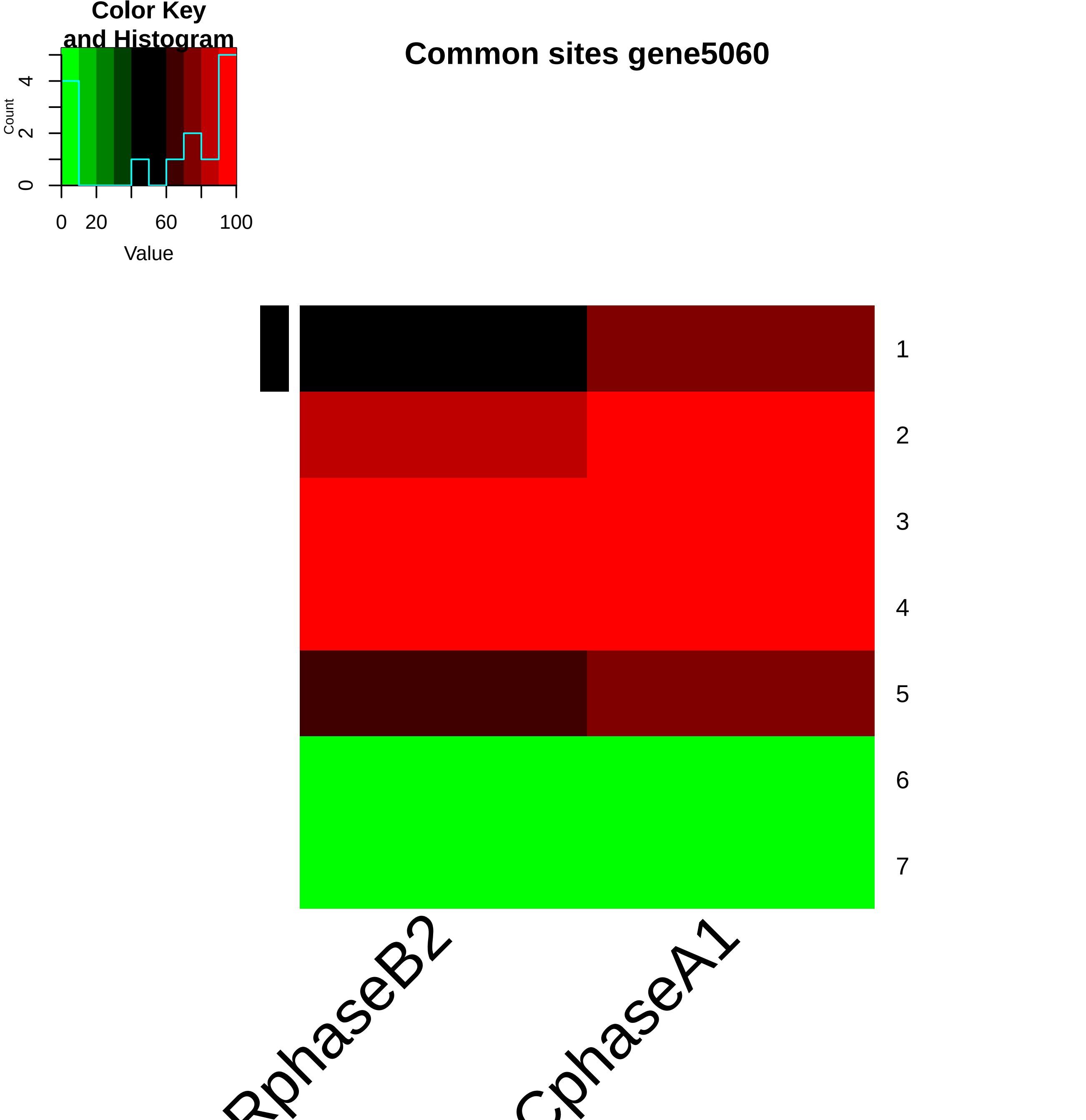
Color Key
and Histogram
Count
0
2
4
0
20
60
100
Value
Common sites gene5060
1
2
3
4
5
6
7
RphaseB2
CphaseA1

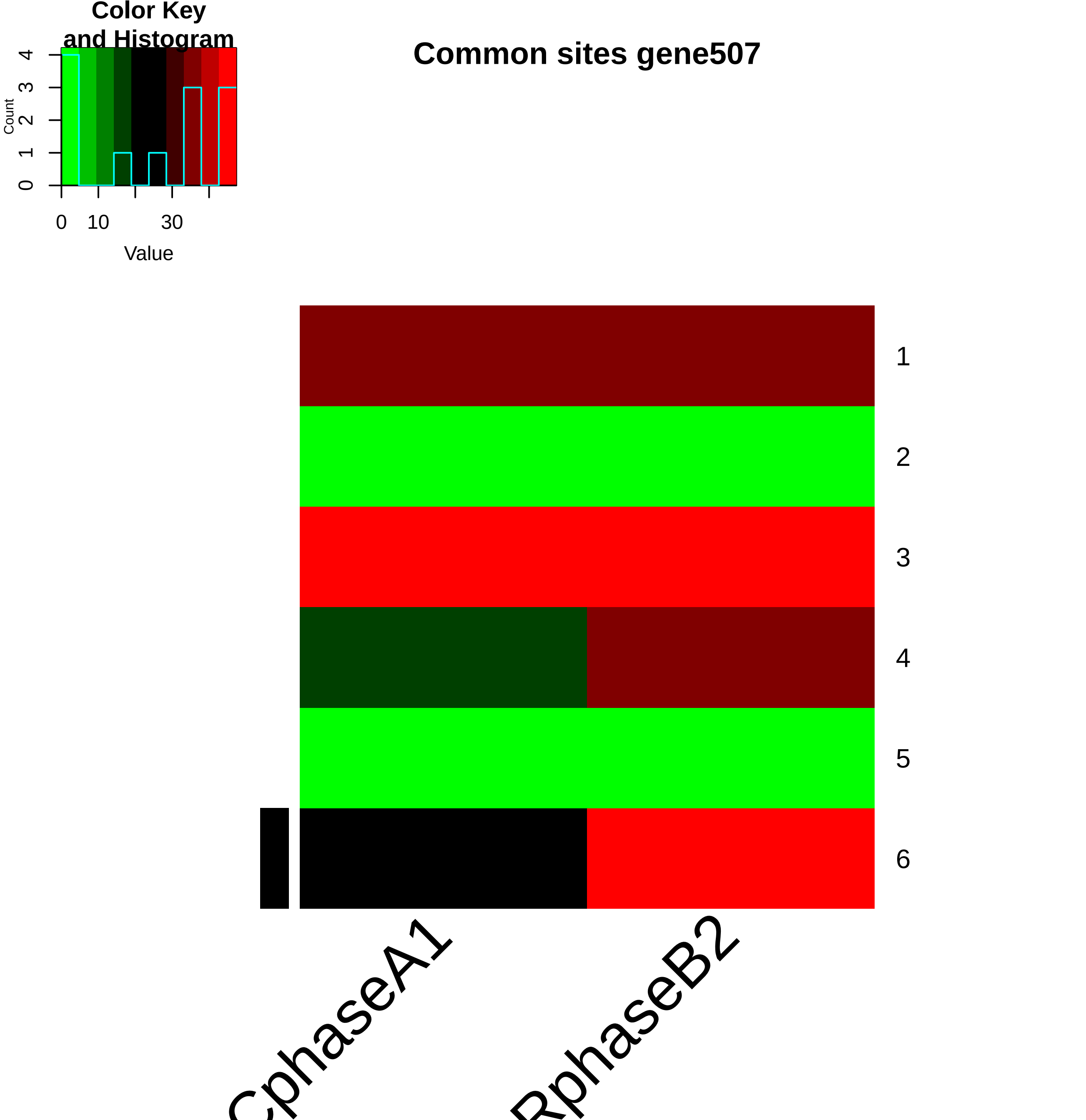
Color Key
and Histogram
Count
0
1
2
3
4
0
10
30
Value
Common sites gene507
1
2
3
4
5
6
CphaseA1
RphaseB2

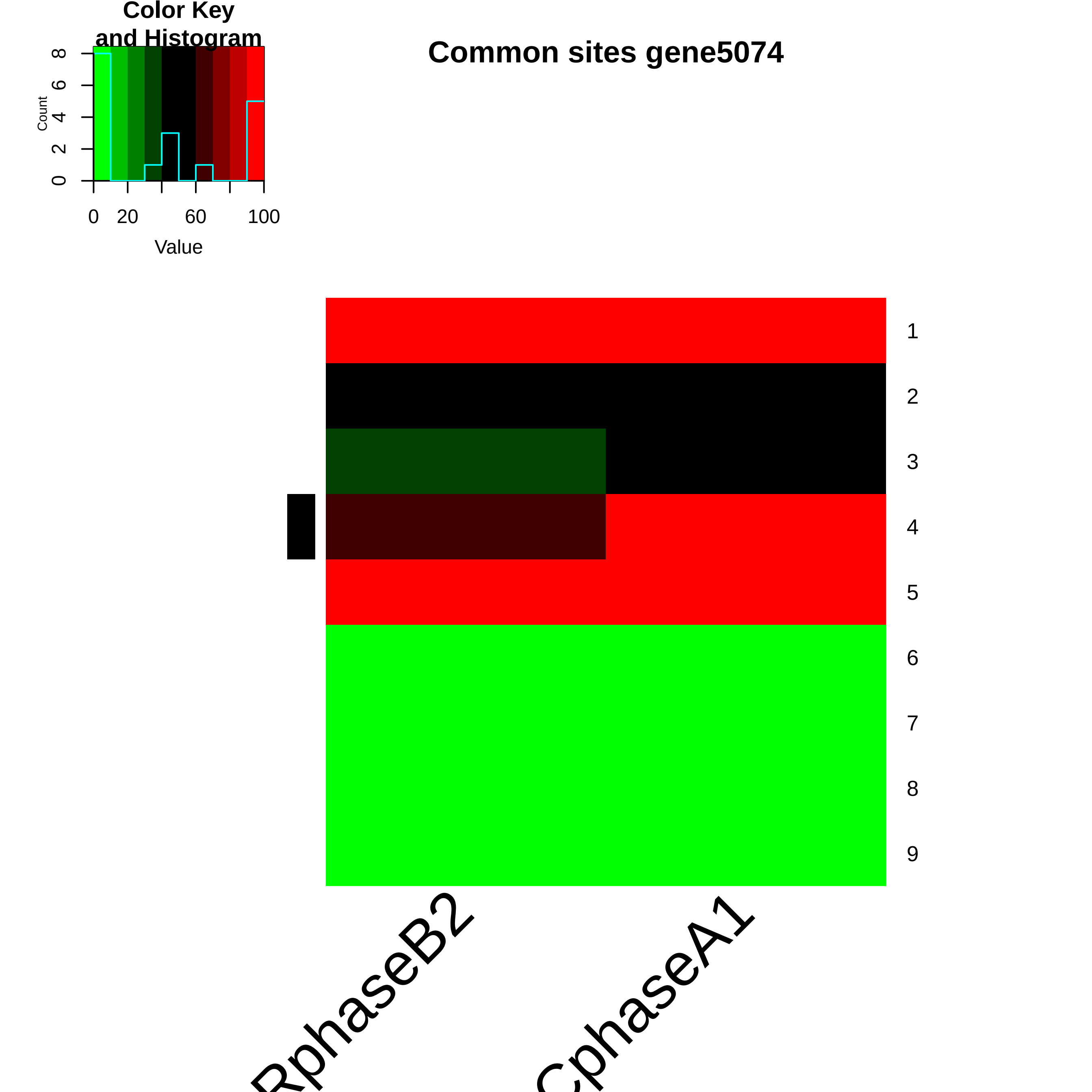
Color Key
and Histogram
Count
0
2
4
6
8
0
20
60
100
Value
Common sites gene5074
1
2
3
4
5
6
7
8
9
RphaseB2
CphaseA1

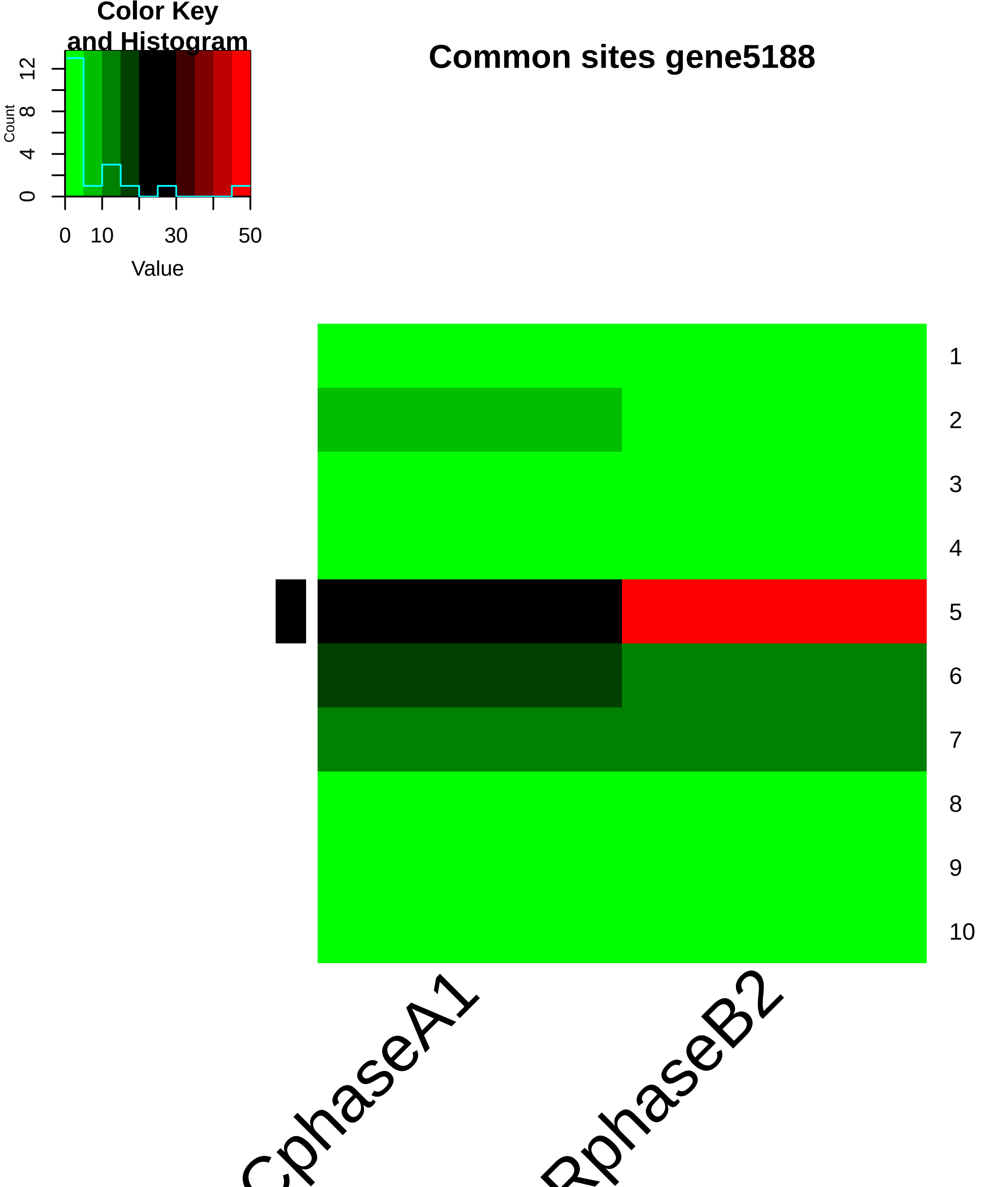

Color Key
and Histogram
Count
0
4
8
12
0
10
30
50
Value
Common sites gene5188
1
2
3
4
5
6
7
8
9
10

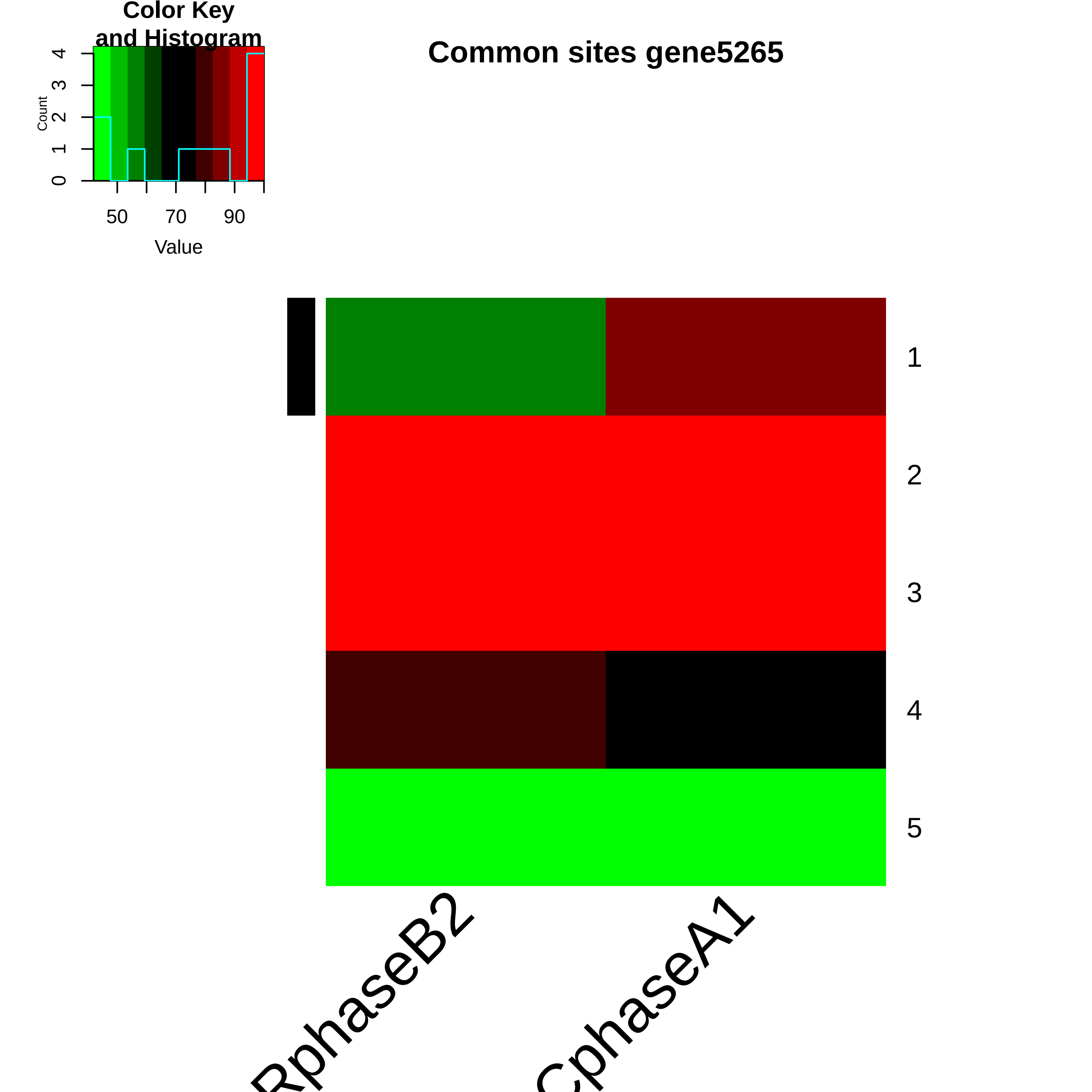

Color Key
and Histogram
Count
0
1
2
3
4
50
70
90
Value
Common sites gene5265
1
2
3
4
5
RphaseB2
CphaseA1

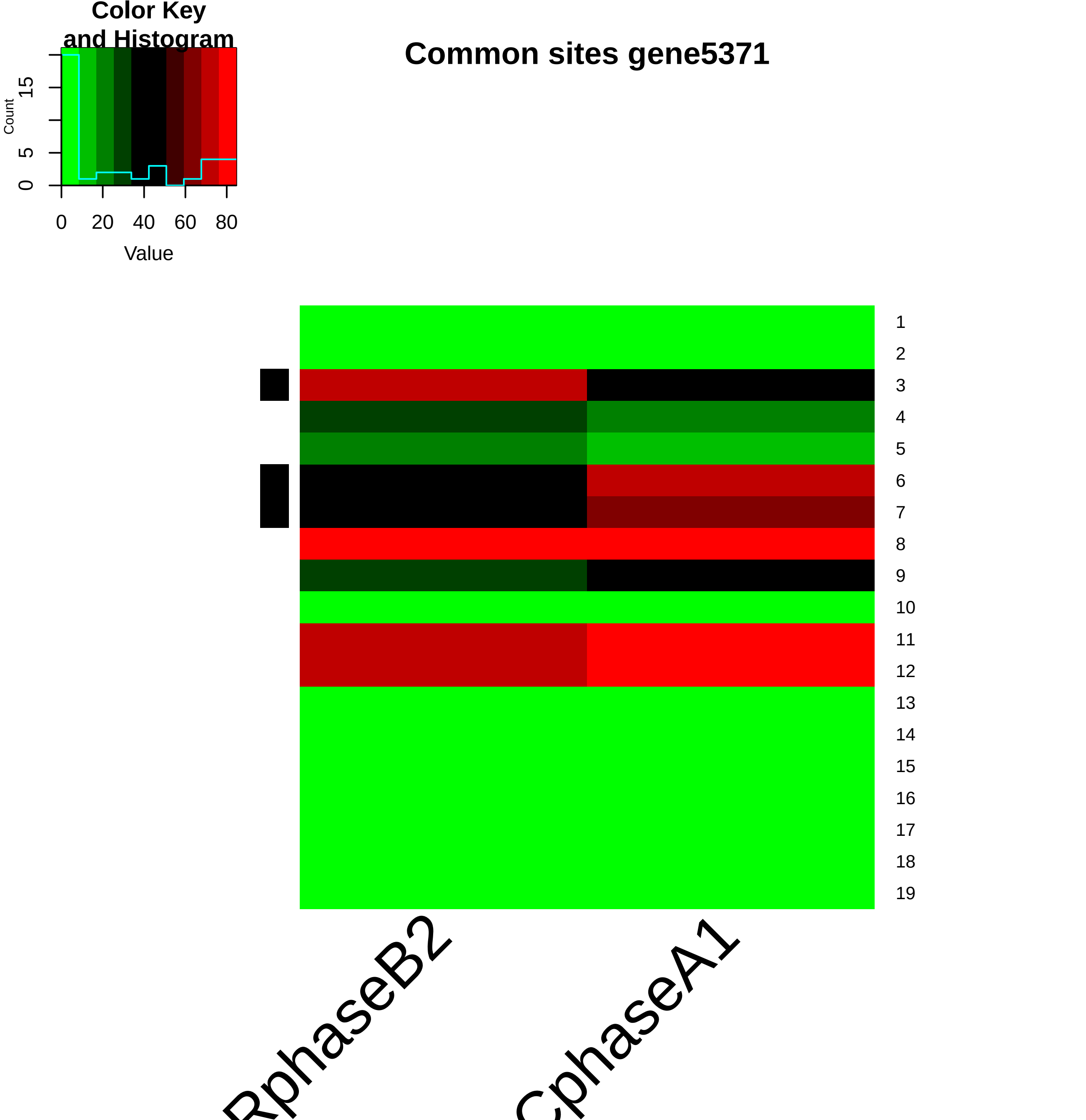

Color Key
and Histogram
Count
0
5
15
0
20
40
60
80
Value
Common sites gene5371
RphaseB2
CphaseA1

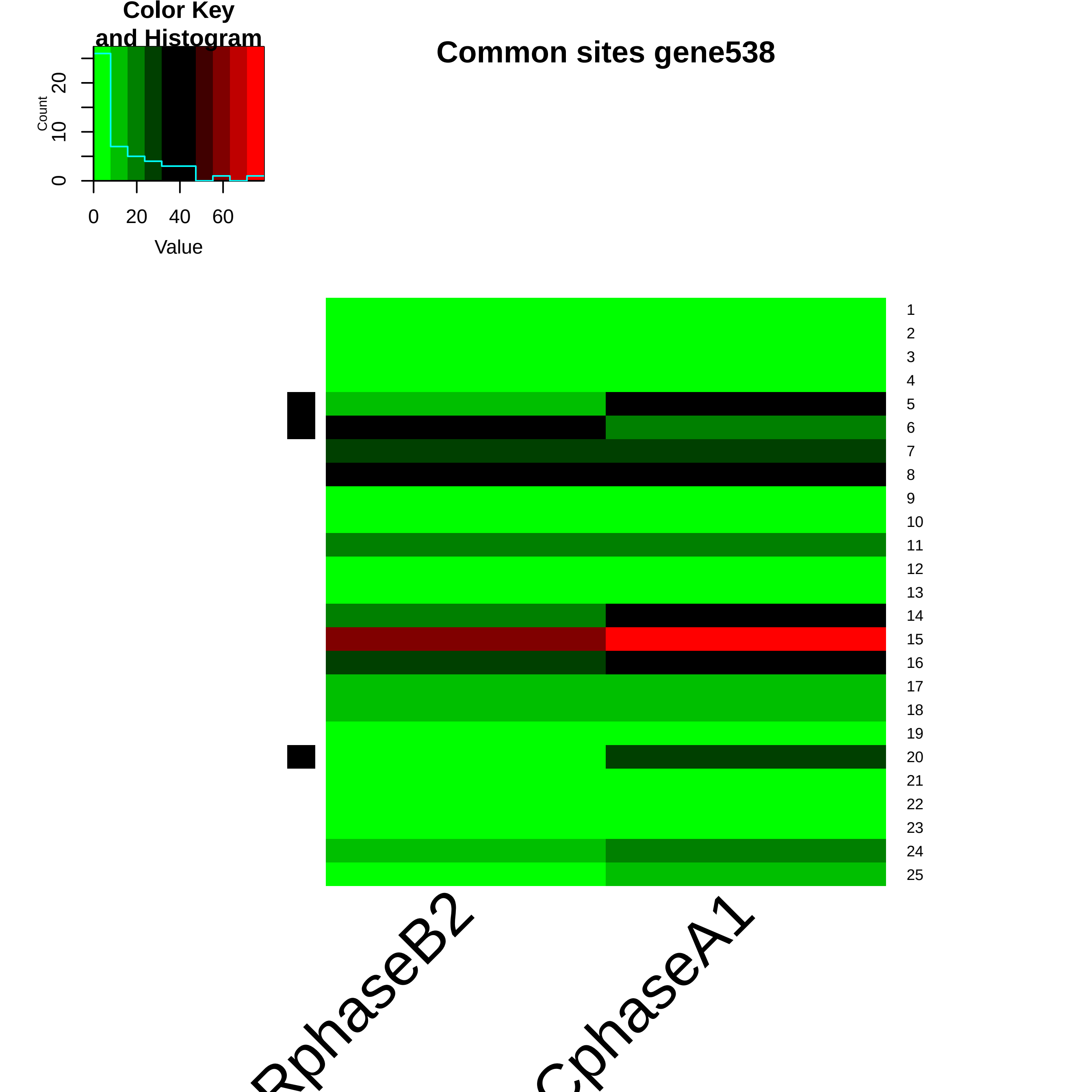
Color Key
and Histogram
Count
0
10
20
0
20
40
60
Value
Common sites gene538
RphaseB2
CphaseA1

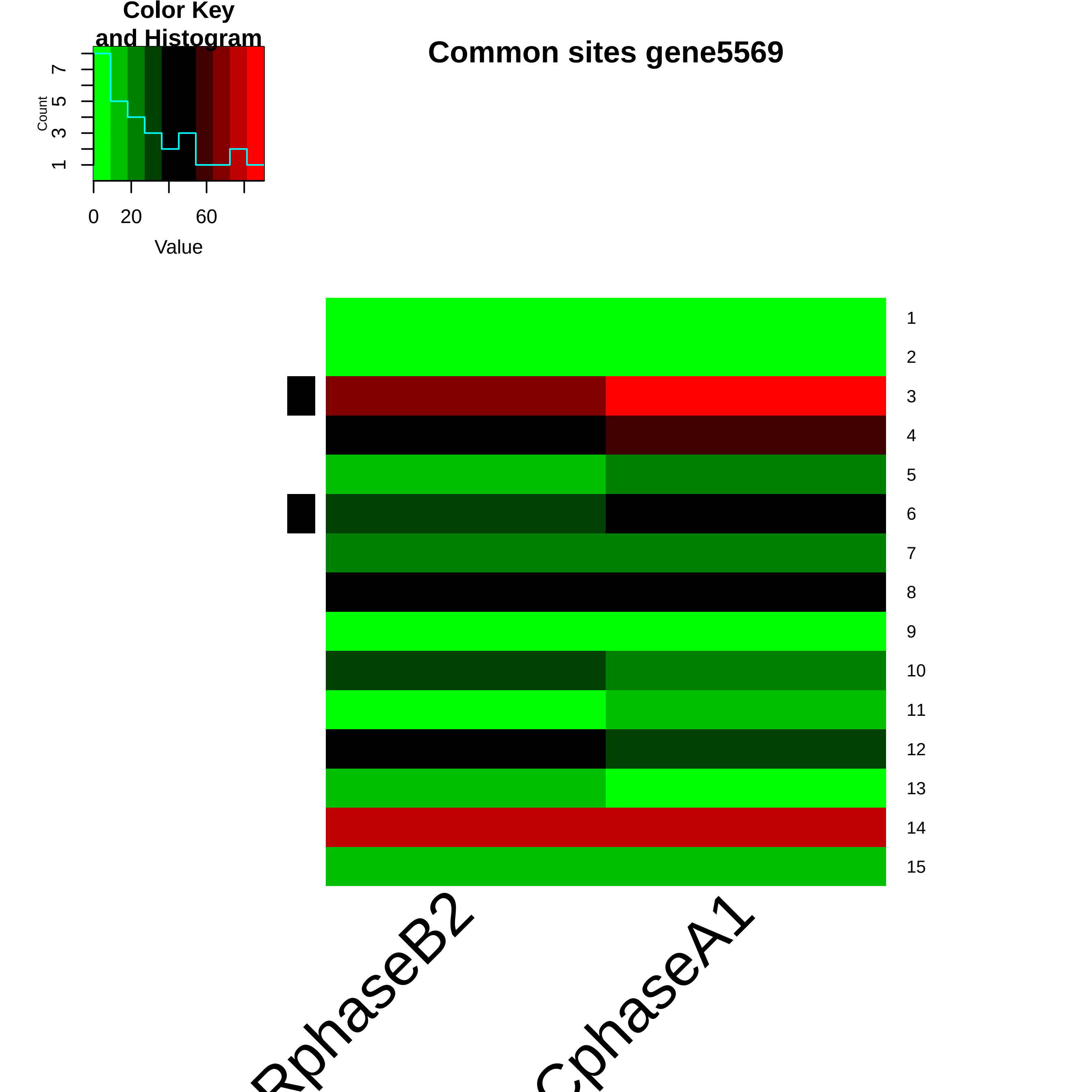
Color Key
and Histogram
Count
1
3
5
7
0
20
60
Value
Common sites gene5569
RphaseB2
CphaseA1

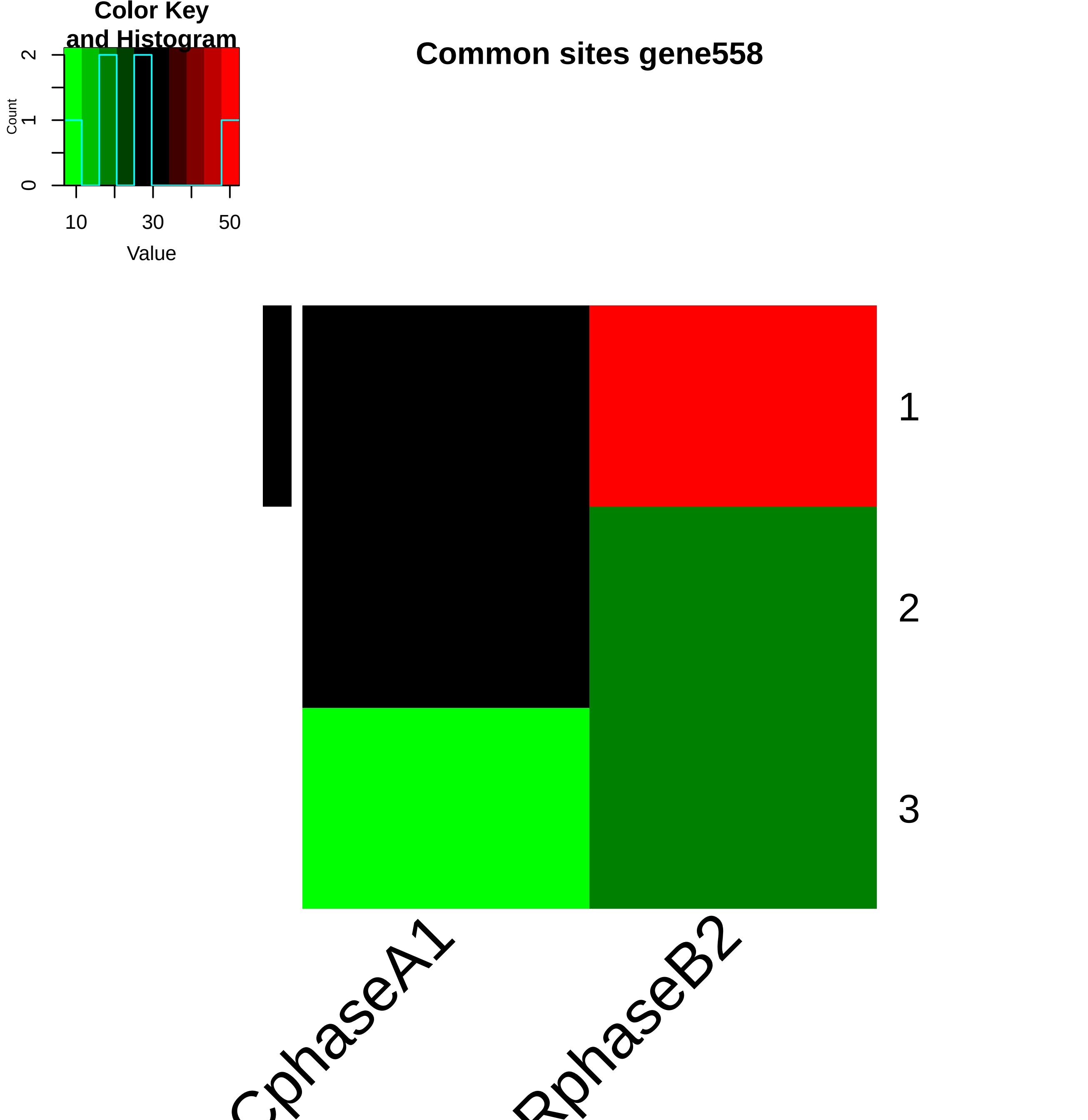
Color Key
and Histogram
Count
0
1
2
10
30
50
Value
Common sites gene558
1
2
3
CphaseA1
RphaseB2

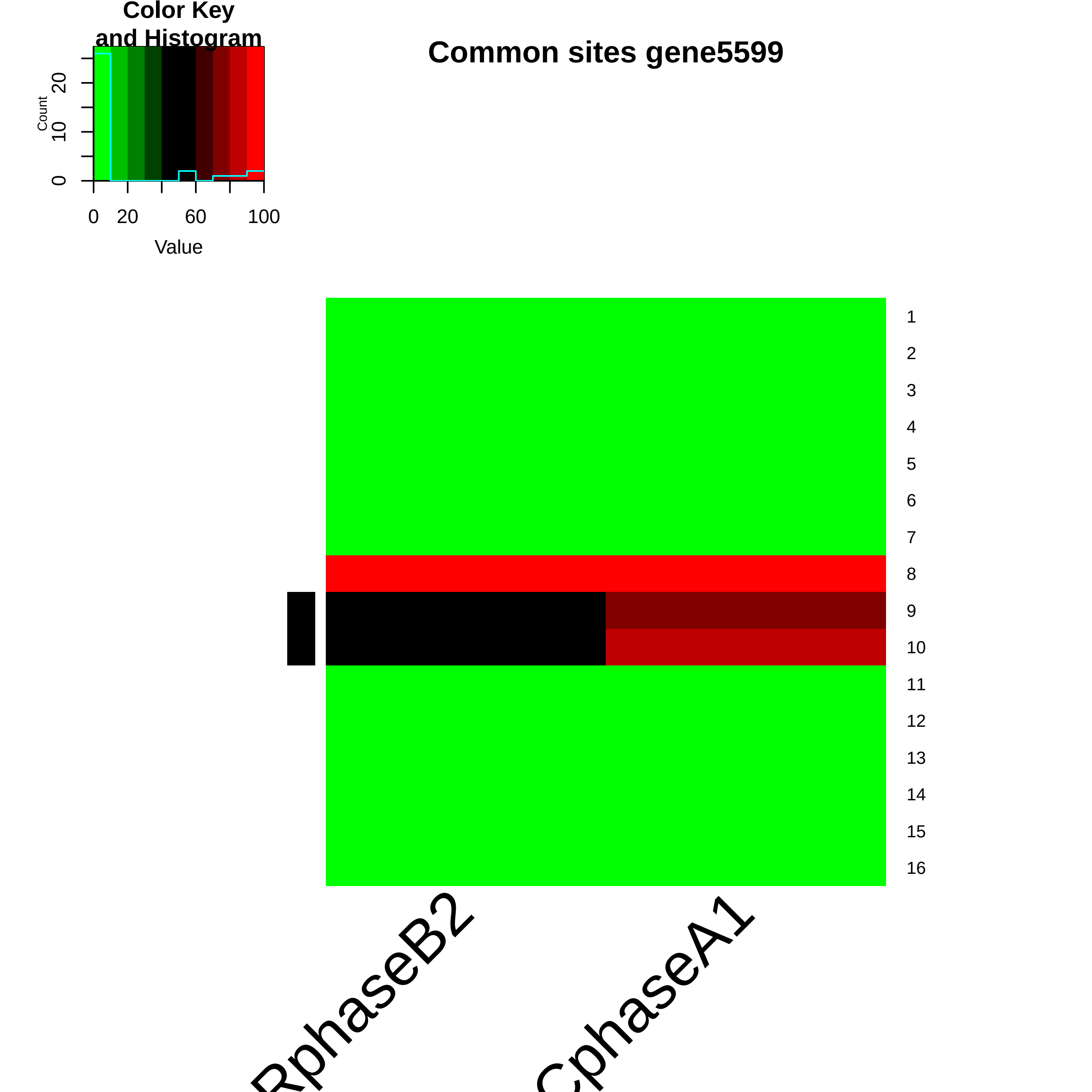

Color Key
and Histogram
Count
0
10
20
0
20
60
100
Value
Common sites gene5599
RphaseB2
CphaseA1

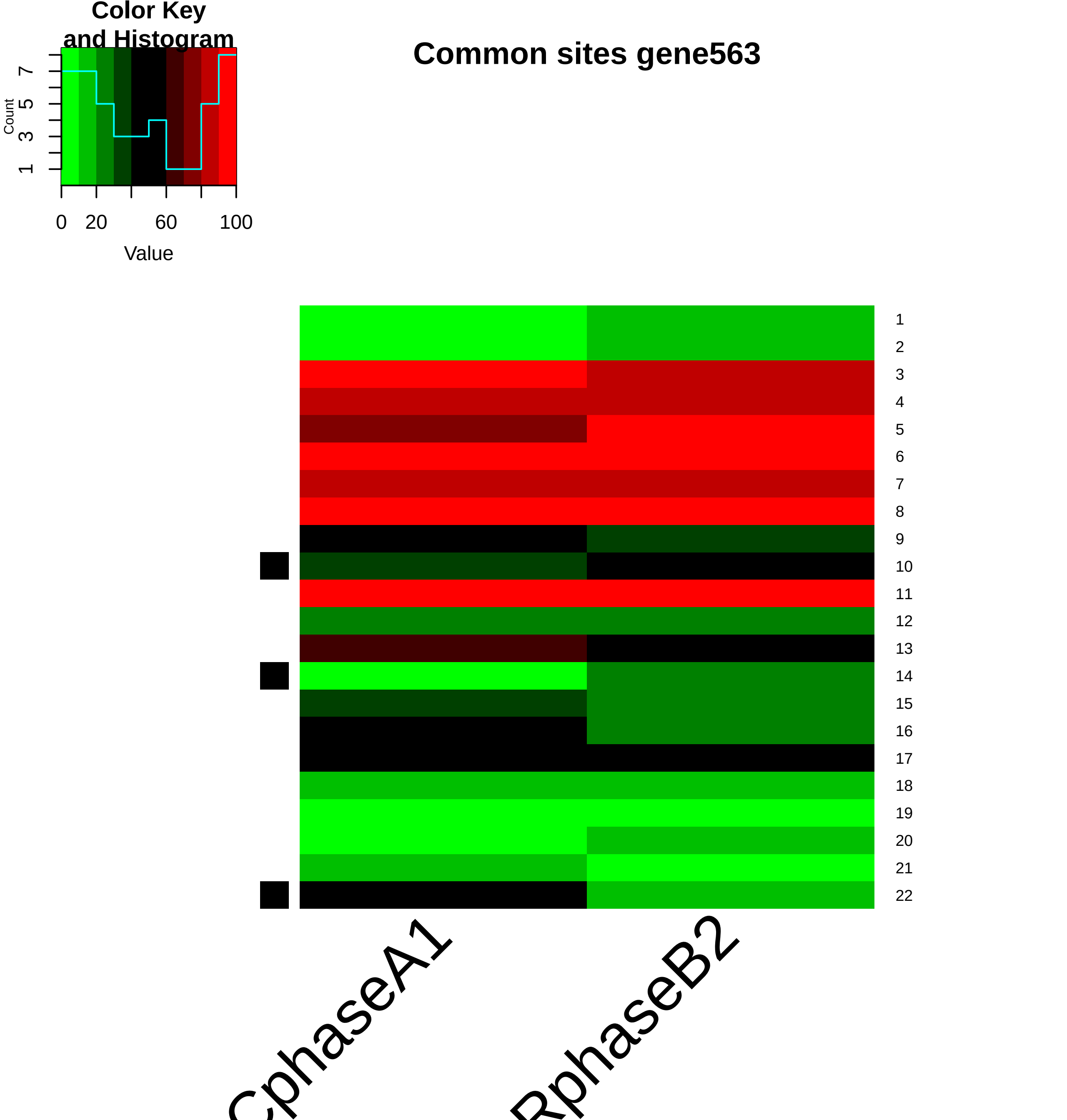

Color Key
and Histogram
Count
1
3
5
7
0
20
60
100
Value
Common sites gene563
CphaseA1
RphaseB2

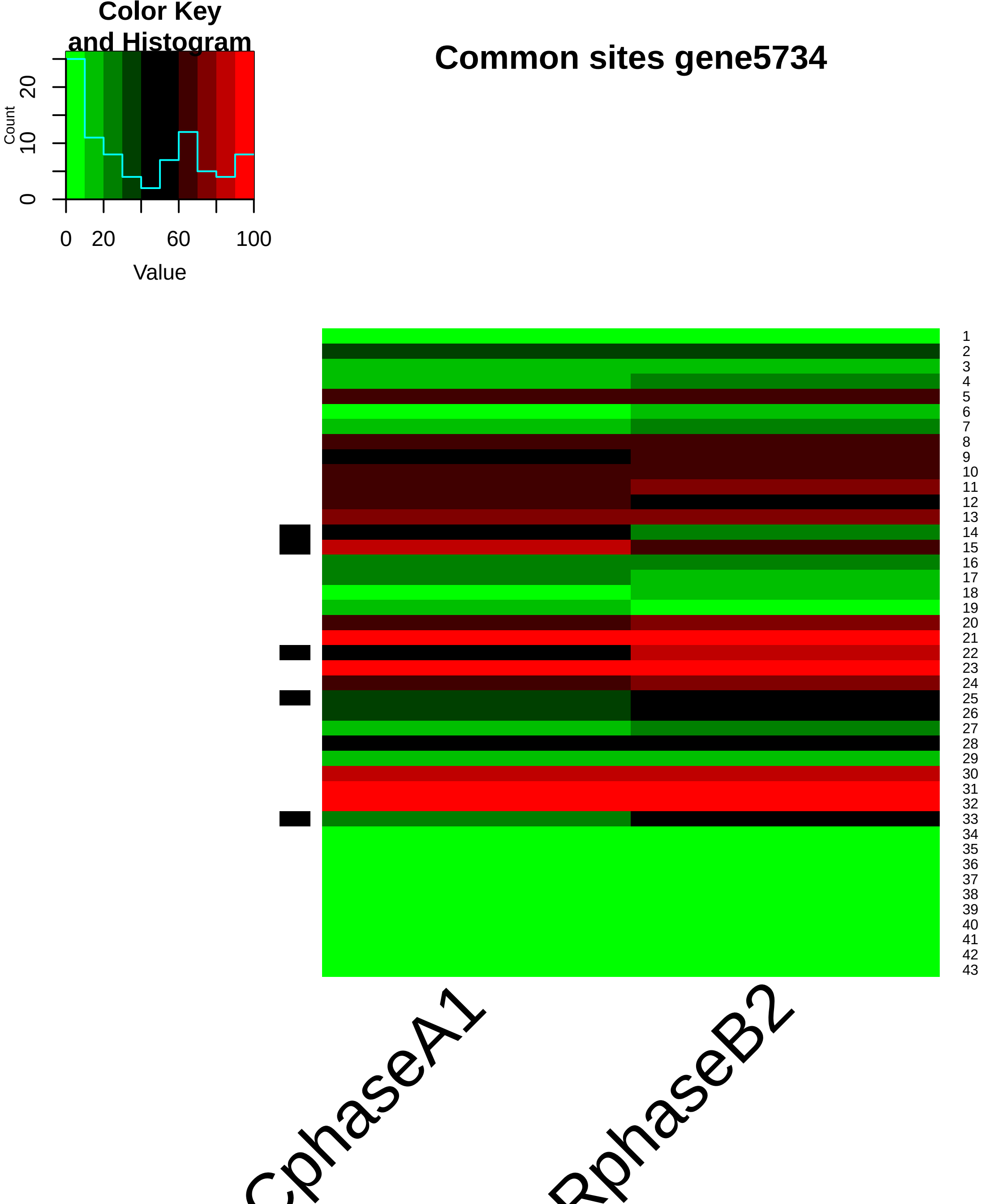

Color Key
and Histogram
Count
0
10
20
0
20
60
100
Value
Common sites gene5734
CphaseA1
RphaseB2

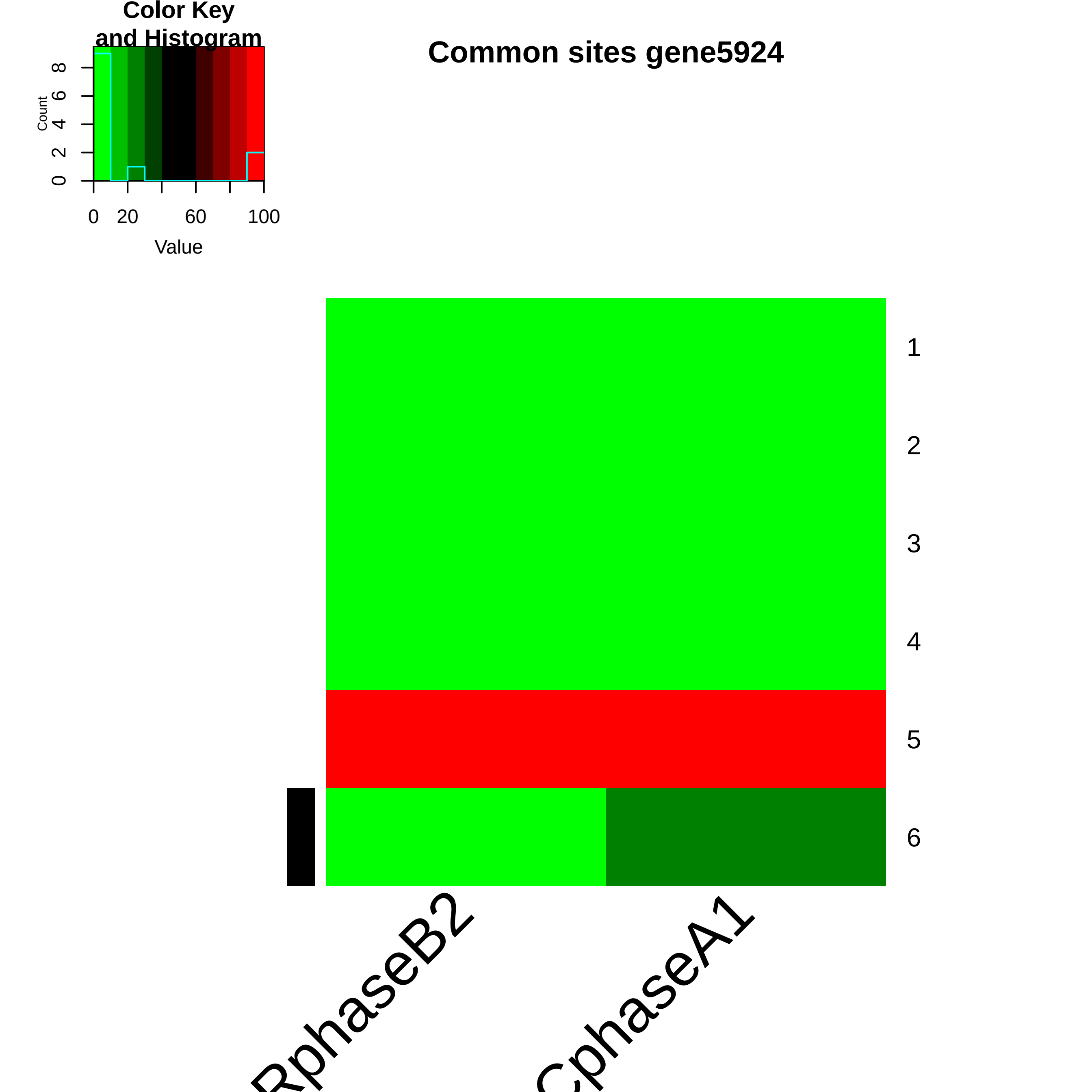

Color Key
and Histogram
Count
0
2
4
6
8
0
20
60
100
Value
Common sites gene5924
1
2
3
4
5
6
RphaseB2
CphaseA1

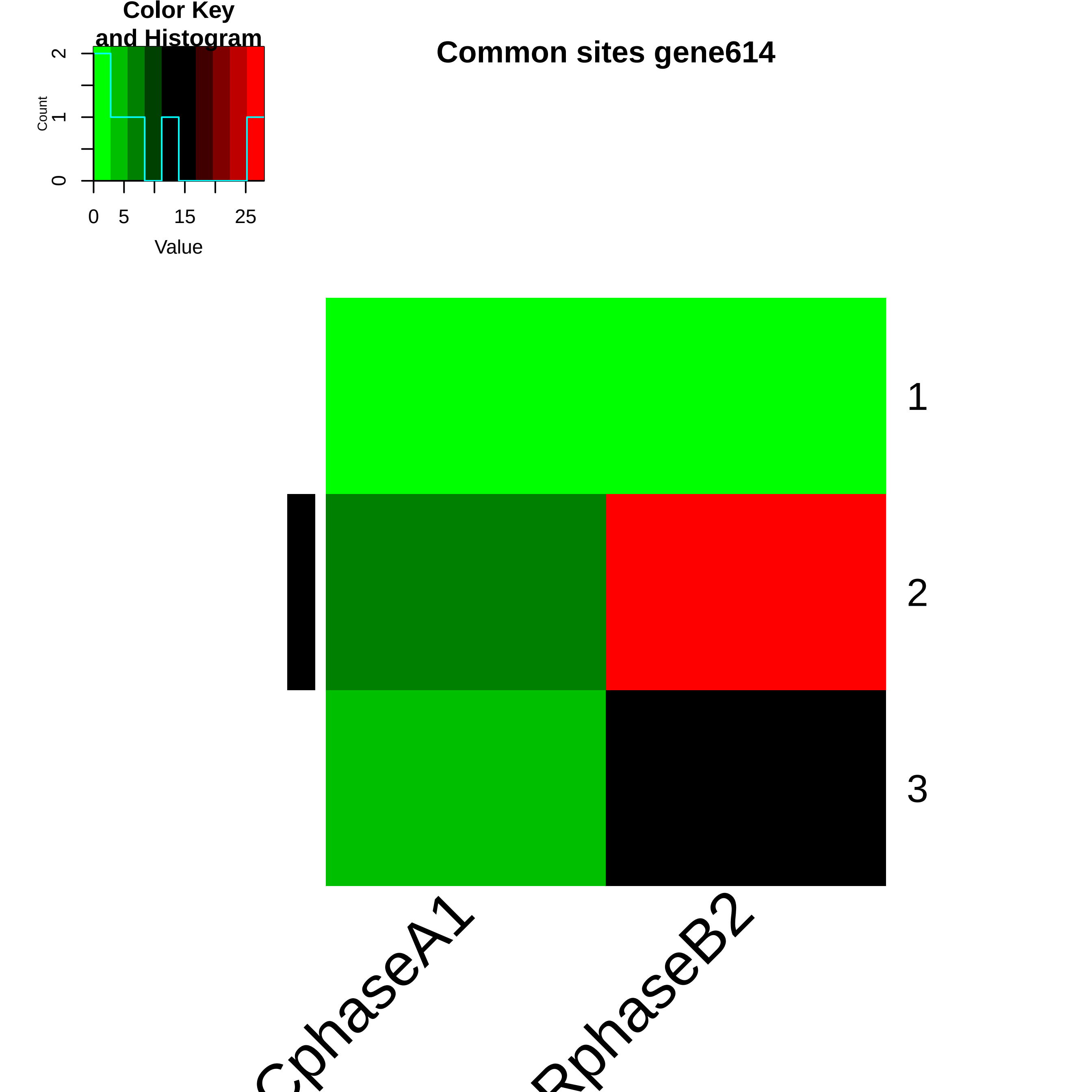

Color Key
and Histogram
Count
0
1
2
0
5
15
25
Value
Common sites gene614
1
2
3
CphaseA1
RphaseB2

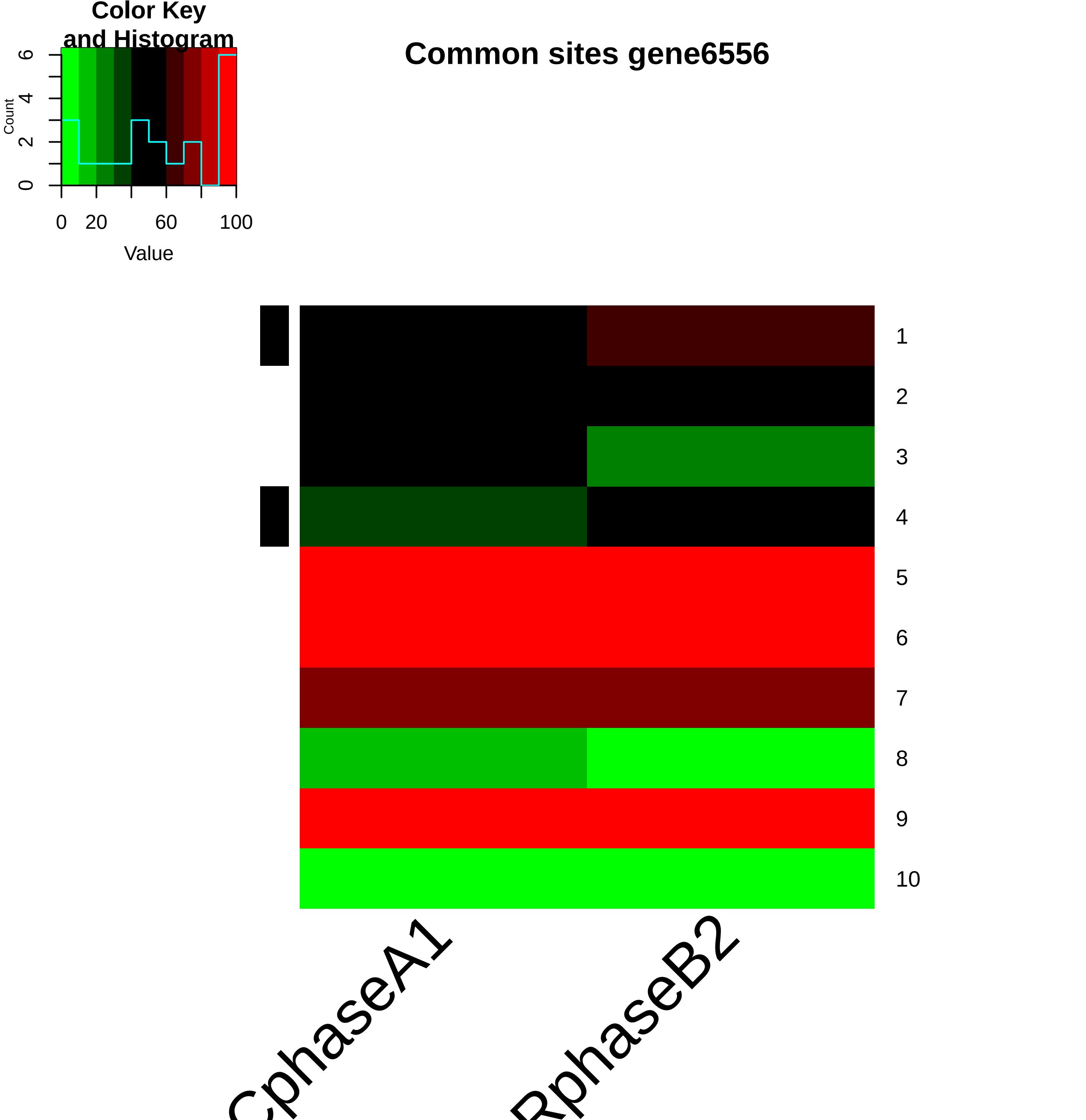

Color Key
and Histogram
Count
0
2
4
6
0
20
60
100
Value
Common sites gene6556
1
2
3
4
5
6
7
8
9
10
CphaseA1
RphaseB2

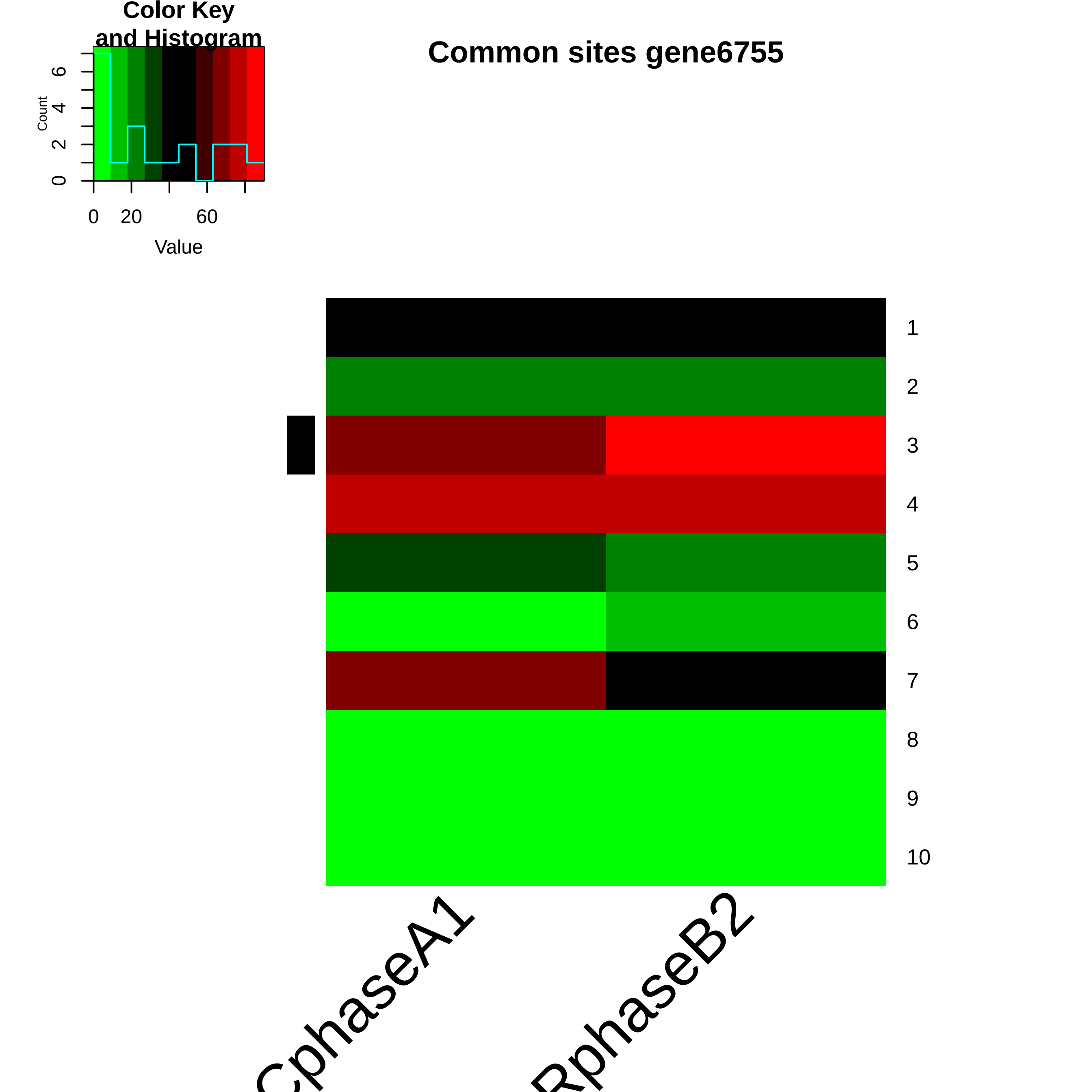

Color Key
and Histogram
Count
0
2
4
6
0
20
60
Value
Common sites gene6755
1
2
3
4
5
6
7
8
9
10
CphaseA1
RphaseB2

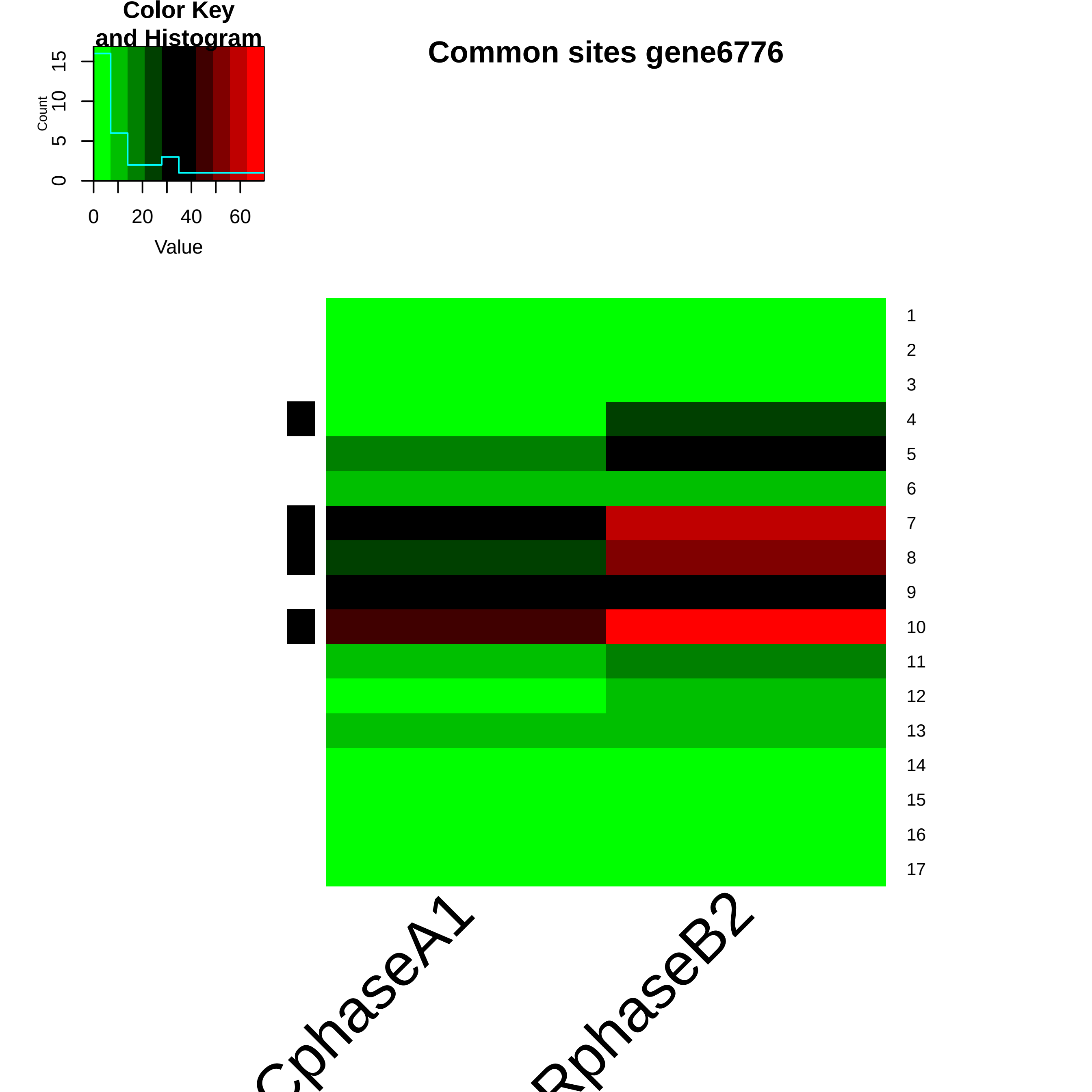
Color Key
and Histogram
Count
0
5
10
15
0
20
40
60
Value
Common sites gene6776
CphaseA1
RphaseB2

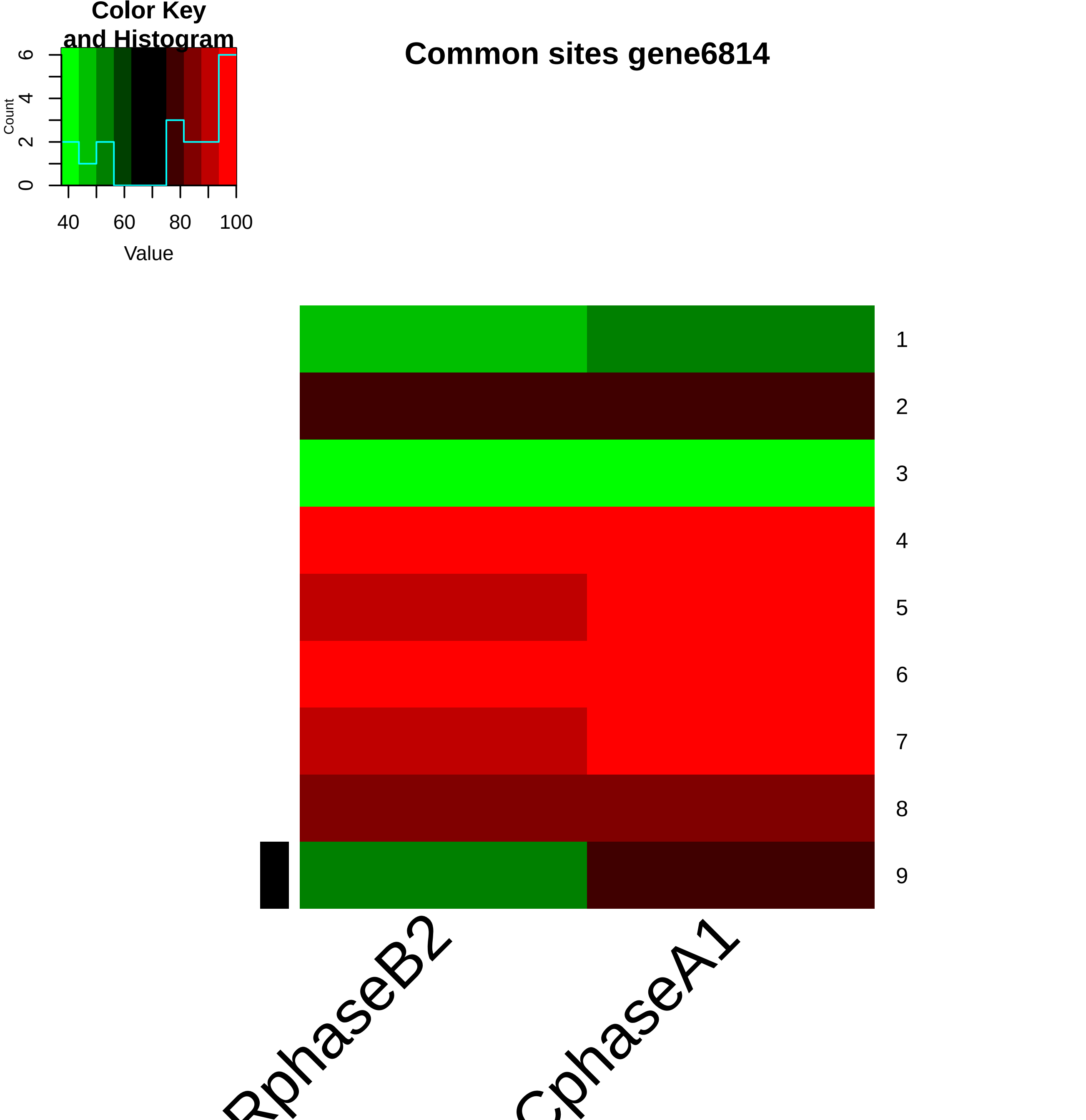
Color Key
and Histogram
Count
0
2
4
6
40
60
80
100
Value
Common sites gene6814
1
2
3
4
5
6
7
8
9
RphaseB2
CphaseA1

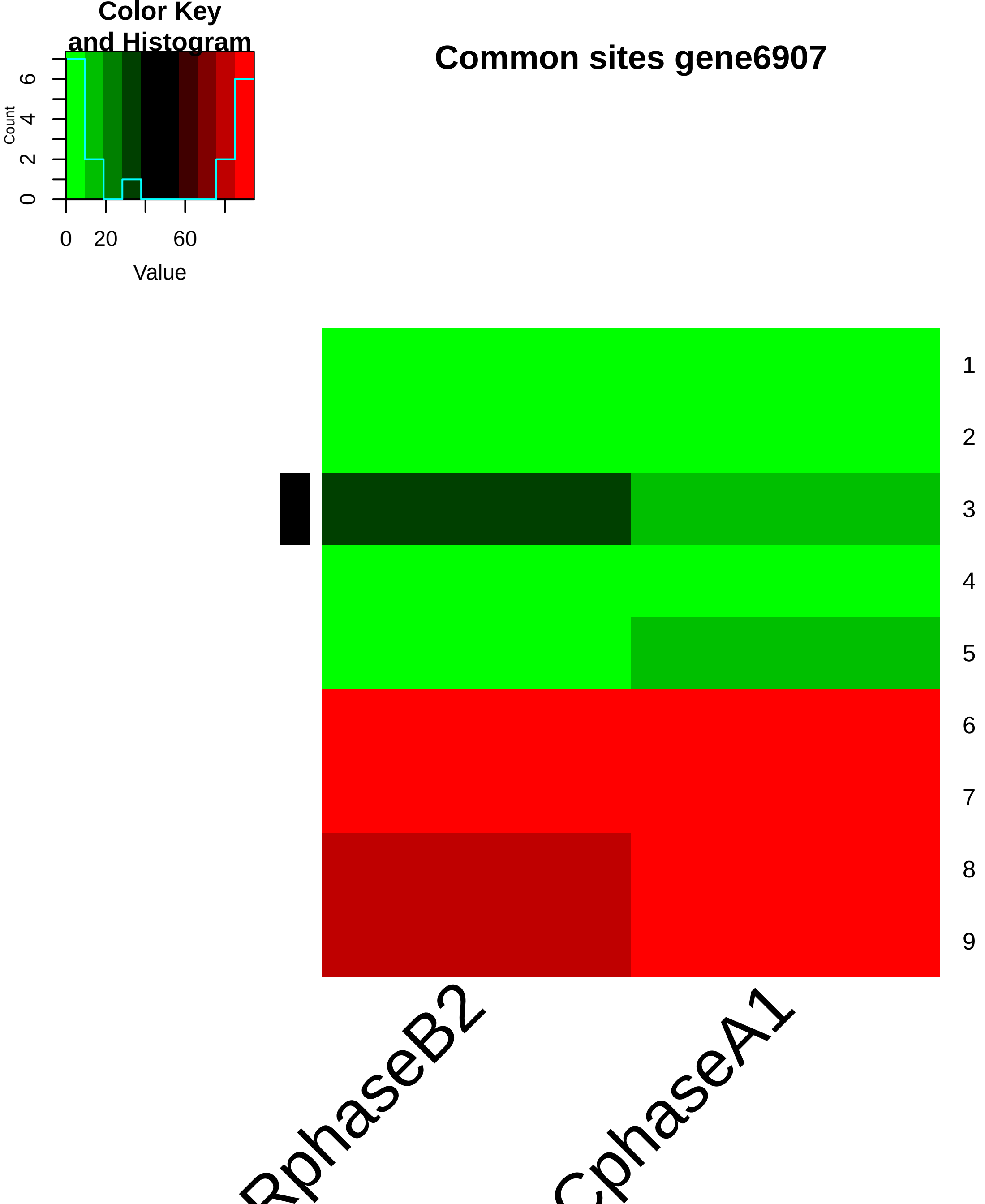
Color Key
and Histogram
Count
0
2
4
6
0
20
60
Value
Common sites gene6907
1
2
3
4
5
6
7
8
9
RphaseB2
CphaseA1

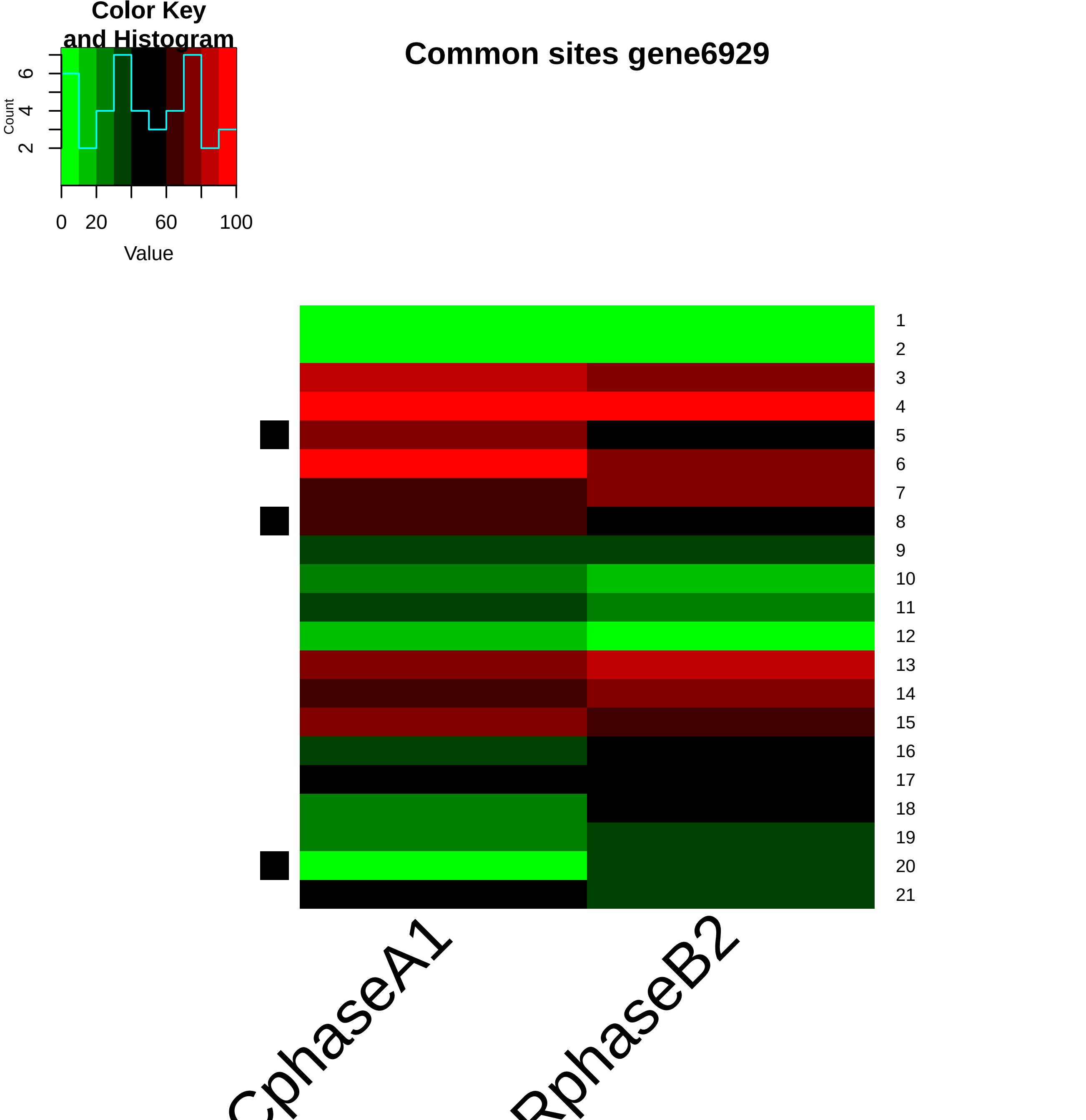
Color Key
and Histogram
Count
2
4
6
0
20
60
100
Value
Common sites gene6929
CphaseA1
RphaseB2

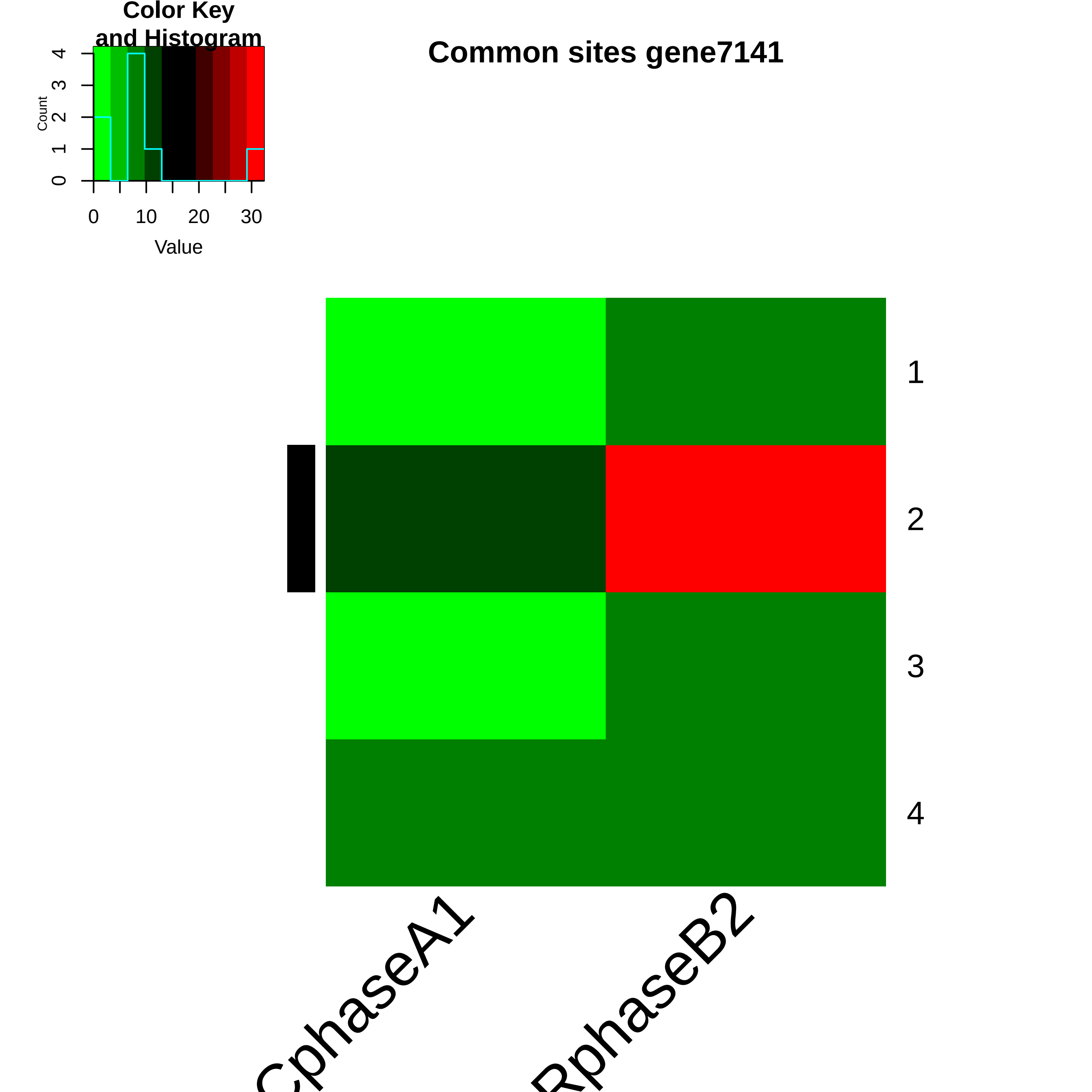

Color Key
and Histogram
Count
Value
0
1
2
3
4
0
10
20
30
Common sites gene7141
1
2
3
4
CphaseA1
RphaseB2

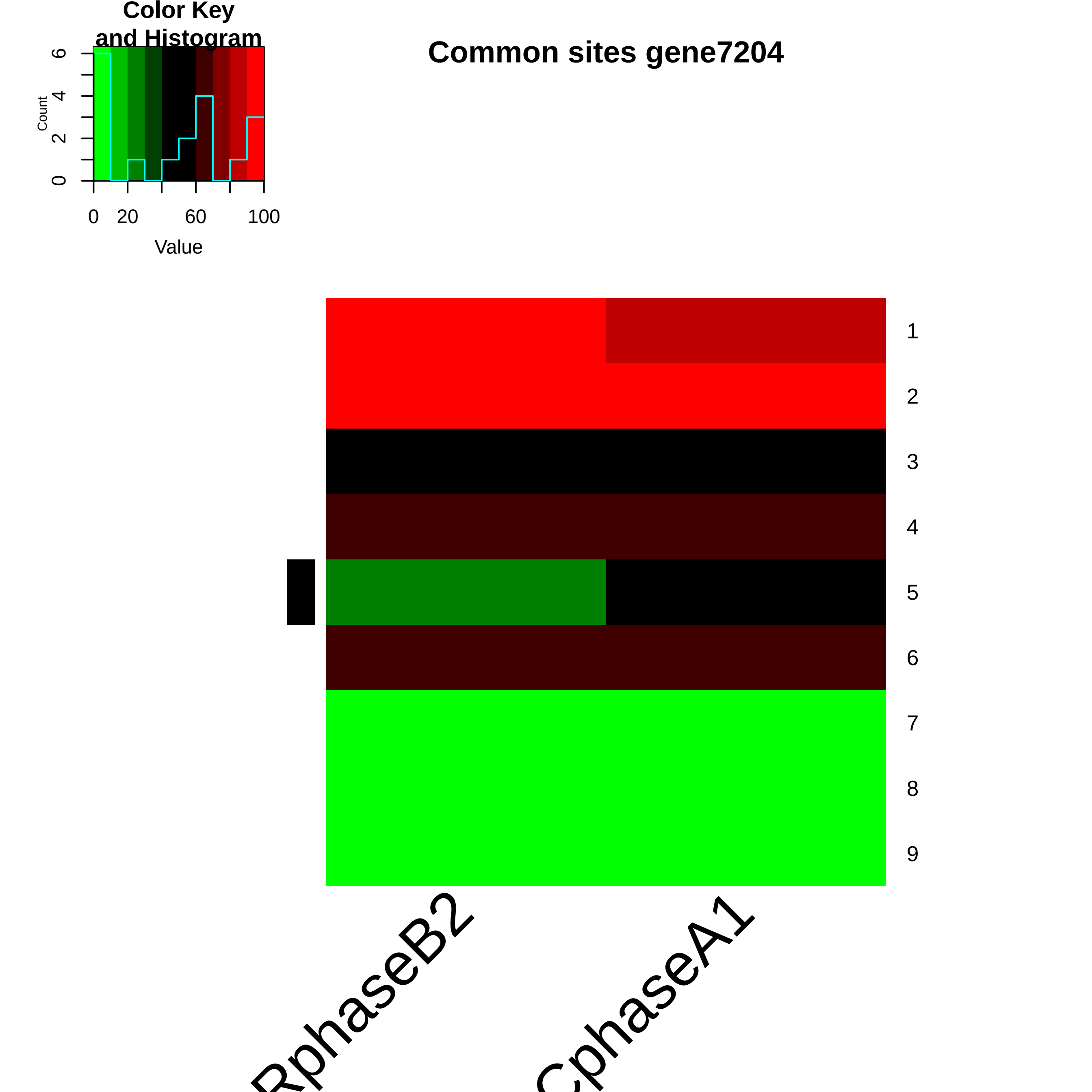
Color Key
and Histogram
Count
0
2
4
6
0
20
60
100
Value
Common sites gene7204
1
2
3
4
5
6
7
8
9
RphaseB2
CphaseA1

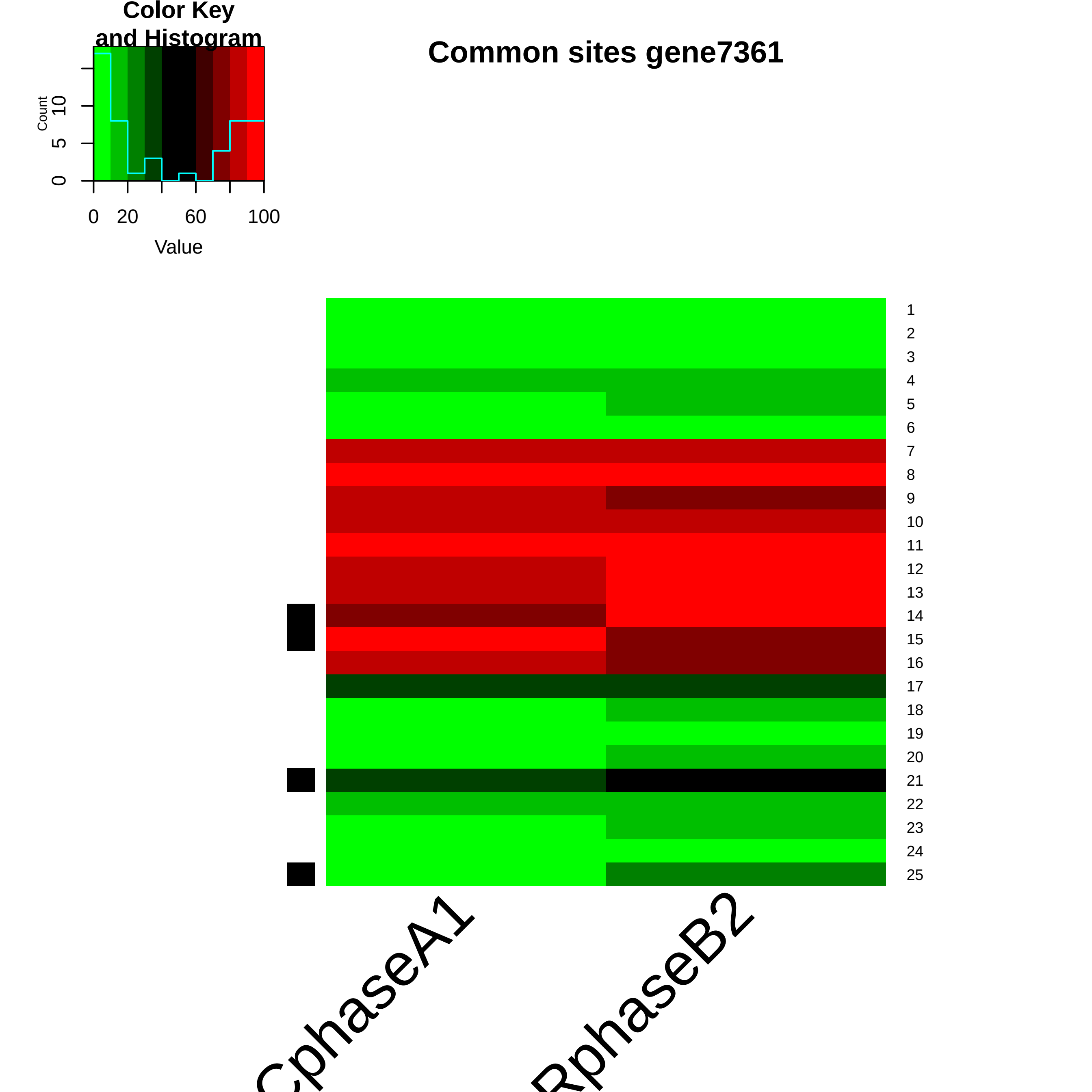
Color Key
and Histogram
Count
0
5
10
0
20
60
100
Value
Common sites gene7361
CphaseA1
RphaseB2

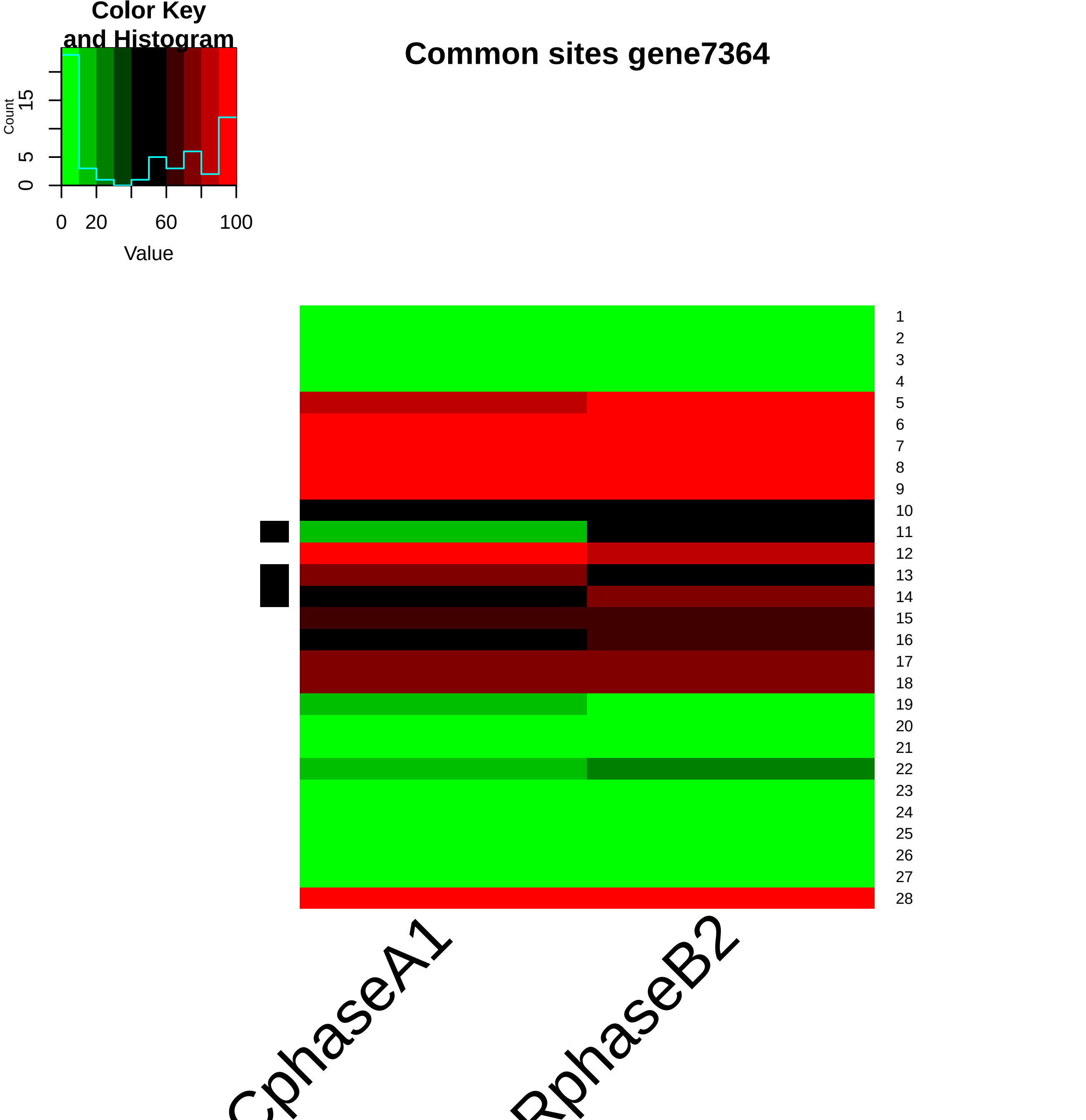

Color Key
and Histogram
Count
0
5
15
0
20
60
100
Value
Common sites gene7364
CphaseA1
RphaseB2

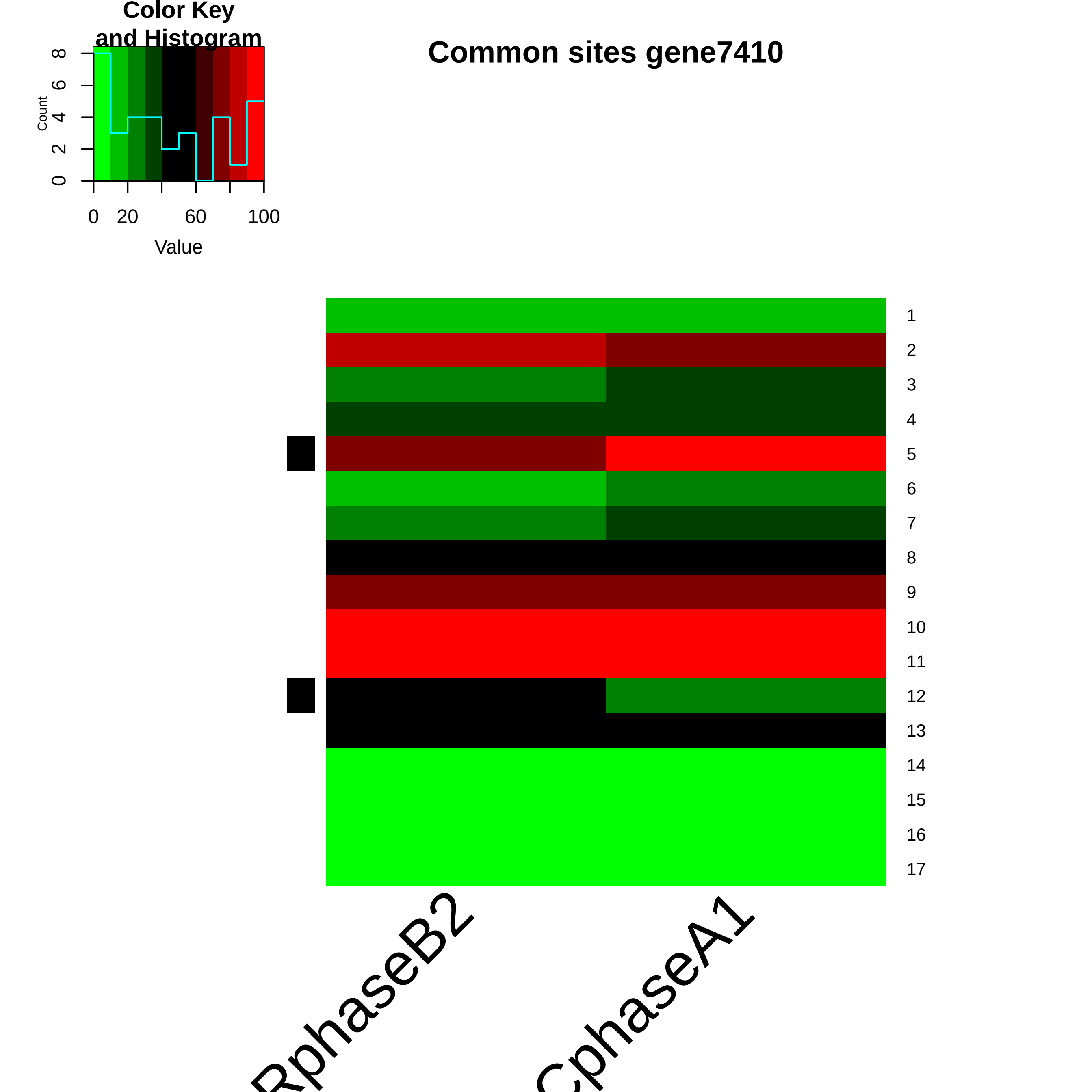

Color Key
and Histogram
Count
0
2
4
6
8
0
20
60
100
Value
Common sites gene7410
RphaseB2
CphaseA1

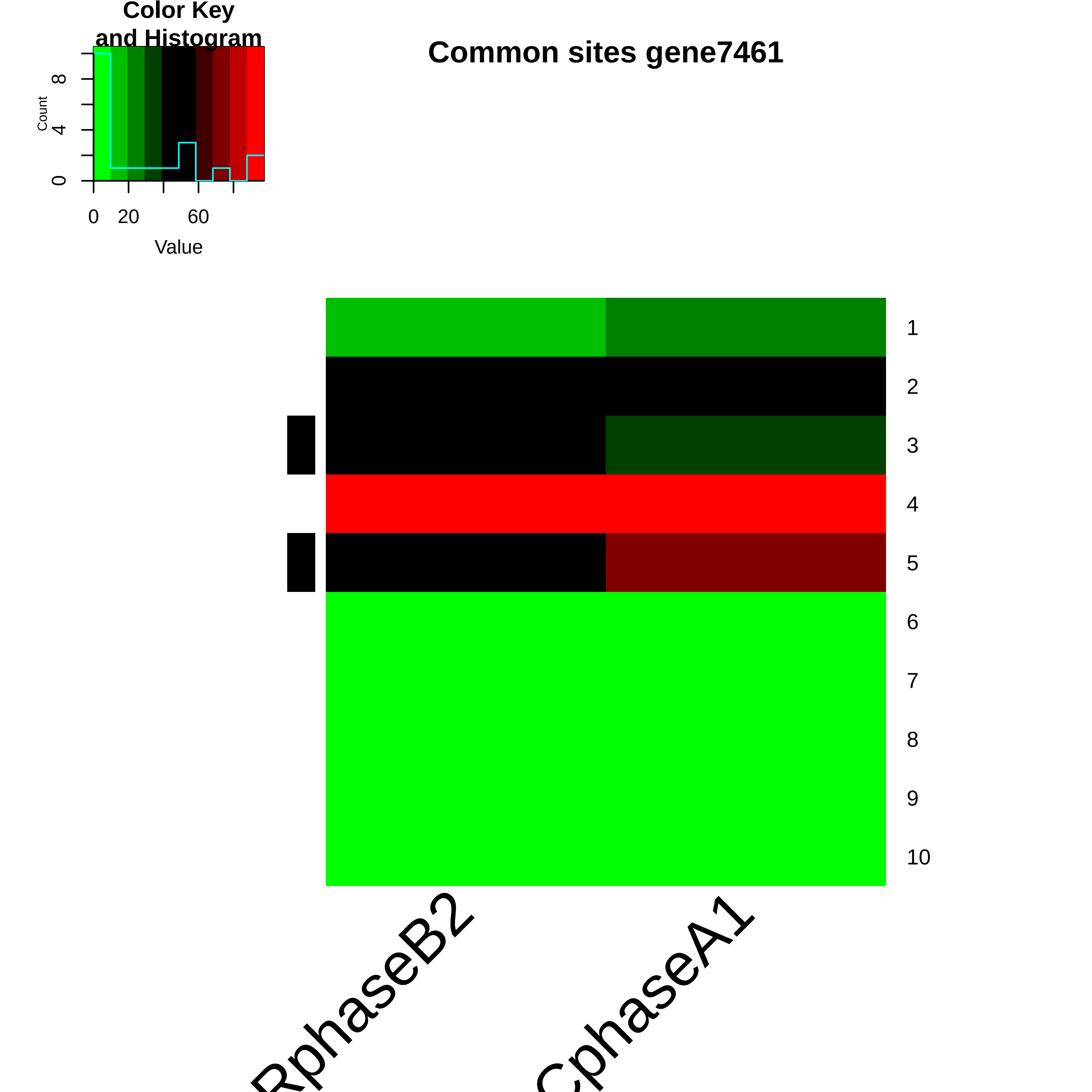

Color Key
and Histogram
Count
0
4
8
0
20
60
Value
Common sites gene7461
1
2
3
4
5
6
7
8
9
10
RphaseB2
CphaseA1

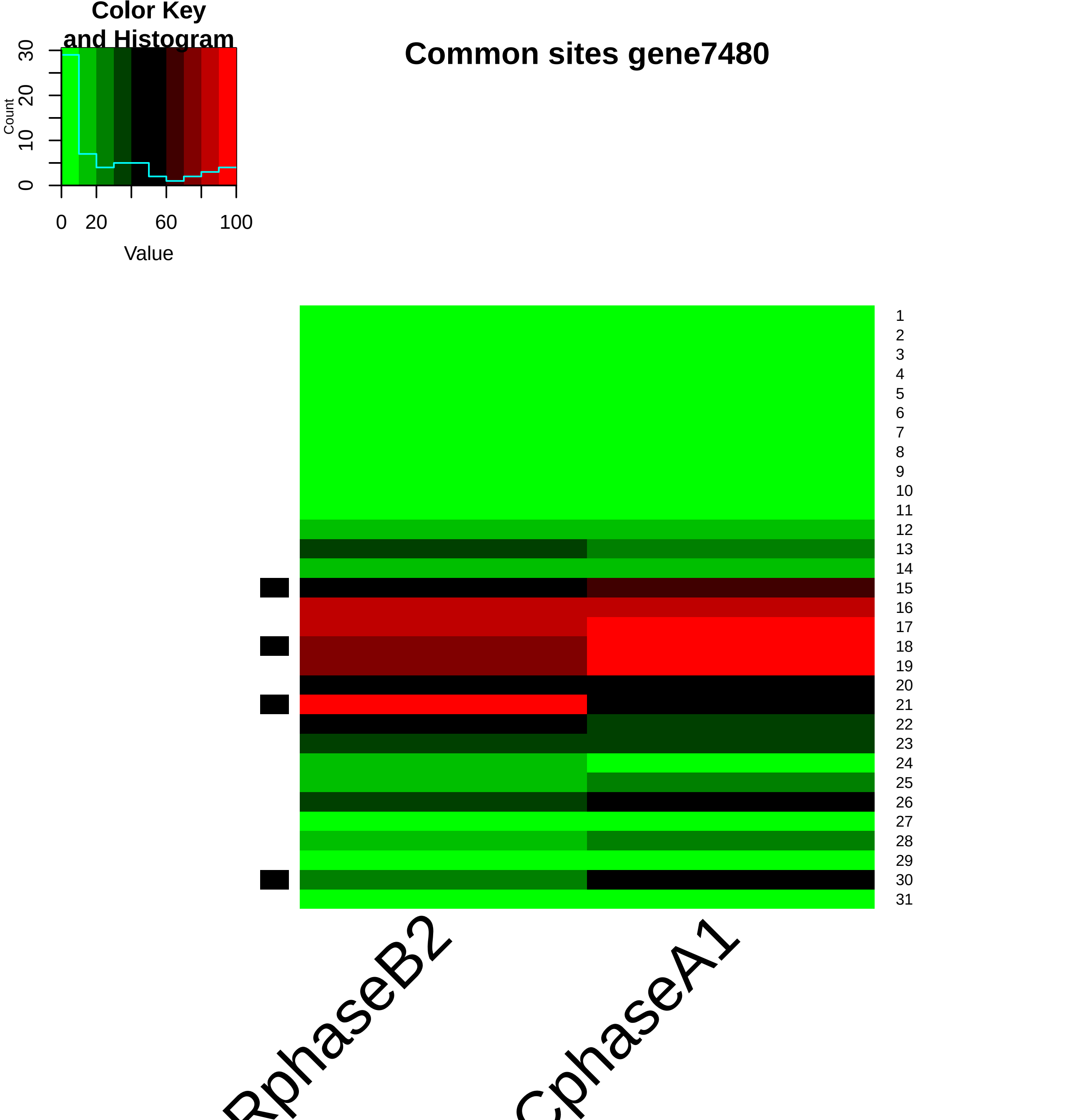
Color Key
and Histogram
Count
0
10
20
30
0
20
60
100
Value
Common sites gene7480
RphaseB2
CphaseA1

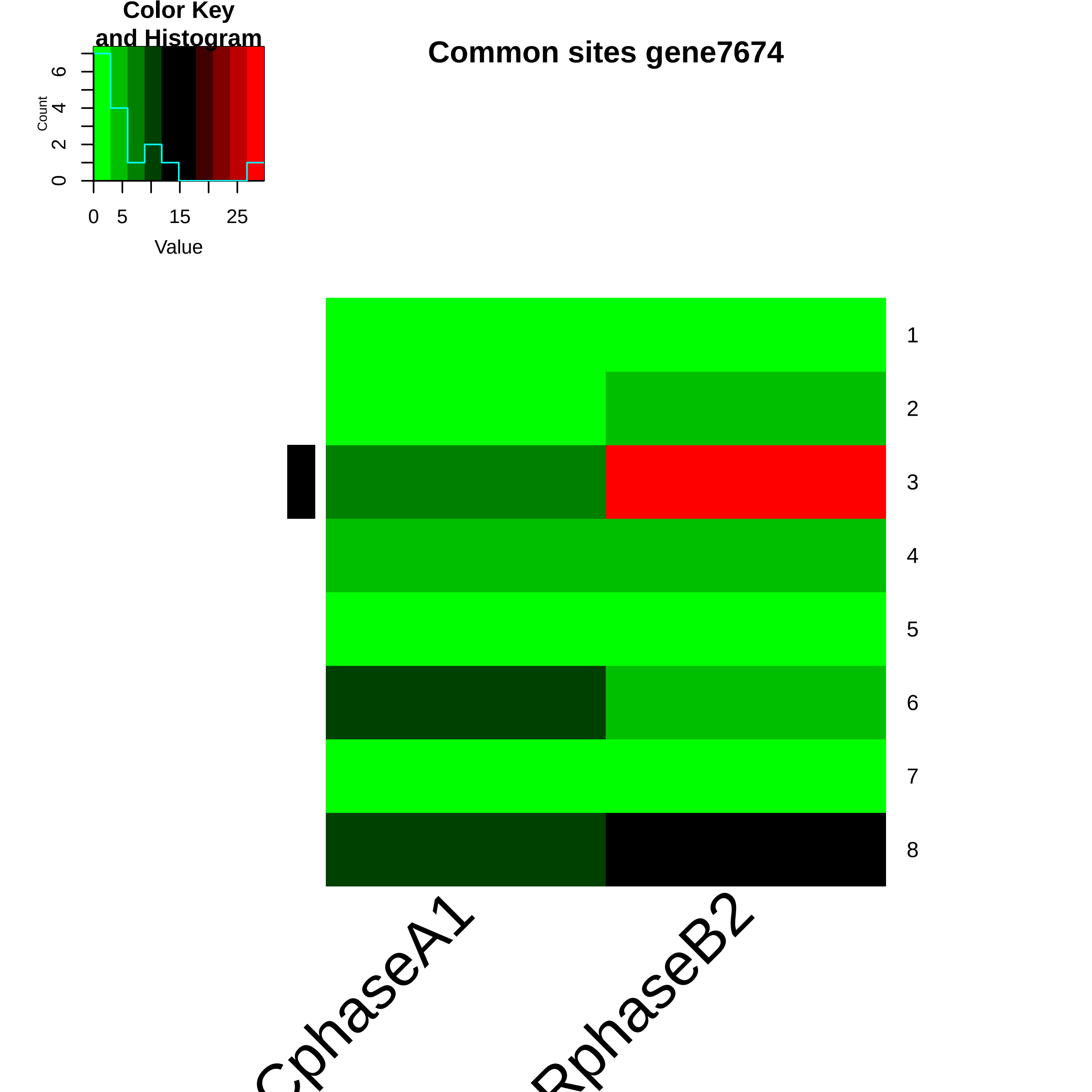

Color Key
and Histogram
Count
0
2
4
6
0
5
15
25
Value
Common sites gene7674
1
2
3
4
5
6
7
8
CphaseA1
RphaseB2

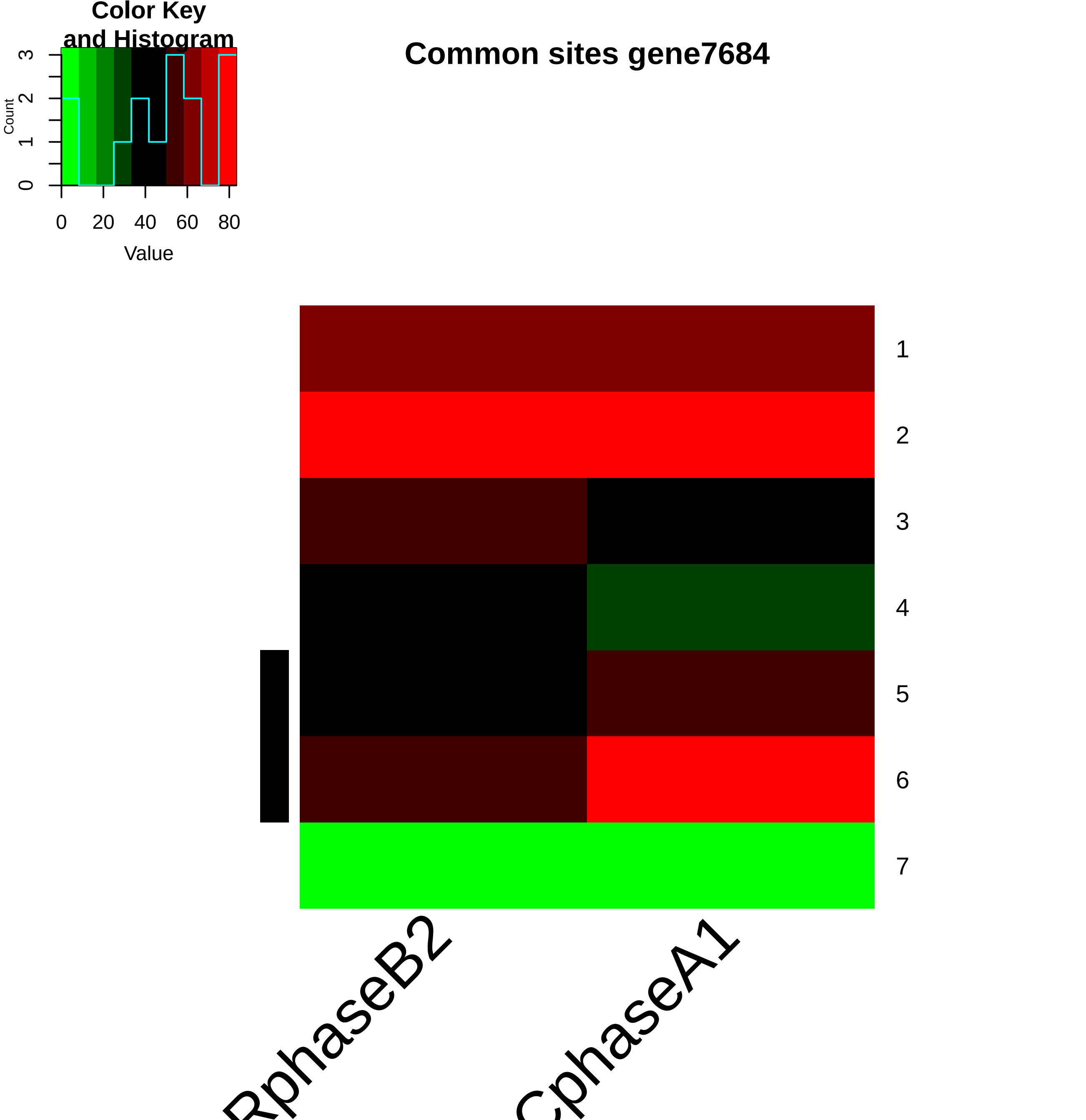
Color Key
and Histogram
Count
0
1
2
3
0
20
40
60
80
Value
Common sites gene7684
1
2
3
4
5
6
7
RphaseB2
CphaseA1

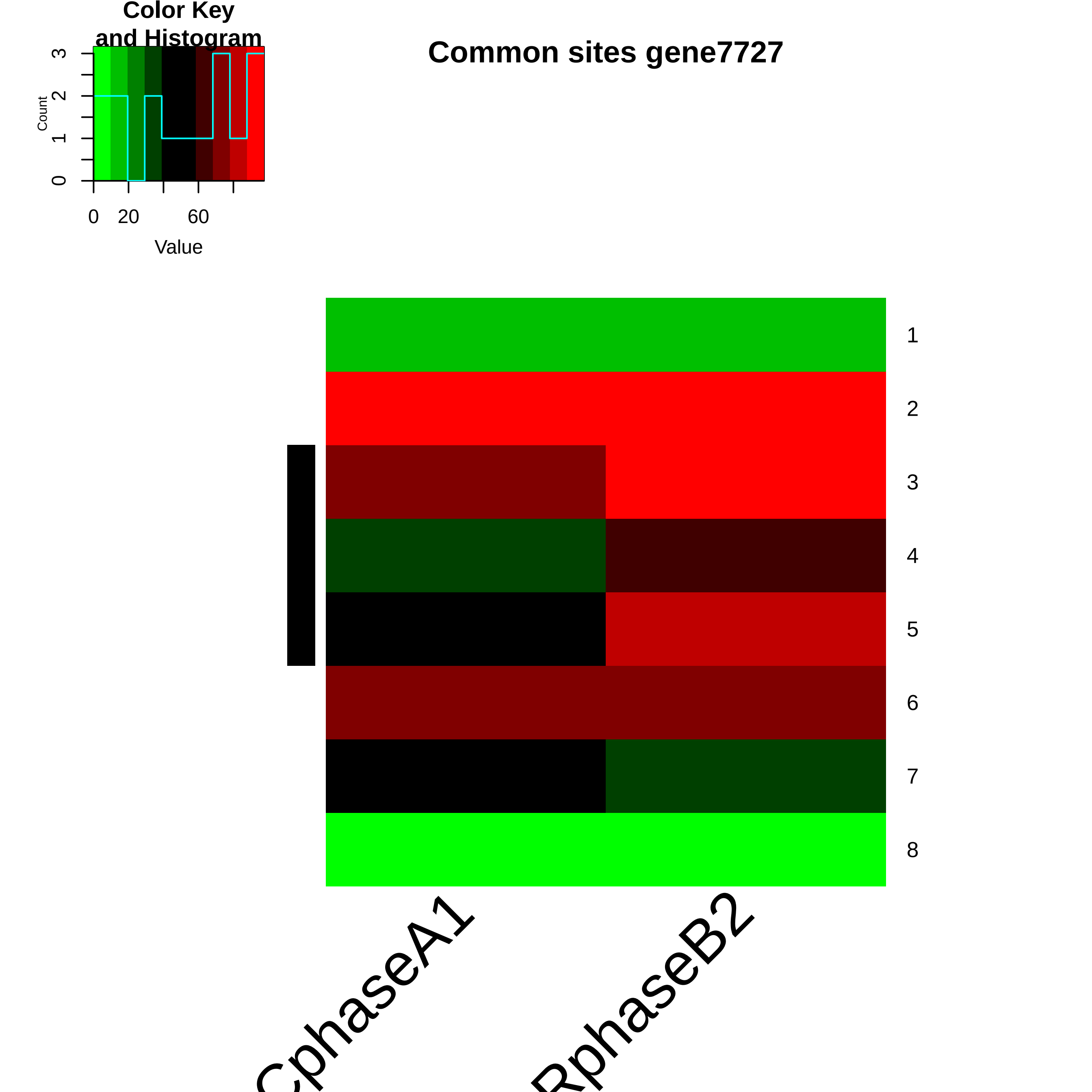

Color Key and Histogram
Common sites gene7727
Count
0
1
2
3
0
20
60
Value
1
2
3
4
5
6
7
8
CphaseA1
RphaseB2

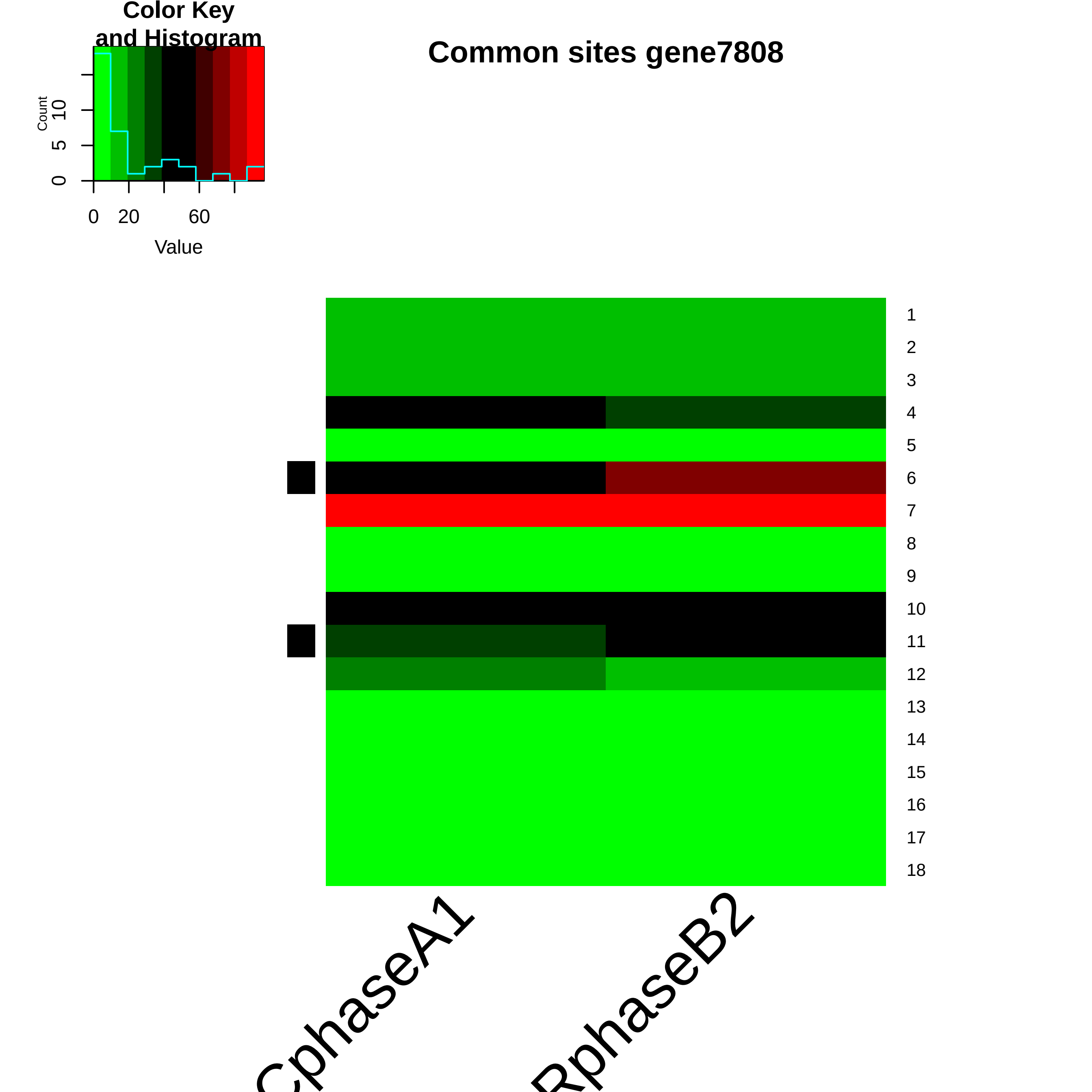
Color Key
and Histogram
Count
0
5
10
0
20
60
Value
Common sites gene7808
CphaseA1
RphaseB2

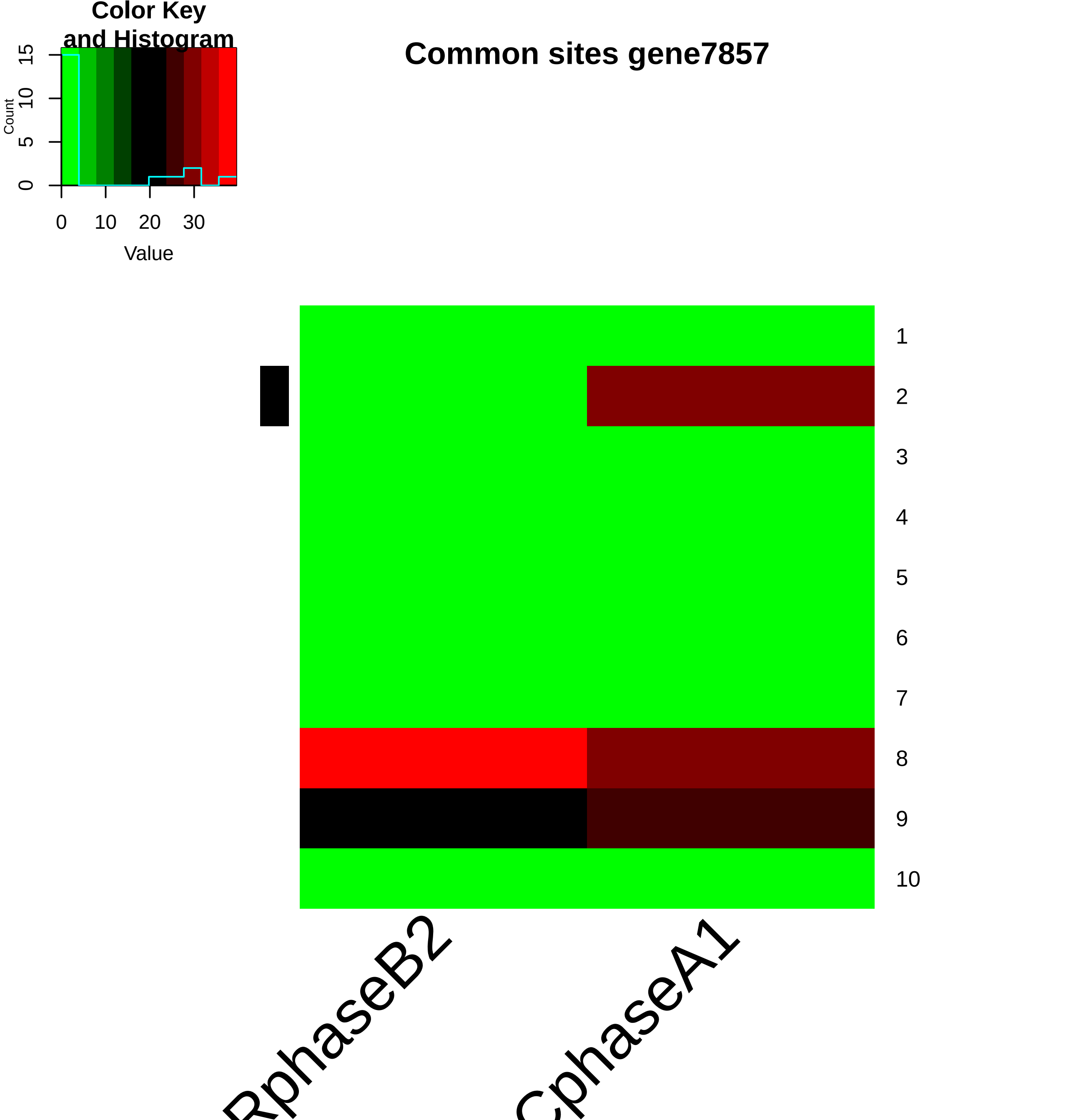

Color Key
and Histogram
Count
0
5
10
15
0
10
20
30
Value
Common sites gene7857
1
2
3
4
5
6
7
8
9
10
RphaseB2
CphaseA1

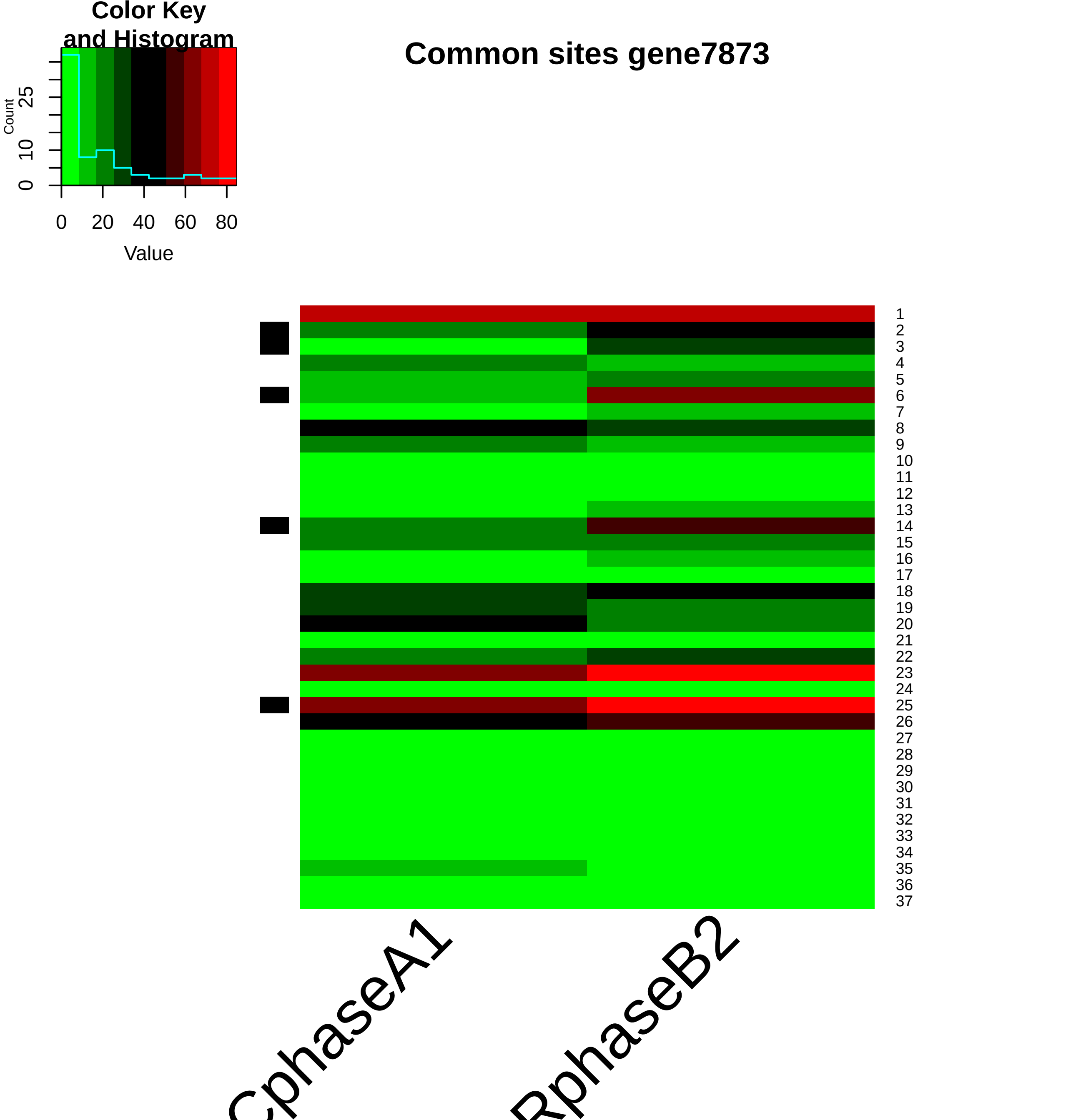

Color Key
and Histogram
Count
0
10
25
0
20
40
60
80
Value
Common sites gene7873
CphaseA1
RphaseB2

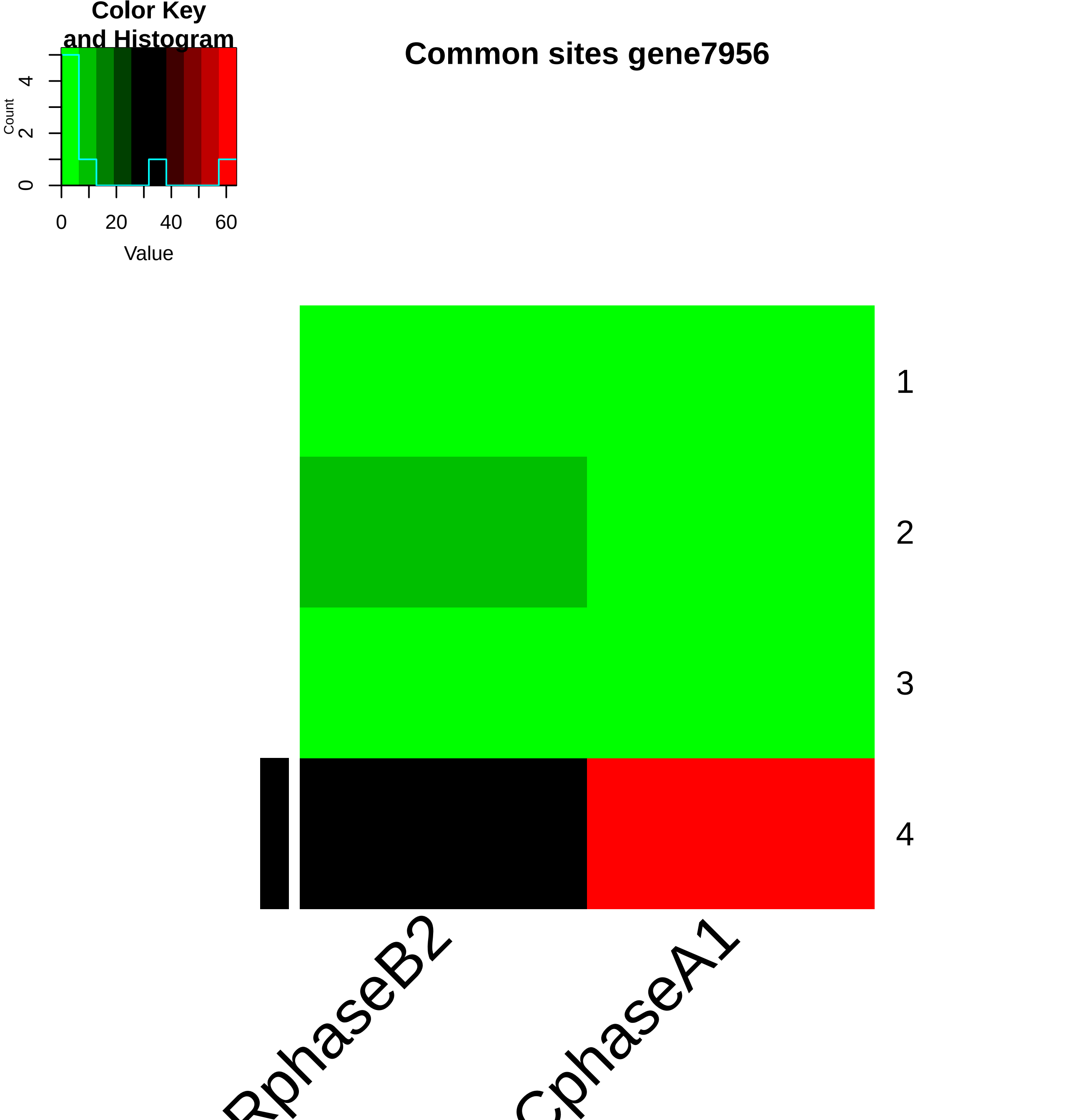

Color Key
and Histogram
Count
0
2
4
0
20
40
60
Value
Common sites gene7956
1
2
3
4
RphaseB2
CphaseA1

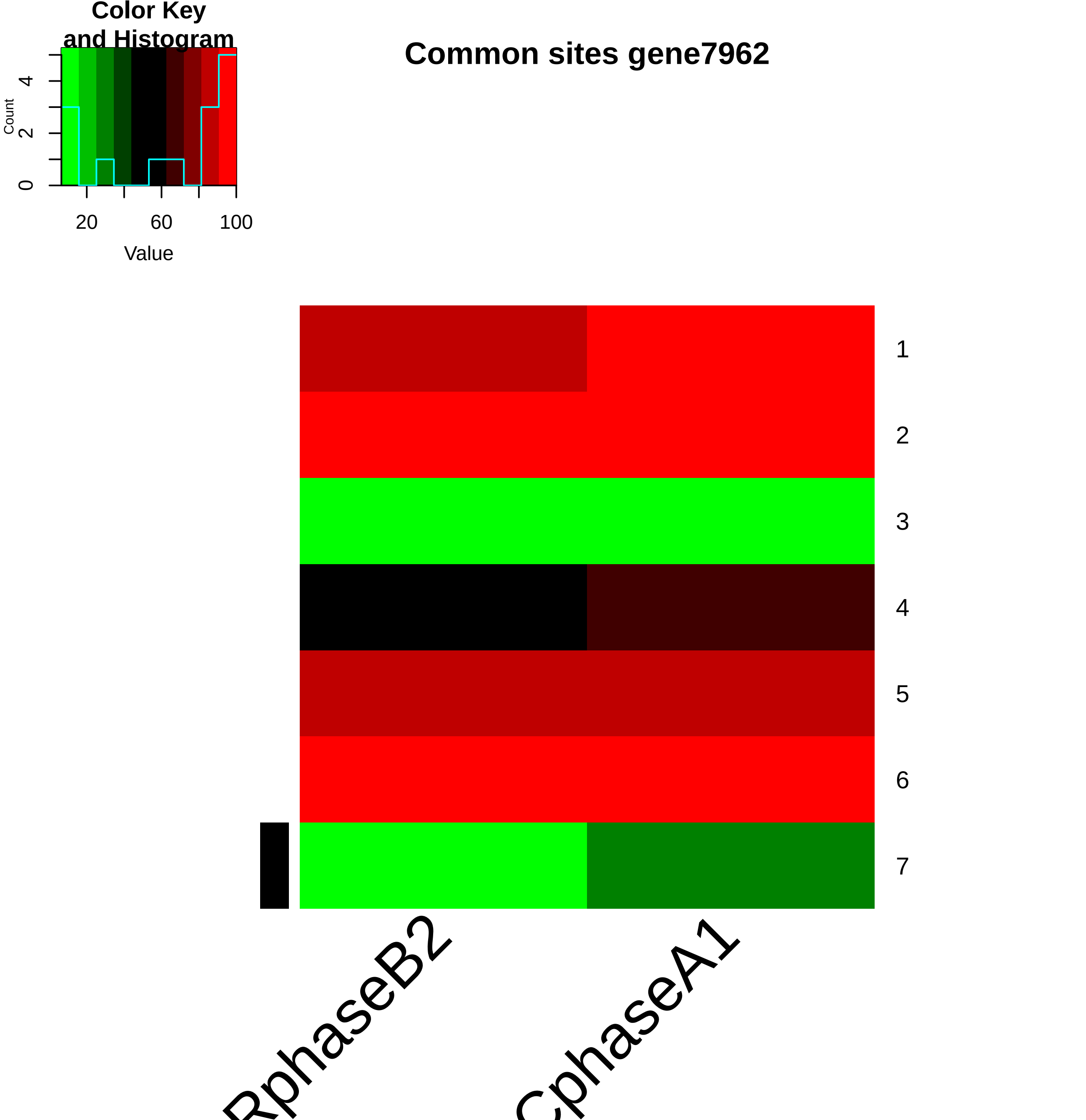
Color Key
and Histogram
Count
0
2
4
20
60
100
Value
Common sites gene7962
1
2
3
4
5
6
7
RphaseB2
CphaseA1

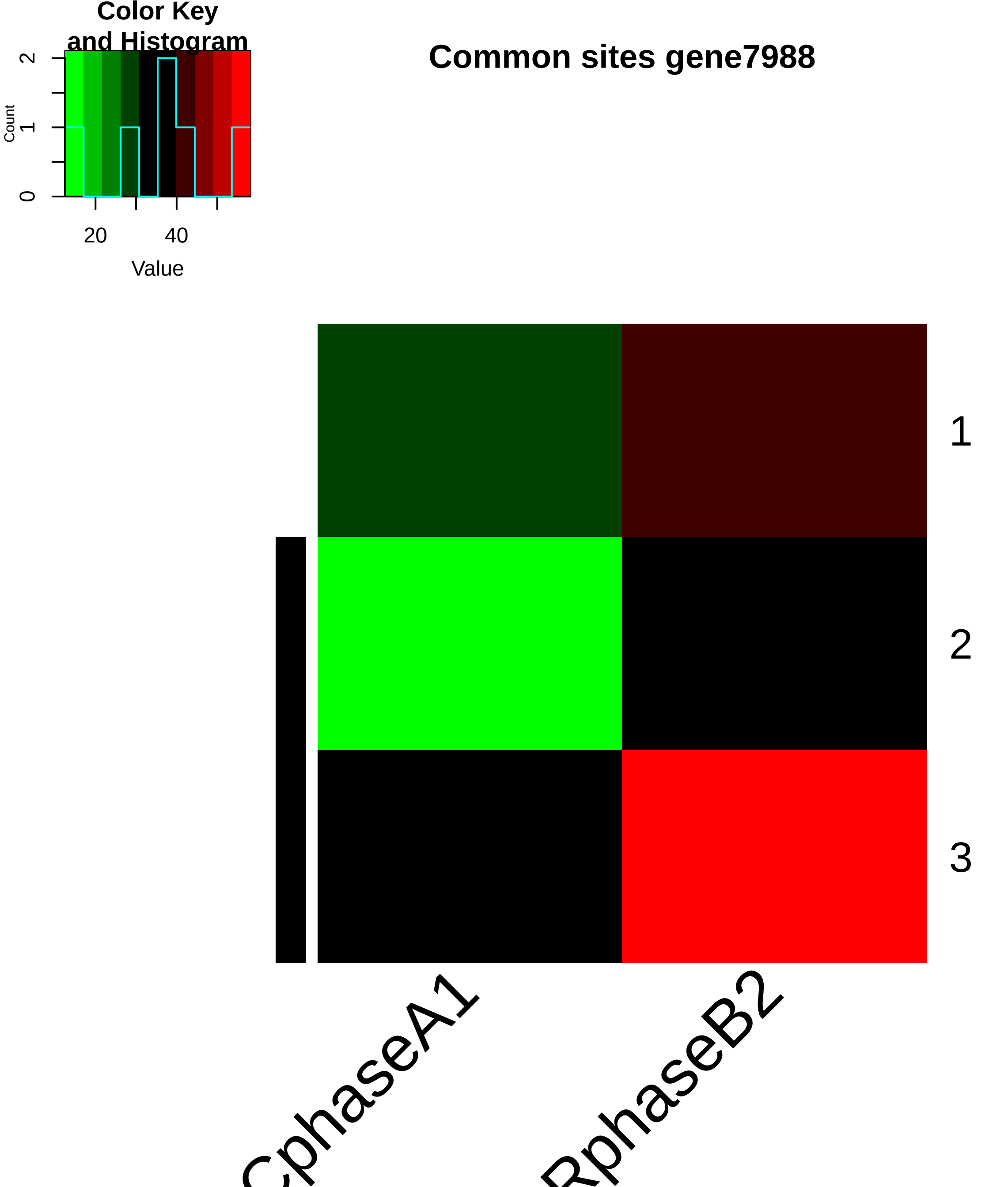

Color Key
and Histogram
Count
0
1
2
20
40
Value
Common sites gene7988
1
2
3
CphaseA1
RphaseB2

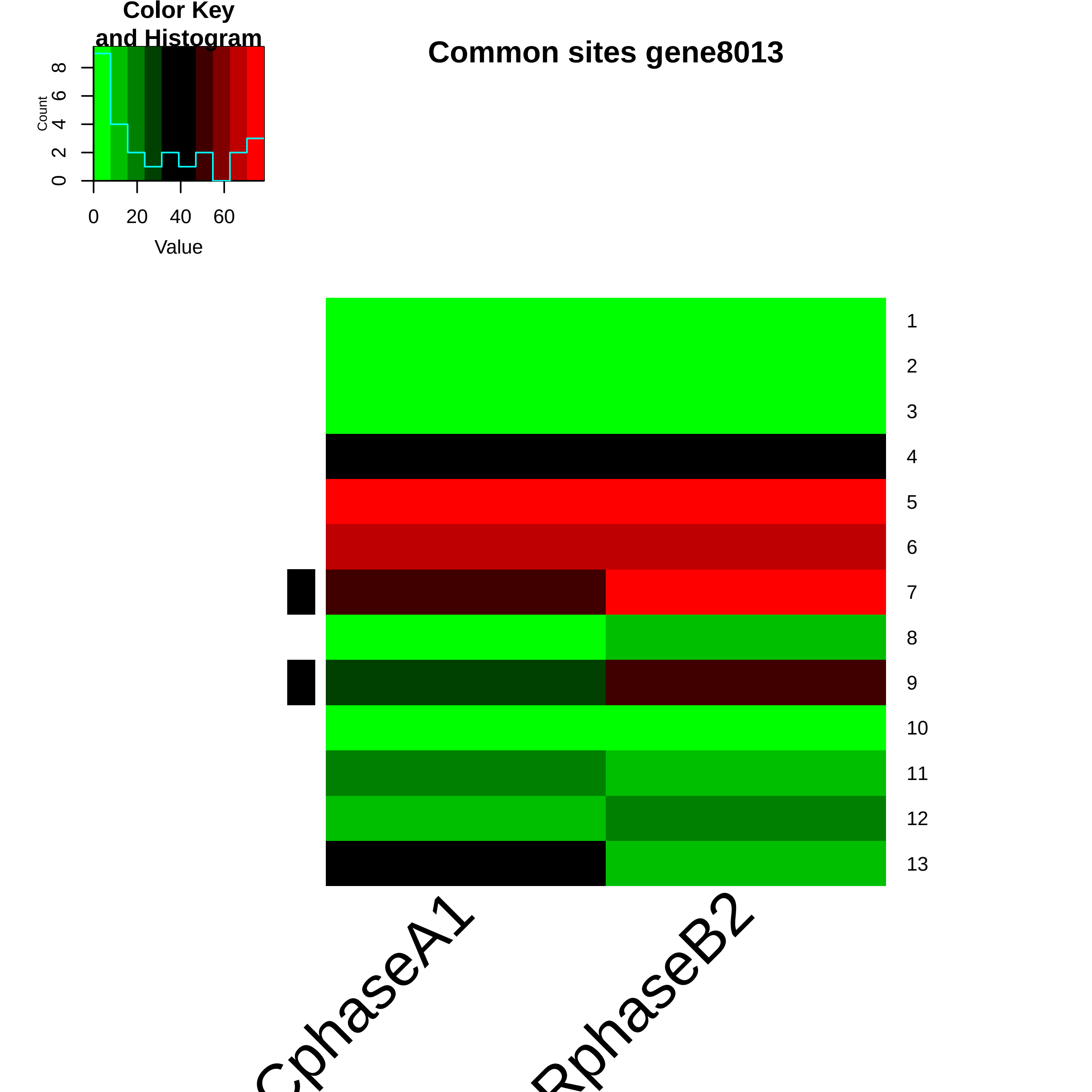

Color Key
and Histogram
Count
0
2
4
6
8
0
20
40
60
Value
Common sites gene8013
1
2
3
4
5
6
7
8
9
10
11
12
13
CphaseA1
RphaseB2

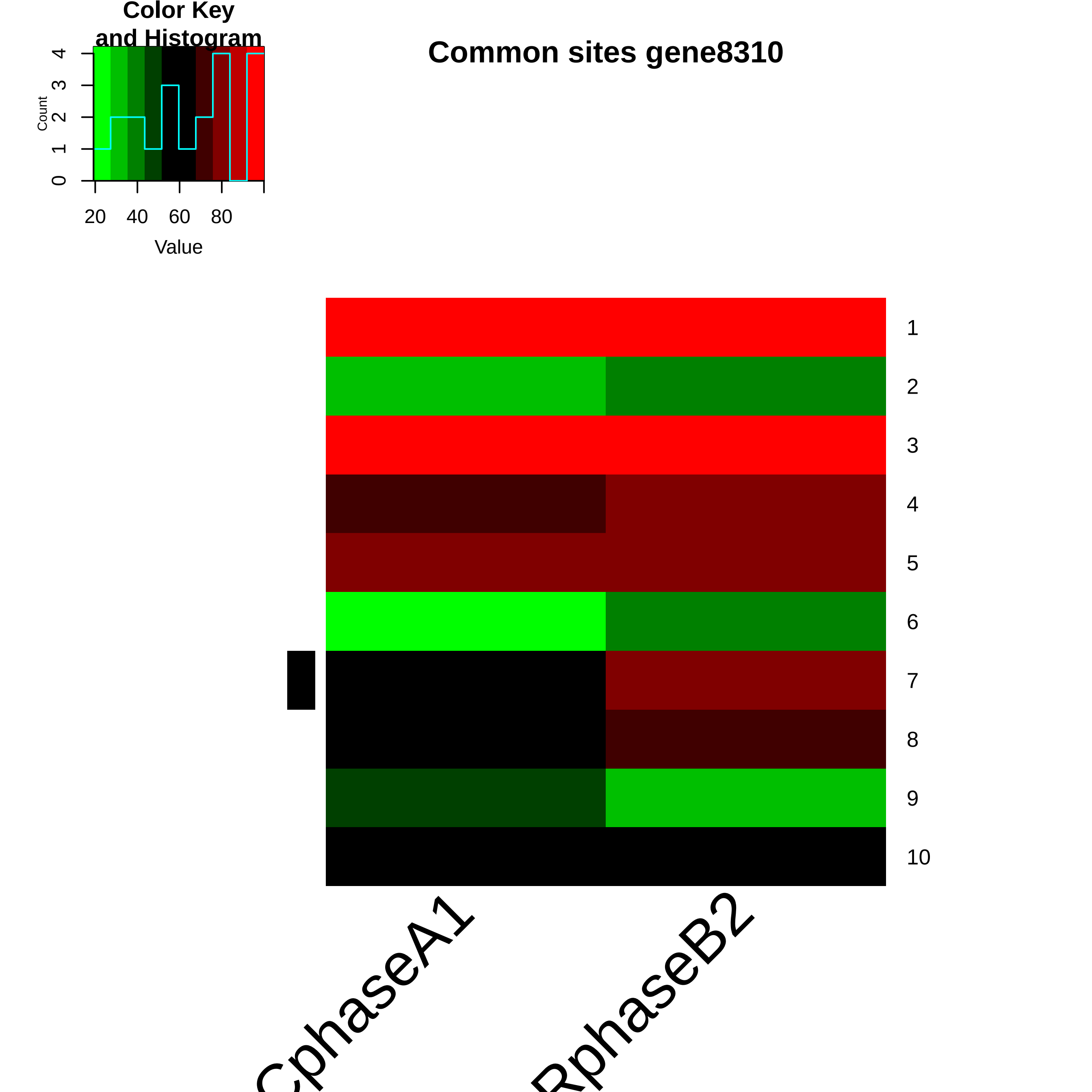

Color Key and Histogram
Count
0 1 2 3 4
20 40 60 80
Value
Common sites gene8310
1
2
3
4
5
6
7
8
9
10
CphaseA1
RphaseB2

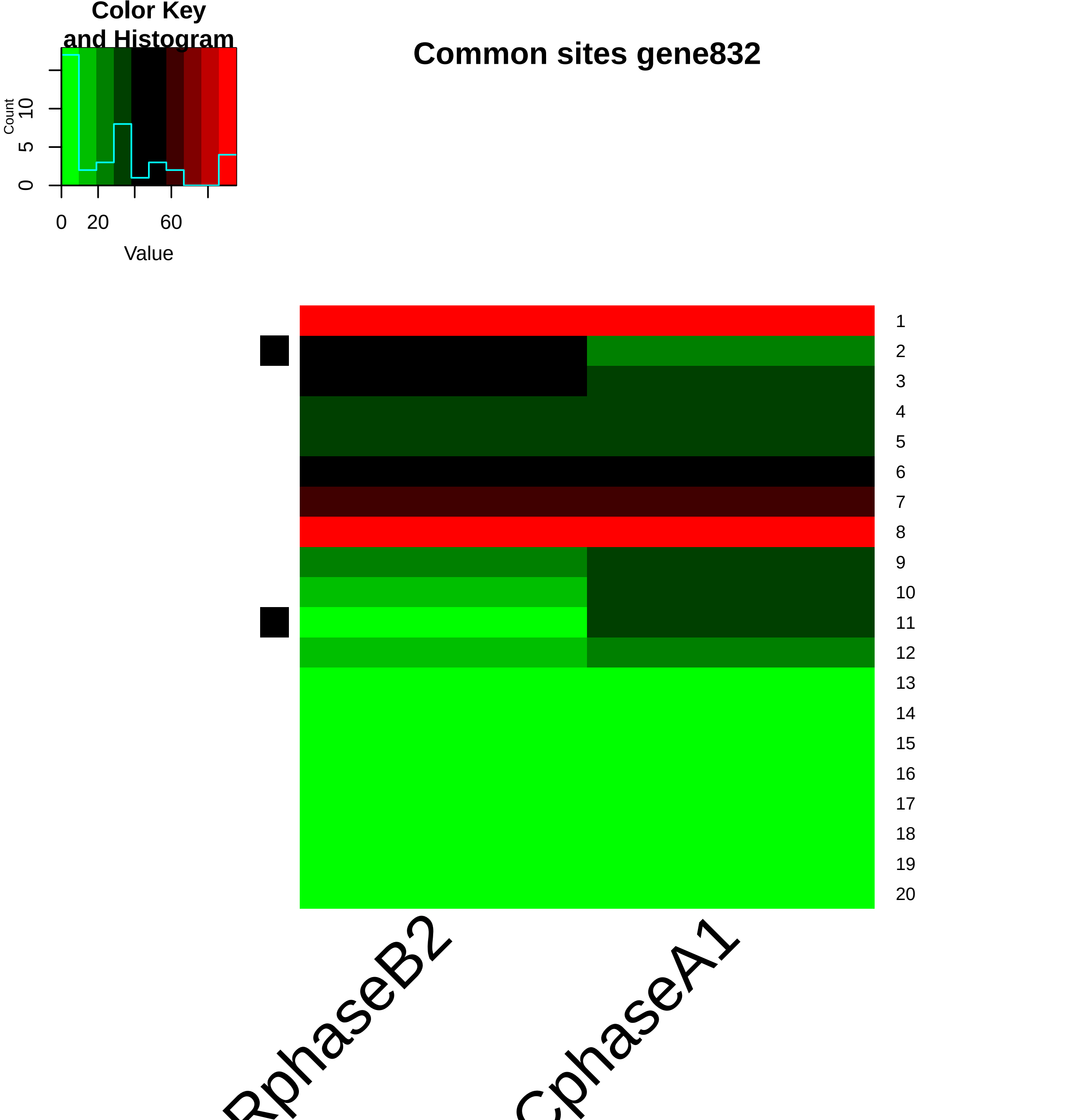

Color Key
and Histogram
Count
0
5
10
0
20
60
Value
Common sites gene832
RphaseB2
CphaseA1

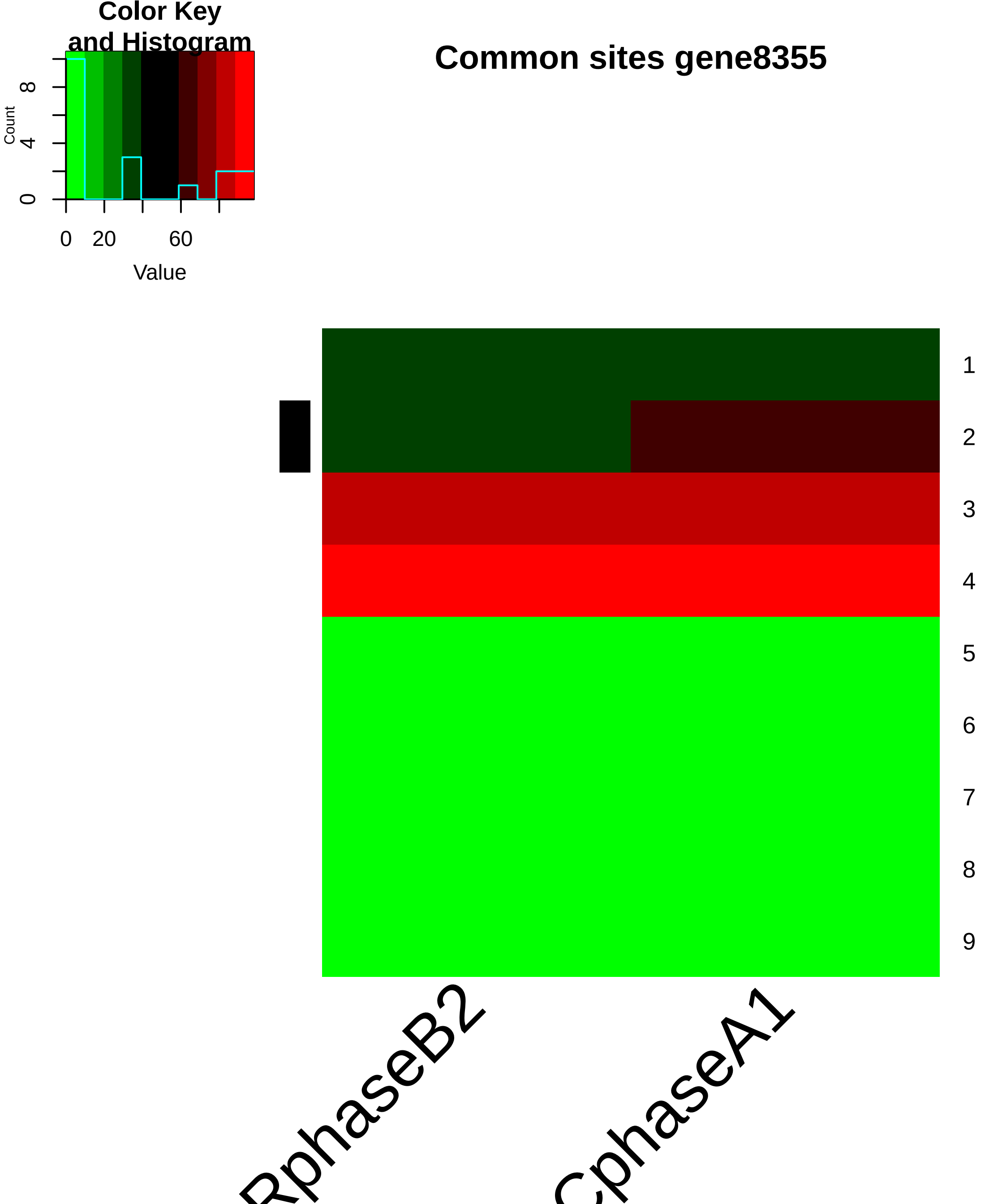
Color Key
and Histogram
Count
0
4
8
0
20
60
Value
Common sites gene8355
1
2
3
4
5
6
7
8
9
RphaseB2
CphaseA1

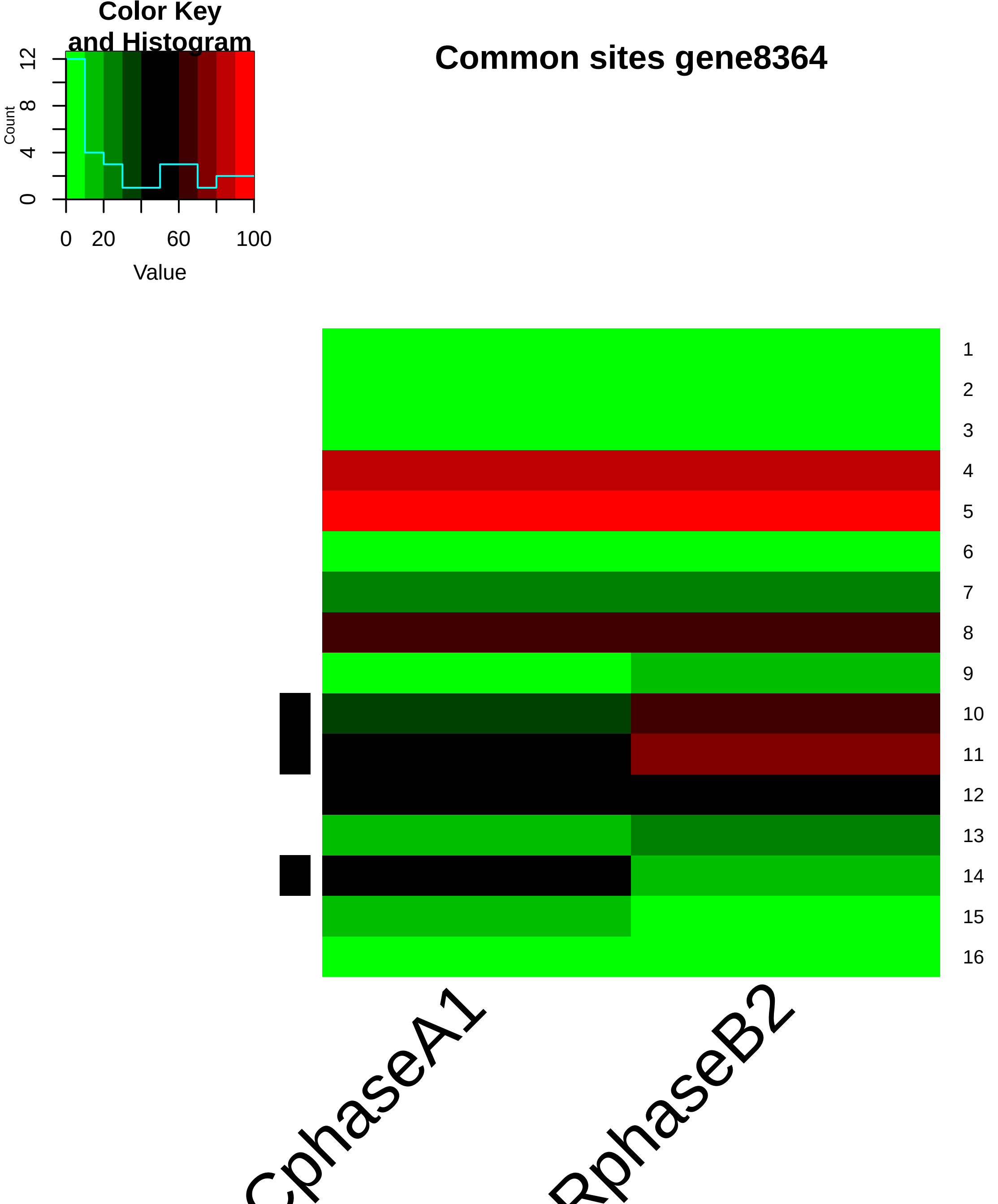
Color Key
and Histogram
Count
0
4
8
12
0
20
60
100
Value
Common sites gene8364
CphaseA1
RphaseB2

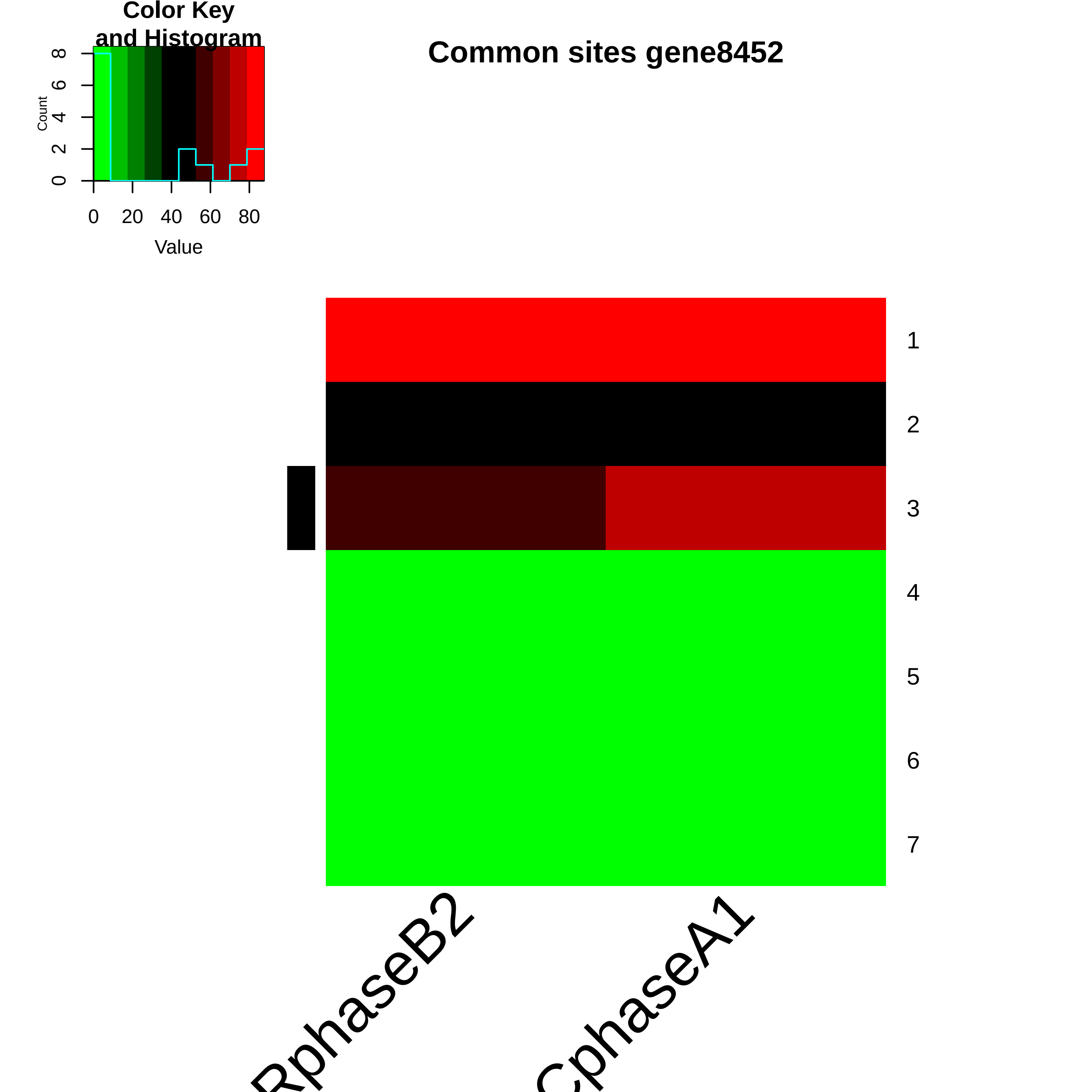

Color Key
and Histogram
Count
0
2
4
6
8
0
20
40
60
80
Value
Common sites gene8452
1
2
3
4
5
6
7
RphaseB2
CphaseA1

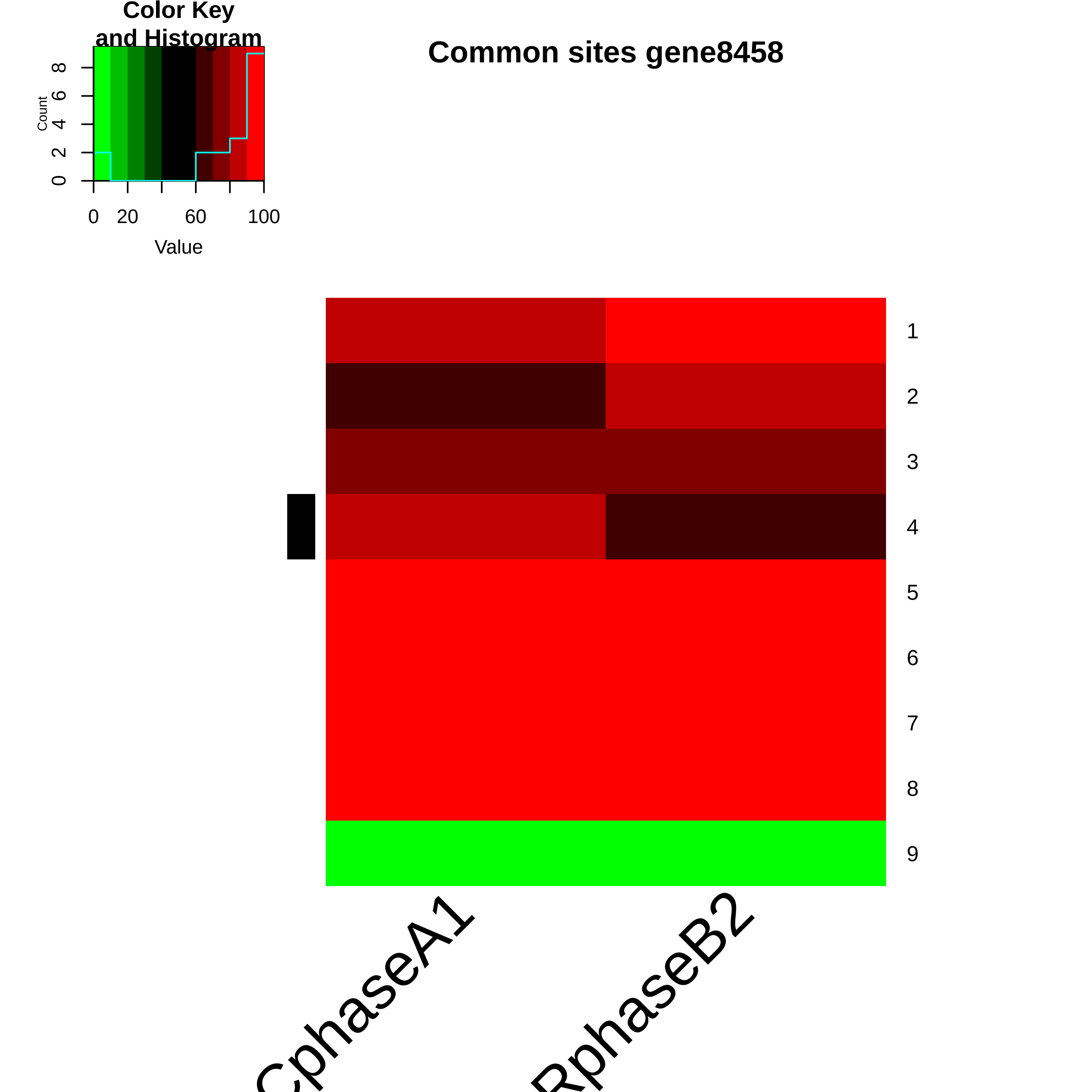

Color Key
and Histogram
Count
0
2
4
6
8
0
20
60
100
Value
Common sites gene8458
1
2
3
4
5
6
7
8
9
CphaseA1
RphaseB2

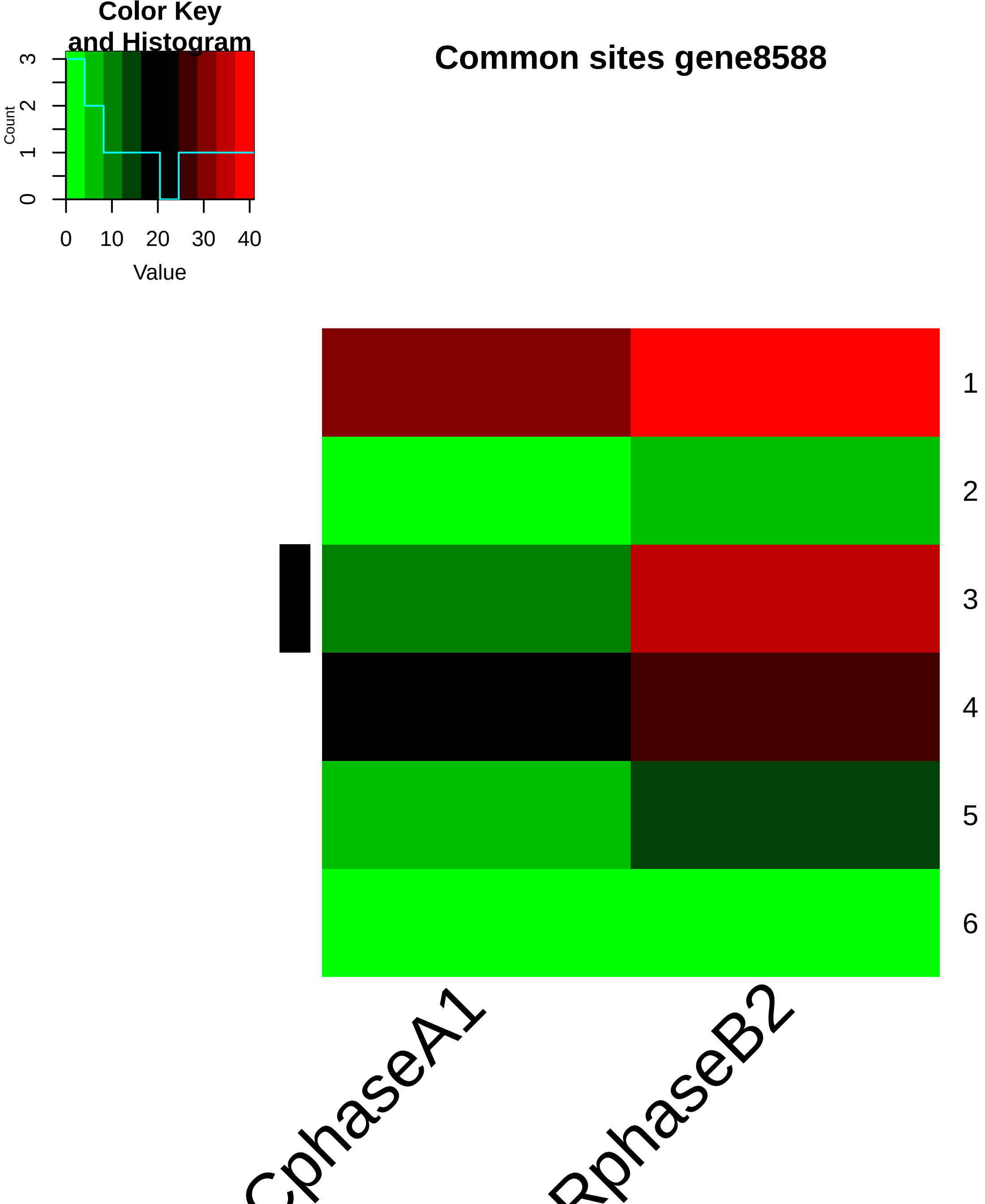
Color Key
and Histogram
Count
Value
0
1
2
3
0
10
20
30
40
Common sites gene8588
1
2
3
4
5
6
CphaseA1
RphaseB2

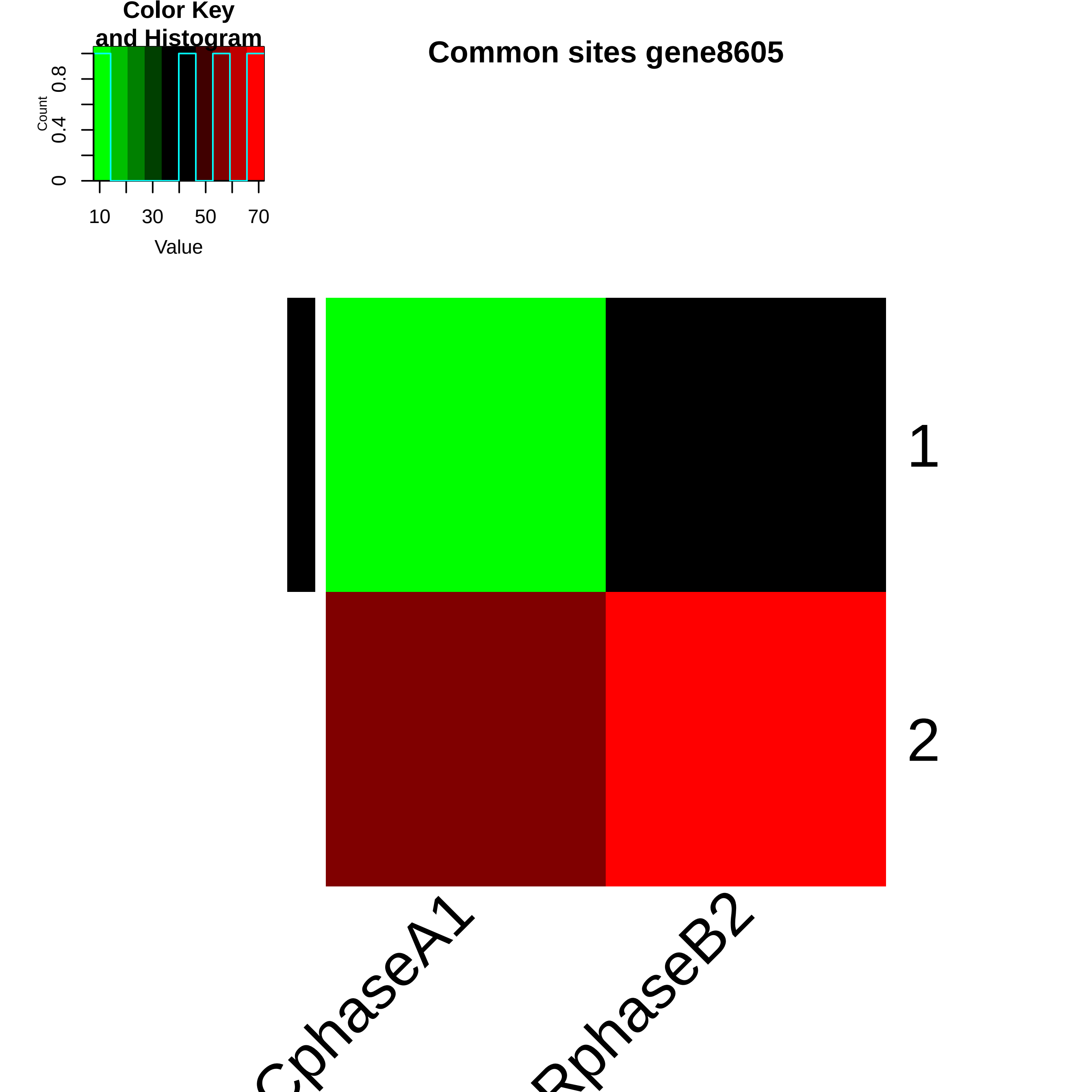

Color Key
and Histogram
Count
0
0.4
0.8
10
30
50
70
Value
Common sites gene8605
1
2
CphaseA1
RphaseB2

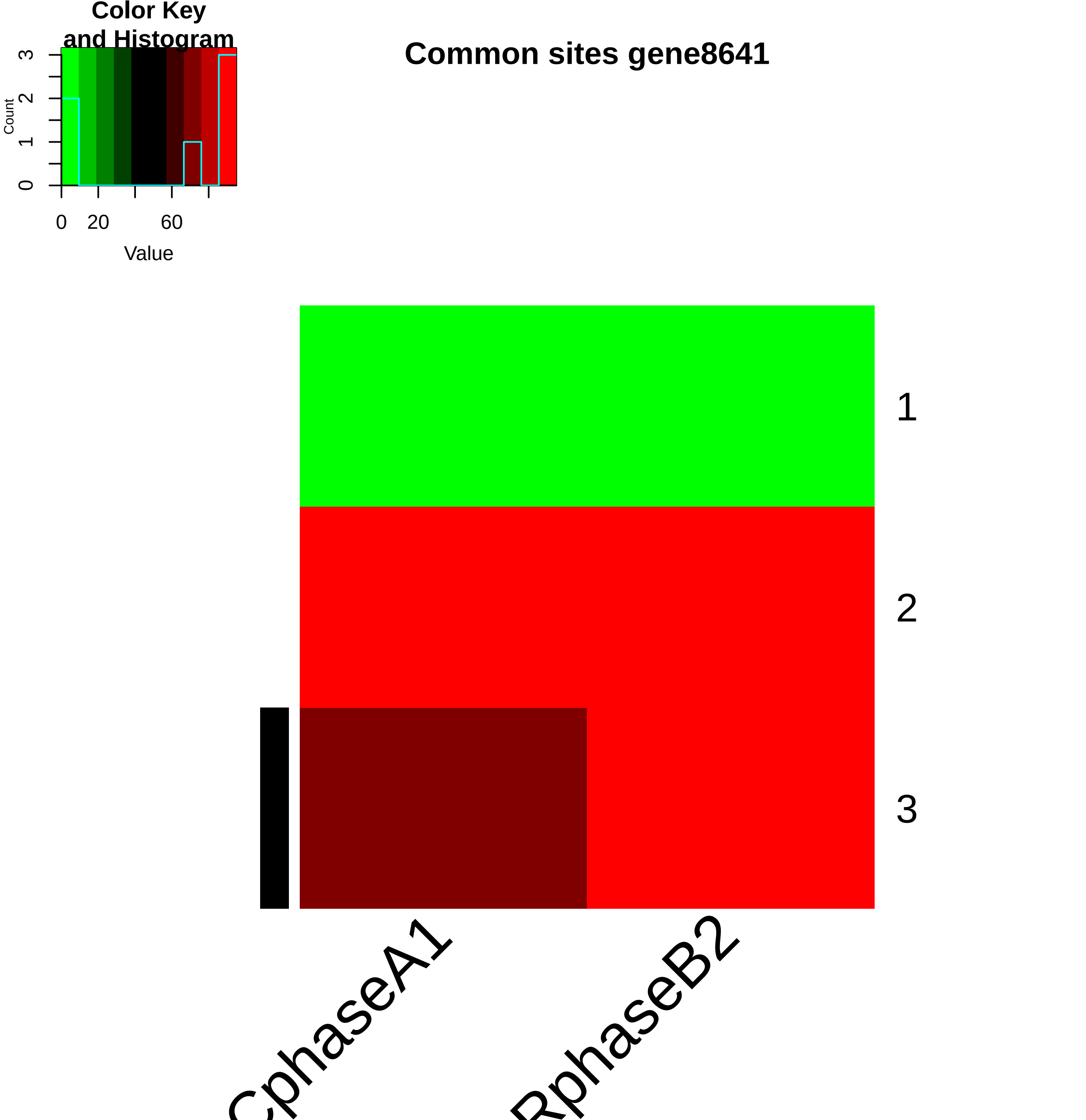
Color Key
and Histogram
Count
0
1
2
3
0
20
60
Value
Common sites gene8641
1
2
3
CphaseA1
RphaseB2

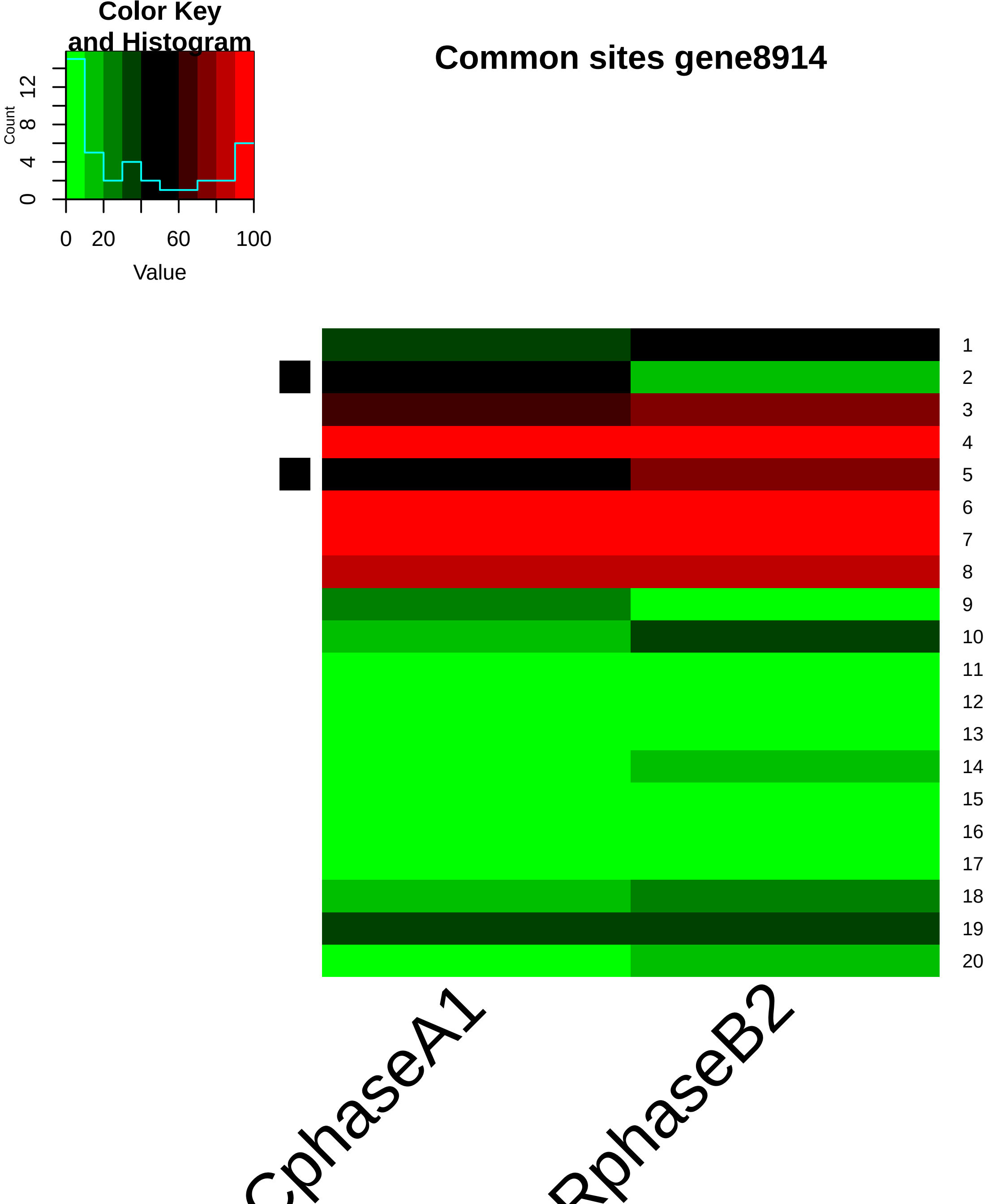
Color Key
and Histogram
Count
0
4
8
12
0
20
60
100
Value
Common sites gene8914
CphaseA1
RphaseB2

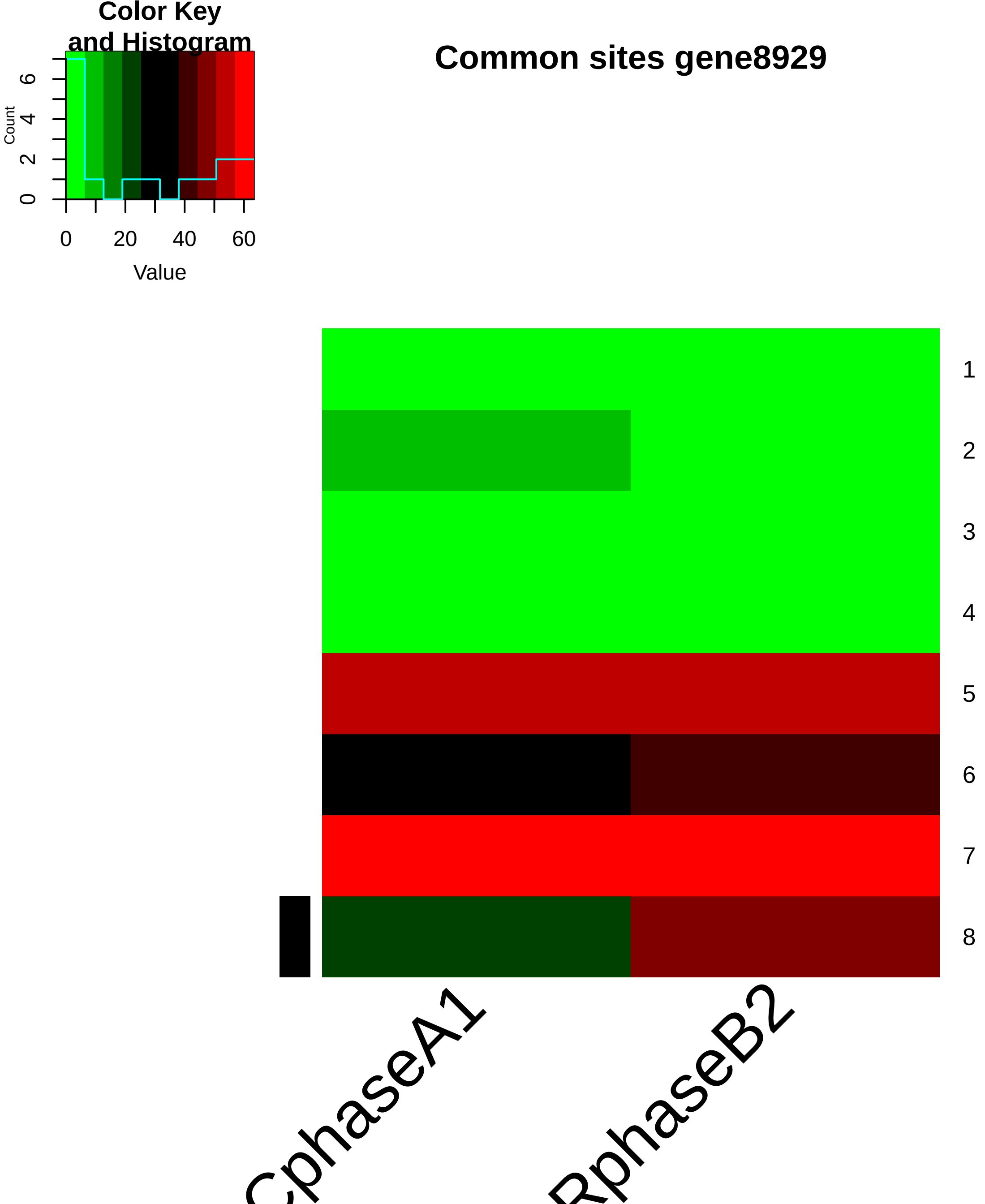

Color Key
and Histogram
Count
0
2
4
6
0
20
40
60
Value
Common sites gene8929
1
2
3
4
5
6
7
8
CphaseA1
RphaseB2

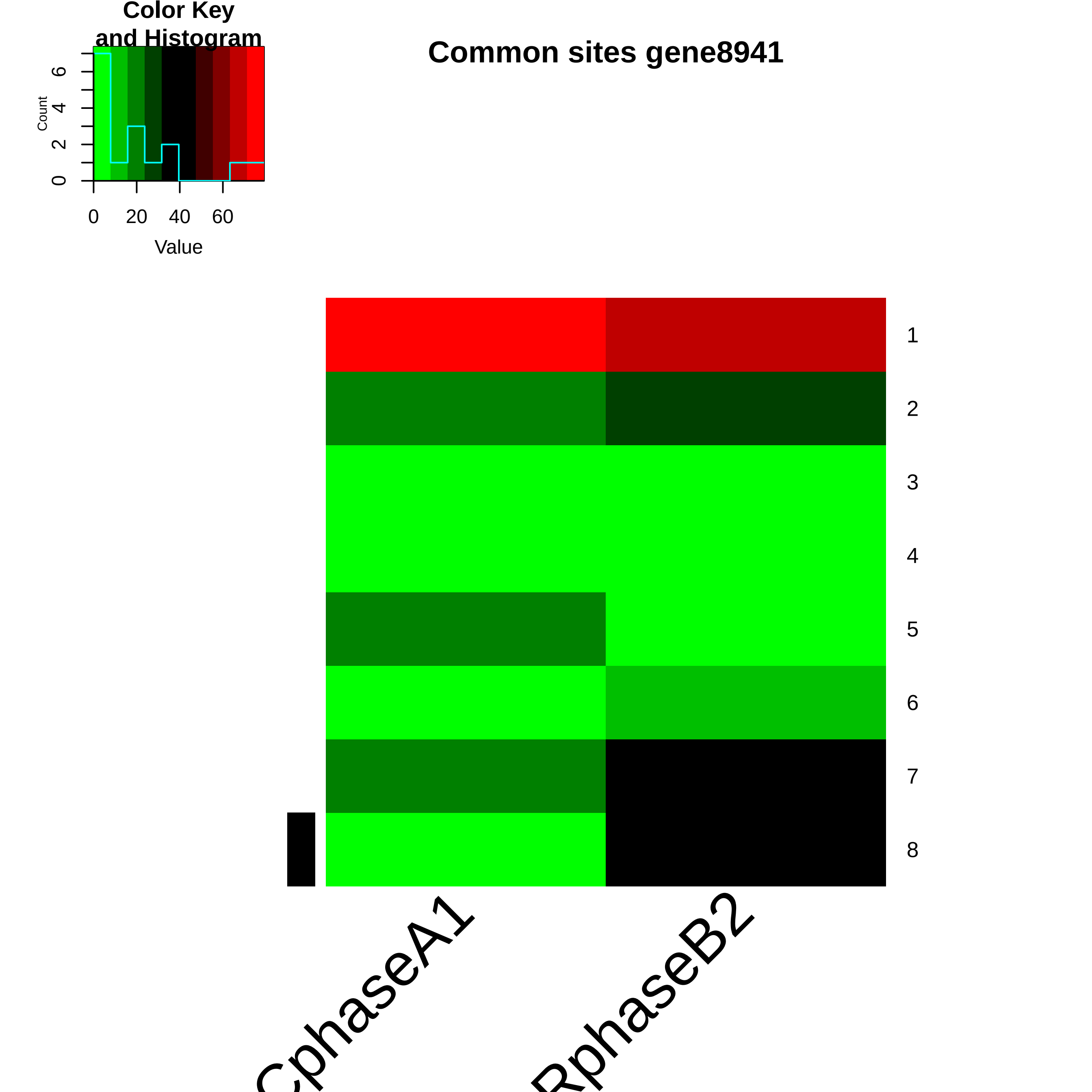

Color Key
and Histogram
Count
0
2
4
6
0
20
40
60
Value
Common sites gene8941
1
2
3
4
5
6
7
8
CphaseA1
RphaseB2

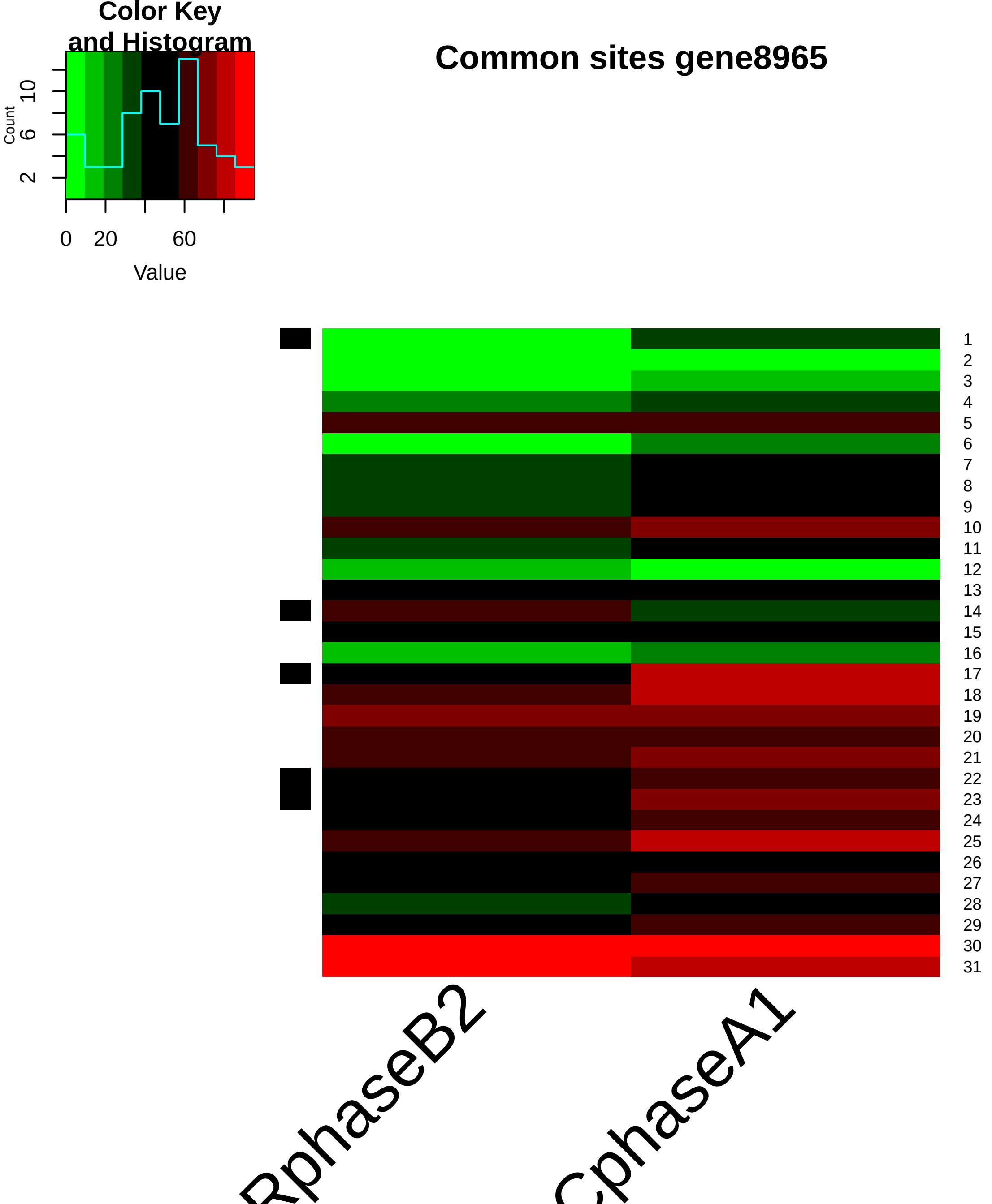
Color Key
and Histogram
Count
2
6
10
0
20
60
Value
Common sites gene8965
RphaseB2
CphaseA1

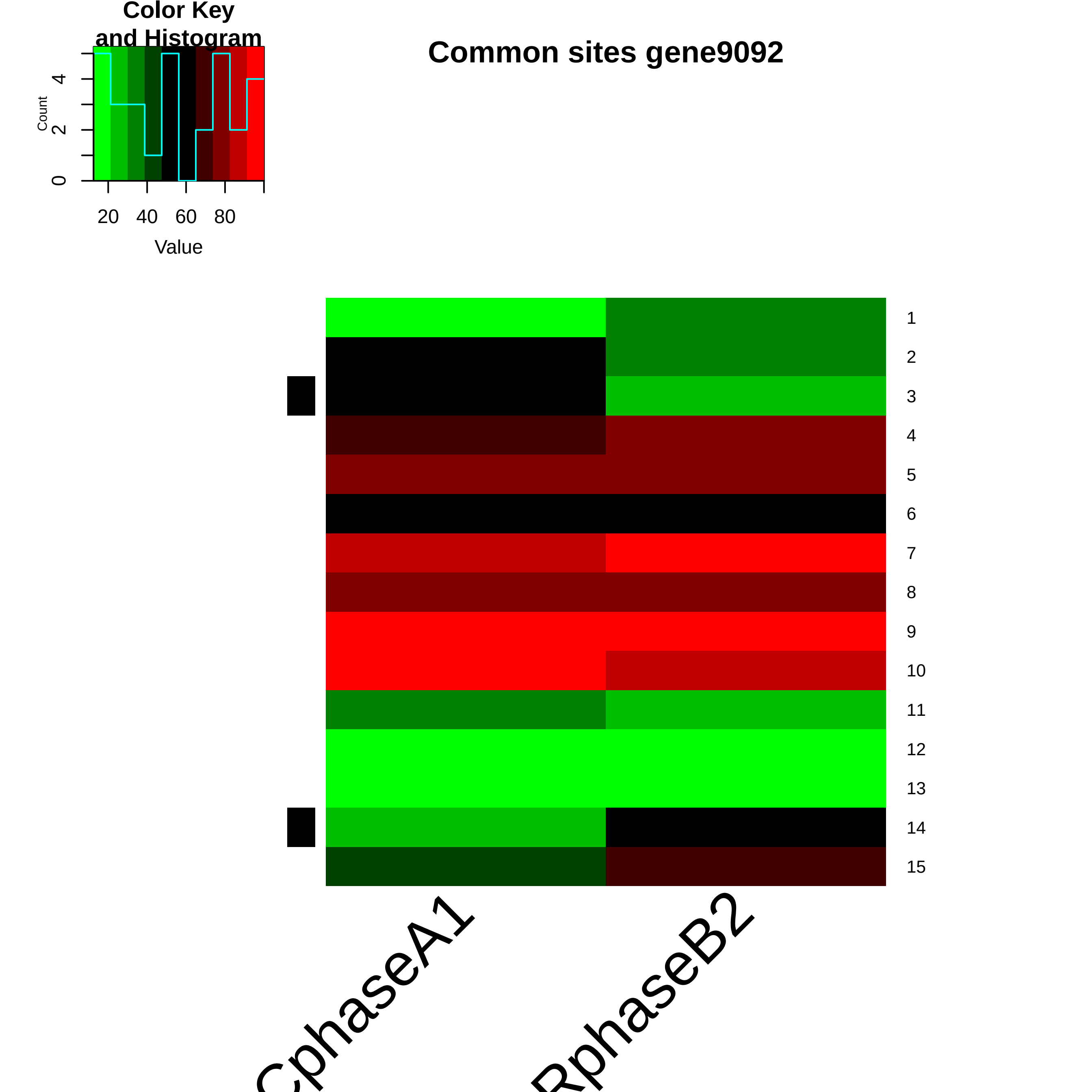
Color Key
and Histogram
Count
0
2
4
20
40
60
80
Value
Common sites gene9092
CphaseA1
RphaseB2

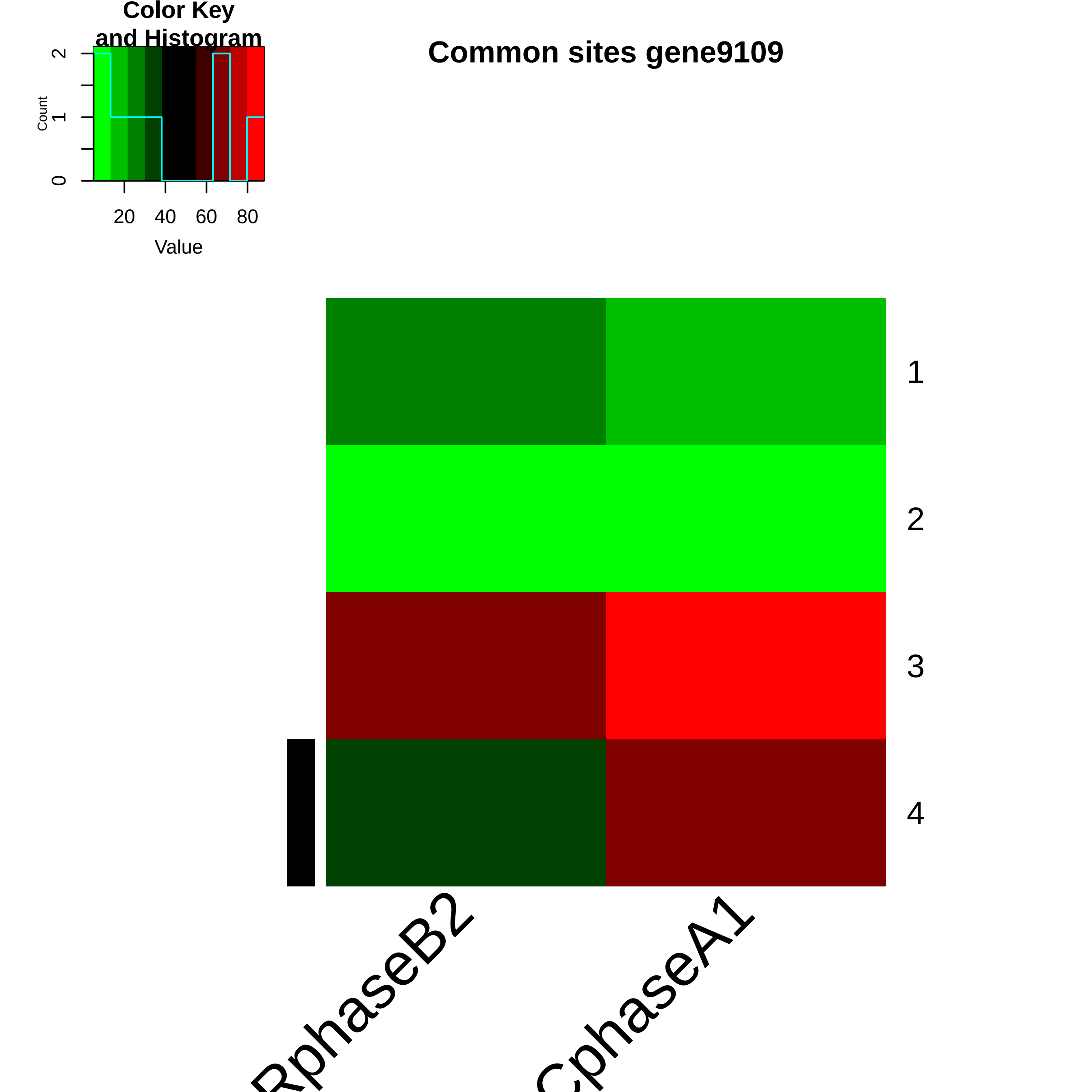
Color Key
and Histogram
Count
0
1
2
20
40
60
80
Value
Common sites gene9109
1
2
3
4
RphaseB2
CphaseA1

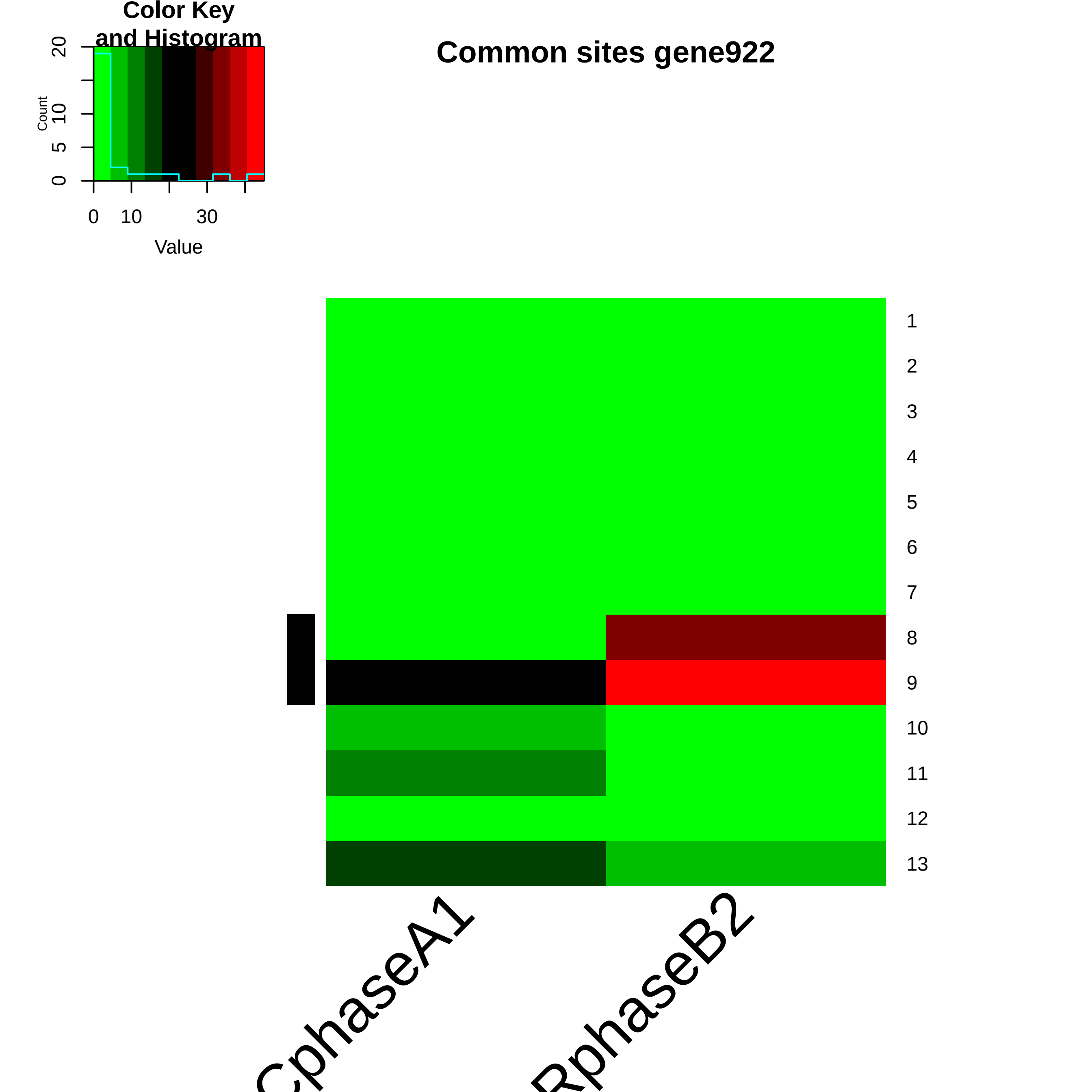

Color Key
and Histogram
Count
0
5
10
20
0
10
30
Value
Common sites gene922
1
2
3
4
5
6
7
8
9
10
11
12
13
CphaseA1
RphaseB2

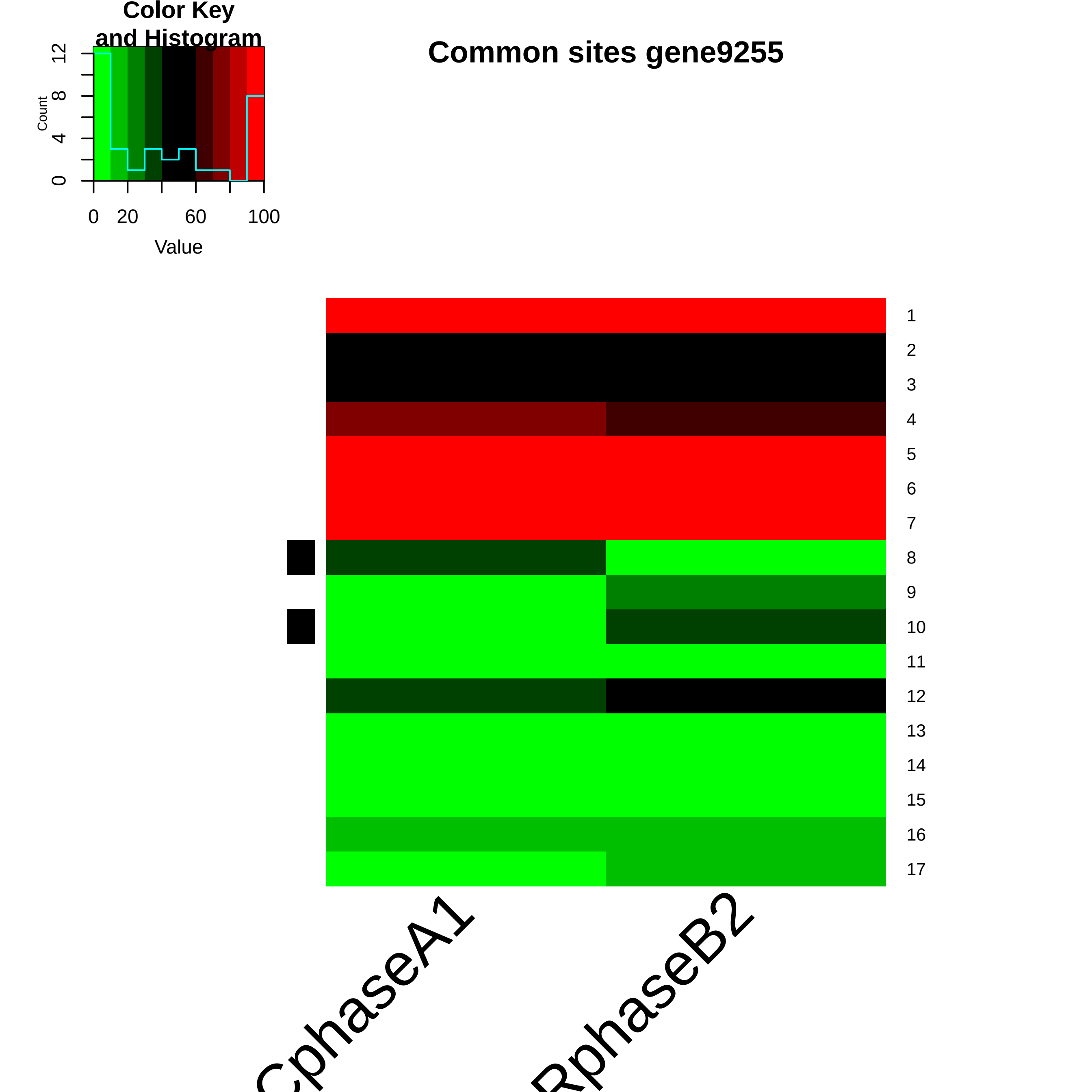
Color Key
and Histogram
Count
0
4
8
12
0
20
60
100
Value
Common sites gene9255
CphaseA1
RphaseB2

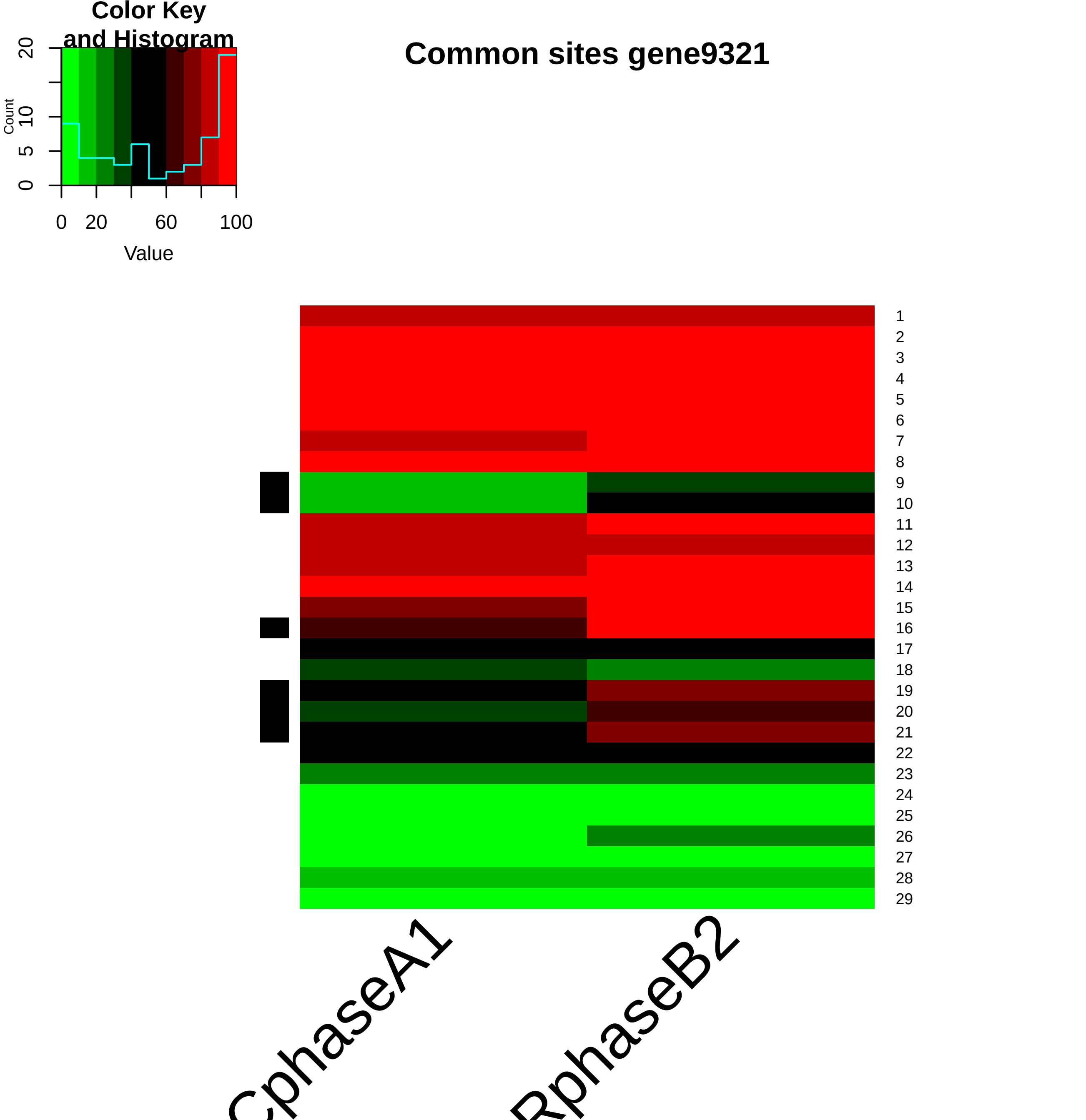
Color Key
and Histogram
Count
0
5
10
20
0
20
60
100
Value
Common sites gene9321
CphaseA1
RphaseB2

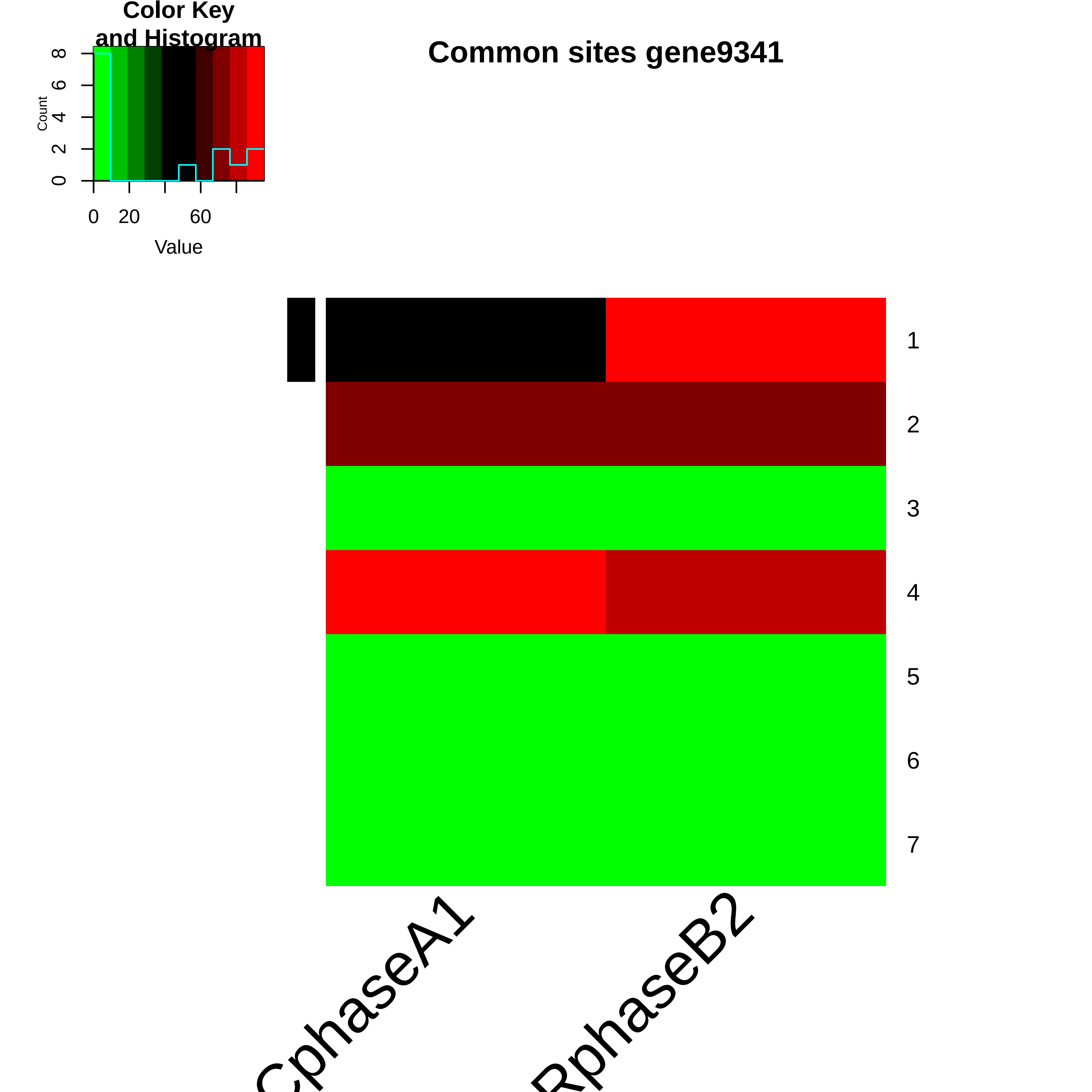

Color Key
and Histogram
Count
0
2
4
6
8
0
20
60
Value
Common sites gene9341
1
2
3
4
5
6
7
CphaseA1
RphaseB2

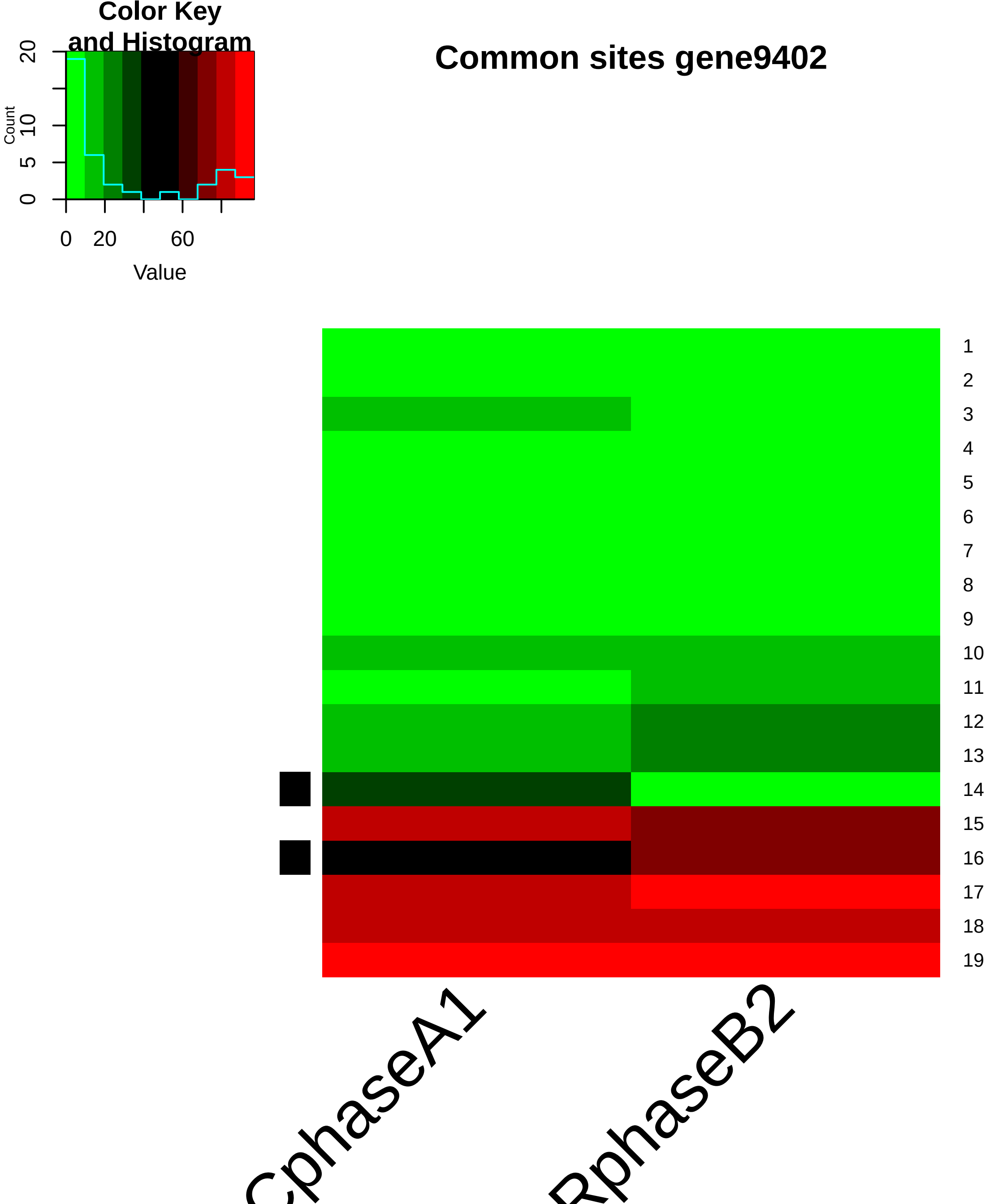
Color Key
and Histogram
Count
0
5
10
20
0
20
60
Value
Common sites gene9402
CphaseA1
RphaseB2

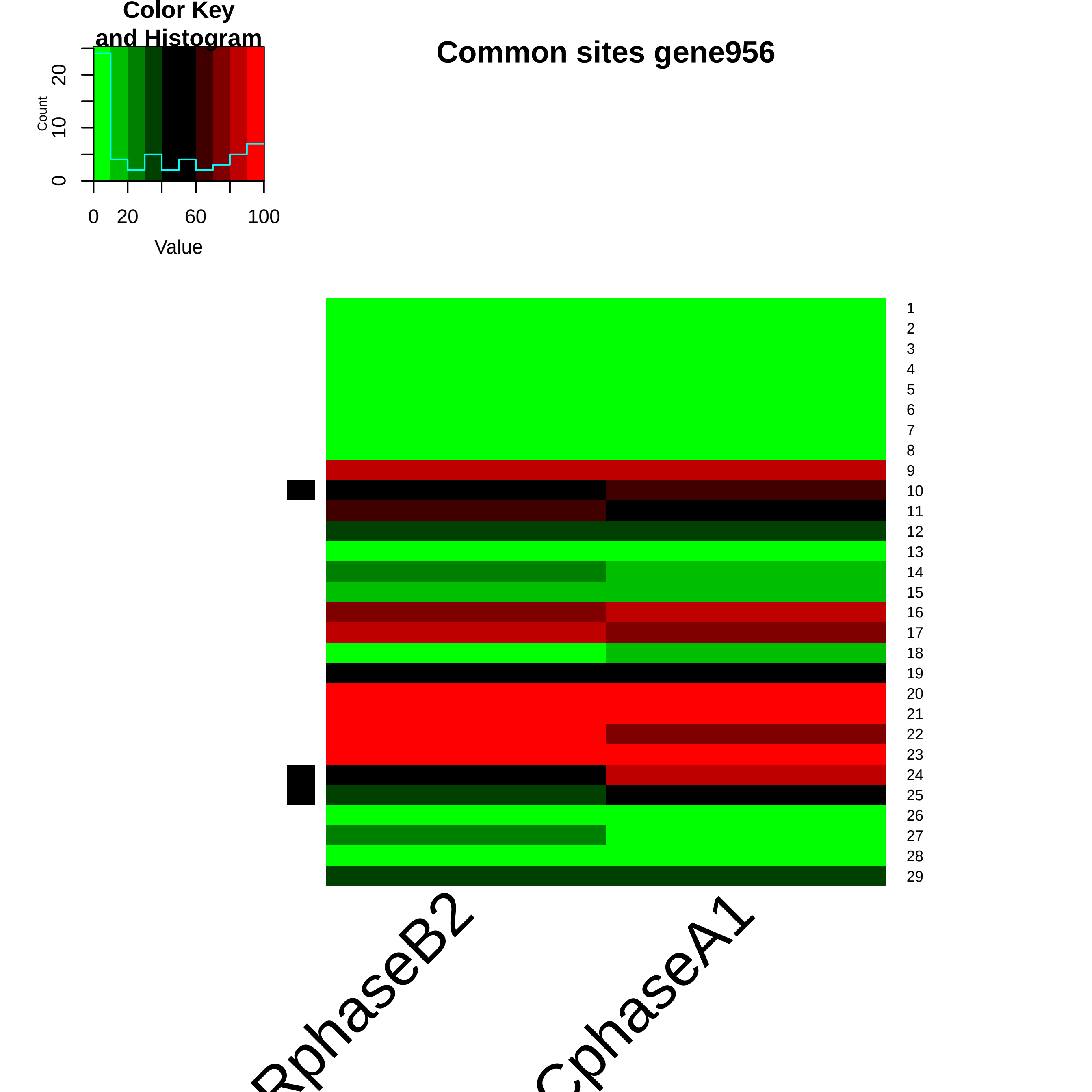
Color Key
and Histogram
Count
0
10
20
0
20
60
100
Value
Common sites gene956
RphaseB2
CphaseA1

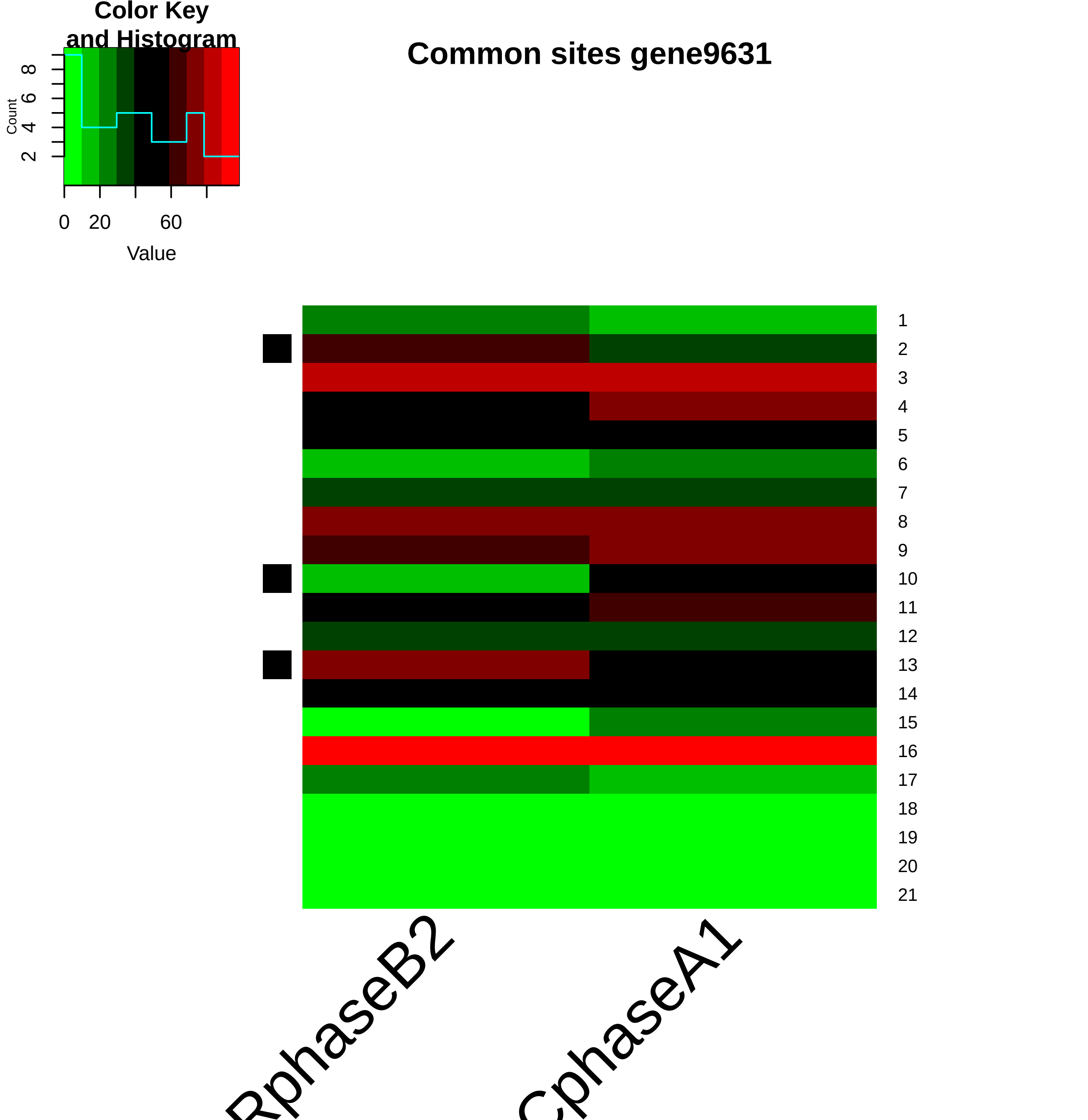
Color Key
and Histogram
Count
2
4
6
8
0
20
60
Value
Common sites gene9631
RphaseB2
CphaseA1

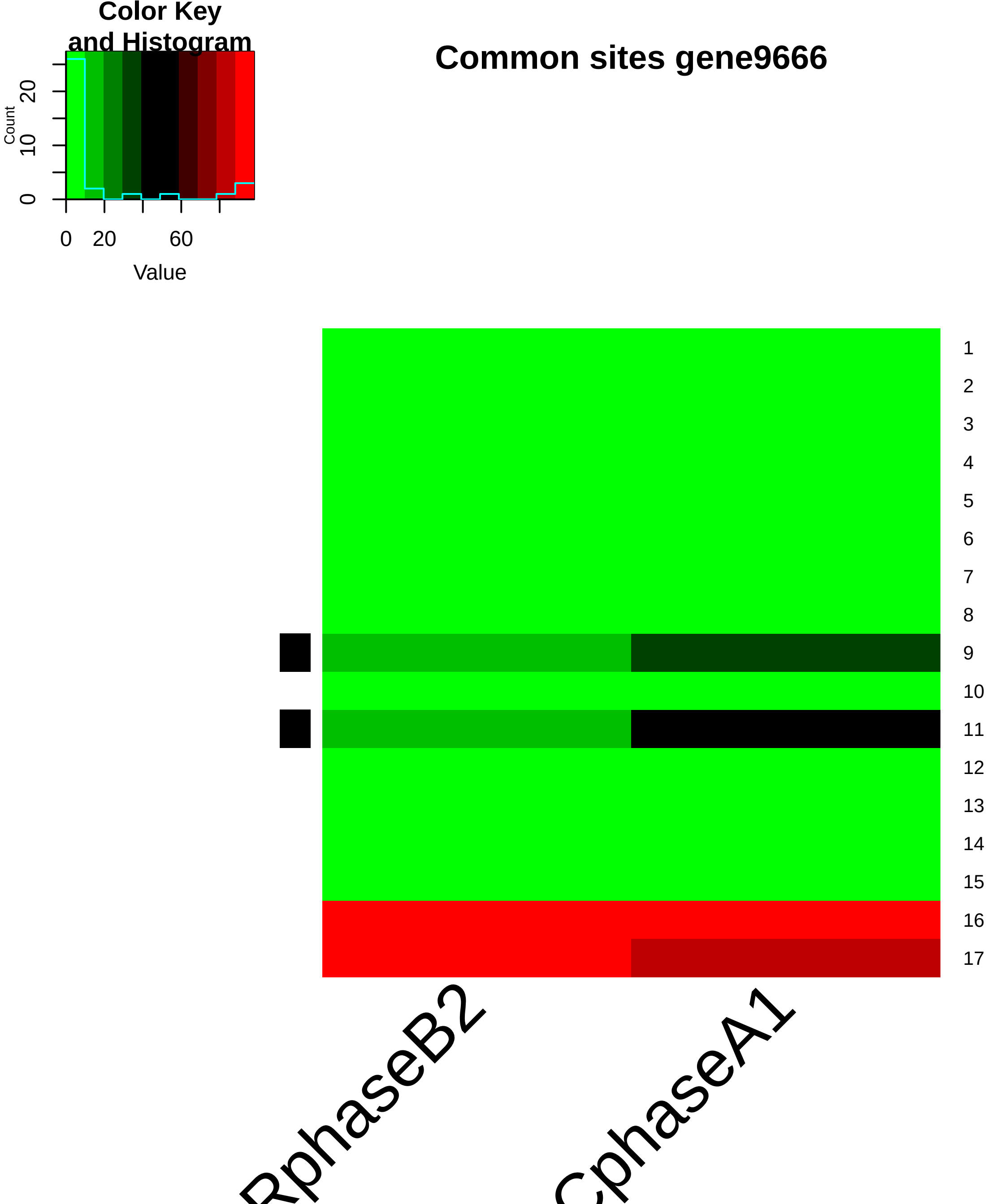

Color Key
and Histogram
Count
0
10
20
0
20
60
Value
Common sites gene9666
RphaseB2
CphaseA1

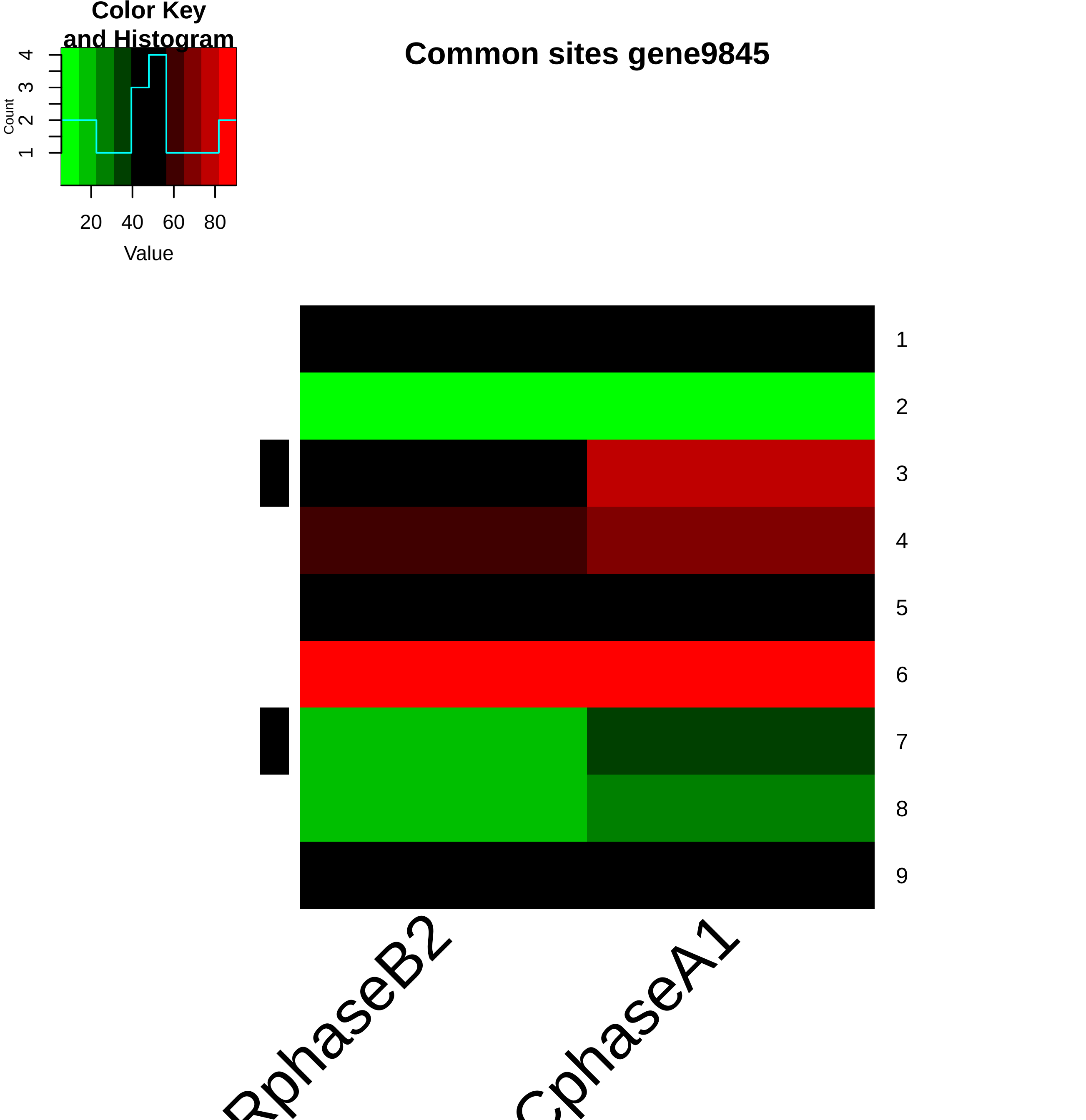

Color Key
and Histogram
Count
1
2
3
4
20
40
60
80
Value
Common sites gene9845
1
2
3
4
5
6
7
8
9
RphaseB2
CphaseA1

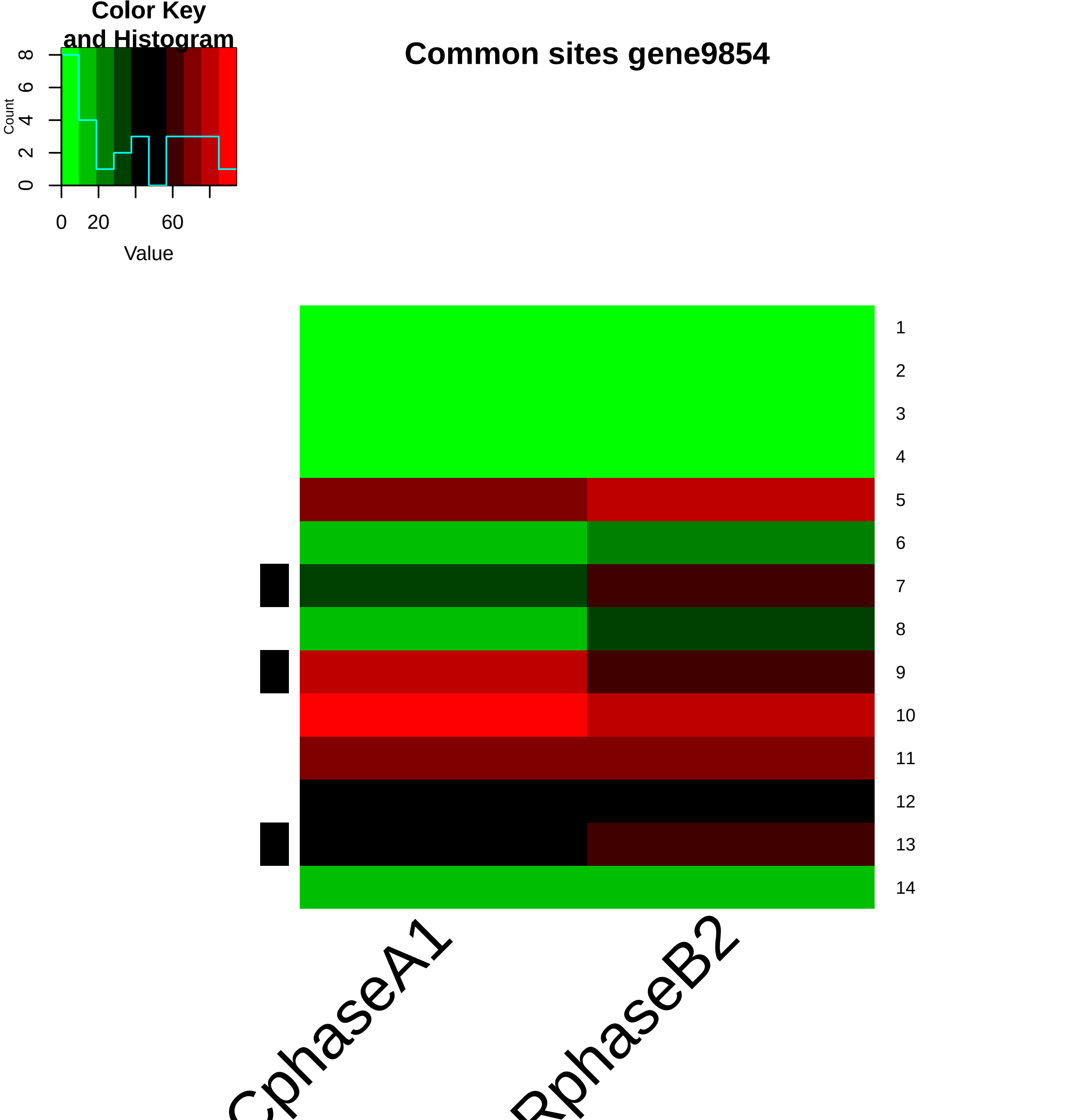
Color Key
and Histogram
Count
0
2
4
6
8
0
20
60
Value
Common sites gene9854
1
2
3
4
5
6
7
8
9
10
11
12
13
14
CphaseA1
RphaseB2

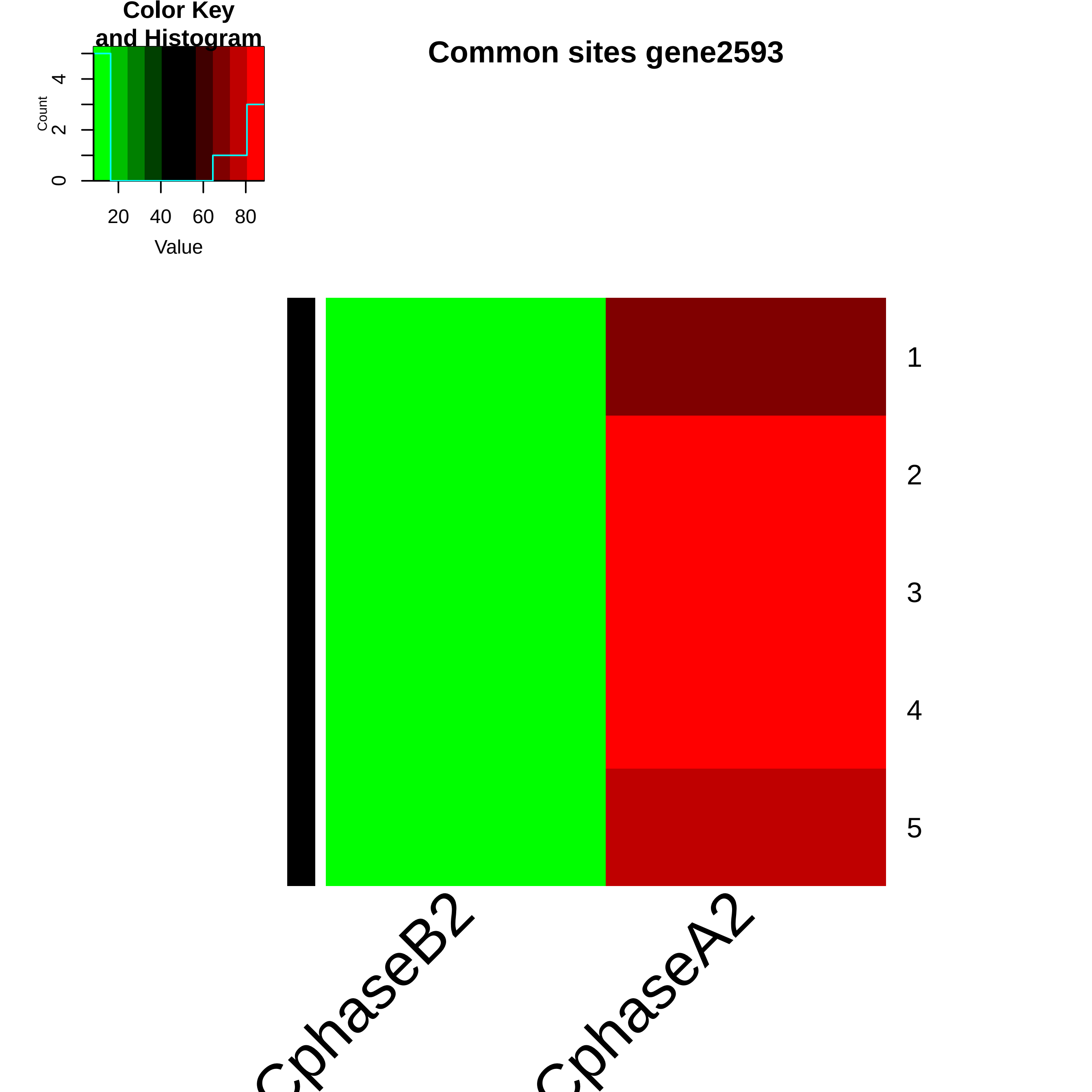

Color Key
and Histogram
Count
0
2
4
20
40
60
80
Value
Common sites gene2593
1
2
3
4
5
CphaseB2
CphaseA2

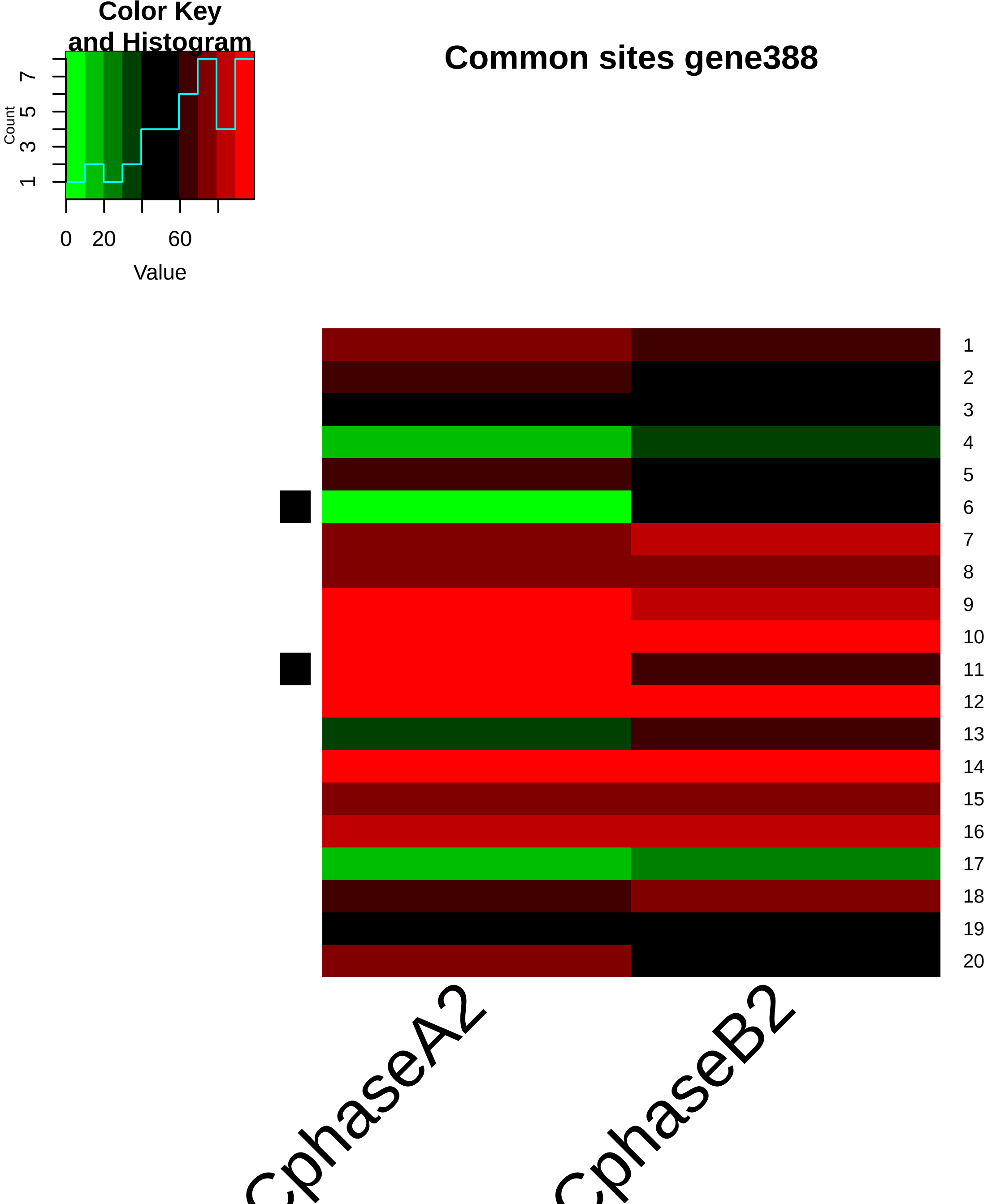

Color Key
and Histogram
Count
1
3
5
7
0
20
60
Value
Common sites gene388
CphaseA2
CphaseB2

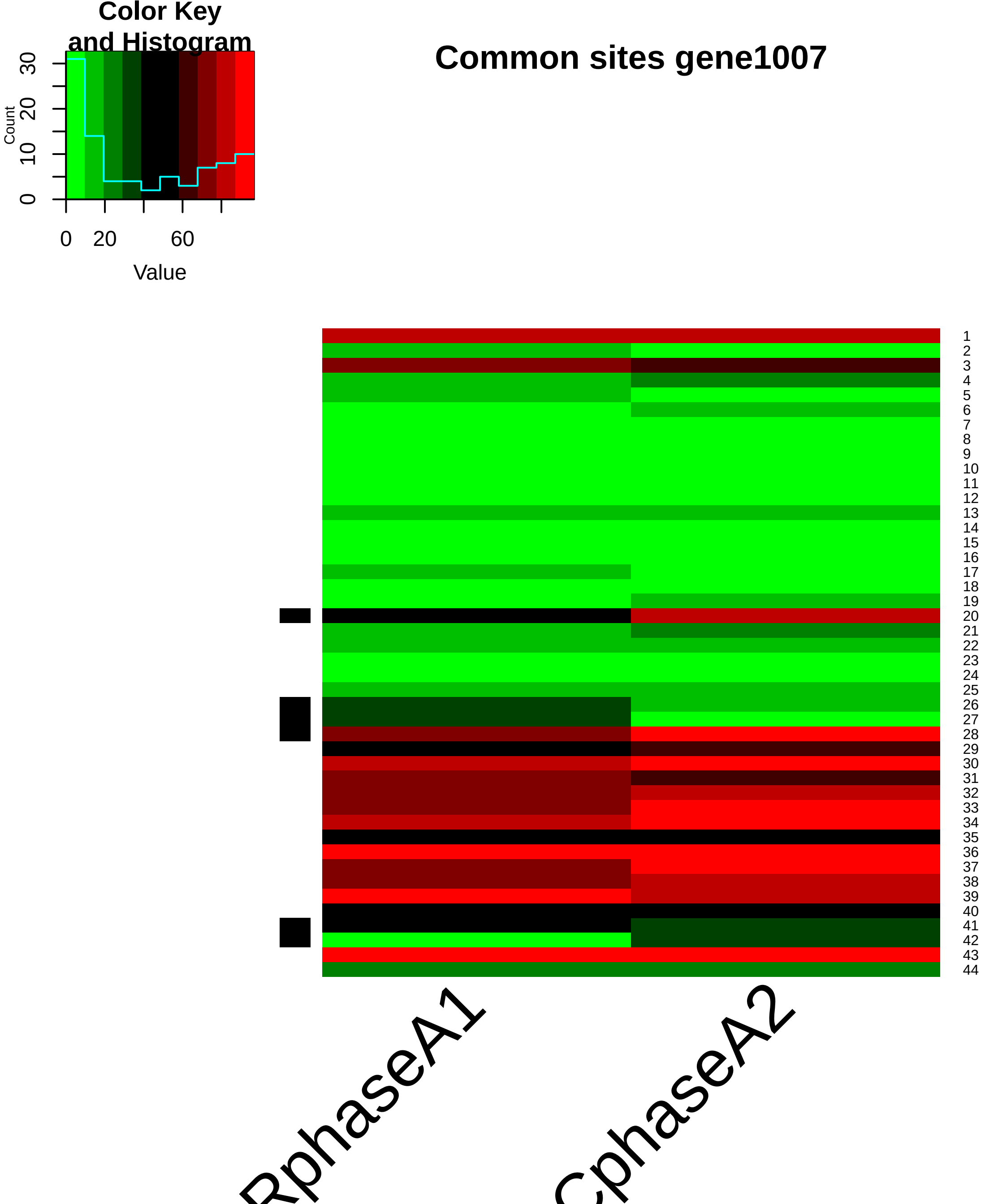
Color Key
and Histogram
Count
0
10
20
30
0
20
60
Value
Common sites gene1007
RphaseA1
CphaseA2

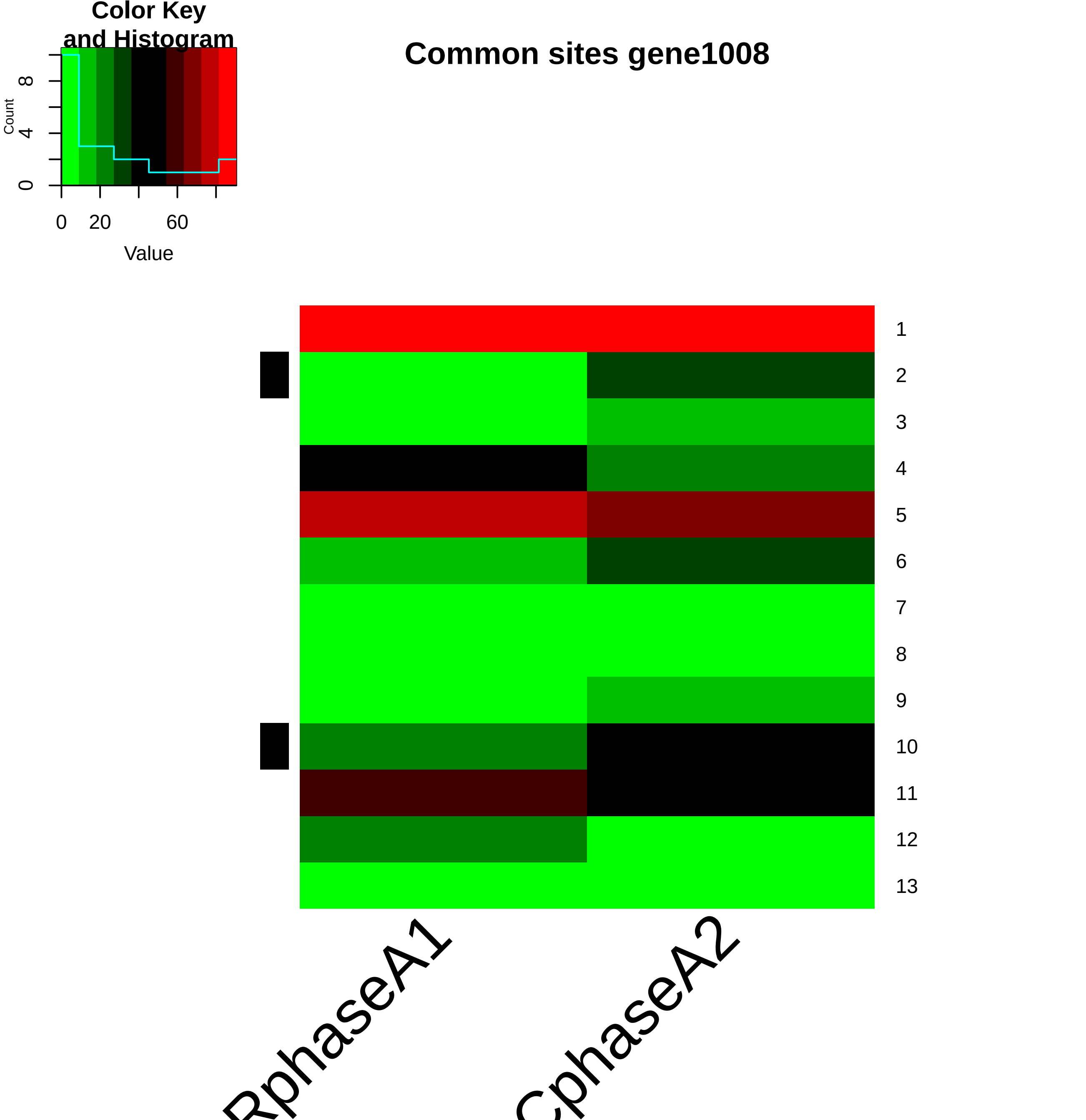
Color Key
and Histogram
Count
0
4
8
0
20
60
Value
Common sites gene1008
1
2
3
4
5
6
7
8
9
10
11
12
13
RphaseA1
CphaseA2

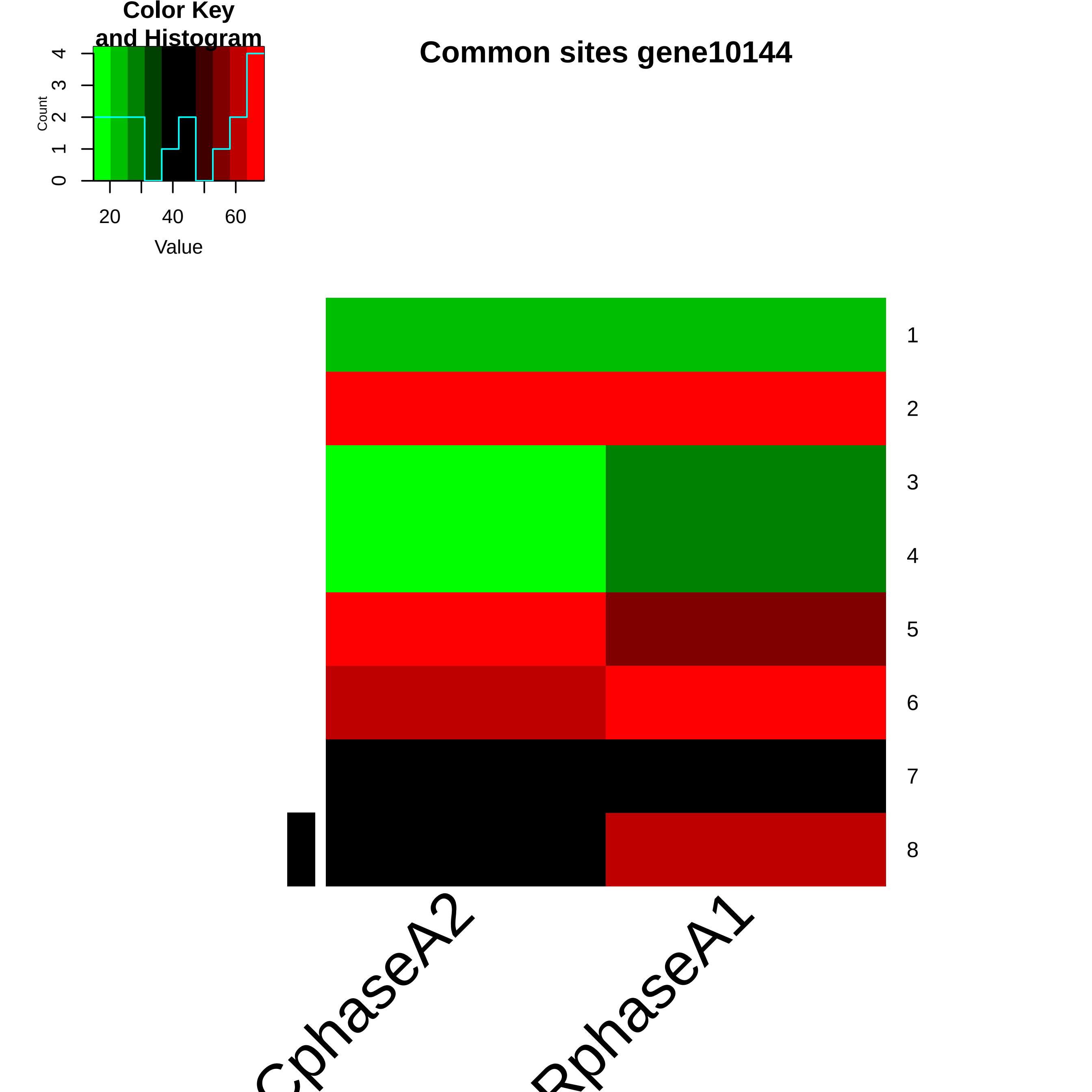
Color Key
and Histogram
Count
0
1
2
3
4
20
40
60
Value
Common sites gene10144
1
2
3
4
5
6
7
8
CphaseA2
RphaseA1

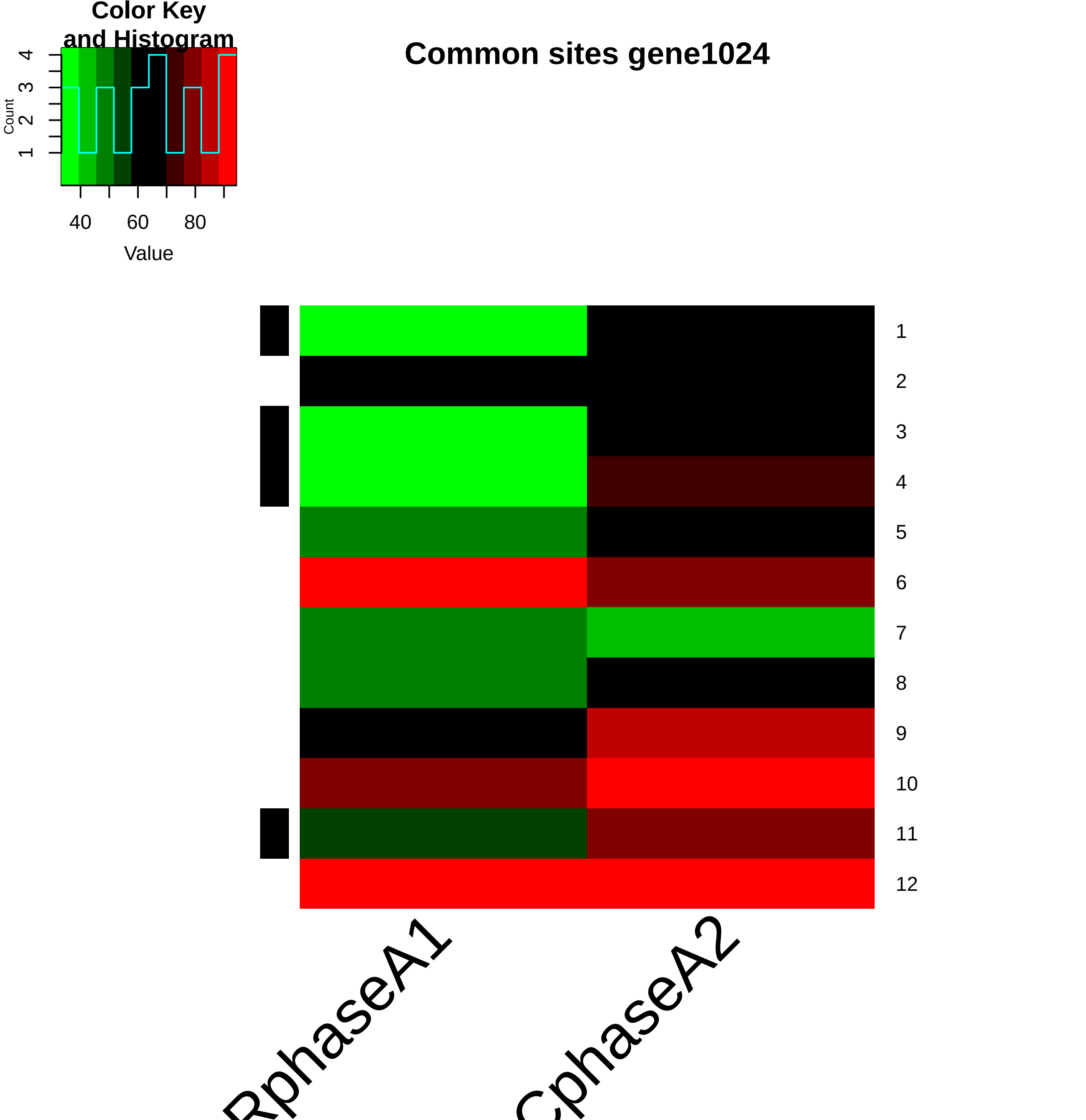
Color Key
and Histogram
Count
1
2
3
4
40
60
80
Value
Common sites gene1024
1
2
3
4
5
6
7
8
9
10
11
12
RphaseA1
CphaseA2

### Common sites gene1025

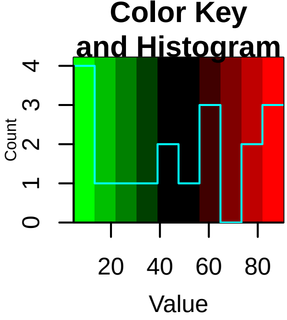

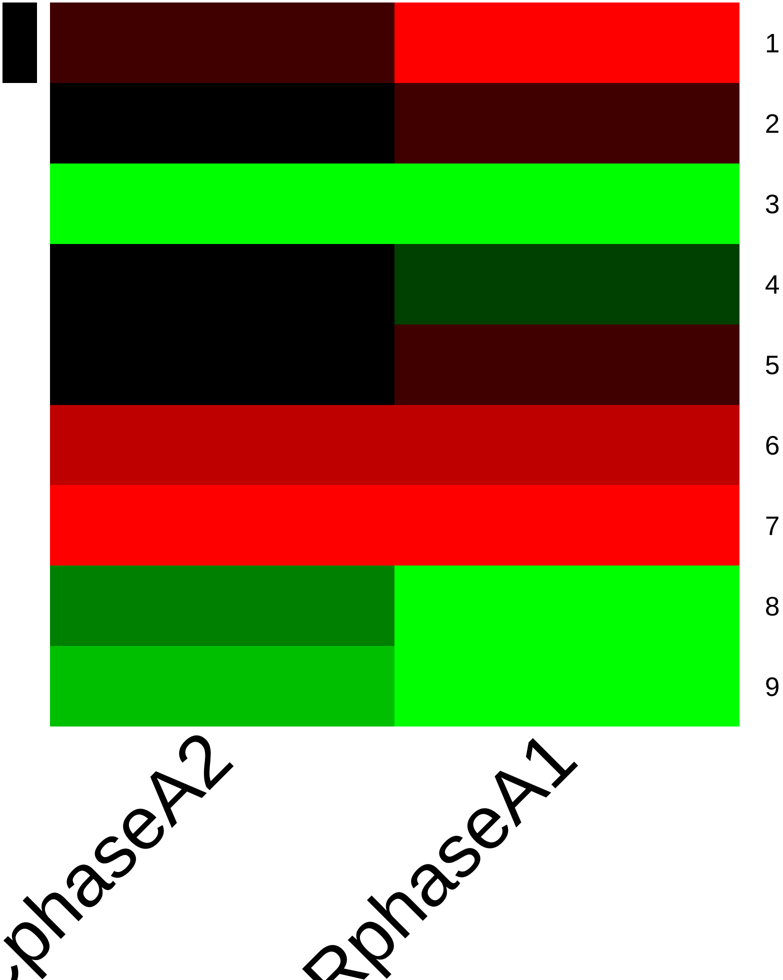

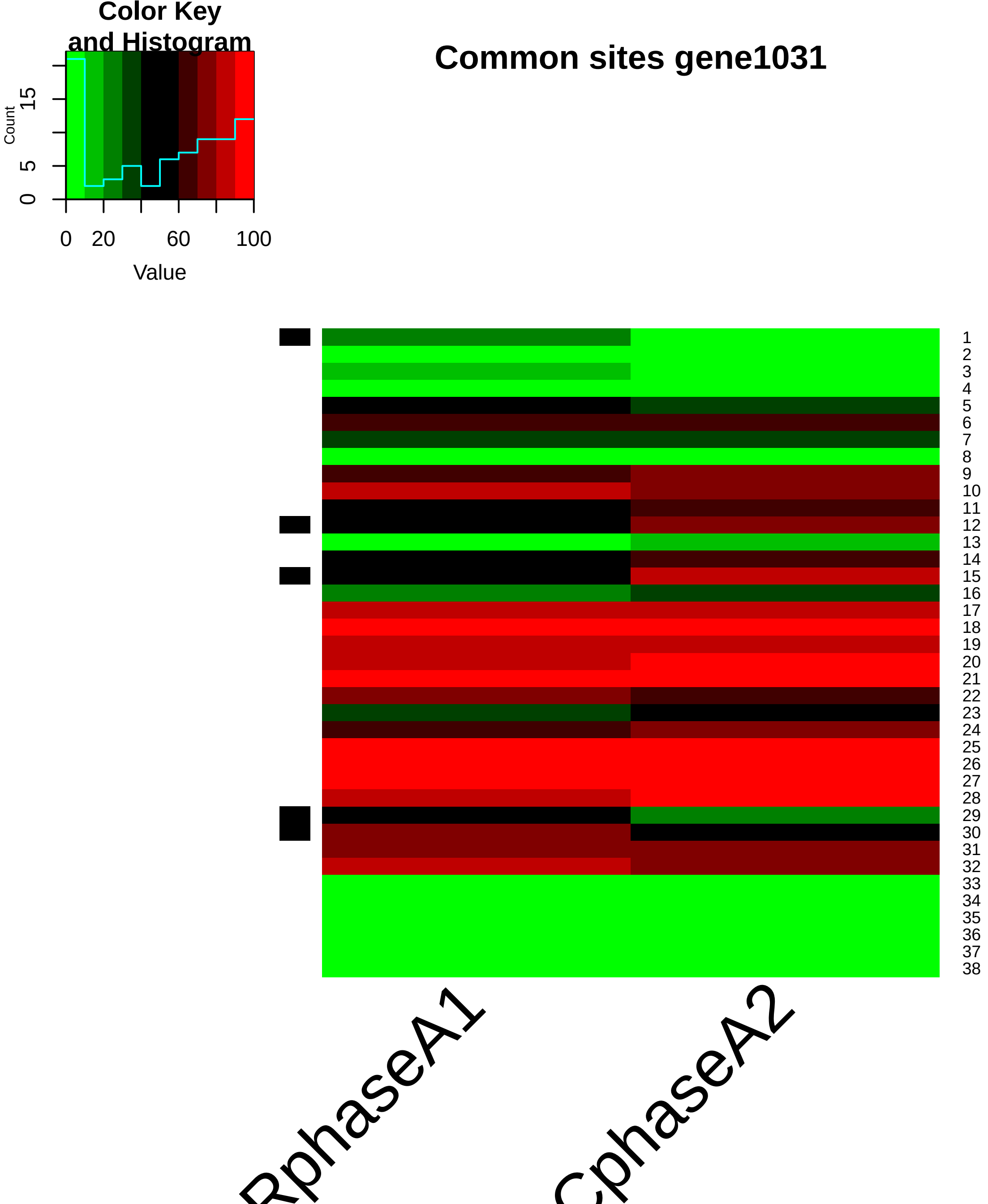

Color Key
and Histogram
Count
0
5
15
0
20
60
100
Value
Common sites gene1031
RphaseA1
CphaseA2

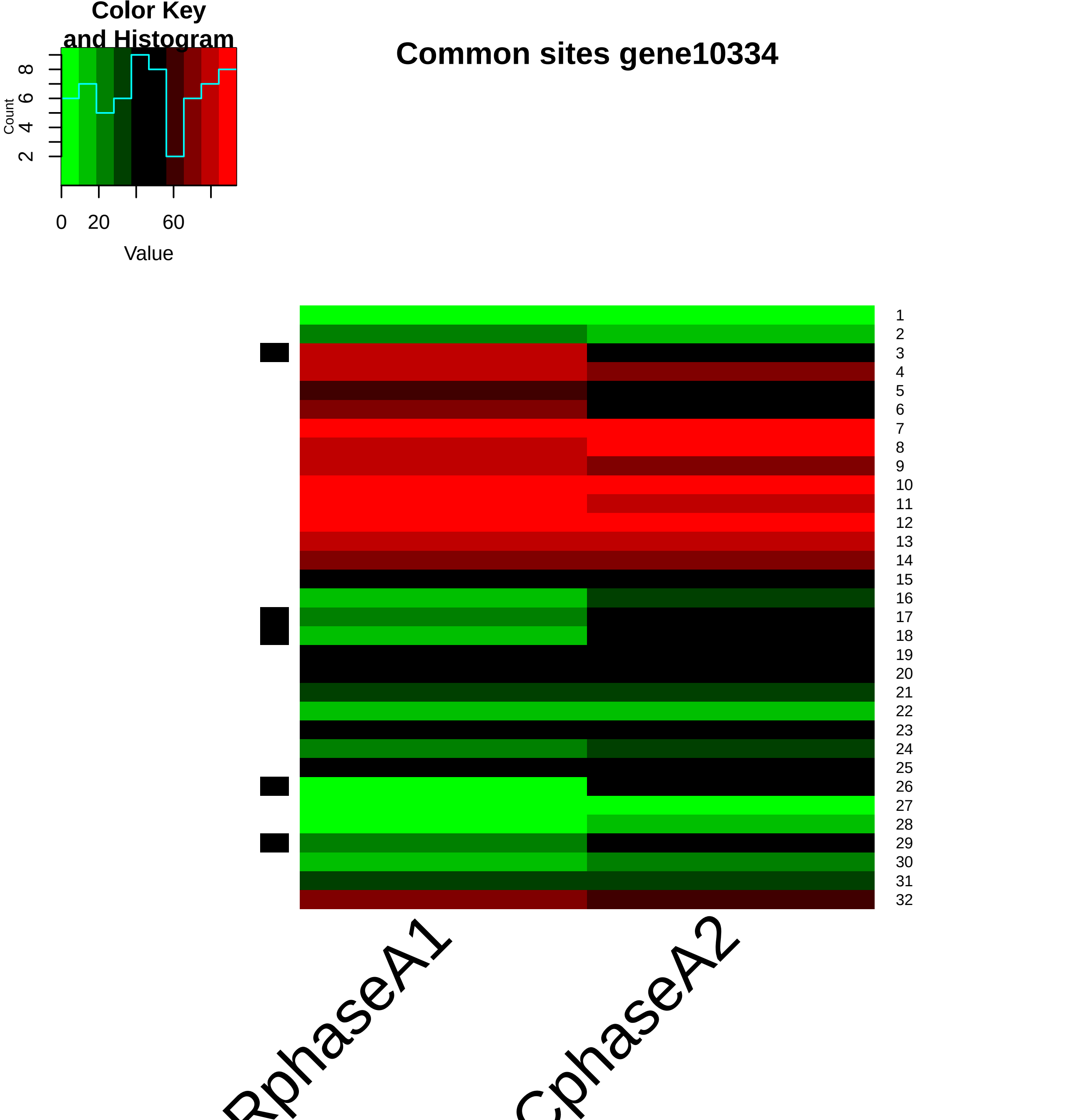
Color Key
and Histogram
Count
2
4
6
8
0
20
60
Value
Common sites gene10334
RphaseA1
CphaseA2

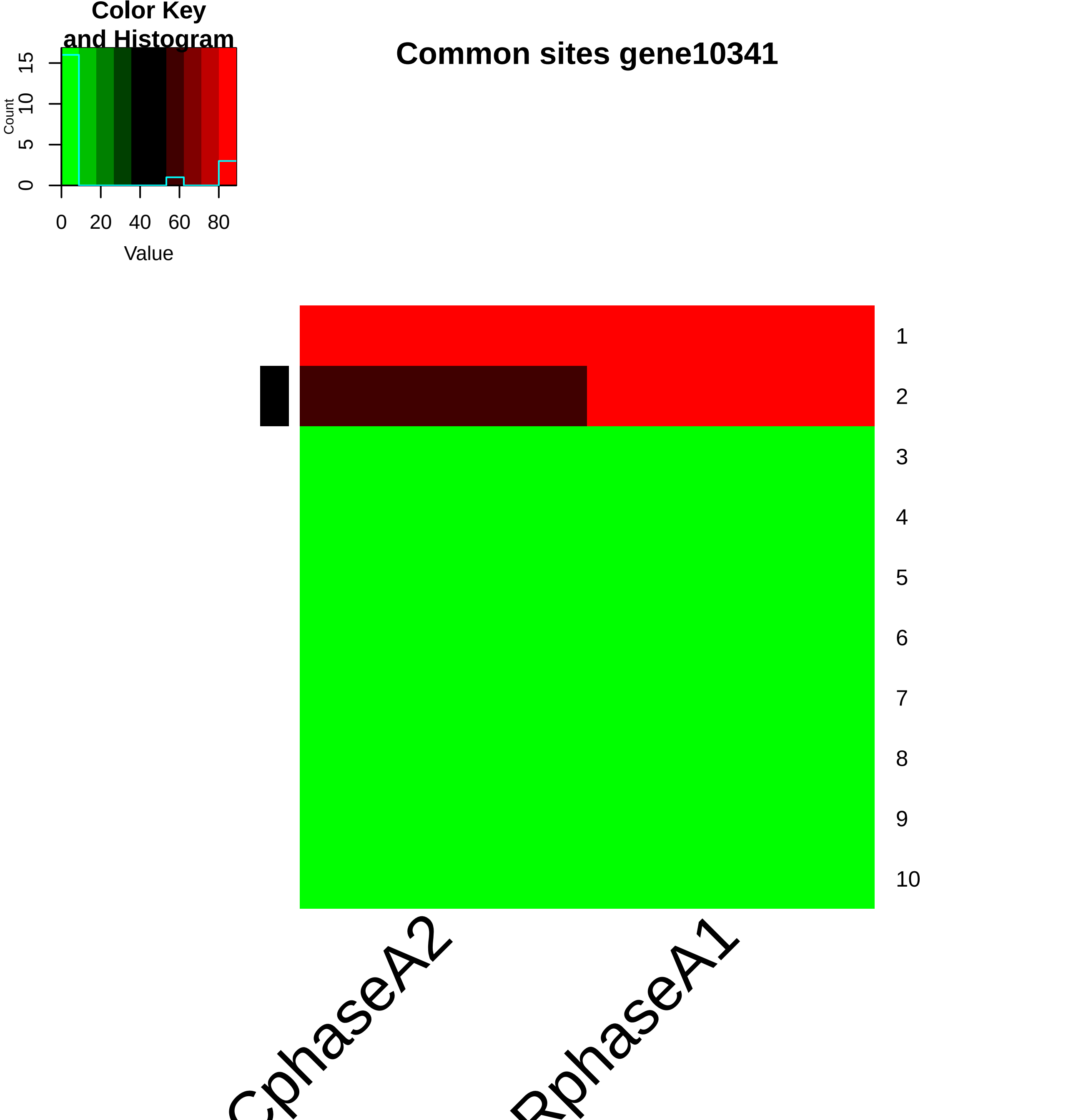

Color Key
and Histogram
Count
0
5
10
15
0
20
40
60
80
Value
Common sites gene10341
1
2
3
4
5
6
7
8
9
10

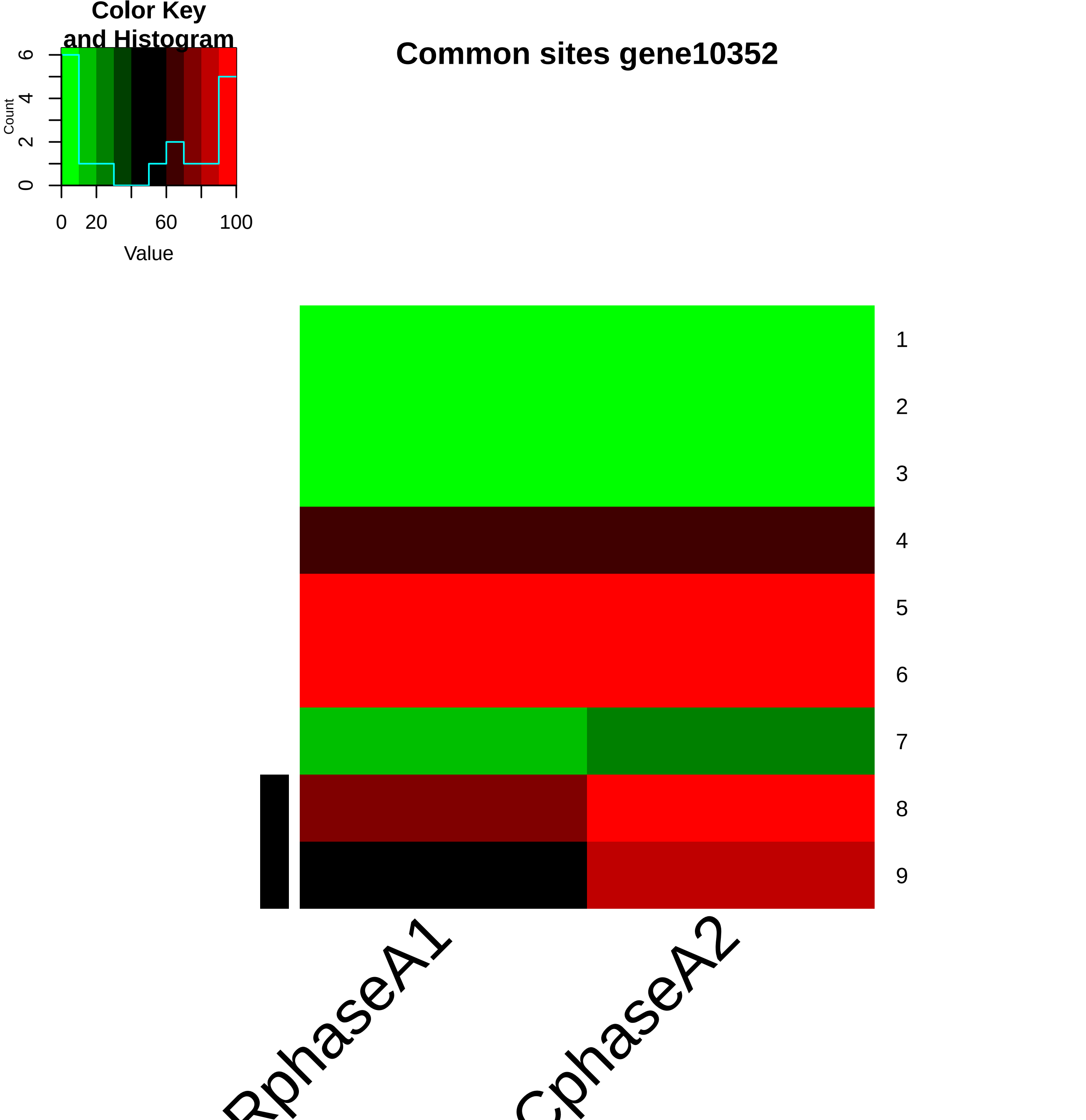
Color Key
and Histogram
Count
0
2
4
6
0
20
60
100
Value
Common sites gene10352
1
2
3
4
5
6
7
8
9
RphaseA1
CphaseA2

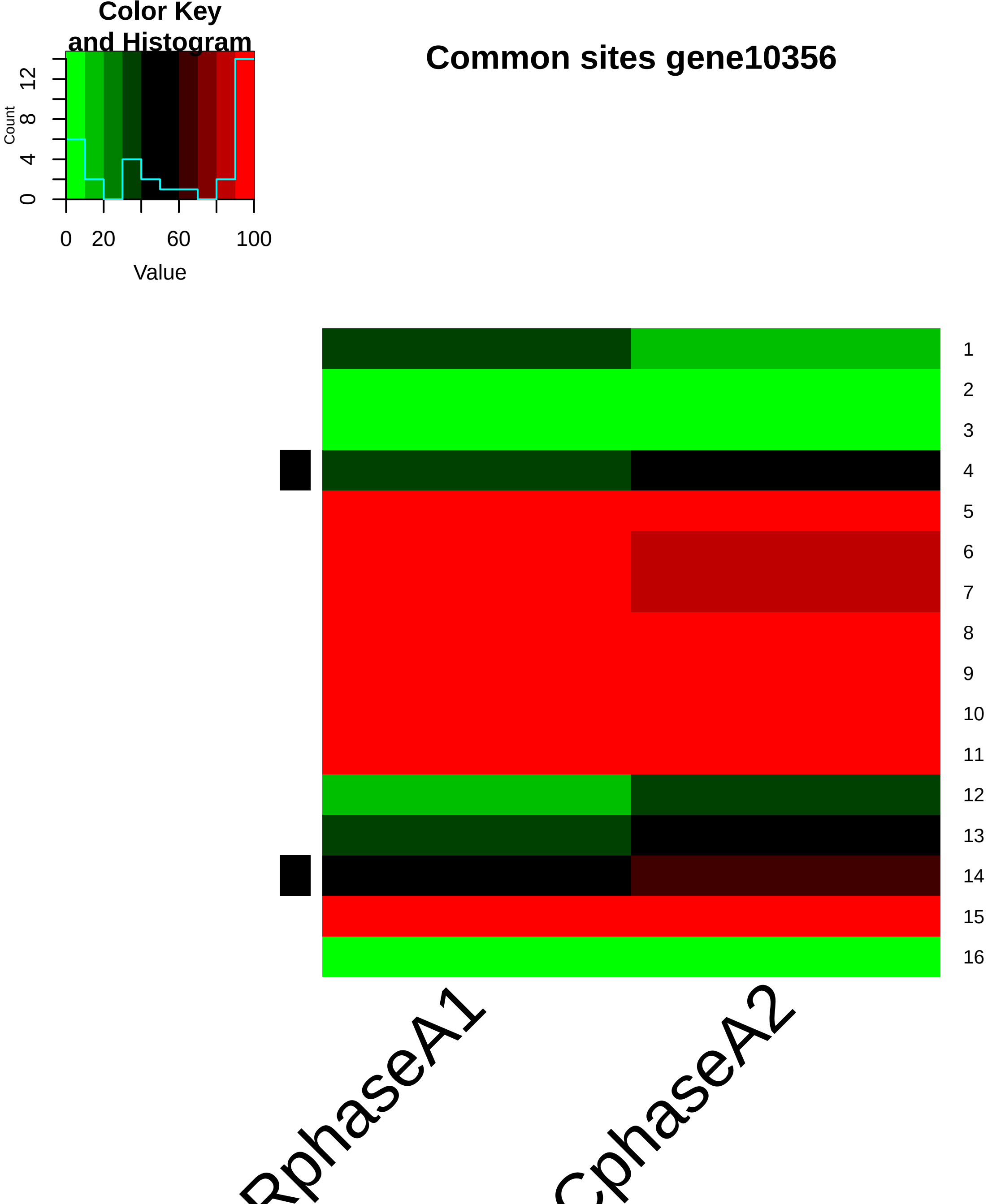
Color Key
and Histogram
Count
0
4
8
12
0
20
60
100
Value
Common sites gene10356
RphaseA1
CphaseA2

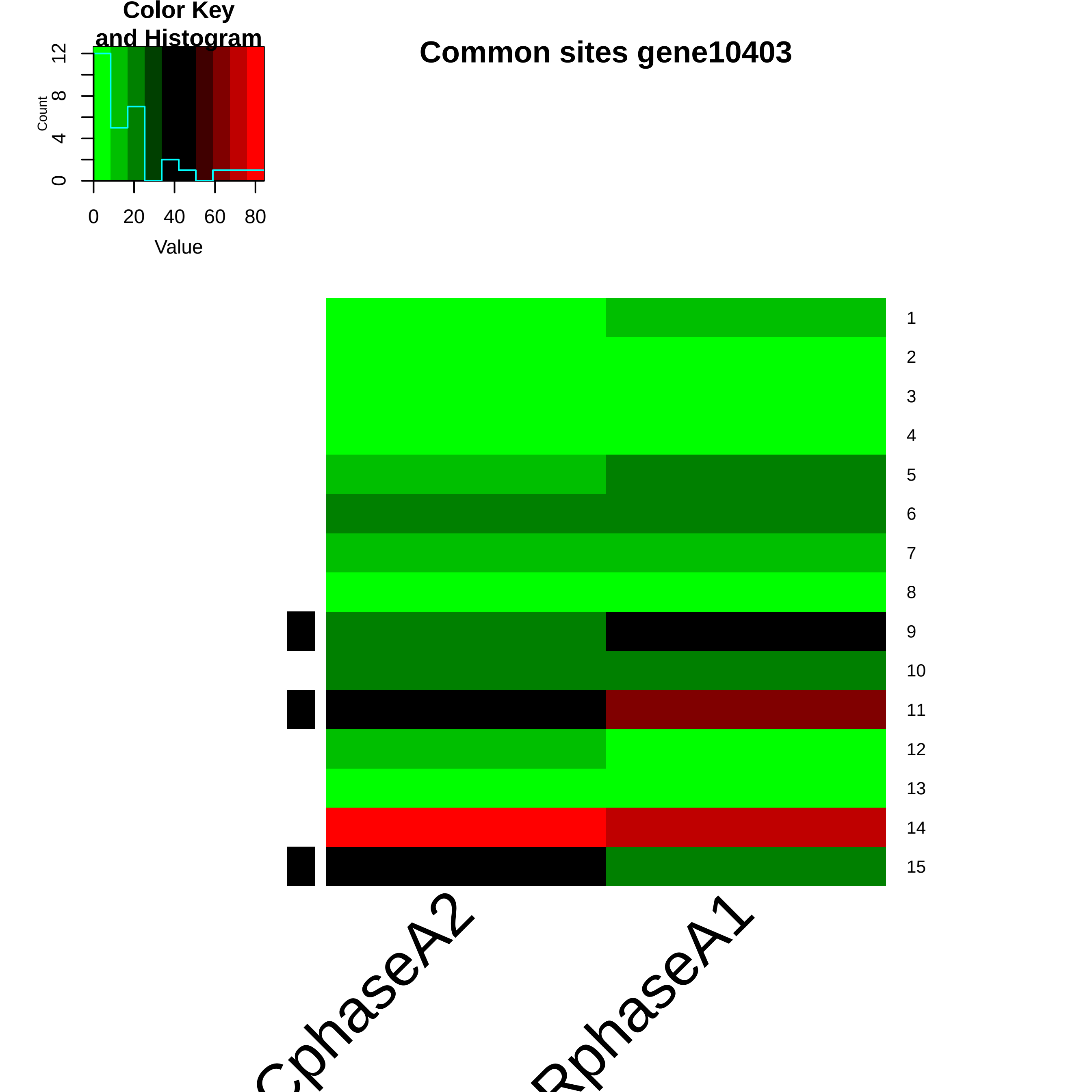
Color Key
and Histogram
Count
0
4
8
12
0
20
40
60
80
Value
Common sites gene10403
CphaseA2
RphaseA1

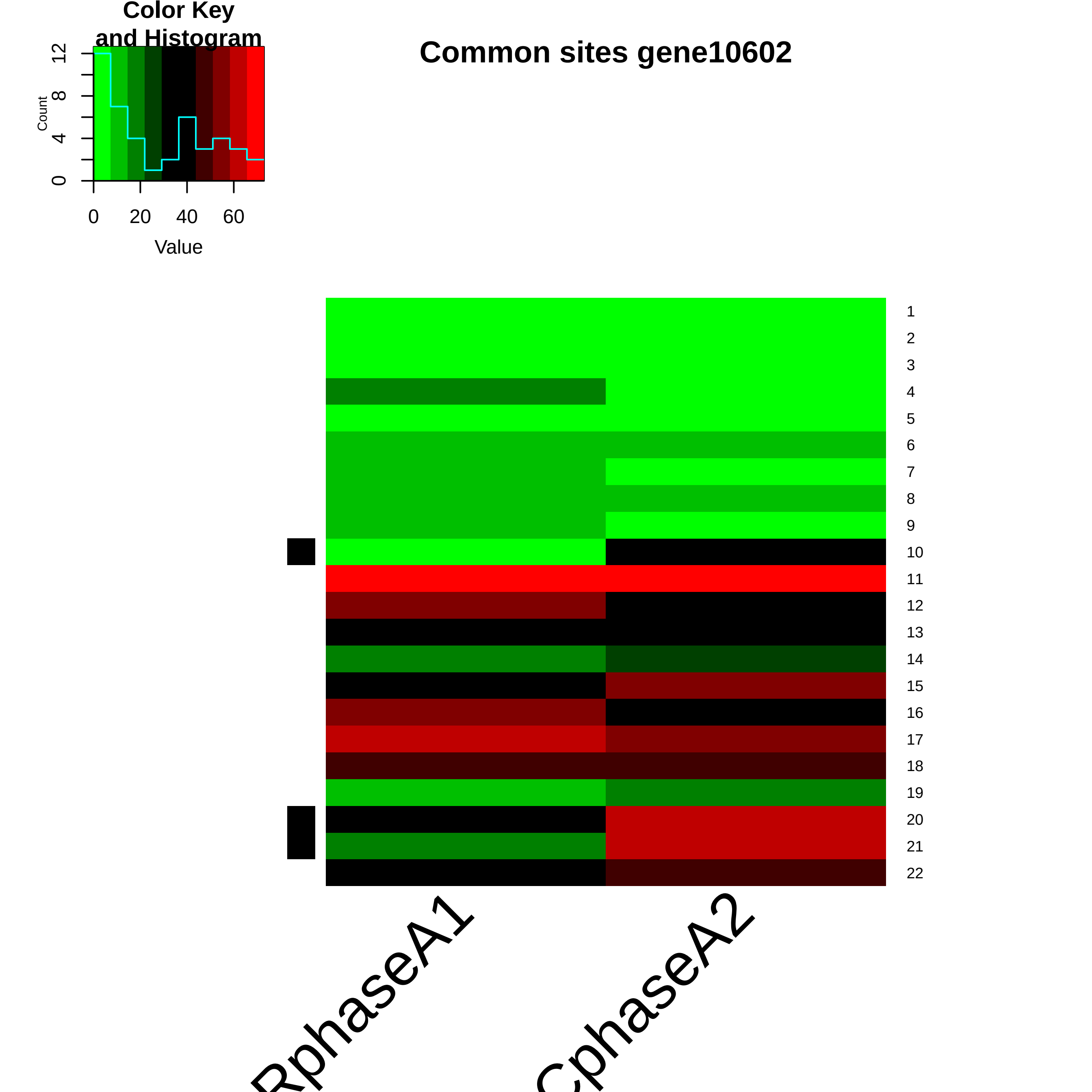
Color Key
and Histogram
Count
0
4
8
12
0
20
40
60
Value
Common sites gene10602

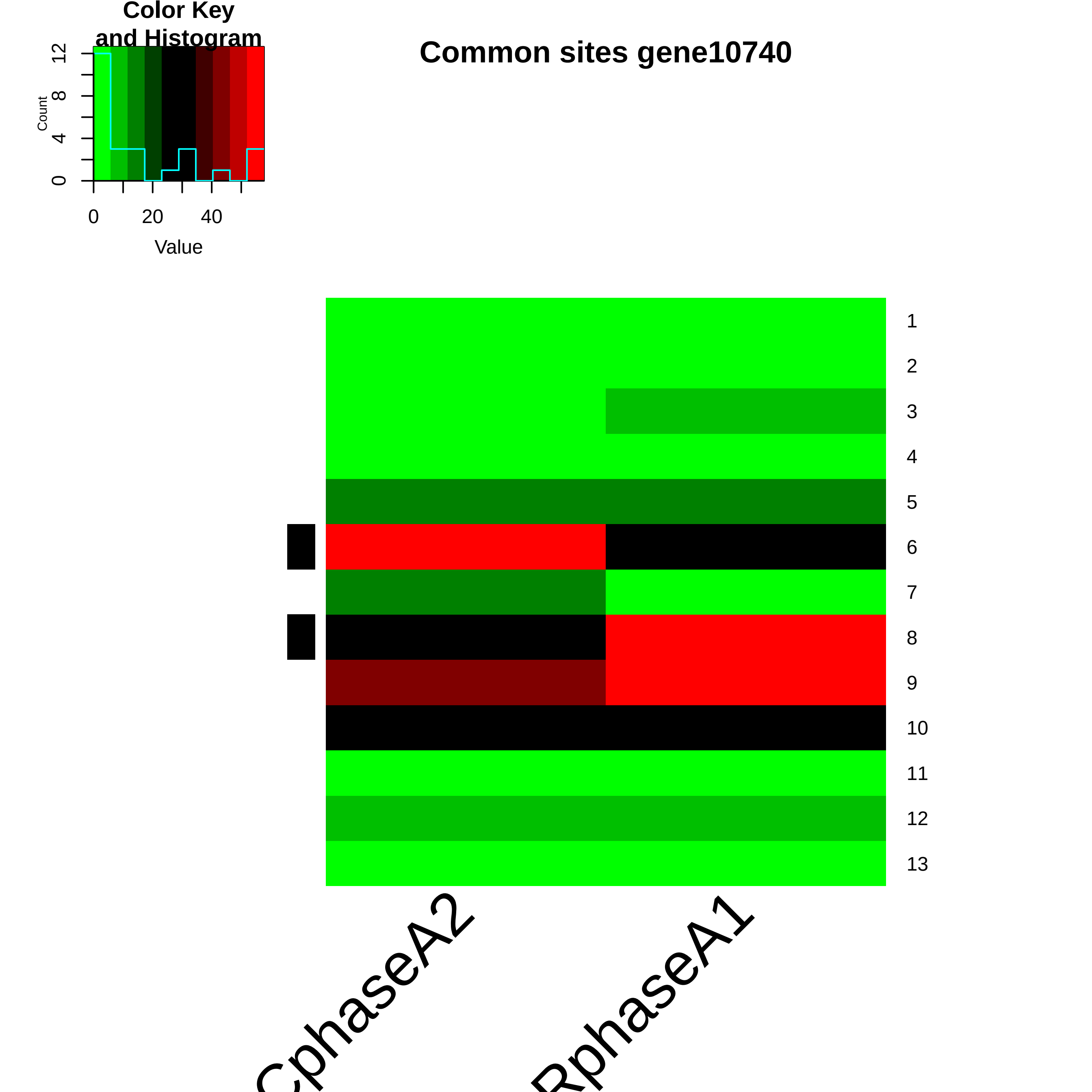

Color Key
and Histogram
Count
0
4
8
12
0
20
40
Value
Common sites gene10740
1
2
3
4
5
6
7
8
9
10
11
12
13
CphaseA2
RphaseA1

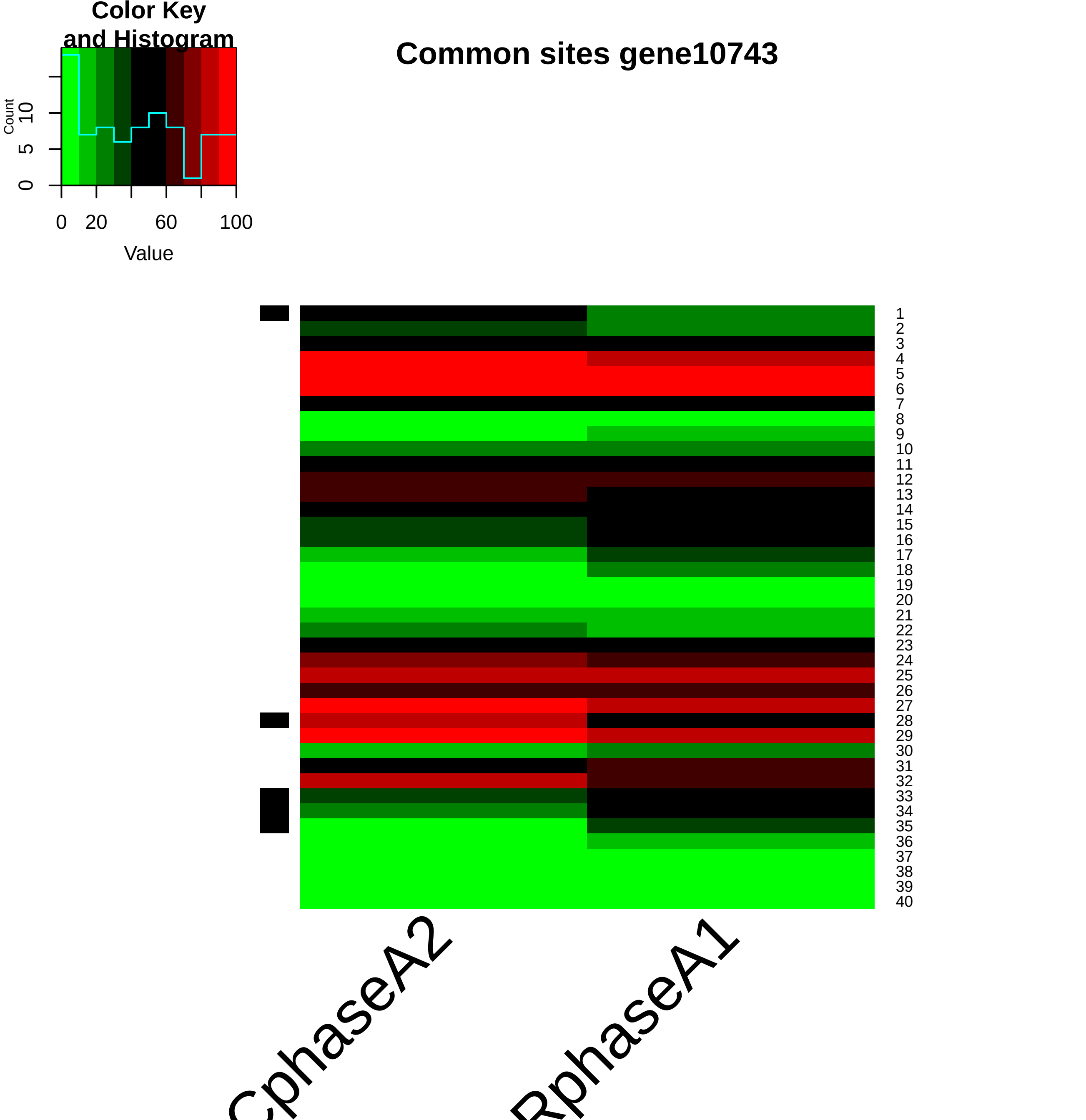

Color Key
and Histogram
Count
0
5
10
0
20
60
100
Value
Common sites gene10743
CphaseA2
RphaseA1

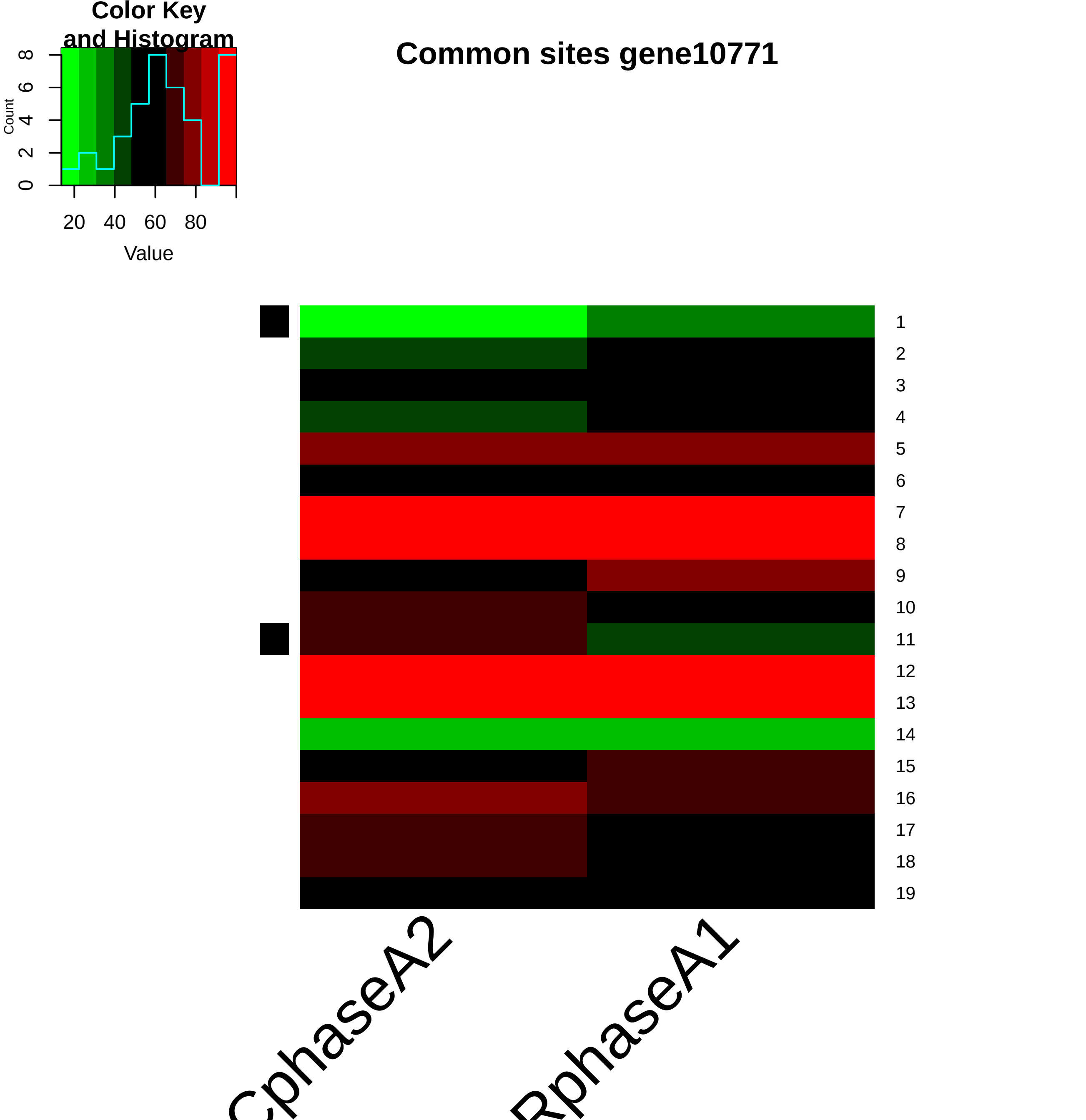

Color Key
and Histogram
Count
0
2
4
6
8
20
40
60
80
Value
Common sites gene10771
CphaseA2
RphaseA1

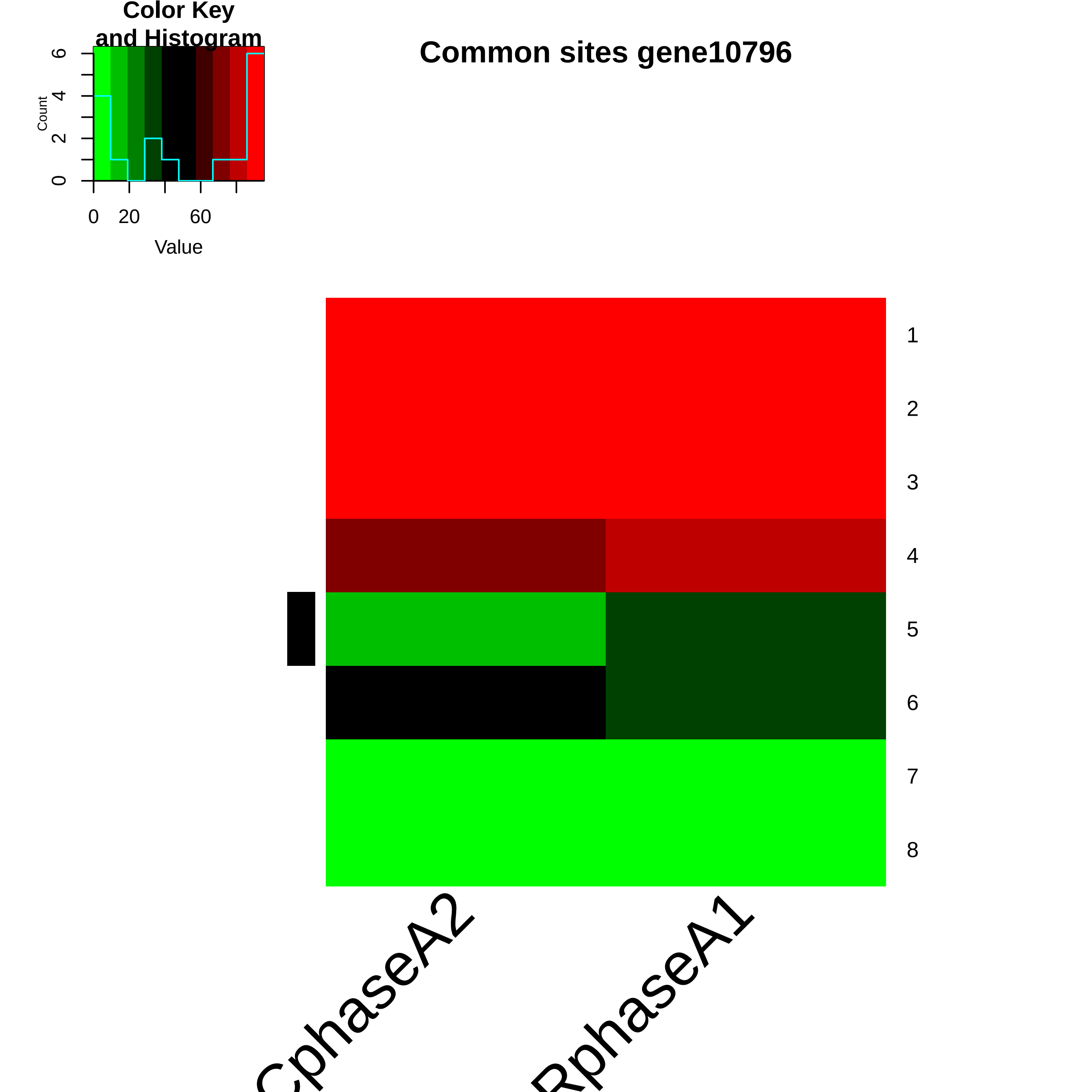
Color Key
and Histogram
Count
0
2
4
6
0
20
60
Value
Common sites gene10796
1
2
3
4
5
6
7
8
CphaseA2
RphaseA1

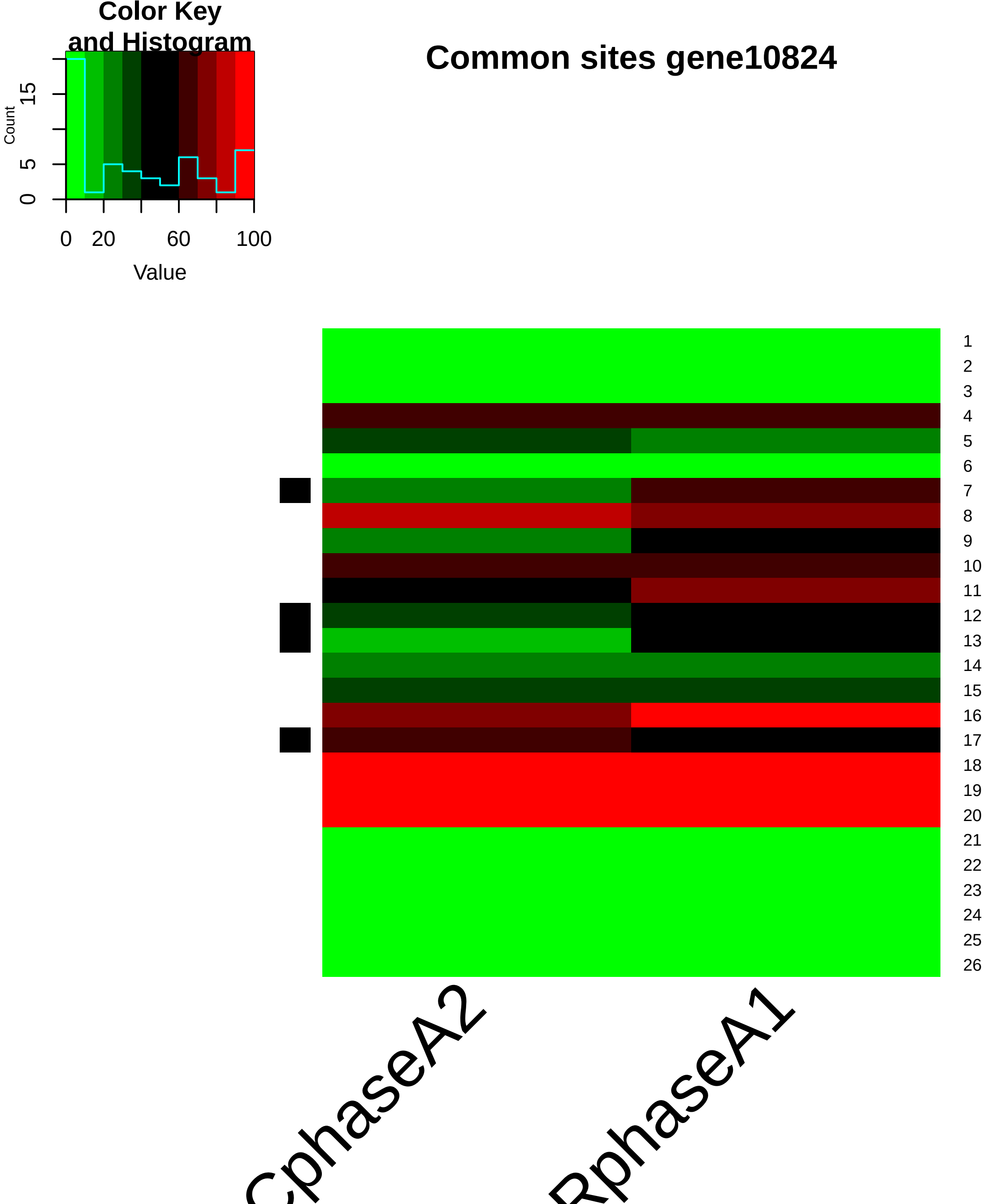
Color Key
and Histogram
Count
0
5
15
0
20
60
100
Value
Common sites gene10824

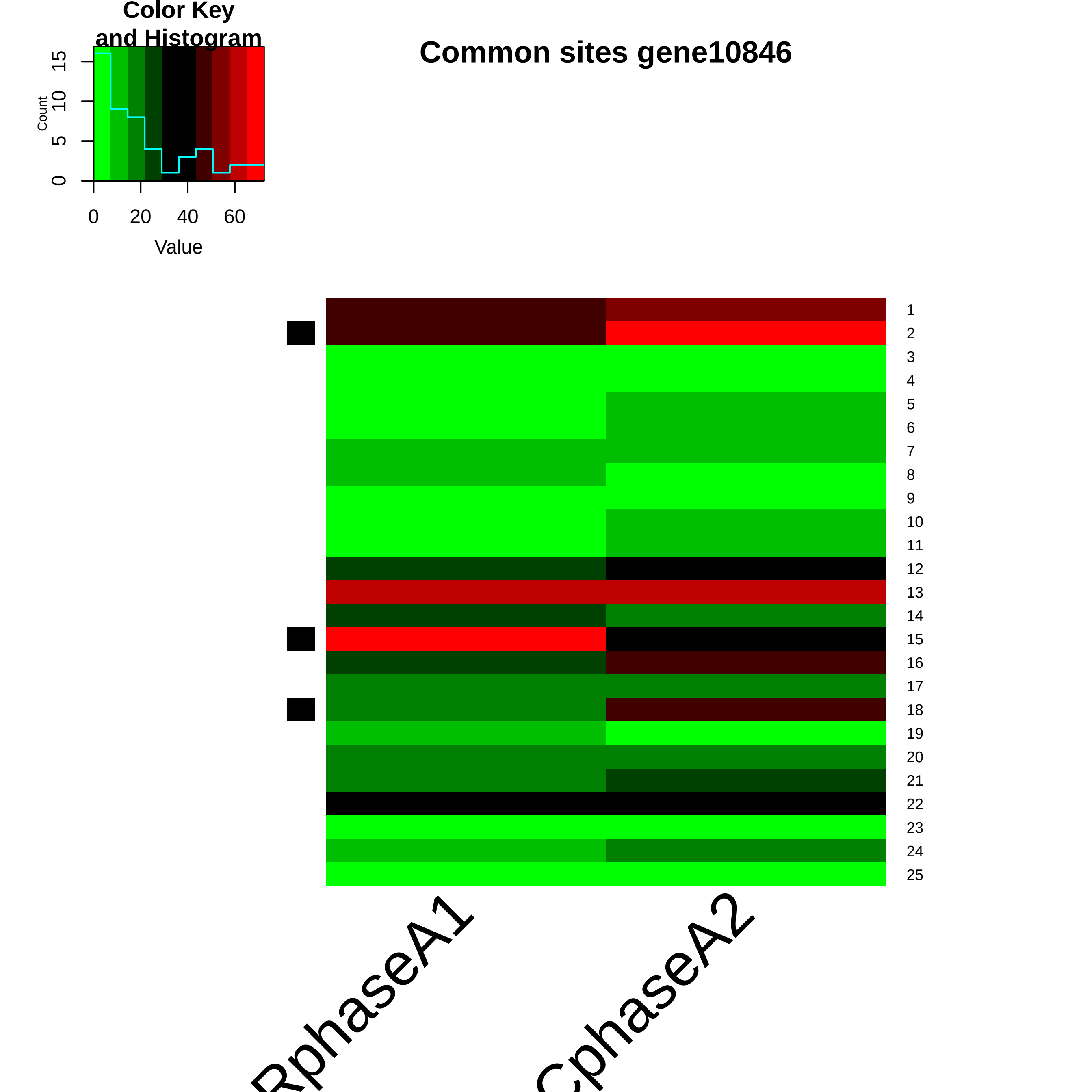

Color Key
and Histogram
Count
Value
0
5
10
15
0
20
40
60
Common sites gene10846
RphaseA1
CphaseA2

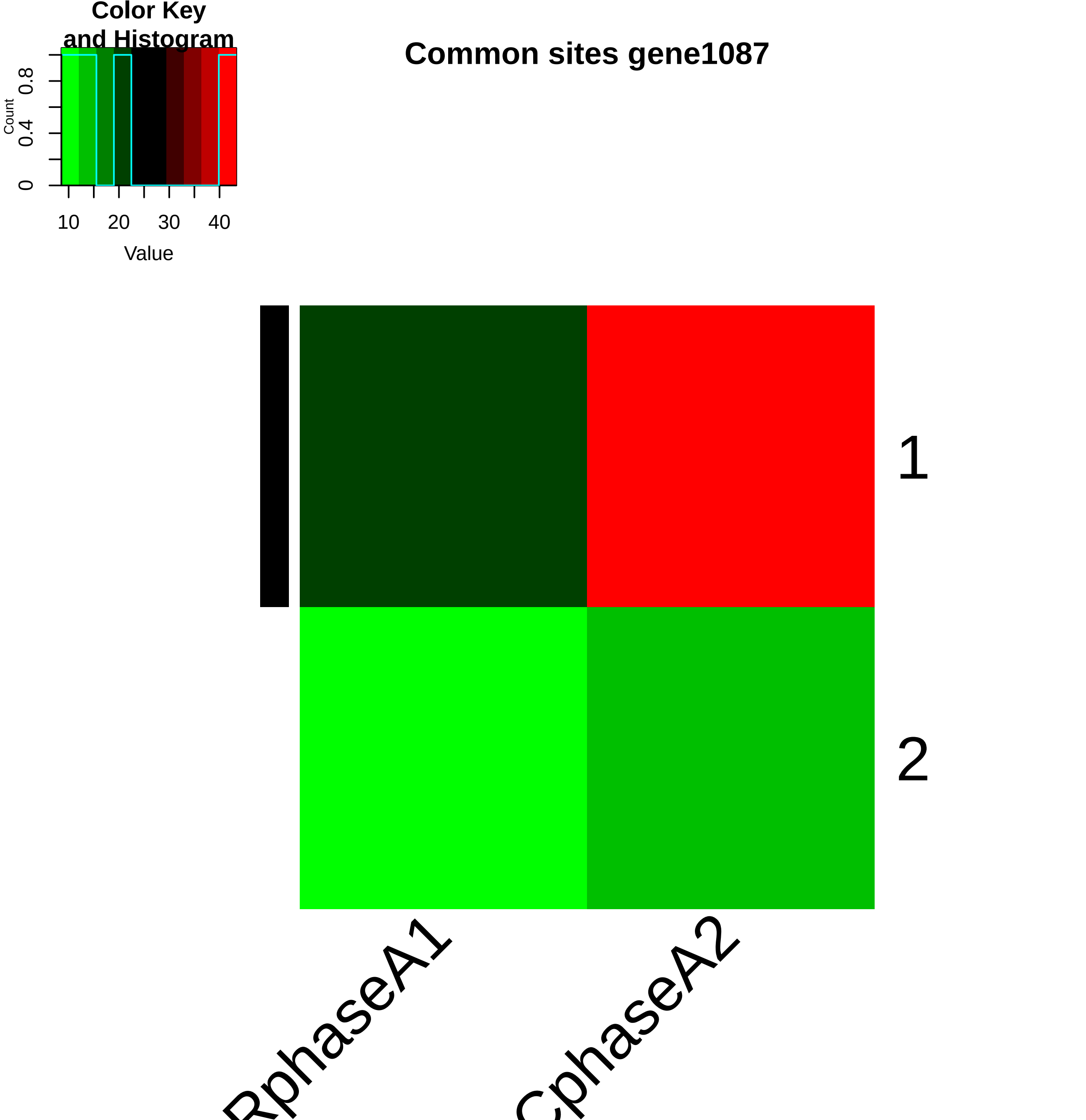
Color Key
and Histogram
Count
0
0.4
0.8
10
20
30
40
Value
Common sites gene1087
1
2
RphaseA1
CphaseA2

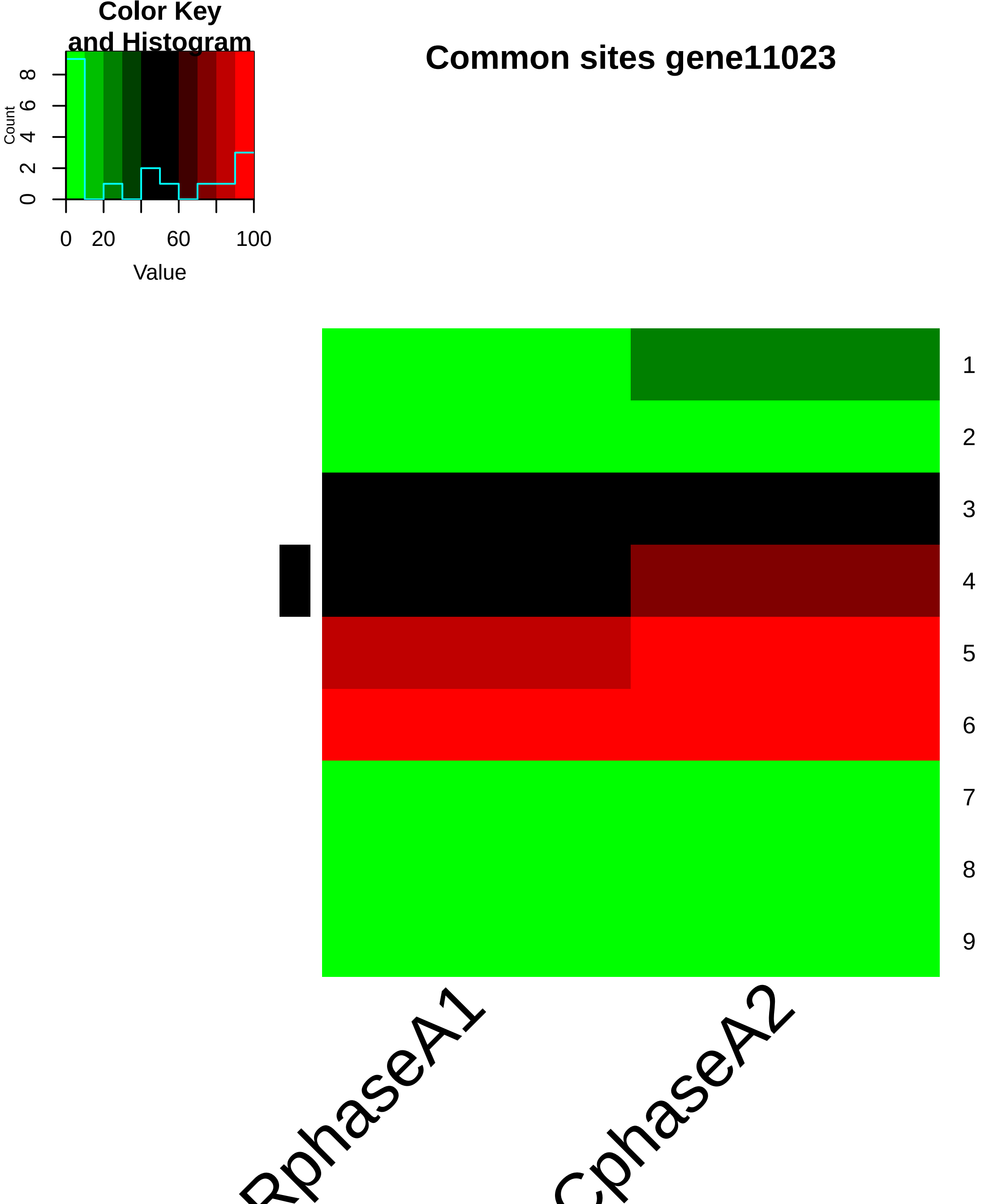
Color Key
and Histogram
Count
0
2
4
6
8
0
20
60
100
Value
Common sites gene11023
1
2
3
4
5
6
7
8
9
RphaseA1
CphaseA2

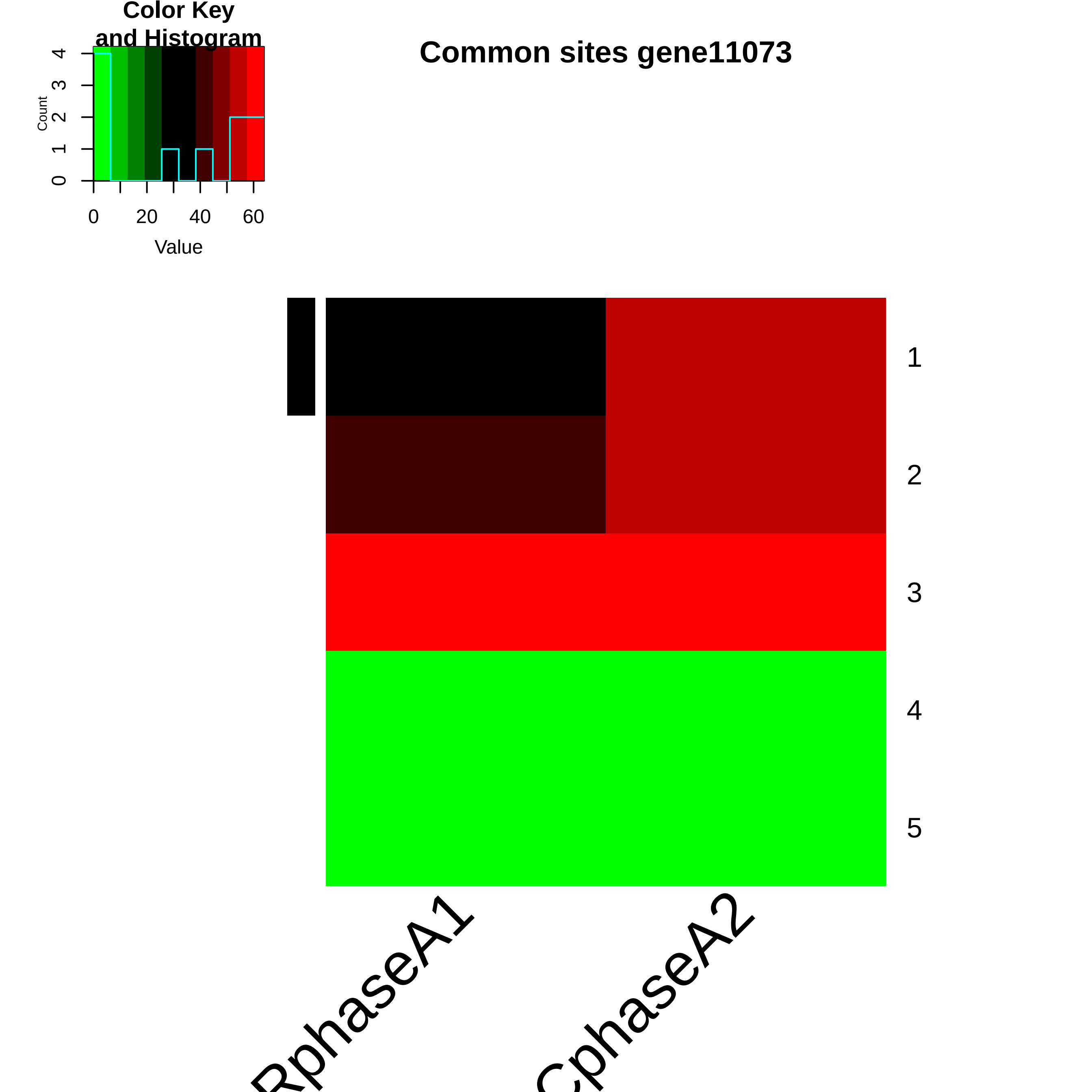
Color Key
and Histogram
Count
0
1
2
3
4
0
20
40
60
Value
Common sites gene11073
1
2
3
4
5
RphaseA1
CphaseA2

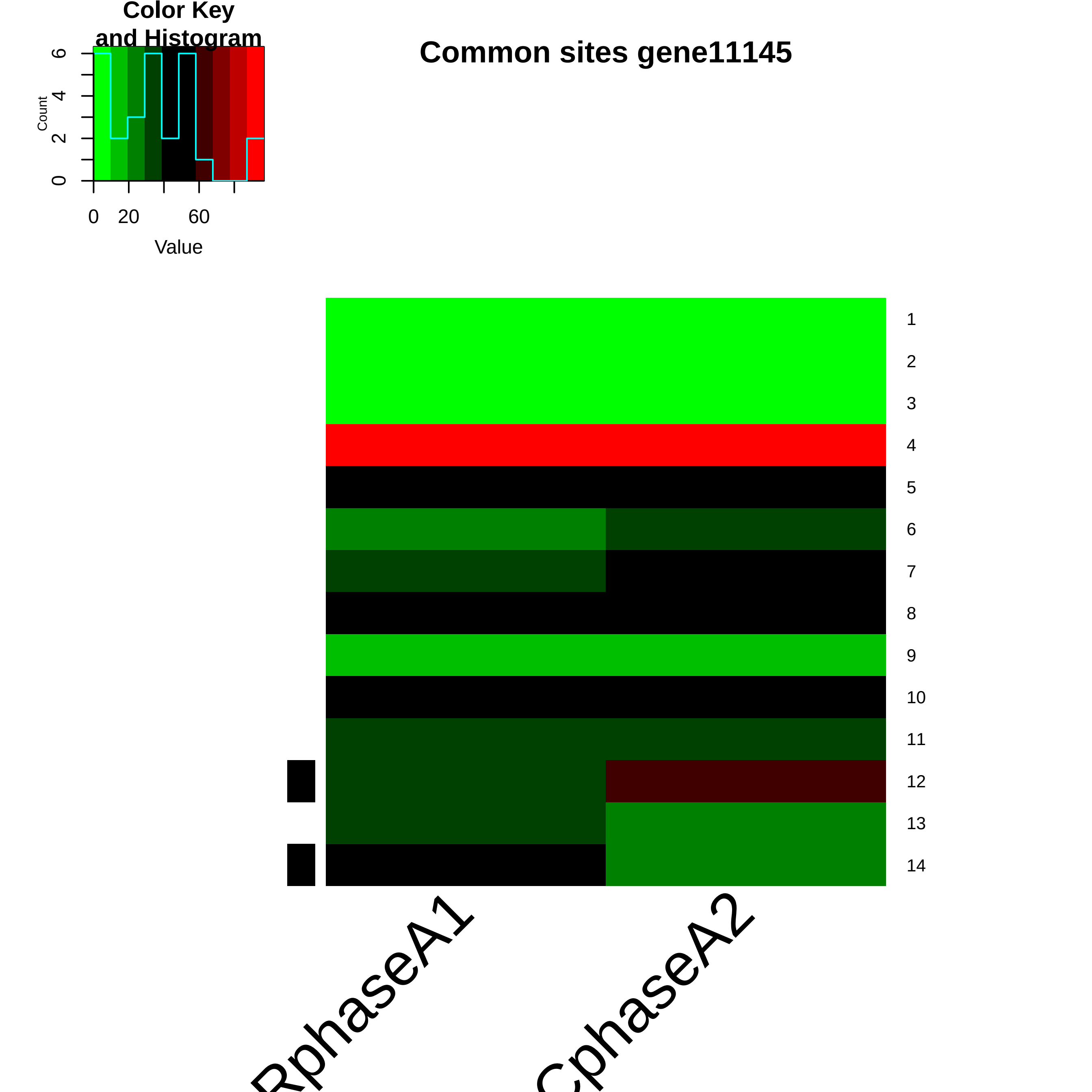

Color Key
and Histogram
Count
0
2
4
6
0
20
60
Value
Common sites gene11145
1
2
3
4
5
6
7
8
9
10
11
12
13
14
RphaseA1
CphaseA2

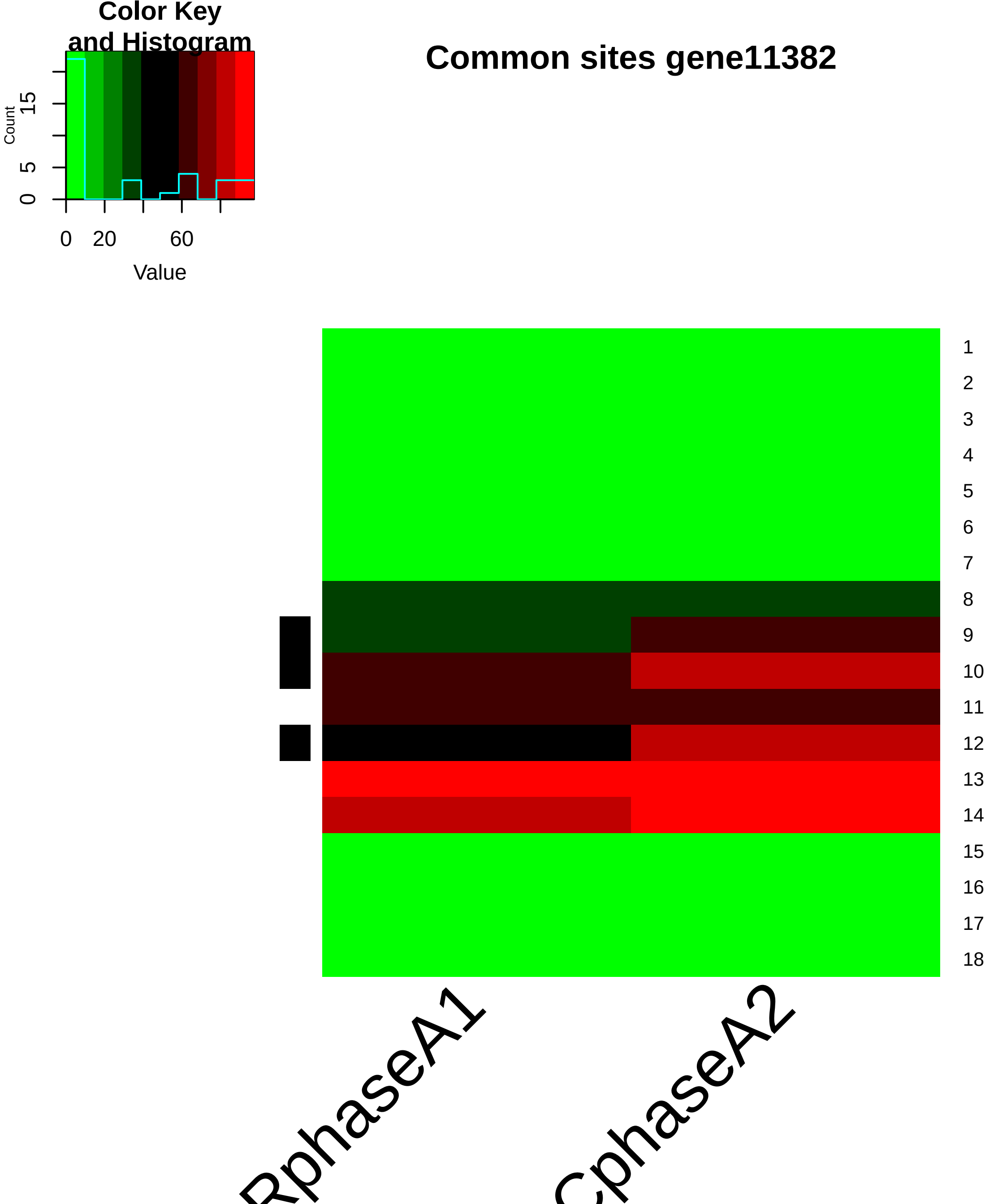

Color Key
and Histogram
Count
0
5
15
0
20
60
Value
Common sites gene11382
RphaseA1
CphaseA2

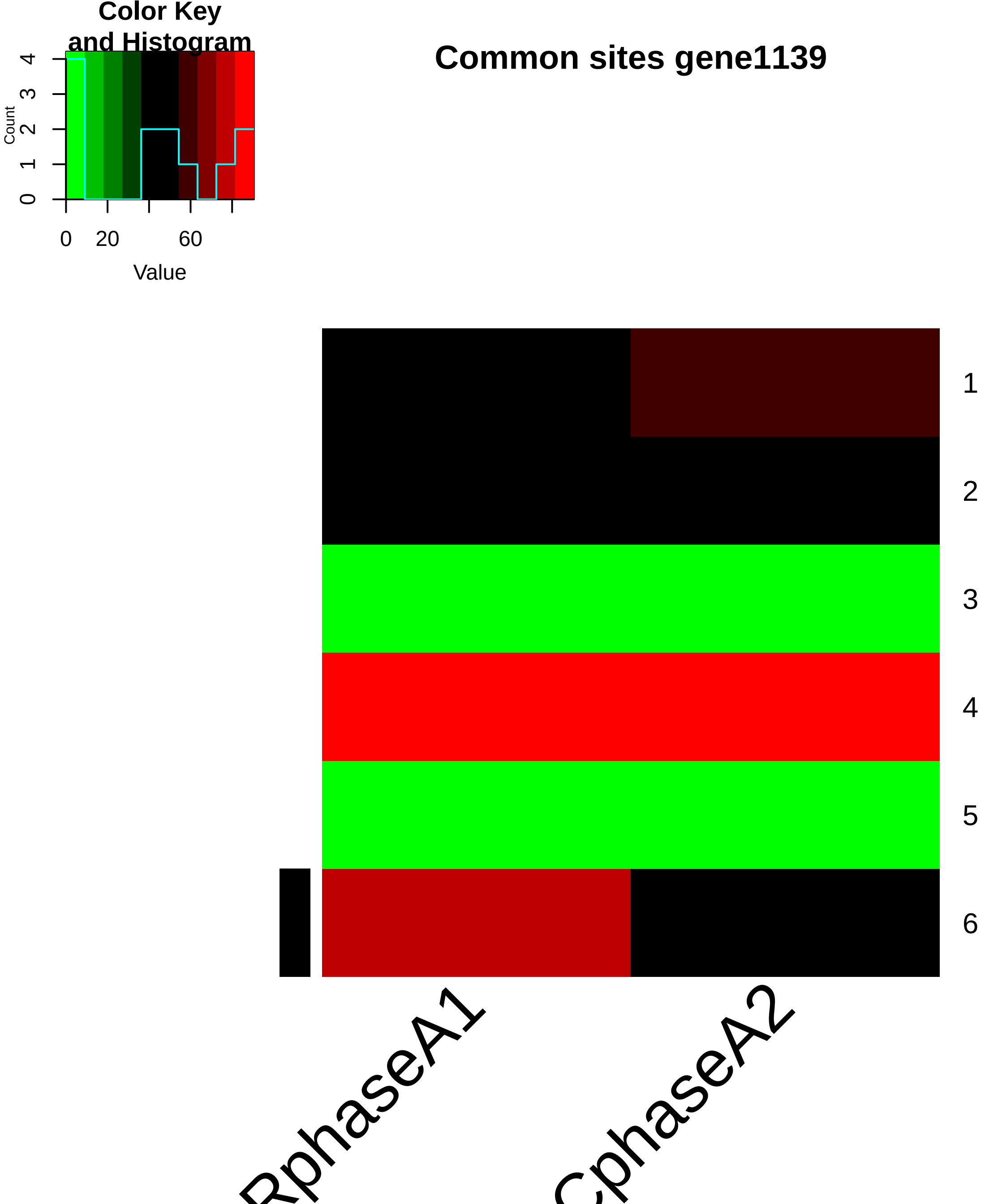
Color Key
and Histogram
Count
0
1
2
3
4
0
20
60
Value
Common sites gene1139
1
2
3
4
5
6
RphaseA1
CphaseA2

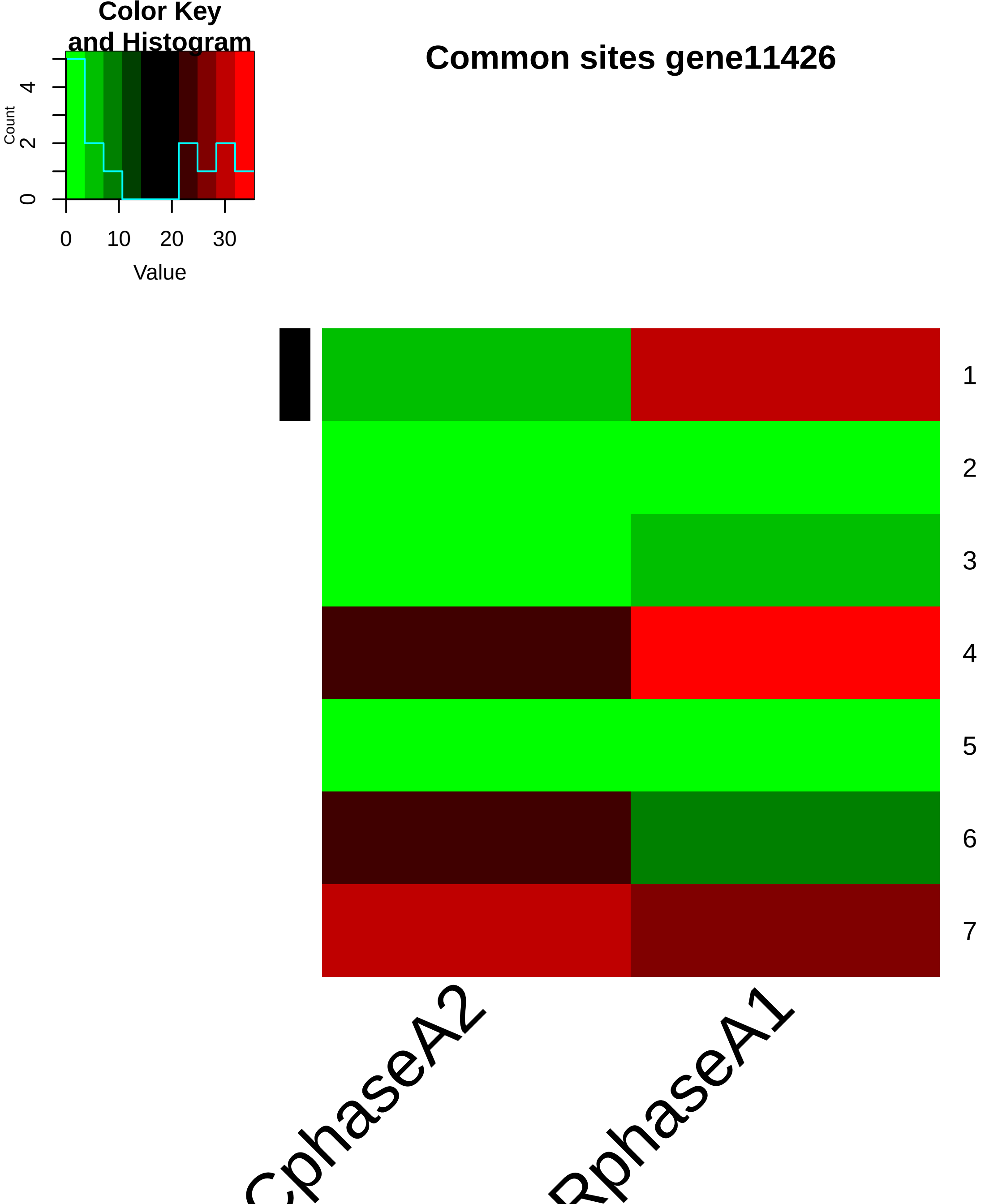
Color Key
and Histogram
Count
0
2
4
0
10
20
30
Value
Common sites gene11426
1
2
3
4
5
6
7
CphaseA2
RphaseA1

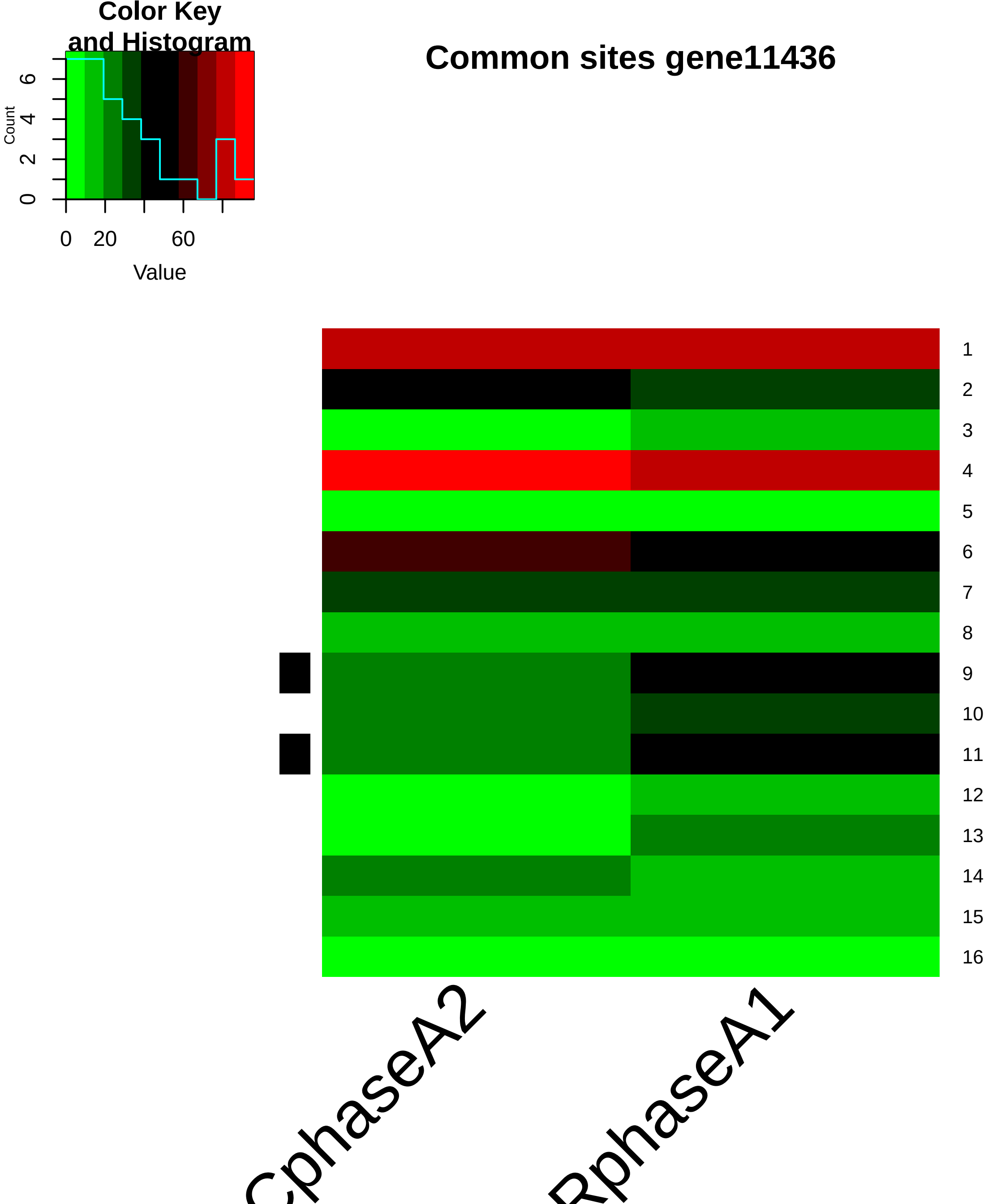

Color Key
and Histogram
Count
0
2
4
6
0
20
60
Value
Common sites gene11436
CphaseA2
RphaseA1

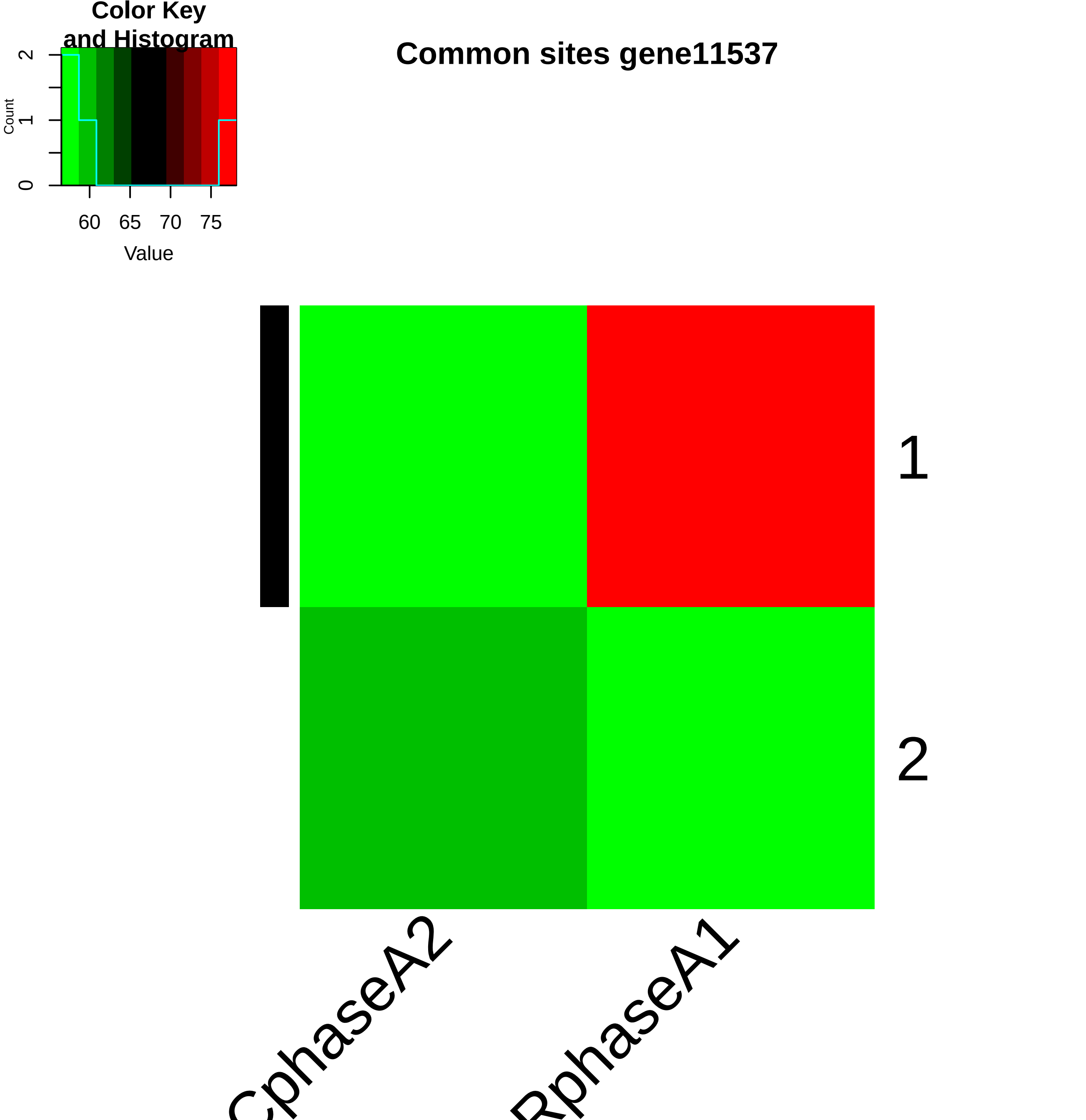
Color Key
and Histogram
Count
0
1
2
60
65
70
75
Value
Common sites gene11537
1
2
CphaseA2
RphaseA1

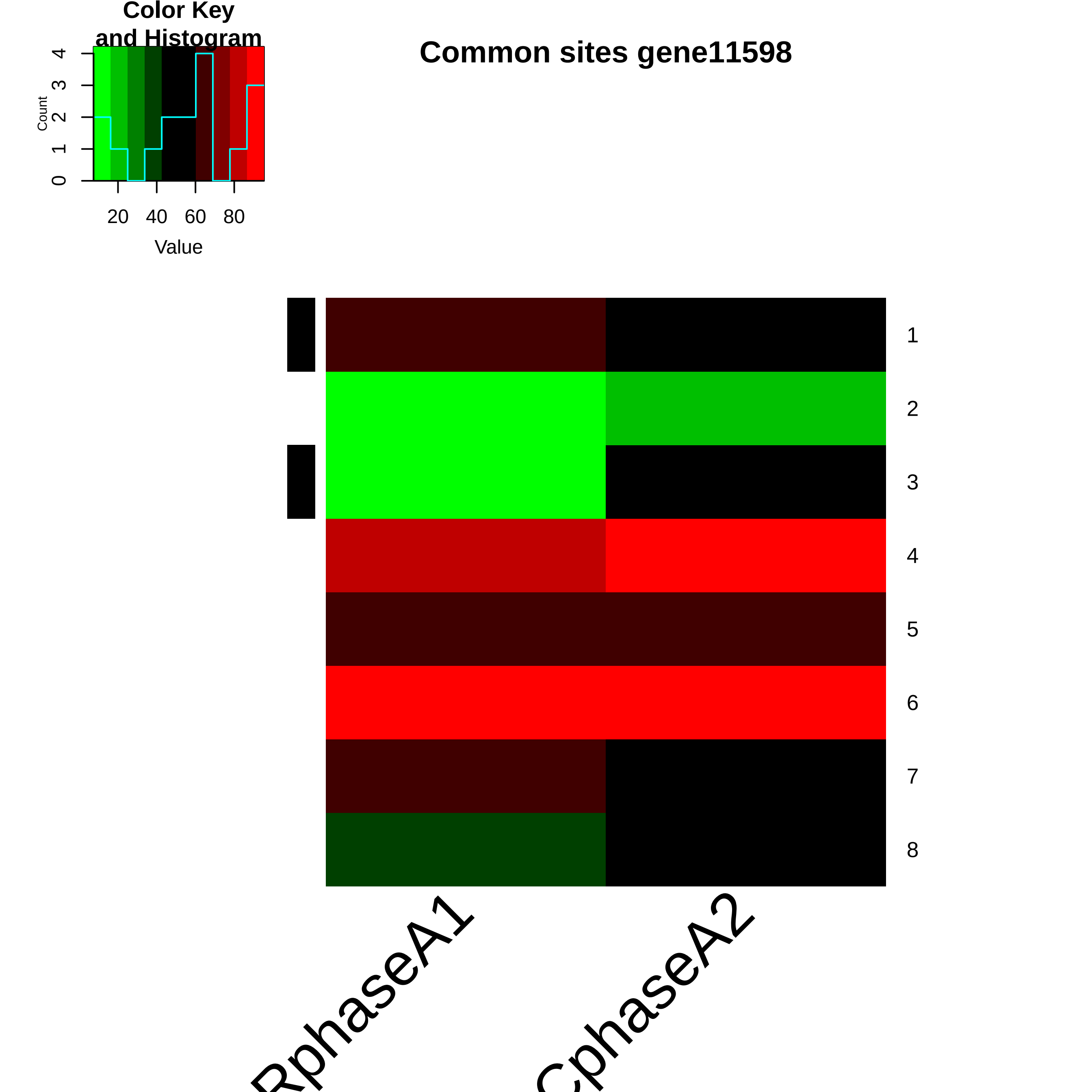

Color Key
and Histogram
Count
0
1
2
3
4
20
40
60
80
Value
Common sites gene11598
1
2
3
4
5
6
7
8
RphaseA1
CphaseA2

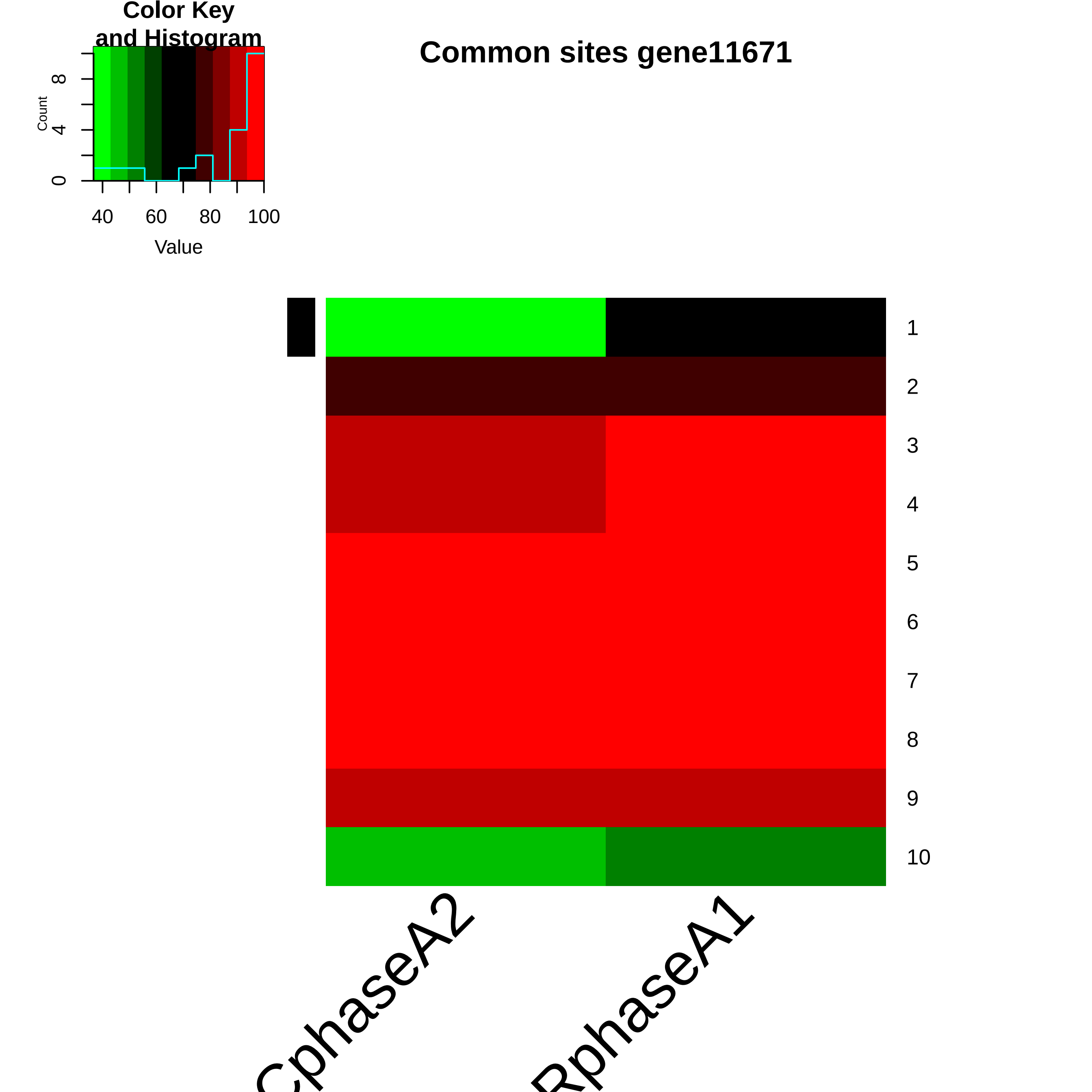

Color Key
and Histogram
Count
0
4
8
40
60
80
100
Value
Common sites gene11671
1
2
3
4
5
6
7
8
9
10
CphaseA2
RphaseA1

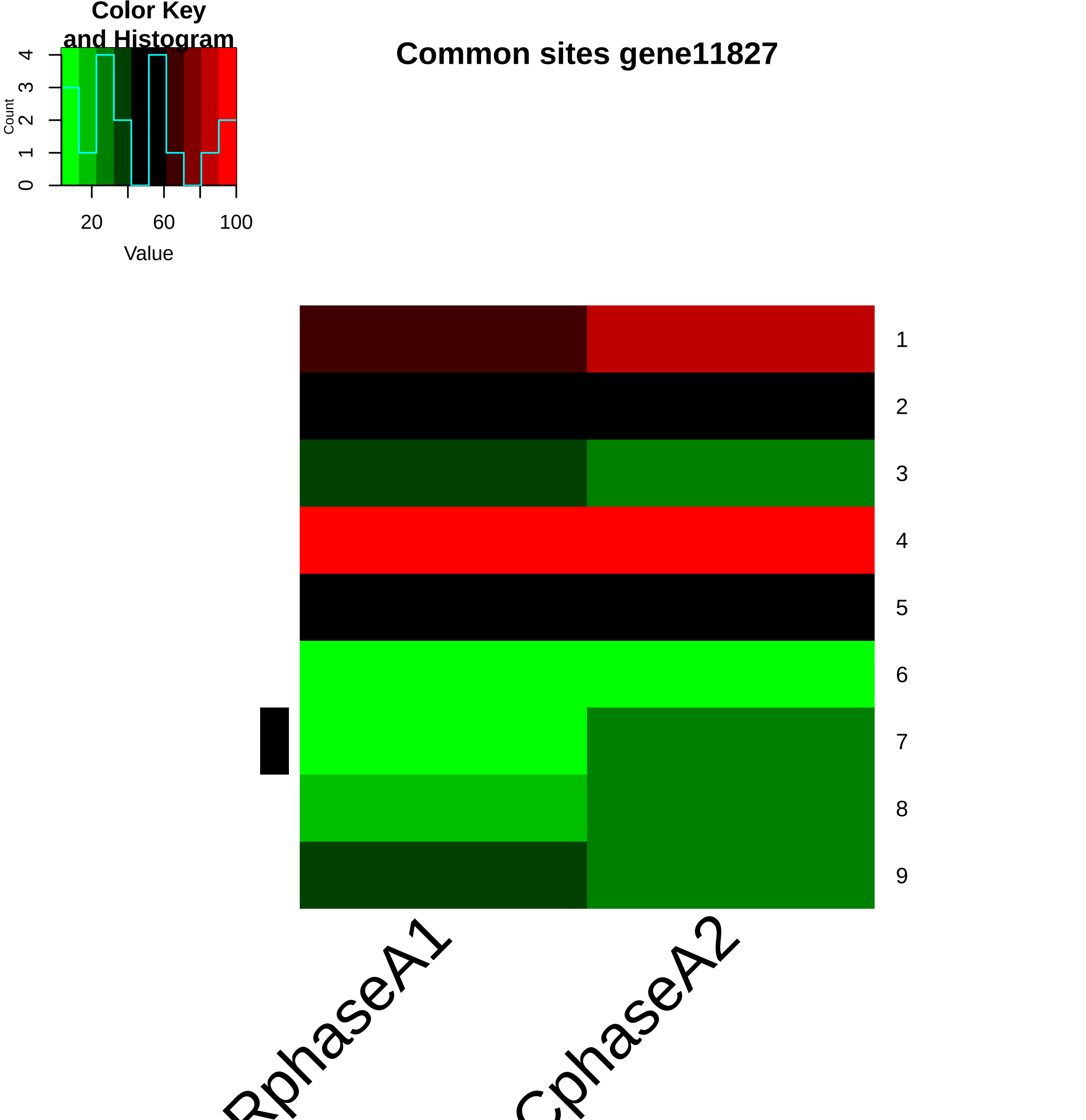
Color Key
and Histogram
Count
0
1
2
3
4
20
60
100
Value
Common sites gene11827
1
2
3
4
5
6
7
8
9
RphaseA1
CphaseA2

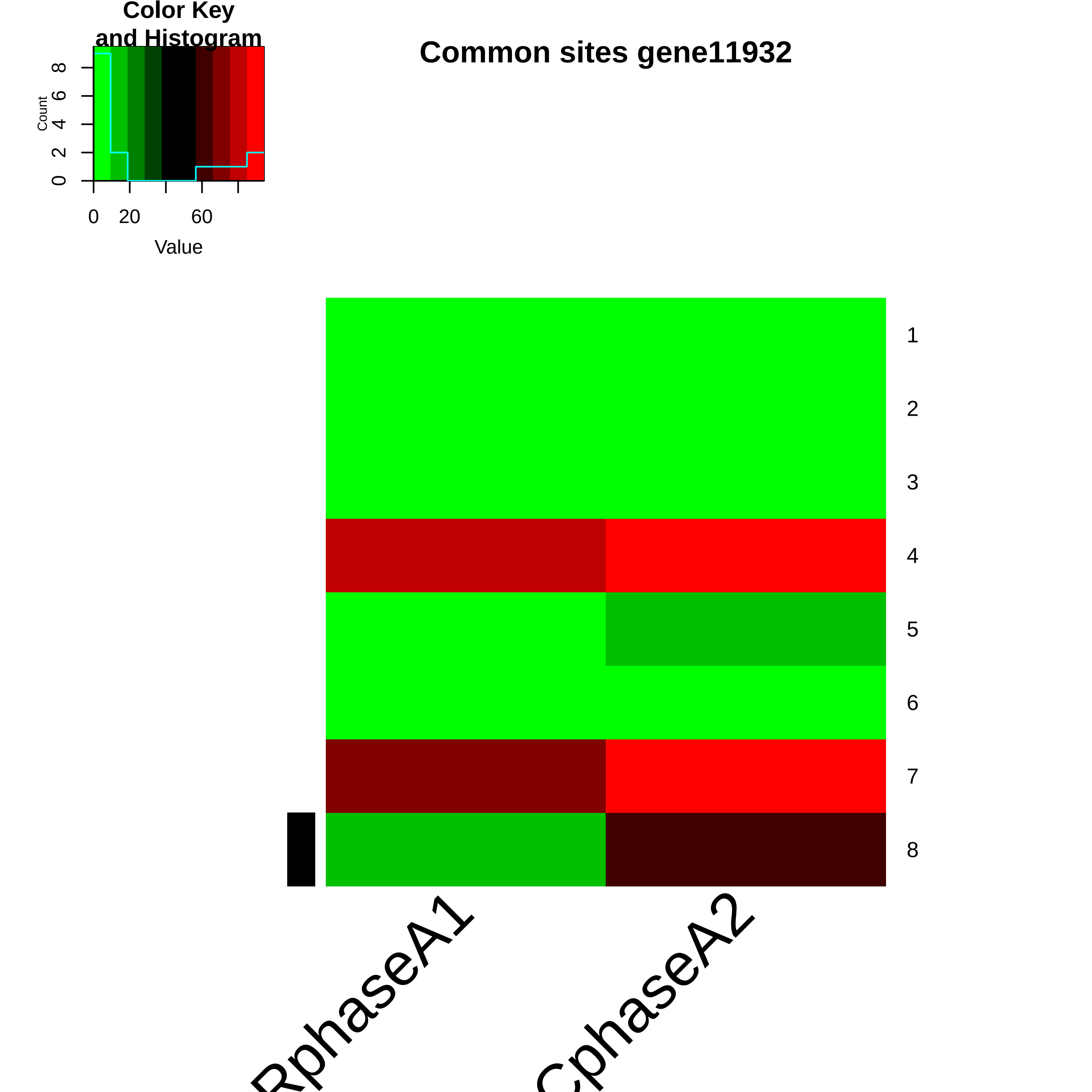
Color Key
and Histogram
Count
0
2
4
6
8
0
20
60
Value
Common sites gene11932
1
2
3
4
5
6
7
8
RphaseA1
CphaseA2

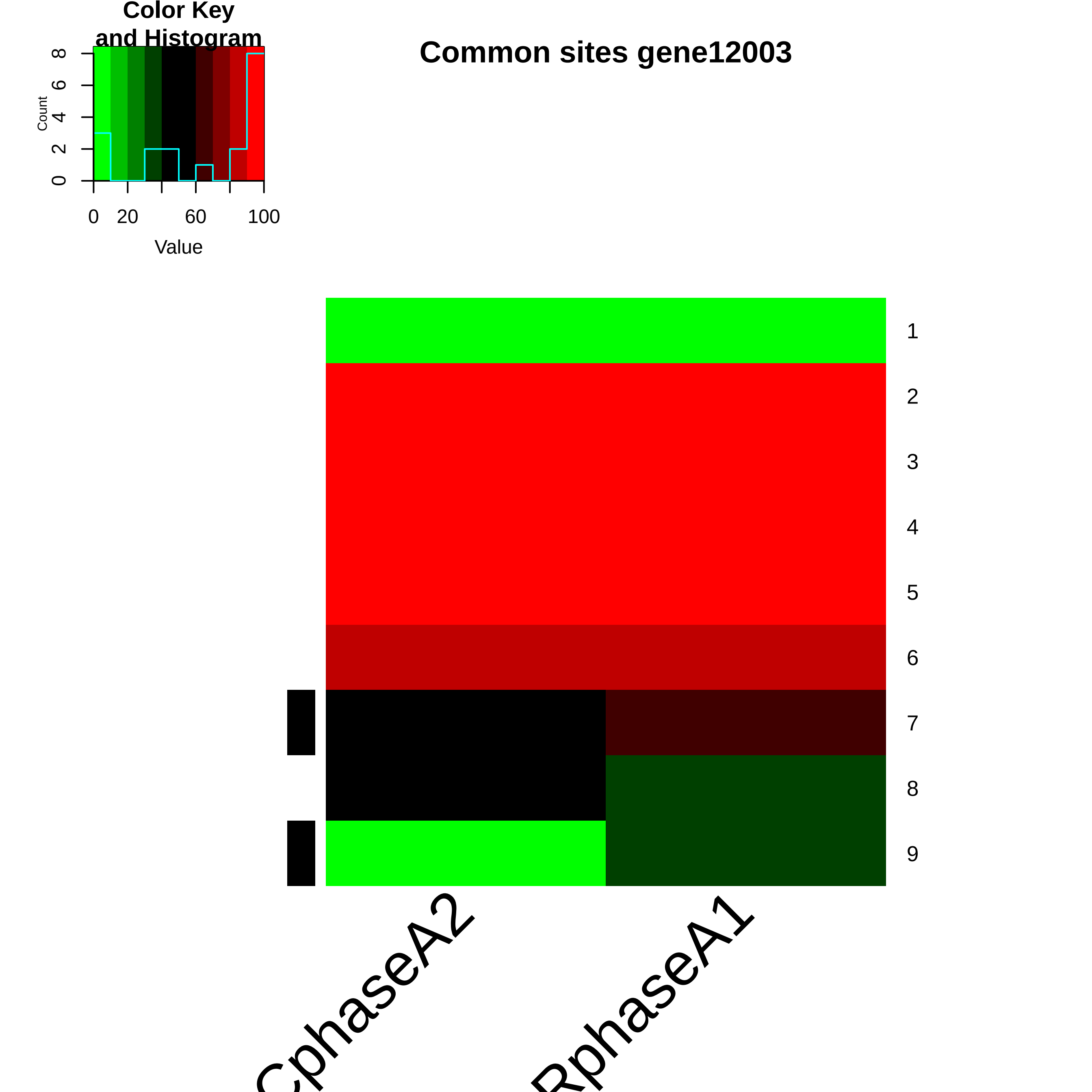
Color Key
and Histogram
Count
0
2
4
6
8
0
20
60
100
Value
Common sites gene12003
1
2
3
4
5
6
7
8
9
CphaseA2
RphaseA1

#### Common sites gene12209

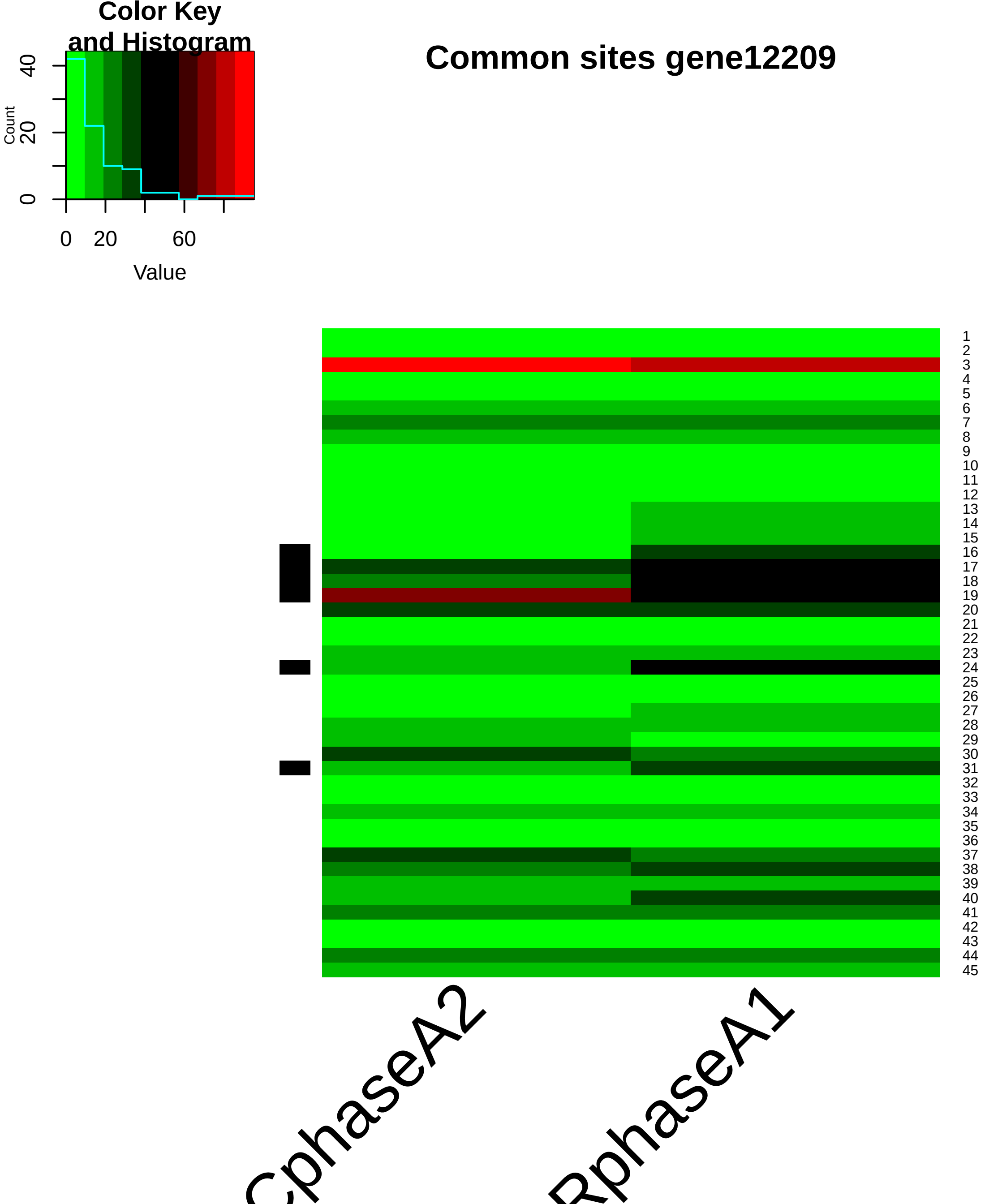

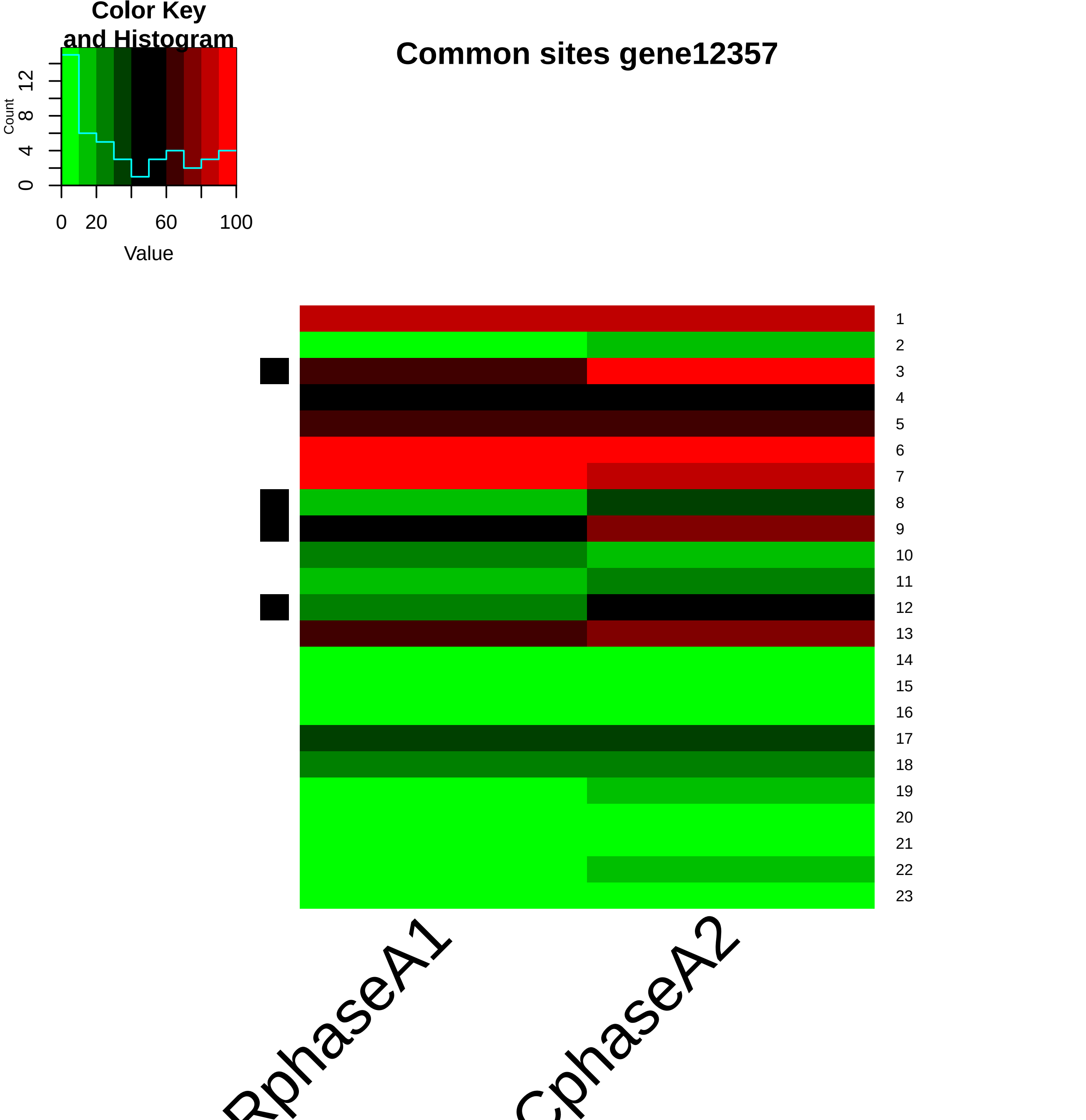
Color Key
and Histogram
Count
0
4
8
12
0
20
60
100
Value
Common sites gene12357
RphaseA1
CphaseA2

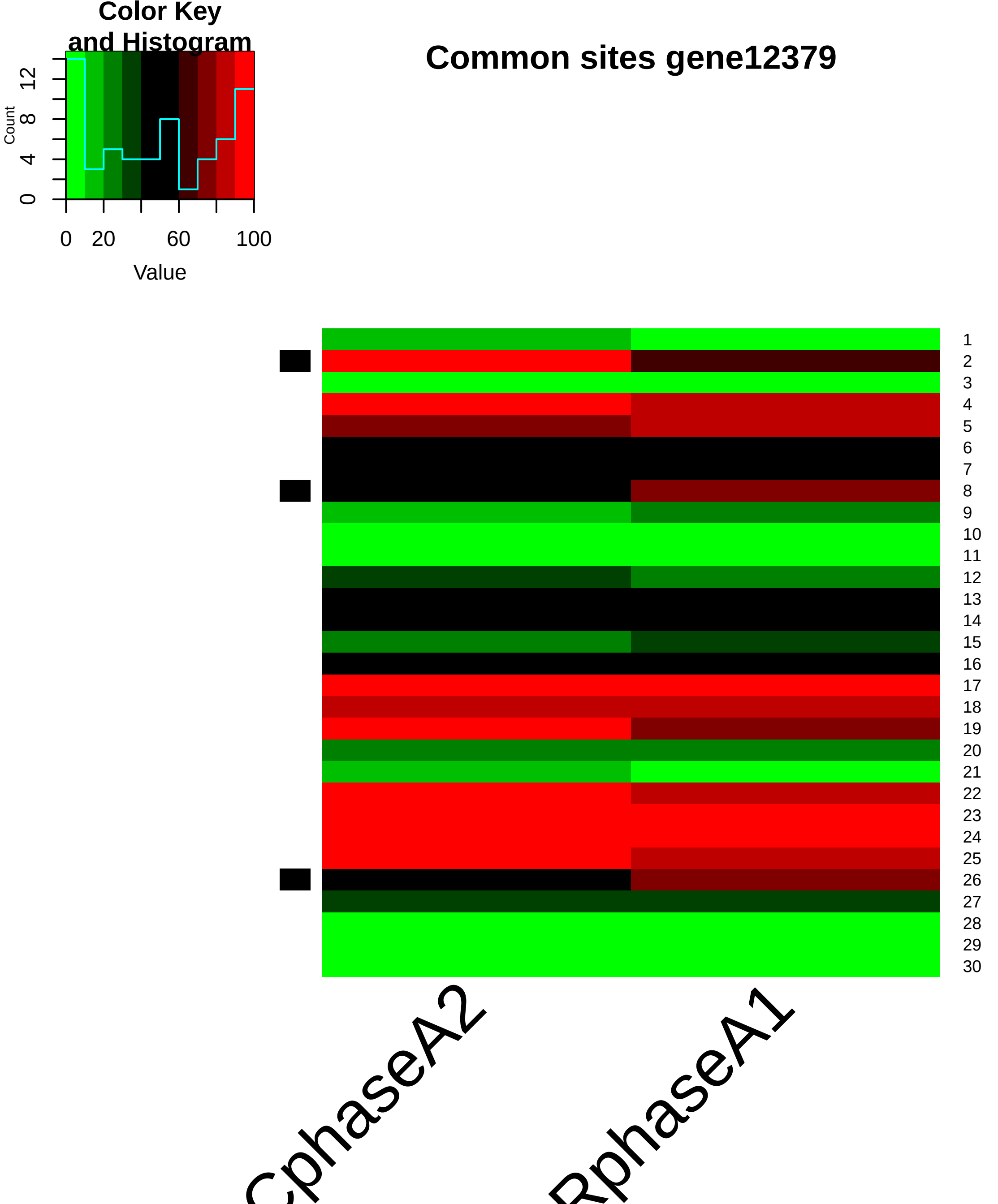

Color Key and Histogram
Count
0
4
8
12
0
20
60
100
Value
Common sites gene12379
CphaseA2
RphaseA1

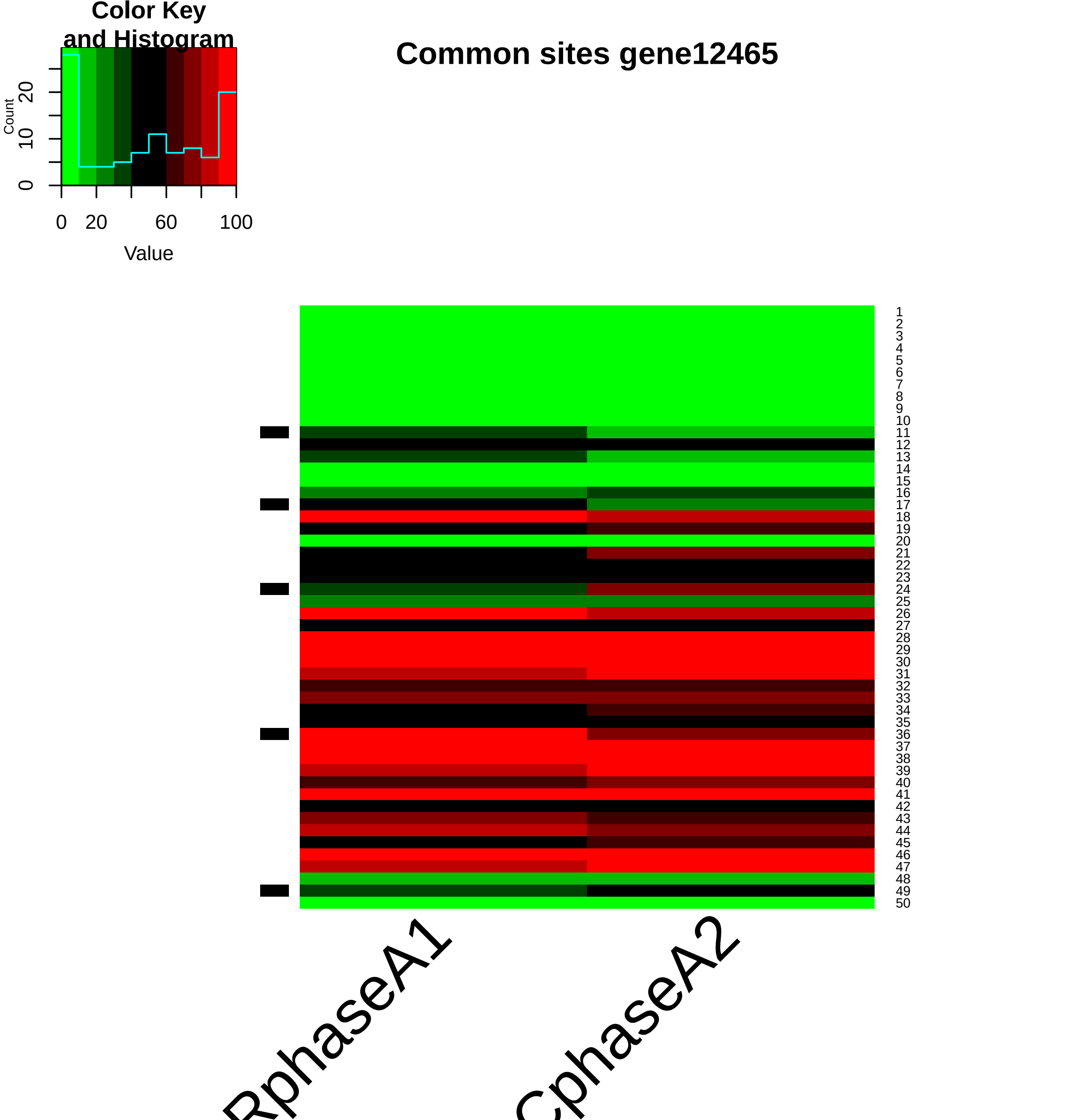
Color Key
and Histogram
Count
0
10
20
0
20
60
100
Value
Common sites gene12465
RphaseA1
CphaseA2

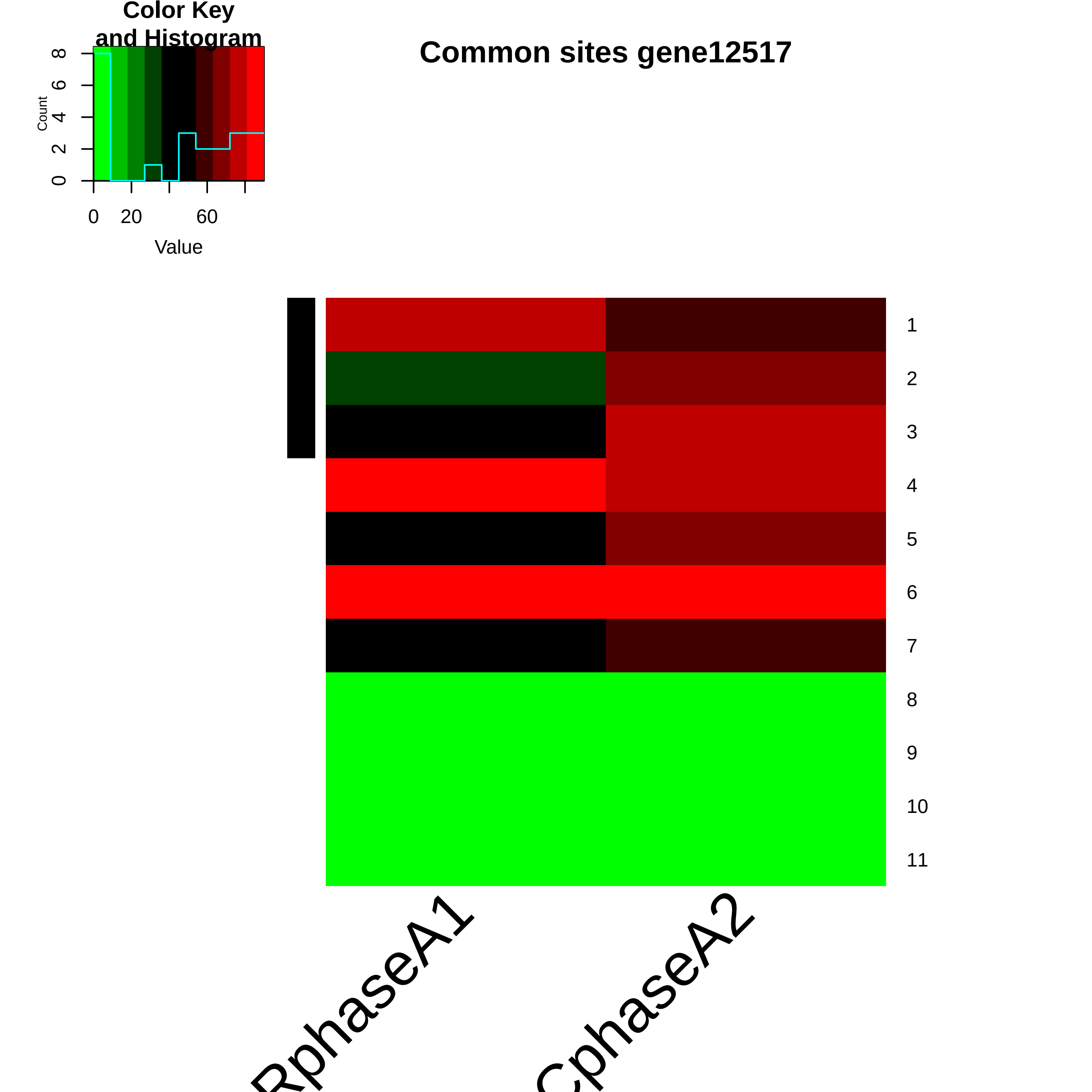

Color Key
and Histogram
Count
0
2
4
6
8
0
20
60
Value
Common sites gene12517
1
2
3
4
5
6
7
8
9
10
11
RphaseA1
CphaseA2

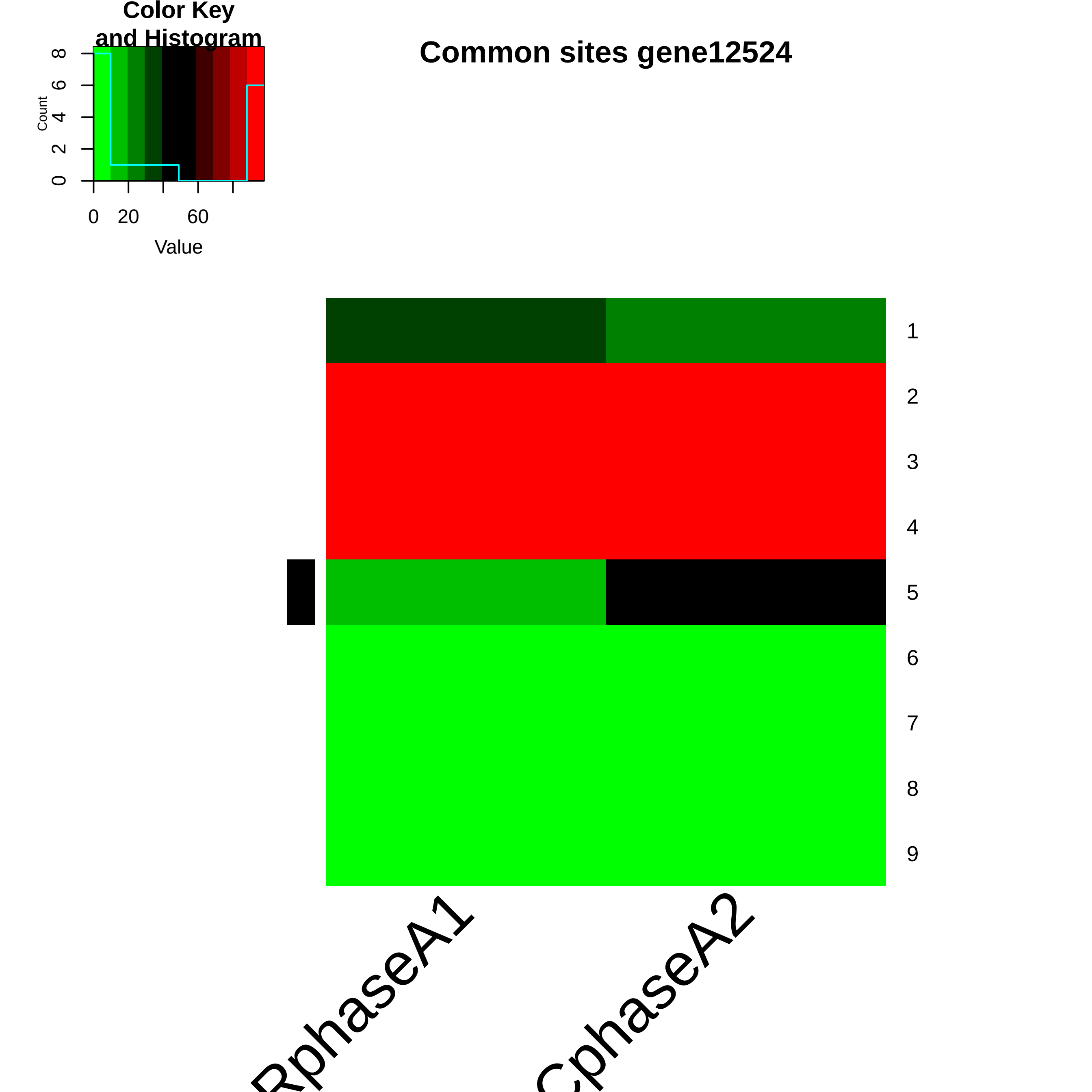
Color Key
and Histogram
Count
0
2
4
6
8
0
20
60
Value
Common sites gene12524
1
2
3
4
5
6
7
8
9
RphaseA1
CphaseA2

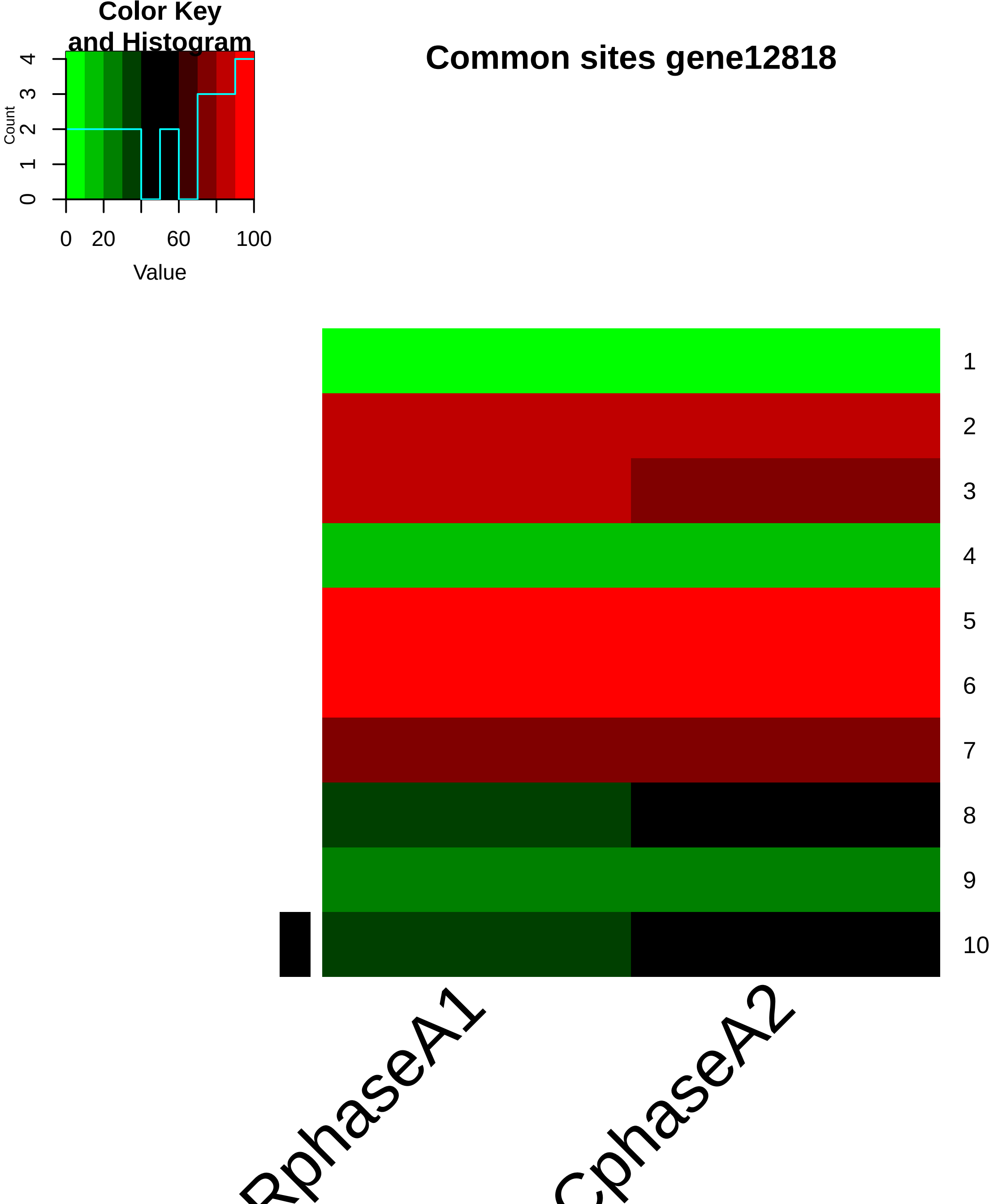
Color Key
and Histogram
Count
0
1
2
3
4
0
20
60
100
Value
Common sites gene12818
1
2
3
4
5
6
7
8
9
10
RphaseA1
CphaseA2

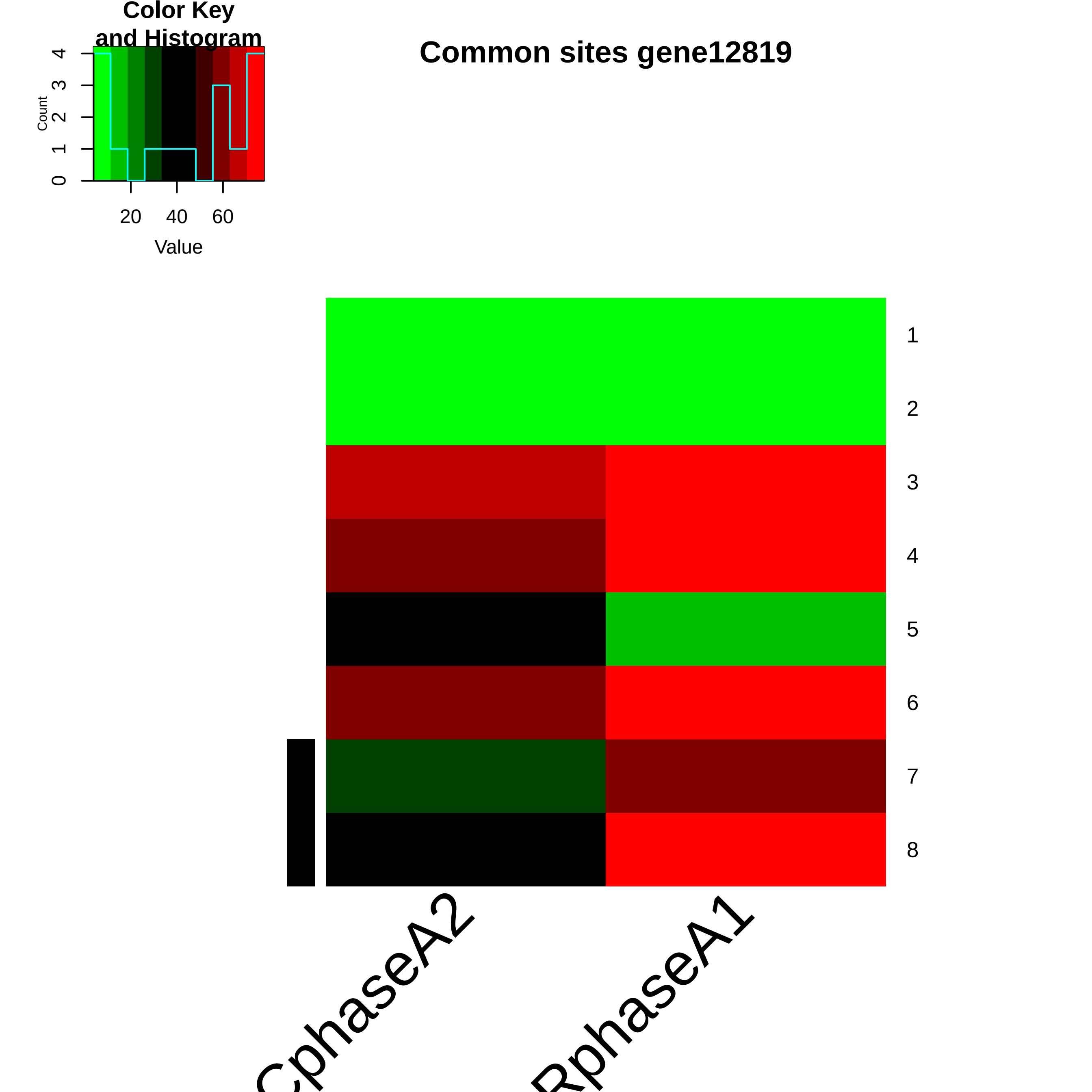
Color Key
and Histogram
Count
0
1
2
3
4
20
40
60
Value
Common sites gene12819
1
2
3
4
5
6
7
8
CphaseA2
RphaseA1

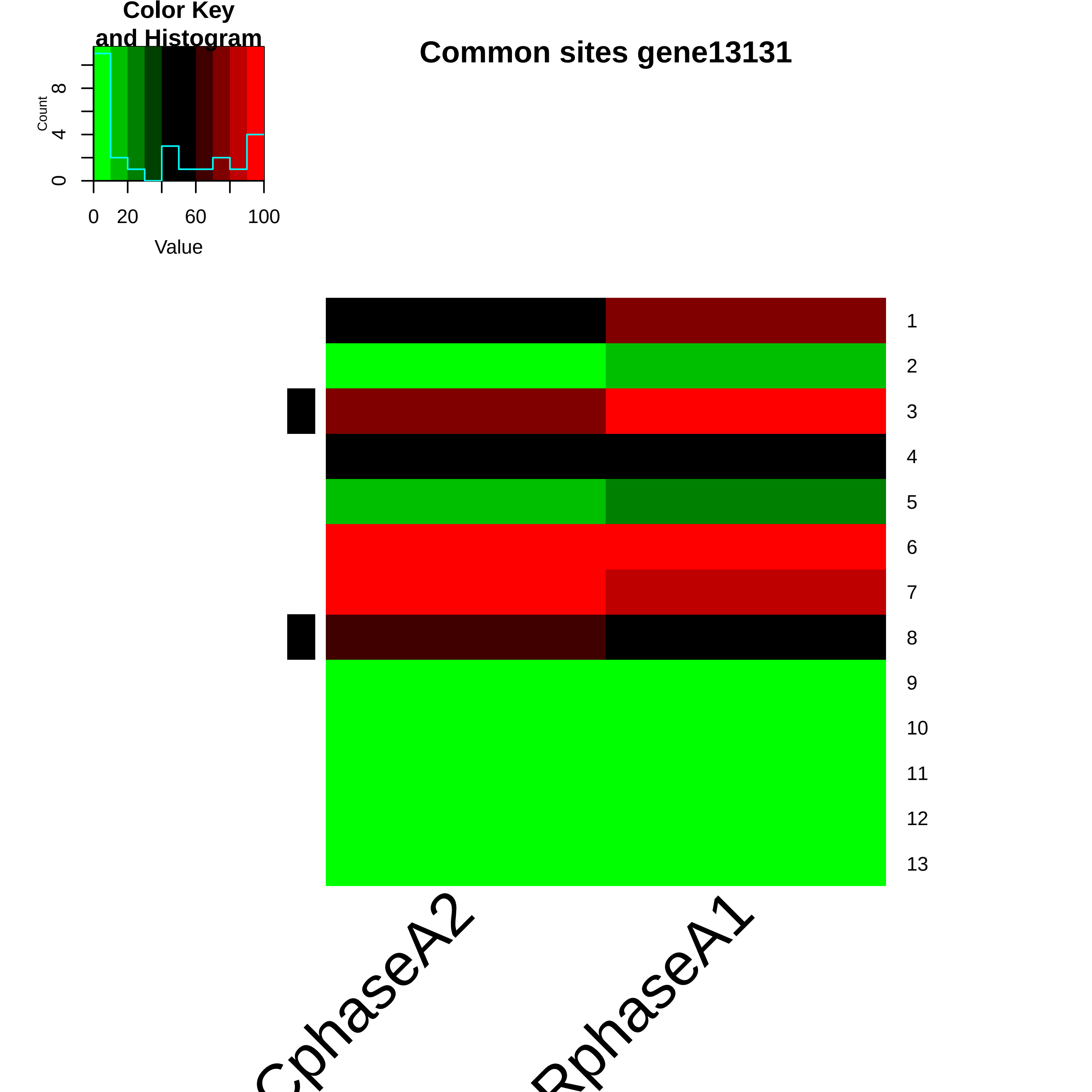

Color Key
and Histogram
Count
0
4
8
0
20
60
100
Value
Common sites gene13131
1
2
3
4
5
6
7
8
9
10
11
12
13
CphaseA2
RphaseA1

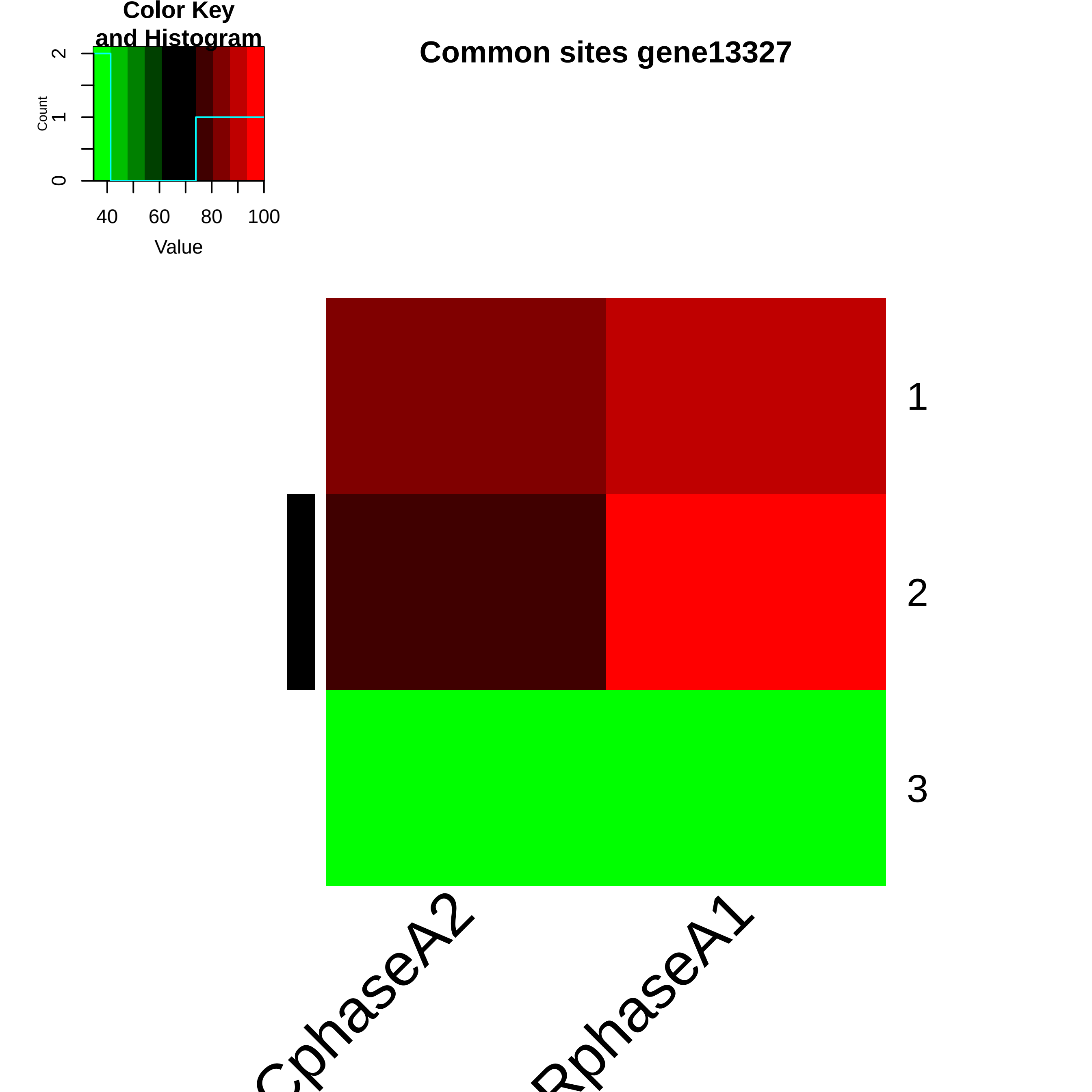

Color Key
and Histogram
Count
0
1
2
40
60
80
100
Value
Common sites gene13327
1
2
3
CphaseA2
RphaseA1

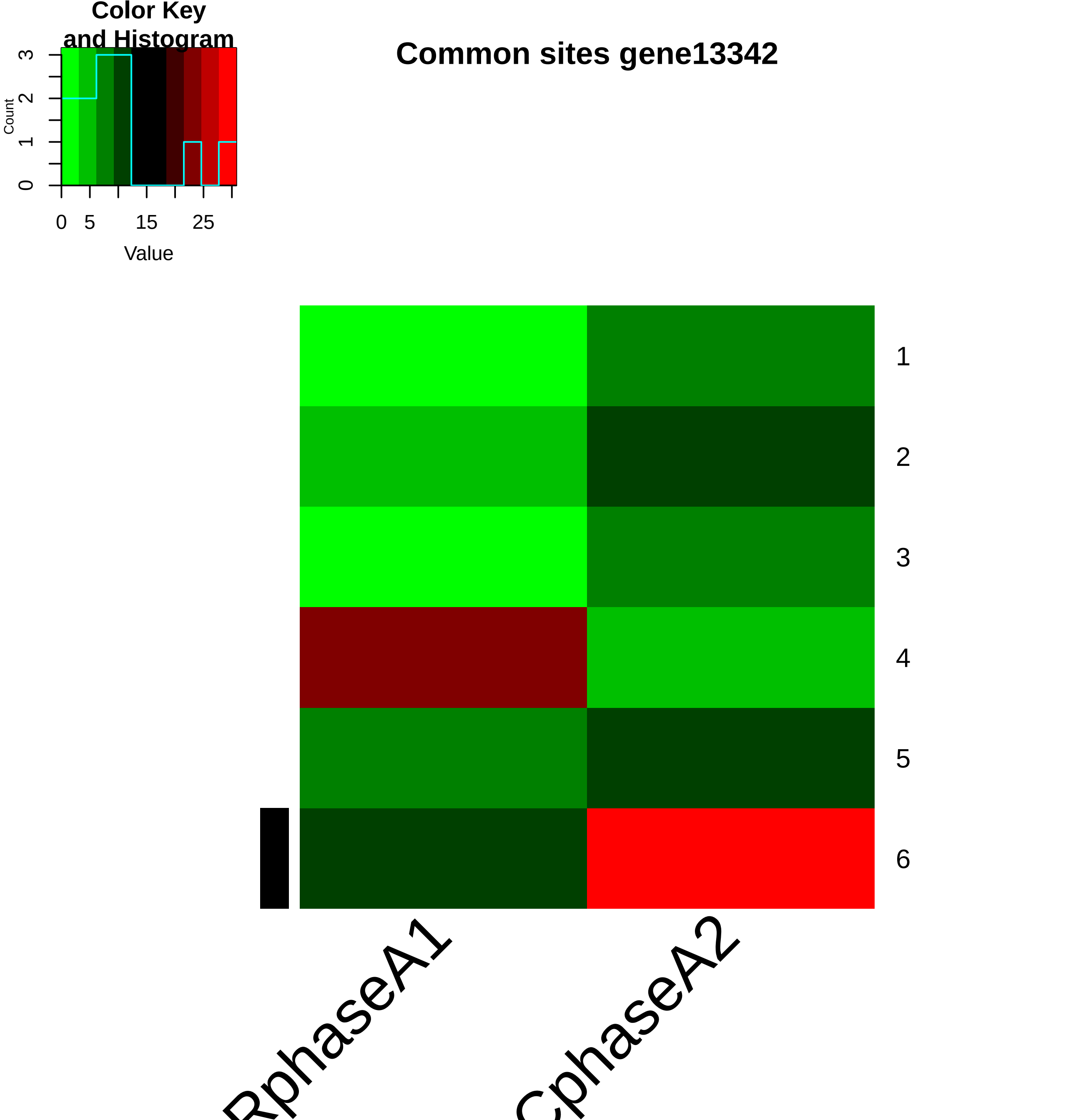
Color Key
and Histogram
Count
0
1
2
3
0
5
15
25
Value
Common sites gene13342
1
2
3
4
5
6

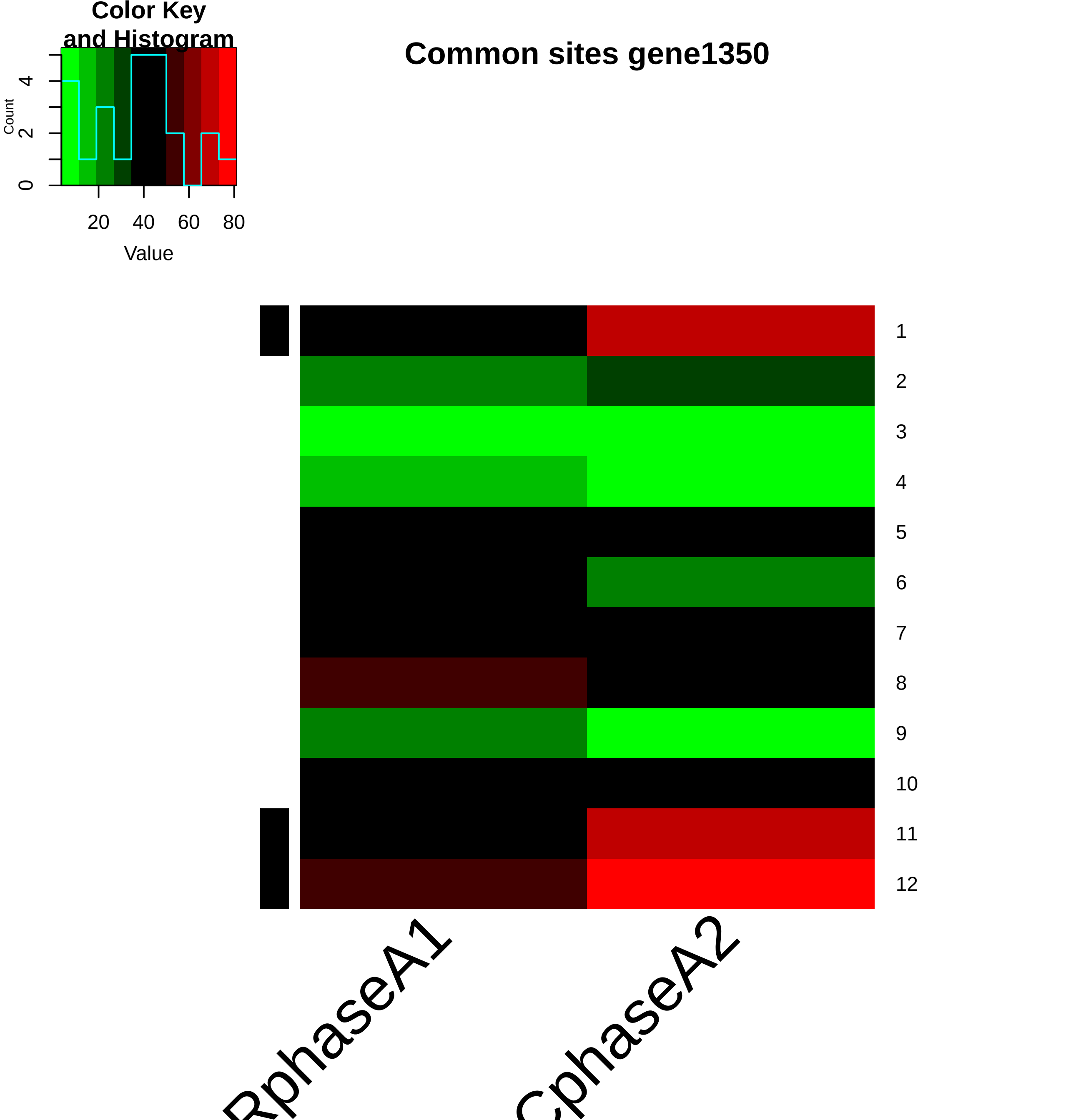
Color Key
and Histogram
Count
0
2
4
20
40
60
80
Value
Common sites gene1350
1
2
3
4
5
6
7
8
9
10
11
12
RphaseA1
CphaseA2

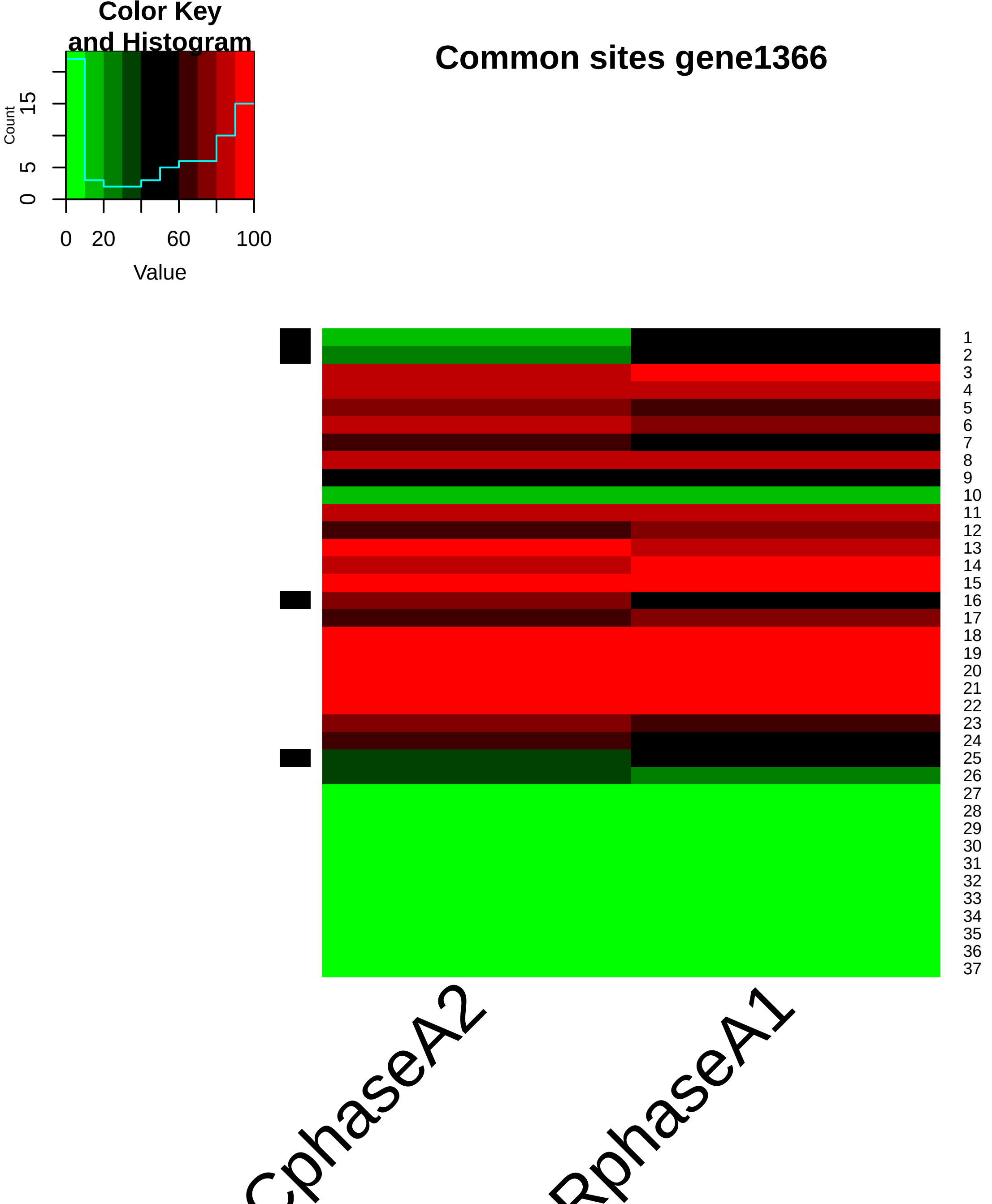

Color Key
and Histogram
Count
0
5
15
0
20
60
100
Value
Common sites gene1366
CphaseA2
RphaseA1

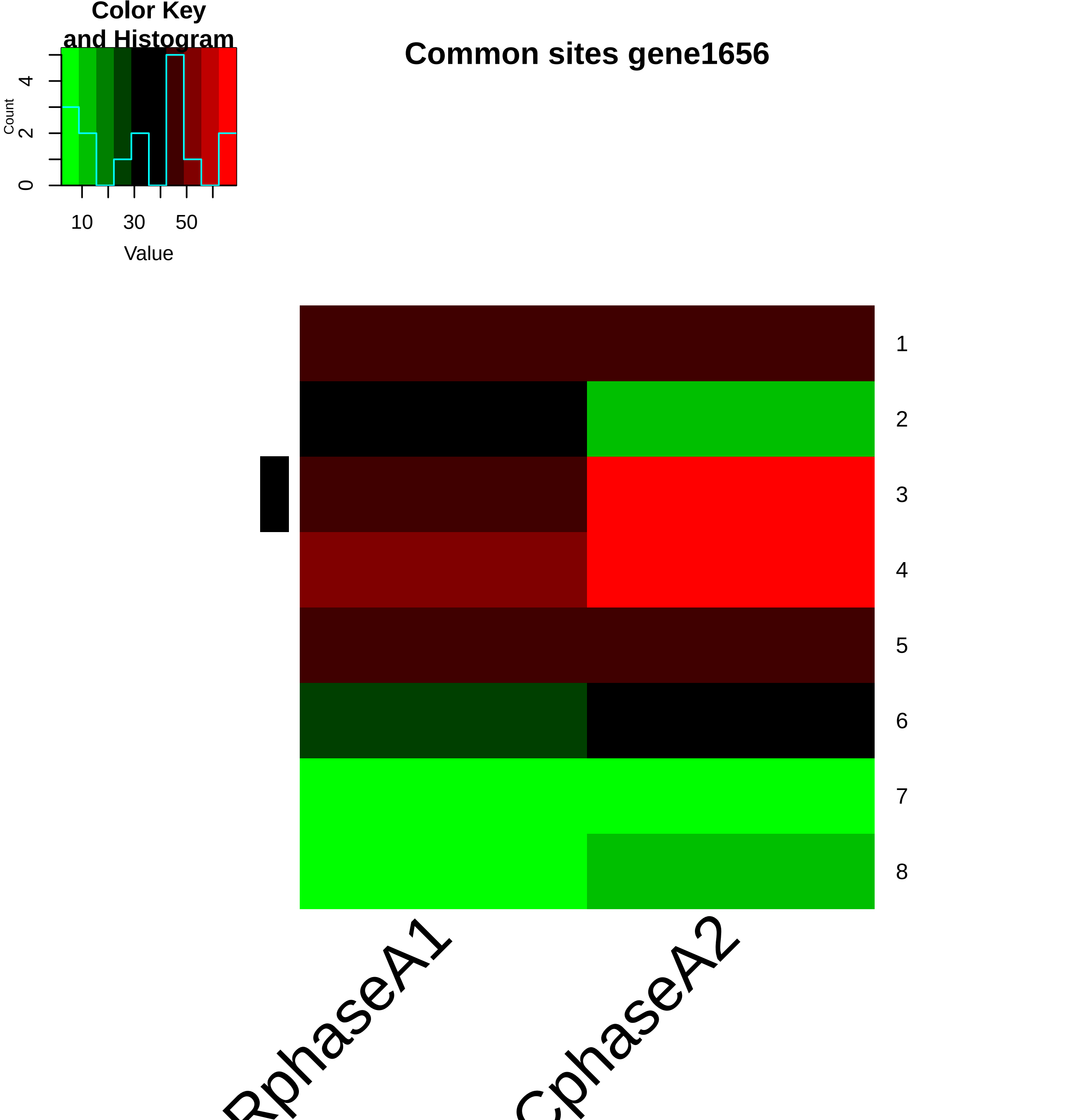
Color Key
and Histogram
Count
0
2
4
10
30
50
Value
Common sites gene1656
1
2
3
4
5
6
7
8
RphaseA1
CphaseA2

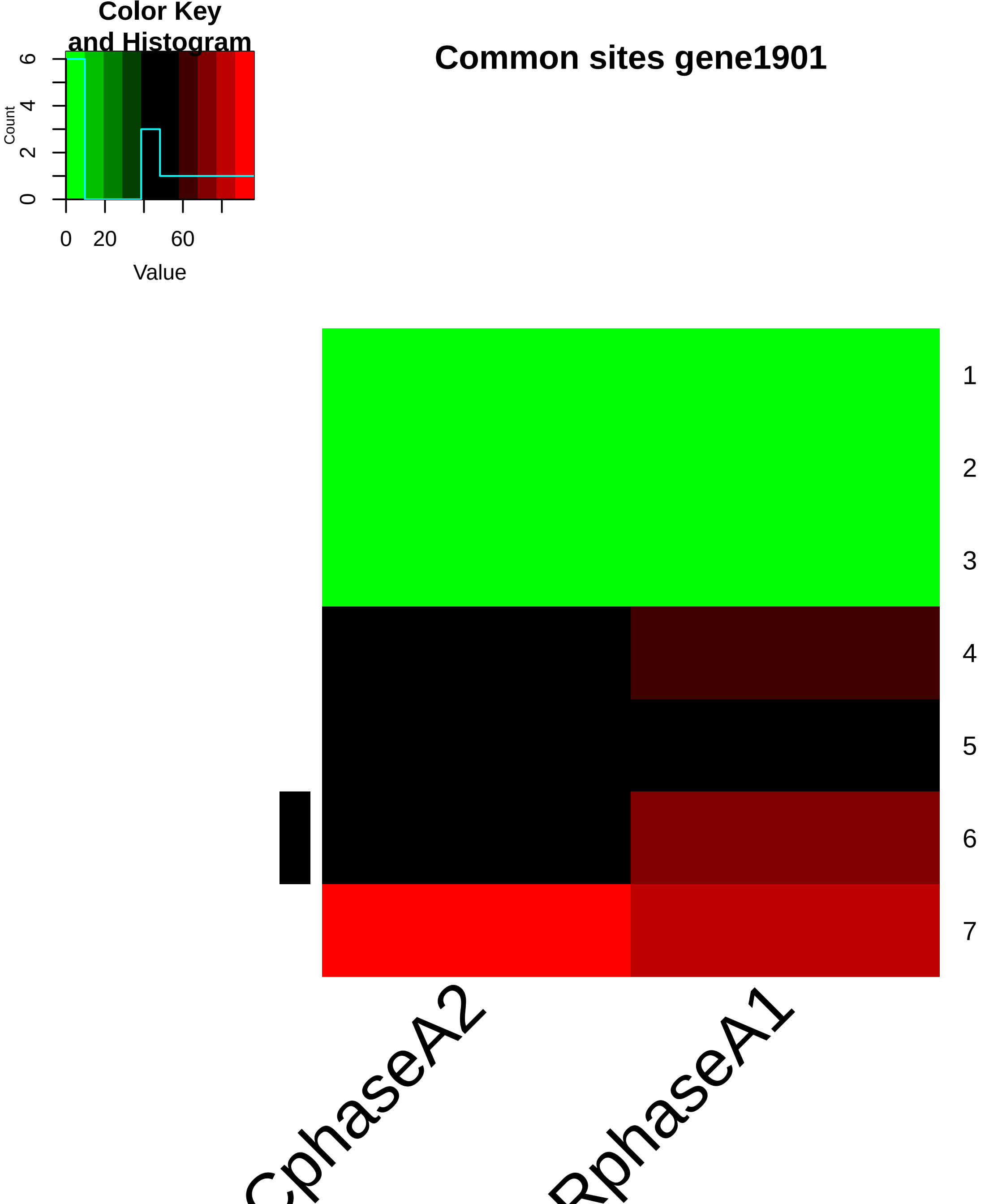
Color Key
and Histogram
Count
0
2
4
6
0
20
60
Value
Common sites gene1901
1
2
3
4
5
6
7
CphaseA2
RphaseA1

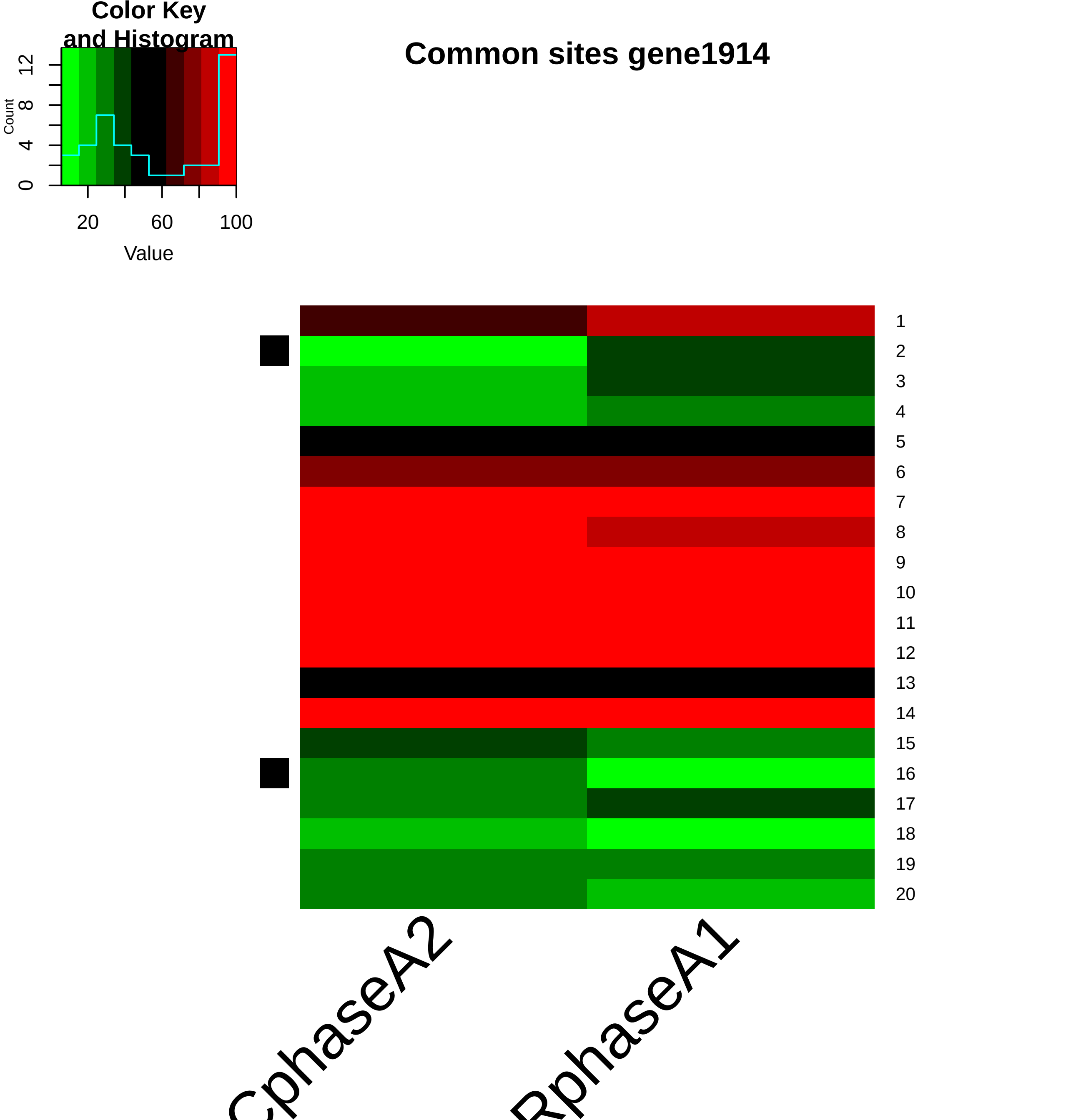
Color Key
and Histogram
Count
0
4
8
12
20
60
100
Value
Common sites gene1914
CphaseA2
RphaseA1

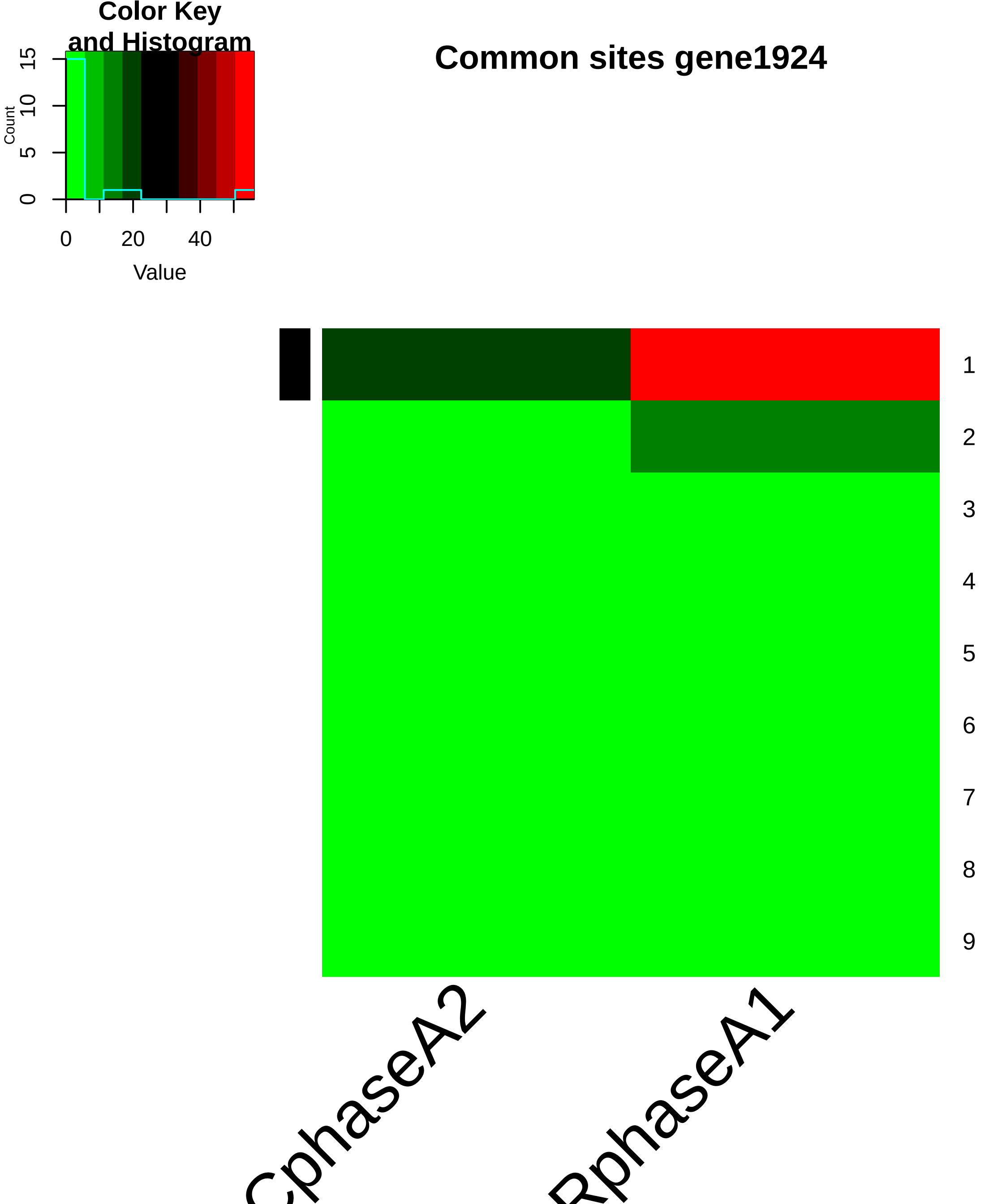
Color Key
and Histogram
Count
Value
0
20
40
0
5
10
15
Common sites gene1924
1
2
3
4
5
6
7
8
9
CphaseA2
RphaseA1

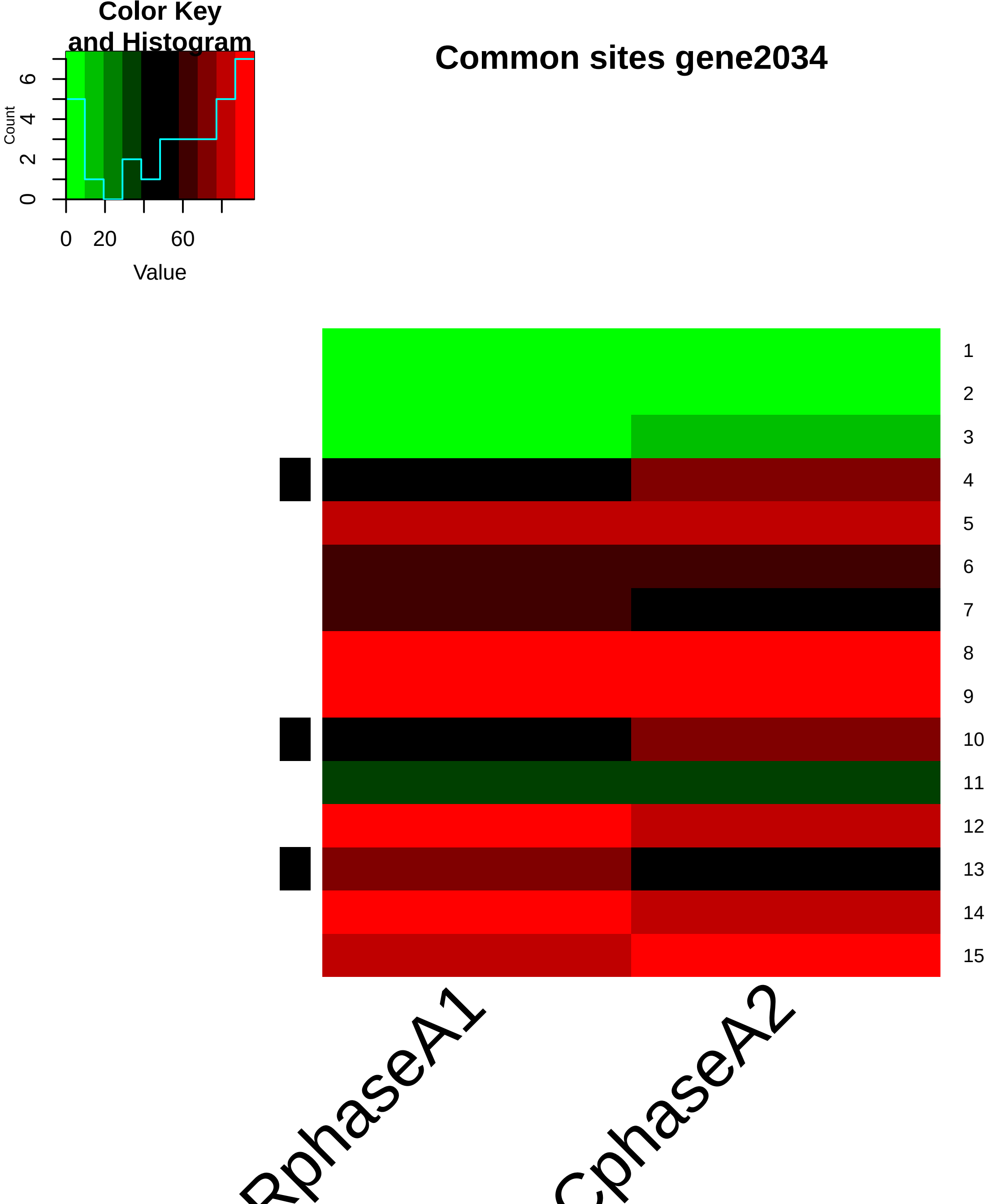

Color Key
and Histogram
Count
0
2
4
6
0
20
60
Value
Common sites gene2034
RphaseA1
CphaseA2

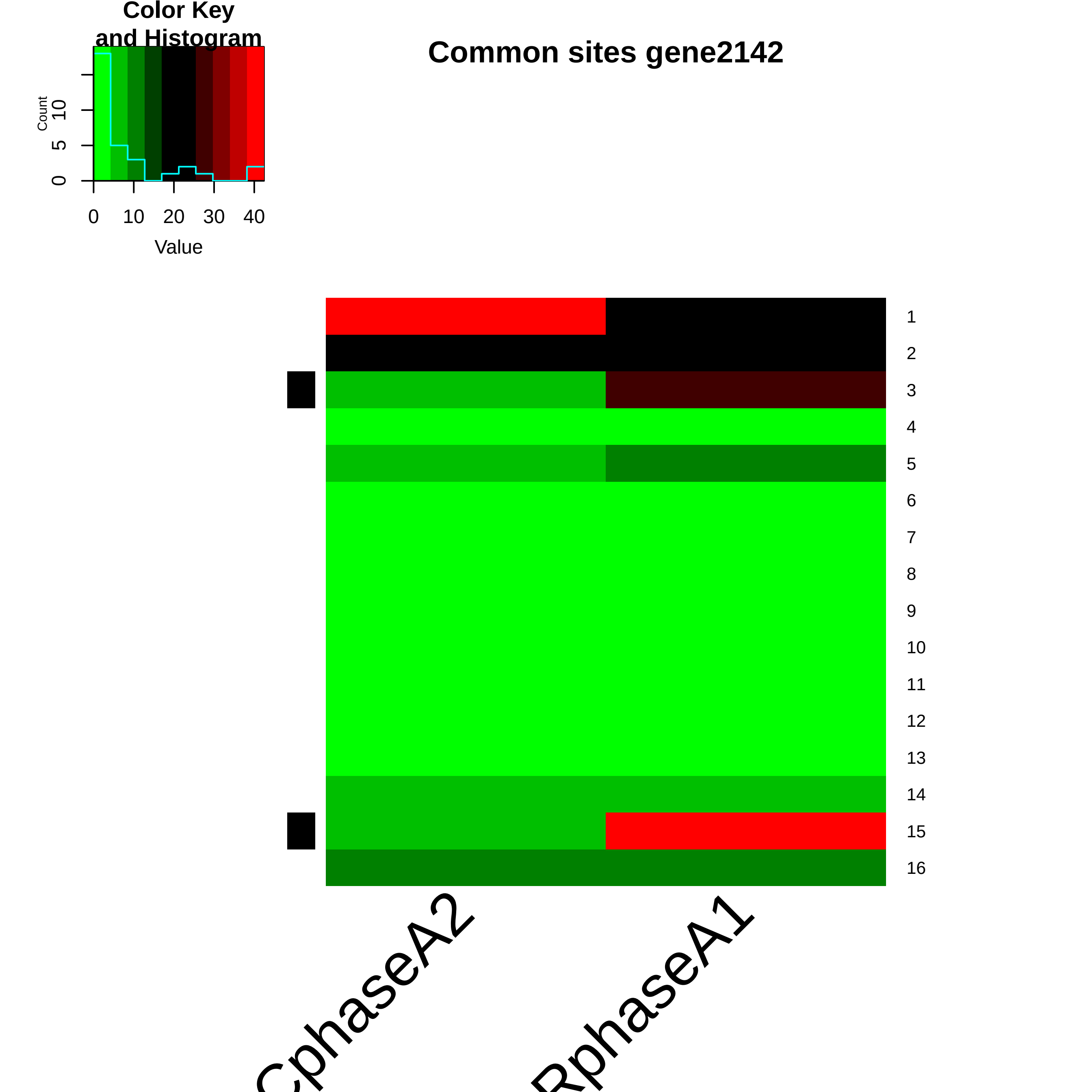

Color Key
and Histogram
Count
0
5
10
0
10
20
30
40
Value
Common sites gene2142
CphaseA2
RphaseA1

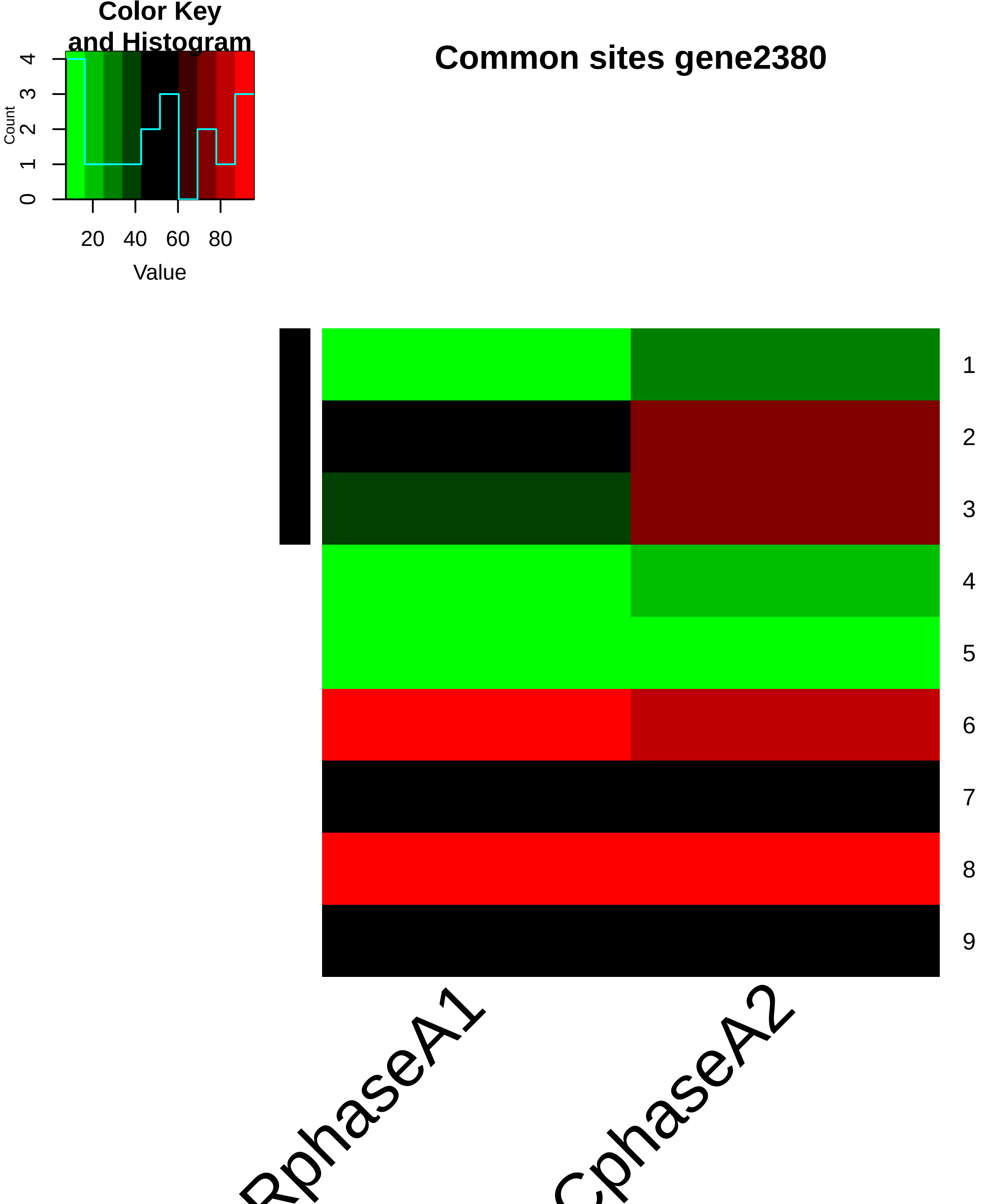

Color Key
and Histogram
Count
0
1
2
3
4
20
40
60
80
Value
Common sites gene2380
1
2
3
4
5
6
7
8
9
RphaseA1
CphaseA2

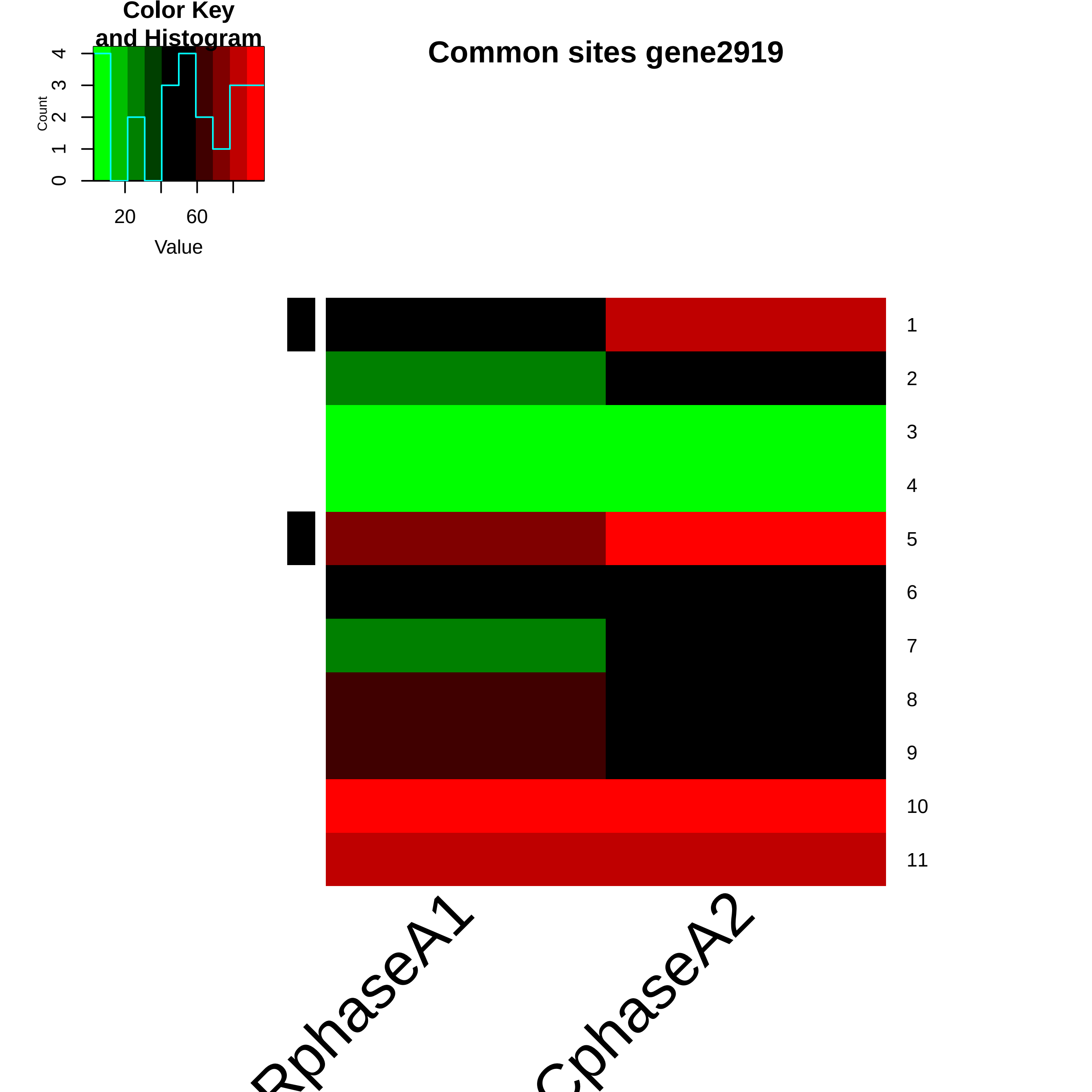
Color Key
and Histogram
Count
0
1
2
3
4
20
60
Value
Common sites gene2919
1
2
3
4
5
6
7
8
9
10
11
RphaseA1
CphaseA2

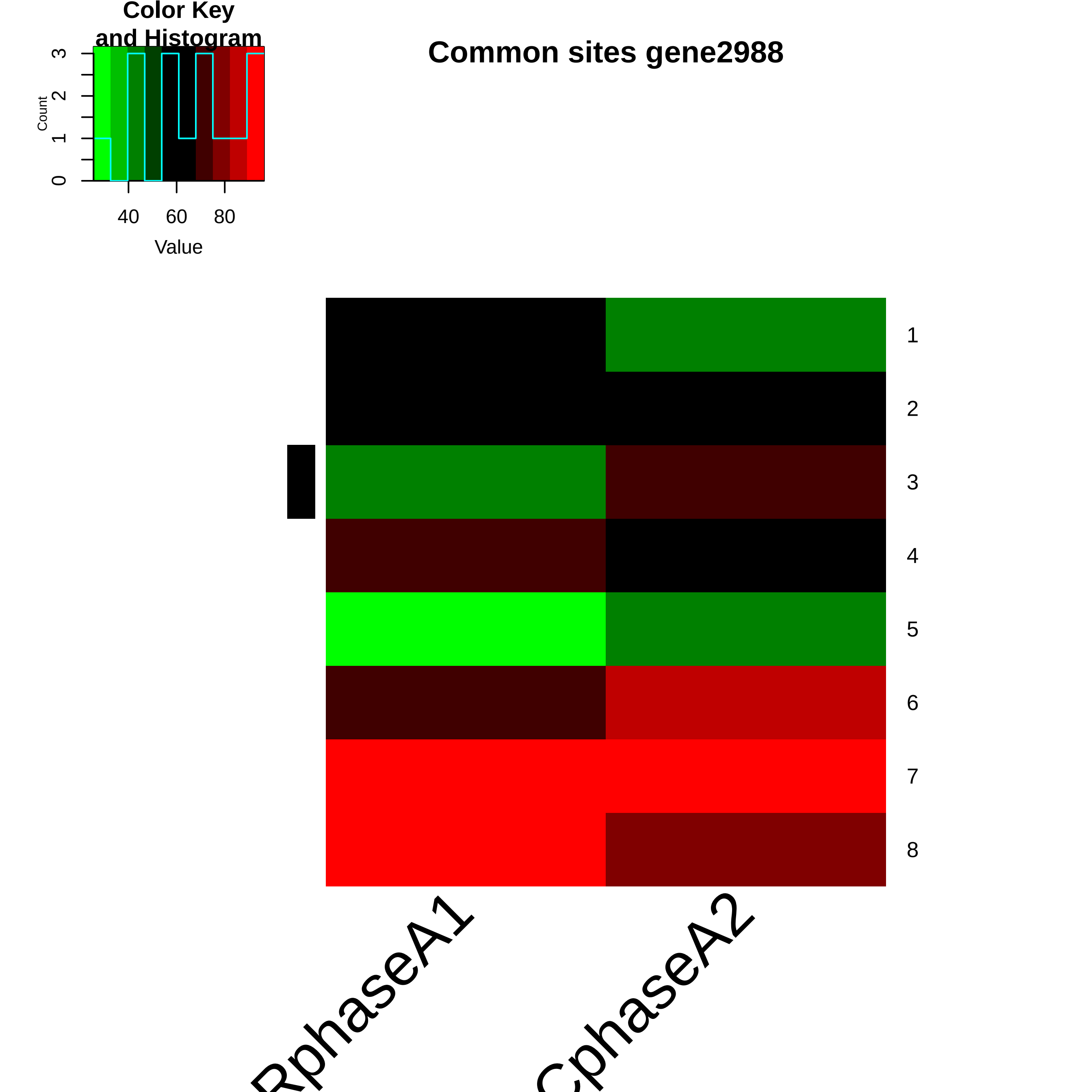

Color Key
and Histogram
Count
0
1
2
3
40
60
80
Value
Common sites gene2988
1
2
3
4
5
6
7
8
RphaseA1
CphaseA2

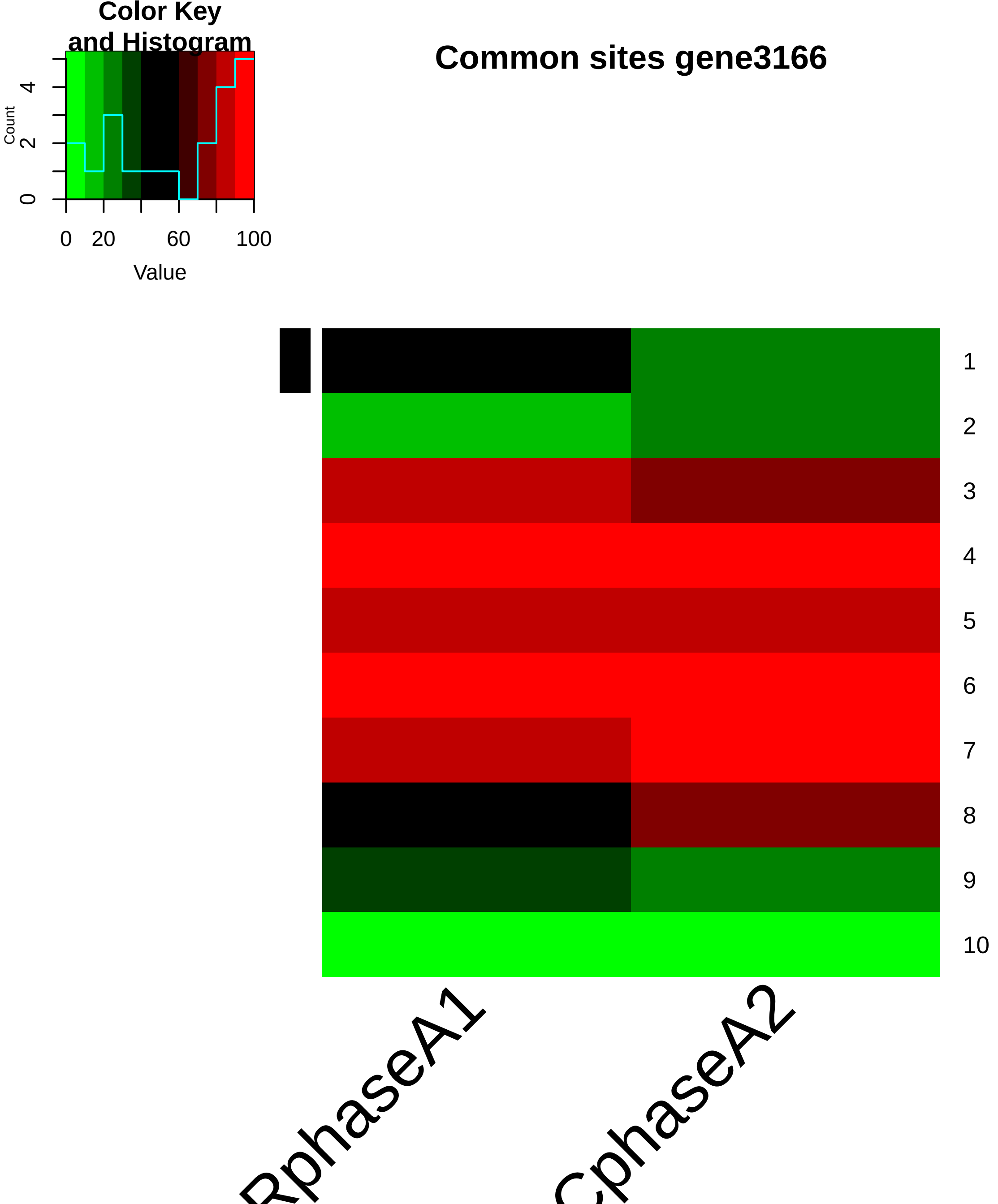
Color Key
and Histogram
Count
0
2
4
0
20
60
100
Value
Common sites gene3166
1
2
3
4
5
6
7
8
9
10
RphaseA1
CphaseA2

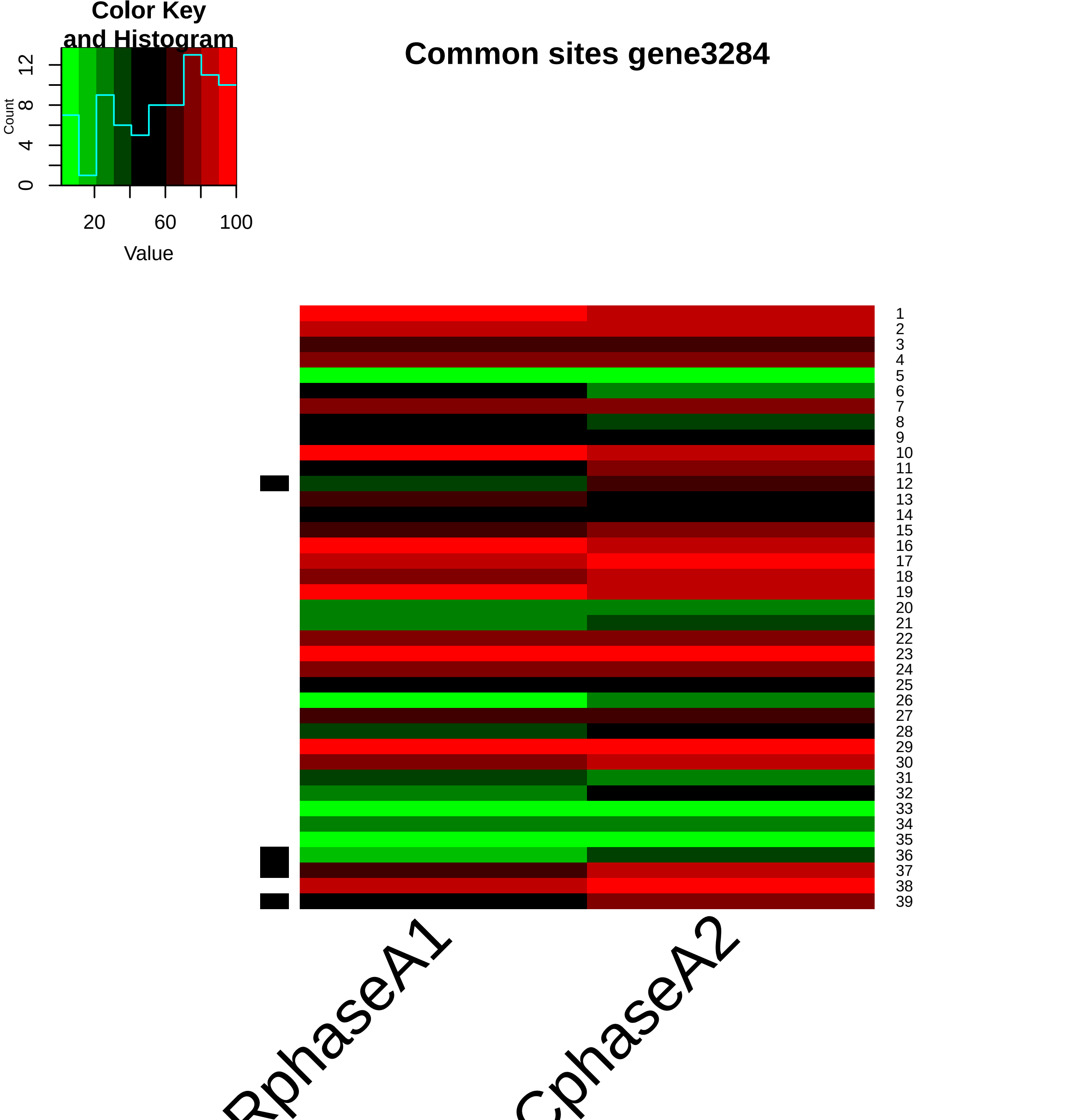
Color Key
and Histogram
Count
0
4
8
12
20
60
100
Value
Common sites gene3284
RphaseA1
CphaseA2

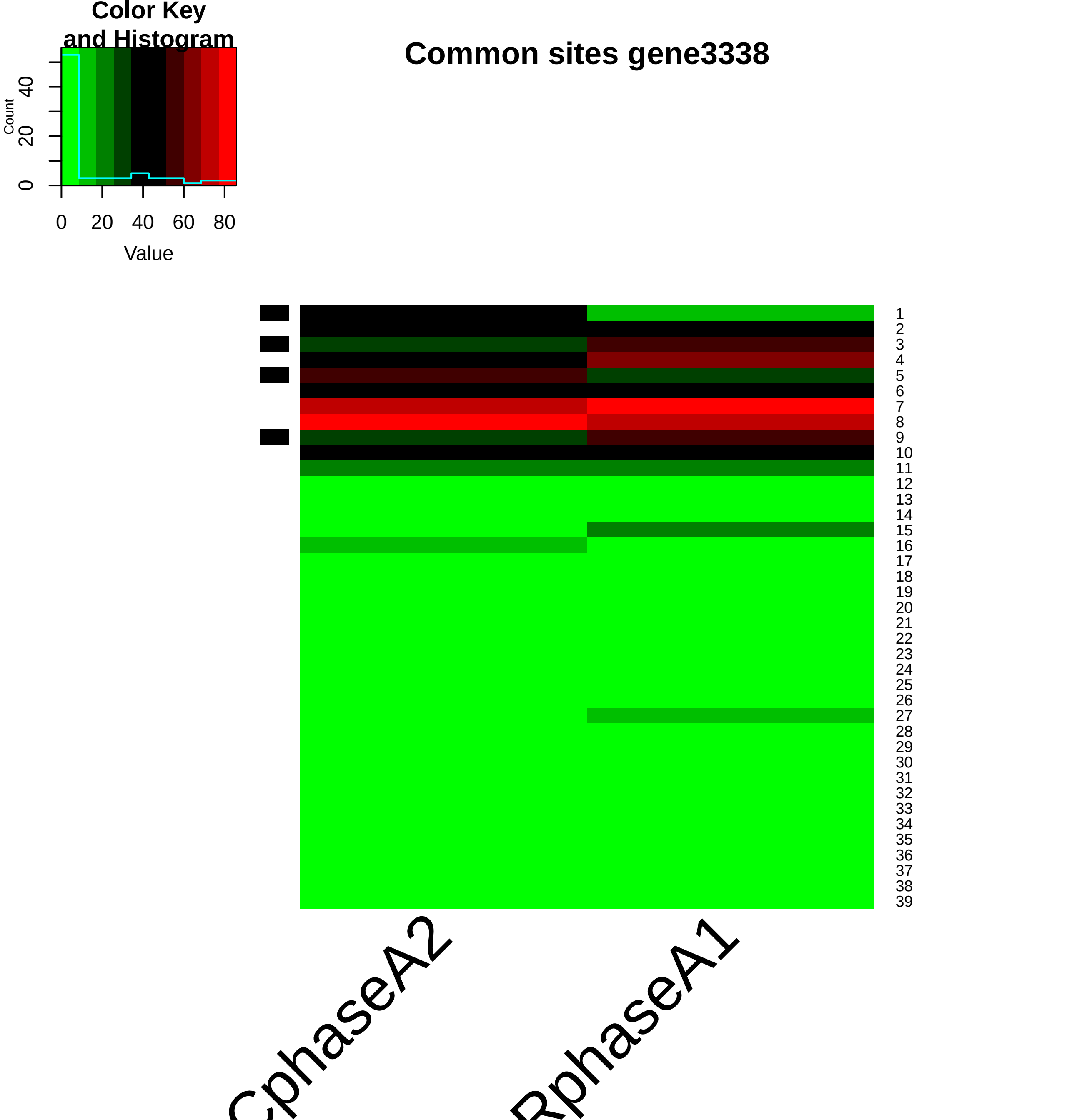
Color Key
and Histogram
Count
0
20
40
0
20
40
60
80
Value
Common sites gene3338
CphaseA2
RphaseA1

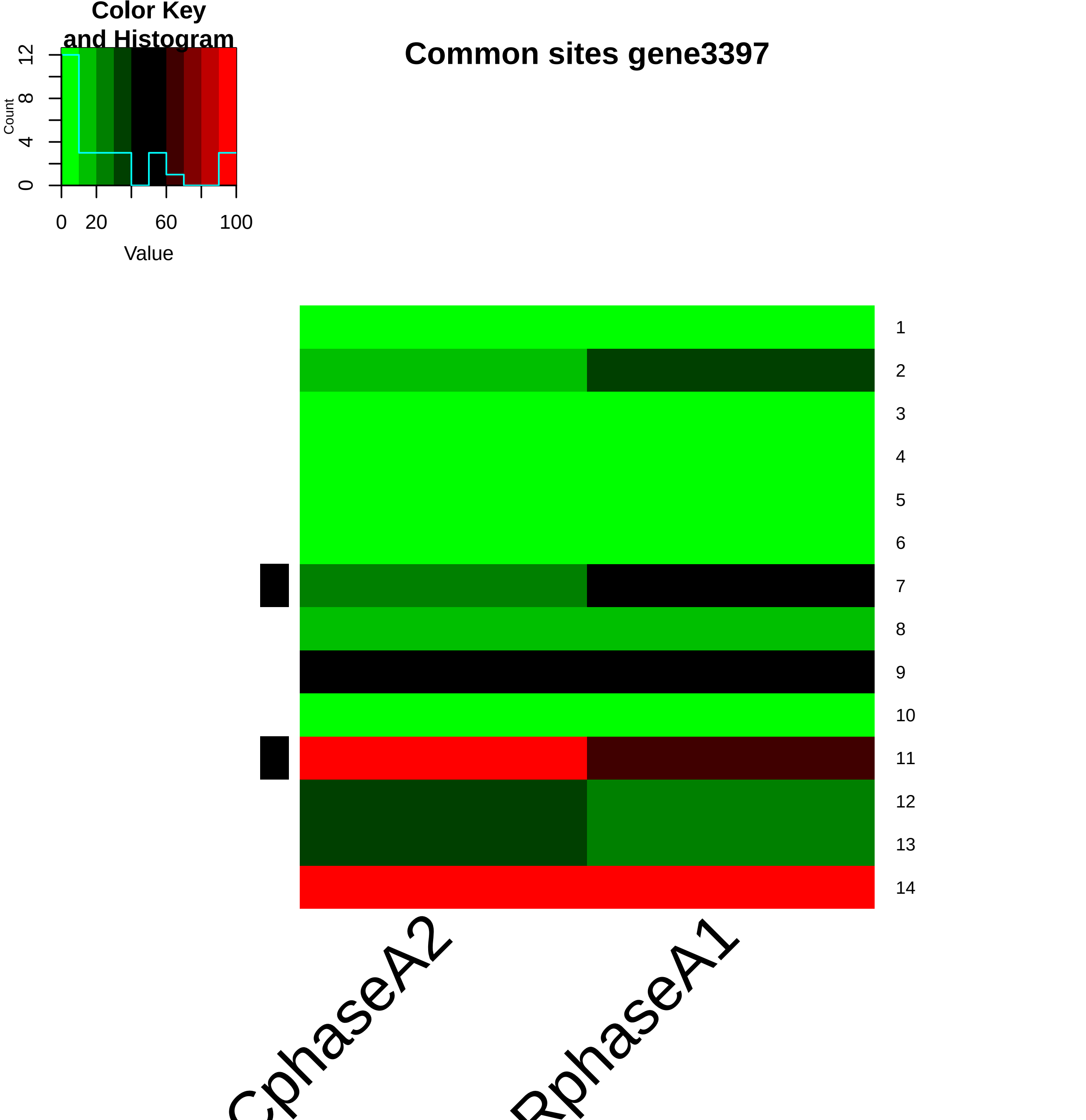

Color Key
and Histogram
Count
Value
0
20
60
100
0
4
8
12
Common sites gene3397
1
2
3
4
5
6
7
8
9
10
11
12
13
14

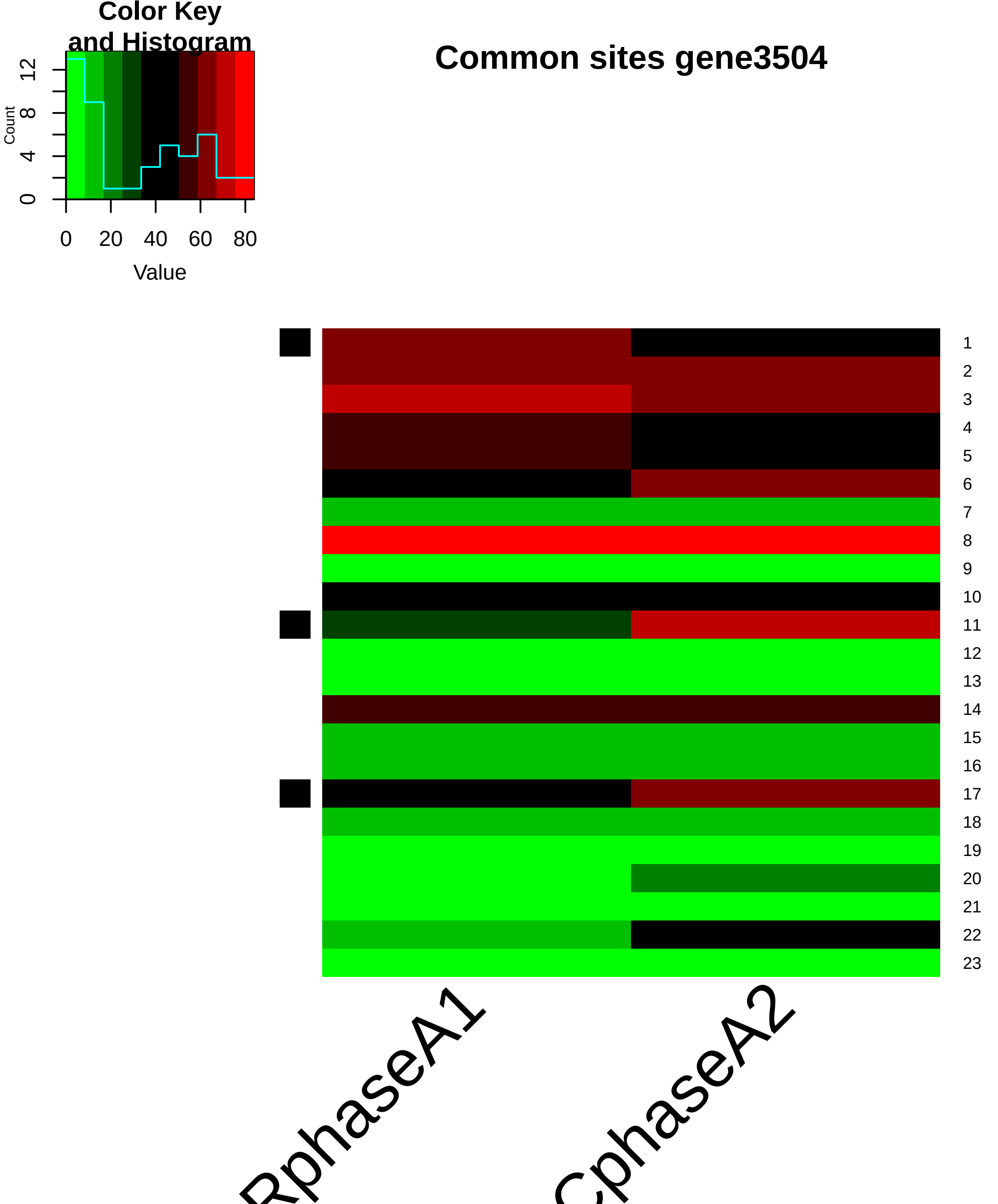
Color Key
and Histogram
Count
0
4
8
12
0
20
40
60
80
Value
Common sites gene3504
RphaseA1
CphaseA2

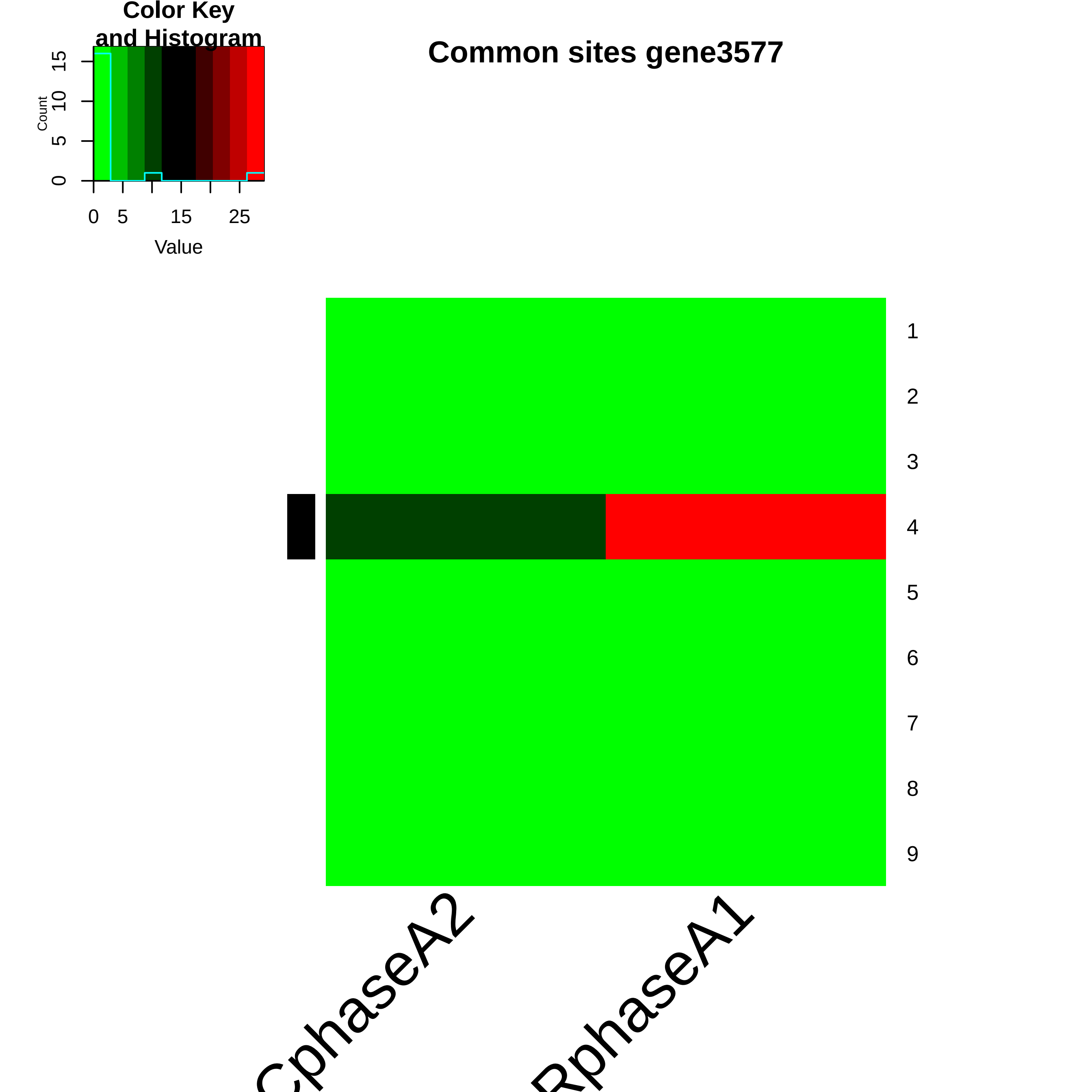

Color Key
and Histogram
Count
0
5
10
15
0
5
15
25
Value
Common sites gene3577
1
2
3
4
5
6
7
8
9
CphaseA2
RphaseA1

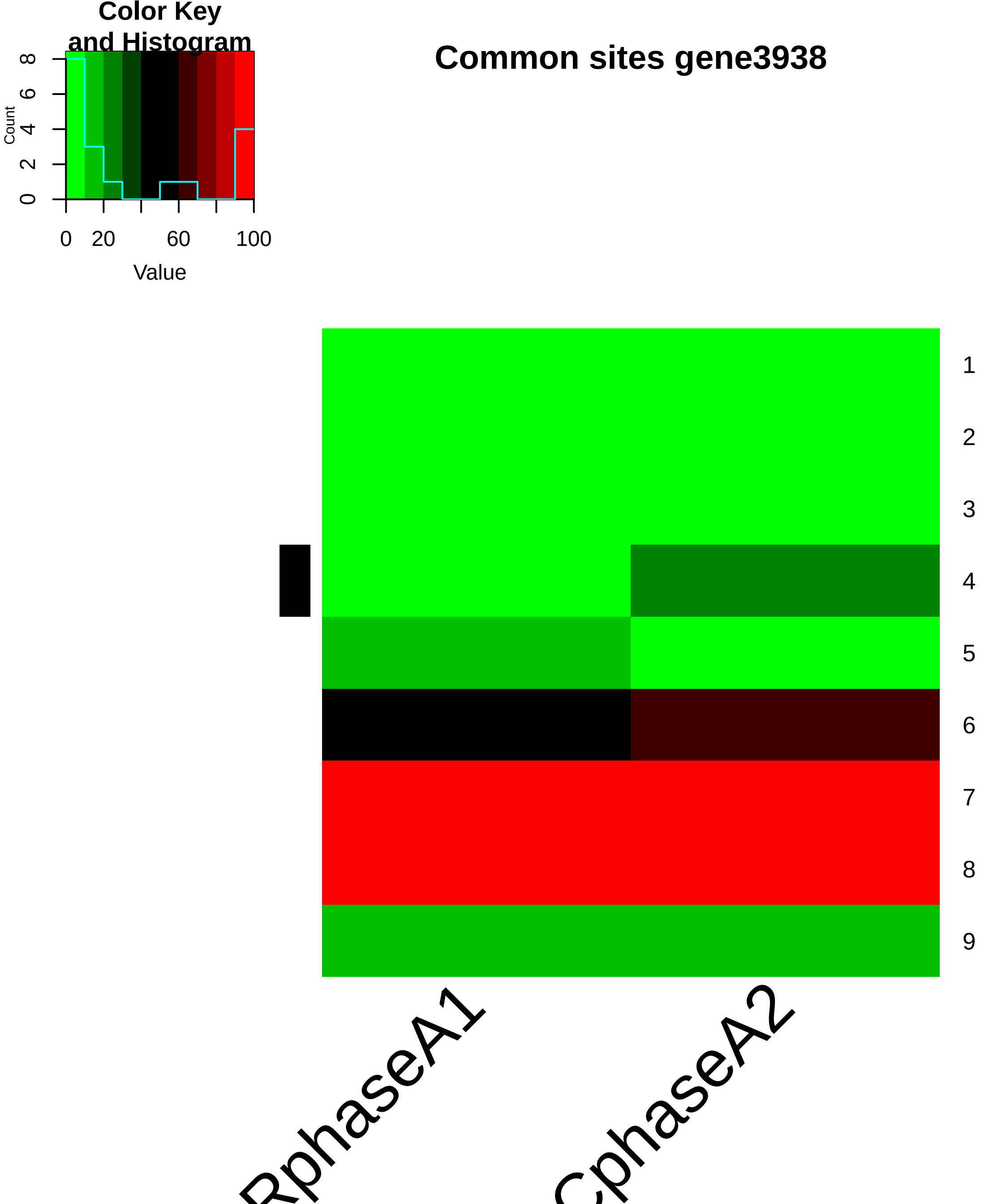
Color Key
and Histogram
Count
0
2
4
6
8
0
20
60
100
Value
Common sites gene3938
1
2
3
4
5
6
7
8
9
RphaseA1
CphaseA2

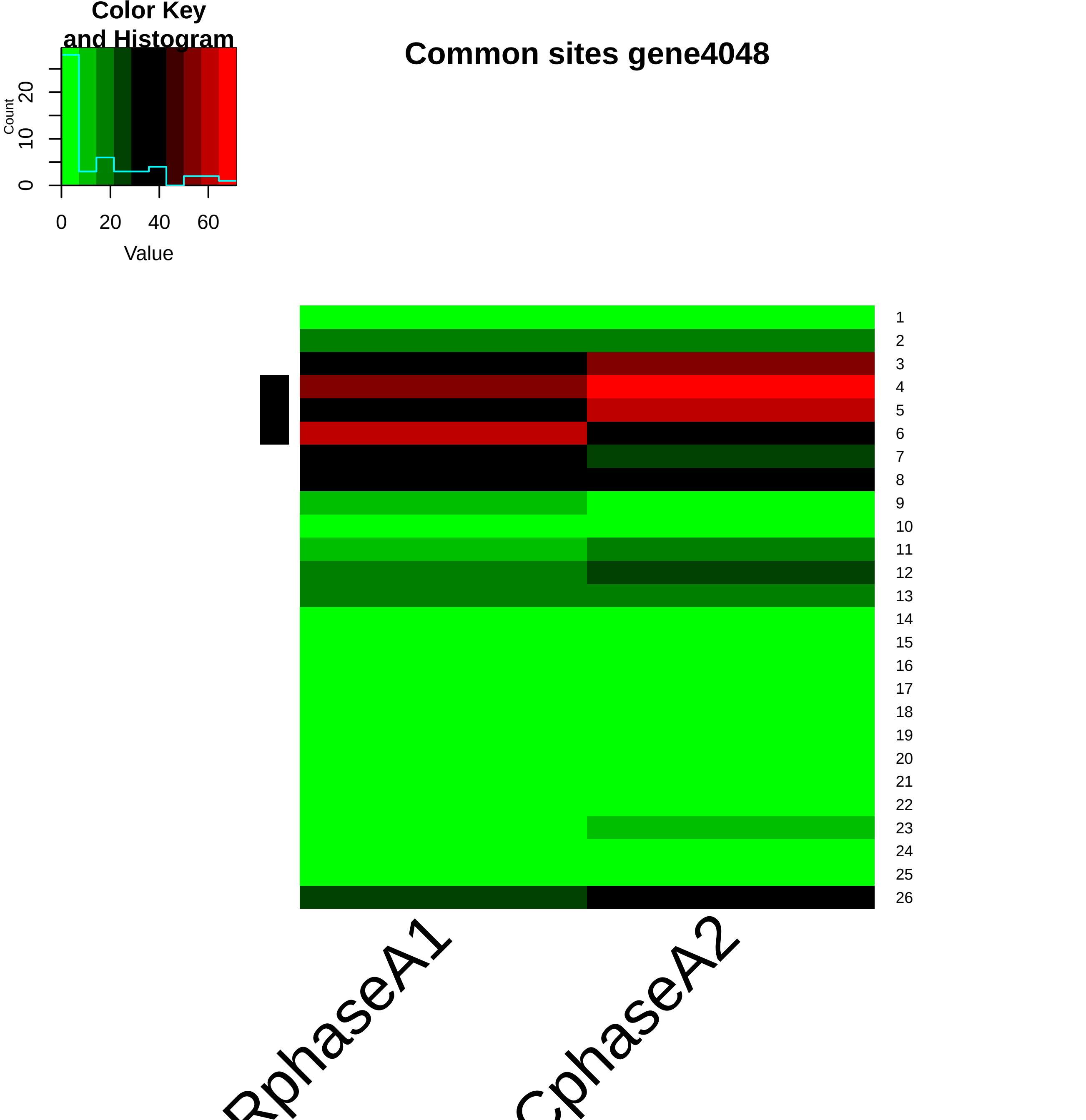

Color Key
and Histogram
Count
0
10
20
0
20
40
60
Value
Common sites gene4048
RphaseA1
CphaseA2

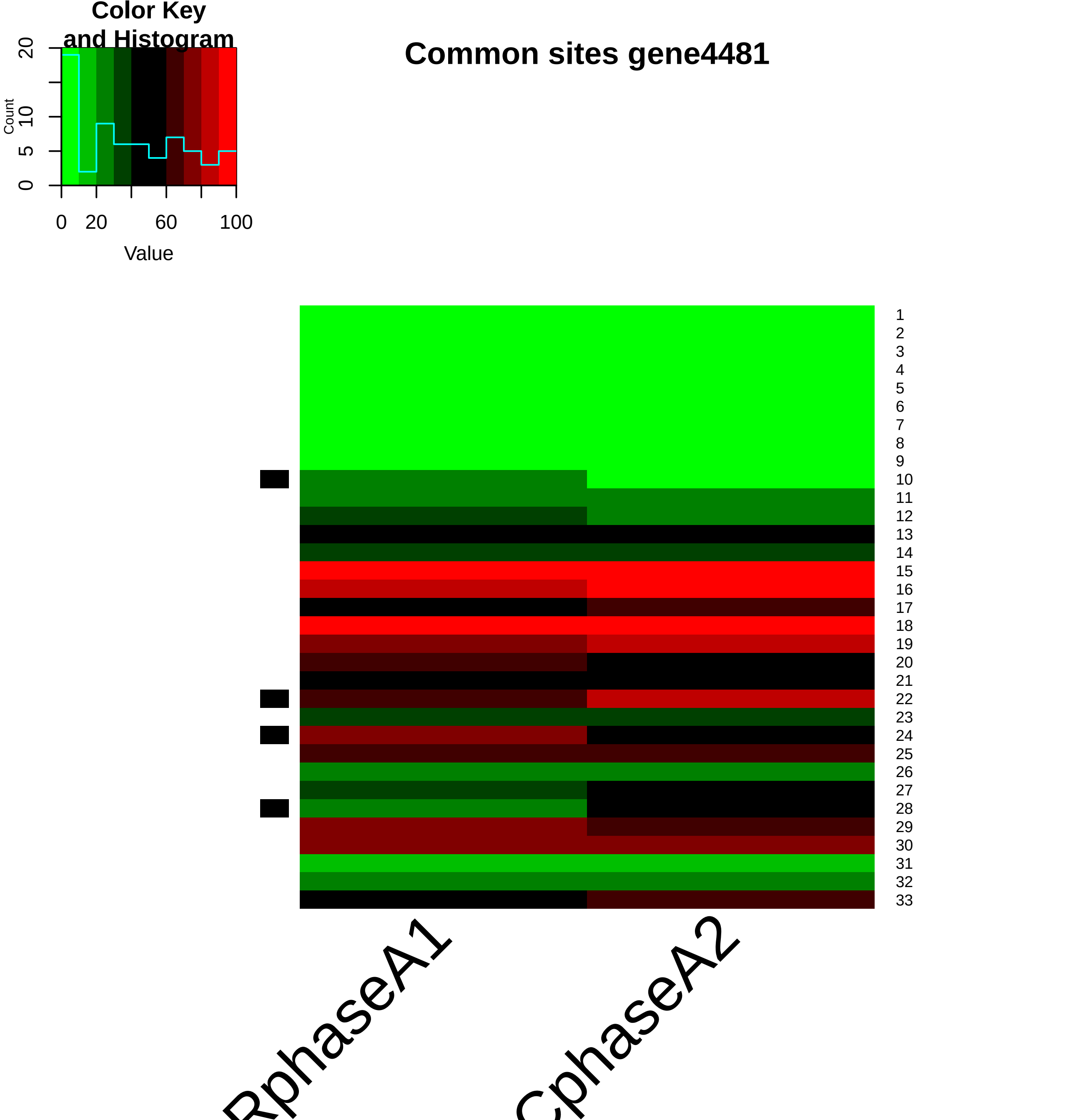
Color Key
and Histogram
Count
0
5
10
20
0
20
60
100
Value
Common sites gene4481
RphaseA1
CphaseA2

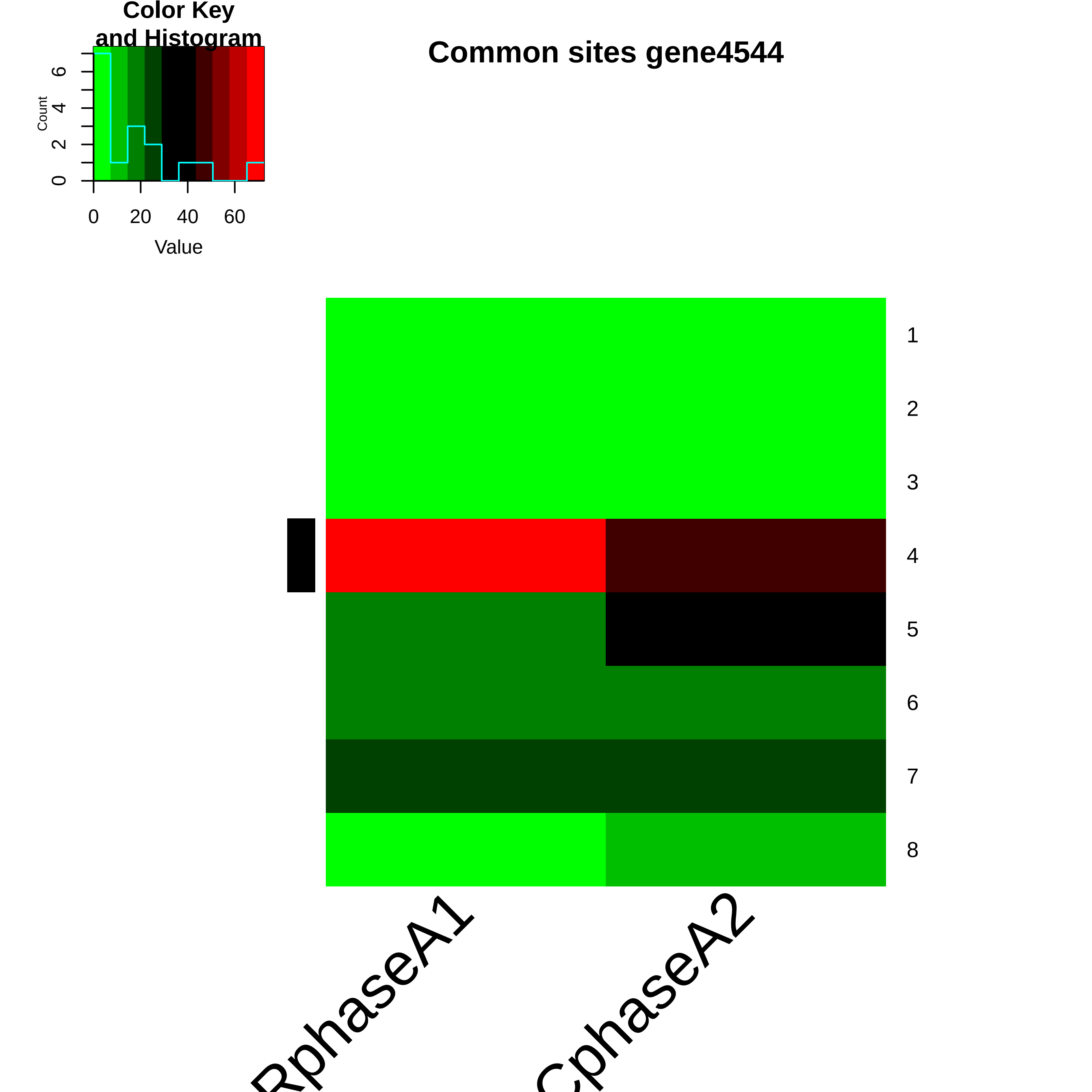
Color Key
and Histogram
Count
0
2
4
6
0
20
40
60
Value
Common sites gene4544
1
2
3
4
5
6
7
8
RphaseA1
CphaseA2

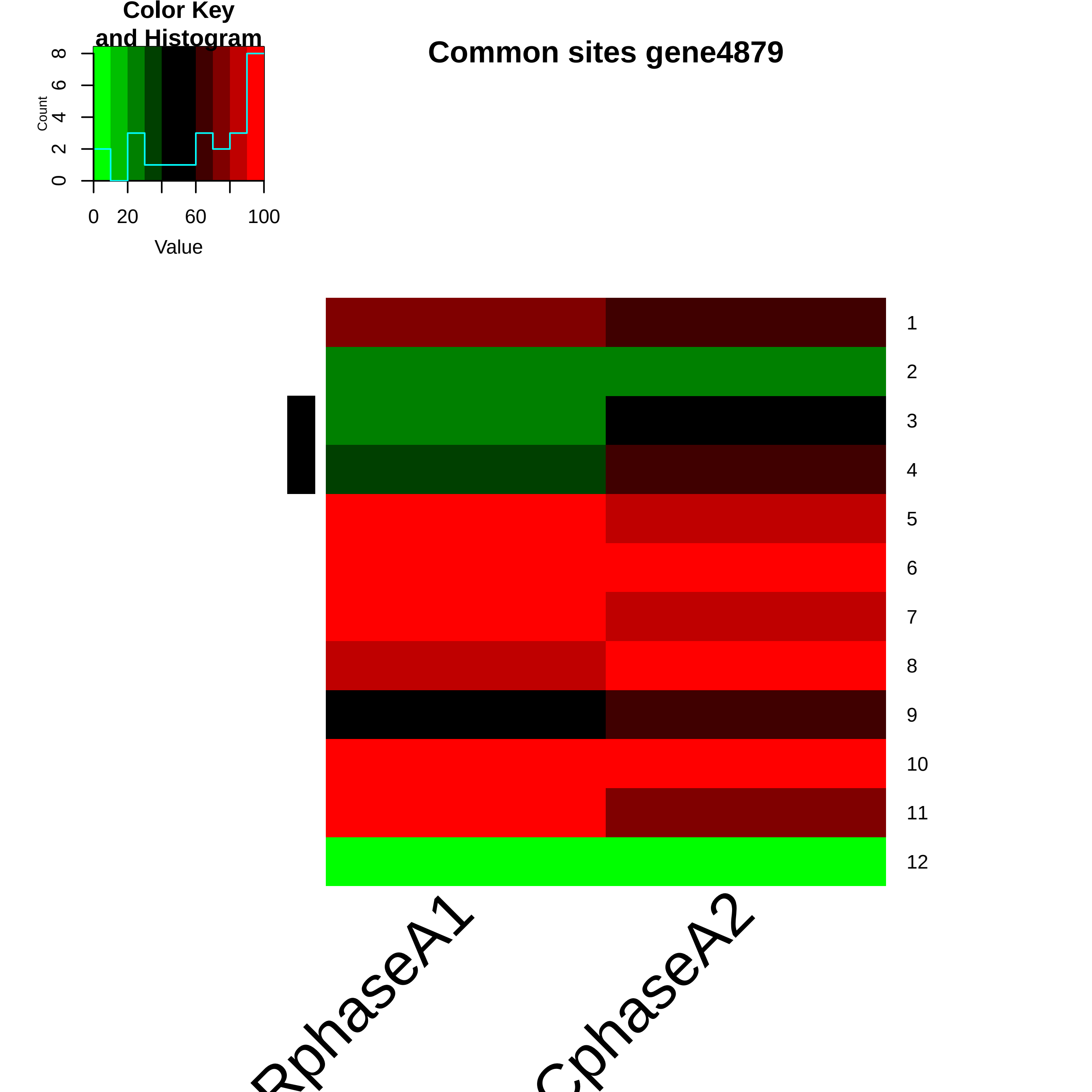

Color Key
and Histogram
Count
0
2
4
6
8
0
20
60
100
Value
Common sites gene4879
1
2
3
4
5
6
7
8
9
10
11
12
RphaseA1
CphaseA2

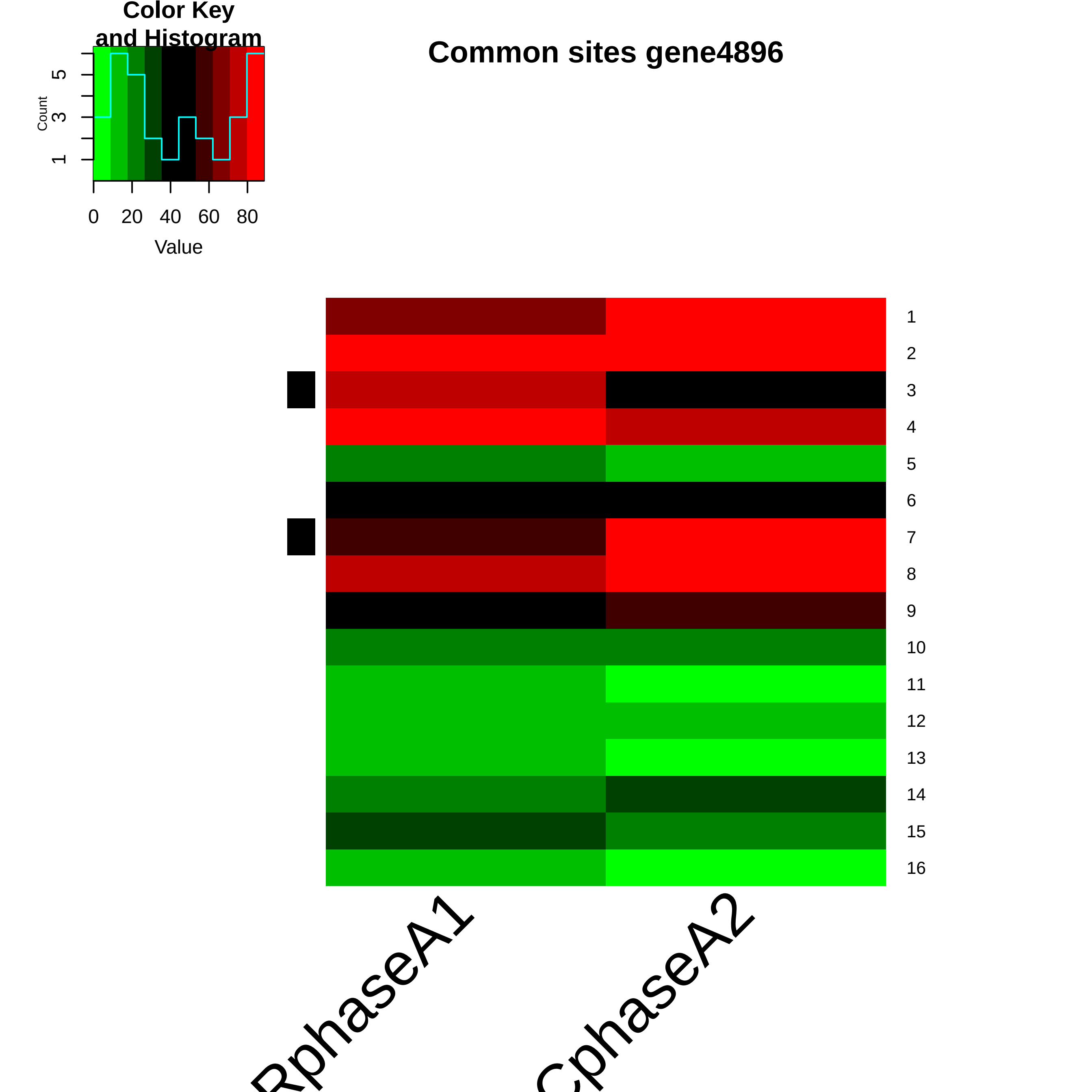
Color Key
and Histogram
Count
1
3
5
0
20
40
60
80
Value
Common sites gene4896
RphaseA1
CphaseA2

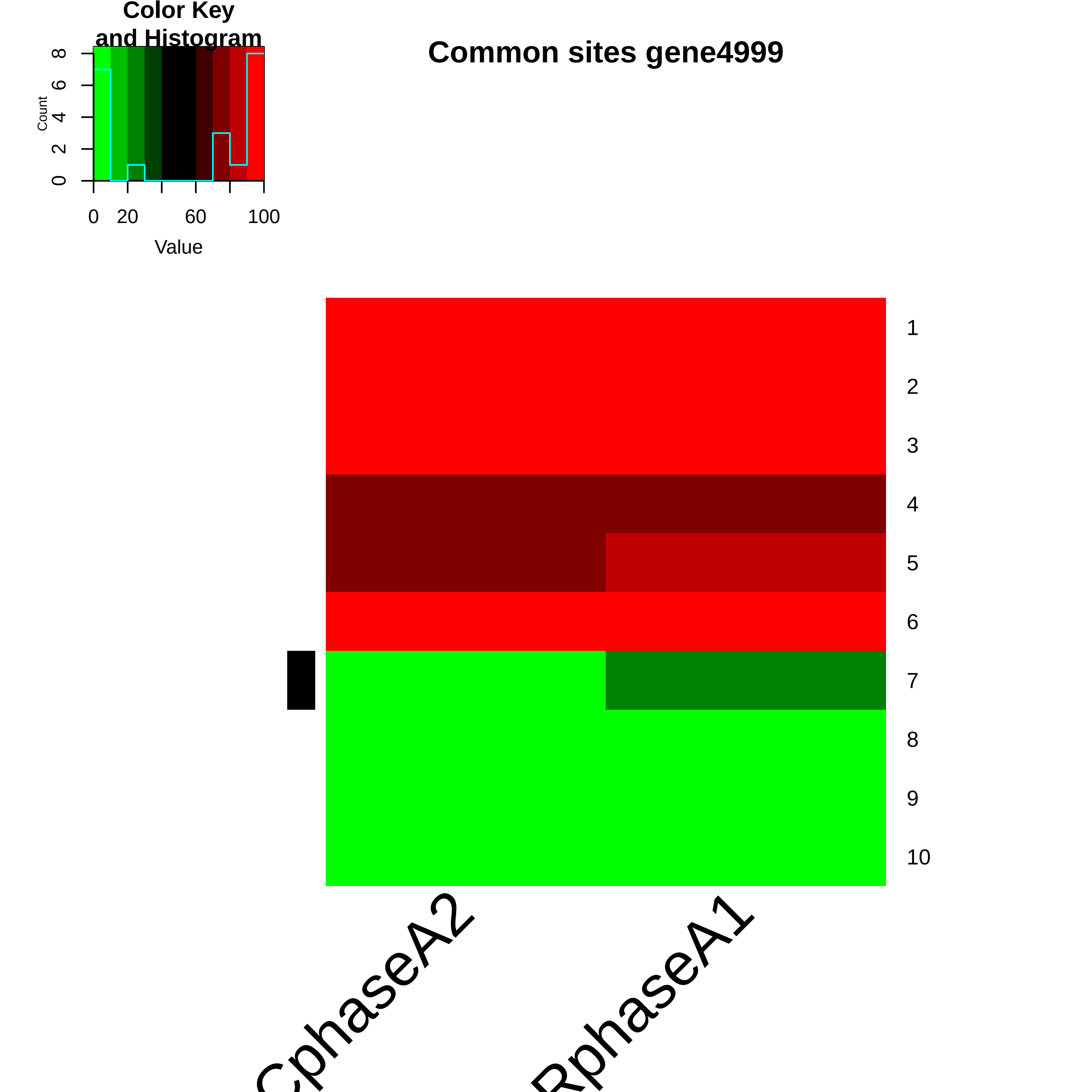

Color Key
and Histogram
Count
0
2
4
6
8
0
20
60
100
Value
Common sites gene4999
1
2
3
4
5
6
7
8
9
10
CphaseA2
RphaseA1

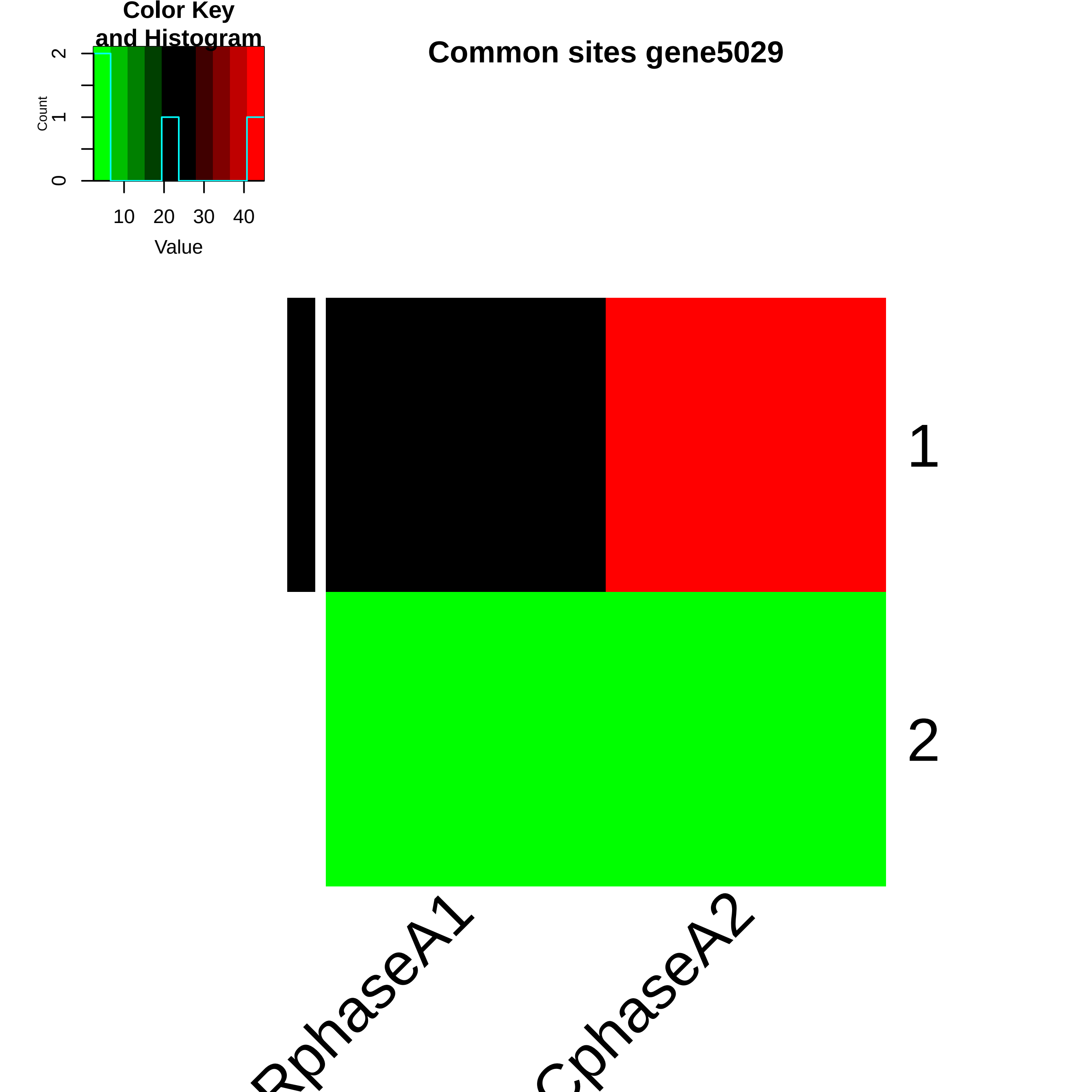
Color Key
and Histogram
Count
0
1
2
10
20
30
40
Value
Common sites gene5029
1
2
RphaseA1
CphaseA2

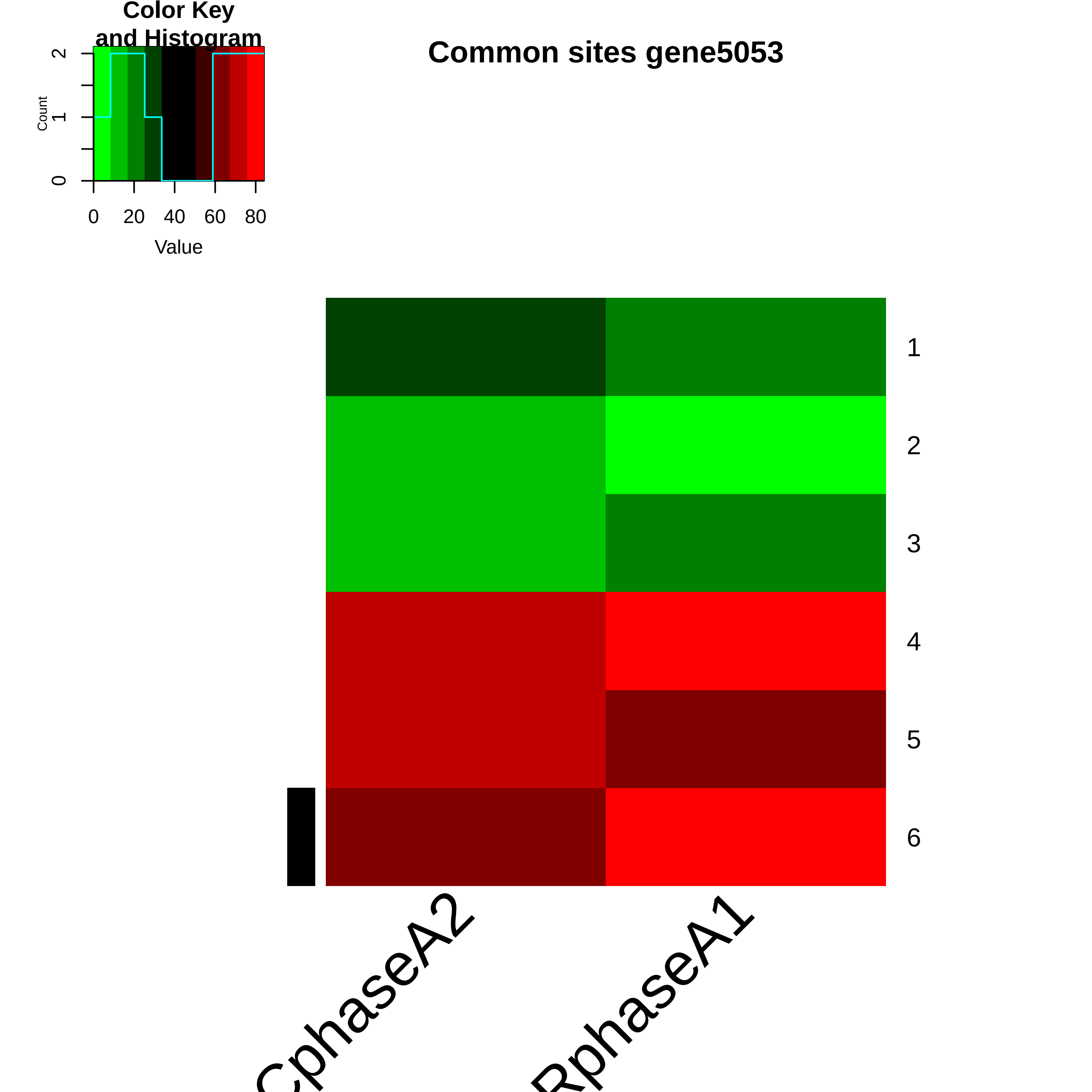

Color Key
and Histogram
Count
0
1
2
0
20
40
60
80
Value
Common sites gene5053
1
2
3
4
5
6
CphaseA2
RphaseA1

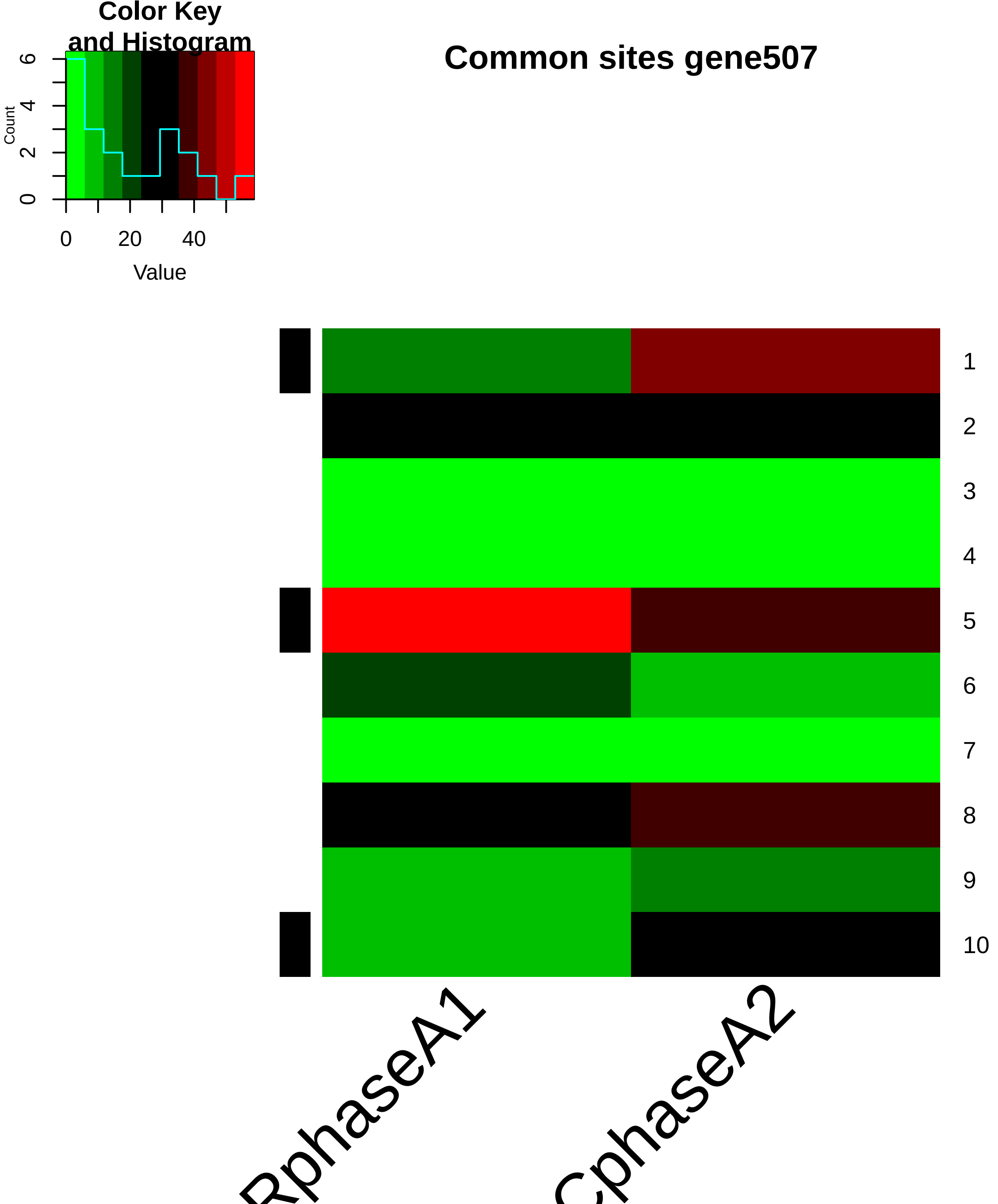

Color Key
and Histogram
Count
Value
0
20
40
0
2
4
6
Common sites gene507
1
2
3
4
5
6
7
8
9
10
RphaseA1
CphaseA2

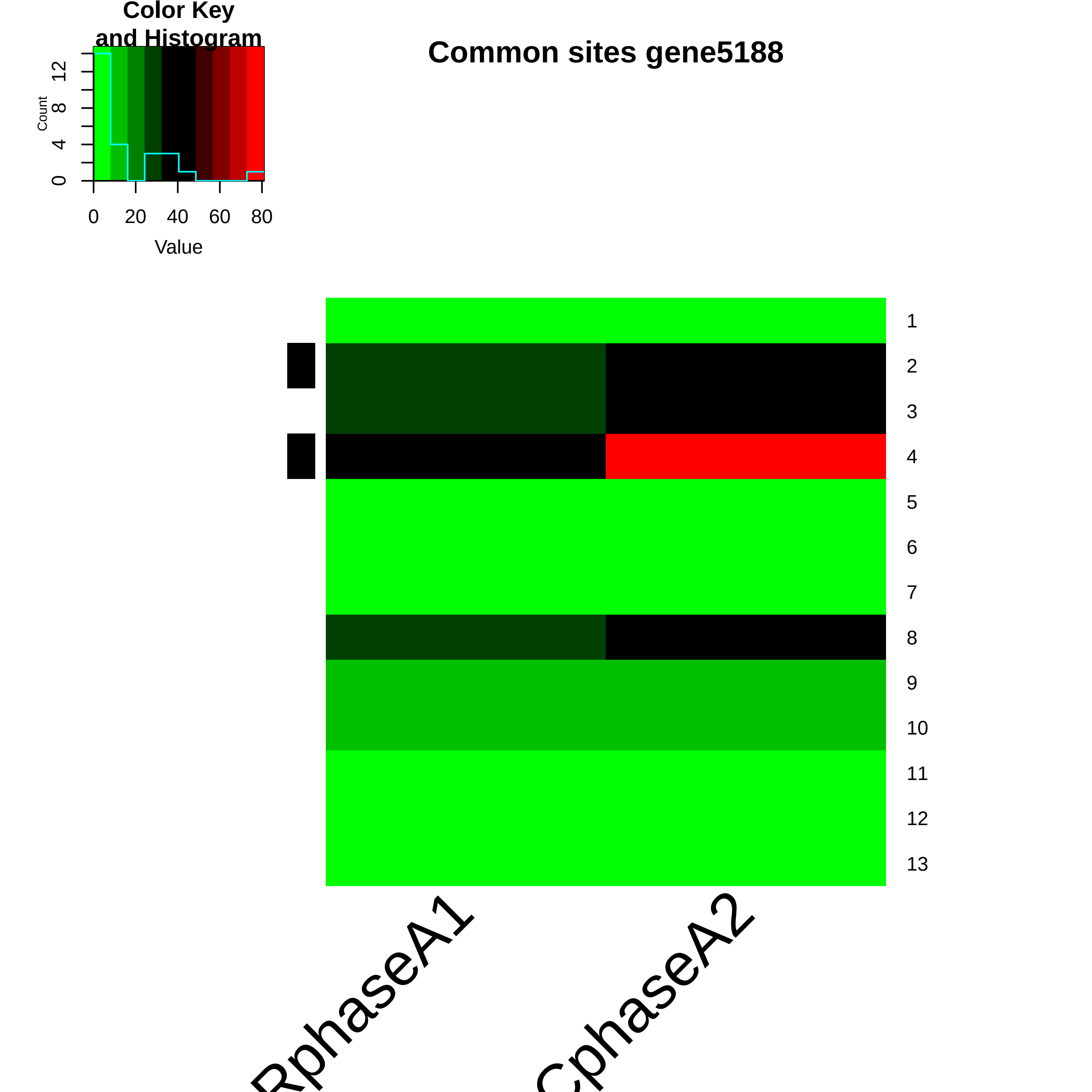
Color Key
and Histogram
Count
0
4
8
12
0
20
40
60
80
Value
Common sites gene5188
1
2
3
4
5
6
7
8
9
10
11
12
13
RphaseA1
CphaseA2

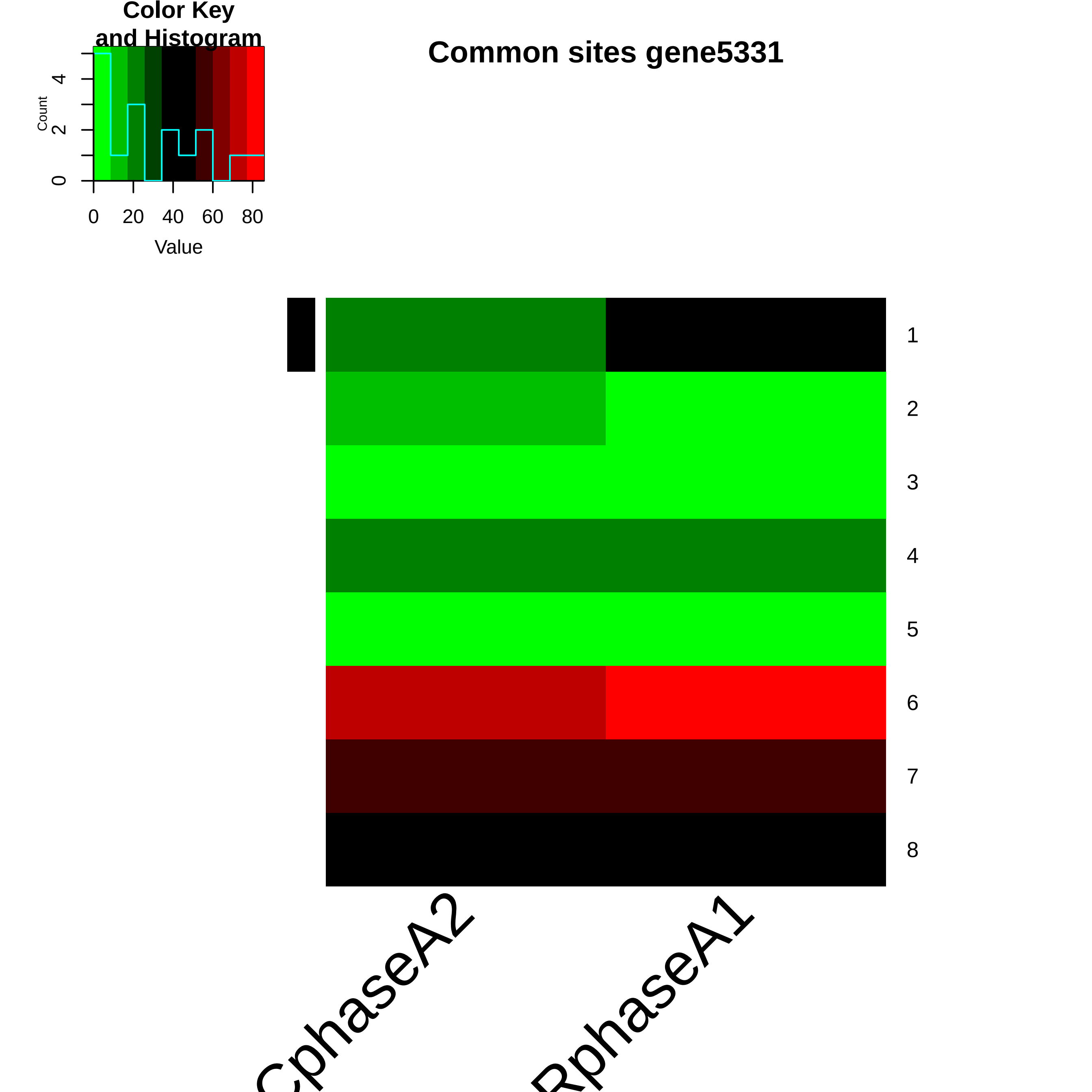

Color Key
and Histogram
Count
0
2
4
0
20
40
60
80
Value
Common sites gene5331
1
2
3
4
5
6
7
8
CphaseA2
RphaseA1

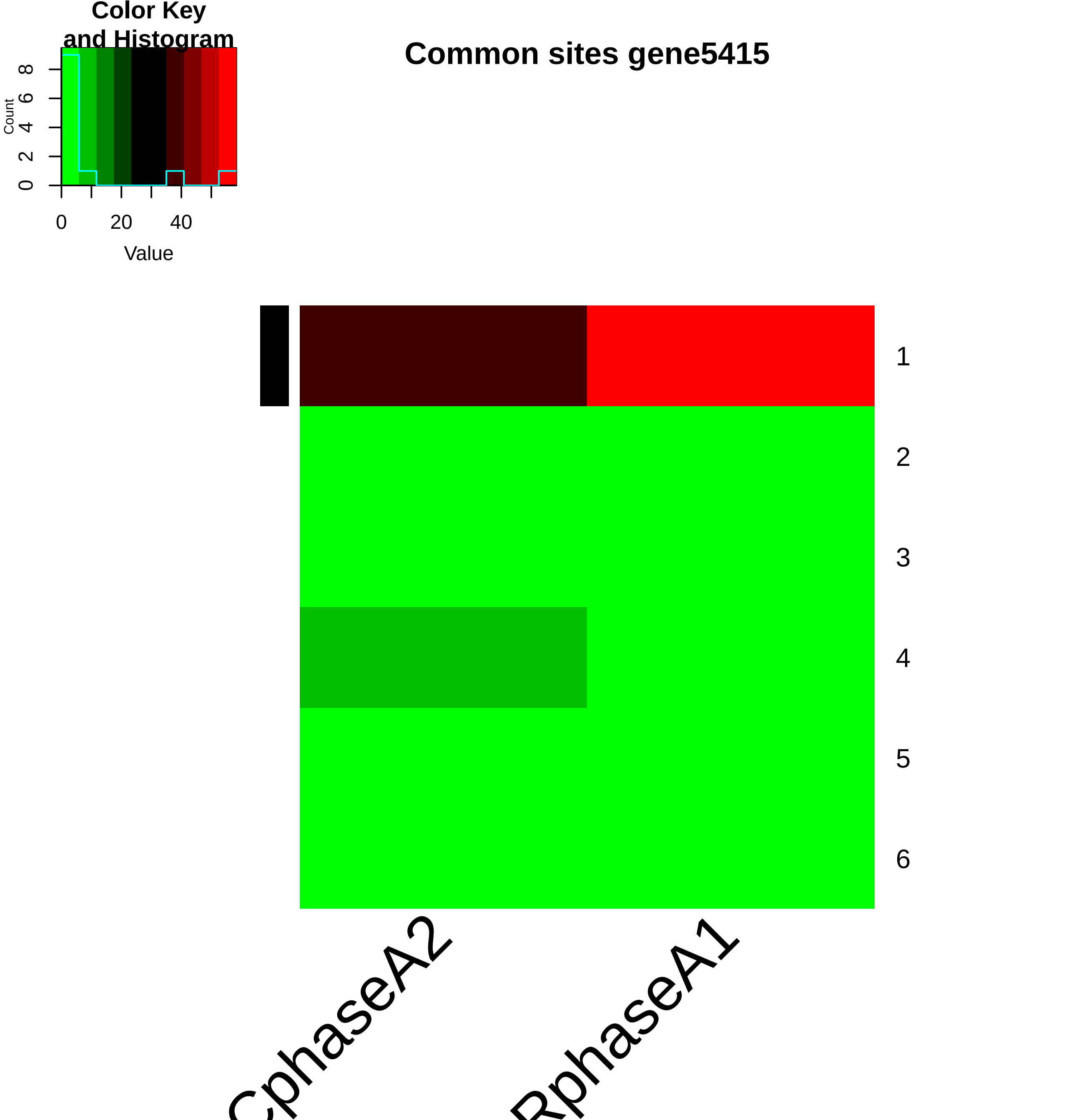
Color Key
and Histogram
Count
0
2
4
6
8
0
20
40
Value
Common sites gene5415
1
2
3
4
5
6
CphaseA2
RphaseA1

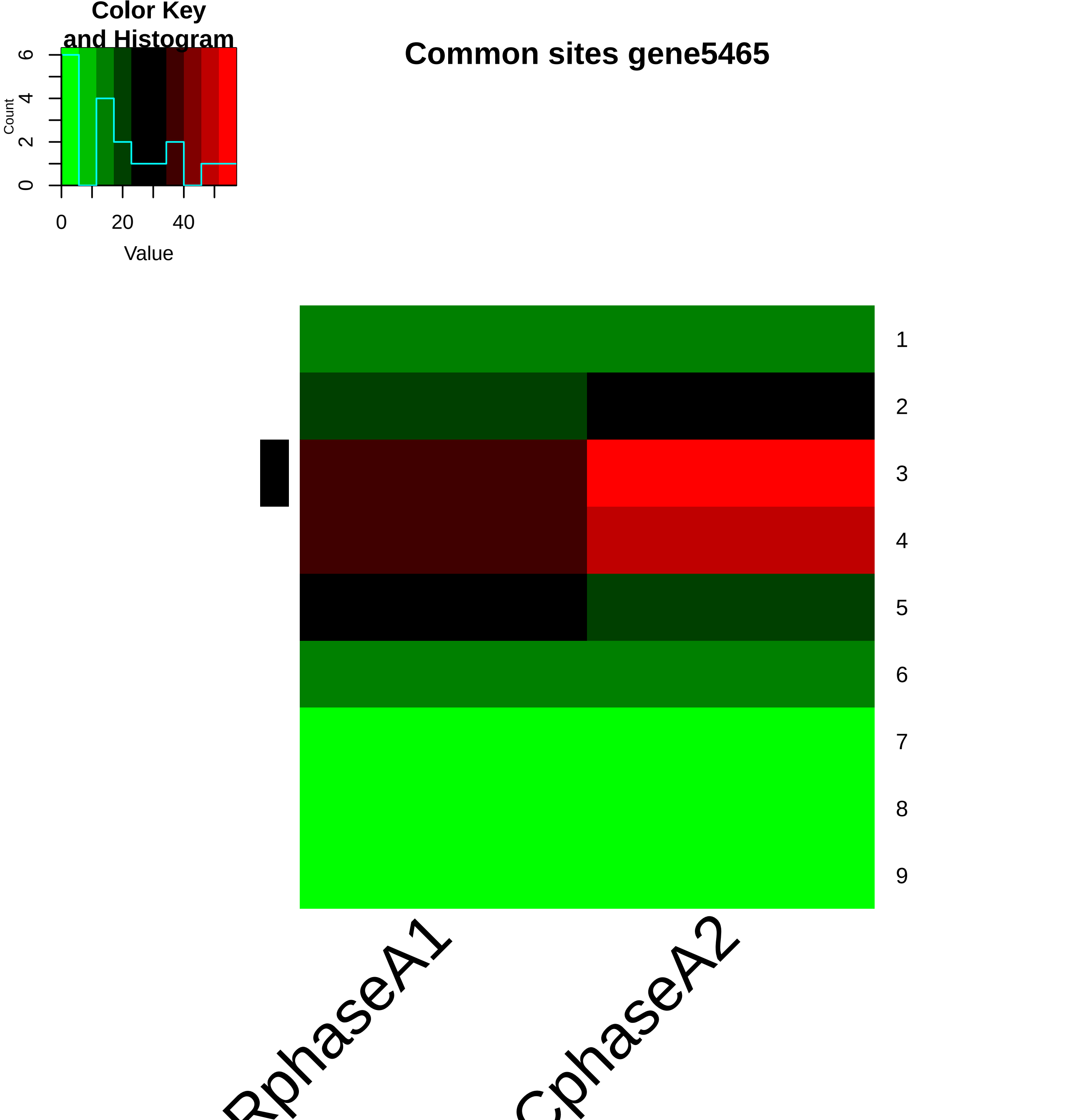
Color Key
and Histogram
Count
0
2
4
6
0
20
40
Value
Common sites gene5465
1
2
3
4
5
6
7
8
9
RphaseA1
CphaseA2

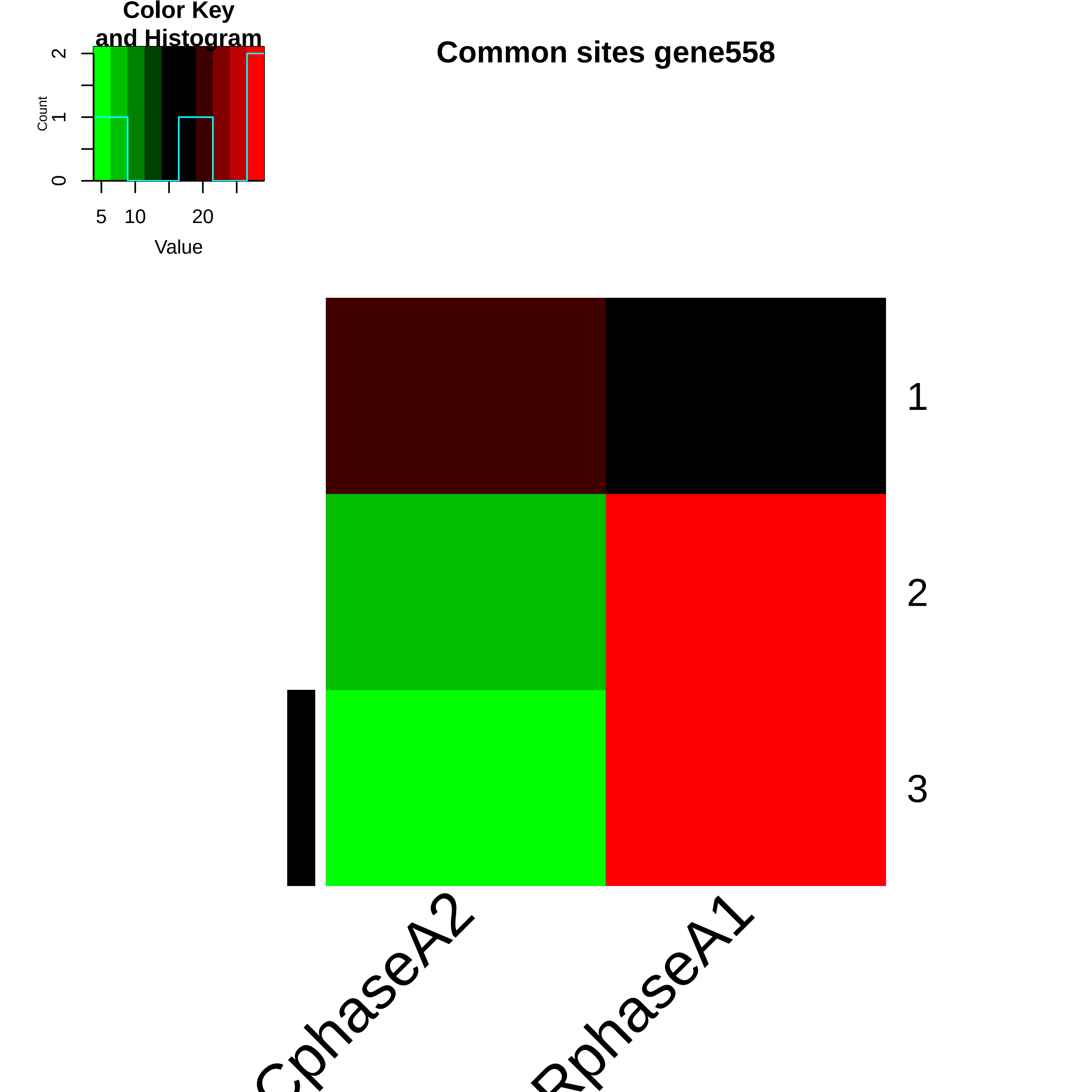
Color Key
and Histogram
Count
0
1
2
5
10
20
Value
Common sites gene558
1
2
3
CphaseA2
RphaseA1

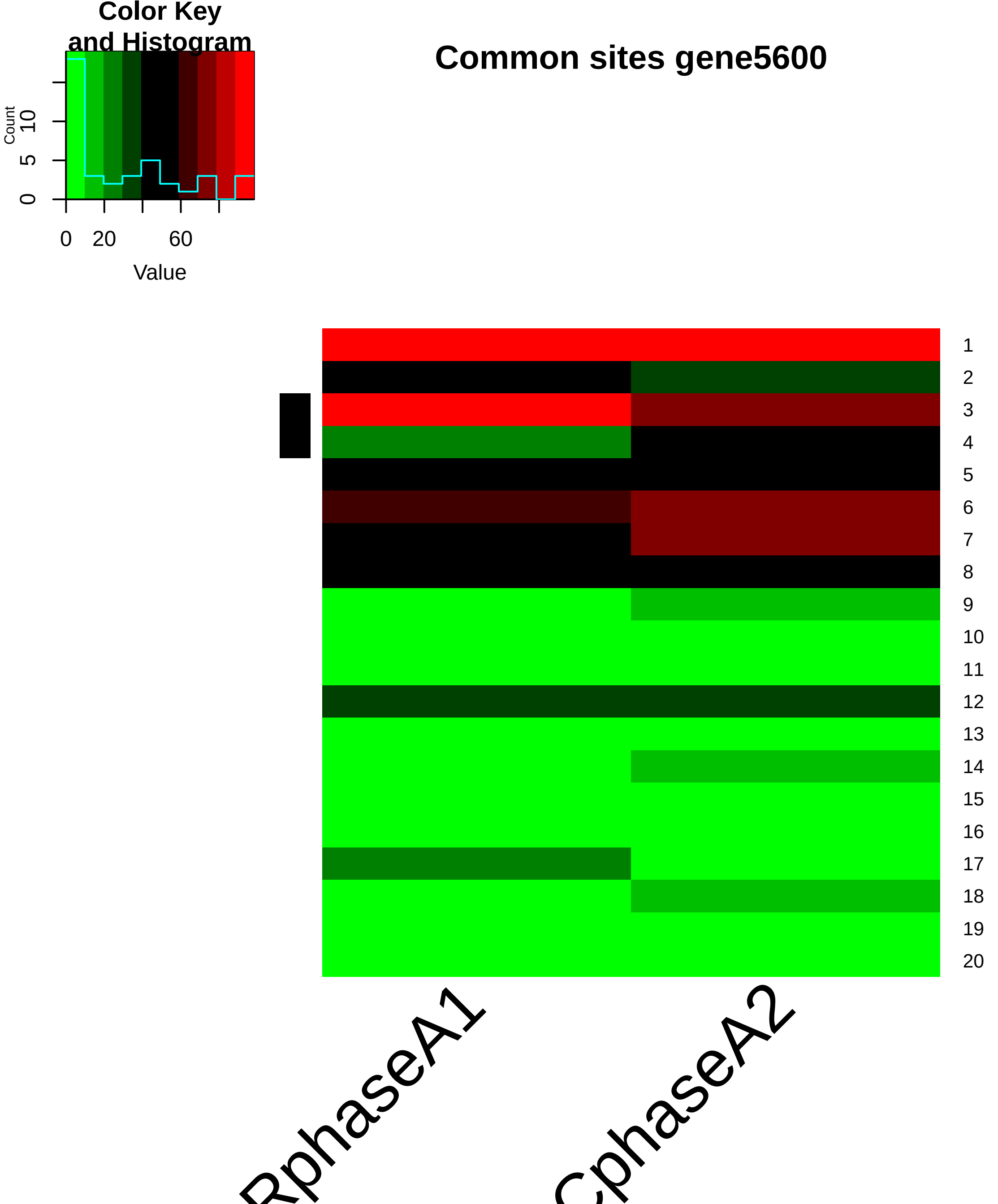
Color Key
and Histogram
Count
0
5
10
0
20
60
Value
Common sites gene5600
RphaseA1
CphaseA2

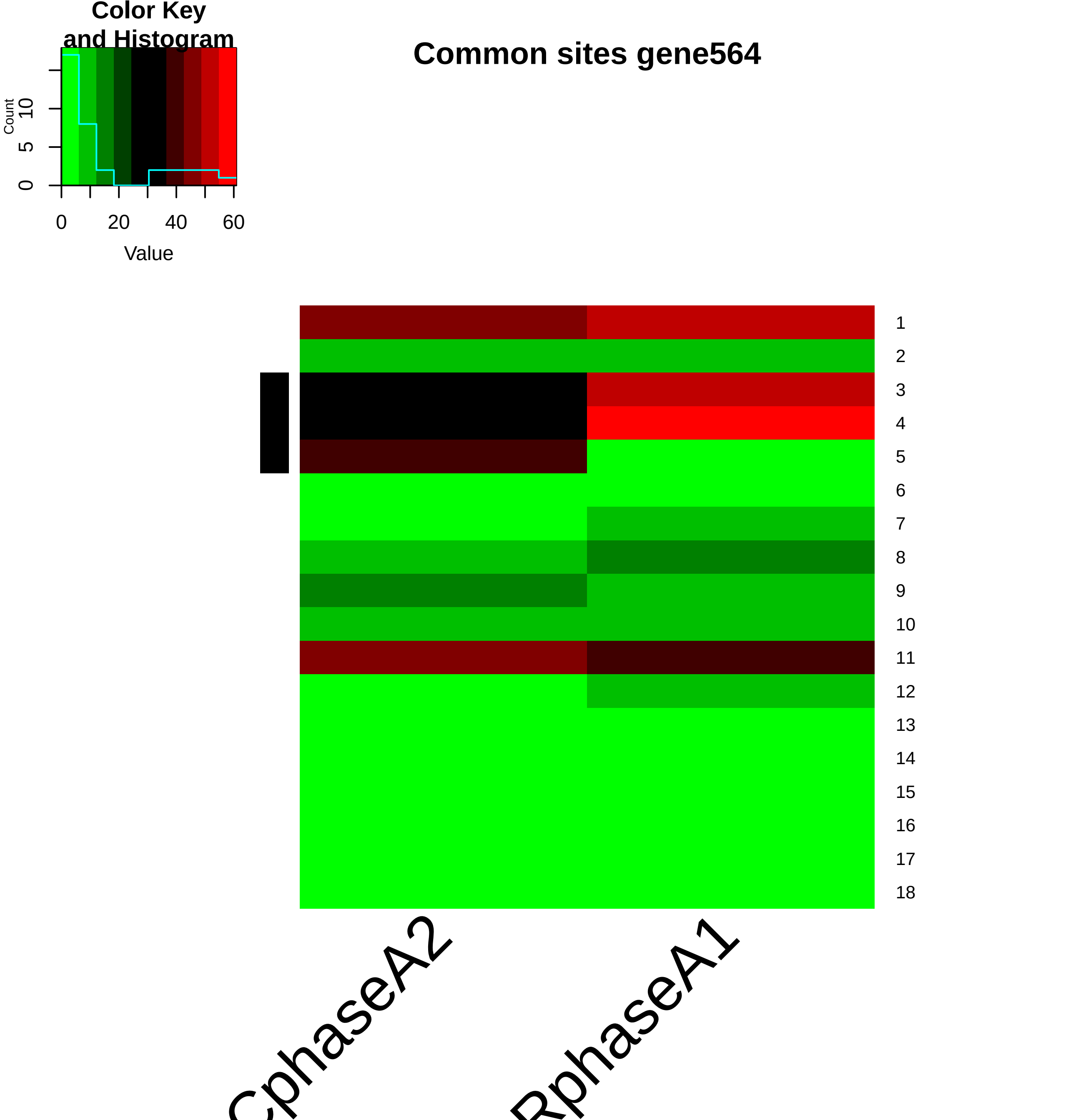
Color Key
and Histogram
Count
0
5
10
0
20
40
60
Value
Common sites gene564
CphaseA2
RphaseA1

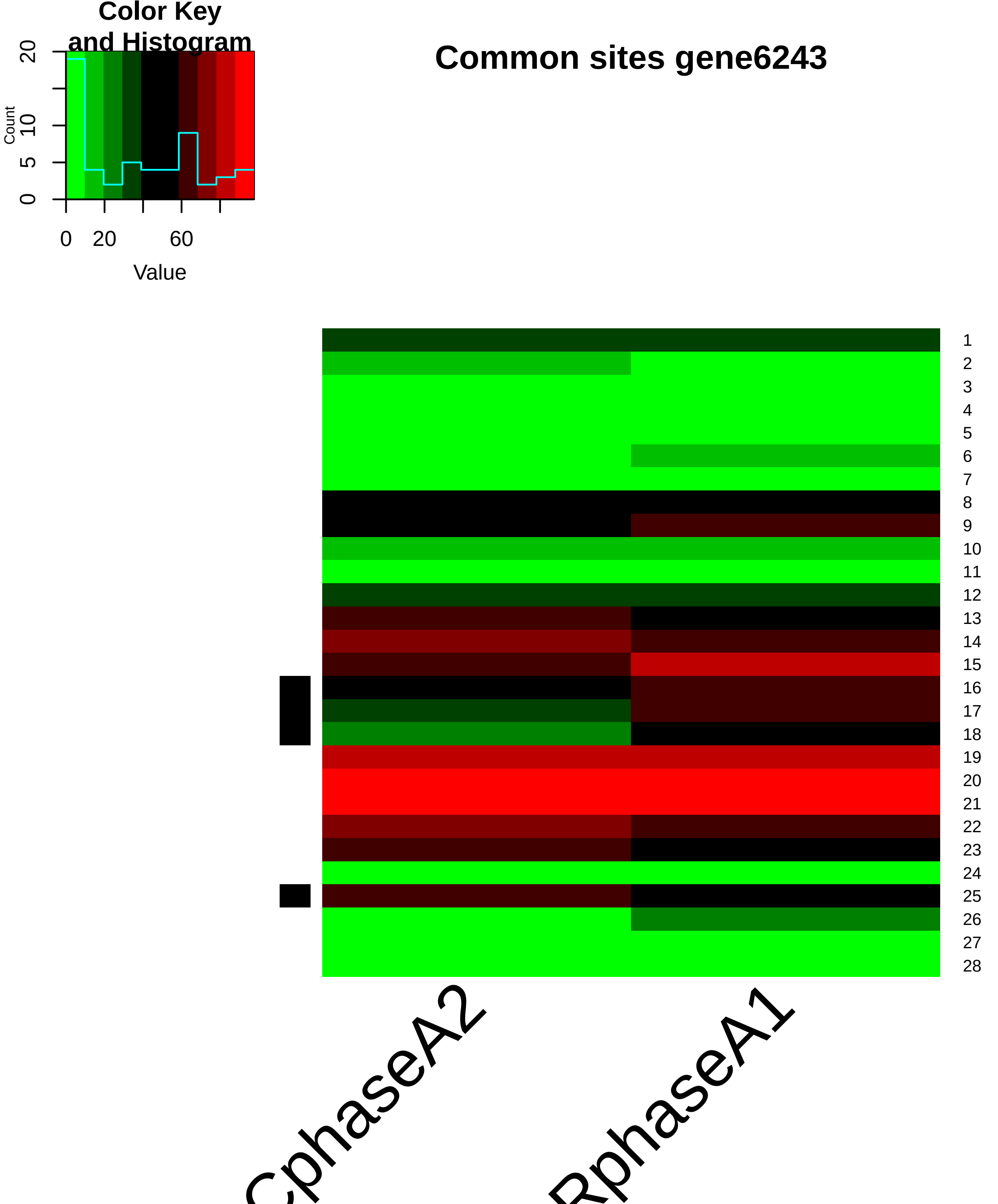

Color Key
and Histogram
Count
0
5
10
20
0
20
60
Value
Common sites gene6243
CphaseA2
RphaseA1

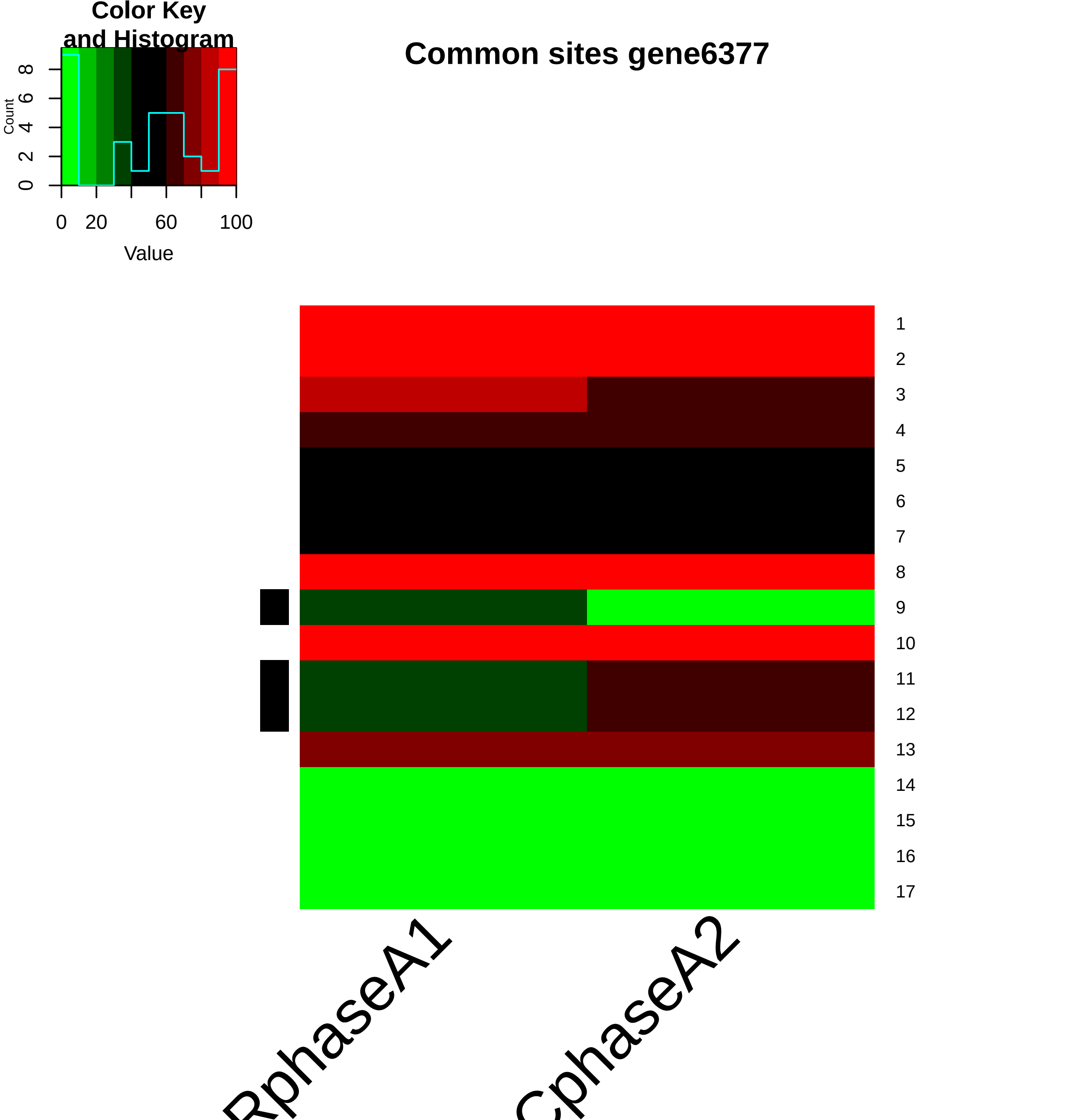
Color Key
and Histogram
Count
0
2
4
6
8
0
20
60
100
Value
Common sites gene6377
RphaseA1
CphaseA2

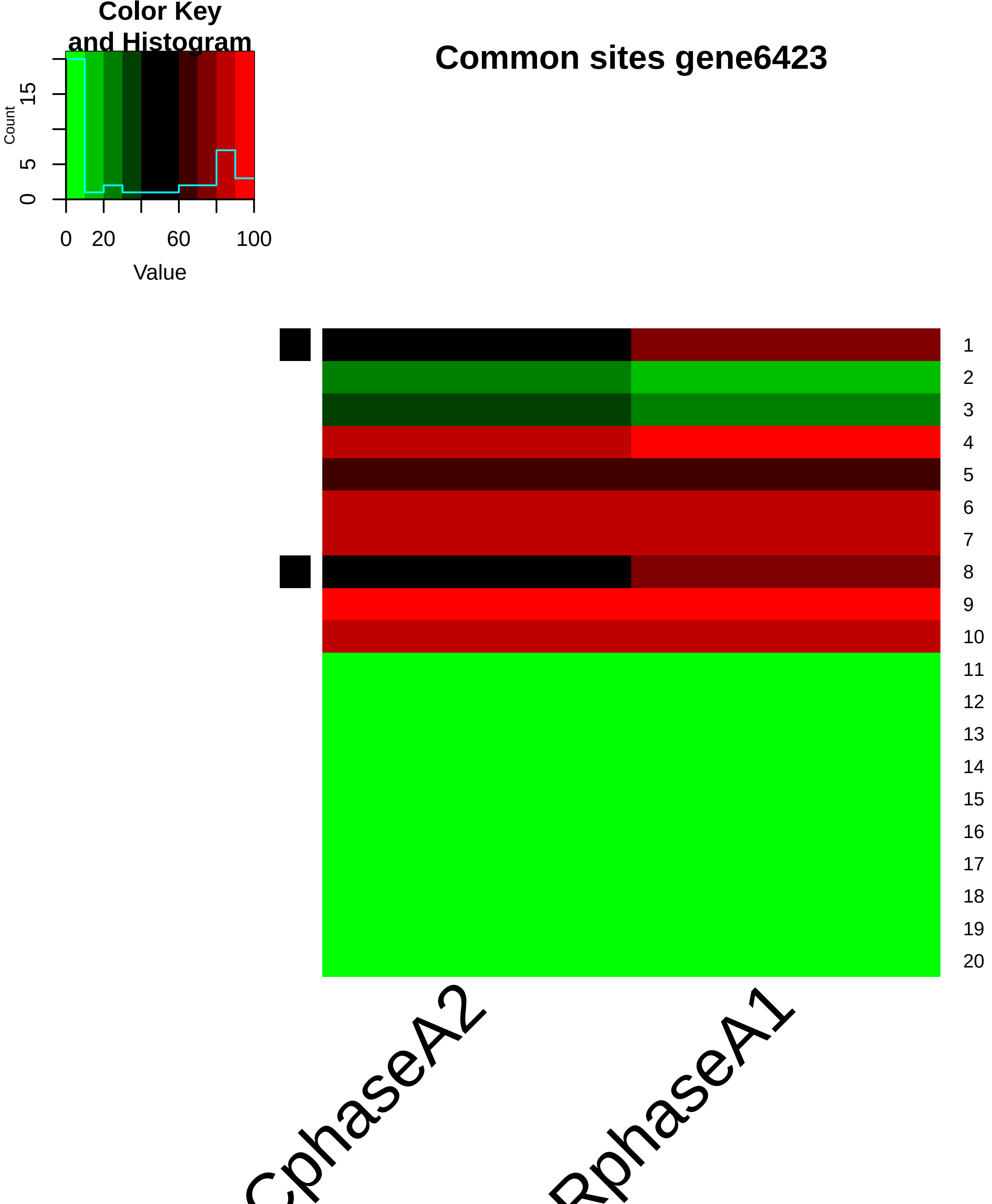
Color Key
and Histogram
Count
0
5
15
0
20
60
100
Value
Common sites gene6423
CphaseA2
RphaseA1

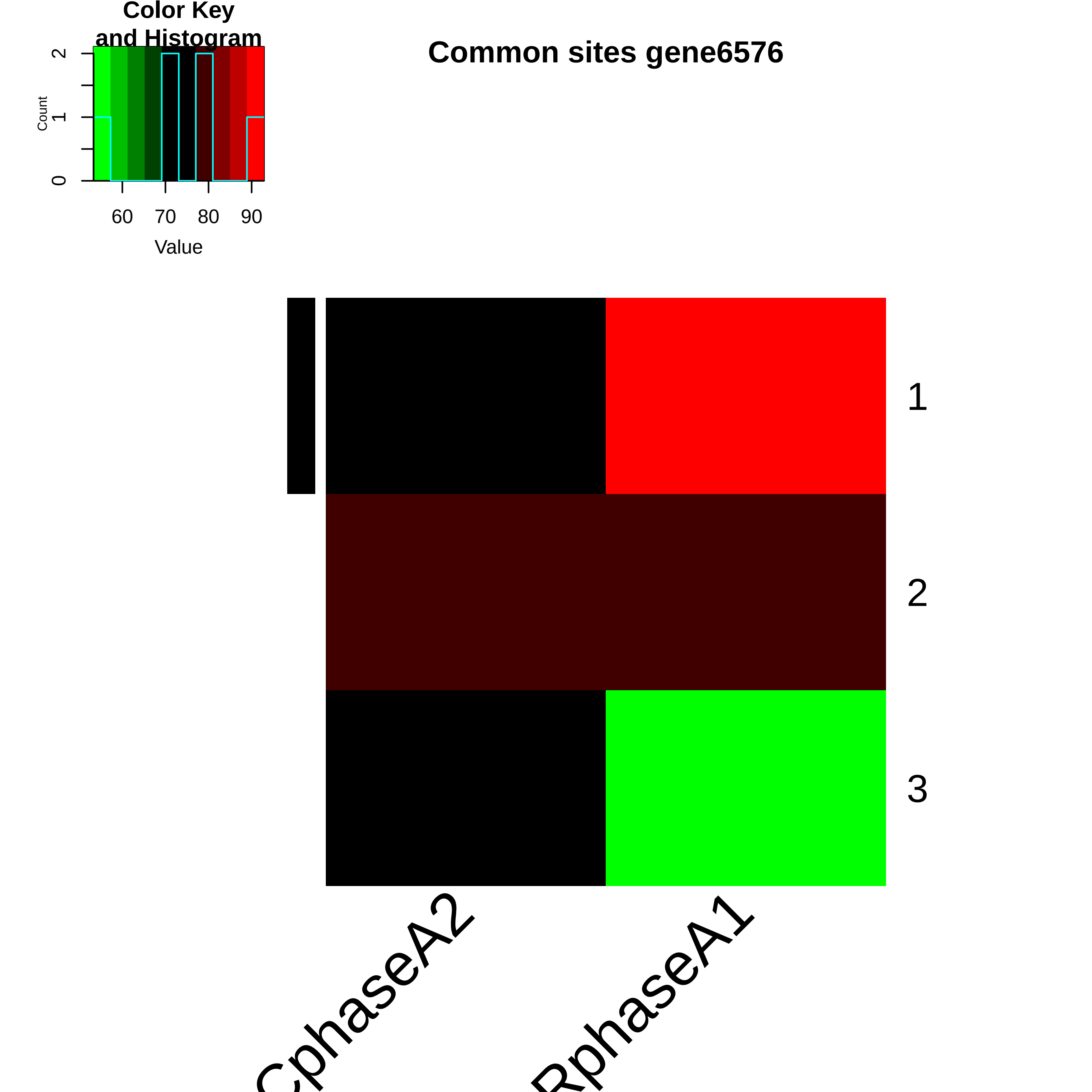

Color Key
and Histogram
Count
0
1
2
60
70
80
90
Value
Common sites gene6576
1
2
3
CphaseA2
RphaseA1

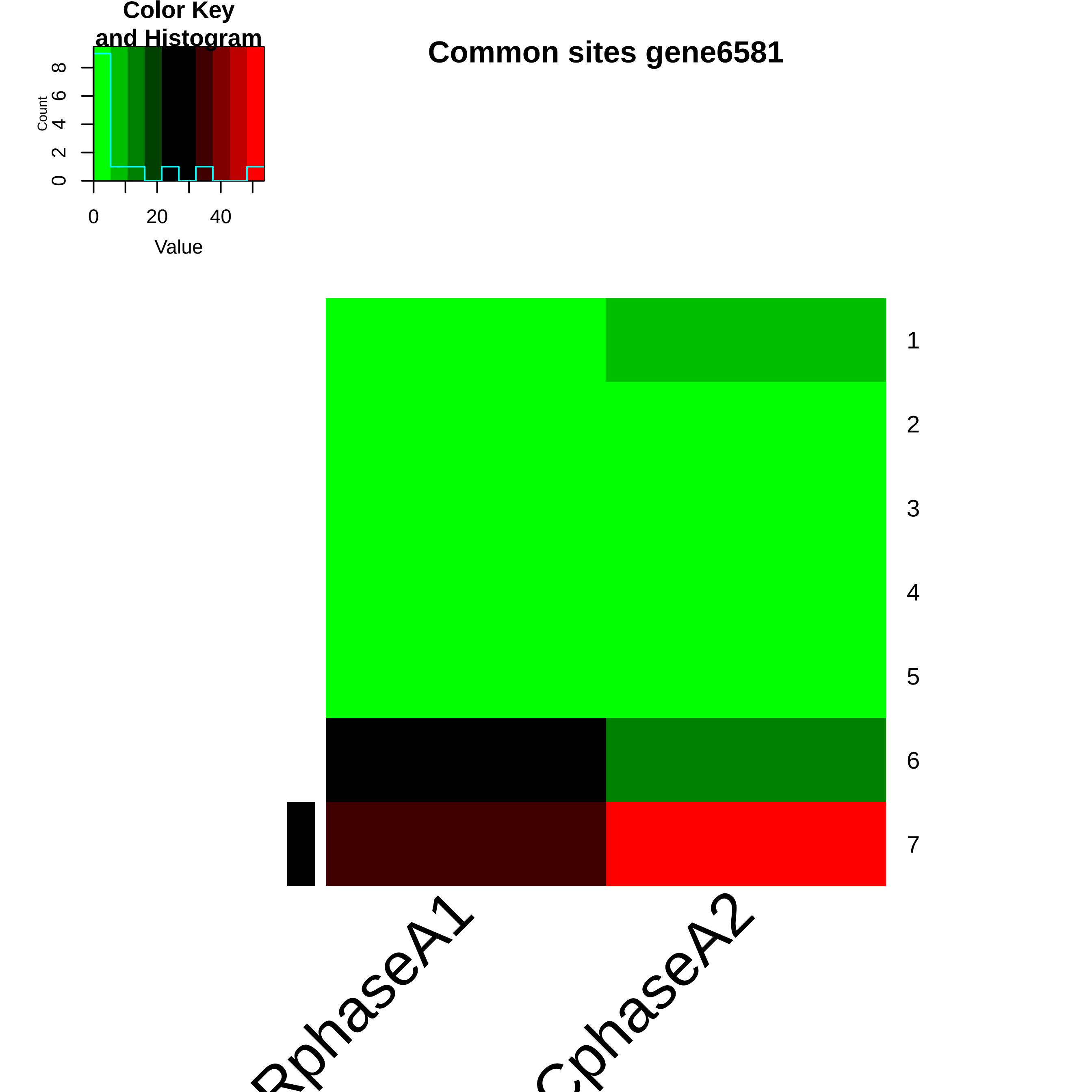

Color Key
and Histogram
Count
0
2
4
6
8
0
20
40
Value
Common sites gene6581
1
2
3
4
5
6
7
RphaseA1
CphaseA2

### Common sites gene6755

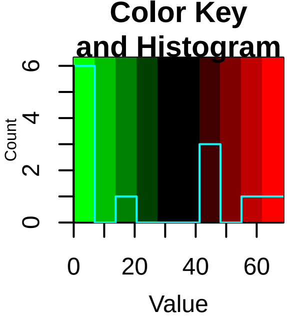

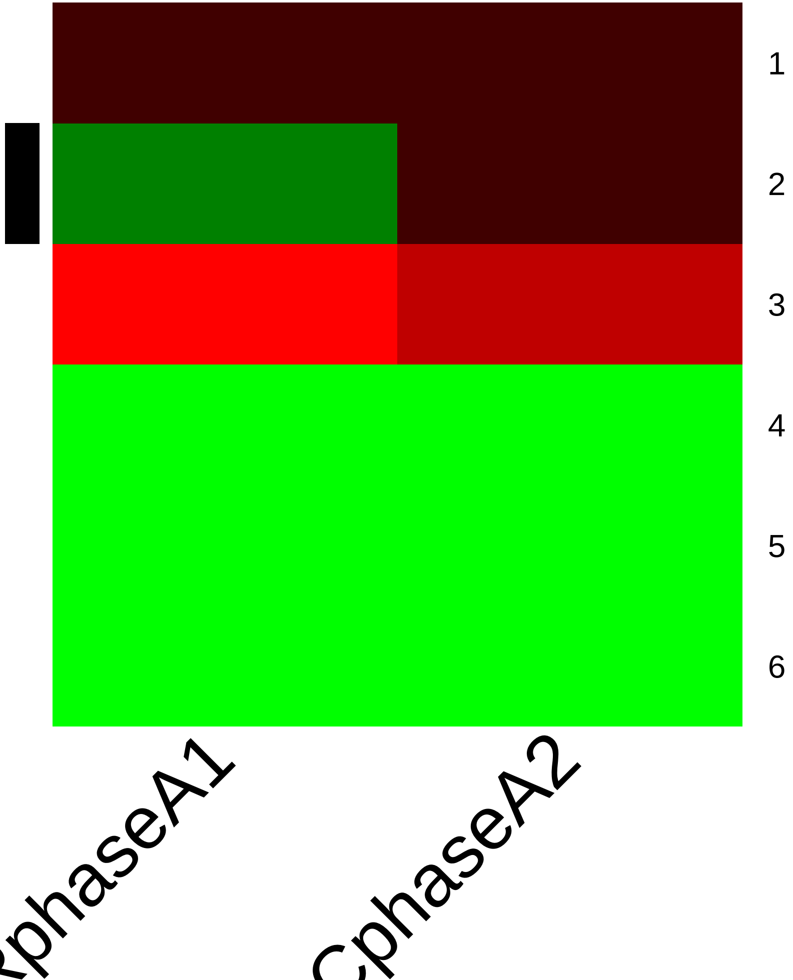

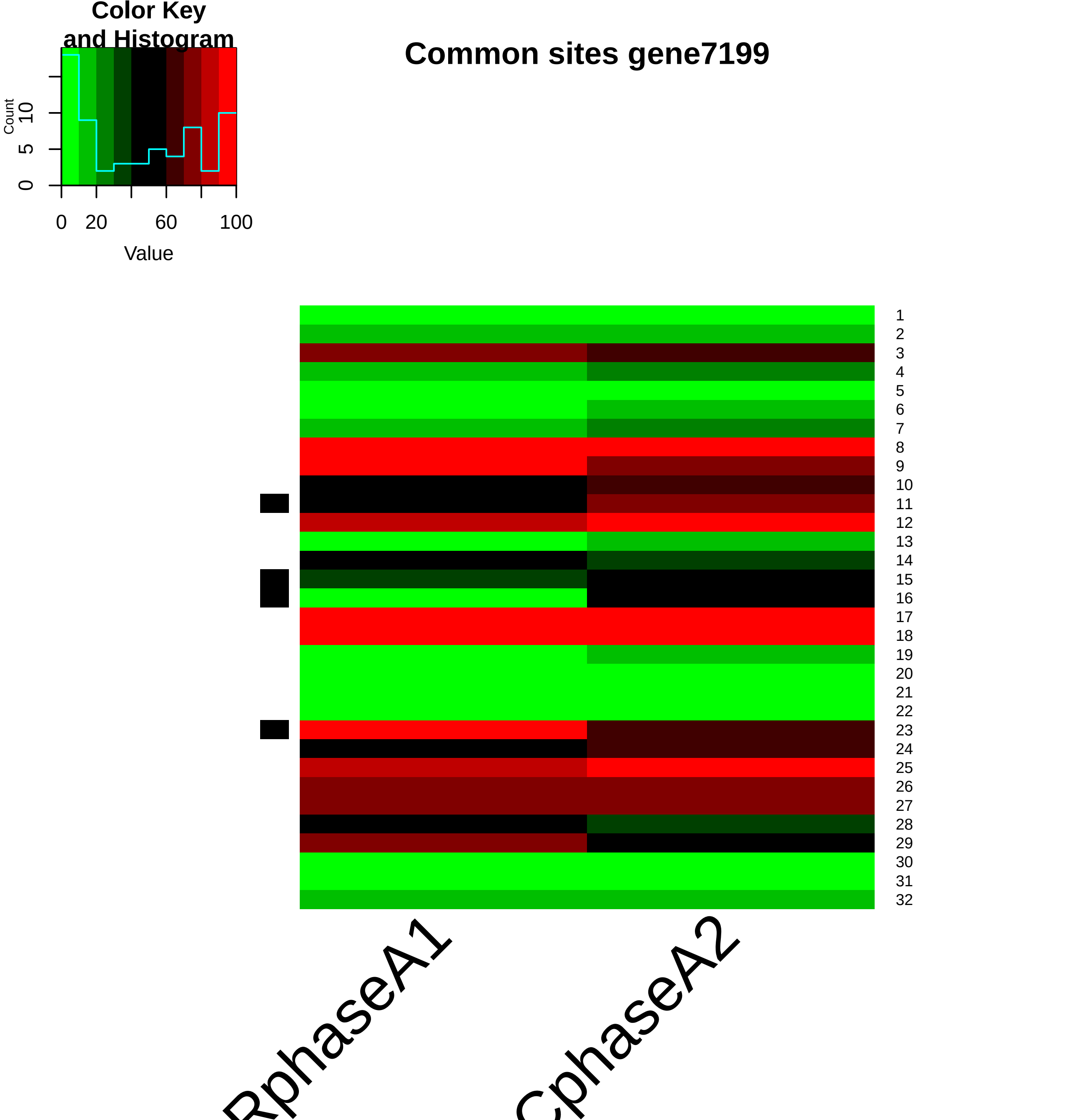

Color Key
and Histogram
Common sites gene7199
Count
0
5
10
0
20
60
100
Value
RphaseA1
CphaseA2

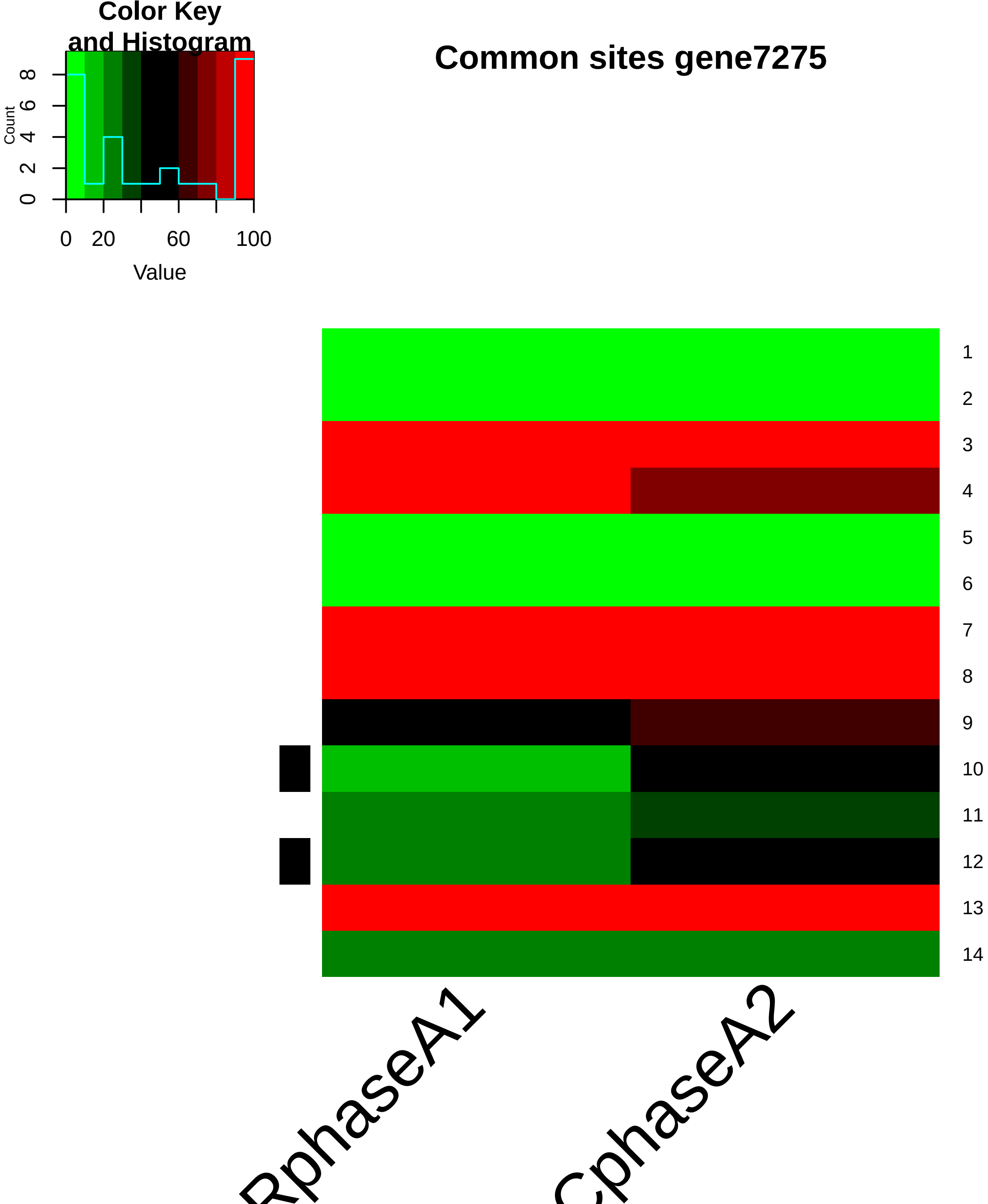
Color Key
and Histogram
Count
0
2
4
6
8
0
20
60
100
Value
Common sites gene7275
1
2
3
4
5
6
7
8
9
10
11
12
13
14
RphaseA1
CphaseA2

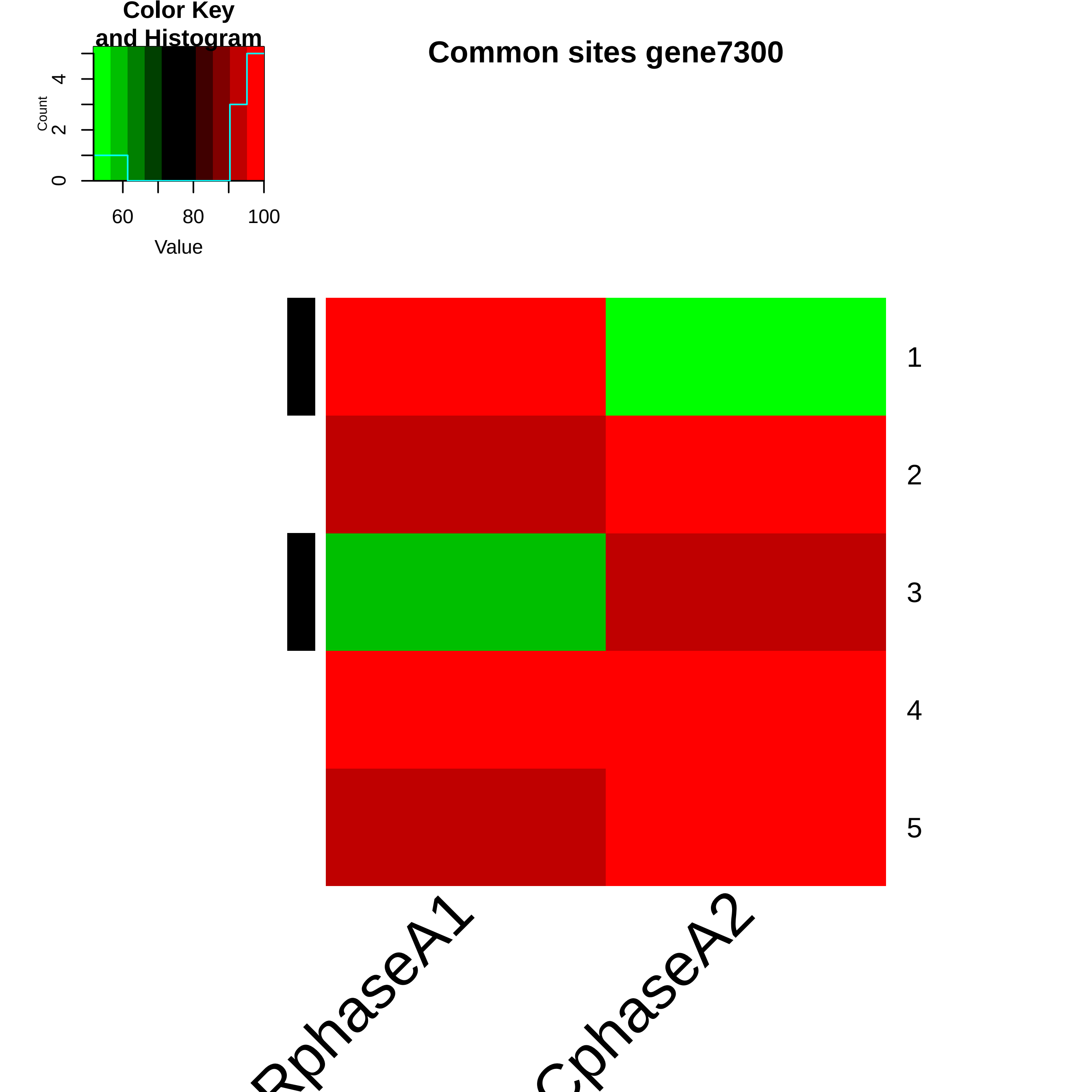
Color Key
and Histogram
Count
0
2
4
60
80
100
Value
Common sites gene7300
1
2
3
4
5
RphaseA1
CphaseA2

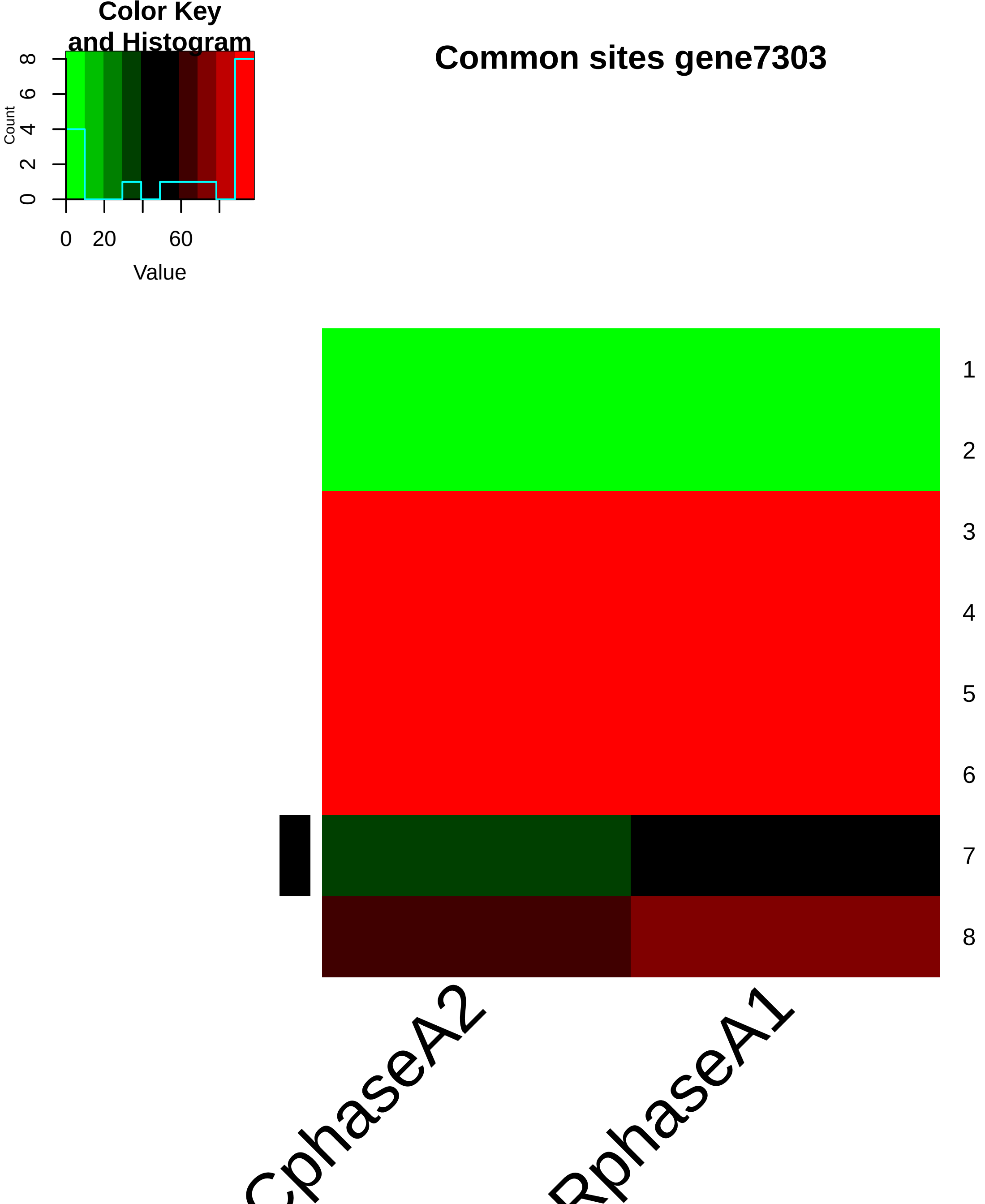

Color Key
and Histogram
Count
0
2
4
6
8
0
20
60
Value
Common sites gene7303
1
2
3
4
5
6
7
8
CphaseA2
RphaseA1

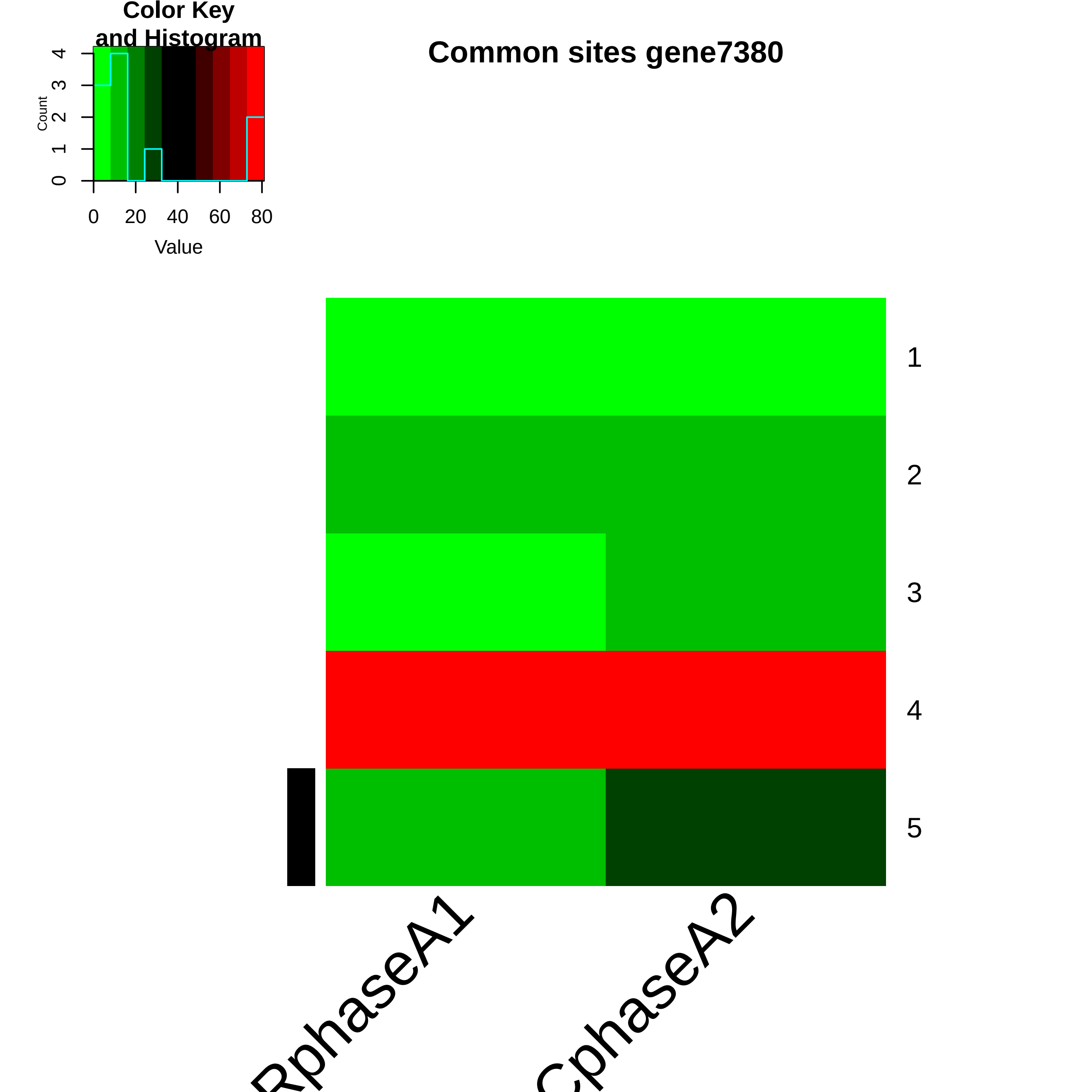
Color Key
and Histogram
Count
0
1
2
3
4
0
20
40
60
80
Value
Common sites gene7380
1
2
3
4
5
RphaseA1
CphaseA2

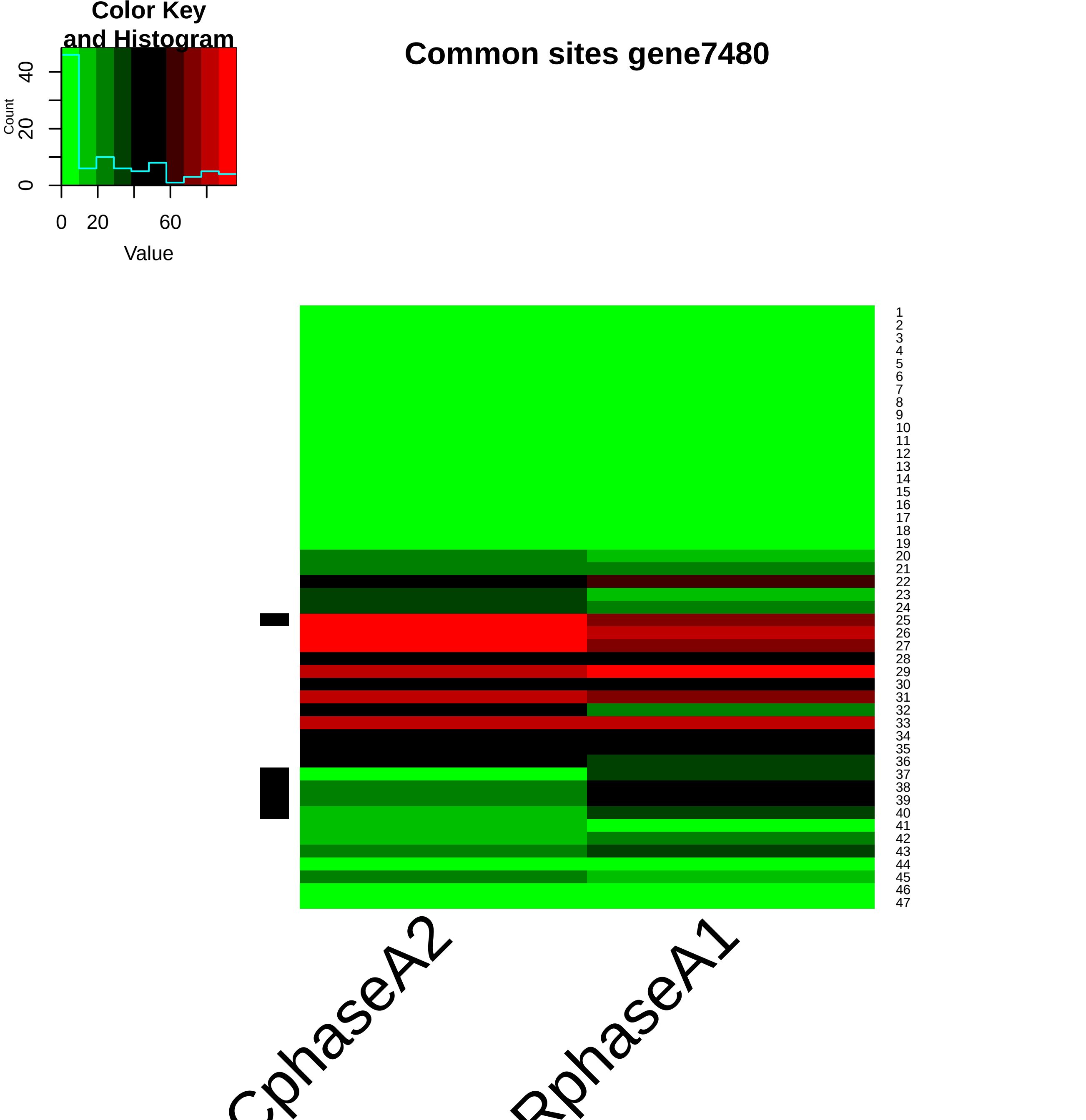
Color Key
and Histogram
Count
0
20
40
0
20
60
Value
Common sites gene7480
CphaseA2
RphaseA1

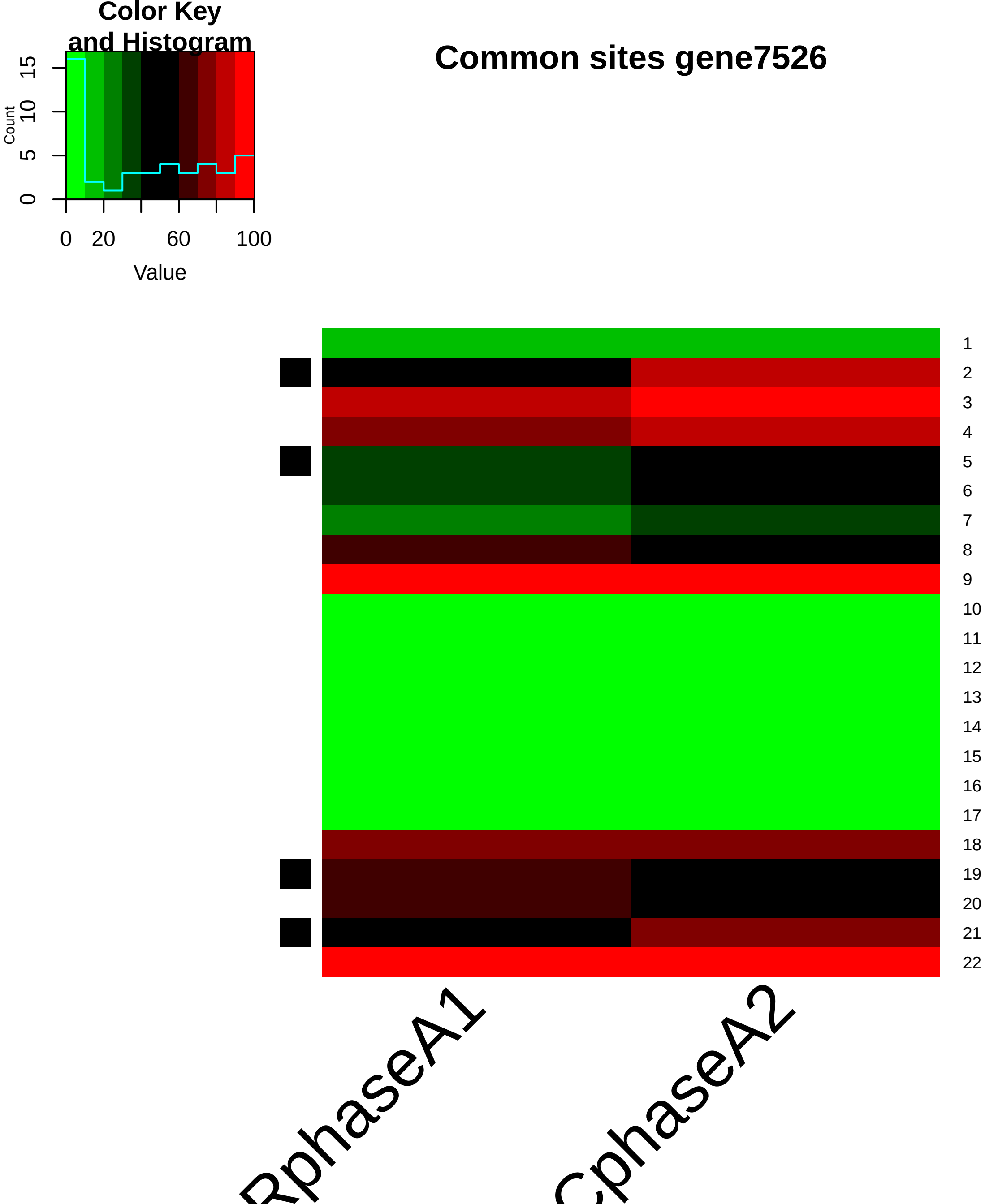

Color Key
and Histogram
Count
0
5
10
15
0
20
60
100
Value
Common sites gene7526
RphaseA1
CphaseA2

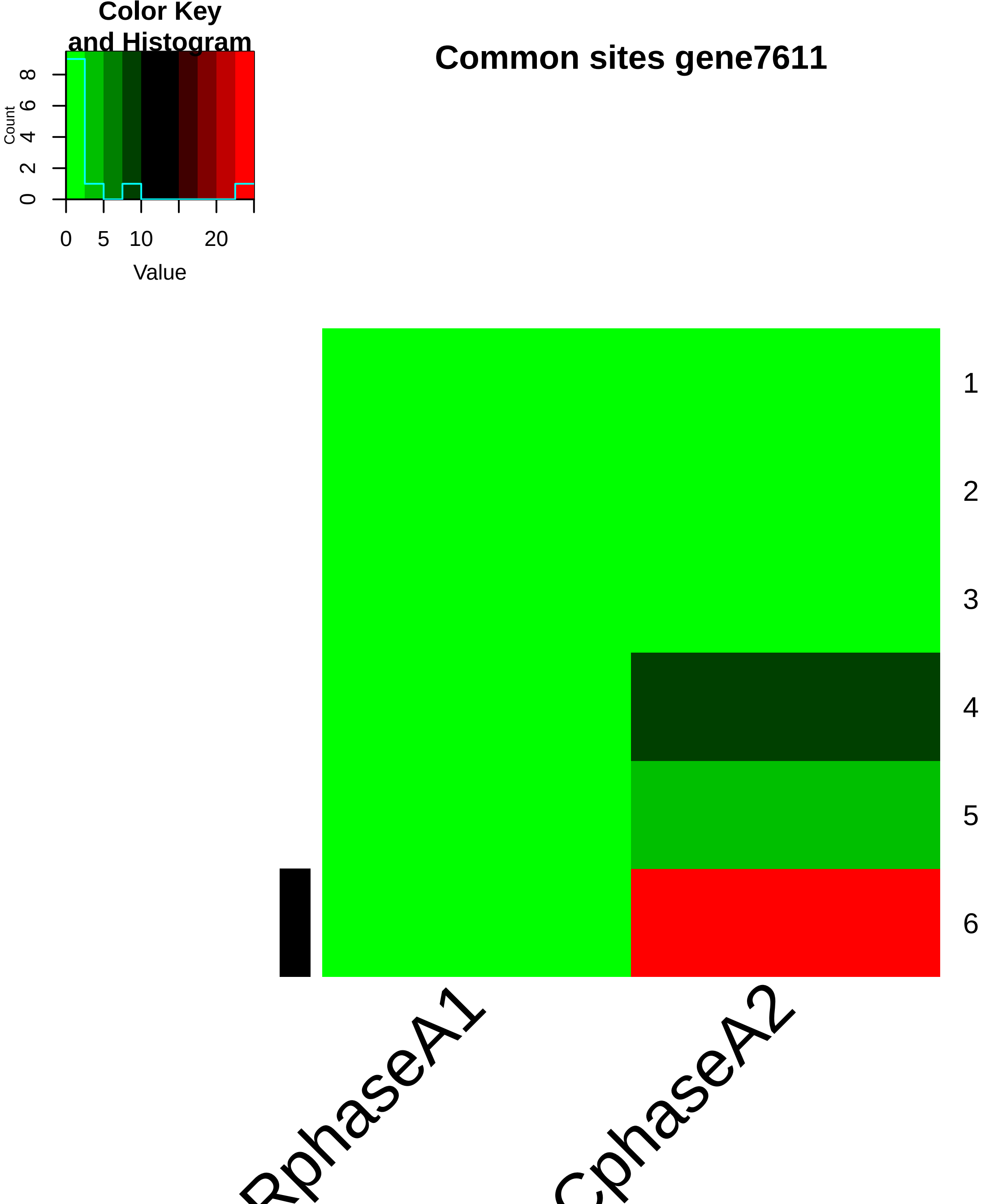
Color Key
and Histogram
Count
0 2 4 6 8
0 5 10 20
Value
Common sites gene7611
1
2
3
4
5
6
RphaseA1
CphaseA2

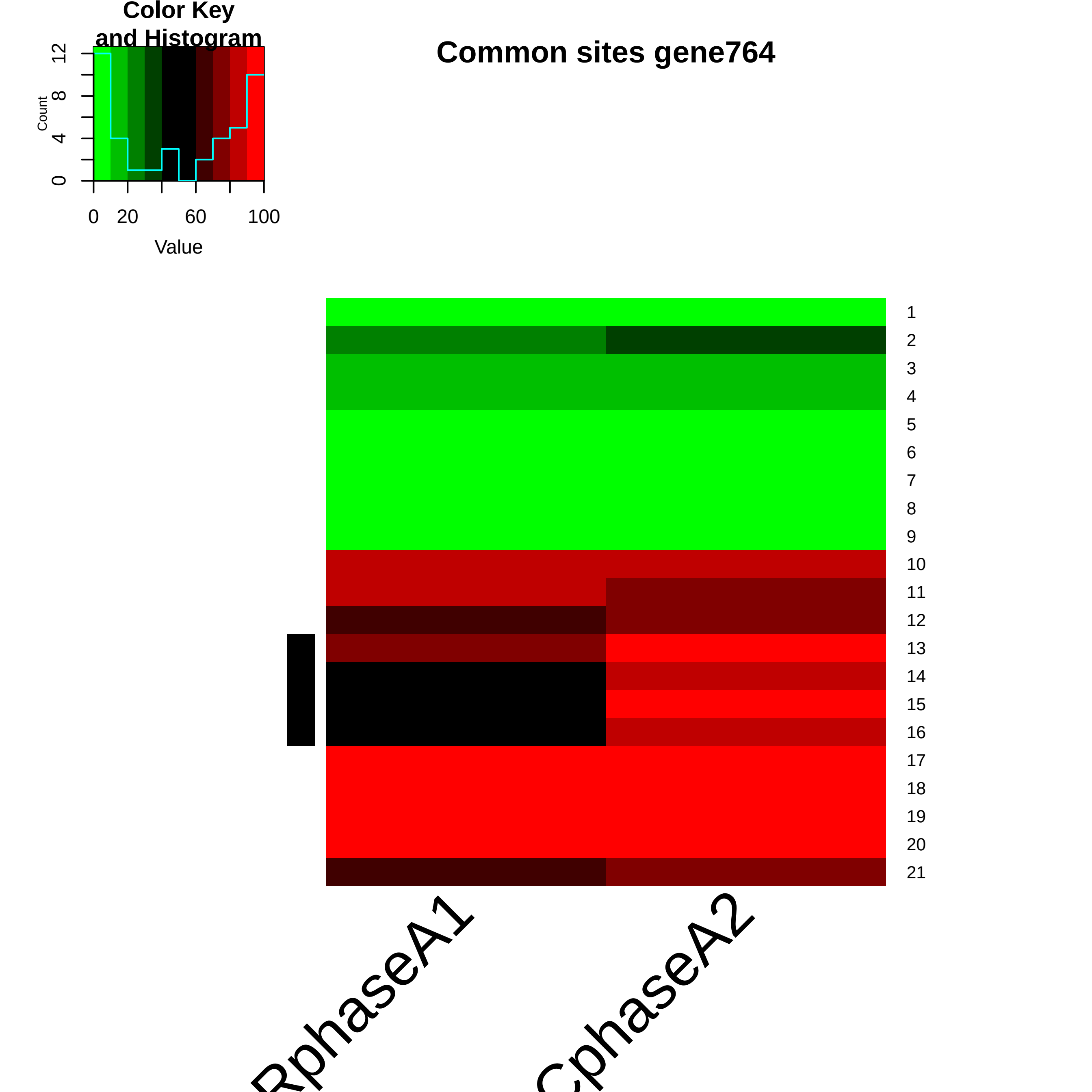

Color Key
and Histogram
Count
0
4
8
12
0
20
60
100
Value
Common sites gene764
RphaseA1
CphaseA2

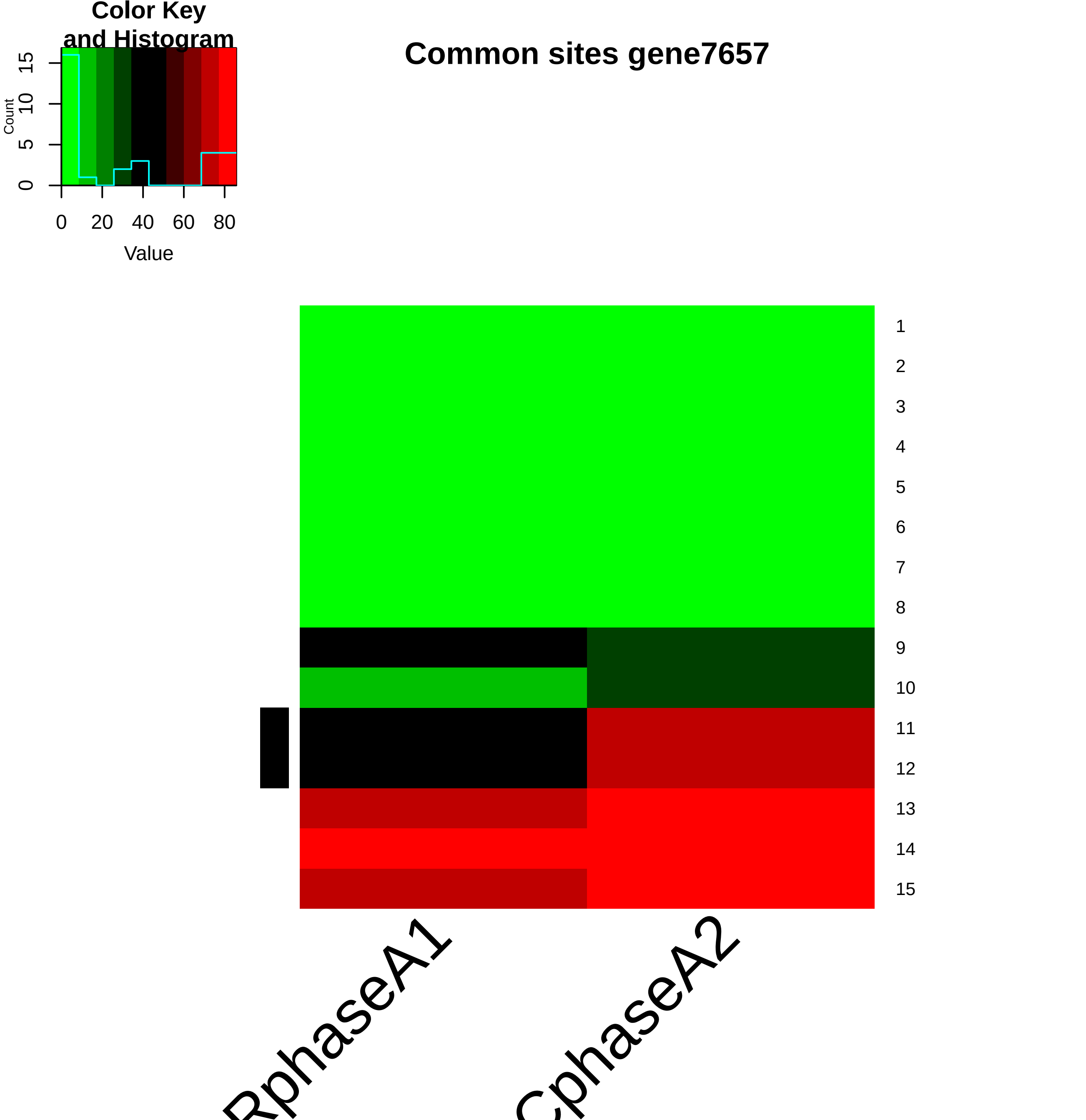
Color Key
and Histogram
Count
0
5
10
15
0
20
40
60
80
Value
Common sites gene7657
RphaseA1
CphaseA2

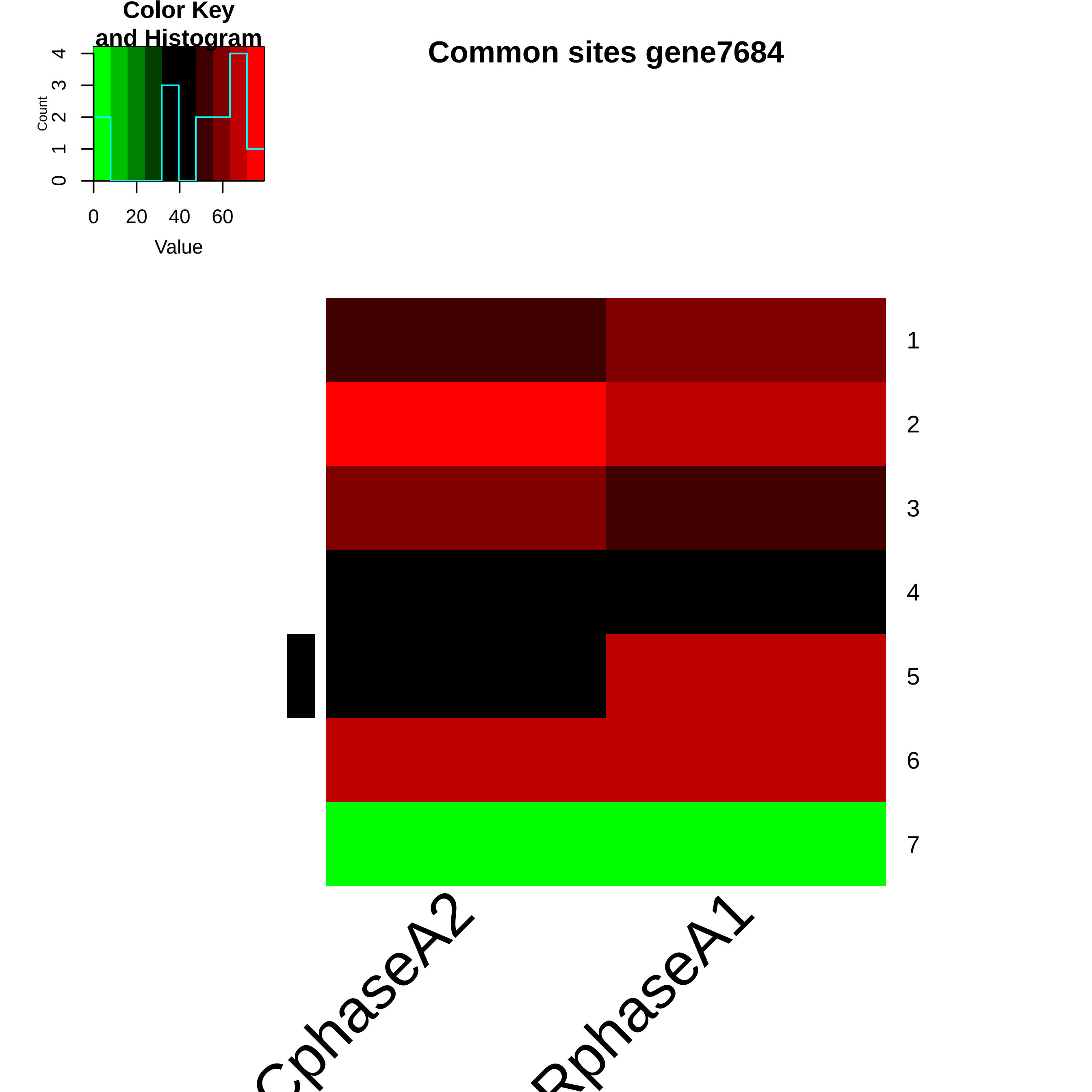
Color Key
and Histogram
Count
0
1
2
3
4
0
20
40
60
Value
Common sites gene7684
1
2
3
4
5
6
7
CphaseA2
RphaseA1

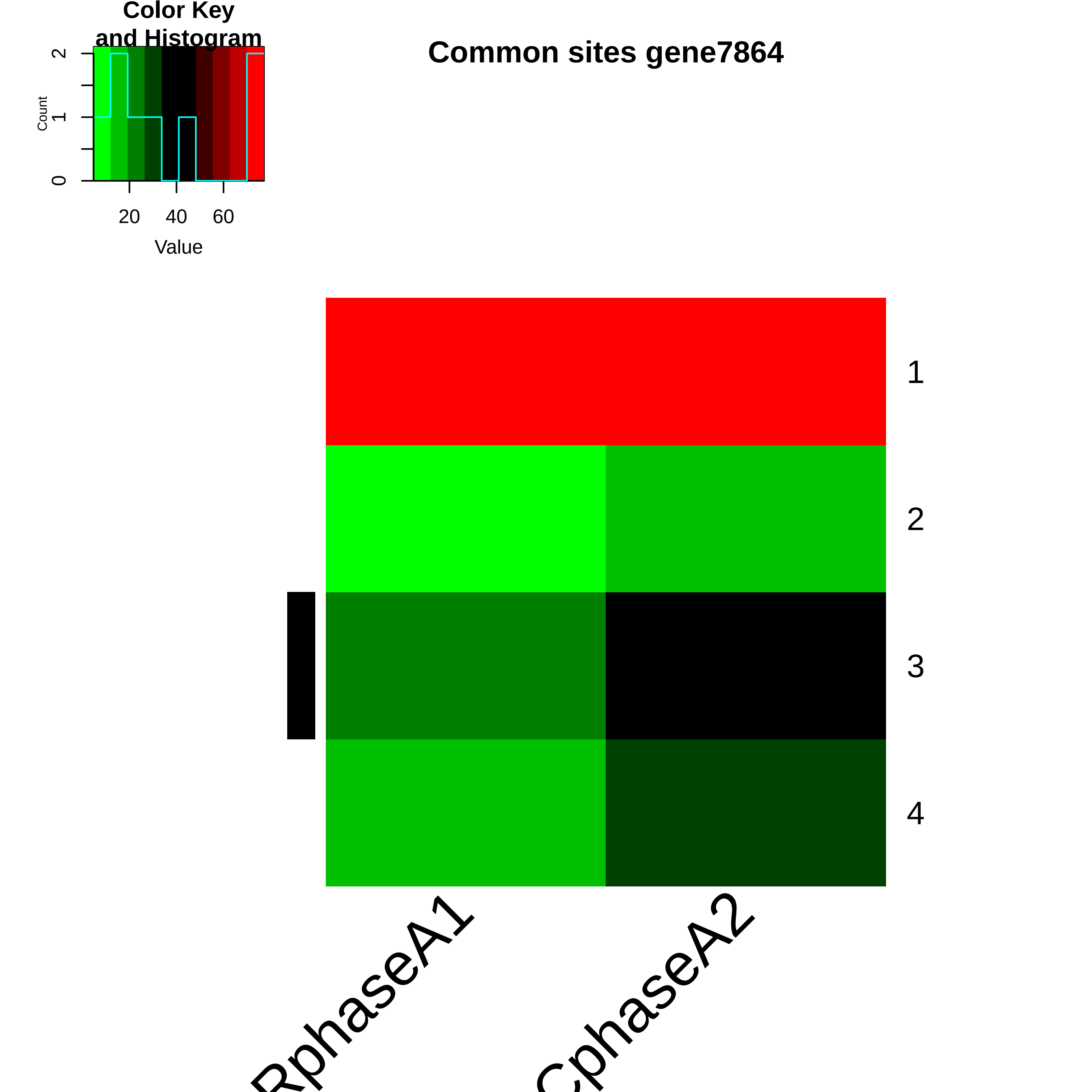
Color Key
and Histogram
Count
0
1
2
20
40
60
Value
Common sites gene7864
1
2
3
4
RphaseA1
CphaseA2

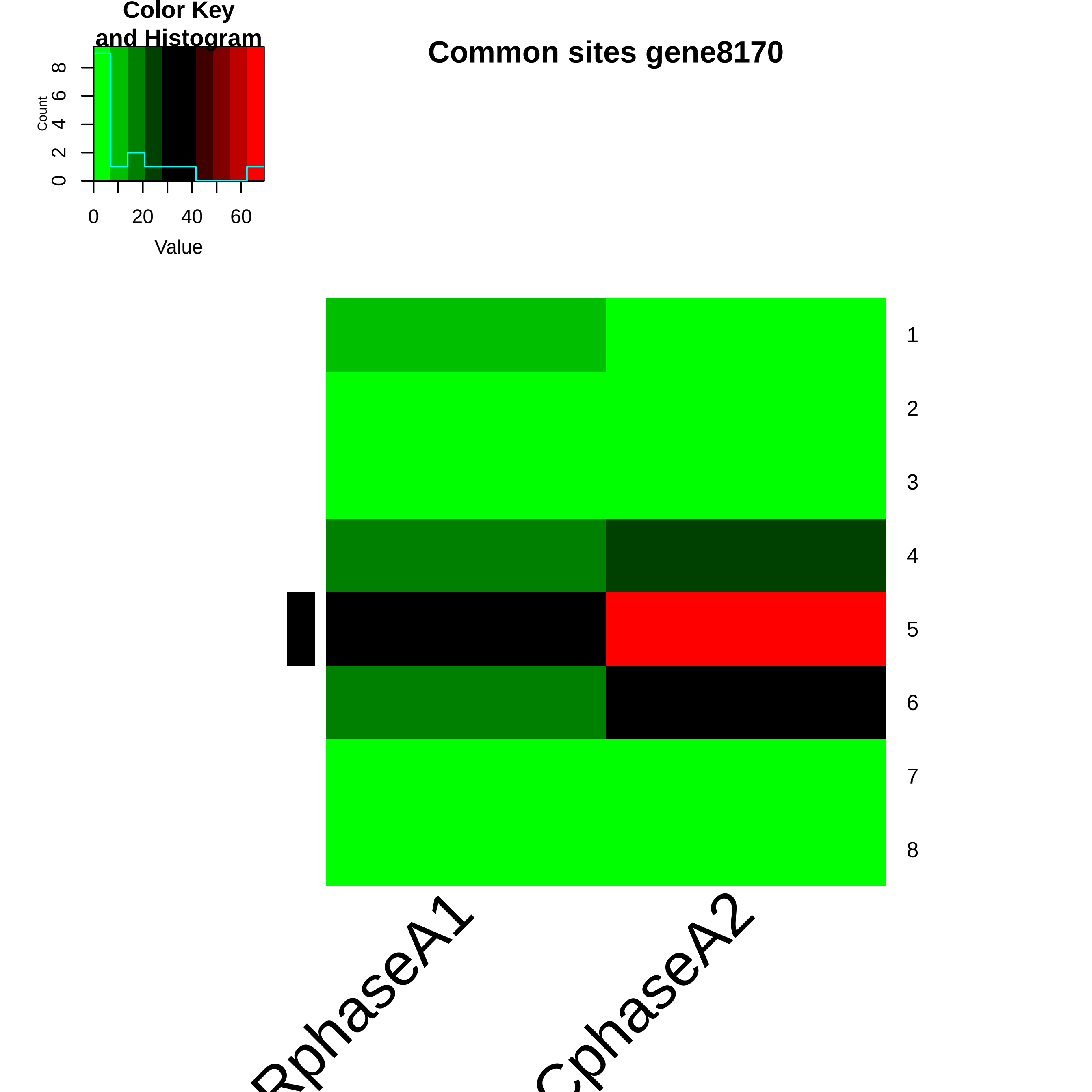
Color Key
and Histogram
Count
0
2
4
6
8
0
20
40
60
Value
Common sites gene8170
1
2
3
4
5
6
7
8
RphaseA1
CphaseA2

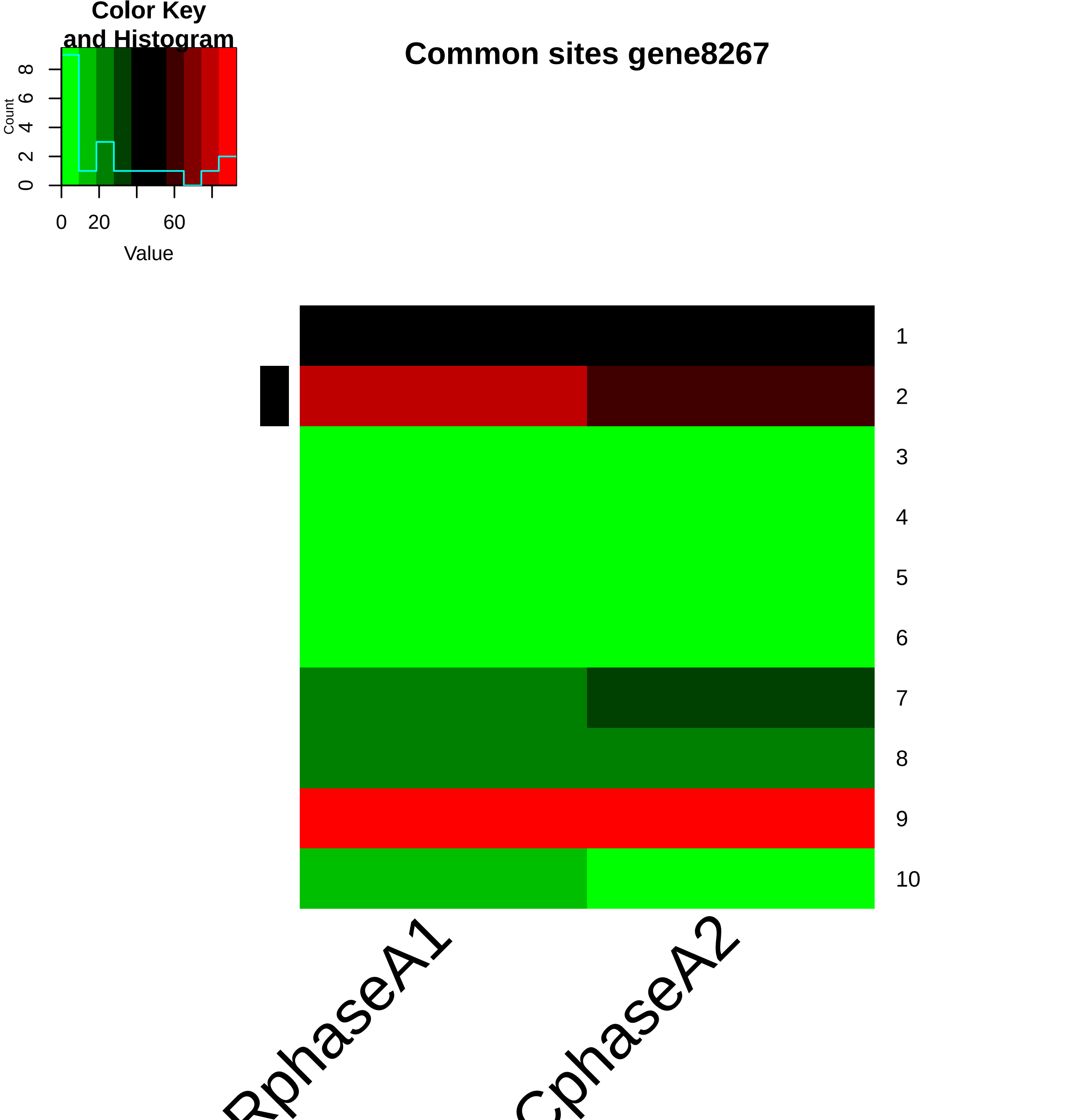

Color Key
and Histogram
Count
0 2 4 6 8
0 20 60
Value
Common sites gene8267
1
2
3
4
5
6
7
8
9
10
RphaseA1
CphaseA2

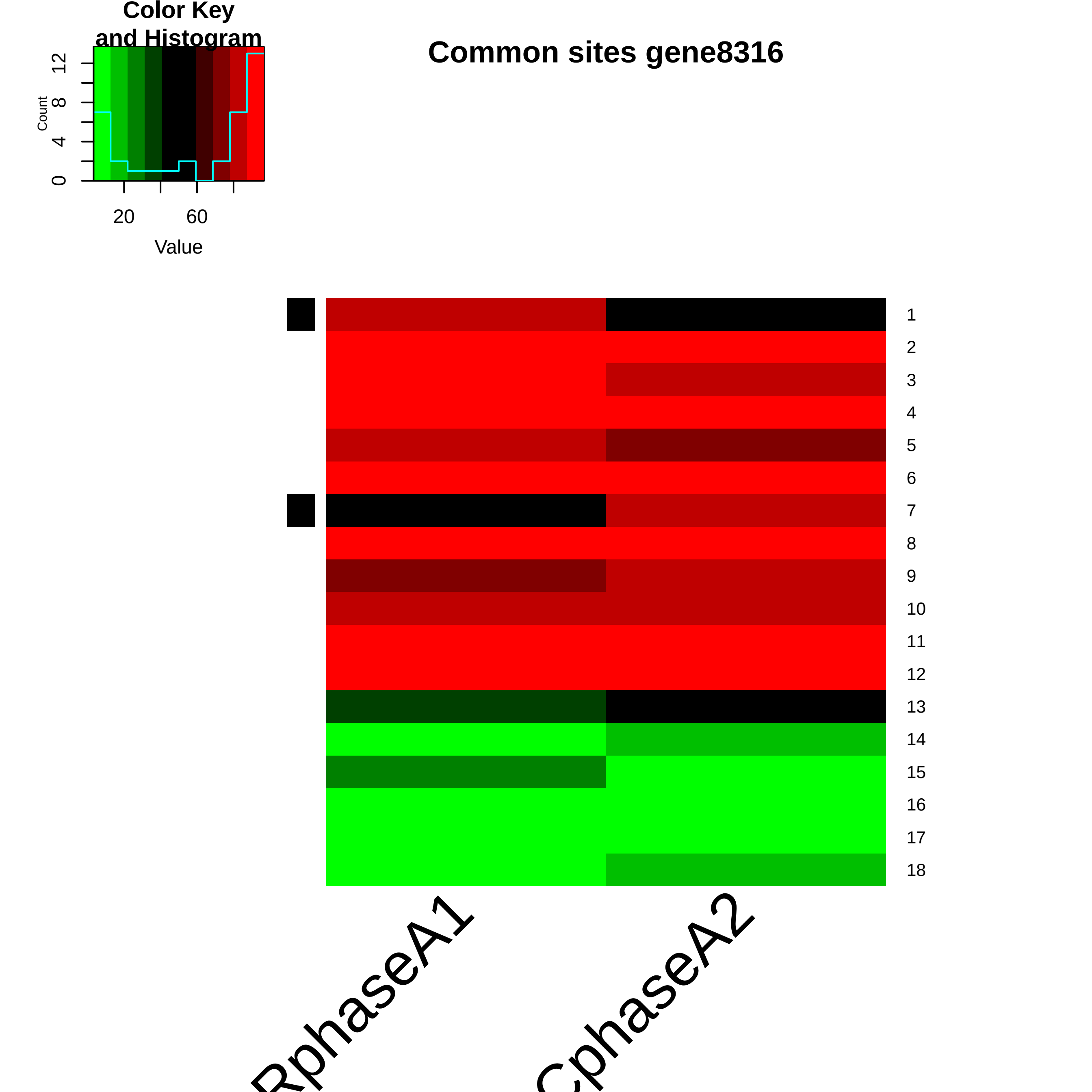
Color Key
and Histogram
Count
0
4
8
12
20
60
Value
Common sites gene8316
RphaseA1
CphaseA2

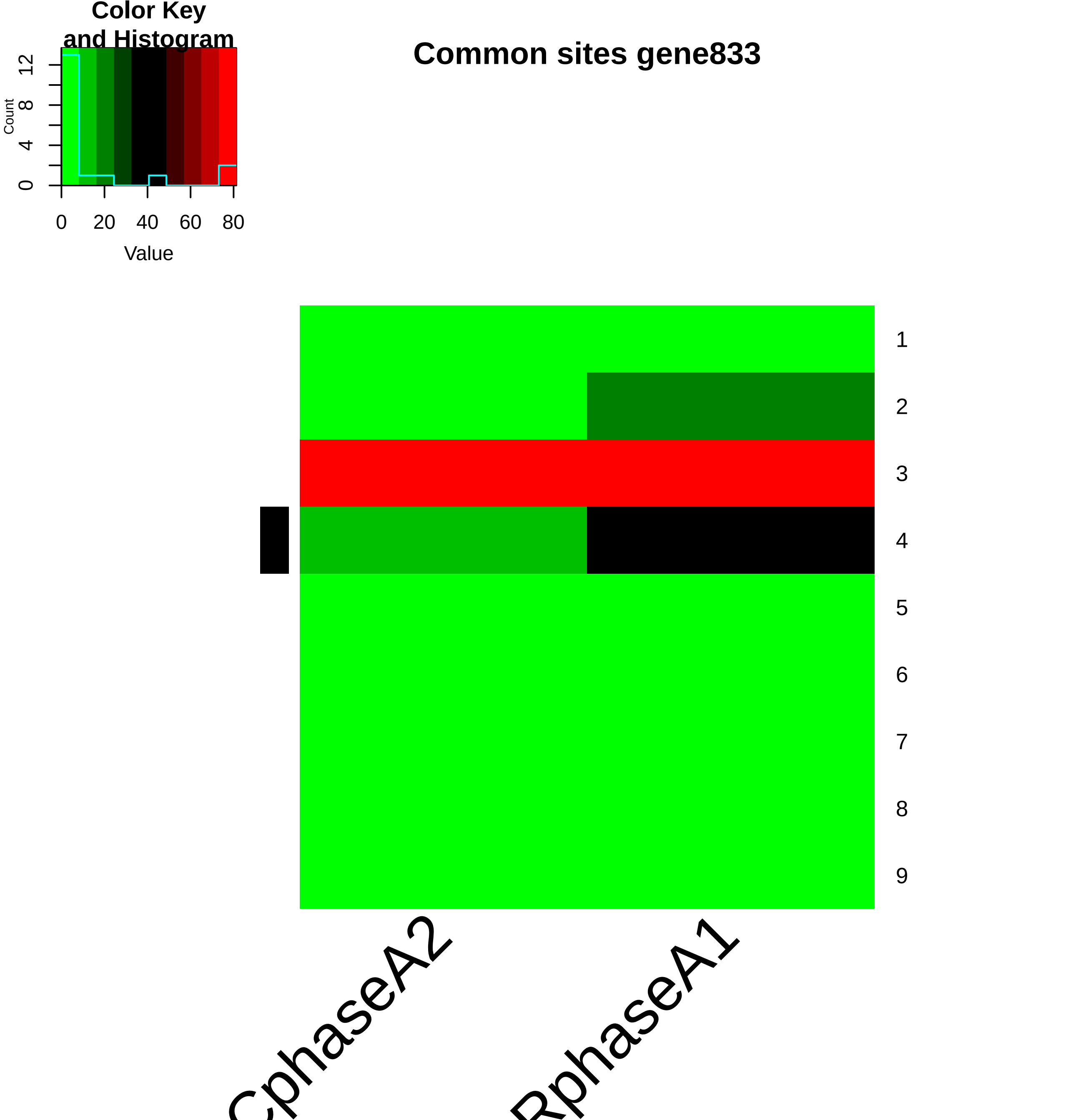

Color Key
and Histogram
Count
0
4
8
12
0
20
40
60
80
Value
Common sites gene833
1
2
3
4
5
6
7
8
9
CphaseA2
RphaseA1

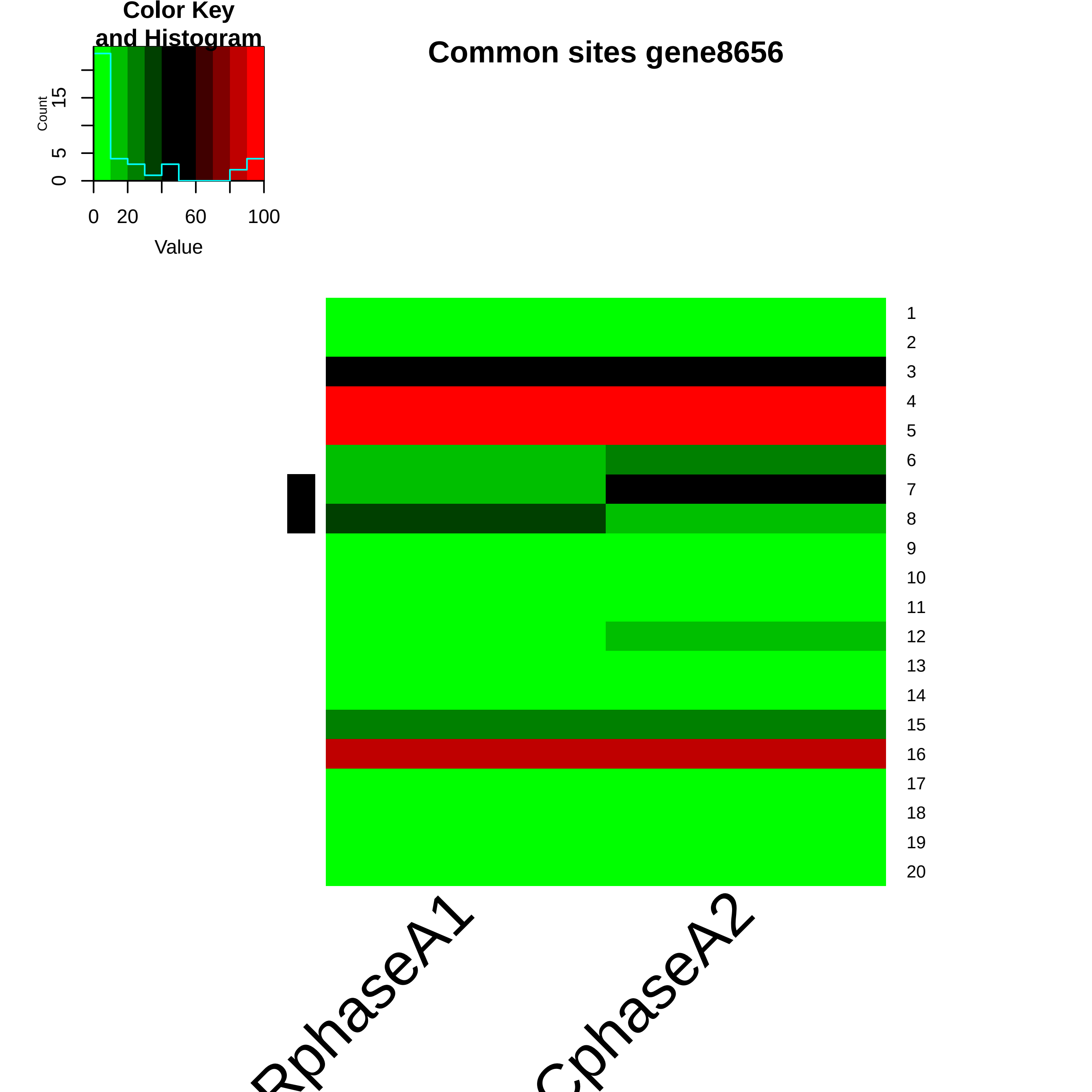
Color Key
and Histogram
Count
0
5
15
0
20
60
100
Value
Common sites gene8656

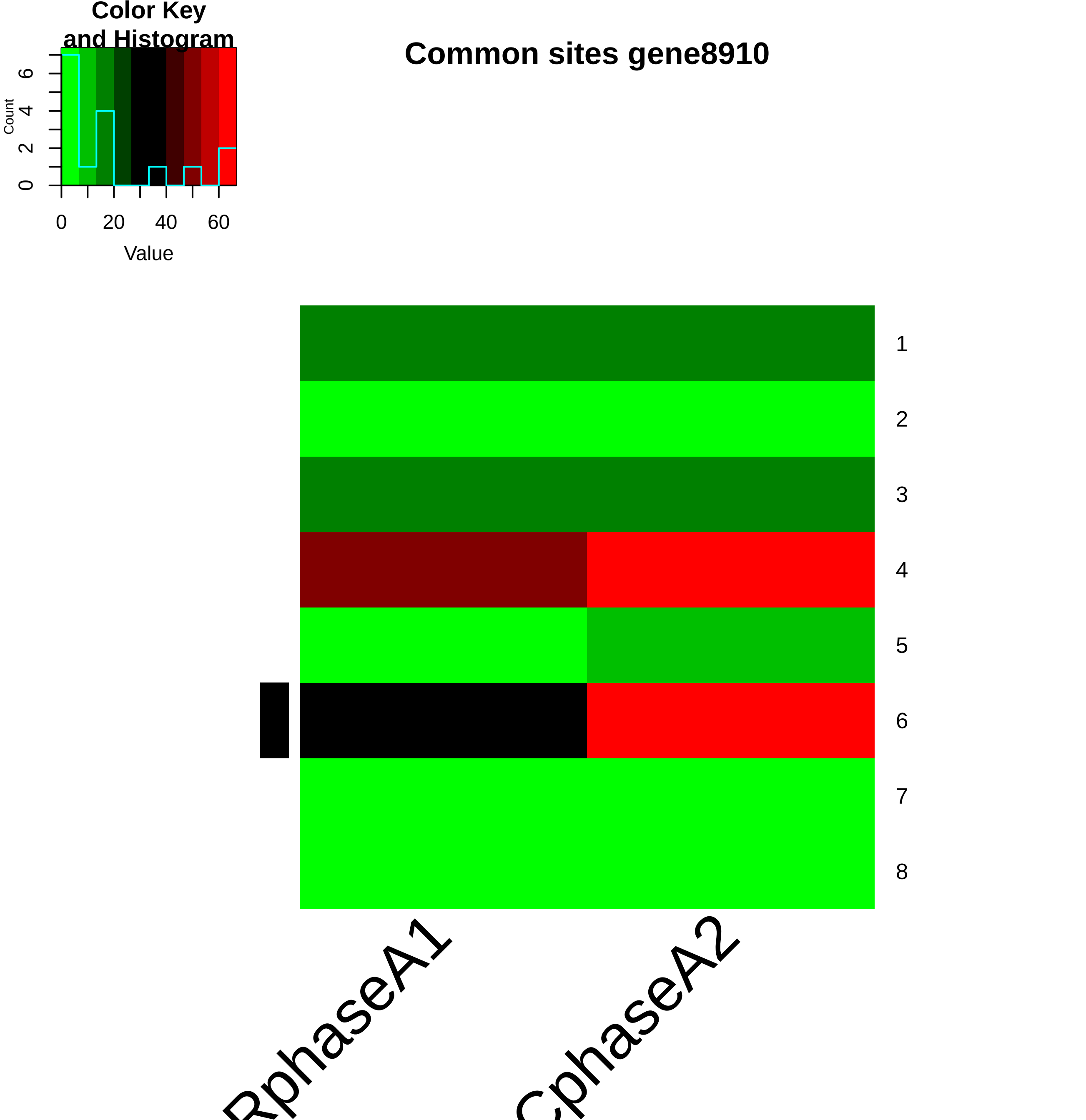
Color Key
and Histogram
Count
0
2
4
6
0
20
40
60
Value
Common sites gene8910
1
2
3
4
5
6
7
8
RphaseA1
CphaseA2

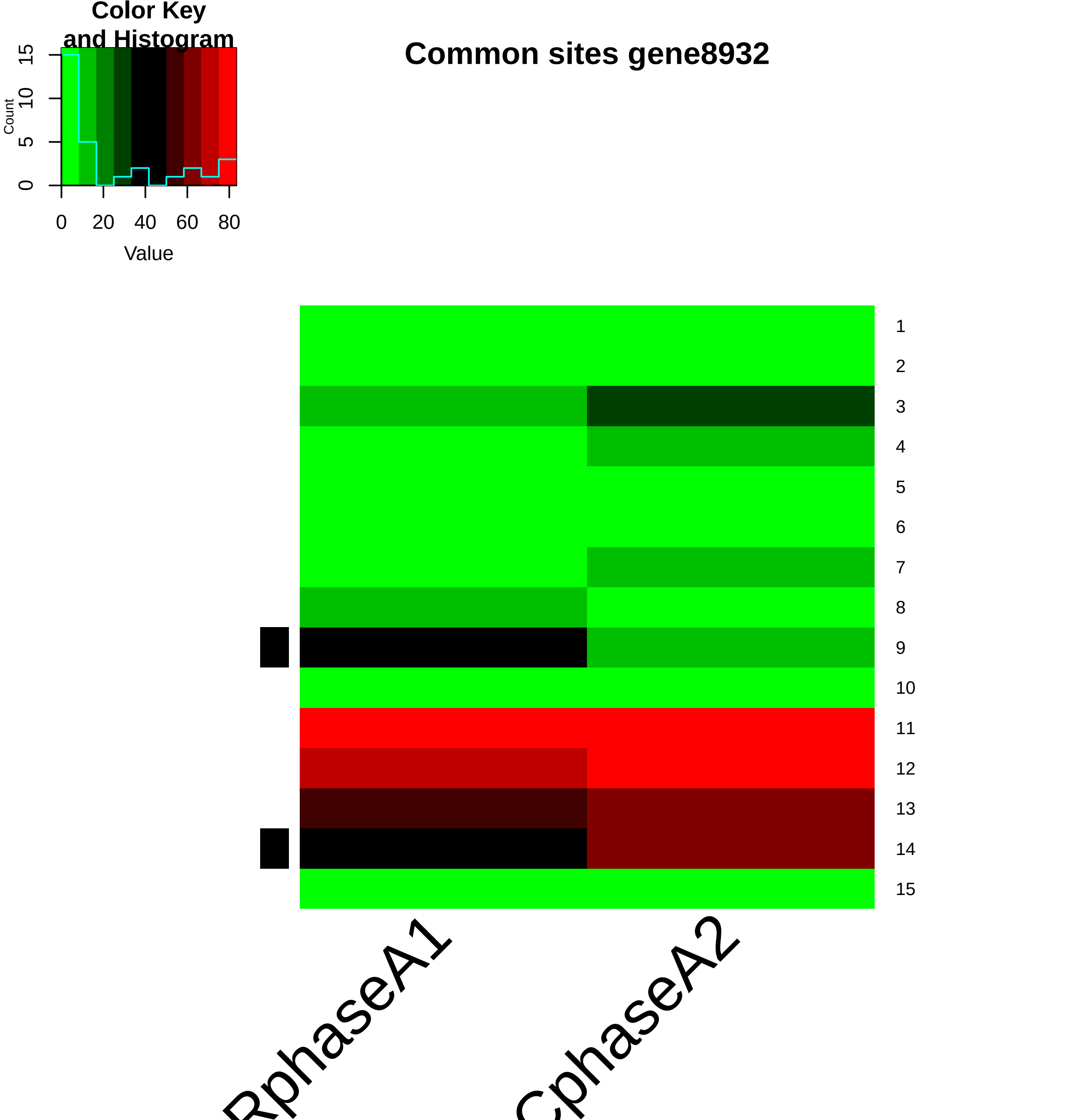
Color Key
and Histogram
Count
0
5
10
15
0
20
40
60
80
Value
Common sites gene8932
RphaseA1
CphaseA2

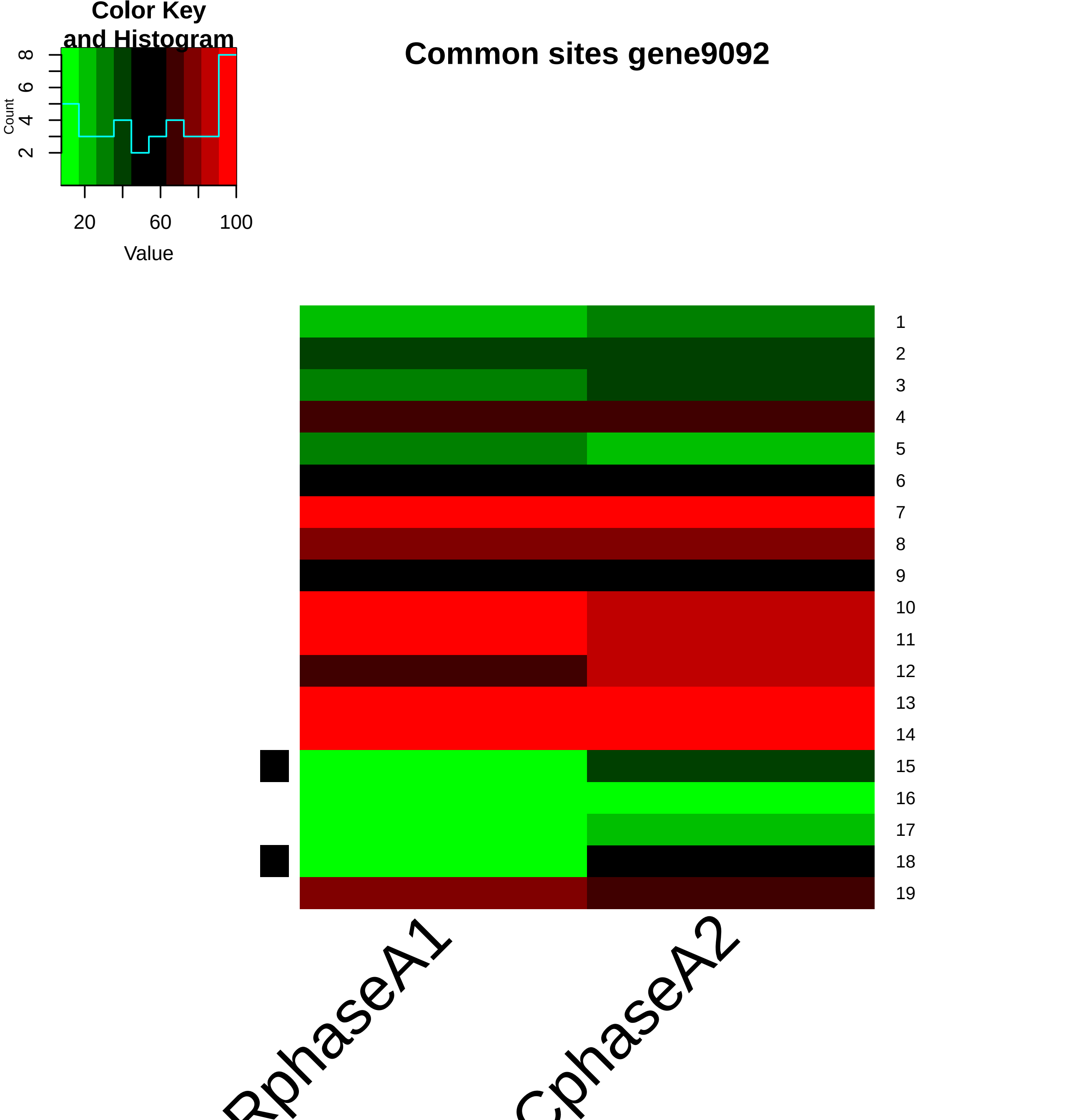
Color Key
and Histogram
Count
2
4
6
8
20
60
100
Value
Common sites gene9092
RphaseA1
CphaseA2

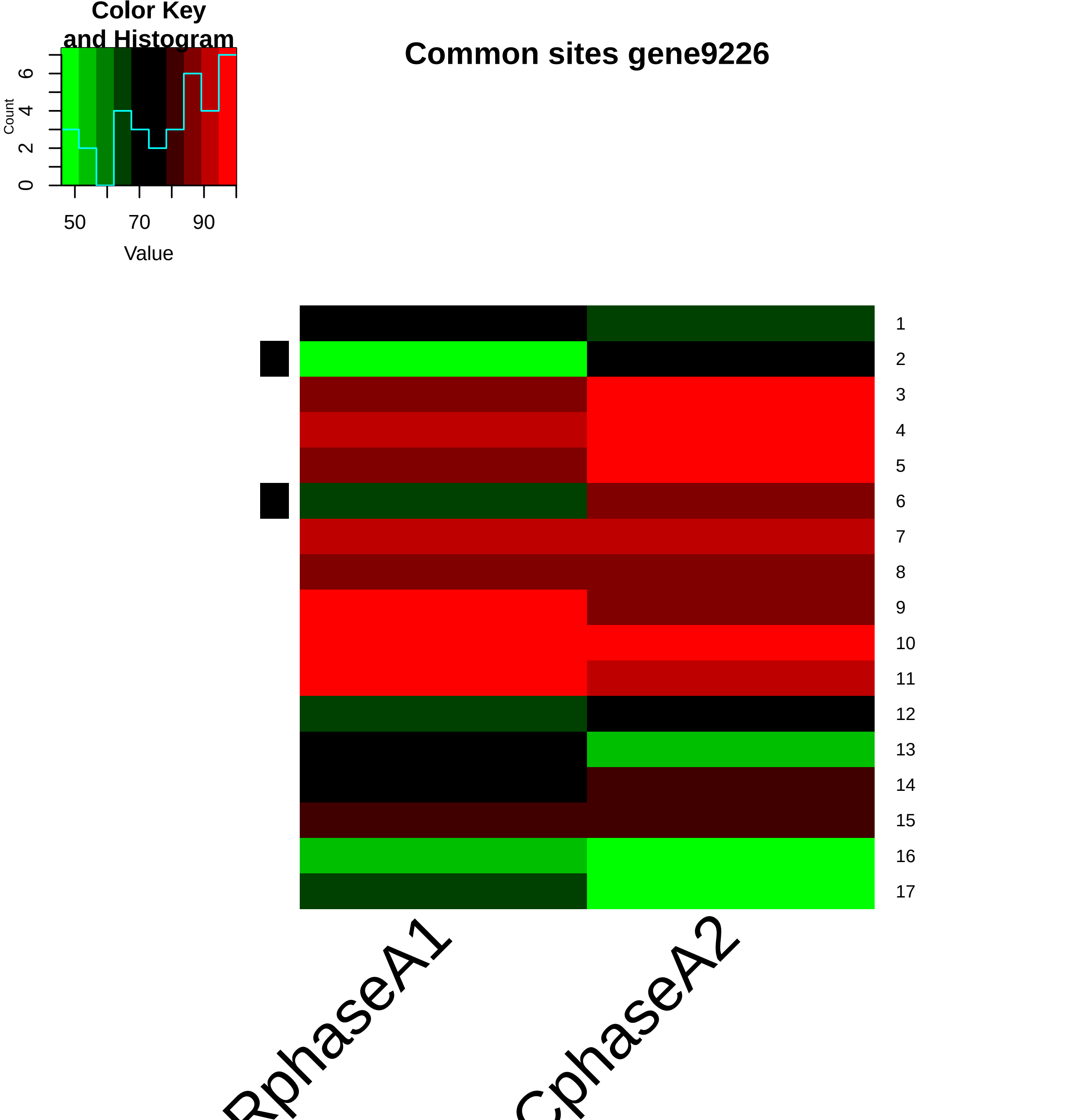

Color Key
and Histogram
Count
0
2
4
6
50
70
90
Value
Common sites gene9226
RphaseA1
CphaseA2

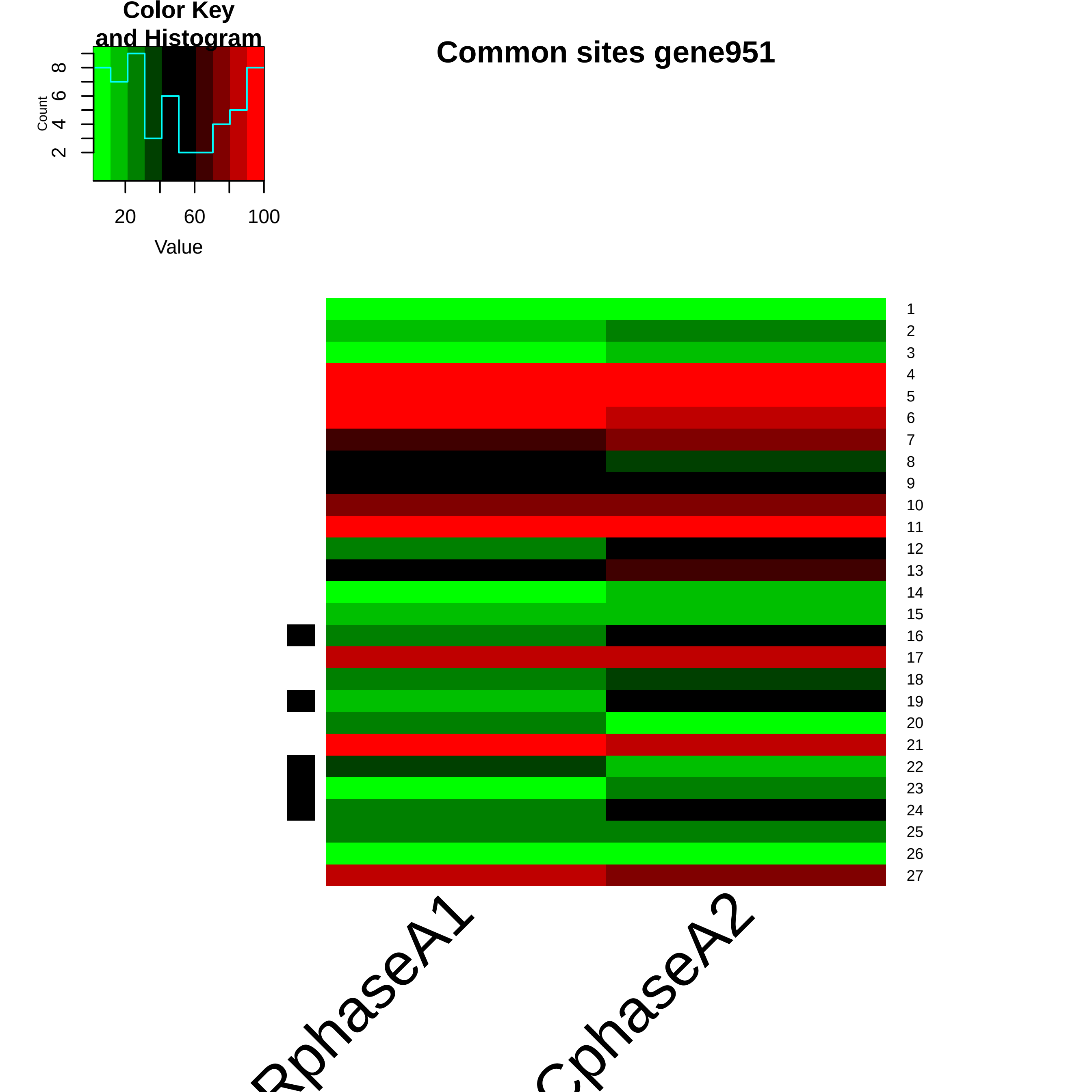
Color Key
and Histogram
Count
2
4
6
8
20
60
100
Value
Common sites gene951
RphaseA1
CphaseA2

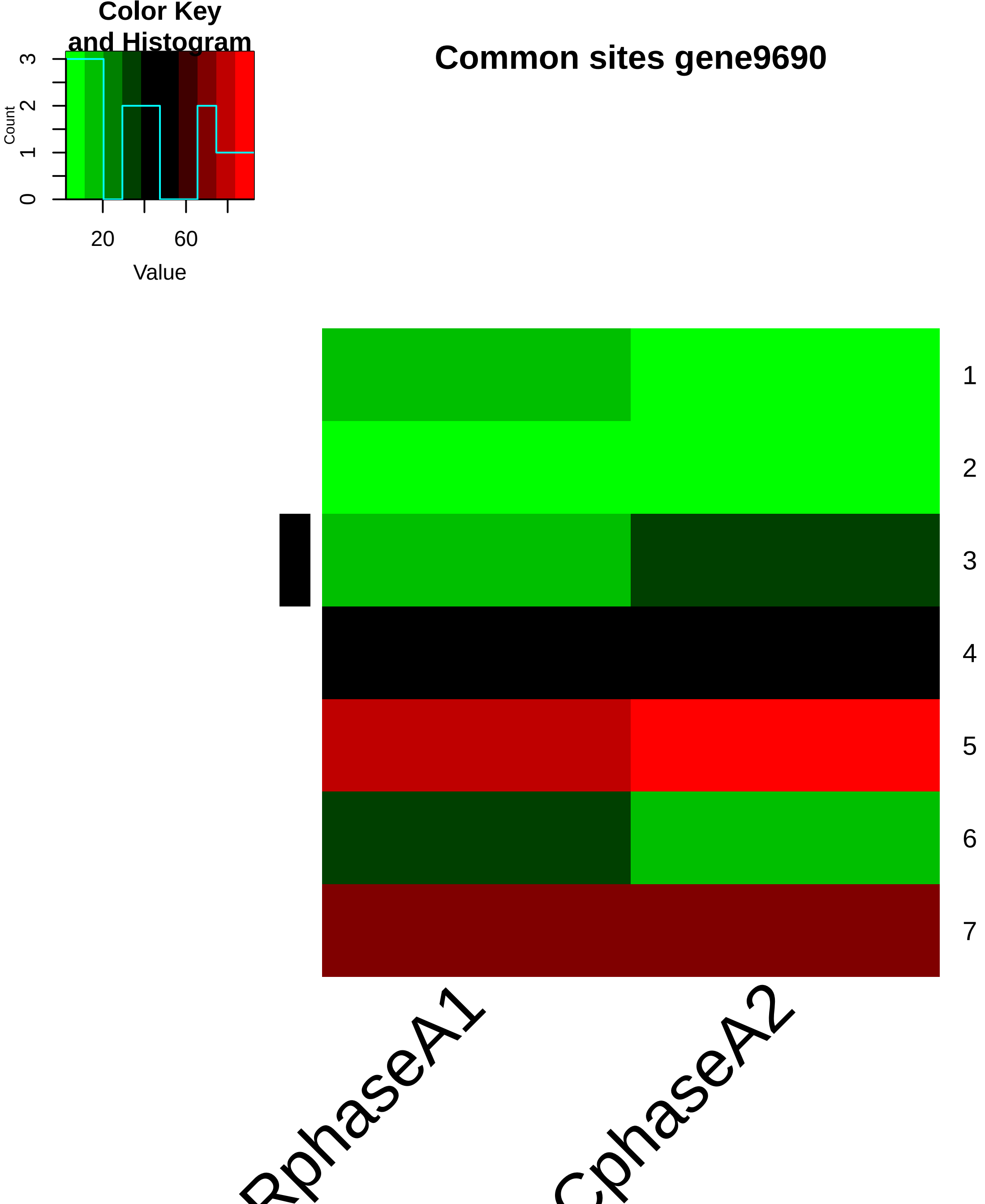

Color Key
and Histogram
Count
0
1
2
3
20
60
Value
Common sites gene9690
1
2
3
4
5
6
7
RphaseA1
CphaseA2

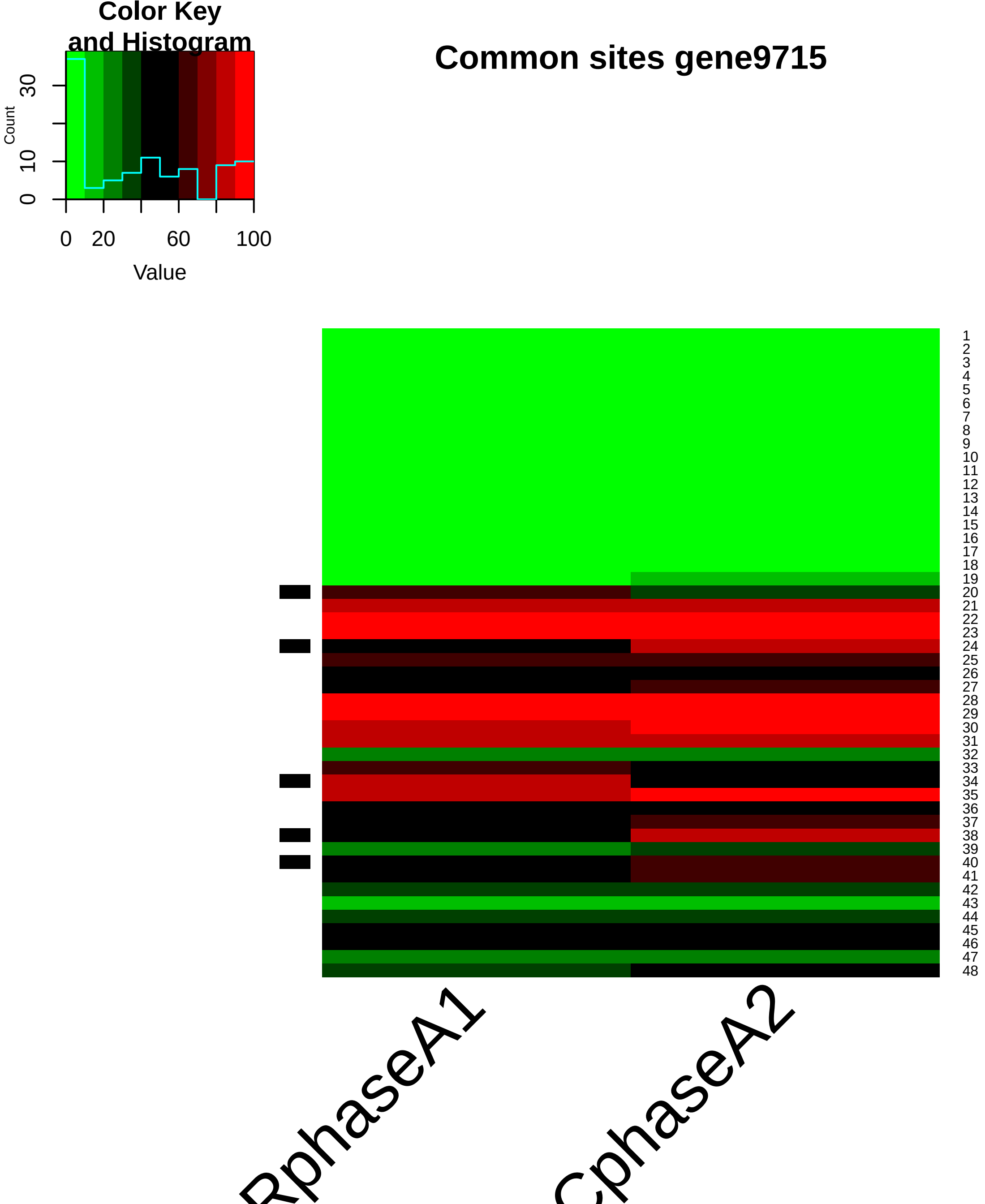
Color Key
and Histogram
Count
0
10
30
0
20
60
100
Value
Common sites gene9715
RphaseA1
CphaseA2

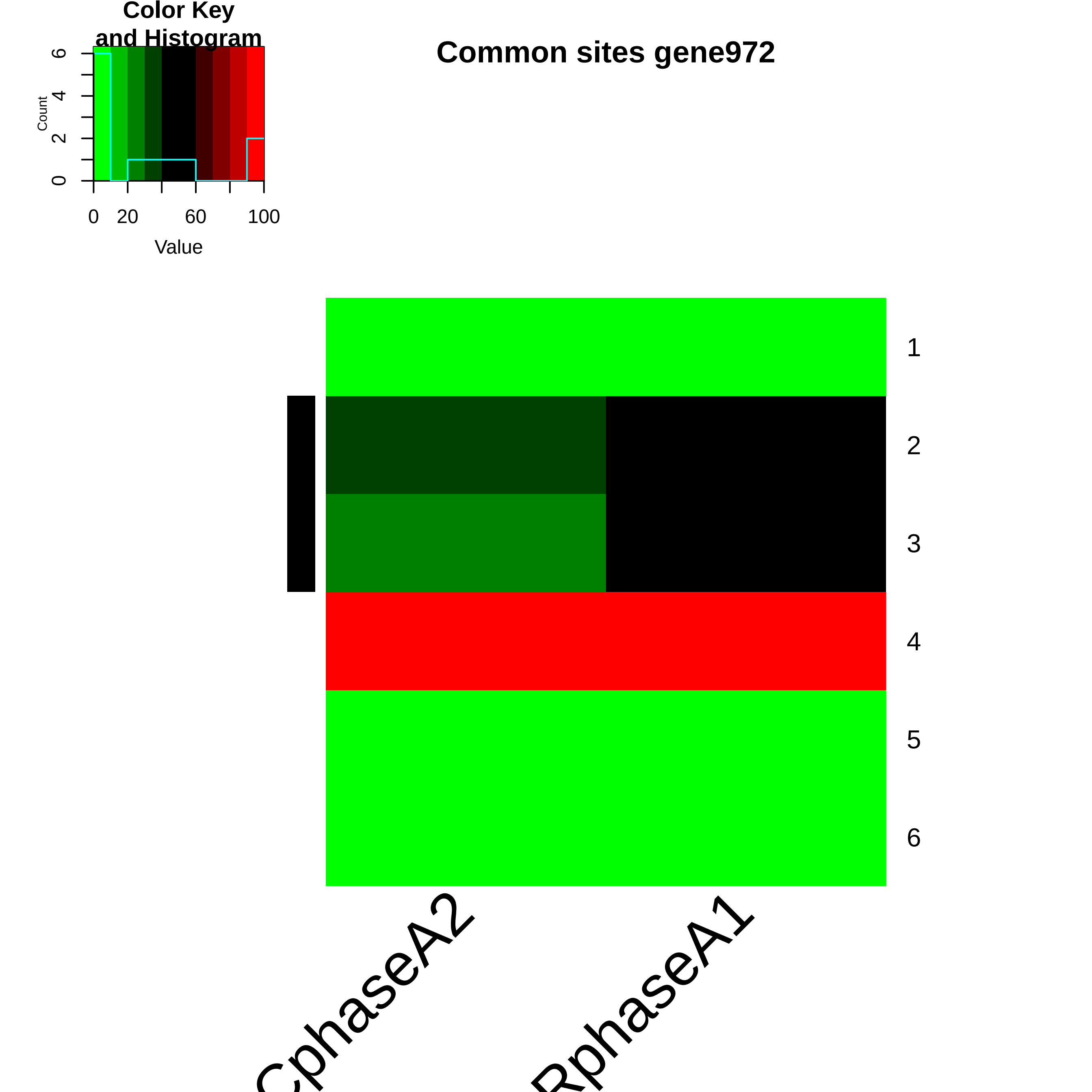
Color Key
and Histogram
Count
0
2
4
6
0
20
60
100
Value
Common sites gene972
1
2
3
4
5
6
CphaseA2
RphaseA1

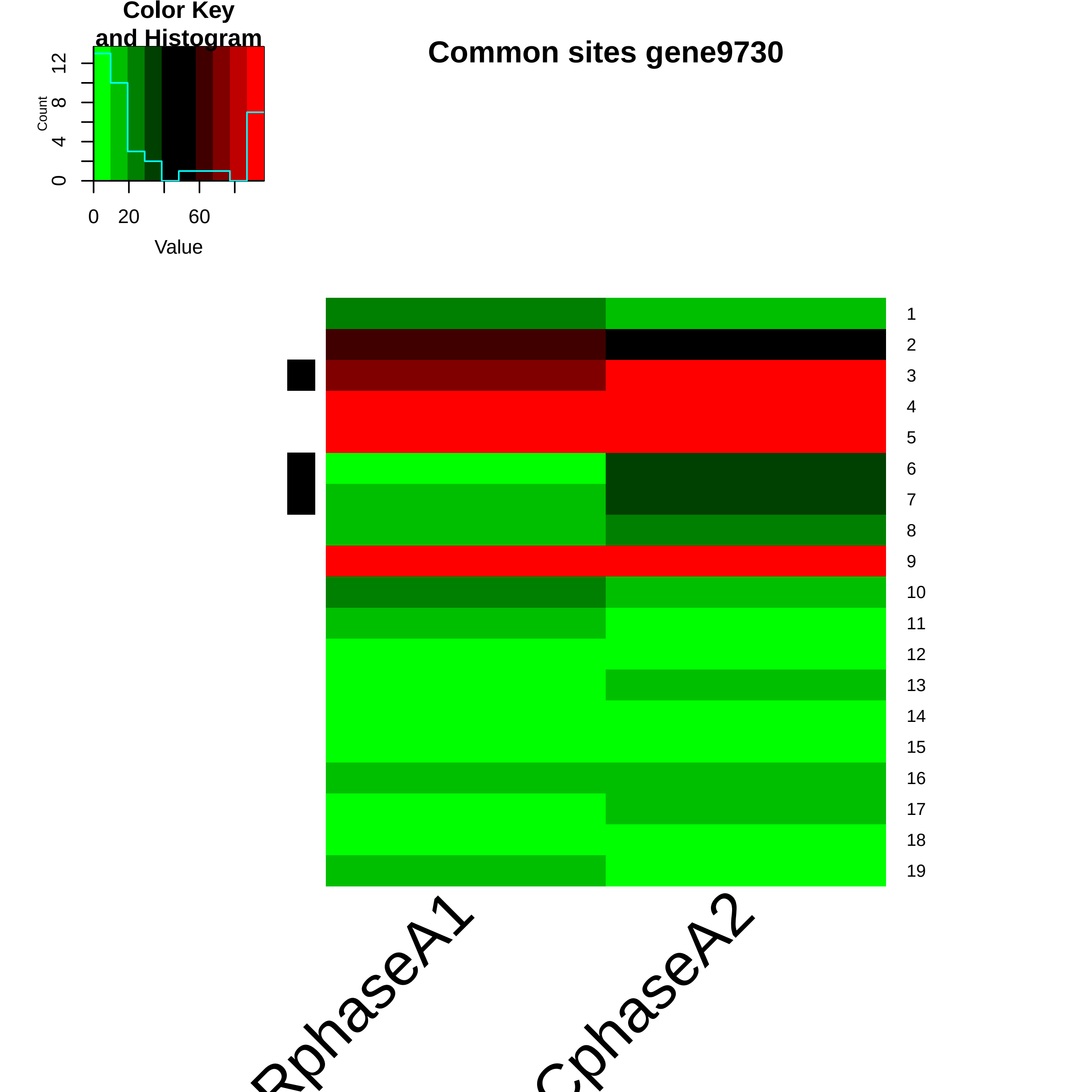
Color Key
and Histogram
Count
0
4
8
12
0
20
60
Value
Common sites gene9730
RphaseA1
CphaseA2

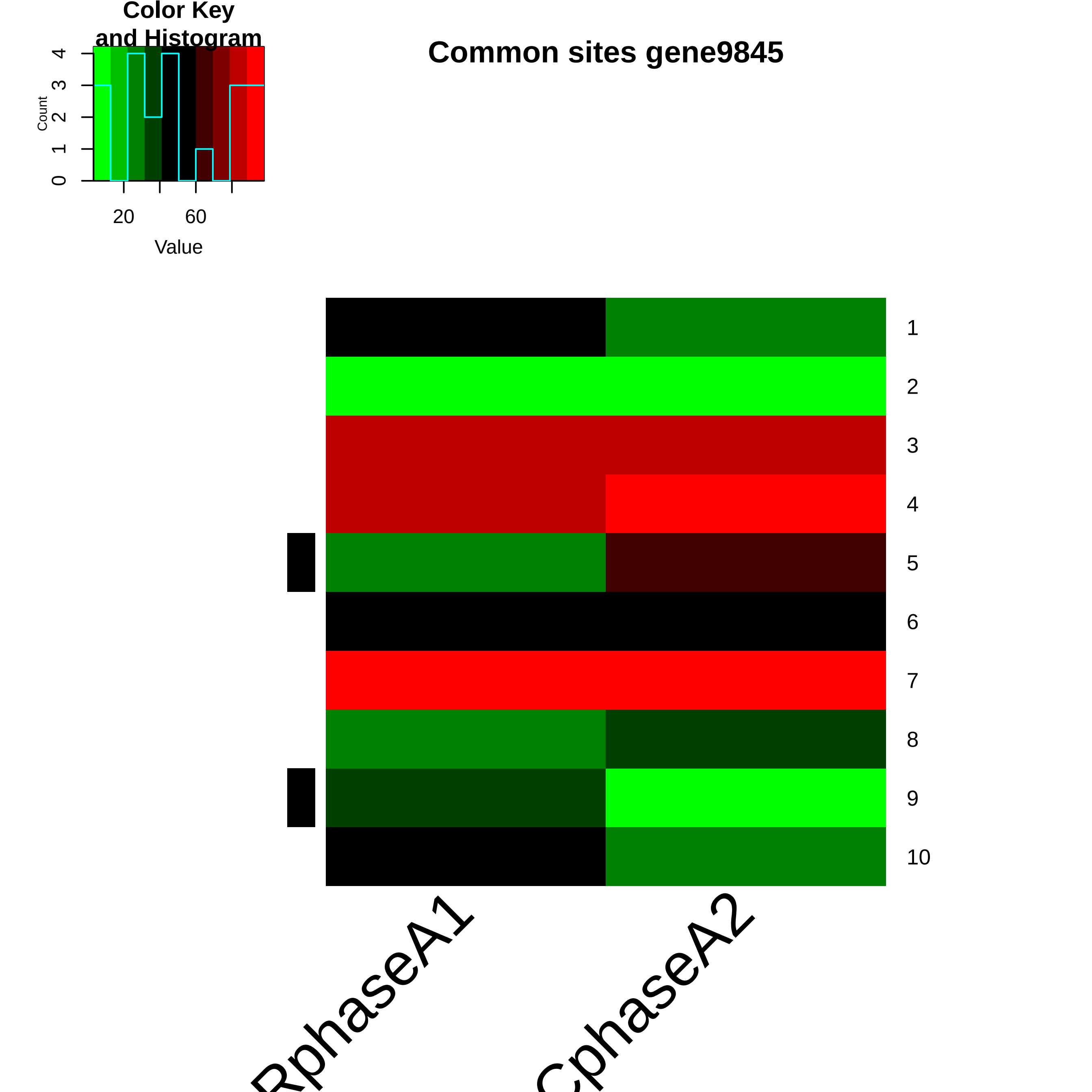

Color Key
and Histogram
Count
0
1
2
3
4
20
60
Value
Common sites gene9845
1
2
3
4
5
6
7
8
9
10
RphaseA1
CphaseA2

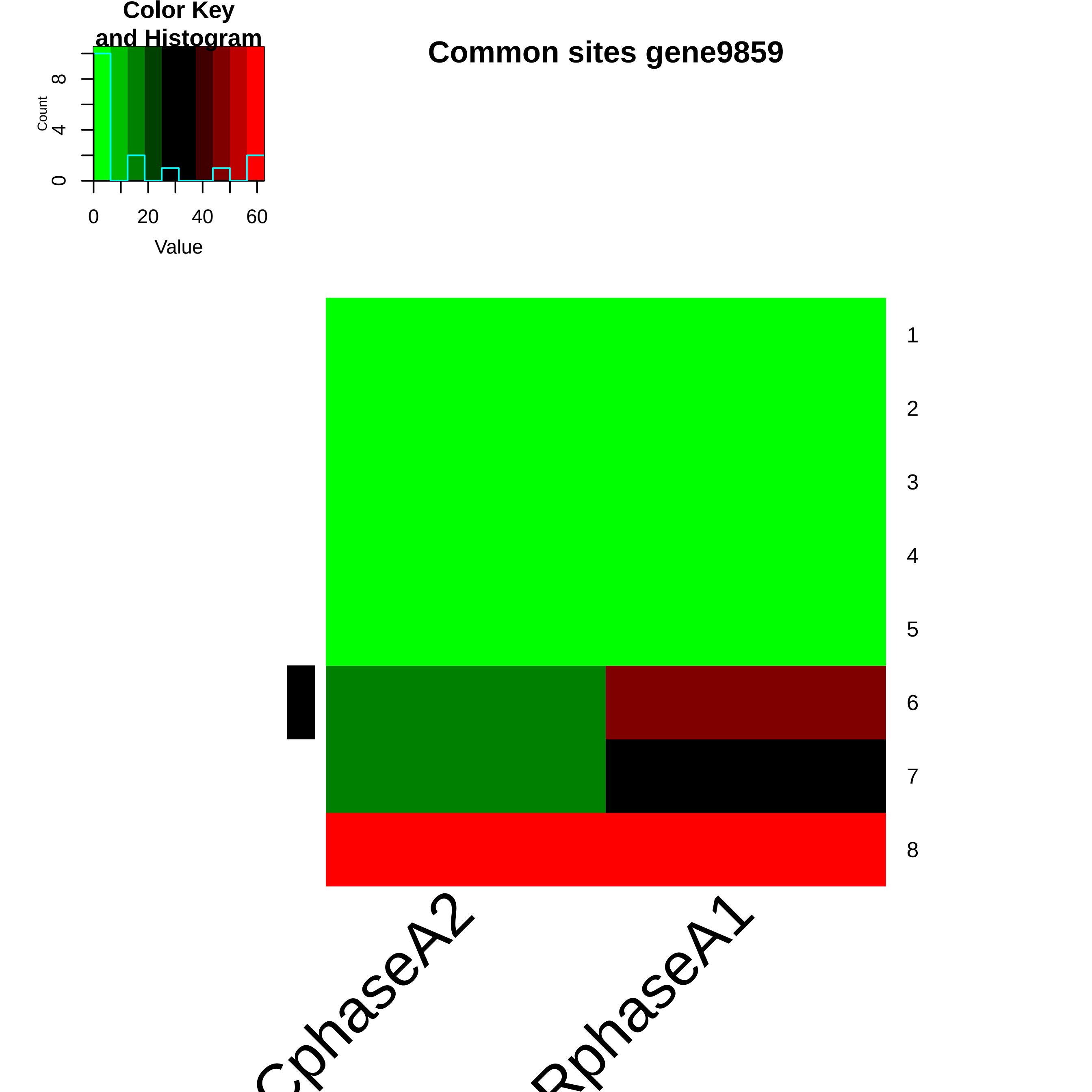
Color Key
and Histogram
Count
0
4
8
0
20
40
60
Value
Common sites gene9859
1
2
3
4
5
6
7
8

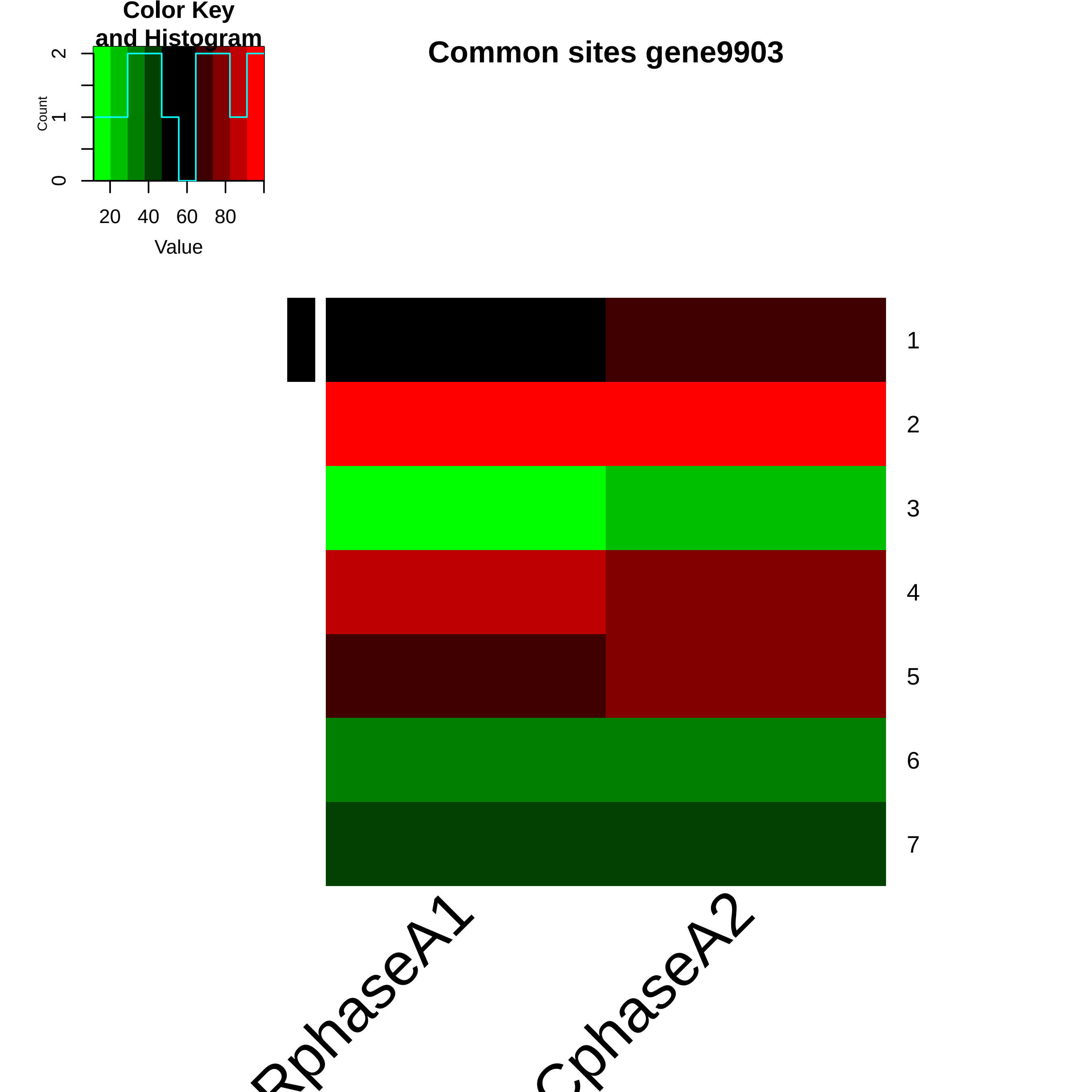
Color Key
and Histogram
Count
0
1
2
20
40
60
80
Value
Common sites gene9903
1
2
3
4
5
6
7
RphaseA1
CphaseA2

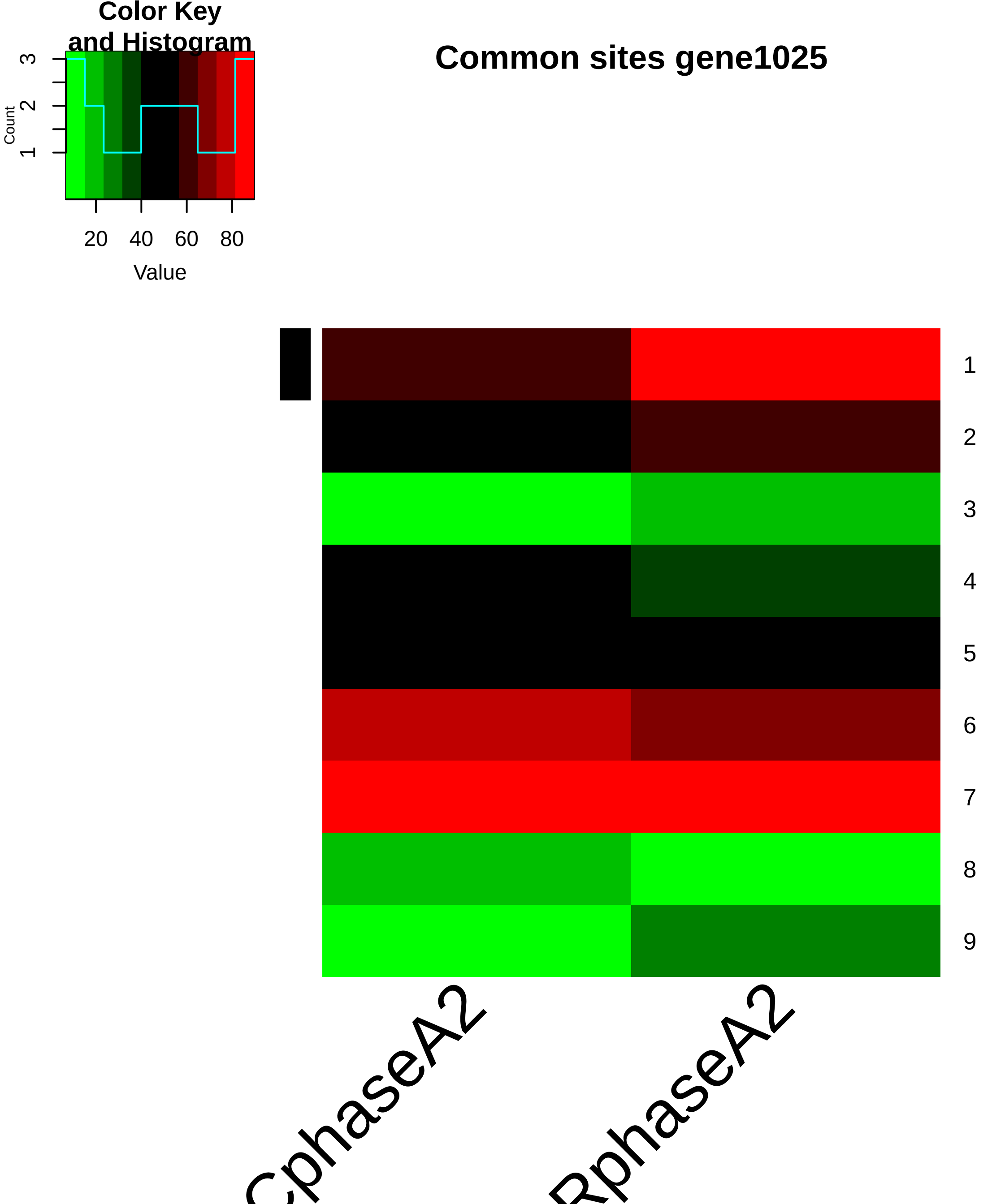

Color Key
and Histogram
Count
3
2
1
20
40
60
80
Value
Common sites gene1025
1
2
3
4
5
6
7
8
9
CphaseA2
RphaseA2

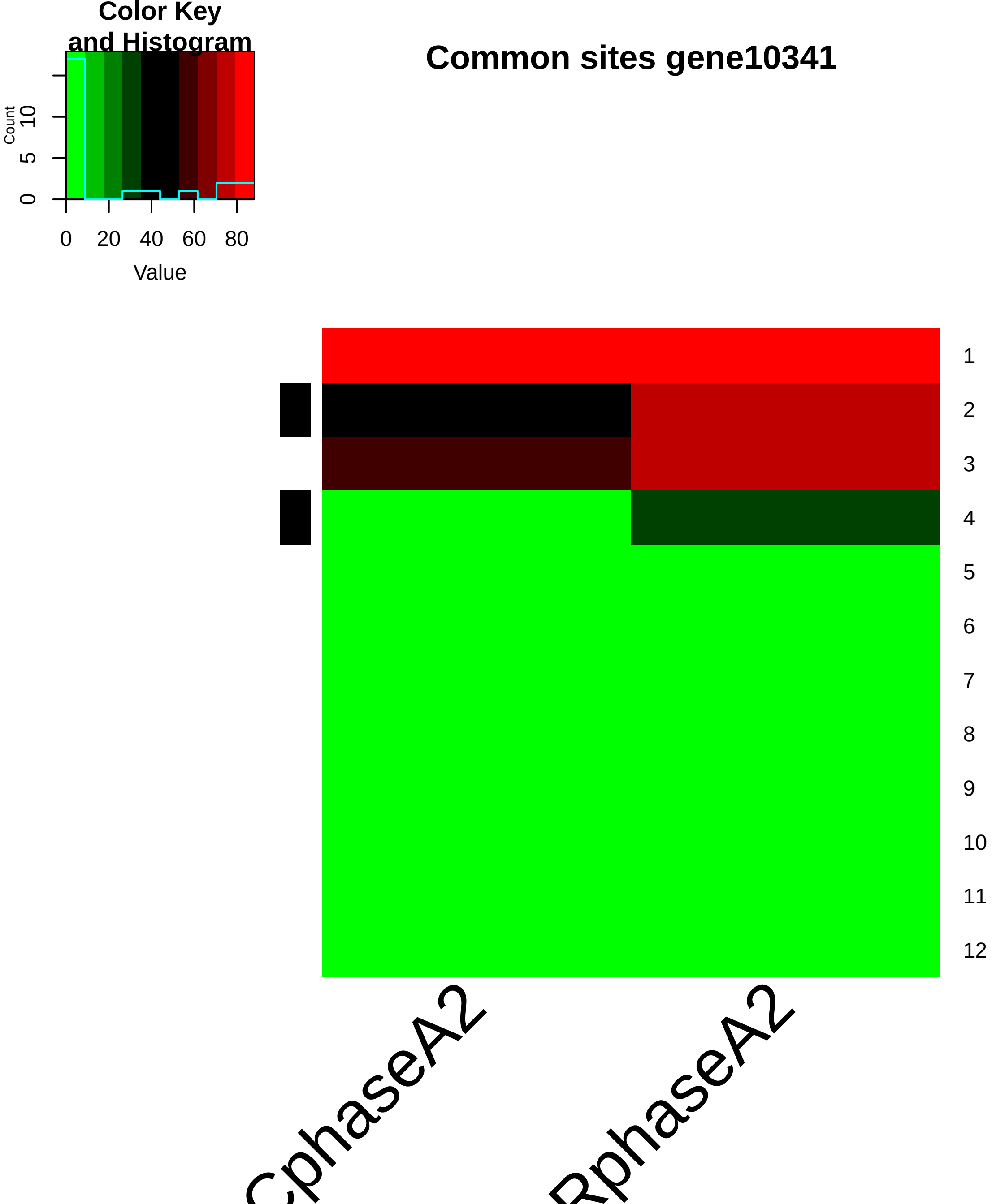
Color Key
and Histogram
Count
0
5
10
0
20
40
60
80
Value
Common sites gene10341
1
2
3
4
5
6
7
8
9
10
11
12

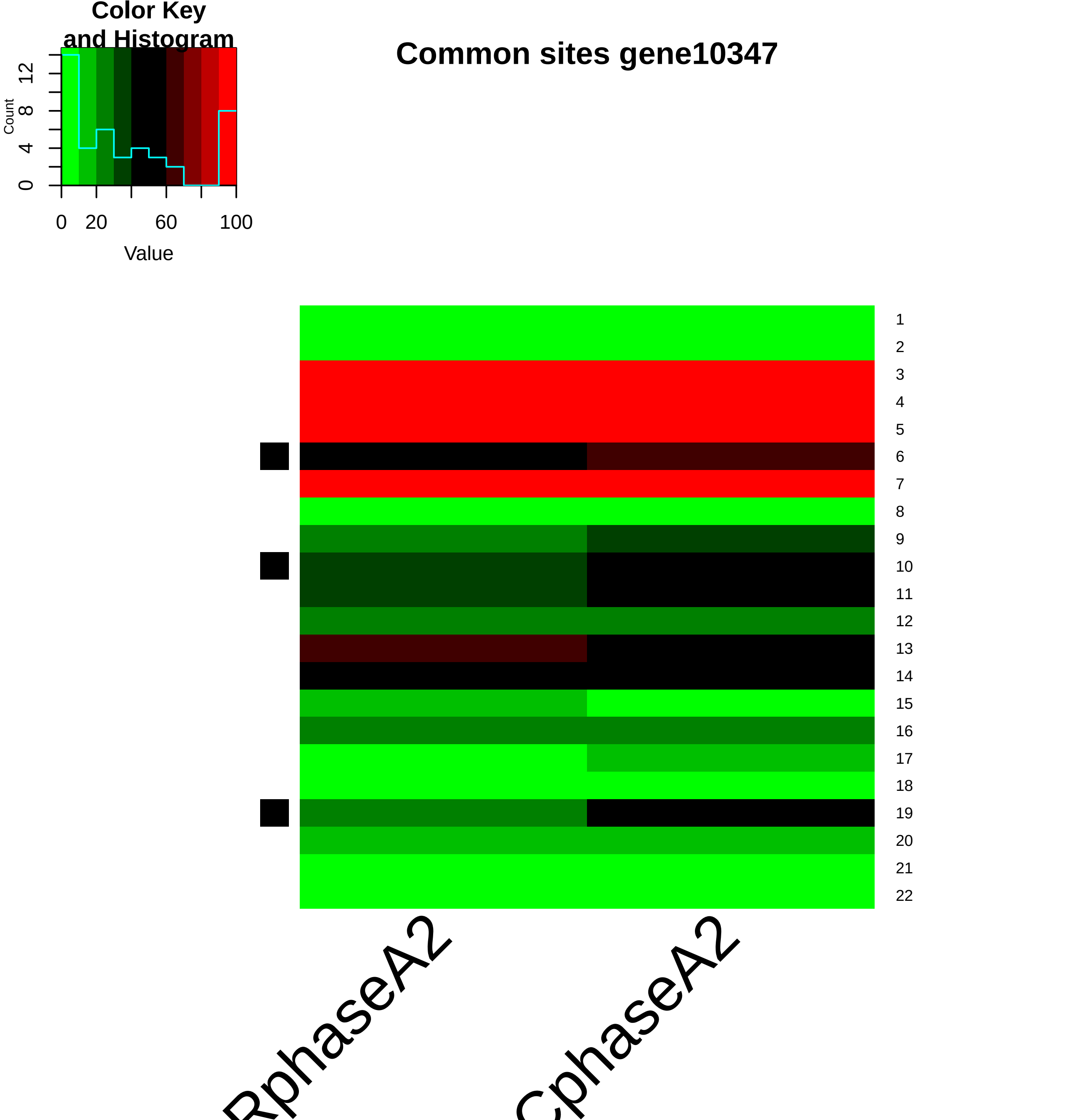

Color Key
and Histogram
Count
0
4
8
12
0
20
60
100
Value
Common sites gene10347
RphaseA2
CphaseA2

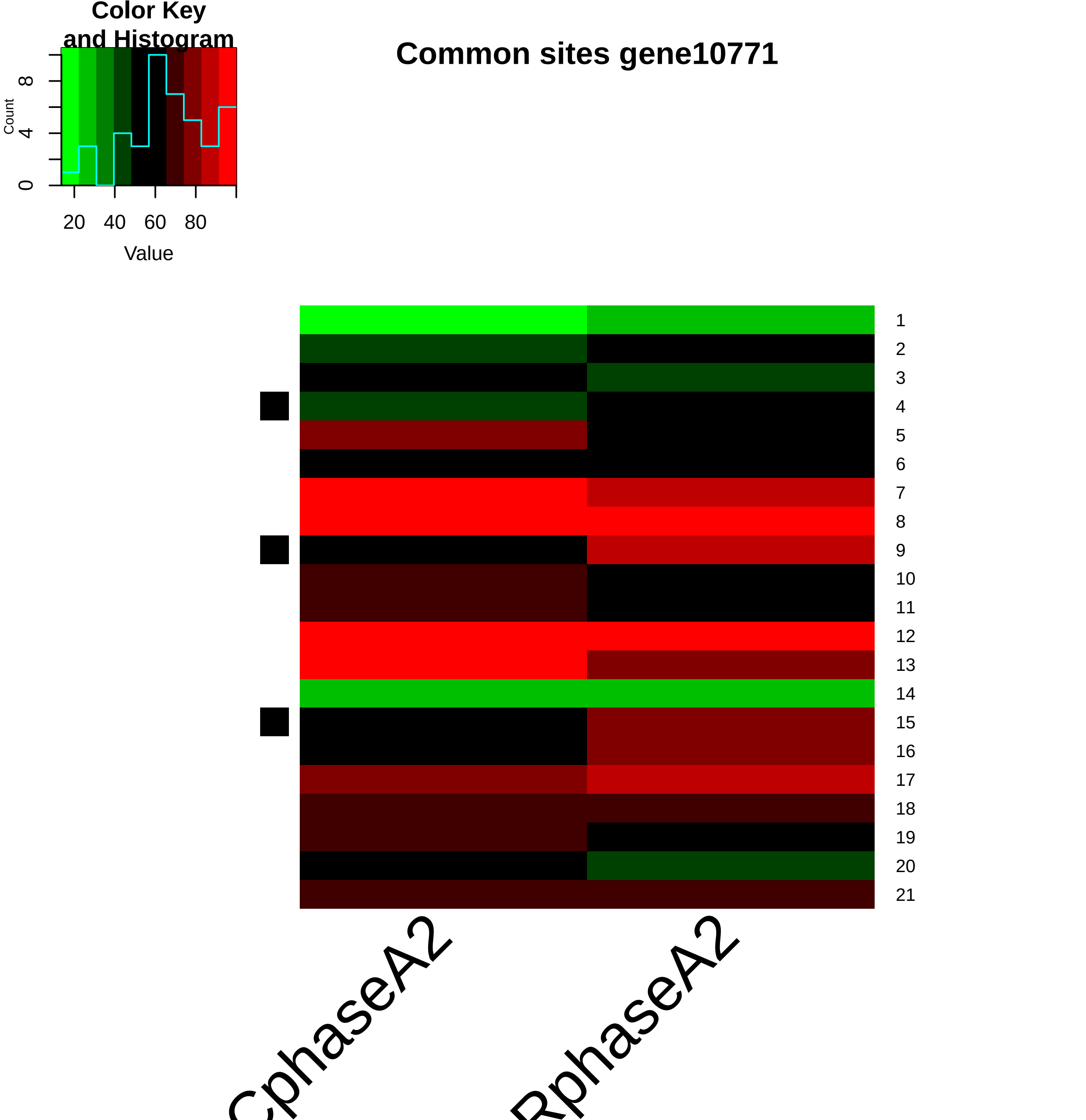

Color Key
and Histogram
Count
0
4
8
20
40
60
80
Value
Common sites gene10771
CphaseA2
RphaseA2

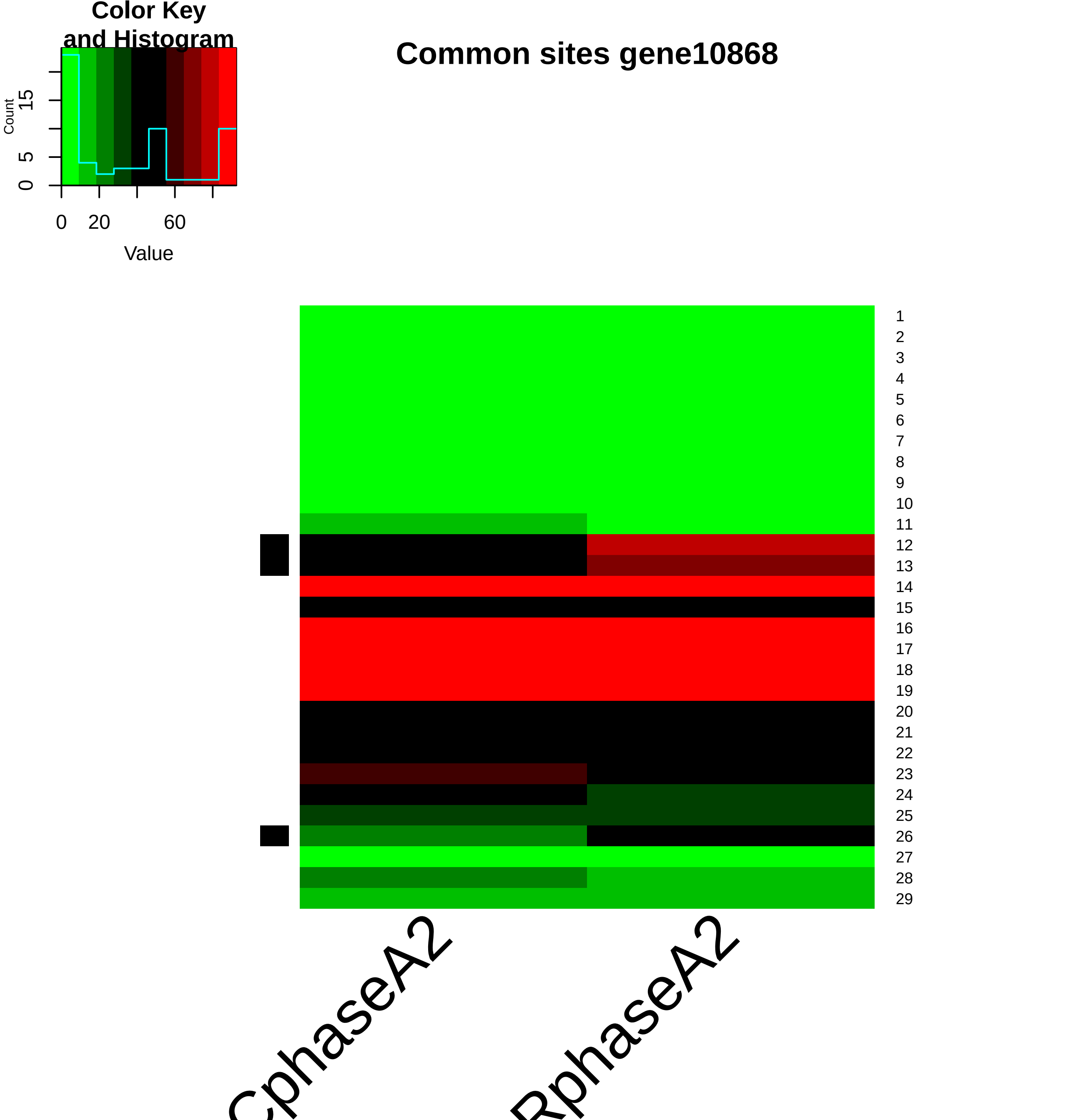

Color Key
and Histogram
Count
0
5
15
0
20
60
Value
Common sites gene10868
CphaseA2
RphaseA2

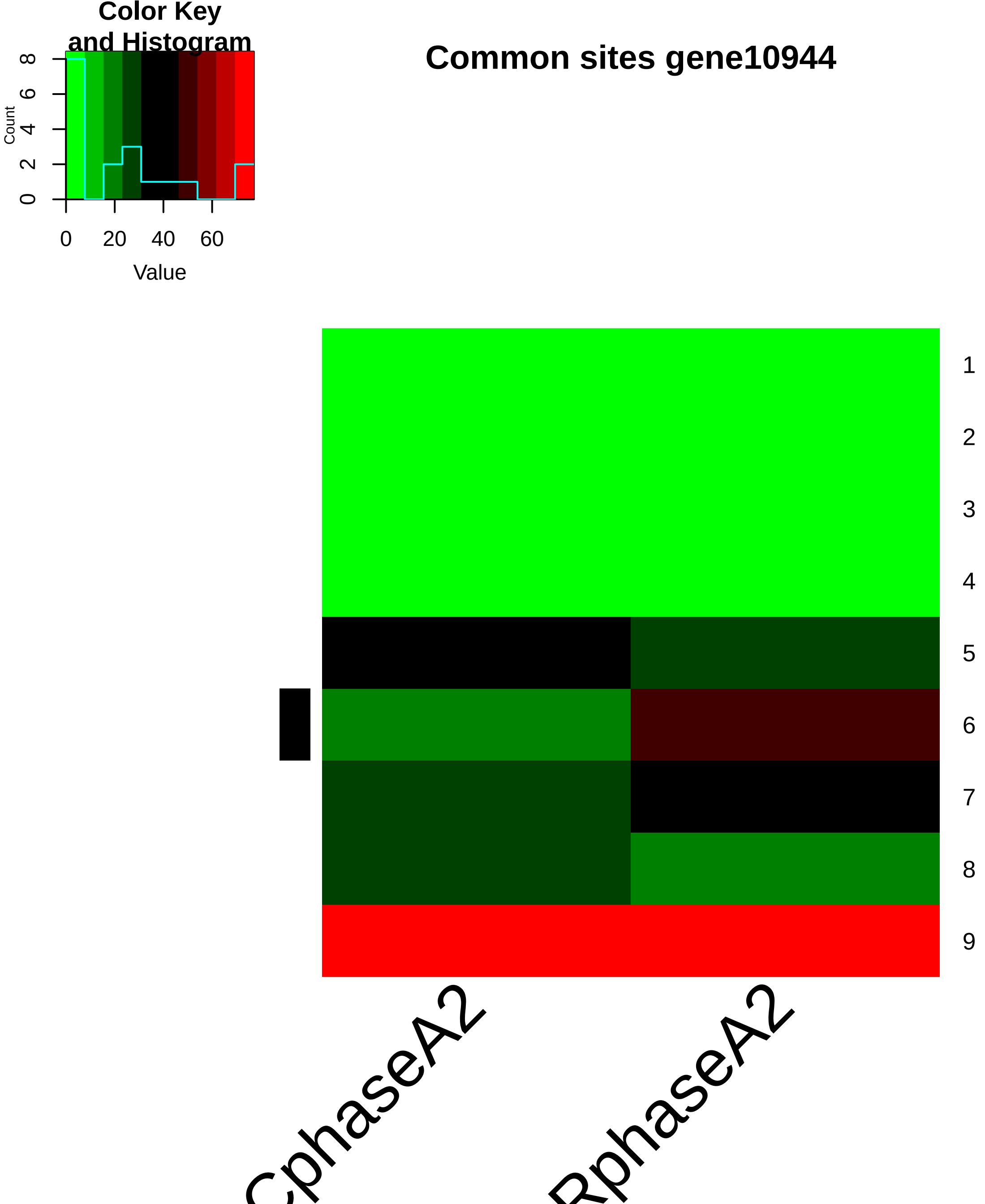
Color Key
and Histogram
Count
0
2
4
6
8
0
20
40
60
Value
Common sites gene10944
1
2
3
4
5
6
7
8
9
CphaseA2
RphaseA2

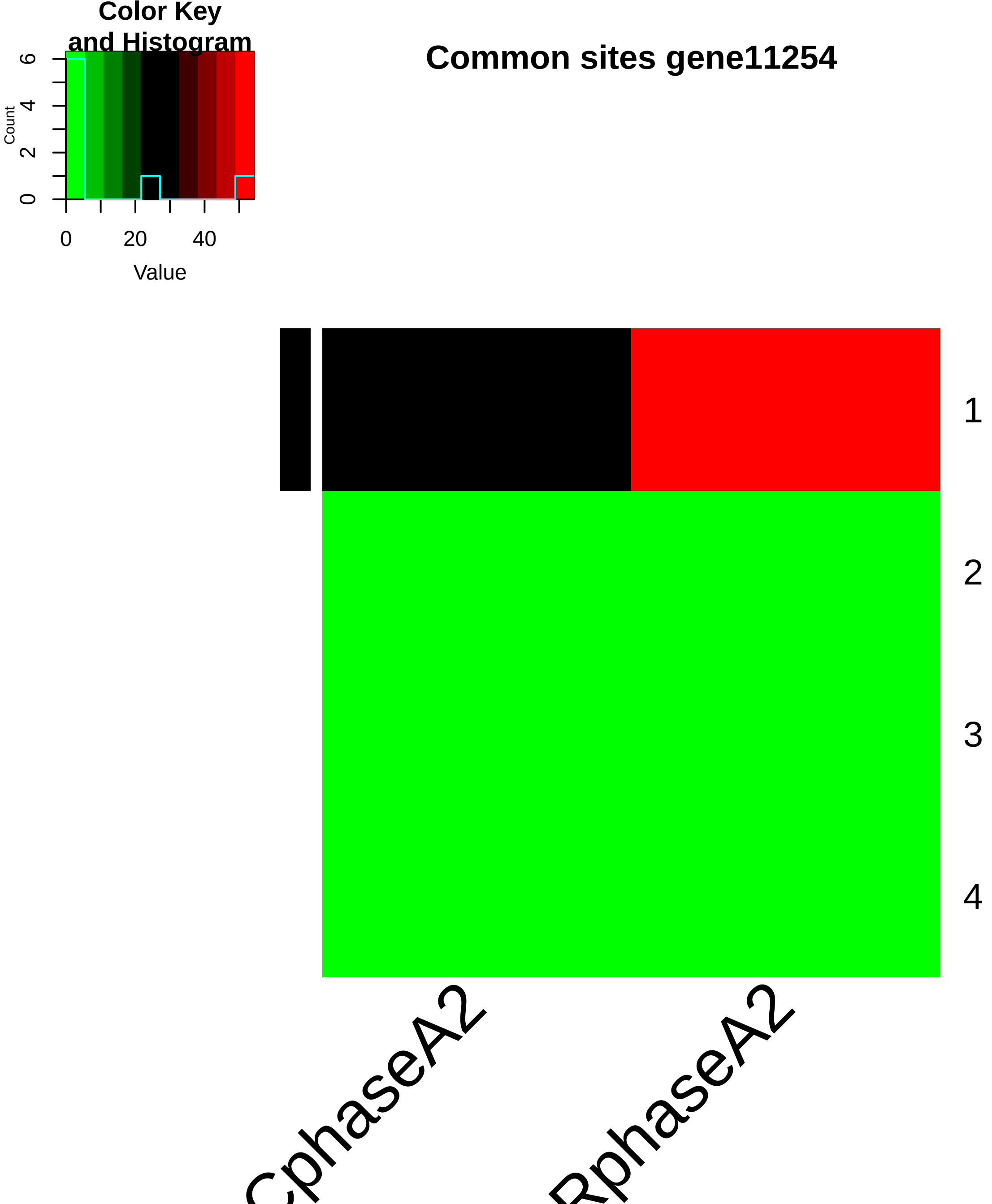
Color Key
and Histogram
Count
0
2
4
6
0
20
40
Value
Common sites gene11254
1
2
3
4
CphaseA2
RphaseA2

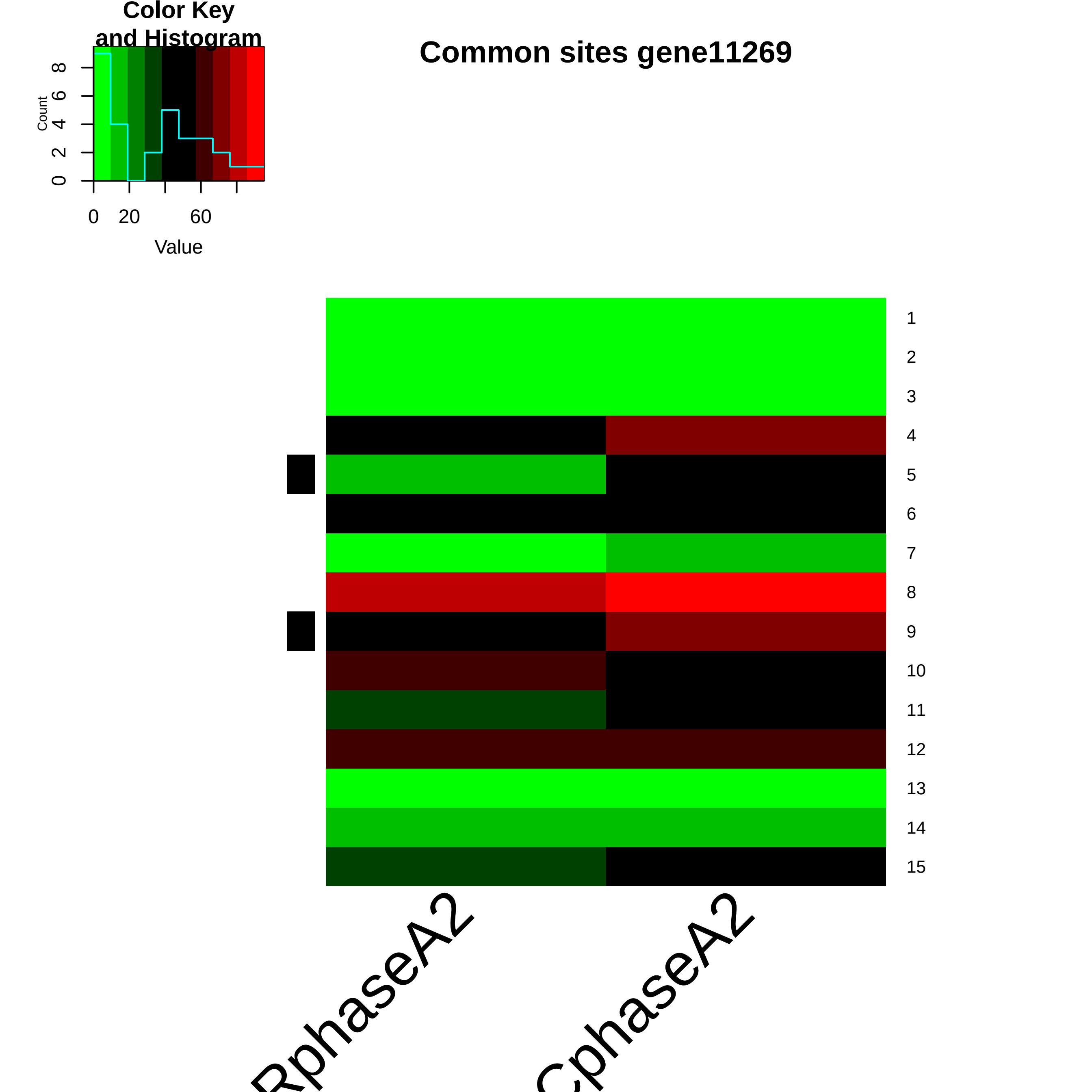

Color Key
and Histogram
Count
0
2
4
6
8
0
20
60
Value
Common sites gene11269
RphaseA2
CphaseA2

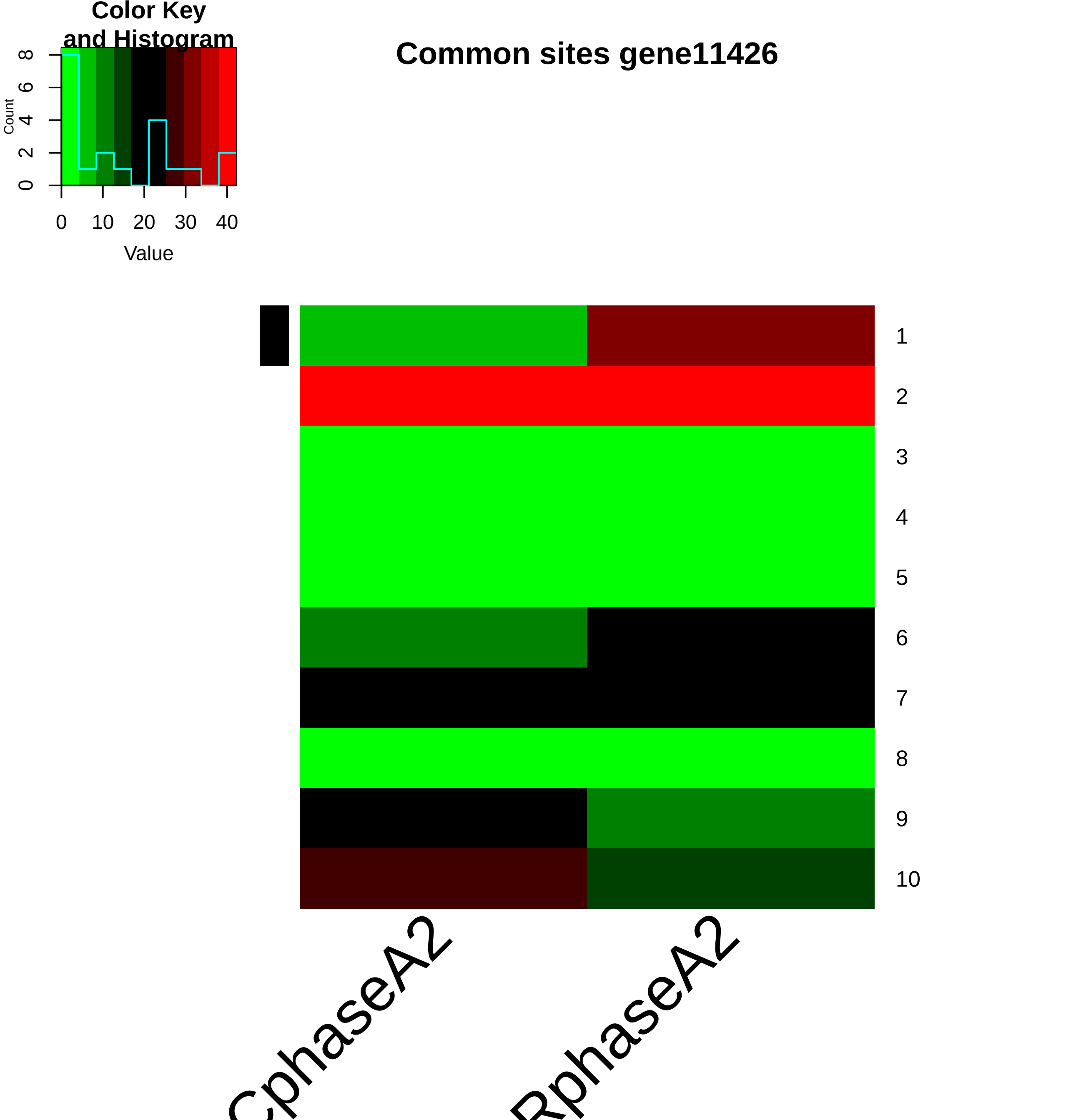

Color Key
and Histogram
Count
0
2
4
6
8
0
10
20
30
40
Value
Common sites gene11426
1
2
3
4
5
6
7
8
9
10
CphaseA2
RphaseA2

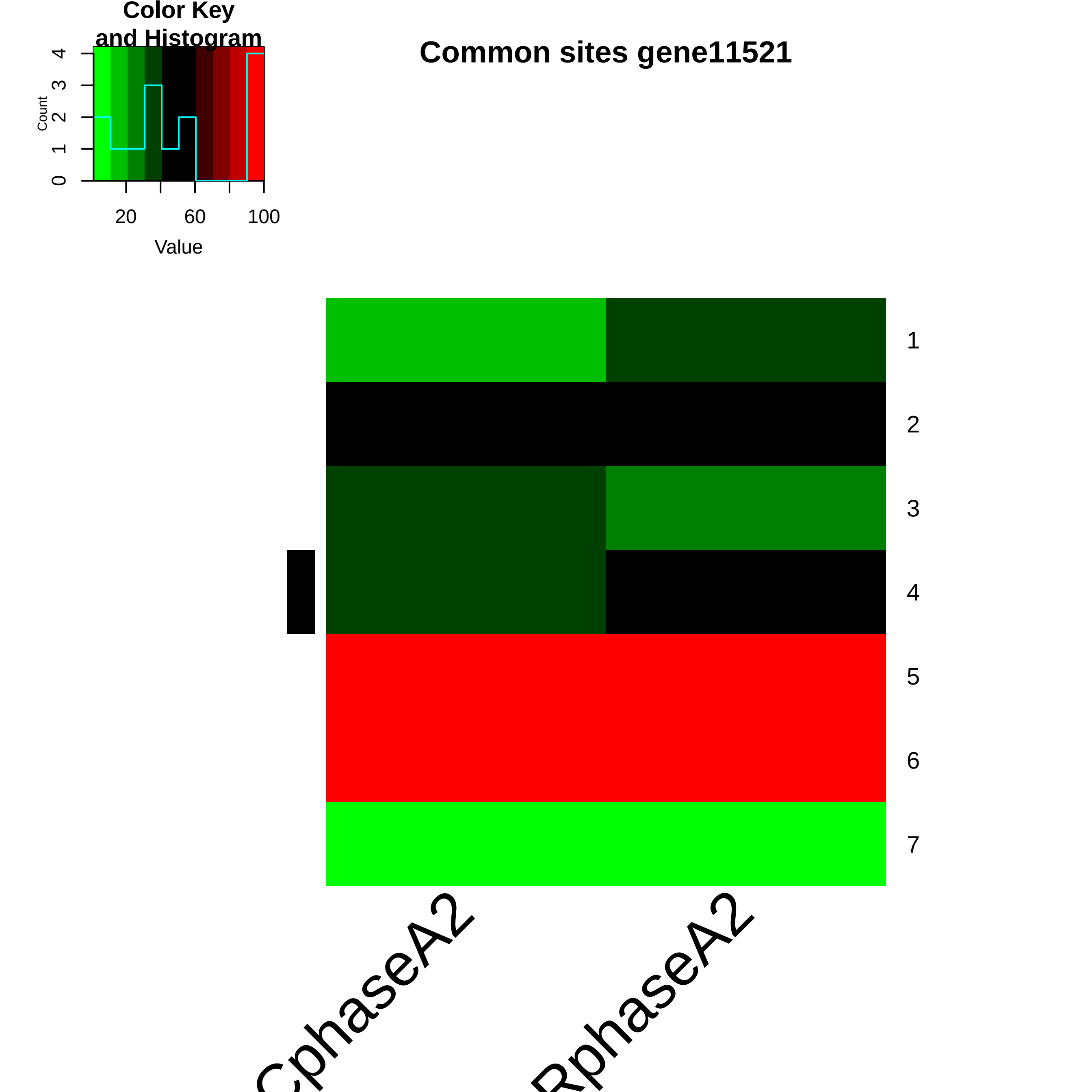

Color Key
and Histogram
Count
0
1
2
3
4
20
60
100
Value
Common sites gene11521
1
2
3
4
5
6
7
CphaseA2
RphaseA2

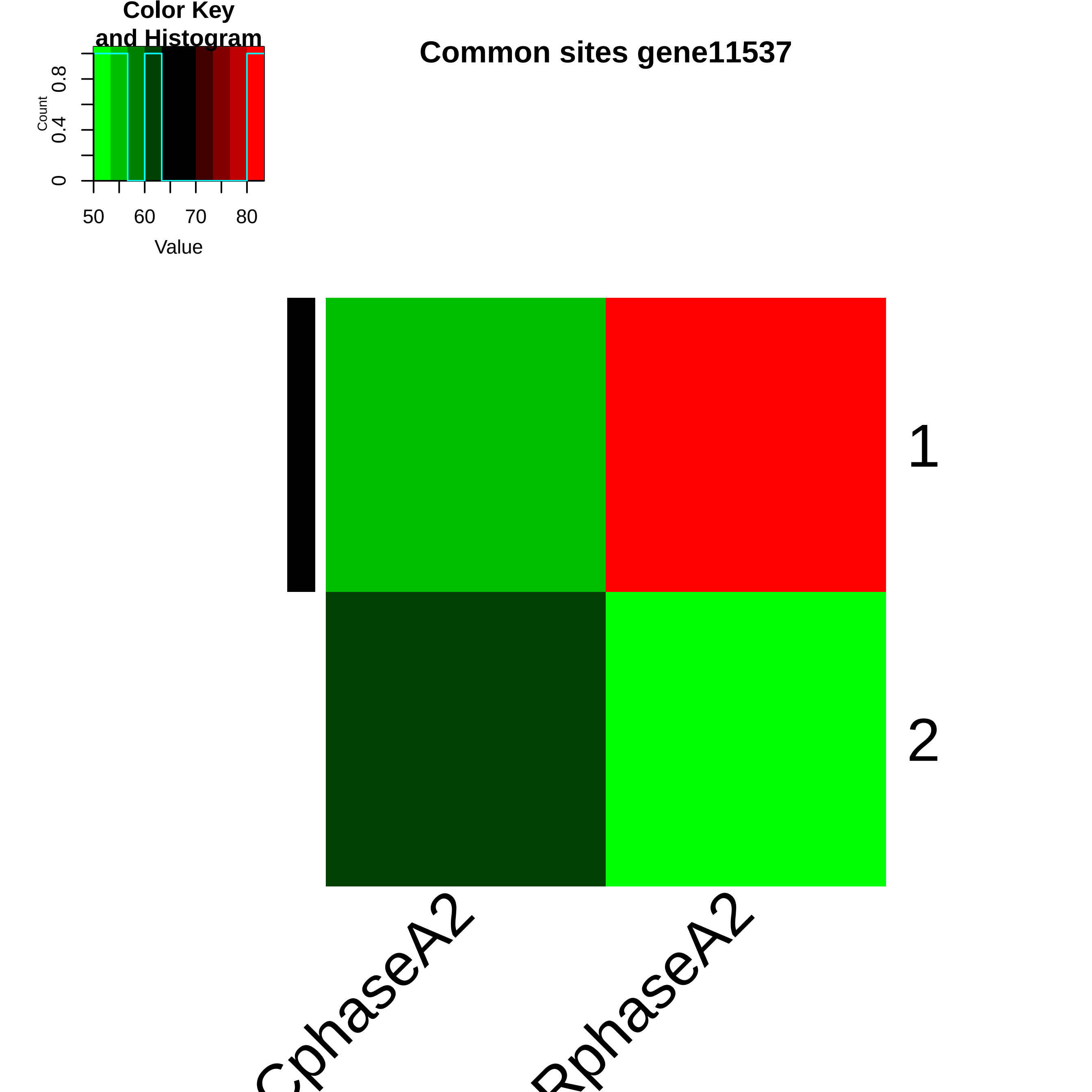

Color Key
and Histogram
Count
0
0.4
0.8
50
60
70
80
Value
Common sites gene11537
1
2
CphaseA2
RphaseA2

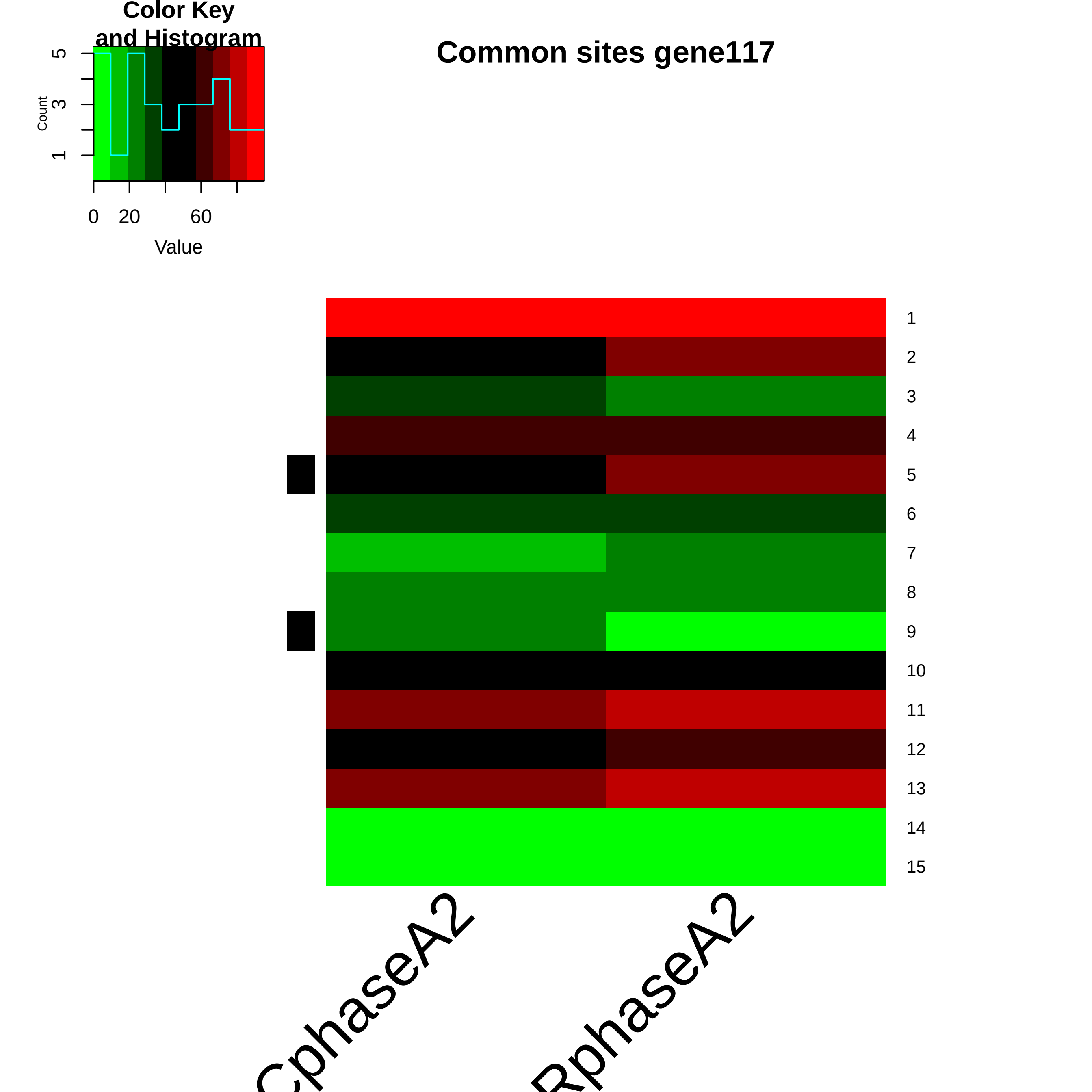

Color Key
and Histogram
Count
1
3
5
0
20
60
Value
Common sites gene117
CphaseA2
RphaseA2

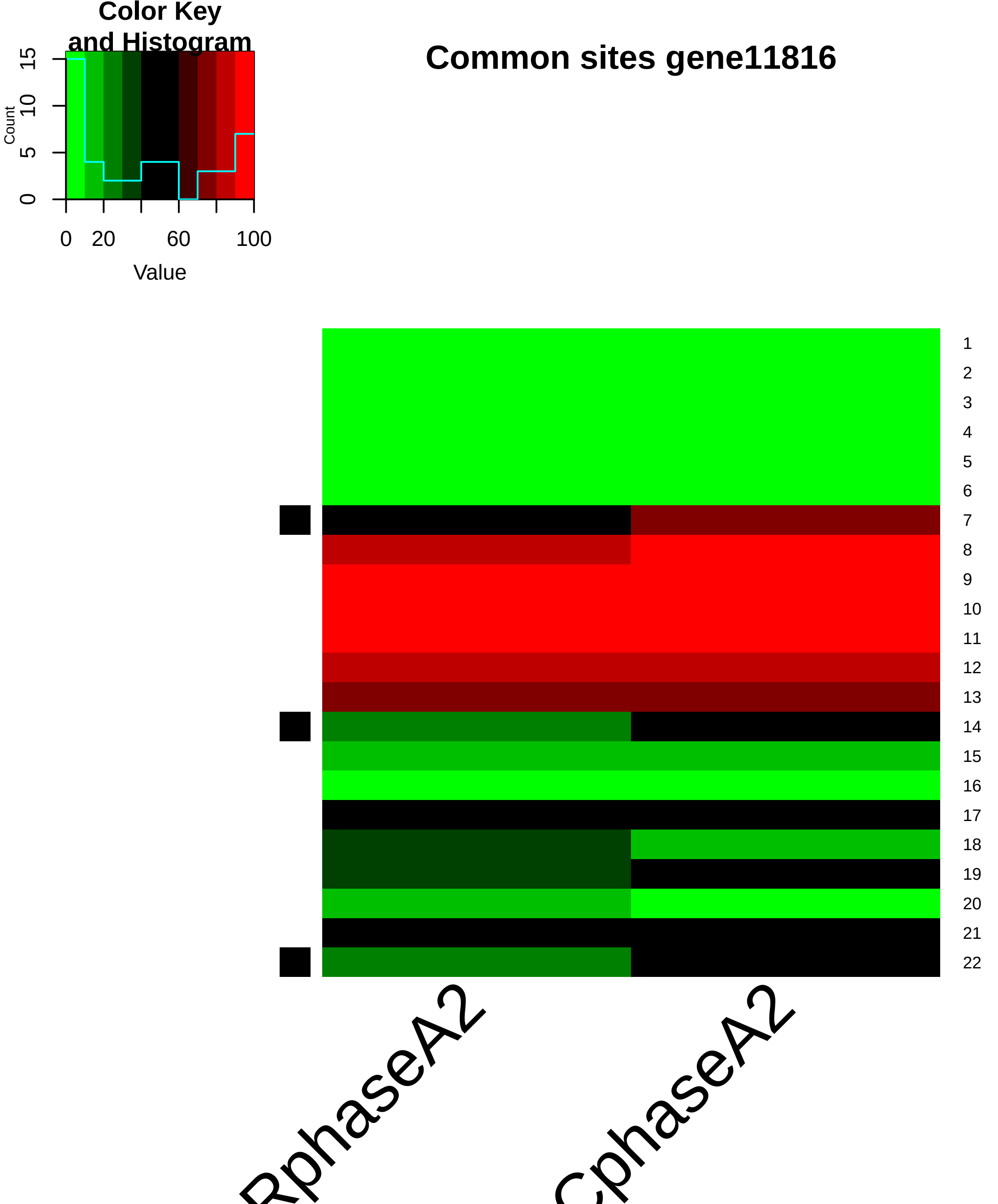
Color Key
and Histogram
Count
0
5
10
15
0
20
60
100
Value
Common sites gene11816
RphaseA2
CphaseA2

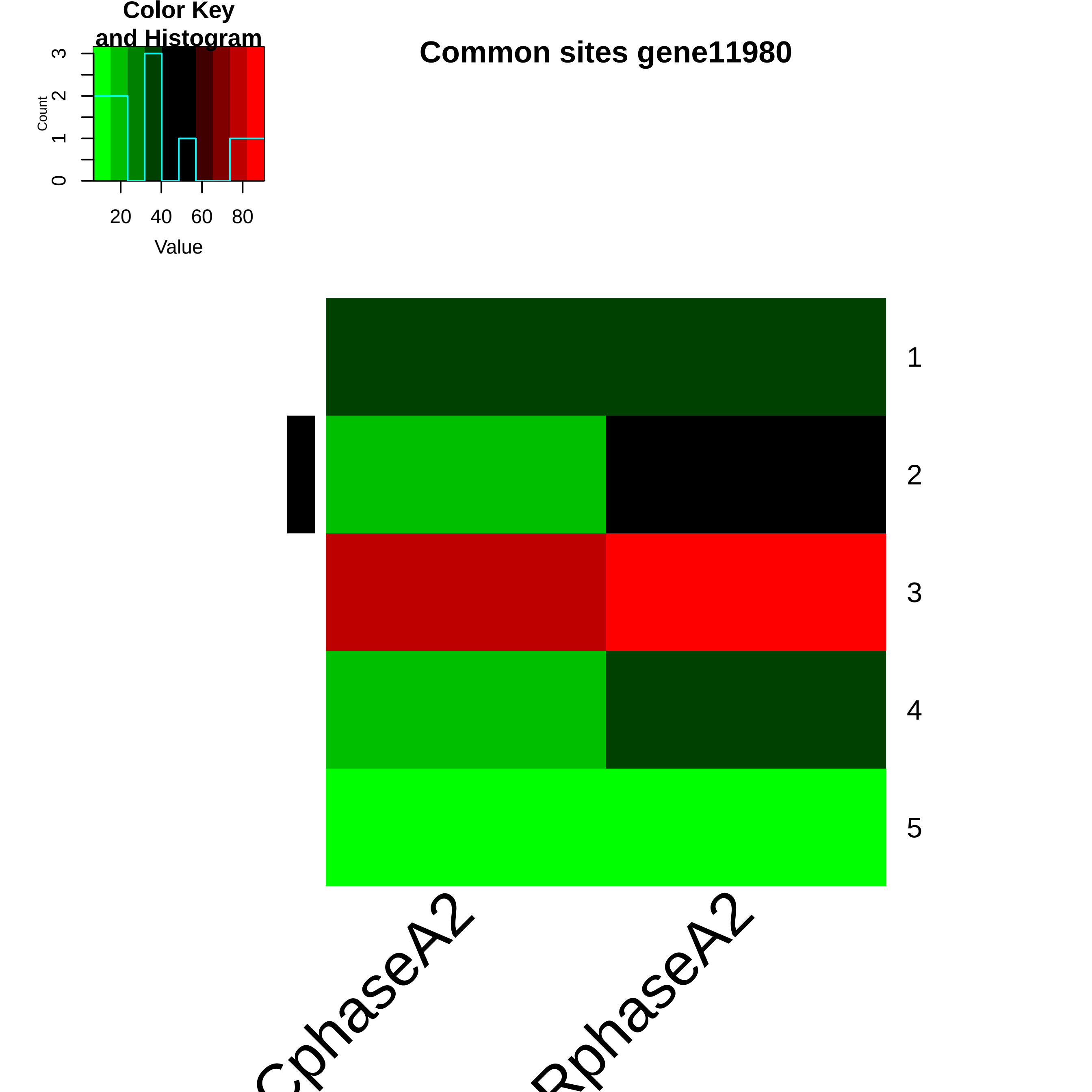

Color Key
and Histogram
Count
0
1
2
3
20
40
60
80
Value
Common sites gene11980
1
2
3
4
5
CphaseA2
RphaseA2

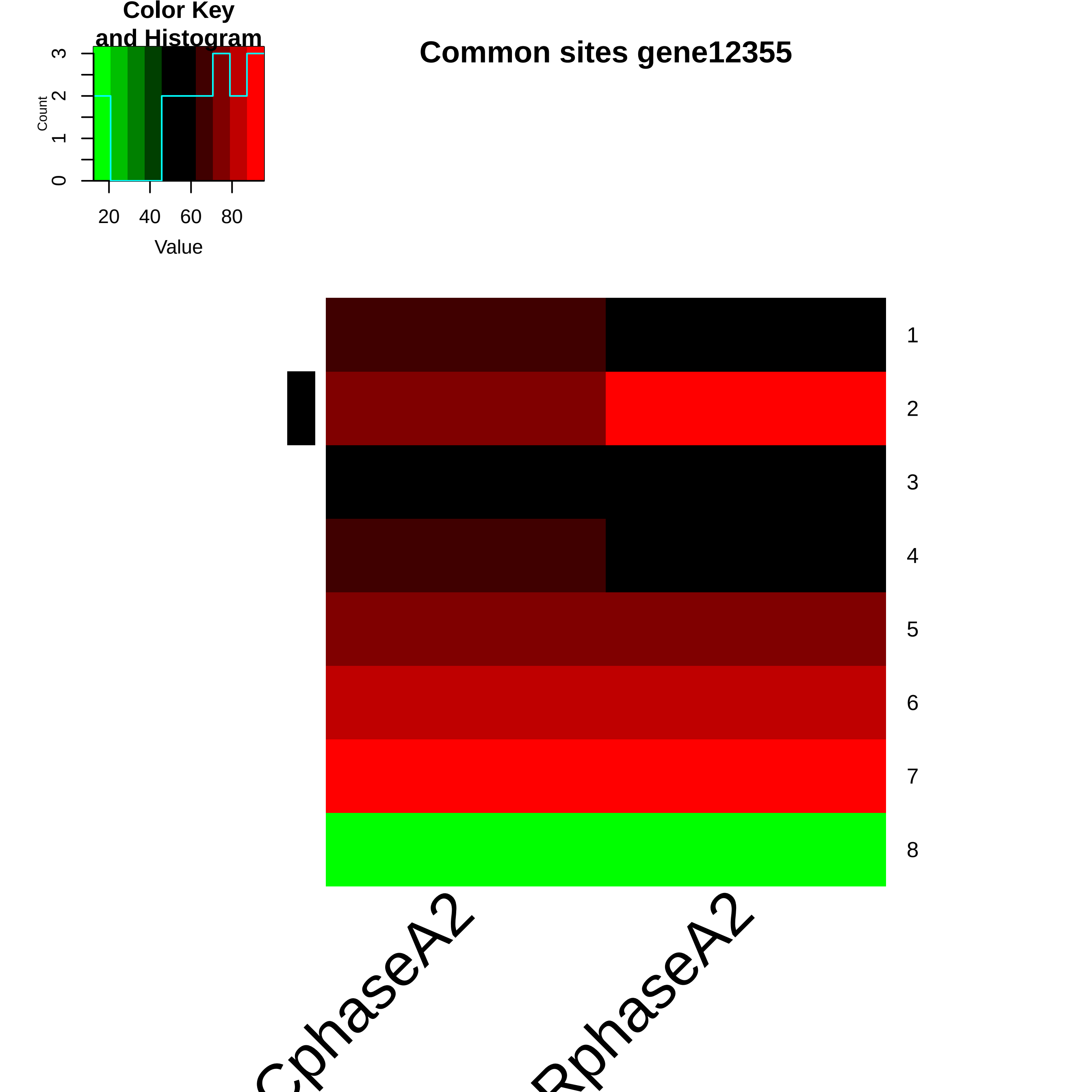
Color Key
and Histogram
Count
0
1
2
3
20
40
60
80
Value
Common sites gene12355
1
2
3
4
5
6
7
8
CphaseA2
RphaseA2

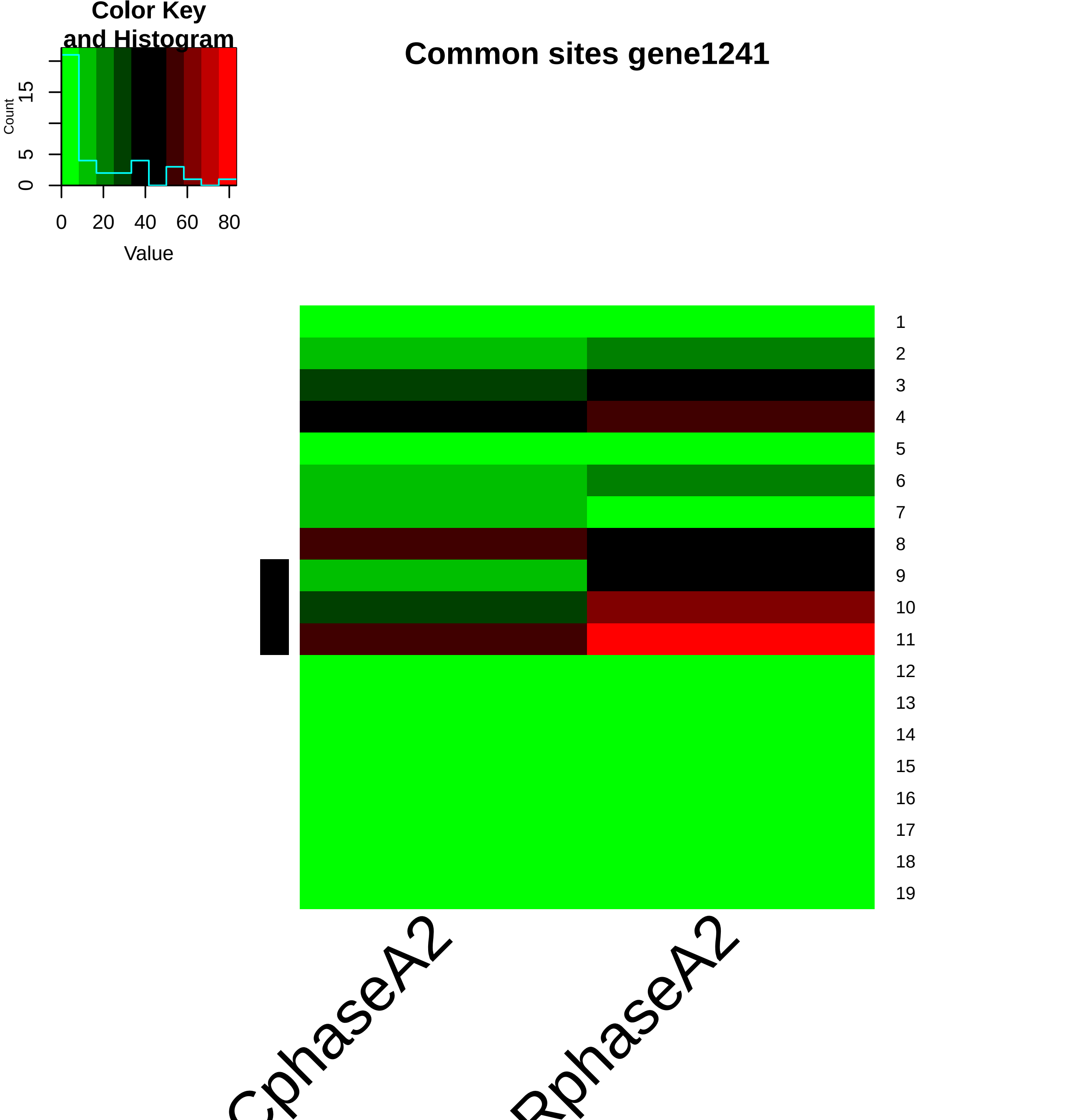
Color Key
and Histogram
Count
0
5
15
0
20
40
60
80
Value
Common sites gene1241
CphaseA2
RphaseA2

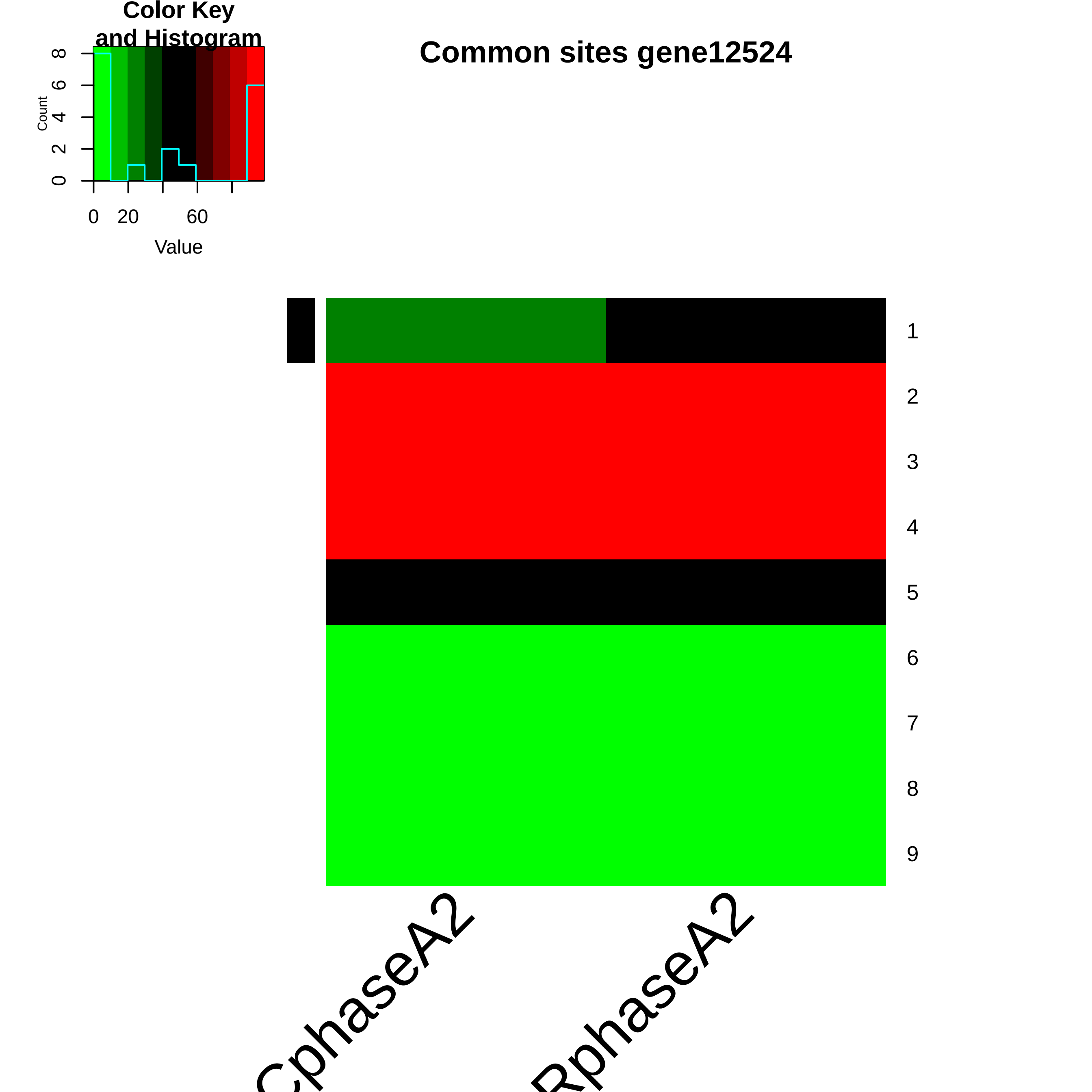
Color Key
and Histogram
Count
0
2
4
6
8
0
20
60
Value
Common sites gene12524
1
2
3
4
5
6
7
8
9
CphaseA2
RphaseA2

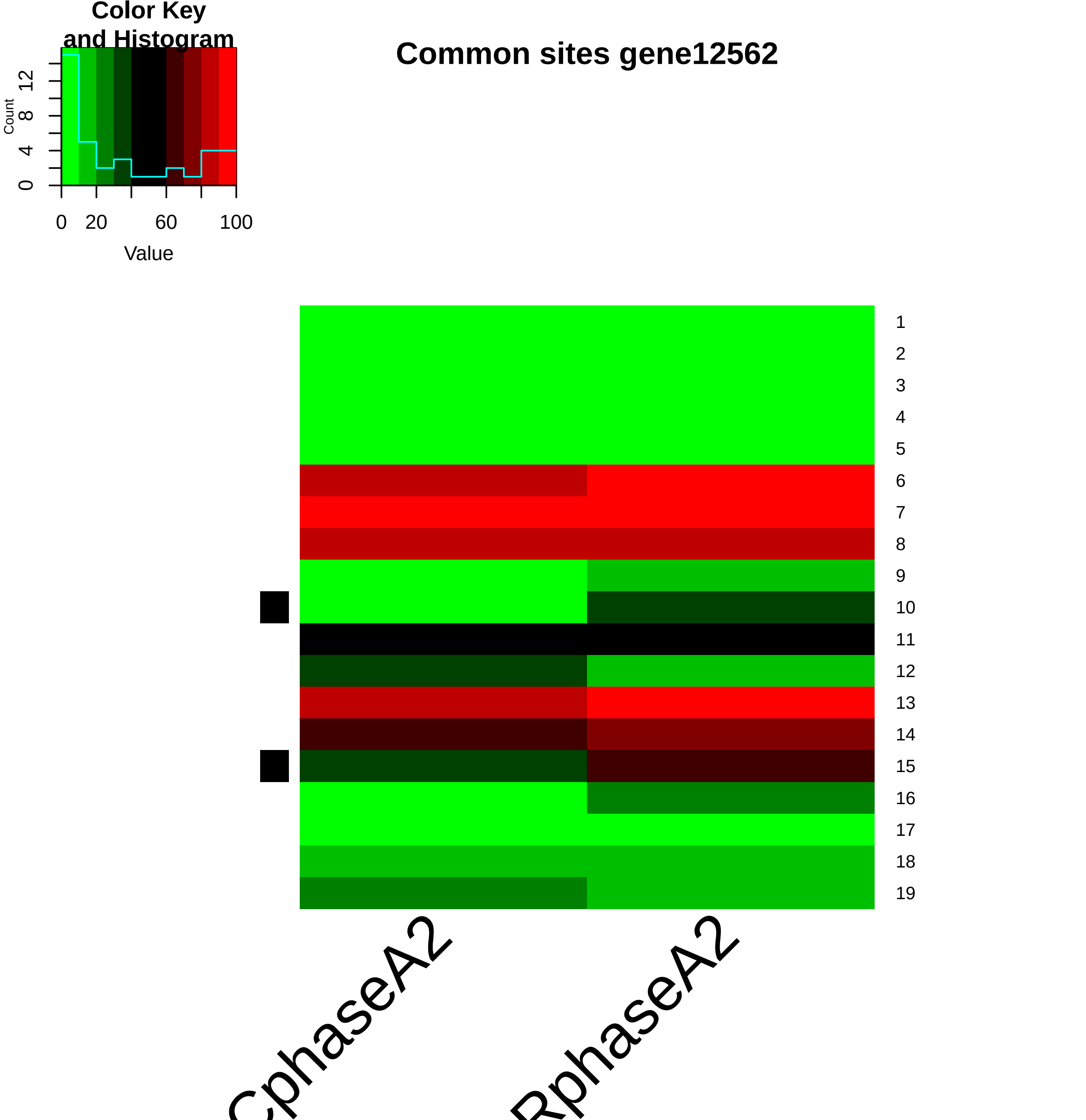

Color Key
and Histogram
Count
0
4
8
12
0
20
60
100
Value
Common sites gene12562
CphaseA2
RphaseA2

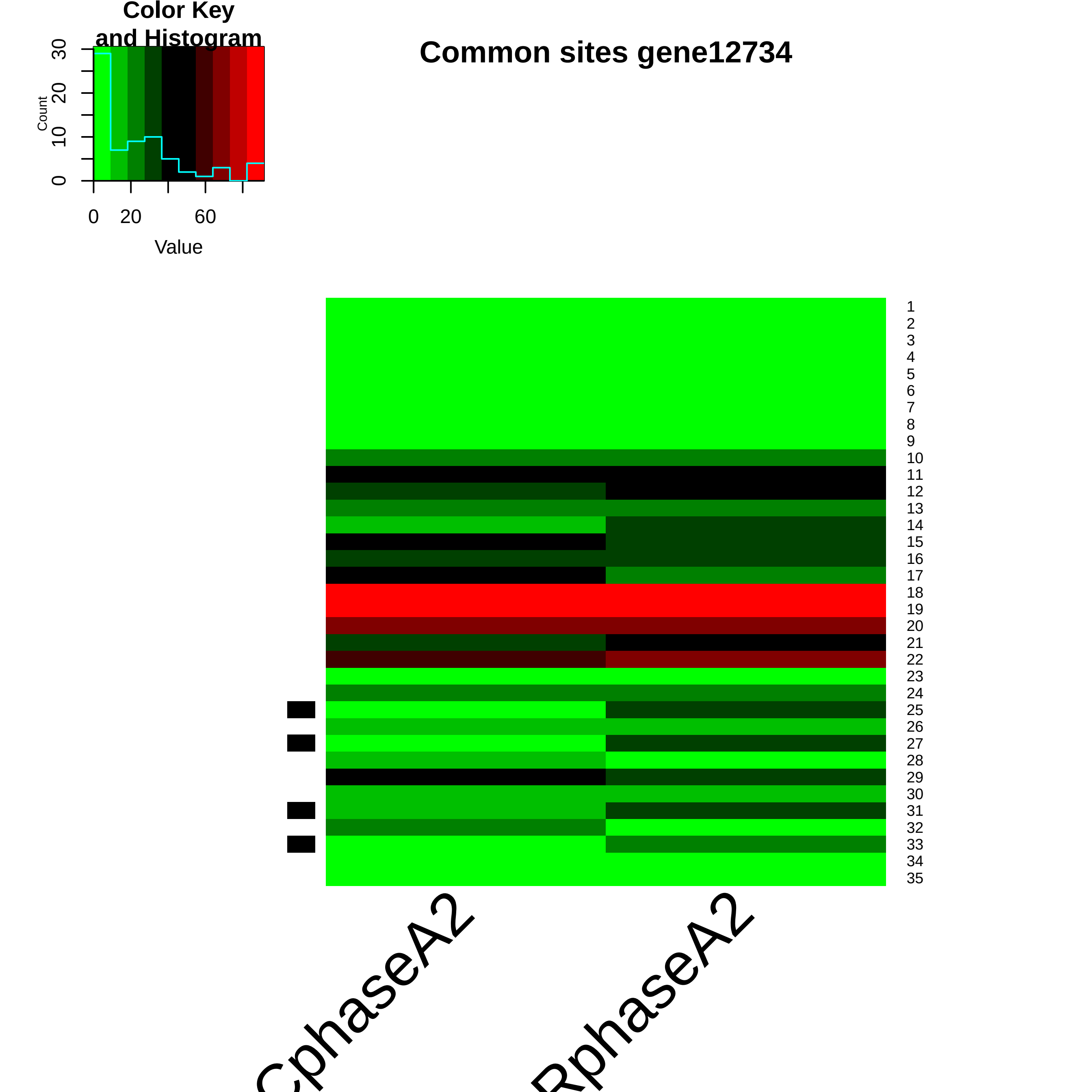
Color Key
and Histogram
Count
0
10
20
30
0
20
60
Value
Common sites gene12734
CphaseA2
RphaseA2

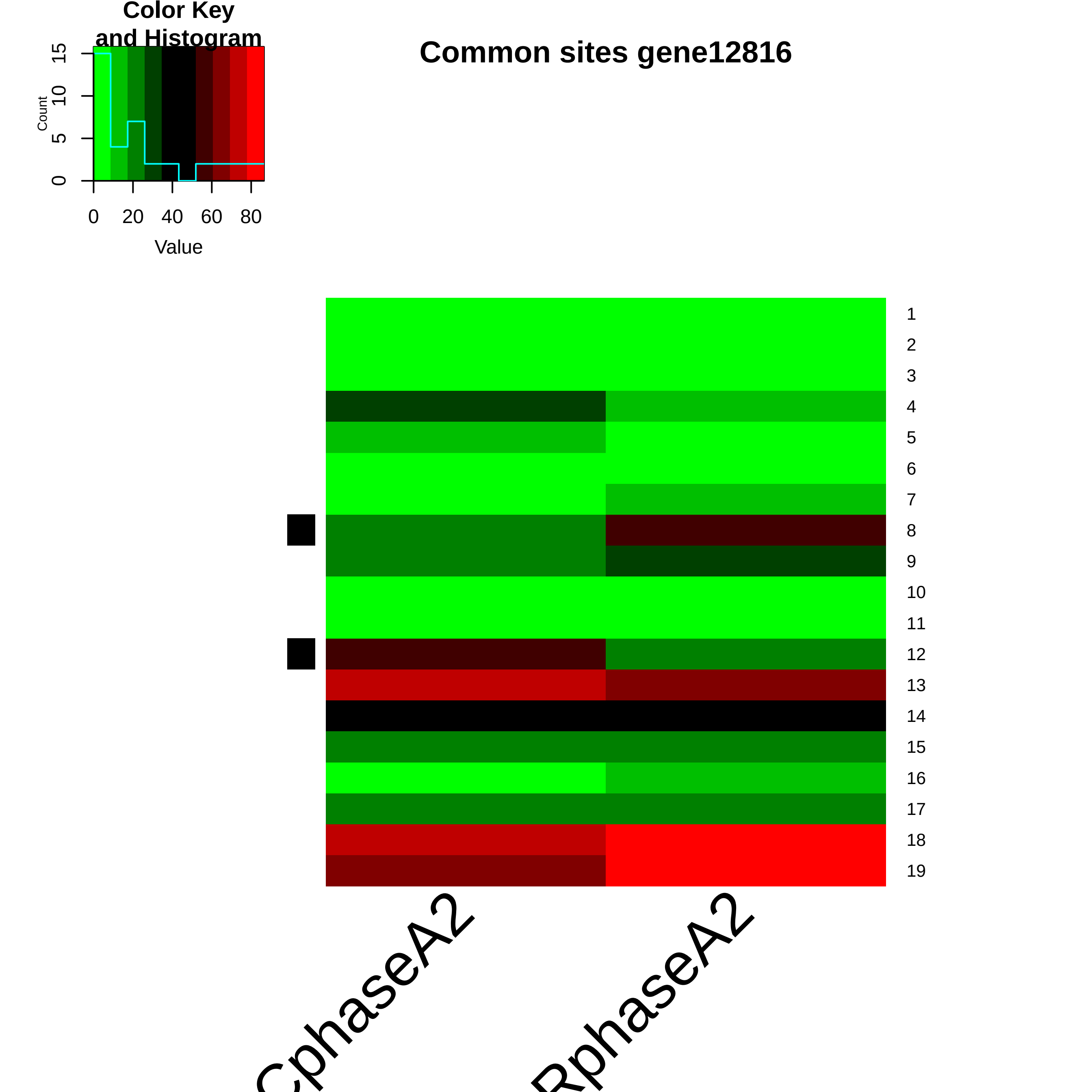

Color Key
and Histogram
Count
0
5
10
15
0
20
40
60
80
Value
Common sites gene12816
CphaseA2
RphaseA2

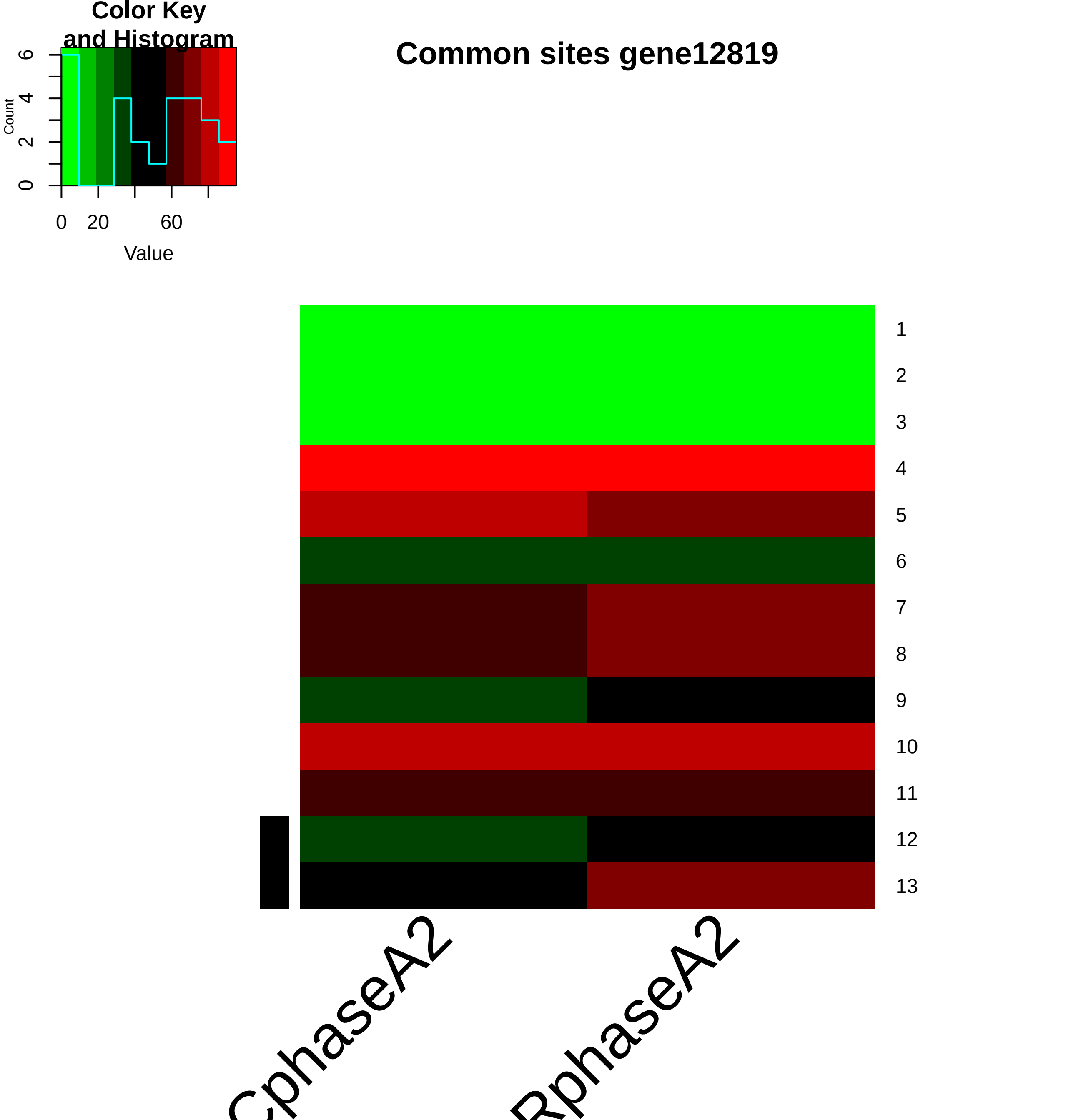

Color Key
and Histogram
Count
0
2
4
6
0
20
60
Value
Common sites gene12819
1
2
3
4
5
6
7
8
9
10
11
12
13
CphaseA2
RphaseA2

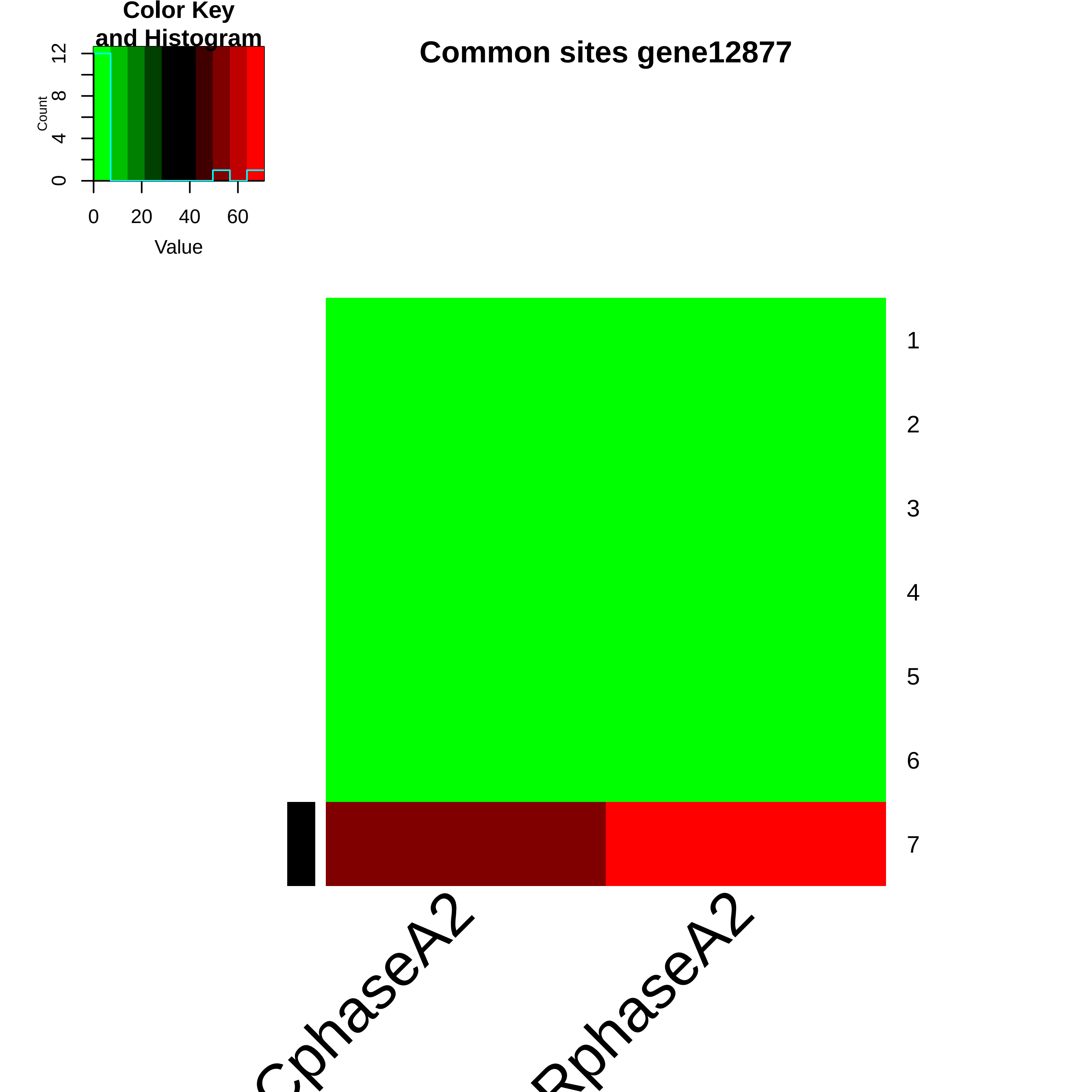

Color Key
and Histogram
Count
0
4
8
12
0
20
40
60
Value
Common sites gene12877
1
2
3
4
5
6
7

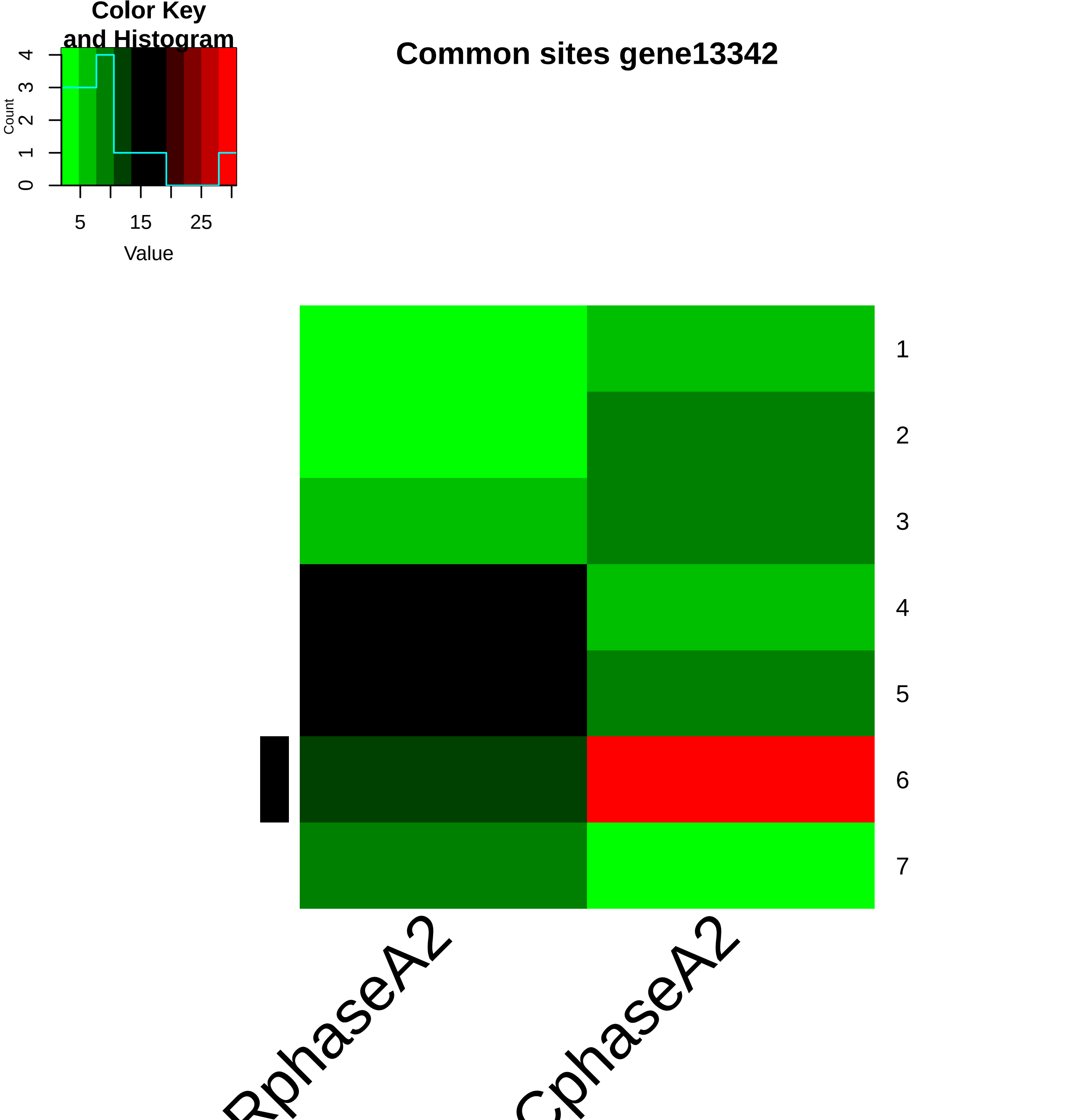
Color Key
and Histogram
Count
0
1
2
3
4
5
15
25
Value
Common sites gene13342
1
2
3
4
5
6
7
RphaseA2
CphaseA2

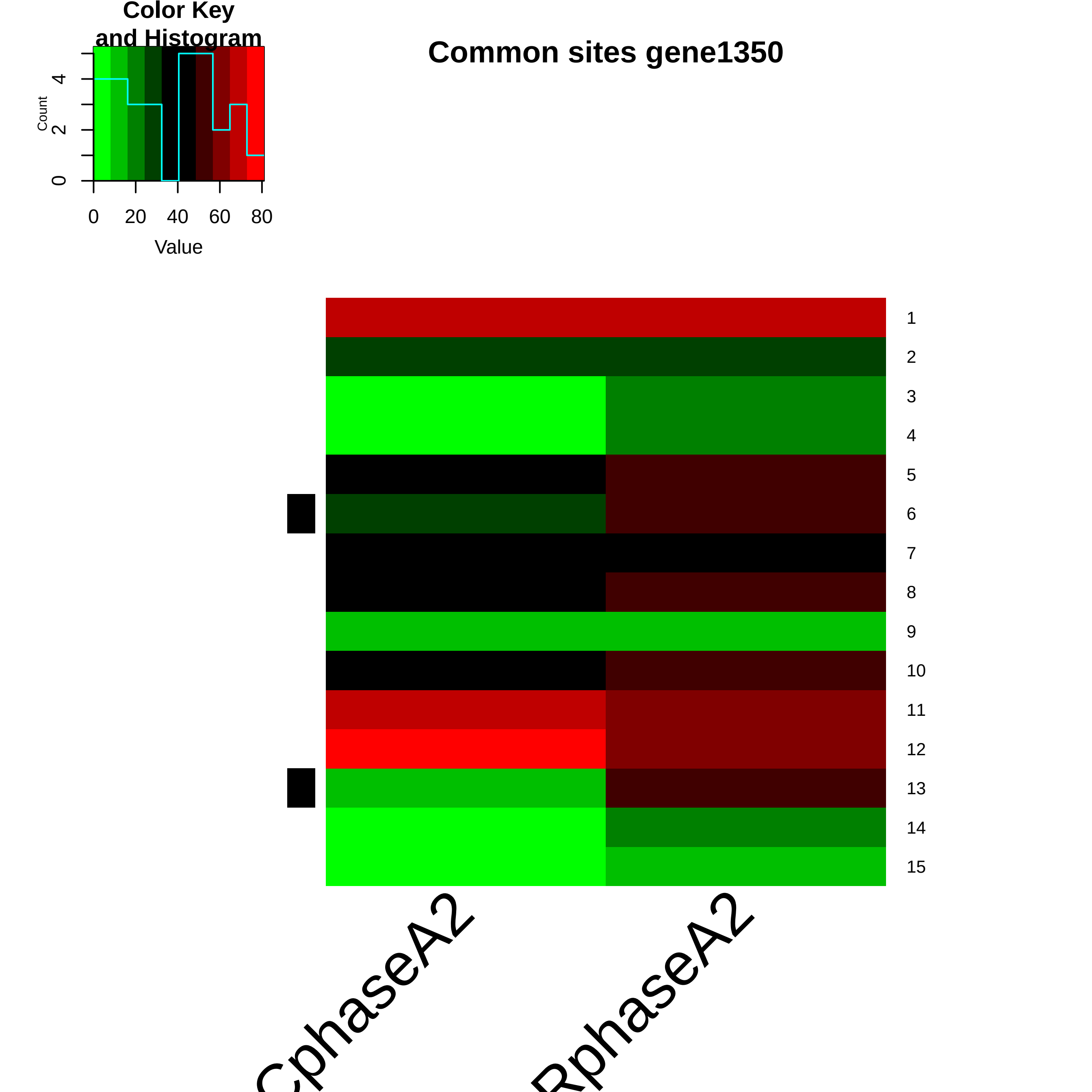

Color Key and Histogram
Count
0
2
4
0
20
40
60
80
Value
Common sites gene1350
CphaseA2
RphaseA2

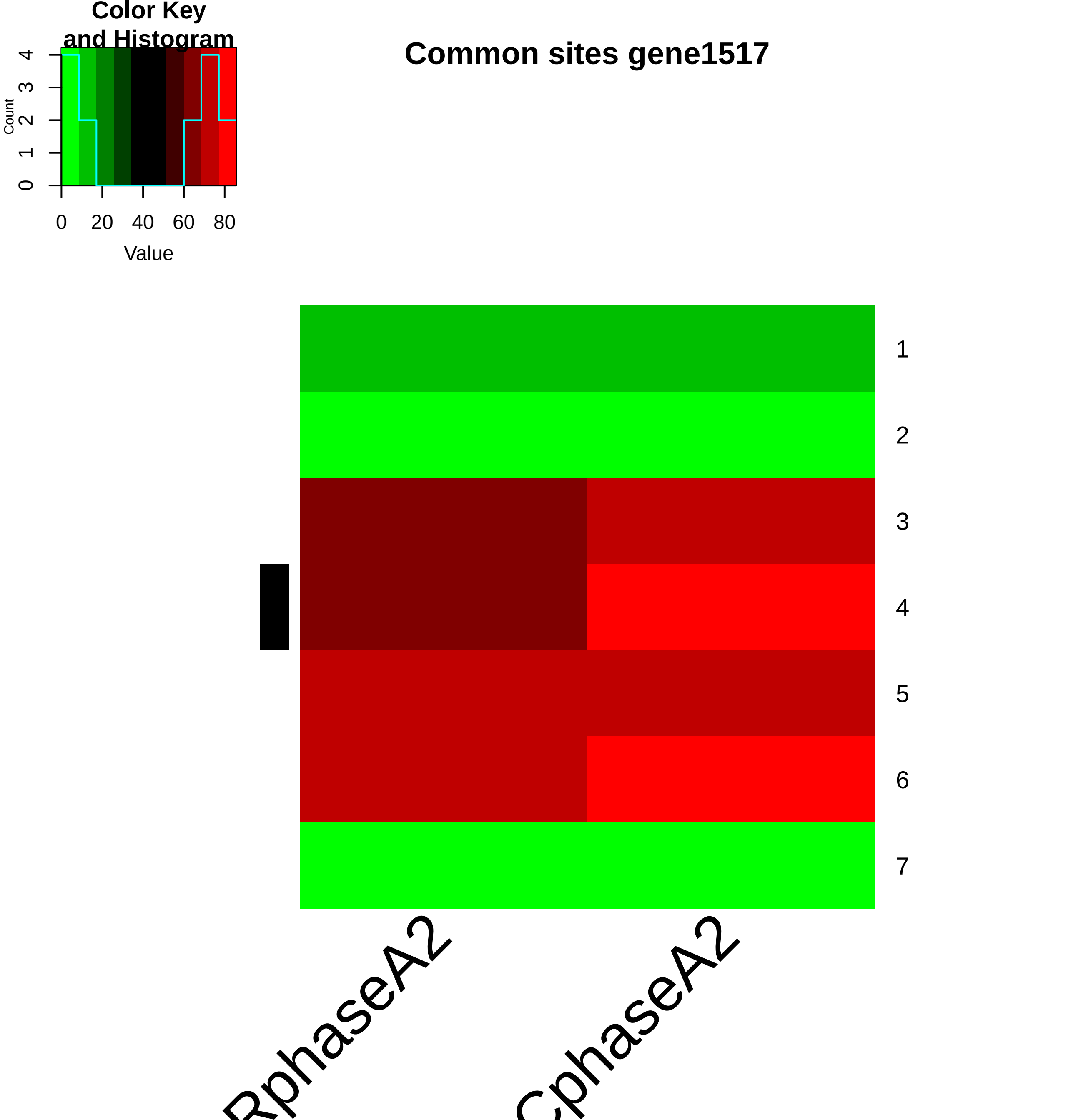
Color Key
and Histogram
Count
0
1
2
3
4
0
20
40
60
80
Value
Common sites gene1517
1
2
3
4
5
6
7
RphaseA2
CphaseA2

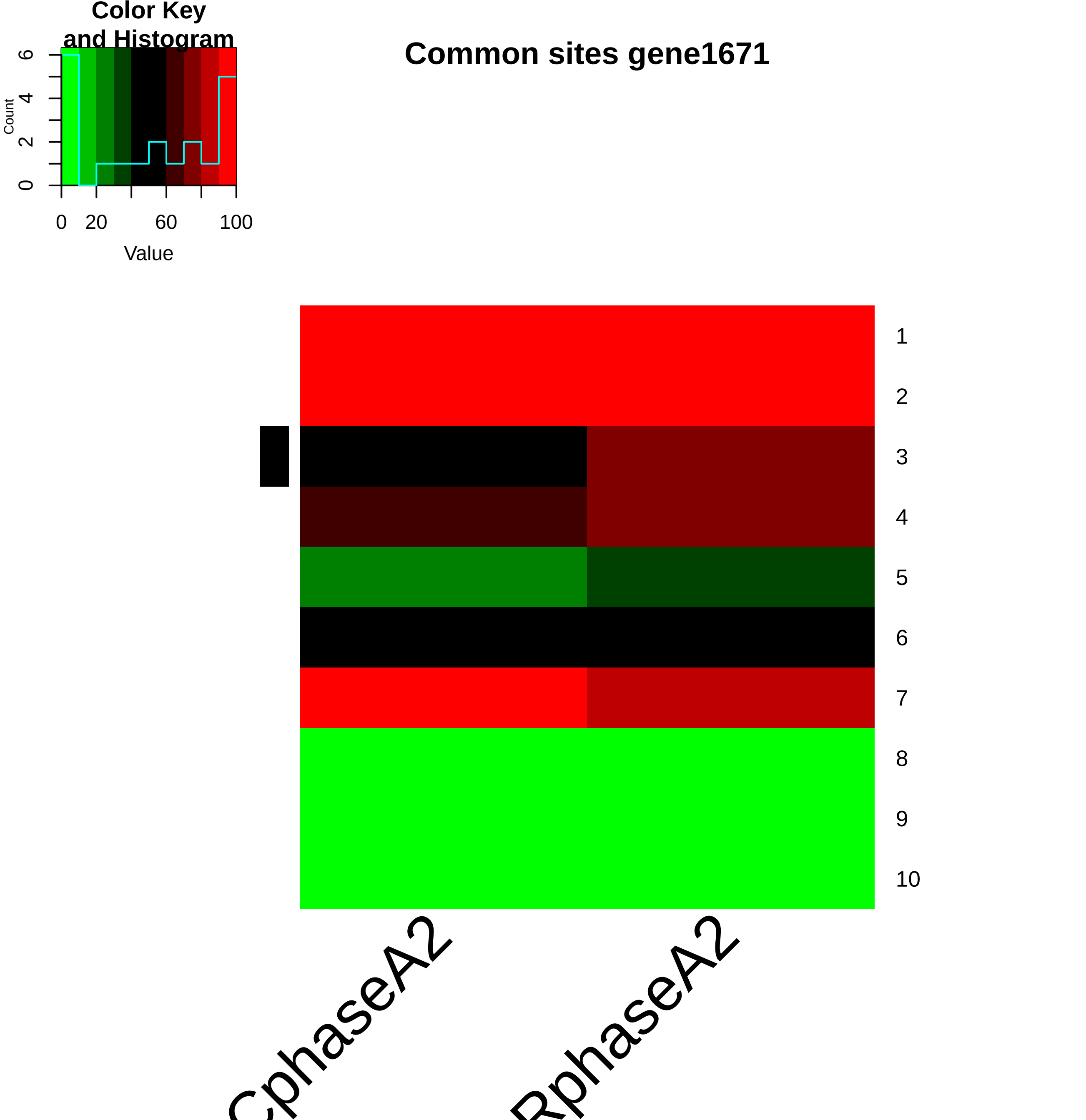

Color Key
and Histogram
Count
0
2
4
6
0
20
60
100
Value
Common sites gene1671
1
2
3
4
5
6
7
8
9
10
CphaseA2
RphaseA2

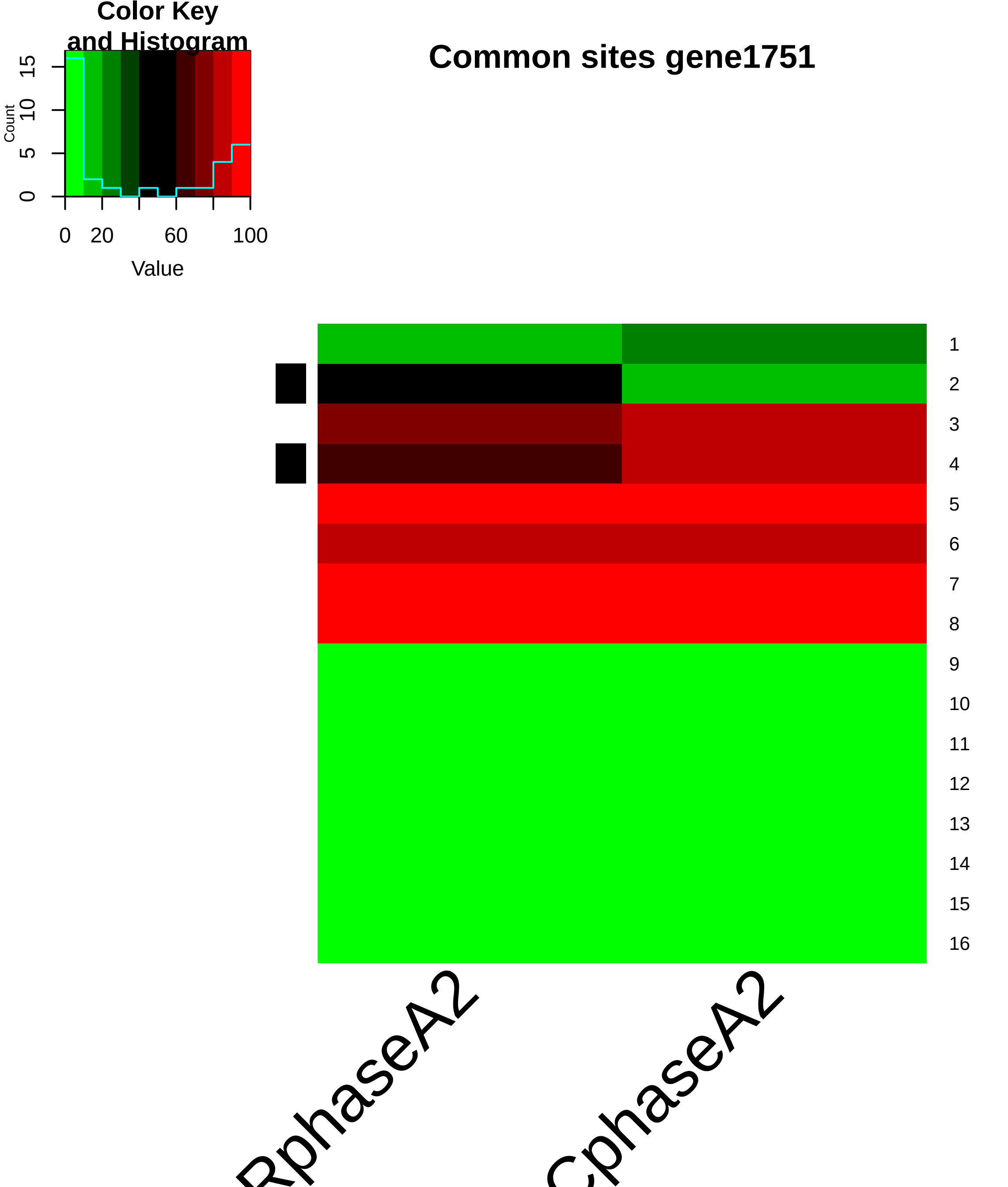

Color Key
and Histogram
Count
0
5
10
15
0
20
60
100
Value
Common sites gene1751
RphaseA2
CphaseA2

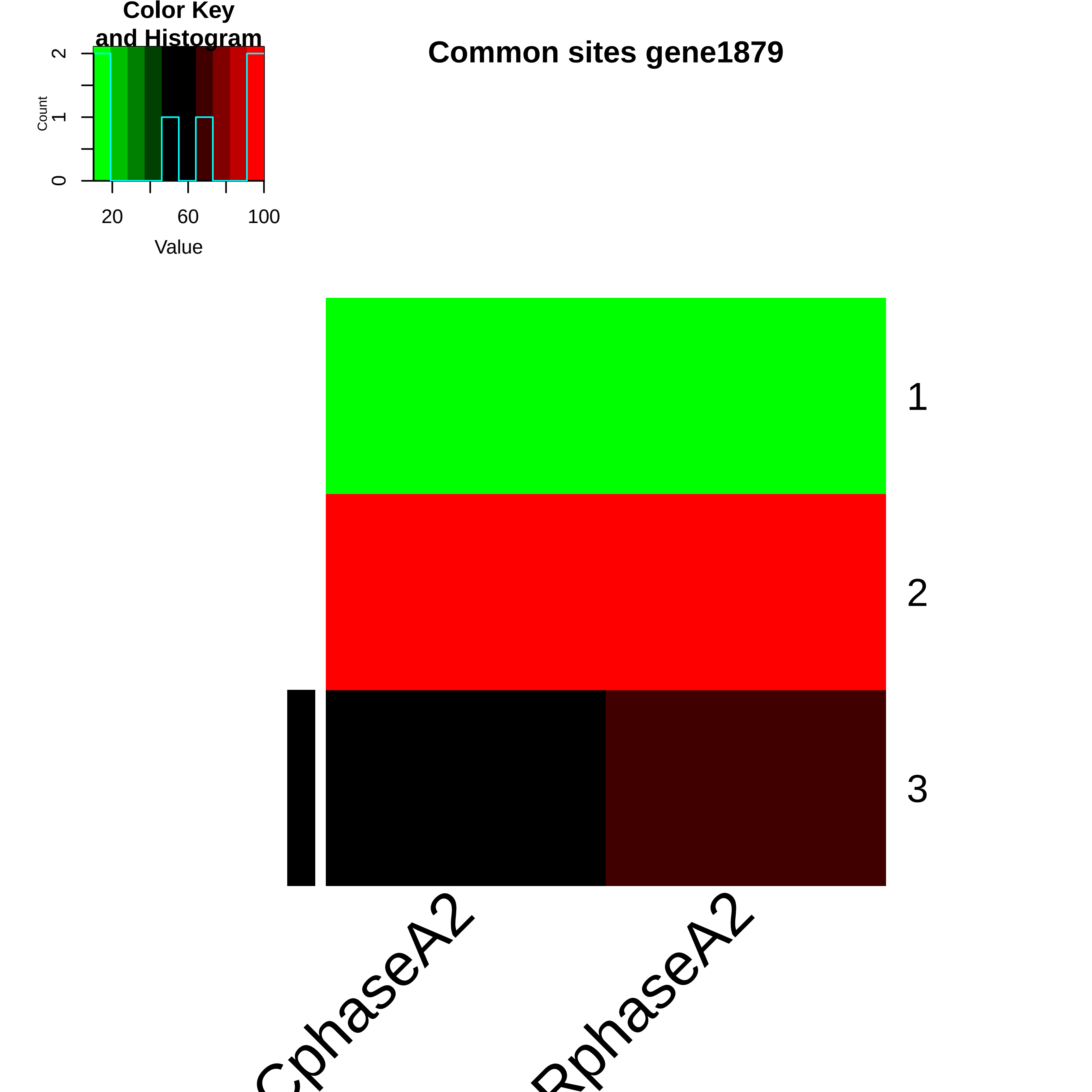

Color Key
and Histogram
Count
0
1
2
20
60
100
Value
Common sites gene1879
1
2
3
CphaseA2
RphaseA2

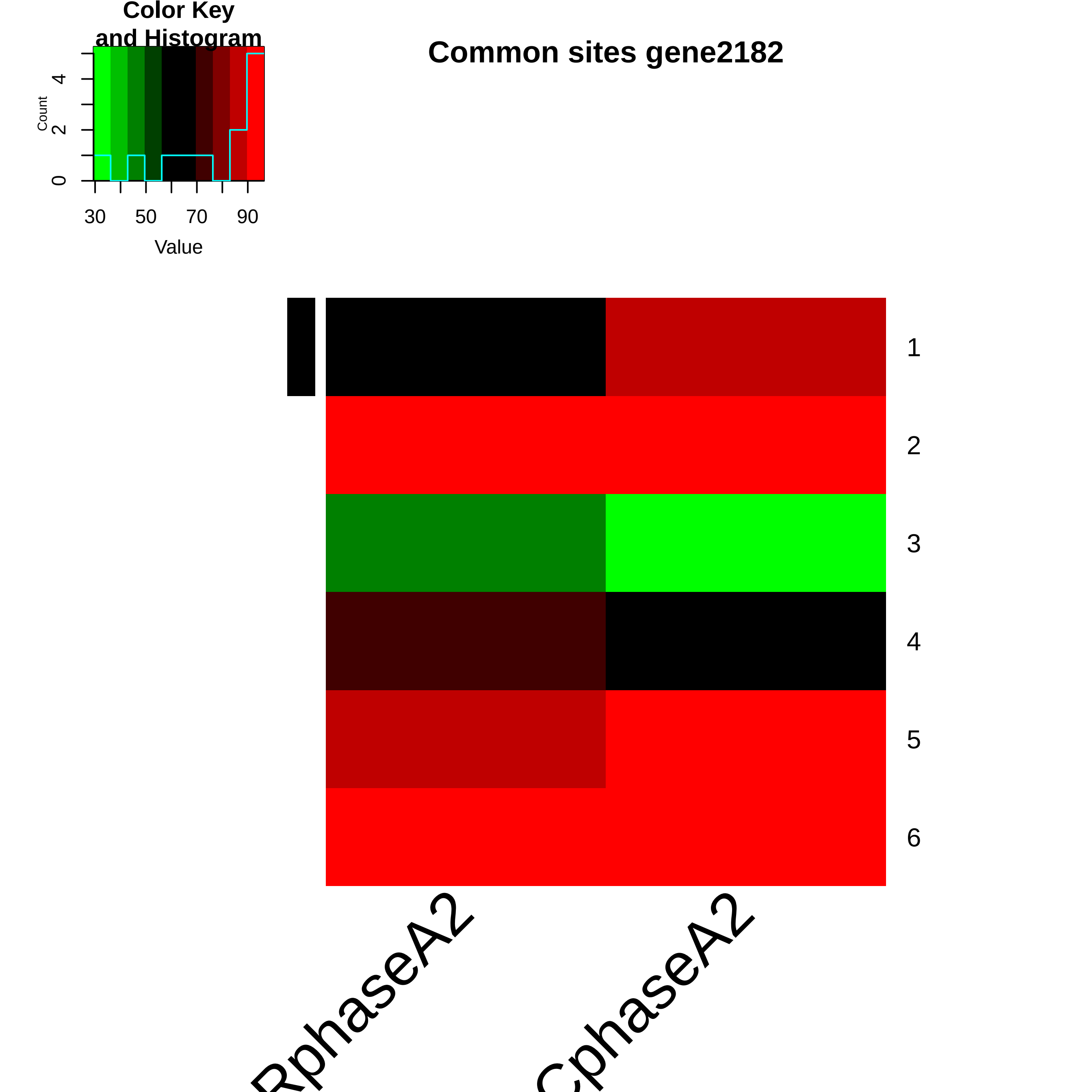

Color Key
and Histogram
Count
0
2
4
30
50
70
90
Value
Common sites gene2182
1
2
3
4
5
6
RphaseA2
CphaseA2

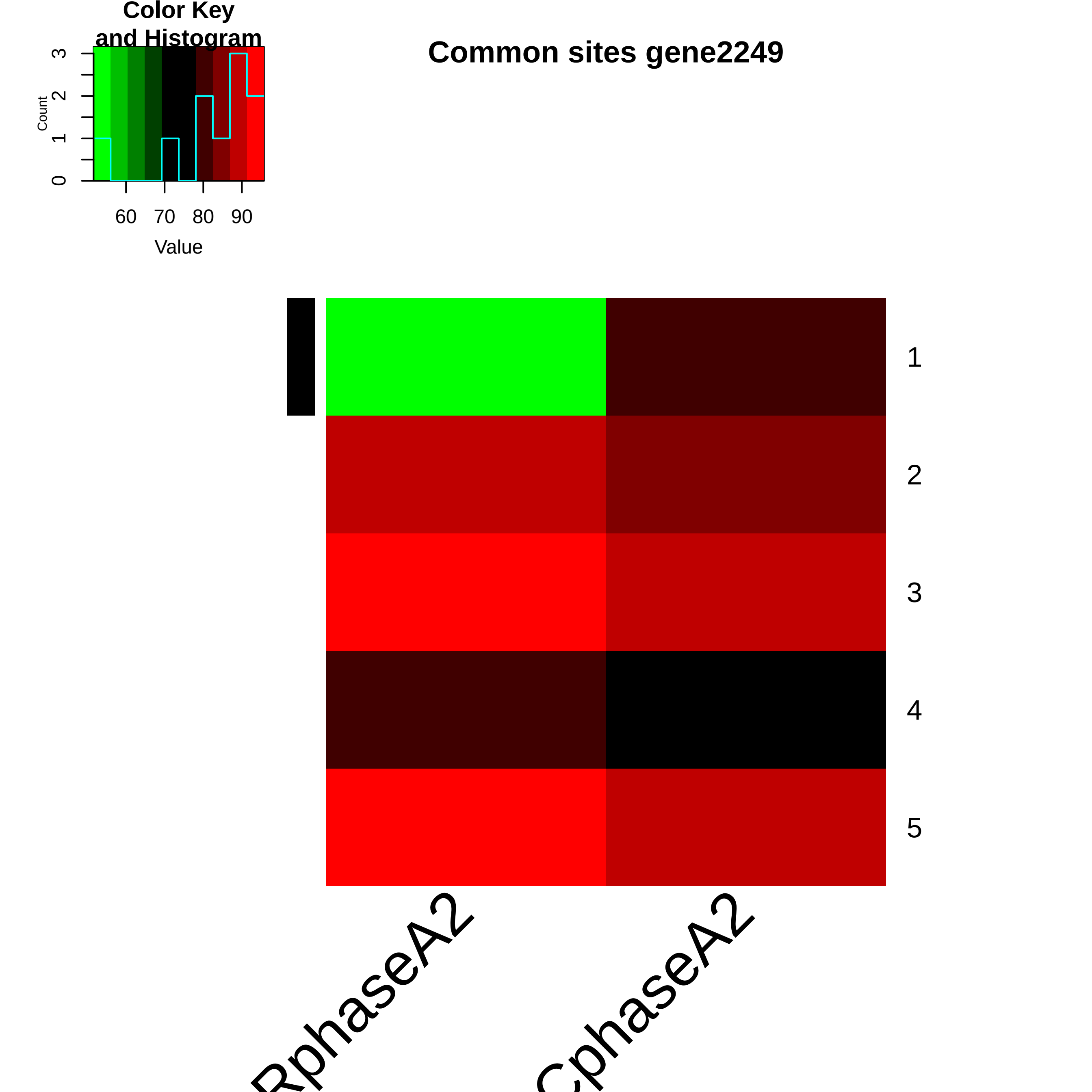
Color Key
and Histogram
Count
0
1
2
3
60
70
80
90
Value
Common sites gene2249
1
2
3
4
5
RphaseA2
CphaseA2

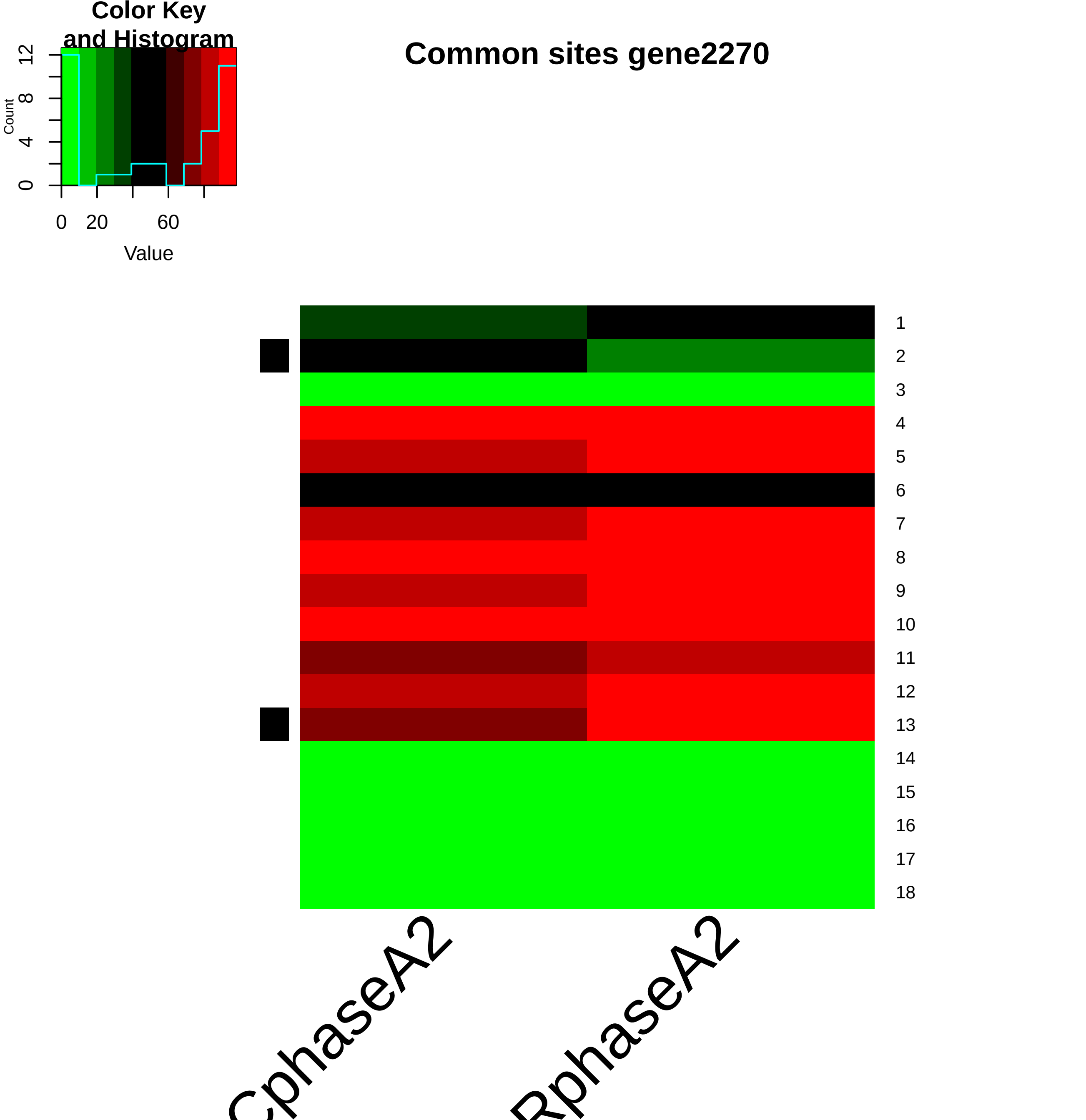
Color Key
and Histogram
Count
0
4
8
12
0
20
60
Value
Common sites gene2270
CphaseA2
RphaseA2

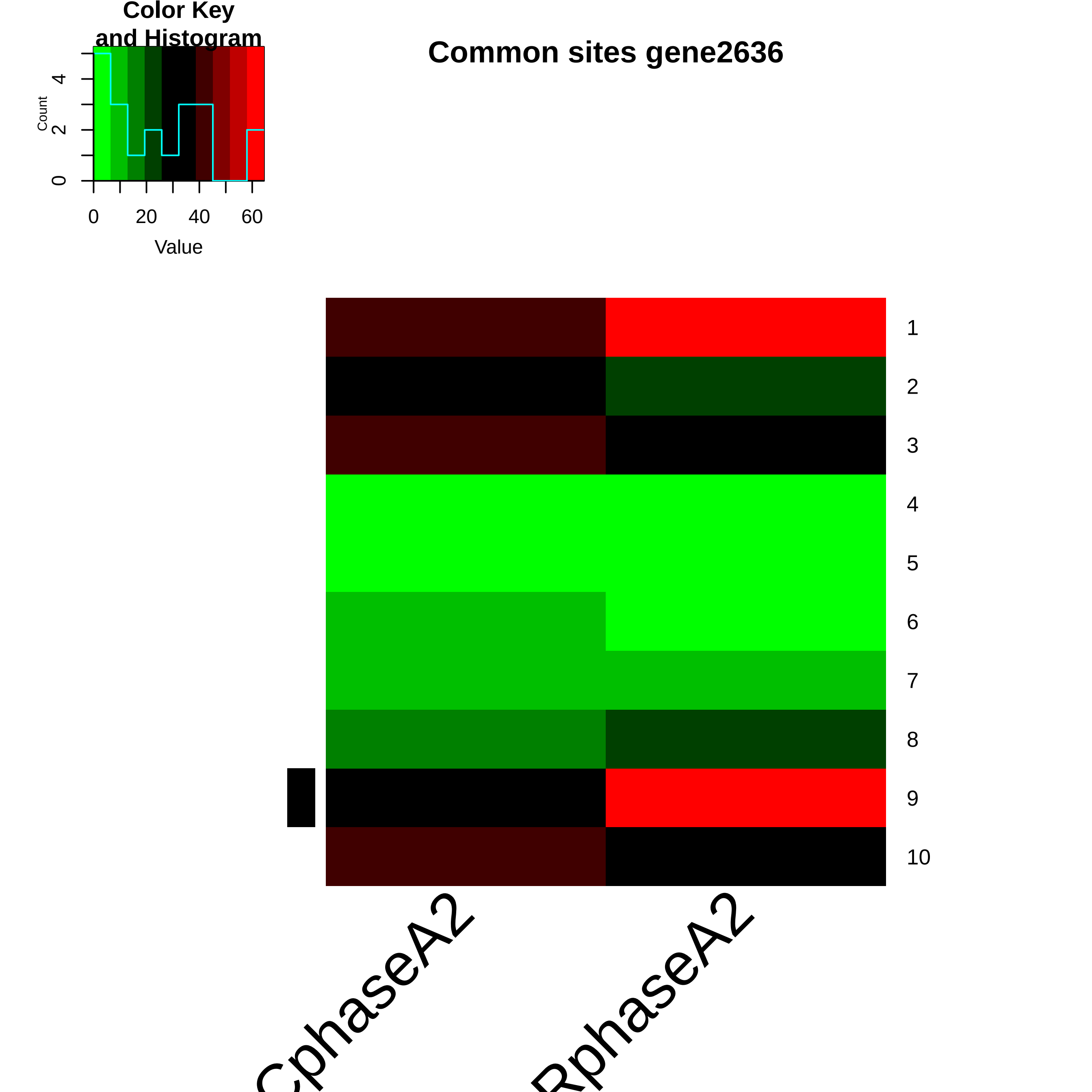

Color Key
and Histogram
Count
0
2
4
0
20
40
60
Value
Common sites gene2636
1
2
3
4
5
6
7
8
9
10
CphaseA2
RphaseA2

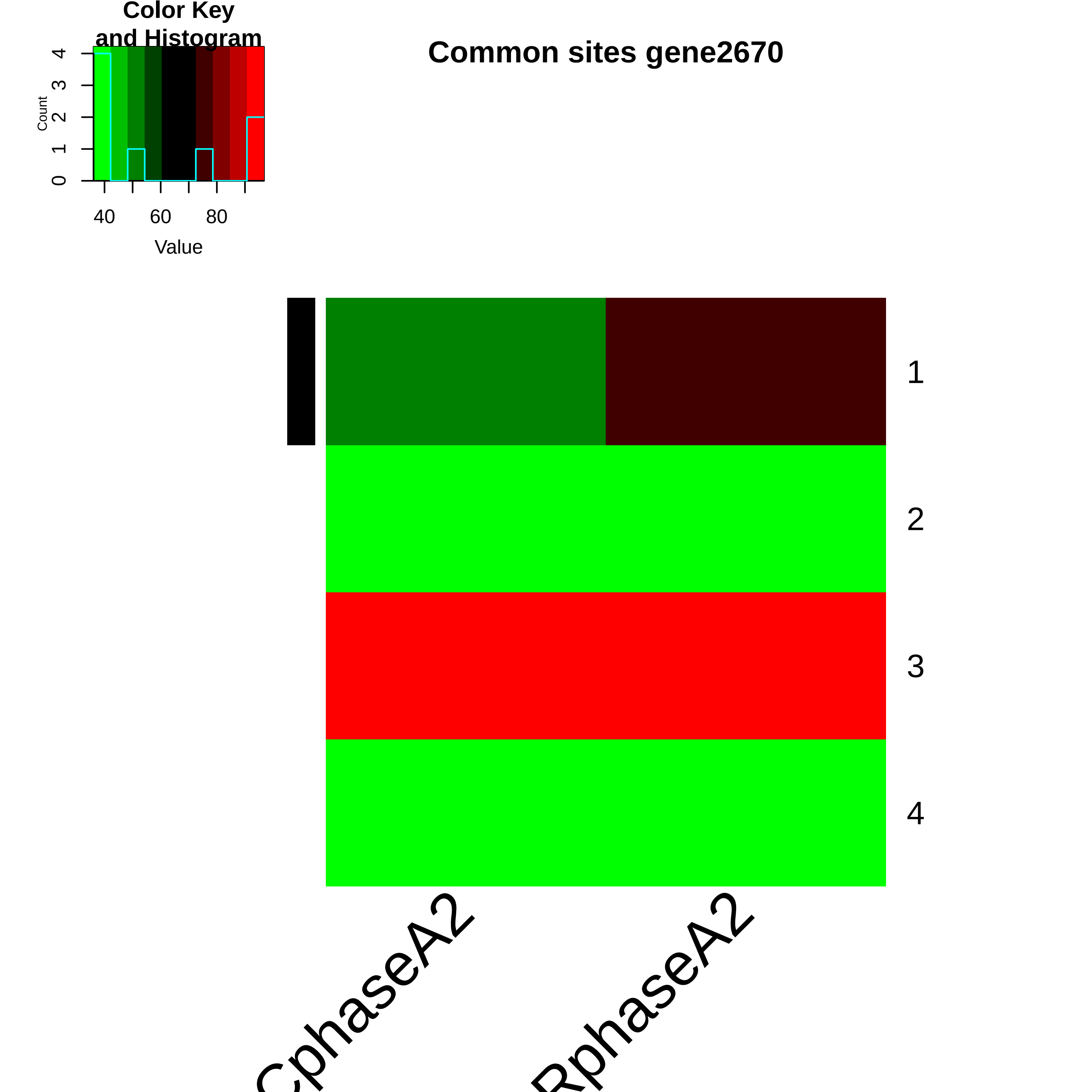
Color Key
and Histogram
Count
0
1
2
3
4
40
60
80
Value
Common sites gene2670
1
2
3
4
CphaseA2
RphaseA2

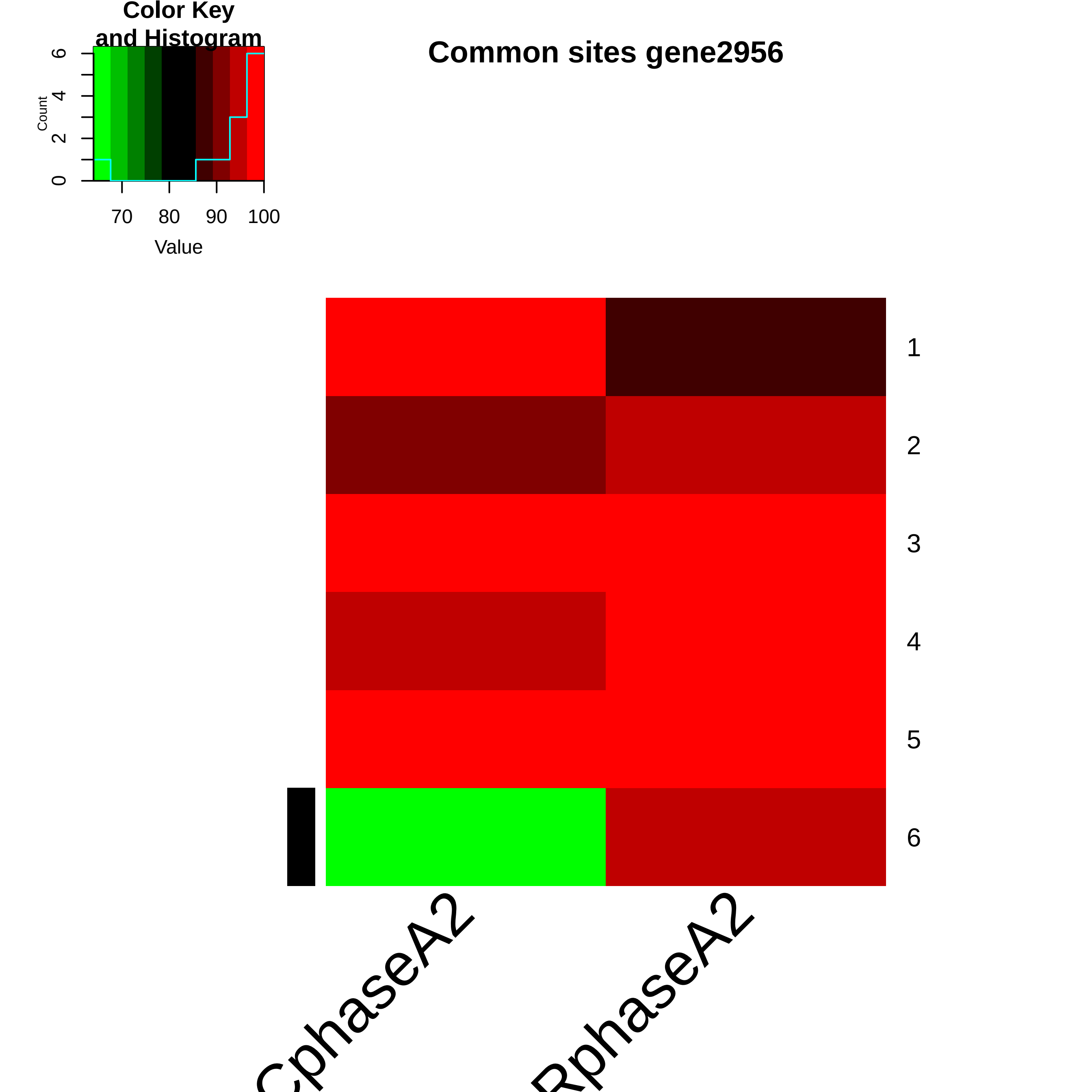

Color Key
and Histogram
Count
0
2
4
6
70
80
90
100
Value
Common sites gene2956
1
2
3
4
5
6
CphaseA2
RphaseA2

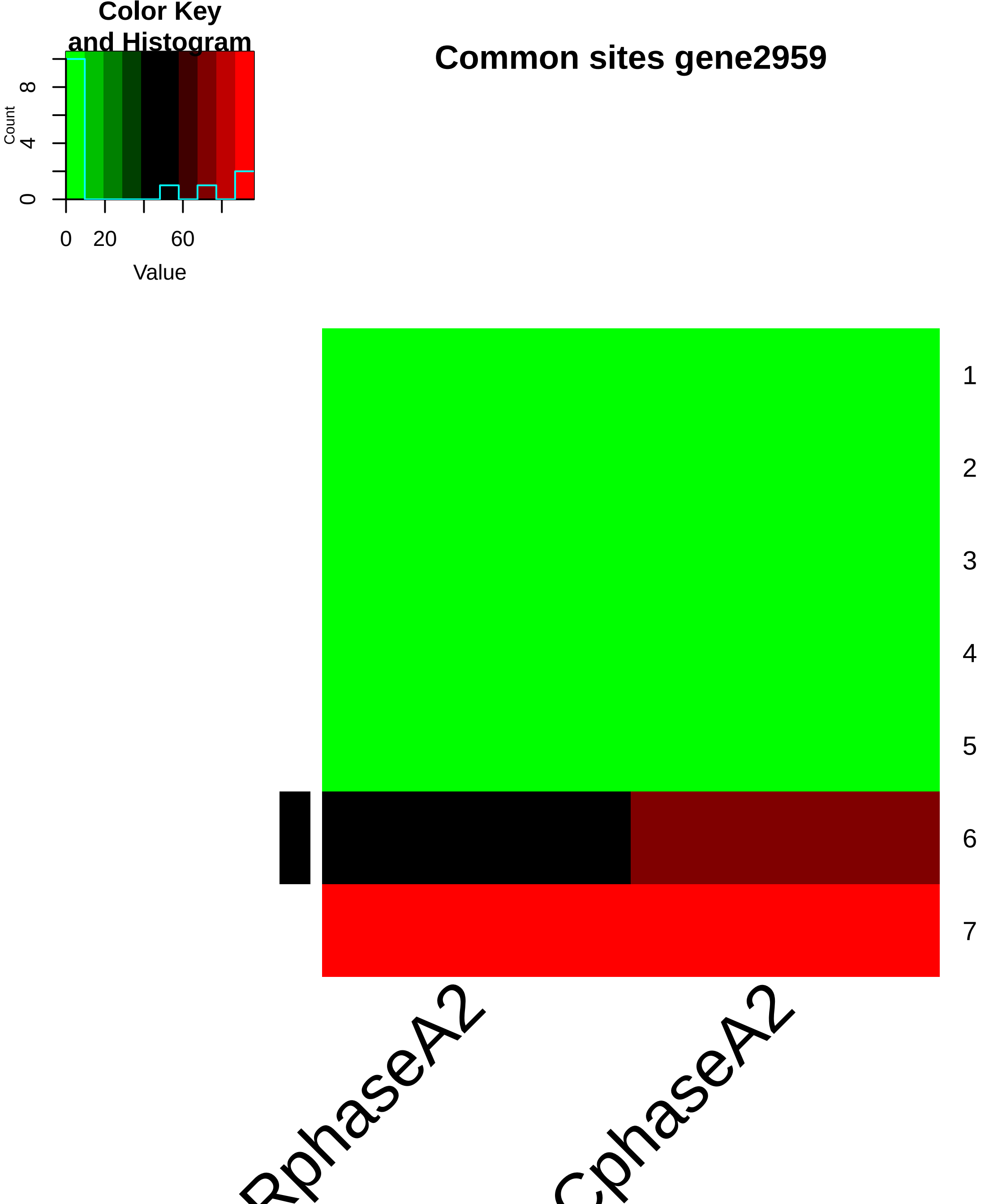

Color Key
and Histogram
Count
0
4
8
0
20
60
Value
Common sites gene2959
1
2
3
4
5
6
7
RphaseA2
CphaseA2

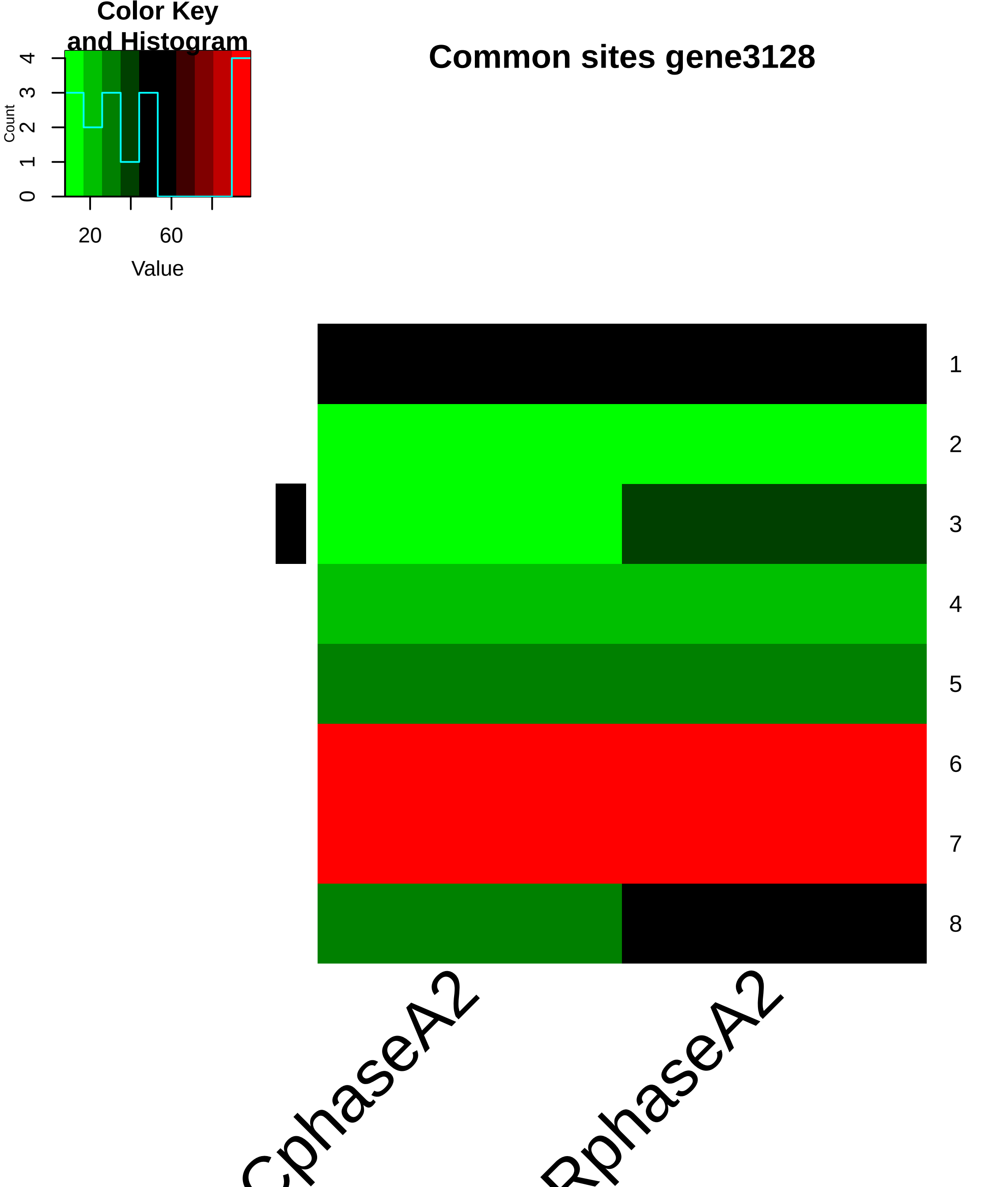

Color Key
and Histogram
Count
0
1
2
3
4
20
60
Value
Common sites gene3128
1
2
3
4
5
6
7
8
CphaseA2
RphaseA2

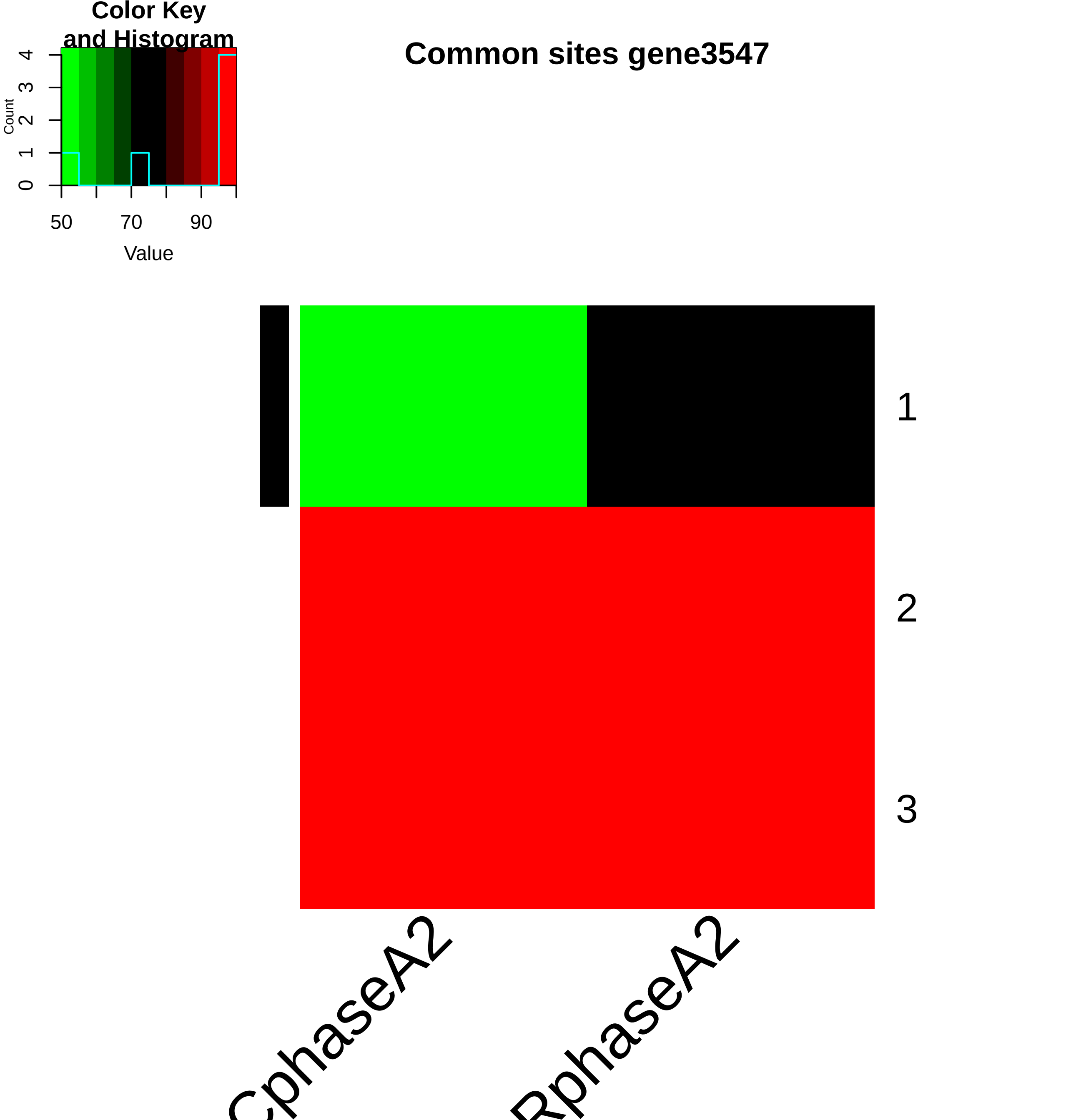
Color Key
and Histogram
Count
0
1
2
3
4
50
70
90
Value
Common sites gene3547
1
2
3
CphaseA2
RphaseA2

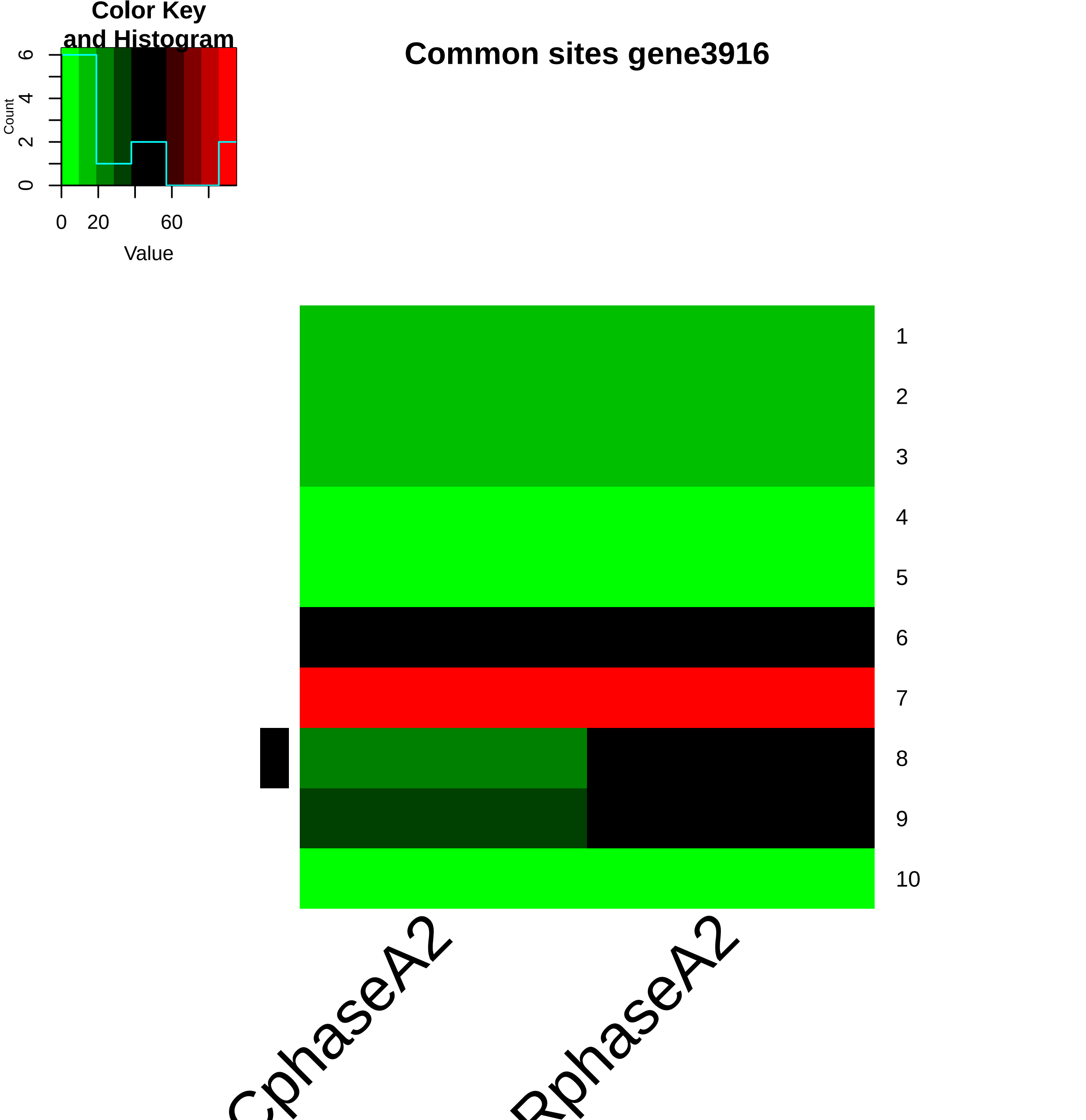
Color Key
and Histogram
Count
0
2
4
6
0
20
60
Value
Common sites gene3916
1
2
3
4
5
6
7
8
9
10
CphaseA2
RphaseA2

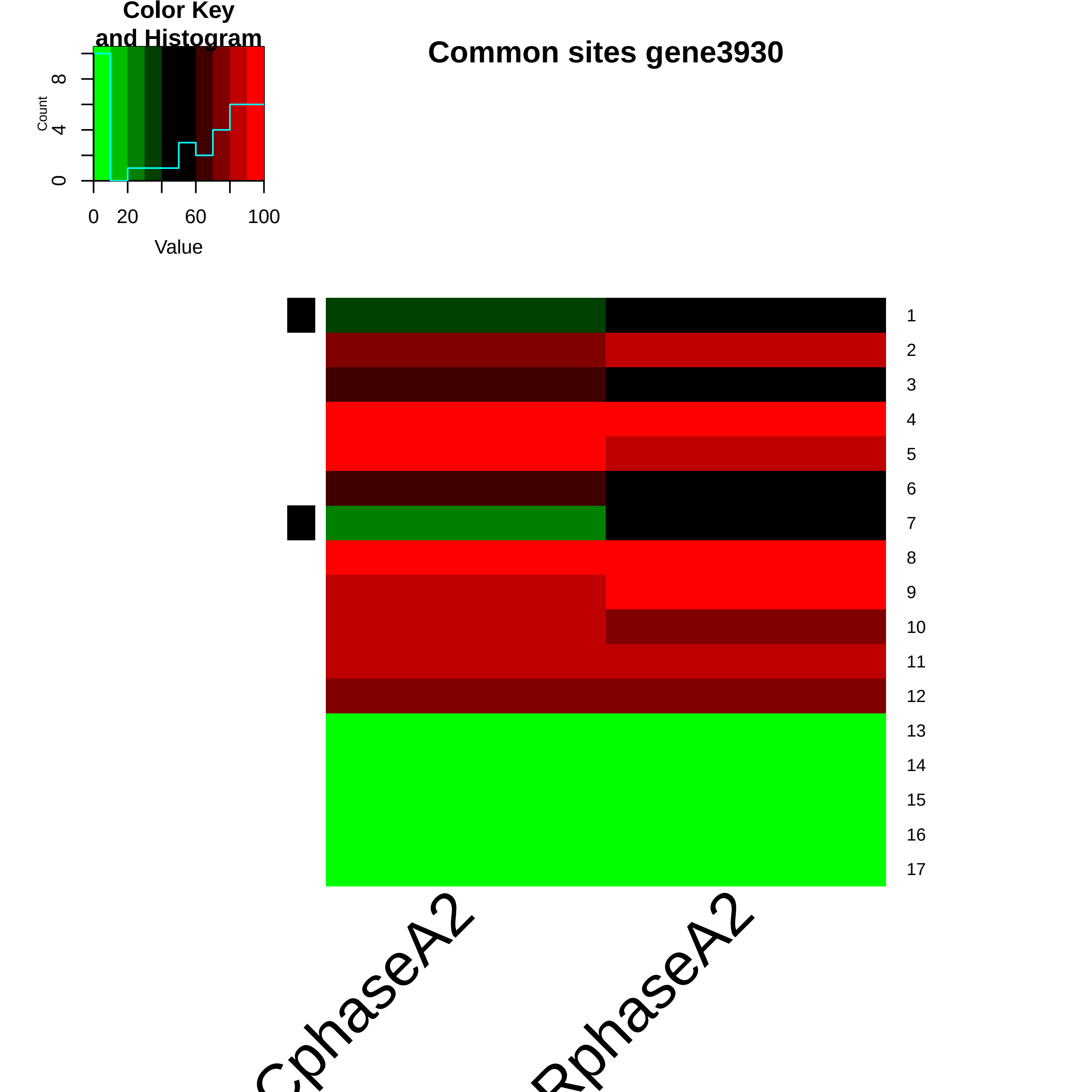

Color Key
and Histogram
Count
0
4
8
0
20
60
100
Value
Common sites gene3930

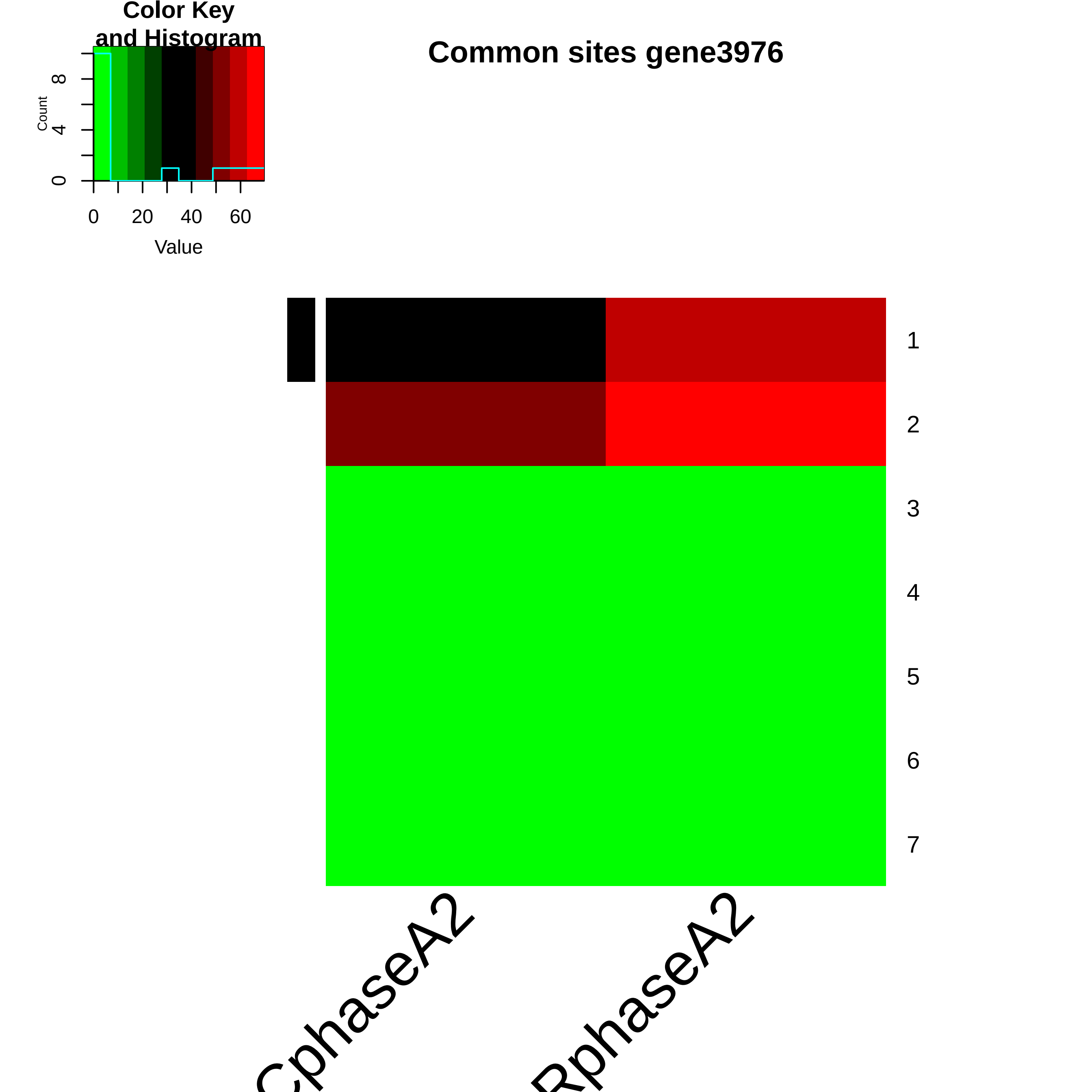
Color Key
and Histogram
Count
0
4
8
0
20
40
60
Value
Common sites gene3976
1
2
3
4
5
6
7
CphaseA2
RphaseA2

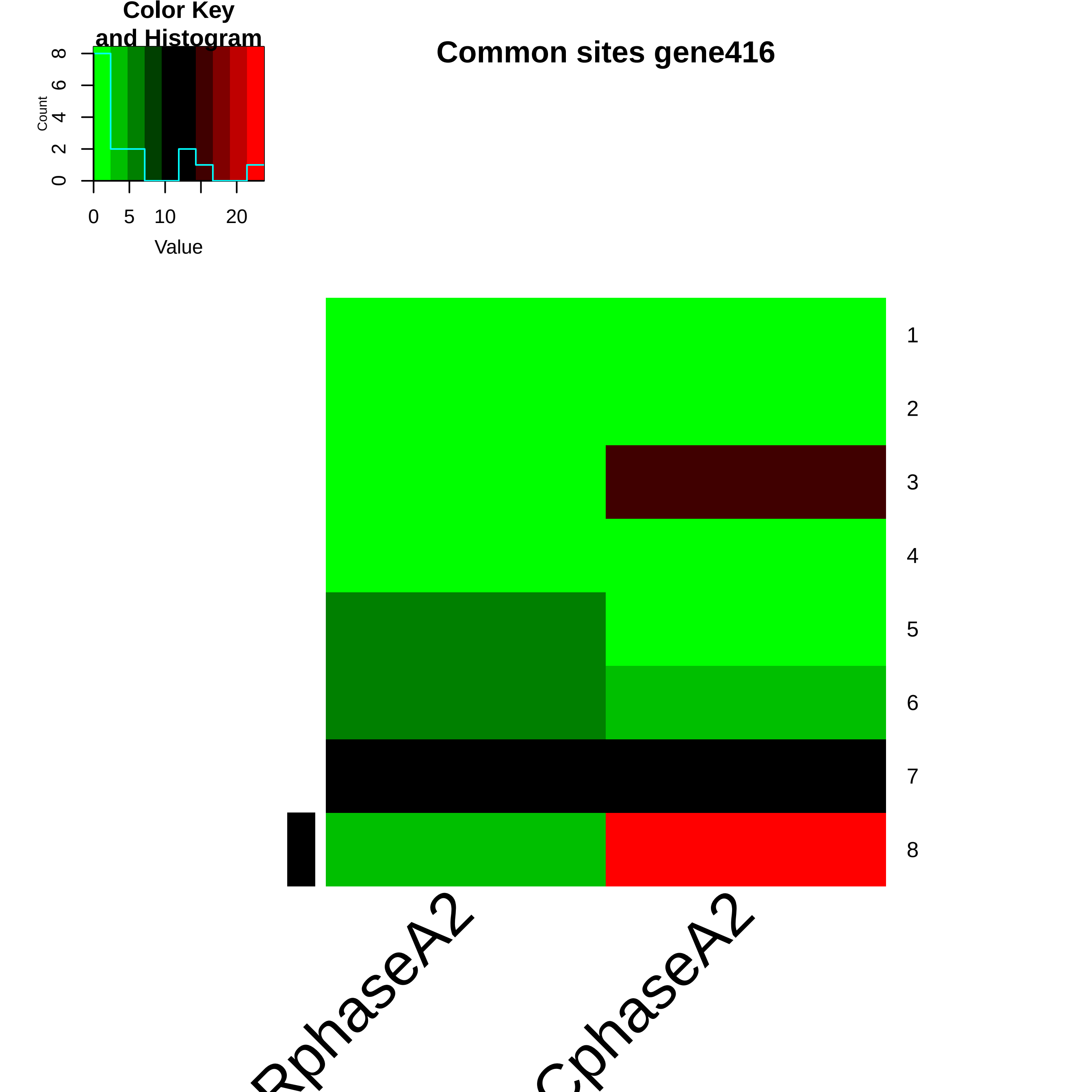
Color Key
and Histogram
Count
0
2
4
6
8
0
5
10
20
Value
Common sites gene416
1
2
3
4
5
6
7
8
RphaseA2
CphaseA2

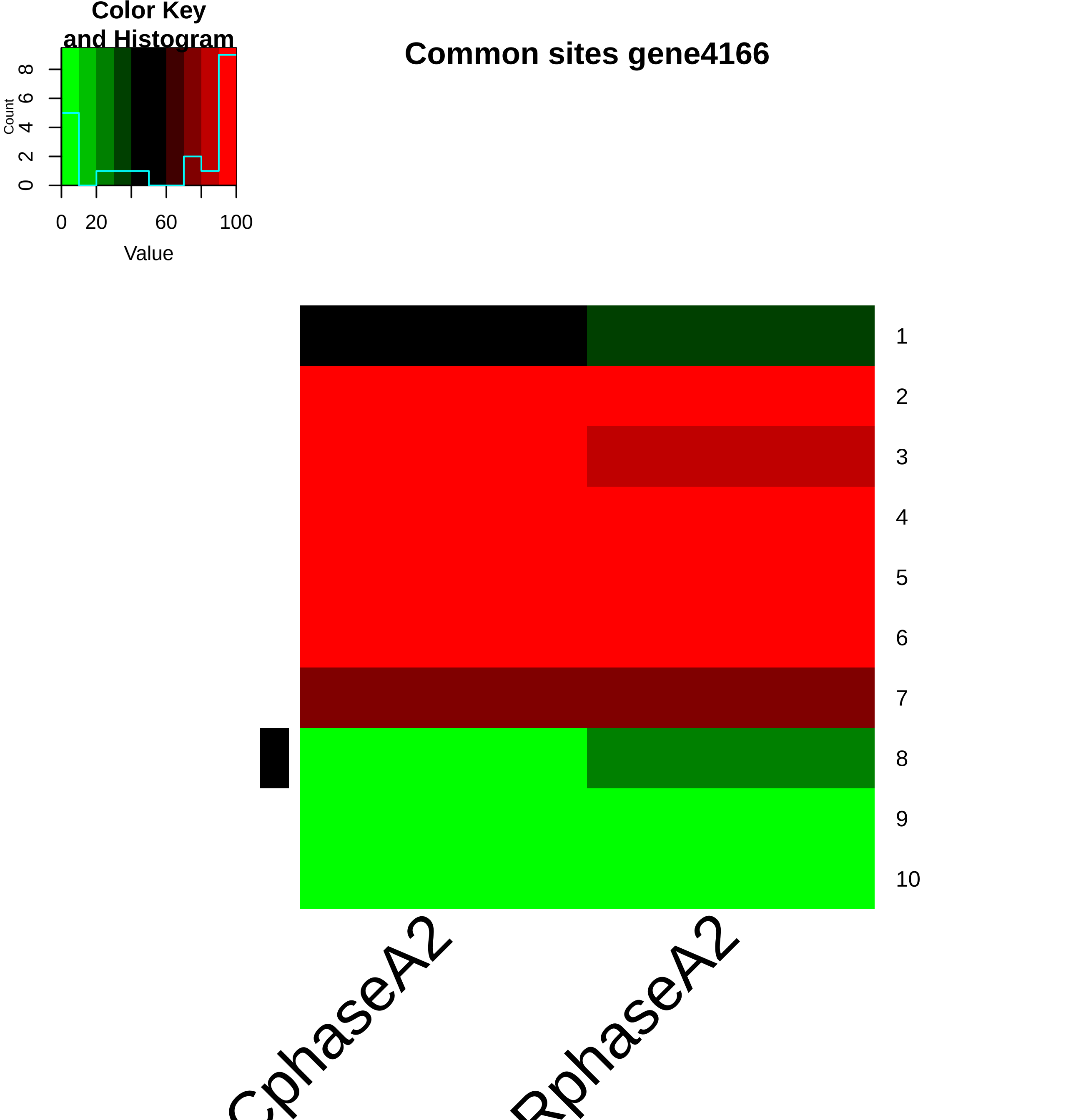
Color Key
and Histogram
Count
0
2
4
6
8
0
20
60
100
Value
Common sites gene4166
1
2
3
4
5
6
7
8
9
10
CphaseA2
RphaseA2

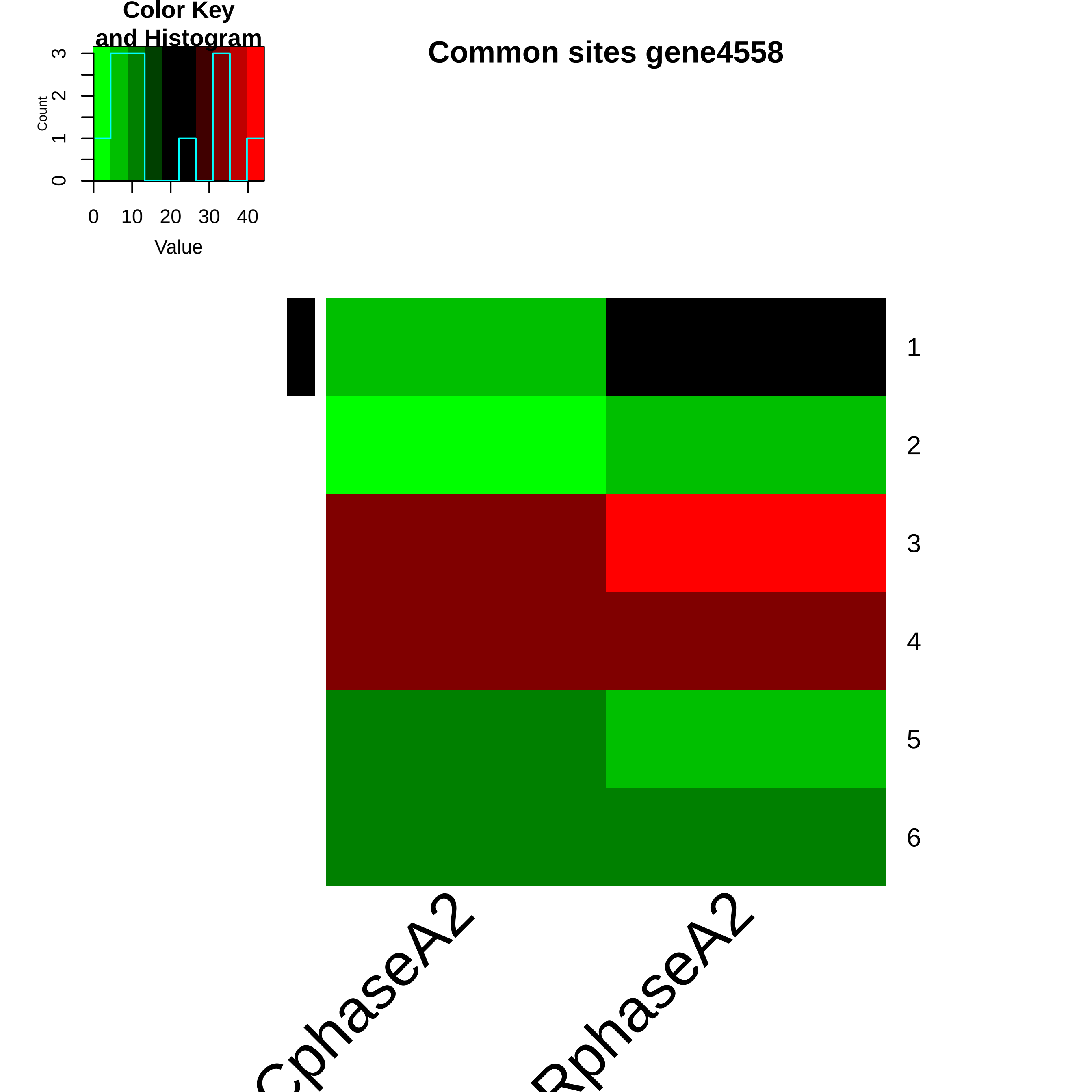
Color Key
and Histogram
Count
Value
0
10
20
30
40
0
1
2
3
Common sites gene4558
1
2
3
4
5
6
CphaseA2
RphaseA2

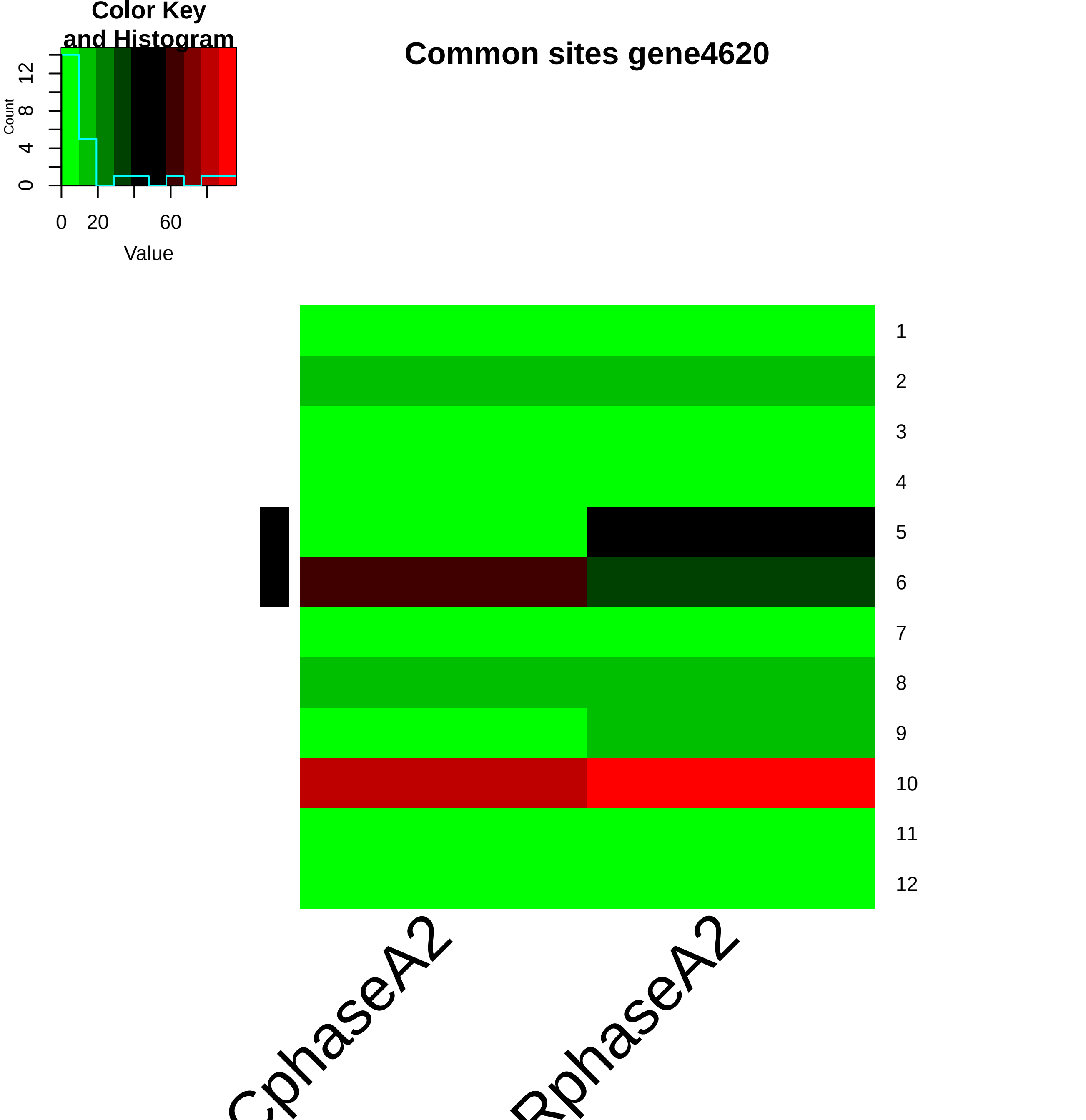
Color Key
and Histogram
Count
0
4
8
12
0
20
60
Value
Common sites gene4620
1
2
3
4
5
6
7
8
9
10
11
12
CphaseA2
RphaseA2

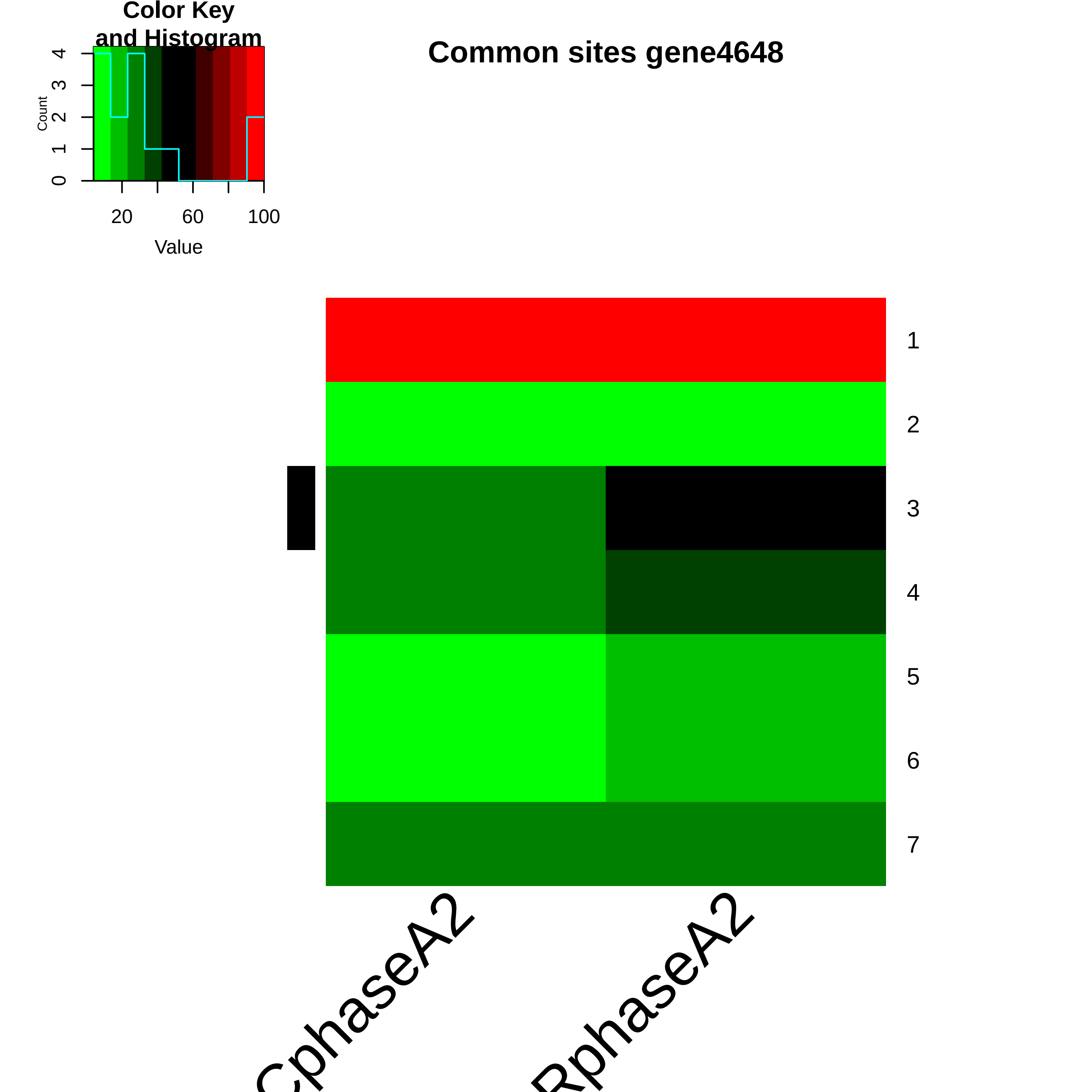
Color Key
and Histogram
Count
0
1
2
3
4
20
60
100
Value
Common sites gene4648
1
2
3
4
5
6
7
CphaseA2
RphaseA2

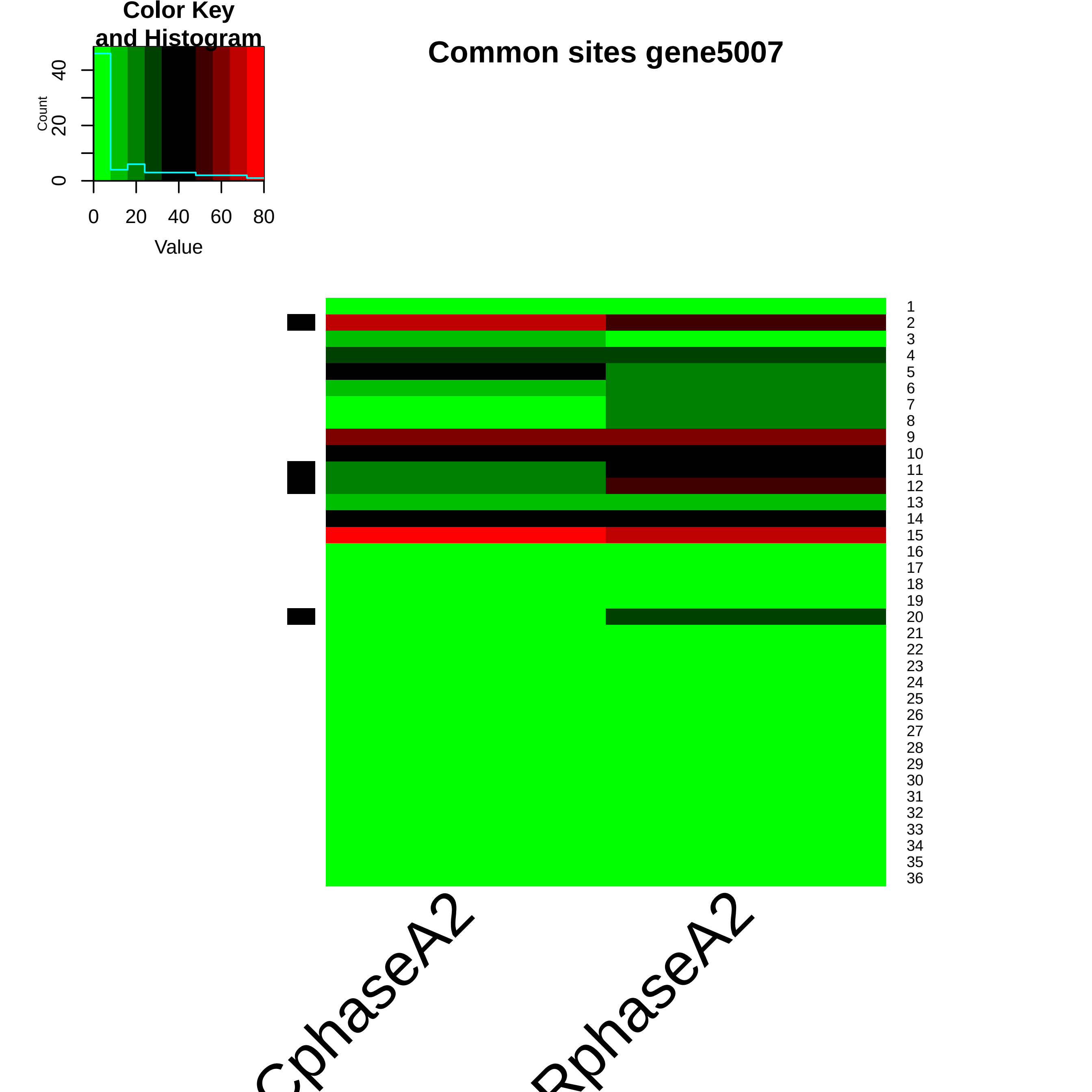

Color Key
and Histogram
Count
0
20
40
0
20
40
60
80
Value
Common sites gene5007
CphaseA2
RphaseA2

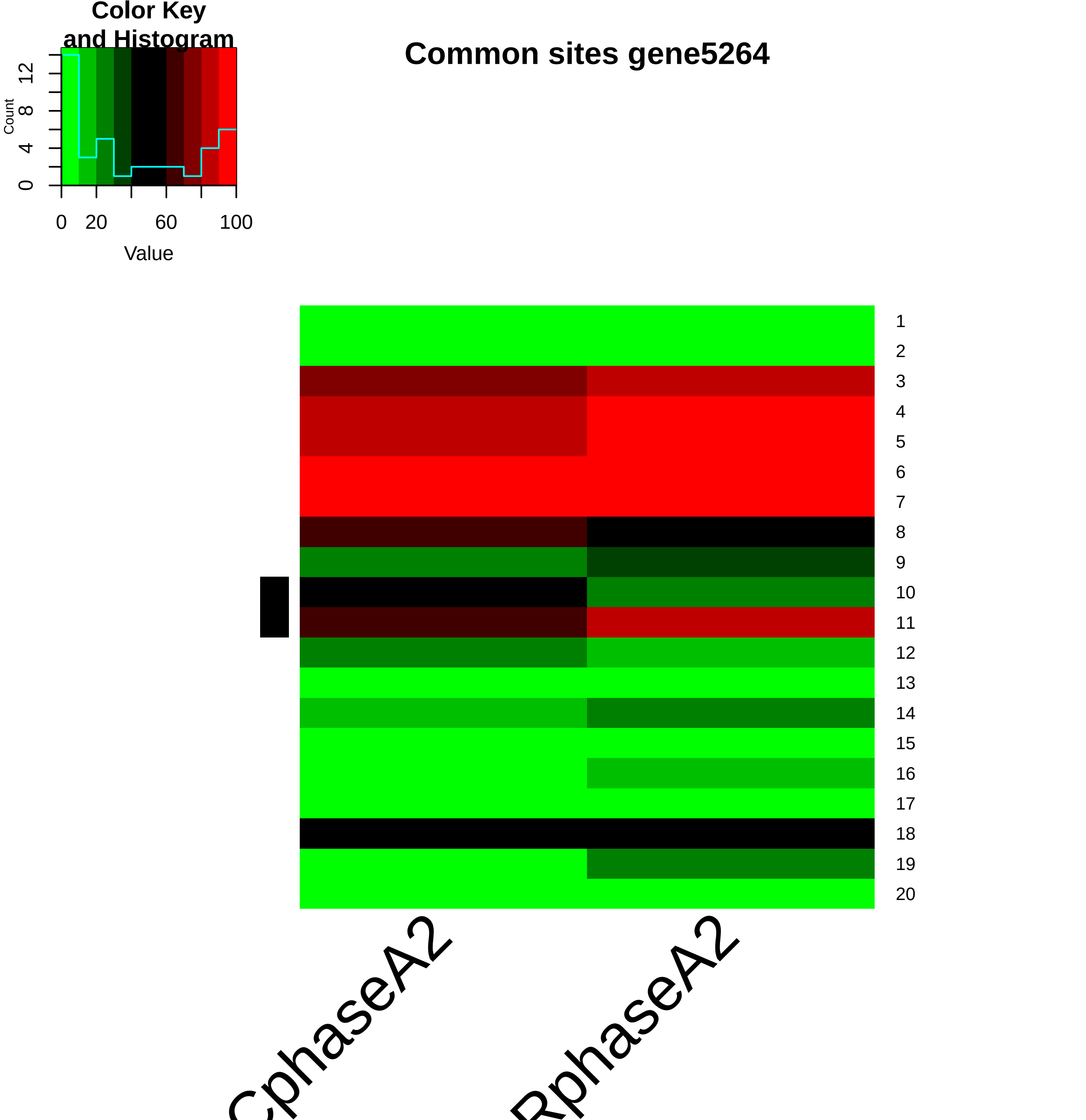
Color Key
and Histogram
Count
0
4
8
12
0
20
60
100
Value
Common sites gene5264
CphaseA2
RphaseA2

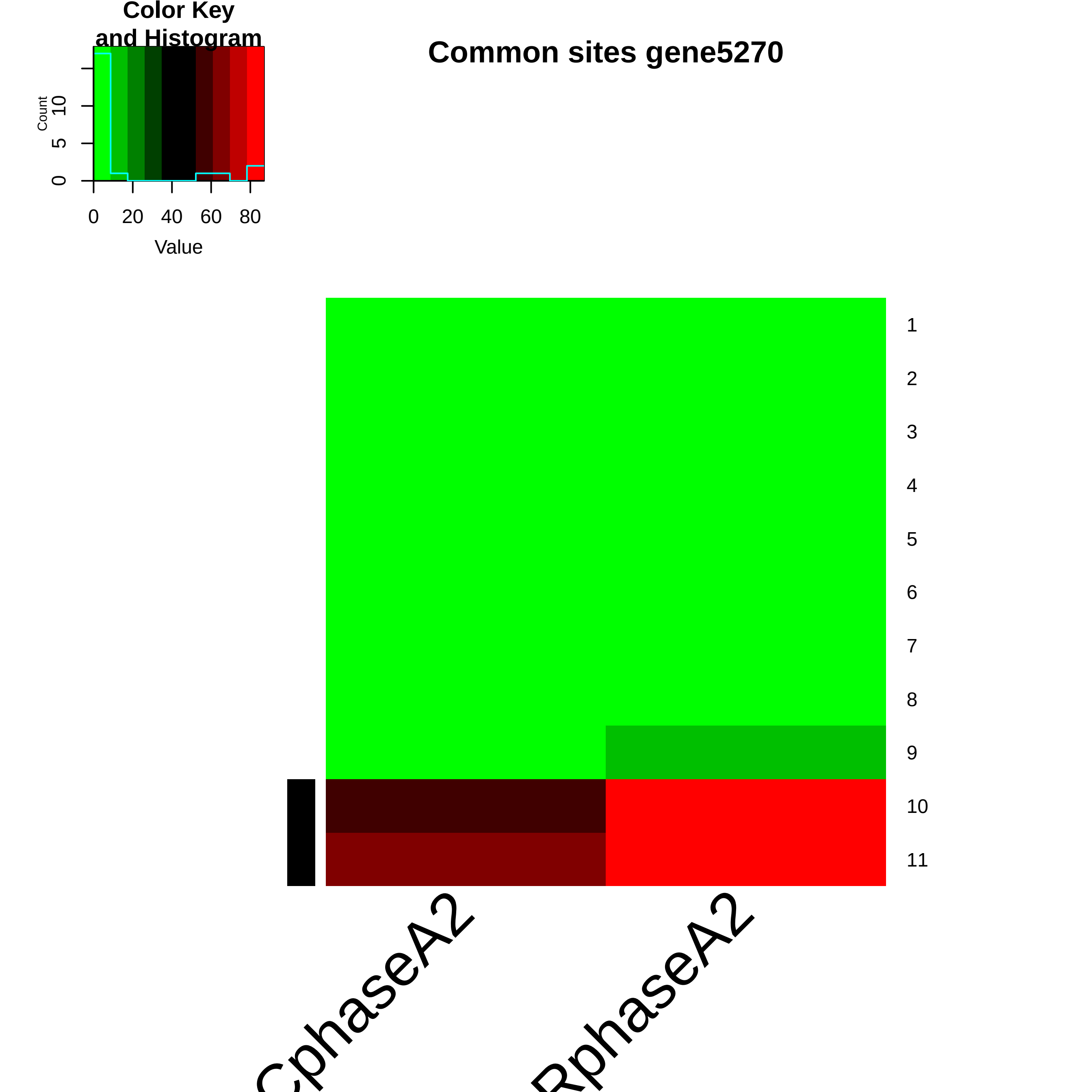

Color Key
and Histogram
Count
0
5
10
0
20
40
60
80
Value
Common sites gene5270
1
2
3
4
5
6
7
8
9
10
11
CphaseA2
RphaseA2

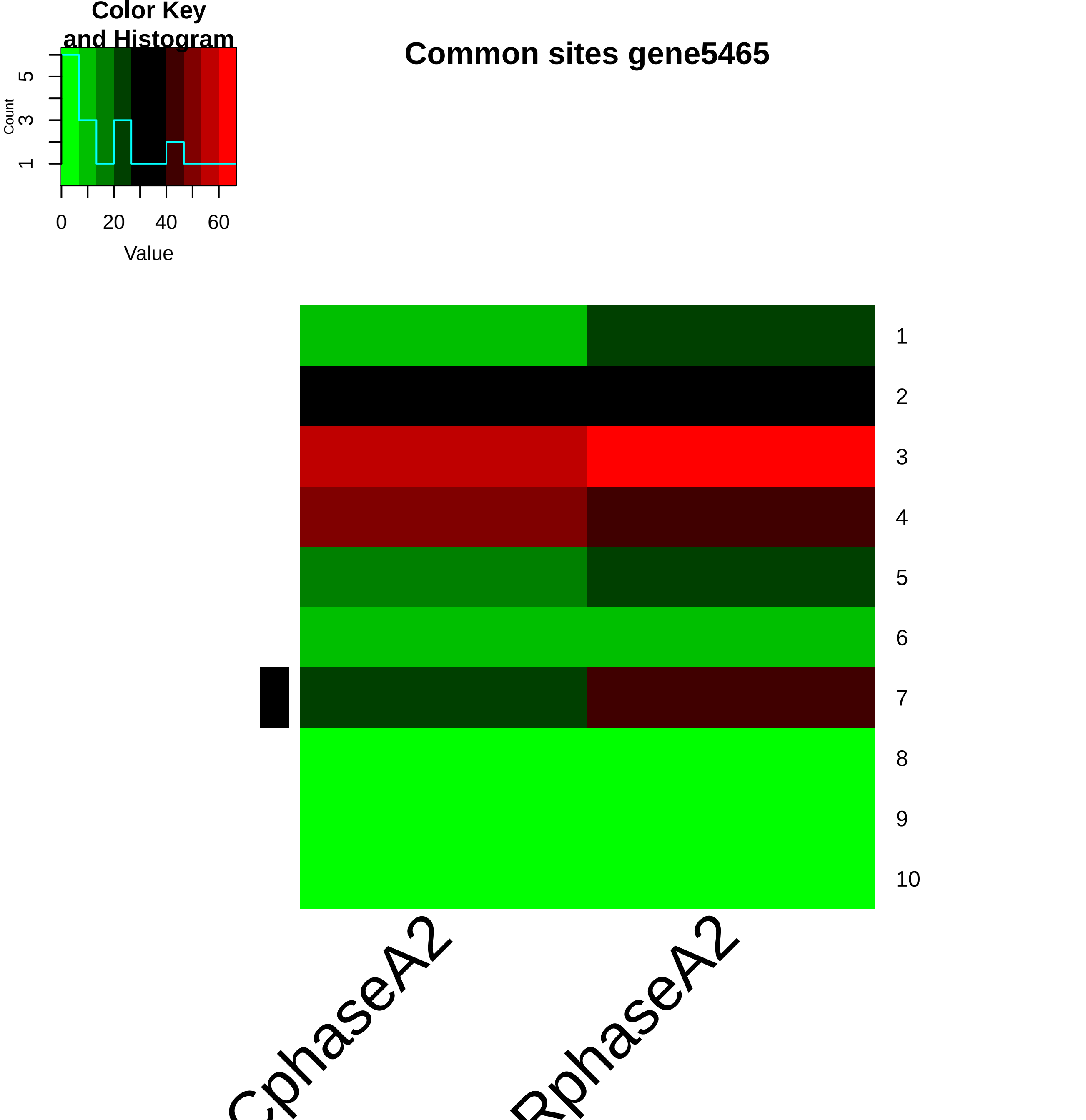

Color Key
and Histogram
Count
1
3
5
0
20
40
60
Value
Common sites gene5465
1
2
3
4
5
6
7
8
9
10
CphaseA2
RphaseA2

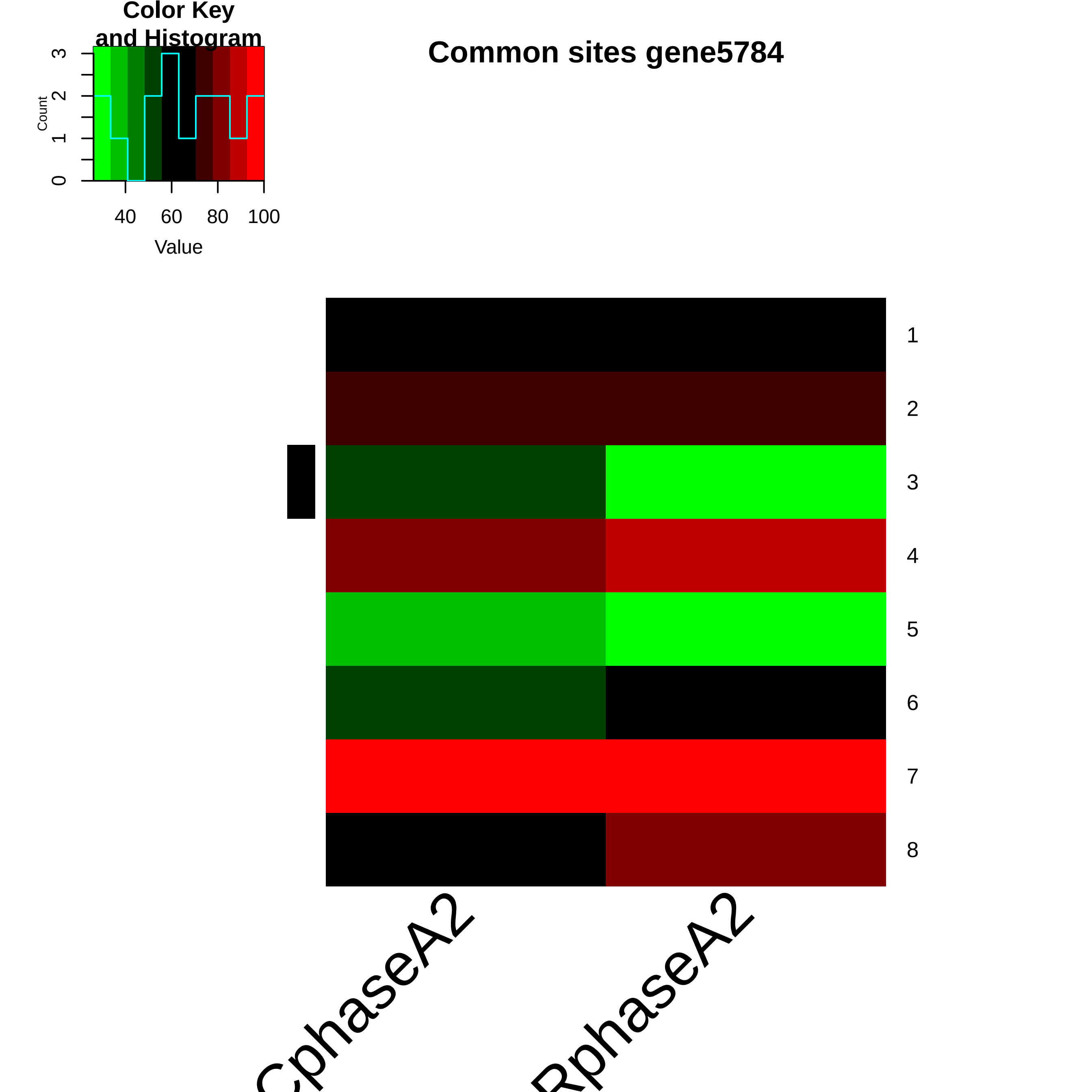
Color Key
and Histogram
Count
0
1
2
3
40
60
80
100
Value
Common sites gene5784
1
2
3
4
5
6
7
8
CphaseA2
RphaseA2

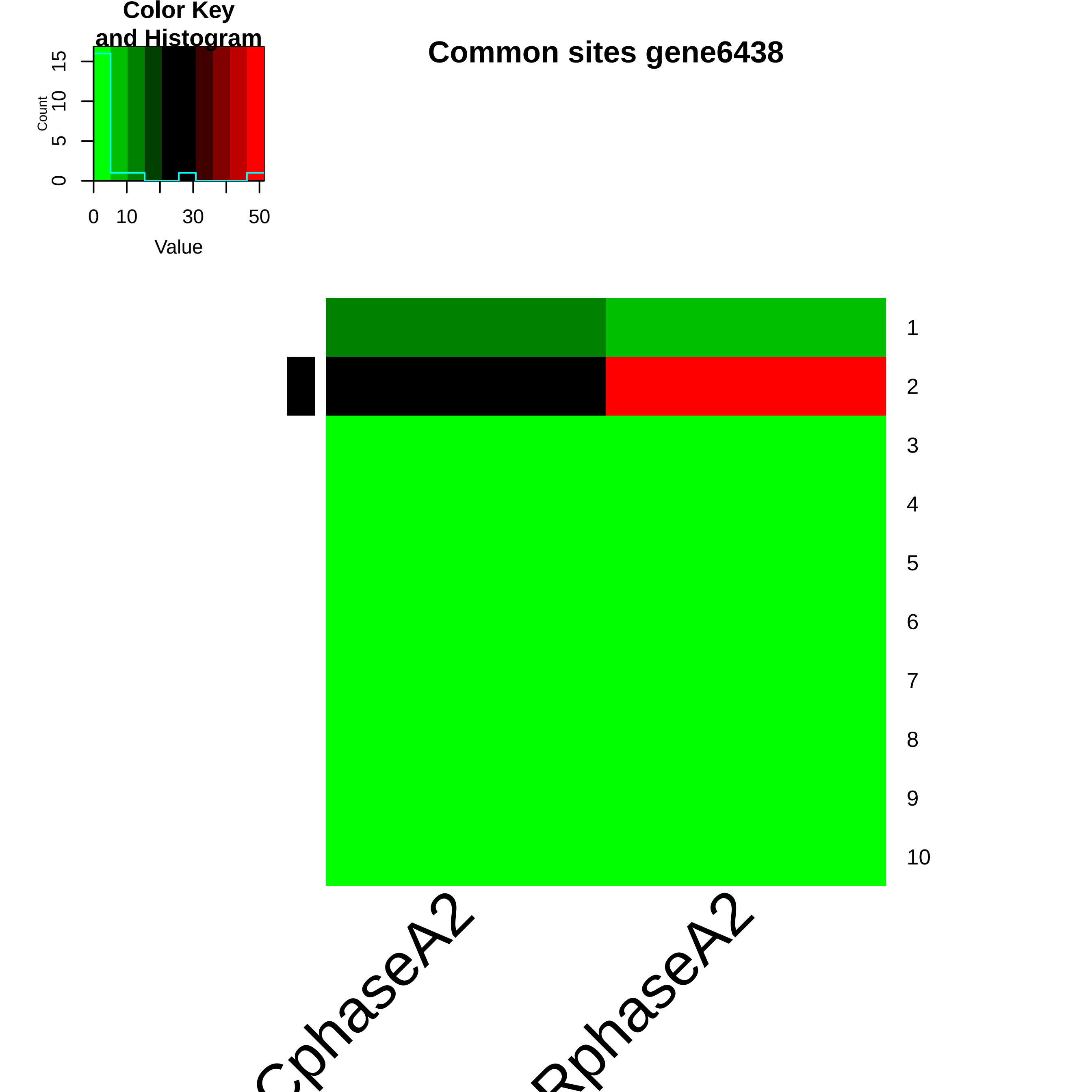
Color Key
and Histogram
Count
0
5
10
15
0
10
30
50
Value
Common sites gene6438
1
2
3
4
5
6
7
8
9
10
CphaseA2
RphaseA2

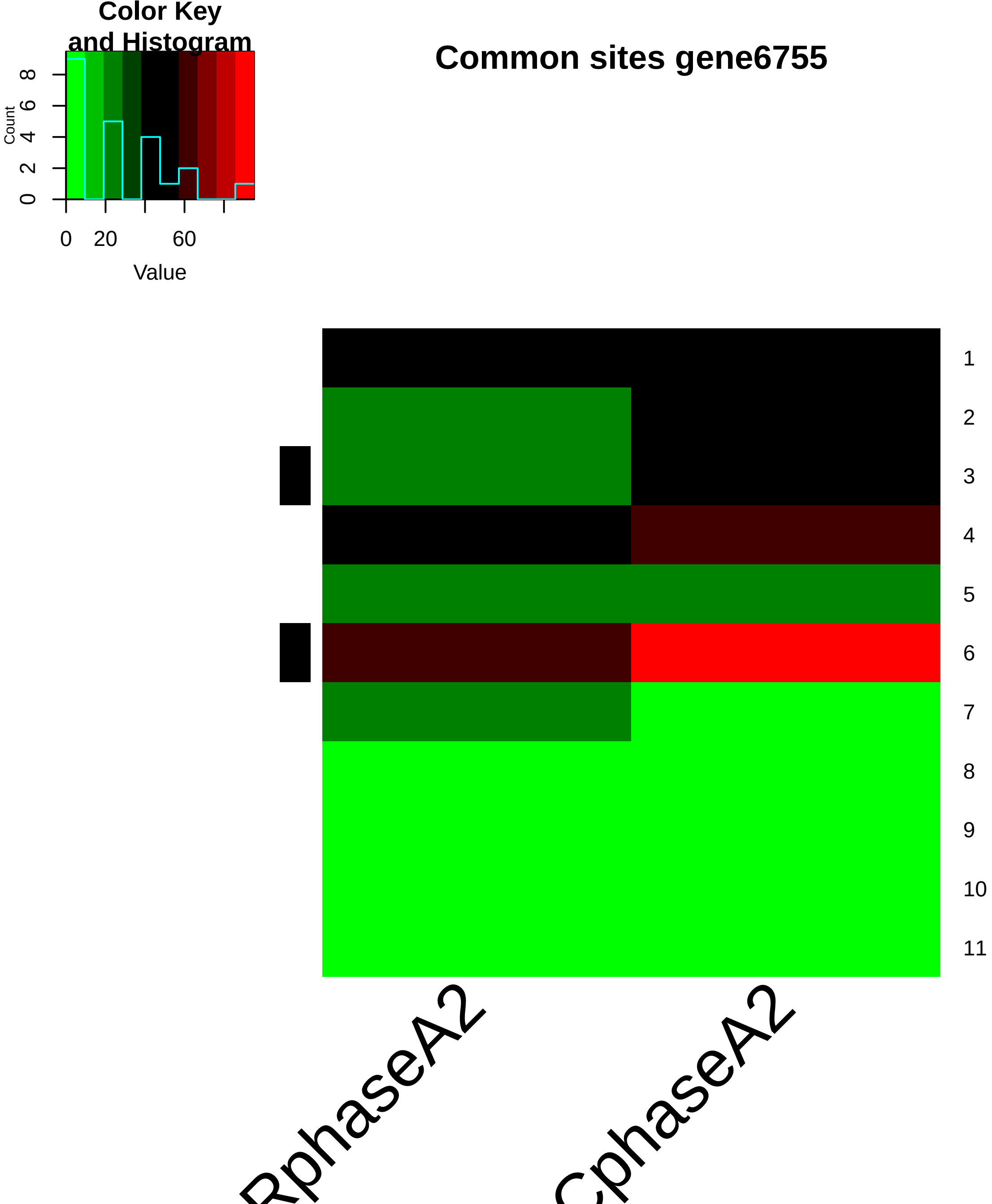
Color Key
and Histogram
Count
0
2
4
6
8
0
20
60
Value
Common sites gene6755
1
2
3
4
5
6
7
8
9
10
11
RphaseA2
CphaseA2

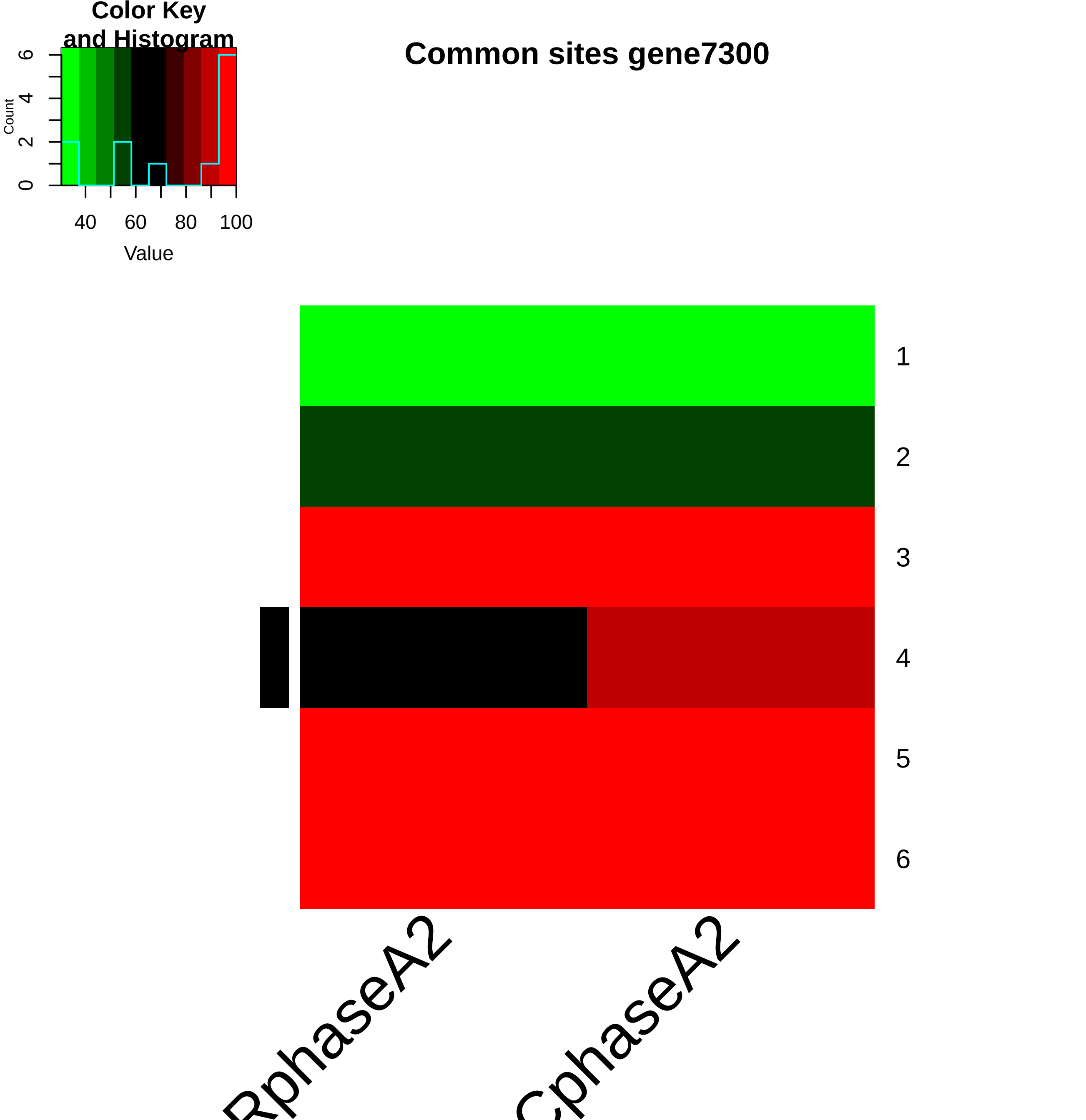
Color Key
and Histogram
Count
0
2
4
6
40
60
80
100
Value
Common sites gene7300
1
2
3
4
5
6
RphaseA2
CphaseA2

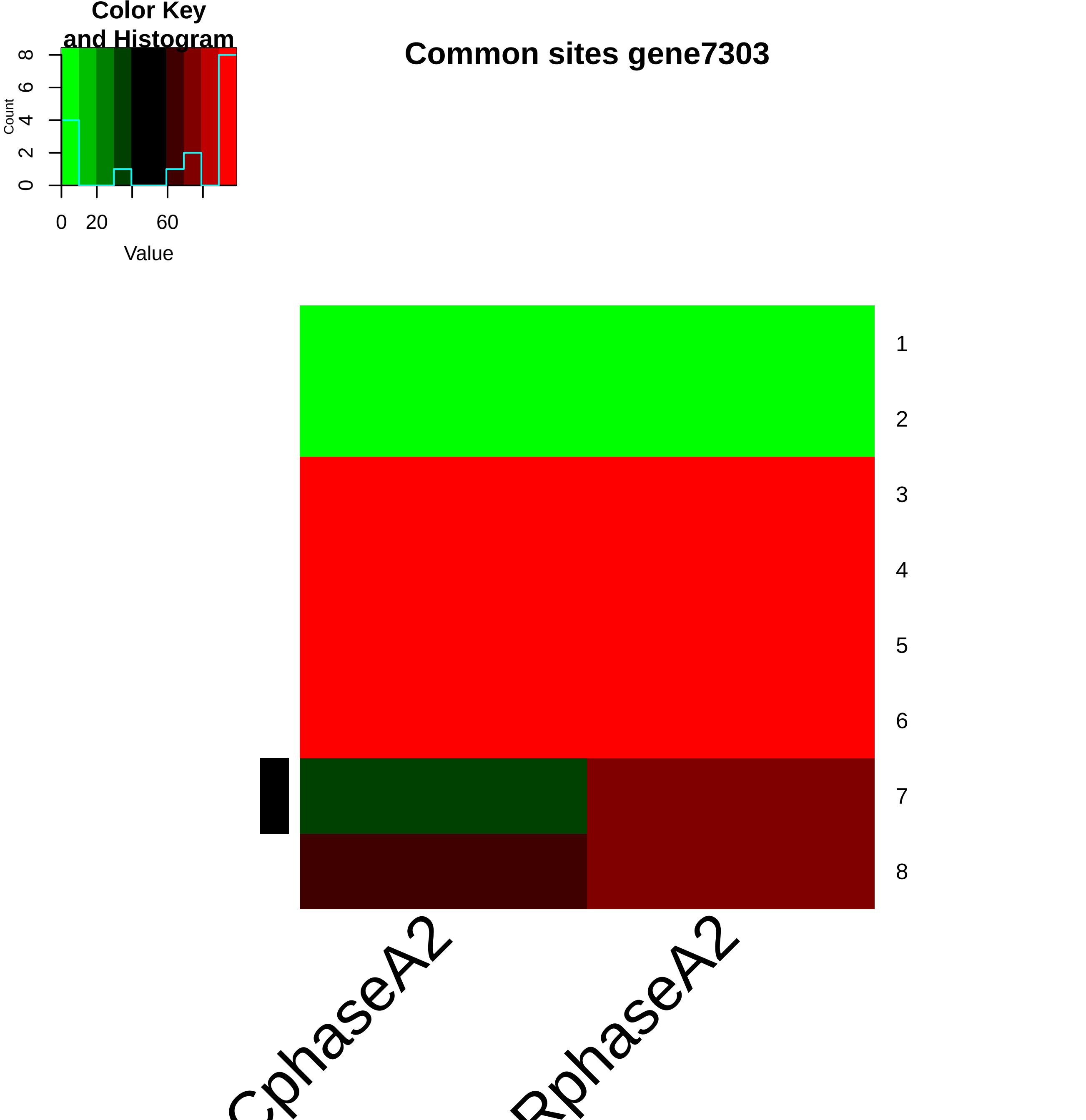
Color Key
and Histogram
Count
0
2
4
6
8
0
20
60
Value
Common sites gene7303
1
2
3
4
5
6
7
8
CphaseA2
RphaseA2

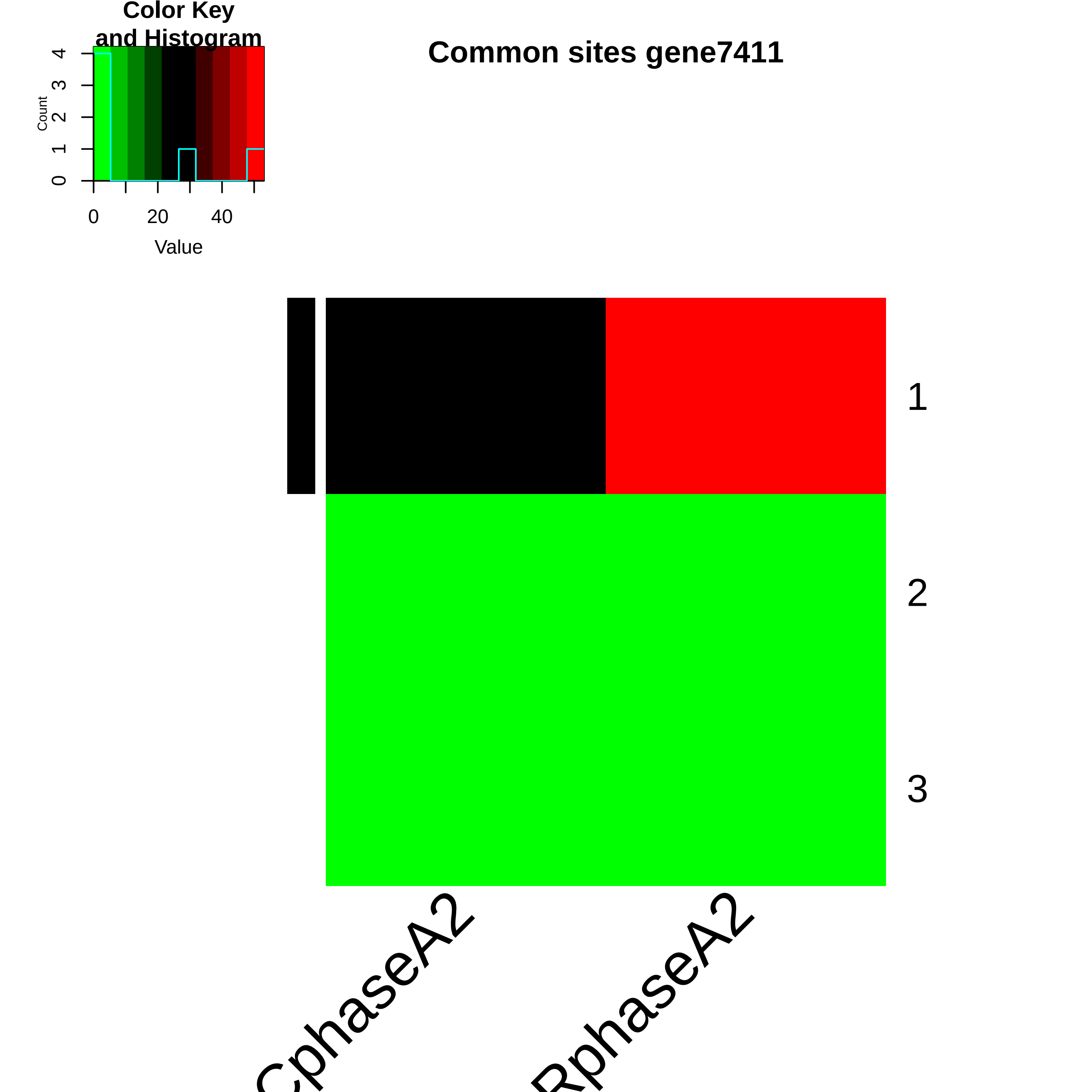
Color Key
and Histogram
Count
0
1
2
3
4
0
20
40
Value
Common sites gene7411
1
2
3
CphaseA2
RphaseA2

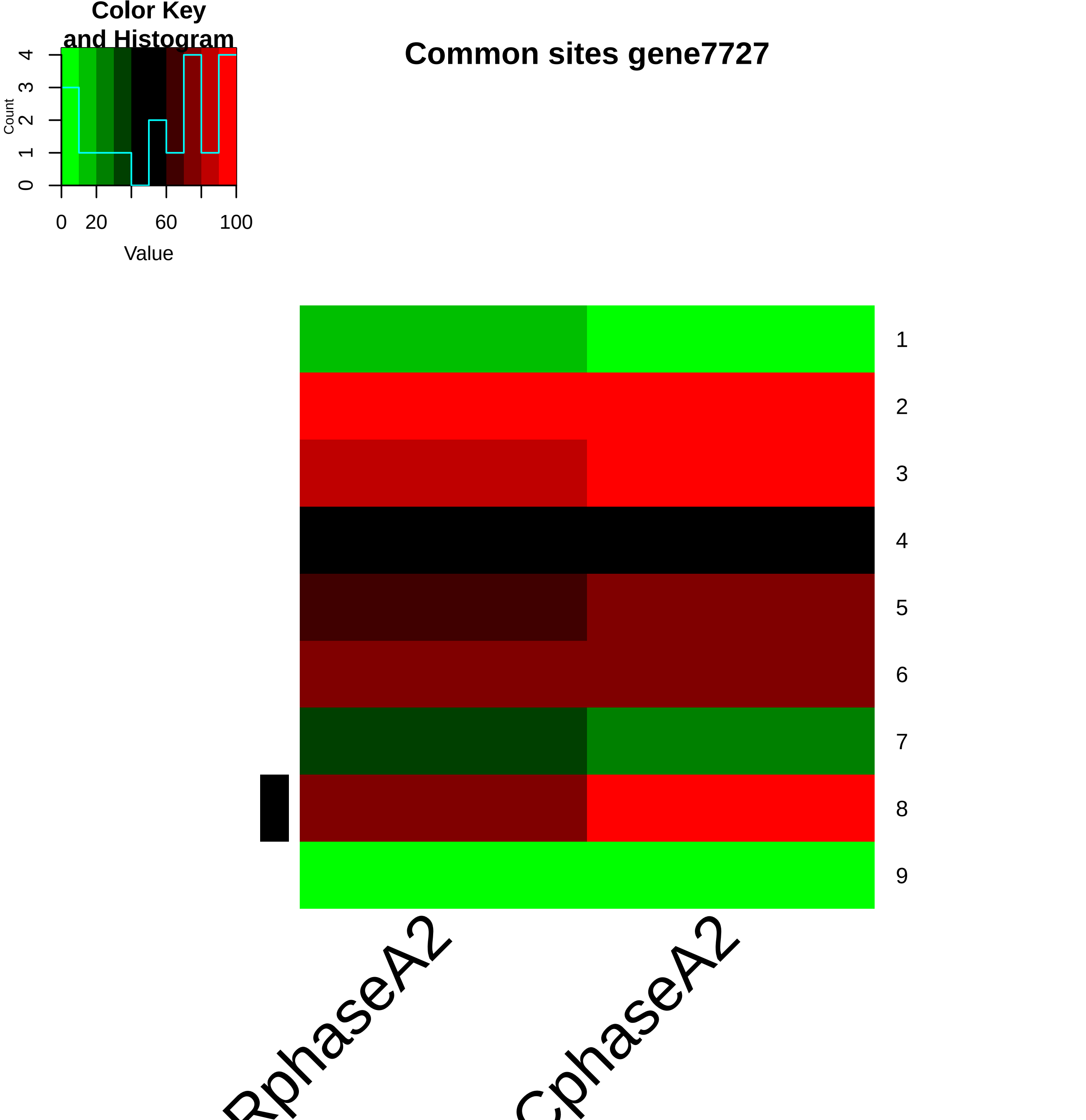
Color Key
and Histogram
Count
0
1
2
3
4
0
20
60
100
Value
Common sites gene7727
1
2
3
4
5
6
7
8
9
RphaseA2
CphaseA2

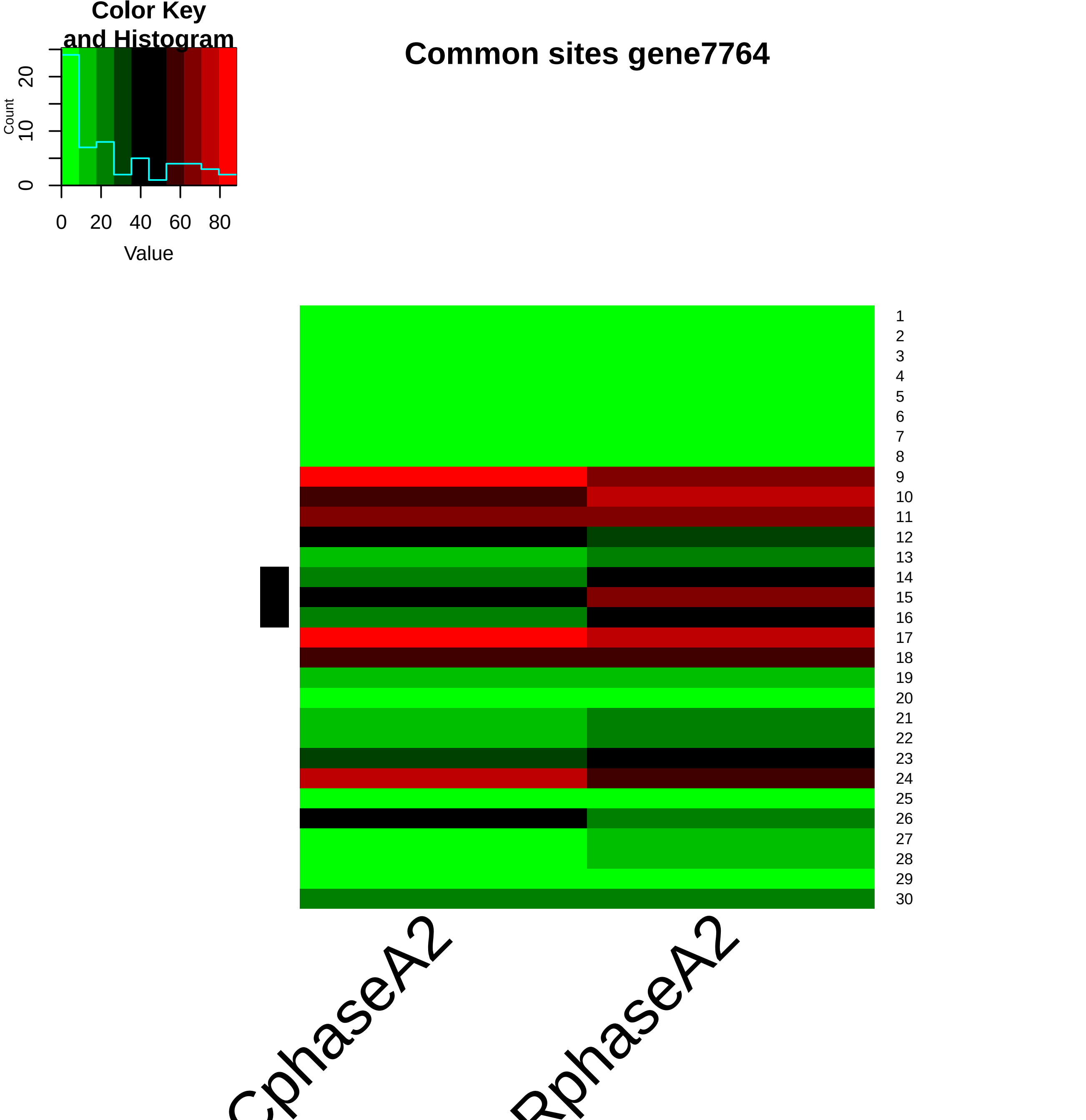

Color Key
and Histogram
Count
0
10
20
0
20
40
60
80
Value
Common sites gene7764
CphaseA2
RphaseA2

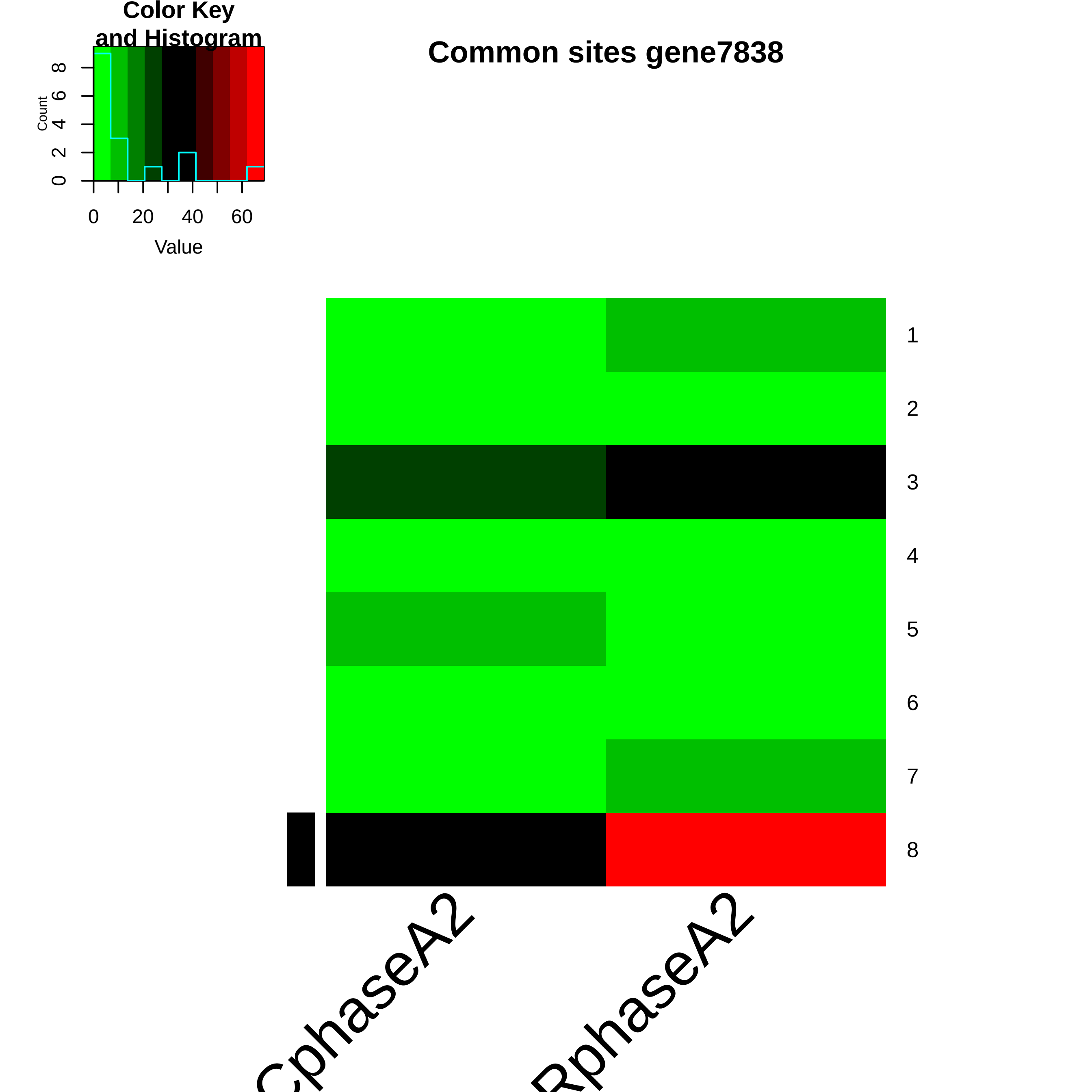
Color Key
and Histogram
Count
0
2
4
6
8
0
20
40
60
Value
Common sites gene7838
1
2
3
4
5
6
7
8
CphaseA2
RphaseA2

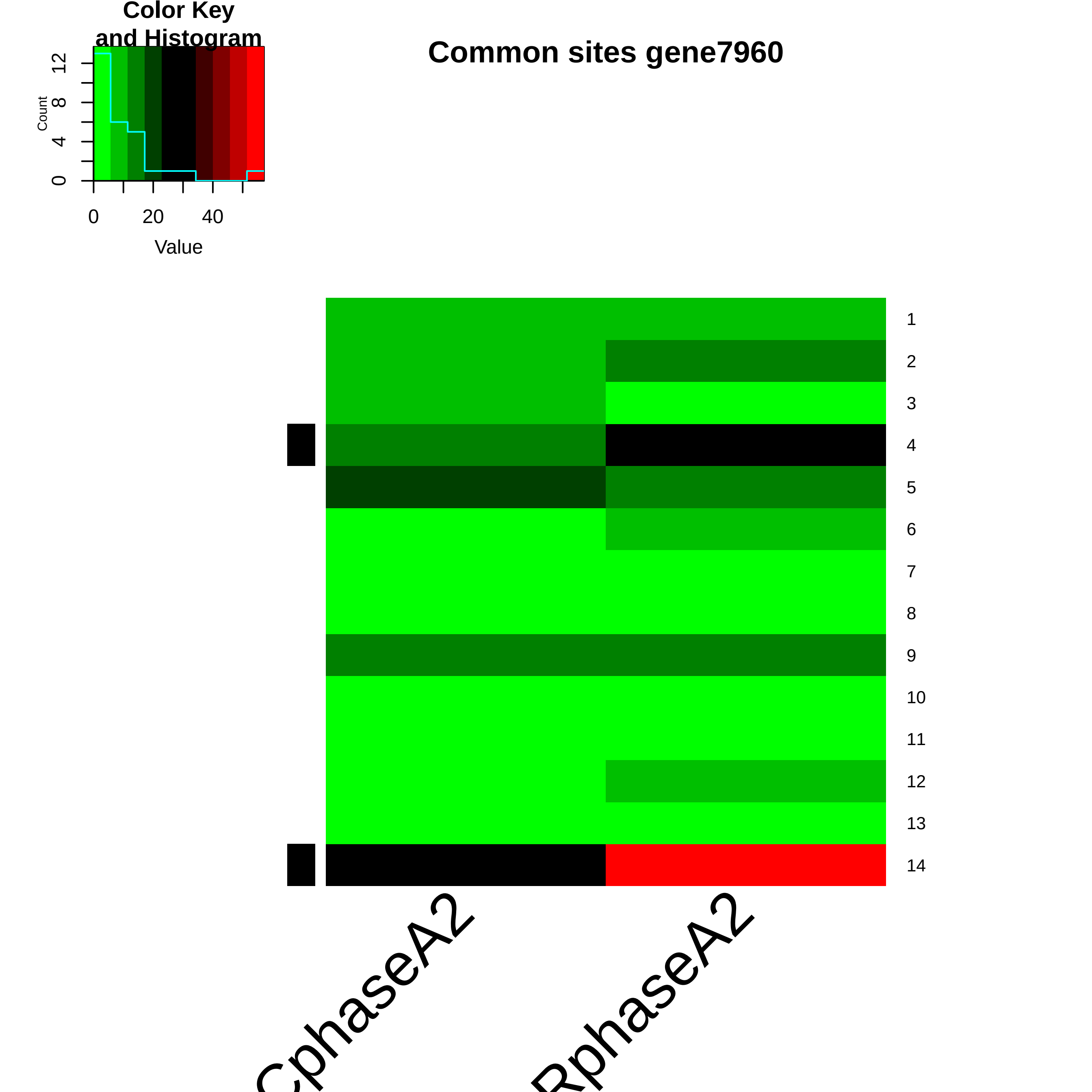
Color Key
and Histogram
Count
0
4
8
12
0
20
40
Value
Common sites gene7960
1
2
3
4
5
6
7
8
9
10
11
12
13
14
CphaseA2
RphaseA2

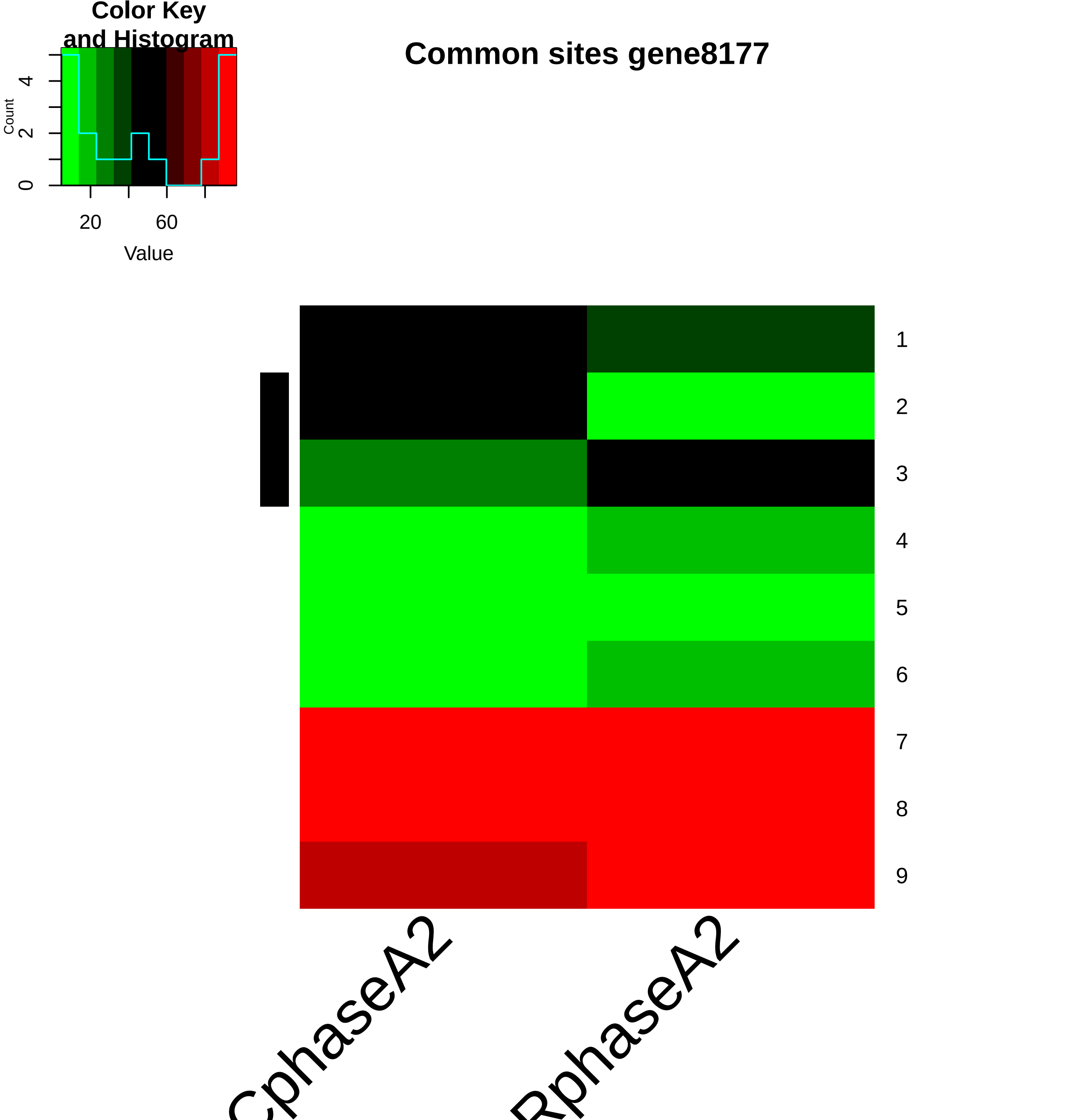

Color Key
and Histogram
Count
0
2
4
20
60
Value
Common sites gene8177
1
2
3
4
5
6
7
8
9
CphaseA2
RphaseA2

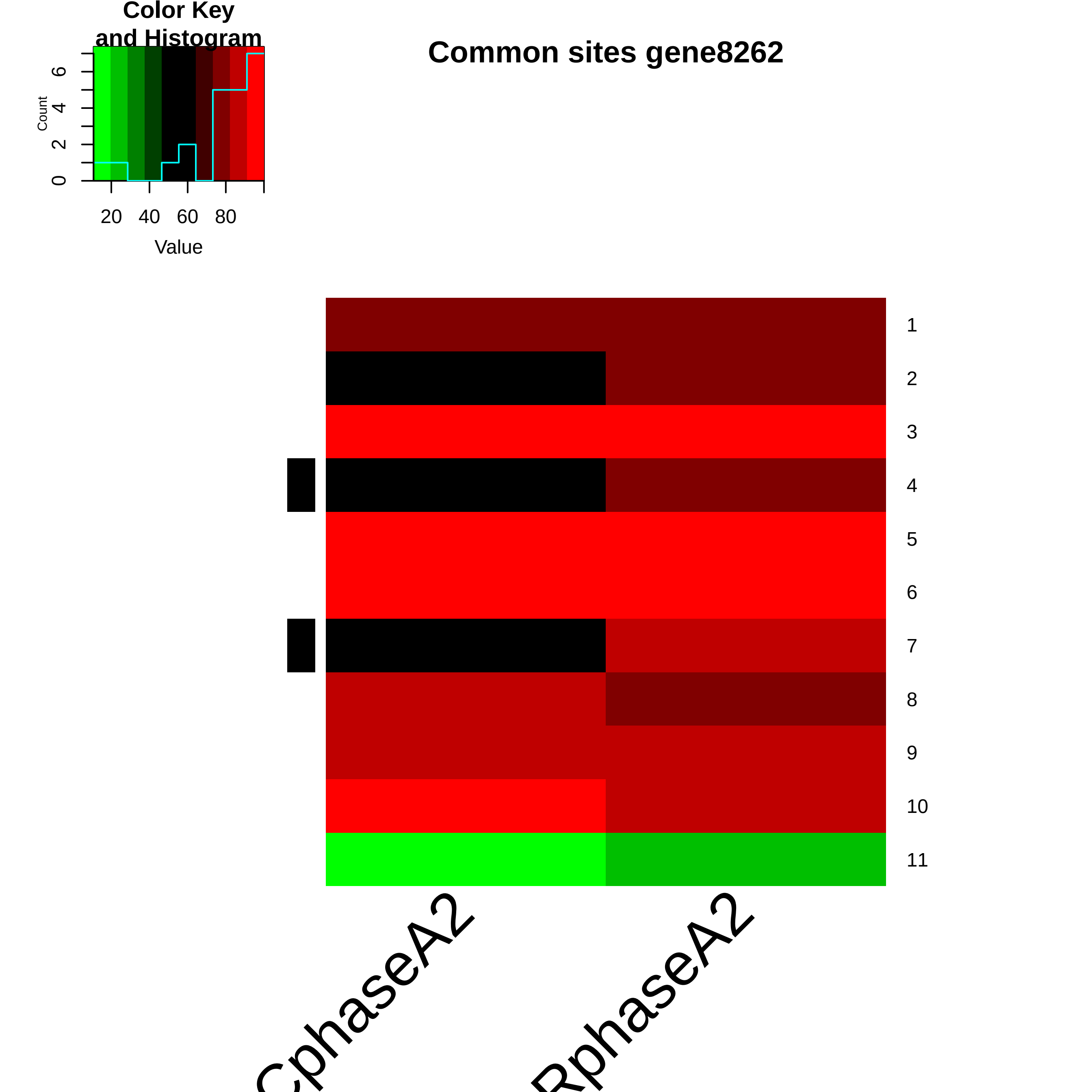
Color Key
and Histogram
Count
0
2
4
6
20
40
60
80
Value
Common sites gene8262
1
2
3
4
5
6
7
8
9
10
11
CphaseA2
RphaseA2

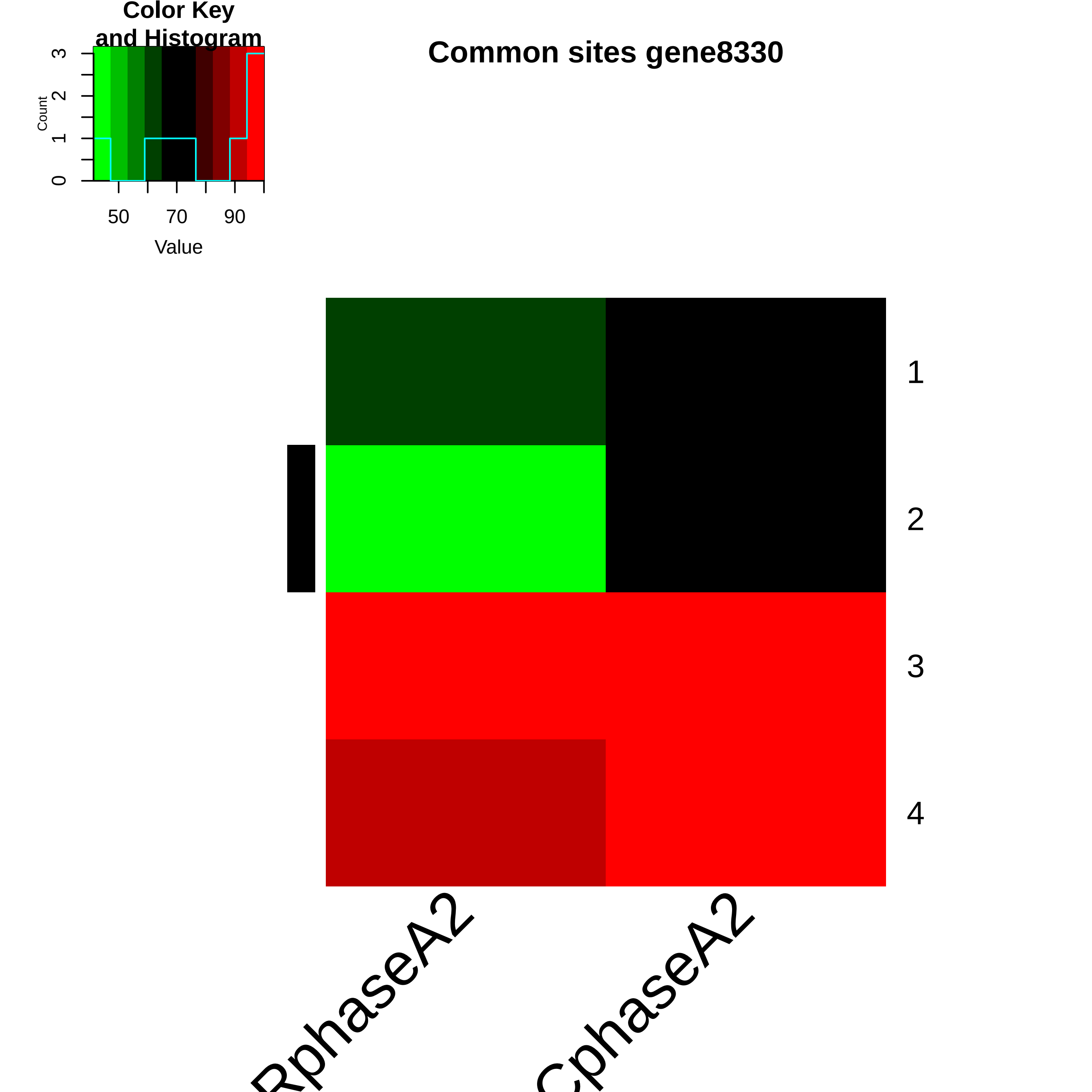
Color Key
and Histogram
Count
0
1
2
3
50
70
90
Value
Common sites gene8330
1
2
3
4
RphaseA2
CphaseA2

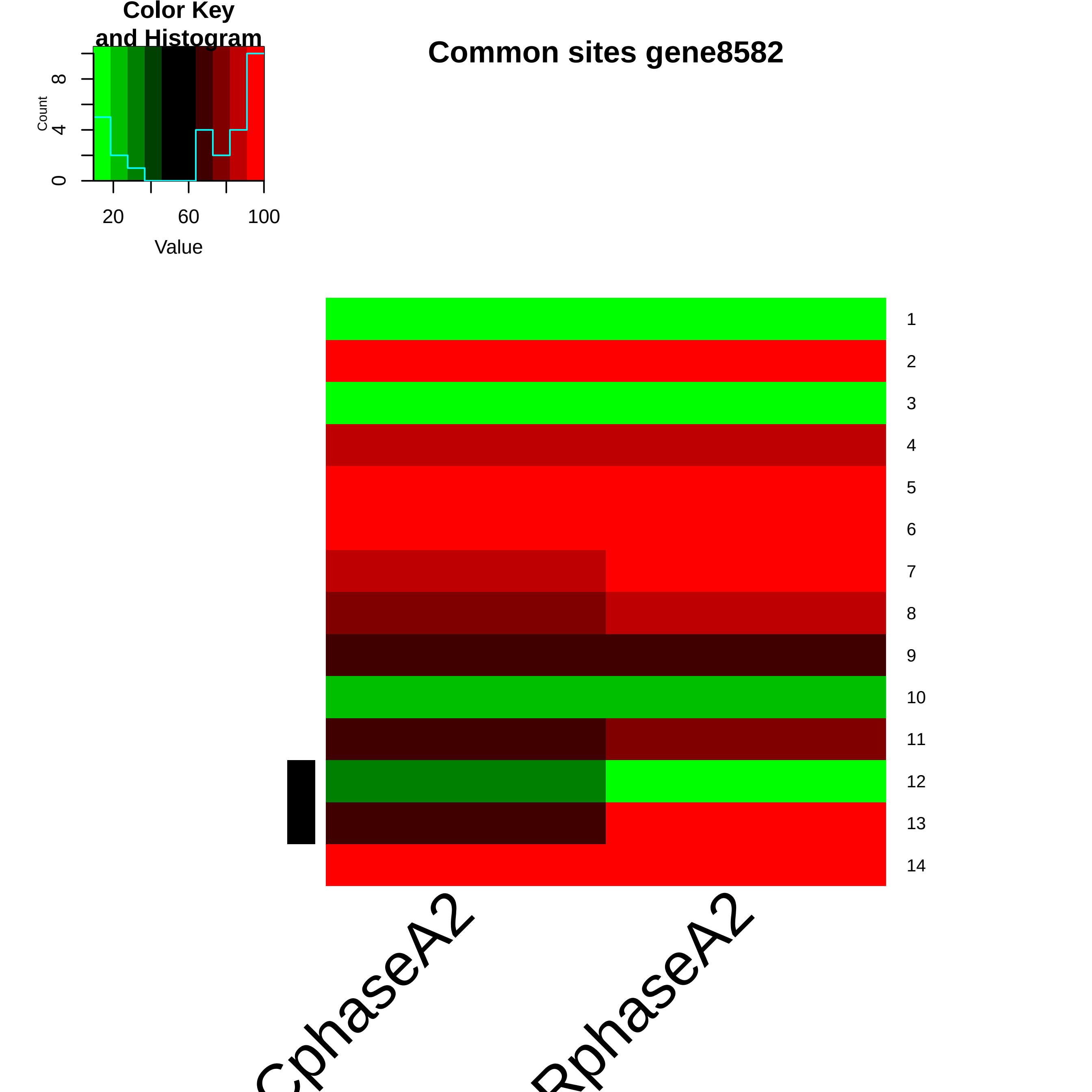
Color Key
and Histogram
Count
0
4
8
20
60
100
Value
Common sites gene8582
1
2
3
4
5
6
7
8
9
10
11
12
13
14
CphaseA2
RphaseA2

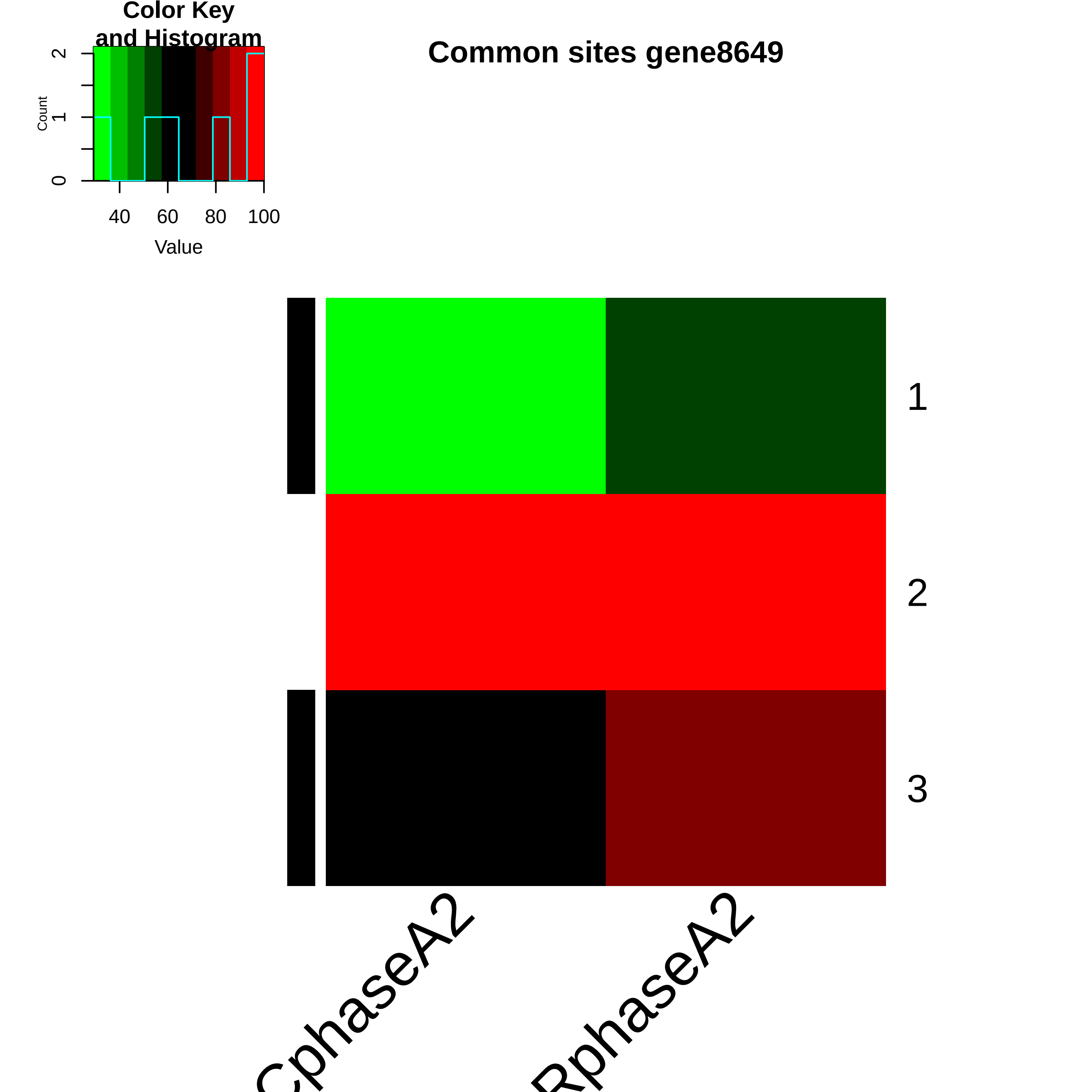

Color Key
and Histogram
Count
0
1
2
40
60
80
100
Value
Common sites gene8649
1
2
3
CphaseA2
RphaseA2

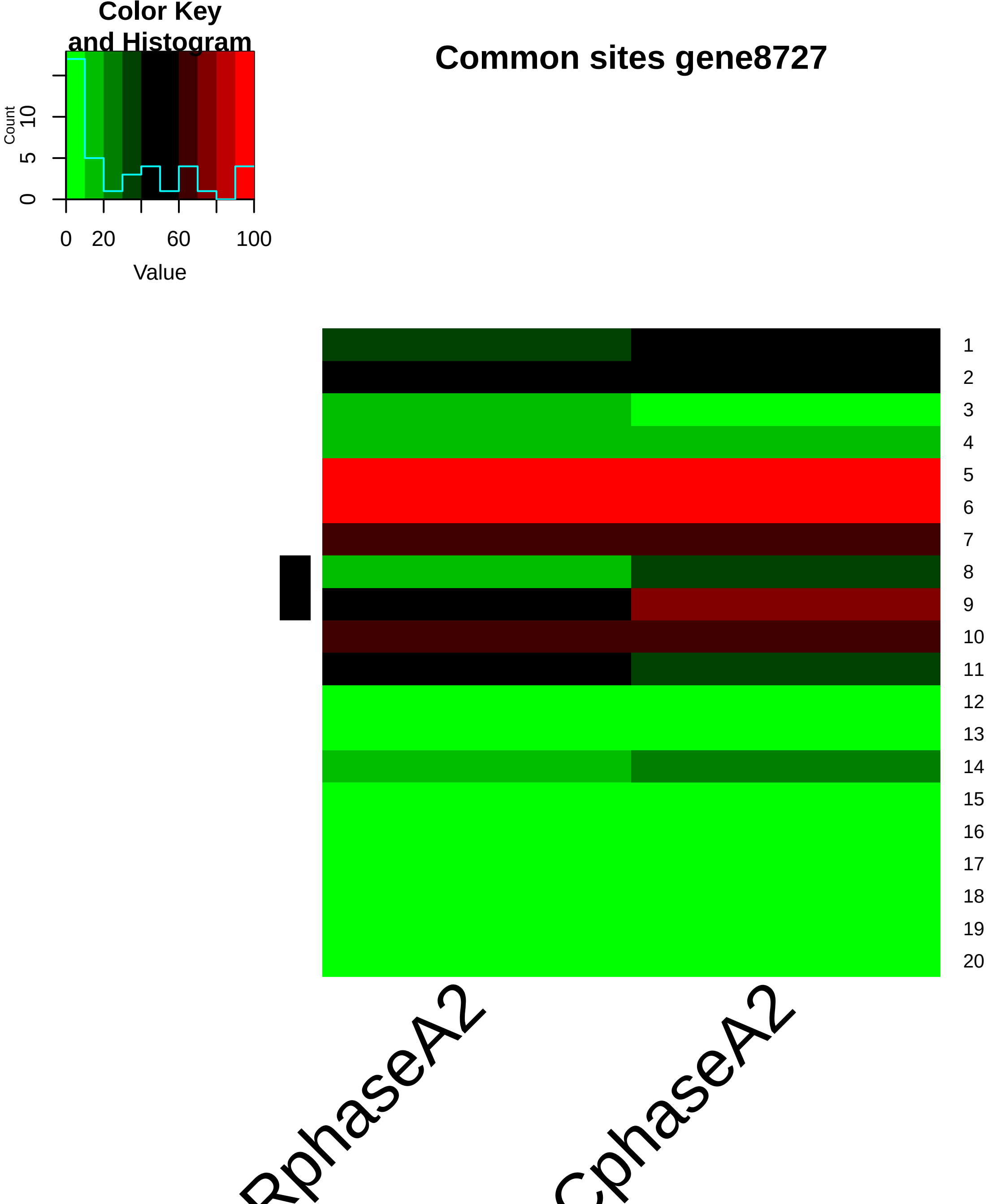
Color Key
and Histogram
Count
0
5
10
0
20
60
100
Value
Common sites gene8727
RphaseA2
CphaseA2

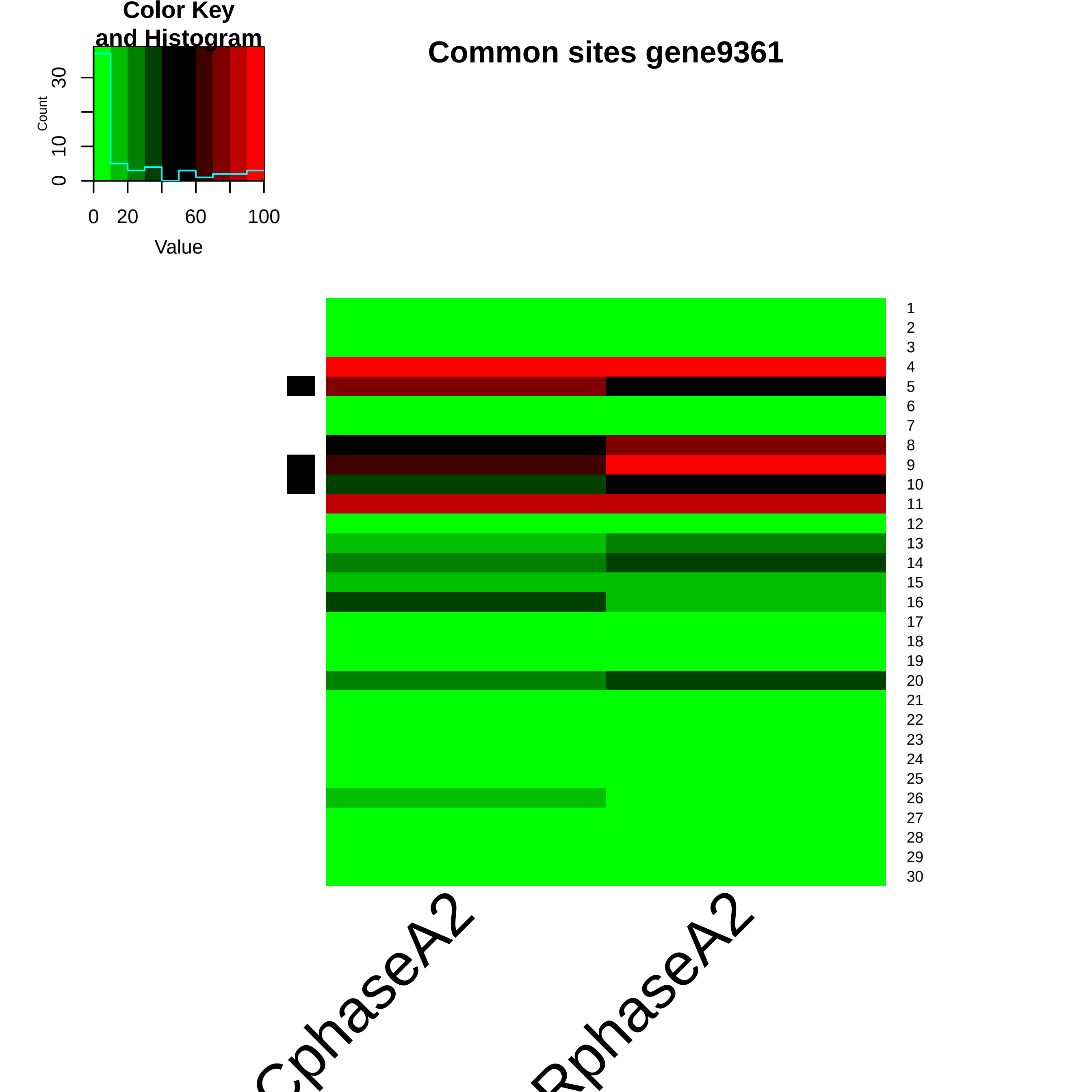
Color Key
and Histogram
Count
0
10
30
0
20
60
100
Value
Common sites gene9361
CphaseA2
RphaseA2

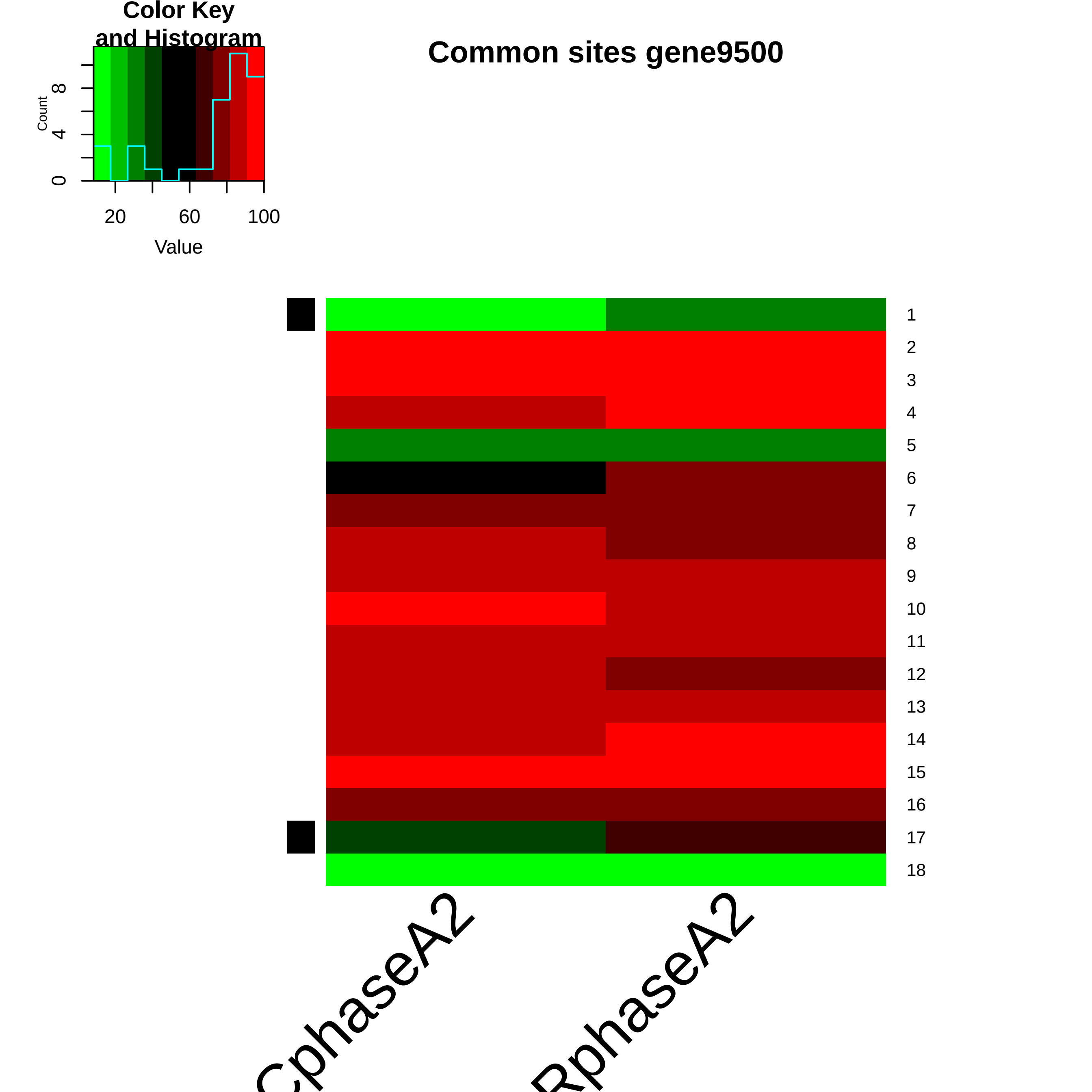

Color Key
and Histogram
Count
0
4
8
20
60
100
Value
Common sites gene9500
CphaseA2
RphaseA2

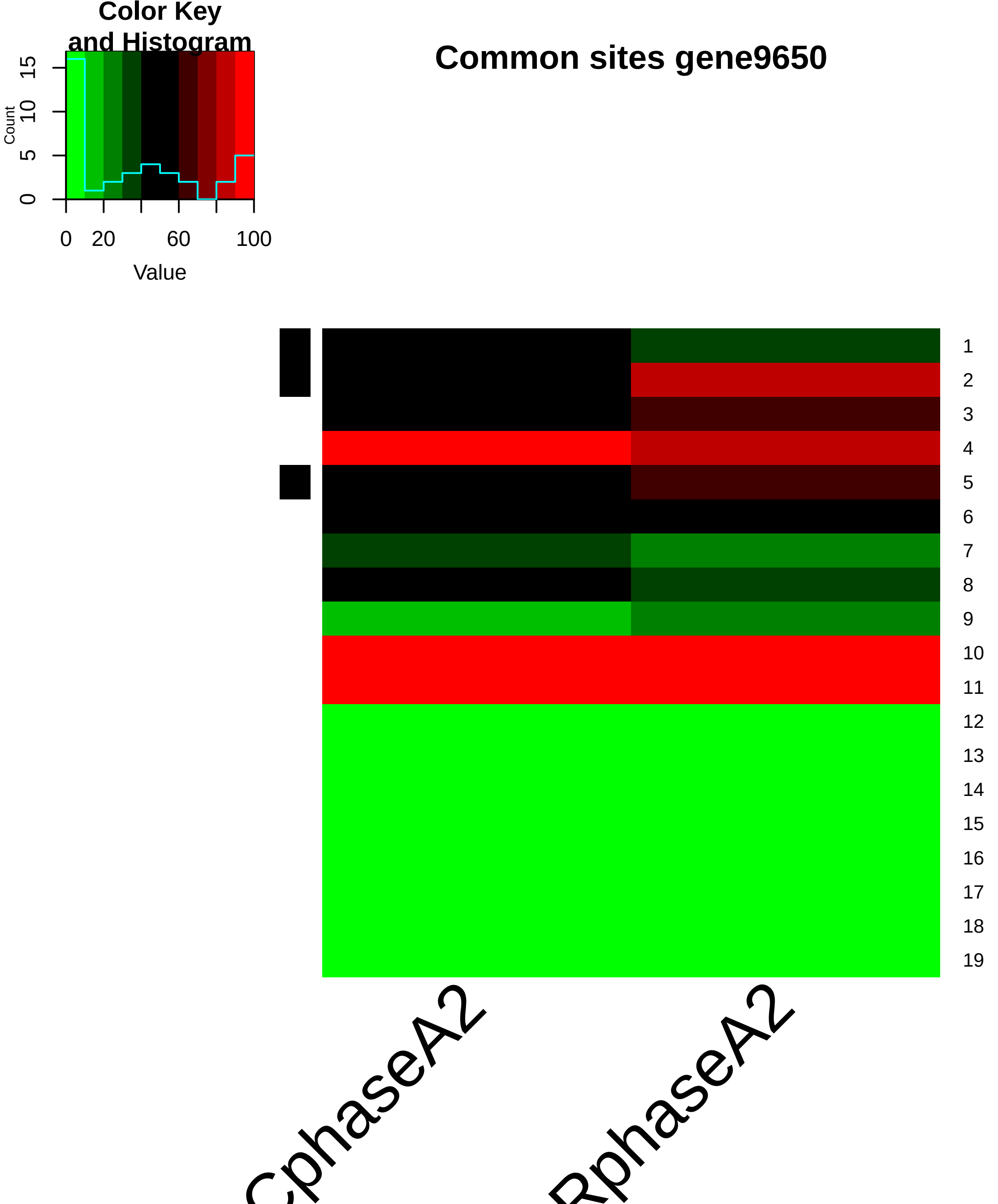
Color Key
and Histogram
Count
0
5
10
15
0
20
60
100
Value
Common sites gene9650
CphaseA2
RphaseA2

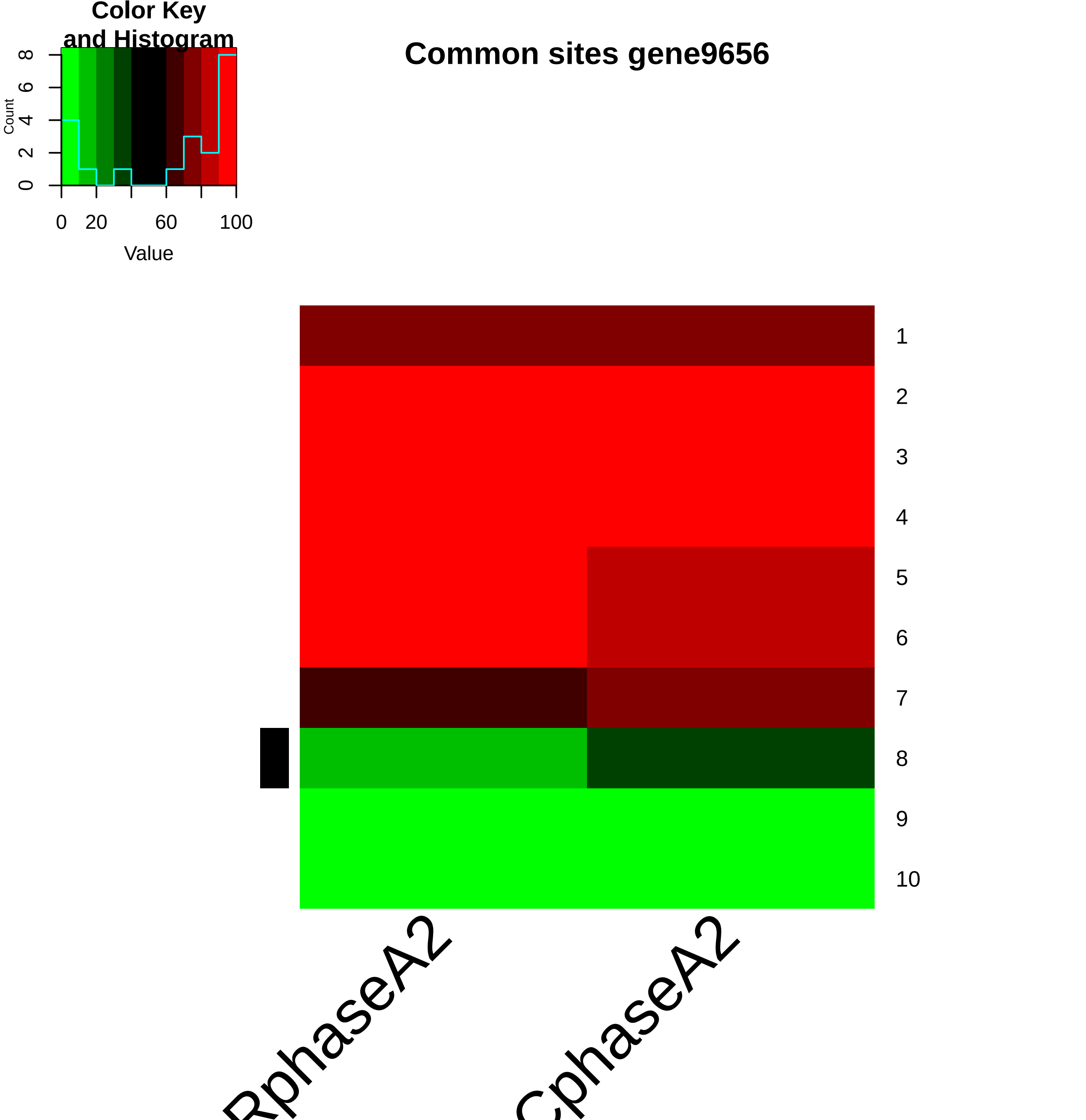
Color Key
and Histogram
Count
0
2
4
6
8
0
20
60
100
Value
Common sites gene9656
1
2
3
4
5
6
7
8
9
10
RphaseA2
CphaseA2

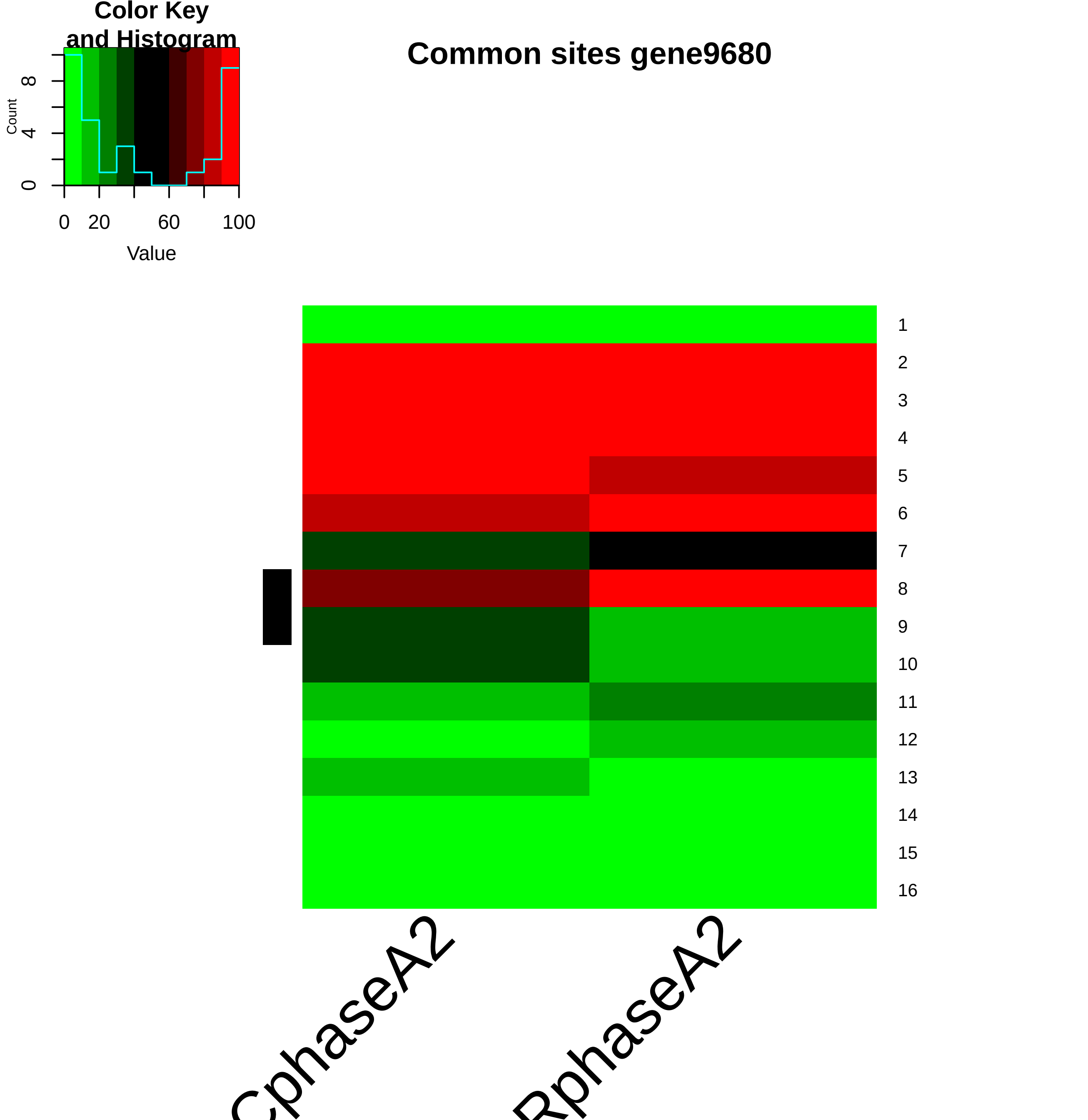
Color Key
and Histogram
Count
0
4
8
0
20
60
100
Value
Common sites gene9680
CphaseA2
RphaseA2

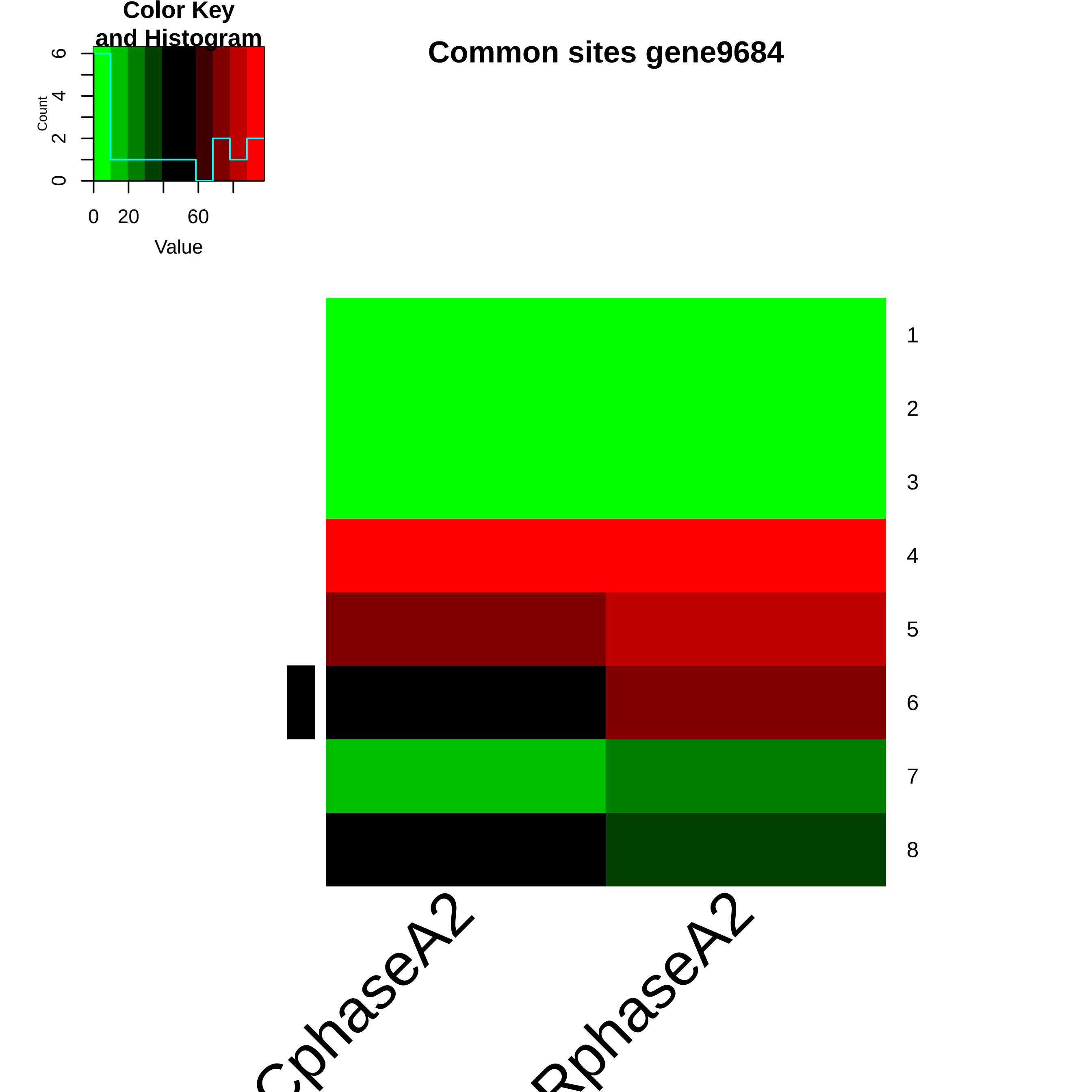

Color Key
and Histogram
Count
0
2
4
6
0
20
60
Value
Common sites gene9684
1
2
3
4
5
6
7
8
CphaseA2
RphaseA2

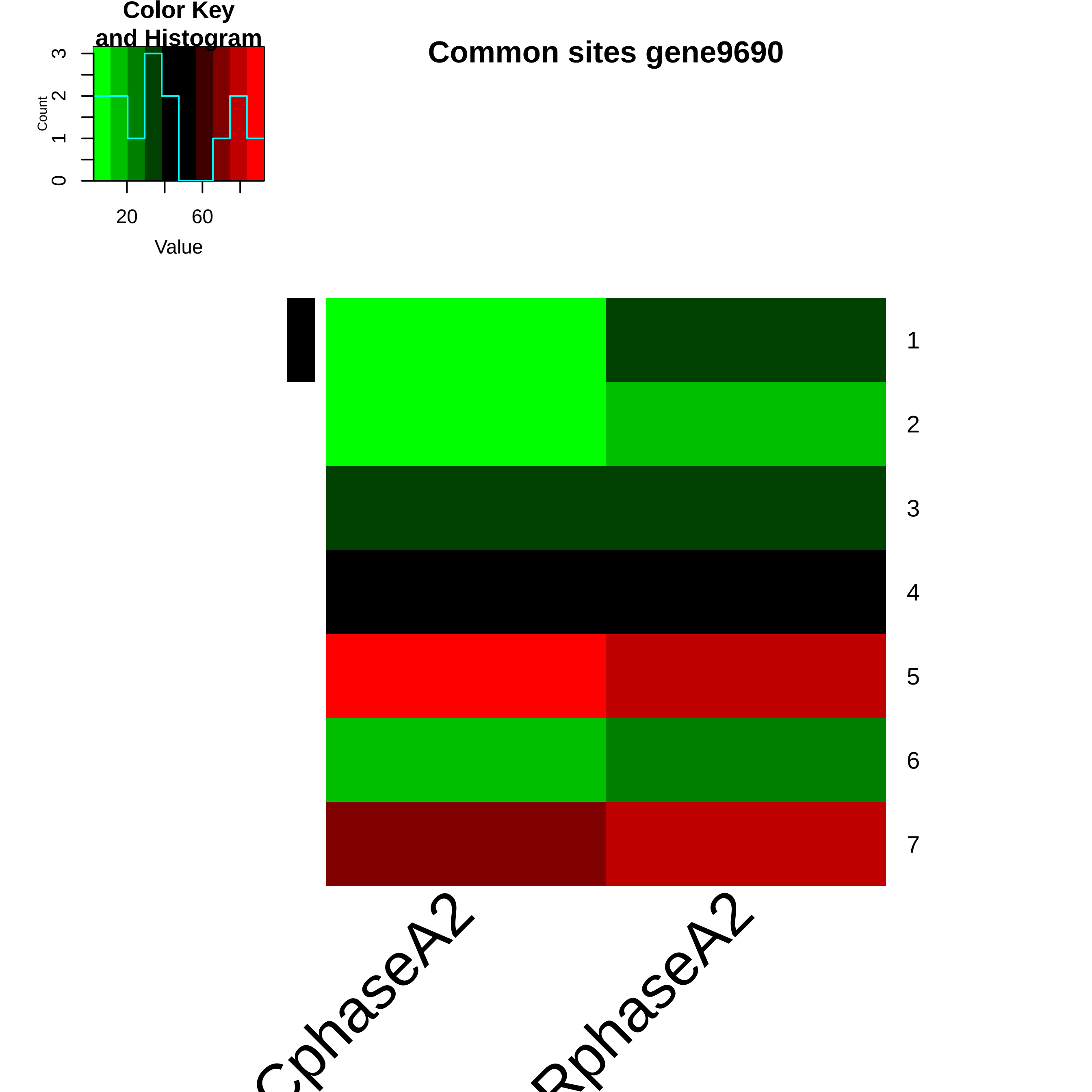

Color Key
and Histogram
Count
0
1
2
3
20
60
Value
Common sites gene9690
1
2
3
4
5
6
7

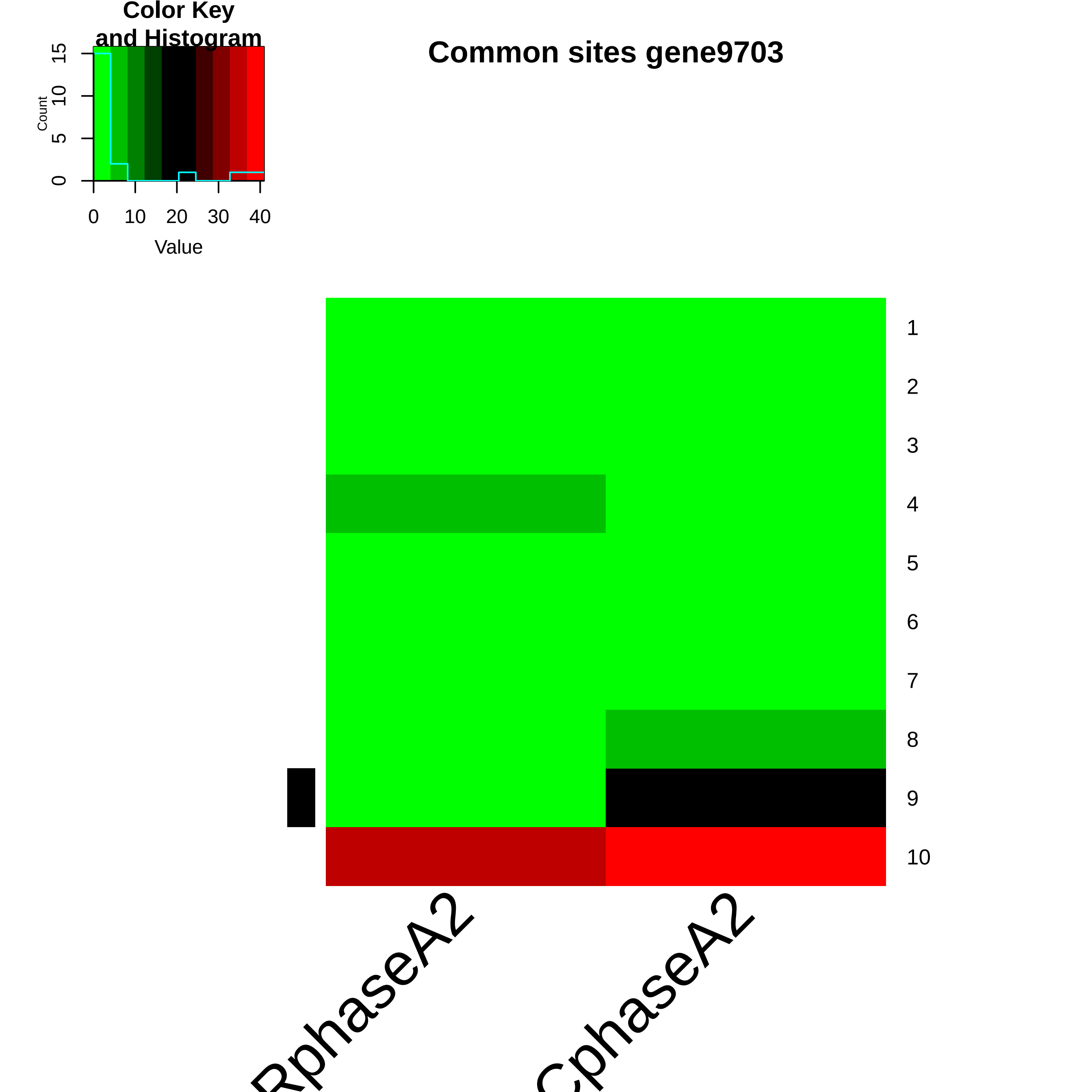
Color Key
and Histogram
Count
0
5
10
15
0
10
20
30
40
Value
Common sites gene9703
1
2
3
4
5
6
7
8
9
10
RphaseA2
CphaseA2

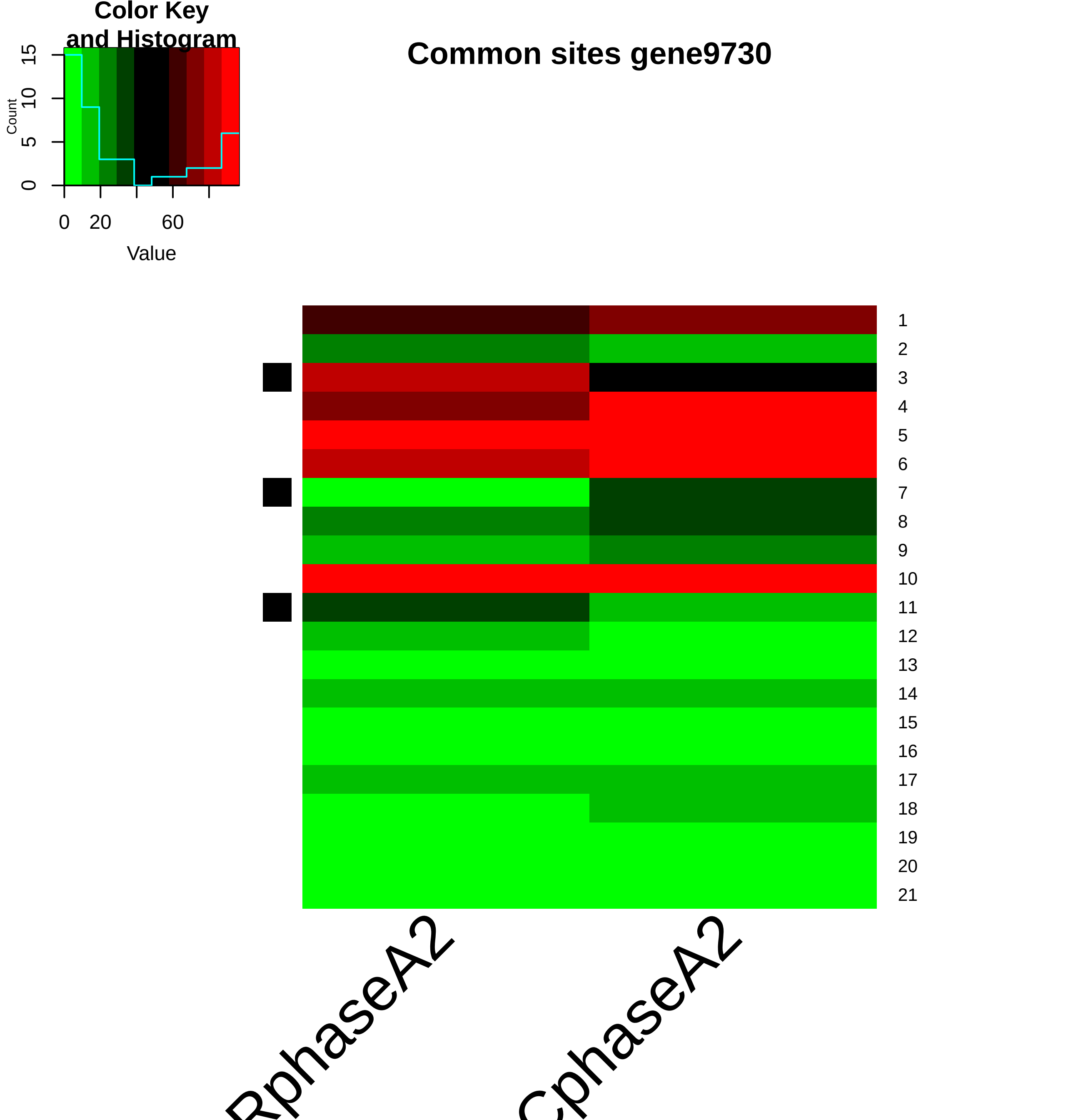

Color Key
and Histogram
Count
0
5
10
15
0
20
60
Value
Common sites gene9730
RphaseA2
CphaseA2

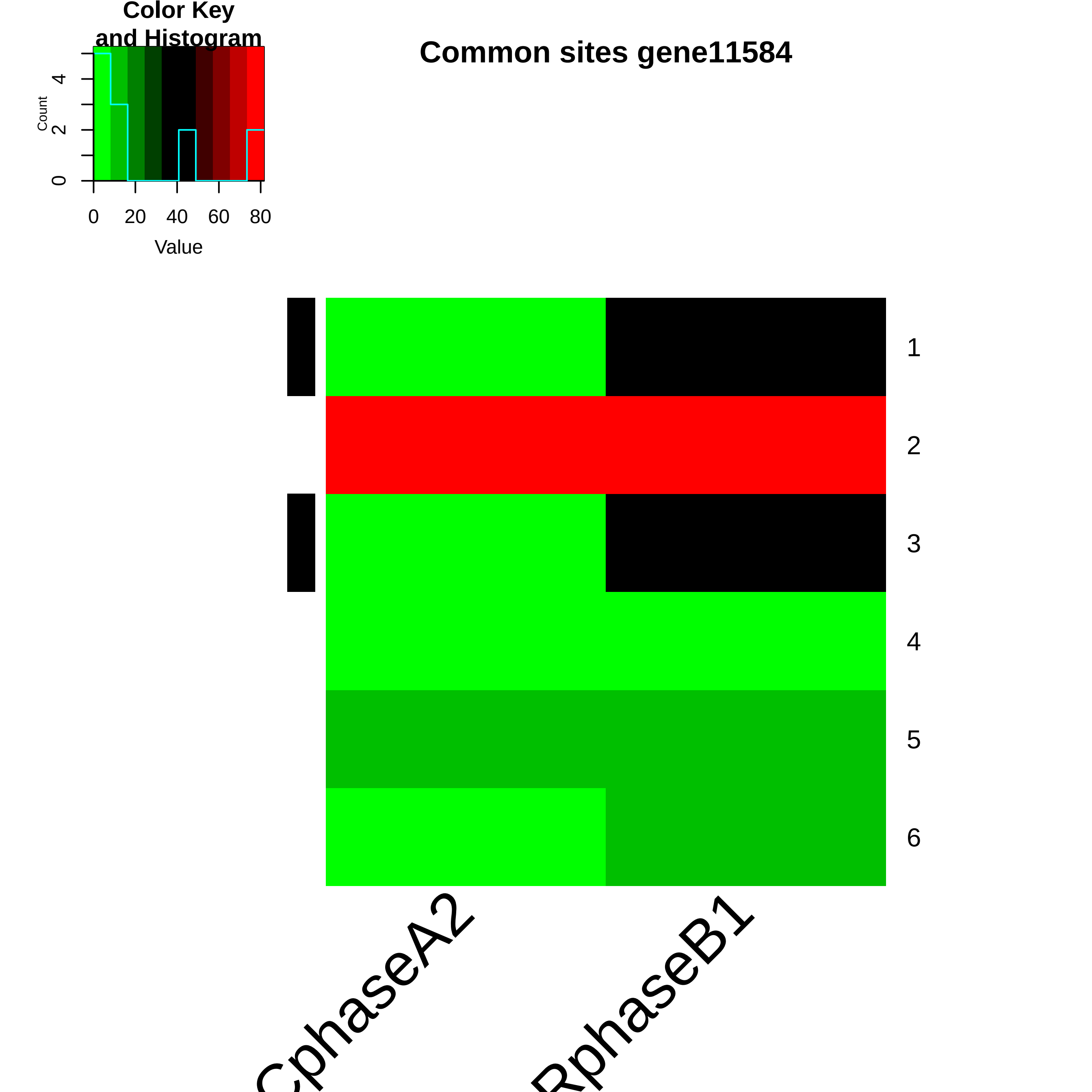

Color Key
and Histogram
Count
0
2
4
0
20
40
60
80
Value
Common sites gene11584
1
2
3
4
5
6
CphaseA2
RphaseB1

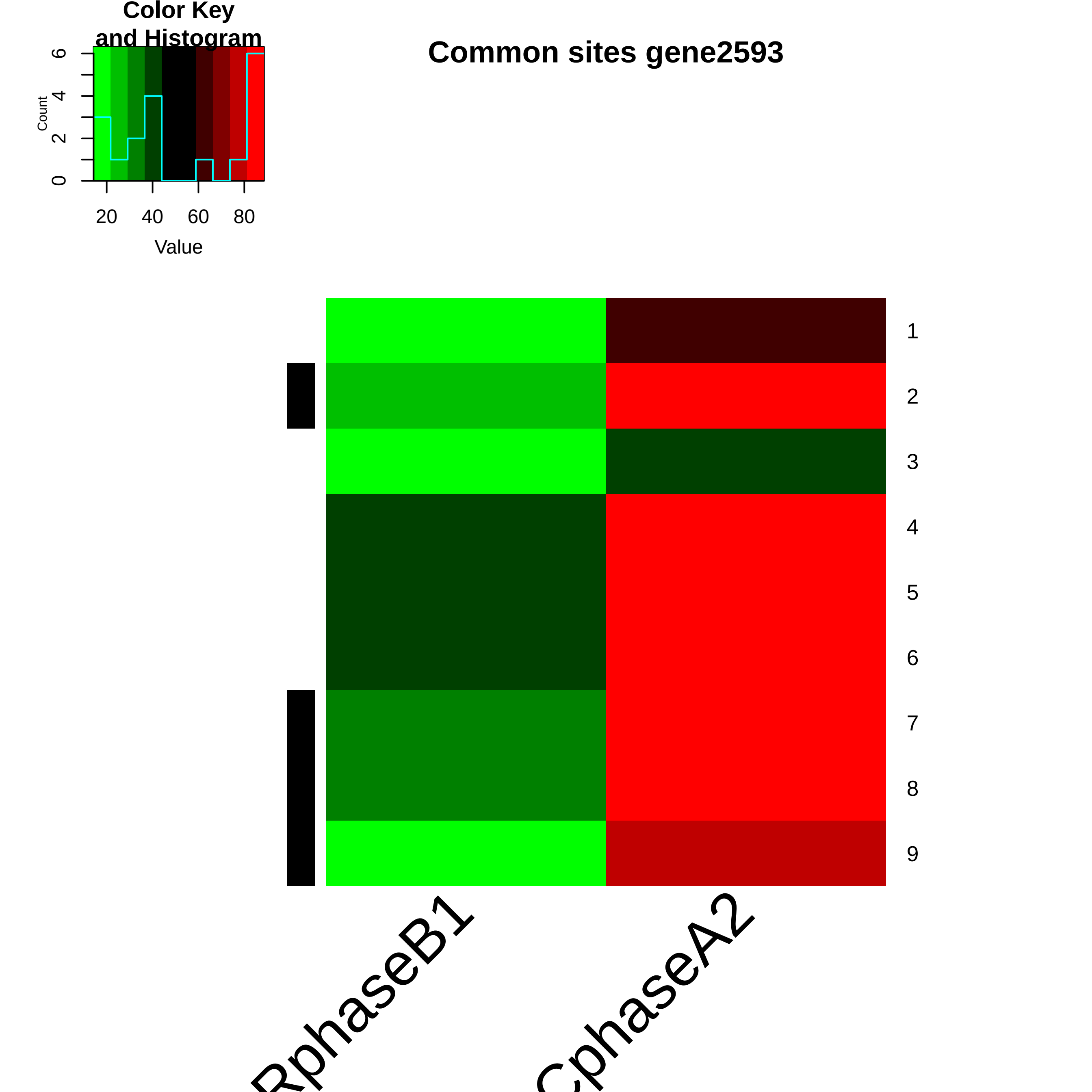

Color Key
and Histogram
Count
Value
0
2
4
6
20
40
60
80
Common sites gene2593
1
2
3
4
5
6
7
8
9
RphaseB1
CphaseA2

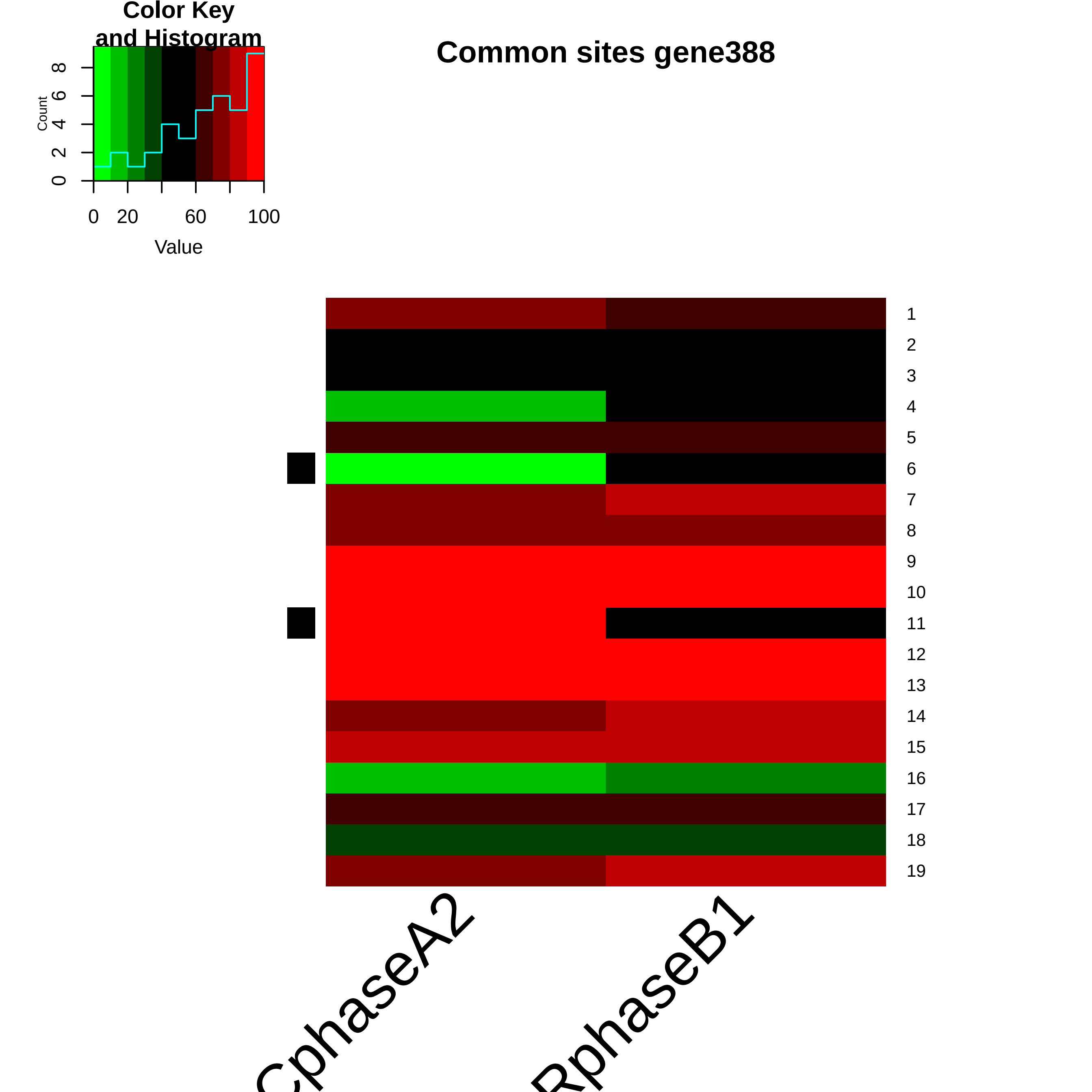

Color Key
and Histogram
Count
0
2
4
6
8
0
20
60
100
Value
Common sites gene388
CphaseA2
RphaseB1

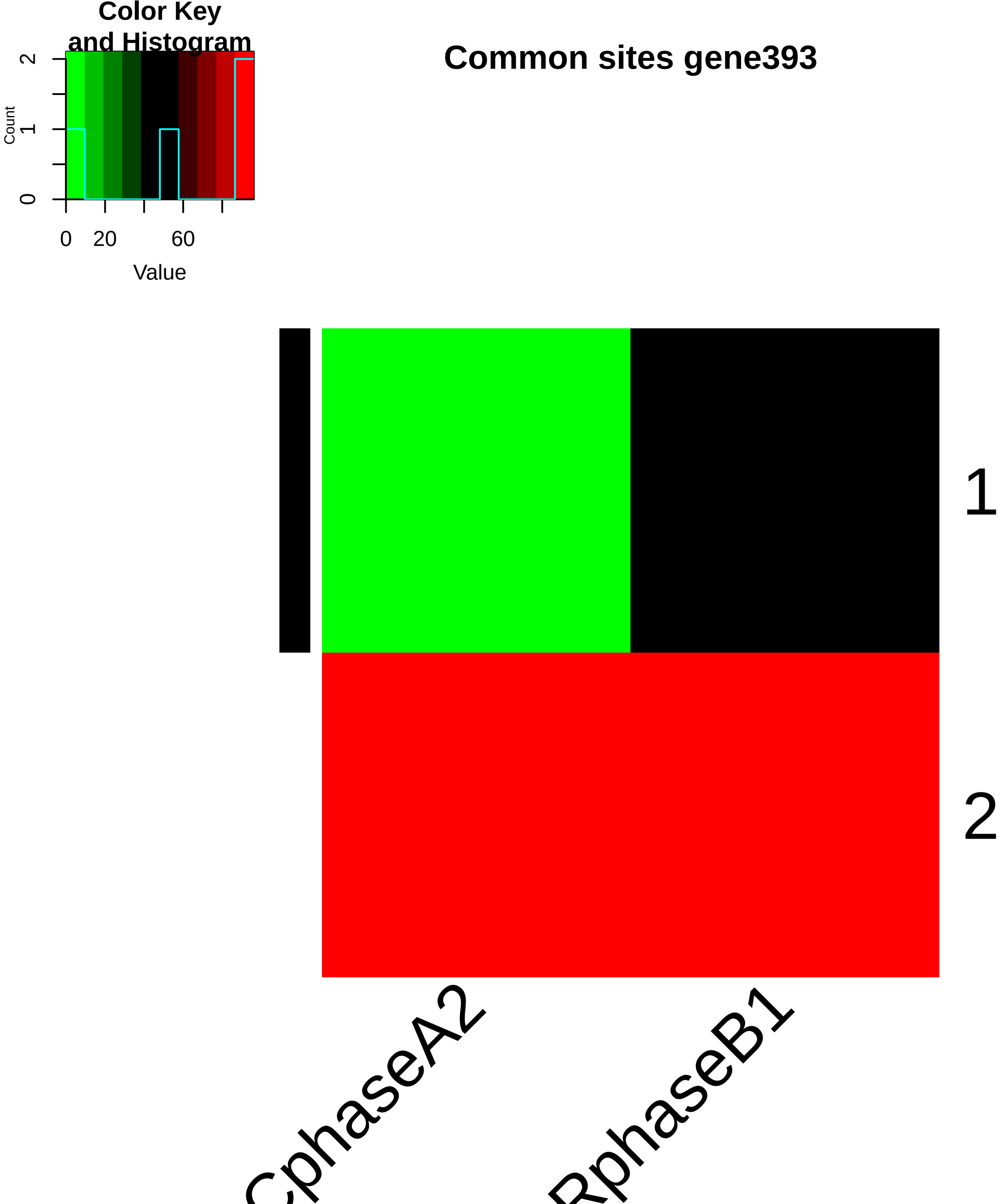

Color Key
and Histogram
Count
0
1
2
0
20
60
Value
Common sites gene393
1
2
CphaseA2
RphaseB1

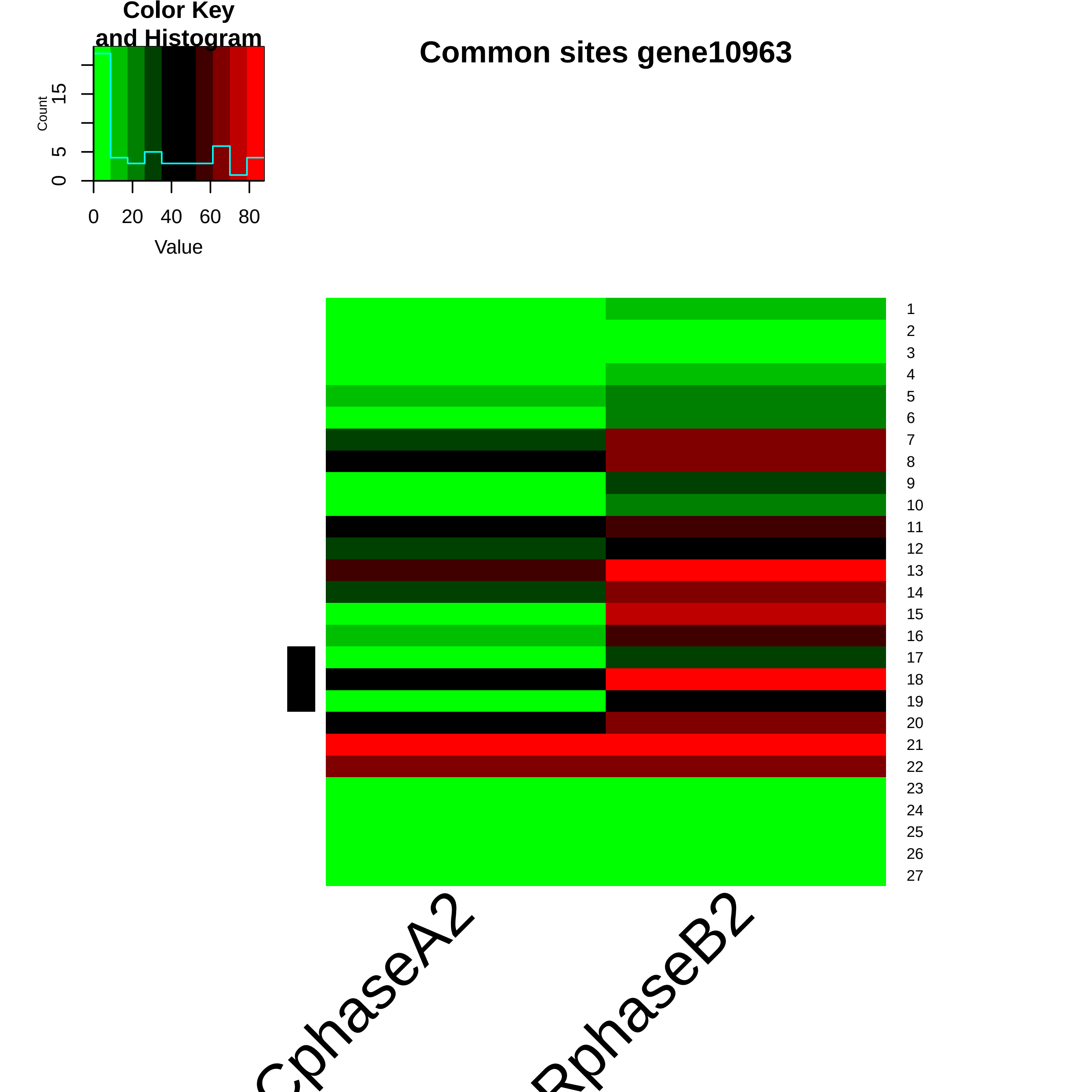

Color Key
and Histogram
Count
0
5
15
0
20
40
60
80
Value
Common sites gene10963
CphaseA2
RphaseB2

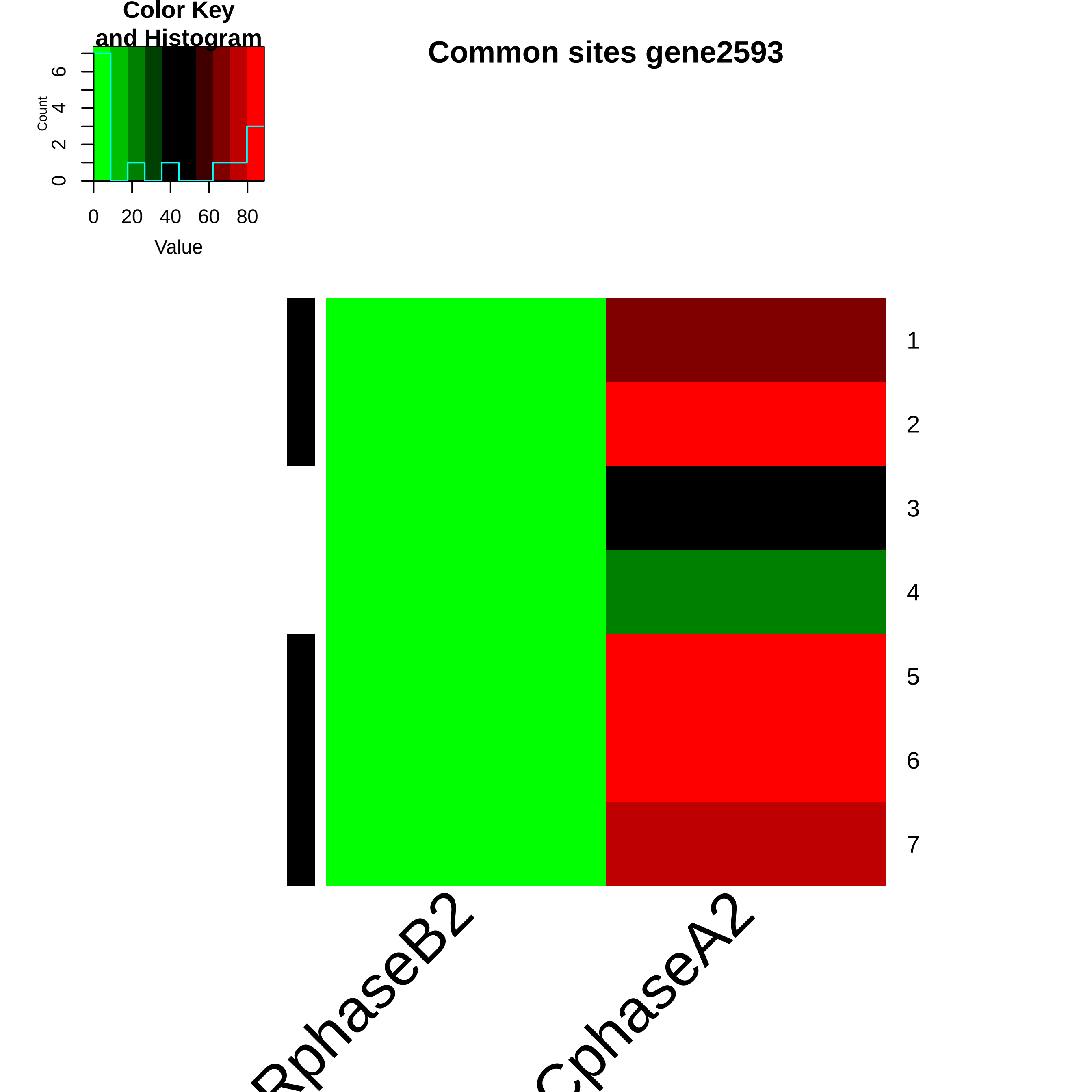

Color Key
and Histogram
Count
0
2
4
6
0
20
40
60
80
Value
Common sites gene2593
1
2
3
4
5
6
7
RphaseB2
CphaseA2

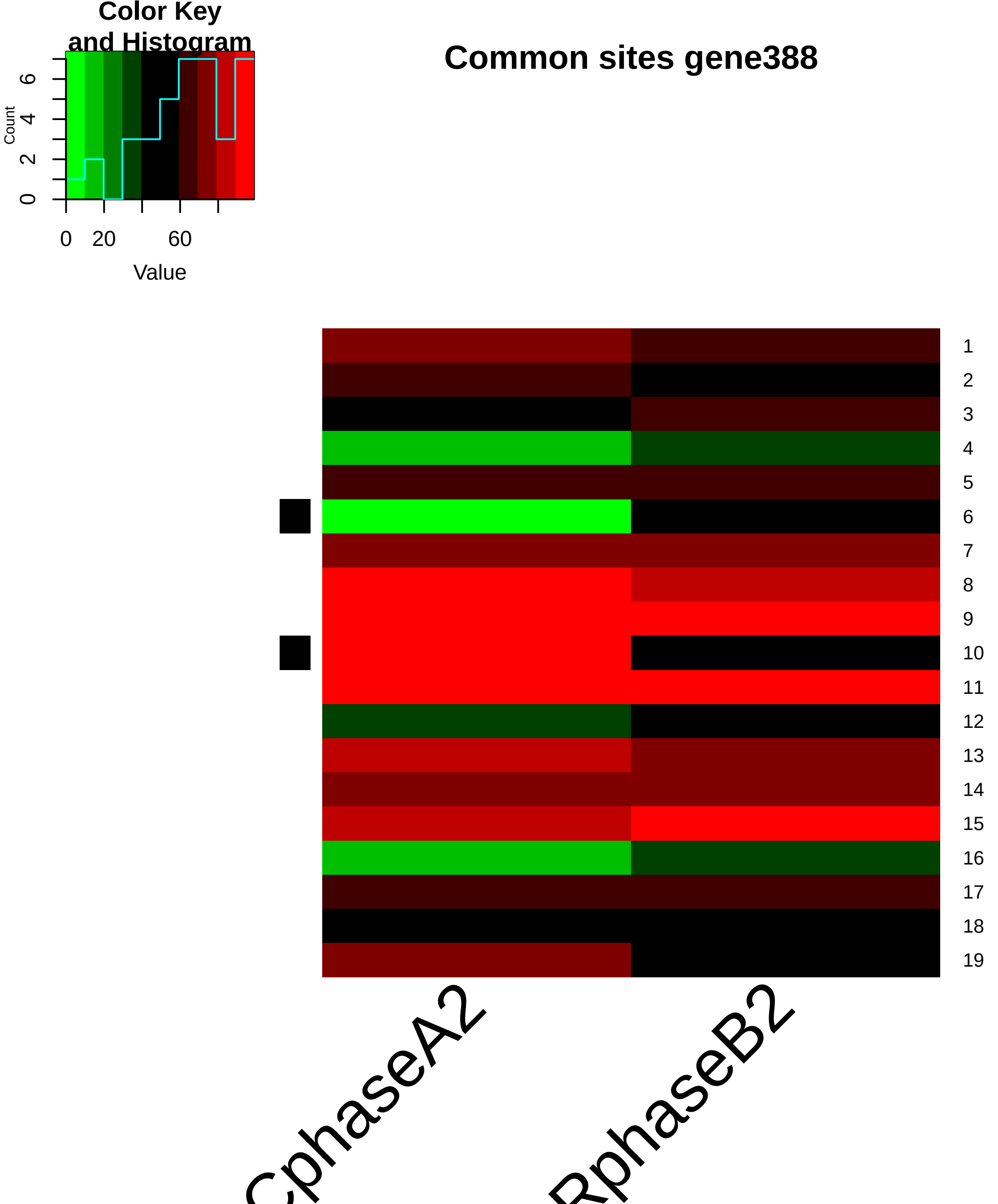
Color Key
and Histogram
Count
0
2
4
6
0
20
60
Value
Common sites gene388
CphaseA2
RphaseB2

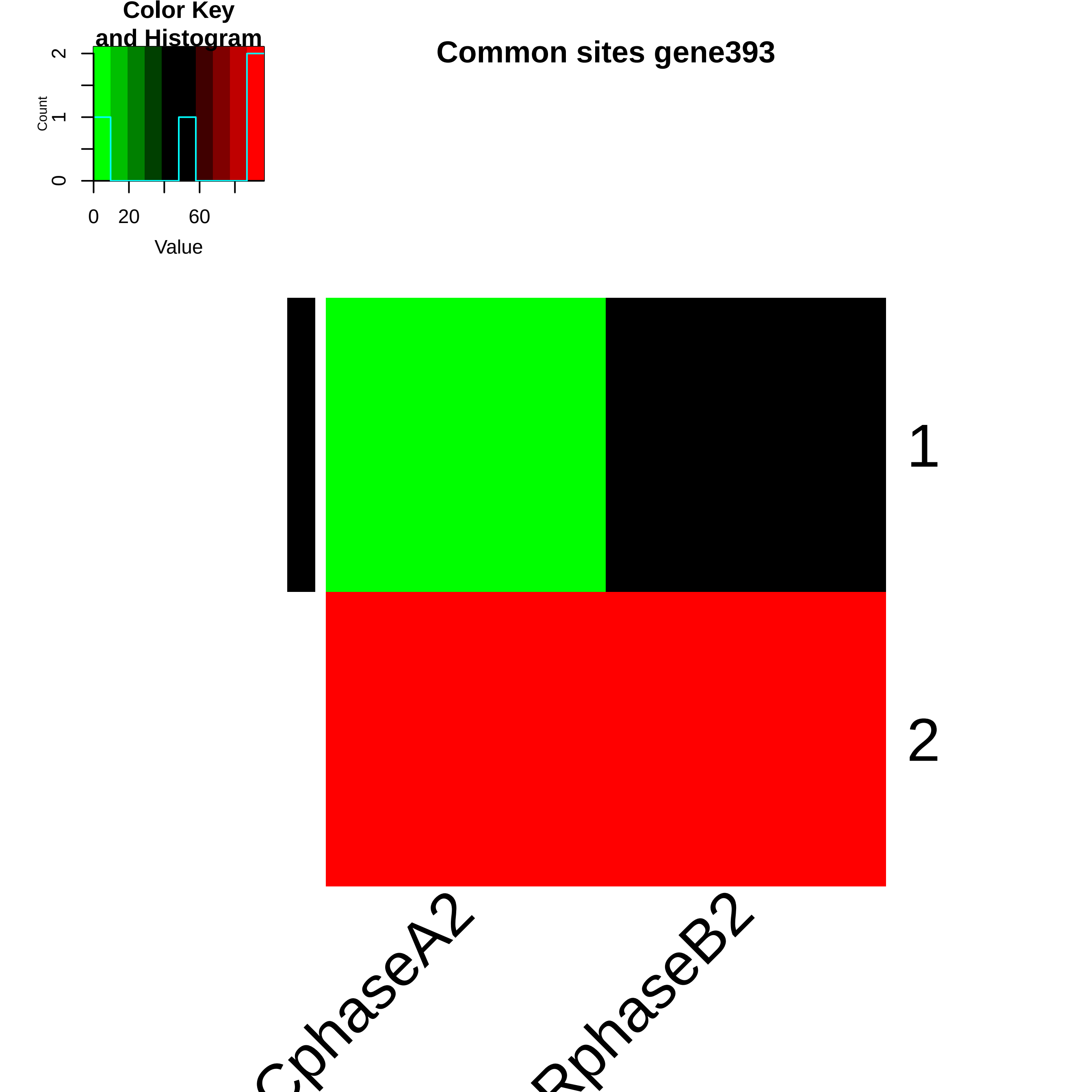
Color Key
and Histogram
Count
0
1
2
0
20
60
Value
Common sites gene393
1
2

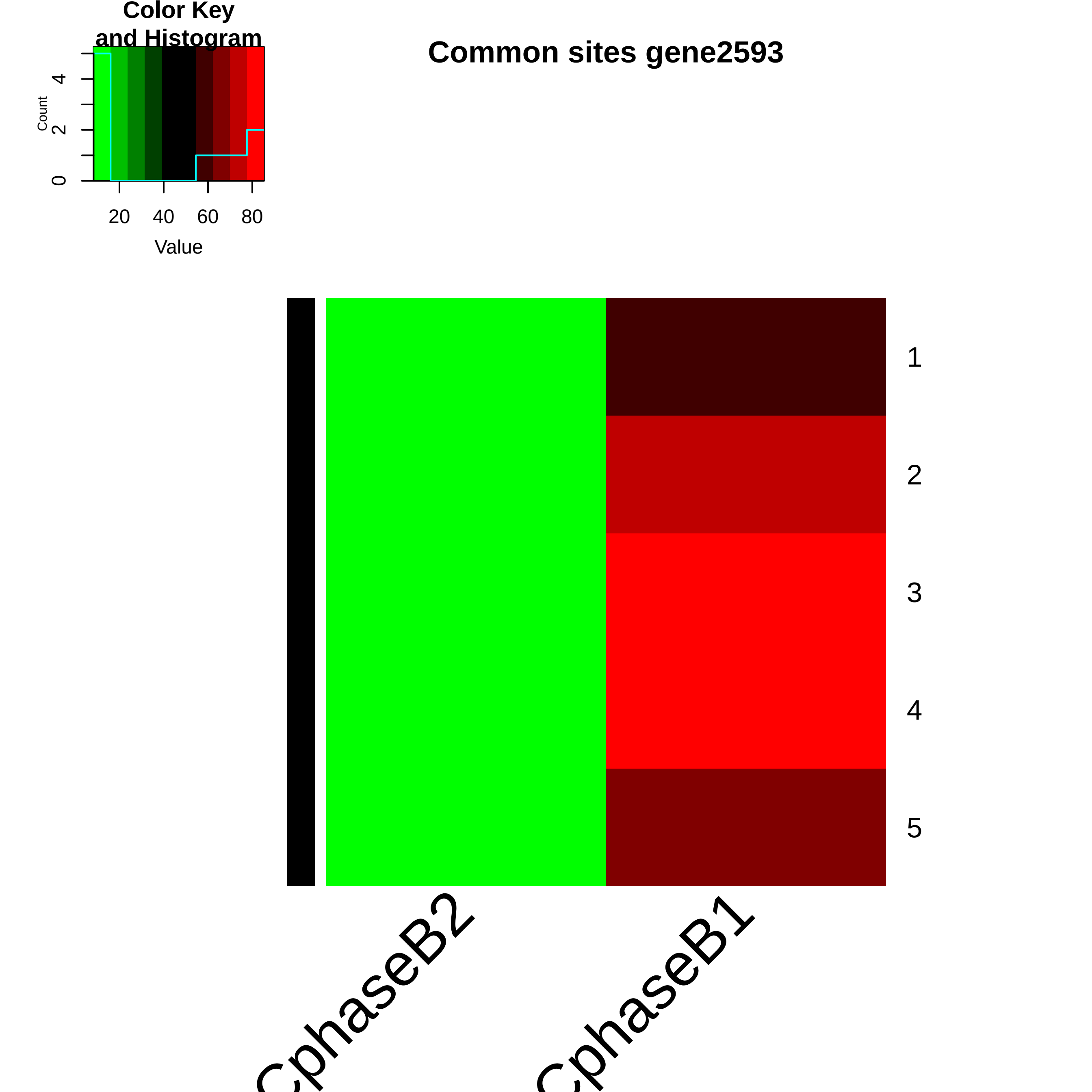

Color Key
and Histogram
Count
0
2
4
20
40
60
80
Value
Common sites gene2593
1
2
3
4
5
CphaseB2
CphaseB1

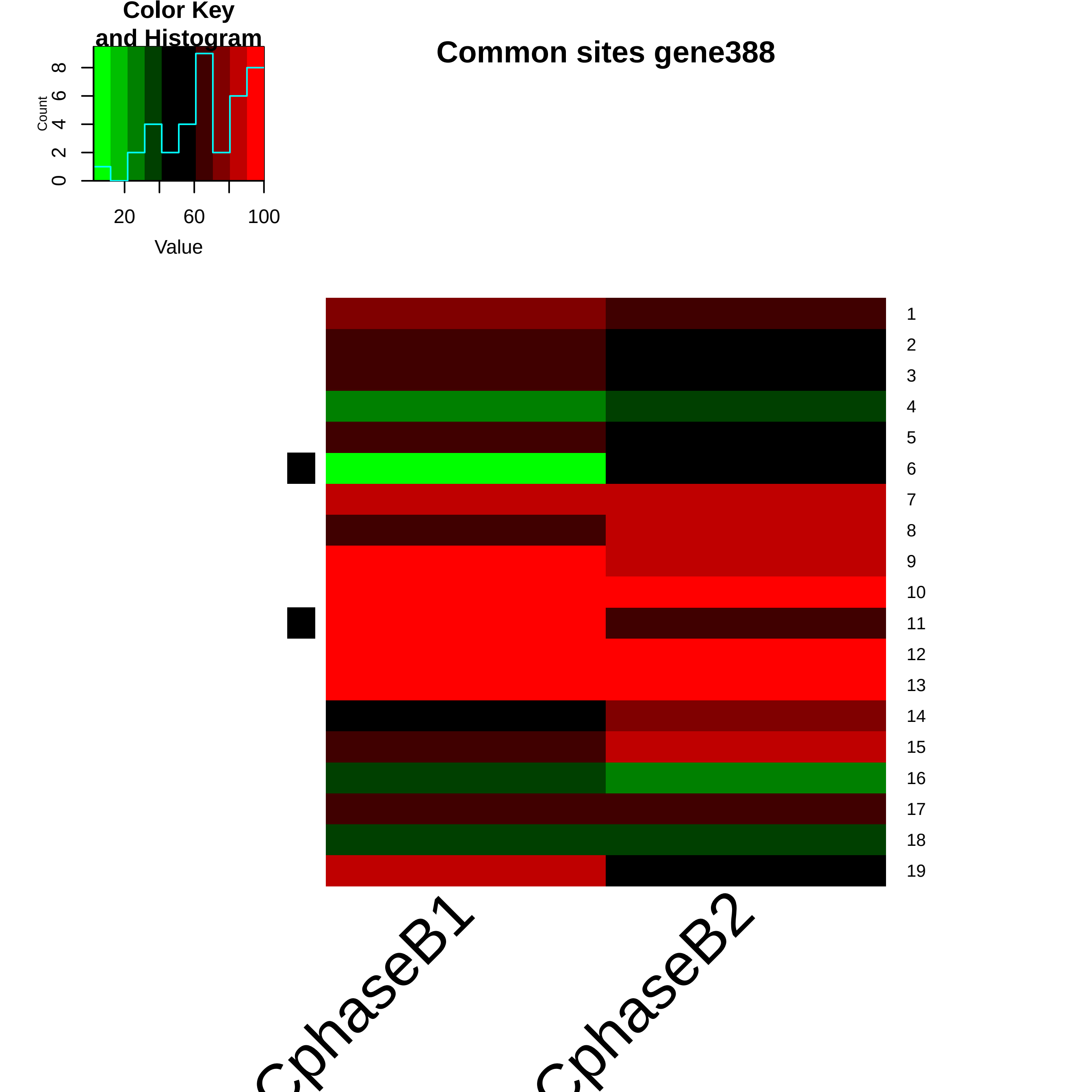

Color Key
and Histogram
Count
0
2
4
6
8
20
60
100
Value
Common sites gene388
CphaseB1
CphaseB2

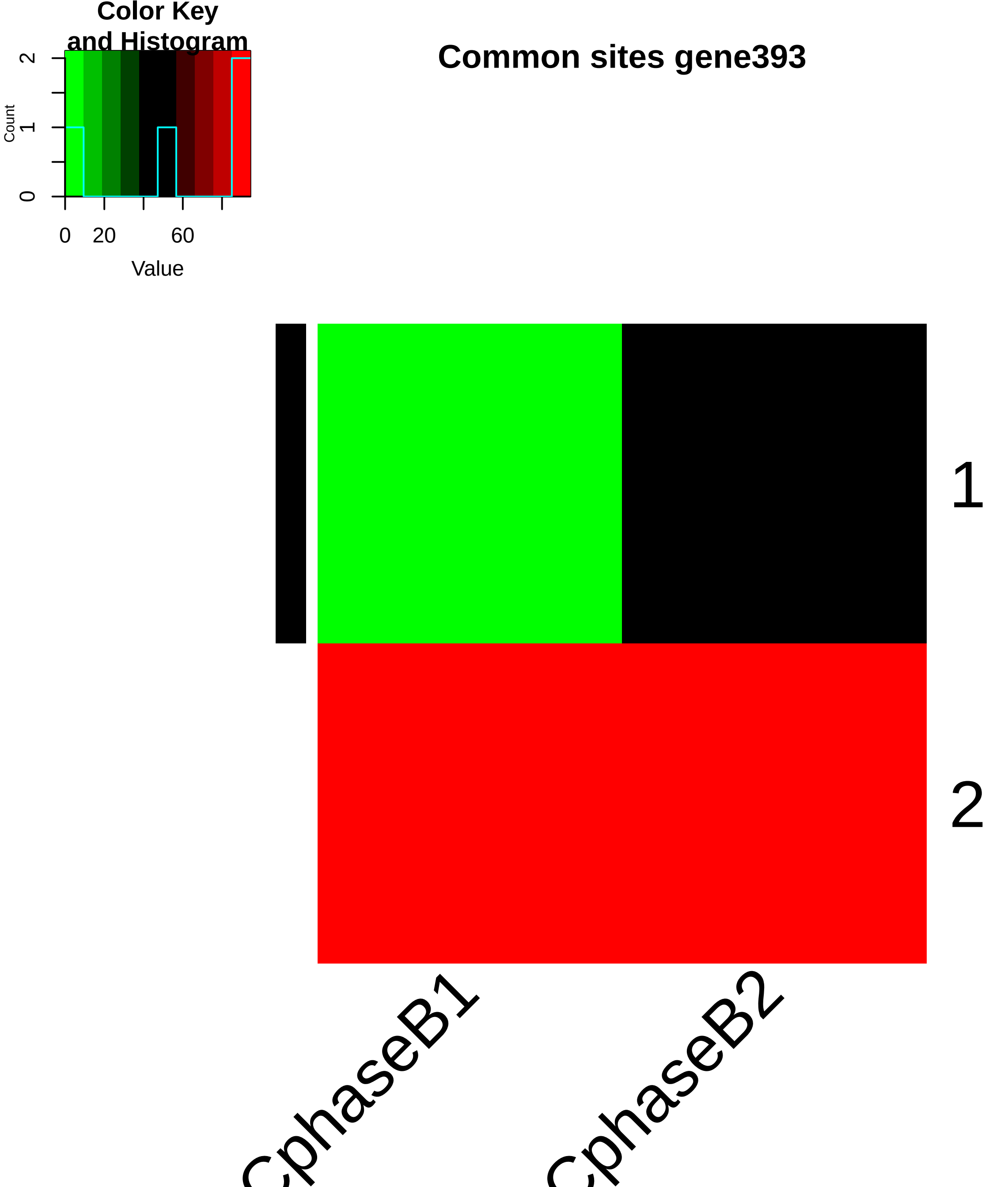
Color Key
and Histogram
Count
0
1
2
0
20
60
Value
Common sites gene393
1
2
CphaseB1
CphaseB2

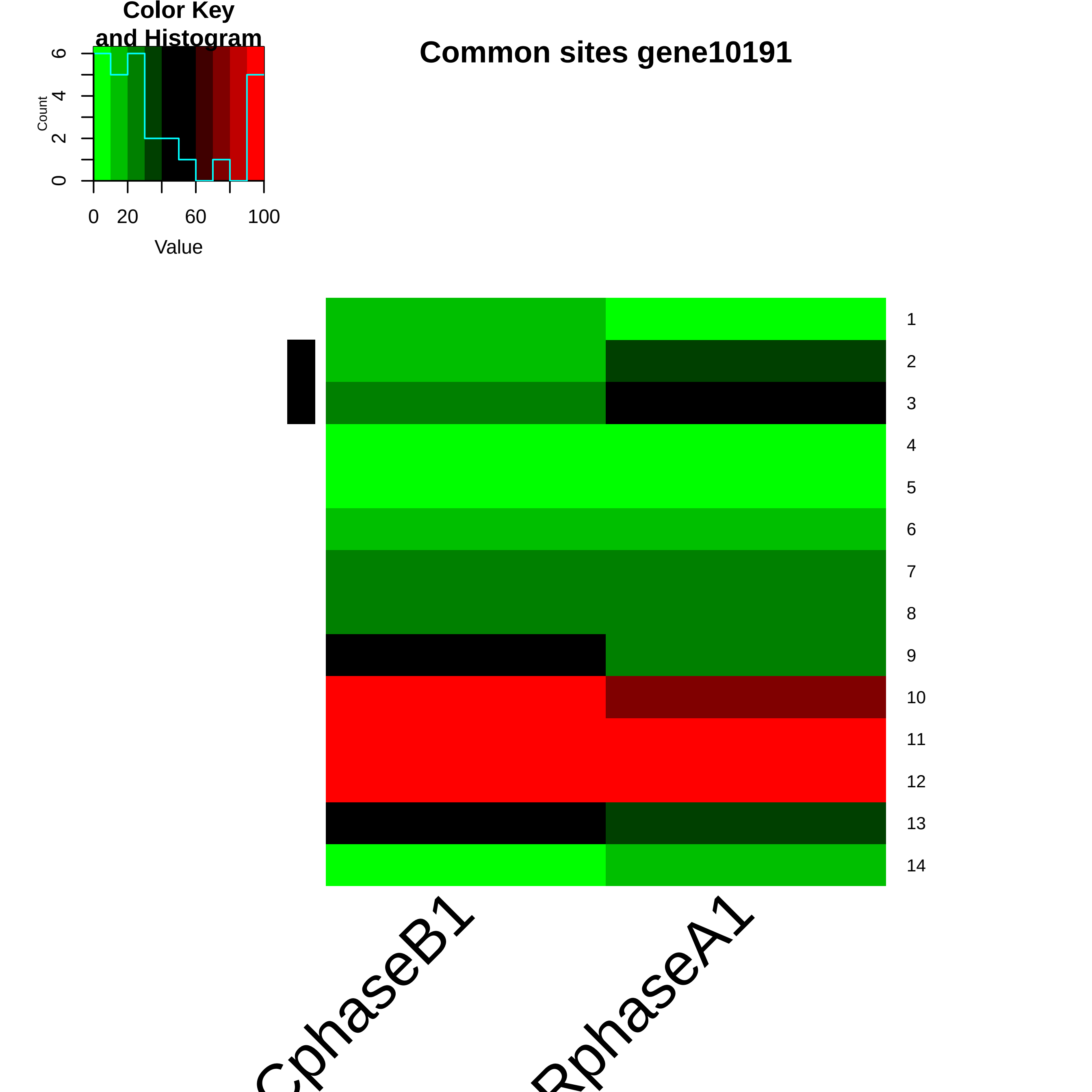
Color Key
and Histogram
Count
0
2
4
6
0
20
60
100
Value
Common sites gene10191
1
2
3
4
5
6
7
8
9
10
11
12
13
14
CphaseB1
RphaseA1

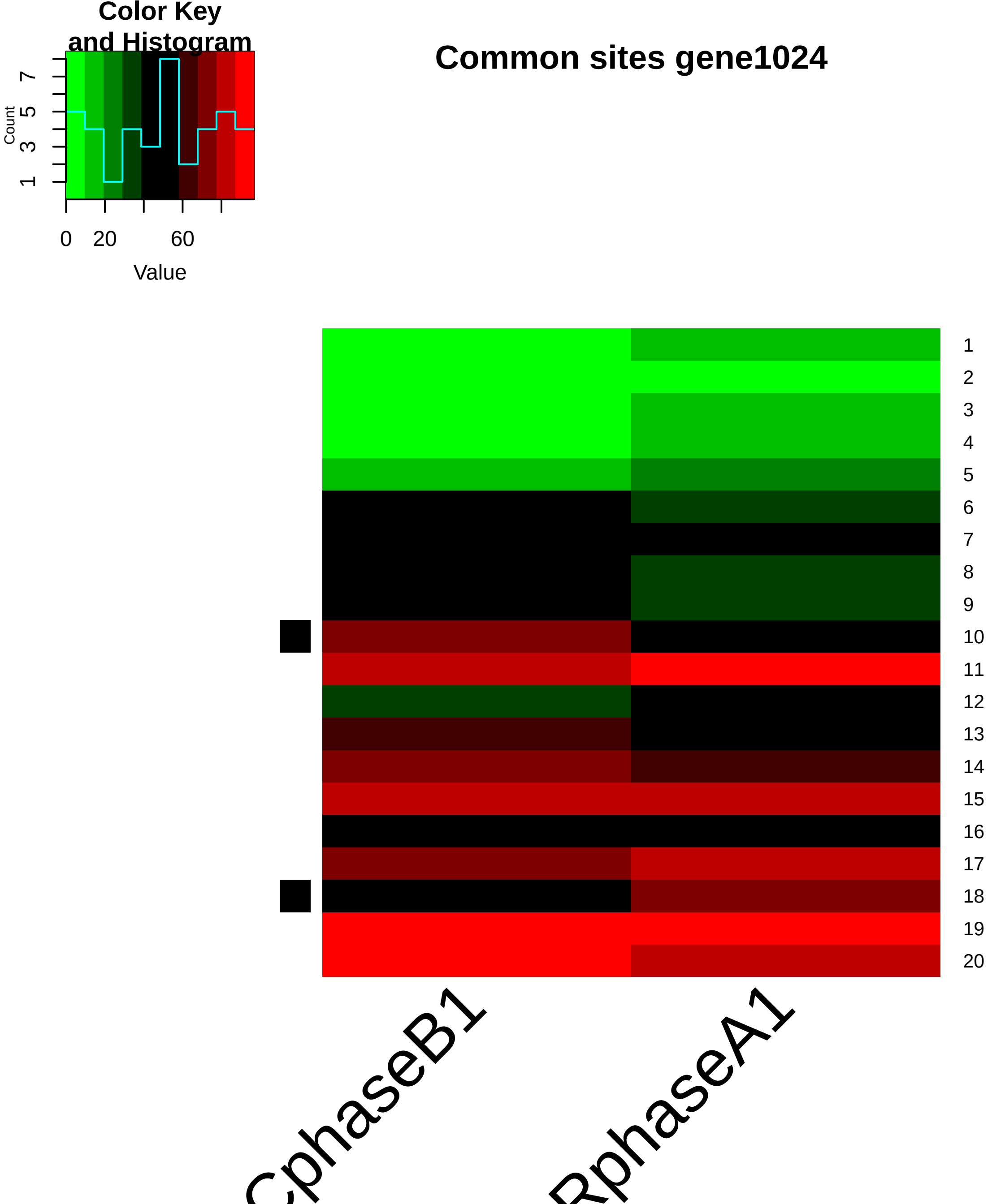
Color Key
and Histogram
Count
1
3
5
7
0
20
60
Value
Common sites gene1024
CphaseB1
RphaseA1

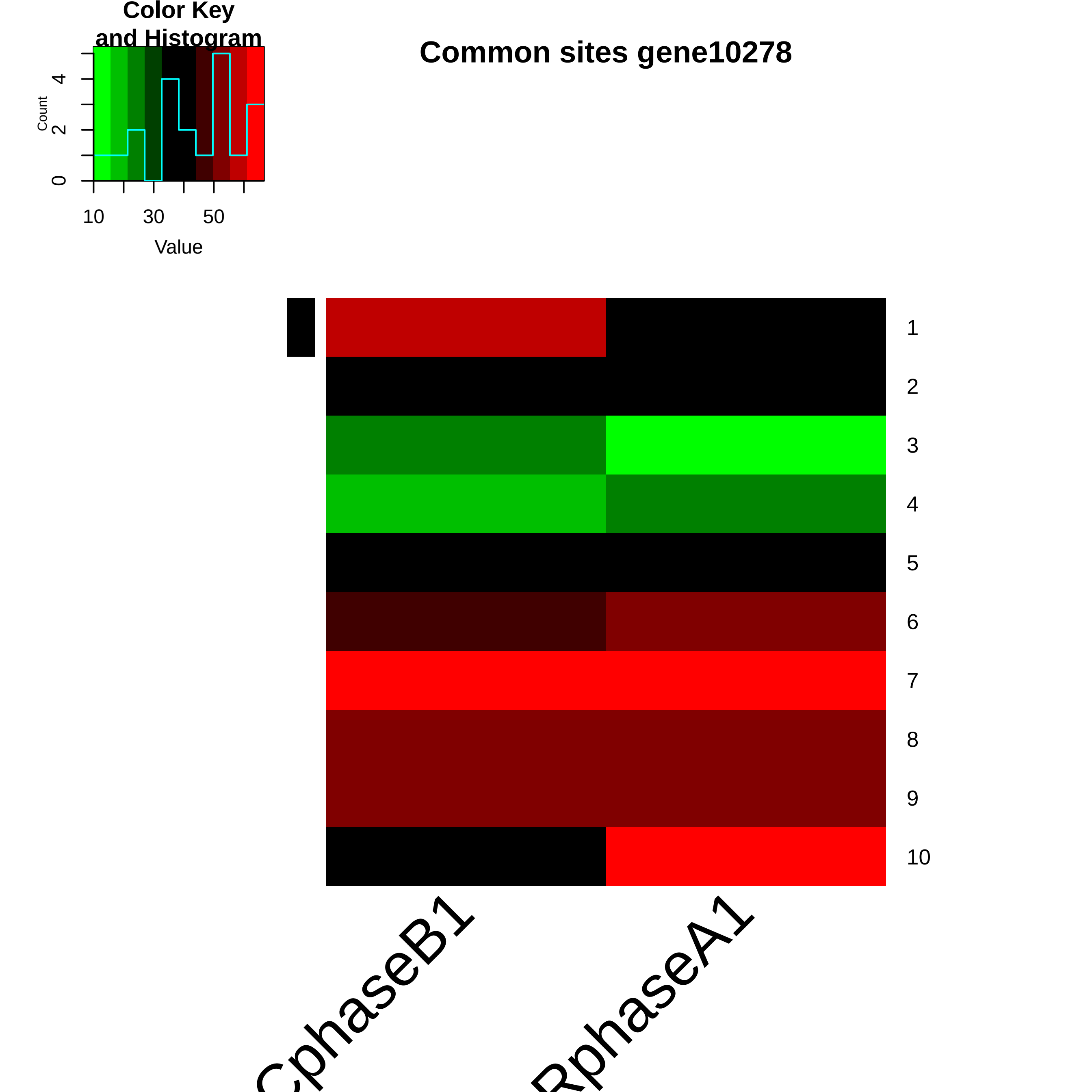
Color Key
and Histogram
Count
0
2
4
10
30
50
Value
Common sites gene10278
1
2
3
4
5
6
7
8
9
10

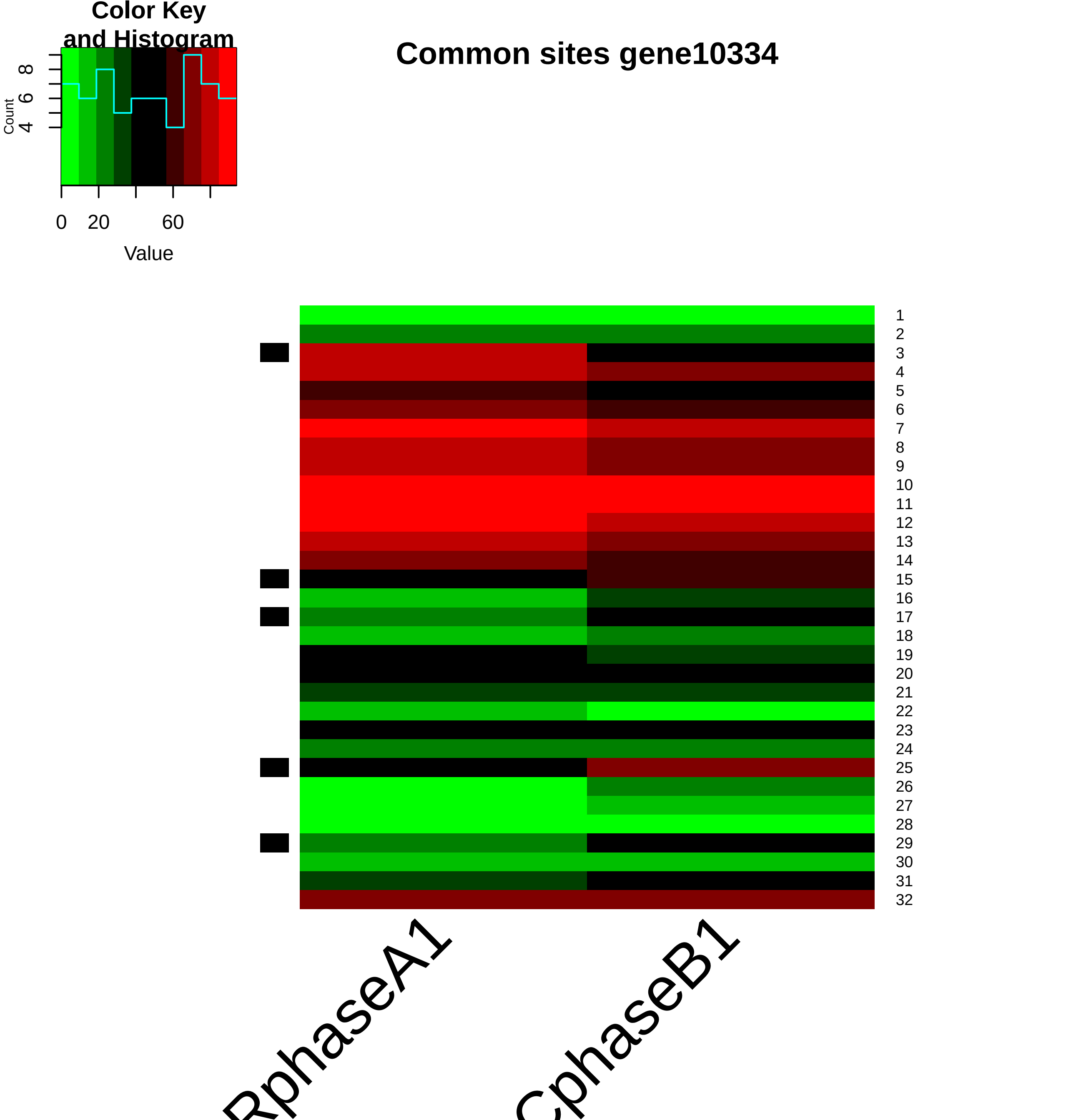
Color Key
and Histogram
Count
4
6
8
0
20
60
Value
Common sites gene10334
RphaseA1
CphaseB1

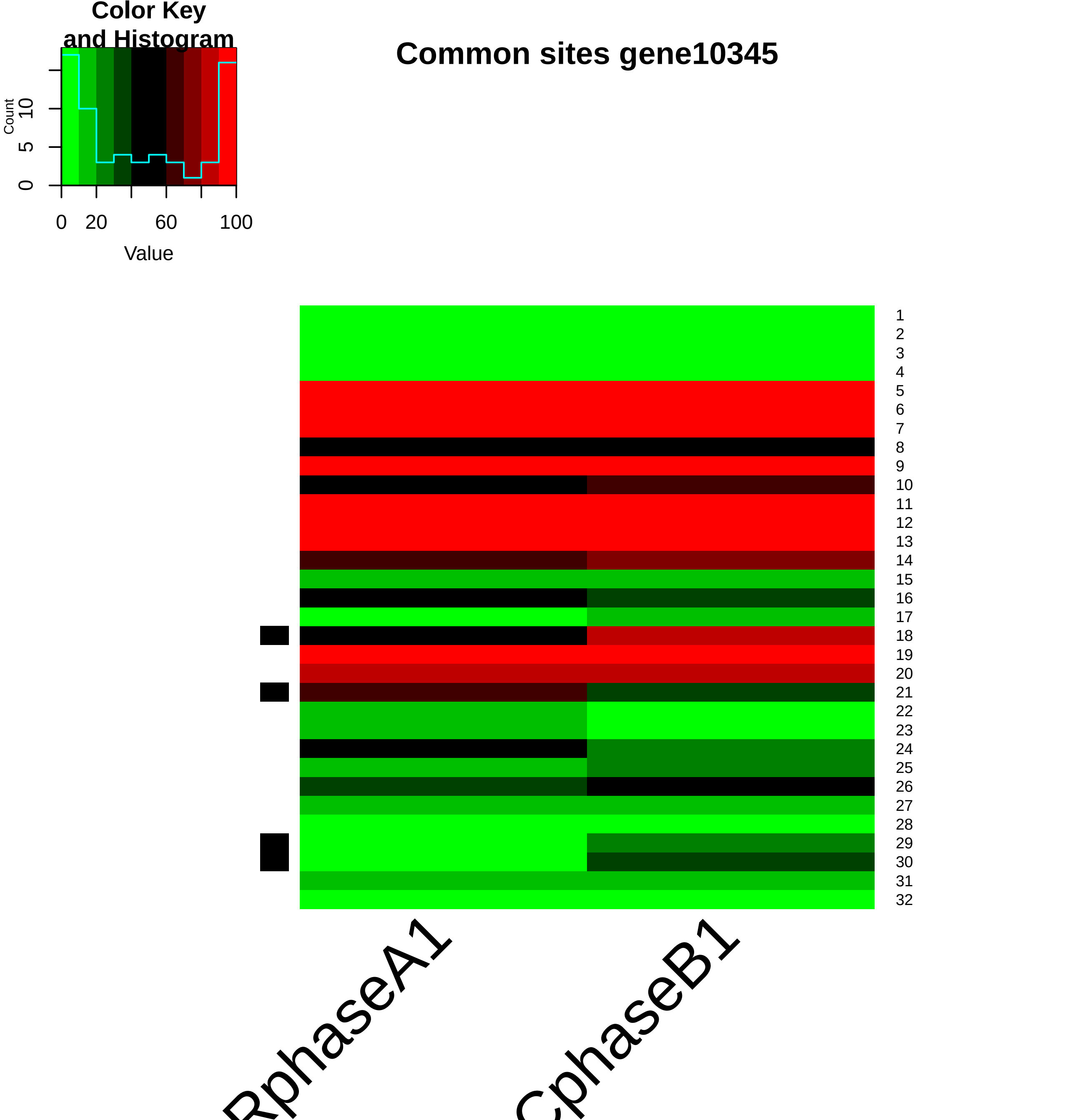

Color Key
and Histogram
Count
0
5
10
0
20
60
100
Value
Common sites gene10345
RphaseA1
CphaseB1

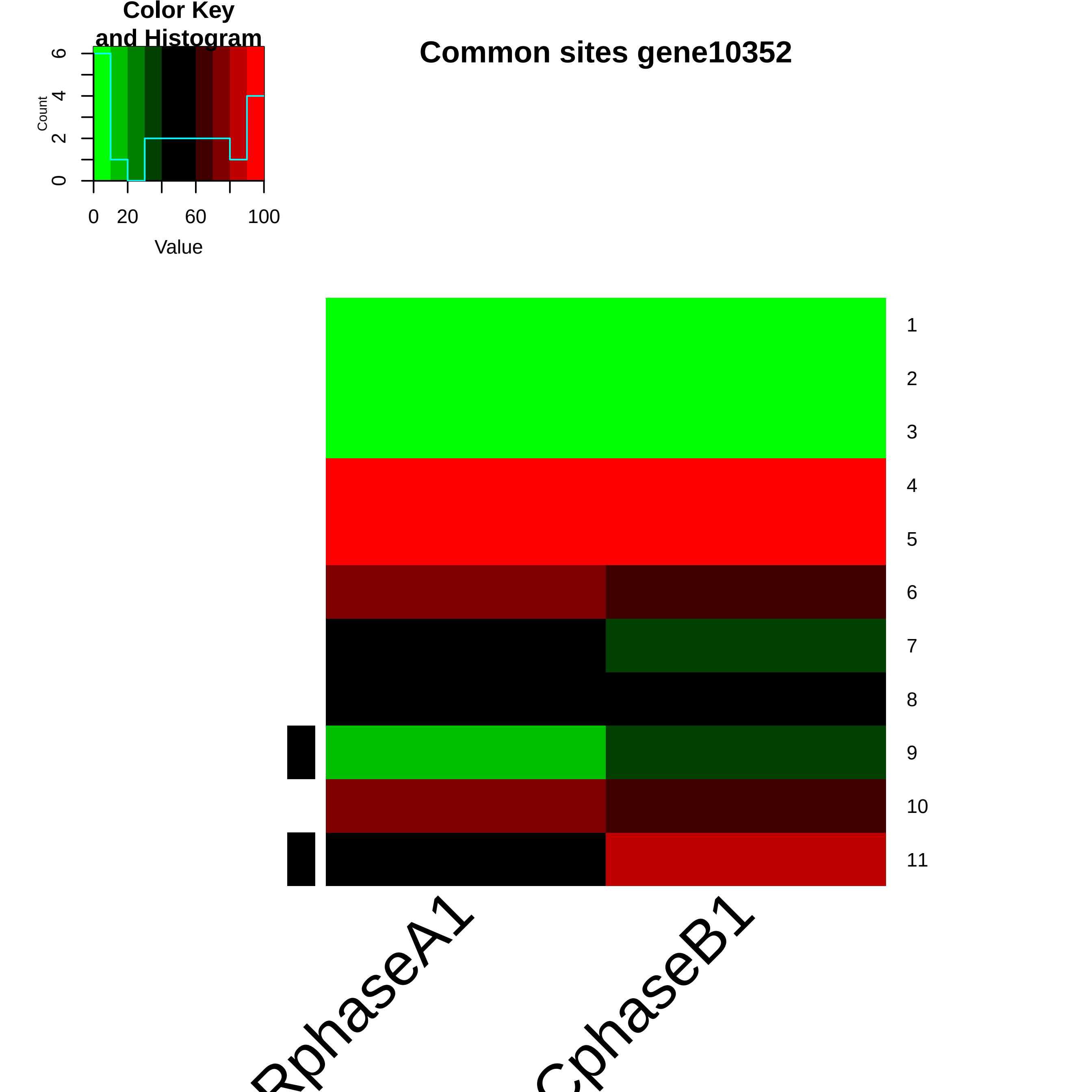
Color Key
and Histogram
Count
0
2
4
6
0
20
60
100
Value
Common sites gene10352
1
2
3
4
5
6
7
8
9
10
11
RphaseA1
CphaseB1

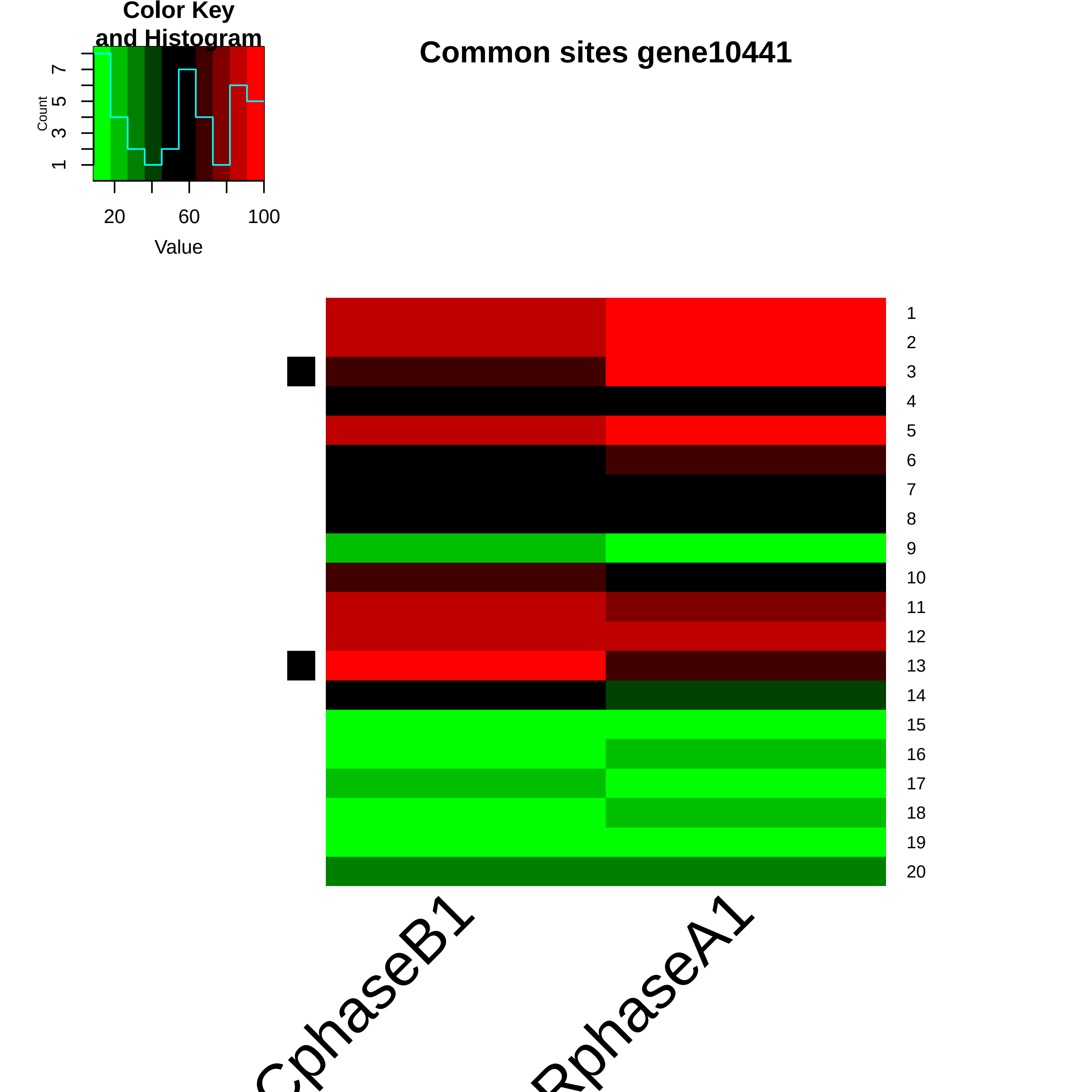
Color Key
and Histogram
Count
1
3
5
7
20
60
100
Value
Common sites gene10441
CphaseB1
RphaseA1

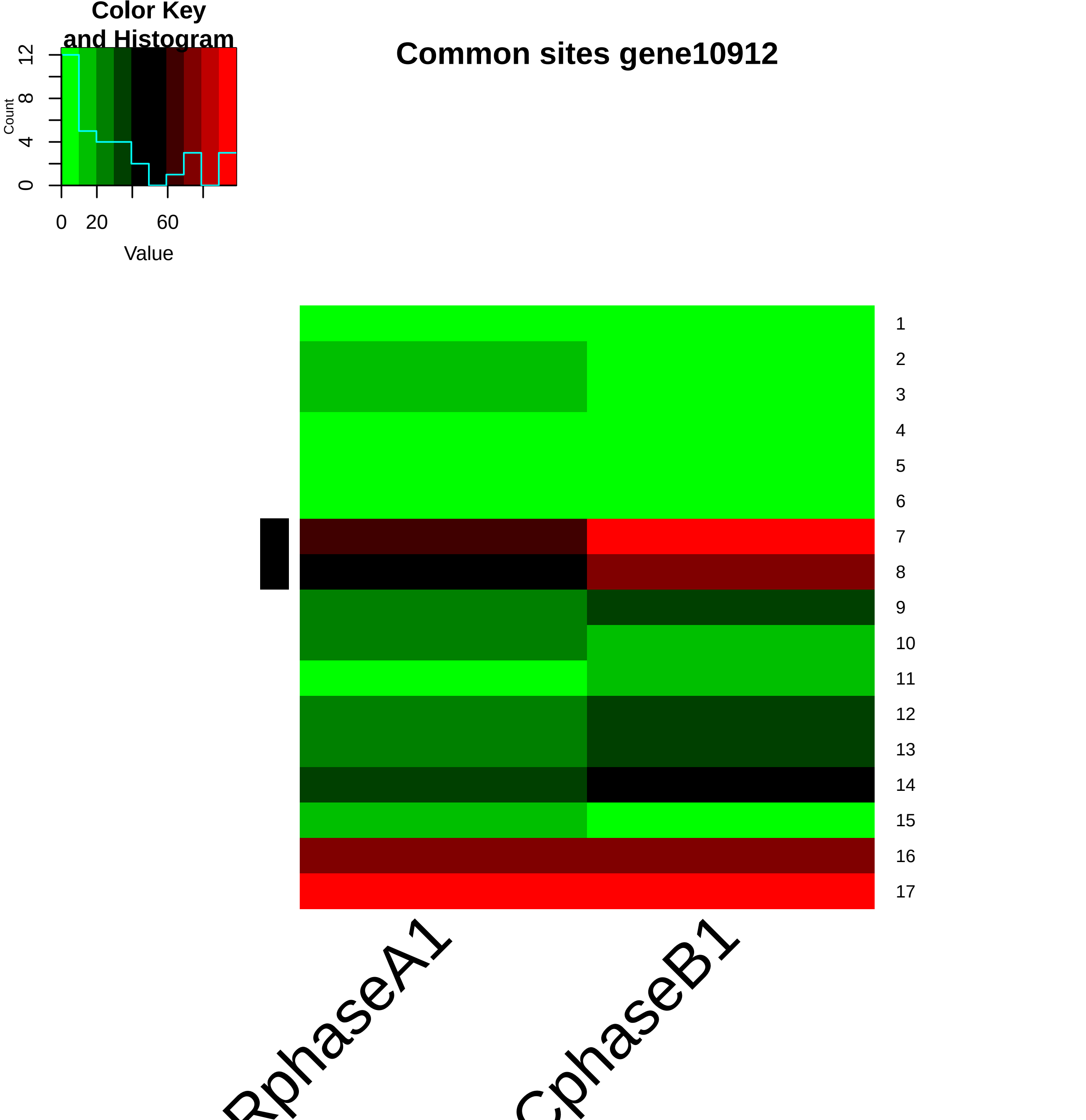

Color Key
and Histogram
Count
0
4
8
12
0
20
60
Value
Common sites gene10912
RphaseA1
CphaseB1

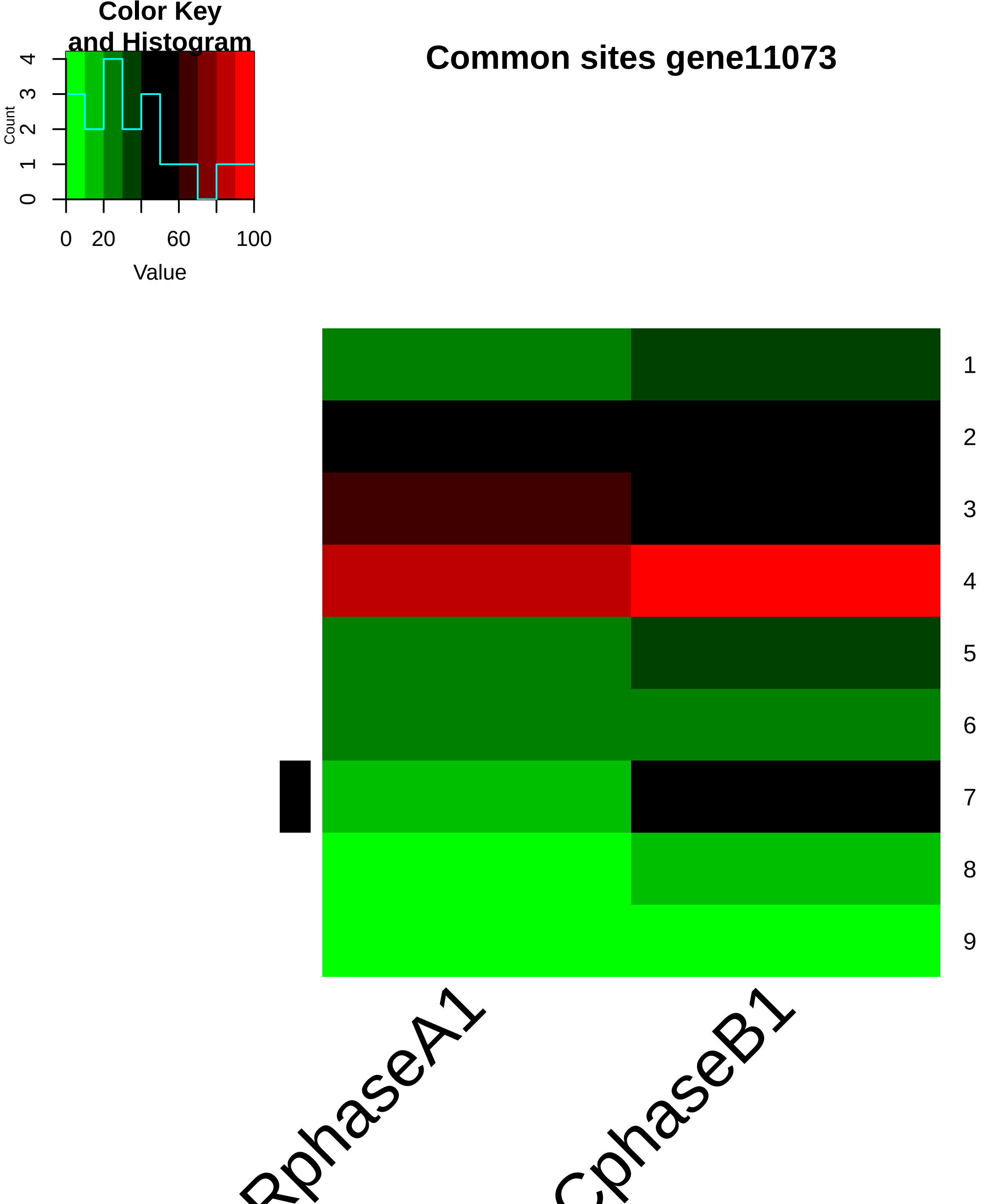

Color Key
and Histogram
Count
0
1
2
3
4
0
20
60
100
Value
Common sites gene11073
1
2
3
4
5
6
7
8
9
RphaseA1
CphaseB1

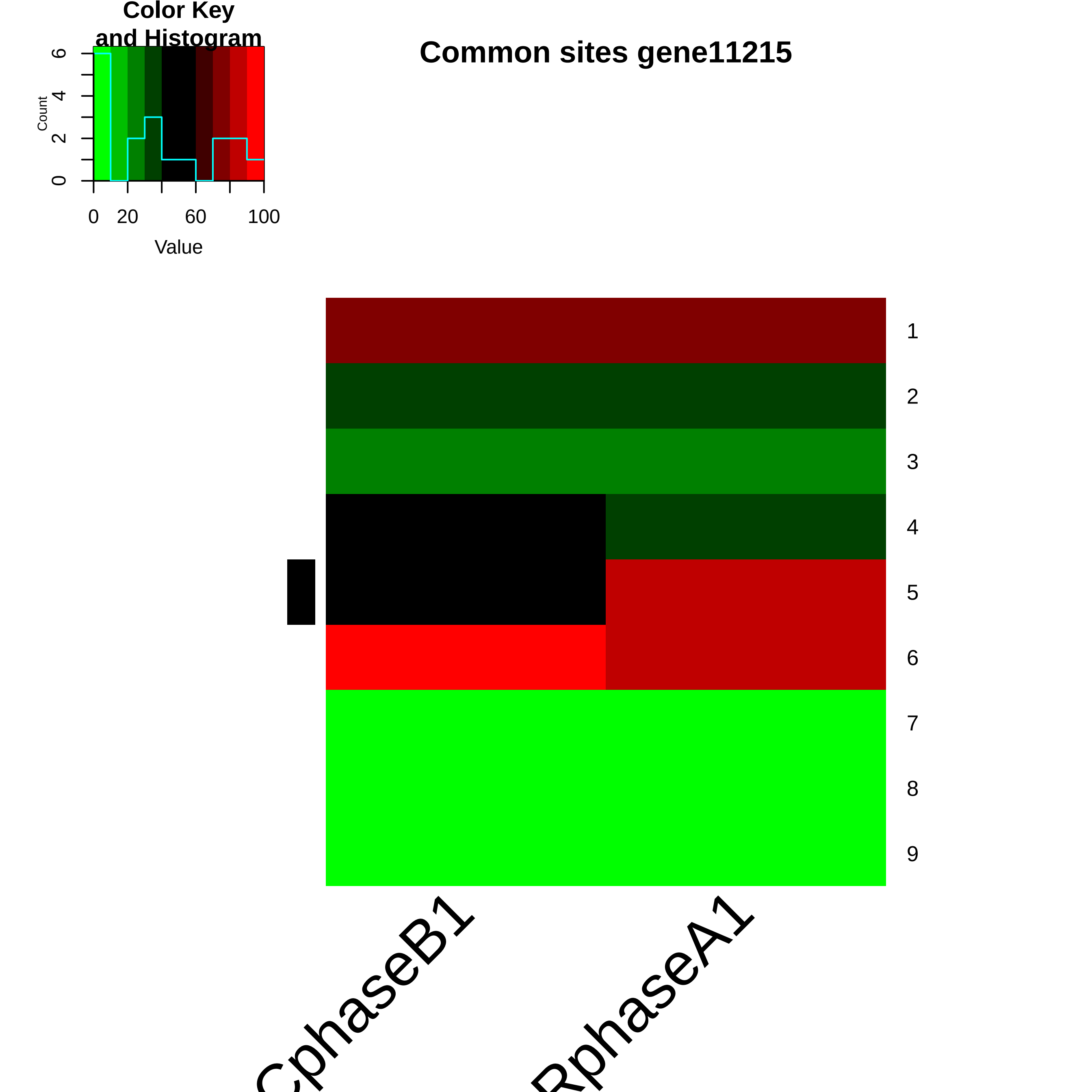
Color Key
and Histogram
Count
0
2
4
6
0
20
60
100
Value
Common sites gene11215
1
2
3
4
5
6
7
8
9
CphaseB1
RphaseA1

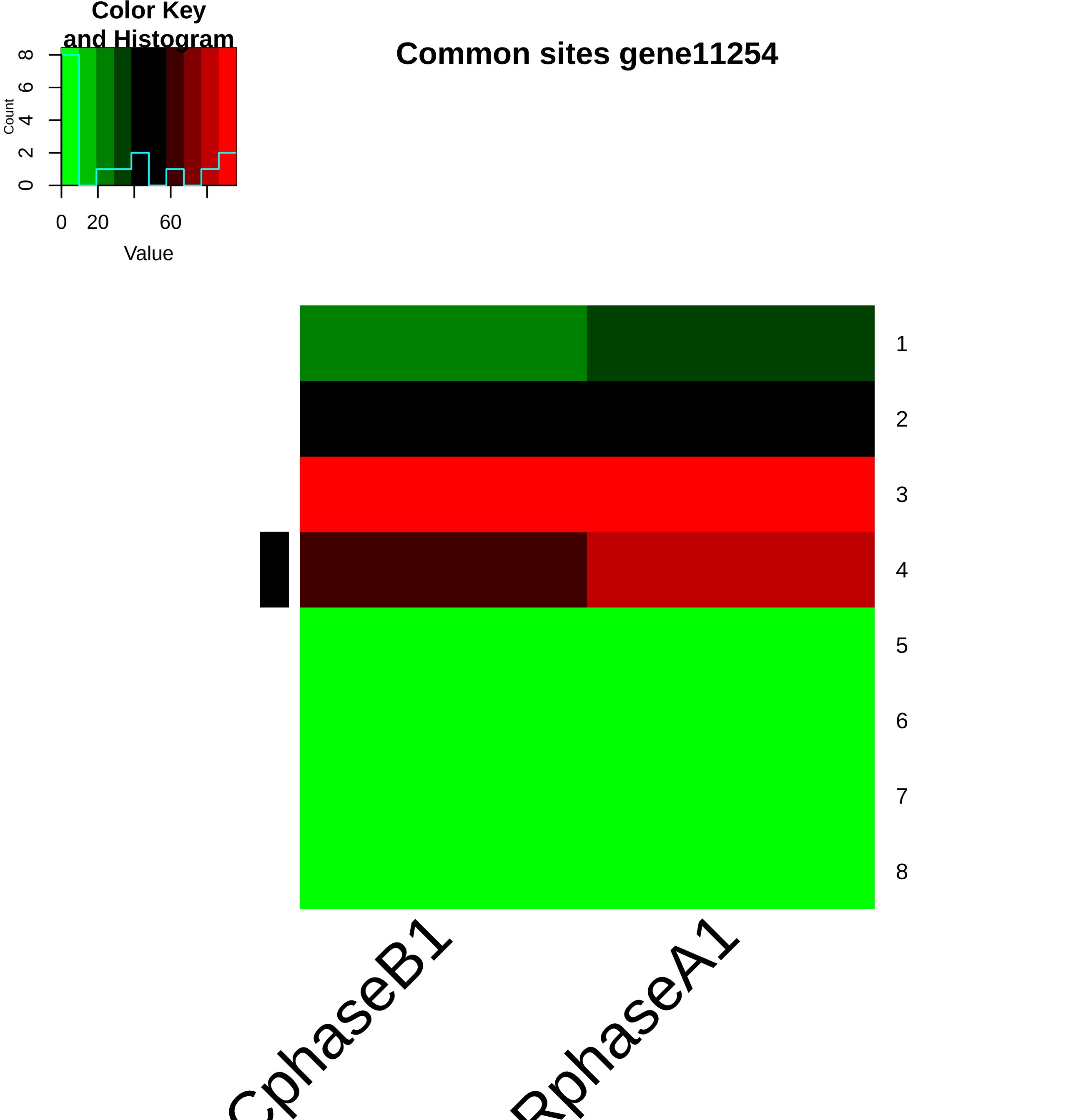

Color Key
and Histogram
Count
0
2
4
6
8
0
20
60
Value
Common sites gene11254
1
2
3
4
5
6
7
8
CphaseB1
RphaseA1

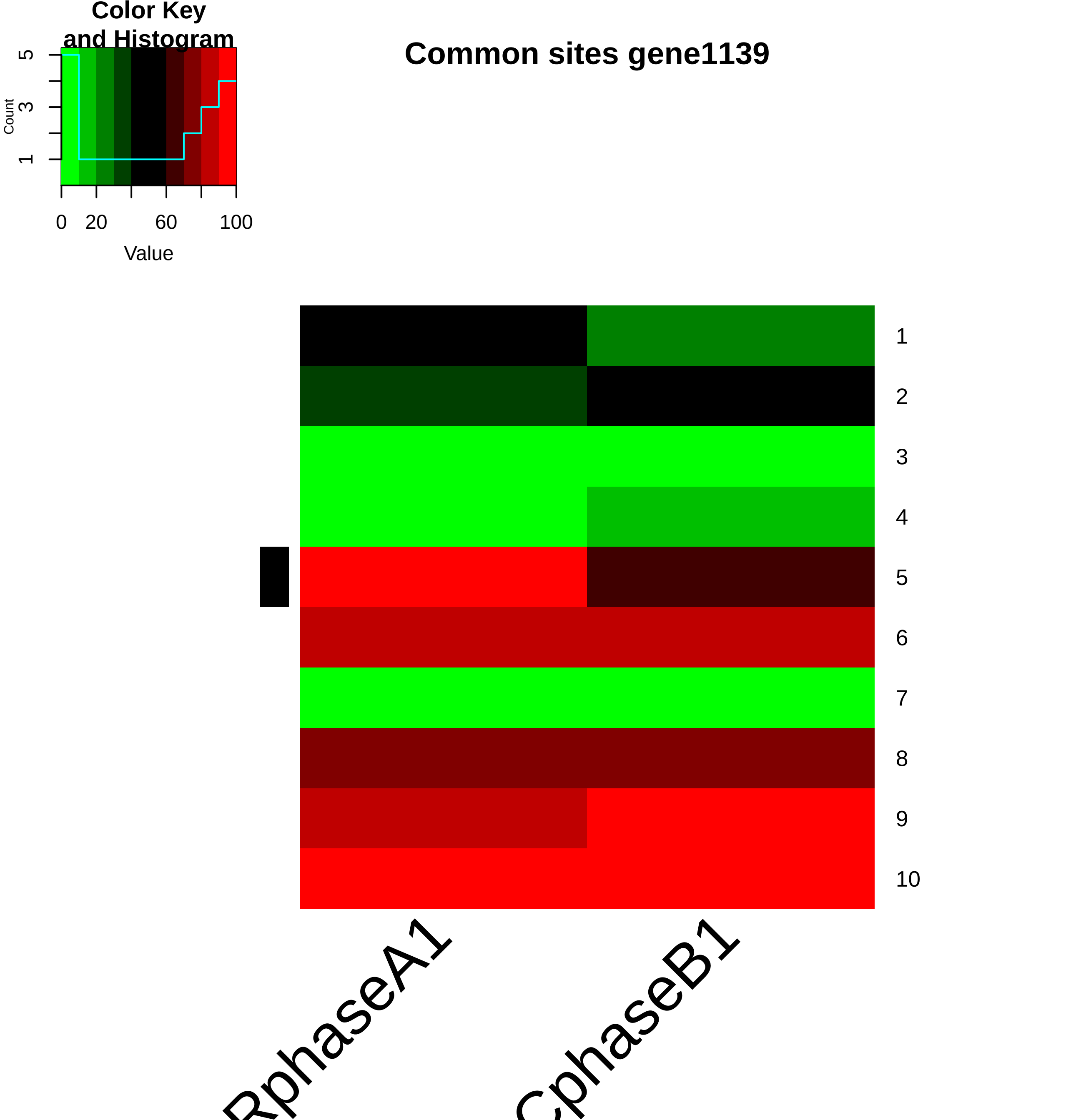

Color Key
and Histogram
Count
5
3
1
0
20
60
100
Value
Common sites gene1139
1
2
3
4
5
6
7
8
9
10
RphaseA1
CphaseB1

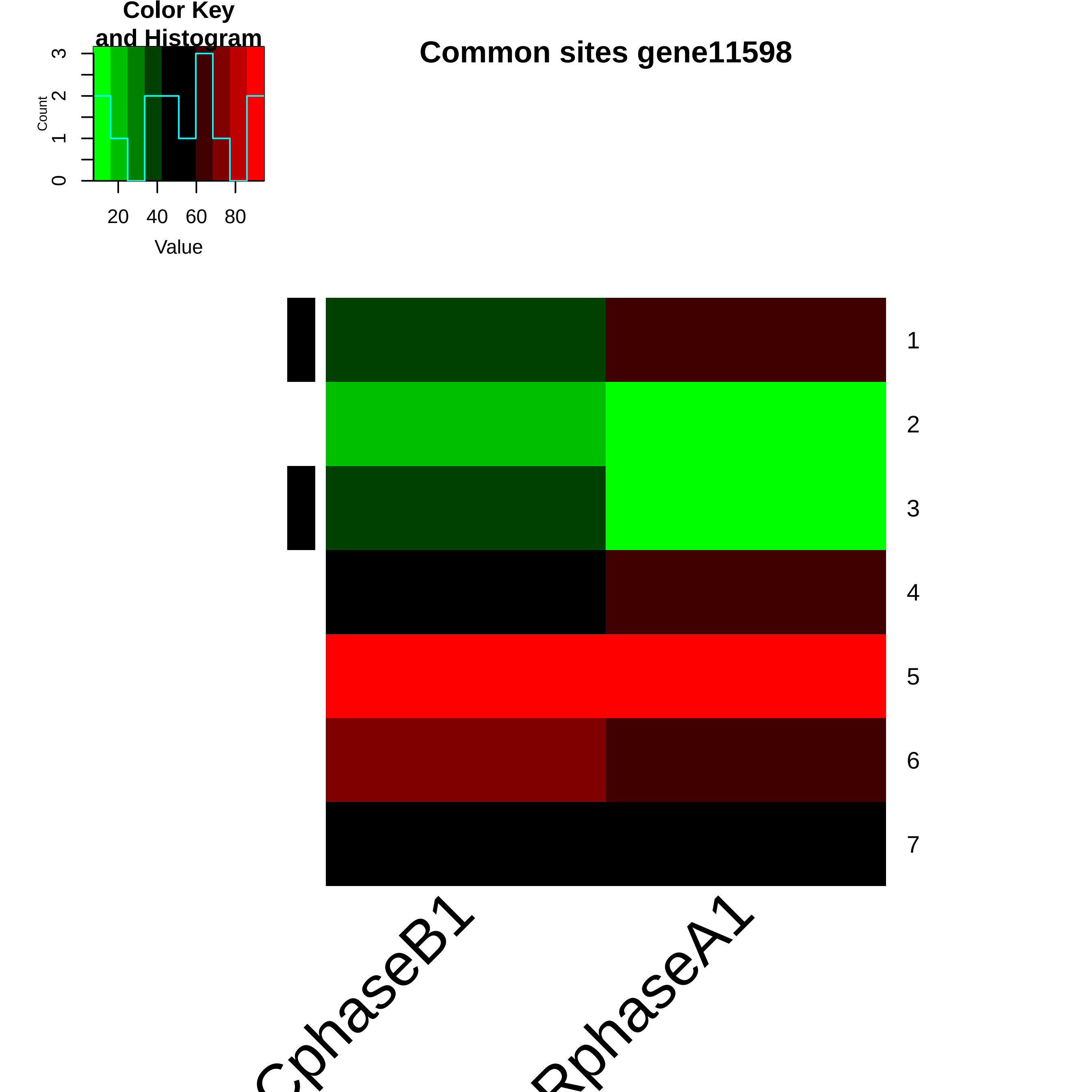

Color Key
and Histogram
Count
0
1
2
3
20
40
60
80
Value
Common sites gene11598
1
2
3
4
5
6
7
CphaseB1
RphaseA1

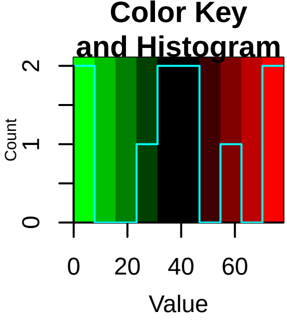
Color Key
and Histogram
Count
0
1
2
0
20
40
60
Value

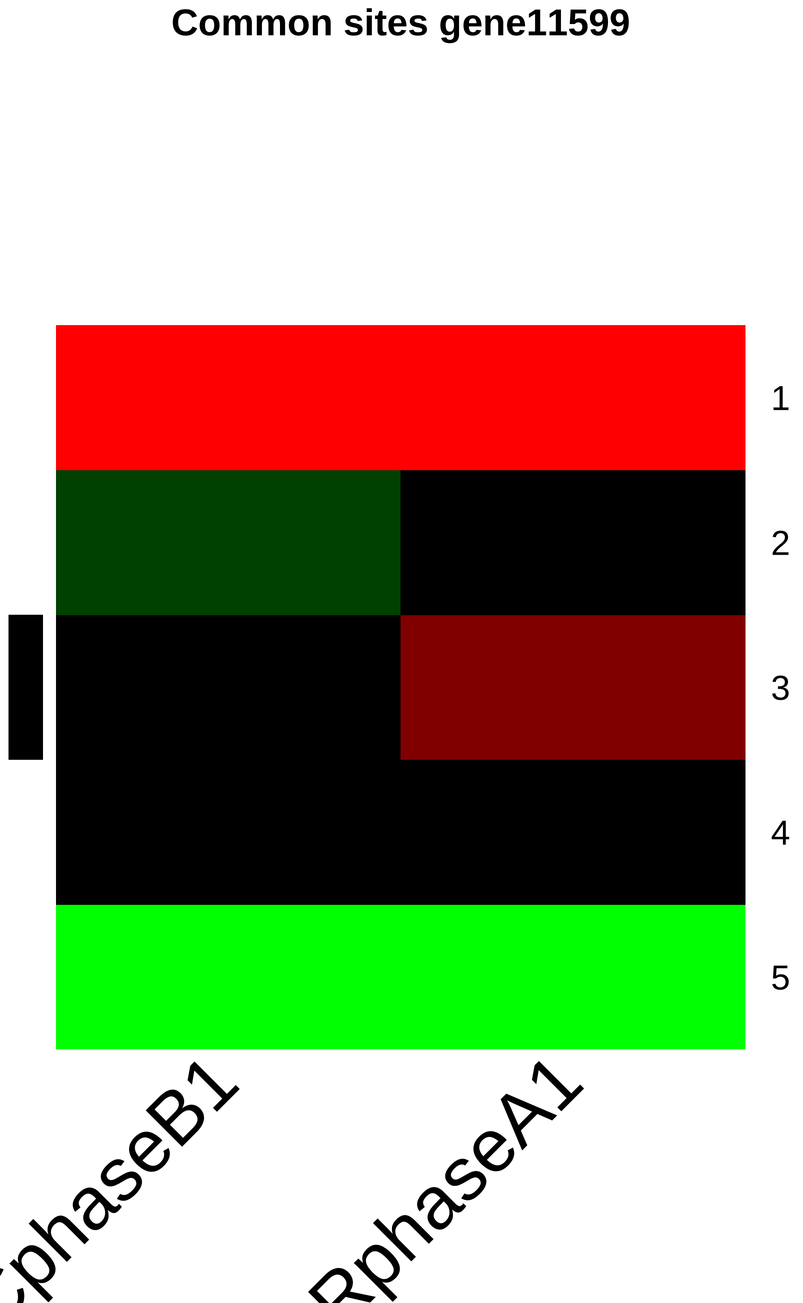
Common sites gene11599
1
2
3
4
5
CphaseB1
RphaseA1

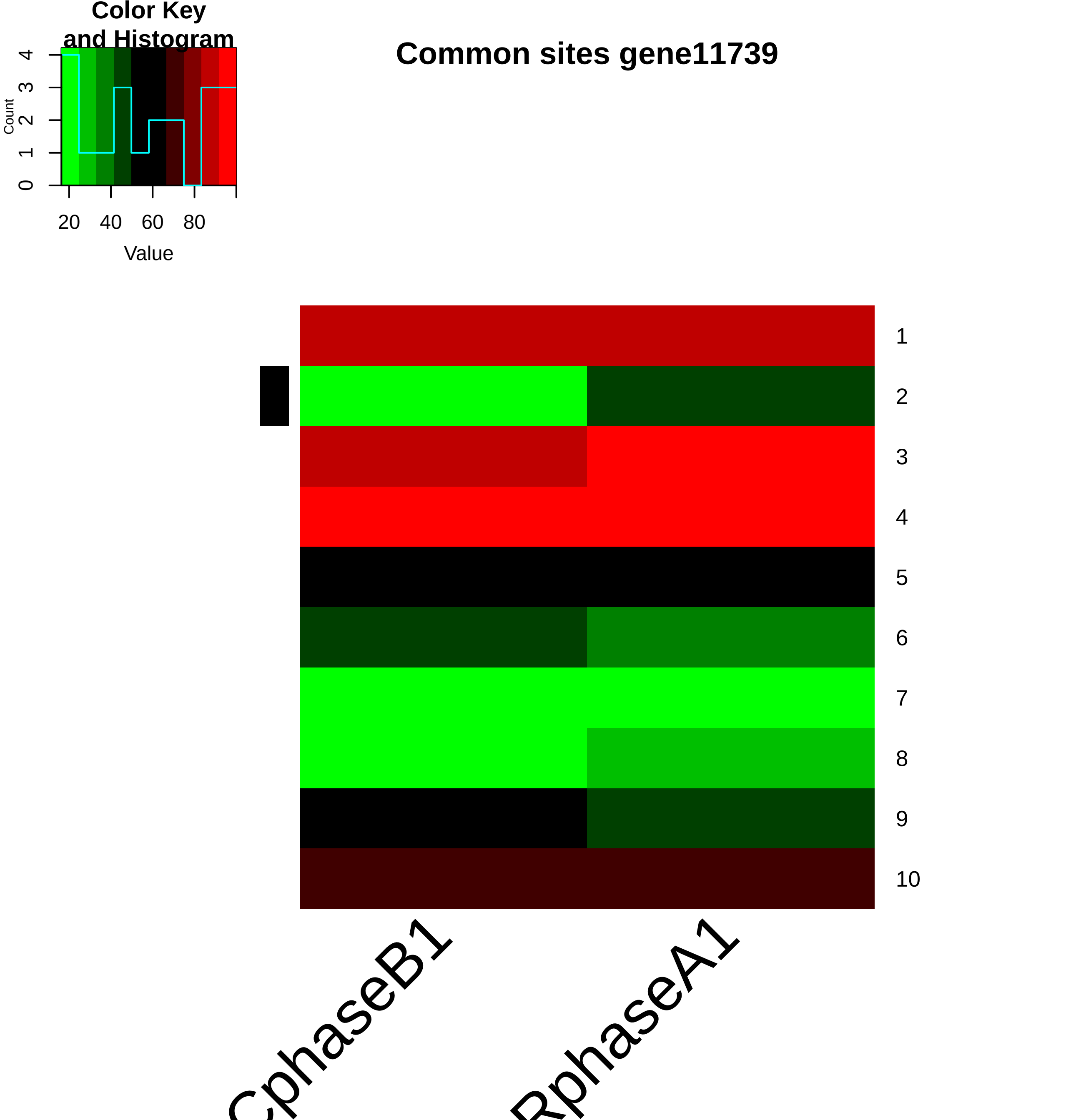

Color Key
and Histogram
Count
0
1
2
3
4
20
40
60
80
Value
Common sites gene11739
1
2
3
4
5
6
7
8
9
10
CphaseB1
RphaseA1

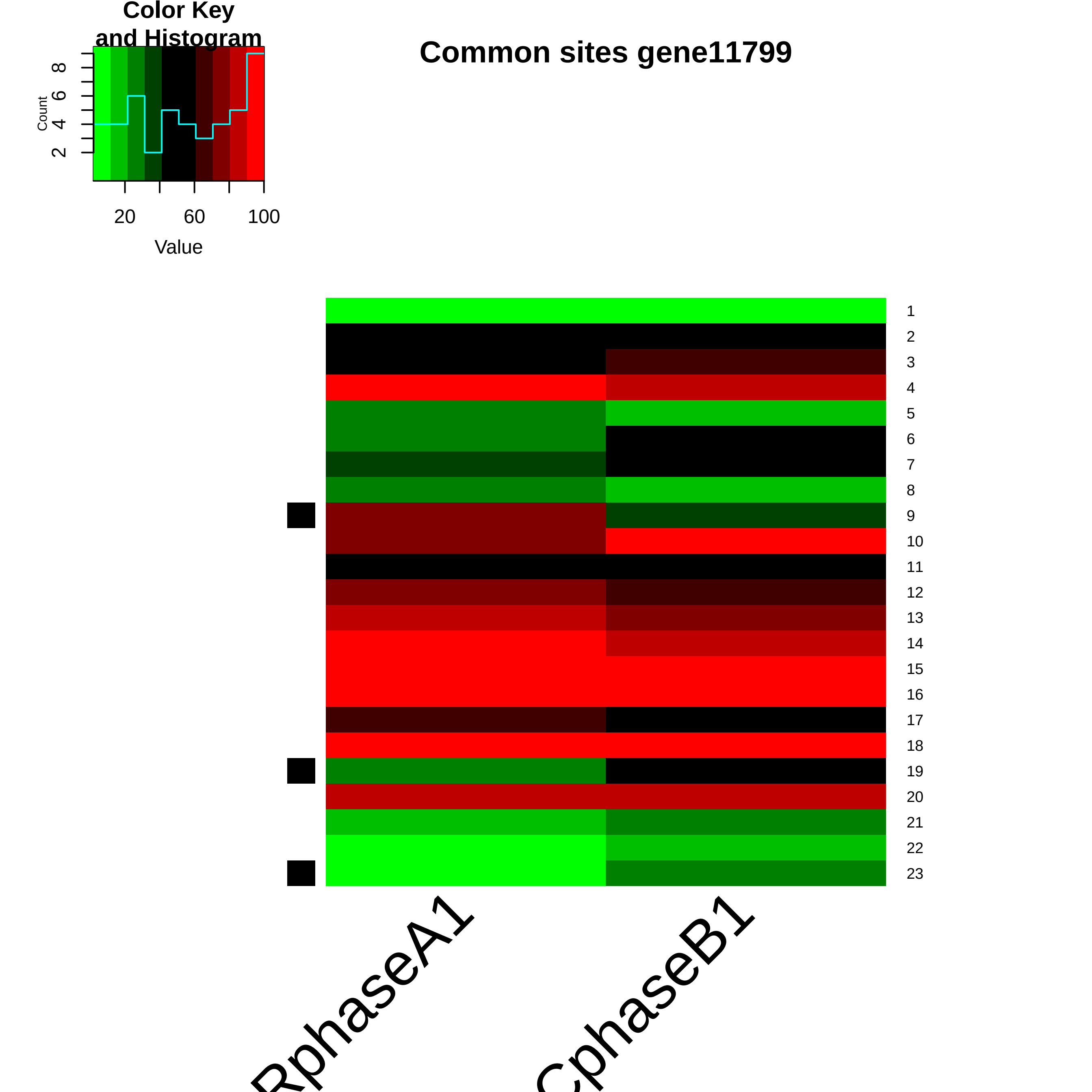
Color Key
and Histogram
Count
2
4
6
8
20
60
100
Value
Common sites gene11799
RphaseA1
CphaseB1

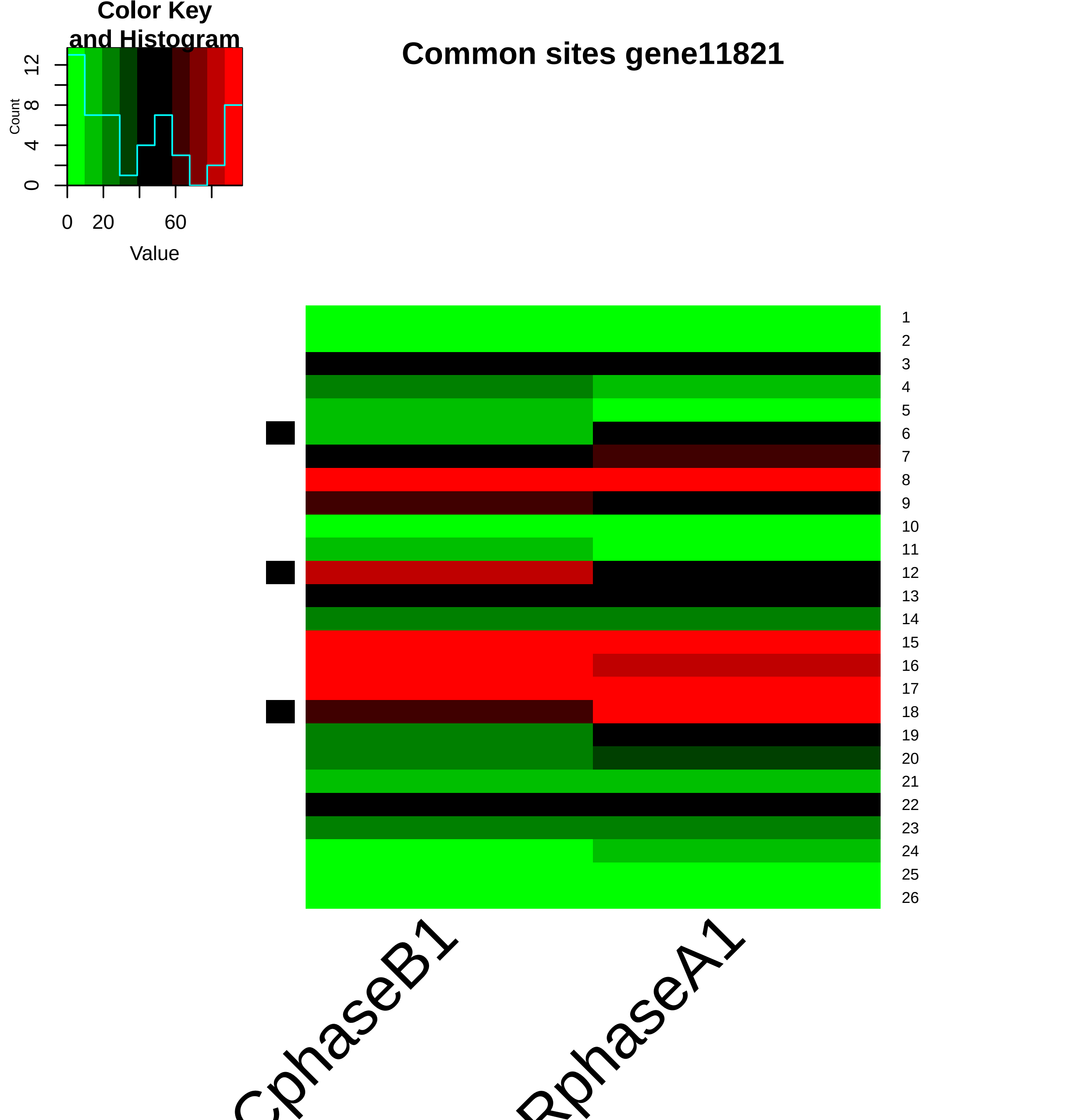
Color Key
and Histogram
Count
0
4
8
12
0
20
60
Value
Common sites gene11821
CphaseB1
RphaseA1

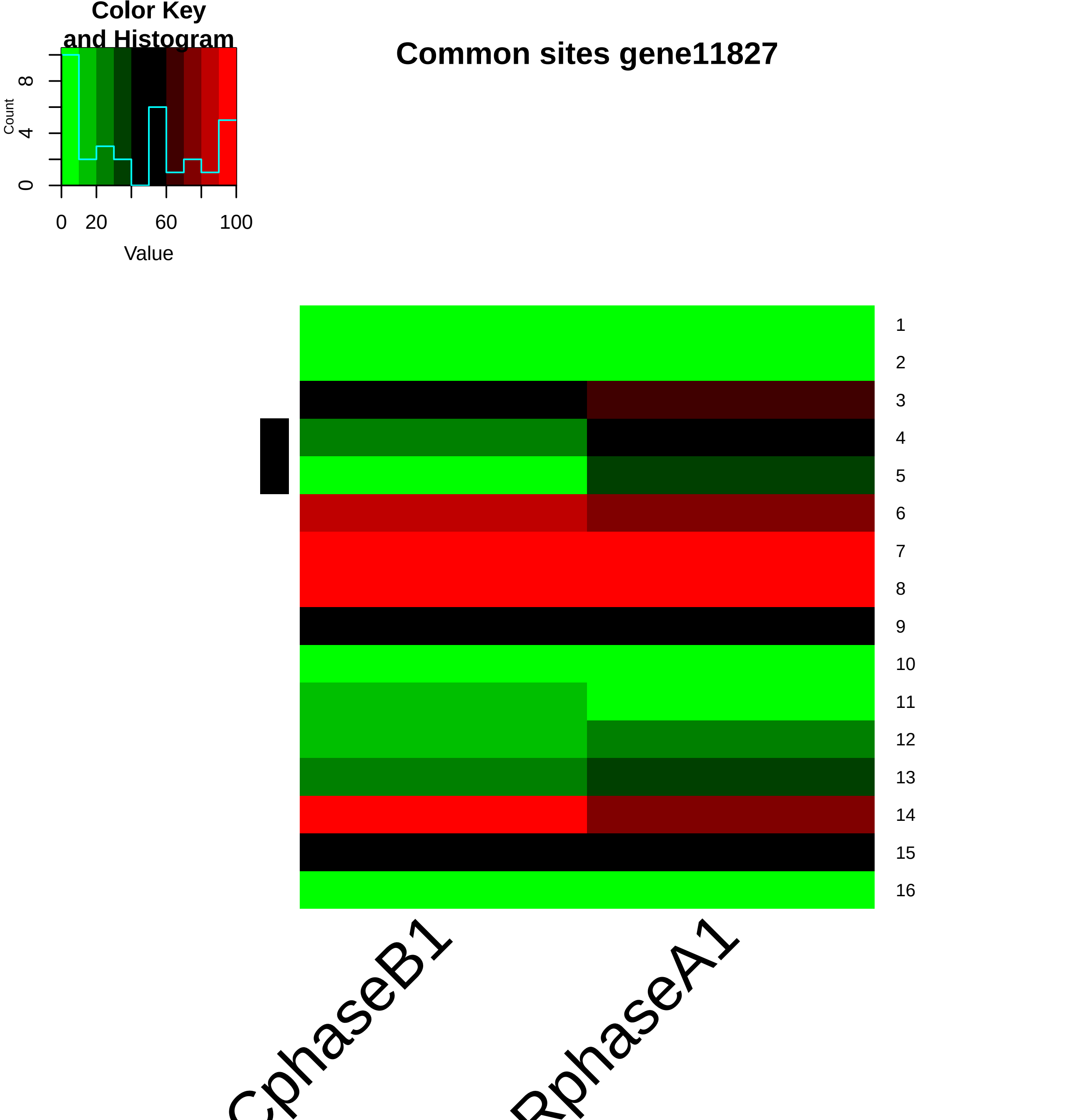
Color Key
and Histogram
Count
0
4
8
0
20
60
100
Value
Common sites gene11827
CphaseB1
RphaseA1

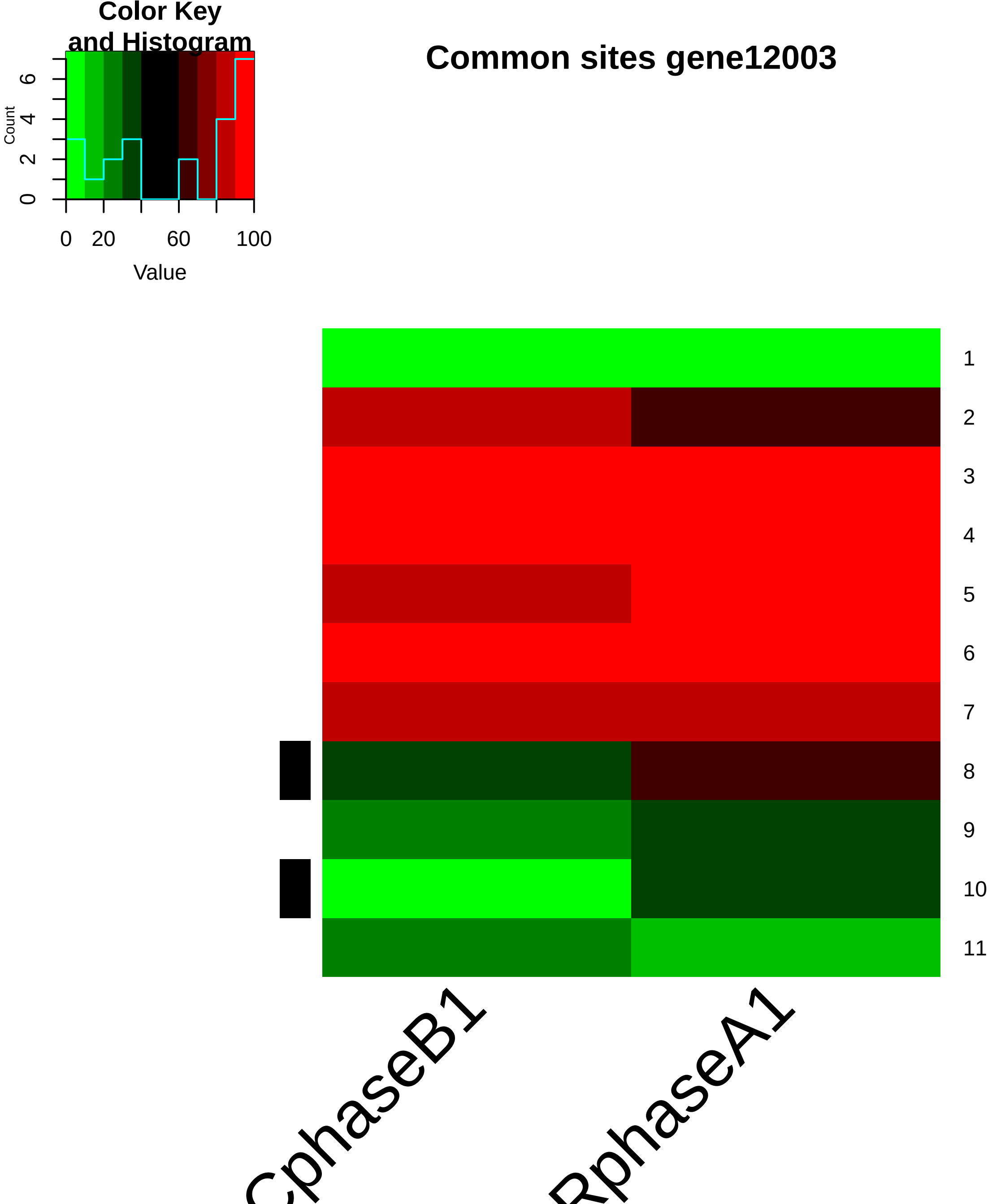
Color Key
and Histogram
Count
0
2
4
6
0
20
60
100
Value
Common sites gene12003
1
2
3
4
5
6
7
8
9
10
11
CphaseB1
RphaseA1

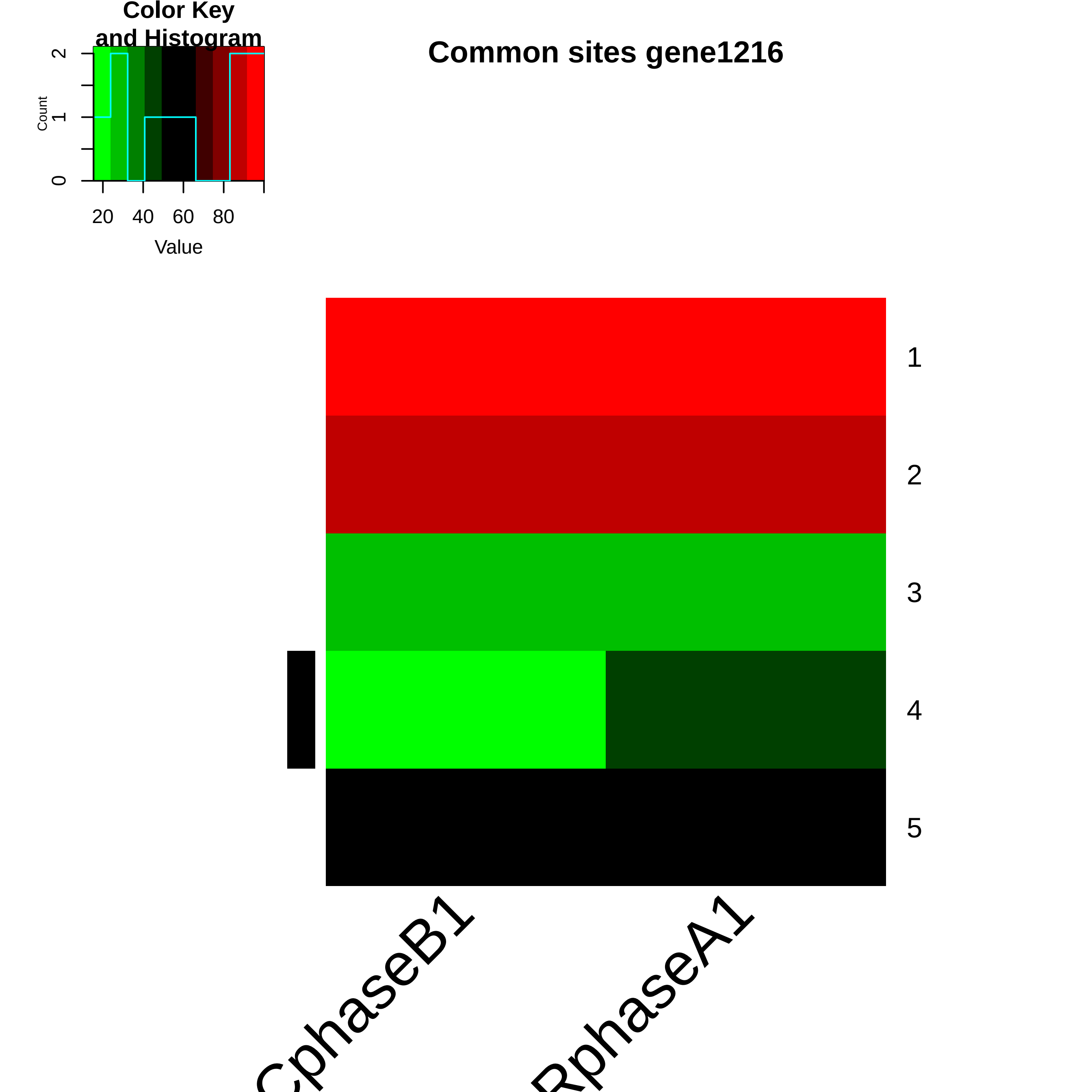

Color Key
and Histogram
Count
0
1
2
20
40
60
80
Value
Common sites gene1216
1
2
3
4
5
CphaseB1
RphaseA1

### Common sites gene12173

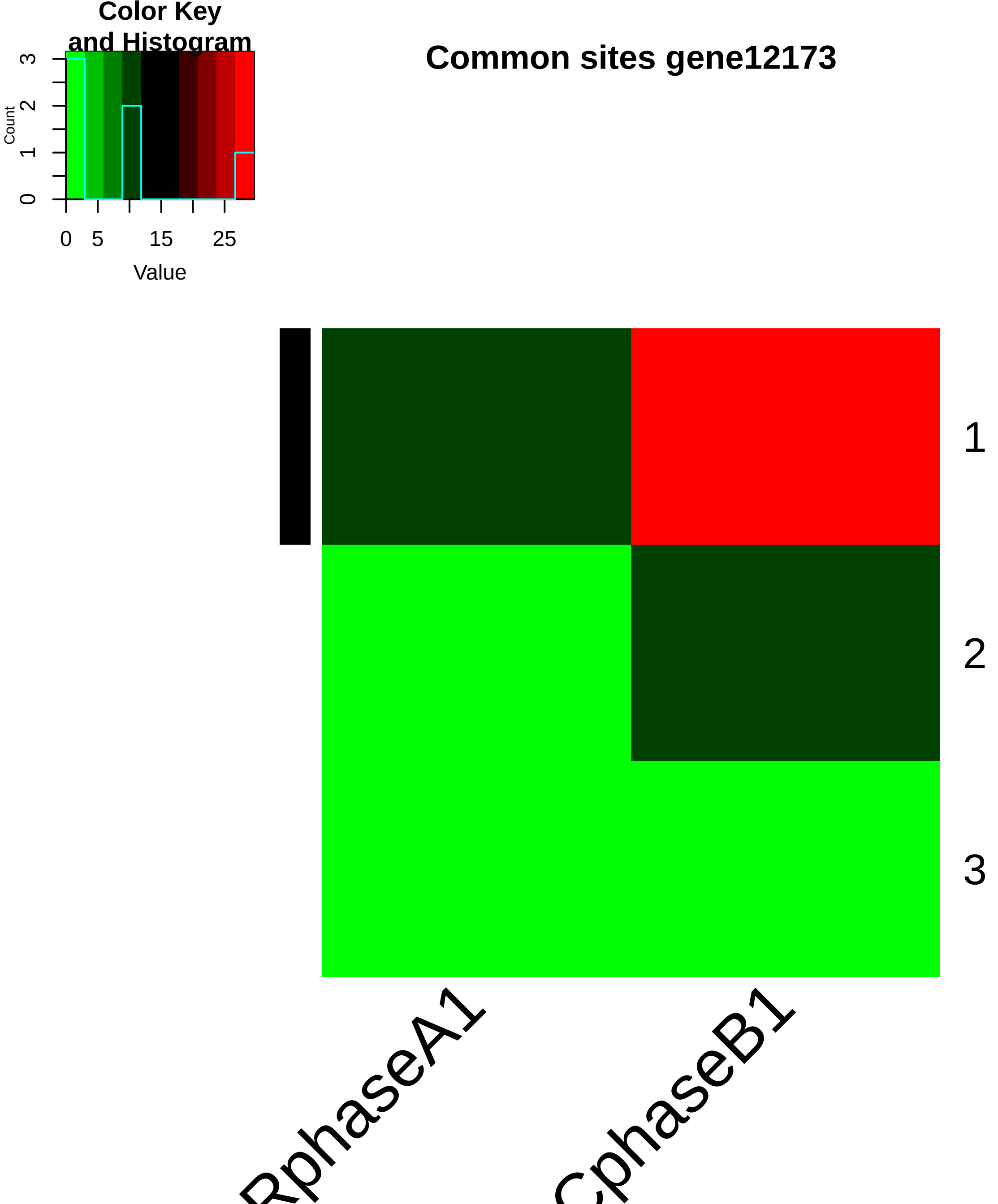

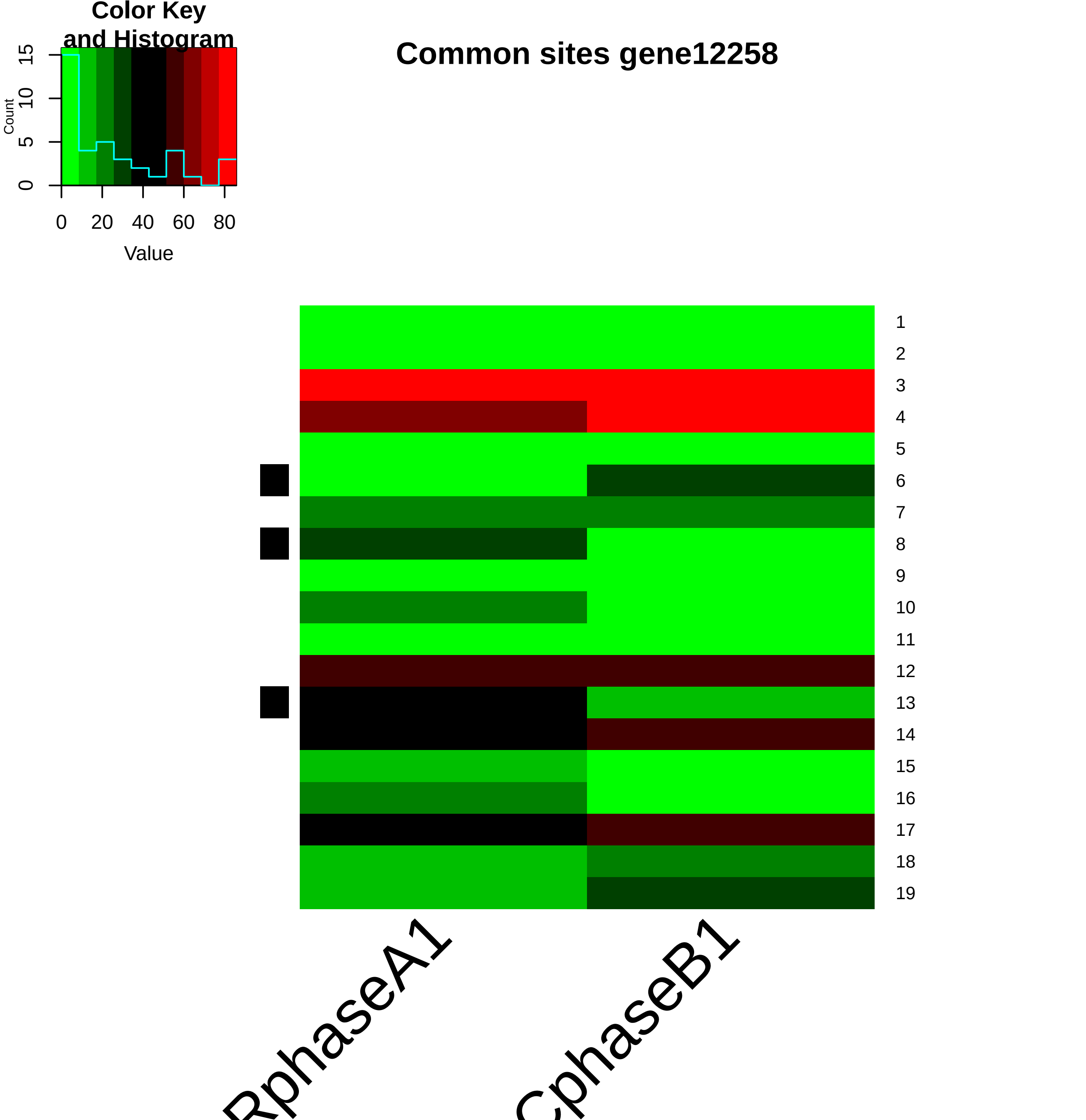
Common sites gene12258
Color Key
and Histogram
Count
0
5
10
15
0
20
40
60
80
Value
RphaseA1
CphaseB1

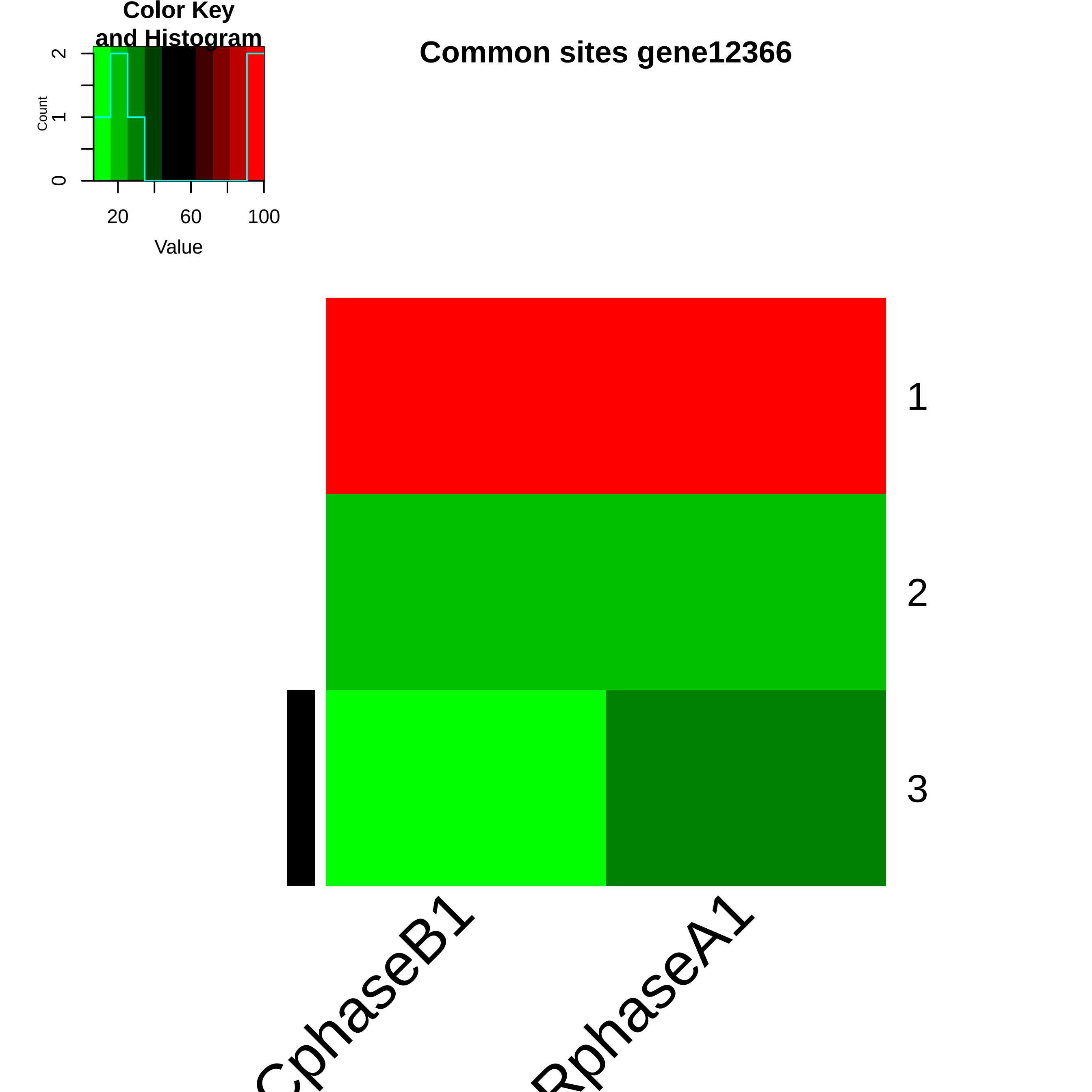

Color Key
and Histogram
Count
0
1
2
20
60
100
Value
Common sites gene12366
1
2
3
CphaseB1
RphaseA1

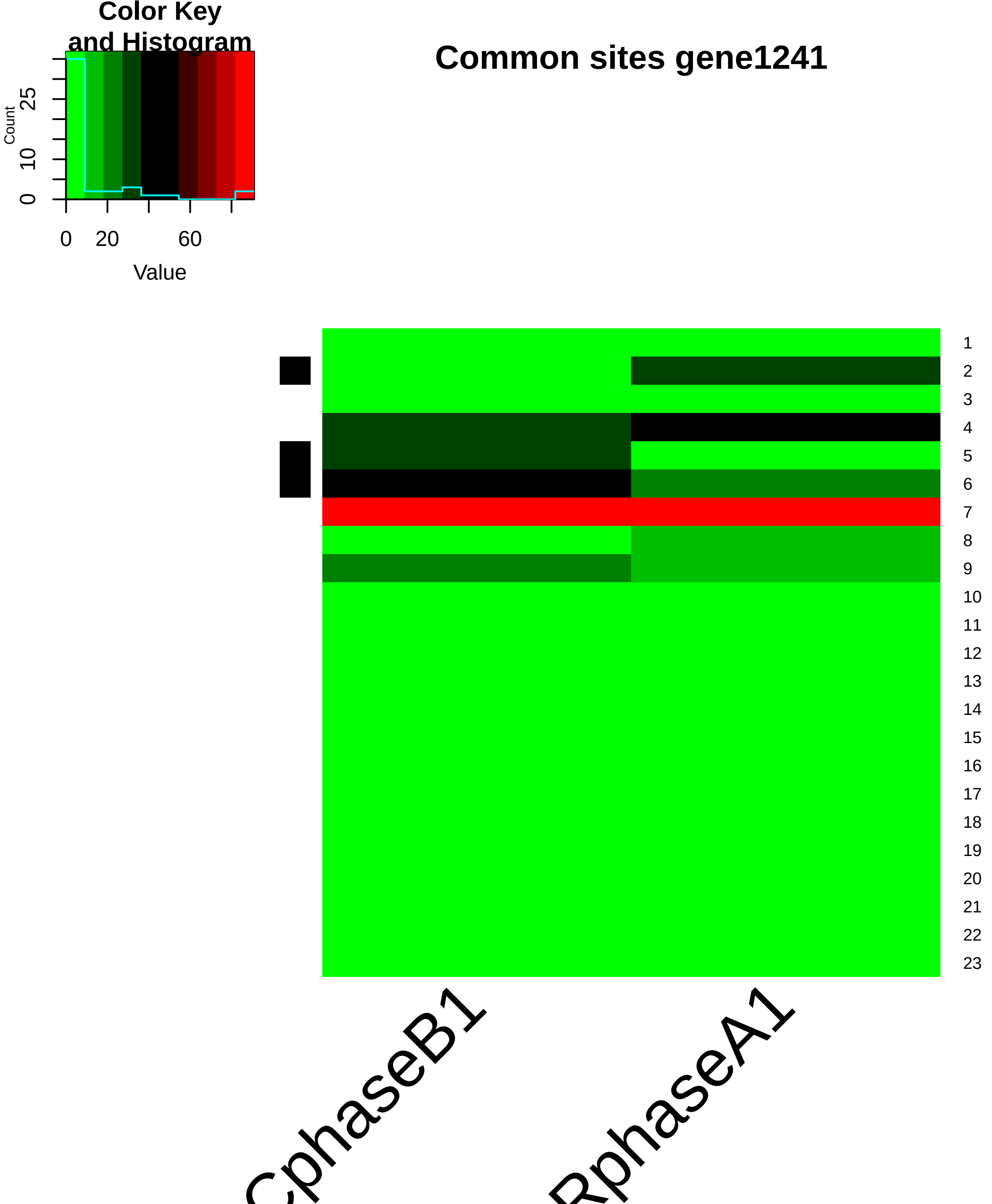

Color Key
and Histogram
Count
0
10
25
0
20
60
Value
Common sites gene1241
CphaseB1
RphaseA1

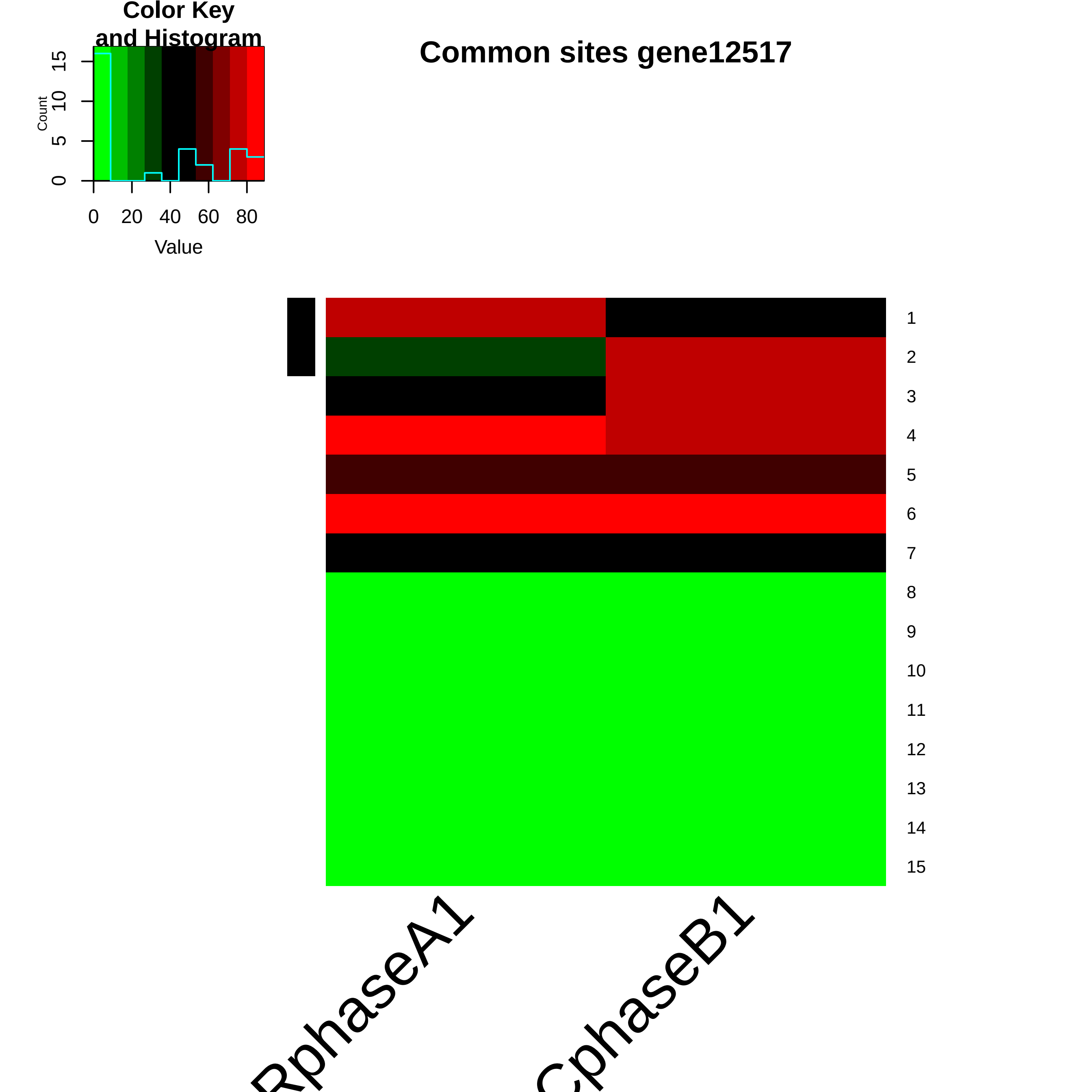
Color Key
and Histogram
Count
0
5
10
15
0
20
40
60
80
Value
Common sites gene12517
RphaseA1
CphaseB1

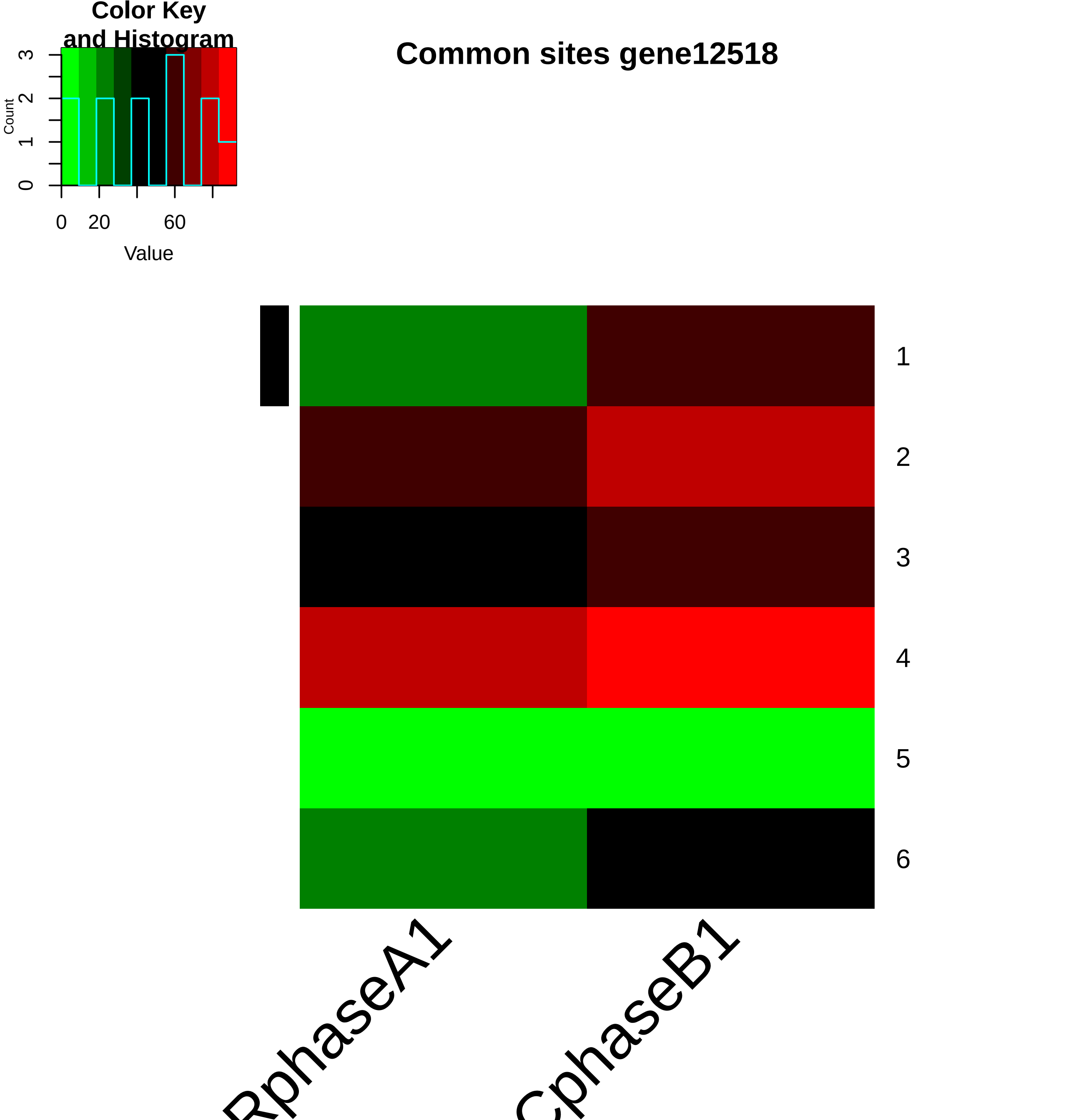
Color Key
and Histogram
Count
0
1
2
3
0
20
60
Value
Common sites gene12518
1
2
3
4
5
6
RphaseA1
CphaseB1

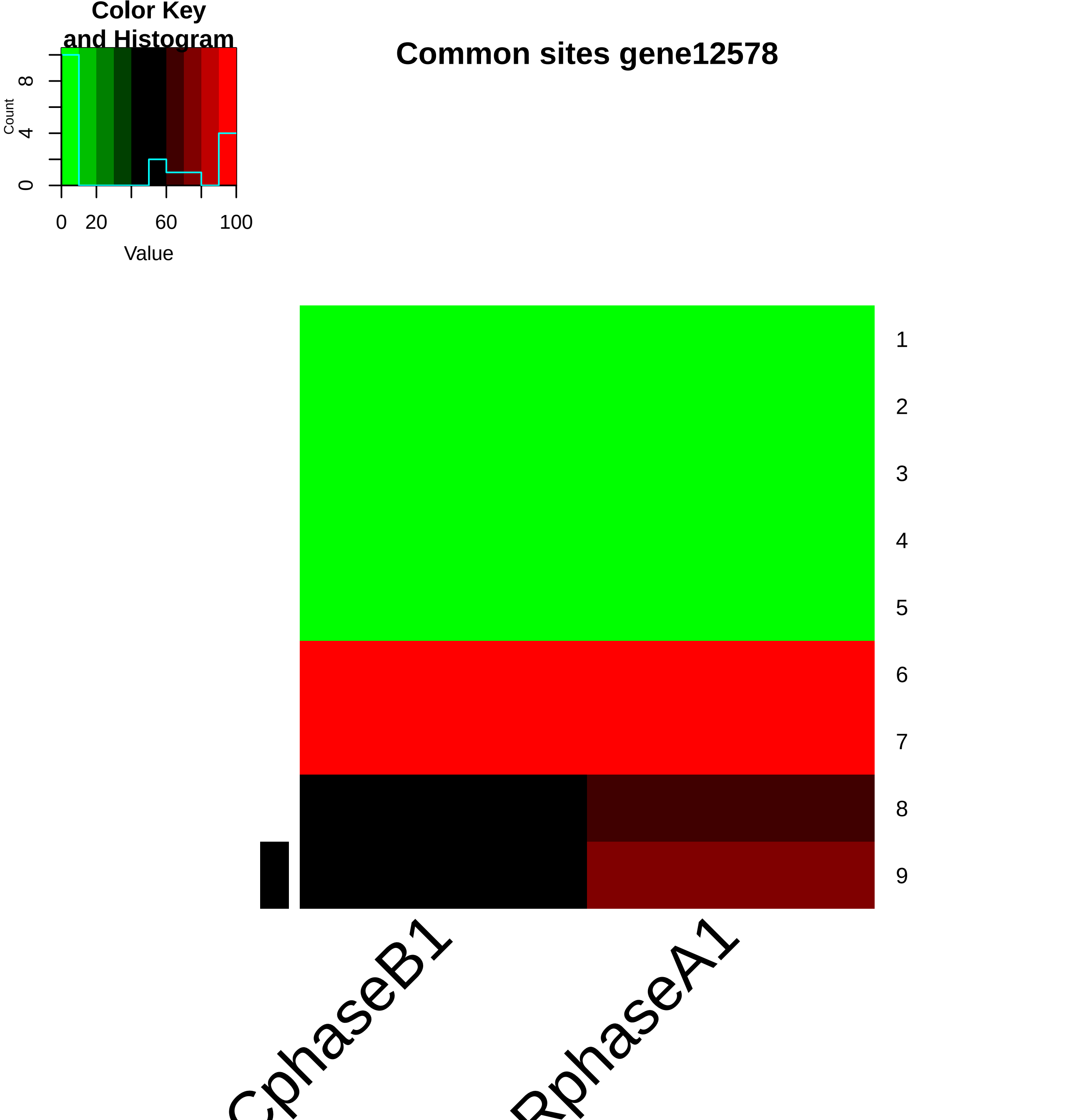

Color Key
and Histogram
Count
0
4
8
0
20
60
100
Value
Common sites gene12578
1
2
3
4
5
6
7
8
9
CphaseB1
RphaseA1

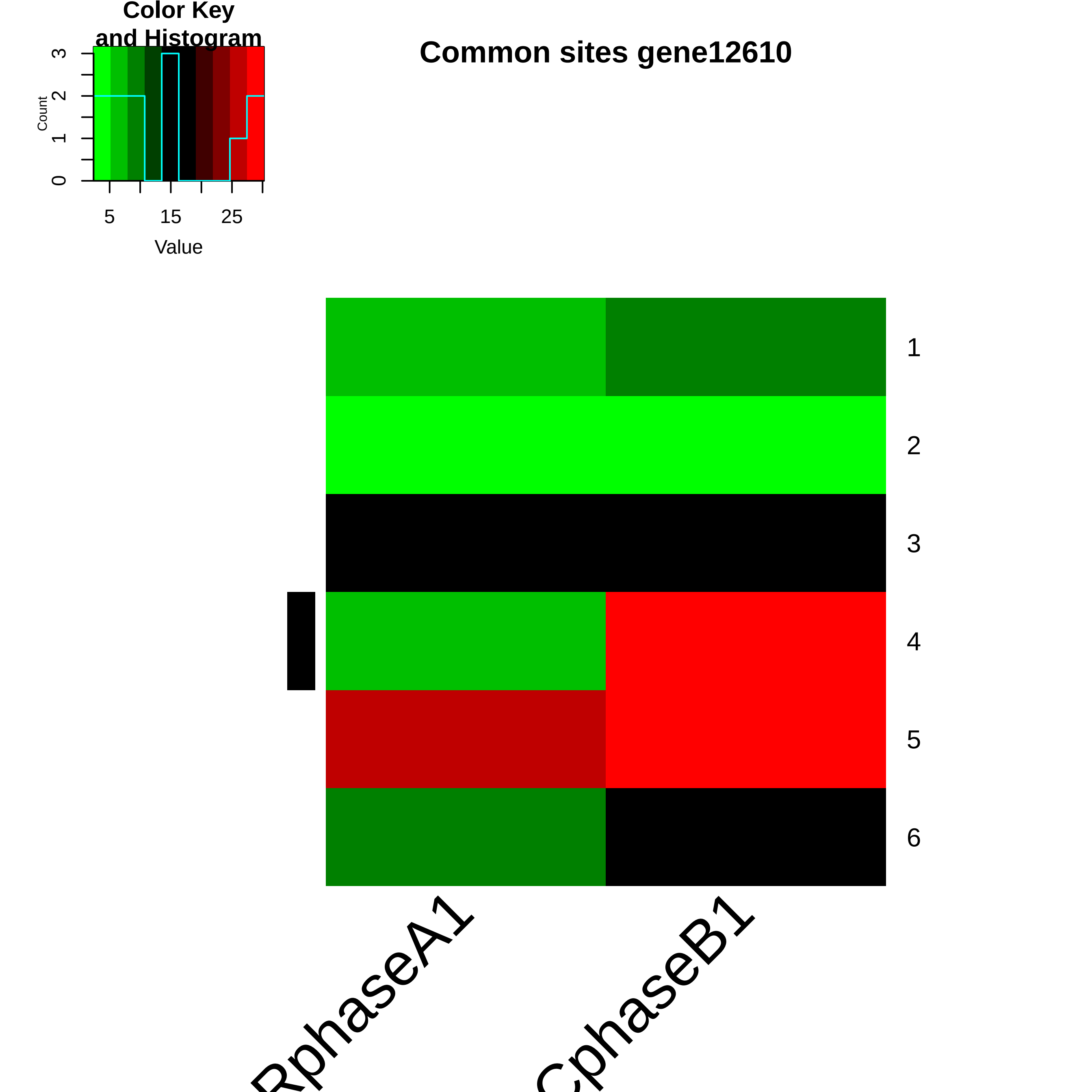
Color Key
and Histogram
Count
0
1
2
3
5
15
25
Value
Common sites gene12610
1
2
3
4
5
6
RphaseA1
CphaseB1

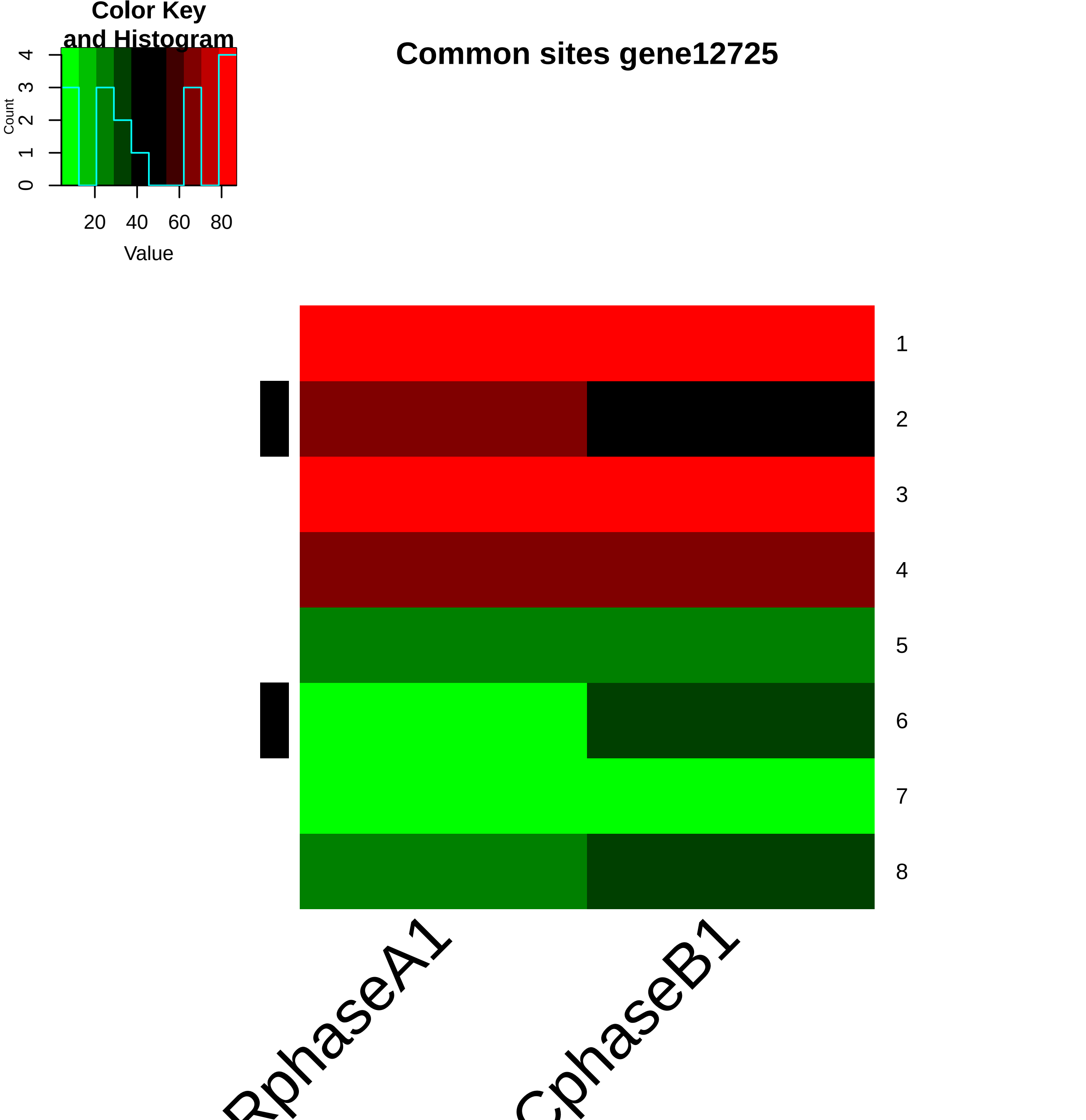
Color Key
and Histogram
Count
0
1
2
3
4
20
40
60
80
Value
Common sites gene12725
1
2
3
4
5
6
7
8
RphaseA1
CphaseB1

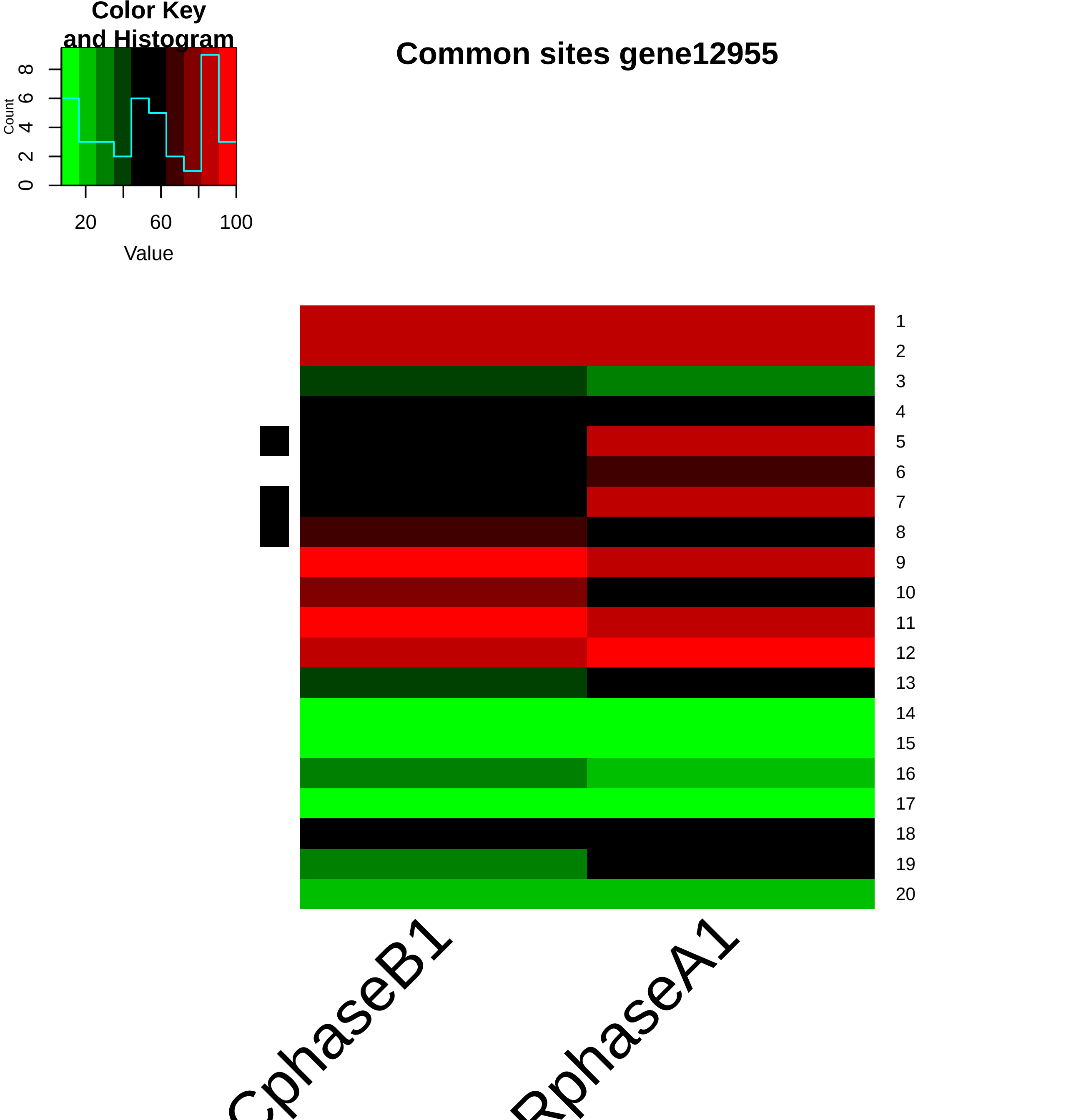
Color Key
and Histogram
Count
0
2
4
6
8
20
60
100
Value
Common sites gene12955
CphaseB1
RphaseA1

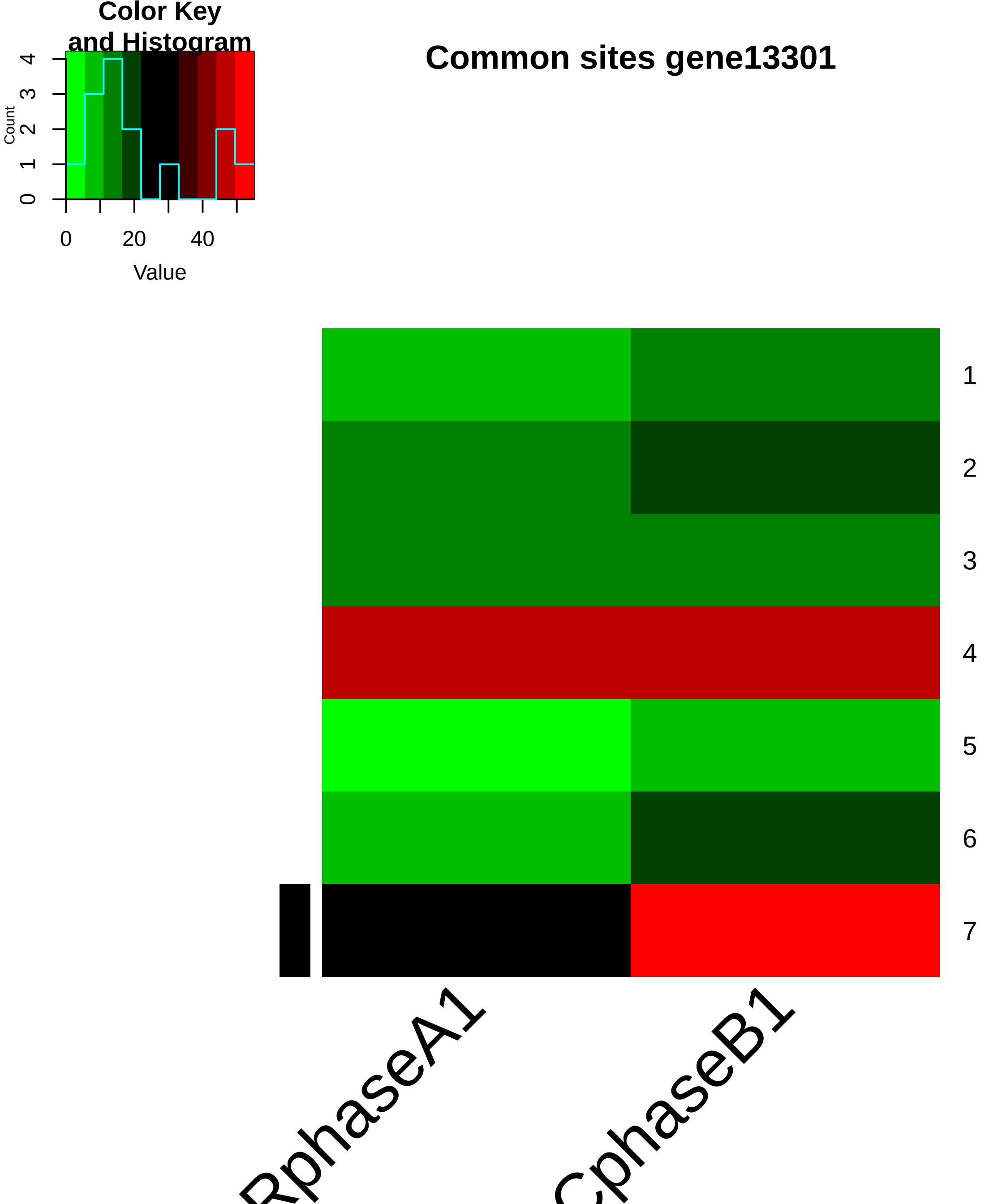

Color Key
and Histogram
Count
0
1
2
3
4
0
20
40
Value
Common sites gene13301
1
2
3
4
5
6
7
RphaseA1
CphaseB1

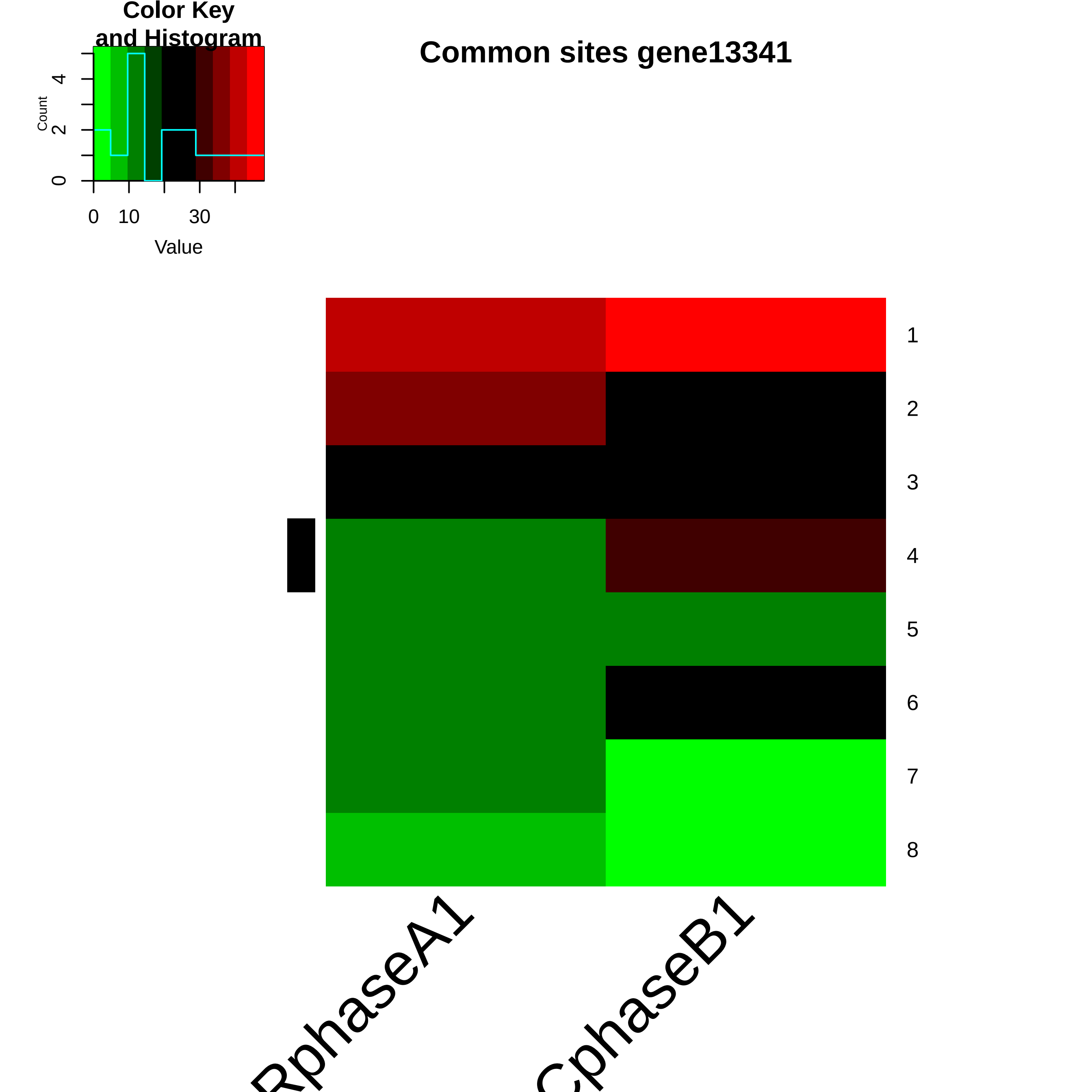

Color Key
and Histogram
Count
0
2
4
0
10
30
Value
Common sites gene13341
1
2
3
4
5
6
7
8
RphaseA1
CphaseB1

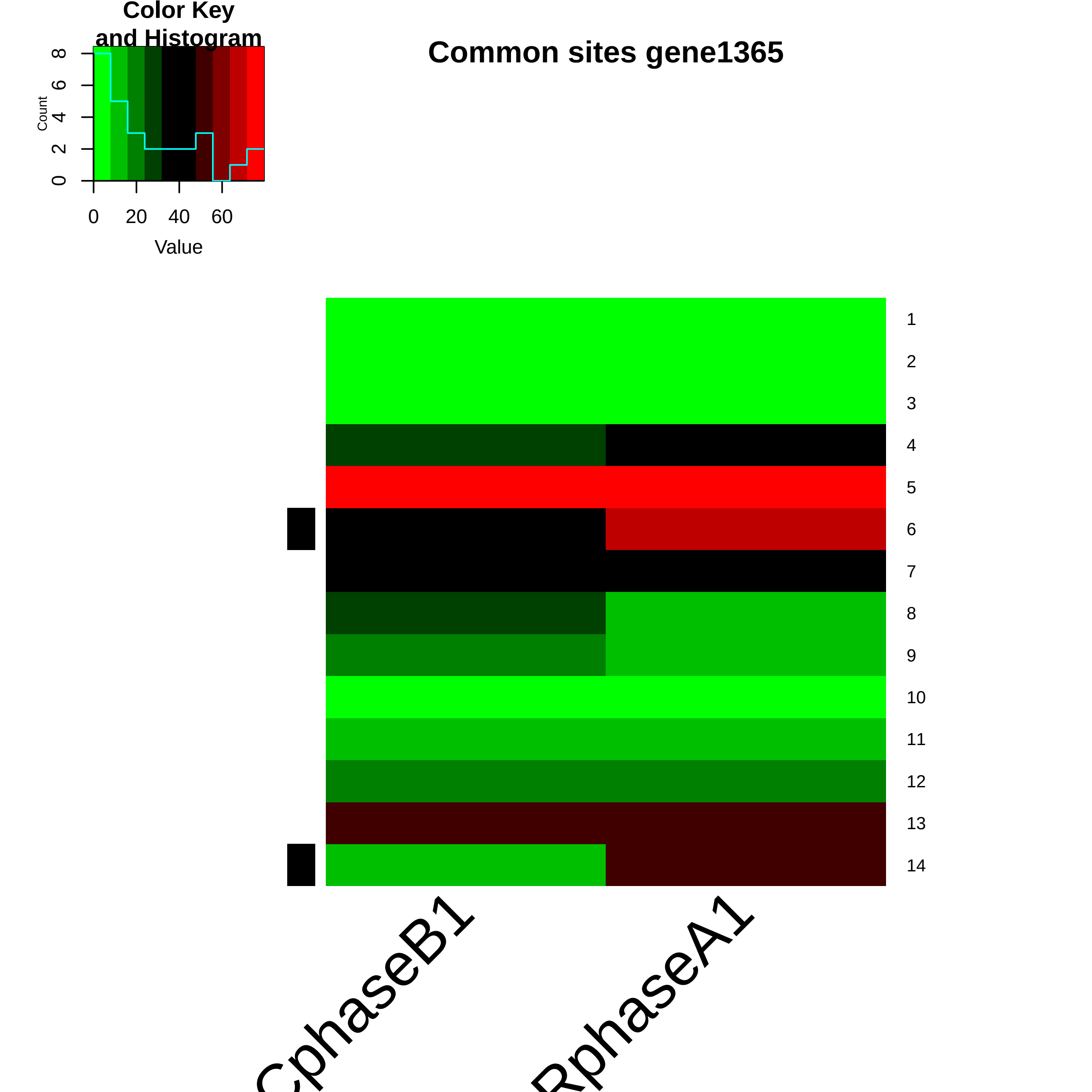

Color Key
and Histogram
Count
0 2 4 6 8
0 20 40 60
Value
Common sites gene1365
1
2
3
4
5
6
7
8
9
10
11
12
13
14
CphaseB1
RphaseA1

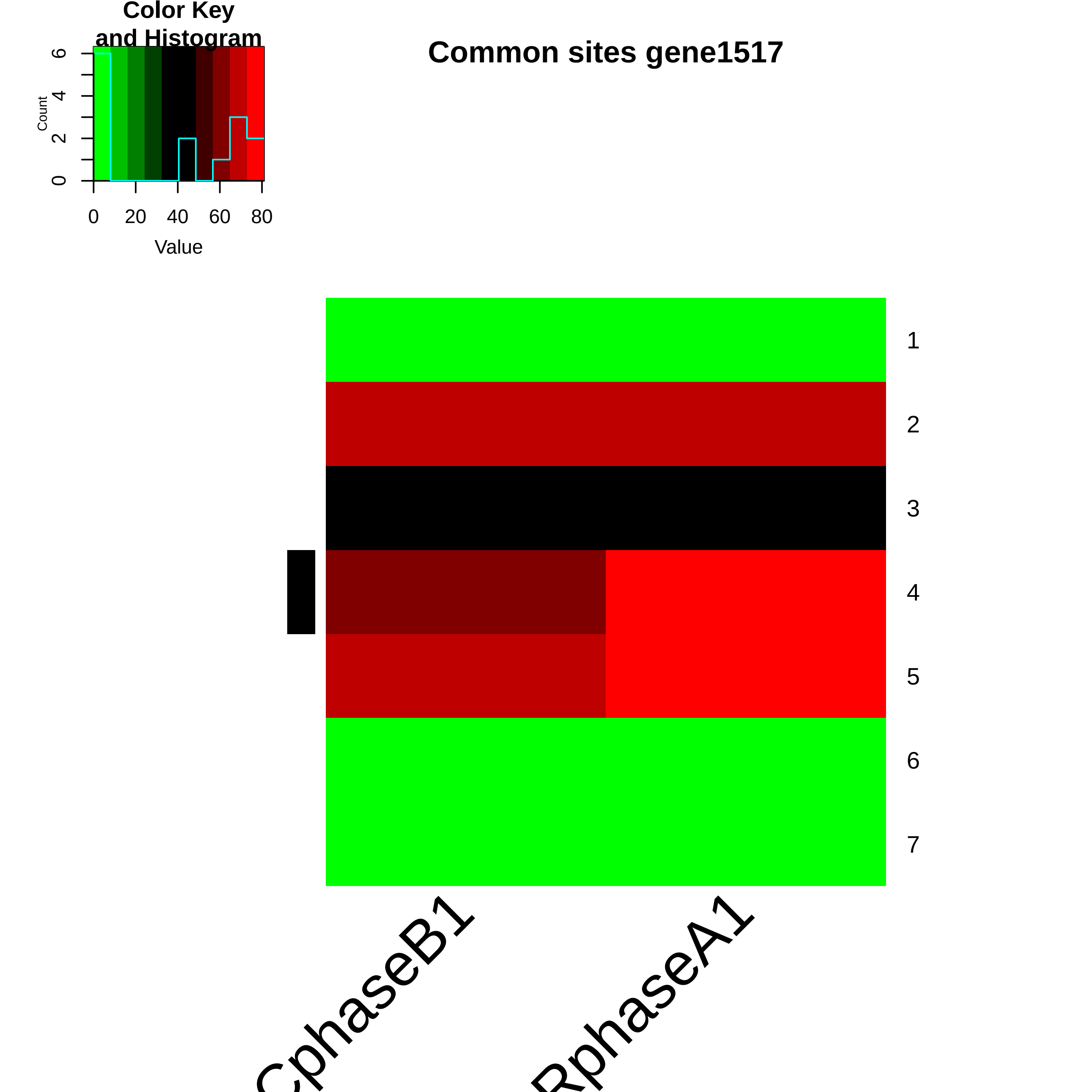
Color Key
and Histogram
Count
0
2
4
6
0
20
40
60
80
Value
Common sites gene1517
1
2
3
4
5
6
7
CphaseB1
RphaseA1

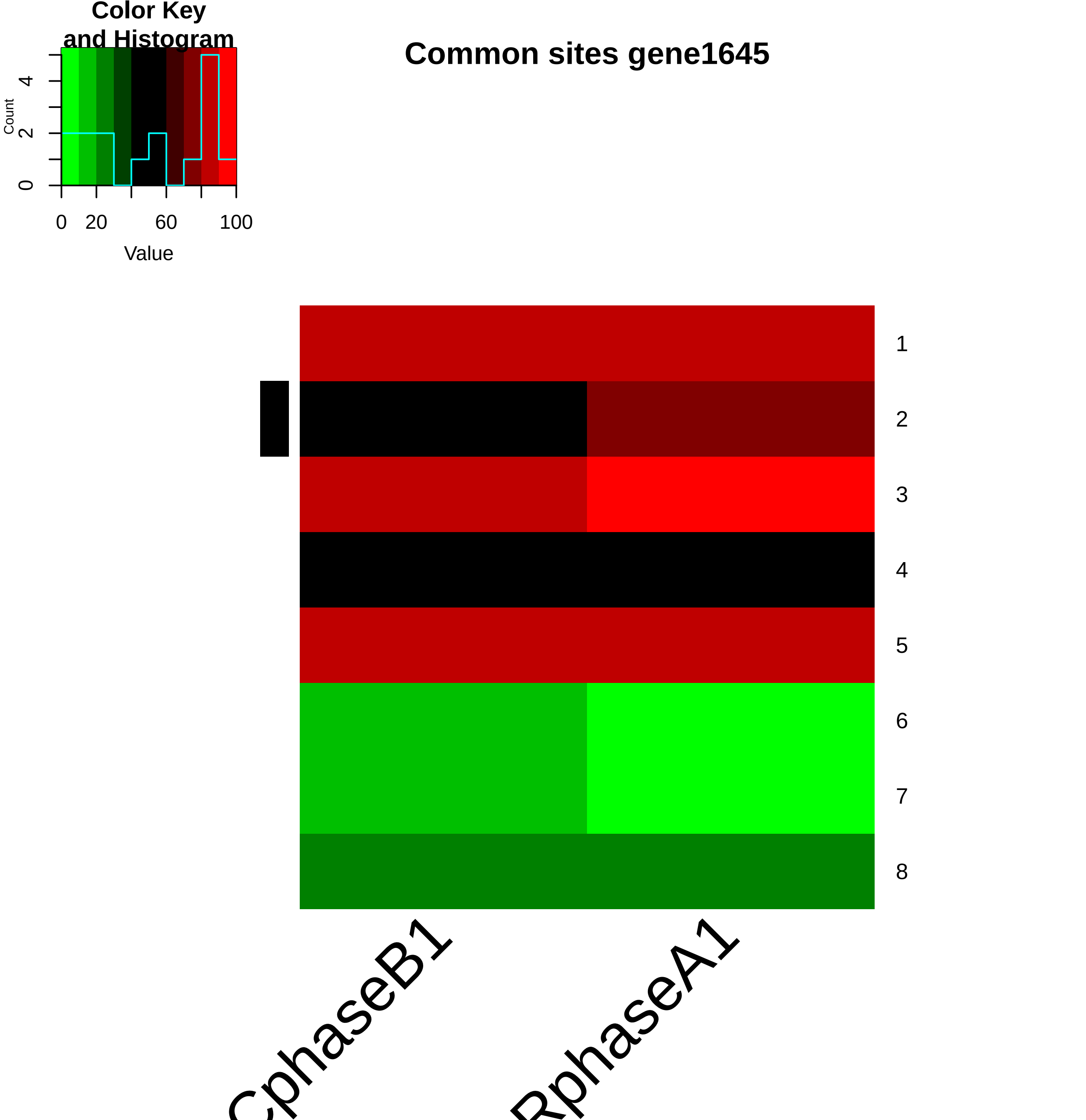
Color Key
and Histogram
Count
0
2
4
0
20
60
100
Value
Common sites gene1645
1
2
3
4
5
6
7
8
CphaseB1
RphaseA1

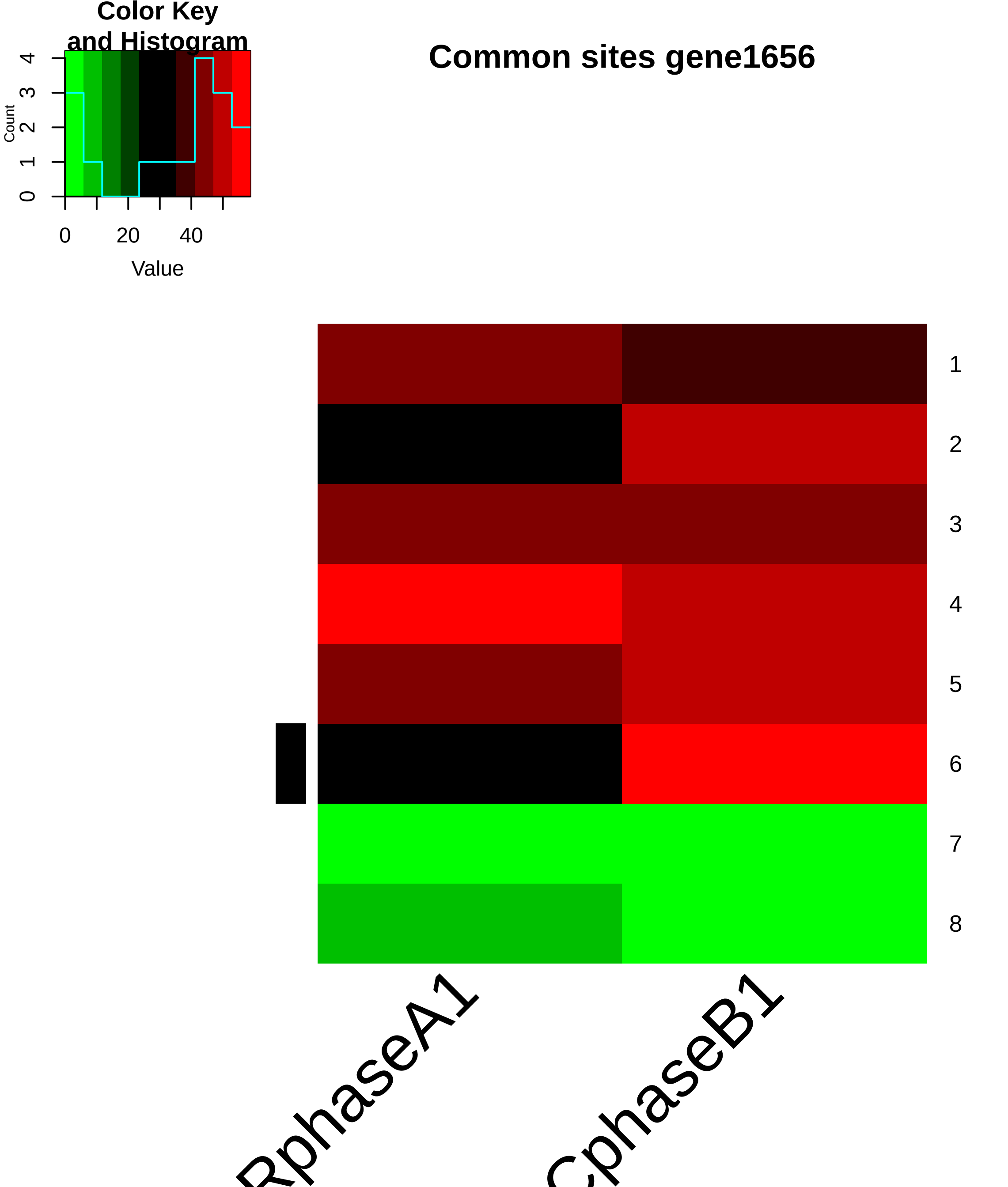
Color Key
and Histogram
Count
0
1
2
3
4
0
20
40
Value
Common sites gene1656
1
2
3
4
5
6
7
8
RphaseA1
CphaseB1

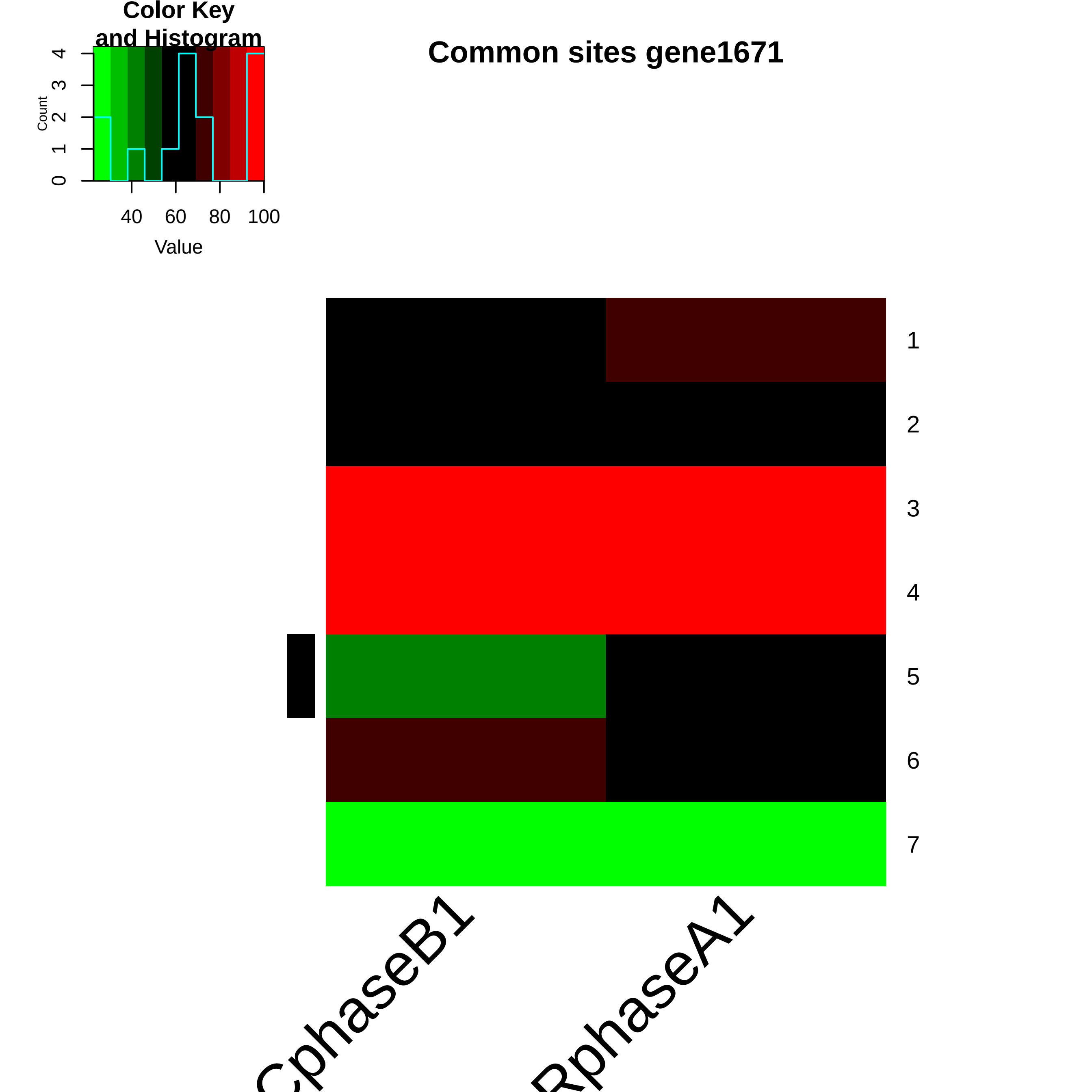
Color Key
and Histogram
Count
0
1
2
3
4
40
60
80
100
Value
Common sites gene1671
1
2
3
4
5
6
7
CphaseB1
RphaseA1

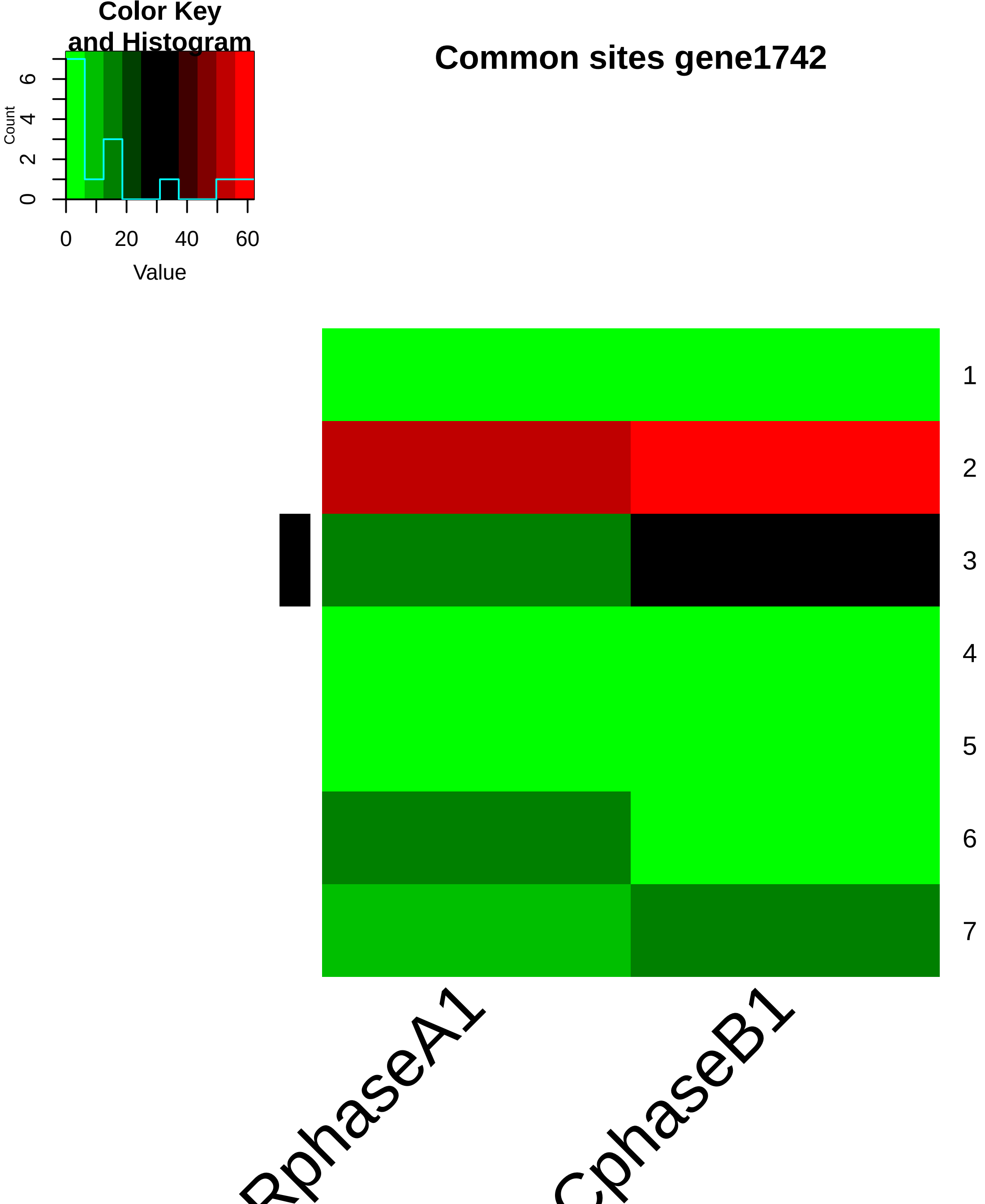
Color Key
and Histogram
Count
Value
0
20
40
60
0
2
4
6
Common sites gene1742
1
2
3
4
5
6
7
RphaseA1
CphaseB1

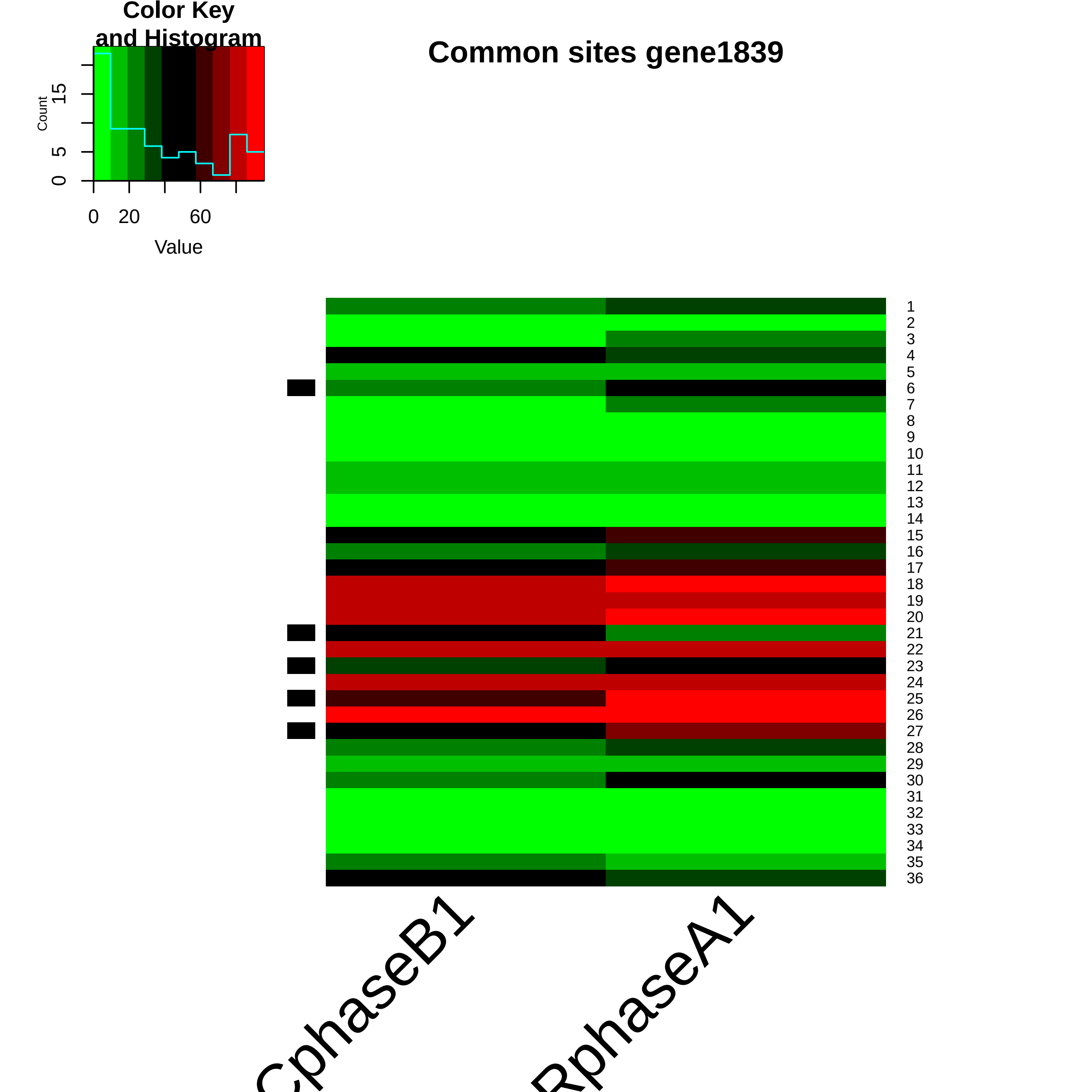

Color Key
and Histogram
Count
0
5
15
0
20
60
Value
Common sites gene1839
CphaseB1
RphaseA1

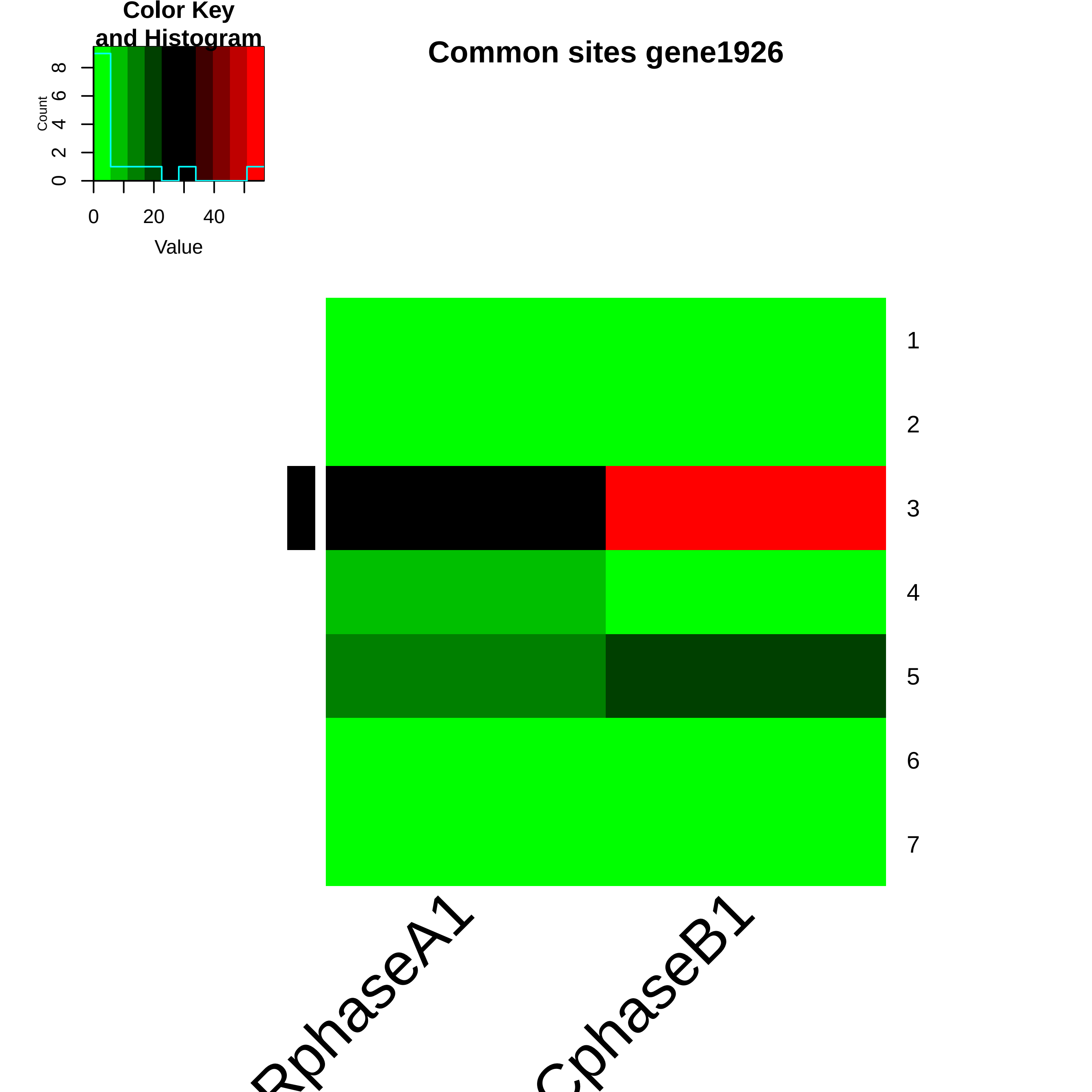

Color Key
and Histogram
Count
0
2
4
6
8
0
20
40
Value
Common sites gene1926
1
2
3
4
5
6
7
RphaseA1
CphaseB1

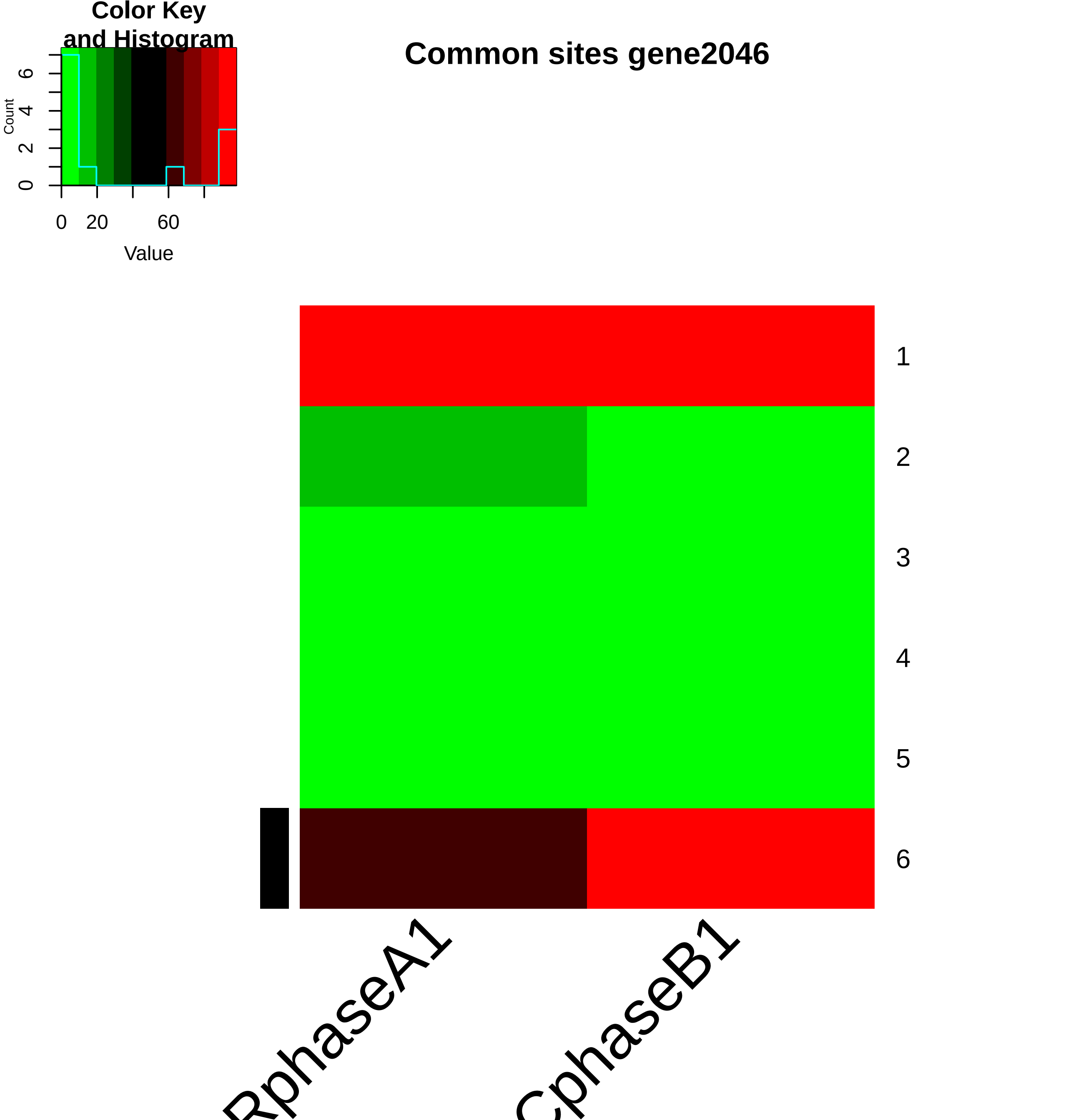
Color Key
and Histogram
Count
0
2
4
6
0
20
60
Value
Common sites gene2046
1
2
3
4
5
6
RphaseA1
CphaseB1

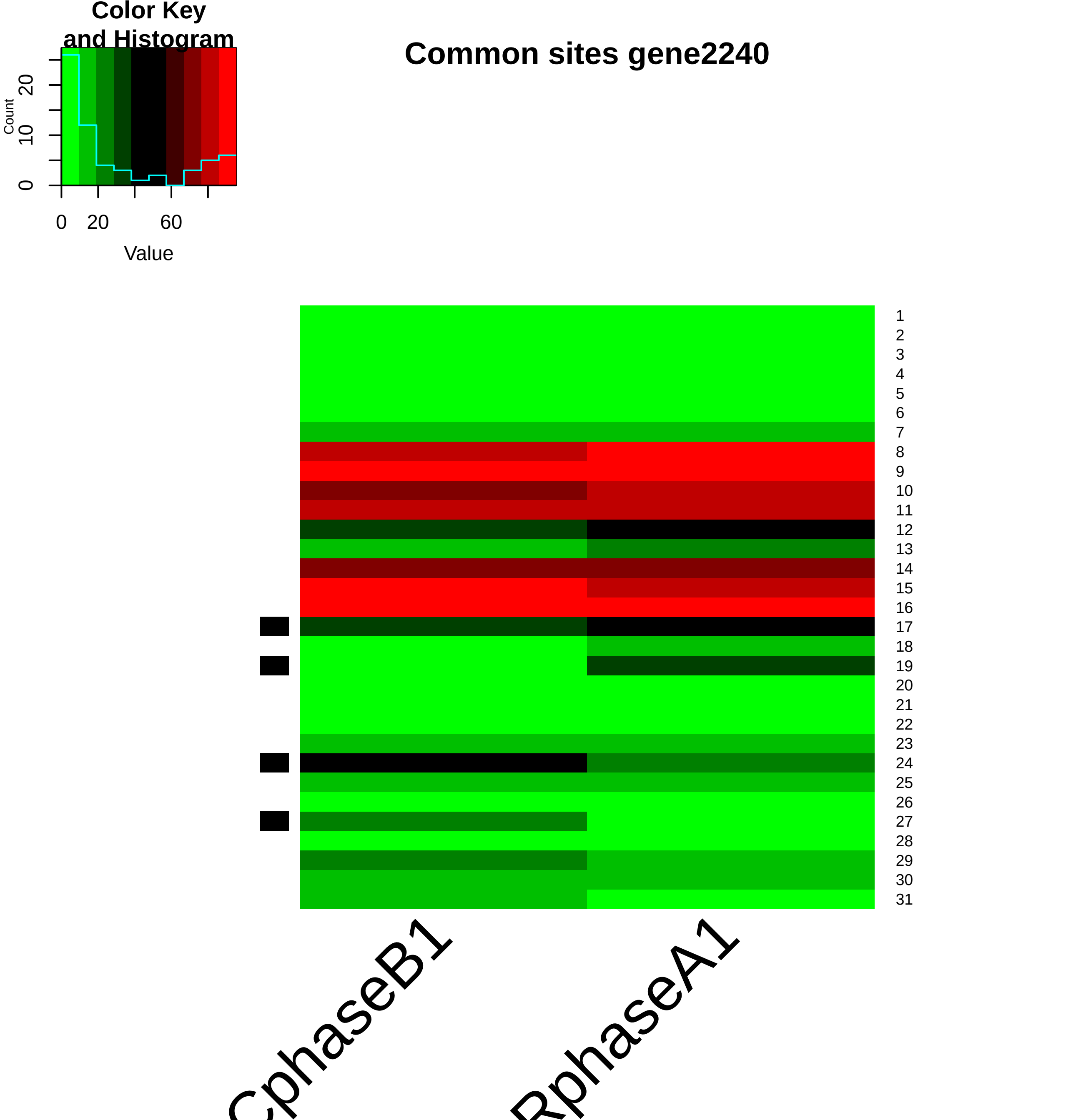
Color Key
and Histogram
Count
0
10
20
0
20
60
Value
Common sites gene2240
CphaseB1
RphaseA1

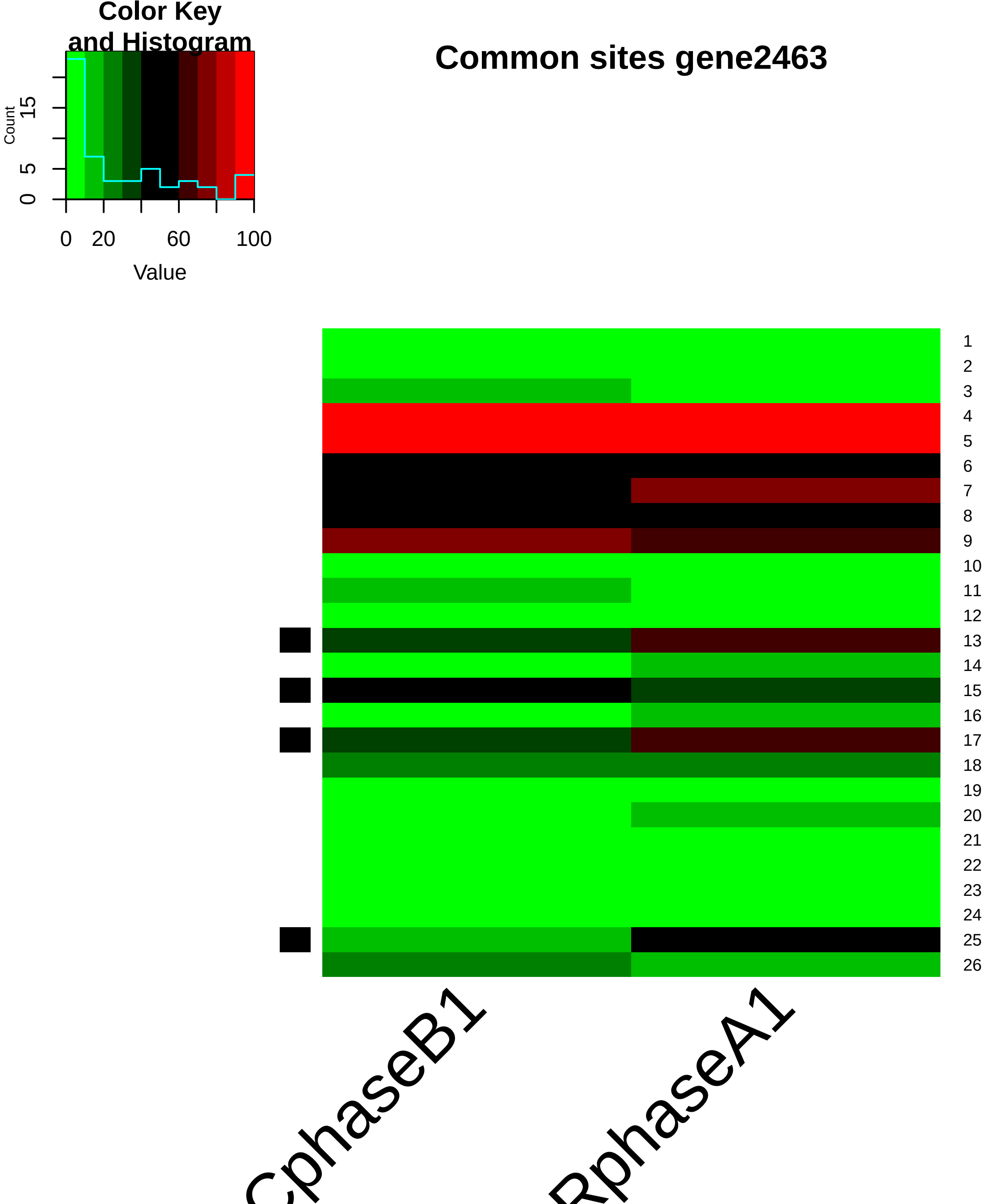

Color Key
and Histogram
Count
0
5
15
0
20
60
100
Value
Common sites gene2463
CphaseB1
RphaseA1

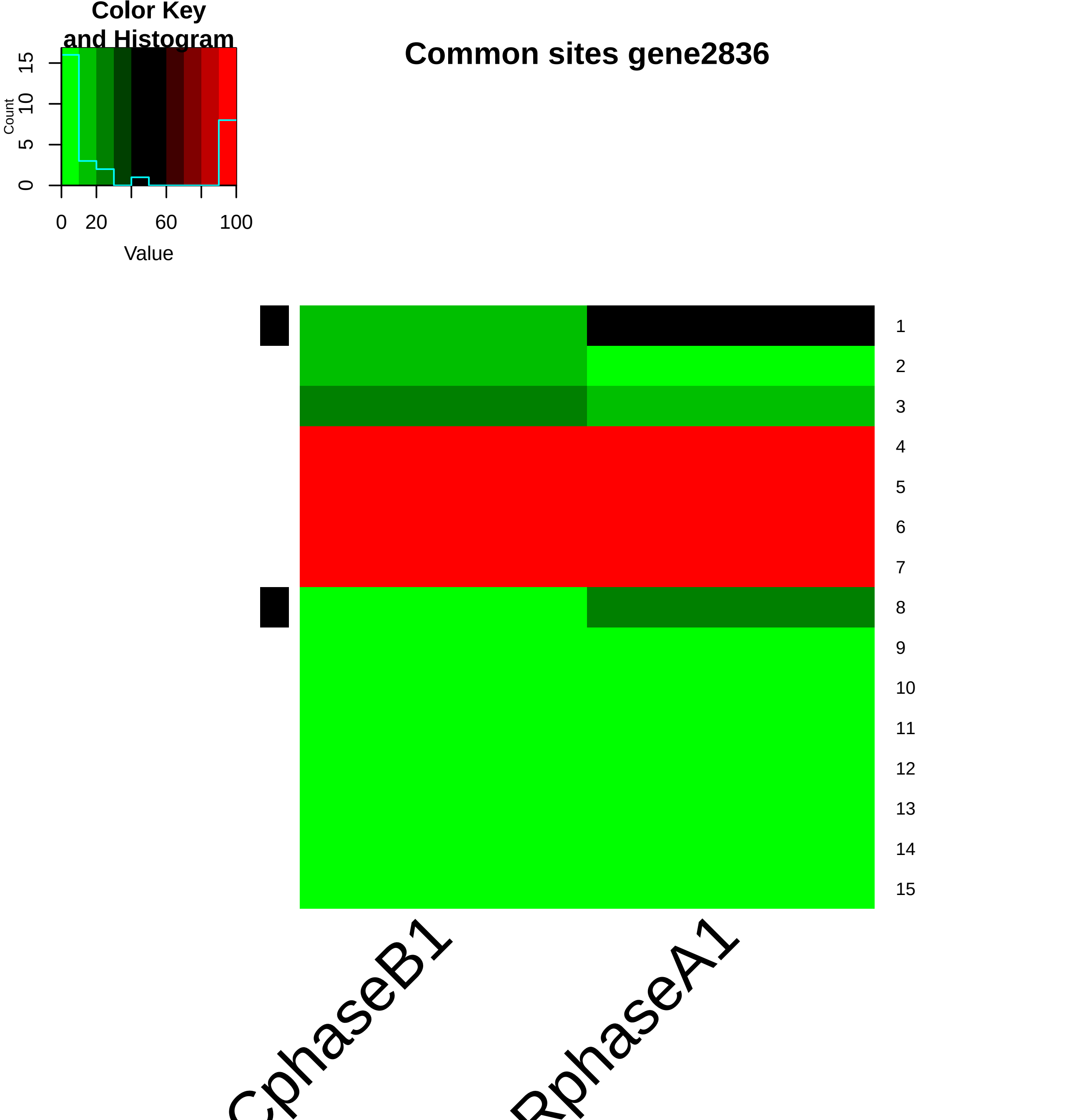
Color Key
and Histogram
Count
0
5
10
15
0
20
60
100
Value
Common sites gene2836
CphaseB1
RphaseA1

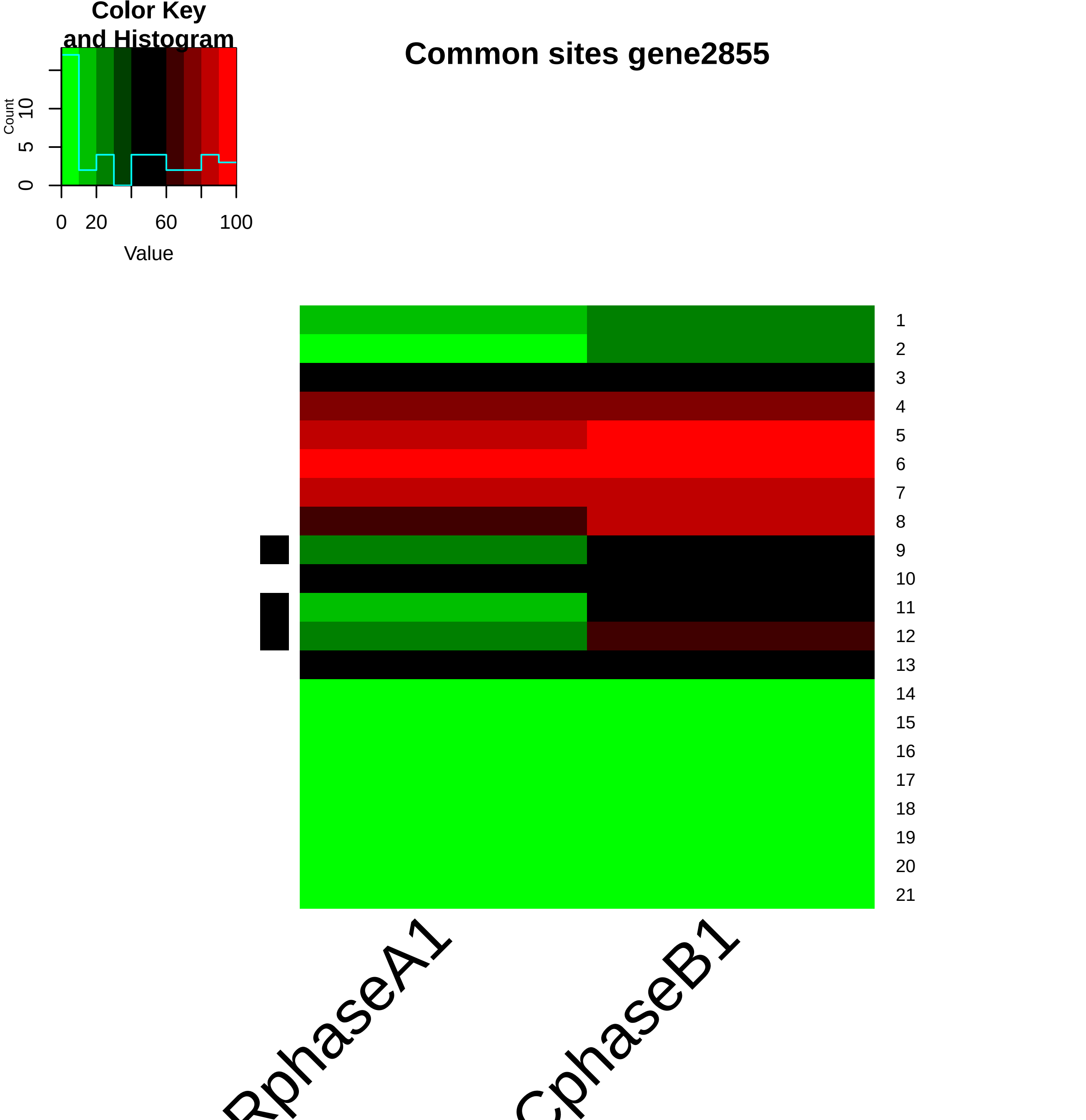

Color Key
and Histogram
Count
0
5
10
0
20
60
100
Value
Common sites gene2855
RphaseA1
CphaseB1

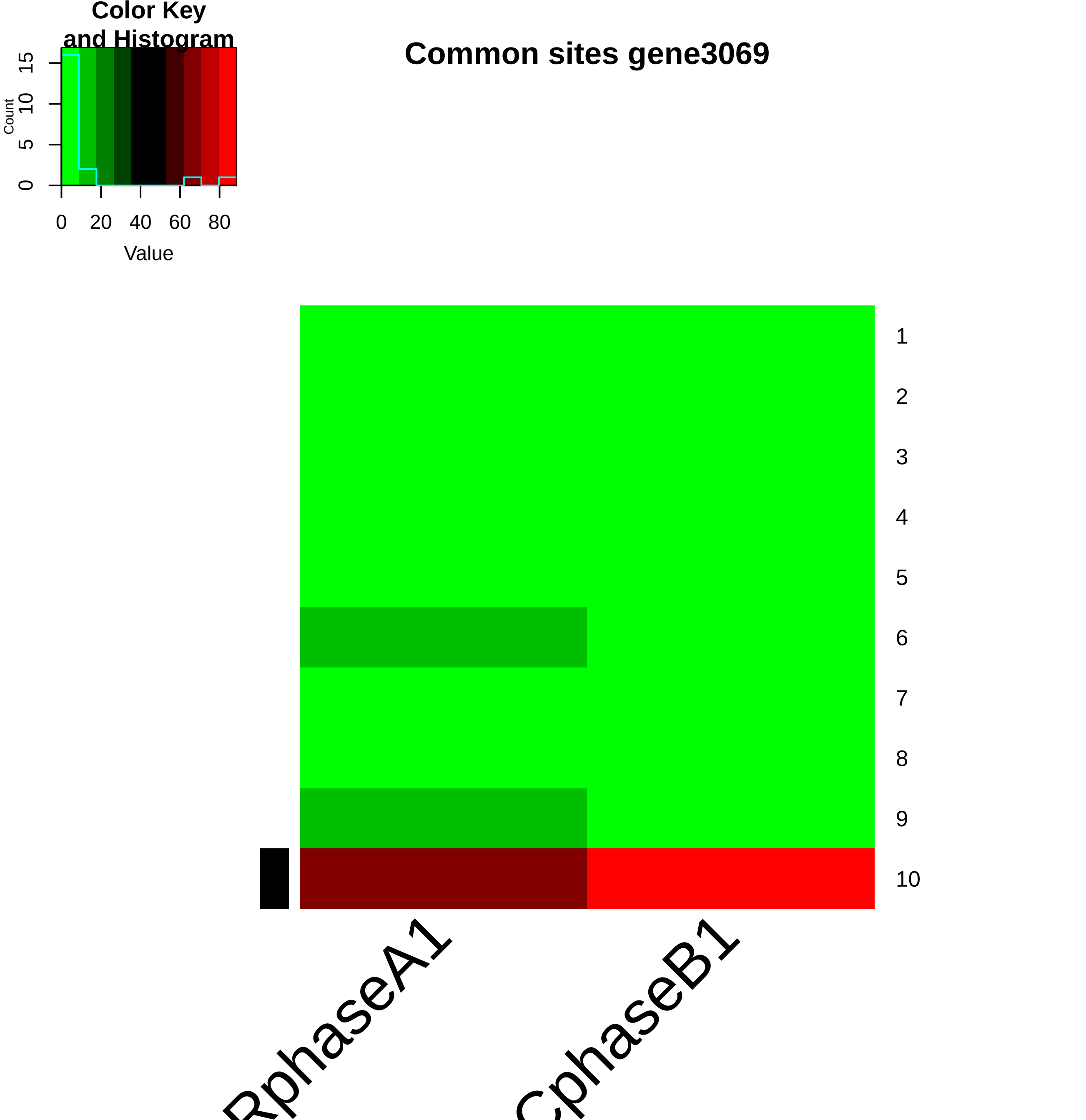
Color Key
and Histogram
Count
0
5
10
15
0
20
40
60
80
Value
Common sites gene3069
1
2
3
4
5
6
7
8
9
10
RphaseA1
CphaseB1

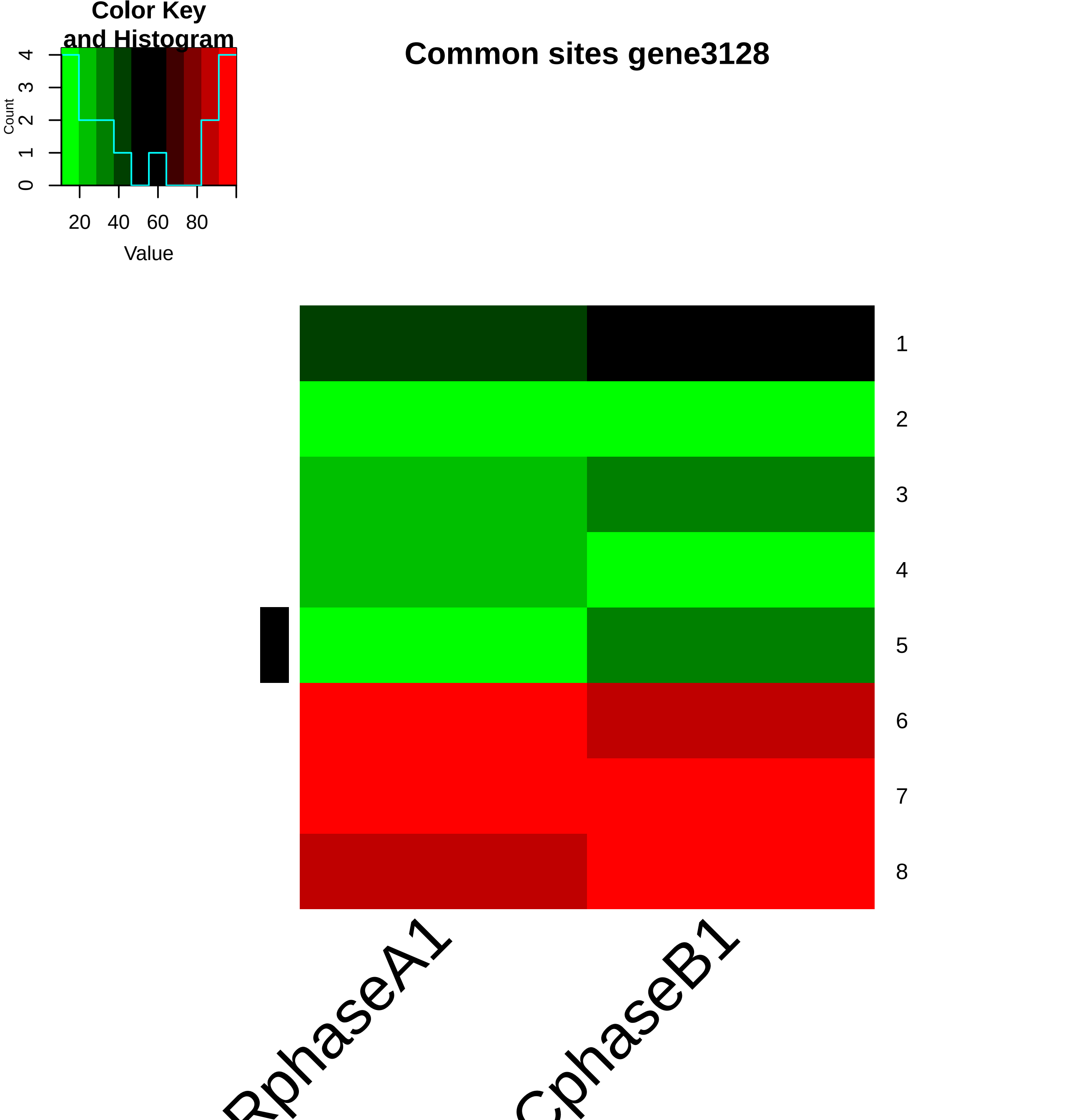
Color Key
and Histogram
Count
0
1
2
3
4
20
40
60
80
Value
Common sites gene3128
1
2
3
4
5
6
7
8
RphaseA1
CphaseB1

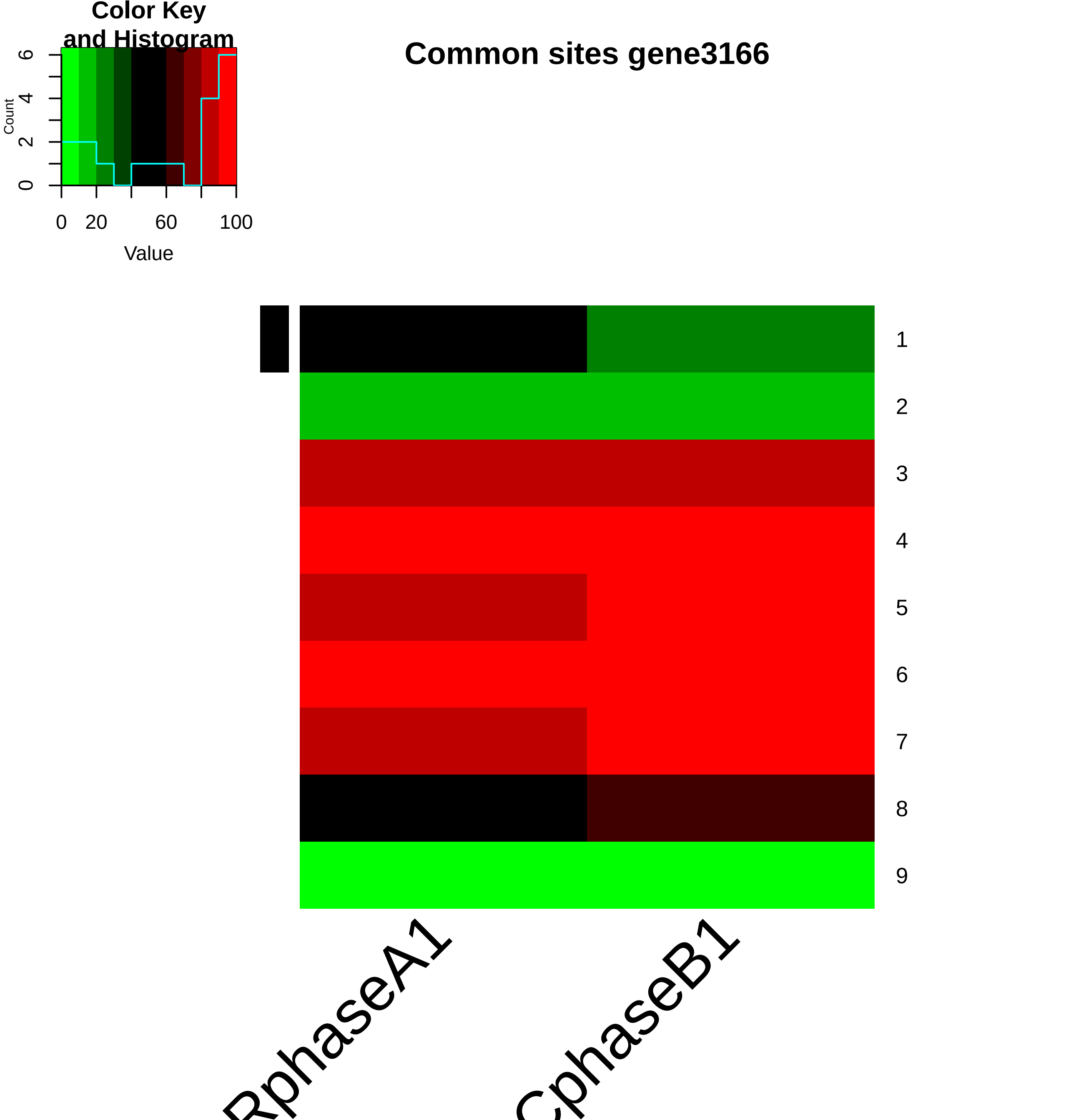

Color Key
and Histogram
Count
0
2
4
6
0
20
60
100
Value
Common sites gene3166
1
2
3
4
5
6
7
8
9
RphaseA1
CphaseB1

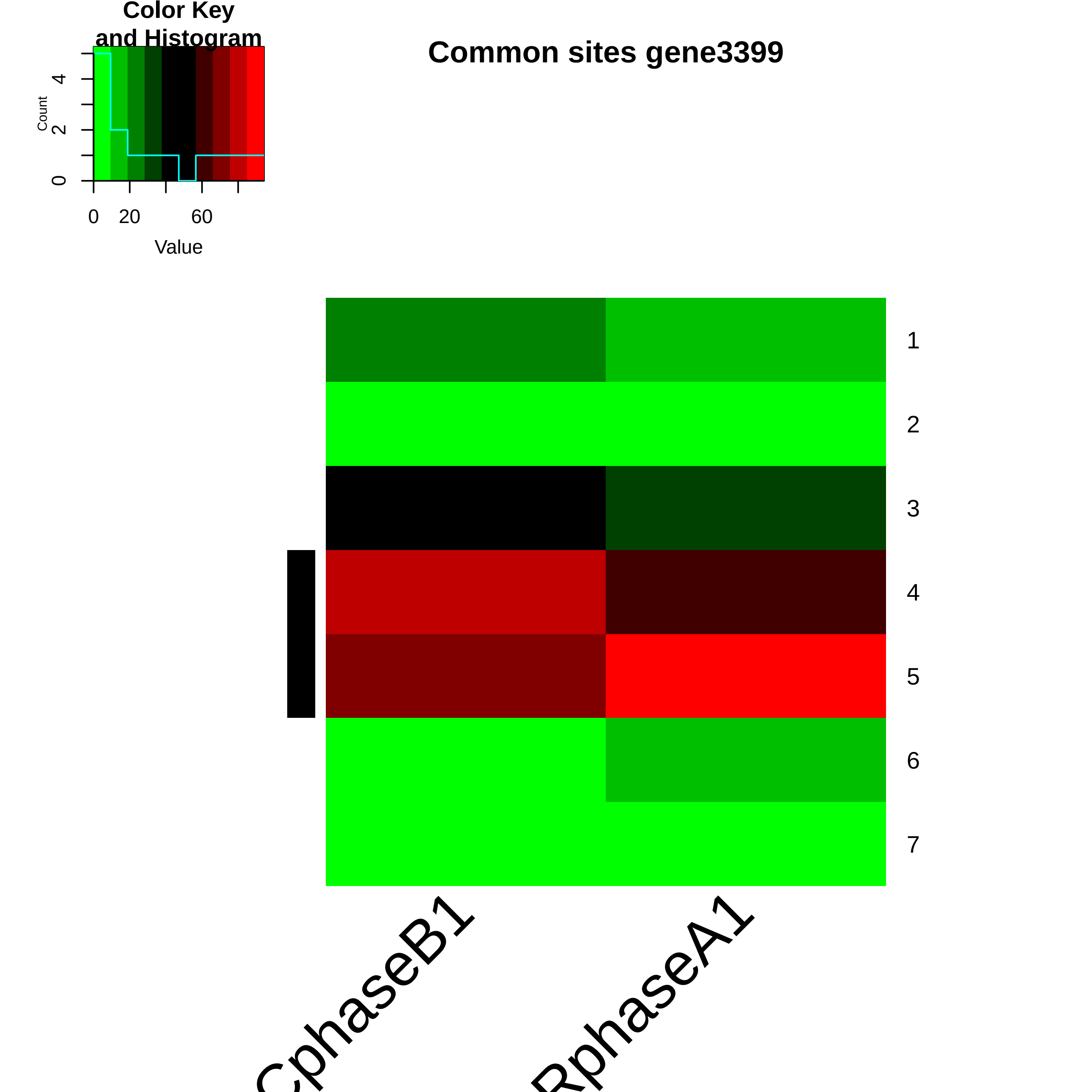
Color Key
and Histogram
Count
0
2
4
0
20
60
Value
Common sites gene3399
1
2
3
4
5
6
7
CphaseB1
RphaseA1

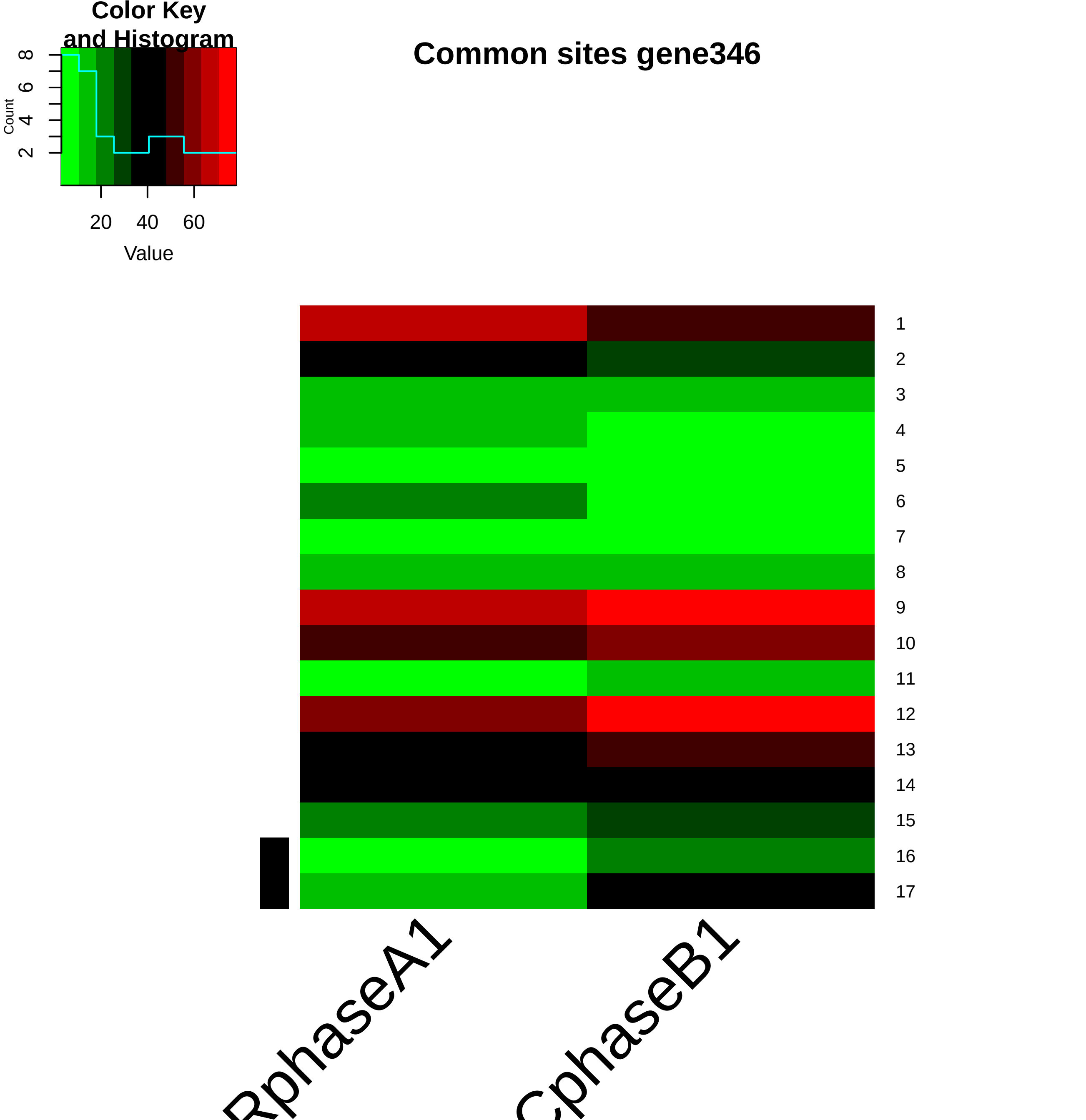
Color Key
and Histogram
Count
2
4
6
8
20
40
60
Value
Common sites gene346
RphaseA1
CphaseB1

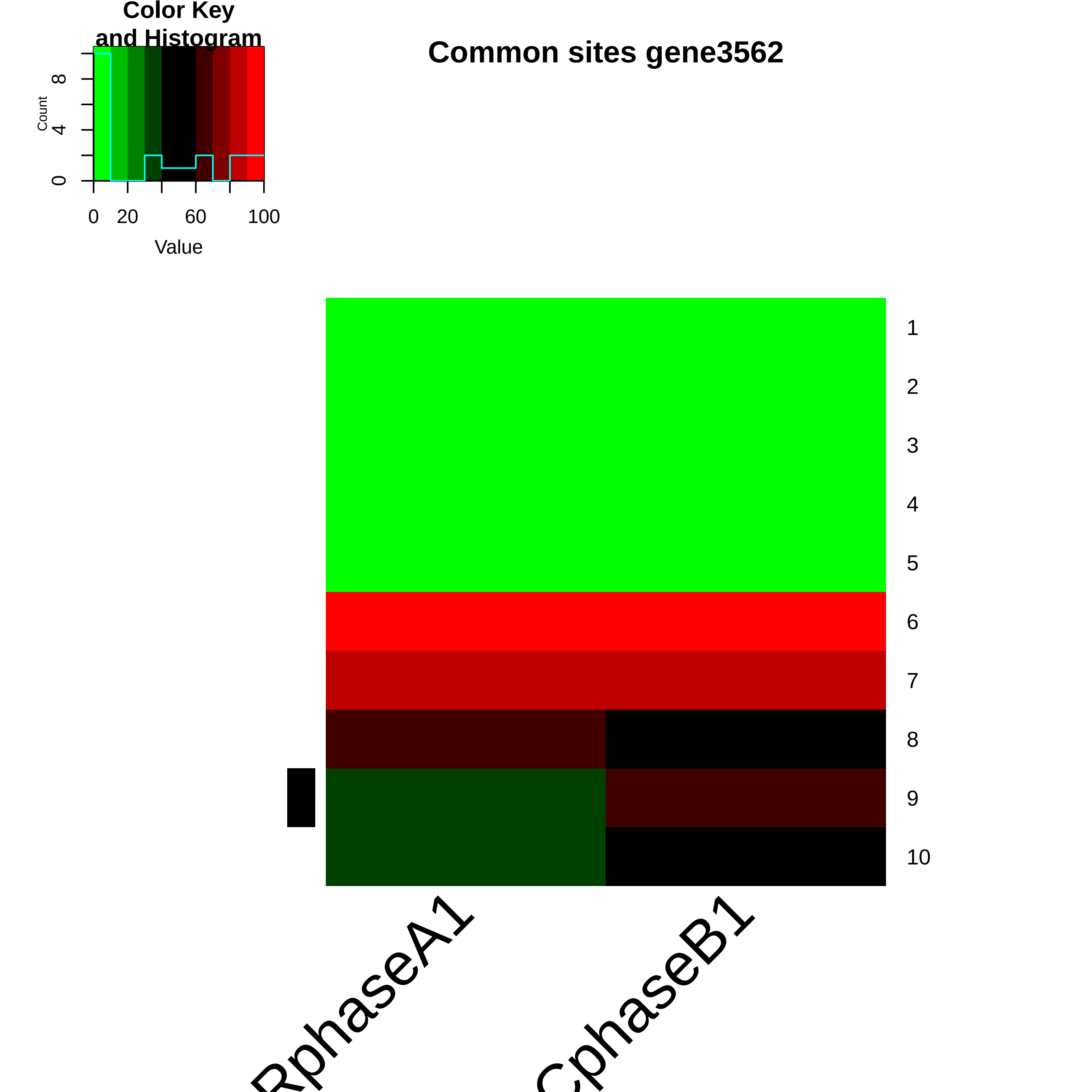
Color Key
and Histogram
Count
0
4
8
0
20
60
100
Value
Common sites gene3562
1
2
3
4
5
6
7
8
9
10
RphaseA1
CphaseB1

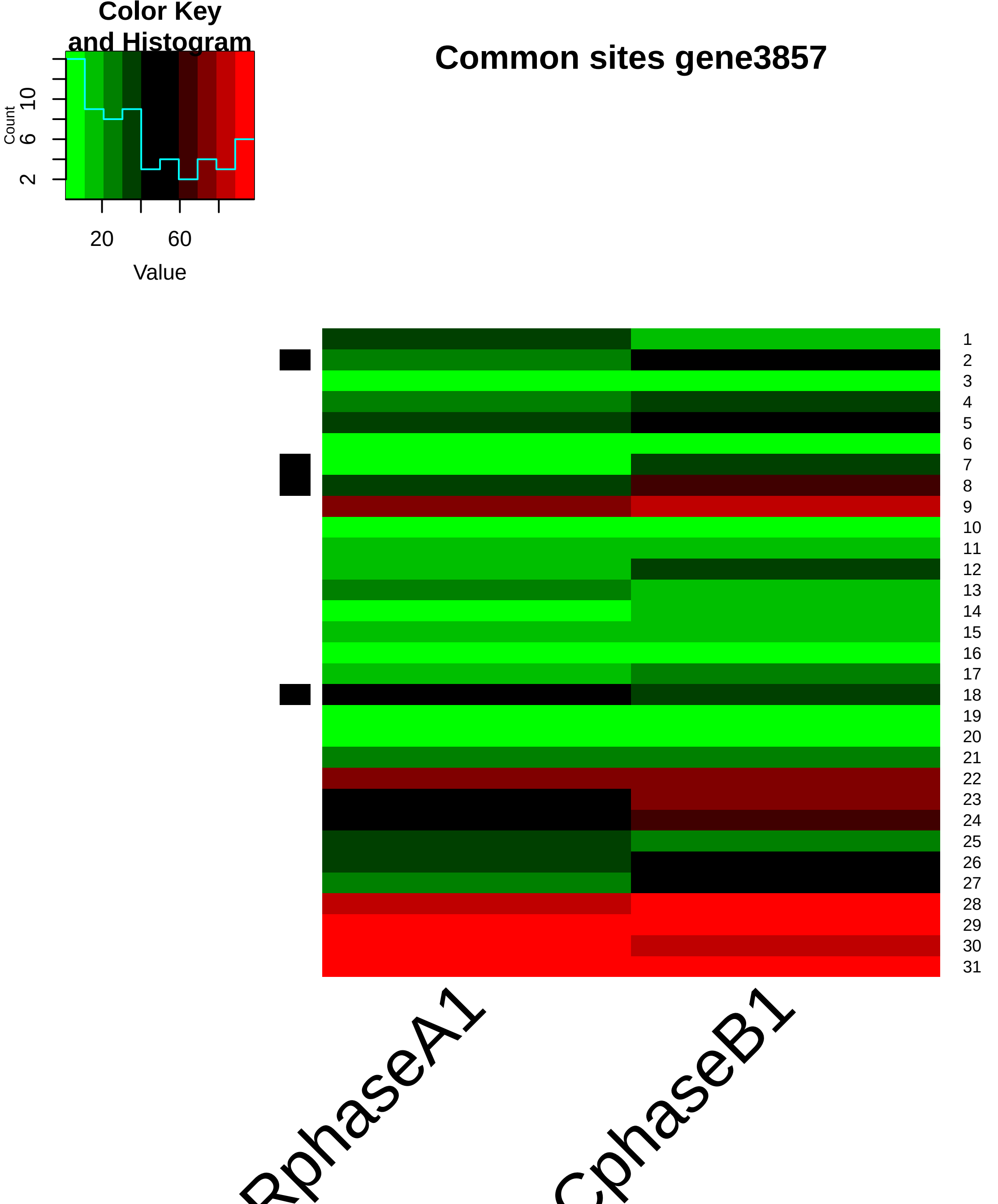

Color Key
and Histogram
Count
2
6
10
20
60
Value
Common sites gene3857
RphaseA1
CphaseB1

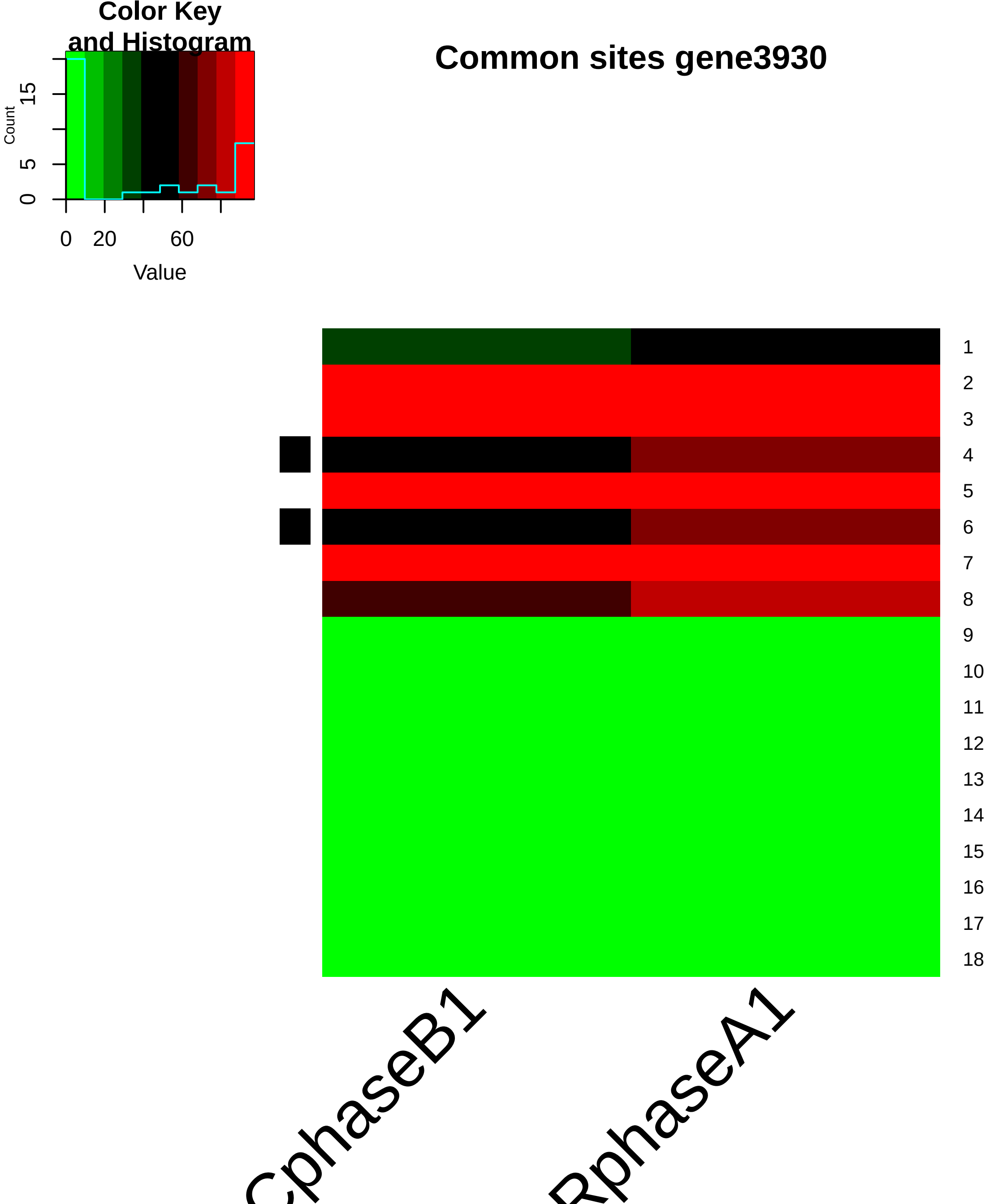
Color Key
and Histogram
Count
0
5
15
0
20
60
Value
Common sites gene3930
CphaseB1
RphaseA1

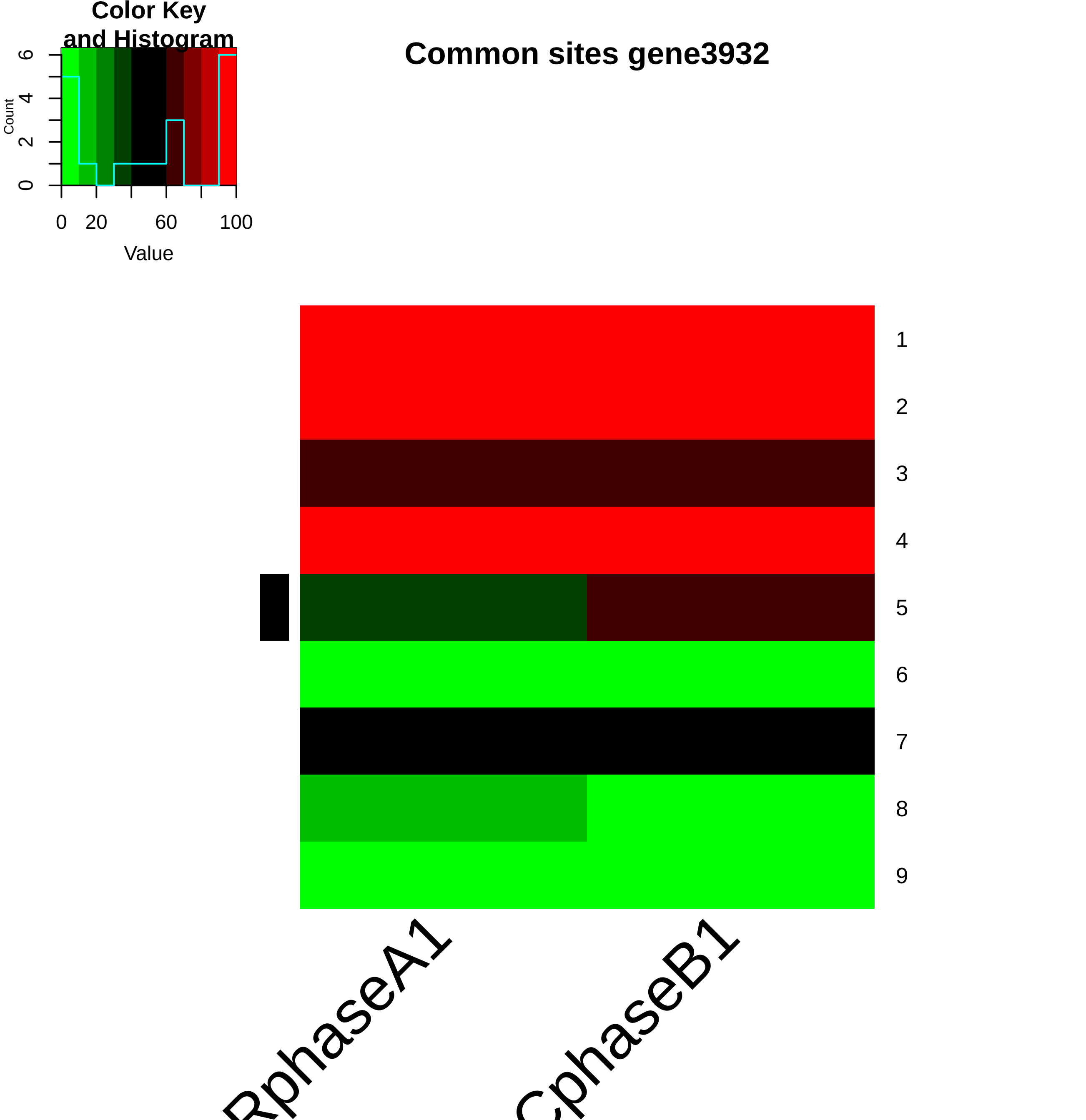
Color Key
and Histogram
Count
0
2
4
6
0
20
60
100
Value
Common sites gene3932
1
2
3
4
5
6
7
8
9
RphaseA1
CphaseB1

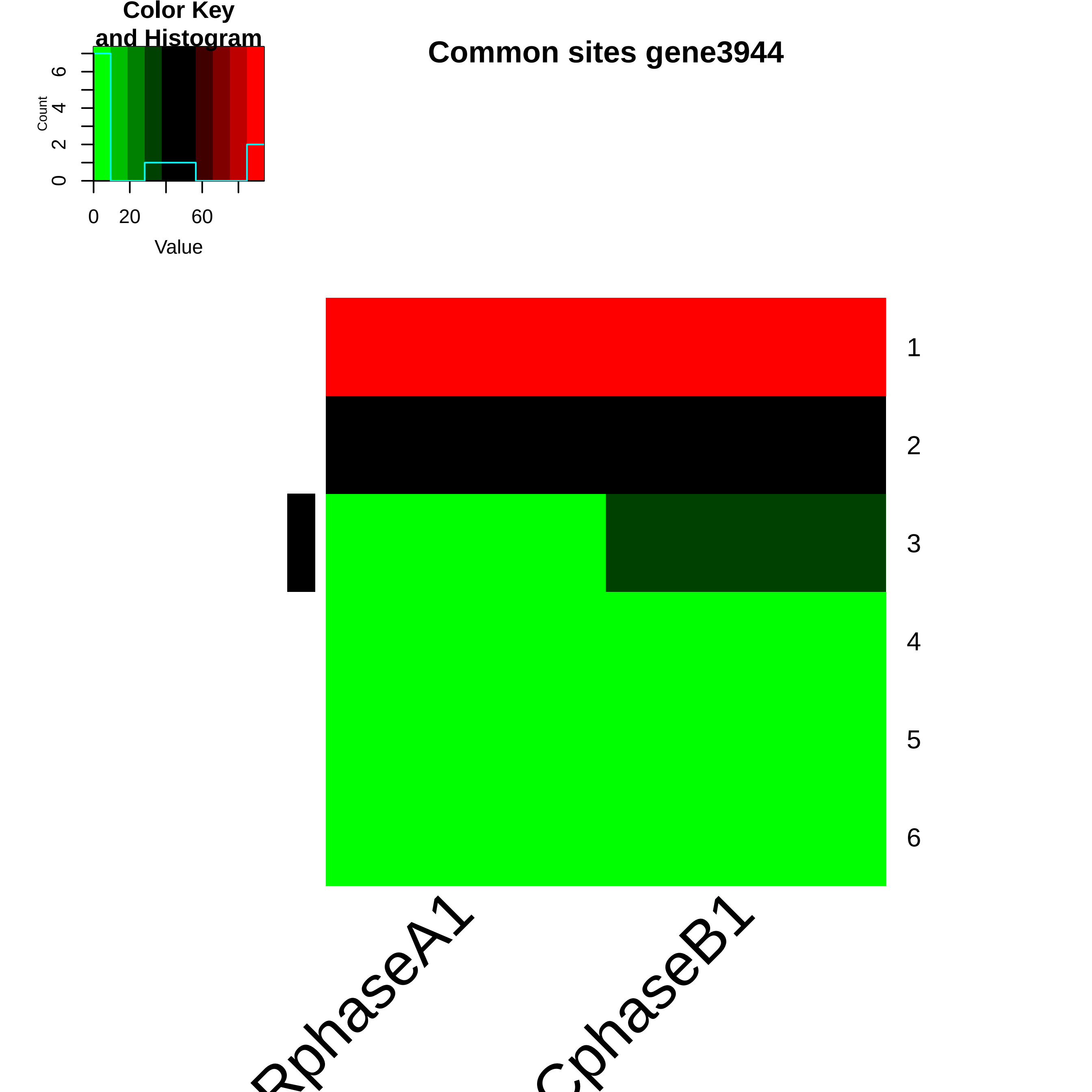
Color Key
and Histogram
Count
0
2
4
6
0
20
60
Value
Common sites gene3944
1
2
3
4
5
6
RphaseA1
CphaseB1

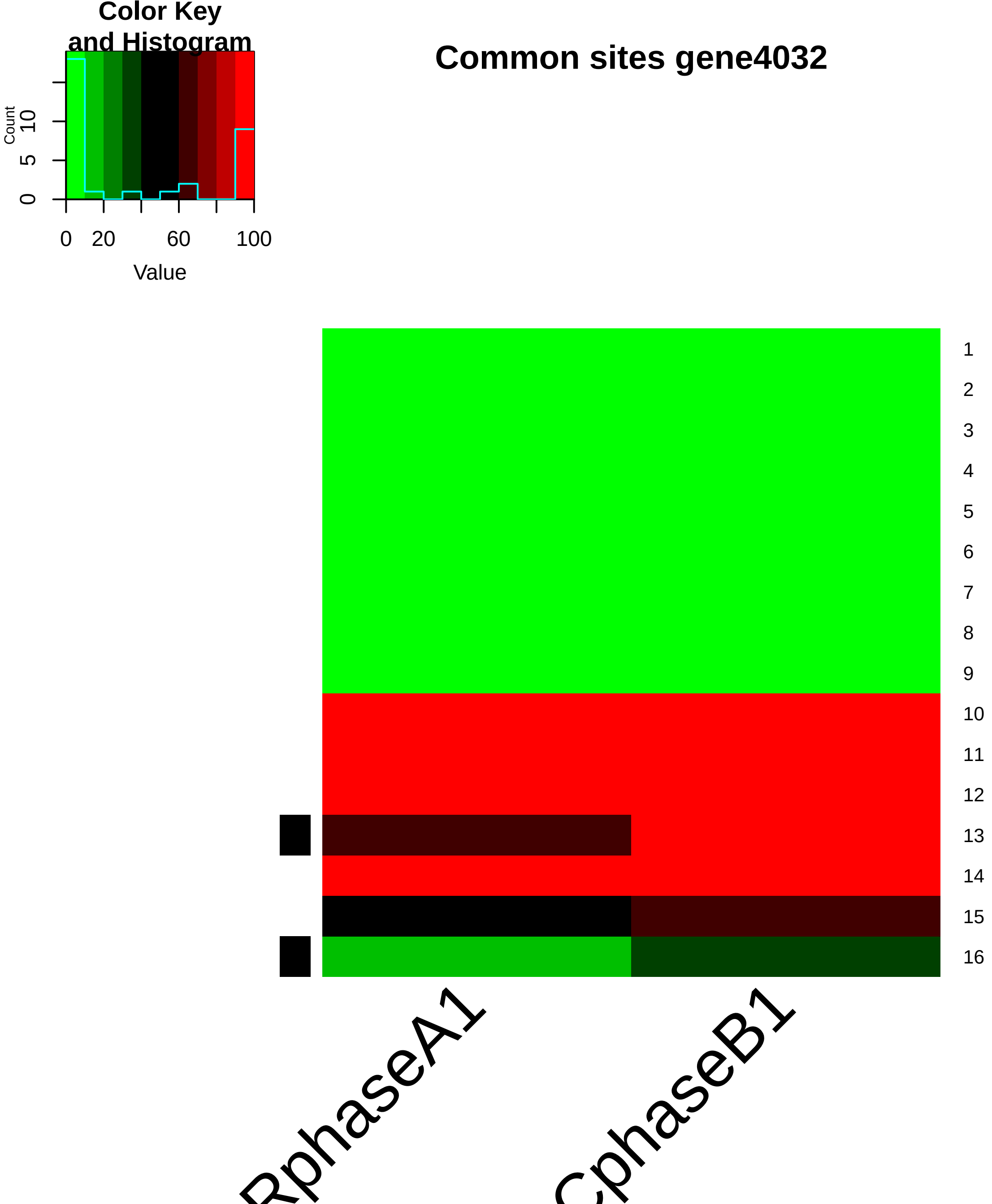
Color Key
and Histogram
Count
0
5
10
0
20
60
100
Value
Common sites gene4032

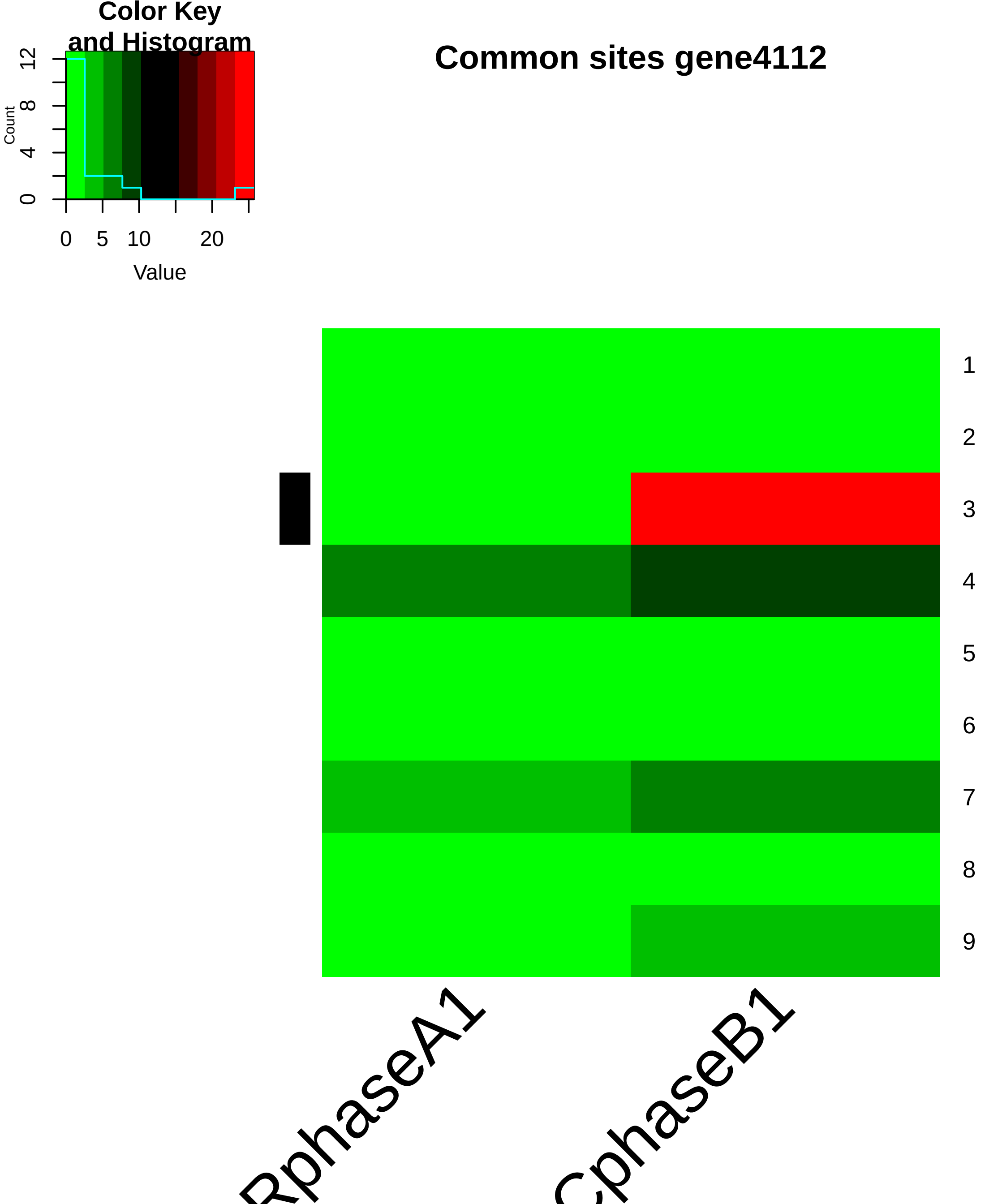
Color Key
and Histogram
Count
0
4
8
12
0
5
10
20
Value
Common sites gene4112
1
2
3
4
5
6
7
8
9
RphaseA1
CphaseB1

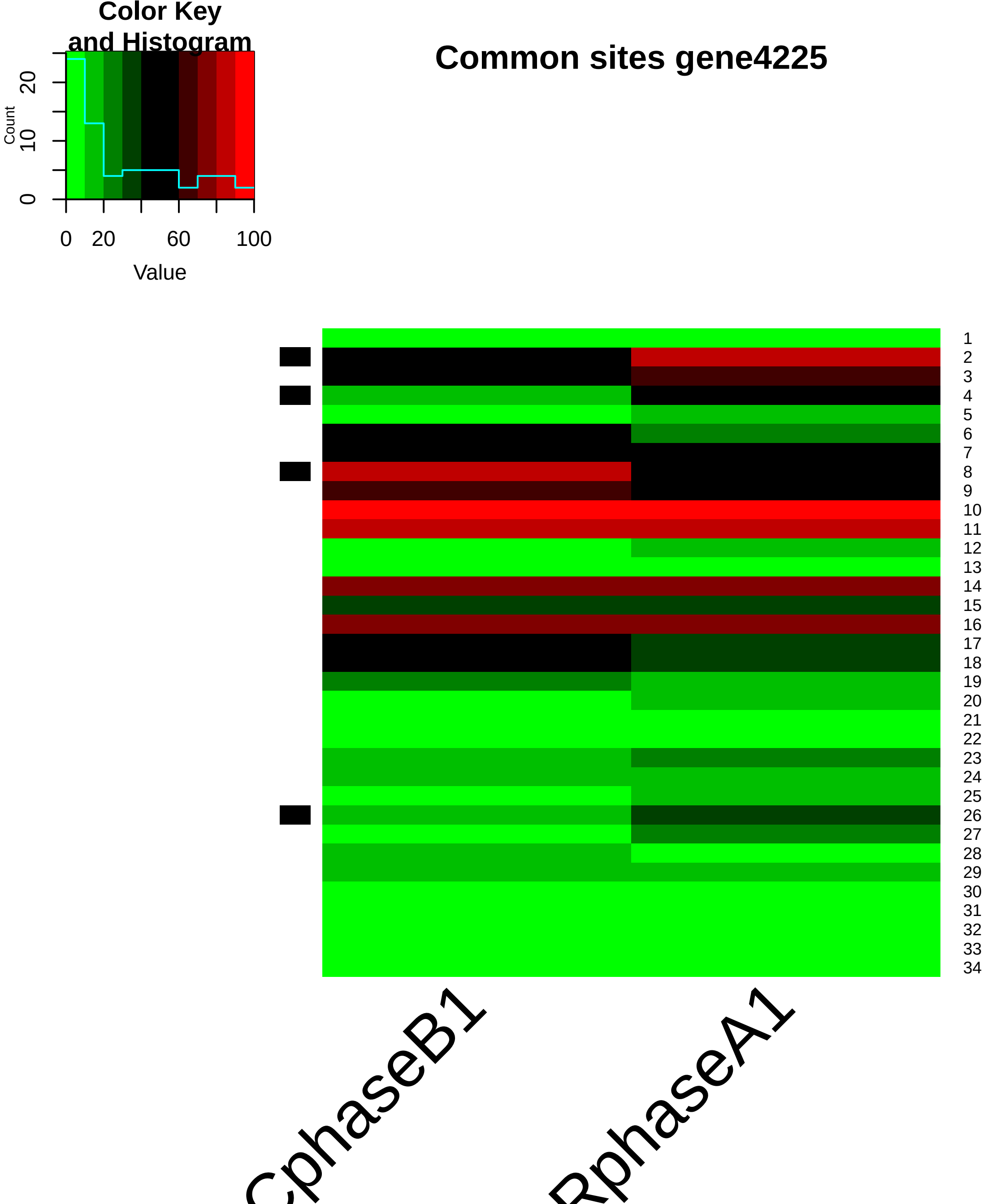

Color Key
and Histogram
Count
0
10
20
0
20
60
100
Value
Common sites gene4225
CphaseB1
RphaseA1

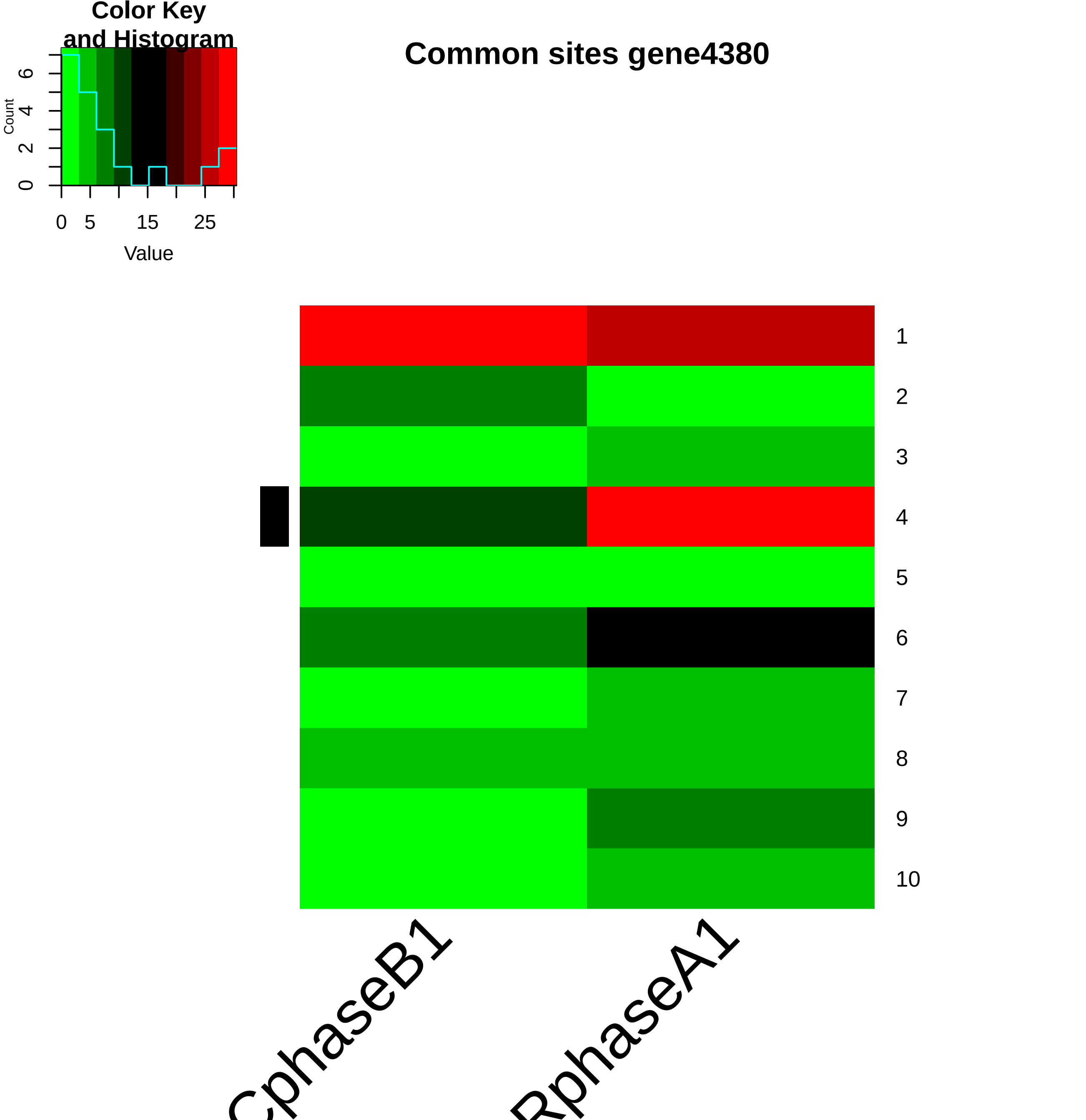
Color Key
and Histogram
Count
0
2
4
6
0
5
15
25
Value
Common sites gene4380
1
2
3
4
5
6
7
8
9
10
CphaseB1
RphaseA1

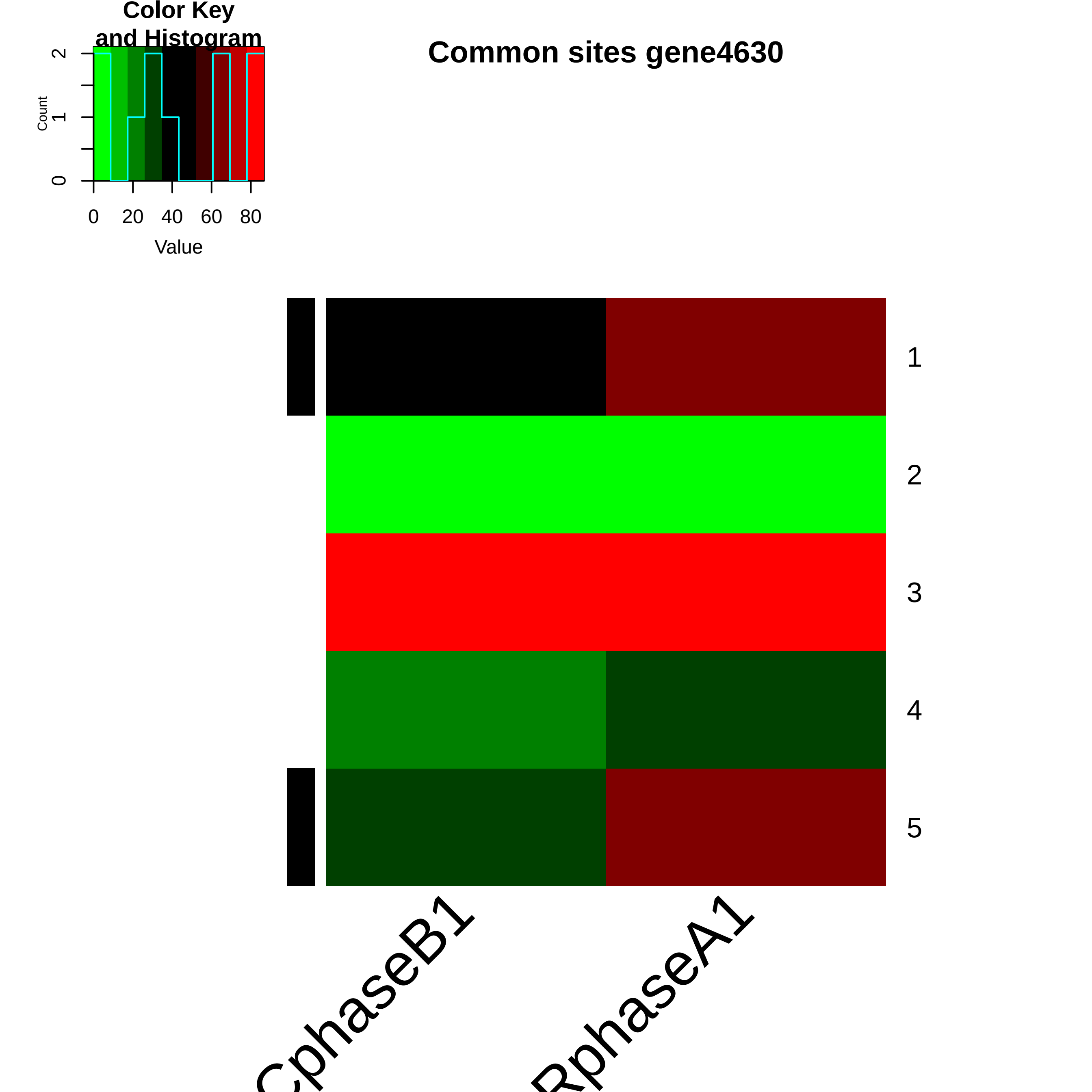
Color Key
and Histogram
Count
Value
0
20
40
60
80
0
1
2
Common sites gene4630
1
2
3
4
5
CphaseB1
RphaseA1

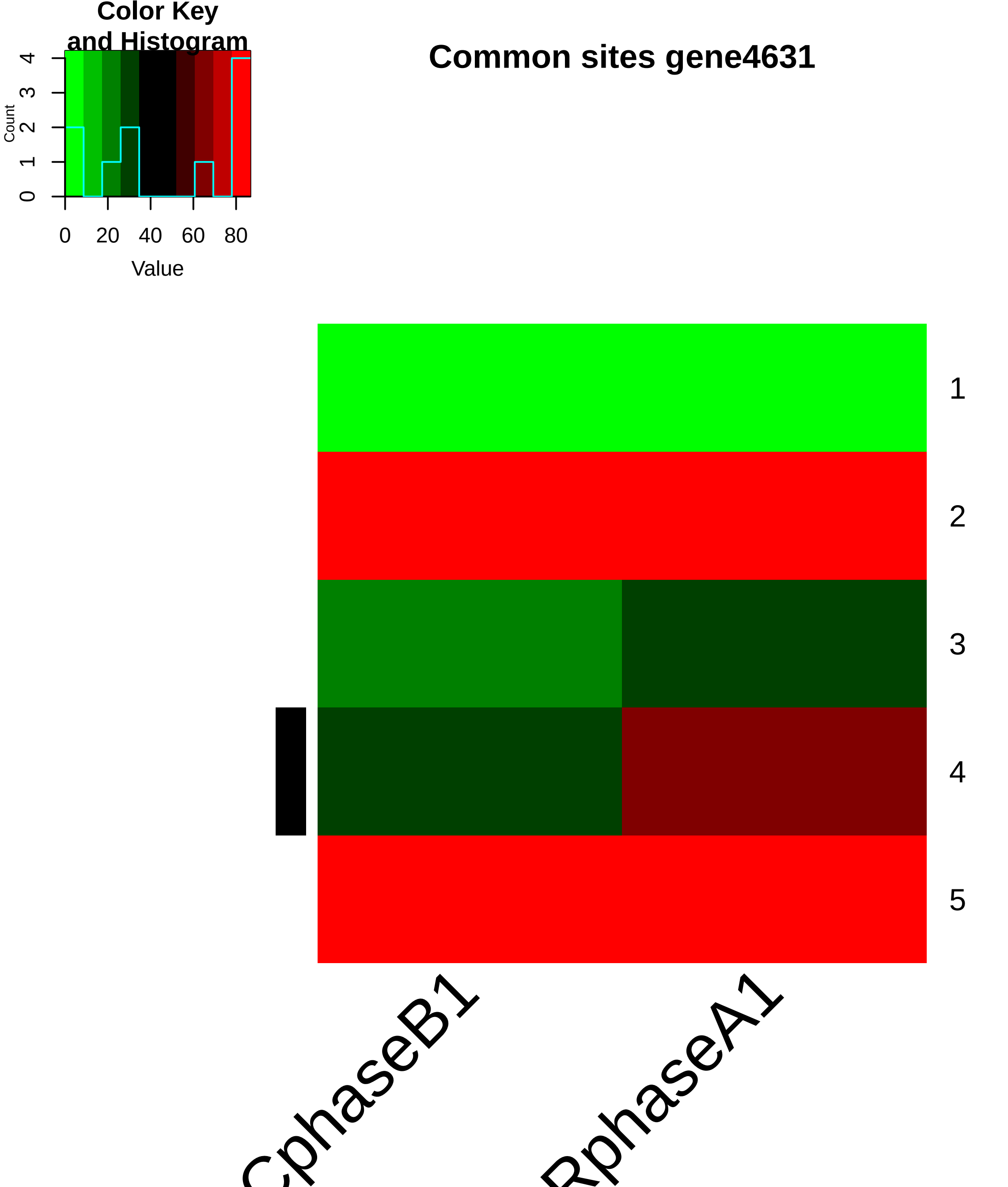
Color Key
and Histogram
Count
4
3
2
1
0
0
20
40
60
80
Value
Common sites gene4631
1
2
3
4
5
CphaseB1
RphaseA1

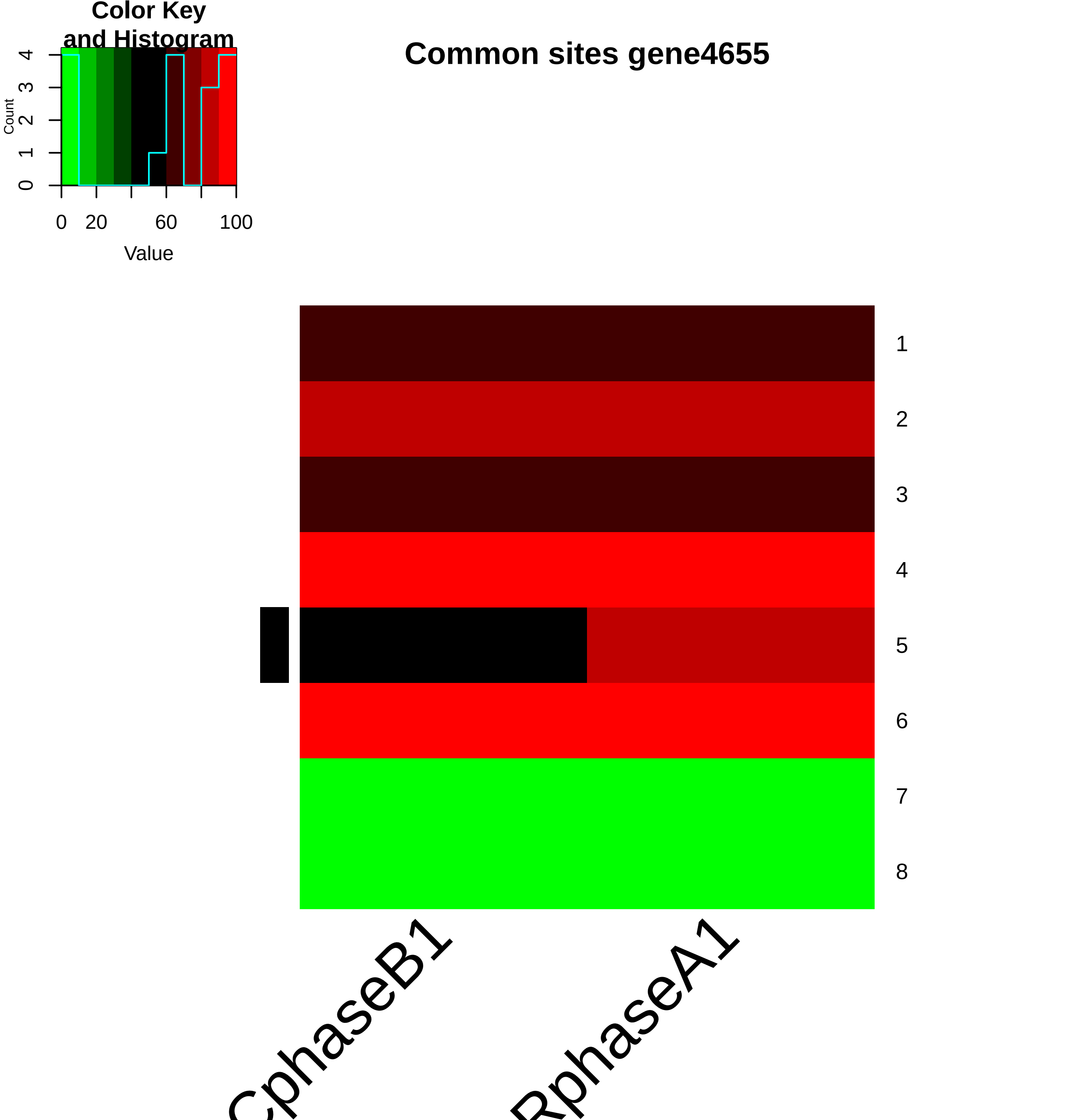
Color Key
and Histogram
Count
0
1
2
3
4
0
20
60
100
Value
Common sites gene4655
1
2
3
4
5
6
7
8
CphaseB1
RphaseA1

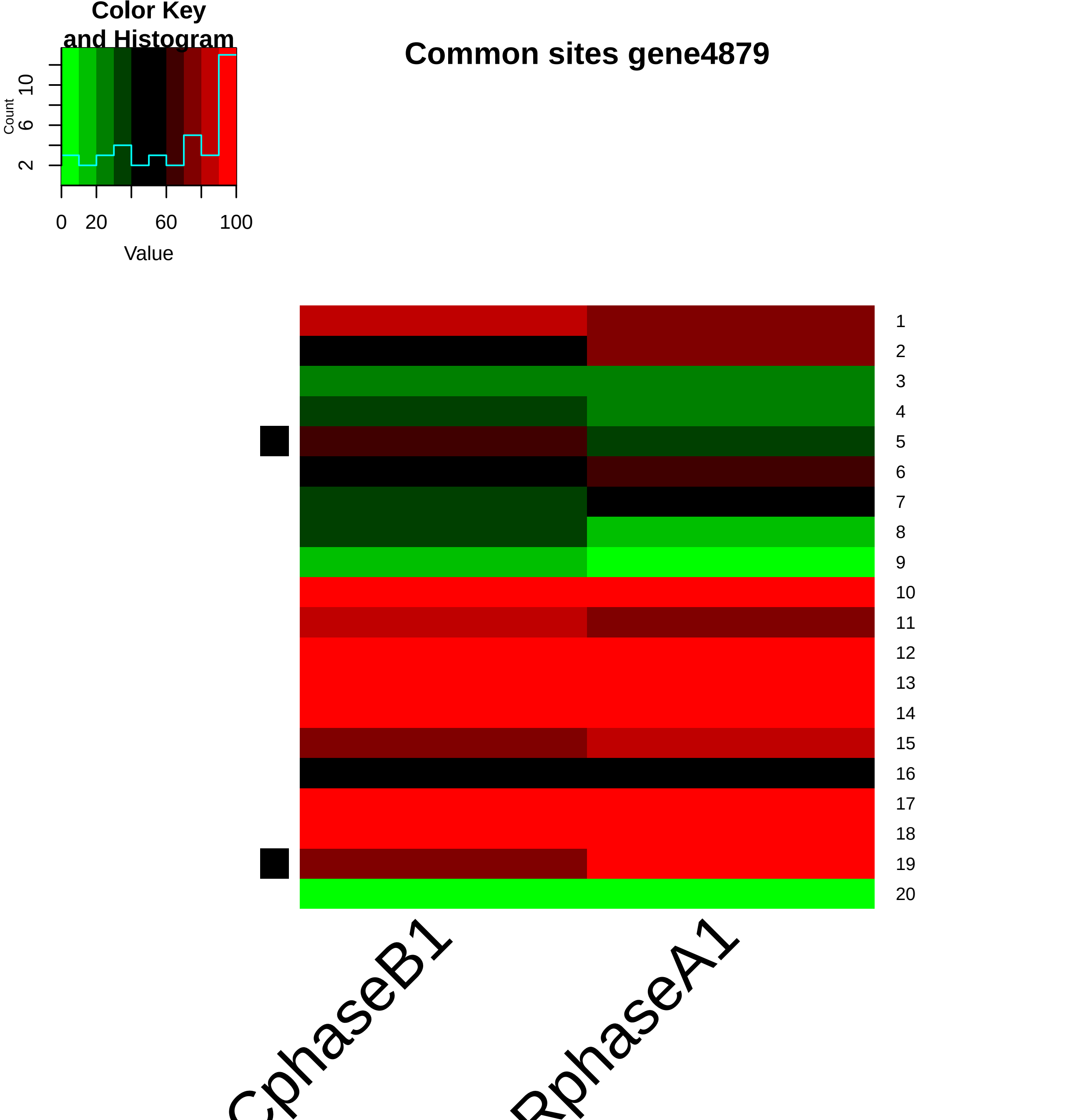
Color Key
and Histogram
Count
2
6
10
0
20
60
100
Value
Common sites gene4879
CphaseB1
RphaseA1

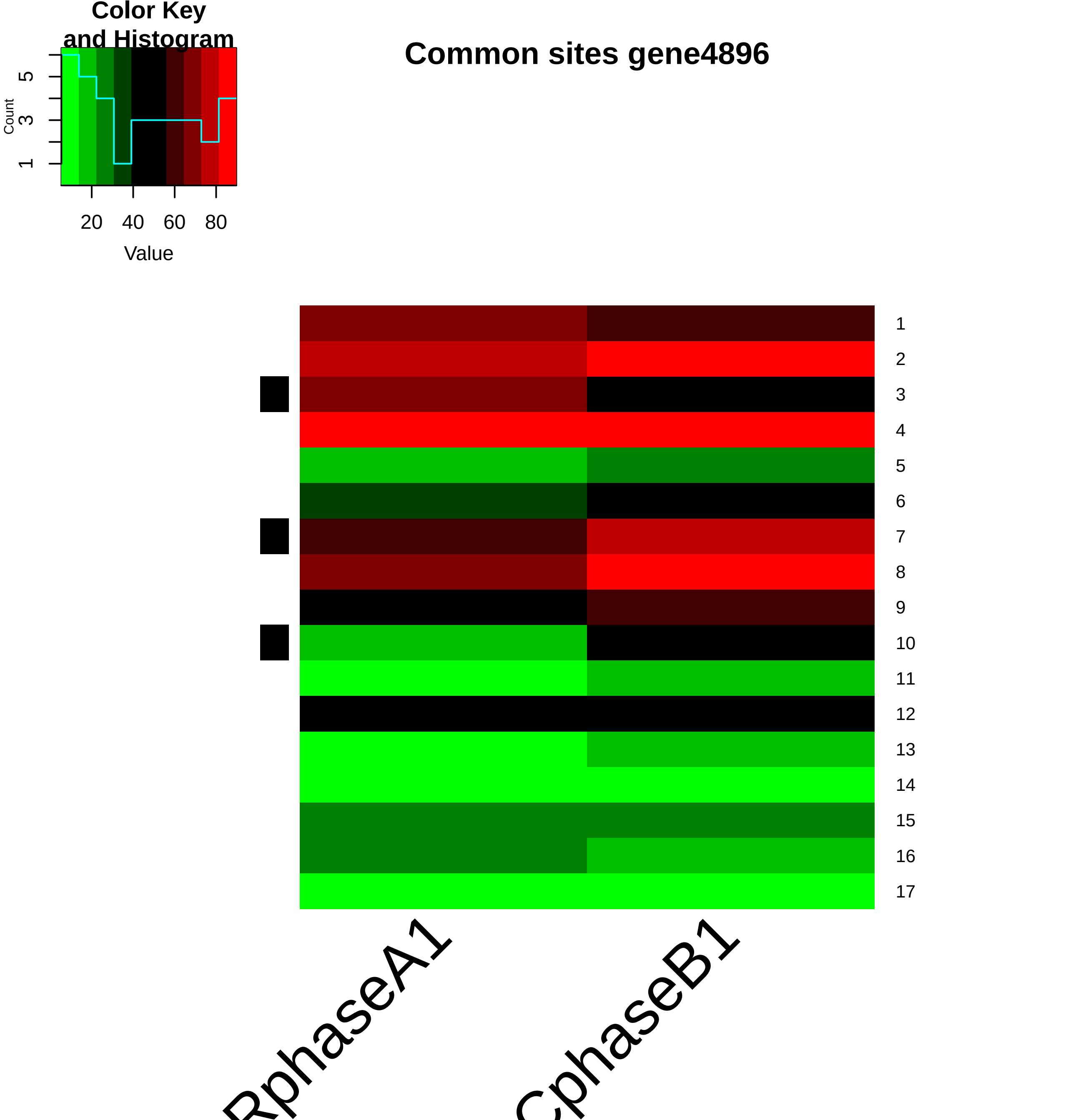

Color Key
and Histogram
Count
1
3
5
20
40
60
80
Value
Common sites gene4896
RphaseA1
CphaseB1

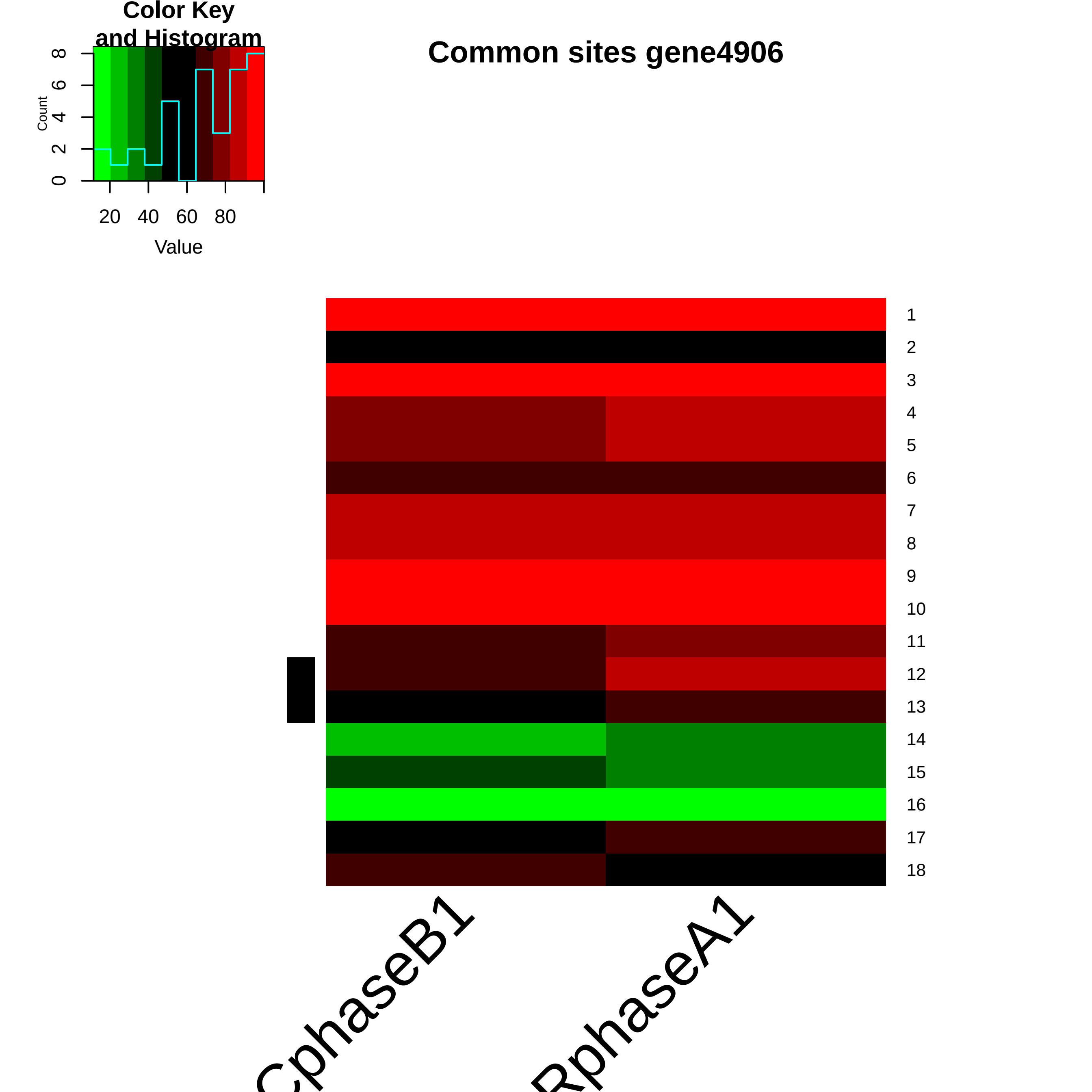
Color Key
and Histogram
Count
0
2
4
6
8
20
40
60
80
Value
Common sites gene4906
CphaseB1
RphaseA1

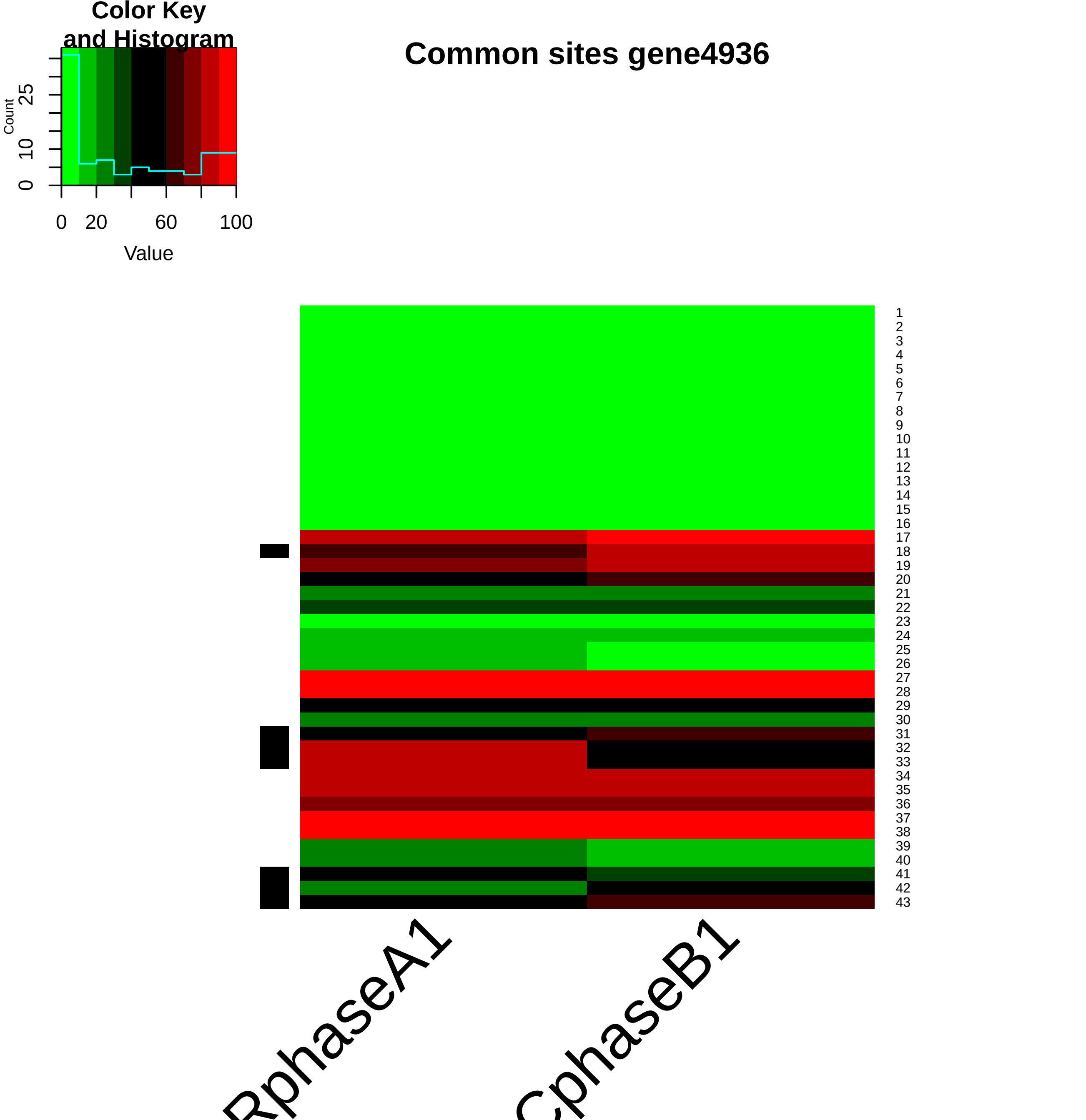

Color Key
and Histogram
Count
0
10
25
0
20
60
100
Value
Common sites gene4936
RphaseA1
CphaseB1

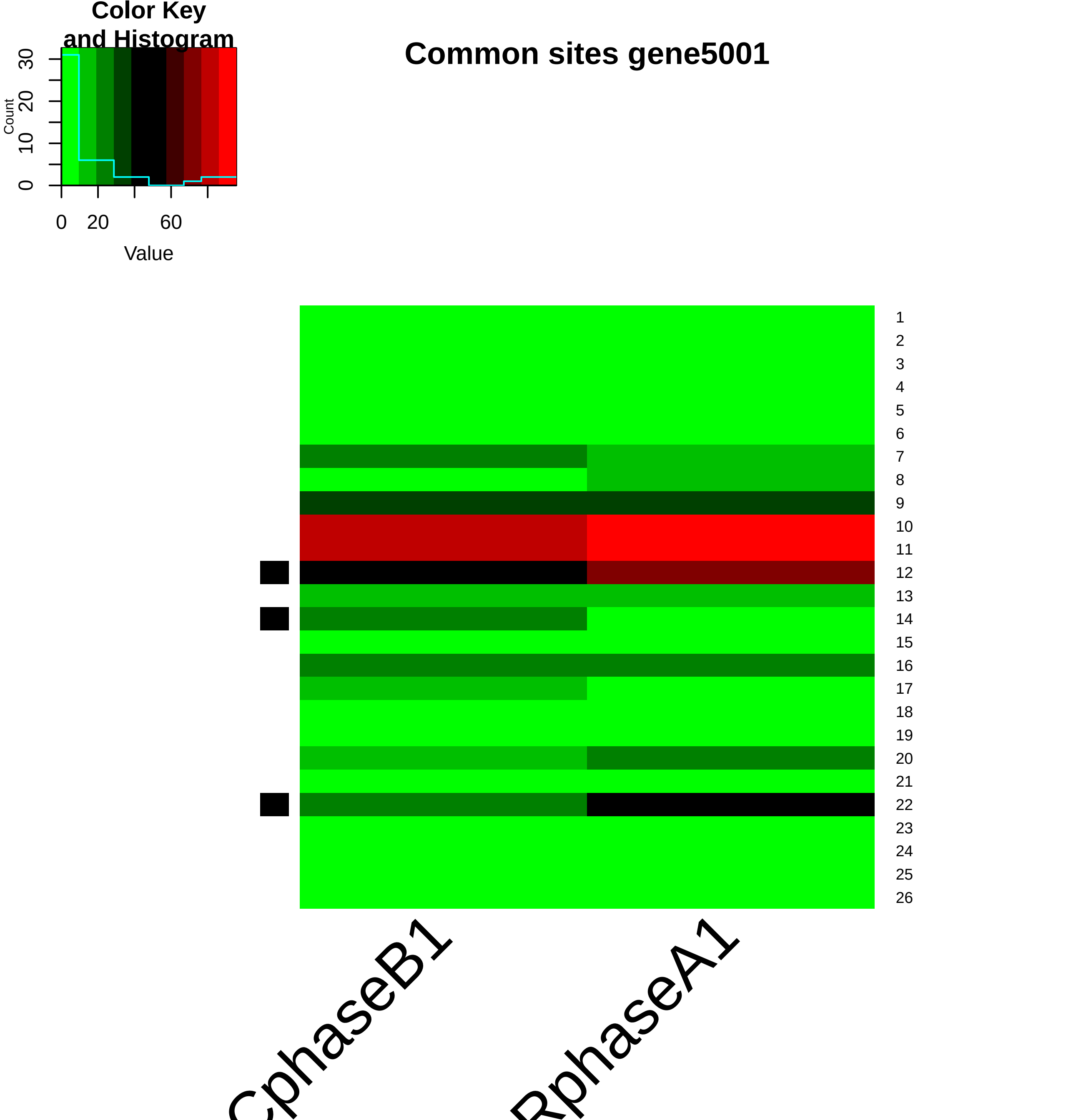
Color Key
and Histogram
Count
0
10
20
30
0
20
60
Value
Common sites gene5001
CphaseB1
RphaseA1

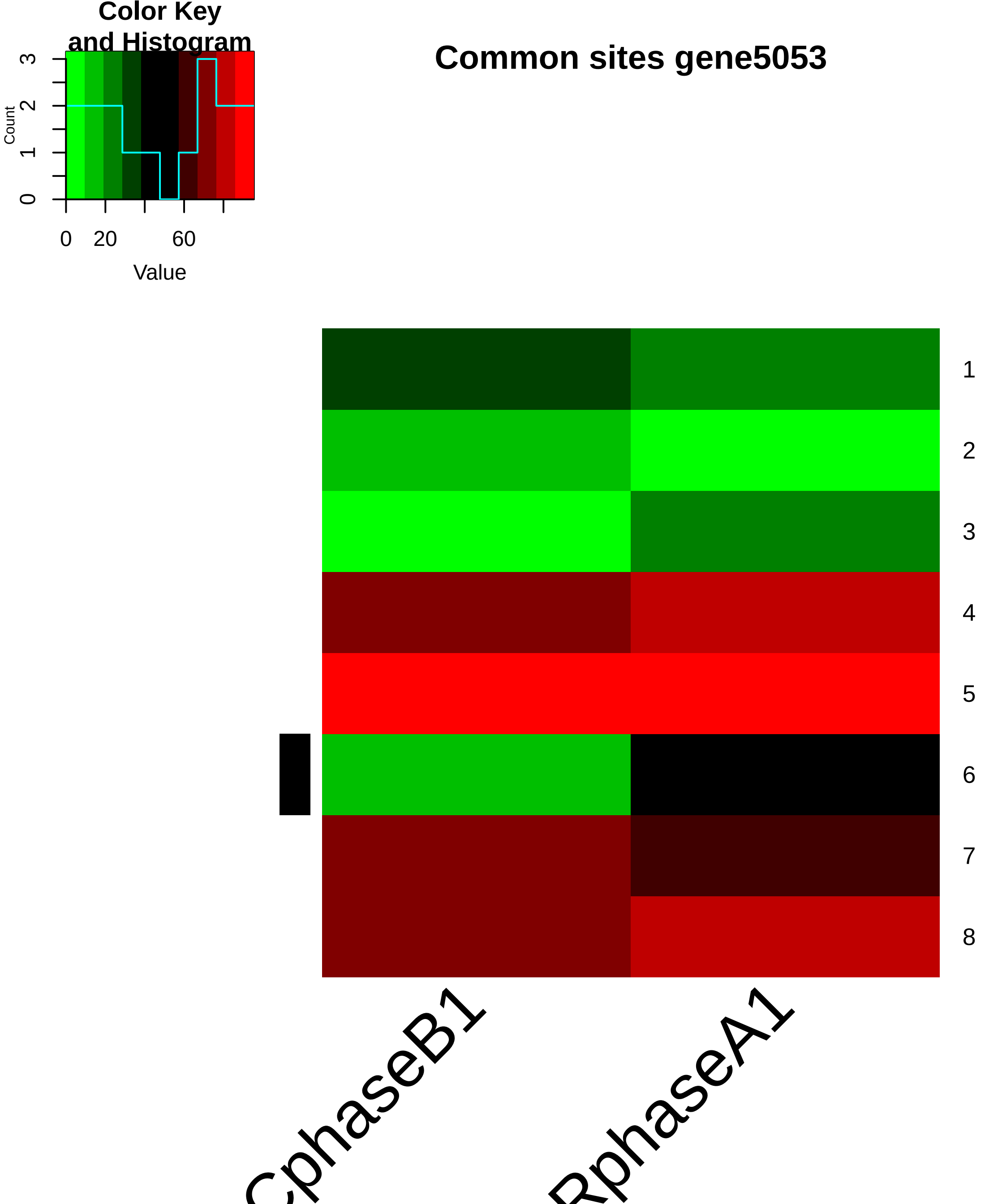

Color Key
and Histogram
Count
0
1
2
3
0
20
60
Value
Common sites gene5053
1
2
3
4
5
6
7
8
CphaseB1
RphaseA1

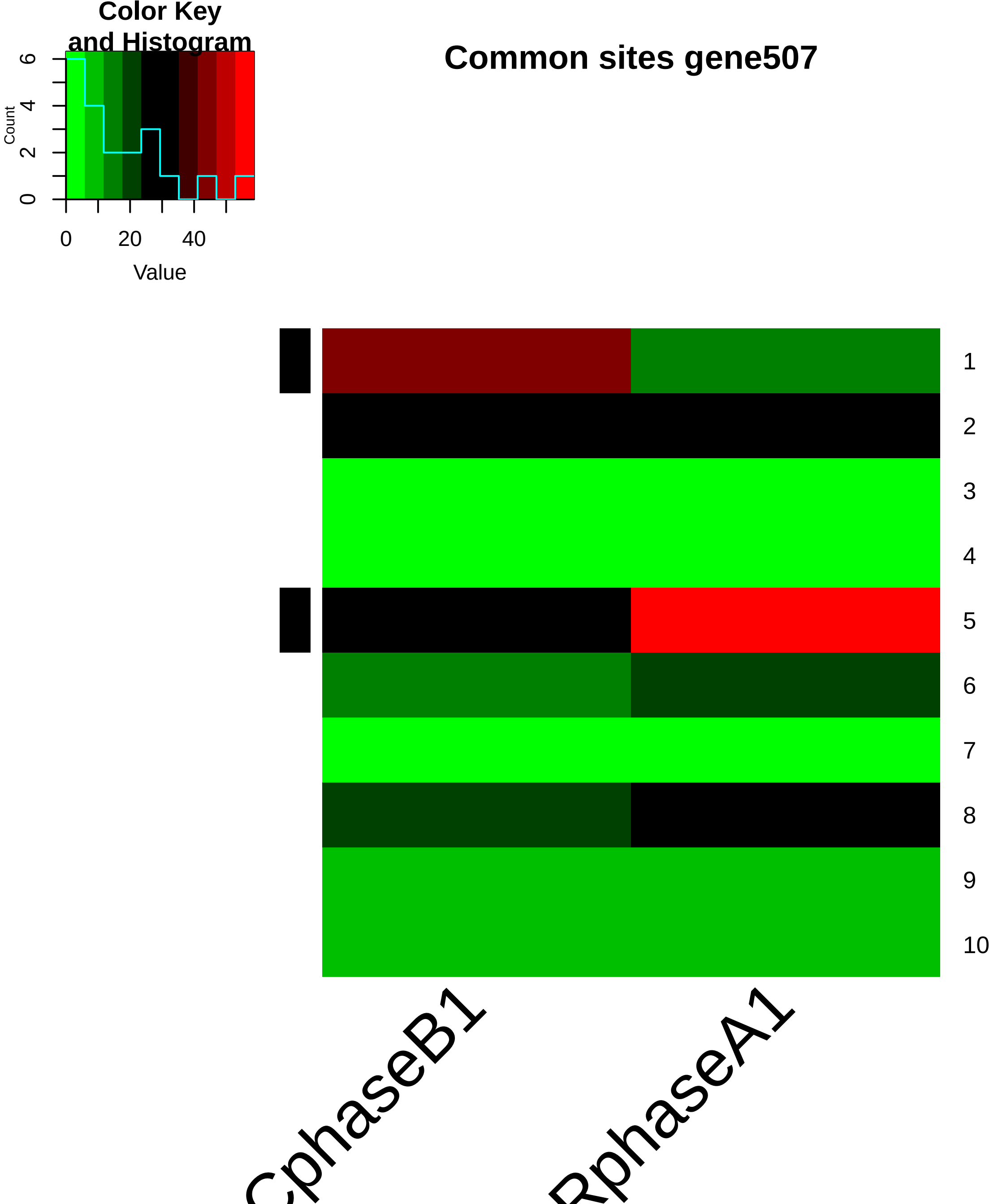
Color Key
and Histogram
Count
0
2
4
6
0
20
40
Value
Common sites gene507
1
2
3
4
5
6
7
8
9
10
CphaseB1
RphaseA1

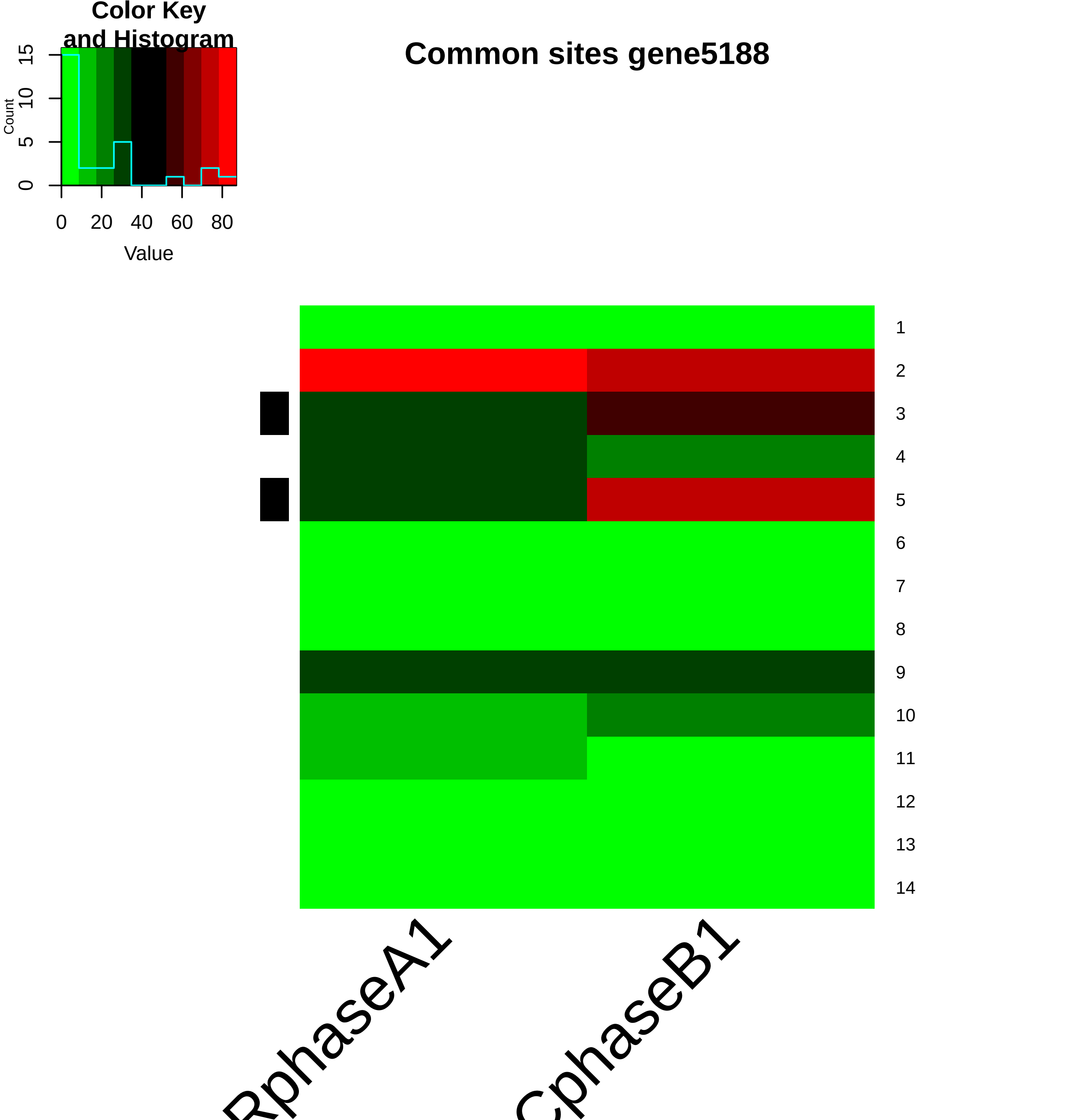
Color Key
and Histogram
Count
0
5
10
15
0
20
40
60
80
Value
Common sites gene5188
1
2
3
4
5
6
7
8
9
10
11
12
13
14
RphaseA1
CphaseB1

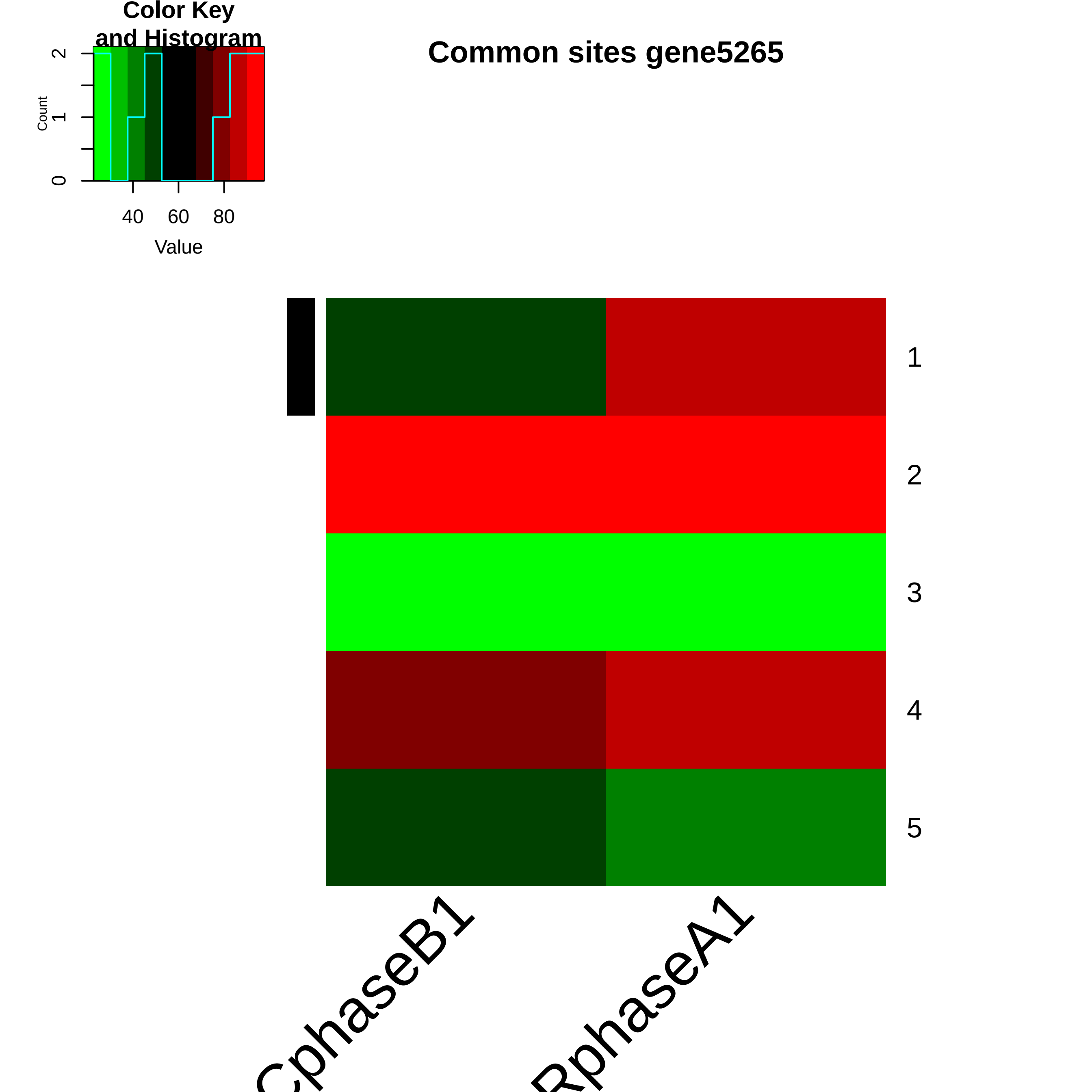

Color Key
and Histogram
Count
0
1
2
40
60
80
Value
Common sites gene5265
1
2
3
4
5
CphaseB1
RphaseA1

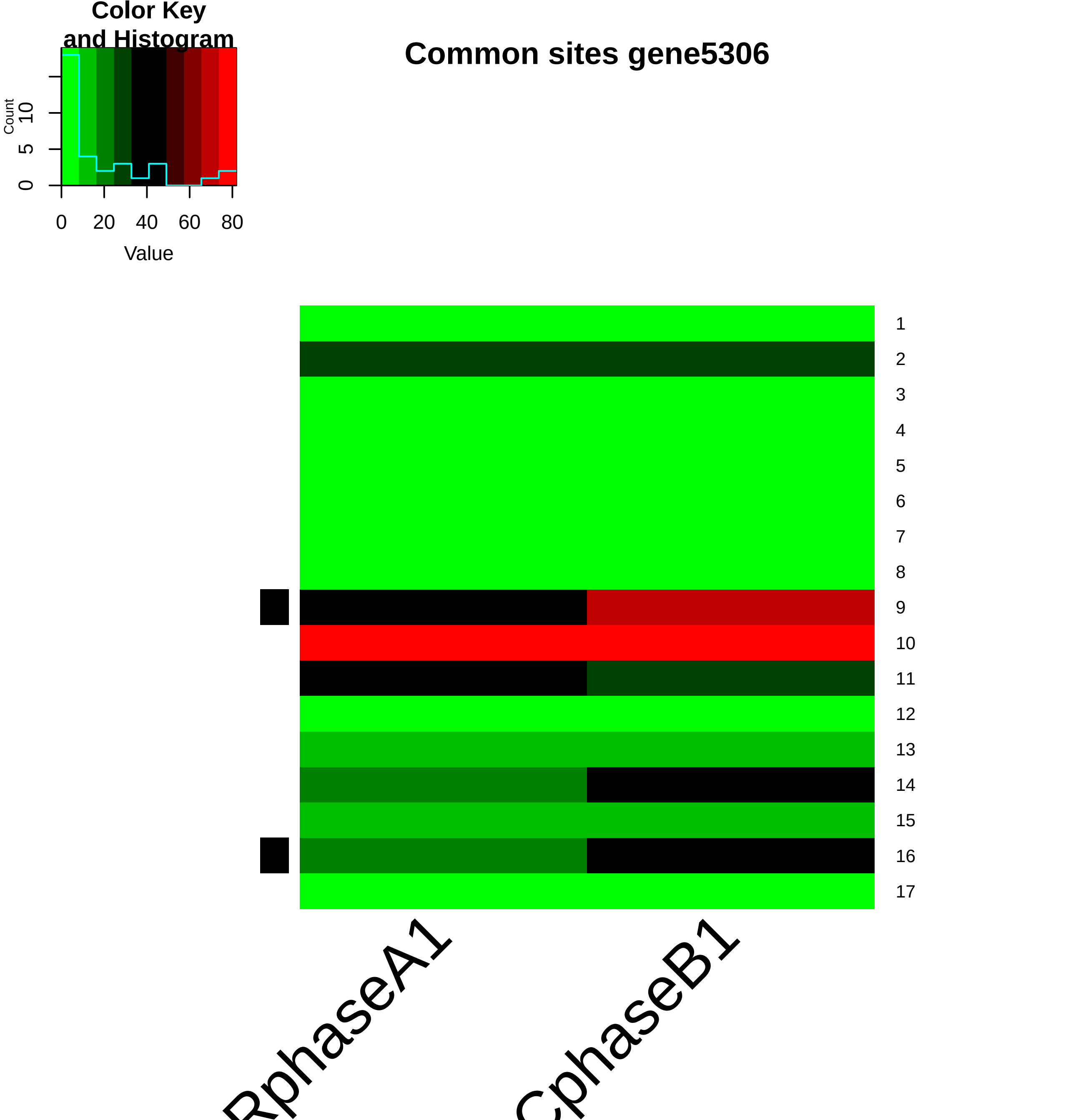
Color Key
and Histogram
Count
0
5
10
0
20
40
60
80
Value
Common sites gene5306
RphaseA1
CphaseB1

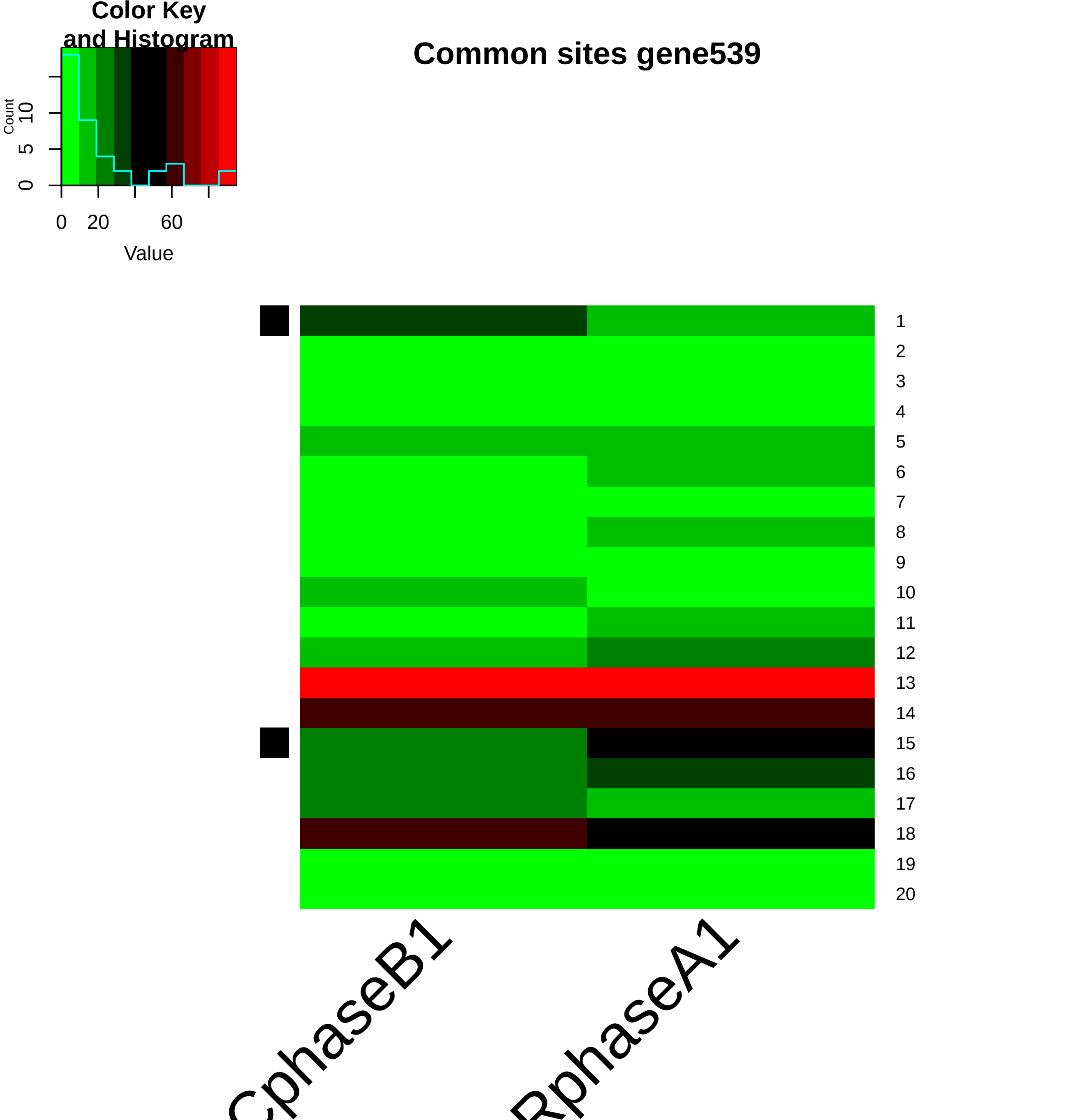

Color Key
and Histogram
Count
0
5
10
0
20
60
Value
Common sites gene539
CphaseB1
RphaseA1

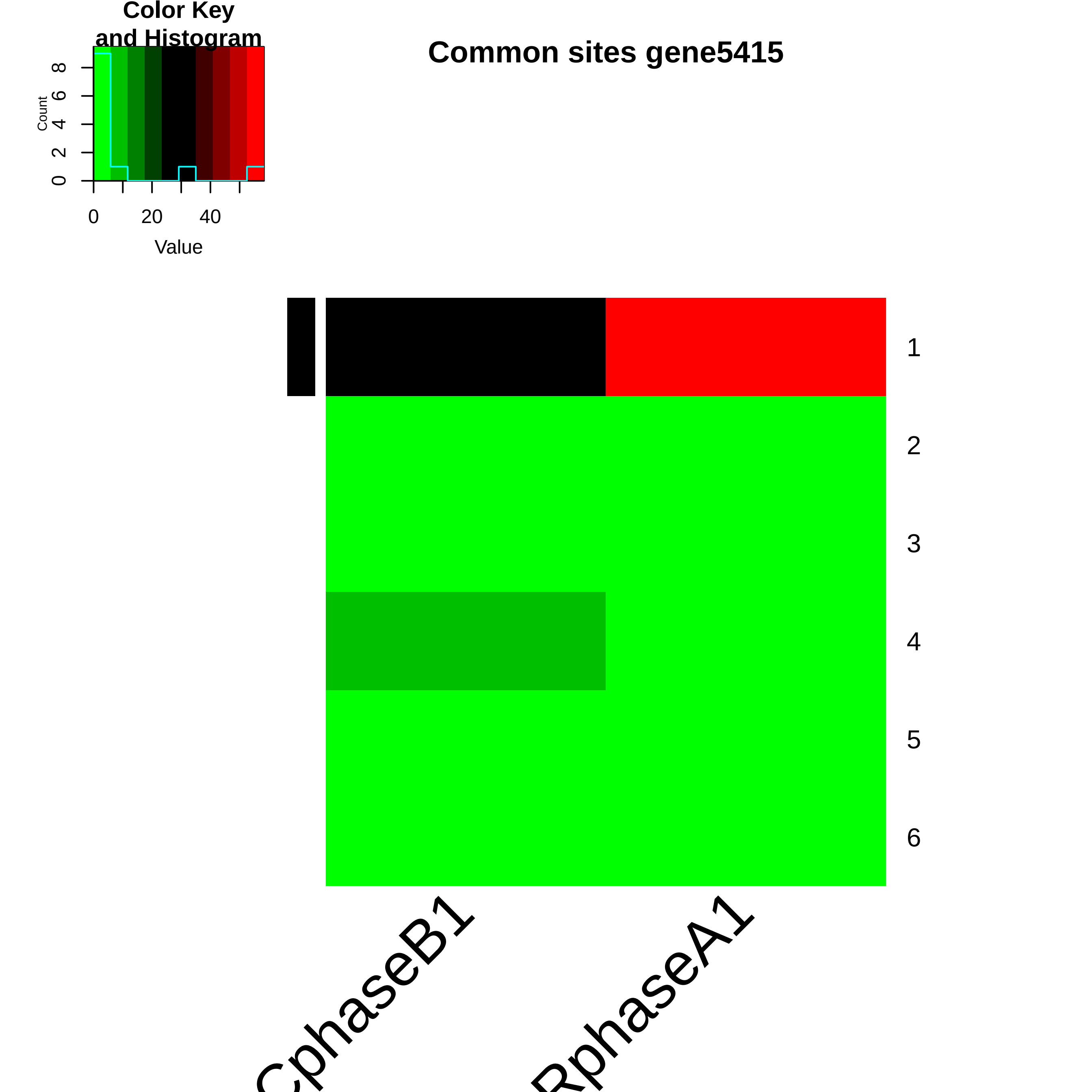

Color Key
and Histogram
Count
0
2
4
6
8
0
20
40
Value
Common sites gene5415
1
2
3
4
5
6
CphaseB1
RphaseA1

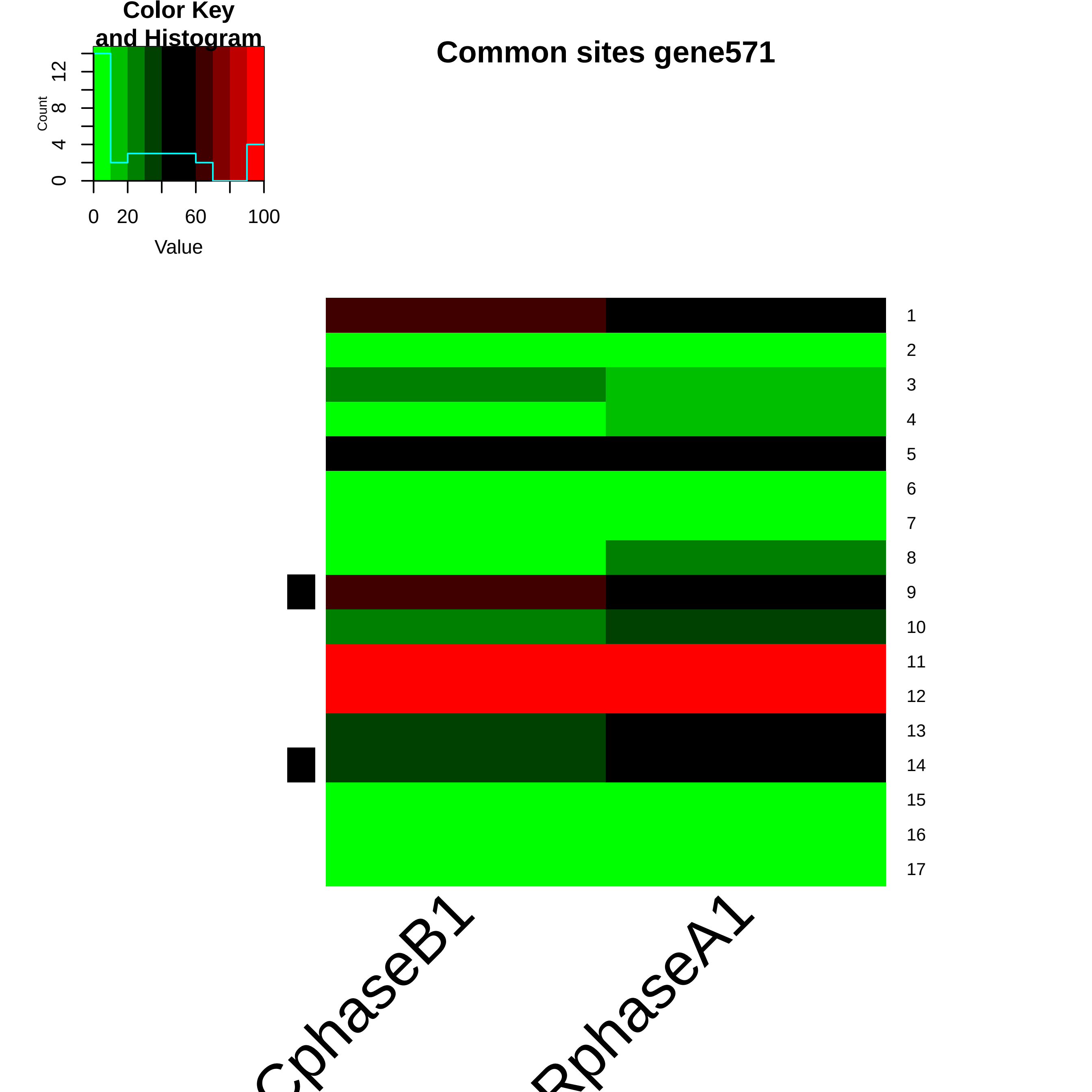
Color Key
and Histogram
Count
0
4
8
12
0
20
60
100
Value
Common sites gene571
CphaseB1
RphaseA1

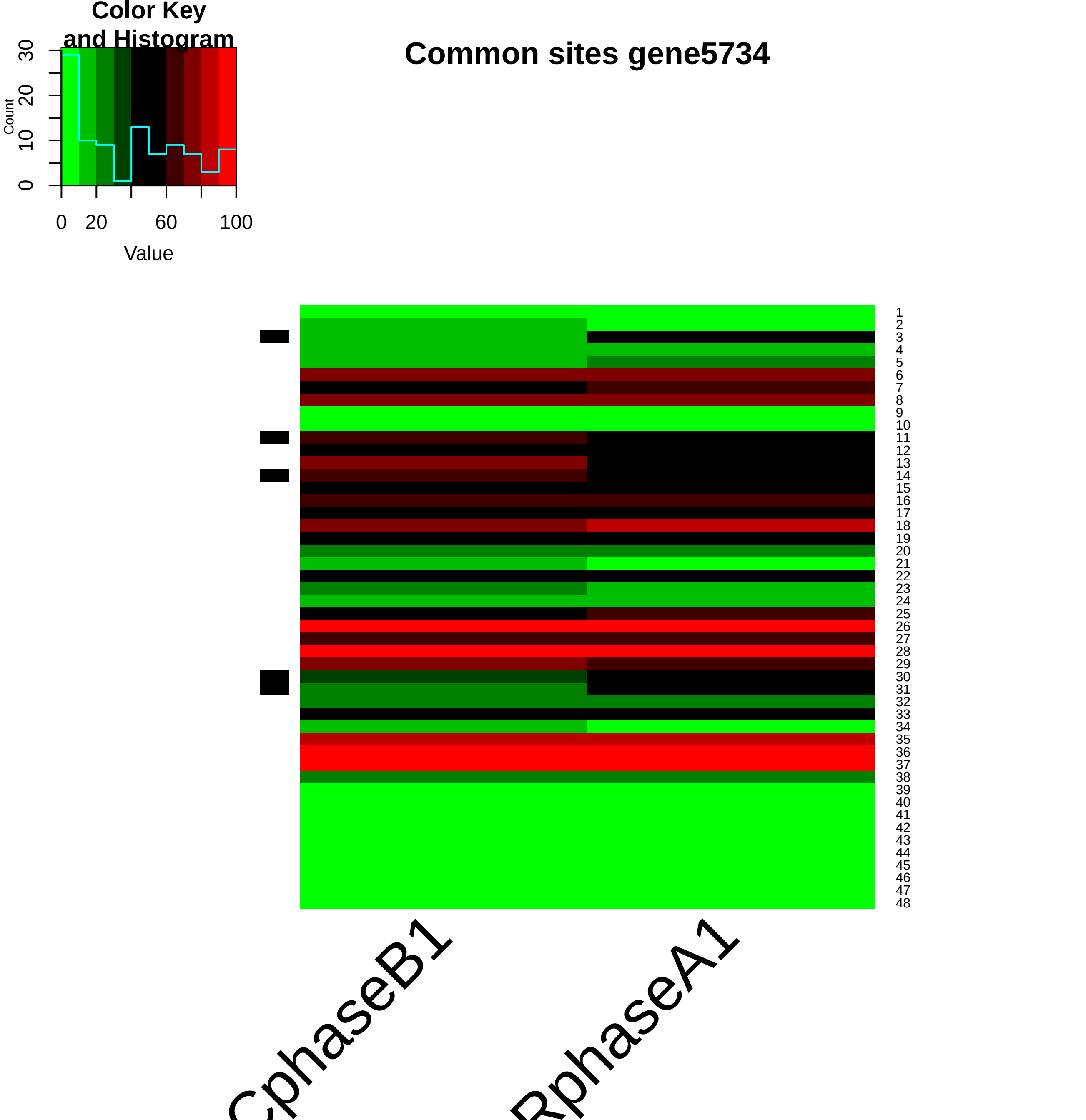
Color Key
and Histogram
Count
0
10
20
30
0
20
60
100
Value
Common sites gene5734

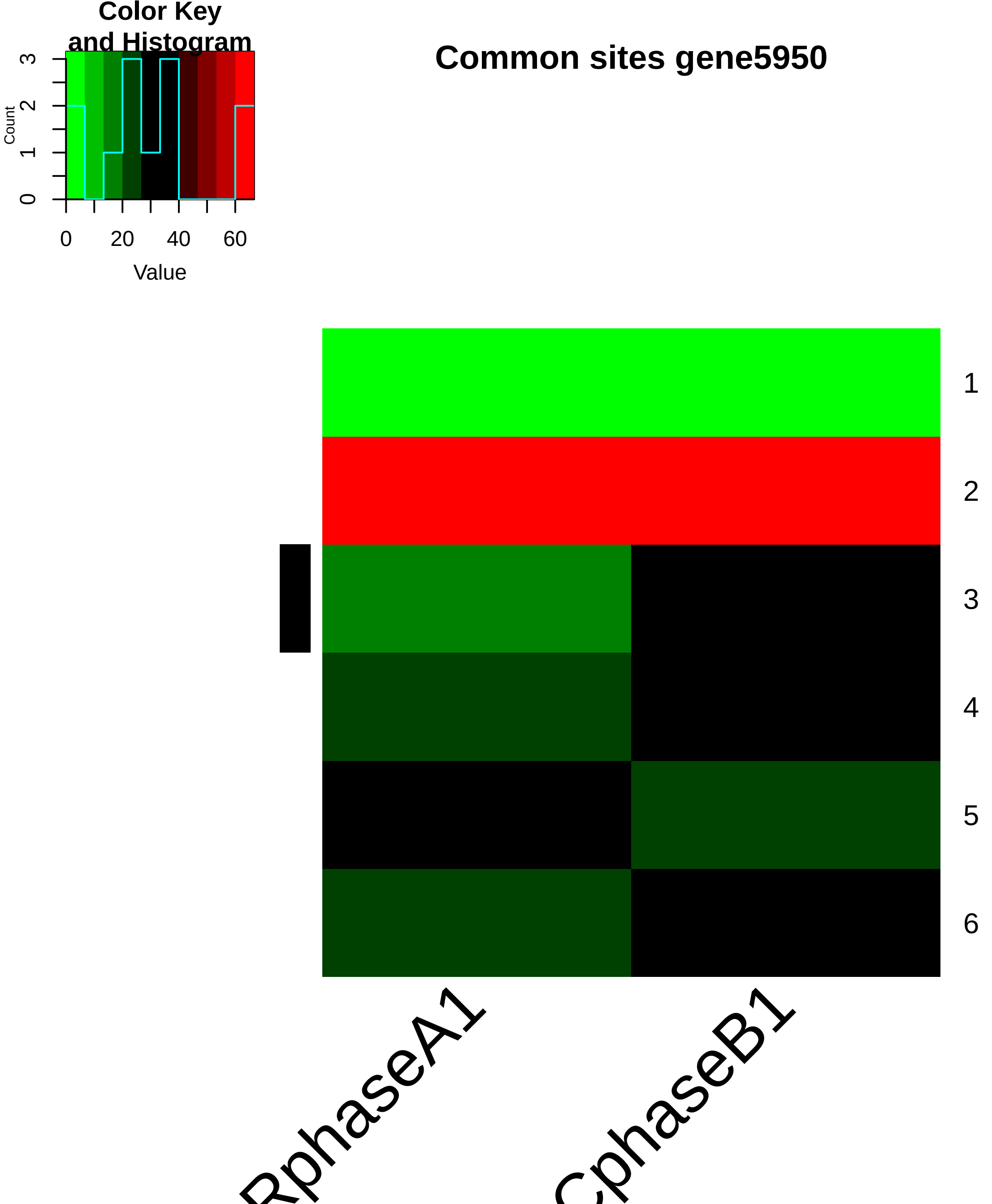
Color Key
and Histogram
Count
0
1
2
3
0
20
40
60
Value
Common sites gene5950
1
2
3
4
5
6
RphaseA1
CphaseB1

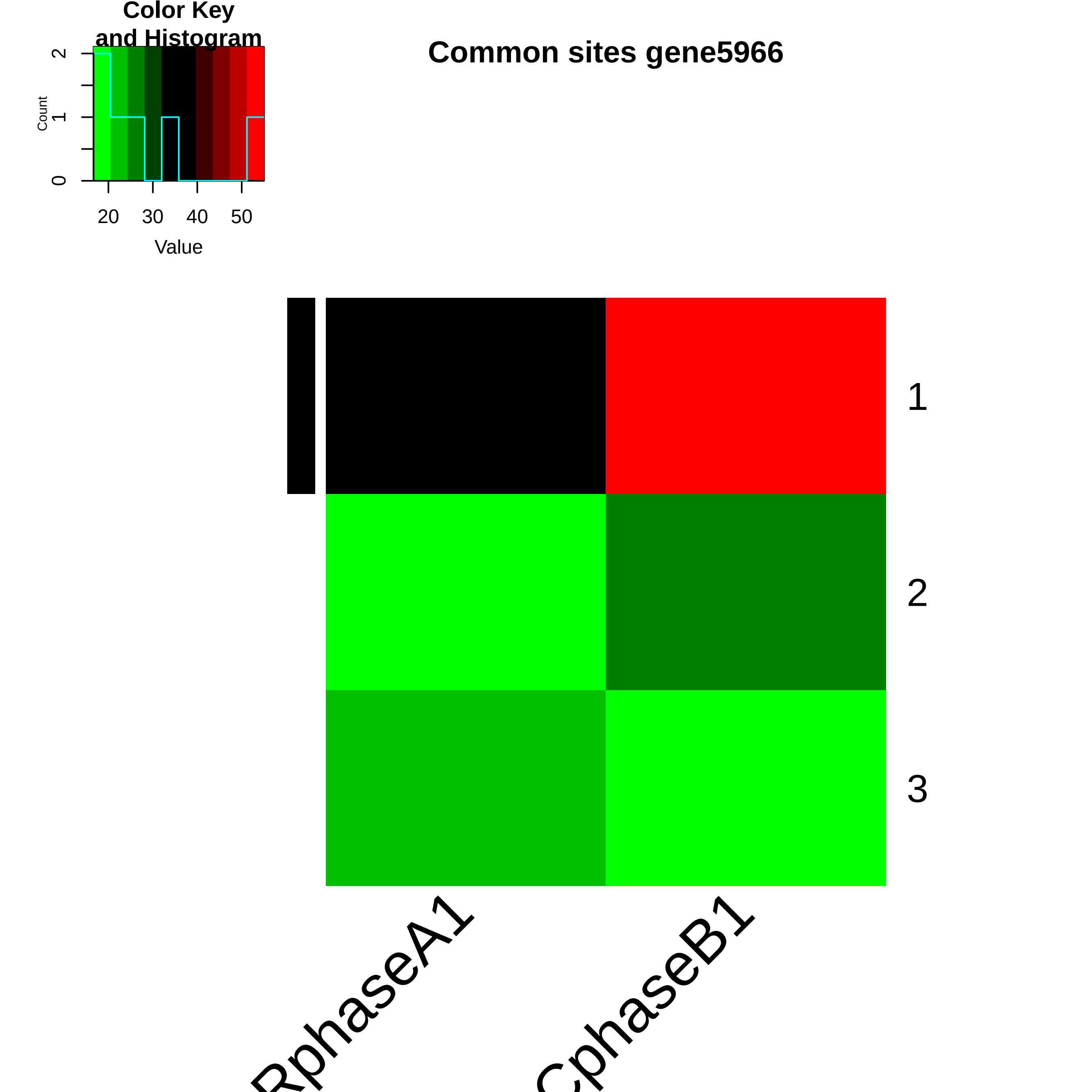

Color Key
and Histogram
Count
Value
0
1
2
20
30
40
50
Common sites gene5966
1
2
3
RphaseA1
CphaseB1

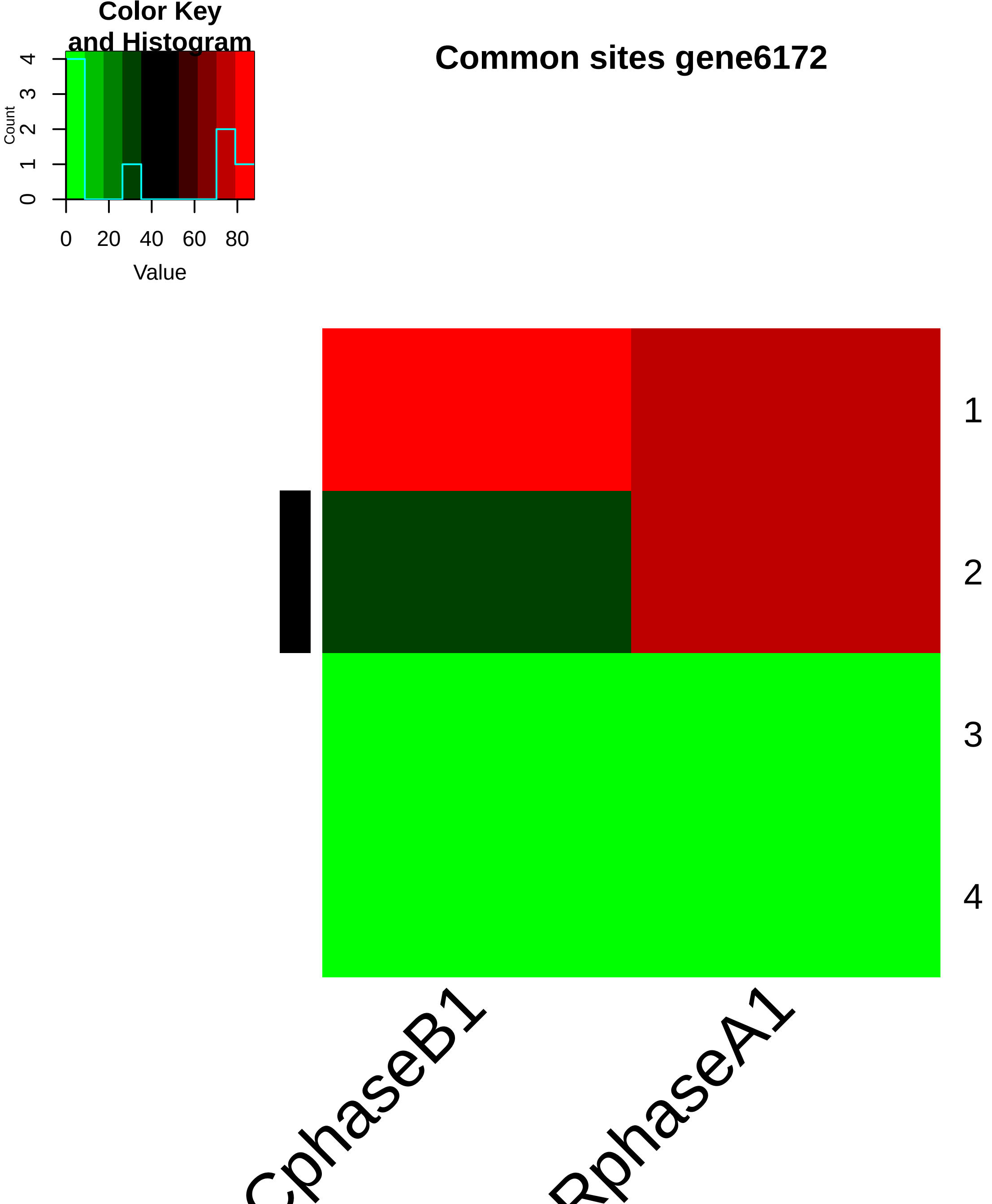
Color Key
and Histogram
Count
Value
0 20 40 60 80
0 1 2 3 4
Common sites gene6172
1
2
3
4
CphaseB1
RphaseA1

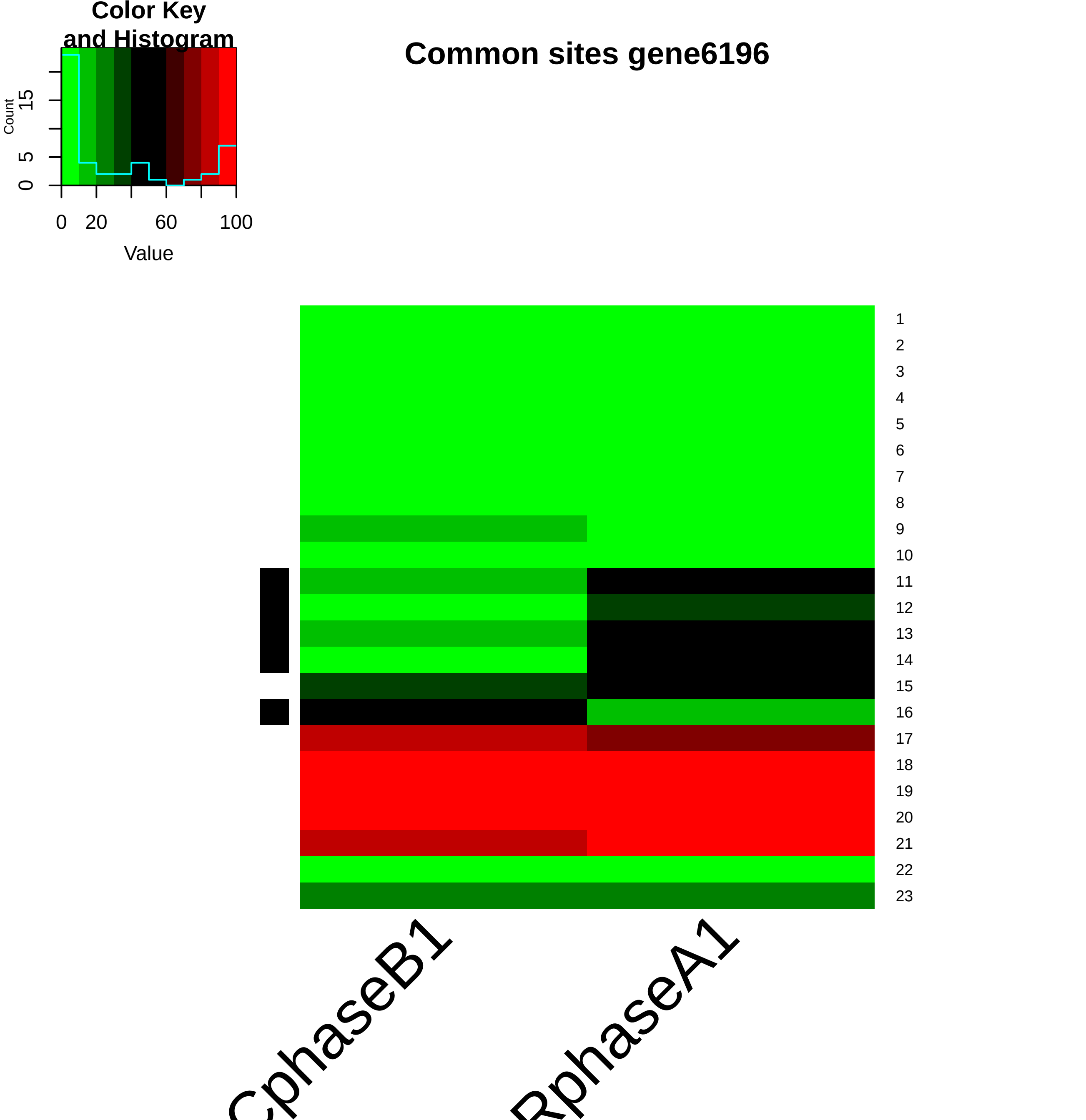

Color Key
and Histogram
Count
0
5
15
0
20
60
100
Value
Common sites gene6196
CphaseB1
RphaseA1

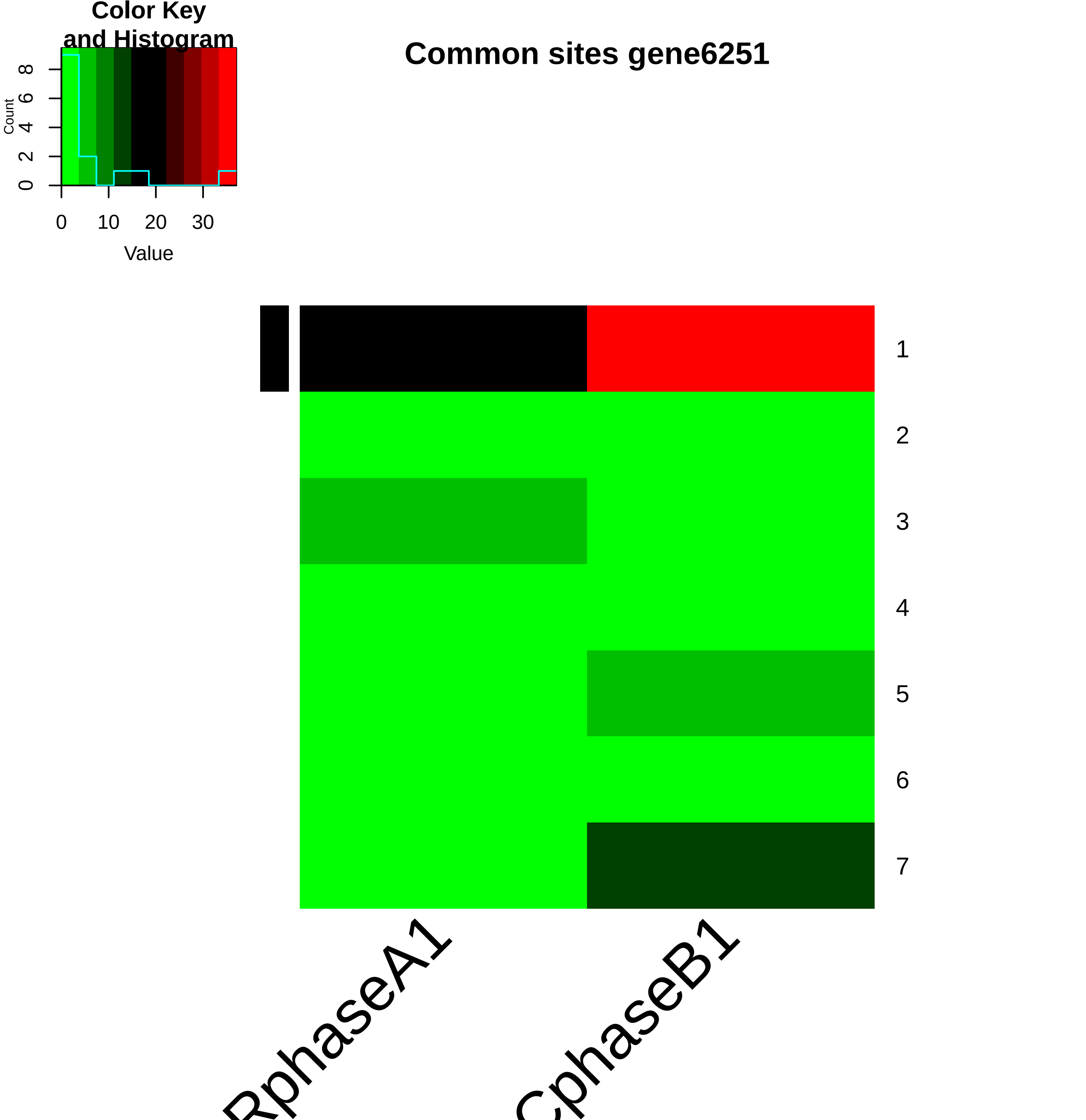

Color Key
and Histogram
Count
0
2
4
6
8
0
10
20
30
Value
Common sites gene6251
1
2
3
4
5
6
7
RphaseA1
CphaseB1

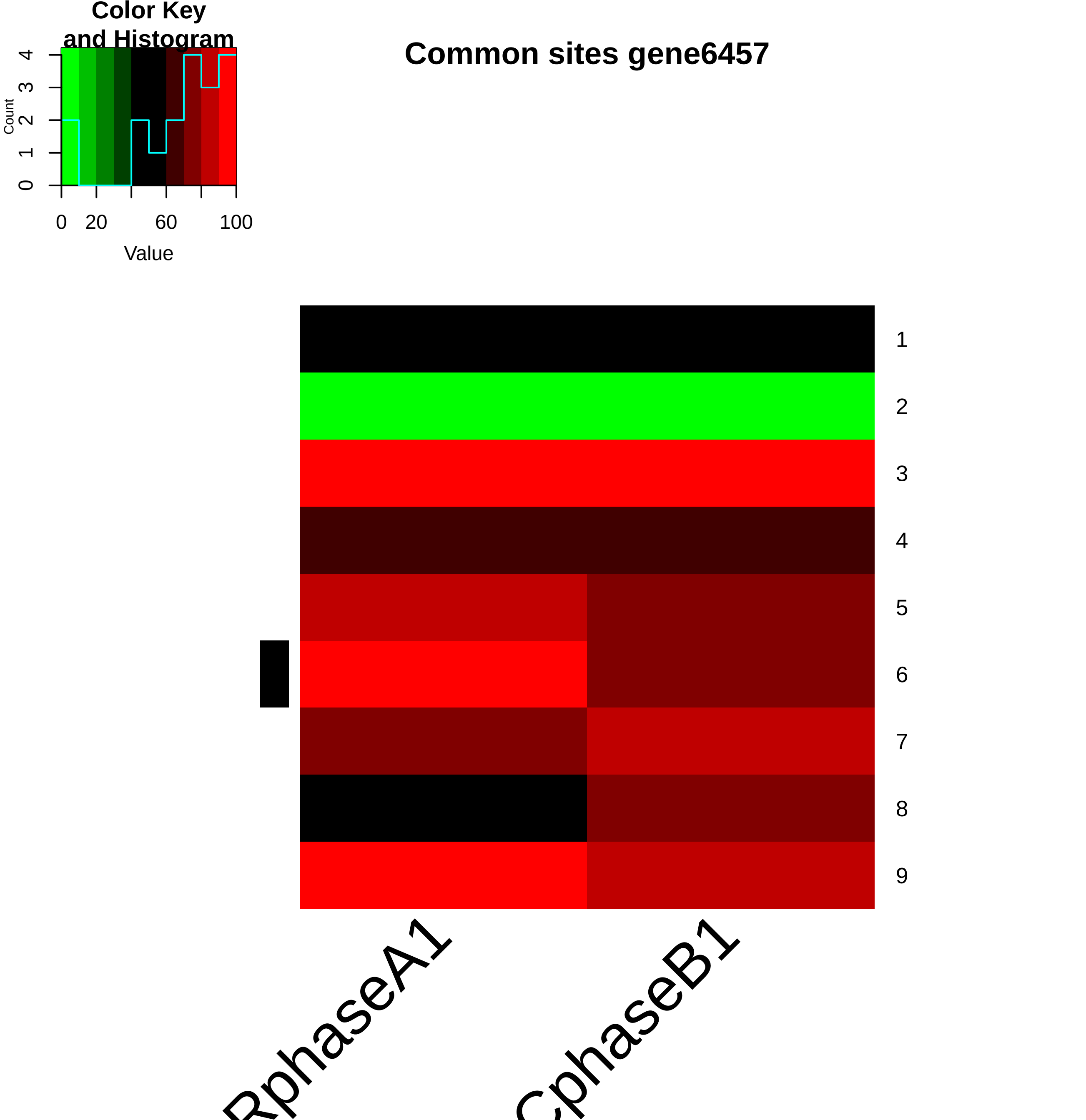

Color Key
and Histogram
Count
0
1
2
3
4
0
20
60
100
Value
Common sites gene6457
1
2
3
4
5
6
7
8
9
RphaseA1
CphaseB1

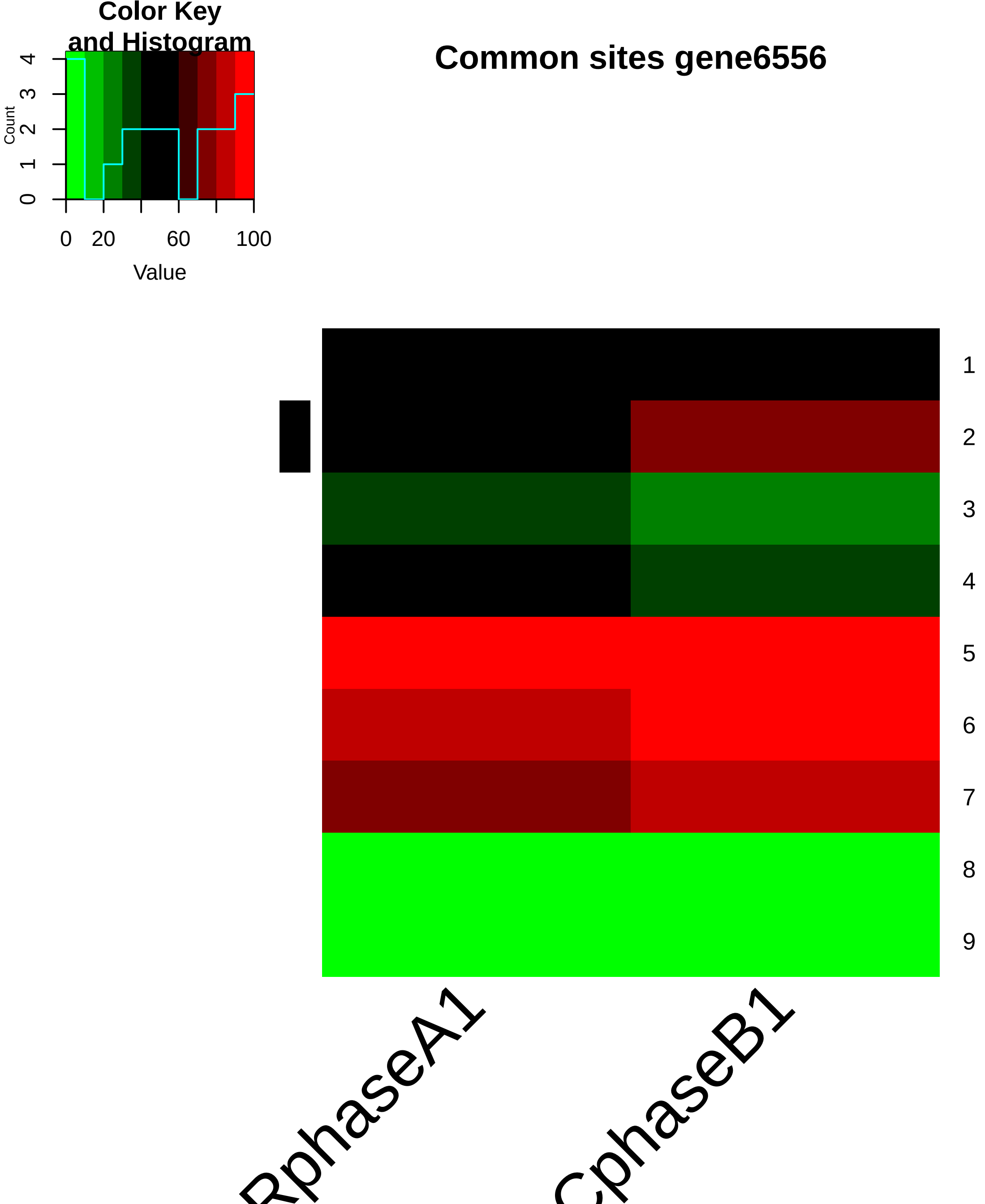

Color Key
and Histogram
Count
0
1
2
3
4
0
20
60
100
Value
Common sites gene6556
1
2
3
4
5
6
7
8
9
RphaseA1
CphaseB1

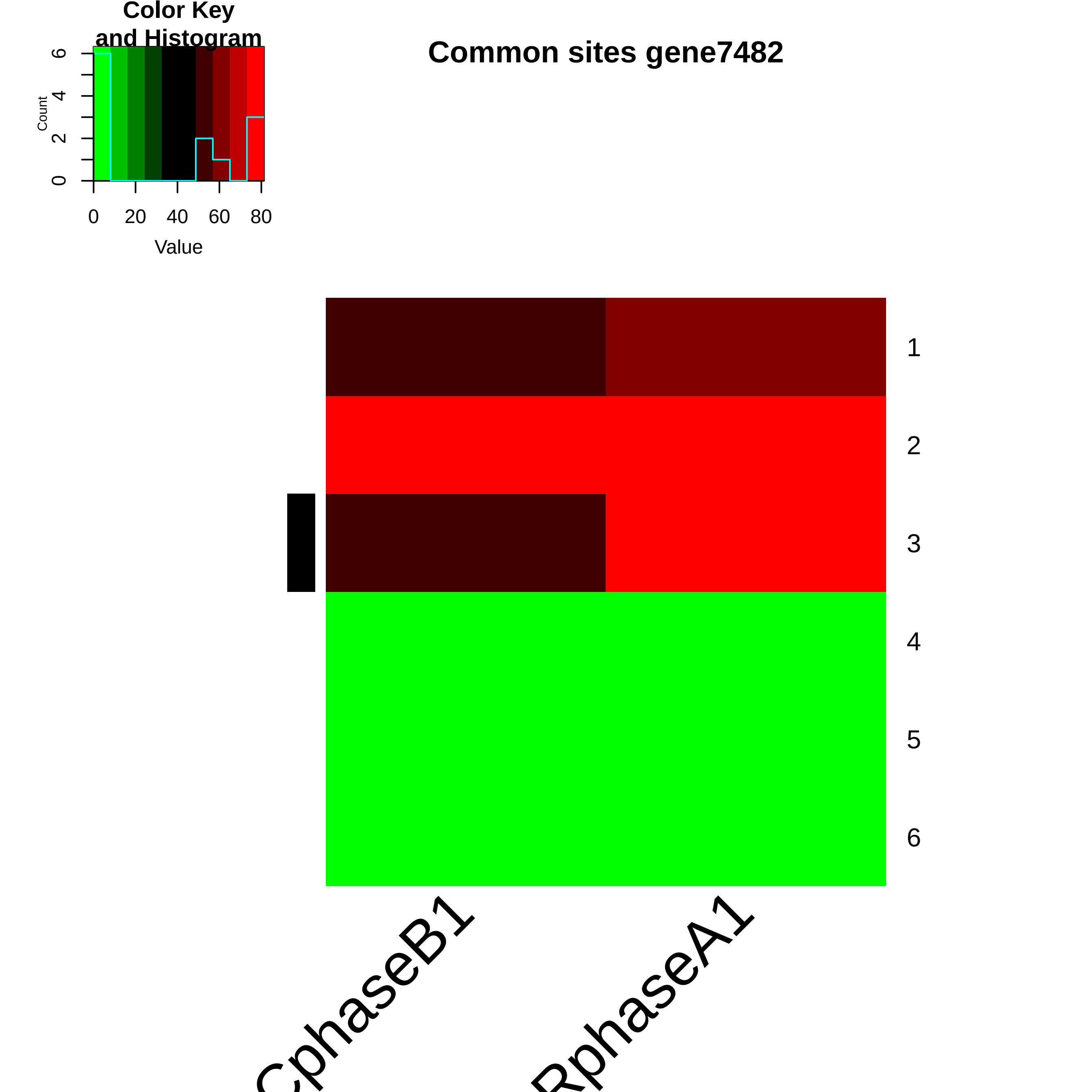
Color Key
and Histogram
Count
0
2
4
6
0
20
40
60
80
Value
Common sites gene7482
1
2
3
4
5
6
CphaseB1
RphaseA1

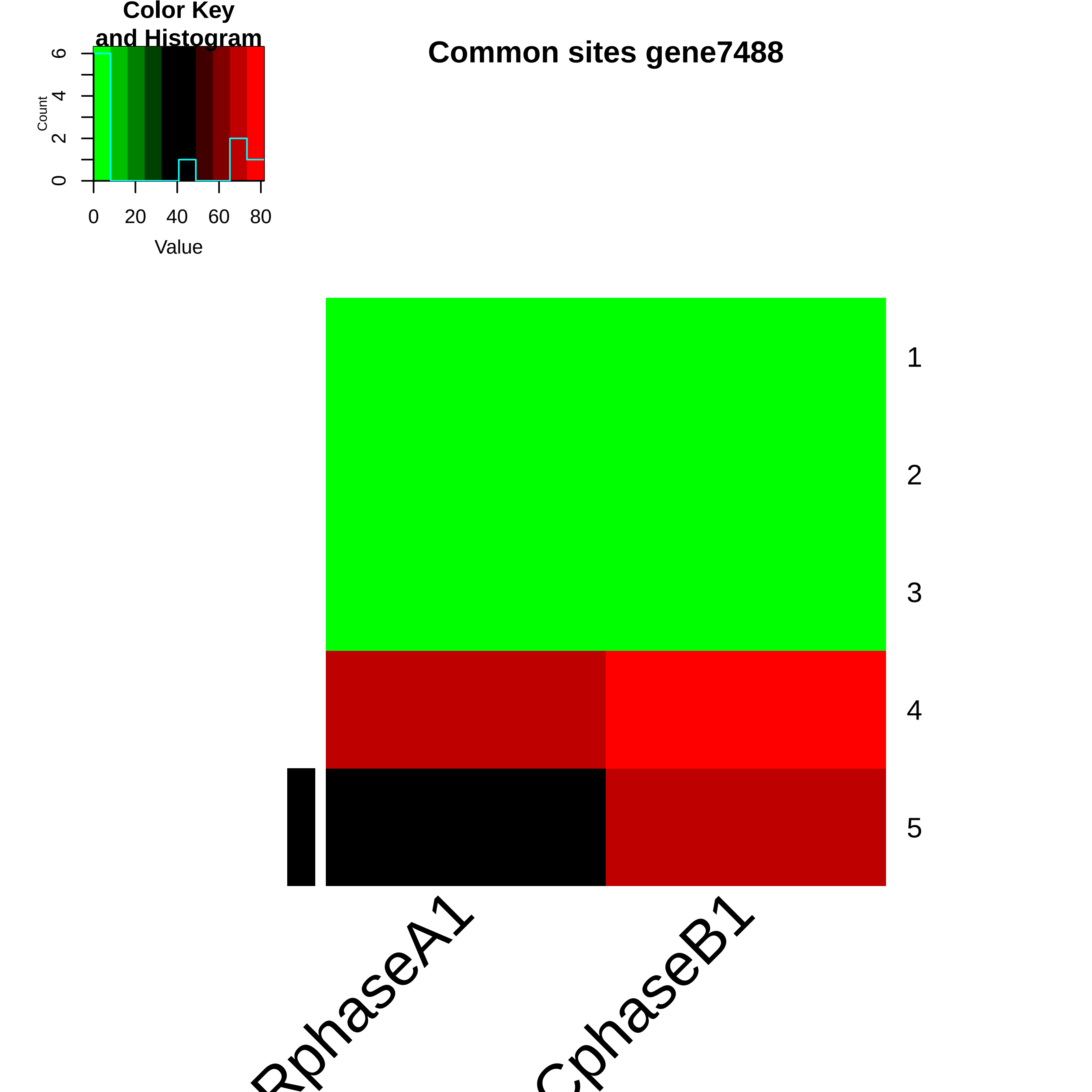

Color Key
and Histogram
Count
0
2
4
6
0
20
40
60
80
Value
Common sites gene7488
1
2
3
4
5
RphaseA1
CphaseB1

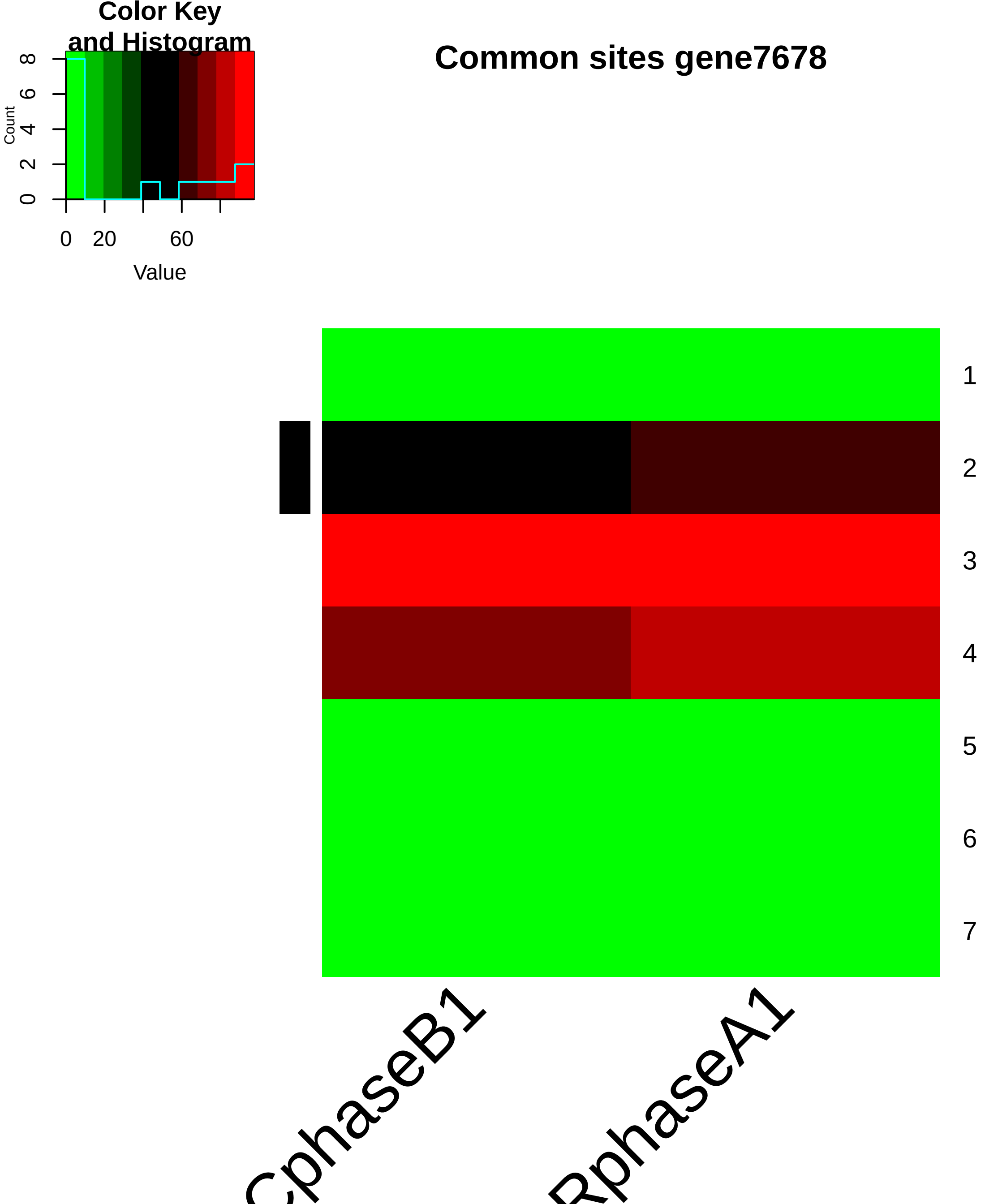
Color Key
and Histogram
Count
0 2 4 6 8
0 20 60
Value
Common sites gene7678
1
2
3
4
5
6
7
CphaseB1
RphaseA1

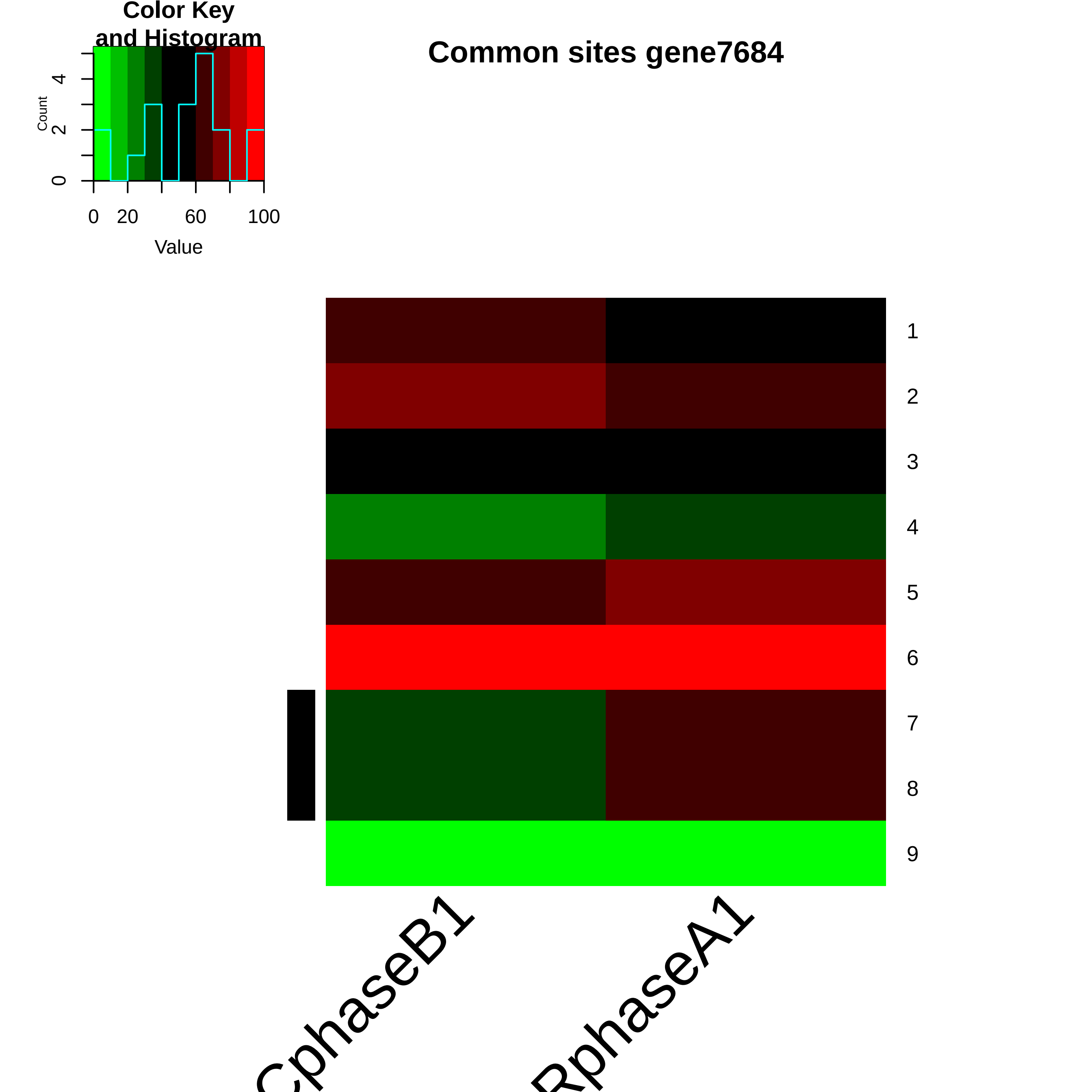

Color Key
and Histogram
Count
0
2
4
0
20
60
100
Value
Common sites gene7684
1
2
3
4
5
6
7
8
9
CphaseB1
RphaseA1

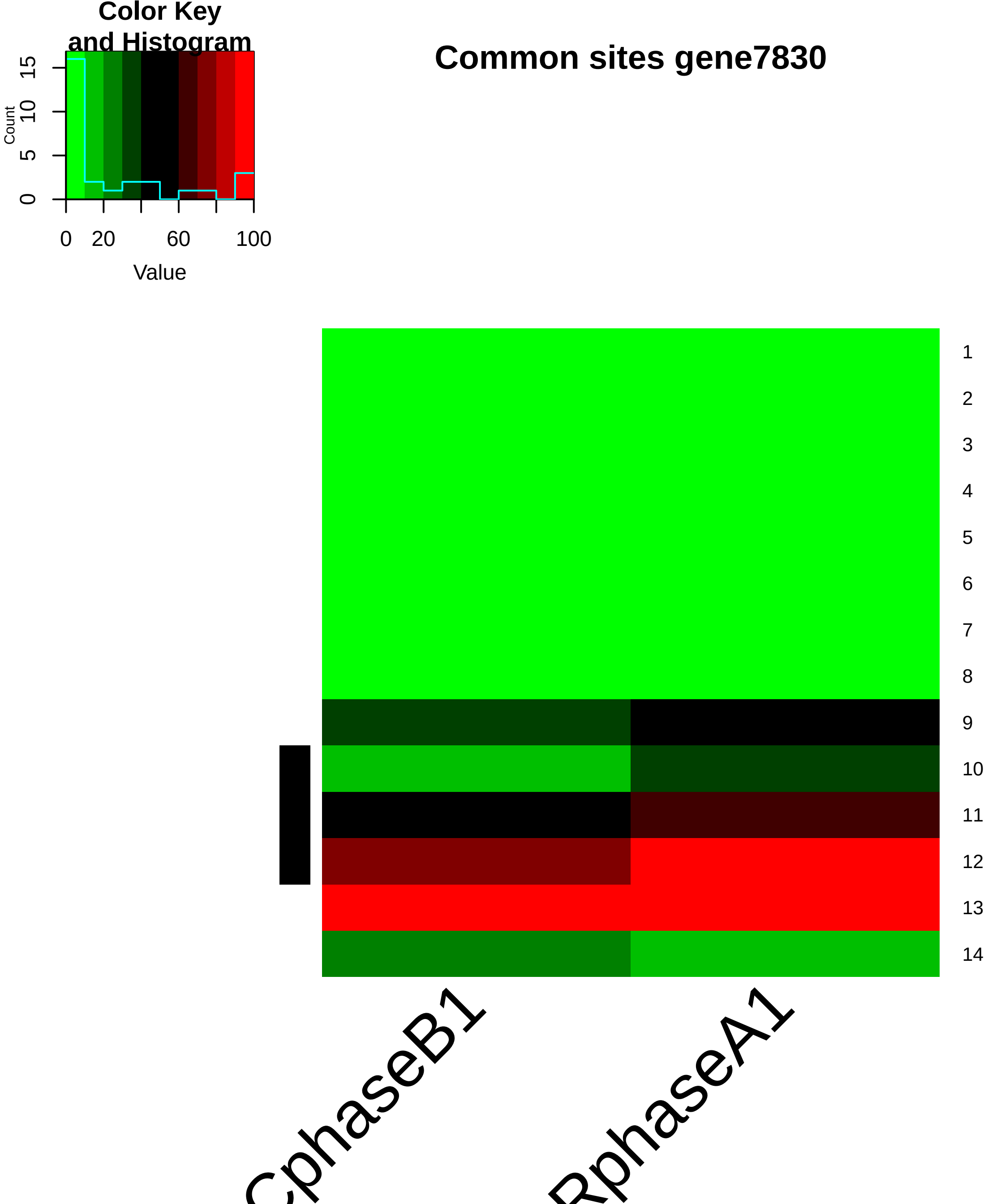
Common sites gene7830
Color Key
and Histogram
Count
0
5
10
15
0
20
60
100
Value
1
2
3
4
5
6
7
8
9
10
11
12
13
14
CphaseB1
RphaseA1

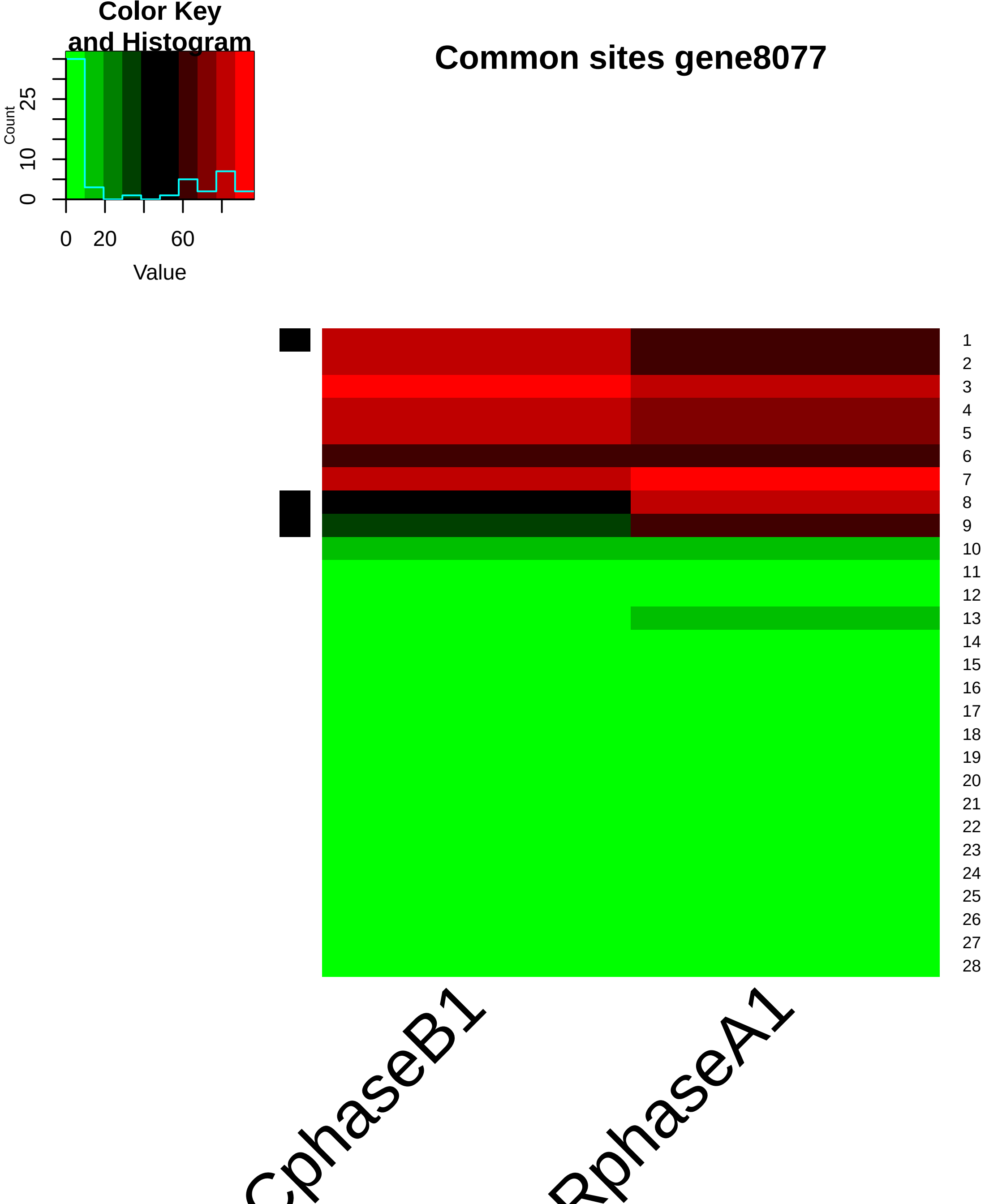
Color Key
and Histogram
Count
0
10
25
0
20
60
Value
Common sites gene8077
CphaseB1
RphaseA1

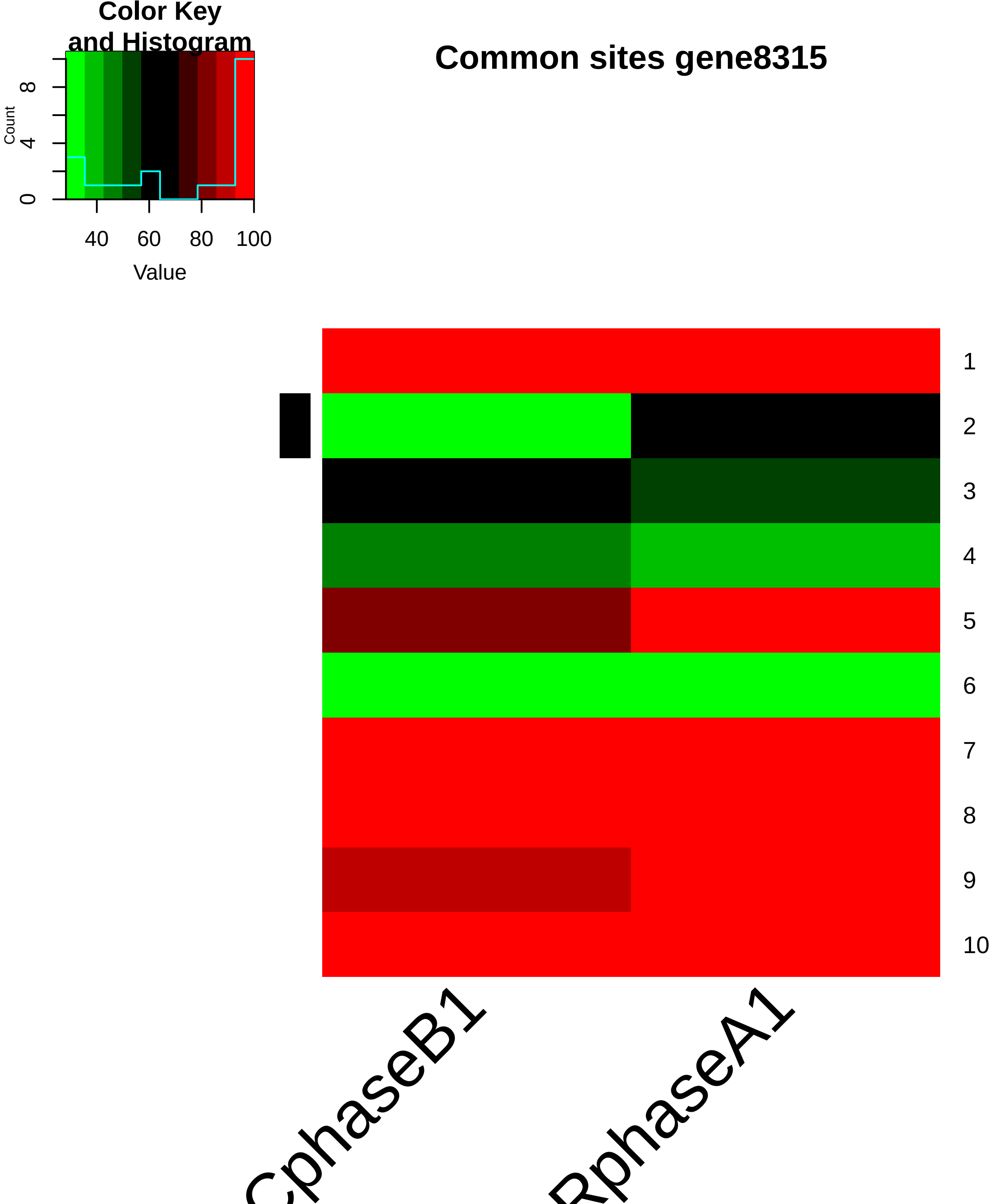
Color Key
and Histogram
Count
0
4
8
40
60
80
100
Value
Common sites gene8315
1
2
3
4
5
6
7
8
9
10
CphaseB1
RphaseA1

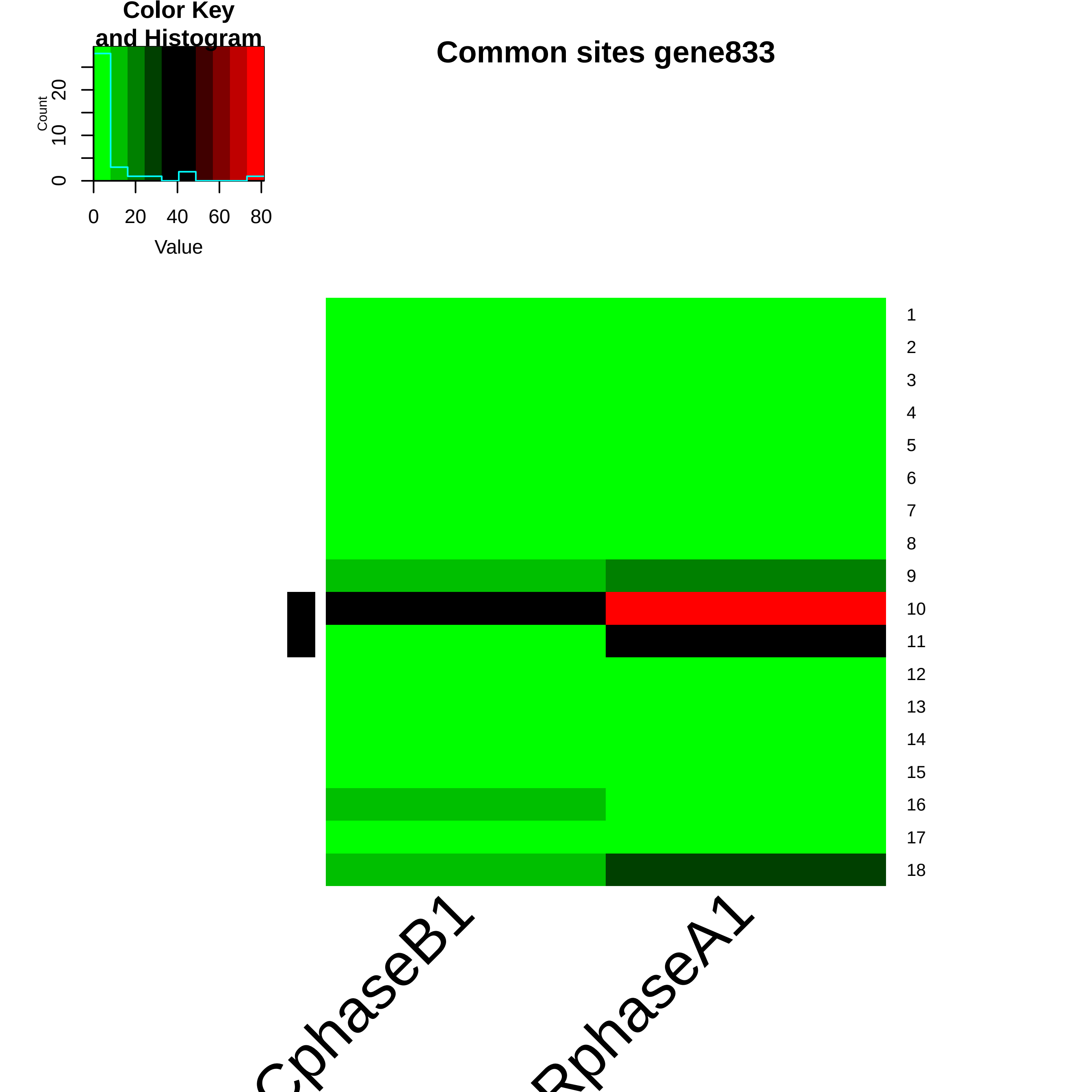
Color Key
and Histogram
Count
0
10
20
0
20
40
60
80
Value
Common sites gene833
CphaseB1
RphaseA1

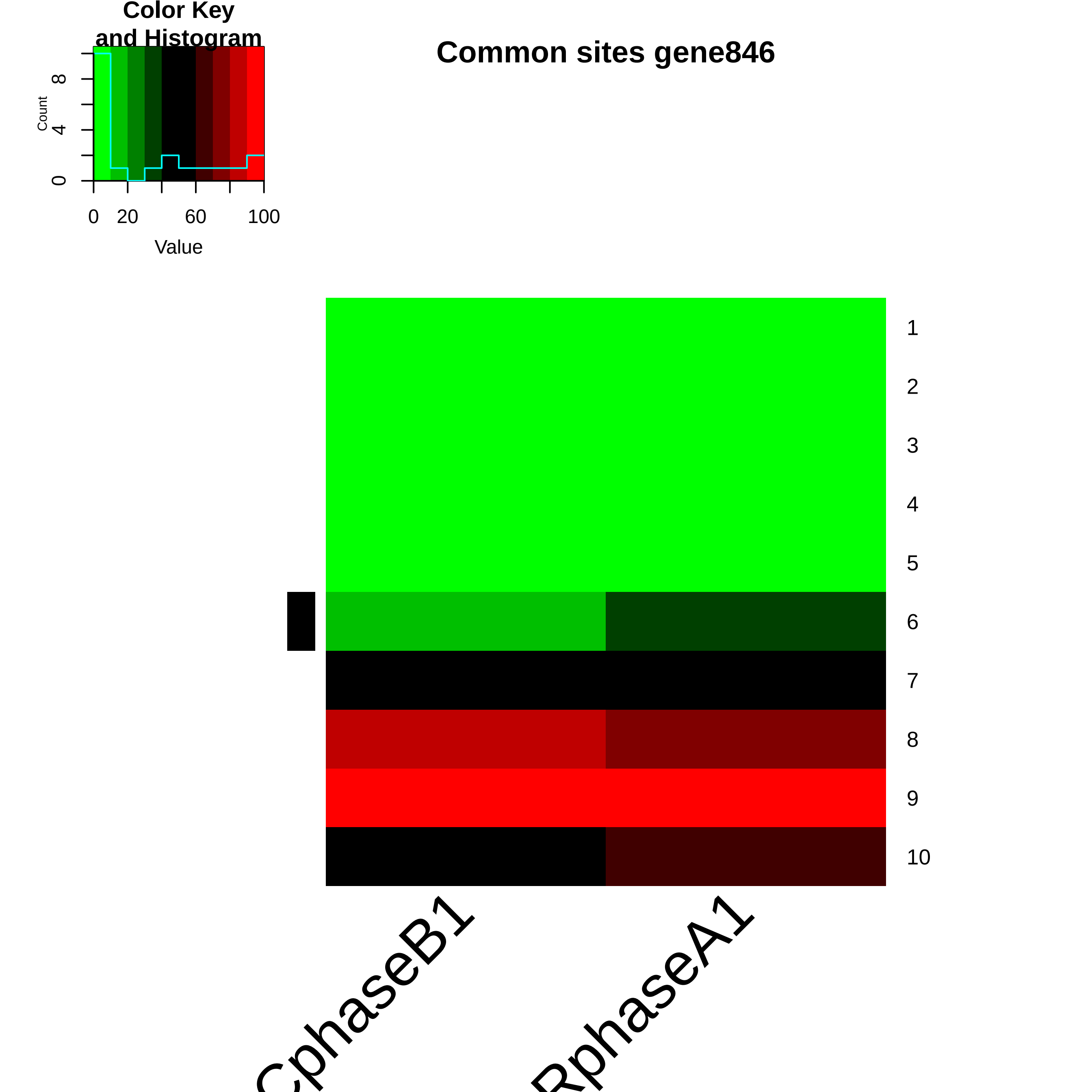
Color Key
and Histogram
Count
0
4
8
0
20
60
100
Value
Common sites gene846
1
2
3
4
5
6
7
8
9
10
CphaseB1
RphaseA1

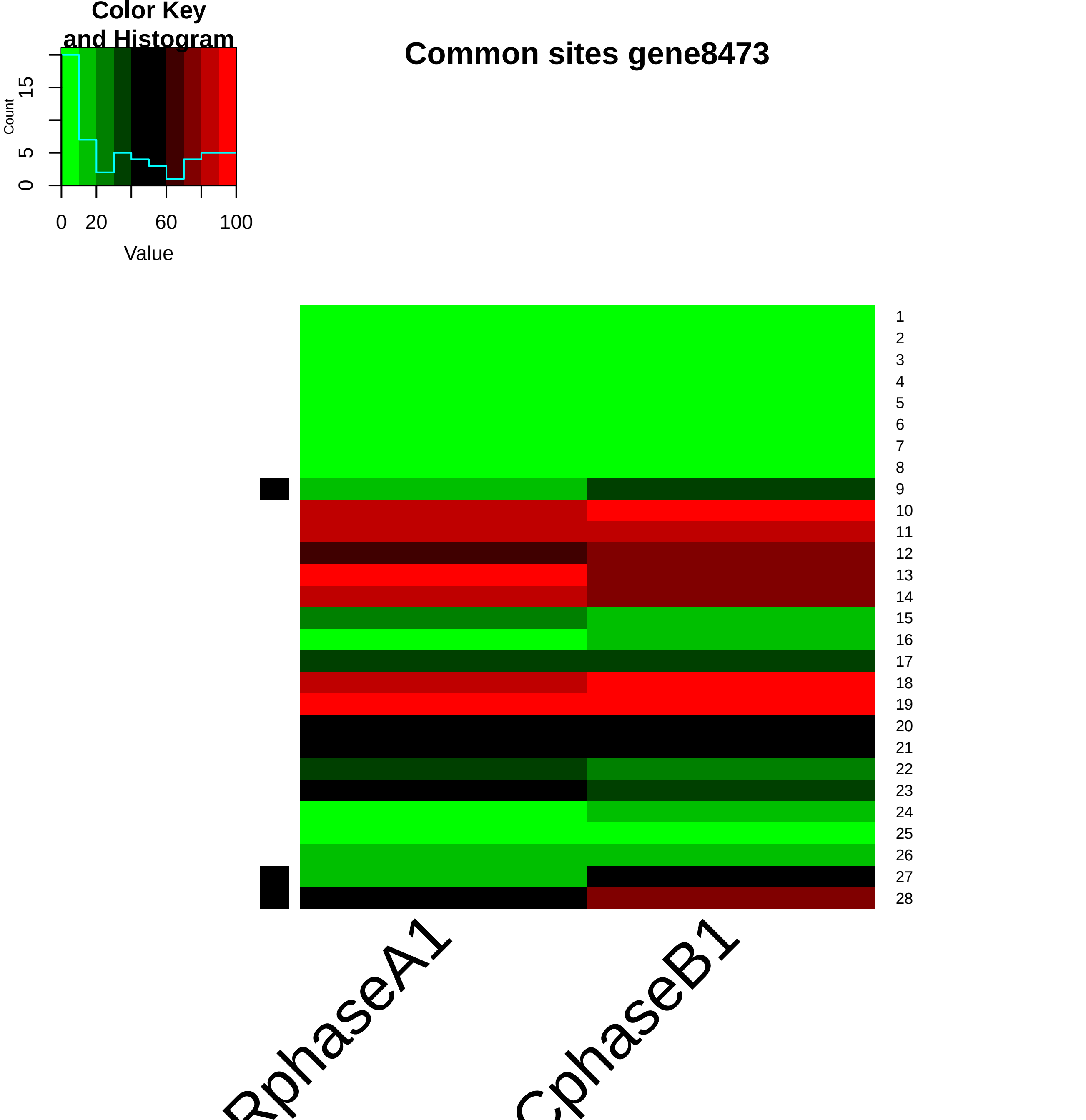
Color Key and Histogram
Count
0
5
15
0
20
60
100
Value
Common sites gene8473
RphaseA1
CphaseB1

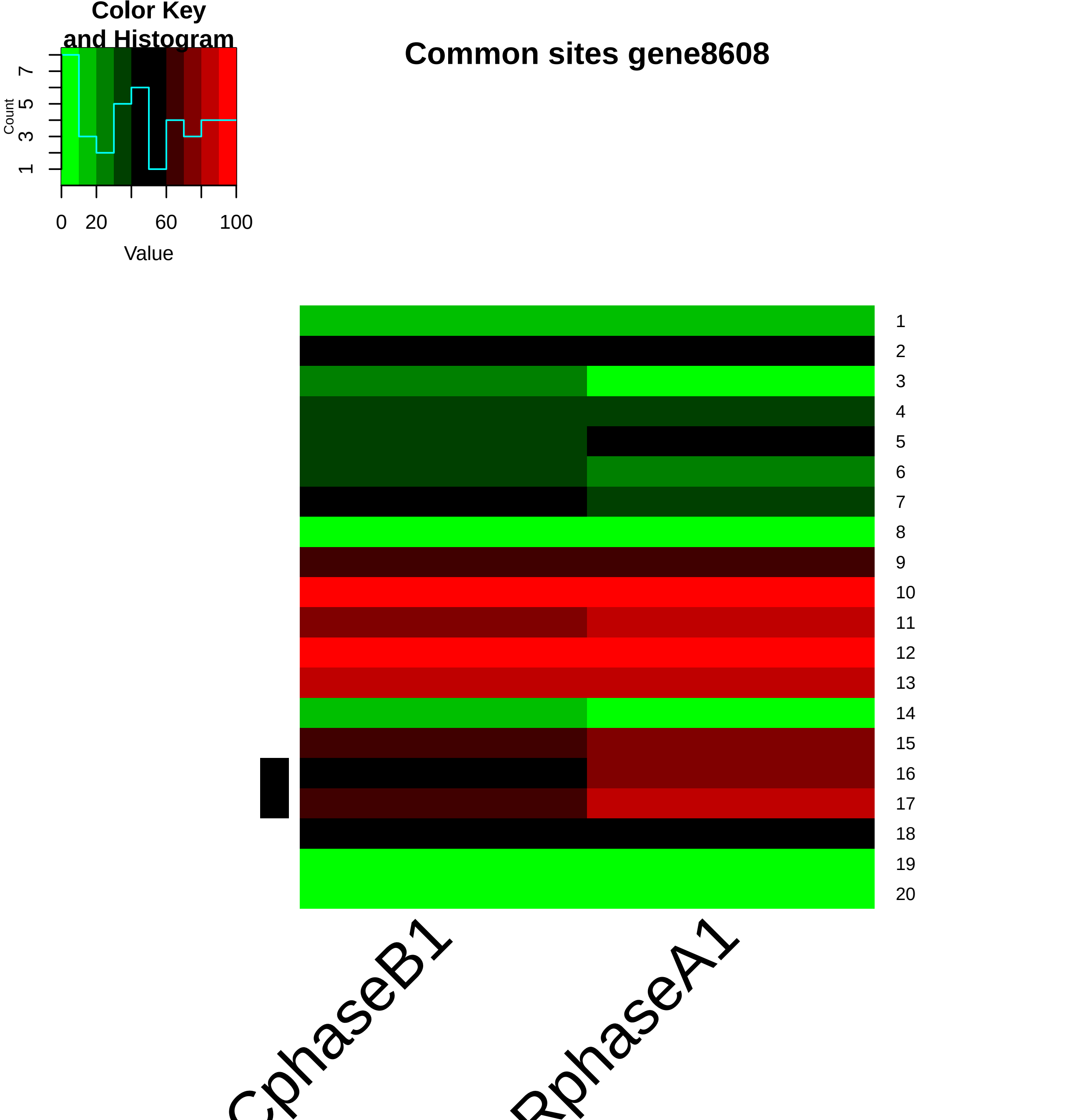
Color Key
and Histogram
Count
1
3
5
7
0
20
60
100
Value
Common sites gene8608
CphaseB1
RphaseA1

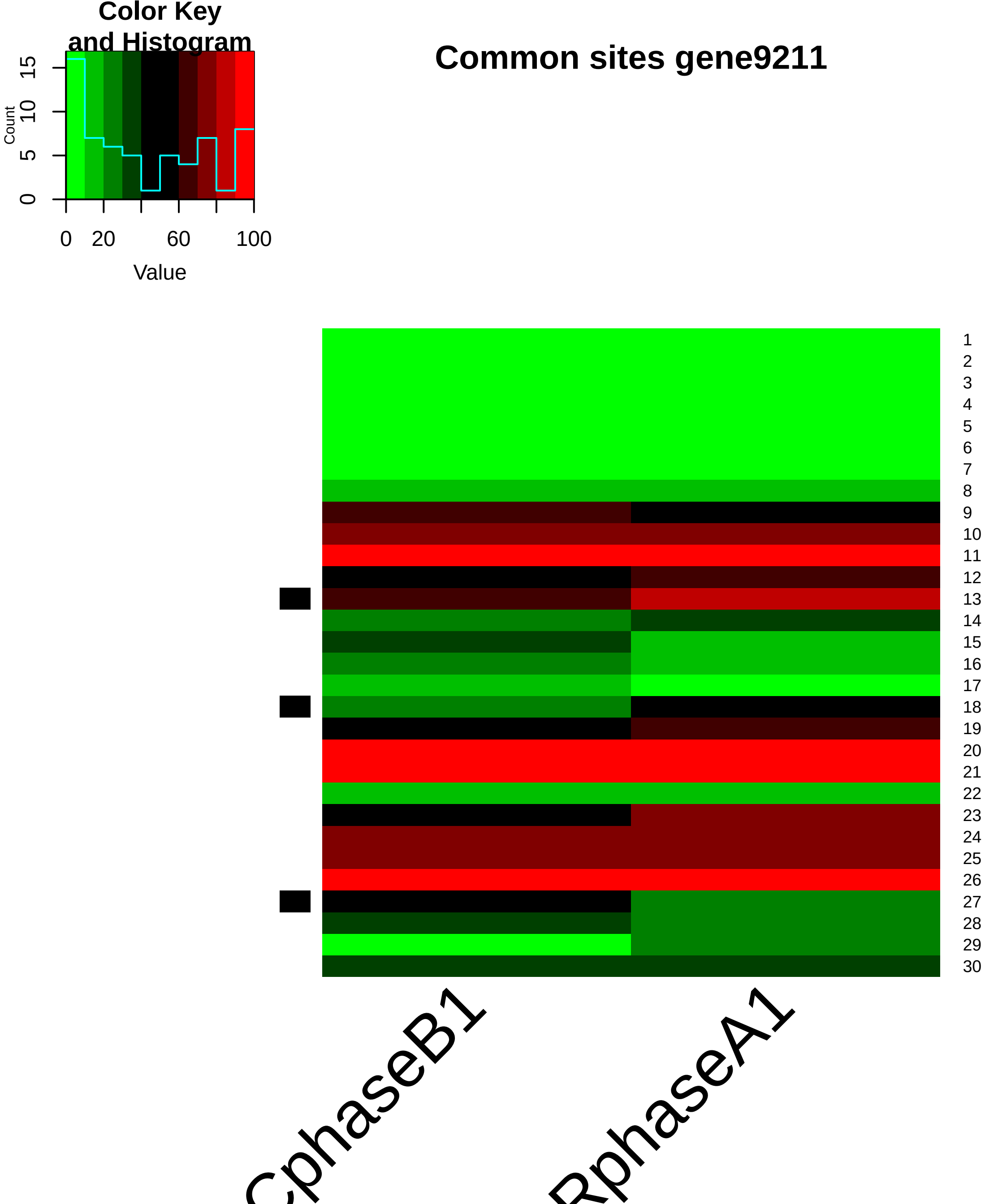
Color Key
and Histogram
Count
0
5
10
15
0
20
60
100
Value
Common sites gene9211
CphaseB1
RphaseA1

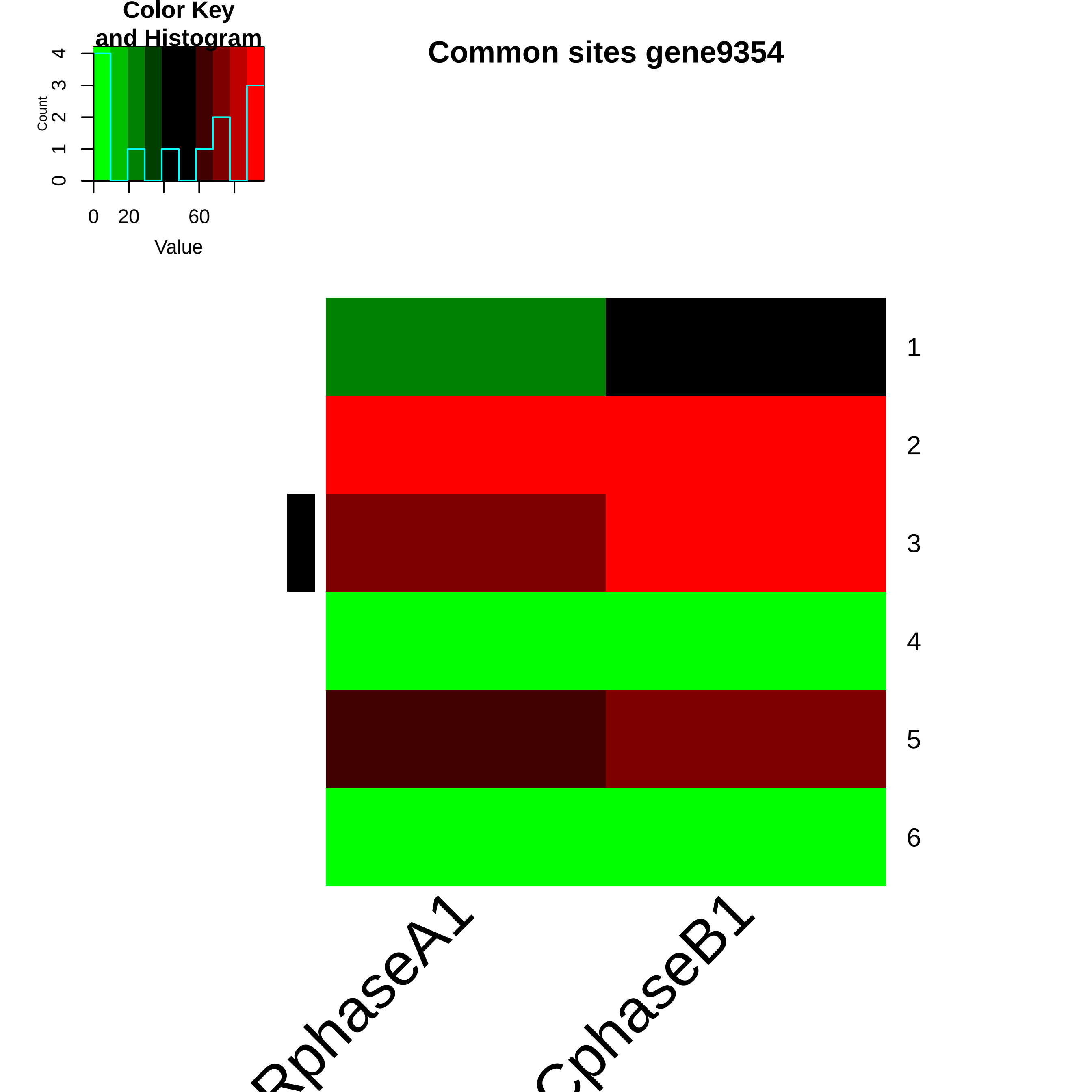
Color Key
and Histogram
Count
0
1
2
3
4
0
20
60
Value
Common sites gene9354
1
2
3
4
5
6
RphaseA1
CphaseB1

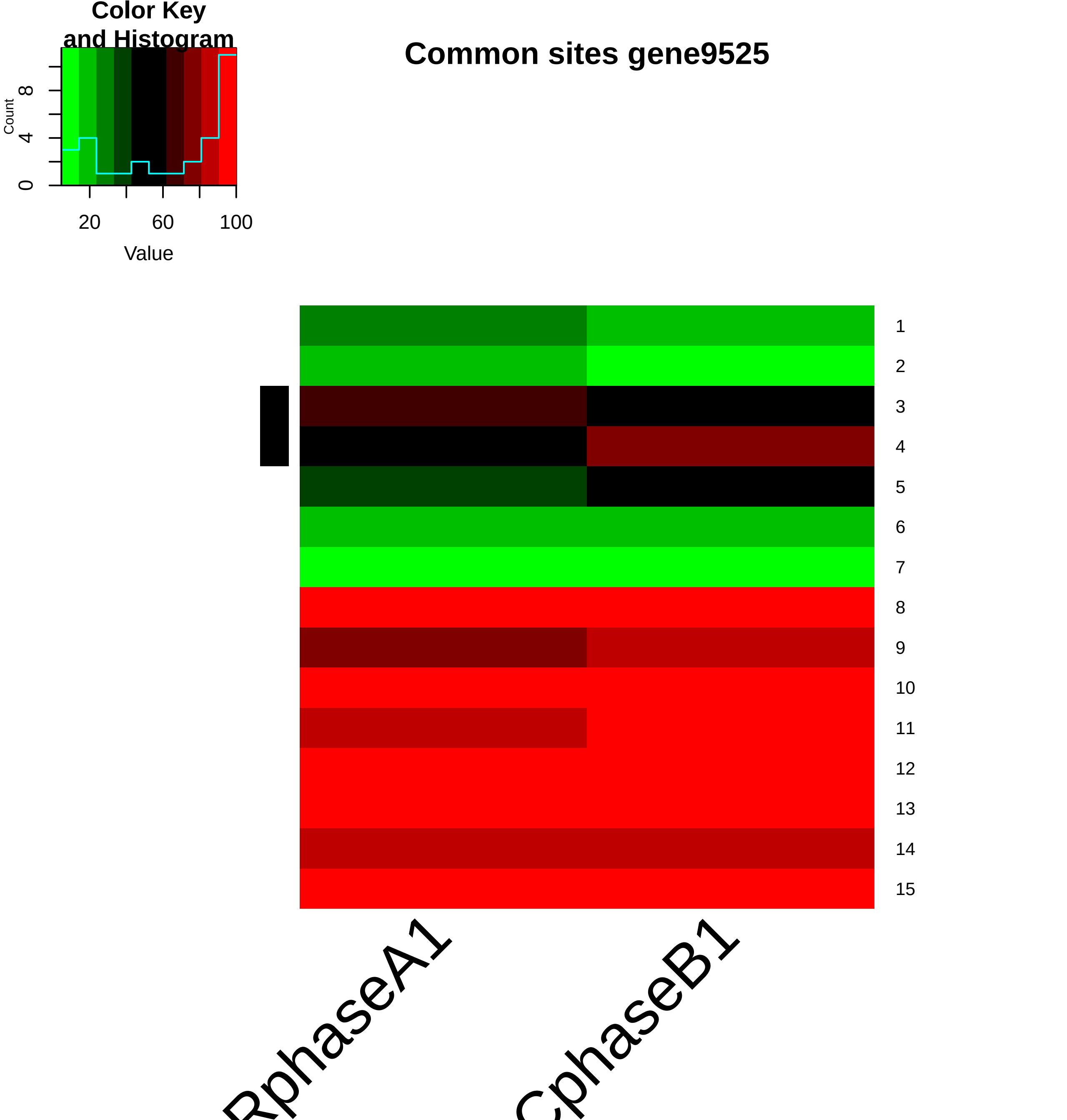
Color Key
and Histogram
Count
0
4
8
20
60
100
Value
Common sites gene9525
RphaseA1
CphaseB1

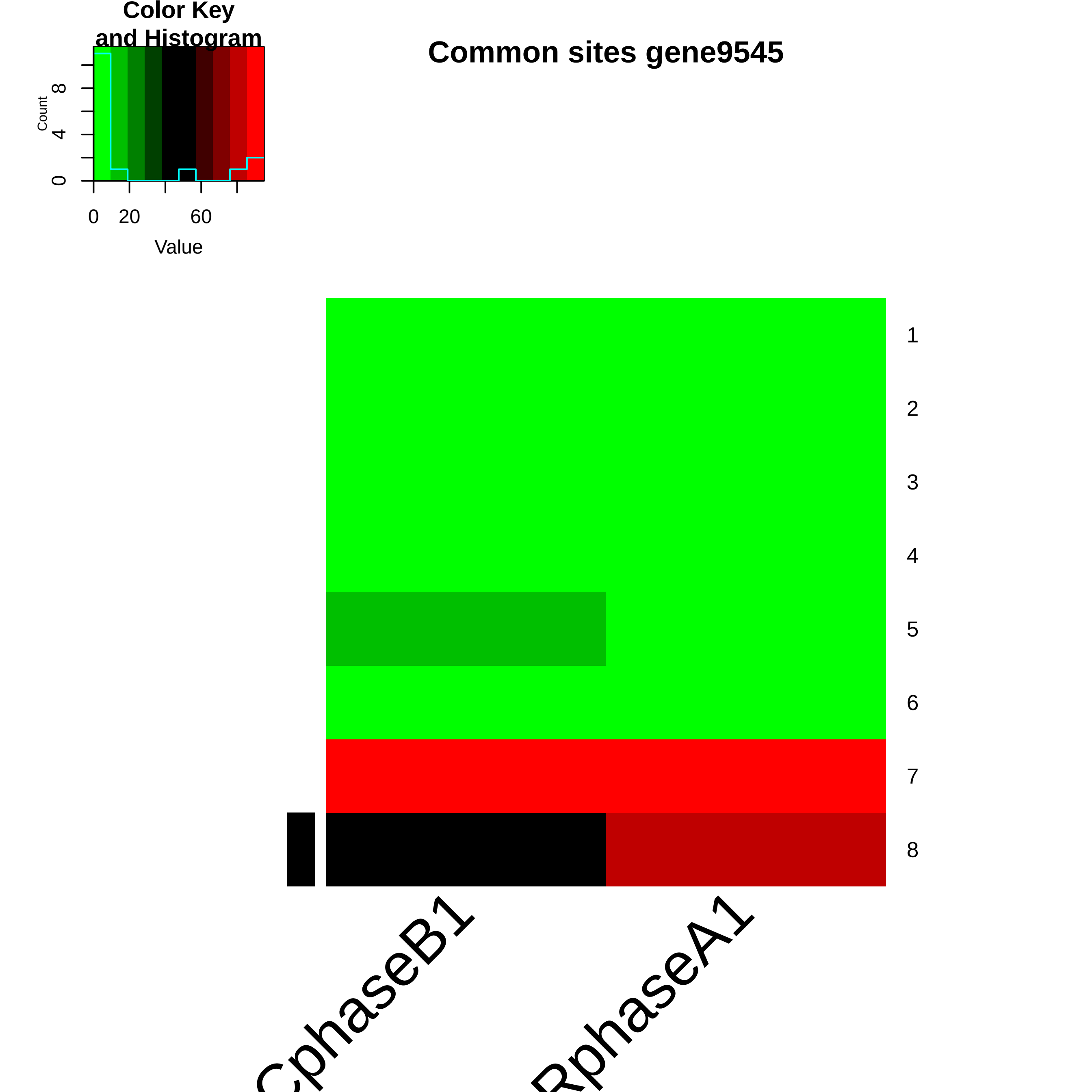
Color Key
and Histogram
Count
0
4
8
0
20
60
Value
Common sites gene9545
1
2
3
4
5
6
7
8
CphaseB1
RphaseA1

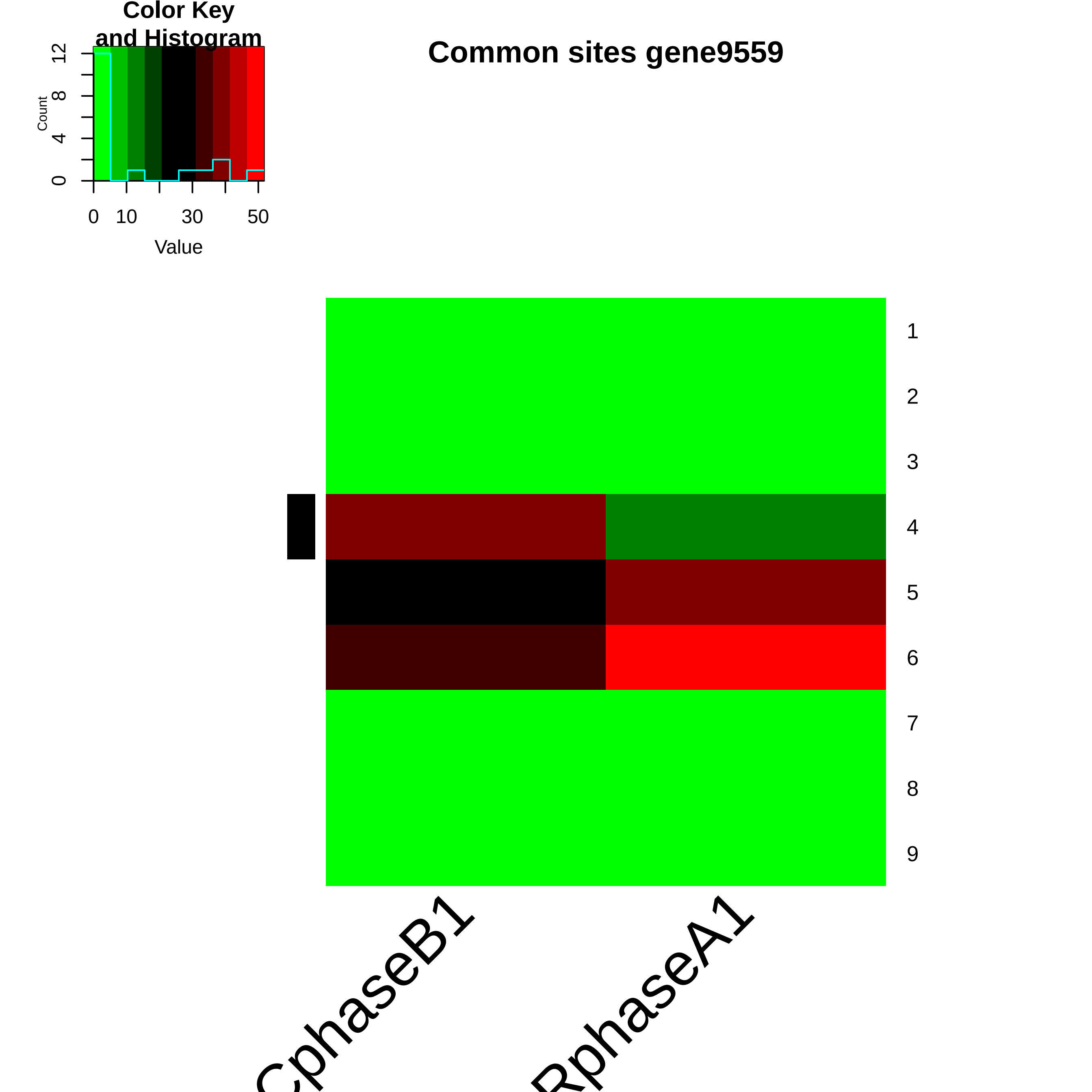

Color Key
and Histogram
Count
0
4
8
12
0
10
30
50
Value
Common sites gene9559
1
2
3
4
5
6
7
8
9
CphaseB1
RphaseA1

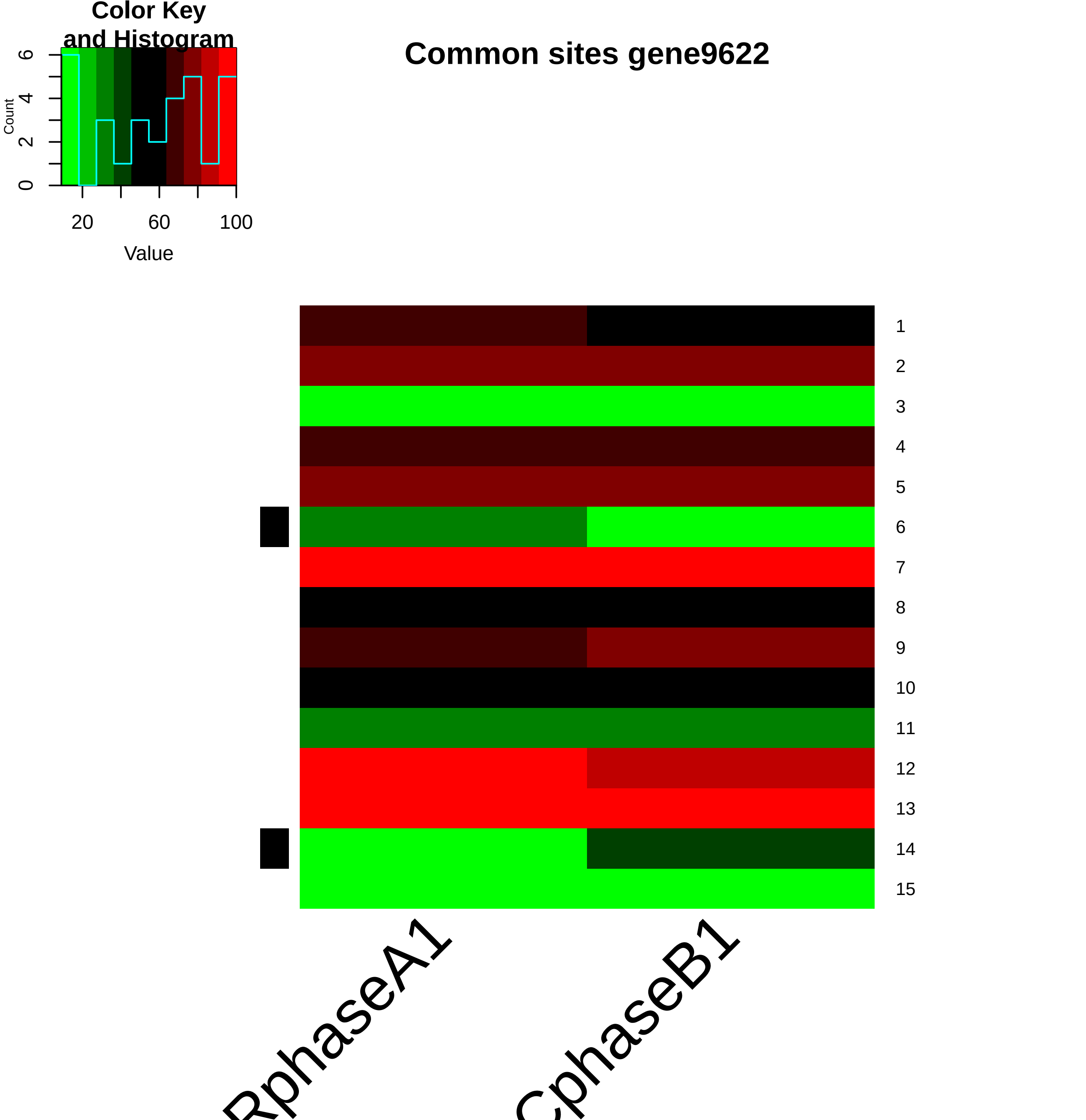

Color Key
and Histogram
Count
0
2
4
6
20
60
100
Value
Common sites gene9622
RphaseA1
CphaseB1

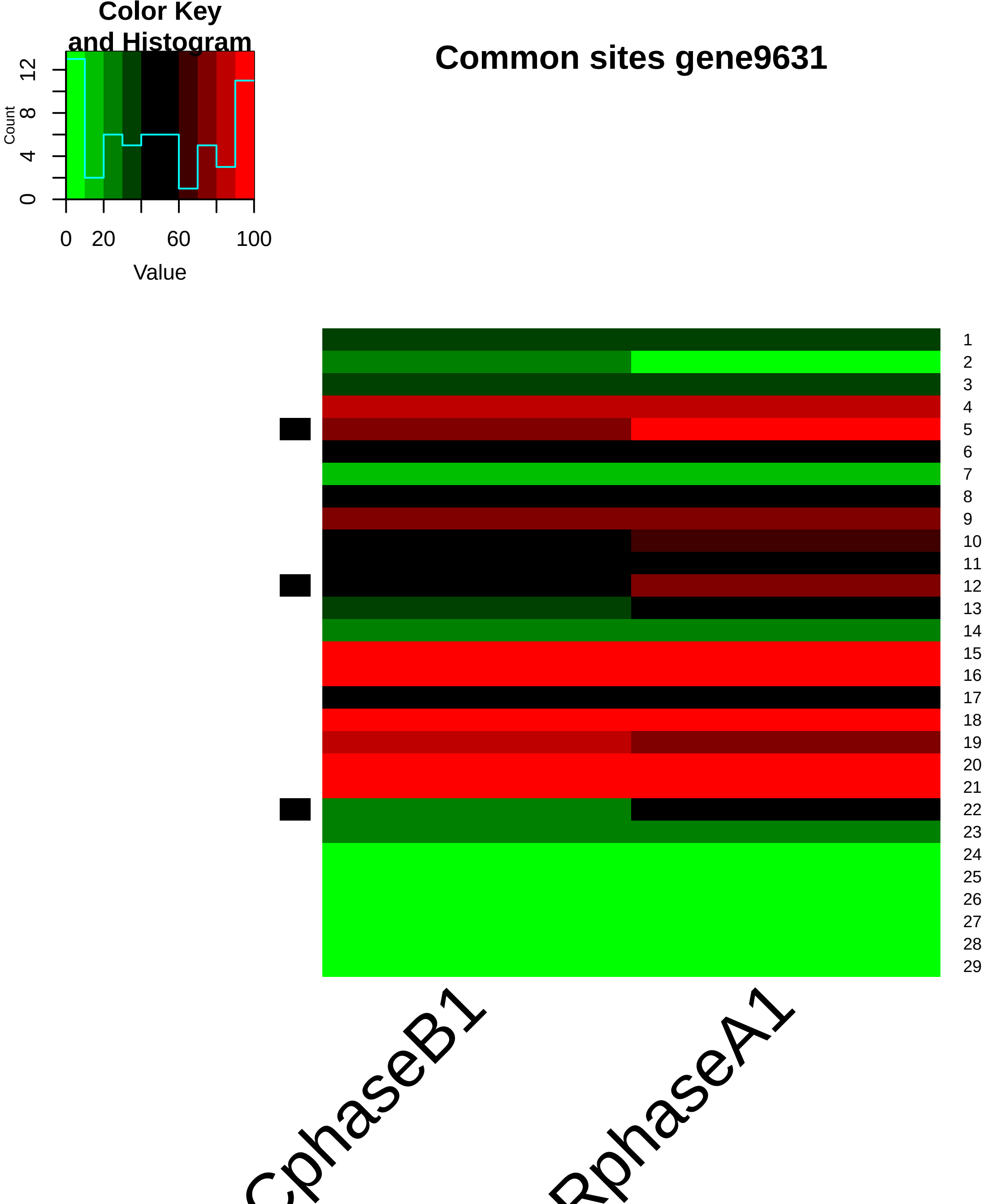
Color Key
and Histogram
Count
0
4
8
12
0
20
60
100
Value
Common sites gene9631
CphaseB1
RphaseA1

### Common sites gene9650

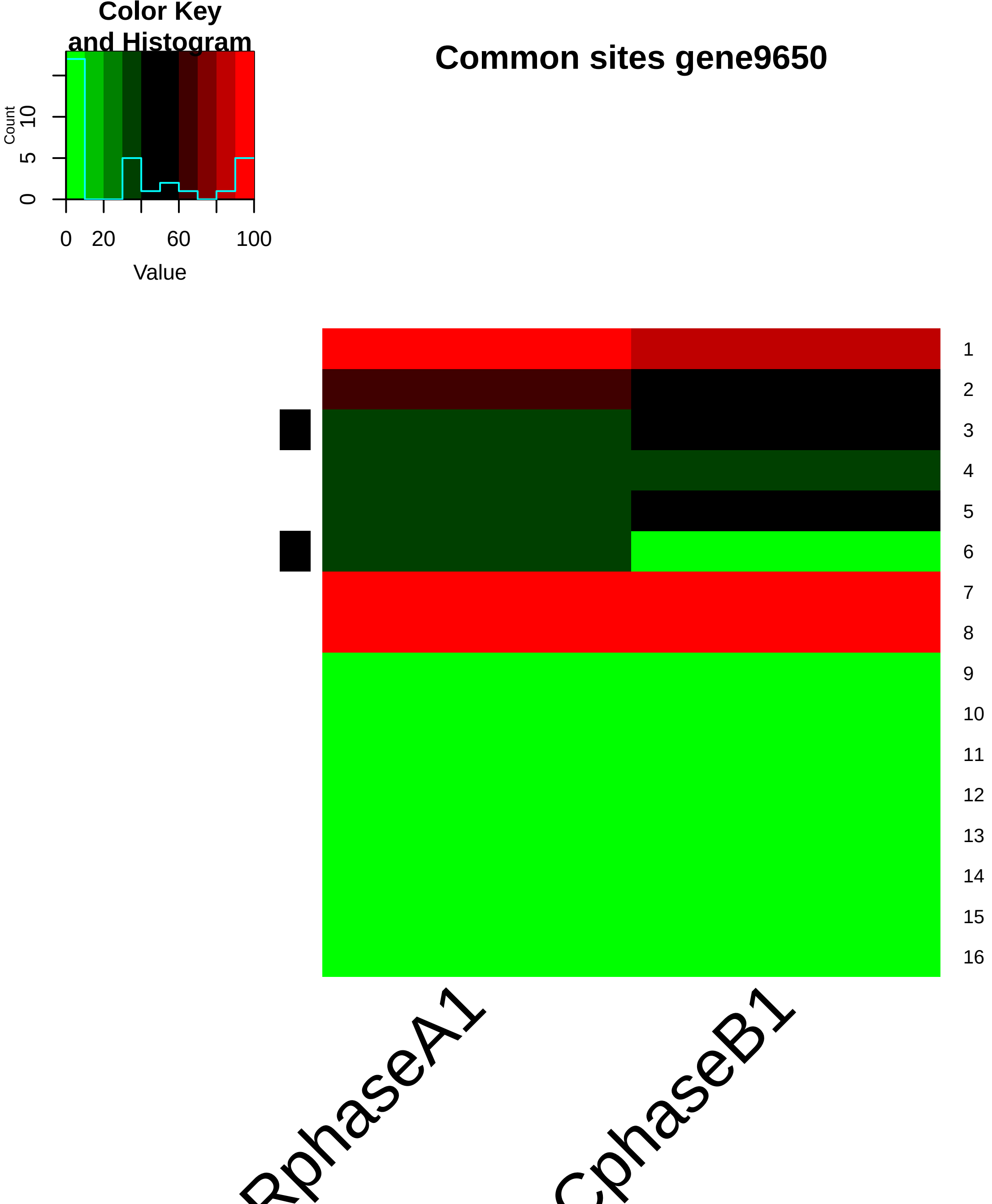

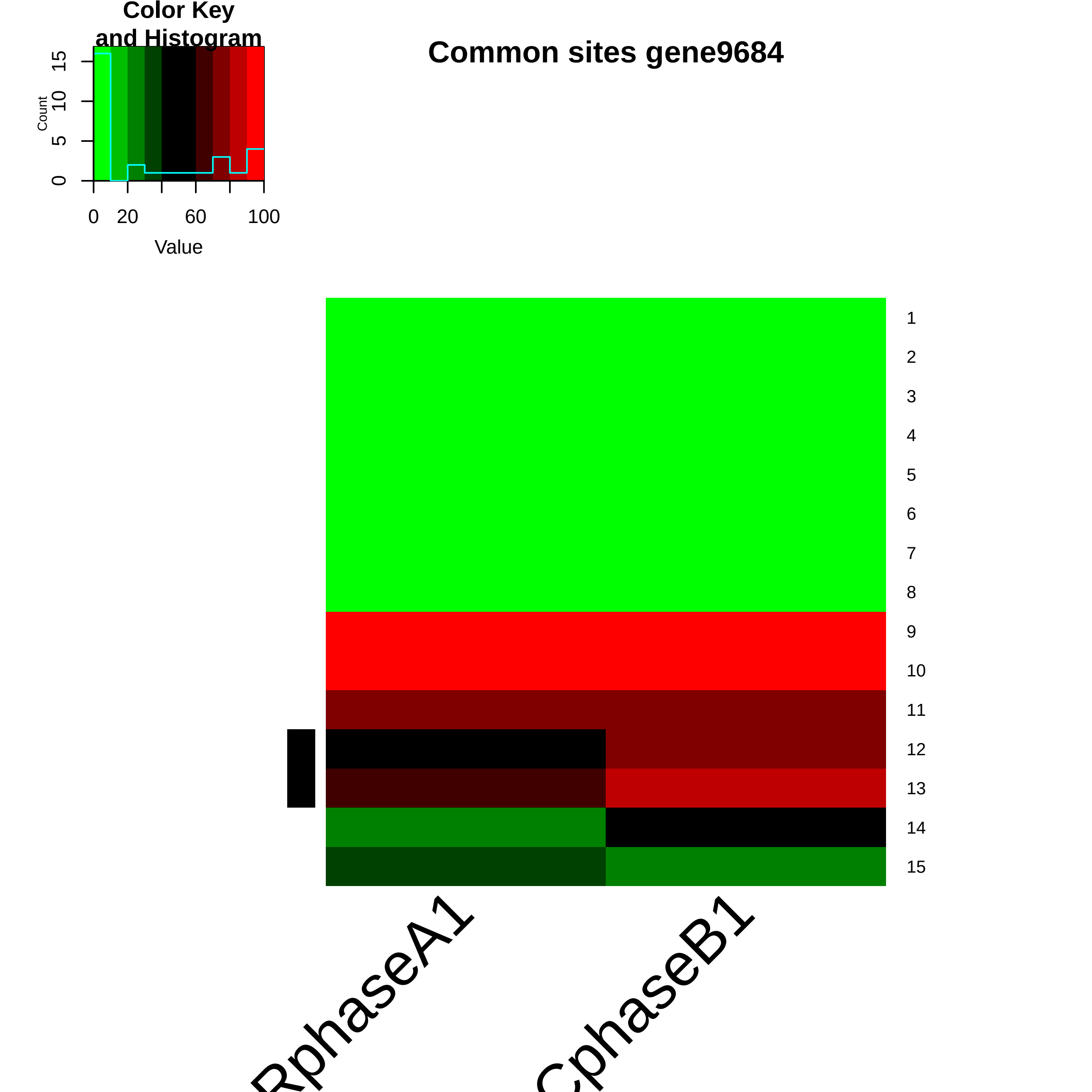
Color Key
and Histogram
Count
0
5
10
15
0
20
60
100
Value
Common sites gene9684
RphaseA1
CphaseB1

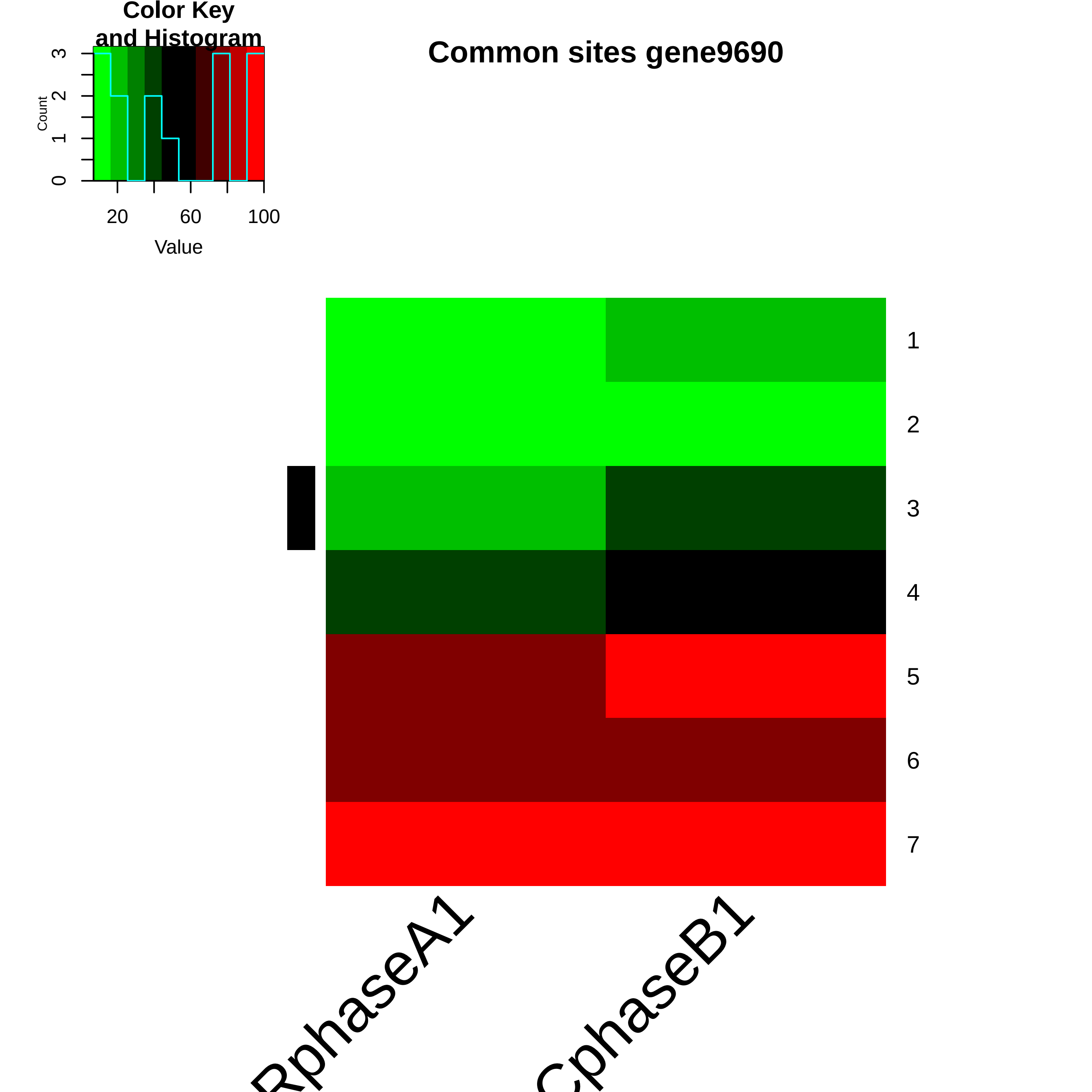

Color Key
and Histogram
Count
0
1
2
3
20
60
100
Value
Common sites gene9690
1
2
3
4
5
6
7
RphaseA1
CphaseB1

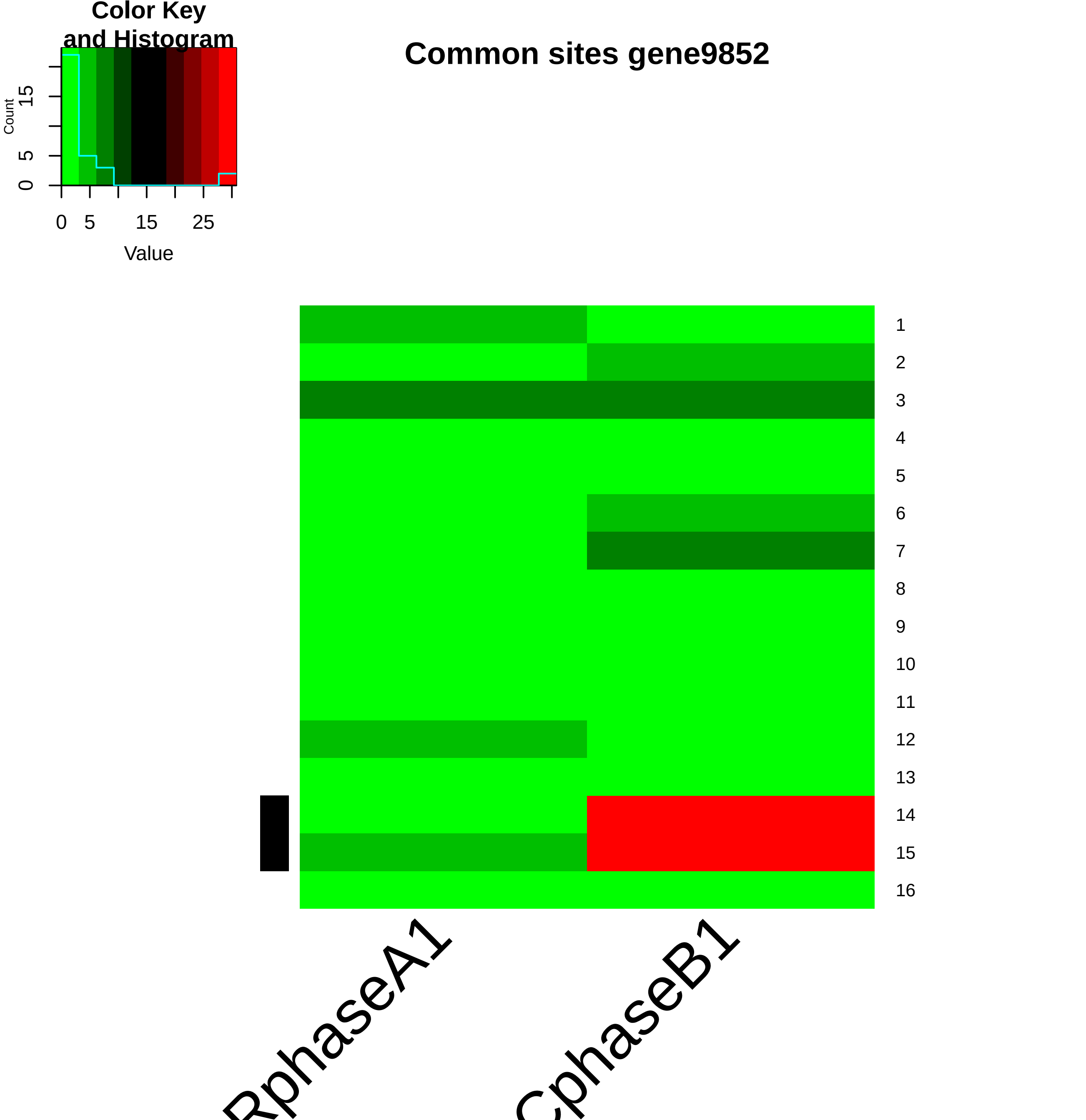
Color Key
and Histogram
Count
0
5
15
0
5
15
25
Value
Common sites gene9852
RphaseA1
CphaseB1

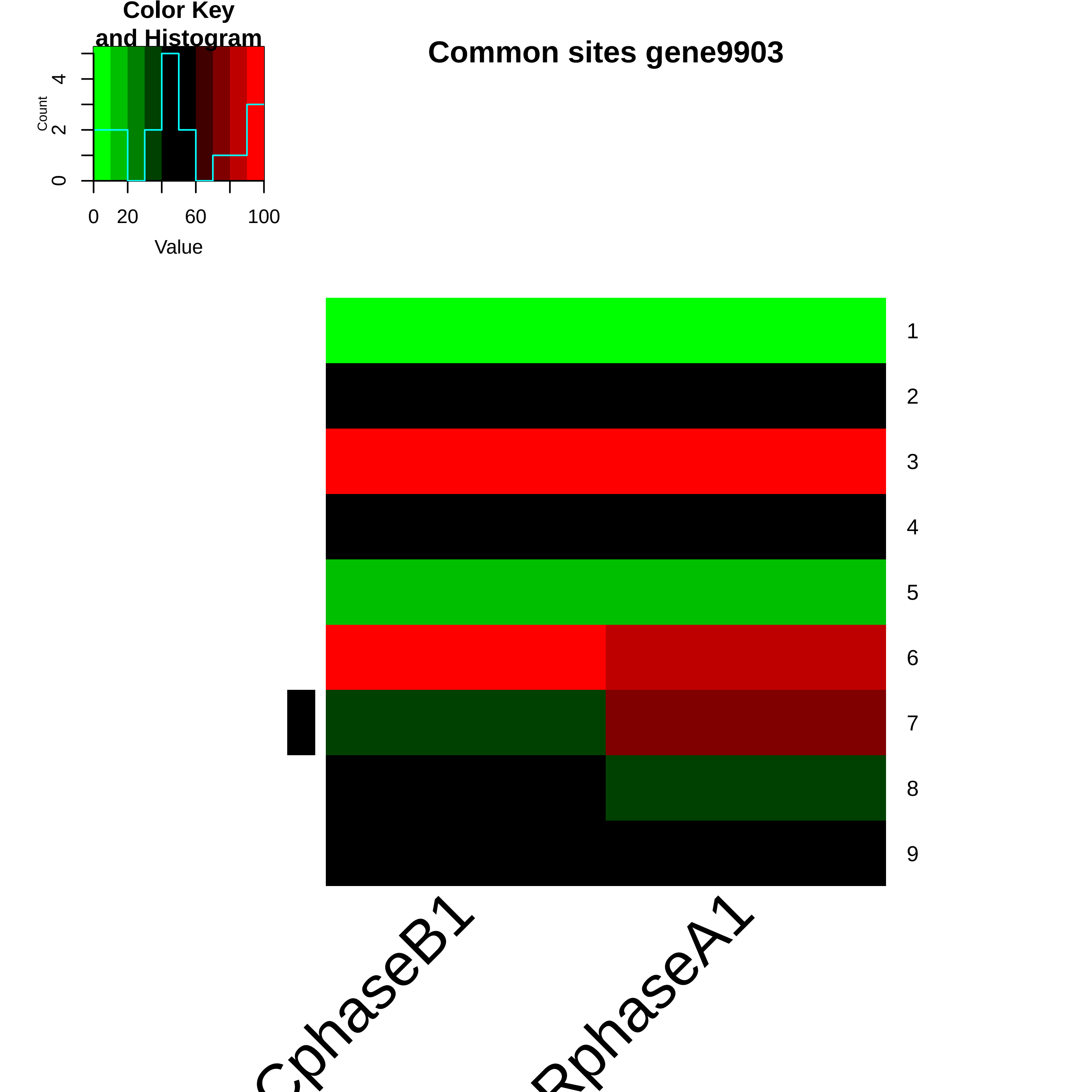

Color Key and Histogram
Count
0
2
4
0
20
60
100
Value
Common sites gene9903
1
2
3
4
5
6
7
8
9
CphaseB1
RphaseA1

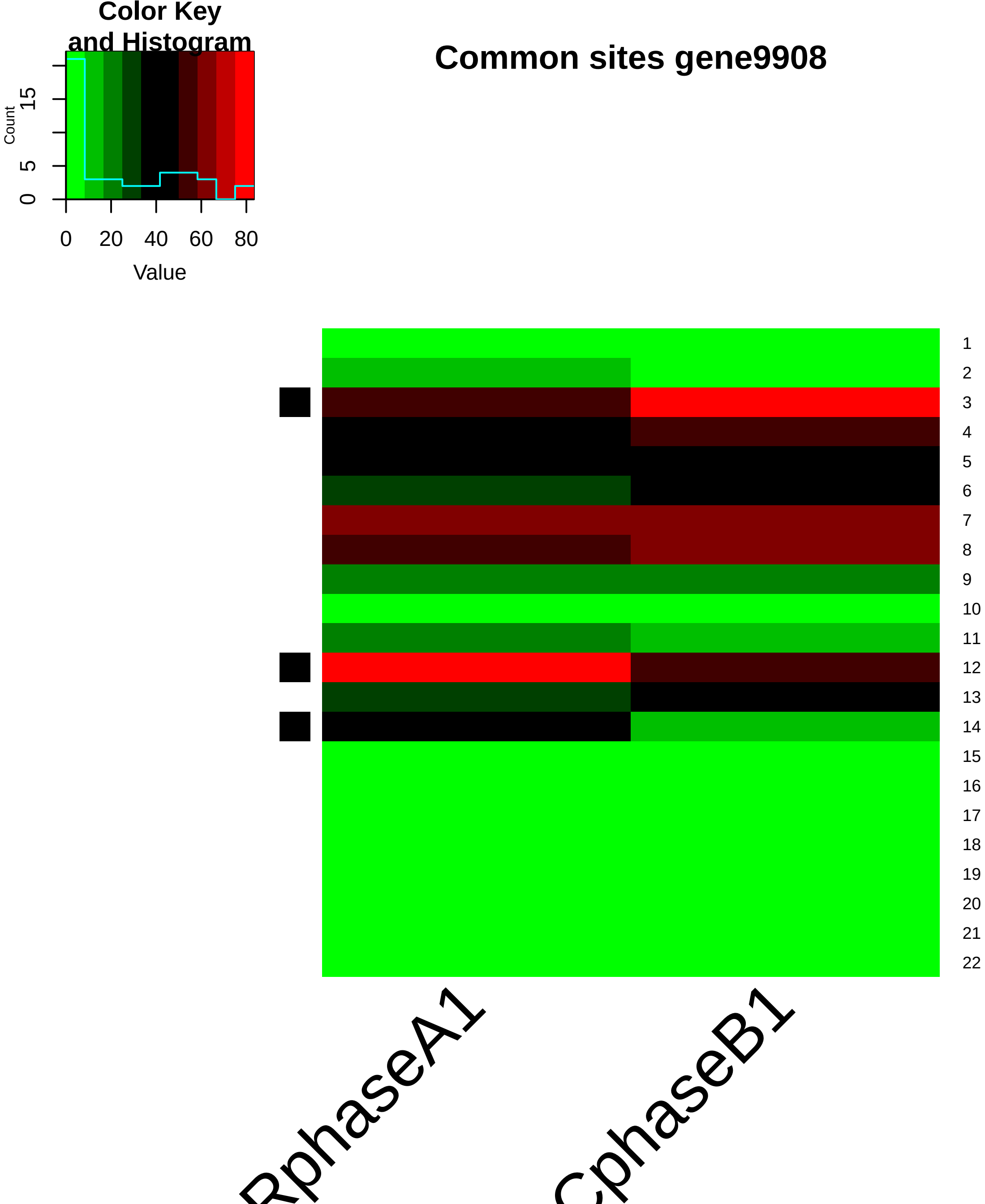
Color Key
and Histogram
Count
0
5
15
0
20
40
60
80
Value
Common sites gene9908
RphaseA1
CphaseB1

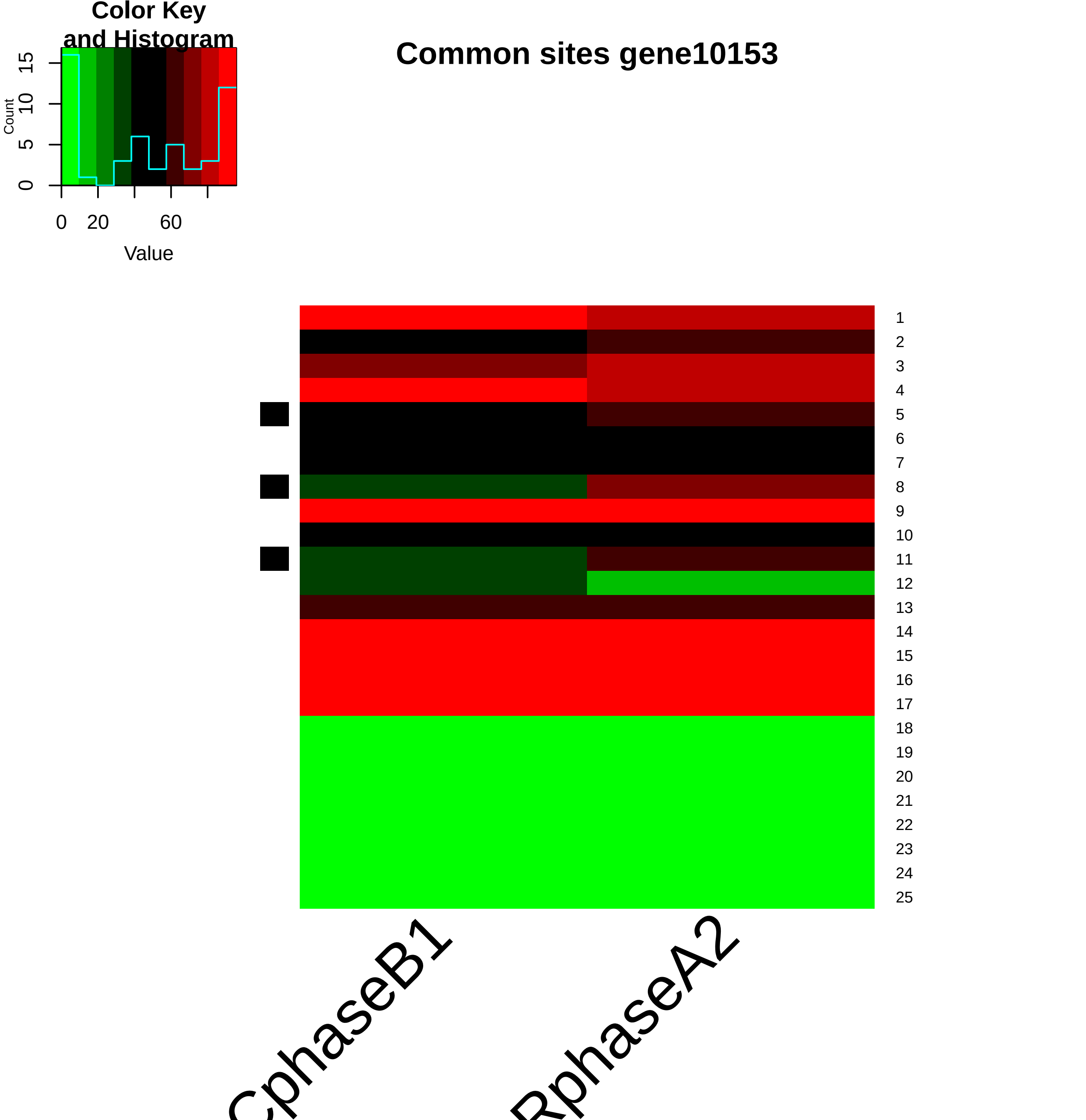
Color Key
and Histogram
Count
0
5
10
15
0
20
60
Value
Common sites gene10153
CphaseB1
RphaseA2

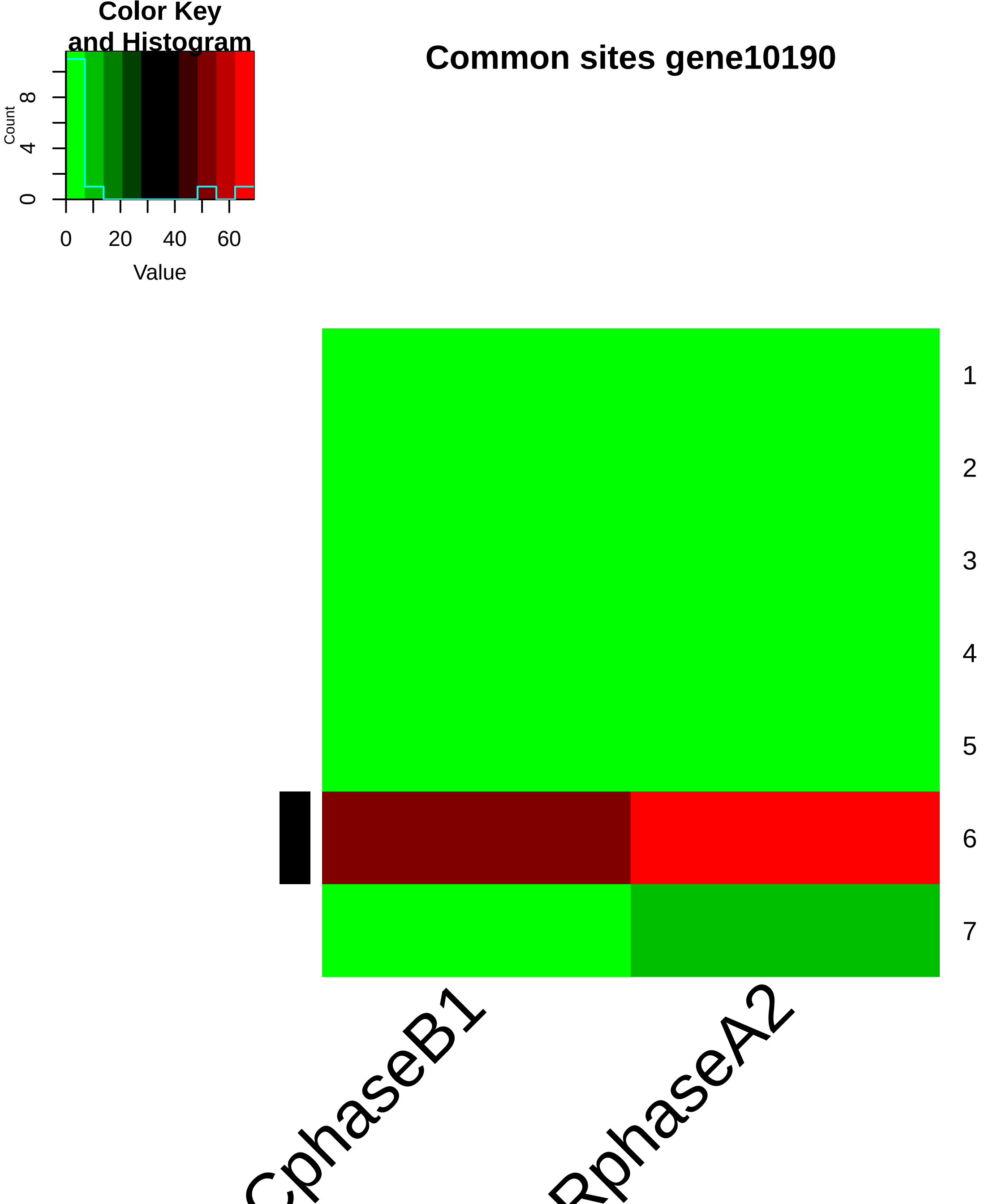

Color Key
and Histogram
Count
0
4
8
0
20
40
60
Value
Common sites gene10190
1
2
3
4
5
6
7
CphaseB1
RphaseA2

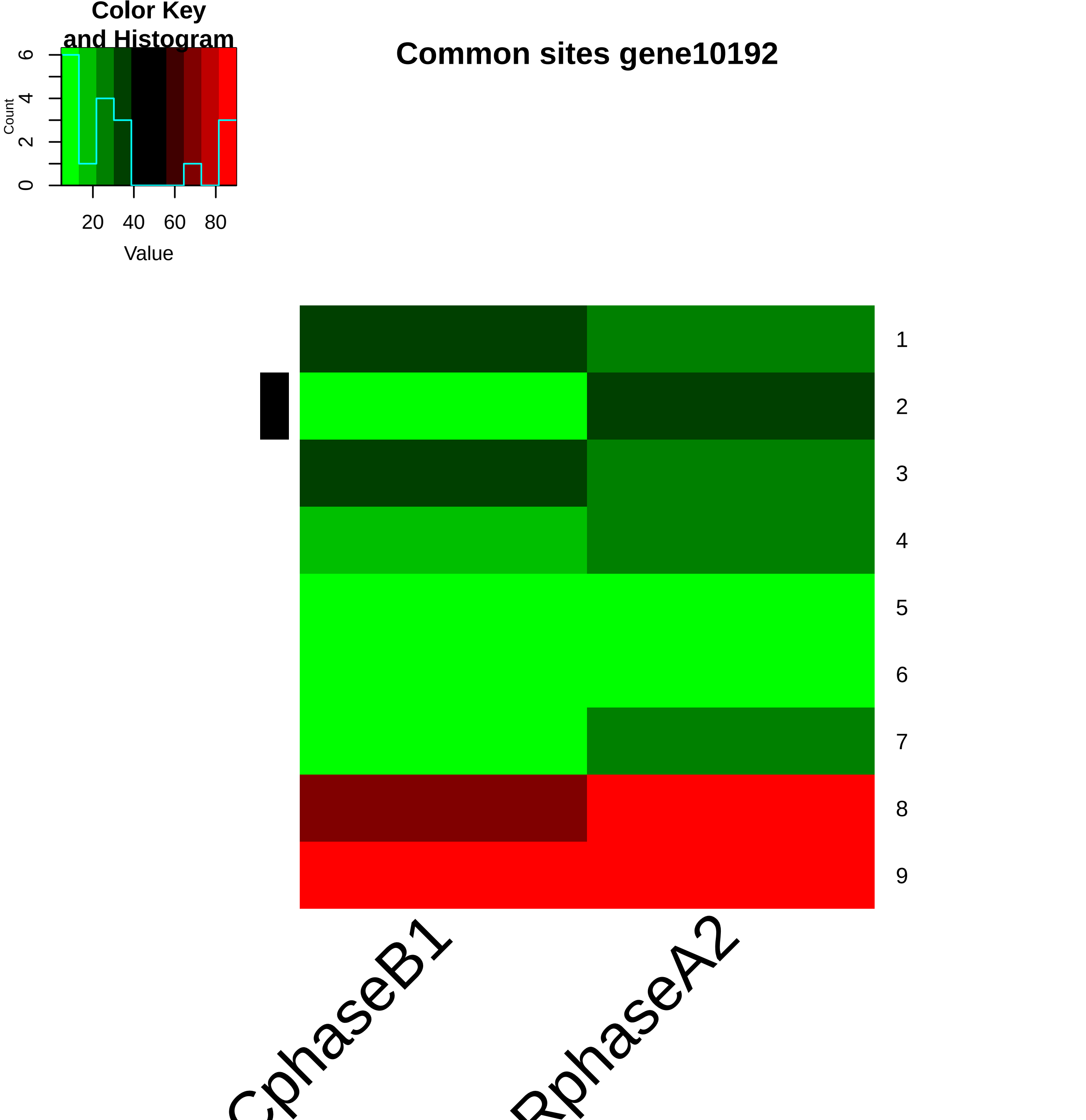
Color Key
and Histogram
Count
0
2
4
6
20
40
60
80
Value
Common sites gene10192
1
2
3
4
5
6
7
8
9
CphaseB1
RphaseA2

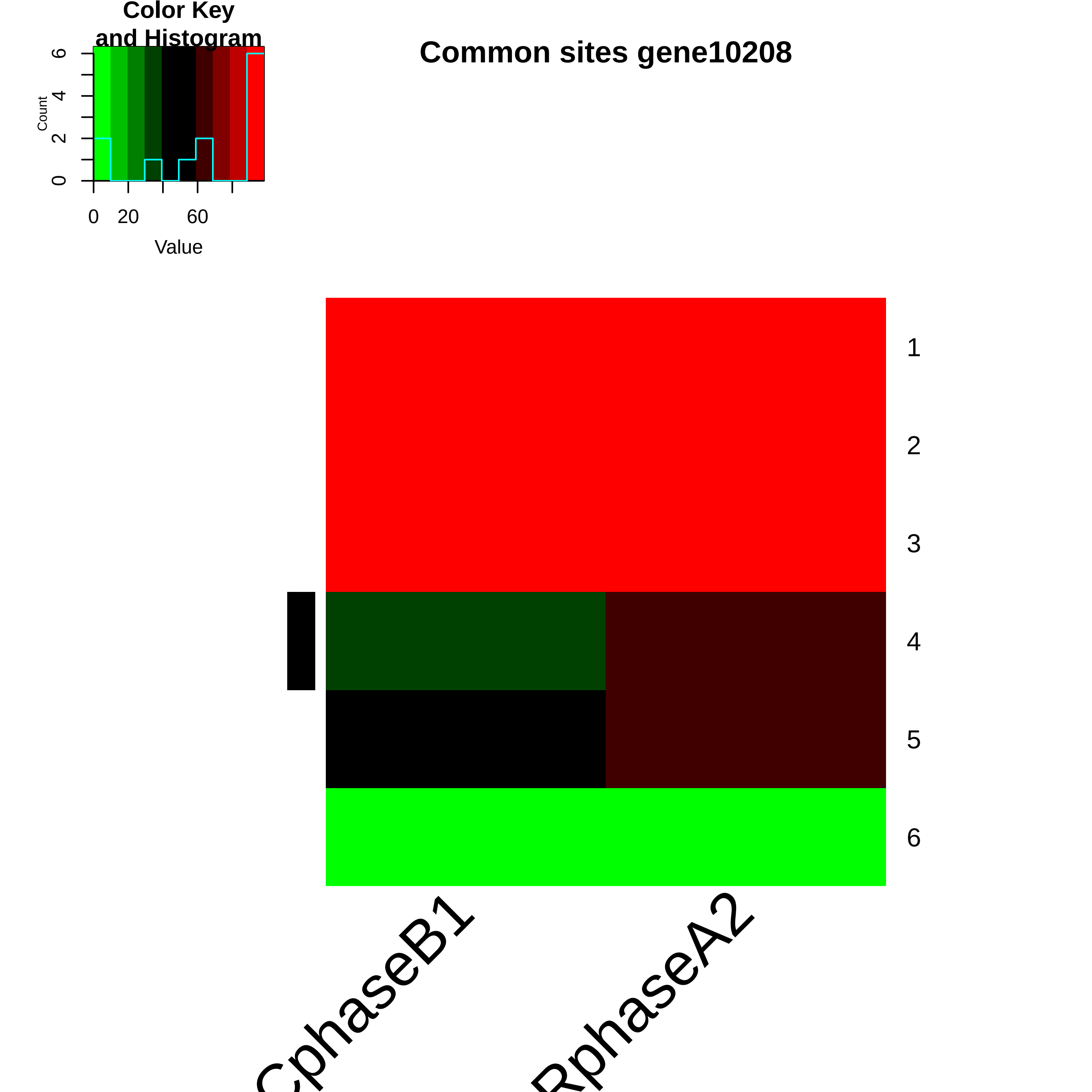
Color Key
and Histogram
Count
0
2
4
6
0
20
60
Value
Common sites gene10208
1
2
3
4
5
6
CphaseB1
RphaseA2

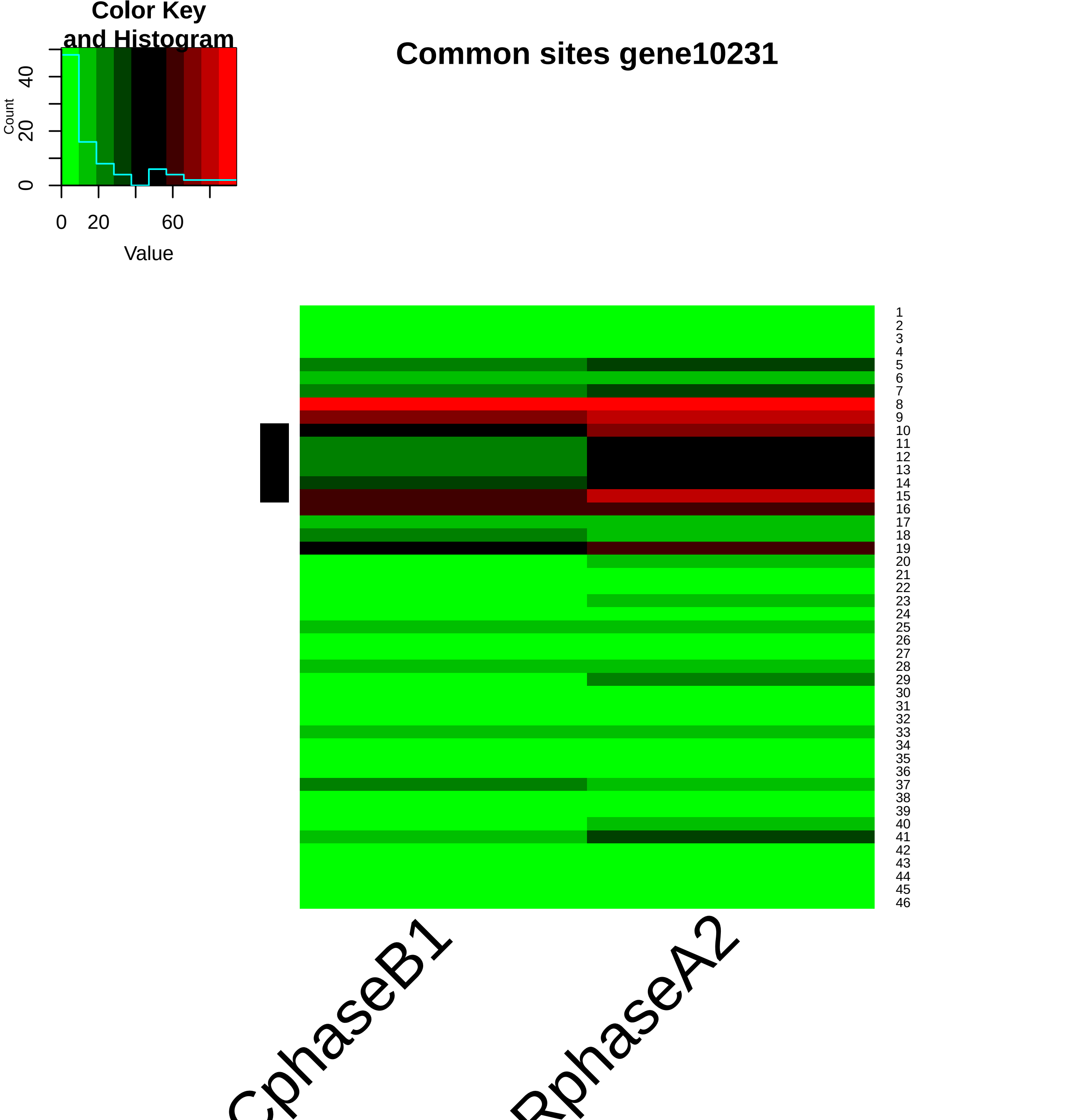

Color Key
and Histogram
Count
0
20
40
0
20
60
Value
Common sites gene10231
CphaseB1
RphaseA2

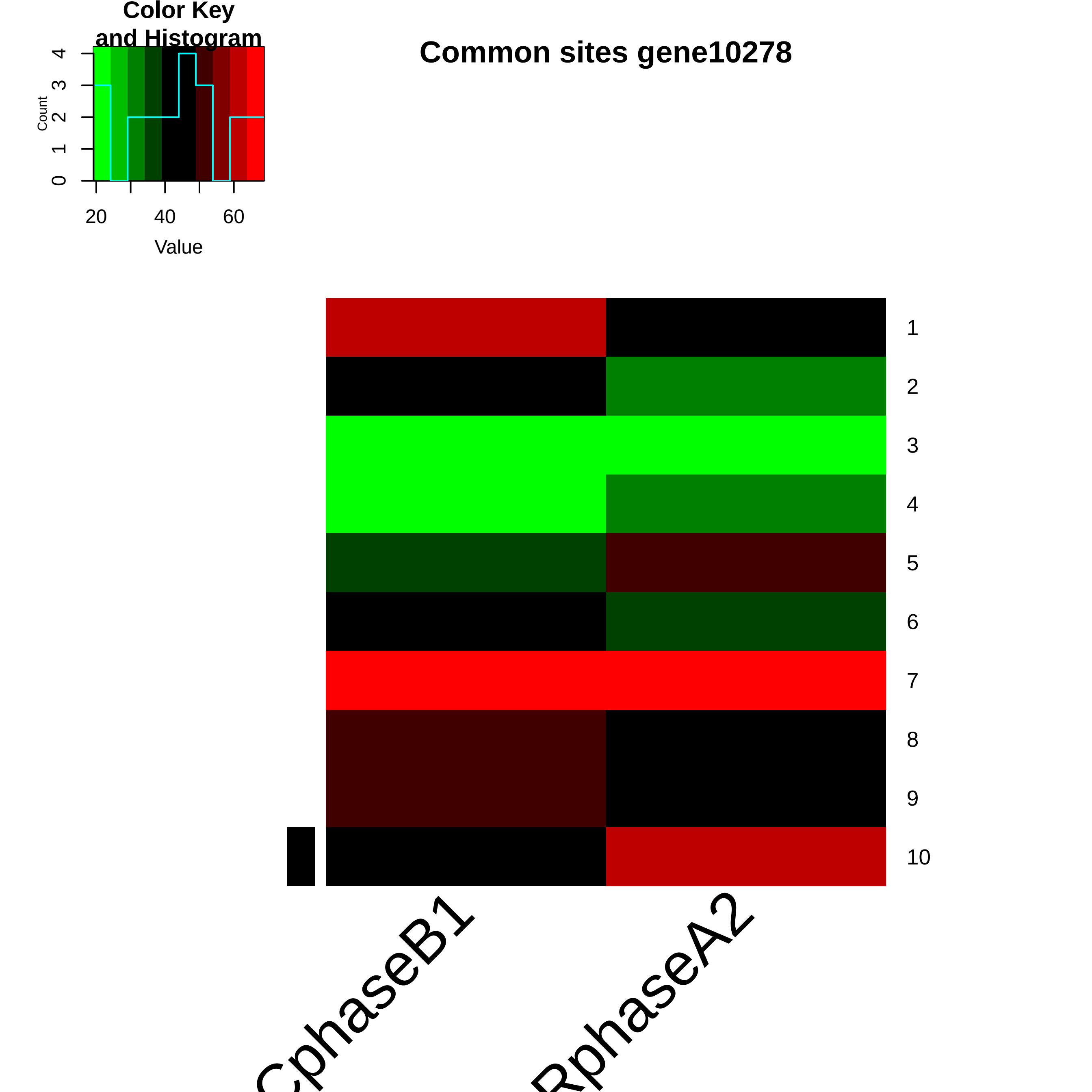
Color Key
and Histogram
Count
0
1
2
3
4
20
40
60
Value
Common sites gene10278
1
2
3
4
5
6
7
8
9
10
CphaseB1
RphaseA2

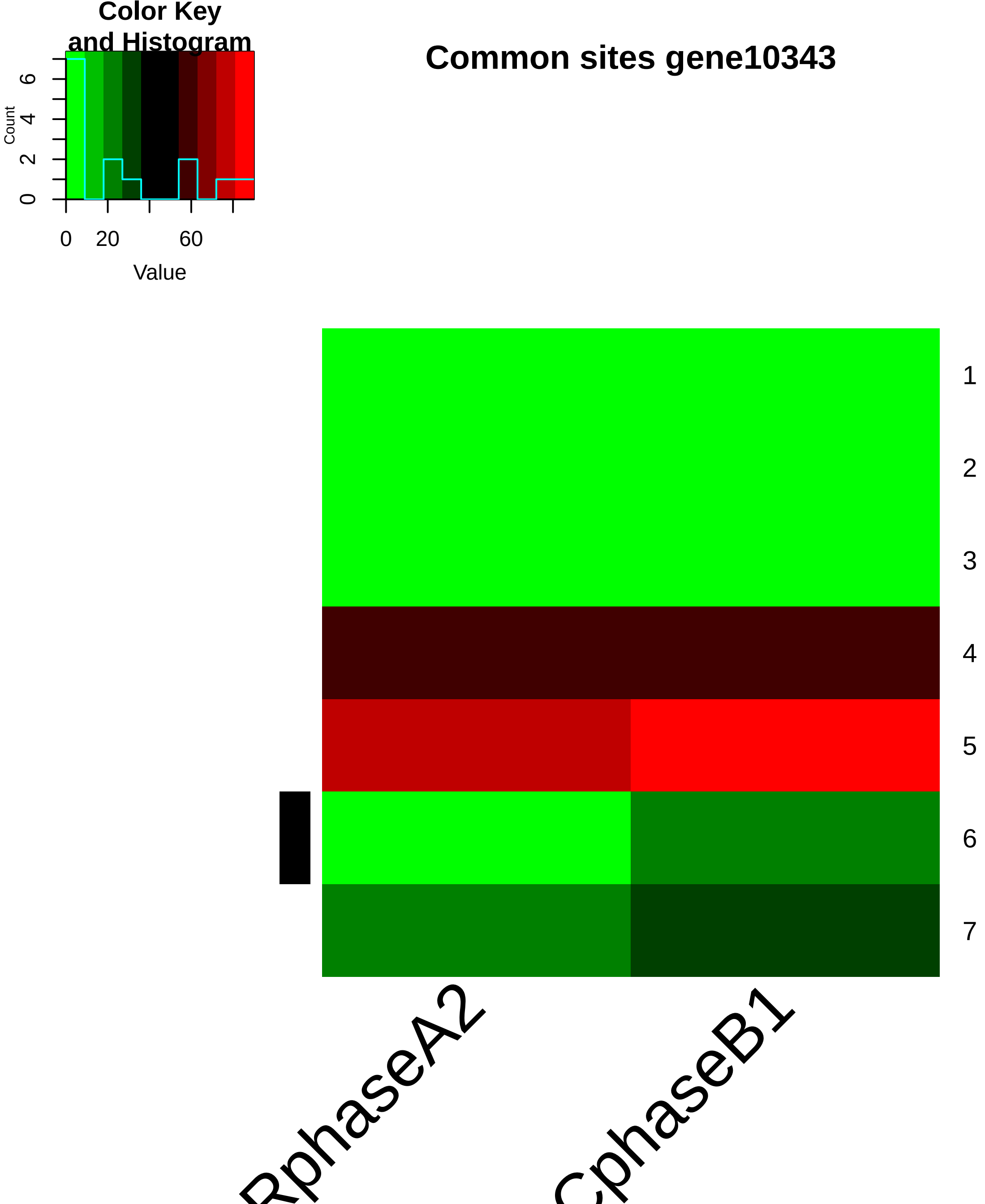
Color Key
and Histogram
Count
0
2
4
6
0
20
60
Value
Common sites gene10343
1
2
3
4
5
6
7
RphaseA2
CphaseB1

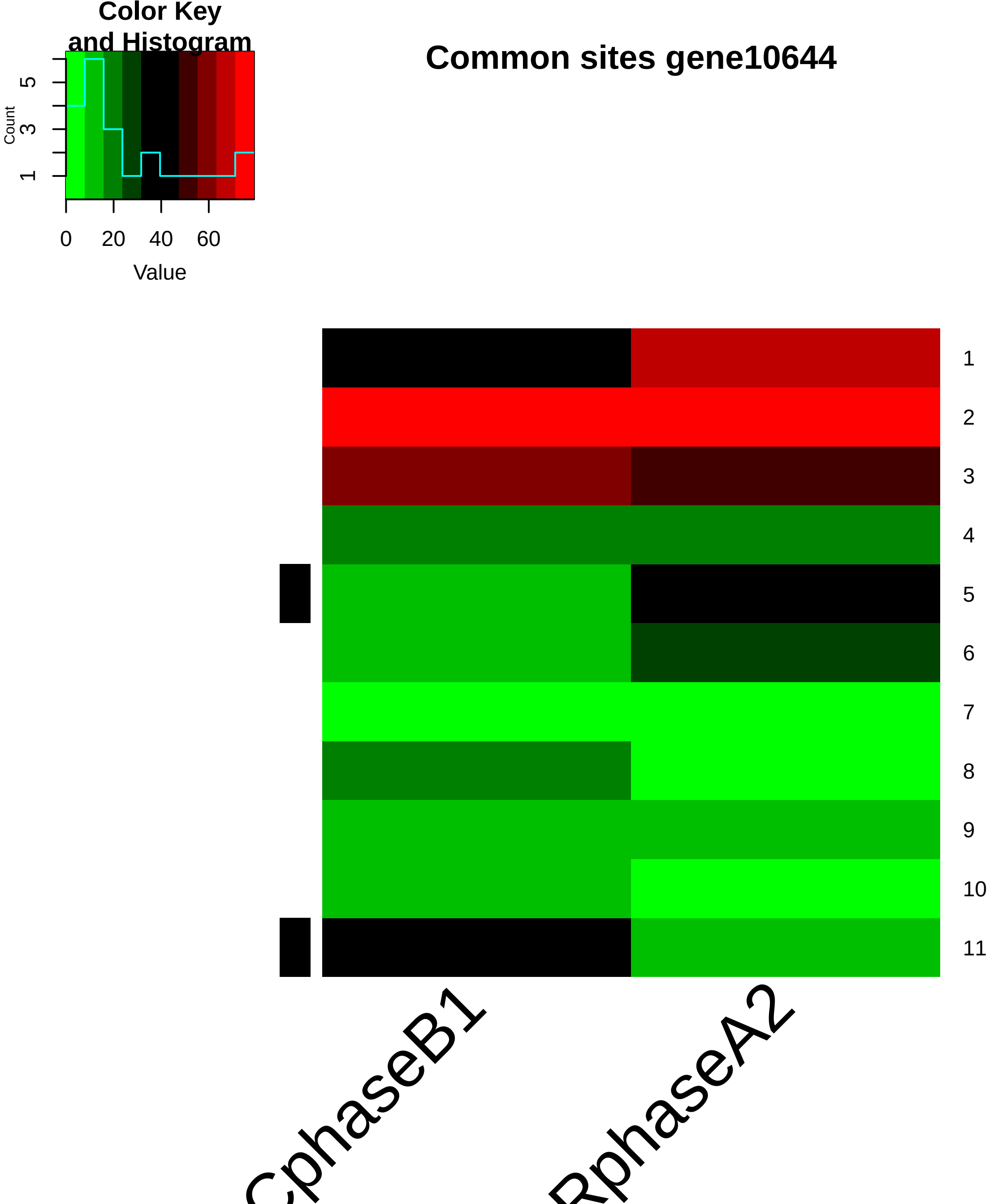
Color Key
and Histogram
Count
1
3
5
0
20
40
60
Value
Common sites gene10644
1
2
3
4
5
6
7
8
9
10
11
CphaseB1
RphaseA2

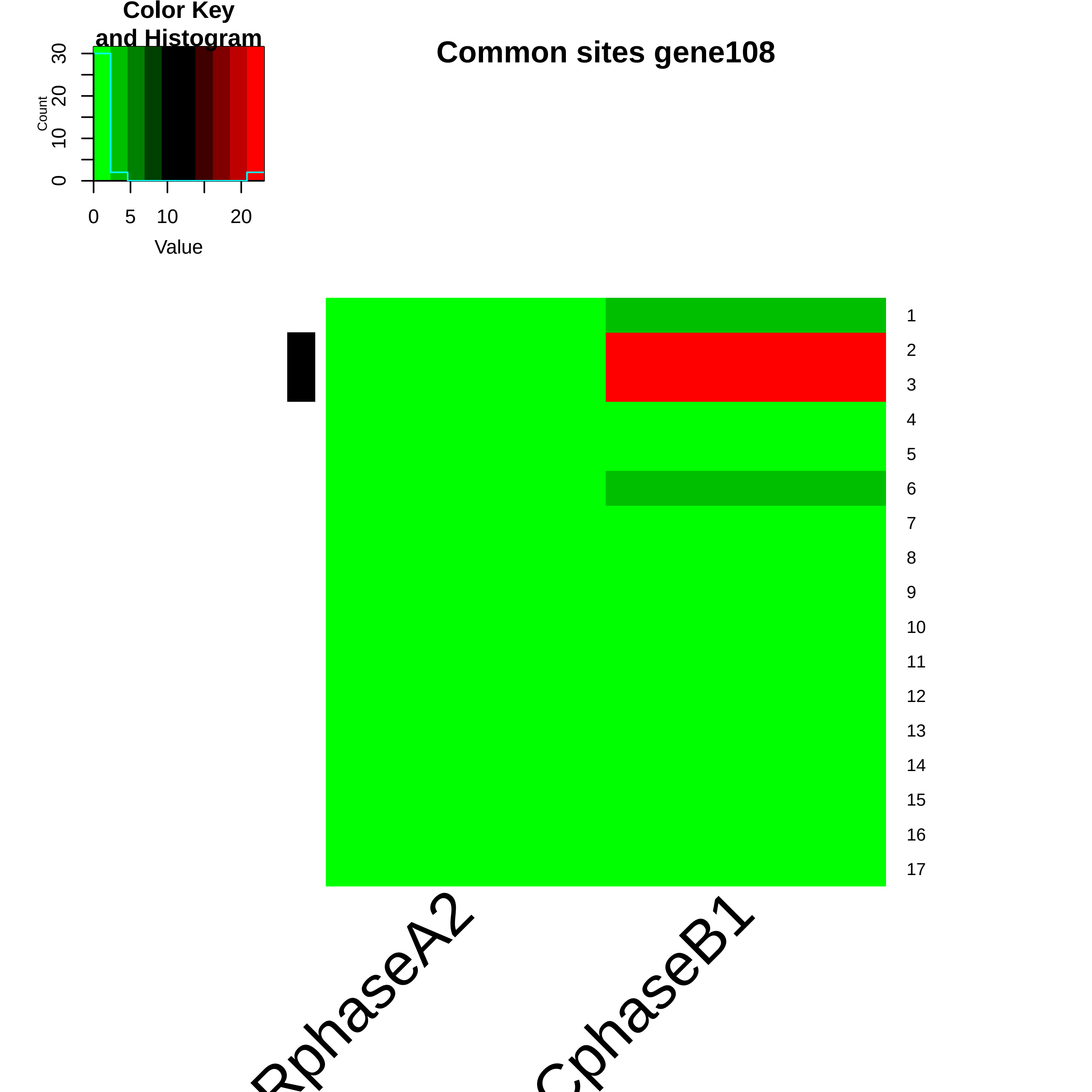
Color Key
and Histogram
Count
0
10
20
30
0
5
10
20
Value
Common sites gene108
RphaseA2
CphaseB1

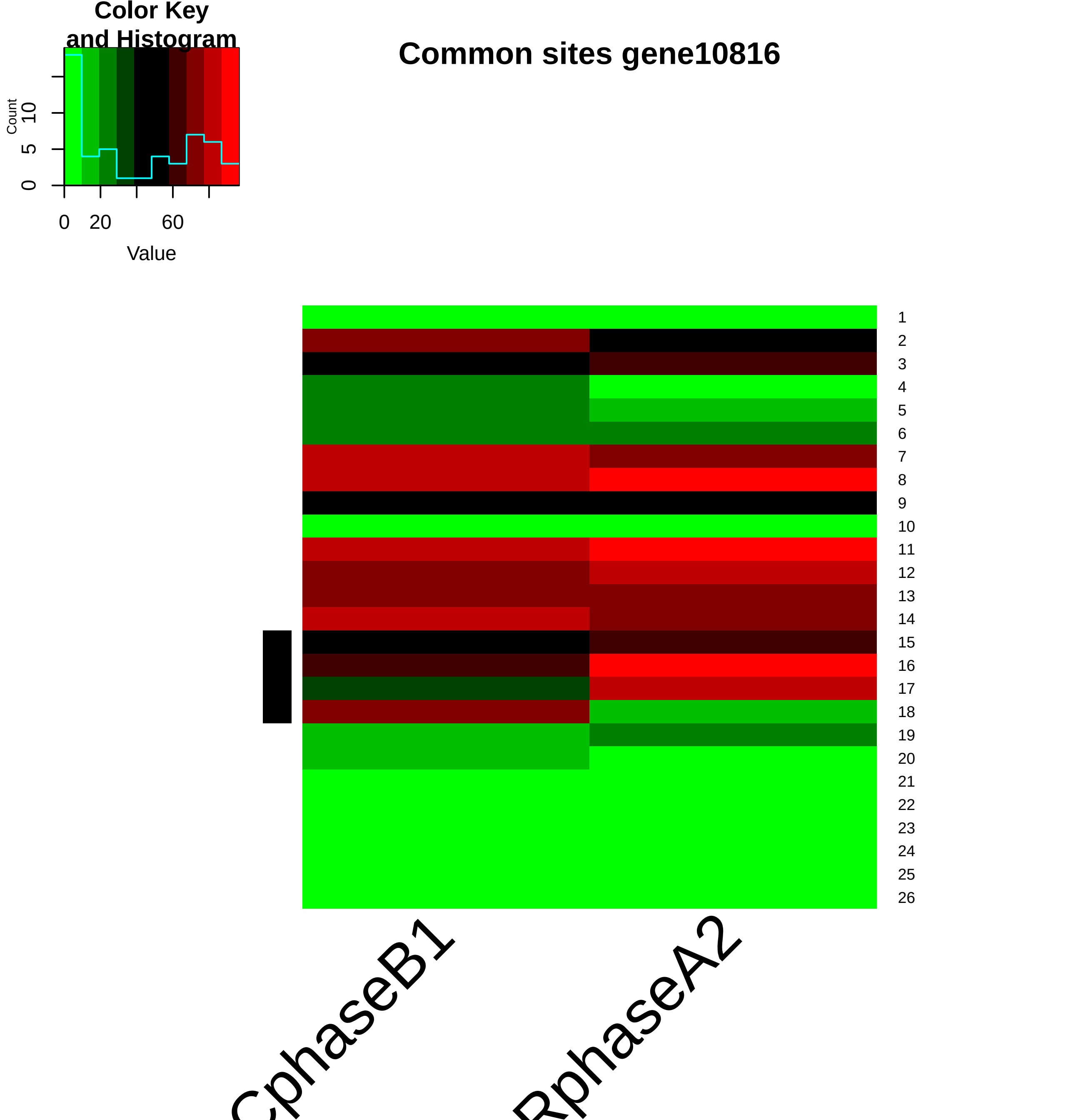
Color Key
and Histogram
Count
0
5
10
0
20
60
Value
Common sites gene10816
CphaseB1
RphaseA2

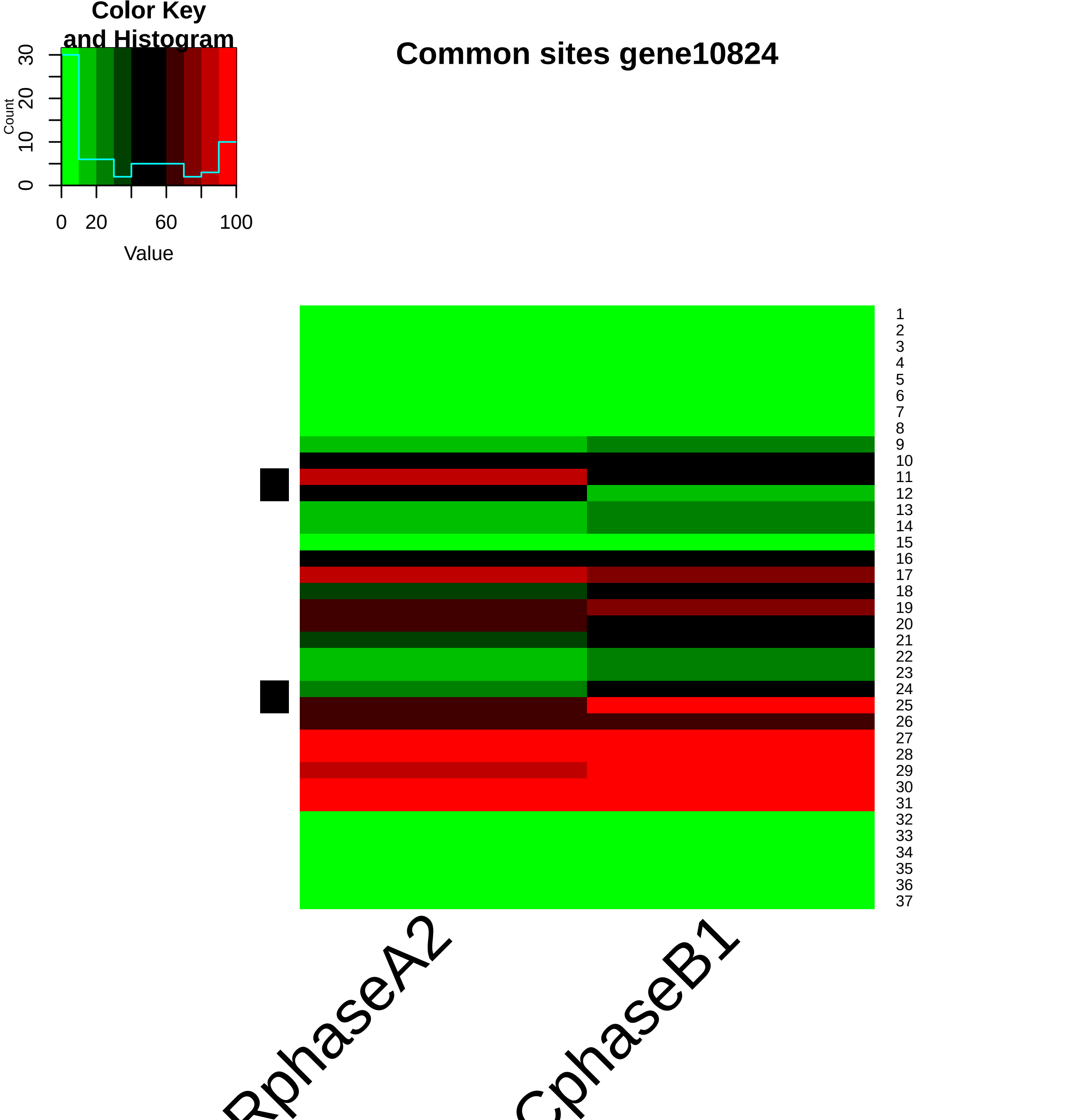
Color Key
and Histogram
Count
0
10
20
30
0
20
60
100
Value
Common sites gene10824
RphaseA2
CphaseB1

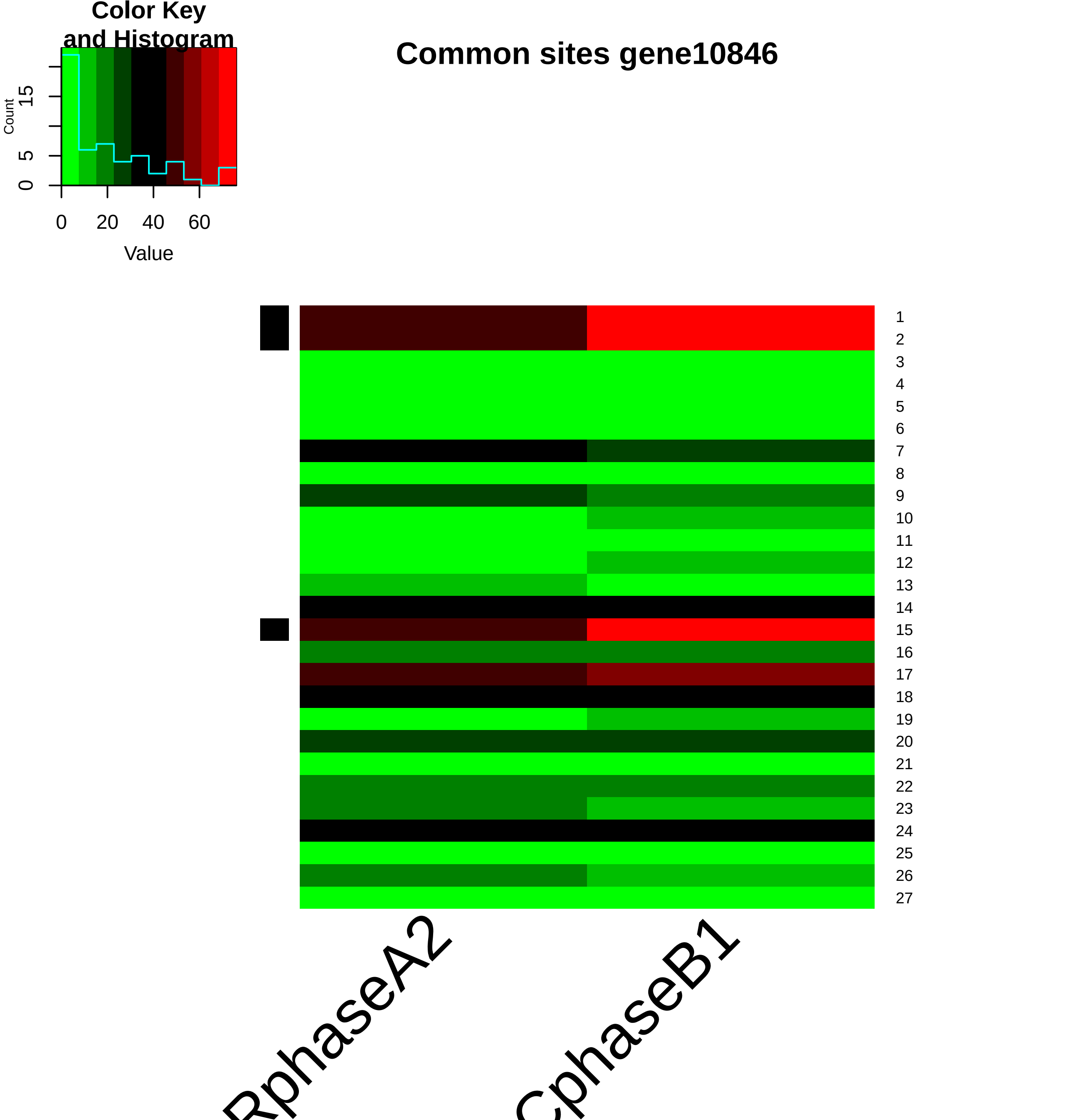

Color Key and Histogram
Count
0
5
15
0
20
40
60
Value
Common sites gene10846
RphaseA2
CphaseB1

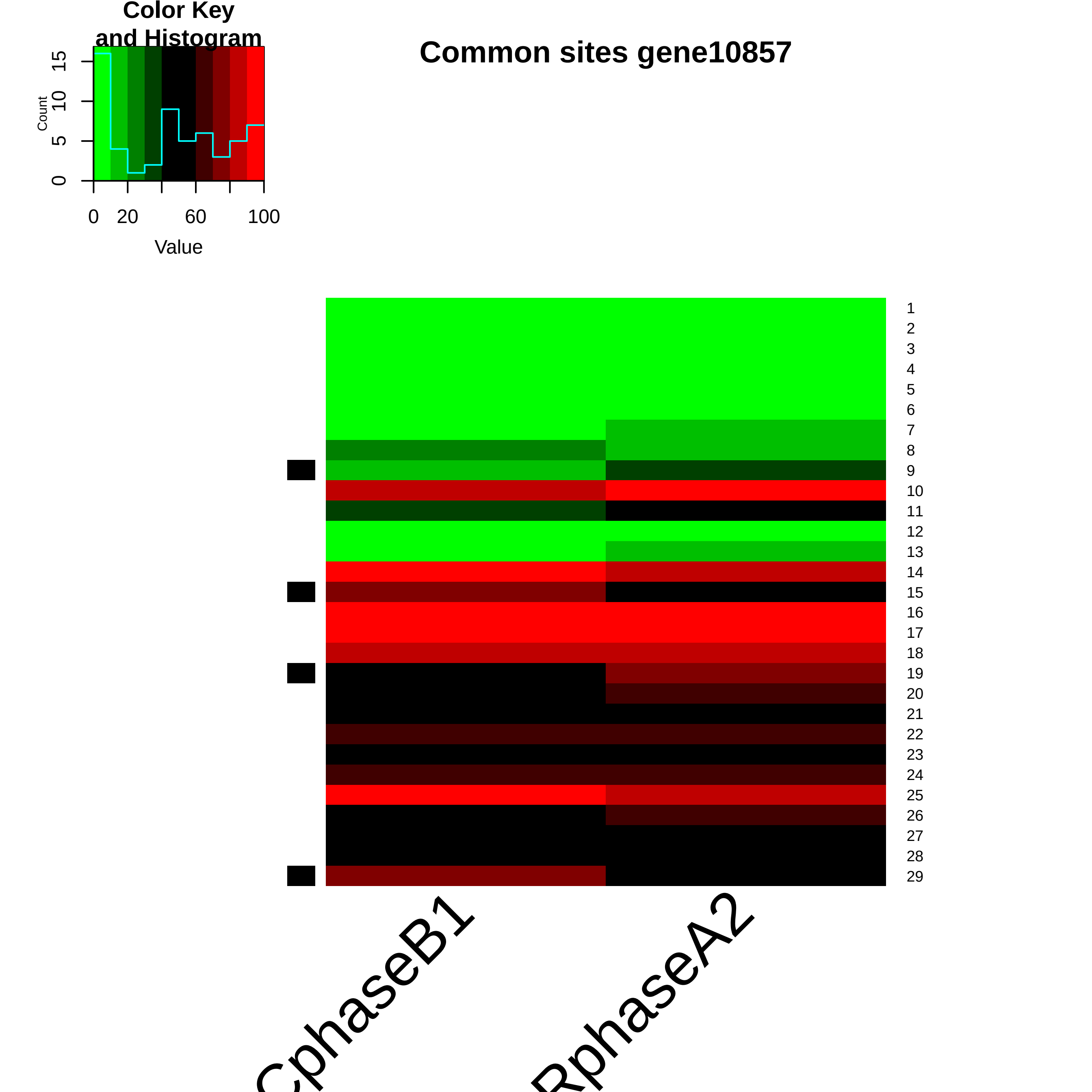
Color Key
and Histogram
Count
0
5
10
15
0
20
60
100
Value
Common sites gene10857
CphaseB1
RphaseA2

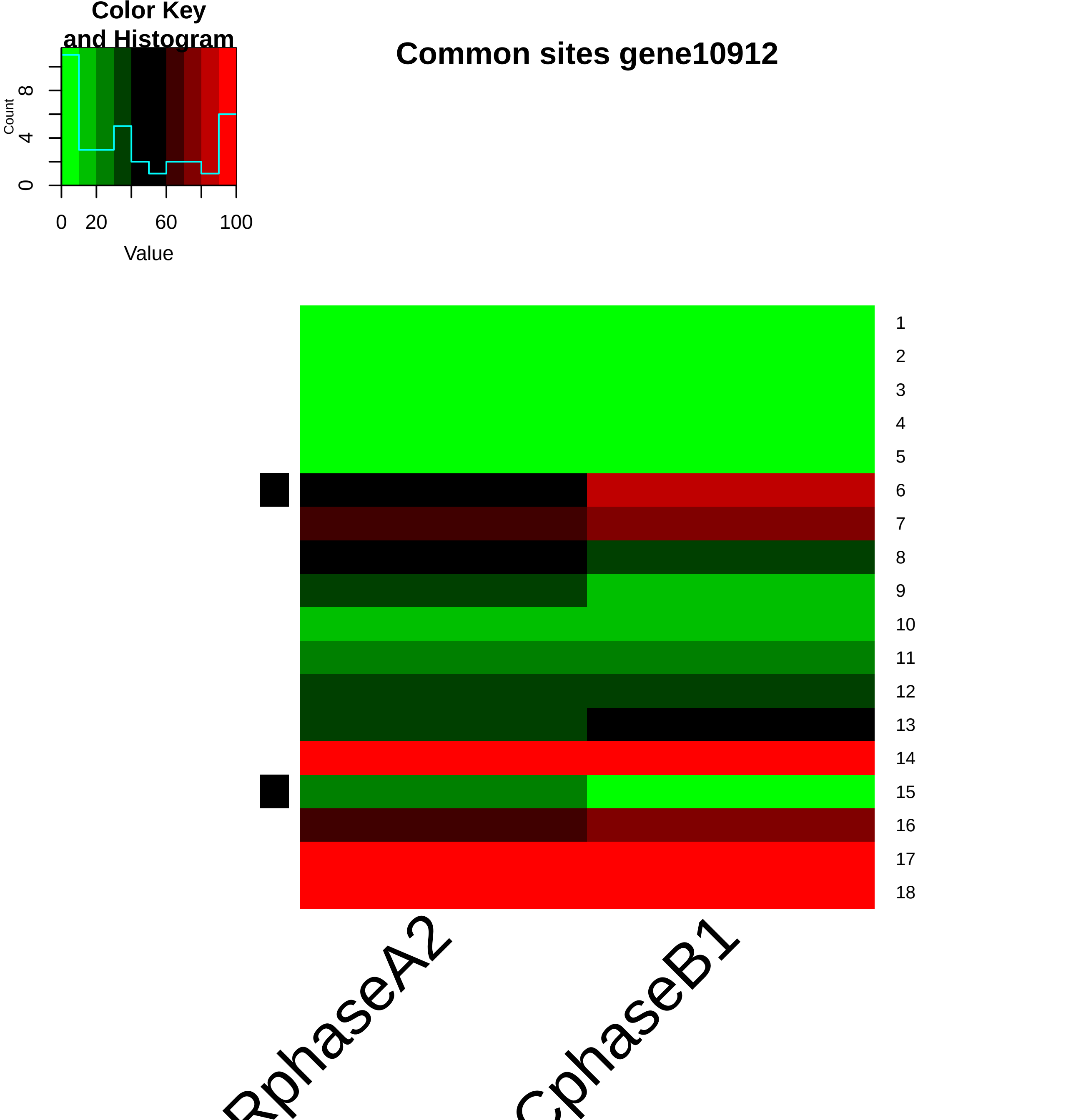
Color Key
and Histogram
Count
0
4
8
0
20
60
100
Value
Common sites gene10912
RphaseA2
CphaseB1

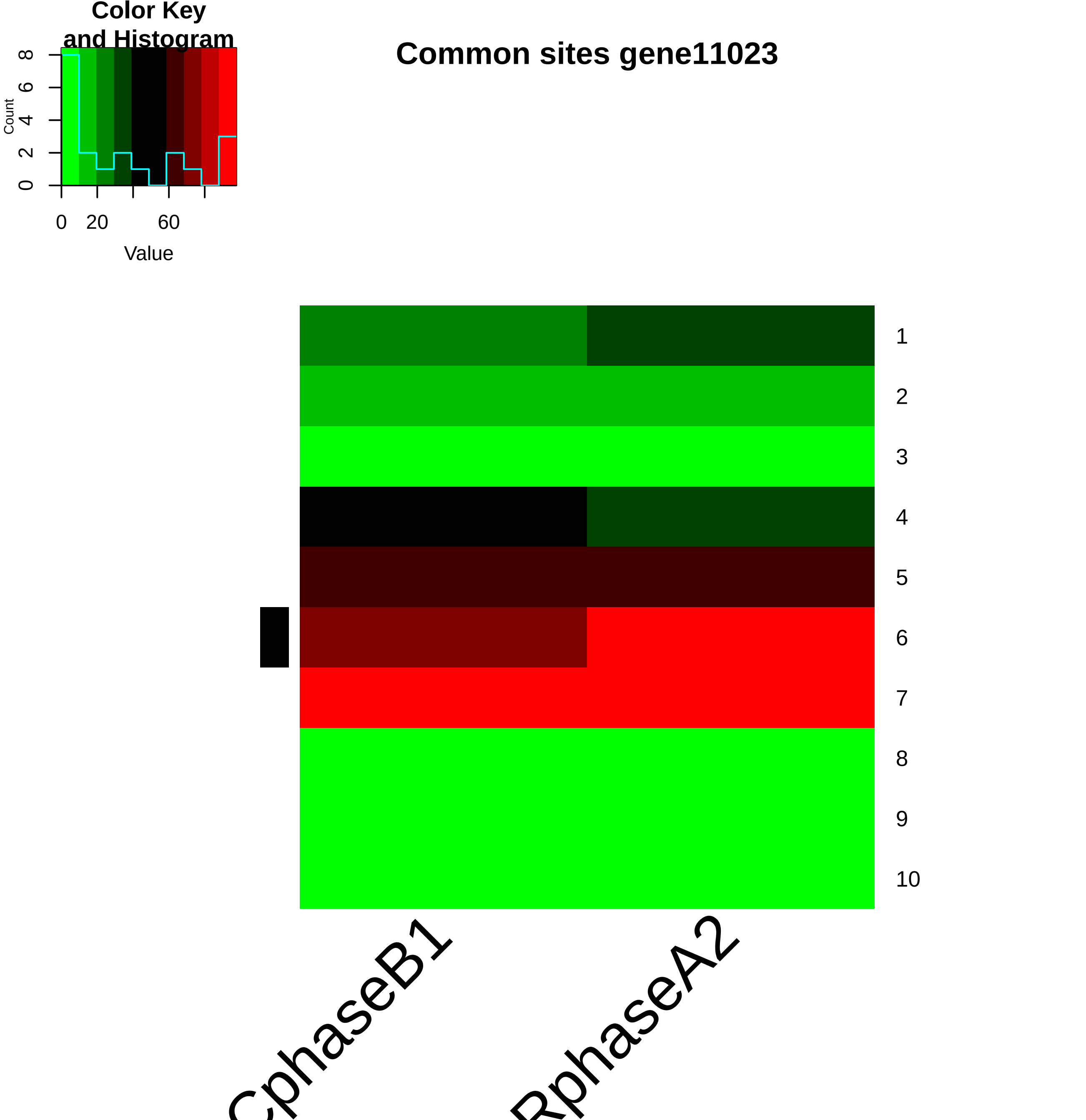
Color Key
and Histogram
Count
0
2
4
6
8
0
20
60
Value
Common sites gene11023
1
2
3
4
5
6
7
8
9
10
CphaseB1
RphaseA2

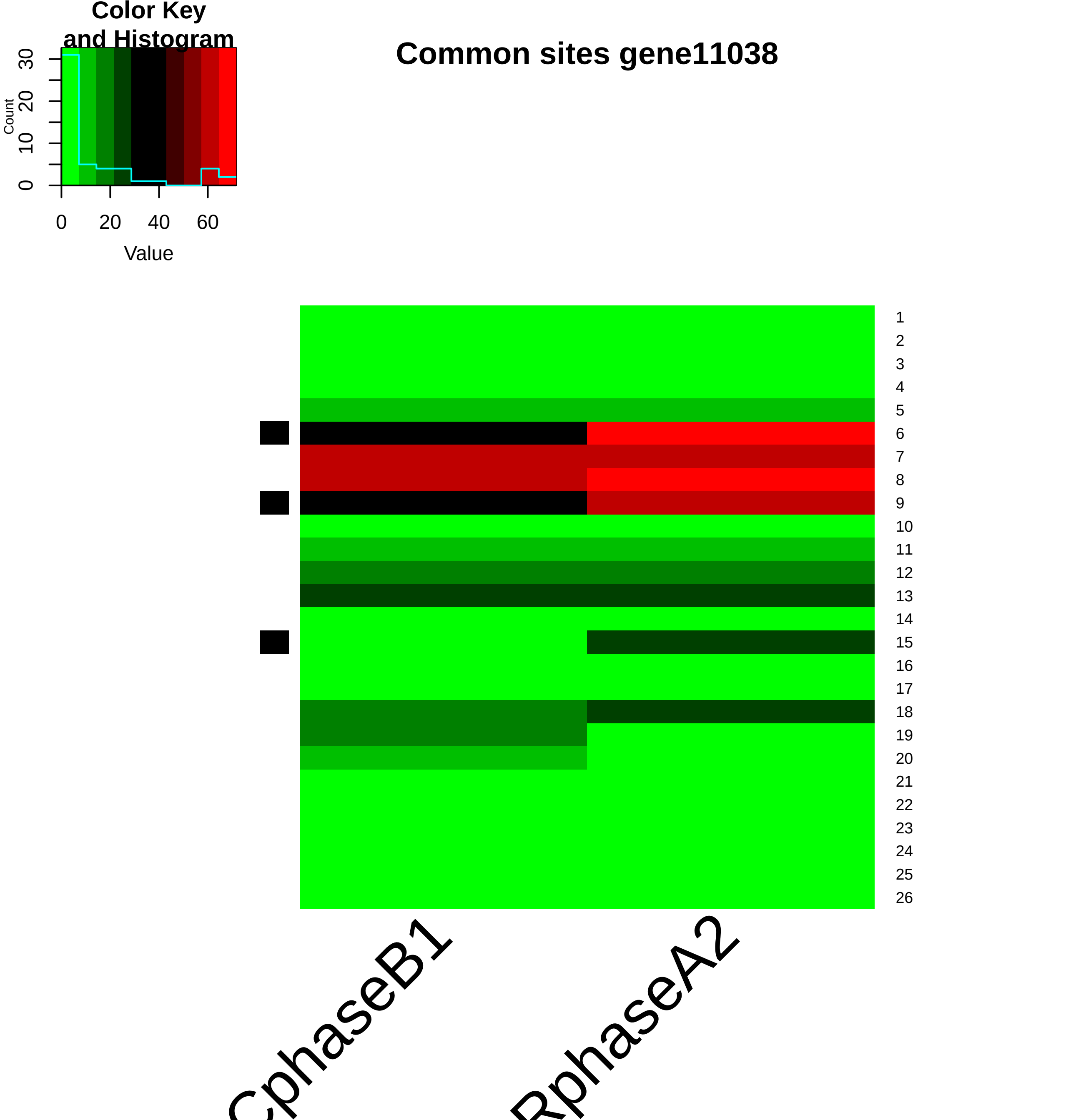
Color Key
and Histogram
Count
0
10
20
30
0
20
40
60
Value
Common sites gene11038
CphaseB1
RphaseA2

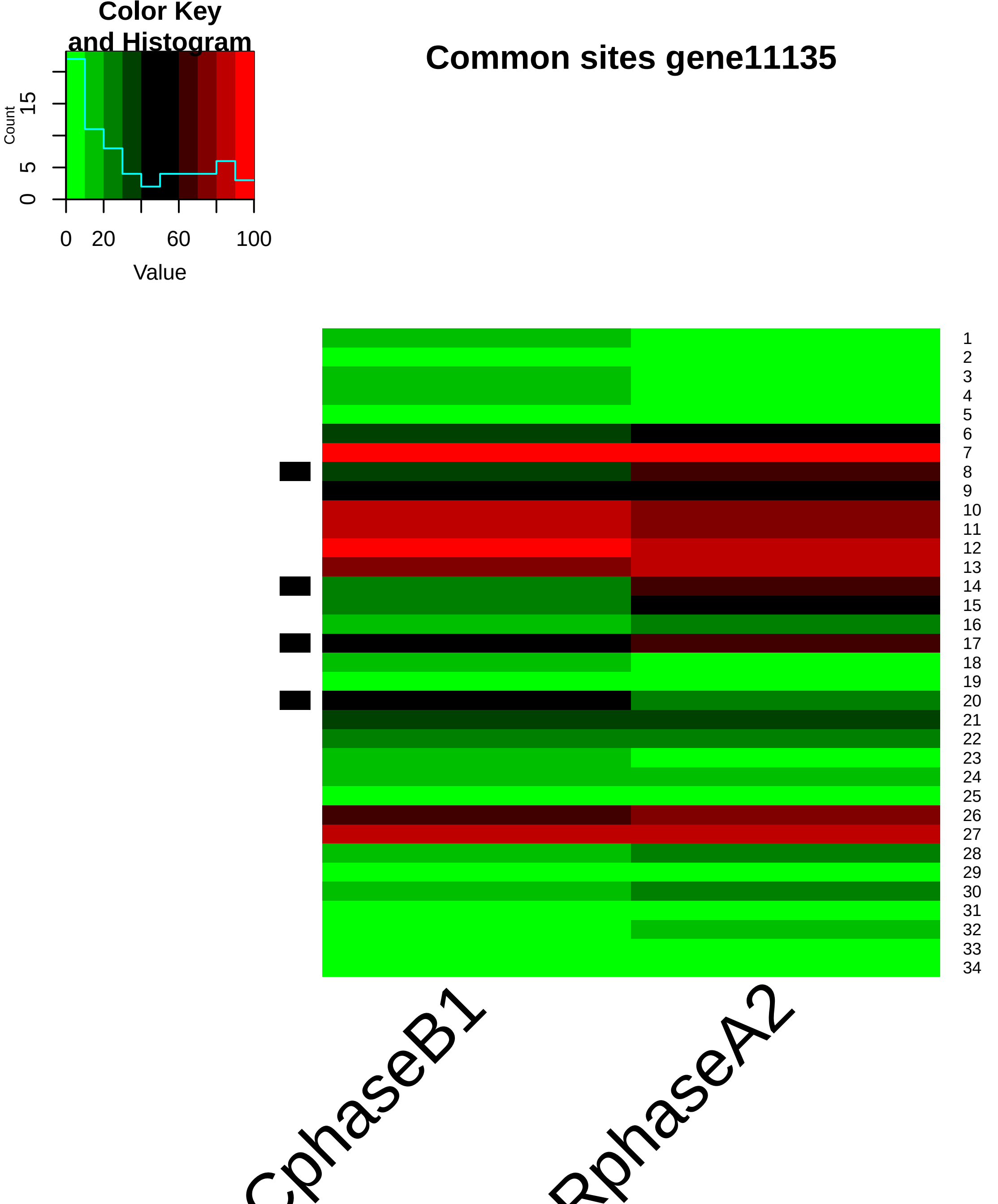
Color Key
and Histogram
Count
0
5
15
0
20
60
100
Value
Common sites gene11135
CphaseB1
RphaseA2

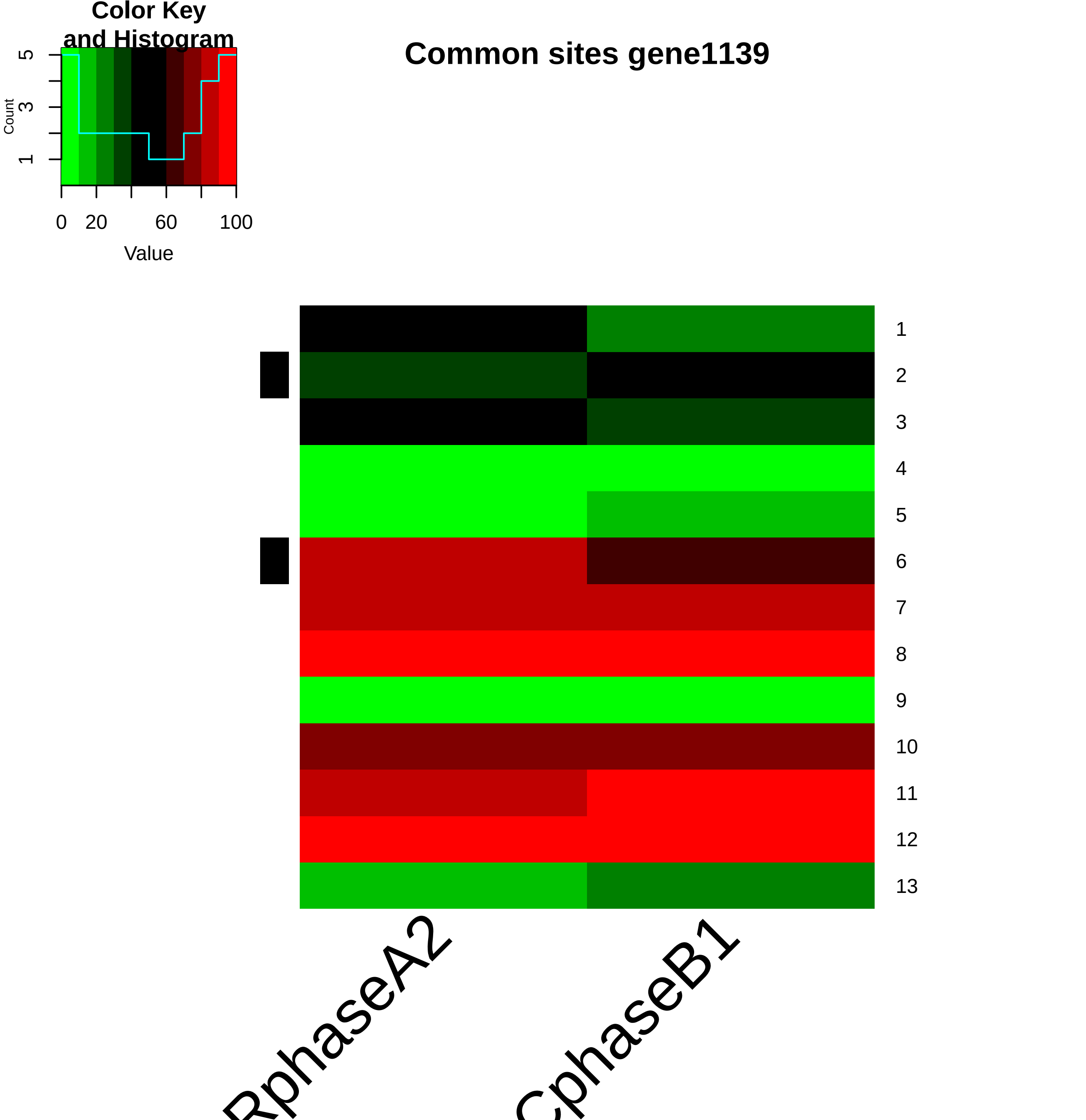

Color Key
and Histogram
Count
1
3
5
0
20
60
100
Value
Common sites gene1139
1
2
3
4
5
6
7
8
9
10
11
12
13
RphaseA2
CphaseB1

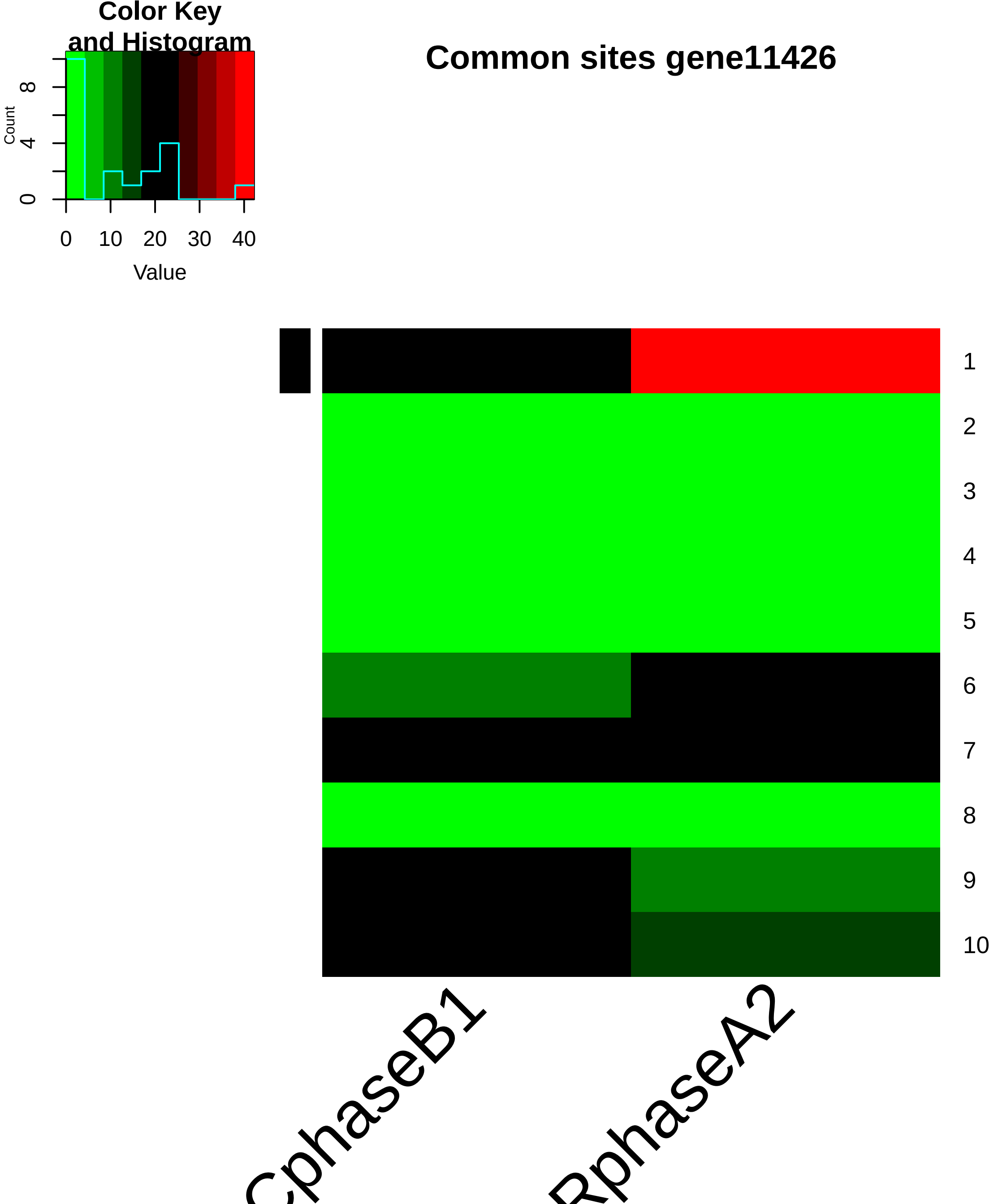
Color Key
and Histogram
Count
0
4
8
0
10
20
30
40
Value
Common sites gene11426
1
2
3
4
5
6
7
8
9
10
CphaseB1
RphaseA2

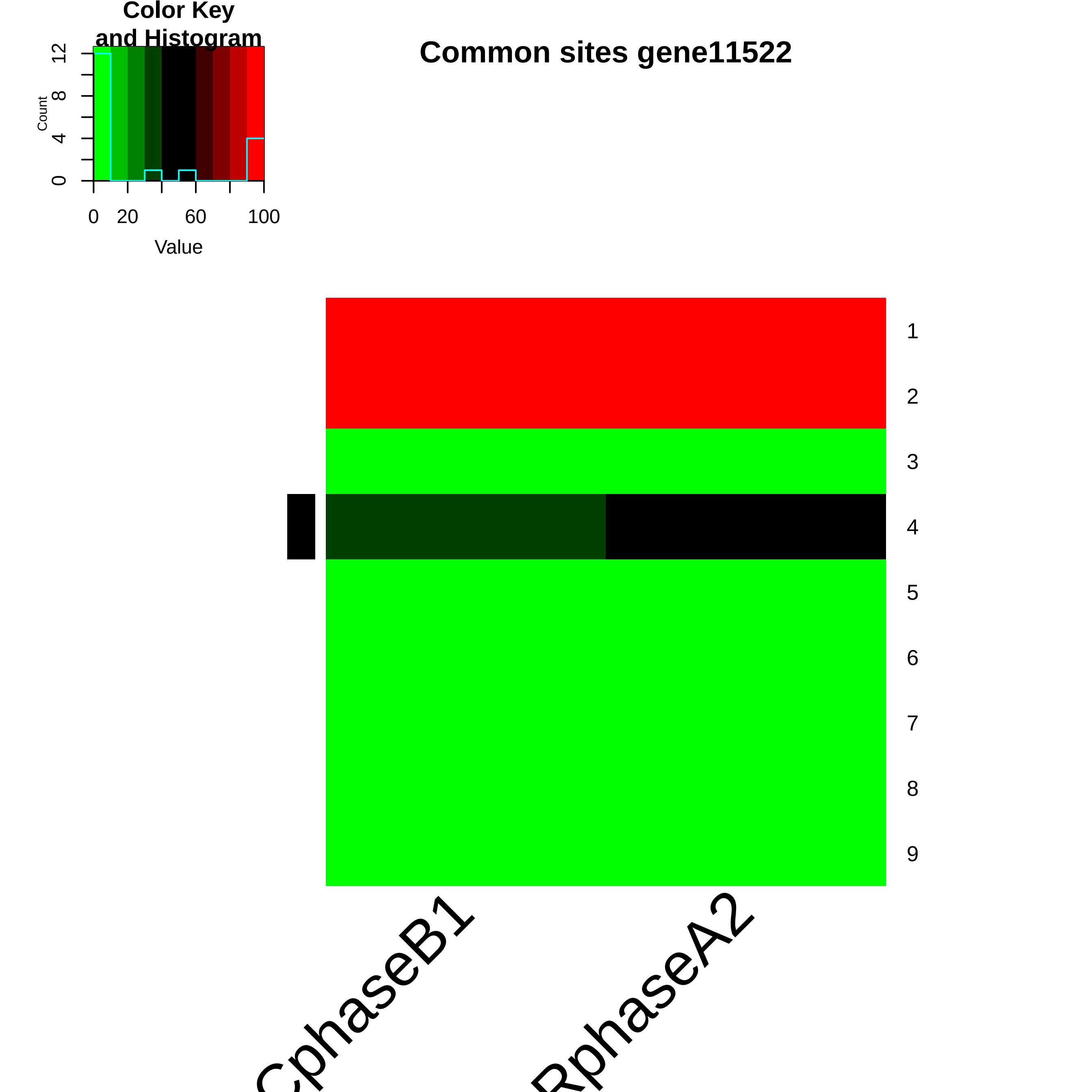

Color Key
and Histogram
Count
0
4
8
12
0
20
60
100
Value
Common sites gene11522
1
2
3
4
5
6
7
8
9
CphaseB1
RphaseA2

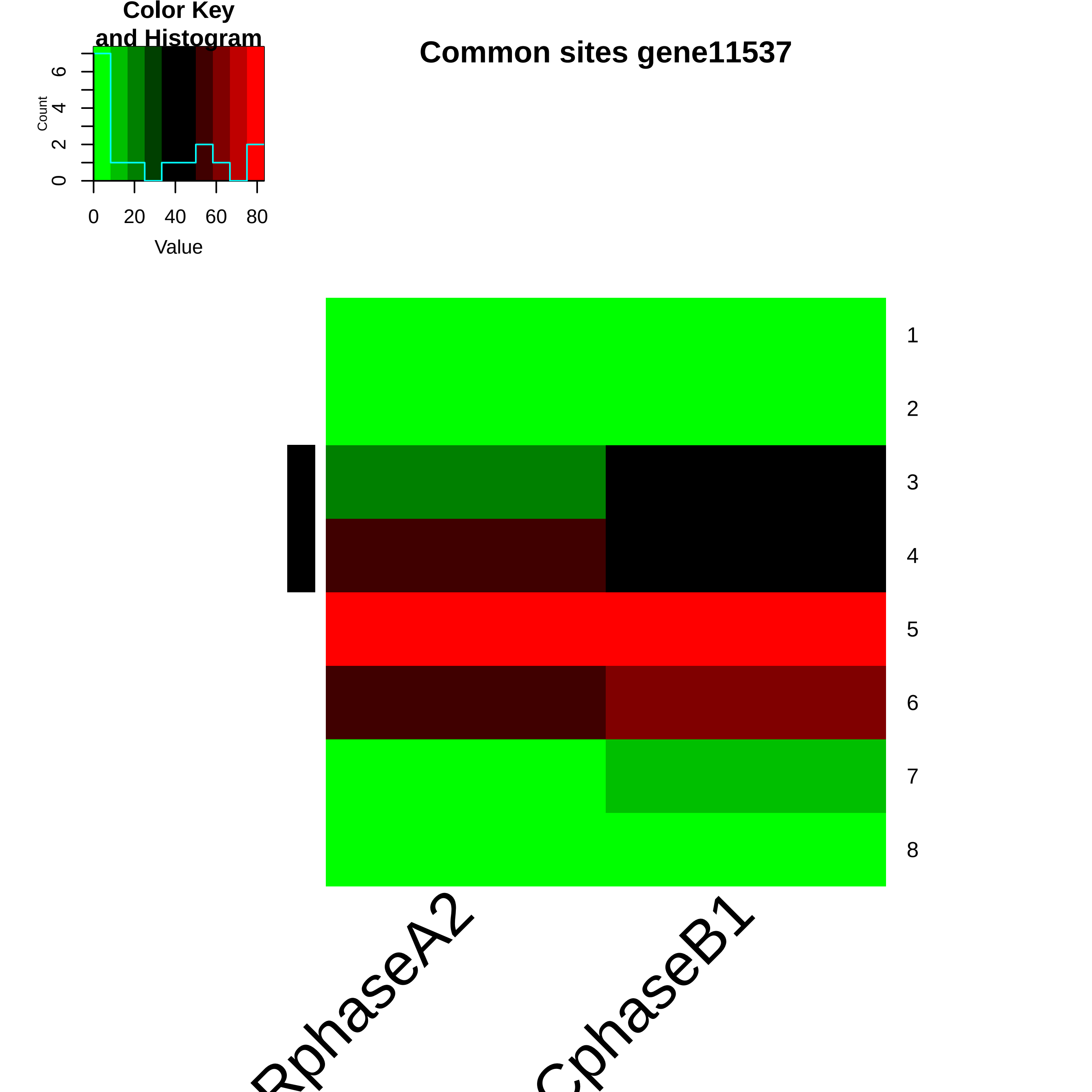
Color Key
and Histogram
Count
Value
0
20
40
60
80
0
2
4
6
Common sites gene11537
1
2
3
4
5
6
7
8
RphaseA2
CphaseB1

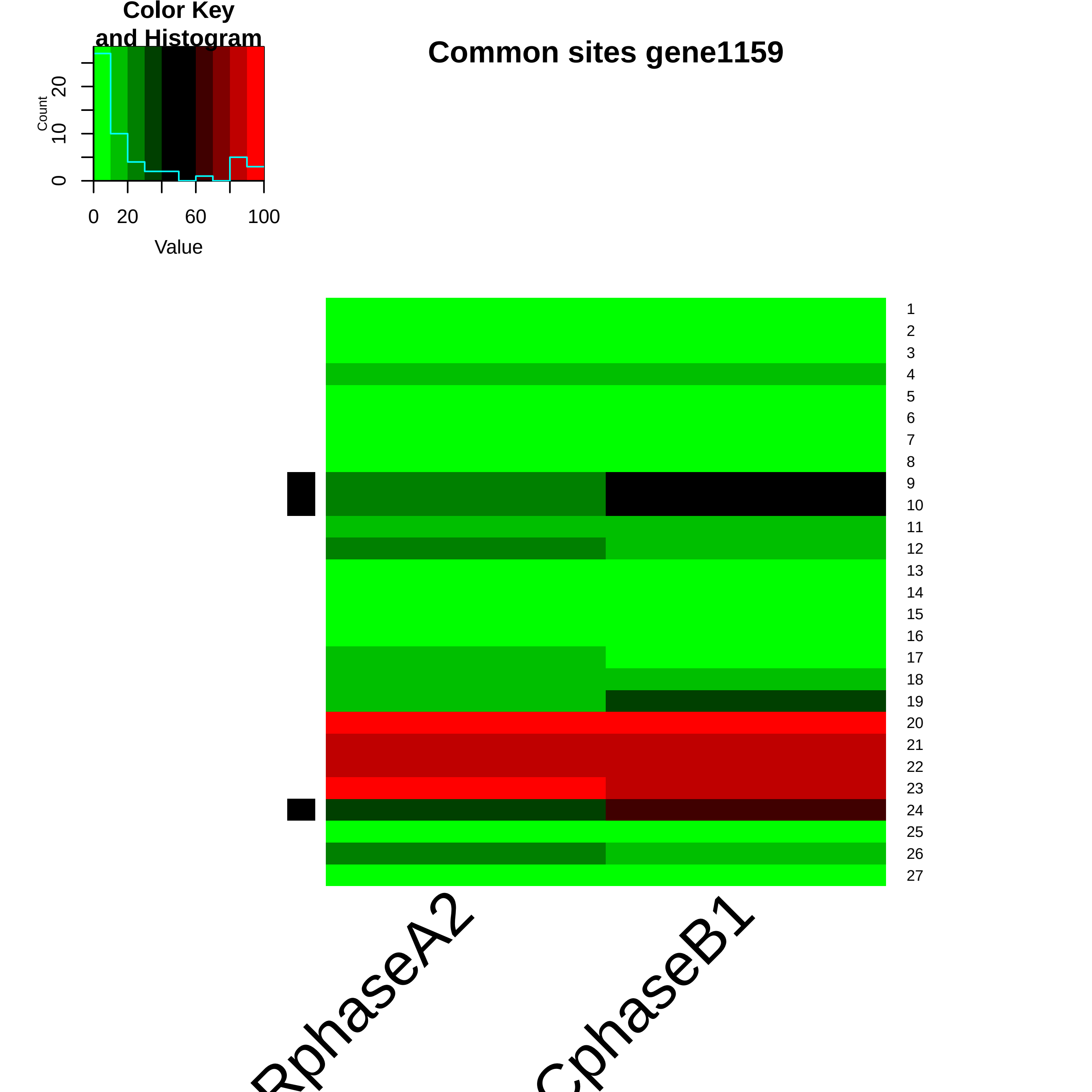

Color Key
and Histogram
Count
0
10
20
0
20
60
100
Value
Common sites gene1159
RphaseA2
CphaseB1

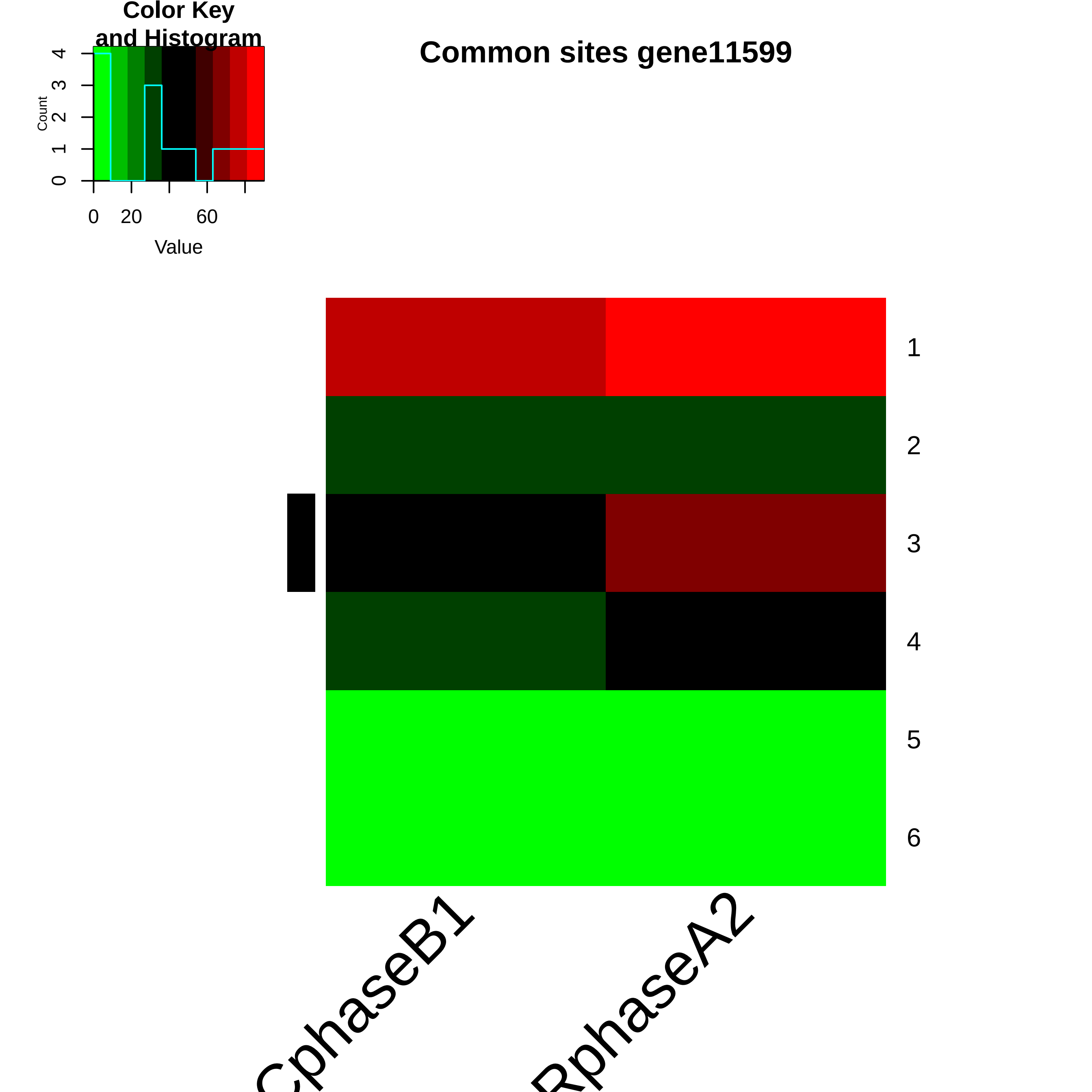

Color Key
and Histogram
Count
Value
0
20
60
0
1
2
3
4
Common sites gene11599
1
2
3
4
5
6
CphaseB1
RphaseA2

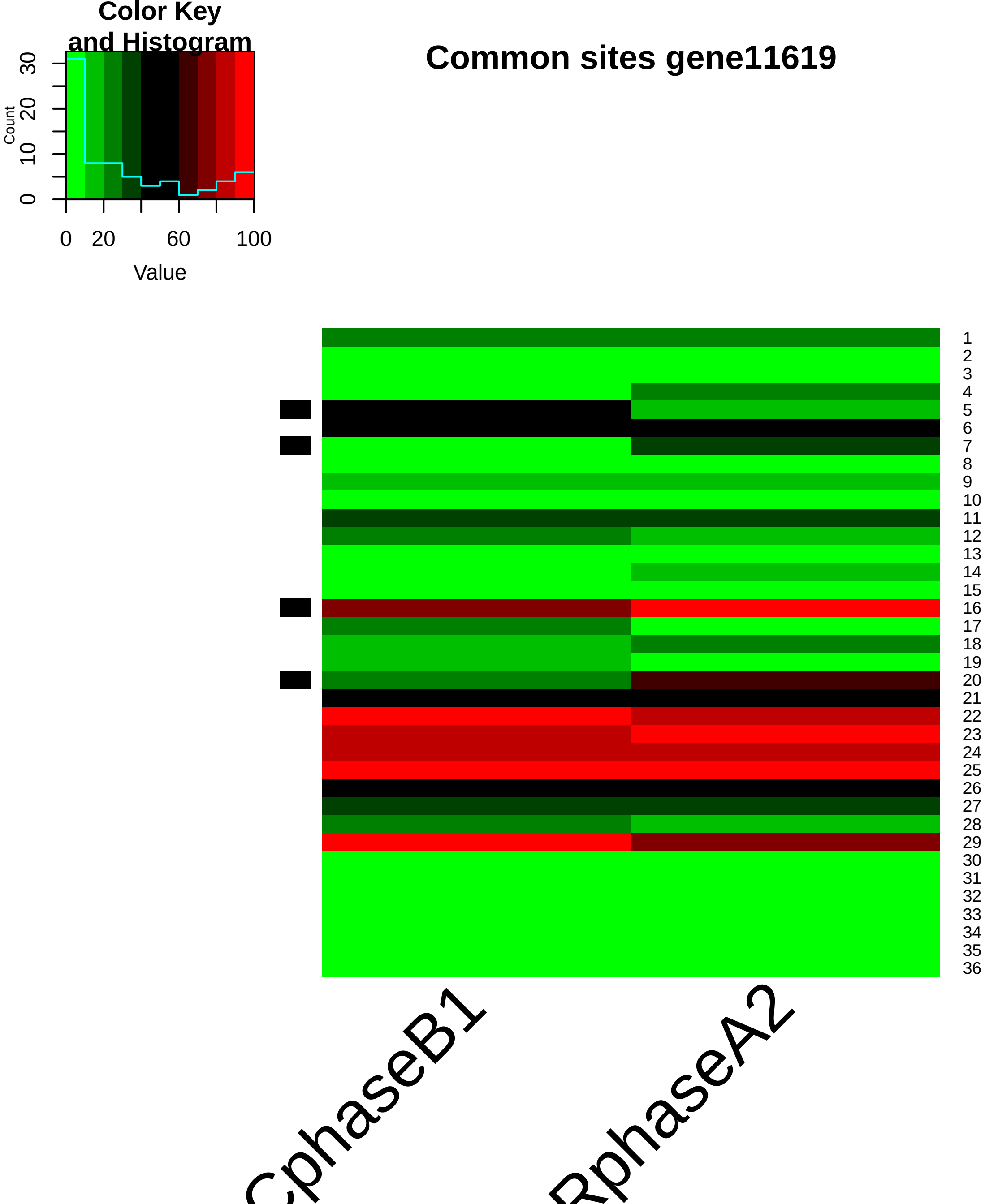
Color Key
and Histogram
Count
0
10
20
30
0
20
60
100
Value
Common sites gene11619
CphaseB1
RphaseA2

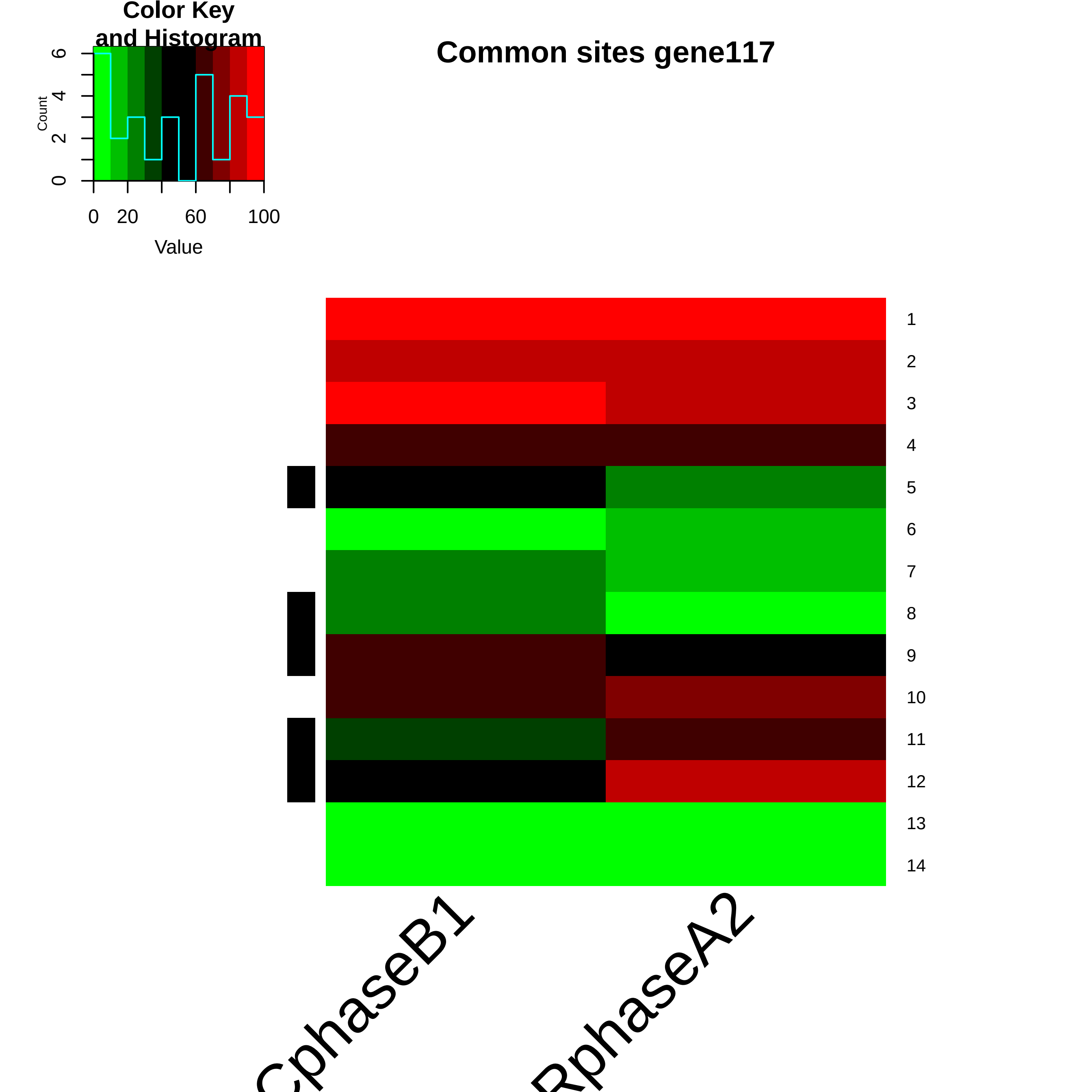
Color Key
and Histogram
Count
Value
0
20
60
100
0
2
4
6
Common sites gene117
1
2
3
4
5
6
7
8
9
10
11
12
13
14
CphaseB1
RphaseA2

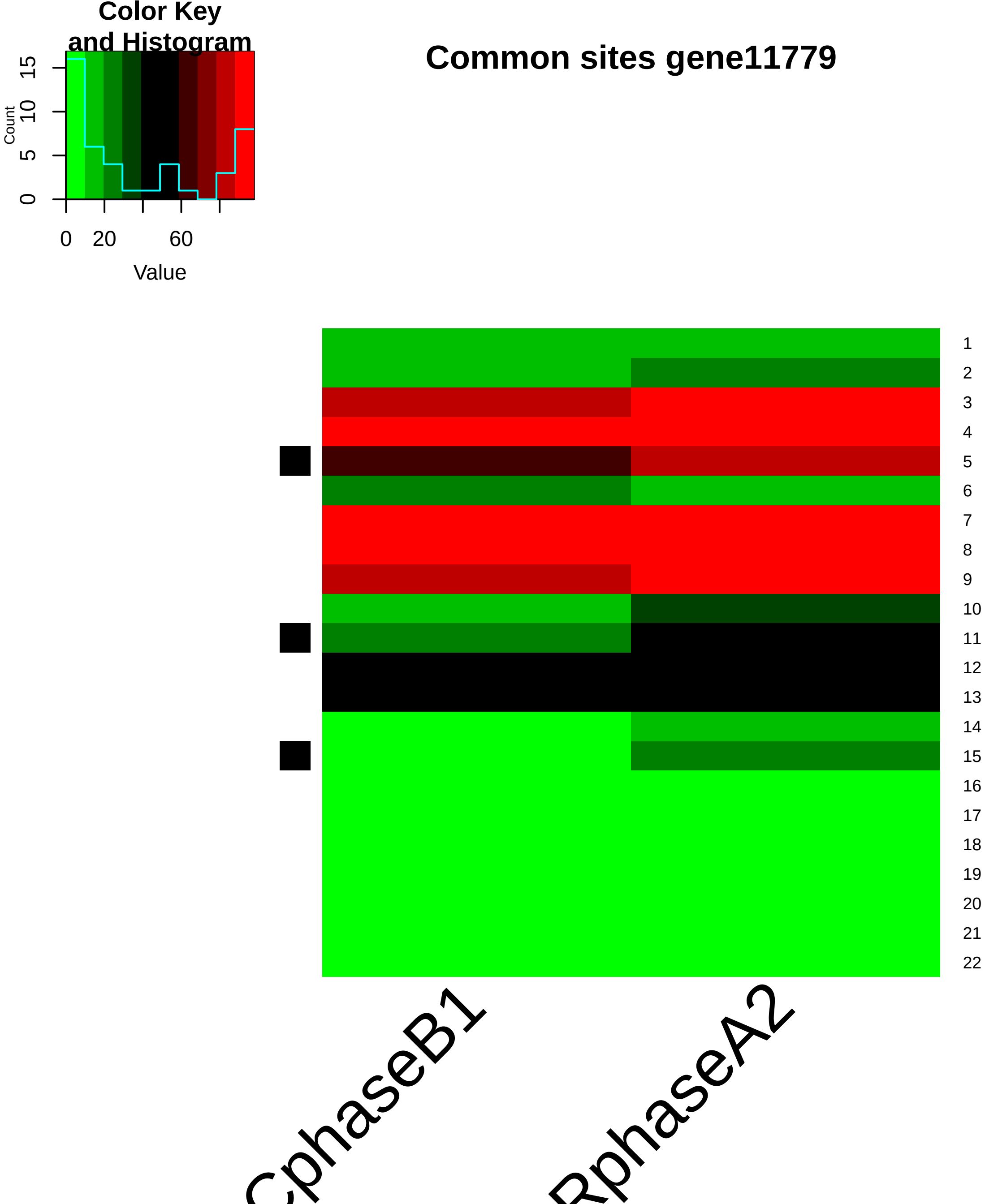

Color Key
and Histogram
Count
0
5
10
15
0
20
60
Value
Common sites gene11779
CphaseB1
RphaseA2

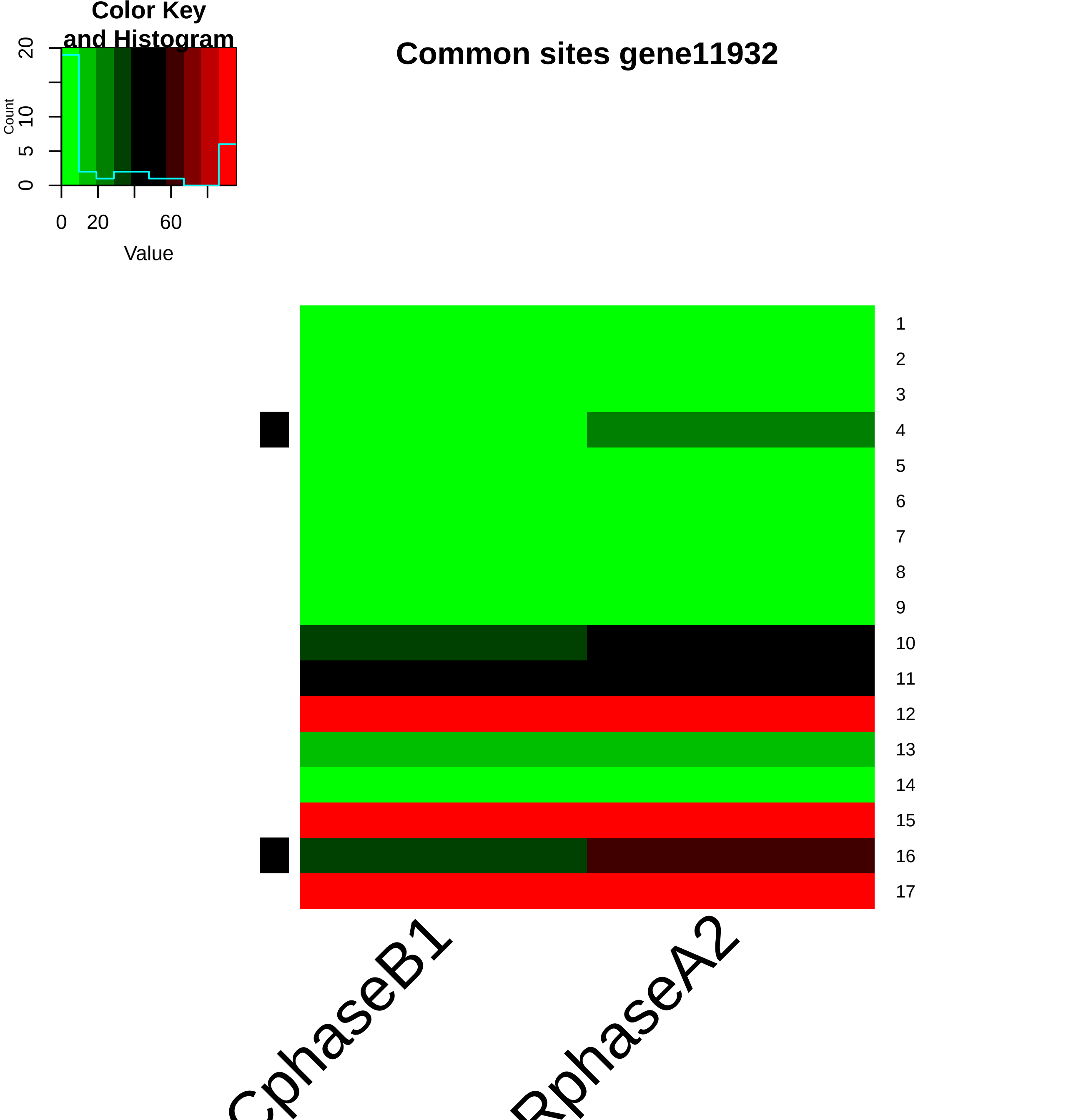
Color Key
and Histogram
Count
0
5
10
20
0
20
60
Value
Common sites gene11932
CphaseB1
RphaseA2

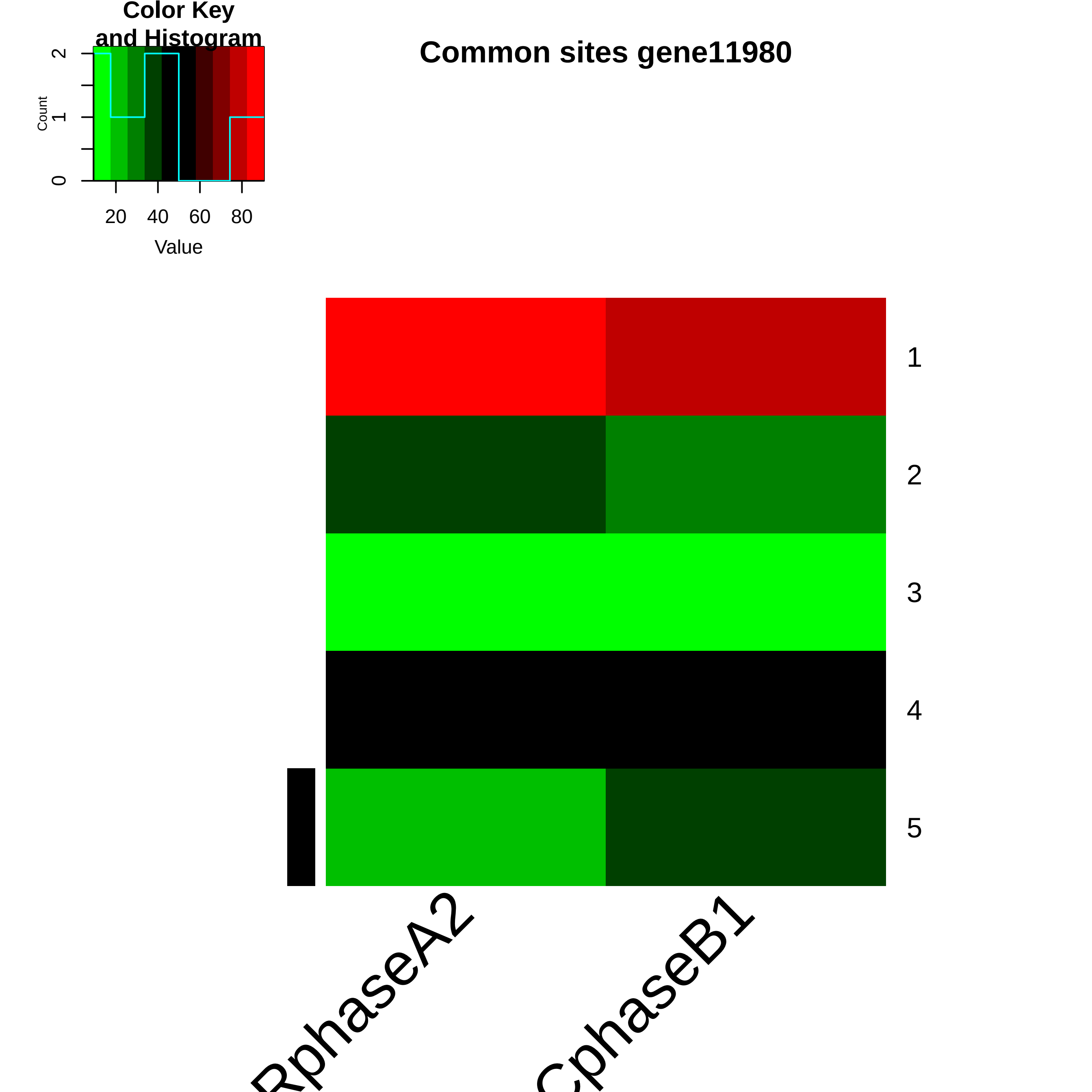

Color Key
and Histogram
Count
0
1
2
20
40
60
80
Value
Common sites gene11980
1
2
3
4
5
RphaseA2
CphaseB1

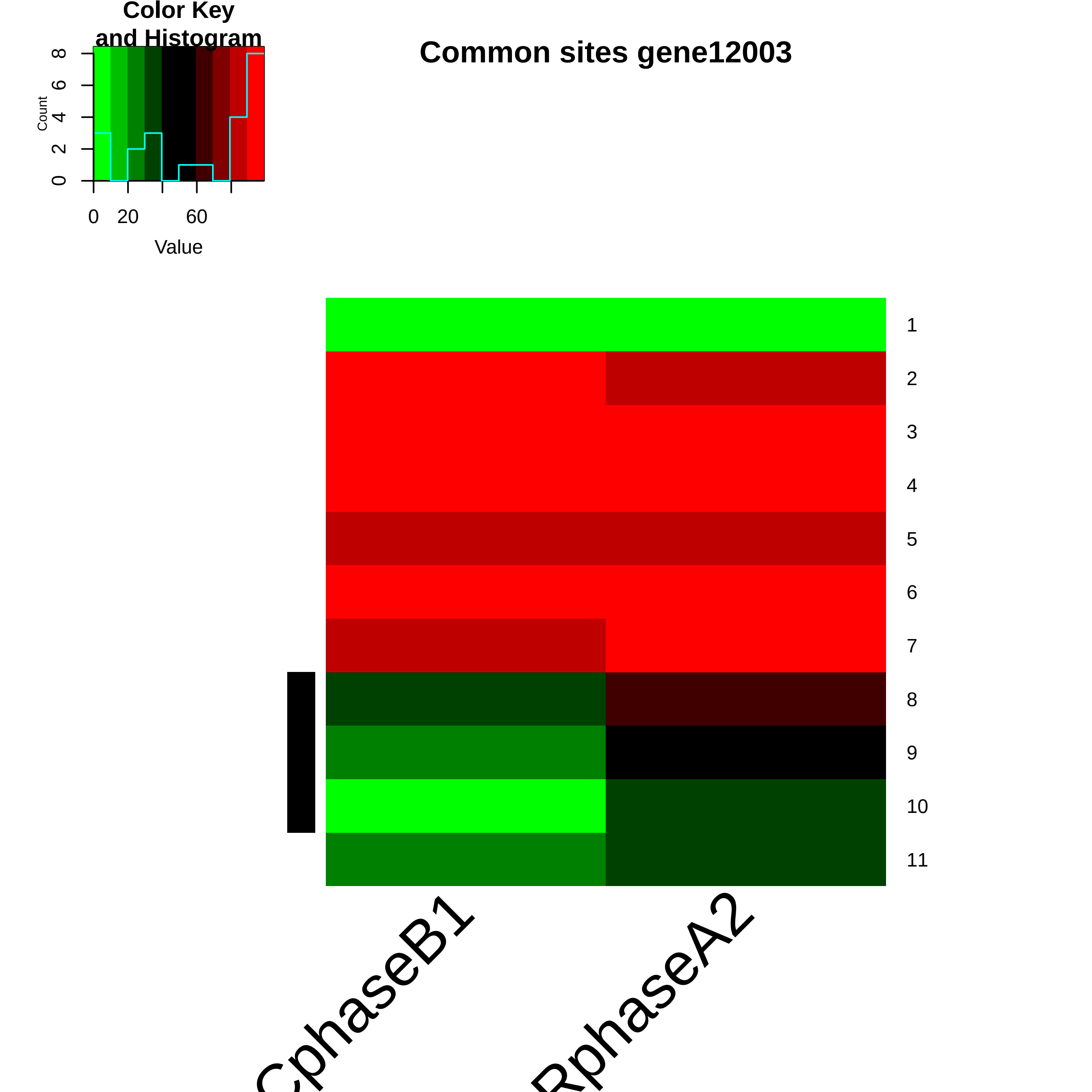

Color Key
and Histogram
Count
0
2
4
6
8
0
20
60
Value
Common sites gene12003
1
2
3
4
5
6
7
8
9
10
11
CphaseB1
RphaseA2

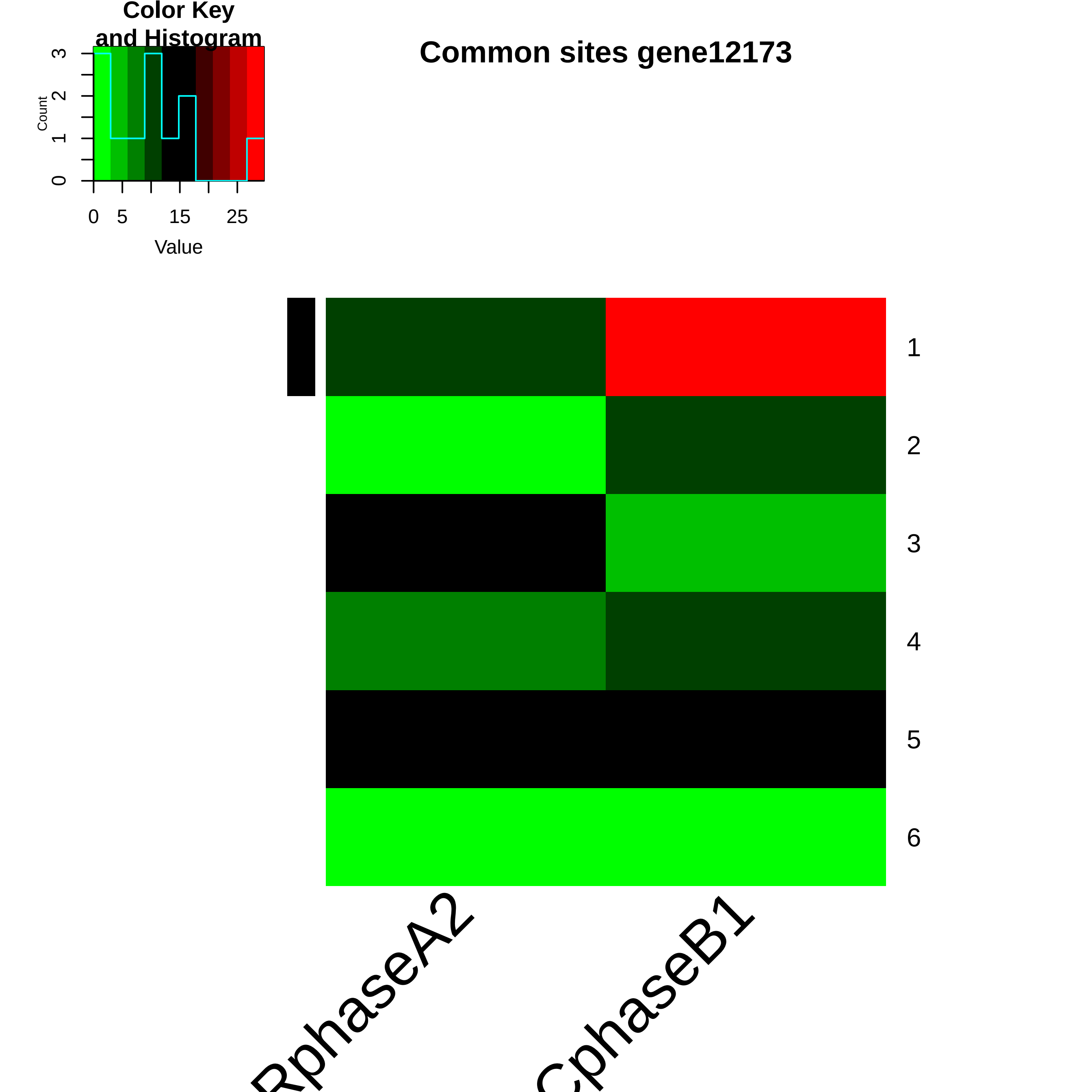

Color Key
and Histogram
Count
0
1
2
3
0
5
15
25
Value
Common sites gene12173
1
2
3
4
5
6
RphaseA2
CphaseB1

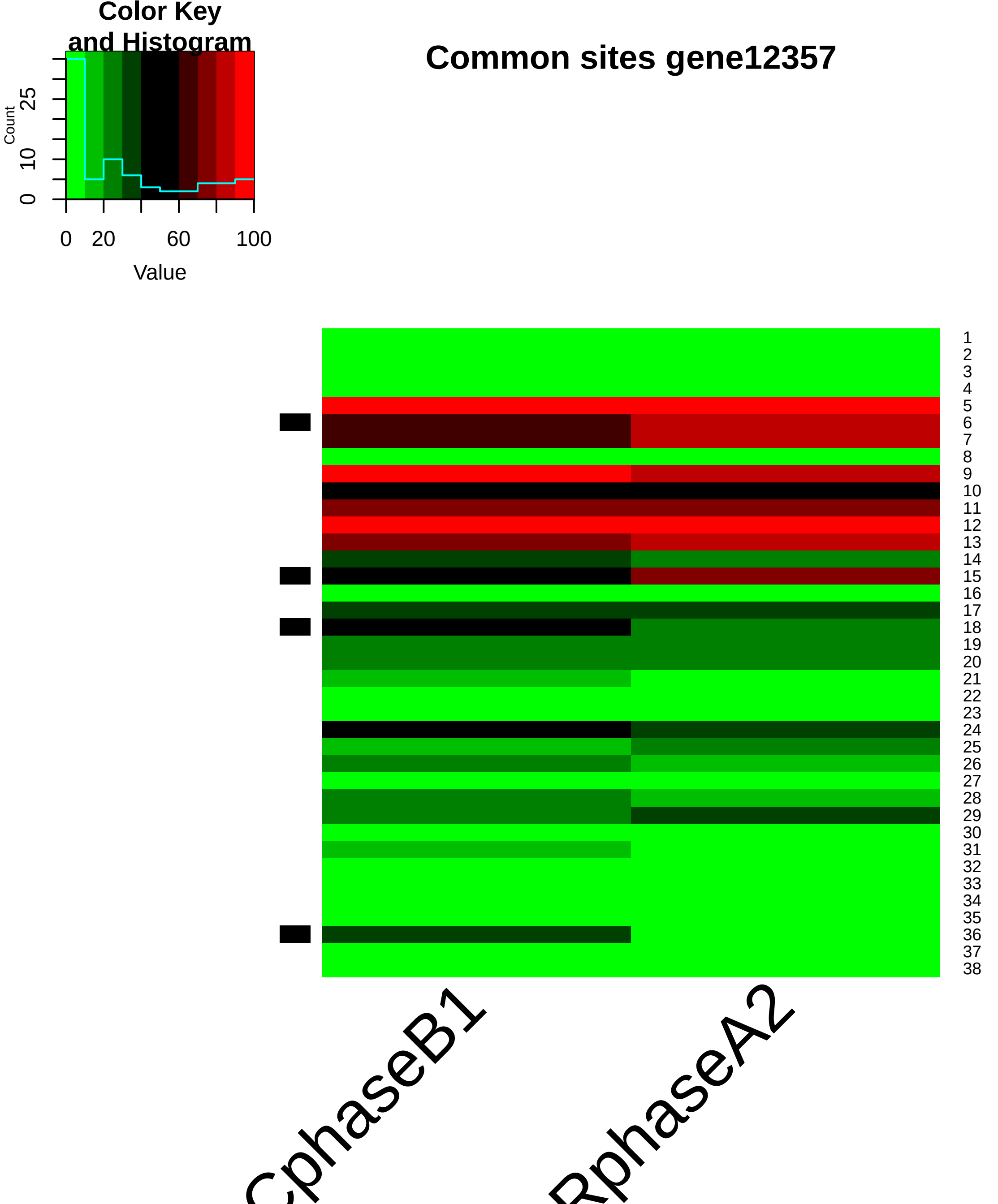
Color Key
and Histogram
Count
0
10
25
0
20
60
100
Value
Common sites gene12357
CphaseB1
RphaseA2

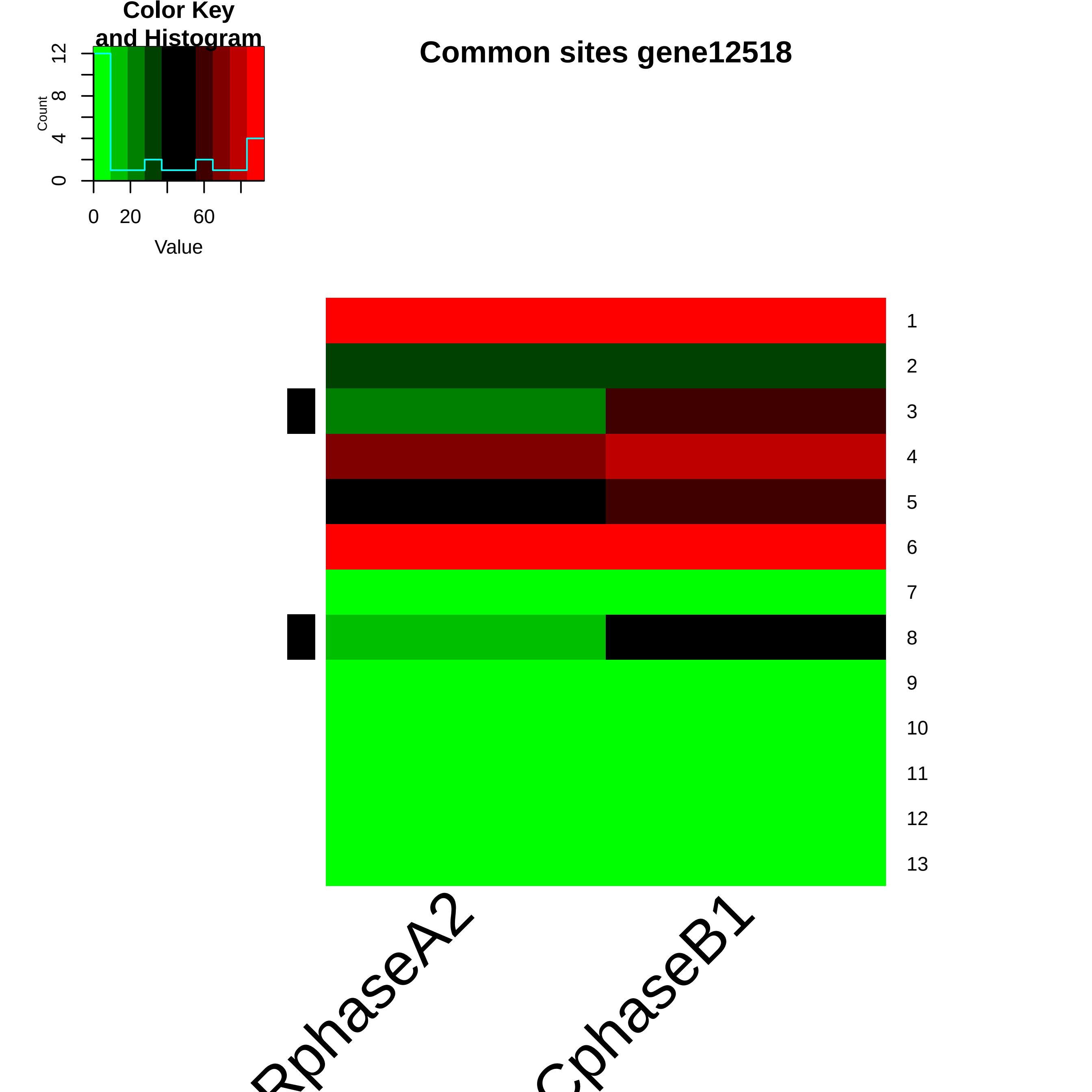
Color Key
and Histogram
Count
0
4
8
12
0
20
60
Value
Common sites gene12518
1
2
3
4
5
6
7
8
9
10
11
12
13
RphaseA2
CphaseB1

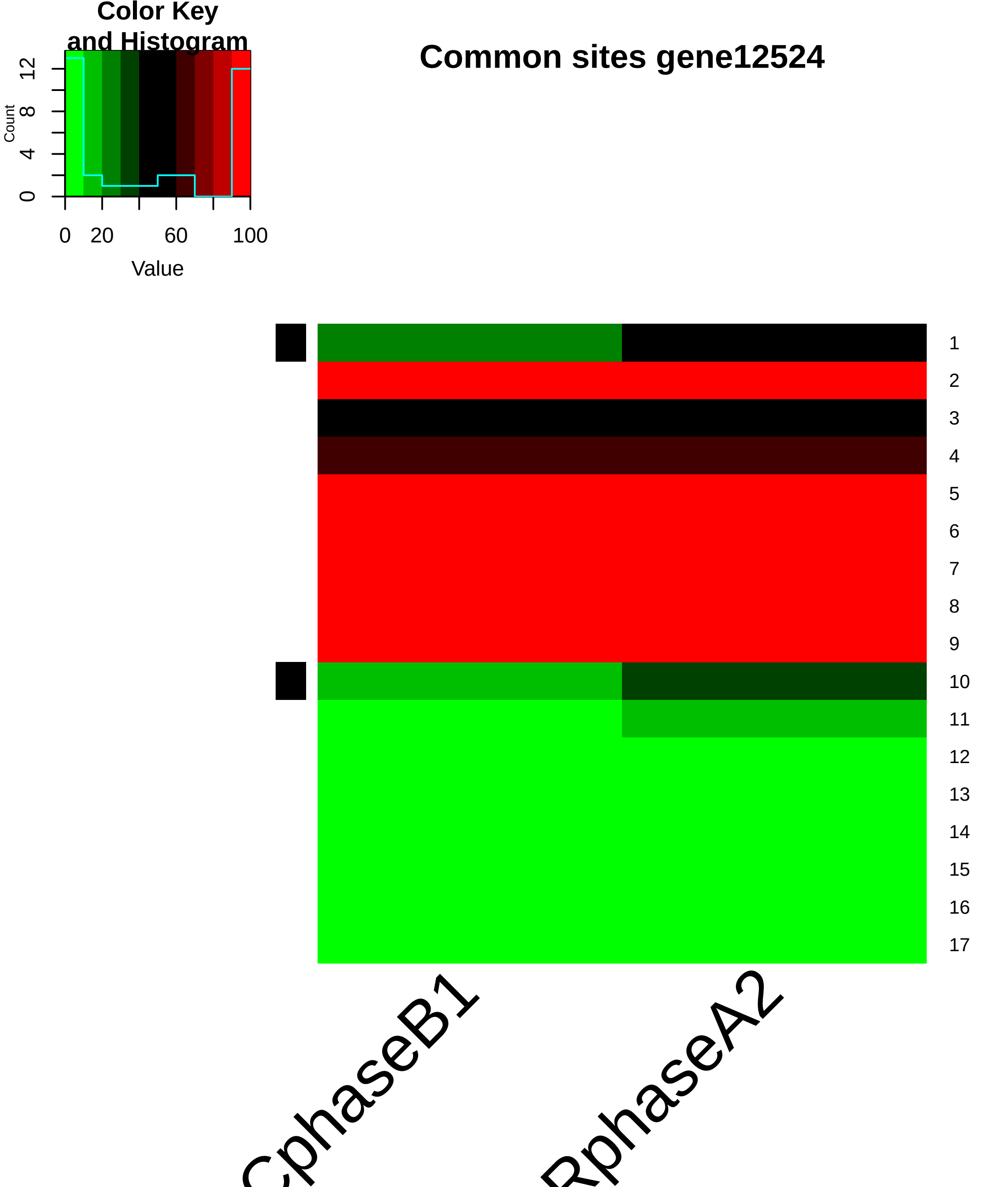
Color Key
and Histogram
Count
0
4
8
12
0
20
60
100
Value
Common sites gene12524
CphaseB1
RphaseA2

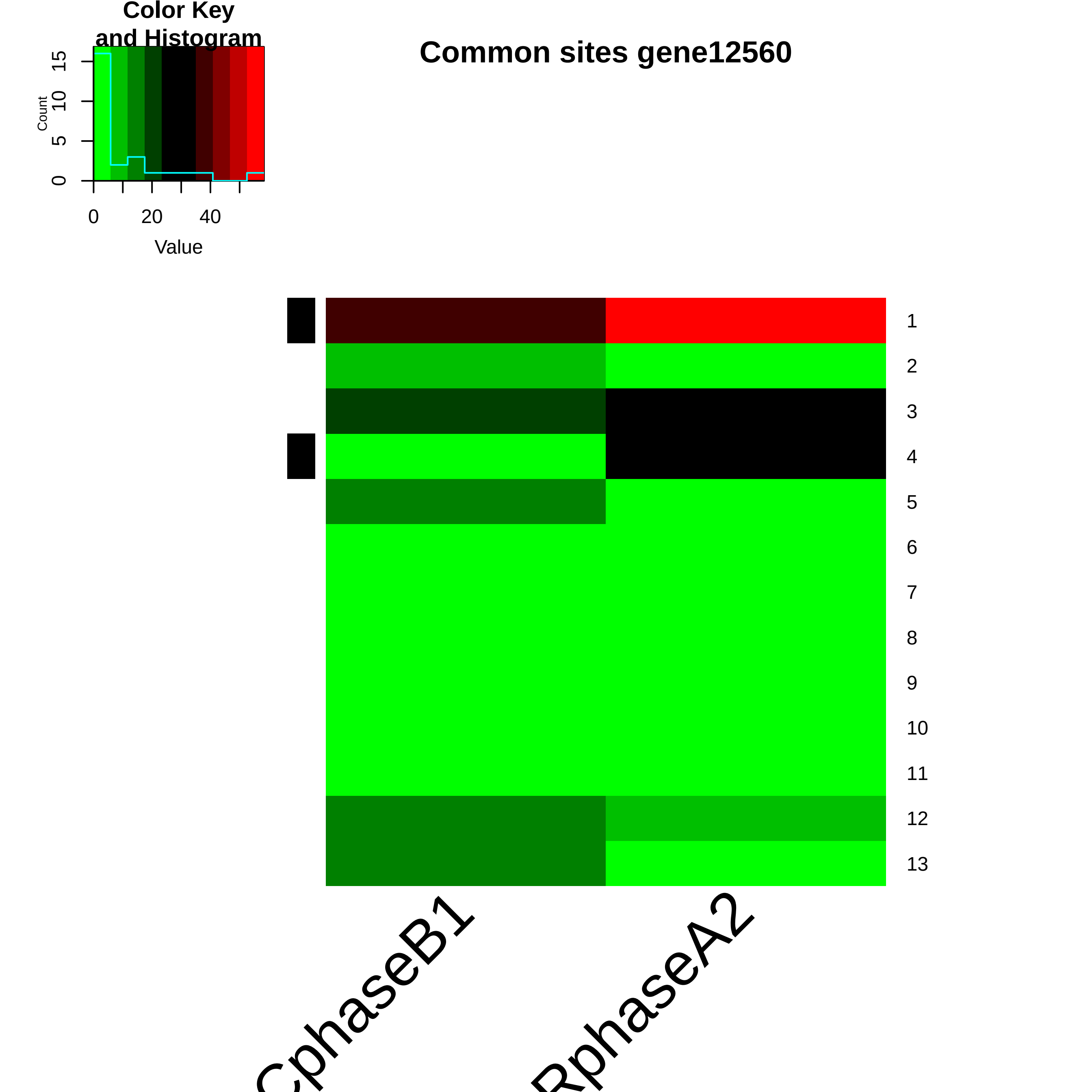

Color Key
and Histogram
Count
0
5
10
15
0
20
40
Value
Common sites gene12560
1
2
3
4
5
6
7
8
9
10
11
12
13
CphaseB1
RphaseA2

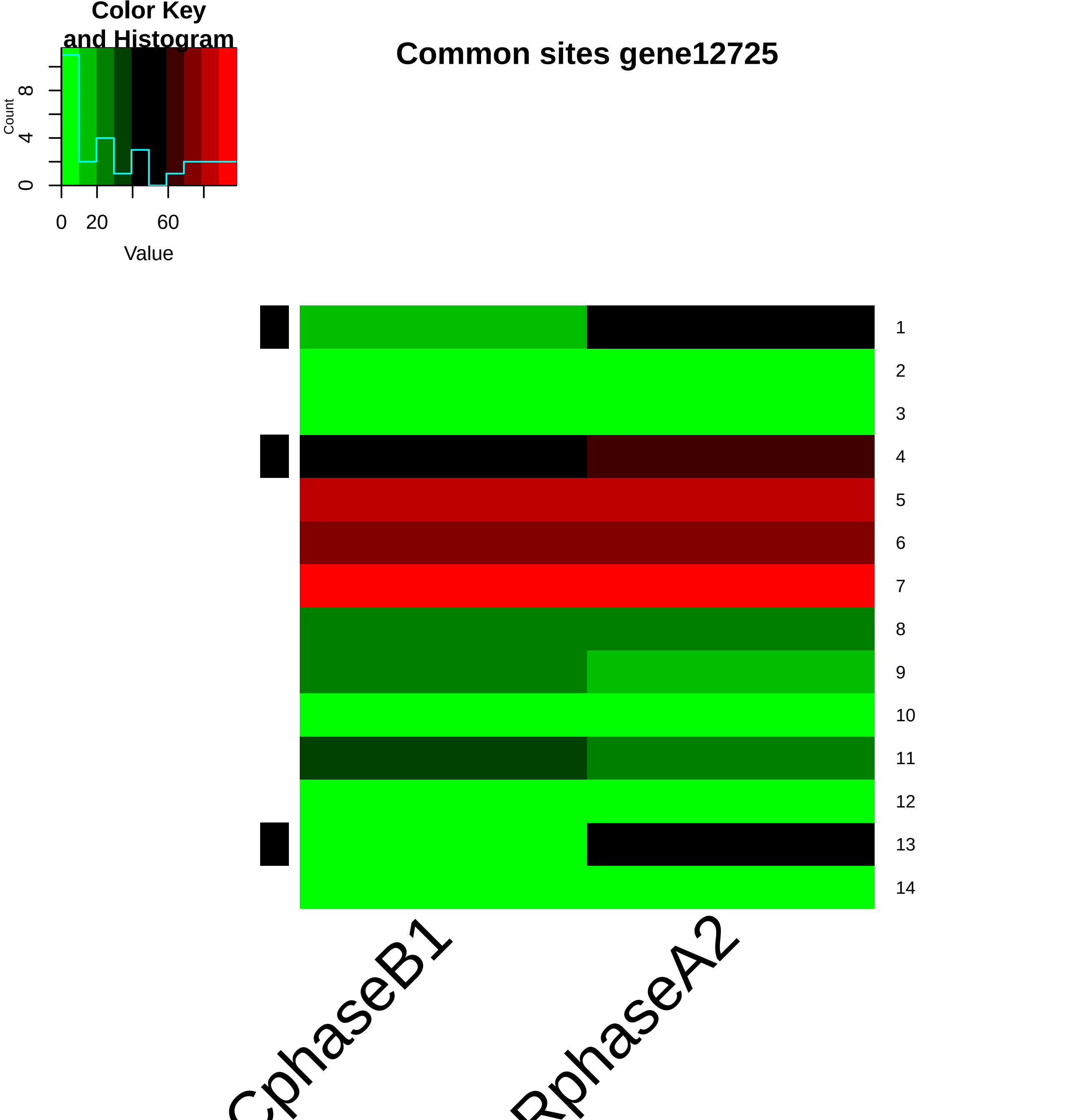

Color Key
and Histogram
Count
0
4
8
0
20
60
Value
Common sites gene12725
1
2
3
4
5
6
7
8
9
10
11
12
13
14
CphaseB1
RphaseA2

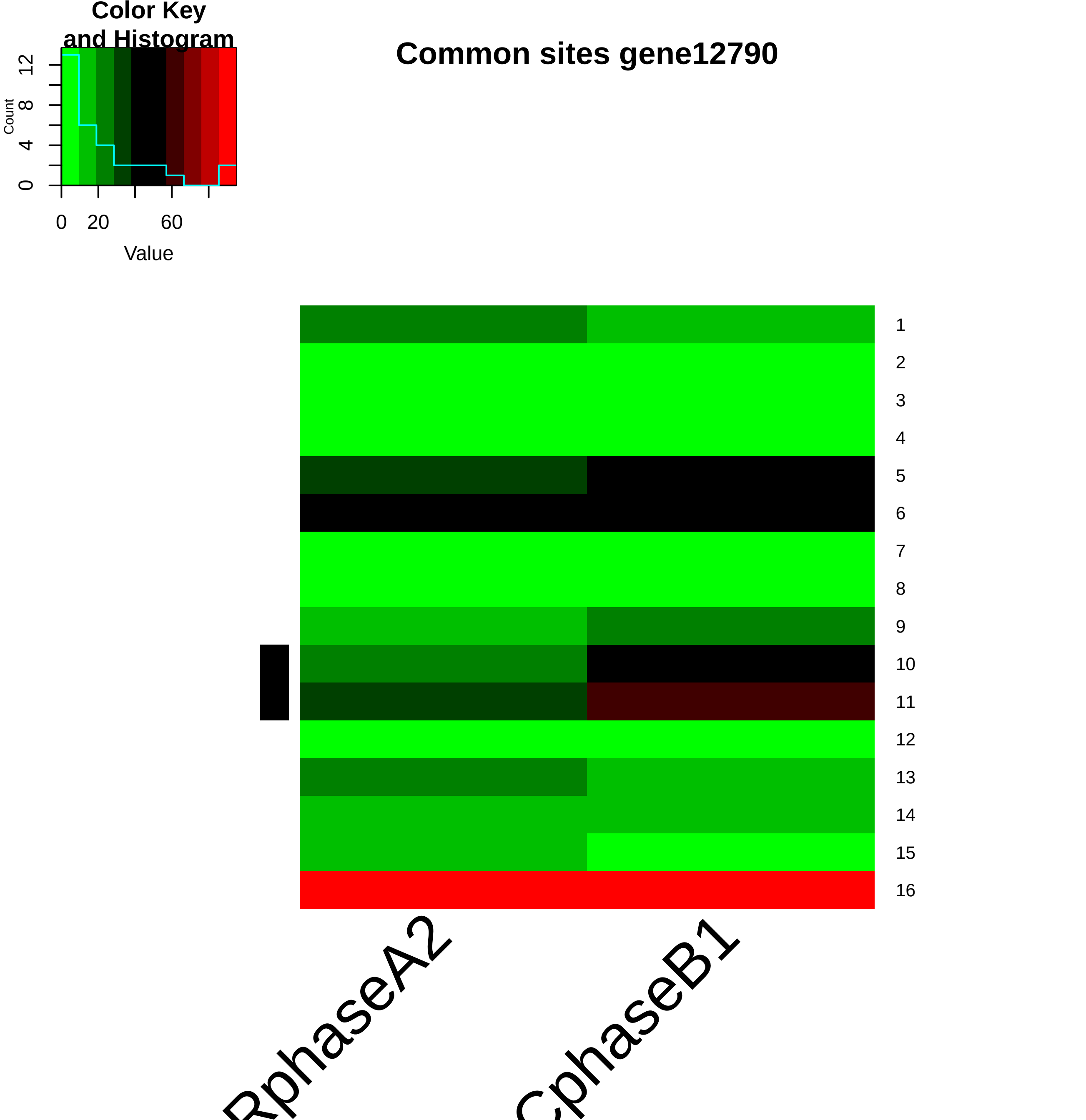

Color Key
and Histogram
Count
0
4
8
12
0
20
60
Value
Common sites gene12790
RphaseA2
CphaseB1

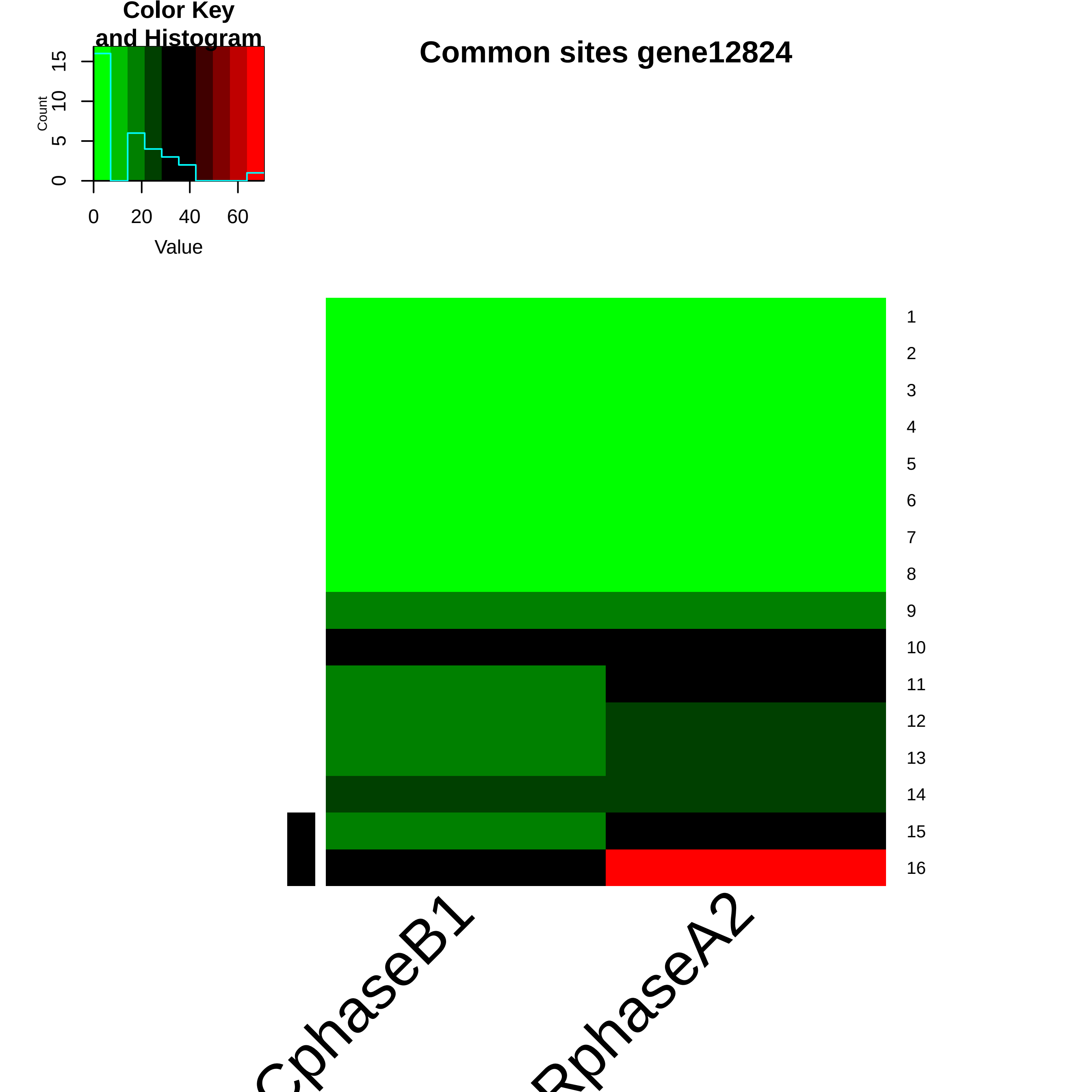

Color Key
and Histogram
Count
0
5
10
15
0
20
40
60
Value
Common sites gene12824
CphaseB1
RphaseA2

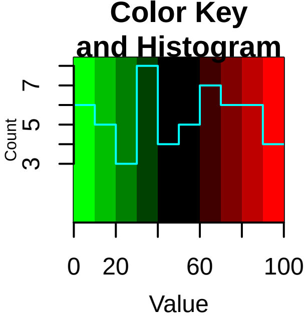

### Common sites gene12955

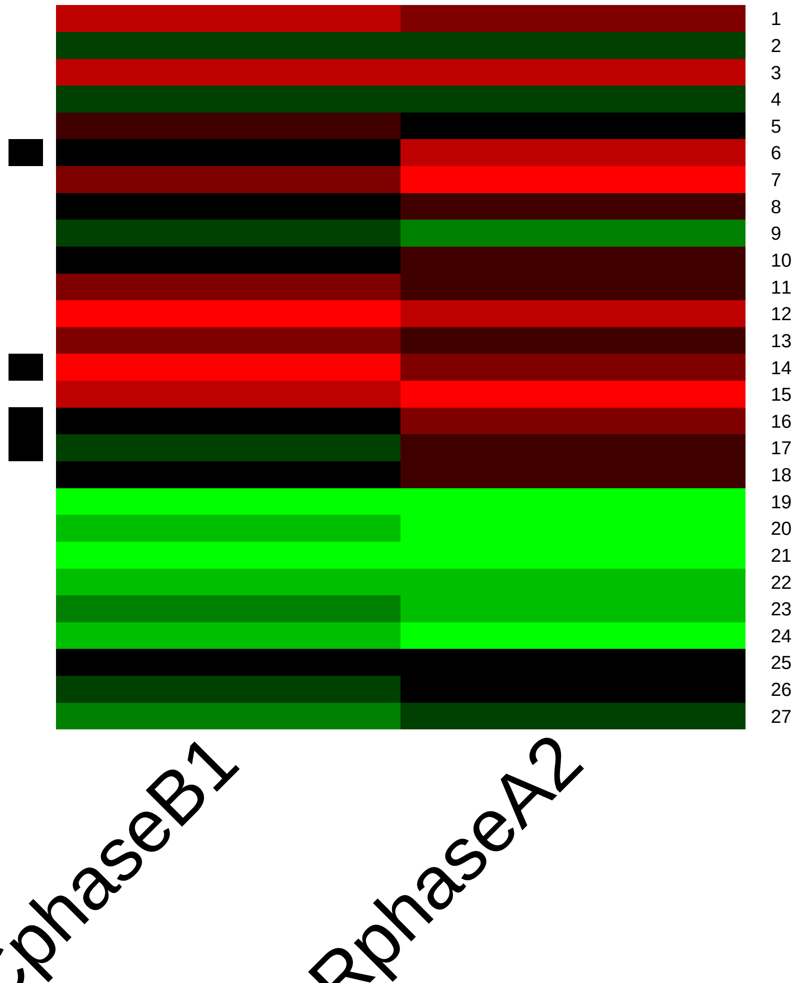

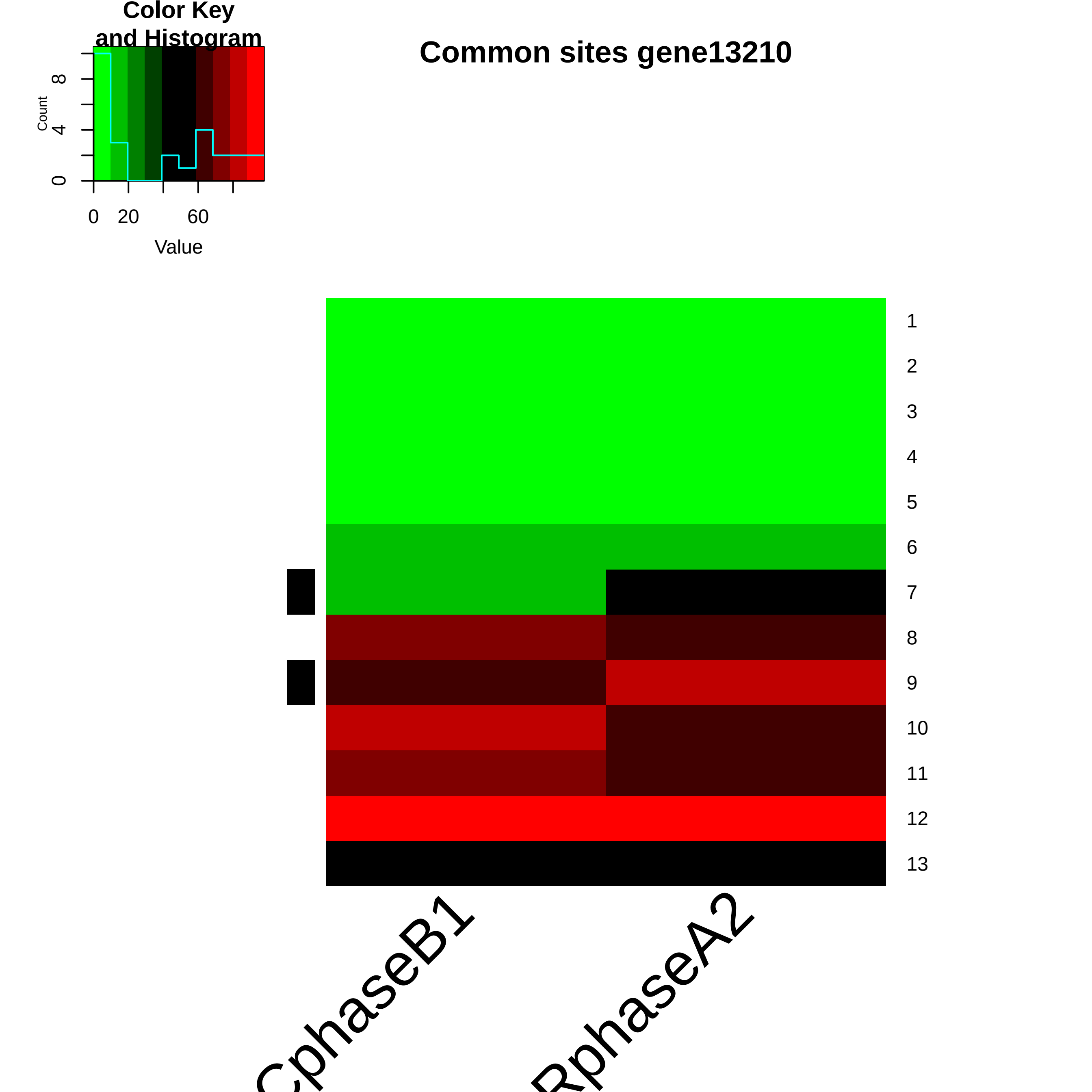

Color Key
and Histogram
Count
0
4
8
0
20
60
Value
Common sites gene13210
1
2
3
4
5
6
7
8
9
10
11
12
13
CphaseB1
RphaseA2

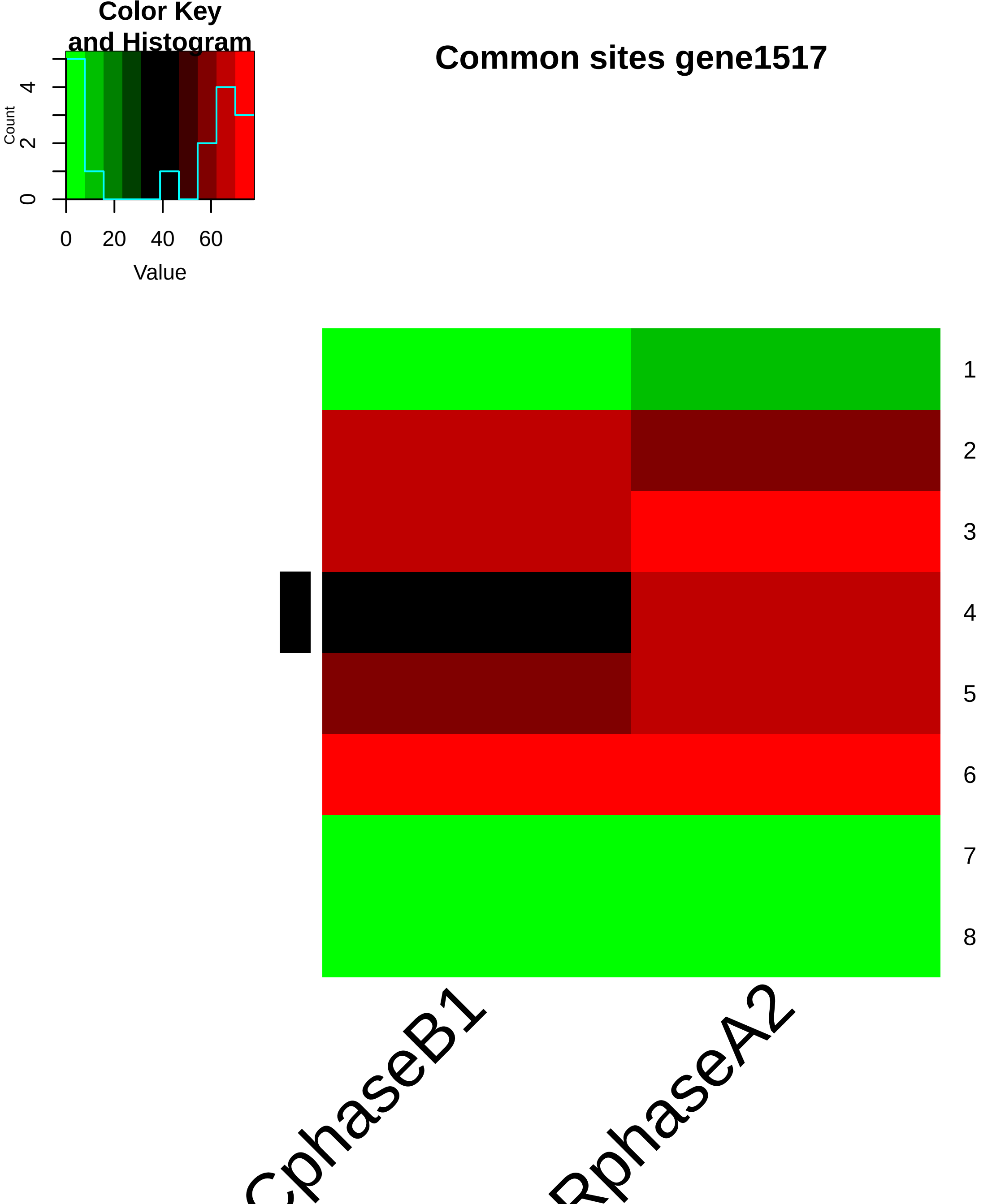
Color Key
and Histogram
Count
0
2
4
0
20
40
60
Value
Common sites gene1517
1
2
3
4
5
6
7
8
CphaseB1
RphaseA2

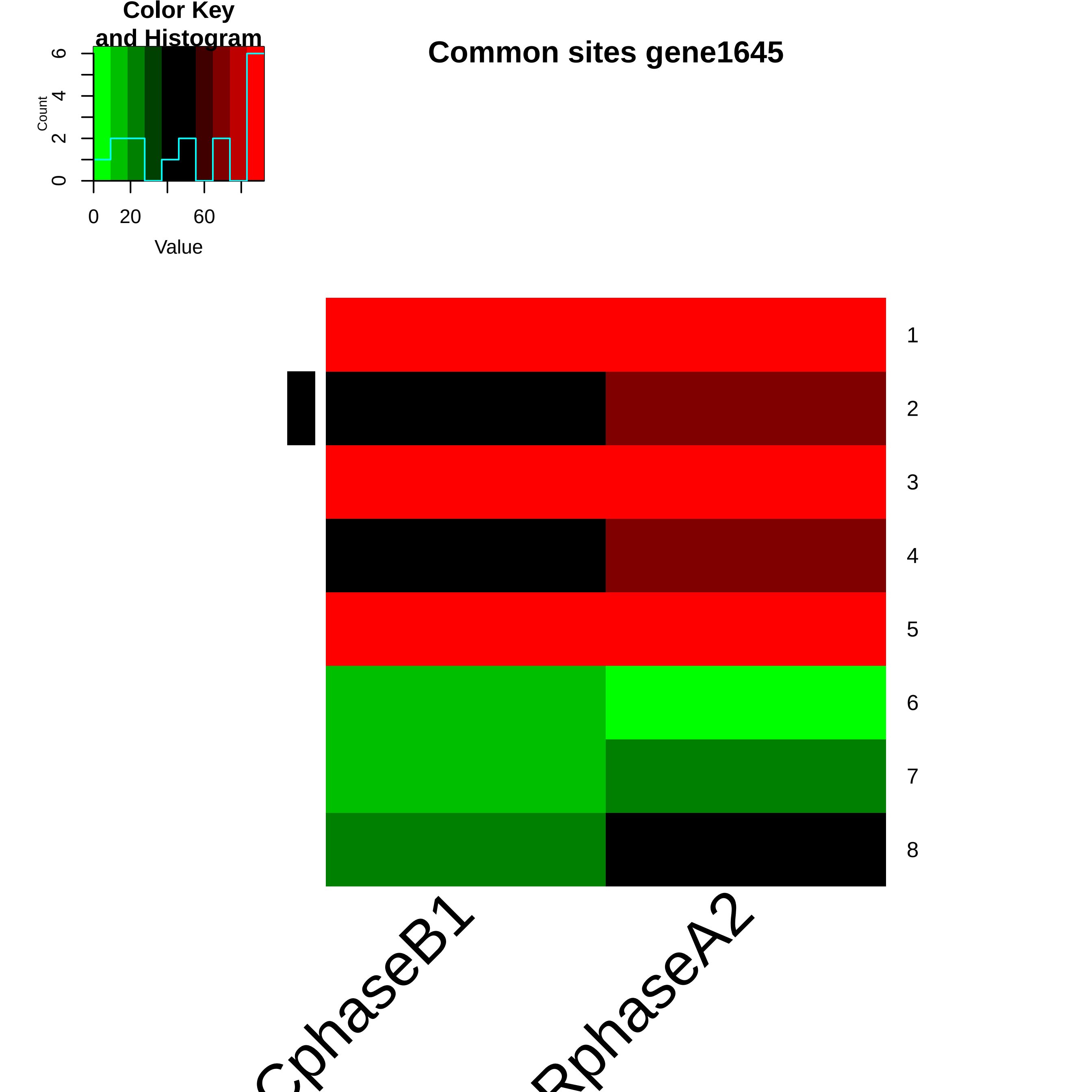
Color Key
and Histogram
Count
0
2
4
6
0
20
60
Value
Common sites gene1645
1
2
3
4
5
6
7
8
CphaseB1
RphaseA2

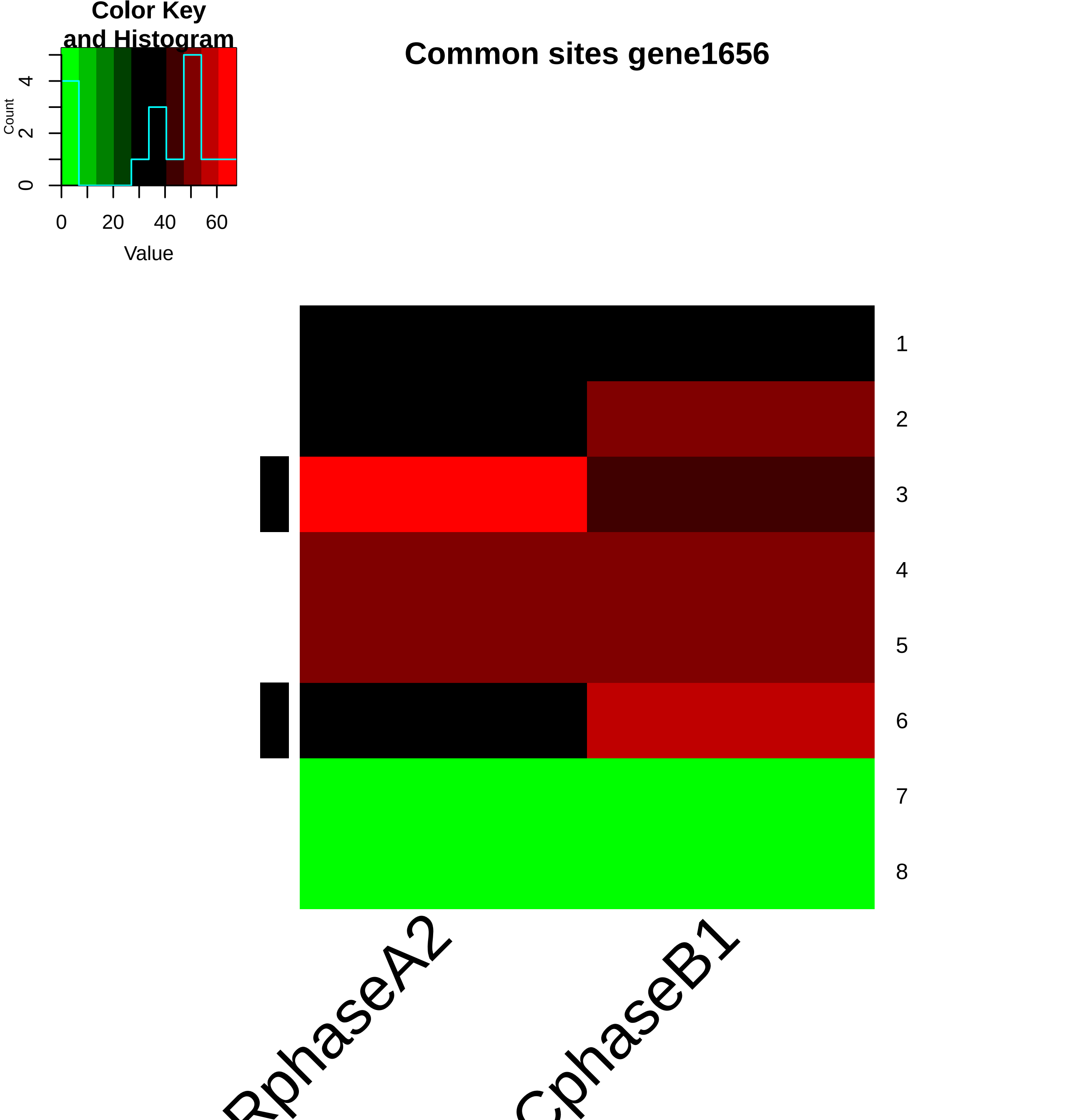
Color Key
and Histogram
Count
0
2
4
0
20
40
60
Value
Common sites gene1656
1
2
3
4
5
6
7
8
RphaseA2
CphaseB1

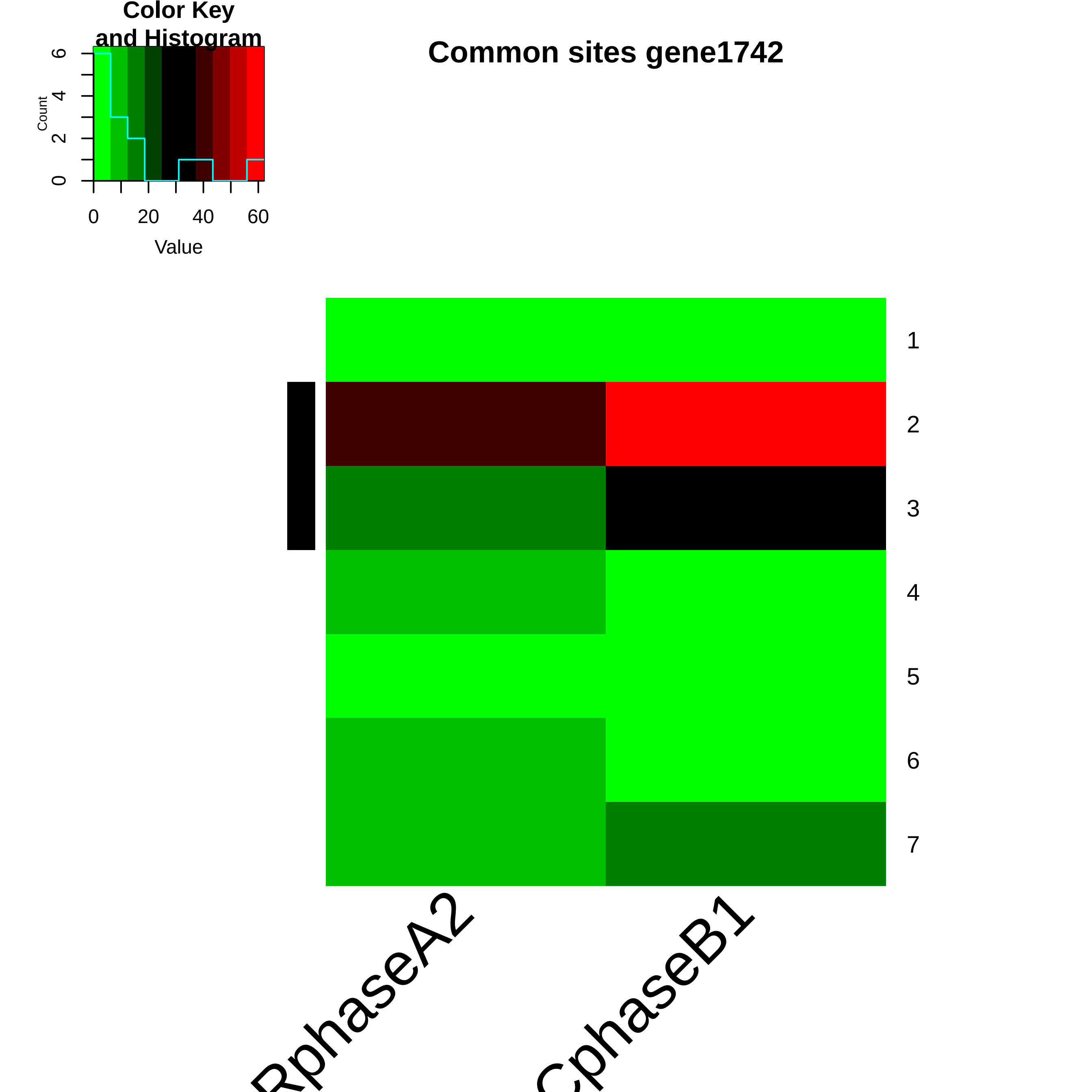
Color Key
and Histogram
Count
0
2
4
6
0
20
40
60
Value
Common sites gene1742
1
2
3
4
5
6
7
RphaseA2
CphaseB1

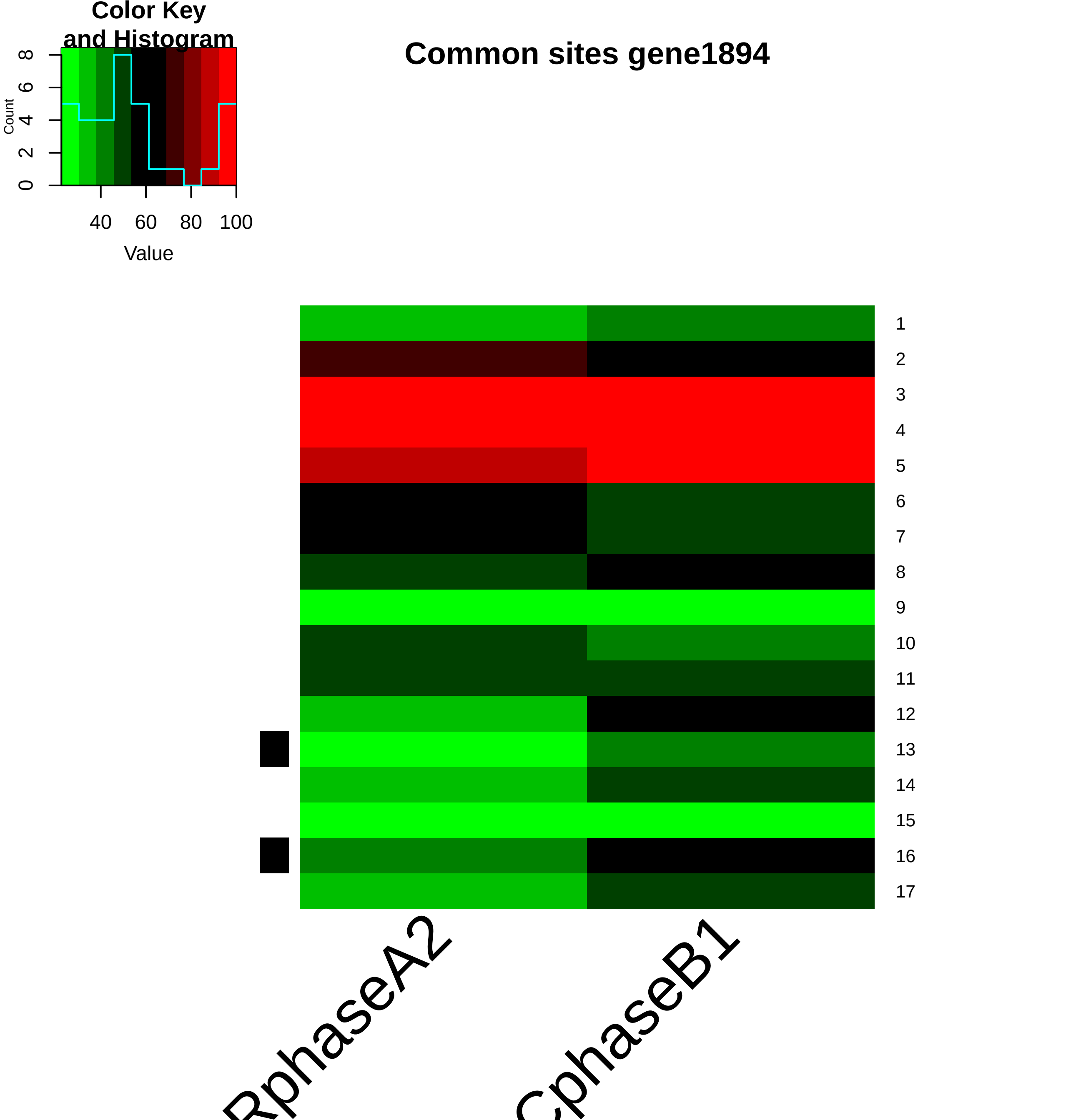
Color Key
and Histogram
Count
0
2
4
6
8
40
60
80
100
Value
Common sites gene1894
RphaseA2
CphaseB1

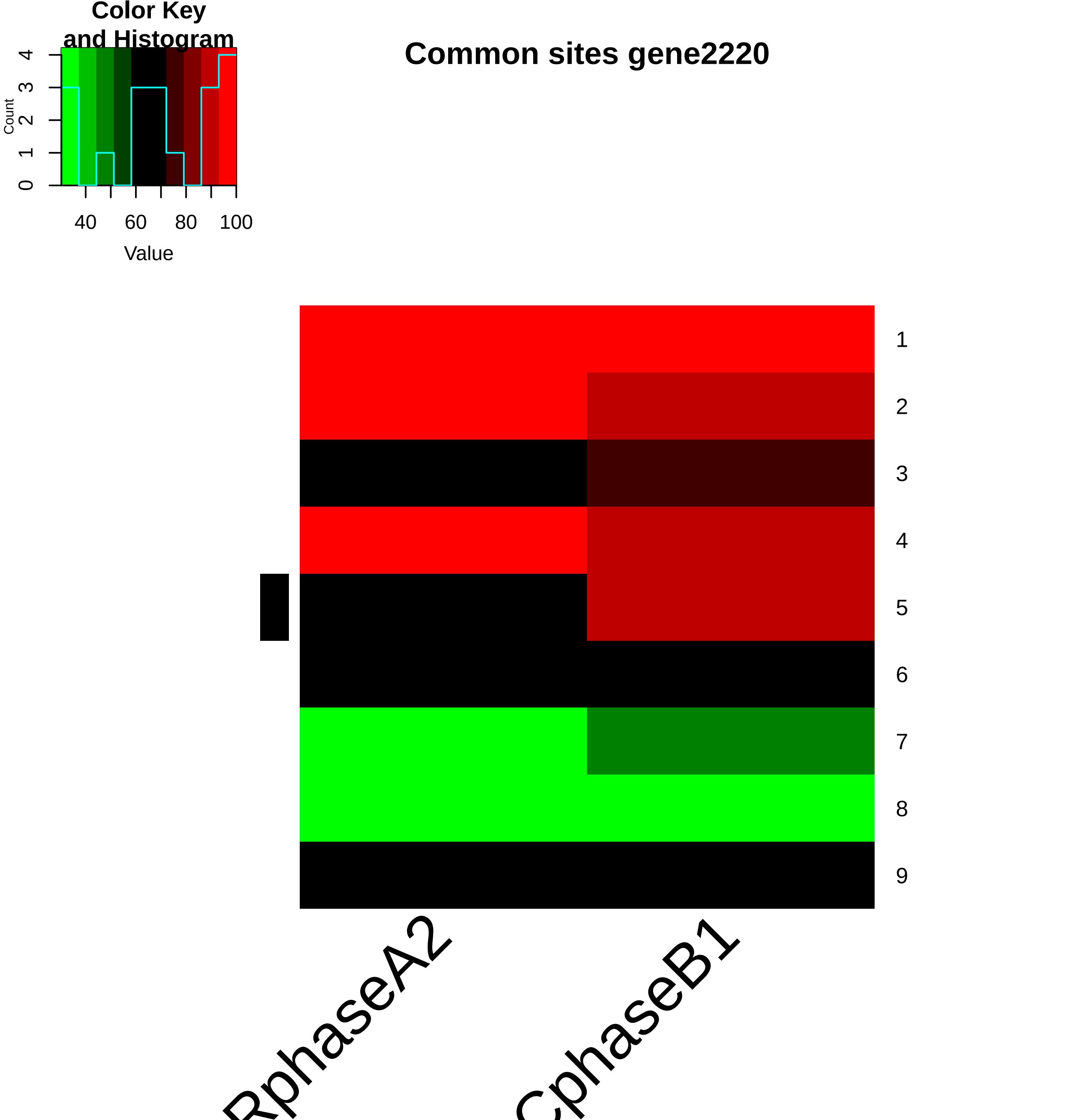
Color Key
and Histogram
Count
0
1
2
3
4
40
60
80
100
Value
Common sites gene2220
1
2
3
4
5
6
7
8
9
RphaseA2
CphaseB1

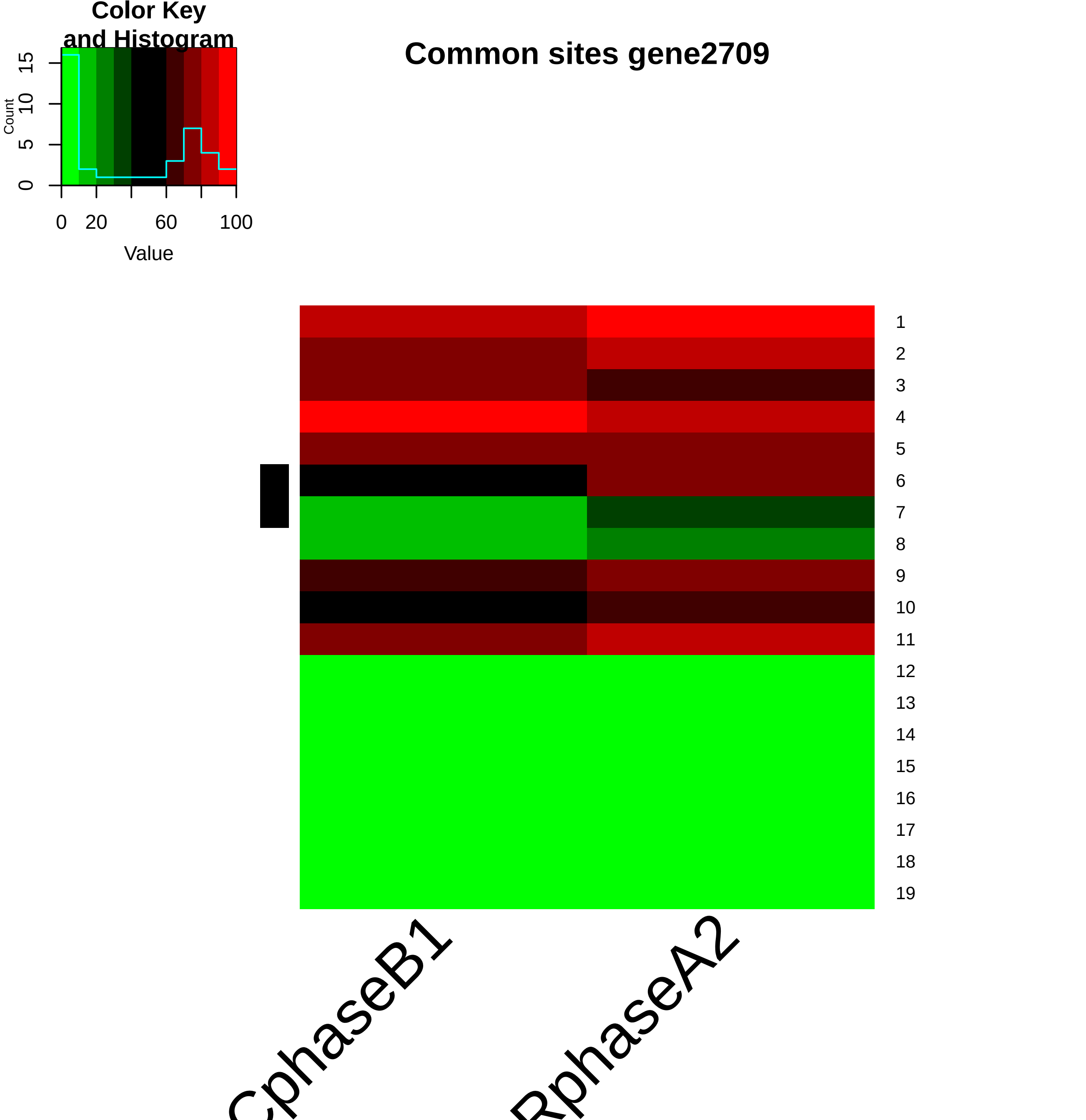

Color Key
and Histogram
Count
0
5
10
15
0
20
60
100
Value
Common sites gene2709
CphaseB1
RphaseA2

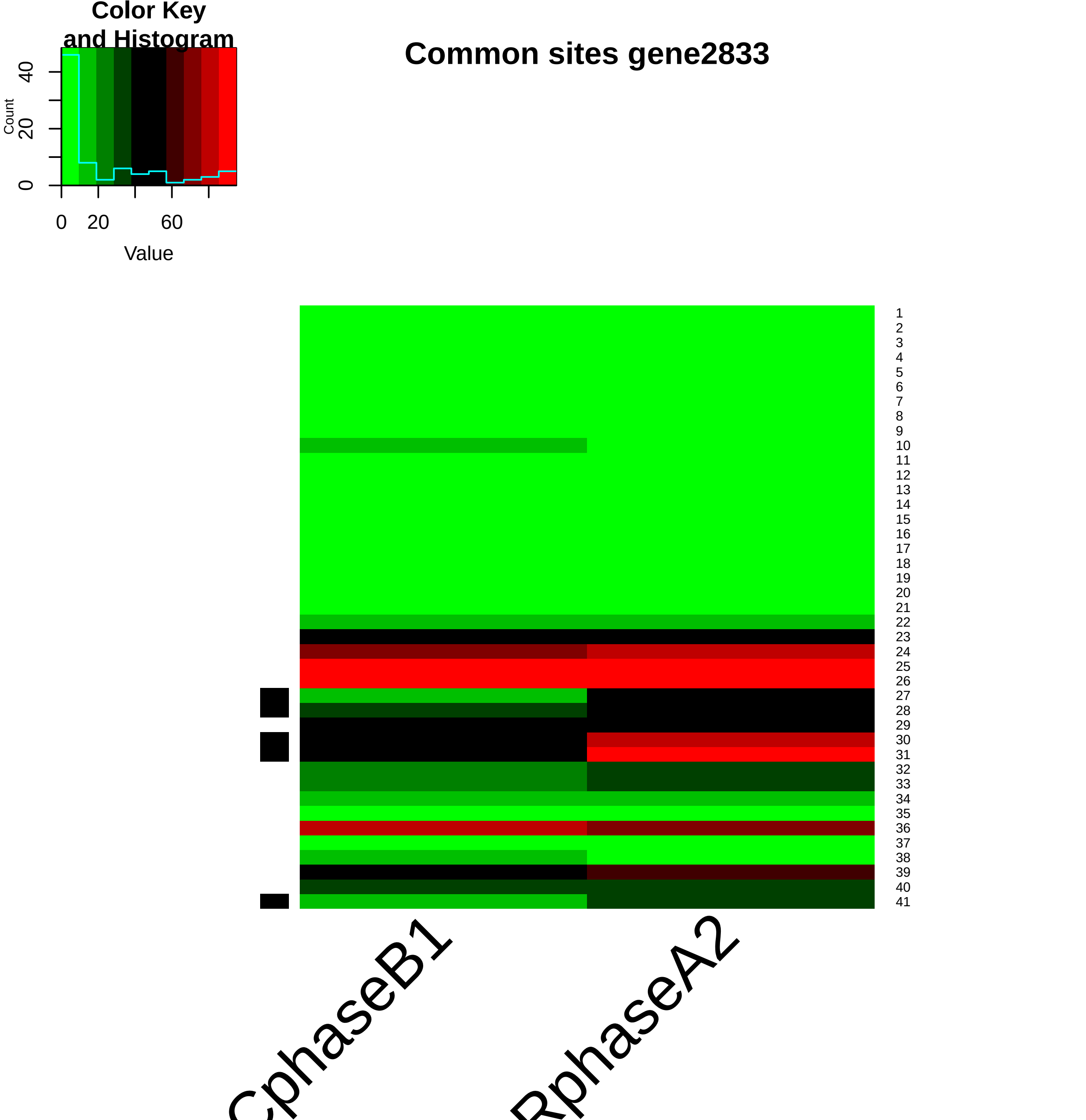
Color Key
and Histogram
Count
0
20
40
0
20
60
Value
Common sites gene2833
CphaseB1
RphaseA2

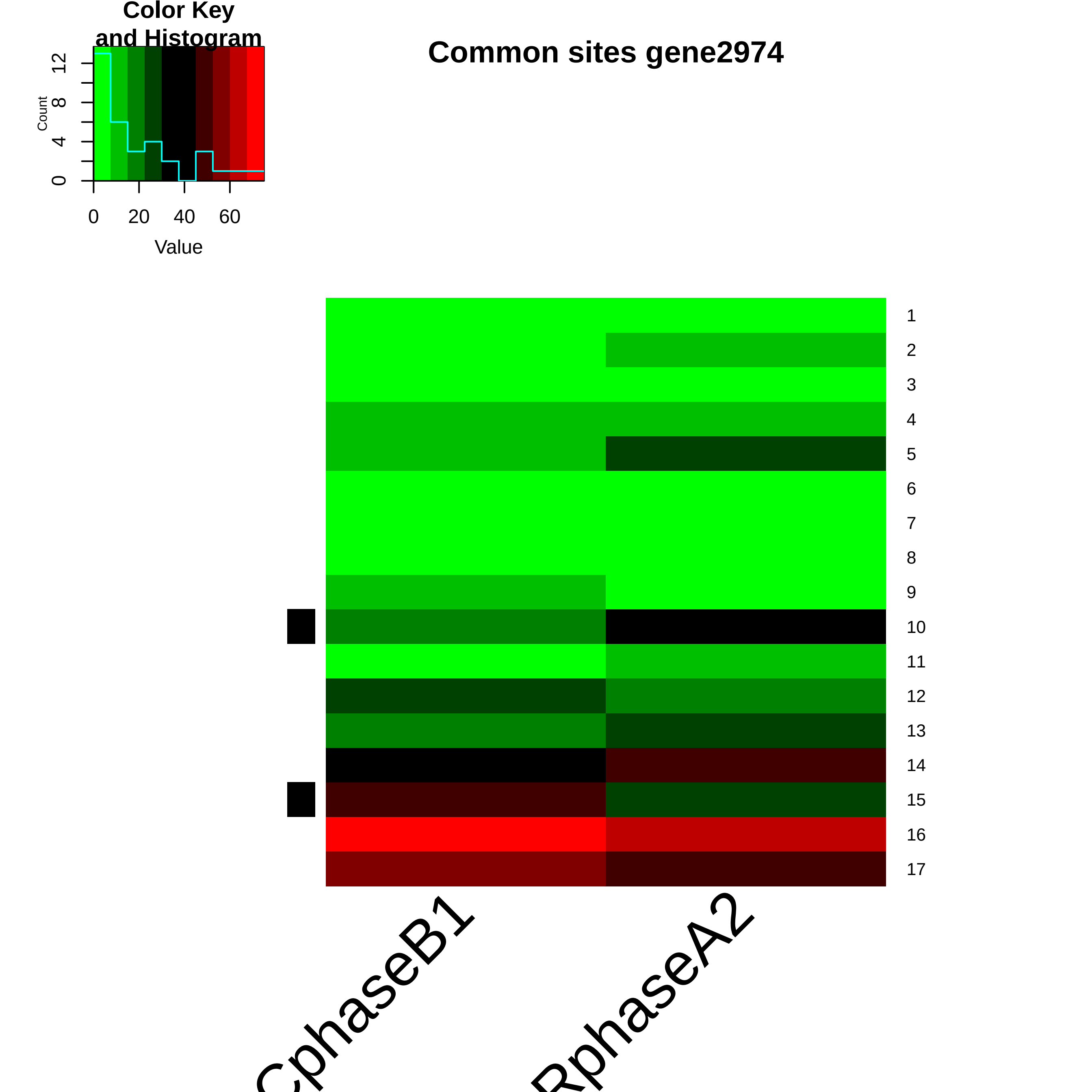

Color Key
and Histogram
Count
0
4
8
12
0
20
40
60
Value
Common sites gene2974
CphaseB1
RphaseA2

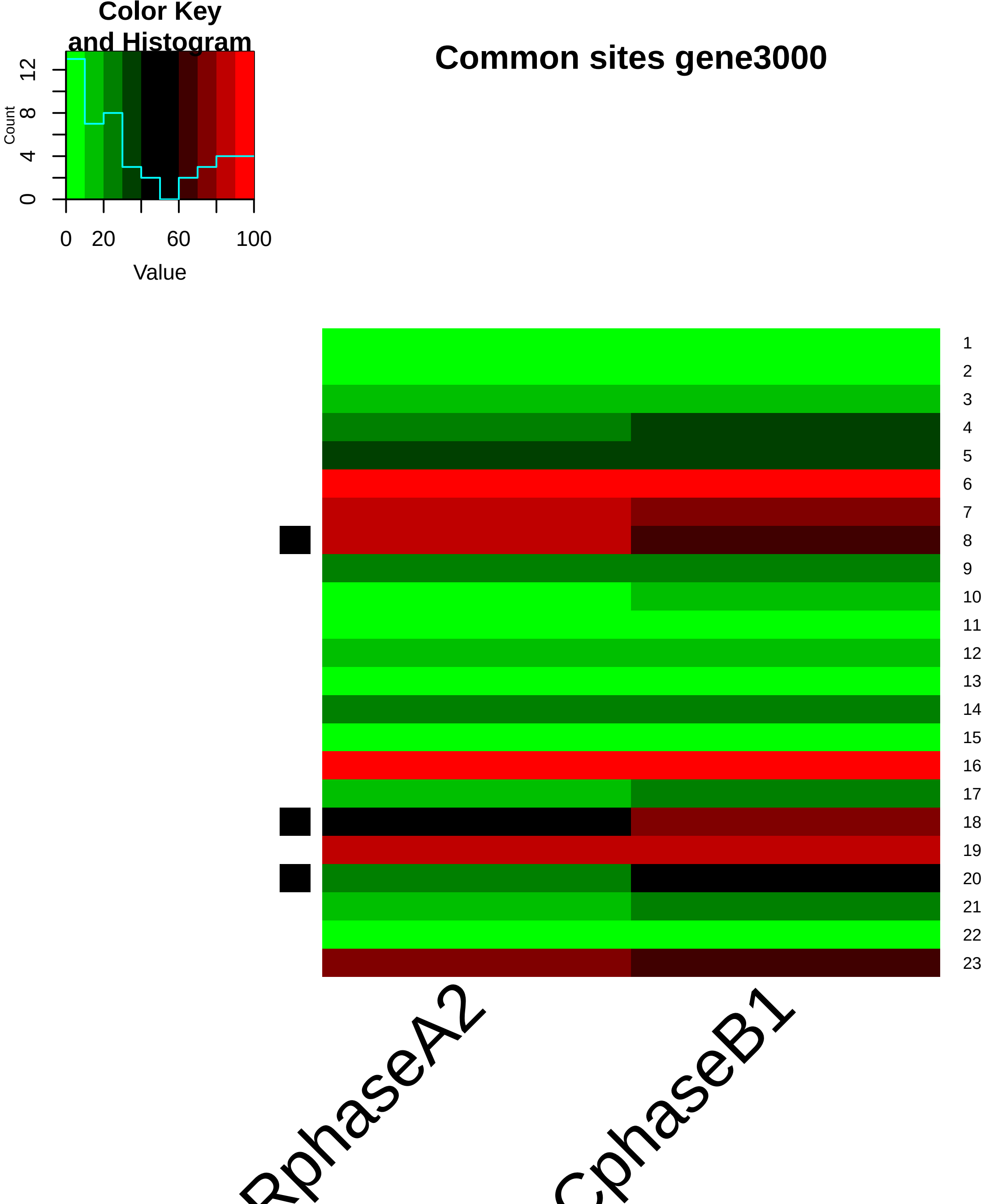
Color Key
and Histogram
Count
0
4
8
12
0
20
60
100
Value
Common sites gene3000
RphaseA2
CphaseB1

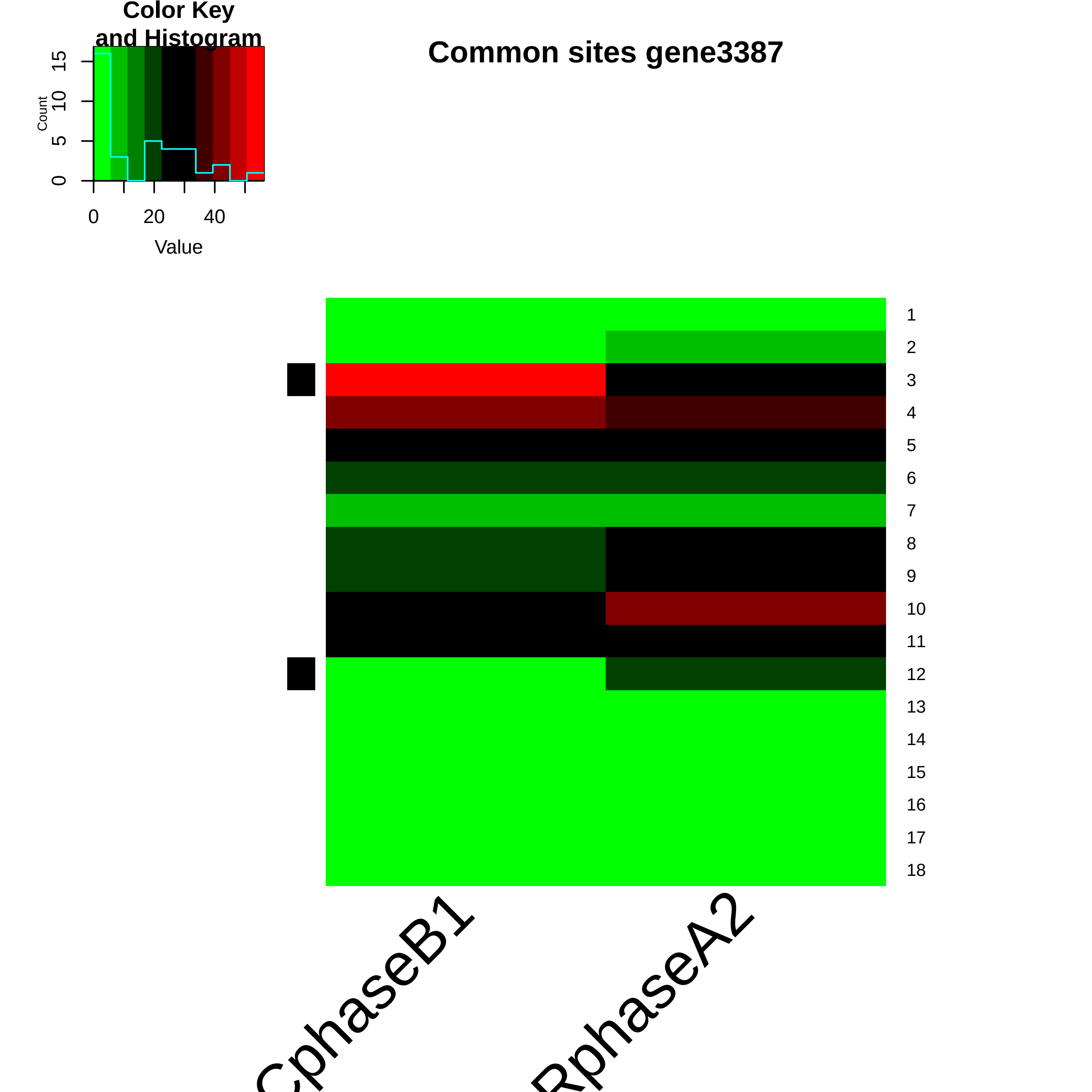

Color Key
and Histogram
Count
0
5
10
15
0
20
40
Value
Common sites gene3387
CphaseB1
RphaseA2

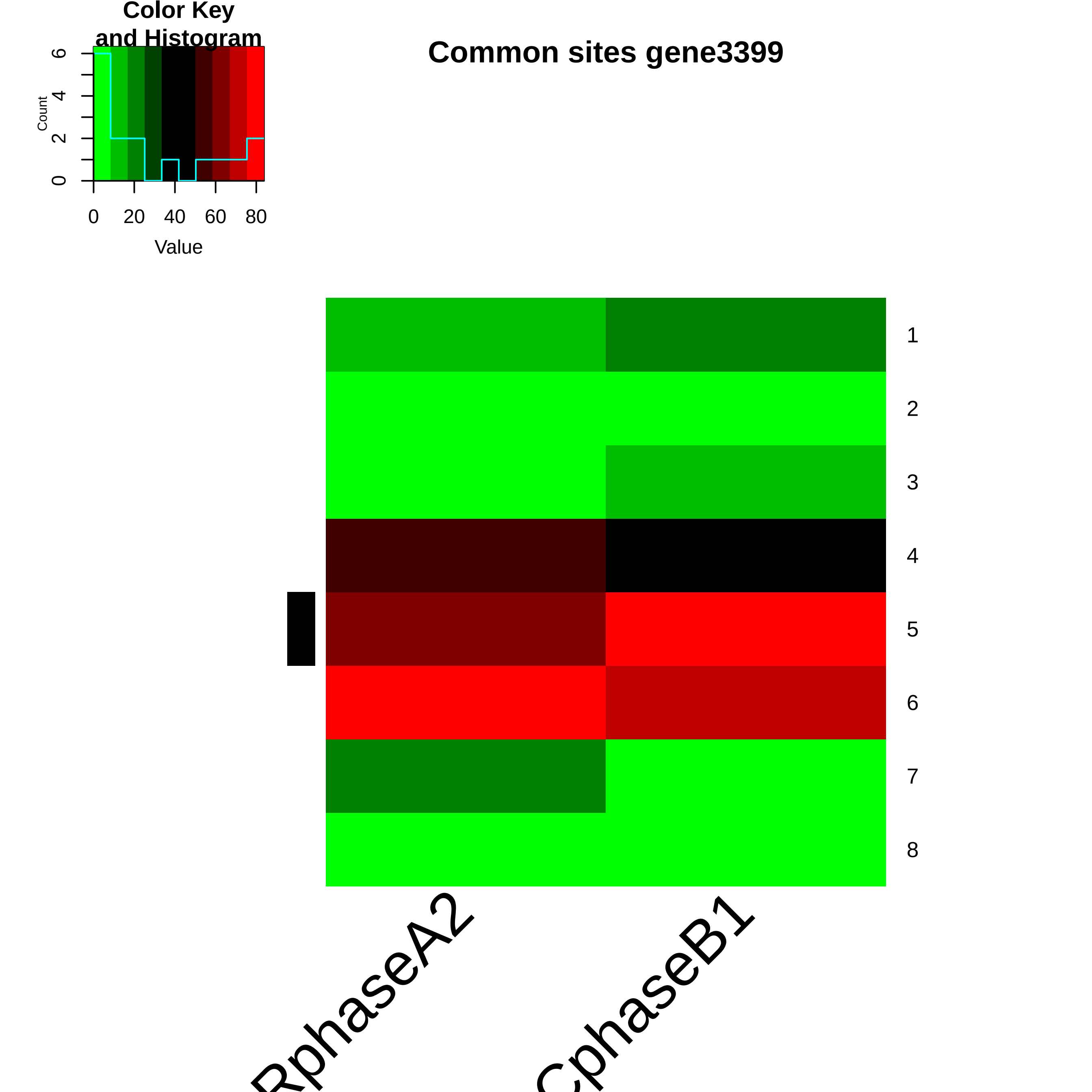
Color Key
and Histogram
Count
0
2
4
6
0
20
40
60
80
Value
Common sites gene3399
1
2
3
4
5
6
7
8
RphaseA2
CphaseB1

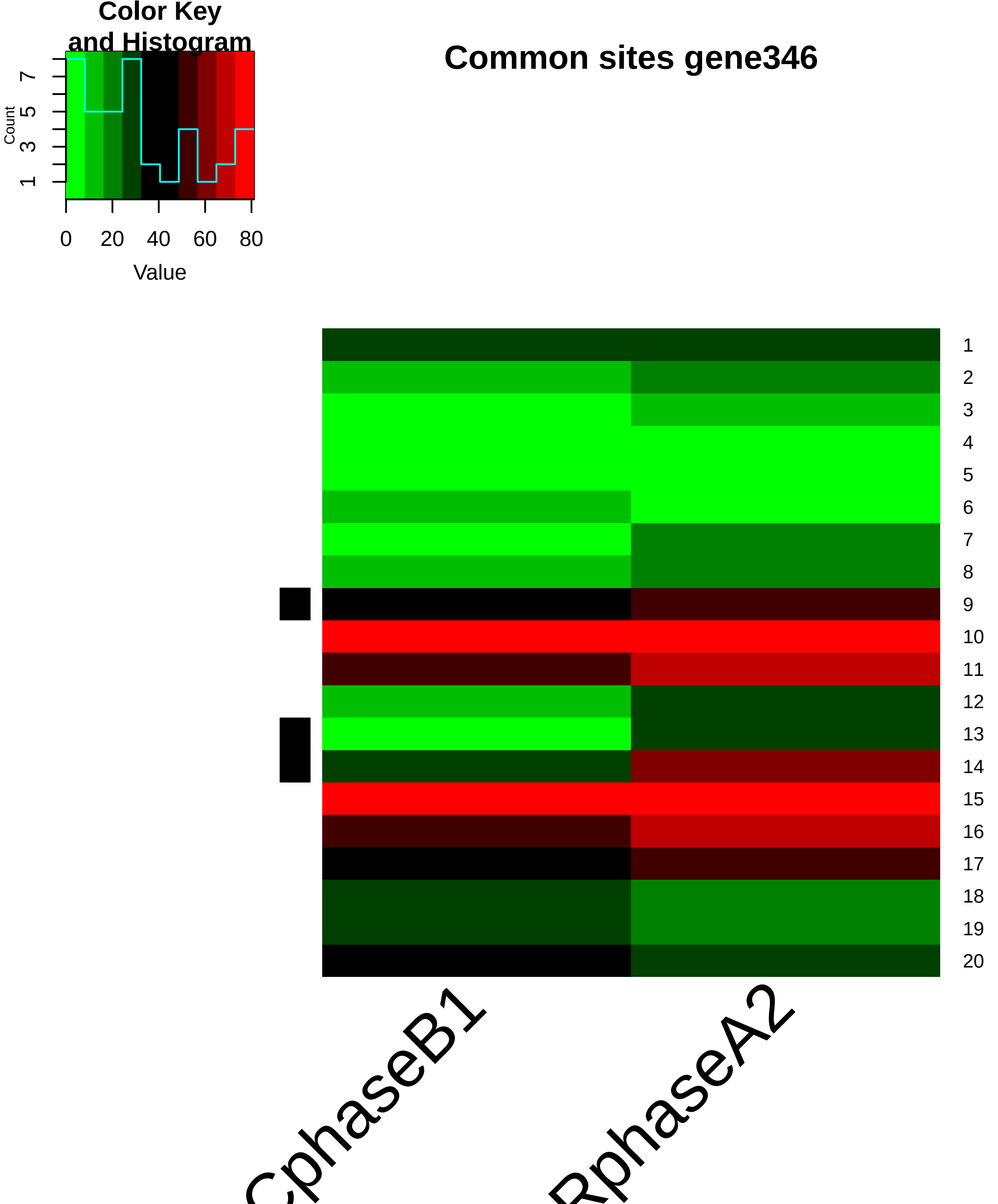
Color Key
and Histogram
Count
1
3
5
7
0
20
40
60
80
Value
Common sites gene346
CphaseB1
RphaseA2

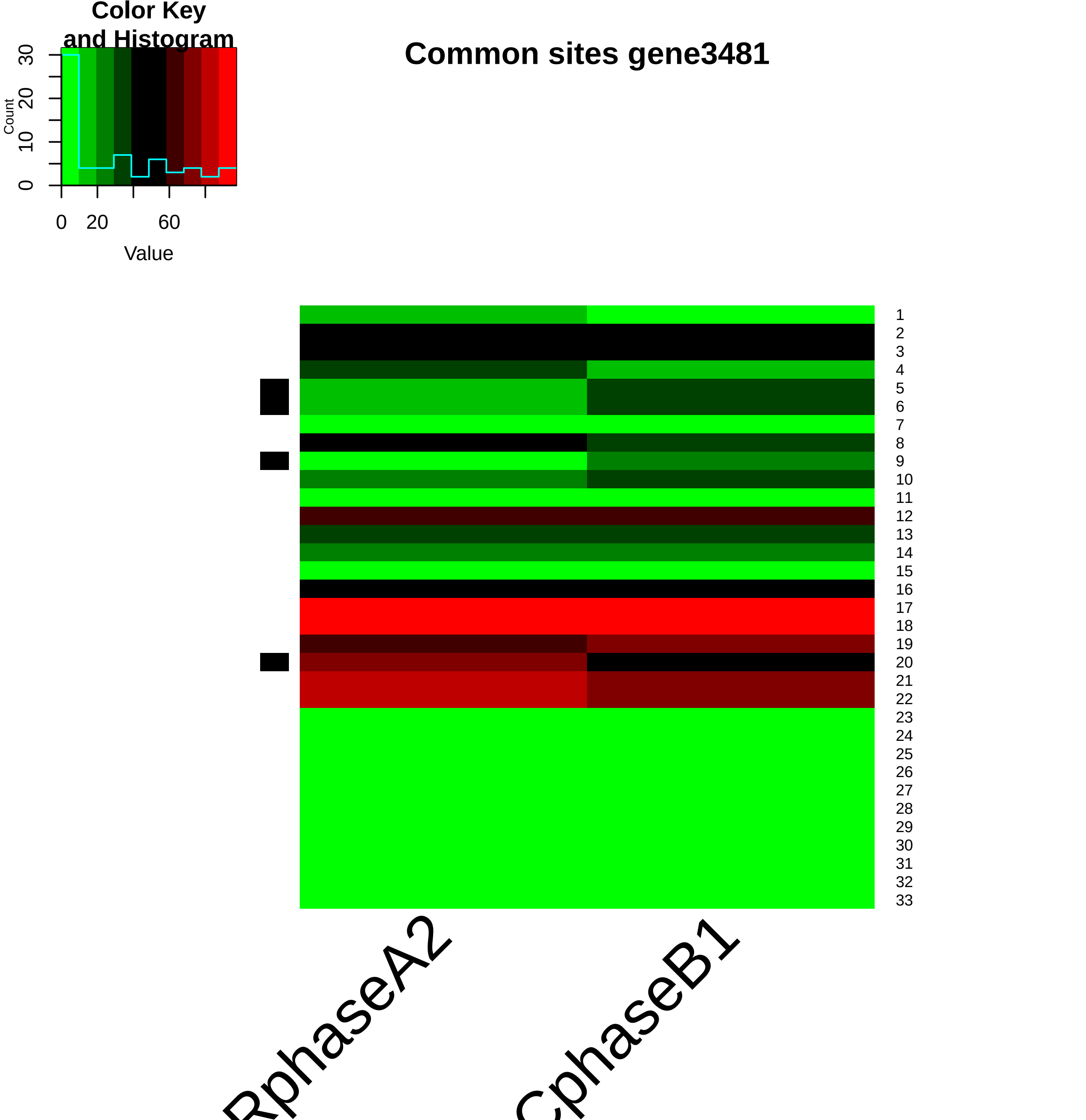

Color Key
and Histogram
Count
0
10
20
30
0
20
60
Value
Common sites gene3481
RphaseA2
CphaseB1

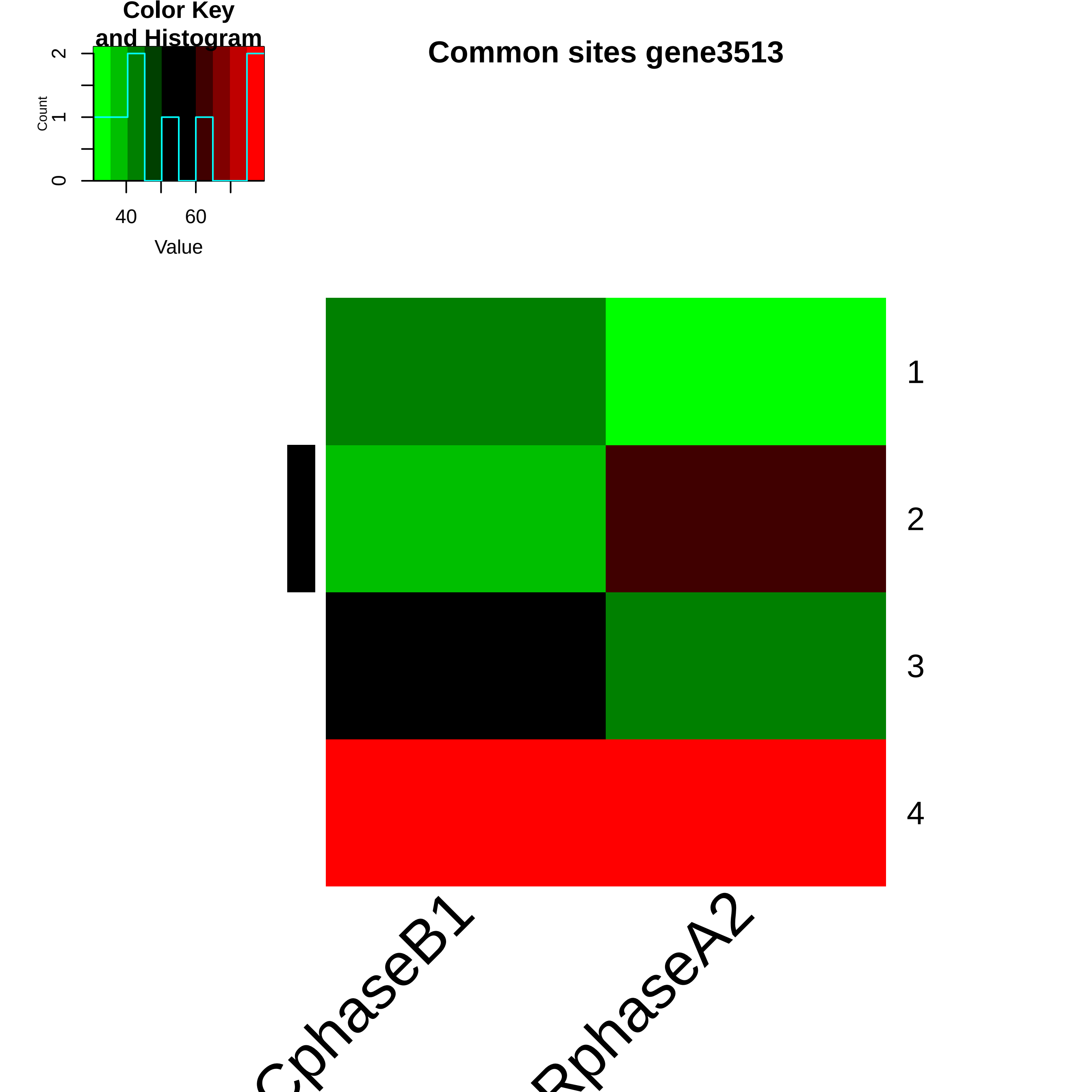
Color Key
and Histogram
Count
0
1
2
40
60
Value
Common sites gene3513
1
2
3
4
CphaseB1
RphaseA2

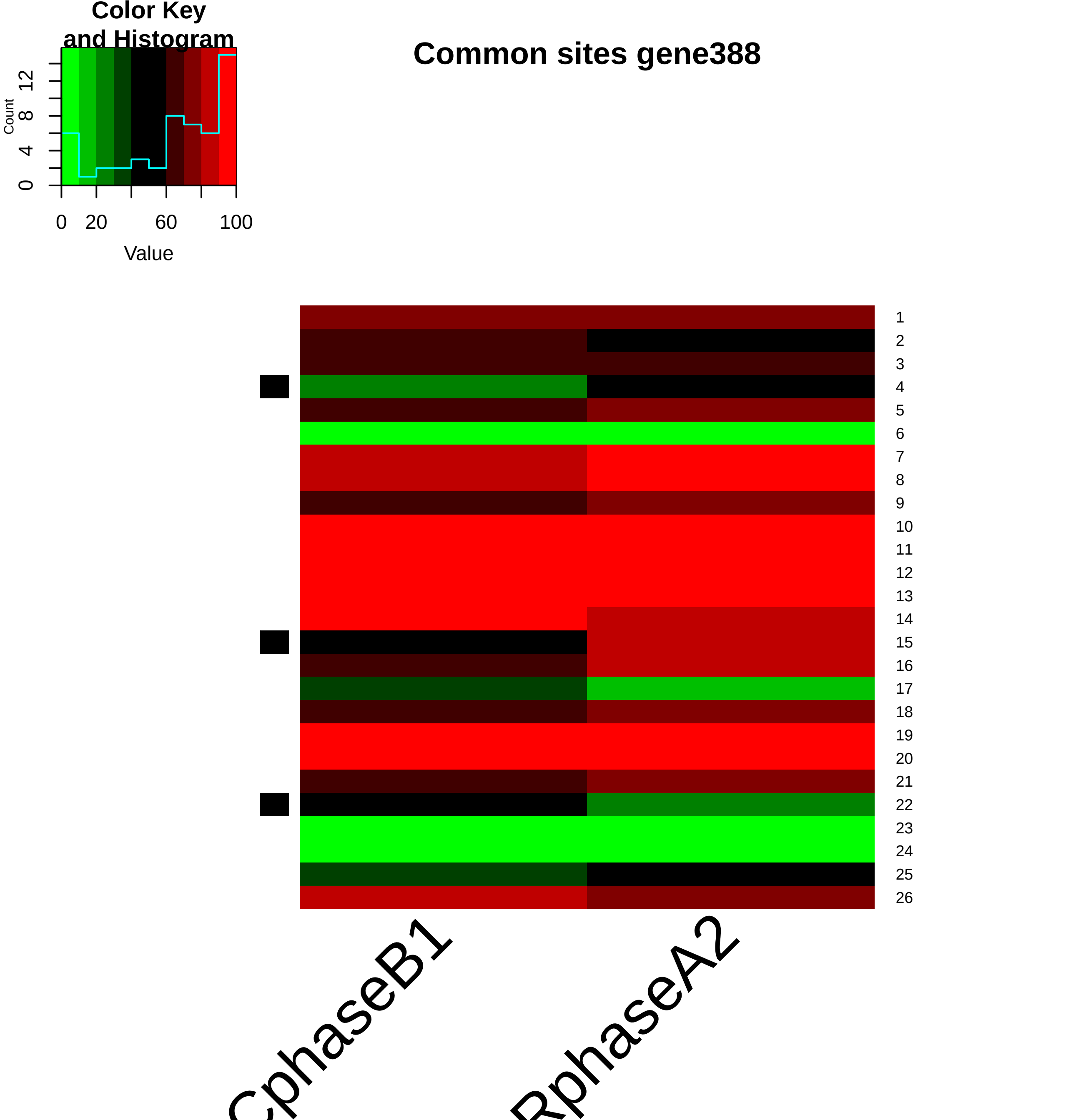

Color Key
and Histogram
Count
0
4
8
12
0
20
60
100
Value
Common sites gene388
CphaseB1
RphaseA2

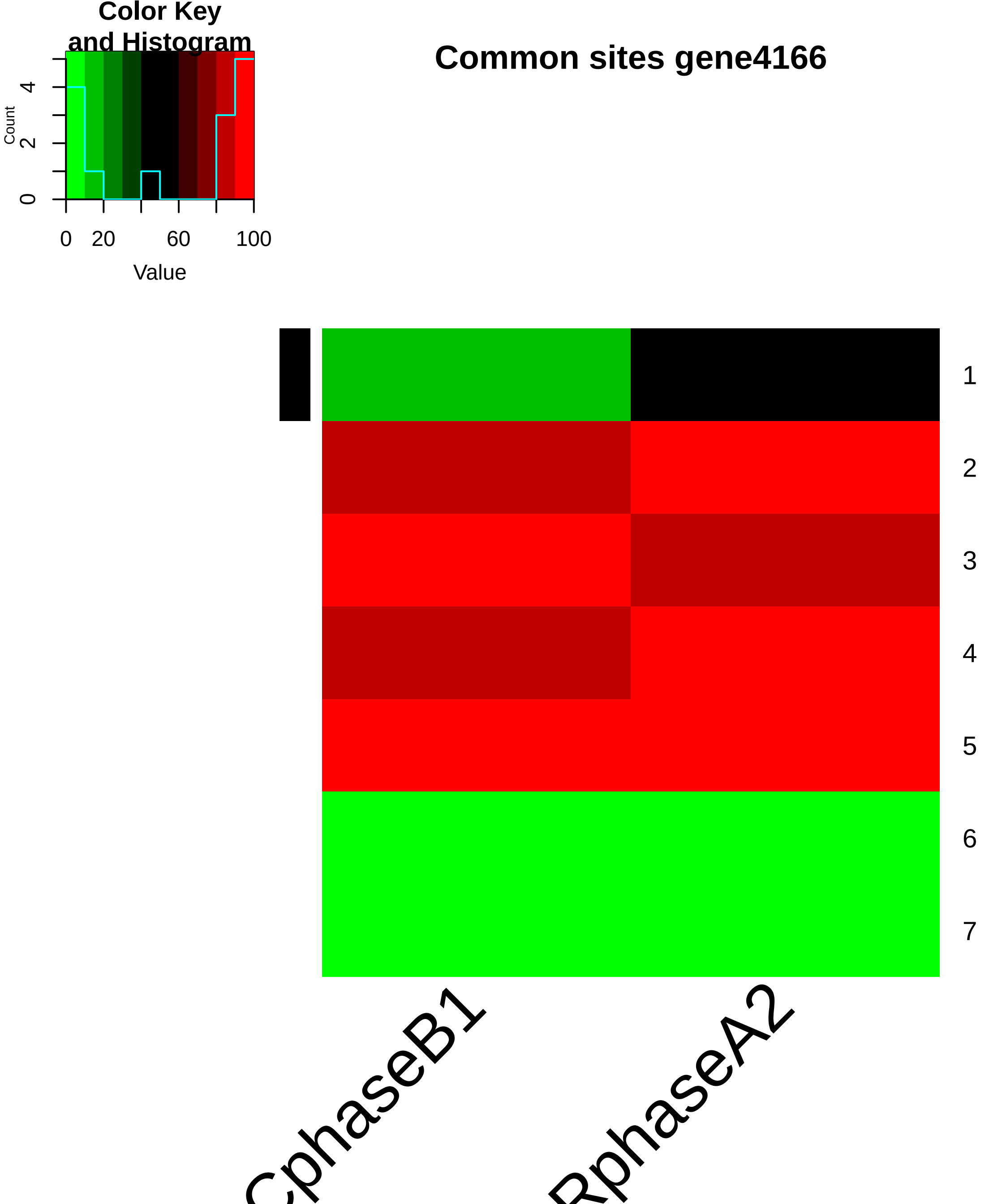

Color Key
and Histogram
Count
0
2
4
0
20
60
100
Value
Common sites gene4166
1
2
3
4
5
6
7

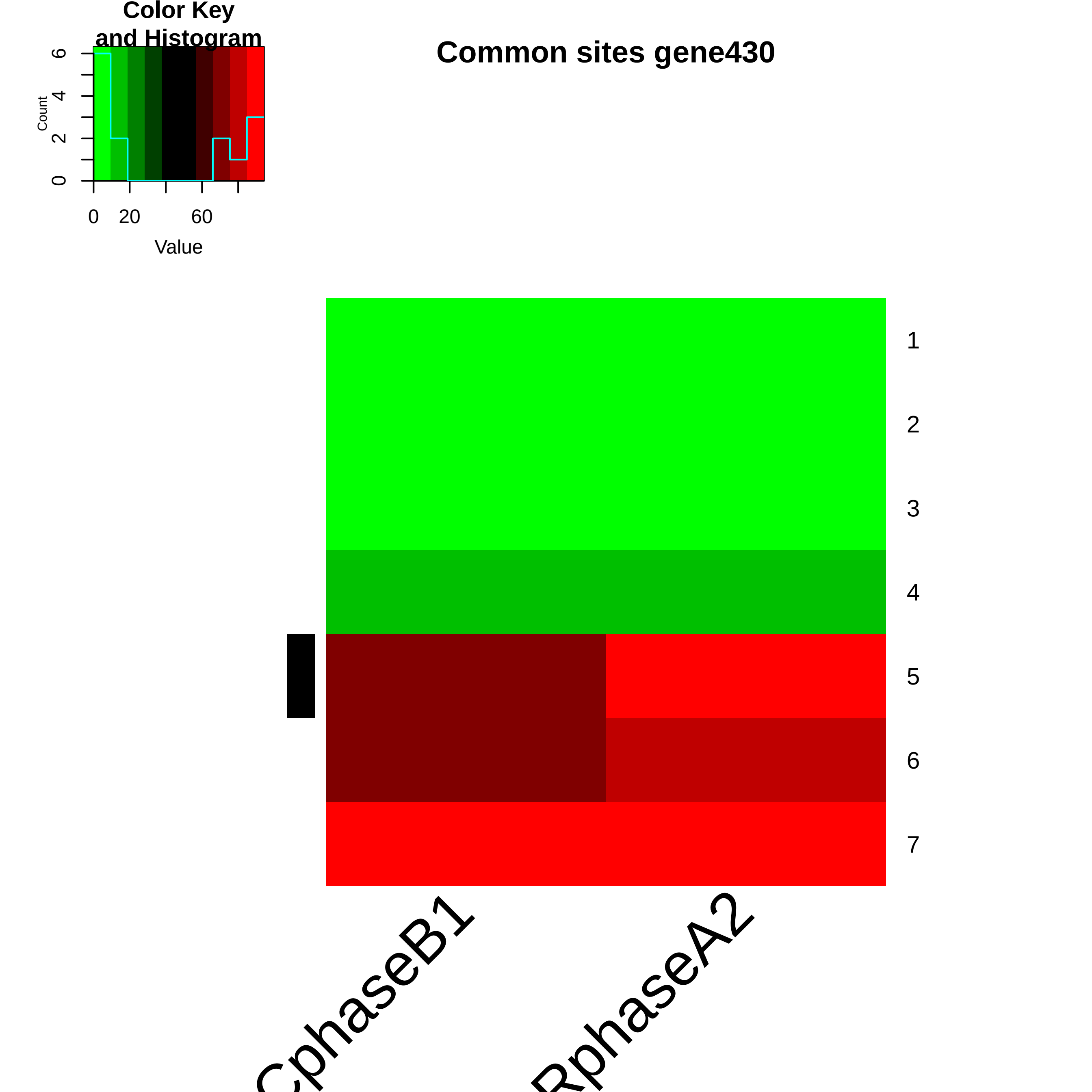

Color Key
and Histogram
Count
0
2
4
6
0
20
60
Value
Common sites gene430
1
2
3
4
5
6
7
CphaseB1
RphaseA2

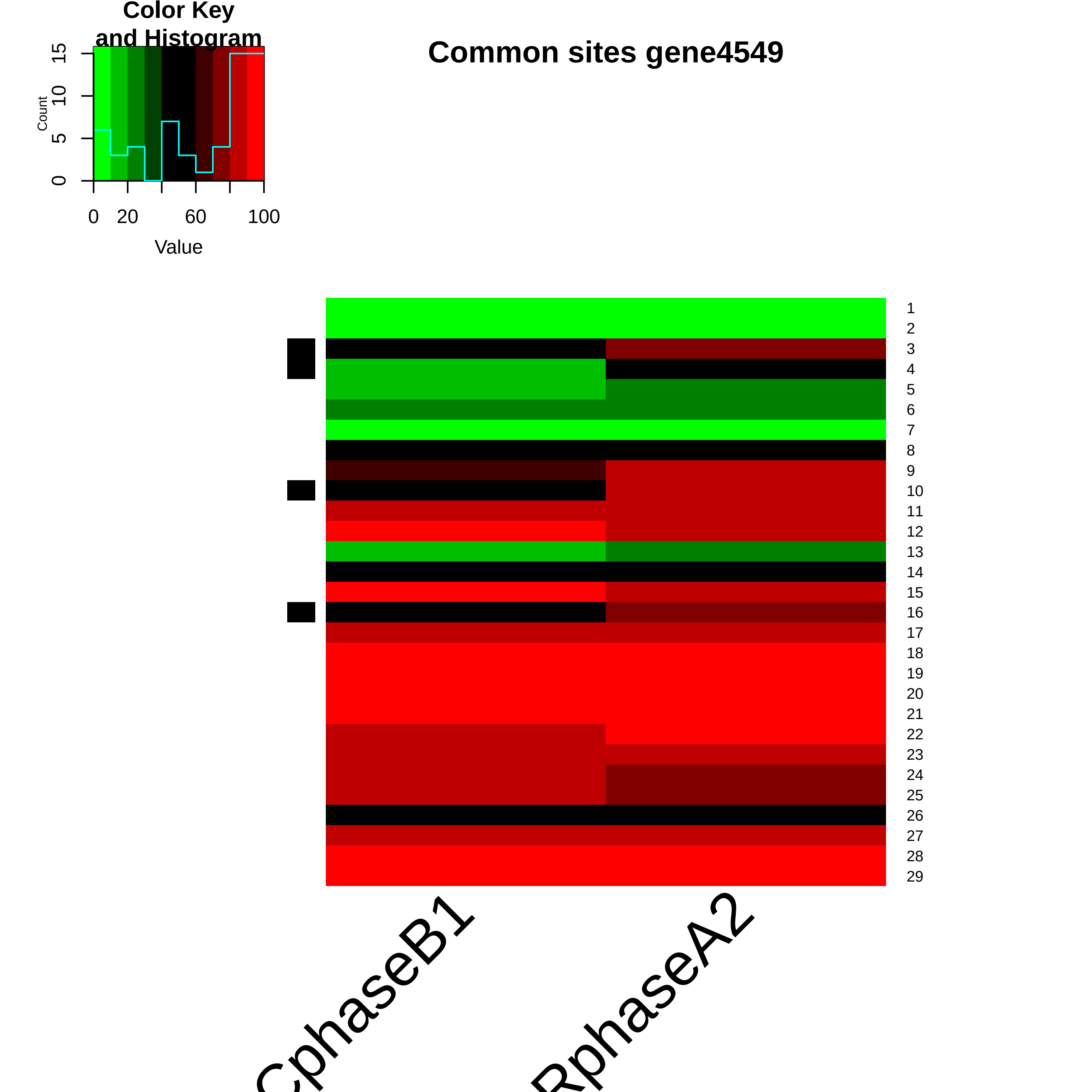
Color Key
and Histogram
Count
0
5
10
15
0
20
60
100
Value
Common sites gene4549
CphaseB1
RphaseA2

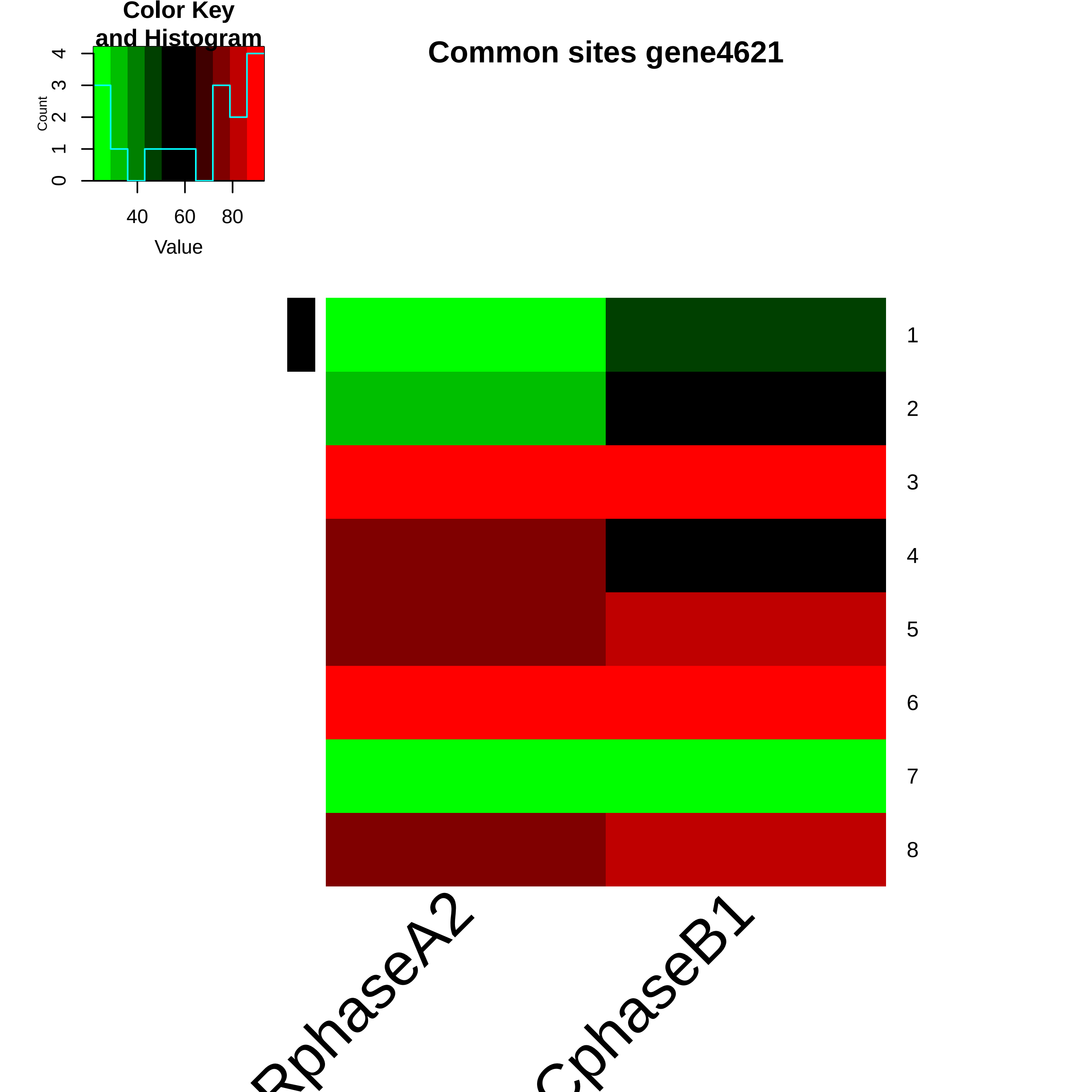
Color Key
and Histogram
Count
0
1
2
3
4
40
60
80
Value
Common sites gene4621
1
2
3
4
5
6
7
8
RphaseA2
CphaseB1

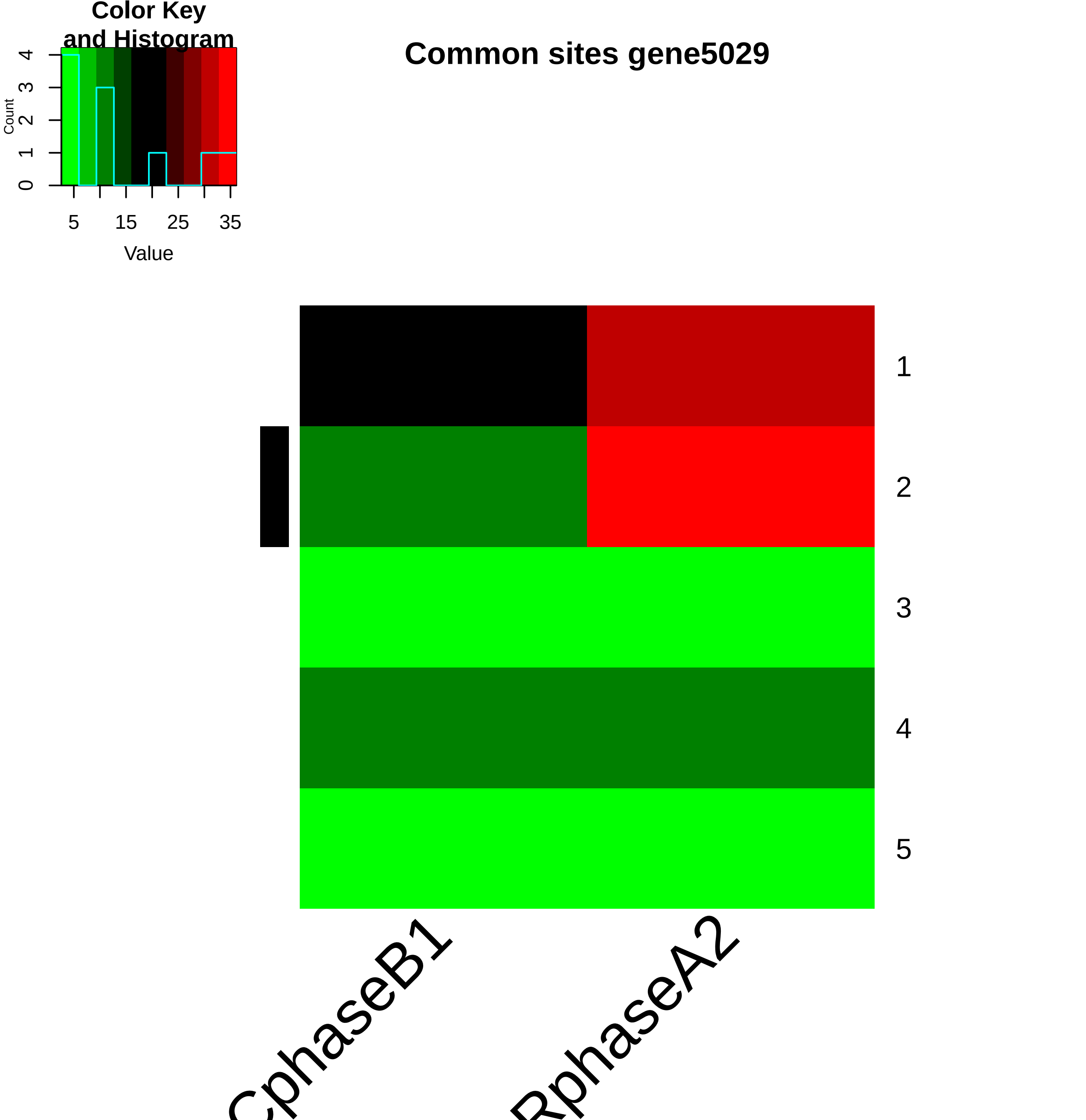
Color Key
and Histogram
Count
Value
5 15 25 35
0 1 2 3 4
Common sites gene5029
1
2
3
4
5
CphaseB1
RphaseA2

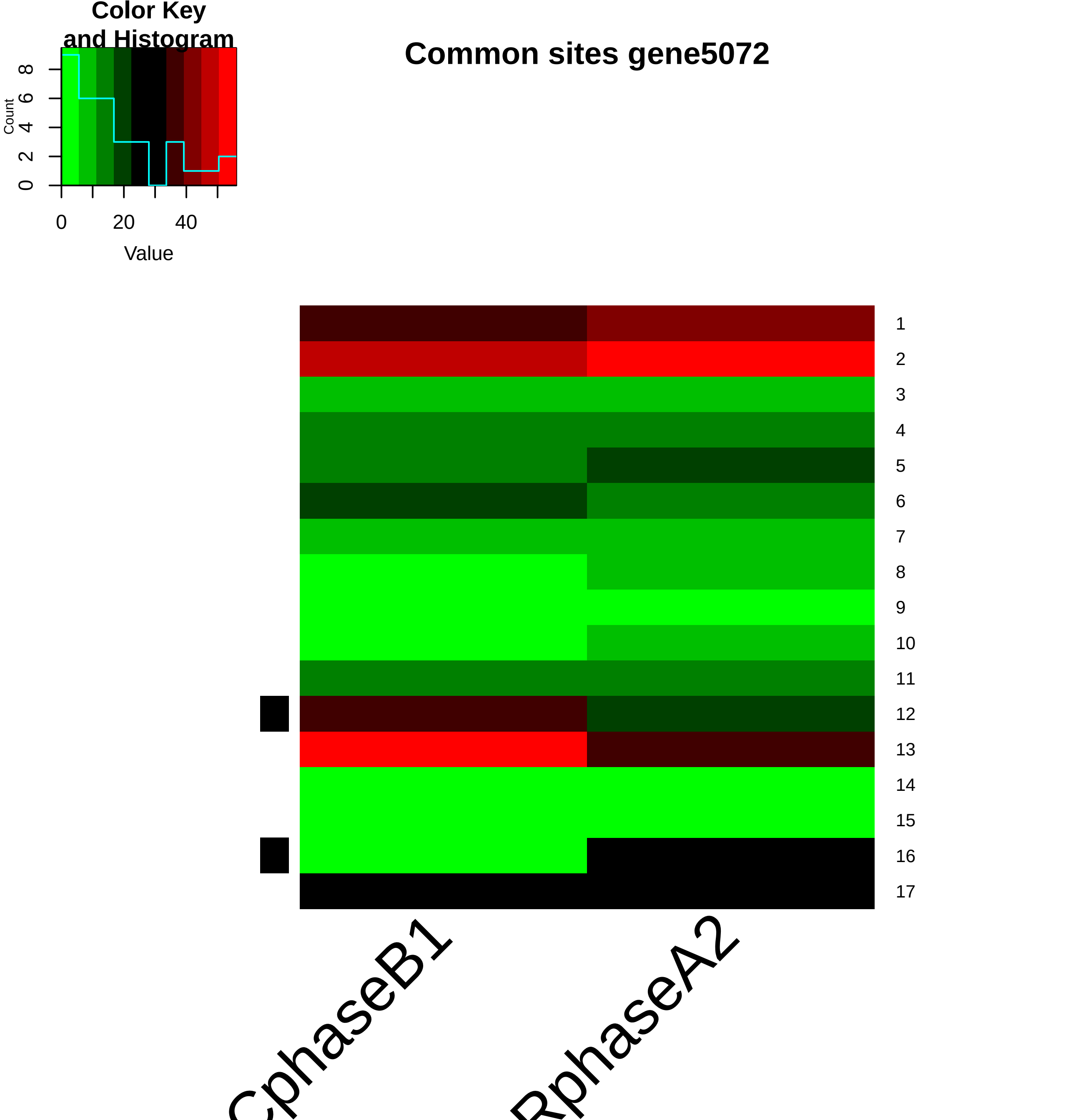
Color Key
and Histogram
Count
Value
0
20
40
0
2
4
6
8
Common sites gene5072
CphaseB1
RphaseA2

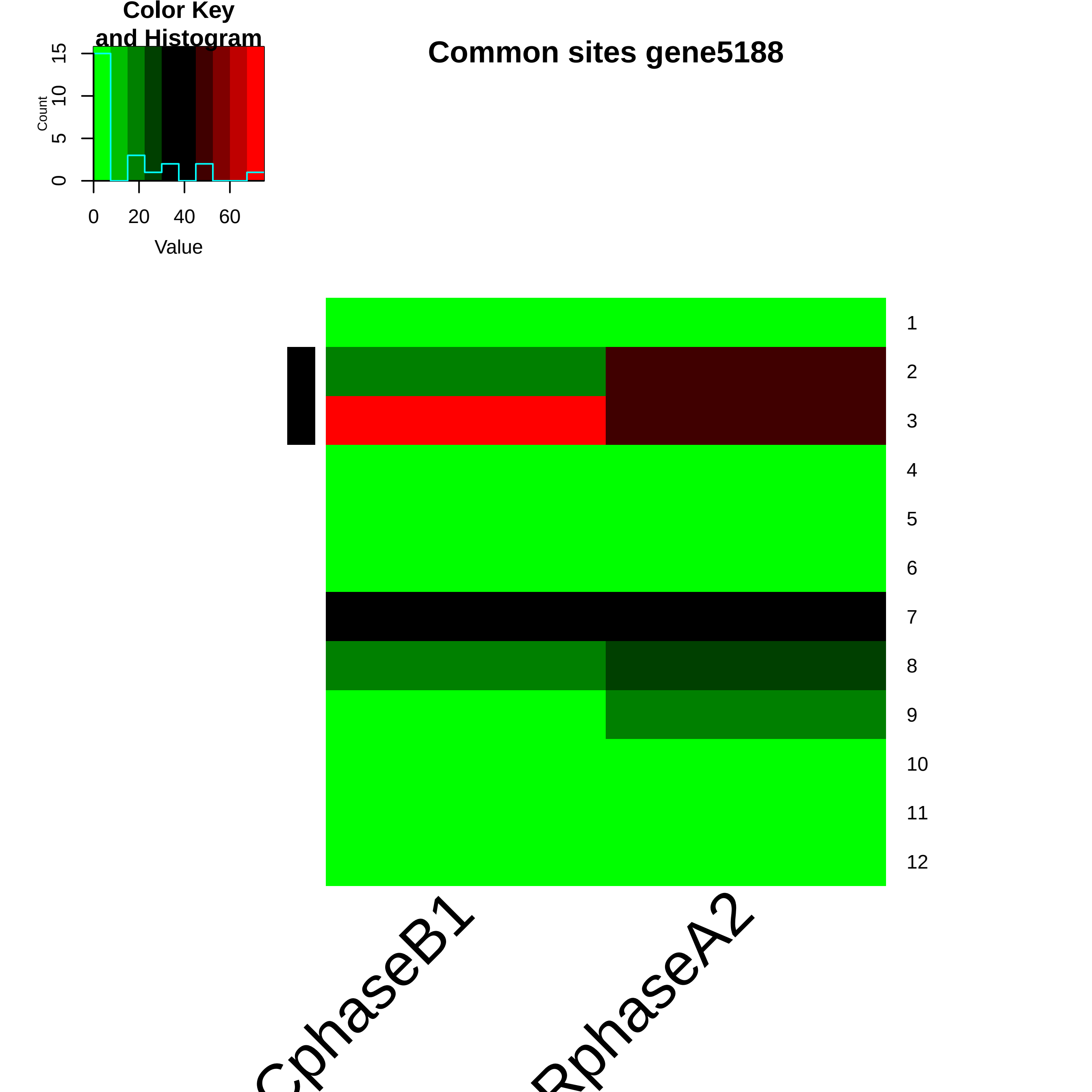
Color Key
and Histogram
Count
0
5
10
15
0
20
40
60
Value
Common sites gene5188
1
2
3
4
5
6
7
8
9
10
11
12
CphaseB1
RphaseA2

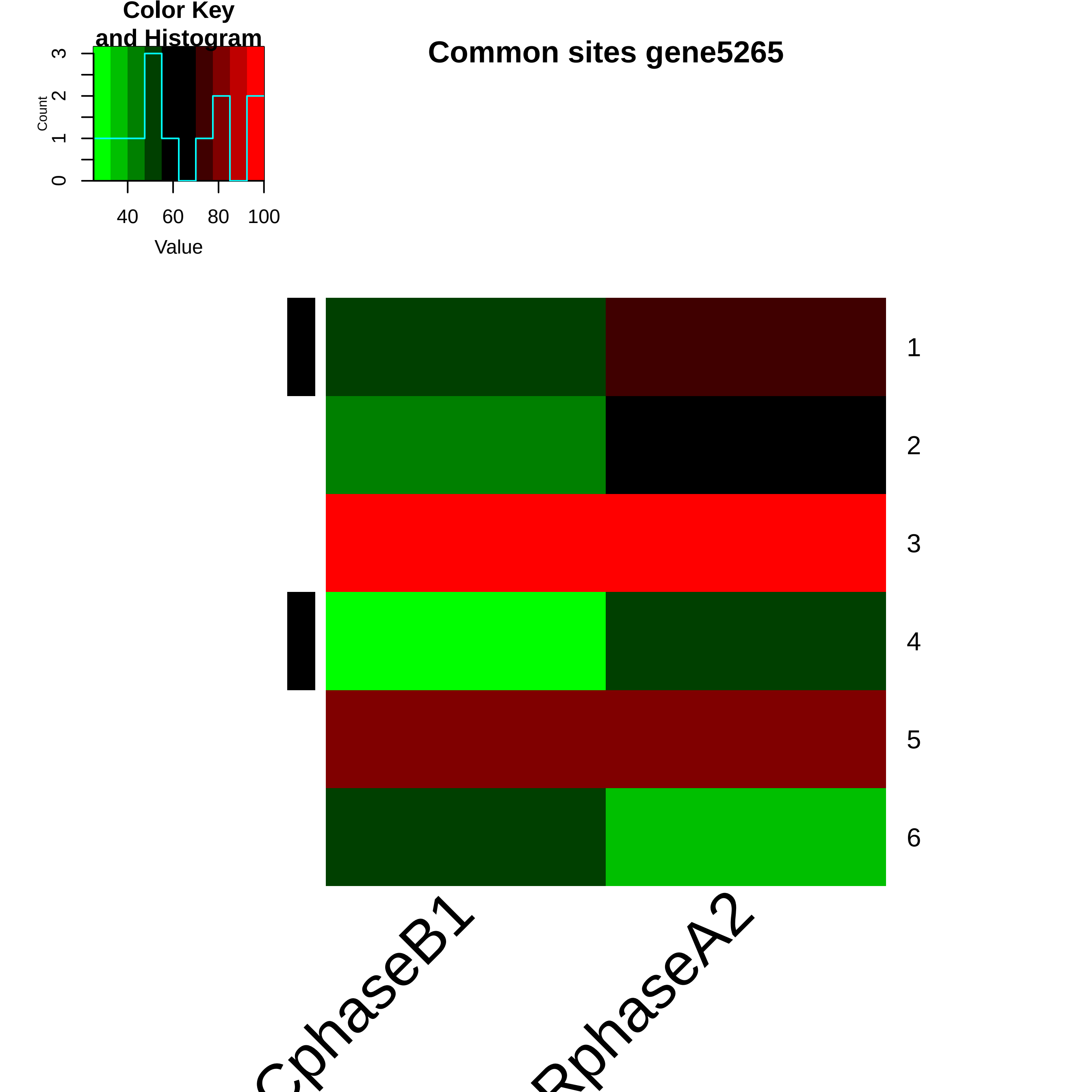
Color Key
and Histogram
Count
0
1
2
3
40
60
80
100
Value
Common sites gene5265
1
2
3
4
5
6
CphaseB1
RphaseA2

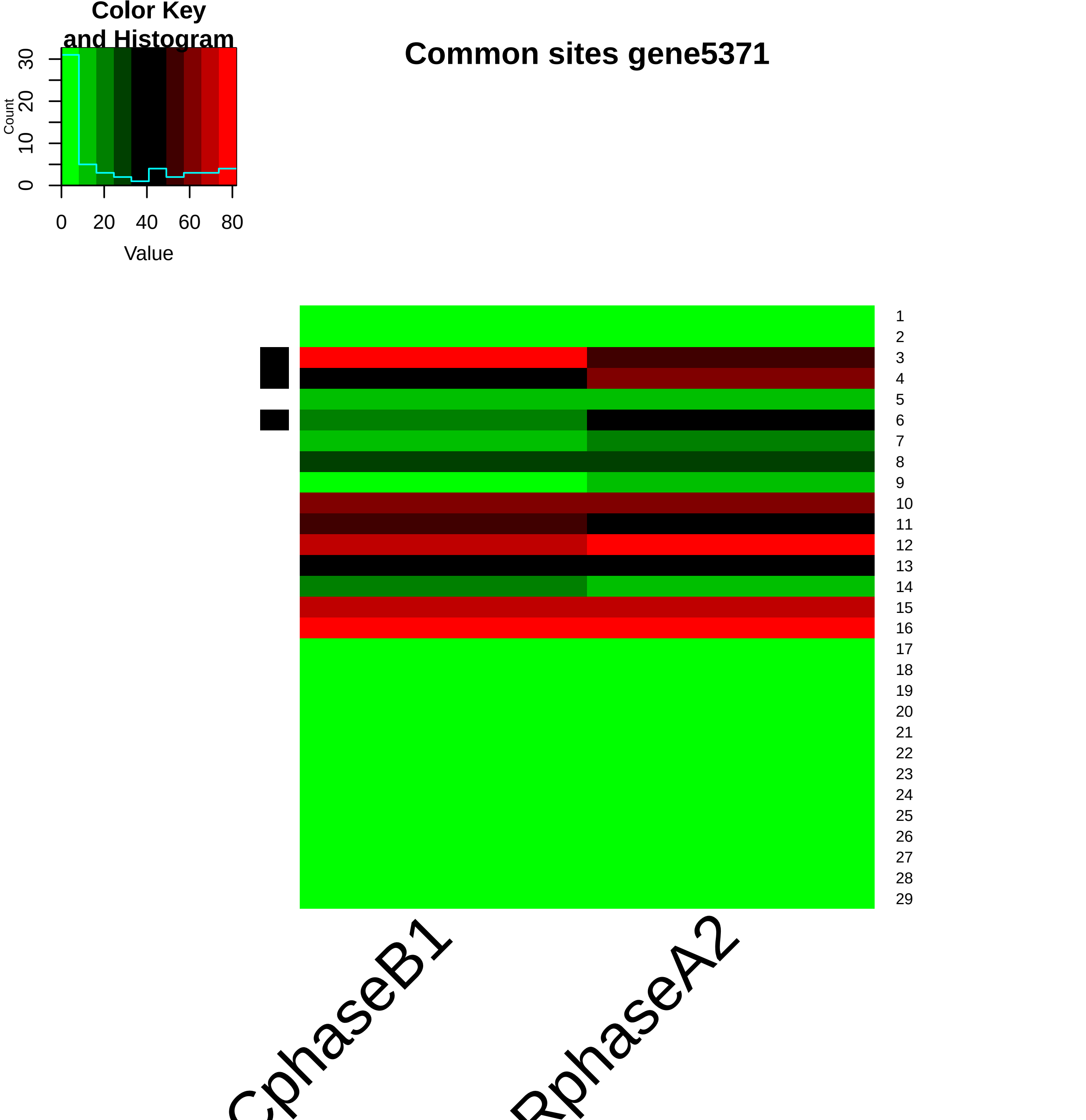
Color Key
and Histogram
Count
0
10
20
30
0
20
40
60
80
Value
Common sites gene5371
CphaseB1
RphaseA2

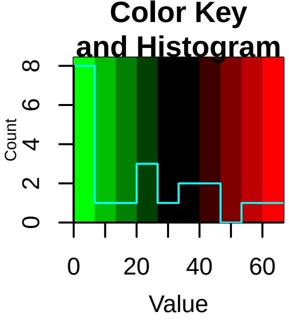

Color Key
and Histogram
Count
0
2
4
6
8
0
20
40
60
Value

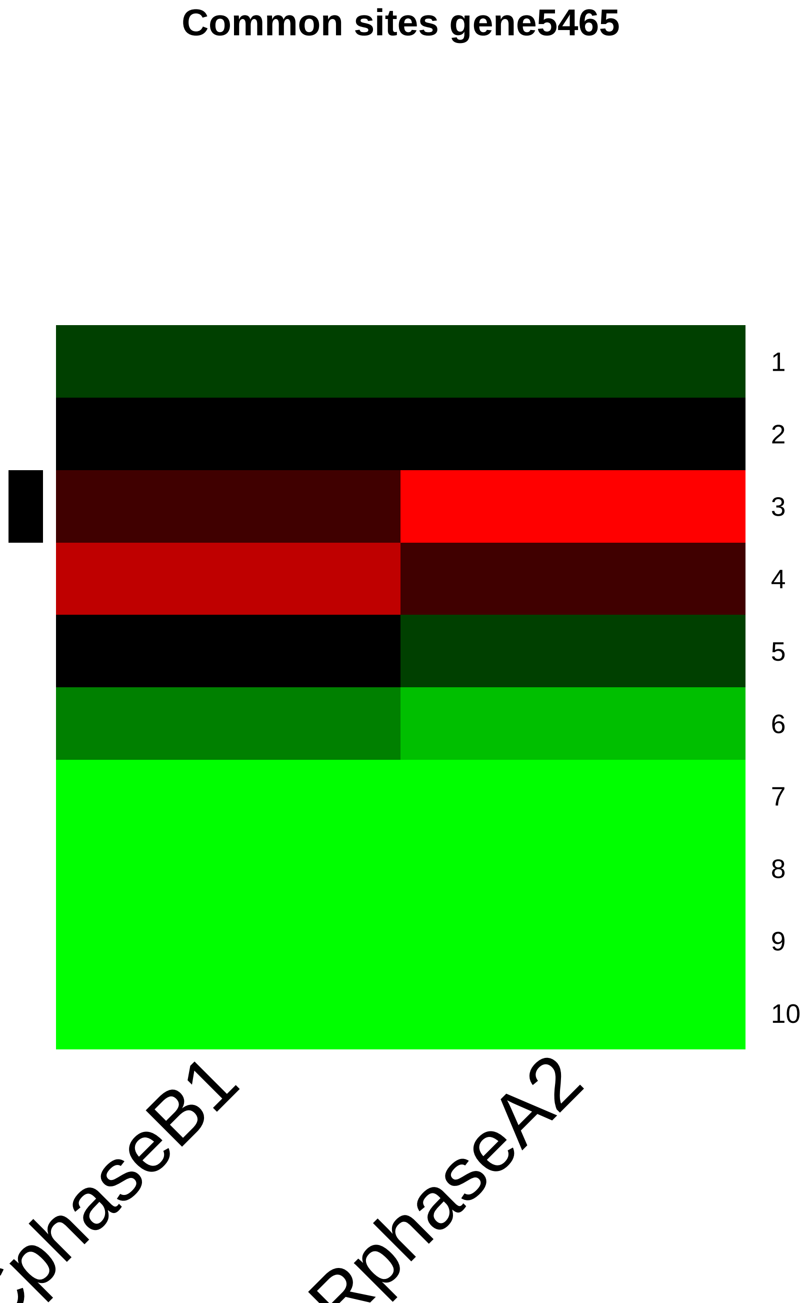

Common sites gene5465
1
2
3
4
5
6
7
8
9
10
CphaseB1
RphaseA2

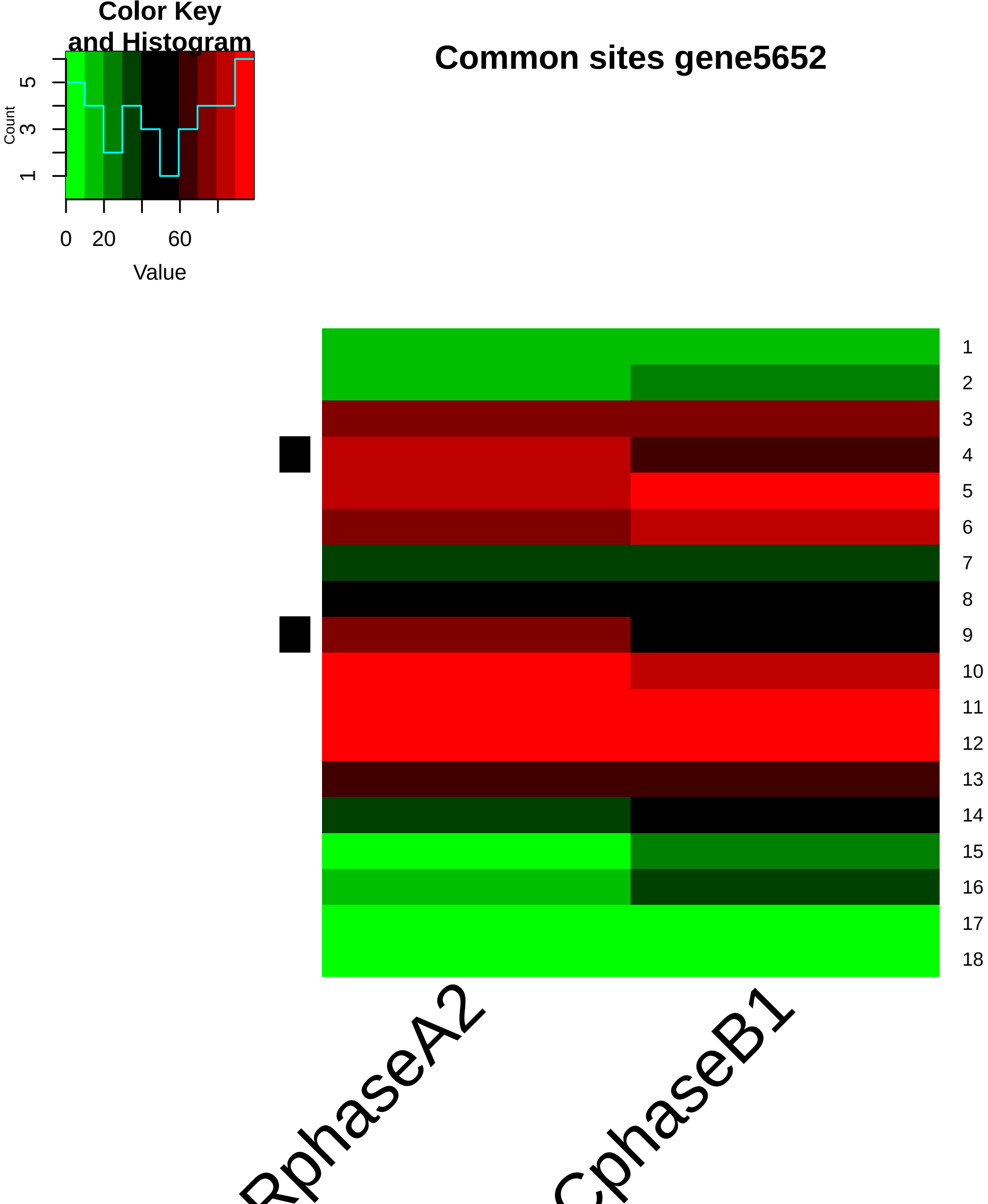
Color Key
and Histogram
Count
1
3
5
0
20
60
Value
Common sites gene5652
RphaseA2
CphaseB1

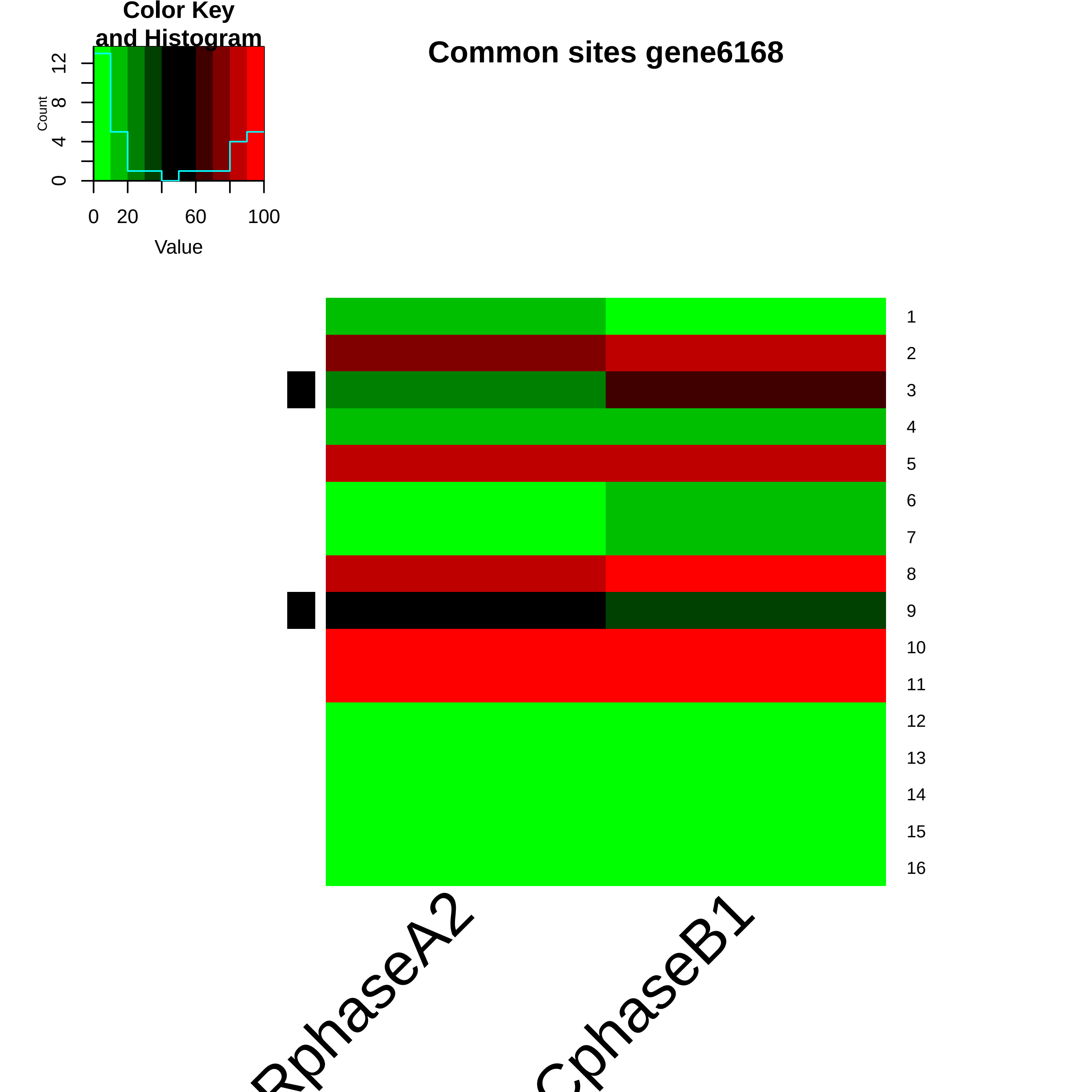
Color Key
and Histogram
Count
0
4
8
12
0
20
60
100
Value
Common sites gene6168
RphaseA2
CphaseB1

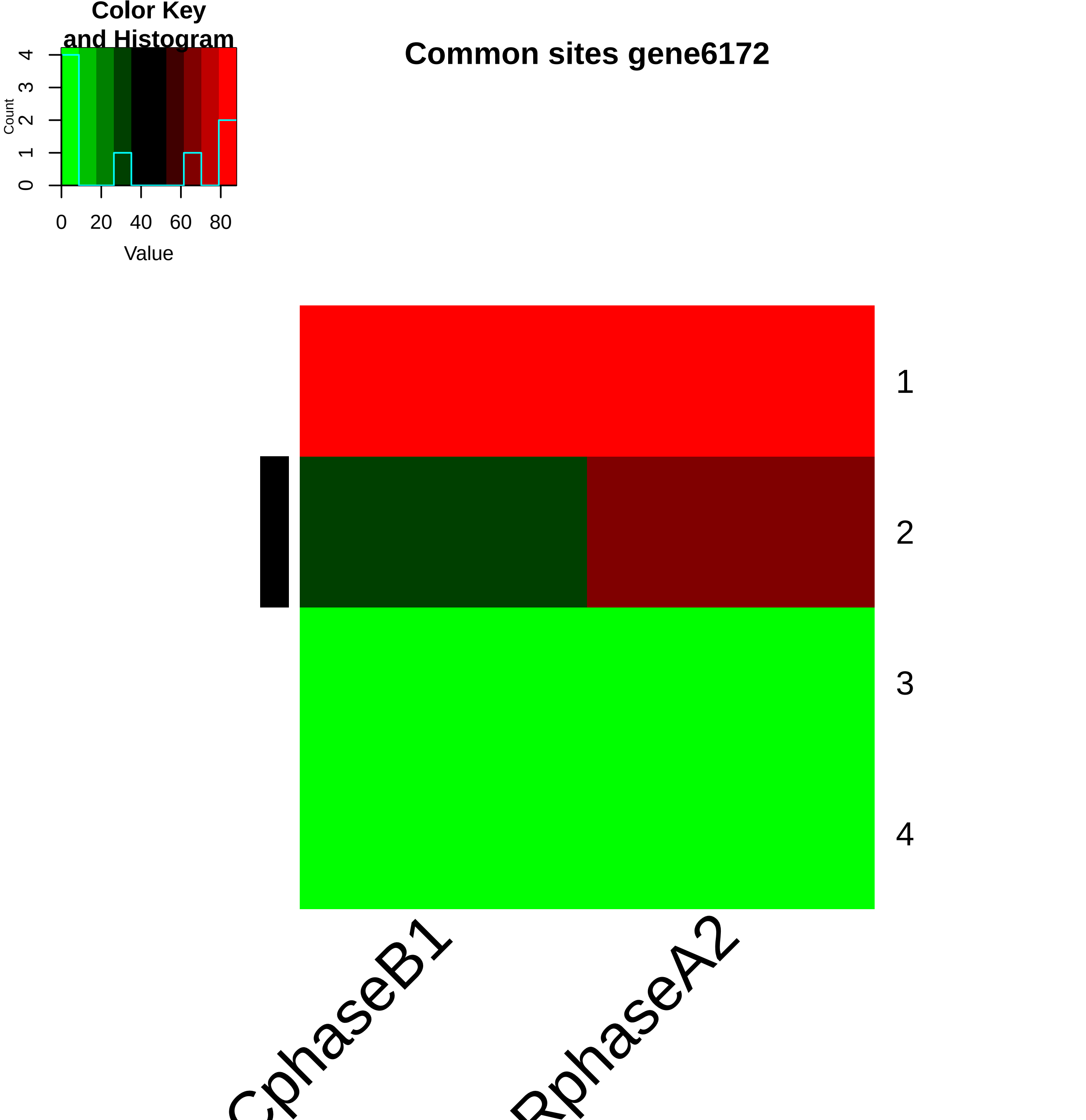
Color Key
and Histogram
Count
0
1
2
3
4
0
20
40
60
80
Value
Common sites gene6172
1
2
3
4
CphaseB1
RphaseA2

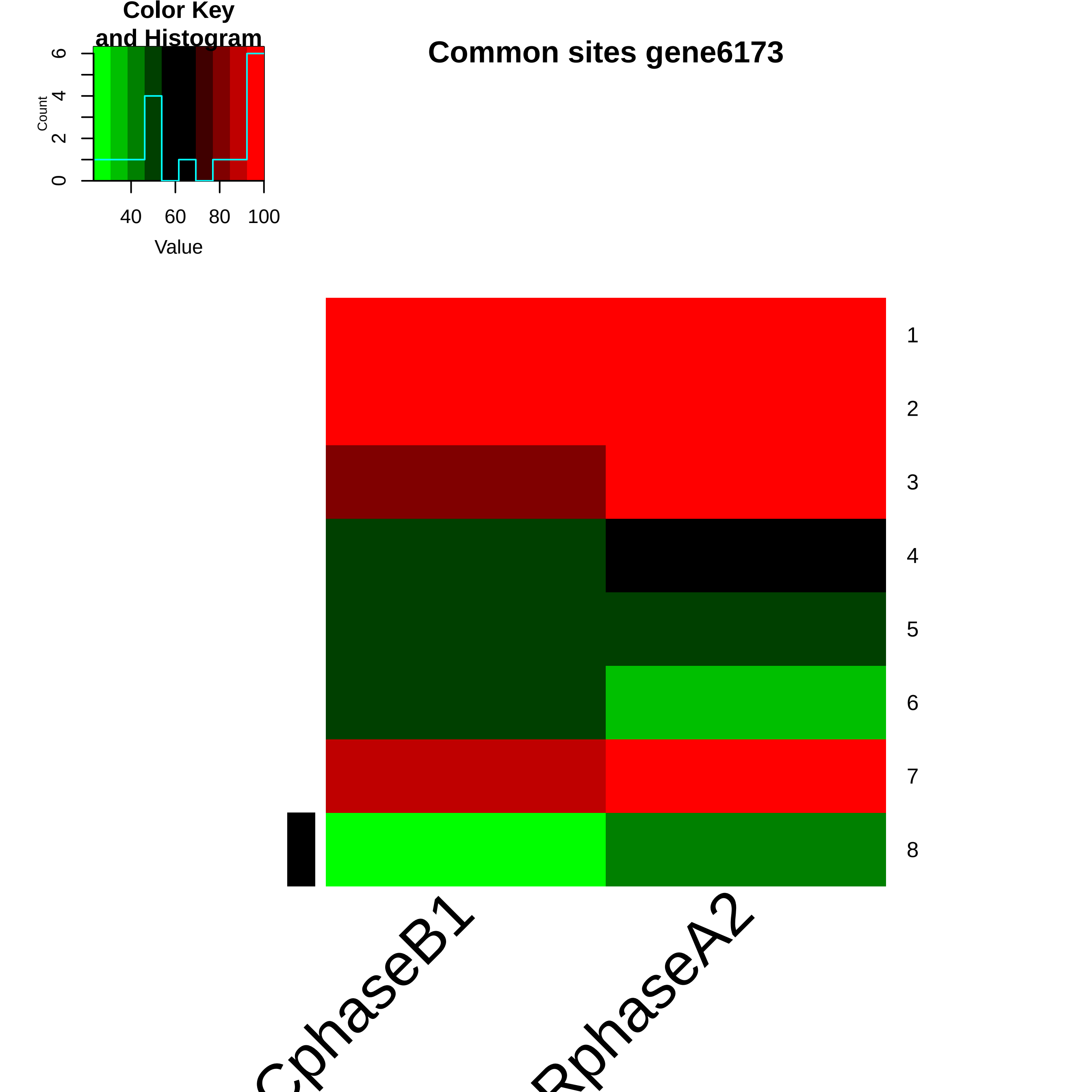

Color Key
and Histogram
Count
0
2
4
6
40
60
80
100
Value
Common sites gene6173
1
2
3
4
5
6
7
8
CphaseB1
RphaseA2

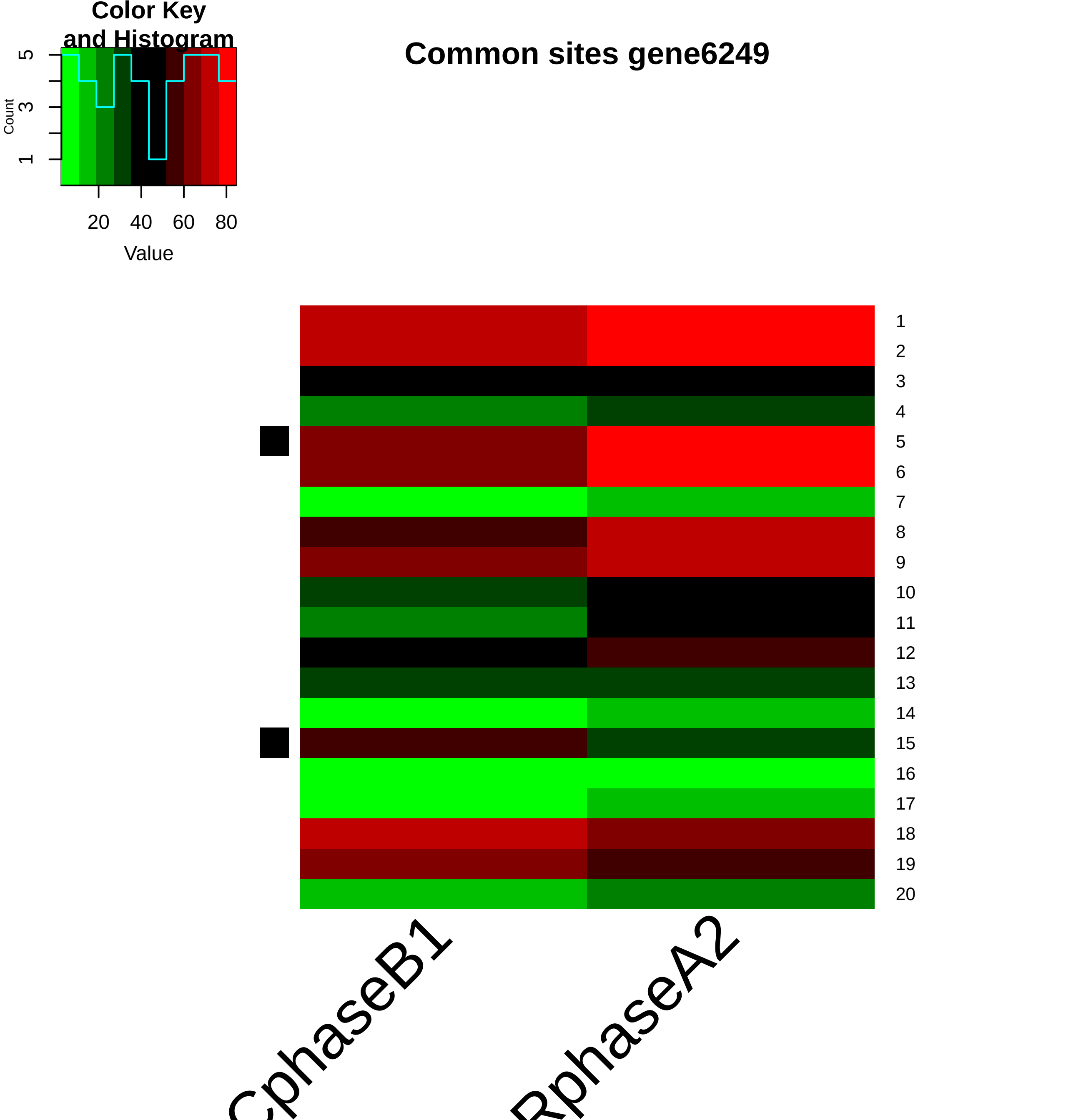
Color Key
and Histogram
Count
1
3
5
20
40
60
80
Value
Common sites gene6249
CphaseB1
RphaseA2

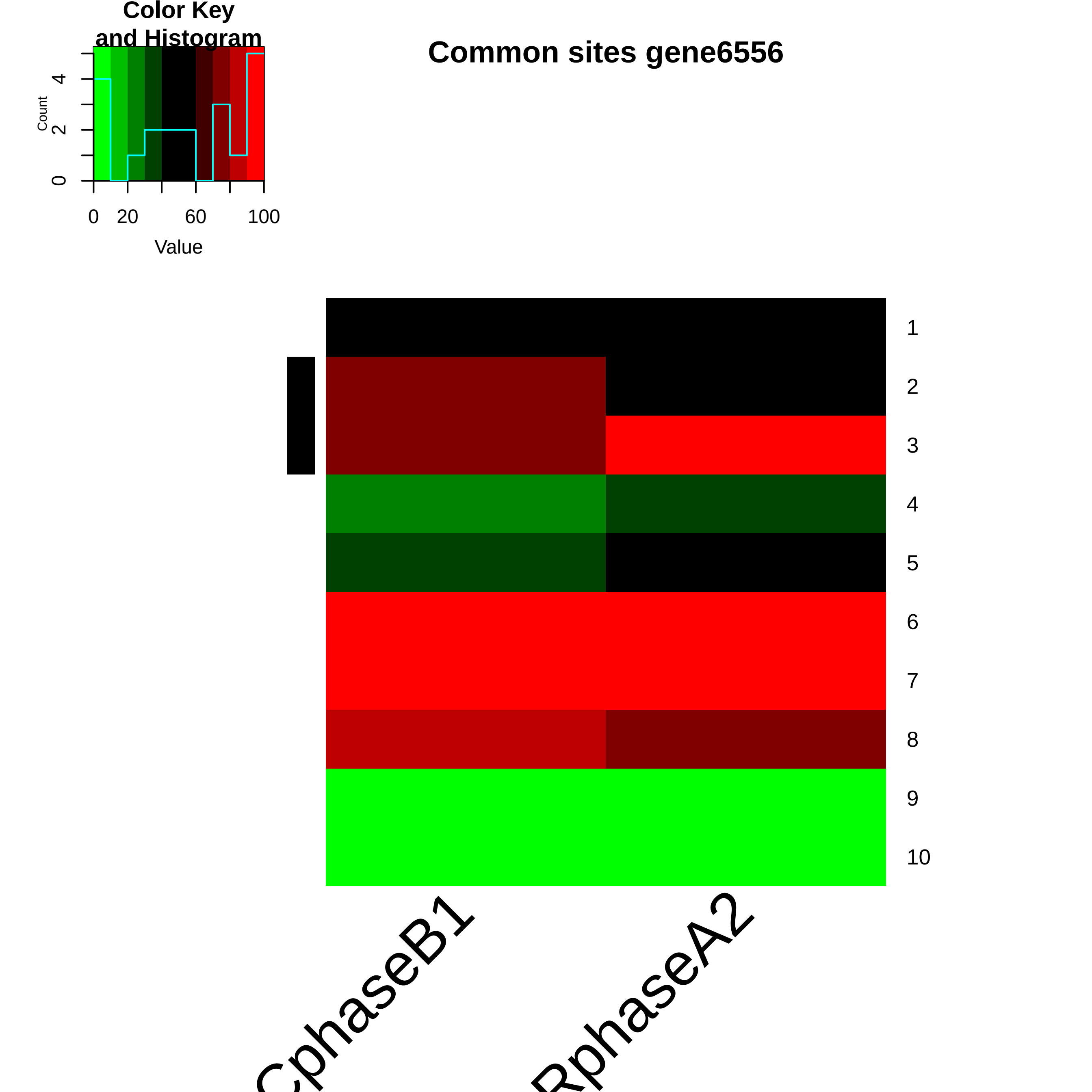
Color Key
and Histogram
Count
0
2
4
0
20
60
100
Value
Common sites gene6556
1
2
3
4
5
6
7
8
9
10
CphaseB1
RphaseA2

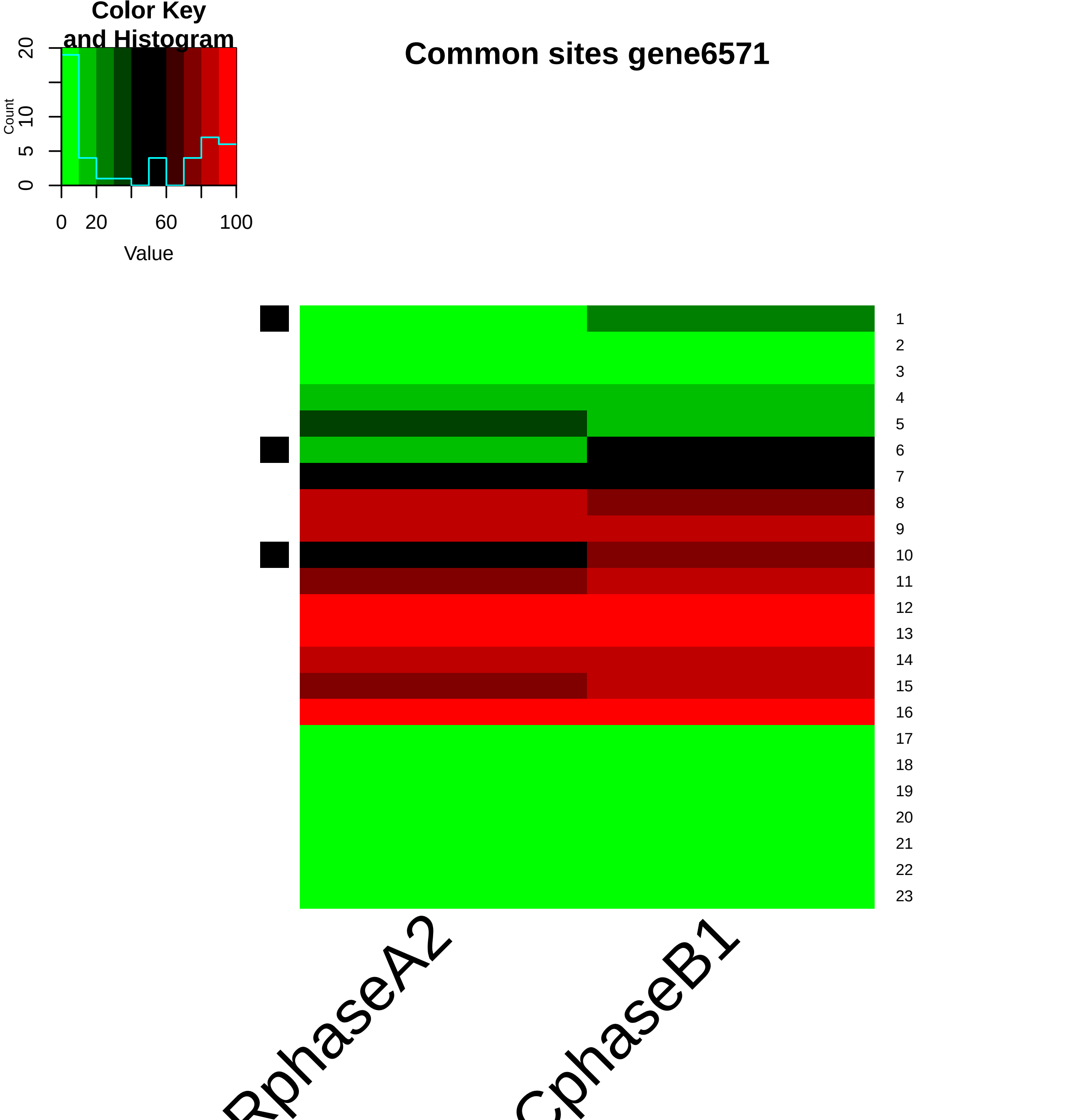

Color Key
and Histogram
Count
Value
0
5
10
20
0
20
60
100
Common sites gene6571
RphaseA2
CphaseB1

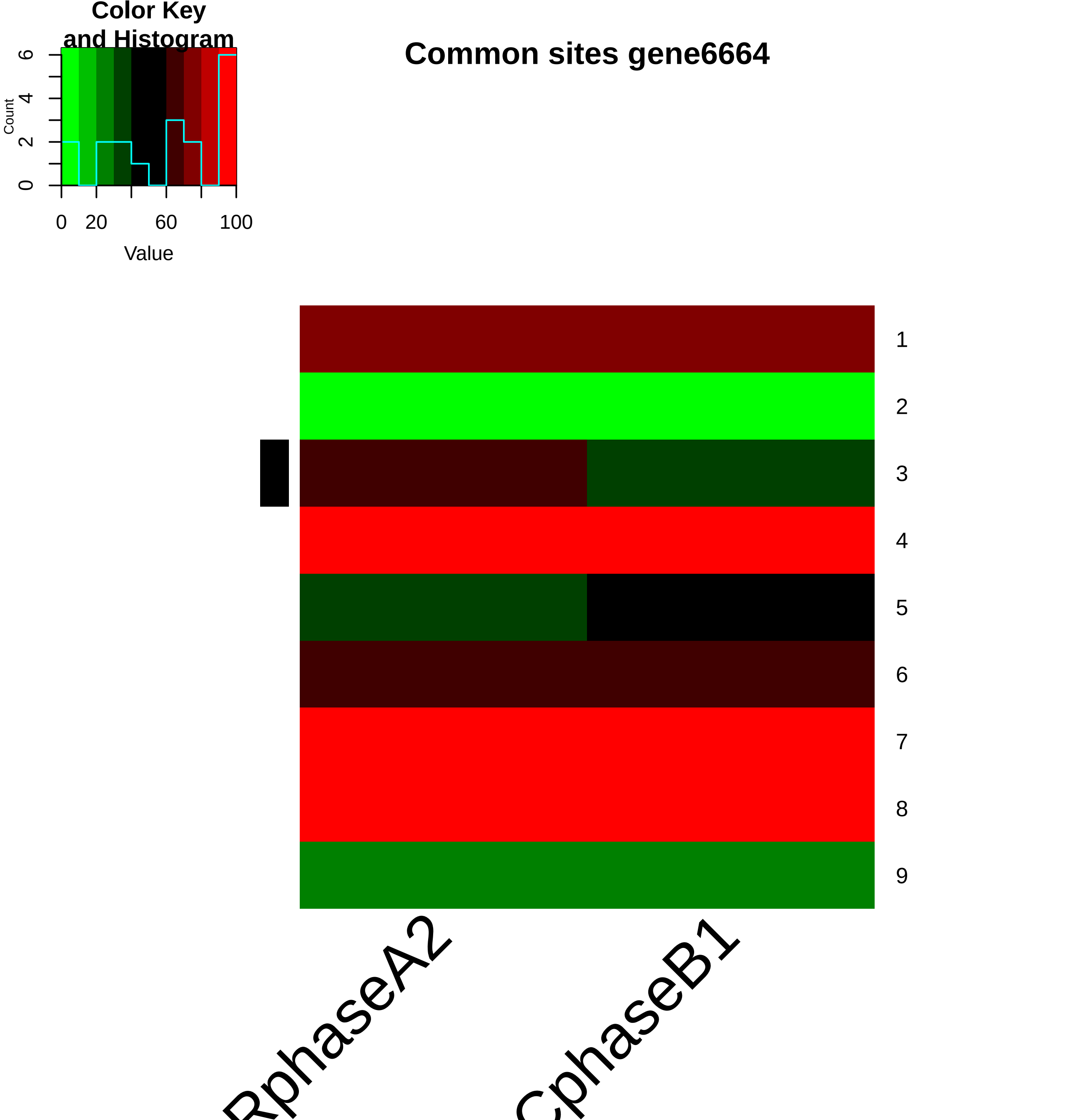
Color Key
and Histogram
Count
0
2
4
6
0
20
60
100
Value
Common sites gene6664
1
2
3
4
5
6
7
8
9
RphaseA2
CphaseB1

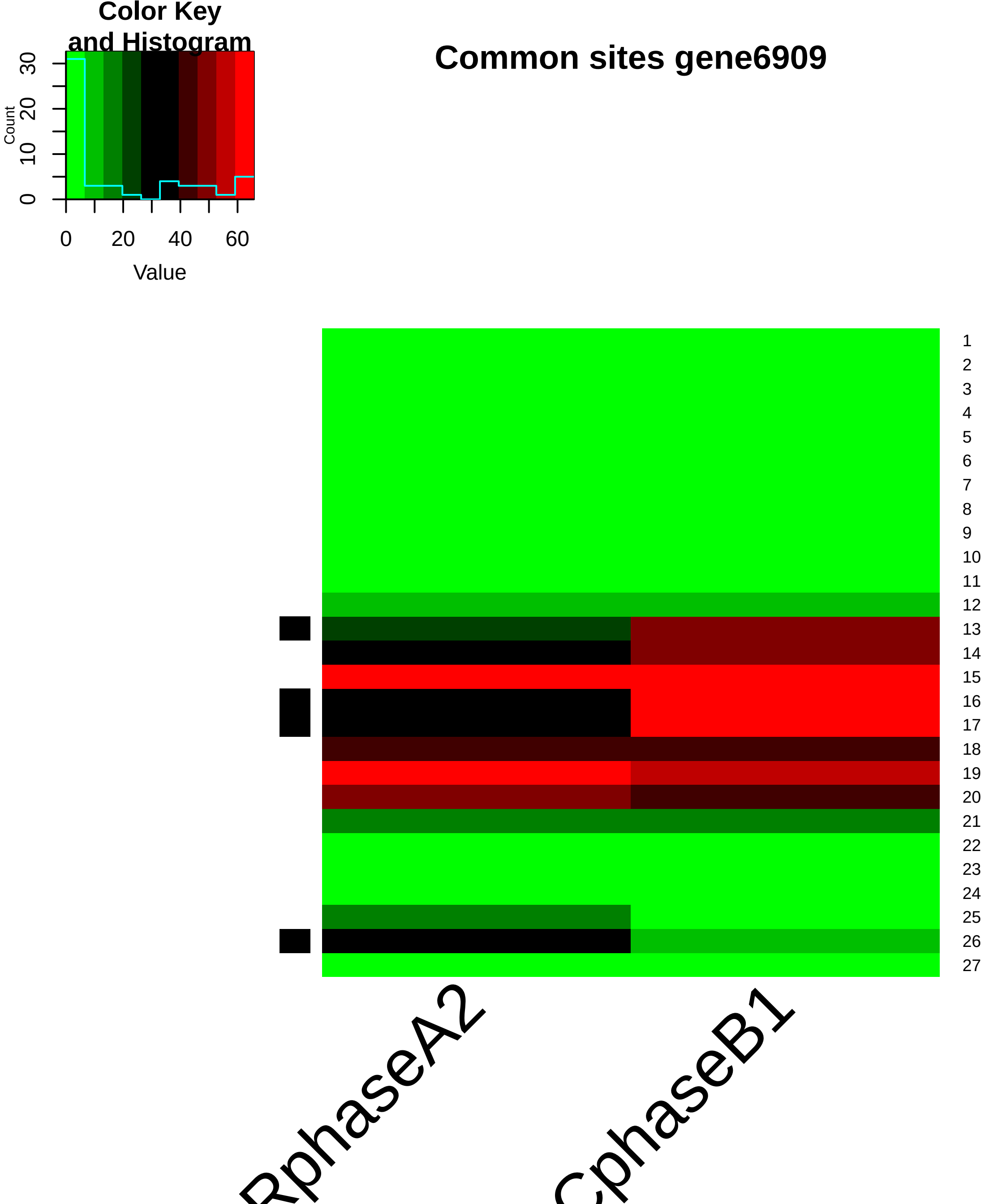
Color Key
and Histogram
Count
0
10
20
30
0
20
40
60
Value
Common sites gene6909
RphaseA2
CphaseB1

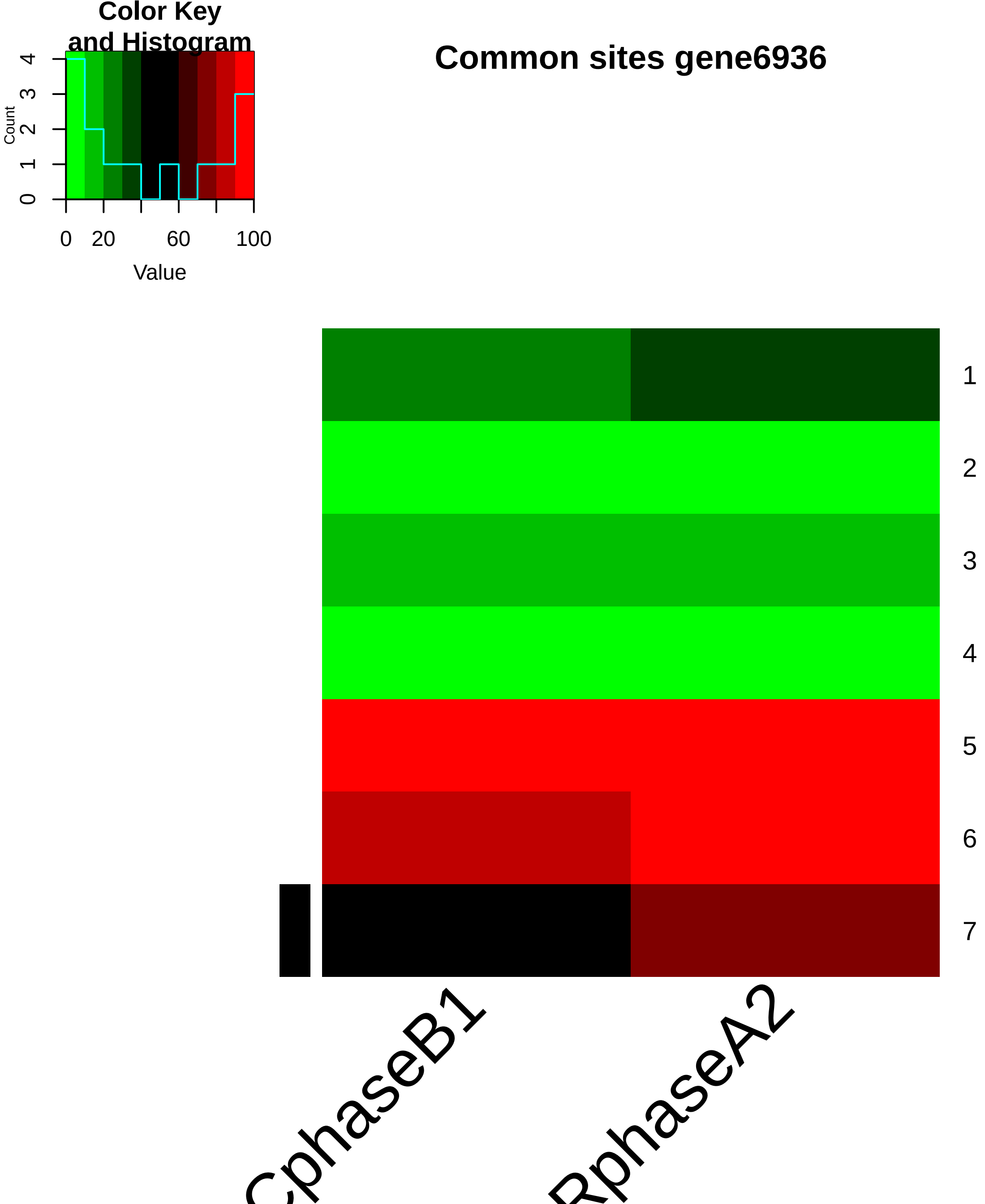
Color Key
and Histogram
Count
Value
0
20
60
100
0
1
2
3
4
Common sites gene6936
1
2
3
4
5
6
7
RphaseA2

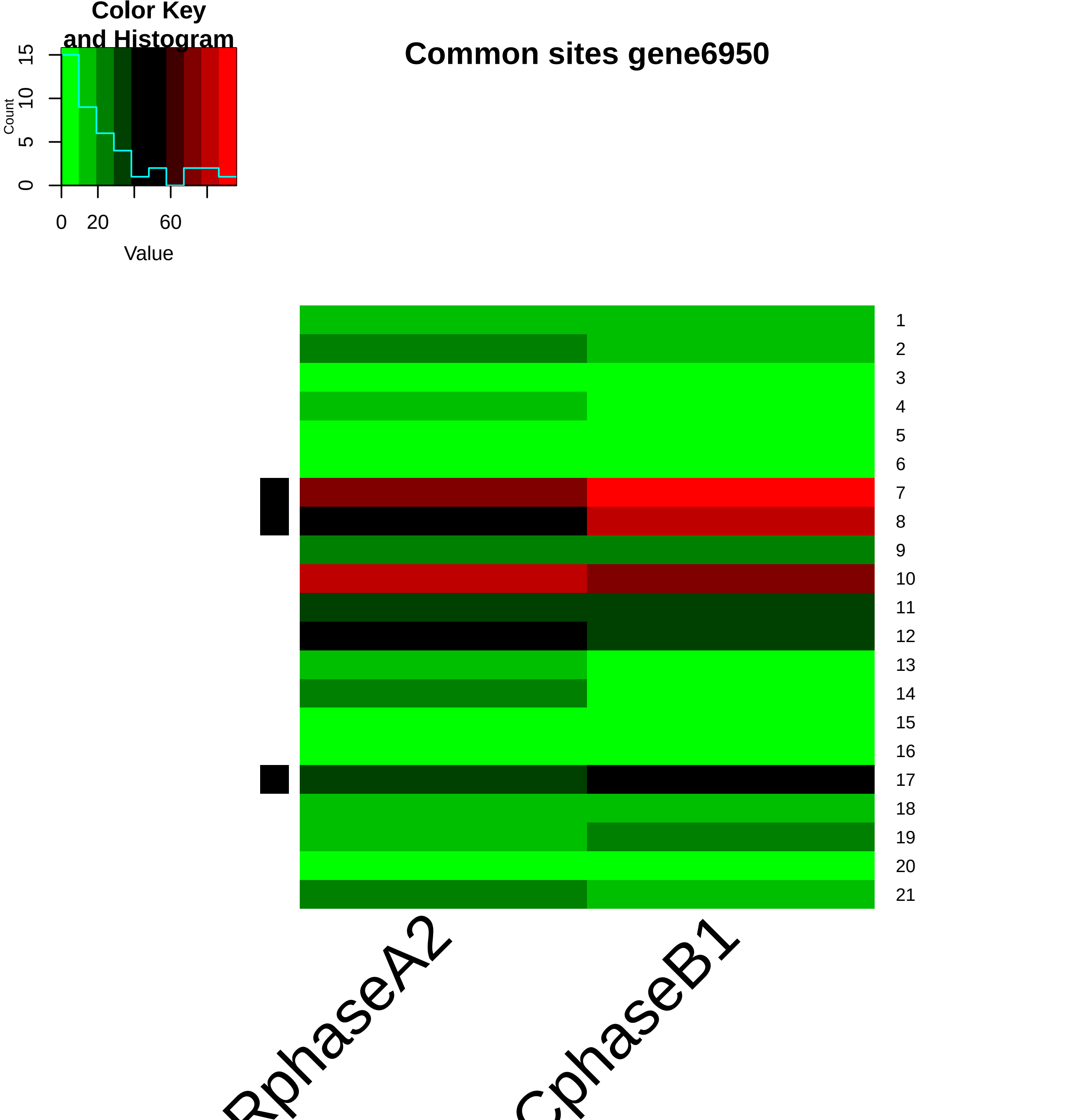

Color Key
and Histogram
Count
Value
0
20
60
0
5
10
15
Common sites gene6950
RphaseA2
CphaseB1

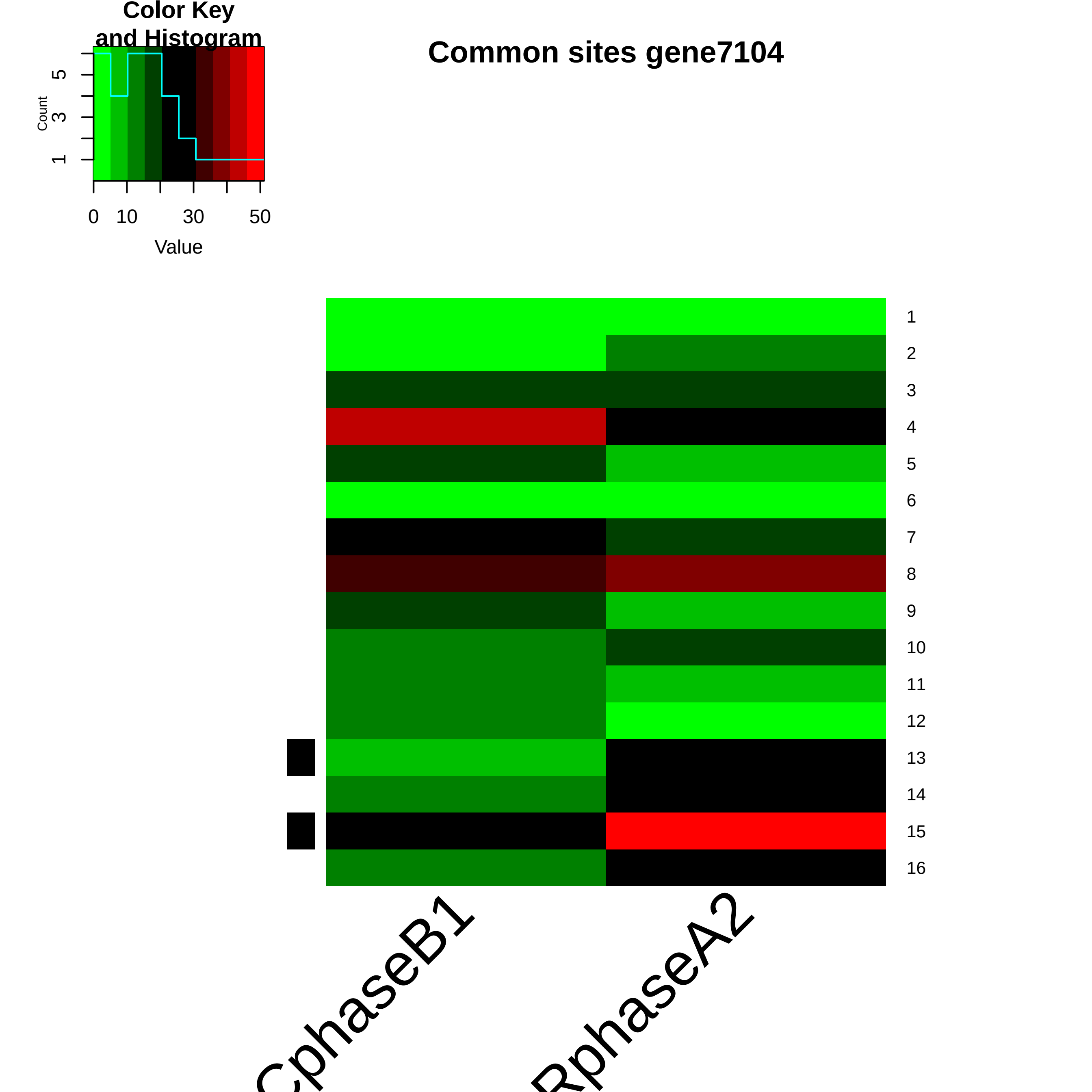

Color Key
and Histogram
Count
1
3
5
0
10
30
50
Value
Common sites gene7104
CphaseB1
RphaseA2

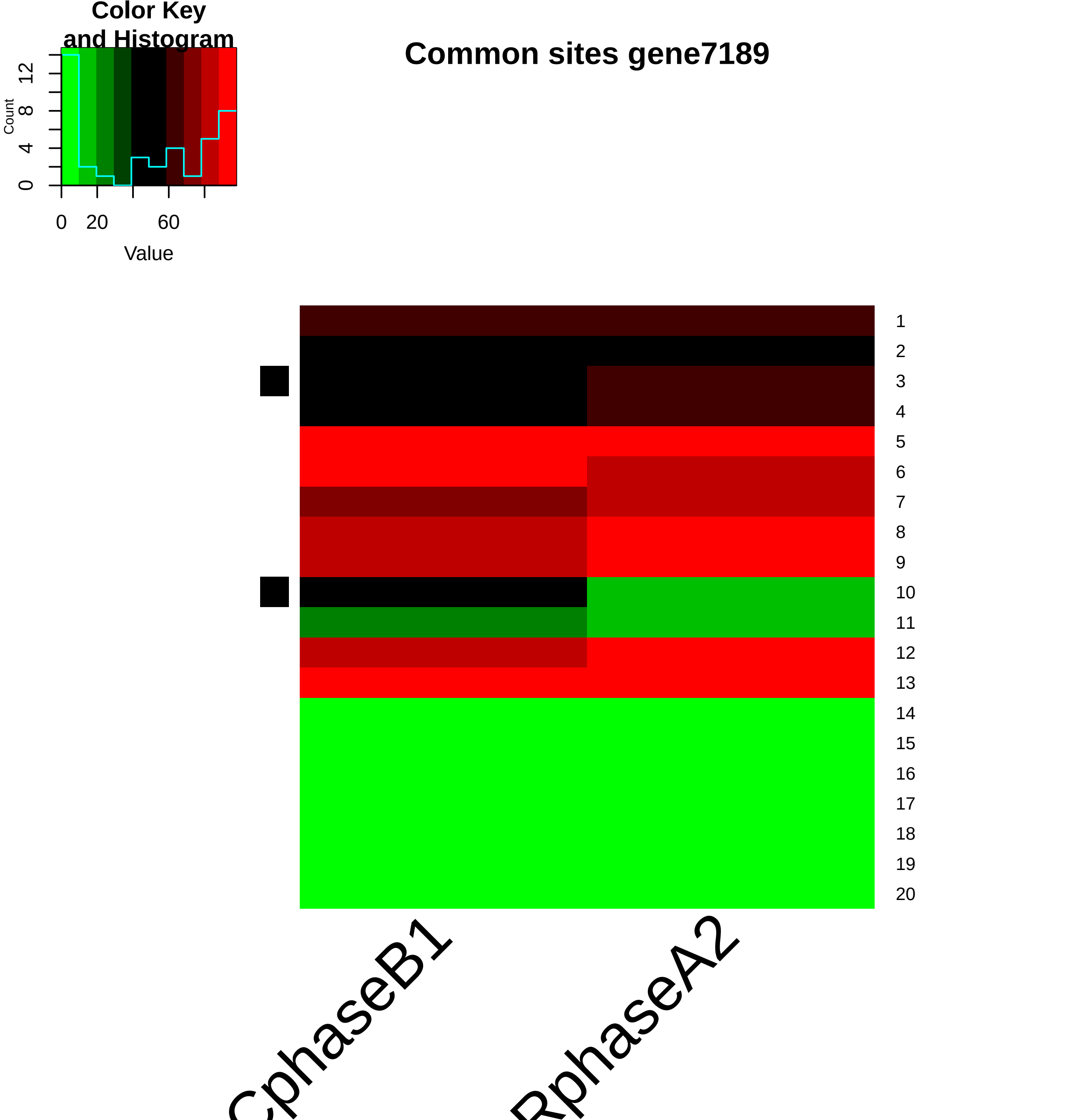
Color Key
and Histogram
Count
0
4
8
12
0
20
60
Value
Common sites gene7189
CphaseB1
RphaseA2

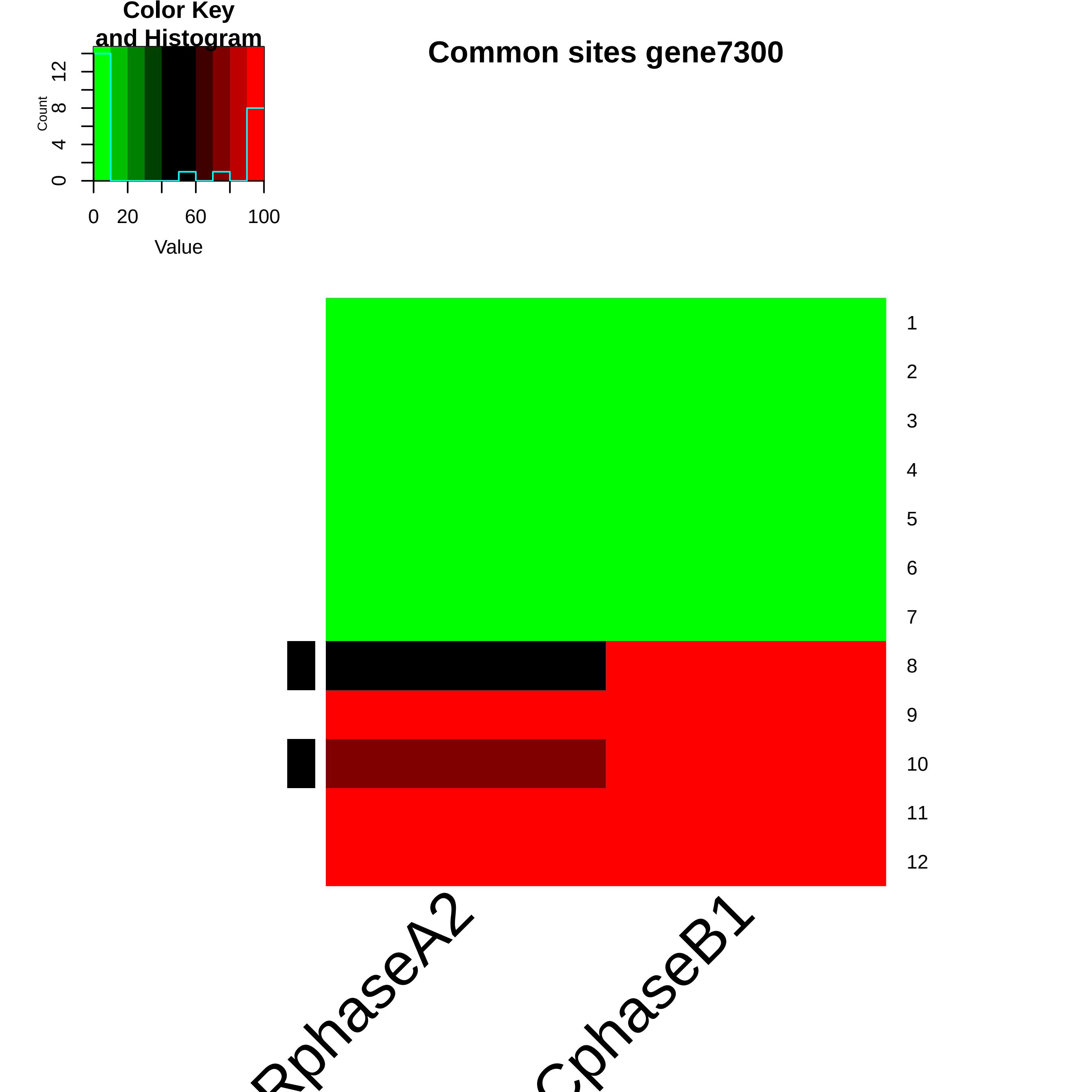
Color Key
and Histogram
Common sites gene7300
Count
0
4
8
12
0
20
60
100
Value
1
2
3
4
5
6
7
8
9
10
11
12
RphaseA2
CphaseB1

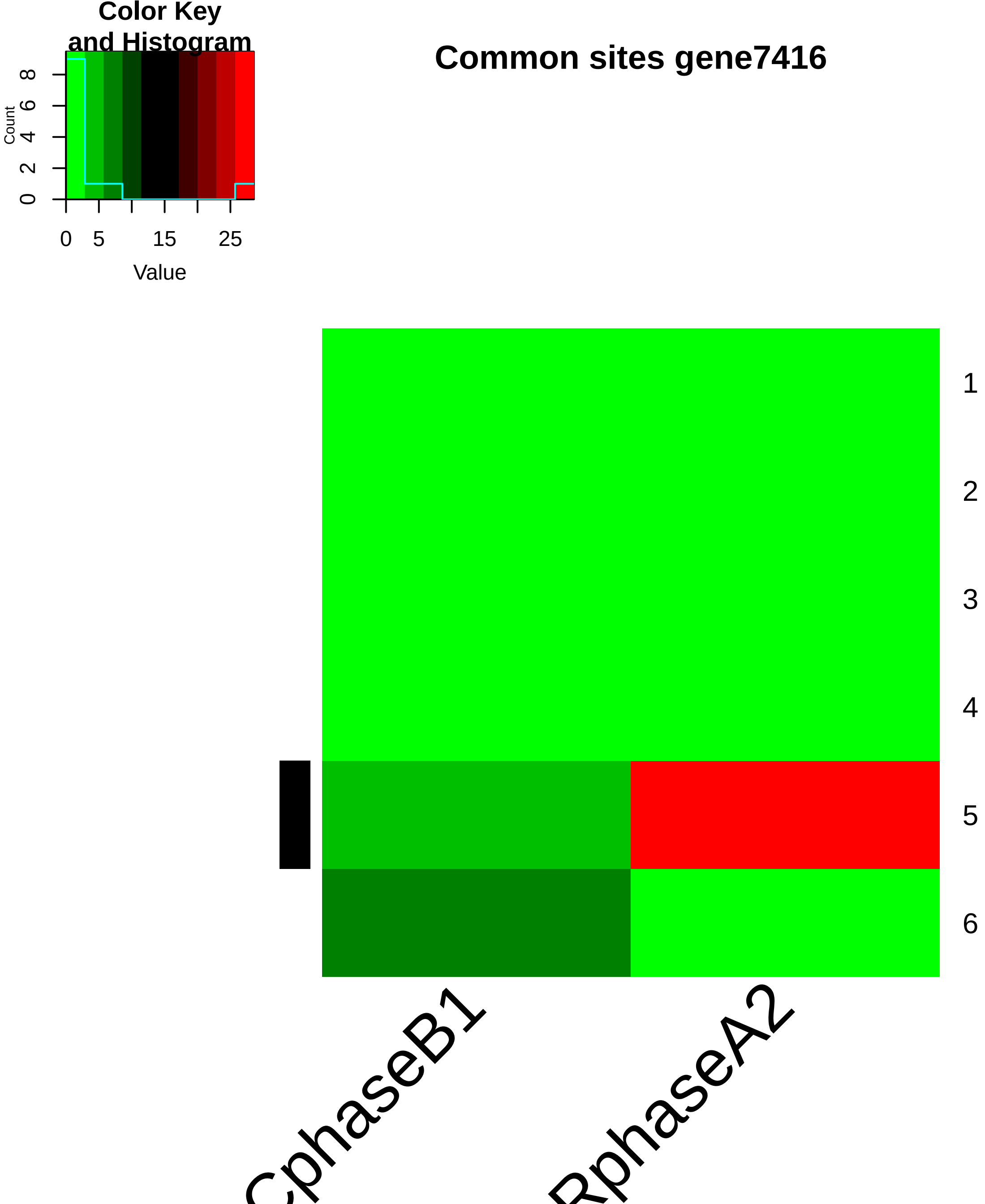
Color Key
and Histogram
Count
0
2
4
6
8
0
5
15
25
Value
Common sites gene7416
1
2
3
4
5
6
CphaseB1
RphaseA2

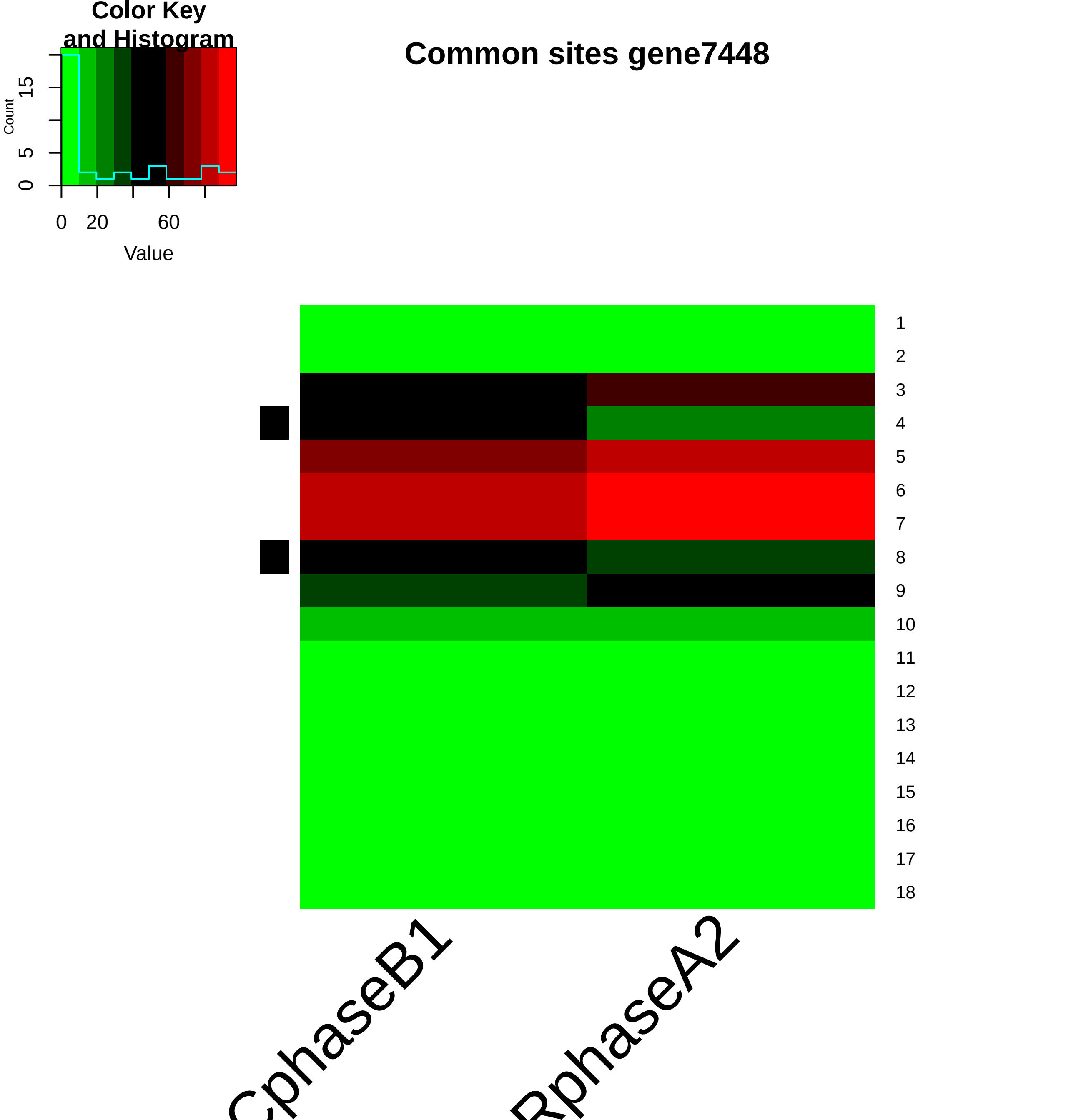

Color Key
and Histogram
Count
0
5
15
0
20
60
Value
Common sites gene7448
CphaseB1
RphaseA2

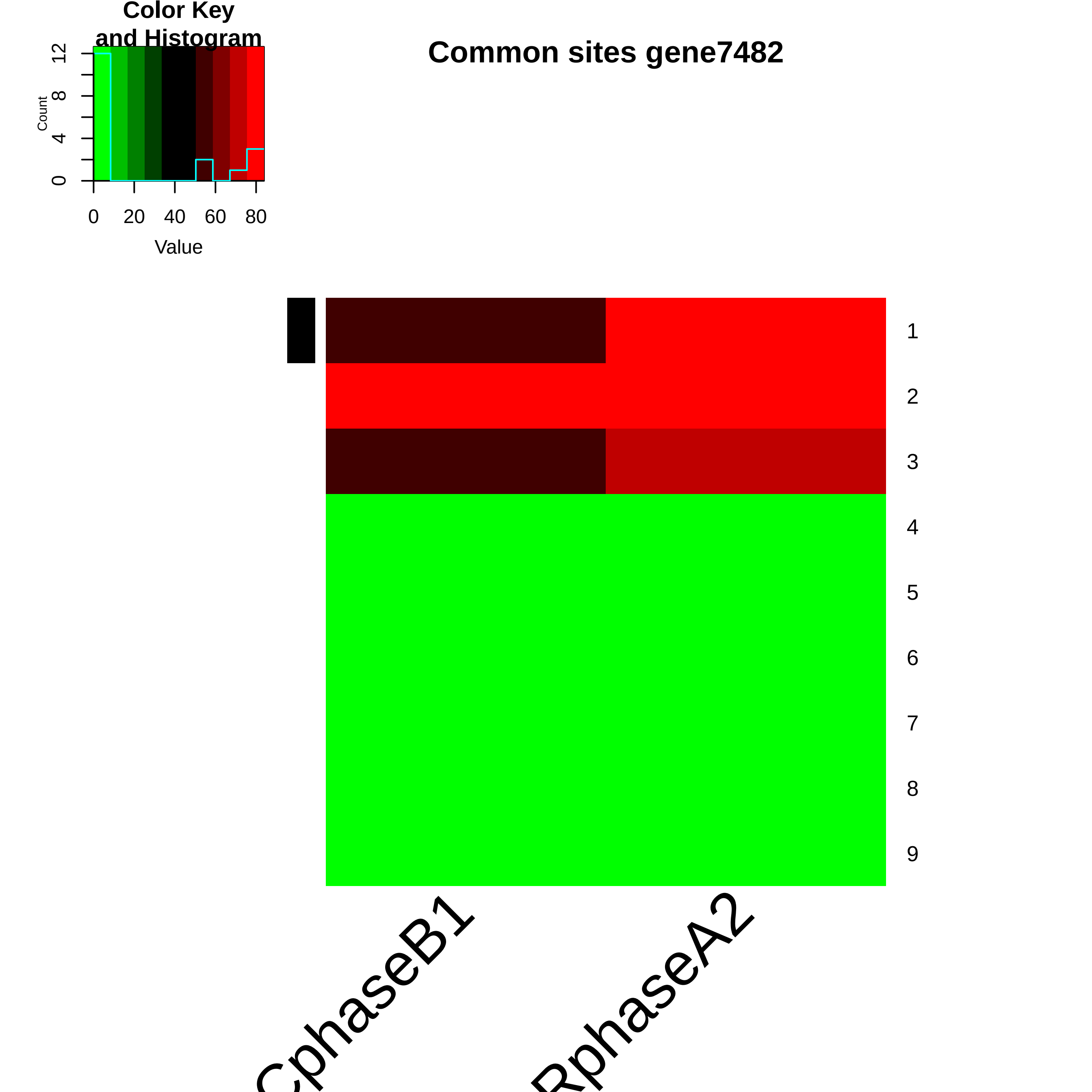
Color Key
and Histogram
Count
0
4
8
12
0
20
40
60
80
Value
Common sites gene7482
1
2
3
4
5
6
7
8
9
CphaseB1
RphaseA2

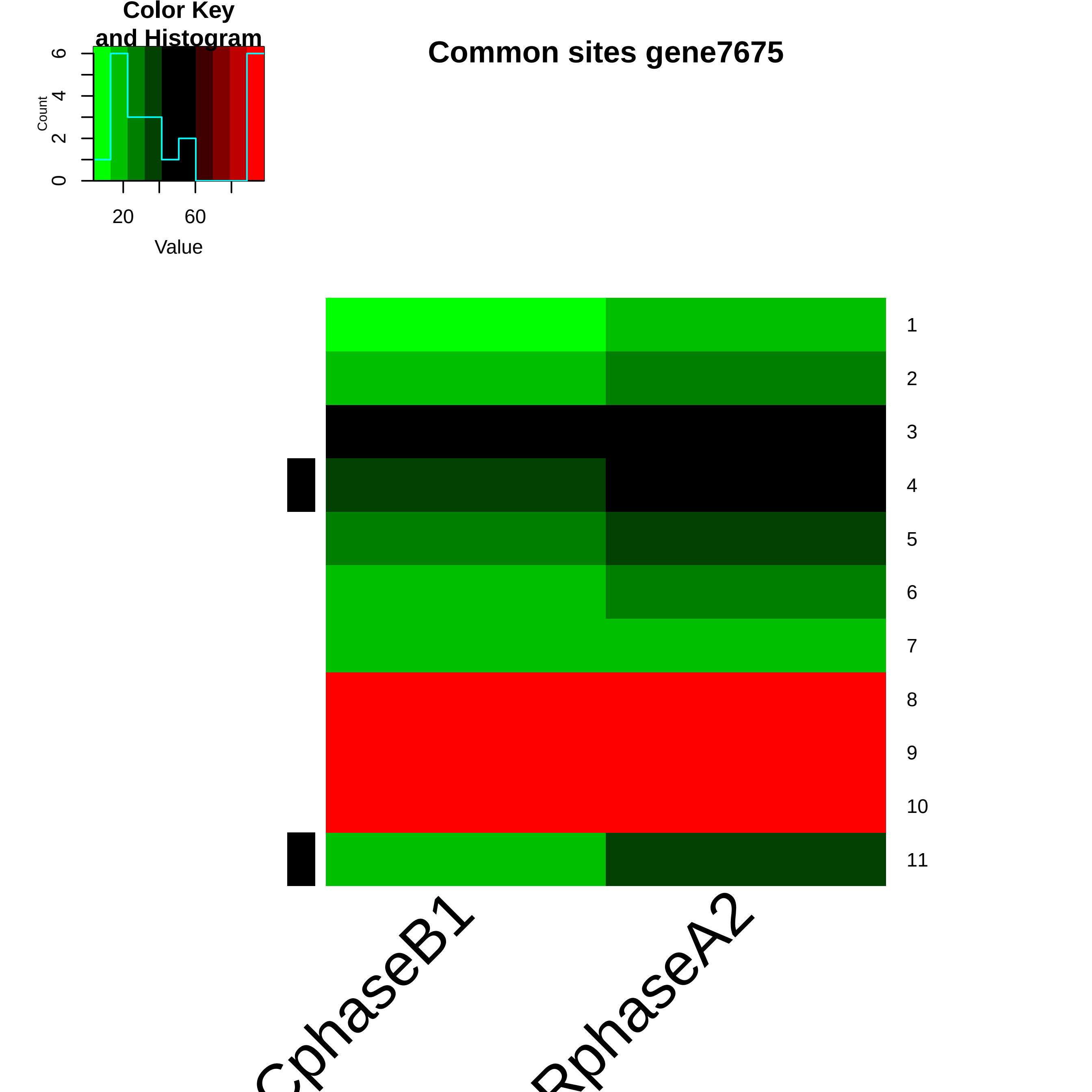

Color Key
and Histogram
Count
0
2
4
6
20
60
Value
Common sites gene7675
1
2
3
4
5
6
7
8
9
10
11
CphaseB1
RphaseA2

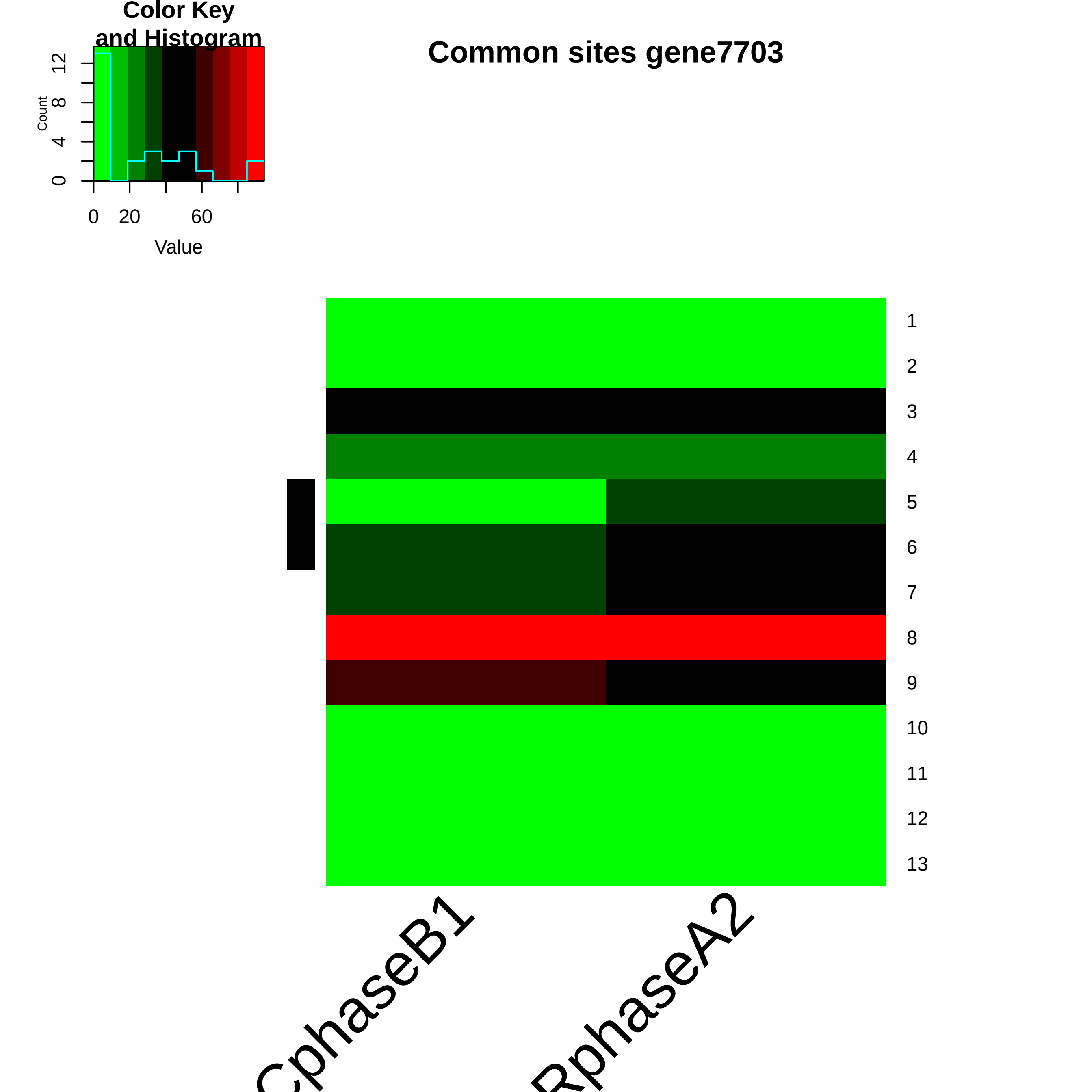

Color Key
and Histogram
Count
0
4
8
12
0
20
60
Value
Common sites gene7703
1
2
3
4
5
6
7
8
9
10
11
12
13

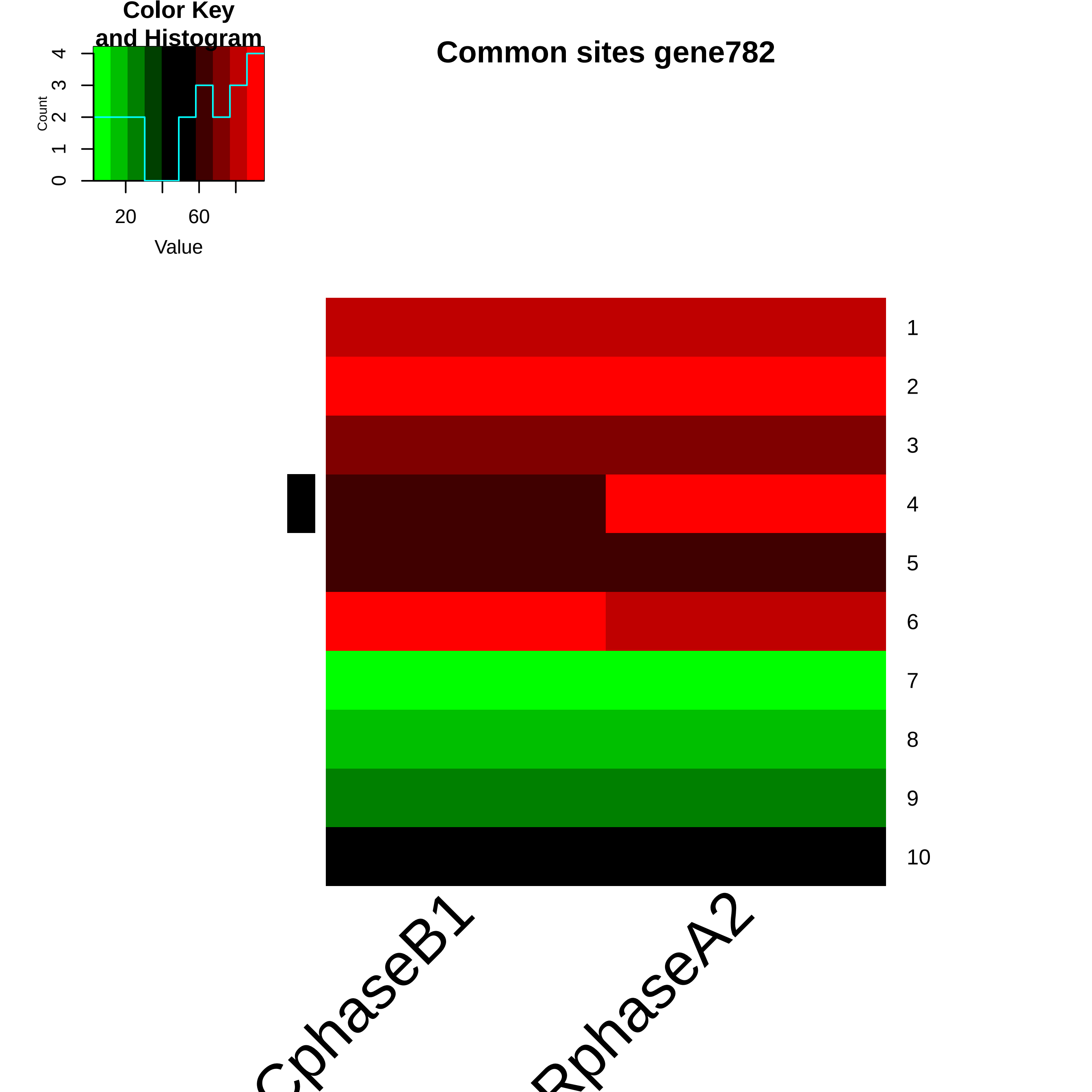
Color Key
and Histogram
Count
0
1
2
3
4
20
60
Value
Common sites gene782
1
2
3
4
5
6
7
8
9
10
CphaseB1
RphaseA2

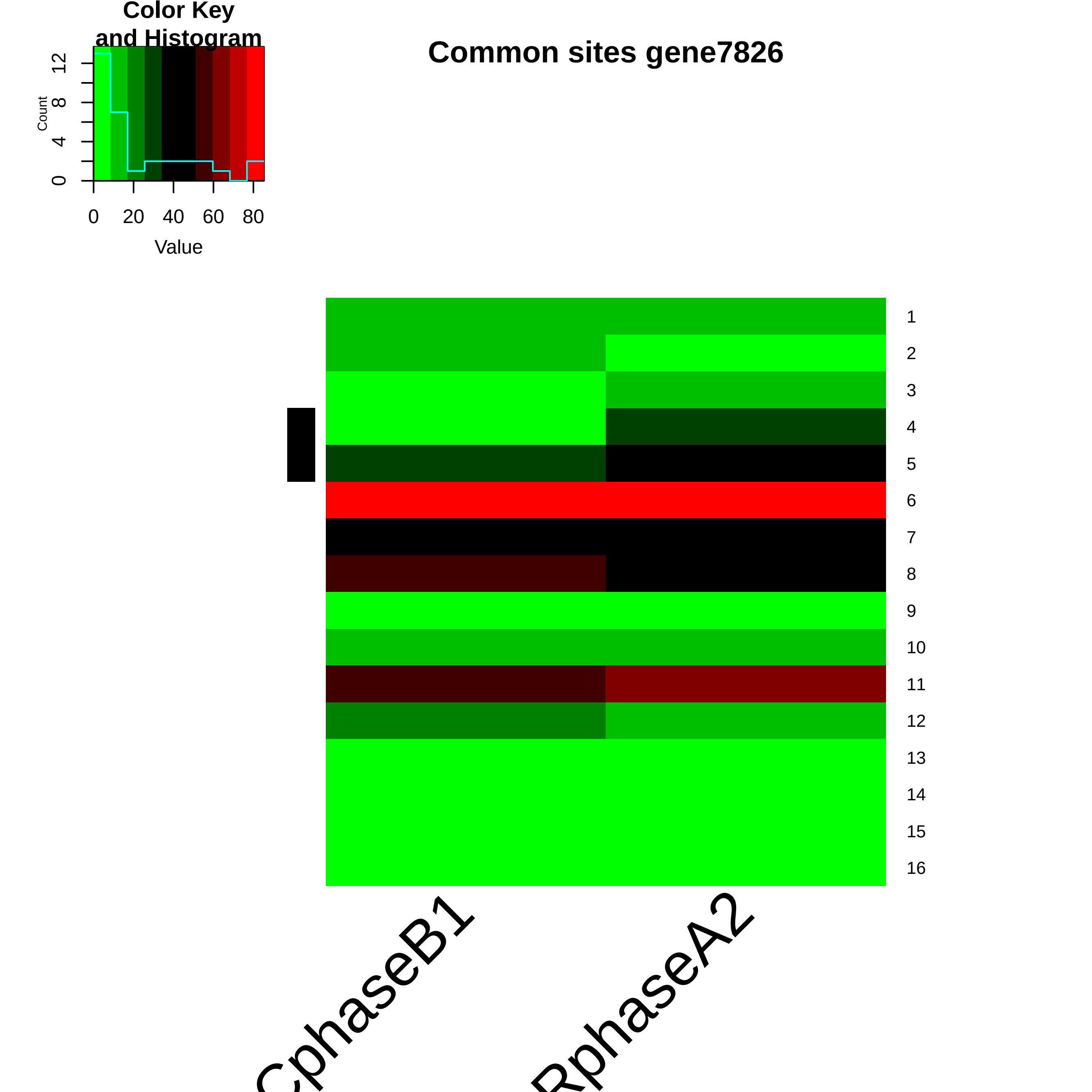

Color Key and Histogram
Count
0
4
8
12
0
20
40
60
80
Value
Common sites gene7826
CphaseB1
RphaseA2

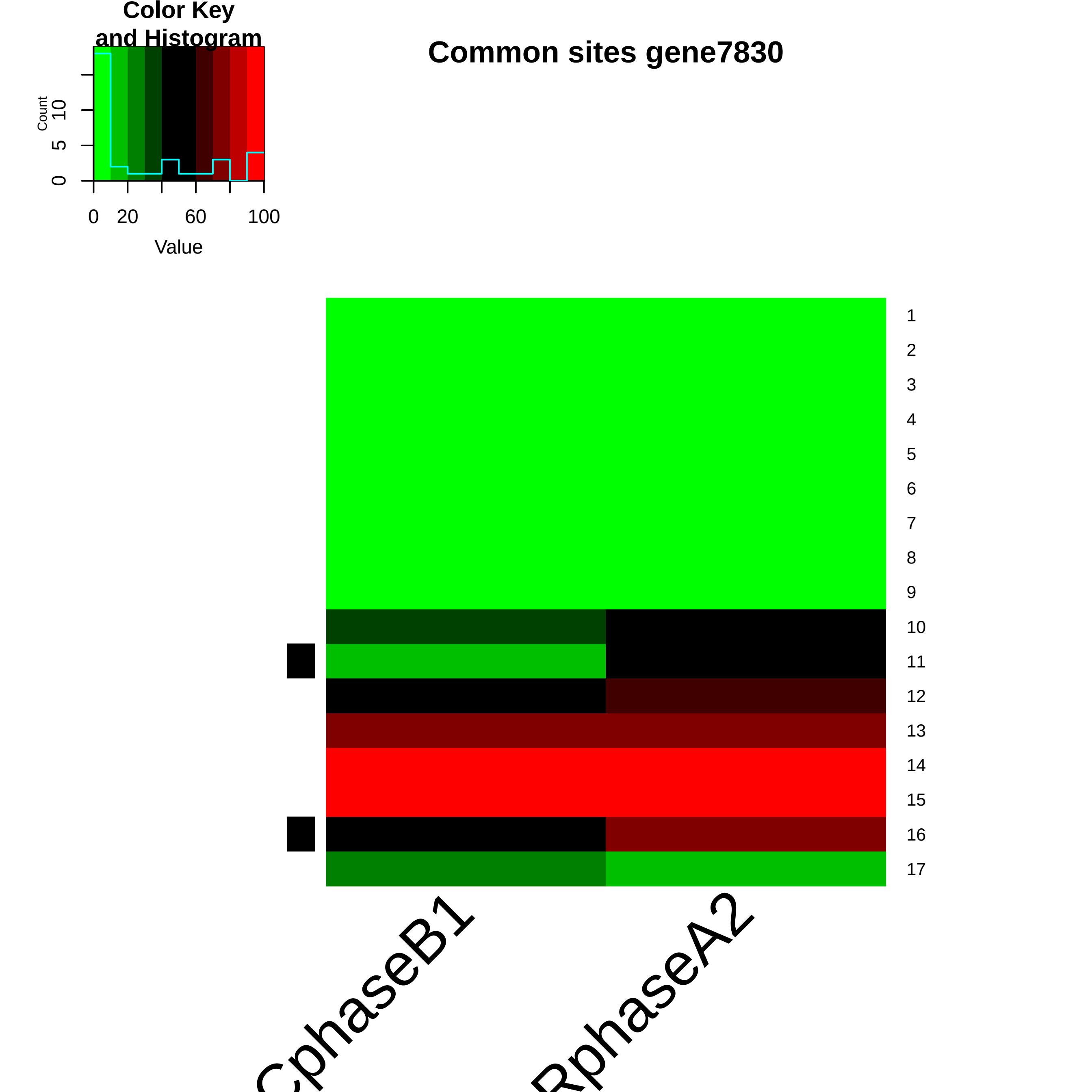
Color Key
and Histogram
Count
0
5
10
0
20
60
100
Value
Common sites gene7830
CphaseB1
RphaseA2

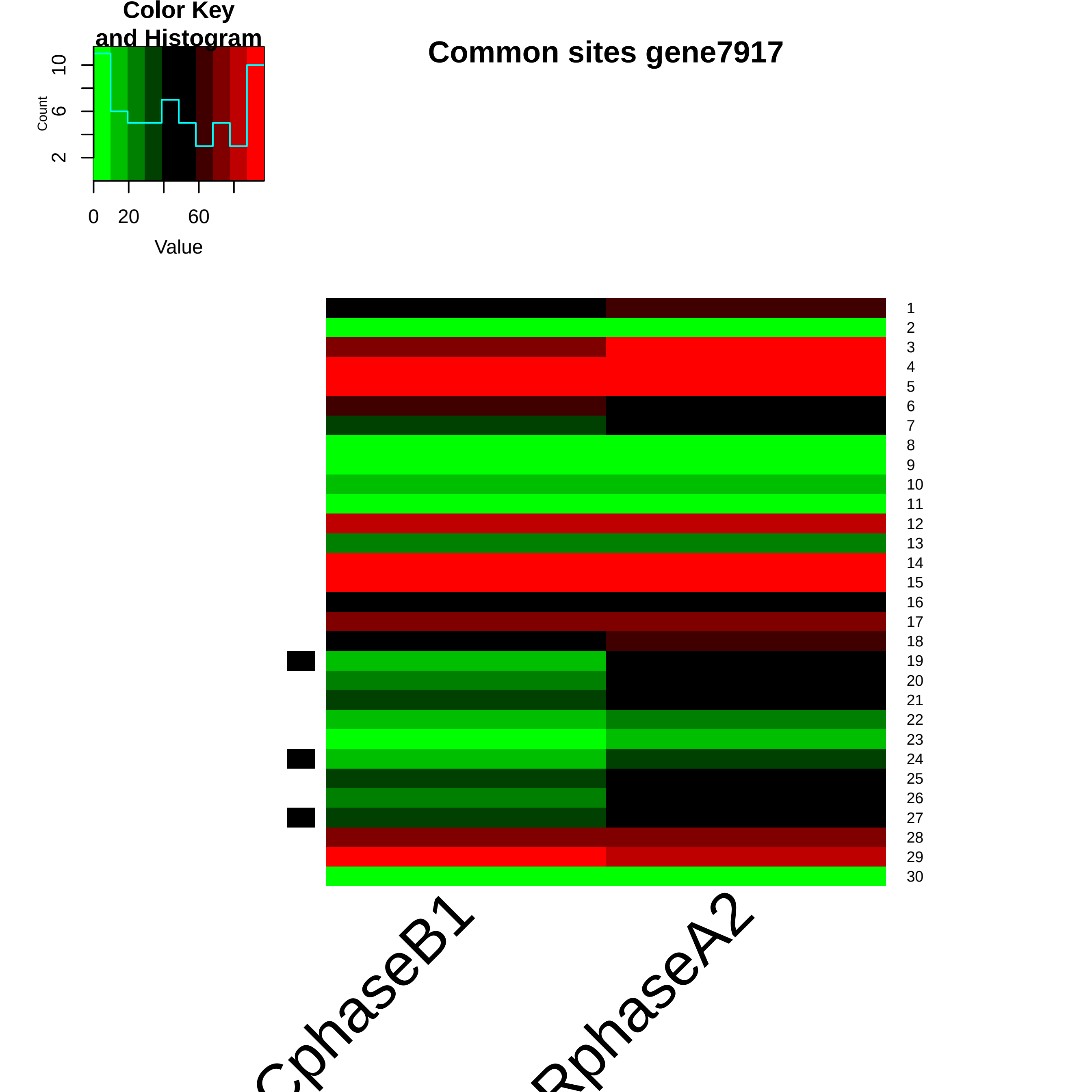
Color Key
and Histogram
Count
10
6
2
0
20
60
Value
Common sites gene7917
CphaseB1
RphaseA2

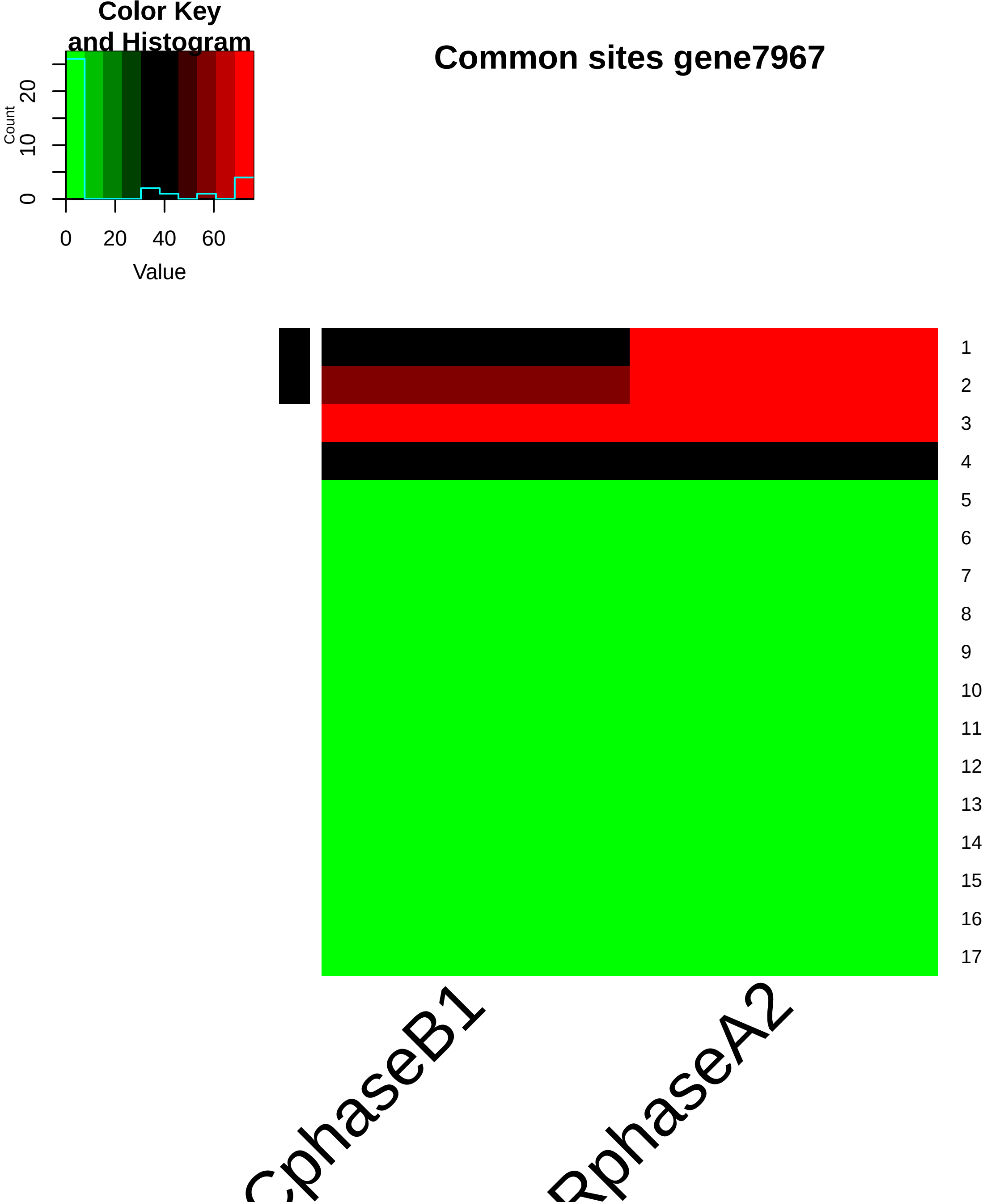
Color Key
and Histogram
Count
0
10
20
0
20
40
60
Value
Common sites gene7967
CphaseB1
RphaseA2

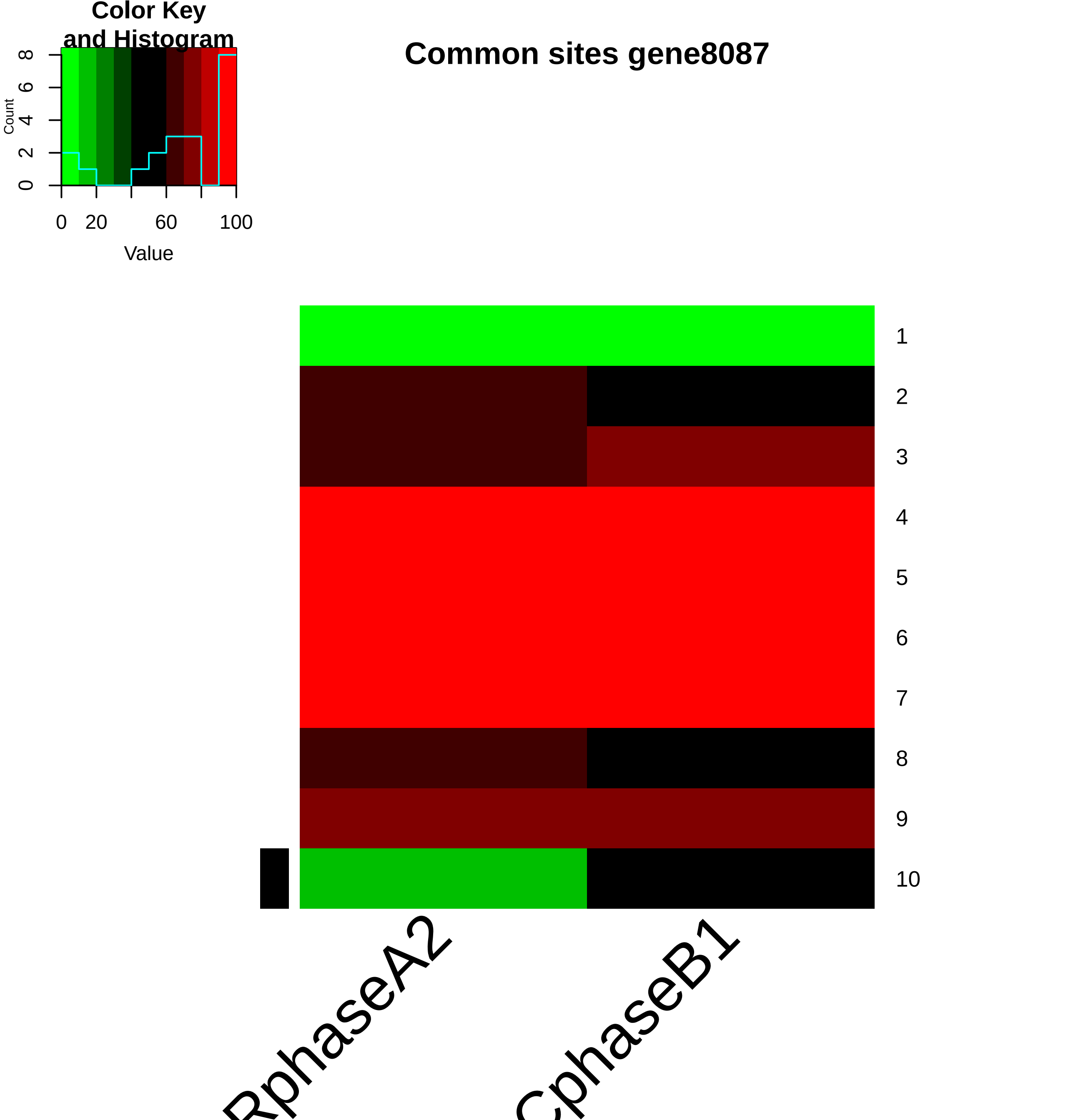
Color Key
and Histogram
Count
0
2
4
6
8
0
20
60
100
Value
Common sites gene8087
1
2
3
4
5
6
7
8
9
10
RphaseA2
CphaseB1

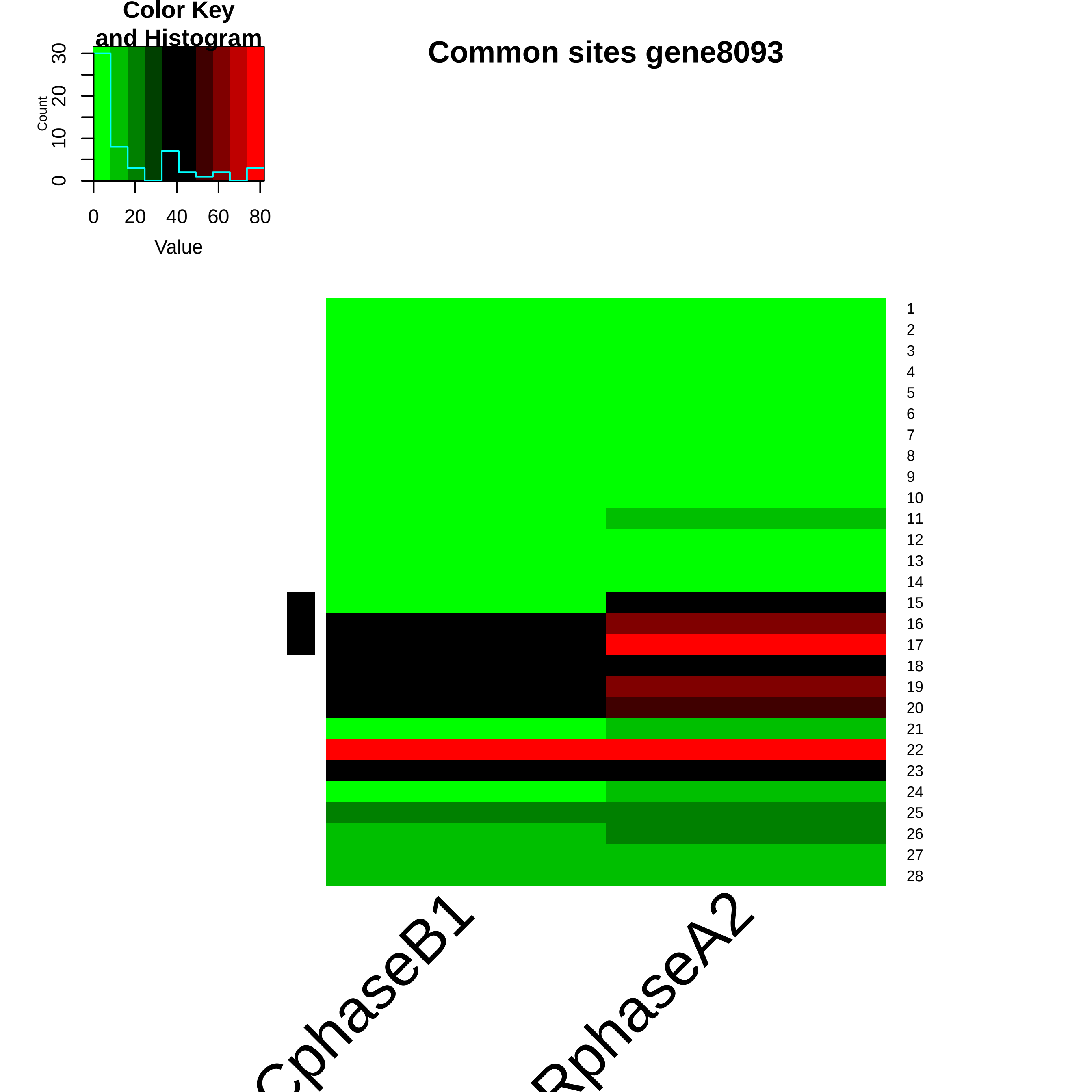

Color Key
and Histogram
Count
0
10
20
30
0
20
40
60
80
Value
Common sites gene8093
CphaseB1
RphaseA2

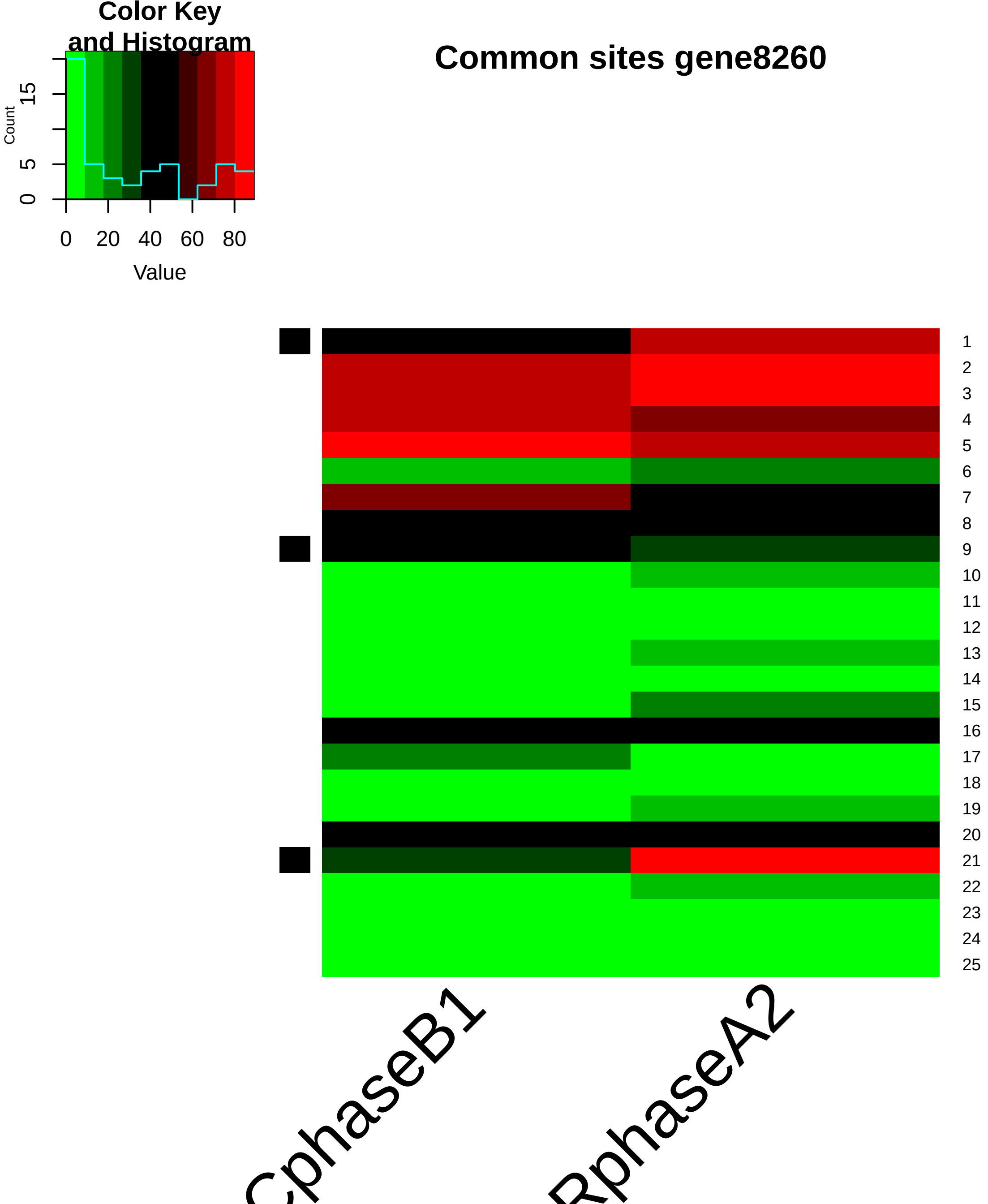
Color Key
and Histogram
Count
0
5
15
0
20
40
60
80
Value
Common sites gene8260
CphaseB1
RphaseA2

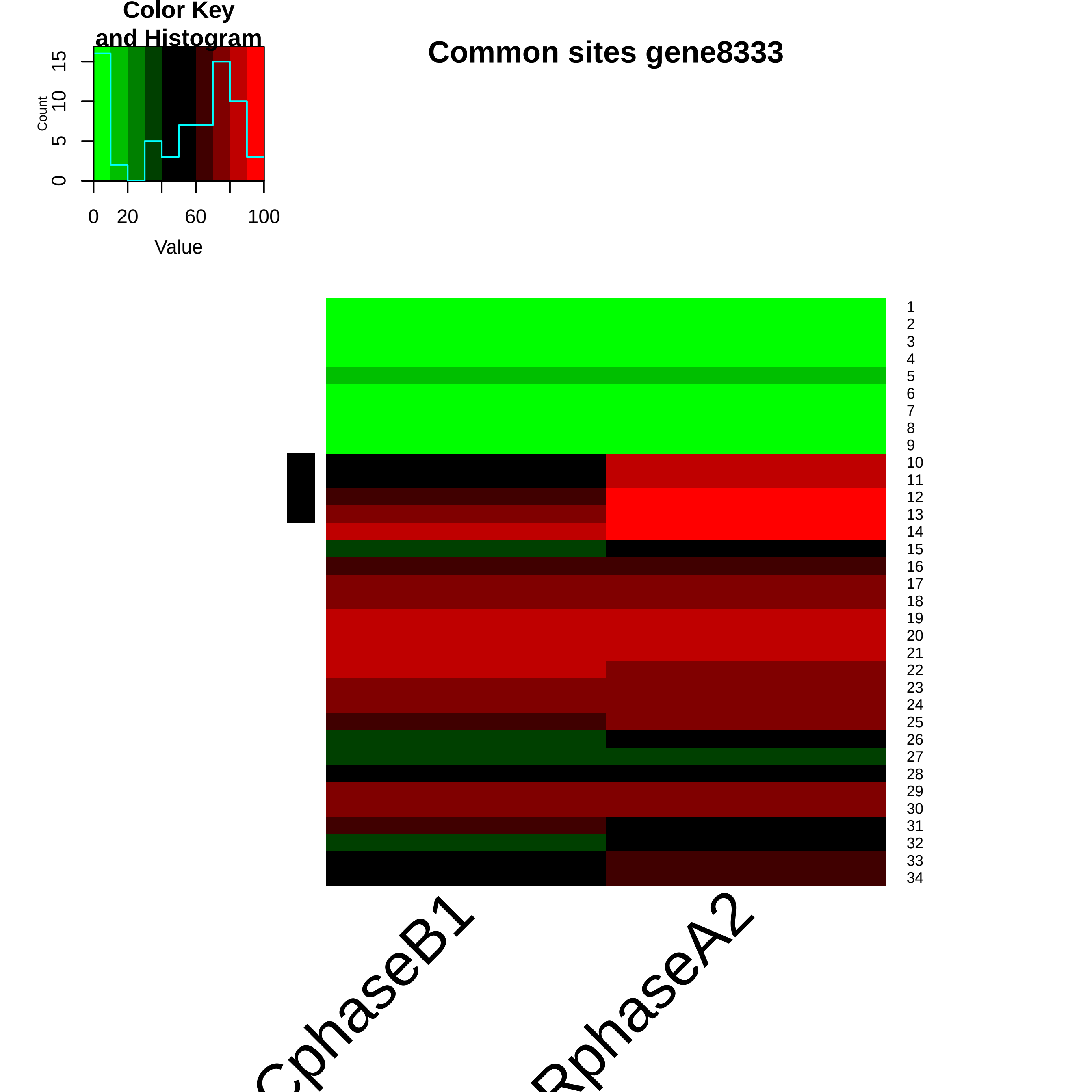
Color Key
and Histogram
Count
0
5
10
15
0
20
60
100
Value
Common sites gene8333
CphaseB1
RphaseA2

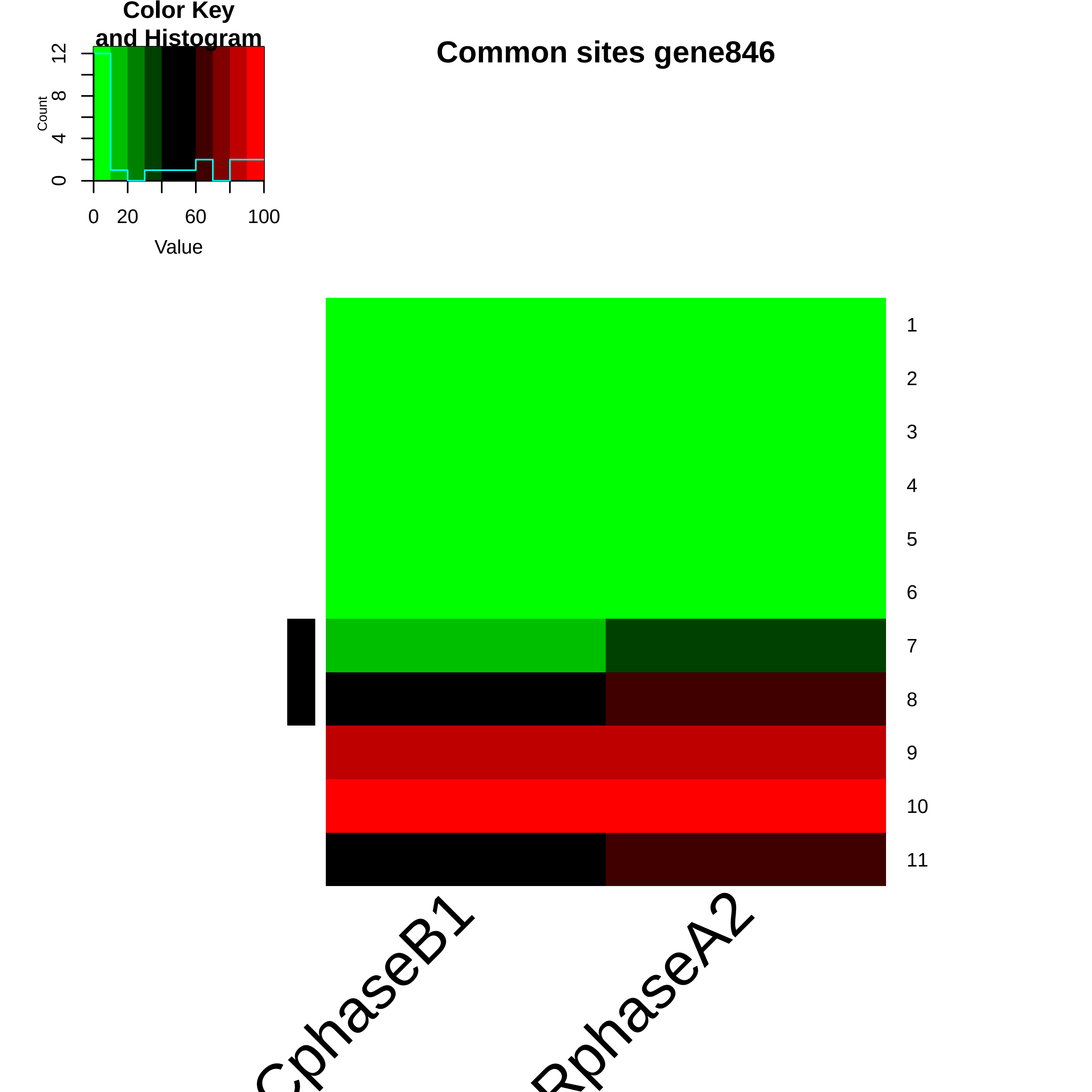
Color Key
and Histogram
Count
0
4
8
12
0
20
60
100
Value
Common sites gene846
1
2
3
4
5
6
7
8
9
10
11
CphaseB1
RphaseA2

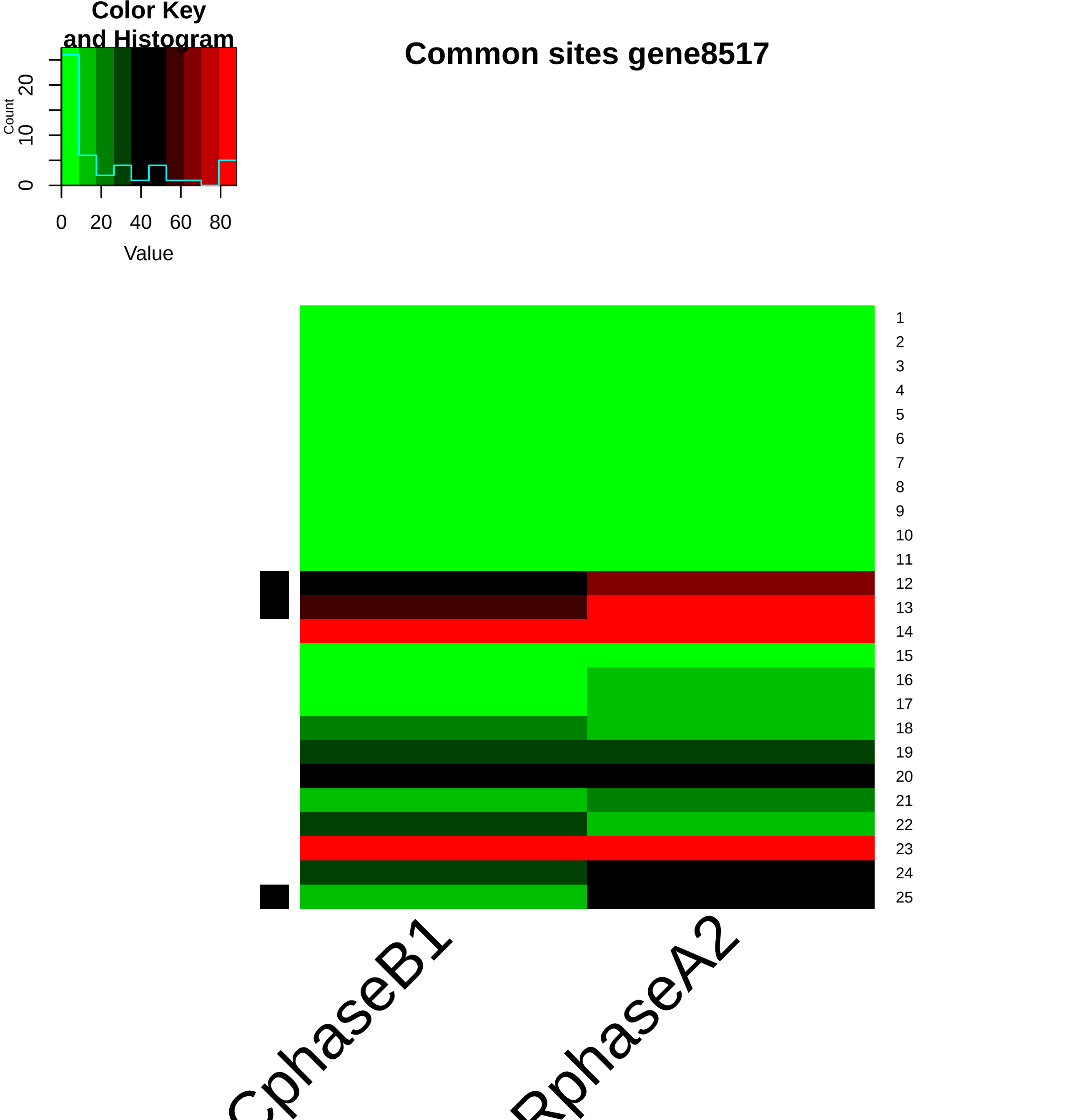
Color Key
and Histogram
Count
0
10
20
0
20
40
60
80
Value
Common sites gene8517
CphaseB1
RphaseA2

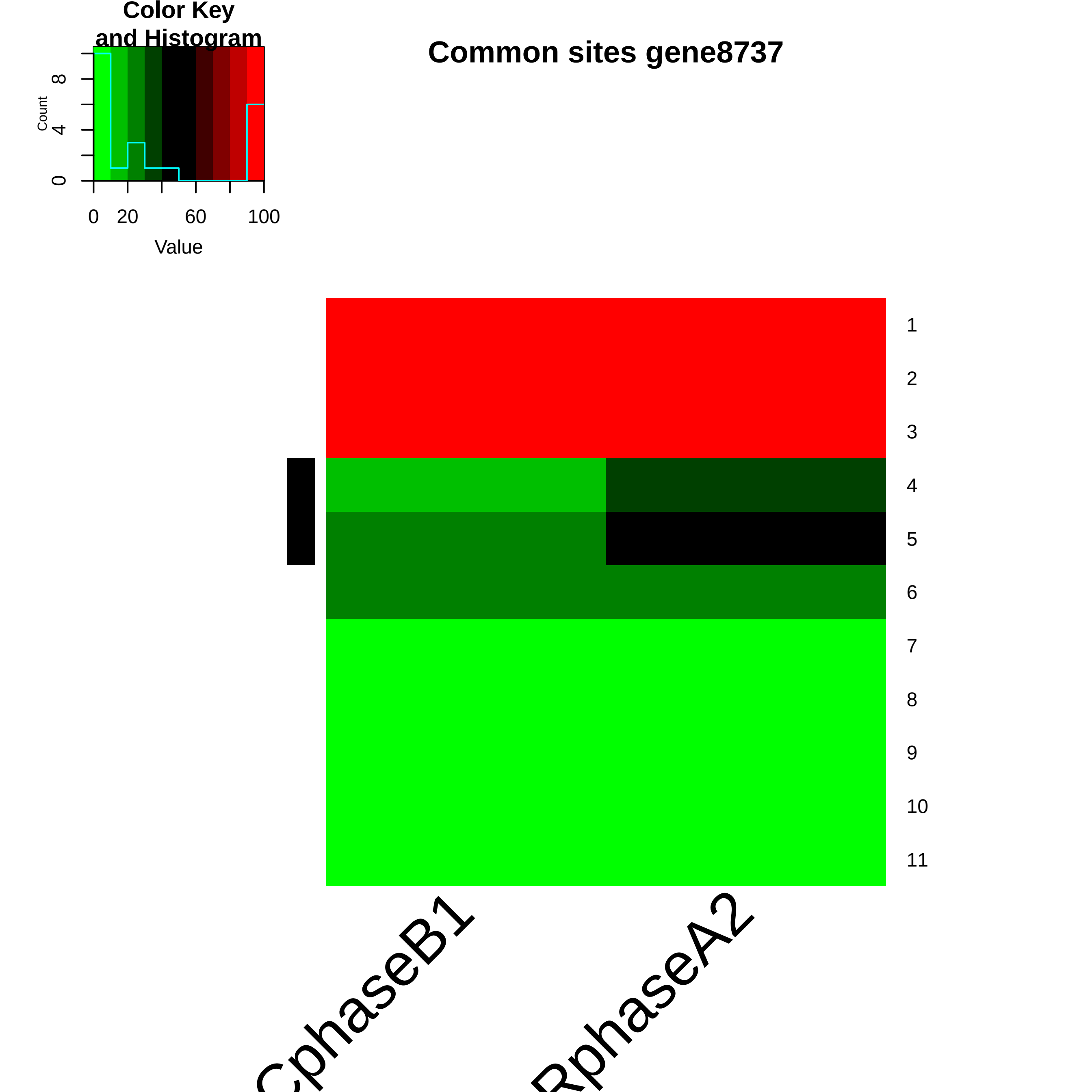
Color Key
and Histogram
Count
0
4
8
0
20
60
100
Value
Common sites gene8737
1
2
3
4
5
6
7
8
9
10
11
CphaseB1
RphaseA2

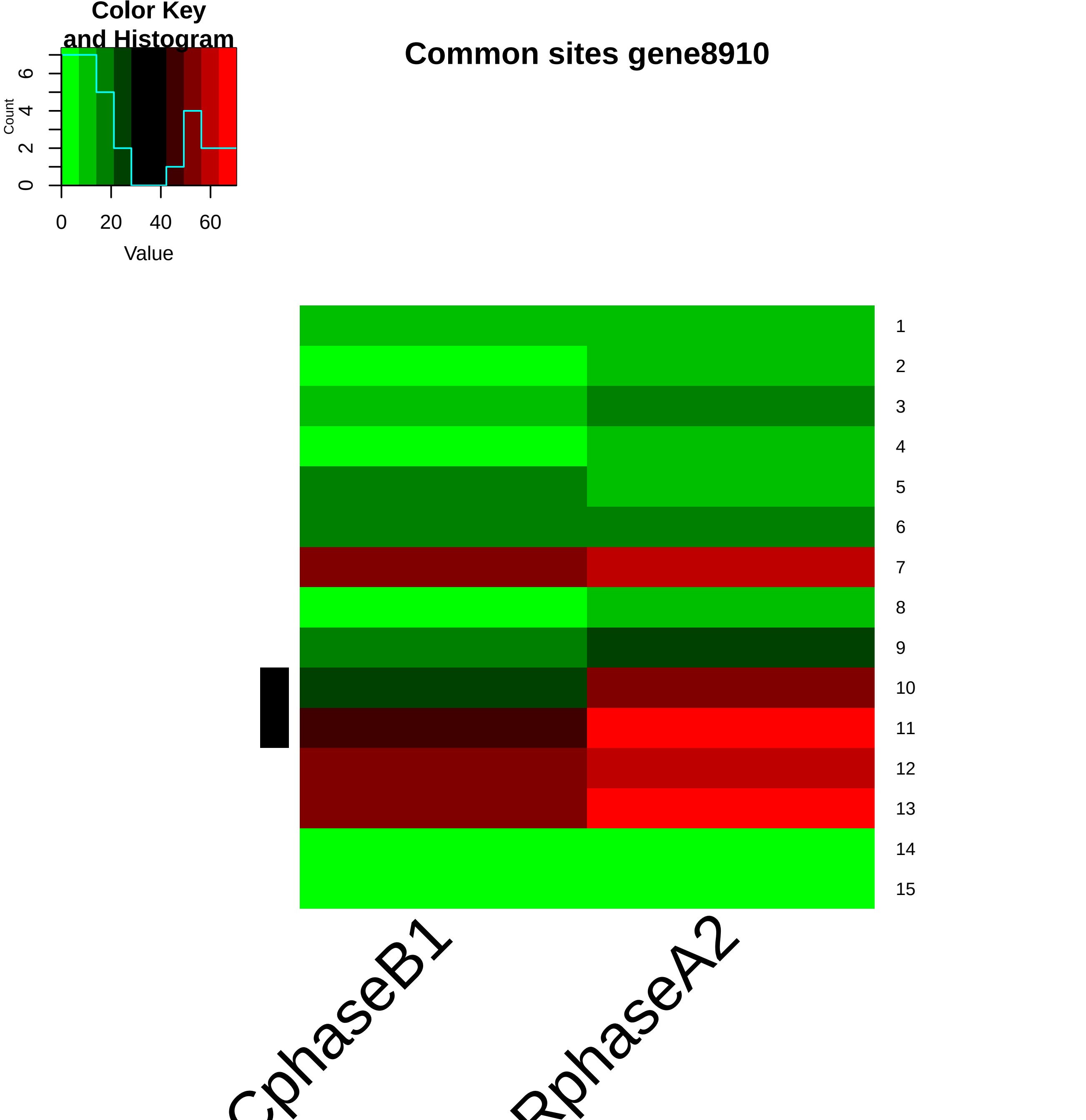

Color Key
and Histogram
Count
0
2
4
6
0
20
40
60
Value
Common sites gene8910
CphaseB1
RphaseA2

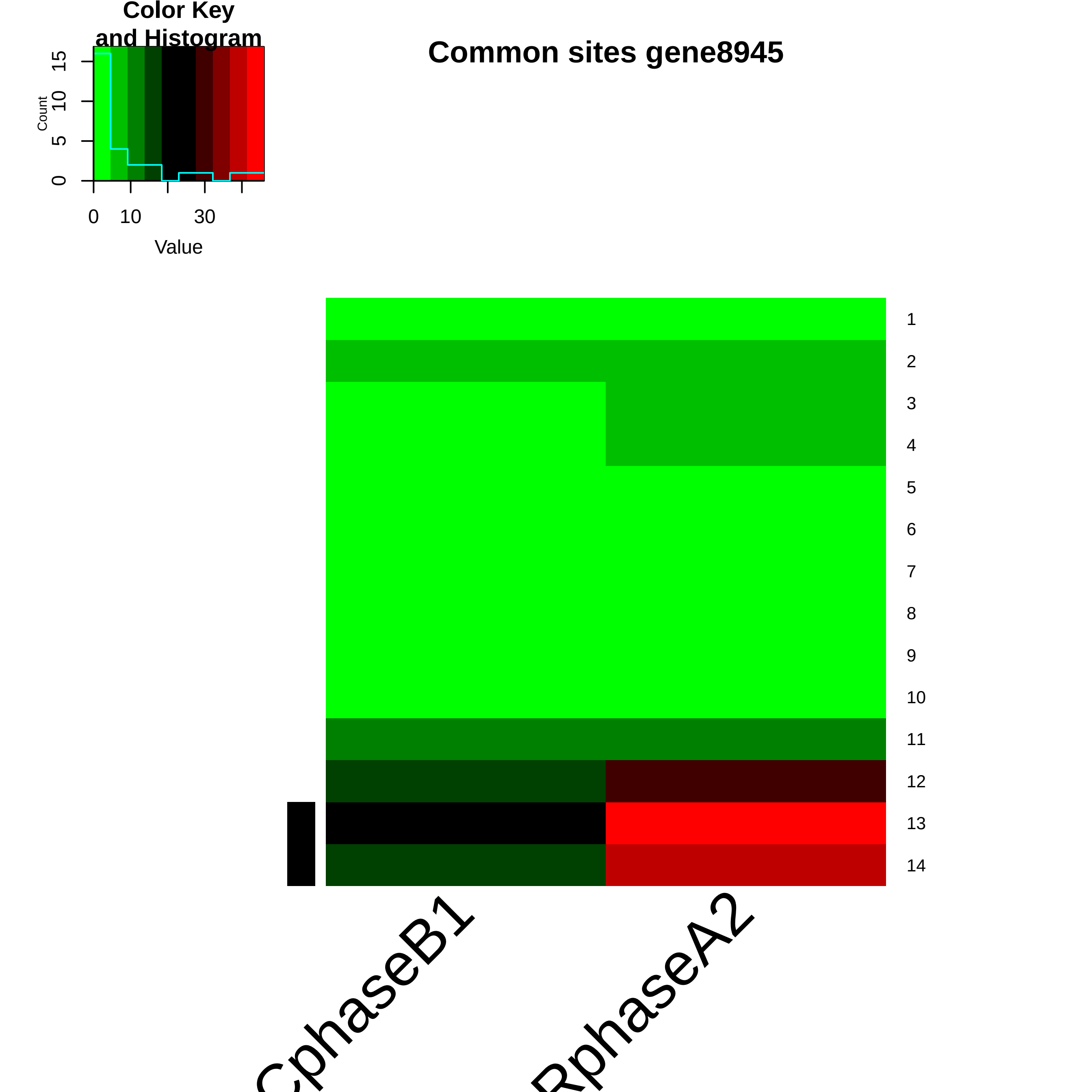

Color Key
and Histogram
Count
0
5
10
15
0
10
30
Value
Common sites gene8945
1
2
3
4
5
6
7
8
9
10
11
12
13
14
CphaseB1
RphaseA2

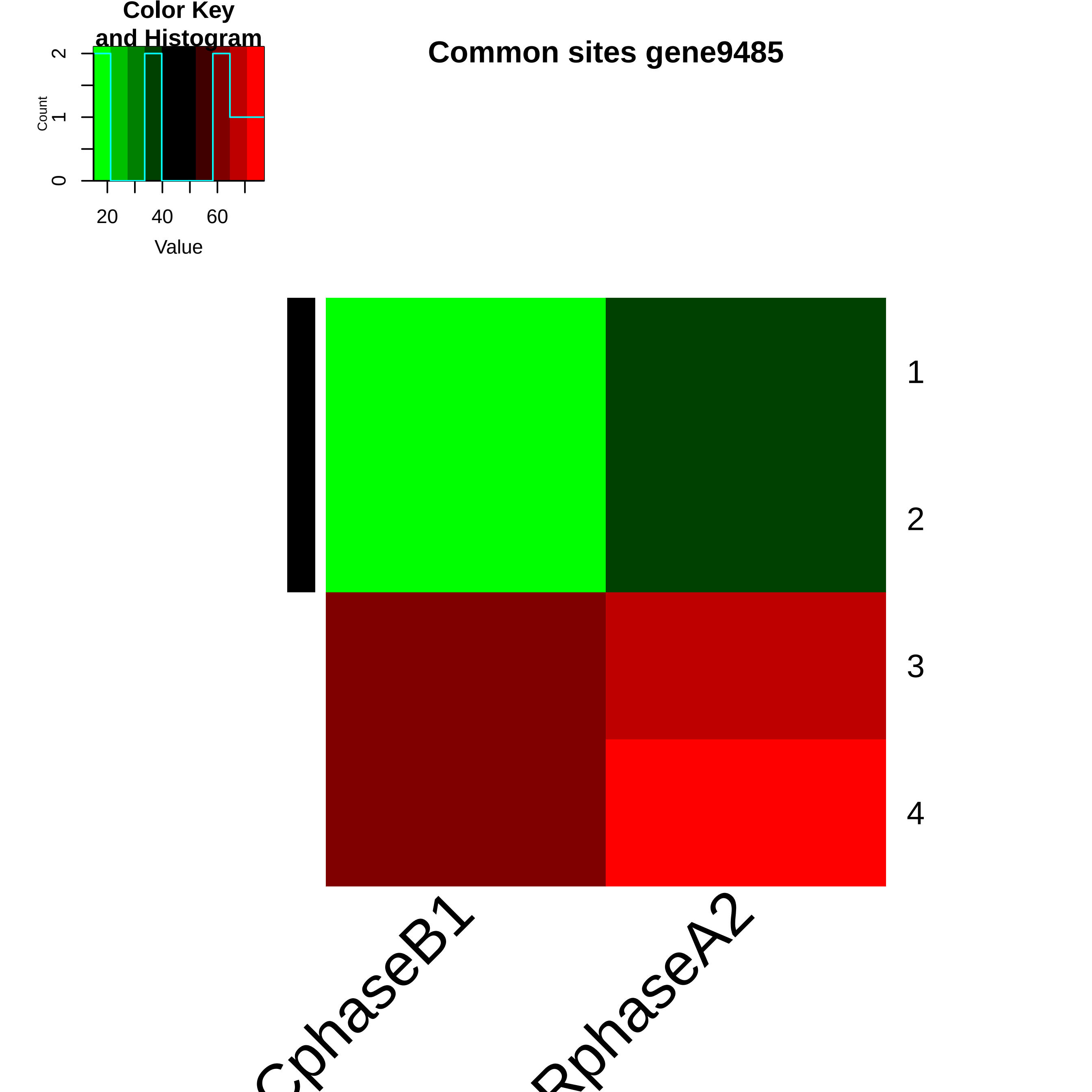
Color Key
and Histogram
Count
0
1
2
20
40
60
Value
Common sites gene9485
1
2
3
4
CphaseB1
RphaseA2

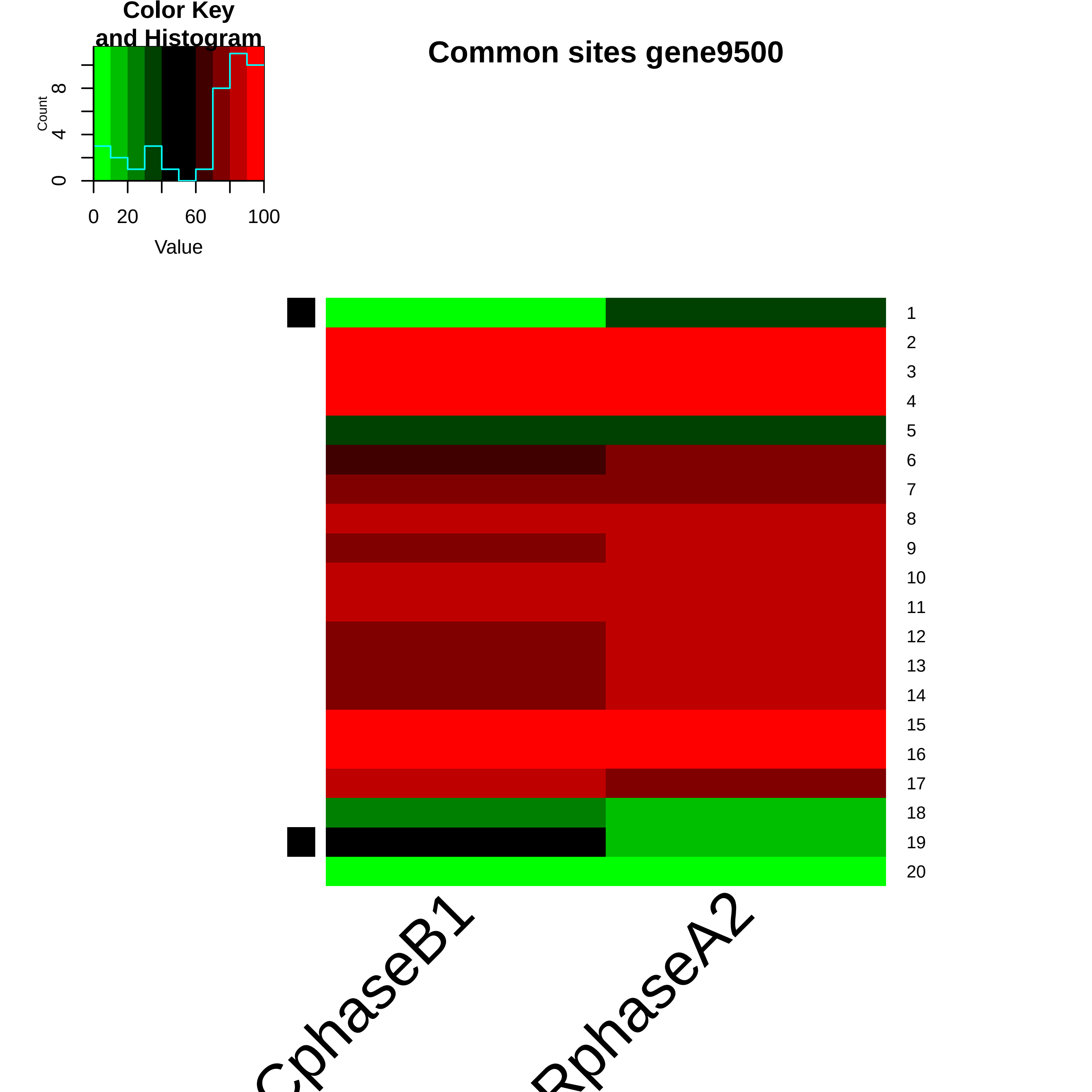

Color Key and Histogram
Count
0
4
8
0
20
60
100
Value
Common sites gene9500
CphaseB1
RphaseA2

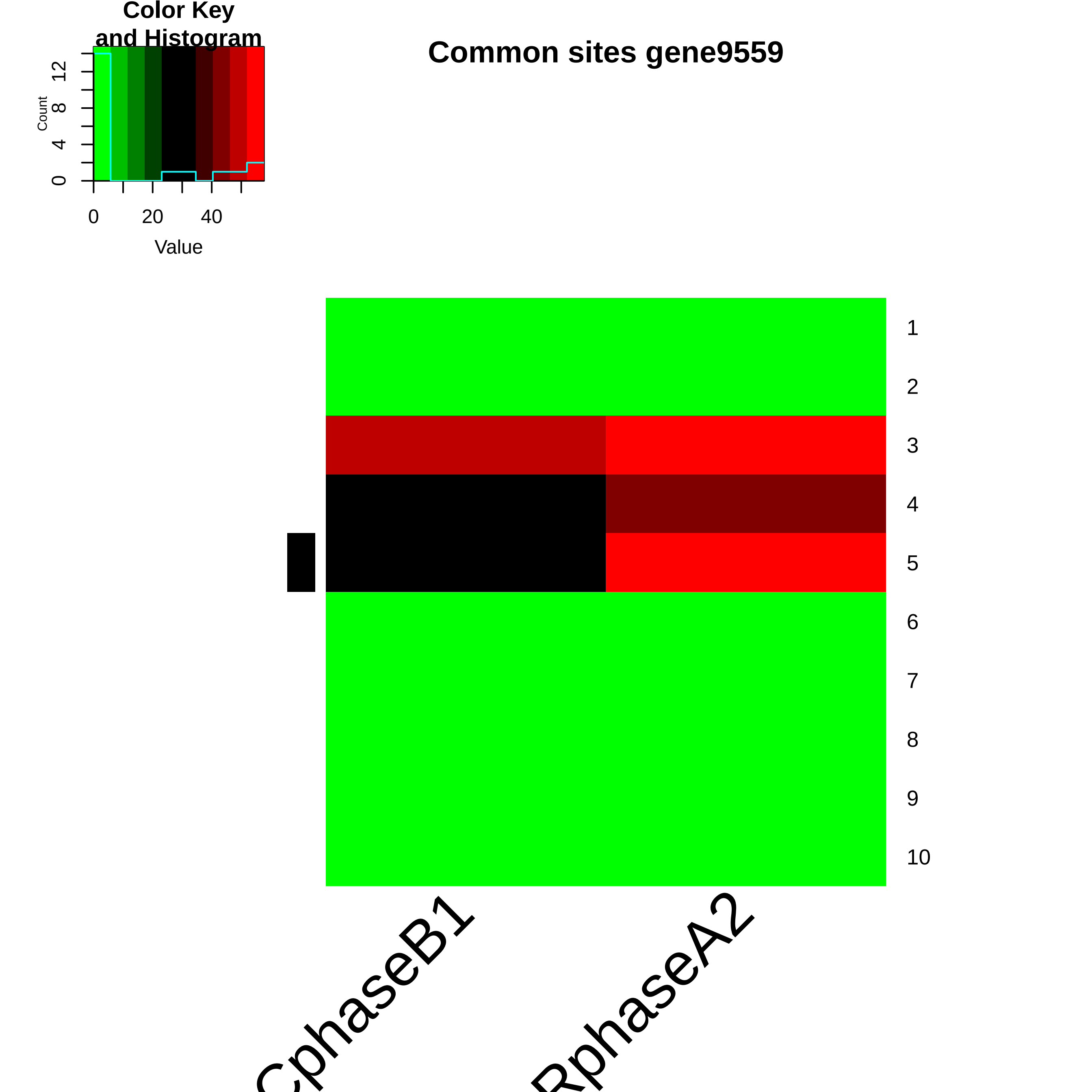

Color Key
and Histogram
Count
0
4
8
12
0
20
40
Value
Common sites gene9559
1
2
3
4
5
6
7
8
9
10
CphaseB1
RphaseA2

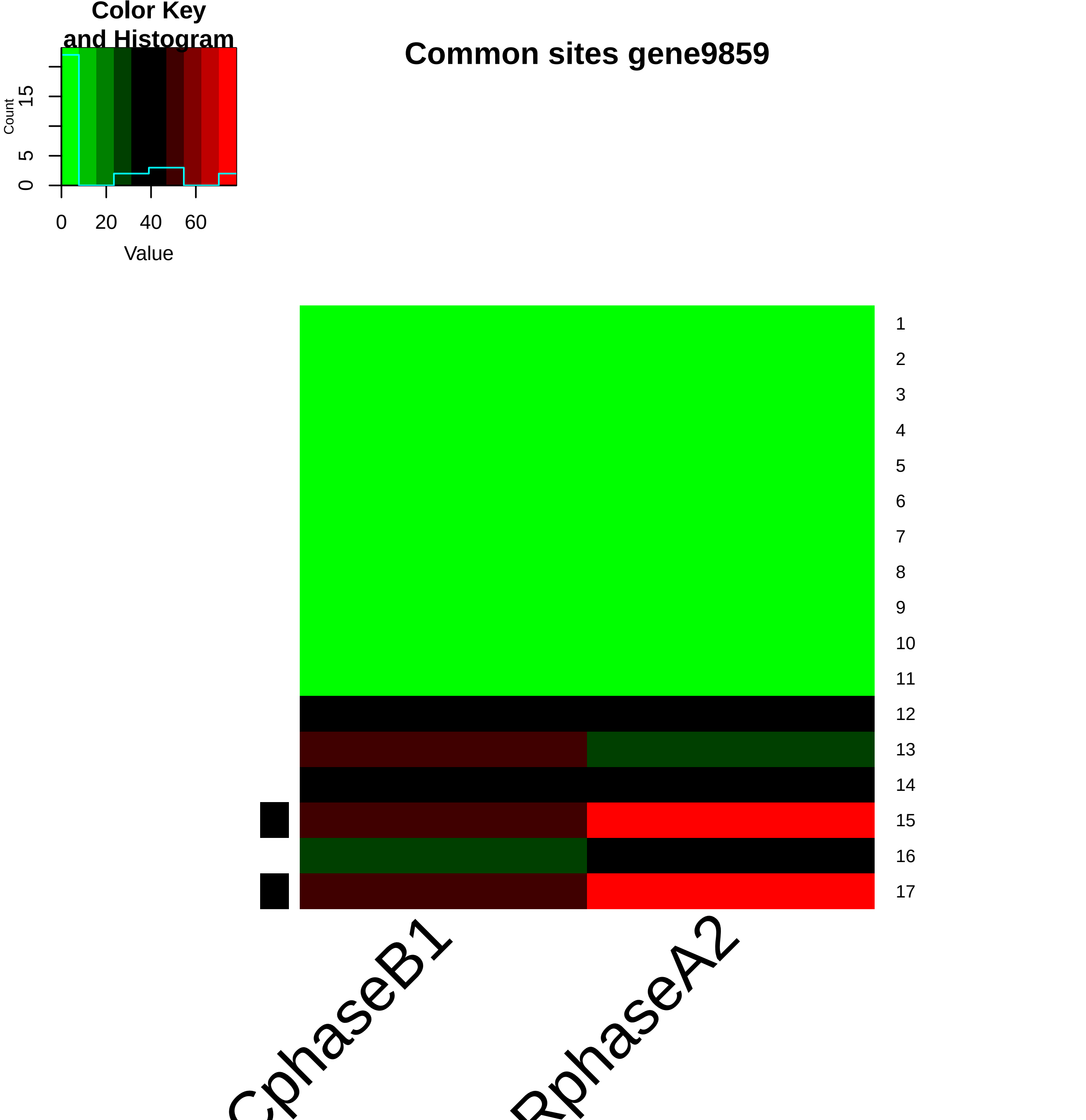

Color Key
and Histogram
Count
0
5
15
0
20
40
60
Value
Common sites gene9859
CphaseB1
RphaseA2

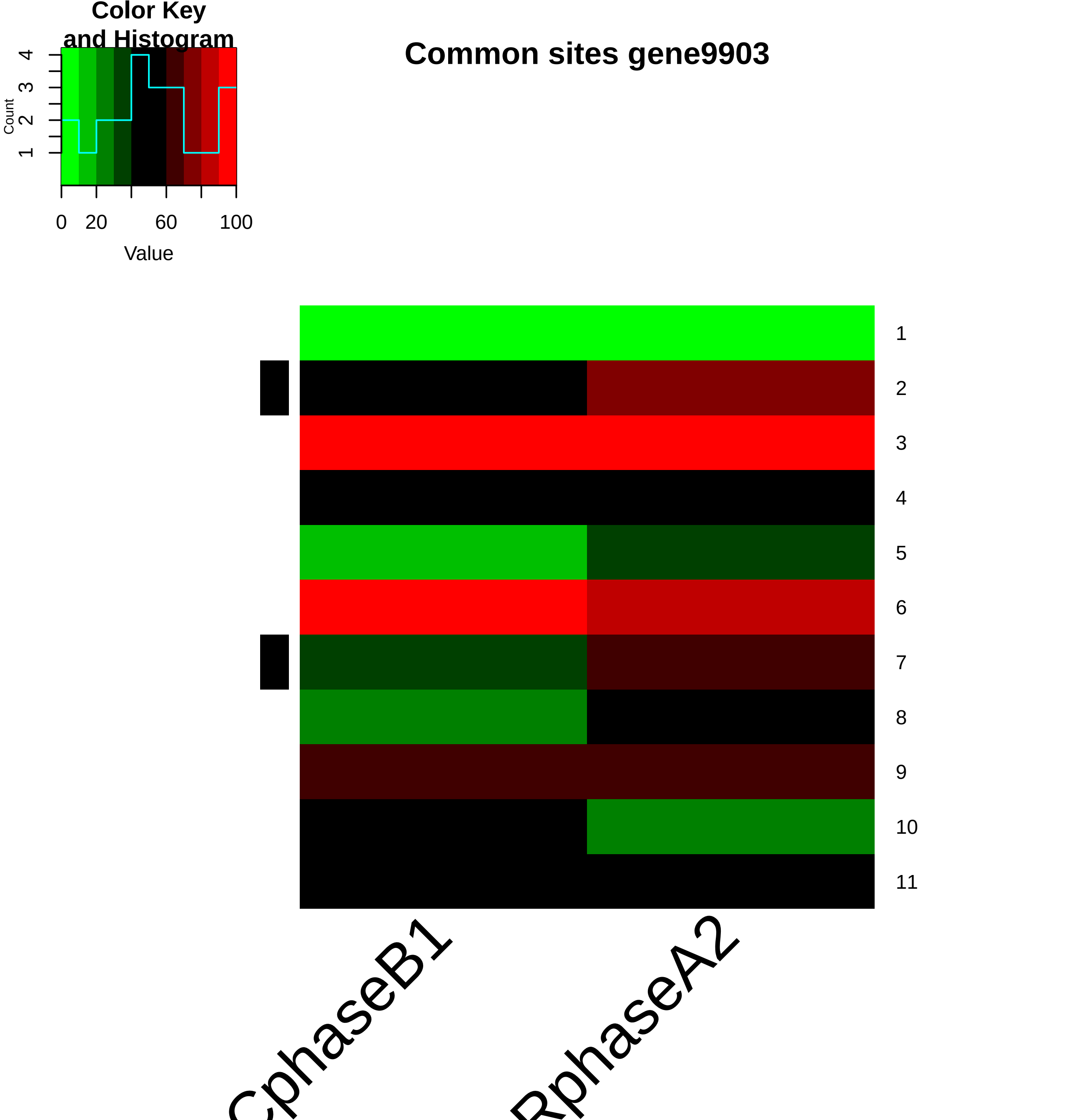

Color Key
and Histogram
Count
1
2
3
4
0
20
60
100
Value
Common sites gene9903
1
2
3
4
5
6
7
8
9
10
11
CphaseB1
RphaseA2

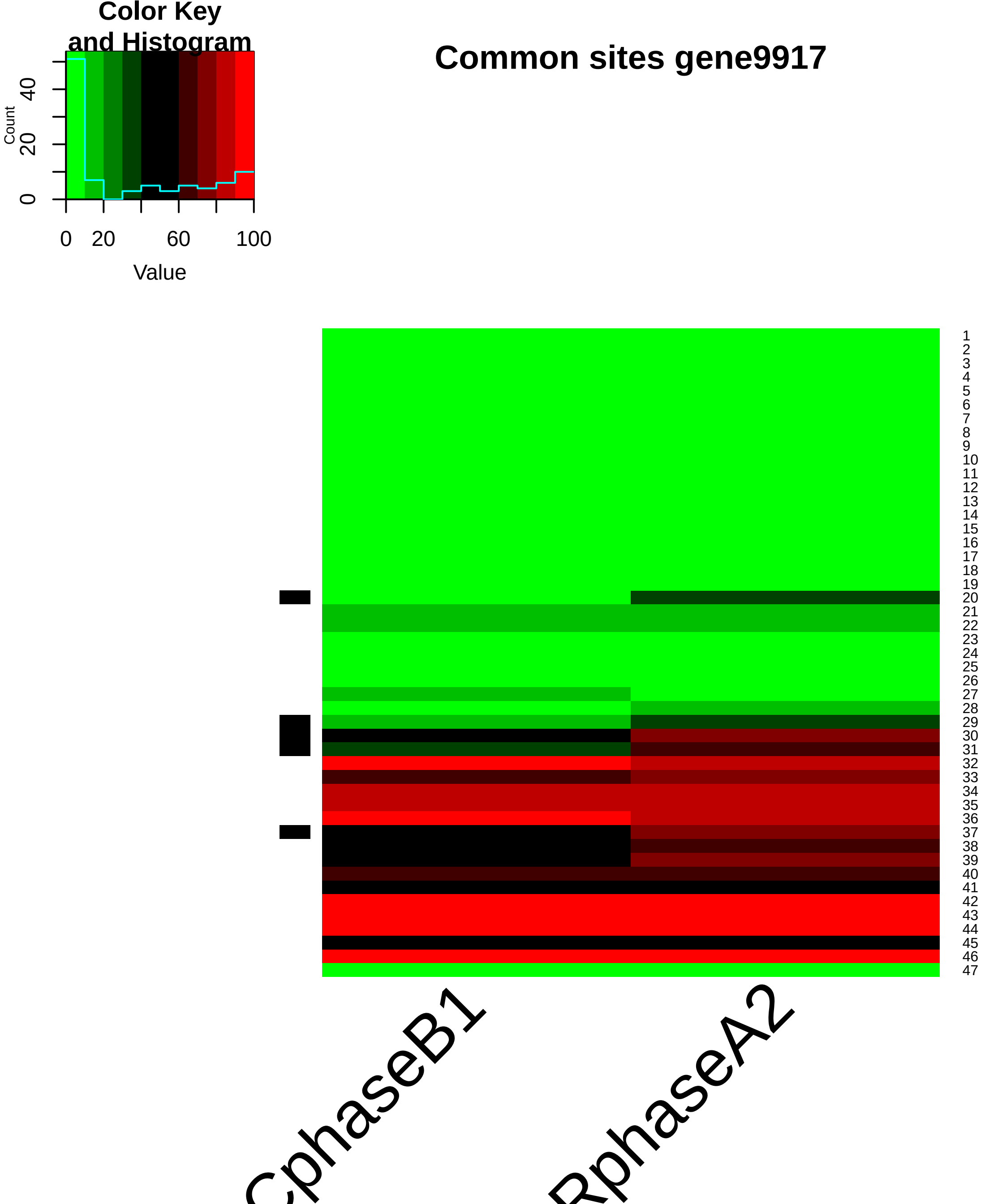
Color Key
and Histogram
Count
Value
0
20
40
60
100
Common sites gene9917
CphaseB1
RphaseA2

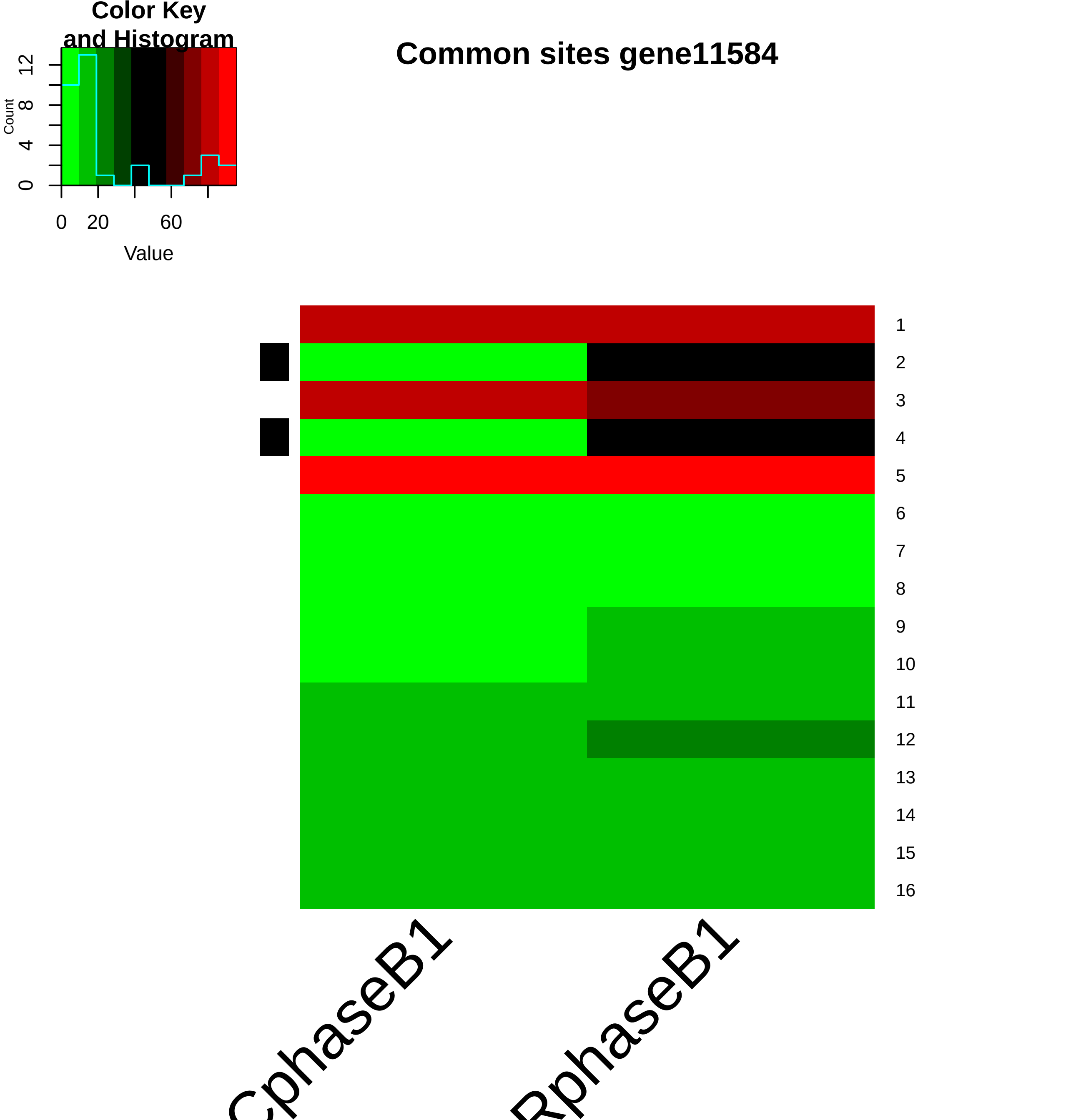
Color Key
and Histogram
Count
0
4
8
12
0
20
60
Value
Common sites gene11584
CphaseB1
RphaseB1

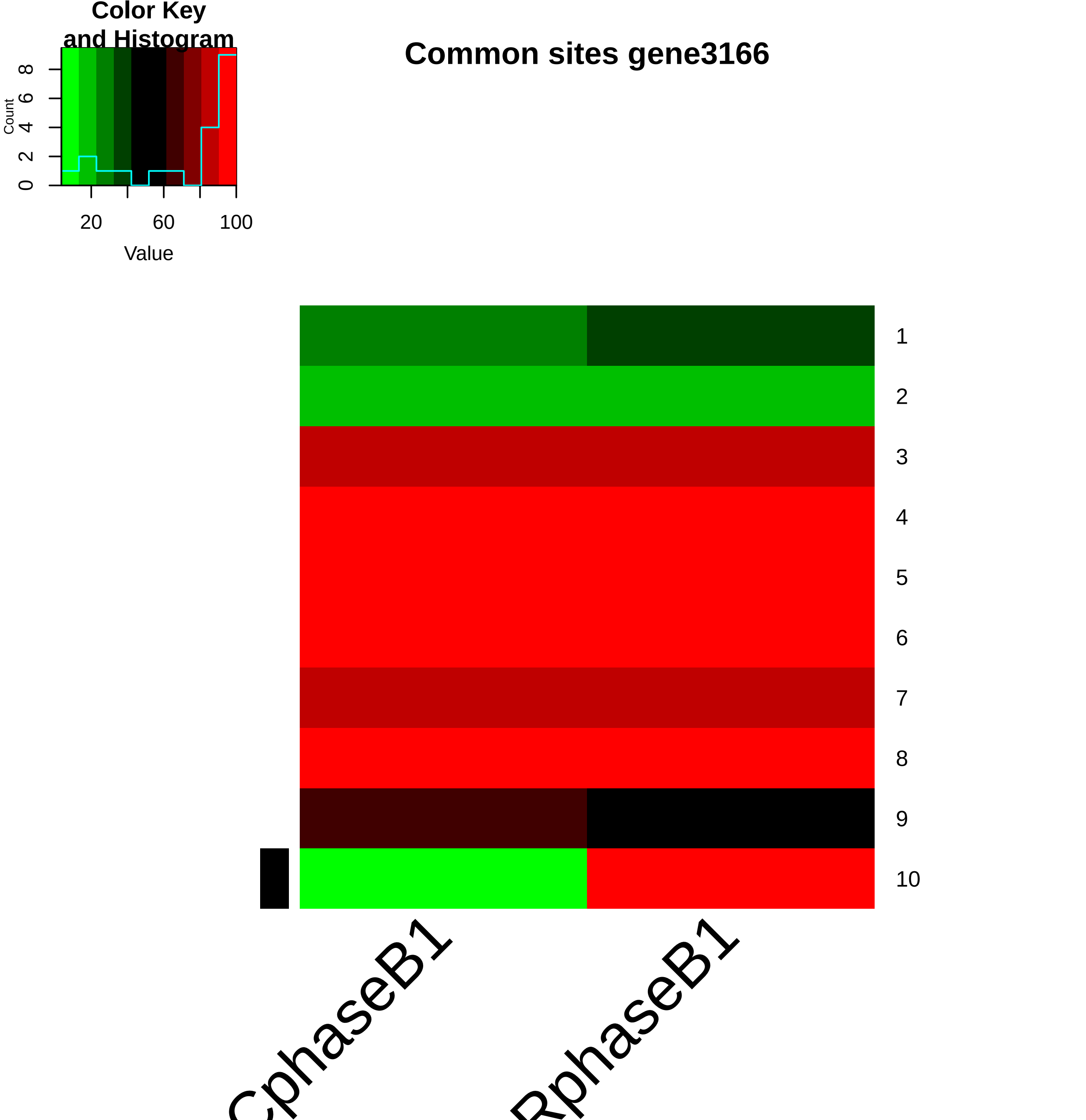
Color Key
and Histogram
Count
0
2
4
6
8
20
60
100
Value
Common sites gene3166
1
2
3
4
5
6
7
8
9
10
CphaseB1
RphaseB1

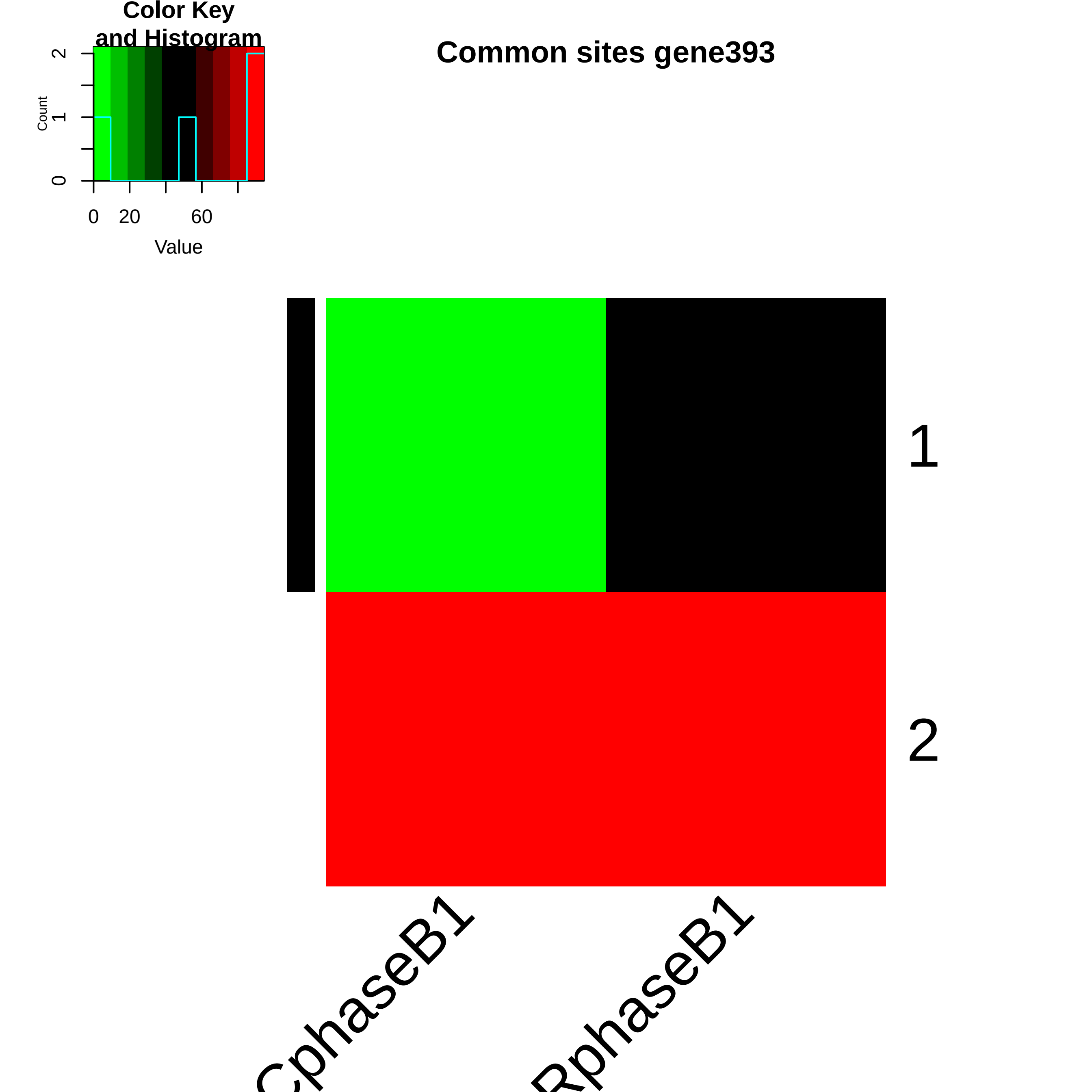
Color Key
and Histogram
Count
0
1
2
0
20
60
Value
Common sites gene393
1
2
CphaseB1
RphaseB1

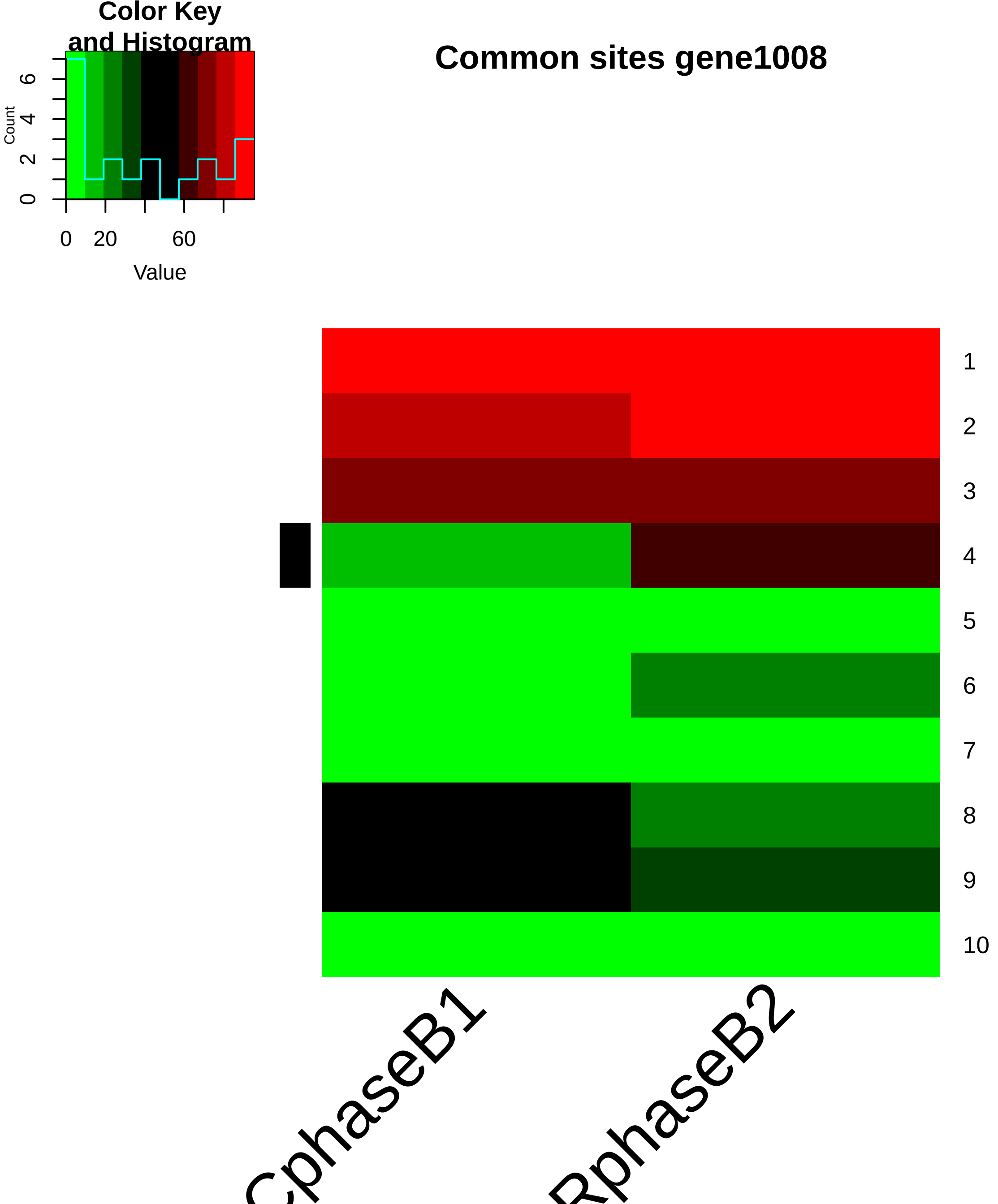
Color Key
and Histogram
Count
0
2
4
6
0
20
60
Value
Common sites gene1008
1
2
3
4
5
6
7
8
9
10
CphaseB1
RphaseB2

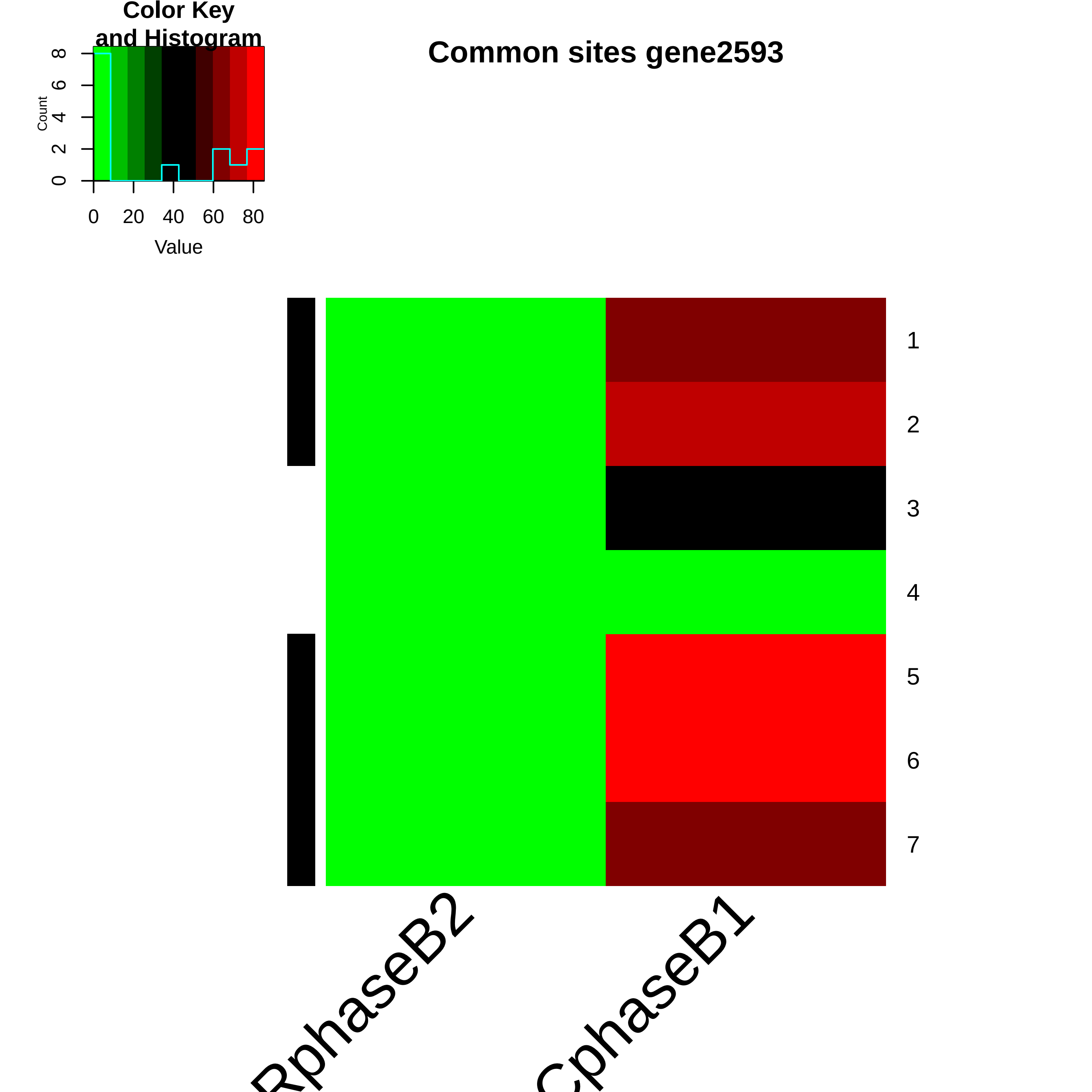
Color Key
and Histogram
Count
0
2
4
6
8
0
20
40
60
80
Value
Common sites gene2593
1
2
3
4
5
6
7
RphaseB2
CphaseB1

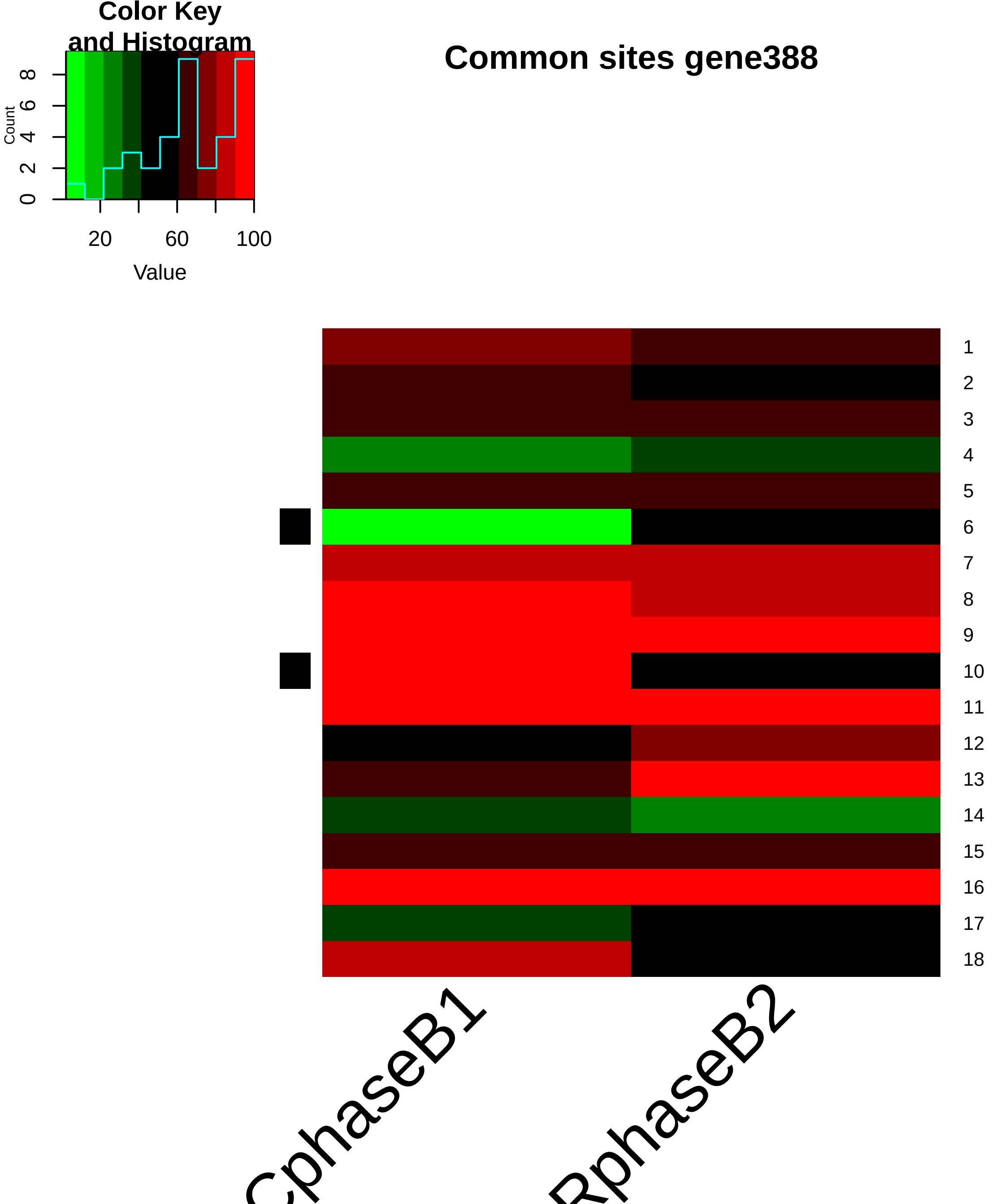
Color Key
and Histogram
Count
0 2 4 6 8
20 60 100
Value
Common sites gene388
CphaseB1
RphaseB2

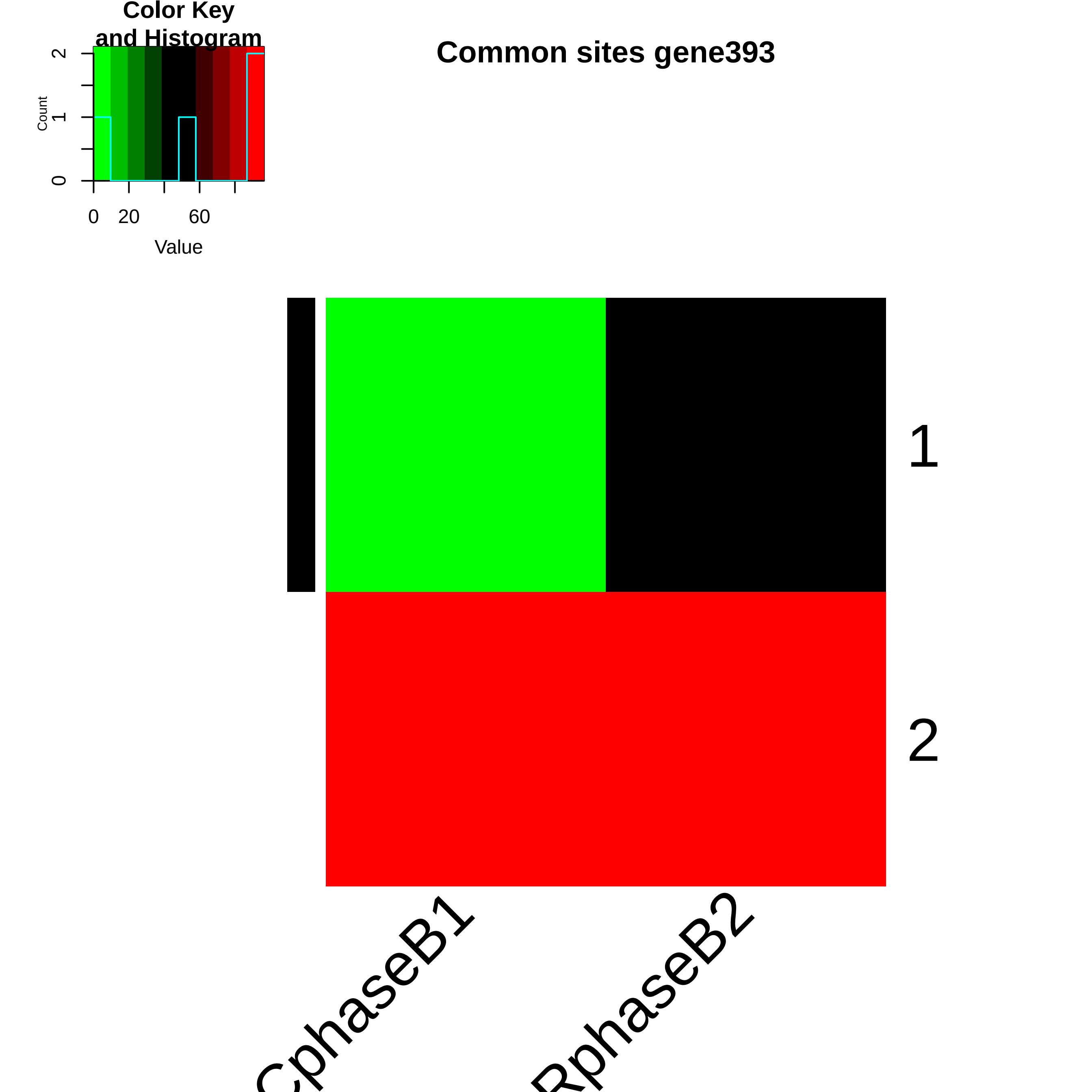

Color Key
and Histogram
Count
0
1
2
0
20
60
Value
Common sites gene393
1
2
CphaseB1
RphaseB2

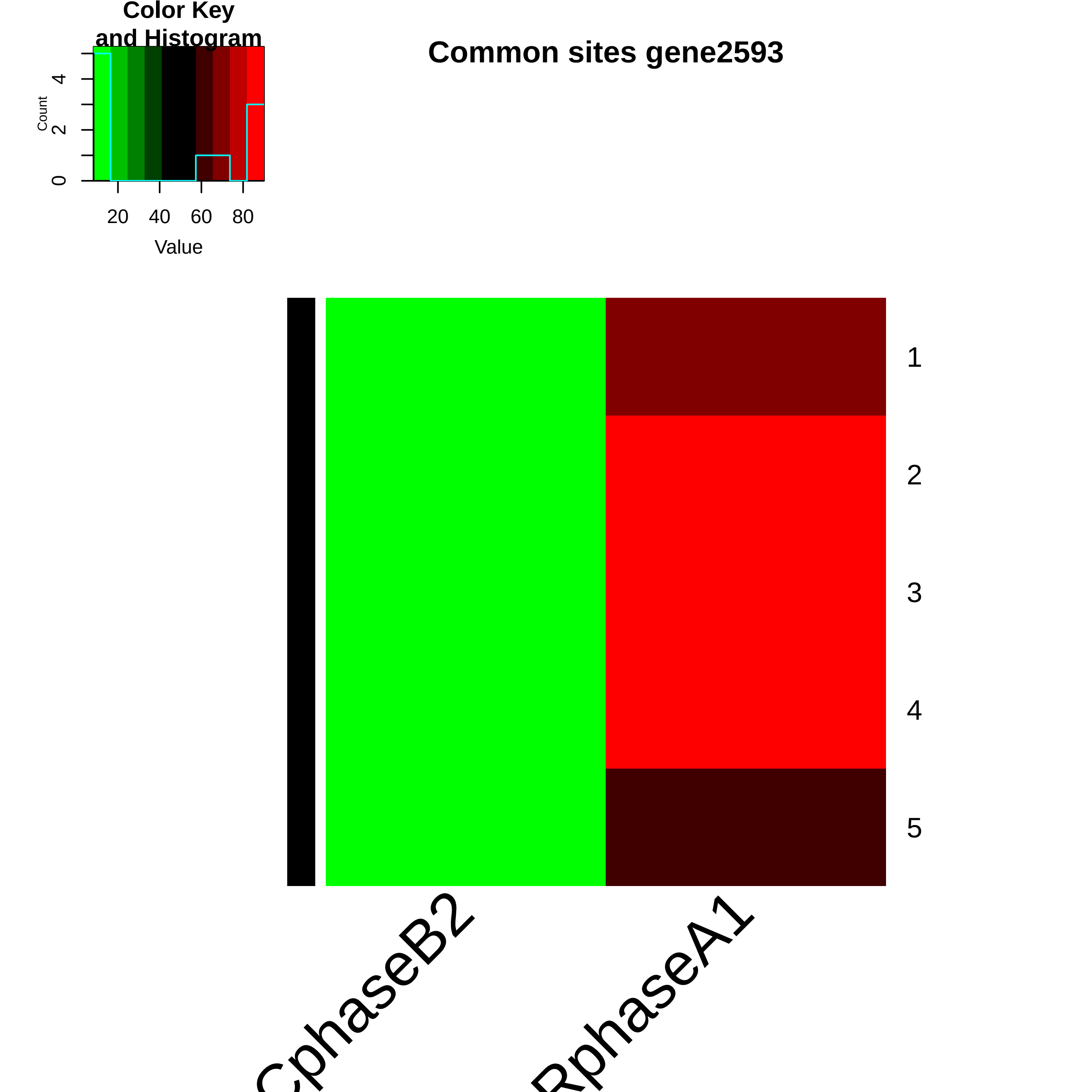

Color Key
and Histogram
Count
0
2
4
20
40
60
80
Value
Common sites gene2593
1
2
3
4
5
RphaseA1

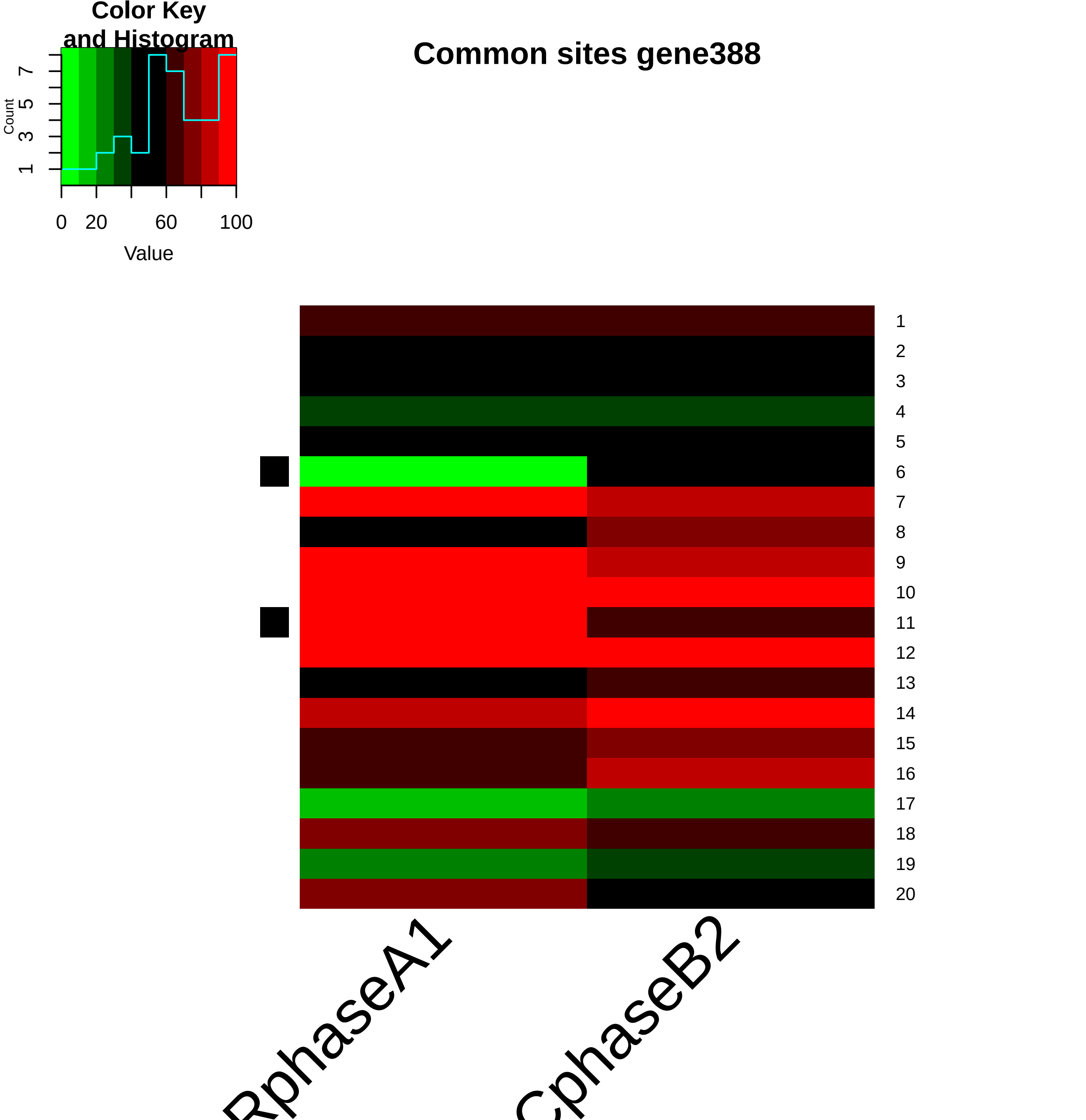
Color Key
and Histogram
Count
1 3 5 7
0 20 60 100
Value
Common sites gene388
RphaseA1
CphaseB2

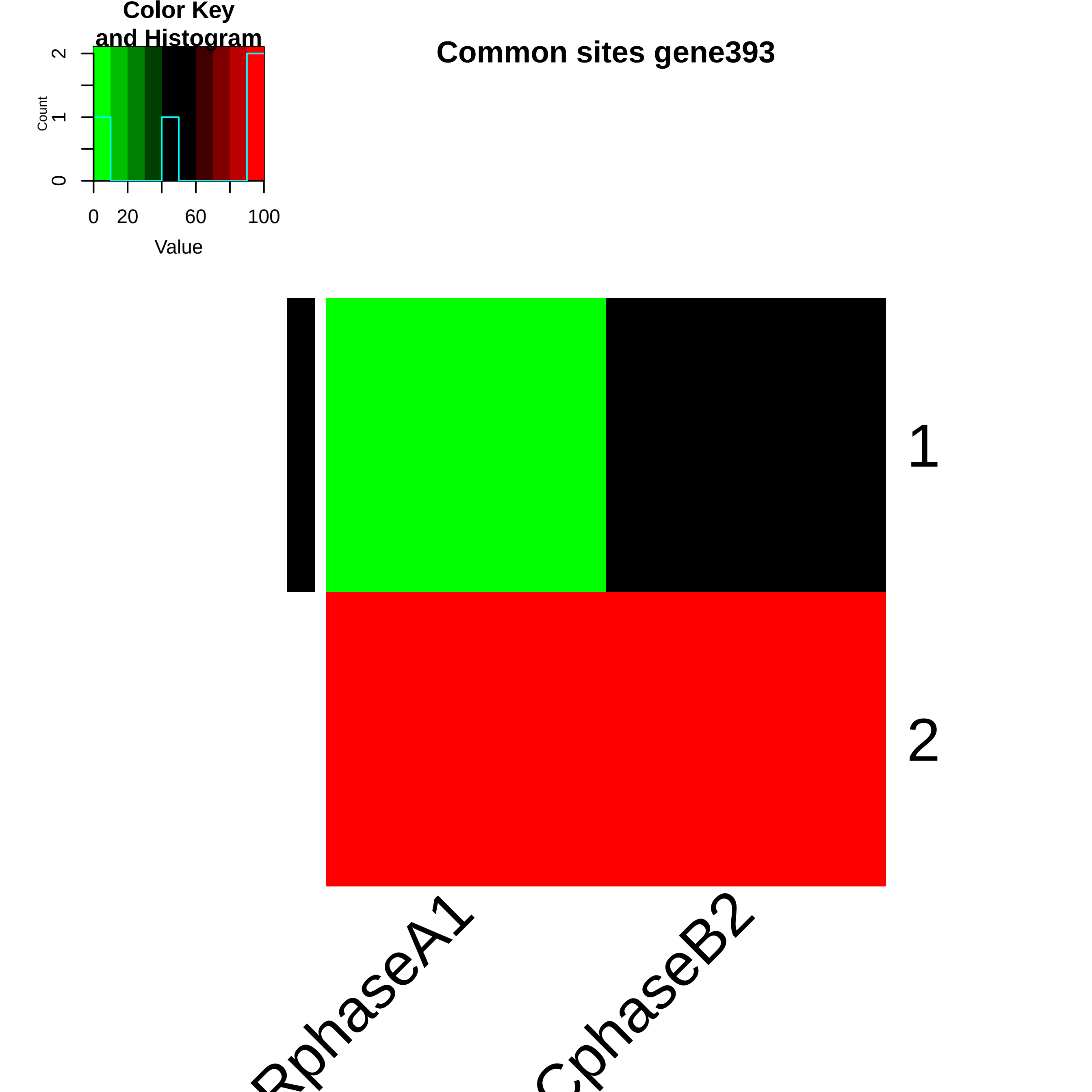

Color Key
and Histogram
Count
0
1
2
0
20
60
100
Value
Common sites gene393
1
2
RphaseA1
CphaseB2

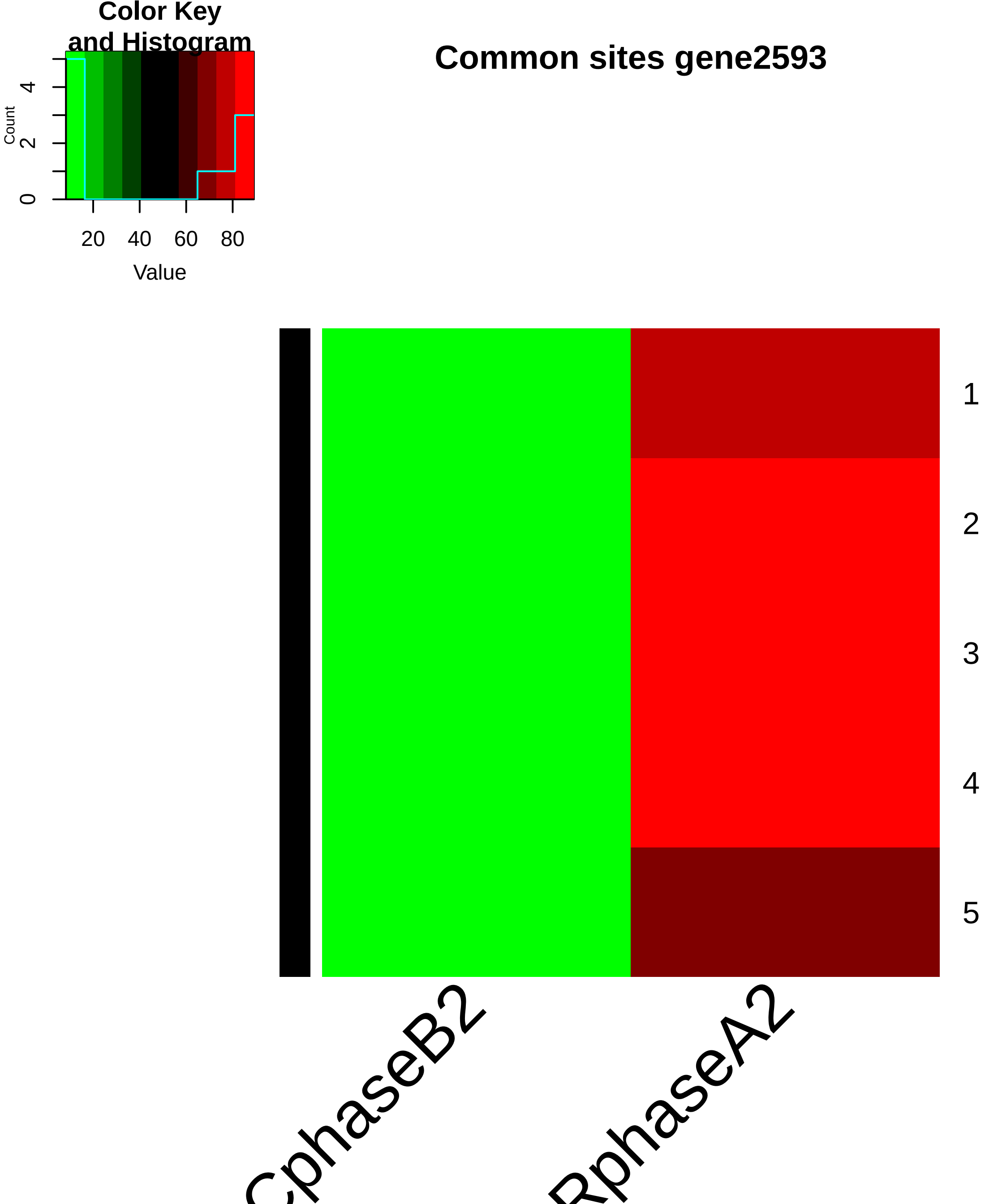
Color Key
and Histogram
Count
0
2
4
20
40
60
80
Value
Common sites gene2593
1
2
3
4
5
CphaseB2
RphaseA2

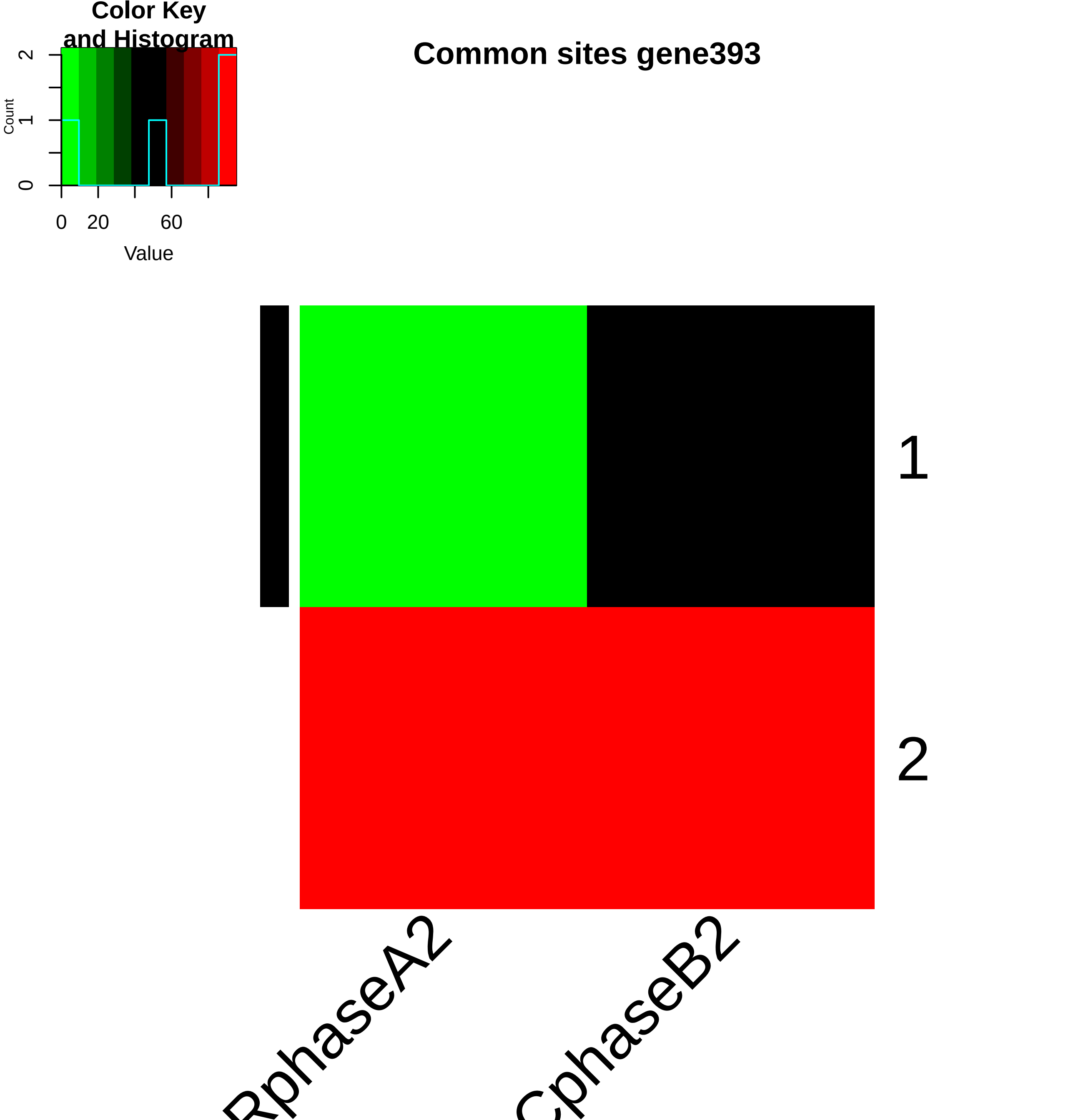

Color Key
and Histogram
Count
Value
0
20
60
0
1
2
Common sites gene393
1
2
RphaseA2
CphaseB2

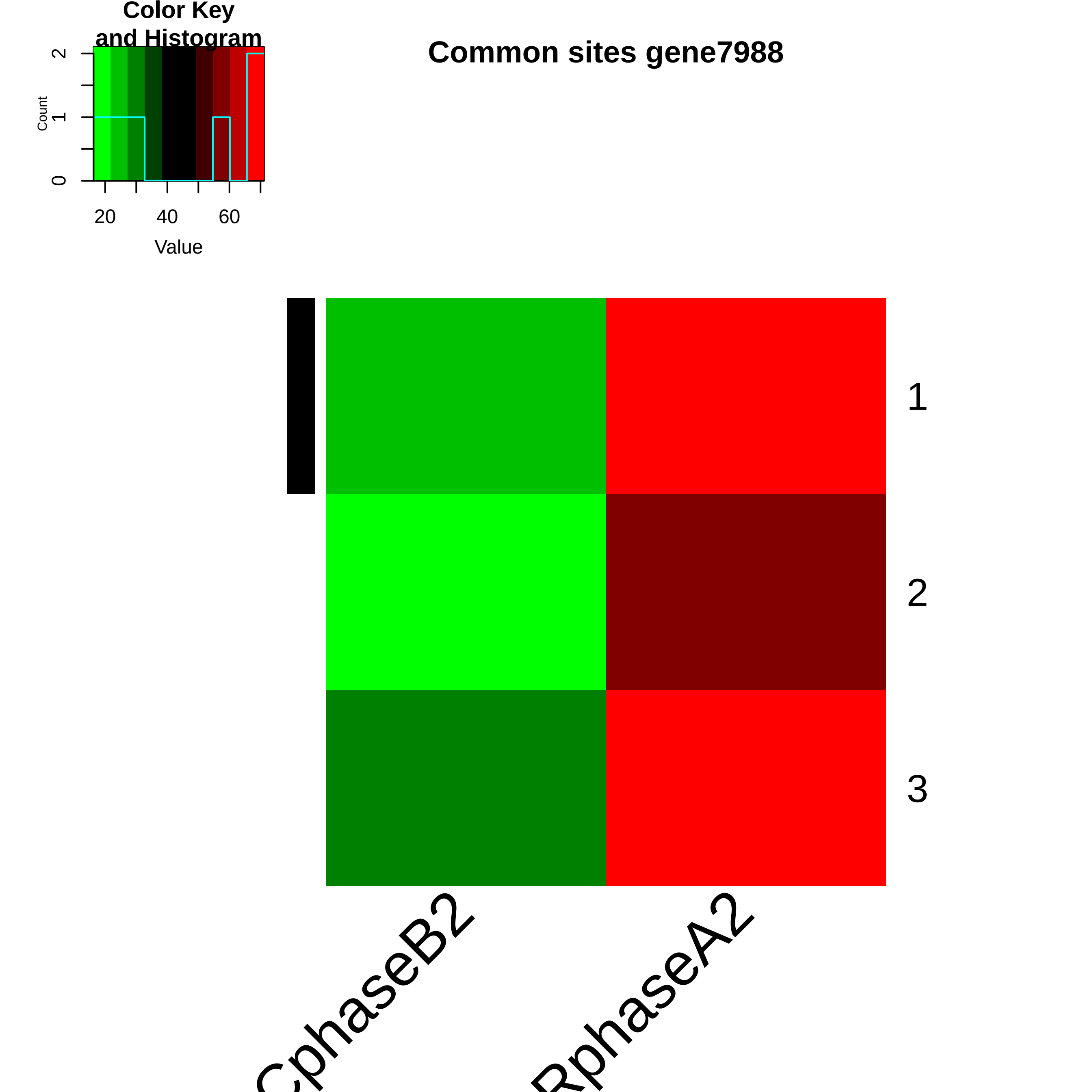
Color Key
and Histogram
Count
0
1
2
20
40
60
Value
Common sites gene7988
1
2
3
CphaseB2
RphaseA2

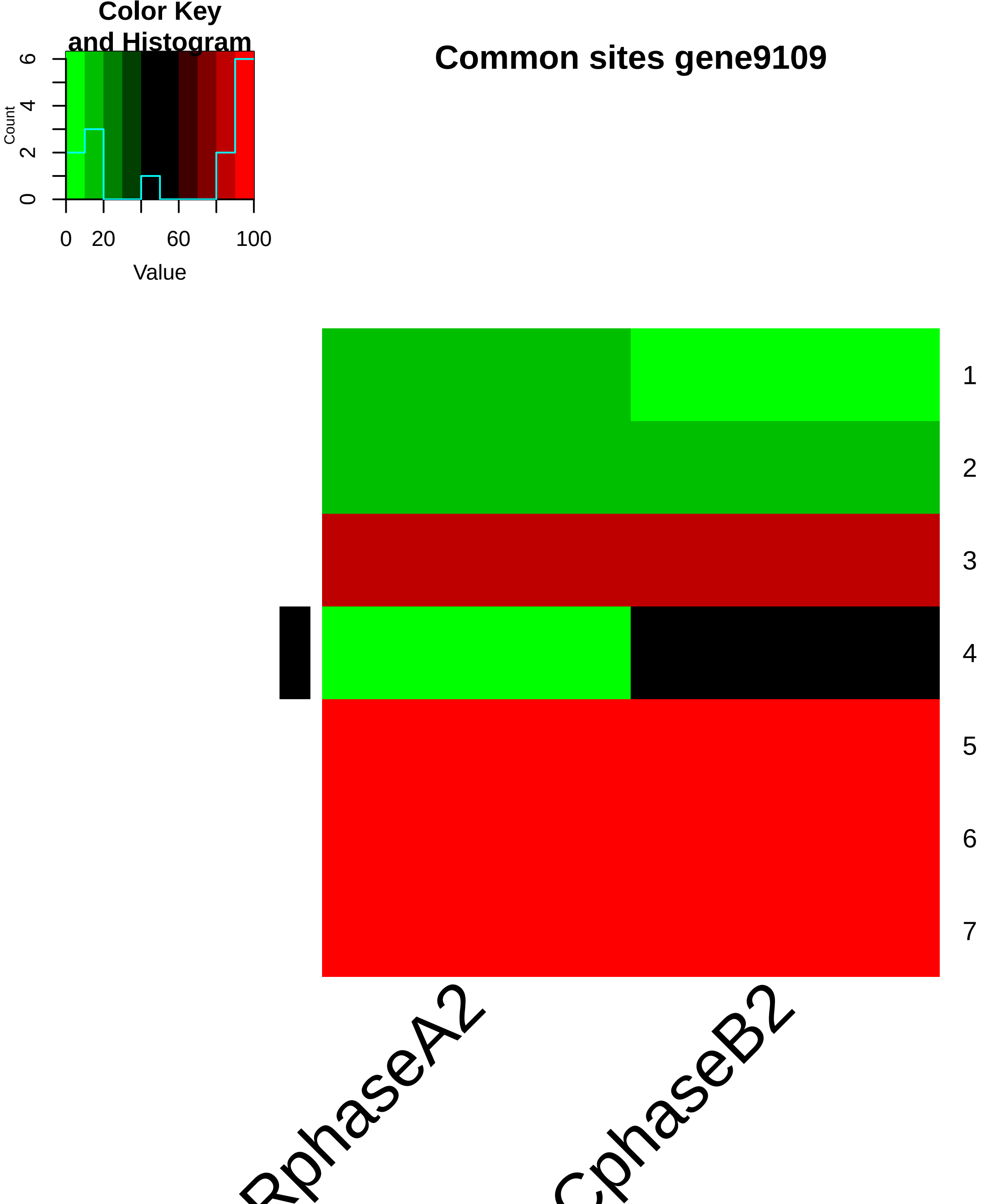
Color Key
and Histogram
Count
0
2
4
6
0
20
60
100
Value
Common sites gene9109
1
2
3
4
5
6
7
RphaseA2
CphaseB2

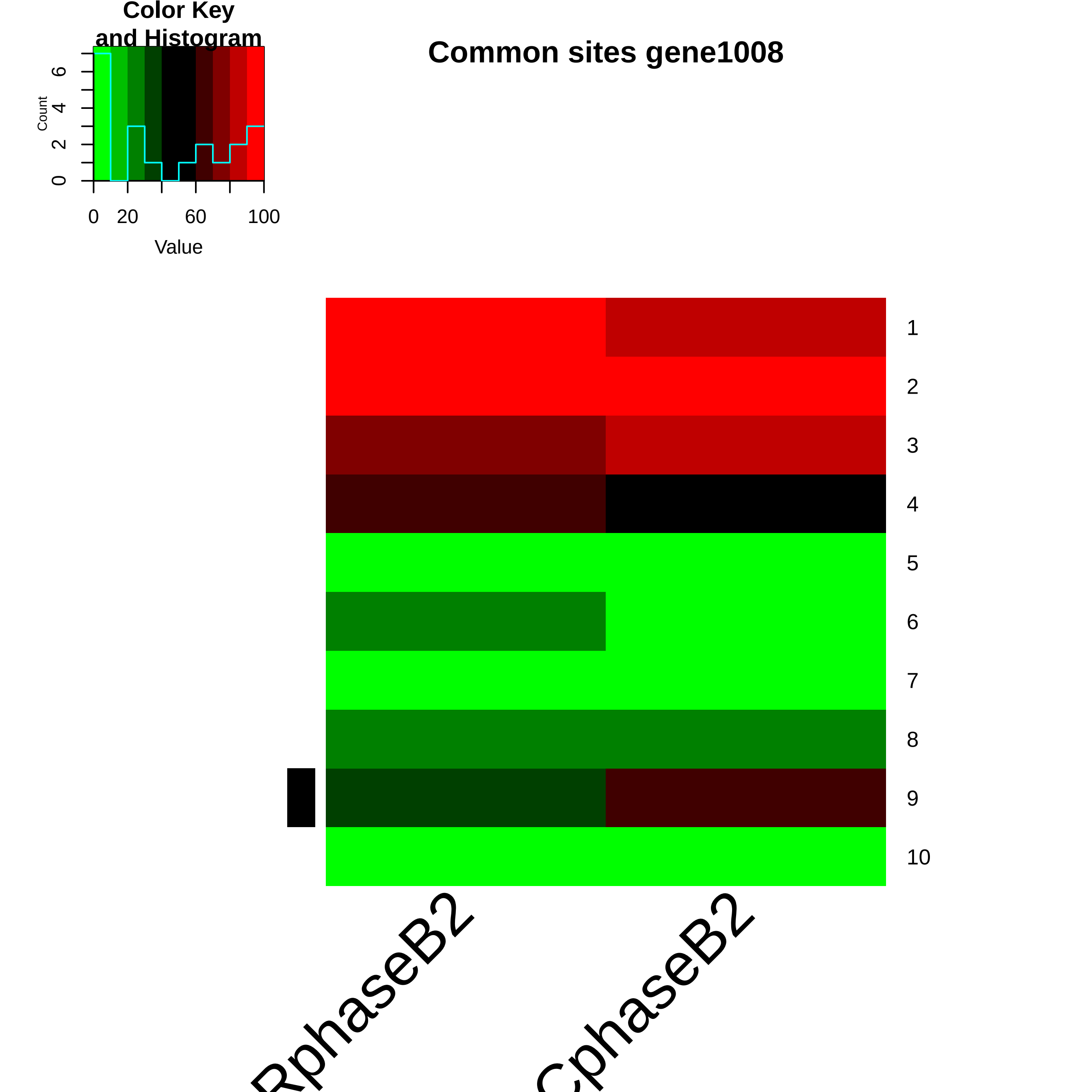
Color Key
and Histogram
Count
0
2
4
6
0
20
60
100
Value
Common sites gene1008
1
2
3
4
5
6
7
8
9
10
RphaseB2
CphaseB2

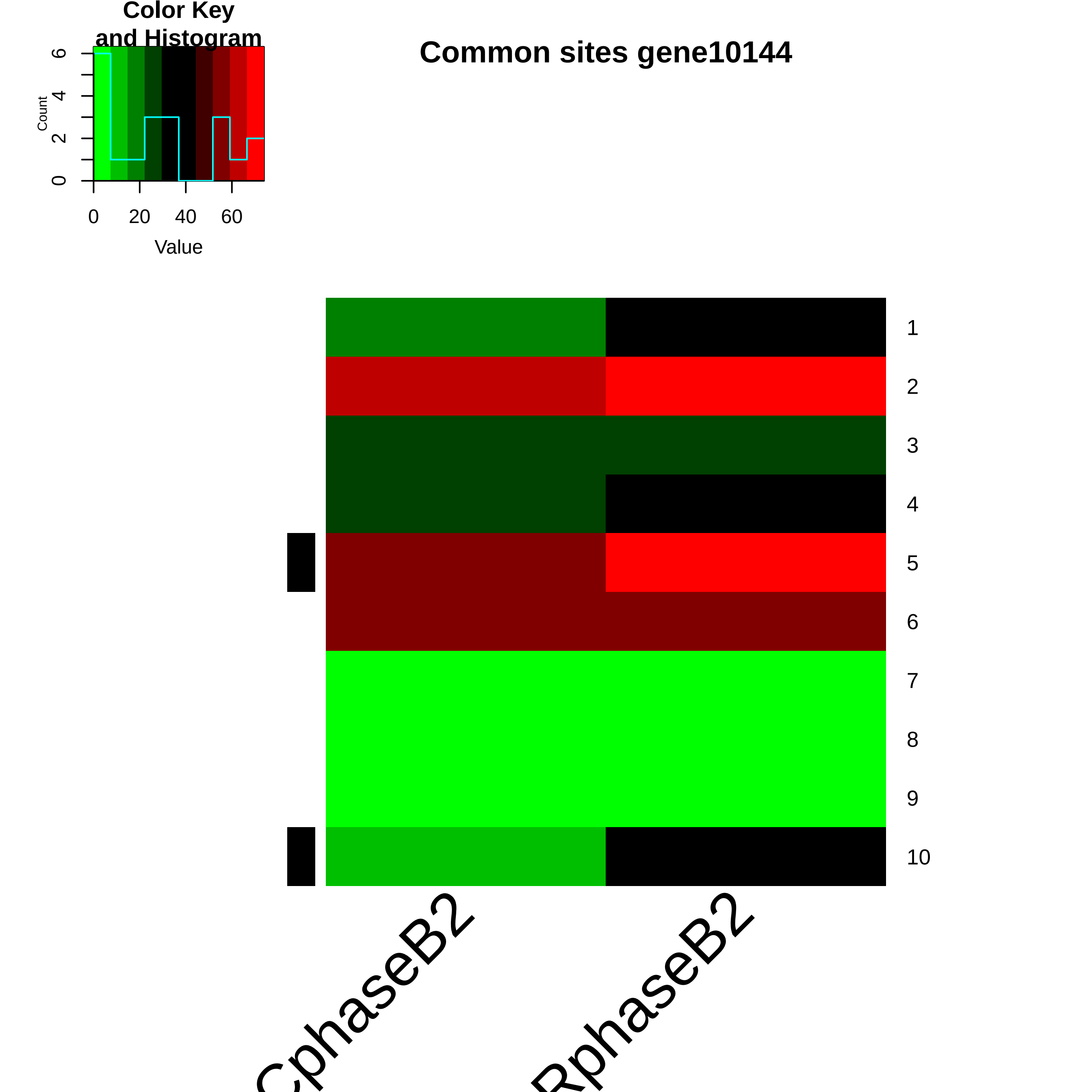
Color Key
and Histogram
Count
0
2
4
6
0
20
40
60
Value
Common sites gene10144
1
2
3
4
5
6
7
8
9
10

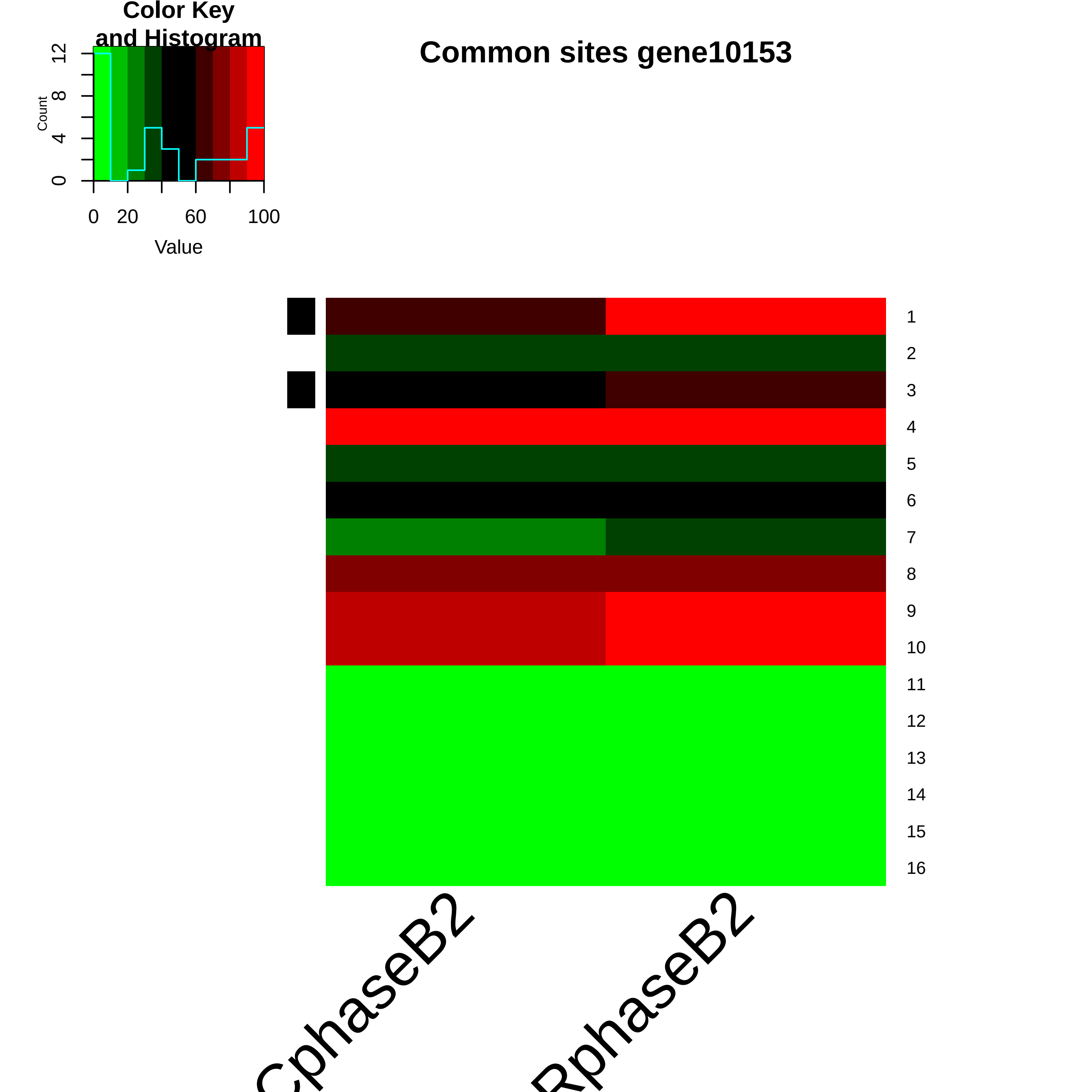

Color Key
and Histogram
Count
0
4
8
12
0
20
60
100
Value
Common sites gene10153
CphaseB2
RphaseB2

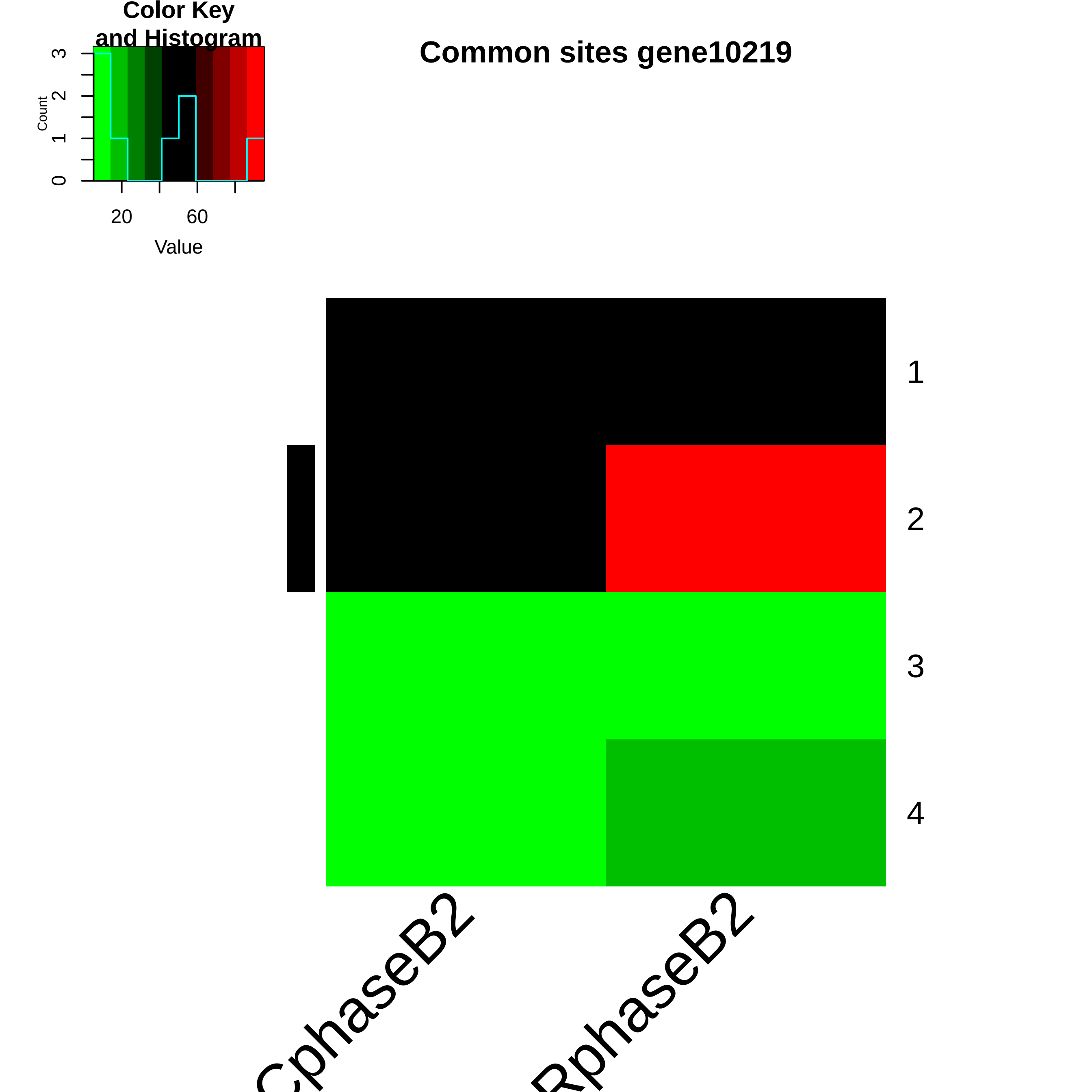

Color Key
and Histogram
Count
Value
0
1
2
3
20
60
Common sites gene10219
1
2
3
4
CphaseB2
RphaseB2

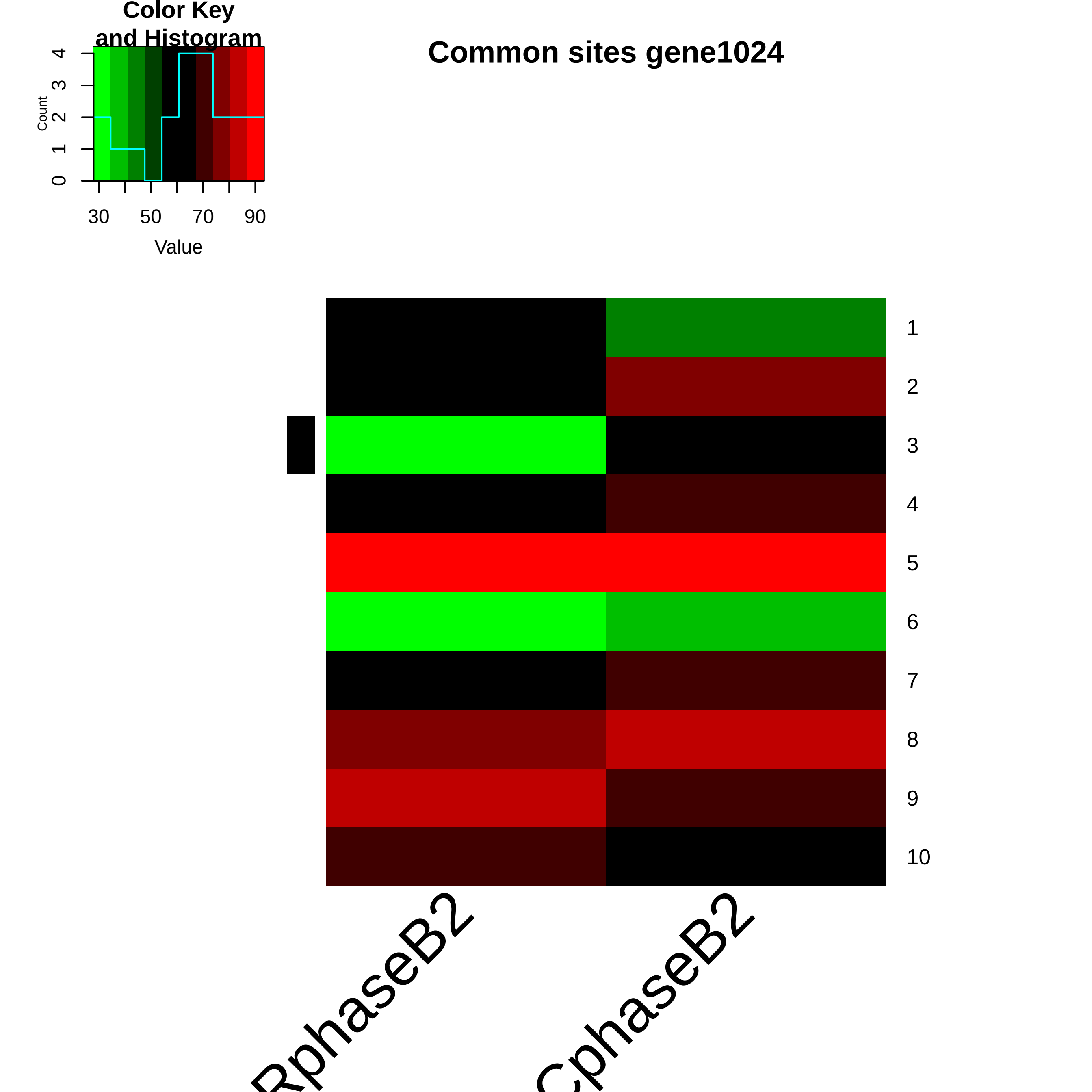
Color Key
and Histogram
Count
0
1
2
3
4
30
50
70
90
Value
Common sites gene1024
1
2
3
4
5
6
7
8
9
10
RphaseB2
CphaseB2

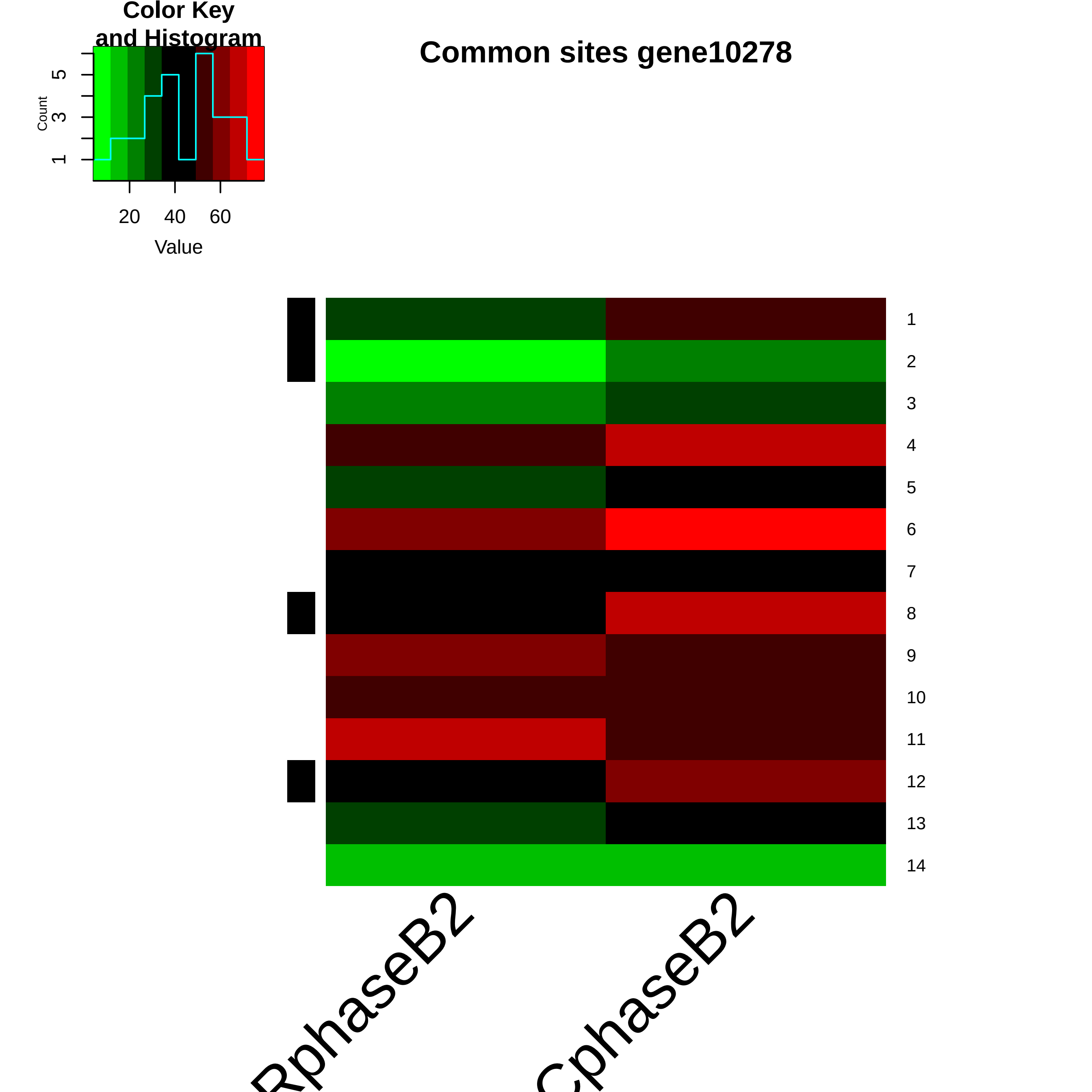

Color Key
and Histogram
Count
1
3
5
20
40
60
Value
Common sites gene10278
1
2
3
4
5
6
7
8
9
10
11
12
13
14
RphaseB2
CphaseB2

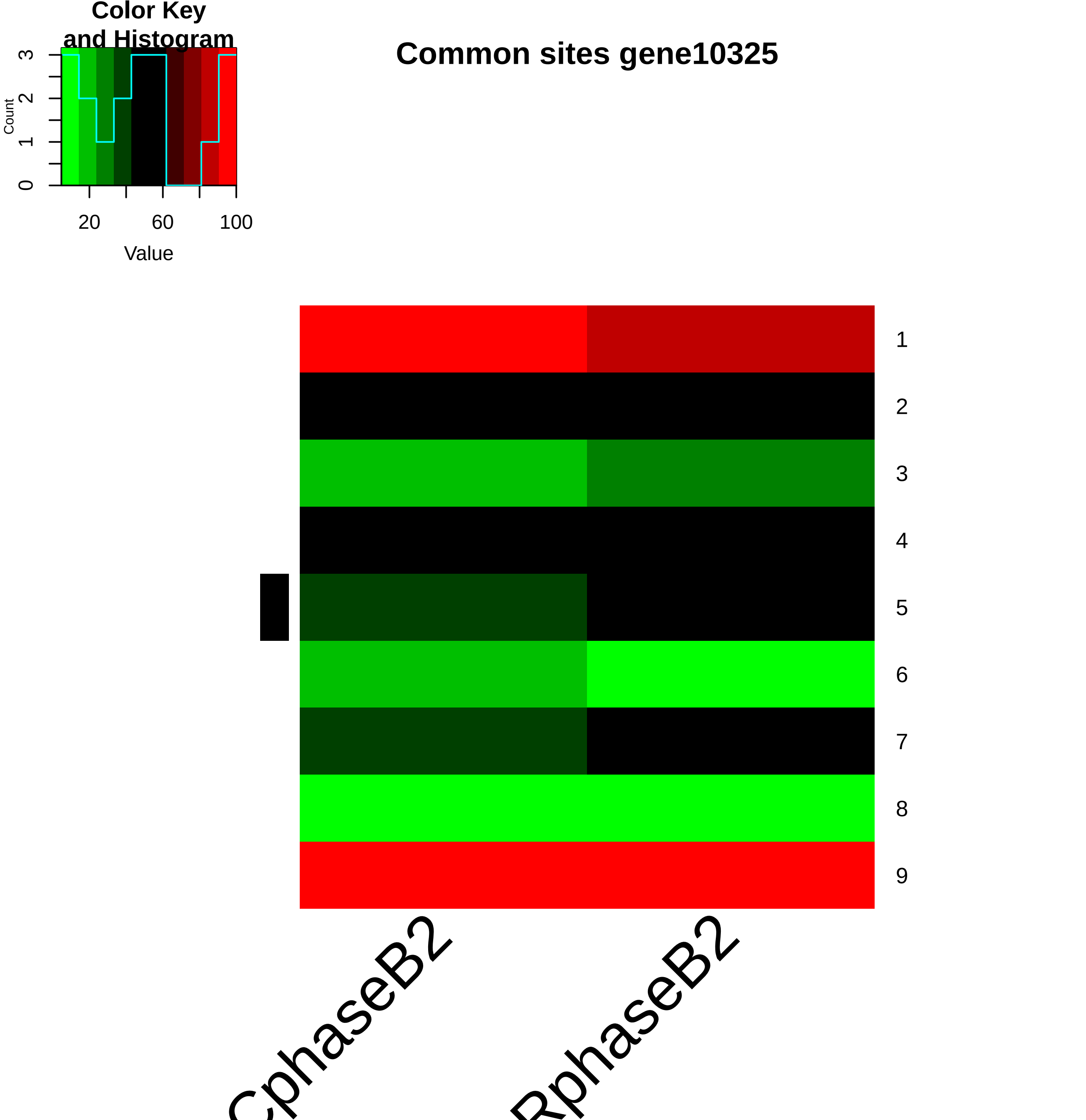

Color Key
and Histogram
Count
0
1
2
3
20
60
100
Value
Common sites gene10325
1
2
3
4
5
6
7
8
9
CphaseB2
RphaseB2

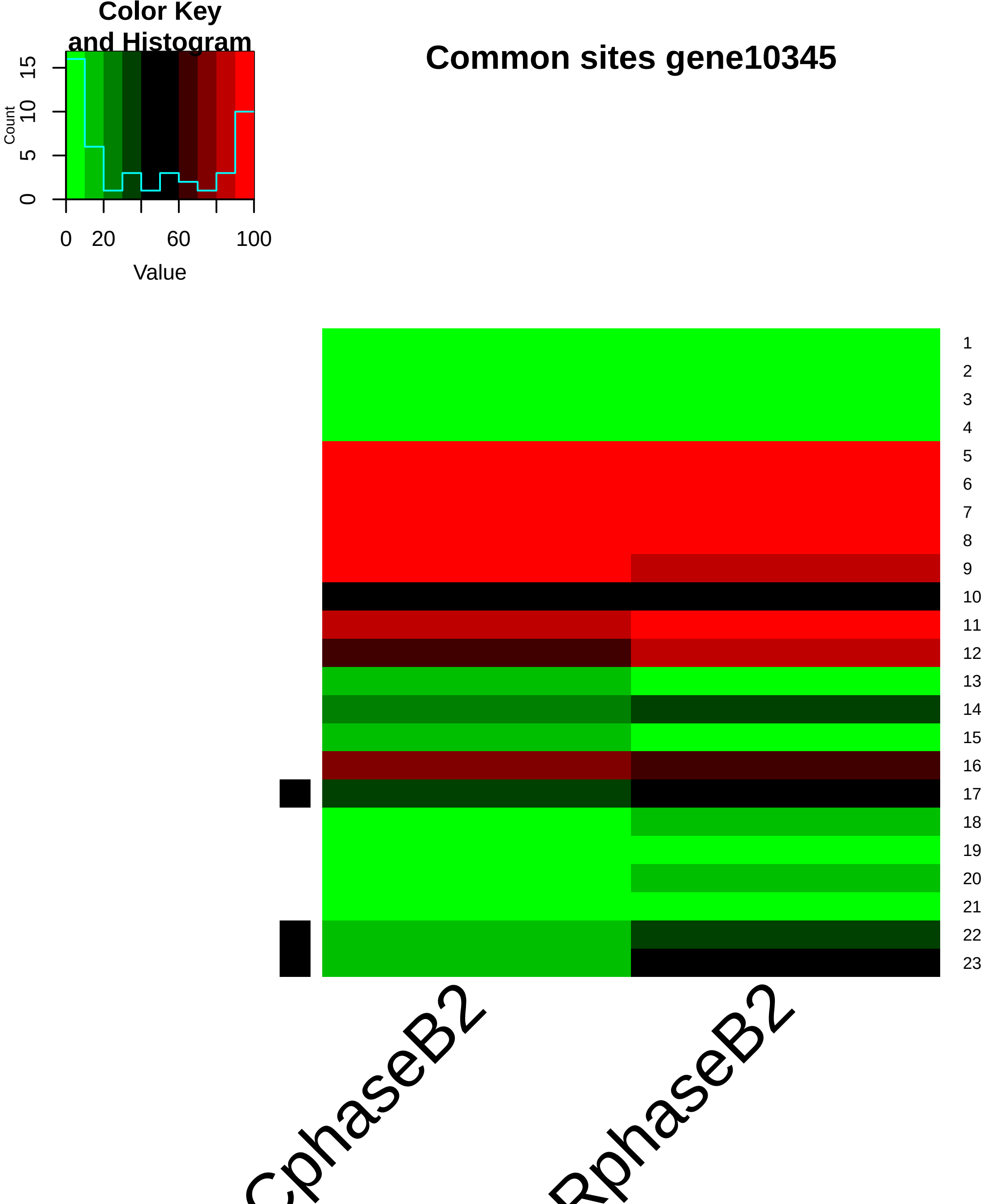

Color Key
and Histogram
Count
0
5
10
15
0
20
60
100
Value
Common sites gene10345
CphaseB2
RphaseB2

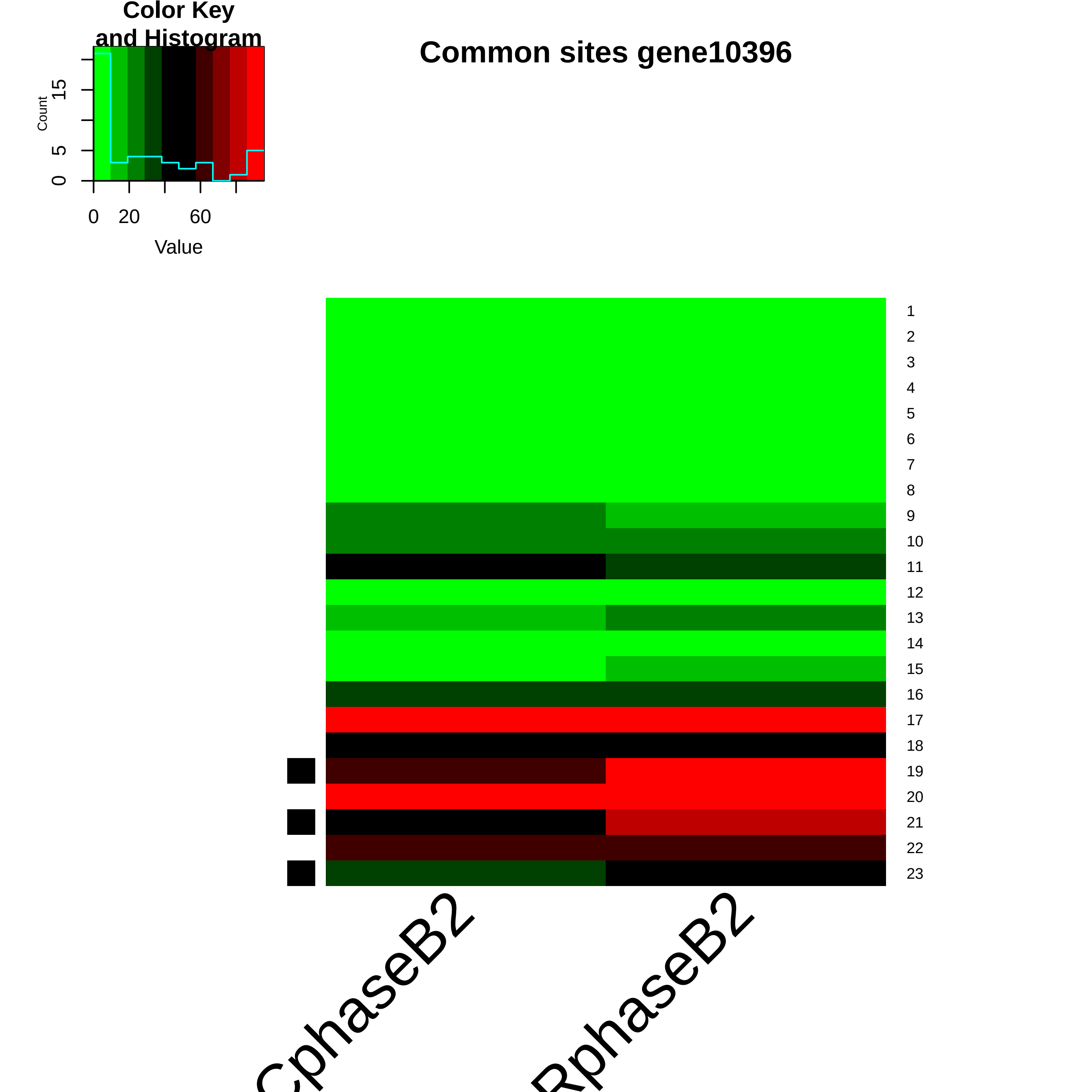

Color Key
and Histogram
Count
0
5
15
0
20
60
Value
Common sites gene10396
CphaseB2
RphaseB2

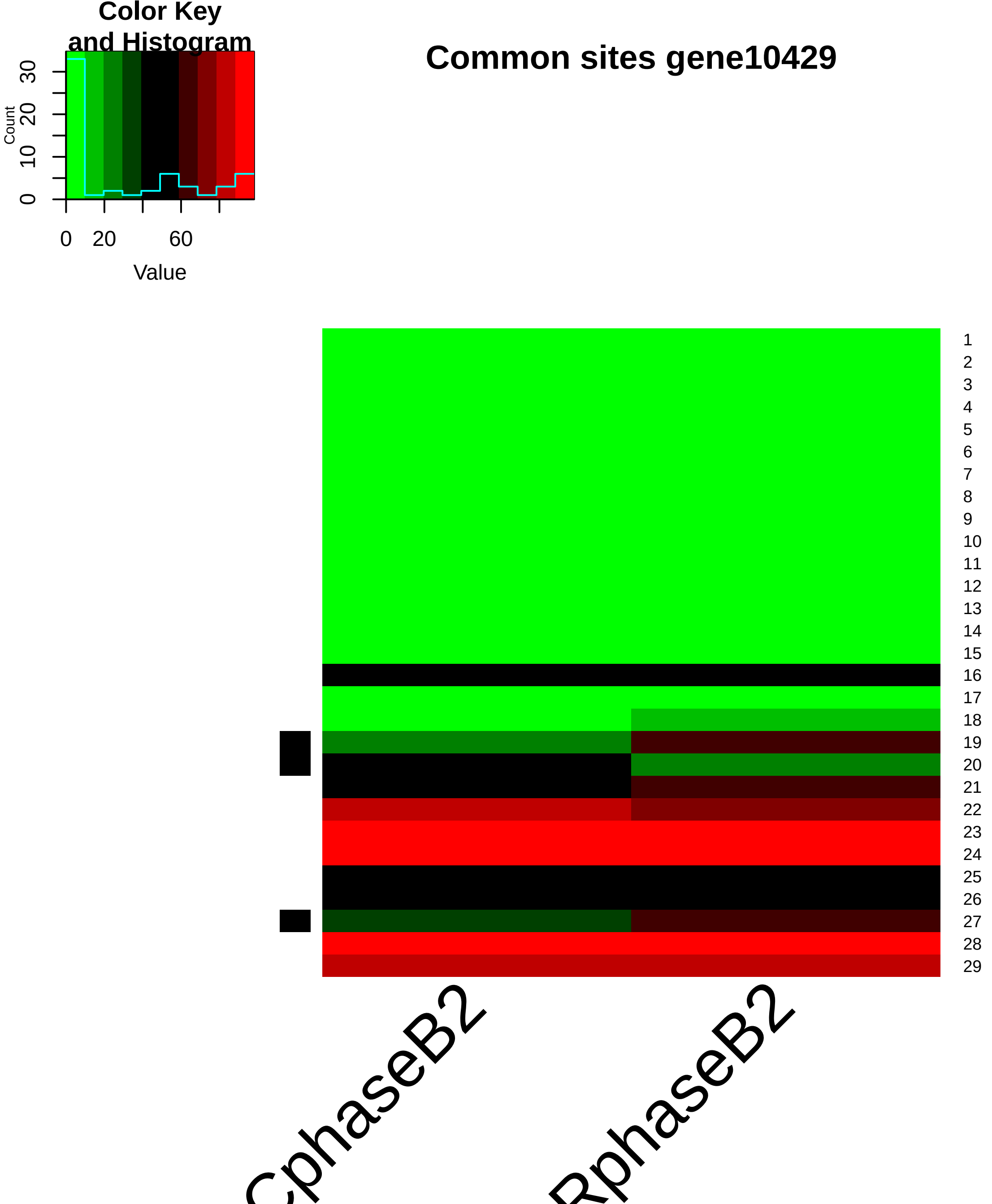
Color Key
and Histogram
Count
0
10
20
30
0
20
60
Value
Common sites gene10429
CphaseB2
RphaseB2

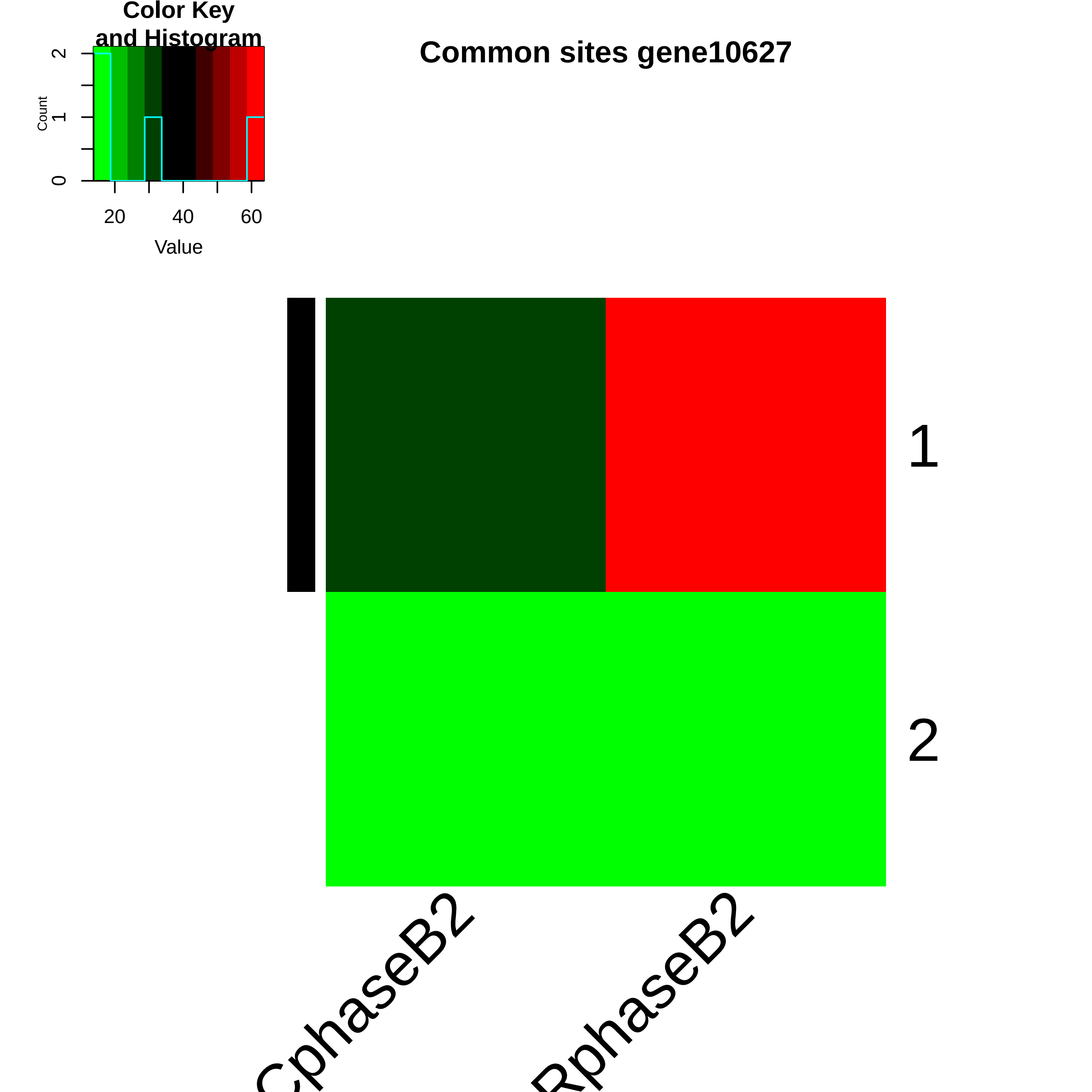

Color Key
and Histogram
Count
0
1
2
20
40
60
Value
Common sites gene10627
1
2
CphaseB2
RphaseB2

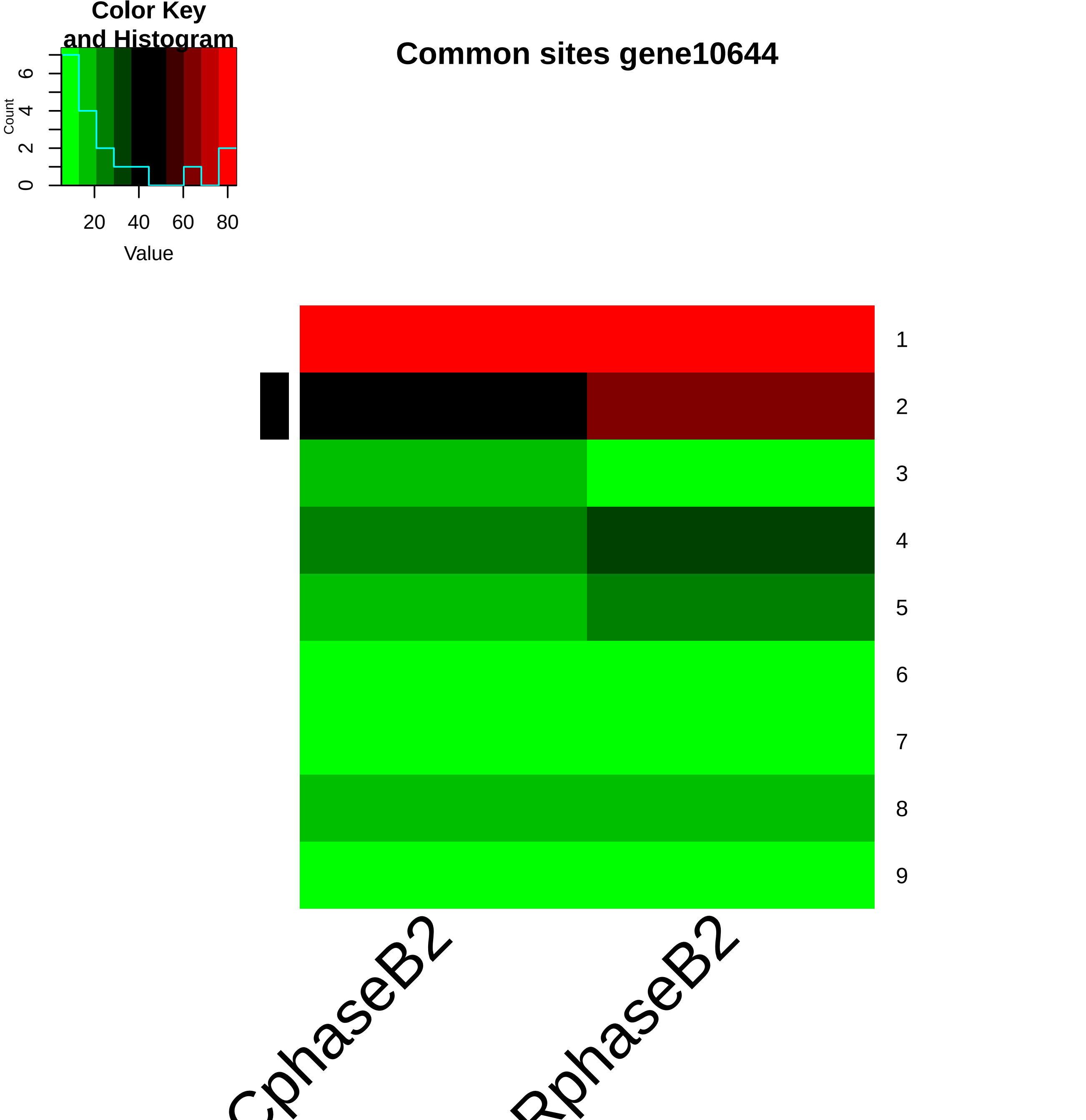

Color Key
and Histogram
Count
0
2
4
6
20
40
60
80
Value
Common sites gene10644
1
2
3
4
5
6
7
8
9
CphaseB2
RphaseB2

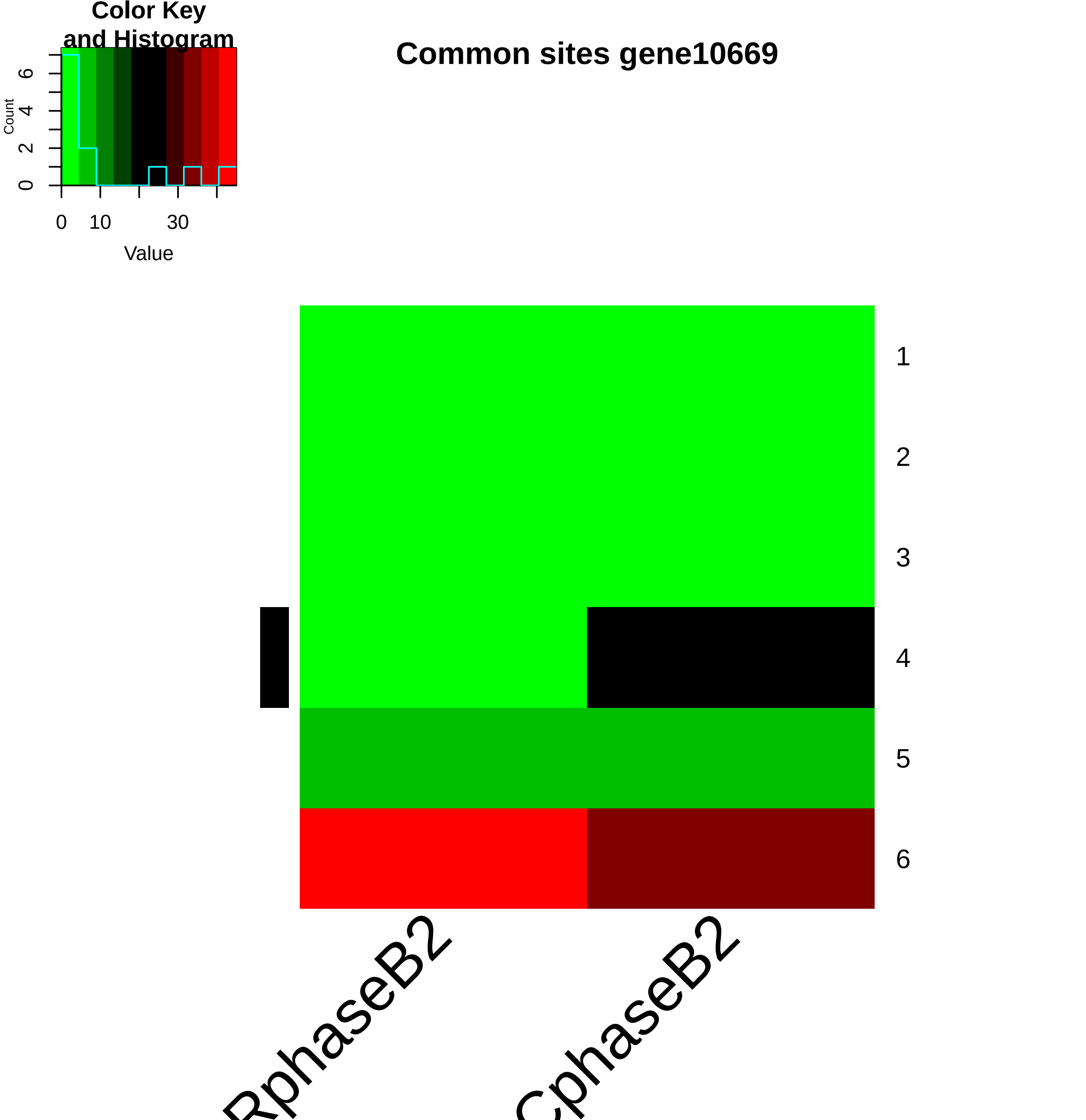

Color Key
and Histogram
Count
0
2
4
6
0
10
30
Value
Common sites gene10669
1
2
3
4
5
6
RphaseB2
CphaseB2

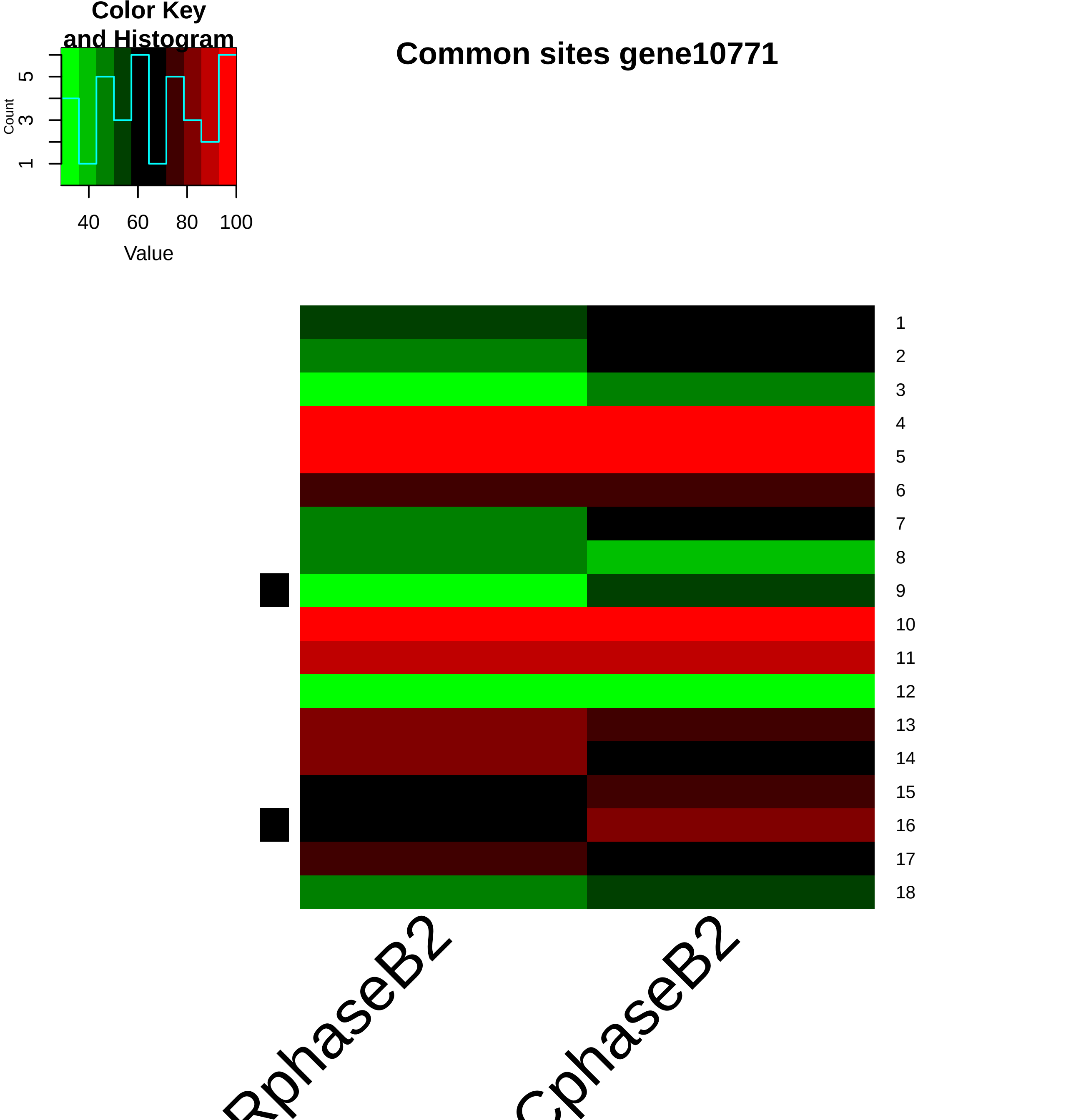

Color Key
and Histogram
Count
1
3
5
40
60
80
100
Value
Common sites gene10771
RphaseB2
CphaseB2

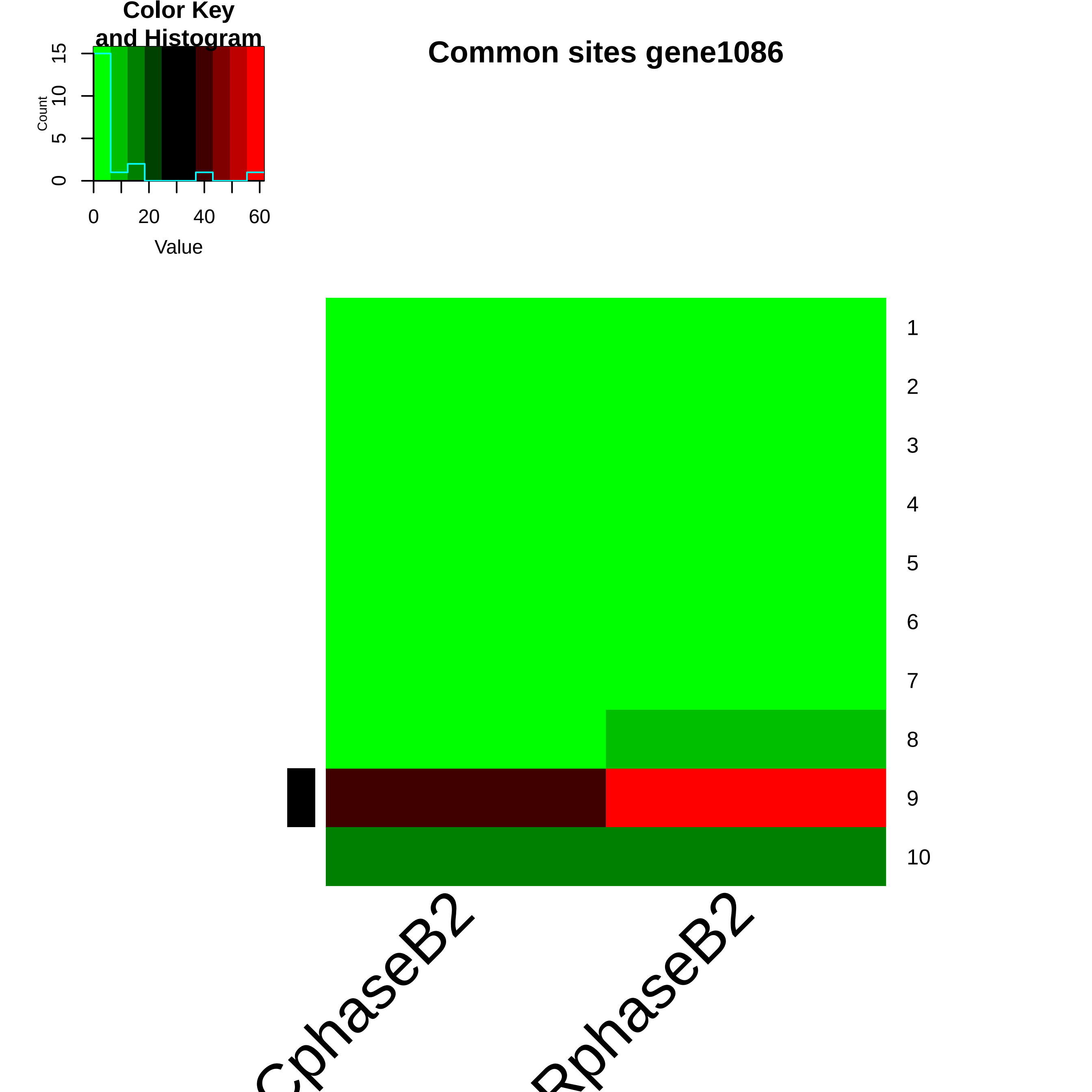

Color Key
and Histogram
Count
0
5
10
15
0
20
40
60
Value
Common sites gene1086
1
2
3
4
5
6
7
8
9
10
CphaseB2
RphaseB2

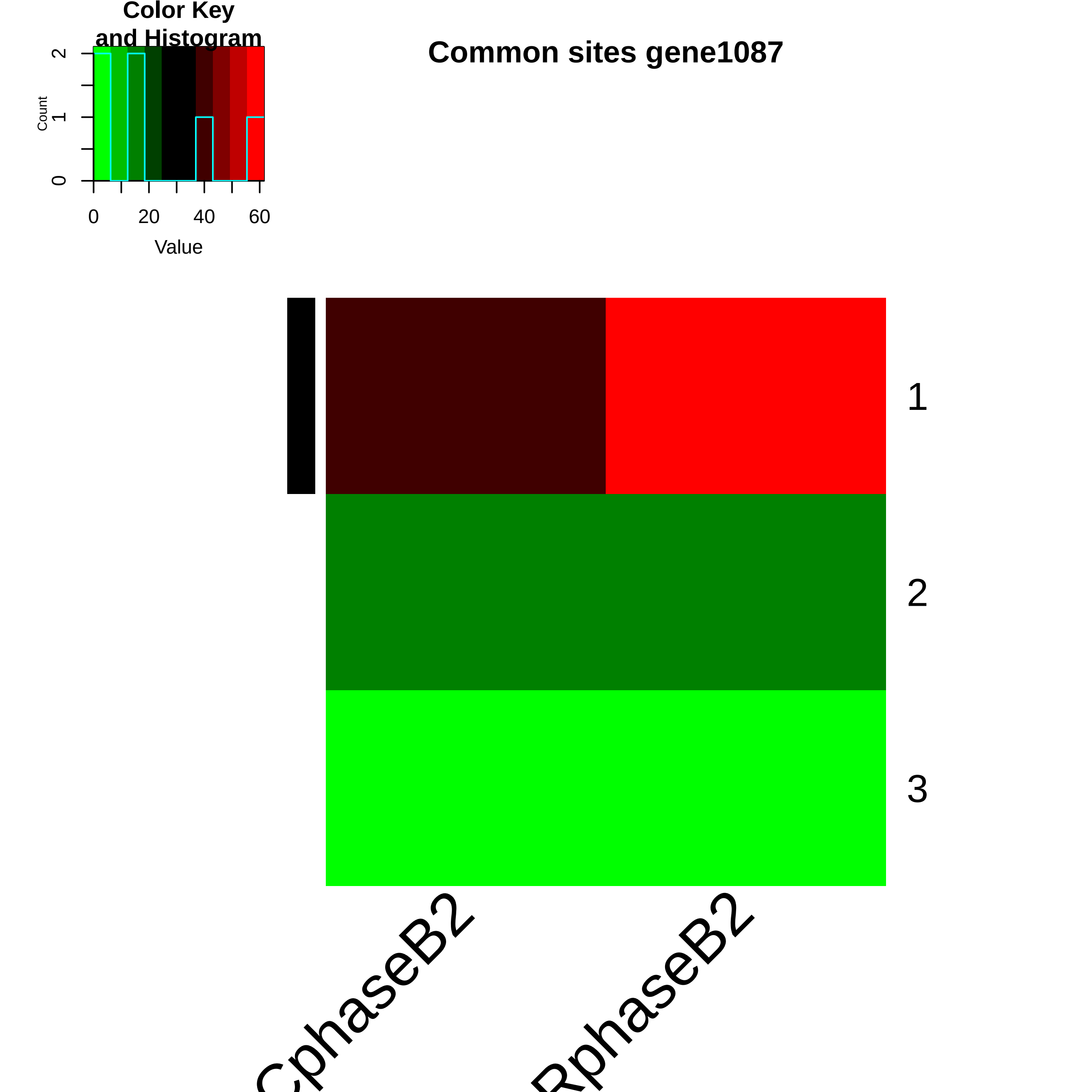

Color Key
and Histogram
Count
0
1
2
0
20
40
60
Value
Common sites gene1087
1
2
3
CphaseB2
RphaseB2

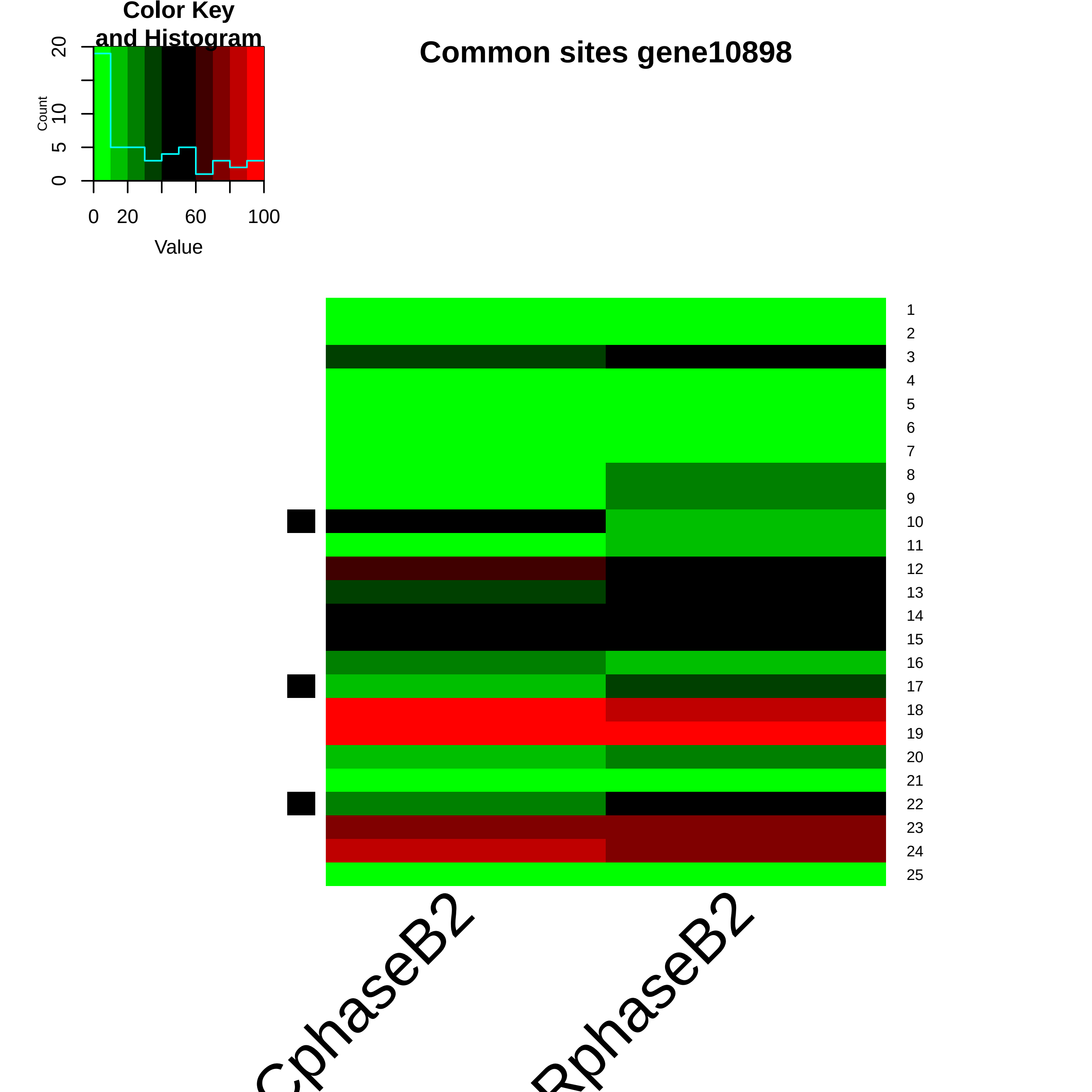
Color Key
and Histogram
Count
0
5
10
20
0
20
60
100
Value
Common sites gene10898
CphaseB2
RphaseB2

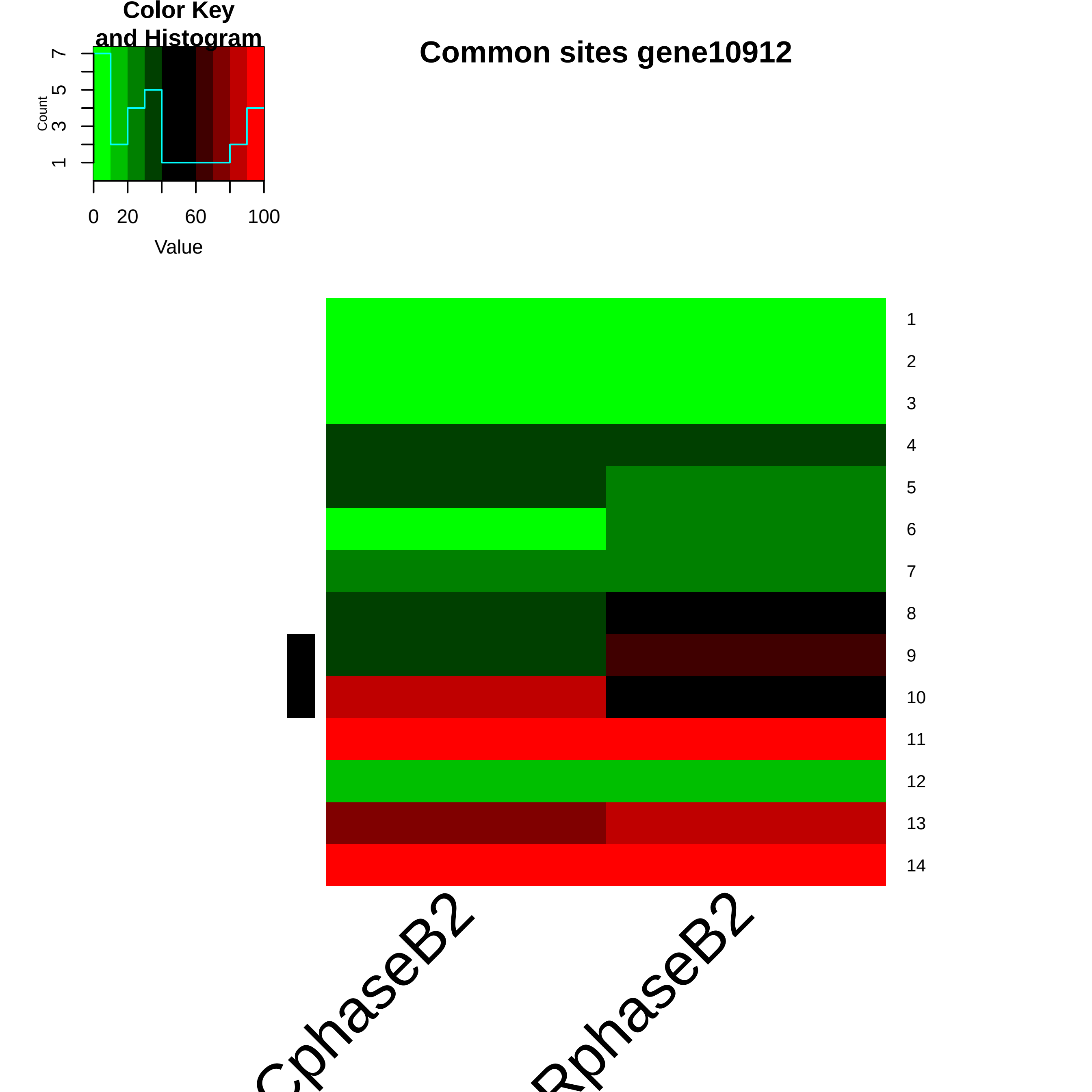
Color Key
and Histogram
Count
1
3
5
7
0
20
60
100
Value
Common sites gene10912
1
2
3
4
5
6
7
8
9
10
11
12
13
14
CphaseB2
RphaseB2

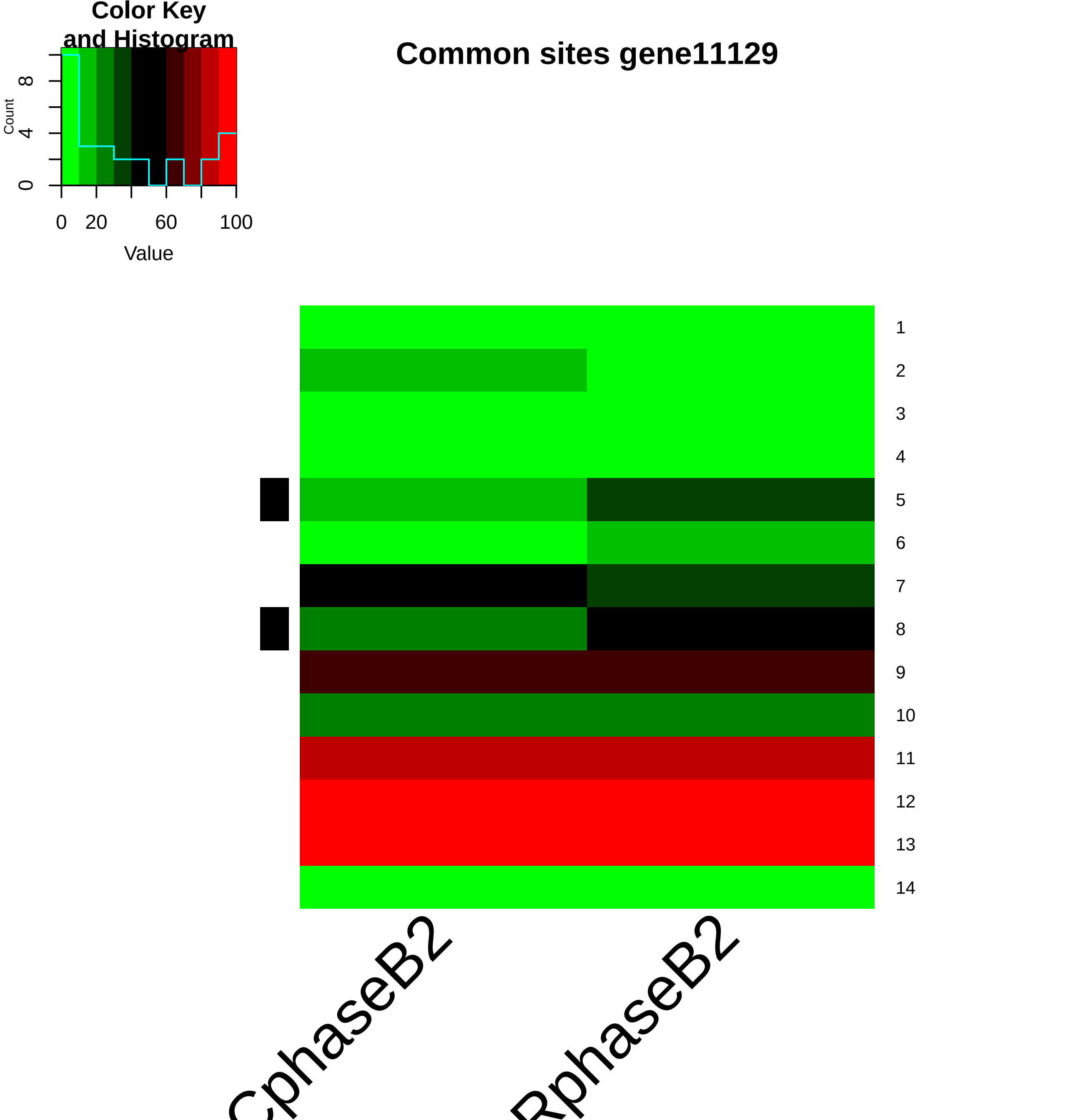
Color Key
and Histogram
Count
0
4
8
0
20
60
100
Value
Common sites gene11129
1
2
3
4
5
6
7
8
9
10
11
12
13
14
CphaseB2
RphaseB2

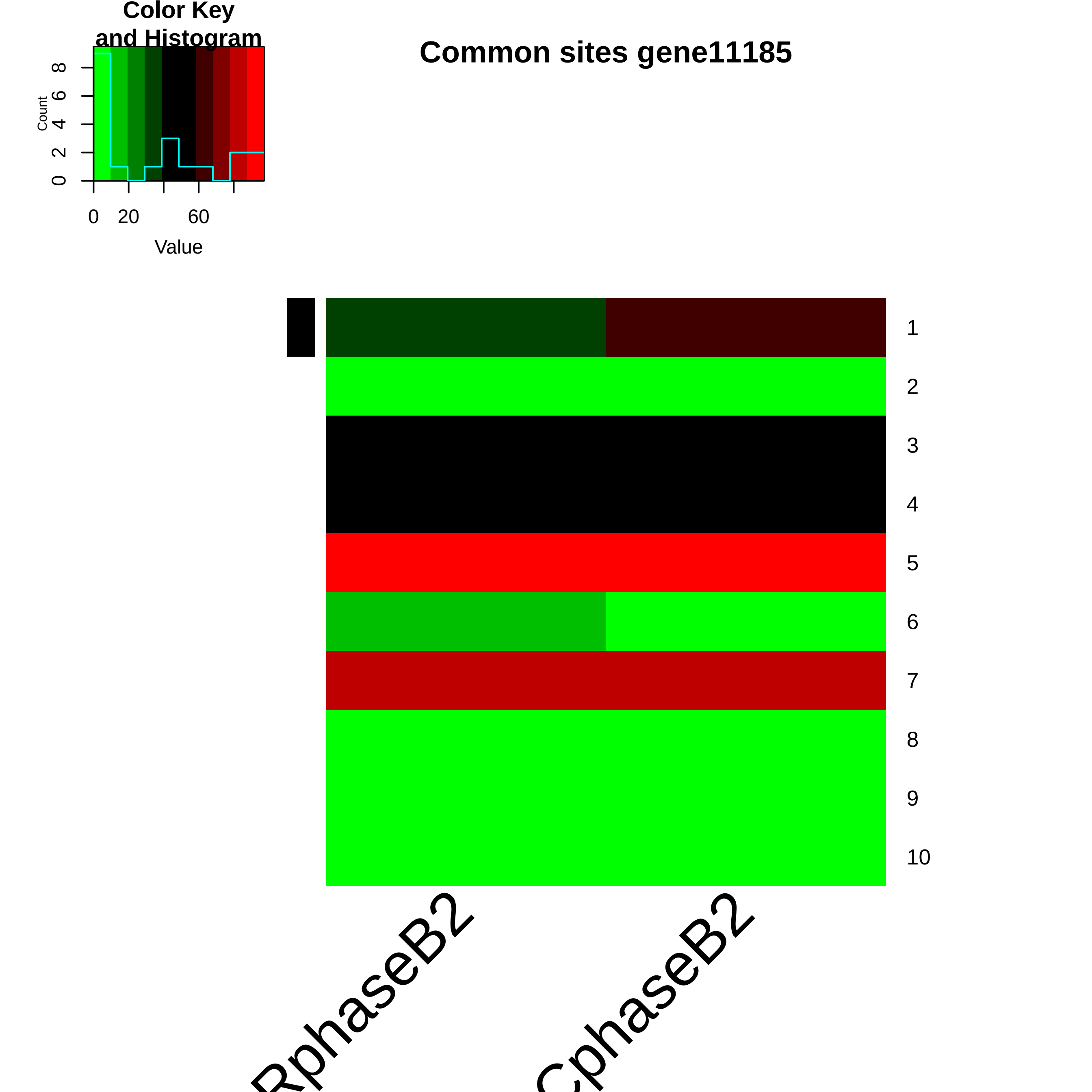

Color Key
and Histogram
Count
0
2
4
6
8
0
20
60
Value
Common sites gene11185
1
2
3
4
5
6
7
8
9
10
RphaseB2
CphaseB2

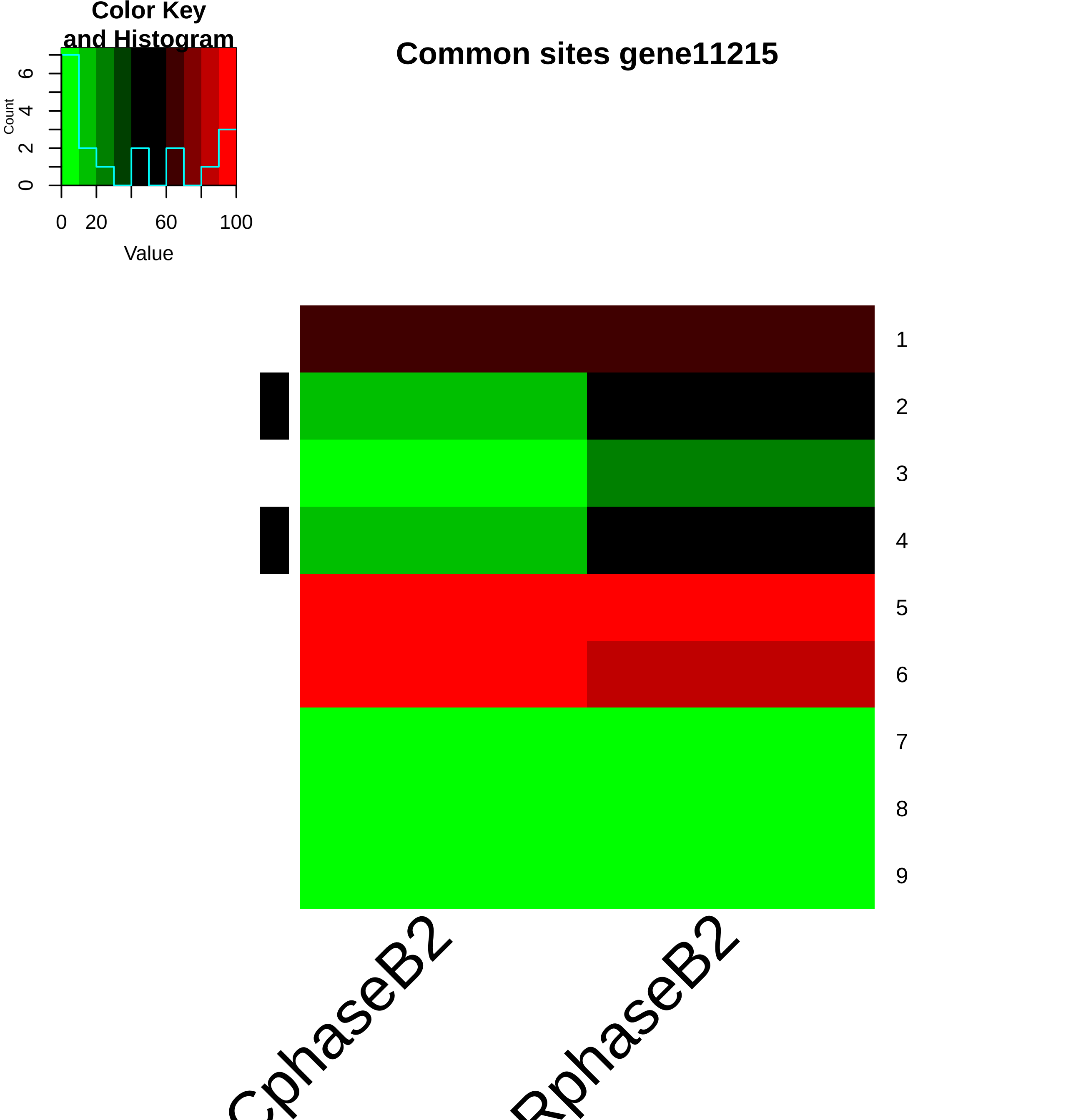
Color Key
and Histogram
Count
0
2
4
6
0
20
60
100
Value
Common sites gene11215
1
2
3
4
5
6
7
8
9
CphaseB2
RphaseB2

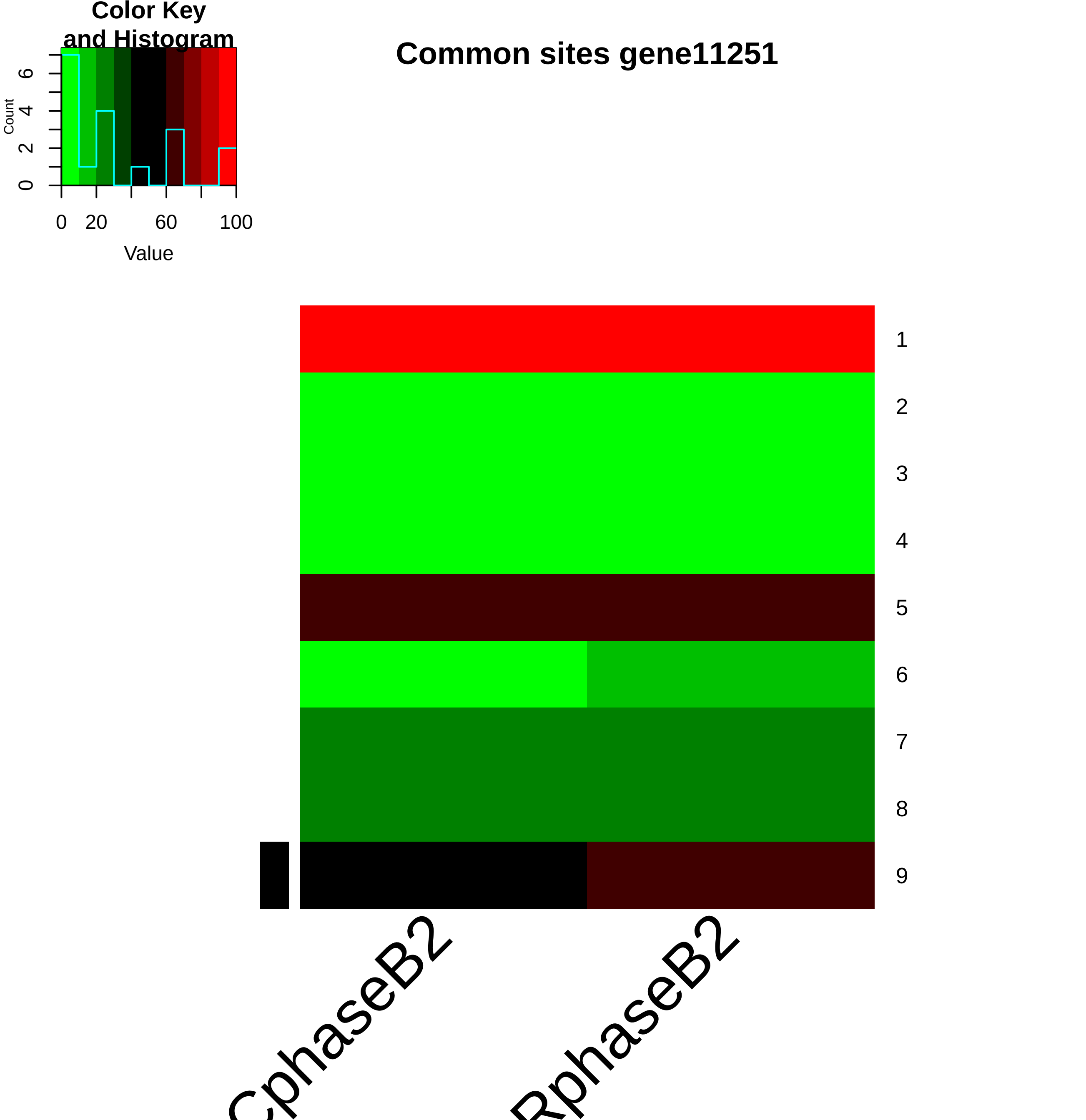
Color Key
and Histogram
Count
0
2
4
6
0
20
60
100
Value
Common sites gene11251
1
2
3
4
5
6
7
8
9
CphaseB2
RphaseB2

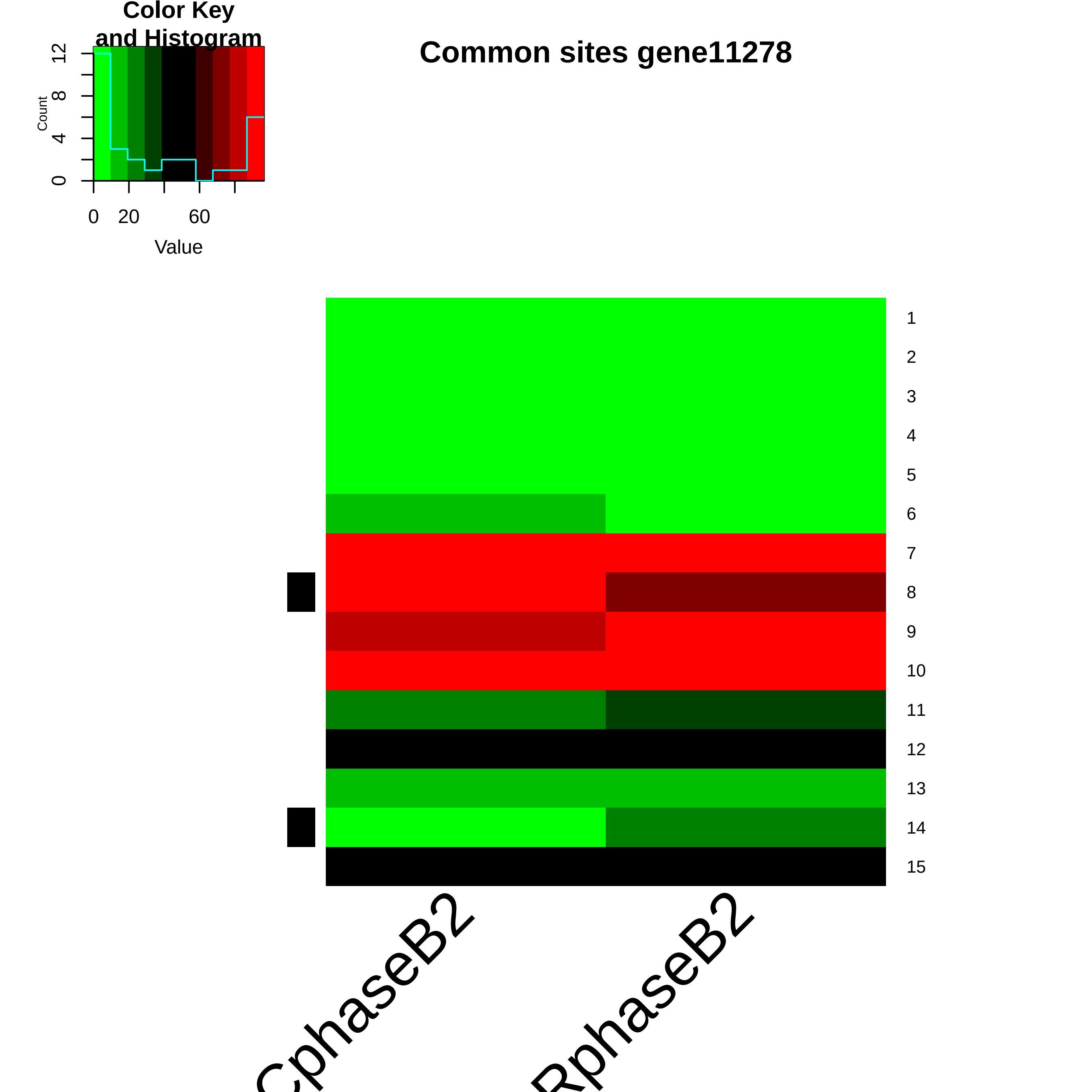
Color Key
and Histogram
Count
0
4
8
12
0
20
60
Value
Common sites gene11278
CphaseB2
RphaseB2

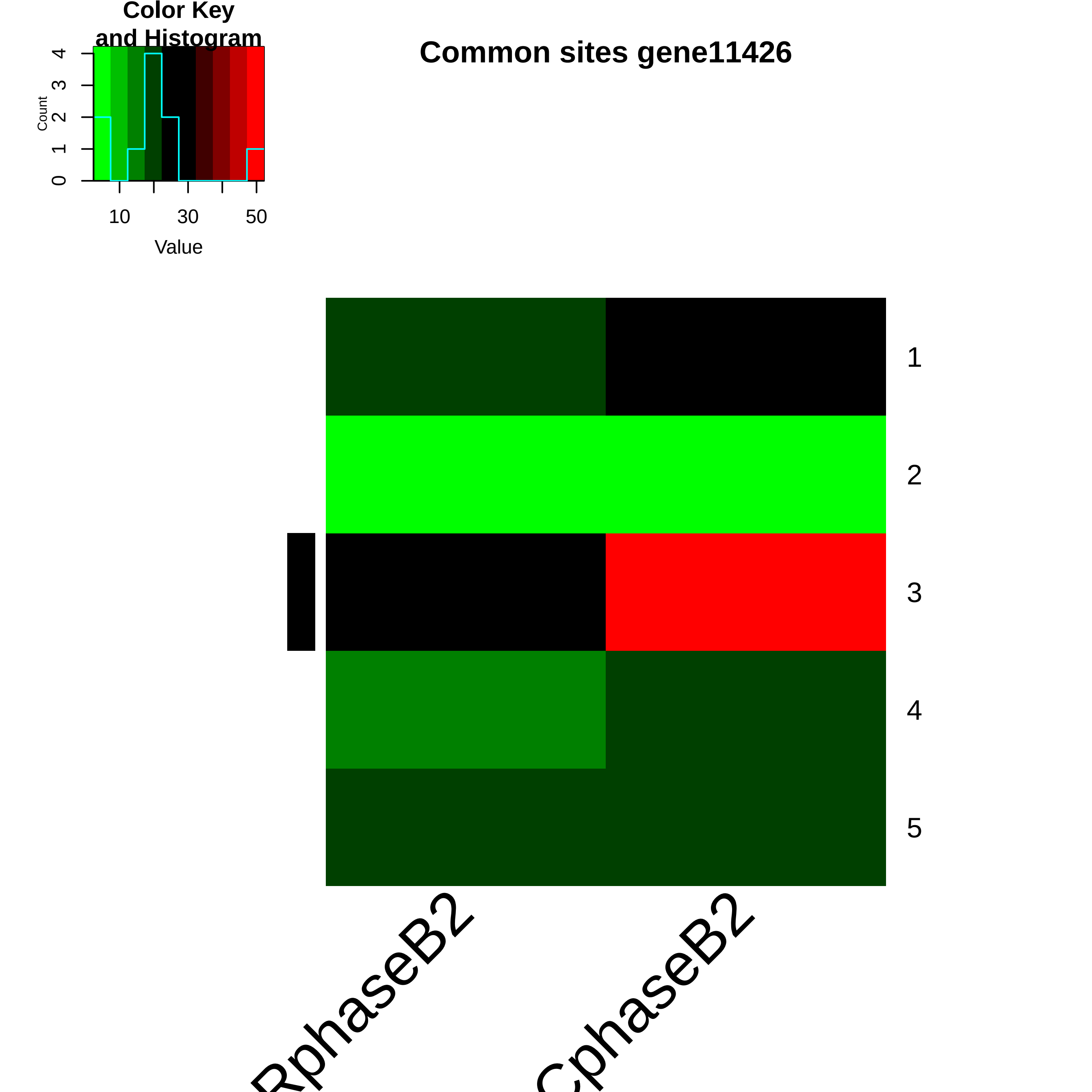

Color Key
and Histogram
Count
Value
10
30
50
0
1
2
3
4
Common sites gene11426
1
2
3
4
5
RphaseB2
CphaseB2

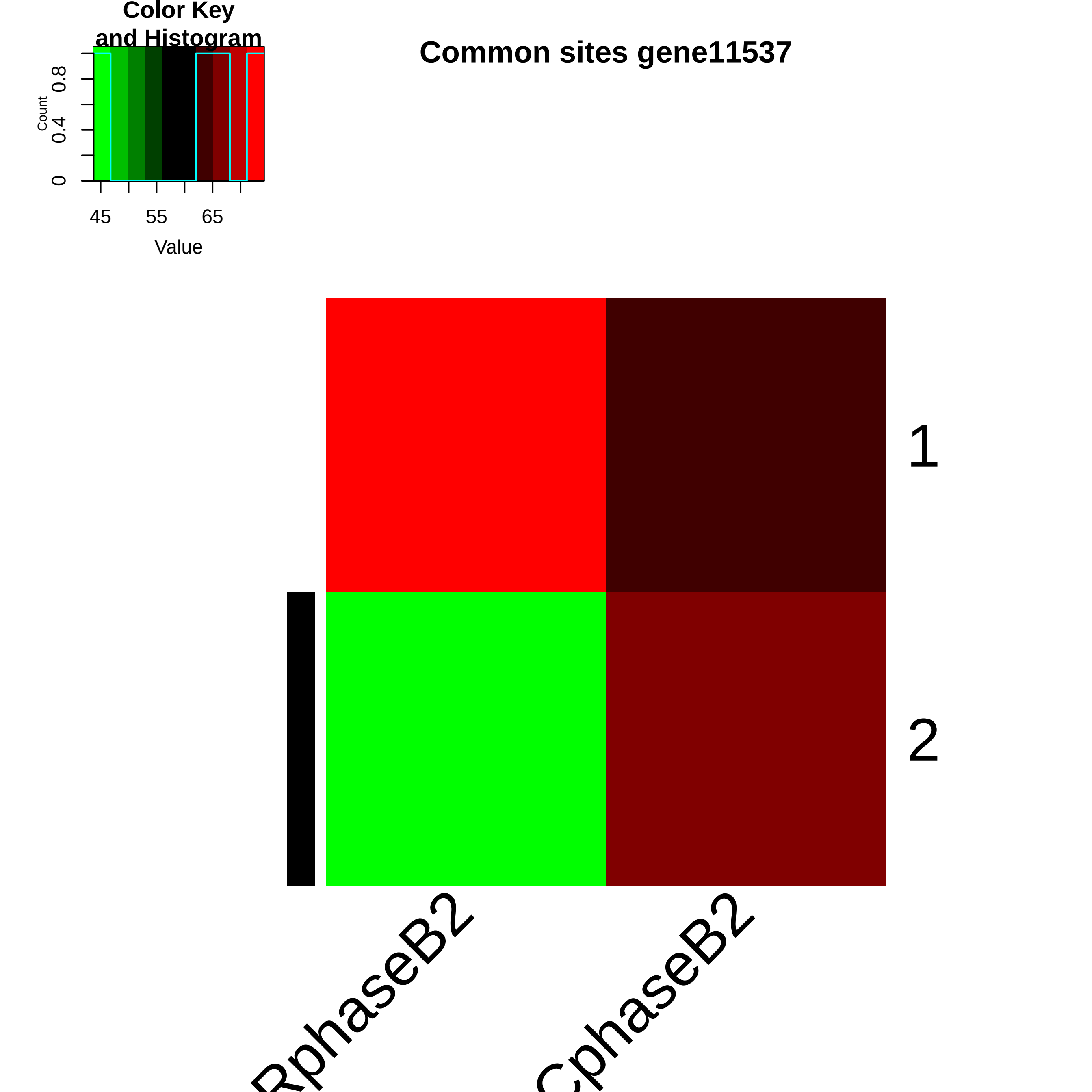
Color Key
and Histogram
Count
0
0.4
0.8
45
55
65
Value
Common sites gene11537
1
2
RphaseB2
CphaseB2

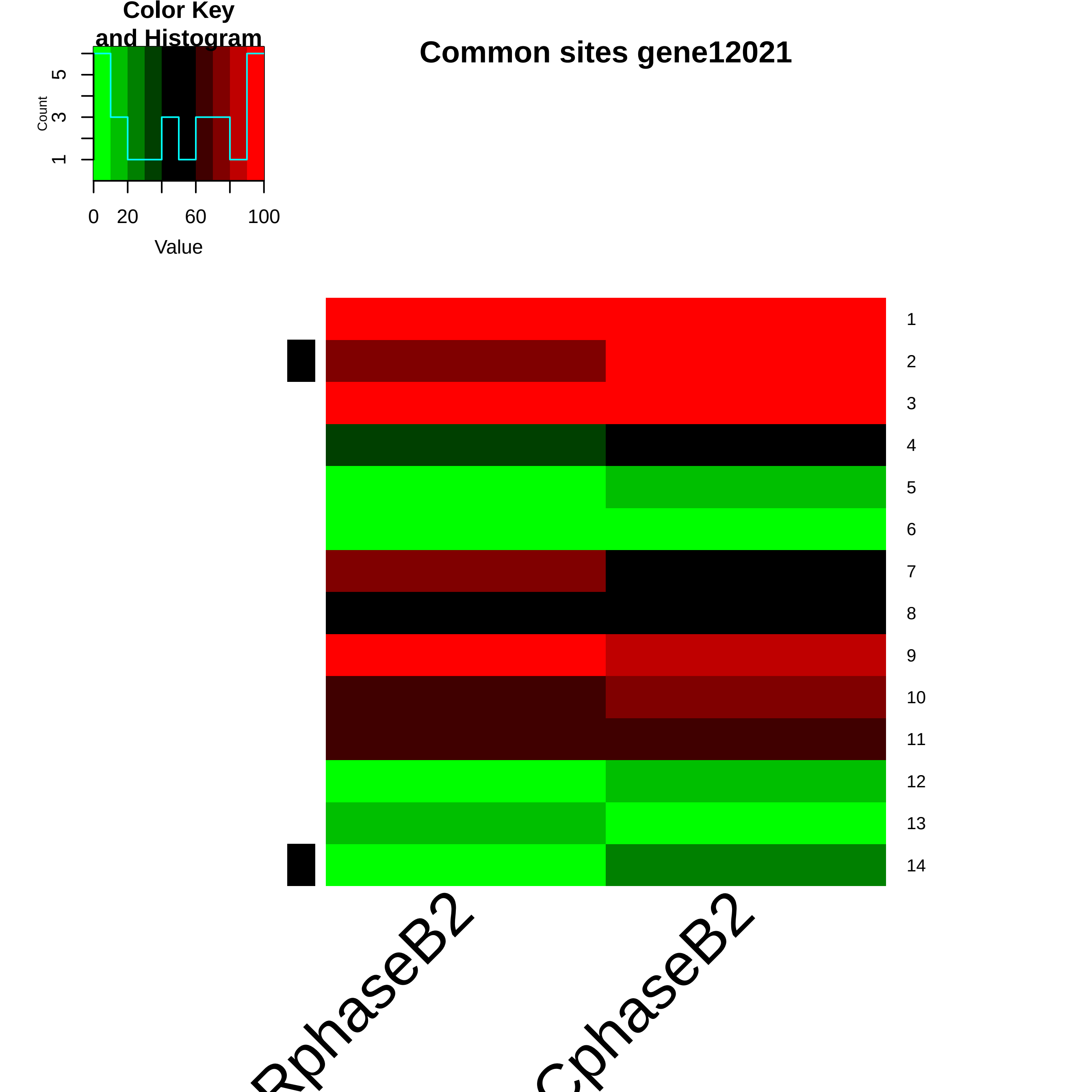

Color Key
and Histogram
Count
1
3
5
0
20
60
100
Value
Common sites gene12021
1
2
3
4
5
6
7
8
9
10
11
12
13
14
RphaseB2
CphaseB2

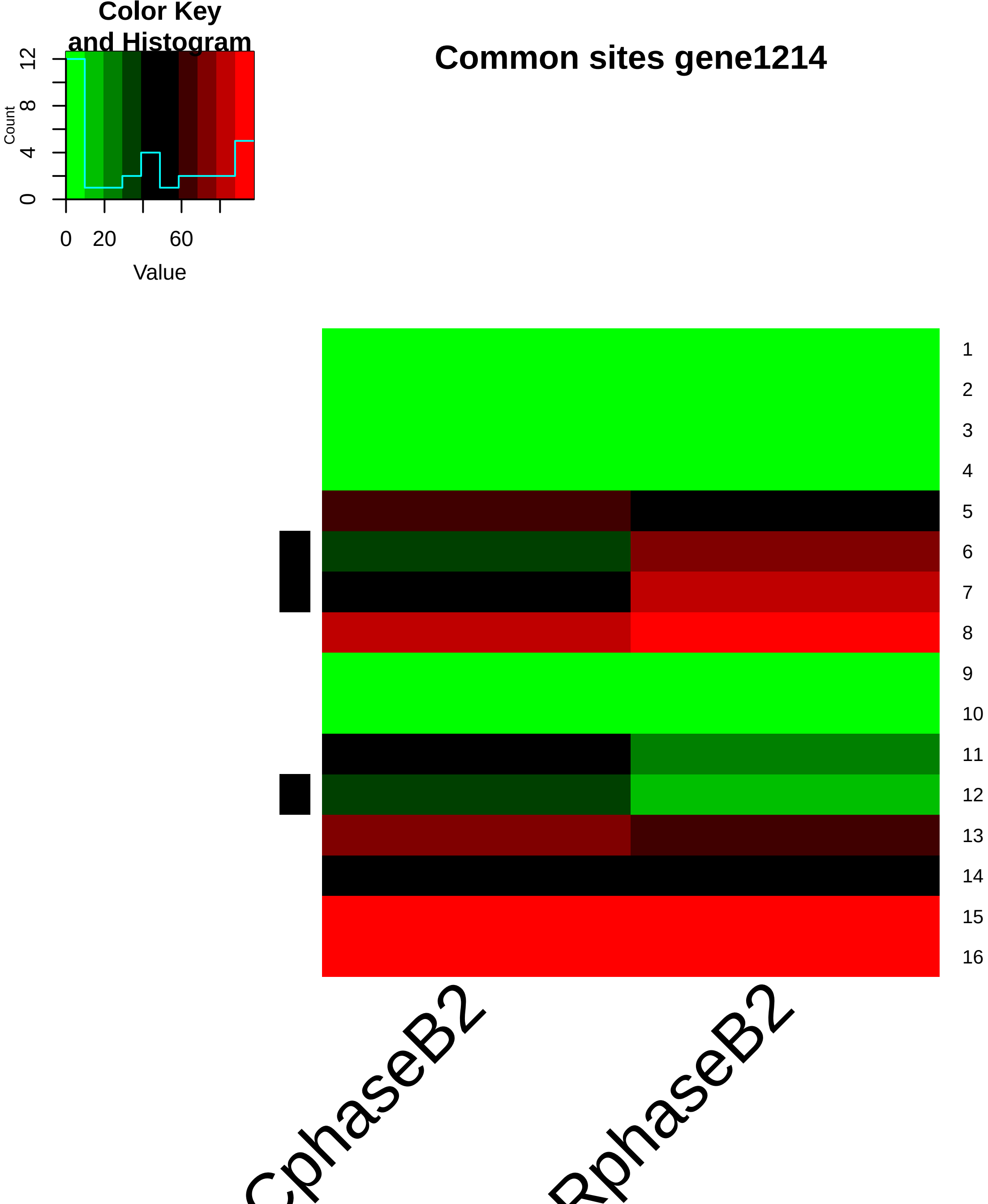

Color Key
and Histogram
Count
0
4
8
12
0
20
60
Value
Common sites gene1214
CphaseB2
RphaseB2

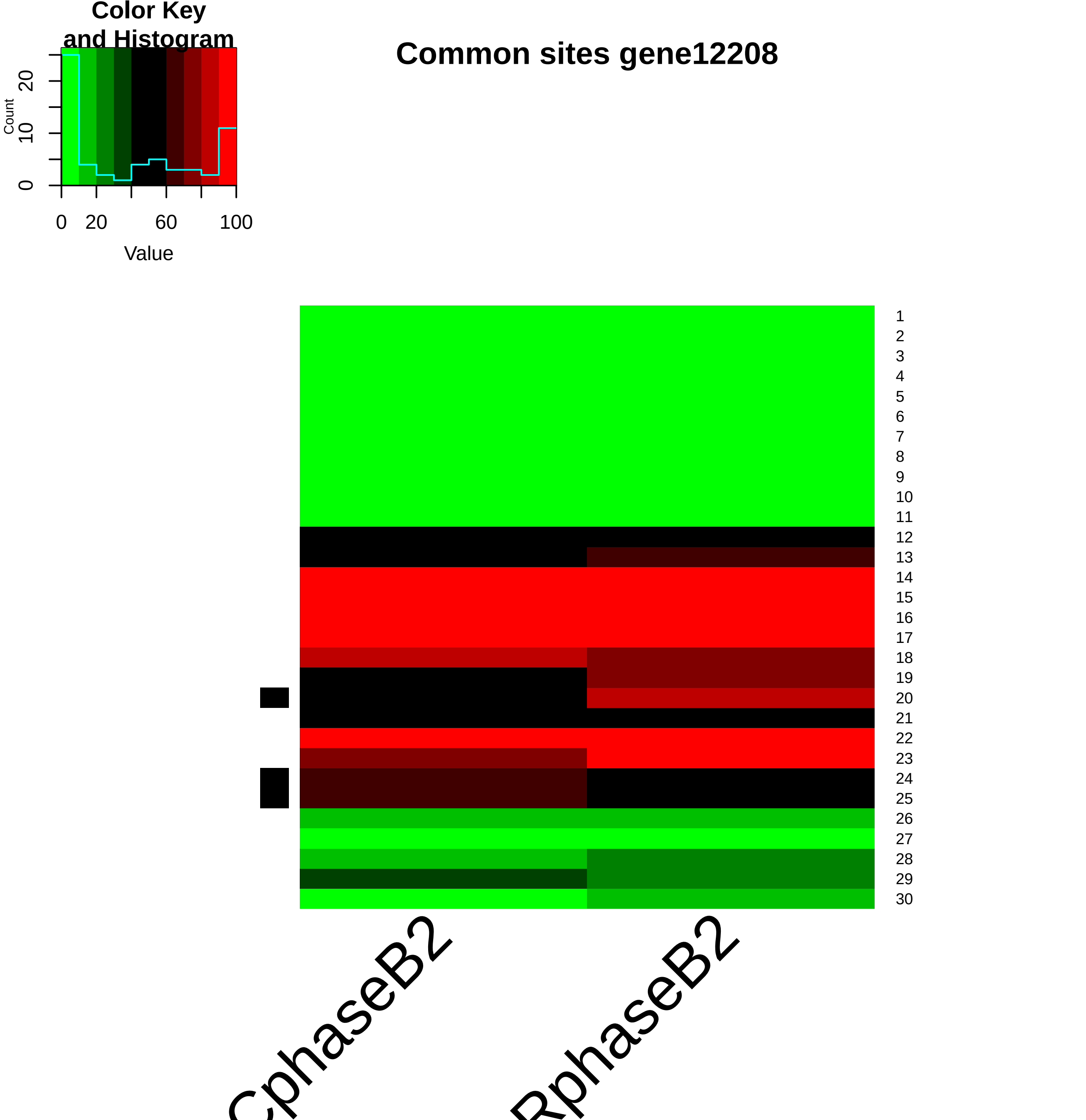
Color Key
and Histogram
Count
0
10
20
0
20
60
100
Value
Common sites gene12208
CphaseB2
RphaseB2

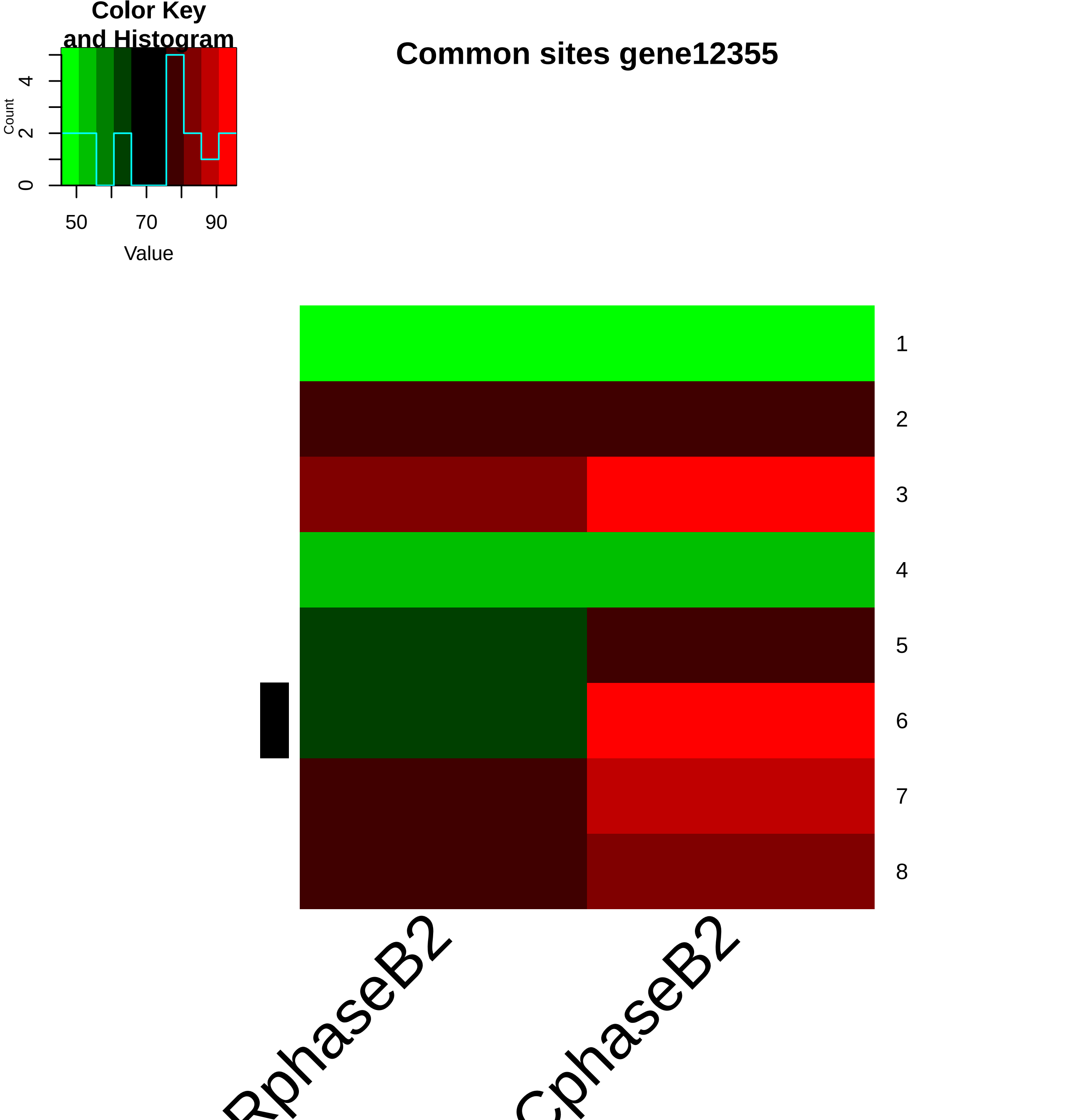
Color Key
and Histogram
Count
0
2
4
50
70
90
Value
Common sites gene12355
1
2
3
4
5
6
7
8
RphaseB2
CphaseB2

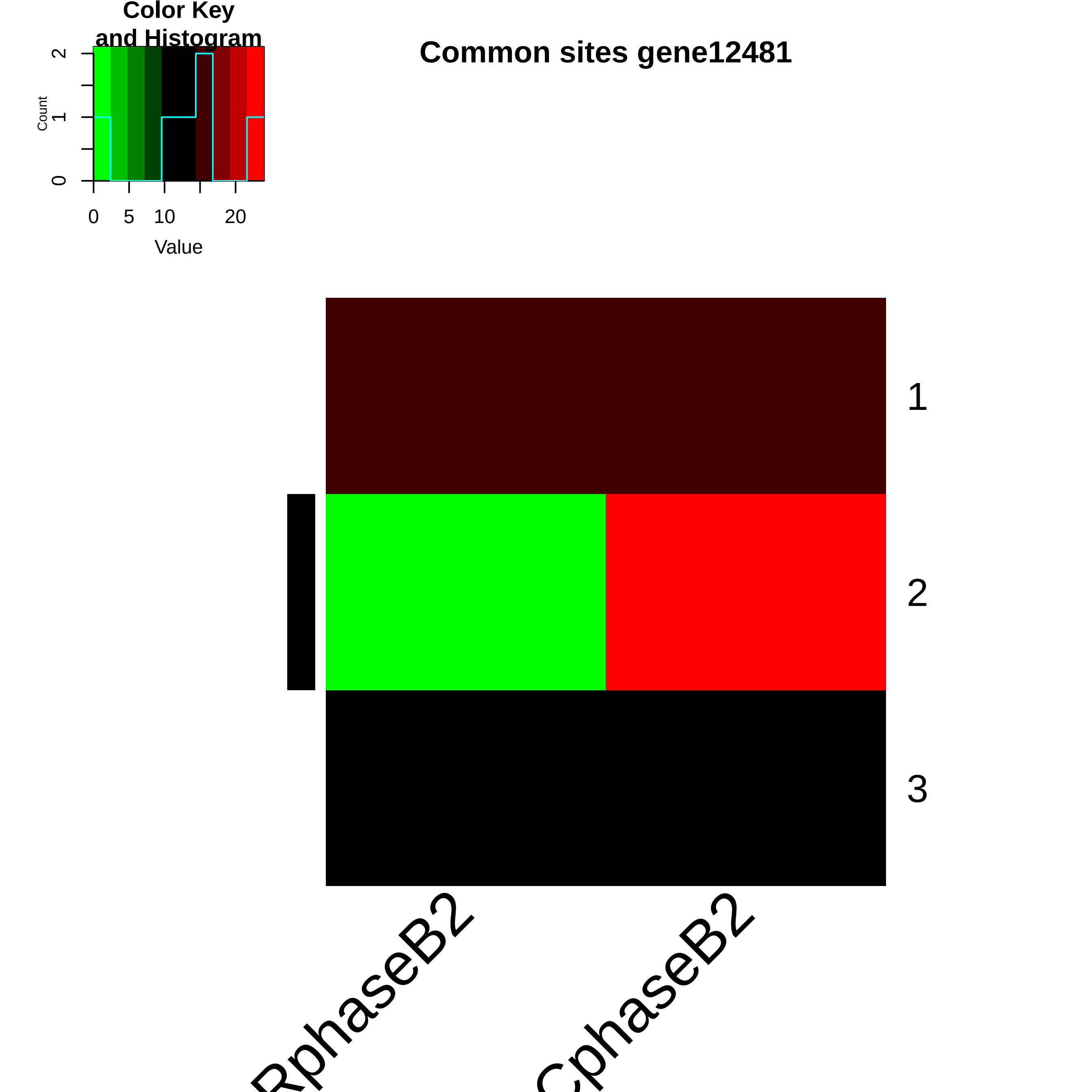

Color Key
and Histogram
Count
0
1
2
0
5
10
20
Value
Common sites gene12481
1
2
3
RphaseB2
CphaseB2

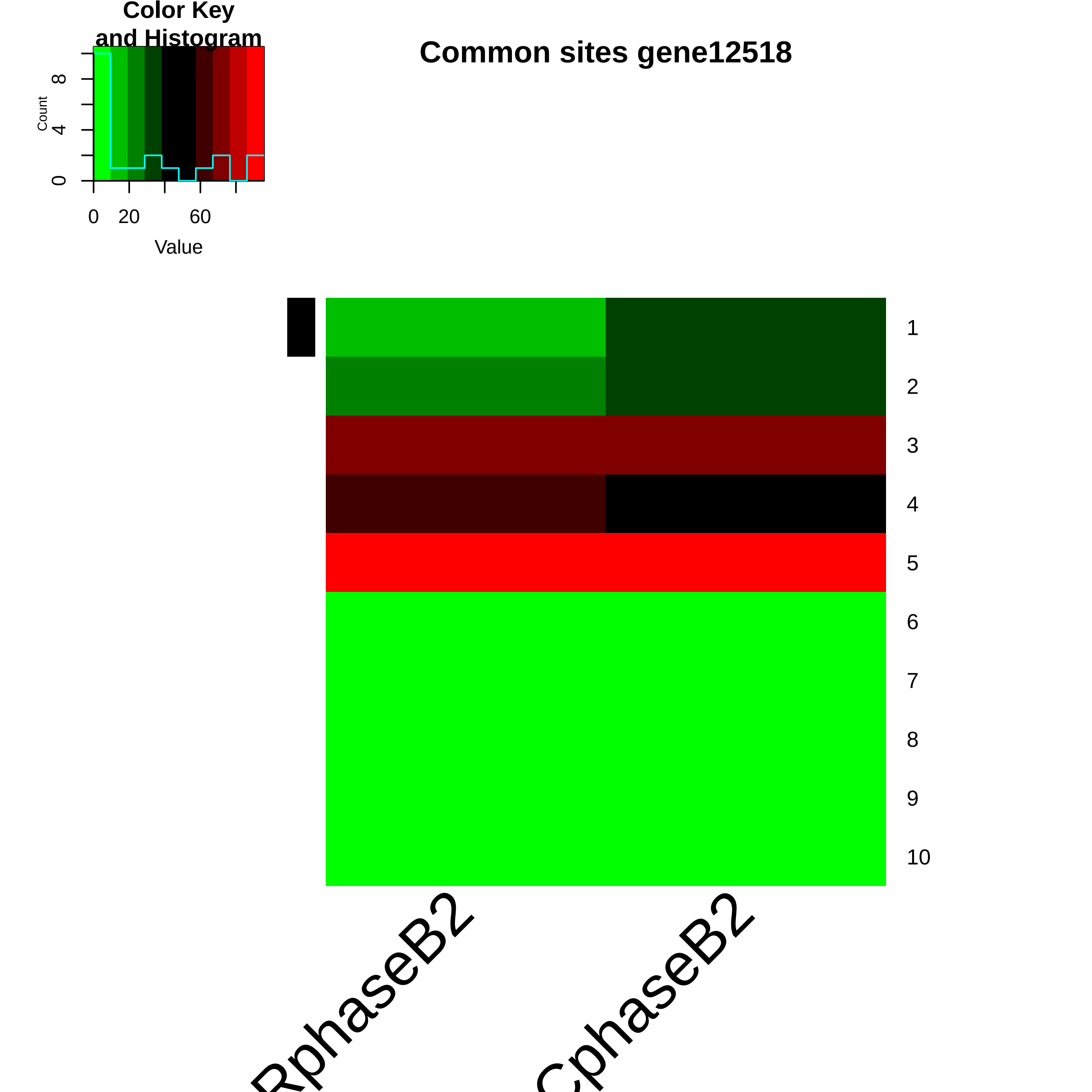
Color Key
and Histogram
Count
0
4
8
0
20
60
Value
Common sites gene12518
1
2
3
4
5
6
7
8
9
10
RphaseB2
CphaseB2

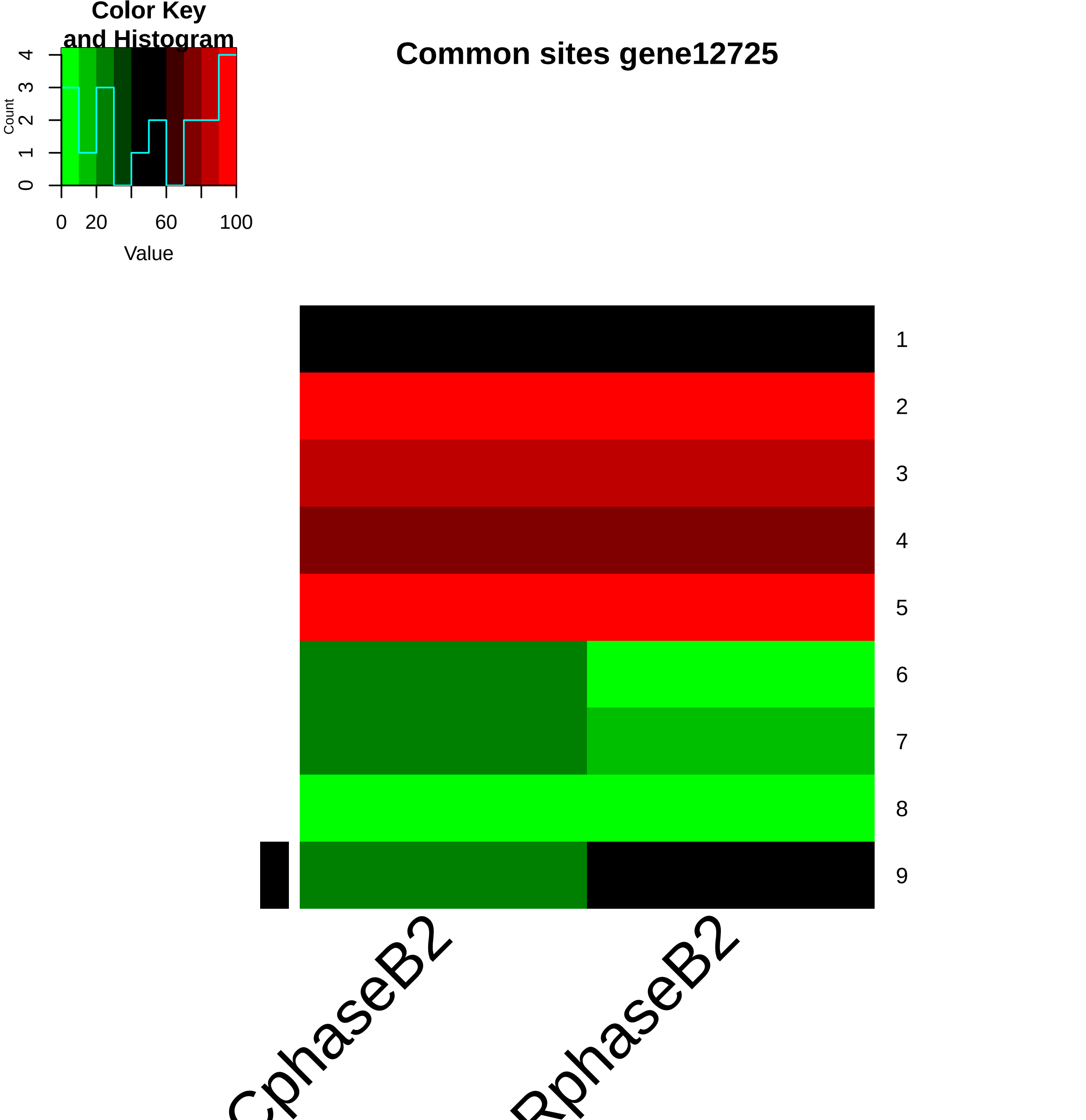

Color Key
and Histogram
Count
0
1
2
3
4
0
20
60
100
Value
Common sites gene12725
1
2
3
4
5
6
7
8
9
CphaseB2
RphaseB2

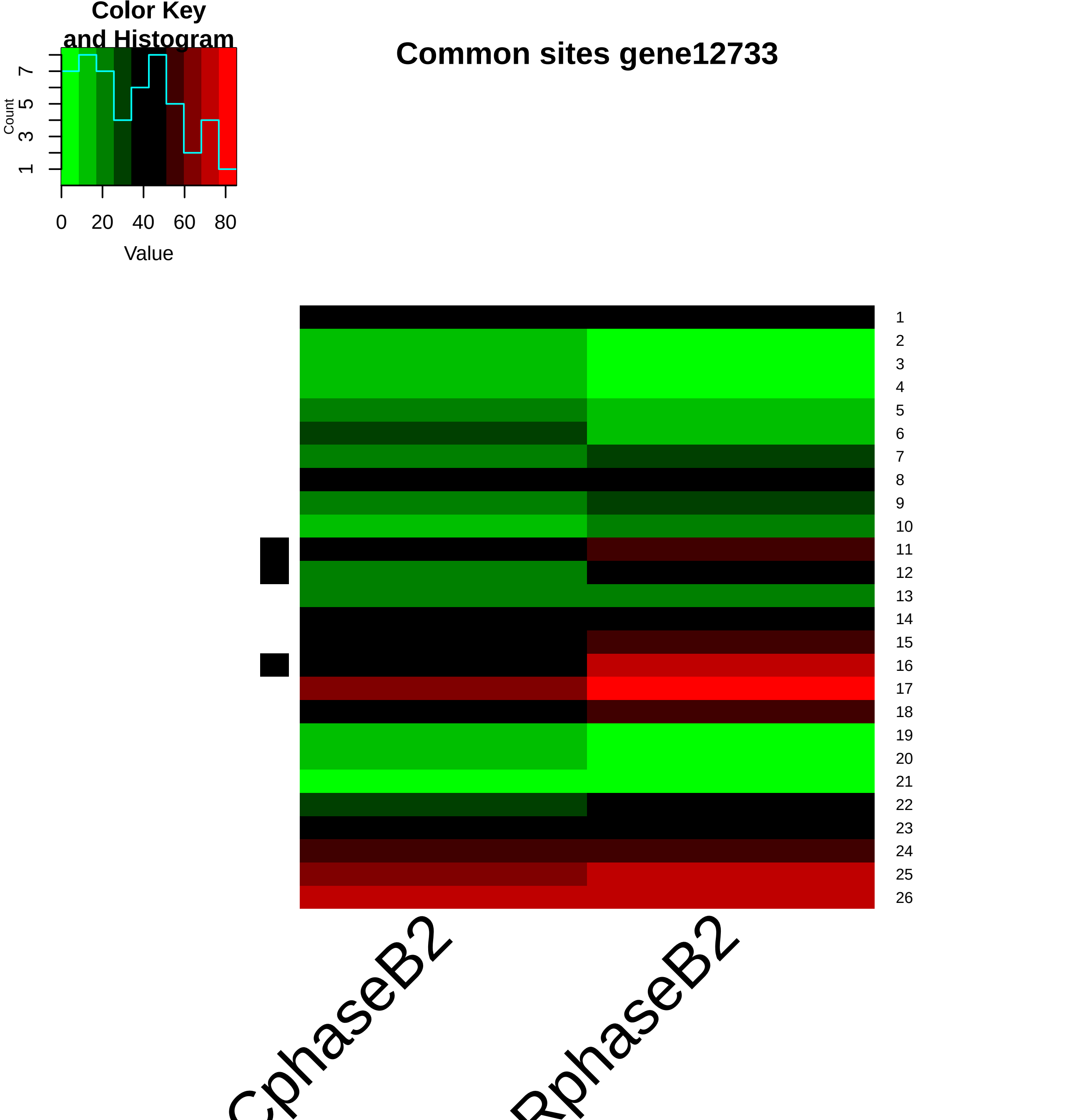

Color Key
and Histogram
Count
1
3
5
7
0
20
40
60
80
Value
Common sites gene12733
CphaseB2
RphaseB2

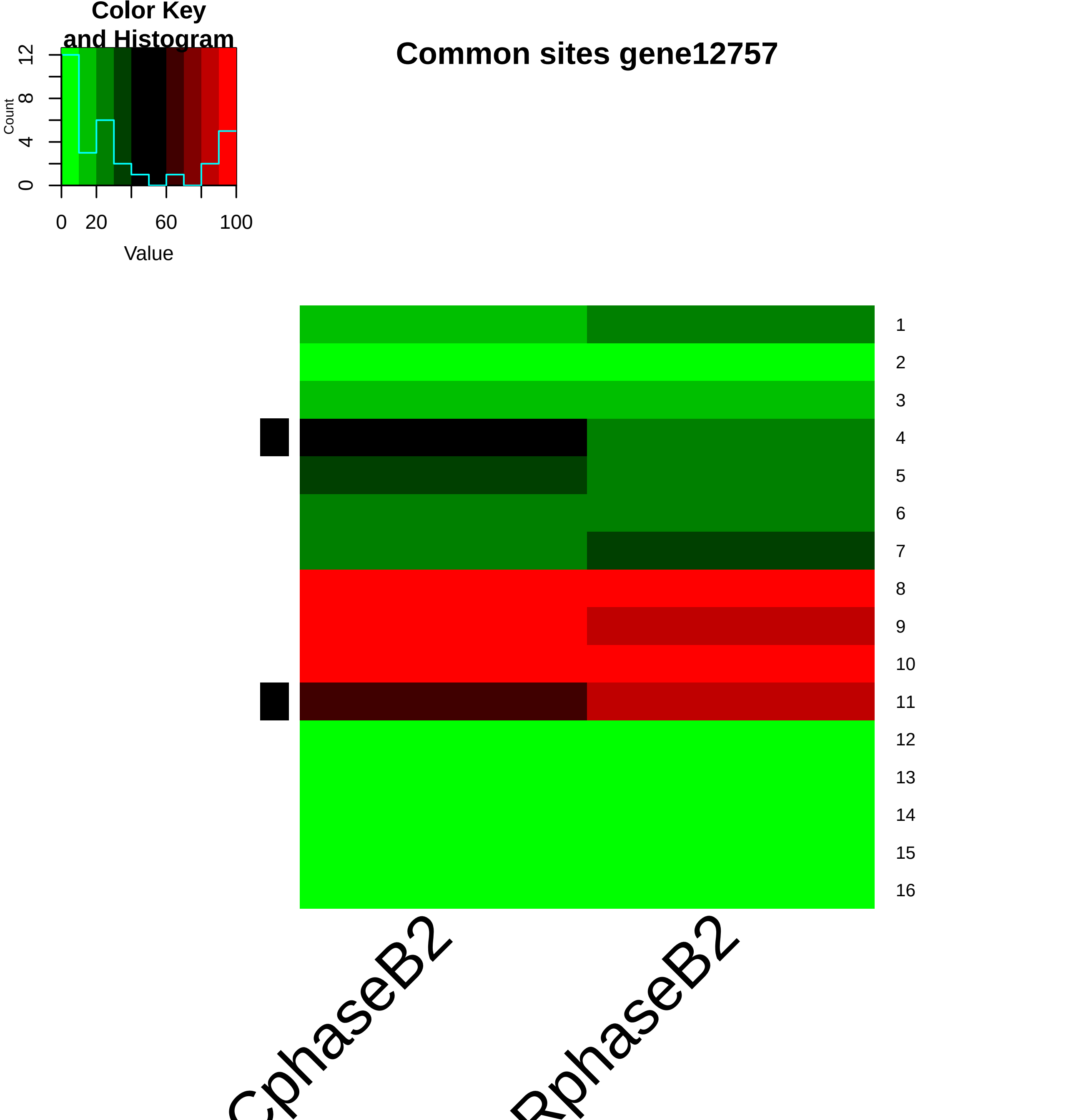
Color Key
and Histogram
Count
0
4
8
12
0
20
60
100
Value
Common sites gene12757
CphaseB2
RphaseB2

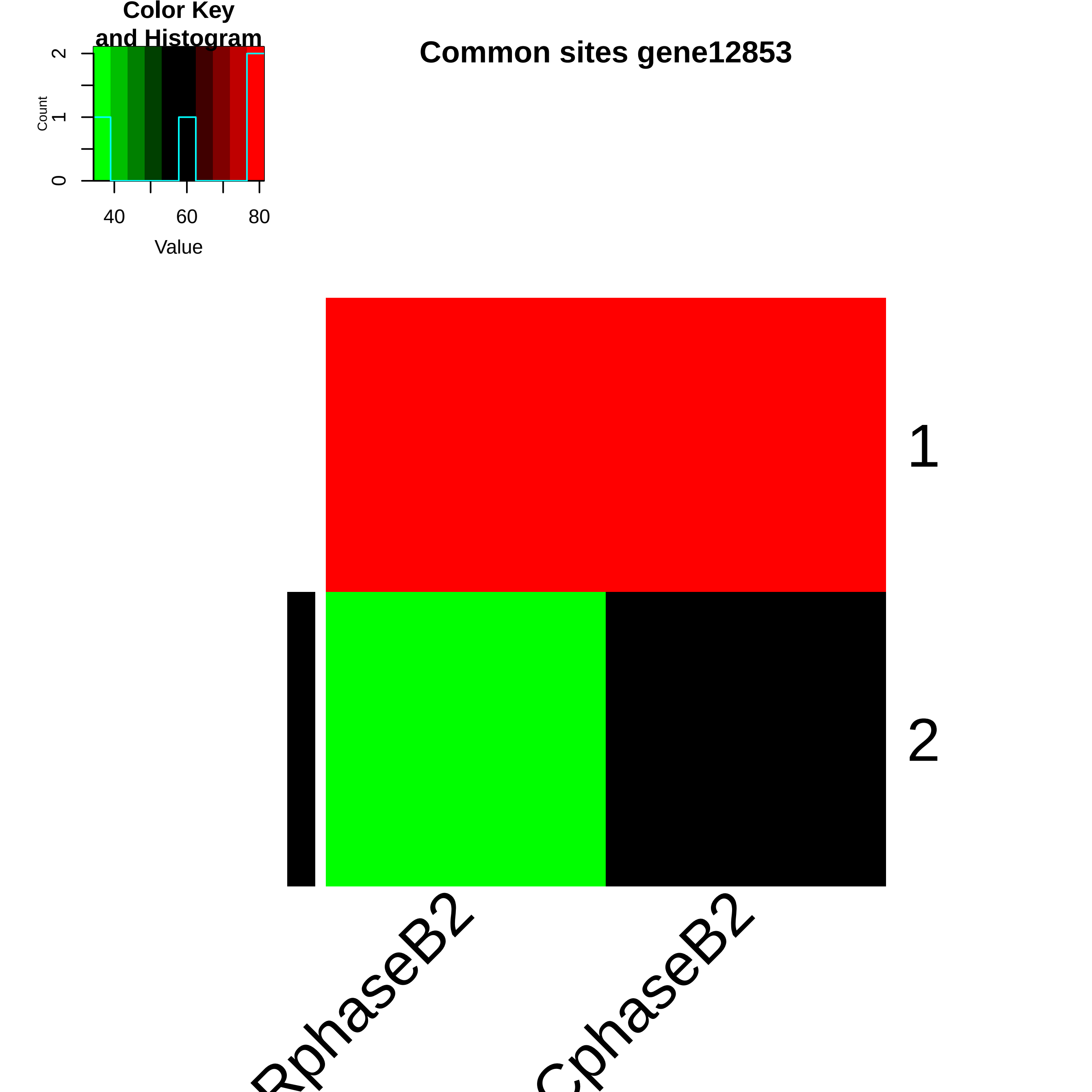
Color Key
and Histogram
Count
0
1
2
40
60
80
Value
Common sites gene12853
1
2
RphaseB2
CphaseB2

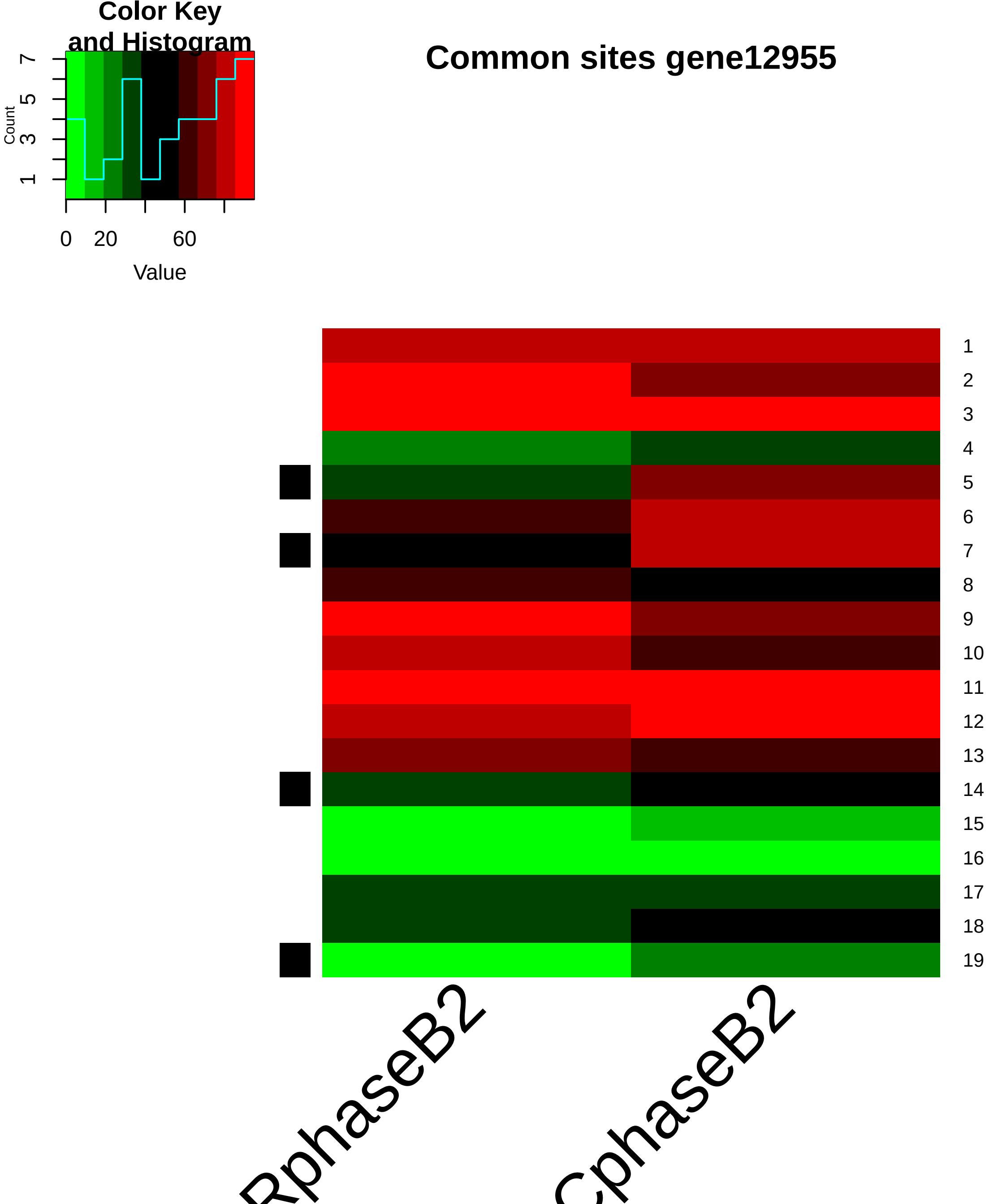

Color Key
and Histogram
Count
1
3
5
7
0
20
60
Value
Common sites gene12955
RphaseB2
CphaseB2

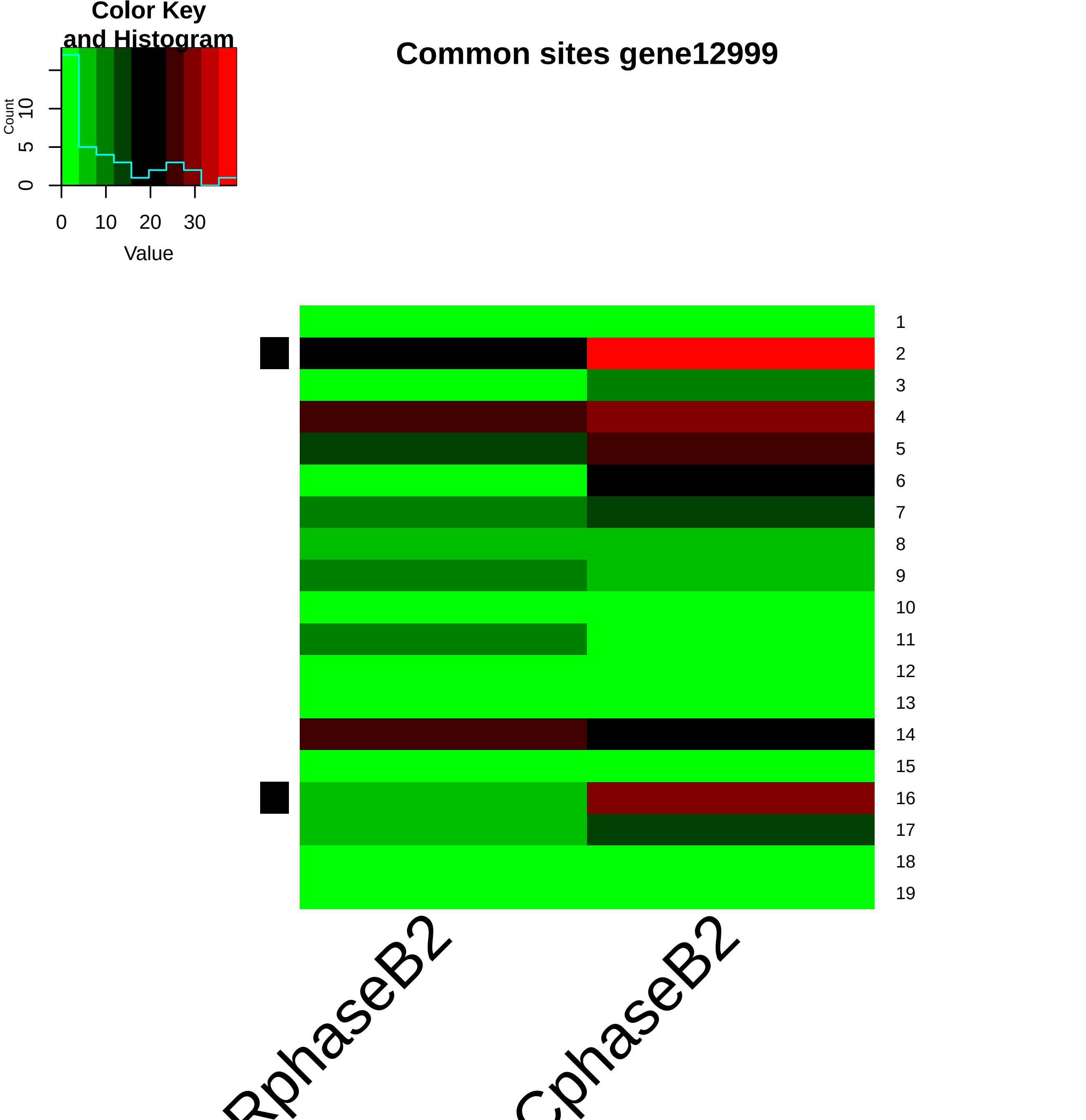
Color Key
and Histogram
Count
0
5
10
0
10
20
30
Value
Common sites gene12999
RphaseB2
CphaseB2

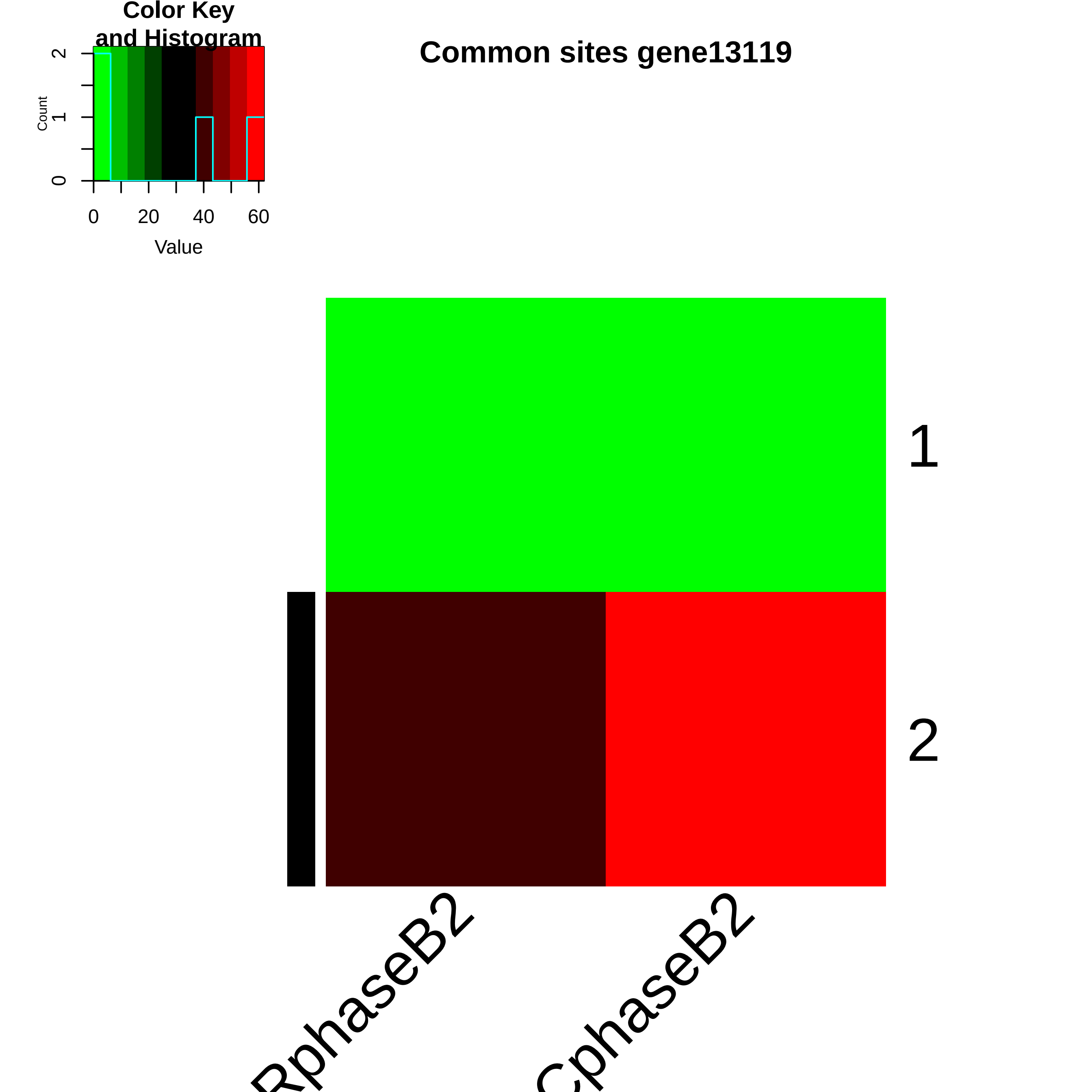

Color Key
and Histogram
Count
0
1
2
0
20
40
60
Value
Common sites gene13119
1
2
RphaseB2
CphaseB2

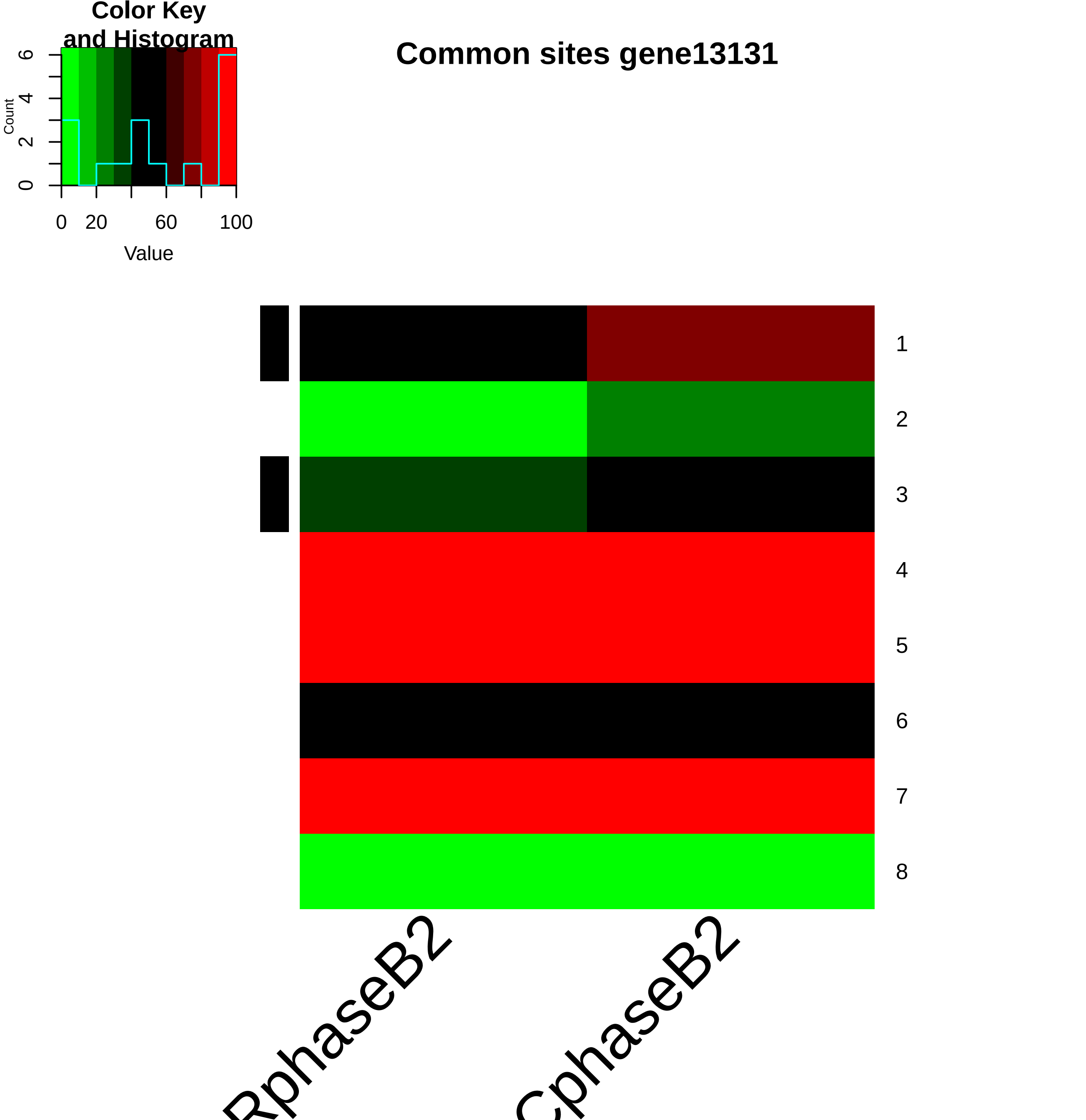
Color Key
and Histogram
Count
0
2
4
6
0
20
60
100
Value
Common sites gene13131
1
2
3
4
5
6
7
8
RphaseB2
CphaseB2

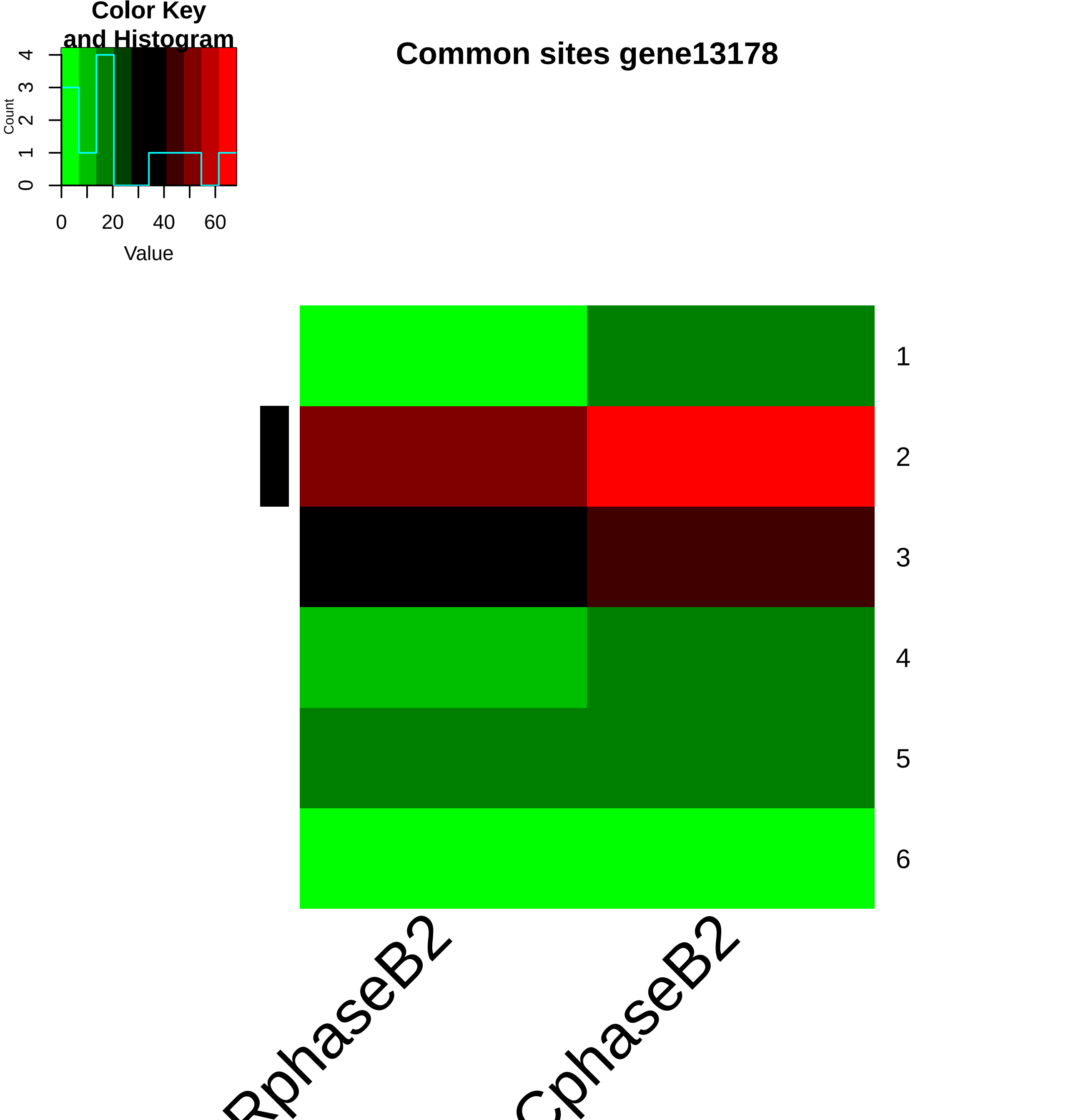

Color Key
and Histogram
Common sites gene13178
Count
0
1
2
3
4
0
20
40
60
Value
1
2
3
4
5
6
RphaseB2
CphaseB2

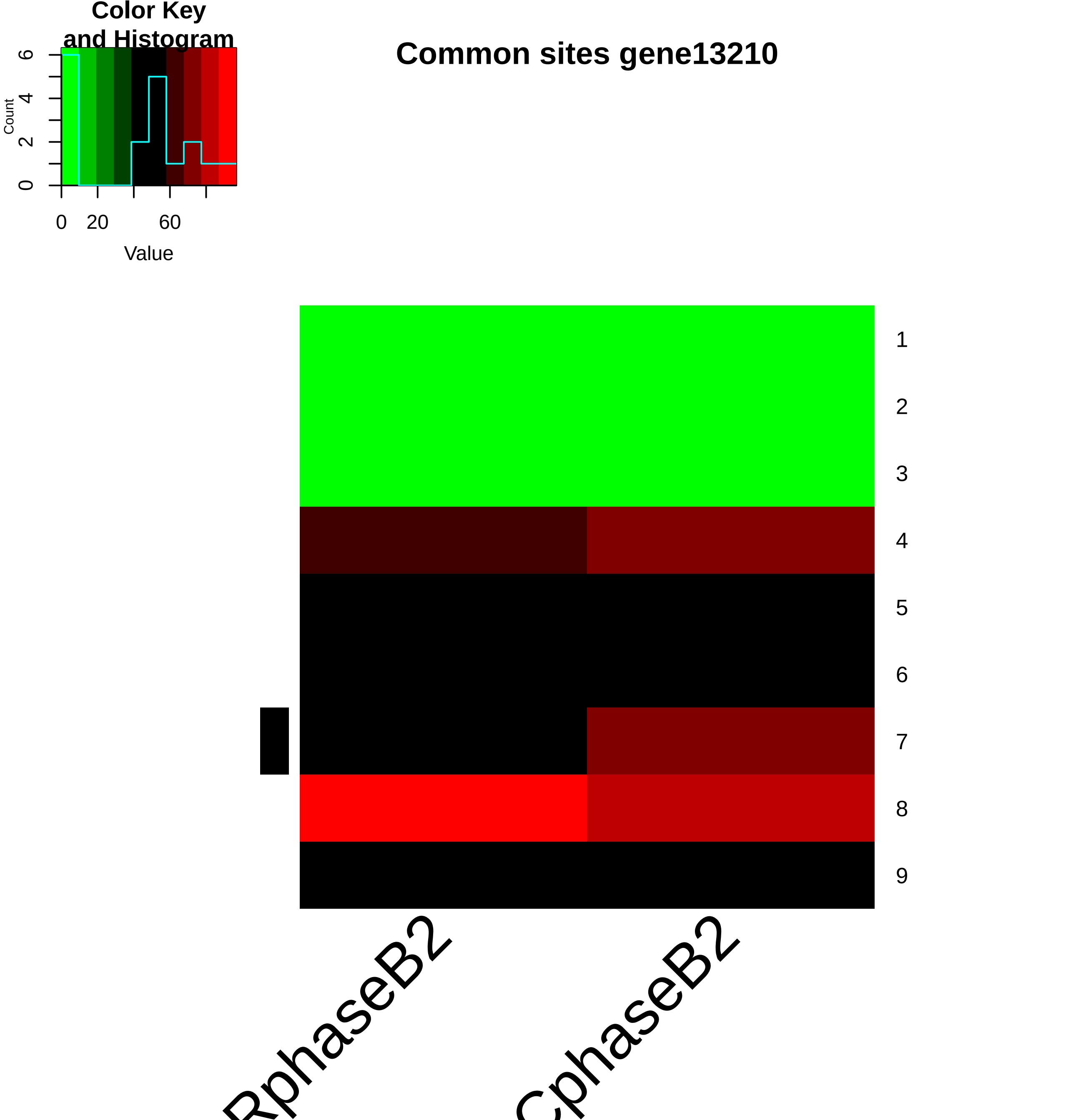
Color Key
and Histogram
Count
0
2
4
6
0
20
60
Value
Common sites gene13210
1
2
3
4
5
6
7
8
9
RphaseB2
CphaseB2

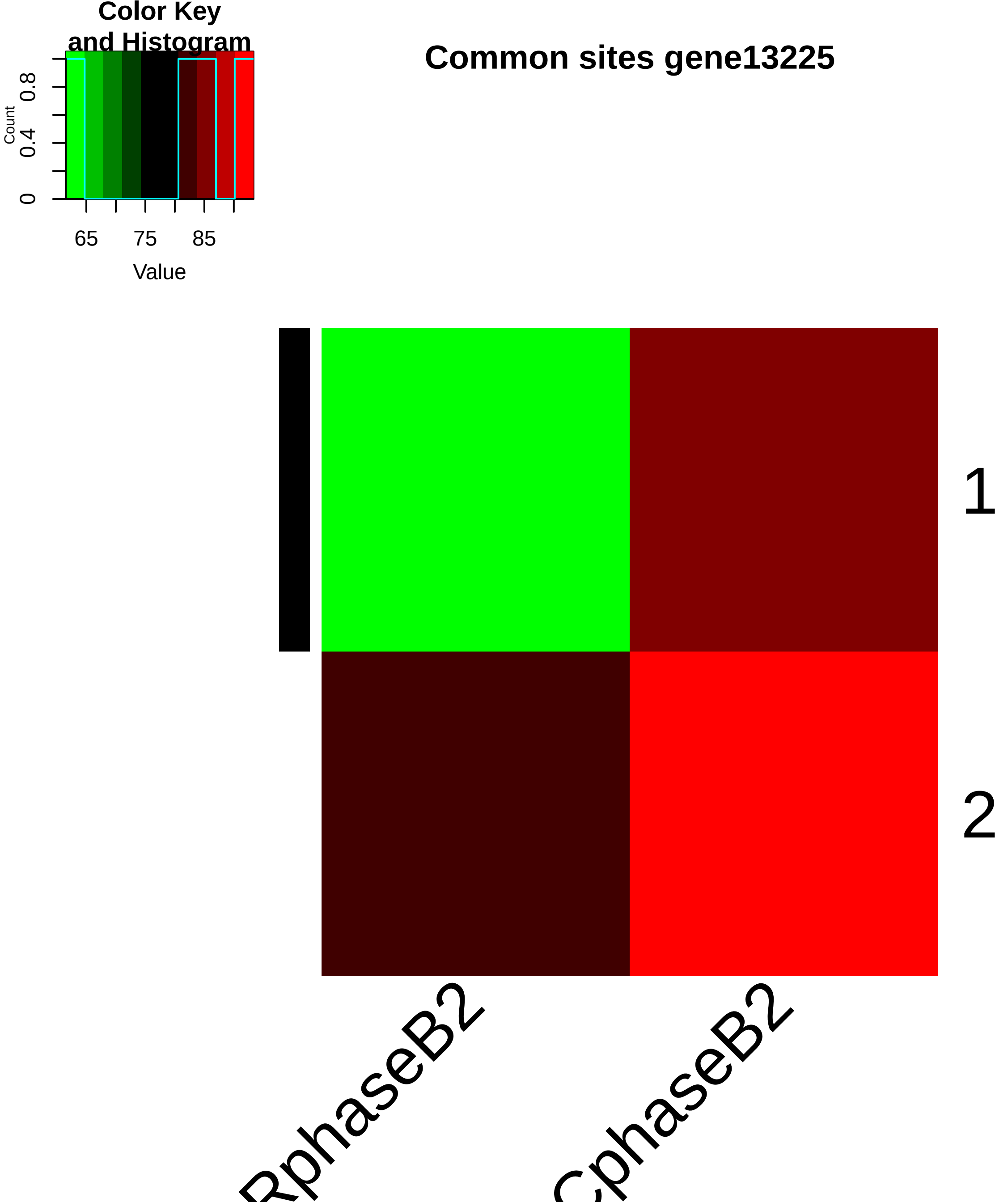

Color Key
and Histogram
Count
0
0.4
0.8
65
75
85
Value
Common sites gene13225
1
2
RphaseB2
CphaseB2

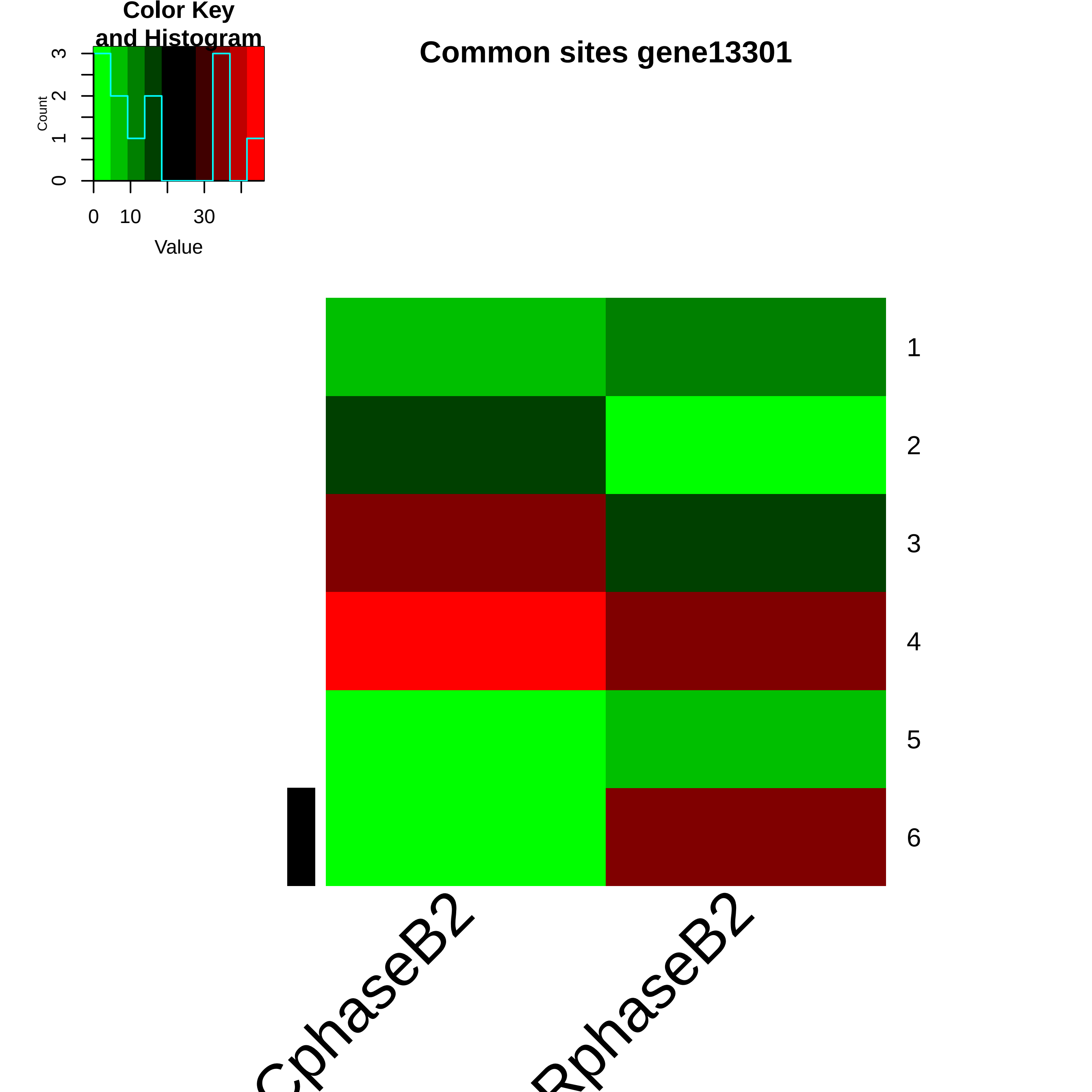

Color Key
and Histogram
Count
Value
0
10
30
0
1
2
3
Common sites gene13301
1
2
3
4
5
6
CphaseB2
RphaseB2

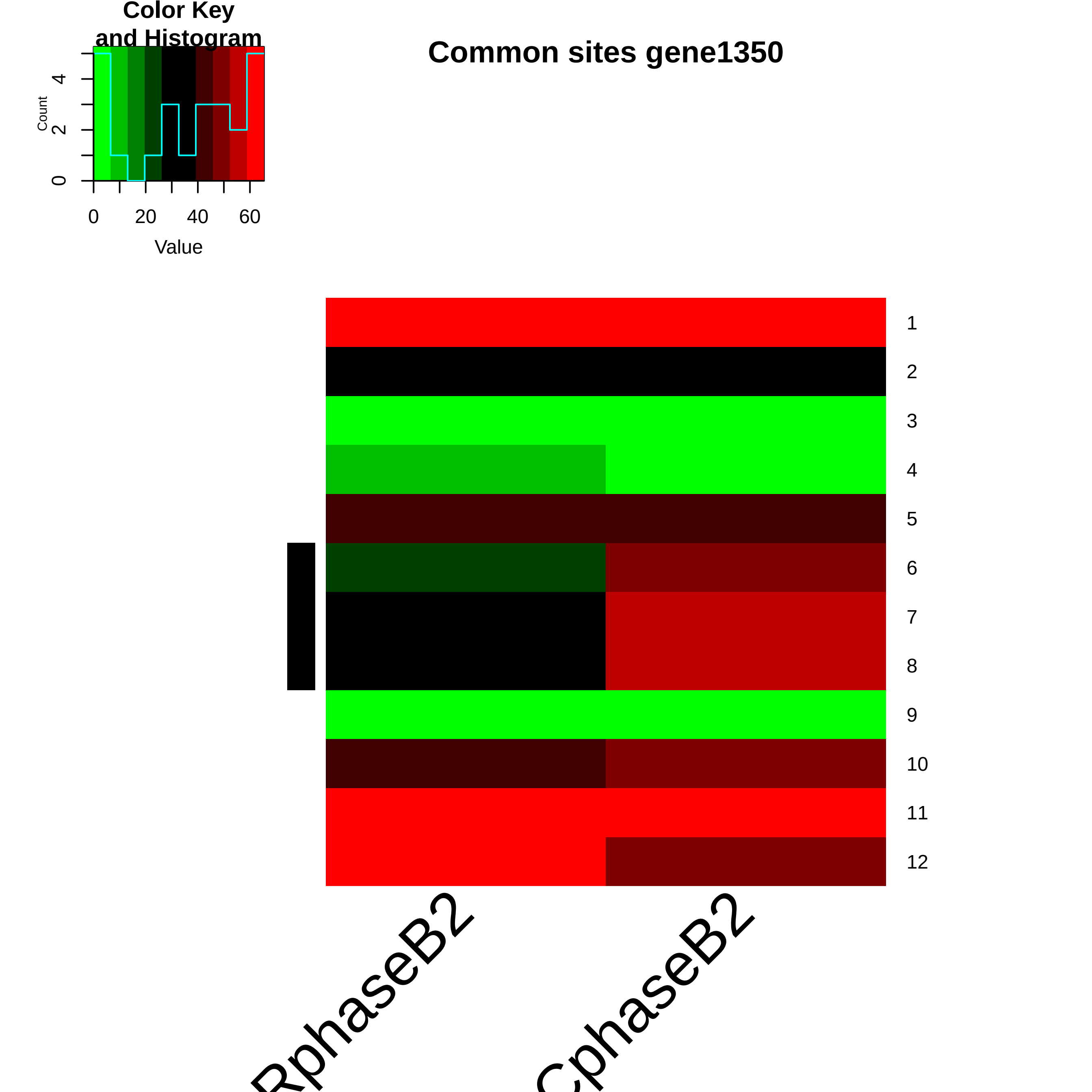

Color Key
and Histogram
Count
0
2
4
0
20
40
60
Value
Common sites gene1350
1
2
3
4
5
6
7
8
9
10
11
12
RphaseB2
CphaseB2

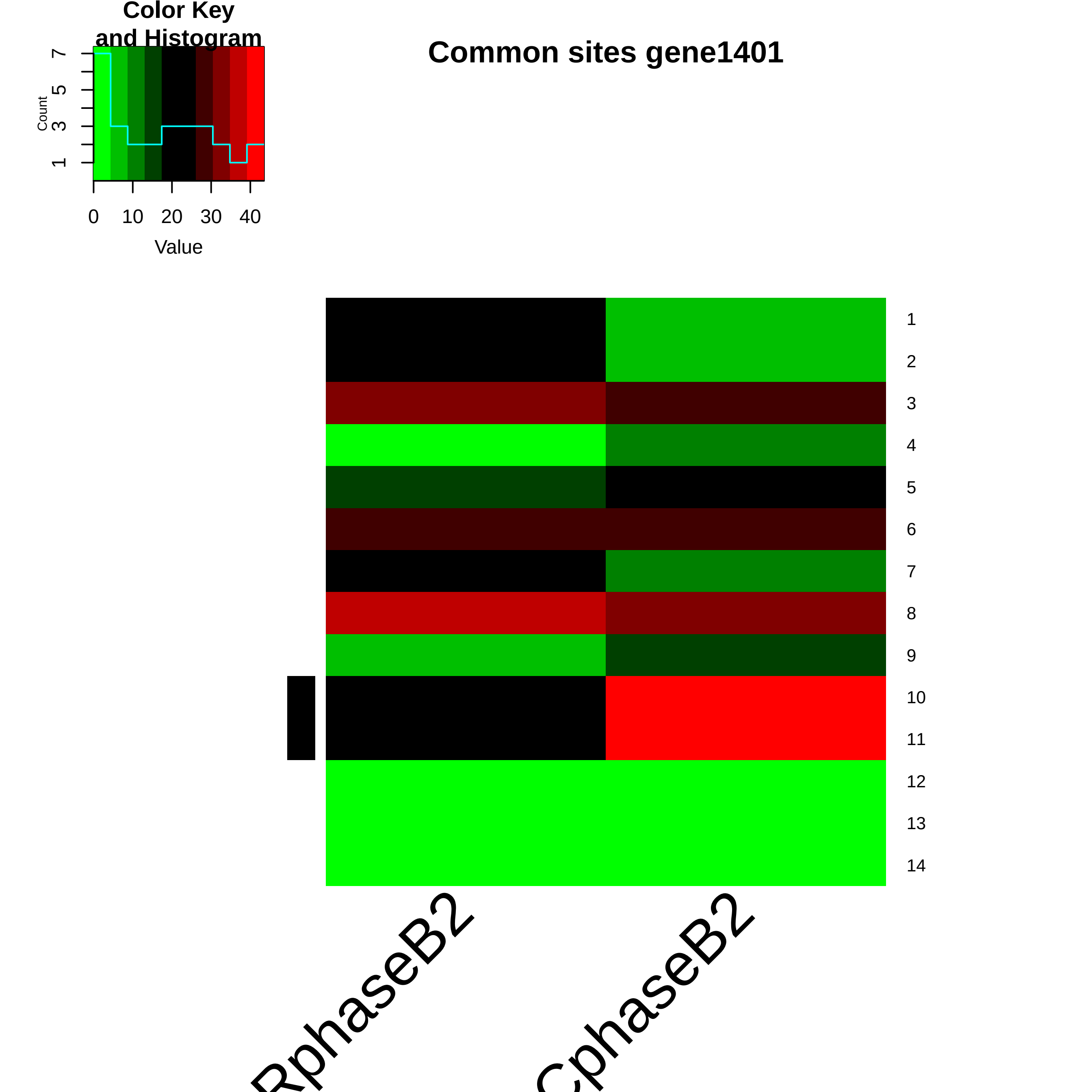
Color Key
and Histogram
Count
Value
Common sites gene1401
1
2
3
4
5
6
7
8
9
10
11
12
13
14
RphaseB2
CphaseB2

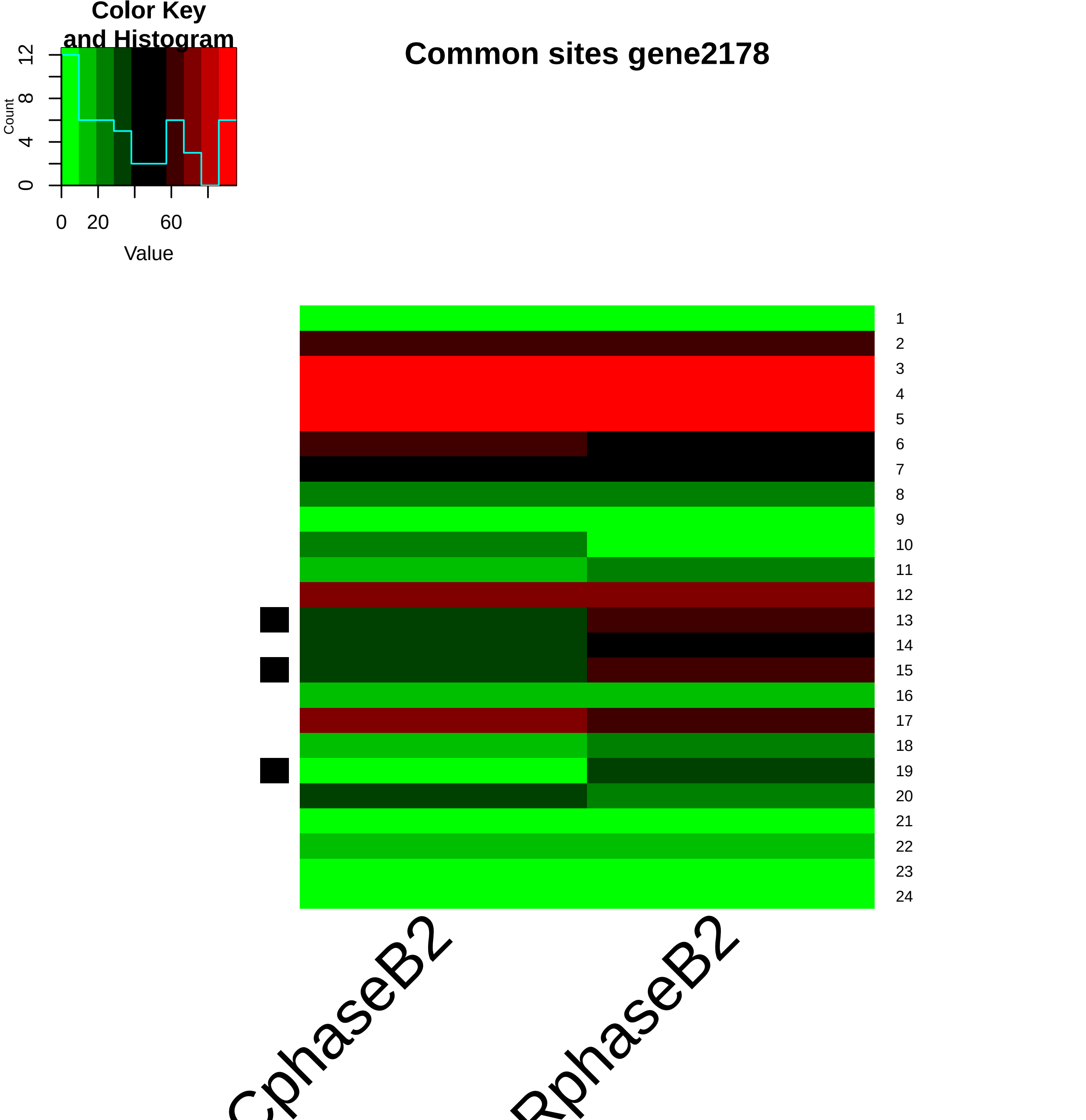
Color Key
and Histogram
Count
0
4
8
12
0
20
60
Value
Common sites gene2178
CphaseB2
RphaseB2

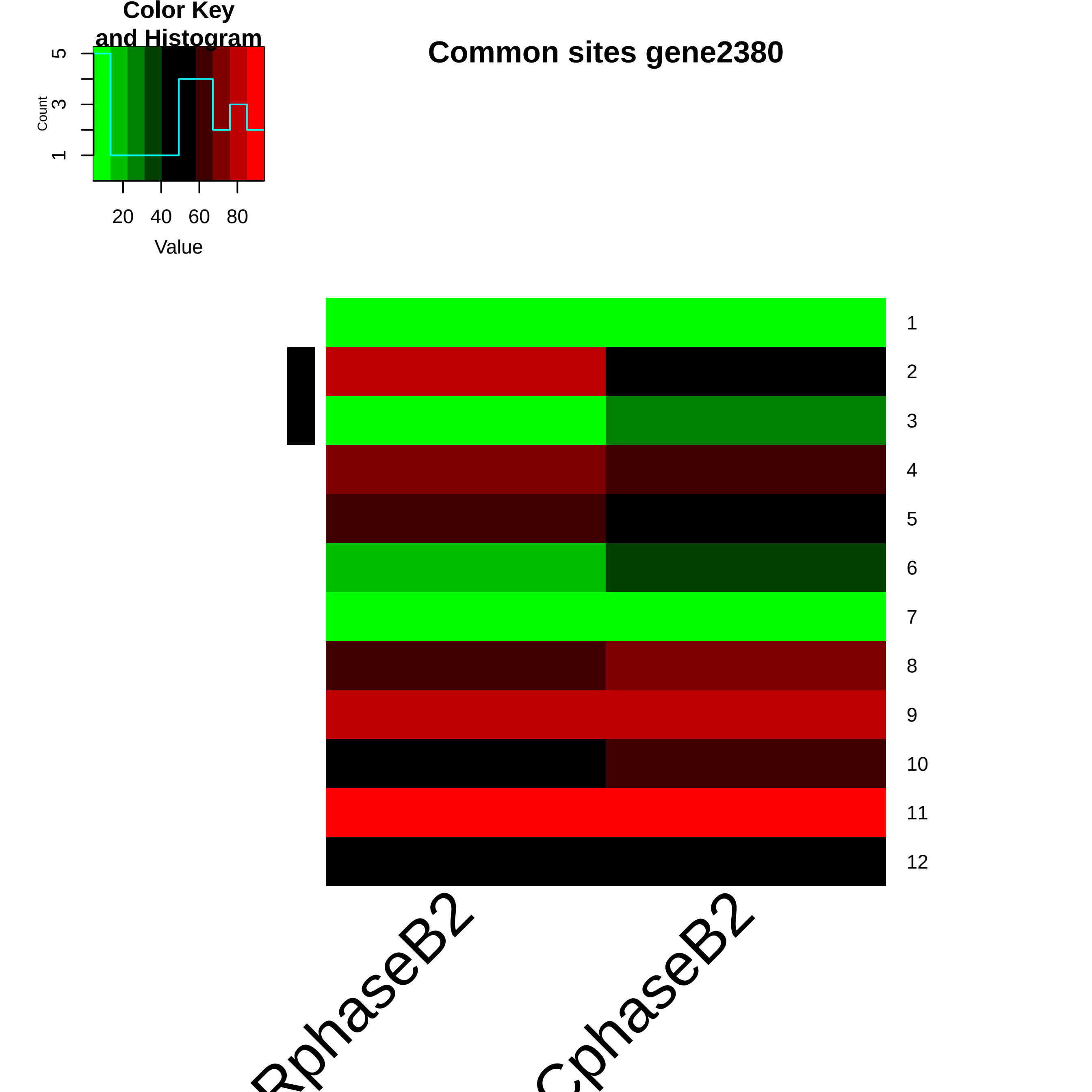
Color Key
and Histogram
Count
1
3
5
20
40
60
80
Value
Common sites gene2380
1
2
3
4
5
6
7
8
9
10
11
12

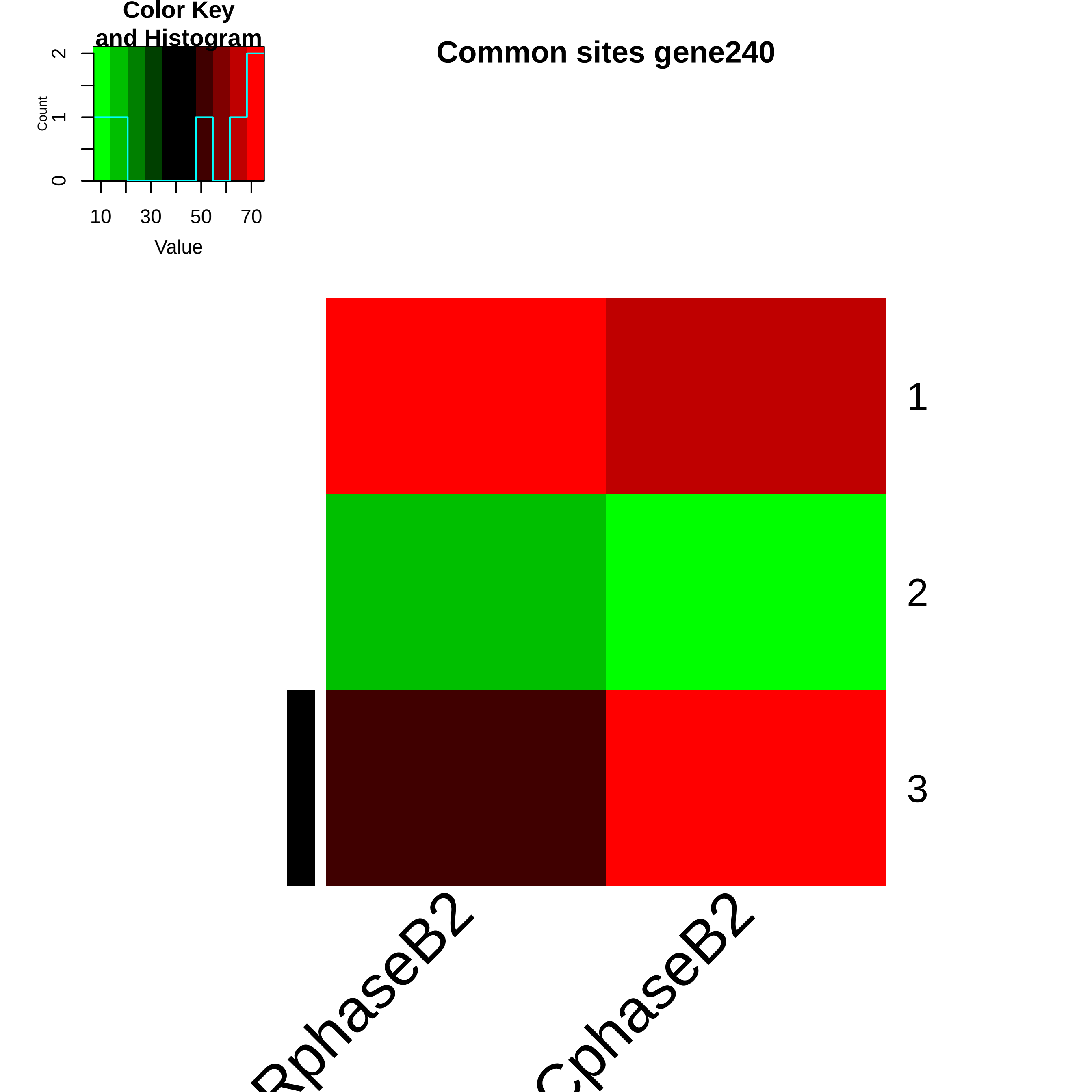

Color Key
and Histogram
Count
0
1
2
10
30
50
70
Value
Common sites gene240
1
2
3
RphaseB2
CphaseB2

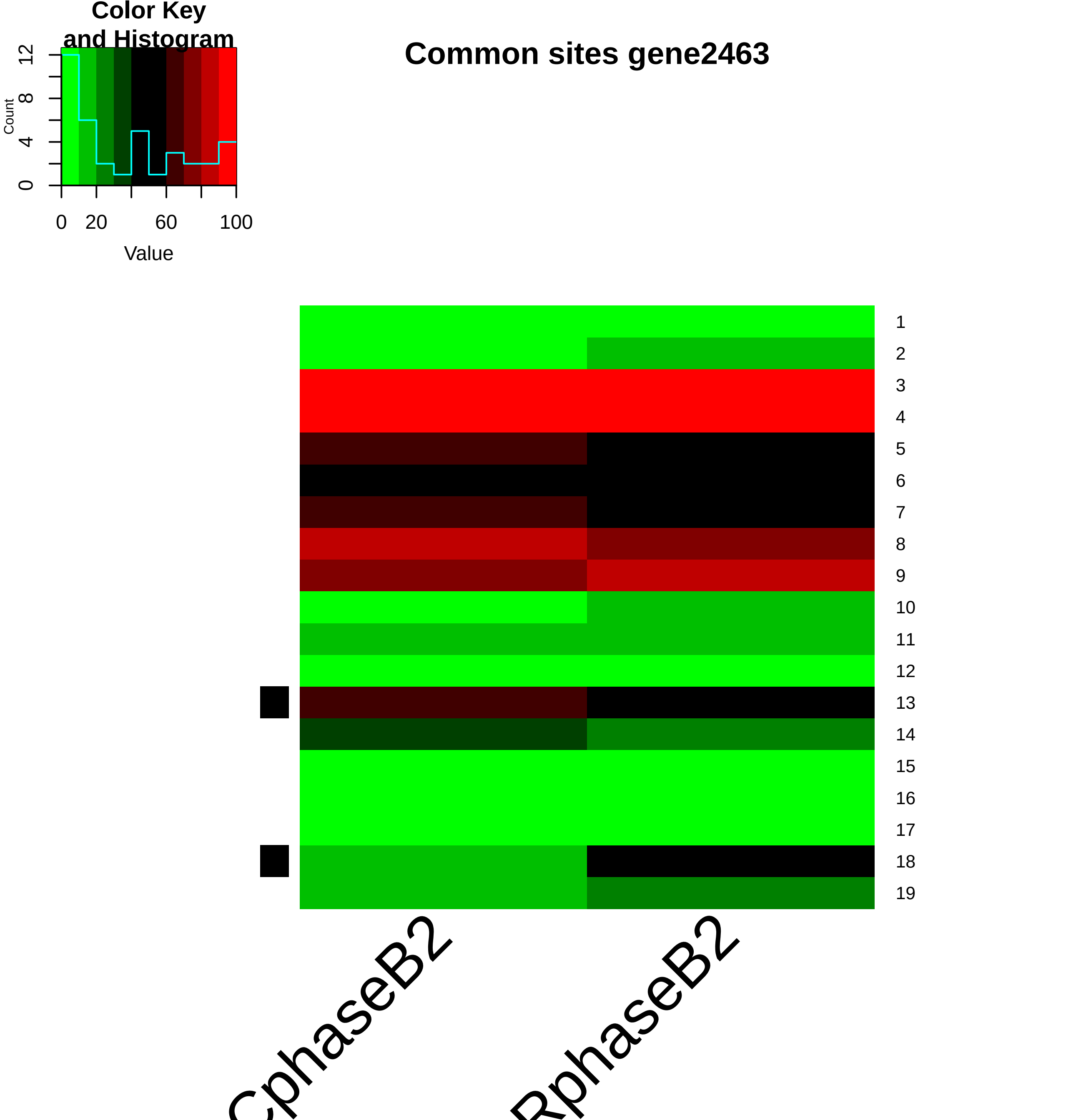

Color Key
and Histogram
Count
0
4
8
12
0
20
60
100
Value
Common sites gene2463
CphaseB2
RphaseB2

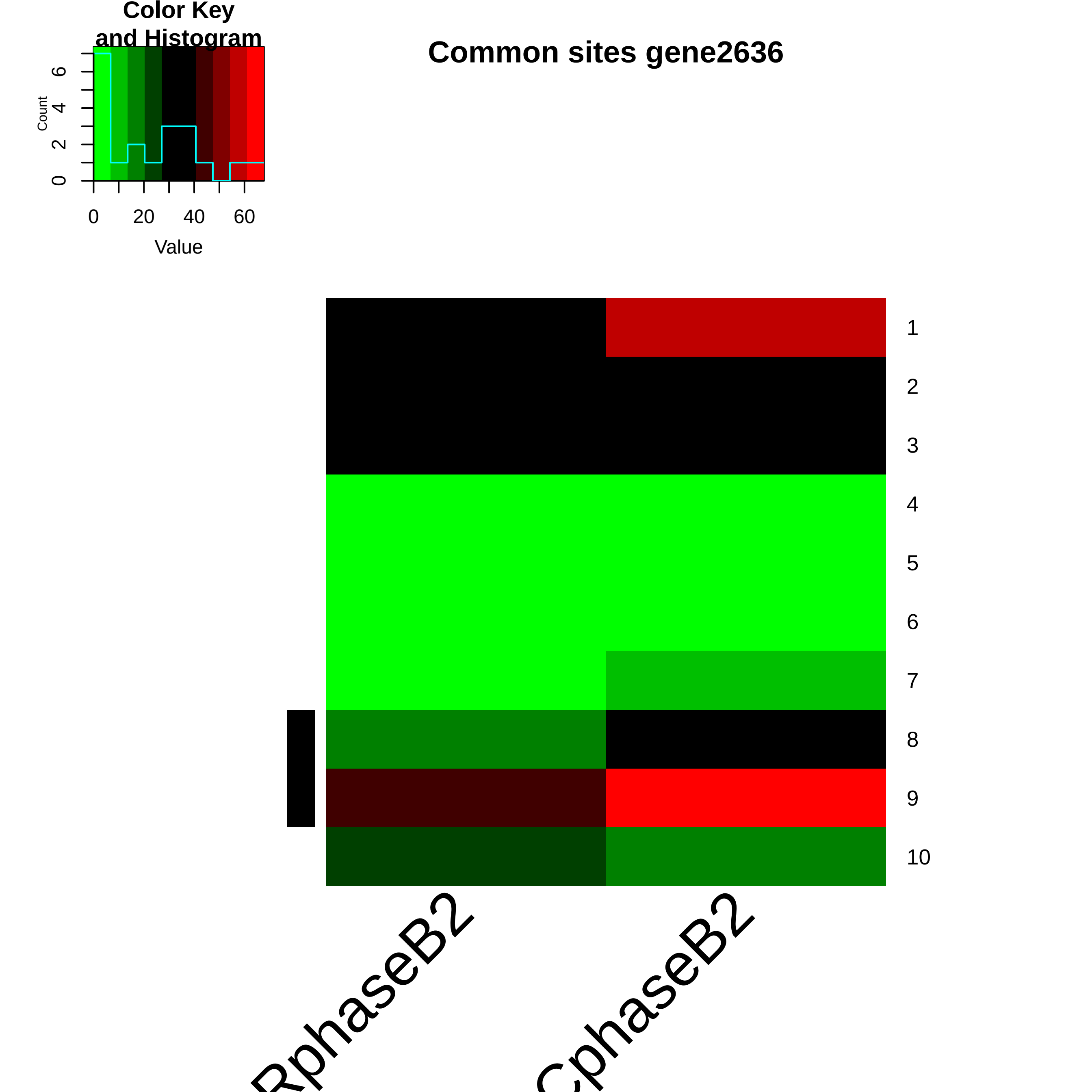

Color Key
and Histogram
Count
0
2
4
6
0
20
40
60
Value
Common sites gene2636
1
2
3
4
5
6
7
8
9
10
RphaseB2
CphaseB2

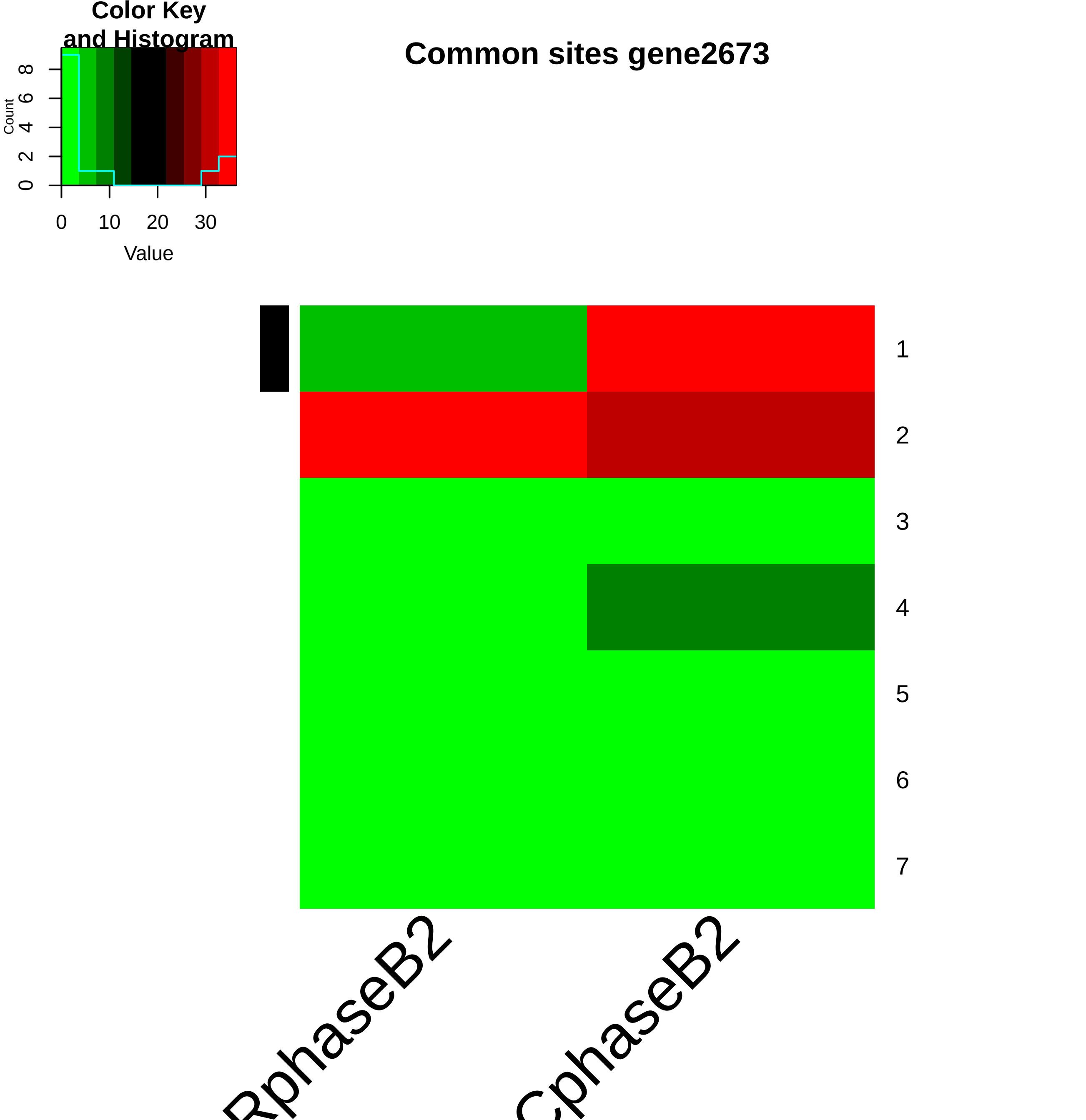

Color Key
and Histogram
Count
0 2 4 6 8
0 10 20 30
Value
Common sites gene2673
1
2
3
4
5
6
7
RphaseB2
CphaseB2

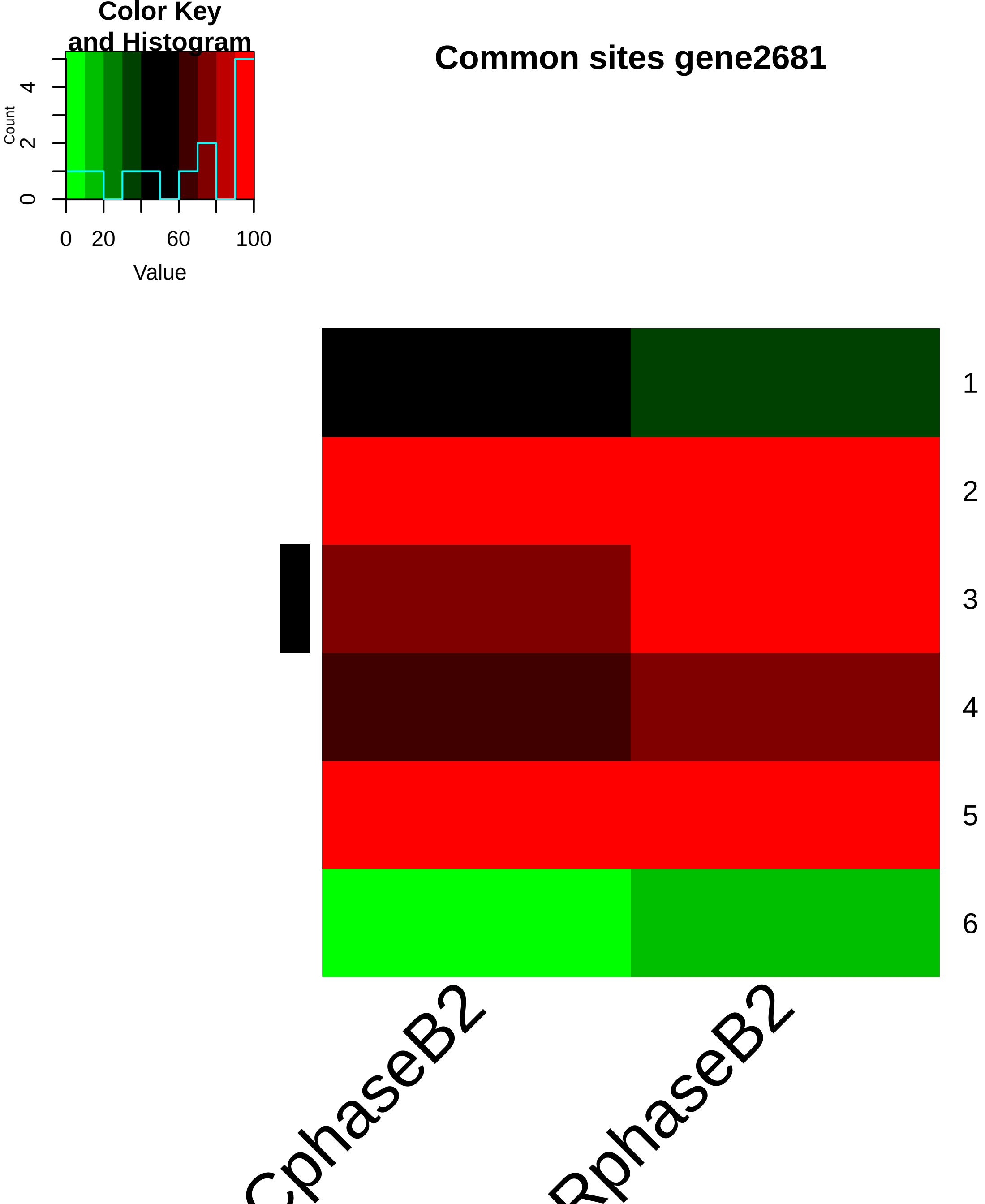

Color Key
and Histogram
Count
0
2
4
0
20
60
100
Value
Common sites gene2681
1
2
3
4
5
6
CphaseB2
RphaseB2

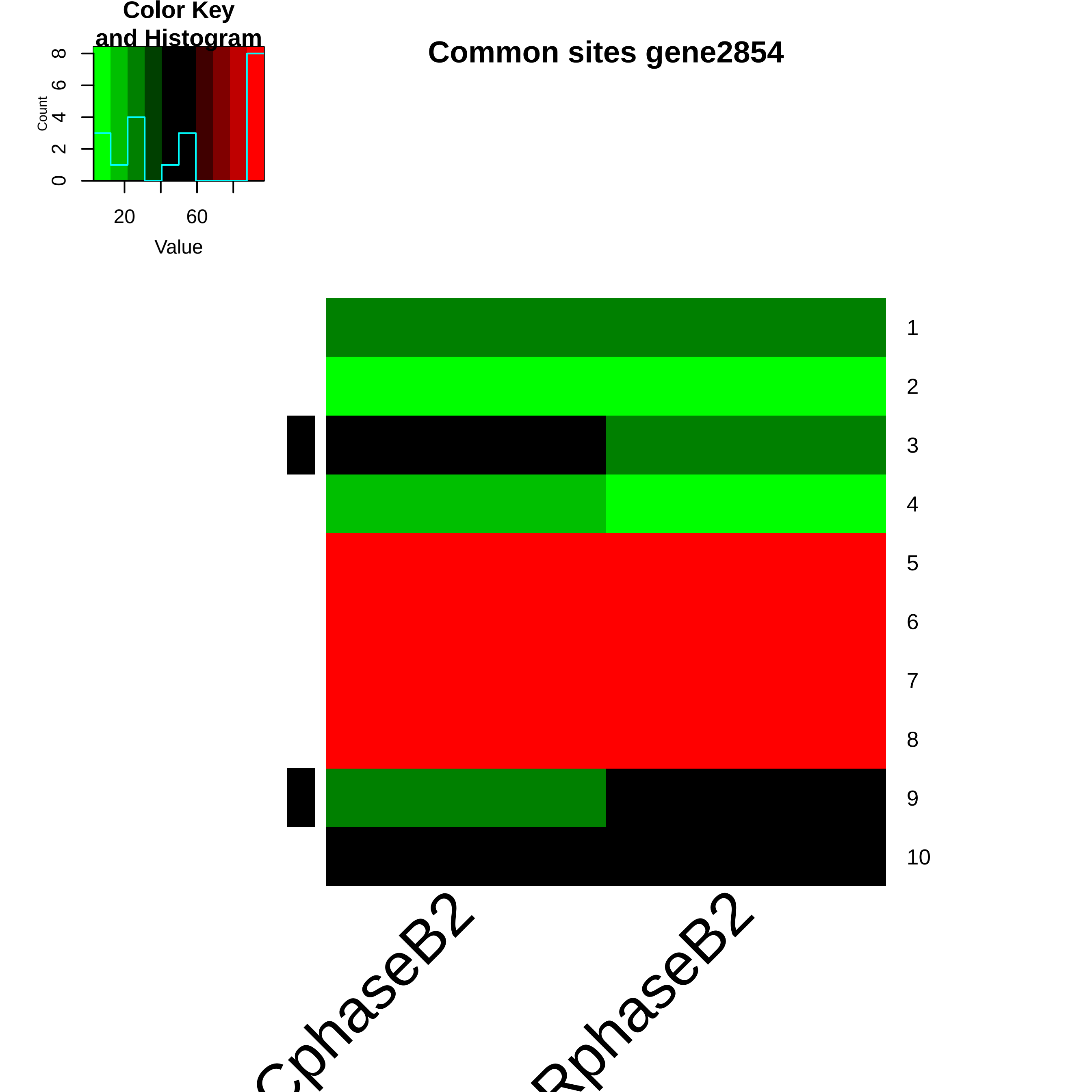
Color Key
and Histogram
Count
0
2
4
6
8
20
60
Value
Common sites gene2854
1
2
3
4
5
6
7
8
9
10
CphaseB2
RphaseB2

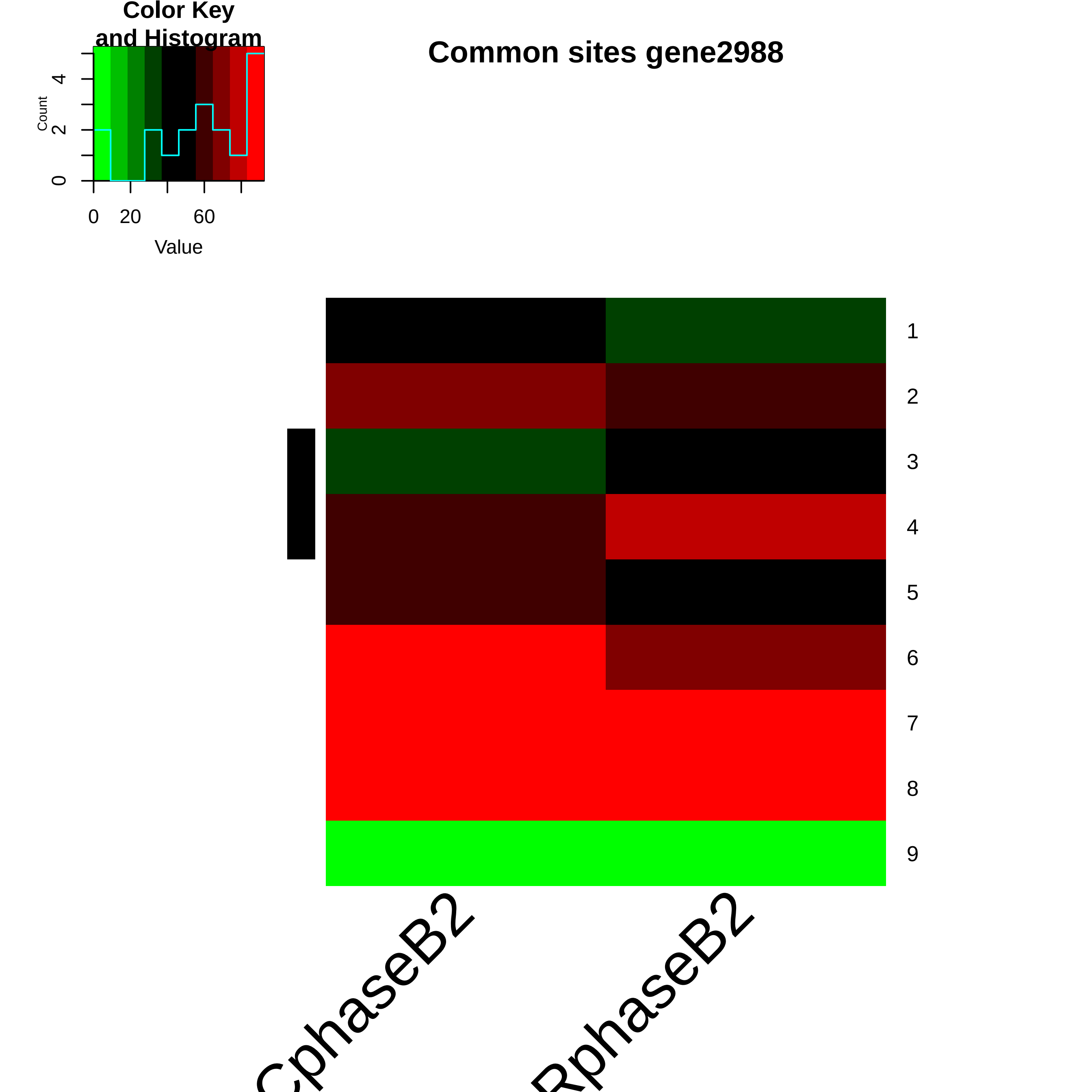

Color Key
and Histogram
Count
0
2
4
0
20
60
Value
Common sites gene2988
1
2
3
4
5
6
7
8
9
CphaseB2
RphaseB2

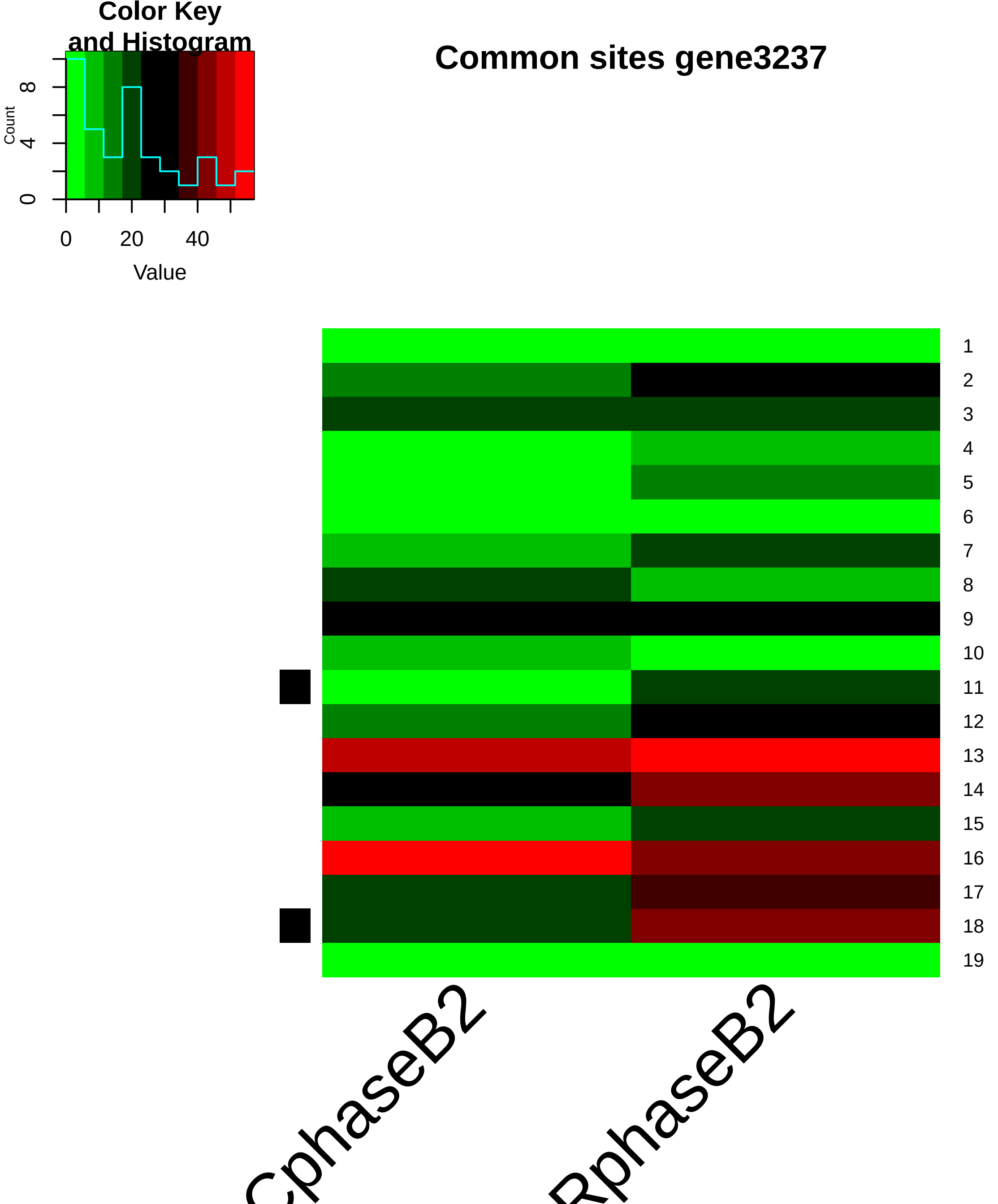
Color Key
and Histogram
Count
0
4
8
0
20
40
Value
Common sites gene3237
CphaseB2
RphaseB2

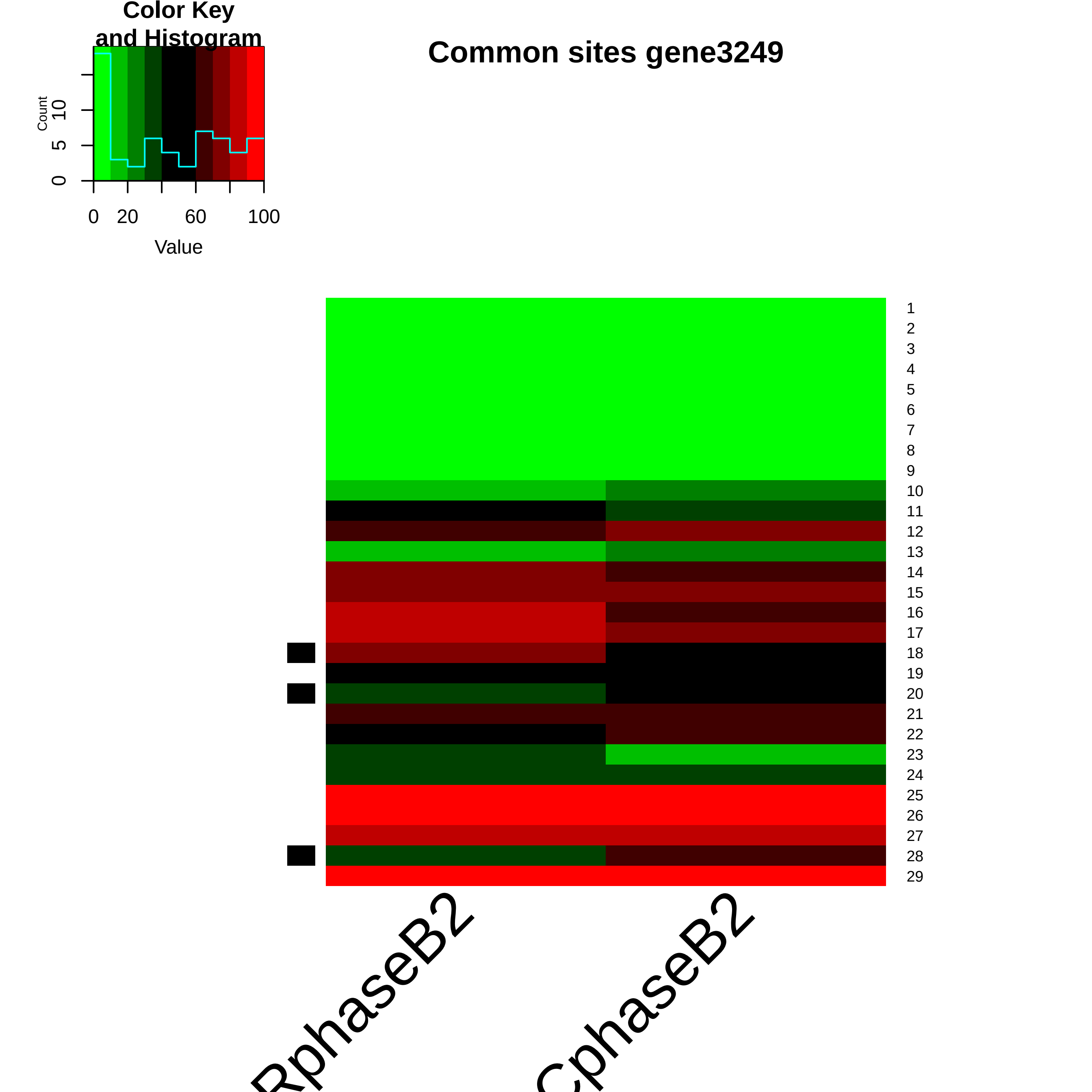
Color Key
and Histogram
Count
0
5
10
0
20
60
100
Value
Common sites gene3249
RphaseB2
CphaseB2

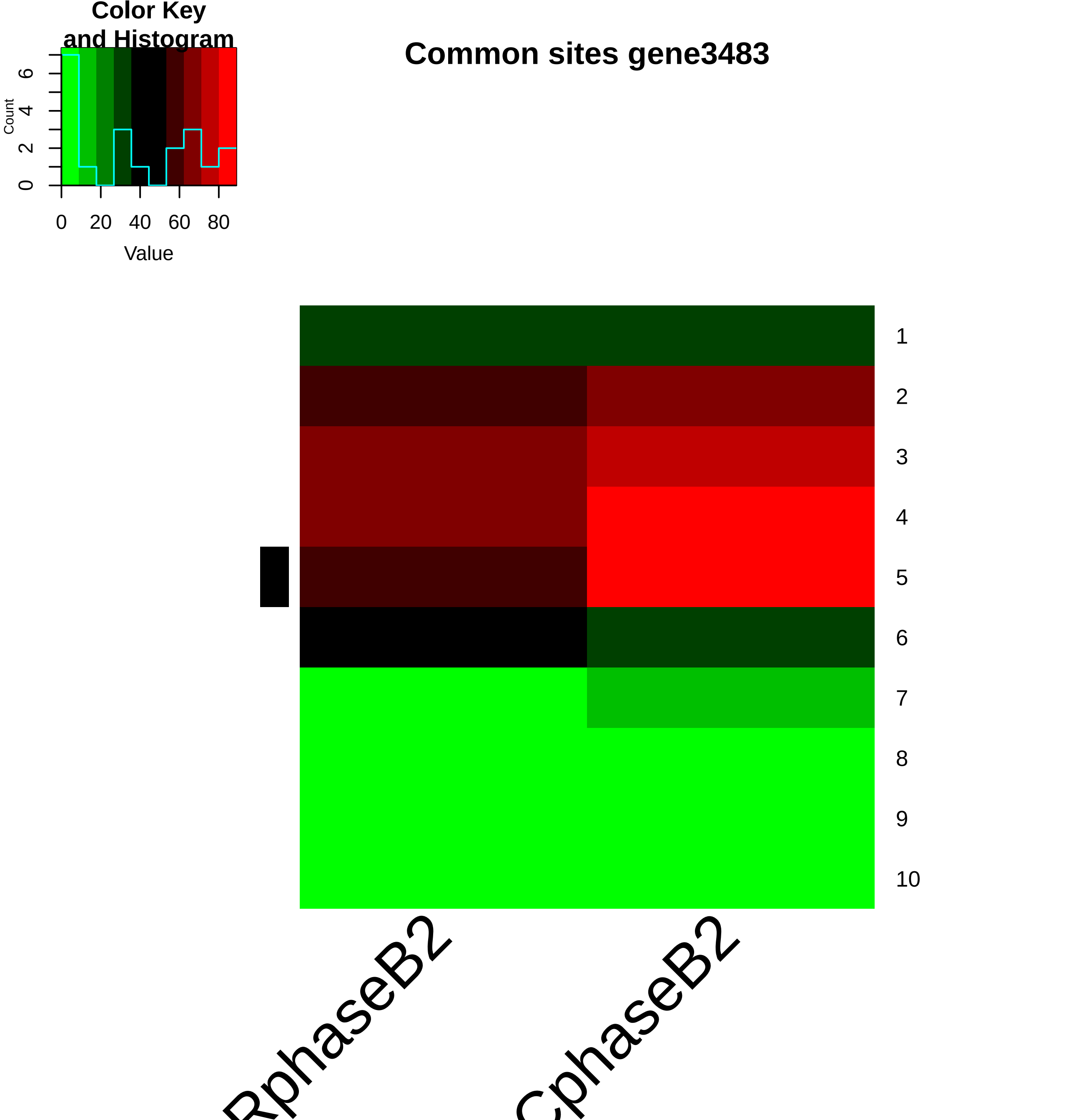
Color Key
and Histogram
Count
0
2
4
6
0
20
40
60
80
Value
Common sites gene3483
1
2
3
4
5
6
7
8
9
10
RphaseB2
CphaseB2

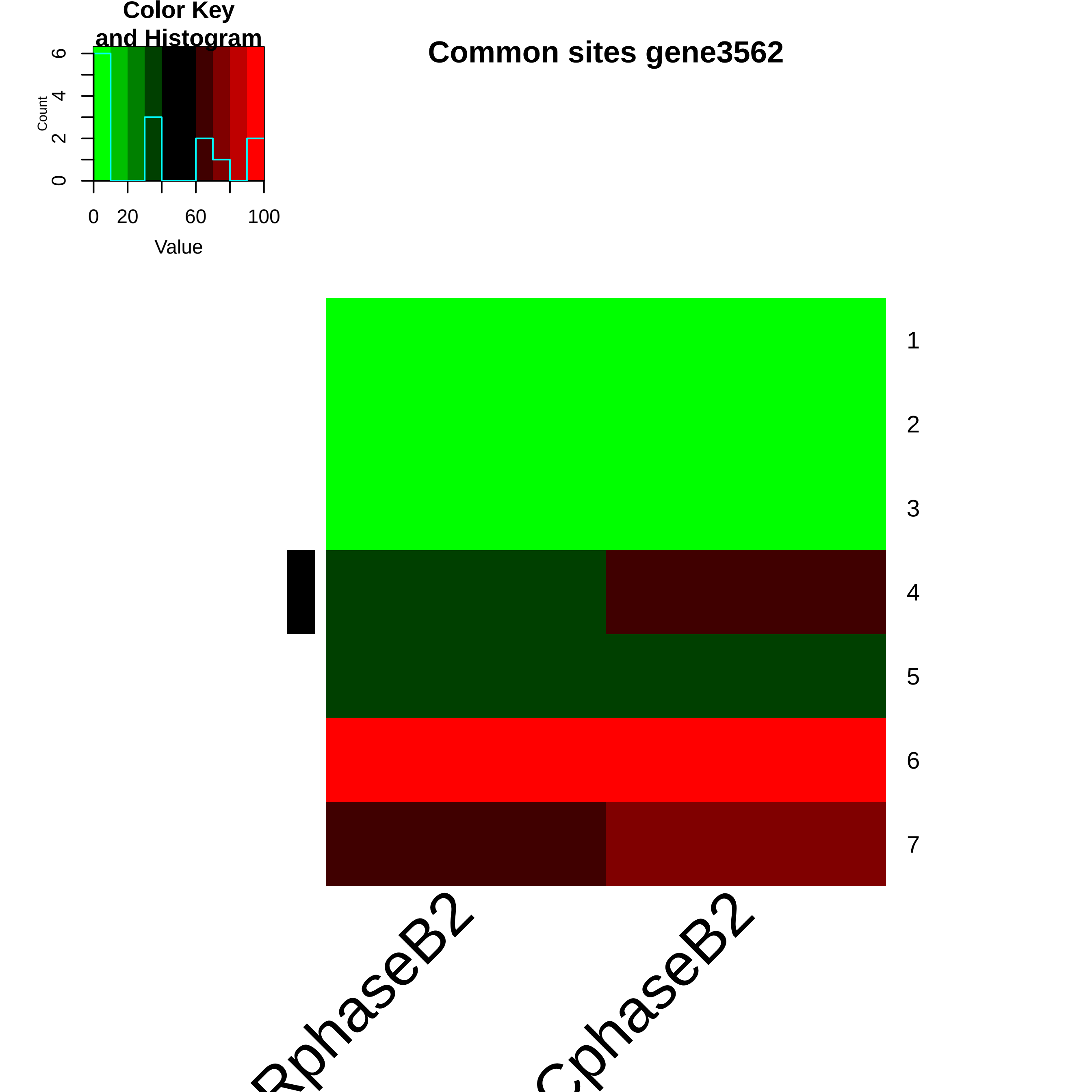

Color Key
and Histogram
Count
Value
0
20
60
100
0
2
4
6
Common sites gene3562
1
2
3
4
5
6
7
RphaseB2
CphaseB2

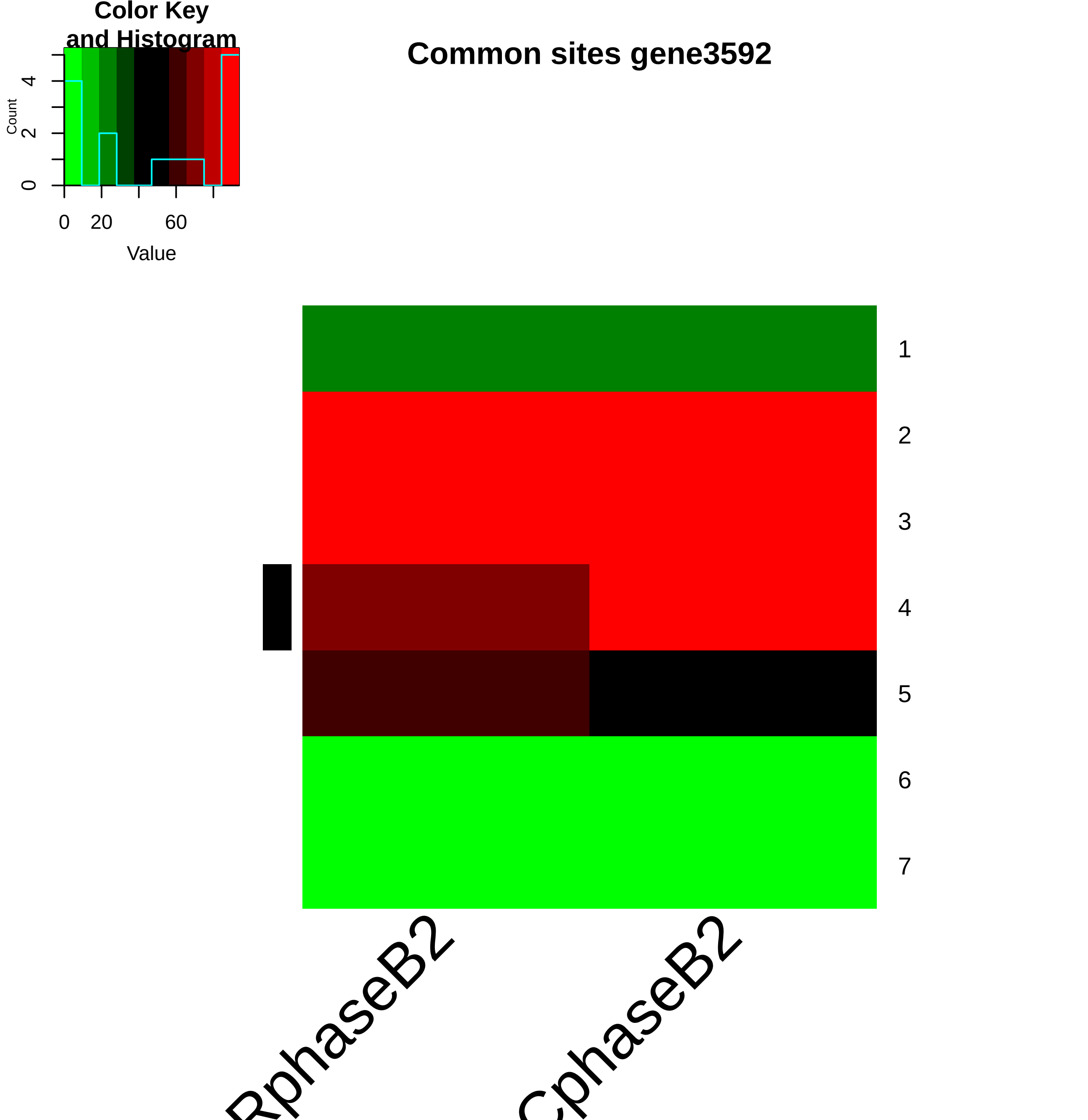
Color Key
and Histogram
Count
0
2
4
0
20
60
Value
Common sites gene3592
1
2
3
4
5
6
7
RphaseB2
CphaseB2

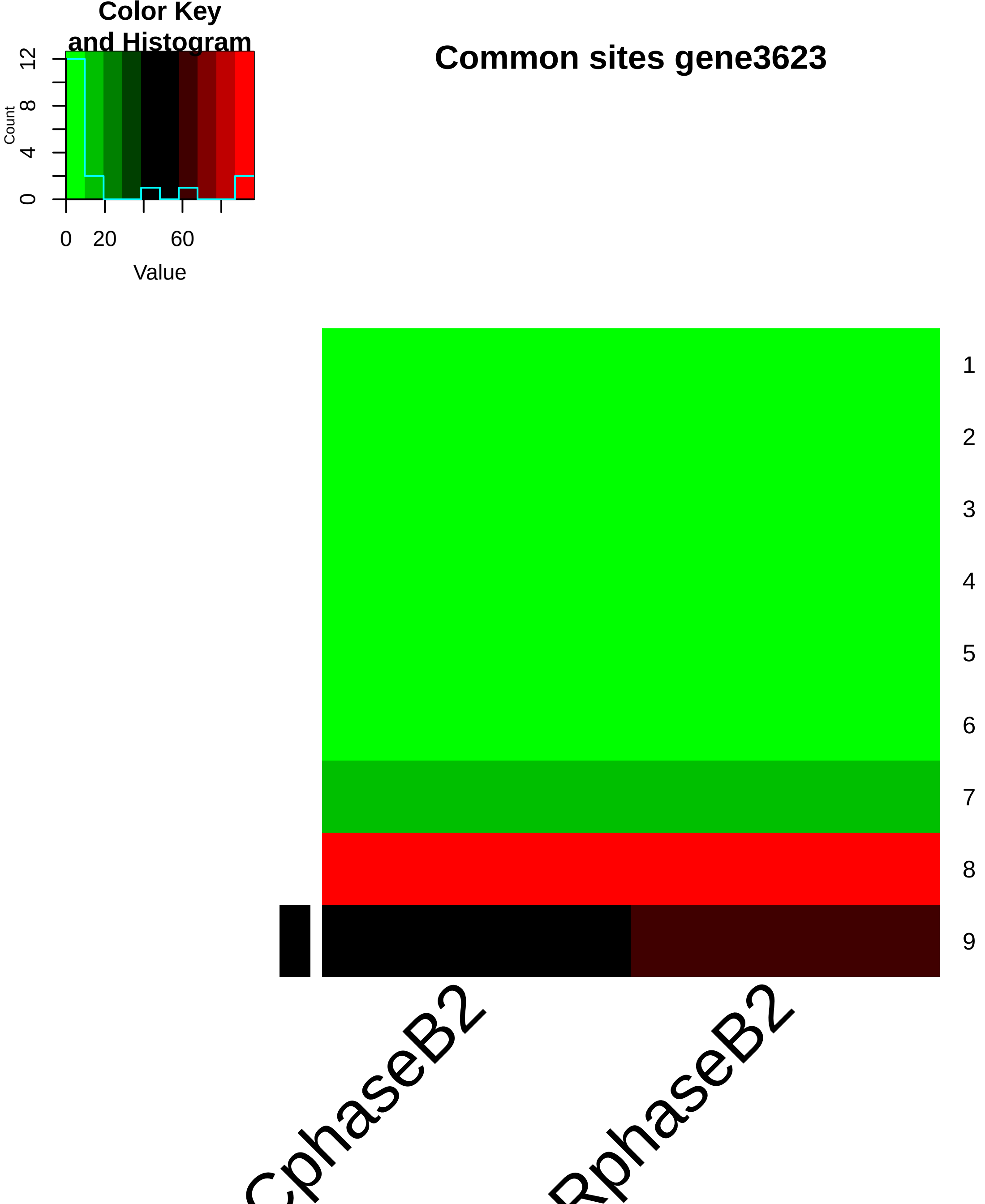

Color Key
and Histogram
Count
0
4
8
12
0
20
60
Value
Common sites gene3623
1
2
3
4
5
6
7
8
9
CphaseB2
RphaseB2

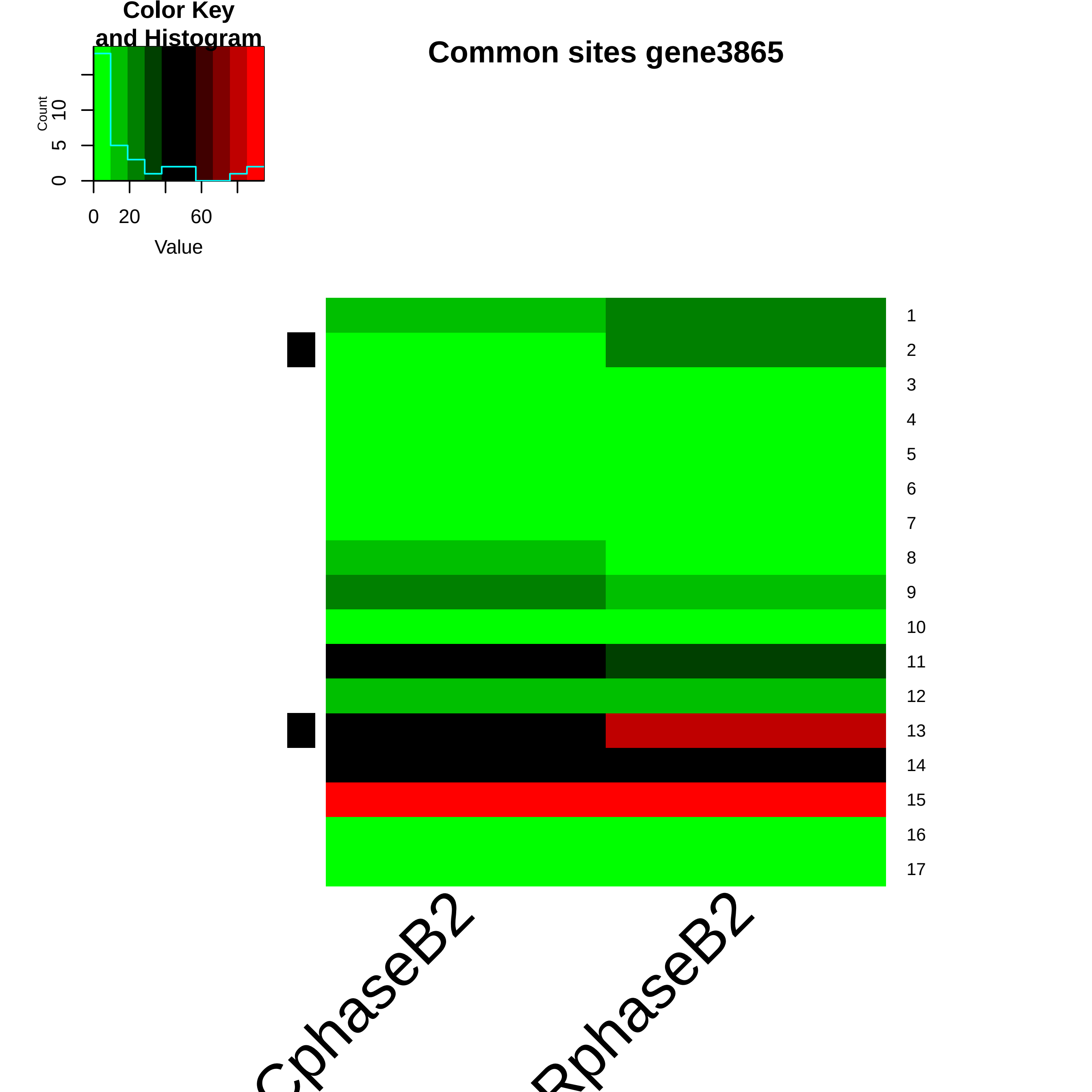
Color Key
and Histogram
Count
0
5
10
0
20
60
Value
Common sites gene3865
CphaseB2
RphaseB2

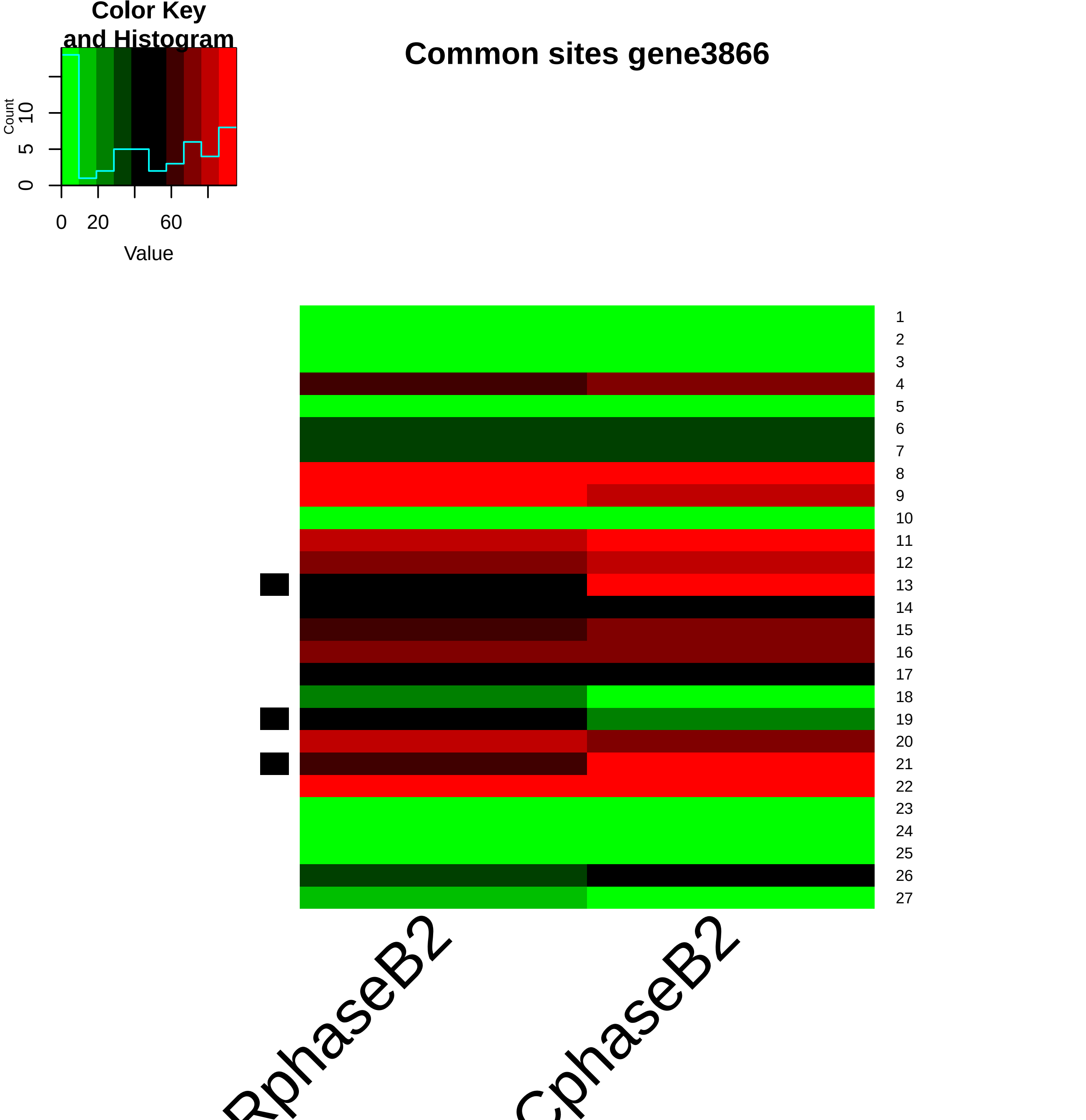
Color Key
and Histogram
Count
0
5
10
0
20
60
Value
Common sites gene3866
RphaseB2
CphaseB2

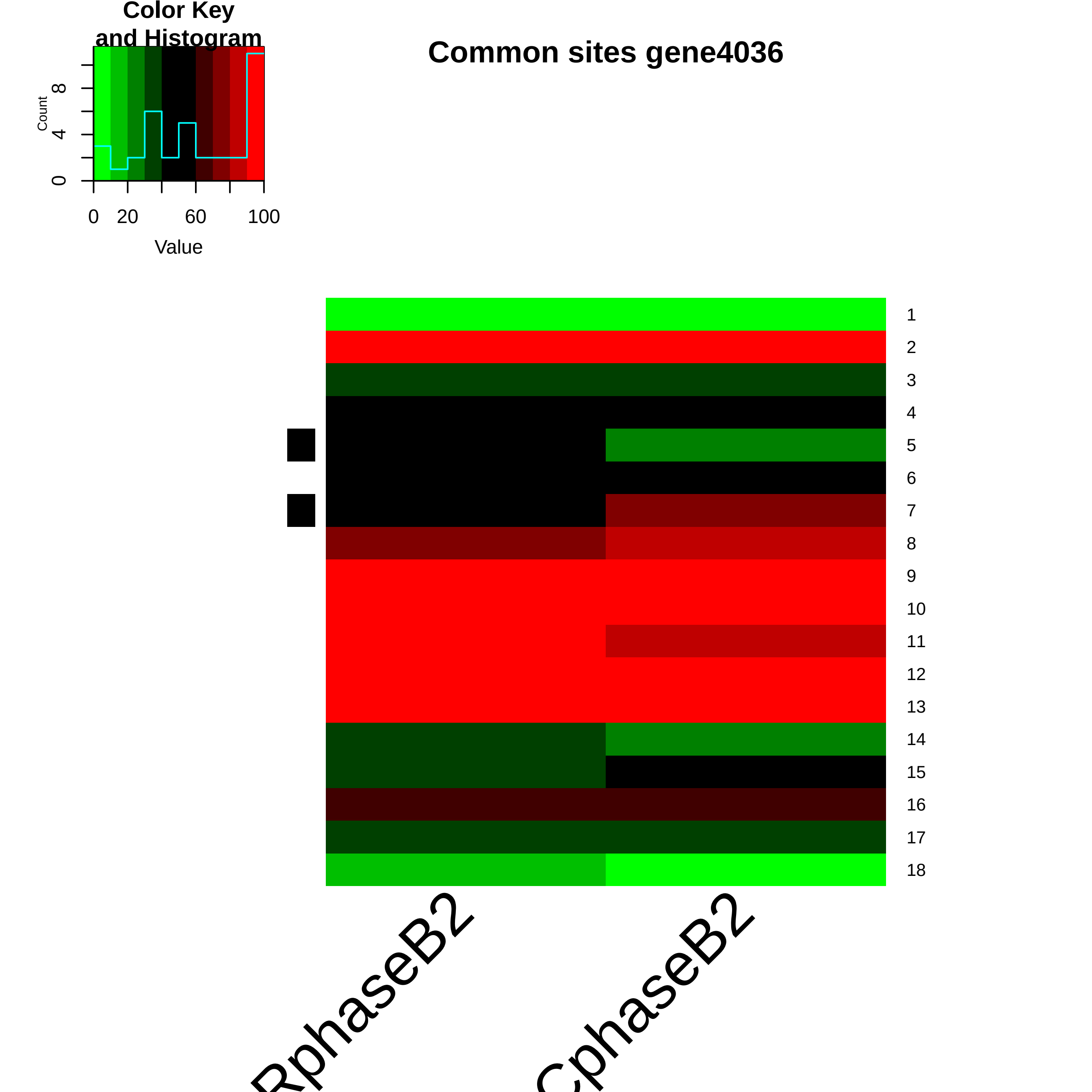

Color Key
and Histogram
Count
0
4
8
0
20
60
100
Value
Common sites gene4036
RphaseB2
CphaseB2

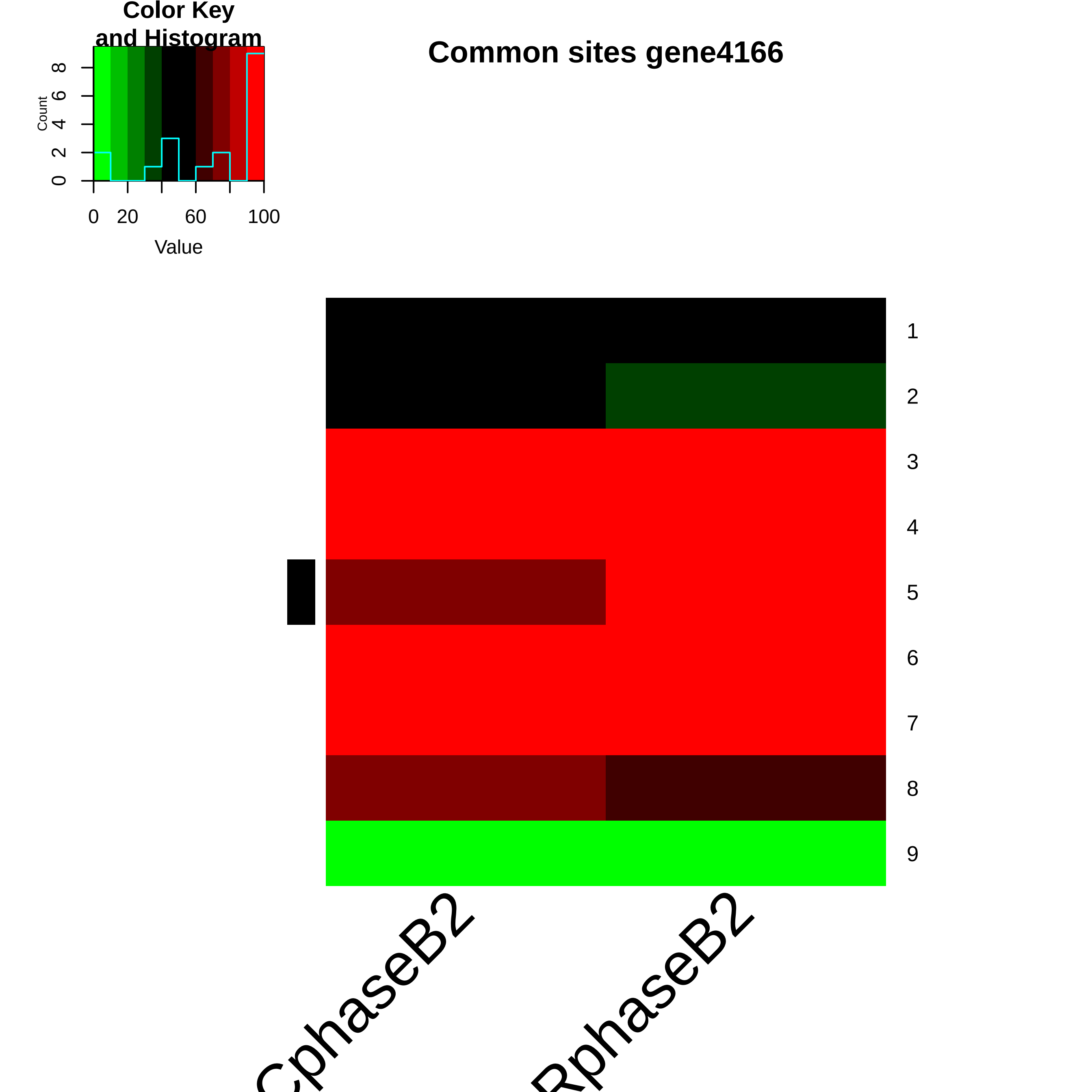
Color Key
and Histogram
Count
0
2
4
6
8
0
20
60
100
Value
Common sites gene4166
1
2
3
4
5
6
7
8
9
CphaseB2
RphaseB2

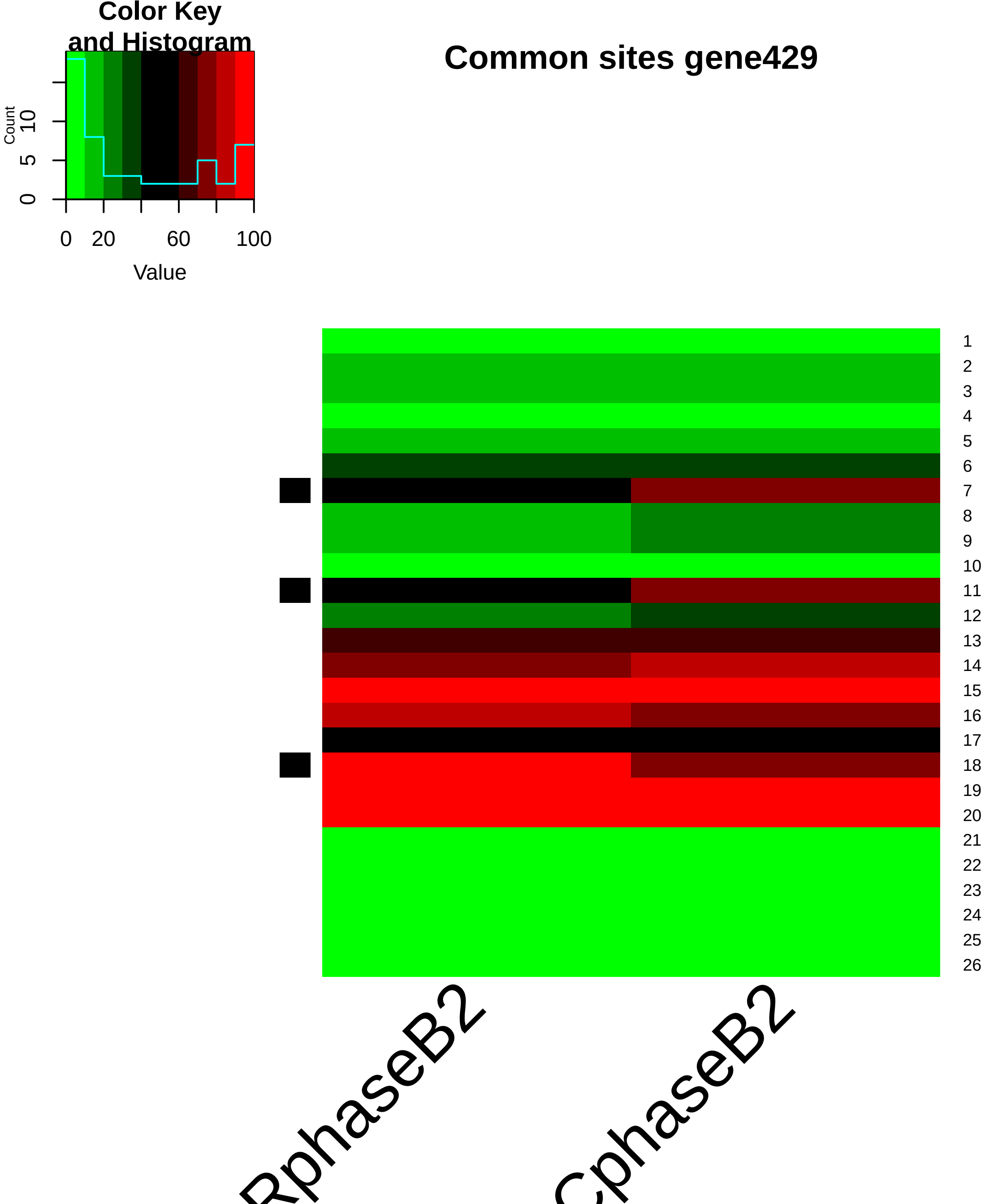
Color Key
and Histogram
Count
0
5
10
0
20
60
100
Value
Common sites gene429
RphaseB2
CphaseB2

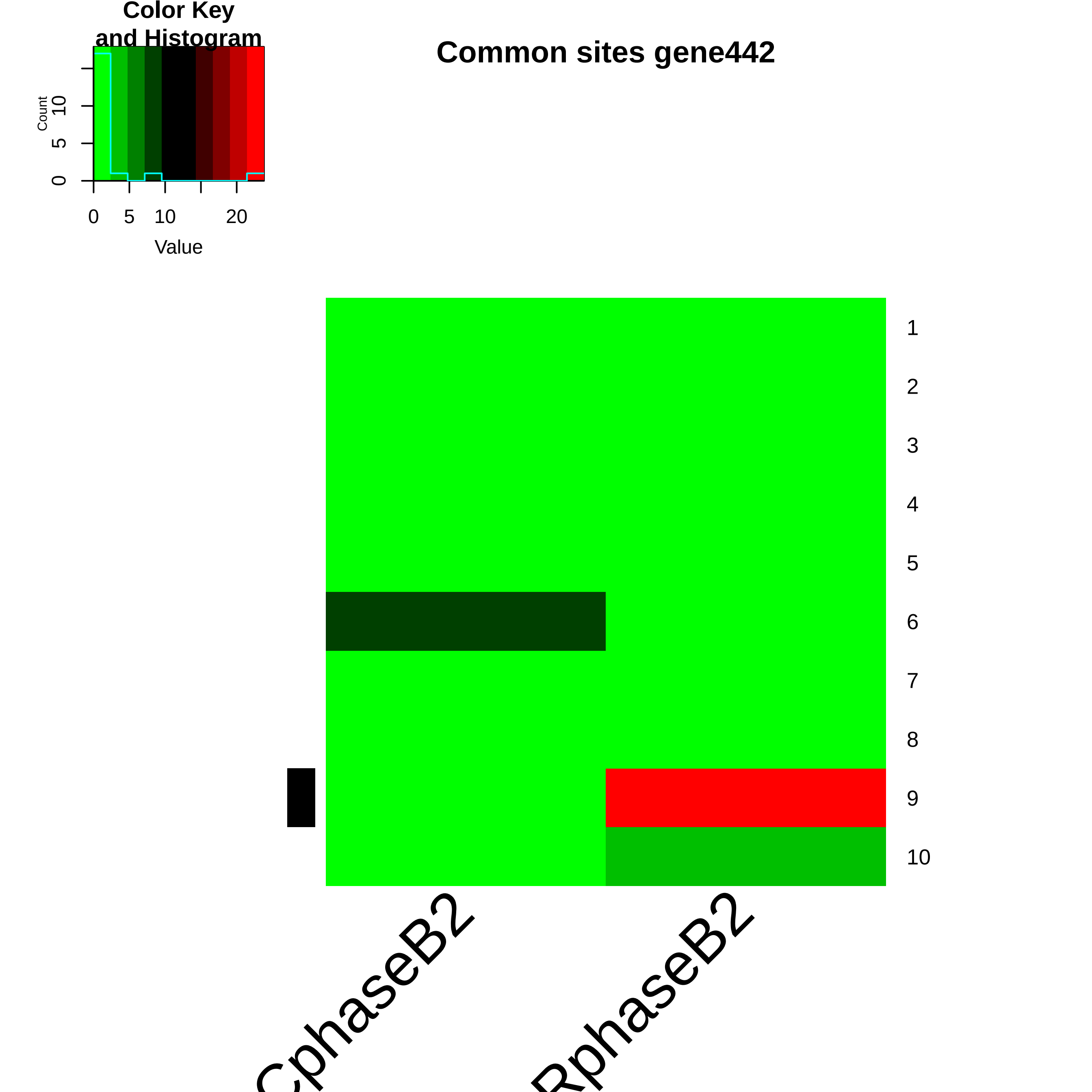
Color Key
and Histogram
Count
0
5
10
0
5
10
20
Value
Common sites gene442
1
2
3
4
5
6
7
8
9
10
CphaseB2
RphaseB2

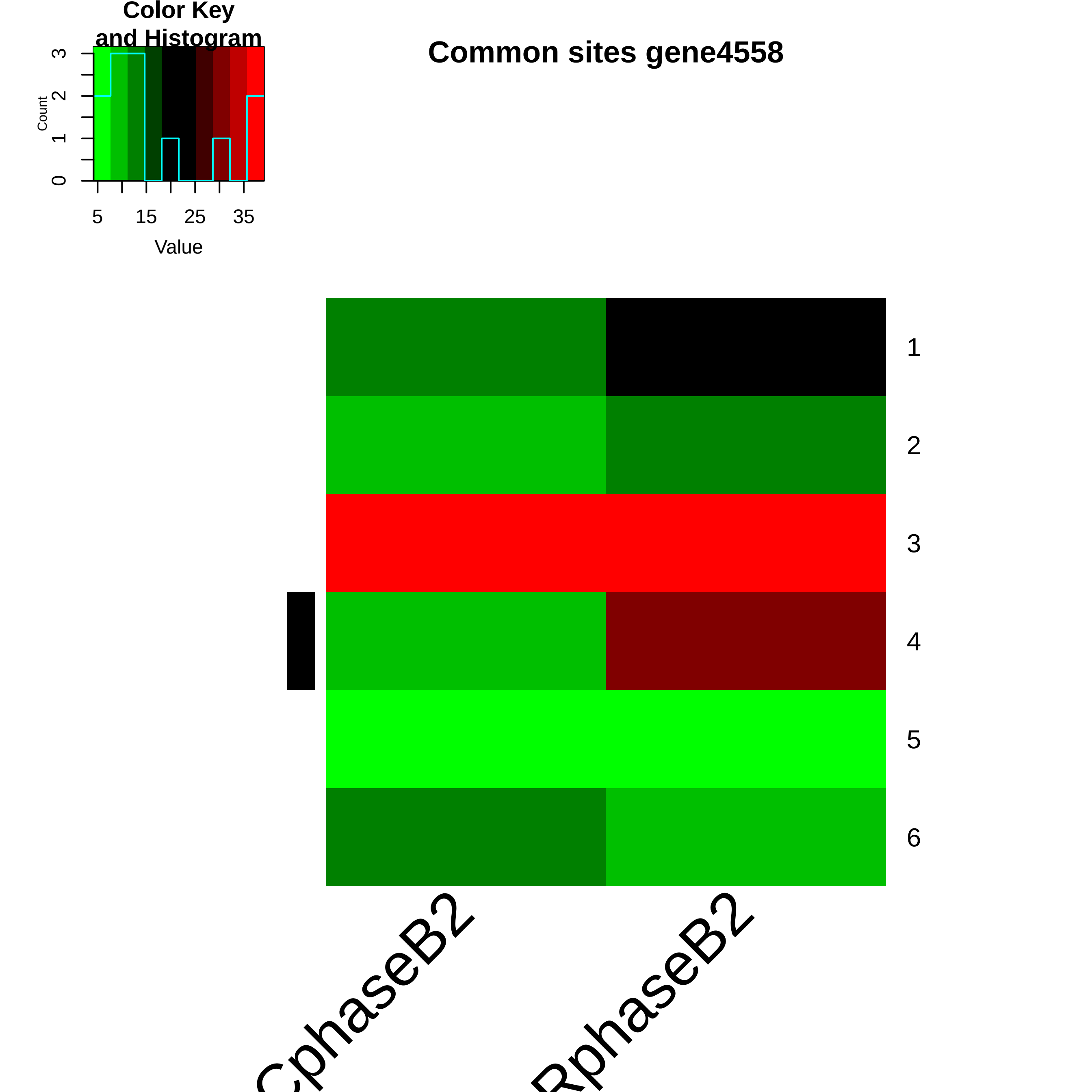
Color Key
and Histogram
Count
0
1
2
3
5
15
25
35
Value
Common sites gene4558
1
2
3
4
5
6
CphaseB2
RphaseB2

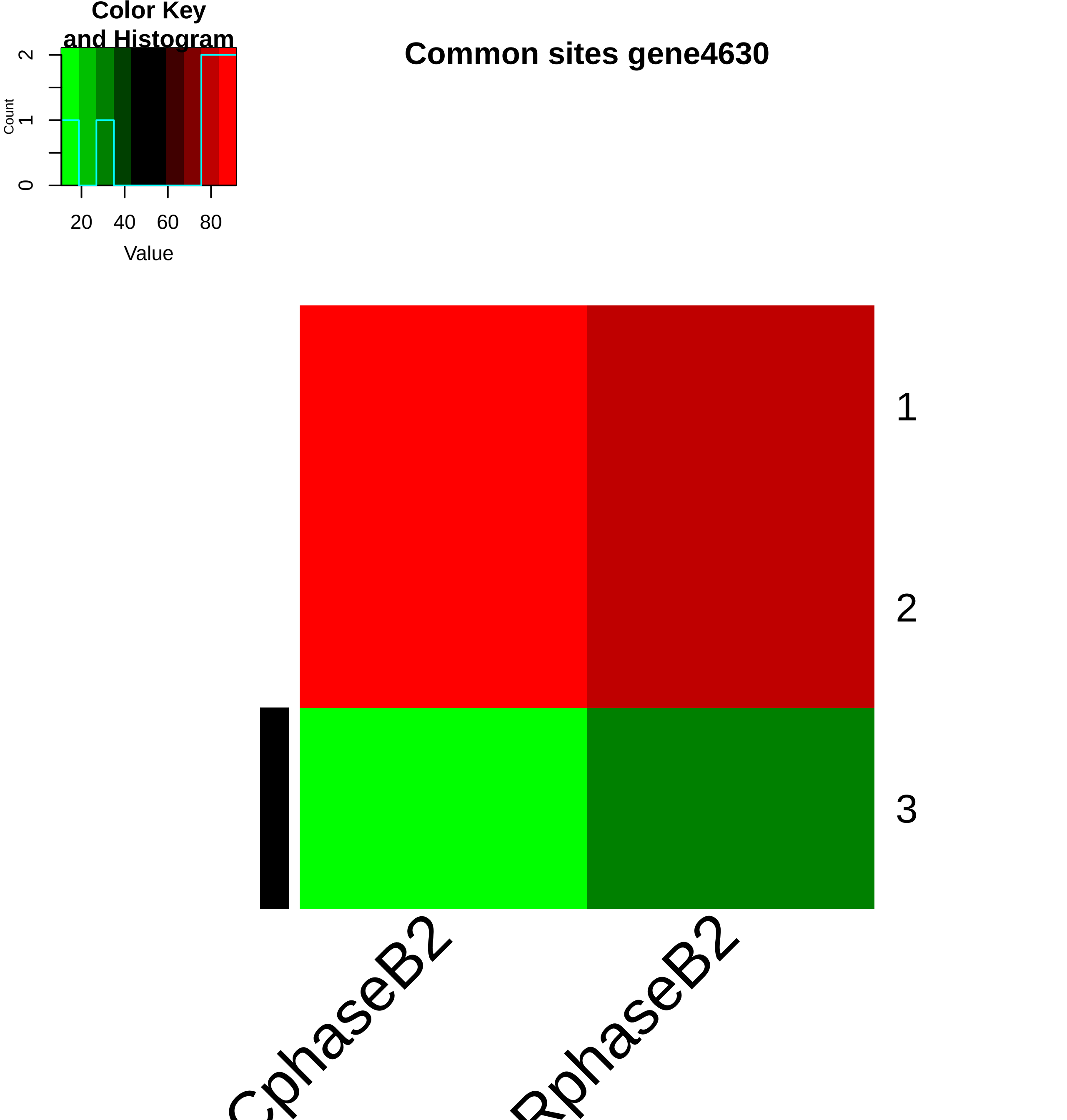
Color Key
and Histogram
Count
Value
0
1
2
20
40
60
80
Common sites gene4630
1
2
3
CphaseB2
RphaseB2

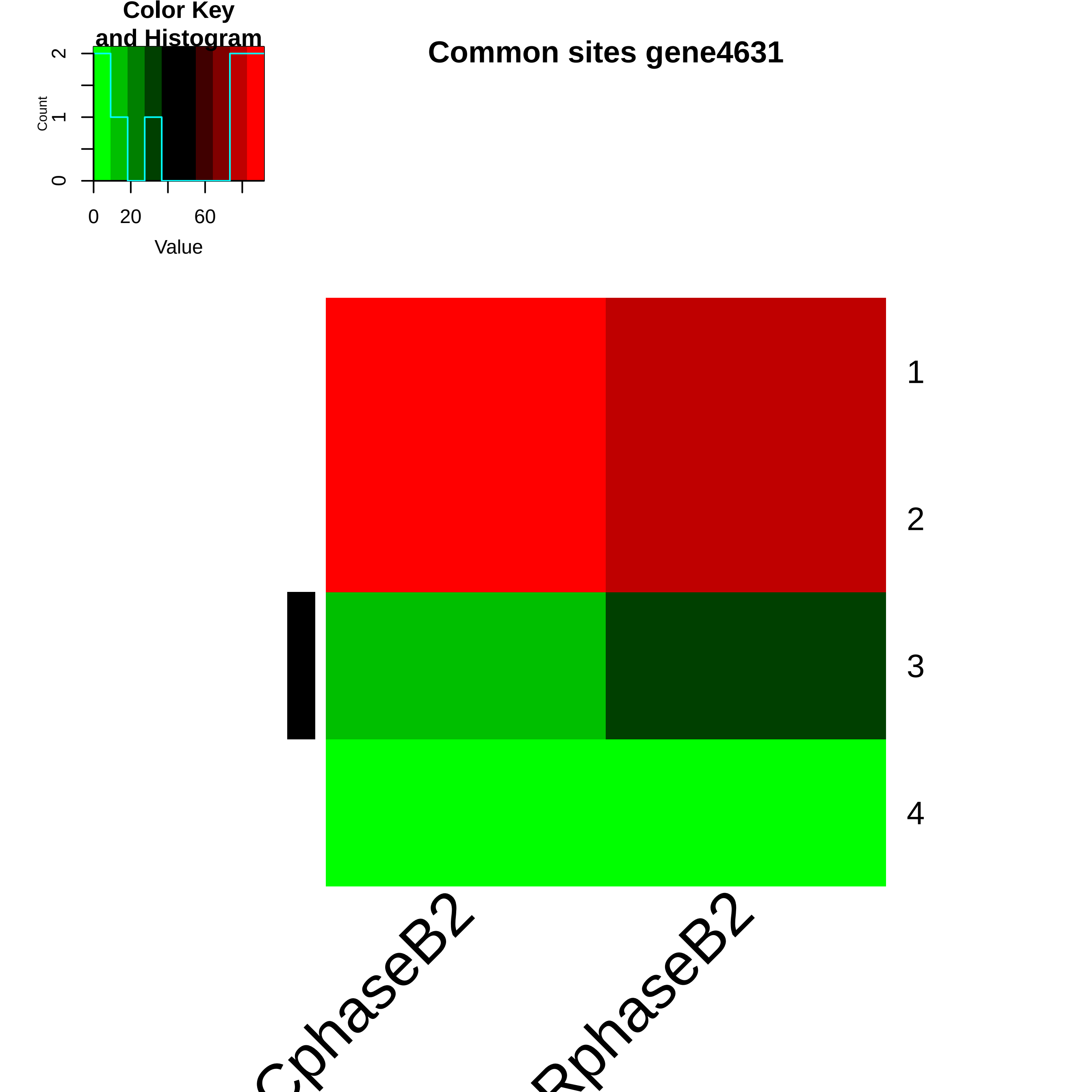
Color Key
and Histogram
Count
Value
0
20
60
0
1
2
Common sites gene4631
1
2
3
4

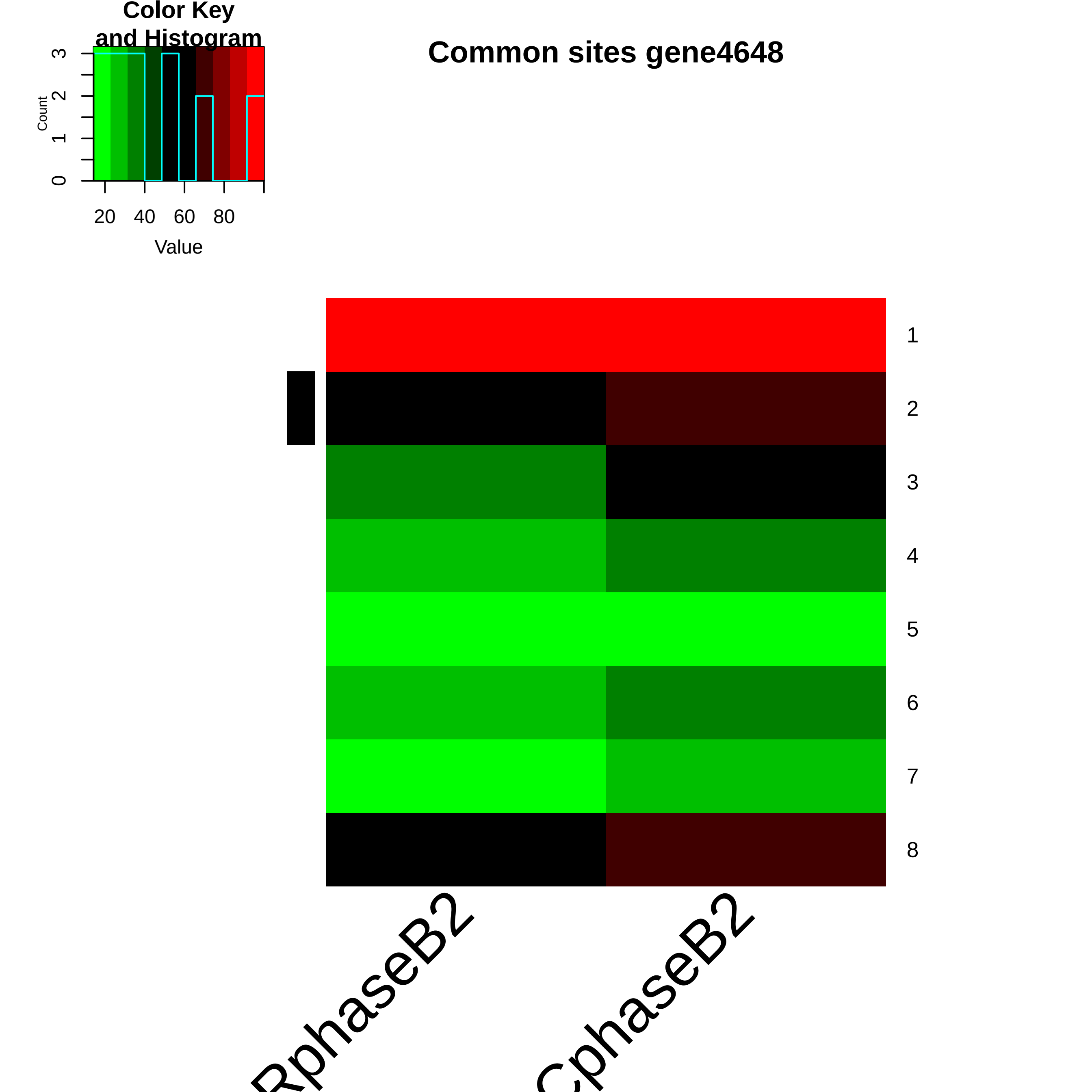
Color Key
and Histogram
Count
0
1
2
3
20
40
60
80
Value
Common sites gene4648
1
2
3
4
5
6
7
8
RphaseB2
CphaseB2

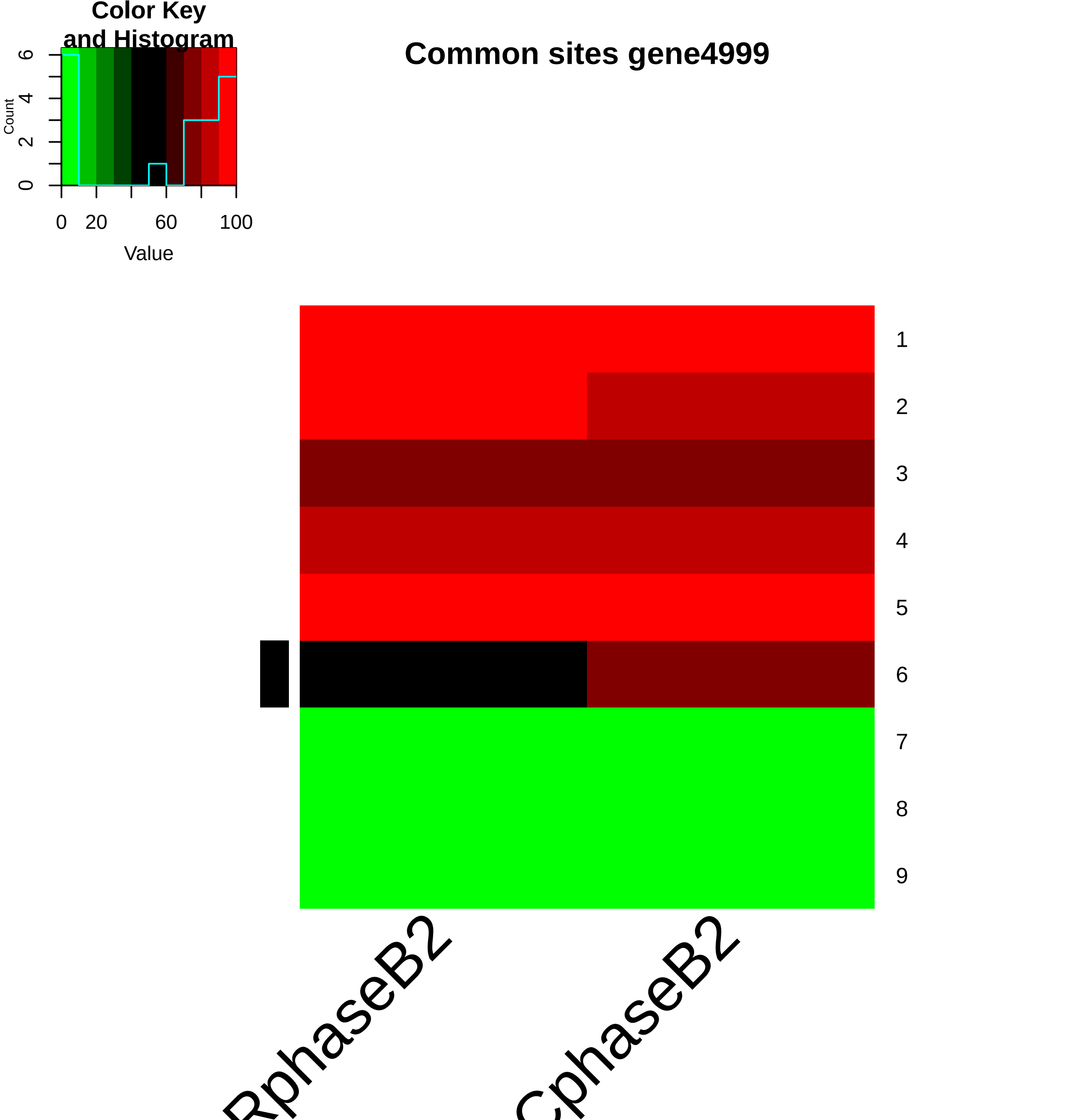
Color Key
and Histogram
Count
0
2
4
6
0
20
60
100
Value
Common sites gene4999
1
2
3
4
5
6
7
8
9
RphaseB2
CphaseB2

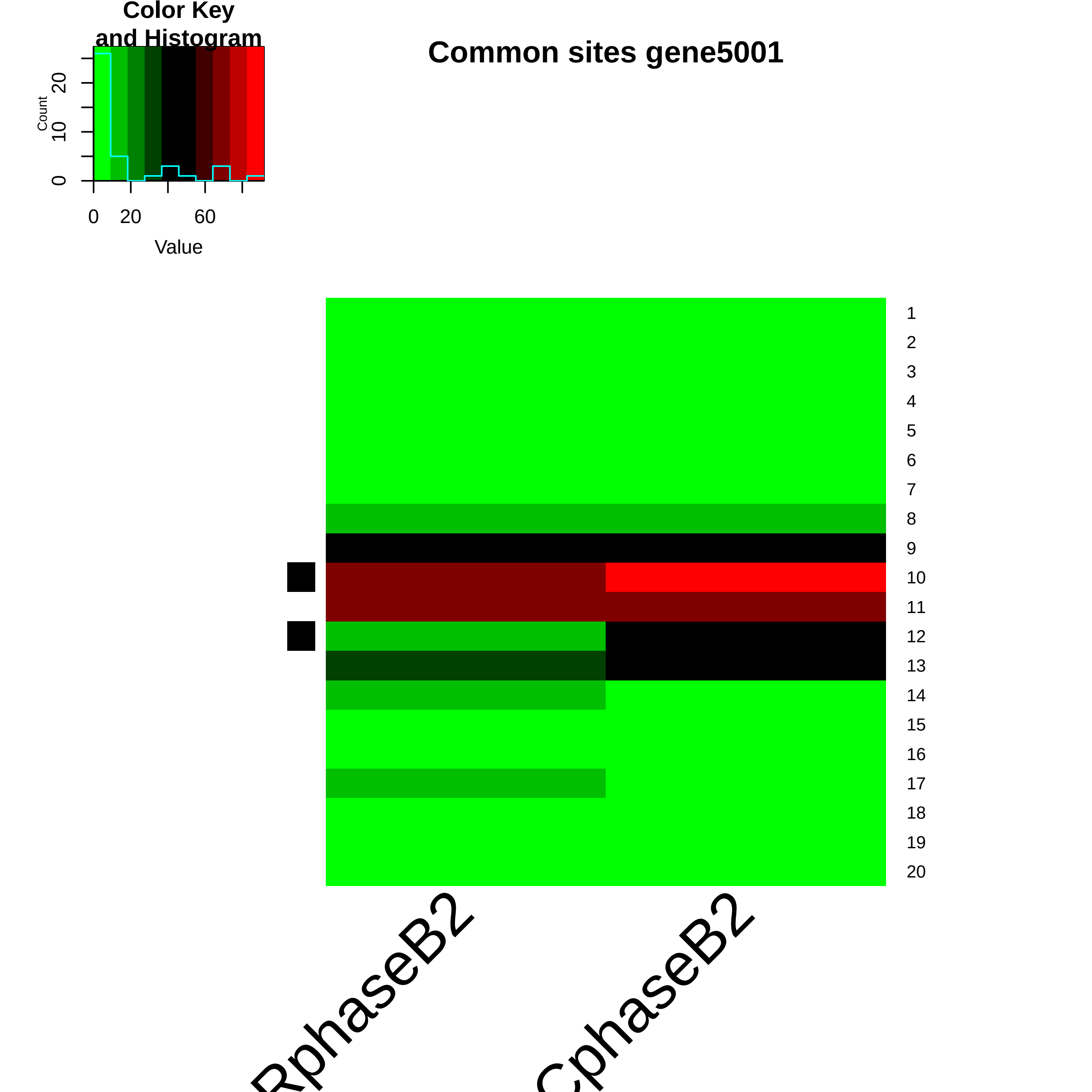
Color Key
and Histogram
Count
0
10
20
0
20
60
Value
Common sites gene5001
RphaseB2
CphaseB2

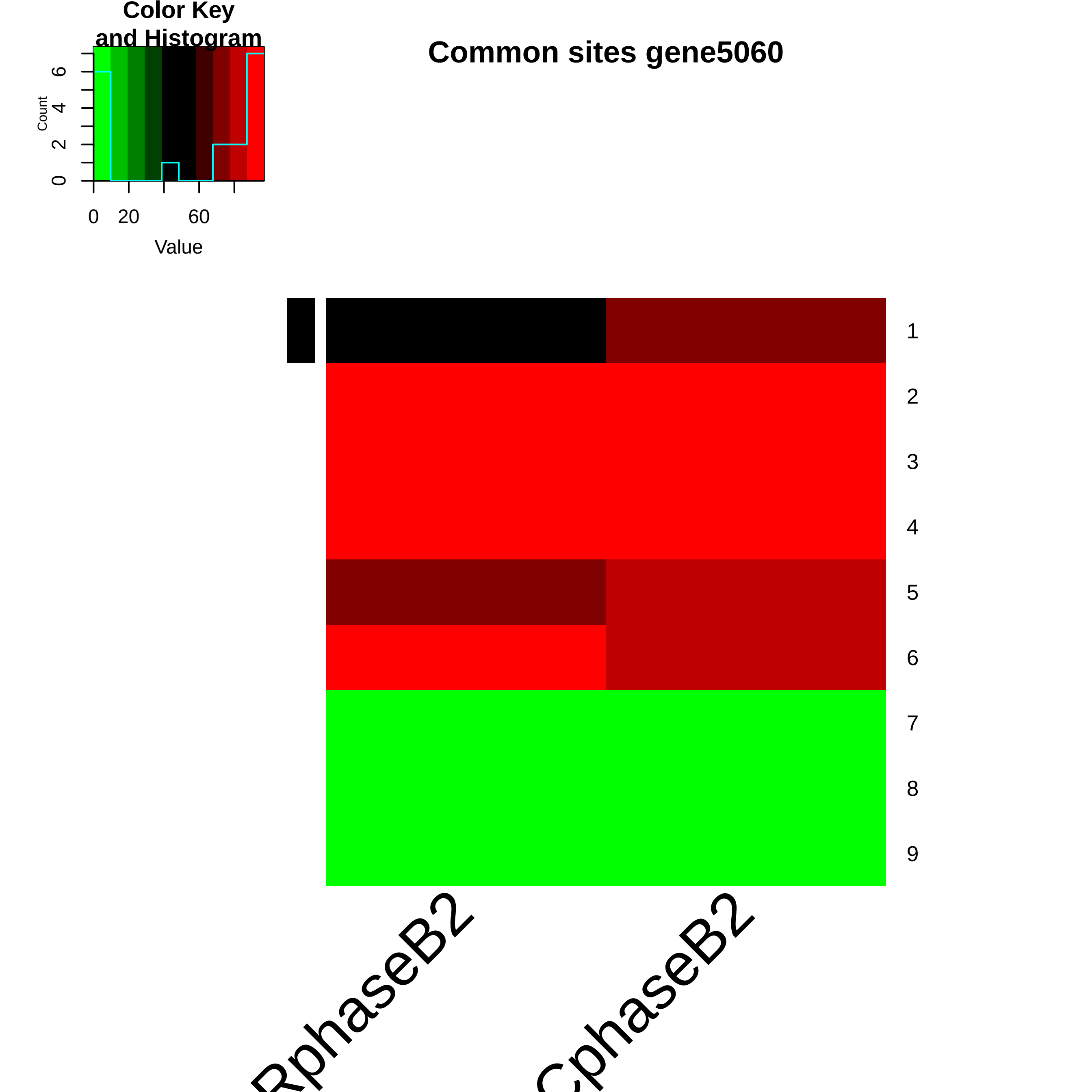

Color Key
and Histogram
Count
0
2
4
6
0
20
60
Value
Common sites gene5060
1
2
3
4
5
6
7
8
9
RphaseB2
CphaseB2

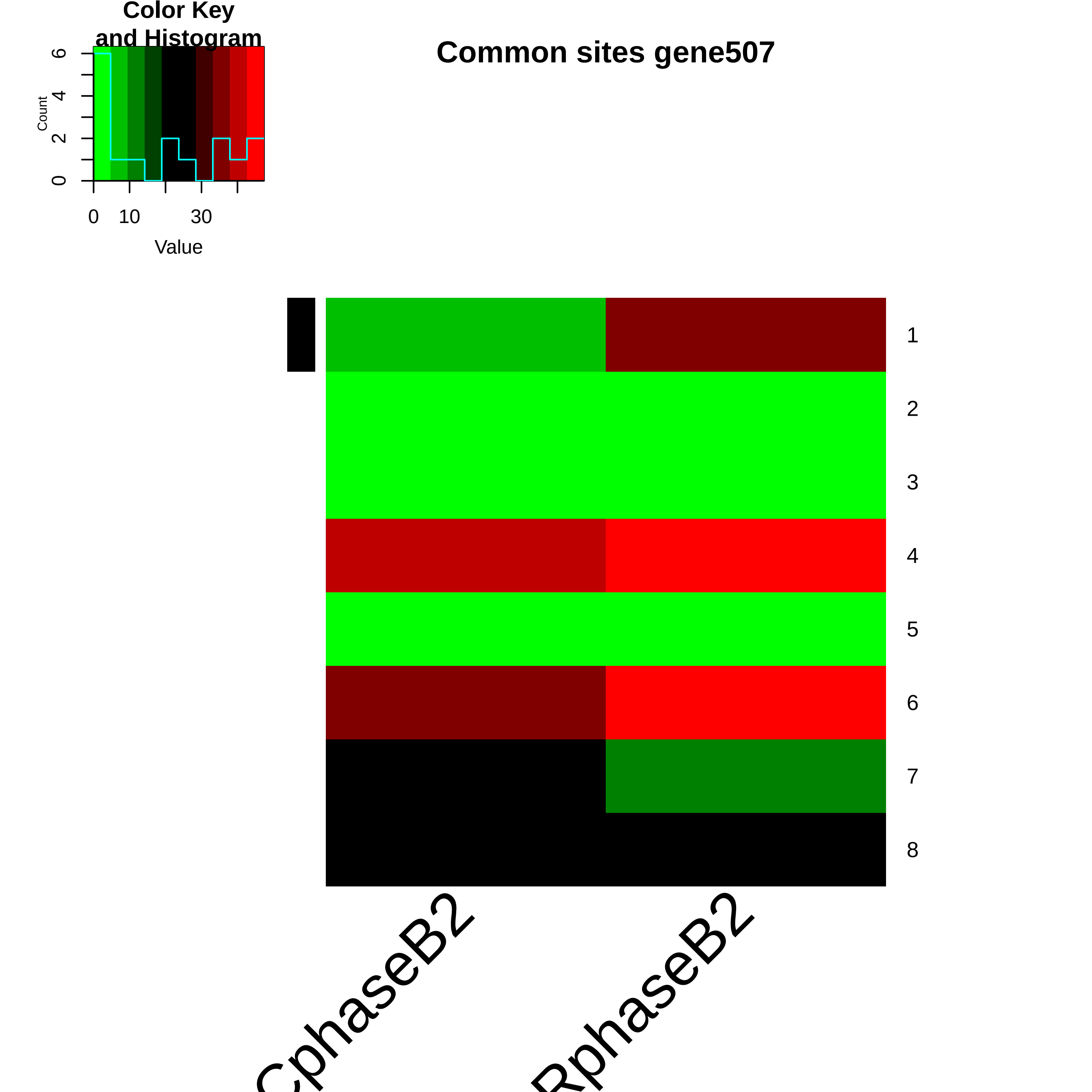
Color Key
and Histogram
Count
Value
Common sites gene507
1
2
3
4
5
6
7
8
CphaseB2
RphaseB2

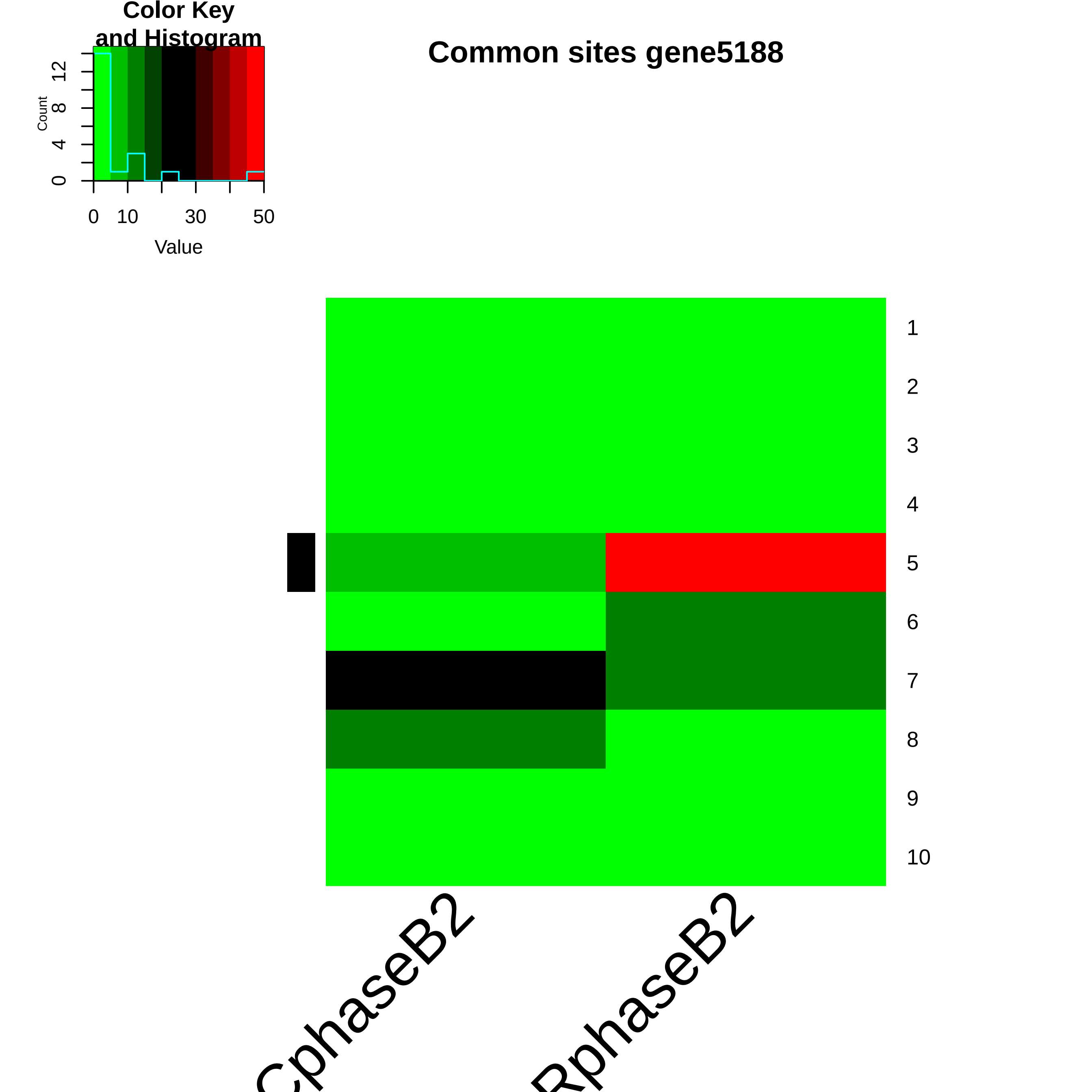
Color Key
and Histogram
Count
0
4
8
12
0
10
30
50
Value
Common sites gene5188
1
2
3
4
5
6
7
8
9
10
CphaseB2
RphaseB2

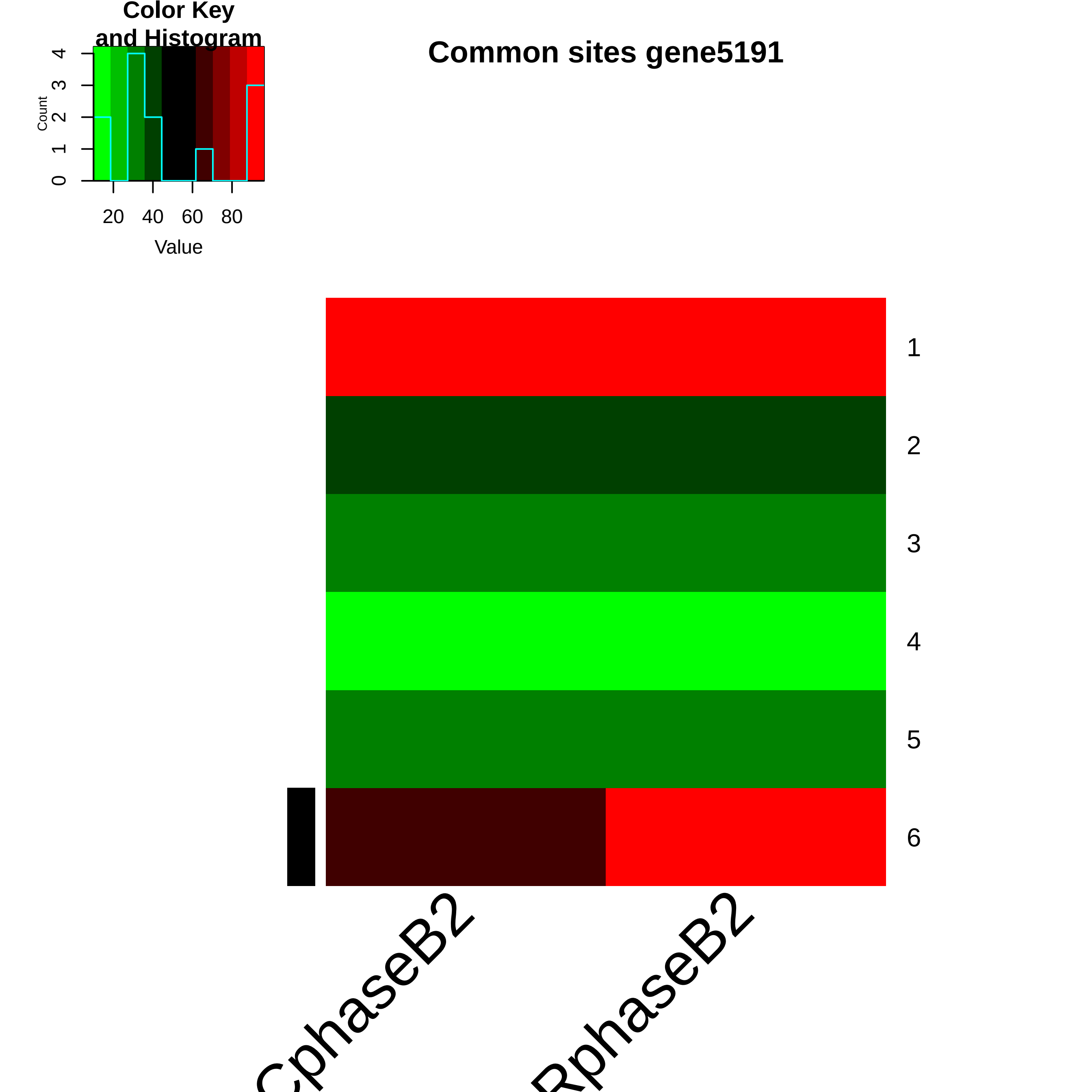
Color Key
and Histogram
Count
0
1
2
3
4
20
40
60
80
Value
Common sites gene5191
1
2
3
4
5
6
CphaseB2
RphaseB2

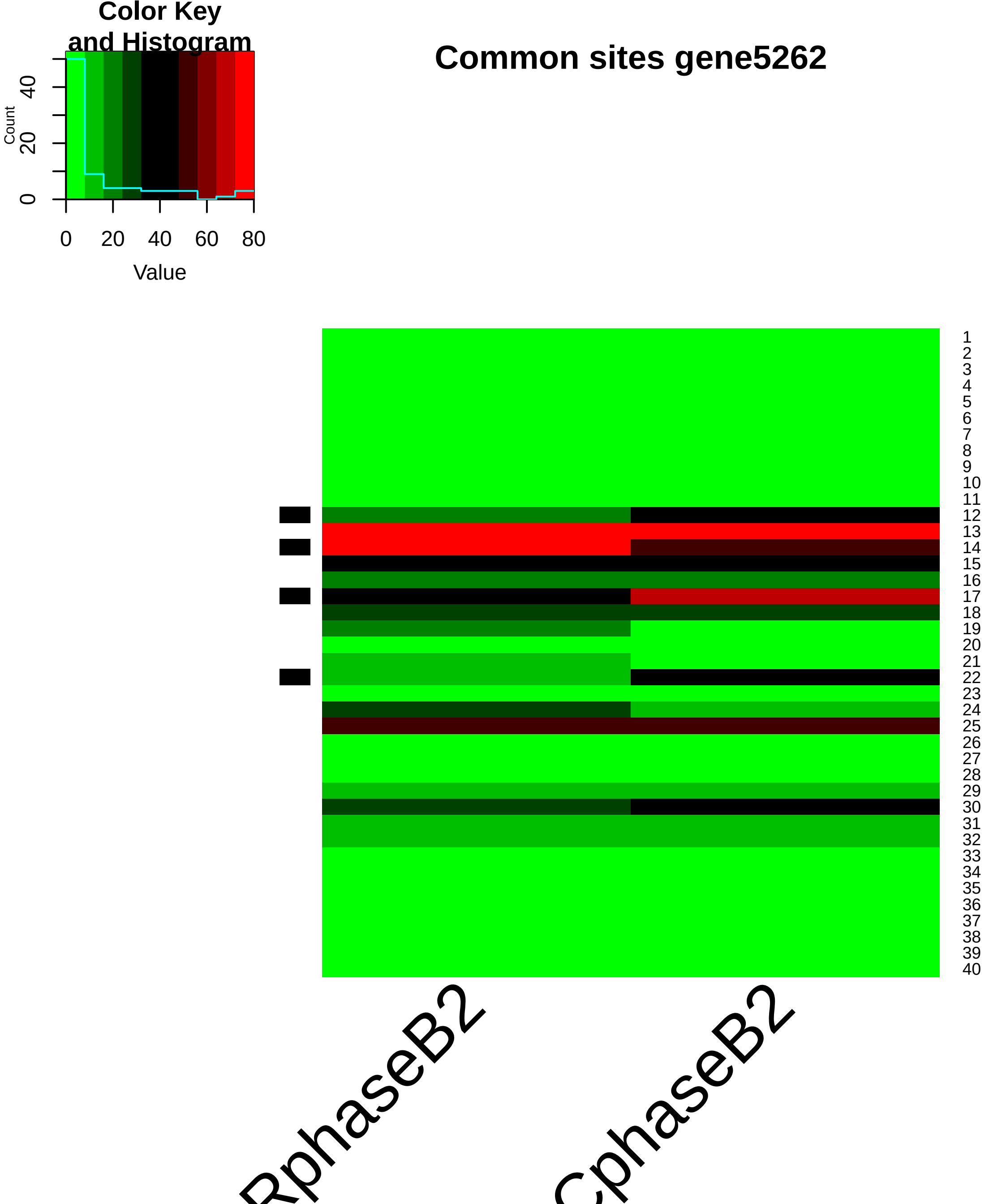
Color Key
and Histogram
Count
0
20
40
0
20
40
60
80
Value
Common sites gene5262
RphaseB2
CphaseB2

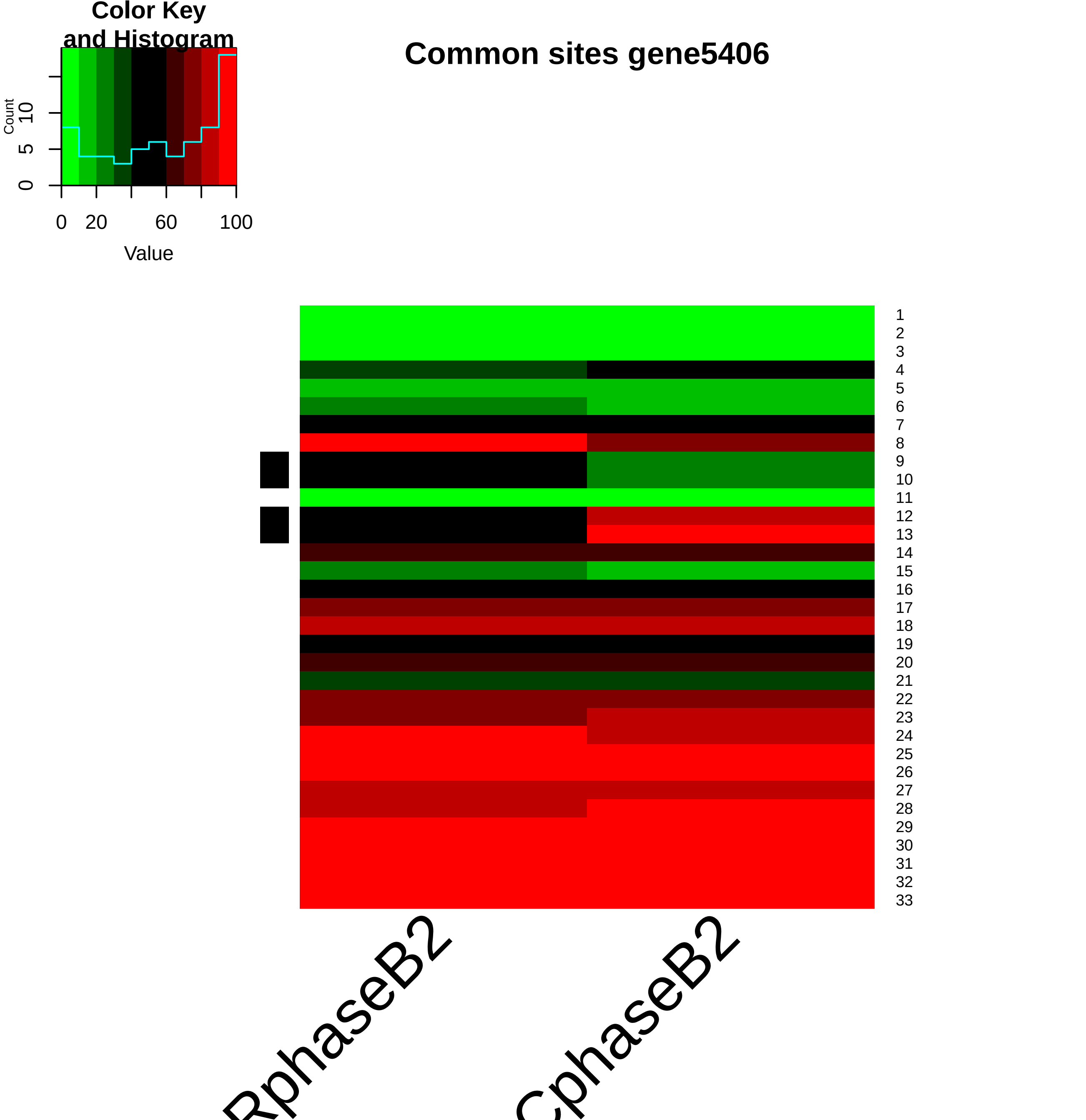

Color Key
and Histogram
Count
0
5
10
0
20
60
100
Value
Common sites gene5406
RphaseB2
CphaseB2

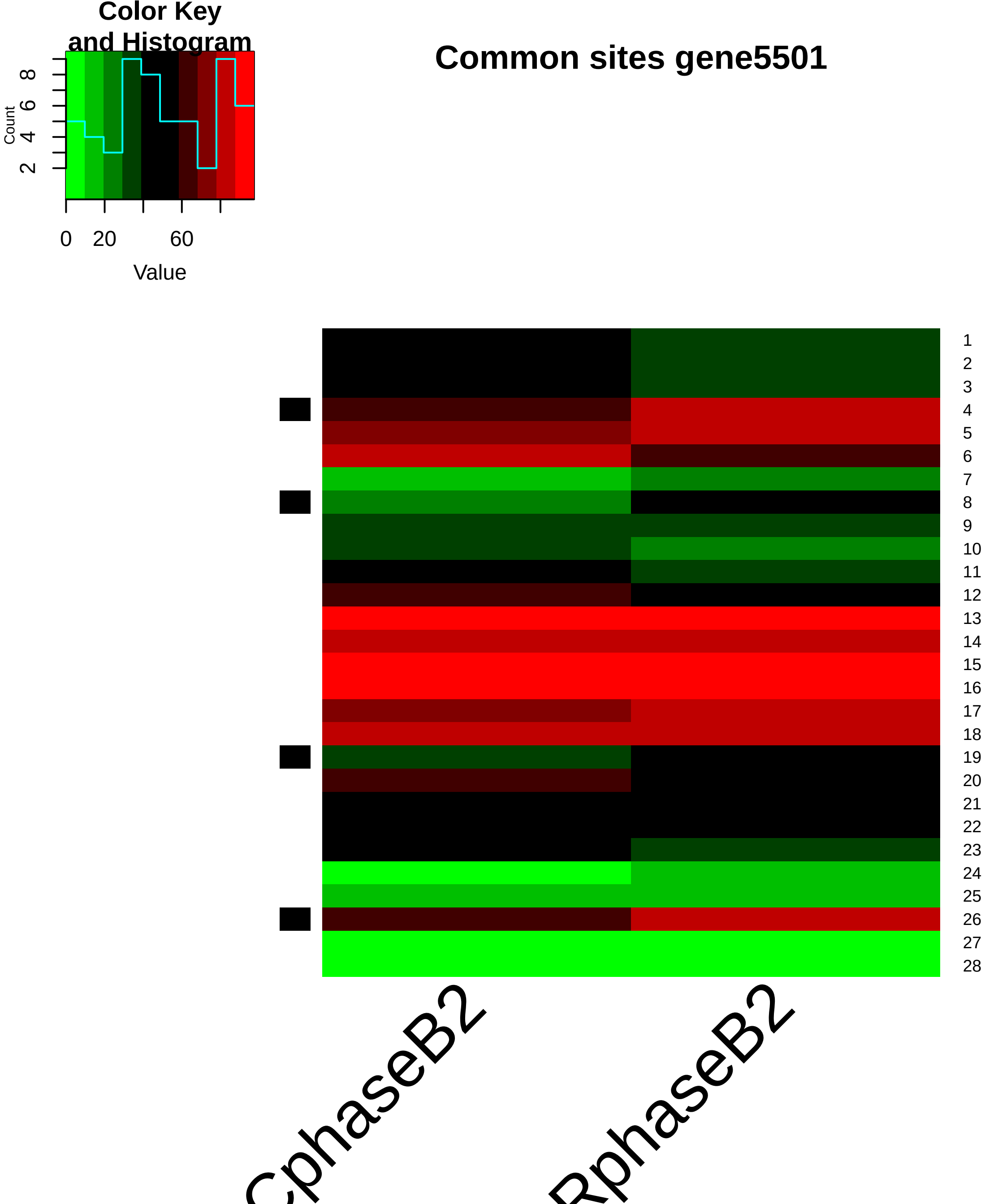
Color Key
and Histogram
Count
2
4
6
8
0
20
60
Value
Common sites gene5501
CphaseB2
RphaseB2

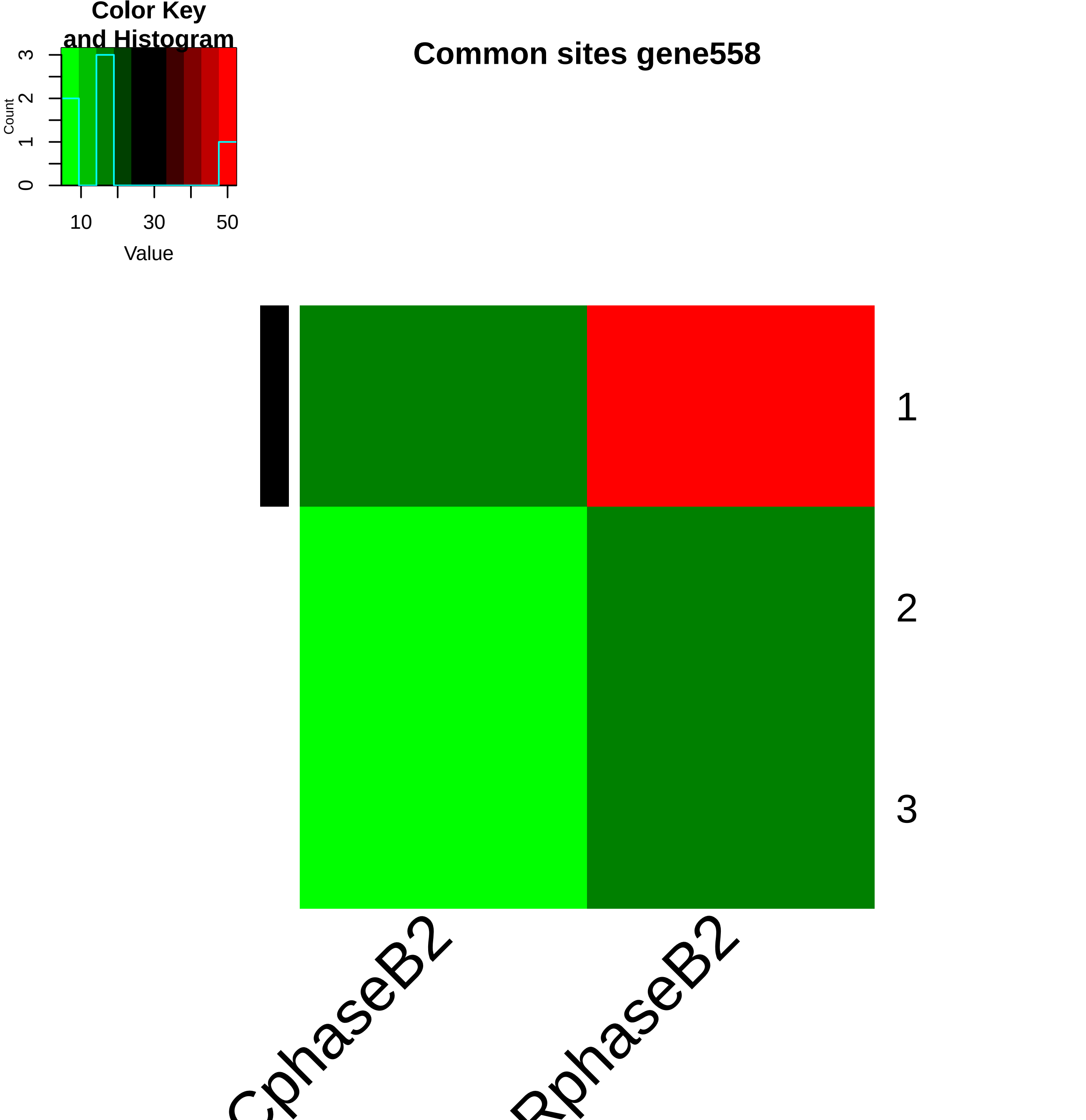
Color Key
and Histogram
Count
0
1
2
3
10
30
50
Value
Common sites gene558
1
2
3
CphaseB2
RphaseB2

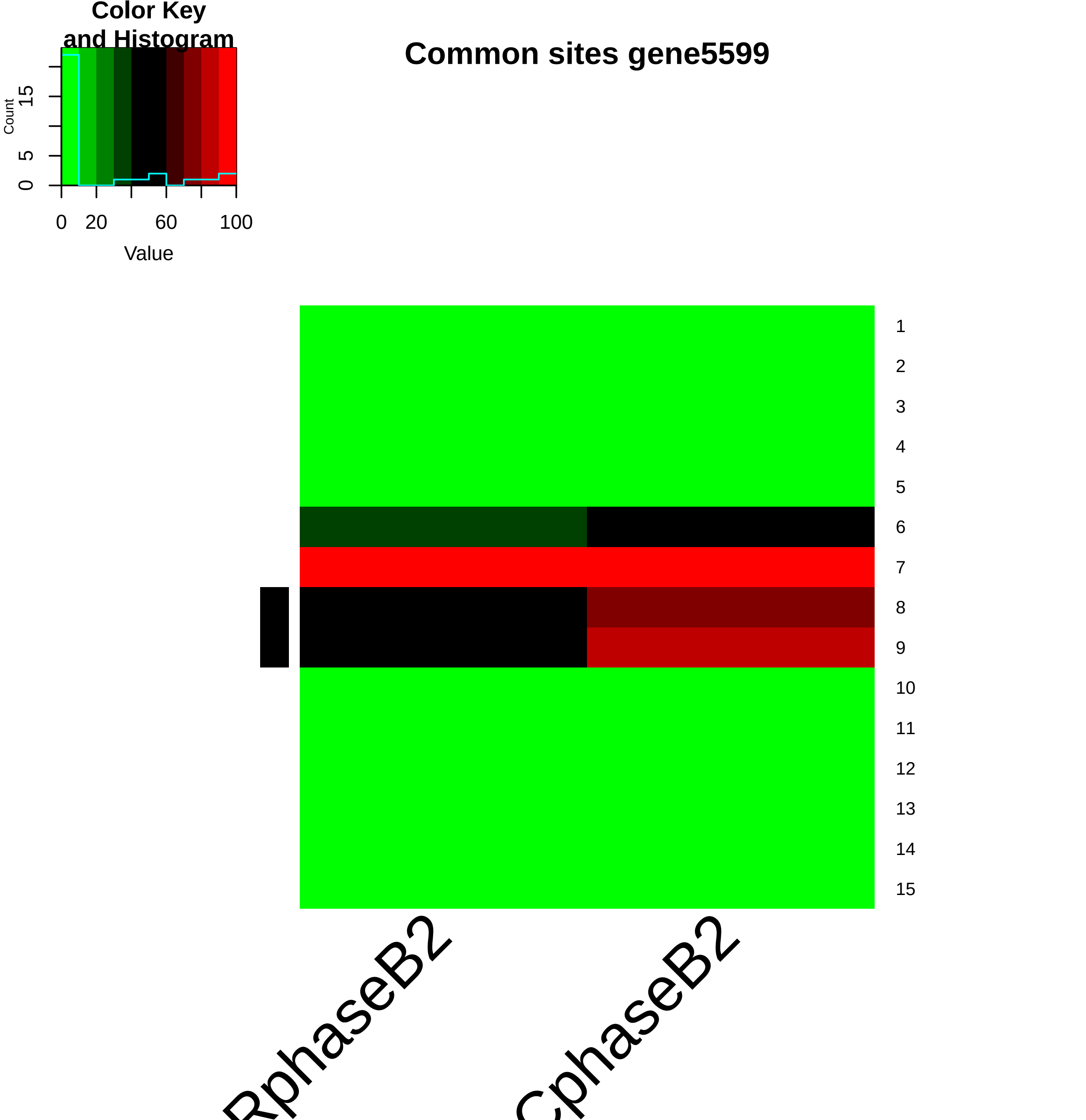

Color Key
and Histogram
Count
0
5
15
0
20
60
100
Value
Common sites gene5599
RphaseB2
CphaseB2

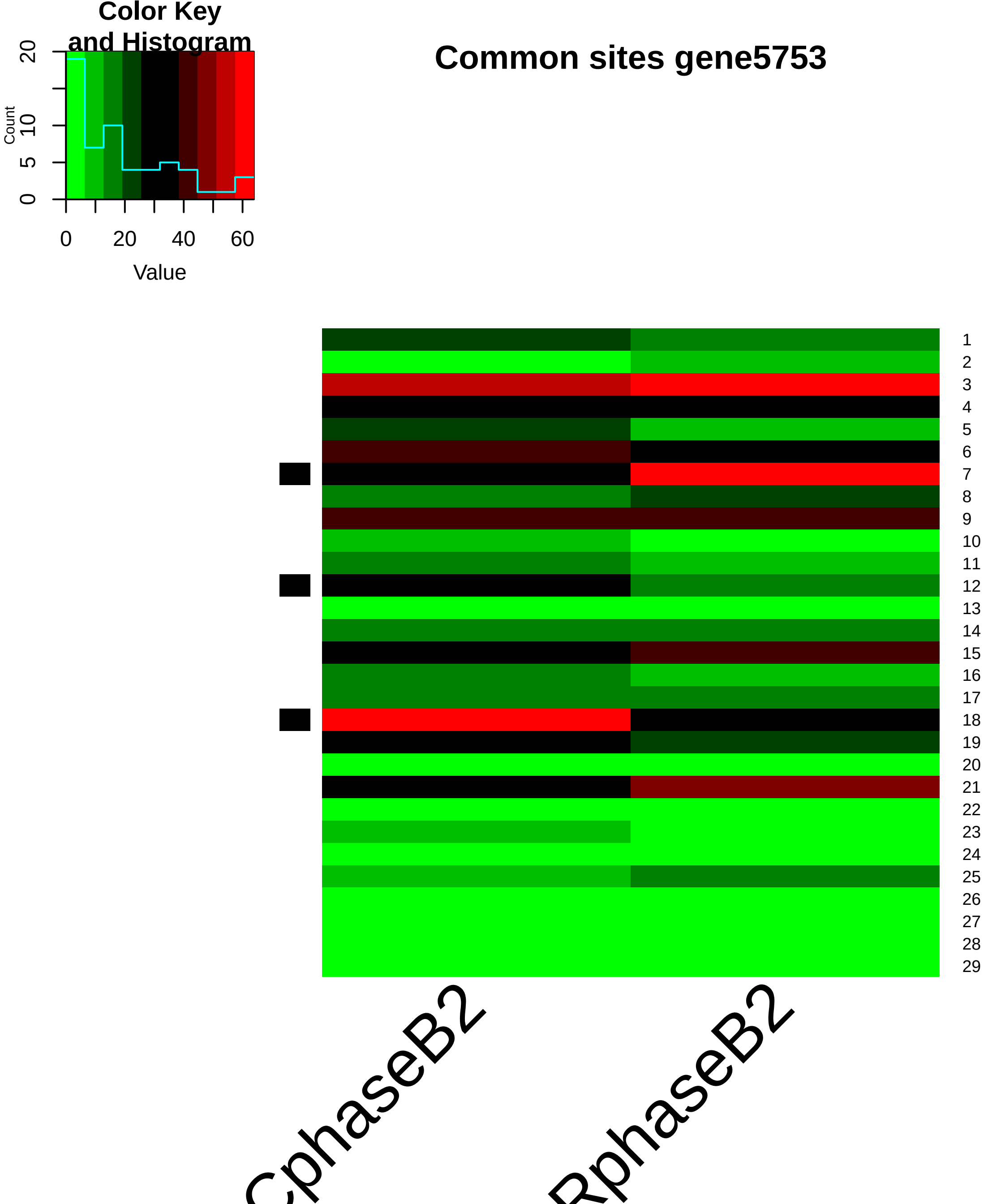
Color Key
and Histogram
Count
Value
Common sites gene5753
CphaseB2
RphaseB2

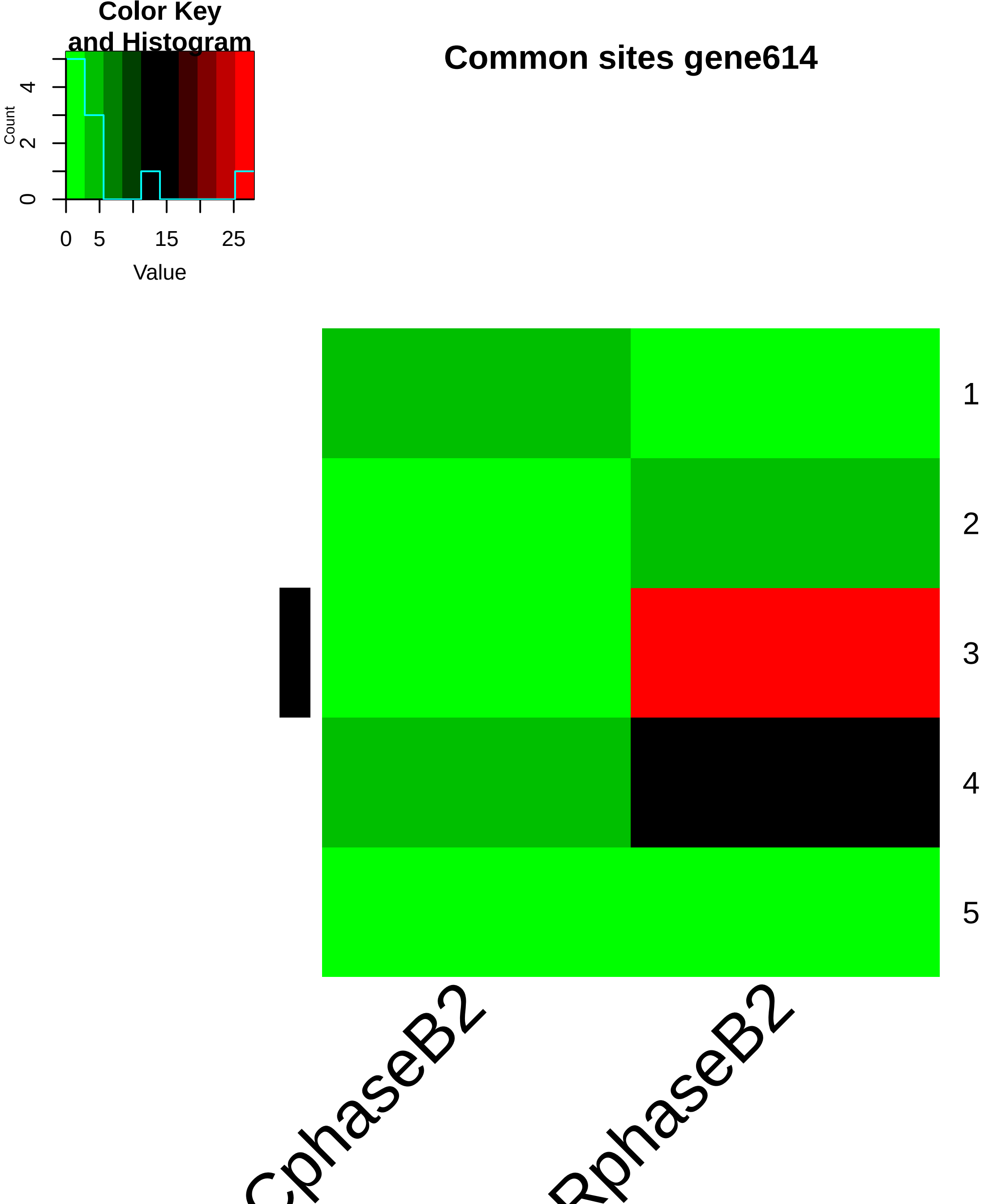
Color Key
and Histogram
Count
0
2
4
0
5
15
25
Value
Common sites gene614
1
2
3
4
5
CphaseB2
RphaseB2

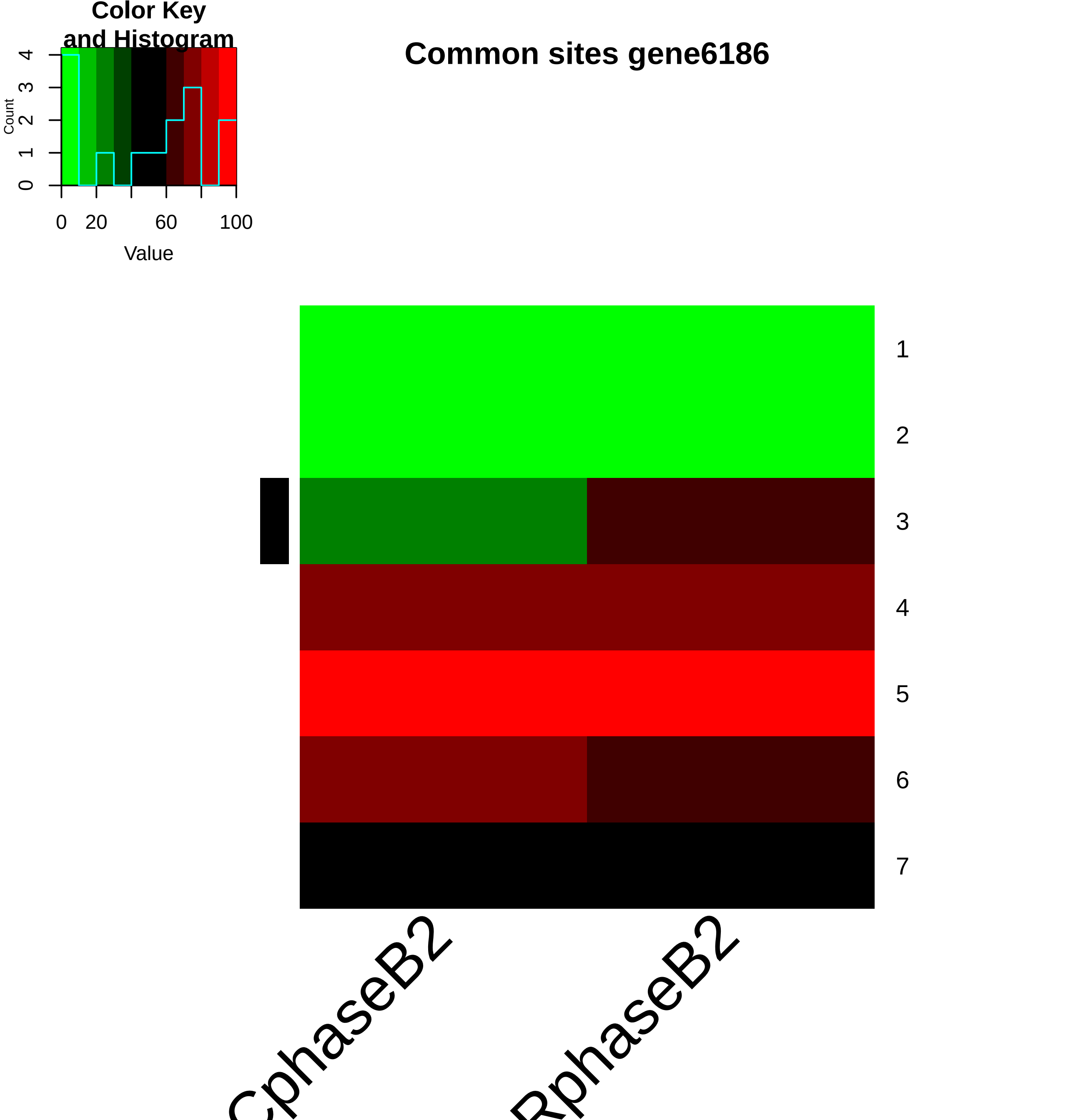

Color Key
and Histogram
Count
0
1
2
3
4
0
20
60
100
Value
Common sites gene6186
1
2
3
4
5
6
7
CphaseB2
RphaseB2

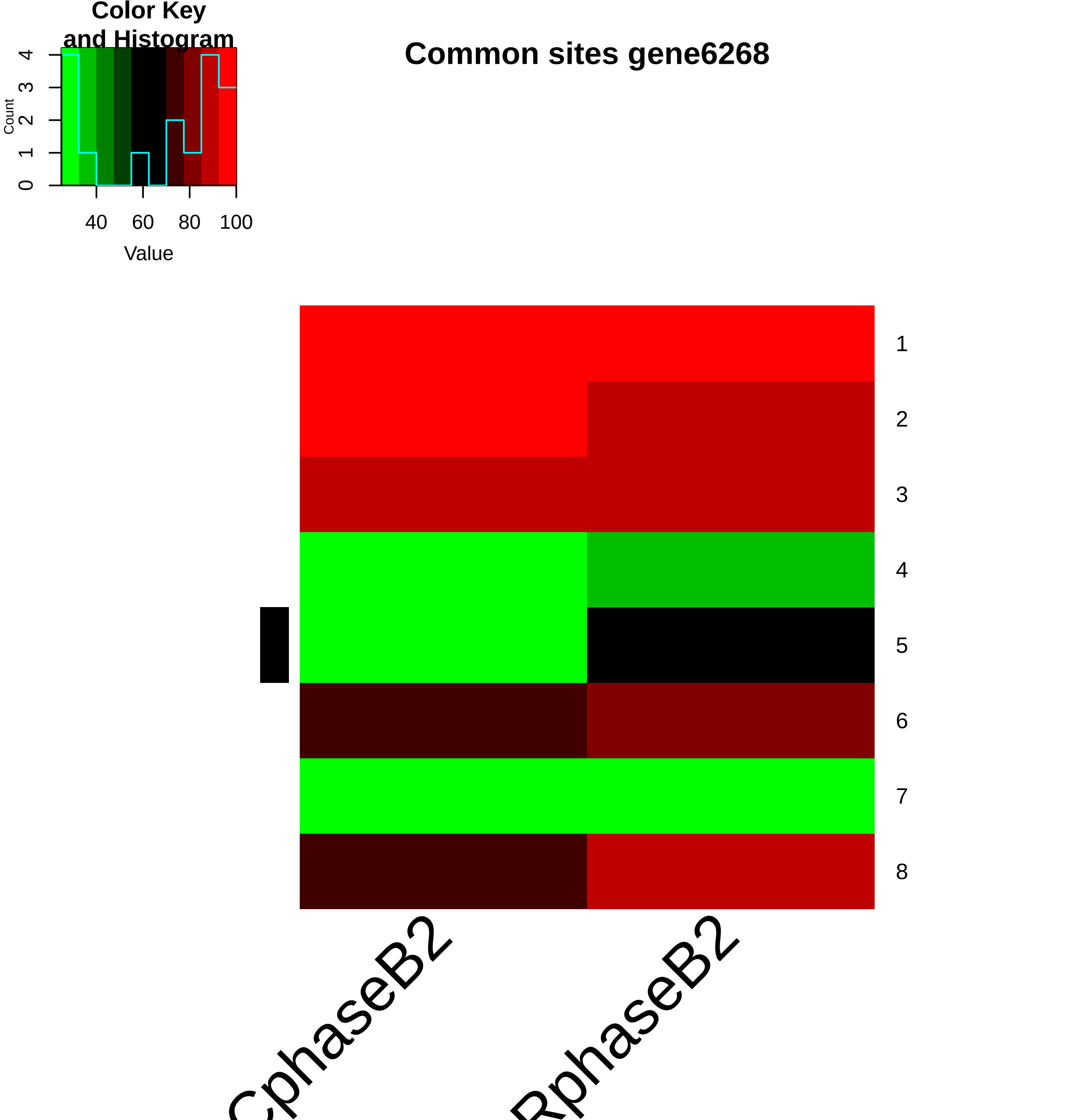

Color Key
and Histogram
Count
0
1
2
3
4
40
60
80
100
Value
Common sites gene6268
1
2
3
4
5
6
7
8
CphaseB2
RphaseB2

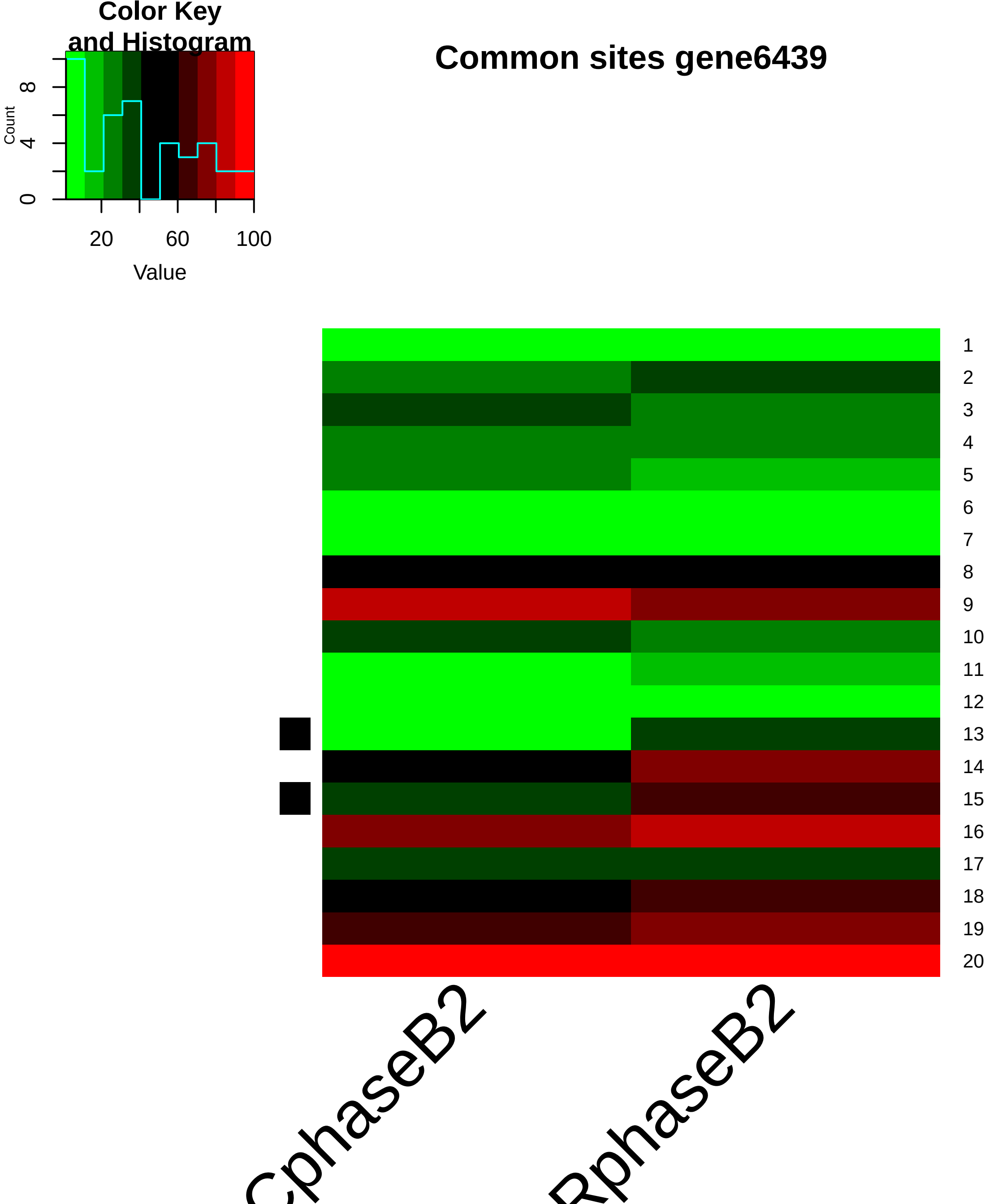
Color Key
and Histogram
Count
0
4
8
20
60
100
Value
Common sites gene6439
CphaseB2
RphaseB2

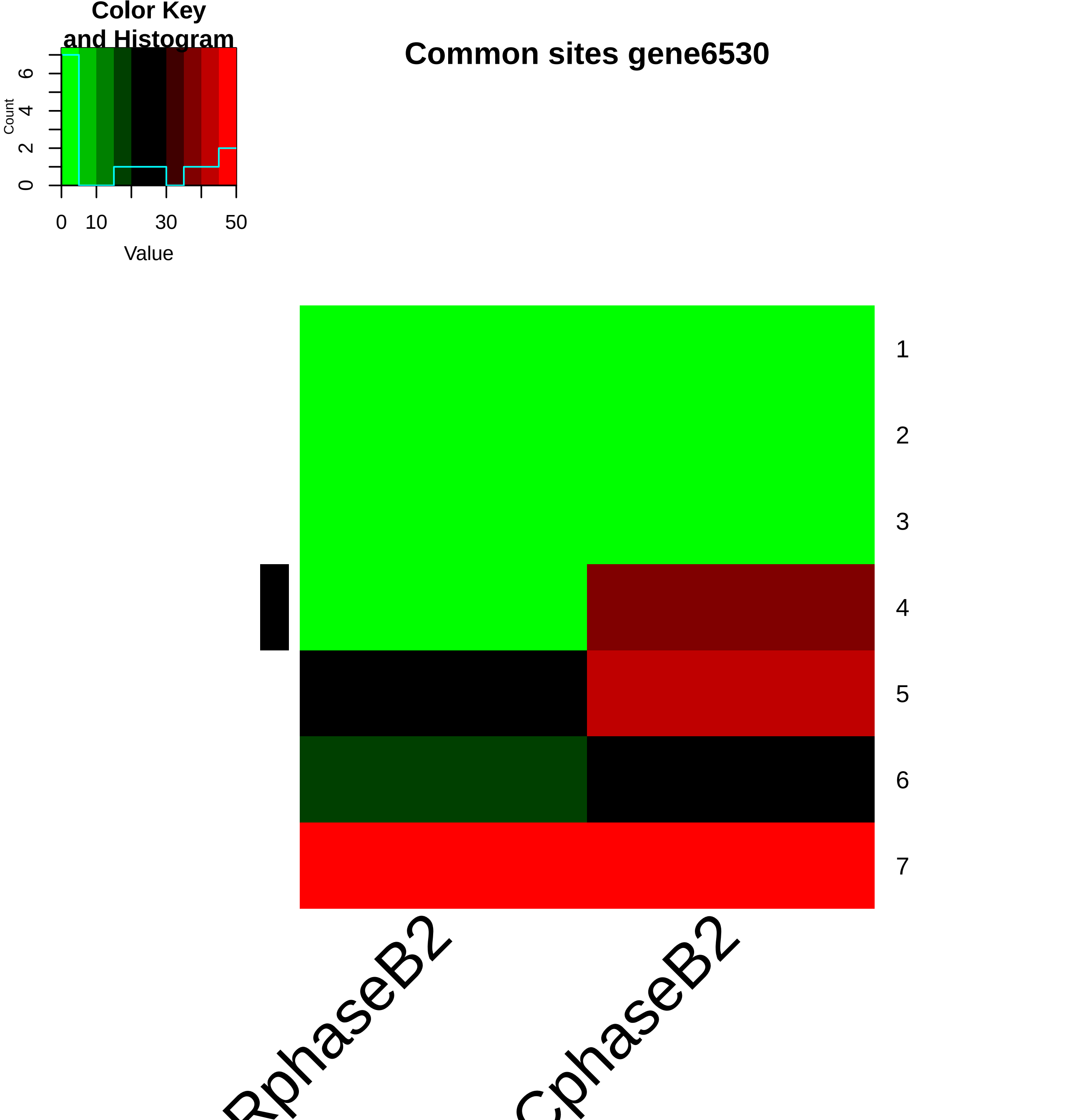

Color Key
and Histogram
Count
0
2
4
6
0
10
30
50
Value
Common sites gene6530
1
2
3
4
5
6
7
RphaseB2
CphaseB2

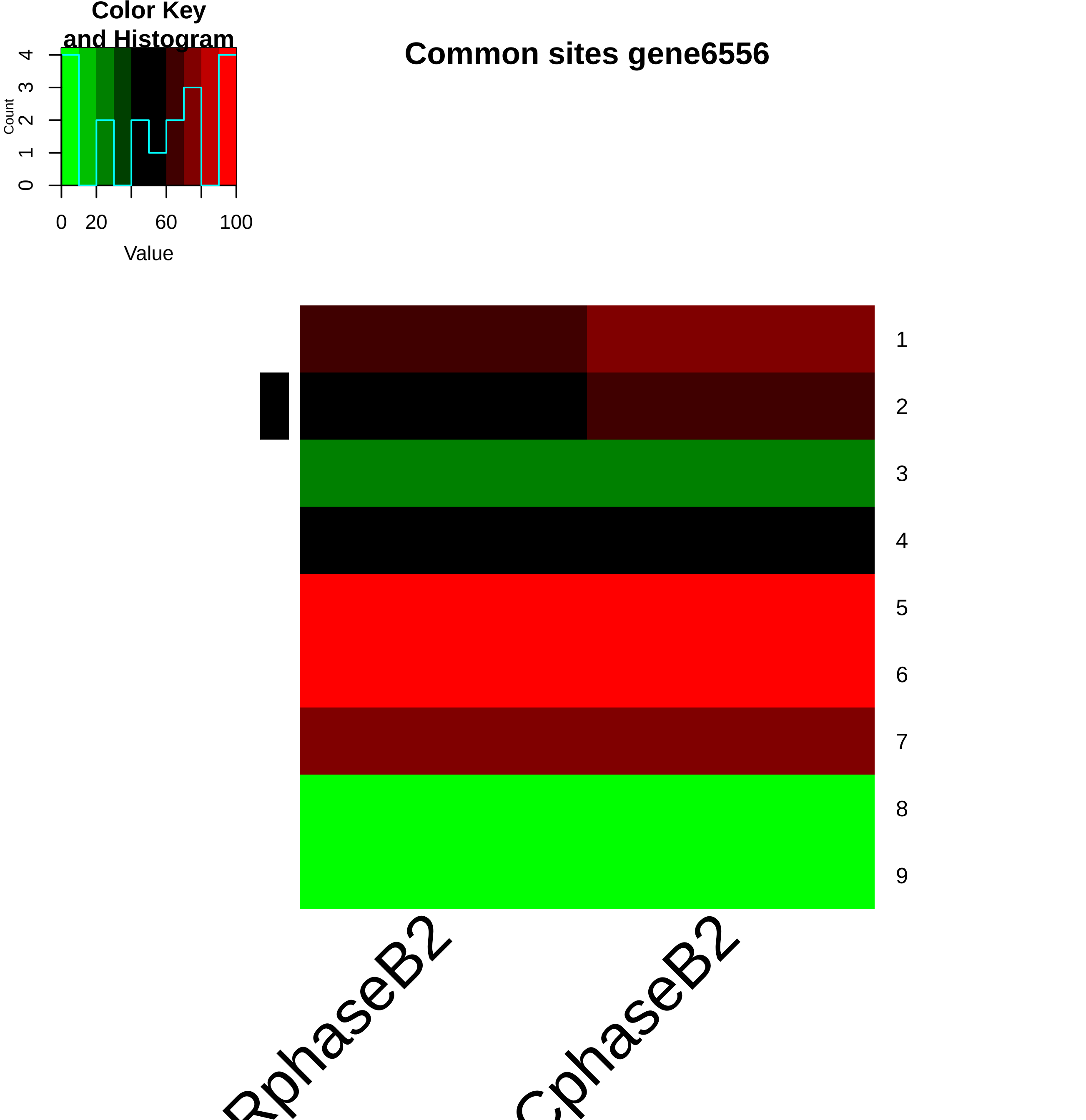
Color Key
and Histogram
Count
0
1
2
3
4
0
20
60
100
Value
Common sites gene6556
1
2
3
4
5
6
7
8
9
RphaseB2
CphaseB2

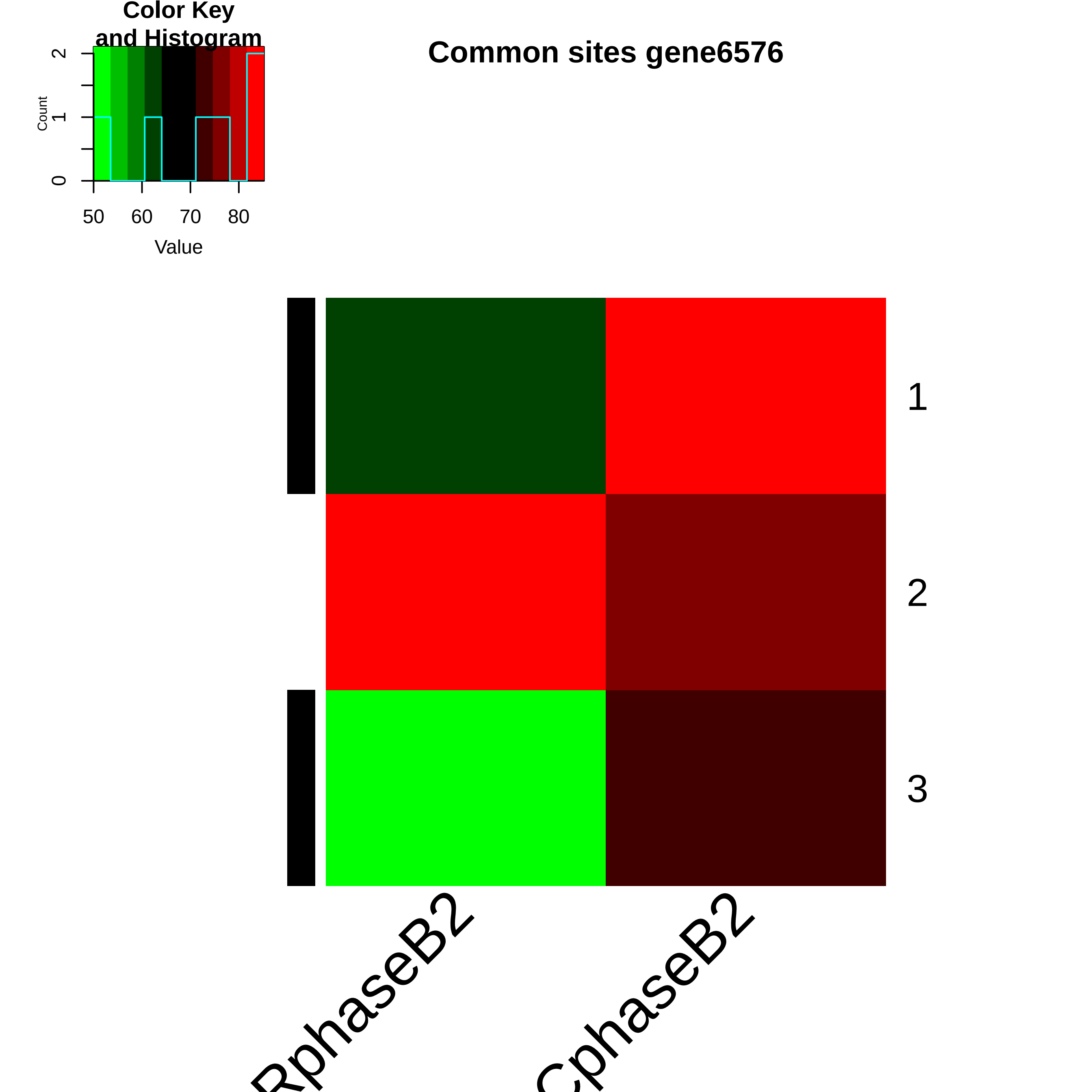
Color Key
and Histogram
Count
0
1
2
50
60
70
80
Value
Common sites gene6576
1
2
3
RphaseB2
CphaseB2

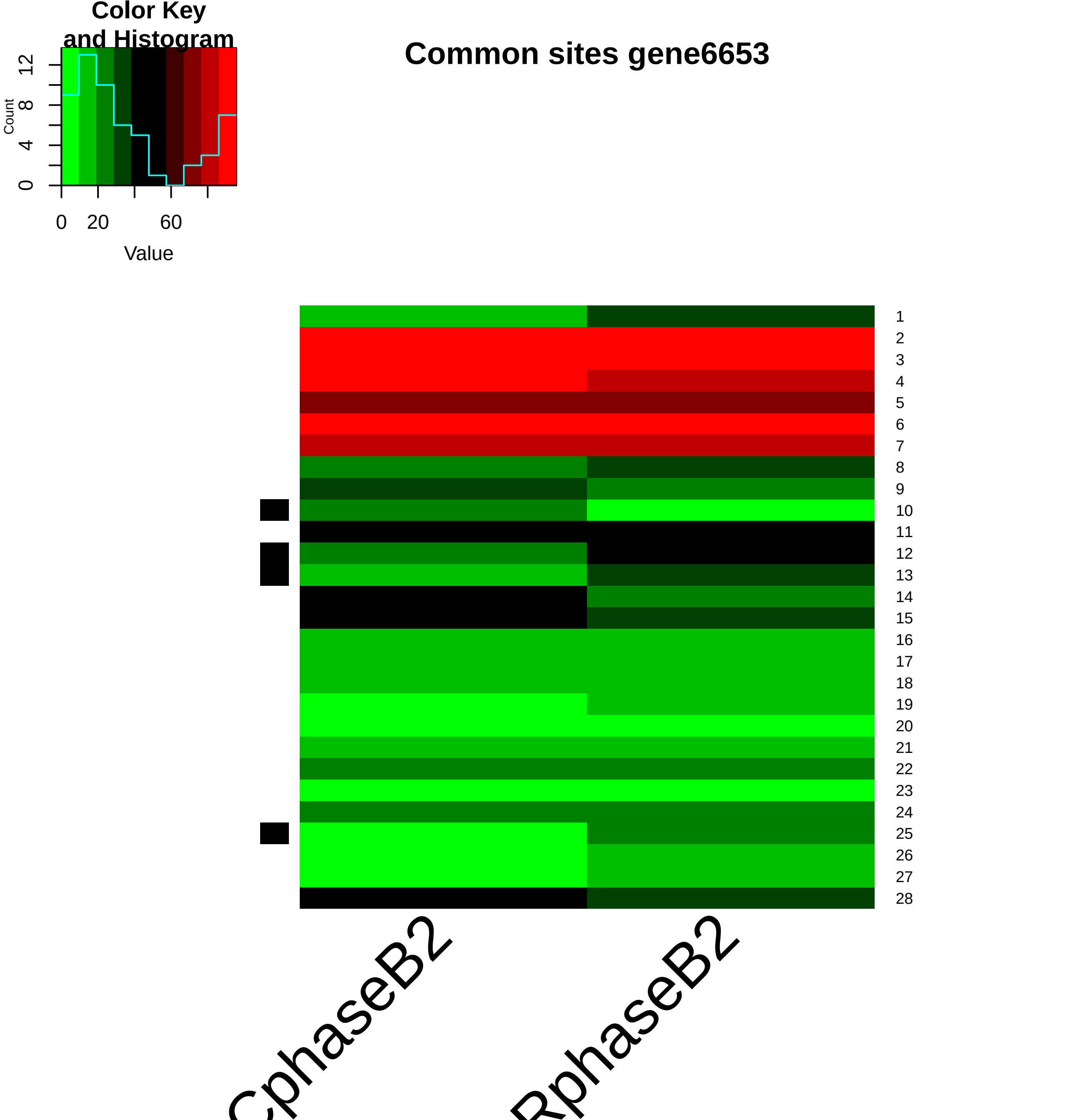

Color Key
and Histogram
Count
0
4
8
12
0
20
60
Value
Common sites gene6653
CphaseB2
RphaseB2

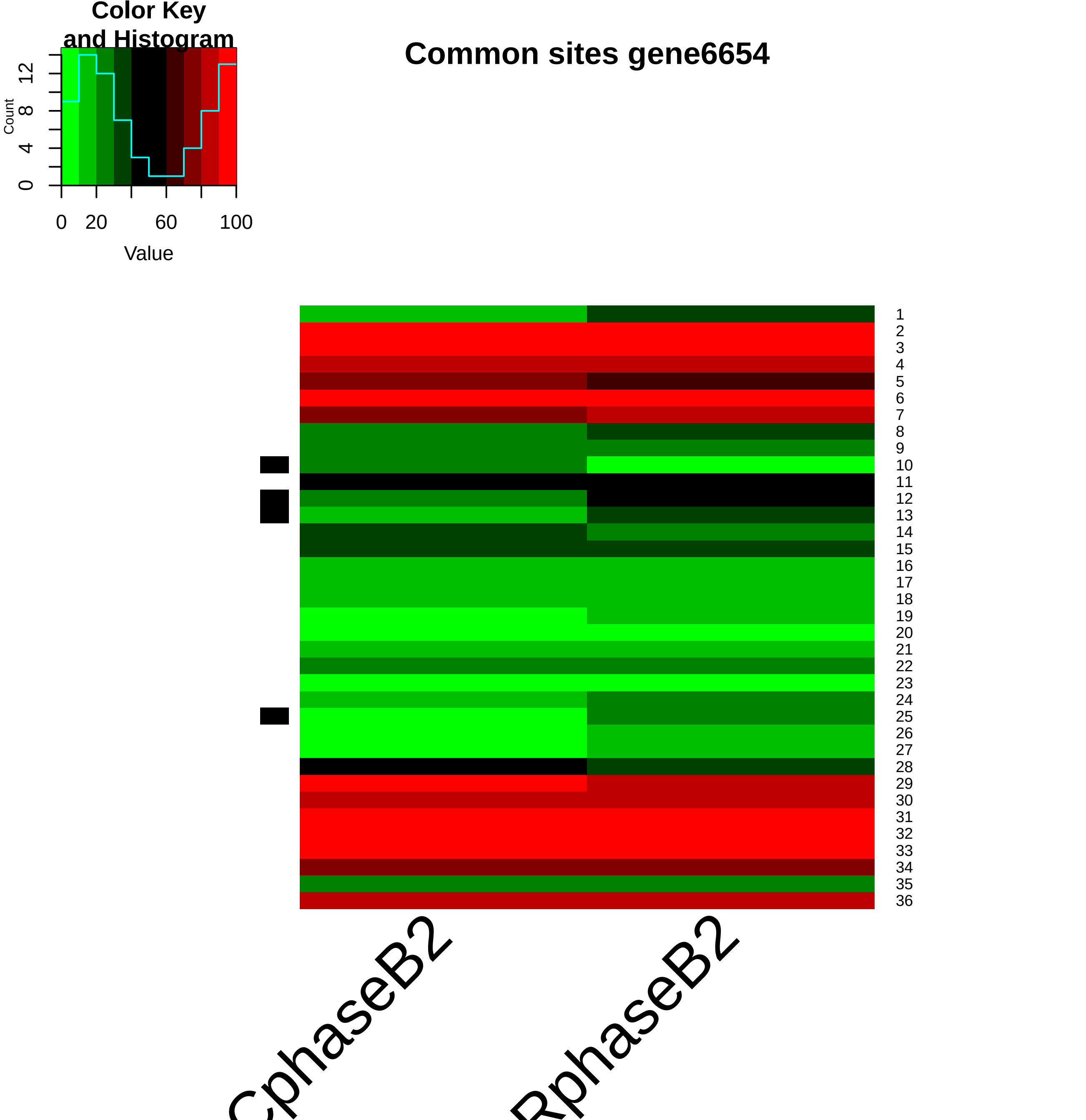

Color Key
and Histogram
Count
0
4
8
12
0
20
60
100
Value
Common sites gene6654
CphaseB2
RphaseB2

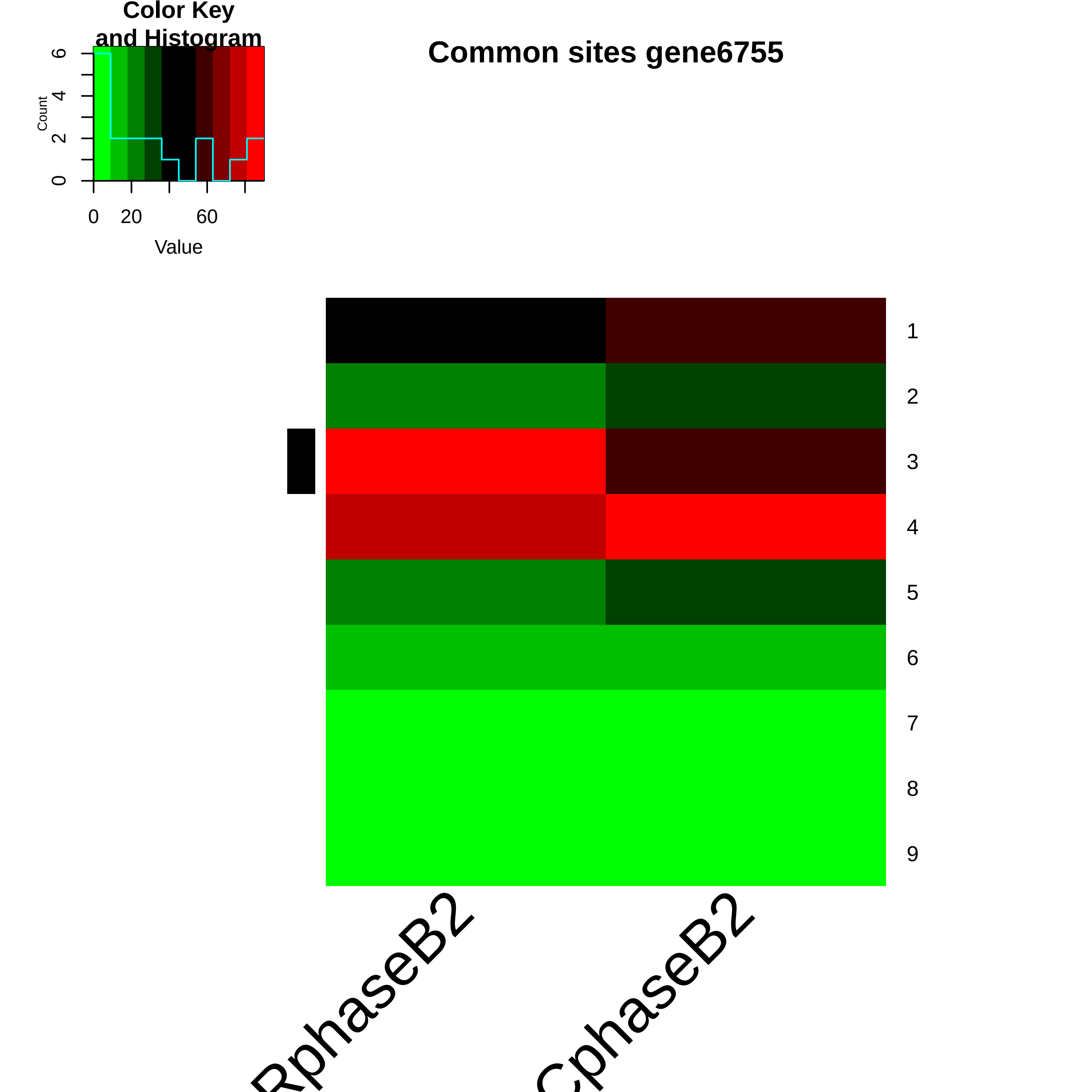

Color Key
and Histogram
Count
0
2
4
6
0
20
60
Value
Common sites gene6755
1
2
3
4
5
6
7
8
9
RphaseB2
CphaseB2

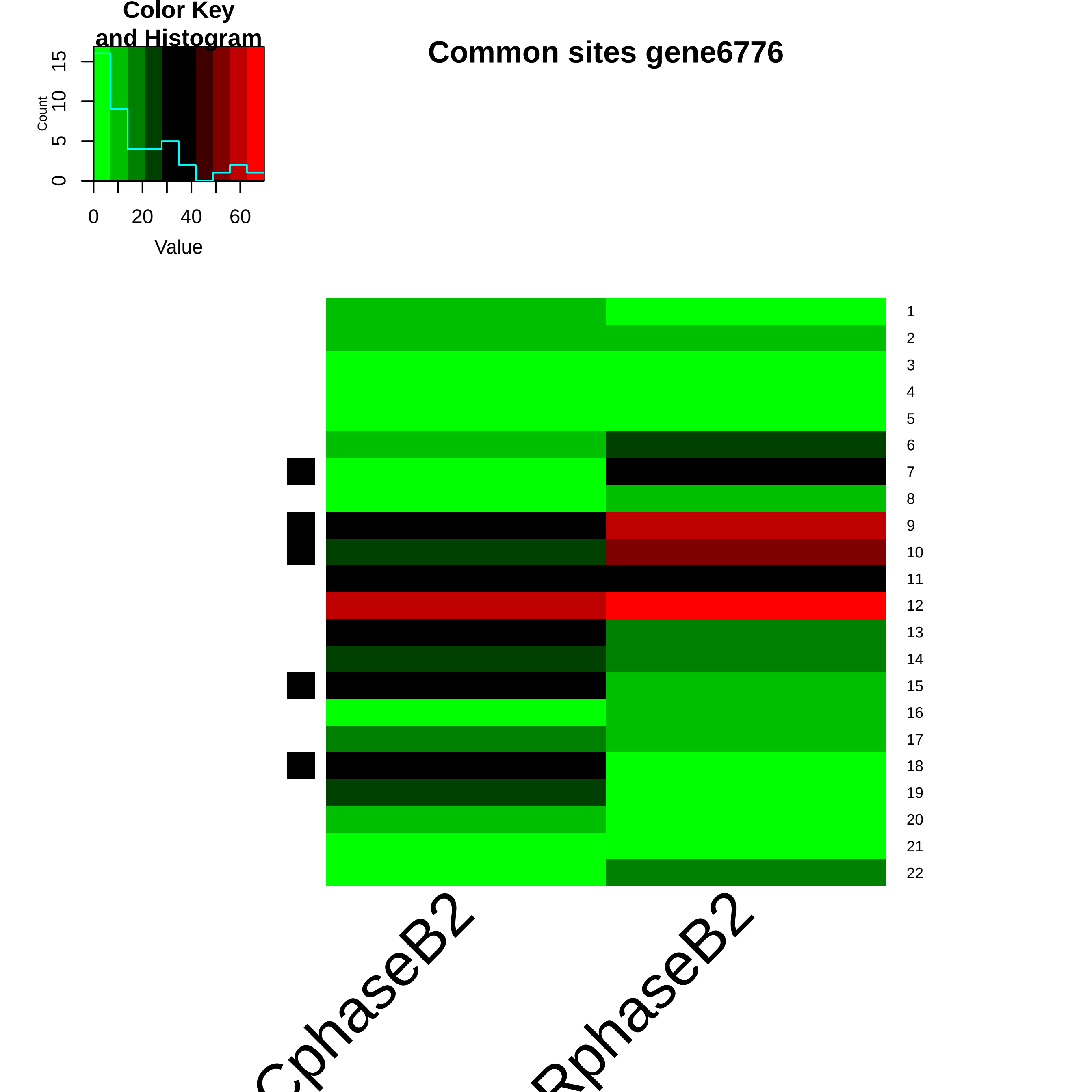

Color Key and Histogram
Count
0
5
10
15
0
20
40
60
Value
Common sites gene6776
CphaseB2
RphaseB2

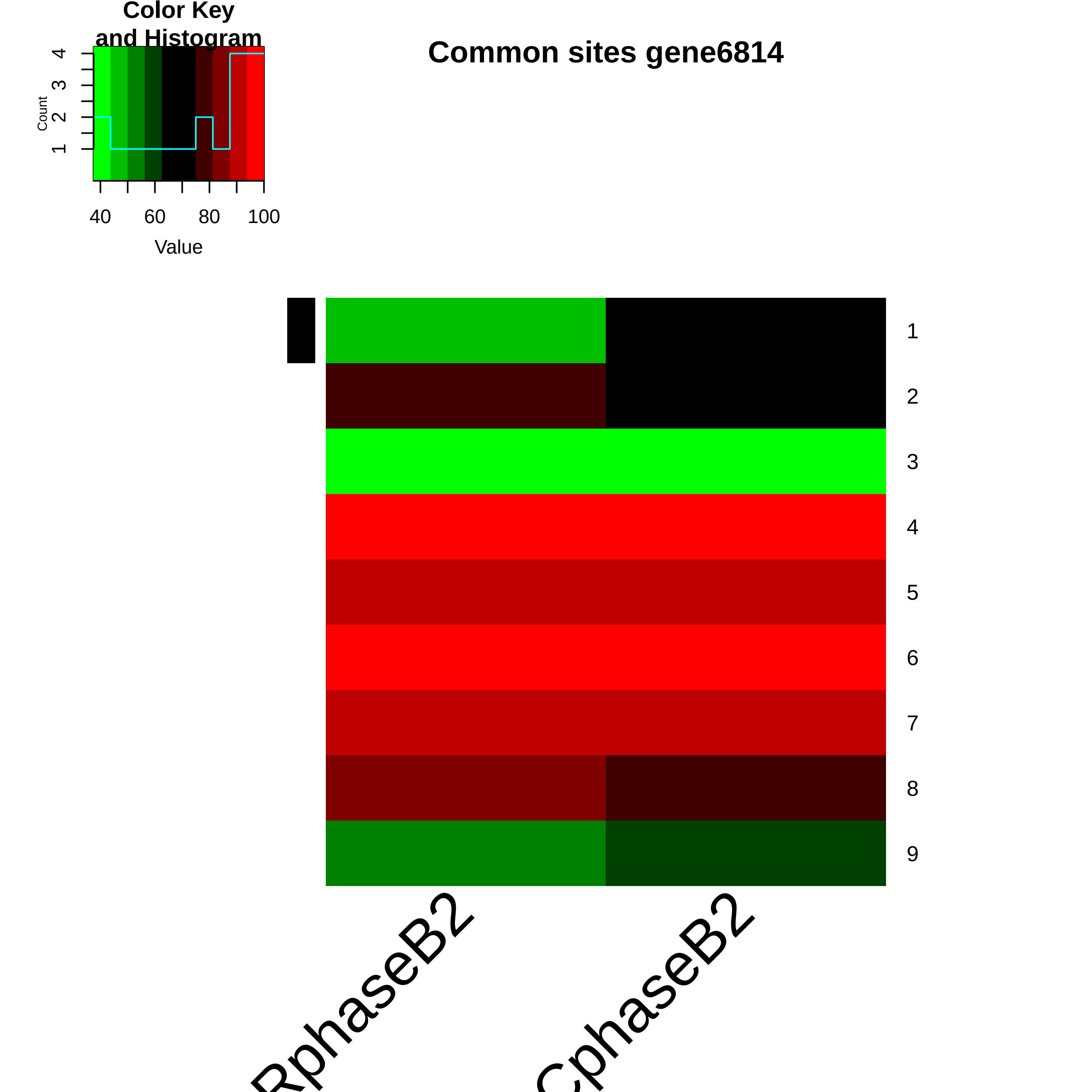
Color Key
and Histogram
Count
1
2
3
4
40
60
80
100
Value
Common sites gene6814
1
2
3
4
5
6
7
8
9

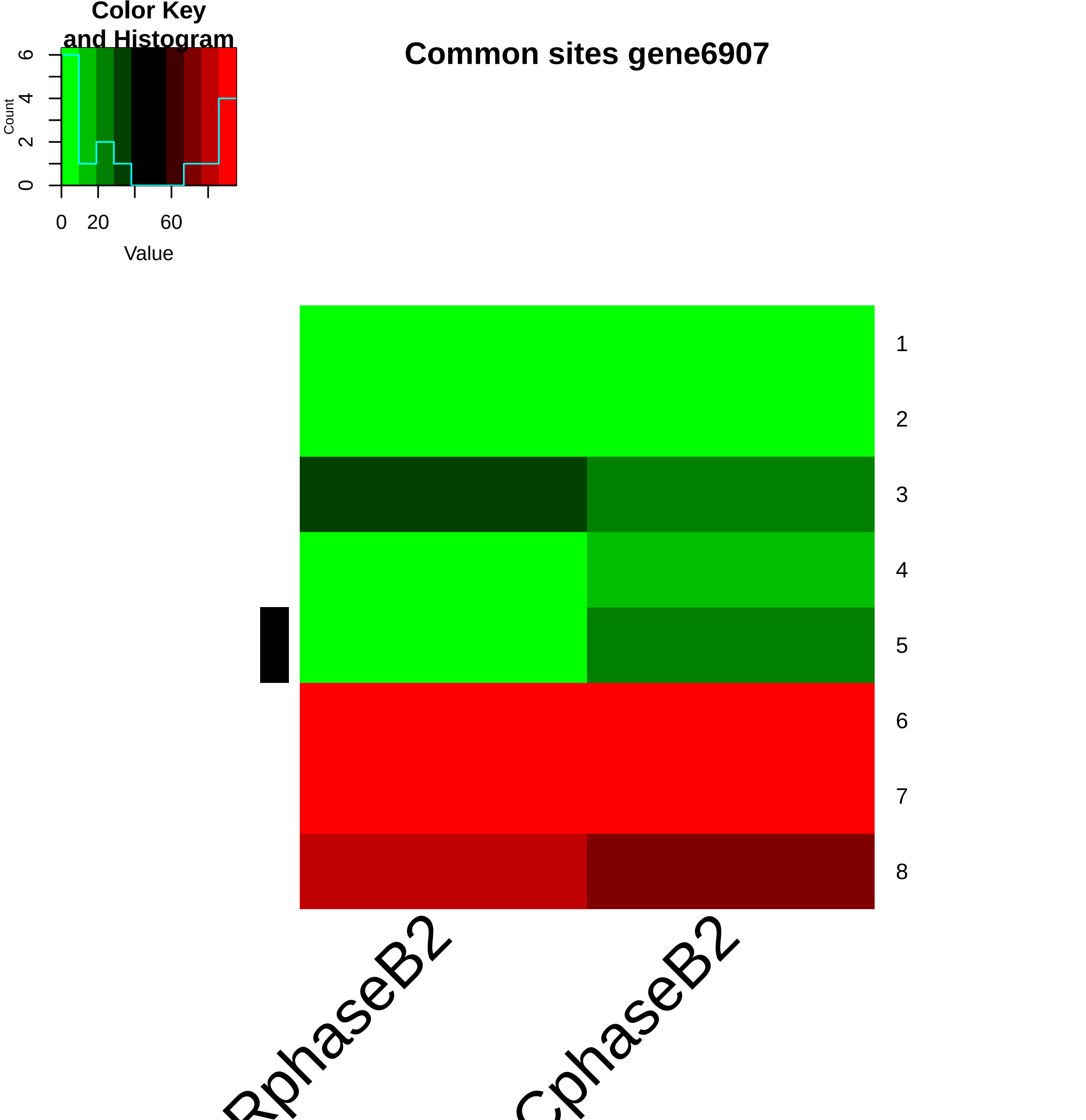
Color Key
and Histogram
Count
0
2
4
6
0
20
60
Value
Common sites gene6907
1
2
3
4
5
6
7
8
RphaseB2
CphaseB2

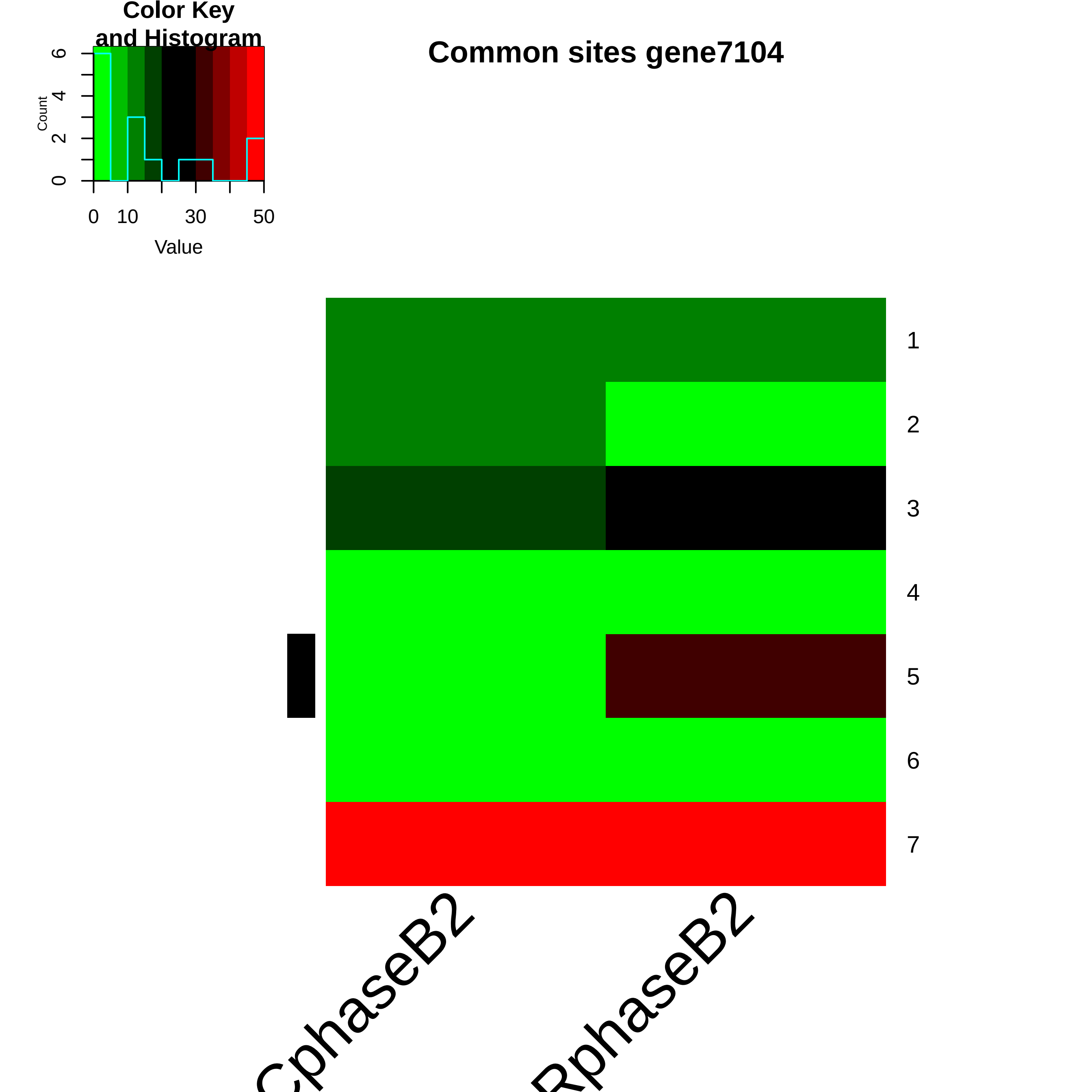

Color Key
and Histogram
Count
0
2
4
6
0
10
30
50
Value
Common sites gene7104
1
2
3
4
5
6
7
CphaseB2
RphaseB2

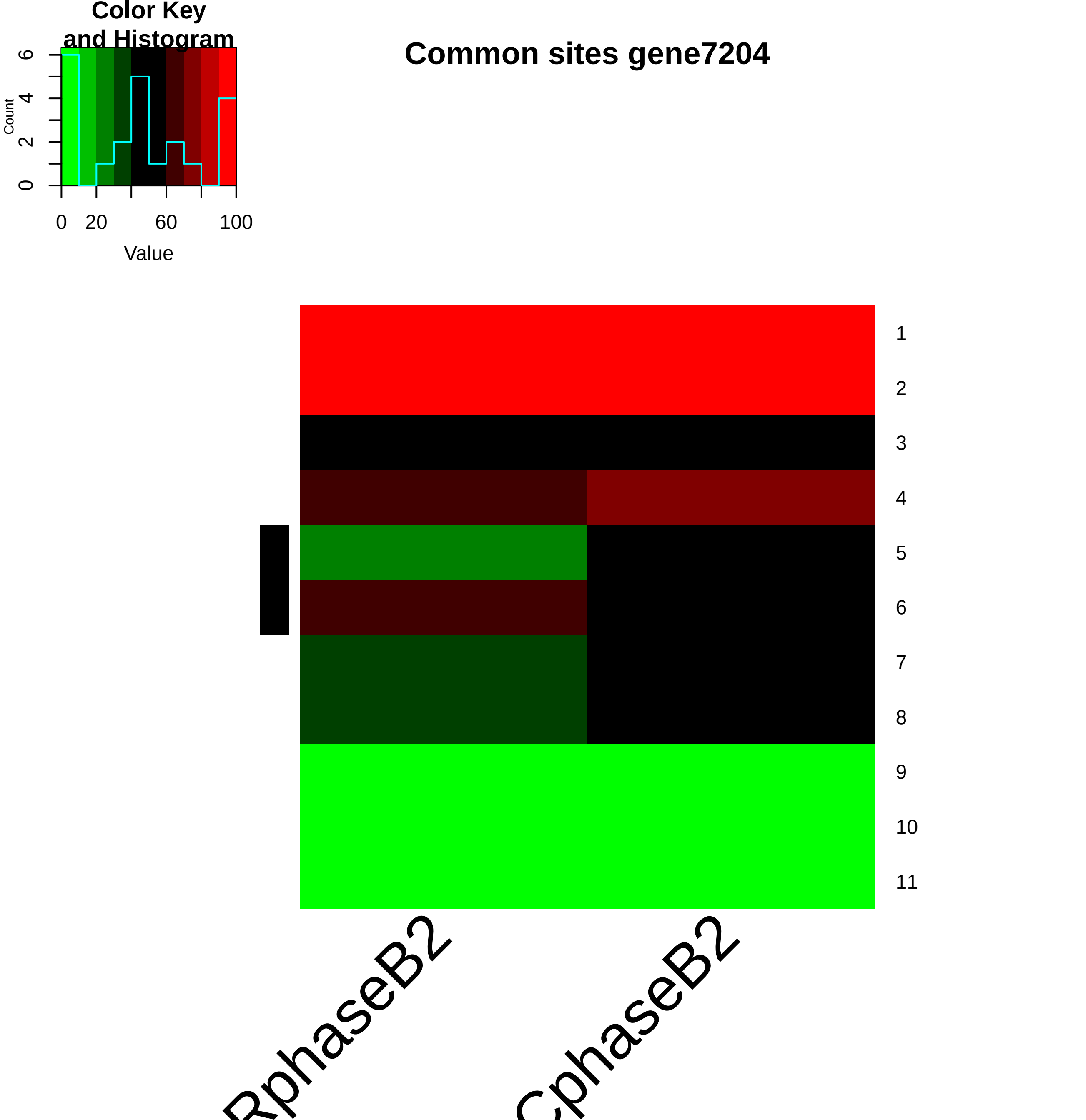

Color Key
and Histogram
Count
0
2
4
6
0
20
60
100
Value
Common sites gene7204
1
2
3
4
5
6
7
8
9
10
11
RphaseB2
CphaseB2

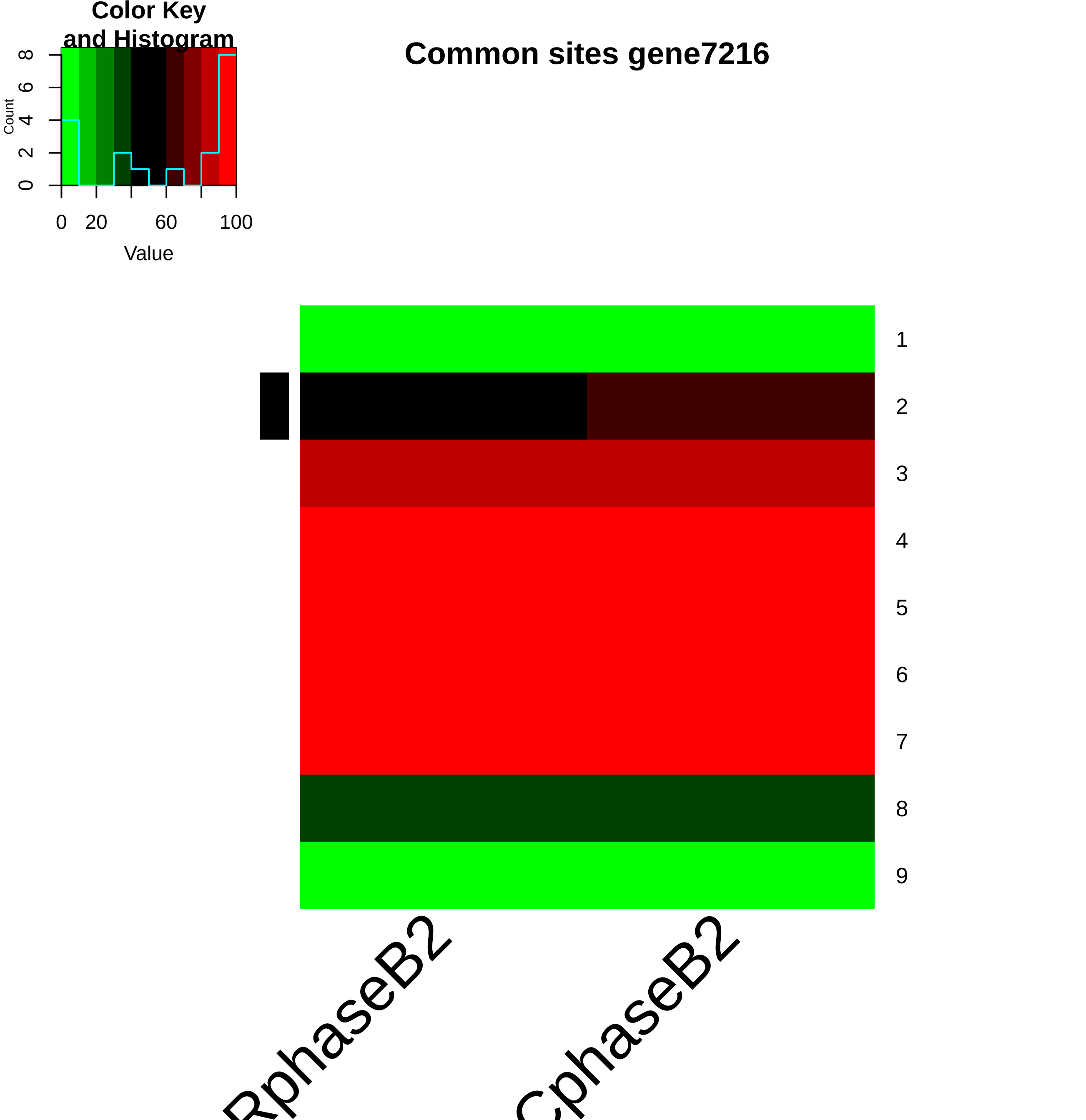

Color Key
and Histogram
Count
0
2
4
6
8
0
20
60
100
Value
Common sites gene7216
1
2
3
4
5
6
7
8
9
RphaseB2
CphaseB2

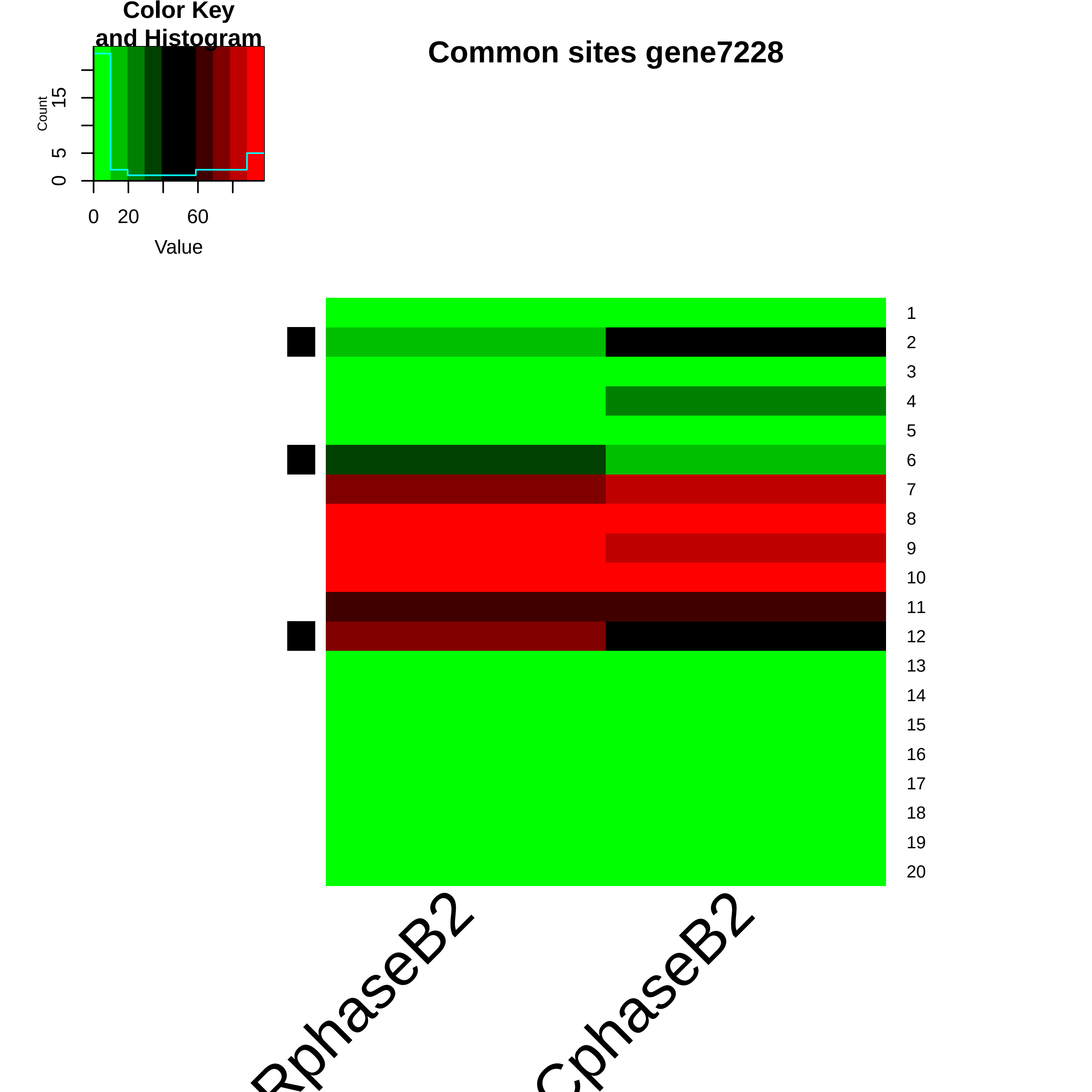
Color Key
and Histogram
Count
0
5
15
0
20
60
Value
Common sites gene7228
RphaseB2
CphaseB2

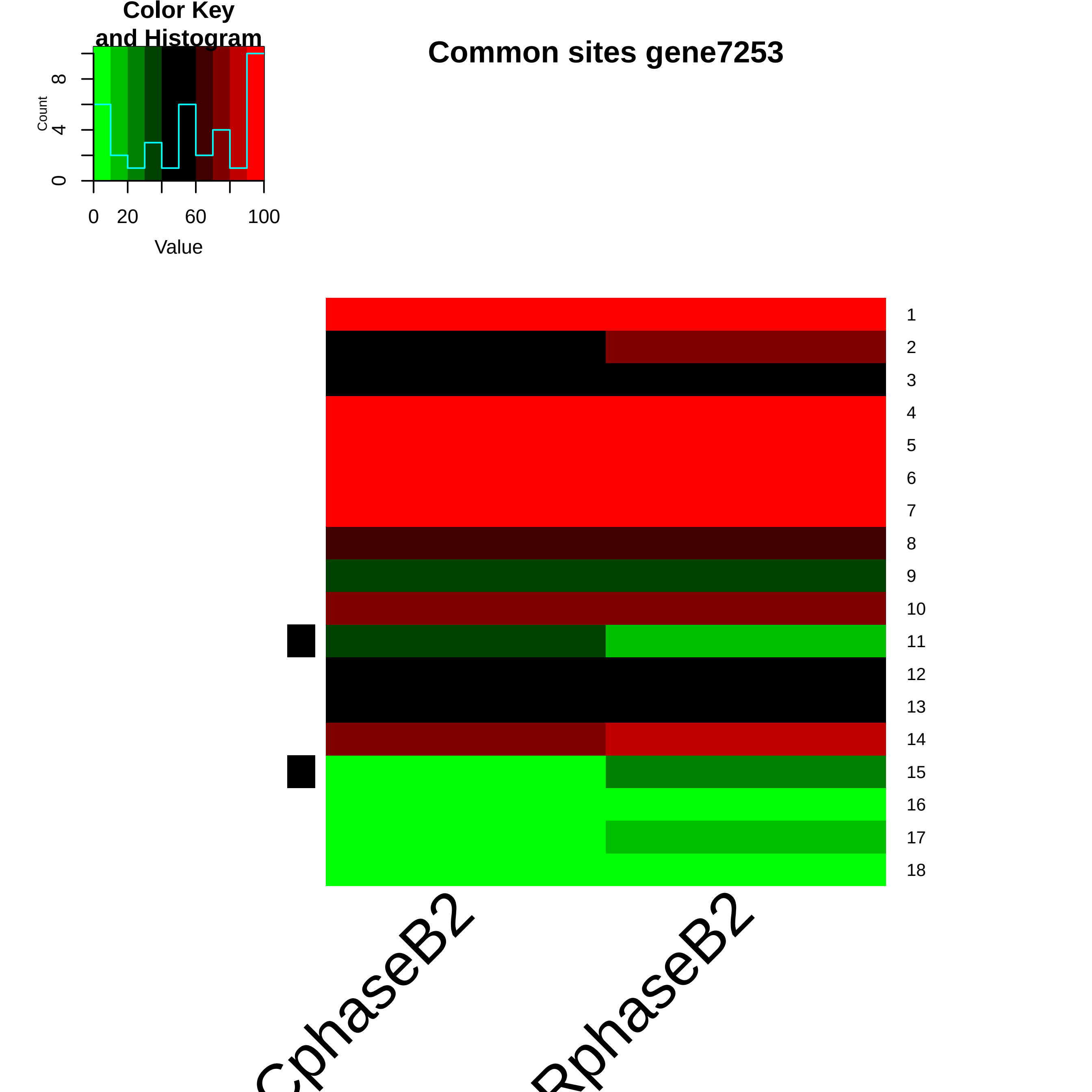

Color Key
and Histogram
Count
0
4
8
0
20
60
100
Value
Common sites gene7253
CphaseB2
RphaseB2

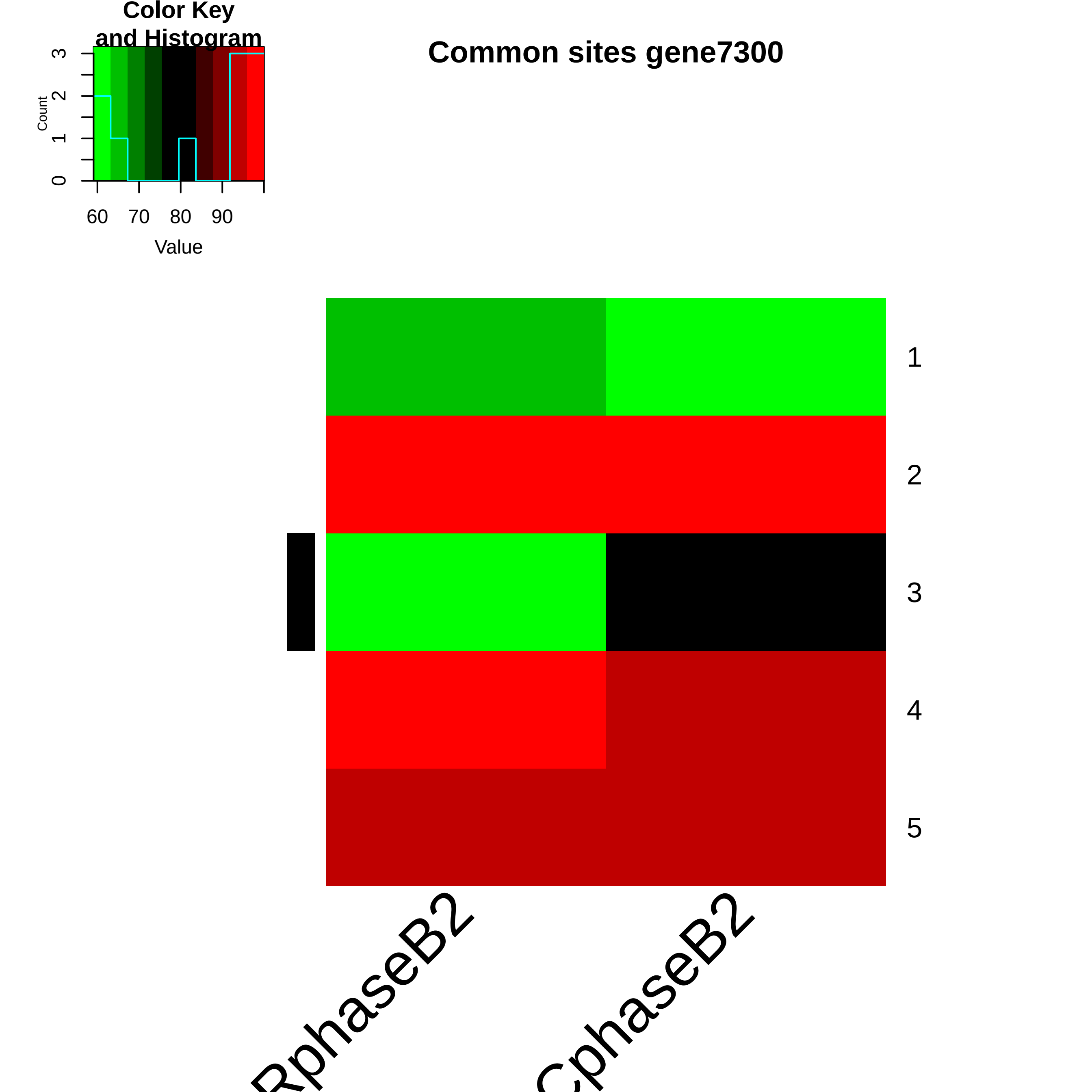
Color Key
and Histogram
Count
0
1
2
3
60
70
80
90
Value
Common sites gene7300
1
2
3
4
5
RphaseB2
CphaseB2

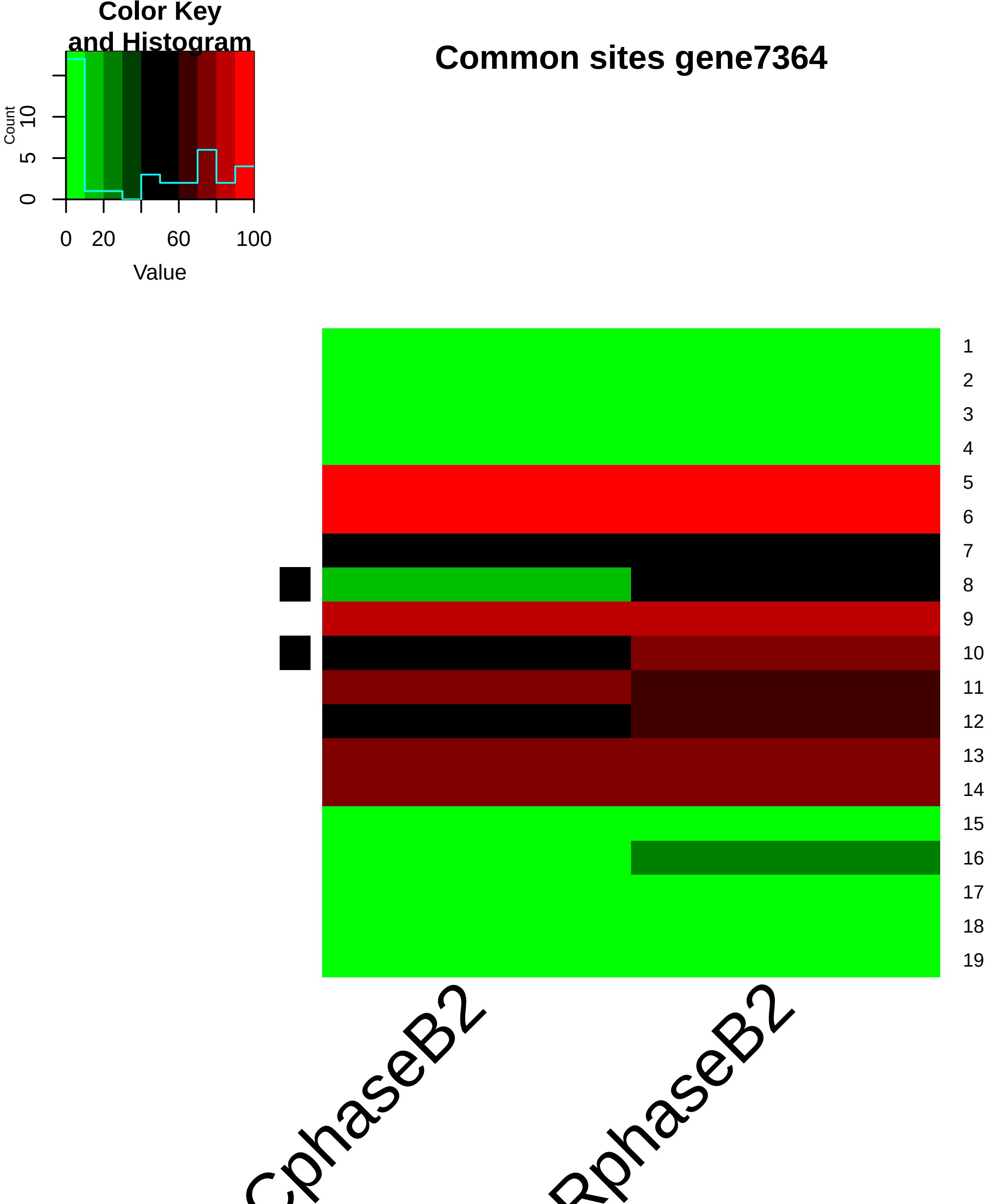
Color Key
and Histogram
Count
0
5
10
0
20
60
100
Value
Common sites gene7364
CphaseB2
RphaseB2

#### Common sites gene7367

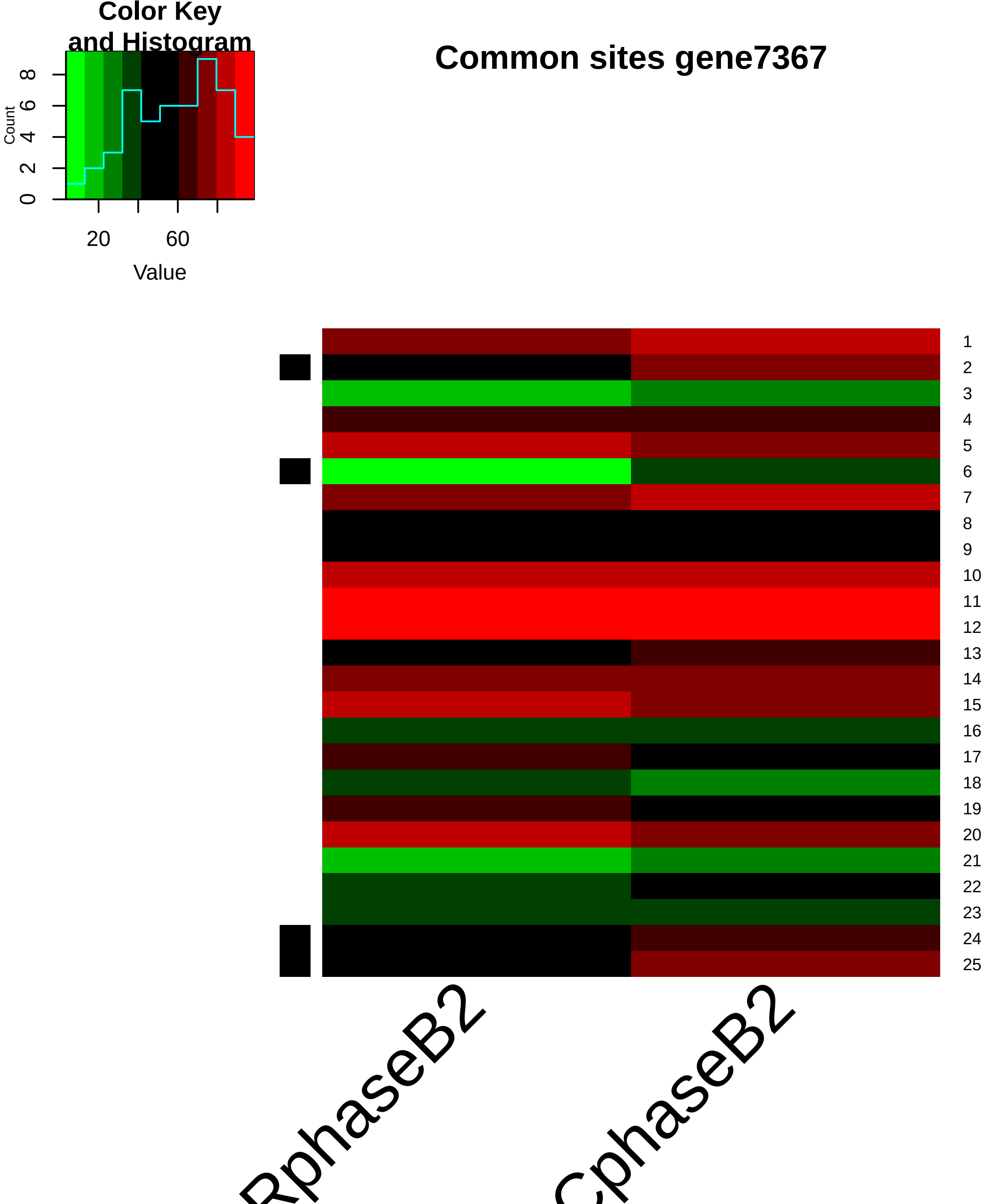

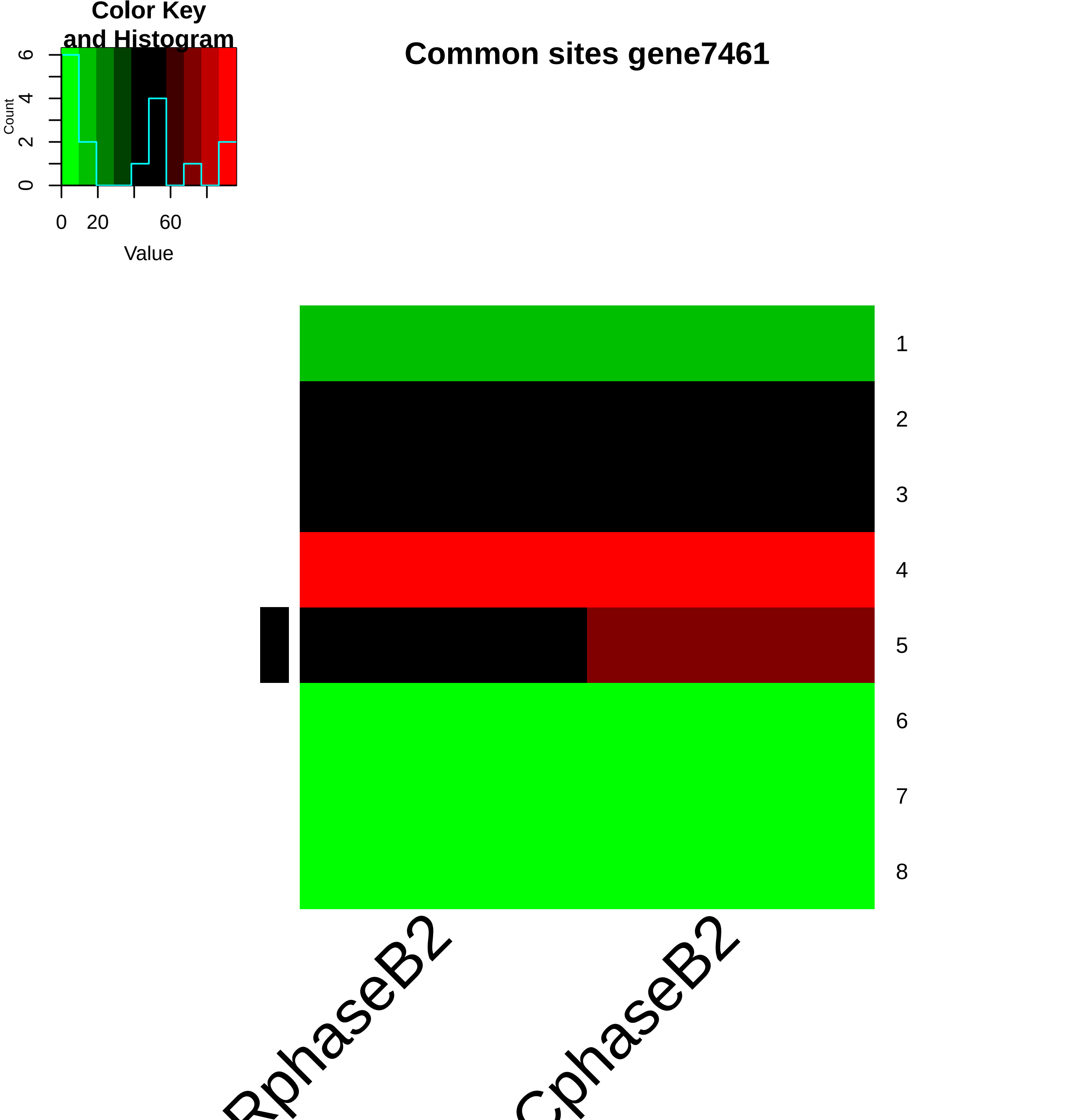
Color Key
and Histogram
Count
0
2
4
6
0
20
60
Value
Common sites gene7461
1
2
3
4
5
6
7
8
RphaseB2
CphaseB2

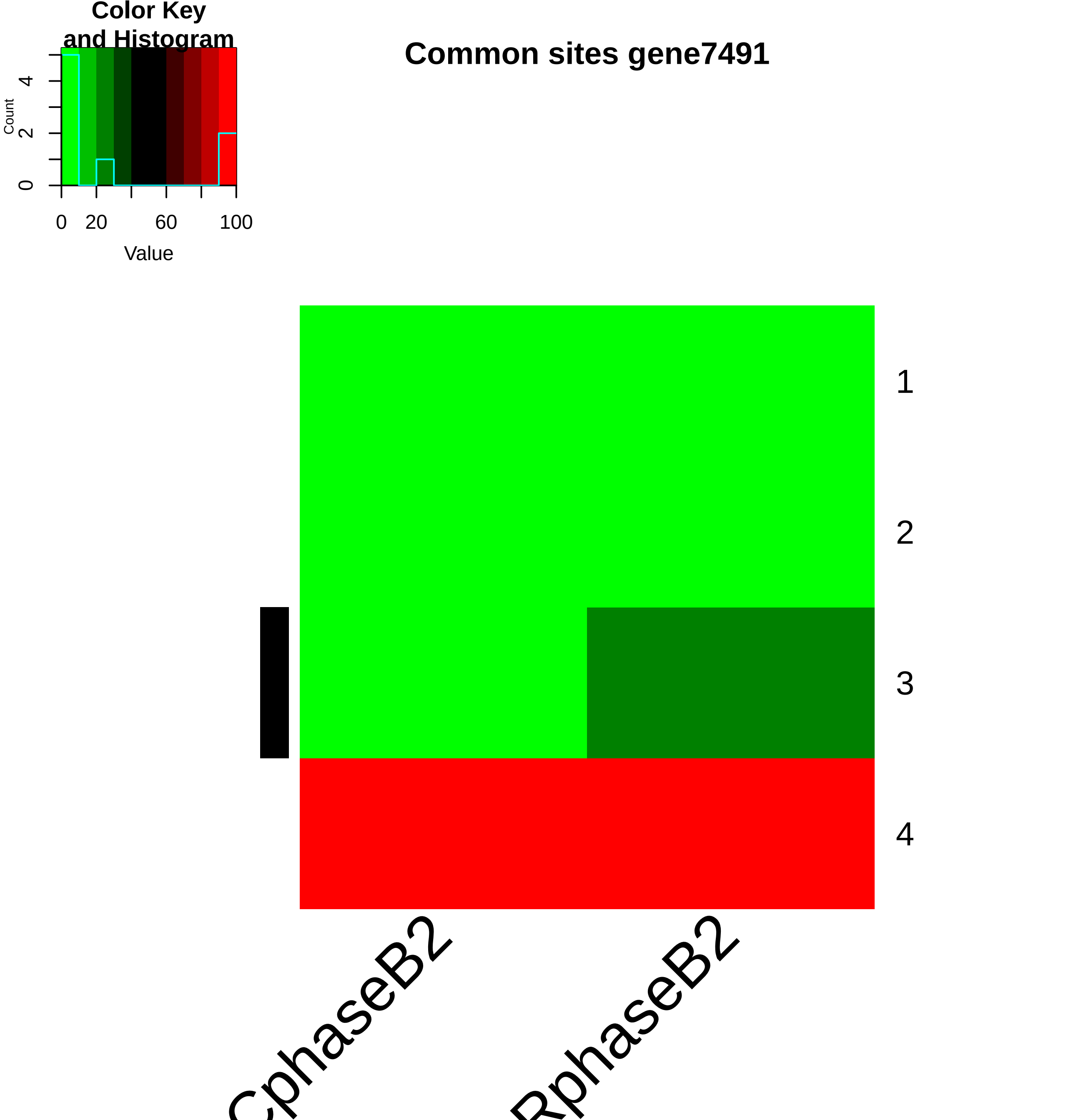
Color Key
and Histogram
Count
0
2
4
0
20
60
100
Value
Common sites gene7491
1
2
3
4
CphaseB2
RphaseB2

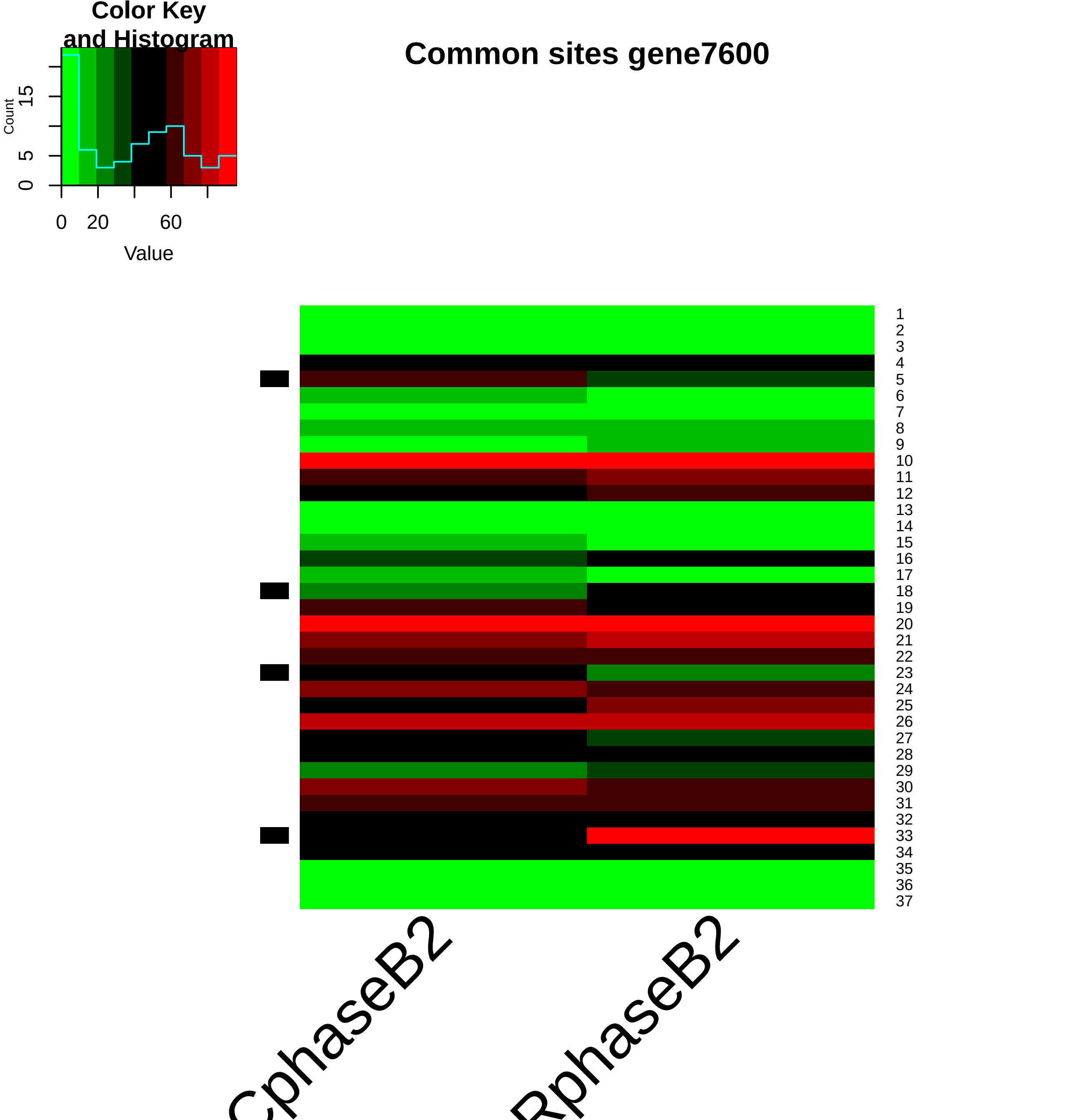
Color Key
and Histogram
Count
0
5
15
0
20
60
Value
Common sites gene7600
CphaseB2
RphaseB2

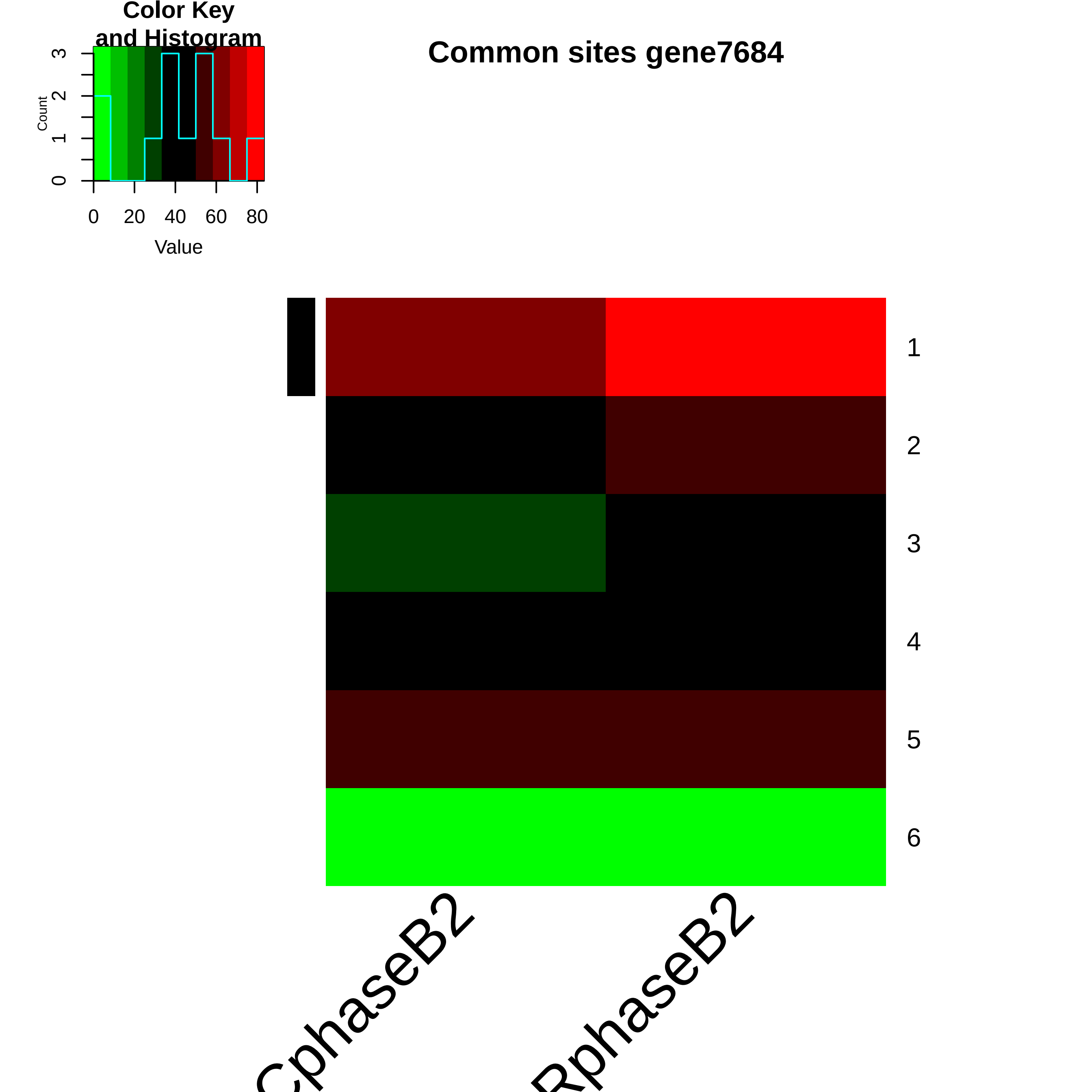
Color Key
and Histogram
Count
0
1
2
3
0
20
40
60
80
Value
Common sites gene7684
1
2
3
4
5
6
CphaseB2
RphaseB2

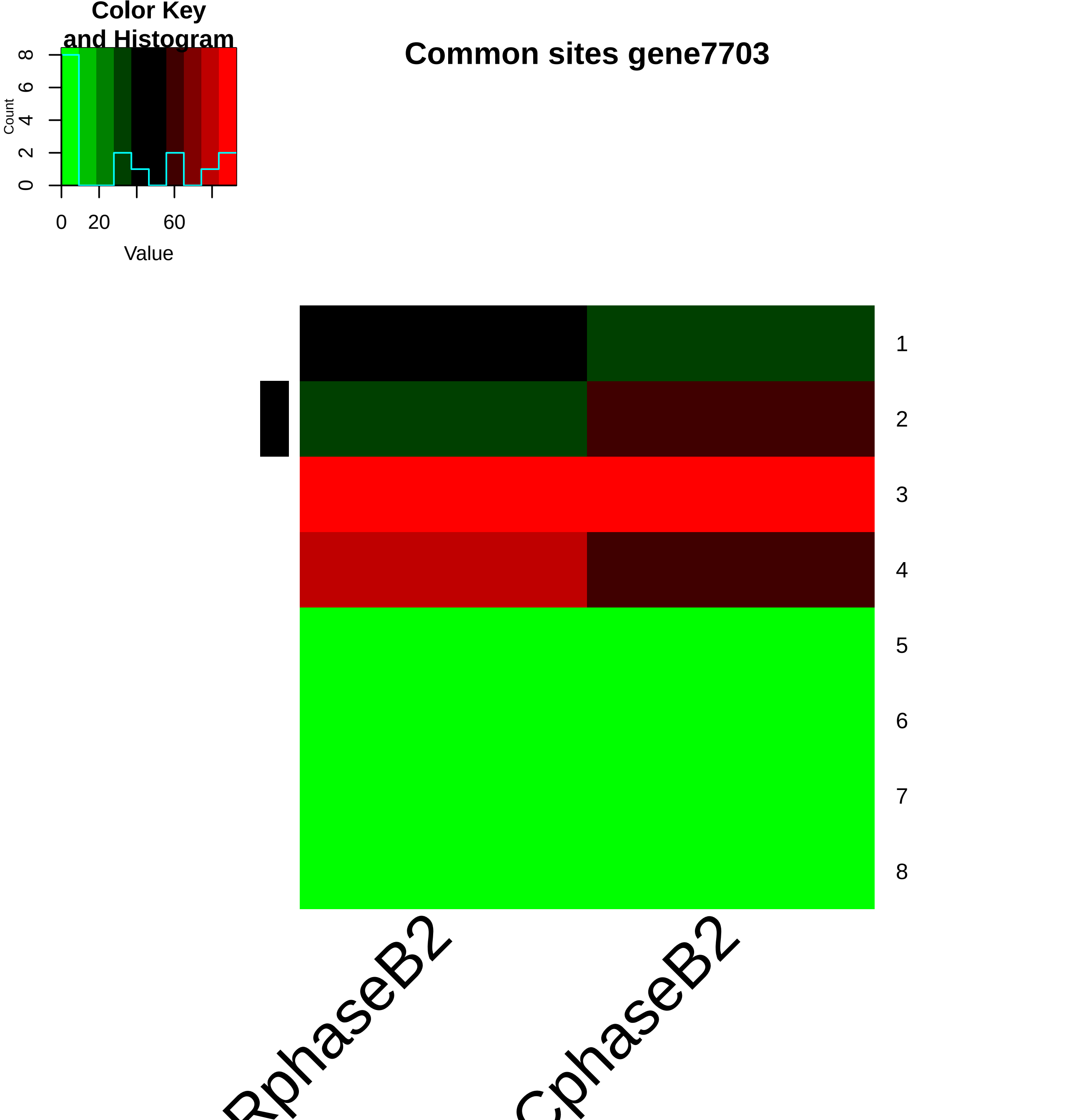
Color Key
and Histogram
Count
0
2
4
6
8
0
20
60
Value
Common sites gene7703
1
2
3
4
5
6
7
8
RphaseB2
CphaseB2

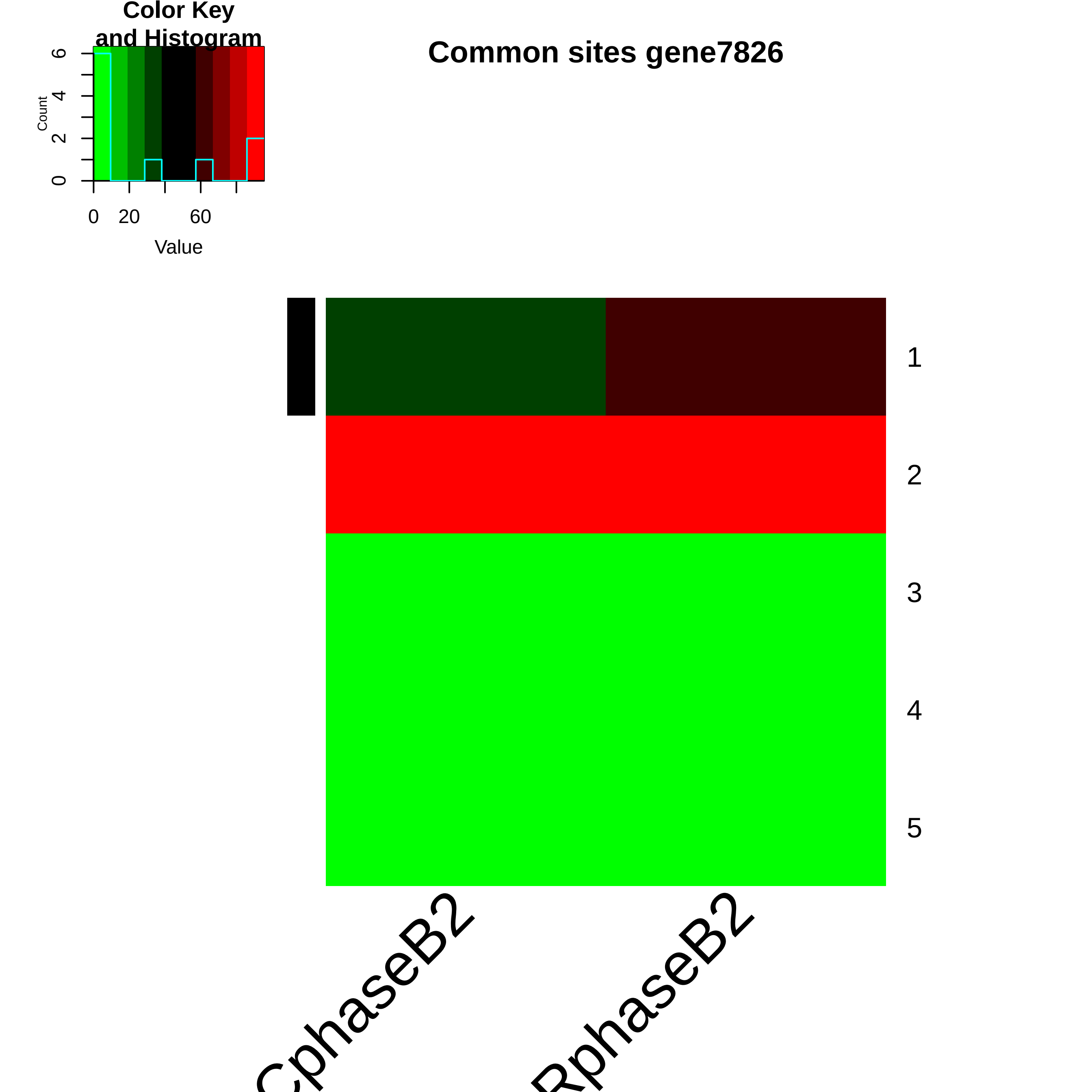
Color Key
and Histogram
Count
Value
0
20
60
0
2
4
6
Common sites gene7826
1
2
3
4
5
CphaseB2
RphaseB2

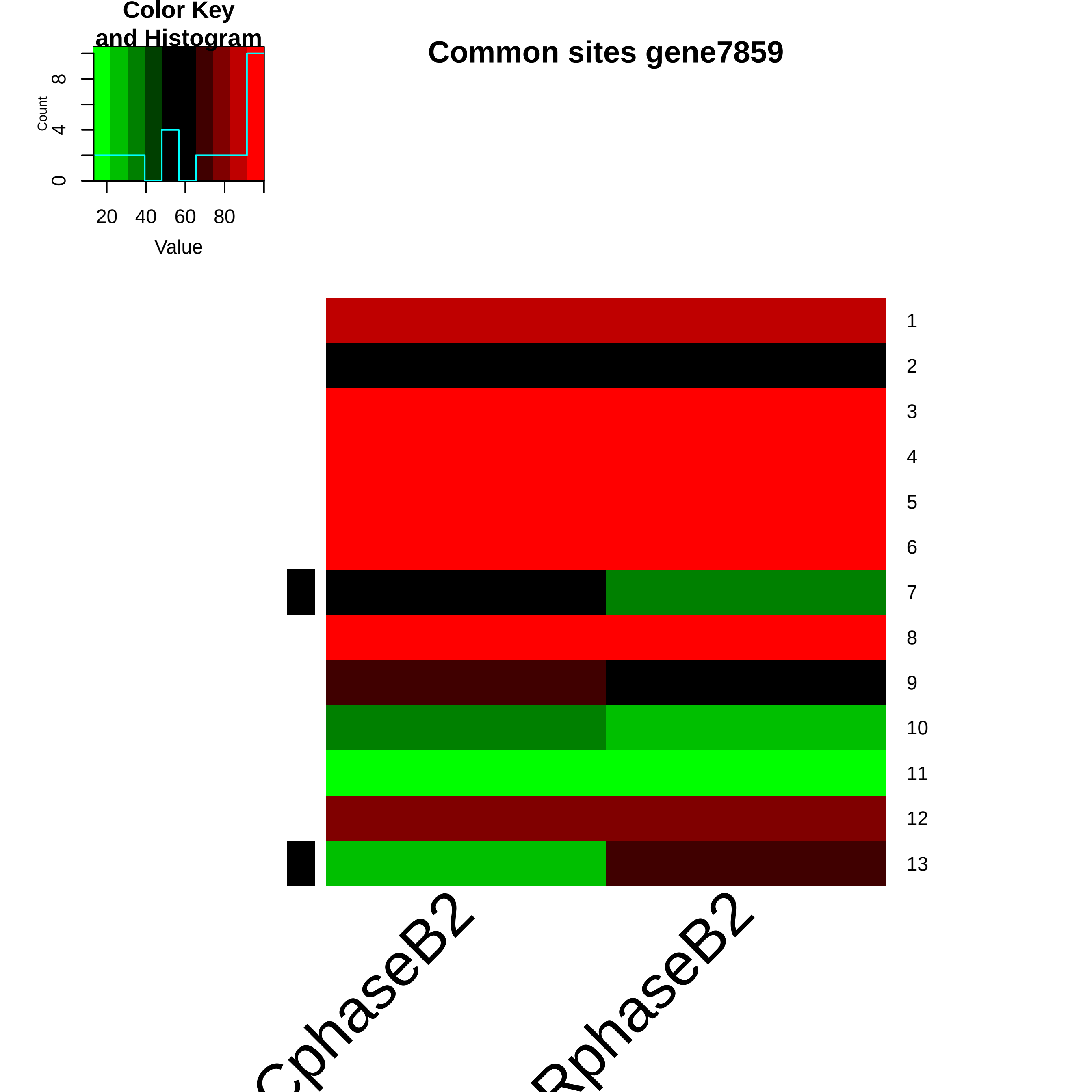

Color Key
and Histogram
Count
0
4
8
20
40
60
80
Value
Common sites gene7859
1
2
3
4
5
6
7
8
9
10
11
12
13
CphaseB2
RphaseB2

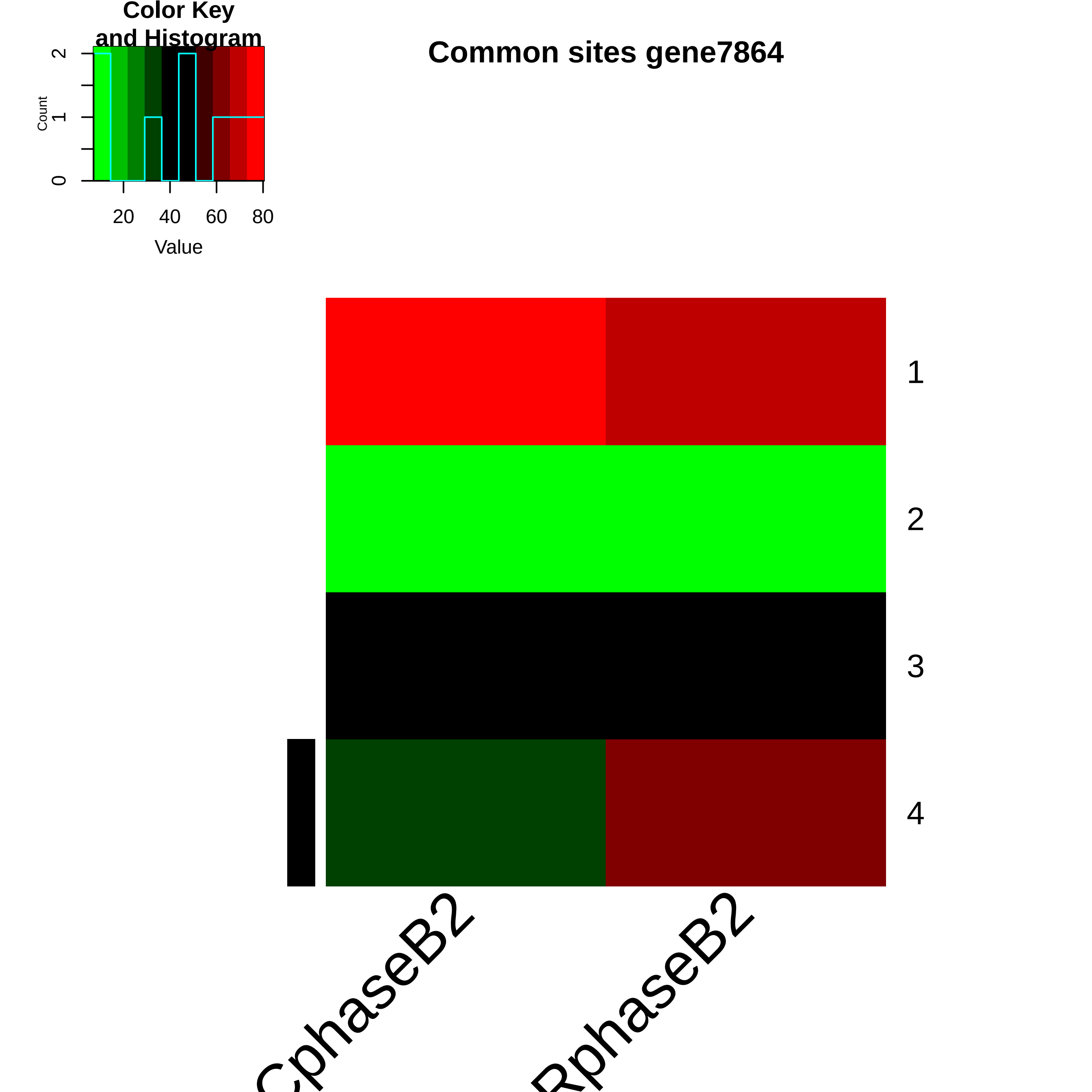

Color Key
and Histogram
Count
0
1
2
20
40
60
80
Value
Common sites gene7864
1
2
3
4
CphaseB2
RphaseB2

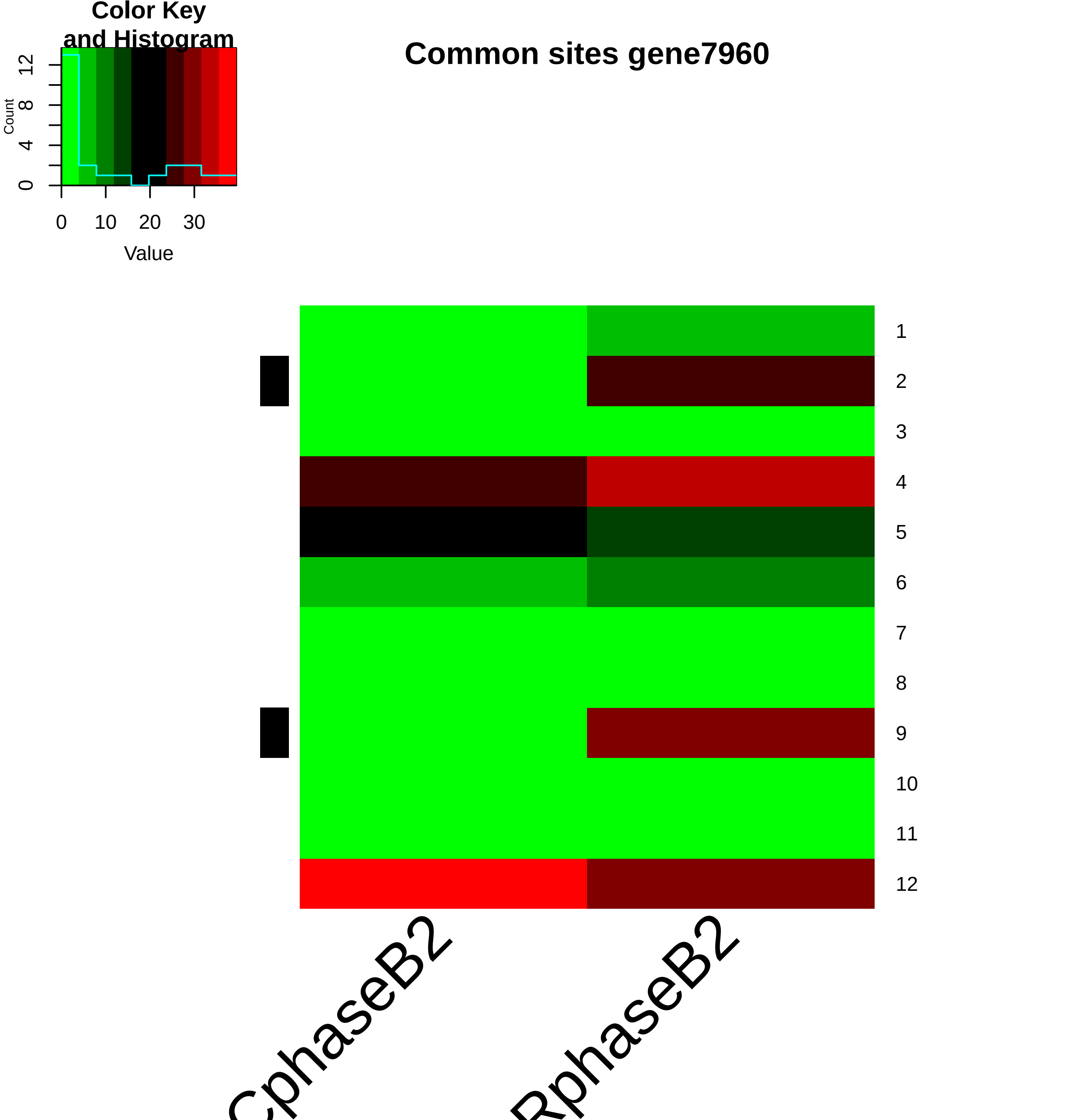
Color Key
and Histogram
Count
Value
0
4
8
12
0
10
20
30
Common sites gene7960
1
2
3
4
5
6
7
8
9
10
11
12
CphaseB2
RphaseB2

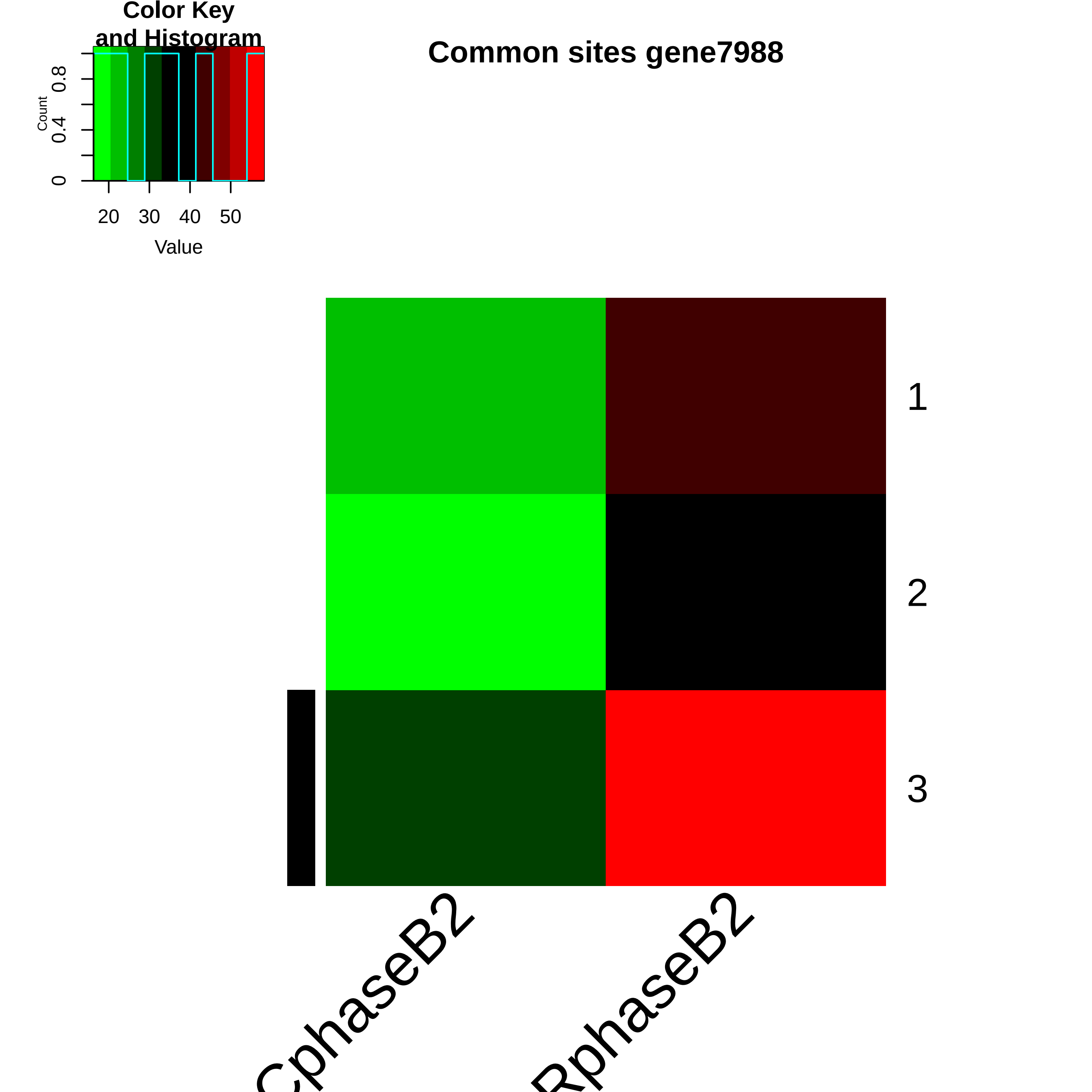

Color Key
and Histogram
Count
0
0.4
0.8
20
30
40
50
Value
Common sites gene7988
1
2
3
CphaseB2
RphaseB2

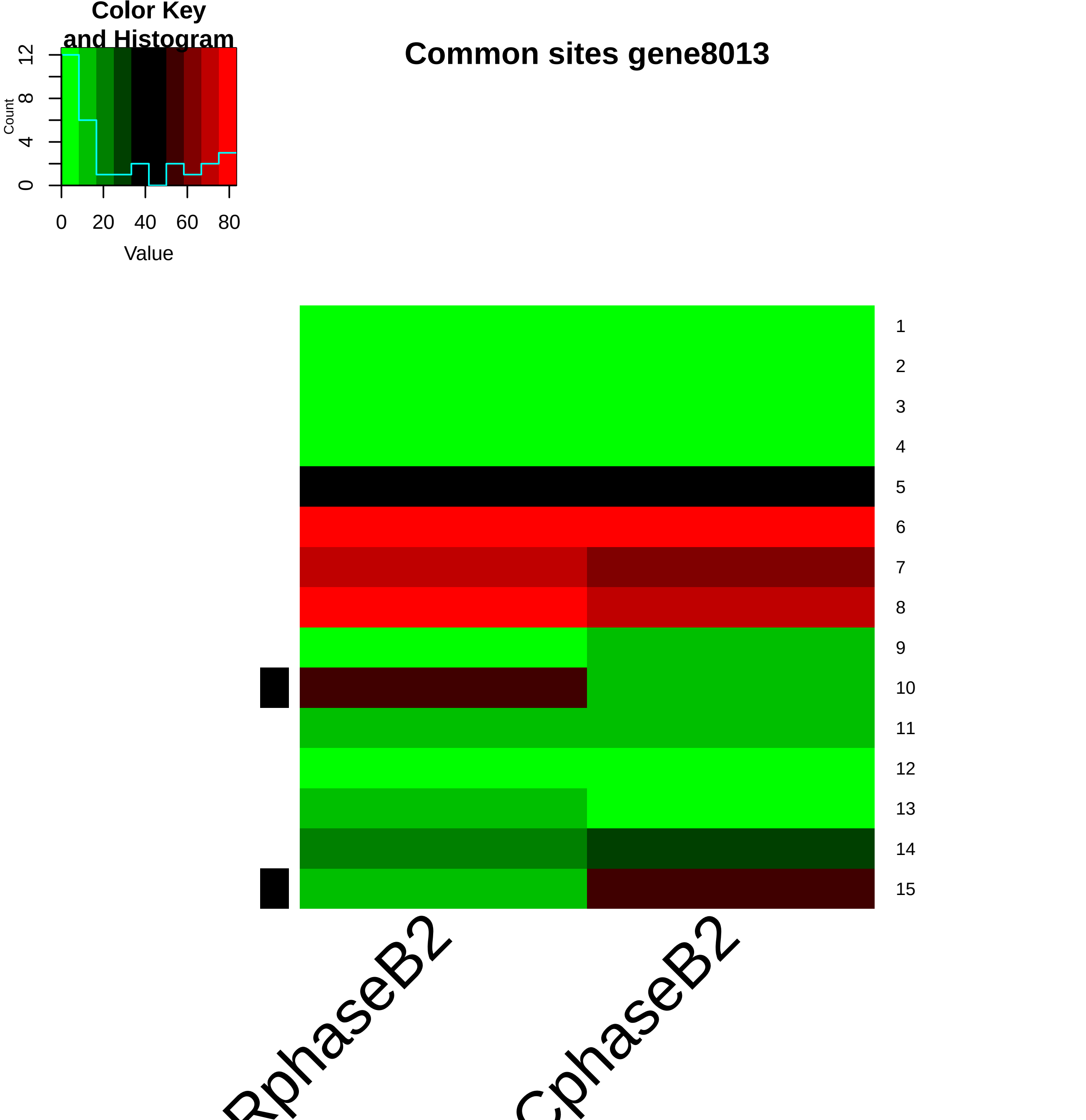
Color Key
and Histogram
Count
0
4
8
12
0
20
40
60
80
Value
Common sites gene8013
RphaseB2
CphaseB2

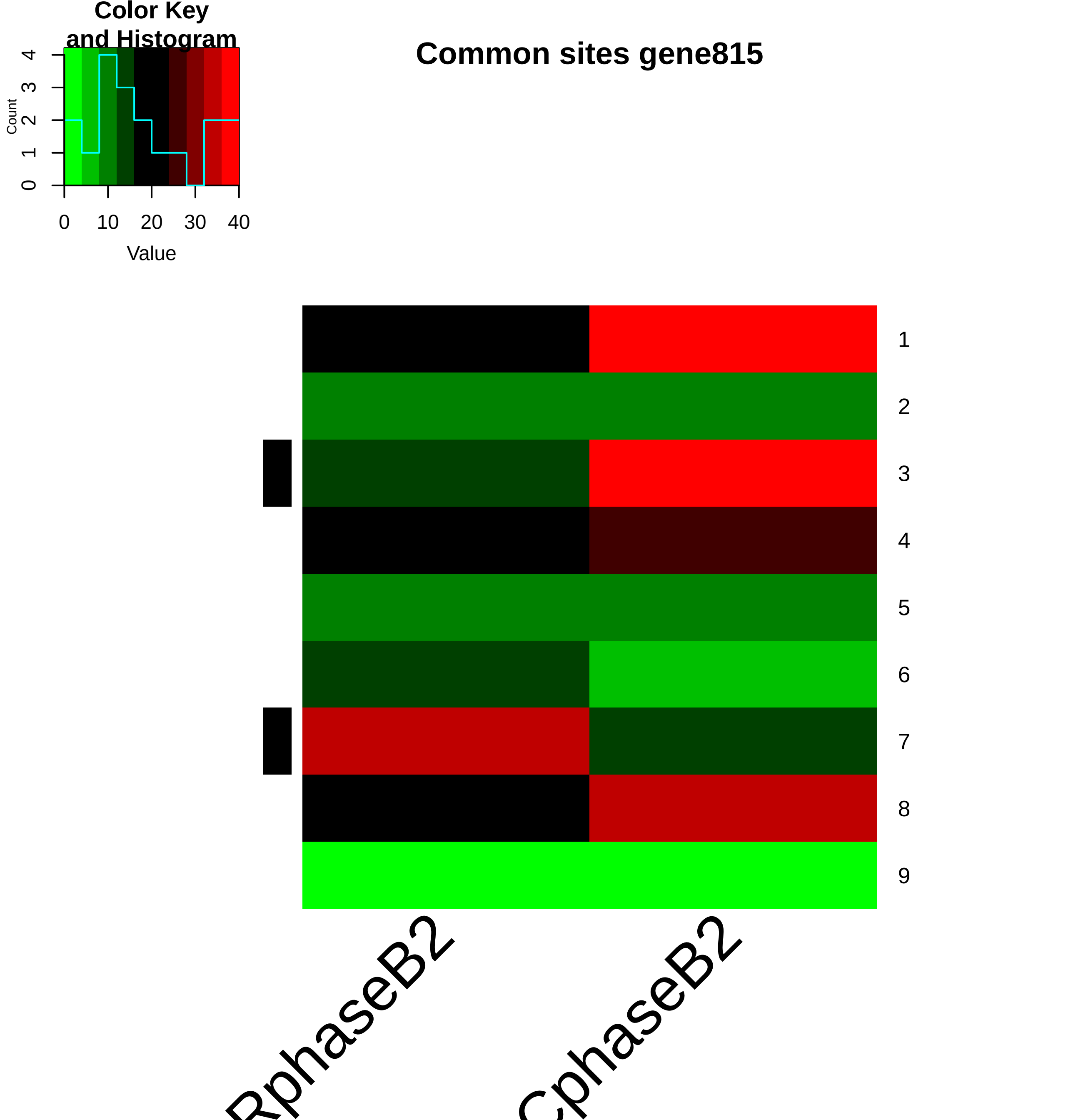
Color Key
and Histogram
Count
0
1
2
3
4
0
10
20
30
40
Value
Common sites gene815
1
2
3
4
5
6
7
8
9
RphaseB2
CphaseB2

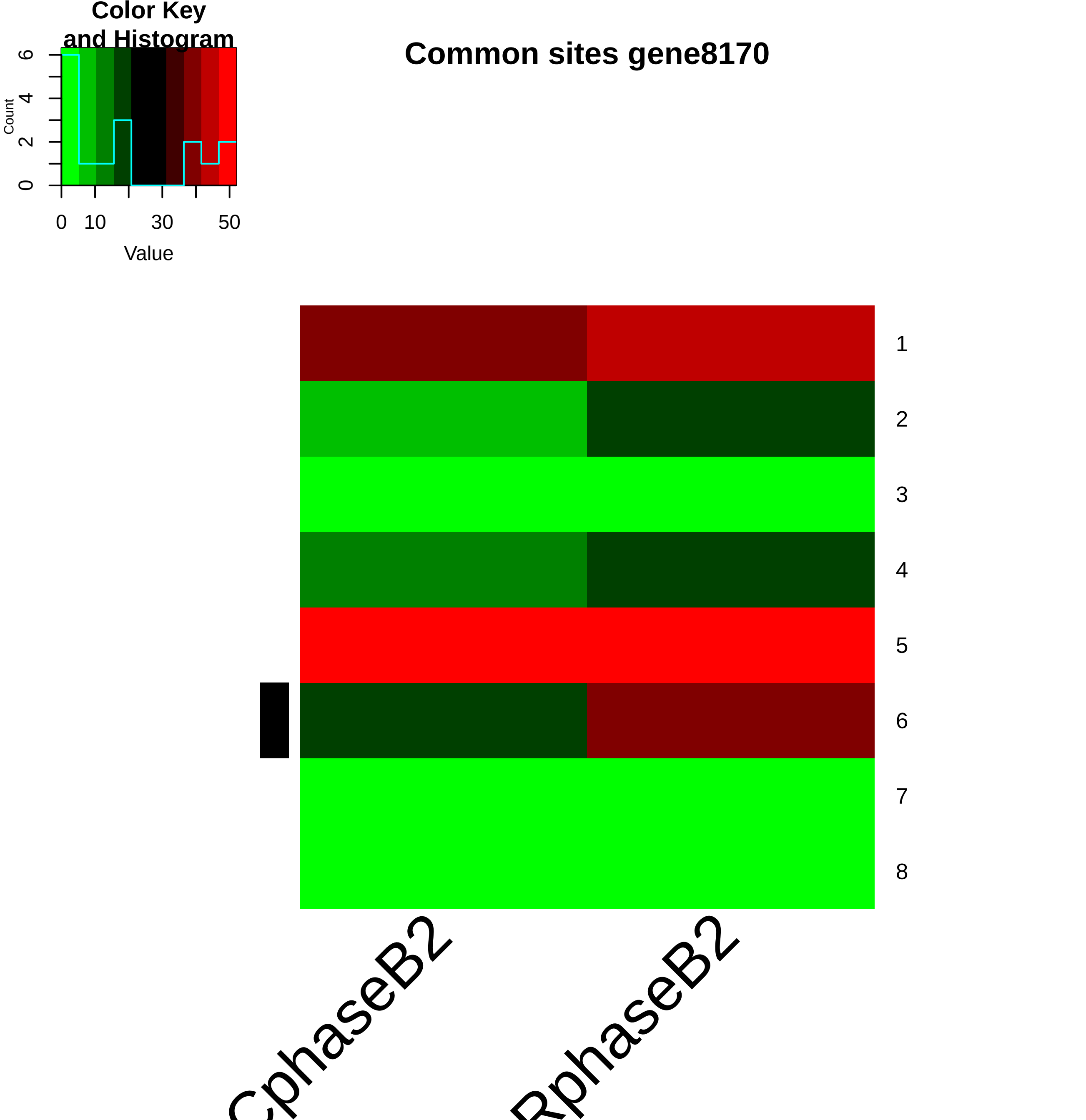
Color Key
and Histogram
Count
0
2
4
6
0
10
30
50
Value
Common sites gene8170
1
2
3
4
5
6
7
8
CphaseB2
RphaseB2

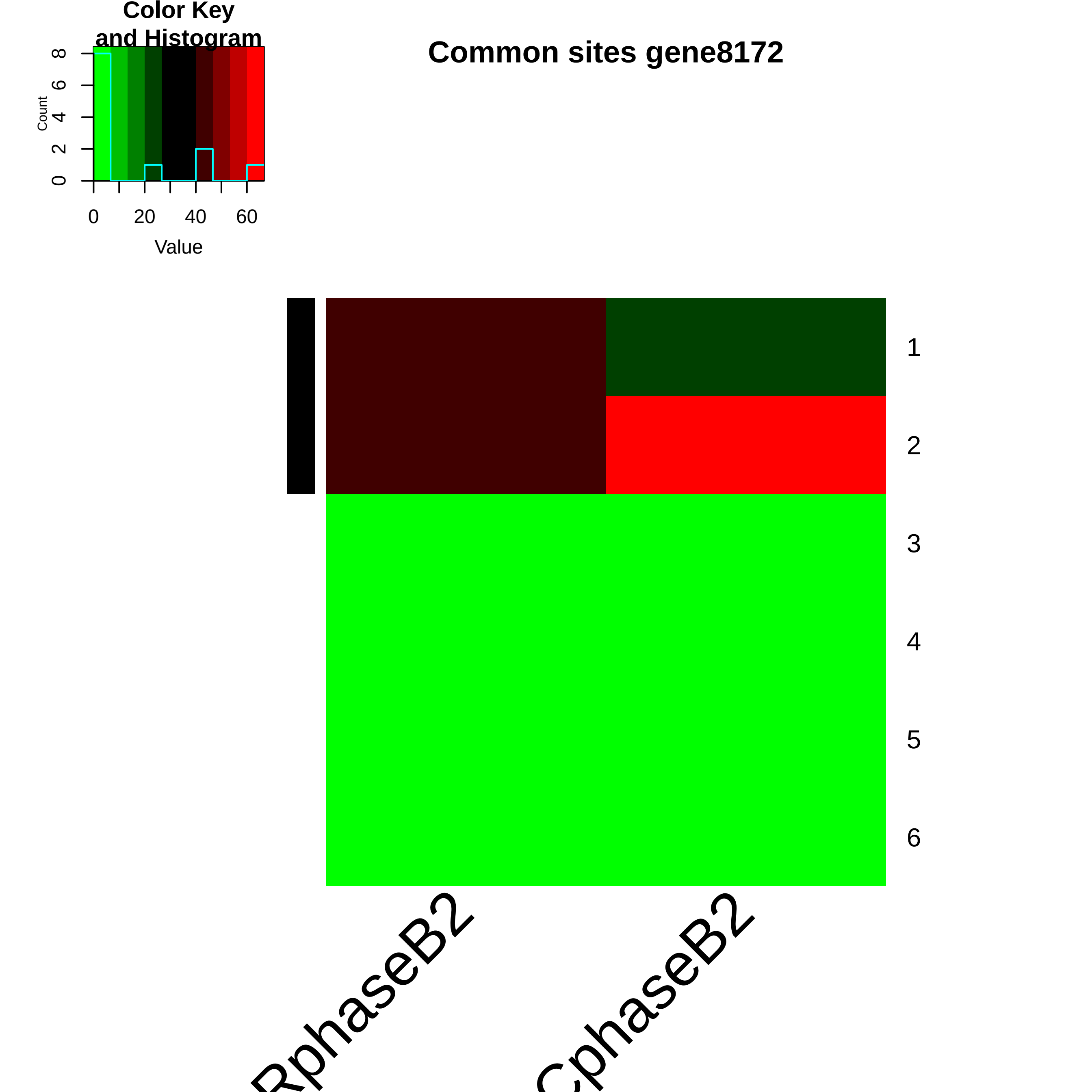
Color Key
and Histogram
Count
0
2
4
6
8
0
20
40
60
Value
Common sites gene8172
1
2
3
4
5
6
RphaseB2
CphaseB2

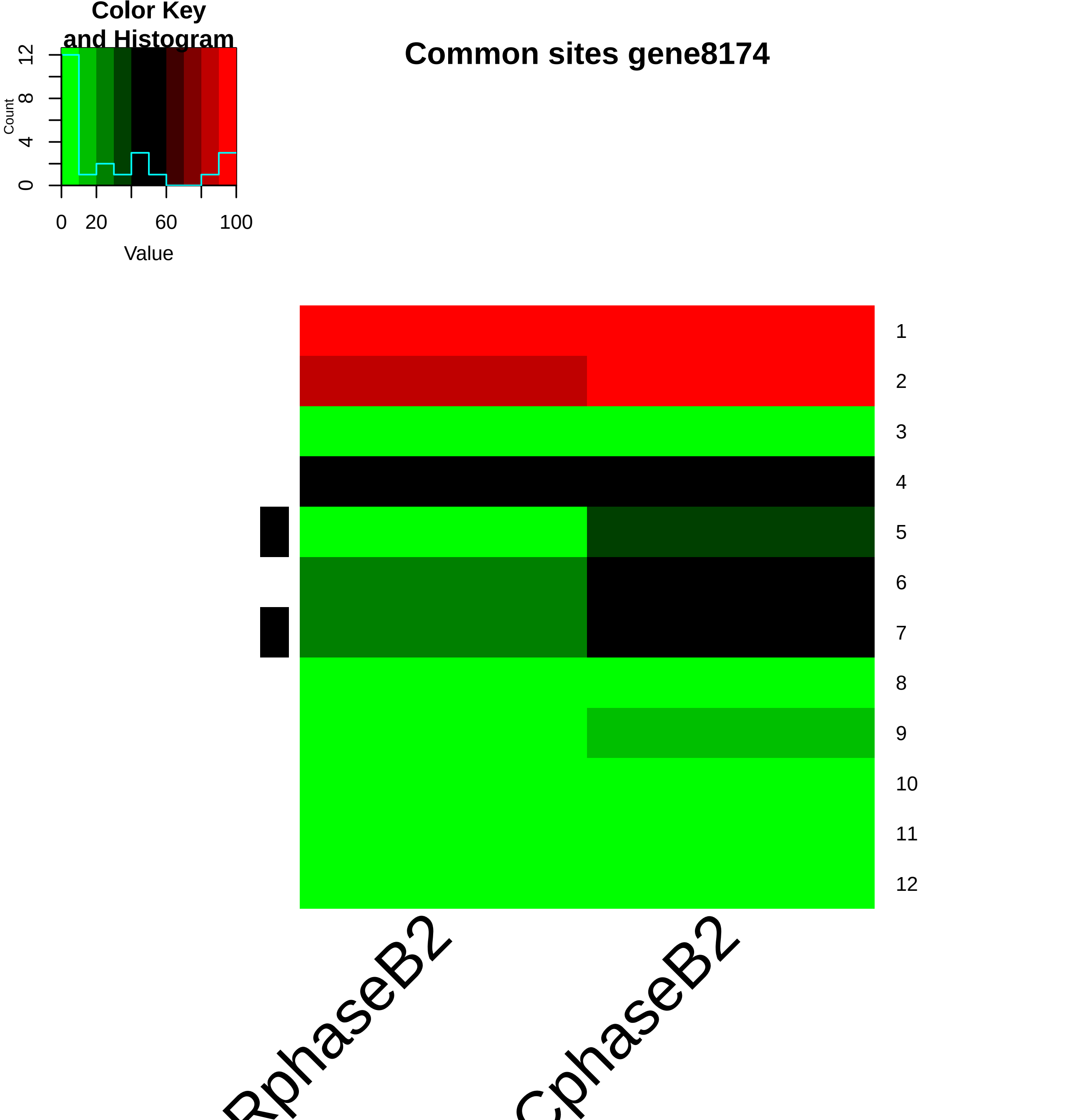
Color Key
and Histogram
Count
0
4
8
12
0
20
60
100
Value
Common sites gene8174
1
2
3
4
5
6
7
8
9
10
11
12
RphaseB2
CphaseB2

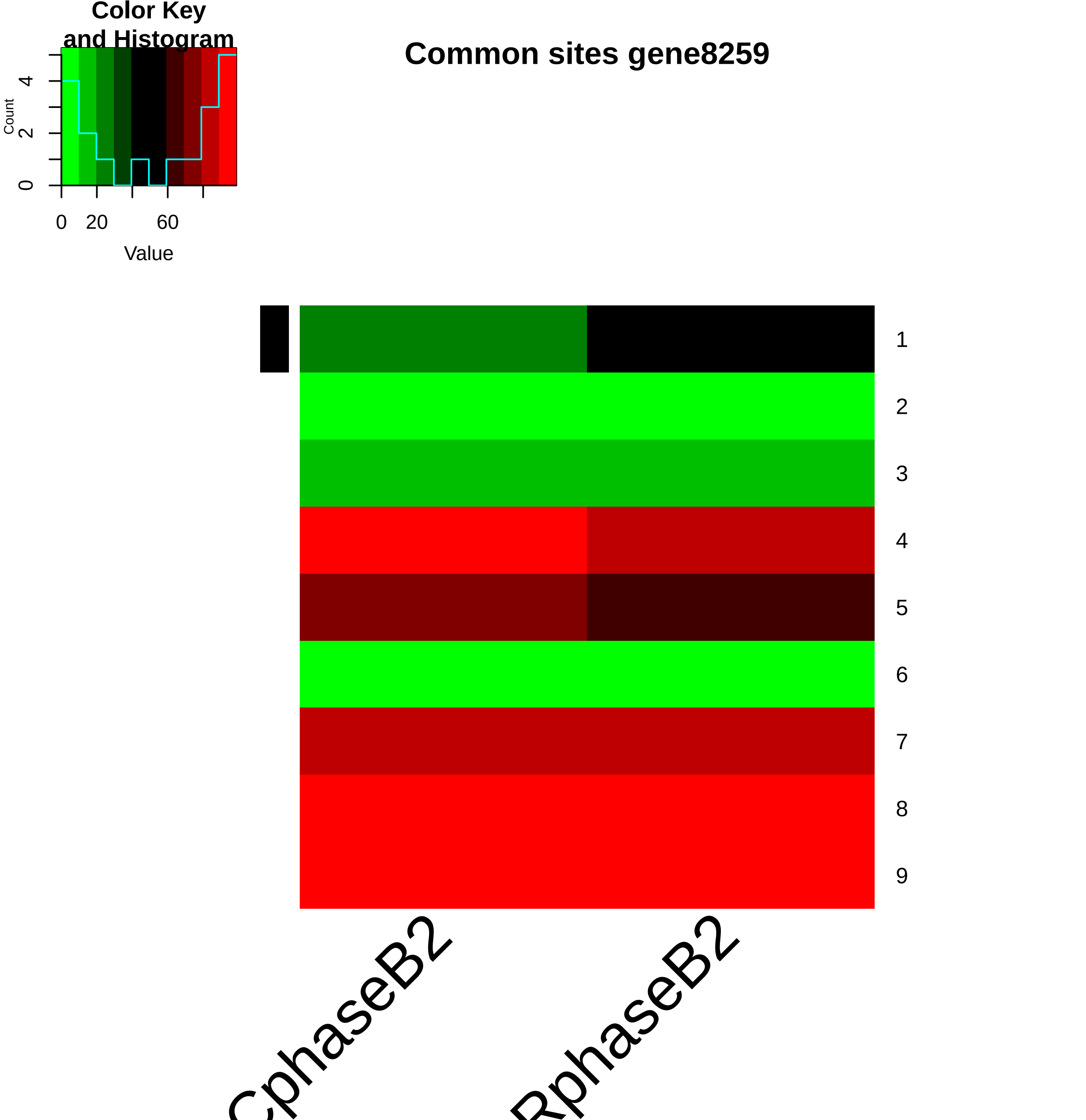
Color Key
and Histogram
Count
0
2
4
0
20
60
Value
Common sites gene8259
1
2
3
4
5
6
7
8
9
CphaseB2
RphaseB2

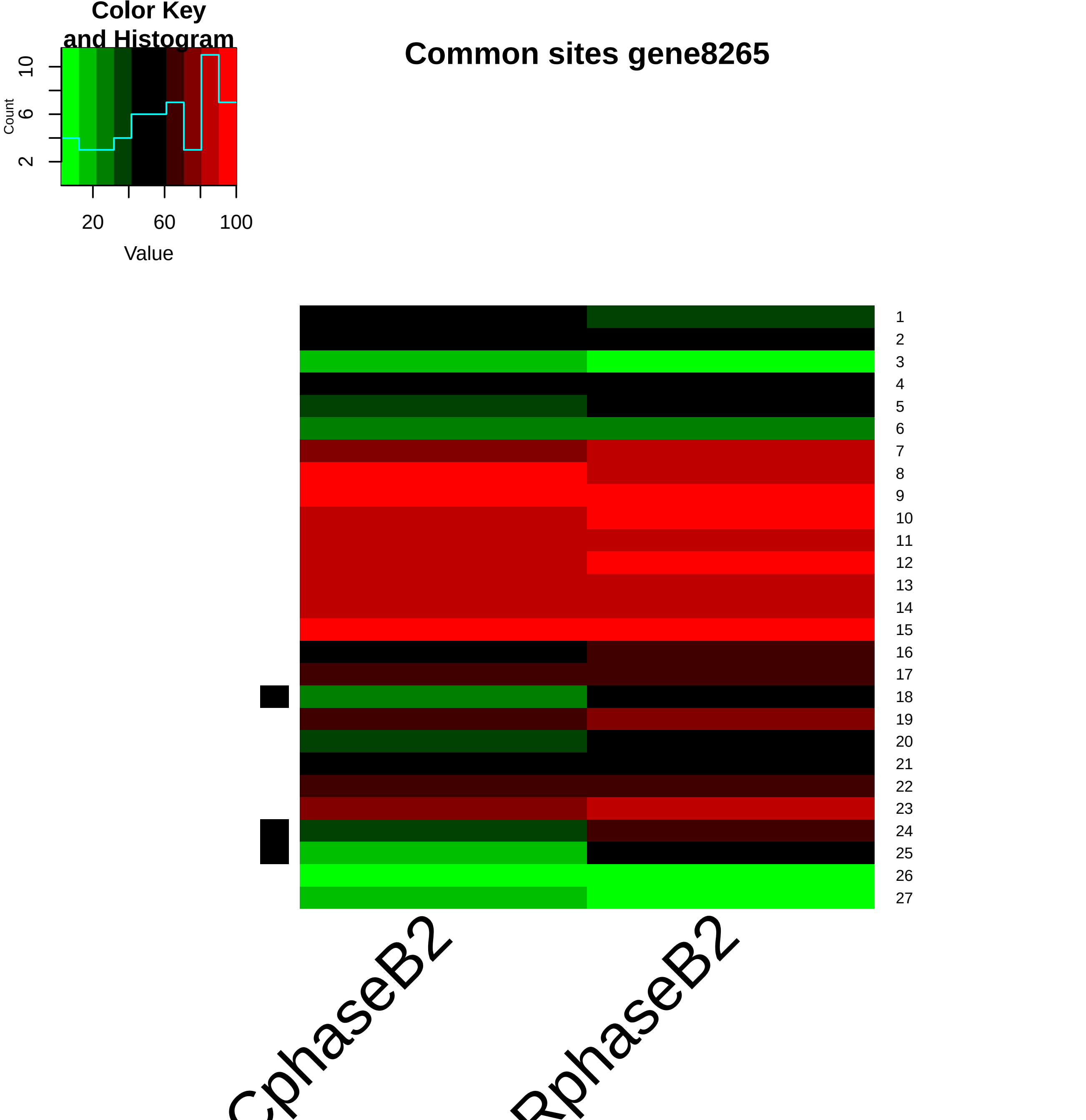

Color Key
and Histogram
Count
10
6
2
20
60
100
Value
Common sites gene8265
CphaseB2
RphaseB2

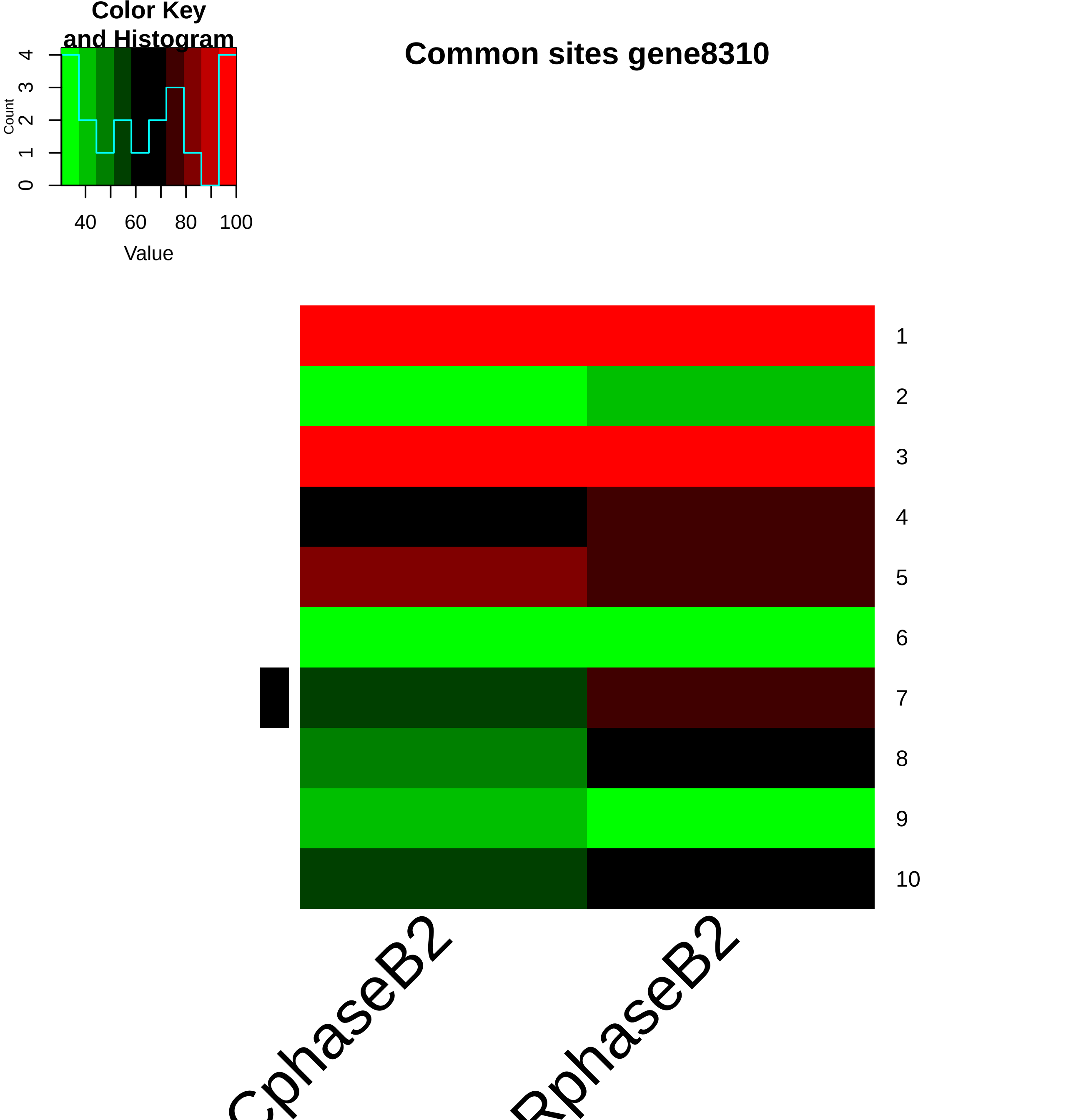

Color Key
and Histogram
Count
0
1
2
3
4
40
60
80
100
Value
Common sites gene8310
1
2
3
4
5
6
7
8
9
10
CphaseB2
RphaseB2

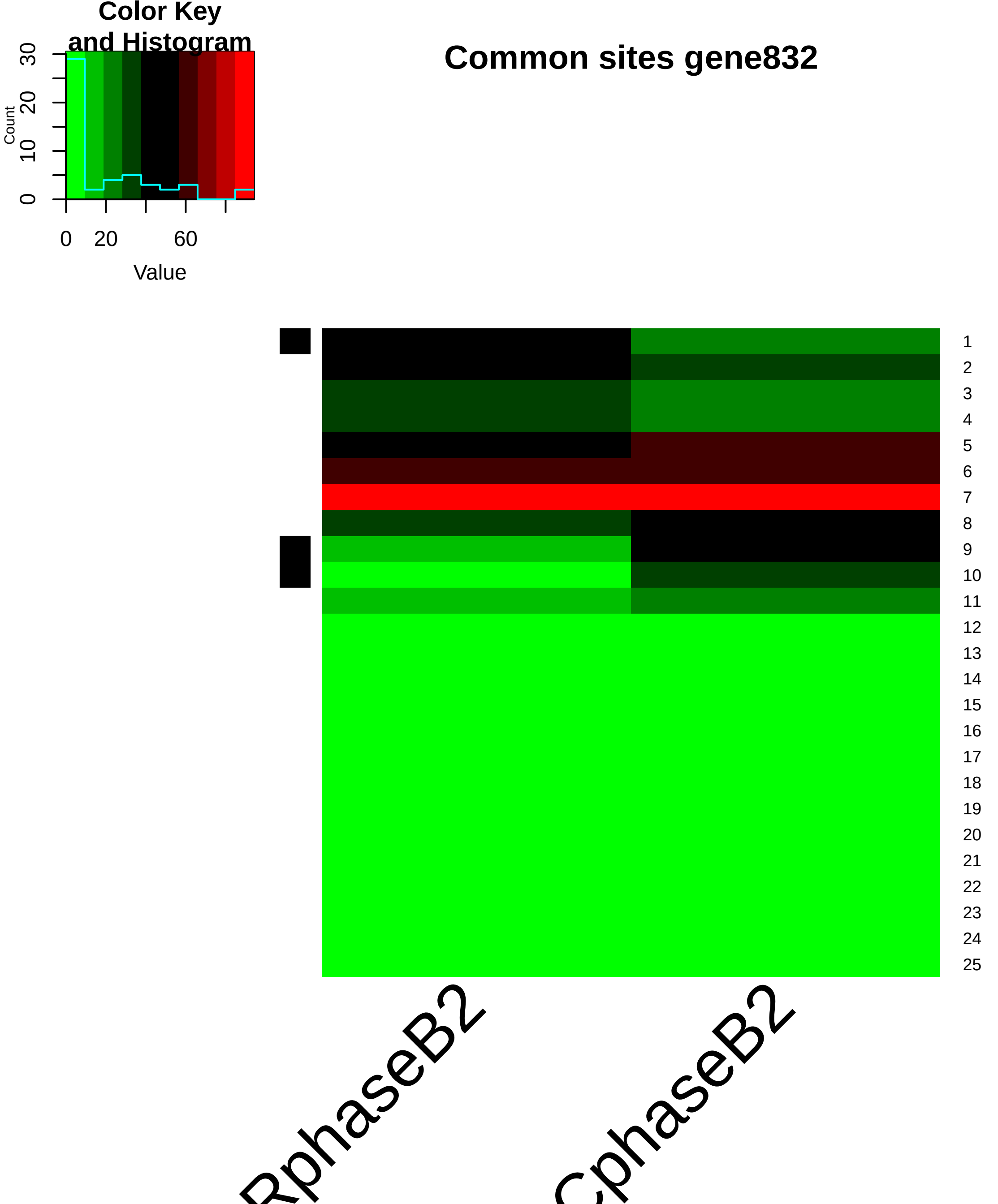

Color Key
and Histogram
Count
0
10
20
30
0
20
60
Value
Common sites gene832
RphaseB2
CphaseB2

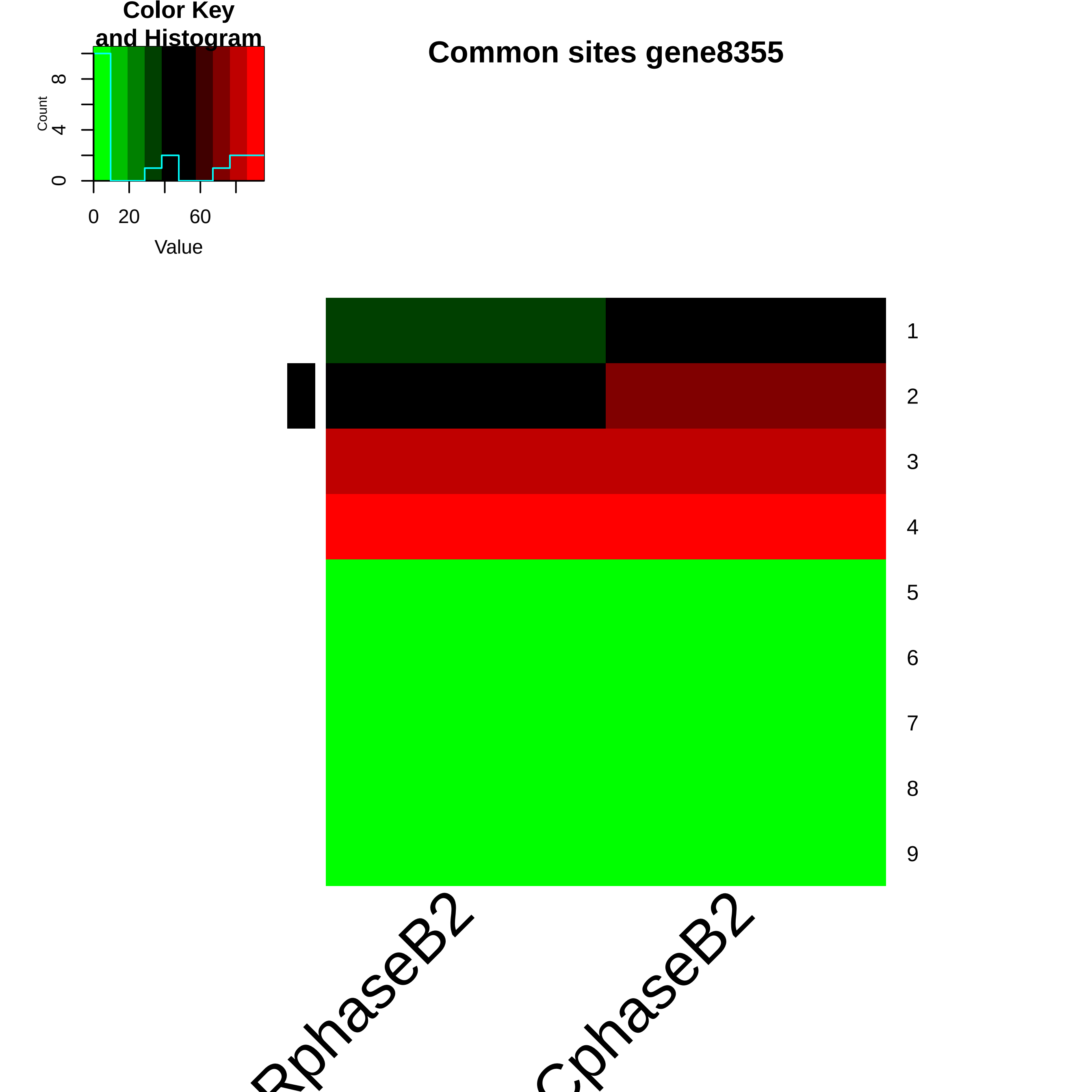

Color Key
and Histogram
Count
0
4
8
0
20
60
Value
Common sites gene8355
1
2
3
4
5
6
7
8
9
RphaseB2
CphaseB2

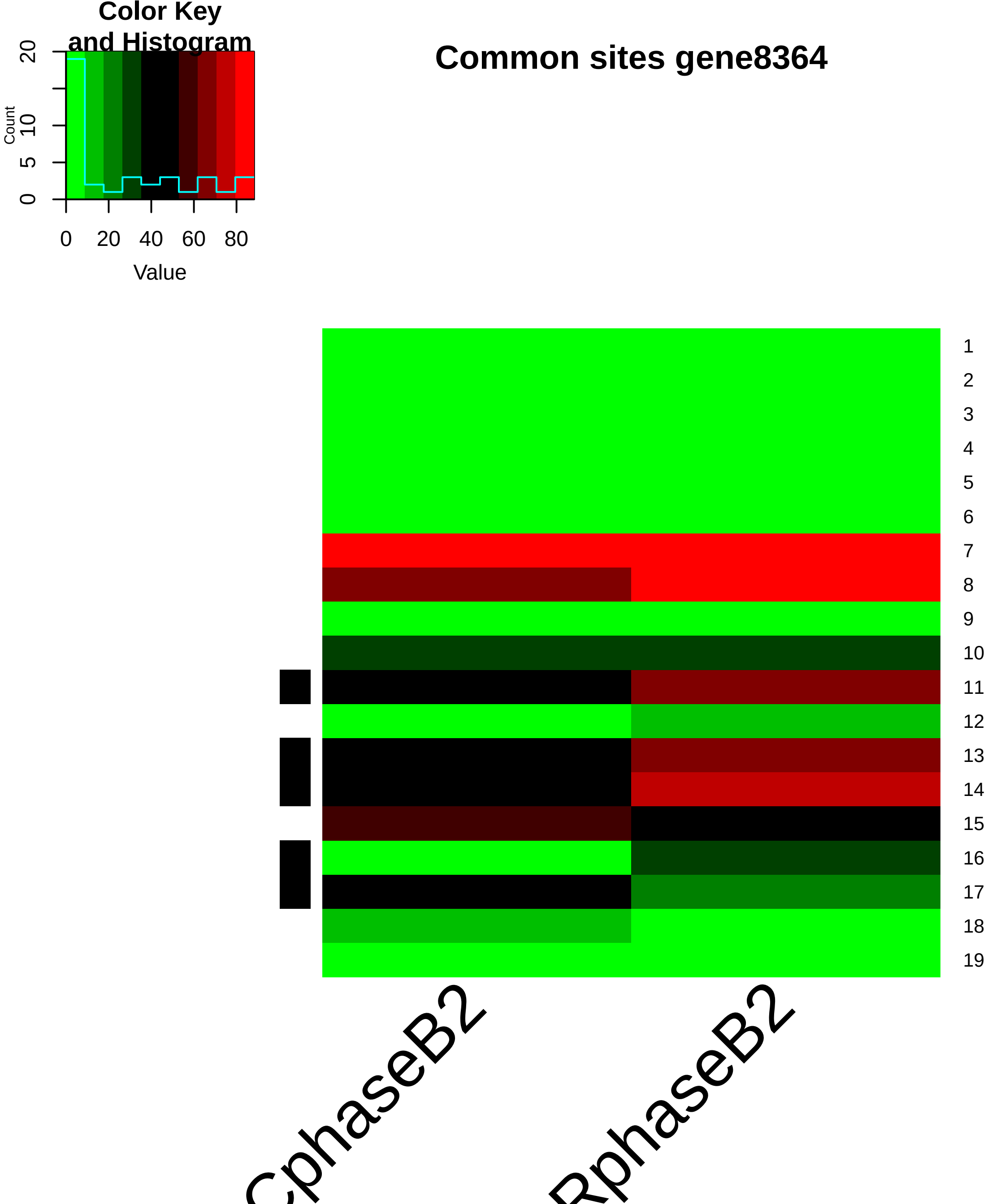

Color Key
and Histogram
Count
0
5
10
20
0
20
40
60
80
Value
Common sites gene8364
CphaseB2
RphaseB2

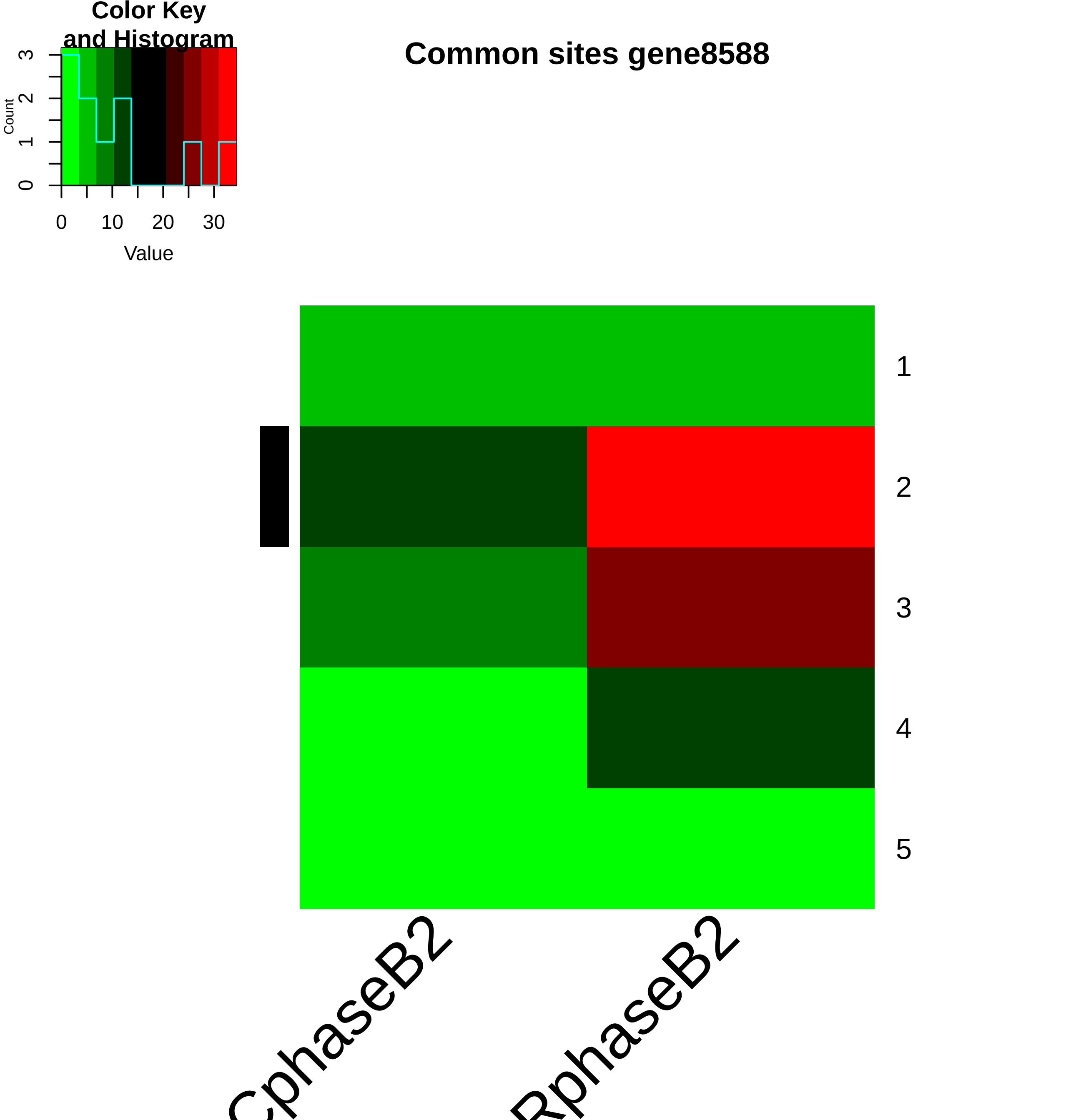
Color Key
and Histogram
Count
0
1
2
3
0
10
20
30
Value
Common sites gene8588
1
2
3
4
5
CphaseB2
RphaseB2

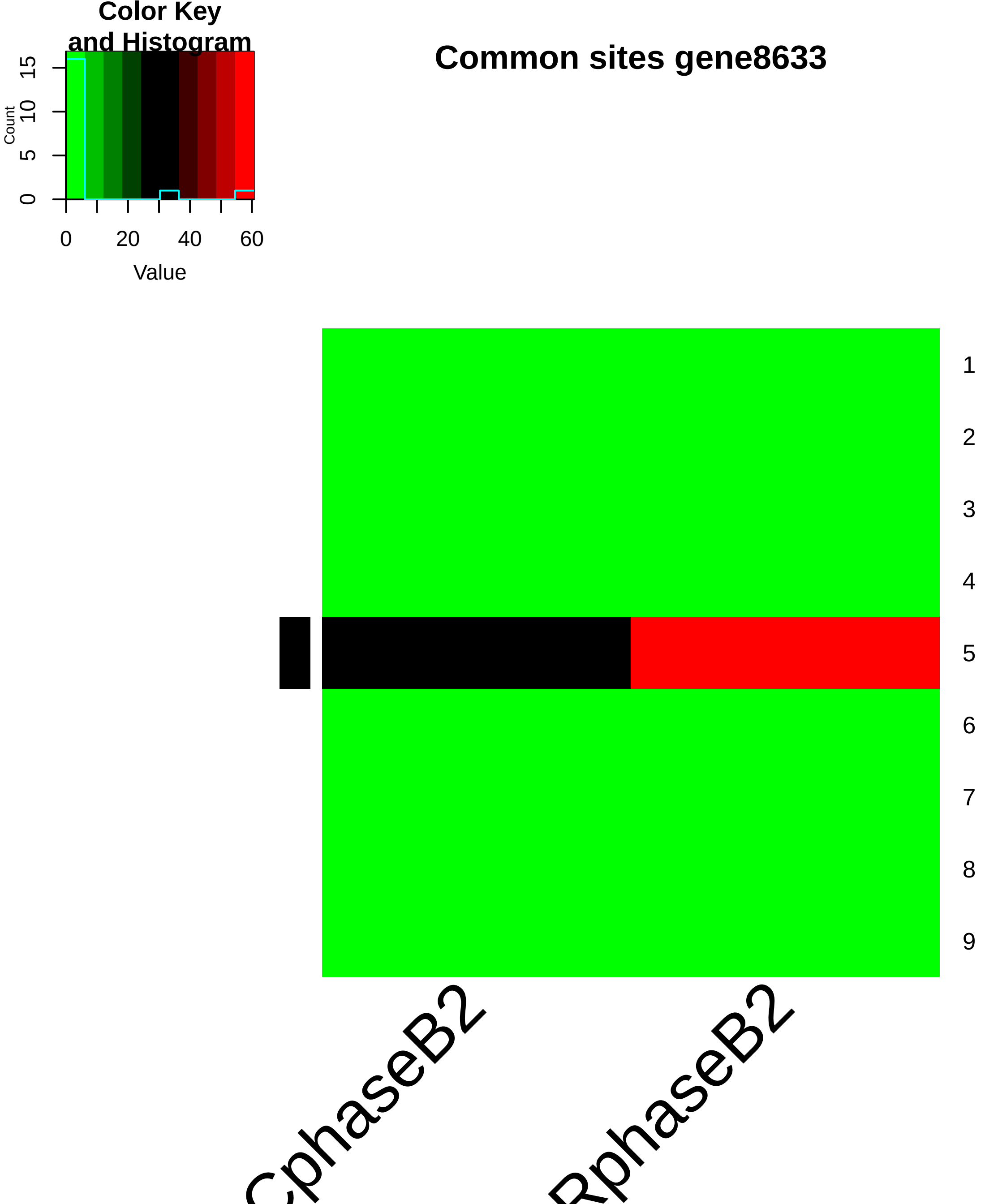
Color Key
and Histogram
Count
0
5
10
15
0
20
40
60
Value
Common sites gene8633
1
2
3
4
5
6
7
8
9
CphaseB2
RphaseB2

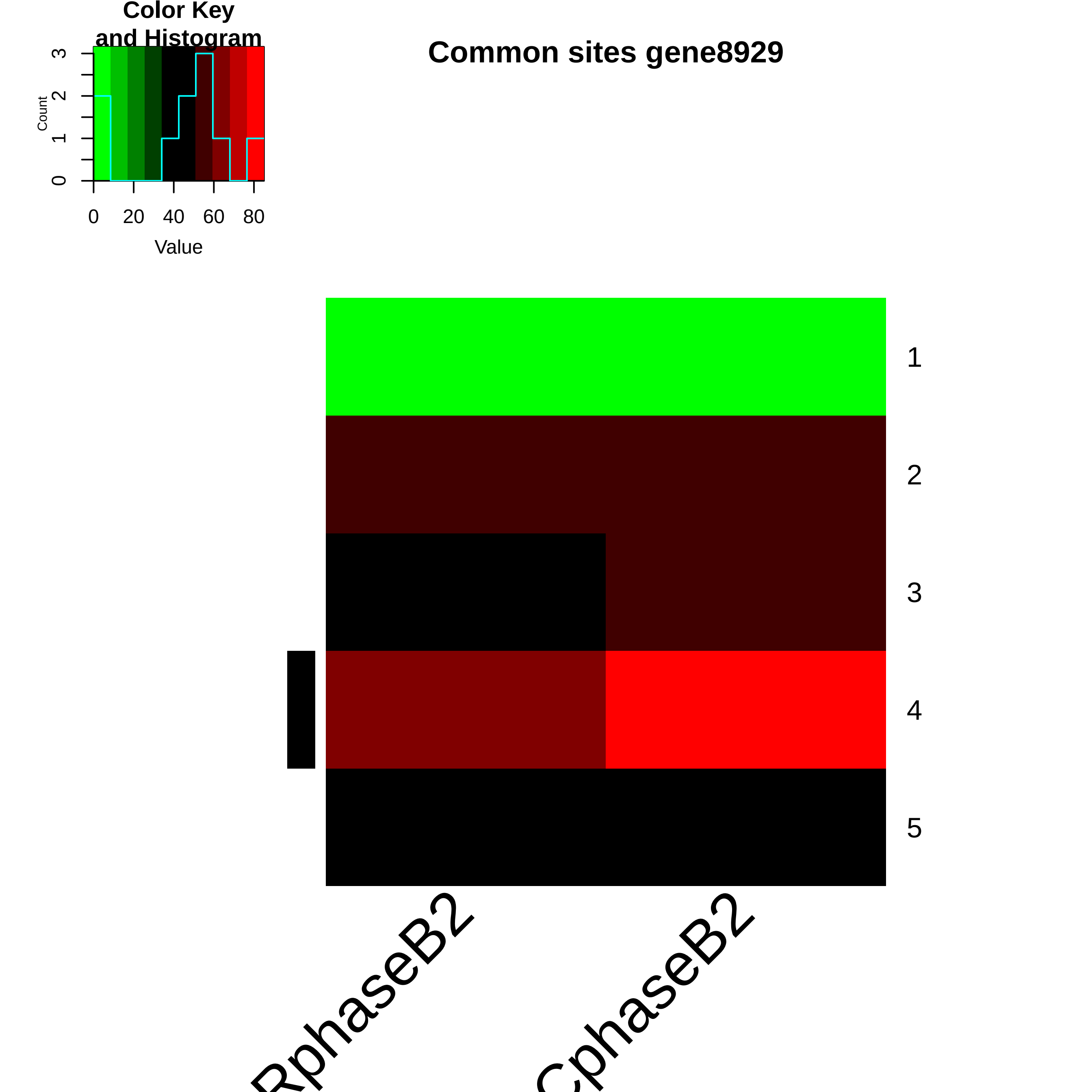

Color Key
and Histogram
Count
0
1
2
3
0
20
40
60
80
Value
Common sites gene8929
1
2
3
4
5
RphaseB2
CphaseB2

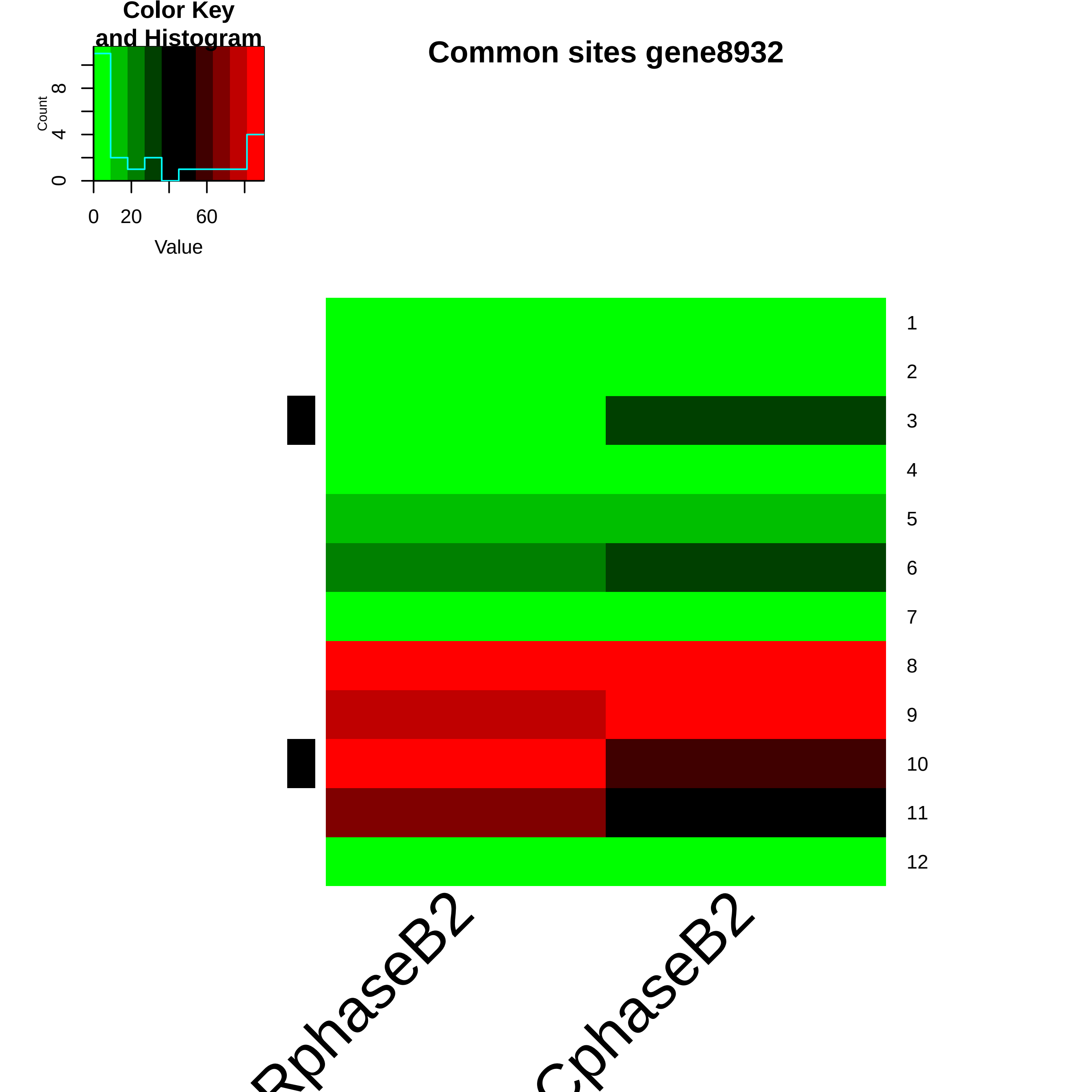

Color Key
and Histogram
Count
0
4
8
0
20
60
Value
Common sites gene8932
1
2
3
4
5
6
7
8
9
10
11
12
RphaseB2
CphaseB2

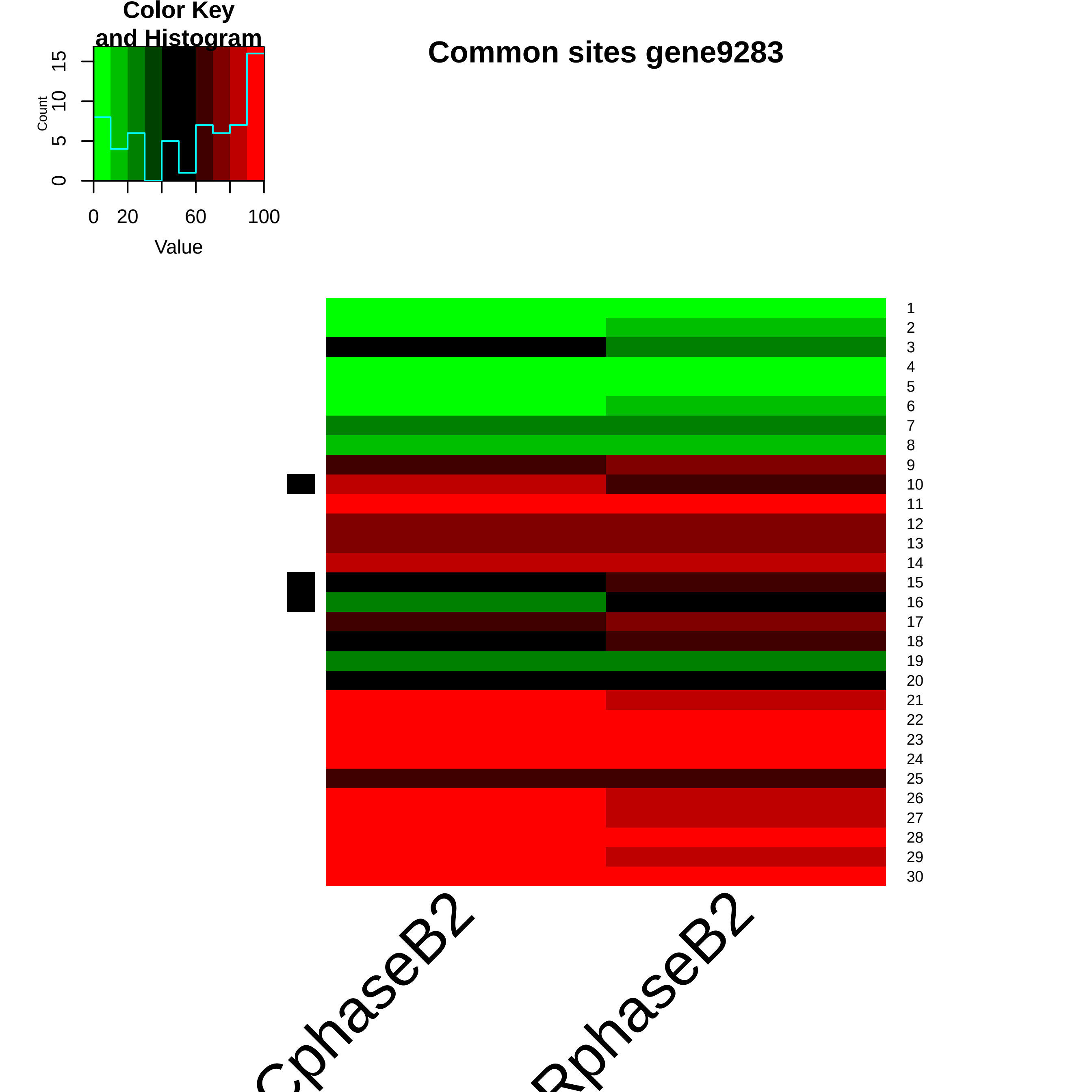

Color Key
and Histogram
Count
0
5
10
15
0
20
60
100
Value
Common sites gene9283
CphaseB2
RphaseB2

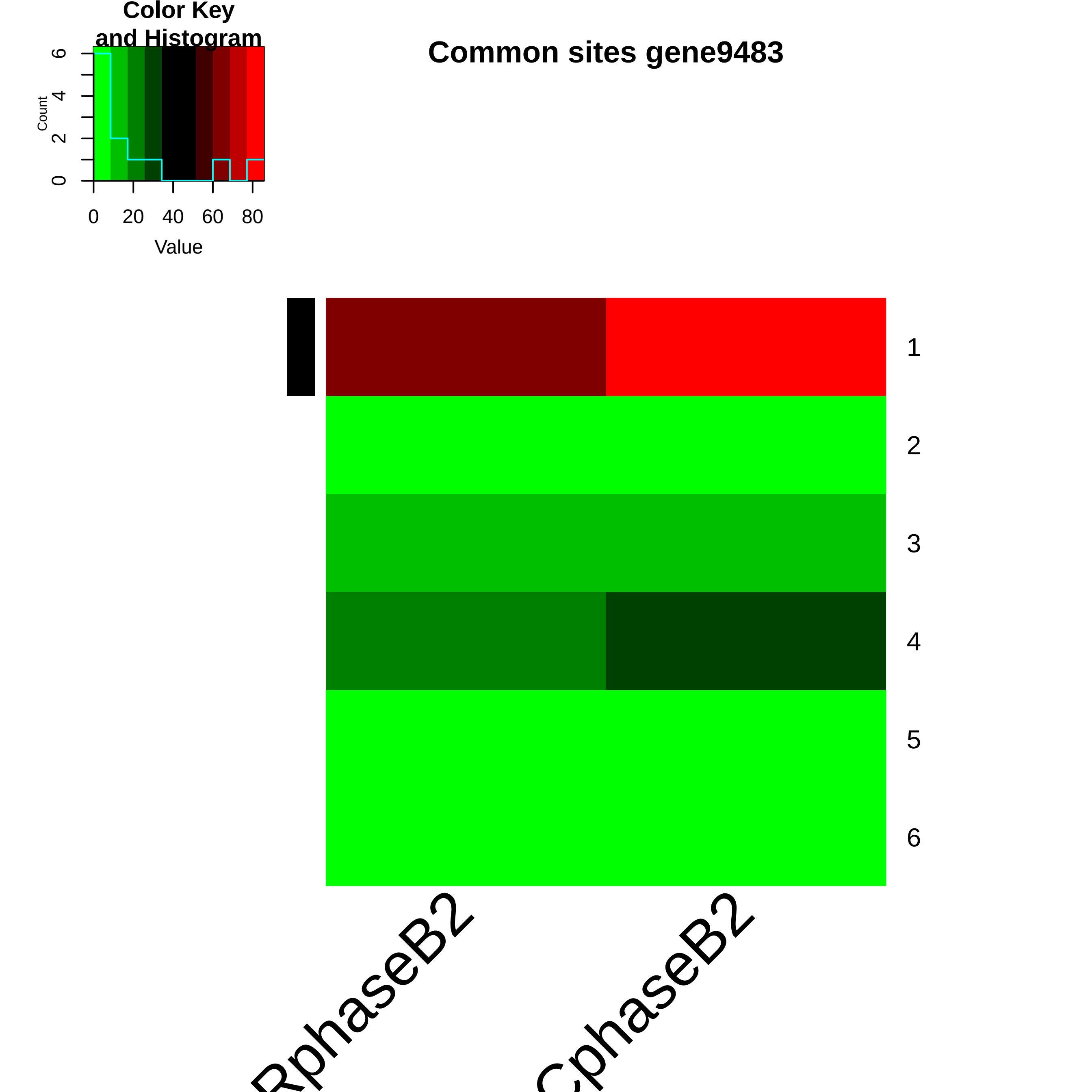
Color Key
and Histogram
Count
0
2
4
6
0
20
40
60
80
Value
Common sites gene9483
1
2
3
4
5
6
RphaseB2
CphaseB2

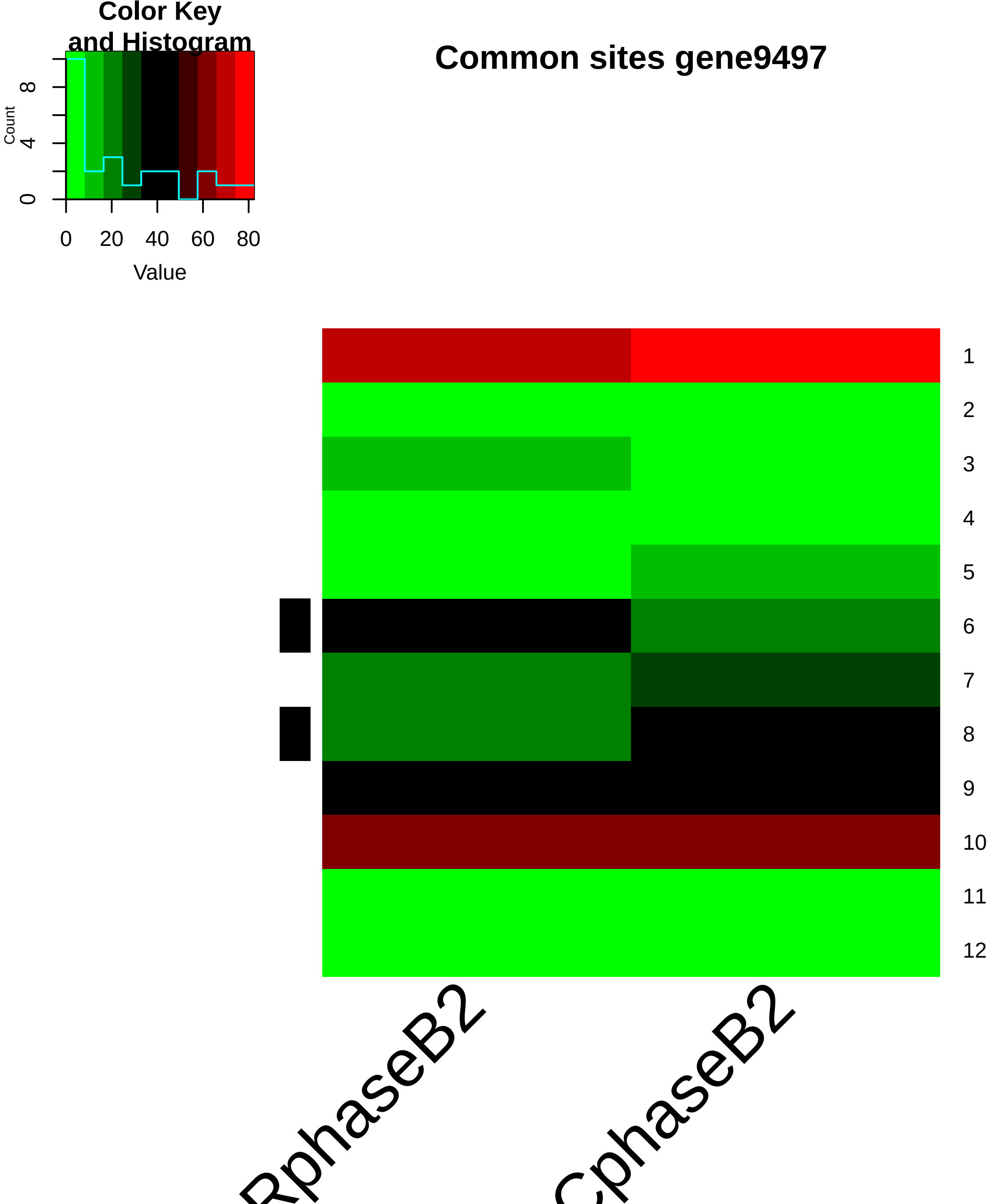

Color Key
and Histogram
Count
0
4
8
0
20
40
60
80
Value
Common sites gene9497
1
2
3
4
5
6
7
8
9
10
11
12
RphaseB2
CphaseB2

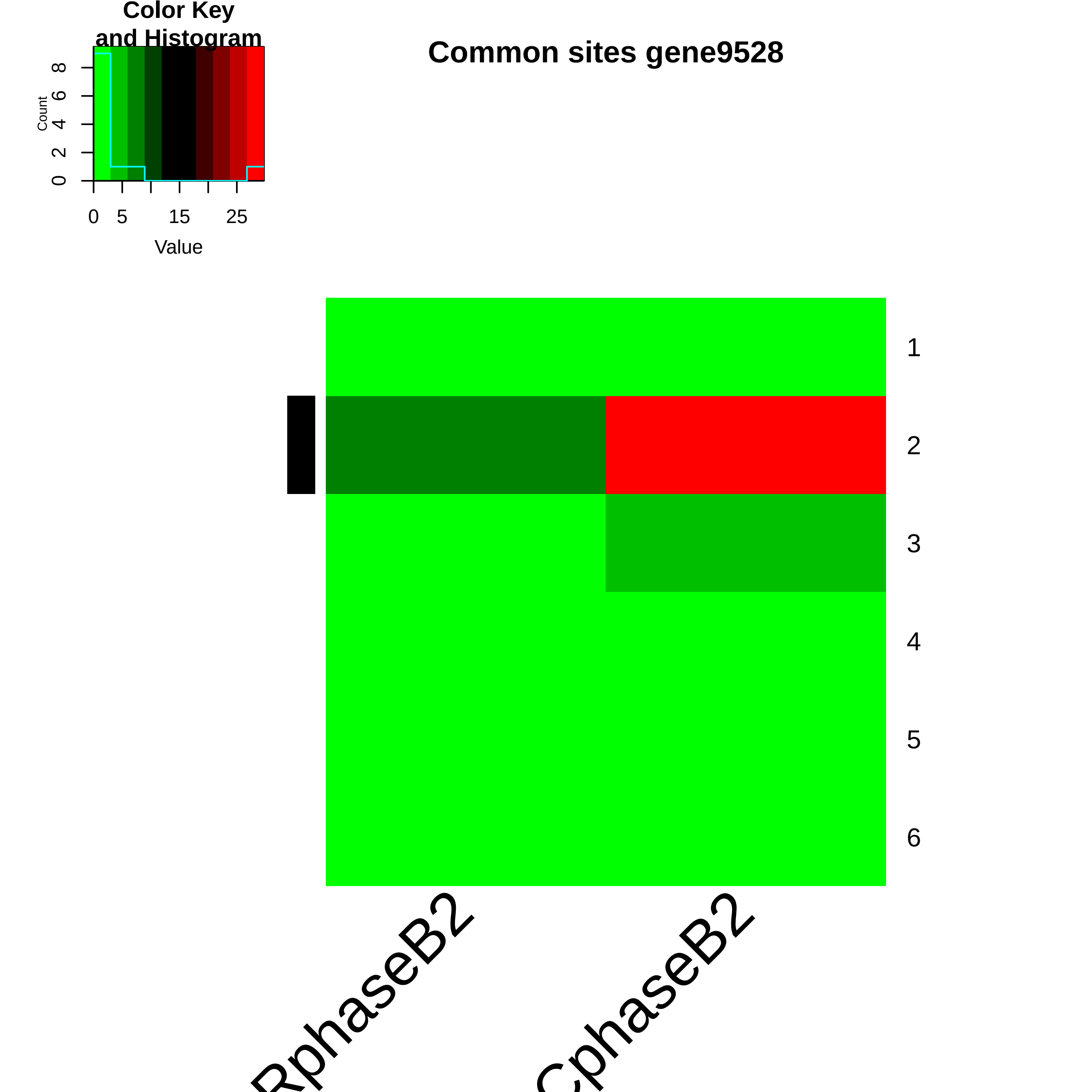

Color Key
and Histogram
Count
0
2
4
6
8
0
5
15
25
Value
Common sites gene9528
1
2
3
4
5
6
RphaseB2
CphaseB2

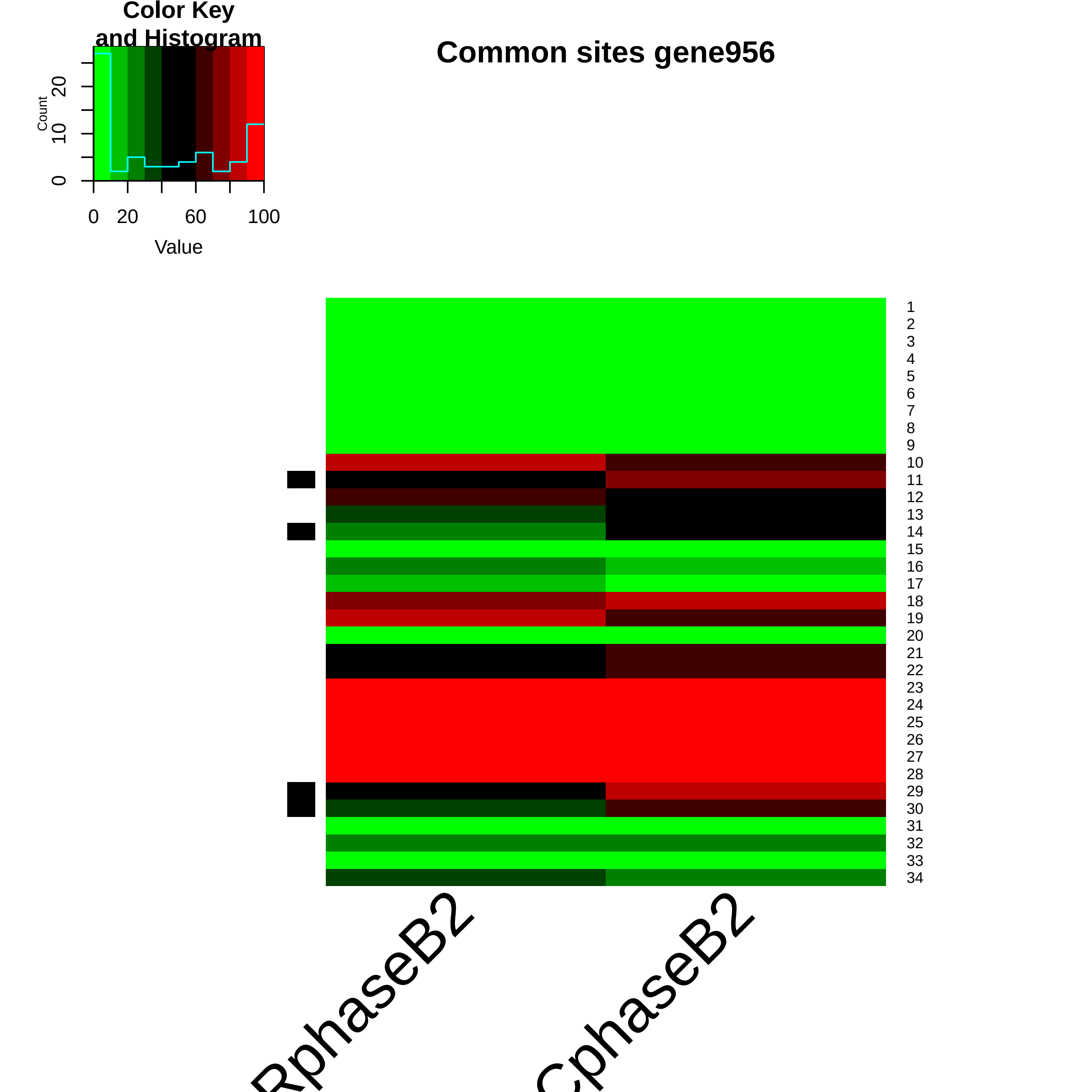
Color Key
and Histogram
Count
0
10
20
0
20
60
100
Value
Common sites gene956
RphaseB2
CphaseB2

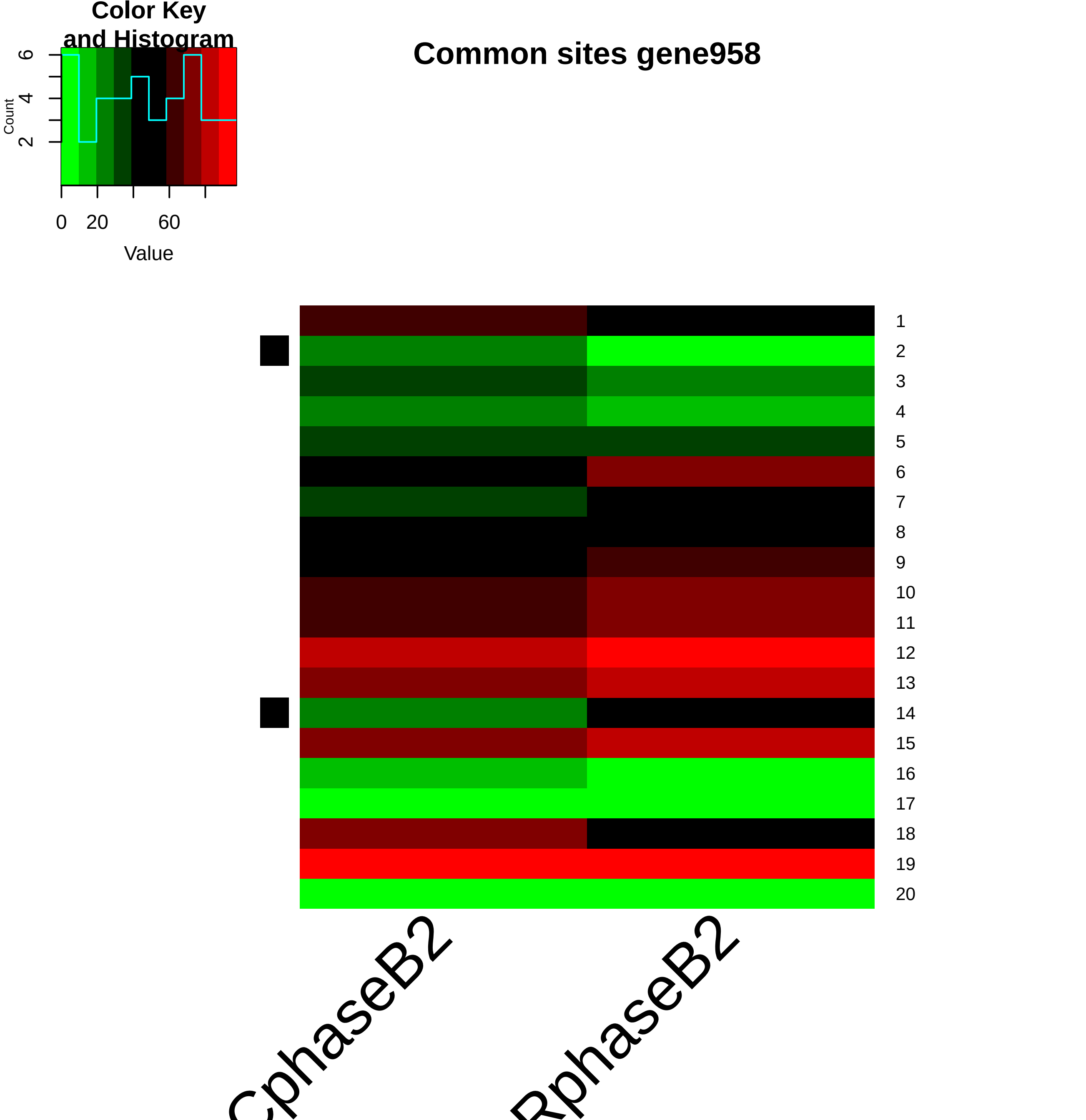

Color Key
and Histogram
Count
Value
0
20
60
2
4
6
Common sites gene958
CphaseB2
RphaseB2

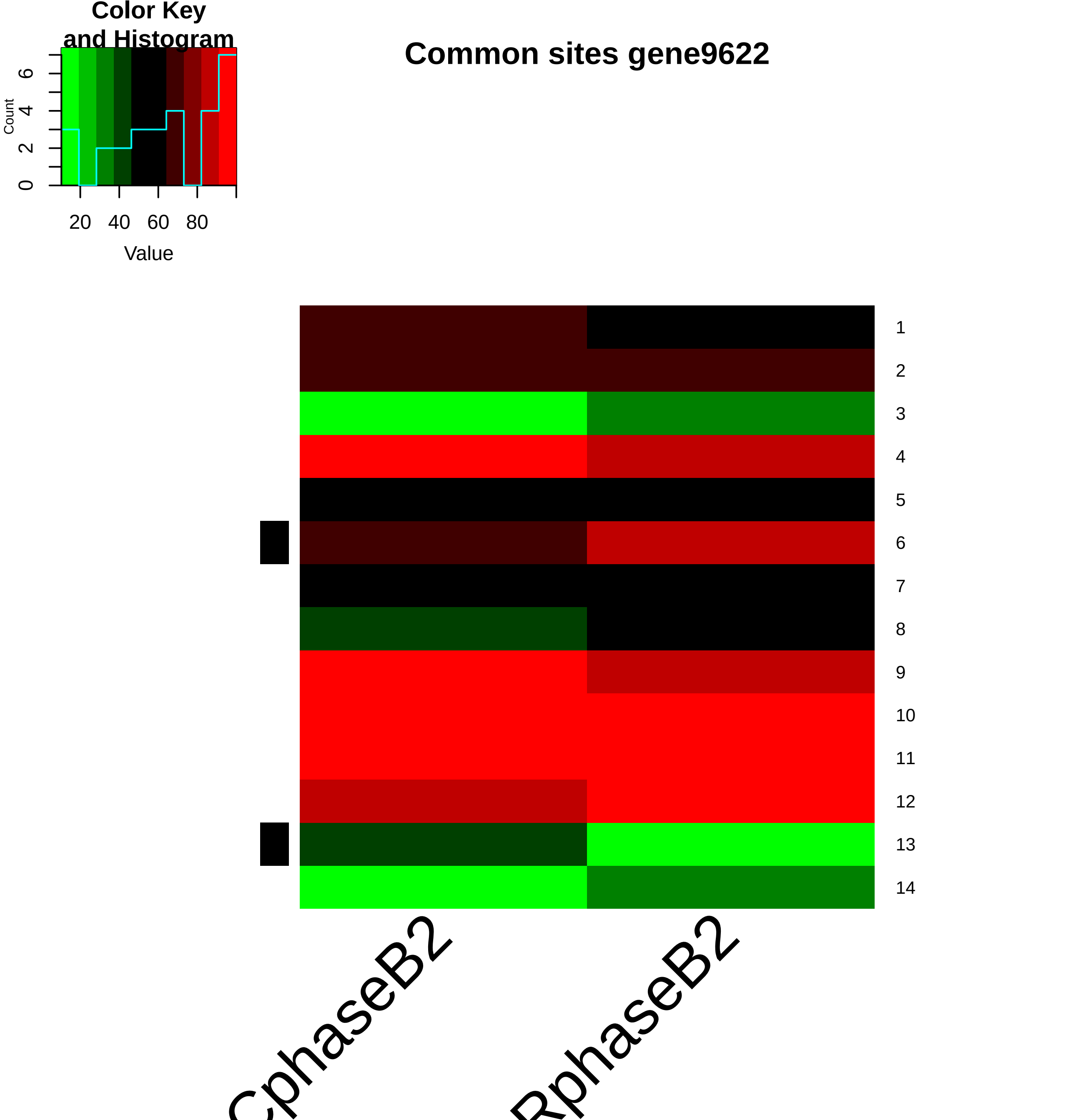
Color Key
and Histogram
Count
0
2
4
6
20
40
60
80
Value
Common sites gene9622
1
2
3
4
5
6
7
8
9
10
11
12
13
14
CphaseB2
RphaseB2

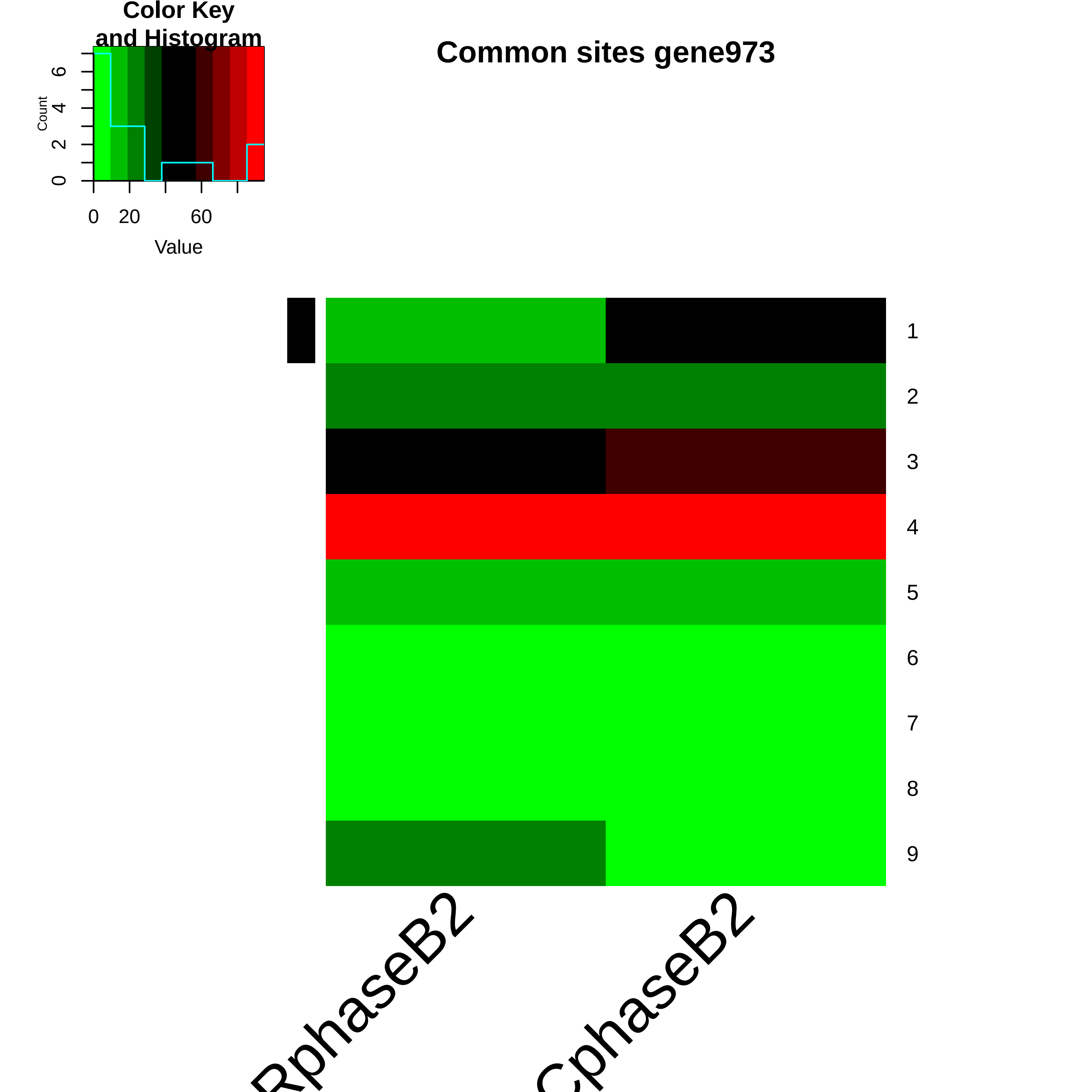
Color Key
and Histogram
Count
0
2
4
6
0
20
60
Value
Common sites gene973
1
2
3
4
5
6
7
8
9
RphaseB2
CphaseB2

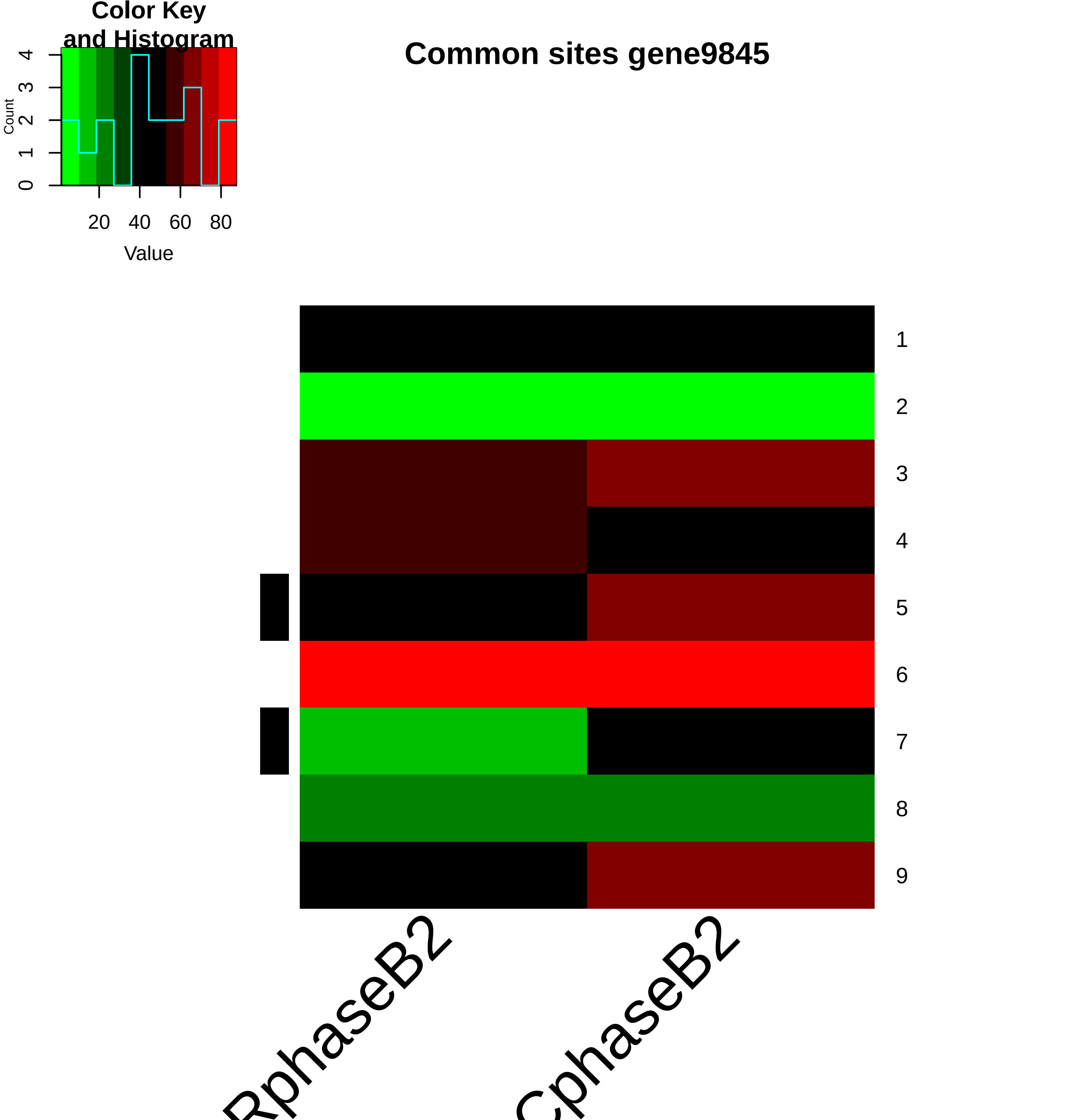

Color Key
and Histogram
Count
0
1
2
3
4
20
40
60
80
Value
Common sites gene9845
1
2
3
4
5
6
7
8
9
RphaseB2
CphaseB2

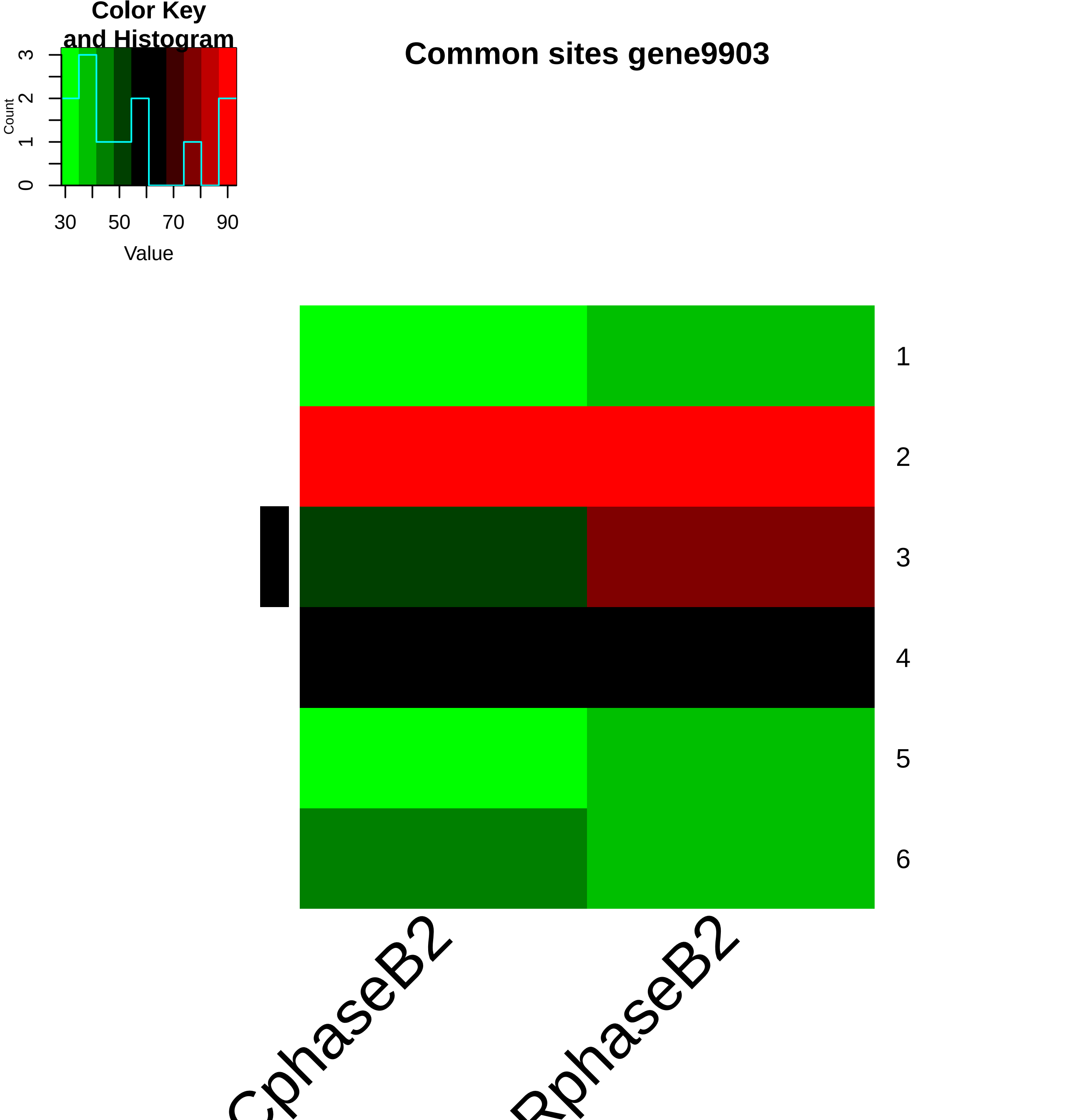

Color Key
and Histogram
Count
0
1
2
3
30
50
70
90
Value
Common sites gene9903
1
2
3
4
5
6
CphaseB2
RphaseB2

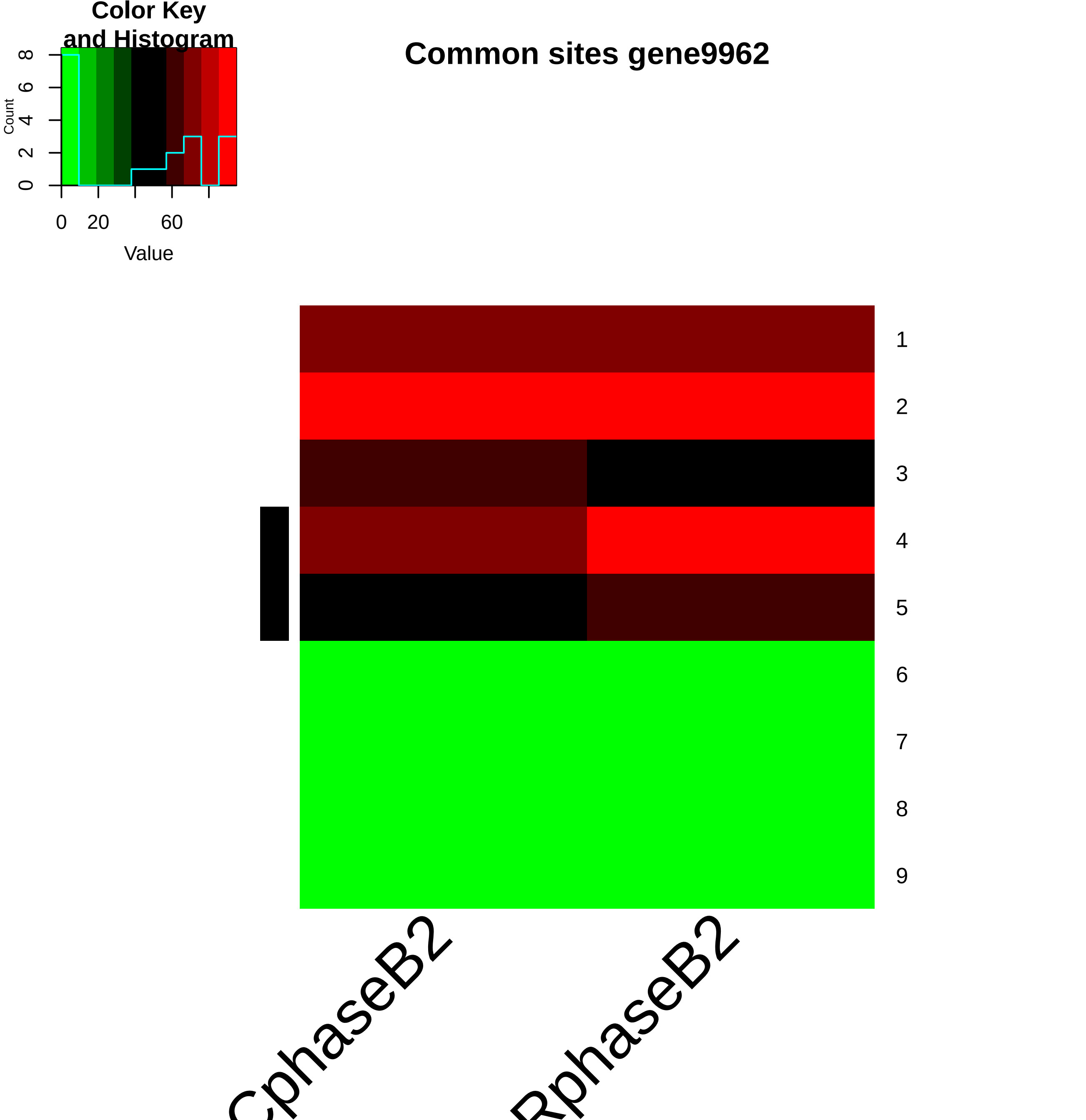
Color Key
and Histogram
Count
0
2
4
6
8
0
20
60
Value
Common sites gene9962
1
2
3
4
5
6
7
8
9

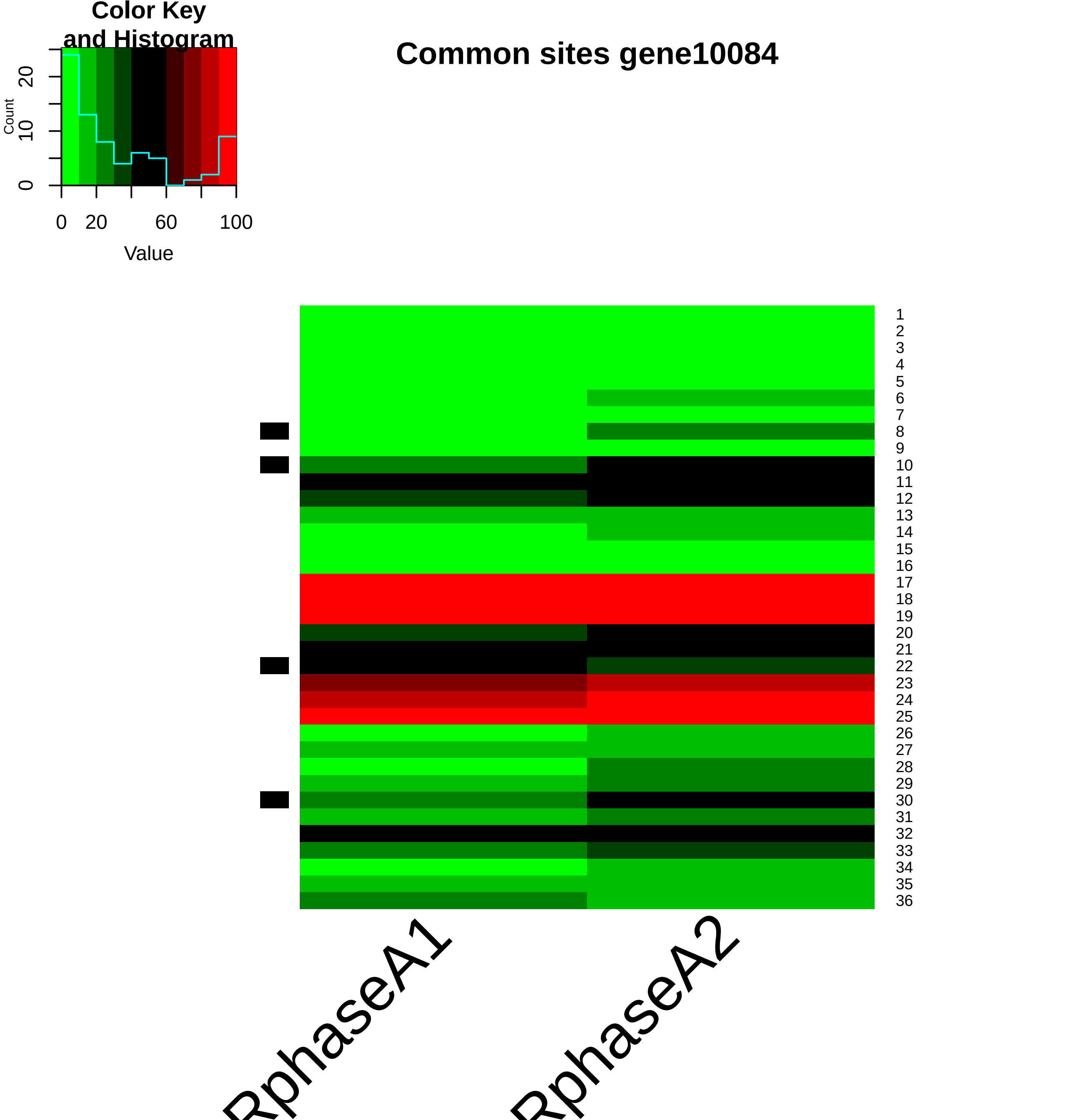
Color Key
and Histogram
Count
0
10
20
0
20
60
100
Value
Common sites gene10084
RphaseA1
RphaseA2

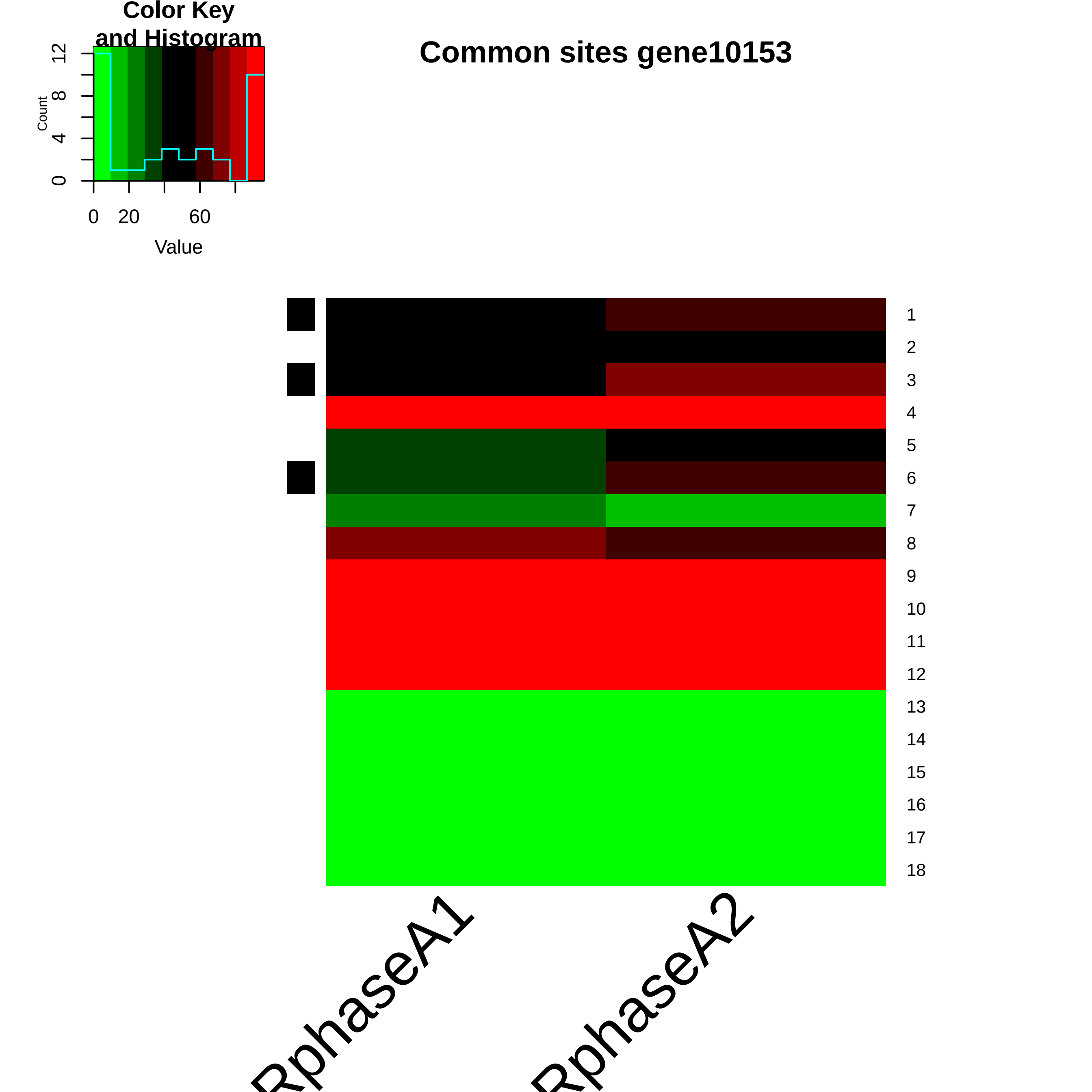
Color Key
and Histogram
Count
0
4
8
12
0
20
60
Value
Common sites gene10153
RphaseA1
RphaseA2

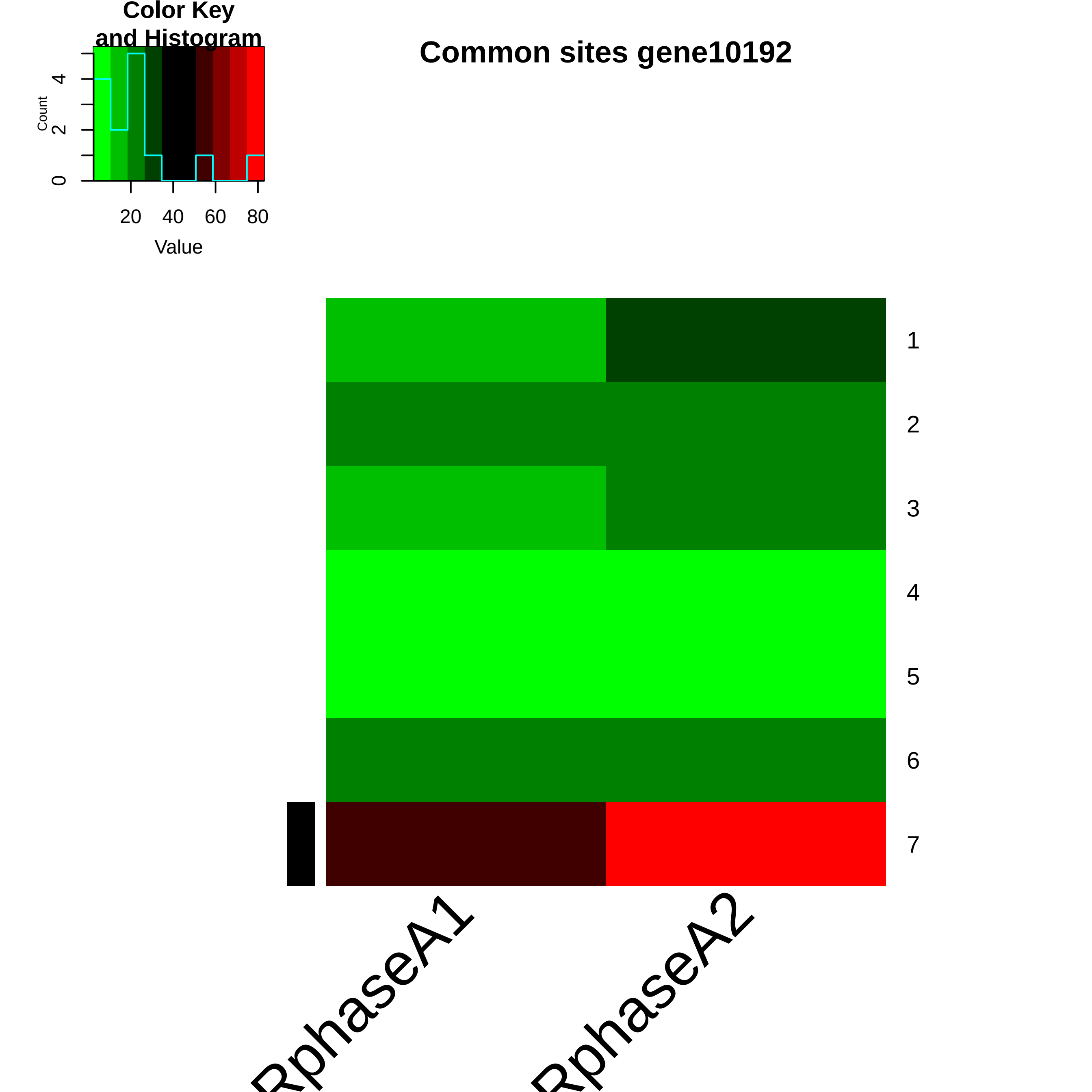
Color Key
and Histogram
Count
0
2
4
20
40
60
80
Value
Common sites gene10192
1
2
3
4
5
6
7
RphaseA1
RphaseA2

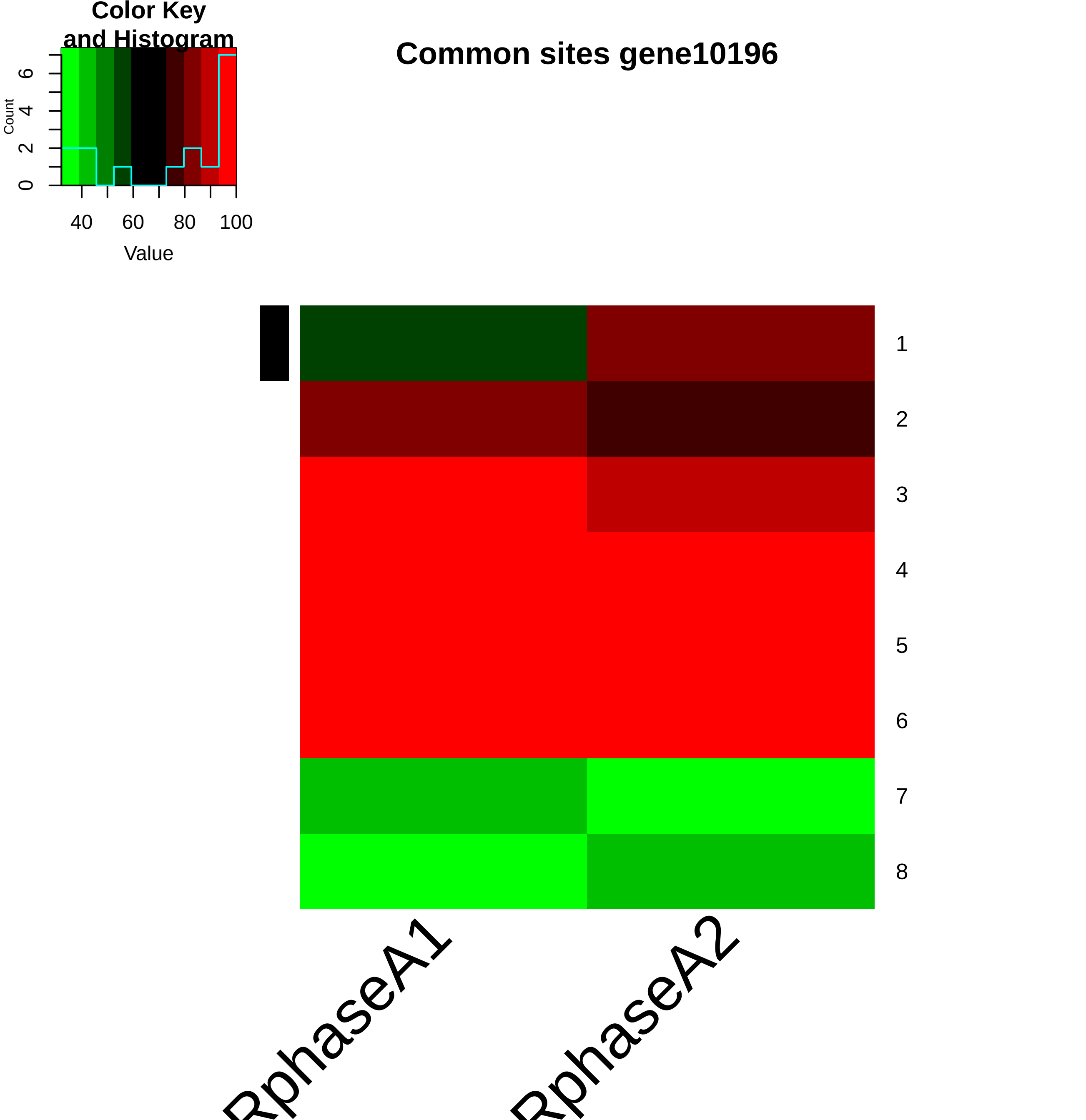

Color Key
and Histogram
Count
0
2
4
6
40
60
80
100
Value
Common sites gene10196
1
2
3
4
5
6
7
8
RphaseA1
RphaseA2

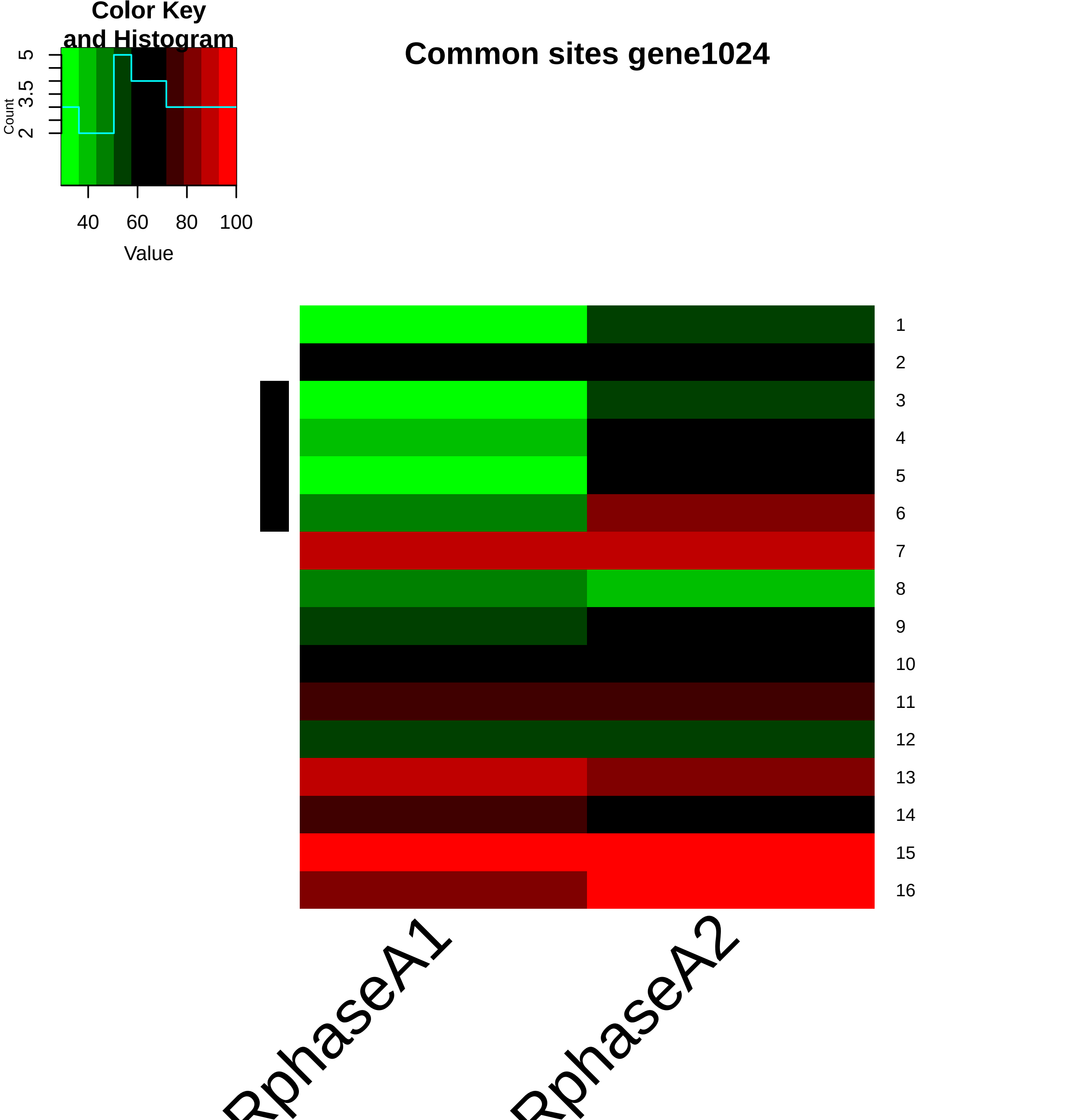

Color Key
and Histogram
Count
2
3.5
5
40
60
80
100
Value
Common sites gene1024
RphaseA1
RphaseA2

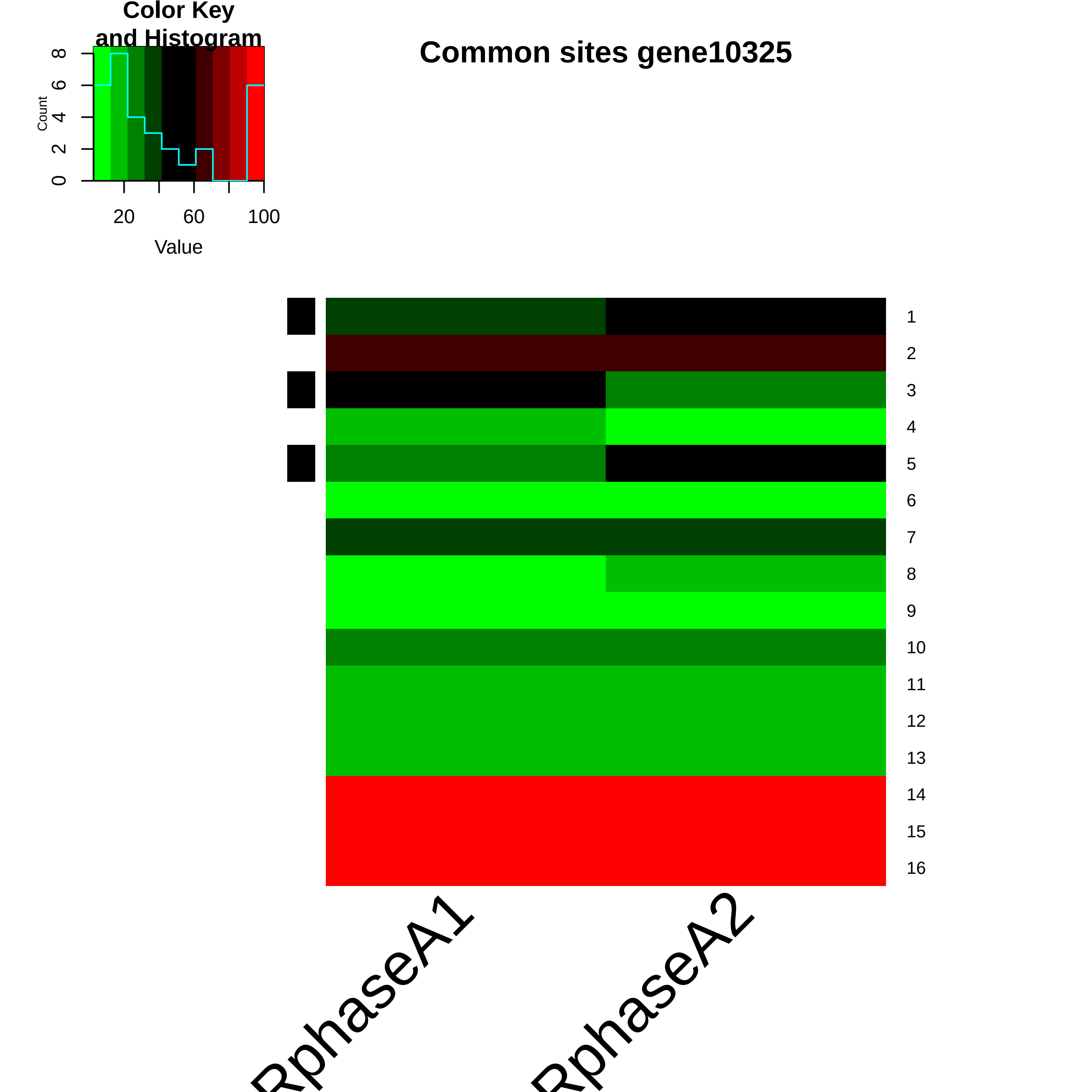
Color Key
and Histogram
Count
Value
0
2
4
6
8
20
60
100
Common sites gene10325
RphaseA1
RphaseA2

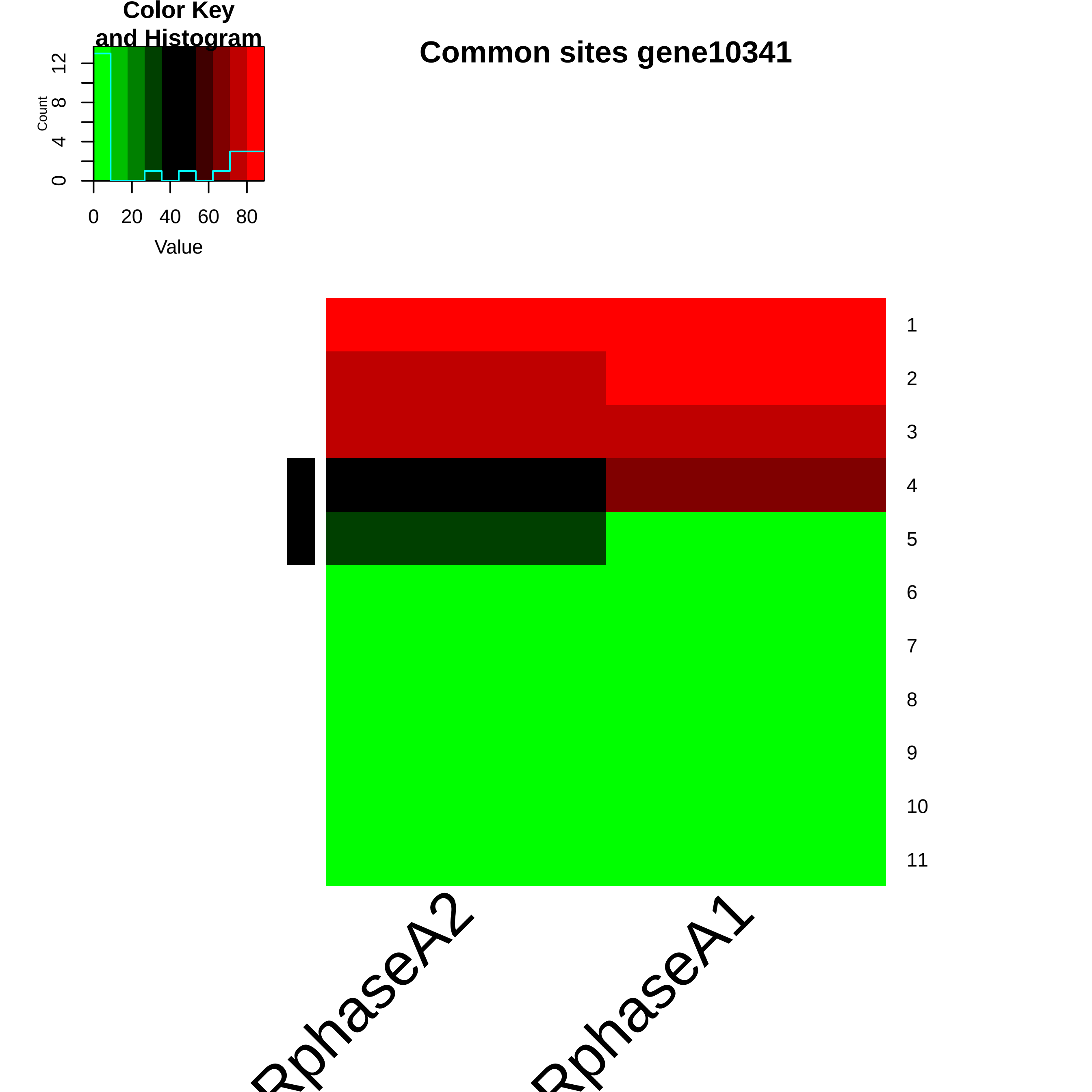
Color Key
and Histogram
Count
0
4
8
12
0
20
40
60
80
Value
Common sites gene10341
1
2
3
4
5
6
7
8
9
10
11
RphaseA2
RphaseA1

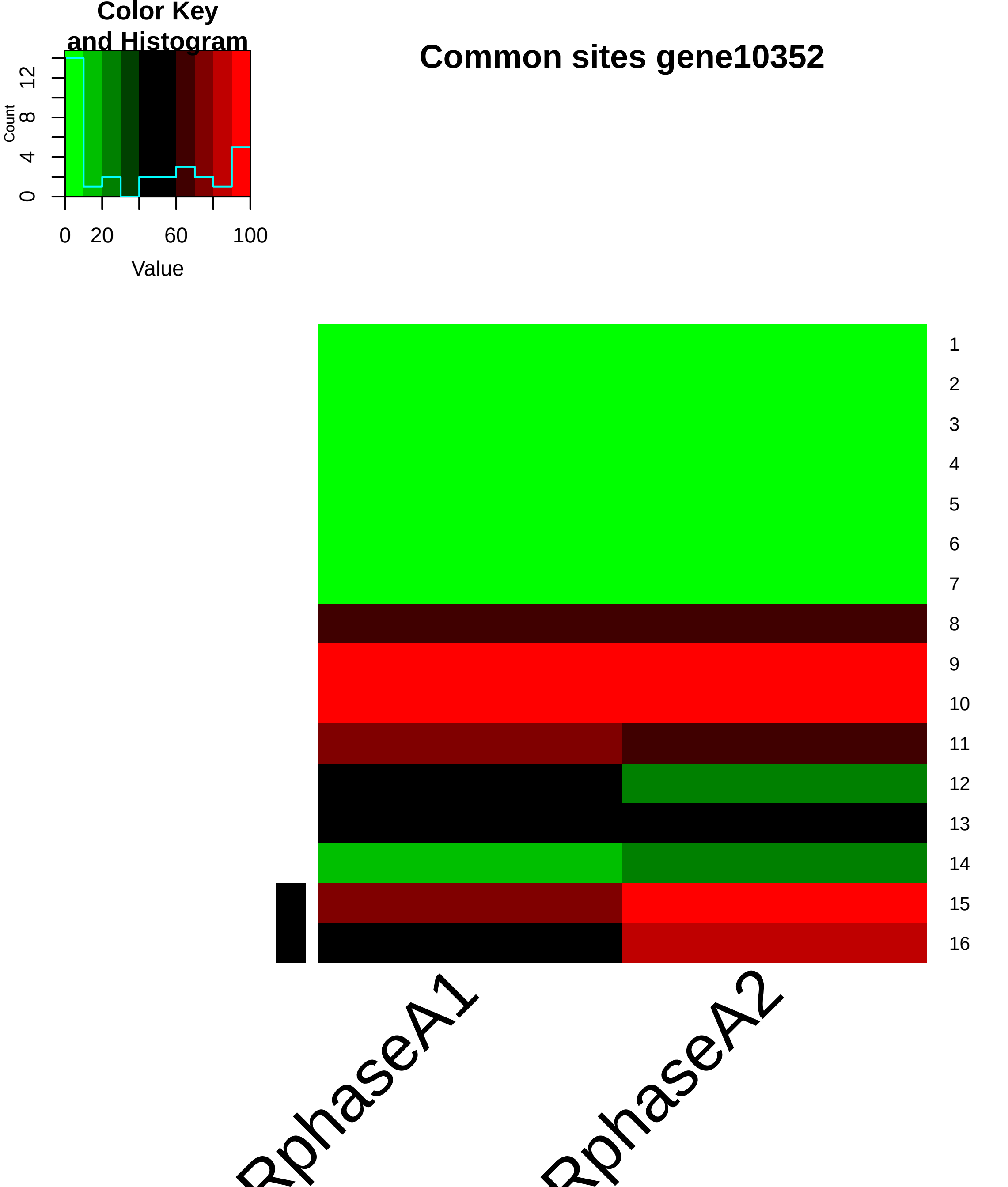

Color Key
and Histogram
Count
0
4
8
12
0
20
60
100
Value
Common sites gene10352
RphaseA1
RphaseA2

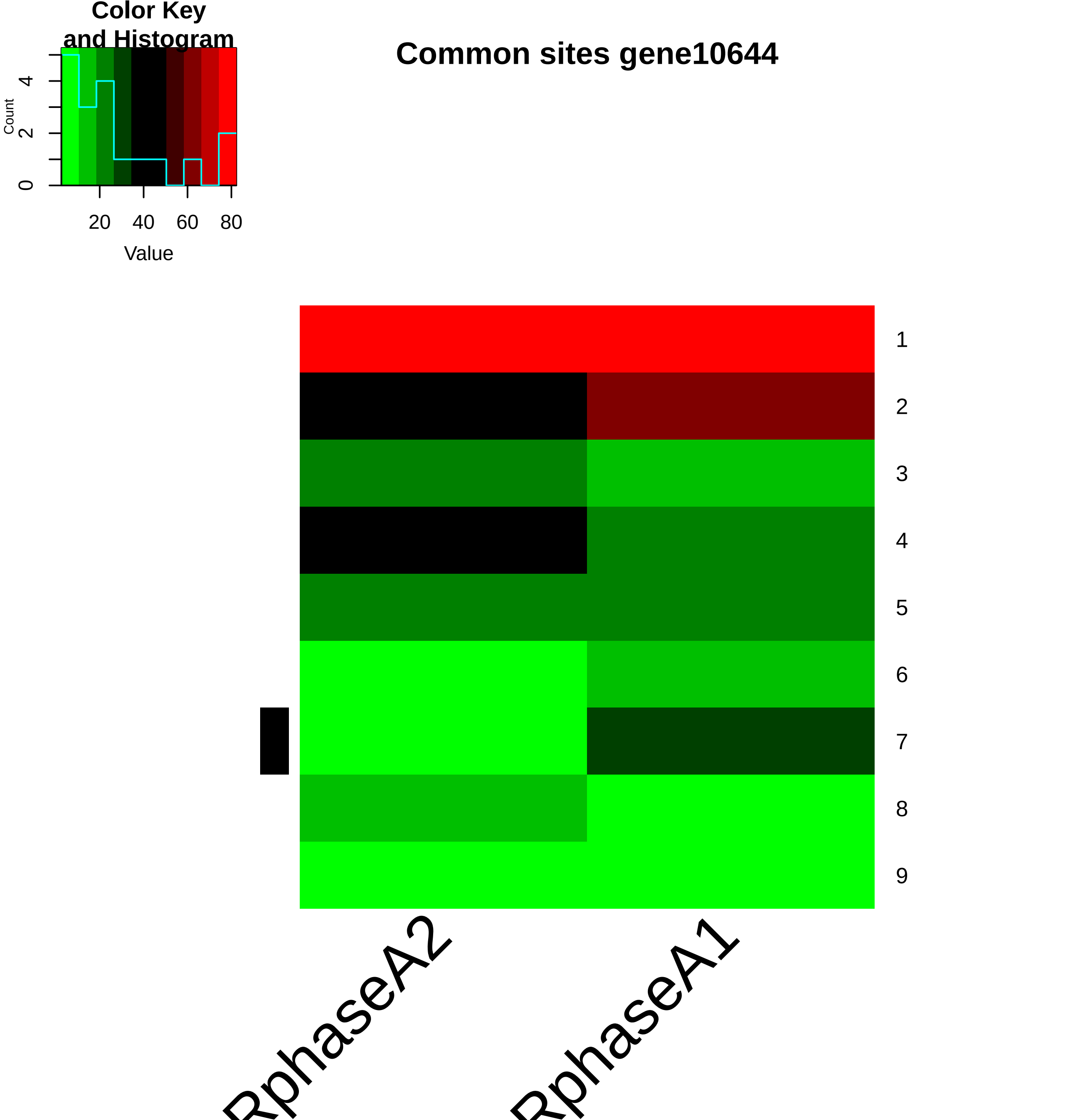

Color Key
and Histogram
Count
0
2
4
20
40
60
80
Value
Common sites gene10644
1
2
3
4
5
6
7
8
9
RphaseA2
RphaseA1

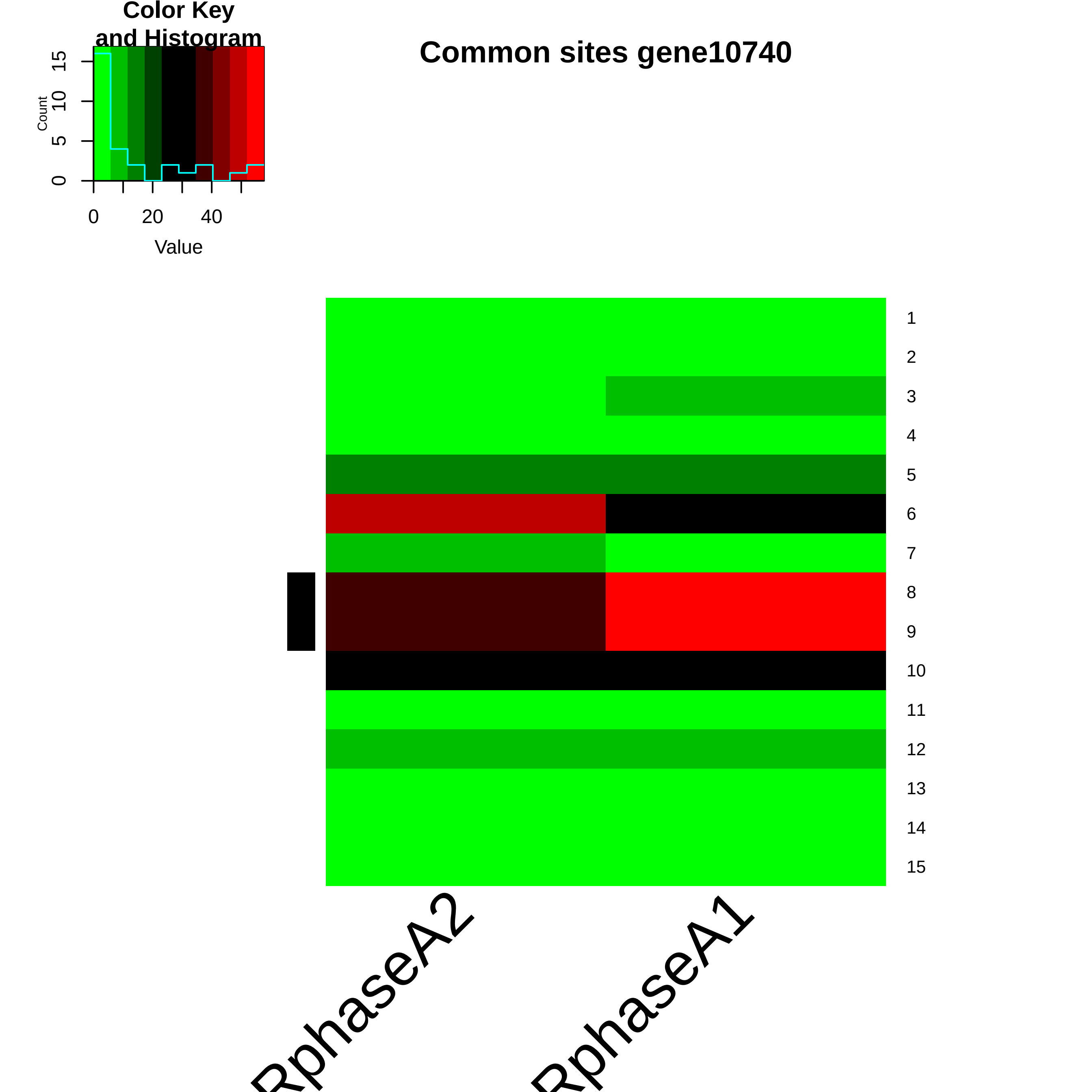

Color Key
and Histogram
Count
0
5
10
15
0
20
40
Value
Common sites gene10740
RphaseA2
RphaseA1

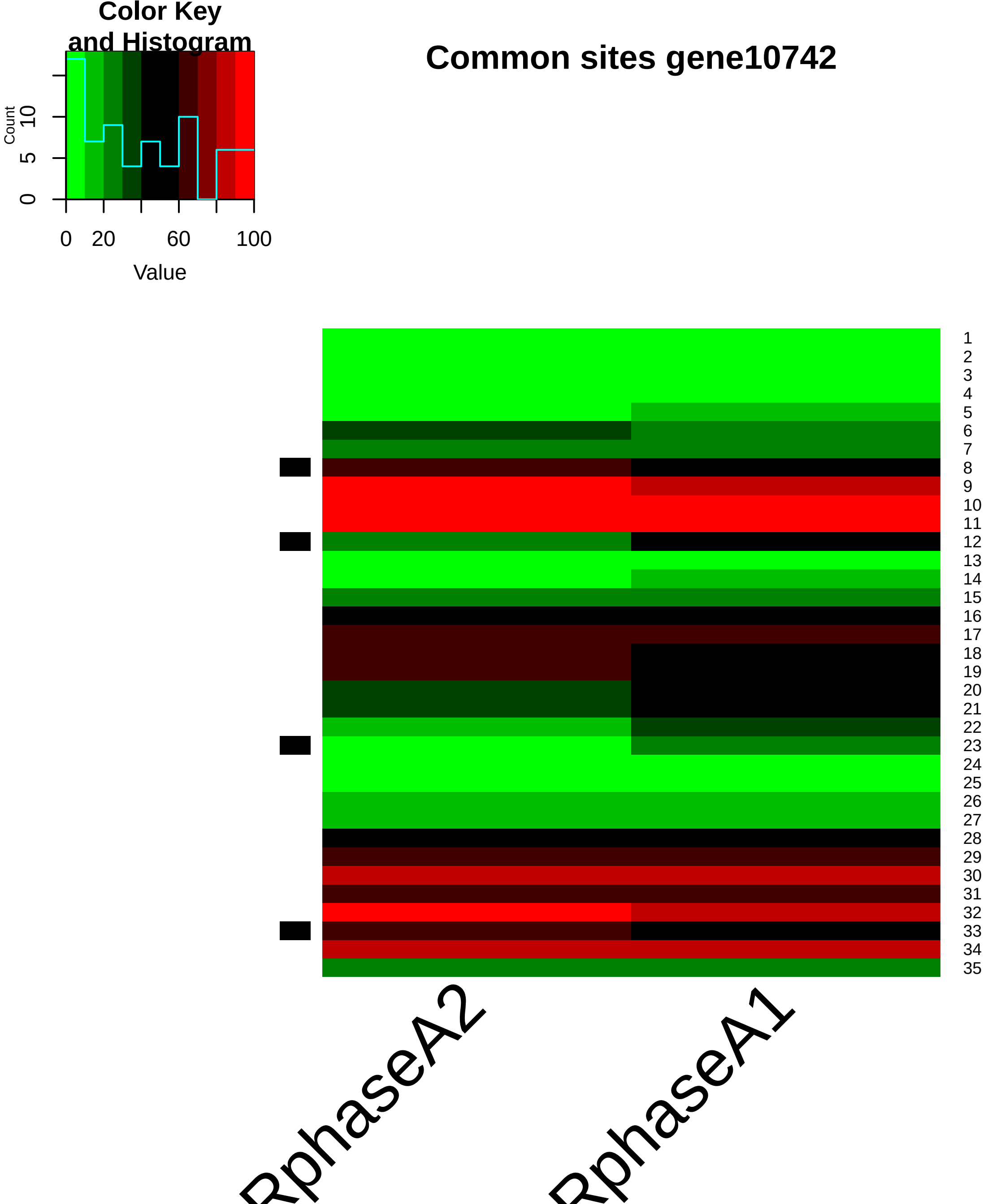

Color Key
and Histogram
Count
0
5
10
0
20
60
100
Value
Common sites gene10742
RphaseA2
RphaseA1

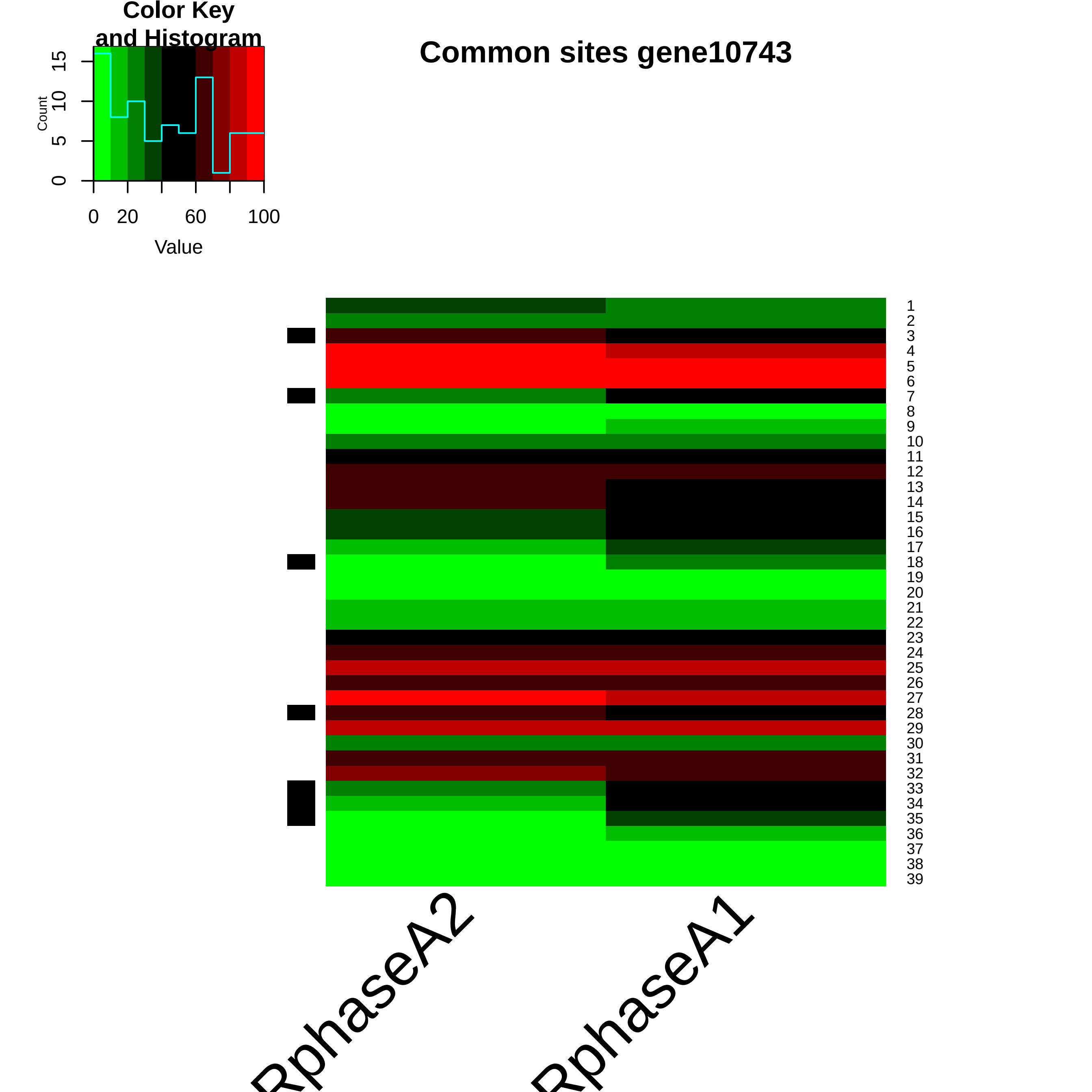

Color Key
and Histogram
Count
0
5
10
15
0
20
60
100
Value
Common sites gene10743
RphaseA2
RphaseA1

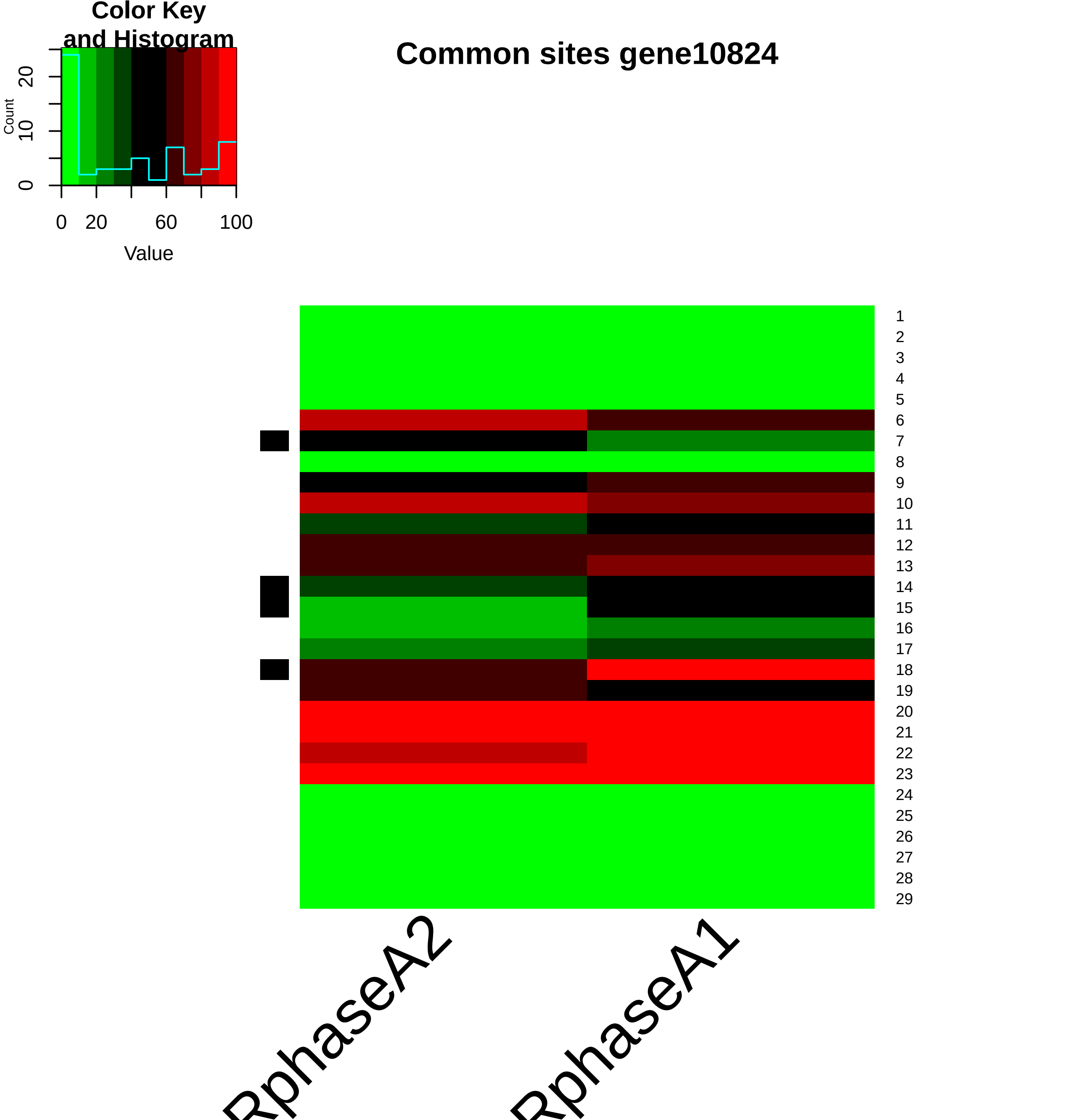
Color Key
and Histogram
Count
0
10
20
0
20
60
100
Value
Common sites gene10824
RphaseA2
RphaseA1

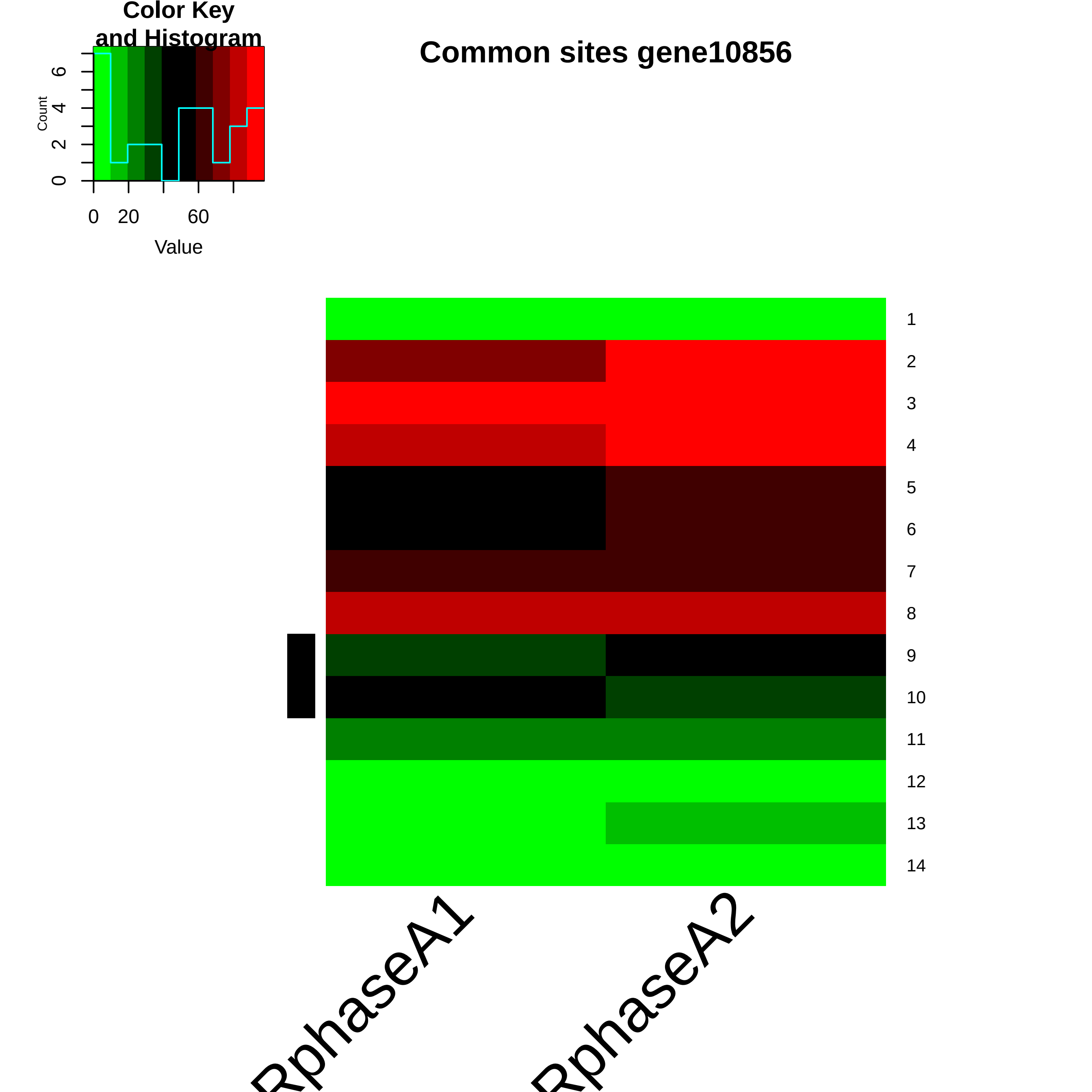
Color Key
and Histogram
Count
0
2
4
6
0
20
60
Value
Common sites gene10856
1
2
3
4
5
6
7
8
9
10
11
12
13
14
RphaseA1
RphaseA2

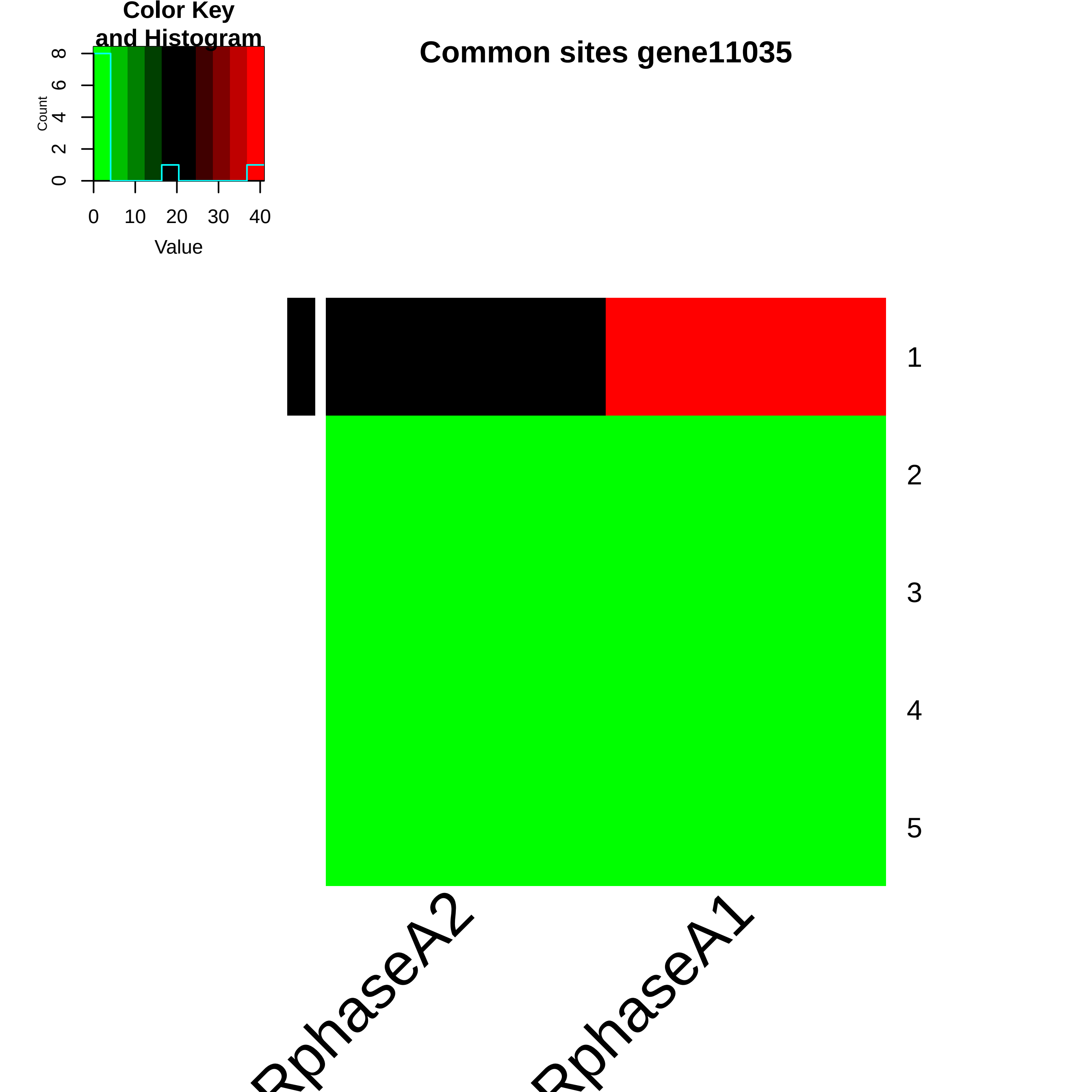
Color Key
and Histogram
Count
Value
0
10
20
30
40
0
2
4
6
8
Common sites gene11035
1
2
3
4
5
RphaseA2
RphaseA1

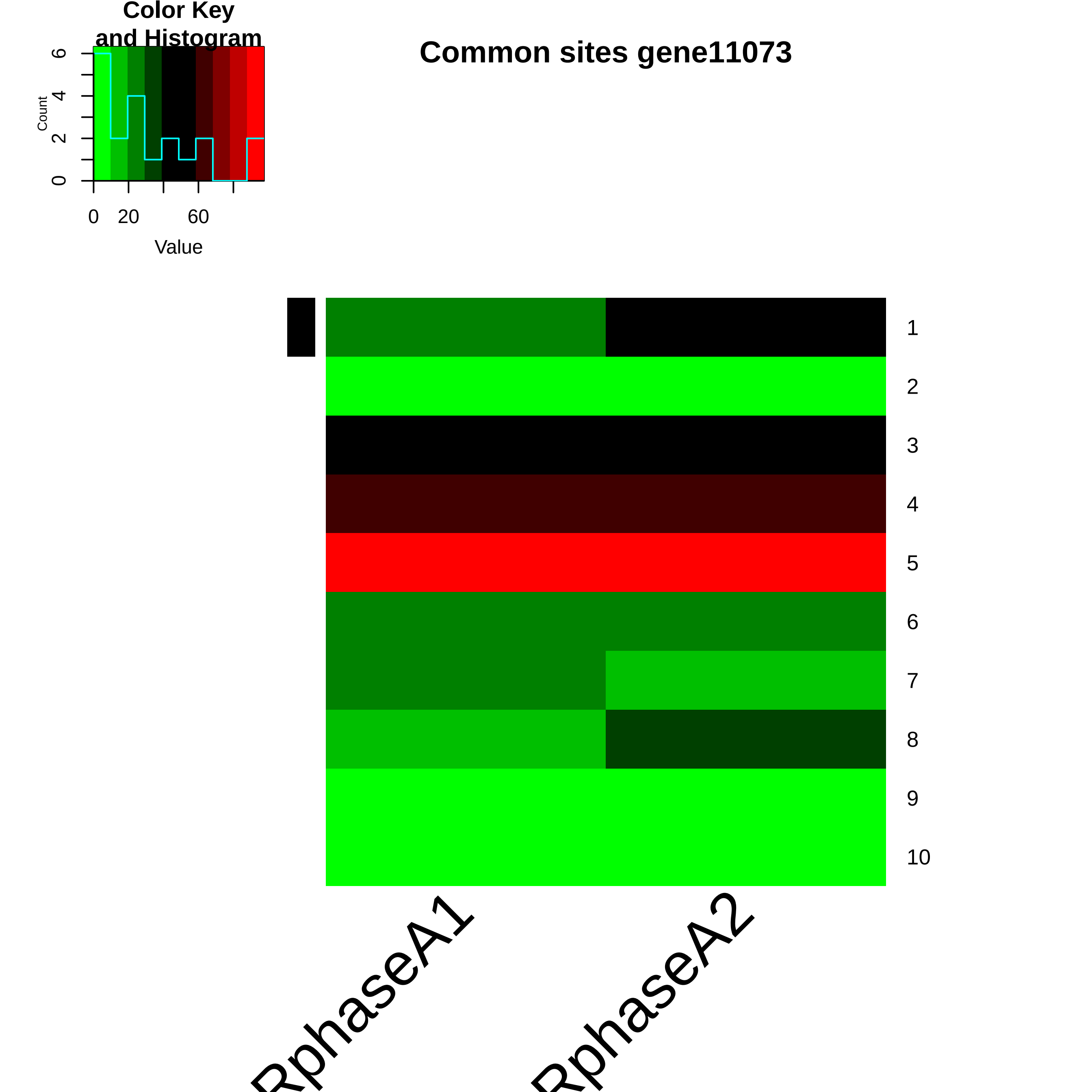

Color Key
and Histogram
Count
0
2
4
6
0
20
60
Value
Common sites gene11073
1
2
3
4
5
6
7
8
9
10
RphaseA1
RphaseA2

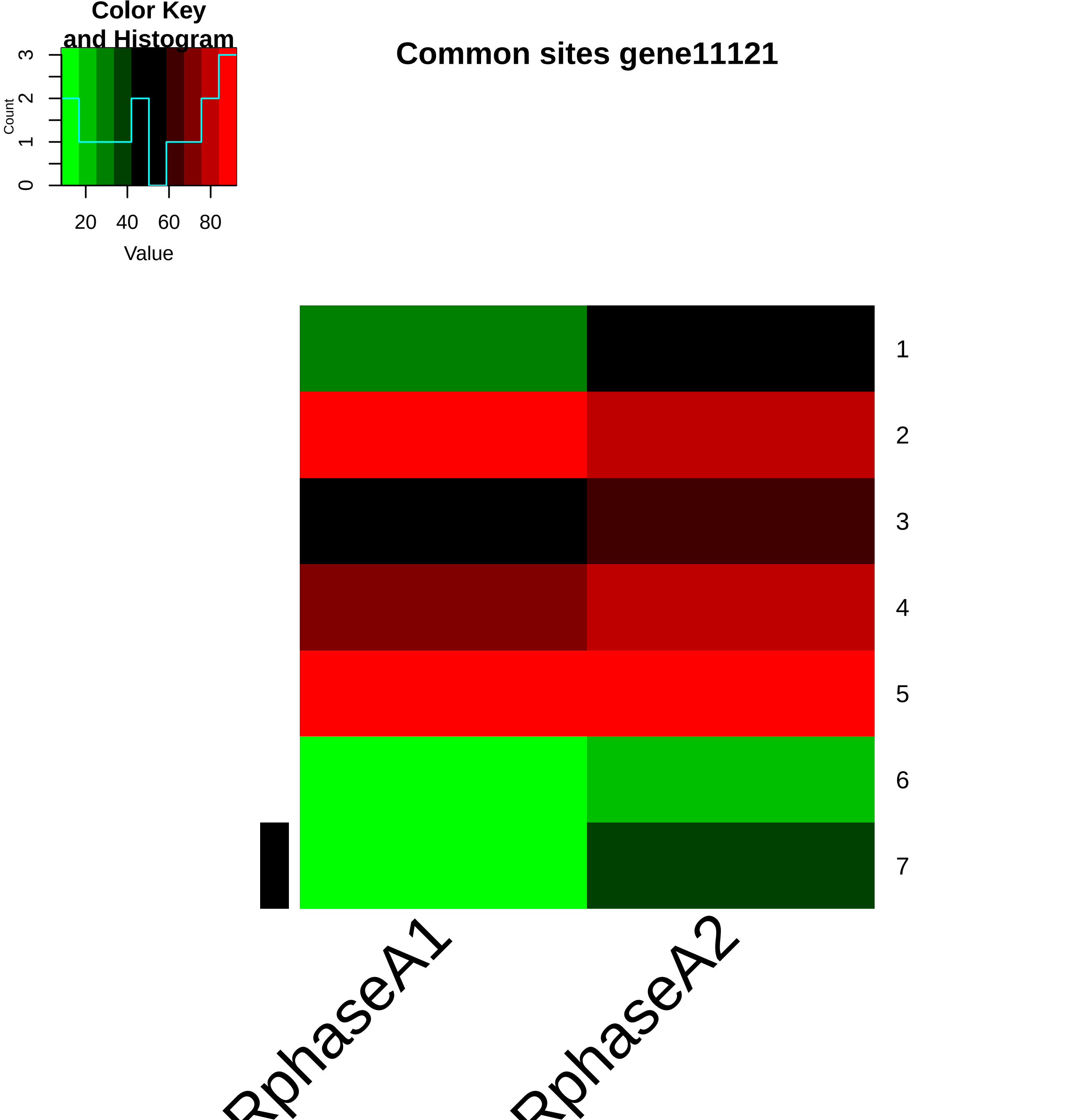
Color Key
and Histogram
Count
0
1
2
3
20
40
60
80
Value
Common sites gene11121
1
2
3
4
5
6
7
RphaseA1
RphaseA2

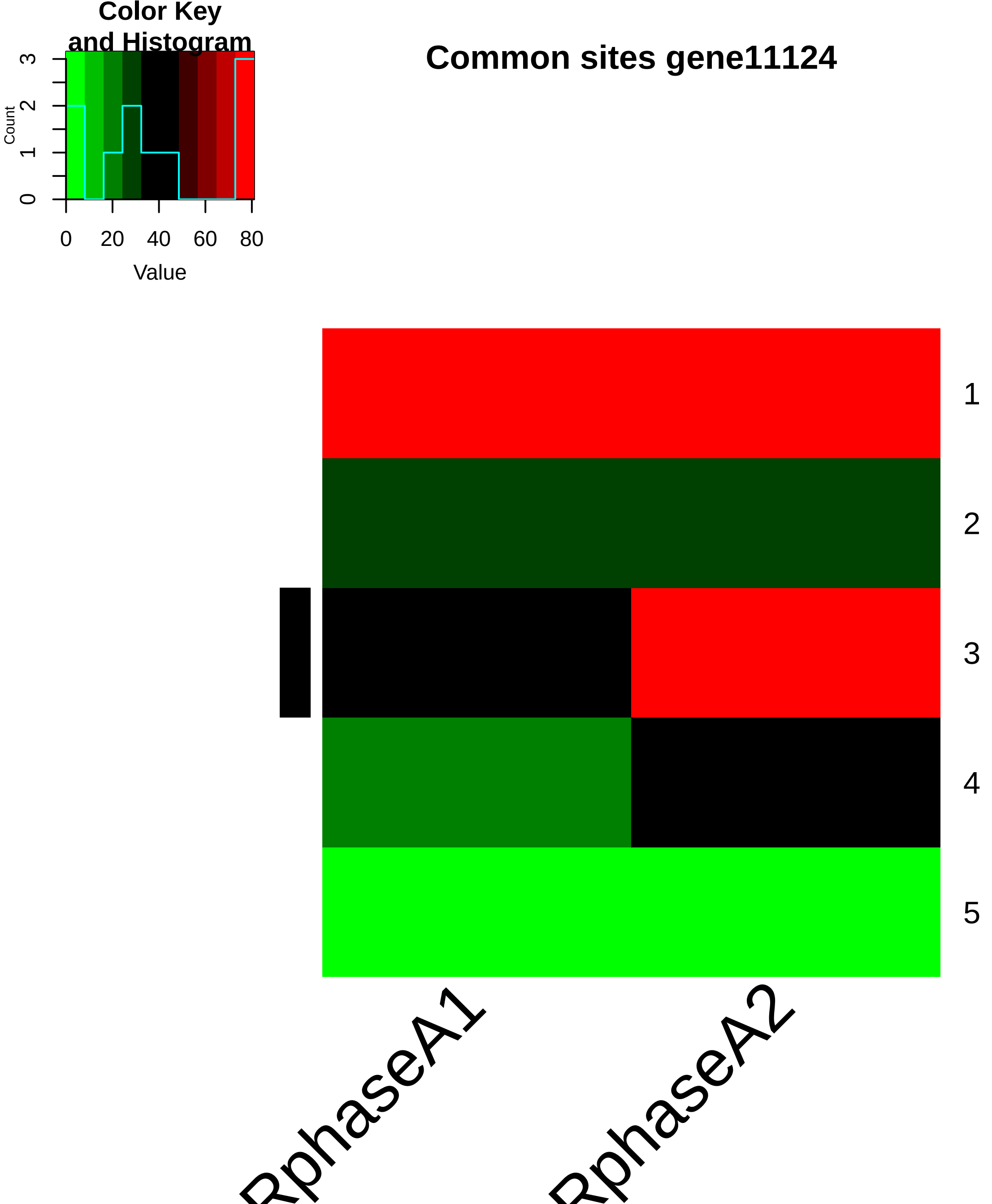
Color Key
and Histogram
Count
0
1
2
3
0
20
40
60
80
Value
Common sites gene11124
1
2
3
4
5
RphaseA1
RphaseA2

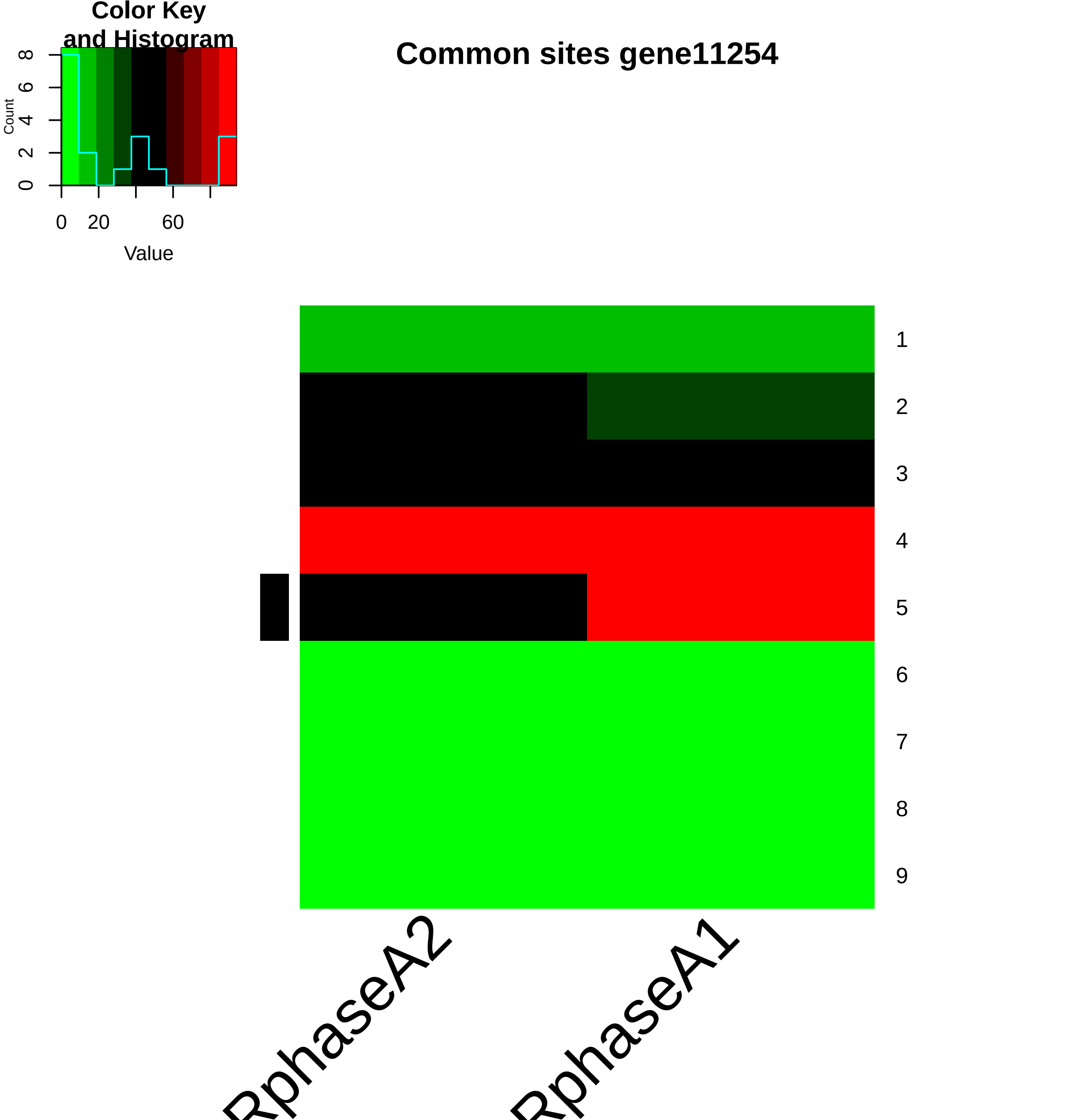
Color Key
and Histogram
Common sites gene11254
Count
0
2
4
6
8
0
20
60
Value
1
2
3
4
5
6
7
8
9
RphaseA2
RphaseA1

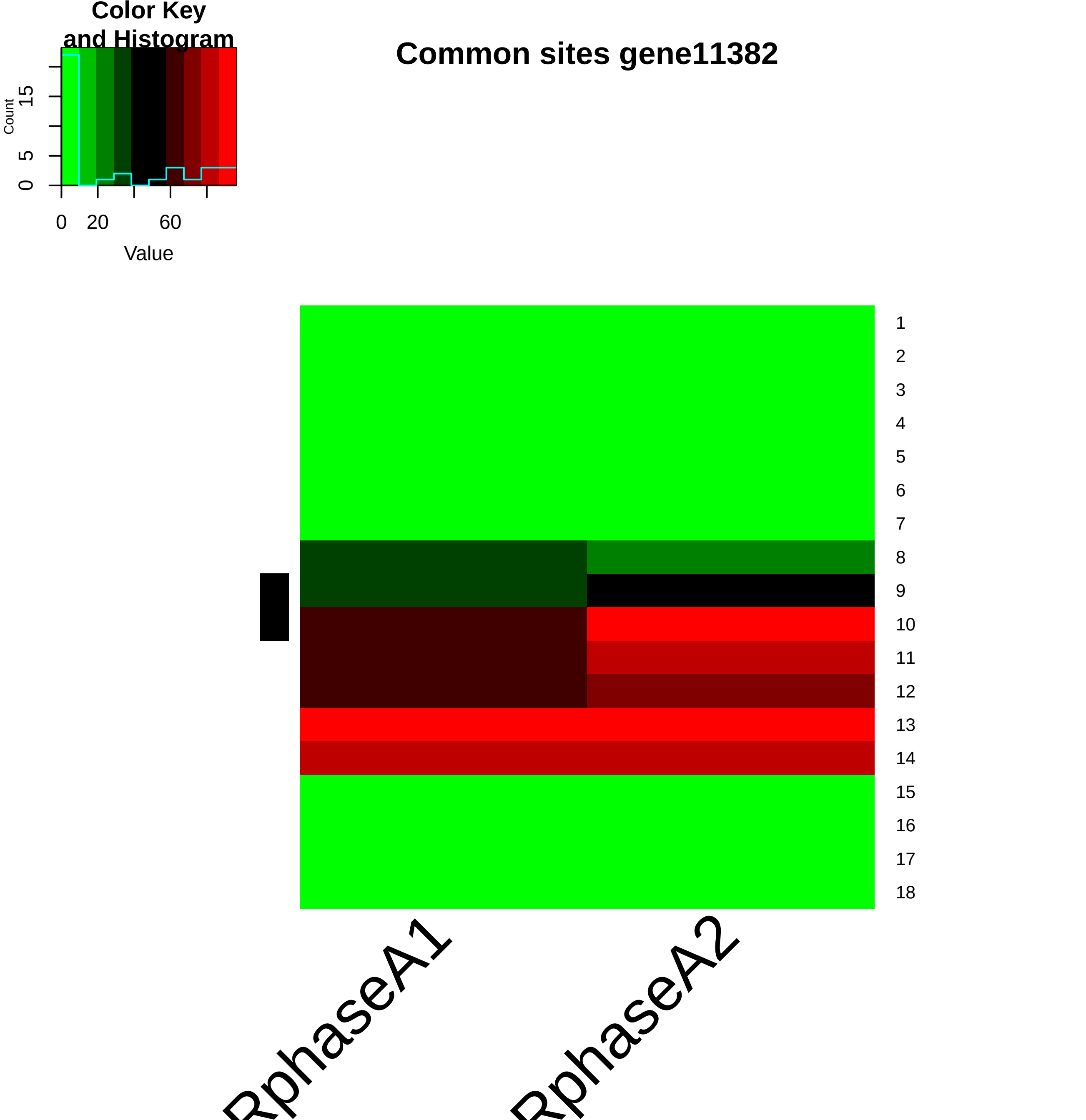

Color Key
and Histogram
Count
0
5
15
0
20
60
Value
Common sites gene11382
RphaseA1
RphaseA2

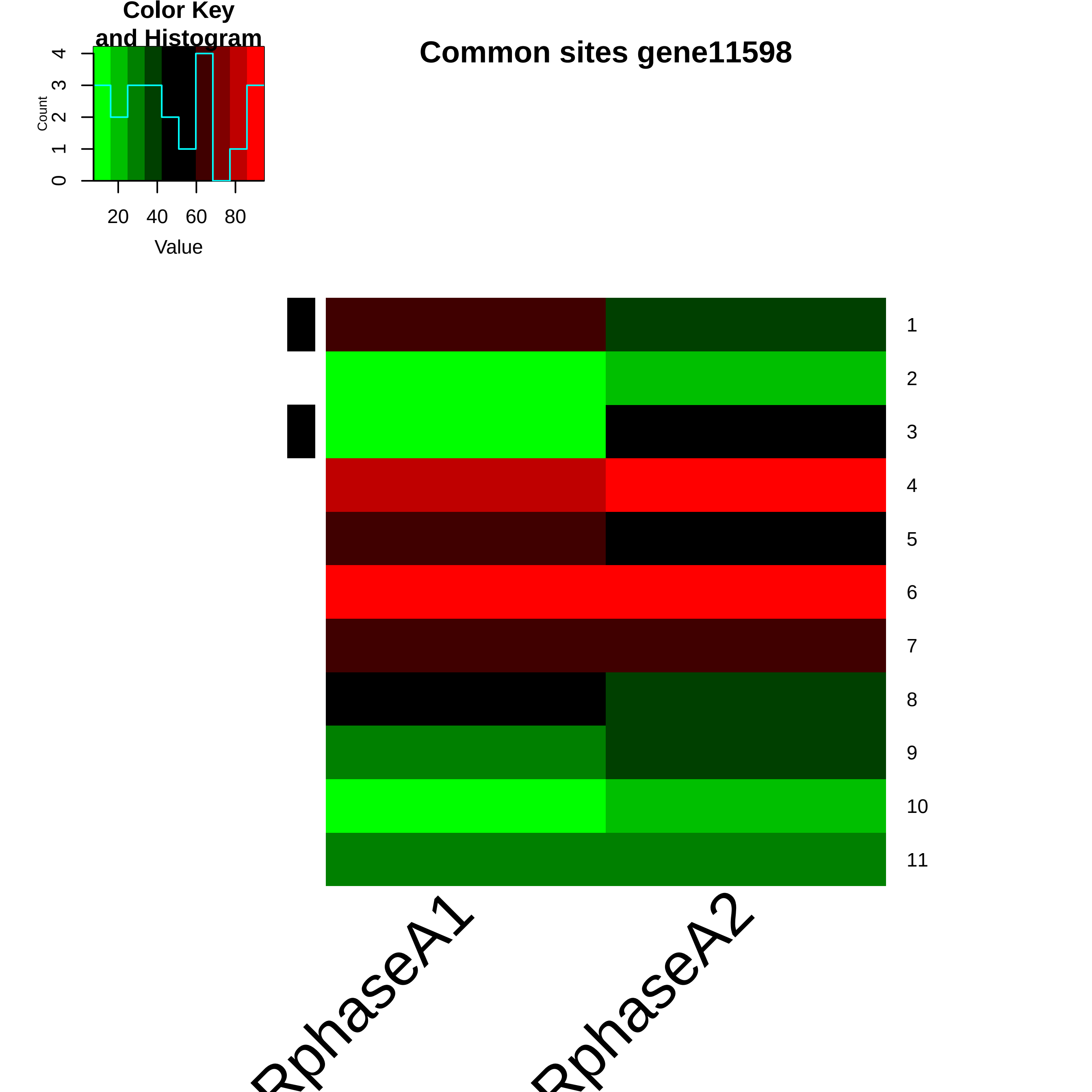
Color Key
and Histogram
Count
0
1
2
3
4
20
40
60
80
Value
Common sites gene11598
1
2
3
4
5
6
7
8
9
10
11
RphaseA1
RphaseA2

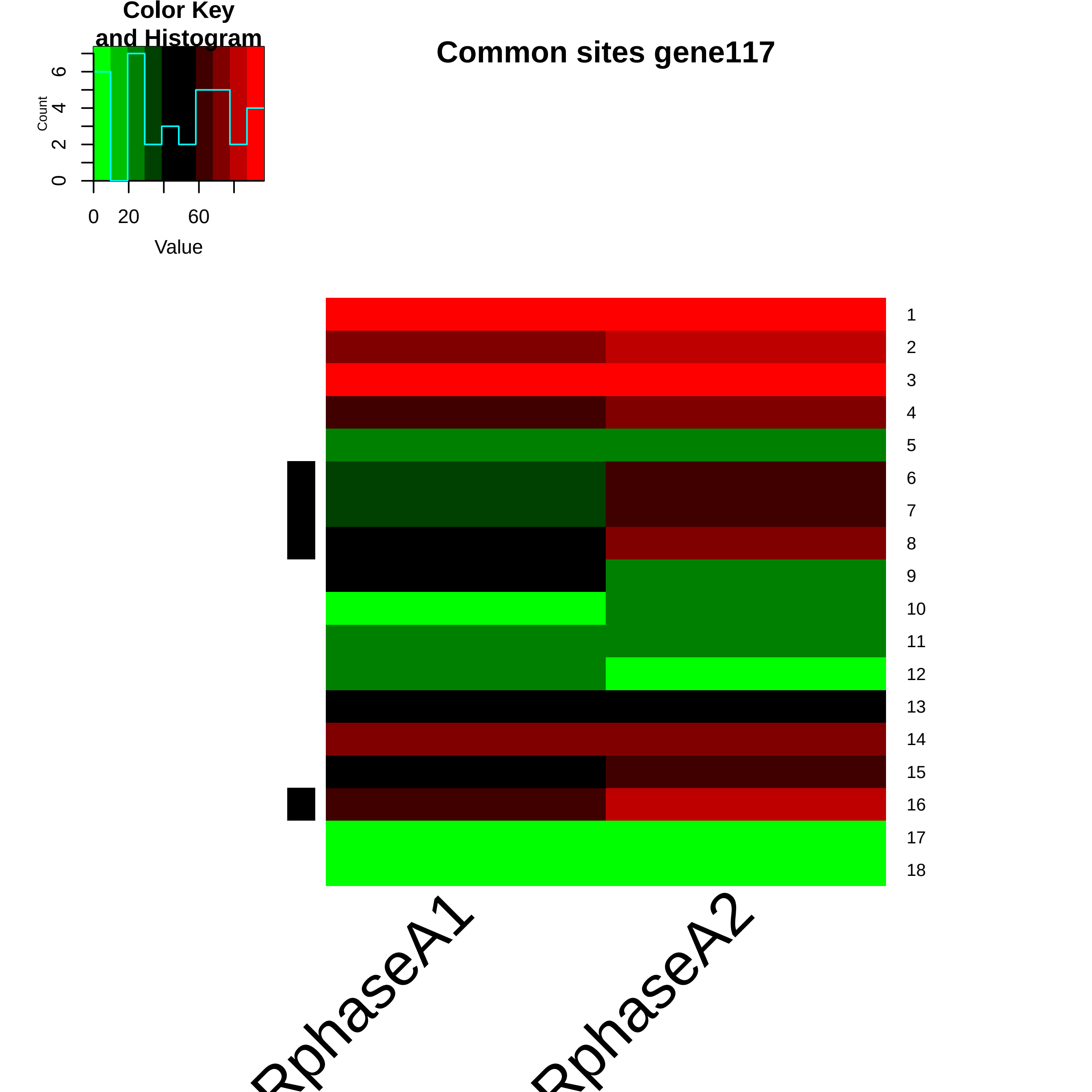

Color Key
and Histogram
Count
0
2
4
6
0
20
60
Value
Common sites gene117
RphaseA1
RphaseA2

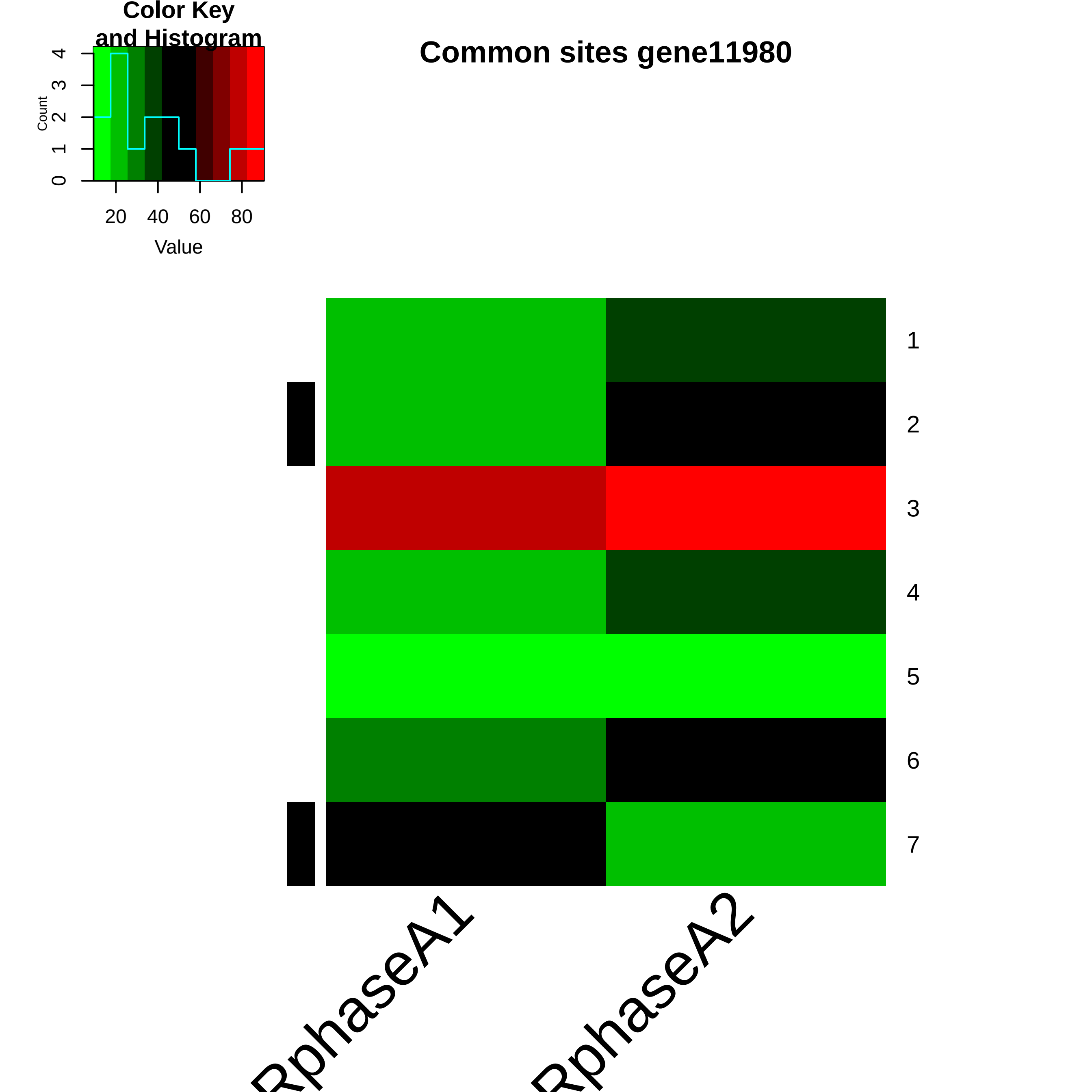

Color Key
and Histogram
Count
Value
20 40 60 80
0 1 2 3 4
Common sites gene11980
1
2
3
4
5
6
7
RphaseA1
RphaseA2

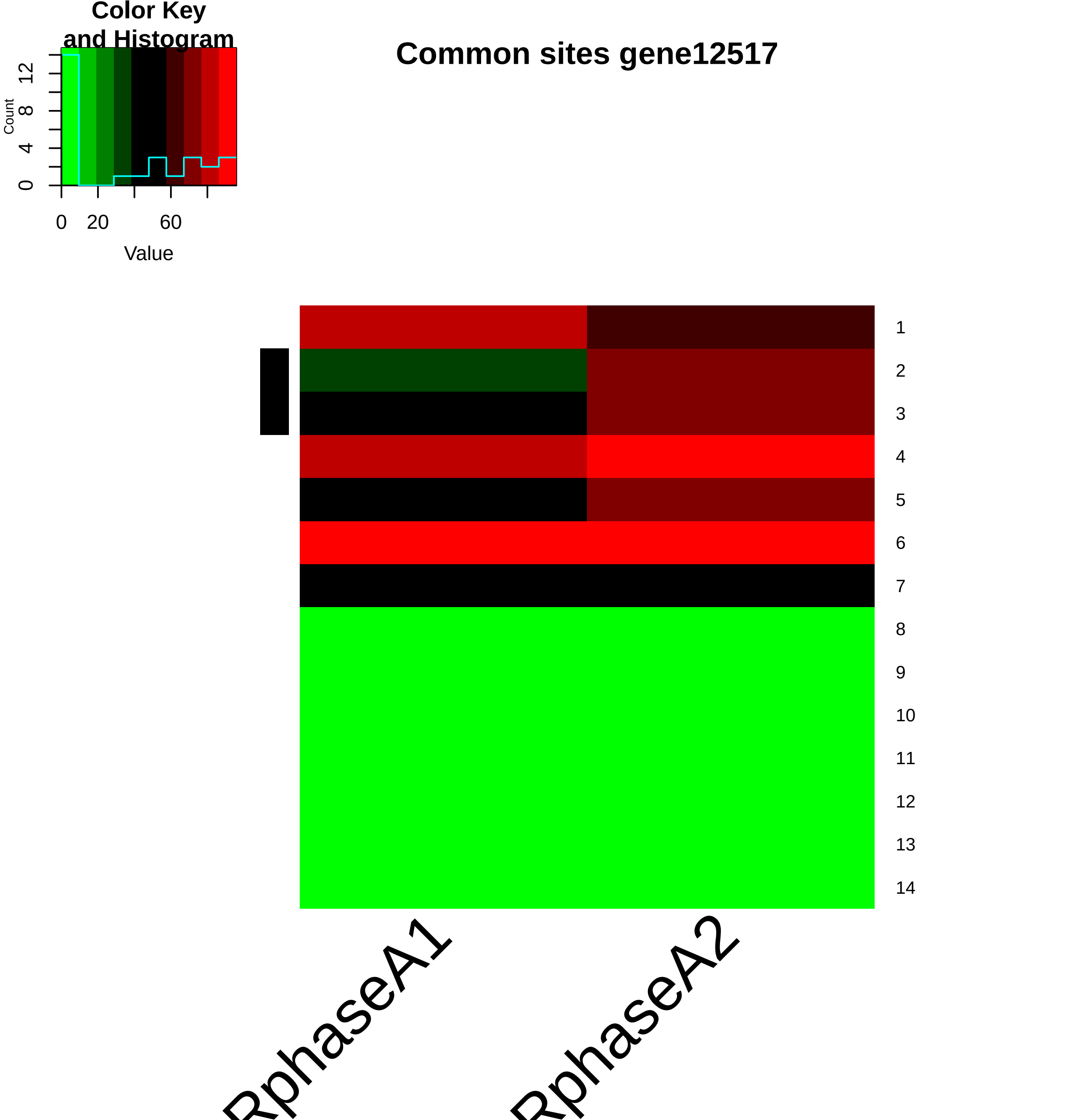
Color Key
and Histogram
Count
0
4
8
12
0
20
60
Value
Common sites gene12517
1
2
3
4
5
6
7
8
9
10
11
12
13
14
RphaseA1
RphaseA2

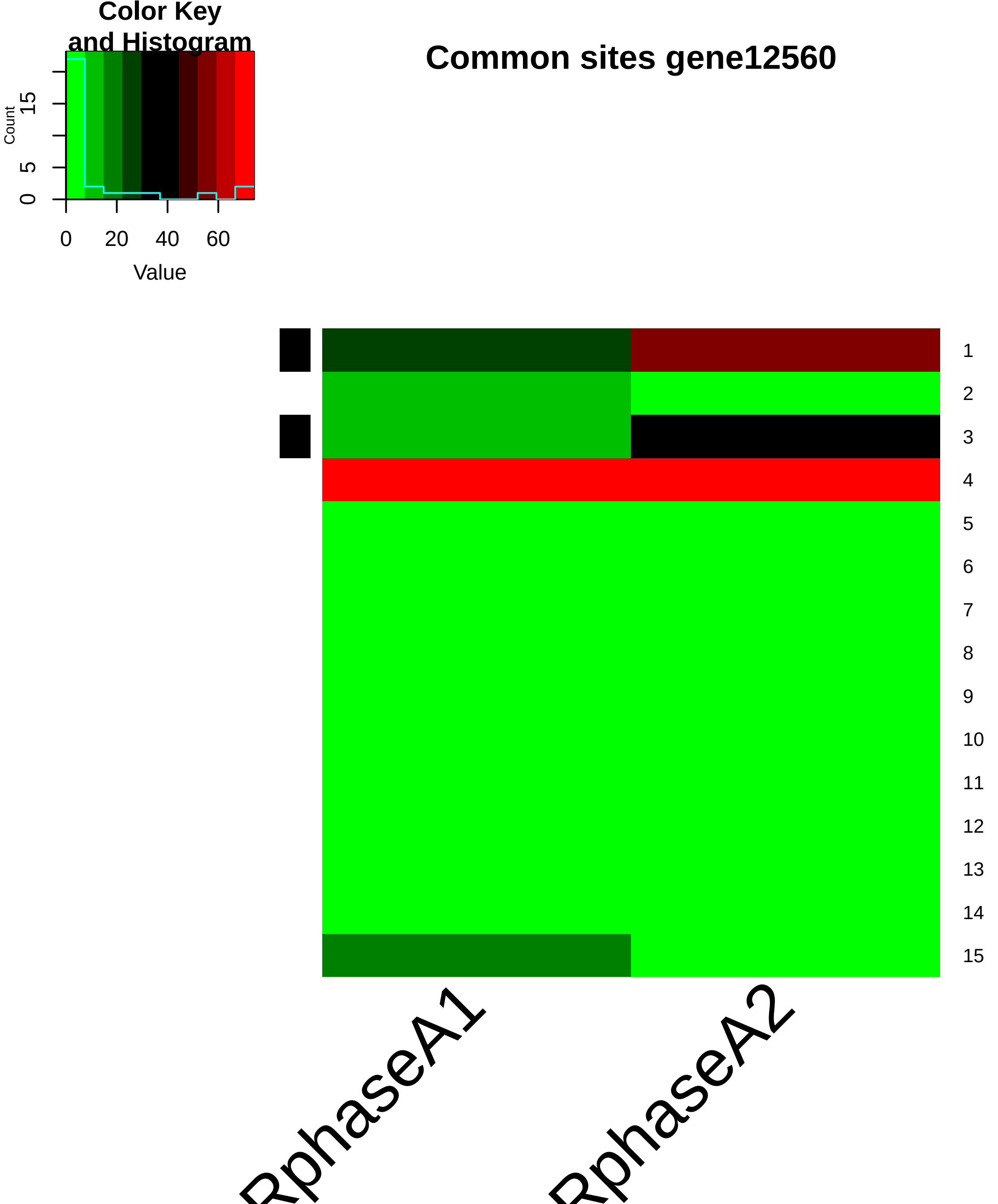
Color Key
and Histogram
Count
0
5
15
0
20
40
60
Value
Common sites gene12560
RphaseA1
RphaseA2

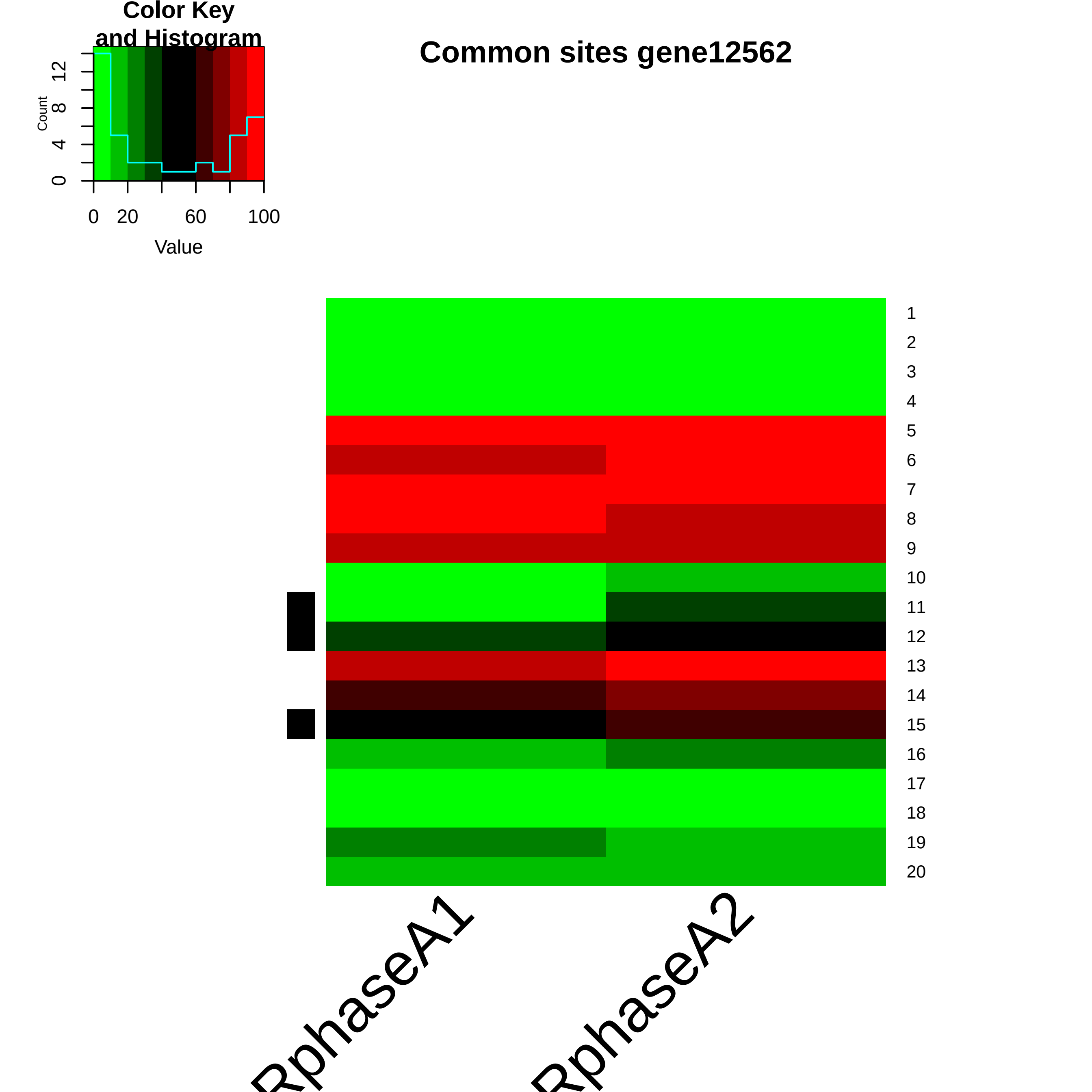

Color Key
and Histogram
Count
0
4
8
12
0
20
60
100
Value
Common sites gene12562
RphaseA1
RphaseA2

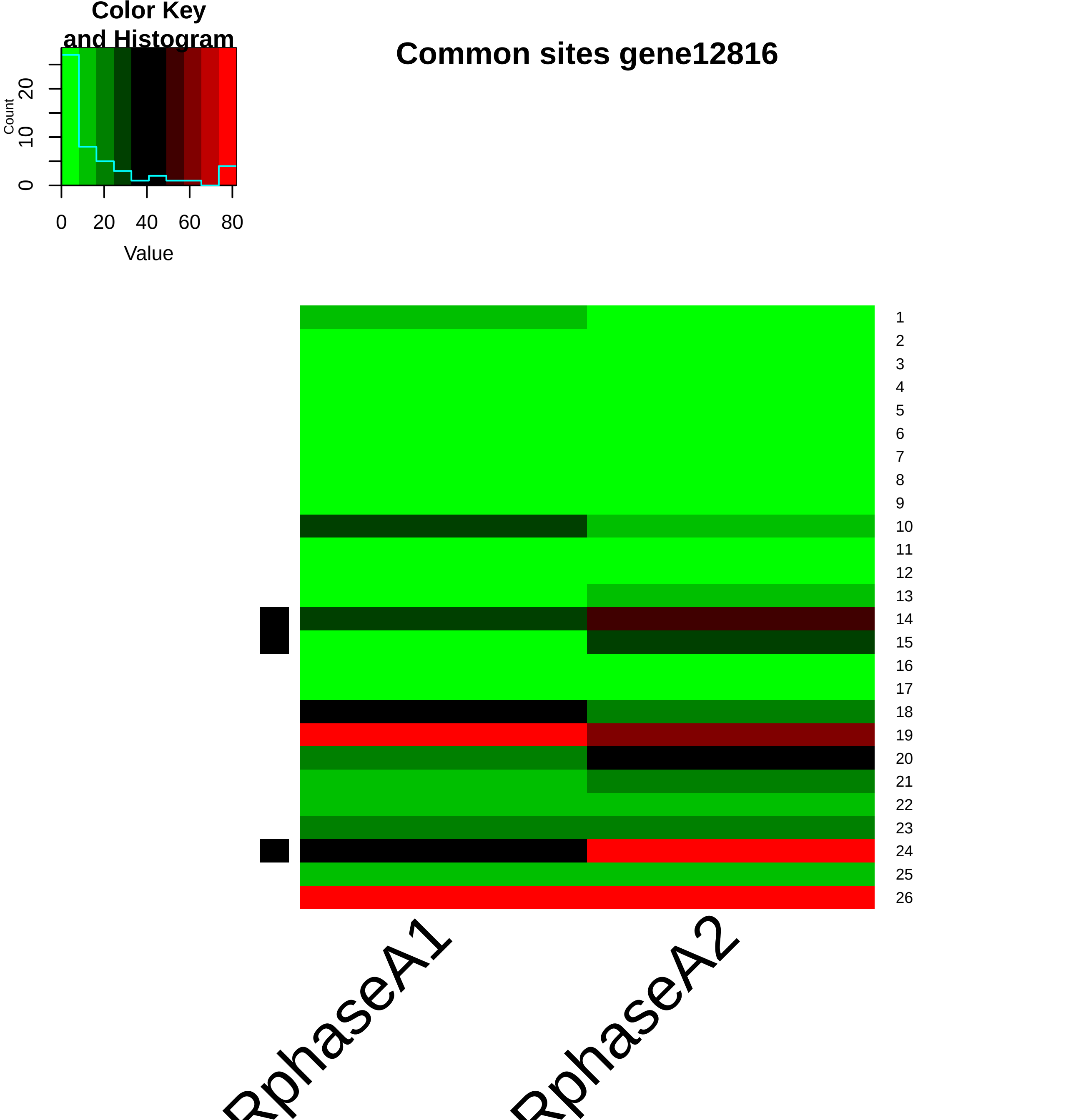
Color Key
and Histogram
Count
0
10
20
0
20
40
60
80
Value
Common sites gene12816
RphaseA1
RphaseA2

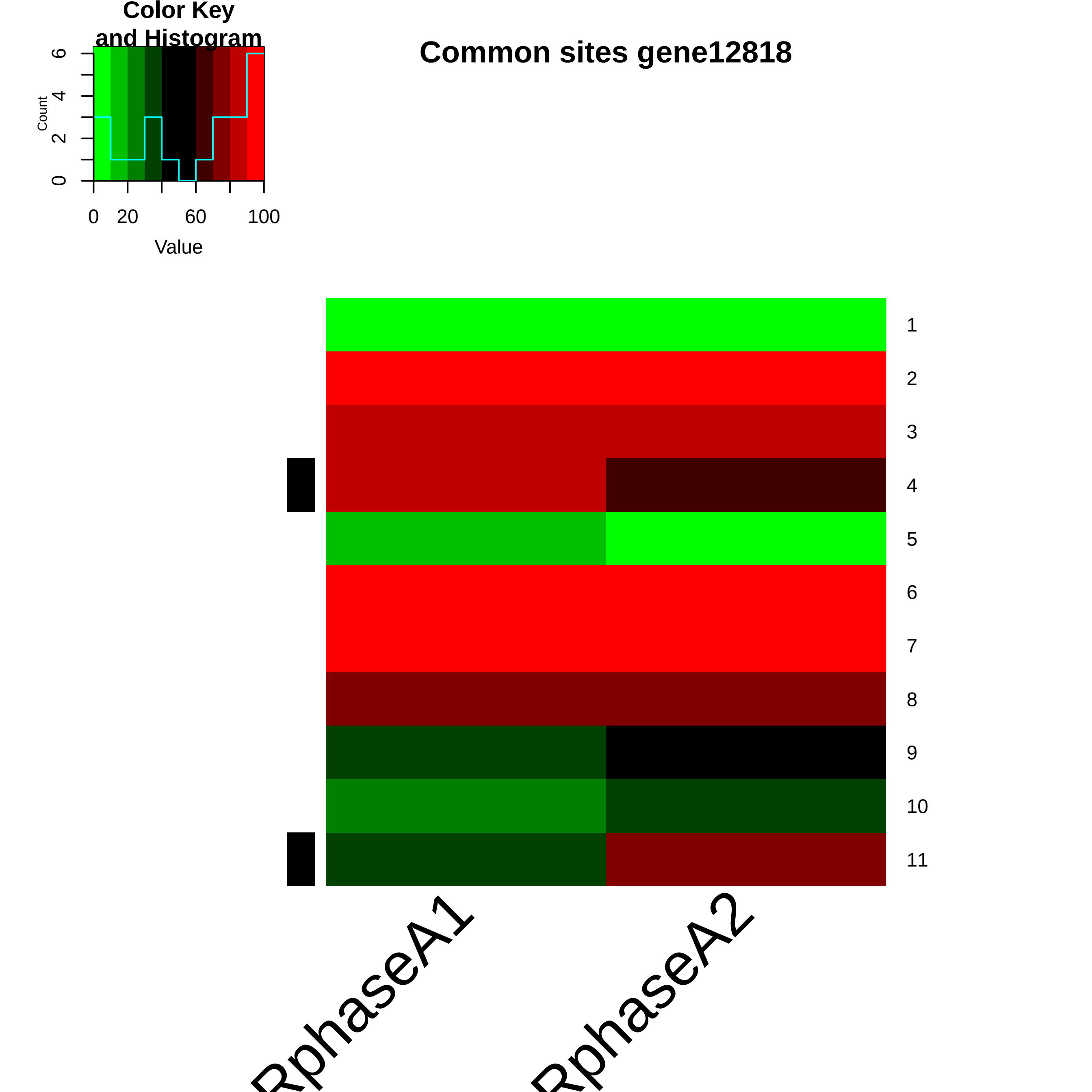
Color Key
and Histogram
Count
Value
Common sites gene12818
1
2
3
4
5
6
7
8
9
10
11
RphaseA1
RphaseA2

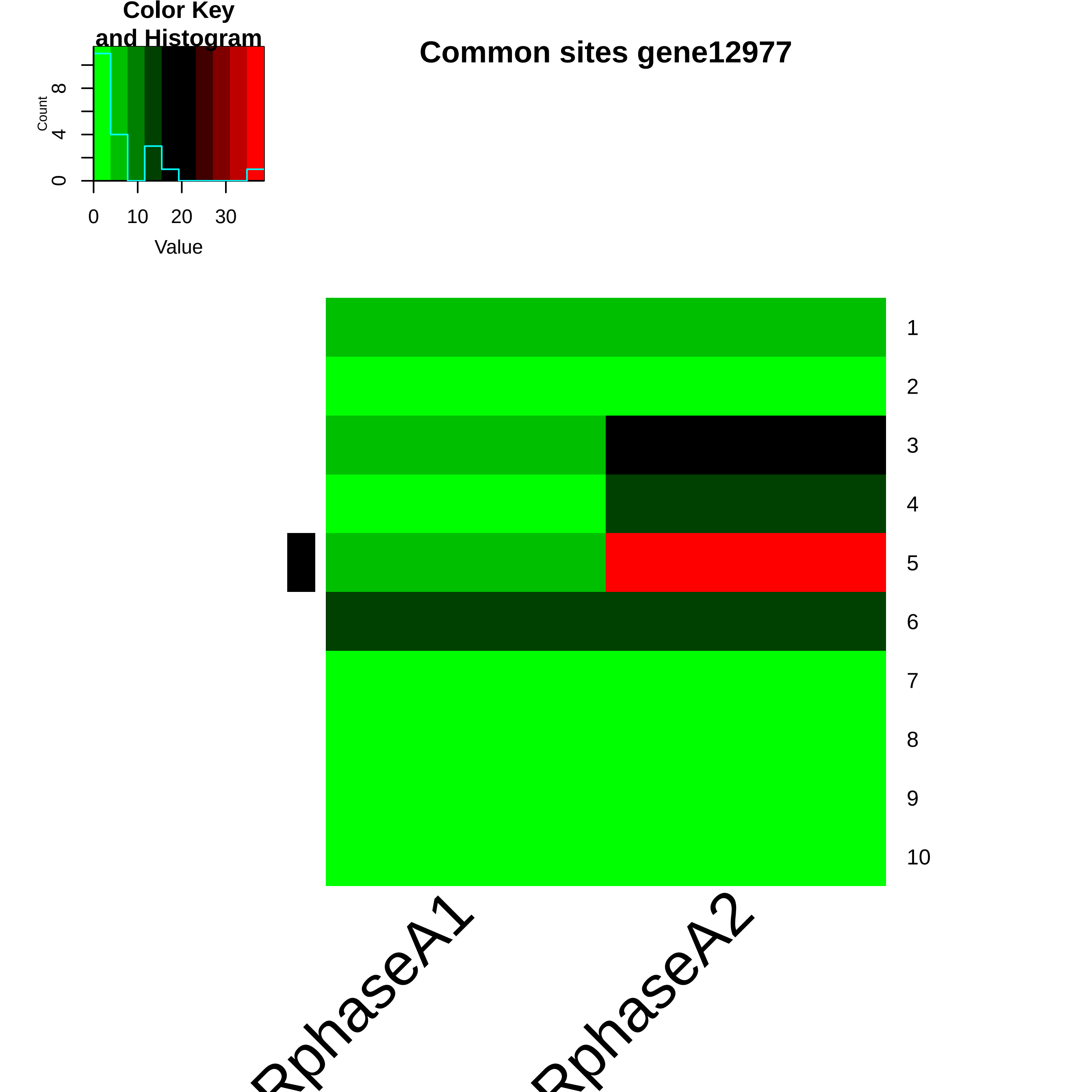
Color Key
and Histogram
Count
0
4
8
0
10
20
30
Value
Common sites gene12977
1
2
3
4
5
6
7
8
9
10
RphaseA1
RphaseA2

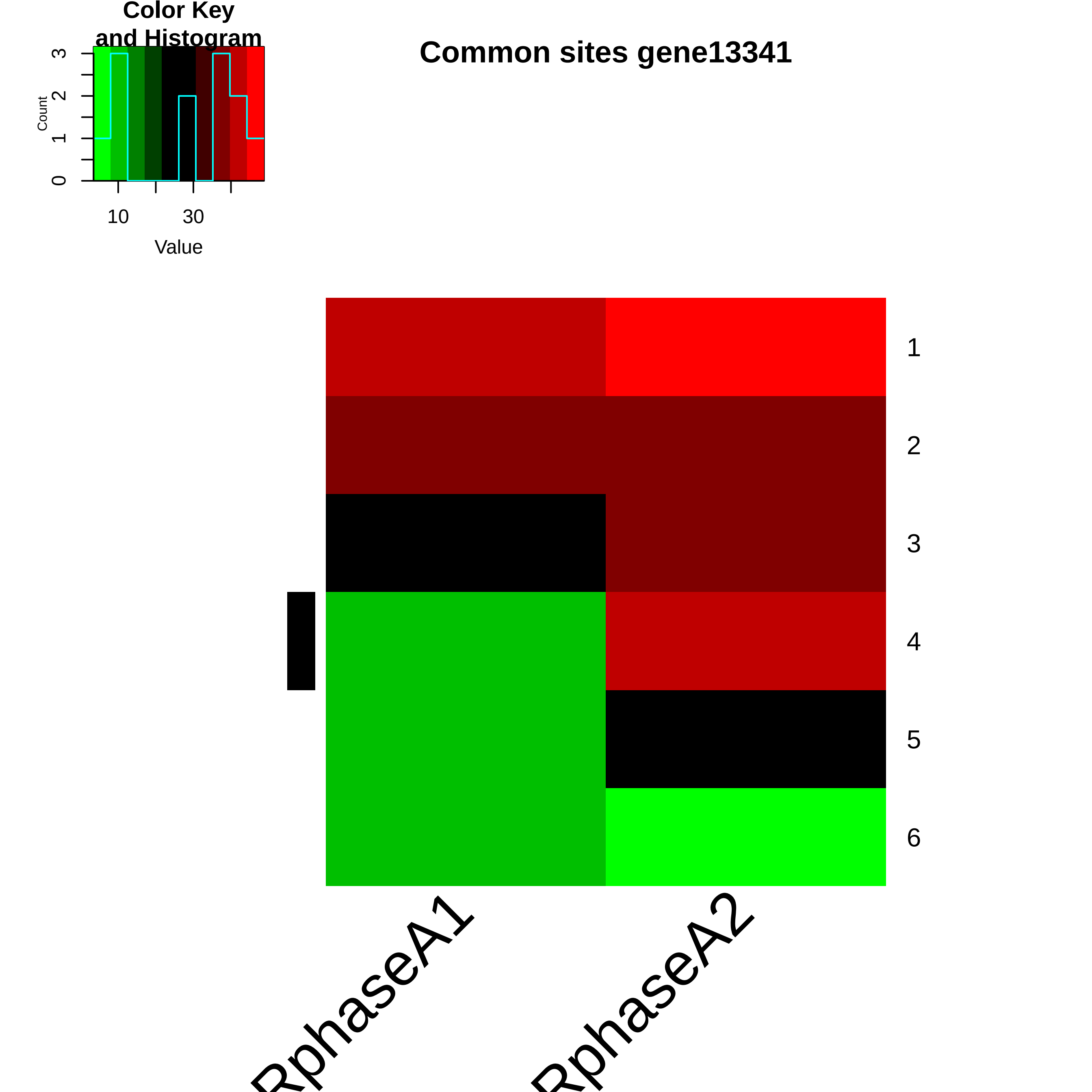
Color Key
and Histogram
Count
0
1
2
3
10
30
Value
Common sites gene13341
1
2
3
4
5
6
RphaseA1
RphaseA2

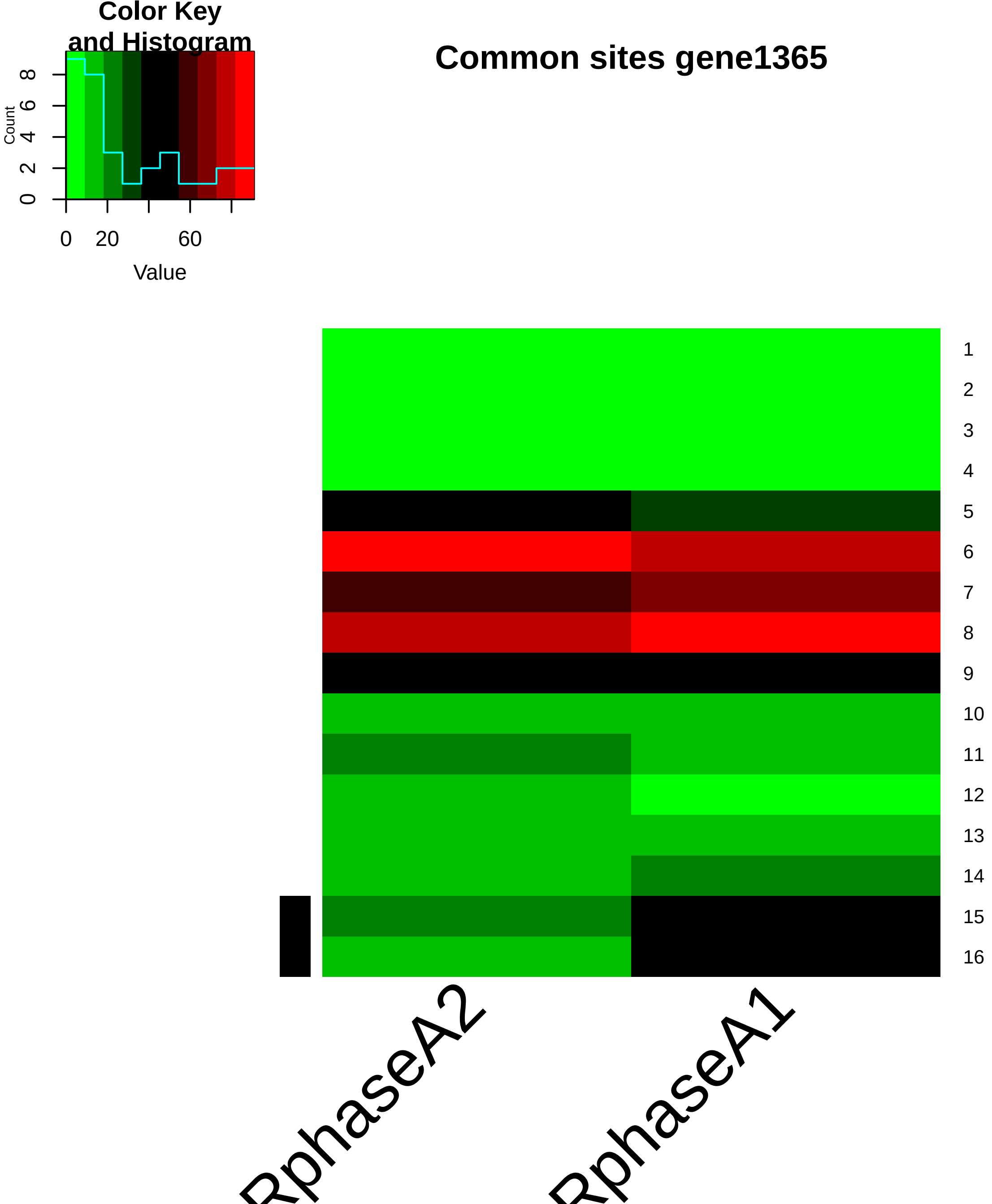
Color Key
and Histogram
Count
0
2
4
6
8
0
20
60
Value
Common sites gene1365
RphaseA2
RphaseA1

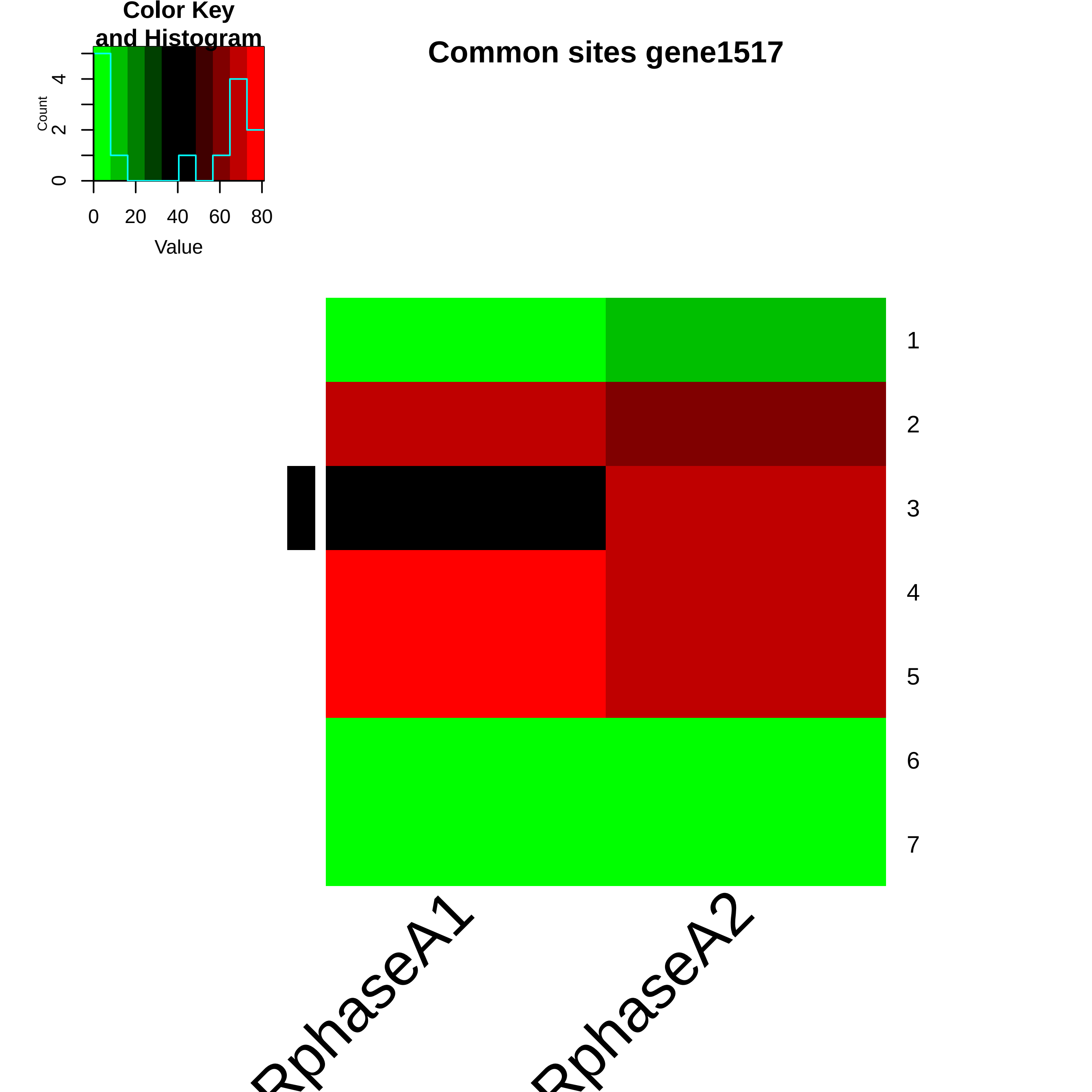
Color Key
and Histogram
Count
Value
0
20
40
60
80
0
2
4
Common sites gene1517
1
2
3
4
5
6
7
RphaseA1
RphaseA2

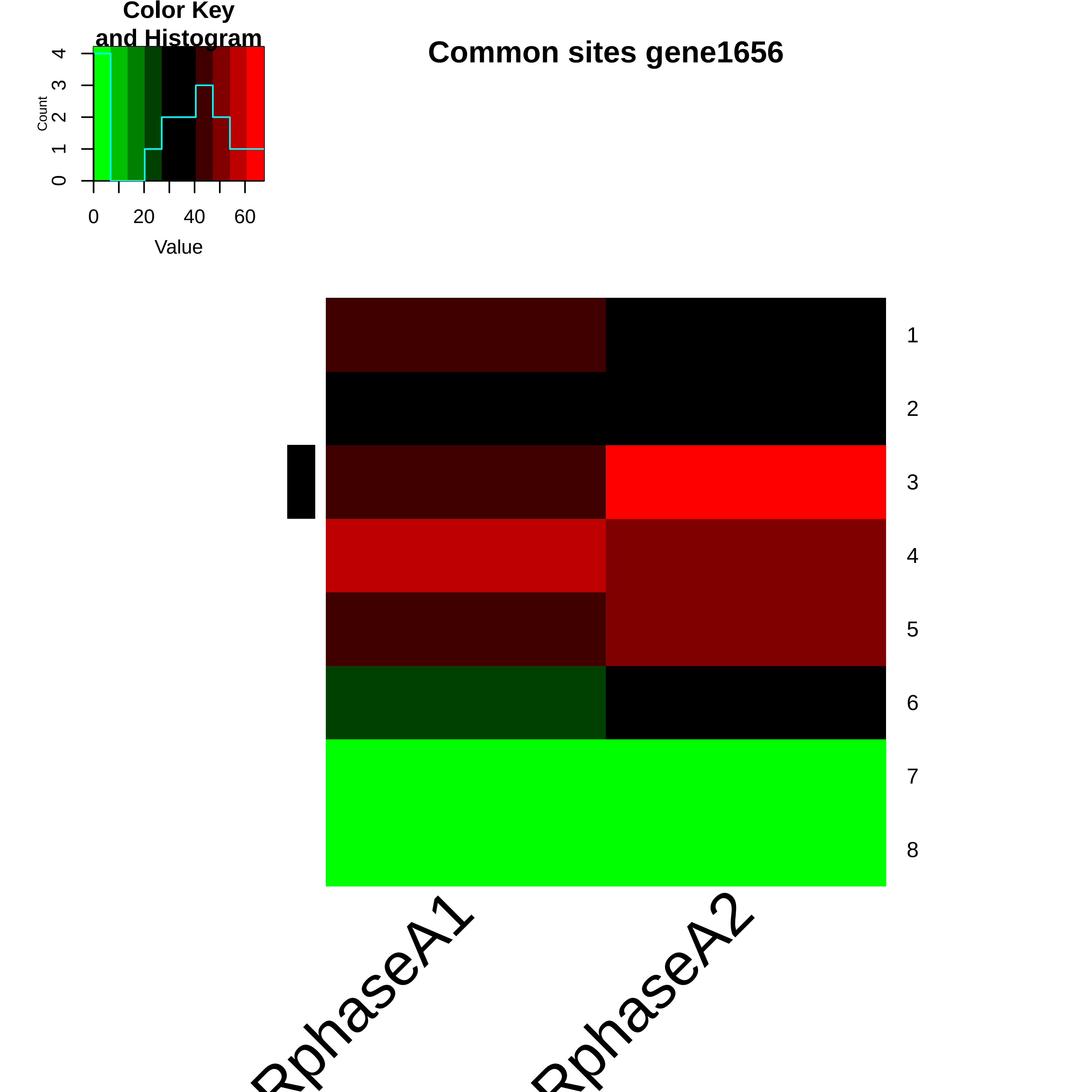
Color Key
and Histogram
Count
0
1
2
3
4
0
20
40
60
Value
Common sites gene1656
1
2
3
4
5
6
7
8
RphaseA1
RphaseA2

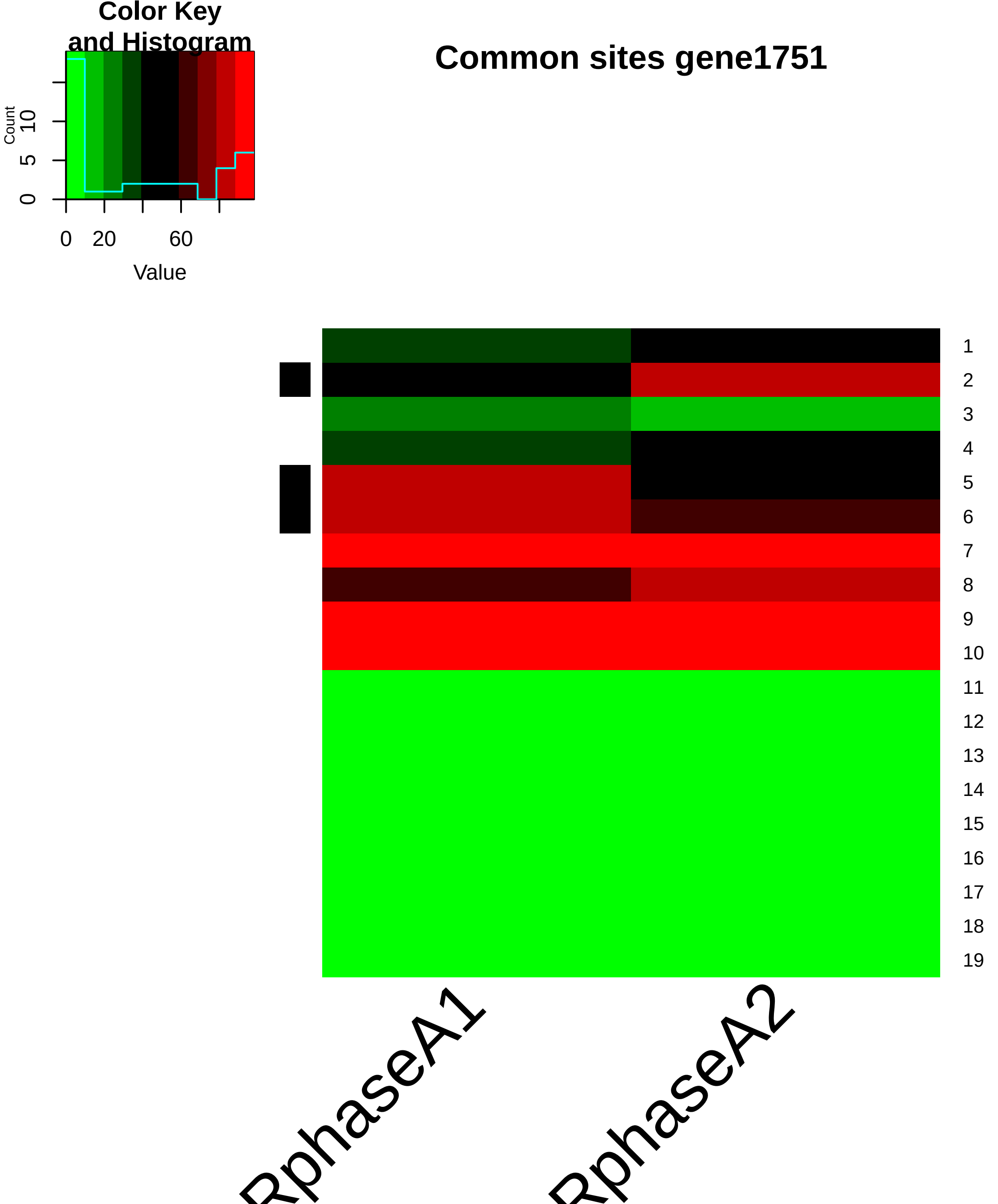

Color Key
and Histogram
Count
0
5
10
0
20
60
Value
Common sites gene1751
RphaseA1
RphaseA2

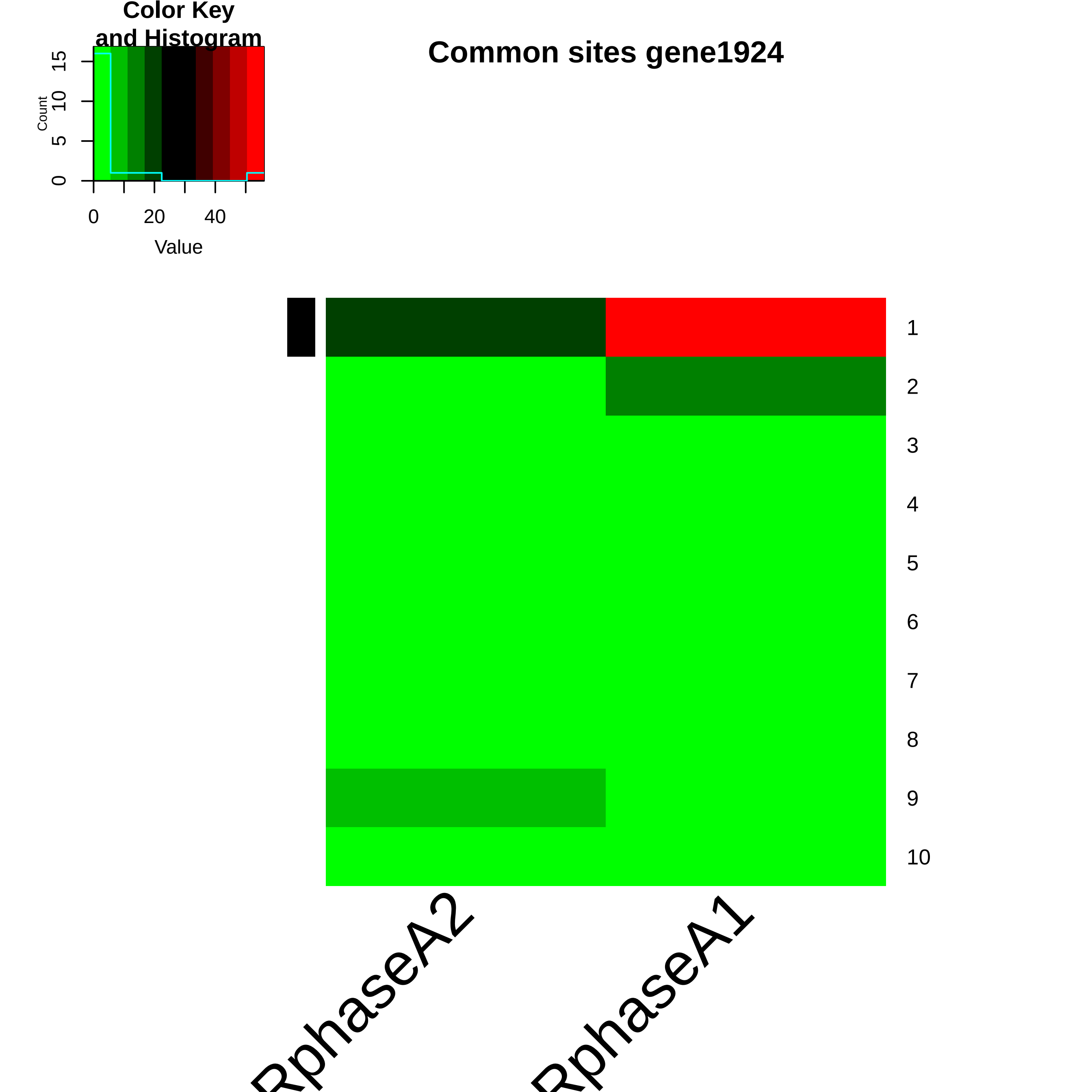

Color Key
and Histogram
Count
0
5
10
15
0
20
40
Value
Common sites gene1924
1
2
3
4
5
6
7
8
9
10
RphaseA2
RphaseA1

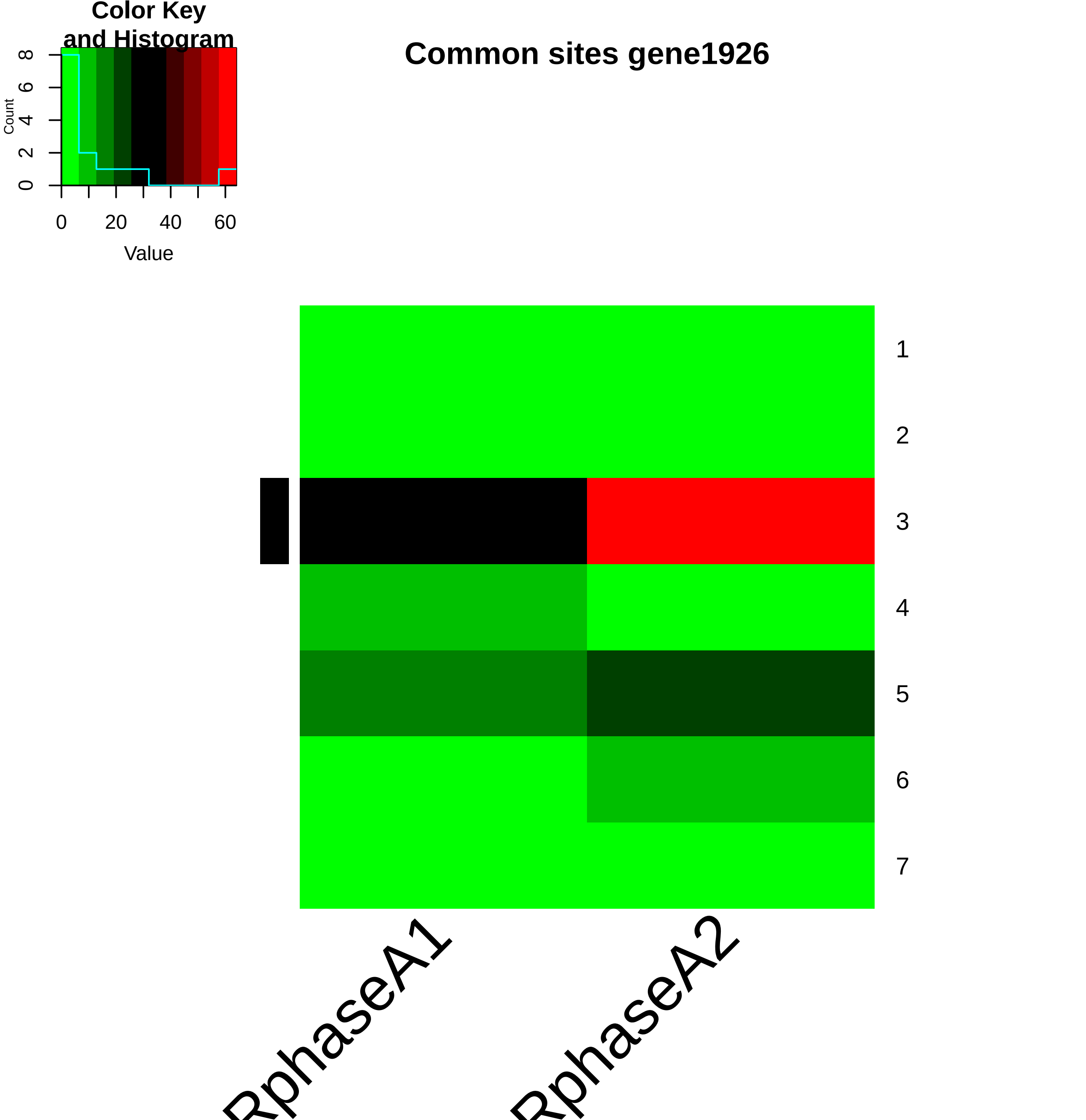
Color Key
and Histogram
Count
Value
0
20
40
60
0
2
4
6
8
Common sites gene1926
1
2
3
4
5
6
7
RphaseA1
RphaseA2

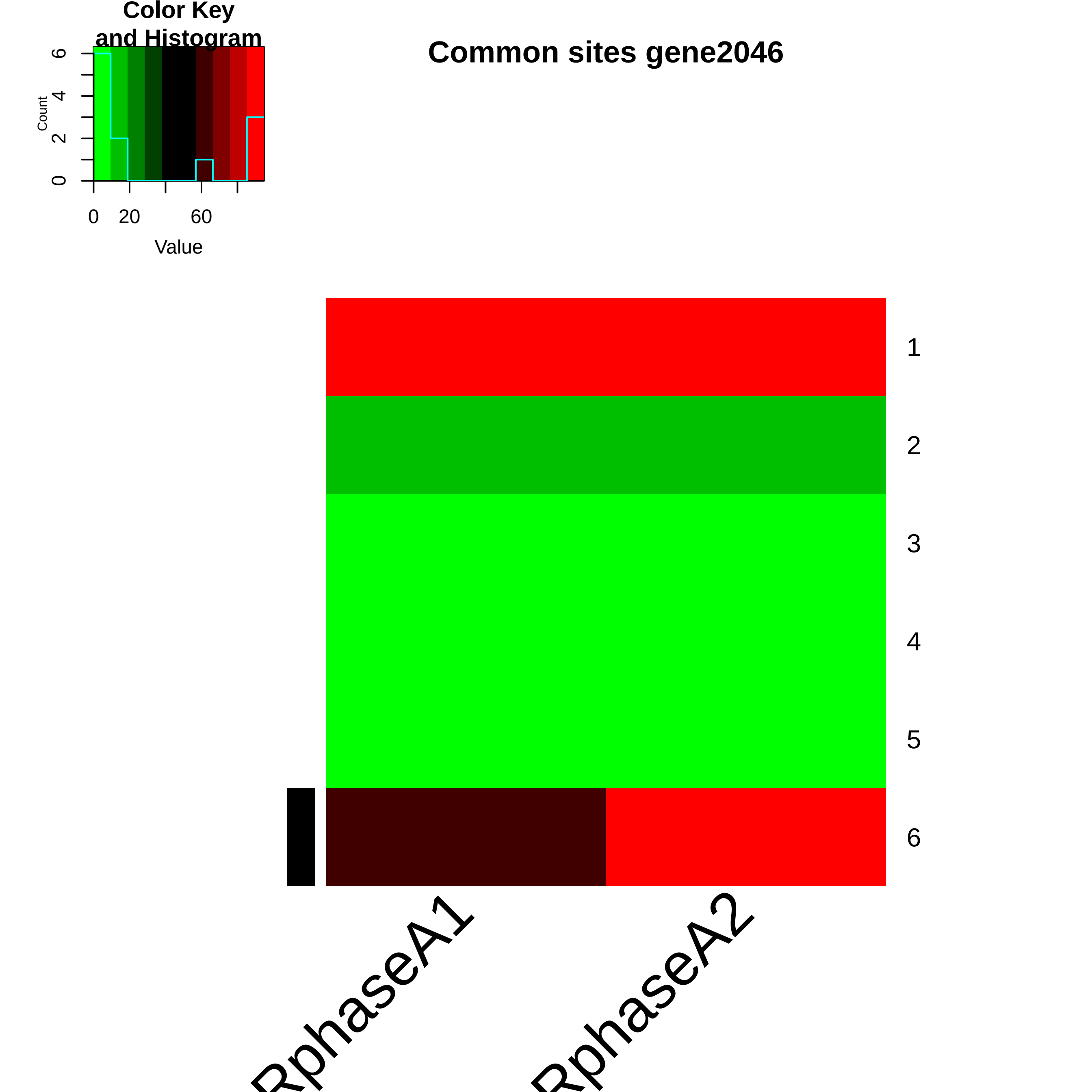

Color Key
and Histogram
Count
0
2
4
6
0
20
60
Value
Common sites gene2046
1
2
3
4
5
6
RphaseA1
RphaseA2

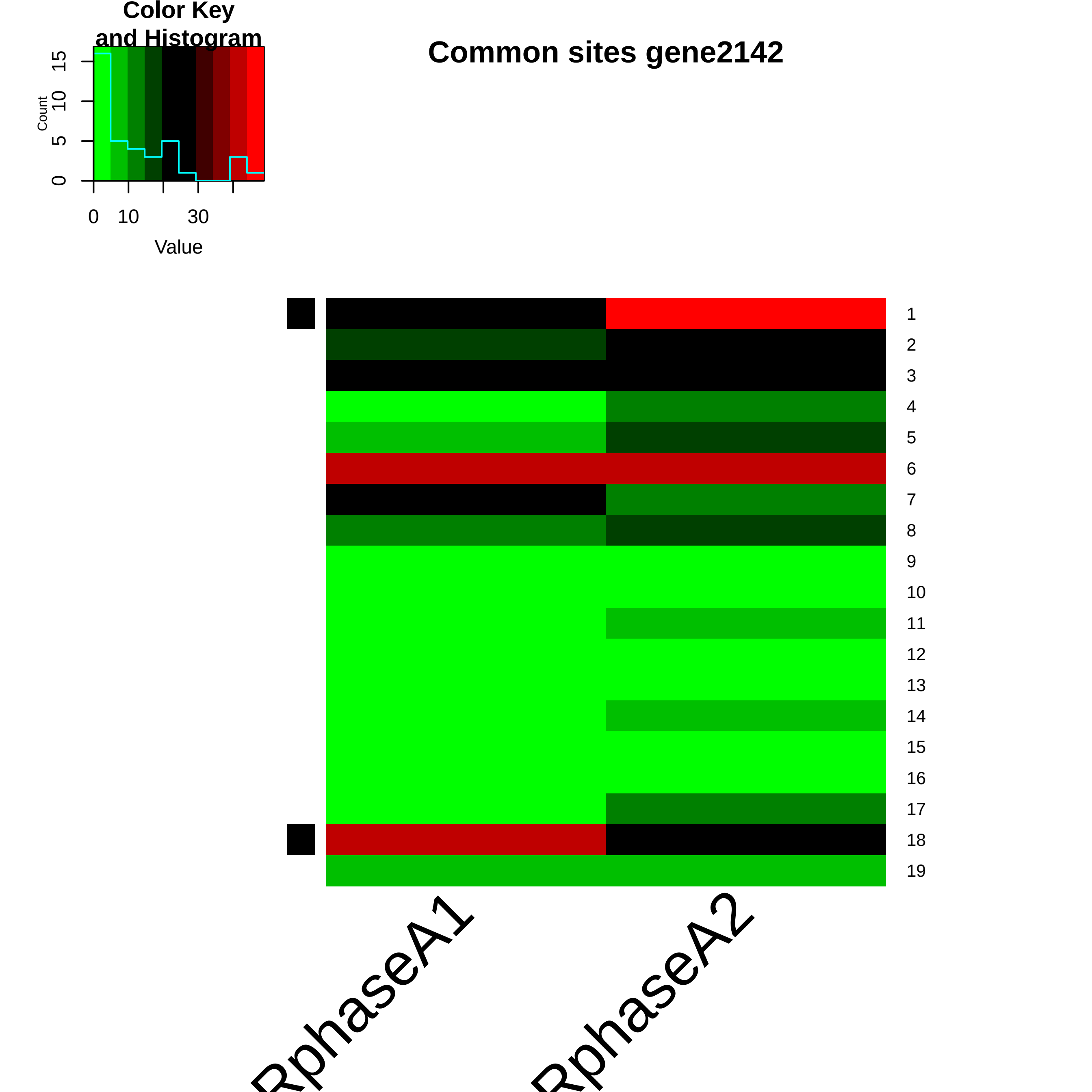

Color Key and Histogram
Count
0
5
10
15
0
10
30
Value
Common sites gene2142
RphaseA1
RphaseA2

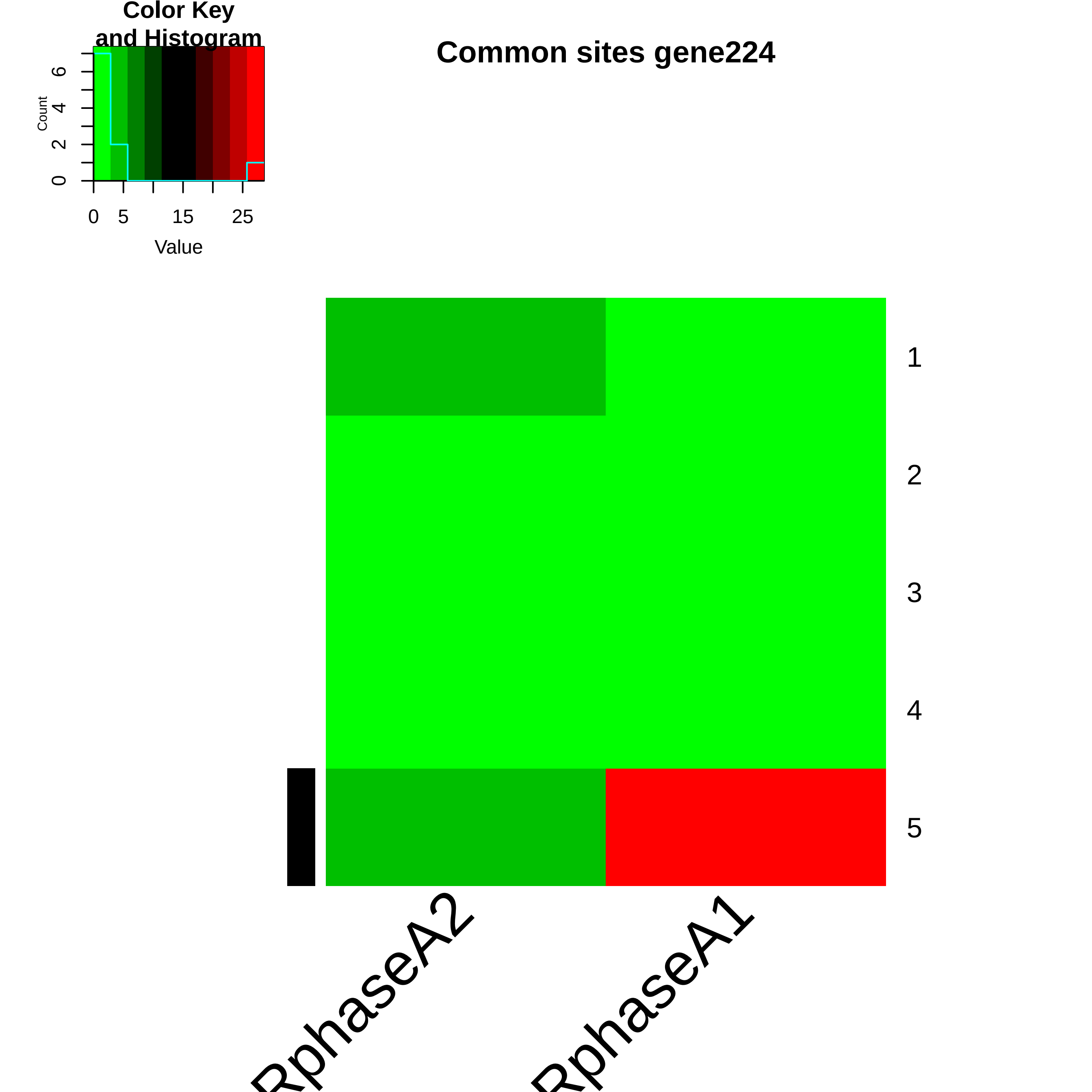
Color Key
and Histogram
Count
0
2
4
6
0
5
15
25
Value
Common sites gene224
1
2
3
4
5
RphaseA2
RphaseA1

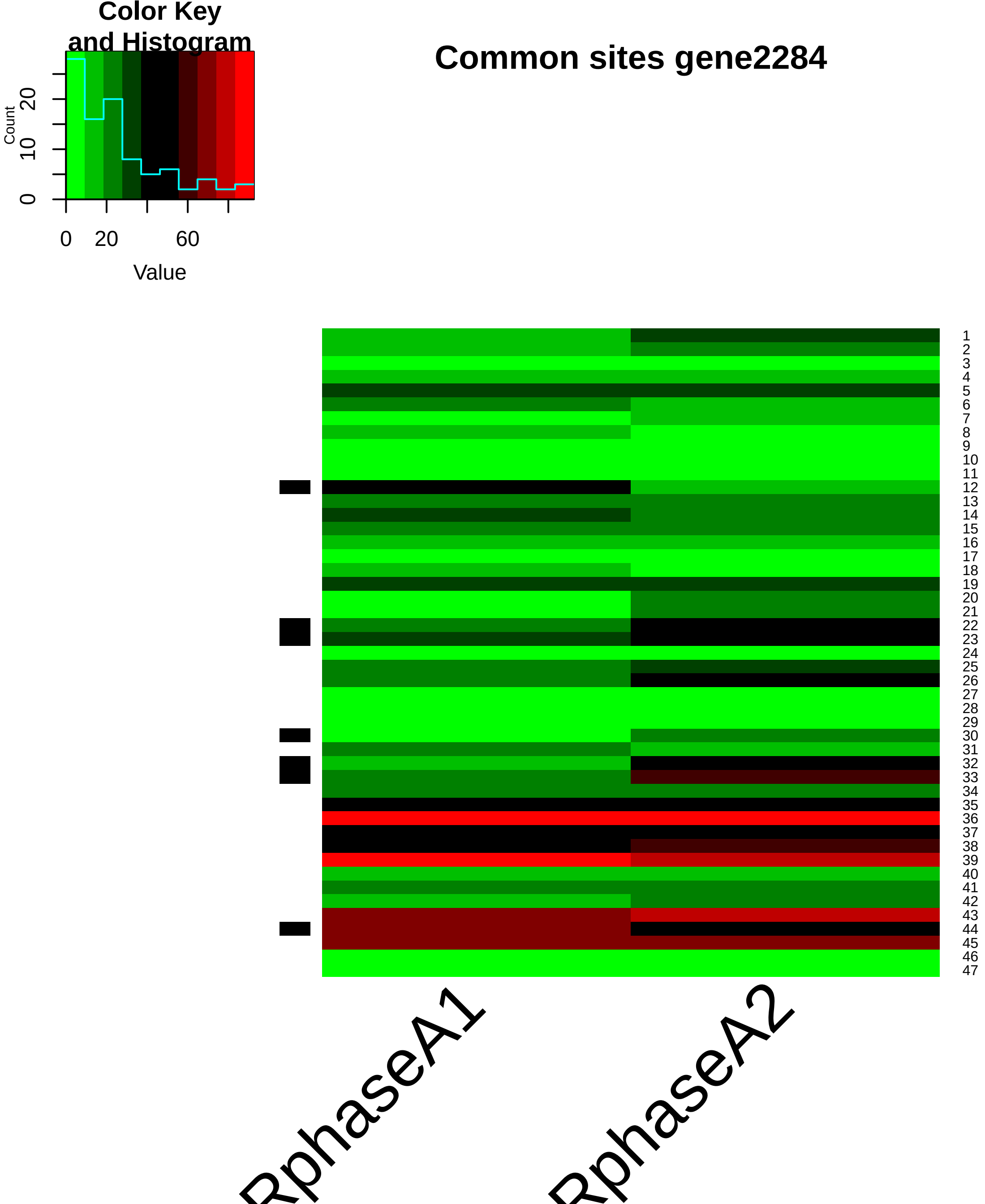
Color Key
and Histogram
Count
0
10
20
0
20
60
Value
Common sites gene2284
RphaseA1
RphaseA2

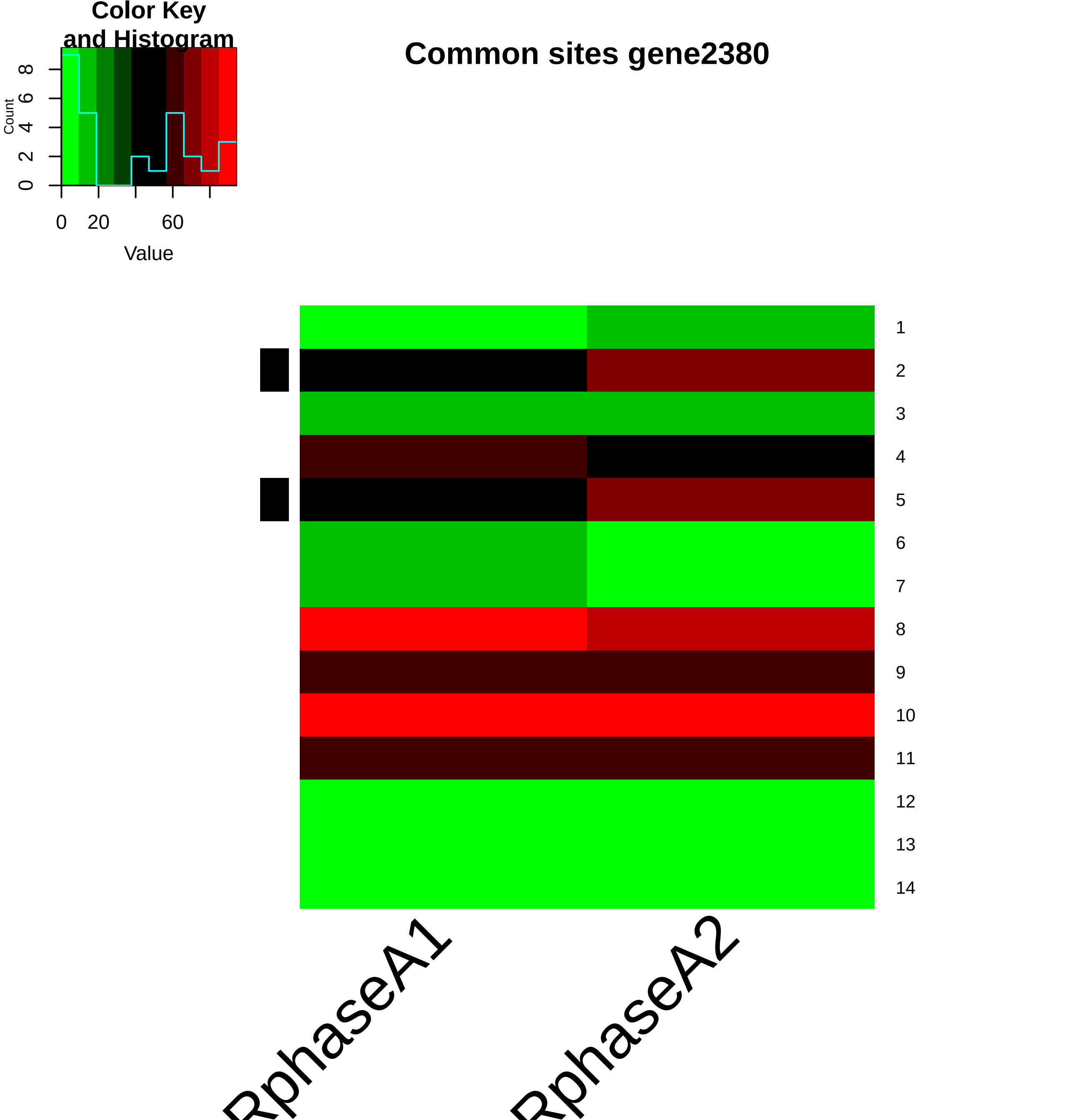
Color Key
and Histogram
Count
0
2
4
6
8
0
20
60
Value
Common sites gene2380
1
2
3
4
5
6
7
8
9
10
11
12
13
14
RphaseA1
RphaseA2

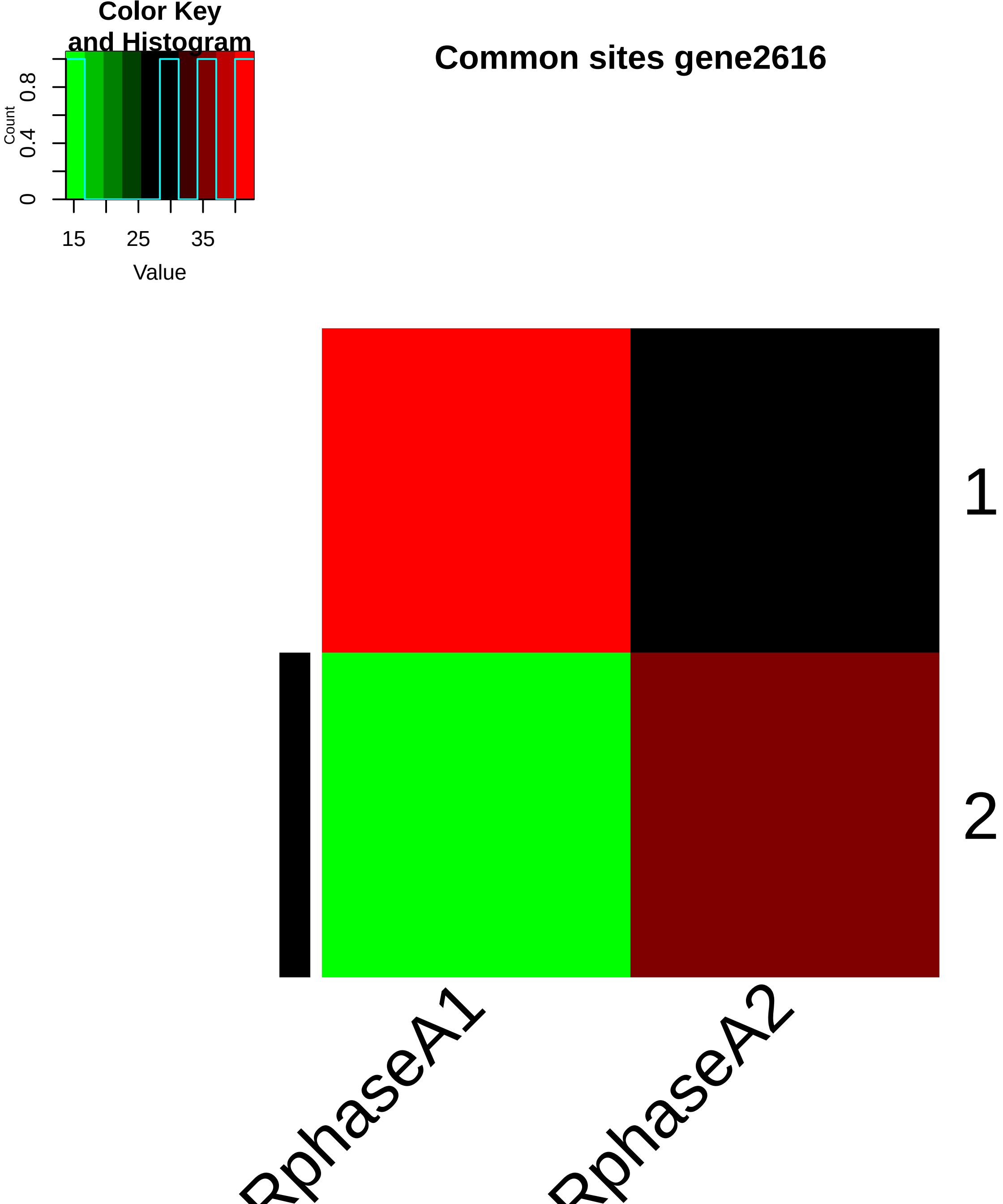
Color Key
and Histogram
Count
0
0.4
0.8
15
25
35
Value
Common sites gene2616
1
2
RphaseA1
RphaseA2

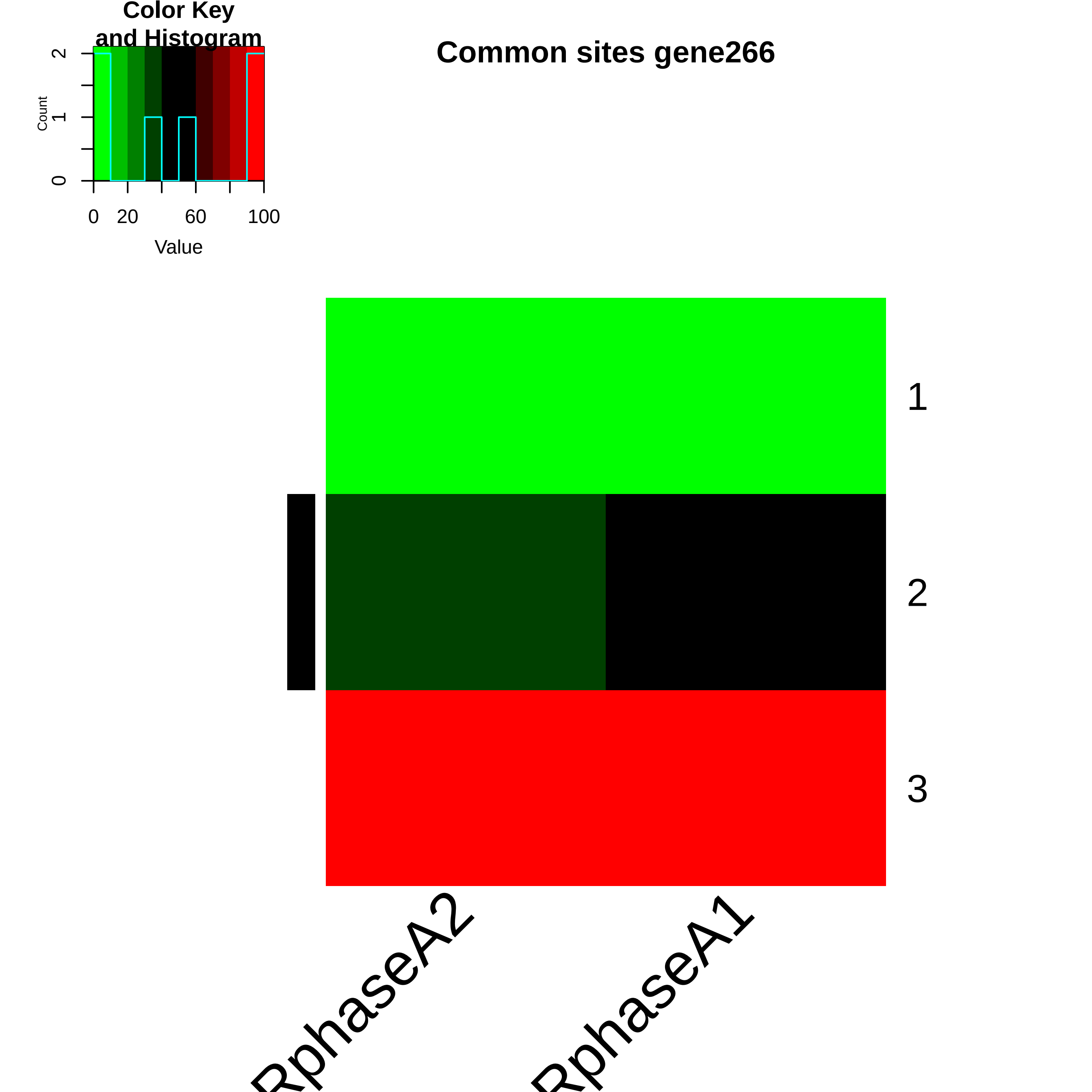
Color Key
and Histogram
Count
0
1
2
0
20
60
100
Value
Common sites gene266
1
2
3
RphaseA2
RphaseA1

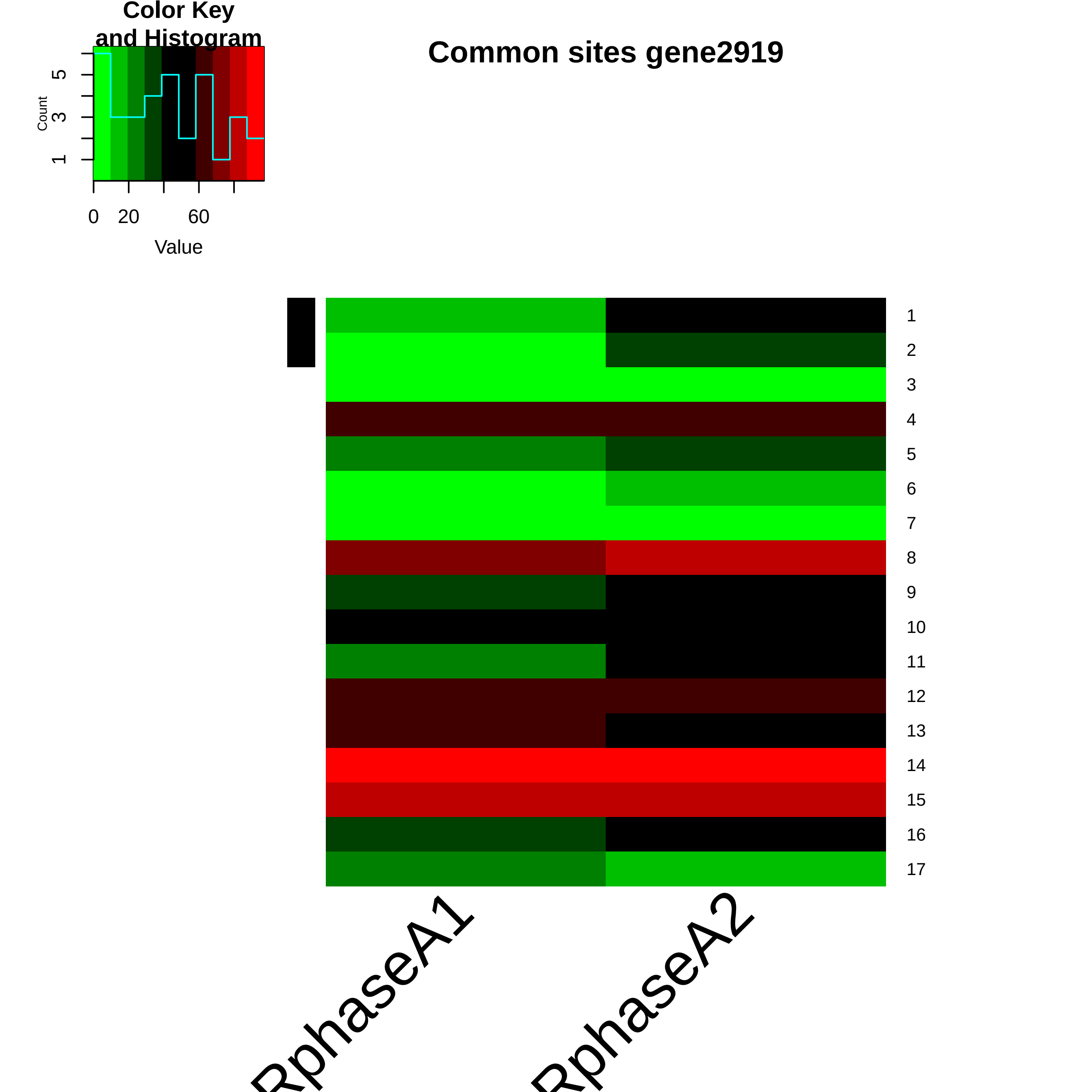
Color Key
and Histogram
Count
1
3
5
0
20
60
Value
Common sites gene2919
RphaseA1
RphaseA2

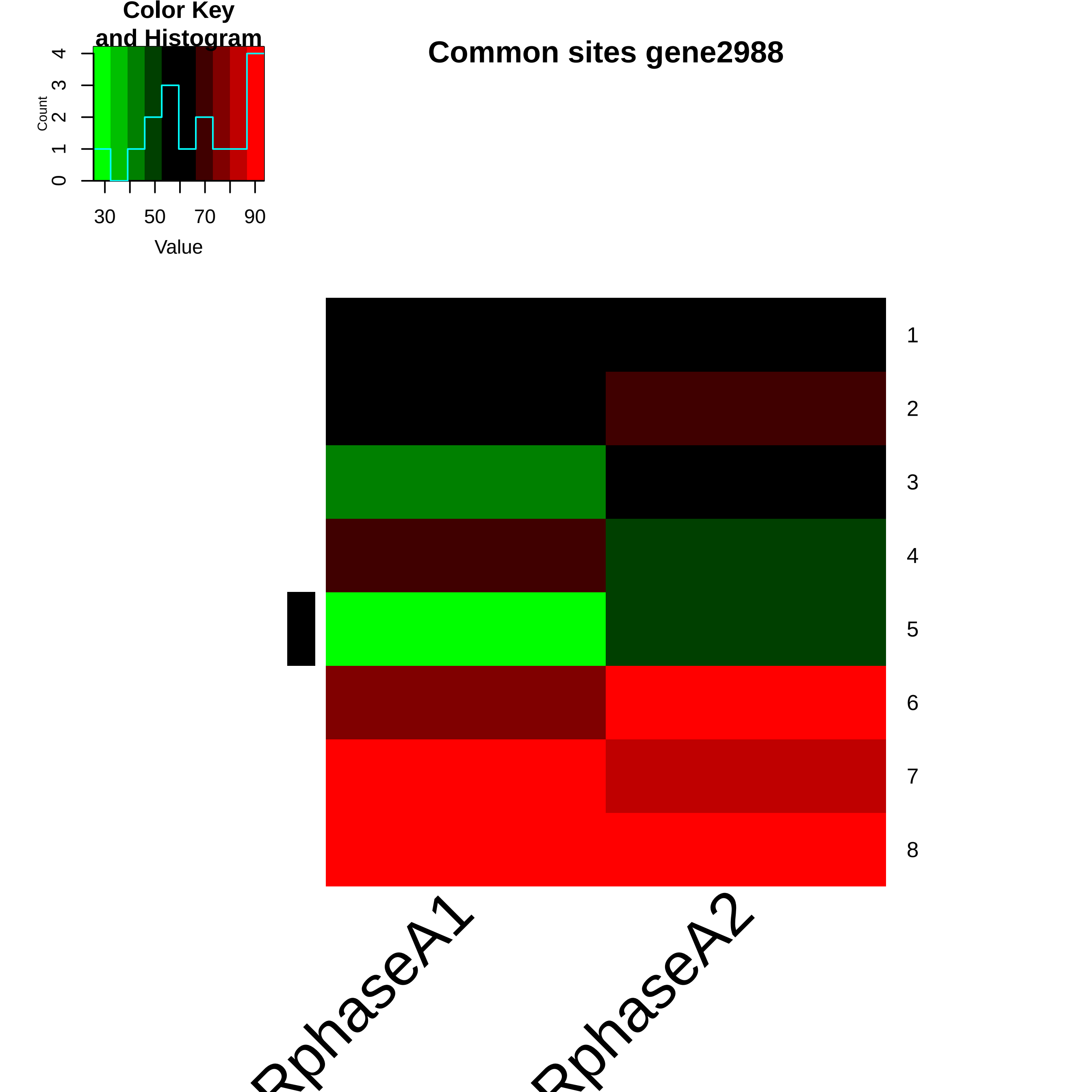
Color Key
and Histogram
Count
0
1
2
3
4
30
50
70
90
Value
Common sites gene2988
1
2
3
4
5
6
7
8
RphaseA1
RphaseA2

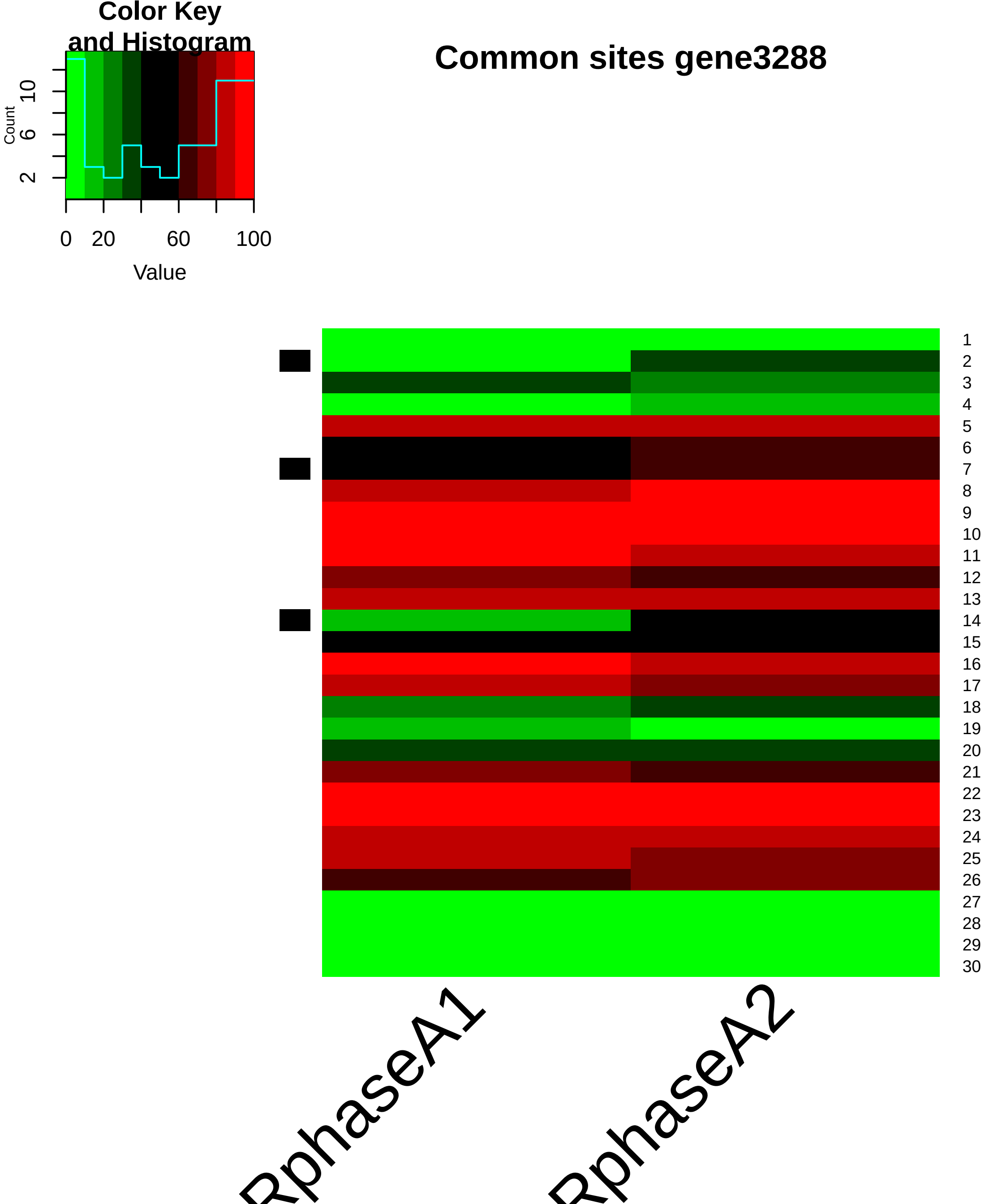
Color Key
and Histogram
Count
2
6
10
0
20
60
100
Value
Common sites gene3288
RphaseA1
RphaseA2

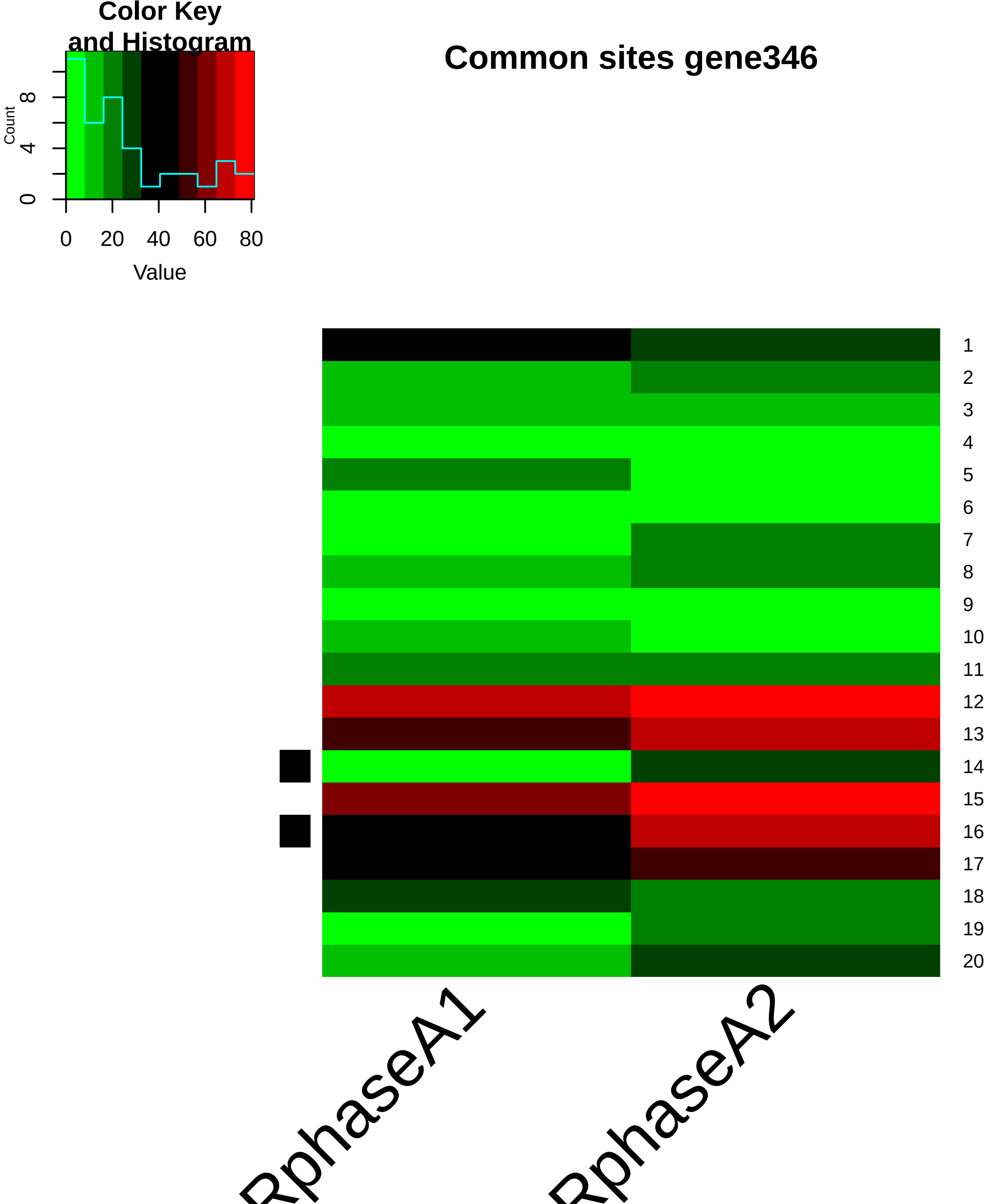
Color Key
and Histogram
Count
0
4
8
0
20
40
60
80
Value
Common sites gene346
RphaseA1
RphaseA2

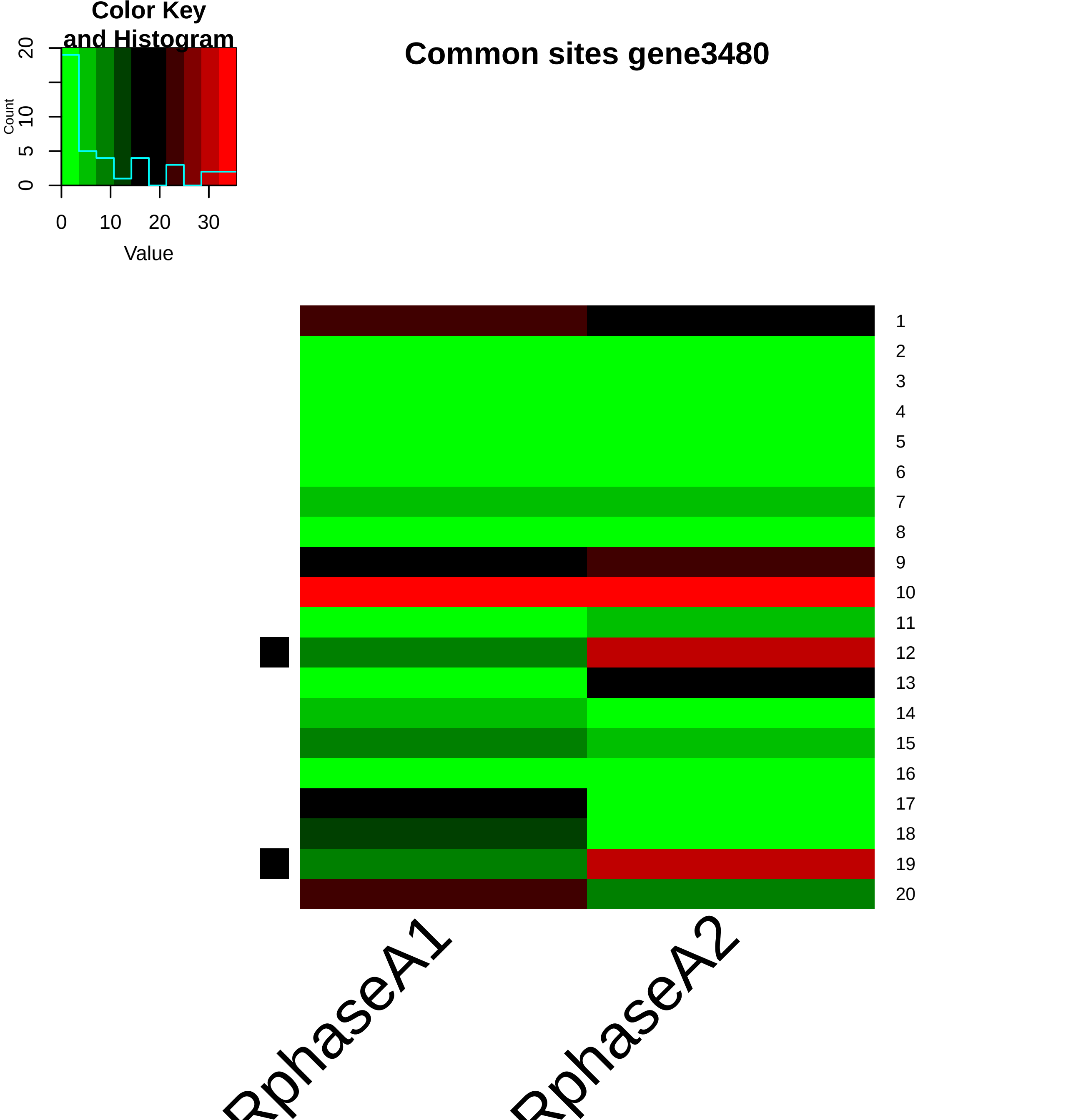

Color Key
and Histogram
Count
0
5
10
20
0
10
20
30
Value
Common sites gene3480
RphaseA1
RphaseA2

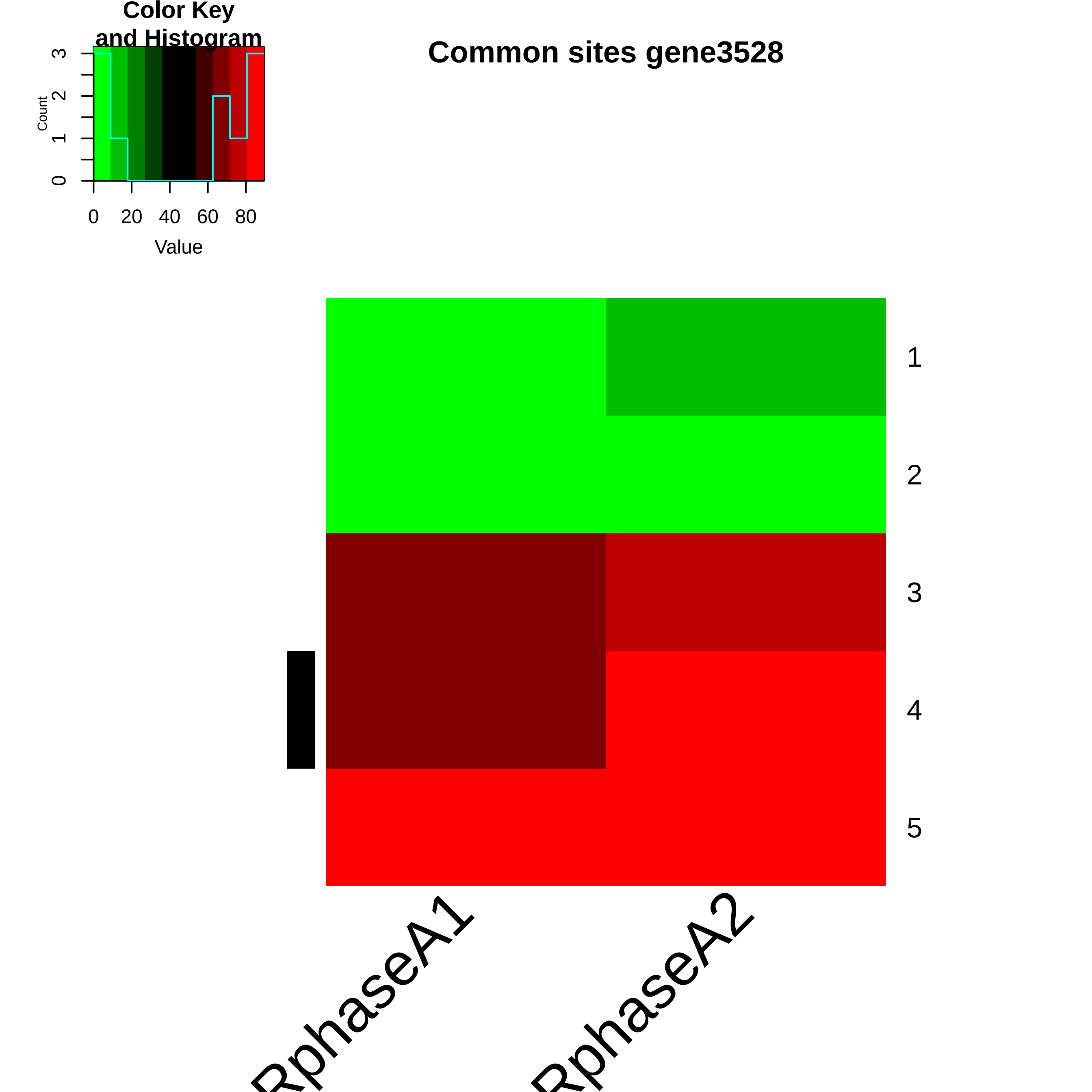

Color Key
and Histogram
Count
0
1
2
3
0
20
40
60
80
Value
Common sites gene3528
1
2
3
4
5
RphaseA1
RphaseA2

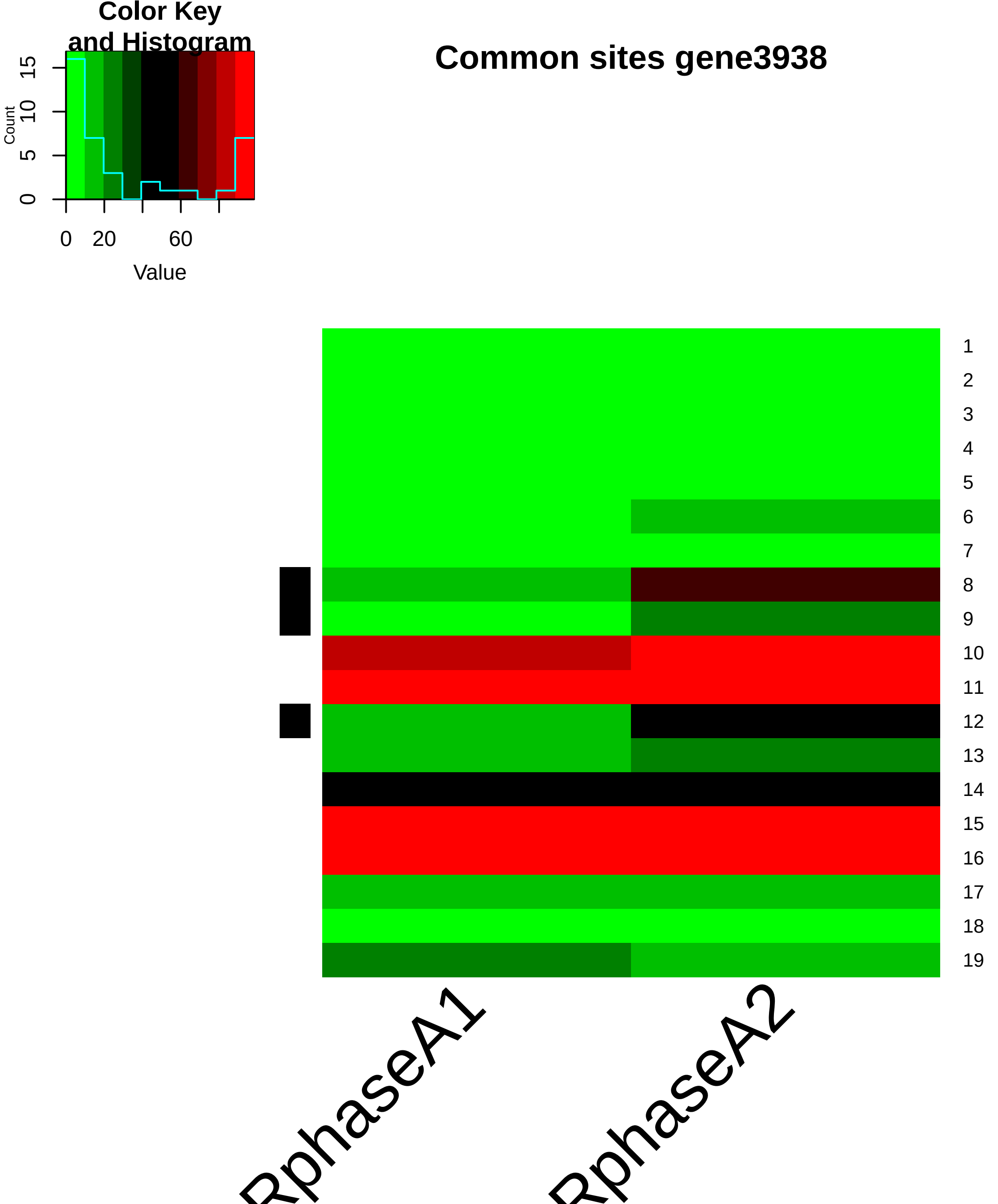

Color Key and Histogram
Count
0
5
10
15
0
20
60
Value
Common sites gene3938
RphaseA1
RphaseA2

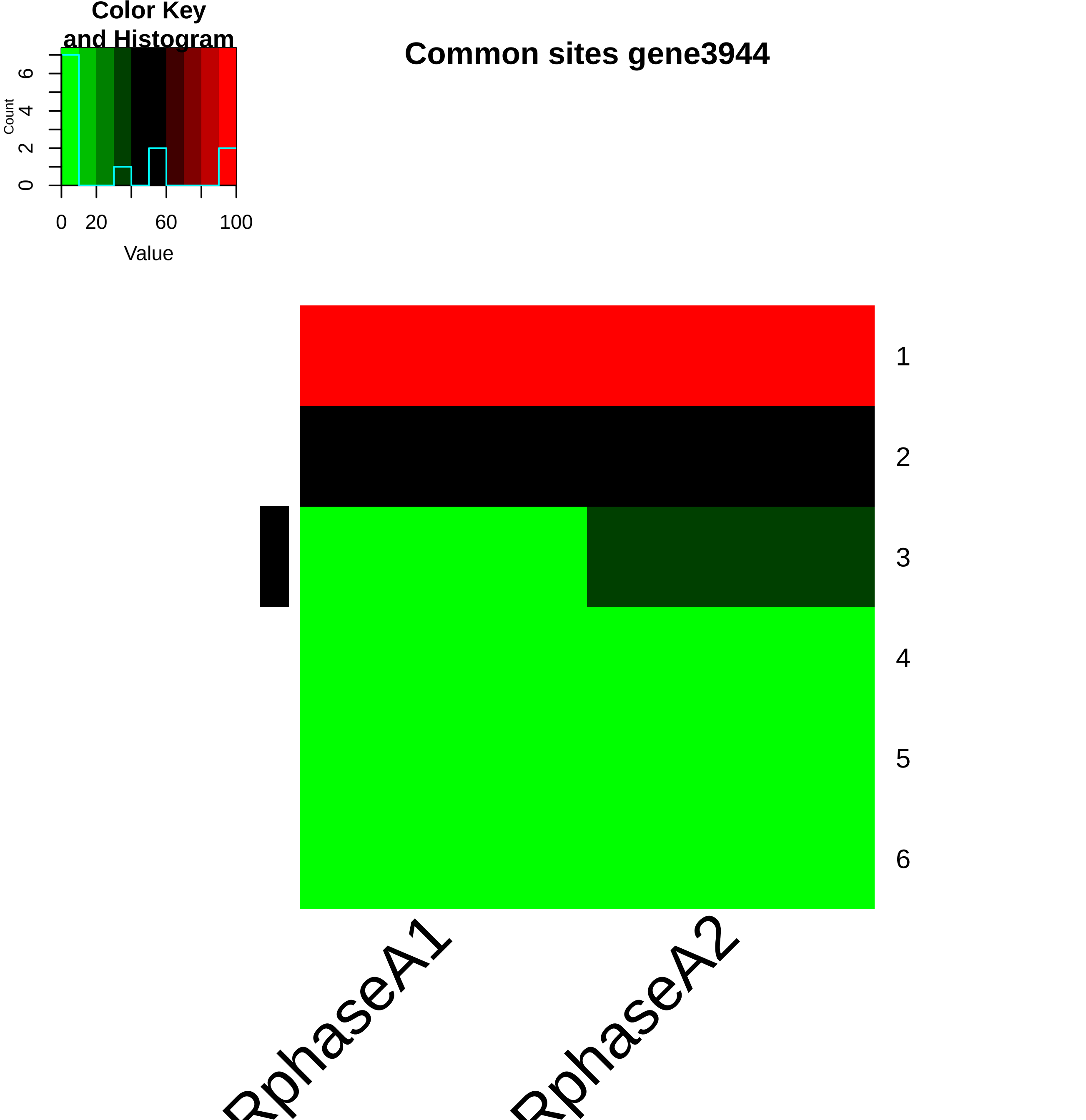
Color Key
and Histogram
Count
0
2
4
6
0
20
60
100
Value
Common sites gene3944
1
2
3
4
5
6
RphaseA1
RphaseA2

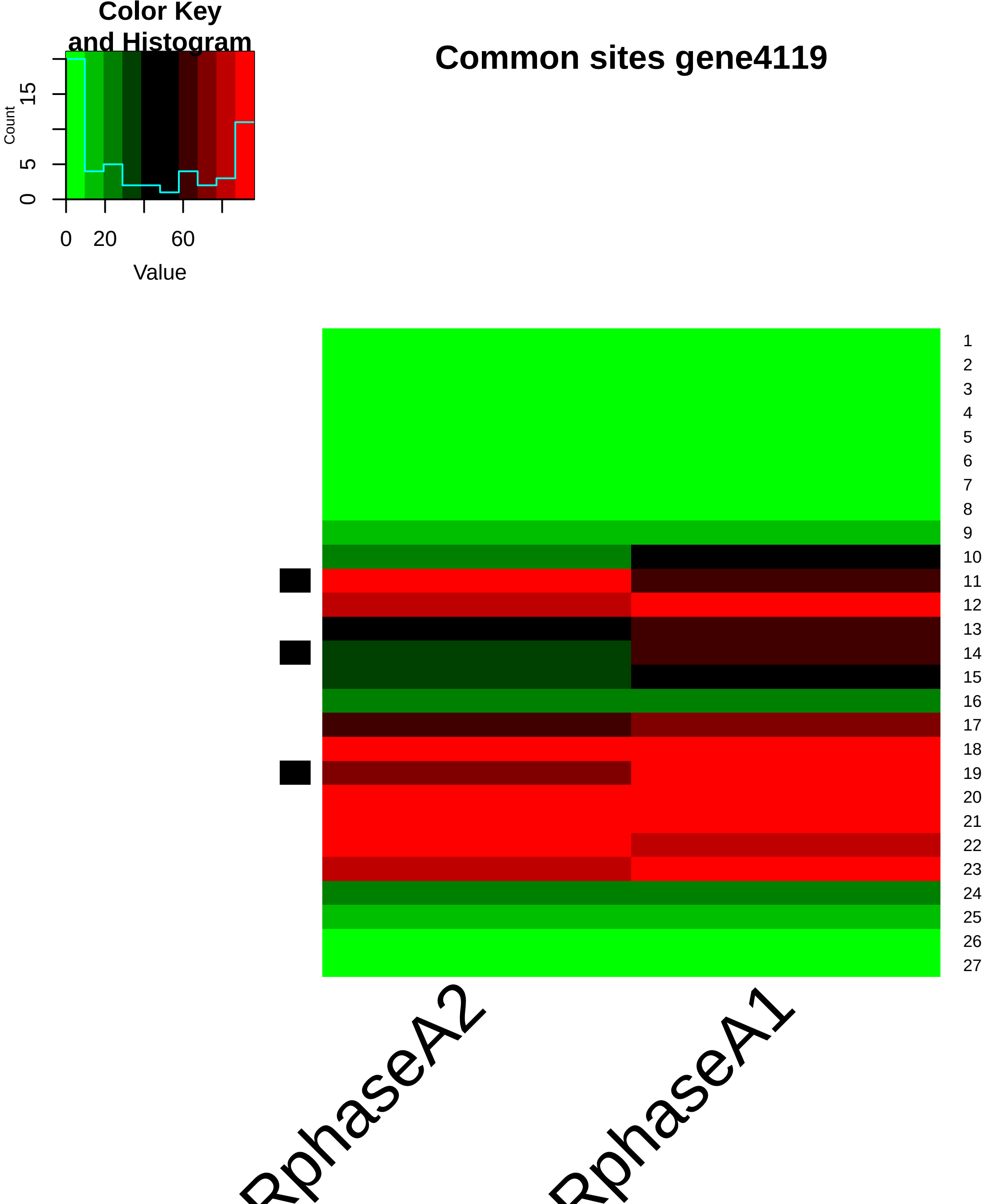
Color Key
and Histogram
Count
0
5
15
0
20
60
Value
Common sites gene4119
RphaseA2
RphaseA1

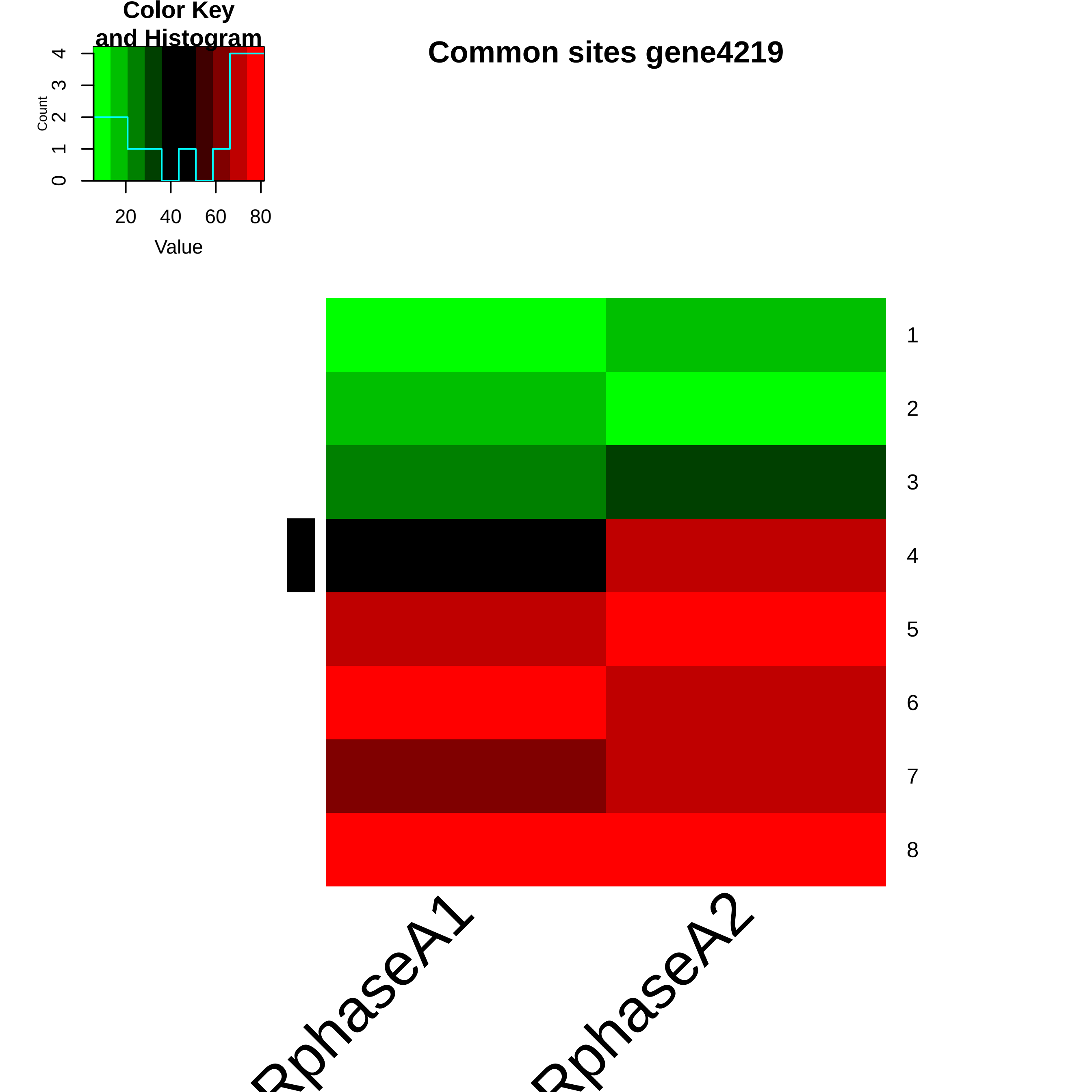

Color Key
and Histogram
Count
0
1
2
3
4
20
40
60
80
Value
Common sites gene4219
1
2
3
4
5
6
7
8
RphaseA1
RphaseA2

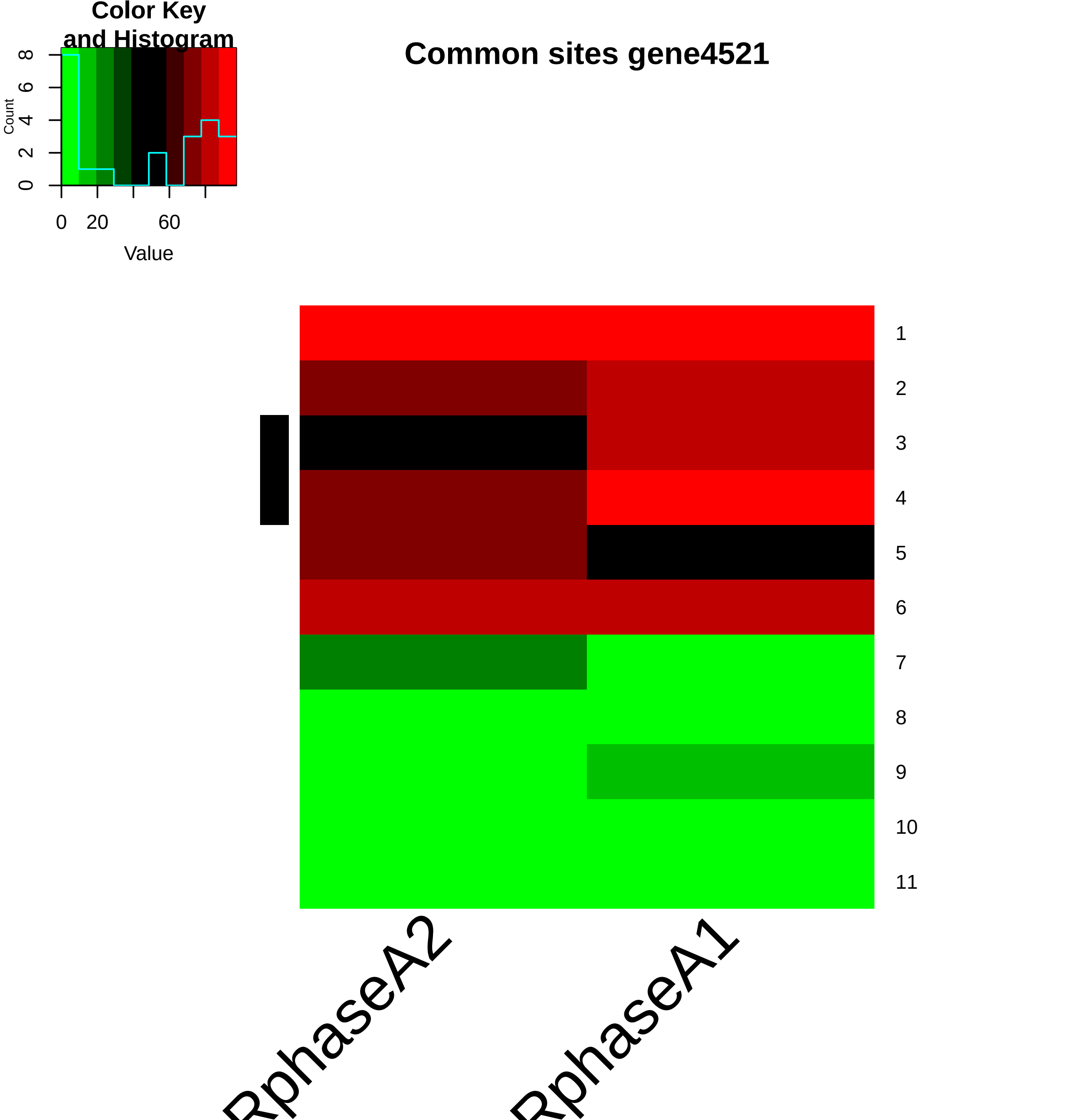
Color Key
and Histogram
Count
0
2
4
6
8
0
20
60
Value
Common sites gene4521
1
2
3
4
5
6
7
8
9
10
11
RphaseA2
RphaseA1

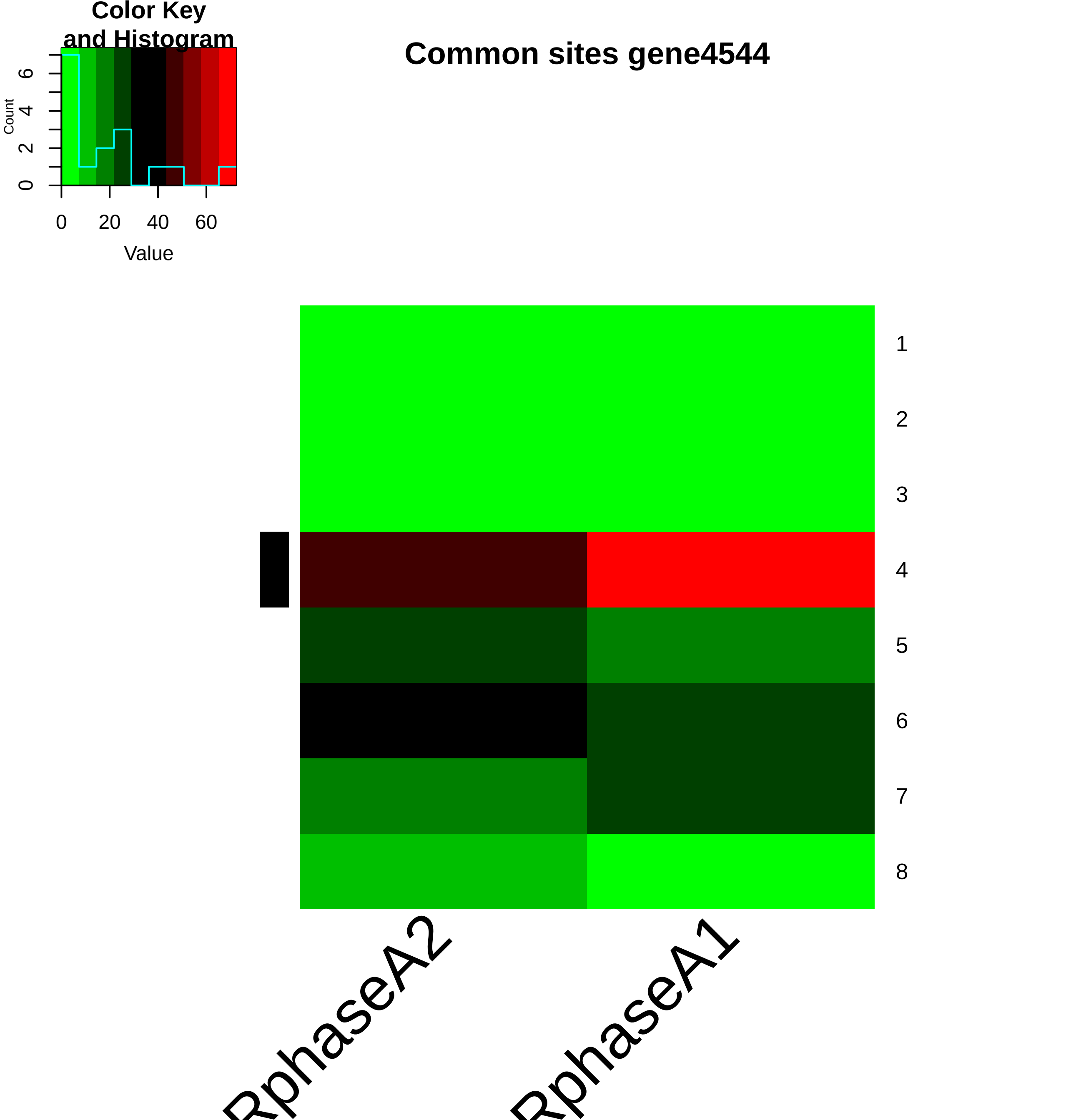
Color Key
and Histogram
Common sites gene4544
Count
0
2
4
6
0
20
40
60
Value
1
2
3
4
5
6
7
8
RphaseA2
RphaseA1

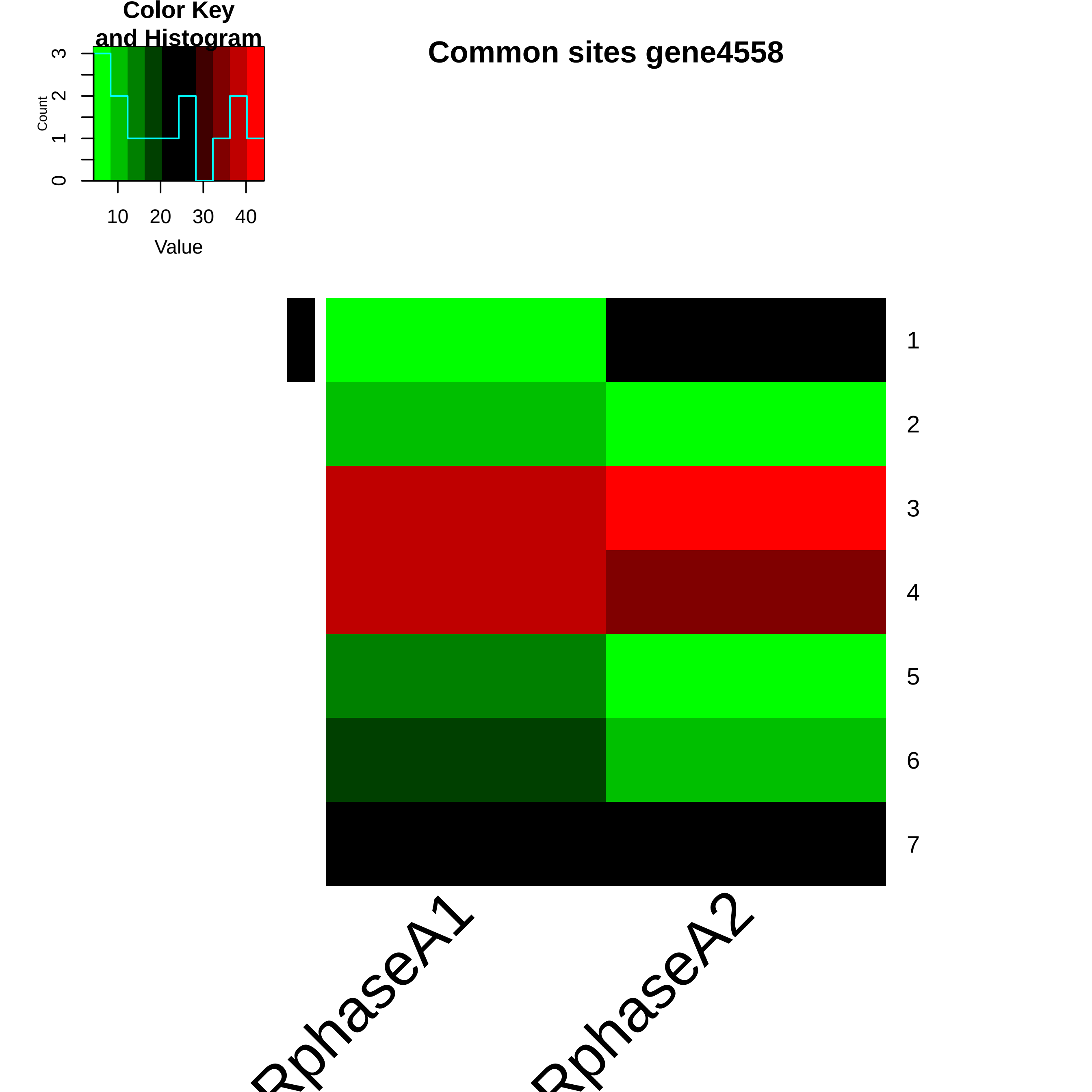
Color Key
and Histogram
Count
Value
Common sites gene4558
1
2
3
4
5
6
7
RphaseA1
RphaseA2

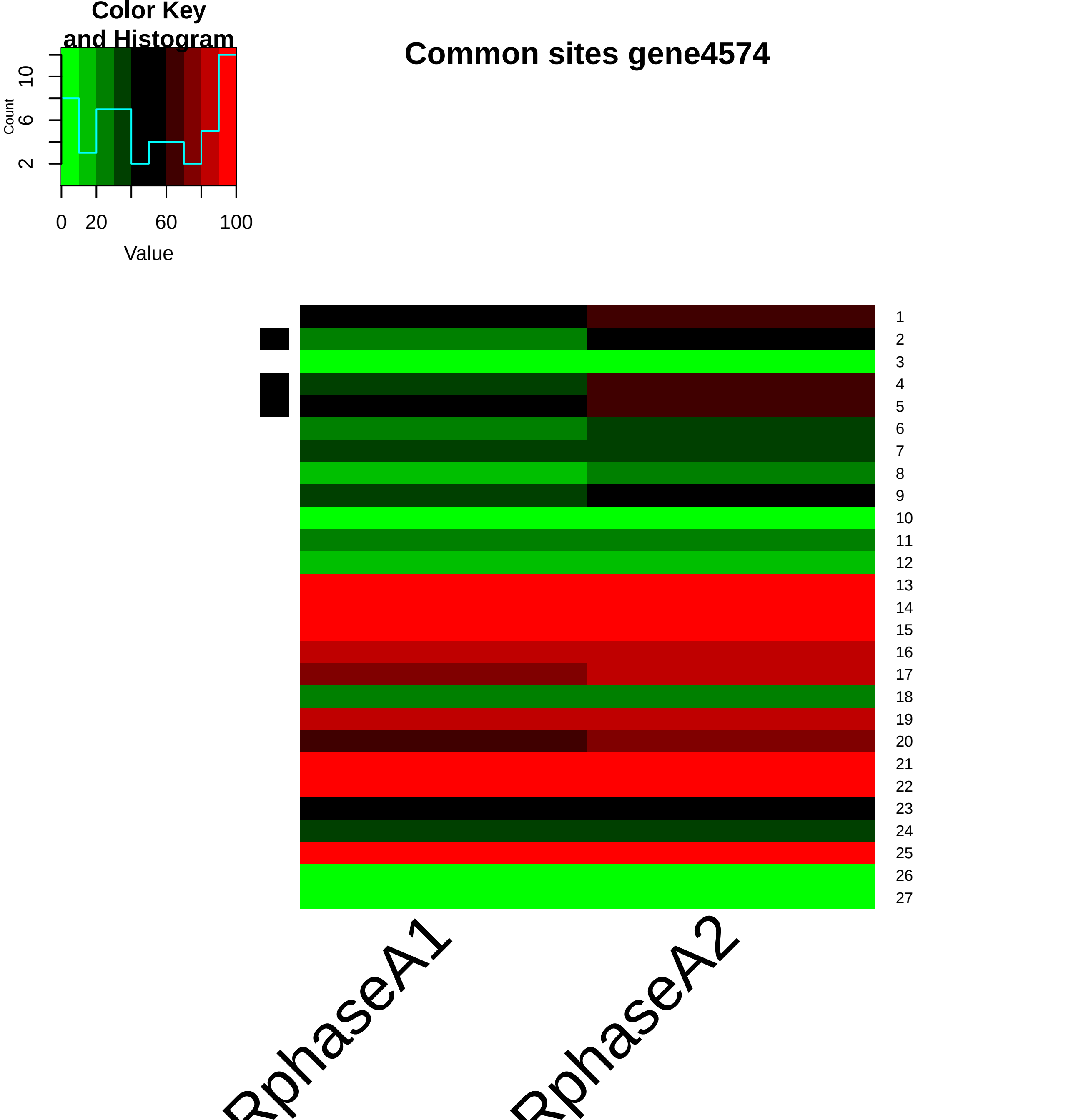
Color Key
and Histogram
Count
2
6
10
0
20
60
100
Value
Common sites gene4574
RphaseA1
RphaseA2

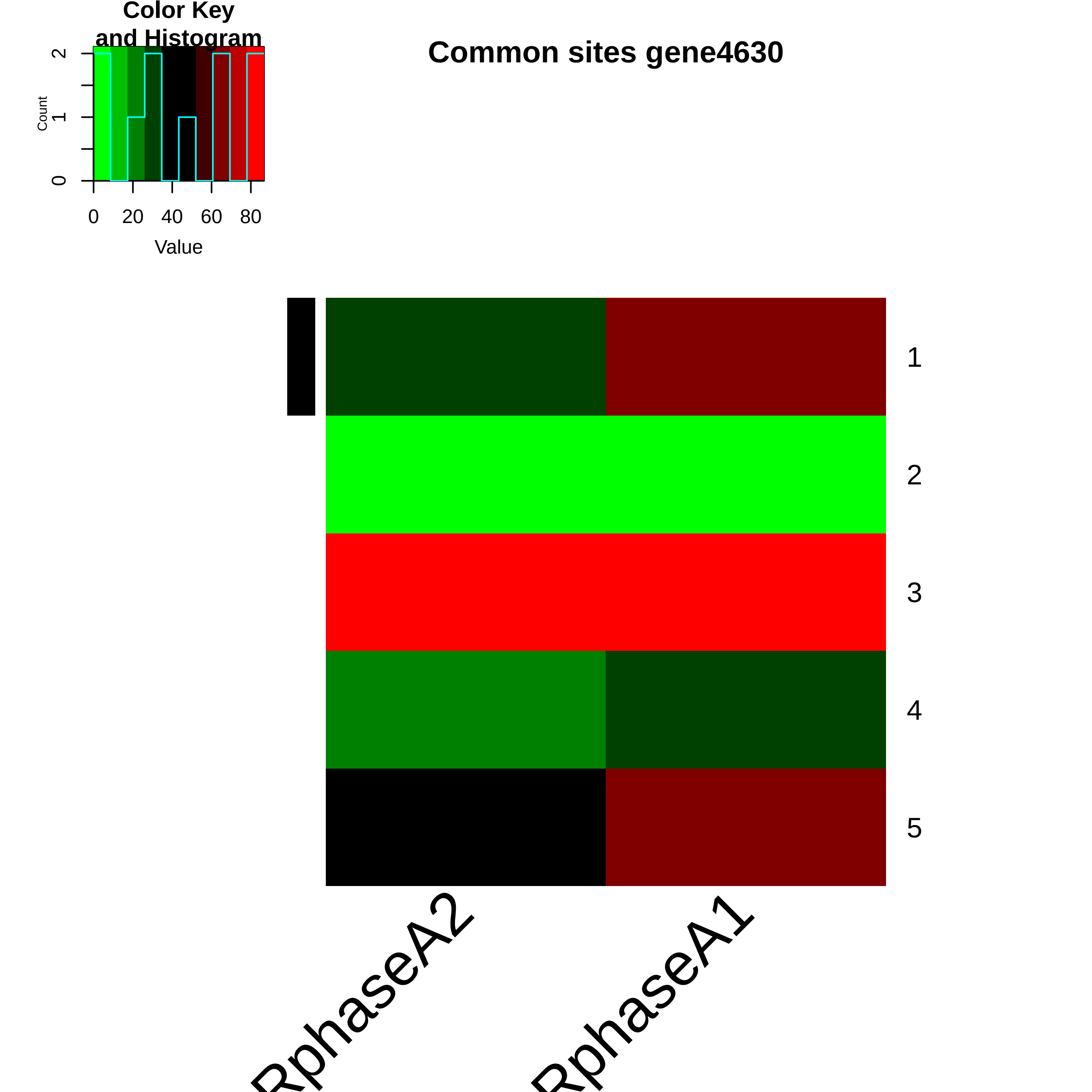

Color Key
and Histogram
Count
0
1
2
0
20
40
60
80
Value
Common sites gene4630
1
2
3
4
5
RphaseA2
RphaseA1

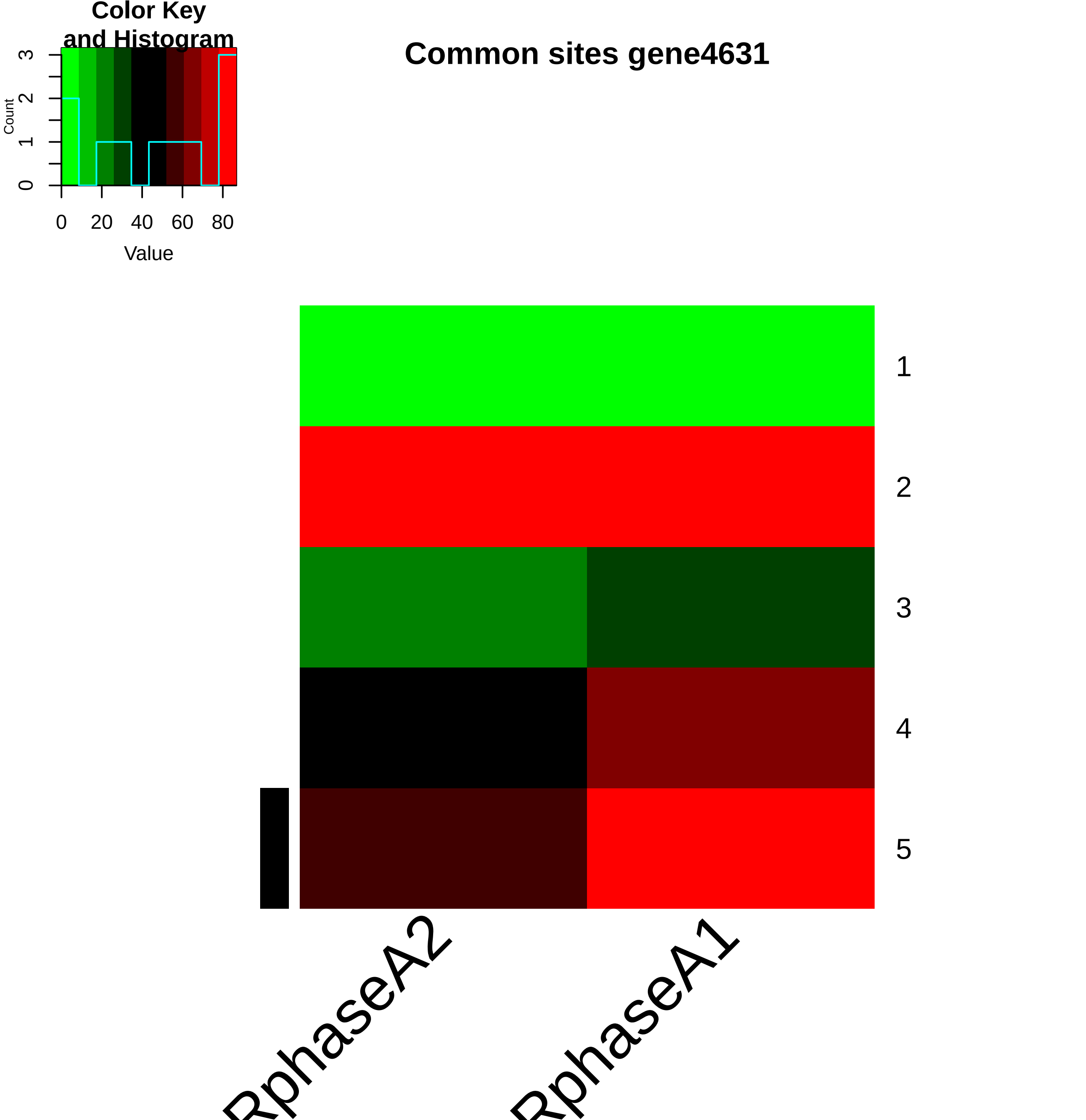
Color Key
and Histogram
Count
0
1
2
3
0
20
40
60
80
Value
Common sites gene4631
1
2
3
4
5
RphaseA2
RphaseA1

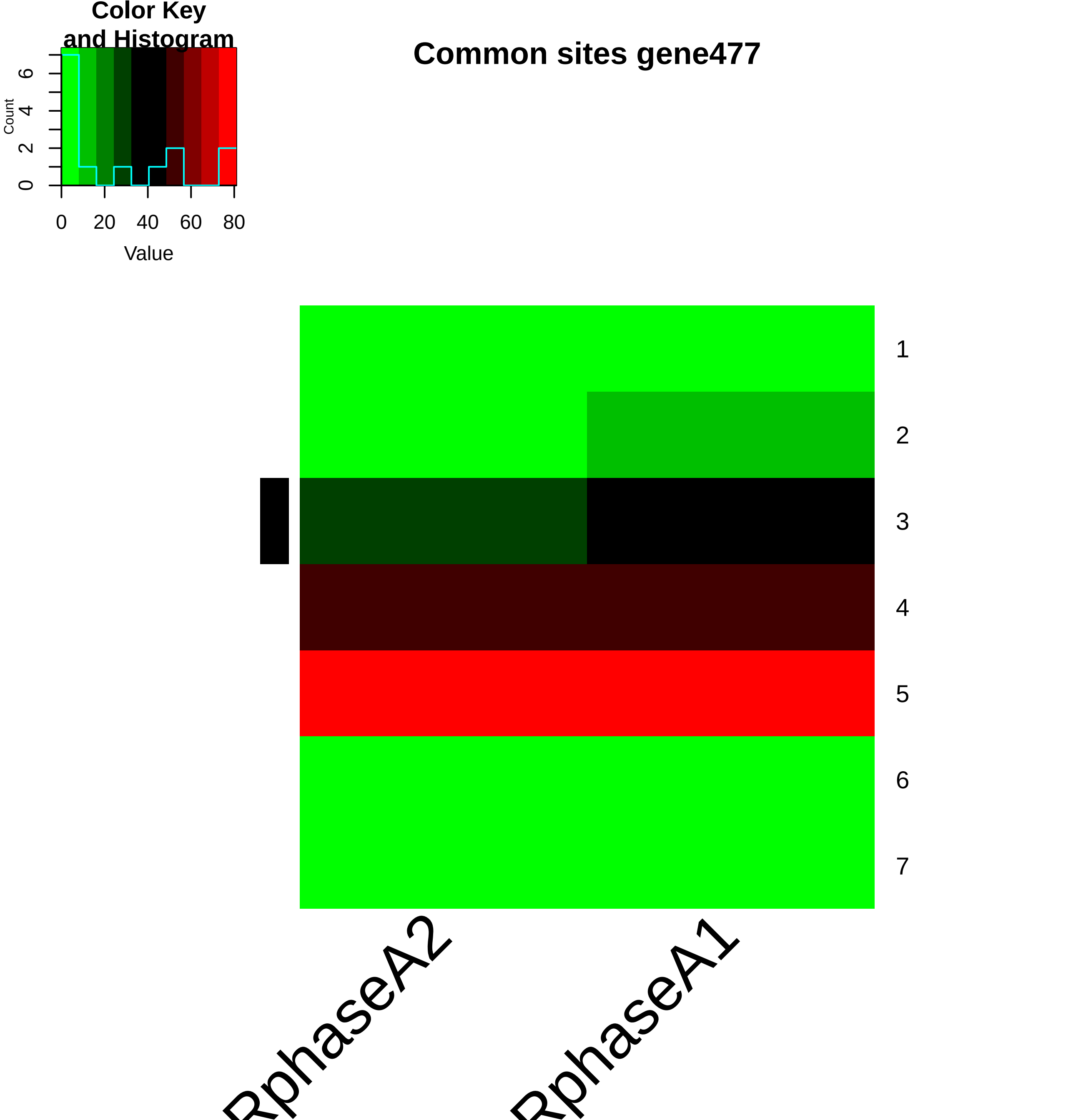

Color Key
and Histogram
Count
0
2
4
6
0
20
40
60
80
Value
Common sites gene477
1
2
3
4
5
6
7
RphaseA2
RphaseA1

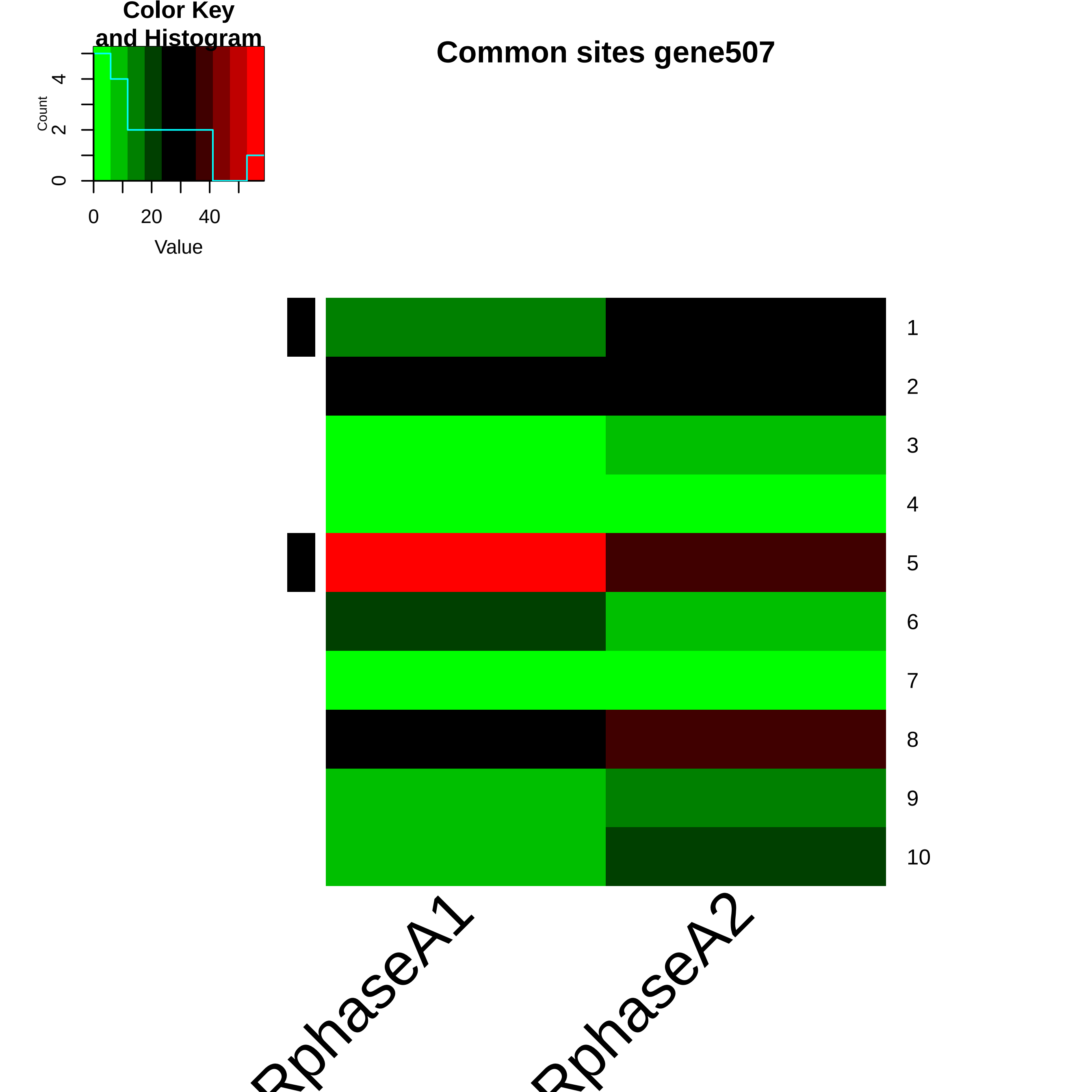

Color Key
and Histogram
Count
0
2
4
0
20
40
Value
Common sites gene507
1
2
3
4
5
6
7
8
9
10
RphaseA1
RphaseA2

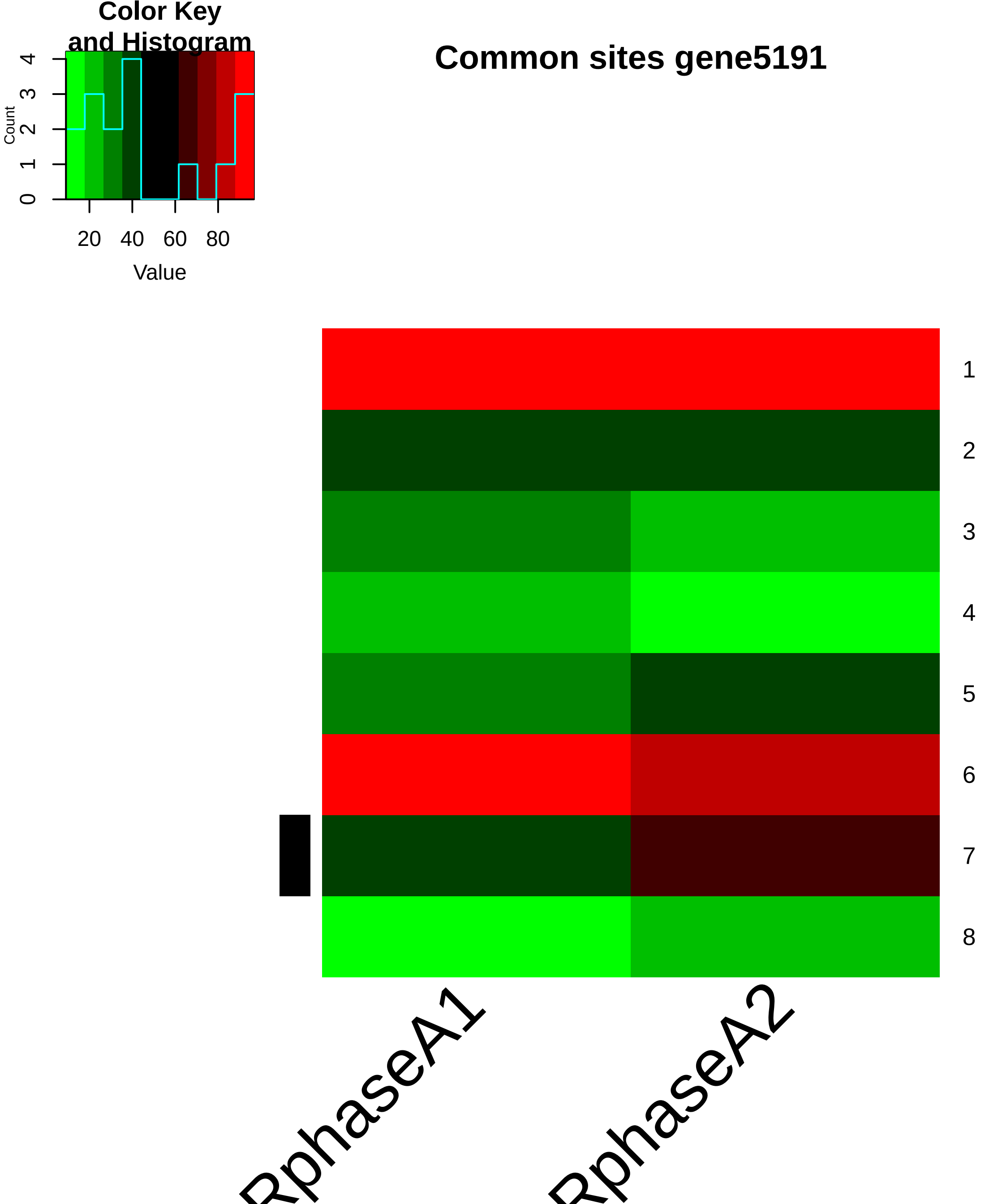
Color Key
and Histogram
Count
0
1
2
3
4
20
40
60
80
Value
Common sites gene5191
1
2
3
4
5
6
7
8
RphaseA1
RphaseA2

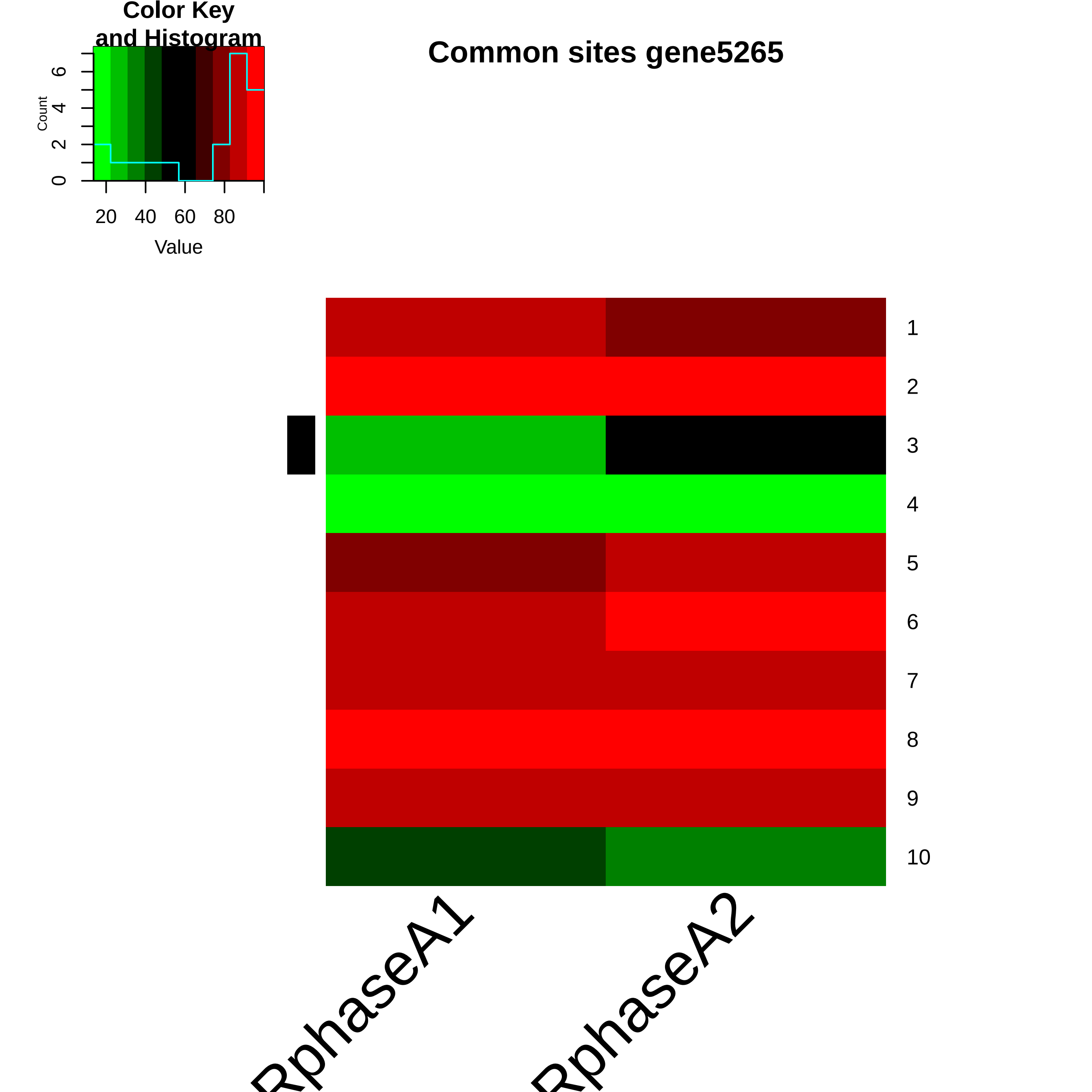
Color Key
and Histogram
Count
0
2
4
6
20
40
60
80
Value
Common sites gene5265
1
2
3
4
5
6
7
8
9
10
RphaseA1
RphaseA2

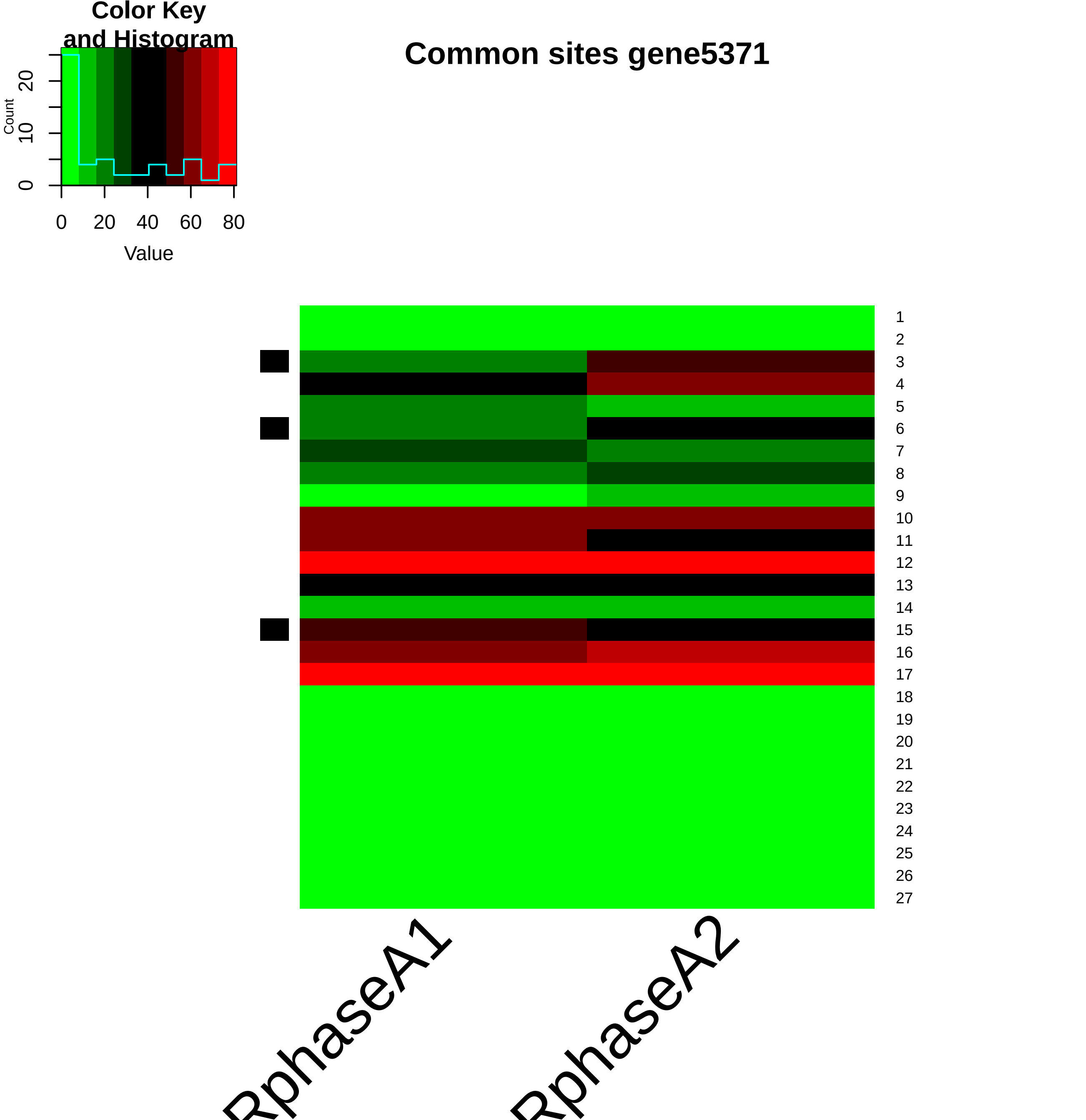

Color Key
and Histogram
Count
0
10
20
0
20
40
60
80
Value
Common sites gene5371
RphaseA1
RphaseA2

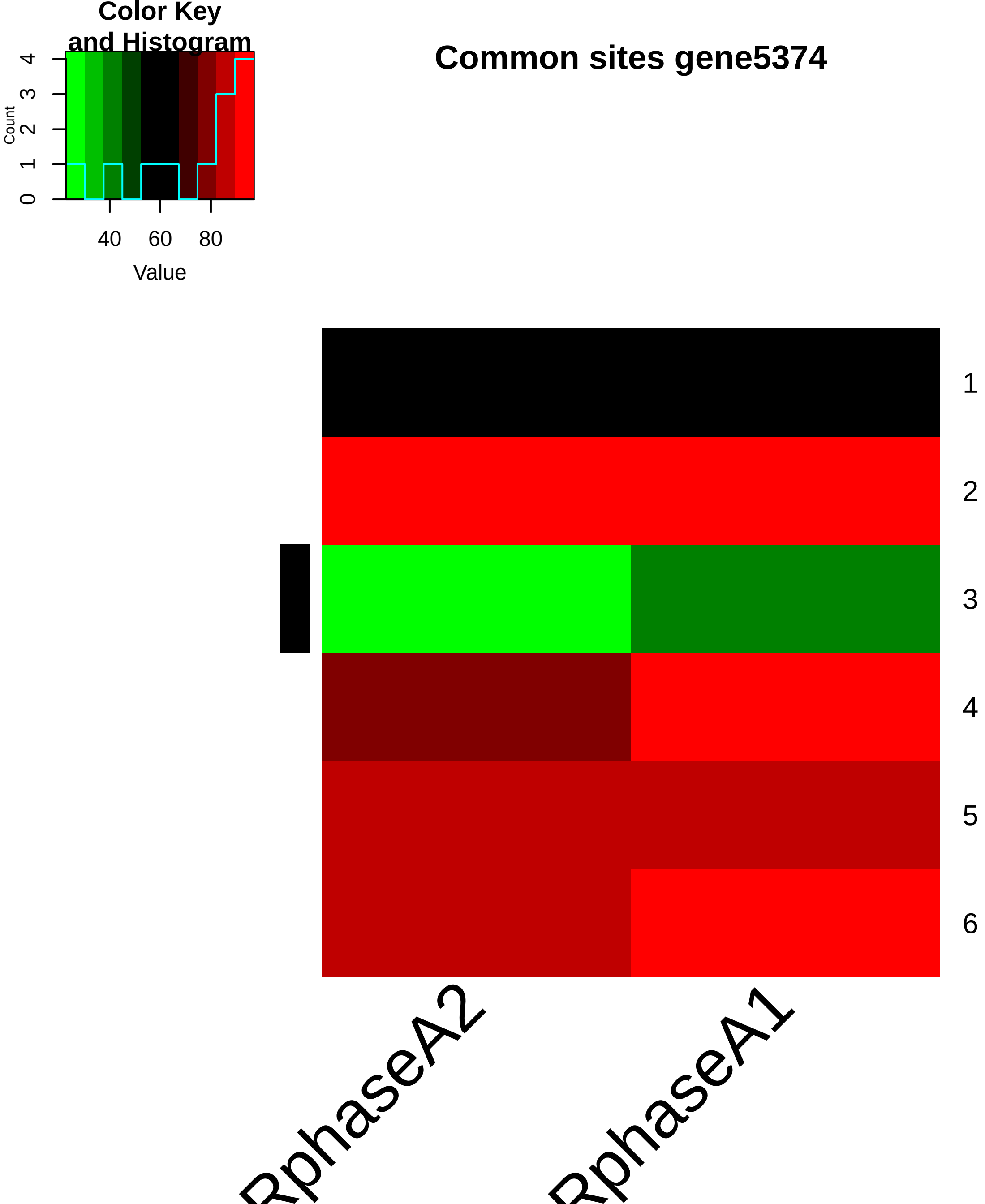
Color Key
and Histogram
Count
0
1
2
3
4
40
60
80
Value
Common sites gene5374
1
2
3
4
5
6
RphaseA2
RphaseA1

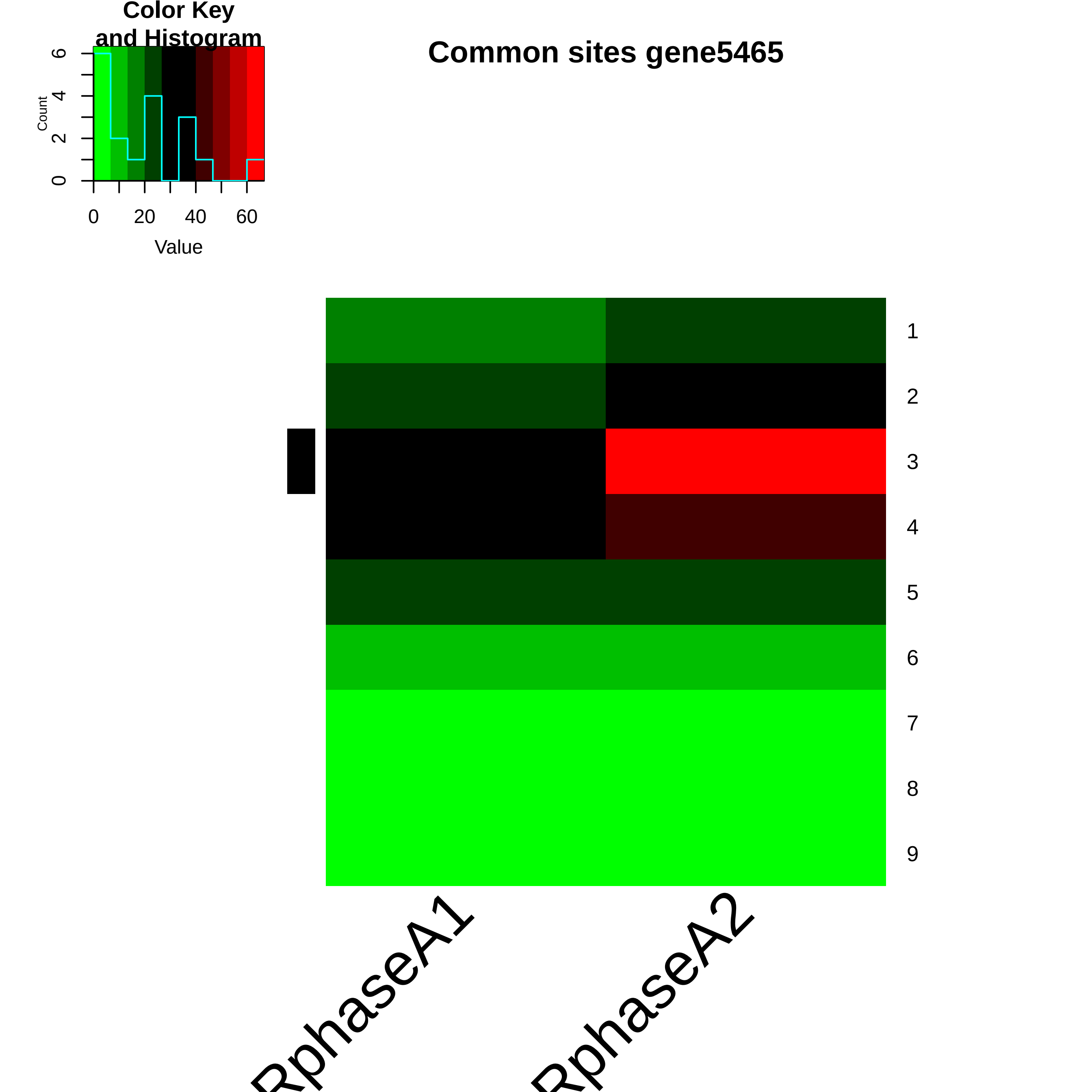

Color Key
and Histogram
Count
0
2
4
6
0
20
40
60
Value
Common sites gene5465
1
2
3
4
5
6
7
8
9
RphaseA1
RphaseA2

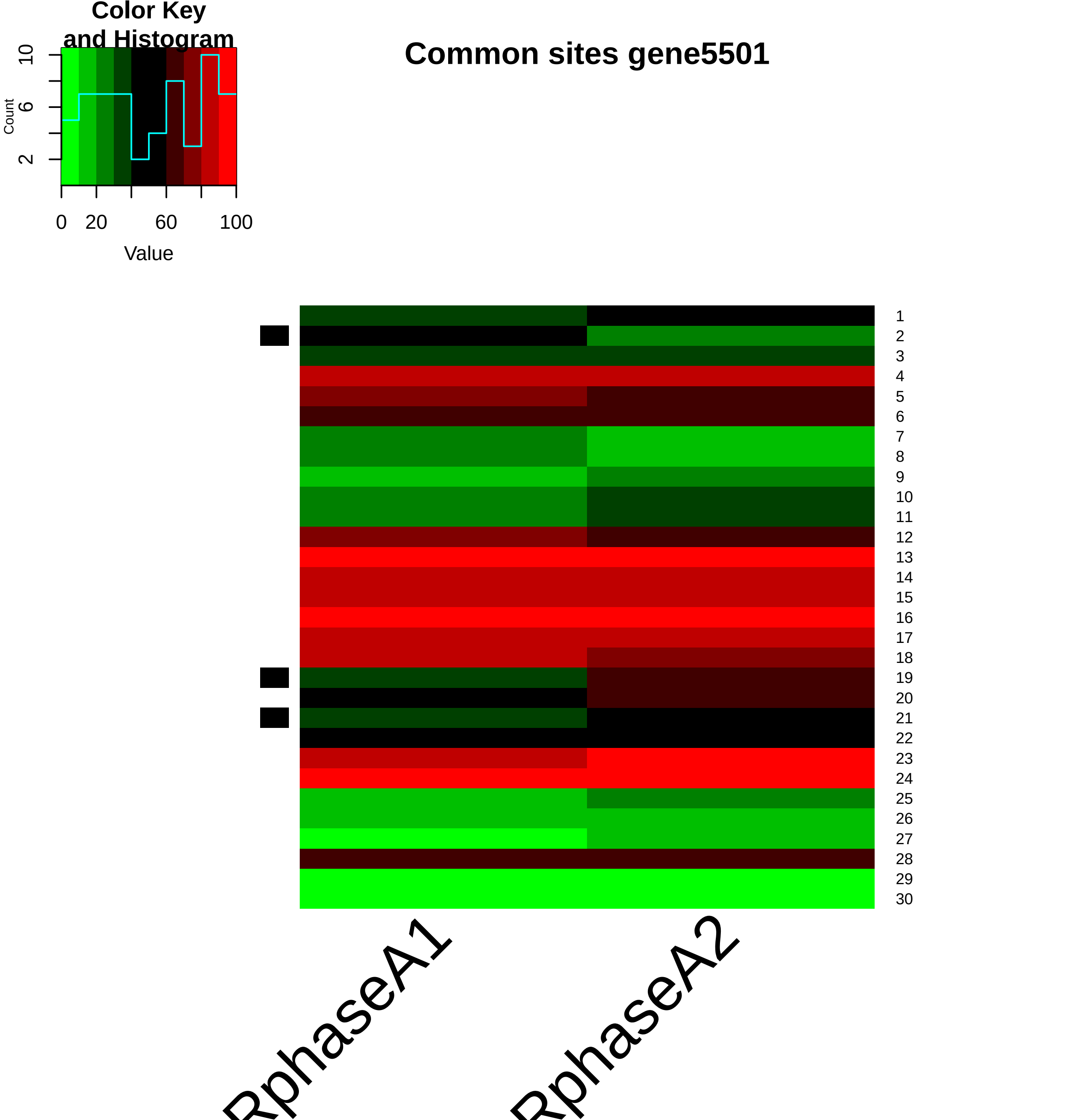
Color Key
and Histogram
Count
2
6
10
0
20
60
100
Value
Common sites gene5501
RphaseA1
RphaseA2

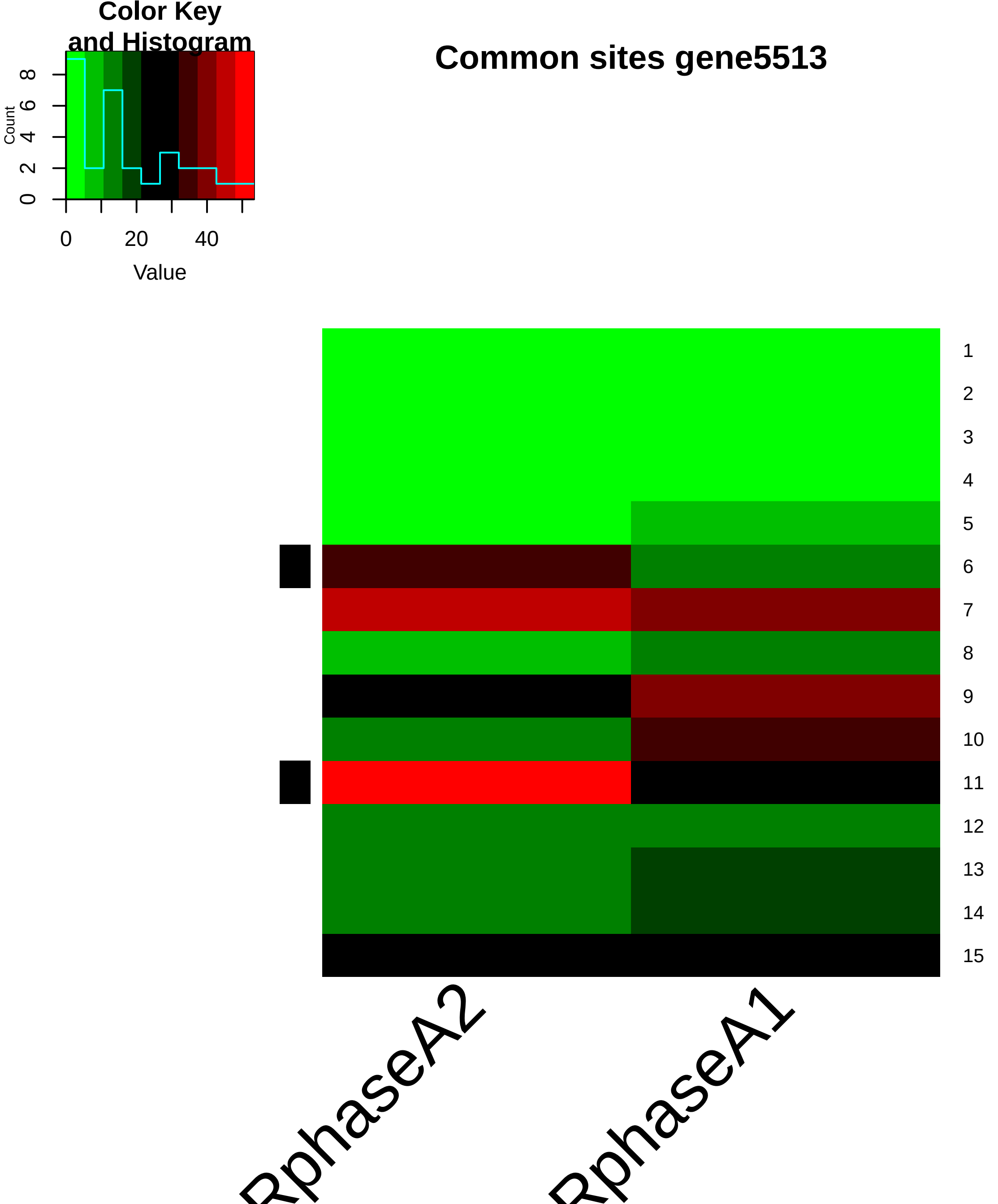

Color Key
and Histogram
Count
0
2
4
6
8
0
20
40
Value
Common sites gene5513
RphaseA2
RphaseA1

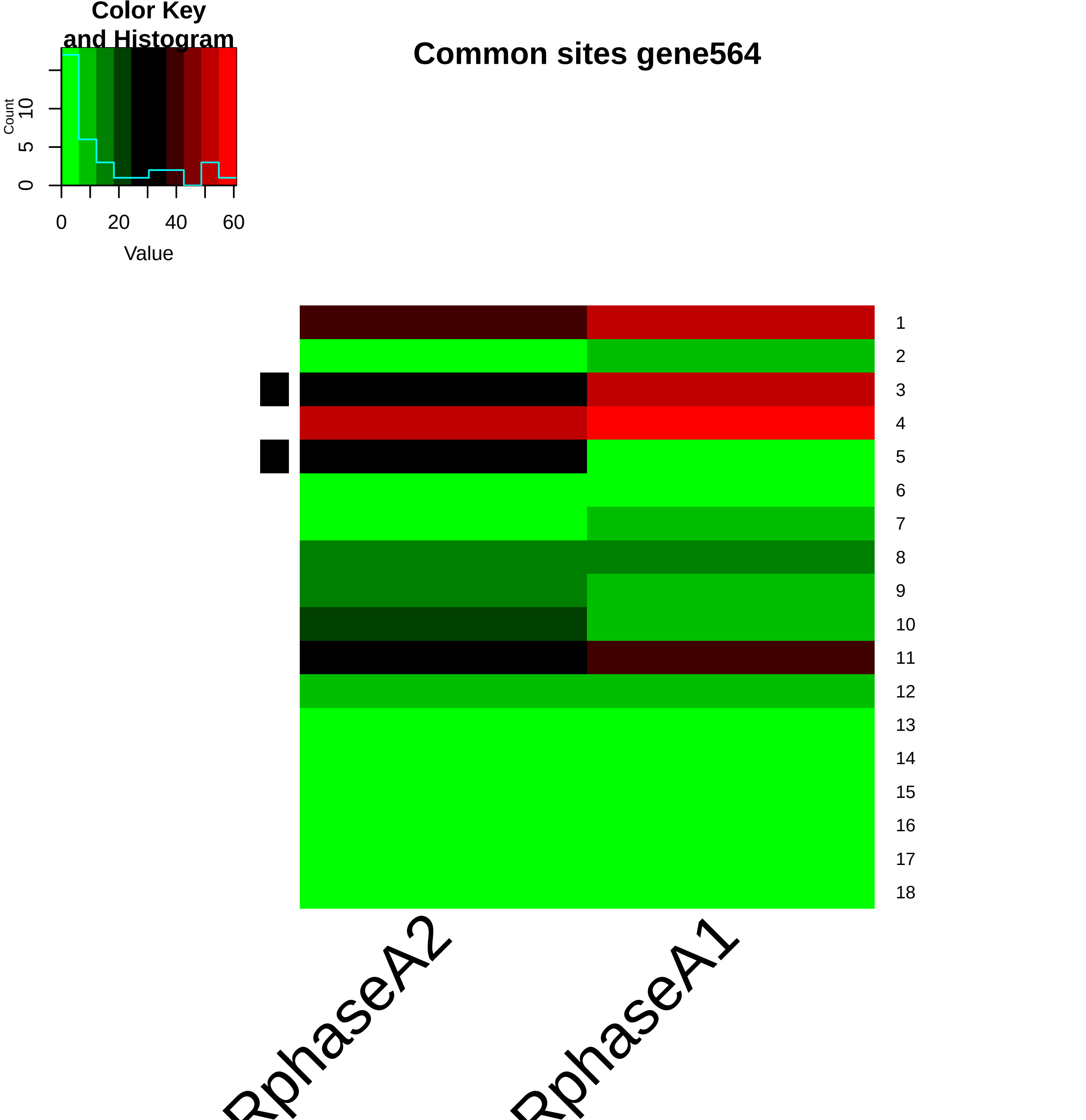

Color Key
and Histogram
Count
0
5
10
0
20
40
60
Value
Common sites gene564
RphaseA2
RphaseA1

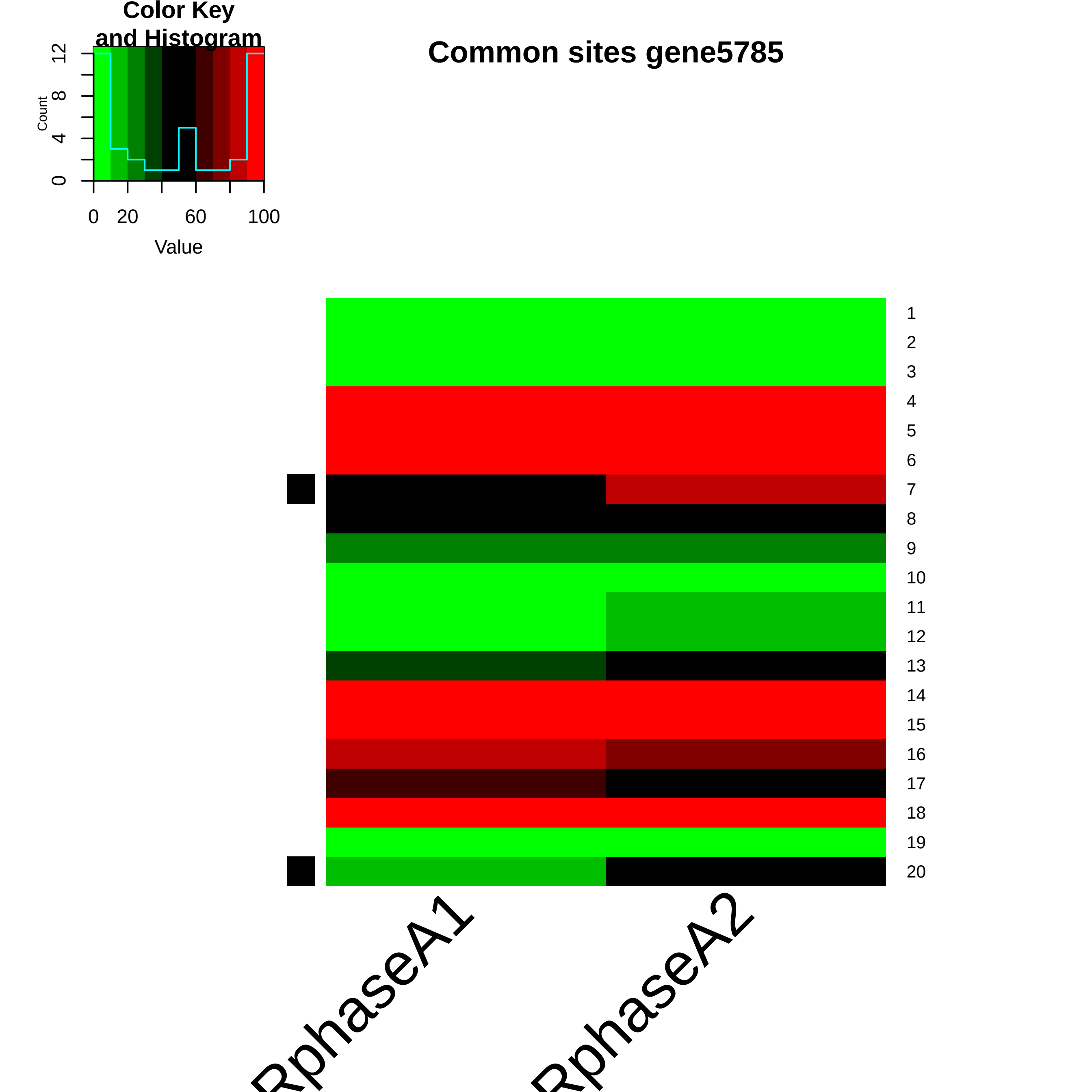
Color Key
and Histogram
Common sites gene5785
Count
0
4
8
12
0
20
60
100
Value
RphaseA1
RphaseA2

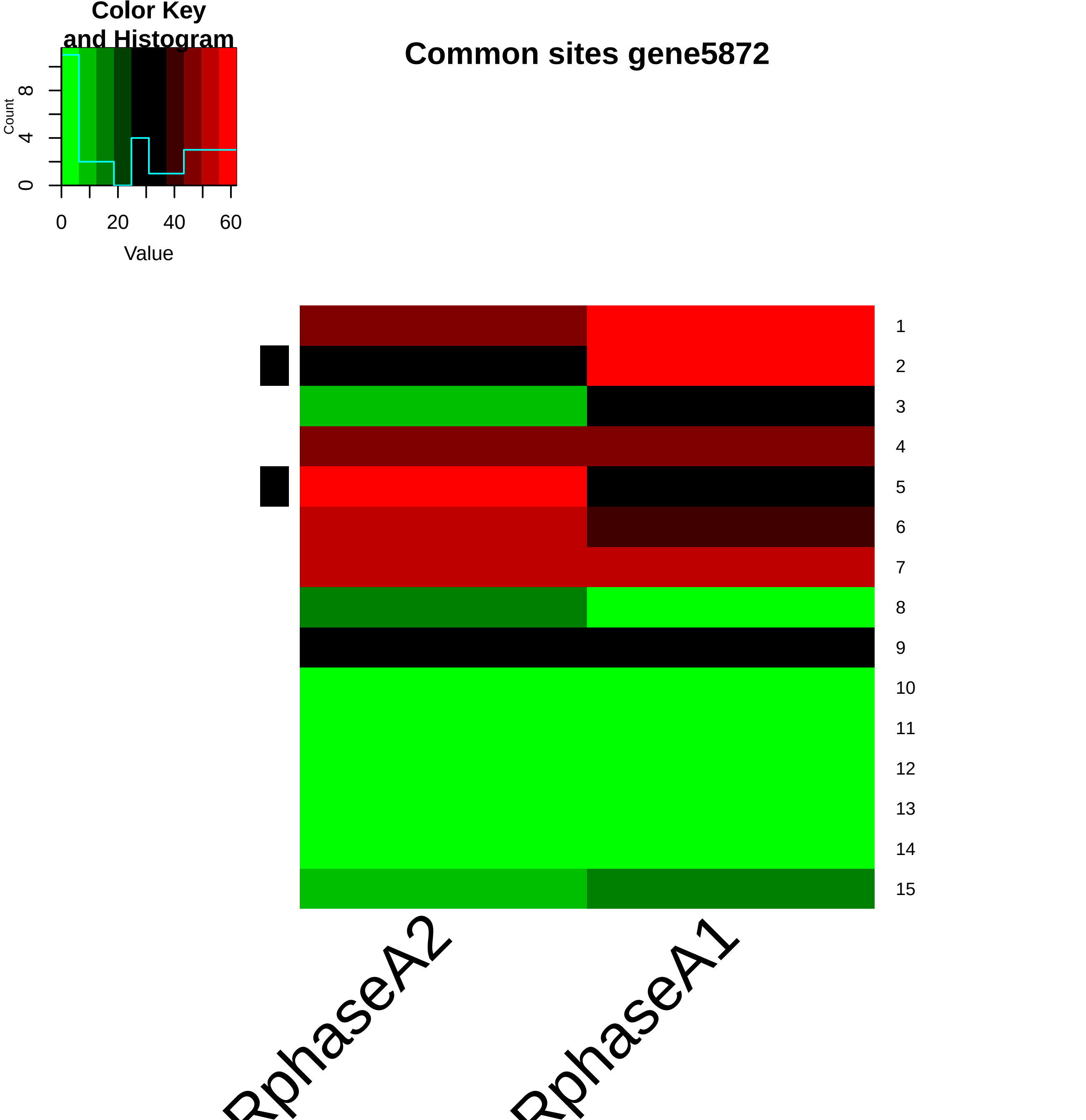
Color Key
and Histogram
Count
0
4
8
0
20
40
60
Value
Common sites gene5872
RphaseA2
RphaseA1

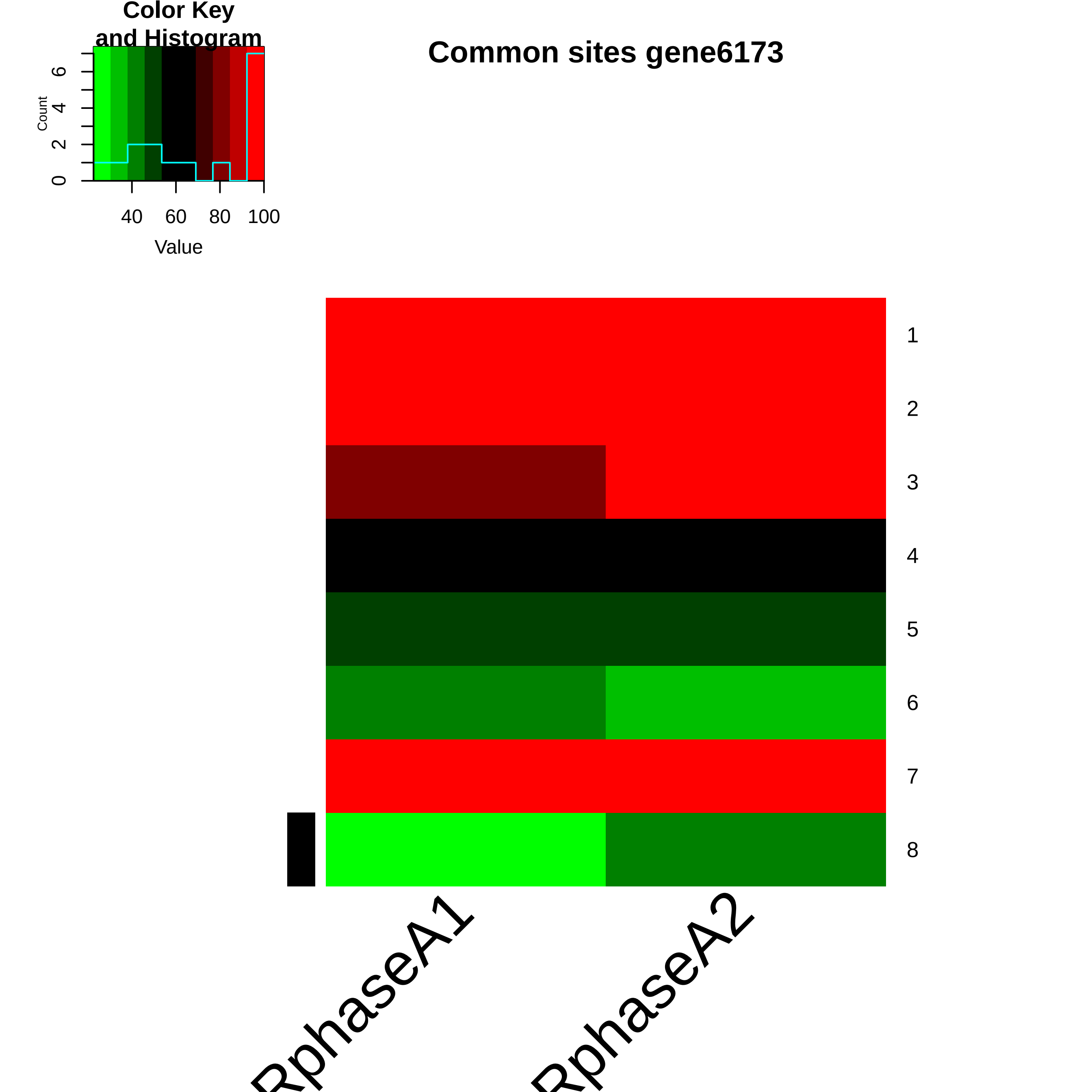
Color Key
and Histogram
Count
0
2
4
6
40
60
80
100
Value
Common sites gene6173
1
2
3
4
5
6
7
8
RphaseA1
RphaseA2

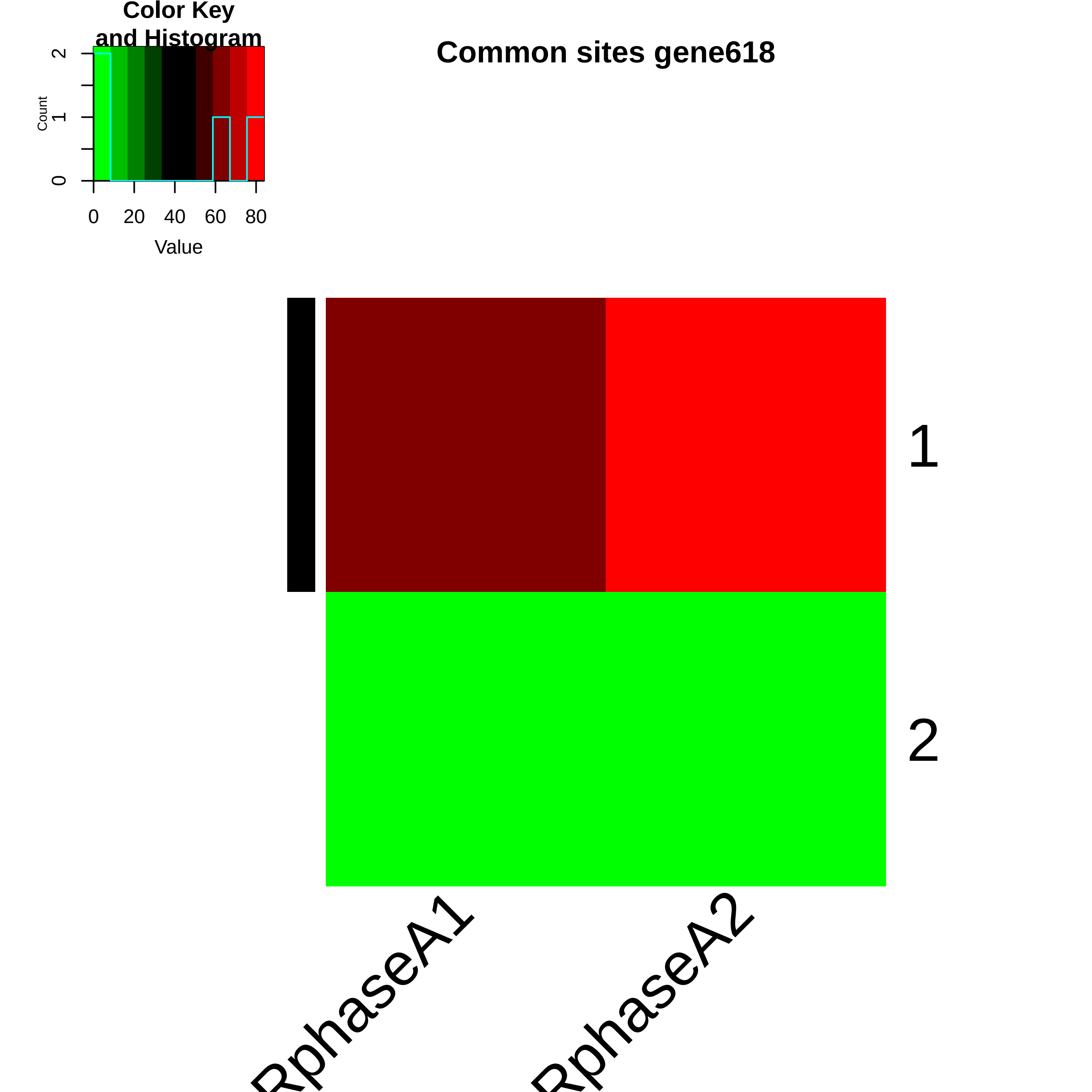
Color Key
and Histogram
Count
0
1
2
0
20
40
60
80
Value
Common sites gene618
1
2
RphaseA1
RphaseA2

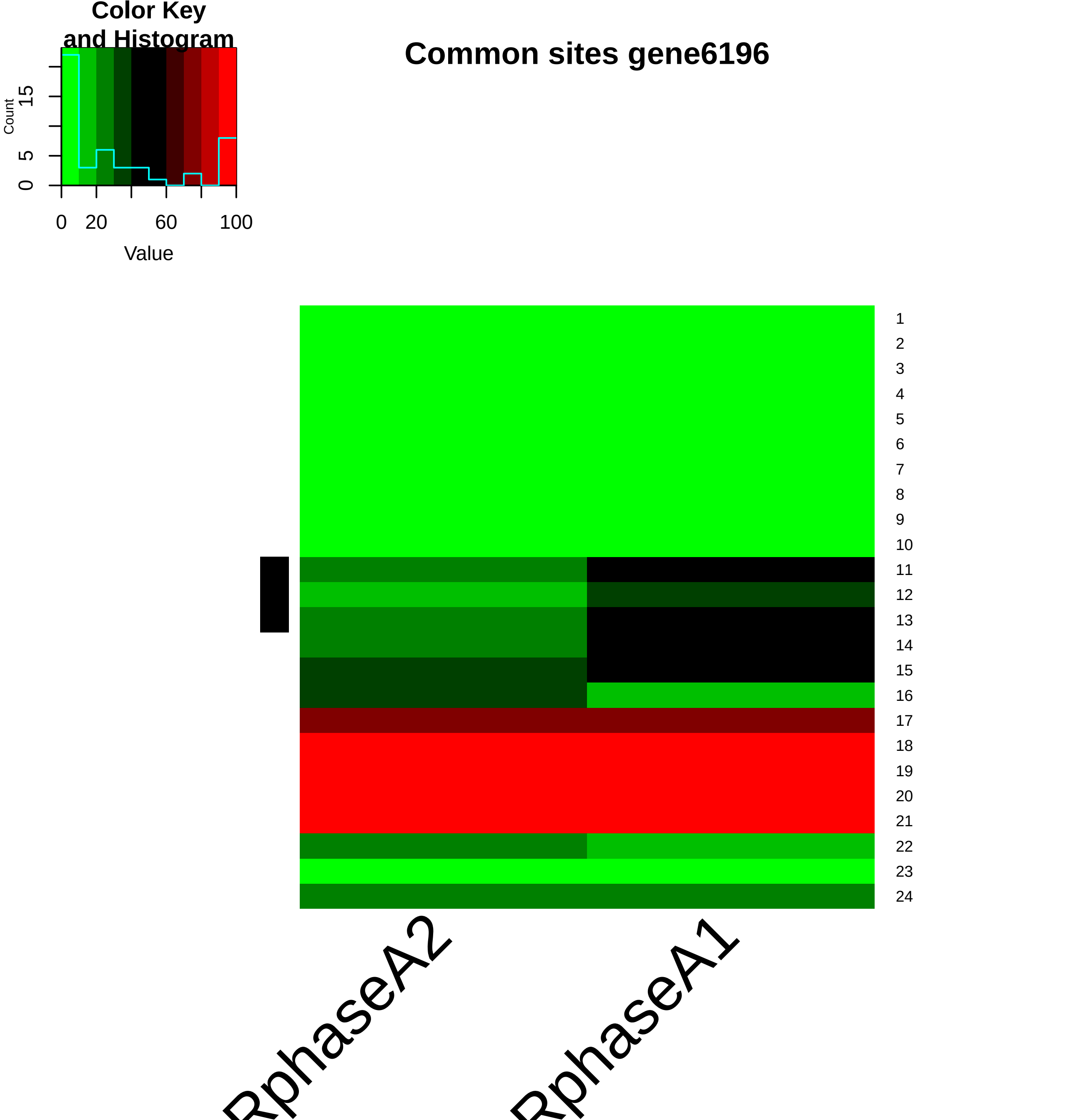

Color Key and Histogram
Count
0
5
15
0
20
60
100
Value
Common sites gene6196
RphaseA2
RphaseA1

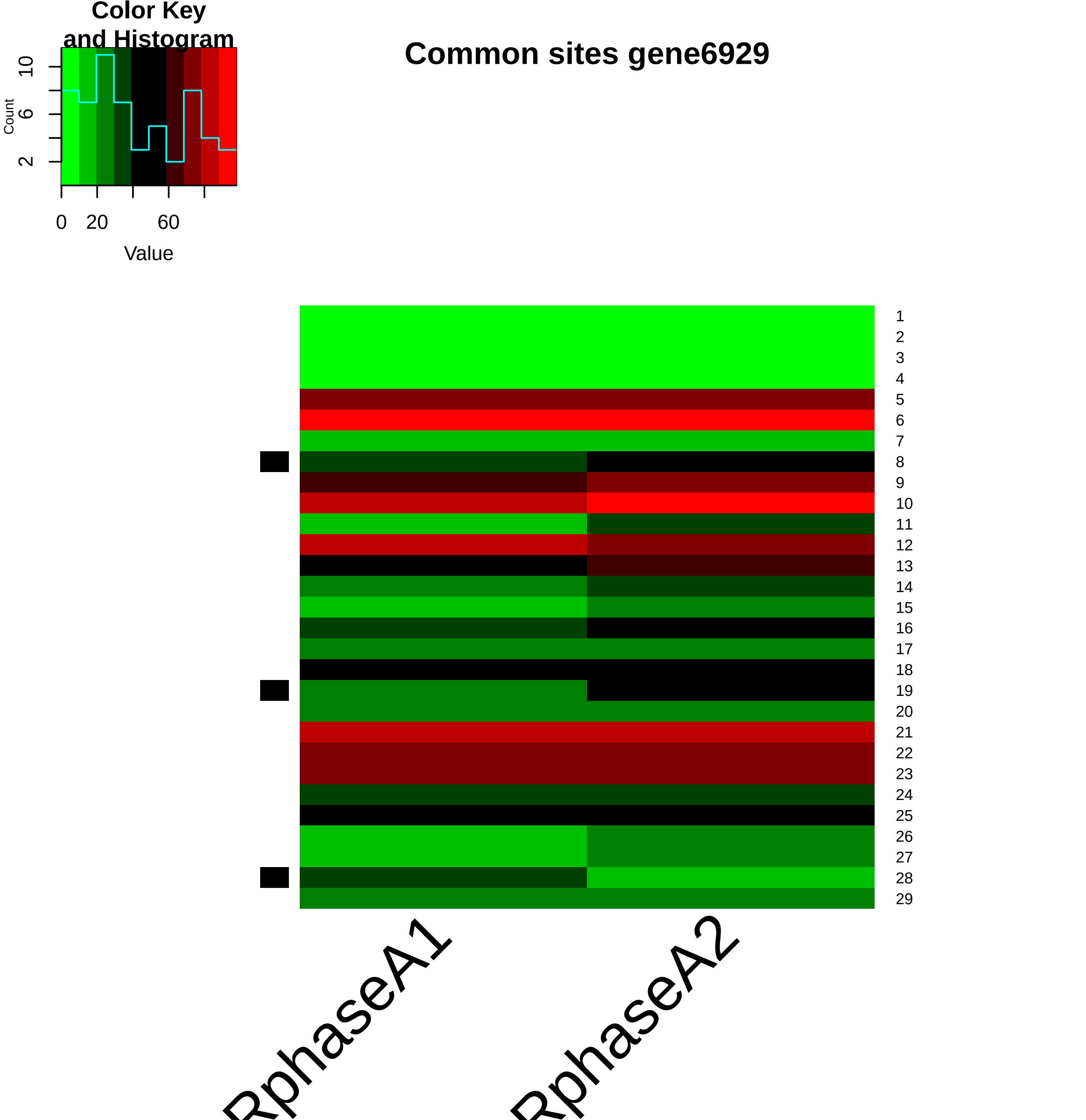
Color Key
and Histogram
Count
2
6
10
0
20
60
Value
Common sites gene6929
RphaseA1
RphaseA2

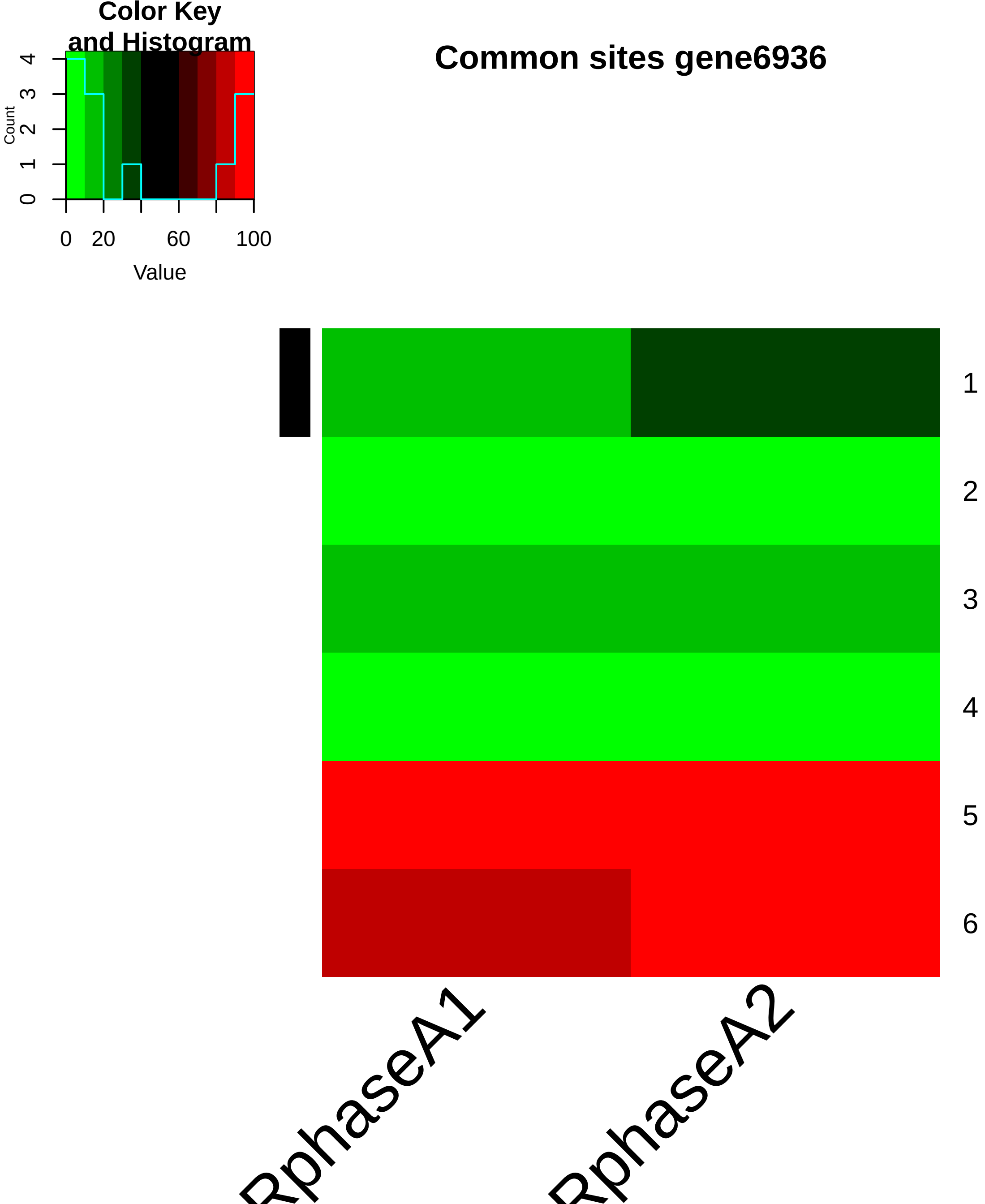
Color Key
and Histogram
Count
0
1
2
3
4
0
20
60
100
Value
Common sites gene6936
1
2
3
4
5
6
RphaseA1
RphaseA2

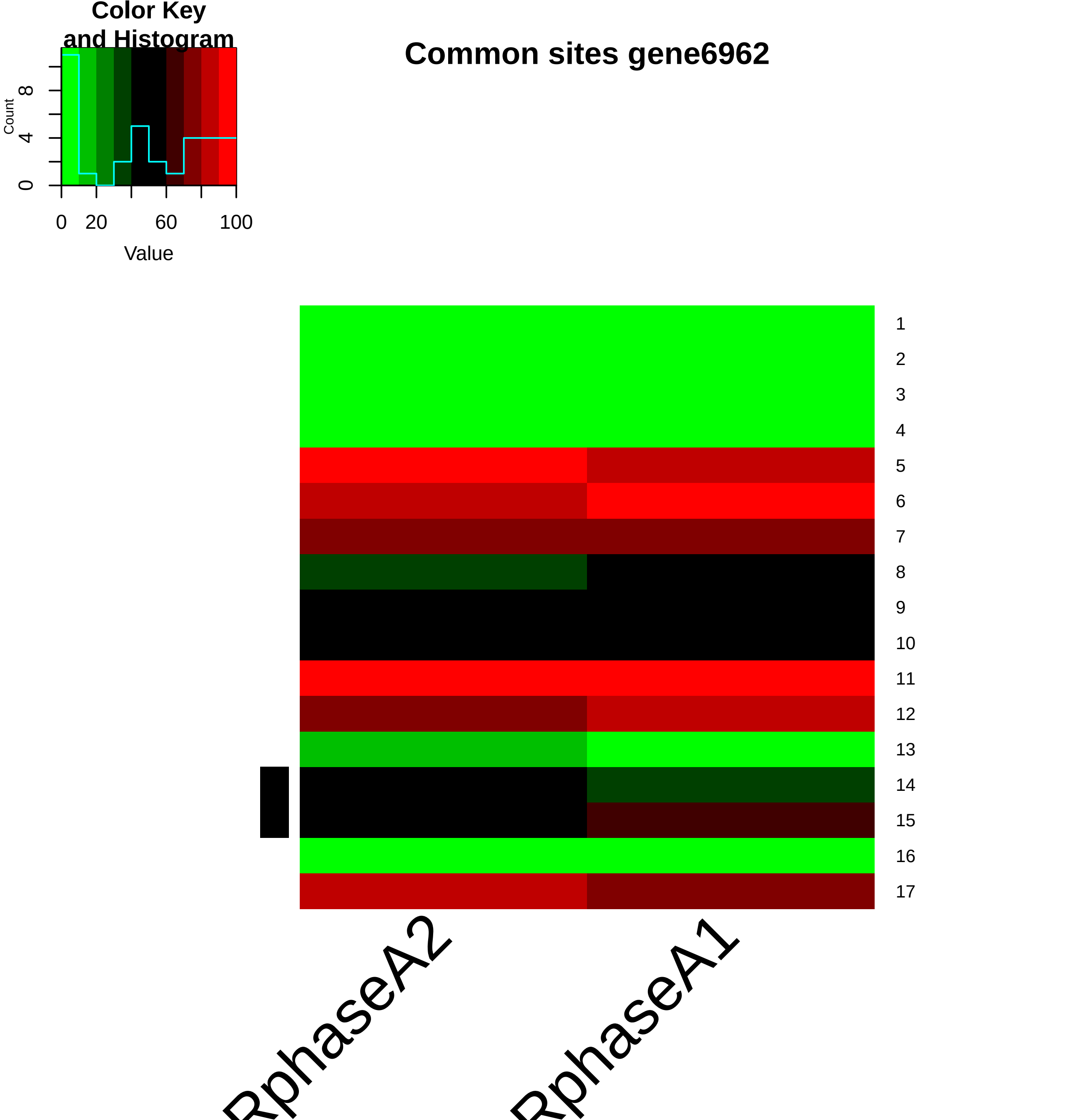

Color Key
and Histogram
Count
0
4
8
0
20
60
100
Value
Common sites gene6962
RphaseA2
RphaseA1

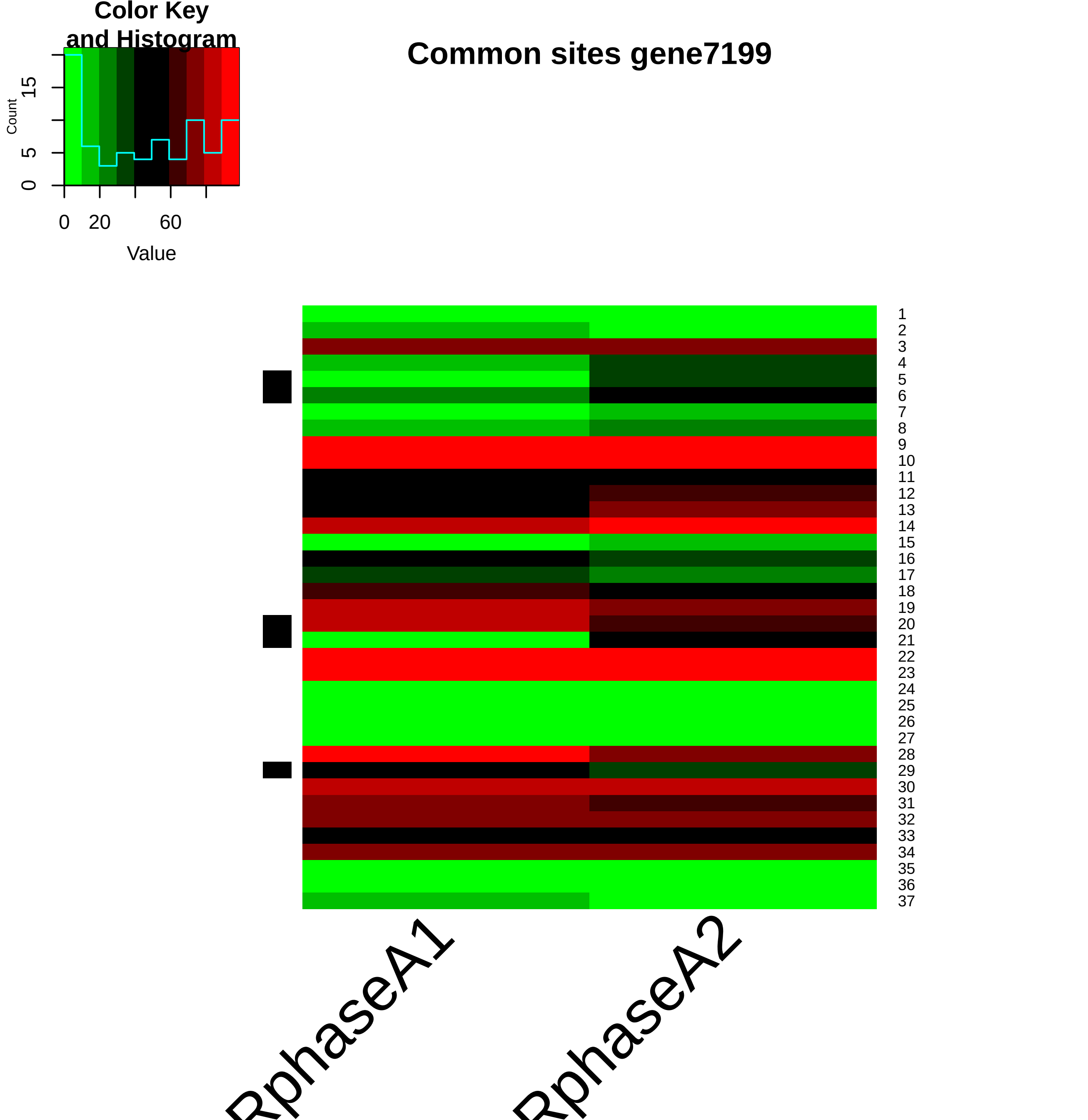

Color Key
and Histogram
Count
0
5
15
0
20
60
Value
Common sites gene7199
RphaseA1
RphaseA2

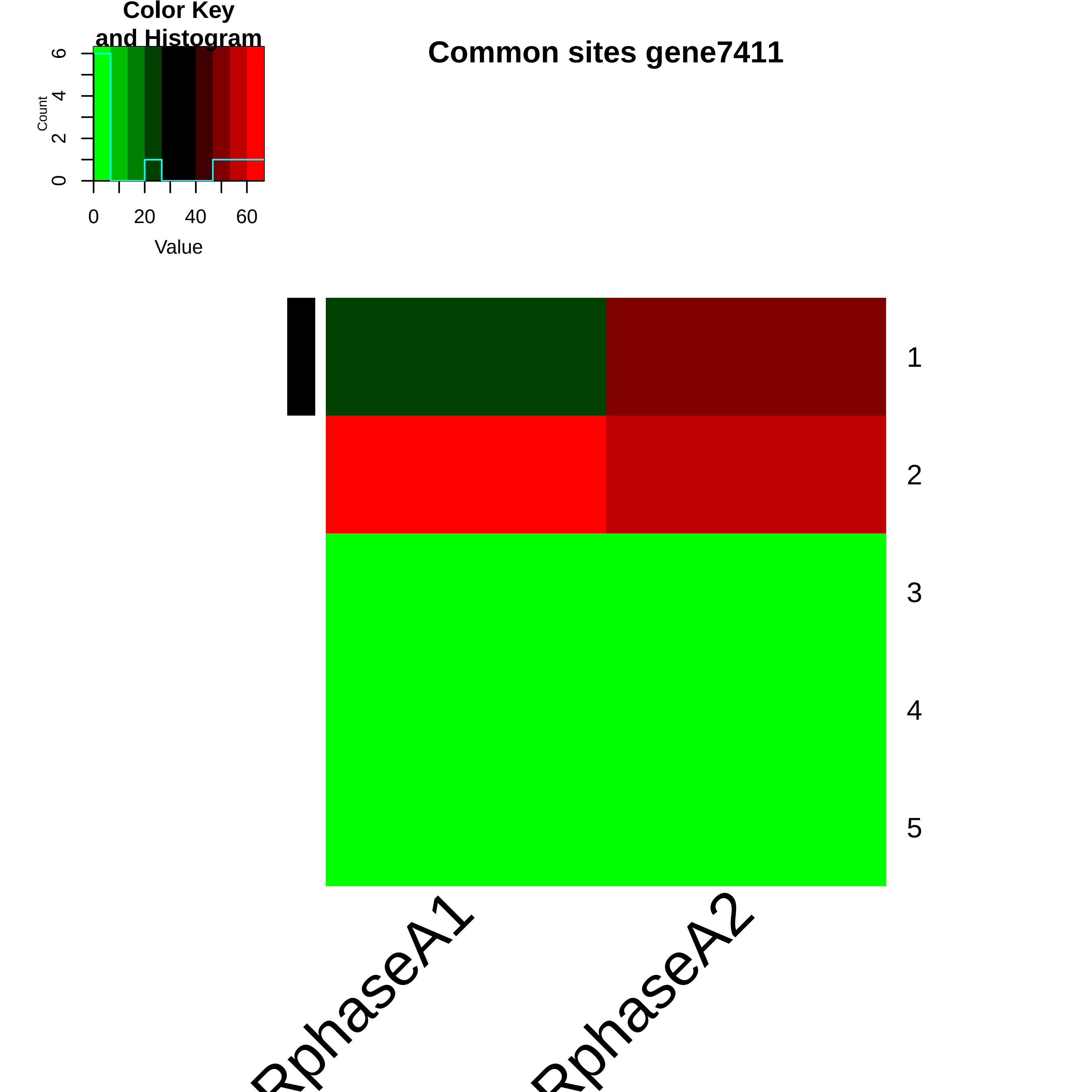
Color Key
and Histogram
Count
0
2
4
6
0
20
40
60
Value
Common sites gene7411
1
2
3
4
5
RphaseA1
RphaseA2

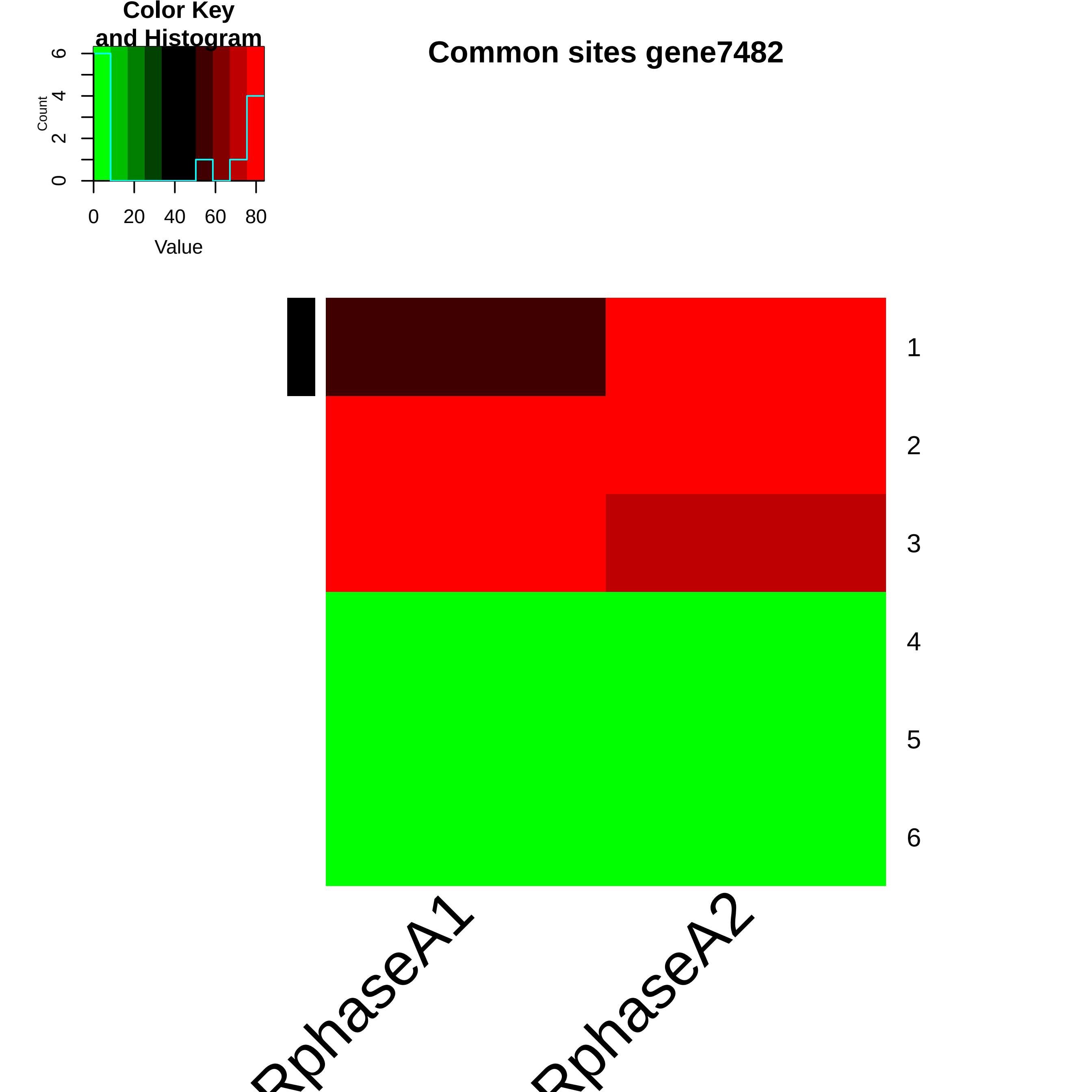

Color Key
and Histogram
Count
0
2
4
6
0
20
40
60
80
Value
Common sites gene7482
1
2
3
4
5
6
RphaseA1
RphaseA2

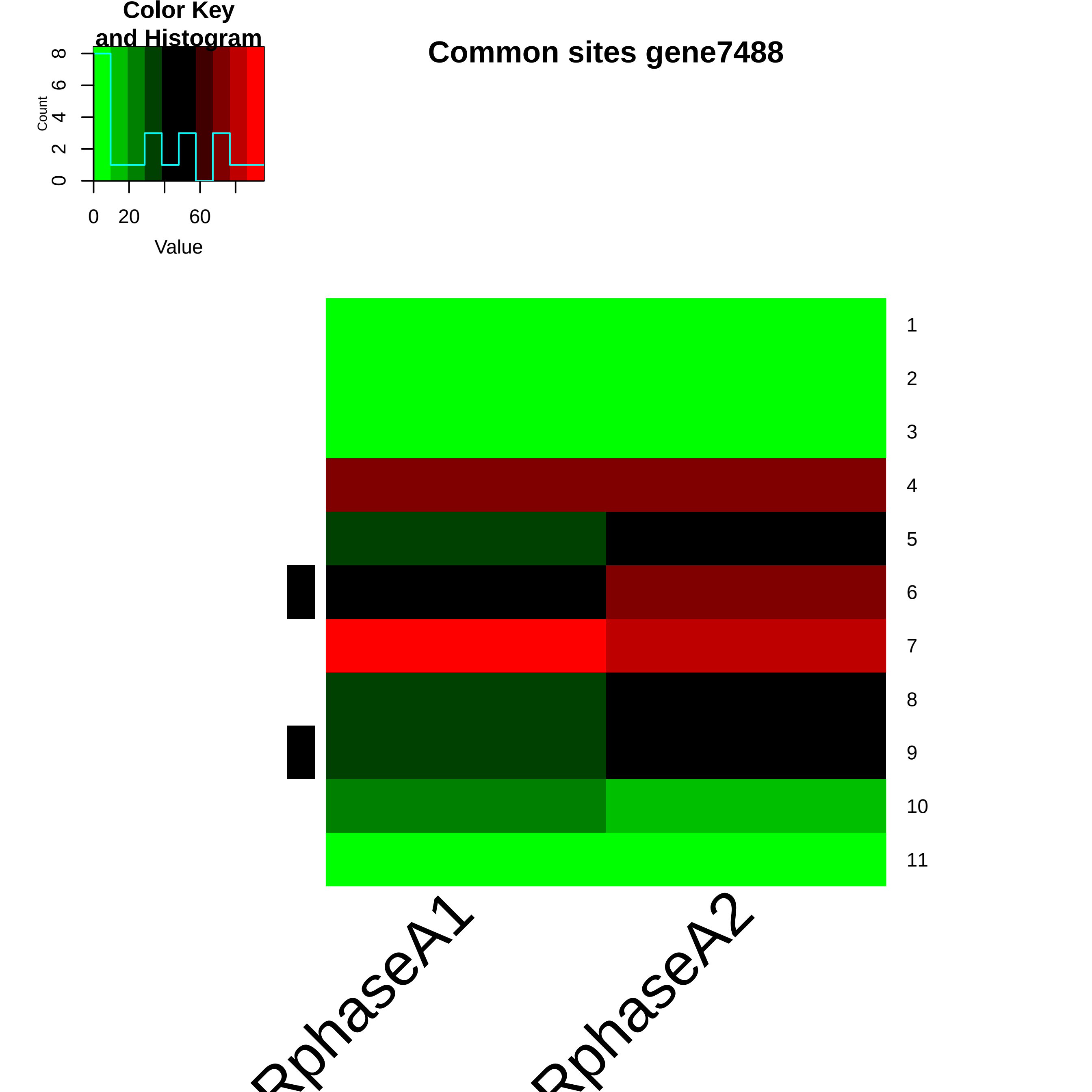

Color Key
and Histogram
Count
0
2
4
6
8
0
20
60
Value
Common sites gene7488
1
2
3
4
5
6
7
8
9
10
11
RphaseA1
RphaseA2

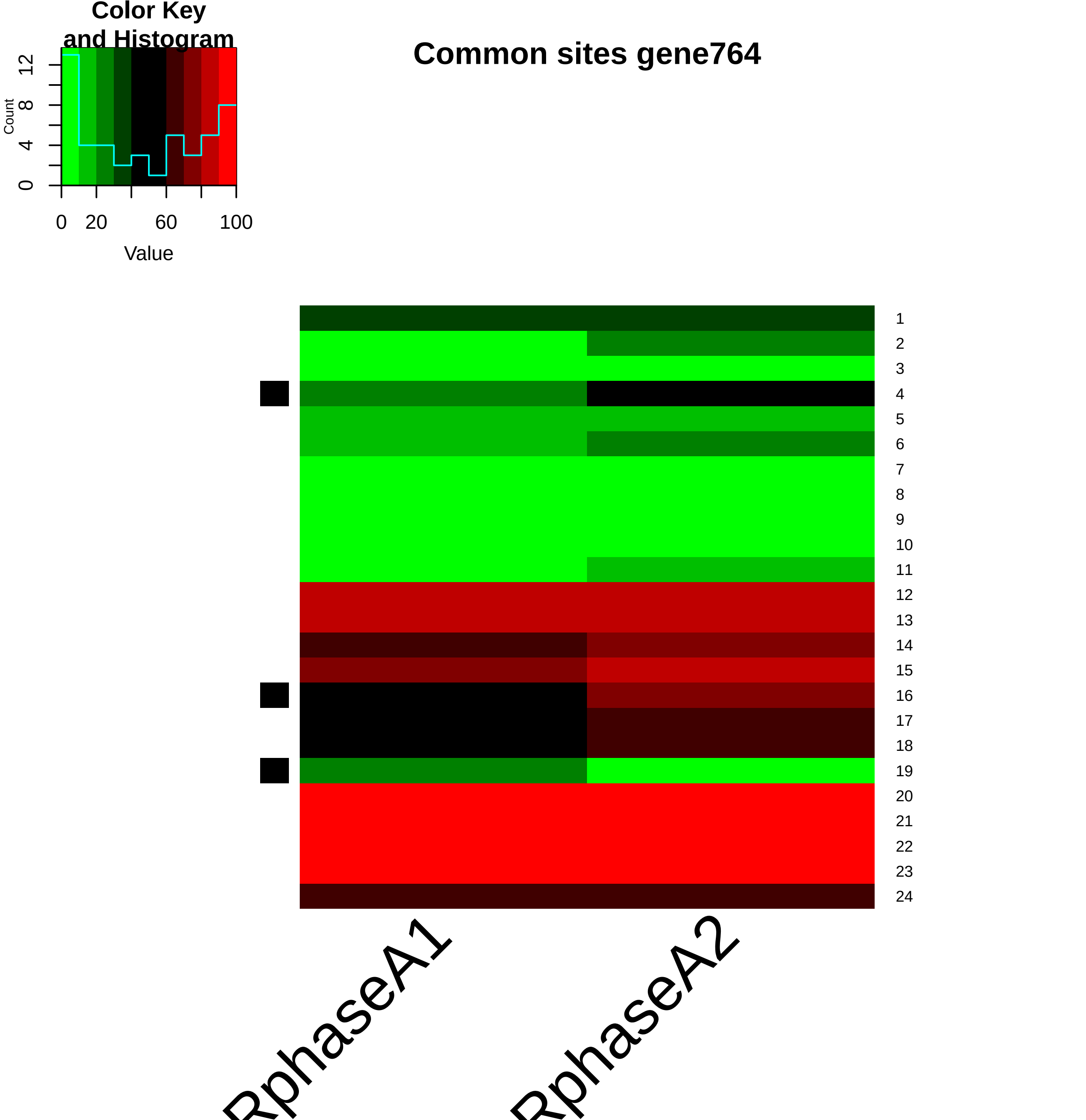
Color Key
and Histogram
Count
0
4
8
12
0
20
60
100
Value
Common sites gene764
RphaseA1
RphaseA2

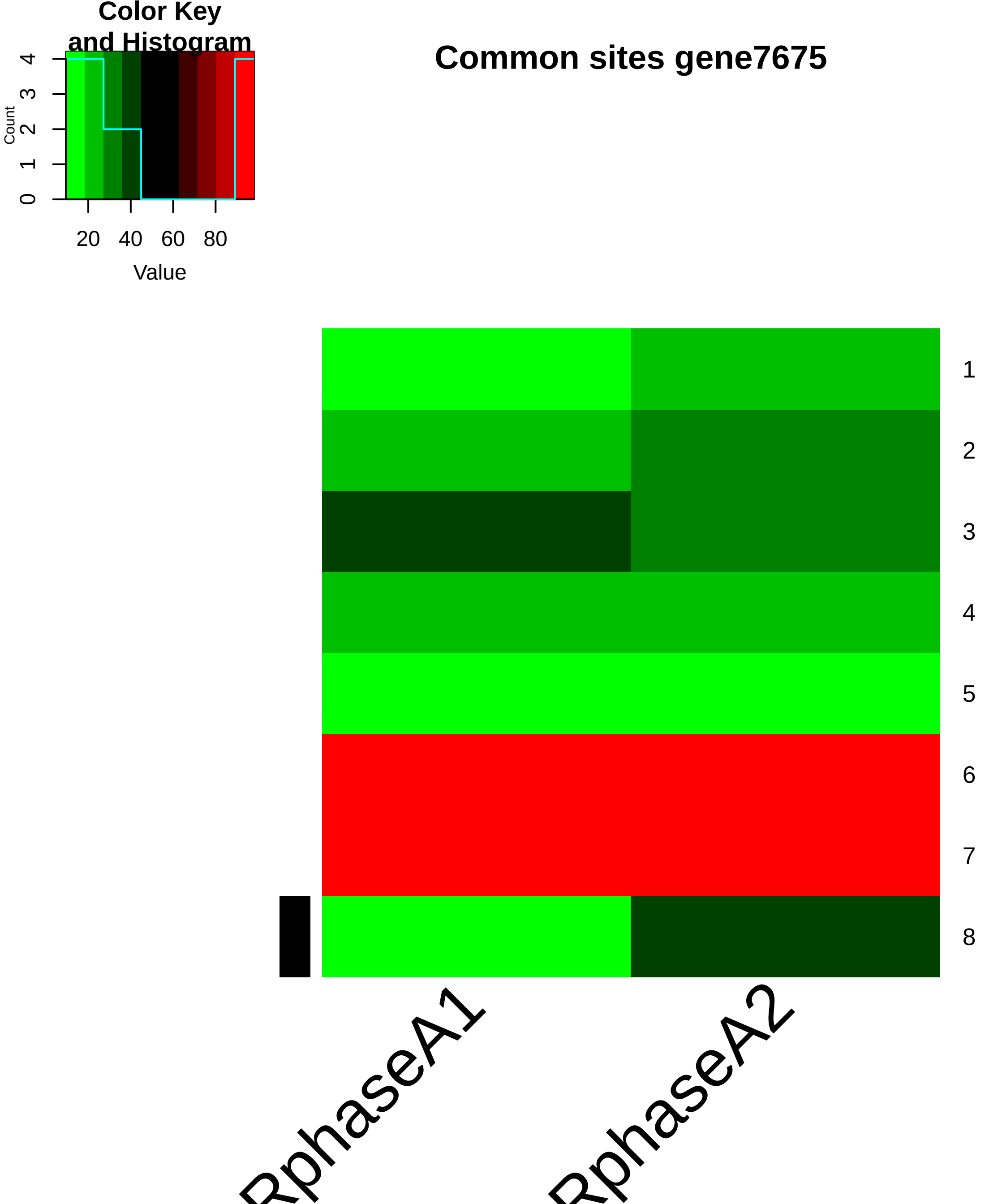
Color Key
and Histogram
Count
0
1
2
3
4
20
40
60
80
Value
Common sites gene7675
1
2
3
4
5
6
7
8
RphaseA1
RphaseA2

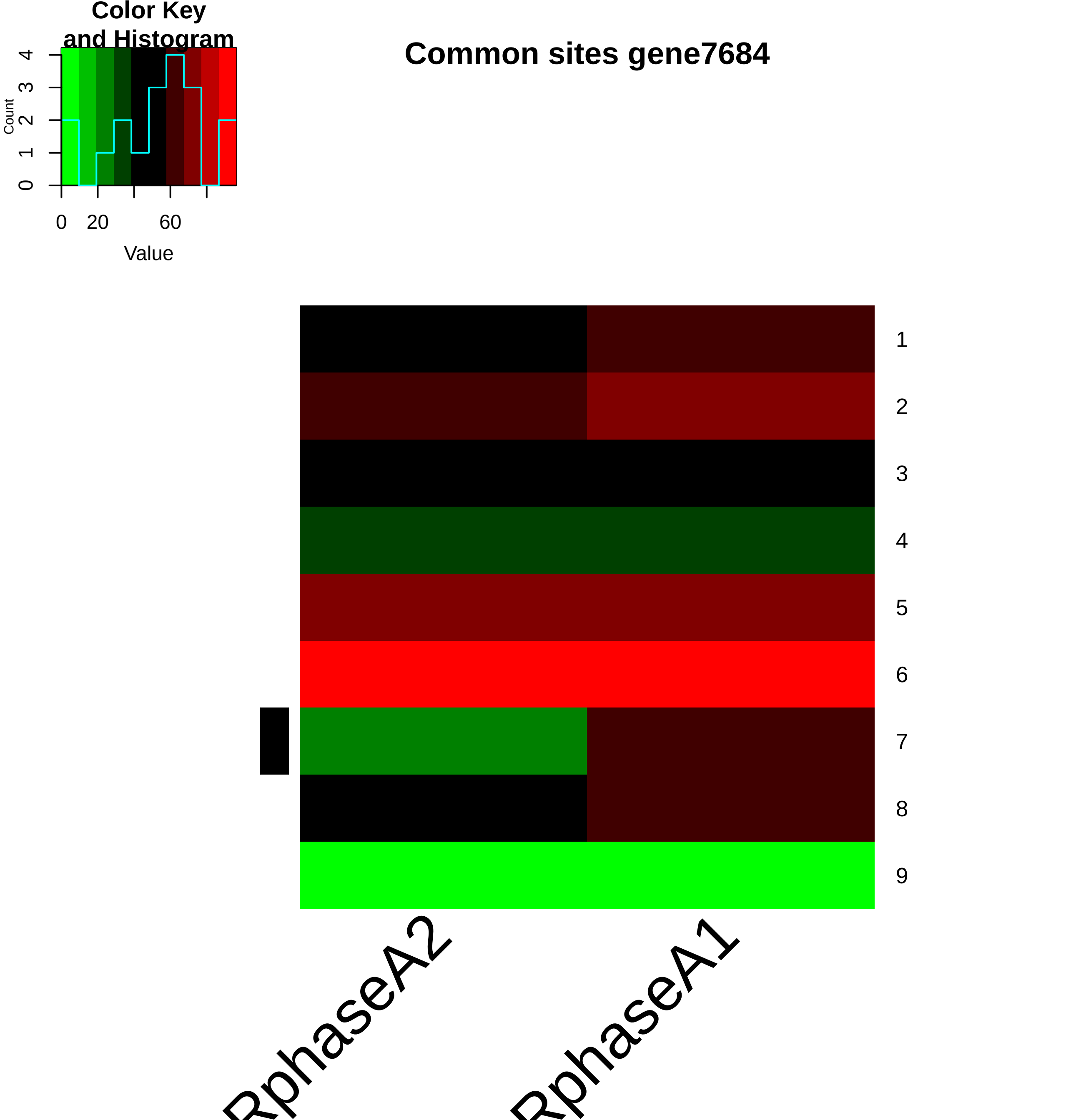

Color Key
and Histogram
Count
0
1
2
3
4
0
20
60
Value
Common sites gene7684
1
2
3
4
5
6
7
8
9
RphaseA2
RphaseA1

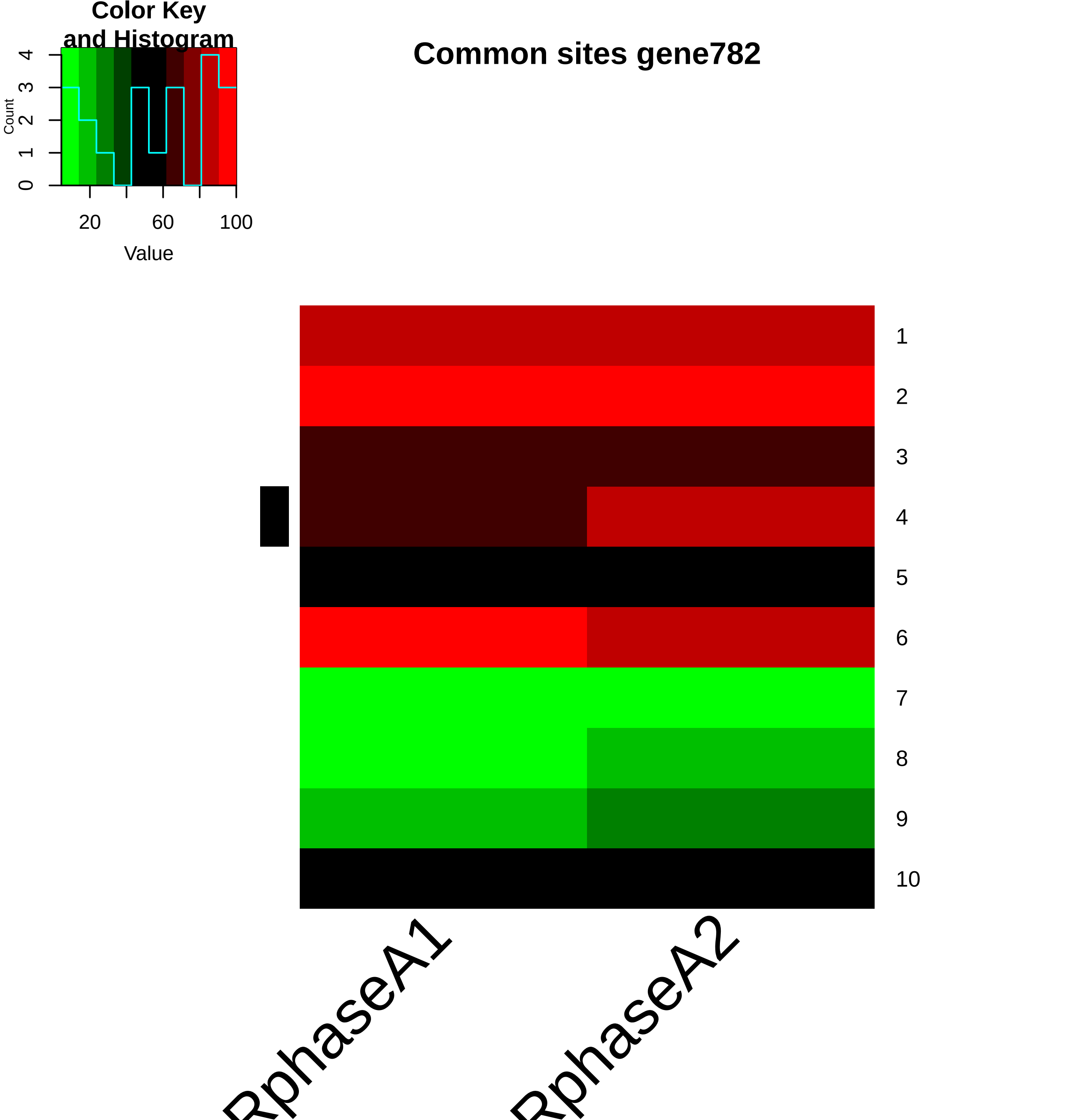
Color Key
and Histogram
Count
0
1
2
3
4
20
60
100
Value
Common sites gene782
1
2
3
4
5
6
7
8
9
10
RphaseA1
RphaseA2

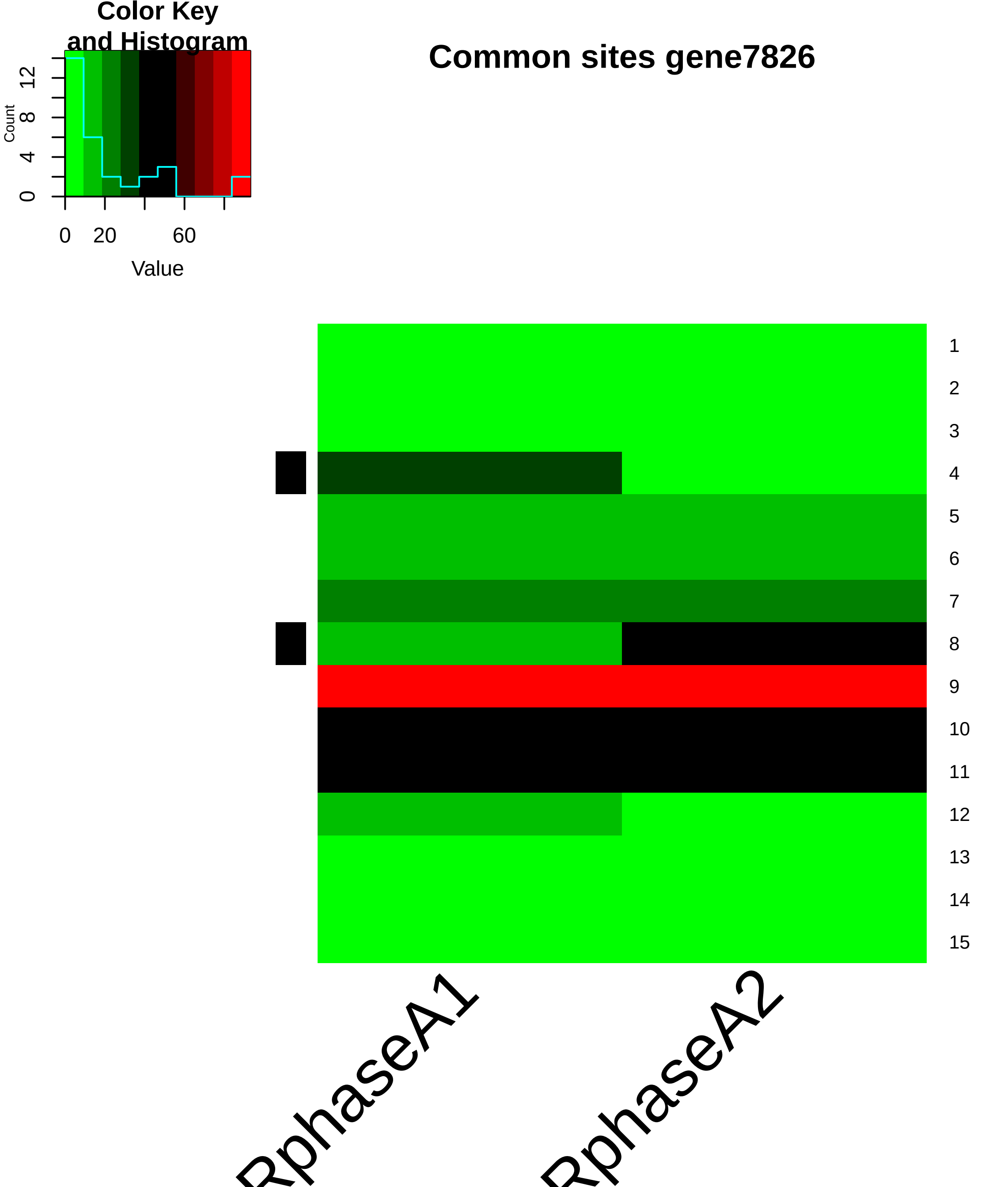
Color Key
and Histogram
Count
0
4
8
12
0
20
60
Value
Common sites gene7826
RphaseA1
RphaseA2

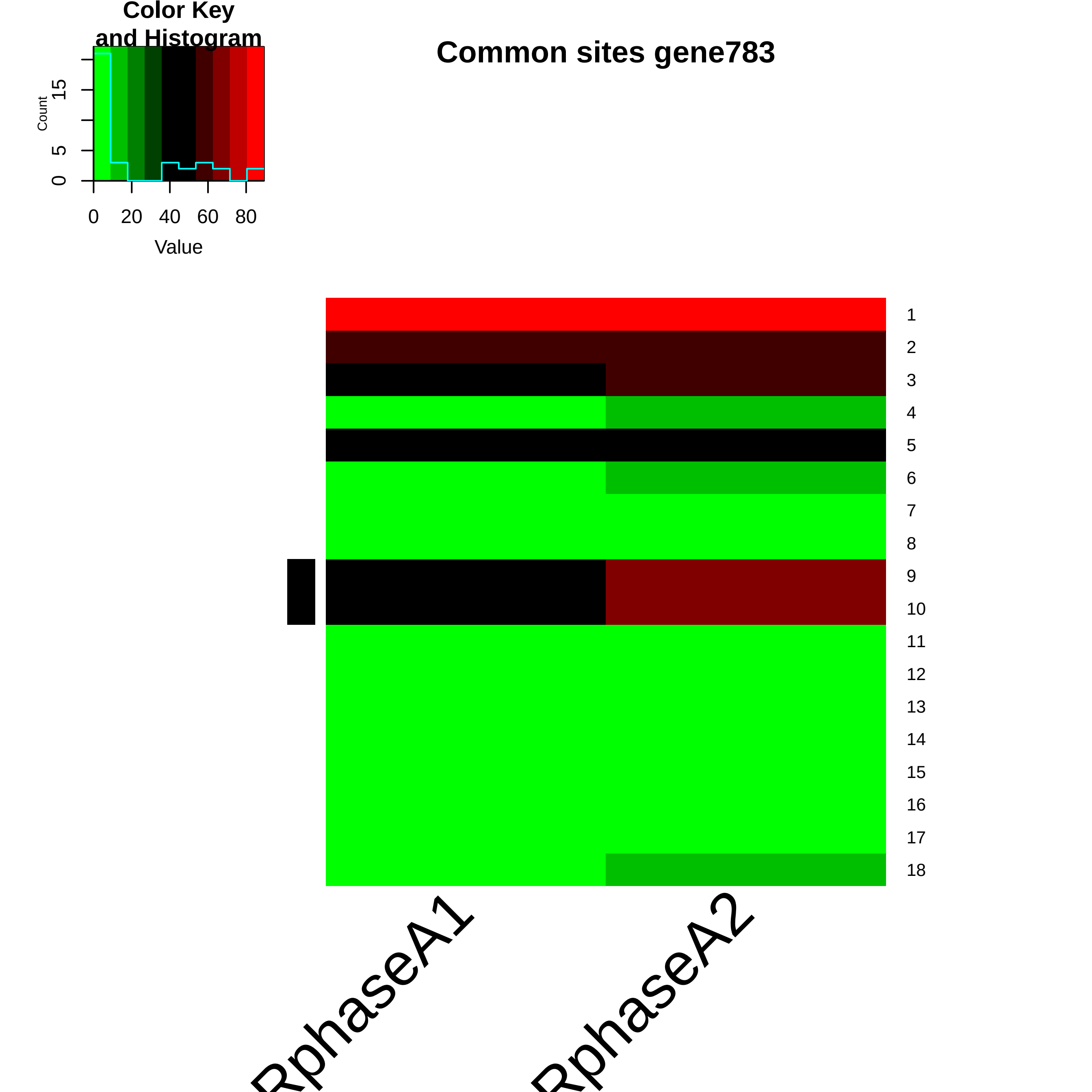
Color Key
and Histogram
Count
0
5
15
0
20
40
60
80
Value
Common sites gene783
RphaseA1
RphaseA2

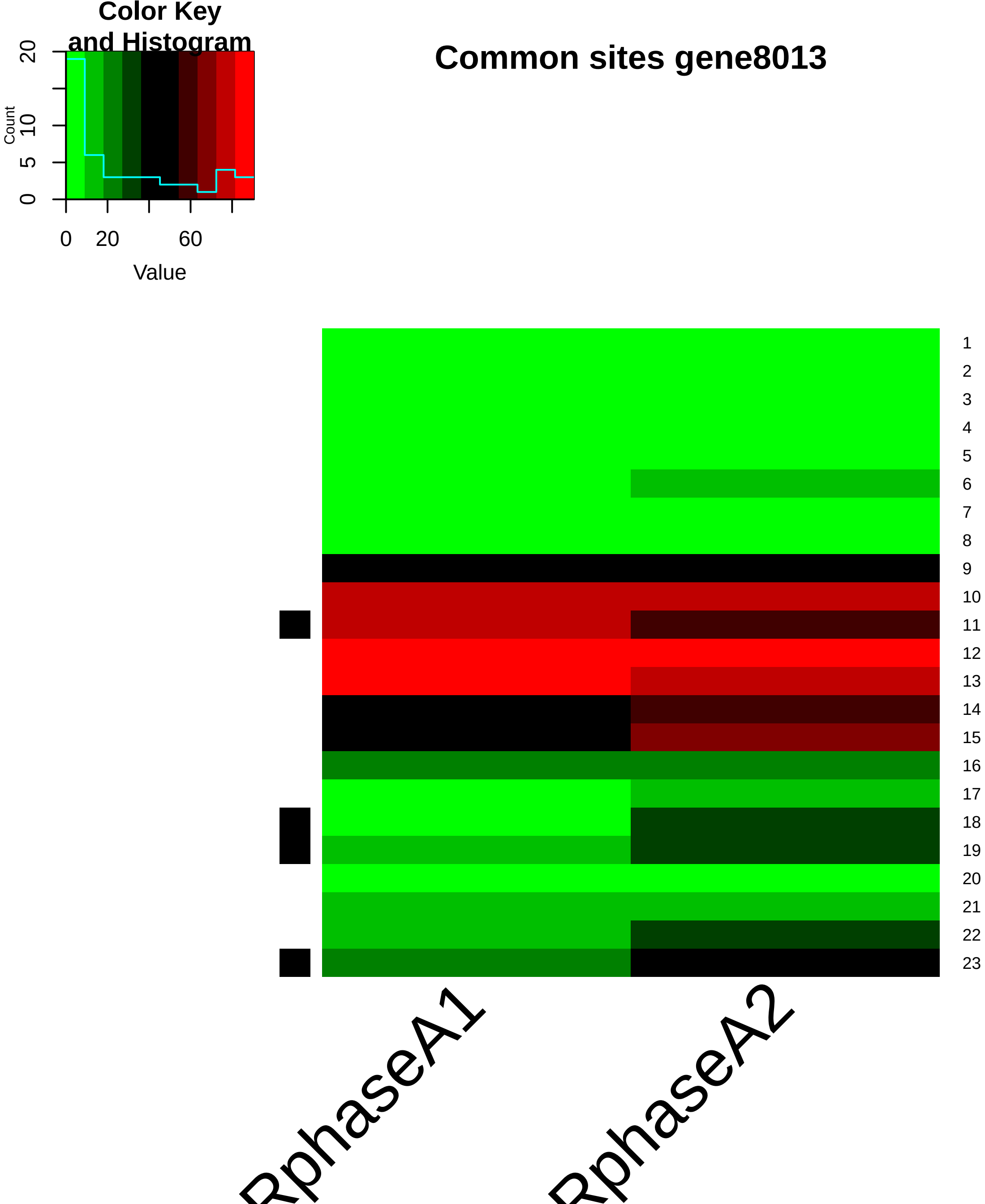
Color Key
and Histogram
Count
0
5
10
20
0
20
60
Value
Common sites gene8013
RphaseA1
RphaseA2

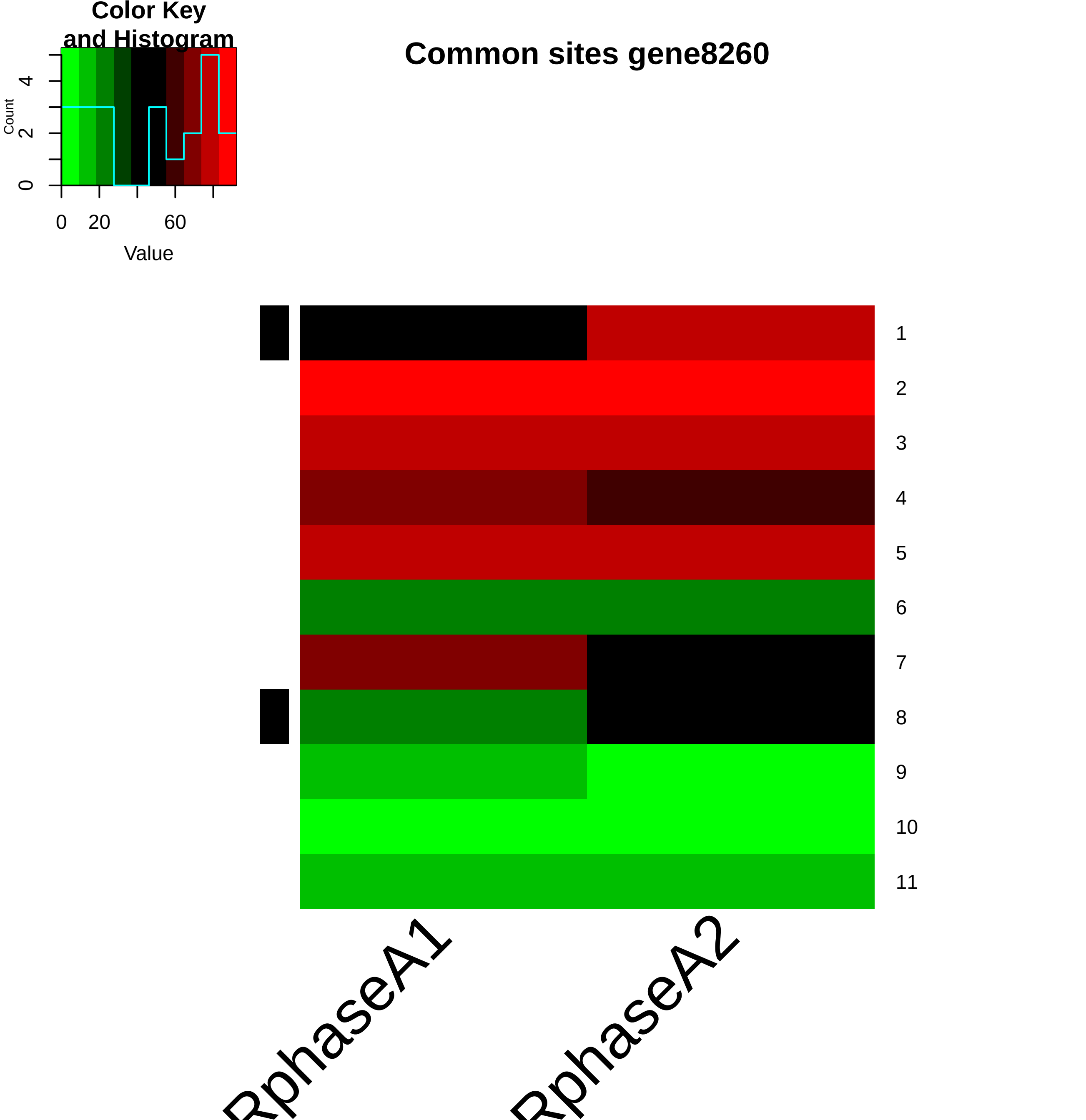
Color Key
and Histogram
Count
0
2
4
0
20
60
Value
Common sites gene8260
1
2
3
4
5
6
7
8
9
10
11
RphaseA1
RphaseA2

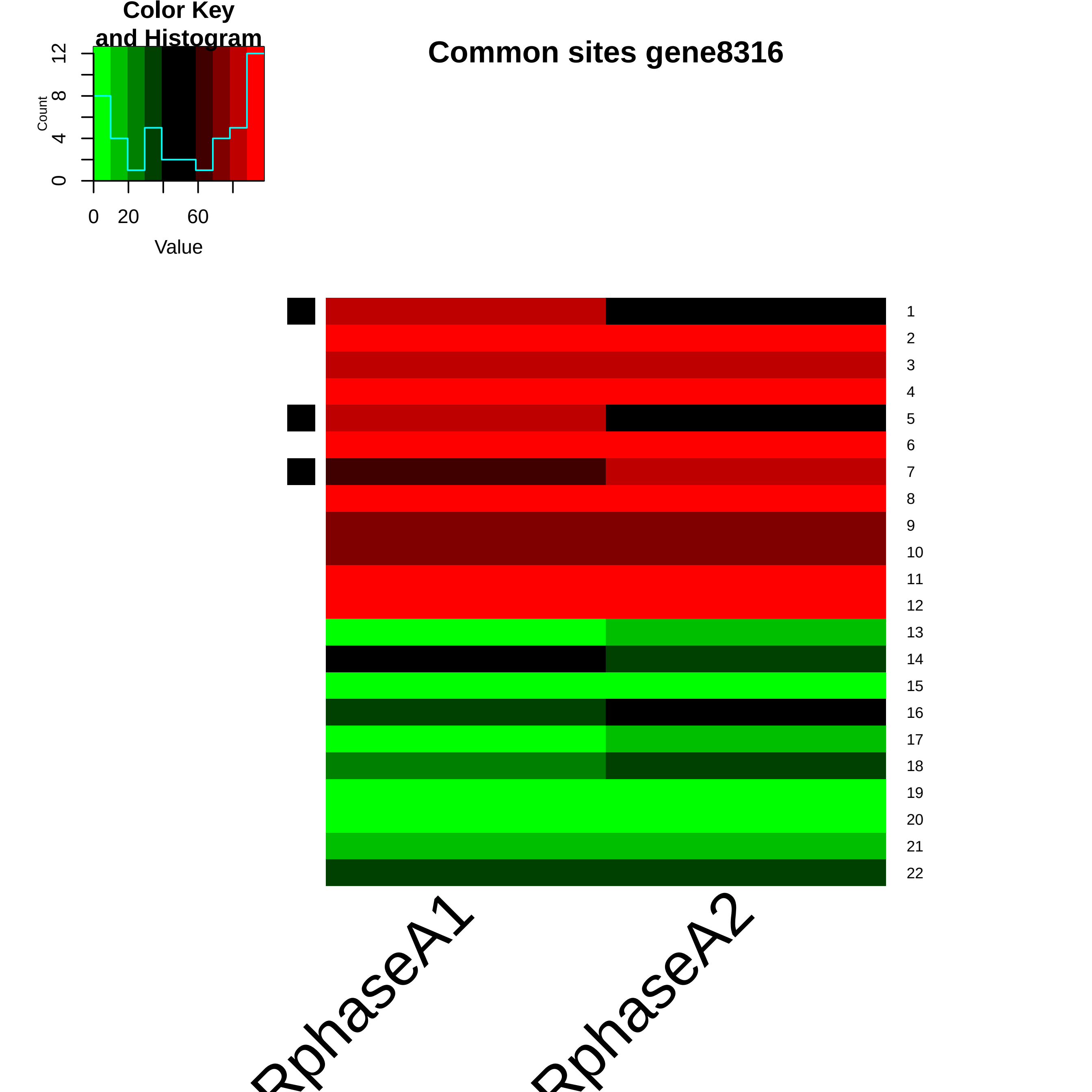

Color Key
and Histogram
Count
0
4
8
12
0
20
60
Value
Common sites gene8316
RphaseA1
RphaseA2

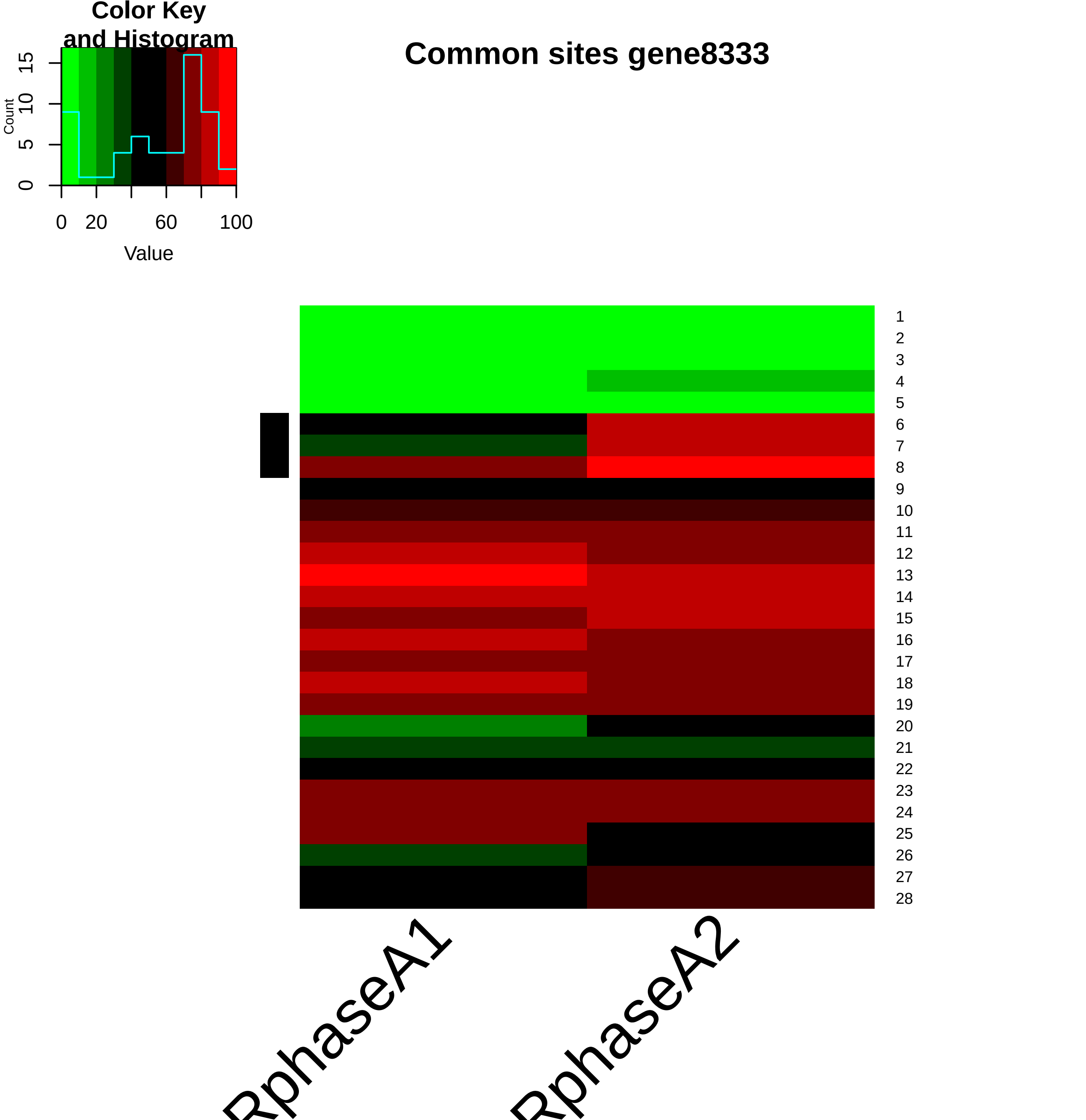
Color Key
and Histogram
Count
0
5
10
15
0
20
60
100
Value
Common sites gene8333
RphaseA1
RphaseA2

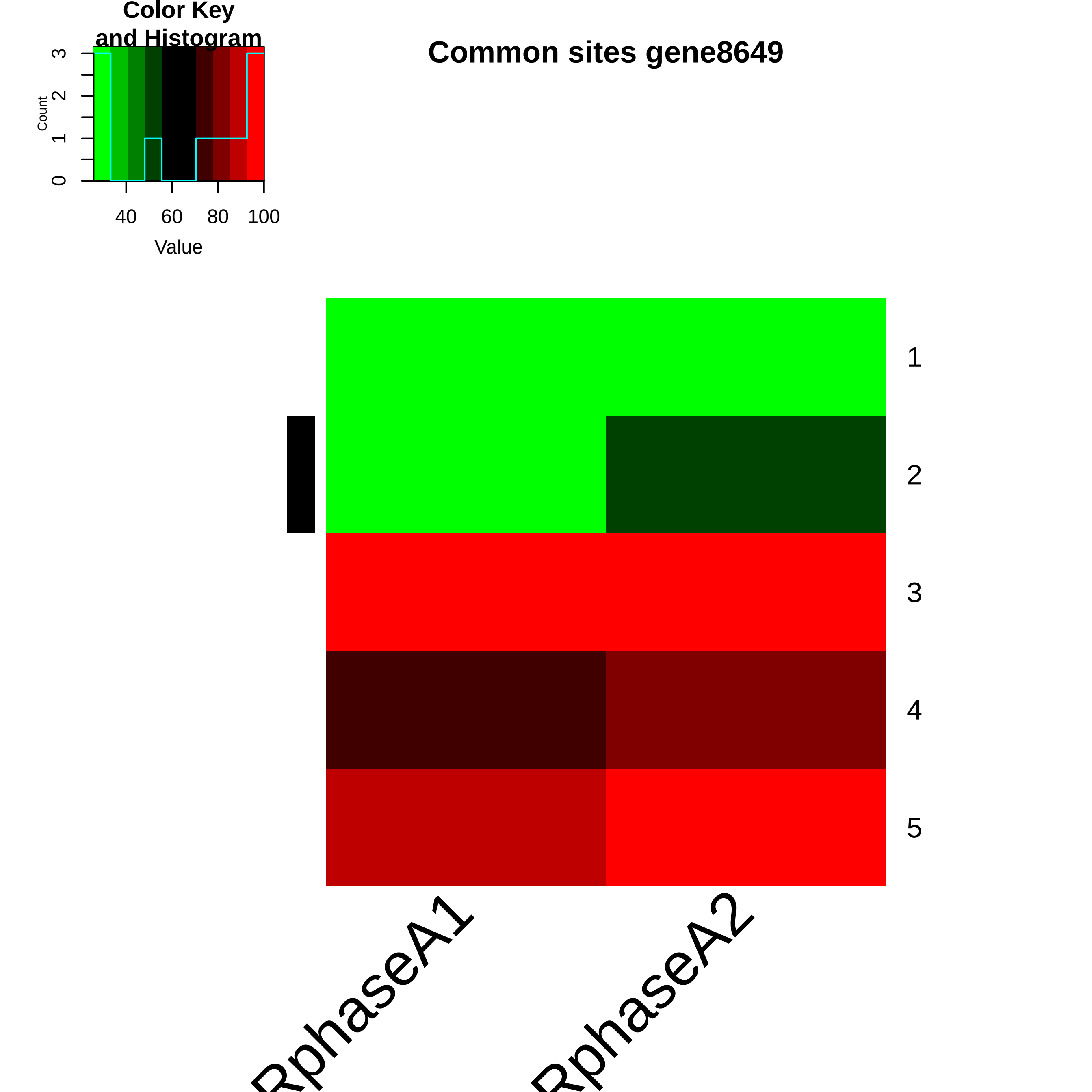

Color Key
and Histogram
Count
0
1
2
3
40
60
80
100
Value
Common sites gene8649
1
2
3
4
5
RphaseA1
RphaseA2

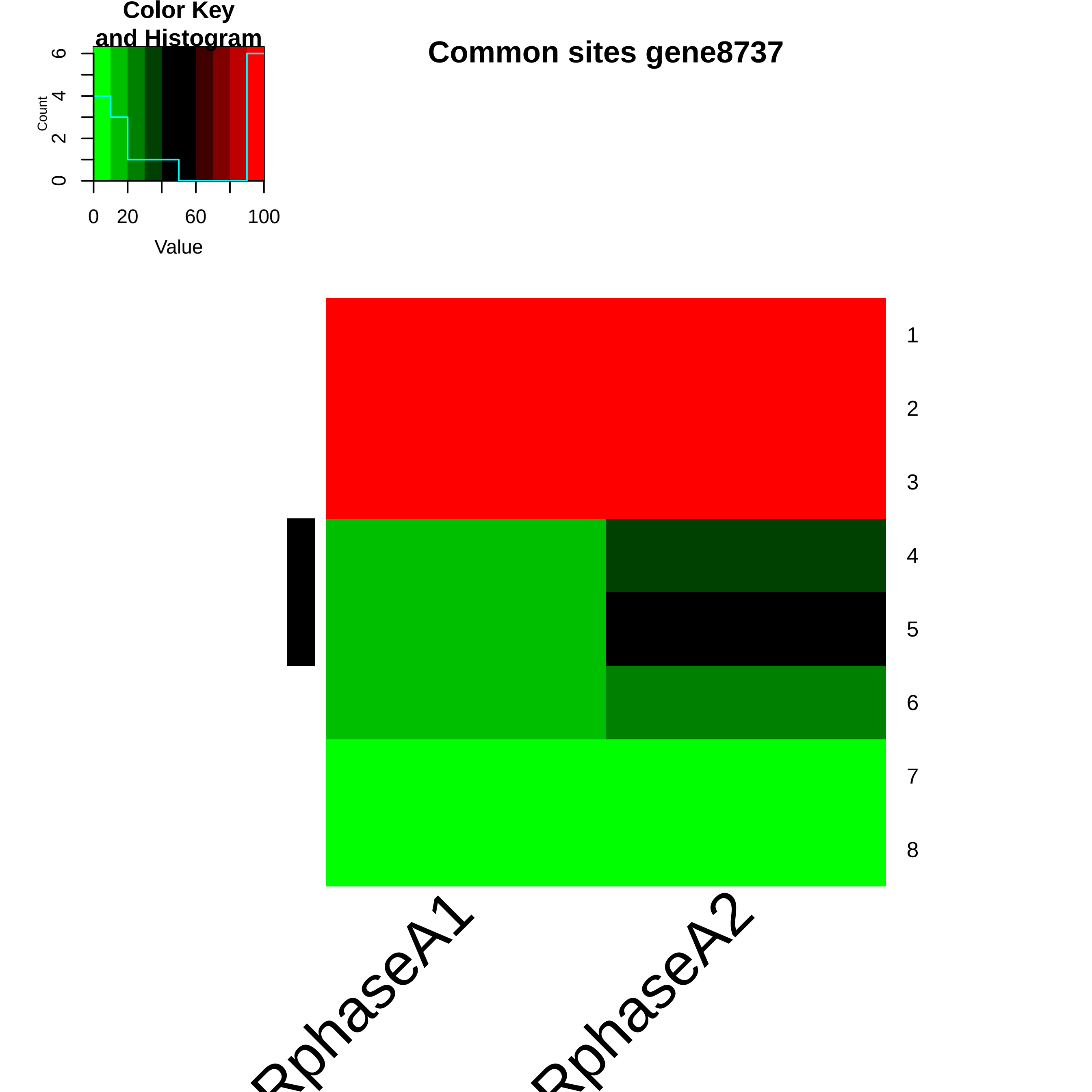
Color Key
and Histogram
Count
0
2
4
6
0
20
60
100
Value
Common sites gene8737
1
2
3
4
5
6
7
8
RphaseA1
RphaseA2

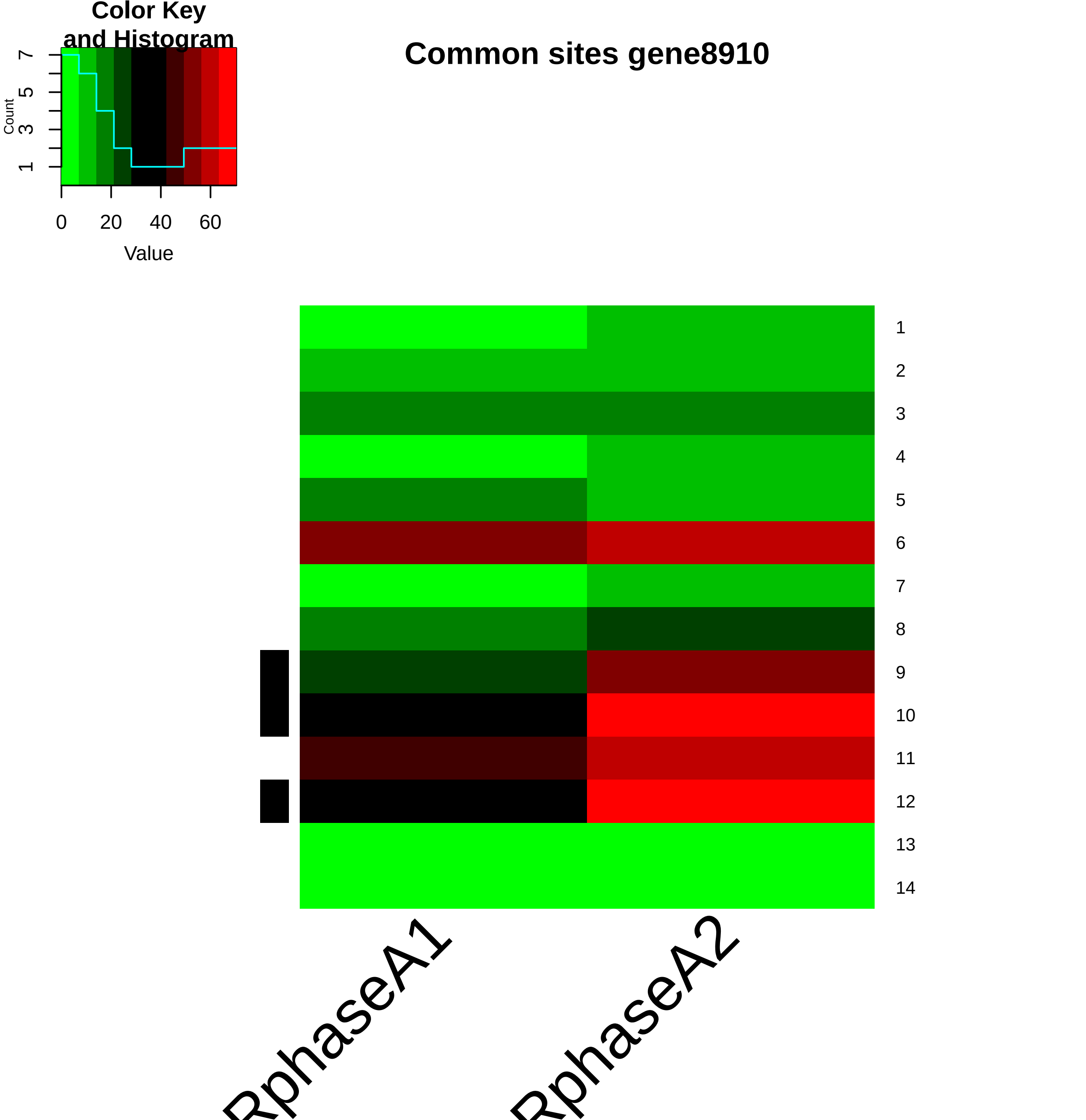

Color Key
and Histogram
Count
Value
Common sites gene8910
1
2
3
4
5
6
7
8
9
10
11
12
13
14
RphaseA1
RphaseA2

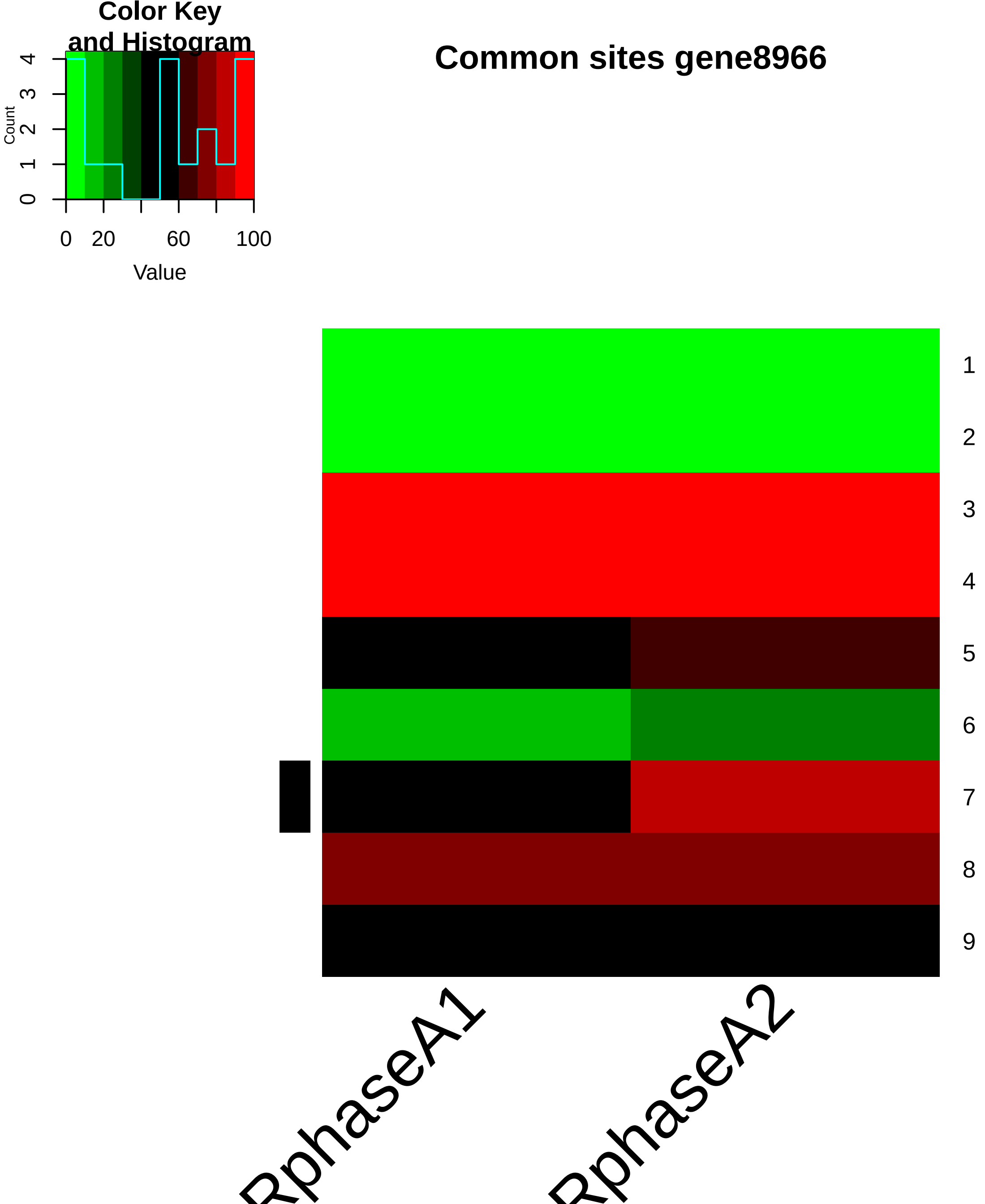
Color Key
and Histogram
Count
Value
0 20 60 100
0 1 2 3 4
Common sites gene8966
1
2
3
4
5
6
7
8
9
RphaseA1
RphaseA2

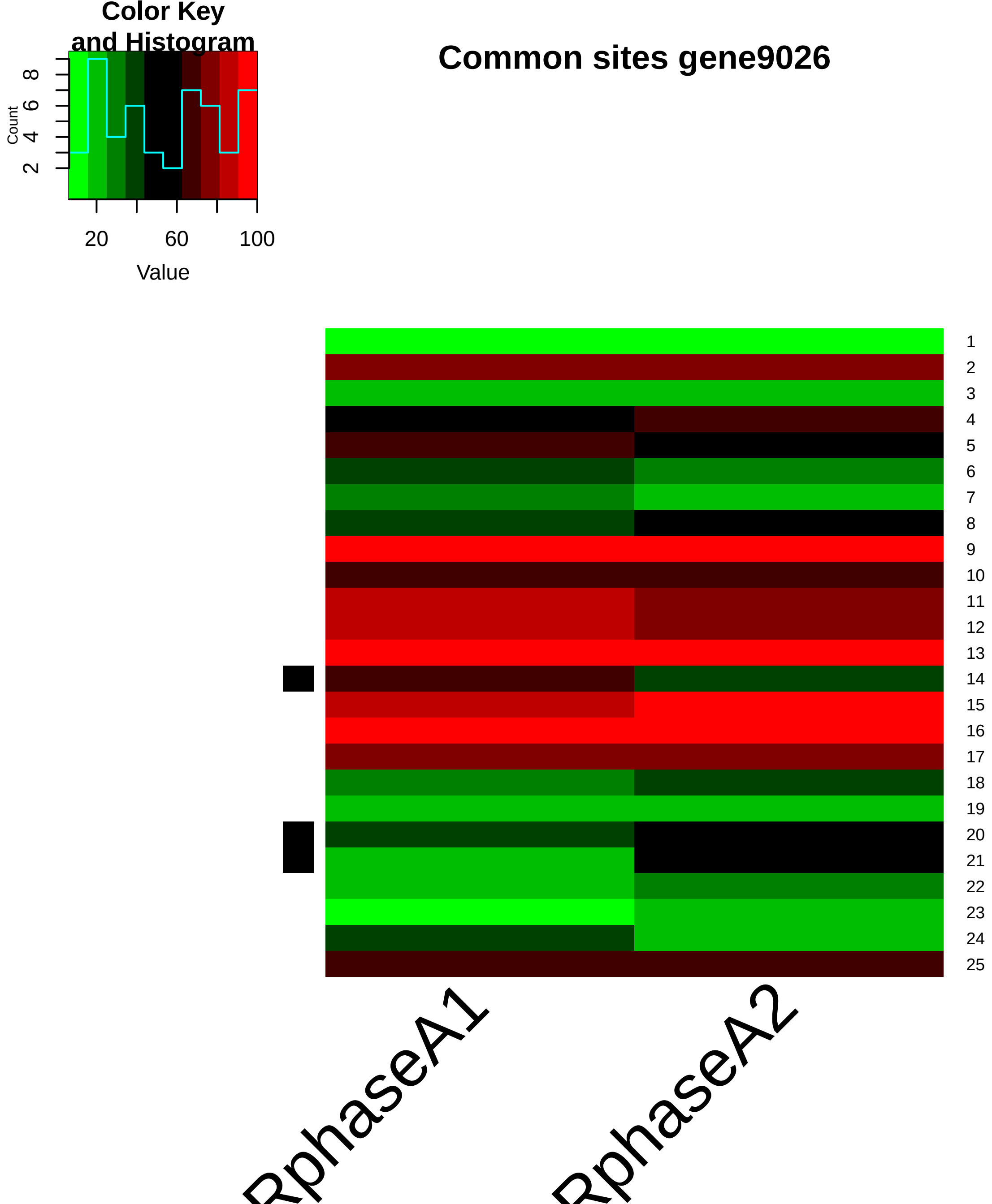

Color Key
and Histogram
Count
2
4
6
8
20
60
100
Value
Common sites gene9026
RphaseA1
RphaseA2

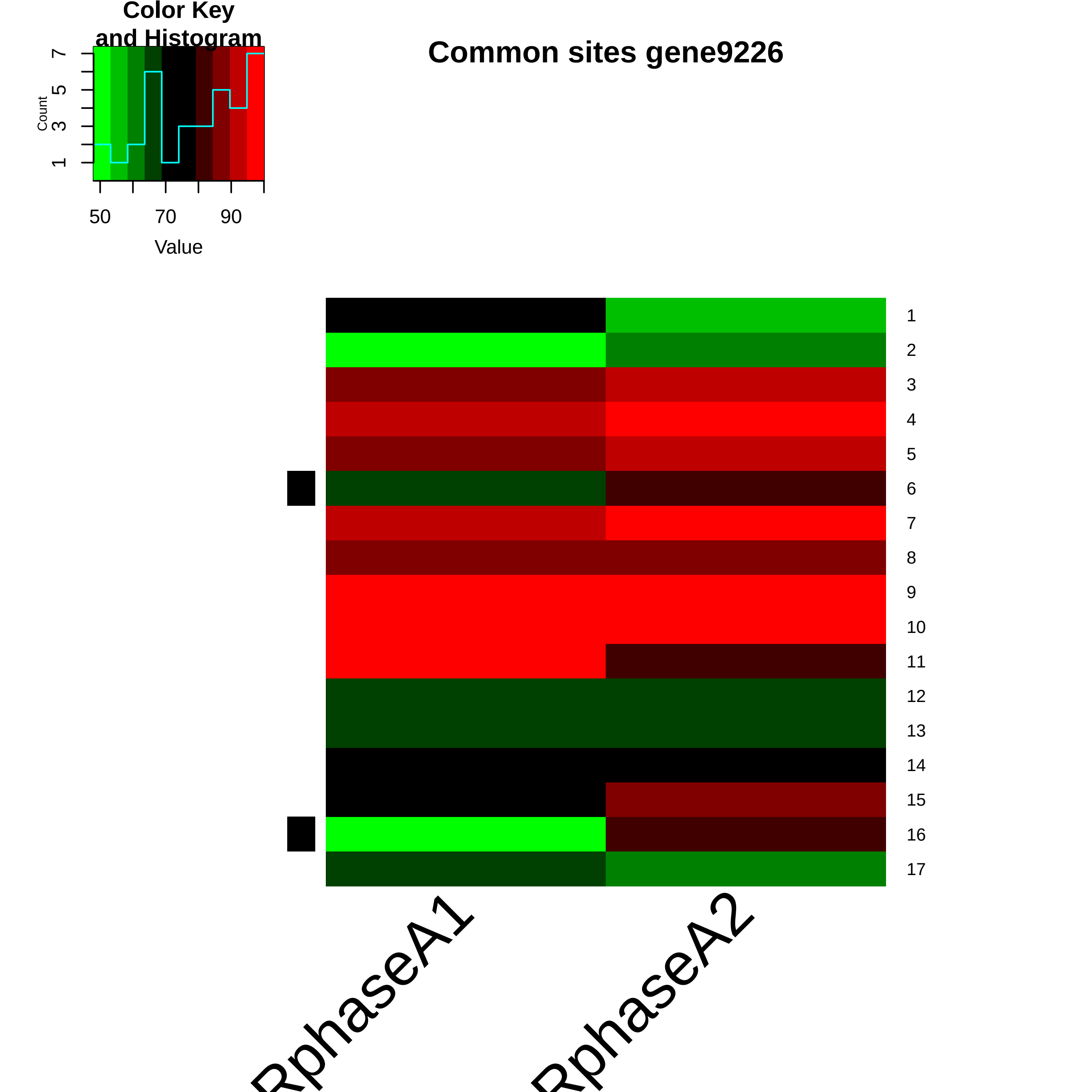

Color Key
and Histogram
Count
1
3
5
7
50
70
90
Value
Common sites gene9226
RphaseA1
RphaseA2

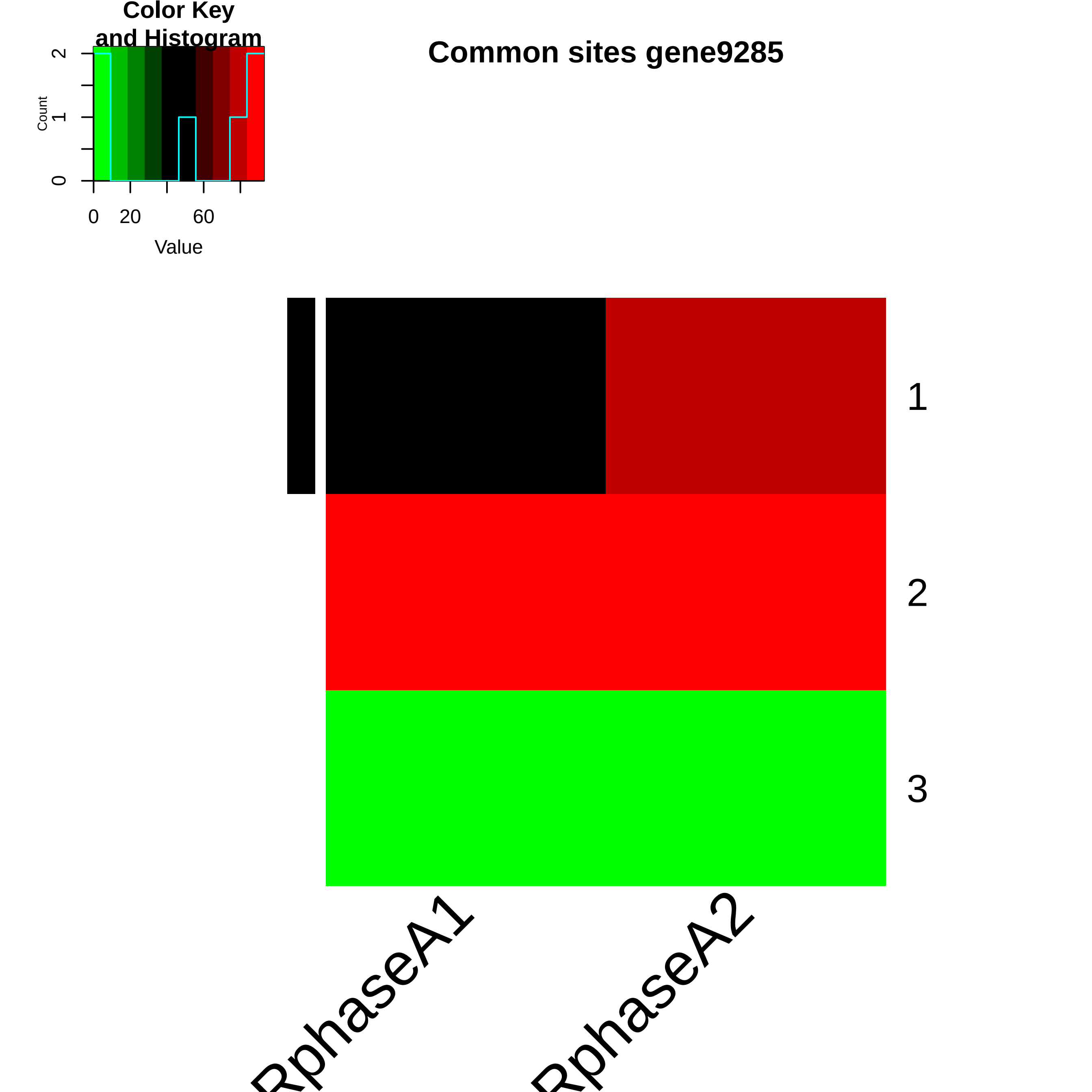

Color Key
and Histogram
Count
0
1
2
0
20
60
Value
Common sites gene9285
1
2
3
RphaseA1
RphaseA2

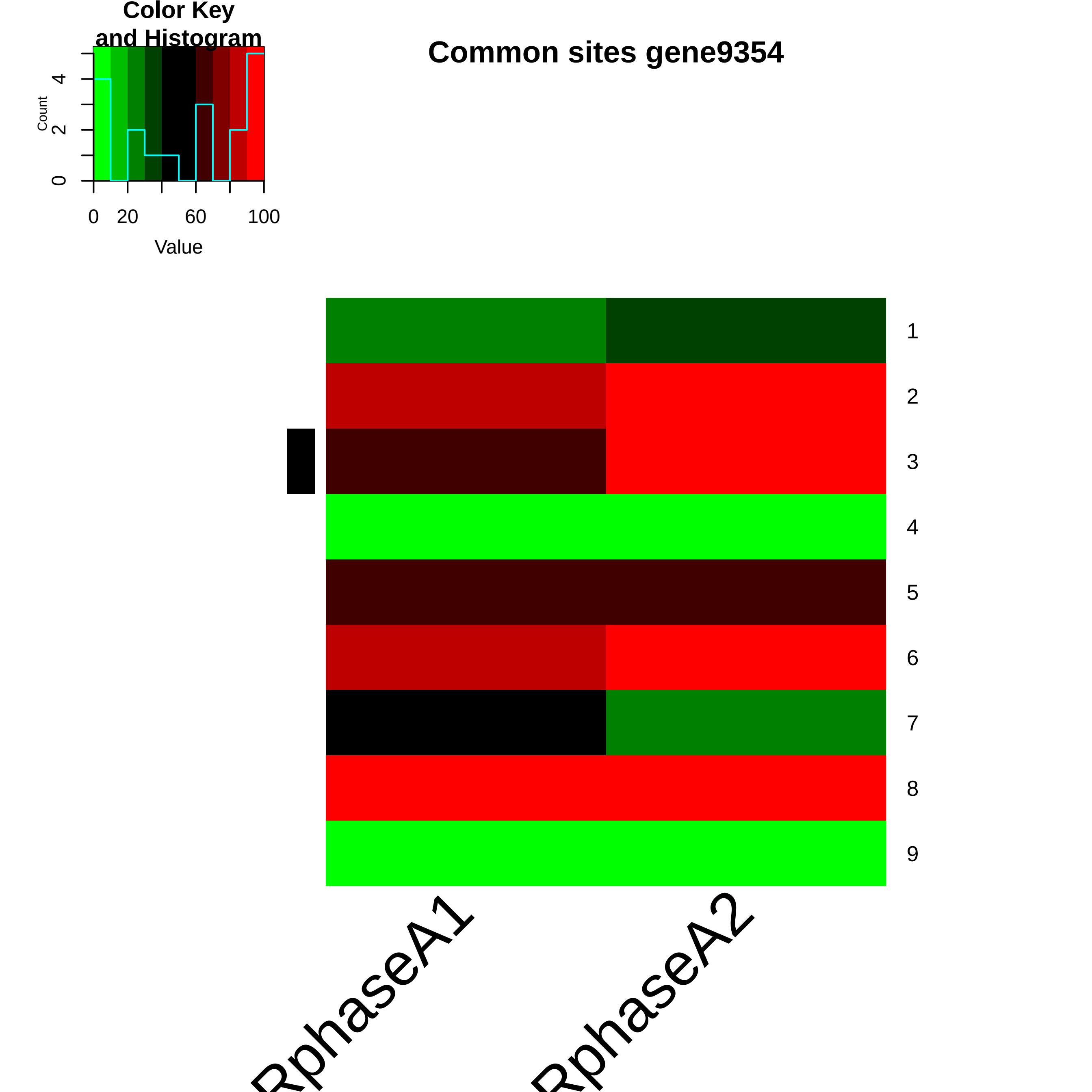
Color Key
and Histogram
Count
0
2
4
0
20
60
100
Value
Common sites gene9354
1
2
3
4
5
6
7
8
9
RphaseA1
RphaseA2

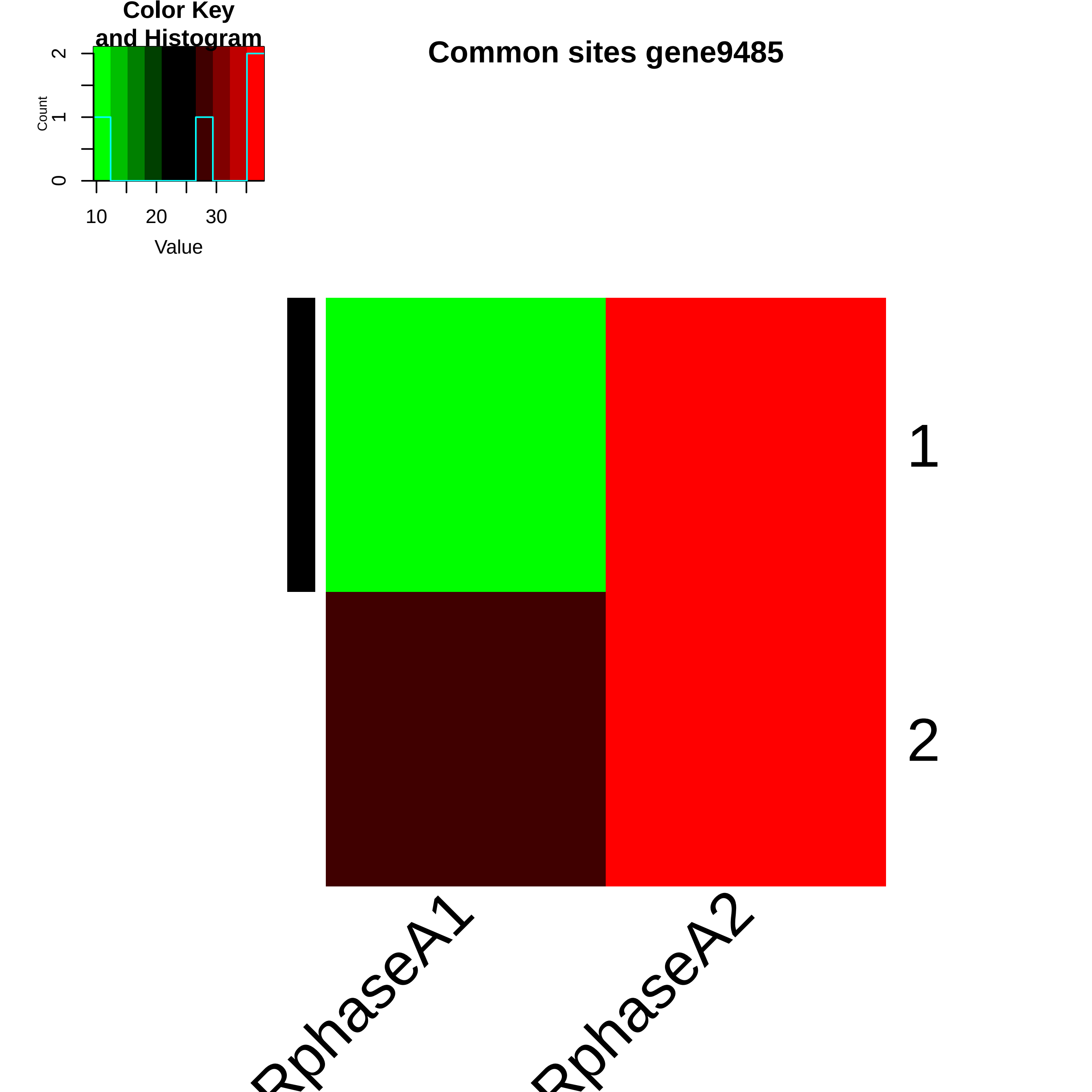

Color Key
and Histogram
Common sites gene9485
Count
0
1
2
10
20
30
Value
1
2
RphaseA1
RphaseA2

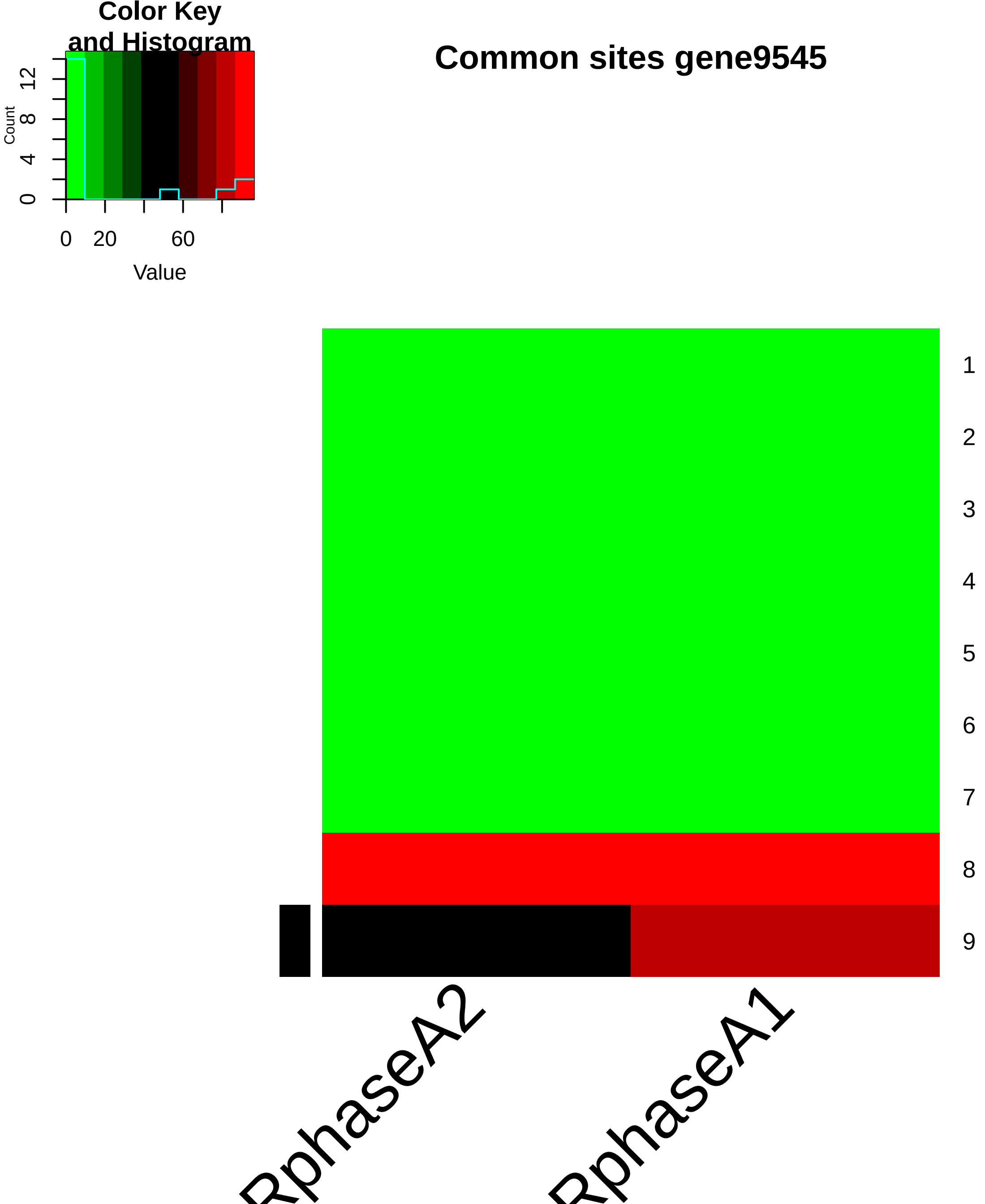

Color Key
and Histogram
Count
0
4
8
12
0
20
60
Value
Common sites gene9545
1
2
3
4
5
6
7
8
9
RphaseA2
RphaseA1

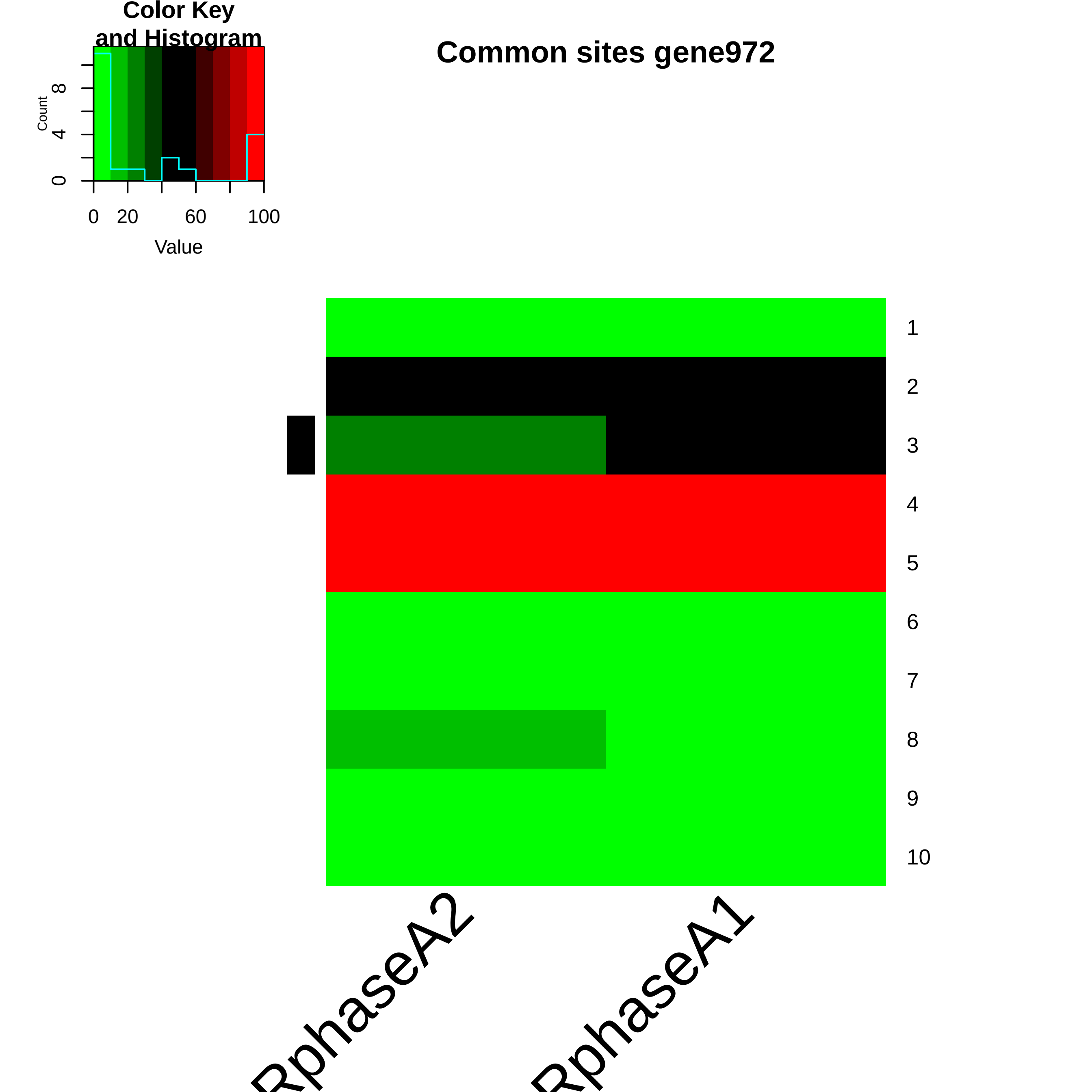
Color Key
and Histogram
Count
0
4
8
0
20
60
100
Value
Common sites gene972
1
2
3
4
5
6
7
8
9
10
RphaseA2
RphaseA1

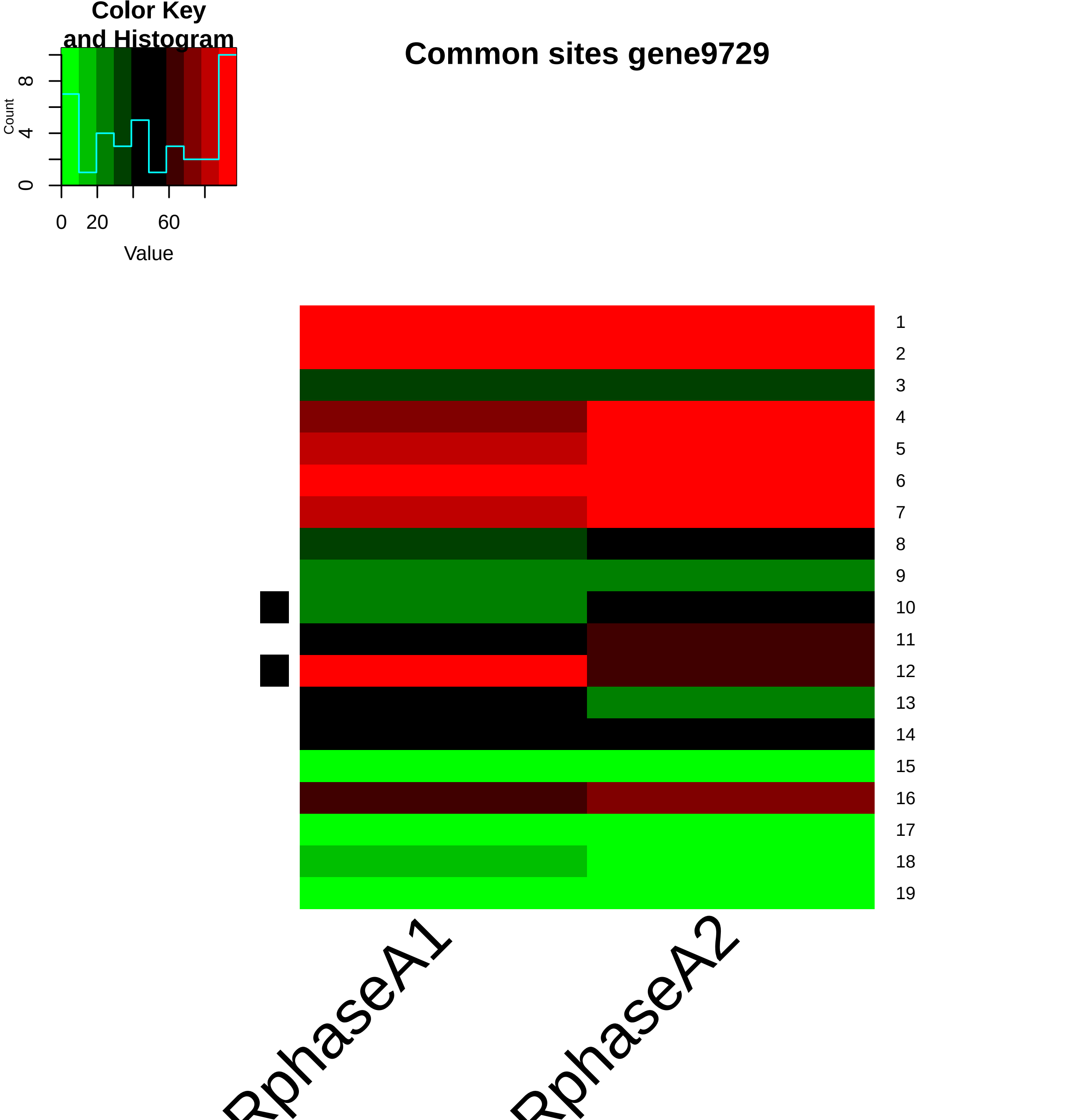
Color Key
and Histogram
Count
0
4
8
0
20
60
Value
Common sites gene9729
RphaseA1
RphaseA2

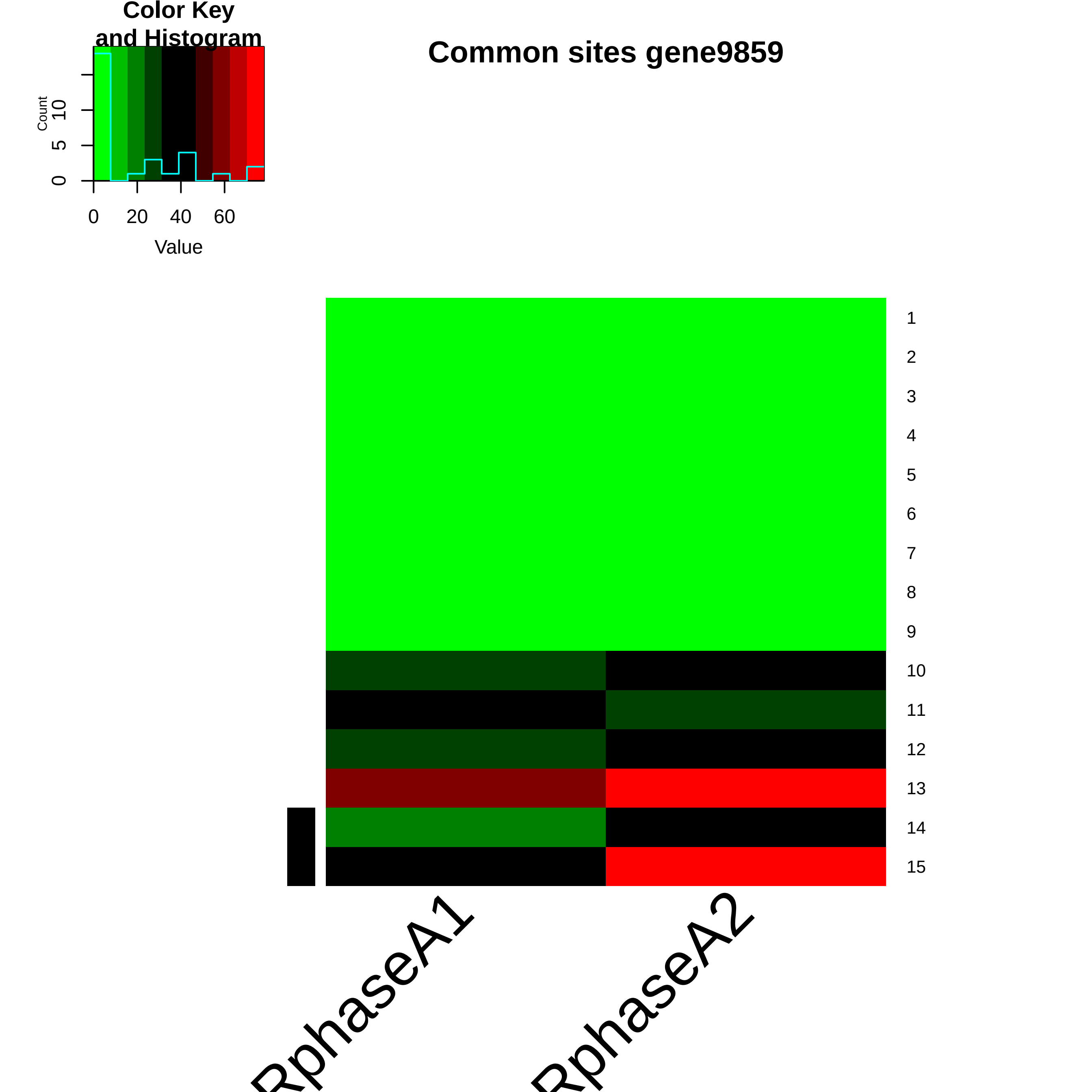
Color Key
and Histogram
Count
0
5
10
0
20
40
60
Value
Common sites gene9859
RphaseA1
RphaseA2

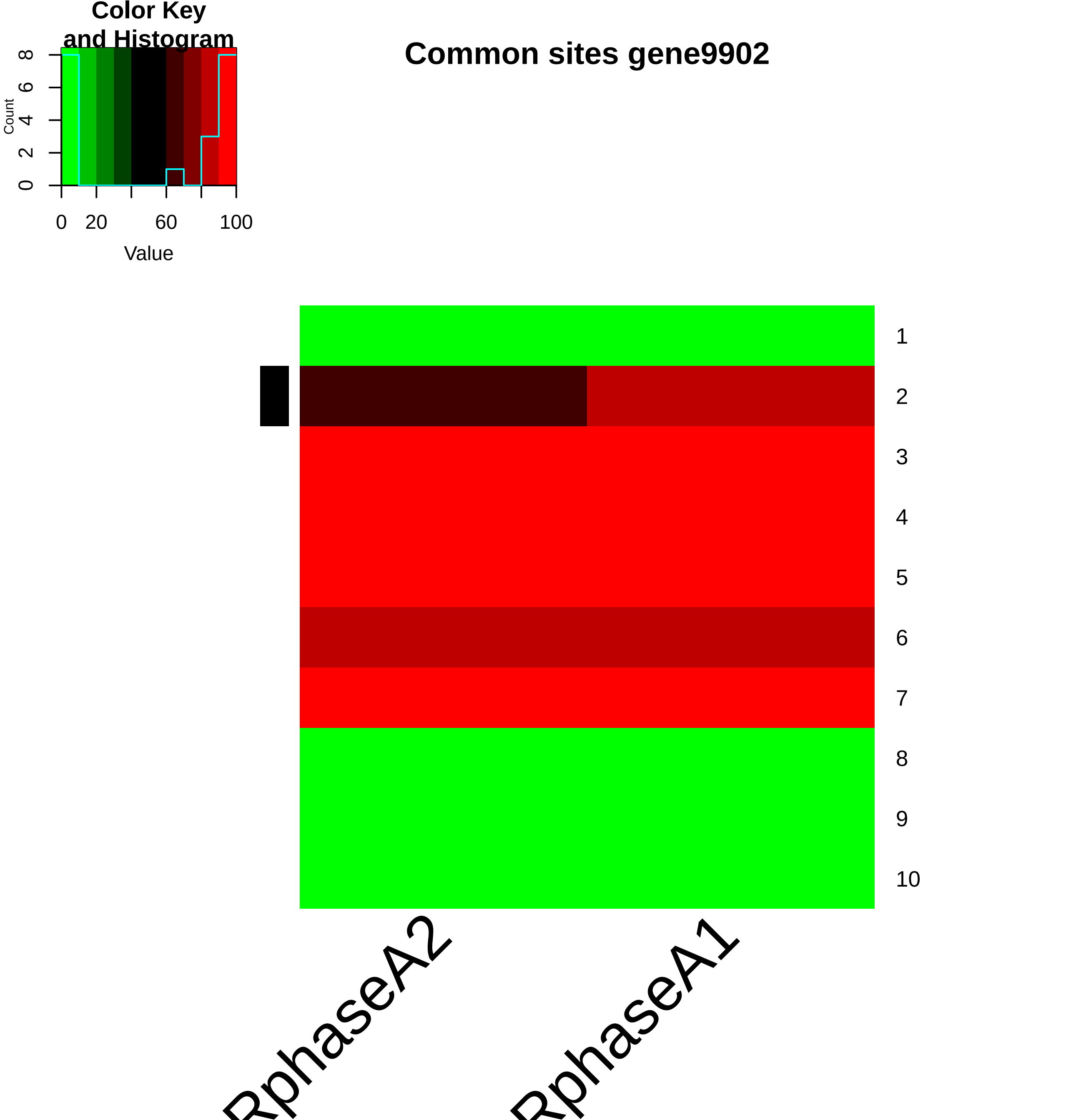
Color Key
and Histogram
Count
0
2
4
6
8
0
20
60
100
Value
Common sites gene9902
1
2
3
4
5
6
7
8
9
10
RphaseA2
RphaseA1

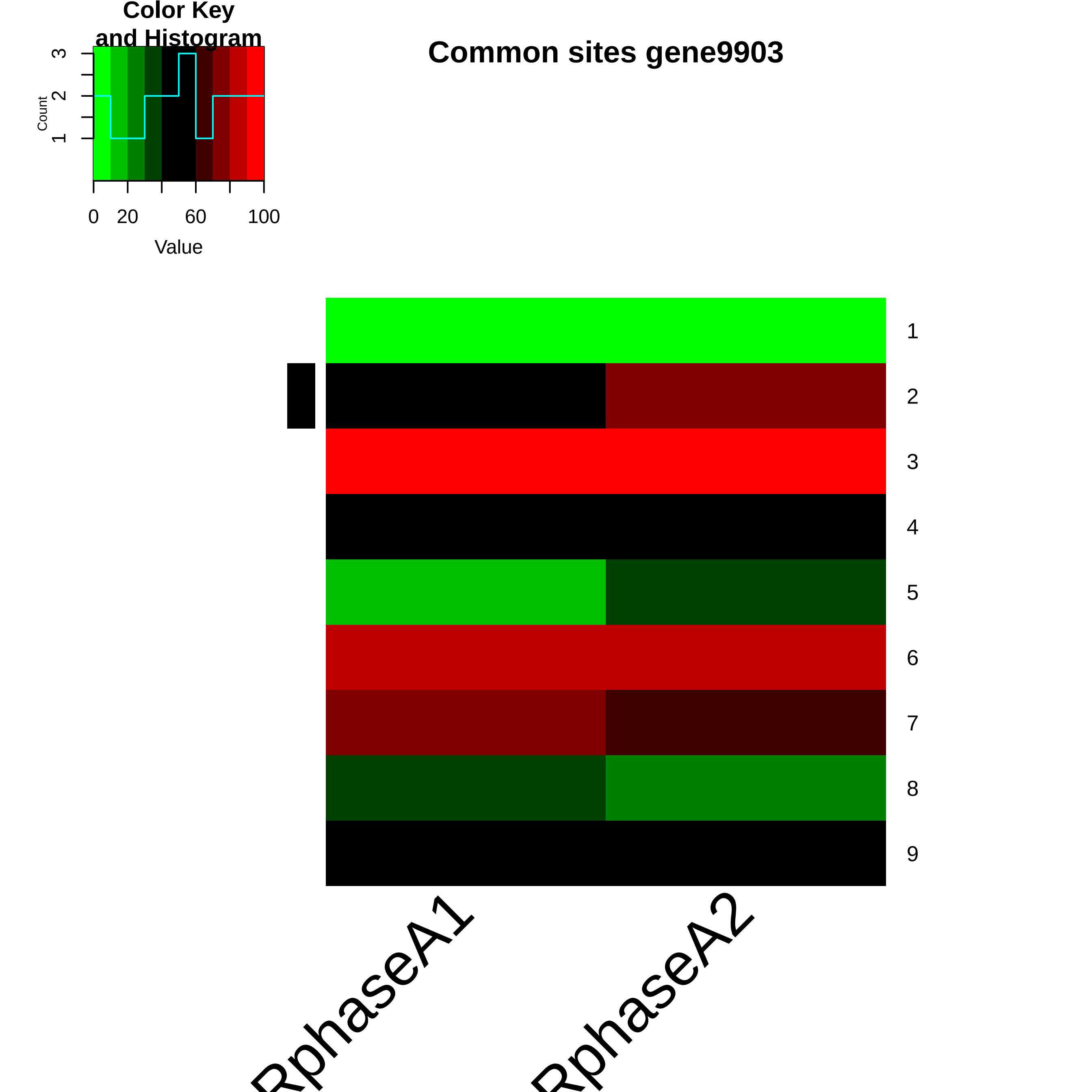

Color Key
and Histogram
Count
3
2
1
0
20
60
100
Value
Common sites gene9903
1
2
3
4
5
6
7
8
9
RphaseA1
RphaseA2

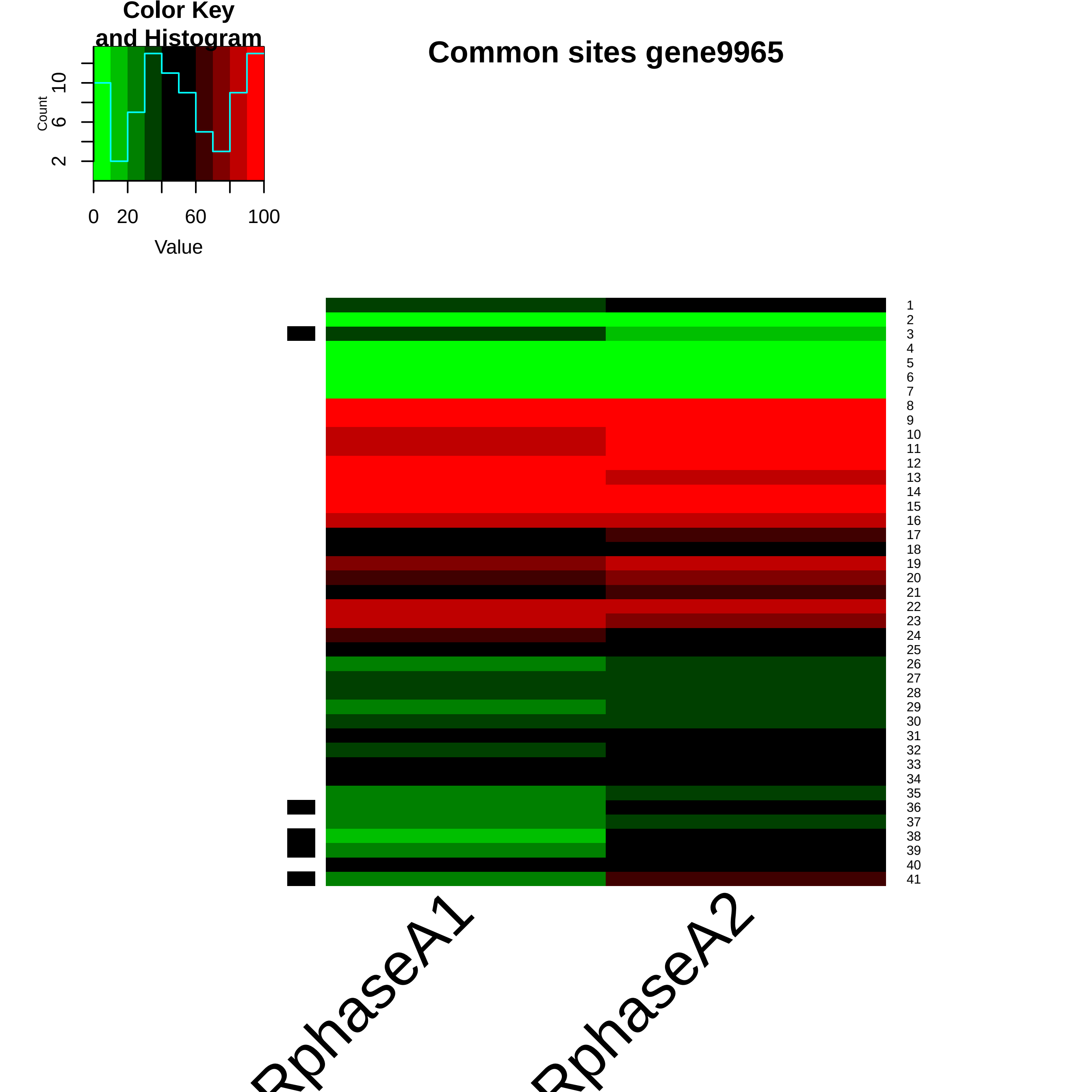

Color Key and Histogram
Count
2
6
10
0
20
60
100
Value
Common sites gene9965
RphaseA1
RphaseA2

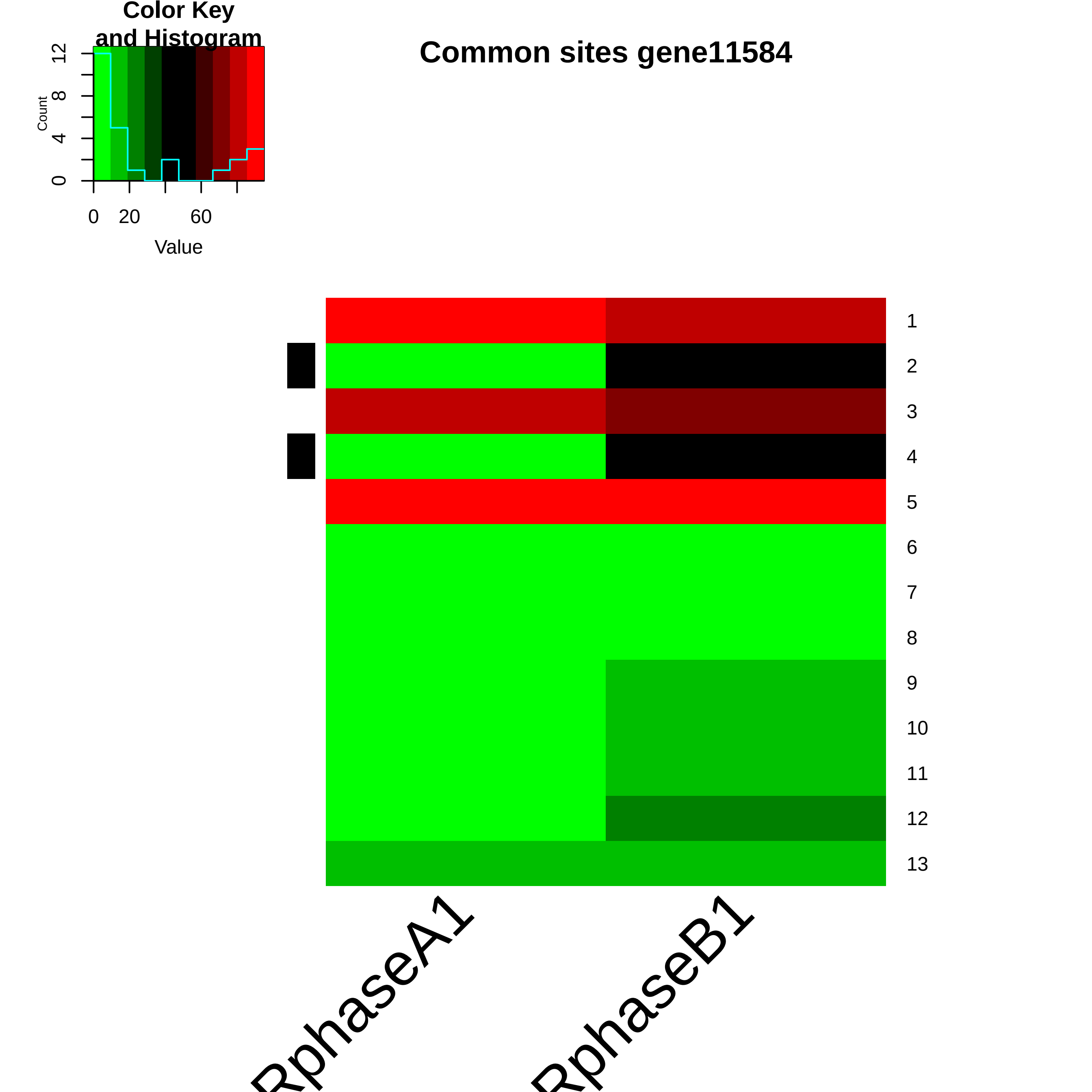
Color Key
and Histogram
Count
0
4
8
12
0
20
60
Value
Common sites gene11584
1
2
3
4
5
6
7
8
9
10
11
12
13
RphaseA1
RphaseB1

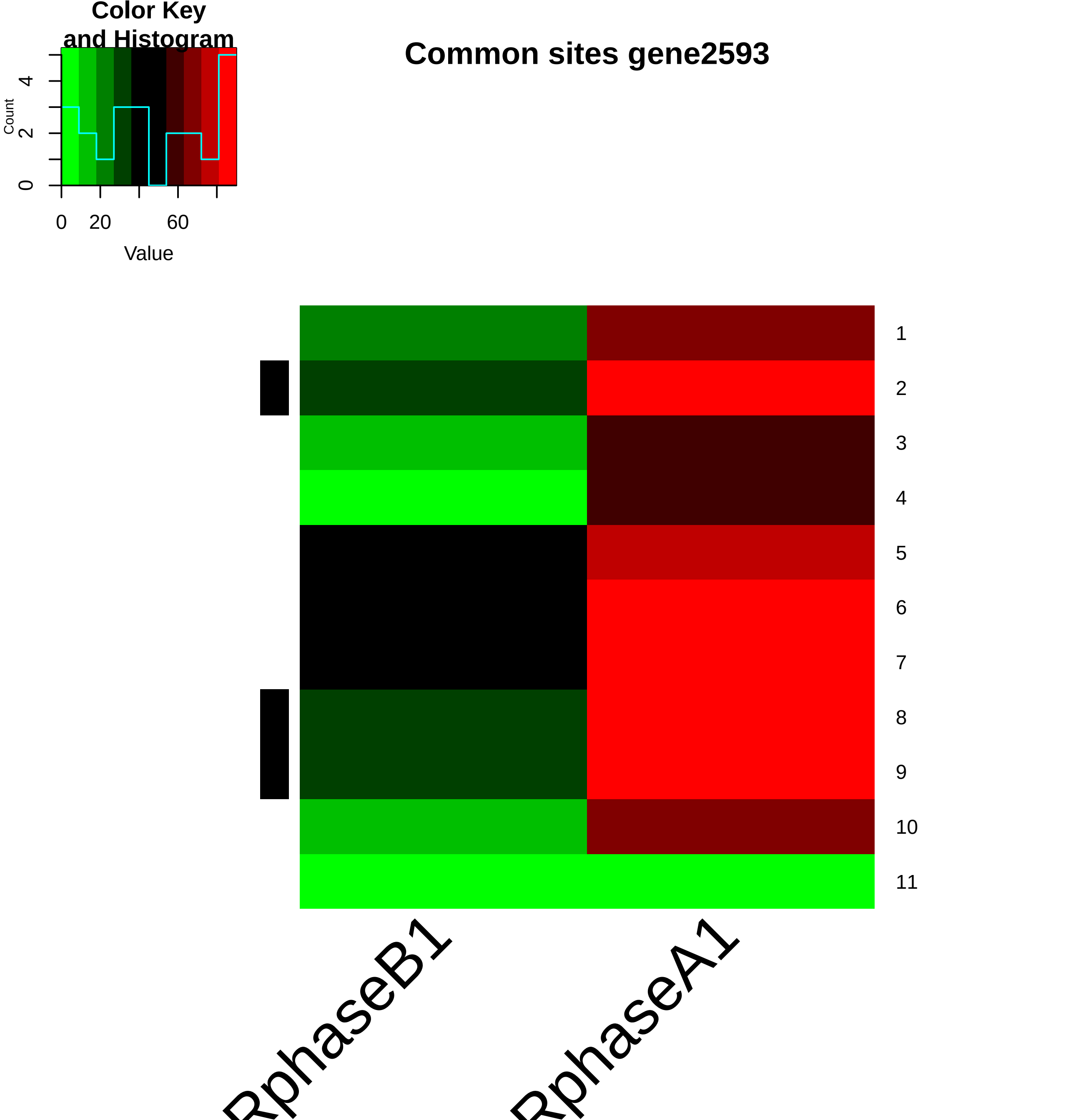
Color Key
and Histogram
Count
0
2
4
0
20
60
Value
Common sites gene2593
1
2
3
4
5
6
7
8
9
10
11
RphaseB1
RphaseA1

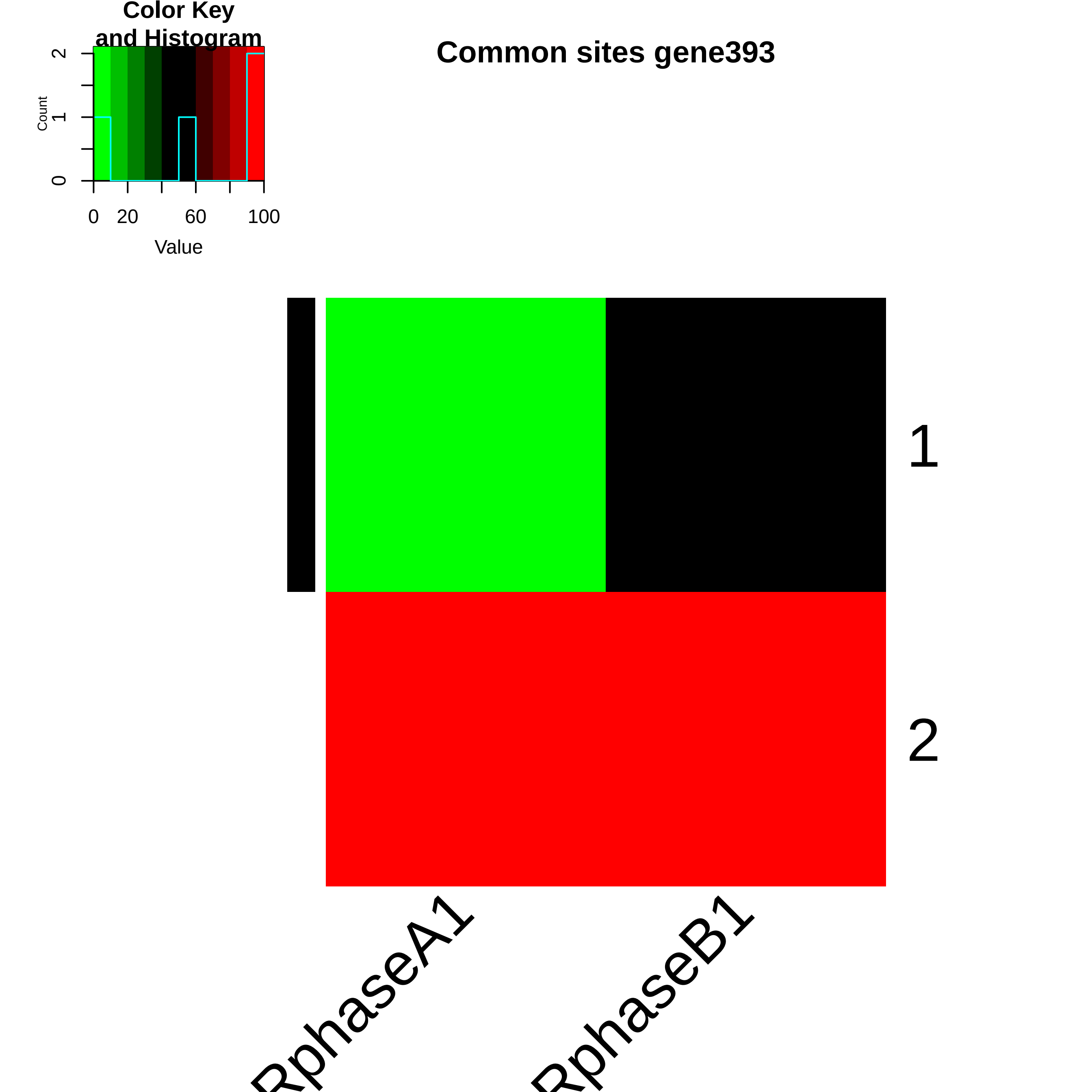
Color Key
and Histogram
Count
0
1
2
0
20
60
100
Value
Common sites gene393
1
2
RphaseA1
RphaseB1

### Common sites gene1008

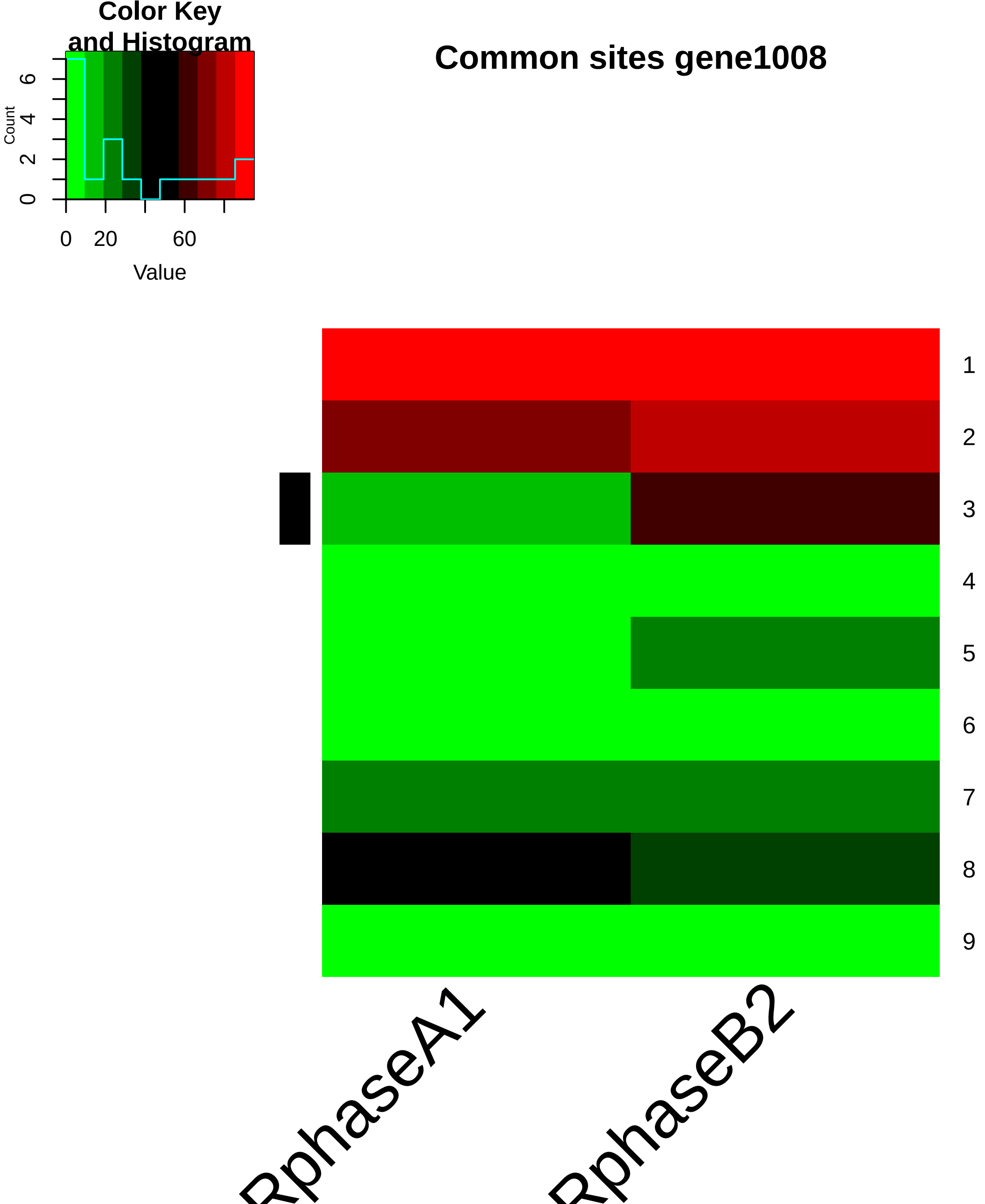

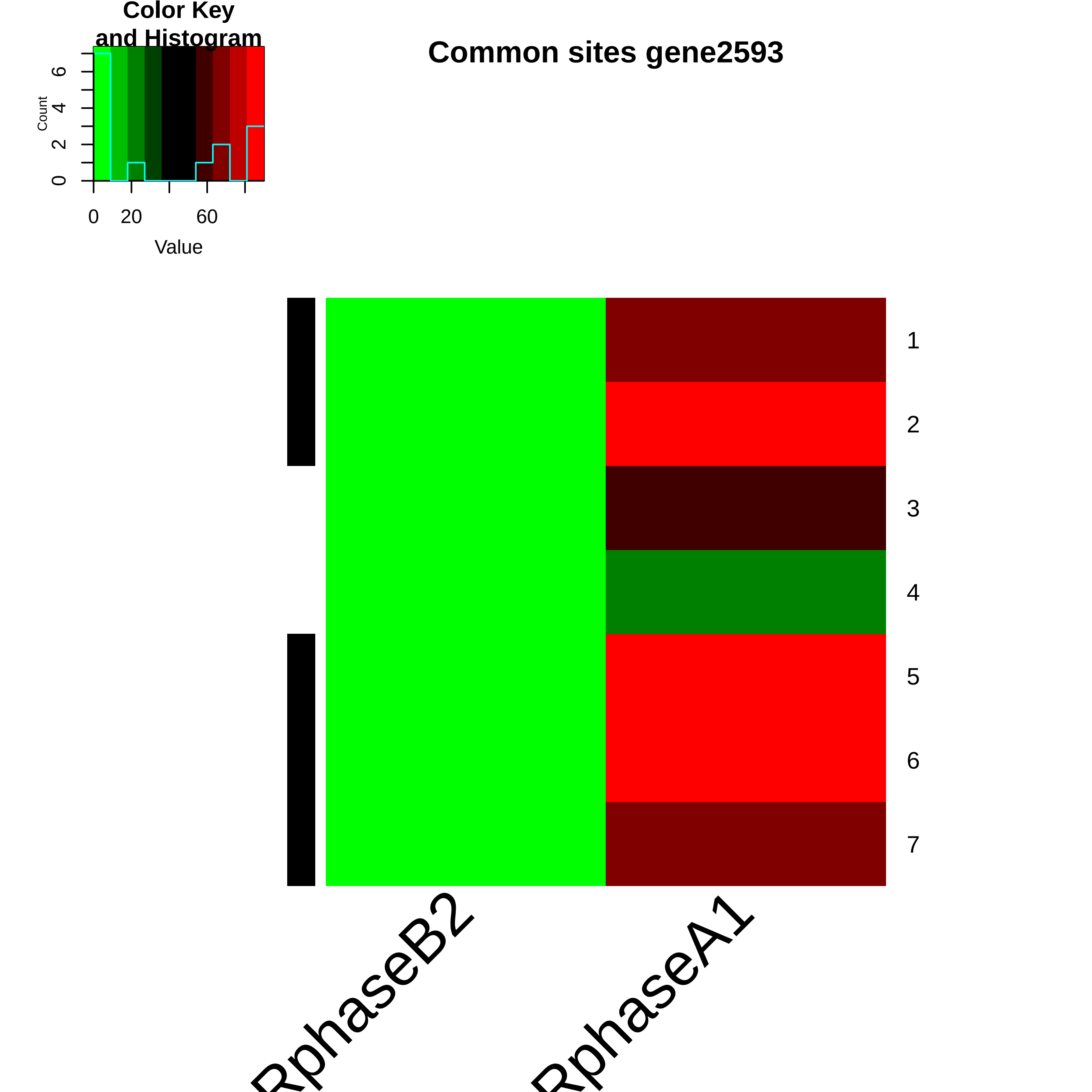
Color Key
and Histogram
Count
0
2
4
6
0
20
60
Value
Common sites gene2593
1
2
3
4
5
6
7
RphaseB2
RphaseA1

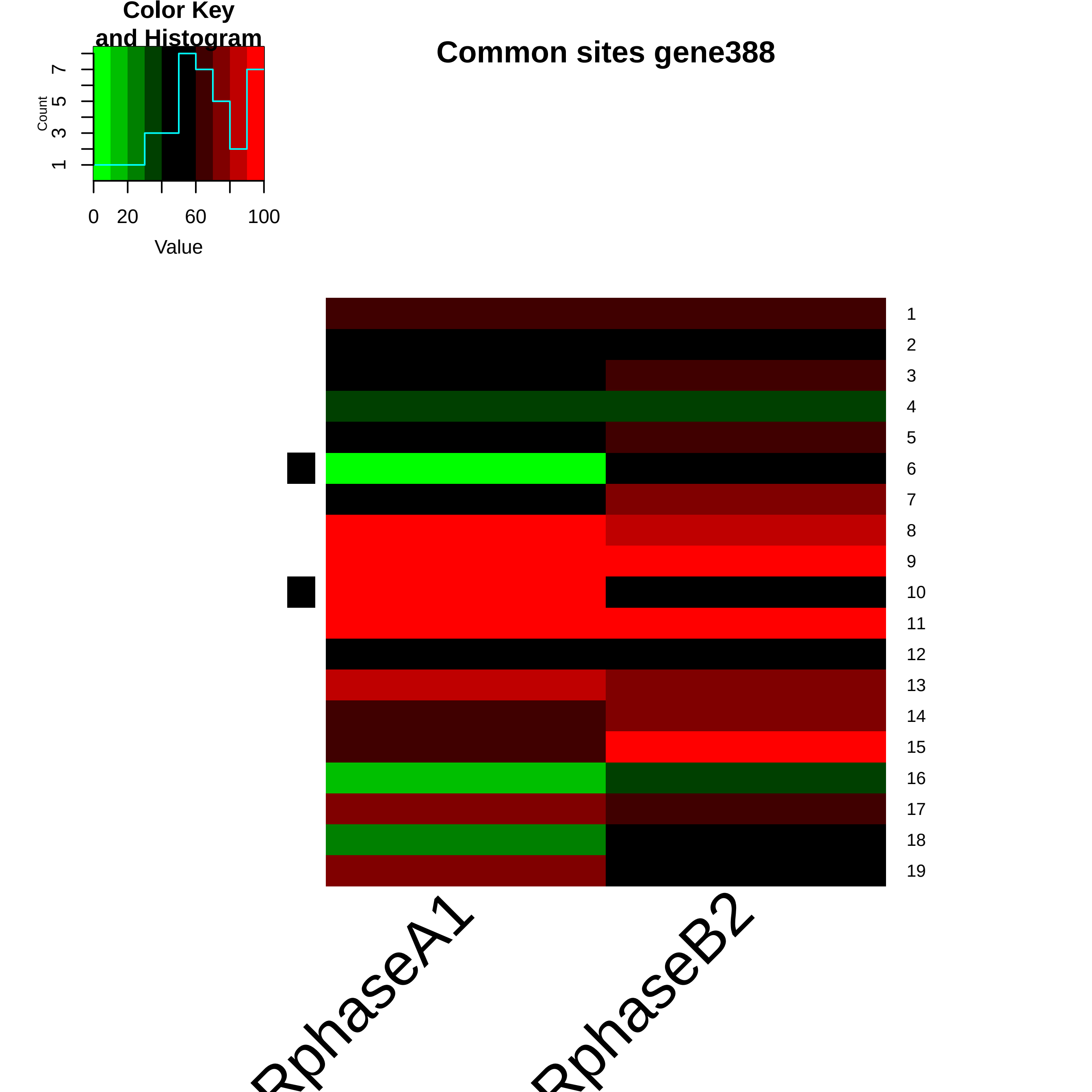
Color Key
and Histogram
Count
1
3
5
7
0
20
60
100
Value
Common sites gene388
RphaseA1
RphaseB2

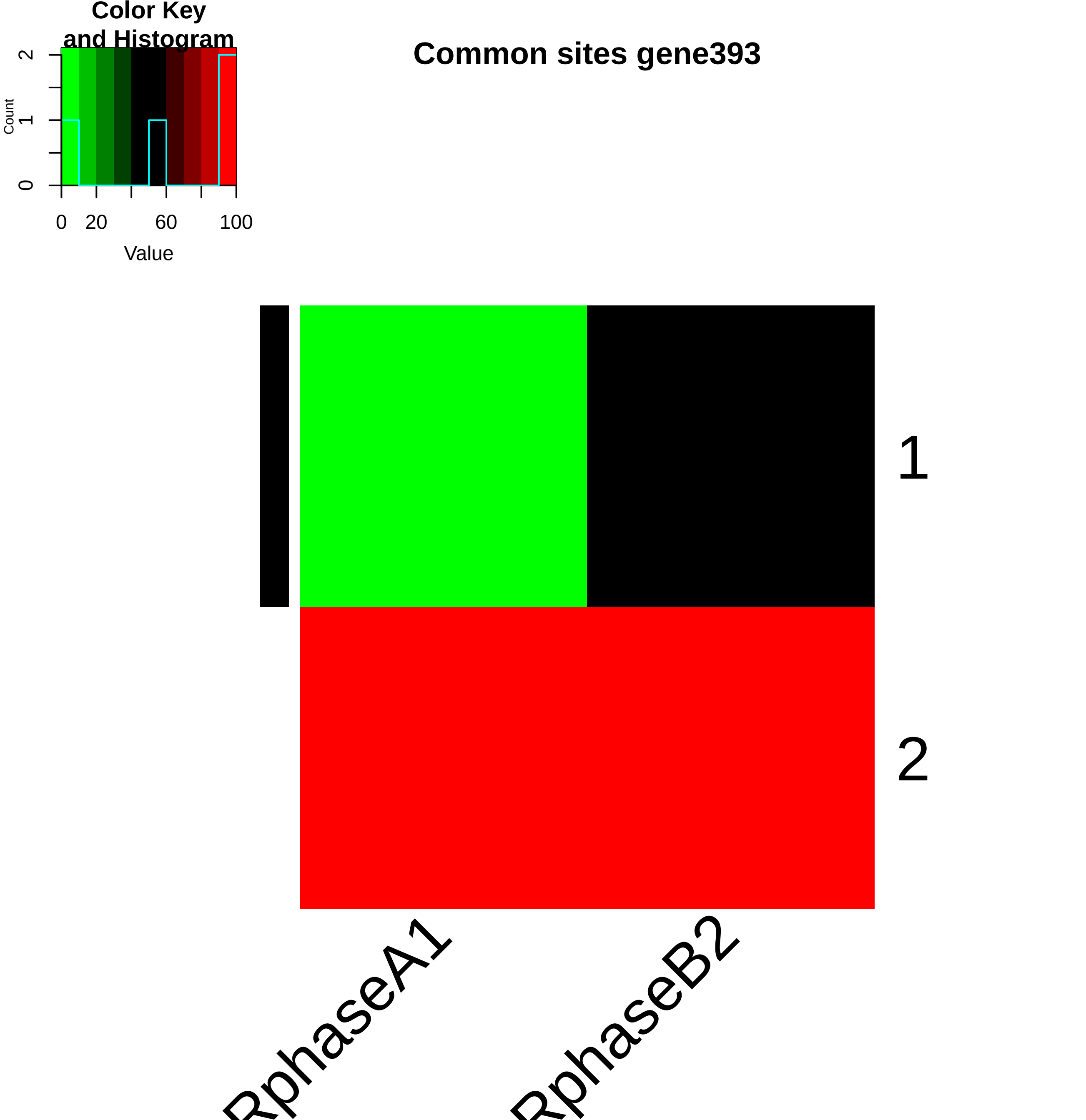
Color Key
and Histogram
Count
0
1
2
0
20
60
100
Value
Common sites gene393
1
2
RphaseA1
RphaseB2

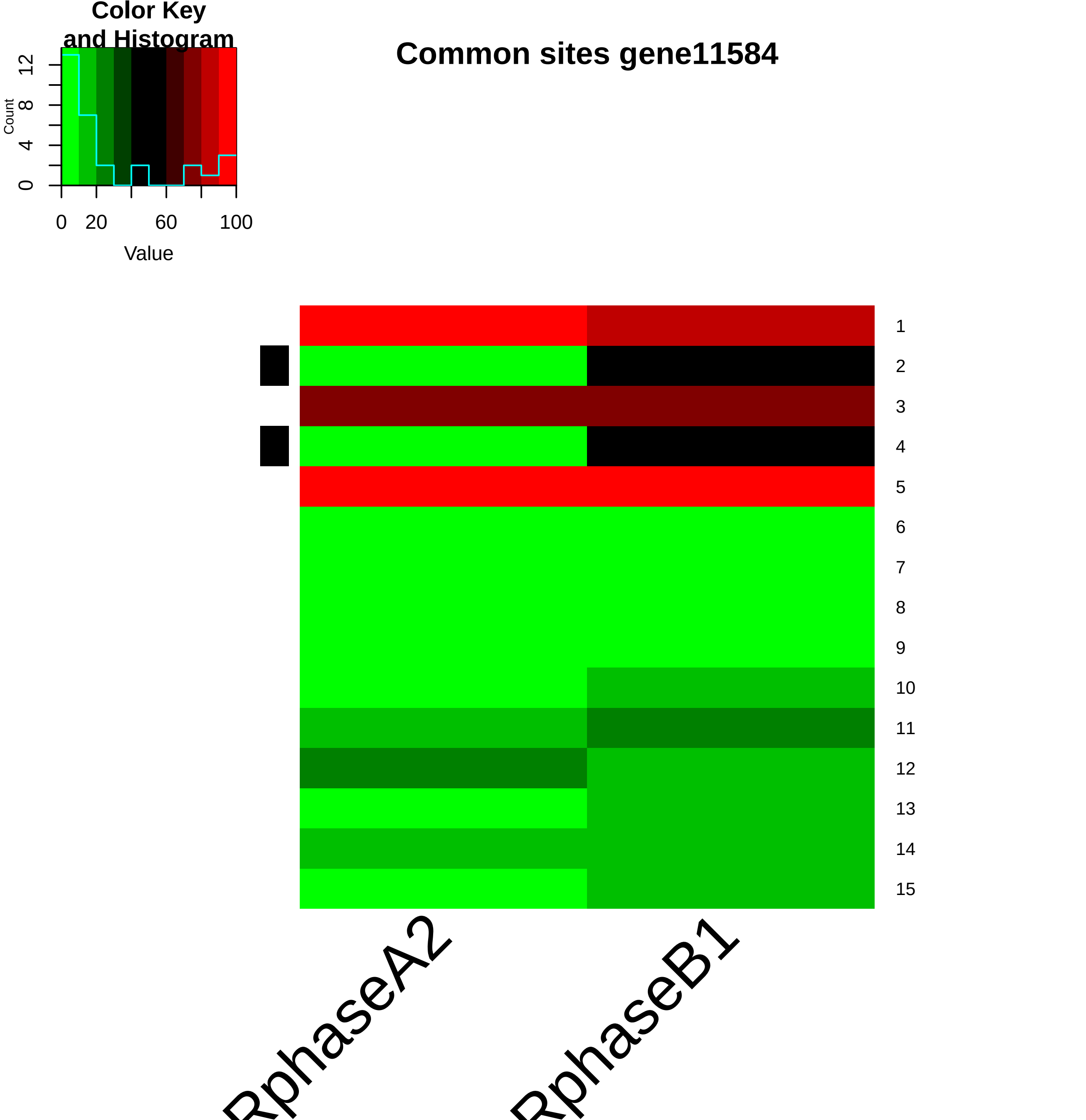
Color Key
and Histogram
Count
0
4
8
12
0
20
60
100
Value
Common sites gene11584
RphaseA2
RphaseB1

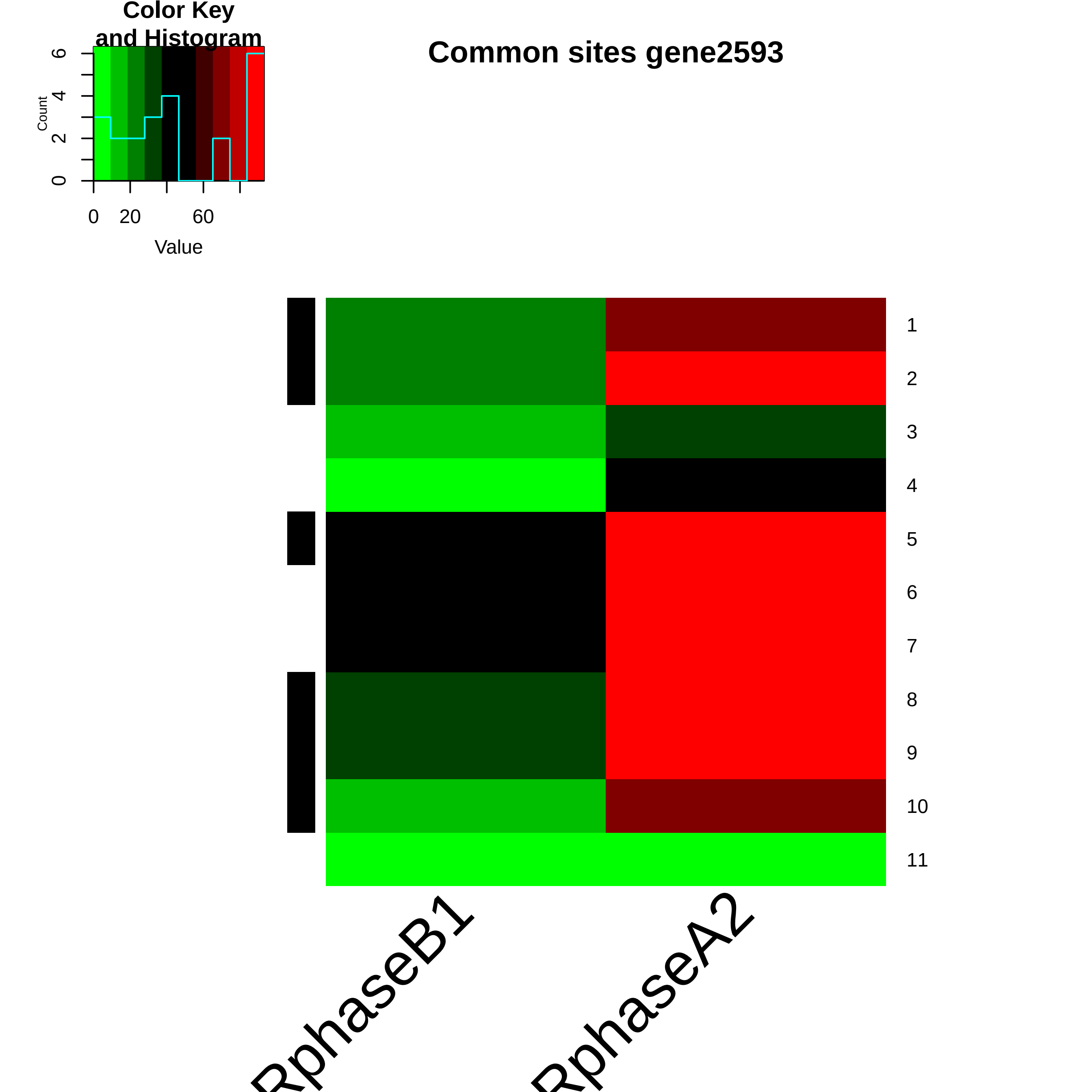

Color Key
and Histogram
Count
Value
0
2
4
6
0
20
60
Common sites gene2593
1
2
3
4
5
6
7
8
9
10
11
RphaseB1
RphaseA2

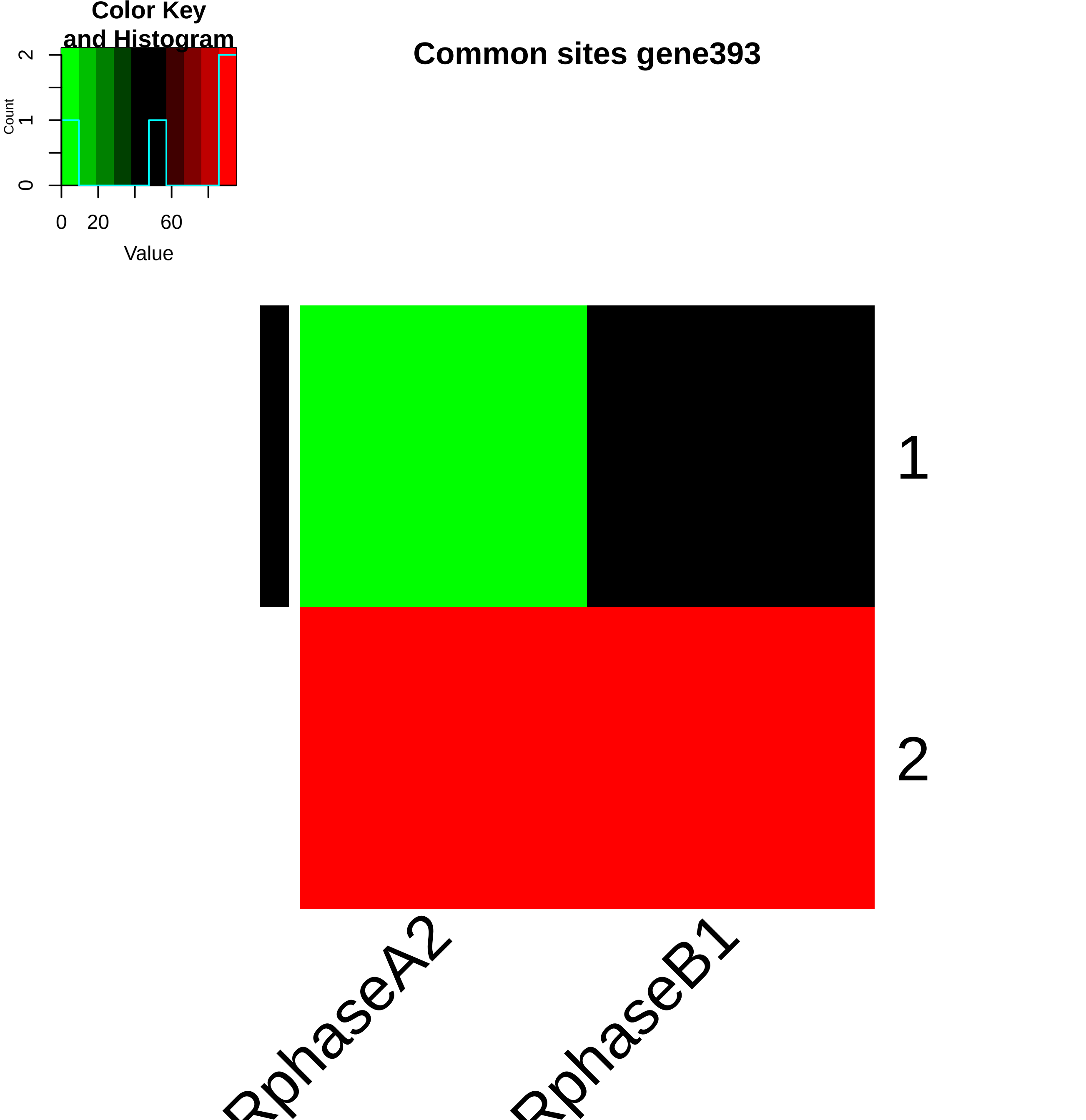
Color Key
and Histogram
Count
0
1
2
0
20
60
Value
Common sites gene393
1
2
RphaseA2
RphaseB1

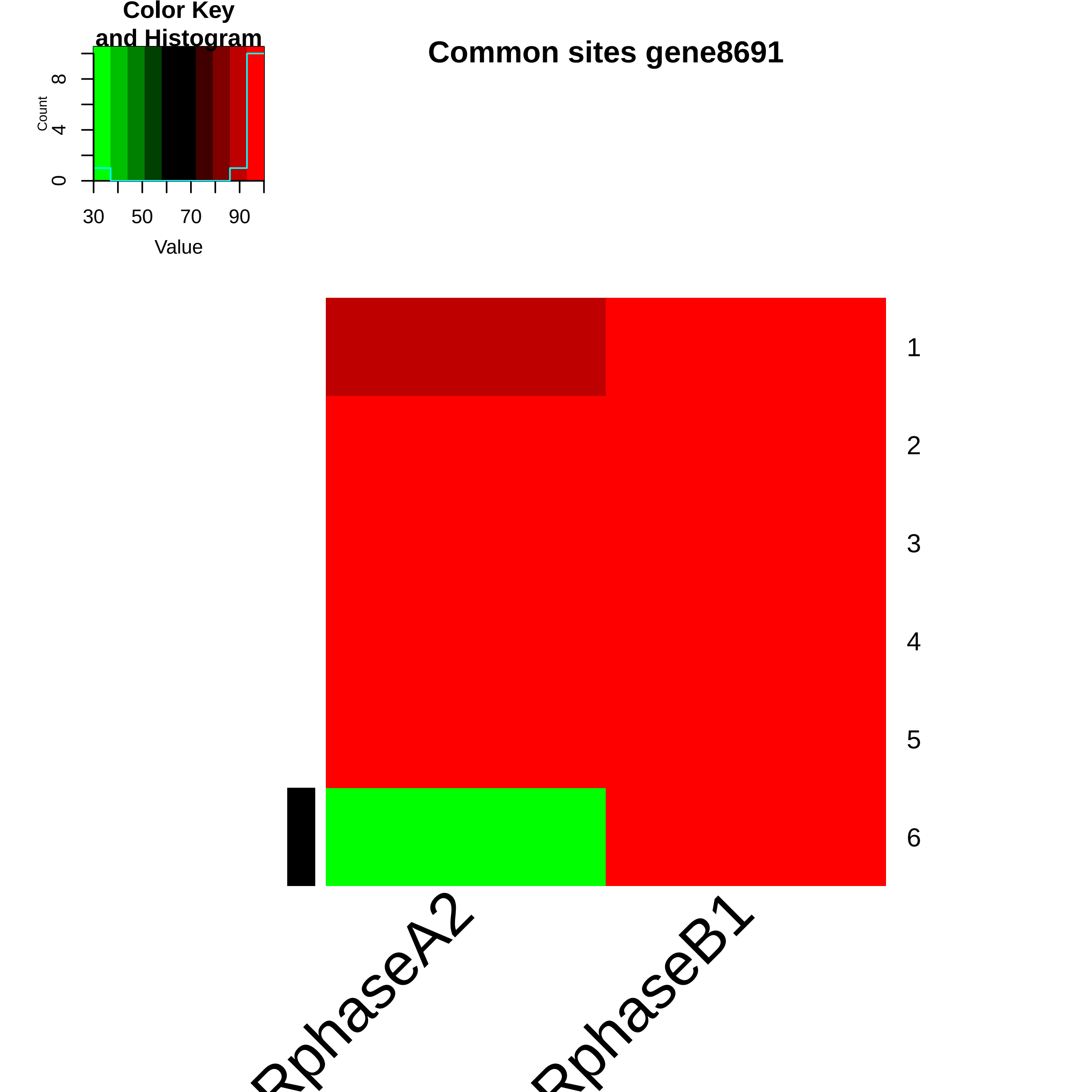

Color Key
and Histogram
Common sites gene8691
Count
0
4
8
30
50
70
90
Value
1
2
3
4
5
6
RphaseA2
RphaseB1

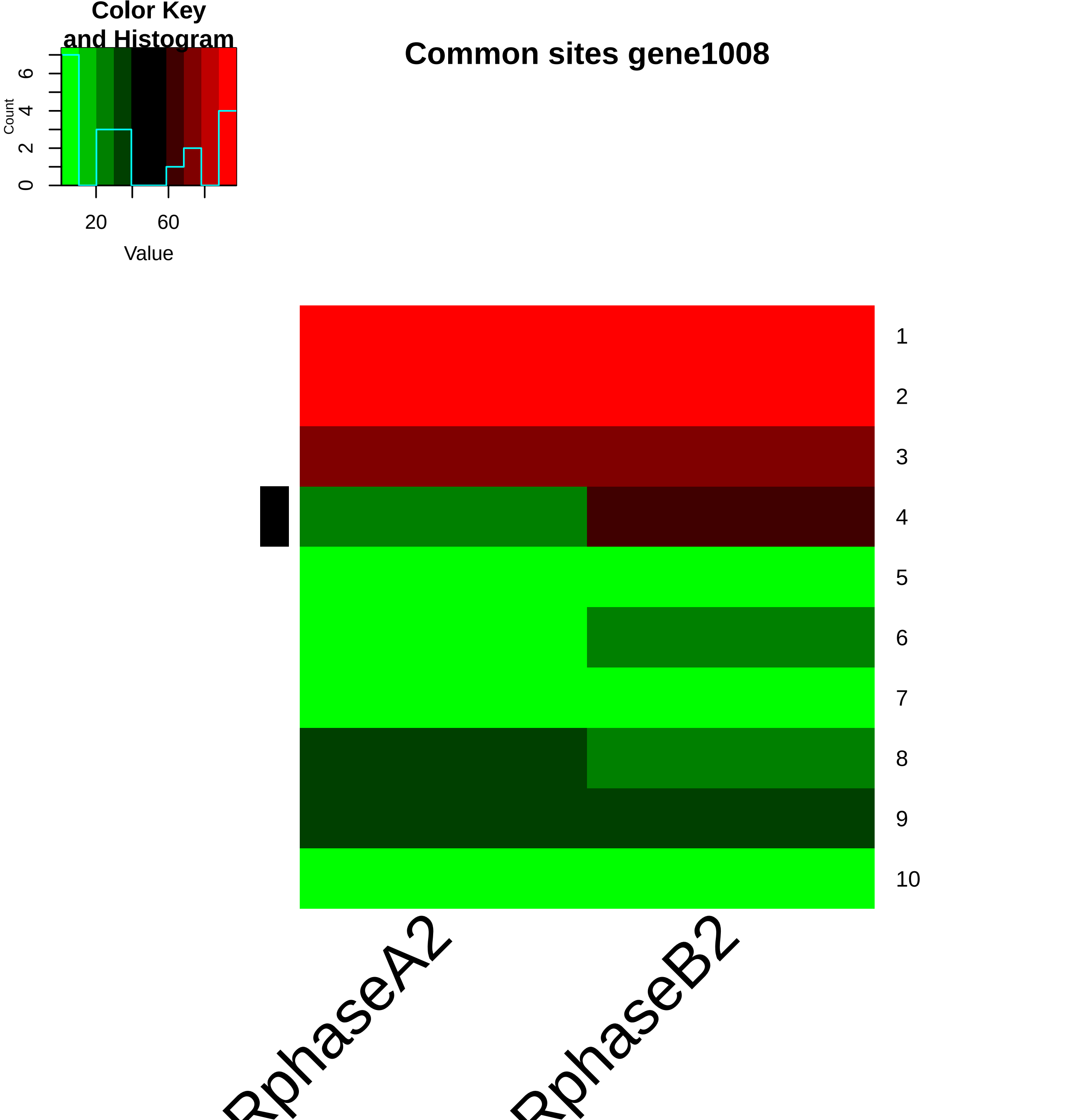

Color Key
and Histogram
Count
0
2
4
6
20
60
Value
Common sites gene1008
1
2
3
4
5
6
7
8
9
10
RphaseA2
RphaseB2

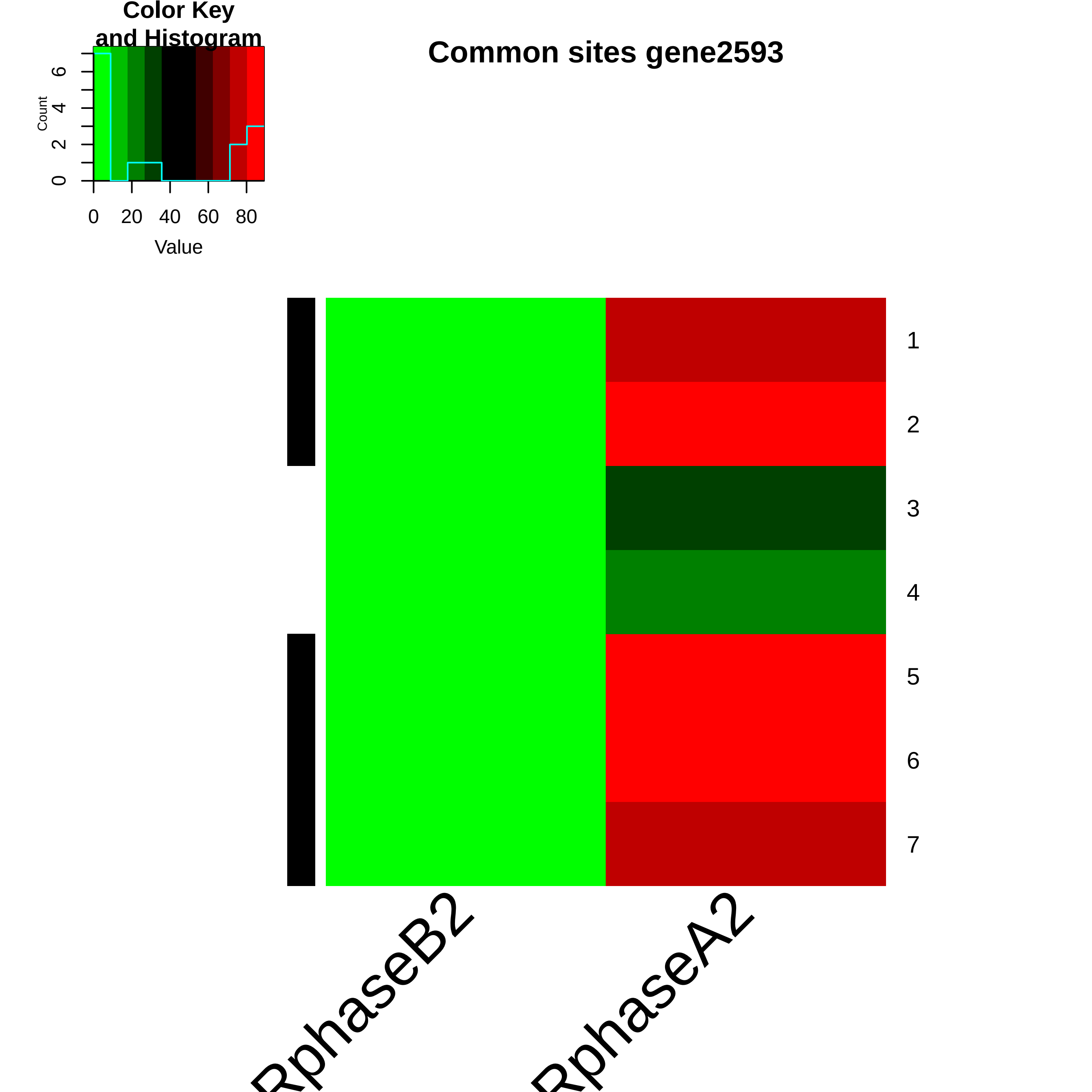

Color Key
and Histogram
Count
0
2
4
6
0
20
40
60
80
Value
Common sites gene2593
1
2
3
4
5
6
7
RphaseB2
RphaseA2

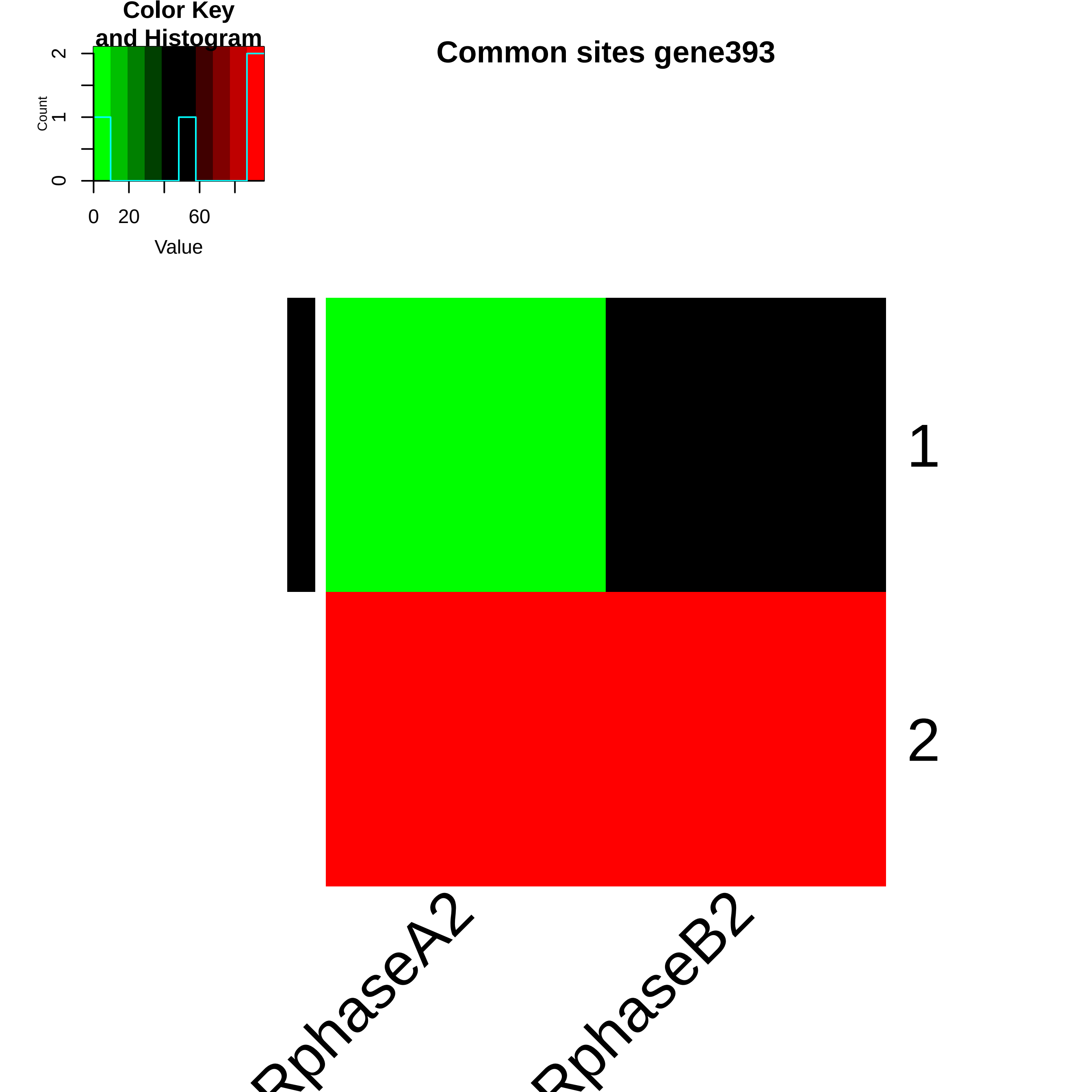
Color Key
and Histogram
Count
0
1
2
0
20
60
Value
Common sites gene393
1
2
RphaseA2
RphaseB2

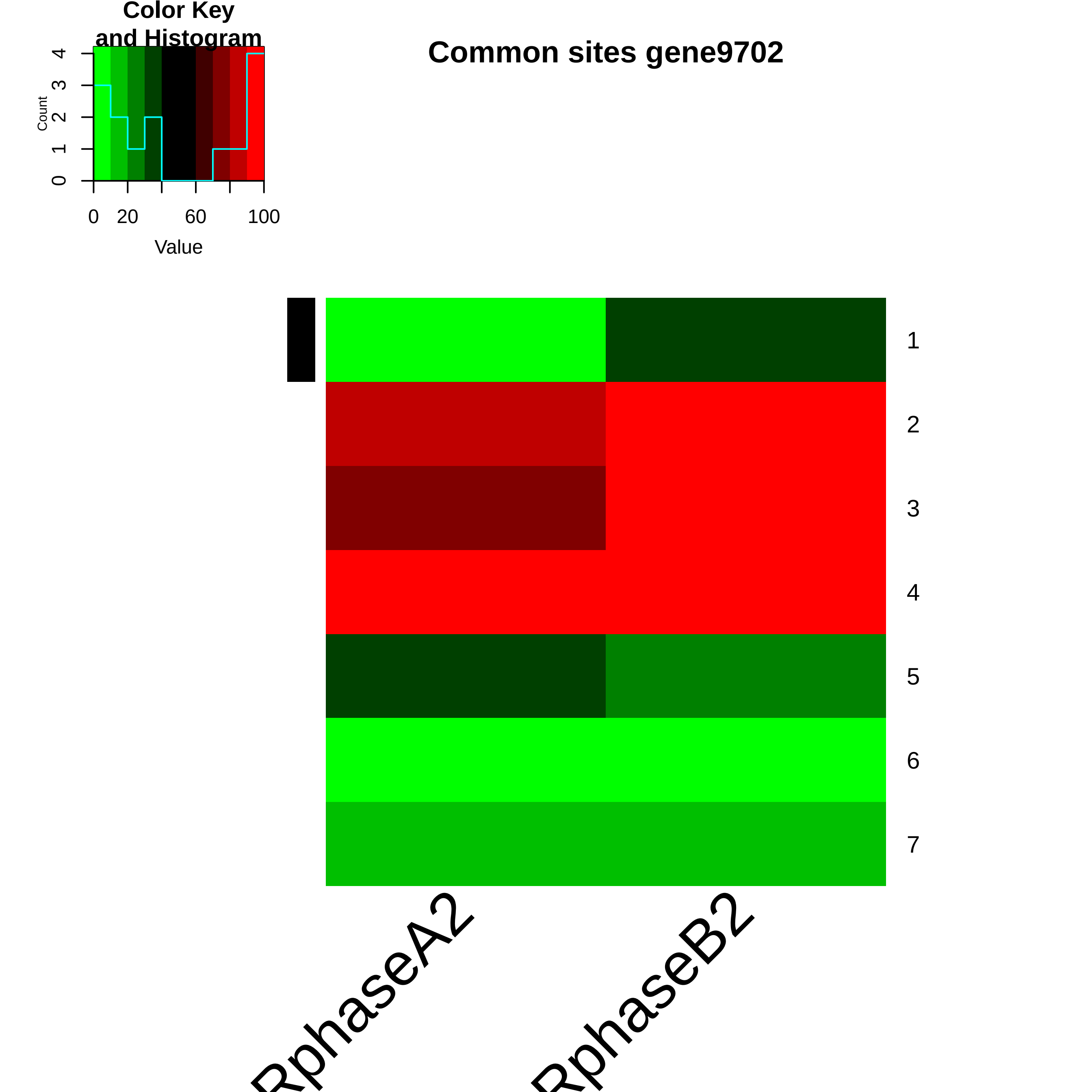
Color Key
and Histogram
Count
0
1
2
3
4
0
20
60
100
Value
Common sites gene9702
1
2
3
4
5
6
7
RphaseA2
RphaseB2

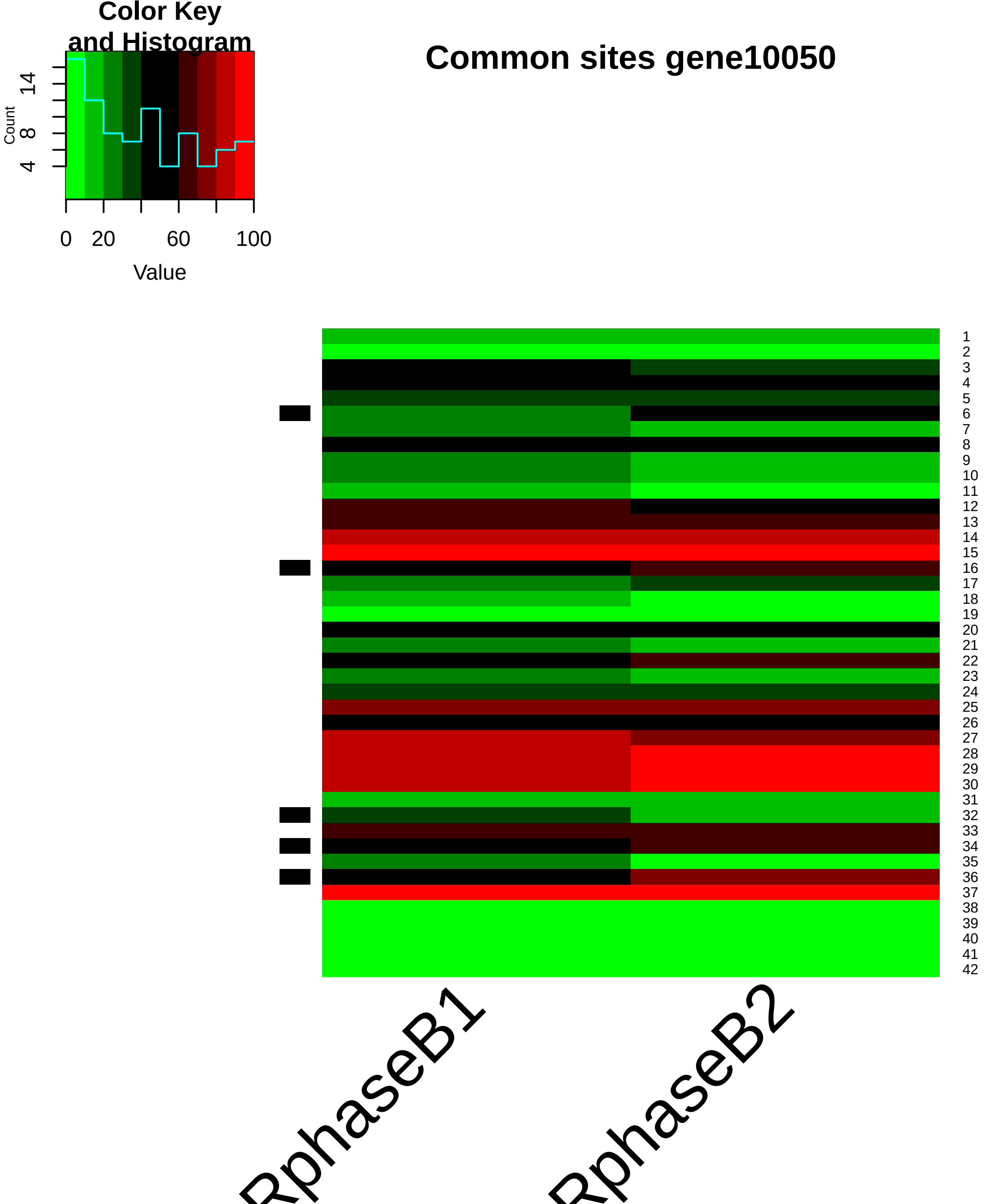
Color Key
and Histogram
Count
14
8
4
0
20
60
100
Value
Common sites gene10050
RphaseB1
RphaseB2

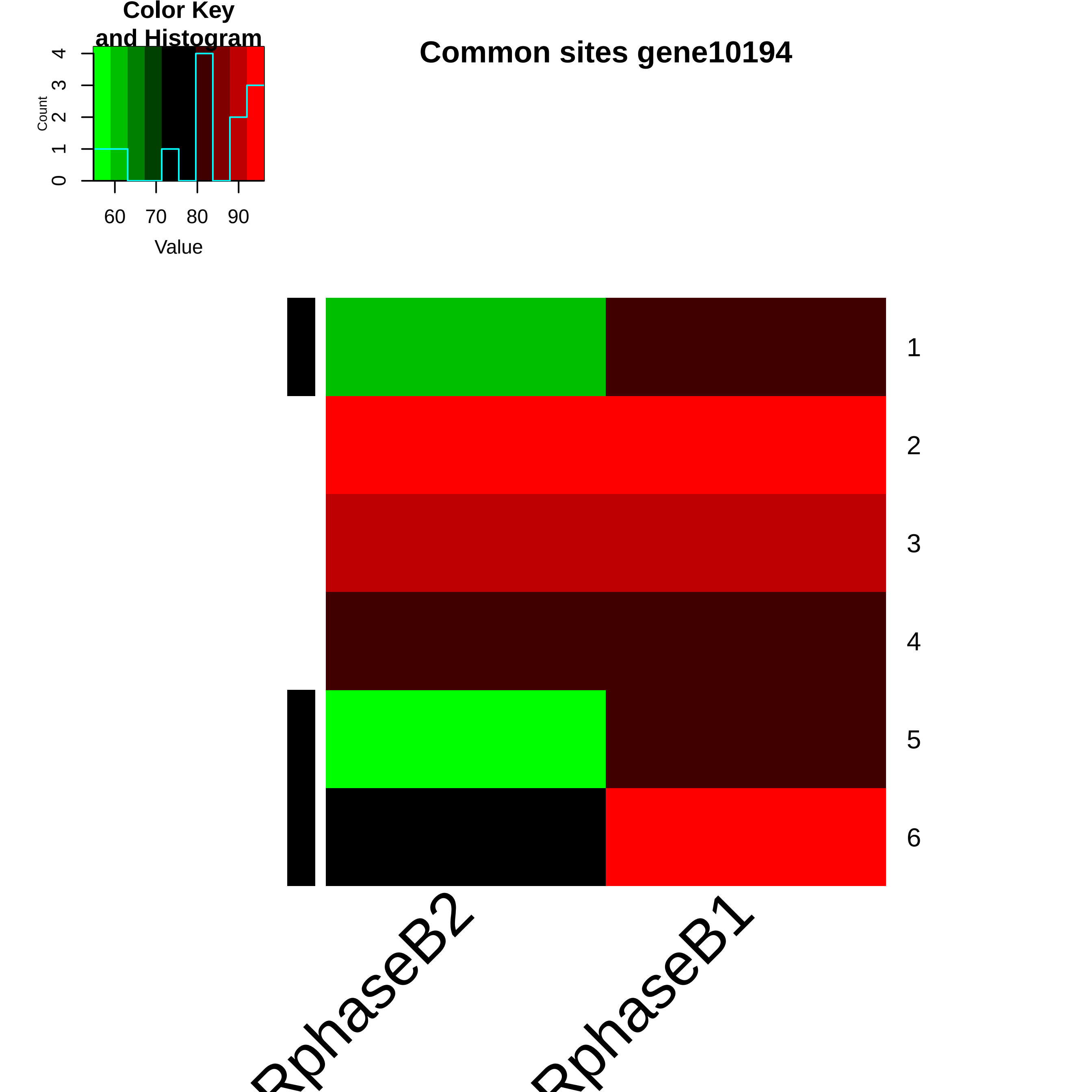

Color Key
and Histogram
Count
Value
60
70
80
90
0
1
2
3
4
Common sites gene10194
1
2
3
4
5
6
RphaseB2
RphaseB1

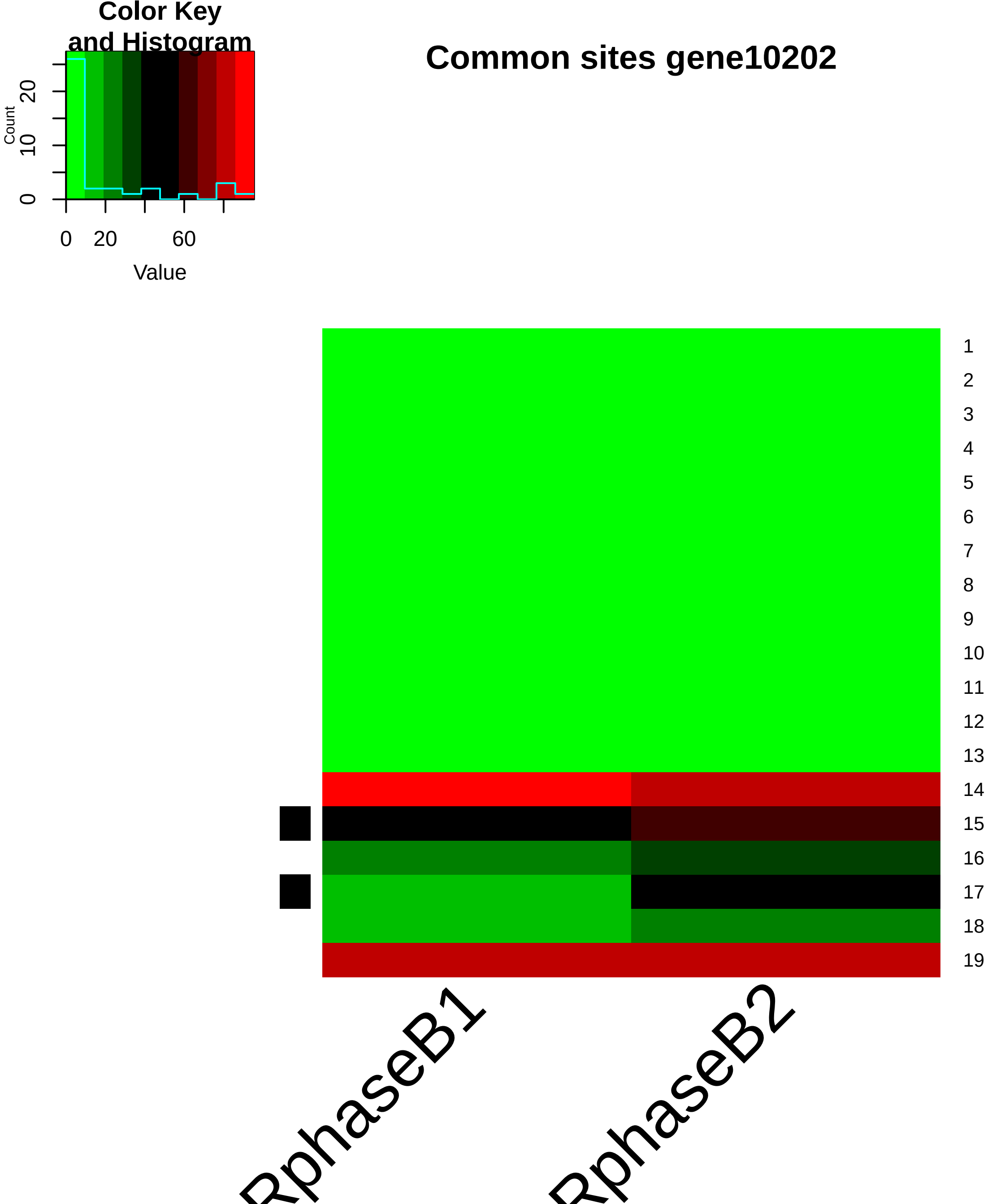

Color Key and Histogram
Count
0
10
20
0
20
60
Value
Common sites gene10202
RphaseB1
RphaseB2

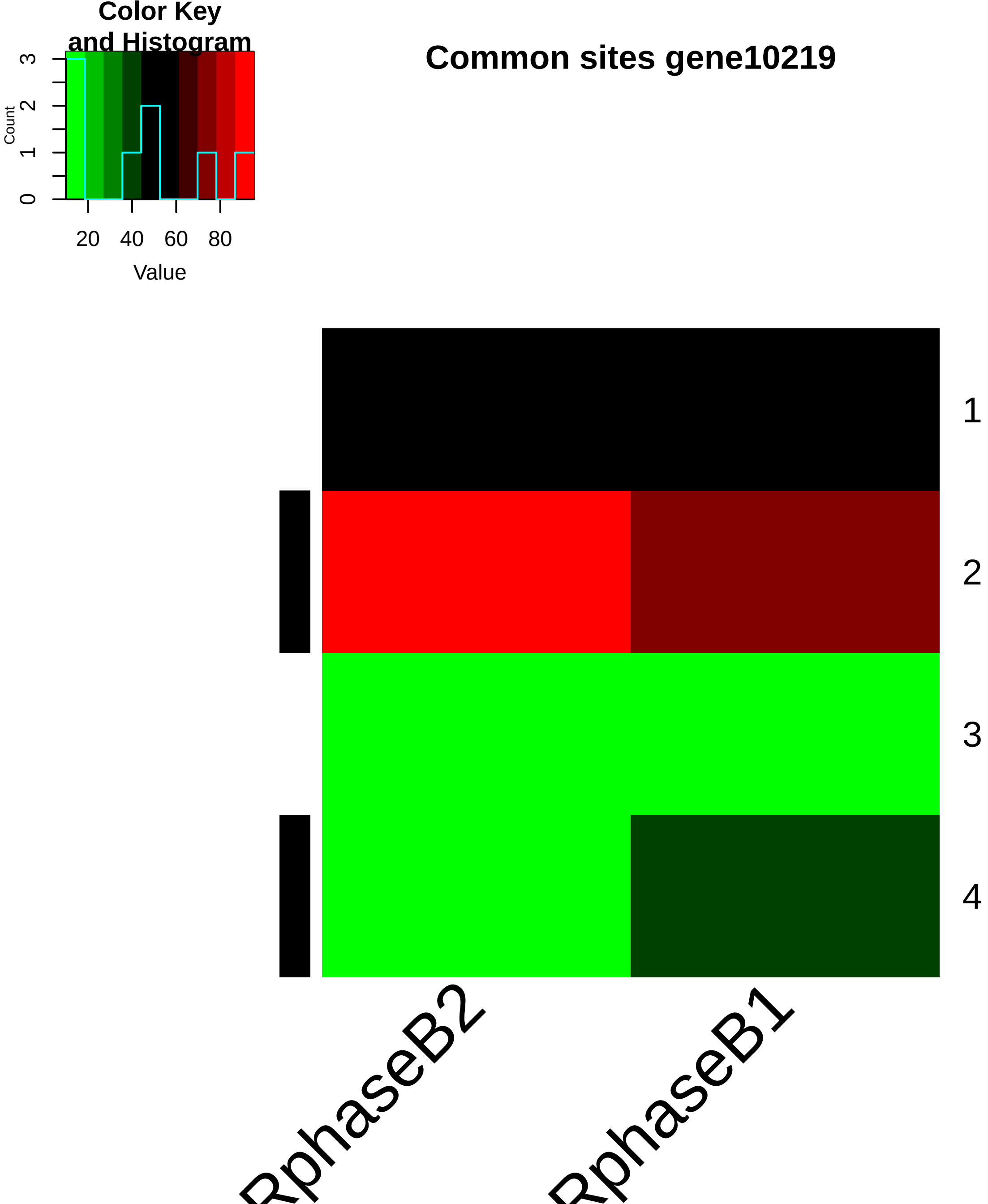
Color Key
and Histogram
Count
0
1
2
3
20
40
60
80
Value
Common sites gene10219
1
2
3
4
RphaseB2
RphaseB1

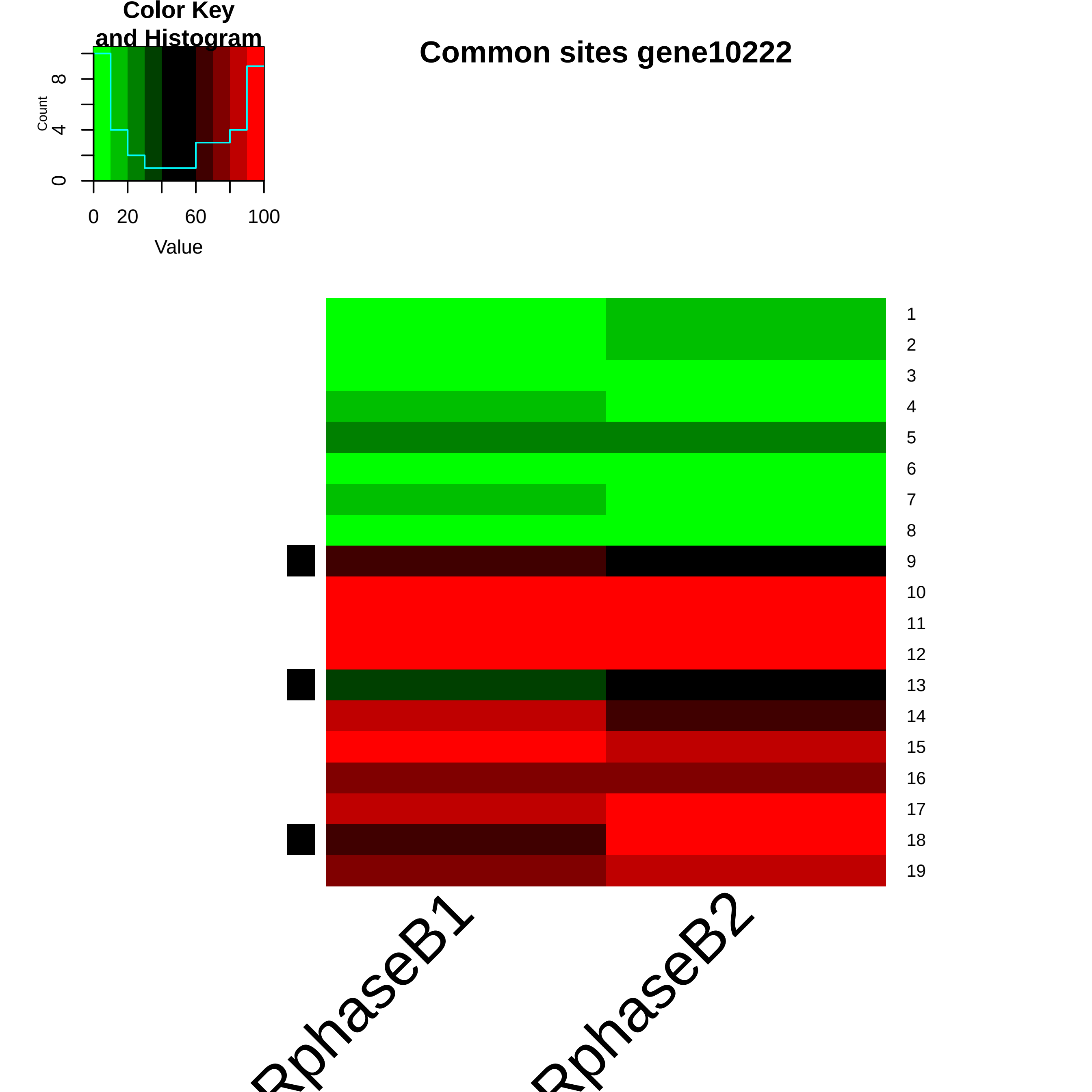

Color Key
and Histogram
Count
0
4
8
0
20
60
100
Value
Common sites gene10222
RphaseB1
RphaseB2

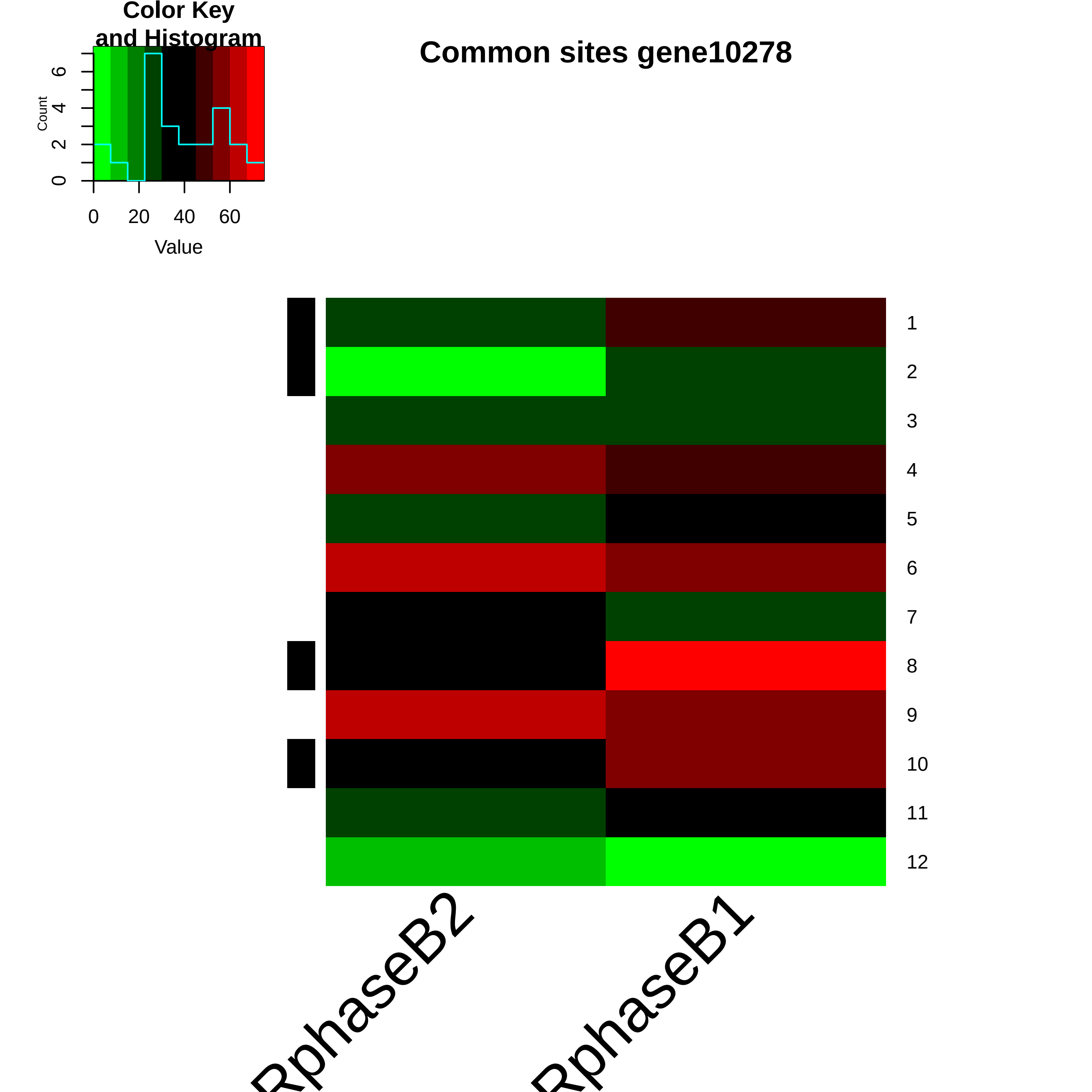
Color Key
and Histogram
Count
0
2
4
6
0
20
40
60
Value
Common sites gene10278
1
2
3
4
5
6
7
8
9
10
11
12
RphaseB2
RphaseB1

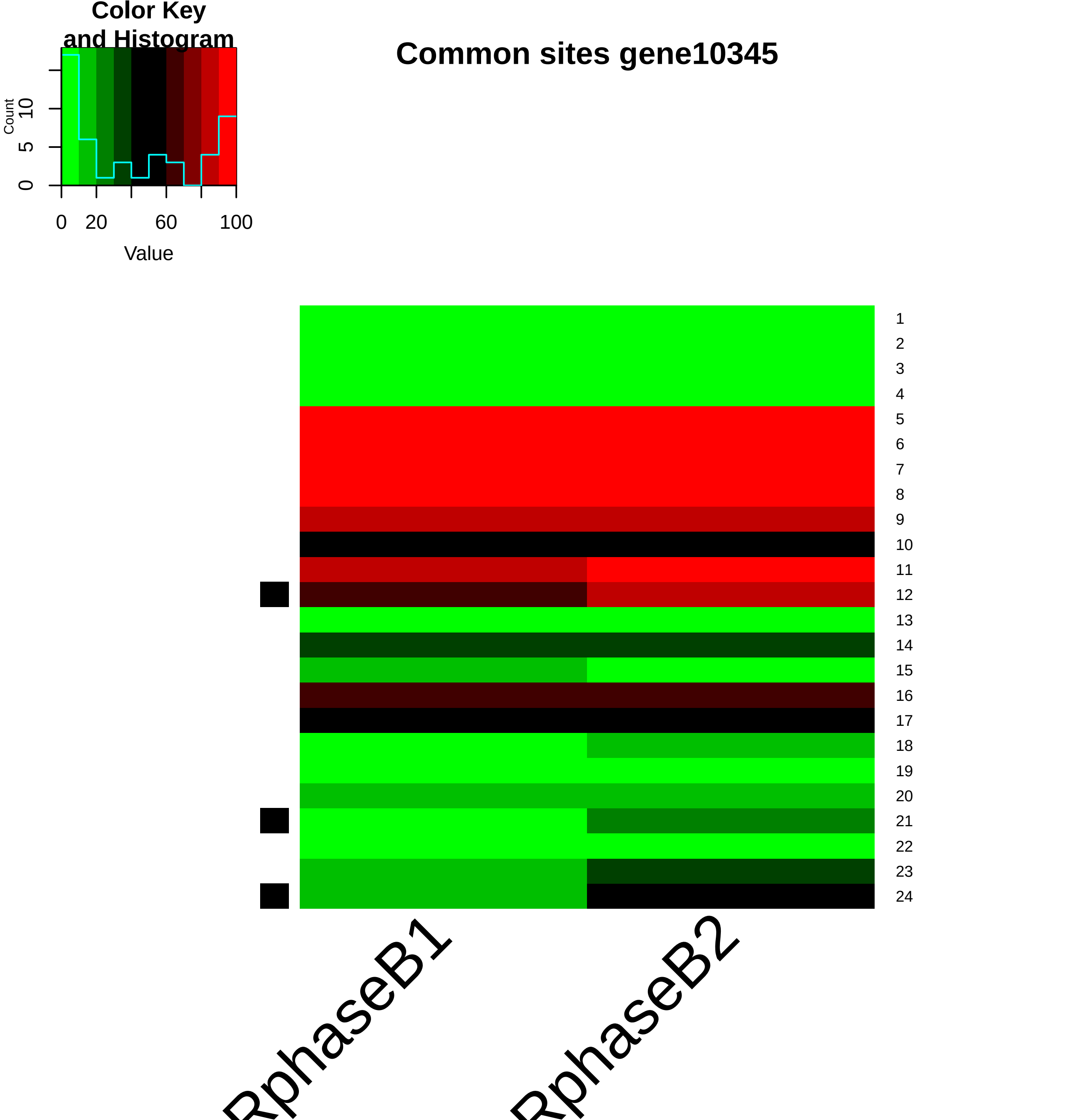
Color Key
and Histogram
Count
0
5
10
0
20
60
100
Value
Common sites gene10345
RphaseB1
RphaseB2

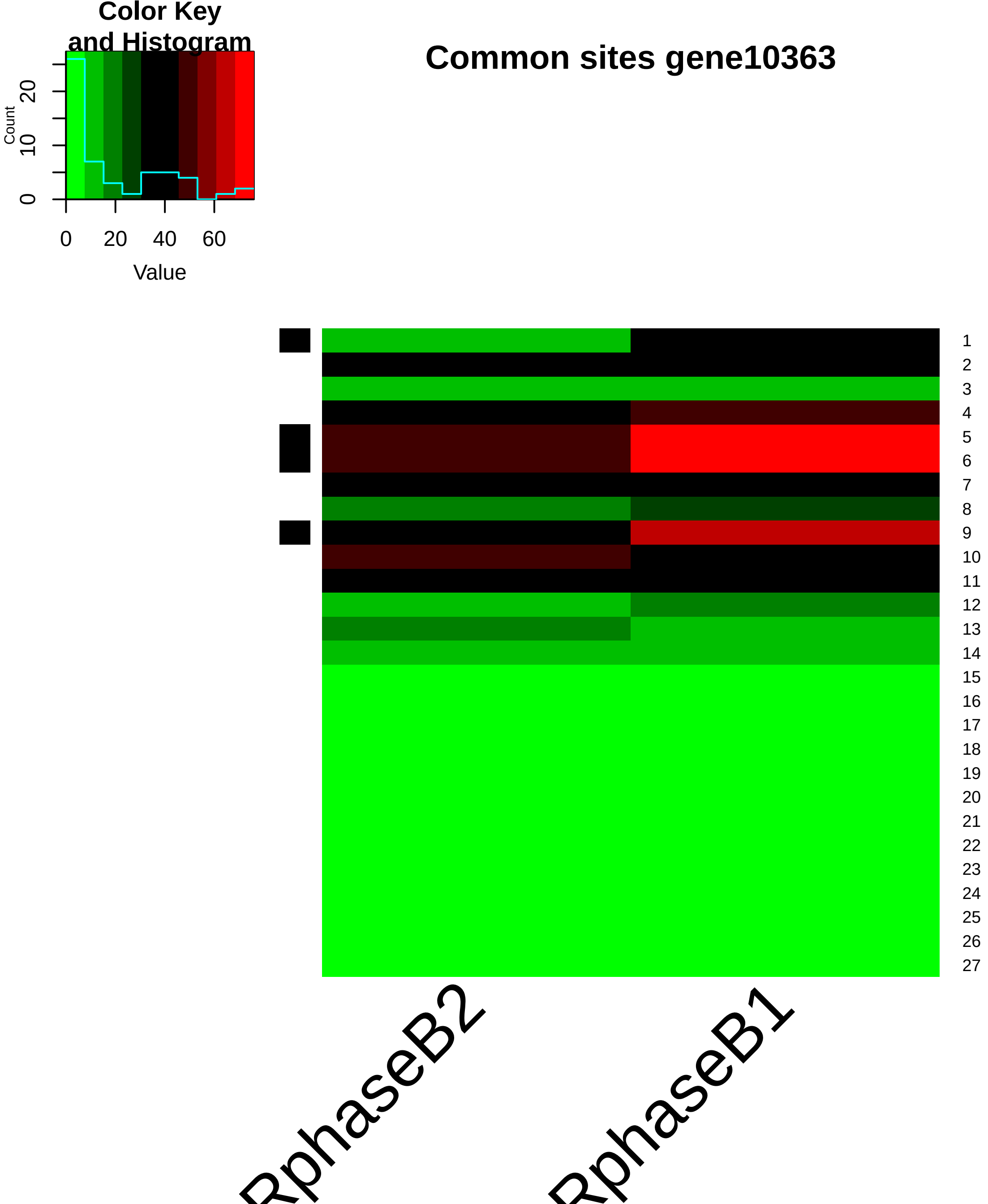

Color Key
and Histogram
Count
0
10
20
0
20
40
60
Value
Common sites gene10363
RphaseB2
RphaseB1

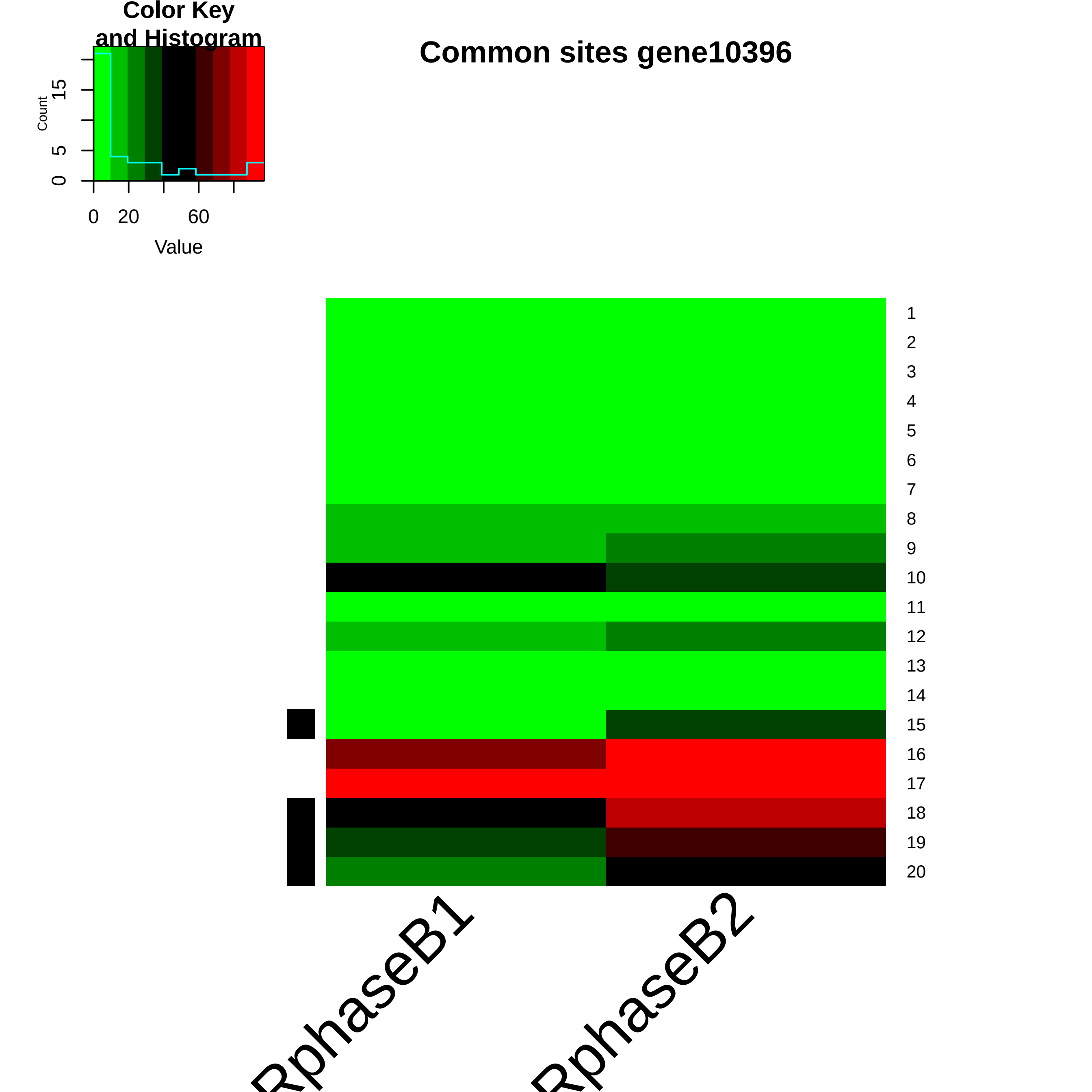

Color Key and Histogram
Count
0
5
15
0
20
60
Value
Common sites gene10396
RphaseB1
RphaseB2

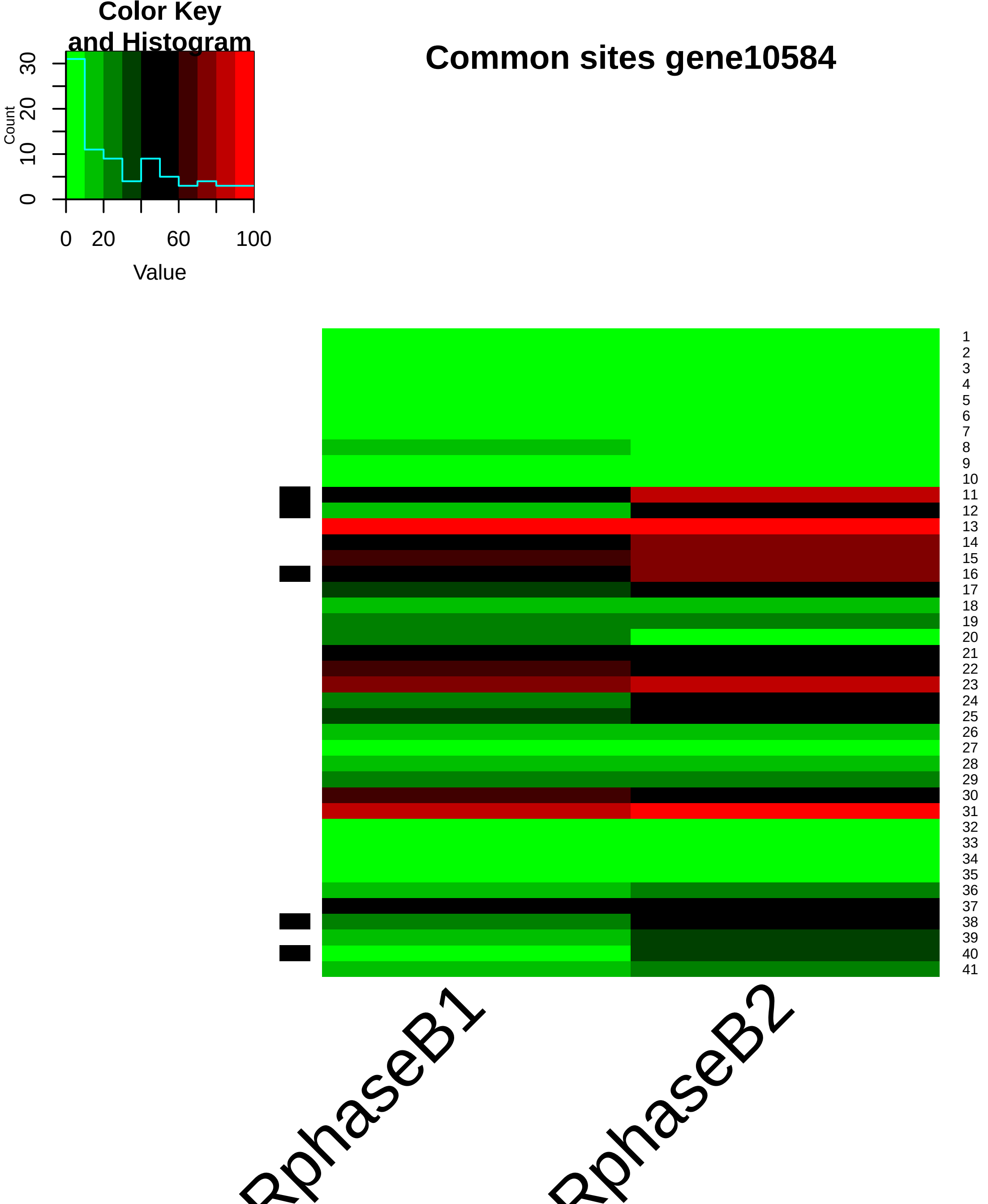
Color Key
and Histogram
Count
0
10
20
30
0
20
60
100
Value
Common sites gene10584
RphaseB1
RphaseB2

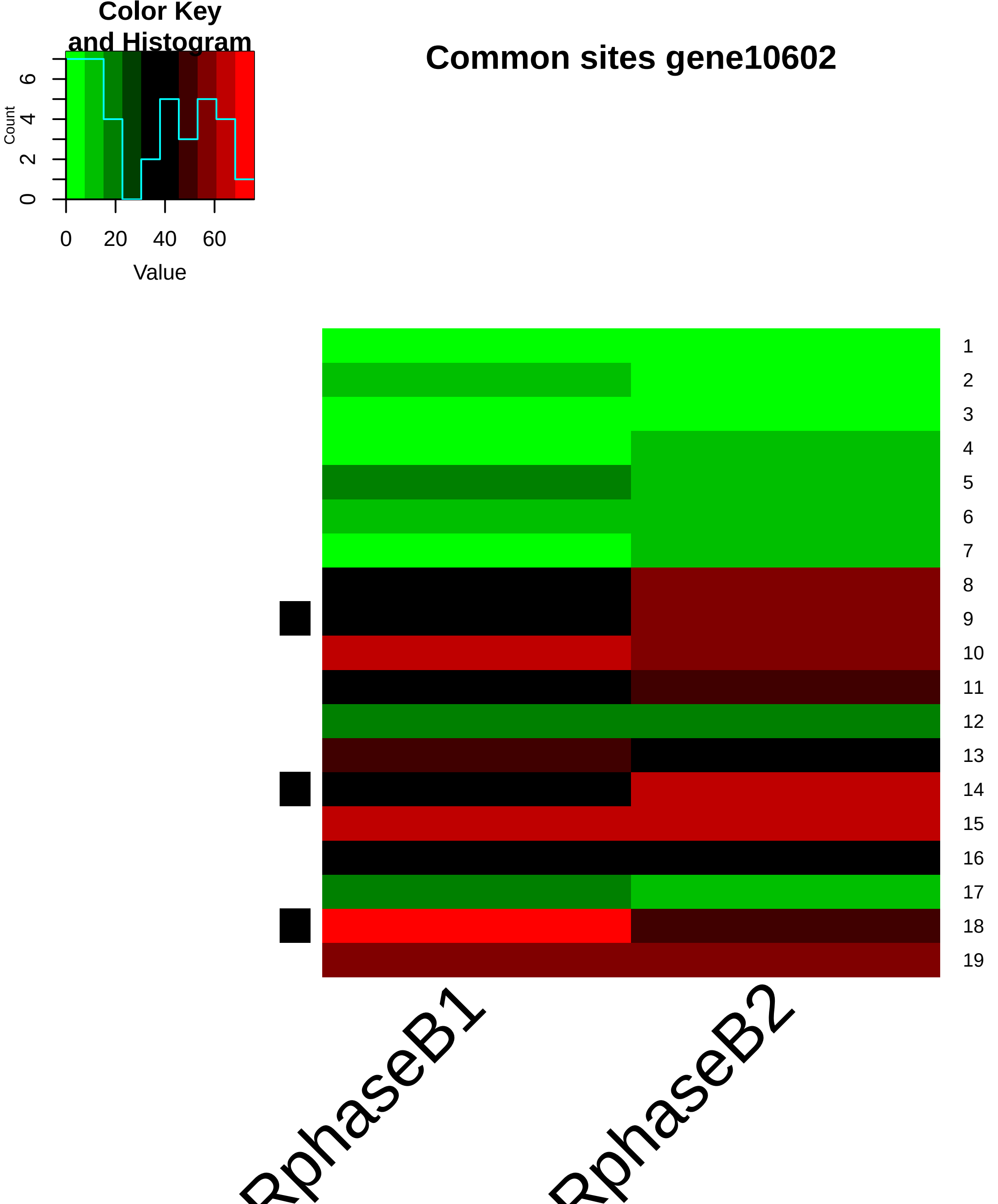
Color Key
and Histogram
Count
0
2
4
6
0
20
40
60
Value
Common sites gene10602
RphaseB1
RphaseB2

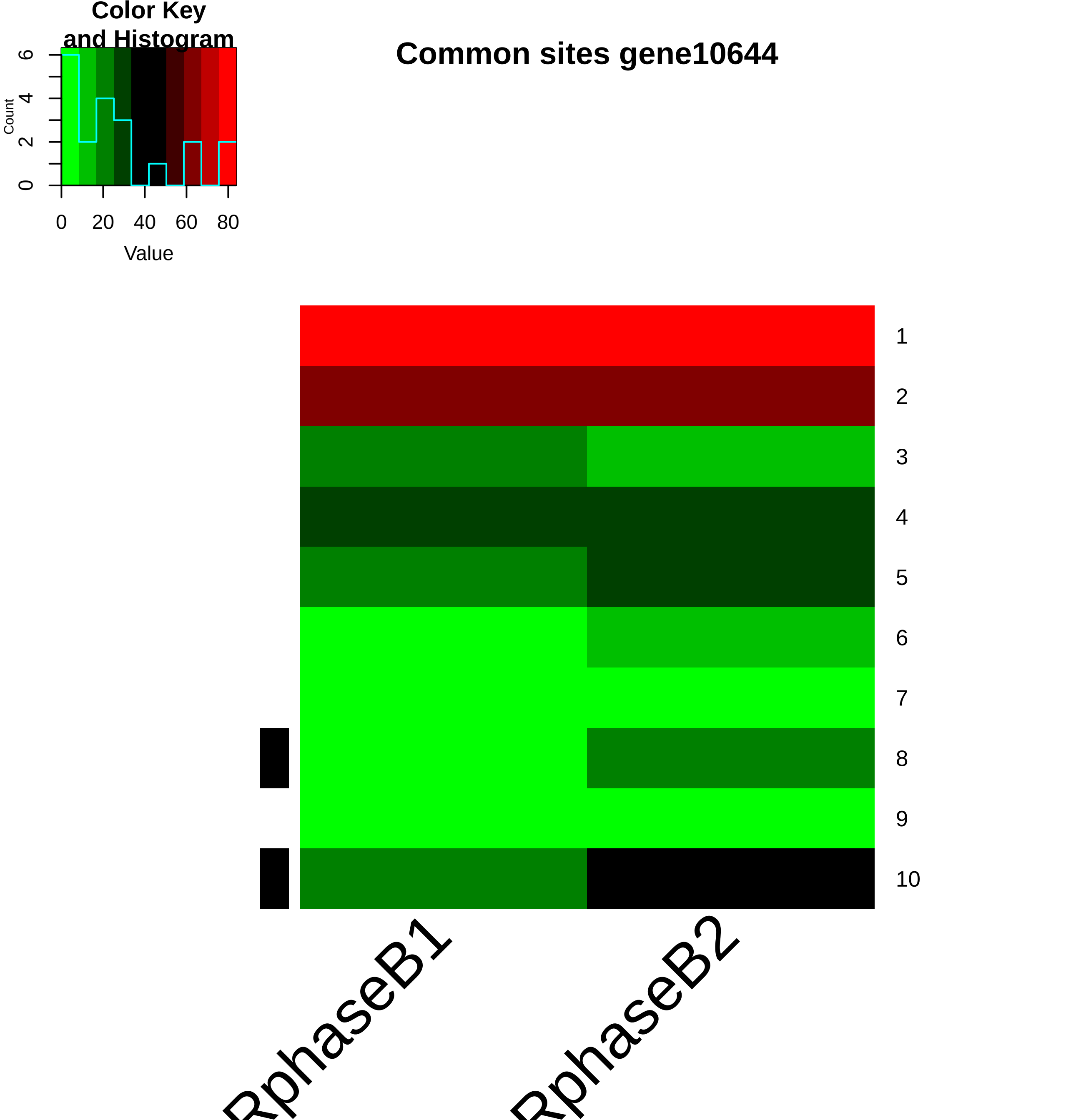

Color Key
and Histogram
Count
Value
0
20
40
60
80
0
2
4
6
Common sites gene10644
1
2
3
4
5
6
7
8
9
10
RphaseB1
RphaseB2

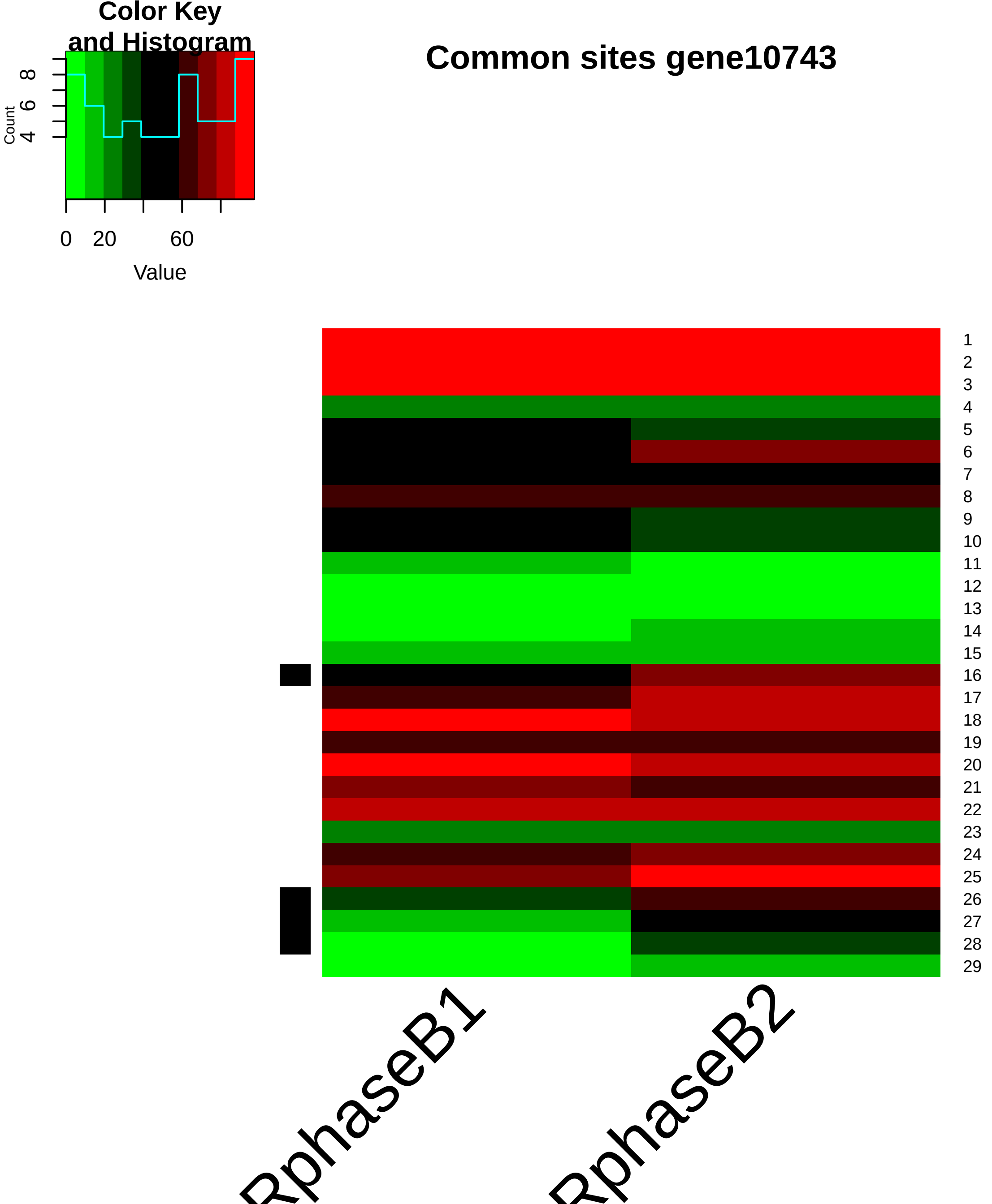
Color Key
and Histogram
Count
4
6
8
0
20
60
Value
Common sites gene10743
RphaseB1
RphaseB2

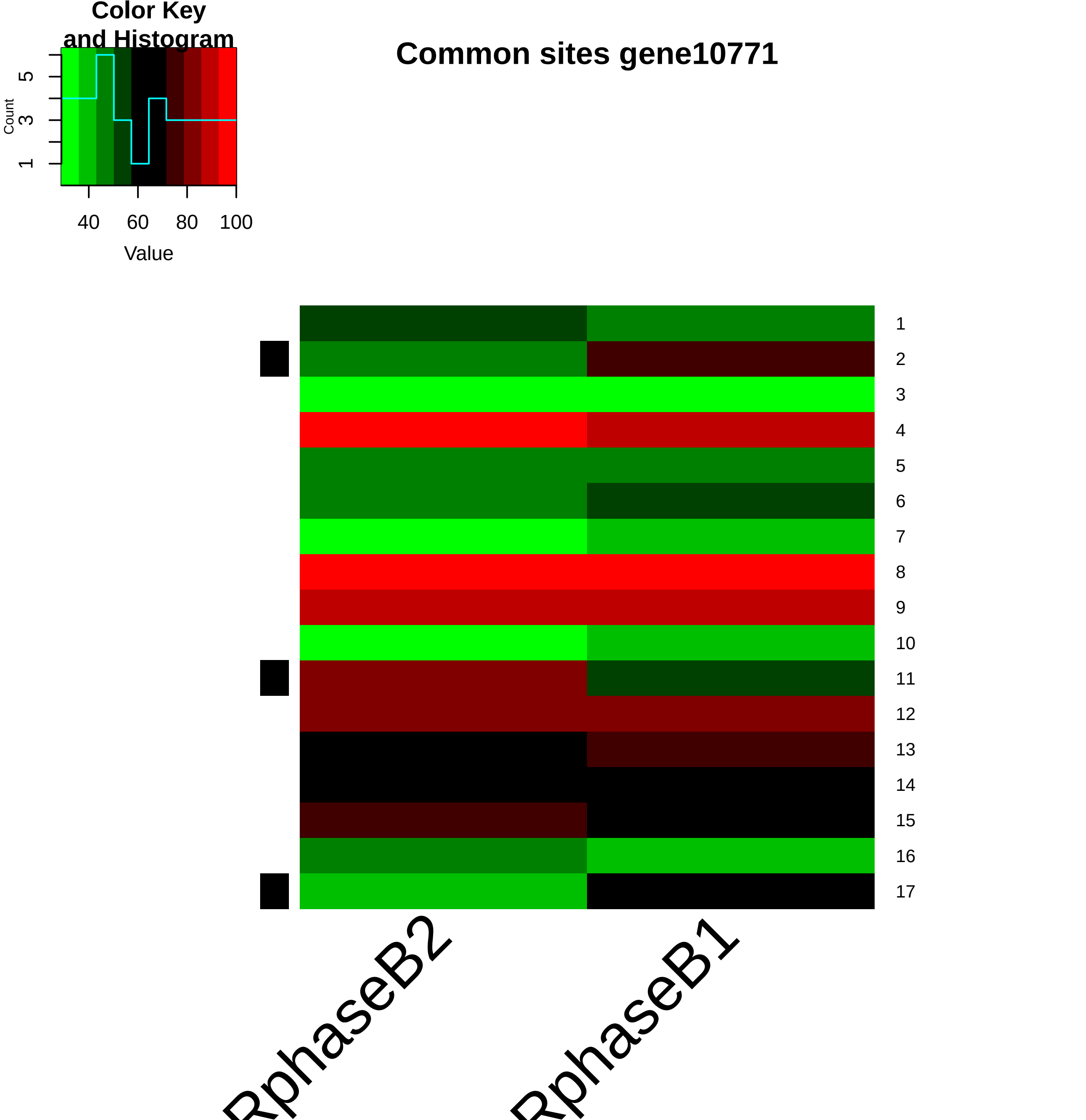

Color Key and Histogram
Count
1
3
5
40
60
80
100
Value
Common sites gene10771
RphaseB2
RphaseB1

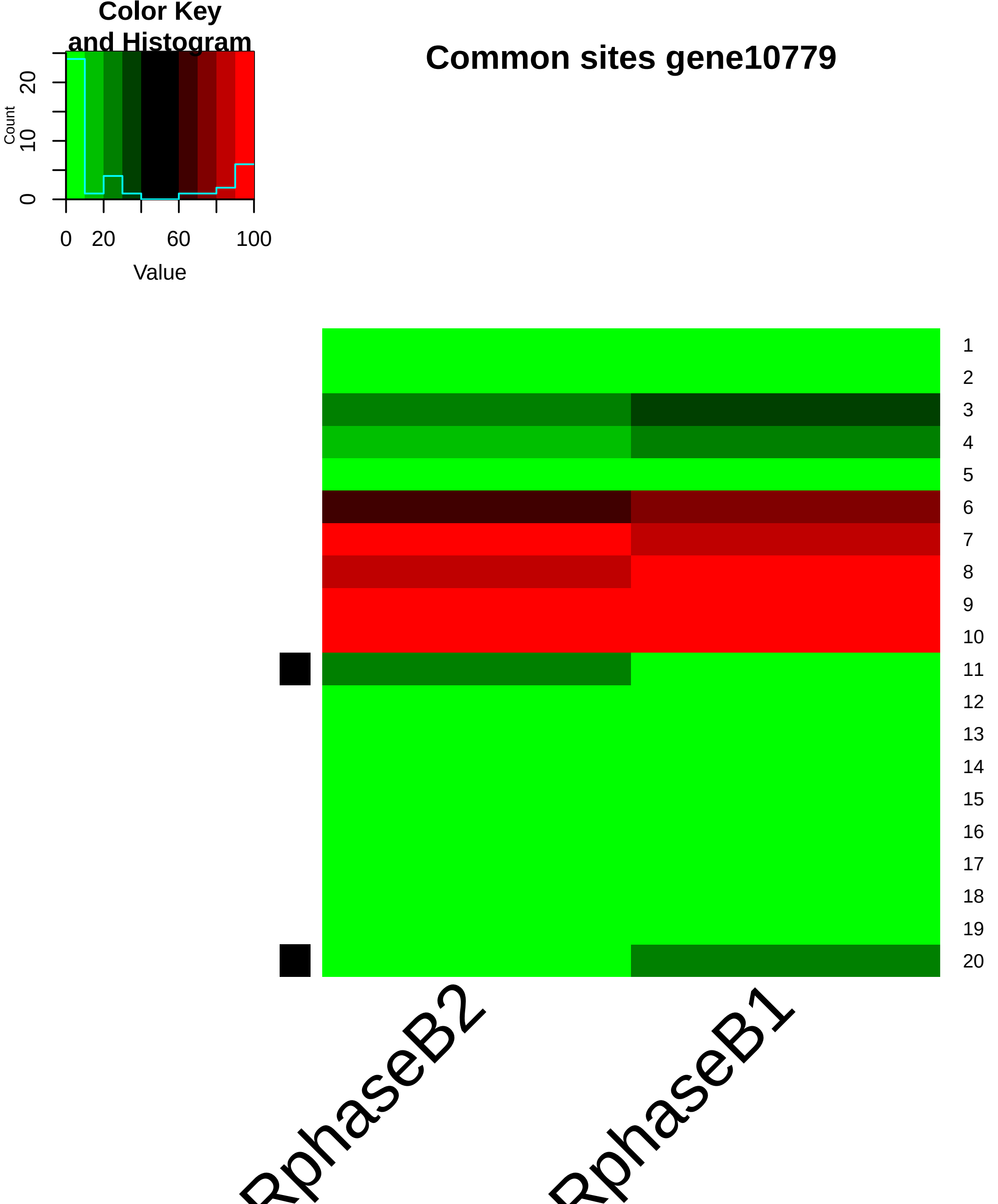

Color Key
and Histogram
Count
0
10
20
0
20
60
100
Value
Common sites gene10779
RphaseB2
RphaseB1

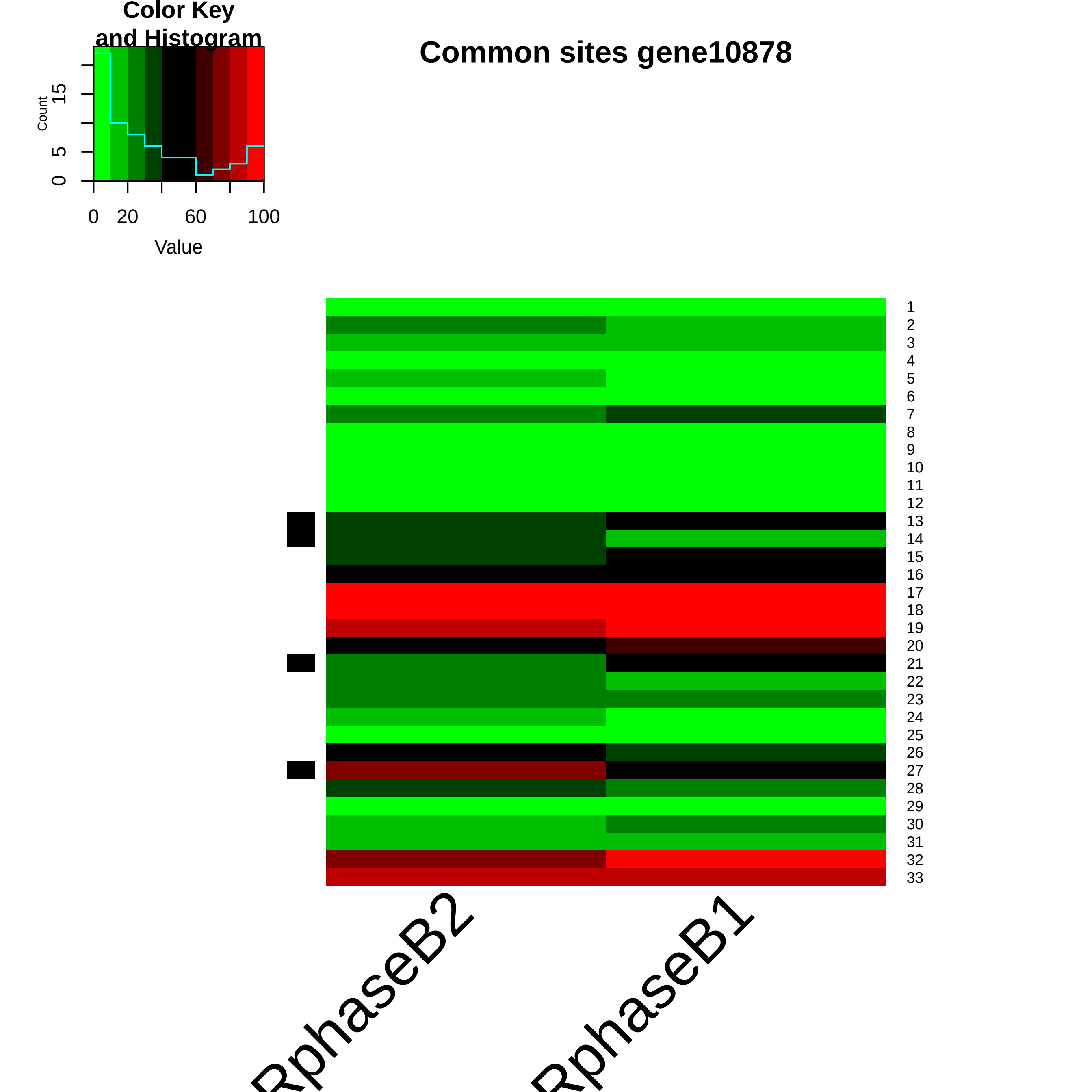
Color Key
and Histogram
Count
0
5
15
0
20
60
100
Value
Common sites gene10878
RphaseB2
RphaseB1

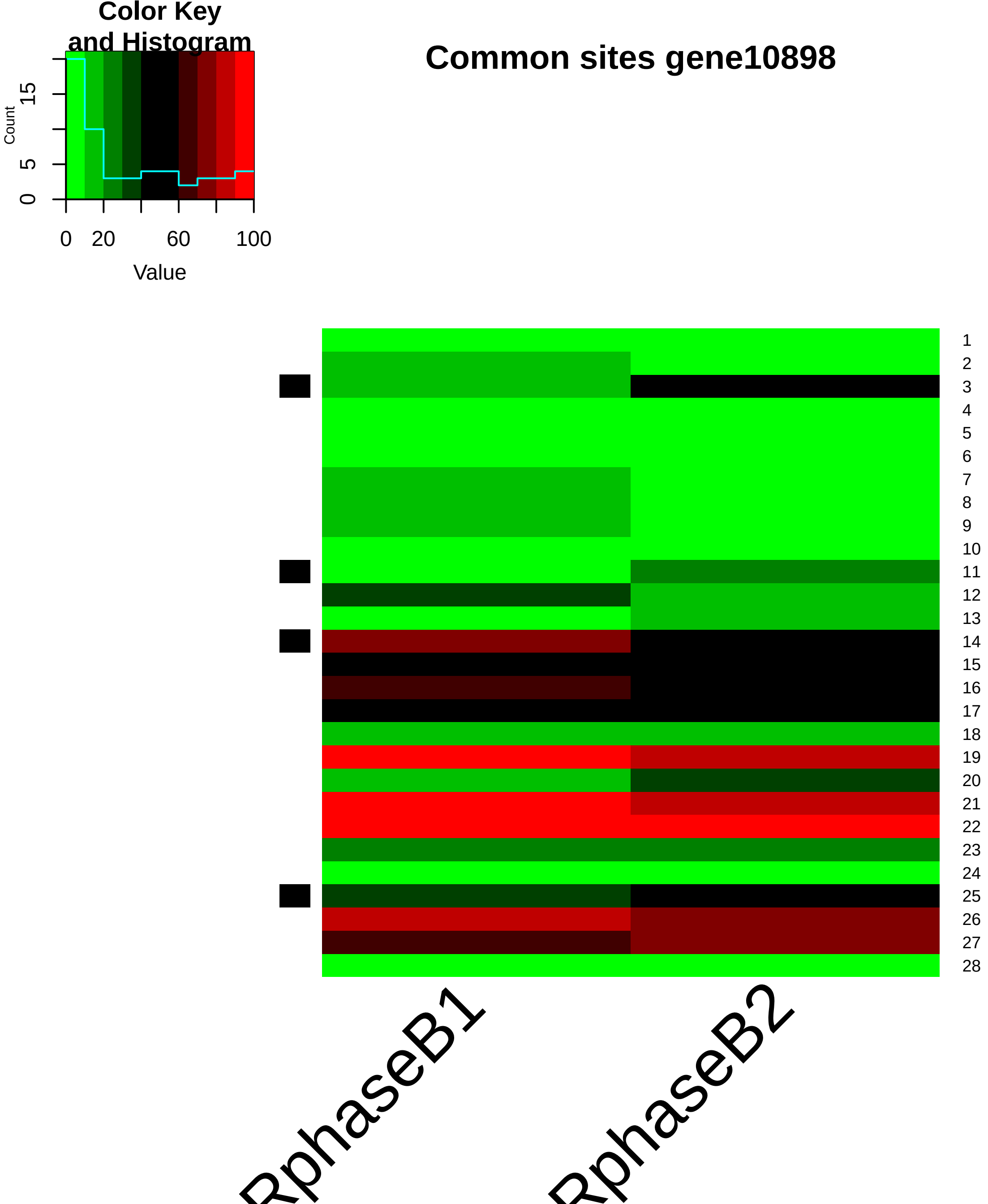
Color Key
and Histogram
Count
0
5
15
0
20
60
100
Value
Common sites gene10898
RphaseB1
RphaseB2

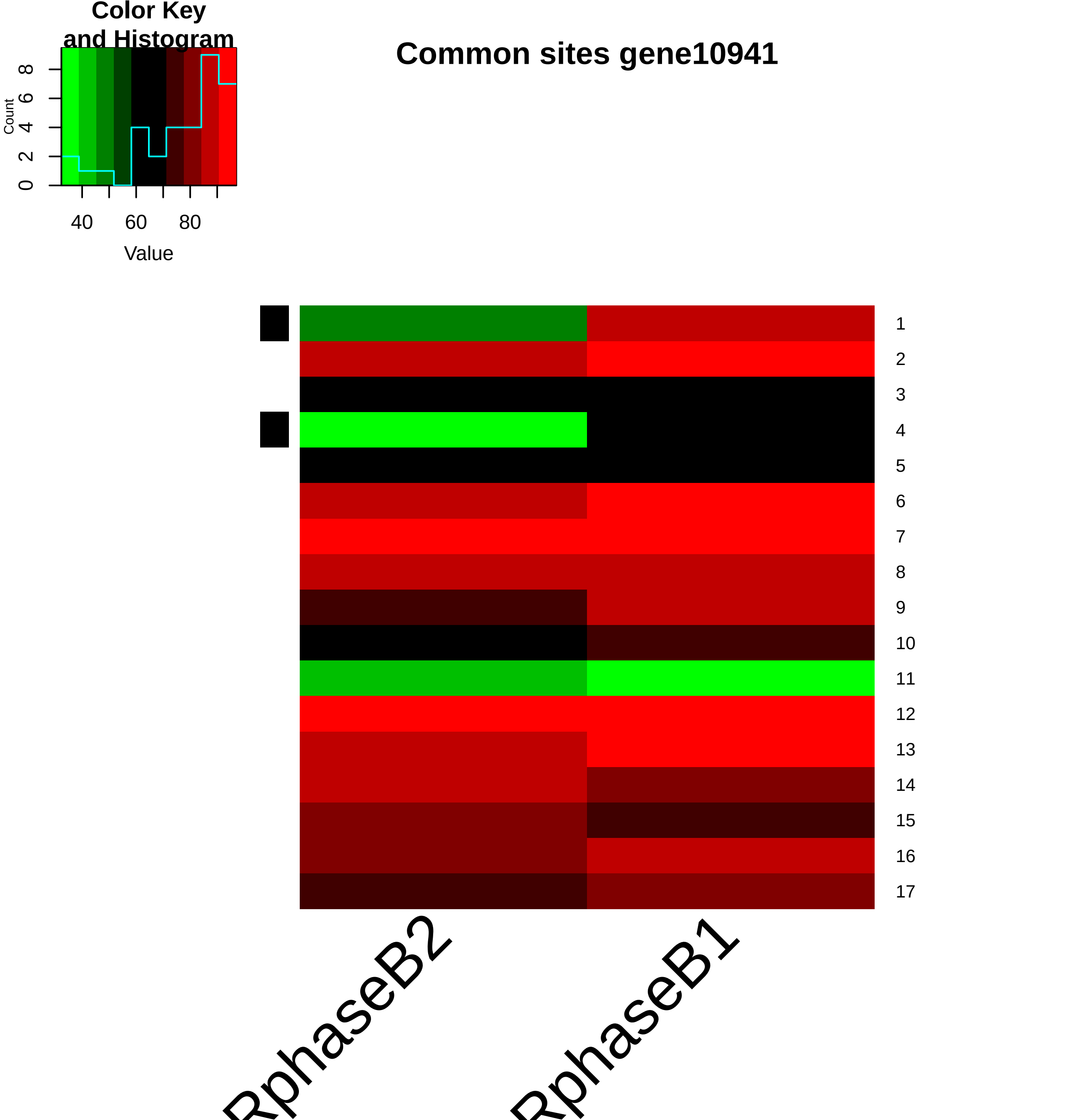
Color Key
and Histogram
Count
0 2 4 6 8
40 60 80
Value
Common sites gene10941
RphaseB2
RphaseB1

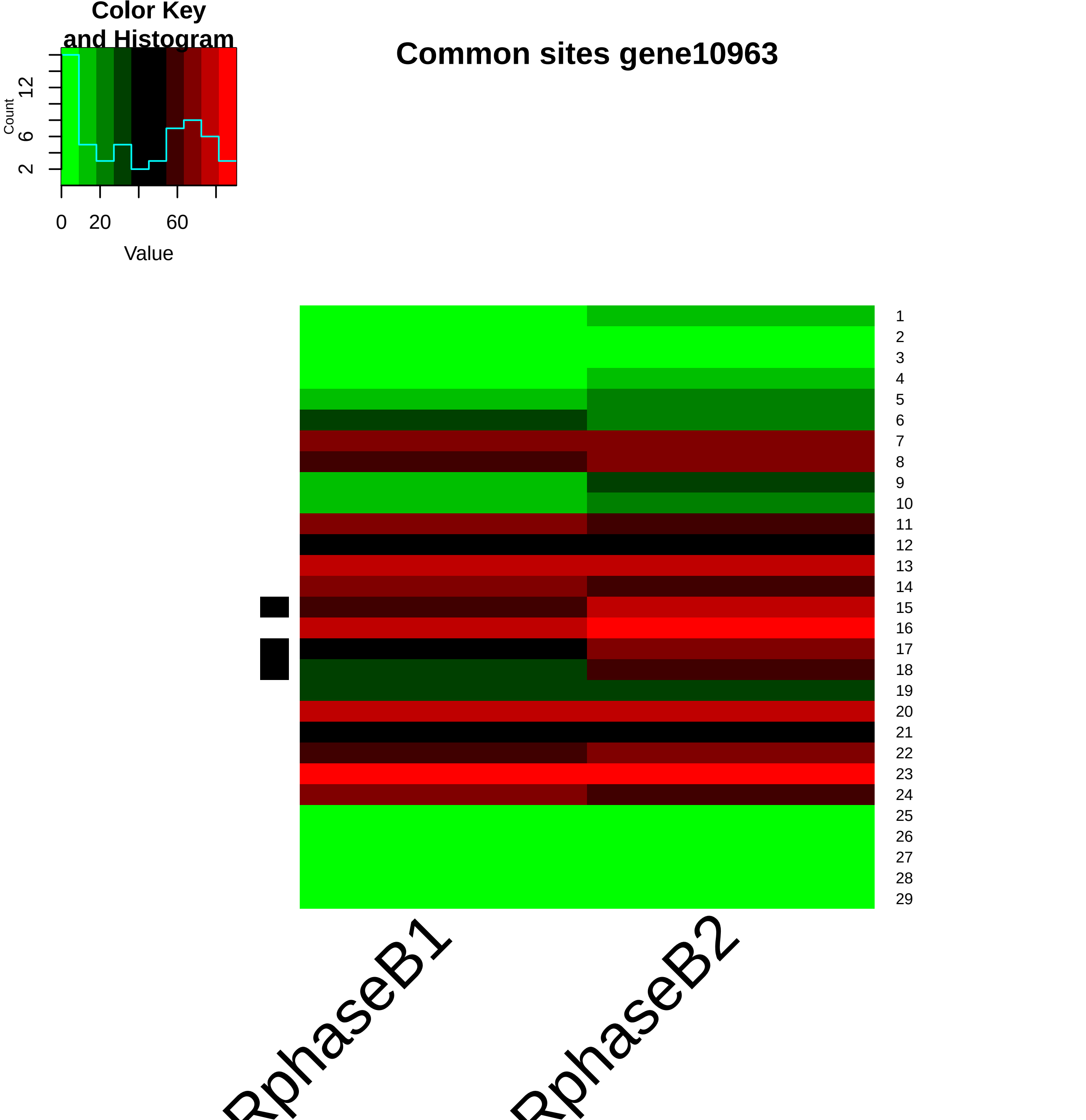
Color Key
and Histogram
Count
2
6
12
0
20
60
Value
Common sites gene10963
RphaseB1
RphaseB2

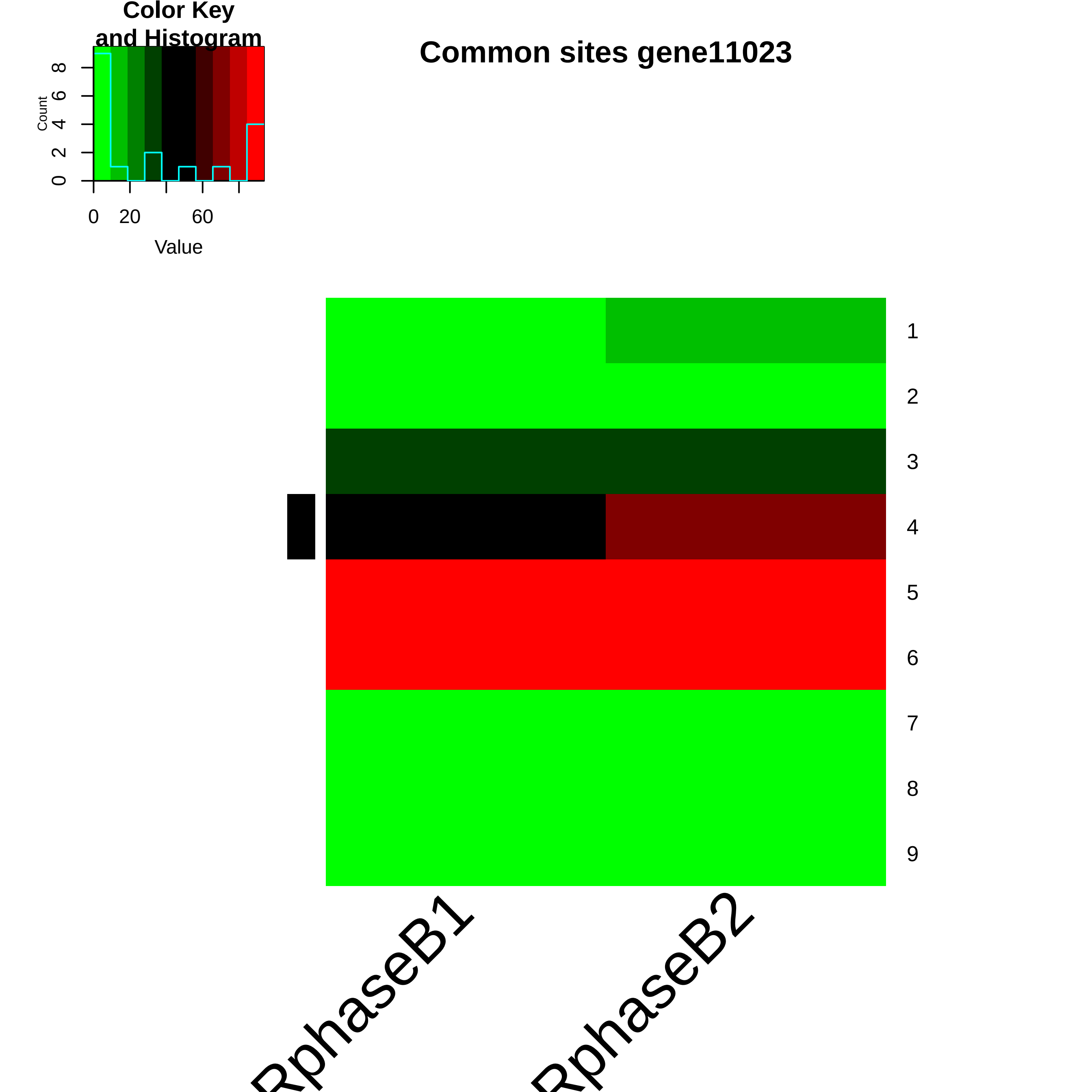

Color Key
and Histogram
Count
0
2
4
6
8
0
20
60
Value
Common sites gene11023
1
2
3
4
5
6
7
8
9
RphaseB1
RphaseB2

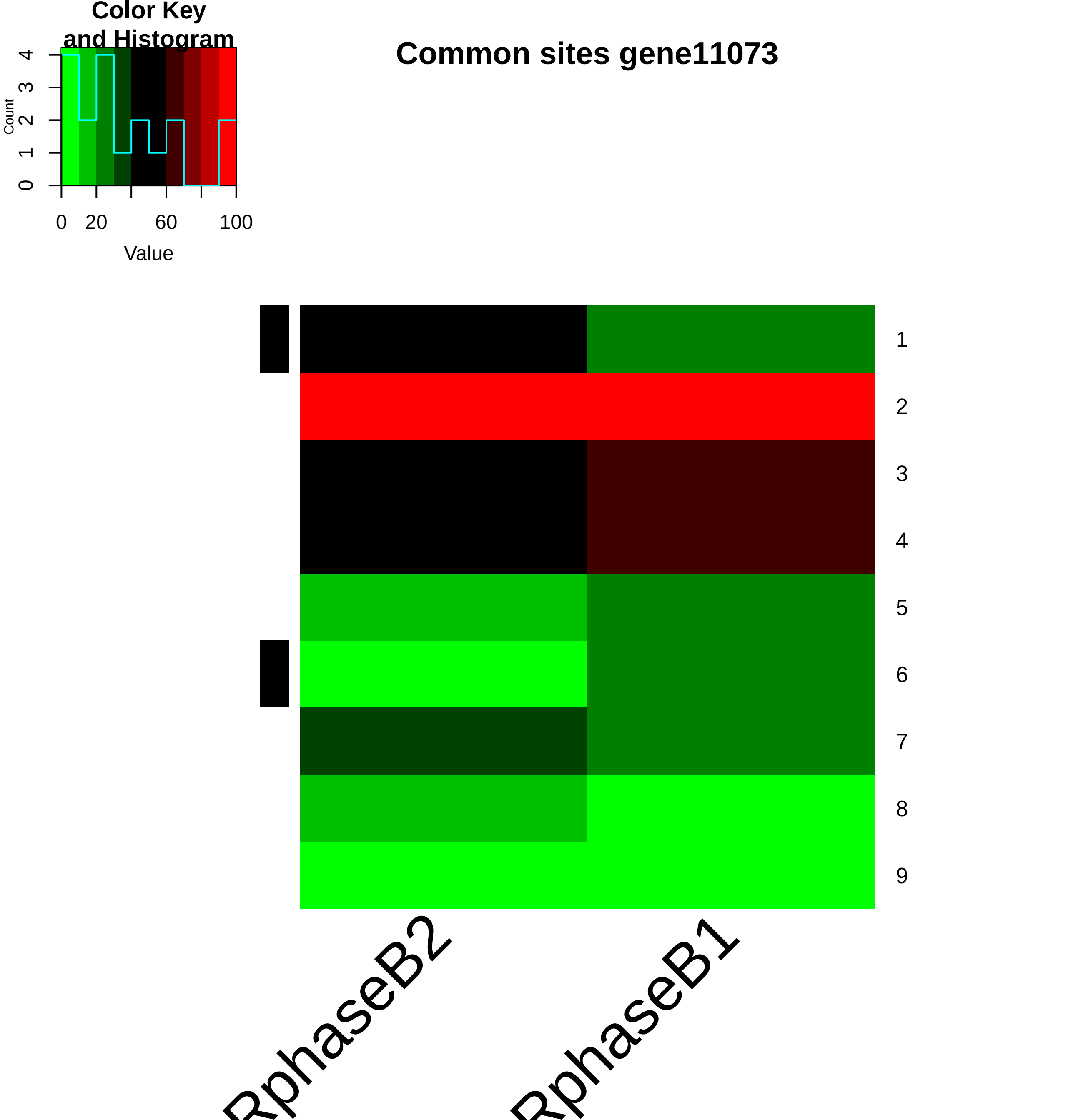

Color Key
and Histogram
Count
0
1
2
3
4
0
20
60
100
Value
Common sites gene11073
1
2
3
4
5
6
7
8
9
RphaseB2
RphaseB1

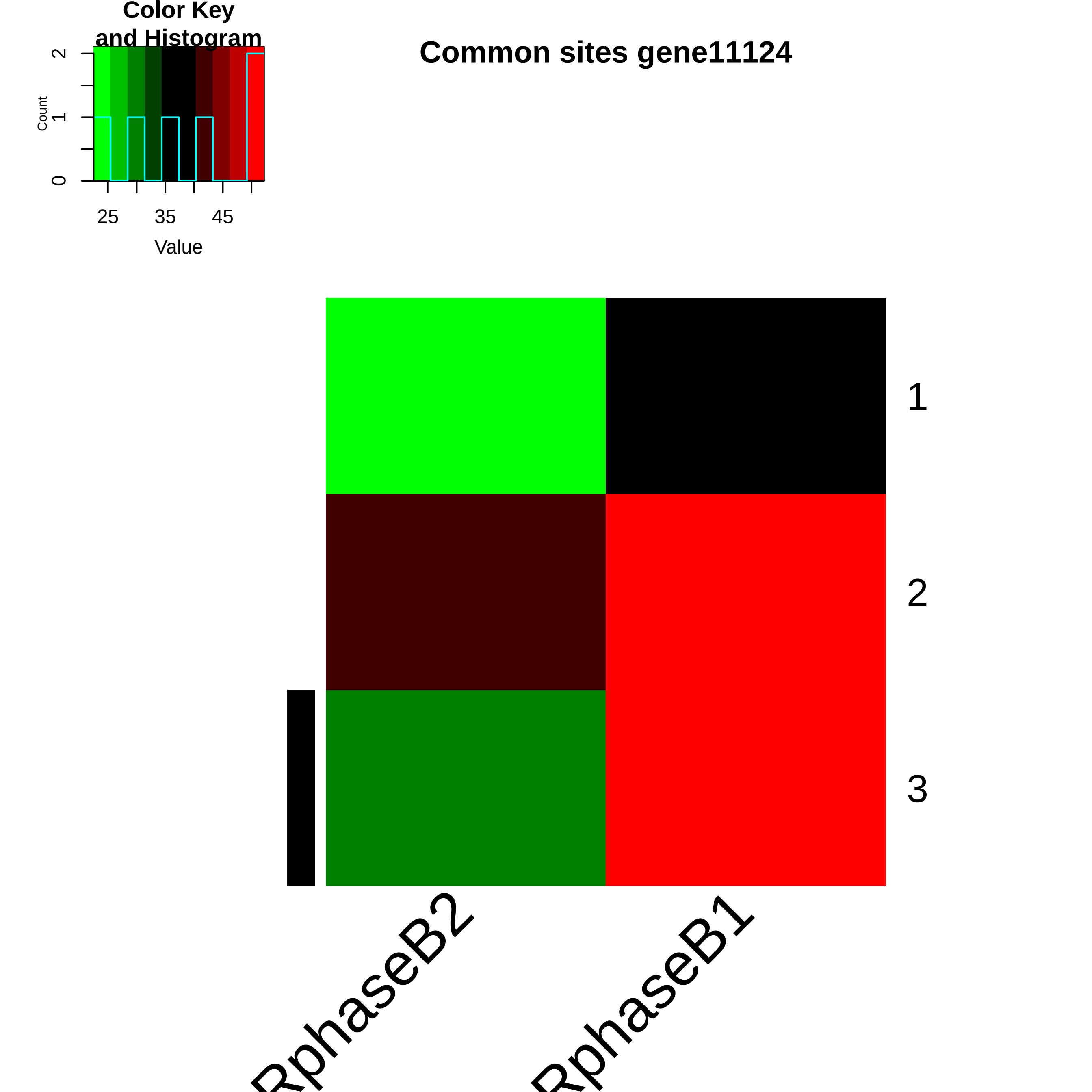

Color Key
and Histogram
Count
Value
25
35
45
0
1
2
Common sites gene11124
1
2
3
RphaseB2
RphaseB1

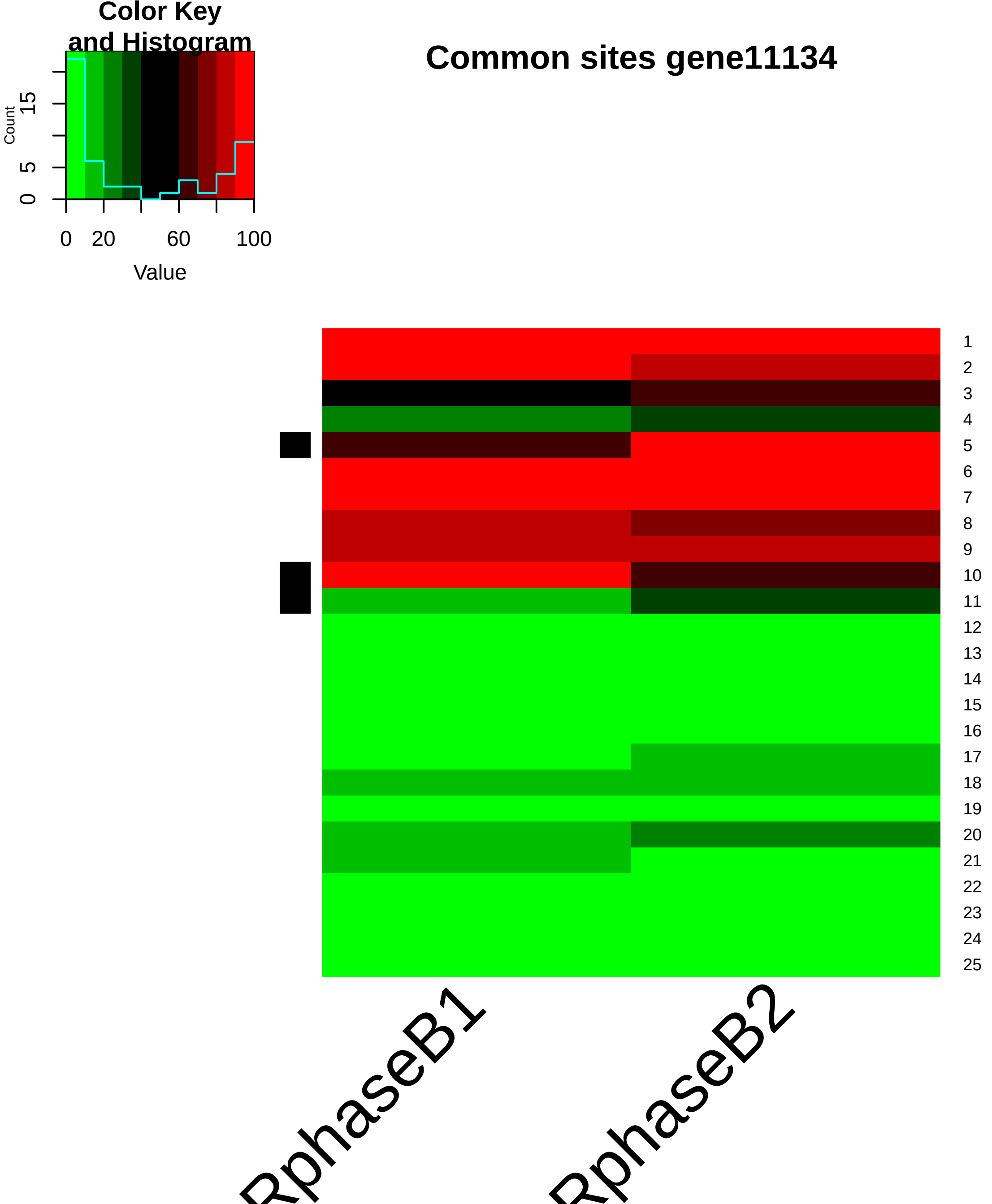
Color Key
and Histogram
Common sites gene11134
Count
0
5
15
0
20
60
100
Value
RphaseB1
RphaseB2

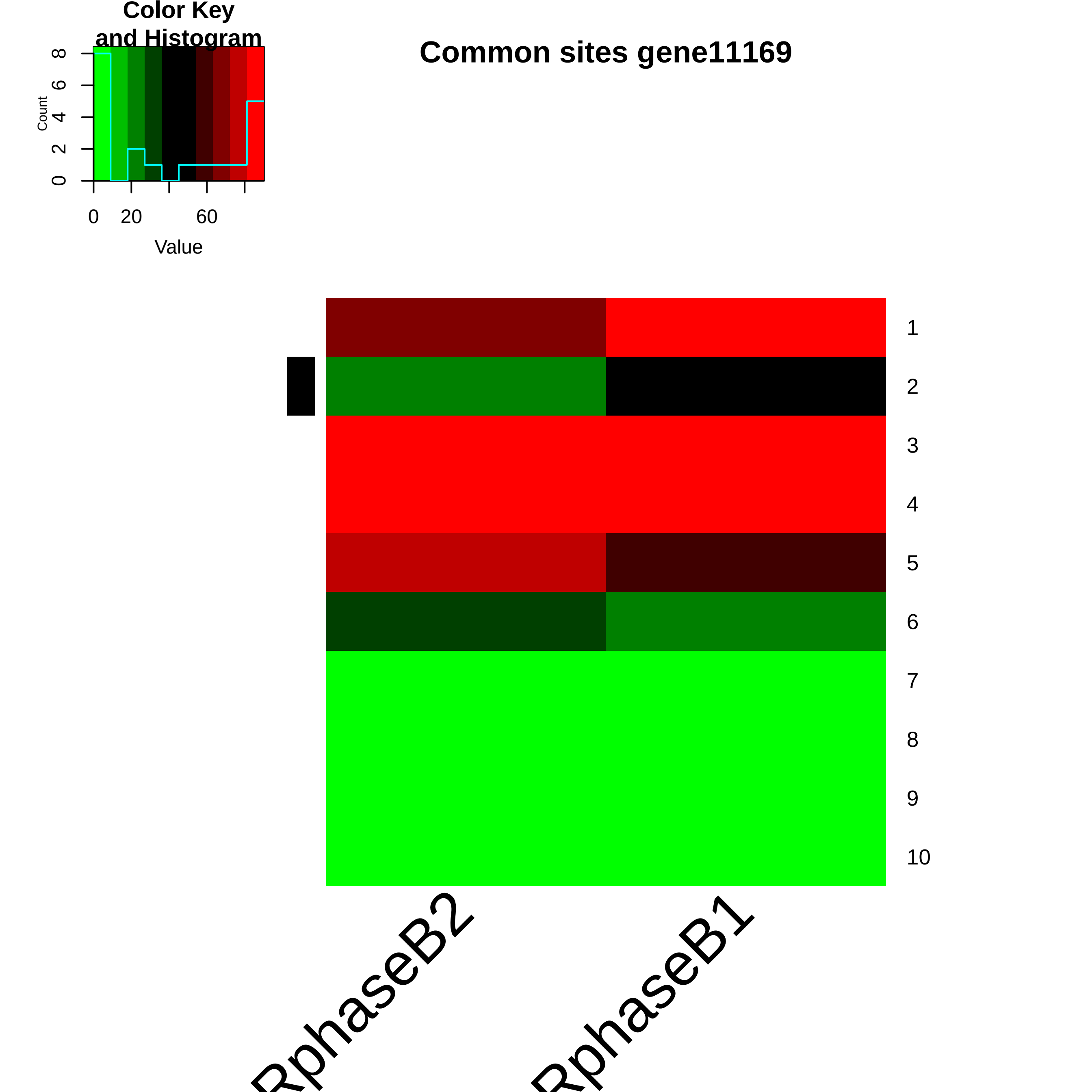
Color Key
and Histogram
Count
0
2
4
6
8
0
20
60
Value
Common sites gene11169
1
2
3
4
5
6
7
8
9
10
RphaseB2
RphaseB1

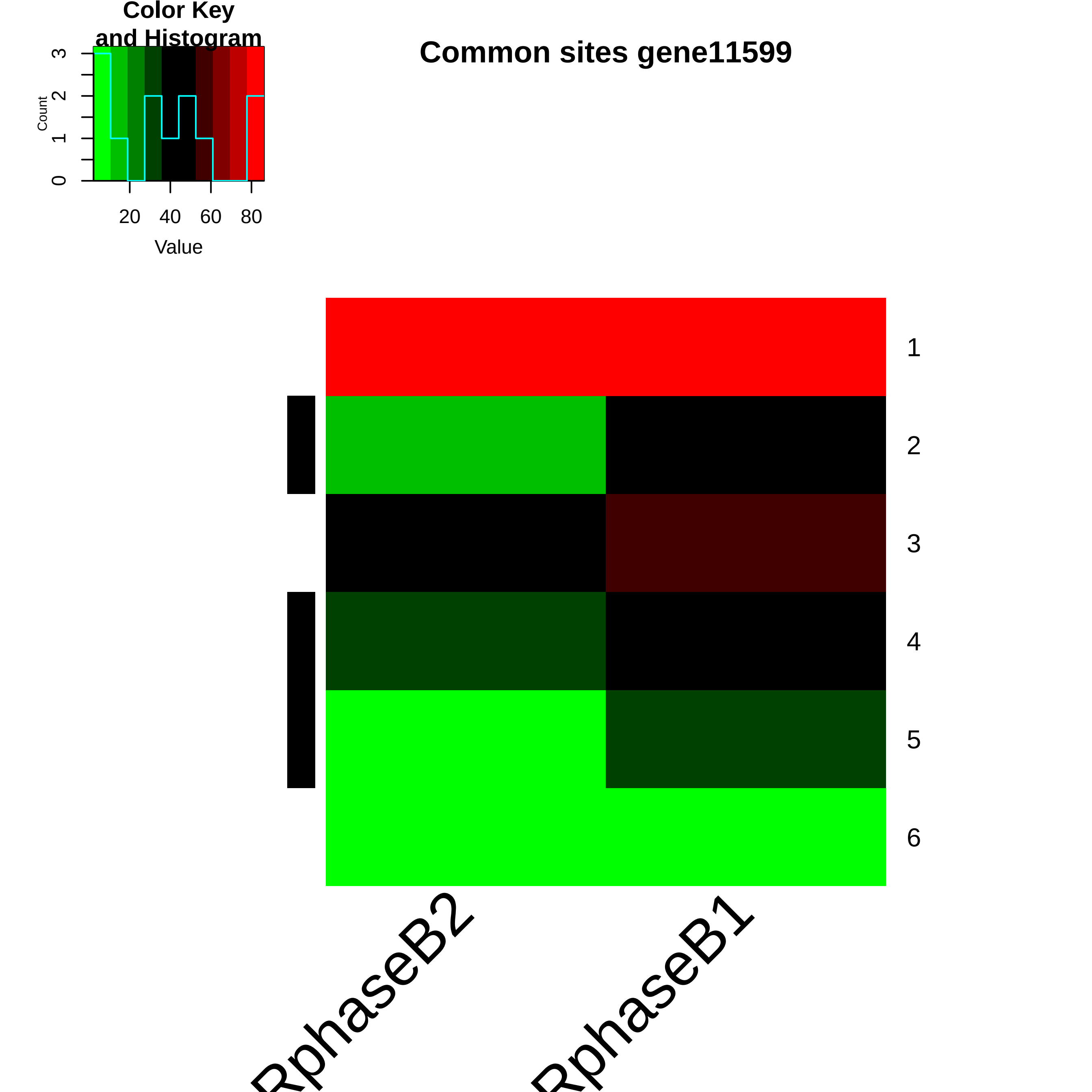

Color Key
and Histogram
Count
0
1
2
3
20
40
60
80
Value
Common sites gene11599
1
2
3
4
5
6
RphaseB2
RphaseB1

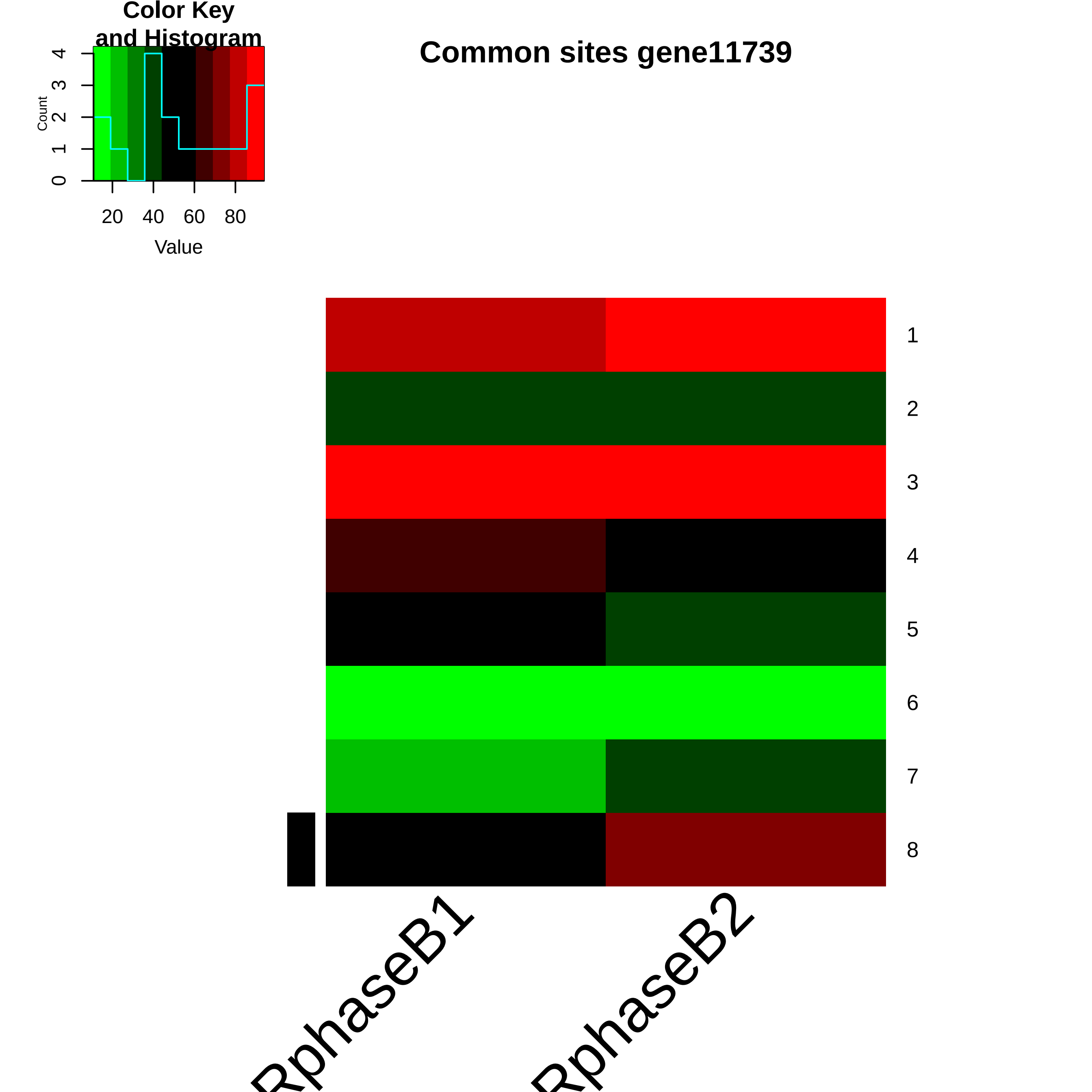
Color Key
and Histogram
Count
0
1
2
3
4
20
40
60
80
Value
Common sites gene11739
1
2
3
4
5
6
7
8
RphaseB1
RphaseB2

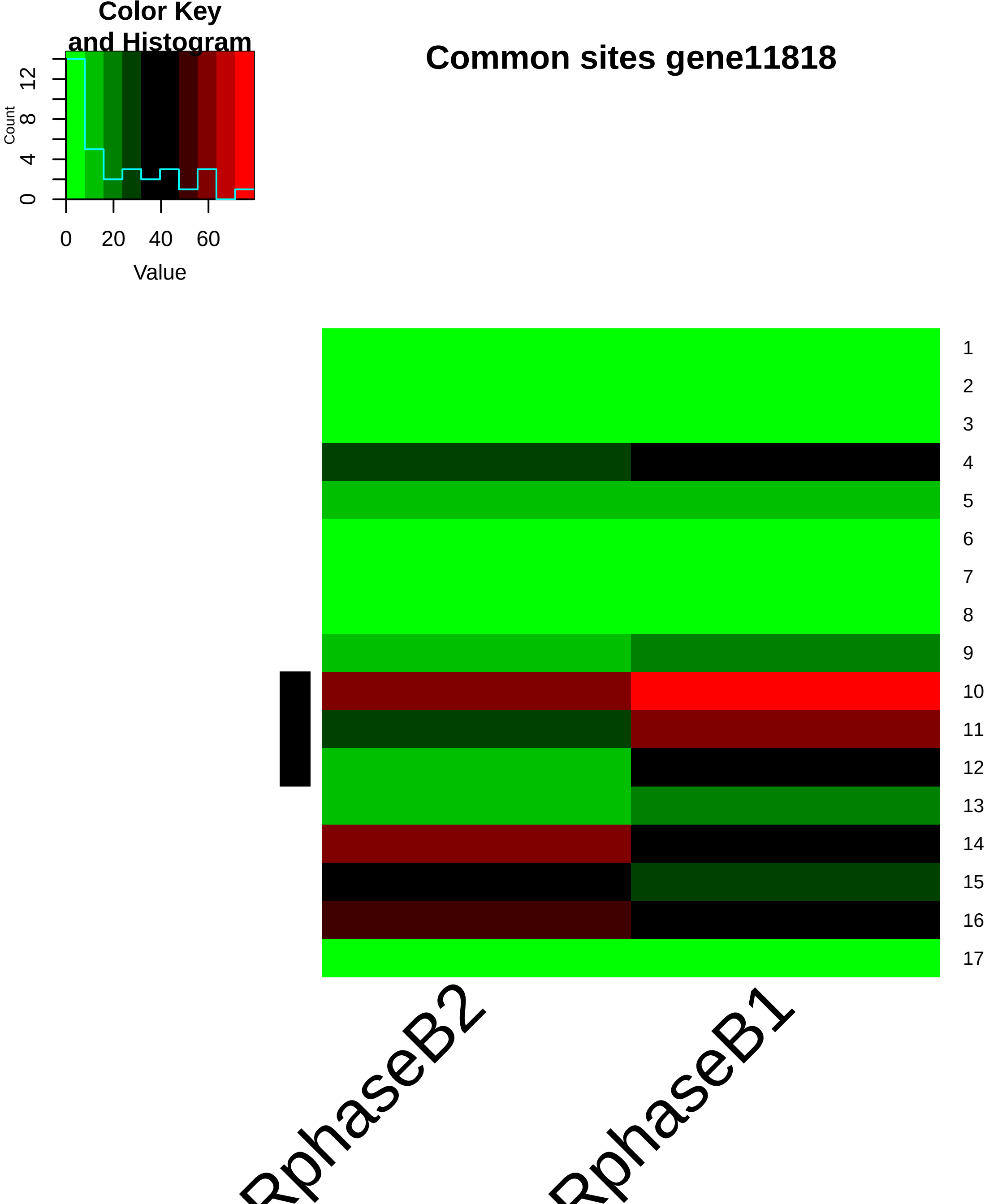
Color Key
and Histogram
Count
0
4
8
12
0
20
40
60
Value
Common sites gene11818
RphaseB2
RphaseB1

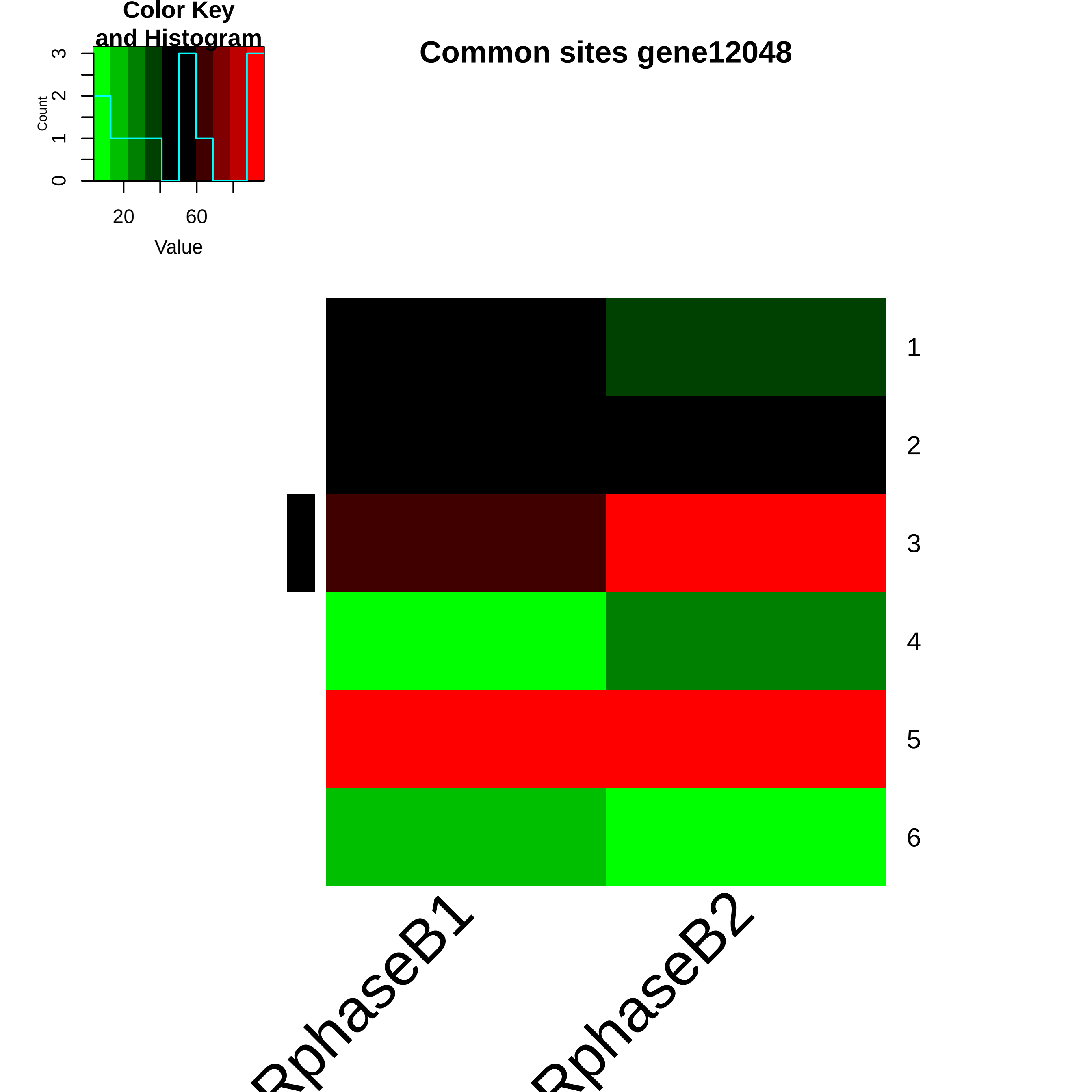

Color Key
and Histogram
Count
0
1
2
3
20
60
Value
Common sites gene12048
1
2
3
4
5
6
RphaseB1
RphaseB2

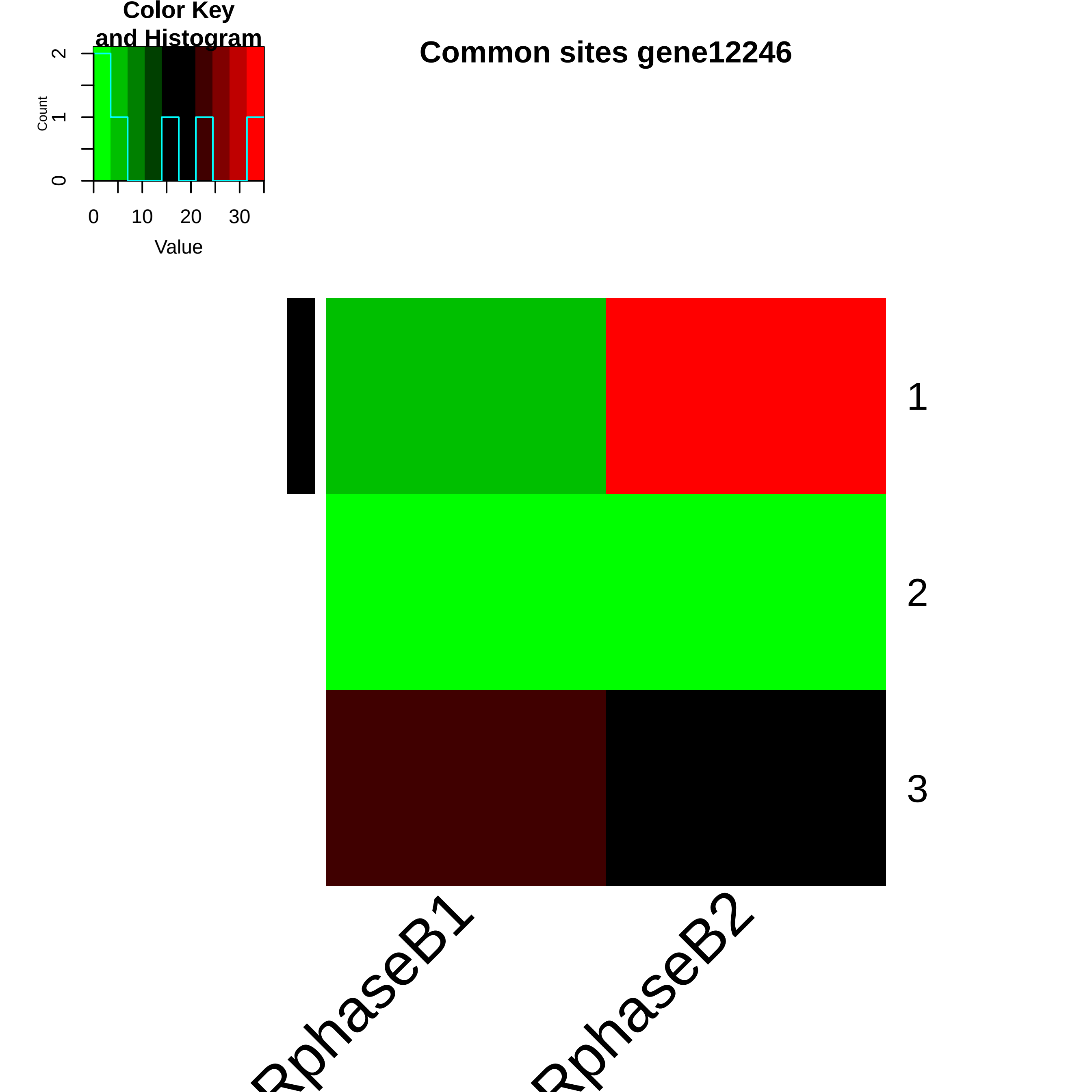

Color Key
and Histogram
Count
0
1
2
0
10
20
30
Value
Common sites gene12246
1
2
3
RphaseB1
RphaseB2

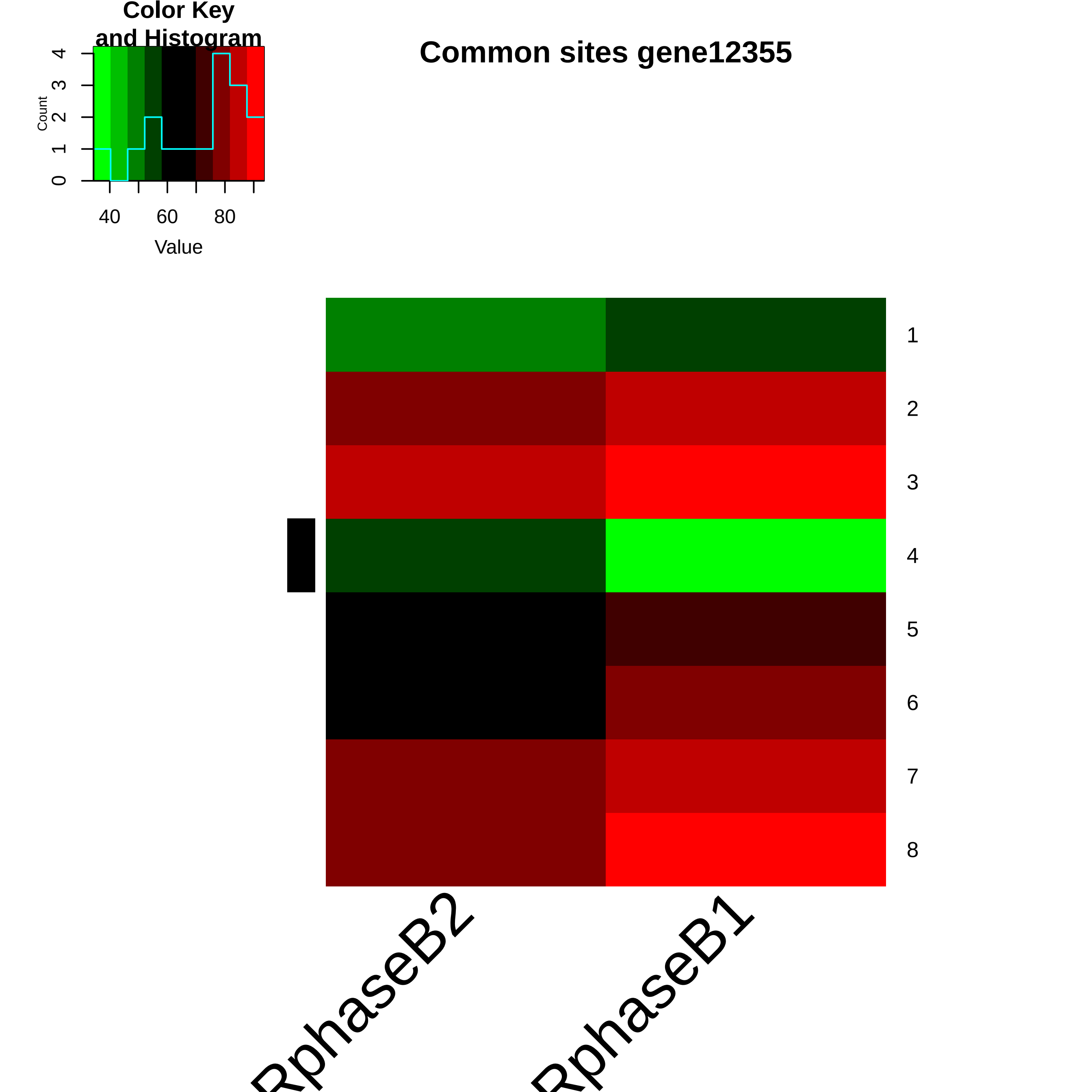
Color Key
and Histogram
Count
0
1
2
3
4
40
60
80
Value
Common sites gene12355
1
2
3
4
5
6
7
8
RphaseB2
RphaseB1

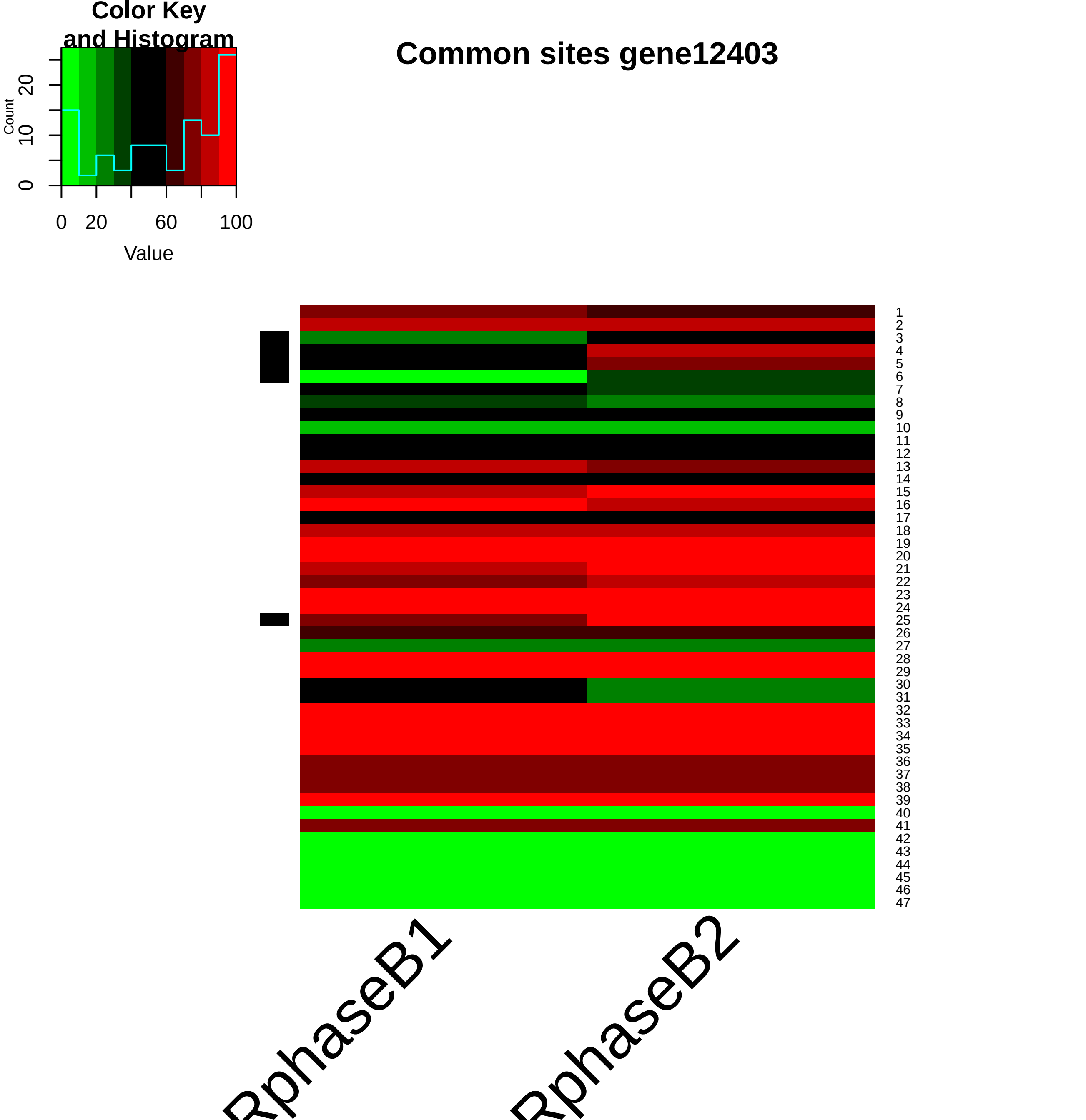
Color Key
and Histogram
Count
0
10
20
0
20
60
100
Value
Common sites gene12403
RphaseB1
RphaseB2

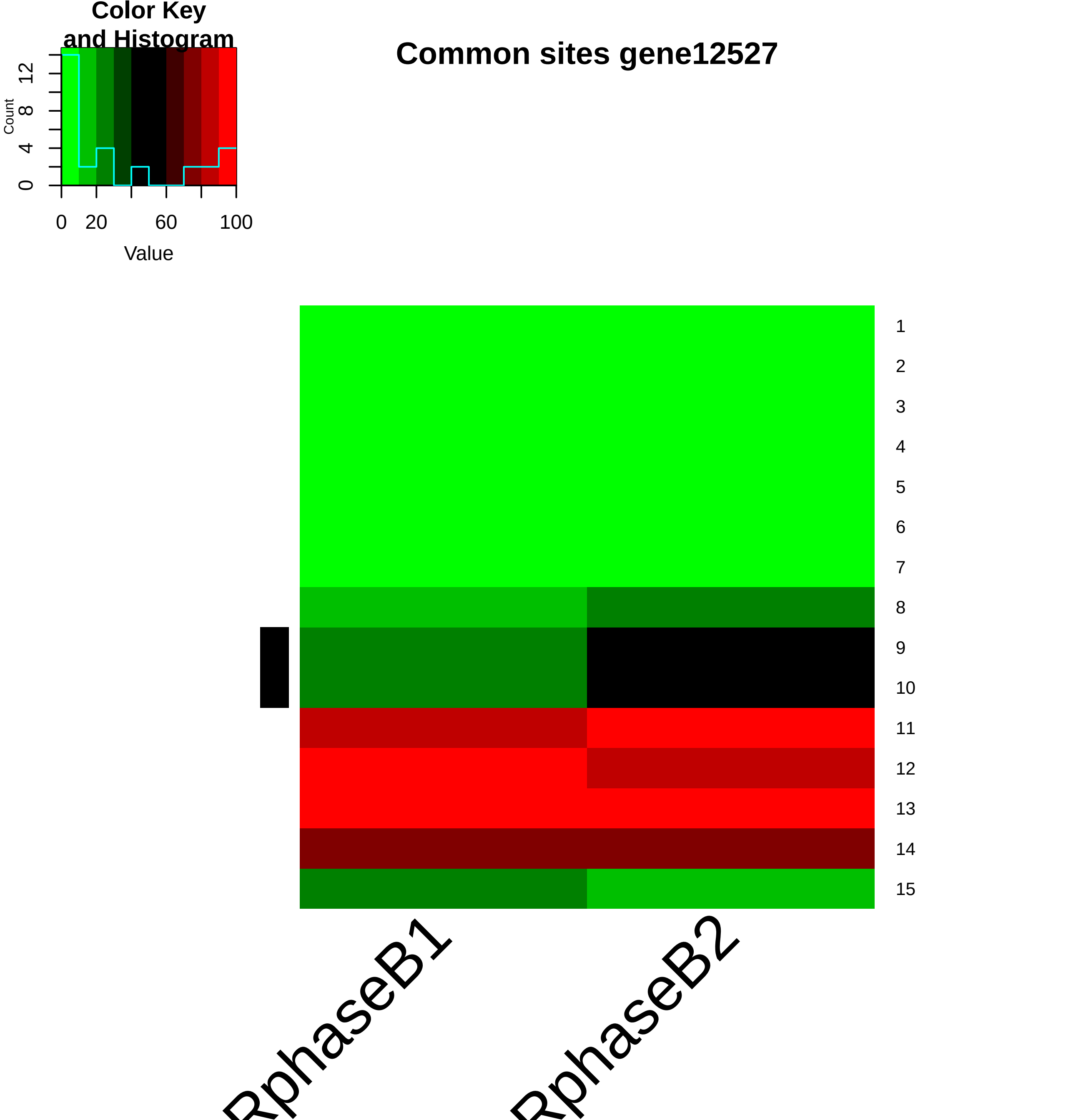
Color Key
and Histogram
Count
0
4
8
12
0
20
60
100
Value
Common sites gene12527
RphaseB1
RphaseB2

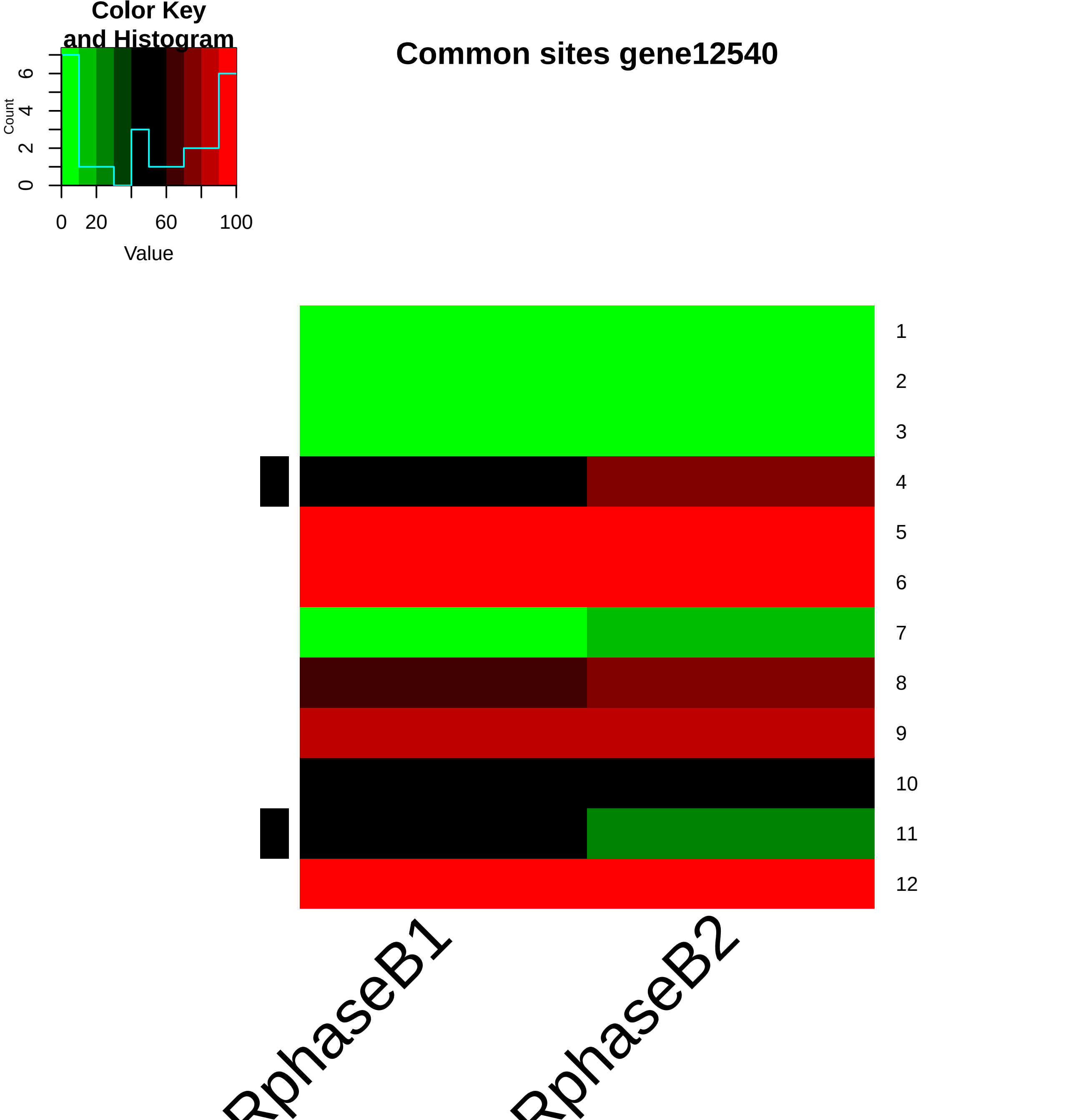
Color Key
and Histogram
Count
0
2
4
6
0
20
60
100
Value
Common sites gene12540
1
2
3
4
5
6
7
8
9
10
11
12
RphaseB1
RphaseB2

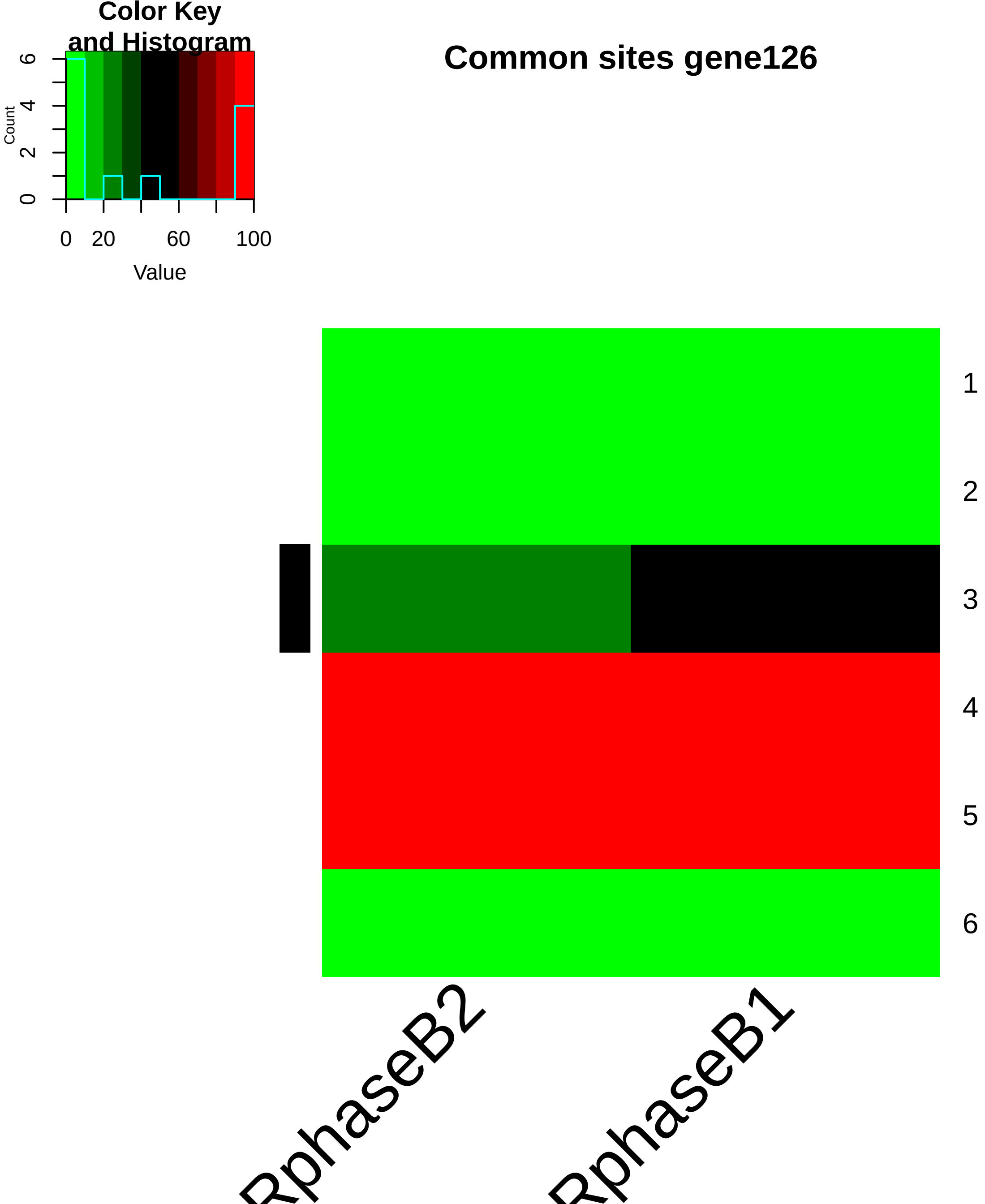
Color Key
and Histogram
Count
0
2
4
6
0
20
60
100
Value
Common sites gene126
1
2
3
4
5
6
RphaseB2
RphaseB1

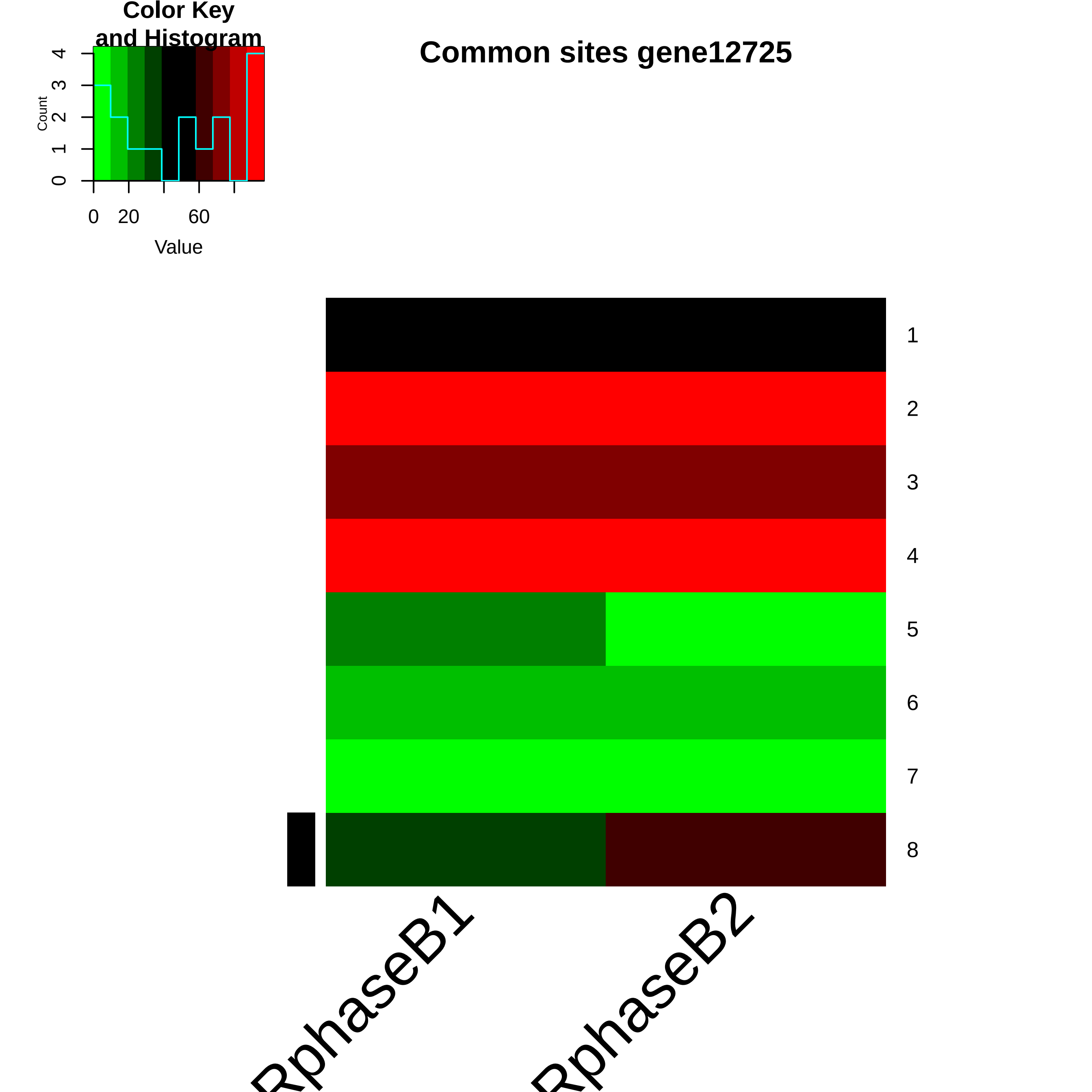
Color Key
and Histogram
Count
0
1
2
3
4
0
20
60
Value
Common sites gene12725
1
2
3
4
5
6
7
8
RphaseB1
RphaseB2

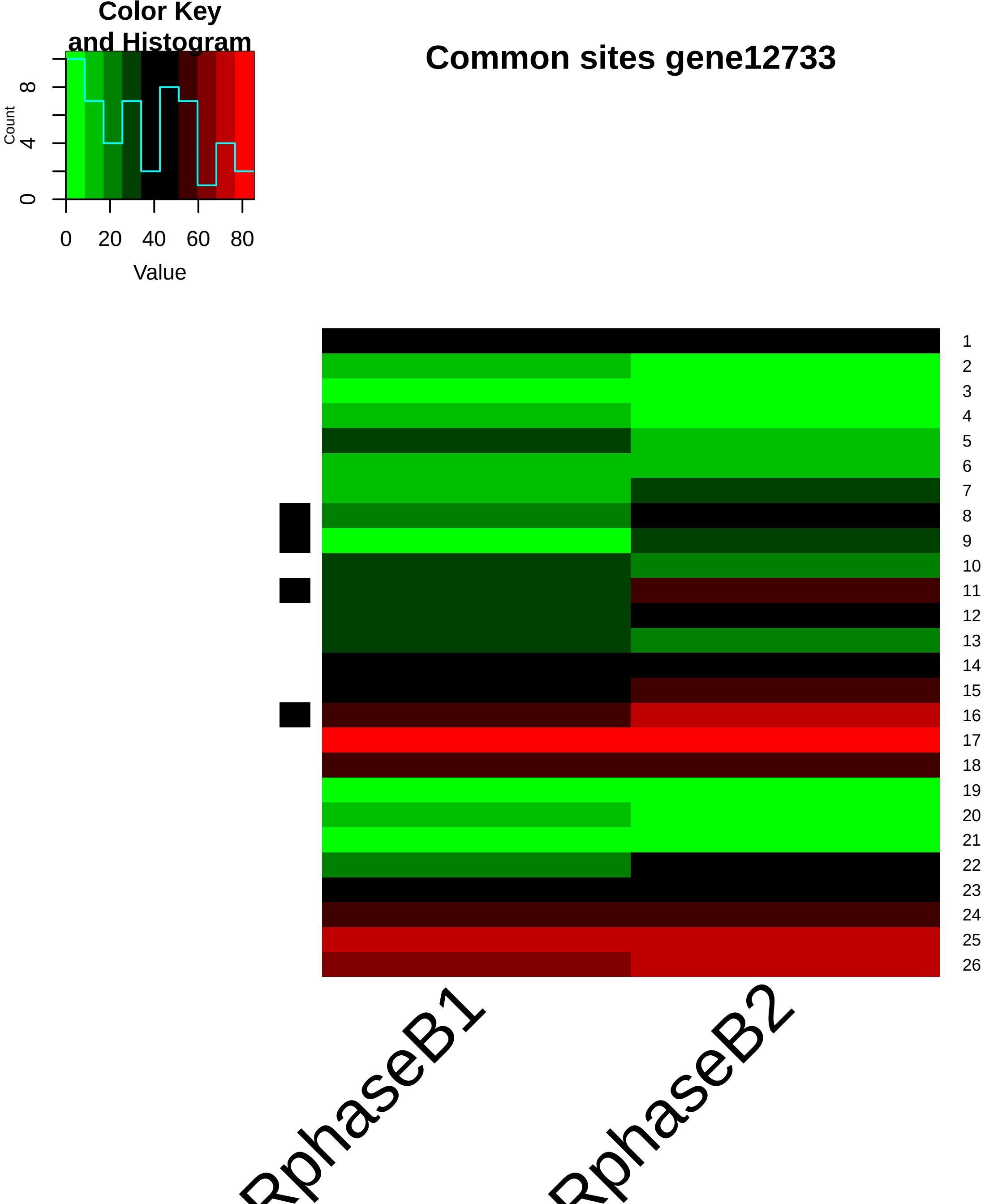

Color Key
and Histogram
Count
0
4
8
0
20
40
60
80
Value
Common sites gene12733
RphaseB1
RphaseB2

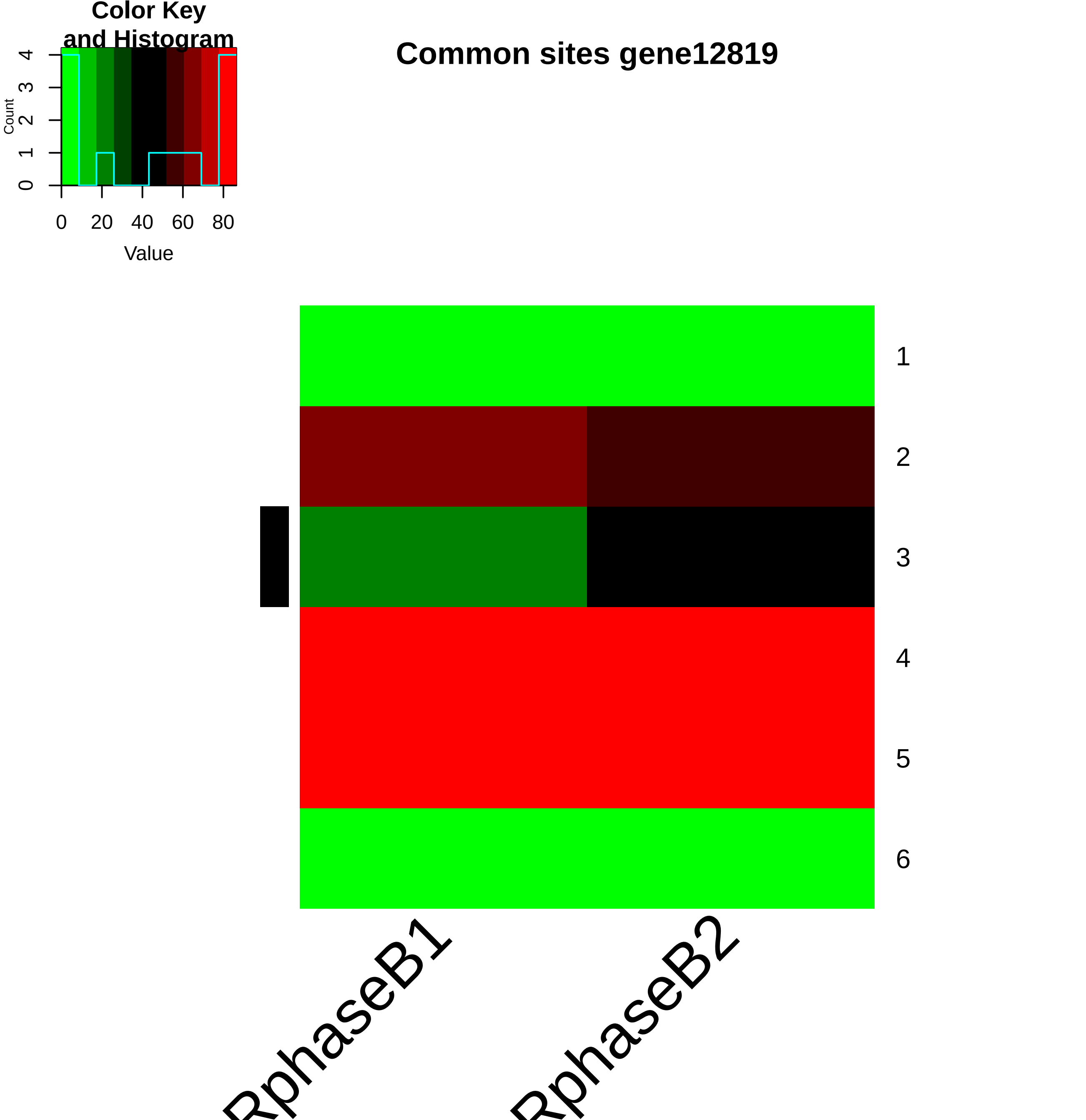

Color Key
and Histogram
Count
0
1
2
3
4
0
20
40
60
80
Value
Common sites gene12819
1
2
3
4
5
6
RphaseB1
RphaseB2

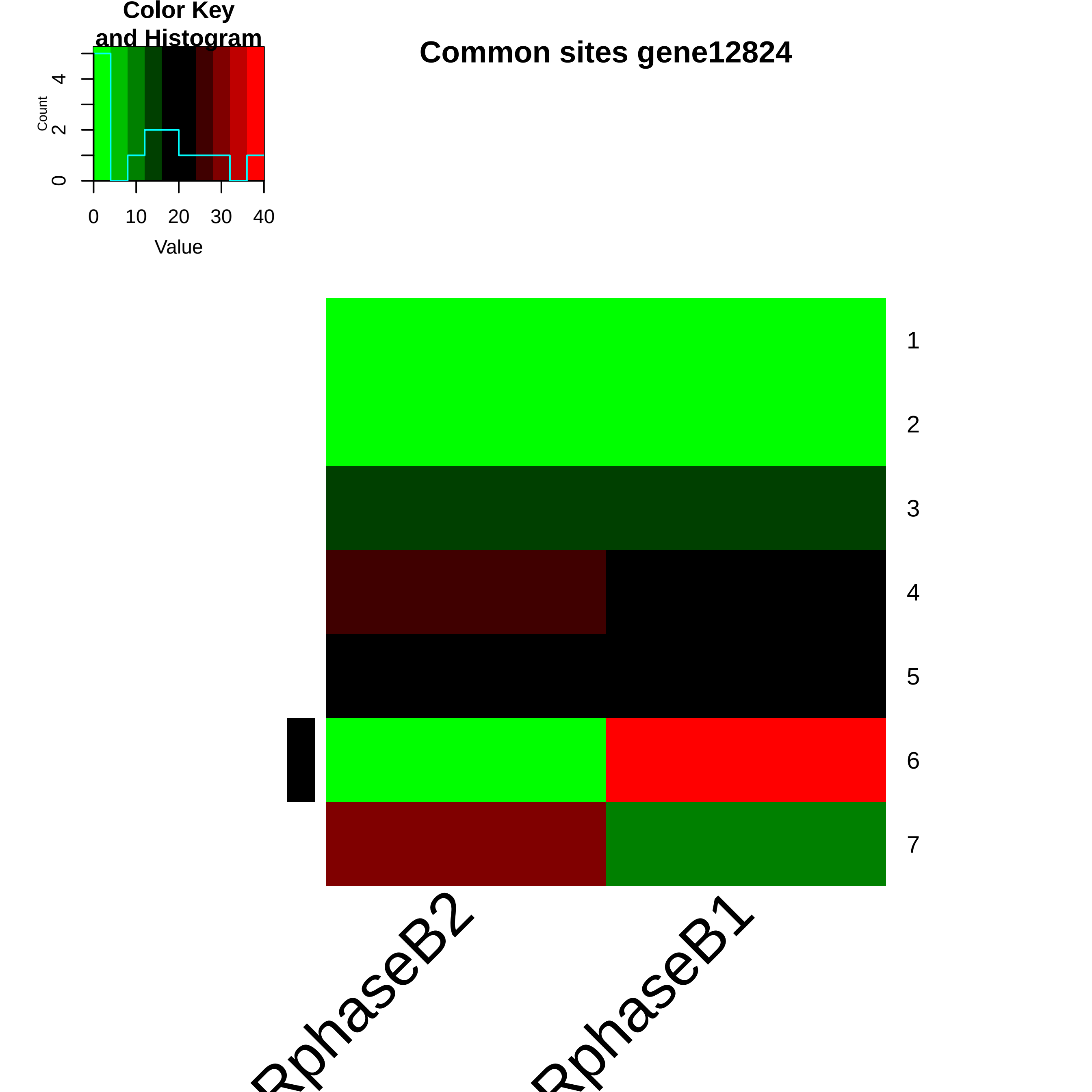
Color Key
and Histogram
Count
0
2
4
0
10
20
30
40
Value
Common sites gene12824
1
2
3
4
5
6
7
RphaseB2
RphaseB1

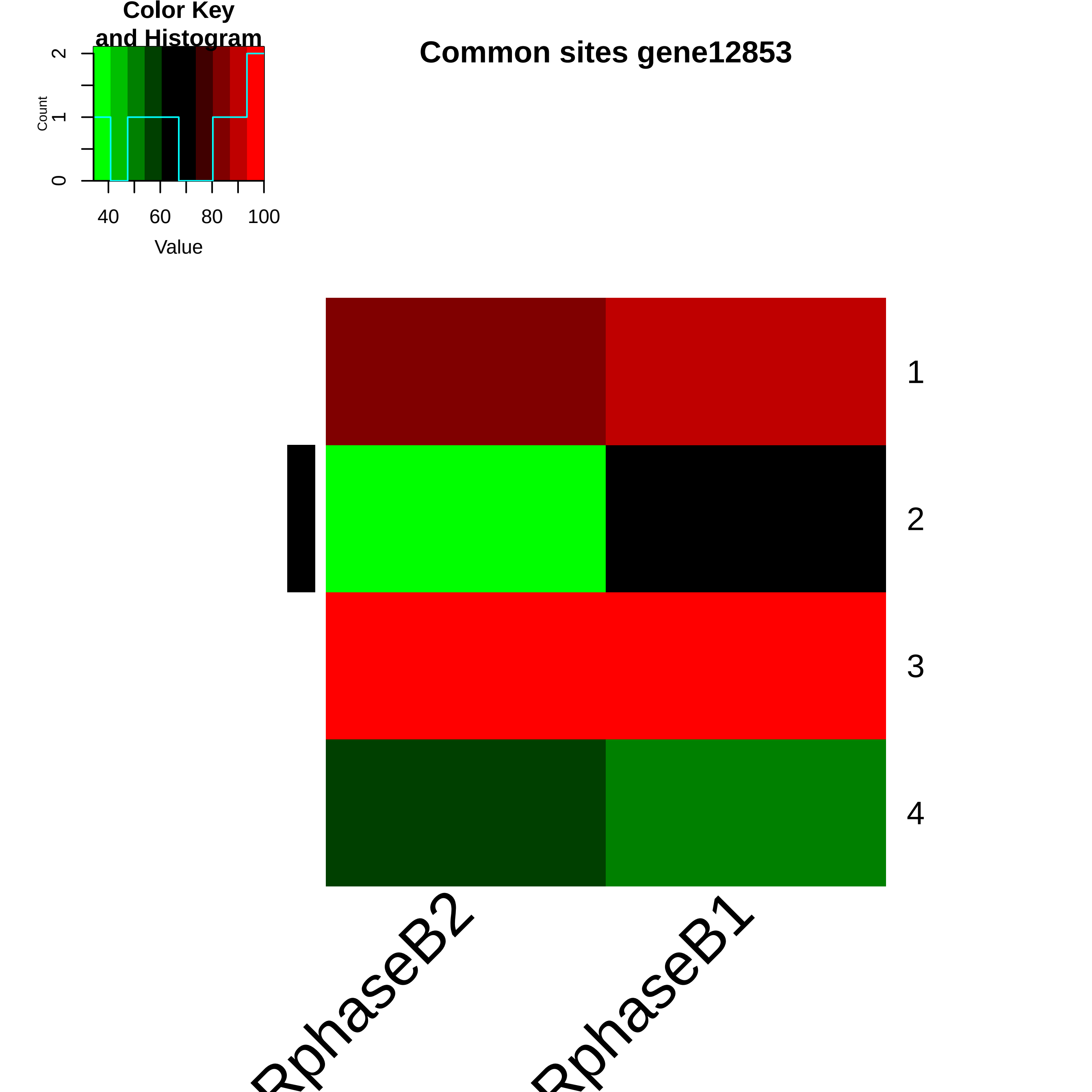

Color Key
and Histogram
Count
0
1
2
40
60
80
100
Value
Common sites gene12853
1
2
3
4
RphaseB2
RphaseB1

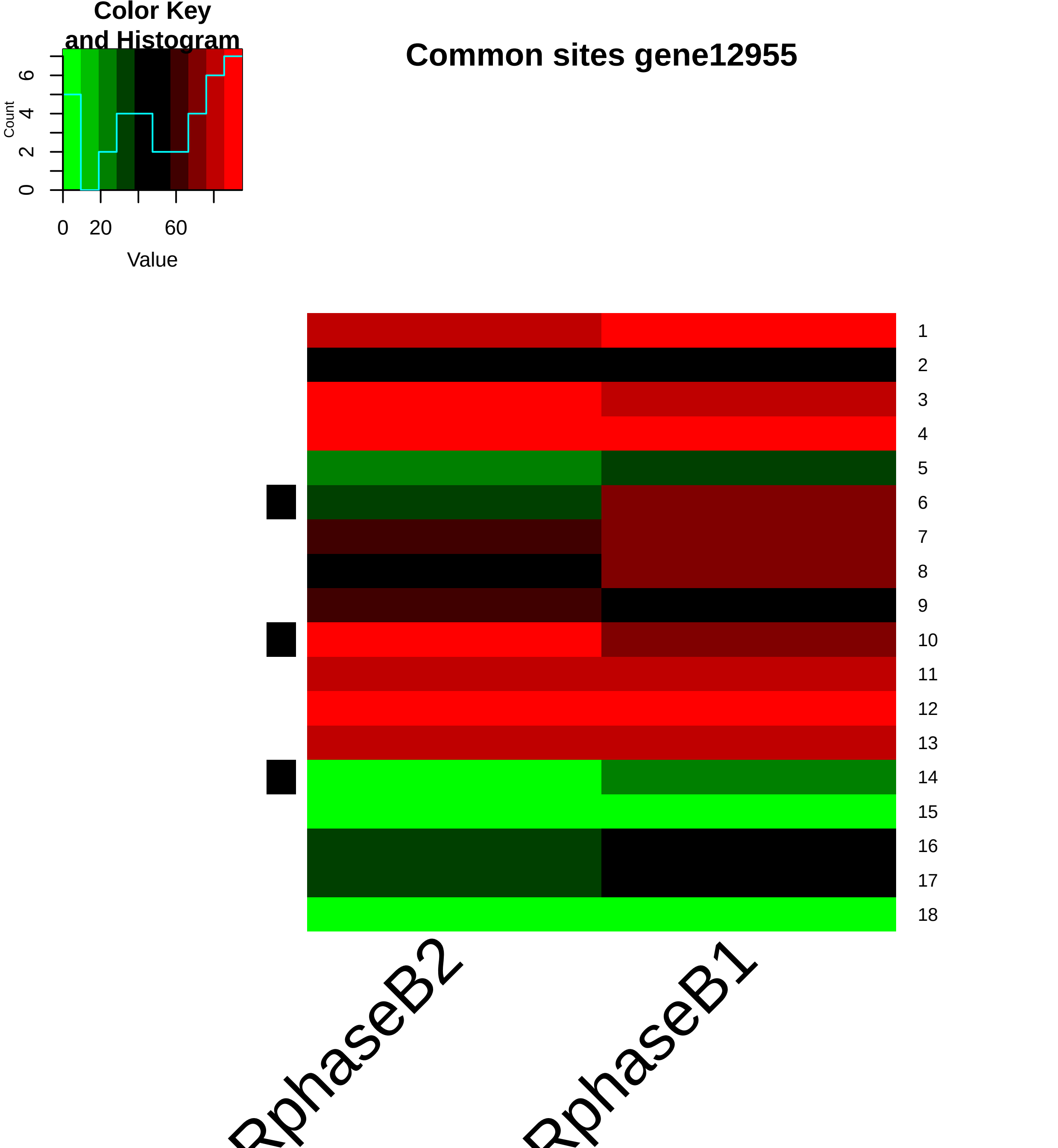
Color Key
and Histogram
Count
0
2
4
6
0
20
60
Value
Common sites gene12955
RphaseB2
RphaseB1

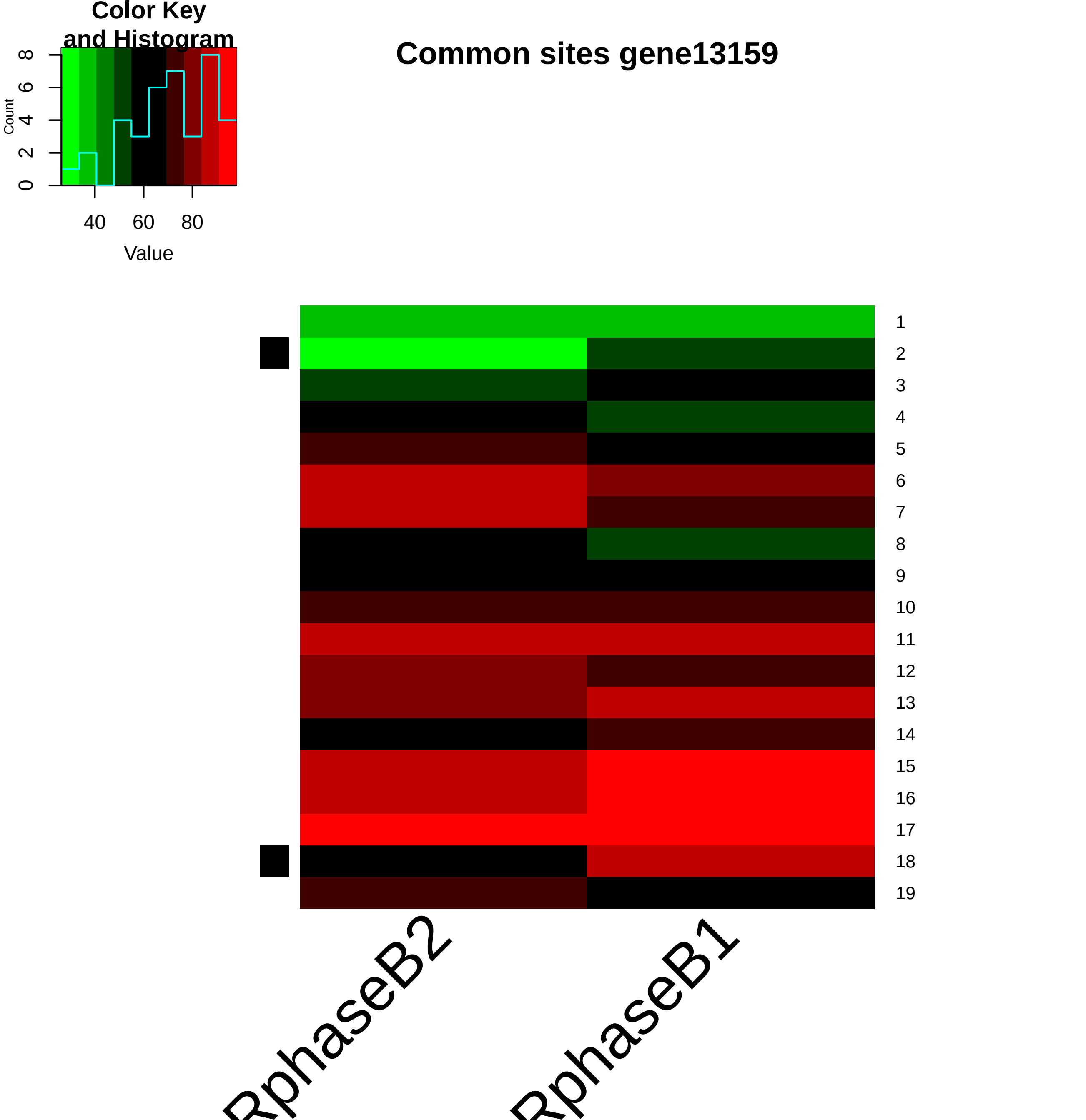

Color Key
and Histogram
Count
0
2
4
6
8
40
60
80
Value
Common sites gene13159
RphaseB2
RphaseB1

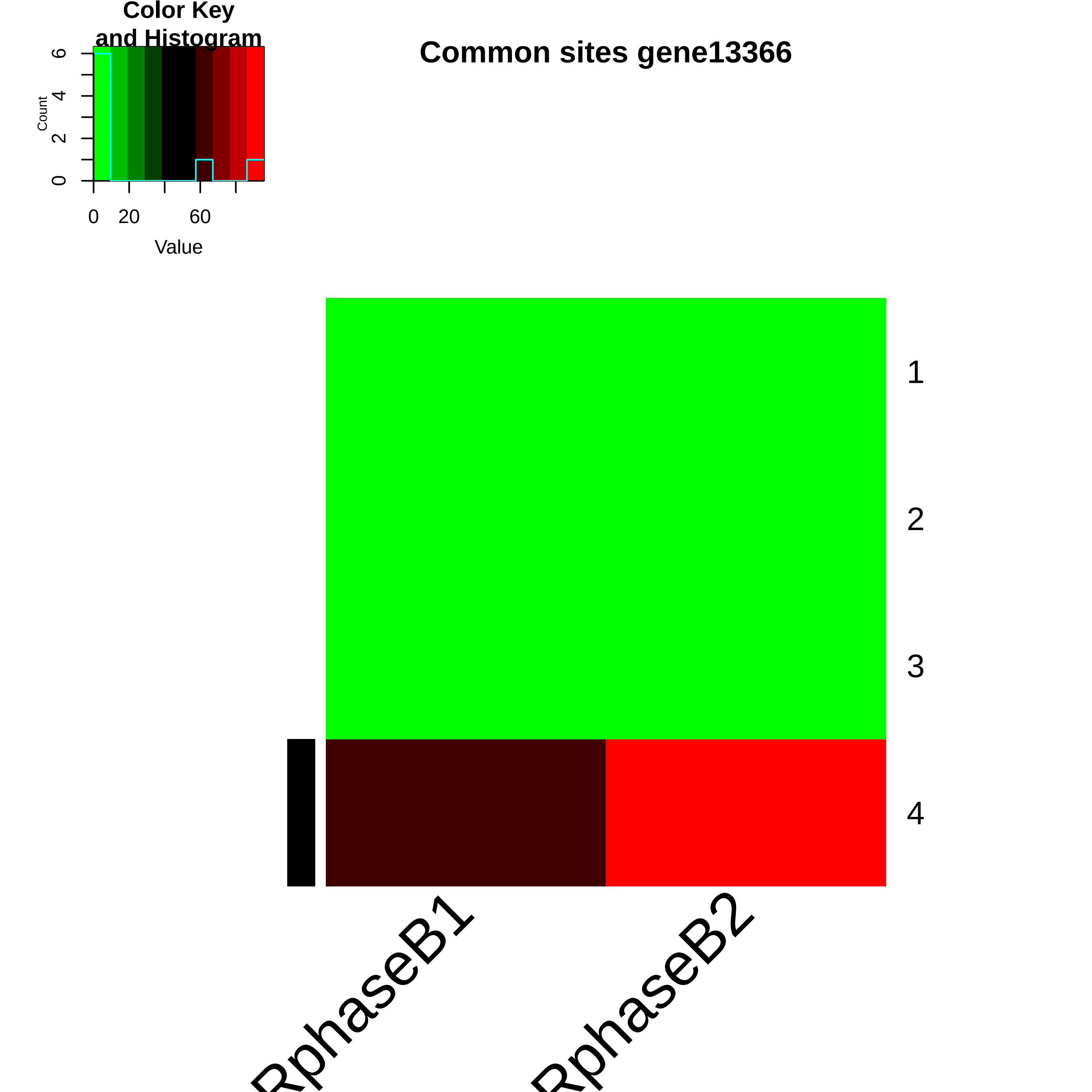

Color Key
and Histogram
Count
Value
Common sites gene13366
1
2
3
4
RphaseB1
RphaseB2

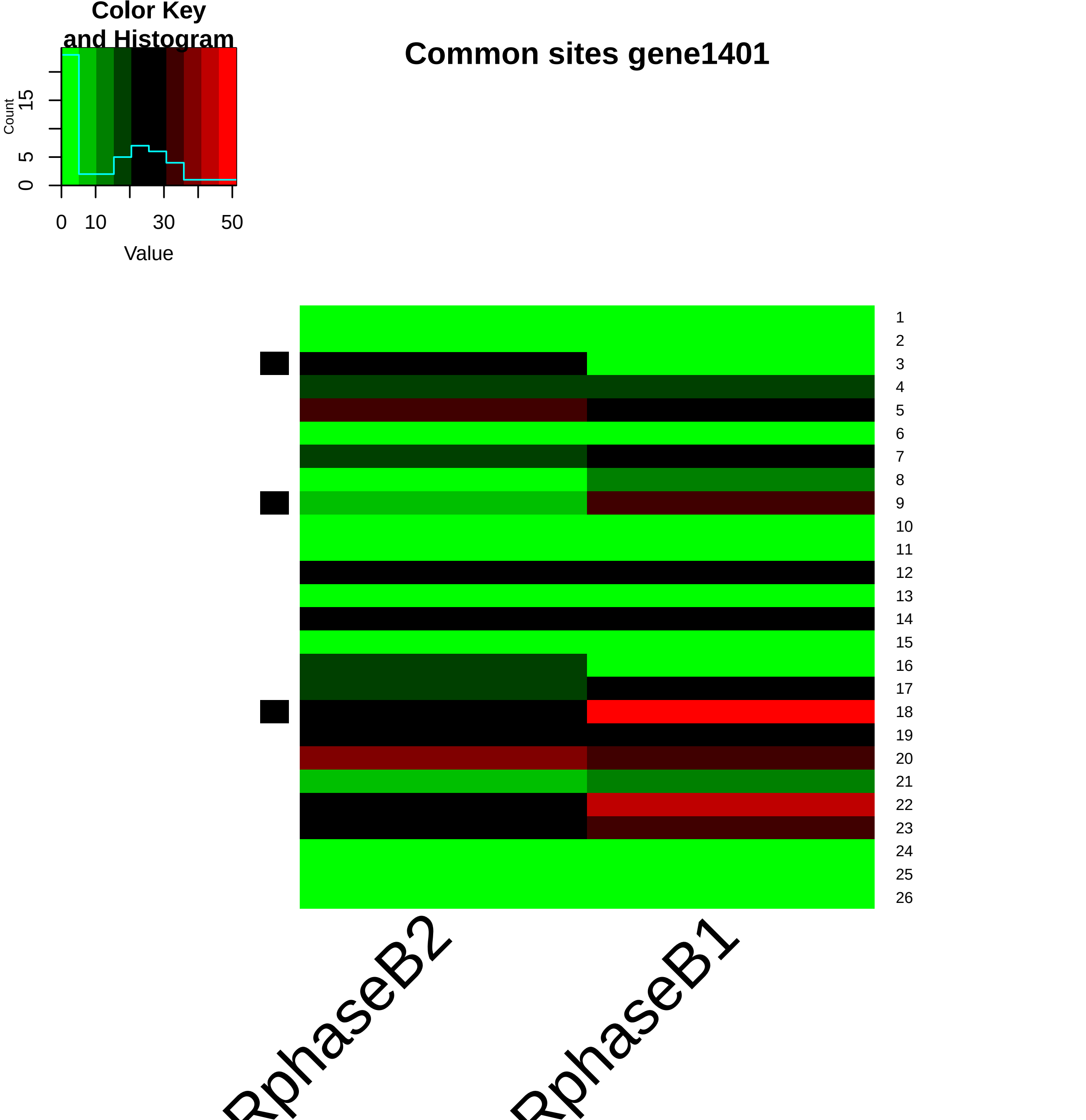
Color Key
and Histogram
Count
0
5
15
0
10
30
50
Value
Common sites gene1401
RphaseB2
RphaseB1

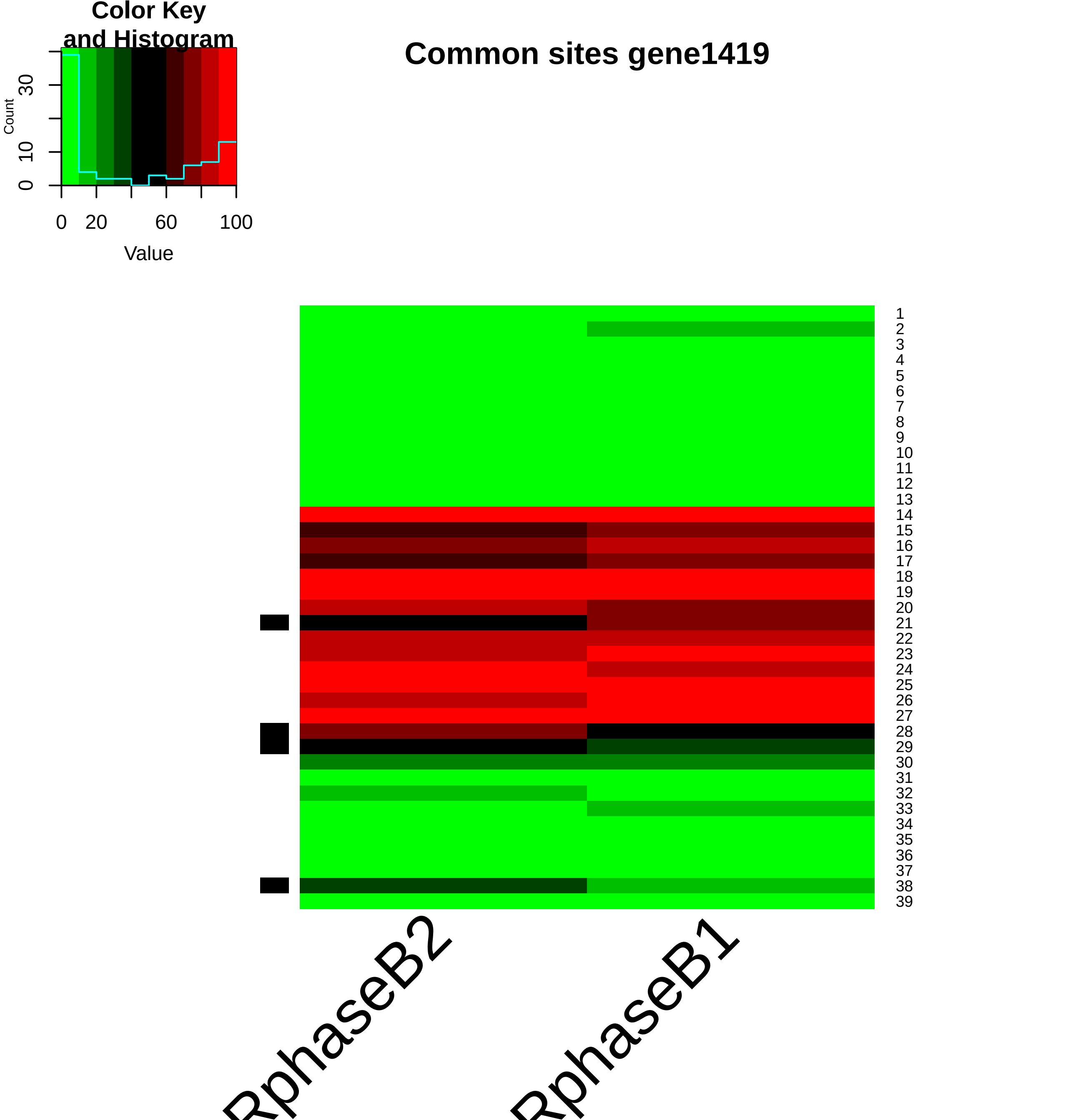
Color Key
and Histogram
Count
0
10
30
0
20
60
100
Value
Common sites gene1419
RphaseB2
RphaseB1

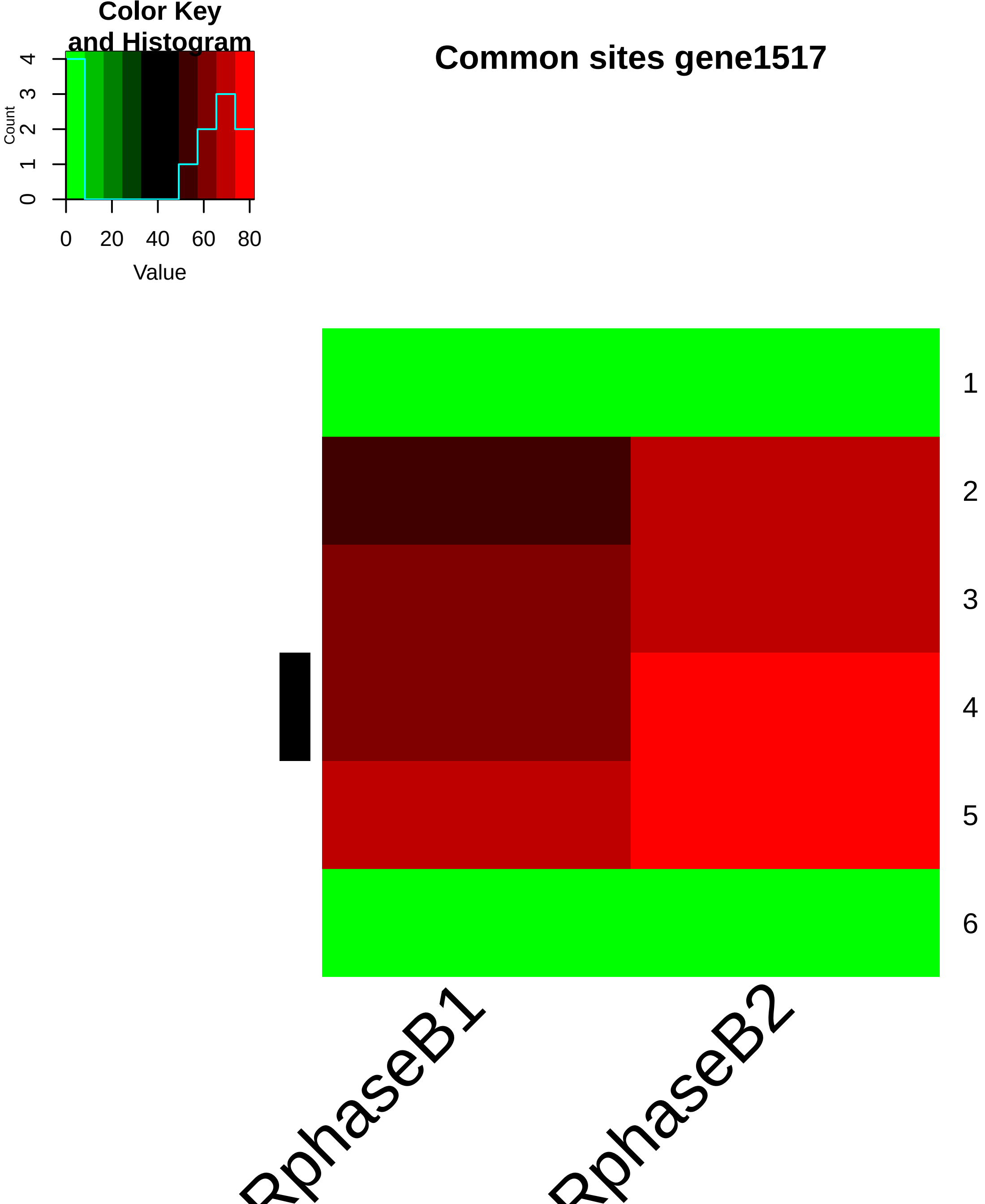

Color Key
and Histogram
Count
Value
0
20
40
60
80
0
1
2
3
4
Common sites gene1517
1
2
3
4
5
6
RphaseB1
RphaseB2

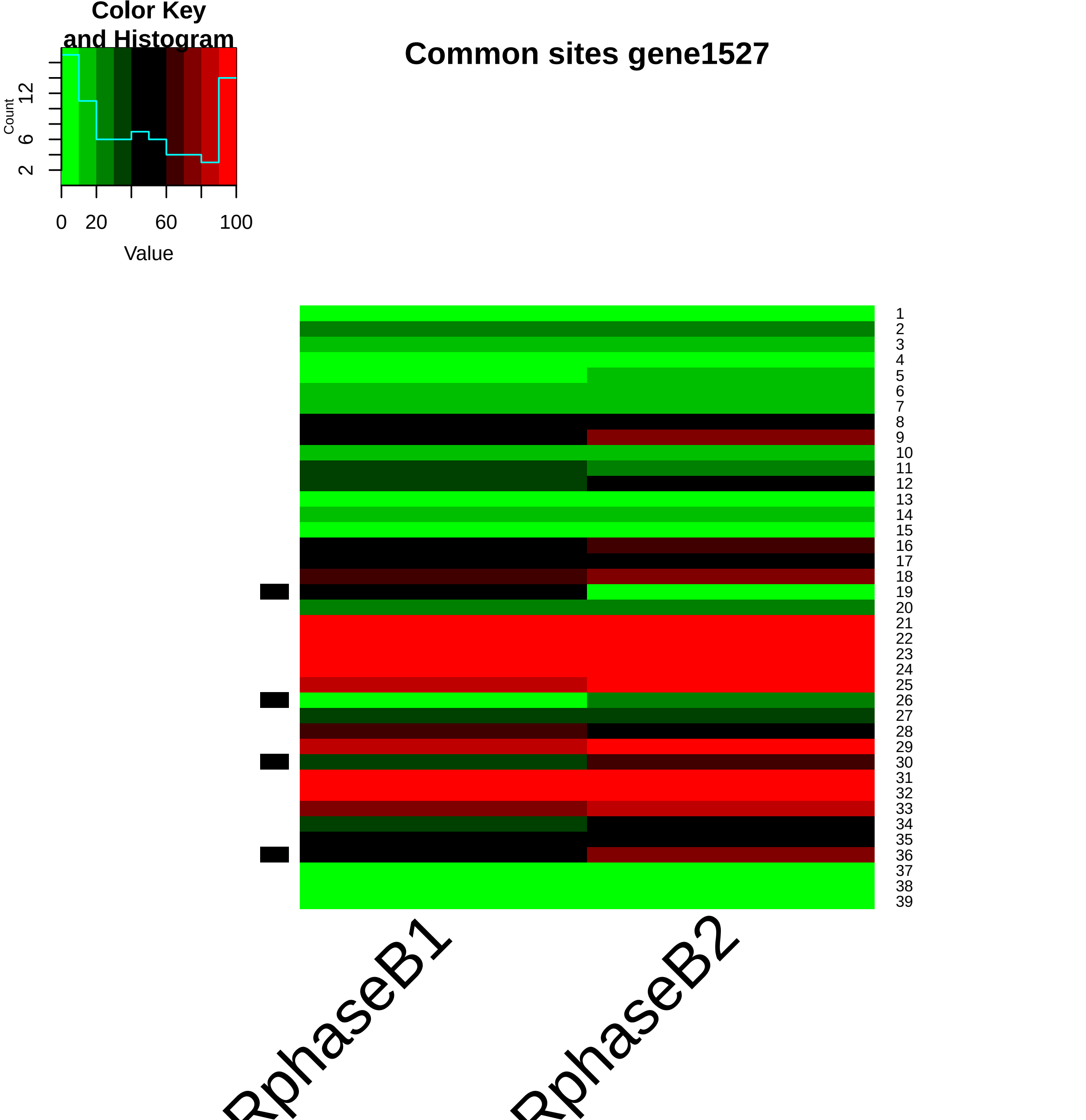

Color Key
and Histogram
Count
2
6
12
0
20
60
100
Value
Common sites gene1527
RphaseB1
RphaseB2

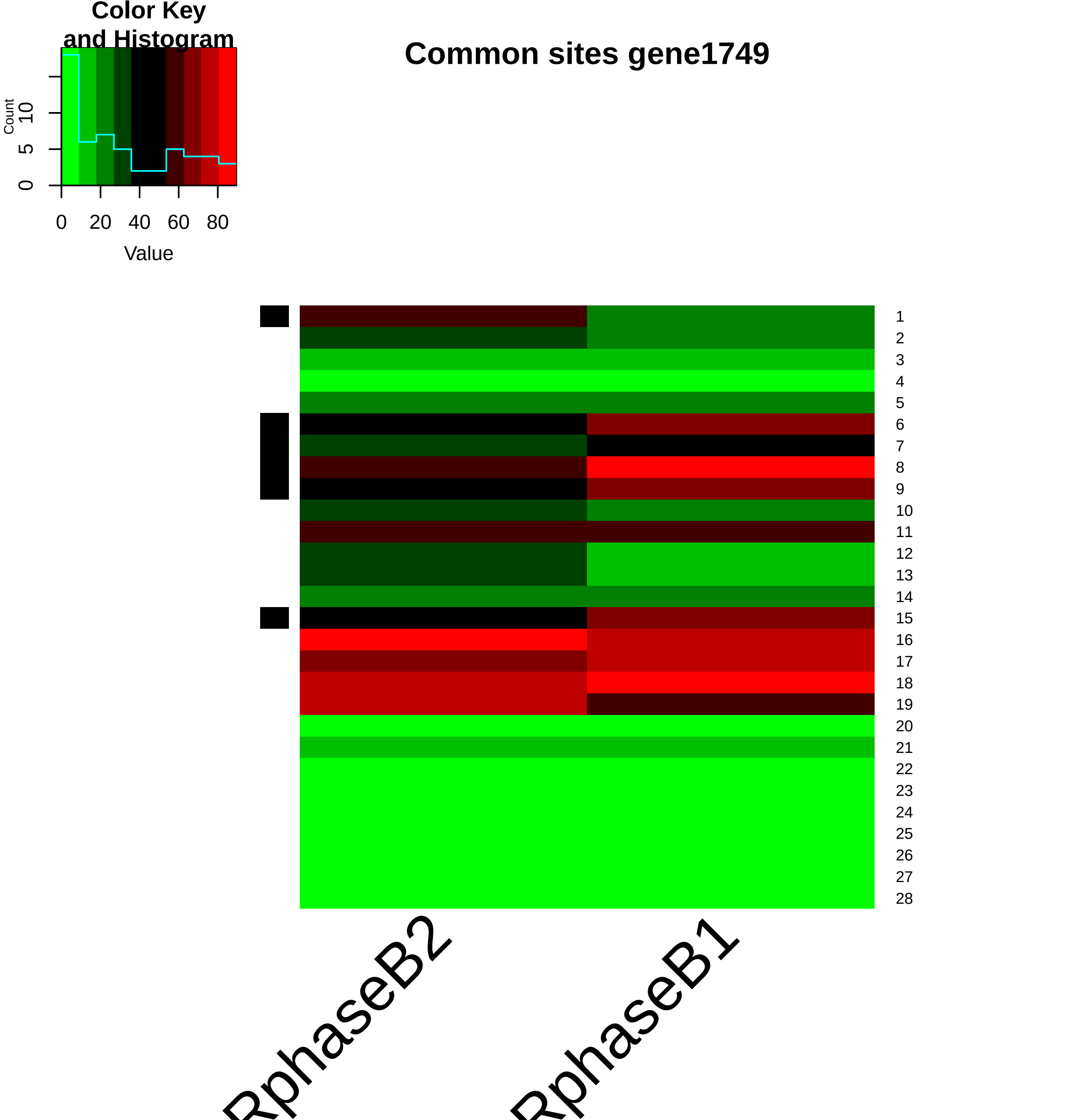
Color Key
and Histogram
Count
0
5
10
0
20
40
60
80
Value
Common sites gene1749
RphaseB2
RphaseB1

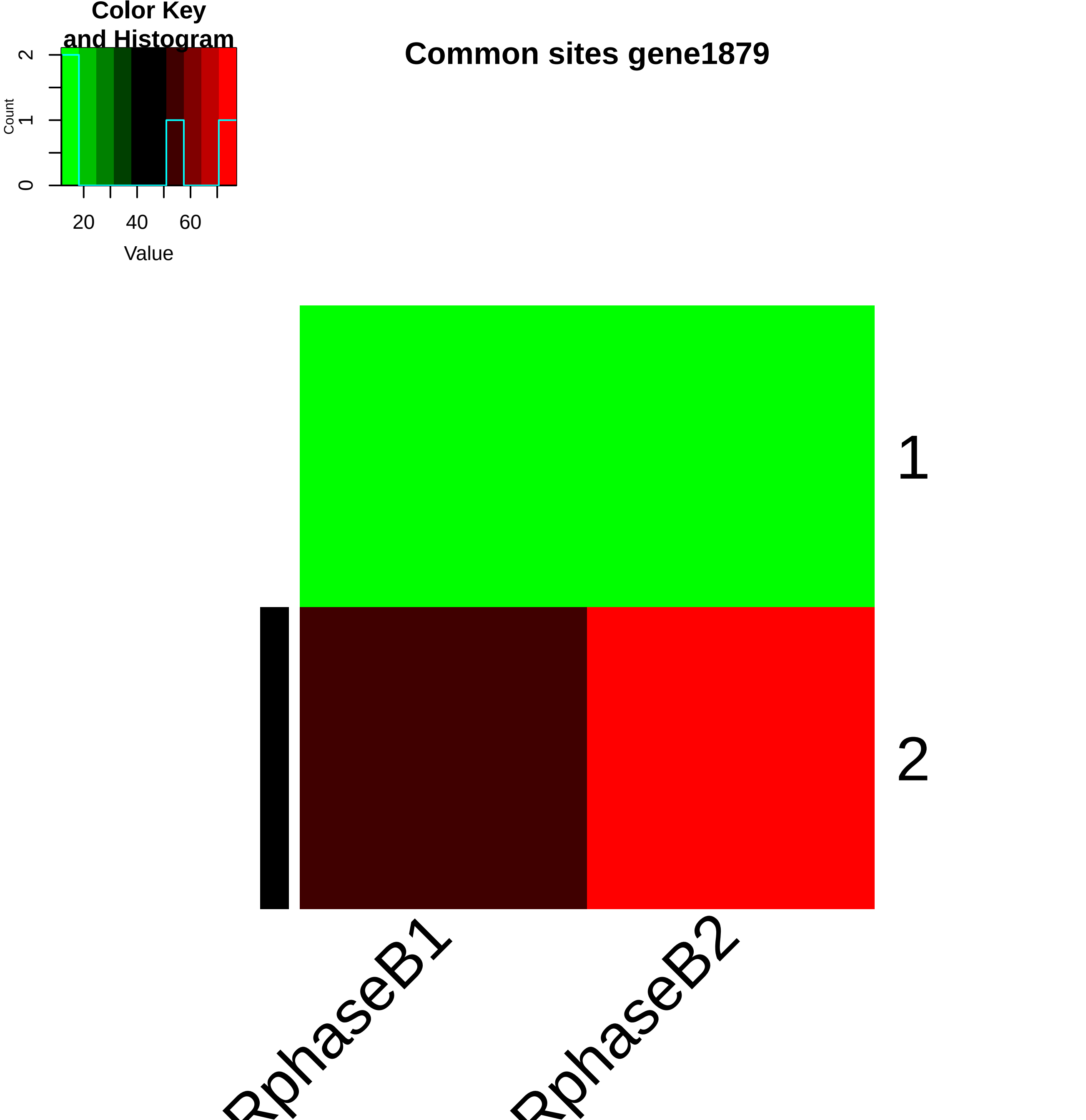
Color Key
and Histogram
Count
0
1
2
20
40
60
Value
Common sites gene1879
1
2
RphaseB1
RphaseB2

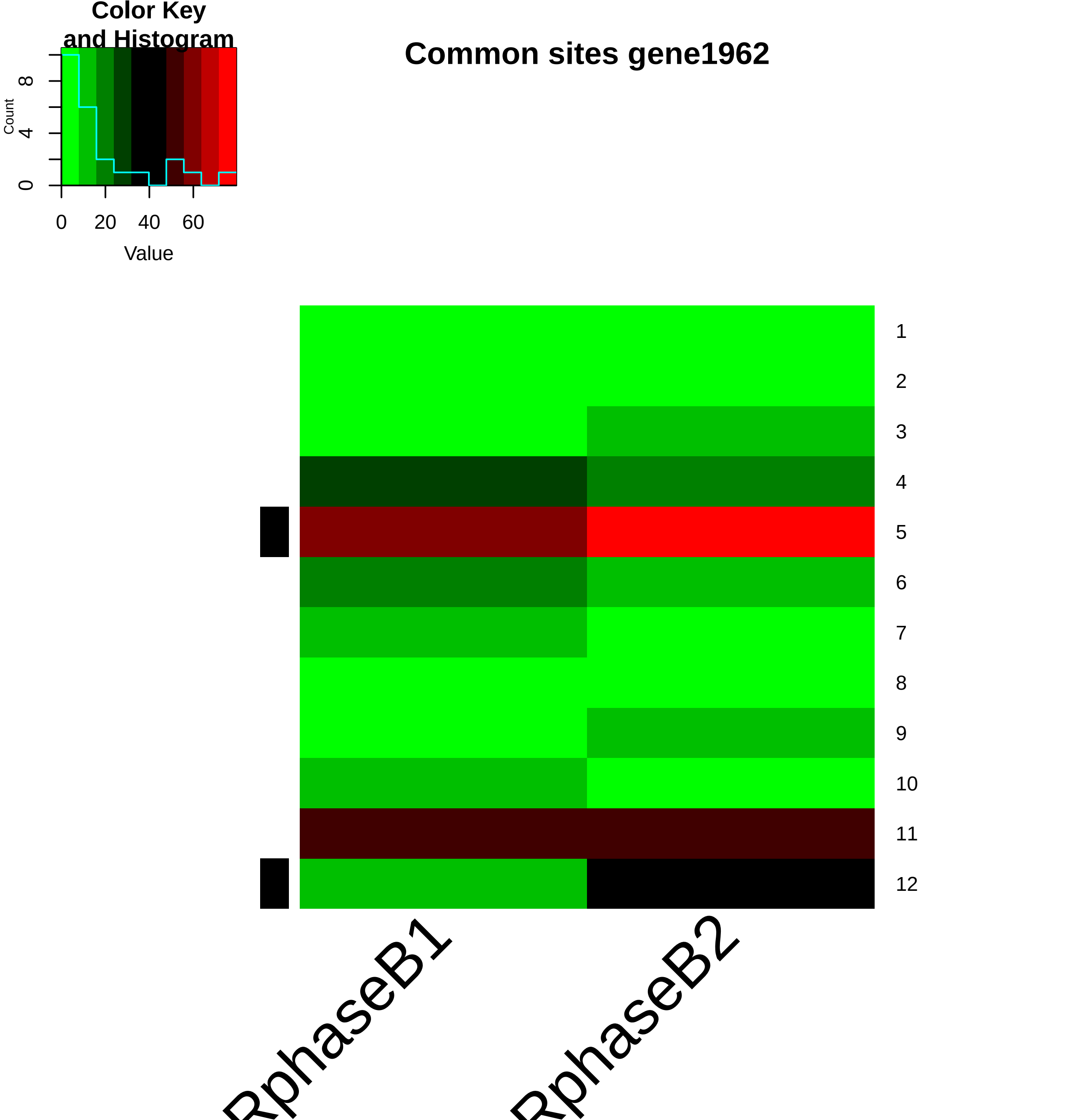
Color Key
and Histogram
Count
0
4
8
0
20
40
60
Value
Common sites gene1962
1
2
3
4
5
6
7
8
9
10
11
12
RphaseB1
RphaseB2

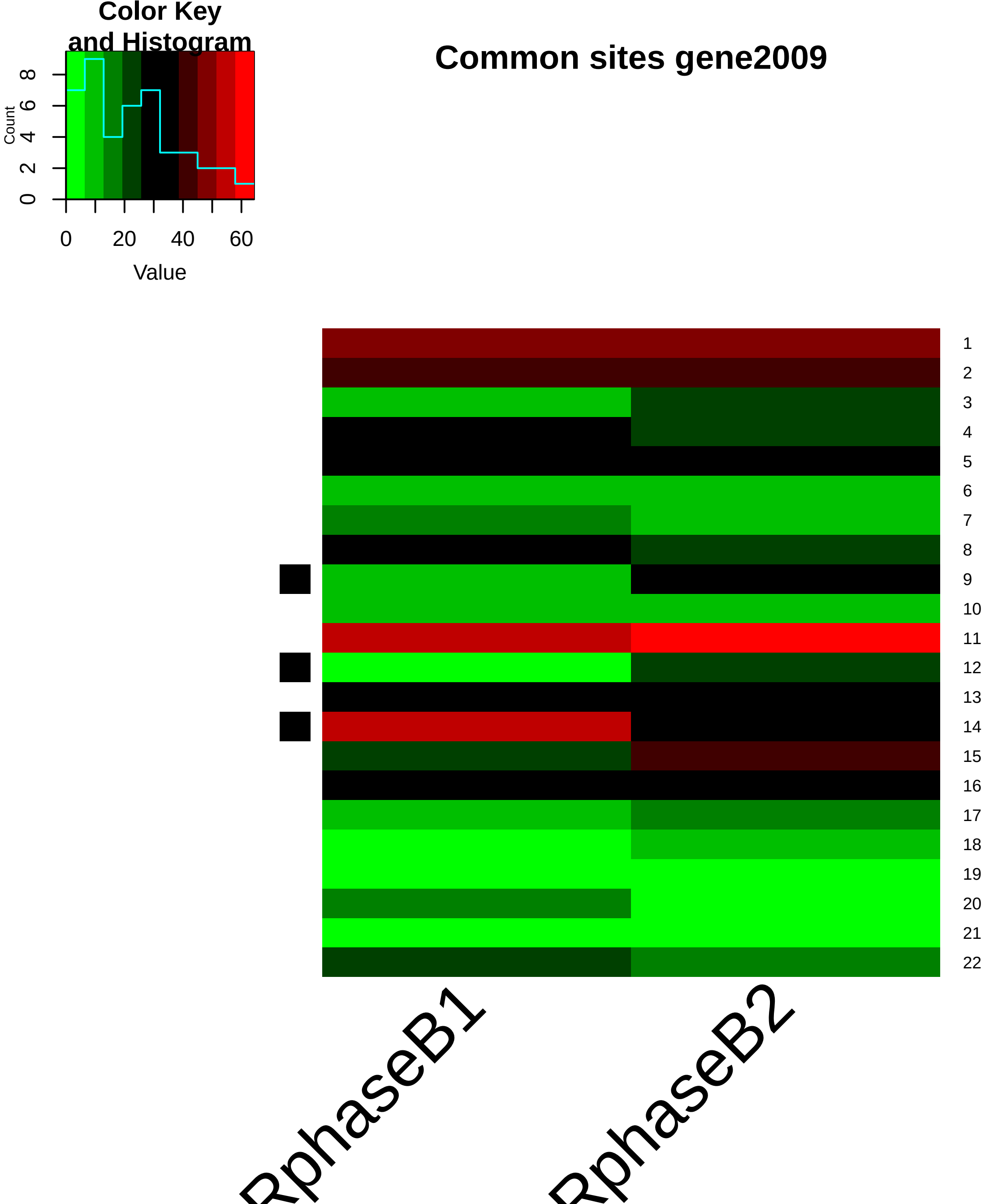
Color Key
and Histogram
Count
0 2 4 6 8
0 20 40 60
Value
Common sites gene2009
RphaseB1
RphaseB2

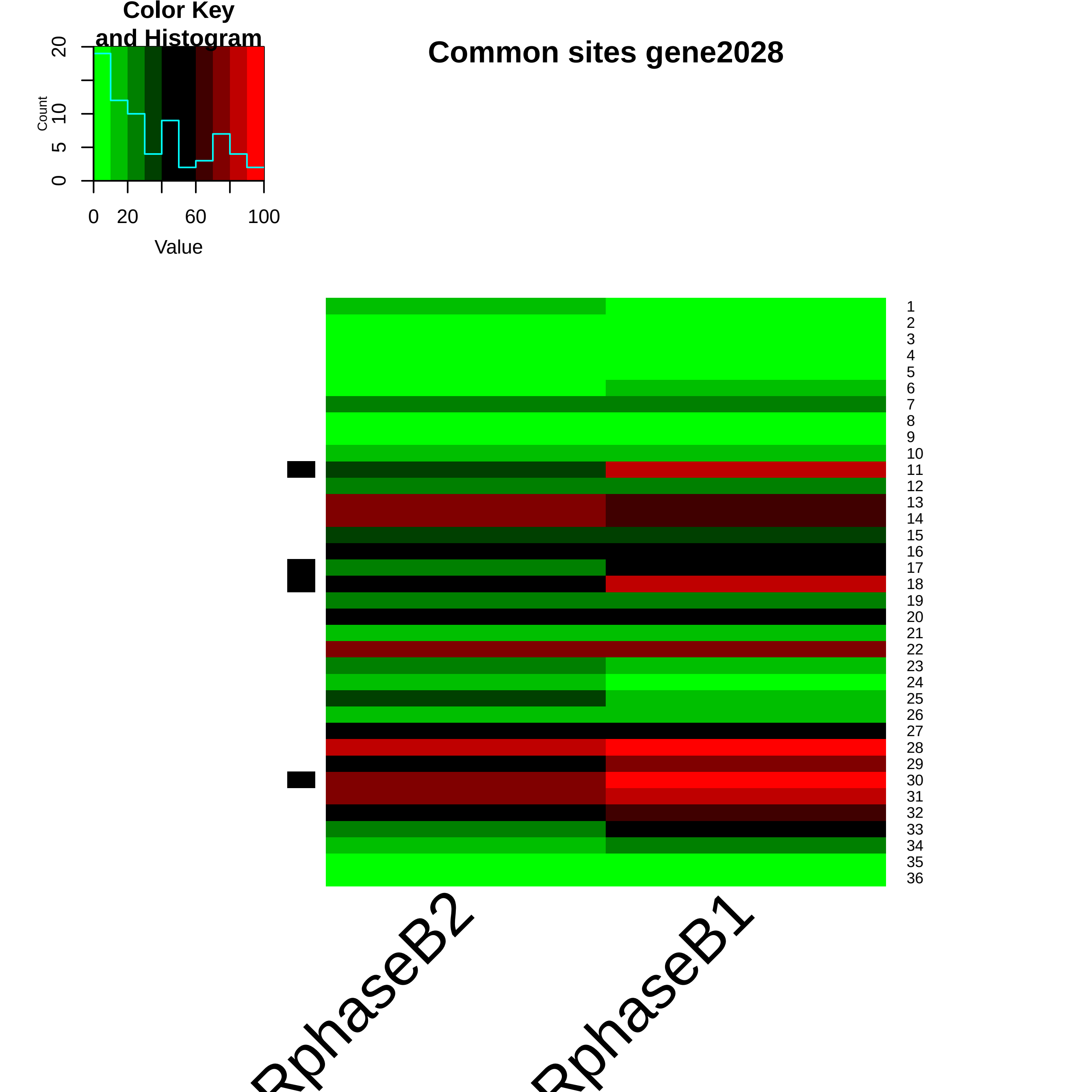

Color Key
and Histogram
Count
Value
0
20
60
100
0
5
10
20
Common sites gene2028
RphaseB2
RphaseB1

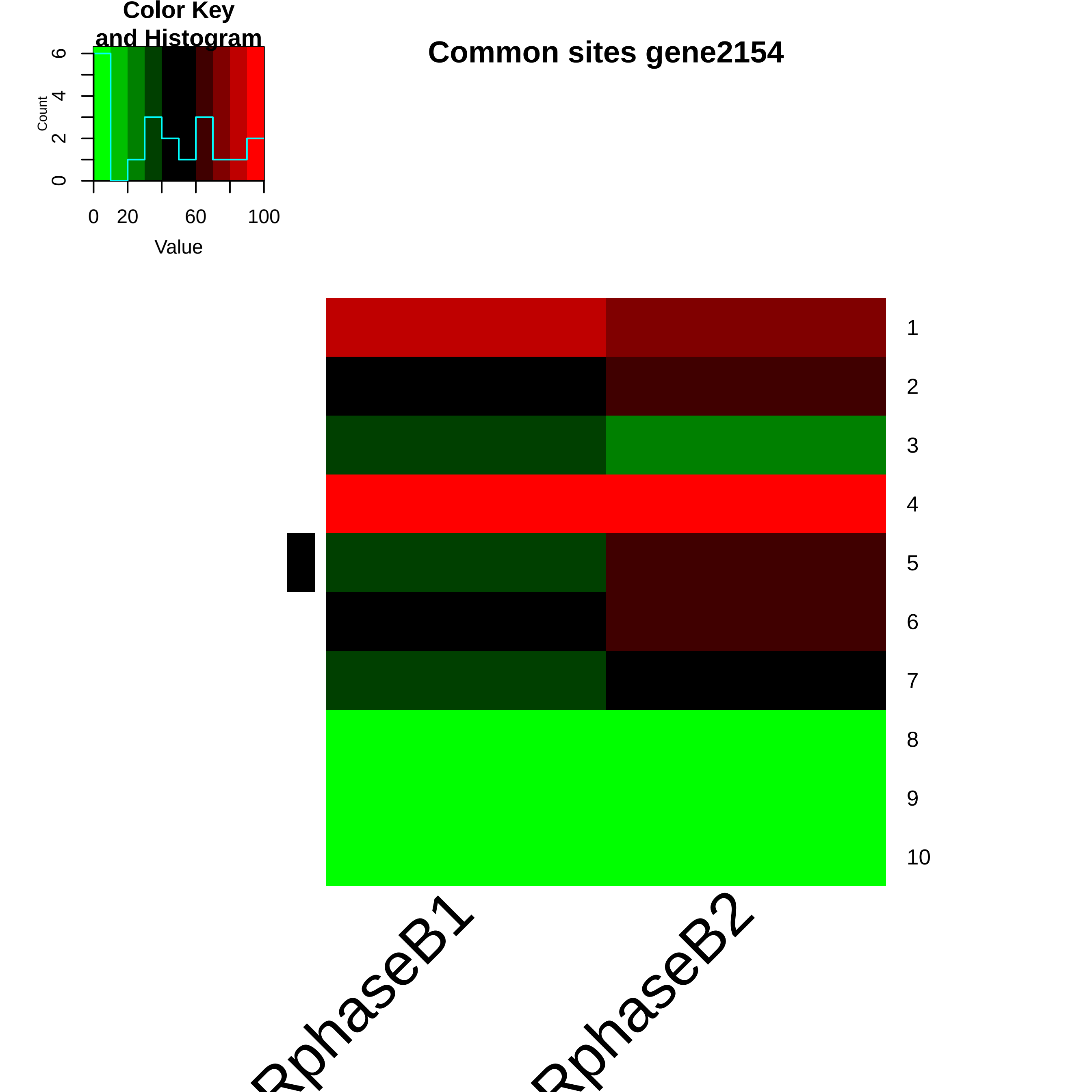
Color Key
and Histogram
Count
0
2
4
6
0
20
60
100
Value
Common sites gene2154
1
2
3
4
5
6
7
8
9
10
RphaseB1
RphaseB2

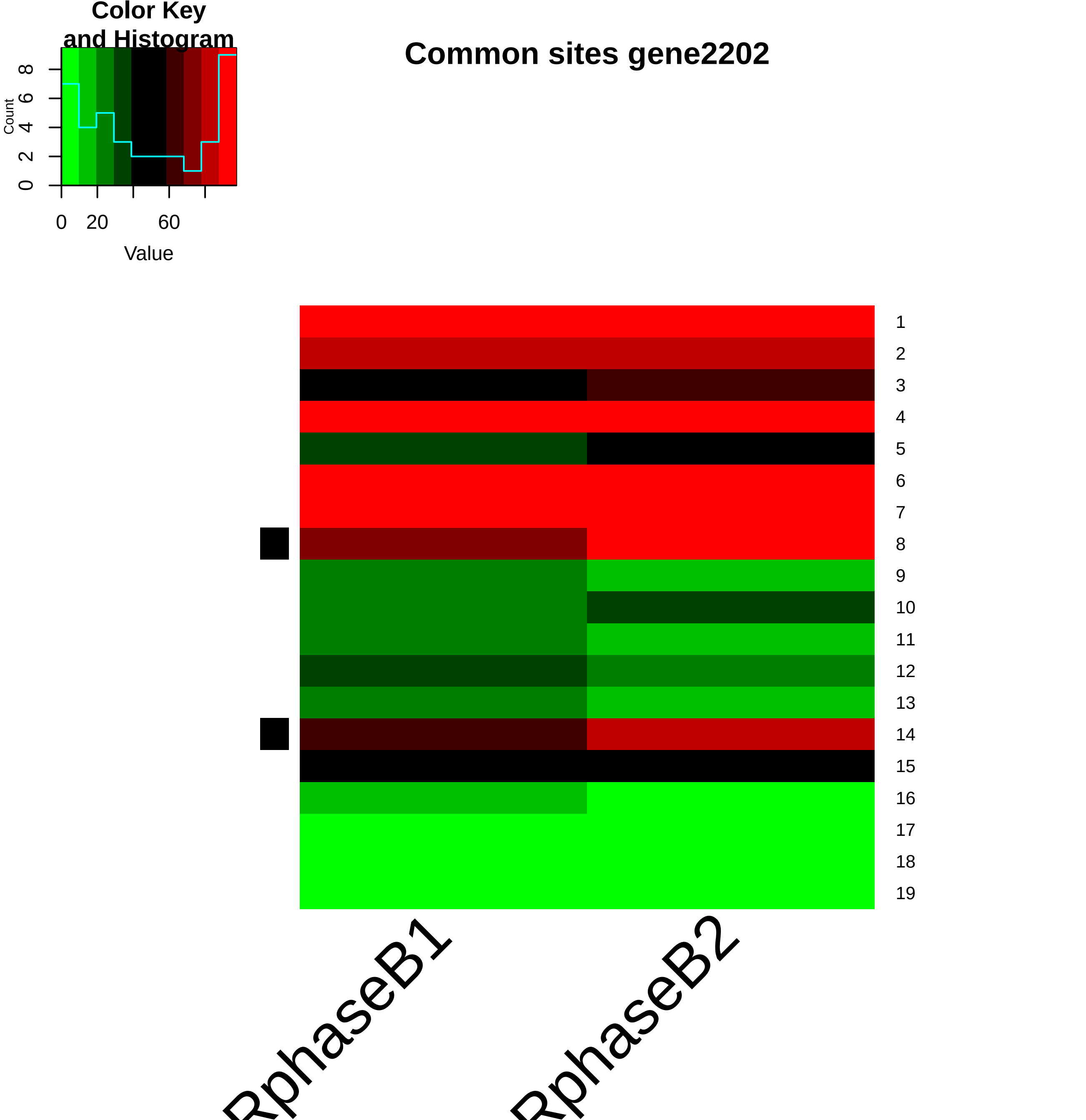

Color Key and Histogram
Count
0
2
4
6
8
0
20
60
Value
Common sites gene2202
RphaseB1
RphaseB2

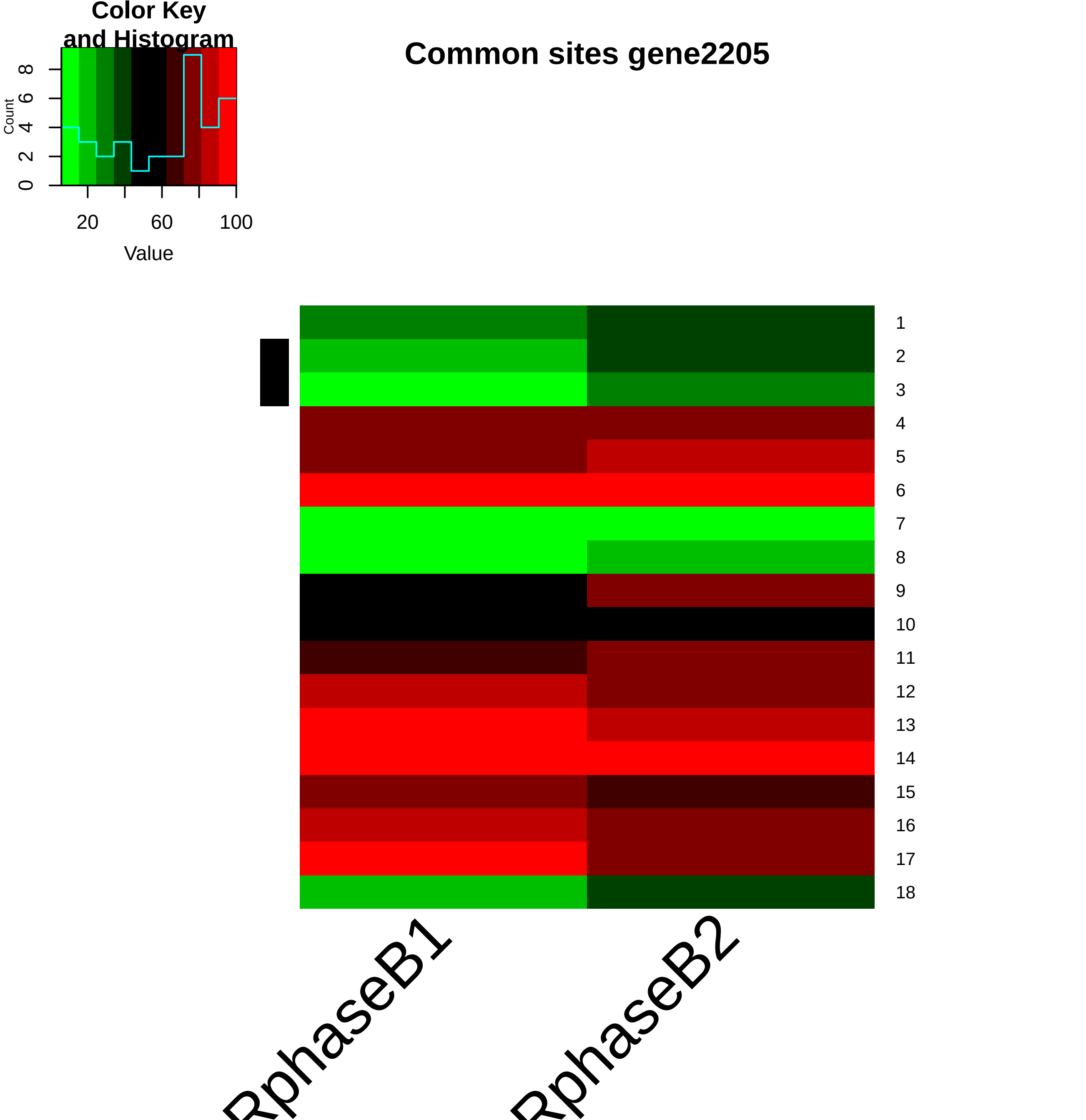
Color Key
and Histogram
Count
0
2
4
6
8
20
60
100
Value
Common sites gene2205
RphaseB1
RphaseB2

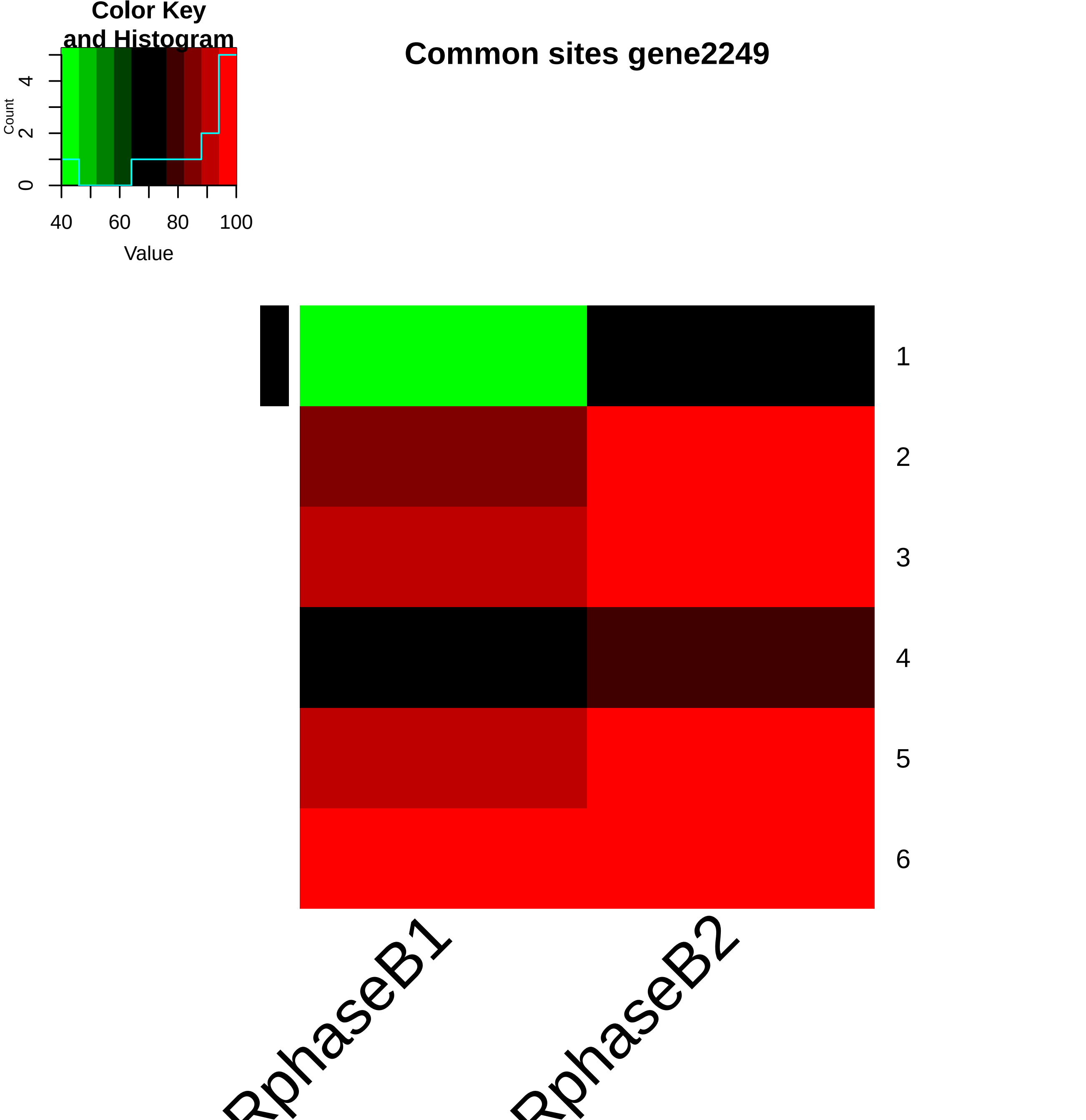

Color Key
and Histogram
Count
0
2
4
40
60
80
100
Value
Common sites gene2249
1
2
3
4
5
6
RphaseB1
RphaseB2

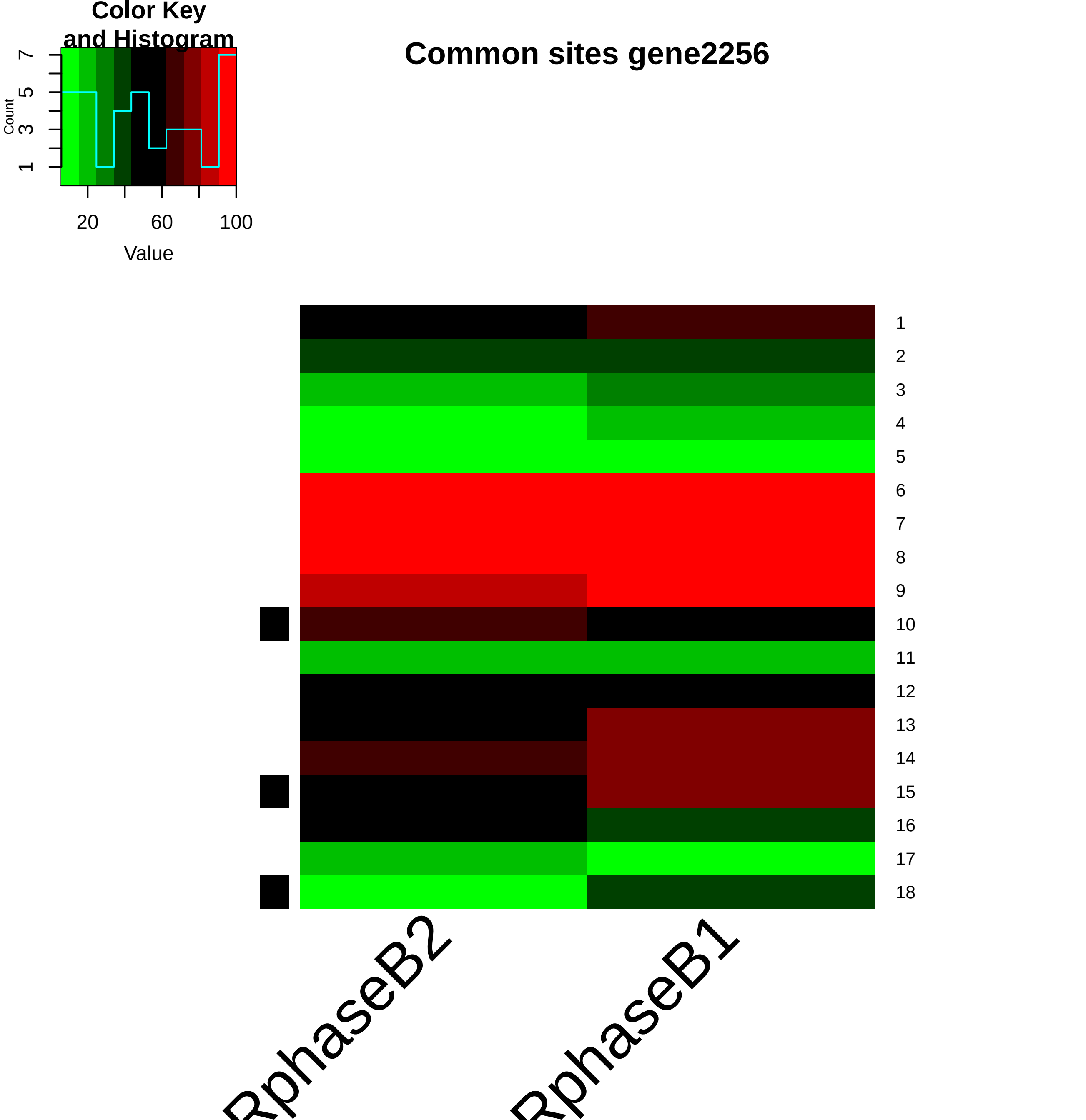

Color Key
and Histogram
Count
1
3
5
7
20
60
100
Value
Common sites gene2256
RphaseB2
RphaseB1

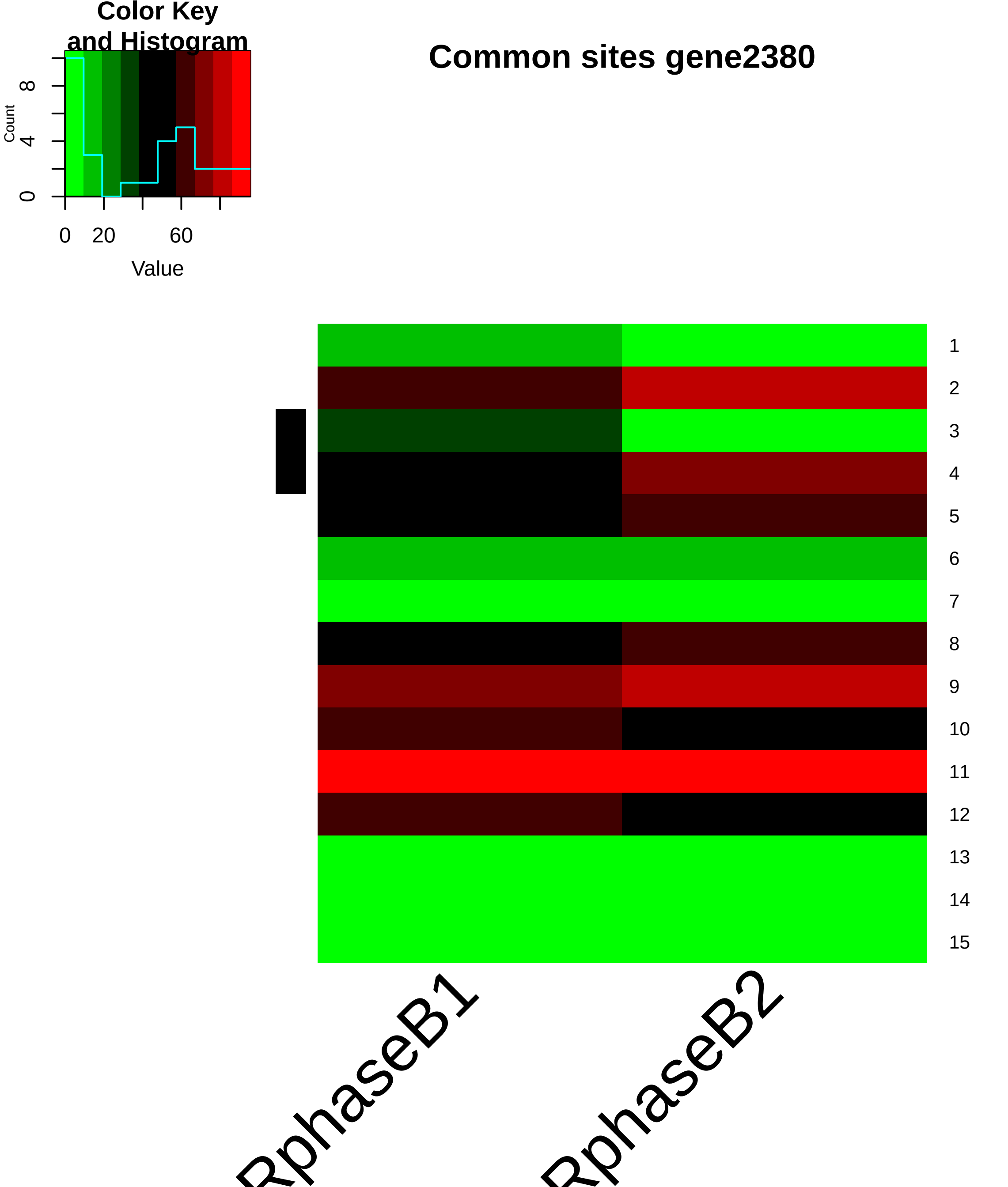
Color Key
and Histogram
Count
0
4
8
0
20
60
Value
Common sites gene2380
RphaseB1
RphaseB2

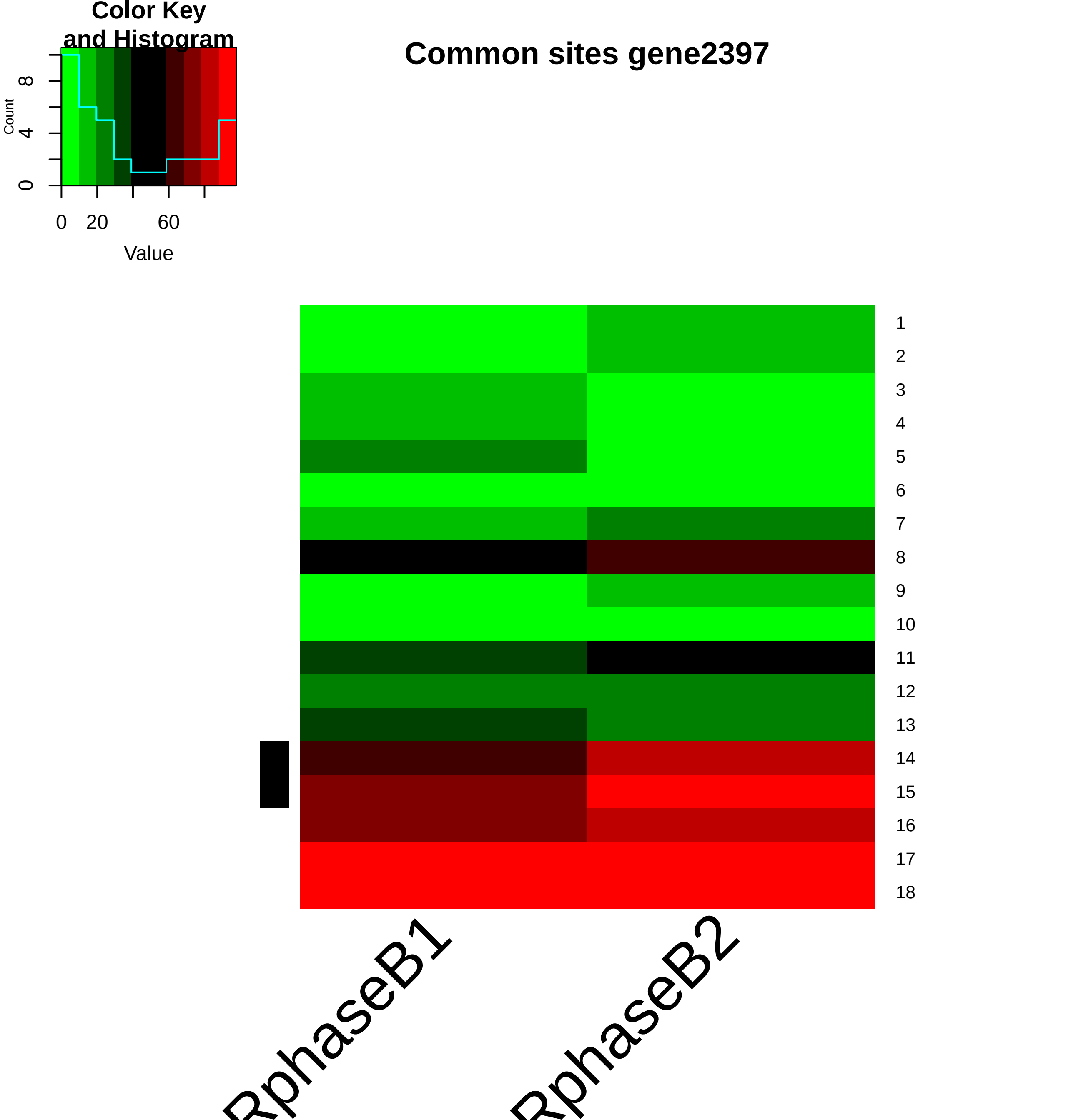

Color Key
and Histogram
Count
0
4
8
0
20
60
Value
Common sites gene2397
RphaseB1
RphaseB2

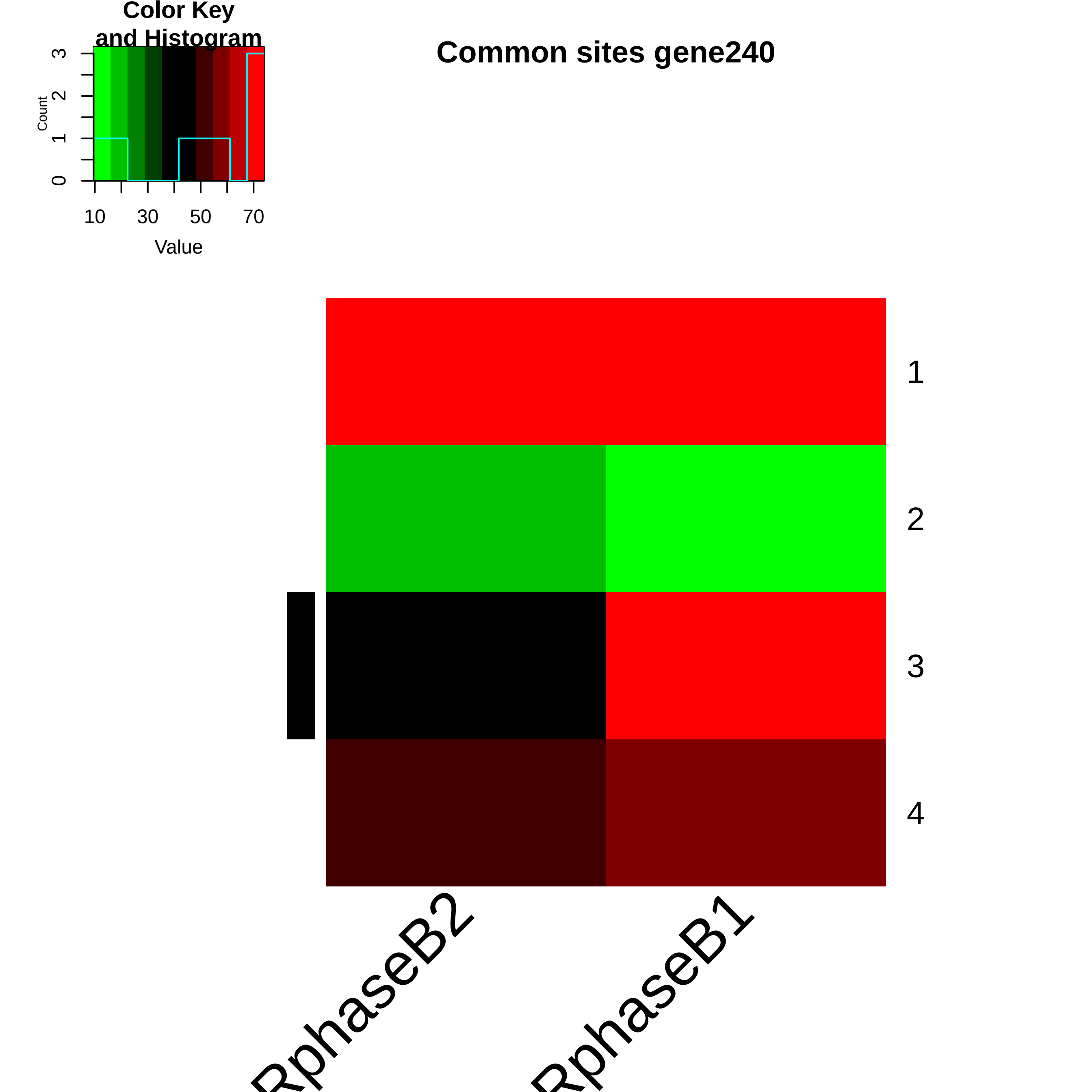
Color Key
and Histogram
Count
0
1
2
3
10
30
50
70
Value
Common sites gene240
1
2
3
4
RphaseB2
RphaseB1

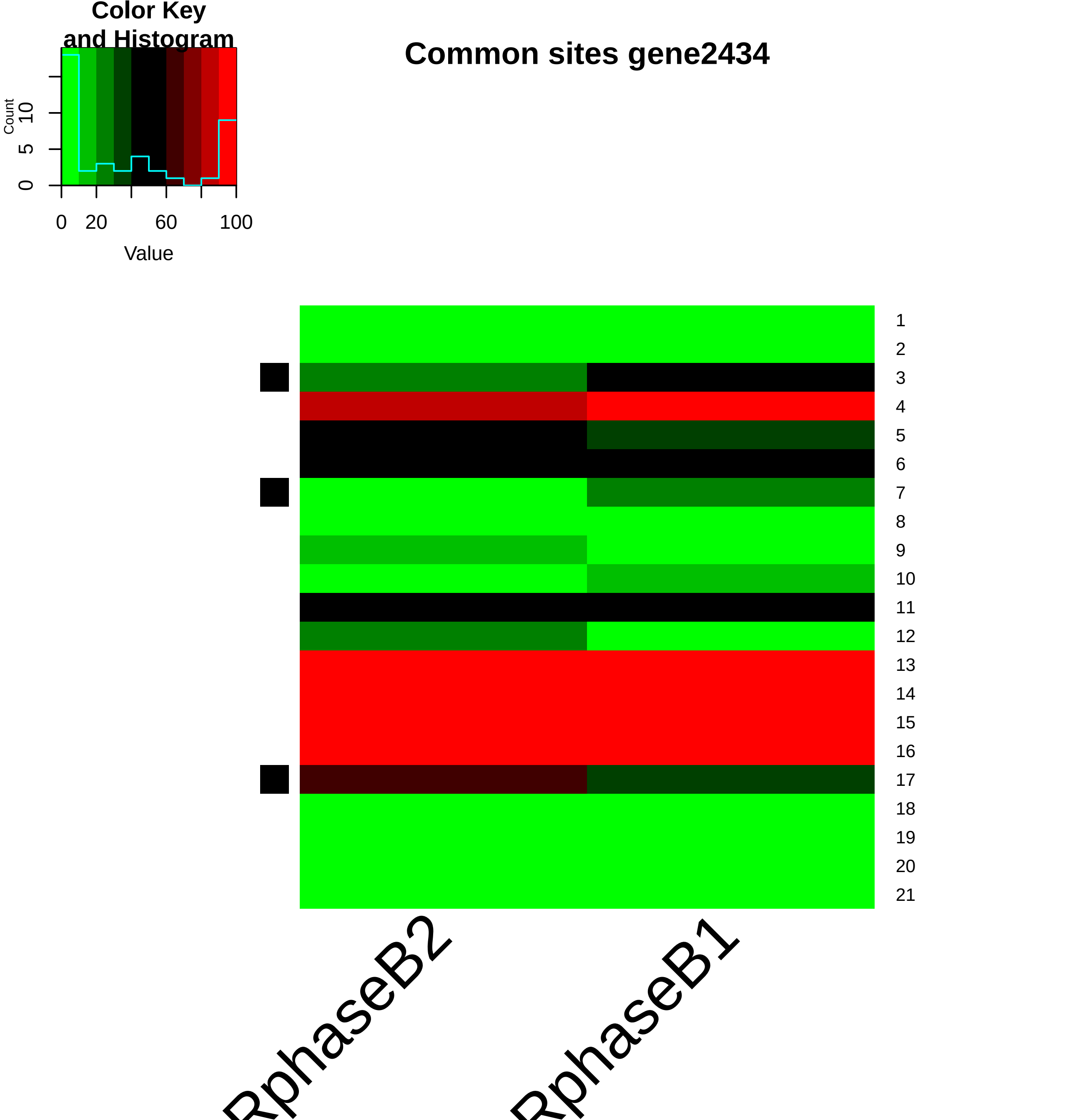

Color Key
and Histogram
Count
0
5
10
0
20
60
100
Value
Common sites gene2434
RphaseB2
RphaseB1

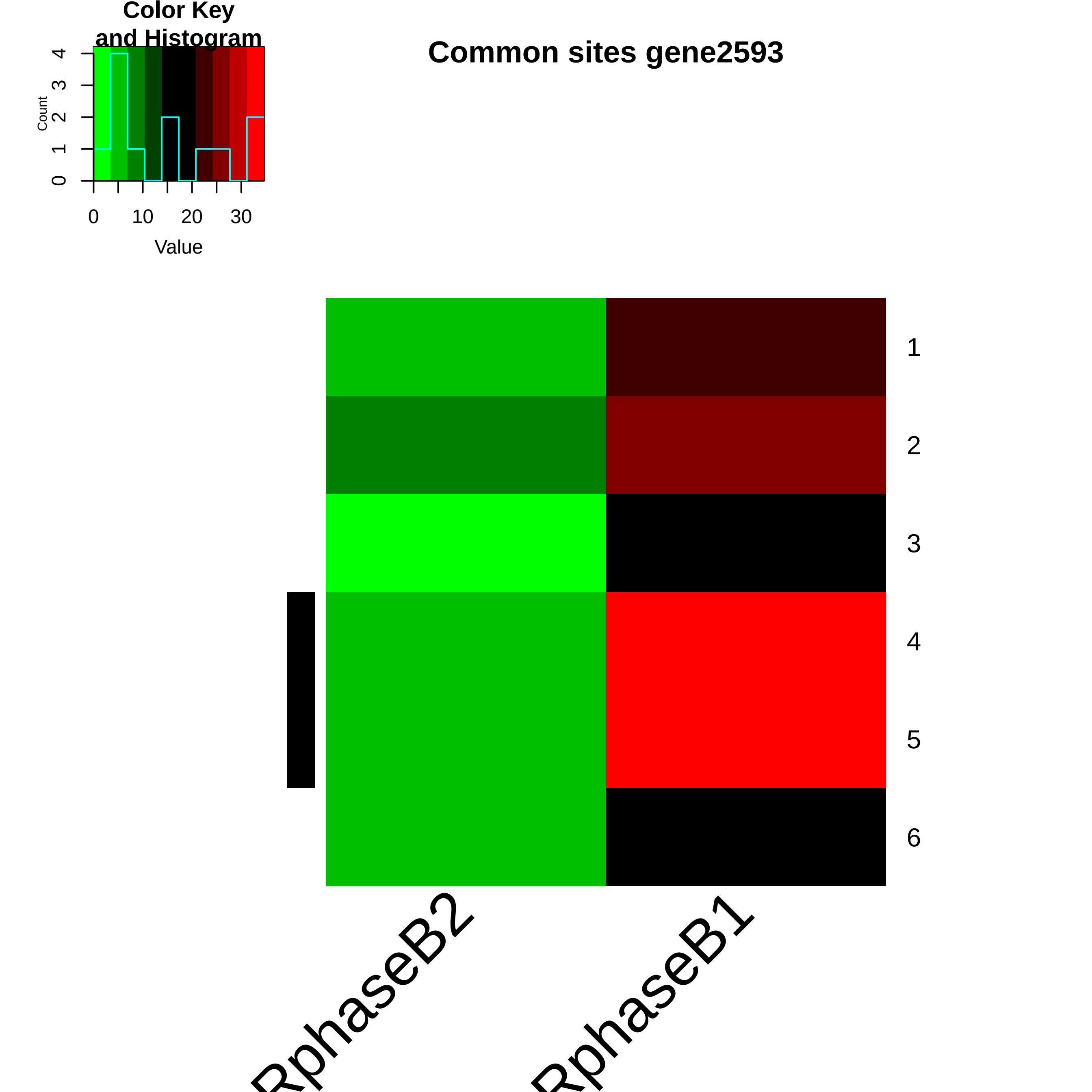

Color Key
and Histogram
Count
0
1
2
3
4
0
10
20
30
Value
Common sites gene2593
1
2
3
4
5
6
RphaseB2
RphaseB1

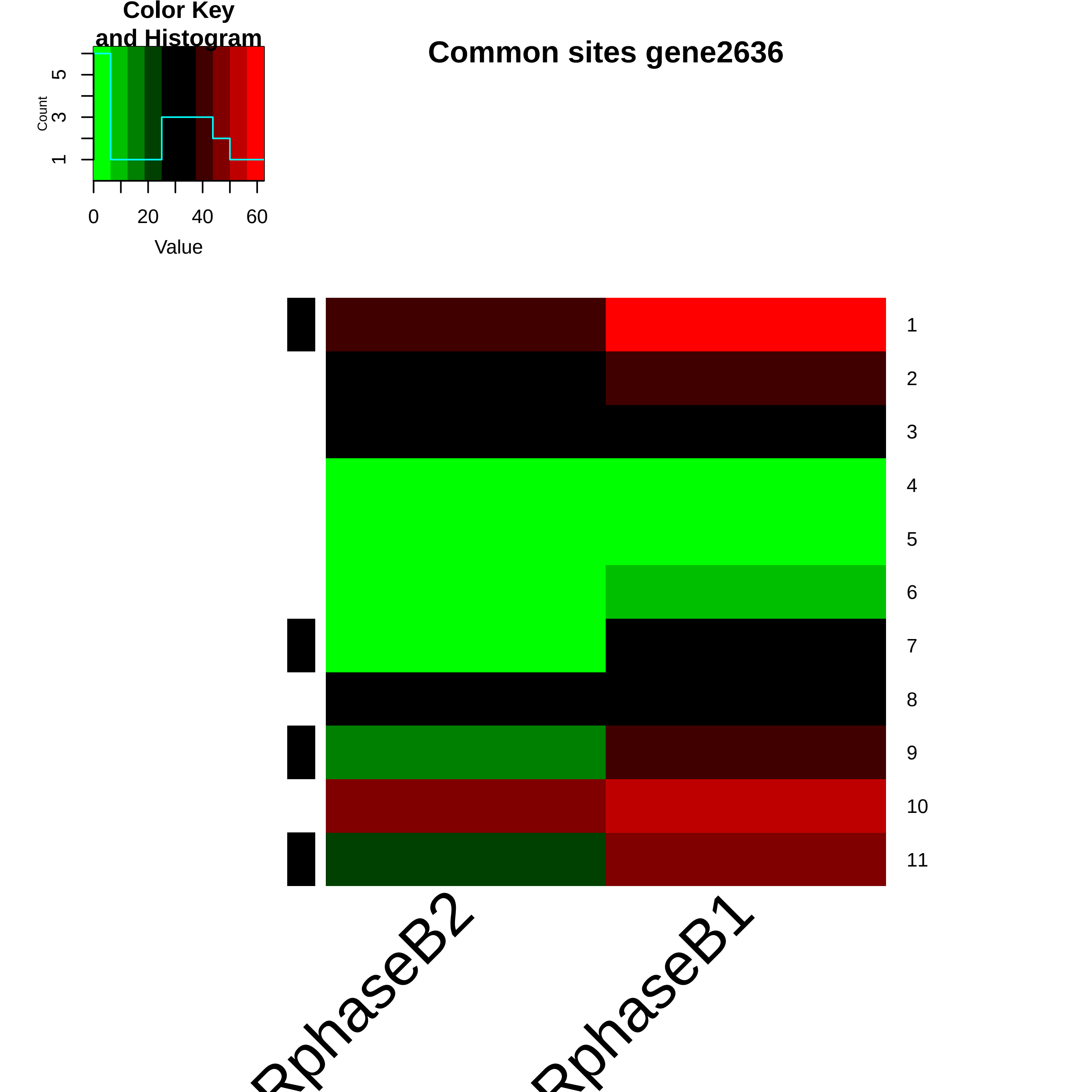

Color Key
and Histogram
Count
1
3
5
0
20
40
60
Value
Common sites gene2636
1
2
3
4
5
6
7
8
9
10
11
RphaseB2
RphaseB1

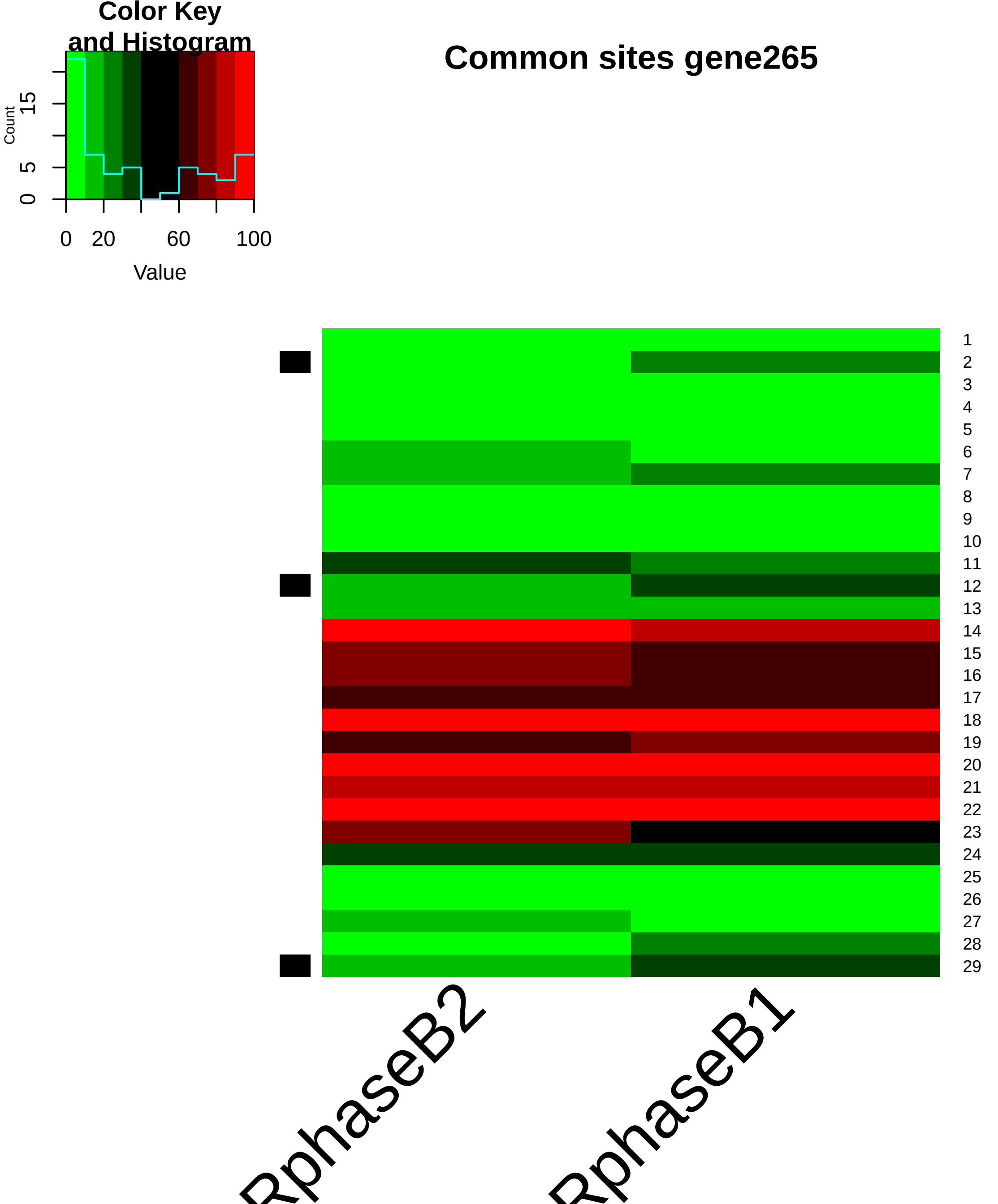
Color Key
and Histogram
Count
0
5
15
0
20
60
100
Value
Common sites gene265
RphaseB2
RphaseB1

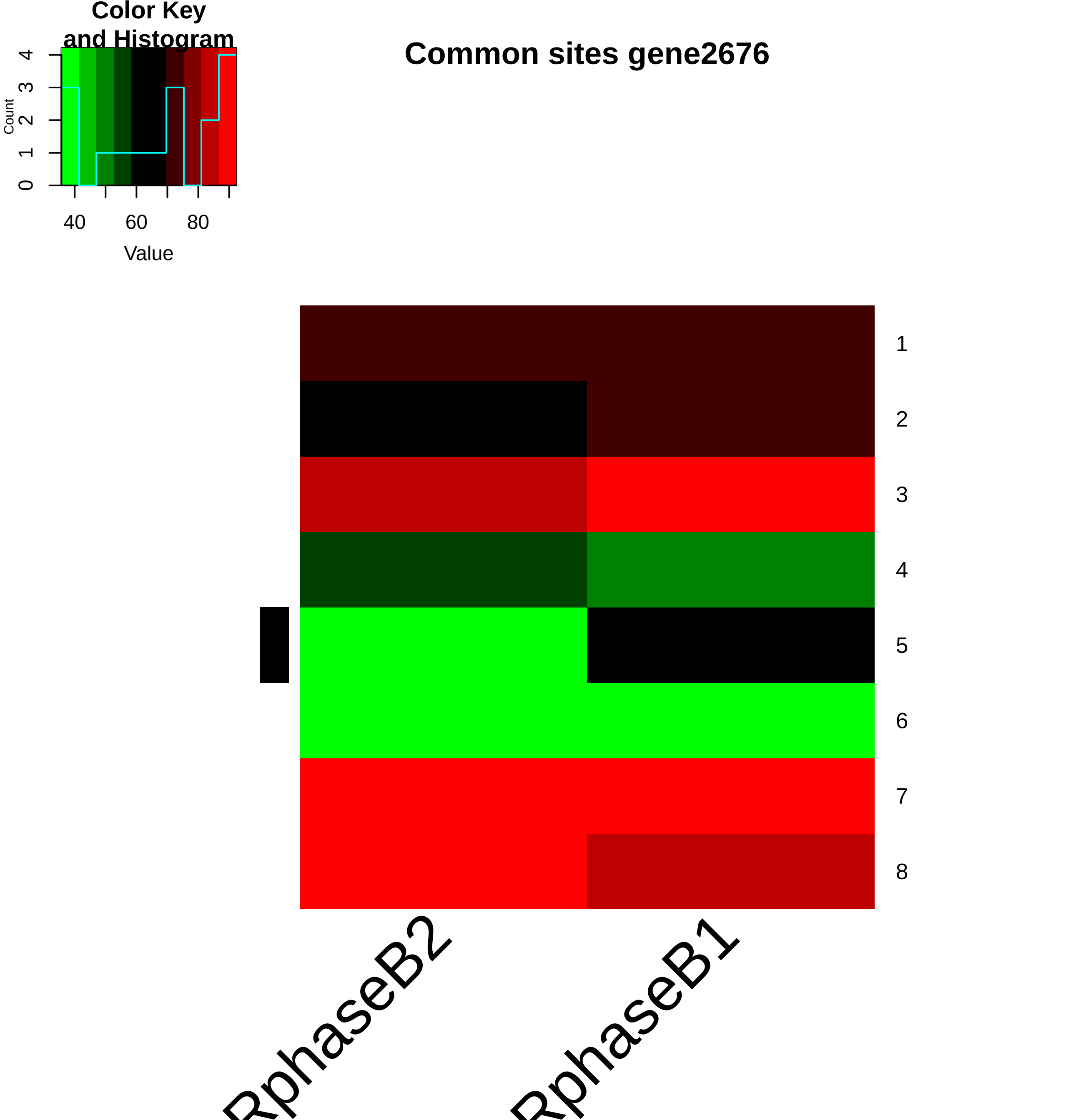
Color Key
and Histogram
Count
0
1
2
3
4
40
60
80
Value
Common sites gene2676
1
2
3
4
5
6
7
8
RphaseB2
RphaseB1

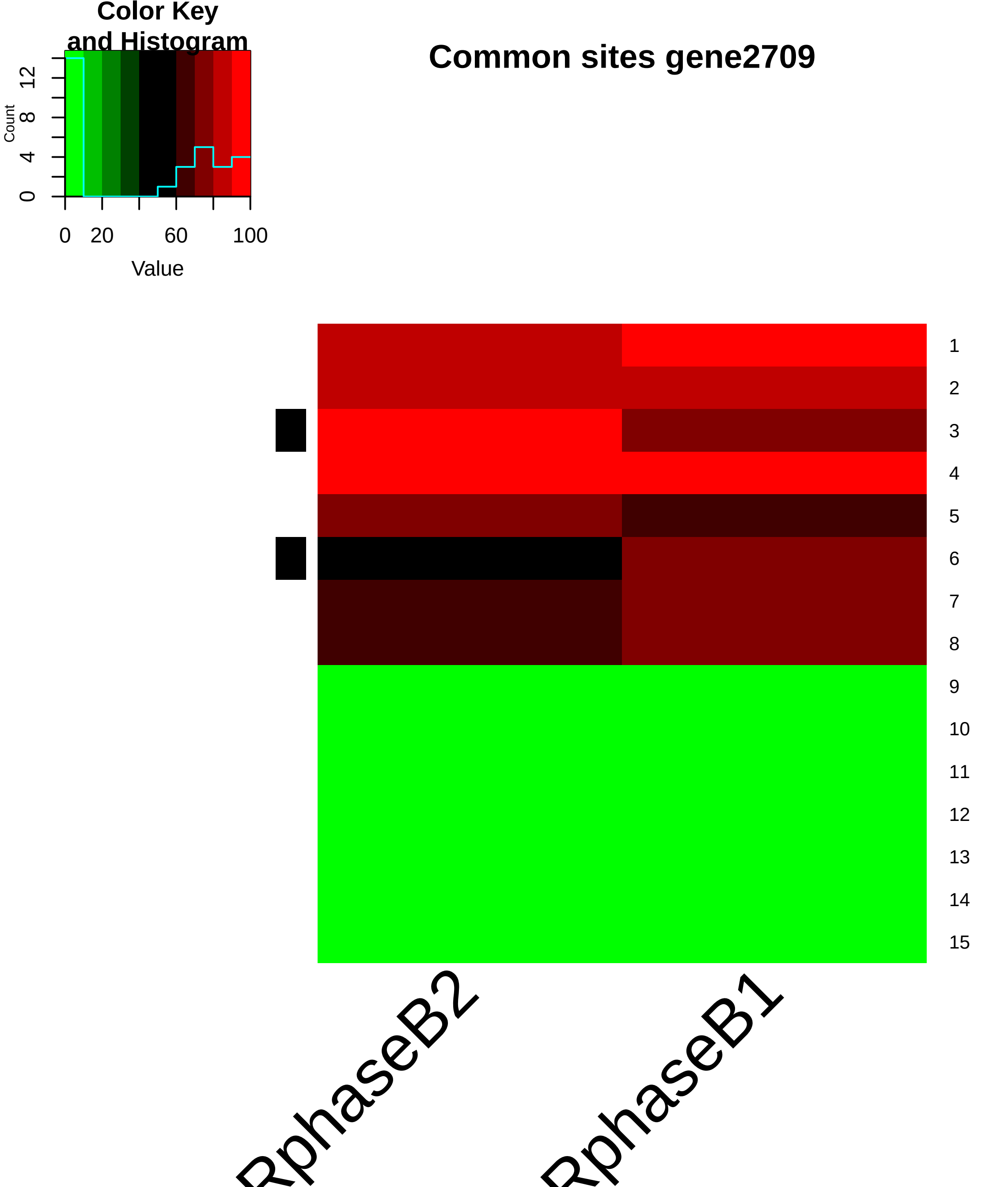

Color Key
and Histogram
Count
0
4
8
12
0
20
60
100
Value
Common sites gene2709
RphaseB2
RphaseB1

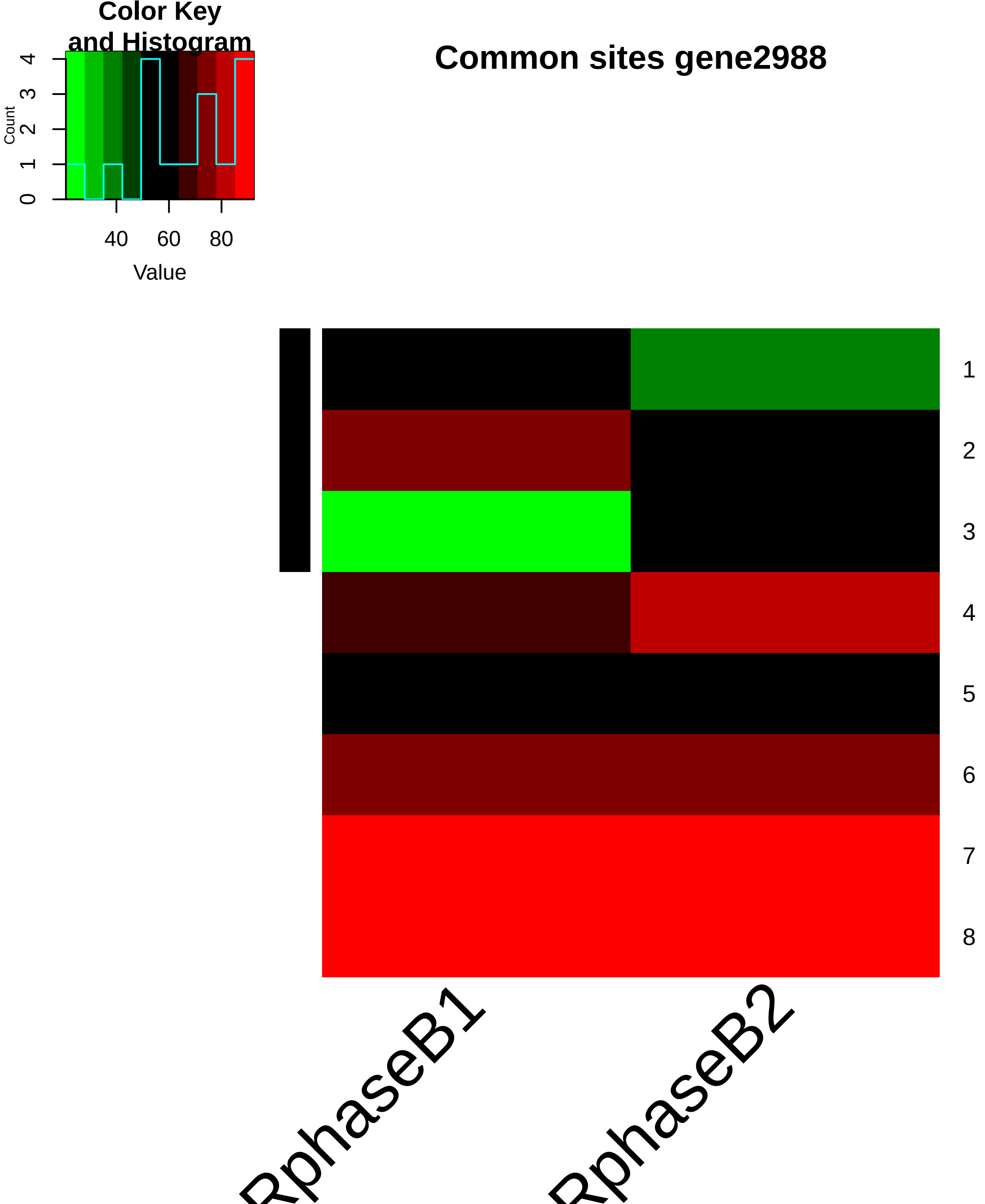

Color Key
and Histogram
Count
0
1
2
3
4
40
60
80
Value
Common sites gene2988
1
2
3
4
5
6
7
8
RphaseB1
RphaseB2

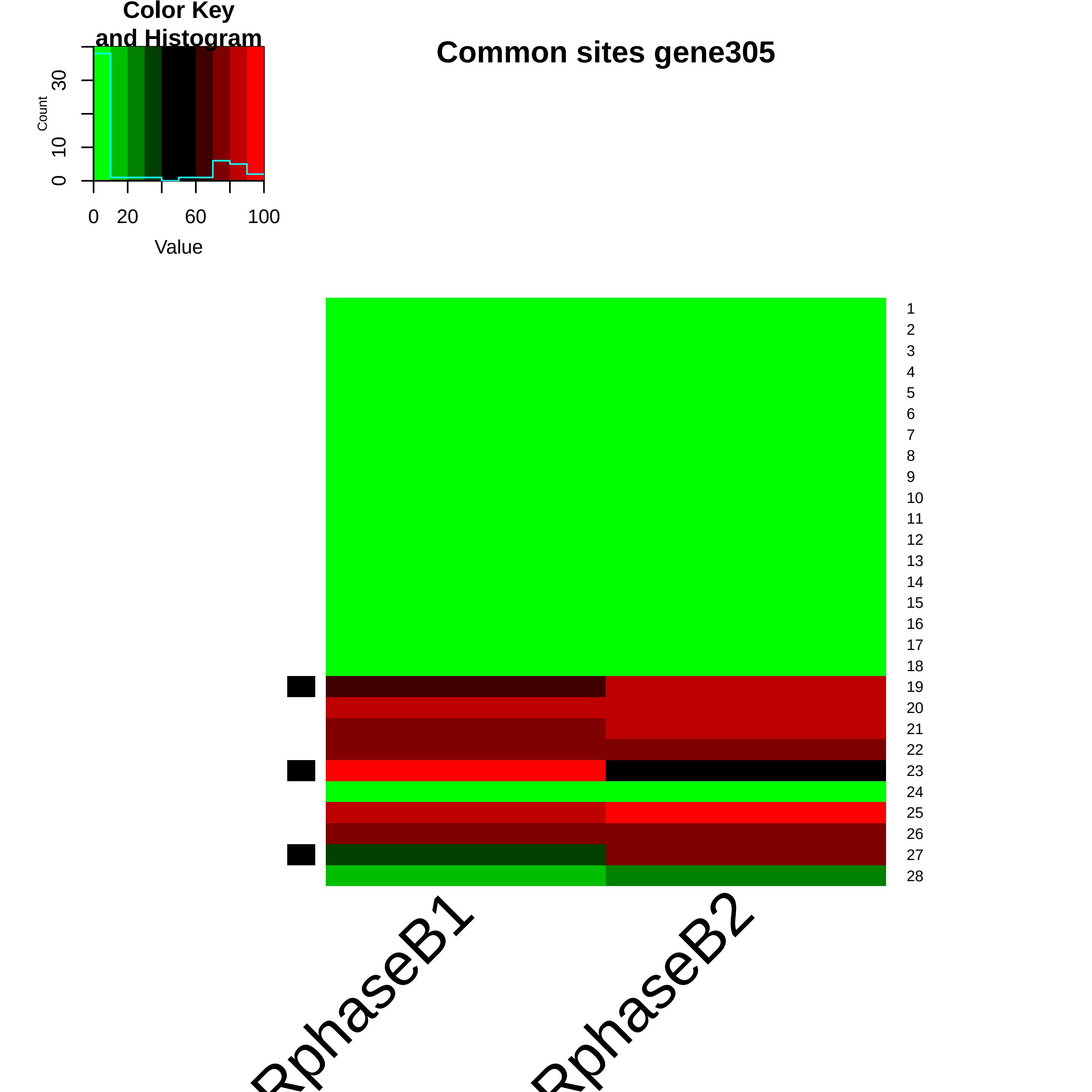

Color Key
and Histogram
Count
0
10
30
0
20
60
100
Value
Common sites gene305
RphaseB1
RphaseB2

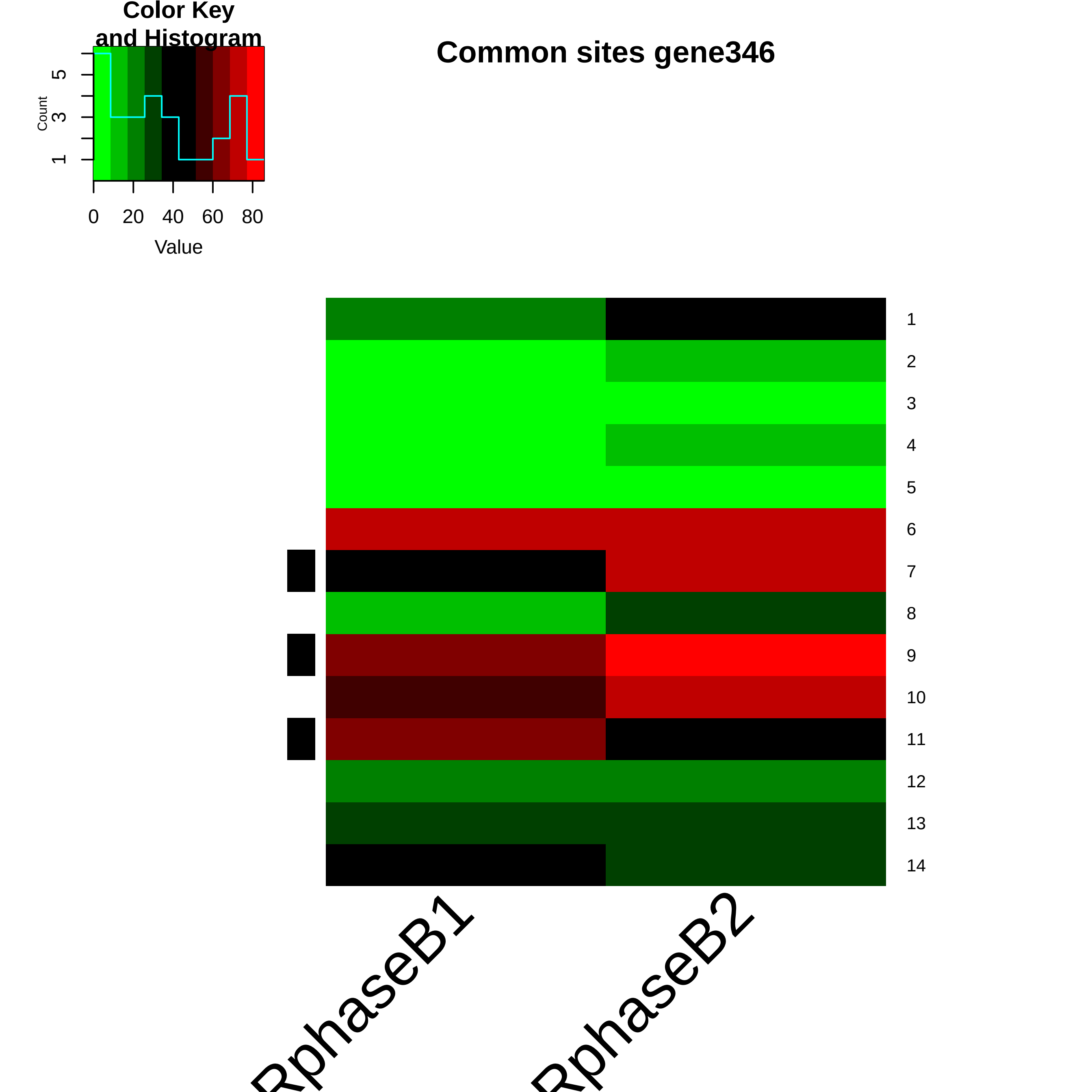
Color Key
and Histogram
Count
1
3
5
0
20
40
60
80
Value
Common sites gene346
1
2
3
4
5
6
7
8
9
10
11
12
13
14
RphaseB1
RphaseB2

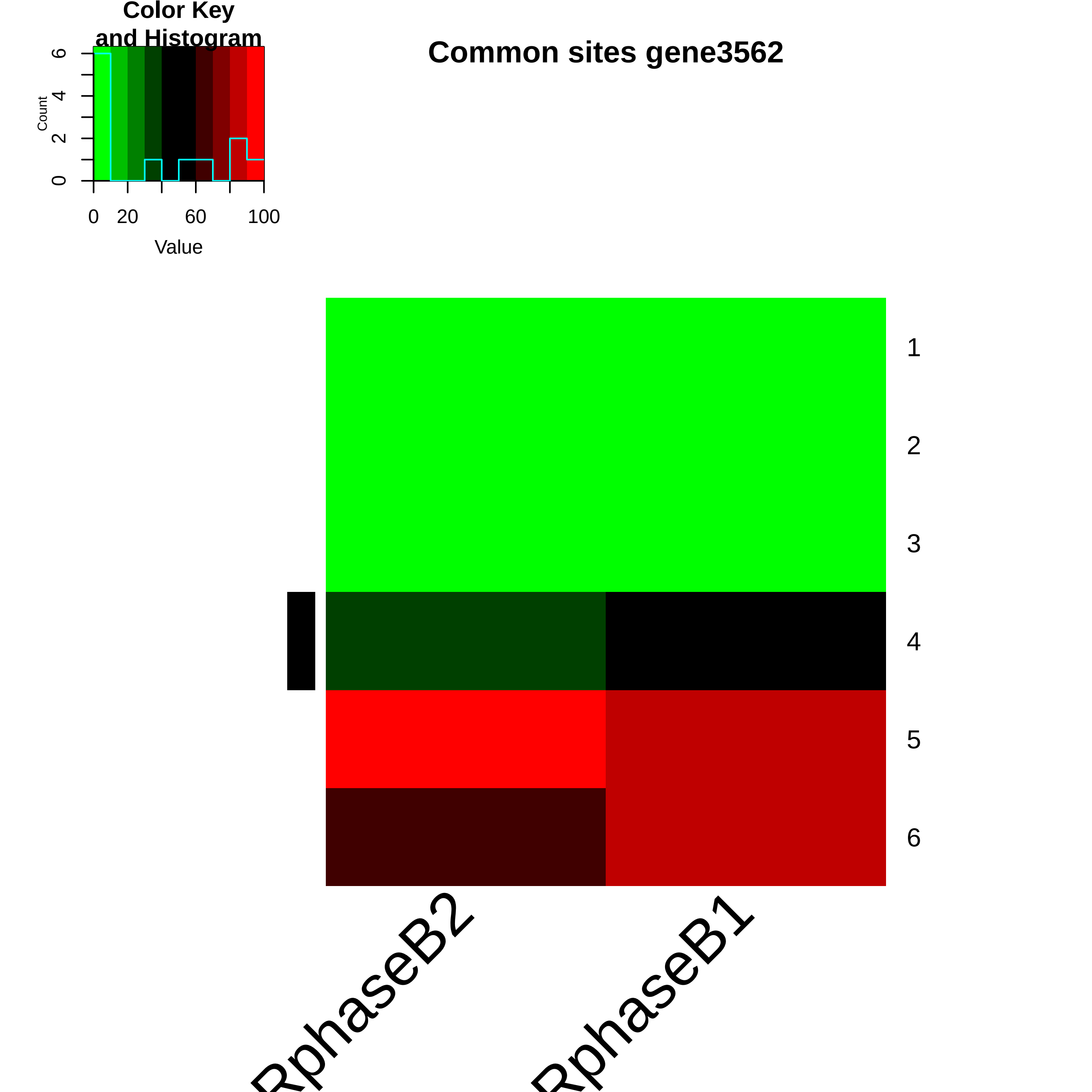
Color Key
and Histogram
Count
Value
0
20
60
100
0
2
4
6
Common sites gene3562
1
2
3
4
5
6
RphaseB2
RphaseB1

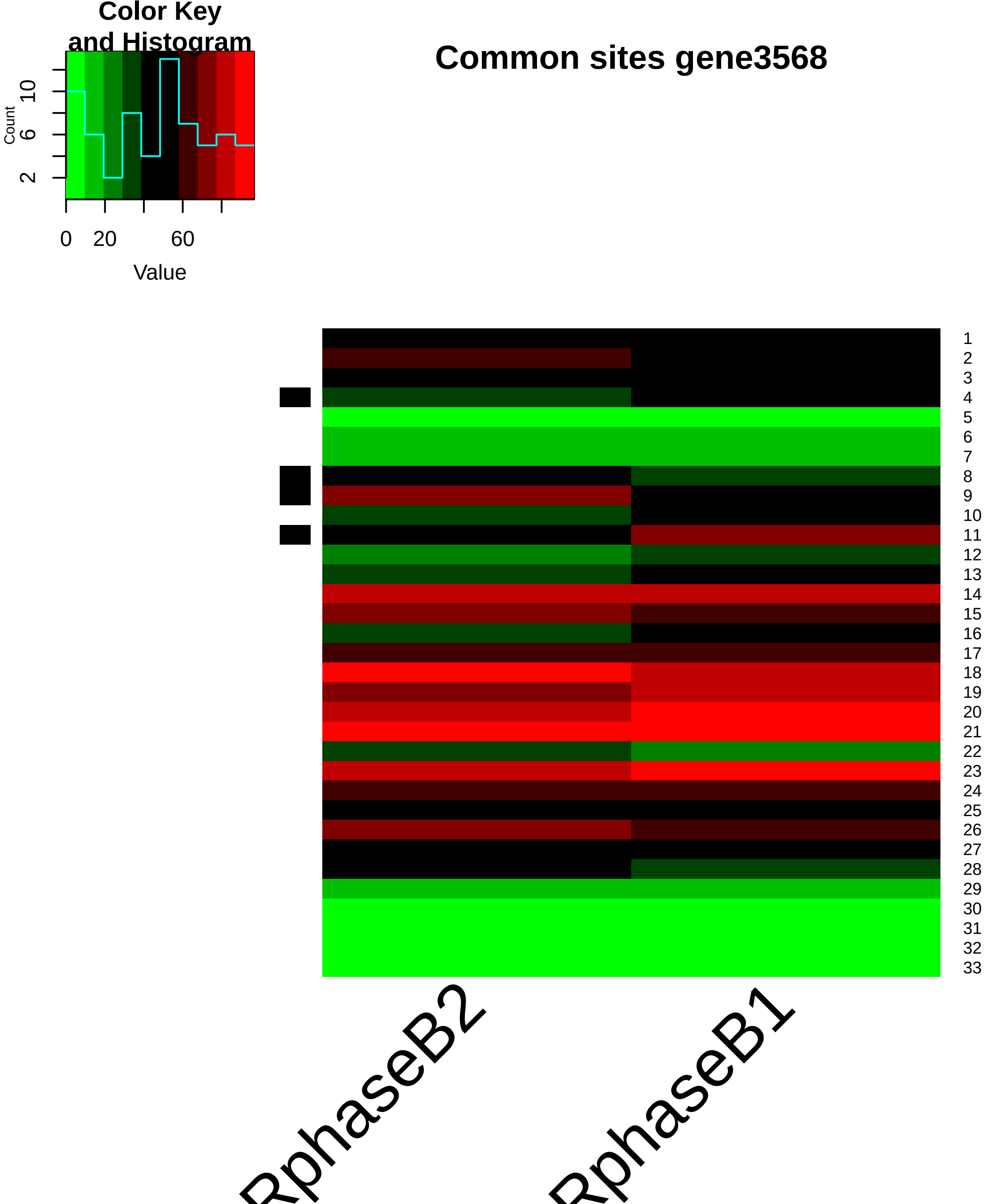

Color Key
and Histogram
Count
2
6
10
0
20
60
Value
Common sites gene3568
RphaseB2
RphaseB1

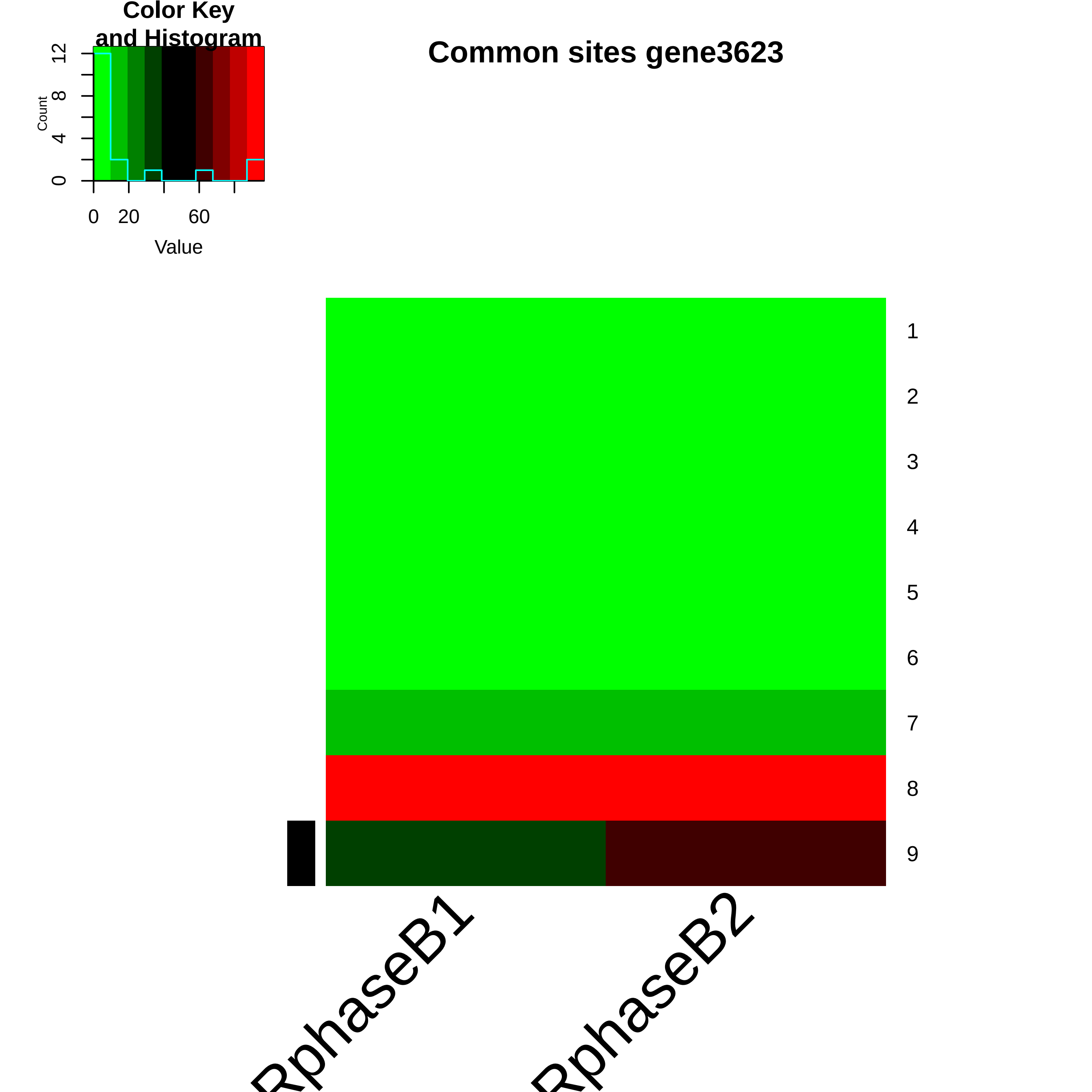

Color Key
and Histogram
Count
0
4
8
12
0
20
60
Value
Common sites gene3623
1
2
3
4
5
6
7
8
9
RphaseB1
RphaseB2

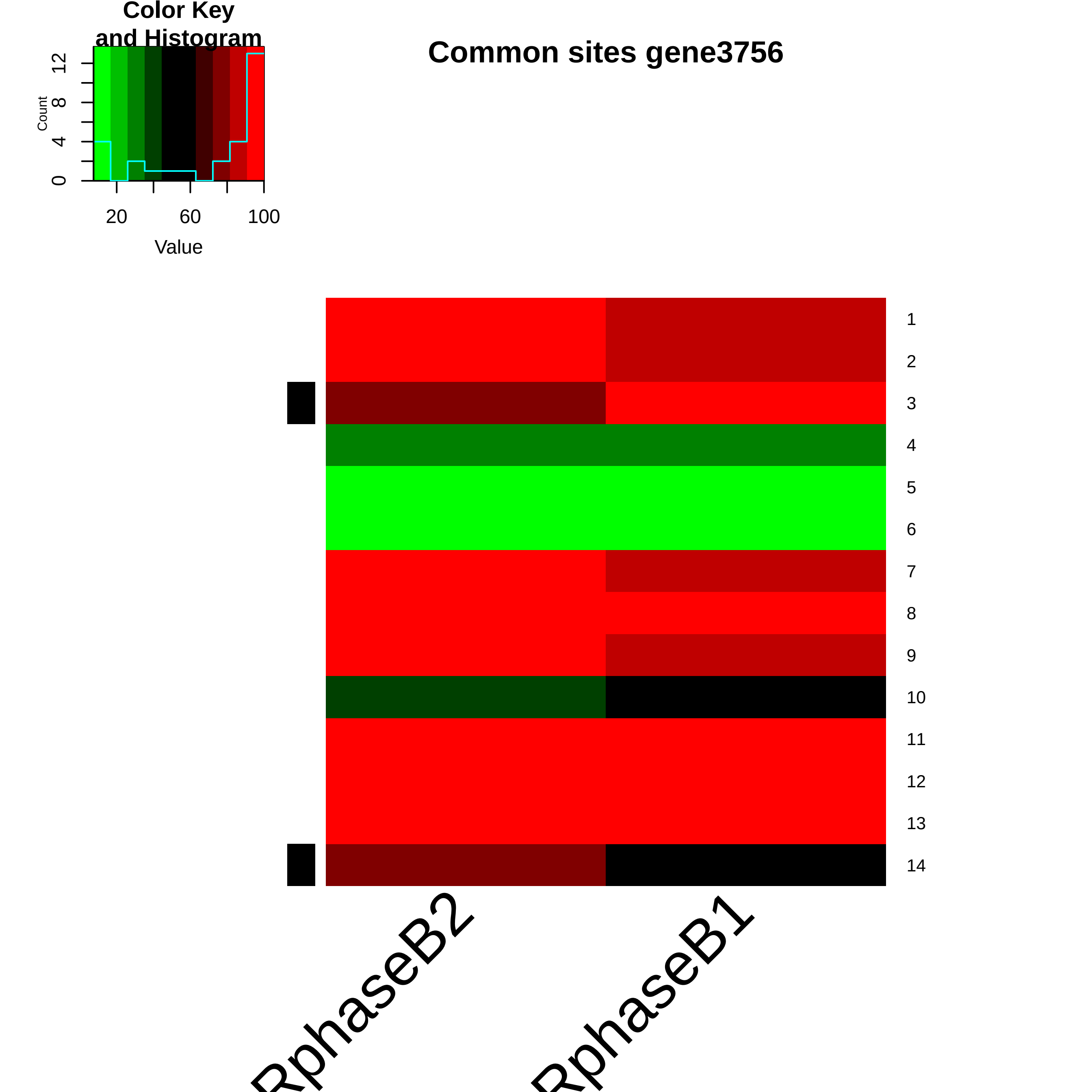
Color Key
and Histogram
Count
0
4
8
12
20
60
100
Value
Common sites gene3756
1
2
3
4
5
6
7
8
9
10
11
12
13
14
RphaseB2
RphaseB1

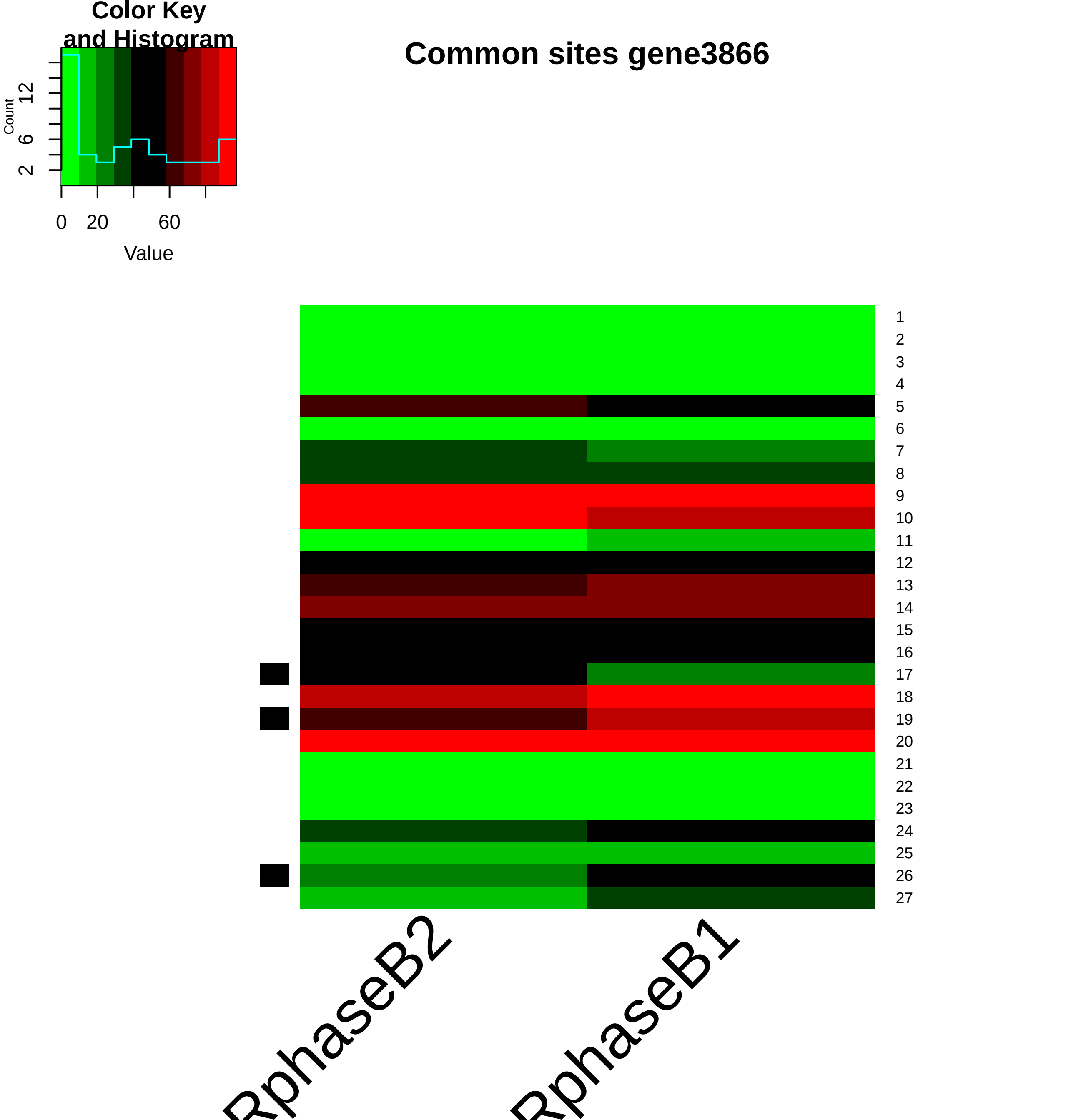

Color Key
and Histogram
Count
2
6
12
0
20
60
Value
Common sites gene3866
RphaseB2
RphaseB1

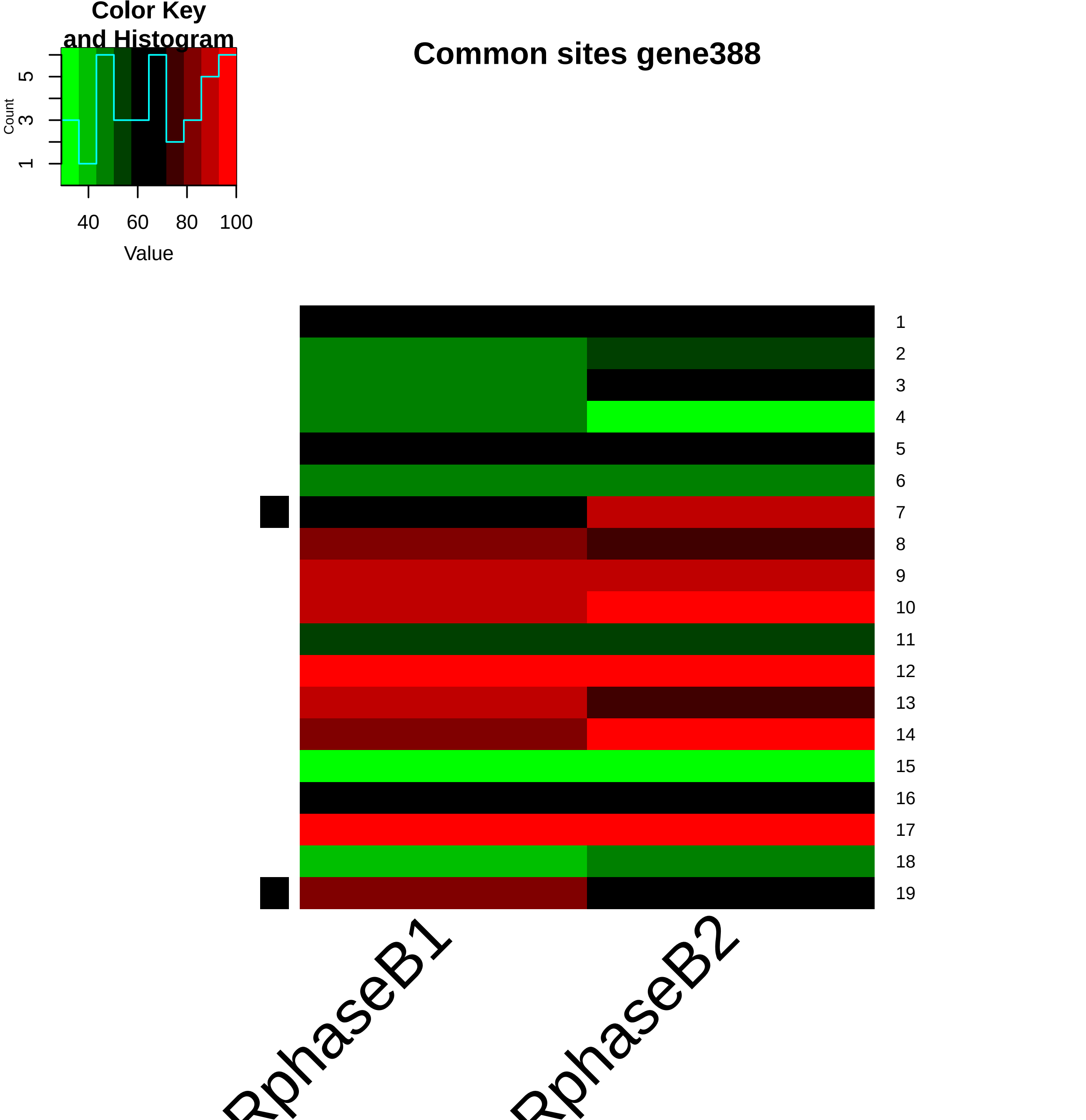

Color Key and Histogram
Count
1
3
5
40
60
80
100
Value
Common sites gene388
RphaseB1
RphaseB2

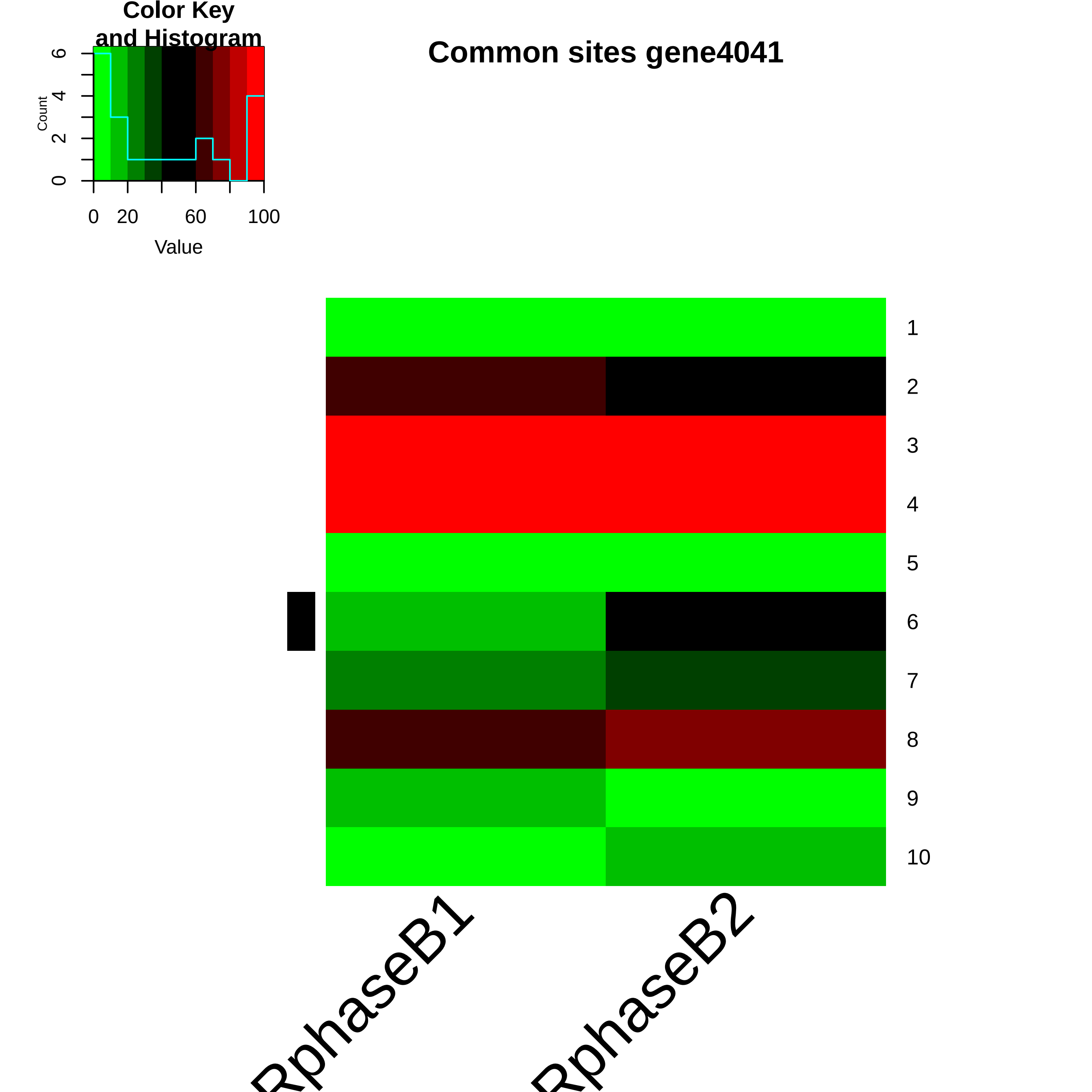
Color Key
and Histogram
Count
0
2
4
6
0
20
60
100
Value
Common sites gene4041
1
2
3
4
5
6
7
8
9
10
RphaseB1
RphaseB2

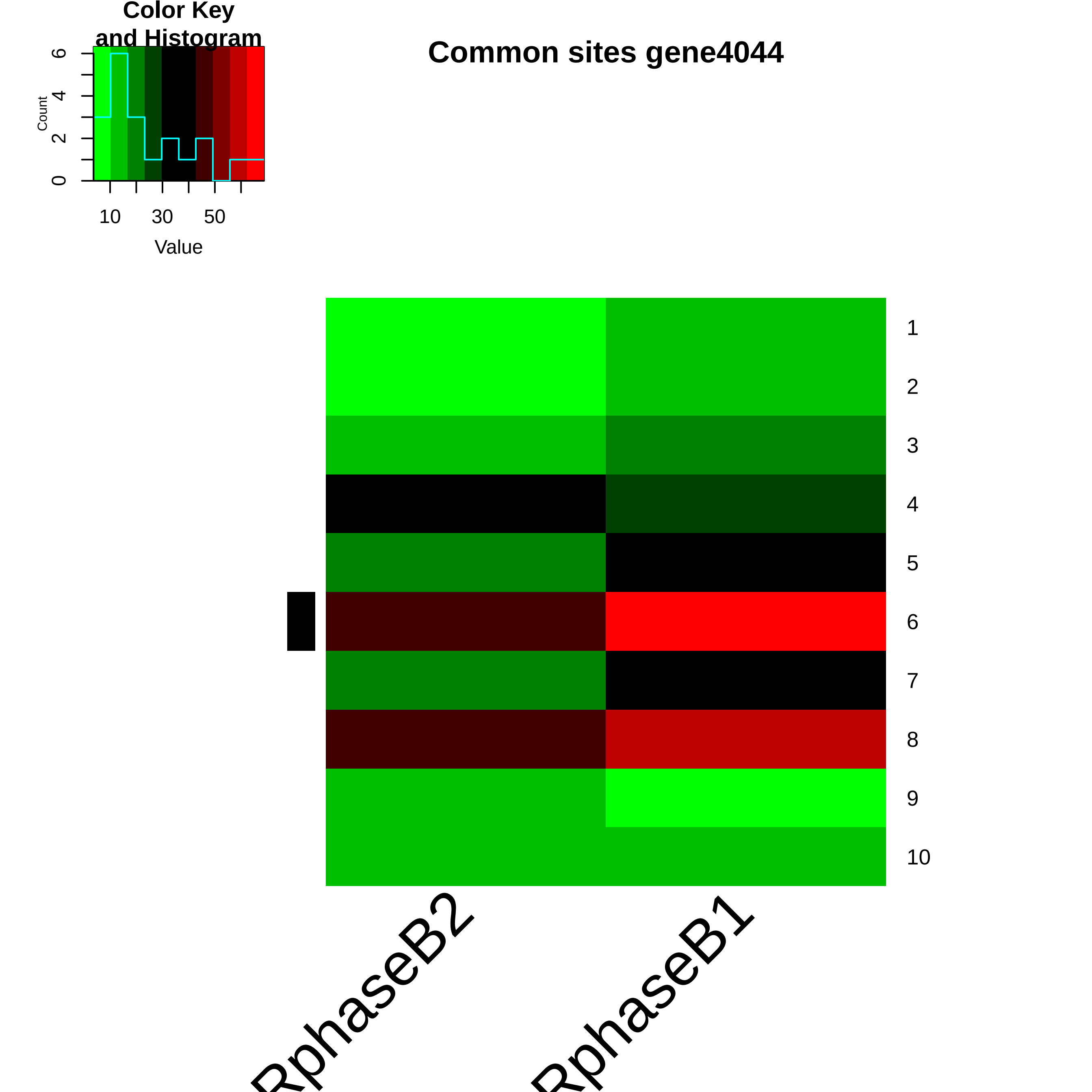
Color Key
and Histogram
Count
0
2
4
6
10
30
50
Value
Common sites gene4044
1
2
3
4
5
6
7
8
9
10
RphaseB2
RphaseB1

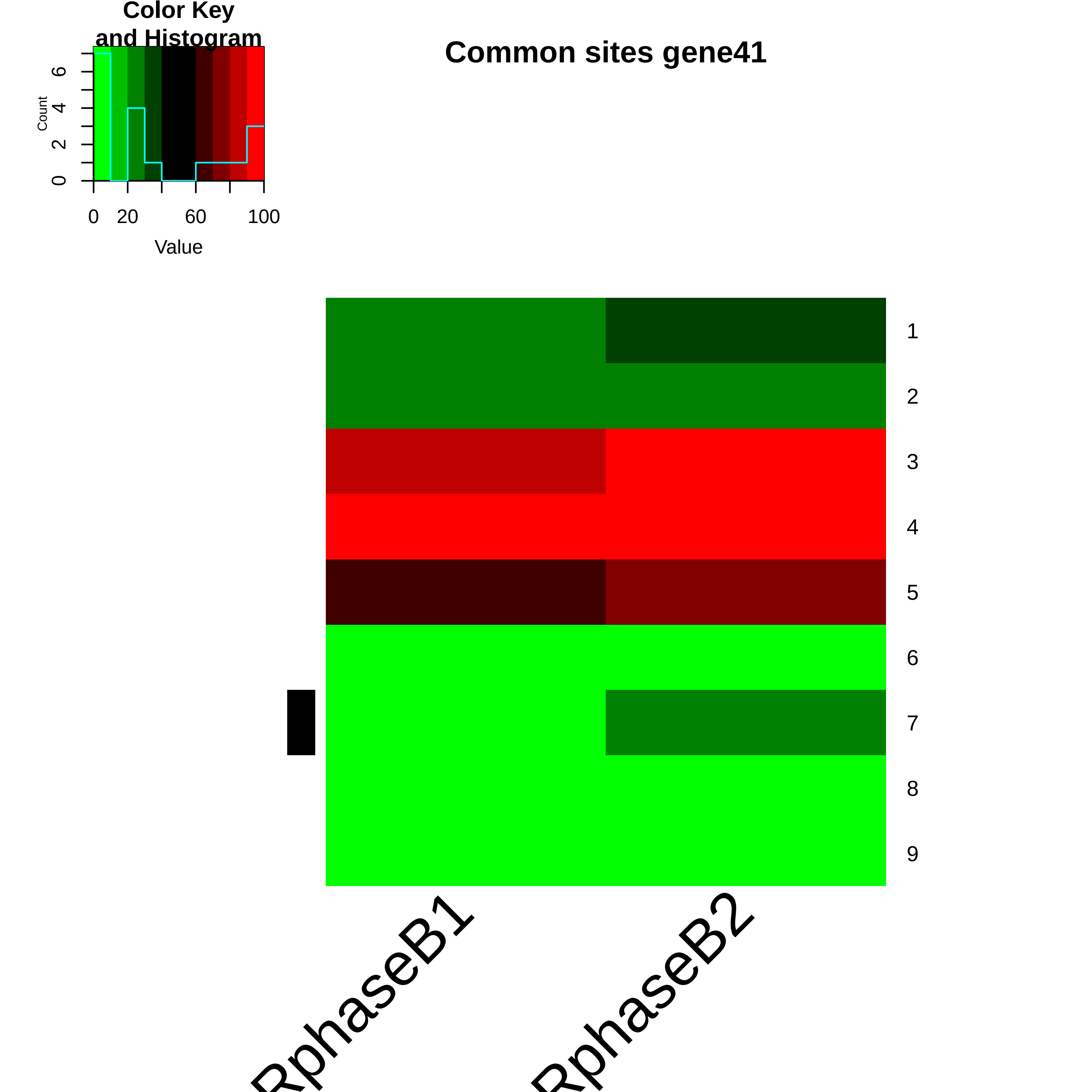
Color Key
and Histogram
Count
0
2
4
6
0
20
60
100
Value
Common sites gene41
1
2
3
4
5
6
7
8
9
RphaseB1
RphaseB2

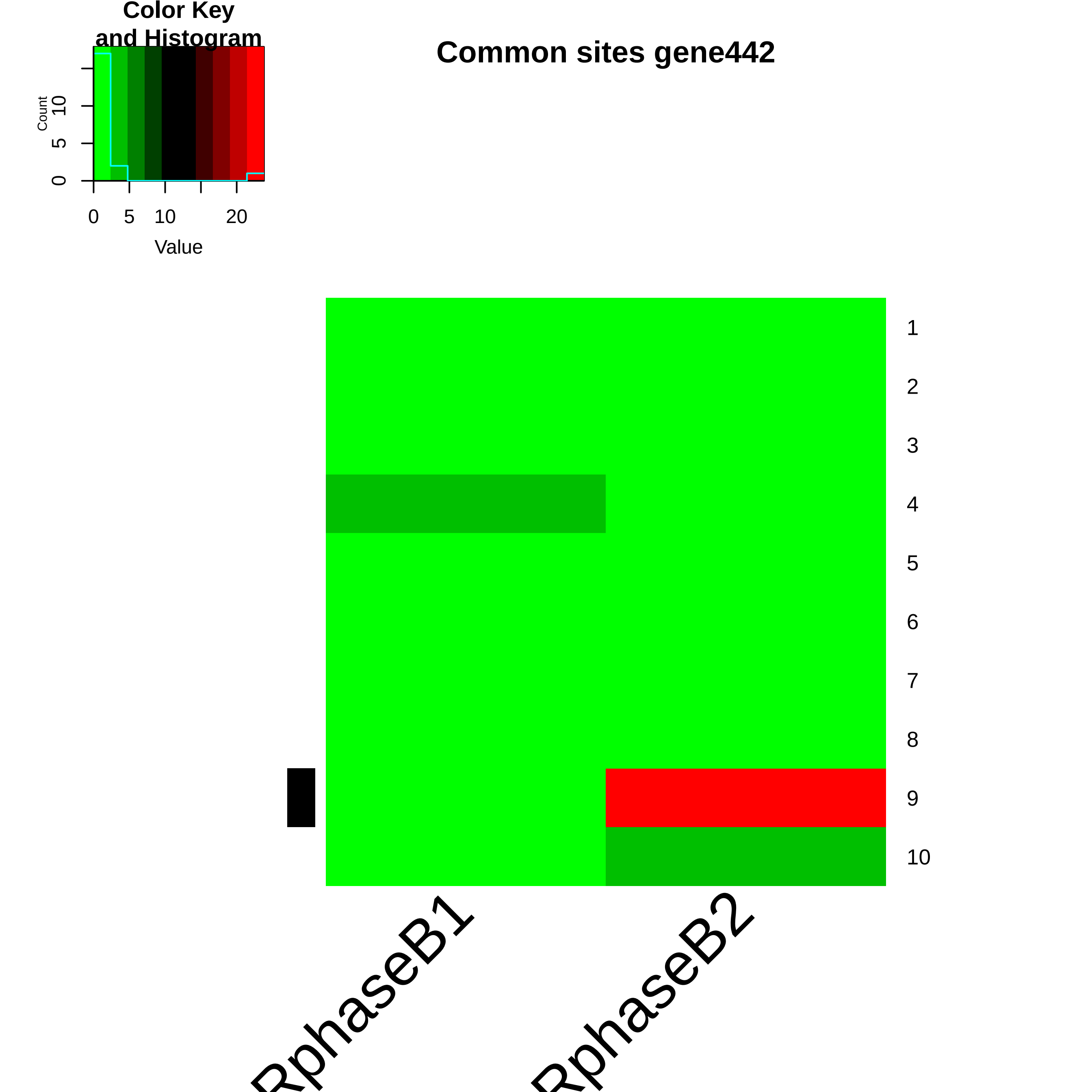
Color Key
and Histogram
Count
0
5
10
0
5
10
20
Value
Common sites gene442
1
2
3
4
5
6
7
8
9
10
RphaseB1
RphaseB2

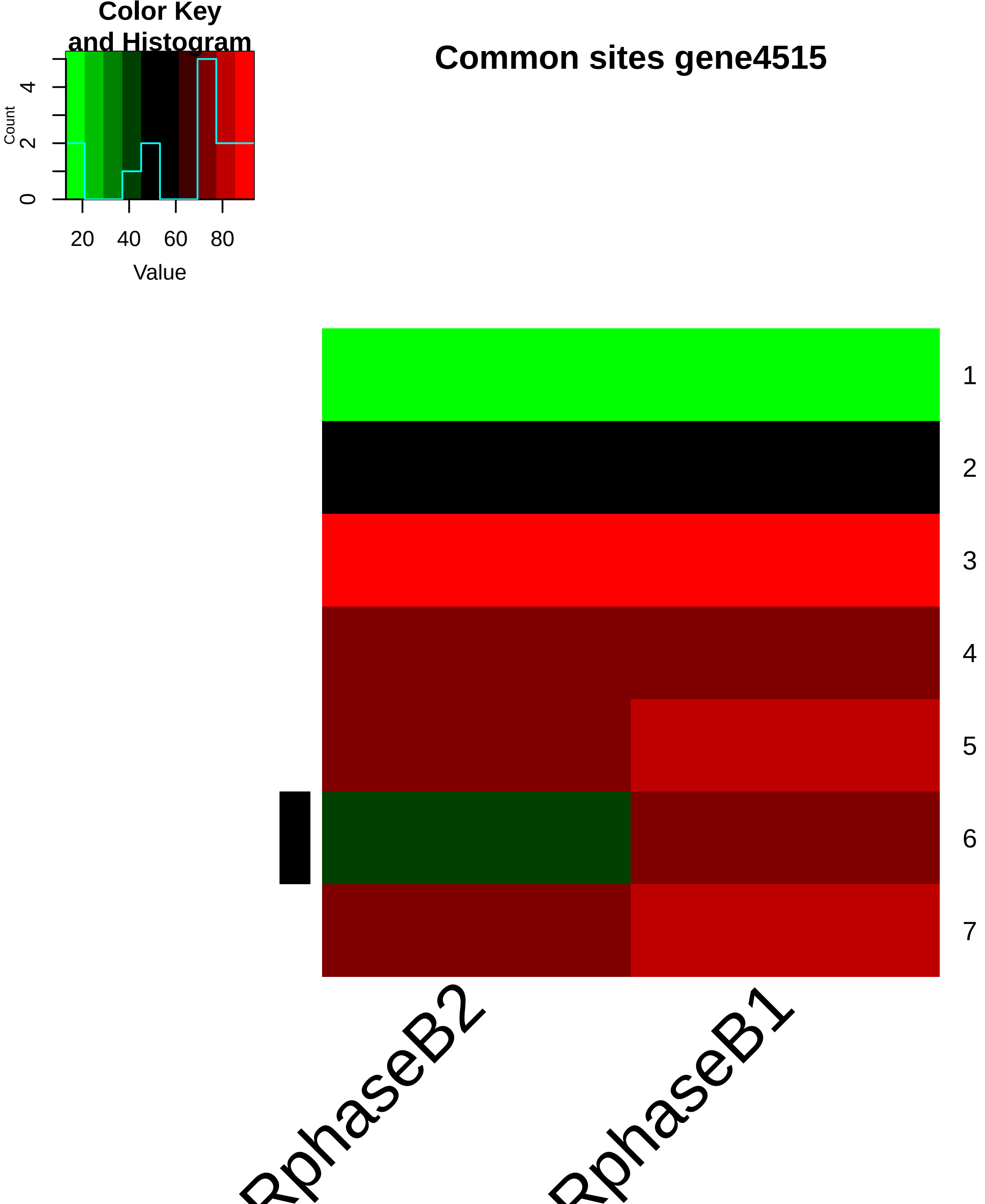

Color Key
and Histogram
Count
0
2
4
20
40
60
80
Value
Common sites gene4515
1
2
3
4
5
6
7
RphaseB2
RphaseB1

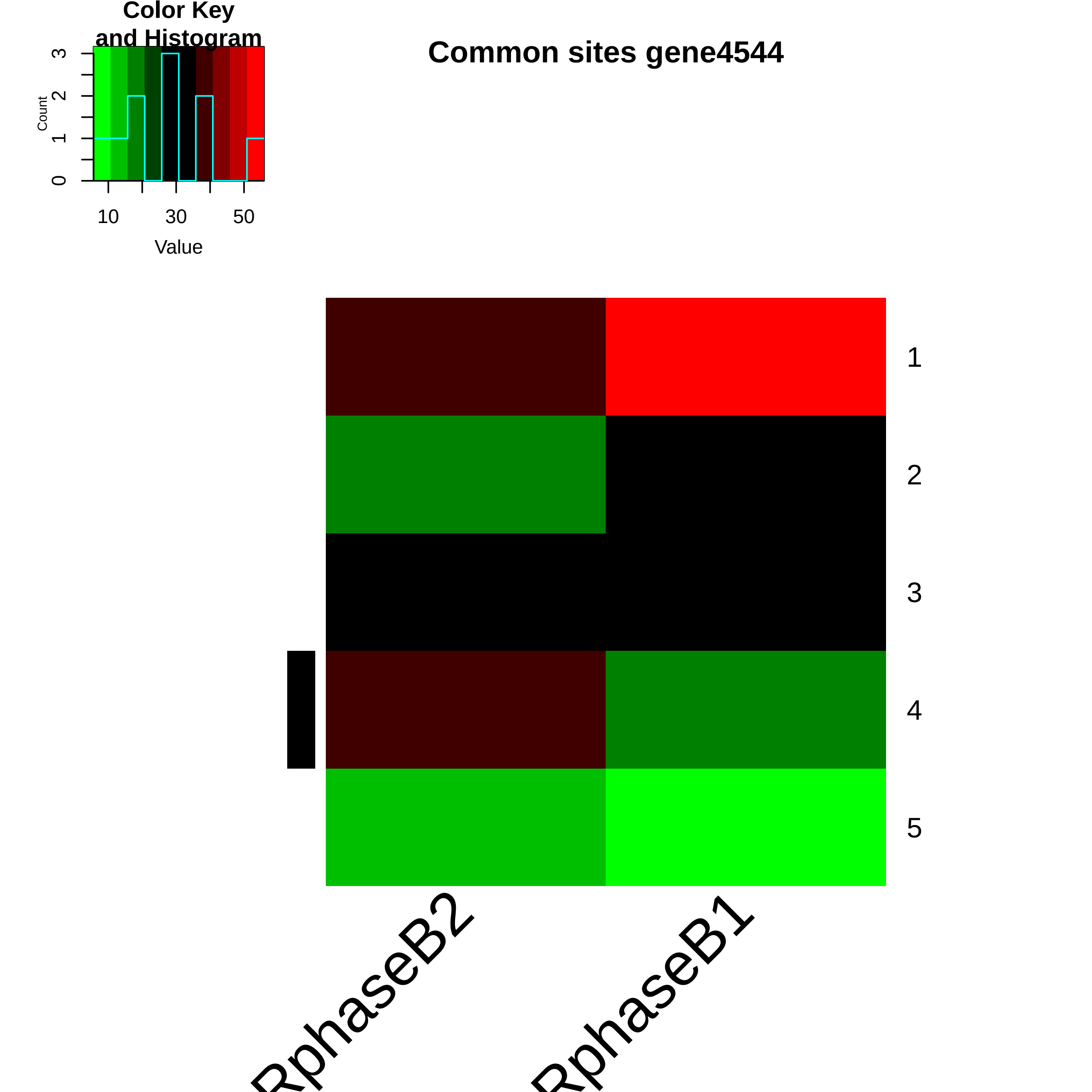

Color Key
and Histogram
Count
0
1
2
3
10
30
50
Value
Common sites gene4544
1
2
3
4
5
RphaseB2
RphaseB1

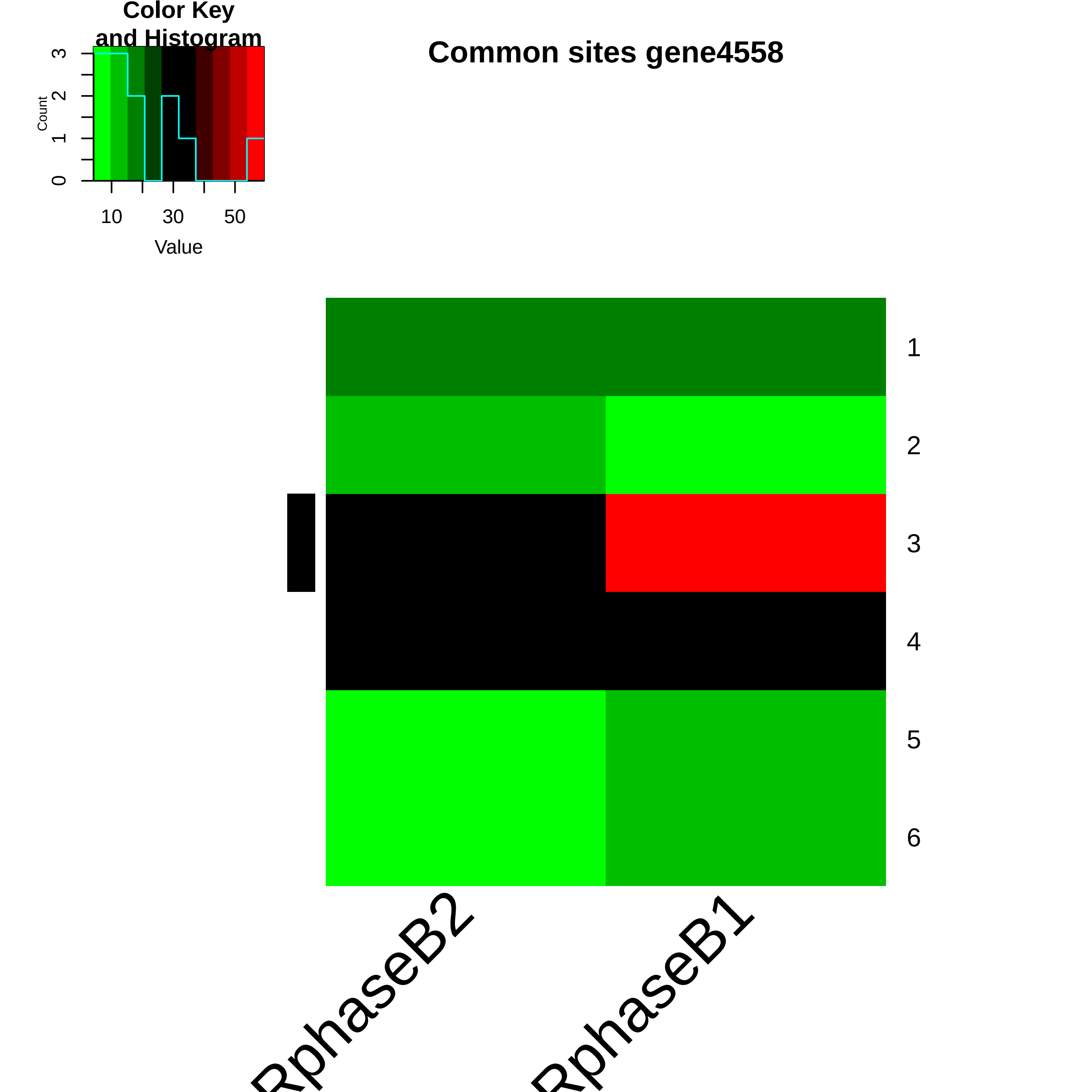
Color Key
and Histogram
Count
0
1
2
3
10
30
50
Value
Common sites gene4558
1
2
3
4
5
6
RphaseB2
RphaseB1

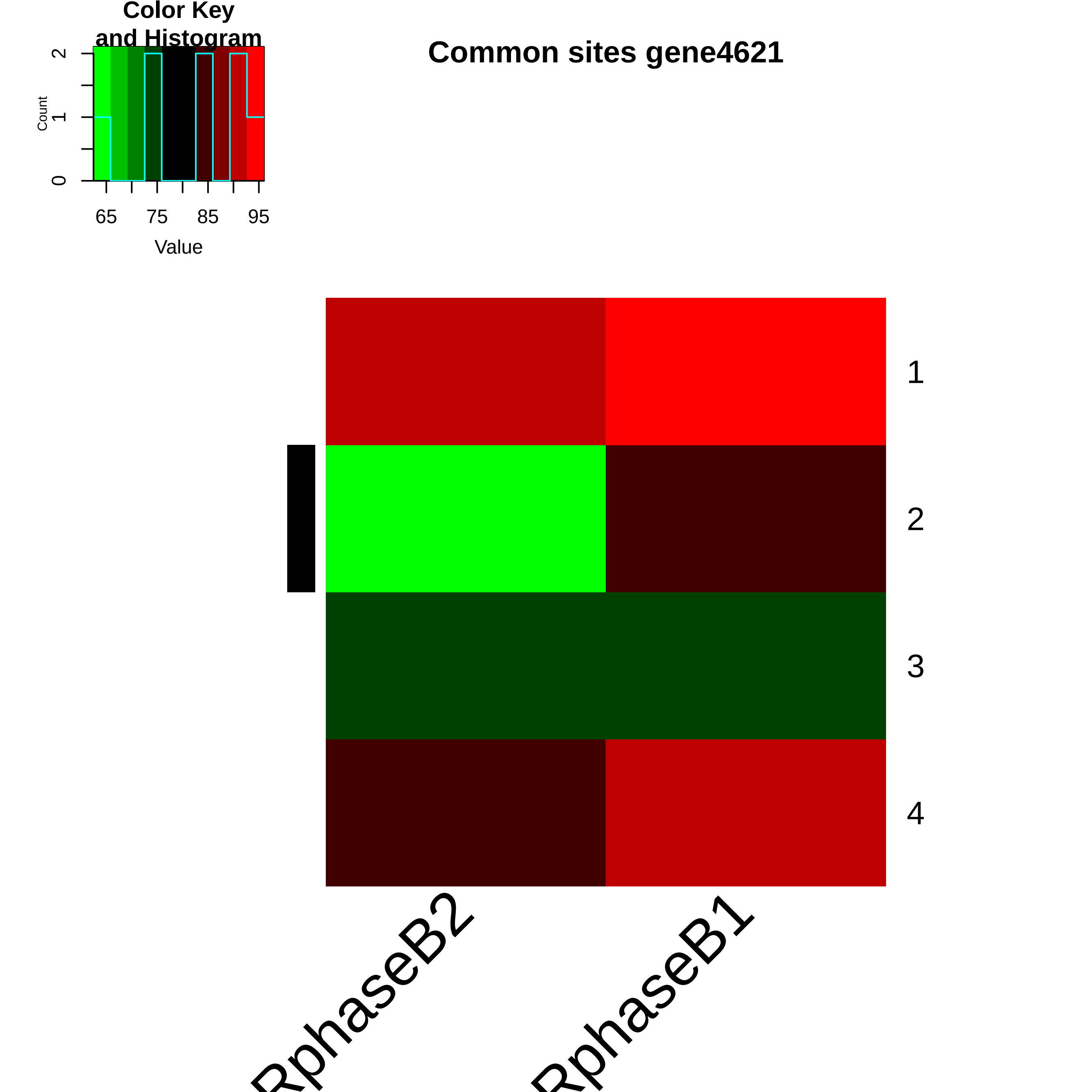
Color Key
and Histogram
Count
0
1
2
65
75
85
95
Value
Common sites gene4621
1
2
3
4
RphaseB2
RphaseB1

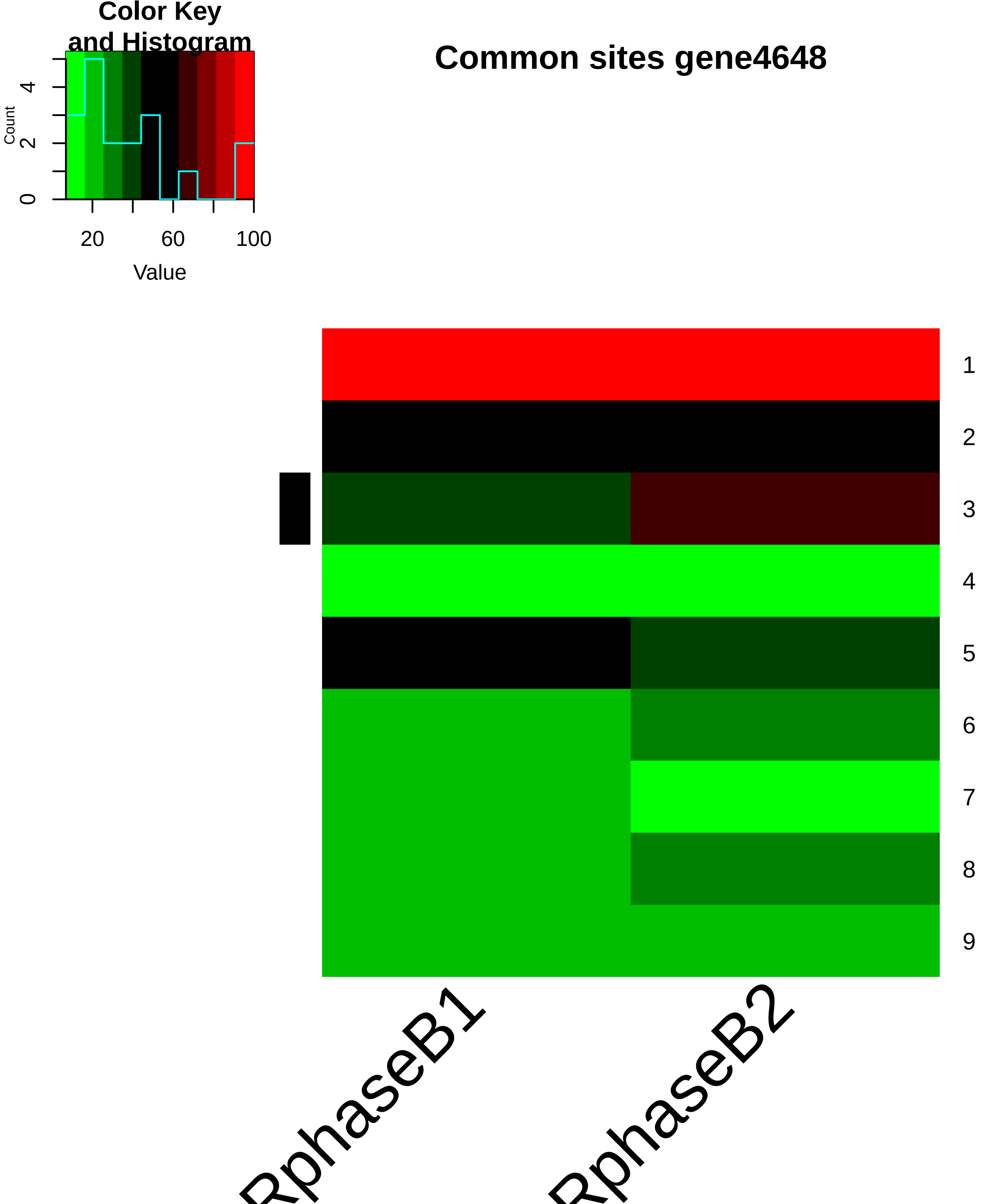

Color Key
and Histogram
Count
0
2
4
20
60
100
Value
Common sites gene4648
1
2
3
4
5
6
7
8
9
RphaseB1
RphaseB2

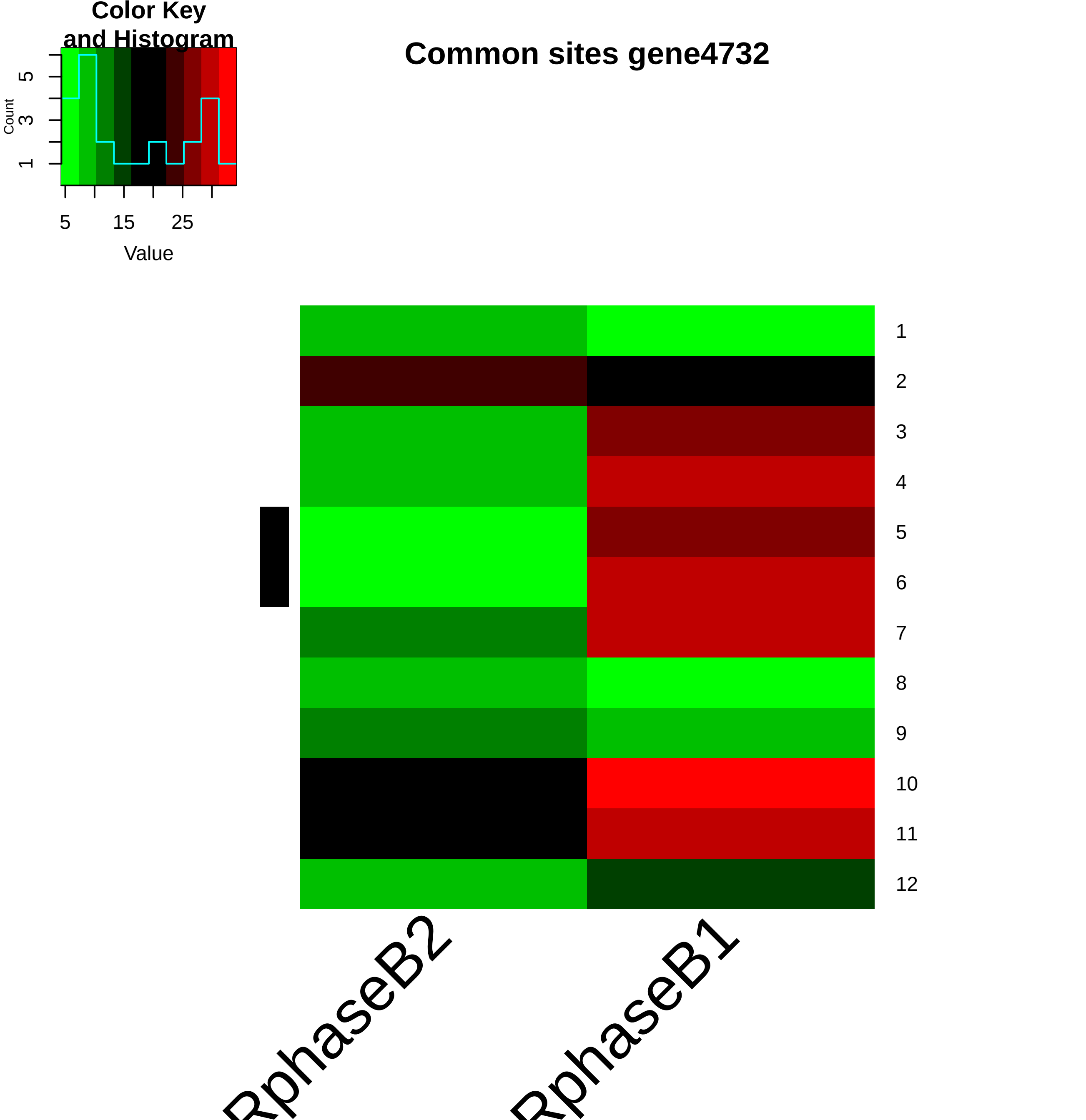
Color Key
and Histogram
Count
1
3
5
5
15
25
Value
Common sites gene4732
1
2
3
4
5
6
7
8
9
10
11
12
RphaseB2
RphaseB1

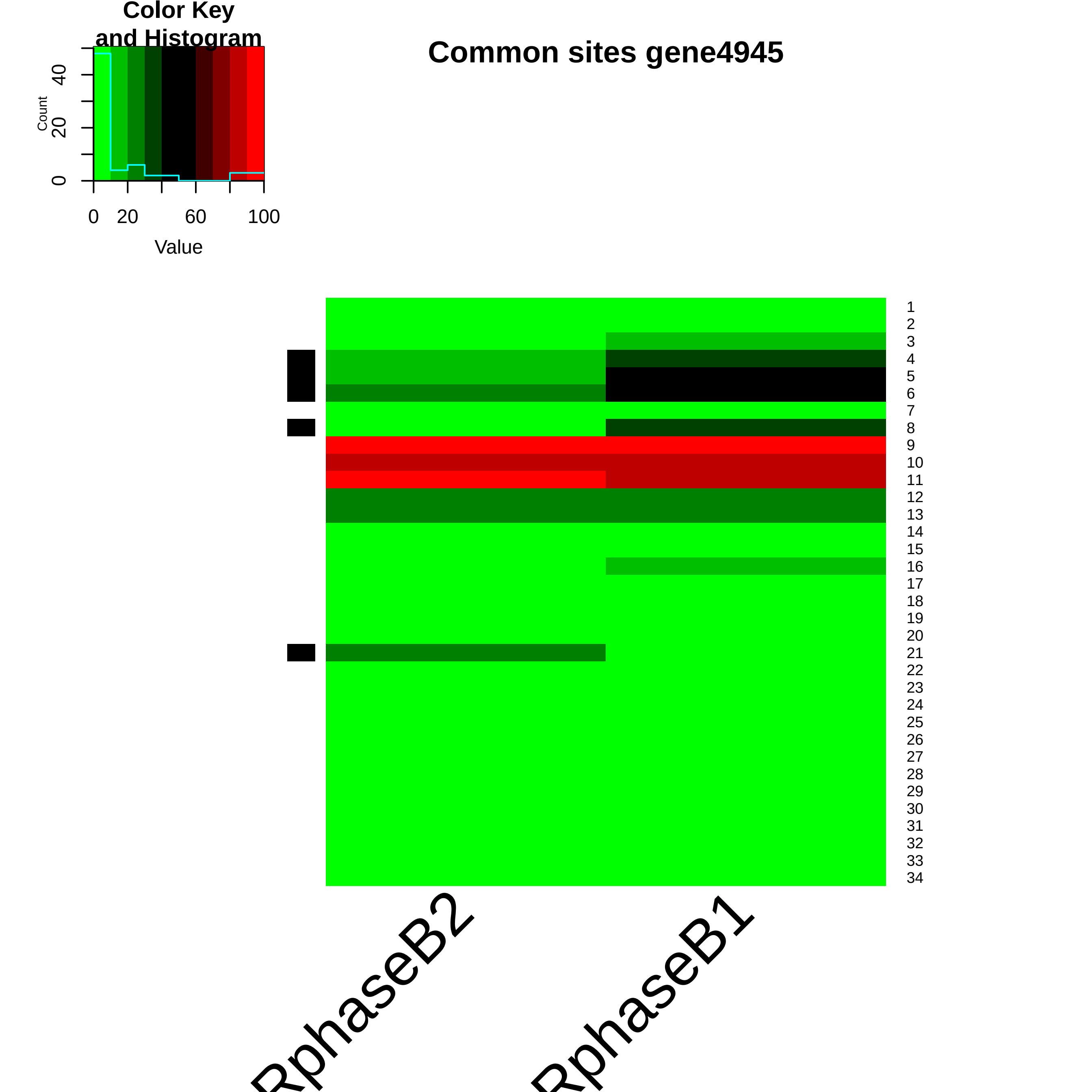

Color Key
and Histogram
Count
0
20
40
0
20
60
100
Value
Common sites gene4945
RphaseB2
RphaseB1

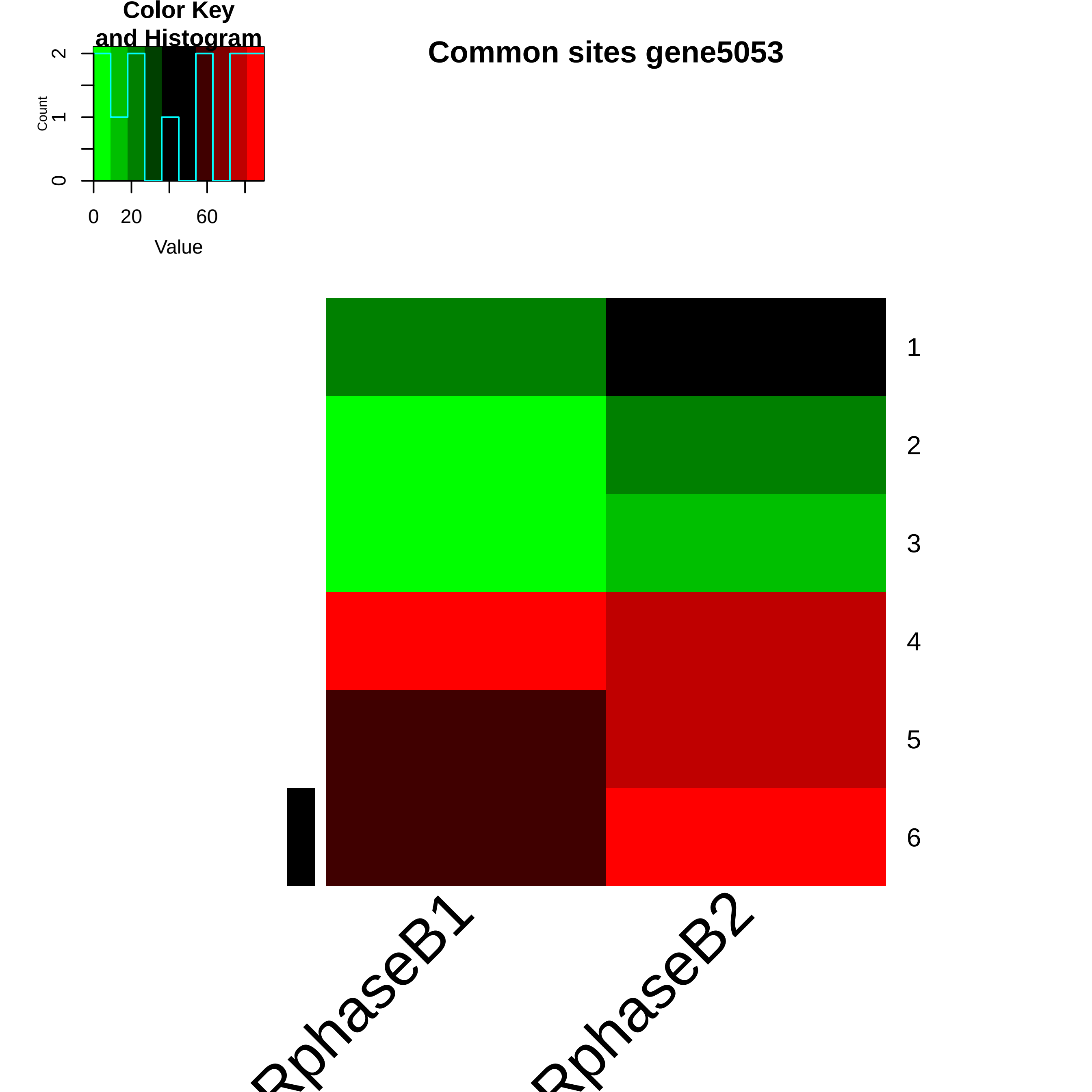
Color Key
and Histogram
Count
0
1
2
0
20
60
Value
Common sites gene5053
1
2
3
4
5
6
RphaseB1
RphaseB2

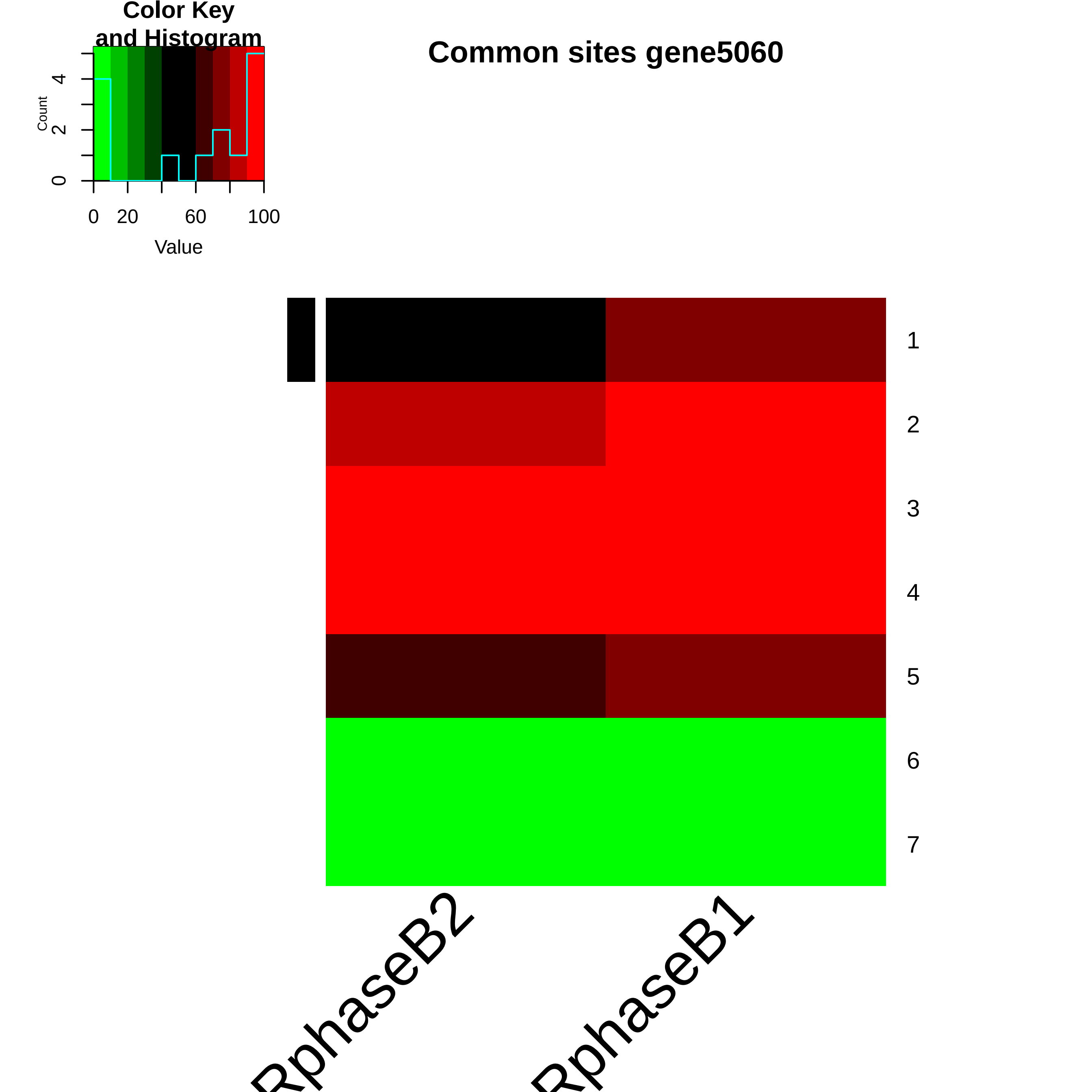
Color Key
and Histogram
Count
0
2
4
0
20
60
100
Value
Common sites gene5060
1
2
3
4
5
6
7
RphaseB2
RphaseB1

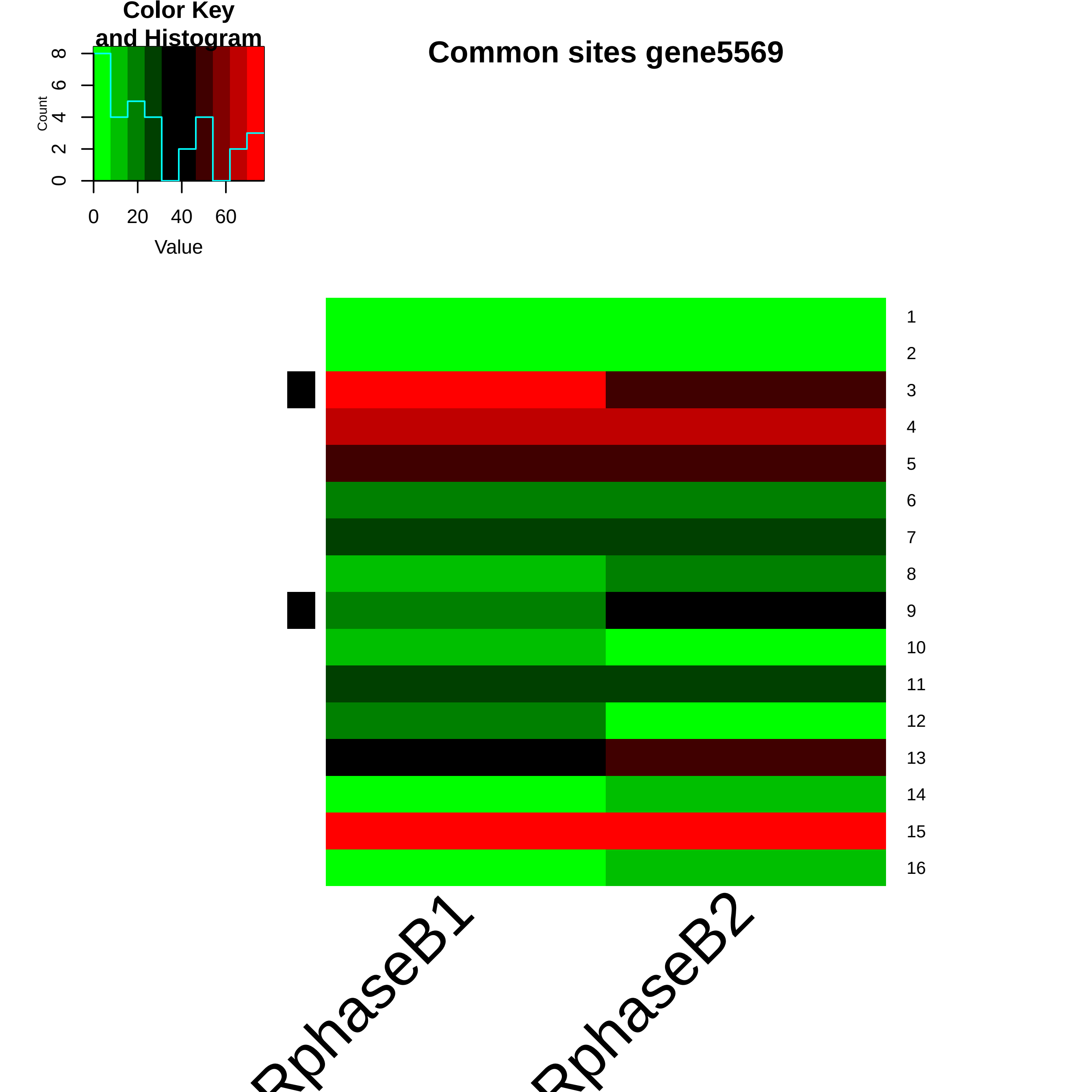
Color Key
and Histogram
Count
0
2
4
6
8
0
20
40
60
Value
Common sites gene5569
RphaseB1
RphaseB2

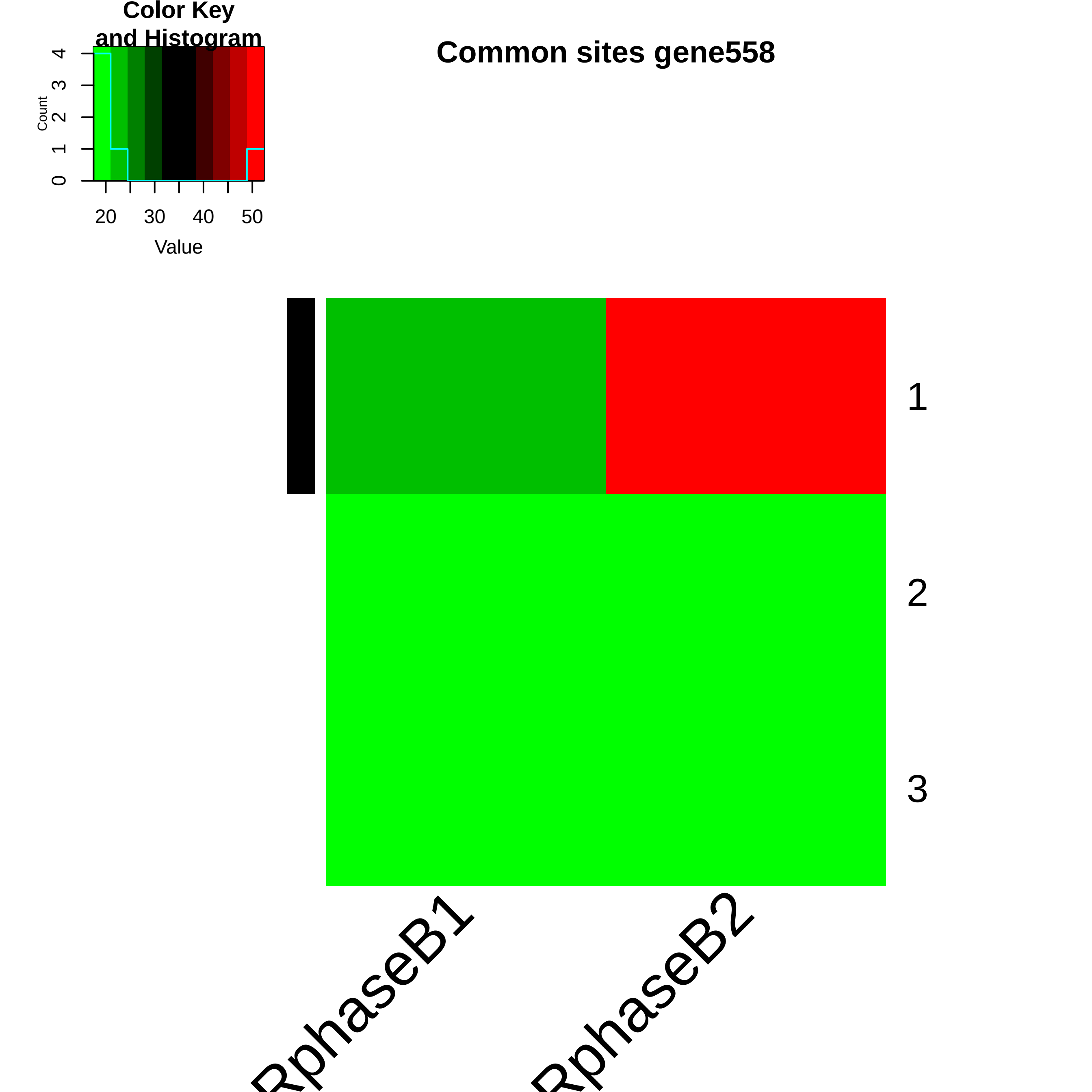

Color Key
and Histogram
Count
Value
0
1
2
3
4
20
30
40
50
Common sites gene558
1
2
3
RphaseB1
RphaseB2

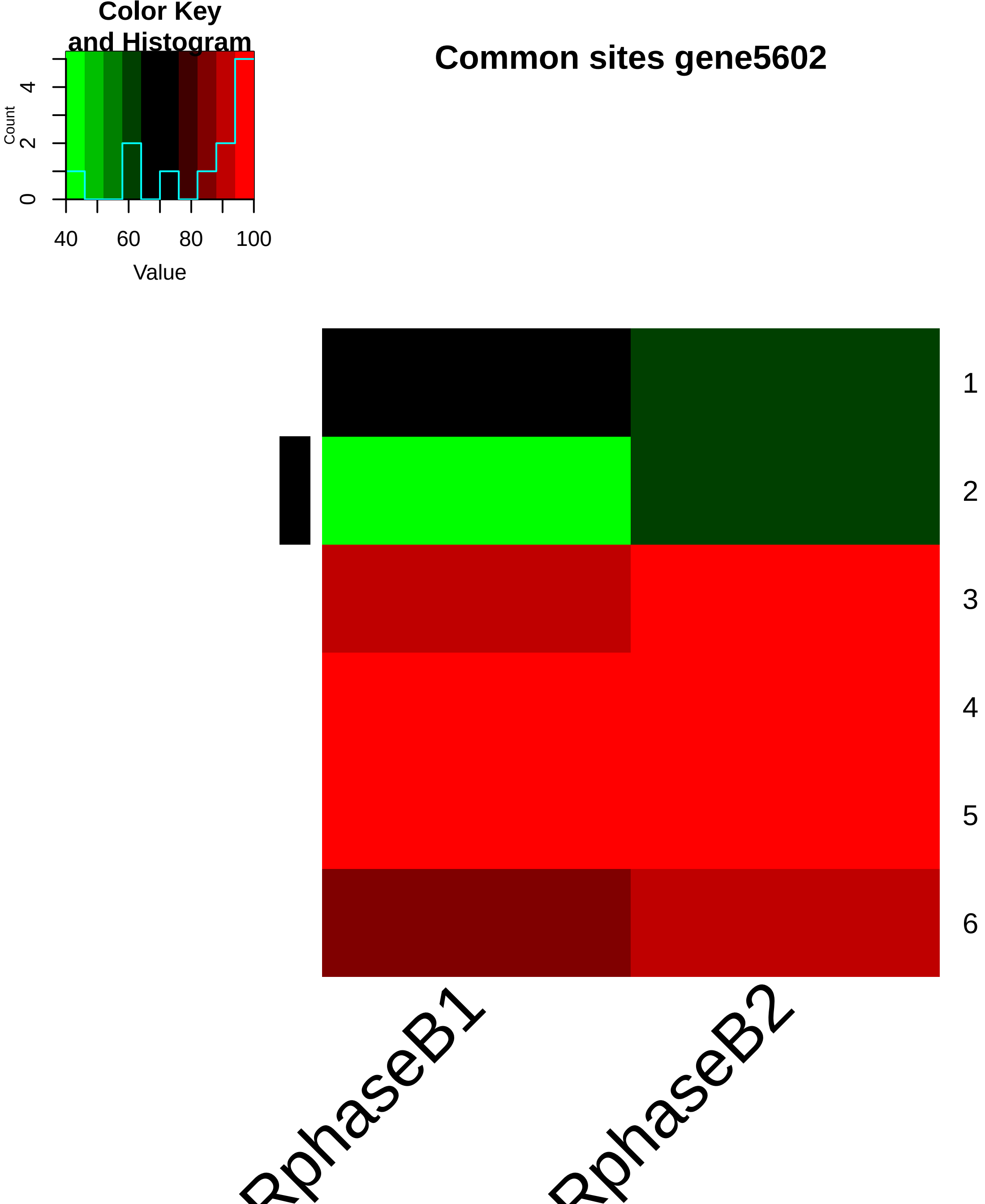
Common sites gene5602
Color Key
and Histogram
Count
0
2
4
40
60
80
100
Value
1
2
3
4
5
6
RphaseB1
RphaseB2

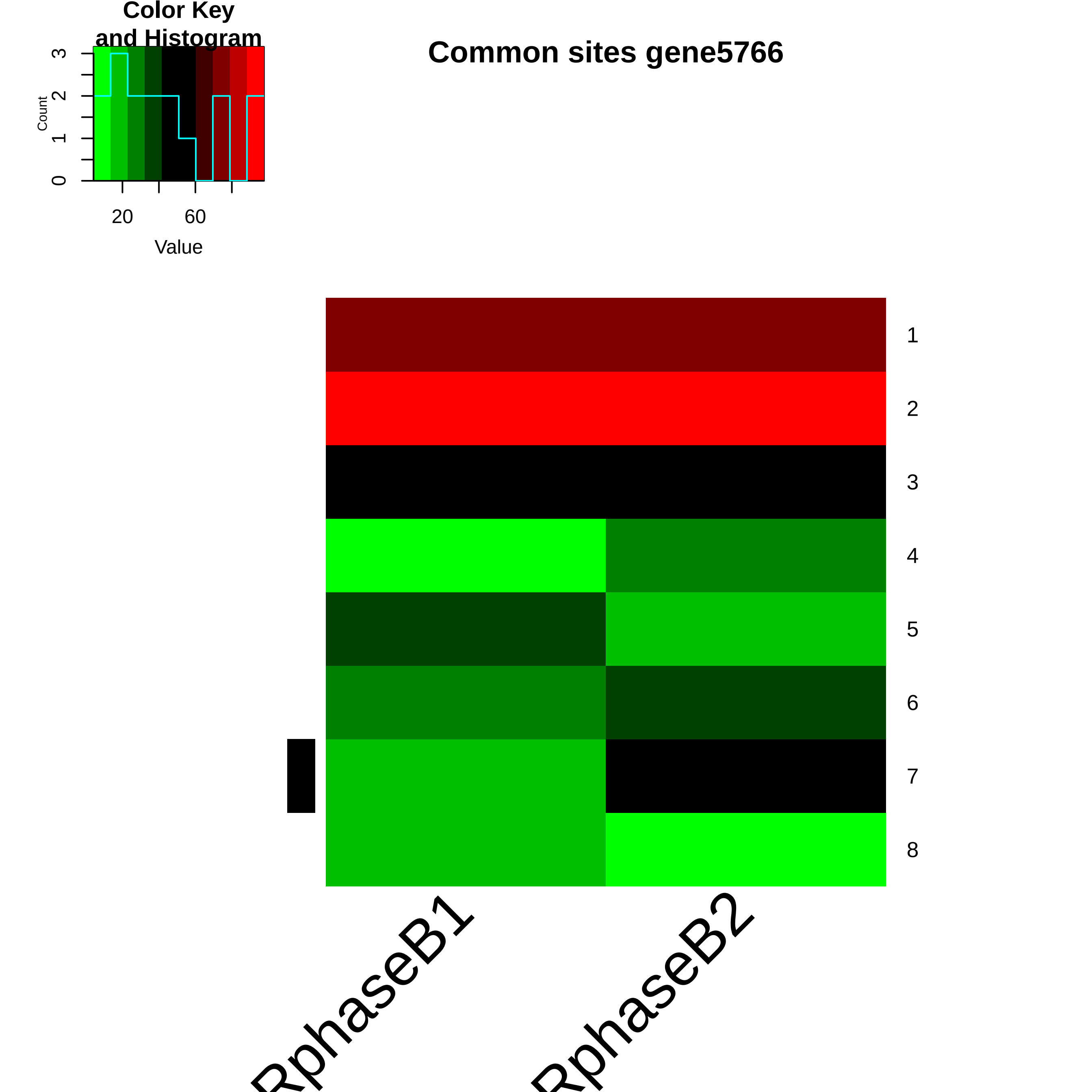
Color Key
and Histogram
Count
0
1
2
3
20
60
Value
Common sites gene5766
1
2
3
4
5
6
7
8
RphaseB1
RphaseB2

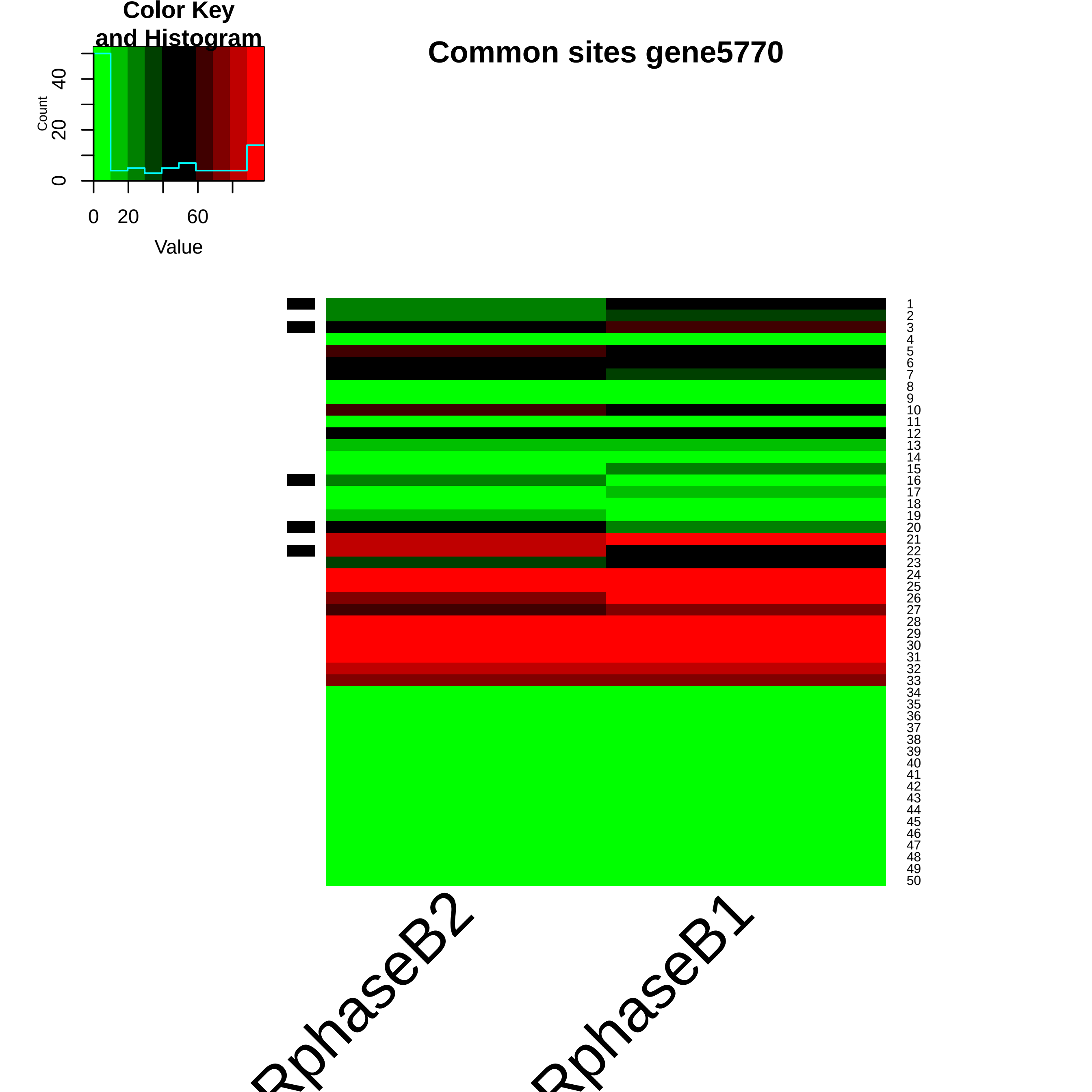

Color Key
and Histogram
Count
0
20
40
0
20
60
Value
Common sites gene5770
RphaseB2
RphaseB1

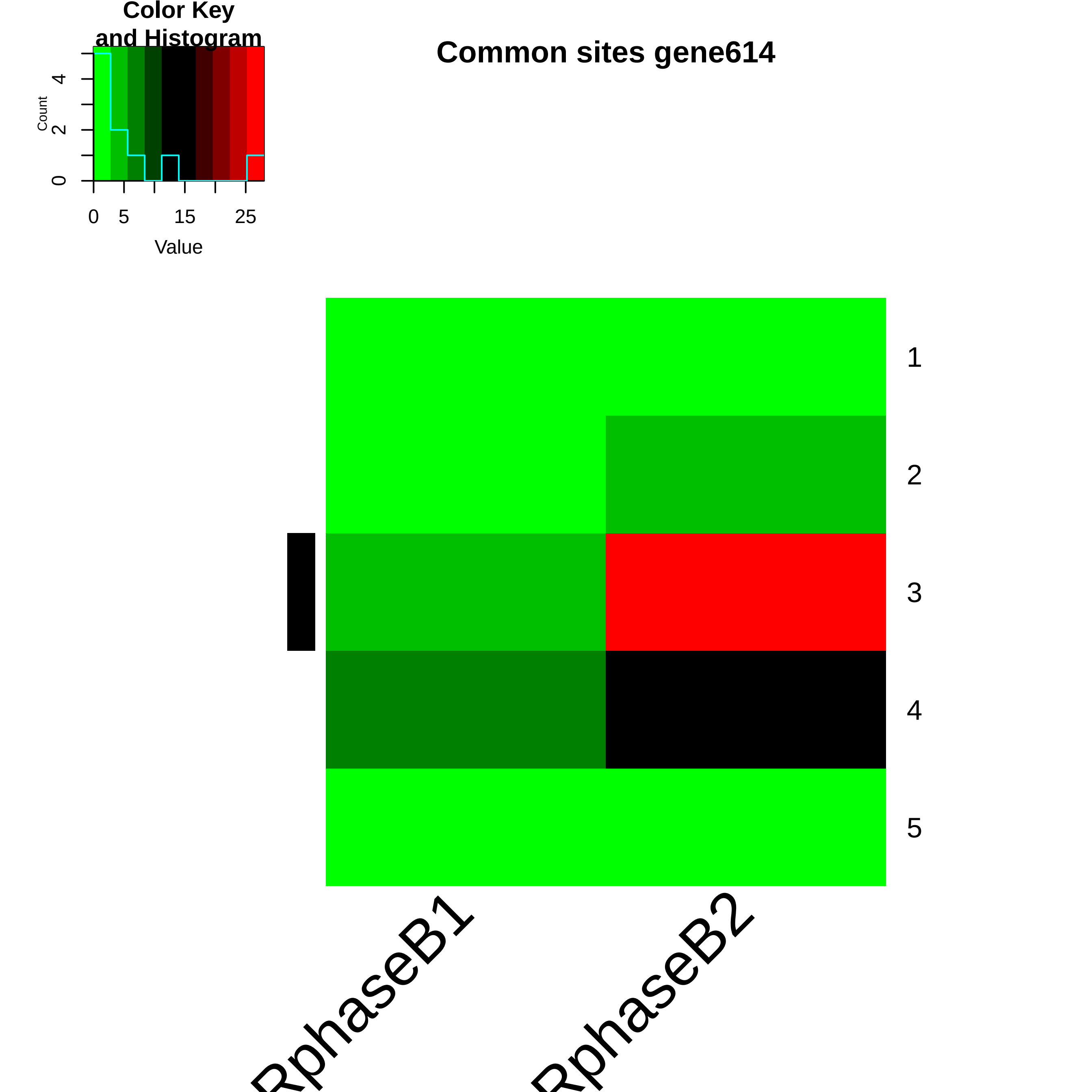
Color Key
and Histogram
Count
0
2
4
0
5
15
25
Value
Common sites gene614
1
2
3
4
5
RphaseB1
RphaseB2

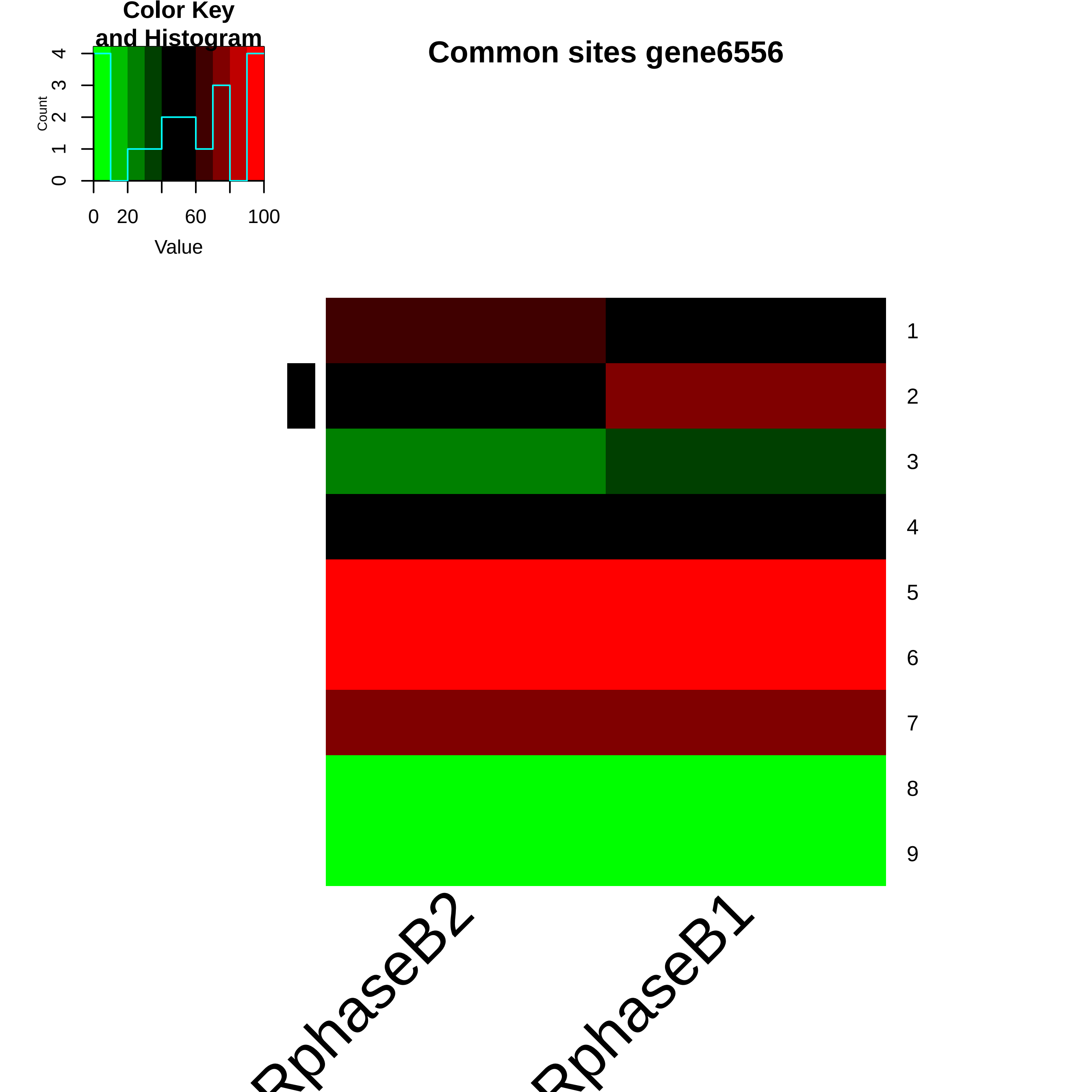

Color Key
and Histogram
Count
0
1
2
3
4
0
20
60
100
Value
Common sites gene6556
1
2
3
4
5
6
7
8
9
RphaseB2
RphaseB1

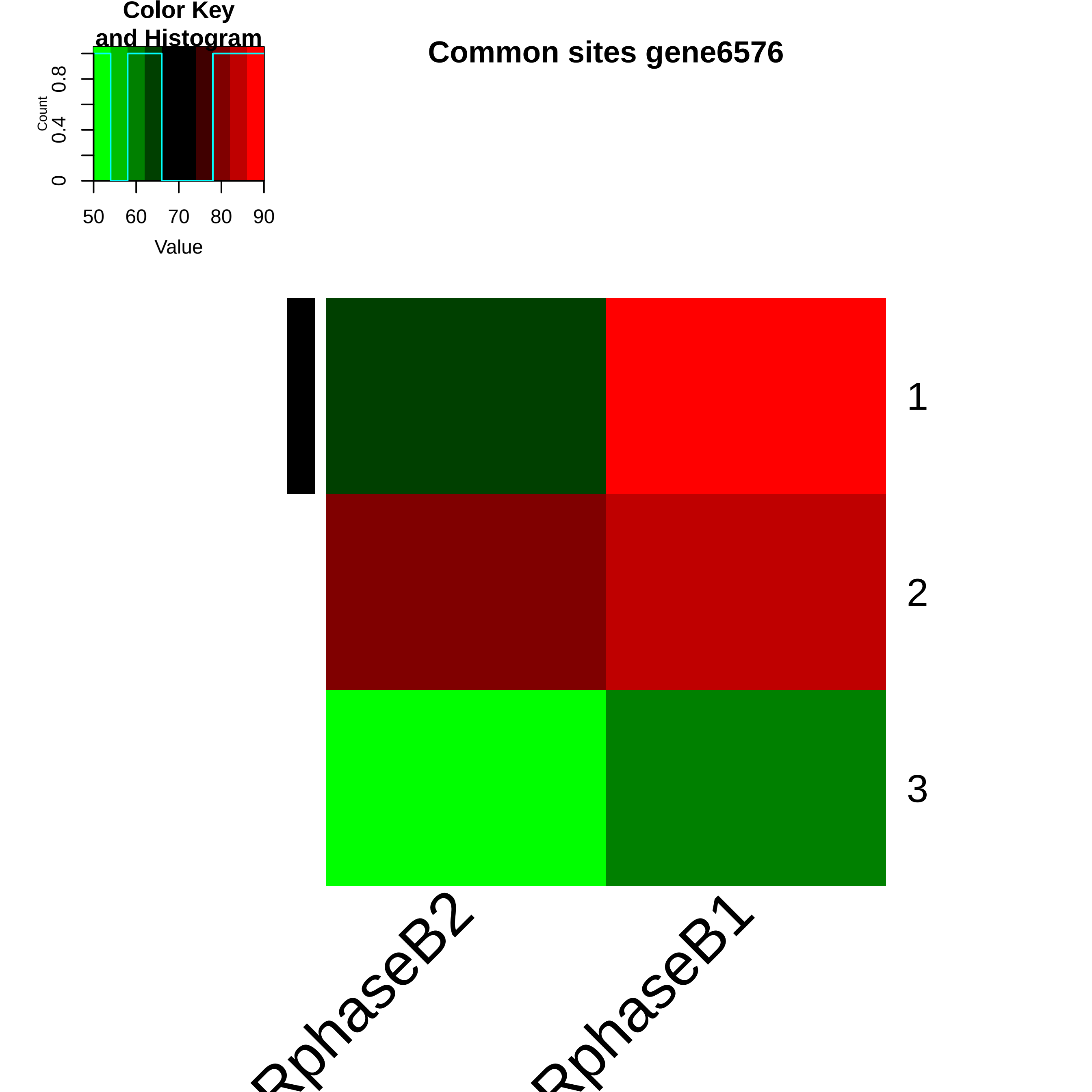
Color Key
and Histogram
Count
0
0.4
0.8
50
60
70
80
90
Value
Common sites gene6576
1
2
3
RphaseB2
RphaseB1

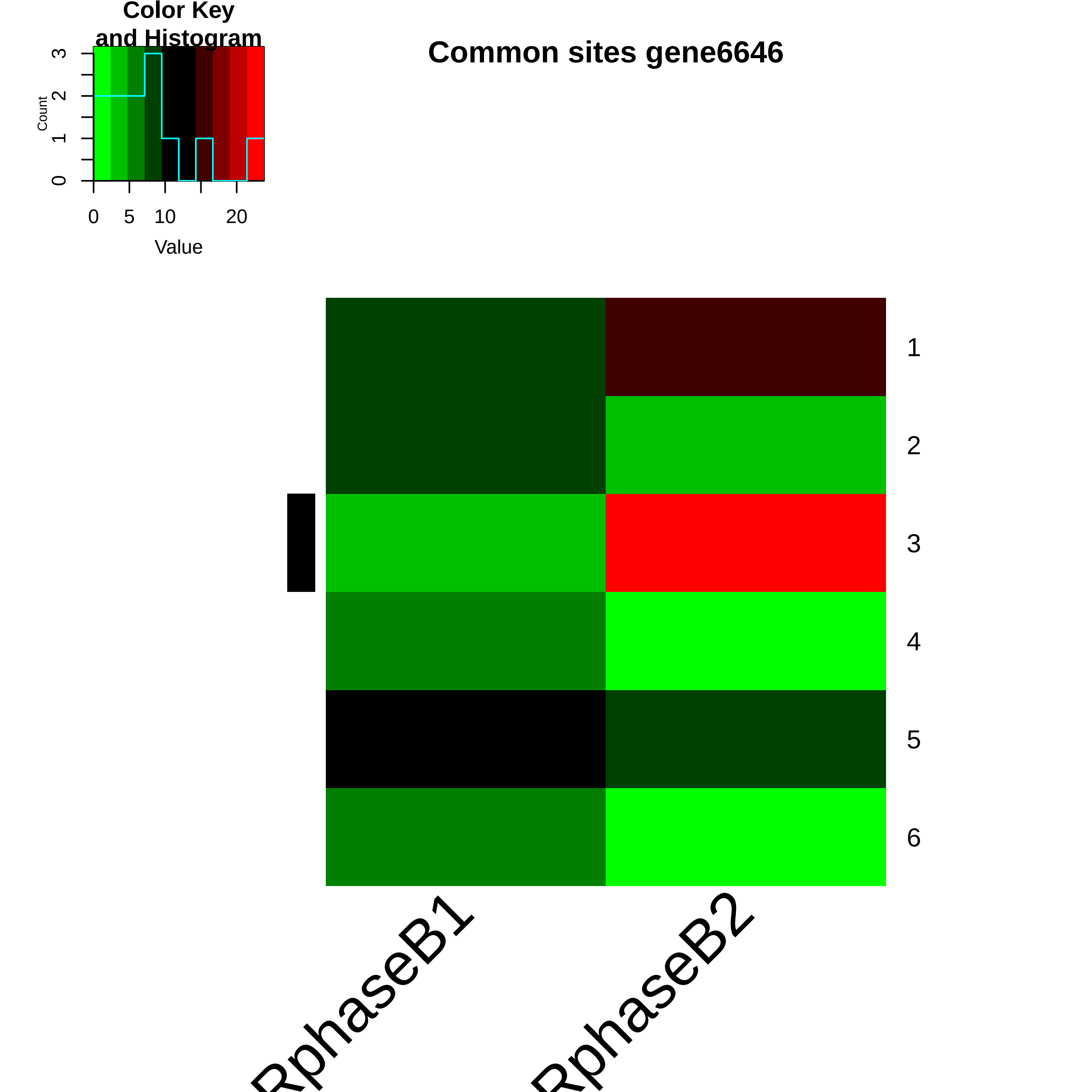
Color Key
and Histogram
Count
0
1
2
3
0
5
10
20
Value
Common sites gene6646
1
2
3
4
5
6
RphaseB1
RphaseB2

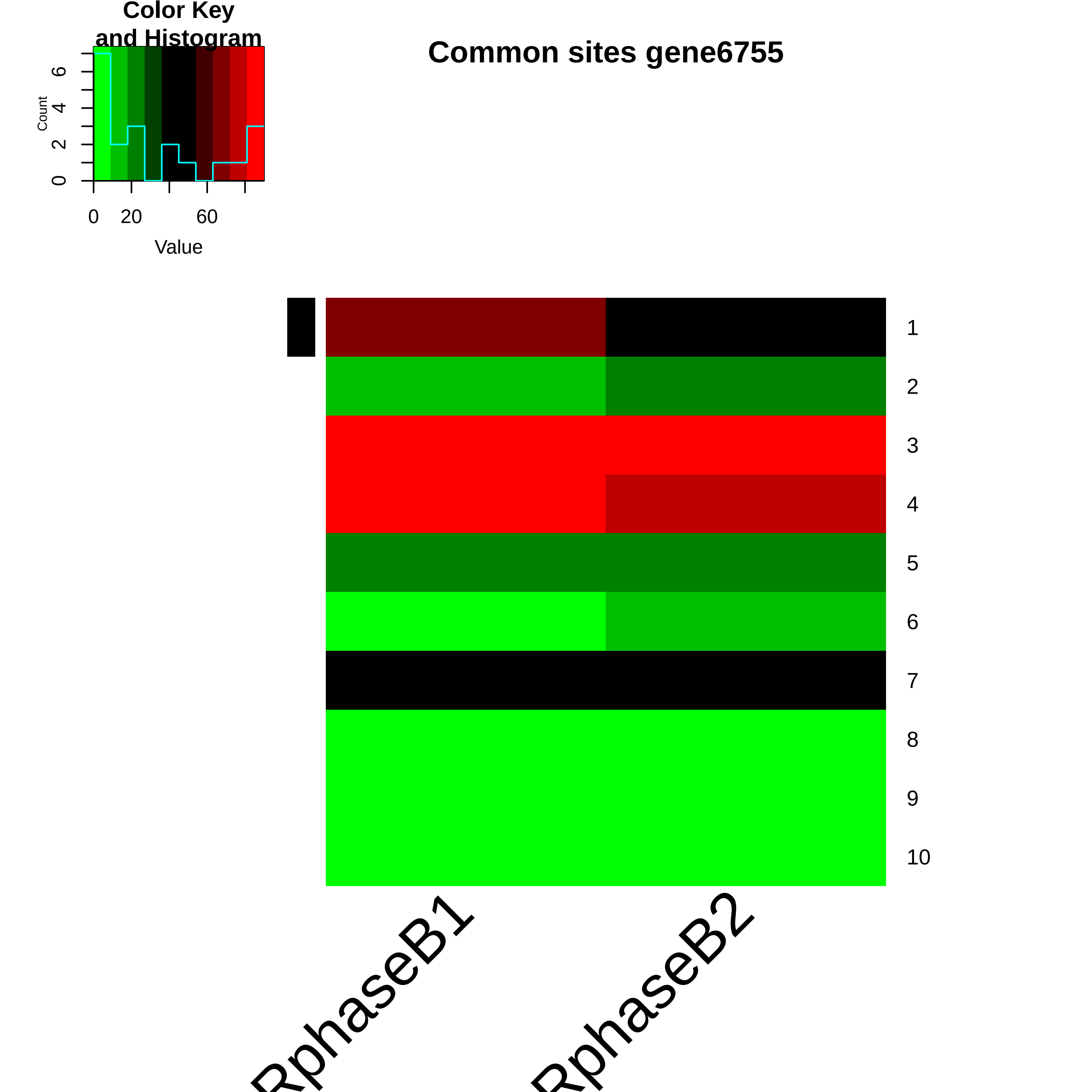

Color Key
and Histogram
Count
0
2
4
6
0
20
60
Value
Common sites gene6755
1
2
3
4
5
6
7
8
9
10
RphaseB1
RphaseB2

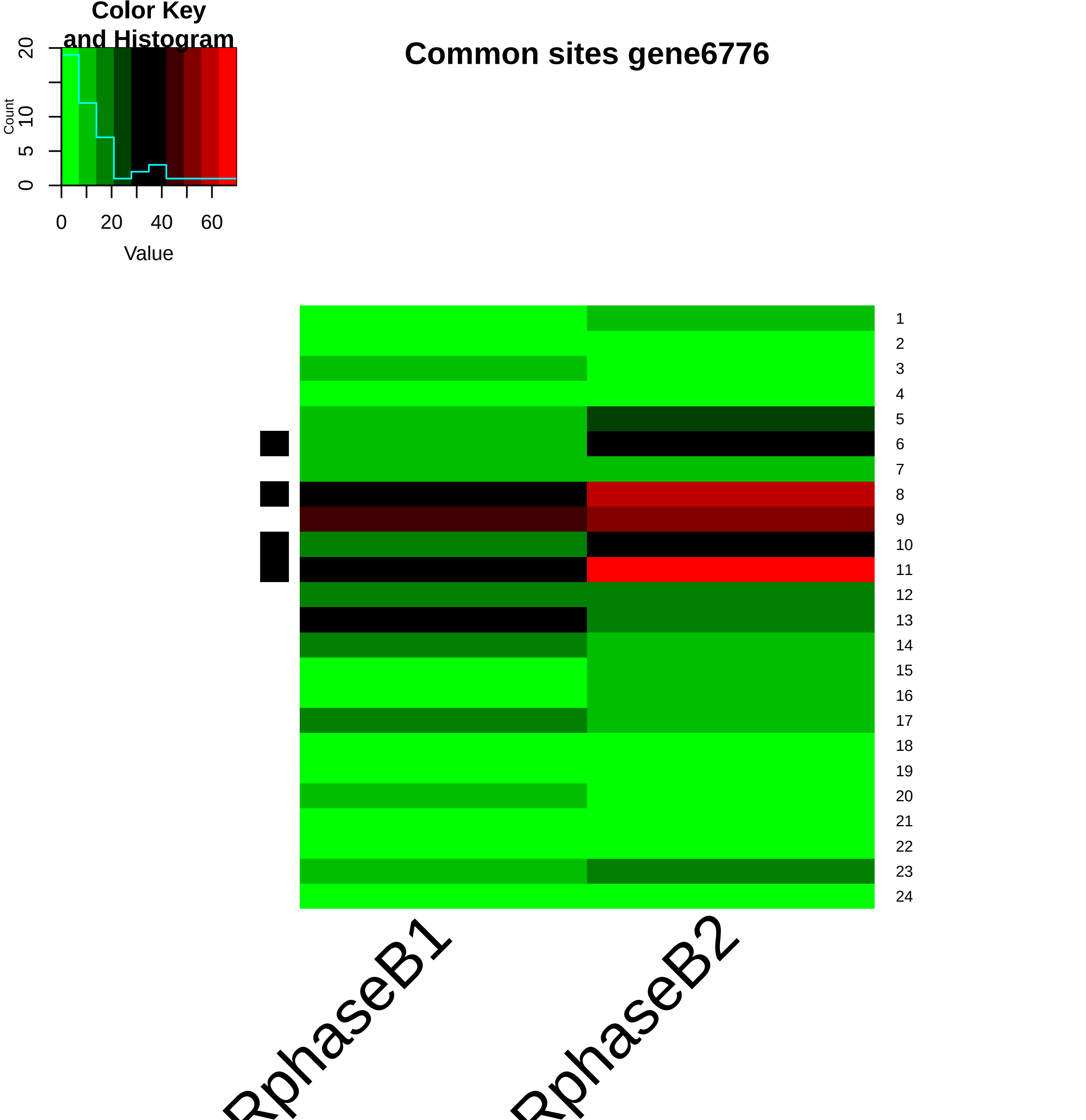
Color Key
and Histogram
Count
0
5
10
20
0
20
40
60
Value
Common sites gene6776
RphaseB1
RphaseB2

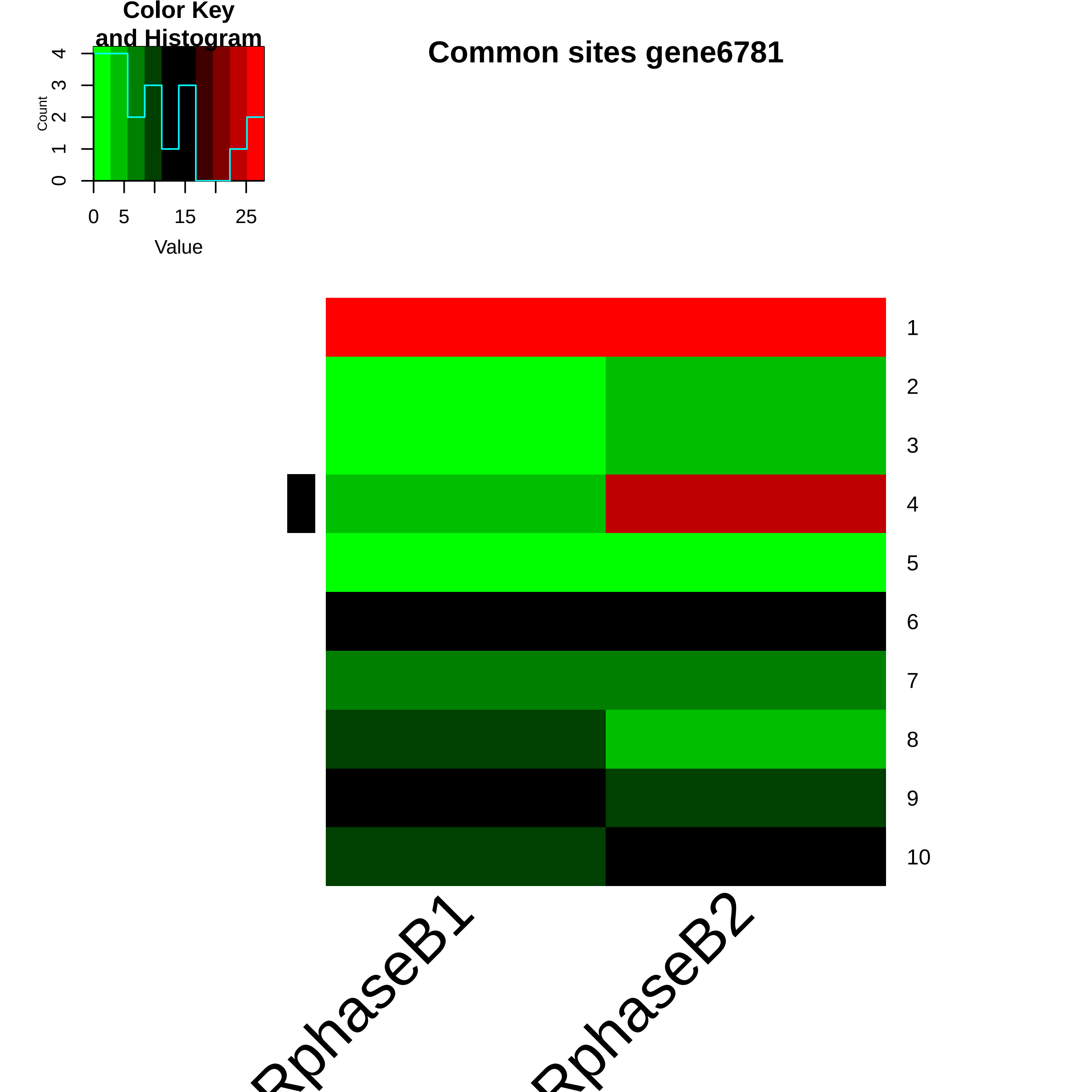
Color Key
and Histogram
Count
0 1 2 3 4
0 5 15 25
Value
Common sites gene6781
1
2
3
4
5
6
7
8
9
10
RphaseB1
RphaseB2

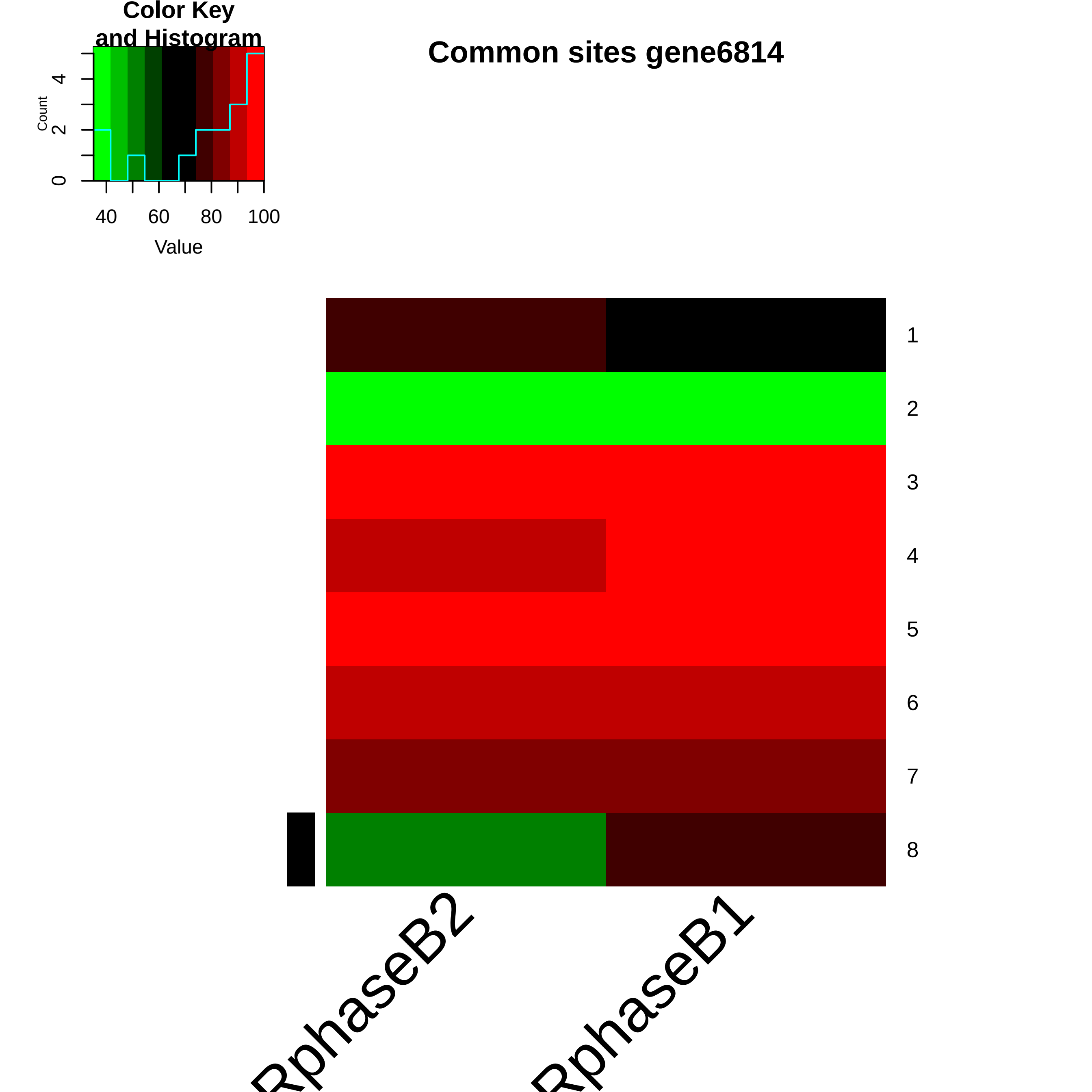

Color Key
and Histogram
Count
0
2
4
40
60
80
100
Value
Common sites gene6814
1
2
3
4
5
6
7
8
RphaseB2
RphaseB1

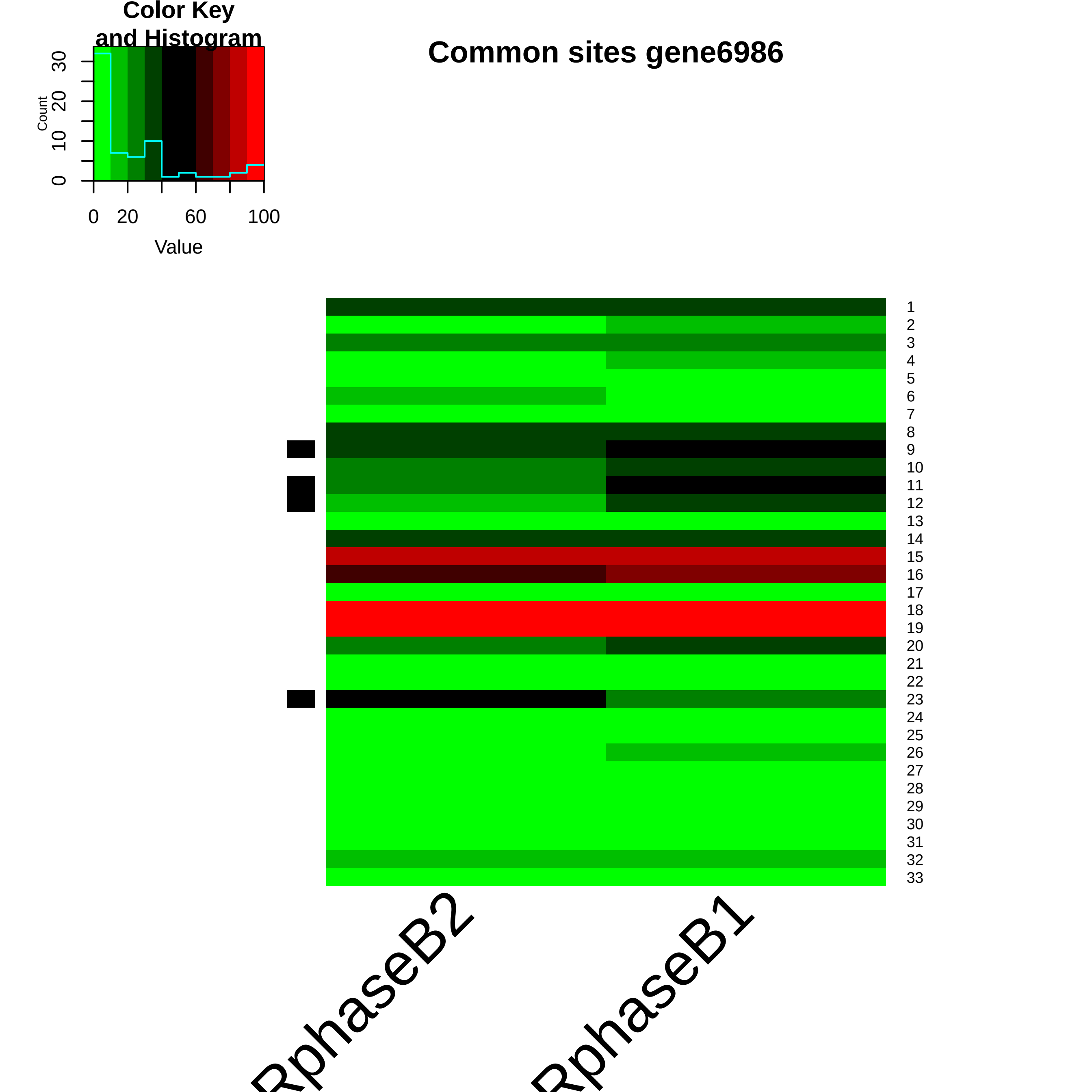
Color Key
and Histogram
Count
0
10
20
30
0
20
60
100
Value
Common sites gene6986
RphaseB2
RphaseB1

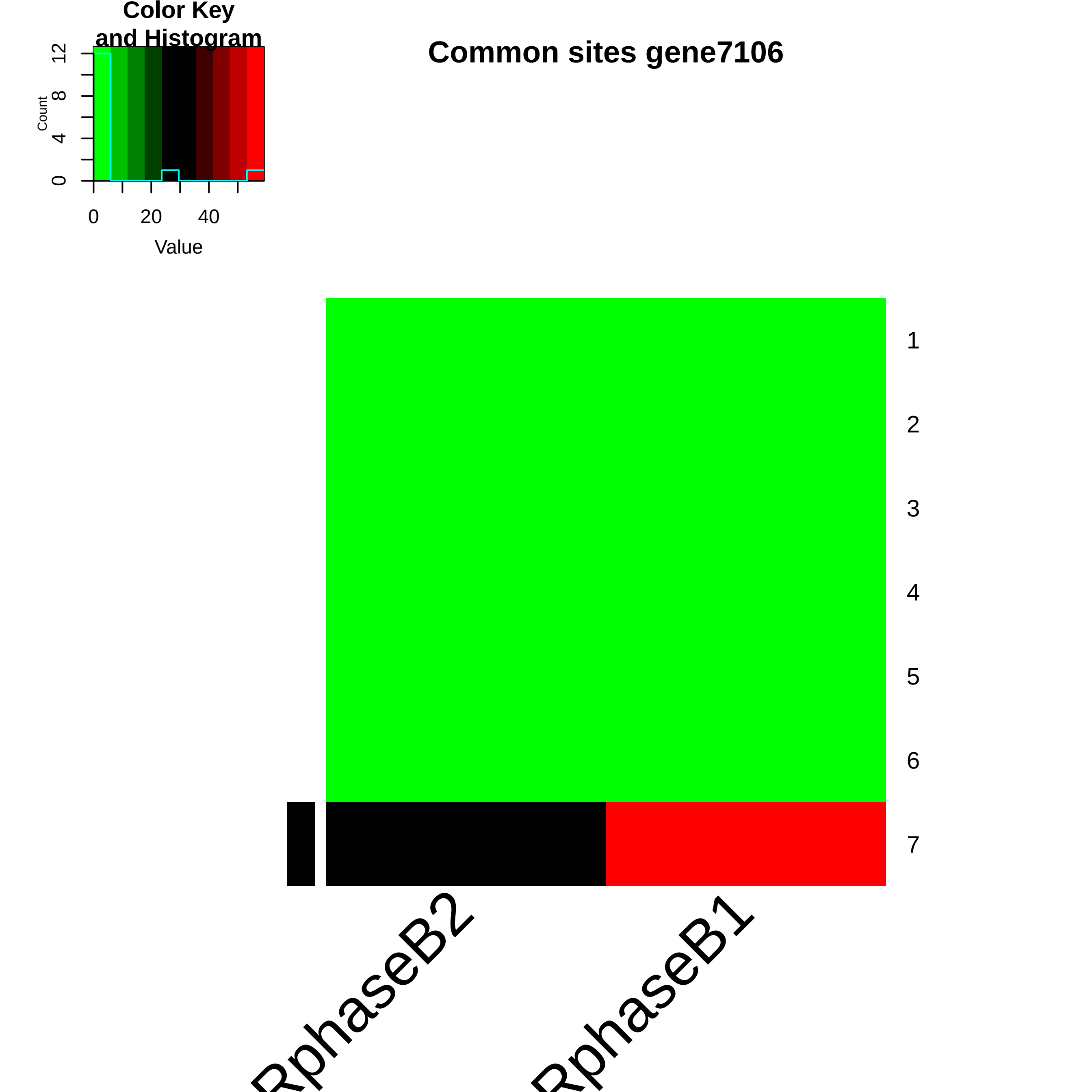
Color Key
and Histogram
Count
0
4
8
12
0
20
40
Value
Common sites gene7106
1
2
3
4
5
6
7
RphaseB2
RphaseB1

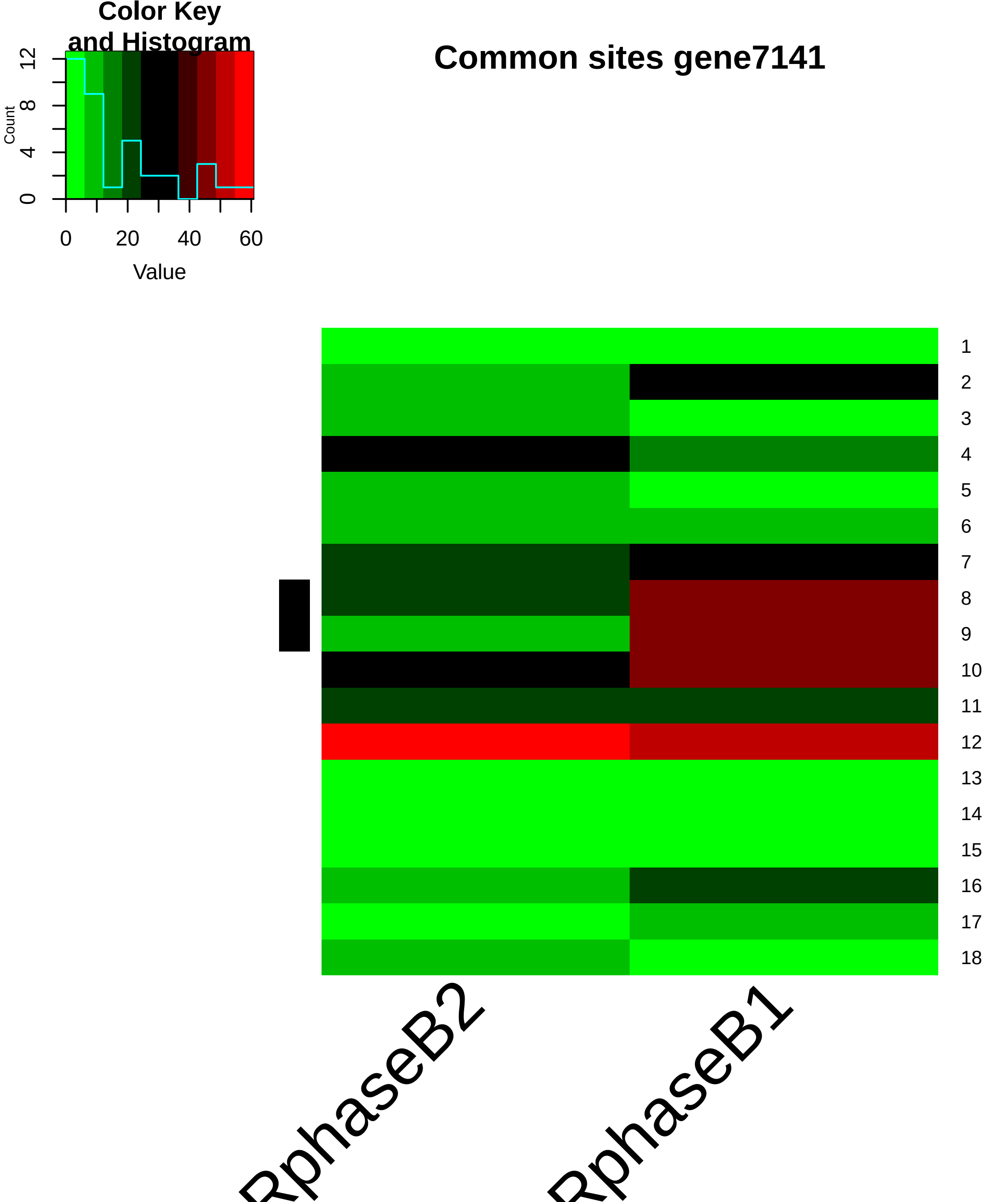
Color Key
and Histogram
Count
0
4
8
12
0
20
40
60
Value
Common sites gene7141
RphaseB2
RphaseB1

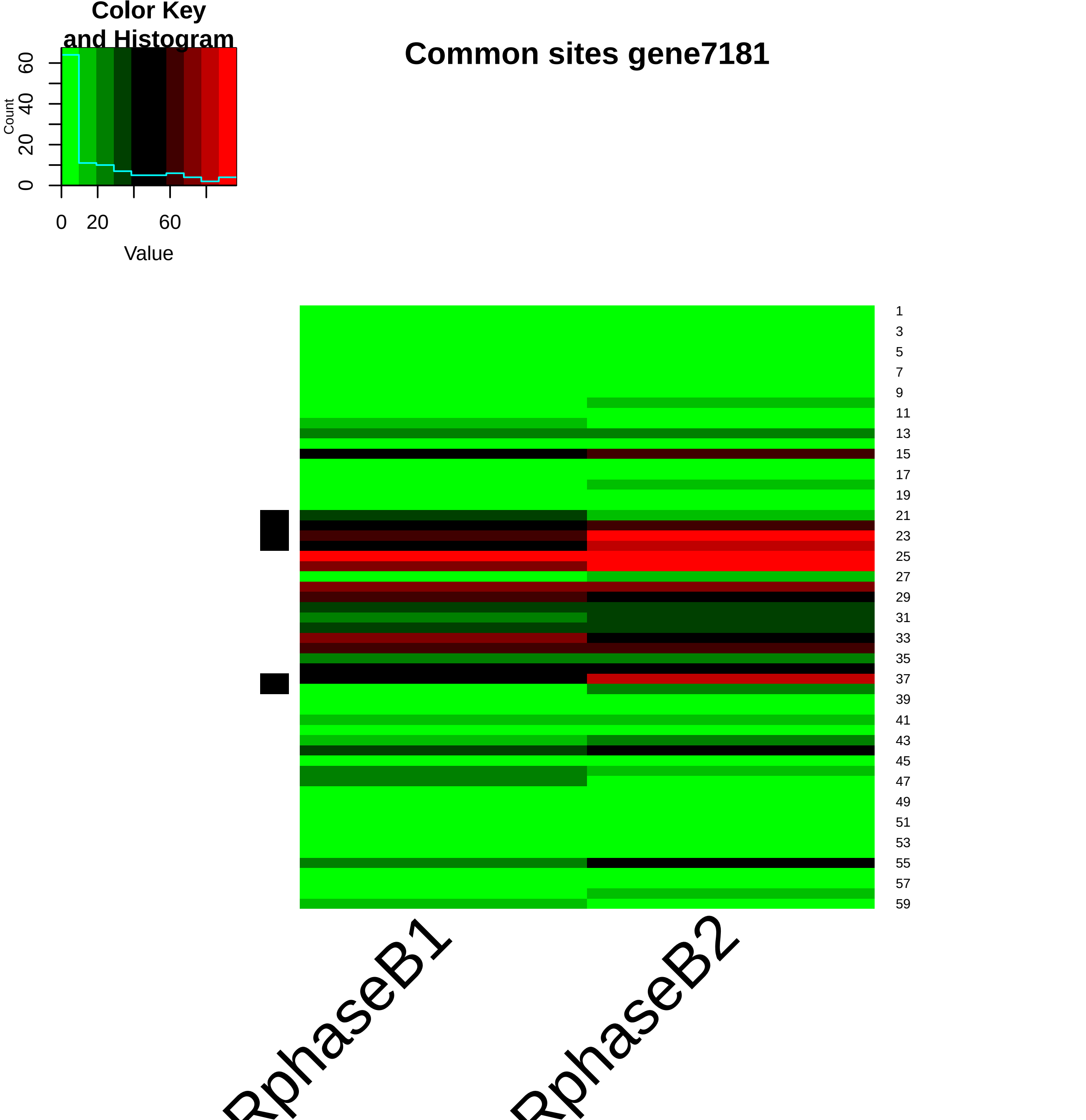
Color Key
and Histogram
Count
0
20
40
60
0
20
60
Value
Common sites gene7181
1
3
5
7
9
11
13
15
17
19
21
23
25
27
29
31
33
35
37
39
41
43
45
47
49
51
53
55
57
59
RphaseB1
RphaseB2

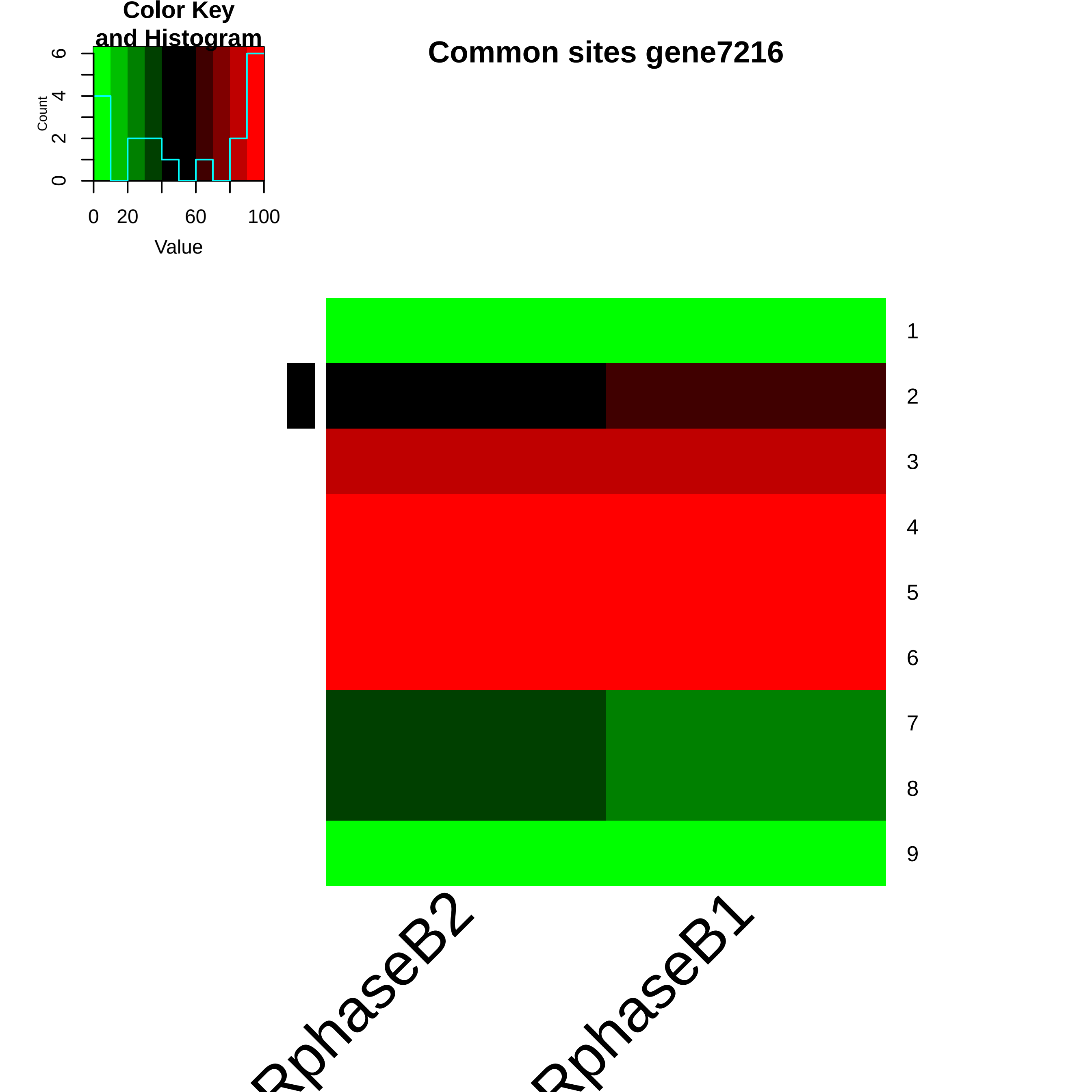
Color Key
and Histogram
Count
Value
0
20
60
100
0
2
4
6
Common sites gene7216
1
2
3
4
5
6
7
8
9
RphaseB2
RphaseB1

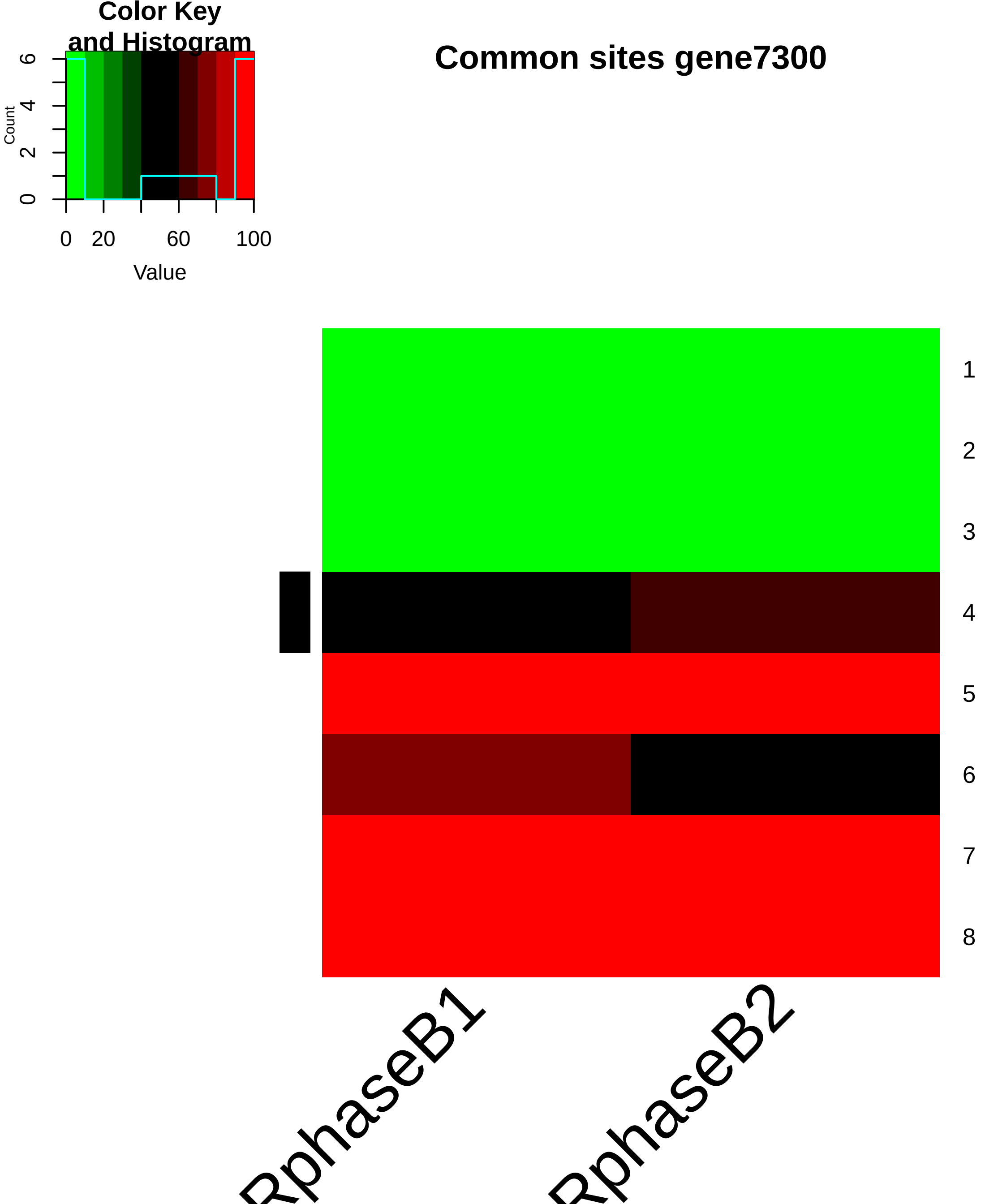
Color Key
and Histogram
Count
0
2
4
6
0
20
60
100
Value
Common sites gene7300
1
2
3
4
5
6
7
8
RphaseB1
RphaseB2

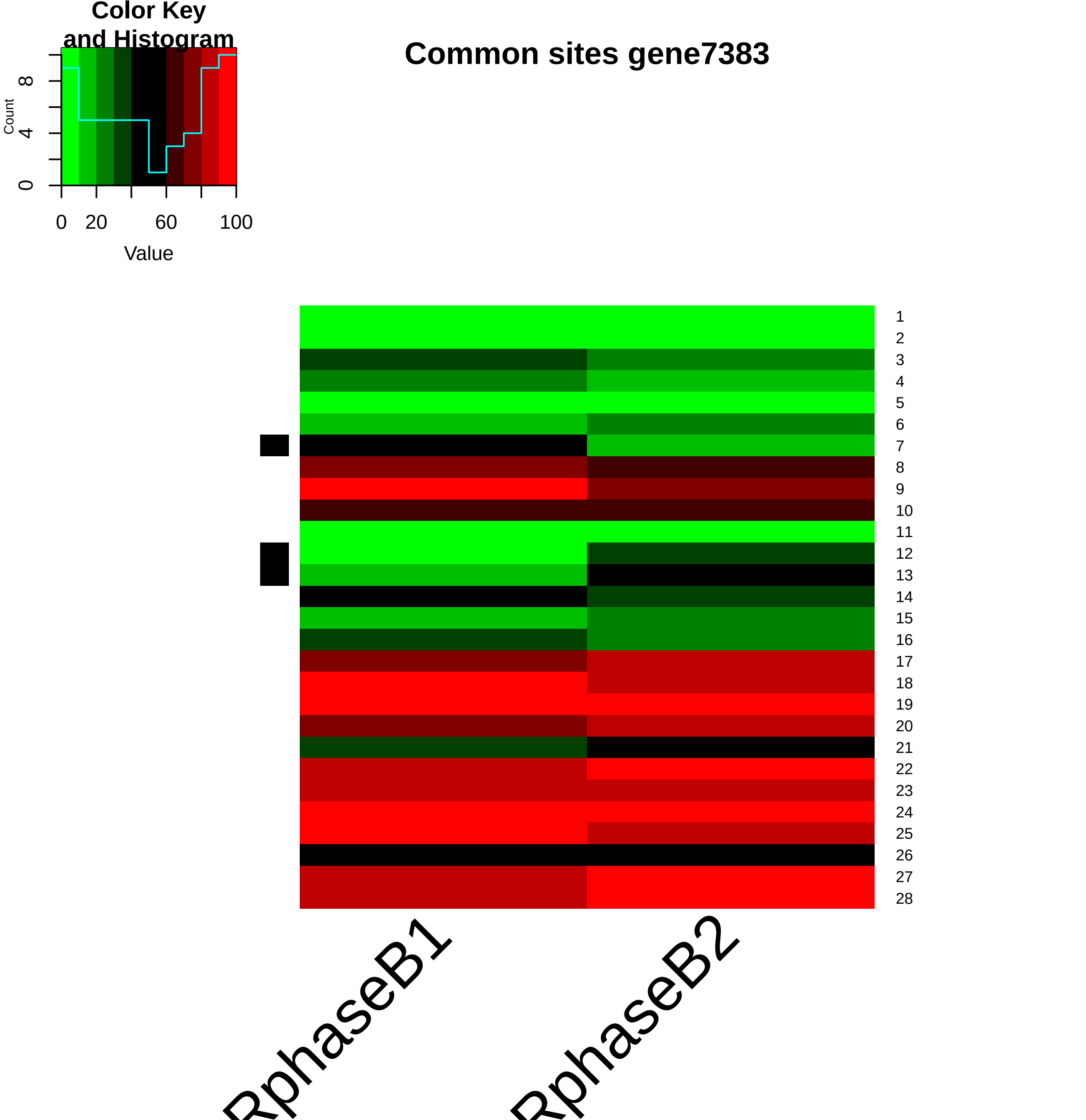
Color Key
and Histogram
Count
0
4
8
0
20
60
100
Value
Common sites gene7383
RphaseB1
RphaseB2

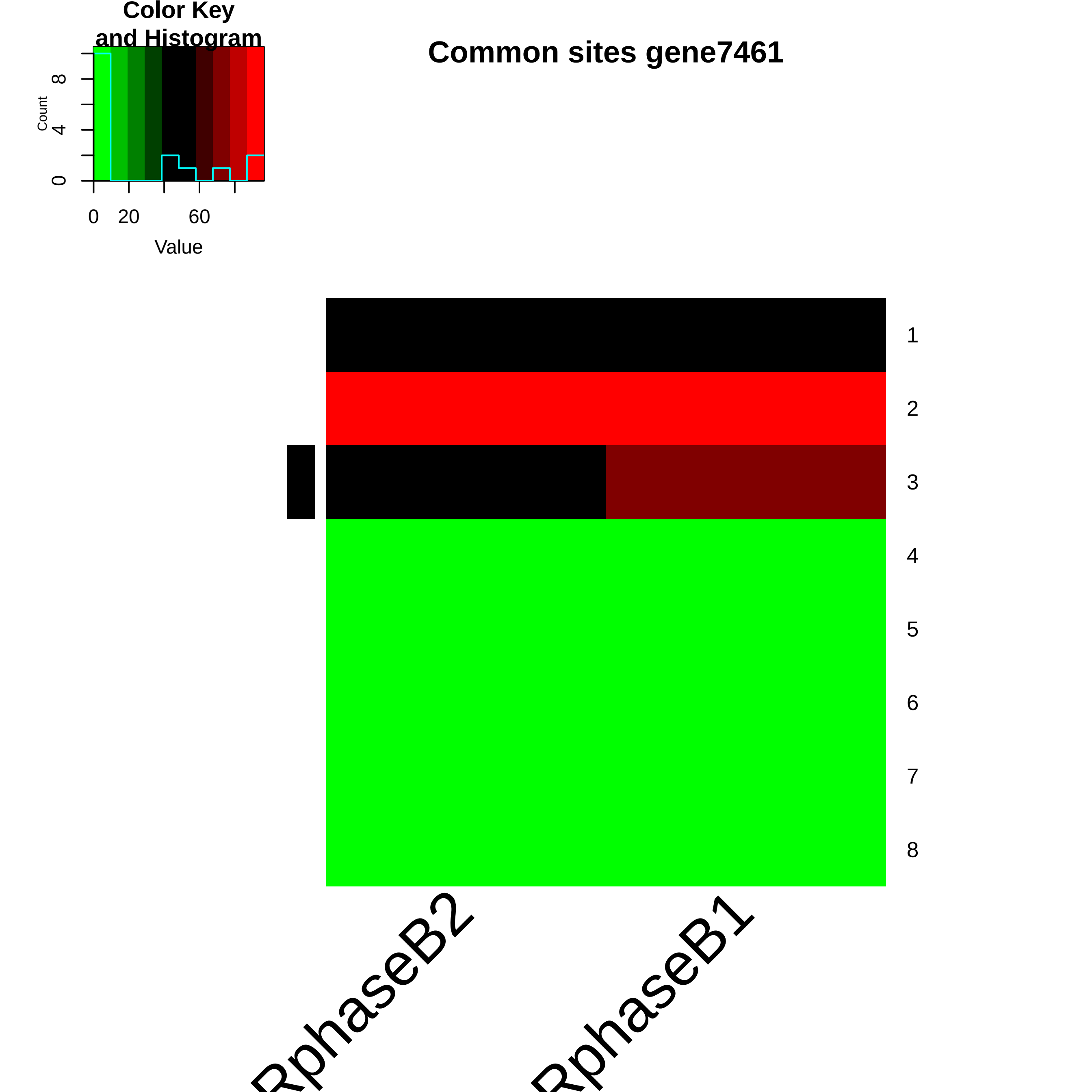

Color Key
and Histogram
Count
0
4
8
0
20
60
Value
Common sites gene7461
1
2
3
4
5
6
7
8
RphaseB2
RphaseB1

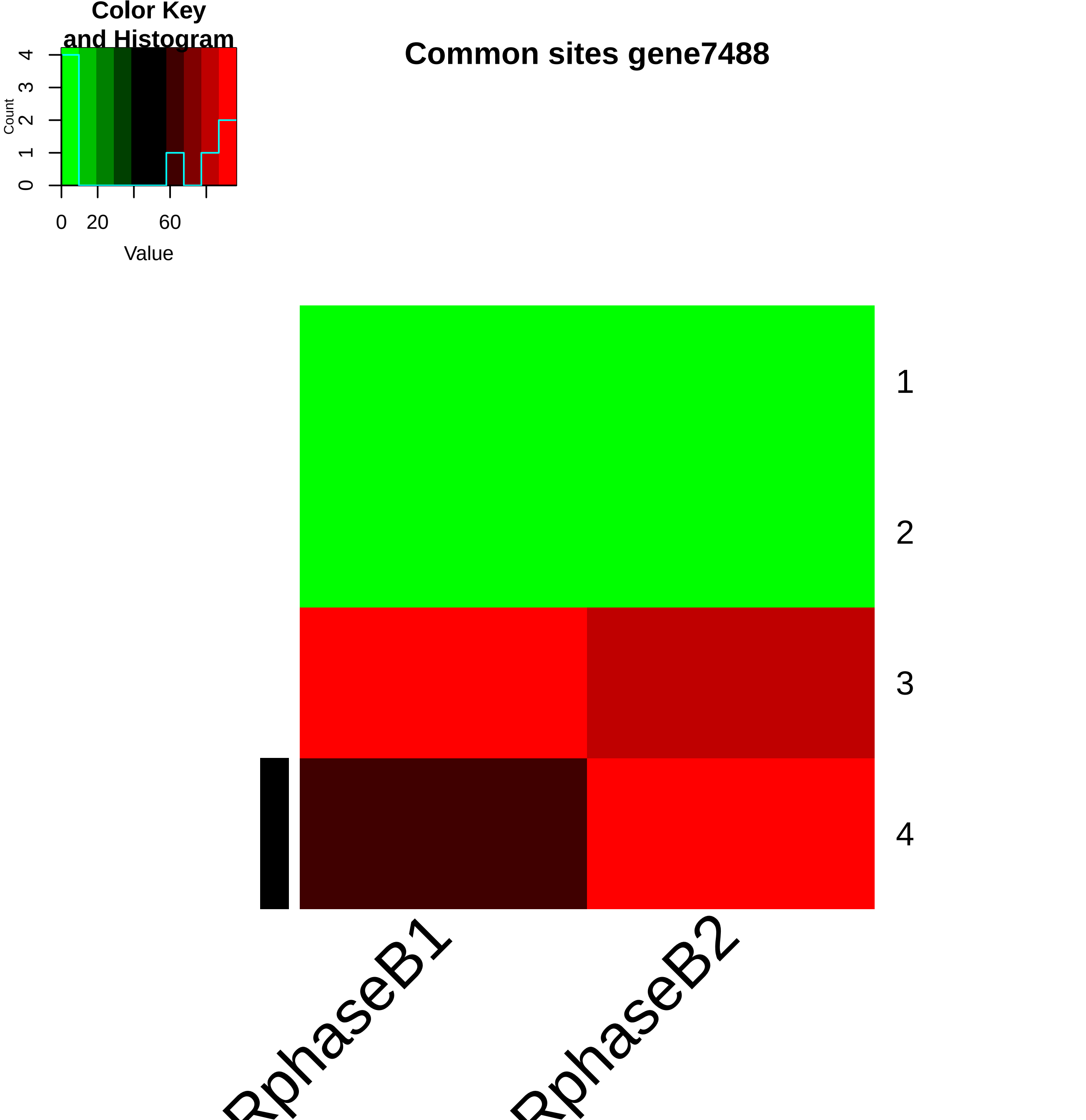
Color Key
and Histogram
Count
0
1
2
3
4
0
20
60
Value
Common sites gene7488
1
2
3
4
RphaseB1
RphaseB2

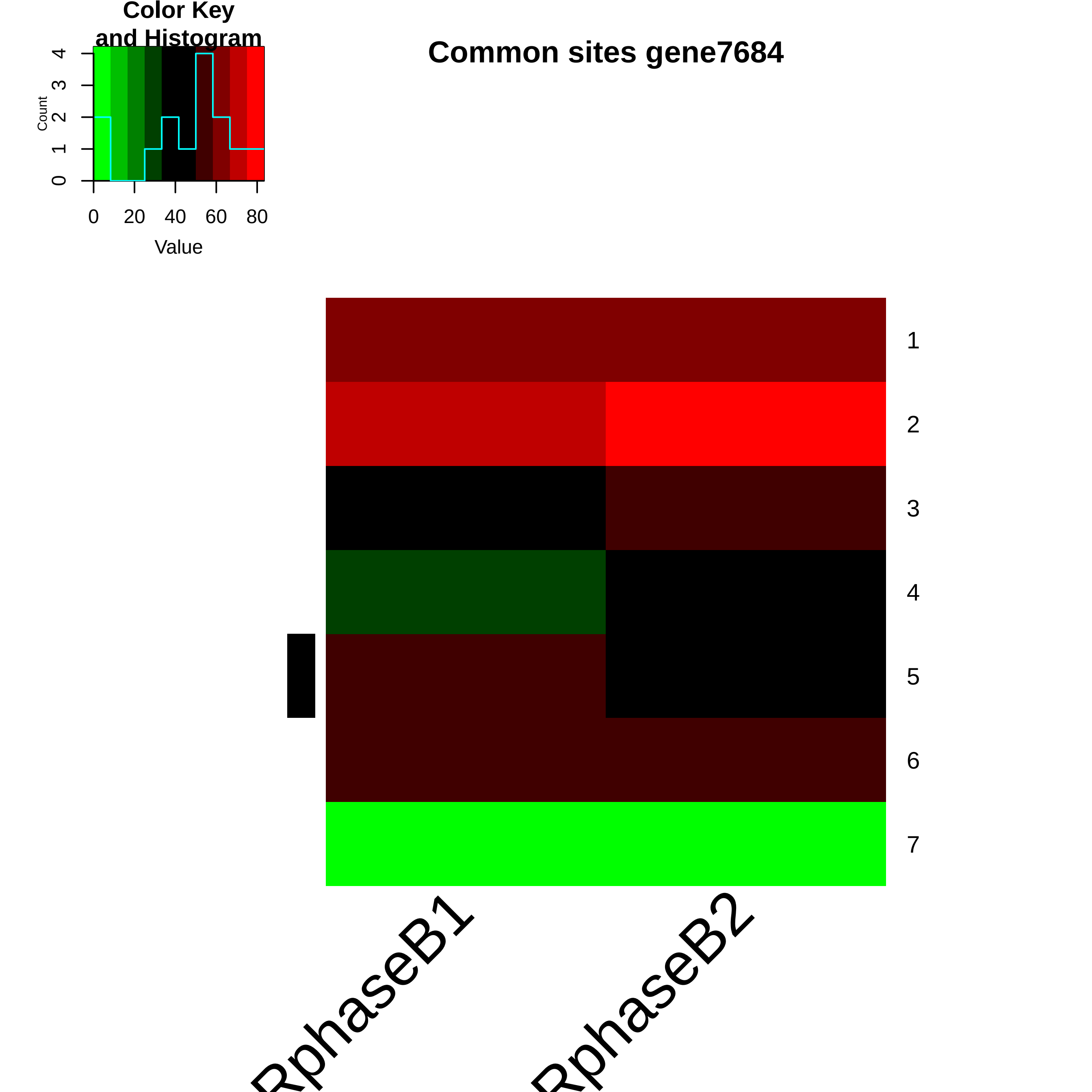
Color Key
and Histogram
Count
0
1
2
3
4
0
20
40
60
80
Value
Common sites gene7684
1
2
3
4
5
6
7
RphaseB1
RphaseB2

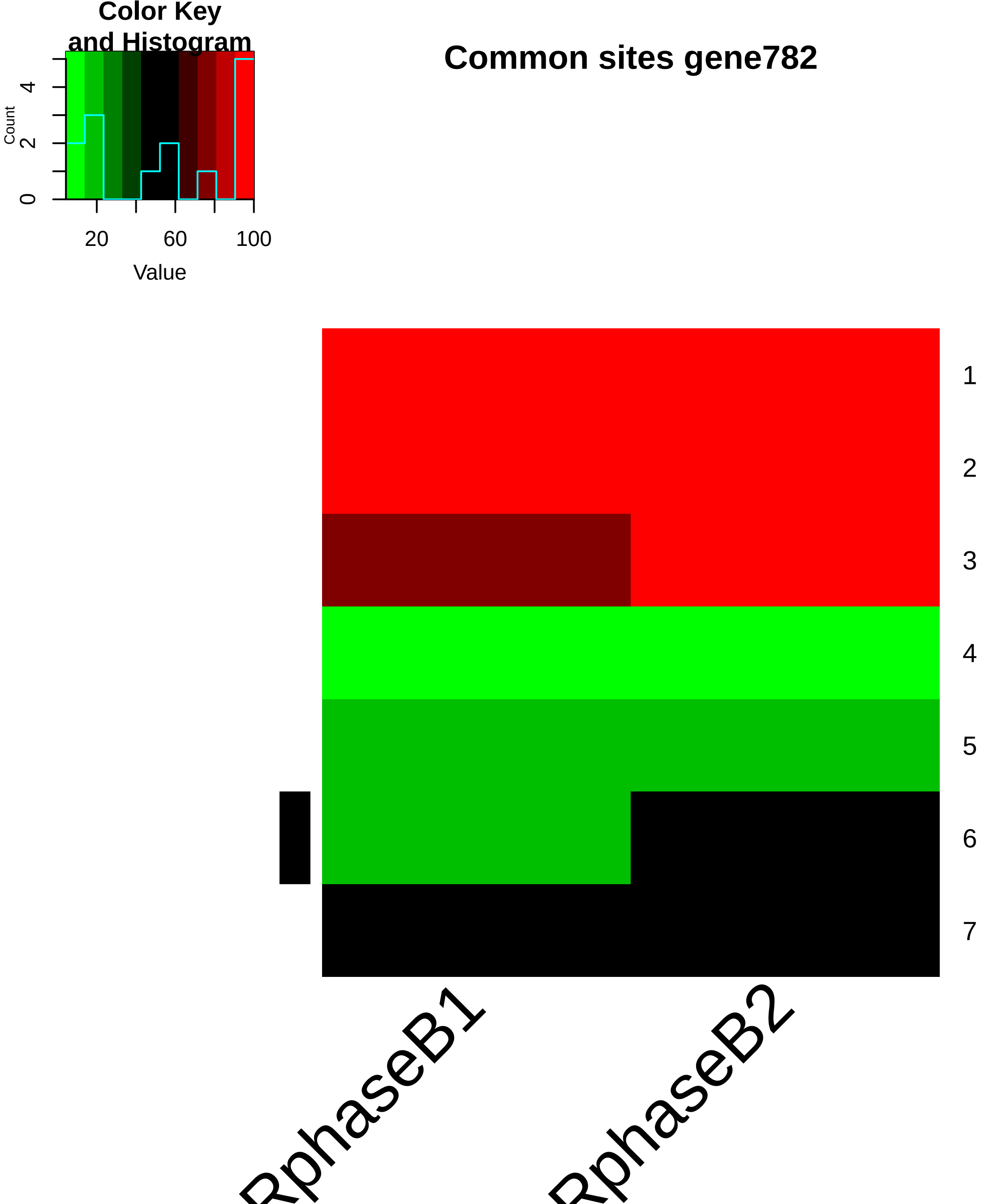

Color Key
and Histogram
Count
0
2
4
20
60
100
Value
Common sites gene782
1
2
3
4
5
6
7
RphaseB1
RphaseB2

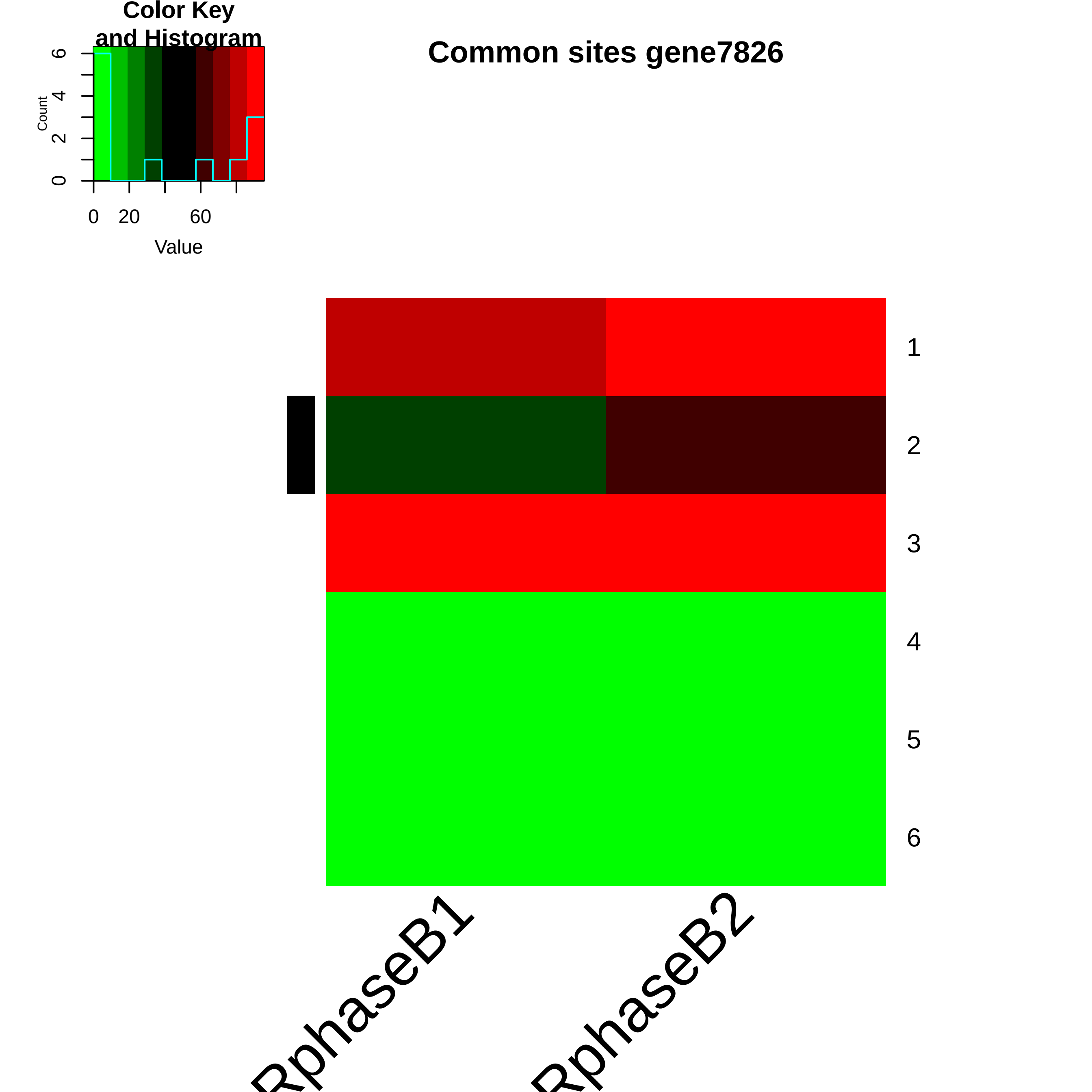
Color Key
and Histogram
Count
0
2
4
6
0
20
60
Value
Common sites gene7826
1
2
3
4
5
6
RphaseB1
RphaseB2

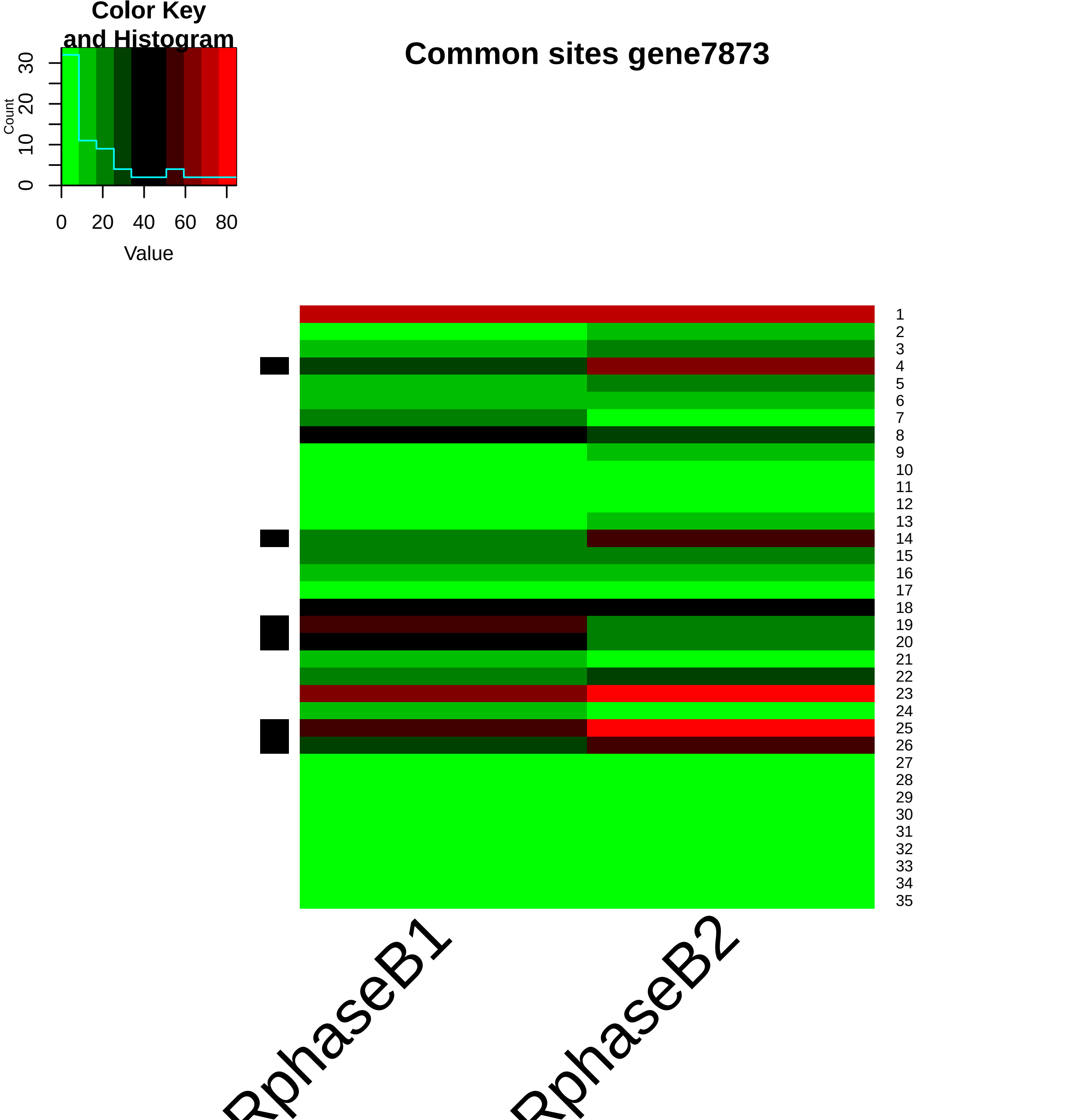

Color Key
and Histogram
Count
0
10
20
30
0
20
40
60
80
Value
Common sites gene7873
RphaseB1
RphaseB2

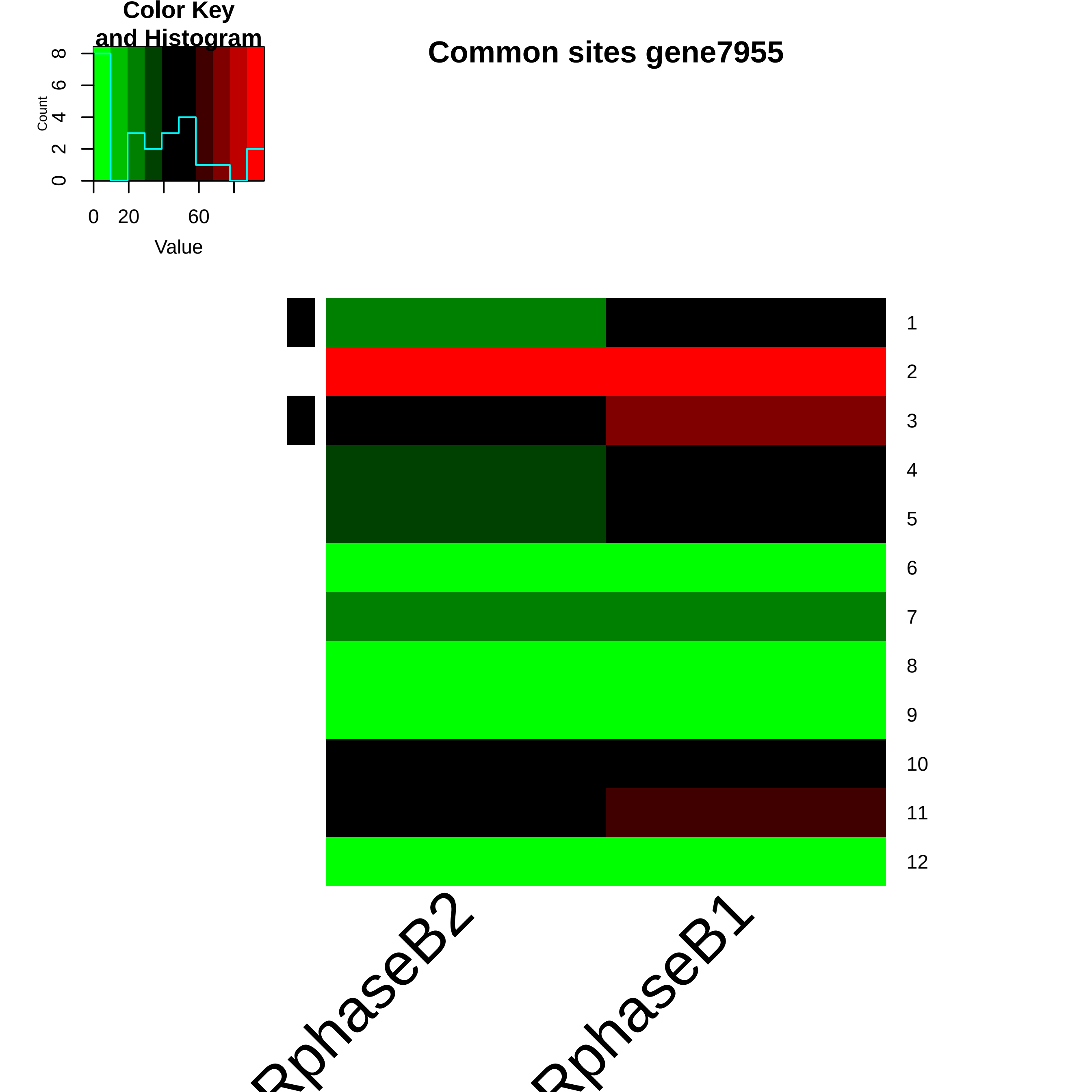
Color Key
and Histogram
Count
0
2
4
6
8
0
20
60
Value
Common sites gene7955
1
2
3
4
5
6
7
8
9
10
11
12
RphaseB2
RphaseB1

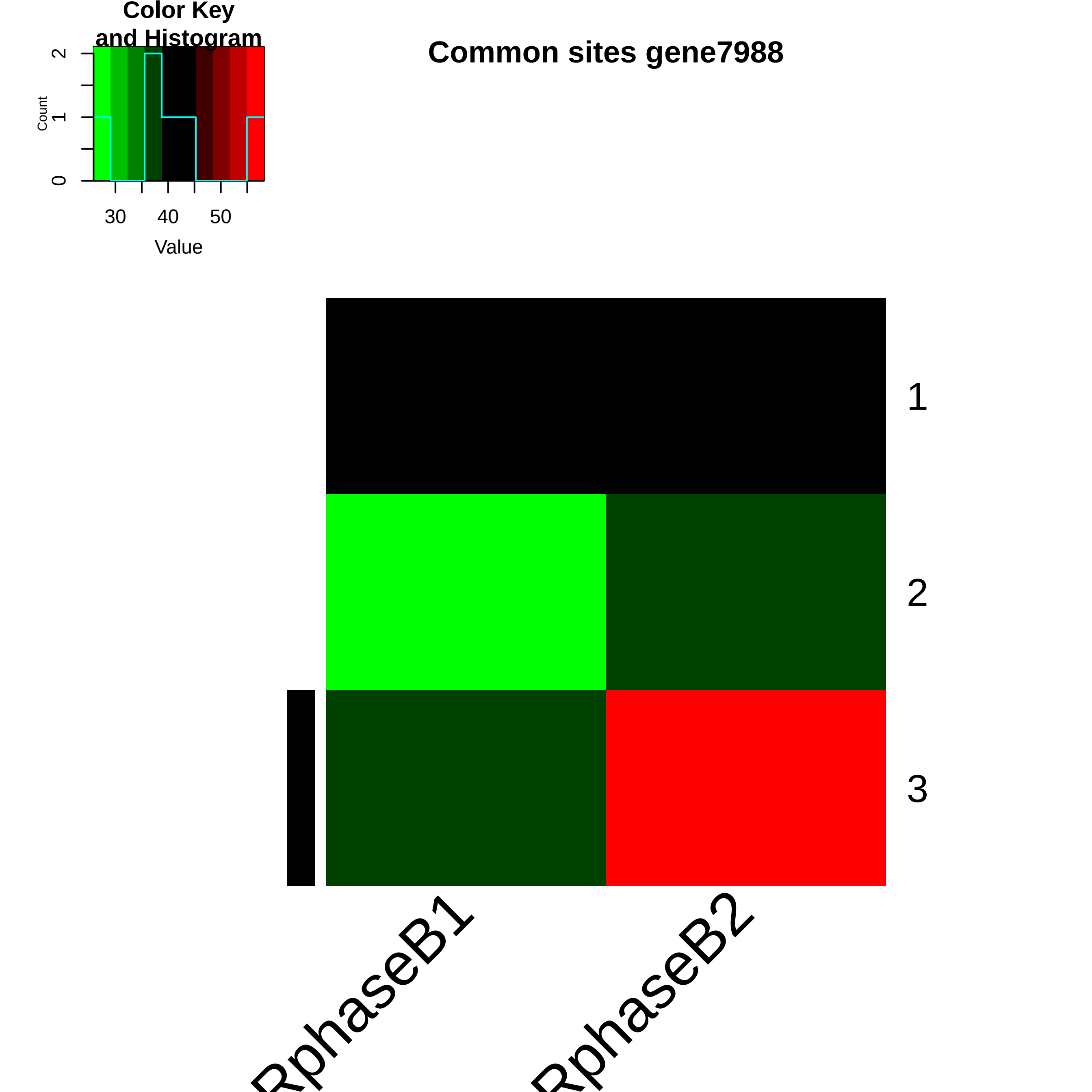

Color Key
and Histogram
Count
0
1
2
30
40
50
Value
Common sites gene7988
1
2
3
RphaseB1
RphaseB2

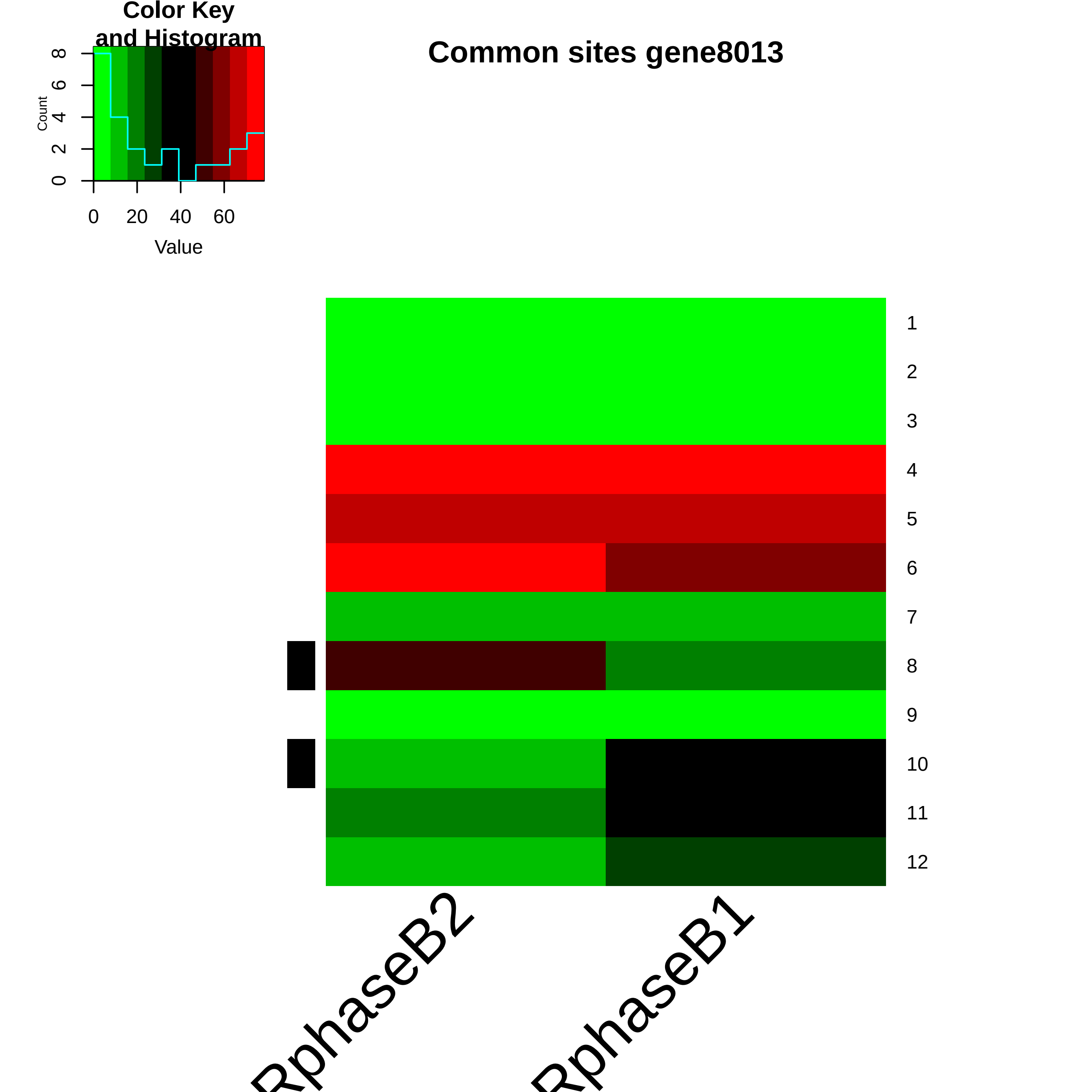

Color Key
and Histogram
Count
Value
0
20
40
60
0
2
4
6
8
Common sites gene8013
1
2
3
4
5
6
7
8
9
10
11
12
RphaseB2
RphaseB1

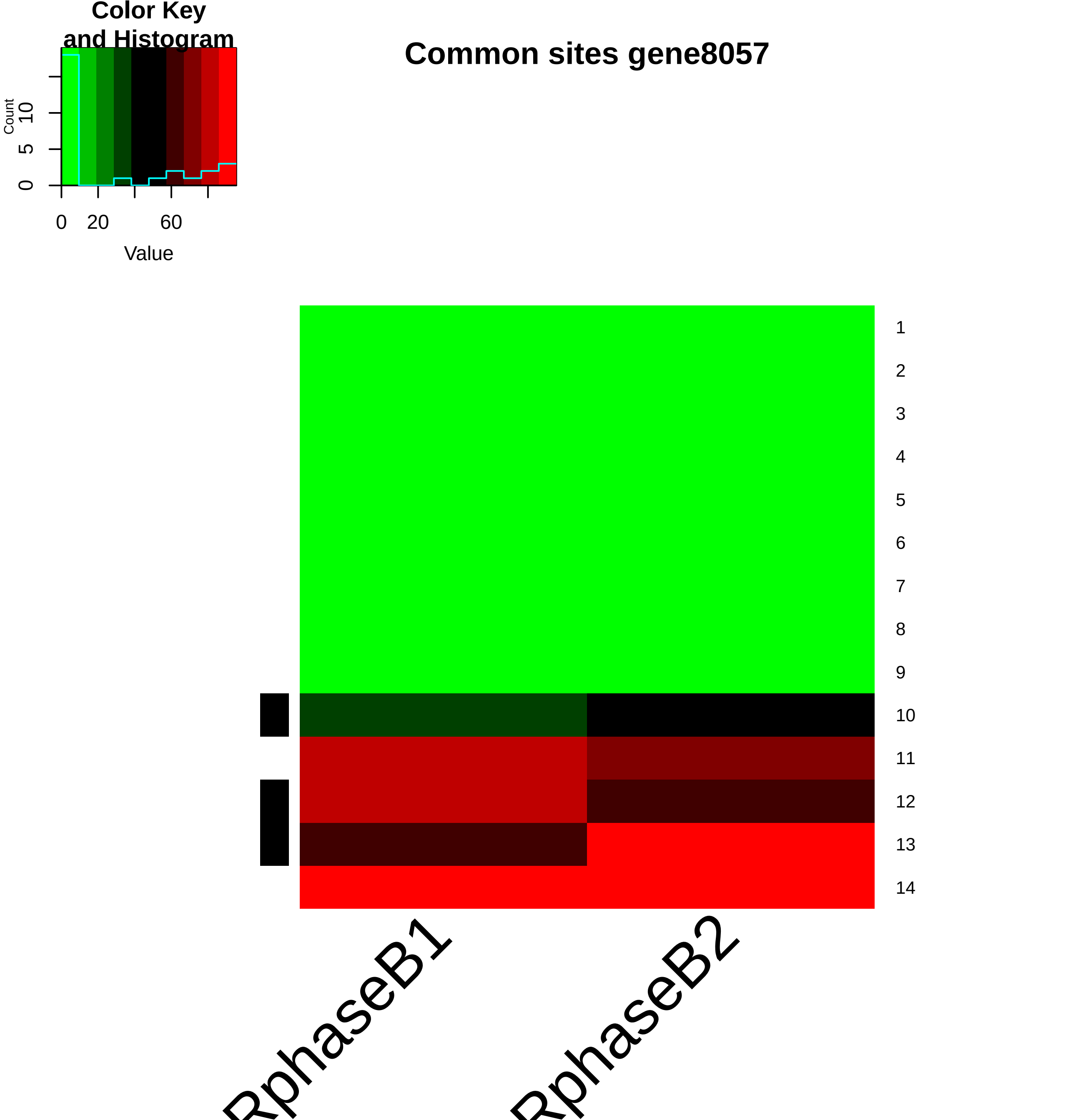
Color Key
and Histogram
Count
0
5
10
0
20
60
Value
Common sites gene8057
1
2
3
4
5
6
7
8
9
10
11
12
13
14
RphaseB1
RphaseB2

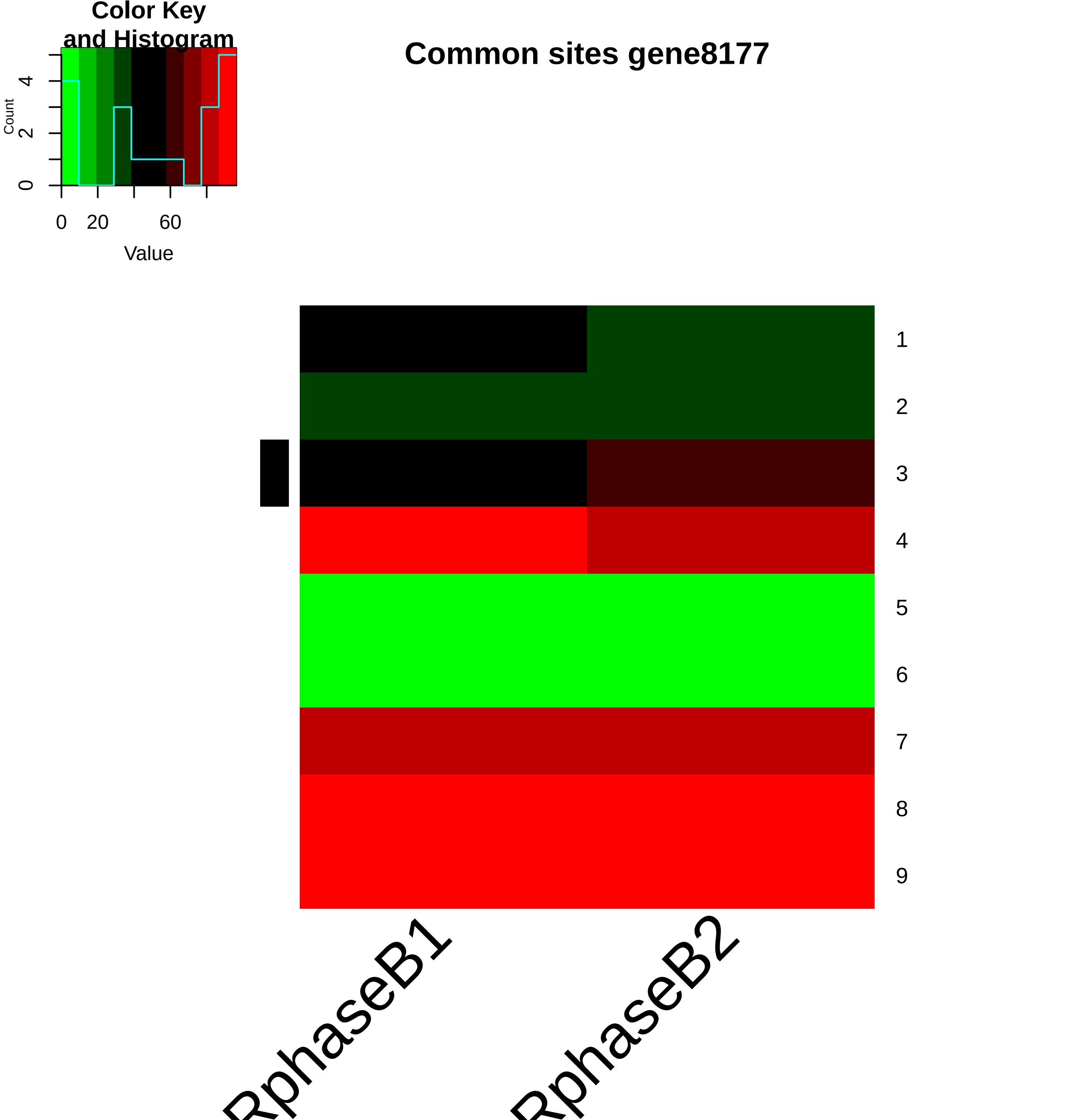
Color Key
and Histogram
Count
0
2
4
0
20
60
Value
Common sites gene8177
1
2
3
4
5
6
7
8
9
RphaseB1
RphaseB2

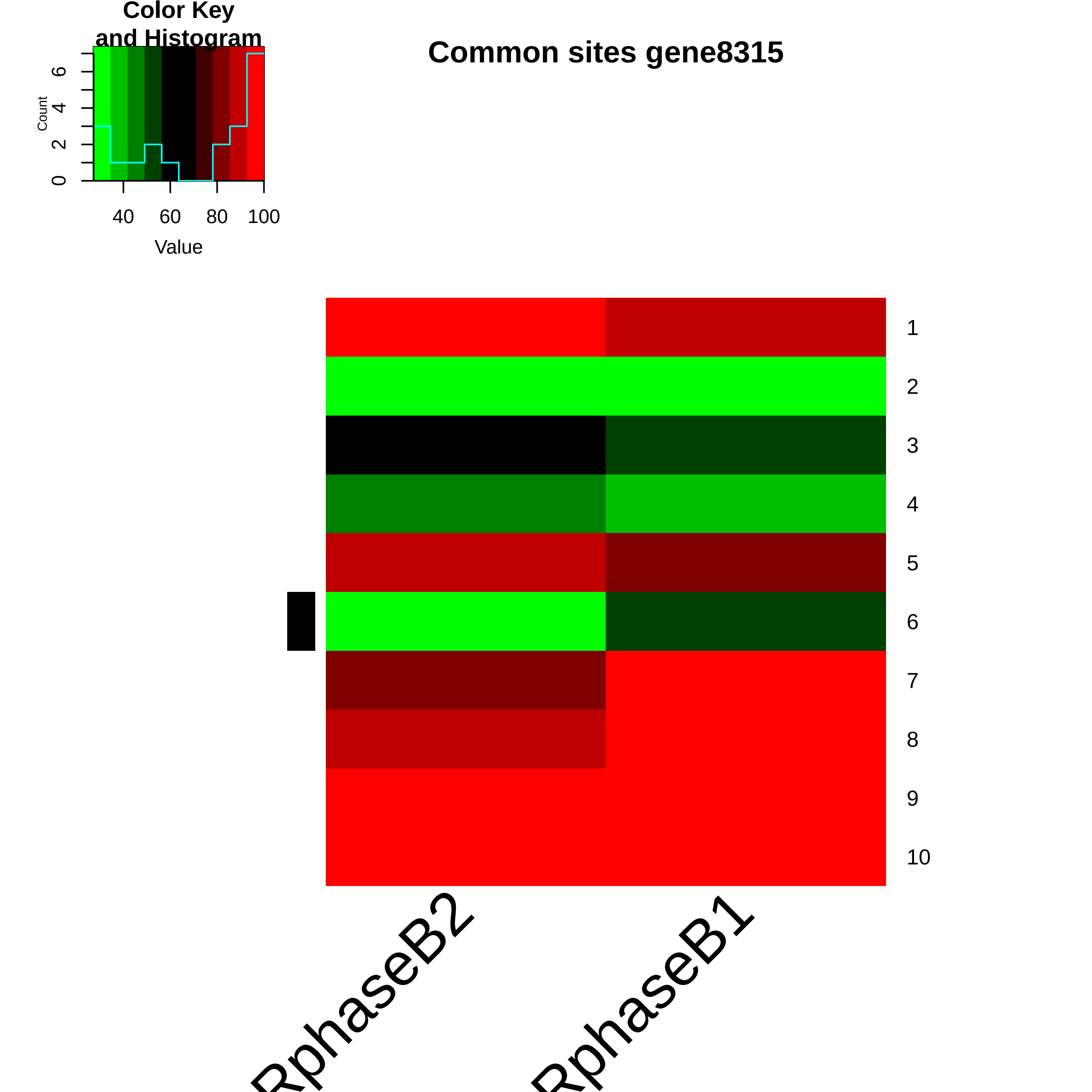
Color Key
and Histogram
Count
0
2
4
6
40
60
80
100
Value
Common sites gene8315
1
2
3
4
5
6
7
8
9
10
RphaseB2
RphaseB1

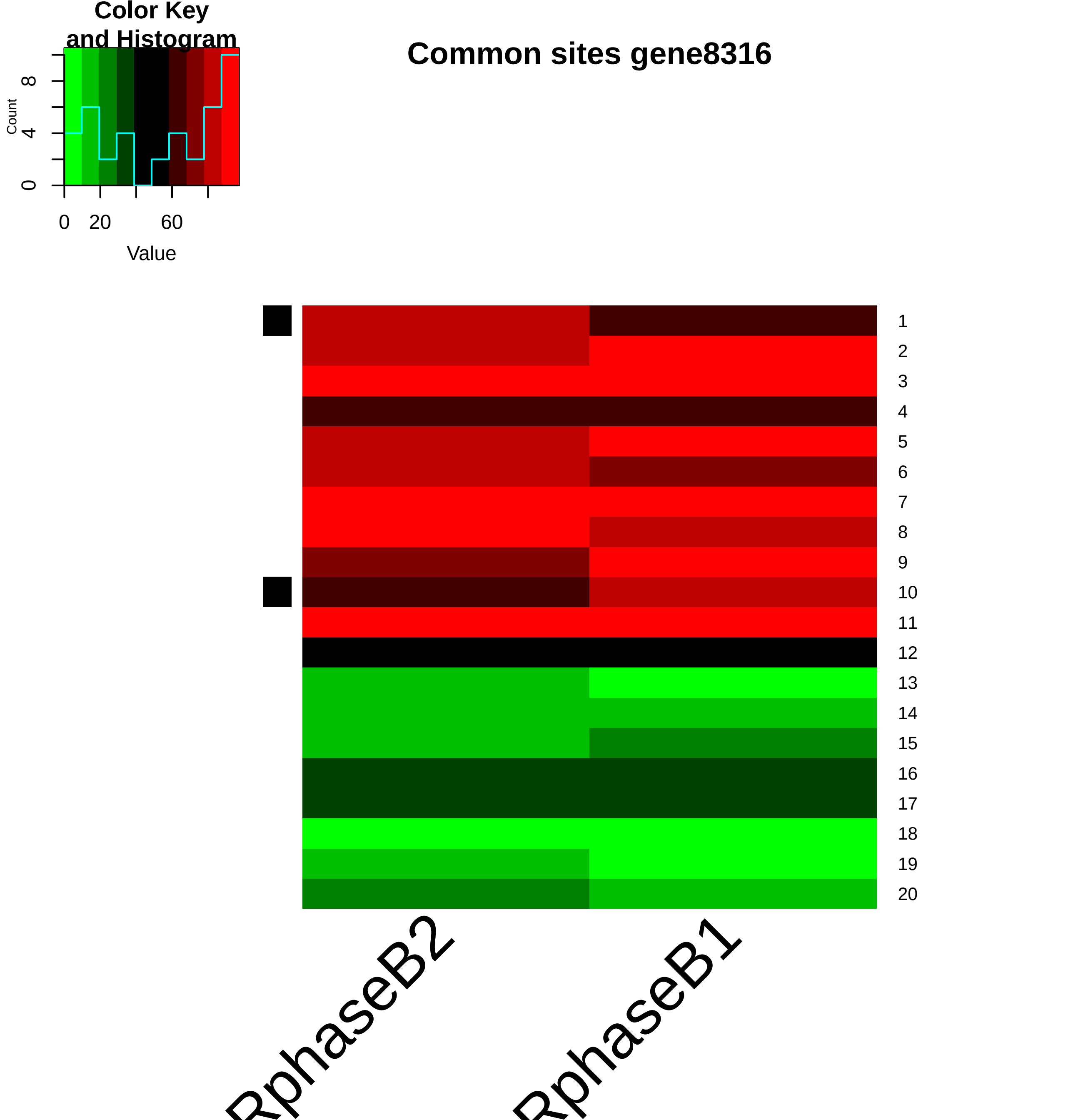
Color Key
and Histogram
Count
0
4
8
0
20
60
Value
Common sites gene8316
RphaseB2
RphaseB1

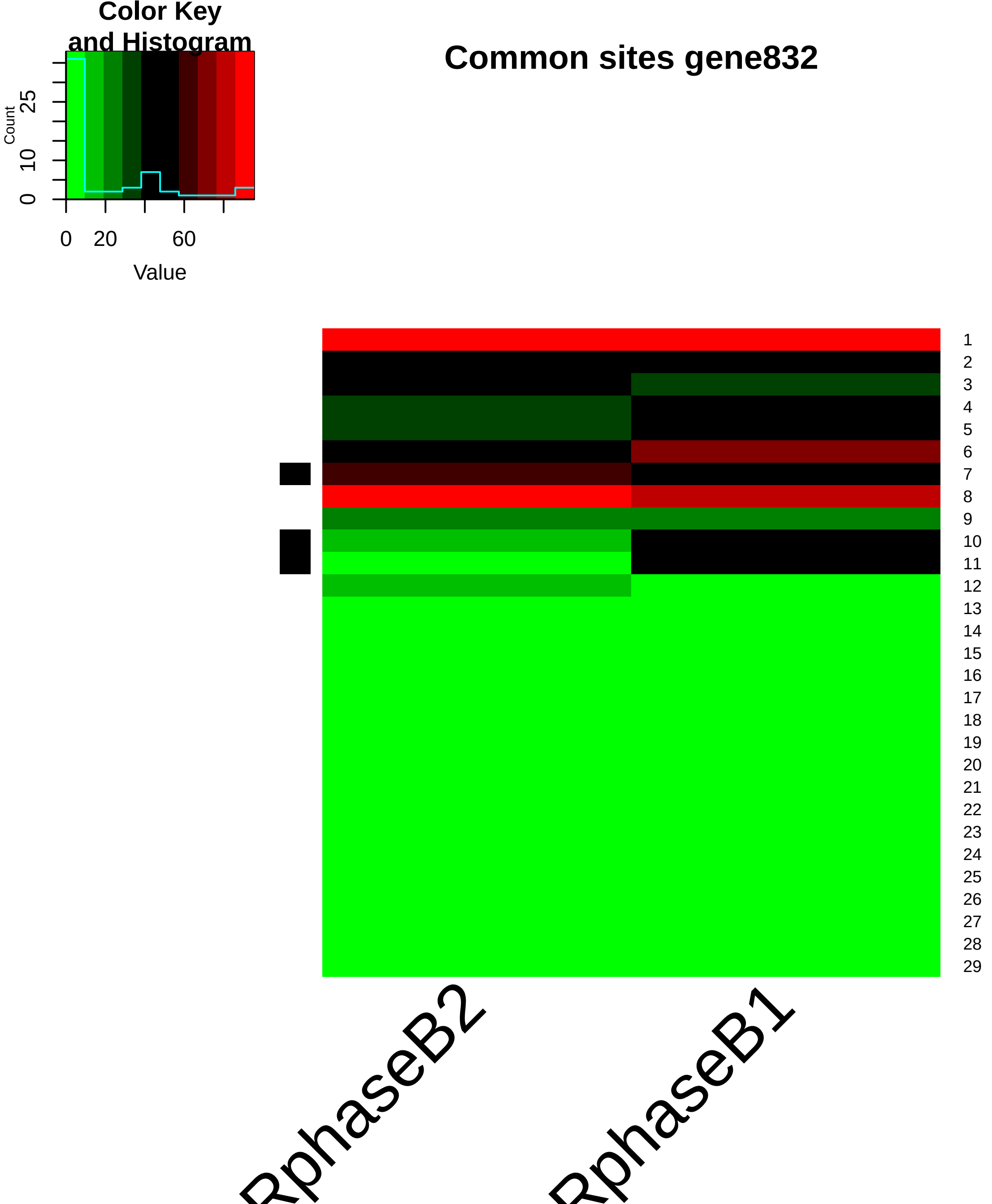
Color Key
and Histogram
Count
0
10
25
0
20
60
Value
Common sites gene832
RphaseB2
RphaseB1

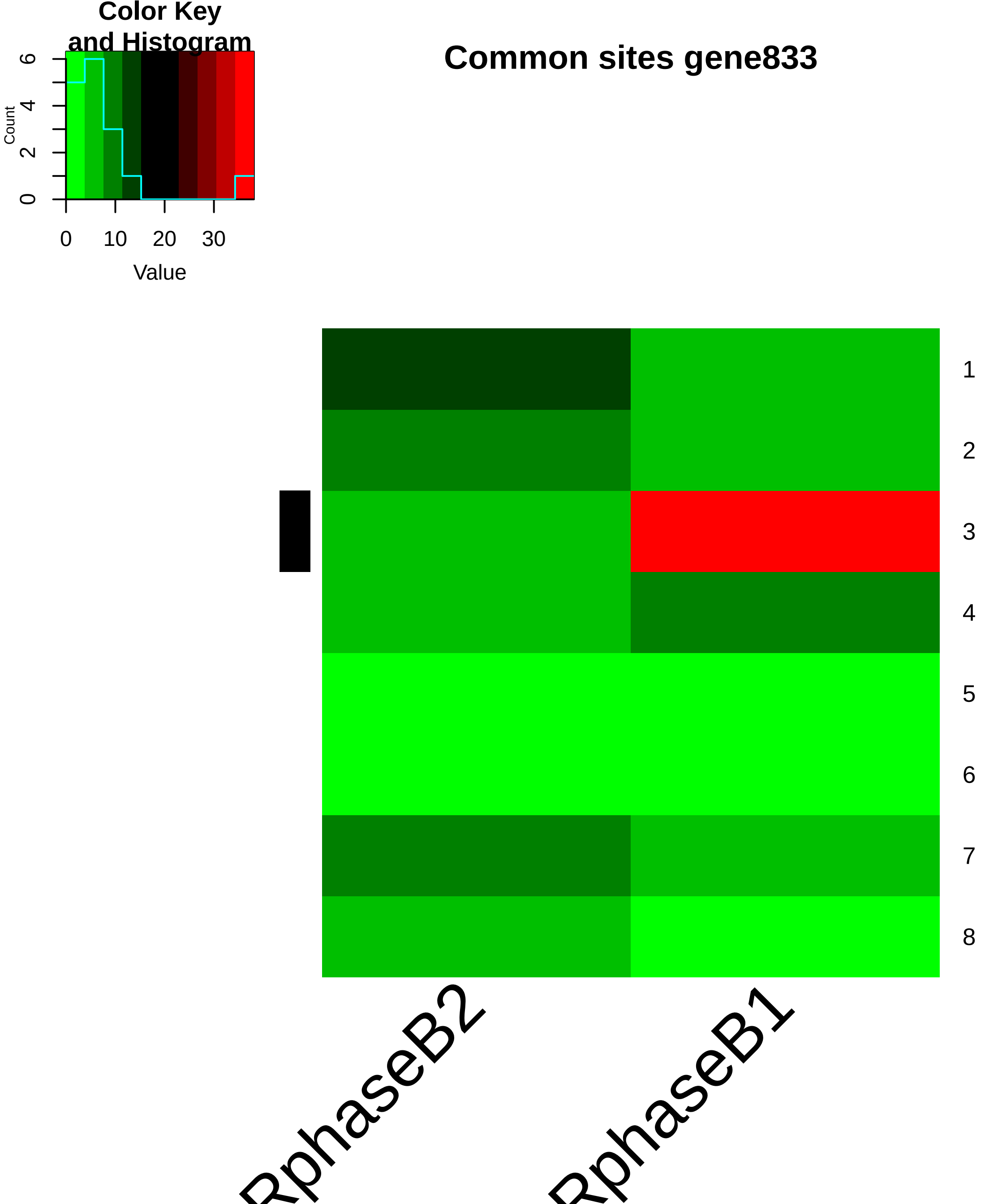

Color Key
and Histogram
Count
Value
0
2
4
6
0
10
20
30
Common sites gene833
1
2
3
4
5
6
7
8
RphaseB2
RphaseB1

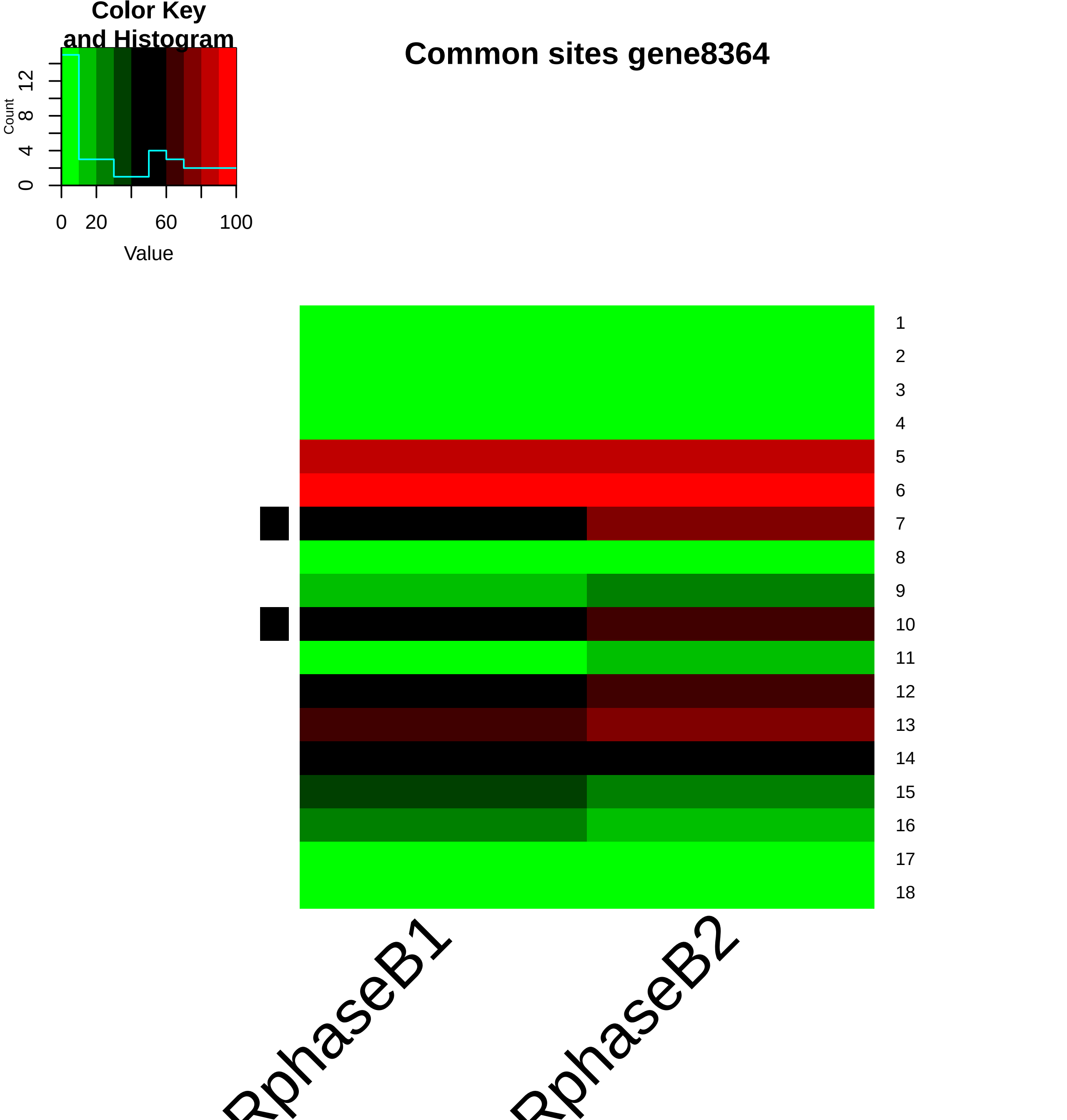
Color Key
and Histogram
Count
0
4
8
12
0
20
60
100
Value
Common sites gene8364
RphaseB1
RphaseB2

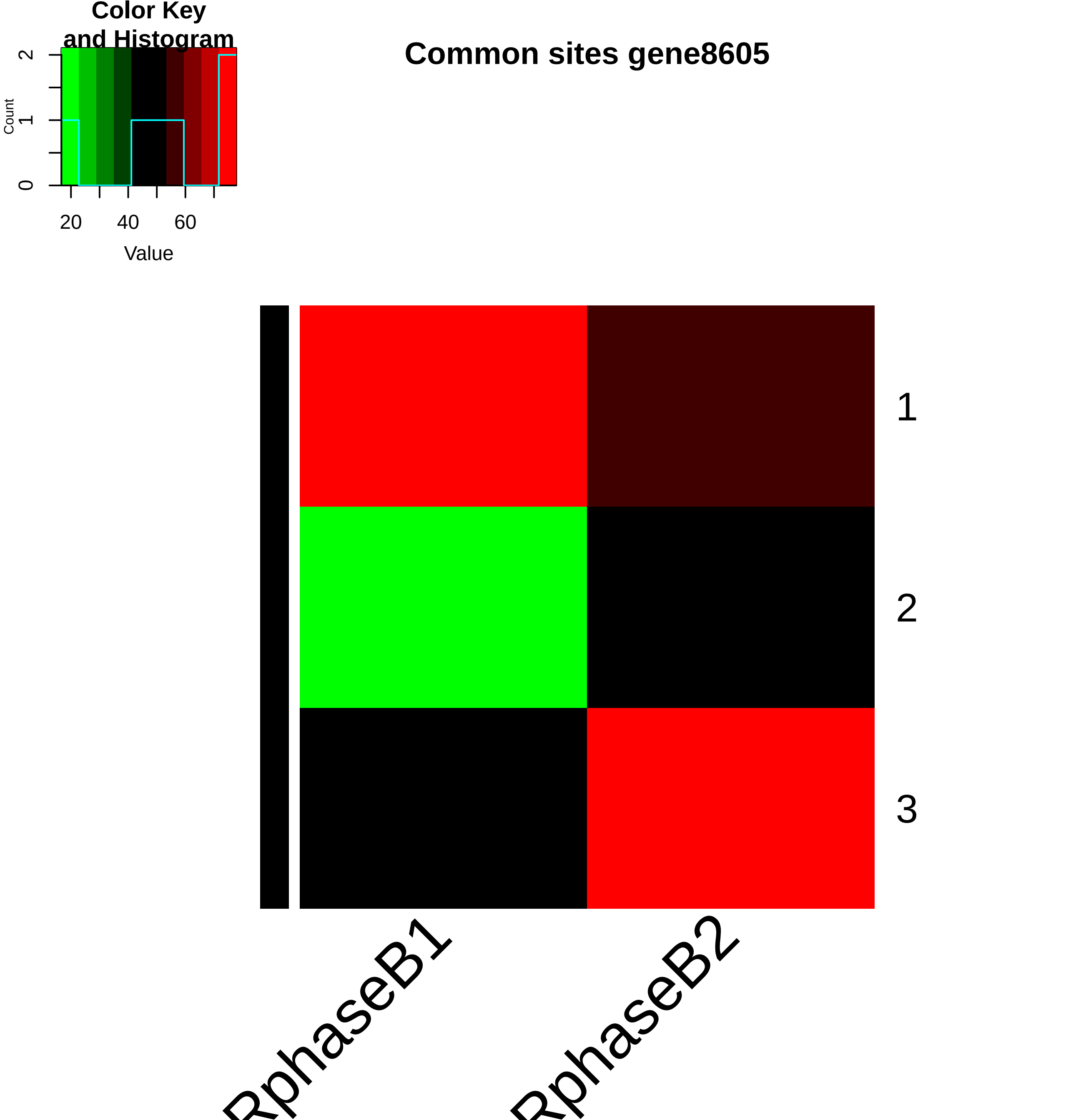

Color Key
and Histogram
Count
0
1
2
20
40
60
Value
Common sites gene8605
1
2
3
RphaseB1
RphaseB2

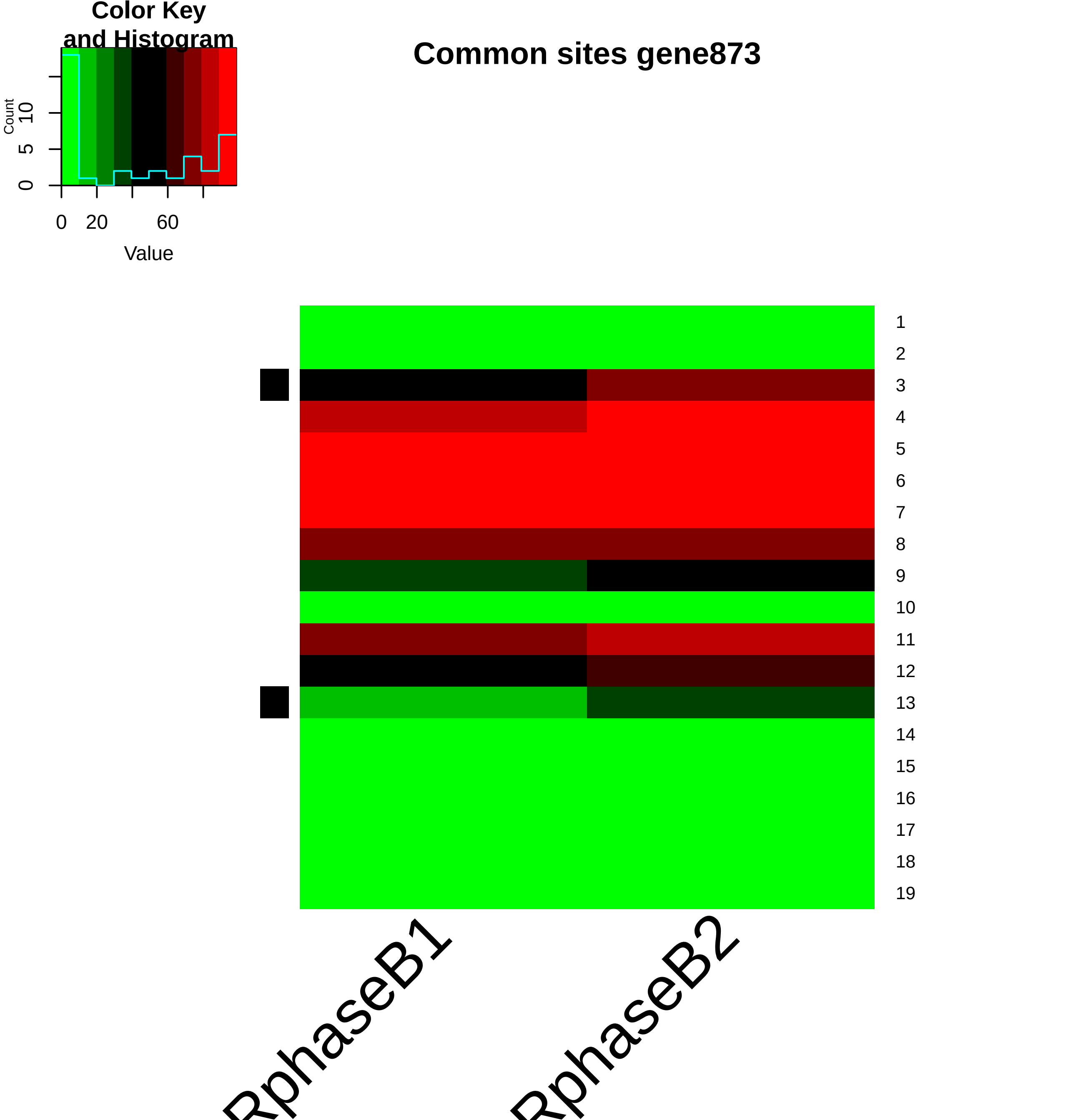

Color Key
and Histogram
Count
0
5
10
0
20
60
Value
Common sites gene873
RphaseB1
RphaseB2

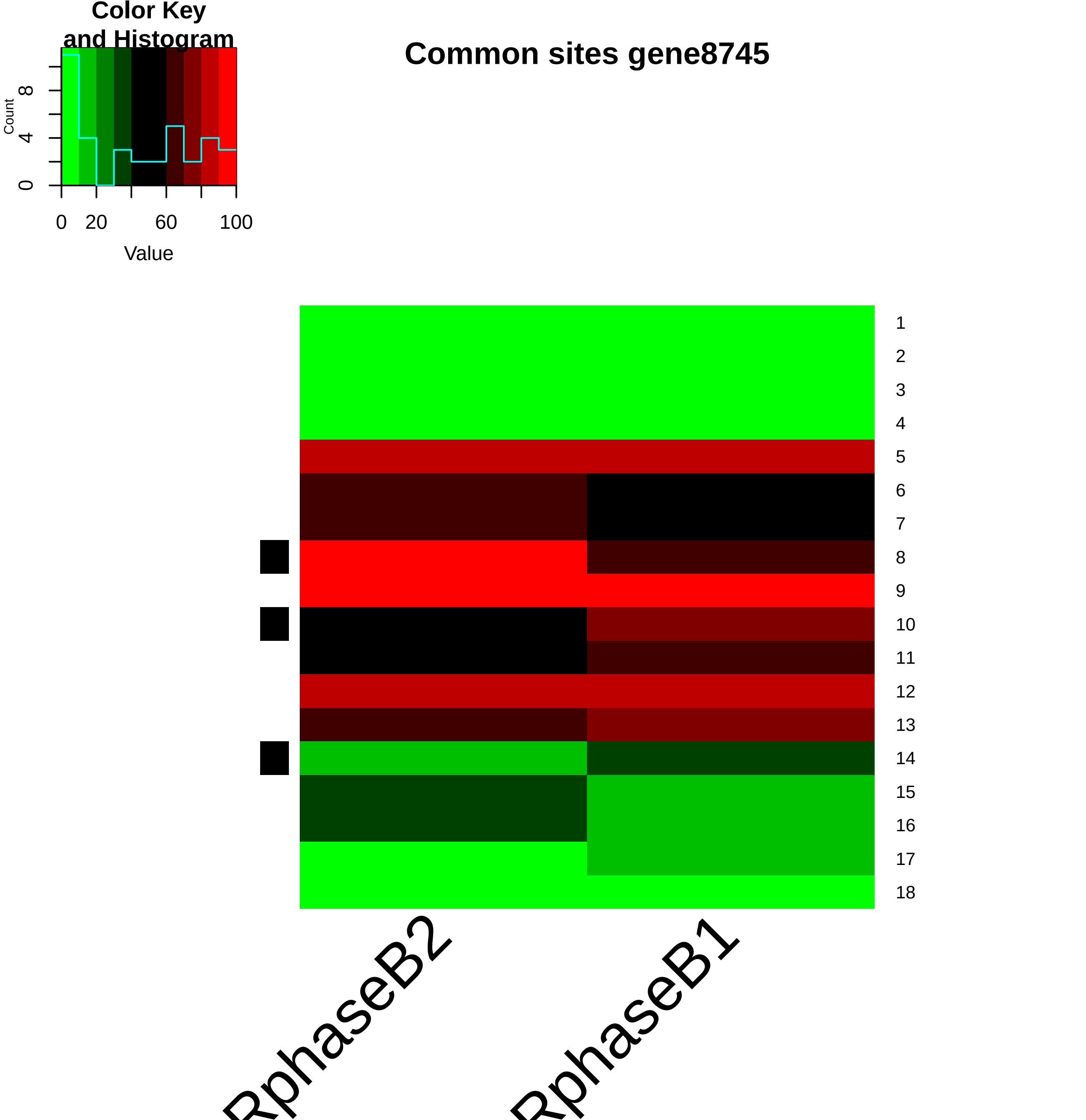

Color Key
and Histogram
Count
0
4
8
0
20
60
100
Value
Common sites gene8745
RphaseB2
RphaseB1

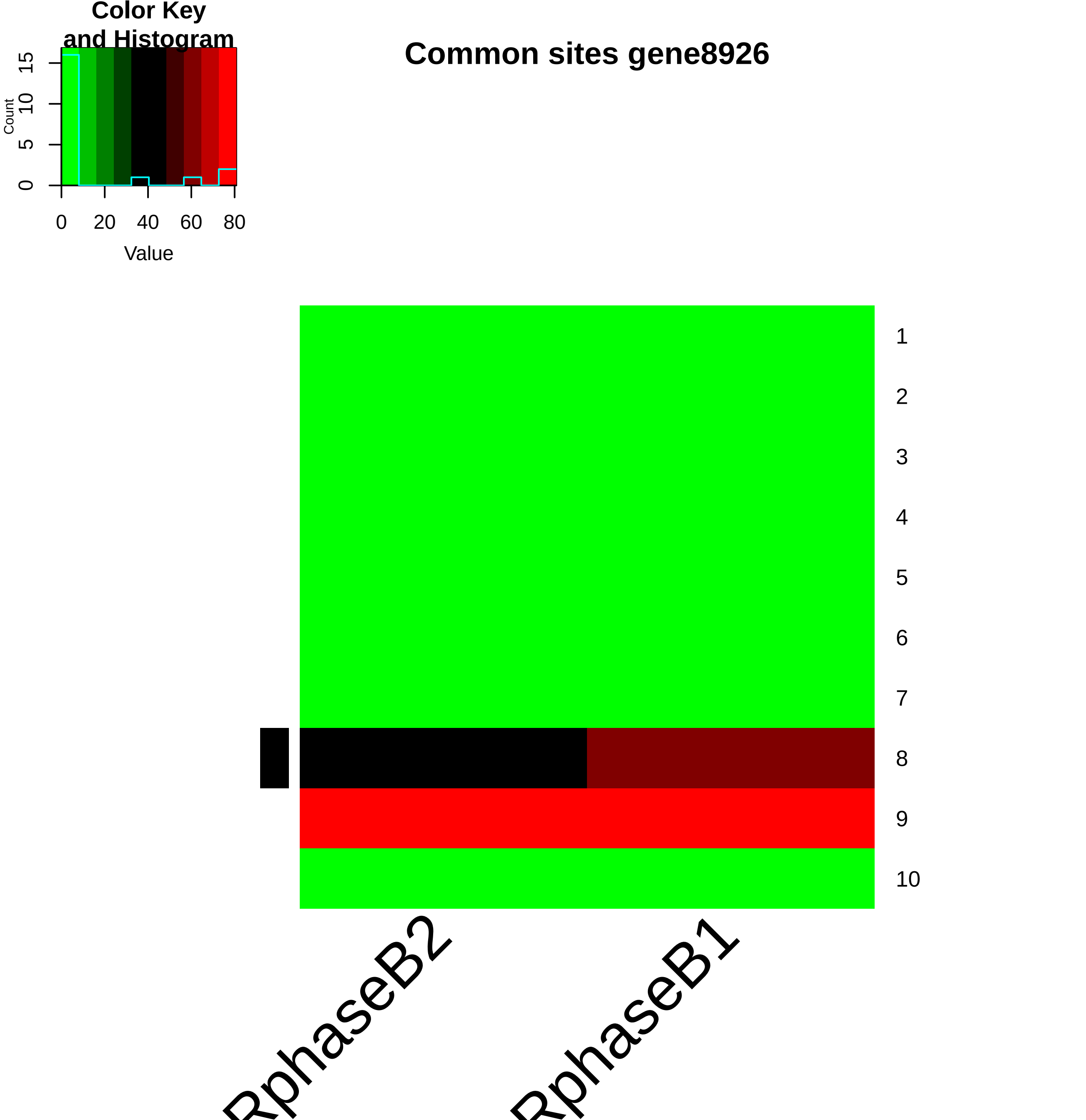
Color Key
and Histogram
Count
0
5
10
15
0
20
40
60
80
Value
Common sites gene8926
1
2
3
4
5
6
7
8
9
10
RphaseB2
RphaseB1

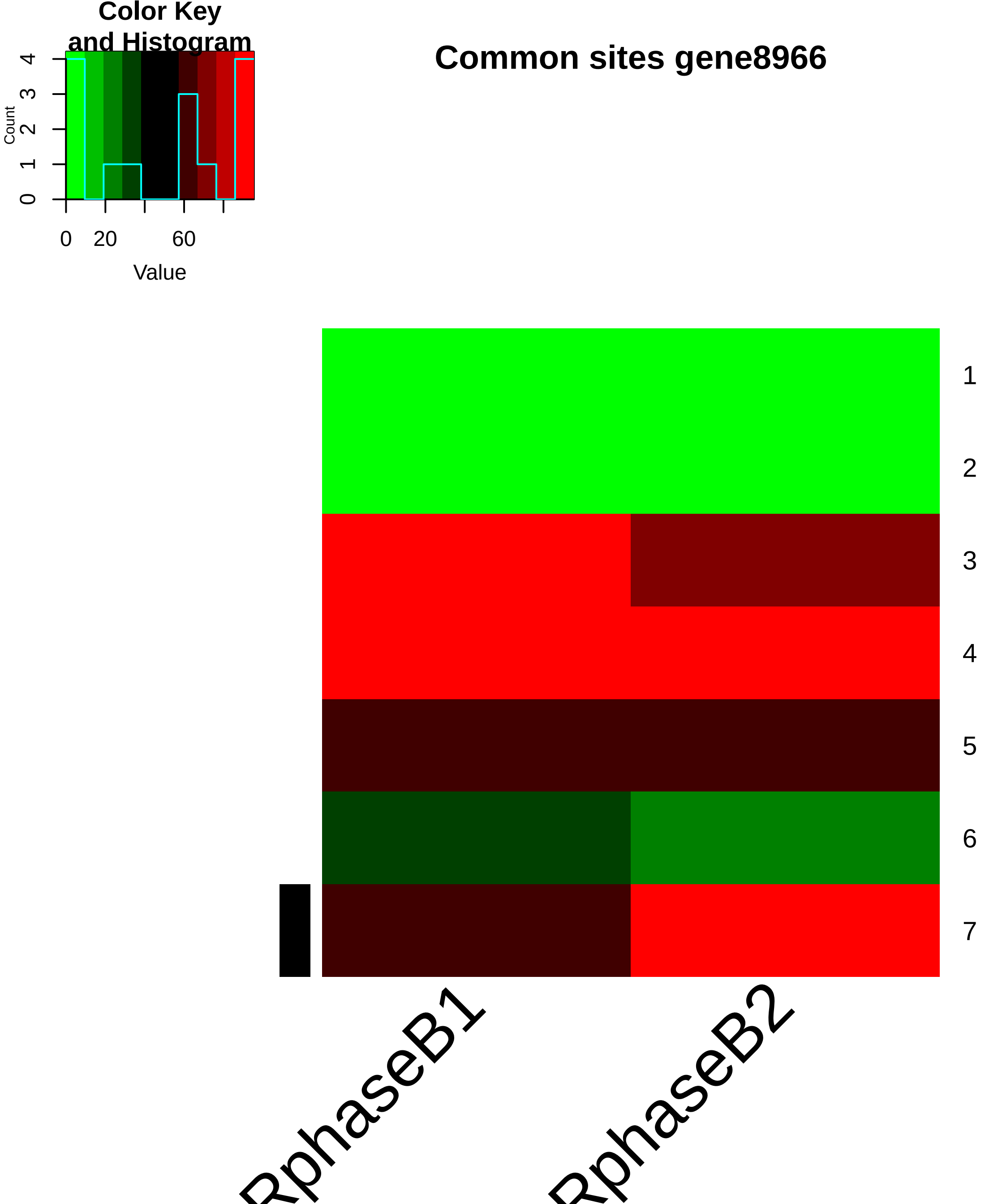
Color Key
and Histogram
Count
0
1
2
3
4
0
20
60
Value
Common sites gene8966
1
2
3
4
5
6
7
RphaseB1
RphaseB2

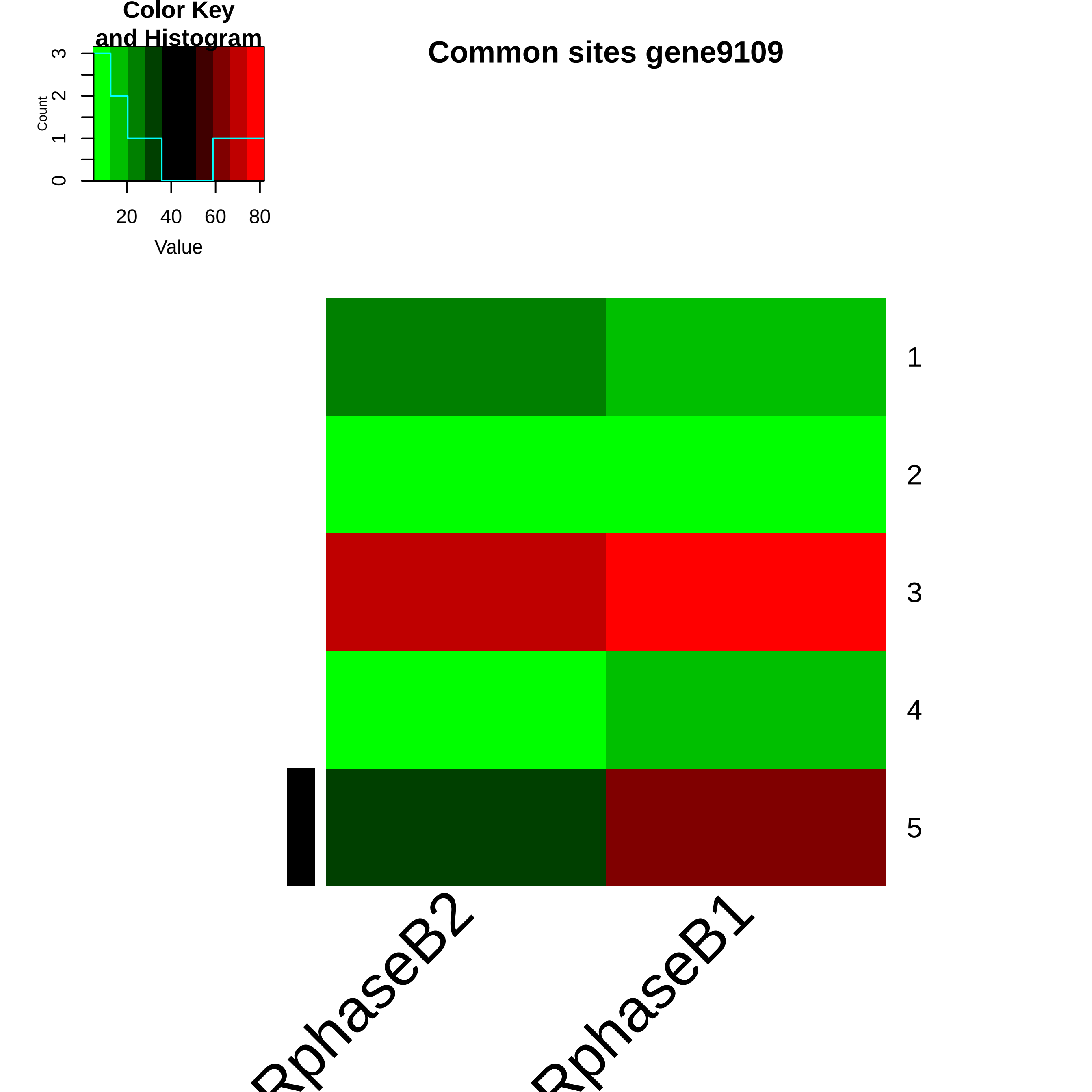
Color Key
and Histogram
Count
3
2
1
0
20
40
60
80
Value
Common sites gene9109
1
2
3
4
5
RphaseB2
RphaseB1

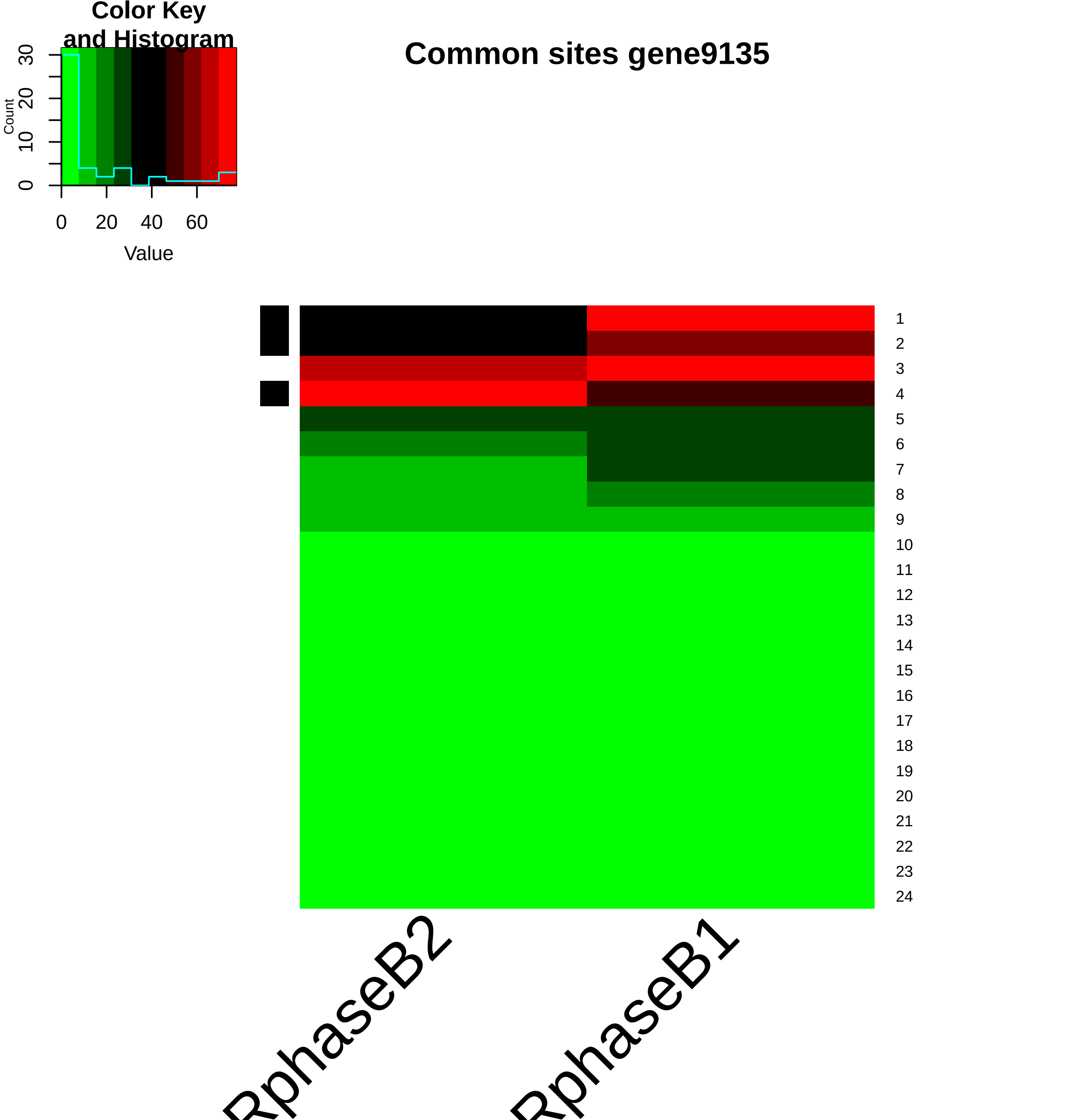
Color Key
and Histogram
Count
0
10
20
30
0
20
40
60
Value
Common sites gene9135
RphaseB2
RphaseB1

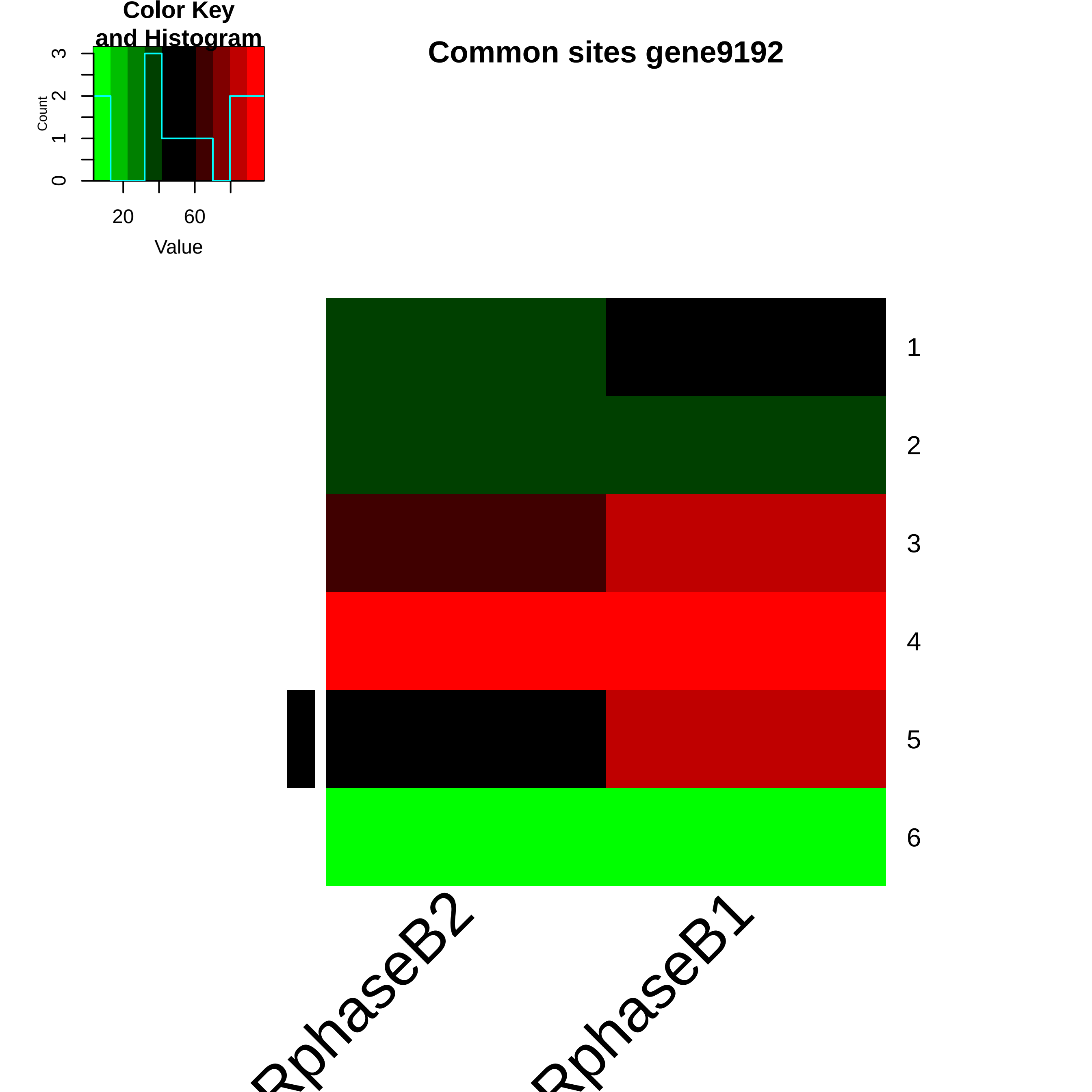
Color Key
and Histogram
Count
0
1
2
3
20
60
Value
Common sites gene9192
1
2
3
4
5
6
RphaseB2
RphaseB1

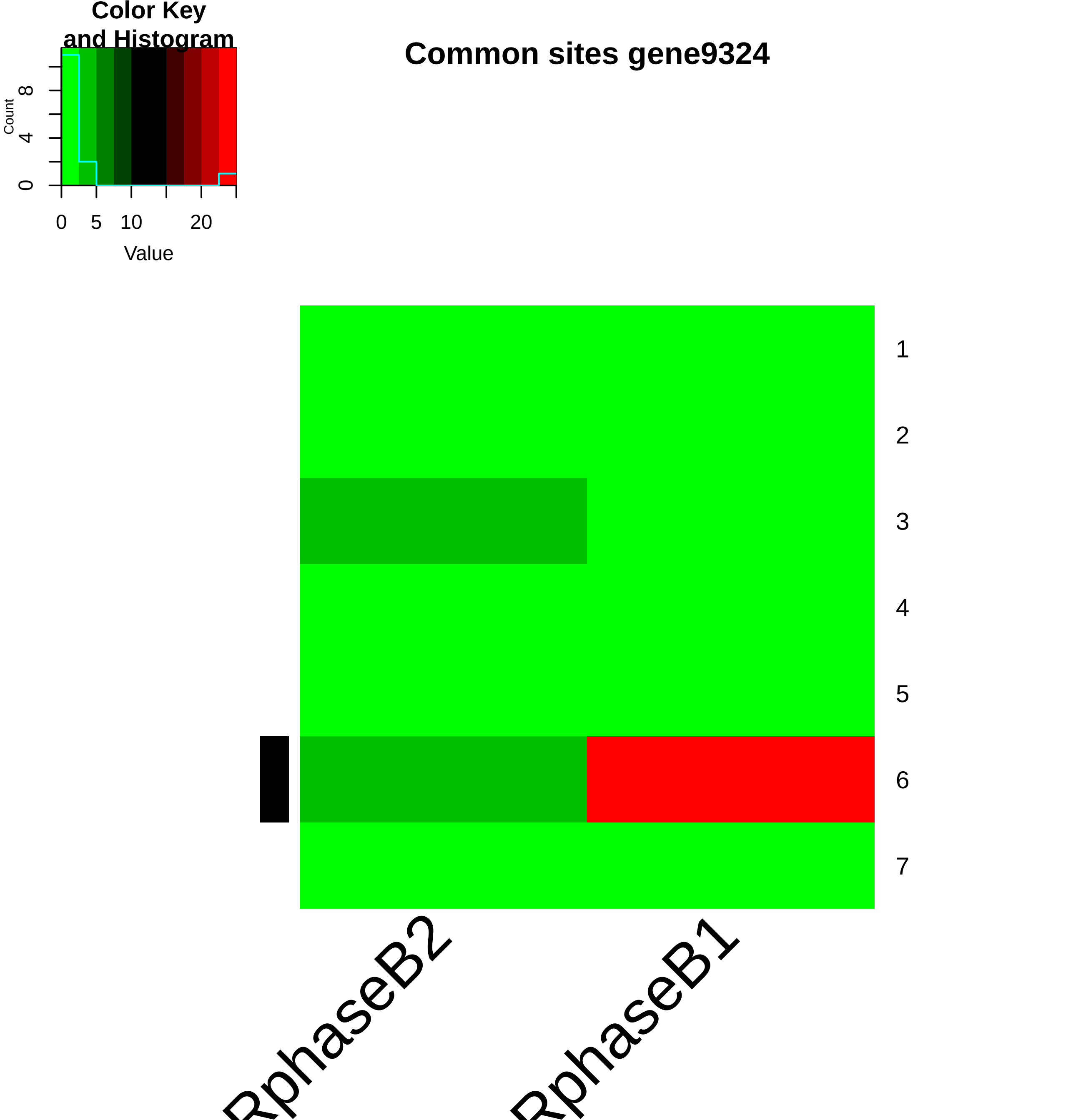

Color Key
and Histogram
Count
0
4
8
0
5
10
20
Value
Common sites gene9324
1
2
3
4
5
6
7
RphaseB2
RphaseB1

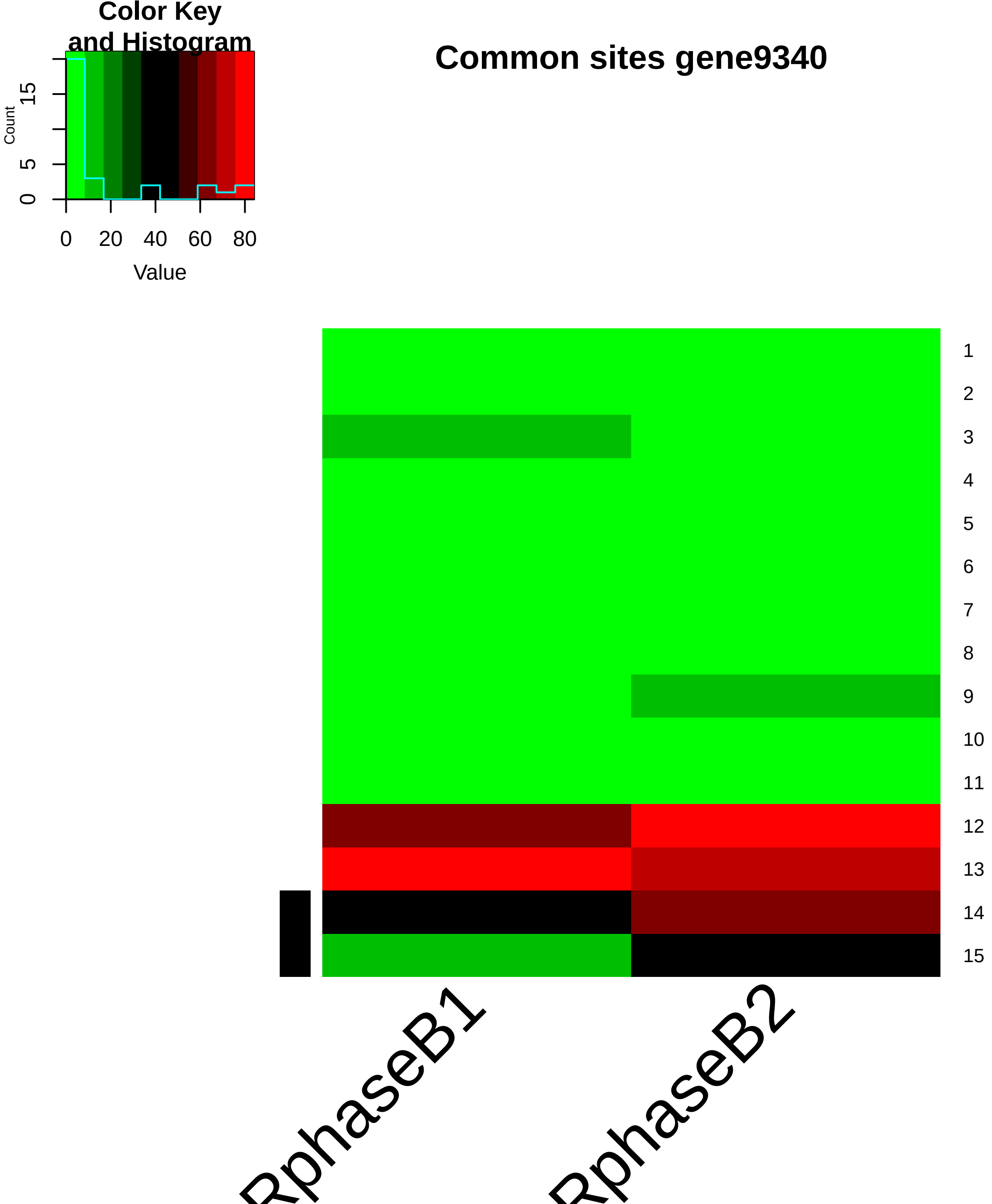
Color Key
and Histogram
Count
0
5
15
0
20
40
60
80
Value
Common sites gene9340
RphaseB1
RphaseB2

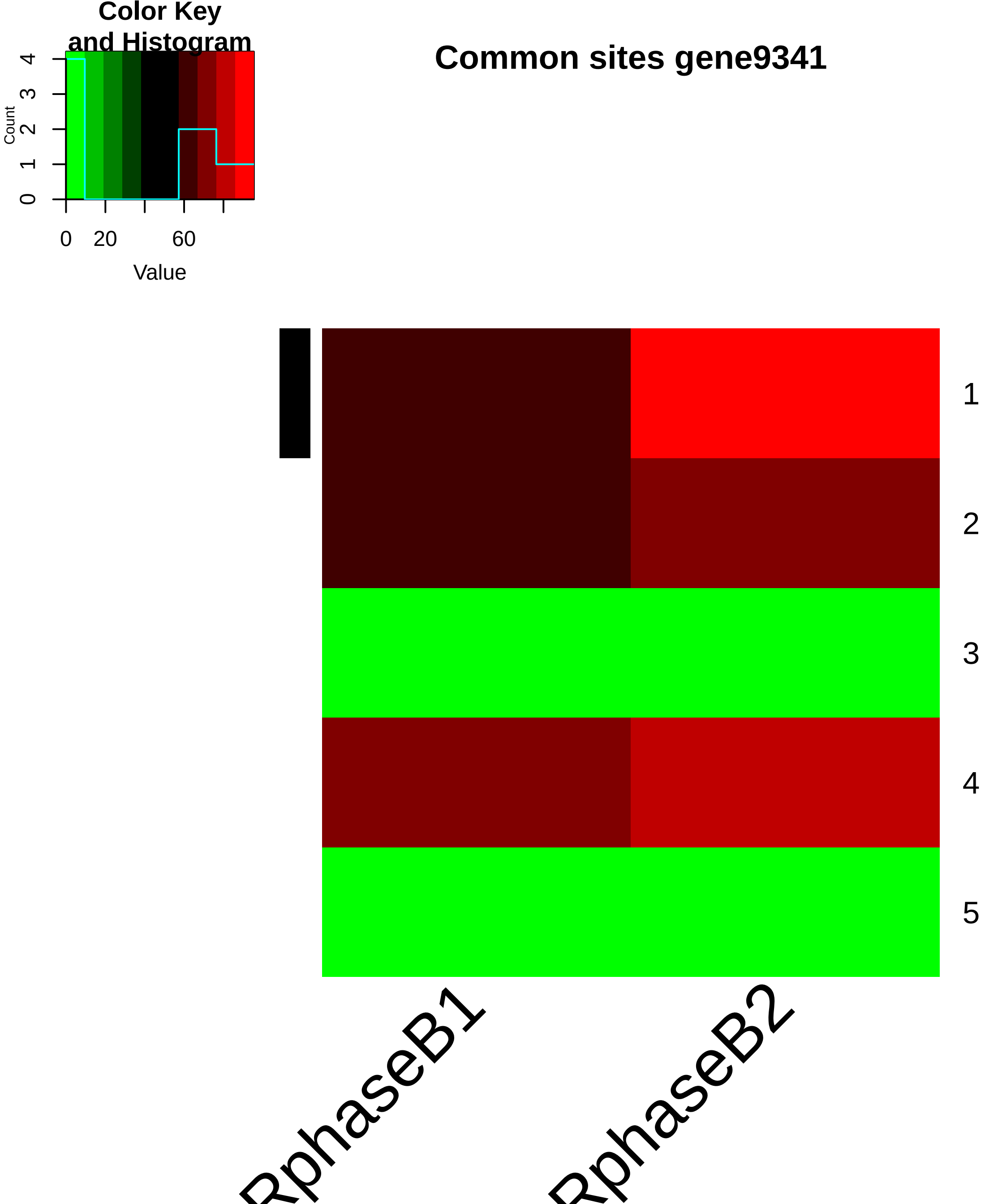

Color Key
and Histogram
Count
0
1
2
3
4
0
20
60
Value
Common sites gene9341
1
2
3
4
5
RphaseB1
RphaseB2

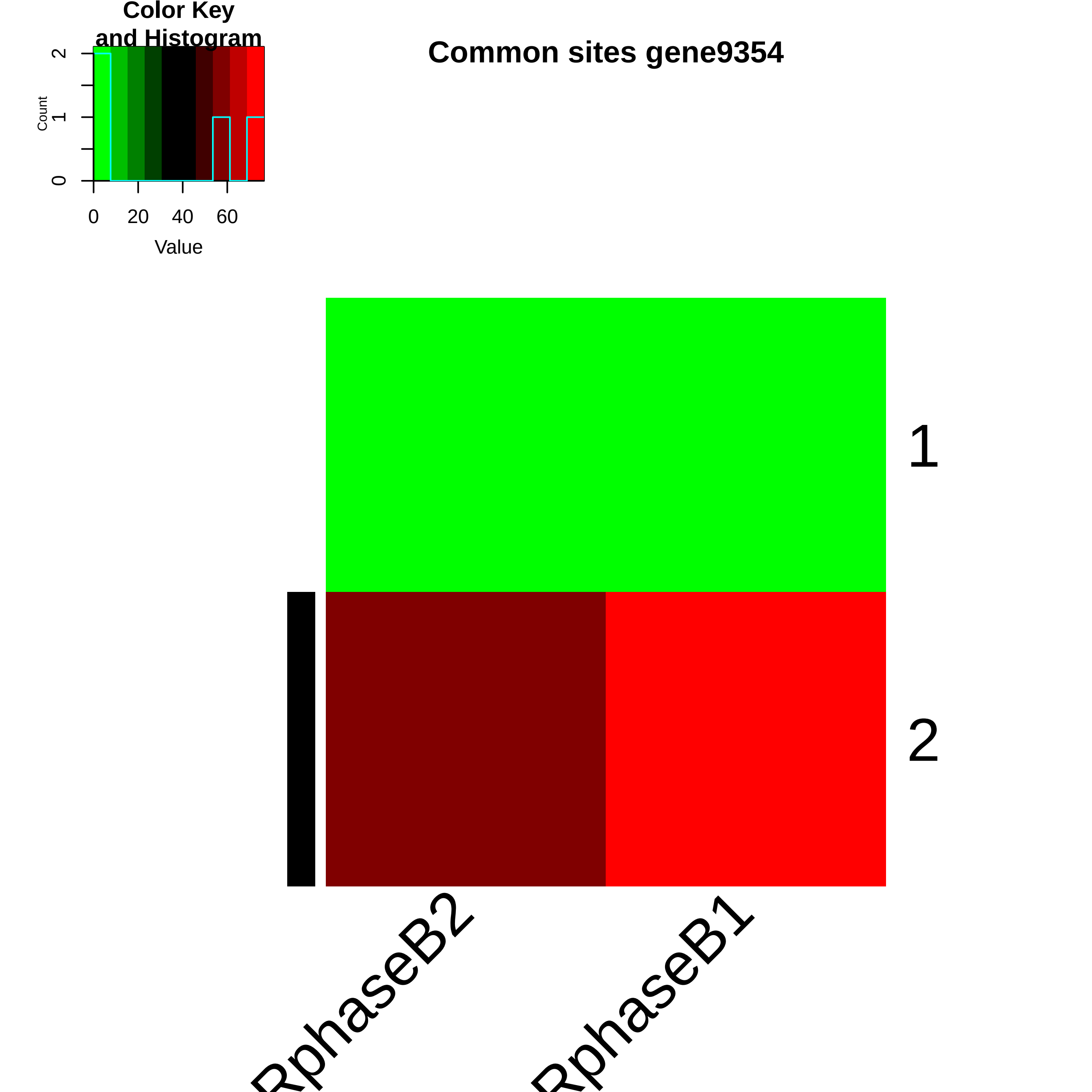
Color Key
and Histogram
Count
0
1
2
0
20
40
60
Value
Common sites gene9354
1
2
RphaseB2
RphaseB1

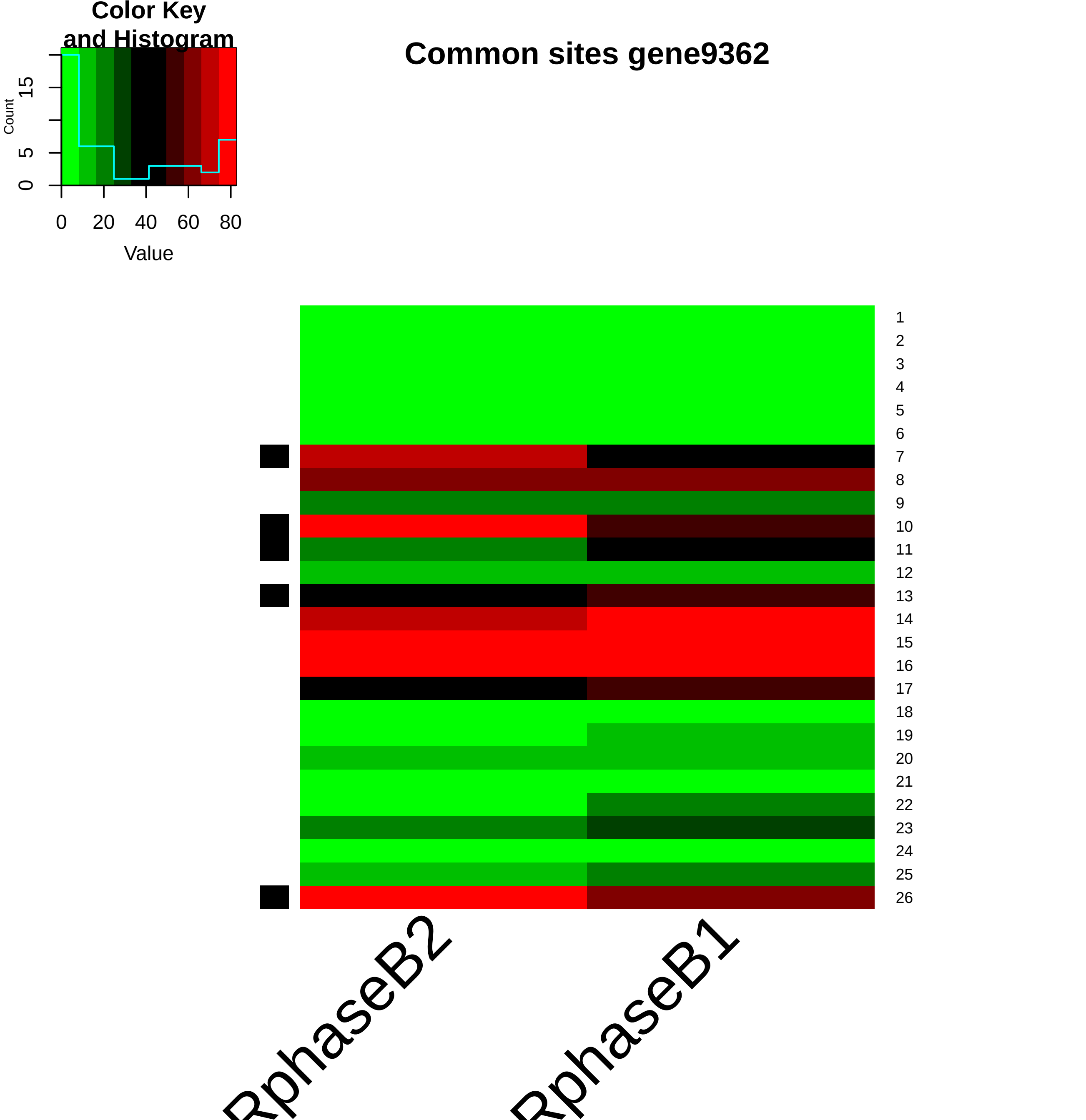
Color Key
and Histogram
Count
0
5
15
0
20
40
60
80
Value
Common sites gene9362
RphaseB2
RphaseB1

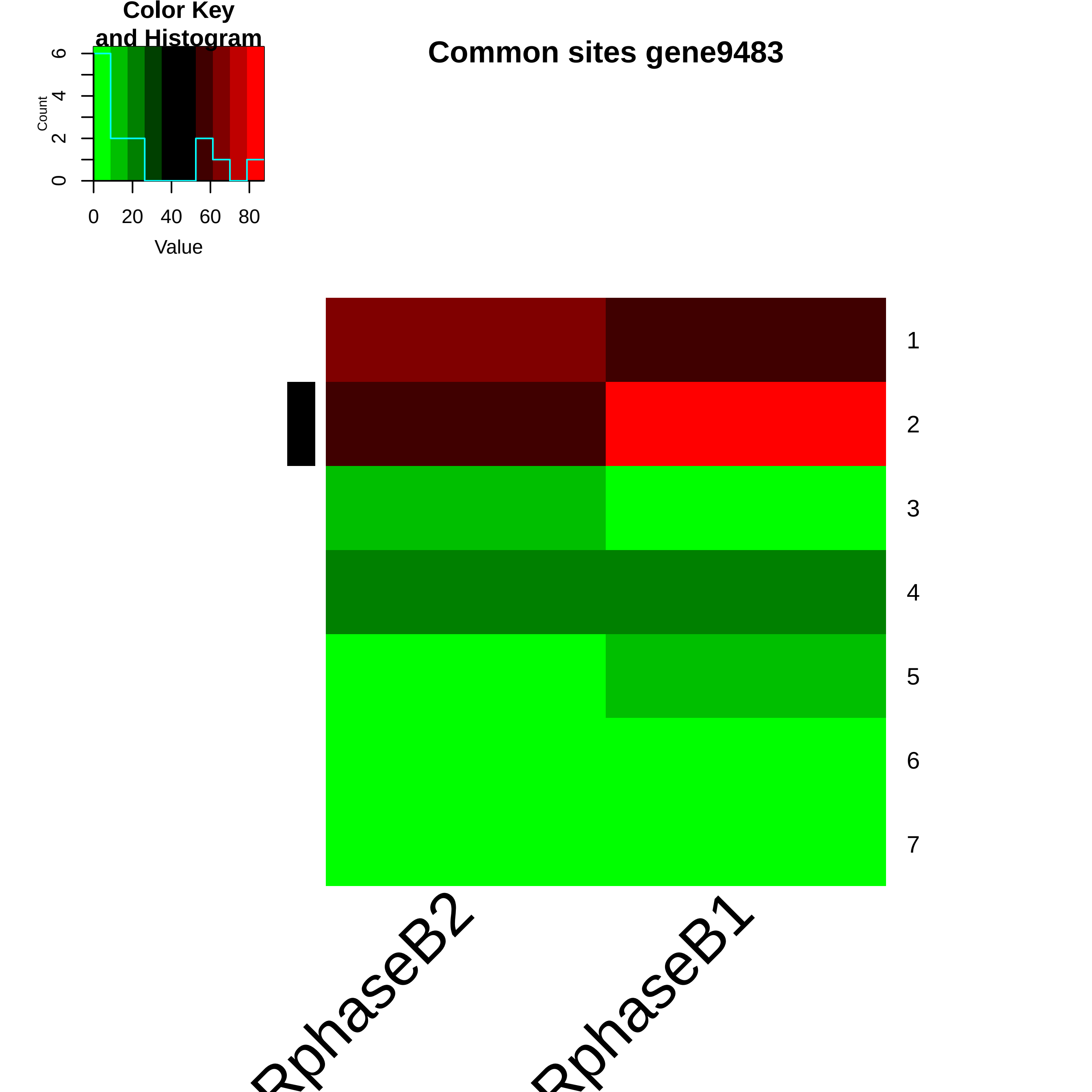

Color Key
and Histogram
Count
0
2
4
6
0
20
40
60
80
Value
Common sites gene9483
1
2
3
4
5
6
7
RphaseB2
RphaseB1

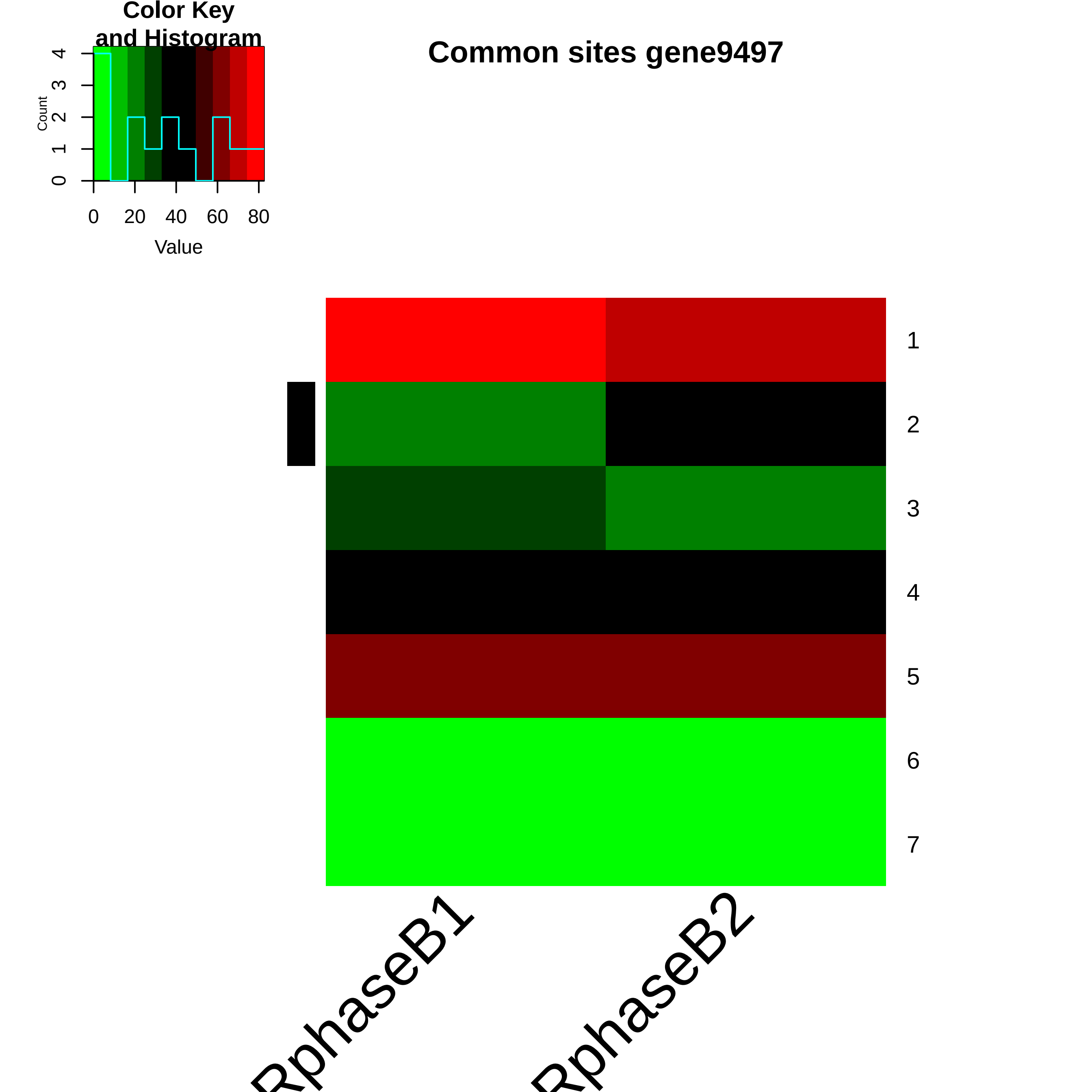

Color Key
and Histogram
Count
0
1
2
3
4
0
20
40
60
80
Value
Common sites gene9497
1
2
3
4
5
6
7
RphaseB1
RphaseB2

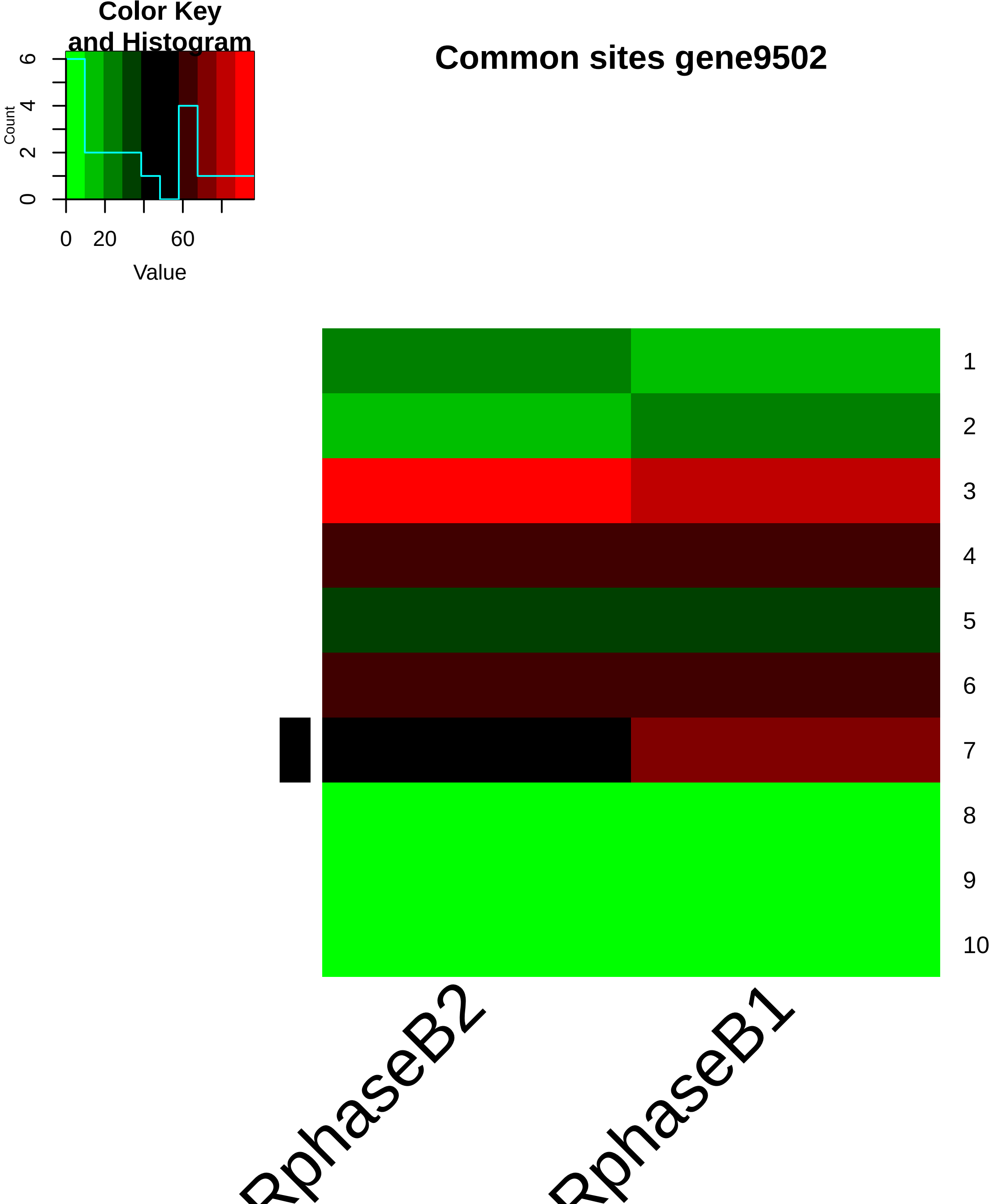
Color Key
and Histogram
Count
0
2
4
6
0
20
60
Value
Common sites gene9502
1
2
3
4
5
6
7
8
9
10
RphaseB2
RphaseB1

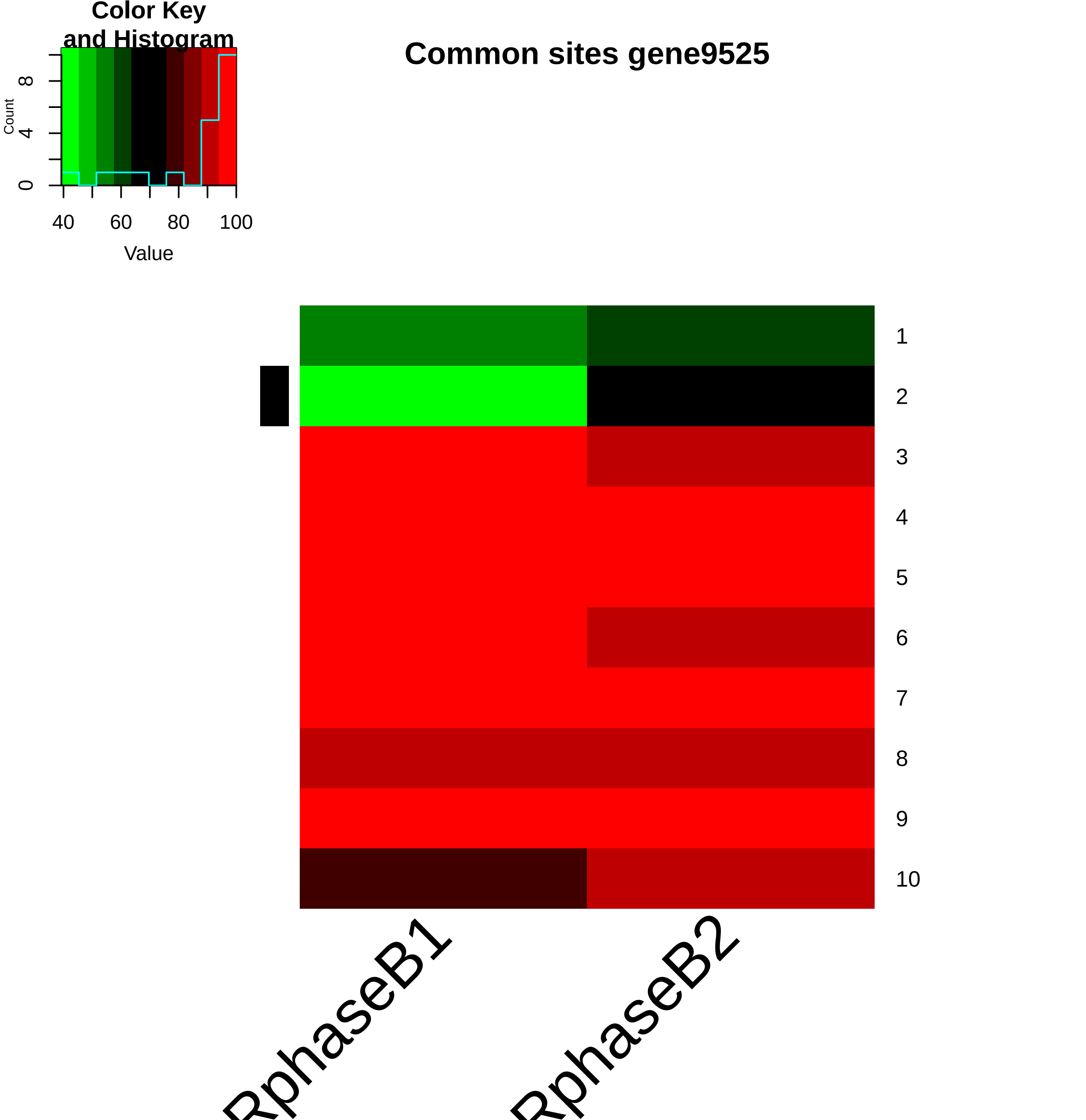
Color Key
and Histogram
Count
0
4
8
40
60
80
100
Value
Common sites gene9525
1
2
3
4
5
6
7
8
9
10
RphaseB1
RphaseB2

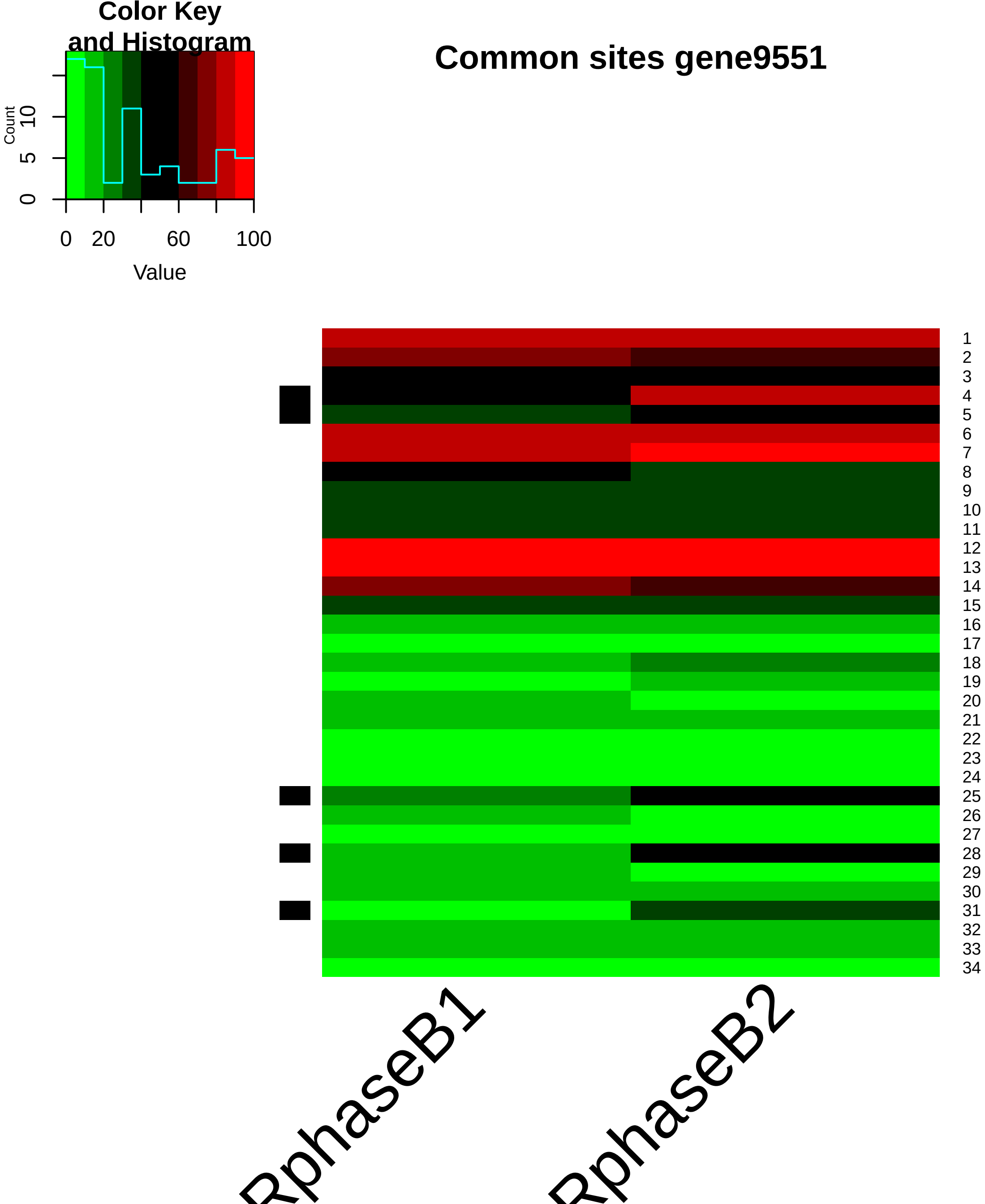

Color Key
and Histogram
Count
0
5
10
0
20
60
100
Value
Common sites gene9551
RphaseB1
RphaseB2

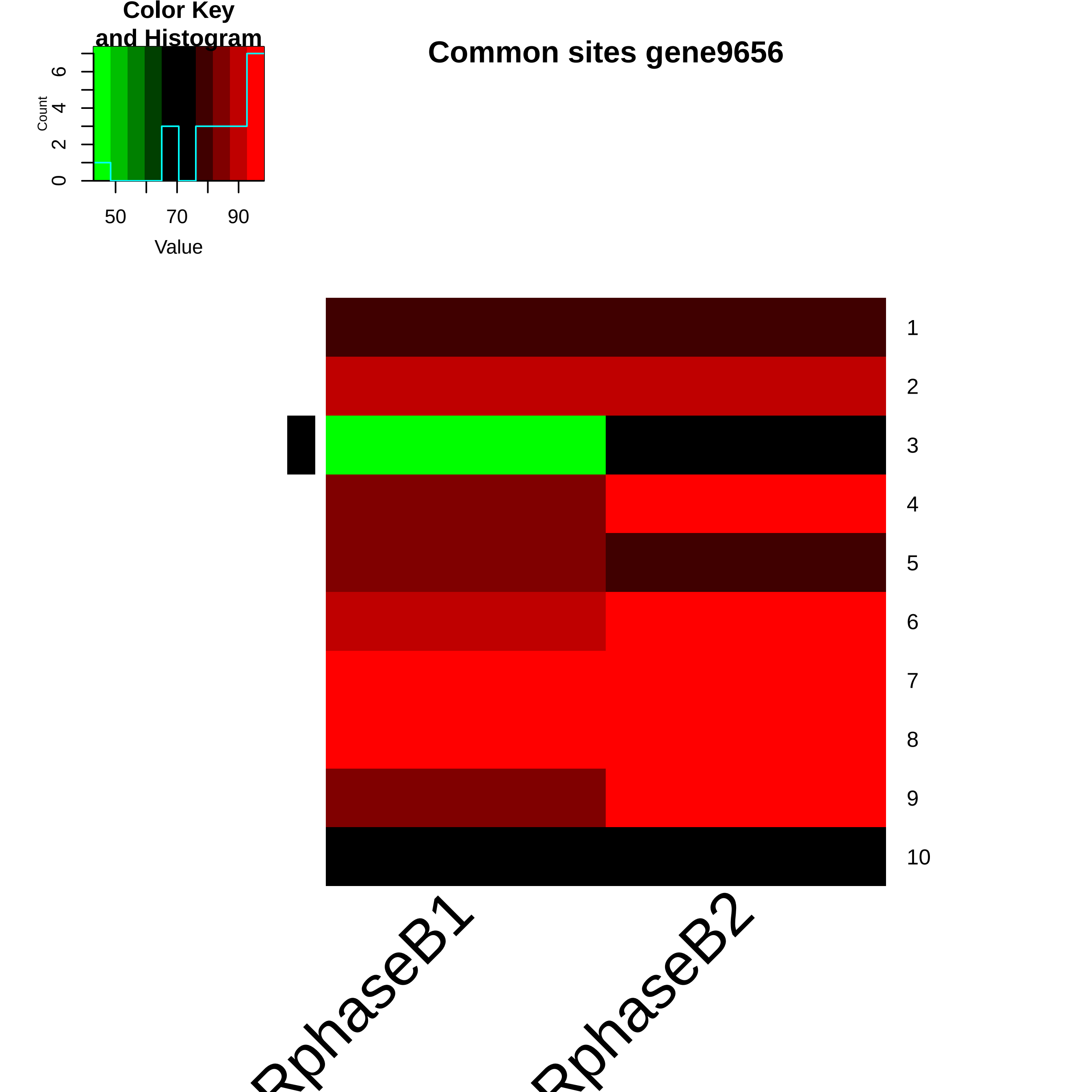
Color Key
and Histogram
Count
0
2
4
6
50
70
90
Value
Common sites gene9656
1
2
3
4
5
6
7
8
9
10
RphaseB1
RphaseB2

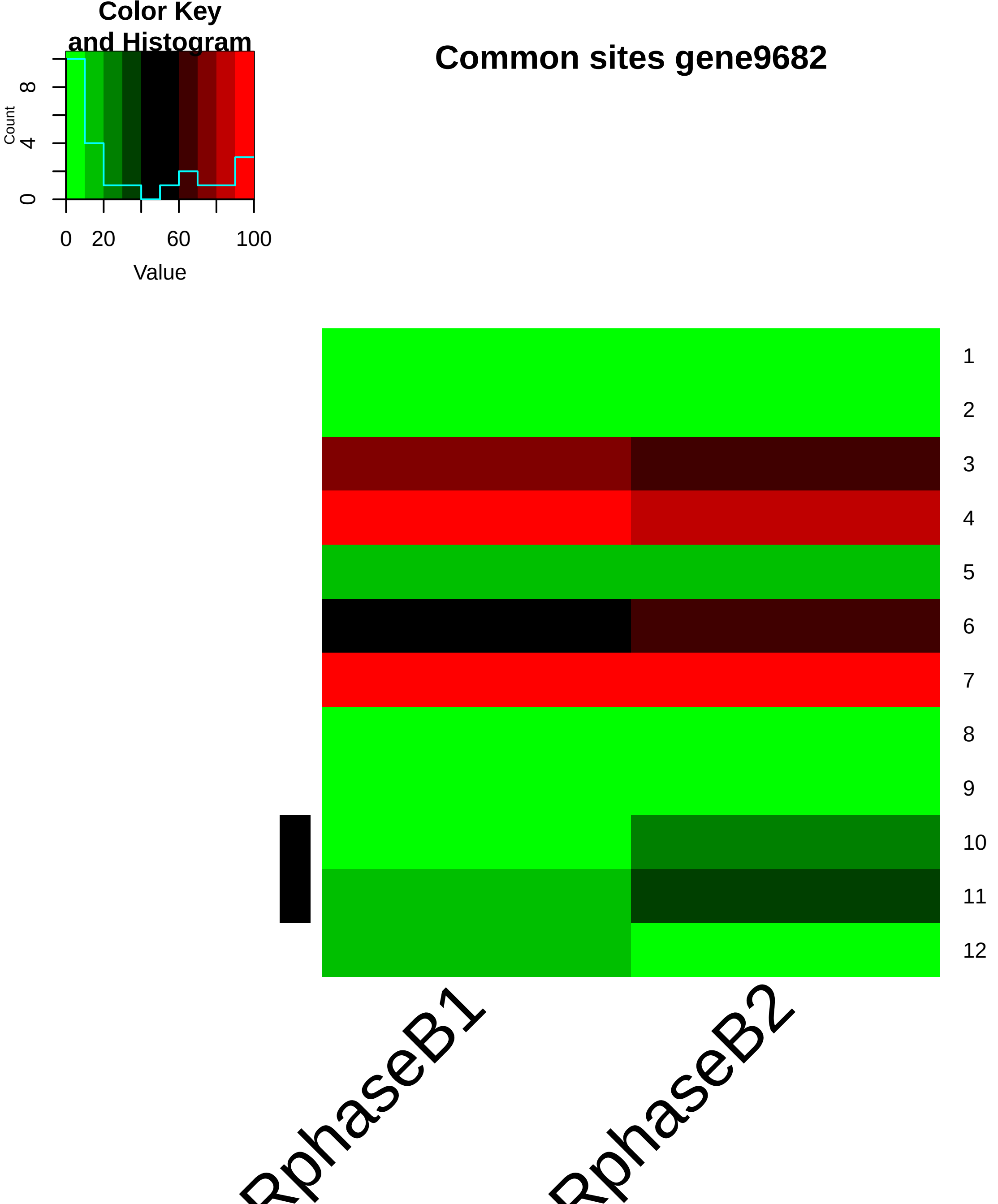

Color Key
and Histogram
Count
0
4
8
0
20
60
100
Value
Common sites gene9682
1
2
3
4
5
6
7
8
9
10
11
12
RphaseB1
RphaseB2

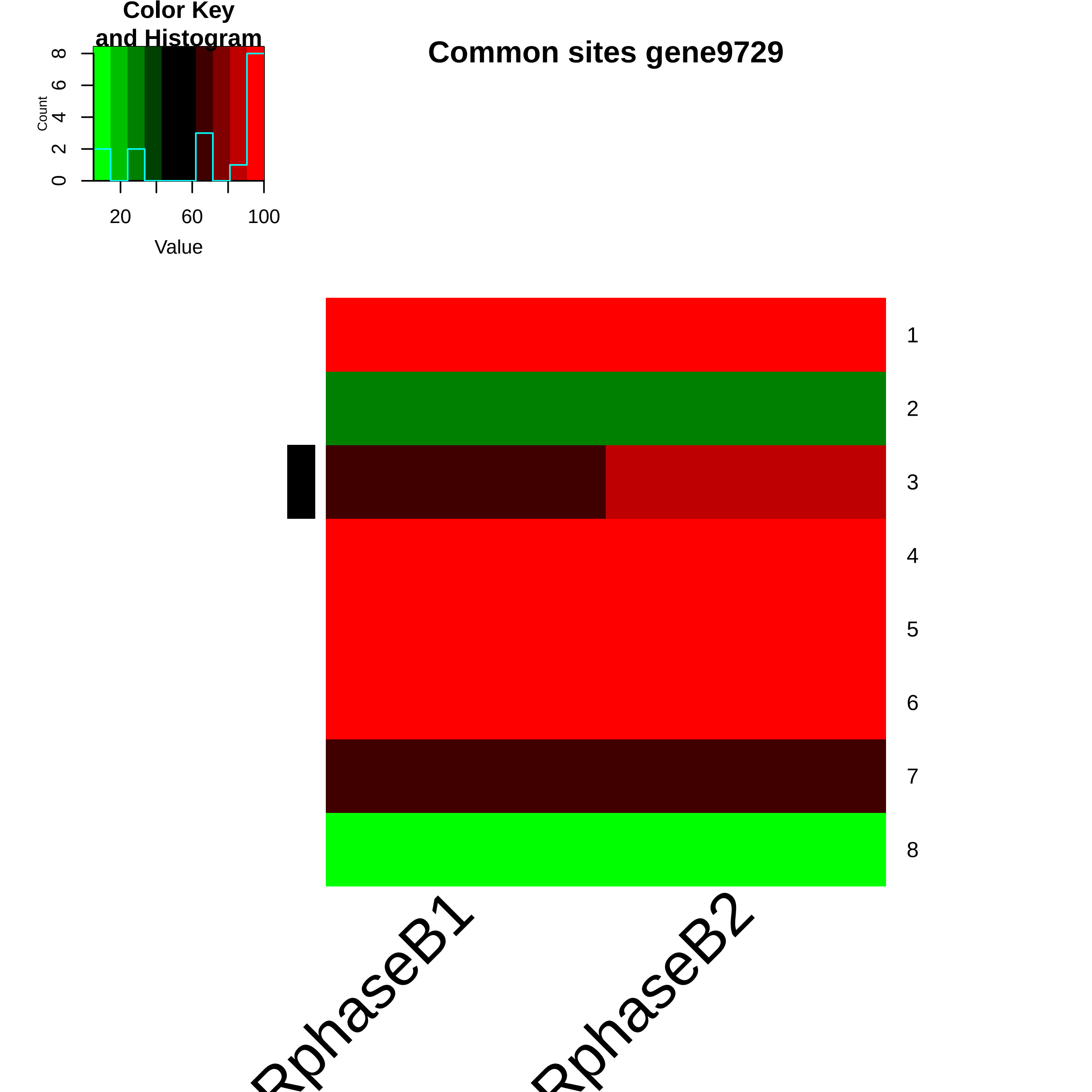

Color Key
and Histogram
Count
0
2
4
6
8
20
60
100
Value
Common sites gene9729
1
2
3
4
5
6
7
8
RphaseB1
RphaseB2

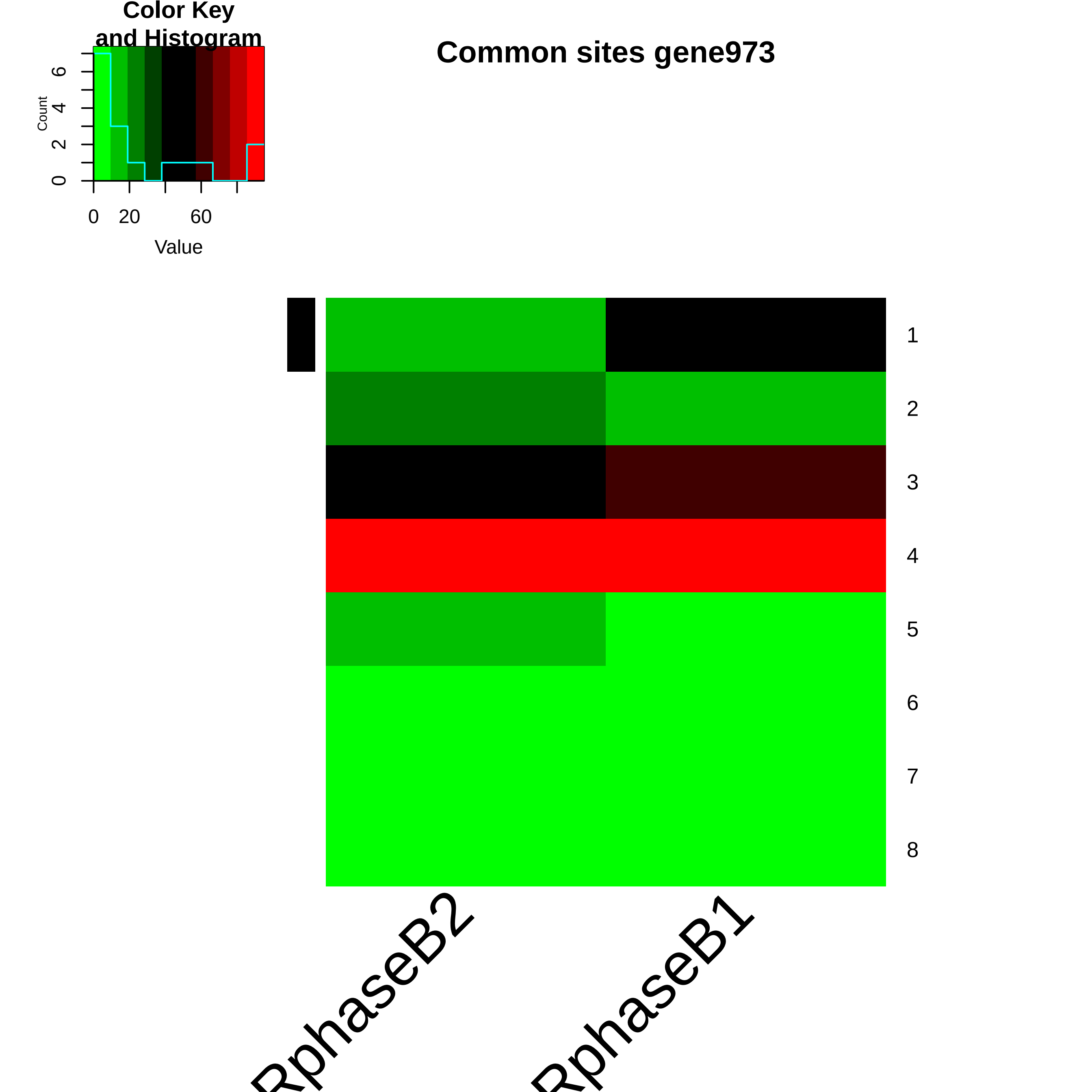
Color Key
and Histogram
Count
0
2
4
6
0
20
60
Value
Common sites gene973
1
2
3
4
5
6
7
8
RphaseB2
RphaseB1

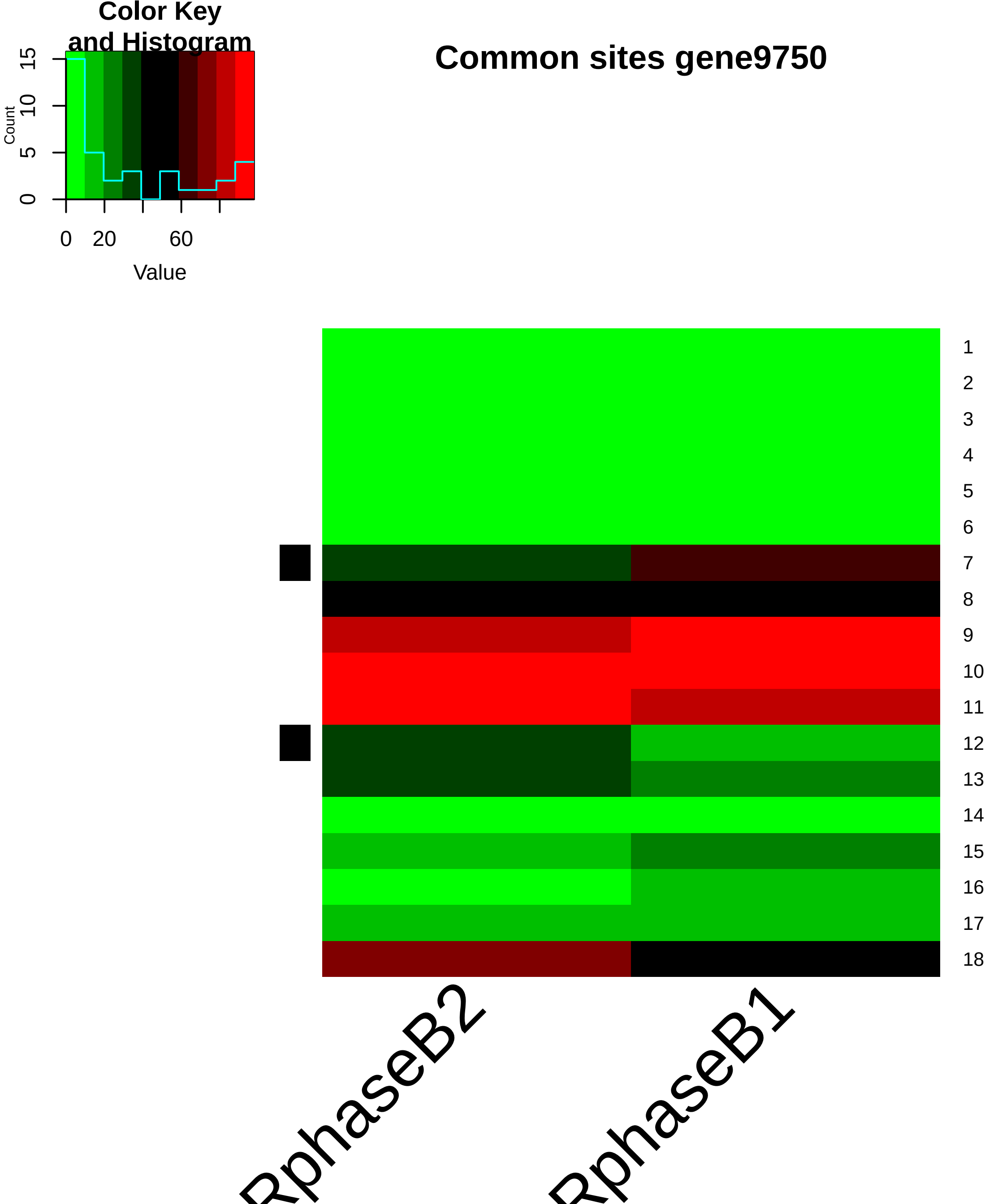
Color Key
and Histogram
Count
0
5
10
15
0
20
60
Value
Common sites gene9750
RphaseB2
RphaseB1

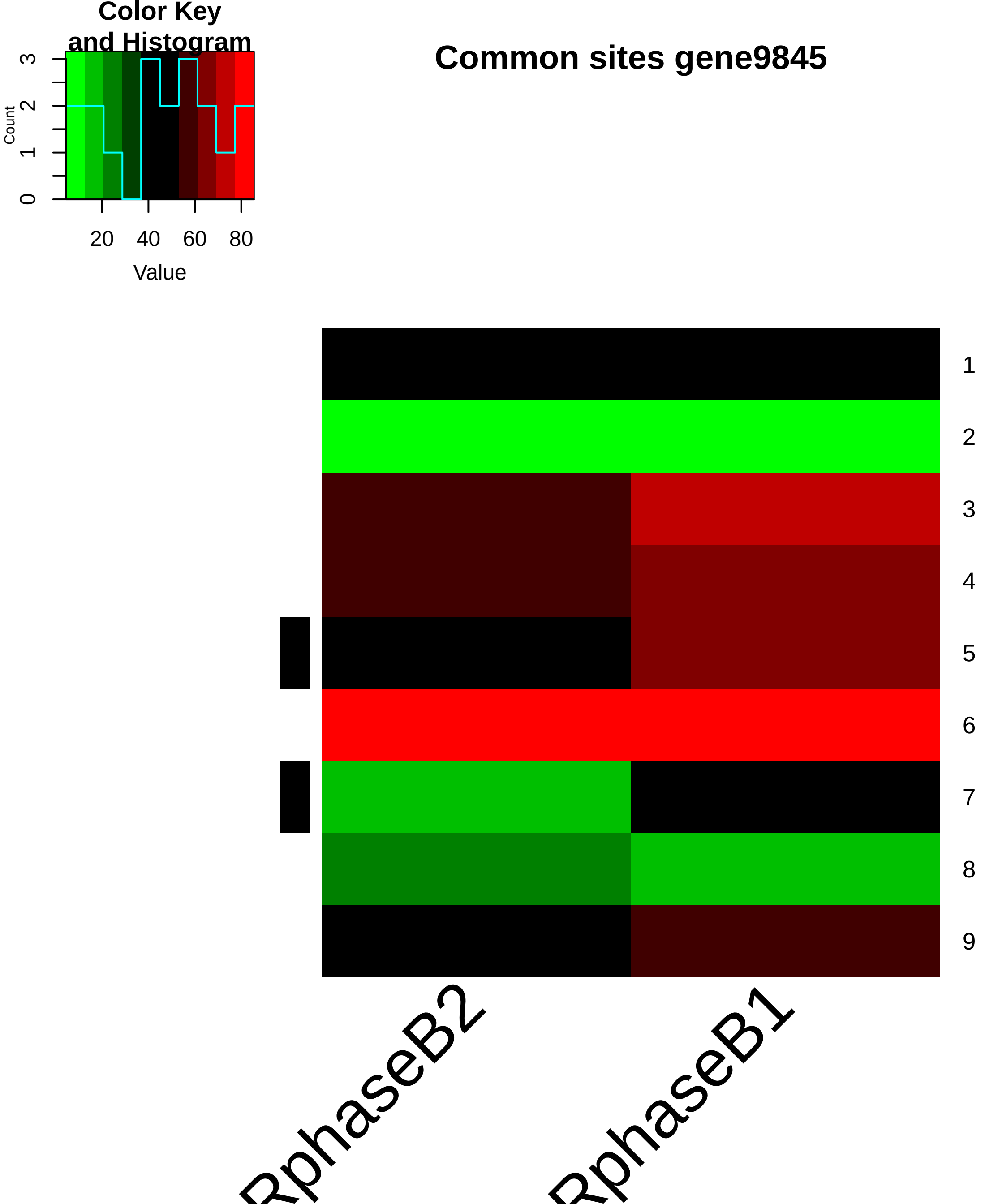
Color Key
and Histogram
Count
0
1
2
3
20
40
60
80
Value
Common sites gene9845
1
2
3
4
5
6
7
8
9
RphaseB2
RphaseB1

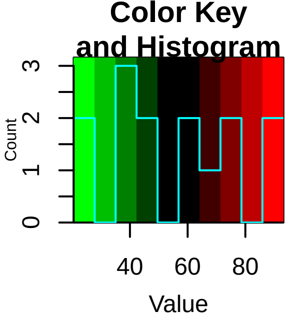

### Common sites gene9903

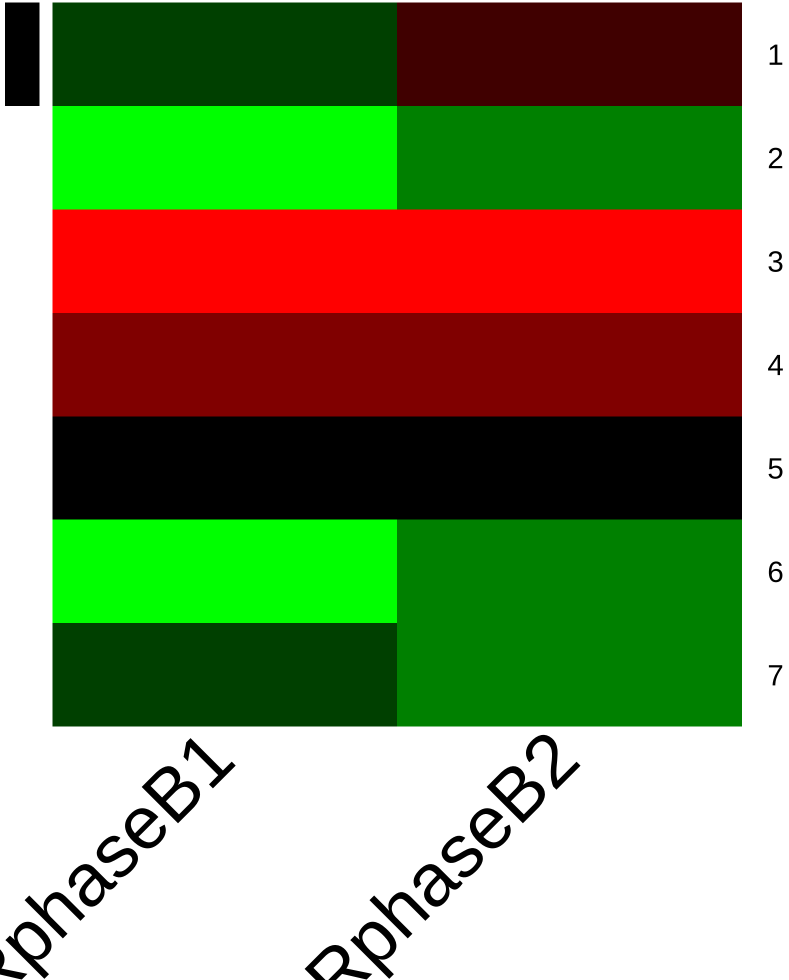

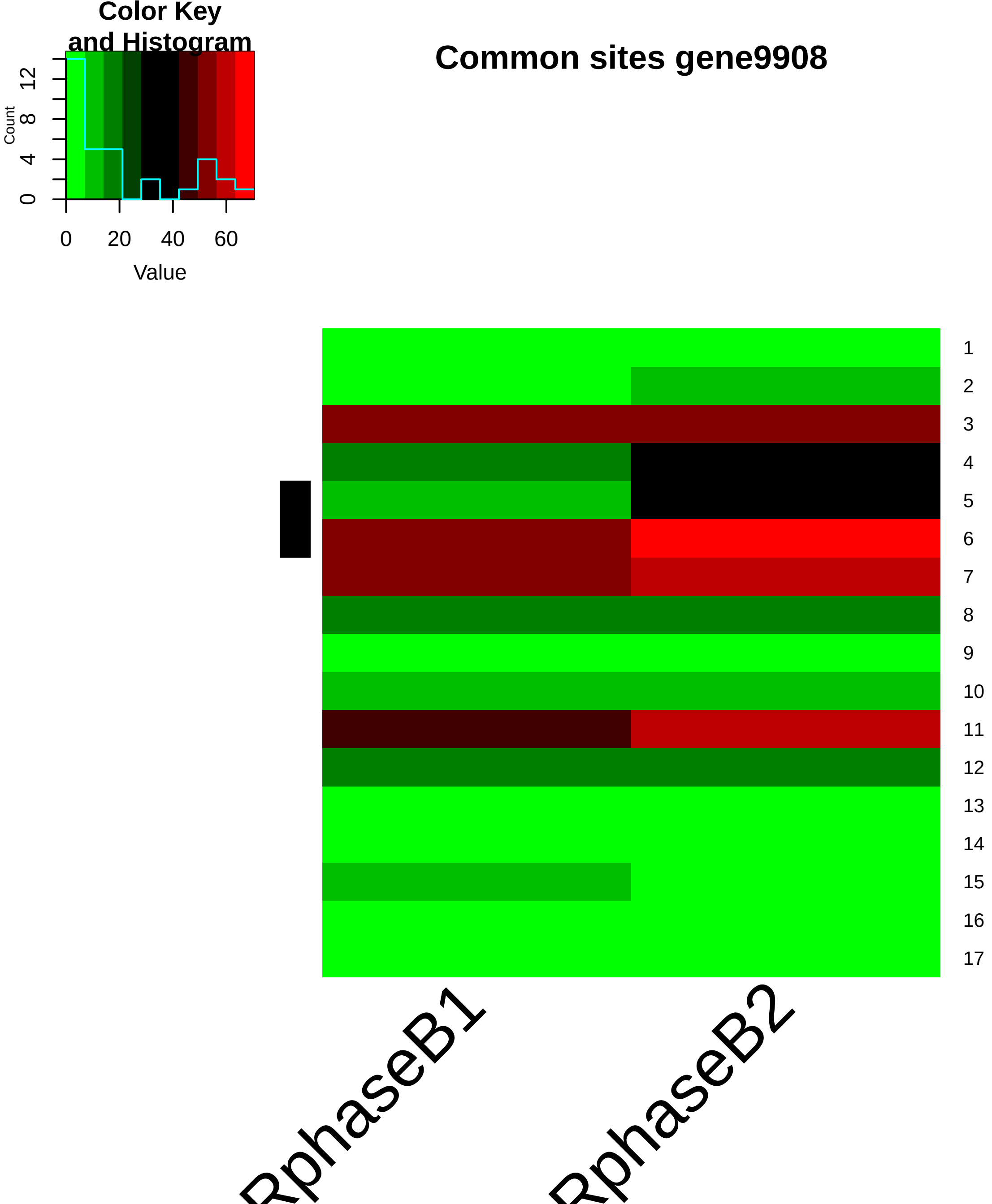
Color Key
and Histogram
Count
0
4
8
12
0
20
40
60
Value
Common sites gene9908
RphaseB1
RphaseB2

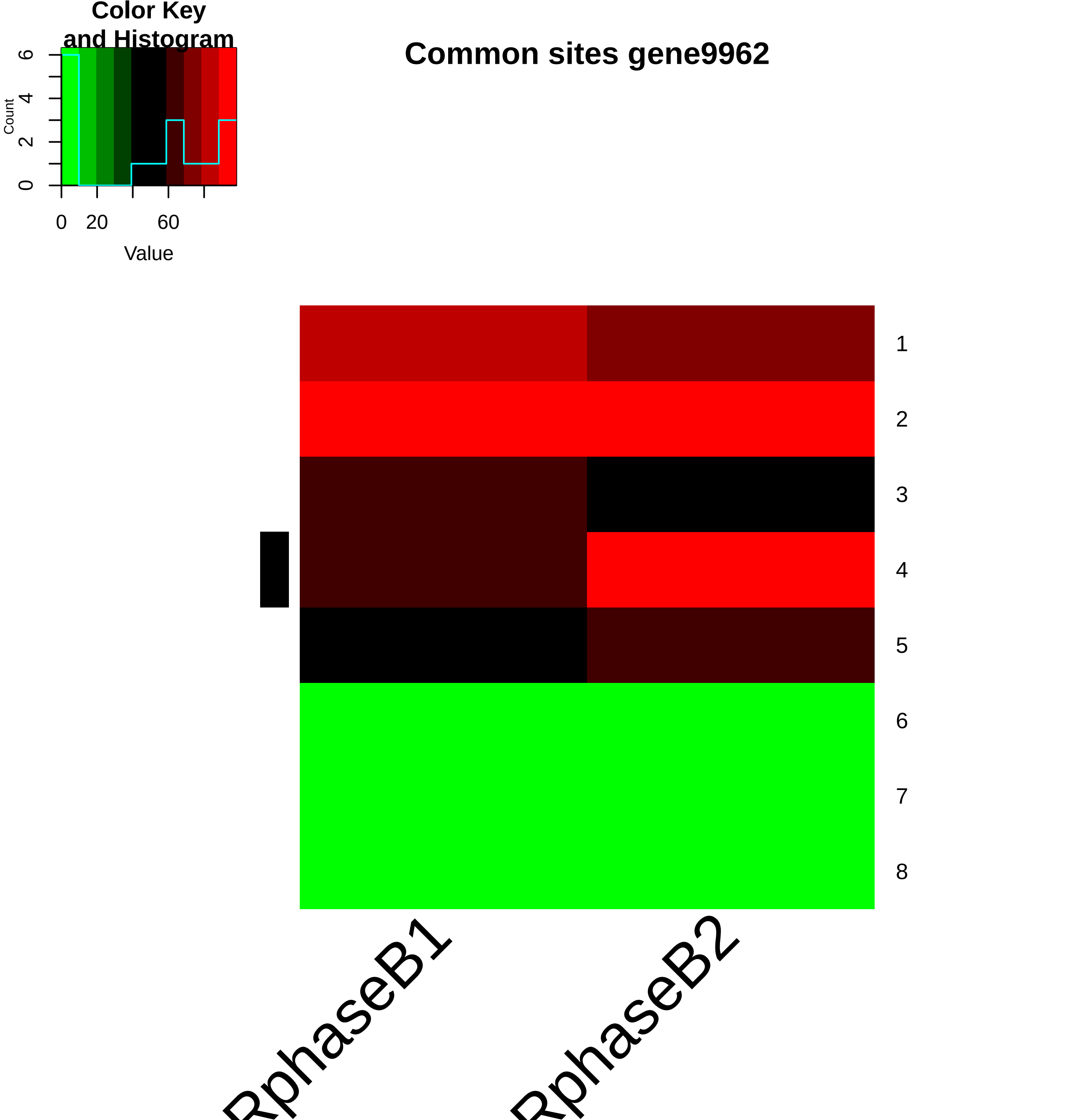
Color Key
and Histogram
Count
0
2
4
6
0
20
60
Value
Common sites gene9962
1
2
3
4
5
6
7
8
RphaseB1
RphaseB2

```
Directory: OGL File: 0READMEogl

OGL - Ordered gene lists.

Input: studyhc annotation   maxgwidth minnbrdmsites
       dmgprp (= output of BWASPR:show_dmsg())

Output: files ogl-*.txt rnk-dmg-*.txt
         ogl-<study>_<sample>.txt ogl-<study>_<sample1>.vs.<sample2>.txt
         rnk-dmg--<study>_<sample1>.vs.<sample2>.txt rnk-dmg--<study>_<sample1>.vs.<sample2>.pdf
         wrt-<study>.txt

Notes: Output files ogl-<study>_<sample>.txt give tables for each sample with columns

        gene_ID gwidth #Sites #per10Kb %perSite %pNucl

       ordered by %pNucl.  If available, a link to an NCBI entry of gene_ID is inserted as second column.
       Abbreviations used: %perSite, percent methylation per site
                           %perNucl, percent methylation per nucleotide of the gene

       Output files ogl-<study>_<sample1>.vs.<sample2>.txt give tables for each comparison with columns

        gene_ID gwidth #Sites #per10Kb #dmSites #dmsp10kb %dmSites %pSite1 %pSite2 DMpSite ADMpSite DMpNucl ADMpNucl

       ordered by DMpSite.  If available, a link to an NCBI entry of gene_ID is inserted as second column.
       Abbreviations used: %dmSites, percent sites that are differentially methylated
                           %pSite1, average per site % methylation for sample1
                           %pSite2, average per site % methylation for sample2
                           DMpSite, average per site difference in % methylation between sample1 and sample2
                           ADMpSite, absolute value of DMpSite
                           DMpNucl, average per nucleotide difference in % methylation between sample1 and sample2
                           ADMpNucl, absolute value of DMpNucl

       Output files rnk-dmg-<study>_<sample1>.vs.<sample2>.txt are equivalent to files
        ogl-<study>_<sample1>.vs.<sample2>.txt but ordered by ADMpNucl.
       Output files rnk-dmg-<study>_<sample1>.vs.<sample2>.pdf provide visualization of the distribution
        of ADMpNucl values.

       Output file wrt-<study>.txt gives results of the Wilcoxon signed rank test comparing
        the %pSite1 and %pSite2 vectors

       In output directory OGL, tables are restricted to genes with gwidth <= maxgwidth and, for
        pairwise comparisons, #dmSites >= minnbrdmsites

       If either maxgwidth or minnbrdmsites is set, then output directory OGLall will show the full tables
        with all genes (for reference).
```

Directory: OGL File: ogl-Cb_LI_BCphaseA1.txt

| gene_ID | gene_link | gwidth | #Sites | #per10Kb | %perSite | %pNucl |
|---|---|---|---|---|---|---|
| gene5911 | https://www.ncbi.nlm.nih.gov/gene/?term=LOC105278844 | 943 | 105 | 1113.47 | 31.24 | 3.48 |
| gene10219 | https://www.ncbi.nlm.nih.gov/gene/?term=LOC105283830 | 235 | 21 | 893.62 | 38.08 | 3.4 |
| gene13384 | https://www.ncbi.nlm.nih.gov/gene/?term=LOC105287447 | 1877 | 216 | 1150.77 | 28.75 | 3.31 |
| gene12538 | https://www.ncbi.nlm.nih.gov/gene/?term=LOC105286531 | 372 | 35 | 940.86 | 34.82 | 3.28 |
| gene5309 | https://www.ncbi.nlm.nih.gov/gene/?term=LOC105278160 | 1704 | 148 | 868.54 | 36.82 | 3.2 |
| gene211 | https://www.ncbi.nlm.nih.gov/gene/?term=LOC105274534 | 1280 | 86 | 671.88 | 45.78 | 3.08 |
| gene12746 | https://www.ncbi.nlm.nih.gov/gene/?term=LOC105286737 | 2688 | 310 | 1153.27 | 25.77 | 2.97 |
| gene3719 | https://www.ncbi.nlm.nih.gov/gene/?term=LOC105276360 | 1471 | 132 | 897.35 | 33.01 | 2.96 |
| gene5917 | https://www.ncbi.nlm.nih.gov/gene/?term=LOC105278849 | 823 | 99 | 1202.92 | 24.3 | 2.92 |
| gene1654 | https://www.ncbi.nlm.nih.gov/gene/?term=LOC105287811 | 2285 | 134 | 586.43 | 48.38 | 2.84 |
| gene3731 | https://www.ncbi.nlm.nih.gov/gene/?term=LOC105276371 | 865 | 55 | 635.84 | 44.05 | 2.8 |
| gene12330 | https://www.ncbi.nlm.nih.gov/gene/?term=LOC105286235 | 1501 | 97 | 646.24 | 43.04 | 2.78 |
| gene1562 | https://www.ncbi.nlm.nih.gov/gene/?term=LOC105287692 | 1491 | 120 | 804.83 | 33.36 | 2.68 |
| gene2797 | https://www.ncbi.nlm.nih.gov/gene/?term=LOC105275314 | 1489 | 79 | 530.56 | 50.1 | 2.66 |
| gene12364 | https://www.ncbi.nlm.nih.gov/gene/?term=LOC105286269 | 1403 | 68 | 484.68 | 54.55 | 2.64 |
| gene458 | https://www.ncbi.nlm.nih.gov/gene/?term=LOC105277254 | 1378 | 109 | 791 | 33.34 | 2.64 |
| gene5301 | https://www.ncbi.nlm.nih.gov/gene/?term=LOC105278148 | 1881 | 182 | 967.57 | 26.39 | 2.55 |
| gene9471 | https://www.ncbi.nlm.nih.gov/gene/?term=LOC105282986 | 1605 | 124 | 772.59 | 32.83 | 2.54 |
| gene7527 | https://www.ncbi.nlm.nih.gov/gene/?term=LOC105280739 | 2686 | 131 | 487.71 | 51.92 | 2.53 |
| gene12652 | https://www.ncbi.nlm.nih.gov/gene/?term=LOC109610640 | 1122 | 55 | 490.2 | 51.51 | 2.52 |
| gene3806 | https://www.ncbi.nlm.nih.gov/gene/?term=LOC105276439 | 2385 | 143 | 599.58 | 42.03 | 2.52 |
| gene2129 | https://www.ncbi.nlm.nih.gov/gene/?term=LOC105274543 | 1631 | 104 | 637.65 | 38.35 | 2.45 |
| gene11569 | https://www.ncbi.nlm.nih.gov/gene/?term=LOC105285394 | 1311 | 46 | 350.88 | 69.57 | 2.44 |
| gene5825 | https://www.ncbi.nlm.nih.gov/gene/?term=LOC105278753 | 1754 | 100 | 570.13 | 42.54 | 2.43 |

...
... only first 25 lines shown ...

Directory: OGL File: ogl-Cb_LI_BCphaseA1.vs.BCphaseA2.txt

| gene_ID | gene_link | gwidth | #Sites | #per10Kb | #dmSites | #dmsp10kb | %dmSites | %pSite1 | %pSite2 | DMpSite | ADMpSite | DMpNucl | ADMpNucl |
|---|---|---|---|---|---|---|---|---|---|---|---|---|---|
| gene393 | https://www.ncbi.nlm.nih.gov/gene/?term=LOC105276566 | 1123 | 2 | 17.81 | 1 | 8.9 | 50 | 77.65 | 48.08 | 29.58 | 29.58 | 0.05 | 0.05 |
| gene9109 | https://www.ncbi.nlm.nih.gov/gene/?term=LOC105282561 | 2400 | 10 | 41.67 | 1 | 4.17 | 10 | 26.72 | 17.84 | 8.89 | 8.89 | 0.04 | 0.04 |
| gene7952 | https://www.ncbi.nlm.nih.gov/gene/?term=LOC105281213 | 2963 | 31 | 104.62 | 1 | 3.37 | 3.23 | 42.96 | 39.57 | 3.39 | 3.39 | 0.04 | 0.04 |
| gene7915 | https://www.ncbi.nlm.nih.gov/gene/?term=LOC105281173 | 1597 | 25 | 156.54 | 1 | 6.26 | 4 | 64.87 | 61.62 | 3.25 | 3.25 | 0.05 | 0.05 |
| gene7368 | https://www.ncbi.nlm.nih.gov/gene/?term=LOC105280556 | 4105 | 40 | 97.44 | 1 | 2.44 | 2.5 | 14.82 | 11.87 | 2.94 | 2.94 | 0.03 | 0.03 |
| gene10213 | https://www.ncbi.nlm.nih.gov/gene/?term=LOC105283826 | 1503 | 53 | 352.63 | 1 | 6.65 | 1.89 | 16.45 | 13.62 | 2.82 | 2.82 | 0.1 | 0.1 |
| gene3168 | https://www.ncbi.nlm.nih.gov/gene/?term=LOC105275711 | 6150 | 71 | 115.45 | 1 | 1.63 | 1.41 | 14.71 | 11.97 | 2.74 | 2.74 | 0.03 | 0.03 |
| gene5490 | https://www.ncbi.nlm.nih.gov/gene/?term=LOC105278358 | 2095 | 25 | 119.33 | 1 | 4.77 | 4 | 36.86 | 34.23 | 2.63 | 2.63 | 0.03 | 0.03 |
| gene388 | https://www.ncbi.nlm.nih.gov/gene/?term=LOC105276538 | 1558 | 19 | 121.95 | 2 | 12.84 | 10.53 | 69.41 | 66.83 | 2.58 | 2.58 | 0.03 | 0.03 |
| gene12981 | https://www.ncbi.nlm.nih.gov/gene/?term=LOC105286999 | 2063 | 39 | 189.05 | 1 | 4.85 | 2.56 | 33.12 | 30.76 | 2.36 | 2.36 | 0.04 | 0.04 |
| gene2404 | https://www.ncbi.nlm.nih.gov/gene/?term=LOC105274853 | 1767 | 33 | 186.76 | 1 | 5.66 | 3.03 | 31.53 | 29.22 | 2.31 | 2.31 | 0.04 | 0.04 |
| gene920 | https://www.ncbi.nlm.nih.gov/gene/?term=LOC105282518 | 1197 | 40 | 334.17 | 1 | 8.35 | 2.5 | 39.94 | 37.69 | 2.25 | 2.25 | 0.08 | 0.08 |
| gene6422 | https://www.ncbi.nlm.nih.gov/gene/?term=LOC105279458 | 2419 | 33 | 136.42 | 1 | 4.13 | 3.03 | 55.1 | 53.04 | 2.06 | 2.06 | 0.03 | 0.03 |
| gene1220 | https://www.ncbi.nlm.nih.gov/gene/?term=LOC105285742 | 3453 | 90 | 260.64 | 1 | 2.9 | 1.11 | 27.79 | 25.88 | 1.92 | 1.92 | 0.05 | 0.05 |
| gene11401 | https://www.ncbi.nlm.nih.gov/gene/?term=LOC105285202 | 1841 | 47 | 255.3 | 1 | 5.43 | 2.13 | 22.36 | 20.6 | 1.77 | 1.77 | 0.05 | 0.05 |
| gene10391 | https://www.ncbi.nlm.nih.gov/gene/?term=LOC109611283 | 1994 | 41 | 205.62 | 1 | 5.02 | 2.44 | 35.16 | 33.4 | 1.76 | 1.76 | 0.04 | 0.04 |
| gene6966 | https://www.ncbi.nlm.nih.gov/gene/?term=LOC105280104 | 3747 | 116 | 309.58 | 1 | 2.67 | 0.86 | 15.51 | 13.86 | 1.65 | 1.65 | 0.05 | 0.05 |
| gene11454 | https://www.ncbi.nlm.nih.gov/gene/?term=LOC105285256 | 1906 | 48 | 251.84 | 1 | 5.25 | 2.08 | 46.56 | 44.93 | 1.63 | 1.63 | 0.04 | 0.04 |
| gene1199 | https://www.ncbi.nlm.nih.gov/gene/?term=LOC105285498 | 2220 | 23 | 103.6 | 1 | 4.5 | 4.35 | 35.45 | 33.87 | 1.58 | 1.58 | 0.02 | 0.02 |
| gene12021 | https://www.ncbi.nlm.nih.gov/gene/?term=LOC105285905 | 1399 | 21 | 150.11 | 1 | 7.15 | 4.76 | 41.18 | 39.62 | 1.55 | 1.55 | 0.02 | 0.02 |
| gene2748 | https://www.ncbi.nlm.nih.gov/gene/?term=LOC105275261 | 3845 | 54 | 140.44 | 1 | 2.6 | 1.85 | 38.15 | 36.73 | 1.42 | 1.42 | 0.02 | 0.02 |
| gene11447 | https://www.ncbi.nlm.nih.gov/gene/?term=LOC105285252 | 7437 | 119 | 160.01 | 1 | 1.34 | 0.84 | 14.91 | 13.69 | 1.22 | 1.22 | 0.02 | 0.02 |
| gene3107 | https://www.ncbi.nlm.nih.gov/gene/?term=LOC105275650 | 5103 | 143 | 280.23 | 1 | 1.96 | 0.7 | 16.9 | 15.68 | 1.21 | 1.21 | 0.03 | 0.03 |
| gene10819 | https://www.ncbi.nlm.nih.gov/gene/?term=LOC105284507 | 5767 | 69 | 119.65 | 1 | 1.73 | 1.45 | 22.66 | 21.47 | 1.19 | 1.19 | 0.01 | 0.01 |
| gene5375 | https://www.ncbi.nlm.nih.gov/gene/?term=LOC105278228 | 8640 | 143 | 165.51 | 1 | 1.16 | 0.7 | 24.34 | 23.23 | 1.11 | 1.11 | 0.02 | 0.02 |
| gene322 | https://www.ncbi.nlm.nih.gov/gene/?term=LOC105275777 | 1606 | 21 | 130.76 | 1 | 6.23 | 4.76 | 49.88 | 48.85 | 1.04 | 1.04 | 0.01 | 0.01 |
| gene5646 | https://www.ncbi.nlm.nih.gov/gene/?term=LOC105278534 | 7475 | 166 | 222.07 | 1 | 1.34 | 0.6 | 18.54 | 17.53 | 1.02 | 1.02 | 0.02 | 0.02 |
| gene10155 | https://www.ncbi.nlm.nih.gov/gene/?term=LOC105283756 | 7030 | 86 | 122.33 | 1 | 1.42 | 1.16 | 7.44 | 6.49 | 0.95 | 0.95 | 0.01 | 0.01 |
| gene4190 | https://www.ncbi.nlm.nih.gov/gene/?term=LOC105276875 | 3150 | 104 | 330.16 | 1 | 3.17 | 0.96 | 25.99 | 25.04 | 0.95 | 0.95 | 0.03 | 0.03 |
| gene10345 | https://www.ncbi.nlm.nih.gov/gene/?term=LOC105283989 | 4444 | 21 | 47.25 | 1 | 2.25 | 4.76 | 48.99 | 48.08 | 0.91 | 0.91 | 0 | 0 |
| gene3286 | https://www.ncbi.nlm.nih.gov/gene/?term=LOC105275848 | 3744 | 121 | 323.18 | 1 | 2.67 | 0.83 | 22.9 | 22 | 0.9 | 0.9 | 0.03 | 0.03 |
| gene5298 | https://www.ncbi.nlm.nih.gov/gene/?term=LOC105278145 | 4060 | 245 | 603.45 | 2 | 4.93 | 0.82 | 11.36 | 10.5 | 0.86 | 0.86 | 0.05 | 0.05 |
| gene3671 | https://www.ncbi.nlm.nih.gov/gene/?term=LOC105276272 | 1966 | 25 | 127.16 | 1 | 5.09 | 4 | 43.89 | 43.09 | 0.8 | 0.8 | 0.01 | 0.01 |
| gene1993 | https://www.ncbi.nlm.nih.gov/gene/?term=LOC105288193 | 1449 | 62 | 427.88 | 1 | 6.9 | 1.61 | 35.55 | 34.76 | 0.78 | 0.78 | 0.03 | 0.03 |
| gene3848 | https://www.ncbi.nlm.nih.gov/gene/?term=LOC105276486 | 5899 | 107 | 181.39 | 1 | 1.7 | 0.93 | 25.75 | 24.98 | 0.77 | 0.77 | 0.01 | 0.01 |
| gene2683 | https://www.ncbi.nlm.nih.gov/gene/?term=LOC105275187 | 5492 | 66 | 120.17 | 1 | 1.82 | 1.52 | 14.03 | 13.27 | 0.76 | 0.76 | 0.01 | 0.01 |
| gene4376 | https://www.ncbi.nlm.nih.gov/gene/?term=LOC105277083 | 4307 | 92 | 213.61 | 1 | 2.32 | 1.09 | 26.82 | 26.06 | 0.76 | 0.76 | 0.02 | 0.02 |
| gene11340 | https://www.ncbi.nlm.nih.gov/gene/?term=LOC105285131 | 2918 | 93 | 318.71 | 1 | 3.43 | 1.08 | 12.16 | 11.4 | 0.75 | 0.75 | 0.02 | 0.02 |
| gene474 | https://www.ncbi.nlm.nih.gov/gene/?term=LOC105277435 | 3625 | 150 | 413.79 | 1 | 2.76 | 0.67 | 12.65 | 11.92 | 0.73 | 0.73 | 0.03 | 0.03 |
| gene1147 | https://www.ncbi.nlm.nih.gov/gene/?term=LOC105284954 | 4022 | 75 | 186.47 | 1 | 2.49 | 1.33 | 14.93 | 14.22 | 0.71 | 0.71 | 0.01 | 0.01 |
| gene2271 | https://www.ncbi.nlm.nih.gov/gene/?term=LOC105274706 | 1448 | 52 | 359.12 | 1 | 6.91 | 1.92 | 37.3 | 36.65 | 0.64 | 0.64 | 0.02 | 0.02 |
| gene7207 | https://www.ncbi.nlm.nih.gov/gene/?term=LOC105280376 | 4659 | 98 | 210.35 | 1 | 2.15 | 1.02 | 30.66 | 30.04 | 0.62 | 0.62 | 0.01 | 0.01 |
| gene657 | https://www.ncbi.nlm.nih.gov/gene/?term=LOC105279667 | 6421 | 64 | 99.67 | 1 | 1.56 | 1.56 | 3.41 | 2.8 | 0.61 | 0.61 | 0.01 | 0.01 |
| gene12573 | https://www.ncbi.nlm.nih.gov/gene/?term=LOC105286515 | 3647 | 106 | 290.65 | 1 | 2.74 | 0.94 | 10.61 | 10.01 | 0.6 | 0.6 | 0.02 | 0.02 |
| gene9645 | https://www.ncbi.nlm.nih.gov/gene/?term=LOC105283181 | 3875 | 118 | 304.52 | 1 | 2.58 | 0.85 | 12.99 | 12.4 | 0.59 | 0.59 | 0.02 | 0.02 |
| gene13036 | https://www.ncbi.nlm.nih.gov/gene/?term=LOC105287059 | 4959 | 150 | 302.48 | 1 | 2.02 | 0.67 | 17.57 | 17 | 0.57 | 0.57 | 0.02 | 0.02 |
| gene3599 | https://www.ncbi.nlm.nih.gov/gene/?term=LOC105276196 | 2955 | 40 | 135.36 | 1 | 3.38 | 2.5 | 52.66 | 52.1 | 0.56 | 0.56 | 0.01 | 0.01 |
| gene5831 | https://www.ncbi.nlm.nih.gov/gene/?term=LOC105278761 | 2506 | 100 | 399.04 | 2 | 7.98 | 2 | 24.32 | 23.77 | 0.54 | 0.54 | 0.02 | 0.02 |
| gene12307 | https://www.ncbi.nlm.nih.gov/gene/?term=LOC105286216 | 10313 | 146 | 141.57 | 1 | 0.97 | 0.68 | 17.27 | 16.76 | 0.51 | 0.51 | 0.01 | 0.01 |
| gene8661 | https://www.ncbi.nlm.nih.gov/gene/?term=LOC105282063 | 1926 | 39 | 202.49 | 1 | 5.19 | 2.56 | 23.72 | 23.24 | 0.48 | 0.48 | 0.01 | 0.01 |
| gene10326 | https://www.ncbi.nlm.nih.gov/gene/?term=LOC105283962 | 3042 | 92 | 302.43 | 1 | 3.29 | 1.09 | 15.29 | 14.85 | 0.44 | 0.44 | 0.01 | 0.01 |
| gene13044 | https://www.ncbi.nlm.nih.gov/gene/?term=LOC105287070 | 5094 | 159 | 312.13 | 1 | 1.96 | 0.63 | 5.69 | 5.29 | 0.4 | 0.4 | 0.01 | 0.01 |
| gene13114 | https://www.ncbi.nlm.nih.gov/gene/?term=LOC105287147 | 7609 | 360 | 473.12 | 1 | 1.31 | 0.28 | 10.34 | 9.95 | 0.39 | 0.39 | 0.02 | 0.02 |
| gene3817 | https://www.ncbi.nlm.nih.gov/gene/?term=LOC105276449 | 1230 | 39 | 317.07 | 1 | 8.13 | 2.56 | 32.77 | 32.39 | 0.38 | 0.38 | 0.01 | 0.01 |
| gene7568 | https://www.ncbi.nlm.nih.gov/gene/?term=LOC105280786 | 2525 | 107 | 423.76 | 1 | 3.96 | 0.93 | 33.06 | 32.68 | 0.38 | 0.38 | 0.02 | 0.02 |
| gene3206 | https://www.ncbi.nlm.nih.gov/gene/?term=LOC105275752 | 3474 | 117 | 336.79 | 1 | 2.88 | 0.85 | 29.75 | 29.38 | 0.37 | 0.37 | 0.01 | 0.01 |
| gene6583 | https://www.ncbi.nlm.nih.gov/gene/?term=LOC105279649 | 7444 | 308 | 413.76 | 1 | 1.34 | 0.32 | 11.75 | 11.41 | 0.34 | 0.34 | 0.01 | 0.01 |
| gene10392 | https://www.ncbi.nlm.nih.gov/gene/?term=LOC105284036 | 3216 | 76 | 236.32 | 1 | 3.11 | 1.32 | 25.16 | 24.85 | 0.31 | 0.31 | 0.01 | 0.01 |
| gene12674 | https://www.ncbi.nlm.nih.gov/gene/?term=LOC105286649 | 18283 | 638 | 348.96 | 1 | 0.55 | 0.16 | 1.31 | 1.05 | 0.26 | 0.26 | 0.01 | 0.01 |
| gene10549 | https://www.ncbi.nlm.nih.gov/gene/?term=LOC105284210 | 7145 | 298 | 417.07 | 1 | 1.4 | 0.34 | 11.27 | 11.01 | 0.25 | 0.25 | 0.01 | 0.01 |

| | | | | | | | | | | | | | |
|---|---|---|---|---|---|---|---|---|---|---|---|---|---|
| gene10600 | https://www.ncbi.nlm.nih.gov/gene/?term=LOC105284270 | 4837 | 97 | 200.54 | 1 | 2.07 | 1.03 | 13.04 | 12.82 | 0.22 | 0.22 | 0 | 0 |
| gene7249 | https://www.ncbi.nlm.nih.gov/gene/?term=LOC105280422 | 1775 | 58 | 326.76 | 1 | 5.63 | 1.72 | 15.27 | 15.08 | 0.19 | 0.19 | 0.01 | 0.01 |
| gene4962 | https://www.ncbi.nlm.nih.gov/gene/?term=LOC105277761 | 4645 | 140 | 301.4 1 | 2.15 | 0.71 | 14.56 | 14.4 | 0.16 | 0.16 | 0 | 0 | |
| gene5967 | https://www.ncbi.nlm.nih.gov/gene/?term=LOC105278918 | 4393 | 121 | 275.44 | 1 | 2.28 | 0.83 | 16.38 | 16.32 | 0.06 | 0.06 | 0 | 0 |
| gene6031 | https://www.ncbi.nlm.nih.gov/gene/?term=LOC105279000 | 5691 | 174 | 305.75 | 1 | 1.76 | 0.57 | 14.03 | 13.97 | 0.06 | 0.06 | 0 | 0 |
| gene11583 | https://www.ncbi.nlm.nih.gov/gene/?term=LOC105285414 | 8192 | 168 | 205.08 | 1 | 1.22 | 0.6 | 8.3 | 8.25 | 0.05 | 0.05 | 0 | 0 |
| gene940 | https://www.ncbi.nlm.nih.gov/gene/?term=LOC105282794 | 3578 | 82 | 229.18 | 1 | 2.79 | 1.22 | 16.49 | 16.45 | 0.04 | 0.04 | 0 | 0 |
| gene5524 | https://www.ncbi.nlm.nih.gov/gene/?term=LOC105278397 | 5177 | 200 | 386.32 | 1 | 1.93 | 0.5 | 24.39 | 24.36 | 0.03 | 0.03 | 0 | 0 |
| gene5746 | https://www.ncbi.nlm.nih.gov/gene/?term=LOC105278666 | 10343 | 387 | 374.17 | 1 | 0.97 | 0.26 | 7.75 | 7.72 | 0.03 | 0.03 | 0 | 0 |
| gene6639 | https://www.ncbi.nlm.nih.gov/gene/?term=LOC105279705 | 7656 | 58 | 75.76 1 | 1.31 | 1.72 | 24.54 | 24.54 | 0 | 0 | 0 | 0 | |
| gene12802 | https://www.ncbi.nlm.nih.gov/gene/?term=LOC105286799 | 3183 | 165 | 518.38 | 1 | 3.14 | 0.61 | 16.21 | 16.23 | -0.02 | 0.02 | 0 | 0 |
| gene2806 | https://www.ncbi.nlm.nih.gov/gene/?term=LOC105275327 | 14257 | 414 | 290.38 | 1 | 0.7 | 0.24 | 7.7 | 7.73 | -0.02 | 0.02 | 0 | 0 |
| gene11579 | https://www.ncbi.nlm.nih.gov/gene/?term=LOC105285406 | 13571 | 248 | 182.74 | 1 | 0.74 | 0.4 | 5.98 | 6.04 | -0.05 | 0.05 | 0 | 0 |
| gene7148 | https://www.ncbi.nlm.nih.gov/gene/?term=LOC105280304 | 8920 | 90 | 100.9 1 | 1.12 | 1.11 | 7.28 | 7.34 | -0.05 | 0.05 | 0 | 0 | |
| gene53 | https://www.ncbi.nlm.nih.gov/gene/?term=LOC105277148 | 19283 | 236 | 122.39 | 1 | 0.52 | 0.42 | 4.4 | 4.47 | -0.07 | 0.07 | 0 | 0 |
| gene5525 | https://www.ncbi.nlm.nih.gov/gene/?term=LOC105278396 | 3090 | 186 | 601.94 | 1 | 3.24 | 0.54 | 24.51 | 24.57 | -0.07 | 0.07 | 0 | 0 |
| gene6620 | https://www.ncbi.nlm.nih.gov/gene/?term=LOC105279687 | 17998 | 294 | 163.35 | 1 | 0.56 | 0.34 | 5.33 | 5.42 | -0.1 | 0.1 | 0 | 0 |
| gene4246 | https://www.ncbi.nlm.nih.gov/gene/?term=LOC105276941 | 16412 | 446 | 271.75 | 1 | 0.61 | 0.22 | 5.36 | 5.47 | -0.11 | 0.11 | 0 | 0 |
| gene9330 | https://www.ncbi.nlm.nih.gov/gene/?term=LOC105282825 | 3936 | 80 | 203.25 | 1 | 2.54 | 1.25 | 13.05 | 13.17 | -0.11 | 0.11 | 0 | 0 |
| gene10018 | https://www.ncbi.nlm.nih.gov/gene/?term=LOC105283607 | 4414 | 181 | 410.06 | 1 | 2.27 | 0.55 | 16.23 | 16.36 | -0.13 | 0.13 | -0.01 | 0.01 |
| gene12203 | https://www.ncbi.nlm.nih.gov/gene/?term=LOC105286099 | 5967 | 70 | 117.31 | 1 | 1.68 | 1.43 | 11.64 | 11.77 | -0.13 | 0.13 | 0 | 0 |
| gene1470 | https://www.ncbi.nlm.nih.gov/gene/?term=LOC105287597 | 2974 | 188 | 632.15 | 1 | 3.36 | 0.53 | 17.15 | 17.35 | -0.19 | 0.19 | -0.01 | 0.01 |
| gene3885 | https://www.ncbi.nlm.nih.gov/gene/?term=LOC105276527 | 7928 | 323 | 407.42 | 1 | 1.26 | 0.31 | 8.78 | 9 | -0.22 | 0.22 | -0.01 | 0.01 |
| gene11568 | https://www.ncbi.nlm.nih.gov/gene/?term=LOC105285395 | 8086 | 160 | 197.87 | 1 | 1.24 | 0.62 | 17.75 | 17.97 | -0.23 | 0.23 | 0 | 0 |
| gene10540 | https://www.ncbi.nlm.nih.gov/gene/?term=LOC105284200 | 5267 | 183 | 347.45 | 1 | 1.9 | 0.55 | 14.63 | 14.93 | -0.3 | 0.3 | -0.01 | 0.01 |
| gene6286 | https://www.ncbi.nlm.nih.gov/gene/?term=LOC105279298 | 2861 | 128 | 447.4 2 | 6.99 | 1.56 | 23.41 | 23.73 | -0.32 | 0.32 | -0.01 | 0.01 | |
| gene3856 | https://www.ncbi.nlm.nih.gov/gene/?term=LOC105276499 | 5236 | 178 | 339.95 | 1 | 1.91 | 0.56 | 9.73 | 10.06 | -0.33 | 0.33 | -0.01 | 0.01 |
| gene4892 | https://www.ncbi.nlm.nih.gov/gene/?term=LOC105277667 | 3898 | 119 | 305.28 | 1 | 2.57 | 0.84 | 27.91 | 28.29 | -0.38 | 0.38 | -0.01 | 0.01 |
| gene12317 | https://www.ncbi.nlm.nih.gov/gene/?term=LOC105286223 | 2235 | 79 | 353.47 | 1 | 4.47 | 1.27 | 30.56 | 30.96 | -0.4 | 0.4 | -0.01 | 0.01 |
| gene1884 | https://www.ncbi.nlm.nih.gov/gene/?term=LOC105288072 | 4169 | 76 | 182.3 1 | 2.4 | 1.32 | 23.55 | 23.95 | -0.4 | 0.4 | -0.01 | 0.01 | |
| gene2056 | https://www.ncbi.nlm.nih.gov/gene/?term=LOC105274452 | 6584 | 206 | 312.88 | 1 | 1.52 | 0.49 | 14.33 | 14.74 | -0.41 | 0.41 | -0.01 | 0.01 |
| gene4641 | https://www.ncbi.nlm.nih.gov/gene/?term=LOC105277398 | 1937 | 68 | 351.06 | 1 | 5.16 | 1.47 | 34.8 | 35.21 | -0.41 | 0.41 | -0.01 | 0.01 |
| gene10606 | https://www.ncbi.nlm.nih.gov/gene/?term=LOC105284273 | 6119 | 227 | 370.98 | 1 | 1.63 | 0.44 | 21.11 | 21.54 | -0.43 | 0.43 | -0.02 | 0.02 |
| gene5649 | https://www.ncbi.nlm.nih.gov/gene/?term=LOC105278538 | 2140 | 77 | 359.81 | 1 | 4.67 | 1.3 | 17.91 | 18.36 | -0.46 | 0.46 | -0.02 | 0.02 |
| gene9755 | https://www.ncbi.nlm.nih.gov/gene/?term=LOC105283296 | 2177 | 79 | 362.88 | 1 | 4.59 | 1.27 | 20.77 | 21.25 | -0.48 | 0.48 | -0.02 | 0.02 |
| gene303 | https://www.ncbi.nlm.nih.gov/gene/?term=LOC105275584 | 3081 | 71 | 230.44 | 1 | 3.25 | 1.41 | 26.68 | 27.17 | -0.49 | 0.49 | -0.01 | 0.01 |
| gene10678 | https://www.ncbi.nlm.nih.gov/gene/?term=LOC105284355 | 4031 | 152 | 377.08 | 1 | 2.48 | 0.66 | 20.4 | 20.91 | -0.51 | 0.51 | -0.02 | 0.02 |
| gene7888 | https://www.ncbi.nlm.nih.gov/gene/?term=LOC105281141 | 4357 | 153 | 351.16 | 1 | 2.3 | 0.65 | 30.21 | 30.79 | -0.57 | 0.57 | -0.02 | 0.02 |
| gene2844 | https://www.ncbi.nlm.nih.gov/gene/?term=LOC105275363 | 4330 | 179 | 413.39 | 1 | 2.31 | 0.56 | 14.53 | 15.15 | -0.61 | 0.61 | -0.03 | 0.03 |
| gene13033 | https://www.ncbi.nlm.nih.gov/gene/?term=LOC105287063 | 1877 | 45 | 239.74 | 1 | 5.33 | 2.22 | 34.91 | 35.54 | -0.62 | 0.62 | -0.01 | 0.01 |
| gene8600 | https://www.ncbi.nlm.nih.gov/gene/?term=LOC105282001 | 8513 | 126 | 148.01 | 1 | 1.17 | 0.79 | 11.25 | 11.9 | -0.65 | 0.65 | -0.01 | 0.01 |
| gene7245 | https://www.ncbi.nlm.nih.gov/gene/?term=LOC105280410 | 12478 | 234 | 187.53 | 1 | 0.8 | 0.43 | 20.56 | 21.23 | -0.67 | 0.67 | -0.01 | 0.01 |
| gene8343 | https://www.ncbi.nlm.nih.gov/gene/?term=LOC105281700 | 2172 | 40 | 184.16 | 1 | 4.6 | 2.5 | 41.18 | 41.86 | -0.68 | 0.68 | -0.01 | 0.01 |
| gene9336 | https://www.ncbi.nlm.nih.gov/gene/?term=LOC105282834 | 5993 | 106 | 176.87 | 1 | 1.67 | 0.94 | 15.44 | 16.15 | -0.71 | 0.71 | -0.01 | 0.01 |
| gene11275 | https://www.ncbi.nlm.nih.gov/gene/?term=LOC105285055 | 4119 | 196 | 475.84 | 1 | 2.43 | 0.51 | 13.91 | 14.65 | -0.74 | 0.74 | -0.04 | 0.04 |
| gene10344 | https://www.ncbi.nlm.nih.gov/gene/?term=LOC105283987 | 4841 | 64 | 132.2 1 | 2.07 | 1.56 | 32.11 | 32.92 | -0.81 | 0.81 | -0.01 | 0.01 | |
| gene292 | https://www.ncbi.nlm.nih.gov/gene/?term=LOC105275398 | 2825 | 77 | 272.57 | 1 | 3.54 | 1.3 | 19.76 | 20.59 | -0.82 | 0.82 | -0.02 | 0.02 |
| gene8729 | https://www.ncbi.nlm.nih.gov/gene/?term=LOC105282137 | 5005 | 195 | 389.61 | 1 | 2 | 0.51 | 16.47 | 17.35 | -0.89 | 0.89 | -0.03 | 0.03 |
| gene298 | https://www.ncbi.nlm.nih.gov/gene/?term=LOC105275460 | 2473 | 107 | 432.67 | 1 | 4.04 | 0.93 | 13.52 | 14.43 | -0.91 | 0.91 | -0.04 | 0.04 |
| gene7166 | https://www.ncbi.nlm.nih.gov/gene/?term=LOC105280326 | 6117 | 195 | 318.78 | 1 | 1.63 | 0.51 | 16.86 | 17.78 | -0.92 | 0.92 | -0.03 | 0.03 |
| gene9101 | https://www.ncbi.nlm.nih.gov/gene/?term=LOC105282553 | 3068 | 104 | 338.98 | 1 | 3.26 | 0.96 | 20.36 | 21.29 | -0.93 | 0.93 | -0.03 | 0.03 |
| gene10531 | https://www.ncbi.nlm.nih.gov/gene/?term=LOC105284190 | 6881 | 164 | 238.34 | 1 | 1.45 | 0.61 | 10.48 | 11.52 | -1.03 | 1.03 | -0.02 | 0.02 |
| gene11819 | https://www.ncbi.nlm.nih.gov/gene/?term=LOC105285671 | 2366 | 91 | 384.62 | 1 | 4.23 | 1.1 | 28.12 | 29.18 | -1.06 | 1.06 | -0.04 | 0.04 |
| gene2173 | https://www.ncbi.nlm.nih.gov/gene/?term=LOC105274590 | 4387 | 64 | 145.89 | 1 | 2.28 | 1.56 | 26 | 27.13 | -1.13 | 1.13 | -0.02 | 0.02 |
| gene6528 | https://www.ncbi.nlm.nih.gov/gene/?term=LOC105279591 | 7222 | 67 | 92.77 1 | 1.38 | 1.49 | 22.84 | 24.15 | -1.32 | 1.32 | -0.01 | 0.01 | |
| gene4023 | https://www.ncbi.nlm.nih.gov/gene/?term=LOC105276692 | 1759 | 35 | 198.98 | 1 | 5.69 | 2.86 | 23.93 | 25.35 | -1.42 | 1.42 | -0.03 | 0.03 |
| gene10984 | https://www.ncbi.nlm.nih.gov/gene/?term=LOC105284704 | 1987 | 61 | 307 1 | 5.03 | 1.64 | 25.94 | 27.43 | -1.49 | 1.49 | -0.05 | 0.05 | |
| gene2970 | https://www.ncbi.nlm.nih.gov/gene/?term=LOC105275518 | 2340 | 18 | 76.92 1 | 4.27 | 5.56 | 47.45 | 48.96 | -1.51 | 1.51 | -0.01 | 0.01 | |
| gene1310 | https://www.ncbi.nlm.nih.gov/gene/?term=LOC105286661 | 2582 | 65 | 251.74 | 1 | 3.87 | 1.54 | 19.96 | 21.58 | -1.62 | 1.62 | -0.04 | 0.04 |
| gene7556 | https://www.ncbi.nlm.nih.gov/gene/?term=LOC105280774 | 2792 | 98 | 351 1 | 3.58 | 1.02 | 31.47 | 33.13 | -1.66 | 1.66 | -0.06 | 0.06 | |
| gene6455 | https://www.ncbi.nlm.nih.gov/gene/?term=LOC105279500 | 2612 | 31 | 118.68 | 1 | 3.83 | 3.23 | 51.05 | 52.76 | -1.71 | 1.71 | -0.02 | 0.02 |
| gene10520 | https://www.ncbi.nlm.nih.gov/gene/?term=LOC105284184 | 2451 | 67 | 273.36 | 1 | 4.08 | 1.49 | 41.63 | 43.45 | -1.81 | 1.81 | -0.05 | 0.05 |
| gene274 | https://www.ncbi.nlm.nih.gov/gene/?term=LOC105275190 | 3488 | 66 | 189.22 | 1 | 2.87 | 1.52 | 20.22 | 22.09 | -1.87 | 1.87 | -0.04 | 0.04 |
| gene3727 | https://www.ncbi.nlm.nih.gov/gene/?term=LOC105276367 | 2484 | 54 | 217.39 | 1 | 4.03 | 1.85 | 48.38 | 50.33 | -1.95 | 1.95 | -0.04 | 0.04 |

| | | | | | | | | | | | | | |
|---|---|---|---|---|---|---|---|---|---|---|---|---|---|
| gene5877 | https://www.ncbi.nlm.nih.gov/gene/?term=LOC105278807 | 1719 | 50 | 290.87 | 1 | 5.82 | 2 | 21.66 | 23.62 | -1.95 | 1.95 | -0.06 | 0.06 |
| gene12516 | https://www.ncbi.nlm.nih.gov/gene/?term=LOC105286452 | 1529 | 35 | 228.91 | 1 | 6.54 | 2.86 | 51.01 | 53.03 | -2.03 | 2.03 | -0.05 | 0.05 |
| gene7183 | https://www.ncbi.nlm.nih.gov/gene/?term=LOC105280348 | 3925 | 51 | 129.94 | 1 | 2.55 | 1.96 | 25.44 | 27.74 | -2.3 | 2.3 | -0.03 | 0.03 |
| gene10107 | https://www.ncbi.nlm.nih.gov/gene/?term=LOC105283712 | 1883 | 23 | 122.15 | 1 | 5.31 | 4.35 | 27.23 | 29.73 | -2.49 | 2.49 | -0.03 | 0.03 |
| gene12406 | https://www.ncbi.nlm.nih.gov/gene/?term=LOC105286316 | 1386 | 29 | 209.24 | 1 | 7.22 | 3.45 | 41.65 | 44.38 | -2.73 | 2.73 | -0.06 | 0.06 |
| gene7398 | https://www.ncbi.nlm.nih.gov/gene/?term=LOC105280589 | 3452 | 41 | 118.77 | 1 | 2.9 | 2.44 | 25.07 | 28.12 | -3.05 | 3.05 | -0.04 | 0.04 |
| gene6334 | https://www.ncbi.nlm.nih.gov/gene/?term=LOC105279356 | 1471 | 40 | 271.92 | 1 | 6.8 | 2.5 | 29.12 | 32.21 | -3.09 | 3.09 | -0.08 | 0.08 |
| gene7734 | https://www.ncbi.nlm.nih.gov/gene/?term=LOC105280972 | 1528 | 26 | 170.16 | 1 | 6.54 | 3.85 | 20.04 | 23.88 | -3.84 | 3.84 | -0.07 | 0.07 |
| gene2824 | https://www.ncbi.nlm.nih.gov/gene/?term=LOC105275337 | 1772 | 58 | 327.31 | 1 | 5.64 | 1.72 | 31.86 | 38.08 | -6.22 | 6.22 | -0.2 | 0.2 |
| gene5714 | https://www.ncbi.nlm.nih.gov/gene/?term=LOC105278729 | 710 | 14 | 197.18 | 1 | 14.08 | 7.14 | 51.67 | 57.98 | -6.31 | 6.31 | -0.12 | 0.12 |
| gene2676 | https://www.ncbi.nlm.nih.gov/gene/?term=LOC105275181 | 1353 | 15 | 110.86 | 1 | 7.39 | 6.67 | 67.39 | 74.05 | -6.66 | 6.66 | -0.07 | 0.07 |
| gene3165 | https://www.ncbi.nlm.nih.gov/gene/?term=LOC105275706 | 2374 | 65 | 273.8 | 6 | 25.27 | 9.23 | 24.65 | 38.08 | -13.43 | 13.43 | -0.37 | 0.37 |
| gene2593 | https://www.ncbi.nlm.nih.gov/gene/?term=LOC105275070 | 1774 | 5 | 28.18 | 2 | 11.27 | 40 | 19.71 | 71.28 | -51.57 | 51.57 | -0.15 | 0.15 |

Directory: OGL File: ogl-Cb_LI_BCphaseA1.vs.BCphaseB1.txt

| gene_ID | gene_link | gwidth | #Sites | #per10Kb | #dmSites | #dmsp10kb | %dmSites | %pSite1 | %pSite2 | DMpSite | ADMpSite | DMpNucl | ADMpNucl |
|---|---|---|---|---|---|---|---|---|---|---|---|---|---|
| gene393 | https://www.ncbi.nlm.nih.gov/gene/?term=LOC105276566 | 1123 | 2 | 17.81 | 1 | 8.9 | 50 | 77.65 | 47.22 | 30.43 | 30.43 | 0.05 | 0.05 |
| gene9109 | https://www.ncbi.nlm.nih.gov/gene/?term=LOC105282561 | 2400 | 12 | 50 | 1 | 4.17 | 8.33 | 20.55 | 12.93 | 7.62 | 7.62 | 0.04 | 0.04 |
| gene10602 | https://www.ncbi.nlm.nih.gov/gene/?term=LOC105284268 | 2122 | 20 | 94.25 | 1 | 4.71 | 5 | 32.63 | 25.61 | 7.02 | 7.02 | 0.07 | 0.07 |
| gene7199 | https://www.ncbi.nlm.nih.gov/gene/?term=LOC105280366 | 1414 | 24 | 169.73 | 1 | 7.07 | 4.17 | 45.18 | 38.6 | 6.59 | 6.59 | 0.11 | 0.11 |
| gene9464 | https://www.ncbi.nlm.nih.gov/gene/?term=LOC105282975 | 1450 | 24 | 165.52 | 1 | 6.9 | 4.17 | 42.63 | 39.26 | 3.37 | 3.37 | 0.06 | 0.06 |
| gene3168 | https://www.ncbi.nlm.nih.gov/gene/?term=LOC105275711 | 6150 | 79 | 128.46 | 1 | 1.63 | 1.27 | 14.18 | 11.09 | 3.09 | 3.09 | 0.04 | 0.04 |
| gene6422 | https://www.ncbi.nlm.nih.gov/gene/?term=LOC105279458 | 2419 | 36 | 148.82 | 1 | 4.13 | 2.78 | 53.83 | 51.06 | 2.78 | 2.78 | 0.04 | 0.04 |
| gene7915 | https://www.ncbi.nlm.nih.gov/gene/?term=LOC105281173 | 1597 | 25 | 156.54 | 1 | 6.26 | 4 | 64.87 | 62.27 | 2.61 | 2.61 | 0.04 | 0.04 |
| gene9740 | https://www.ncbi.nlm.nih.gov/gene/?term=LOC105283277 | 2887 | 138 | 478 | 1 | 3.46 | 0.72 | 23.22 | 20.78 | 2.44 | 2.44 | 0.12 | 0.12 |
| gene7368 | https://www.ncbi.nlm.nih.gov/gene/?term=LOC105280556 | 4105 | 49 | 119.37 | 1 | 2.44 | 2.04 | 12.67 | 10.47 | 2.2 | 2.2 | 0.03 | 0.03 |
| gene652 | https://www.ncbi.nlm.nih.gov/gene/?term=LOC105279528 | 5806 | 42 | 72.34 | 1 | 1.72 | 2.38 | 16.48 | 14.36 | 2.12 | 2.12 | 0.02 | 0.02 |
| gene11401 | https://www.ncbi.nlm.nih.gov/gene/?term=LOC105285202 | 1841 | 50 | 271.59 | 1 | 5.43 | 2 | 21.11 | 19.1 | 2.01 | 2.01 | 0.05 | 0.05 |
| gene1199 | https://www.ncbi.nlm.nih.gov/gene/?term=LOC105285498 | 2220 | 26 | 117.12 | 1 | 4.5 | 3.85 | 33.11 | 31.11 | 2 | 2 | 0.02 | 0.02 |
| gene388 | https://www.ncbi.nlm.nih.gov/gene/?term=LOC105276538 | 1558 | 19 | 121.95 | 1 | 6.42 | 5.26 | 69.93 | 68.17 | 1.77 | 1.77 | 0.02 | 0.02 |
| gene10213 | https://www.ncbi.nlm.nih.gov/gene/?term=LOC105283826 | 1503 | 54 | 359.28 | 1 | 6.65 | 1.85 | 16.14 | 14.48 | 1.66 | 1.66 | 0.06 | 0.06 |
| gene10326 | https://www.ncbi.nlm.nih.gov/gene/?term=LOC105283962 | 3042 | 93 | 305.72 | 1 | 3.29 | 1.08 | 11.63 | 10.03 | 1.6 | 1.6 | 0.05 | 0.05 |
| gene3727 | https://www.ncbi.nlm.nih.gov/gene/?term=LOC105276367 | 2484 | 50 | 201.29 | 1 | 4.03 | 2 | 50.02 | 48.5 | 1.52 | 1.52 | 0.03 | 0.03 |
| gene7345 | https://www.ncbi.nlm.nih.gov/gene/?term=LOC105280529 | 3833 | 87 | 226.98 | 1 | 2.61 | 1.15 | 18.24 | 16.79 | 1.44 | 1.44 | 0.03 | 0.03 |
| gene322 | https://www.ncbi.nlm.nih.gov/gene/?term=LOC105275777 | 1606 | 22 | 136.99 | 1 | 6.23 | 4.55 | 47.62 | 46.19 | 1.43 | 1.43 | 0.02 | 0.02 |
| gene3599 | https://www.ncbi.nlm.nih.gov/gene/?term=LOC105276196 | 2955 | 54 | 182.74 | 1 | 3.38 | 1.85 | 40.6 | 39.18 | 1.42 | 1.42 | 0.03 | 0.03 |
| gene11340 | https://www.ncbi.nlm.nih.gov/gene/?term=LOC105285131 | 2918 | 97 | 332.42 | 1 | 3.43 | 1.03 | 14.75 | 13.34 | 1.41 | 1.41 | 0.05 | 0.05 |
| gene7248 | https://www.ncbi.nlm.nih.gov/gene/?term=LOC105280408 | 5175 | 44 | 85.02 | 1 | 1.93 | 2.27 | 21.35 | 20.08 | 1.27 | 1.27 | 0.01 | 0.01 |
| gene4190 | https://www.ncbi.nlm.nih.gov/gene/?term=LOC105276875 | 3150 | 130 | 412.7 | 1 | 3.17 | 0.77 | 21.07 | 19.84 | 1.23 | 1.23 | 0.05 | 0.05 |
| gene5298 | https://www.ncbi.nlm.nih.gov/gene/?term=LOC105278145 | 4060 | 250 | 615.76 | 2 | 4.93 | 0.8 | 10.79 | 9.61 | 1.17 | 1.17 | 0.07 | 0.07 |
| gene10606 | https://www.ncbi.nlm.nih.gov/gene/?term=LOC105284273 | 6119 | 249 | 406.93 | 2 | 3.27 | 0.8 | 20.32 | 19.15 | 1.16 | 1.16 | 0.05 | 0.05 |
| gene920 | https://www.ncbi.nlm.nih.gov/gene/?term=LOC105282518 | 1197 | 43 | 359.23 | 1 | 8.35 | 2.33 | 37.5 | 36.41 | 1.09 | 1.09 | 0.04 | 0.04 |
| gene2271 | https://www.ncbi.nlm.nih.gov/gene/?term=LOC105274706 | 1448 | 47 | 324.59 | 1 | 6.91 | 2.13 | 41.16 | 40.14 | 1.03 | 1.03 | 0.03 | 0.03 |
| gene7247 | https://www.ncbi.nlm.nih.gov/gene/?term=LOC105280407 | 10486 | 205 | 195.5 | 1 | 0.95 | 0.49 | 17 | 15.99 | 1.01 | 1.01 | 0.02 | 0.02 |
| gene4376 | https://www.ncbi.nlm.nih.gov/gene/?term=LOC105277083 | 4307 | 92 | 213.61 | 1 | 2.32 | 1.09 | 26.68 | 25.69 | 0.99 | 0.99 | 0.02 | 0.02 |
| gene564 | https://www.ncbi.nlm.nih.gov/gene/?term=LOC105278478 | 2802 | 13 | 46.4 | 1 | 3.57 | 7.69 | 25.47 | 24.48 | 0.99 | 0.99 | 0 | 0 |
| gene1220 | https://www.ncbi.nlm.nih.gov/gene/?term=LOC105285742 | 3453 | 94 | 272.23 | 1 | 2.9 | 1.06 | 26.72 | 25.74 | 0.97 | 0.97 | 0.03 | 0.03 |
| gene6031 | https://www.ncbi.nlm.nih.gov/gene/?term=LOC105279000 | 5691 | 180 | 316.29 | 1 | 1.76 | 0.56 | 14.05 | 13.11 | 0.94 | 0.94 | 0.03 | 0.03 |
| gene4962 | https://www.ncbi.nlm.nih.gov/gene/?term=LOC105277761 | 4645 | 140 | 301.4 | 1 | 2.15 | 0.71 | 15.27 | 14.38 | 0.89 | 0.89 | 0.03 | 0.03 |
| gene7249 | https://www.ncbi.nlm.nih.gov/gene/?term=LOC105280422 | 1775 | 61 | 343.66 | 1 | 5.63 | 1.64 | 15.16 | 14.36 | 0.81 | 0.81 | 0.03 | 0.03 |
| gene10392 | https://www.ncbi.nlm.nih.gov/gene/?term=LOC105284036 | 3216 | 84 | 261.19 | 2 | 6.22 | 2.38 | 23.38 | 22.62 | 0.76 | 0.76 | 0.02 | 0.02 |
| gene657 | https://www.ncbi.nlm.nih.gov/gene/?term=LOC105279667 | 6421 | 78 | 121.48 | 1 | 1.56 | 1.28 | 4.46 | 3.72 | 0.74 | 0.74 | 0.01 | 0.01 |
| gene10844 | https://www.ncbi.nlm.nih.gov/gene/?term=LOC105284535 | 9086 | 220 | 242.13 | 1 | 1.1 | 0.45 | 16.85 | 16.12 | 0.73 | 0.73 | 0.02 | 0.02 |
| gene6966 | https://www.ncbi.nlm.nih.gov/gene/?term=LOC105280104 | 3747 | 124 | 330.93 | 1 | 2.67 | 0.81 | 15.19 | 14.51 | 0.68 | 0.68 | 0.02 | 0.02 |
| gene12203 | https://www.ncbi.nlm.nih.gov/gene/?term=LOC105286099 | 5967 | 78 | 130.72 | 1 | 1.68 | 1.28 | 11.28 | 10.61 | 0.67 | 0.67 | 0.01 | 0.01 |
| gene3793 | https://www.ncbi.nlm.nih.gov/gene/?term=LOC105276429 | 2901 | 112 | 386.07 | 1 | 3.45 | 0.89 | 26.42 | 25.76 | 0.65 | 0.65 | 0.03 | 0.03 |
| gene1470 | https://www.ncbi.nlm.nih.gov/gene/?term=LOC105287597 | 2974 | 180 | 605.25 | 1 | 3.36 | 0.56 | 17.88 | 17.25 | 0.63 | 0.63 | 0.04 | 0.04 |
| gene11454 | https://www.ncbi.nlm.nih.gov/gene/?term=LOC105285256 | 1906 | 52 | 272.82 | 1 | 5.25 | 1.92 | 42.98 | 42.36 | 0.62 | 0.62 | 0.02 | 0.02 |
| gene1147 | https://www.ncbi.nlm.nih.gov/gene/?term=LOC105284954 | 4022 | 87 | 216.31 | 1 | 2.49 | 1.15 | 15.12 | 14.51 | 0.61 | 0.61 | 0.01 | 0.01 |
| gene12021 | https://www.ncbi.nlm.nih.gov/gene/?term=LOC105285905 | 1399 | 21 | 150.11 | 1 | 7.15 | 4.76 | 44.76 | 44.21 | 0.55 | 0.55 | 0.01 | 0.01 |
| gene2683 | https://www.ncbi.nlm.nih.gov/gene/?term=LOC105275187 | 5492 | 75 | 136.56 | 1 | 1.82 | 1.33 | 14.15 | 13.62 | 0.53 | 0.53 | 0.01 | 0.01 |
| gene11583 | https://www.ncbi.nlm.nih.gov/gene/?term=LOC105285414 | 8192 | 179 | 218.51 | 1 | 1.22 | 0.56 | 9.12 | 8.6 | 0.52 | 0.52 | 0.01 | 0.01 |
| gene2748 | https://www.ncbi.nlm.nih.gov/gene/?term=LOC105275261 | 3845 | 54 | 140.44 | 1 | 2.6 | 1.85 | 38.38 | 37.86 | 0.52 | 0.52 | 0.01 | 0.01 |
| gene7568 | https://www.ncbi.nlm.nih.gov/gene/?term=LOC105280786 | 2525 | 121 | 479.21 | 1 | 3.96 | 0.83 | 29.4 | 28.89 | 0.52 | 0.52 | 0.02 | 0.02 |
| gene13036 | https://www.ncbi.nlm.nih.gov/gene/?term=LOC105287059 | 4959 | 166 | 334.74 | 1 | 2.02 | 0.6 | 17.36 | 16.86 | 0.5 | 0.5 | 0.02 | 0.02 |
| gene7801 | https://www.ncbi.nlm.nih.gov/gene/?term=LOC105281047 | 5137 | 113 | 219.97 | 1 | 1.95 | 0.88 | 10.88 | 10.38 | 0.5 | 0.5 | 0.01 | 0.01 |
| gene474 | https://www.ncbi.nlm.nih.gov/gene/?term=LOC105277435 | 3625 | 155 | 427.59 | 1 | 2.76 | 0.65 | 12.2 | 11.71 | 0.49 | 0.49 | 0.02 | 0.02 |
| gene5525 | https://www.ncbi.nlm.nih.gov/gene/?term=LOC105278396 | 3090 | 187 | 605.18 | 1 | 3.24 | 0.53 | 24.38 | 23.89 | 0.49 | 0.49 | 0.03 | 0.03 |
| gene10549 | https://www.ncbi.nlm.nih.gov/gene/?term=LOC105284210 | 7145 | 319 | 446.47 | 1 | 1.4 | 0.31 | 10.59 | 10.18 | 0.4 | 0.4 | 0.02 | 0.02 |
| gene6286 | https://www.ncbi.nlm.nih.gov/gene/?term=LOC105279298 | 2861 | 134 | 468.37 | 1 | 3.5 | 0.75 | 23.06 | 22.69 | 0.37 | 0.37 | 0.02 | 0.02 |
| gene5524 | https://www.ncbi.nlm.nih.gov/gene/?term=LOC105278397 | 5177 | 201 | 388.26 | 1 | 1.93 | 0.5 | 24.26 | 23.91 | 0.35 | 0.35 | 0.01 | 0.01 |
| gene927 | https://www.ncbi.nlm.nih.gov/gene/?term=LOC105282600 | 3727 | 57 | 152.94 | 1 | 2.68 | 1.75 | 29.17 | 28.82 | 0.35 | 0.35 | 0.01 | 0.01 |
| gene12573 | https://www.ncbi.nlm.nih.gov/gene/?term=LOC105286515 | 3647 | 102 | 279.68 | 1 | 2.74 | 0.98 | 10.87 | 10.55 | 0.32 | 0.32 | 0.01 | 0.01 |
| gene5746 | https://www.ncbi.nlm.nih.gov/gene/?term=LOC105278666 | 10343 | 405 | 391.57 | 1 | 0.97 | 0.25 | 7.03 | 6.73 | 0.3 | 0.3 | 0.01 | 0.01 |
| gene10155 | https://www.ncbi.nlm.nih.gov/gene/?term=LOC105283756 | 7030 | 97 | 137.98 | 1 | 1.42 | 1.03 | 6.83 | 6.6 | 0.23 | 0.23 | 0 | 0 |
| gene4213 | https://www.ncbi.nlm.nih.gov/gene/?term=LOC105276901 | 3257 | 95 | 291.68 | 1 | 3.07 | 1.05 | 23.09 | 22.87 | 0.22 | 0.22 | 0.01 | 0.01 |

| | | | | | | | | | | | | | |
|---|---|---|---|---|---|---|---|---|---|---|---|---|---|
| gene6620 | https://www.ncbi.nlm.nih.gov/gene/?term=LOC105279687 | 17998 | 363 | 201.69 | 1 | 0.56 | 0.28 | 5.06 | 4.84 | 0.22 | 0.22 | 0 | 0 |
| gene5831 | https://www.ncbi.nlm.nih.gov/gene/?term=LOC105278761 | 2506 | 103 | 411.01 | 2 | 7.98 | 1.94 | 24.82 | 24.61 | 0.21 | 0.21 | 0.01 | 0.01 |
| gene8274 | https://www.ncbi.nlm.nih.gov/gene/?term=LOC105281624 | 5425 | 203 | 374.19 | 1 | 1.84 | 0.49 | 10.66 | 10.47 | 0.2 | 0.2 | 0.01 | 0.01 |
| gene3206 | https://www.ncbi.nlm.nih.gov/gene/?term=LOC105275752 | 3474 | 123 | 354.06 | 1 | 2.88 | 0.81 | 28.92 | 28.73 | 0.19 | 0.19 | 0.01 | 0.01 |
| gene10600 | https://www.ncbi.nlm.nih.gov/gene/?term=LOC105284270 | 4837 | 103 | 212.94 | 1 | 2.07 | 0.97 | 12.92 | 12.79 | 0.13 | 0.13 | 0 | 0 |
| gene5646 | https://www.ncbi.nlm.nih.gov/gene/?term=LOC105278534 | 7475 | 185 | 247.49 | 1 | 1.34 | 0.54 | 17.64 | 17.56 | 0.08 | 0.08 | 0 | 0 |
| gene5967 | https://www.ncbi.nlm.nih.gov/gene/?term=LOC105278918 | 4393 | 132 | 300.48 | 1 | 2.28 | 0.76 | 15.7 | 15.62 | 0.08 | 0.08 | 0 | 0 |
| gene12802 | https://www.ncbi.nlm.nih.gov/gene/?term=LOC105286799 | 3183 | 170 | 534.09 | 1 | 3.14 | 0.59 | 15.91 | 15.85 | 0.06 | 0.06 | 0 | 0 |
| gene3470 | https://www.ncbi.nlm.nih.gov/gene/?term=LOC105276059 | 8716 | 159 | 182.42 | 1 | 1.15 | 0.63 | 9.24 | 9.2 | 0.04 | 0.04 | 0 | 0 |
| gene2806 | https://www.ncbi.nlm.nih.gov/gene/?term=LOC105275327 | 14257 | 479 | 335.98 | 1 | 0.7 | 0.21 | 7.13 | 7.12 | 0.01 | 0.01 | 0 | 0 |
| gene5375 | https://www.ncbi.nlm.nih.gov/gene/?term=LOC105278228 | 8640 | 161 | 186.34 | 1 | 1.16 | 0.62 | 23.31 | 23.3 | 0.01 | 0.01 | 0 | 0 |
| gene2056 | https://www.ncbi.nlm.nih.gov/gene/?term=LOC105274452 | 6584 | 206 | 312.88 | 1 | 1.52 | 0.49 | 14.18 | 14.18 | 0 | 0 | 0 | 0 |
| gene11579 | https://www.ncbi.nlm.nih.gov/gene/?term=LOC105285406 | 13571 | 282 | 207.8 1 | 0.74 | 0.35 | 5.68 | 5.69 | -0.01 | 0.01 | 0 | 0 | |
| gene3107 | https://www.ncbi.nlm.nih.gov/gene/?term=LOC105275650 | 5103 | 150 | 293.94 | 1 | 1.96 | 0.67 | 17.09 | 17.11 | -0.02 | 0.02 | 0 | 0 |
| gene2173 | https://www.ncbi.nlm.nih.gov/gene/?term=LOC105274590 | 4387 | 79 | 180.08 | 1 | 2.28 | 1.27 | 22.08 | 22.11 | -0.03 | 0.03 | 0 | 0 |
| gene13114 | https://www.ncbi.nlm.nih.gov/gene/?term=LOC105287147 | 7609 | 381 | 500.72 | 1 | 1.31 | 0.26 | 9.67 | 9.7 | -0.04 | 0.04 | 0 | 0 |
| gene2668 | https://www.ncbi.nlm.nih.gov/gene/?term=LOC105275170 | 12496 | 336 | 268.89 | 1 | 0.8 | 0.3 | 2.05 | 2.12 | -0.07 | 0.07 | 0 | 0 |
| gene6698 | https://www.ncbi.nlm.nih.gov/gene/?term=LOC105279778 | 4099 | 138 | 336.67 | 1 | 2.44 | 0.72 | 9.41 | 9.48 | -0.07 | 0.07 | 0 | 0 |
| gene53 | https://www.ncbi.nlm.nih.gov/gene/?term=LOC105277148 | 19283 | 270 | 140.02 | 1 | 0.52 | 0.37 | 3.92 | 4.02 | -0.1 | 0.1 | 0 | 0 |
| gene11568 | https://www.ncbi.nlm.nih.gov/gene/?term=LOC105285395 | 8086 | 176 | 217.66 | 1 | 1.24 | 0.57 | 15.78 | 15.89 | -0.11 | 0.11 | 0 | 0 |
| gene10596 | https://www.ncbi.nlm.nih.gov/gene/?term=LOC105284262 | 6520 | 228 | 349.69 | 1 | 1.53 | 0.44 | 3.34 | 3.5 | -0.16 | 0.16 | -0.01 | 0.01 |
| gene10540 | https://www.ncbi.nlm.nih.gov/gene/?term=LOC105284200 | 5267 | 193 | 366.43 | 1 | 1.9 | 0.52 | 13.99 | 14.21 | -0.22 | 0.22 | -0.01 | 0.01 |
| gene303 | https://www.ncbi.nlm.nih.gov/gene/?term=LOC105275584 | 3081 | 86 | 279.13 | 1 | 3.25 | 1.16 | 23.78 | 24 | -0.22 | 0.22 | -0.01 | 0.01 |
| gene5184 | https://www.ncbi.nlm.nih.gov/gene/?term=LOC105277999 | 5528 | 147 | 265.92 | 1 | 1.81 | 0.68 | 12.4 | 12.64 | -0.24 | 0.24 | -0.01 | 0.01 |
| gene7888 | https://www.ncbi.nlm.nih.gov/gene/?term=LOC105281141 | 4357 | 155 | 355.75 | 1 | 2.3 | 0.65 | 29.83 | 30.06 | -0.24 | 0.24 | -0.01 | 0.01 |
| gene7398 | https://www.ncbi.nlm.nih.gov/gene/?term=LOC105280589 | 3452 | 41 | 118.77 | 1 | 2.9 | 2.44 | 25.07 | 25.33 | -0.26 | 0.26 | 0 | 0 |
| gene7166 | https://www.ncbi.nlm.nih.gov/gene/?term=LOC105280326 | 6117 | 208 | 340.04 | 1 | 1.63 | 0.48 | 15.97 | 16.24 | -0.27 | 0.27 | -0.01 | 0.01 |
| gene6528 | https://www.ncbi.nlm.nih.gov/gene/?term=LOC105279591 | 7222 | 67 | 92.77 1 | 1.38 | 1.49 | 23.03 | 23.32 | -0.28 | 0.28 | 0 | 0 | |
| gene4023 | https://www.ncbi.nlm.nih.gov/gene/?term=LOC105276692 | 1759 | 38 | 216.03 | 1 | 5.69 | 2.63 | 22.43 | 22.73 | -0.3 | 0.3 | -0.01 | 0.01 |
| gene6455 | https://www.ncbi.nlm.nih.gov/gene/?term=LOC105279500 | 2612 | 32 | 122.51 | 1 | 3.83 | 3.12 | 49.77 | 50.07 | -0.3 | 0.3 | 0 | 0 |
| gene2443 | https://www.ncbi.nlm.nih.gov/gene/?term=LOC105274895 | 10433 | 345 | 330.68 | 1 | 0.96 | 0.29 | 6.76 | 7.09 | -0.33 | 0.33 | -0.01 | 0.01 |
| gene10520 | https://www.ncbi.nlm.nih.gov/gene/?term=LOC105284184 | 2451 | 82 | 334.56 | 1 | 4.08 | 1.22 | 34.55 | 34.9 | -0.35 | 0.35 | -0.01 | 0.01 |
| gene2404 | https://www.ncbi.nlm.nih.gov/gene/?term=LOC105274853 | 1767 | 37 | 209.39 | 1 | 5.66 | 2.7 | 31.99 | 32.37 | -0.37 | 0.37 | -0.01 | 0.01 |
| gene10531 | https://www.ncbi.nlm.nih.gov/gene/?term=LOC105284190 | 6881 | 192 | 279.03 | 1 | 1.45 | 0.52 | 9.5 | 9.91 | -0.4 | 0.4 | -0.01 | 0.01 |
| gene3856 | https://www.ncbi.nlm.nih.gov/gene/?term=LOC105276499 | 5236 | 197 | 376.24 | 1 | 1.91 | 0.51 | 9.94 | 10.34 | -0.41 | 0.41 | -0.02 | 0.02 |
| gene4892 | https://www.ncbi.nlm.nih.gov/gene/?term=LOC105277667 | 3898 | 120 | 307.85 | 1 | 2.57 | 0.83 | 28.07 | 28.49 | -0.42 | 0.42 | -0.01 | 0.01 |
| gene6583 | https://www.ncbi.nlm.nih.gov/gene/?term=LOC105279649 | 7444 | 317 | 425.85 | 1 | 1.34 | 0.32 | 11.9 | 12.34 | -0.44 | 0.44 | -0.02 | 0.02 |
| gene11275 | https://www.ncbi.nlm.nih.gov/gene/?term=LOC105285055 | 4119 | 213 | 517.12 | 1 | 2.43 | 0.47 | 13.26 | 13.73 | -0.47 | 0.47 | -0.02 | 0.02 |
| gene9411 | https://www.ncbi.nlm.nih.gov/gene/?term=LOC105282914 | 5534 | 41 | 74.09 1 | 1.81 | 2.44 | 49.68 | 50.18 | -0.49 | 0.49 | 0 | 0 | |
| gene10678 | https://www.ncbi.nlm.nih.gov/gene/?term=LOC105284355 | 4031 | 170 | 421.73 | 1 | 2.48 | 0.59 | 18.52 | 19.03 | -0.5 | 0.5 | -0.02 | 0.02 |
| gene8178 | https://www.ncbi.nlm.nih.gov/gene/?term=LOC105281515 | 8647 | 44 | 50.88 1 | 1.16 | 2.27 | 29.36 | 29.86 | -0.5 | 0.5 | 0 | 0 | |
| gene8600 | https://www.ncbi.nlm.nih.gov/gene/?term=LOC105282001 | 8513 | 153 | 179.73 | 1 | 1.17 | 0.65 | 9.37 | 9.87 | -0.5 | 0.5 | -0.01 | 0.01 |
| gene6639 | https://www.ncbi.nlm.nih.gov/gene/?term=LOC105279705 | 7656 | 68 | 88.82 1 | 1.31 | 1.47 | 22.19 | 22.7 | -0.51 | 0.51 | 0 | 0 | |
| gene3885 | https://www.ncbi.nlm.nih.gov/gene/?term=LOC105276527 | 7928 | 346 | 436.43 | 1 | 1.26 | 0.29 | 8.3 | 8.89 | -0.6 | 0.6 | -0.03 | 0.03 |
| gene10018 | https://www.ncbi.nlm.nih.gov/gene/?term=LOC105283607 | 4414 | 204 | 462.17 | 1 | 2.27 | 0.49 | 14.8 | 15.47 | -0.67 | 0.67 | -0.03 | 0.03 |
| gene274 | https://www.ncbi.nlm.nih.gov/gene/?term=LOC105275190 | 3488 | 70 | 200.69 | 1 | 2.87 | 1.43 | 19.35 | 20.02 | -0.67 | 0.67 | -0.01 | 0.01 |
| gene1332 | https://www.ncbi.nlm.nih.gov/gene/?term=LOC105286872 | 2460 | 141 | 573.17 | 1 | 4.07 | 0.71 | 21.92 | 22.68 | -0.76 | 0.76 | -0.04 | 0.04 |
| gene8729 | https://www.ncbi.nlm.nih.gov/gene/?term=LOC105282137 | 5005 | 199 | 397.6 1 | 2 | 0.5 | 16.19 | 16.99 | -0.79 | 0.79 | -0.03 | 0.03 | |
| gene11819 | https://www.ncbi.nlm.nih.gov/gene/?term=LOC105285671 | 2366 | 96 | 405.75 | 1 | 4.23 | 1.04 | 27.73 | 28.57 | -0.84 | 0.84 | -0.03 | 0.03 |
| gene9336 | https://www.ncbi.nlm.nih.gov/gene/?term=LOC105282834 | 5993 | 115 | 191.89 | 1 | 1.67 | 0.87 | 14.71 | 15.58 | -0.87 | 0.87 | -0.02 | 0.02 |
| gene10107 | https://www.ncbi.nlm.nih.gov/gene/?term=LOC105283712 | 1883 | 28 | 148.7 1 | 5.31 | 3.57 | 28.01 | 28.9 | -0.88 | 0.88 | -0.01 | 0.01 | |
| gene1310 | https://www.ncbi.nlm.nih.gov/gene/?term=LOC105286661 | 2582 | 65 | 251.74 | 1 | 3.87 | 1.54 | 19.96 | 20.95 | -0.99 | 0.99 | -0.03 | 0.03 |
| gene2844 | https://www.ncbi.nlm.nih.gov/gene/?term=LOC105275363 | 4330 | 187 | 431.87 | 1 | 2.31 | 0.53 | 14.36 | 15.35 | -0.99 | 0.99 | -0.04 | 0.04 |
| gene292 | https://www.ncbi.nlm.nih.gov/gene/?term=LOC105275398 | 2825 | 76 | 269.03 | 1 | 3.54 | 1.32 | 20.33 | 21.34 | -1.01 | 1.01 | -0.03 | 0.03 |
| gene7183 | https://www.ncbi.nlm.nih.gov/gene/?term=LOC105280348 | 3925 | 52 | 132.48 | 2 | 5.1 | 3.85 | 25.66 | 26.67 | -1.01 | 1.01 | -0.01 | 0.01 |
| gene8343 | https://www.ncbi.nlm.nih.gov/gene/?term=LOC105281700 | 2172 | 44 | 202.58 | 1 | 4.6 | 2.27 | 37.43 | 38.56 | -1.13 | 1.13 | -0.02 | 0.02 |
| gene4034 | https://www.ncbi.nlm.nih.gov/gene/?term=LOC105276704 | 2065 | 84 | 406.78 | 1 | 4.84 | 1.19 | 12.86 | 14.01 | -1.15 | 1.15 | -0.05 | 0.05 |
| gene298 | https://www.ncbi.nlm.nih.gov/gene/?term=LOC105275460 | 2473 | 115 | 465.02 | 1 | 4.04 | 0.87 | 12.68 | 13.89 | -1.21 | 1.21 | -0.06 | 0.06 |
| gene5101 | https://www.ncbi.nlm.nih.gov/gene/?term=LOC105277903 | 1813 | 34 | 187.53 | 1 | 5.52 | 2.94 | 52.57 | 53.85 | -1.27 | 1.27 | -0.02 | 0.02 |
| gene570 | https://www.ncbi.nlm.nih.gov/gene/?term=LOC105278549 | 1673 | 63 | 376.57 | 1 | 5.98 | 1.59 | 40.52 | 41.79 | -1.27 | 1.27 | -0.05 | 0.05 |
| gene2856 | https://www.ncbi.nlm.nih.gov/gene/?term=LOC105275384 | 1987 | 76 | 382.49 | 1 | 5.03 | 1.32 | 32.95 | 34.23 | -1.28 | 1.28 | -0.05 | 0.05 |
| gene9101 | https://www.ncbi.nlm.nih.gov/gene/?term=LOC105282553 | 3068 | 103 | 335.72 | 1 | 3.26 | 0.97 | 20.56 | 21.9 | -1.34 | 1.34 | -0.04 | 0.04 |
| gene4641 | https://www.ncbi.nlm.nih.gov/gene/?term=LOC105277398 | 1937 | 70 | 361.38 | 1 | 5.16 | 1.43 | 35.36 | 36.76 | -1.4 | 1.4 | -0.05 | 0.05 |
| gene5877 | https://www.ncbi.nlm.nih.gov/gene/?term=LOC105278807 | 1719 | 51 | 296.68 | 1 | 5.82 | 1.96 | 22.99 | 24.43 | -1.44 | 1.44 | -0.04 | 0.04 |

| | | | | | | | | | | | | | |
|---|---|---|---|---|---|---|---|---|---|---|---|---|---|
| gene5402 | https://www.ncbi.nlm.nih.gov/gene/?term=LOC105278260 | 3068 | 42 | 136.9 | 1 | 3.26 | 2.38 | 12.8 | 14.3 | -1.5 | 1.5 | -0.02 | 0.02 |
| gene12406 | https://www.ncbi.nlm.nih.gov/gene/?term=LOC105286316 | 1386 | 30 | 216.45 | 1 | 7.22 | 3.33 | 39.2 | 40.74 | -1.55 | 1.55 | -0.03 | 0.03 |
| gene1884 | https://www.ncbi.nlm.nih.gov/gene/?term=LOC105288072 | 4169 | 80 | 191.89 | 1 | 2.4 | 1.25 | 22.74 | 24.44 | -1.7 | 1.7 | -0.03 | 0.03 |
| gene1855 | https://www.ncbi.nlm.nih.gov/gene/?term=LOC105288036 | 2387 | 87 | 364.47 | 1 | 4.19 | 1.15 | 28.59 | 30.36 | -1.76 | 1.76 | -0.06 | 0.06 |
| gene10345 | https://www.ncbi.nlm.nih.gov/gene/?term=LOC105283989 | 4444 | 29 | 65.26 | 1 | 2.25 | 3.45 | 48.19 | 50.2 | -2.01 | 2.01 | -0.01 | 0.01 |
| gene6334 | https://www.ncbi.nlm.nih.gov/gene/?term=LOC105279356 | 1471 | 43 | 292.32 | 1 | 6.8 | 2.33 | 28.25 | 30.28 | -2.03 | 2.03 | -0.06 | 0.06 |
| gene12516 | https://www.ncbi.nlm.nih.gov/gene/?term=LOC105286452 | 1529 | 35 | 228.91 | 1 | 6.54 | 2.86 | 51.01 | 53.47 | -2.46 | 2.46 | -0.06 | 0.06 |
| gene2676 | https://www.ncbi.nlm.nih.gov/gene/?term=LOC105275181 | 1353 | 14 | 103.47 | 1 | 7.39 | 7.14 | 65.95 | 68.91 | -2.96 | 2.96 | -0.03 | 0.03 |
| gene2970 | https://www.ncbi.nlm.nih.gov/gene/?term=LOC105275518 | 2340 | 20 | 85.47 | 1 | 4.27 | 5 | 47.36 | 50.74 | -3.38 | 3.38 | -0.03 | 0.03 |
| gene5714 | https://www.ncbi.nlm.nih.gov/gene/?term=LOC105278729 | 710 | 14 | 197.18 | 1 | 14.08 | 7.14 | 51.67 | 55.07 | -3.41 | 3.41 | -0.07 | 0.07 |
| gene5647 | https://www.ncbi.nlm.nih.gov/gene/?term=LOC105278567 | 3118 | 38 | 121.87 | 1 | 3.21 | 2.63 | 12.56 | 16.06 | -3.5 | 3.5 | -0.04 | 0.04 |
| gene1962 | https://www.ncbi.nlm.nih.gov/gene/?term=LOC105288157 | 1246 | 16 | 128.41 | 1 | 8.03 | 6.25 | 13.95 | 18.55 | -4.6 | 4.6 | -0.06 | 0.06 |
| gene3165 | https://www.ncbi.nlm.nih.gov/gene/?term=LOC105275706 | 2374 | 66 | 278.01 | 4 | 16.85 | 6.06 | 24.58 | 35.31 | -10.73 | 10.73 | -0.3 | 0.3 |
| gene8929 | https://www.ncbi.nlm.nih.gov/gene/?term=LOC105282371 | 1849 | 8 | 43.27 | 1 | 5.41 | 12.5 | 22.25 | 44.14 | -21.89 | 21.89 | -0.09 | 0.09 |
| gene2593 | https://www.ncbi.nlm.nih.gov/gene/?term=LOC105275070 | 1774 | 6 | 33.82 | 1 | 5.64 | 16.67 | 16.42 | 54.02 | -37.6 | 37.6 | -0.13 | 0.13 |

Directory: OGL File: ogl-Cb_LI_BCphaseA1.vs.BCphaseB2.txt

| gene_ID | gene_link | gwidth | #Sites | #per10Kb | #dmSites | #dmsp10kb | %dmSites | %pSite1 | %pSite2 | DMpSite | ADMpSite | DMpNucl | ADMpNucl |
|---|---|---|---|---|---|---|---|---|---|---|---|---|---|
| gene7106 | https://www.ncbi.nlm.nih.gov/gene/?term=LOC105280261 | 970 | 4 | 41.24 | 1 | 10.31 | 25 | 82.16 | 67.27 | 14.89 | 14.89 | 0.06 | 0.06 |
| gene11980 | https://www.ncbi.nlm.nih.gov/gene/?term=LOC109611395 | 571 | 3 | 52.54 | 1 | 17.51 | 33.33 | 22.75 | 11.76 | 10.99 | 10.99 | 0.06 | 0.06 |
| gene558 | https://www.ncbi.nlm.nih.gov/gene/?term=LOC105278458 | 216 | 3 | 138.89 | 1 | 46.3 | 33.33 | 20.87 | 10.44 | 10.43 | 10.43 | 0.14 | 0.14 |
| gene7684 | https://www.ncbi.nlm.nih.gov/gene/?term=LOC105280926 | 1727 | 6 | 34.74 | 2 | 11.58 | 33.33 | 47.53 | 37.53 | 10 | 10 | 0.03 | 0.03 |
| gene10219 | https://www.ncbi.nlm.nih.gov/gene/?term=LOC105283830 | 235 | 12 | 510.64 | 1 | 42.55 | 8.33 | 42.96 | 33.59 | 9.36 | 9.36 | 0.48 | 0.48 |
| gene3395 | https://www.ncbi.nlm.nih.gov/gene/?term=LOC105275978 | 2468 | 5 | 20.26 | 1 | 4.05 | 20 | 19.06 | 10.64 | 8.42 | 8.42 | 0.02 | 0.02 |
| gene12003 | https://www.ncbi.nlm.nih.gov/gene/?term=LOC105285885 | 1336 | 11 | 82.34 | 2 | 14.97 | 18.18 | 63.58 | 55.36 | 8.22 | 8.22 | 0.07 | 0.07 |
| gene1656 | https://www.ncbi.nlm.nih.gov/gene/?term=LOC105287814 | 812 | 7 | 86.21 | 1 | 12.32 | 14.29 | 37.14 | 29.51 | 7.63 | 7.63 | 0.07 | 0.07 |
| gene4643 | https://www.ncbi.nlm.nih.gov/gene/?term=LOC109610847 | 429 | 5 | 116.55 | 1 | 23.31 | 20 | 29.86 | 22.36 | 7.5 | 7.5 | 0.09 | 0.09 |
| gene10325 | https://www.ncbi.nlm.nih.gov/gene/?term=LOC105283963 | 1286 | 9 | 69.98 | 1 | 7.78 | 11.11 | 57.17 | 49.99 | 7.19 | 7.19 | 0.05 | 0.05 |
| gene8172 | https://www.ncbi.nlm.nih.gov/gene/?term=LOC105281507 | 946 | 5 | 52.85 | 1 | 10.57 | 20 | 11.82 | 4.83 | 6.99 | 6.99 | 0.04 | 0.04 |
| gene405 | https://www.ncbi.nlm.nih.gov/gene/?term=LOC105276706 | 1442 | 16 | 110.96 | 1 | 6.93 | 6.25 | 82.68 | 76.42 | 6.27 | 6.27 | 0.07 | 0.07 |
| gene8965 | https://www.ncbi.nlm.nih.gov/gene/?term=LOC105282405 | 1310 | 31 | 236.64 | 3 | 22.9 | 9.68 | 51.28 | 45.18 | 6.1 | 6.1 | 0.14 | 0.14 |
| gene539 | https://www.ncbi.nlm.nih.gov/gene/?term=LOC105278243 | 1952 | 18 | 92.21 | 1 | 5.12 | 5.56 | 20.77 | 14.99 | 5.77 | 5.77 | 0.05 | 0.05 |
| gene10310 | https://www.ncbi.nlm.nih.gov/gene/?term=LOC109611277 | 1730 | 8 | 46.24 | 1 | 5.78 | 12.5 | 12.29 | 6.59 | 5.7 | 5.7 | 0.03 | 0.03 |
| gene11269 | https://www.ncbi.nlm.nih.gov/gene/?term=LOC105285051 | 1363 | 11 | 80.7 | 3 | 22.01 | 27.27 | 46.02 | 40.39 | 5.64 | 5.64 | 0.05 | 0.05 |
| gene8458 | https://www.ncbi.nlm.nih.gov/gene/?term=LOC105281831 | 1053 | 8 | 75.97 | 1 | 9.5 | 12.5 | 87.04 | 81.5 | 5.54 | 5.54 | 0.04 | 0.04 |
| gene7962 | https://www.ncbi.nlm.nih.gov/gene/?term=LOC105281226 | 1280 | 6 | 46.88 | 1 | 7.81 | 16.67 | 62.79 | 57.27 | 5.53 | 5.53 | 0.03 | 0.03 |
| gene10204 | https://www.ncbi.nlm.nih.gov/gene/?term=LOC105283813 | 1312 | 8 | 60.98 | 1 | 7.62 | 12.5 | 29.71 | 24.26 | 5.45 | 5.45 | 0.03 | 0.03 |
| gene10792 | https://www.ncbi.nlm.nih.gov/gene/?term=LOC105284483 | 2910 | 24 | 82.47 | 1 | 3.44 | 4.17 | 22.77 | 17.6 | 5.17 | 5.17 | 0.04 | 0.04 |
| gene10357 | https://www.ncbi.nlm.nih.gov/gene/?term=LOC105284002 | 1605 | 12 | 74.77 | 1 | 6.23 | 8.33 | 44.27 | 39.3 | 4.98 | 4.98 | 0.04 | 0.04 |
| gene1914 | https://www.ncbi.nlm.nih.gov/gene/?term=LOC105288105 | 1289 | 23 | 178.43 | 2 | 15.52 | 8.7 | 60.89 | 56.04 | 4.85 | 4.85 | 0.09 | 0.09 |
| gene3296 | https://www.ncbi.nlm.nih.gov/gene/?term=LOC105275859 | 2340 | 21 | 89.74 | 2 | 8.55 | 9.52 | 53.4 | 48.62 | 4.78 | 4.78 | 0.04 | 0.04 |
| gene939 | https://www.ncbi.nlm.nih.gov/gene/?term=LOC105282729 | 1450 | 11 | 75.86 | 1 | 6.9 | 9.09 | 28.64 | 23.91 | 4.73 | 4.73 | 0.04 | 0.04 |

...
... only first 25 lines shown ...

Directory: OGL File: ogl-Cb_LI_BCphaseA1.vs.RphaseA1.txt

| gene_ID | gene_link | gwidth | #Sites | #per10Kb | #dmSites | #dmsp10kb | %dmSites | %pSite1 | %pSite2 | DMpSite | ADMpSite | DMpNucl | ADMpNucl |
|---|---|---|---|---|---|---|---|---|---|---|---|---|---|
| gene393 | https://www.ncbi.nlm.nih.gov/gene/?term=LOC105276566 | 1123 | 2 | 17.81 | 1 | 8.9 | 50 | 77.65 | 50 | 27.65 | 27.65 | 0.05 | 0.05 |
| gene7199 | https://www.ncbi.nlm.nih.gov/gene/?term=LOC105280366 | 1414 | 26 | 183.88 | 1 | 7.07 | 3.85 | 47.43 | 38.76 | 8.67 | 8.67 | 0.16 | 0.16 |
| gene12517 | https://www.ncbi.nlm.nih.gov/gene/?term=LOC105286451 | 1626 | 10 | 61.5 | 1 | 6.15 | 10 | 52.19 | 44.01 | 8.18 | 8.18 | 0.05 | 0.05 |
| gene388 | https://www.ncbi.nlm.nih.gov/gene/?term=LOC105276538 | 1558 | 19 | 121.95 | 1 | 6.42 | 5.26 | 69.41 | 64.89 | 4.52 | 4.52 | 0.06 | 0.06 |
| gene7915 | https://www.ncbi.nlm.nih.gov/gene/?term=LOC105281173 | 1597 | 25 | 156.54 | 1 | 6.26 | 4 | 64.87 | 60.99 | 3.88 | 3.88 | 0.06 | 0.06 |
| gene3168 | https://www.ncbi.nlm.nih.gov/gene/?term=LOC105275711 | 6150 | 77 | 125.2 | 1 | 1.63 | 1.3 | 14.51 | 10.99 | 3.51 | 3.51 | 0.04 | 0.04 |
| gene6422 | https://www.ncbi.nlm.nih.gov/gene/?term=LOC105279458 | 2419 | 37 | 152.96 | 1 | 4.13 | 2.7 | 54.1 | 51.26 | 2.83 | 2.83 | 0.04 | 0.04 |
| gene10714 | https://www.ncbi.nlm.nih.gov/gene/?term=LOC105284396 | 1966 | 53 | 269.58 | 1 | 5.09 | 1.89 | 40.95 | 38.4 | 2.55 | 2.55 | 0.07 | 0.07 |
| gene322 | https://www.ncbi.nlm.nih.gov/gene/?term=LOC105275777 | 1606 | 23 | 143.21 | 1 | 6.23 | 4.35 | 45.55 | 43.01 | 2.54 | 2.54 | 0.04 | 0.04 |
| gene11401 | https://www.ncbi.nlm.nih.gov/gene/?term=LOC105285202 | 1841 | 49 | 266.16 | 1 | 5.43 | 2.04 | 21.54 | 19.38 | 2.16 | 2.16 | 0.06 | 0.06 |
| gene7368 | https://www.ncbi.nlm.nih.gov/gene/?term=LOC105280556 | 4105 | 46 | 112.06 | 1 | 2.44 | 2.17 | 13.39 | 11.38 | 2 | 2 | 0.02 | 0.02 |
| gene10213 | https://www.ncbi.nlm.nih.gov/gene/?term=LOC105283826 | 1503 | 52 | 345.97 | 1 | 6.65 | 1.92 | 16.76 | 14.79 | 1.97 | 1.97 | 0.07 | 0.07 |
| gene12021 | https://www.ncbi.nlm.nih.gov/gene/?term=LOC105285905 | 1399 | 23 | 164.4 | 1 | 7.15 | 4.35 | 41.24 | 39.35 | 1.89 | 1.89 | 0.03 | 0.03 |
| gene2271 | https://www.ncbi.nlm.nih.gov/gene/?term=LOC105274706 | 1448 | 49 | 338.4 | 1 | 6.91 | 2.04 | 38.87 | 37.01 | 1.86 | 1.86 | 0.06 | 0.06 |
| gene4190 | https://www.ncbi.nlm.nih.gov/gene/?term=LOC105276875 | 3150 | 124 | 393.65 | 1 | 3.17 | 0.81 | 21.46 | 19.89 | 1.58 | 1.58 | 0.06 | 0.06 |
| gene11454 | https://www.ncbi.nlm.nih.gov/gene/?term=LOC105285256 | 1906 | 52 | 272.82 | 1 | 5.25 | 1.92 | 42.98 | 41.47 | 1.51 | 1.51 | 0.04 | 0.04 |
| gene2752 | https://www.ncbi.nlm.nih.gov/gene/?term=LOC105275266 | 2111 | 49 | 232.12 | 1 | 4.74 | 2.04 | 23.87 | 22.43 | 1.44 | 1.44 | 0.03 | 0.03 |
| gene920 | https://www.ncbi.nlm.nih.gov/gene/?term=LOC105282518 | 1197 | 42 | 350.88 | 1 | 8.35 | 2.38 | 38.27 | 36.85 | 1.43 | 1.43 | 0.05 | 0.05 |
| gene5649 | https://www.ncbi.nlm.nih.gov/gene/?term=LOC105278538 | 2140 | 76 | 355.14 | 1 | 4.67 | 1.32 | 18.24 | 16.87 | 1.37 | 1.37 | 0.05 | 0.05 |
| gene9740 | https://www.ncbi.nlm.nih.gov/gene/?term=LOC105283277 | 2887 | 135 | 467.61 | 1 | 3.46 | 0.74 | 21.88 | 20.52 | 1.37 | 1.37 | 0.06 | 0.06 |
| gene2748 | https://www.ncbi.nlm.nih.gov/gene/?term=LOC105275261 | 3845 | 55 | 143.04 | 1 | 2.6 | 1.82 | 38.68 | 37.31 | 1.36 | 1.36 | 0.02 | 0.02 |
| gene10326 | https://www.ncbi.nlm.nih.gov/gene/?term=LOC105283962 | 3042 | 94 | 309.01 | 1 | 3.29 | 1.06 | 12.22 | 10.91 | 1.31 | 1.31 | 0.04 | 0.04 |
| gene6031 | https://www.ncbi.nlm.nih.gov/gene/?term=LOC105279000 | 5691 | 175 | 307.5 | 1 | 1.76 | 0.57 | 14.52 | 13.23 | 1.29 | 1.29 | 0.04 | 0.04 |
| gene10844 | https://www.ncbi.nlm.nih.gov/gene/?term=LOC105284535 | 9086 | 218 | 239.93 | 1 | 1.1 | 0.46 | 16.62 | 15.4 | 1.22 | 1.22 | 0.03 | 0.03 |
| gene4376 | https://www.ncbi.nlm.nih.gov/gene/?term=LOC105277083 | 4307 | 90 | 208.96 | 1 | 2.32 | 1.11 | 26.1 | 24.89 | 1.21 | 1.21 | 0.03 | 0.03 |
| gene3599 | https://www.ncbi.nlm.nih.gov/gene/?term=LOC105276196 | 2955 | 55 | 186.13 | 1 | 3.38 | 1.82 | 40.49 | 39.41 | 1.07 | 1.07 | 0.02 | 0.02 |
| gene5298 | https://www.ncbi.nlm.nih.gov/gene/?term=LOC105278145 | 4060 | 240 | 591.13 | 2 | 4.93 | 0.83 | 11.5 | 10.44 | 1.06 | 1.06 | 0.06 | 0.06 |
| gene7248 | https://www.ncbi.nlm.nih.gov/gene/?term=LOC105280408 | 5175 | 44 | 85.02 | 1 | 1.93 | 2.27 | 22.41 | 21.41 | 0.99 | 0.99 | 0.01 | 0.01 |
| gene6806 | https://www.ncbi.nlm.nih.gov/gene/?term=LOC105279894 | 3649 | 39 | 106.88 | 1 | 2.74 | 2.56 | 35.25 | 34.29 | 0.97 | 0.97 | 0.01 | 0.01 |
| gene10392 | https://www.ncbi.nlm.nih.gov/gene/?term=LOC105284036 | 3216 | 83 | 258.08 | 1 | 3.11 | 1.2 | 24.77 | 23.81 | 0.96 | 0.96 | 0.02 | 0.02 |
| gene12203 | https://www.ncbi.nlm.nih.gov/gene/?term=LOC105286099 | 5967 | 69 | 115.64 | 1 | 1.68 | 1.45 | 11.63 | 10.68 | 0.96 | 0.96 | 0.01 | 0.01 |
| gene4962 | https://www.ncbi.nlm.nih.gov/gene/?term=LOC105277761 | 4645 | 147 | 316.47 | 1 | 2.15 | 0.68 | 13.9 | 13.02 | 0.88 | 0.88 | 0.03 | 0.03 |
| gene12307 | https://www.ncbi.nlm.nih.gov/gene/?term=LOC105286216 | 10313 | 145 | 140.6 | 1 | 0.97 | 0.69 | 17.77 | 16.9 | 0.87 | 0.87 | 0.01 | 0.01 |
| gene6966 | https://www.ncbi.nlm.nih.gov/gene/?term=LOC105280104 | 3747 | 119 | 317.59 | 1 | 2.67 | 0.84 | 15.37 | 14.52 | 0.85 | 0.85 | 0.03 | 0.03 |
| gene6639 | https://www.ncbi.nlm.nih.gov/gene/?term=LOC105279705 | 7656 | 70 | 91.43 | 1 | 1.31 | 1.43 | 22.25 | 21.42 | 0.84 | 0.84 | 0.01 | 0.01 |
| gene7568 | https://www.ncbi.nlm.nih.gov/gene/?term=LOC105280786 | 2525 | 116 | 459.41 | 1 | 3.96 | 0.86 | 28.86 | 28.06 | 0.8 | 0.8 | 0.04 | 0.04 |
| gene7550 | https://www.ncbi.nlm.nih.gov/gene/?term=LOC105280763 | 2173 | 93 | 427.98 | 1 | 4.6 | 1.08 | 27 | 26.23 | 0.77 | 0.77 | 0.03 | 0.03 |
| gene7249 | https://www.ncbi.nlm.nih.gov/gene/?term=LOC105280422 | 1775 | 61 | 343.66 | 1 | 5.63 | 1.64 | 15.16 | 14.42 | 0.75 | 0.75 | 0.03 | 0.03 |
| gene10606 | https://www.ncbi.nlm.nih.gov/gene/?term=LOC105284273 | 6119 | 243 | 397.12 | 2 | 3.27 | 0.82 | 20.74 | 20.03 | 0.71 | 0.71 | 0.03 | 0.03 |
| gene10600 | https://www.ncbi.nlm.nih.gov/gene/?term=LOC105284270 | 4837 | 95 | 196.4 | 1 | 2.07 | 1.05 | 13.92 | 13.23 | 0.69 | 0.69 | 0.01 | 0.01 |
| gene6286 | https://www.ncbi.nlm.nih.gov/gene/?term=LOC105279298 | 2861 | 136 | 475.36 | 1 | 3.5 | 0.74 | 22.78 | 22.14 | 0.64 | 0.64 | 0.03 | 0.03 |
| gene3107 | https://www.ncbi.nlm.nih.gov/gene/?term=LOC105275650 | 5103 | 145 | 284.15 | 1 | 1.96 | 0.69 | 17.36 | 16.74 | 0.62 | 0.62 | 0.02 | 0.02 |
| gene11340 | https://www.ncbi.nlm.nih.gov/gene/?term=LOC105285131 | 2918 | 97 | 332.42 | 1 | 3.43 | 1.03 | 14.71 | 14.15 | 0.56 | 0.56 | 0.02 | 0.02 |
| gene2404 | https://www.ncbi.nlm.nih.gov/gene/?term=LOC105274853 | 1767 | 39 | 220.71 | 1 | 5.66 | 2.56 | 31.07 | 30.51 | 0.56 | 0.56 | 0.01 | 0.01 |
| gene10155 | https://www.ncbi.nlm.nih.gov/gene/?term=LOC105283756 | 7030 | 93 | 132.29 | 1 | 1.42 | 1.08 | 7.03 | 6.49 | 0.54 | 0.54 | 0.01 | 0.01 |
| gene3206 | https://www.ncbi.nlm.nih.gov/gene/?term=LOC105275752 | 3474 | 126 | 362.69 | 1 | 2.88 | 0.79 | 28.38 | 27.85 | 0.53 | 0.53 | 0.02 | 0.02 |
| gene657 | https://www.ncbi.nlm.nih.gov/gene/?term=LOC105279667 | 6421 | 77 | 119.92 | 1 | 1.56 | 1.3 | 5.63 | 5.1 | 0.53 | 0.53 | 0.01 | 0.01 |
| gene7801 | https://www.ncbi.nlm.nih.gov/gene/?term=LOC105281047 | 5137 | 109 | 212.19 | 1 | 1.95 | 0.92 | 11.18 | 10.67 | 0.5 | 0.5 | 0.01 | 0.01 |
| gene7398 | https://www.ncbi.nlm.nih.gov/gene/?term=LOC105280589 | 3452 | 40 | 115.87 | 1 | 2.9 | 2.5 | 25.2 | 24.72 | 0.48 | 0.48 | 0.01 | 0.01 |
| gene1470 | https://www.ncbi.nlm.nih.gov/gene/?term=LOC105287597 | 2974 | 181 | 608.61 | 1 | 3.36 | 0.55 | 17.88 | 17.42 | 0.47 | 0.47 | 0.03 | 0.03 |
| gene5746 | https://www.ncbi.nlm.nih.gov/gene/?term=LOC105278666 | 10343 | 393 | 379.97 | 1 | 0.97 | 0.25 | 7.69 | 7.22 | 0.47 | 0.47 | 0.02 | 0.02 |
| gene940 | https://www.ncbi.nlm.nih.gov/gene/?term=LOC105282794 | 3578 | 84 | 234.77 | 1 | 2.79 | 1.19 | 14.53 | 14.06 | 0.47 | 0.47 | 0.01 | 0.01 |
| gene8729 | https://www.ncbi.nlm.nih.gov/gene/?term=LOC105282137 | 5005 | 200 | 399.6 | 1 | 2 | 0.5 | 15.61 | 15.22 | 0.39 | 0.39 | 0.02 | 0.02 |
| gene5524 | https://www.ncbi.nlm.nih.gov/gene/?term=LOC105278397 | 5177 | 199 | 384.39 | 1 | 1.93 | 0.5 | 24.51 | 24.16 | 0.35 | 0.35 | 0.01 | 0.01 |
| gene5831 | https://www.ncbi.nlm.nih.gov/gene/?term=LOC105278761 | 2506 | 99 | 395.05 | 1 | 3.99 | 1.01 | 25.53 | 25.18 | 0.35 | 0.35 | 0.01 | 0.01 |
| gene7600 | https://www.ncbi.nlm.nih.gov/gene/?term=LOC105280819 | 1615 | 41 | 253.87 | 1 | 6.19 | 2.44 | 40.74 | 40.39 | 0.35 | 0.35 | 0.01 | 0.01 |
| gene13114 | https://www.ncbi.nlm.nih.gov/gene/?term=LOC105287147 | 7609 | 370 | 486.27 | 1 | 1.31 | 0.27 | 9.54 | 9.22 | 0.32 | 0.32 | 0.02 | 0.02 |
| gene1147 | https://www.ncbi.nlm.nih.gov/gene/?term=LOC105284954 | 4022 | 87 | 216.31 | 1 | 2.49 | 1.15 | 14.67 | 14.36 | 0.3 | 0.3 | 0.01 | 0.01 |
| gene2173 | https://www.ncbi.nlm.nih.gov/gene/?term=LOC105274590 | 4387 | 78 | 177.8 | 1 | 2.28 | 1.28 | 22.31 | 22.01 | 0.3 | 0.3 | 0.01 | 0.01 |
| gene9101 | https://www.ncbi.nlm.nih.gov/gene/?term=LOC105282553 | 3068 | 103 | 335.72 | 1 | 3.26 | 0.97 | 20.28 | 19.99 | 0.29 | 0.29 | 0.01 | 0.01 |

| | | | | | | | | | | | | | | |
|---|---|---|---|---|---|---|---|---|---|---|---|---|---|---|
| gene13036 | https://www.ncbi.nlm.nih.gov/gene/?term=LOC105287059 | 4959 | 157 | 316.6 1 | 2.02 | 0.64 | 17.83 | 17.59 | 0.25 | 0.25 | 0.01 | 0.01 | | |
| gene10549 | https://www.ncbi.nlm.nih.gov/gene/?term=LOC105284210 | 7145 | 313 | 438.07 | 1 | 1.4 | 0.32 | 10.77 | 10.55 | 0.22 | 0.22 | 0.01 | 0.01 | |
| gene8274 | https://www.ncbi.nlm.nih.gov/gene/?term=LOC105281624 | 5425 | 189 | 348.39 | 1 | 1.84 | 0.53 | 11.18 | 10.97 | 0.21 | 0.21 | 0.01 | 0.01 | |
| gene12802 | https://www.ncbi.nlm.nih.gov/gene/?term=LOC105286799 | 3183 | 170 | 534.09 | 1 | 3.14 | 0.59 | 15.96 | 15.78 | 0.18 | 0.18 | 0.01 | 0.01 | |
| gene5525 | https://www.ncbi.nlm.nih.gov/gene/?term=LOC105278396 | 3090 | 185 | 598.71 | 1 | 3.24 | 0.54 | 24.64 | 24.46 | 0.18 | 0.18 | 0.01 | 0.01 | |
| gene1220 | https://www.ncbi.nlm.nih.gov/gene/?term=LOC105285742 | 3453 | 86 | 249.06 | 1 | 2.9 | 1.16 | 21.68 | 21.51 | 0.17 | 0.17 | 0 | 0 | |
| gene7247 | https://www.ncbi.nlm.nih.gov/gene/?term=LOC105280407 | 10486 | 206 | 196.45 | 1 | 0.95 | 0.49 | 17.14 | 16.97 | 0.17 | 0.17 | 0 | 0 | |
| gene4213 | https://www.ncbi.nlm.nih.gov/gene/?term=LOC105276901 | 3257 | 96 | 294.75 | 1 | 3.07 | 1.04 | 22.87 | 22.72 | 0.14 | 0.14 | 0 | 0 | |
| gene10107 | https://www.ncbi.nlm.nih.gov/gene/?term=LOC105283712 | 1883 | 29 | 154.01 | 1 | 5.31 | 3.45 | 27.22 | 27.09 | 0.13 | 0.13 | 0 | 0 | |
| gene9645 | https://www.ncbi.nlm.nih.gov/gene/?term=LOC105283181 | 3875 | 134 | 345.81 | 1 | 2.58 | 0.75 | 11.45 | 11.37 | 0.08 | 0.08 | 0 | 0 | |
| gene6620 | https://www.ncbi.nlm.nih.gov/gene/?term=LOC105279687 | 17998 | 354 | 196.69 | 1 | 0.56 | 0.28 | 5.08 | 5.05 | 0.03 | 0.03 | 0 | 0 | |
| gene11583 | https://www.ncbi.nlm.nih.gov/gene/?term=LOC105285414 | 8192 | 175 | 213.62 | 1 | 1.22 | 0.57 | 9.21 | 9.24 | -0.03 | 0.03 | 0 | 0 | |
| gene2806 | https://www.ncbi.nlm.nih.gov/gene/?term=LOC105275327 | 14257 | 462 | 324.05 | 1 | 0.7 | 0.22 | 7.37 | 7.4 | -0.03 | 0.03 | 0 | 0 | |
| gene11579 | https://www.ncbi.nlm.nih.gov/gene/?term=LOC105285406 | 13571 | 260 | 191.58 | 1 | 0.74 | 0.38 | 5.75 | 5.79 | -0.04 | 0.04 | 0 | 0 | |
| gene7166 | https://www.ncbi.nlm.nih.gov/gene/?term=LOC105280326 | 6117 | 208 | 340.04 | 1 | 1.63 | 0.48 | 16.25 | 16.28 | -0.04 | 0.04 | 0 | 0 | |
| gene12406 | https://www.ncbi.nlm.nih.gov/gene/?term=LOC105286316 | 1386 | 30 | 216.45 | 1 | 7.22 | 3.33 | 39.2 | 39.29 | -0.09 | 0.09 | 0 | 0 | |
| gene5375 | https://www.ncbi.nlm.nih.gov/gene/?term=LOC105278228 | 8640 | 148 | 171.3 1 | 1.16 | 0.68 | 24.78 | 24.89 | -0.11 | 0.11 | 0 | 0 | | |
| gene12573 | https://www.ncbi.nlm.nih.gov/gene/?term=LOC105286515 | 3647 | 105 | 287.91 | 1 | 2.74 | 0.95 | 10.51 | 10.65 | -0.14 | 0.14 | 0 | 0 | |
| gene1310 | https://www.ncbi.nlm.nih.gov/gene/?term=LOC105286661 | 2582 | 63 | 244 1 | 3.87 | 1.59 | 19.4 | 19.54 | -0.14 | 0.14 | 0 | 0 | | |
| gene10531 | https://www.ncbi.nlm.nih.gov/gene/?term=LOC105284190 | 6881 | 180 | 261.59 | 1 | 1.45 | 0.56 | 10.06 | 10.22 | -0.16 | 0.16 | 0 | 0 | |
| gene2443 | https://www.ncbi.nlm.nih.gov/gene/?term=LOC105274895 | 10433 | 330 | 316.3 1 | 0.96 | 0.3 | 5.47 | 5.65 | -0.18 | 0.18 | -0.01 | 0.01 | | |
| gene4892 | https://www.ncbi.nlm.nih.gov/gene/?term=LOC105277667 | 3898 | 116 | 297.59 | 1 | 2.57 | 0.86 | 28.81 | 28.99 | -0.19 | 0.19 | -0.01 | 0.01 | |
| gene1332 | https://www.ncbi.nlm.nih.gov/gene/?term=LOC105286872 | 2460 | 142 | 577.24 | 1 | 4.07 | 0.7 | 21.72 | 21.98 | -0.26 | 0.26 | -0.02 | 0.02 | |
| gene1632 | https://www.ncbi.nlm.nih.gov/gene/?term=LOC105287764 | 18061 | 334 | 184.93 | 1 | 0.55 | 0.3 | 2.71 | 2.98 | -0.27 | 0.27 | 0 | 0 | |
| gene7183 | https://www.ncbi.nlm.nih.gov/gene/?term=LOC105280348 | 3925 | 53 | 135.03 | 1 | 2.55 | 1.89 | 25.26 | 25.53 | -0.27 | 0.27 | 0 | 0 | |
| gene6455 | https://www.ncbi.nlm.nih.gov/gene/?term=LOC105279500 | 2612 | 29 | 111.03 | 1 | 3.83 | 3.45 | 48.01 | 48.34 | -0.33 | 0.33 | 0 | 0 | |
| gene3335 | https://www.ncbi.nlm.nih.gov/gene/?term=LOC105275898 | 4665 | 128 | 274.38 | 1 | 2.14 | 0.78 | 13.41 | 13.77 | -0.36 | 0.36 | -0.01 | 0.01 | |
| gene3856 | https://www.ncbi.nlm.nih.gov/gene/?term=LOC105276499 | 5236 | 188 | 359.05 | 1 | 1.91 | 0.53 | 10.14 | 10.5 | -0.36 | 0.36 | -0.01 | 0.01 | |
| gene6583 | https://www.ncbi.nlm.nih.gov/gene/?term=LOC105279649 | 7444 | 308 | 413.76 | 1 | 1.34 | 0.32 | 11.09 | 11.46 | -0.37 | 0.37 | -0.02 | 0.02 | |
| gene10018 | https://www.ncbi.nlm.nih.gov/gene/?term=LOC105283607 | 4414 | 194 | 439.51 | 1 | 2.27 | 0.52 | 15.27 | 15.67 | -0.4 | 0.4 | -0.02 | 0.02 | |
| gene11275 | https://www.ncbi.nlm.nih.gov/gene/?term=LOC105285055 | 4119 | 202 | 490.41 | 1 | 2.43 | 0.5 | 12.81 | 13.23 | -0.42 | 0.42 | -0.02 | 0.02 | |
| gene10243 | https://www.ncbi.nlm.nih.gov/gene/?term=LOC105283862 | 3239 | 38 | 117.32 | 1 | 3.09 | 2.63 | 30.66 | 31.1 | -0.44 | 0.44 | -0.01 | 0.01 | |
| gene12978 | https://www.ncbi.nlm.nih.gov/gene/?term=LOC105286997 | 2858 | 134 | 468.86 | 1 | 3.5 | 0.75 | 11.8 | 12.24 | -0.44 | 0.44 | -0.02 | 0.02 | |
| gene8600 | https://www.ncbi.nlm.nih.gov/gene/?term=LOC105282001 | 8513 | 148 | 173.85 | 1 | 1.17 | 0.68 | 9.45 | 9.9 | -0.45 | 0.45 | -0.01 | 0.01 | |
| gene10540 | https://www.ncbi.nlm.nih.gov/gene/?term=LOC105284200 | 5267 | 195 | 370.23 | 1 | 1.9 | 0.51 | 13.62 | 14.09 | -0.47 | 0.47 | -0.02 | 0.02 | |
| gene10520 | https://www.ncbi.nlm.nih.gov/gene/?term=LOC105284184 | 2451 | 80 | 326.4 1 | 4.08 | 1.25 | 34.85 | 35.33 | -0.48 | 0.48 | -0.02 | 0.02 | | |
| gene13033 | https://www.ncbi.nlm.nih.gov/gene/?term=LOC105287063 | 1877 | 45 | 239.74 | 1 | 5.33 | 2.22 | 34.91 | 35.5 | -0.59 | 0.59 | -0.01 | 0.01 | |
| gene4023 | https://www.ncbi.nlm.nih.gov/gene/?term=LOC105276692 | 1759 | 36 | 204.66 | 1 | 5.69 | 2.78 | 23.26 | 23.9 | -0.64 | 0.64 | -0.01 | 0.01 | |
| gene8343 | https://www.ncbi.nlm.nih.gov/gene/?term=LOC105281700 | 2172 | 45 | 207.18 | 1 | 4.6 | 2.22 | 38.72 | 39.37 | -0.65 | 0.65 | -0.01 | 0.01 | |
| gene2844 | https://www.ncbi.nlm.nih.gov/gene/?term=LOC105275363 | 4330 | 185 | 427.25 | 1 | 2.31 | 0.54 | 14.56 | 15.46 | -0.9 | 0.9 | -0.04 | 0.04 | |
| gene564 | https://www.ncbi.nlm.nih.gov/gene/?term=LOC105278478 | 2802 | 12 | 42.83 1 | 3.57 | 8.33 | 20.51 | 21.62 | -1.11 | 1.11 | 0 | 0 | | |
| gene6528 | https://www.ncbi.nlm.nih.gov/gene/?term=LOC105279591 | 7222 | 68 | 94.16 1 | 1.38 | 1.47 | 22.7 | 23.89 | -1.19 | 1.19 | -0.01 | 0.01 | | |
| gene1855 | https://www.ncbi.nlm.nih.gov/gene/?term=LOC105288036 | 2387 | 86 | 360.28 | 1 | 4.19 | 1.16 | 28.48 | 29.73 | -1.24 | 1.24 | -0.04 | 0.04 | |
| gene8873 | https://www.ncbi.nlm.nih.gov/gene/?term=LOC105282297 | 2055 | 78 | 379.56 | 1 | 4.87 | 1.28 | 33.62 | 34.98 | -1.37 | 1.37 | -0.05 | 0.05 | |
| gene570 | https://www.ncbi.nlm.nih.gov/gene/?term=LOC105278549 | 1673 | 63 | 376.57 | 1 | 5.98 | 1.59 | 41.36 | 42.83 | -1.47 | 1.47 | -0.06 | 0.06 | |
| gene5052 | https://www.ncbi.nlm.nih.gov/gene/?term=LOC105277856 | 3167 | 27 | 85.25 1 | 3.16 | 3.7 | 36.57 | 38.1 | -1.52 | 1.52 | -0.01 | 0.01 | | |
| gene9478 | https://www.ncbi.nlm.nih.gov/gene/?term=LOC105282991 | 3726 | 48 | 128.82 | 1 | 2.68 | 2.08 | 31.5 | 33.03 | -1.53 | 1.53 | -0.02 | 0.02 | |
| gene274 | https://www.ncbi.nlm.nih.gov/gene/?term=LOC105275190 | 3488 | 65 | 186.35 | 1 | 2.87 | 1.54 | 21.06 | 22.74 | -1.69 | 1.69 | -0.03 | 0.03 | |
| gene4034 | https://www.ncbi.nlm.nih.gov/gene/?term=LOC105276704 | 2065 | 80 | 387.41 | 1 | 4.84 | 1.25 | 13.38 | 15.07 | -1.69 | 1.69 | -0.07 | 0.07 | |
| gene1884 | https://www.ncbi.nlm.nih.gov/gene/?term=LOC105288072 | 4169 | 79 | 189.49 | 1 | 2.4 | 1.27 | 22.97 | 24.67 | -1.7 | 1.7 | -0.03 | 0.03 | |
| gene298 | https://www.ncbi.nlm.nih.gov/gene/?term=LOC105275460 | 2473 | 115 | 465.02 | 1 | 4.04 | 0.87 | 12.68 | 14.4 | -1.73 | 1.73 | -0.08 | 0.08 | |
| gene4641 | https://www.ncbi.nlm.nih.gov/gene/?term=LOC105277398 | 1937 | 70 | 361.38 | 1 | 5.16 | 1.43 | 35.36 | 37.39 | -2.03 | 2.03 | -0.07 | 0.07 | |
| gene10345 | https://www.ncbi.nlm.nih.gov/gene/?term=LOC105283989 | 4444 | 27 | 60.76 1 | 2.25 | 3.7 | 48.21 | 50.36 | -2.14 | 2.14 | -0.01 | 0.01 | | |
| gene6334 | https://www.ncbi.nlm.nih.gov/gene/?term=LOC105279356 | 1471 | 43 | 292.32 | 1 | 6.8 | 2.33 | 28.25 | 30.45 | -2.2 | 2.2 | -0.06 | 0.06 | |
| gene11819 | https://www.ncbi.nlm.nih.gov/gene/?term=LOC105285671 | 2366 | 90 | 380.39 | 1 | 4.23 | 1.11 | 28.49 | 30.76 | -2.27 | 2.27 | -0.09 | 0.09 | |
| gene5877 | https://www.ncbi.nlm.nih.gov/gene/?term=LOC105278807 | 1719 | 51 | 296.68 | 1 | 5.82 | 1.96 | 21.68 | 24.24 | -2.55 | 2.55 | -0.08 | 0.08 | |
| gene12516 | https://www.ncbi.nlm.nih.gov/gene/?term=LOC105286452 | 1529 | 35 | 228.91 | 1 | 6.54 | 2.86 | 51.01 | 53.56 | -2.56 | 2.56 | -0.06 | 0.06 | |
| gene2676 | https://www.ncbi.nlm.nih.gov/gene/?term=LOC105275181 | 1353 | 14 | 103.47 | 1 | 7.39 | 7.14 | 65.95 | 69.25 | -3.3 | 3.3 | -0.03 | 0.03 | |
| gene5101 | https://www.ncbi.nlm.nih.gov/gene/?term=LOC105277903 | 1813 | 34 | 187.53 | 1 | 5.52 | 2.94 | 52.57 | 56.73 | -4.16 | 4.16 | -0.08 | 0.08 | |
| gene5647 | https://www.ncbi.nlm.nih.gov/gene/?term=LOC105278567 | 3118 | 32 | 102.63 | 1 | 3.21 | 3.12 | 13.07 | 18.25 | -5.18 | 5.18 | -0.05 | 0.05 | |
| gene3165 | https://www.ncbi.nlm.nih.gov/gene/?term=LOC105275706 | 2374 | 59 | 248.53 | 4 | 16.85 | 6.78 | 23.46 | 34.48 | -11.02 | | 11.02 | -0.27 | 0.27 |
| gene2593 | https://www.ncbi.nlm.nih.gov/gene/?term=LOC105275070 | 1774 | 6 | 33.82 2 | 11.27 | 33.33 | 16.42 | 60.02 | -43.6 | 43.6 | -0.15 | 0.15 | | |

Directory: OGL File: ogl-Cb_LI_BCphaseA1.vs.RphaseA2.txt

| gene_ID | gene_link | gwidth | #Sites | #per10Kb | #dmSites | #dmsp10kb | %dmSites | %pSite1 | %pSite2 | DMpSite | ADMpSite | DMpNucl | ADMpNucl |
|---|---|---|---|---|---|---|---|---|---|---|---|---|---|
| gene393 | https://www.ncbi.nlm.nih.gov/gene/?term=LOC105276566 | 1123 | 2 | 17.81 | 1 | 8.9 | 50 | 77.65 | 47.62 | 30.03 | 30.03 | 0.05 | 0.05 |
| gene4630 | https://www.ncbi.nlm.nih.gov/gene/?term=LOC105277380 | 1893 | 5 | 26.41 | 1 | 5.28 | 20 | 69.42 | 53.65 | 15.77 | 15.77 | 0.04 | 0.04 |
| gene9109 | https://www.ncbi.nlm.nih.gov/gene/?term=LOC105282561 | 2400 | 15 | 62.5 | 1 | 4.17 | 6.67 | 34.05 | 29.4 | 4.65 | 4.65 | 0.03 | 0.03 |
| gene7952 | https://www.ncbi.nlm.nih.gov/gene/?term=LOC105281213 | 2963 | 43 | 145.12 | 1 | 3.37 | 2.33 | 42.11 | 38.78 | 3.33 | 3.33 | 0.05 | 0.05 |
| gene3200 | https://www.ncbi.nlm.nih.gov/gene/?term=LOC105275743 | 1786 | 77 | 431.13 | 1 | 5.6 | 1.3 | 22.67 | 19.88 | 2.79 | 2.79 | 0.12 | 0.12 |
| gene7368 | https://www.ncbi.nlm.nih.gov/gene/?term=LOC105280556 | 4105 | 49 | 119.37 | 1 | 2.44 | 2.04 | 12.57 | 9.91 | 2.66 | 2.66 | 0.03 | 0.03 |
| gene920 | https://www.ncbi.nlm.nih.gov/gene/?term=LOC105282518 | 1197 | 43 | 359.23 | 1 | 8.35 | 2.33 | 37.5 | 34.97 | 2.52 | 2.52 | 0.09 | 0.09 |
| gene7915 | https://www.ncbi.nlm.nih.gov/gene/?term=LOC105281173 | 1597 | 25 | 156.54 | 1 | 6.26 | 4 | 64.87 | 62.52 | 2.35 | 2.35 | 0.04 | 0.04 |
| gene3168 | https://www.ncbi.nlm.nih.gov/gene/?term=LOC105275711 | 6150 | 80 | 130.08 | 1 | 1.63 | 1.25 | 14.39 | 12.36 | 2.03 | 2.03 | 0.03 | 0.03 |
| gene10213 | https://www.ncbi.nlm.nih.gov/gene/?term=LOC105283826 | 1503 | 54 | 359.28 | 1 | 6.65 | 1.85 | 16.14 | 14.22 | 1.92 | 1.92 | 0.07 | 0.07 |
| gene1199 | https://www.ncbi.nlm.nih.gov/gene/?term=LOC105285498 | 2220 | 26 | 117.12 | 1 | 4.5 | 3.85 | 33.11 | 31.3 | 1.81 | 1.81 | 0.02 | 0.02 |
| gene7398 | https://www.ncbi.nlm.nih.gov/gene/?term=LOC105280589 | 3452 | 41 | 118.77 | 1 | 2.9 | 2.44 | 25.07 | 23.32 | 1.75 | 1.75 | 0.02 | 0.02 |
| gene11454 | https://www.ncbi.nlm.nih.gov/gene/?term=LOC105285256 | 1906 | 51 | 267.58 | 1 | 5.25 | 1.96 | 43.83 | 42.42 | 1.4 | 1.4 | 0.04 | 0.04 |
| gene3671 | https://www.ncbi.nlm.nih.gov/gene/?term=LOC105276272 | 1966 | 27 | 137.33 | 1 | 5.09 | 3.7 | 41.96 | 40.71 | 1.25 | 1.25 | 0.02 | 0.02 |
| gene322 | https://www.ncbi.nlm.nih.gov/gene/?term=LOC105275777 | 1606 | 23 | 143.21 | 1 | 6.23 | 4.35 | 45.55 | 44.33 | 1.21 | 1.21 | 0.02 | 0.02 |
| gene6237 | https://www.ncbi.nlm.nih.gov/gene/?term=LOC105279244 | 1484 | 26 | 175.2 | 1 | 6.74 | 3.85 | 46.7 | 45.51 | 1.19 | 1.19 | 0.02 | 0.02 |
| gene10326 | https://www.ncbi.nlm.nih.gov/gene/?term=LOC105283962 | 3042 | 102 | 335.31 | 1 | 3.29 | 0.98 | 14.13 | 13.04 | 1.09 | 1.09 | 0.04 | 0.04 |
| gene4190 | https://www.ncbi.nlm.nih.gov/gene/?term=LOC105276875 | 3150 | 132 | 419.05 | 1 | 3.17 | 0.76 | 21.24 | 20.15 | 1.09 | 1.09 | 0.05 | 0.05 |
| gene7345 | https://www.ncbi.nlm.nih.gov/gene/?term=LOC105280529 | 3833 | 92 | 240.02 | 1 | 2.61 | 1.09 | 17.99 | 16.93 | 1.05 | 1.05 | 0.03 | 0.03 |
| gene10819 | https://www.ncbi.nlm.nih.gov/gene/?term=LOC105284507 | 5767 | 83 | 143.92 | 1 | 1.73 | 1.2 | 21.4 | 20.36 | 1.04 | 1.04 | 0.02 | 0.02 |
| gene3286 | https://www.ncbi.nlm.nih.gov/gene/?term=LOC105275848 | 3744 | 132 | 352.56 | 1 | 2.67 | 0.76 | 21.71 | 20.73 | 0.98 | 0.98 | 0.03 | 0.03 |
| gene13429 | https://www.ncbi.nlm.nih.gov/gene/?term=LOC105287501 | 3714 | 102 | 274.64 | 1 | 2.69 | 0.98 | 22.78 | 21.81 | 0.97 | 0.97 | 0.03 | 0.03 |
| gene814 | https://www.ncbi.nlm.nih.gov/gene/?term=LOC105281296 | 2722 | 43 | 157.97 | 1 | 3.67 | 2.33 | 29.73 | 28.81 | 0.92 | 0.92 | 0.01 | 0.01 |
| gene5298 | https://www.ncbi.nlm.nih.gov/gene/?term=LOC105278145 | 4060 | 257 | 633 | 2 | 4.93 | 0.78 | 10.97 | 10.08 | 0.88 | 0.88 | 0.06 | 0.06 |
| gene11447 | https://www.ncbi.nlm.nih.gov/gene/?term=LOC105285252 | 7437 | 135 | 181.52 | 1 | 1.34 | 0.74 | 13.54 | 12.74 | 0.8 | 0.8 | 0.01 | 0.01 |
| gene1159 | https://www.ncbi.nlm.nih.gov/gene/?term=LOC105285084 | 3282 | 27 | 82.27 | 1 | 3.05 | 3.7 | 22 | 21.23 | 0.77 | 0.77 | 0.01 | 0.01 |
| gene652 | https://www.ncbi.nlm.nih.gov/gene/?term=LOC105279528 | 5806 | 44 | 75.78 | 1 | 1.72 | 2.27 | 16.35 | 15.59 | 0.76 | 0.76 | 0.01 | 0.01 |
| gene6966 | https://www.ncbi.nlm.nih.gov/gene/?term=LOC105280104 | 3747 | 125 | 333.6 | 1 | 2.67 | 0.8 | 14.94 | 14.21 | 0.73 | 0.73 | 0.02 | 0.02 |
| gene9722 | https://www.ncbi.nlm.nih.gov/gene/?term=LOC105283264 | 3951 | 88 | 222.73 | 1 | 2.53 | 1.14 | 24.81 | 24.09 | 0.72 | 0.72 | 0.02 | 0.02 |
| gene11340 | https://www.ncbi.nlm.nih.gov/gene/?term=LOC105285131 | 2918 | 102 | 349.55 | 1 | 3.43 | 0.98 | 14.35 | 13.65 | 0.7 | 0.7 | 0.02 | 0.02 |
| gene9464 | https://www.ncbi.nlm.nih.gov/gene/?term=LOC105282975 | 1450 | 25 | 172.41 | 1 | 6.9 | 4 | 41.13 | 40.46 | 0.67 | 0.67 | 0.01 | 0.01 |
| gene12573 | https://www.ncbi.nlm.nih.gov/gene/?term=LOC105286515 | 3647 | 115 | 315.33 | 1 | 2.74 | 0.87 | 11.99 | 11.33 | 0.66 | 0.66 | 0.02 | 0.02 |
| gene4376 | https://www.ncbi.nlm.nih.gov/gene/?term=LOC105277083 | 4307 | 96 | 222.89 | 1 | 2.32 | 1.04 | 25.73 | 25.08 | 0.66 | 0.66 | 0.01 | 0.01 |
| gene11401 | https://www.ncbi.nlm.nih.gov/gene/?term=LOC105285202 | 1841 | 50 | 271.59 | 1 | 5.43 | 2 | 21.11 | 20.46 | 0.65 | 0.65 | 0.02 | 0.02 |
| gene7600 | https://www.ncbi.nlm.nih.gov/gene/?term=LOC105280819 | 1615 | 41 | 253.87 | 1 | 6.19 | 2.44 | 40.74 | 40.08 | 0.65 | 0.65 | 0.02 | 0.02 |
| gene10155 | https://www.ncbi.nlm.nih.gov/gene/?term=LOC105283756 | 7030 | 97 | 137.98 | 1 | 1.42 | 1.03 | 6.83 | 6.25 | 0.59 | 0.59 | 0.01 | 0.01 |
| gene10714 | https://www.ncbi.nlm.nih.gov/gene/?term=LOC105284396 | 1966 | 57 | 289.93 | 1 | 5.09 | 1.75 | 38.23 | 37.64 | 0.59 | 0.59 | 0.02 | 0.02 |
| gene4962 | https://www.ncbi.nlm.nih.gov/gene/?term=LOC105277761 | 4645 | 150 | 322.93 | 1 | 2.15 | 0.67 | 14.77 | 14.2 | 0.57 | 0.57 | 0.02 | 0.02 |
| gene12203 | https://www.ncbi.nlm.nih.gov/gene/?term=LOC105286099 | 5967 | 78 | 130.72 | 1 | 1.68 | 1.28 | 11.28 | 10.77 | 0.51 | 0.51 | 0.01 | 0.01 |
| gene1809 | https://www.ncbi.nlm.nih.gov/gene/?term=LOC105287989 | 7010 | 73 | 104.14 | 1 | 1.43 | 1.37 | 23.7 | 23.28 | 0.42 | 0.42 | 0 | 0 |
| gene3107 | https://www.ncbi.nlm.nih.gov/gene/?term=LOC105275650 | 5103 | 151 | 295.9 | 1 | 1.96 | 0.66 | 17.08 | 16.7 | 0.38 | 0.38 | 0.01 | 0.01 |
| gene5646 | https://www.ncbi.nlm.nih.gov/gene/?term=LOC105278534 | 7475 | 186 | 248.83 | 1 | 1.34 | 0.54 | 18.03 | 17.67 | 0.36 | 0.36 | 0.01 | 0.01 |
| gene10392 | https://www.ncbi.nlm.nih.gov/gene/?term=LOC105284036 | 3216 | 83 | 258.08 | 2 | 6.22 | 2.41 | 23.78 | 23.42 | 0.35 | 0.35 | 0.01 | 0.01 |
| gene1220 | https://www.ncbi.nlm.nih.gov/gene/?term=LOC105285742 | 3453 | 99 | 286.71 | 1 | 2.9 | 1.01 | 25.65 | 25.32 | 0.33 | 0.33 | 0.01 | 0.01 |
| gene3848 | https://www.ncbi.nlm.nih.gov/gene/?term=LOC105276486 | 5899 | 121 | 205.12 | 1 | 1.7 | 0.83 | 23.22 | 22.92 | 0.3 | 0.3 | 0.01 | 0.01 |
| gene11583 | https://www.ncbi.nlm.nih.gov/gene/?term=LOC105285414 | 8192 | 177 | 216.06 | 1 | 1.22 | 0.56 | 8.82 | 8.53 | 0.29 | 0.29 | 0.01 | 0.01 |
| gene5831 | https://www.ncbi.nlm.nih.gov/gene/?term=LOC105278761 | 2506 | 104 | 415 | 1 | 3.99 | 0.96 | 25.24 | 24.96 | 0.28 | 0.28 | 0.01 | 0.01 |
| gene12307 | https://www.ncbi.nlm.nih.gov/gene/?term=LOC105286216 | 10313 | 167 | 161.93 | 1 | 0.97 | 0.6 | 15.75 | 15.52 | 0.23 | 0.23 | 0 | 0 |
| gene657 | https://www.ncbi.nlm.nih.gov/gene/?term=LOC105279667 | 6421 | 80 | 124.59 | 1 | 1.56 | 1.25 | 3.15 | 2.93 | 0.21 | 0.21 | 0 | 0 |
| gene13114 | https://www.ncbi.nlm.nih.gov/gene/?term=LOC105287147 | 7609 | 389 | 511.24 | 1 | 1.31 | 0.26 | 9.77 | 9.58 | 0.19 | 0.19 | 0.01 | 0.01 |
| gene5746 | https://www.ncbi.nlm.nih.gov/gene/?term=LOC105278666 | 10343 | 409 | 395.44 | 1 | 0.97 | 0.24 | 7.43 | 7.26 | 0.17 | 0.17 | 0.01 | 0.01 |
| gene940 | https://www.ncbi.nlm.nih.gov/gene/?term=LOC105282794 | 3578 | 88 | 245.95 | 1 | 2.79 | 1.14 | 15.65 | 15.49 | 0.16 | 0.16 | 0 | 0 |
| gene12674 | https://www.ncbi.nlm.nih.gov/gene/?term=LOC105286649 | 18283 | 680 | 371.93 | 1 | 0.55 | 0.15 | 1.31 | 1.16 | 0.15 | 0.15 | 0.01 | 0.01 |
| gene6031 | https://www.ncbi.nlm.nih.gov/gene/?term=LOC105279000 | 5691 | 185 | 325.07 | 1 | 1.76 | 0.54 | 13.85 | 13.69 | 0.15 | 0.15 | 0 | 0 |
| gene3599 | https://www.ncbi.nlm.nih.gov/gene/?term=LOC105276196 | 2955 | 56 | 189.51 | 1 | 3.38 | 1.79 | 40.12 | 39.99 | 0.13 | 0.13 | 0 | 0 |
| gene2404 | https://www.ncbi.nlm.nih.gov/gene/?term=LOC105274853 | 1767 | 42 | 237.69 | 1 | 5.66 | 2.38 | 28.85 | 28.79 | 0.07 | 0.07 | 0 | 0 |
| gene2991 | https://www.ncbi.nlm.nih.gov/gene/?term=LOC105275536 | 11224 | 214 | 190.66 | 1 | 0.89 | 0.47 | 5.64 | 5.58 | 0.05 | 0.05 | 0 | 0 |
| gene4527 | https://www.ncbi.nlm.nih.gov/gene/?term=LOC105277273 | 3686 | 151 | 409.66 | 1 | 2.71 | 0.66 | 12.71 | 12.65 | 0.05 | 0.05 | 0 | 0 |
| gene4996 | https://www.ncbi.nlm.nih.gov/gene/?term=LOC105277798 | 5617 | 64 | 113.94 | 1 | 1.78 | 1.56 | 10.86 | 10.81 | 0.05 | 0.05 | 0 | 0 |
| gene2266 | https://www.ncbi.nlm.nih.gov/gene/?term=LOC105274699 | 2391 | 63 | 263.49 | 1 | 4.18 | 1.59 | 23.04 | 22.99 | 0.04 | 0.04 | 0 | 0 |

| | | | | | | | | | | | | | |
|---|---|---|---|---|---|---|---|---|---|---|---|---|---|
| gene7148 | https://www.ncbi.nlm.nih.gov/gene/?term=LOC105280304 | 8920 | 123 | 137.89 | 1 | 1.12 | 0.81 | 5.79 | 5.75 | 0.04 | 0.04 | 0 | 0 |
| gene53 | https://www.ncbi.nlm.nih.gov/gene/?term=LOC105277148 | 19283 | 269 | 139.5 | 1 | 0.52 | 0.37 | 3.9 | 3.86 | 0.03 | 0.03 | 0 | 0 |
| gene9645 | https://www.ncbi.nlm.nih.gov/gene/?term=LOC105283181 | 3875 | 138 | 356.13 | 1 | 2.58 | 0.72 | 11.99 | 11.98 | 0.01 | 0.01 | 0 | 0 |
| gene10549 | https://www.ncbi.nlm.nih.gov/gene/?term=LOC105284210 | 7145 | 323 | 452.06 | 1 | 1.4 | 0.31 | 10.47 | 10.47 | 0 | 0 | 0 | 0 |
| gene1470 | https://www.ncbi.nlm.nih.gov/gene/?term=LOC105287597 | 2974 | 192 | 645.6 | 1 | 3.36 | 0.52 | 16.86 | 16.86 | 0 | 0 | 0 | 0 |
| gene9755 | https://www.ncbi.nlm.nih.gov/gene/?term=LOC105283296 | 2177 | 90 | 413.41 | 1 | 4.59 | 1.11 | 19.36 | 19.36 | 0 | 0 | 0 | 0 |
| gene9411 | https://www.ncbi.nlm.nih.gov/gene/?term=LOC105282914 | 5534 | 41 | 74.09 | 1 | 1.81 | 2.44 | 49.68 | 49.72 | -0.04 | 0.04 | 0 | 0 |
| gene13036 | https://www.ncbi.nlm.nih.gov/gene/?term=LOC105287059 | 4959 | 165 | 332.73 | 1 | 2.02 | 0.61 | 17.35 | 17.4 | -0.05 | 0.05 | 0 | 0 |
| gene7207 | https://www.ncbi.nlm.nih.gov/gene/?term=LOC105280376 | 4659 | 105 | 225.37 | 1 | 2.15 | 0.95 | 28.77 | 28.82 | -0.05 | 0.05 | 0 | 0 |
| gene10606 | https://www.ncbi.nlm.nih.gov/gene/?term=LOC105284273 | 6119 | 252 | 411.83 | 2 | 3.27 | 0.79 | 19.91 | 19.99 | -0.08 | 0.08 | 0 | 0 |
| gene11568 | https://www.ncbi.nlm.nih.gov/gene/?term=LOC105285395 | 8086 | 184 | 227.55 | 1 | 1.24 | 0.54 | 16.42 | 16.51 | -0.08 | 0.08 | 0 | 0 |
| gene3470 | https://www.ncbi.nlm.nih.gov/gene/?term=LOC105276059 | 8716 | 161 | 184.72 | 1 | 1.15 | 0.62 | 9.12 | 9.21 | -0.08 | 0.08 | 0 | 0 |
| gene8274 | https://www.ncbi.nlm.nih.gov/gene/?term=LOC105281624 | 5425 | 208 | 383.41 | 1 | 1.84 | 0.48 | 10.51 | 10.6 | -0.09 | 0.09 | 0 | 0 |
| gene2748 | https://www.ncbi.nlm.nih.gov/gene/?term=LOC105275261 | 3845 | 58 | 150.85 | 1 | 2.6 | 1.72 | 39.98 | 40.08 | -0.1 | 0.1 | 0 | 0 |
| gene1632 | https://www.ncbi.nlm.nih.gov/gene/?term=LOC105287764 | 18061 | 354 | 196 | 1 | 0.55 | 0.28 | 2.69 | 2.8 | -0.11 | 0.11 | 0 | 0 |
| gene10596 | https://www.ncbi.nlm.nih.gov/gene/?term=LOC105284262 | 6520 | 235 | 360.43 | 1 | 1.53 | 0.43 | 3.69 | 3.81 | -0.12 | 0.12 | 0 | 0 |
| gene10864 | https://www.ncbi.nlm.nih.gov/gene/?term=LOC105284557 | 1666 | 78 | 468.19 | 1 | 6 | 1.28 | 30.45 | 30.59 | -0.13 | 0.13 | -0.01 | 0.01 |
| gene5649 | https://www.ncbi.nlm.nih.gov/gene/?term=LOC105278538 | 2140 | 84 | 392.52 | 1 | 4.67 | 1.19 | 18.7 | 18.83 | -0.13 | 0.13 | -0.01 | 0.01 |
| gene2683 | https://www.ncbi.nlm.nih.gov/gene/?term=LOC105275187 | 5492 | 75 | 136.56 | 1 | 1.82 | 1.33 | 14.14 | 14.34 | -0.2 | 0.2 | 0 | 0 |
| gene6583 | https://www.ncbi.nlm.nih.gov/gene/?term=LOC105279649 | 7444 | 330 | 443.31 | 1 | 1.34 | 0.3 | 11.47 | 11.66 | -0.2 | 0.2 | -0.01 | 0.01 |
| gene11579 | https://www.ncbi.nlm.nih.gov/gene/?term=LOC105285406 | 13571 | 293 | 215.9 | 1 | 0.74 | 0.34 | 5.86 | 6.07 | -0.21 | 0.21 | 0 | 0 |
| gene2806 | https://www.ncbi.nlm.nih.gov/gene/?term=LOC105275327 | 14257 | 480 | 336.68 | 1 | 0.7 | 0.21 | 7.1 | 7.32 | -0.22 | 0.22 | -0.01 | 0.01 |
| gene7249 | https://www.ncbi.nlm.nih.gov/gene/?term=LOC105280422 | 1775 | 61 | 343.66 | 1 | 5.63 | 1.64 | 15.16 | 15.4 | -0.24 | 0.24 | -0.01 | 0.01 |
| gene9101 | https://www.ncbi.nlm.nih.gov/gene/?term=LOC105282553 | 3068 | 104 | 338.98 | 1 | 3.26 | 0.96 | 20.36 | 20.64 | -0.27 | 0.27 | -0.01 | 0.01 |
| gene5613 | https://www.ncbi.nlm.nih.gov/gene/?term=LOC105278504 | 8524 | 211 | 247.54 | 1 | 1.17 | 0.47 | 5.46 | 5.74 | -0.28 | 0.28 | -0.01 | 0.01 |
| gene6620 | https://www.ncbi.nlm.nih.gov/gene/?term=LOC105279687 | 17998 | 360 | 200.02 | 1 | 0.56 | 0.28 | 5.06 | 5.34 | -0.28 | 0.28 | -0.01 | 0.01 |
| gene1573 | https://www.ncbi.nlm.nih.gov/gene/?term=LOC105287704 | 3030 | 131 | 432.34 | 1 | 3.3 | 0.76 | 20.77 | 21.09 | -0.32 | 0.32 | -0.01 | 0.01 |
| gene2056 | https://www.ncbi.nlm.nih.gov/gene/?term=LOC105274452 | 6584 | 212 | 321.99 | 1 | 1.52 | 0.47 | 13.94 | 14.26 | -0.33 | 0.33 | -0.01 | 0.01 |
| gene1147 | https://www.ncbi.nlm.nih.gov/gene/?term=LOC105284954 | 4022 | 82 | 203.88 | 1 | 2.49 | 1.22 | 15.94 | 16.29 | -0.34 | 0.34 | -0.01 | 0.01 |
| gene6286 | https://www.ncbi.nlm.nih.gov/gene/?term=LOC105279298 | 2861 | 139 | 485.84 | 2 | 6.99 | 1.44 | 22.43 | 22.79 | -0.36 | 0.36 | -0.02 | 0.02 |
| gene6639 | https://www.ncbi.nlm.nih.gov/gene/?term=LOC105279705 | 7656 | 69 | 90.13 | 1 | 1.31 | 1.45 | 21.96 | 22.32 | -0.36 | 0.36 | 0 | 0 |
| gene10600 | https://www.ncbi.nlm.nih.gov/gene/?term=LOC105284270 | 4837 | 105 | 217.08 | 1 | 2.07 | 0.95 | 12.85 | 13.22 | -0.37 | 0.37 | -0.01 | 0.01 |
| gene3856 | https://www.ncbi.nlm.nih.gov/gene/?term=LOC105276499 | 5236 | 201 | 383.88 | 1 | 1.91 | 0.5 | 10.34 | 10.72 | -0.39 | 0.39 | -0.01 | 0.01 |
| gene7568 | https://www.ncbi.nlm.nih.gov/gene/?term=LOC105280786 | 2525 | 123 | 487.13 | 1 | 3.96 | 0.81 | 28.92 | 29.32 | -0.39 | 0.39 | -0.02 | 0.02 |
| gene10520 | https://www.ncbi.nlm.nih.gov/gene/?term=LOC105284184 | 2451 | 82 | 334.56 | 1 | 4.08 | 1.22 | 34.55 | 34.95 | -0.4 | 0.4 | -0.01 | 0.01 |
| gene7888 | https://www.ncbi.nlm.nih.gov/gene/?term=LOC105281141 | 4357 | 165 | 378.7 | 1 | 2.3 | 0.61 | 28.04 | 28.47 | -0.43 | 0.43 | -0.02 | 0.02 |
| gene5375 | https://www.ncbi.nlm.nih.gov/gene/?term=LOC105278228 | 8640 | 159 | 184.03 | 1 | 1.16 | 0.63 | 24.17 | 24.62 | -0.45 | 0.45 | -0.01 | 0.01 |
| gene4892 | https://www.ncbi.nlm.nih.gov/gene/?term=LOC105277667 | 3898 | 122 | 312.98 | 1 | 2.57 | 0.82 | 27.68 | 28.18 | -0.5 | 0.5 | -0.02 | 0.02 |
| gene7550 | https://www.ncbi.nlm.nih.gov/gene/?term=LOC105280763 | 2173 | 98 | 450.99 | 1 | 4.6 | 1.02 | 26.91 | 27.42 | -0.51 | 0.51 | -0.02 | 0.02 |
| gene3885 | https://www.ncbi.nlm.nih.gov/gene/?term=LOC105276527 | 7928 | 352 | 444 | 1 | 1.26 | 0.28 | 8.21 | 8.74 | -0.52 | 0.52 | -0.02 | 0.02 |
| gene4225 | https://www.ncbi.nlm.nih.gov/gene/?term=LOC105276917 | 2443 | 26 | 106.43 | 1 | 4.09 | 3.85 | 36.73 | 37.26 | -0.53 | 0.53 | -0.01 | 0.01 |
| gene5967 | https://www.ncbi.nlm.nih.gov/gene/?term=LOC105278918 | 4393 | 135 | 307.31 | 1 | 2.28 | 0.74 | 16.02 | 16.55 | -0.53 | 0.53 | -0.02 | 0.02 |
| gene9330 | https://www.ncbi.nlm.nih.gov/gene/?term=LOC105282825 | 3936 | 90 | 228.66 | 1 | 2.54 | 1.11 | 15.33 | 15.86 | -0.53 | 0.53 | -0.01 | 0.01 |
| gene7166 | https://www.ncbi.nlm.nih.gov/gene/?term=LOC105280326 | 6117 | 213 | 348.21 | 1 | 1.63 | 0.47 | 15.89 | 16.43 | -0.54 | 0.54 | -0.02 | 0.02 |
| gene474 | https://www.ncbi.nlm.nih.gov/gene/?term=LOC105277435 | 3625 | 158 | 435.86 | 1 | 2.76 | 0.63 | 12.73 | 13.28 | -0.55 | 0.55 | -0.02 | 0.02 |
| gene8600 | https://www.ncbi.nlm.nih.gov/gene/?term=LOC105282001 | 8513 | 158 | 185.6 | 1 | 1.17 | 0.63 | 9.54 | 10.09 | -0.55 | 0.55 | -0.01 | 0.01 |
| gene7245 | https://www.ncbi.nlm.nih.gov/gene/?term=LOC105280410 | 12478 | 260 | 208.37 | 1 | 0.8 | 0.38 | 19.03 | 19.62 | -0.59 | 0.59 | -0.01 | 0.01 |
| gene10678 | https://www.ncbi.nlm.nih.gov/gene/?term=LOC105284355 | 4031 | 164 | 406.85 | 1 | 2.48 | 0.61 | 19.18 | 19.78 | -0.6 | 0.6 | -0.02 | 0.02 |
| gene3385 | https://www.ncbi.nlm.nih.gov/gene/?term=LOC105275966 | 3305 | 59 | 178.52 | 1 | 3.03 | 1.69 | 50.29 | 50.9 | -0.61 | 0.61 | -0.01 | 0.01 |
| gene10379 | https://www.ncbi.nlm.nih.gov/gene/?term=LOC105284028 | 6974 | 74 | 106.11 | 1 | 1.43 | 1.35 | 8.99 | 9.61 | -0.62 | 0.62 | -0.01 | 0.01 |
| gene11275 | https://www.ncbi.nlm.nih.gov/gene/?term=LOC105285055 | 4119 | 216 | 524.4 | 1 | 2.43 | 0.46 | 13.08 | 13.7 | -0.62 | 0.62 | -0.03 | 0.03 |
| gene12802 | https://www.ncbi.nlm.nih.gov/gene/?term=LOC105286799 | 3183 | 172 | 540.37 | 1 | 3.14 | 0.58 | 15.78 | 16.4 | -0.62 | 0.62 | -0.03 | 0.03 |
| gene388 | https://www.ncbi.nlm.nih.gov/gene/?term=LOC105276538 | 1558 | 20 | 128.37 | 2 | 12.84 | 10 | 70.94 | 71.62 | -0.68 | 0.68 | -0.01 | 0.01 |
| gene3817 | https://www.ncbi.nlm.nih.gov/gene/?term=LOC105276449 | 1230 | 48 | 390.24 | 1 | 8.13 | 2.08 | 26.73 | 27.42 | -0.69 | 0.69 | -0.03 | 0.03 |
| gene10018 | https://www.ncbi.nlm.nih.gov/gene/?term=LOC105283607 | 4414 | 198 | 448.57 | 1 | 2.27 | 0.51 | 15.25 | 15.95 | -0.7 | 0.7 | -0.03 | 0.03 |
| gene2271 | https://www.ncbi.nlm.nih.gov/gene/?term=LOC105274706 | 1448 | 53 | 366.02 | 1 | 6.91 | 1.89 | 36.91 | 37.62 | -0.71 | 0.71 | -0.03 | 0.03 |
| gene8661 | https://www.ncbi.nlm.nih.gov/gene/?term=LOC105282063 | 1926 | 43 | 223.26 | 1 | 5.19 | 2.33 | 23.84 | 24.54 | -0.71 | 0.71 | -0.02 | 0.02 |
| gene3206 | https://www.ncbi.nlm.nih.gov/gene/?term=LOC105275752 | 3474 | 125 | 359.82 | 1 | 2.88 | 0.8 | 28.34 | 29.06 | -0.72 | 0.72 | -0.03 | 0.03 |
| gene927 | https://www.ncbi.nlm.nih.gov/gene/?term=LOC105282600 | 3727 | 65 | 174.4 | 1 | 2.68 | 1.54 | 26.3 | 27.02 | -0.73 | 0.73 | -0.01 | 0.01 |
| gene2844 | https://www.ncbi.nlm.nih.gov/gene/?term=LOC105275363 | 4330 | 192 | 443.42 | 1 | 2.31 | 0.52 | 14.13 | 14.88 | -0.75 | 0.75 | -0.03 | 0.03 |
| gene7556 | https://www.ncbi.nlm.nih.gov/gene/?term=LOC105280774 | 2792 | 117 | 419.05 | 1 | 3.58 | 0.85 | 27.44 | 28.23 | -0.79 | 0.79 | -0.03 | 0.03 |
| gene2173 | https://www.ncbi.nlm.nih.gov/gene/?term=LOC105274590 | 4387 | 84 | 191.47 | 1 | 2.28 | 1.19 | 20.89 | 21.7 | -0.81 | 0.81 | -0.02 | 0.02 |
| gene9598 | https://www.ncbi.nlm.nih.gov/gene/?term=LOC105283132 | 1769 | 54 | 305.26 | 1 | 5.65 | 1.85 | 40.78 | 41.59 | -0.81 | 0.81 | -0.02 | 0.02 |
| gene2330 | https://www.ncbi.nlm.nih.gov/gene/?term=LOC105274773 | 13104 | 172 | 131.26 | 1 | 0.76 | 0.58 | 15.95 | 16.78 | -0.83 | 0.83 | -0.01 | 0.01 |

| | | | | | | | | | | | | | |
|---|---|---|---|---|---|---|---|---|---|---|---|---|---|
| gene8729 | https://www.ncbi.nlm.nih.gov/gene/?term=LOC105282137 | 5005 | 205 | 409.59 | 1 | 2 | 0.49 | 15.76 | 16.6 | -0.84 | 0.84 | -0.03 | 0.03 |
| gene6528 | https://www.ncbi.nlm.nih.gov/gene/?term=LOC105279591 | 7222 | 75 | 103.85 | 1 | 1.38 | 1.33 | 20.71 | 21.6 | -0.89 | 0.89 | -0.01 | 0.01 |
| gene4641 | https://www.ncbi.nlm.nih.gov/gene/?term=LOC105277398 | 1937 | 70 | 361.38 | 1 | 5.16 | 1.43 | 35.36 | 36.26 | -0.9 | 0.9 | -0.03 | 0.03 |
| gene10540 | https://www.ncbi.nlm.nih.gov/gene/?term=LOC105284200 | 5267 | 199 | 377.82 | 1 | 1.9 | 0.5 | 13.8 | 14.75 | -0.95 | 0.95 | -0.04 | 0.04 |
| gene10345 | https://www.ncbi.nlm.nih.gov/gene/?term=LOC105283989 | 4444 | 30 | 67.51 | 1 | 2.25 | 3.33 | 46.58 | 47.61 | -1.03 | 1.03 | -0.01 | 0.01 |
| gene10531 | https://www.ncbi.nlm.nih.gov/gene/?term=LOC105284190 | 6881 | 199 | 289.2 | 1 | 1.45 | 0.5 | 9.56 | 10.66 | -1.1 | 1.1 | -0.03 | 0.03 |
| gene5524 | https://www.ncbi.nlm.nih.gov/gene/?term=LOC105278397 | 5177 | 200 | 386.32 | 1 | 1.93 | 0.5 | 24.39 | 25.66 | -1.27 | 1.27 | -0.05 | 0.05 |
| gene298 | https://www.ncbi.nlm.nih.gov/gene/?term=LOC105275460 | 2473 | 112 | 452.89 | 1 | 4.04 | 0.89 | 12.88 | 14.18 | -1.3 | 1.3 | -0.06 | 0.06 |
| gene4070 | https://www.ncbi.nlm.nih.gov/gene/?term=LOC105276745 | 3957 | 72 | 181.96 | 1 | 2.53 | 1.39 | 20.98 | 22.28 | -1.3 | 1.3 | -0.02 | 0.02 |
| gene9336 | https://www.ncbi.nlm.nih.gov/gene/?term=LOC105282834 | 5993 | 109 | 181.88 | 1 | 1.67 | 0.92 | 15.48 | 16.78 | -1.31 | 1.31 | -0.02 | 0.02 |
| gene1962 | https://www.ncbi.nlm.nih.gov/gene/?term=LOC105288157 | 1246 | 16 | 128.41 | 1 | 8.03 | 6.25 | 13.95 | 15.31 | -1.36 | 1.36 | -0.02 | 0.02 |
| gene13033 | https://www.ncbi.nlm.nih.gov/gene/?term=LOC105287063 | 1877 | 45 | 239.74 | 1 | 5.33 | 2.22 | 34.91 | 36.28 | -1.37 | 1.37 | -0.03 | 0.03 |
| gene7183 | https://www.ncbi.nlm.nih.gov/gene/?term=LOC105280348 | 3925 | 53 | 135.03 | 1 | 2.55 | 1.89 | 25.26 | 26.63 | -1.37 | 1.37 | -0.02 | 0.02 |
| gene8178 | https://www.ncbi.nlm.nih.gov/gene/?term=LOC105281515 | 8647 | 46 | 53.2 | 1 | 1.16 | 2.17 | 28.18 | 29.56 | -1.38 | 1.38 | -0.01 | 0.01 |
| gene1884 | https://www.ncbi.nlm.nih.gov/gene/?term=LOC105288072 | 4169 | 82 | 196.69 | 1 | 2.4 | 1.22 | 22.32 | 23.72 | -1.41 | 1.41 | -0.03 | 0.03 |
| gene2257 | https://www.ncbi.nlm.nih.gov/gene/?term=LOC105274689 | 3658 | 75 | 205.03 | 1 | 2.73 | 1.33 | 12.35 | 13.76 | -1.41 | 1.41 | -0.03 | 0.03 |
| gene5877 | https://www.ncbi.nlm.nih.gov/gene/?term=LOC105278807 | 1719 | 52 | 302.5 | 1 | 5.82 | 1.92 | 22.91 | 24.37 | -1.46 | 1.46 | -0.04 | 0.04 |
| gene8343 | https://www.ncbi.nlm.nih.gov/gene/?term=LOC105281700 | 2172 | 46 | 211.79 | 1 | 4.6 | 2.17 | 39.86 | 41.32 | -1.46 | 1.46 | -0.03 | 0.03 |
| gene5525 | https://www.ncbi.nlm.nih.gov/gene/?term=LOC105278396 | 3090 | 186 | 601.94 | 1 | 3.24 | 0.54 | 24.51 | 25.99 | -1.48 | 1.48 | -0.09 | 0.09 |
| gene6334 | https://www.ncbi.nlm.nih.gov/gene/?term=LOC105279356 | 1471 | 43 | 292.32 | 1 | 6.8 | 2.33 | 28.25 | 29.87 | -1.62 | 1.62 | -0.05 | 0.05 |
| gene1310 | https://www.ncbi.nlm.nih.gov/gene/?term=LOC105286661 | 2582 | 65 | 251.74 | 1 | 3.87 | 1.54 | 19.96 | 21.59 | -1.63 | 1.63 | -0.04 | 0.04 |
| gene4023 | https://www.ncbi.nlm.nih.gov/gene/?term=LOC105276692 | 1759 | 39 | 221.72 | 1 | 5.69 | 2.56 | 19.85 | 21.5 | -1.65 | 1.65 | -0.04 | 0.04 |
| gene6455 | https://www.ncbi.nlm.nih.gov/gene/?term=LOC105279500 | 2612 | 32 | 122.51 | 1 | 3.83 | 3.12 | 49.77 | 51.44 | -1.68 | 1.68 | -0.02 | 0.02 |
| gene11819 | https://www.ncbi.nlm.nih.gov/gene/?term=LOC105285671 | 2366 | 95 | 401.52 | 1 | 4.23 | 1.05 | 27.1 | 28.85 | -1.76 | 1.76 | -0.07 | 0.07 |
| gene5402 | https://www.ncbi.nlm.nih.gov/gene/?term=LOC105278260 | 3068 | 41 | 133.64 | 1 | 3.26 | 2.44 | 13.01 | 14.77 | -1.77 | 1.77 | -0.02 | 0.02 |
| gene10984 | https://www.ncbi.nlm.nih.gov/gene/?term=LOC105284704 | 1987 | 64 | 322.09 | 1 | 5.03 | 1.56 | 25.47 | 27.27 | -1.8 | 1.8 | -0.06 | 0.06 |
| gene12317 | https://www.ncbi.nlm.nih.gov/gene/?term=LOC105286223 | 2235 | 84 | 375.84 | 1 | 4.47 | 1.19 | 29.87 | 31.79 | -1.93 | 1.93 | -0.07 | 0.07 |
| gene10243 | https://www.ncbi.nlm.nih.gov/gene/?term=LOC105283862 | 3239 | 37 | 114.23 | 1 | 3.09 | 2.7 | 31.49 | 33.45 | -1.96 | 1.96 | -0.02 | 0.02 |
| gene3727 | https://www.ncbi.nlm.nih.gov/gene/?term=LOC105276367 | 2484 | 54 | 217.39 | 1 | 4.03 | 1.85 | 48.38 | 50.37 | -1.99 | 1.99 | -0.04 | 0.04 |
| gene2970 | https://www.ncbi.nlm.nih.gov/gene/?term=LOC105275518 | 2340 | 21 | 89.74 | 1 | 4.27 | 4.76 | 45.1 | 47.46 | -2.36 | 2.36 | -0.02 | 0.02 |
| gene12021 | https://www.ncbi.nlm.nih.gov/gene/?term=LOC105285905 | 1399 | 23 | 164.4 | 1 | 7.15 | 4.35 | 41.24 | 43.7 | -2.46 | 2.46 | -0.04 | 0.04 |
| gene12406 | https://www.ncbi.nlm.nih.gov/gene/?term=LOC105286316 | 1386 | 34 | 245.31 | 1 | 7.22 | 2.94 | 39.04 | 41.82 | -2.78 | 2.78 | -0.07 | 0.07 |
| gene4034 | https://www.ncbi.nlm.nih.gov/gene/?term=LOC105276704 | 2065 | 87 | 421.31 | 2 | 9.69 | 2.3 | 12.36 | 15.2 | -2.84 | 2.84 | -0.12 | 0.12 |
| gene12558 | https://www.ncbi.nlm.nih.gov/gene/?term=LOC105286502 | 1285 | 43 | 334.63 | 1 | 7.78 | 2.33 | 24.55 | 27.41 | -2.86 | 2.86 | -0.1 | 0.1 |
| gene5647 | https://www.ncbi.nlm.nih.gov/gene/?term=LOC105278567 | 3118 | 37 | 118.67 | 1 | 3.21 | 2.7 | 11.3 | 14.2 | -2.9 | 2.9 | -0.03 | 0.03 |
| gene12516 | https://www.ncbi.nlm.nih.gov/gene/?term=LOC105286452 | 1529 | 35 | 228.91 | 1 | 6.54 | 2.86 | 51.01 | 54.04 | -3.03 | 3.03 | -0.07 | 0.07 |
| gene10107 | https://www.ncbi.nlm.nih.gov/gene/?term=LOC105283712 | 1883 | 27 | 143.39 | 1 | 5.31 | 3.7 | 26.21 | 29.43 | -3.23 | 3.23 | -0.05 | 0.05 |
| gene2335 | https://www.ncbi.nlm.nih.gov/gene/?term=LOC105274778 | 3945 | 38 | 96.32 | 1 | 2.53 | 2.63 | 45.06 | 48.85 | -3.79 | 3.79 | -0.04 | 0.04 |
| gene274 | https://www.ncbi.nlm.nih.gov/gene/?term=LOC105275190 | 3488 | 73 | 209.29 | 1 | 2.87 | 1.37 | 18.68 | 22.58 | -3.9 | 3.9 | -0.08 | 0.08 |
| gene12961 | https://www.ncbi.nlm.nih.gov/gene/?term=LOC105286983 | 3094 | 70 | 226.24 | 1 | 3.23 | 1.43 | 0.98 | 5.77 | -4.79 | 4.79 | -0.11 | 0.11 |
| gene5052 | https://www.ncbi.nlm.nih.gov/gene/?term=LOC105277856 | 3167 | 28 | 88.41 | 1 | 3.16 | 3.57 | 35.73 | 41.1 | -5.37 | 5.37 | -0.05 | 0.05 |
| gene5714 | https://www.ncbi.nlm.nih.gov/gene/?term=LOC105278729 | 710 | 14 | 197.18 | 1 | 14.08 | 7.14 | 51.67 | 57.2 | -5.53 | 5.53 | -0.11 | 0.11 |
| gene6814 | https://www.ncbi.nlm.nih.gov/gene/?term=LOC105279898 | 1343 | 15 | 111.69 | 1 | 7.45 | 6.67 | 59.84 | 67.04 | -7.2 | 7.2 | -0.08 | 0.08 |
| gene3165 | https://www.ncbi.nlm.nih.gov/gene/?term=LOC105275706 | 2374 | 67 | 282.22 | 4 | 16.85 | 5.97 | 25.11 | 35.96 | -10.84 | 10.84 | -0.31 | 0.31 |
| gene2593 | https://www.ncbi.nlm.nih.gov/gene/?term=LOC105275070 | 1774 | 6 | 33.82 | 3 | 16.91 | 50 | 16.42 | 57.88 | -41.46 | 41.46 | -0.14 | 0.14 |

Directory: OGL File: ogl-Cb_LI_BCphaseA1.vs.RphaseB1.txt

| gene_ID | gene_link | gwidth | #Sites | #per10Kb | #dmSites | #dmsp10kb | %dmSites | %pSite1 | %pSite2 | DMpSite | ADMpSite | DMpNucl | ADMpNucl |
|---|---|---|---|---|---|---|---|---|---|---|---|---|---|
| gene10784 | https://www.ncbi.nlm.nih.gov/gene/?term=LOC105284474 | 481 | 1 | 20.79 | 1 | 20.79 | 100 | 66.67 | 38.1 | 28.57 | 28.57 | 0.06 | 0.06 |
| gene2022 | https://www.ncbi.nlm.nih.gov/gene/?term=LOC105288227 | 1245 | 3 | 24.1 | 1 | 8.03 | 33.33 | 86.21 | 76.94 | 9.26 | 9.26 | 0.02 | 0.02 |
| gene10237 | https://www.ncbi.nlm.nih.gov/gene/?term=LOC105283856 | 1259 | 25 | 198.57 | 5 | 39.71 | 20 | 64.65 | 55.66 | 9 | 9 | 0.18 | 0.18 |
| gene12003 | https://www.ncbi.nlm.nih.gov/gene/?term=LOC105285885 | 1336 | 10 | 74.85 | 1 | 7.49 | 10 | 64.94 | 57.31 | 7.63 | 7.63 | 0.06 | 0.06 |
| gene4643 | https://www.ncbi.nlm.nih.gov/gene/?term=LOC109610847 | 429 | 5 | 116.55 | 1 | 23.31 | 20 | 29.86 | 22.57 | 7.29 | 7.29 | 0.09 | 0.09 |
| gene8965 | https://www.ncbi.nlm.nih.gov/gene/?term=LOC105282405 | 1310 | 31 | 236.64 | 2 | 15.27 | 6.45 | 51.28 | 44.69 | 6.59 | 6.59 | 0.16 | 0.16 |
| gene11436 | https://www.ncbi.nlm.nih.gov/gene/?term=LOC105285241 | 2207 | 14 | 63.43 | 2 | 9.06 | 14.29 | 48.57 | 42.07 | 6.5 | 6.5 | 0.04 | 0.04 |
| gene7962 | https://www.ncbi.nlm.nih.gov/gene/?term=LOC105281226 | 1280 | 10 | 78.12 | 1 | 7.81 | 10 | 77.35 | 71.1 | 6.25 | 6.25 | 0.05 | 0.05 |
| gene12819 | https://www.ncbi.nlm.nih.gov/gene/?term=LOC105286816 | 4989 | 10 | 20.04 | 1 | 2 | 10 | 37.75 | 31.69 | 6.06 | 6.06 | 0.01 | 0.01 |
| gene571 | https://www.ncbi.nlm.nih.gov/gene/?term=LOC105278541 | 1569 | 16 | 101.98 | 1 | 6.37 | 6.25 | 28.82 | 23.23 | 5.58 | 5.58 | 0.06 | 0.06 |
| gene305 | https://www.ncbi.nlm.nih.gov/gene/?term=LOC105275539 | 1277 | 23 | 180.11 | 2 | 15.66 | 8.7 | 20.8 | 15.48 | 5.32 | 5.32 | 0.1 | 0.1 |
| gene8172 | https://www.ncbi.nlm.nih.gov/gene/?term=LOC105281507 | 946 | 5 | 52.85 | 1 | 10.57 | 20 | 11.82 | 6.67 | 5.15 | 5.15 | 0.03 | 0.03 |
| gene10187 | https://www.ncbi.nlm.nih.gov/gene/?term=LOC105283798 | 2511 | 20 | 79.65 | 1 | 3.98 | 5 | 33.53 | 28.81 | 4.72 | 4.72 | 0.04 | 0.04 |
| gene7684 | https://www.ncbi.nlm.nih.gov/gene/?term=LOC105280926 | 1727 | 7 | 40.53 | 1 | 5.79 | 14.29 | 49.96 | 45.26 | 4.69 | 4.69 | 0.02 | 0.02 |
| gene9483 | https://www.ncbi.nlm.nih.gov/gene/?term=LOC105282996 | 3739 | 9 | 24.07 | 2 | 5.35 | 22.22 | 28.24 | 23.6 | 4.64 | 4.64 | 0.01 | 0.01 |
| gene10334 | https://www.ncbi.nlm.nih.gov/gene/?term=LOC105283973 | 1536 | 33 | 214.84 | 4 | 26.04 | 12.12 | 58.47 | 53.86 | 4.61 | 4.61 | 0.1 | 0.1 |
| gene4999 | https://www.ncbi.nlm.nih.gov/gene/?term=LOC105277805 | 6695 | 13 | 19.42 | 1 | 1.49 | 7.69 | 45.92 | 41.43 | 4.49 | 4.49 | 0.01 | 0.01 |
| gene5066 | https://www.ncbi.nlm.nih.gov/gene/?term=LOC105277868 | 2120 | 15 | 70.75 | 2 | 9.43 | 13.33 | 54.5 | 50.07 | 4.44 | 4.44 | 0.03 | 0.03 |
| gene11269 | https://www.ncbi.nlm.nih.gov/gene/?term=LOC105285051 | 1363 | 10 | 73.37 | 1 | 7.34 | 10 | 46.13 | 41.71 | 4.41 | 4.41 | 0.03 | 0.03 |
| gene470 | https://www.ncbi.nlm.nih.gov/gene/?term=LOC105277393 | 799 | 14 | 175.22 | 1 | 12.52 | 7.14 | 32.95 | 28.75 | 4.2 | 4.2 | 0.07 | 0.07 |
| gene4986 | https://www.ncbi.nlm.nih.gov/gene/?term=LOC105277787 | 1212 | 7 | 57.76 | 1 | 8.25 | 14.29 | 24.61 | 20.42 | 4.18 | 4.18 | 0.02 | 0.02 |
| gene10346 | https://www.ncbi.nlm.nih.gov/gene/?term=LOC105283988 | 2518 | 11 | 43.69 | 1 | 3.97 | 9.09 | 45.73 | 41.61 | 4.12 | 4.12 | 0.02 | 0.02 |
| gene9905 | https://www.ncbi.nlm.nih.gov/gene/?term=LOC105283473 | 937 | 9 | 96.05 | 2 | 21.34 | 22.22 | 34.37 | 30.37 | 4 | 4 | 0.04 | 0.04 |
| gene11135 | https://www.ncbi.nlm.nih.gov/gene/?term=LOC105284881 | 2024 | 21 | 103.75 | 2 | 9.88 | 9.52 | 26.62 | 22.75 | 3.88 | 3.88 | 0.04 | 0.04 |

...
... only first 25 lines shown ...

Directory: OGL File: ogl-Cb_LI_BCphaseA1.vs.RphaseB2.txt

| gene_ID | gene_link | gwidth | #Sites | #per10Kb | #dmSites | #dmsp10kb | %dmSites | %pSite1 | %pSite2 | DMpSite | ADMpSite | DMpNucl | ADMpNucl |
|---|---|---|---|---|---|---|---|---|---|---|---|---|---|
| gene2022 | https://www.ncbi.nlm.nih.gov/gene/?term=LOC105288227 | 1245 | 2 | 16.06 | 1 | 8.03 | 50 | 79.31 | 55.55 | 23.76 | 23.76 | 0.04 | 0.04 |
| gene2593 | https://www.ncbi.nlm.nih.gov/gene/?term=LOC105275070 | 1774 | 5 | 28.18 | 1 | 5.64 | 20 | 19.71 | 4.7 | 15.01 | 15.01 | 0.04 | 0.04 |
| gene11599 | https://www.ncbi.nlm.nih.gov/gene/?term=LOC105285431 | 1550 | 5 | 32.26 | 2 | 12.9 | 40 | 49.86 | 35.35 | 14.51 | 14.51 | 0.05 | 0.05 |
| gene1656 | https://www.ncbi.nlm.nih.gov/gene/?term=LOC105287814 | 812 | 5 | 61.58 | 2 | 24.63 | 40 | 35.51 | 23.39 | 12.12 | 12.12 | 0.07 | 0.07 |
| gene9109 | https://www.ncbi.nlm.nih.gov/gene/?term=LOC105282561 | 2400 | 4 | 16.67 | 1 | 4.17 | 25 | 43.84 | 31.75 | 12.09 | 12.09 | 0.02 | 0.02 |
| gene9845 | https://www.ncbi.nlm.nih.gov/gene/?term=LOC105283410 | 1401 | 9 | 64.24 | 2 | 14.28 | 22.22 | 50.57 | 42.07 | 8.5 | 8.5 | 0.05 | 0.05 |
| gene4044 | https://www.ncbi.nlm.nih.gov/gene/?term=LOC105276718 | 1729 | 9 | 52.05 | 3 | 17.35 | 33.33 | 30.94 | 22.72 | 8.22 | 8.22 | 0.04 | 0.04 |
| gene4620 | https://www.ncbi.nlm.nih.gov/gene/?term=LOC105277368 | 1739 | 6 | 34.5 | 1 | 5.75 | 16.67 | 22.3 | 14.79 | 7.5 | 7.5 | 0.03 | 0.03 |
| gene8965 | https://www.ncbi.nlm.nih.gov/gene/?term=LOC105282405 | 1310 | 31 | 236.64 | 5 | 38.17 | 16.13 | 51.28 | 44.76 | 6.52 | 6.52 | 0.15 | 0.15 |
| gene11145 | https://www.ncbi.nlm.nih.gov/gene/?term=LOC105284899 | 1781 | 14 | 78.61 | 1 | 5.61 | 7.14 | 40.04 | 33.58 | 6.46 | 6.46 | 0.05 | 0.05 |
| gene5060 | https://www.ncbi.nlm.nih.gov/gene/?term=LOC105277862 | 1458 | 7 | 48.01 | 1 | 6.86 | 14.29 | 62.13 | 55.84 | 6.3 | 6.3 | 0.03 | 0.03 |
| gene10363 | https://www.ncbi.nlm.nih.gov/gene/?term=LOC105284009 | 1503 | 26 | 172.99 | 3 | 19.96 | 11.54 | 22.01 | 15.79 | 6.22 | 6.22 | 0.11 | 0.11 |
| gene12853 | https://www.ncbi.nlm.nih.gov/gene/?term=LOC105286857 | 1589 | 4 | 25.17 | 1 | 6.29 | 25 | 73.58 | 67.52 | 6.05 | 6.05 | 0.02 | 0.02 |
| gene11598 | https://www.ncbi.nlm.nih.gov/gene/?term=LOC105285429 | 1324 | 8 | 60.42 | 1 | 7.55 | 12.5 | 53.59 | 47.67 | 5.92 | 5.92 | 0.04 | 0.04 |
| gene5006 | https://www.ncbi.nlm.nih.gov/gene/?term=LOC105277810 | 1382 | 21 | 151.95 | 1 | 7.24 | 4.76 | 45.33 | 39.66 | 5.66 | 5.66 | 0.09 | 0.09 |
| gene4944 | https://www.ncbi.nlm.nih.gov/gene/?term=LOC105277745 | 1344 | 8 | 59.52 | 1 | 7.44 | 12.5 | 67.79 | 62.31 | 5.48 | 5.48 | 0.03 | 0.03 |
| gene2226 | https://www.ncbi.nlm.nih.gov/gene/?term=LOC109611583 | 3276 | 11 | 33.58 | 1 | 3.05 | 9.09 | 13.95 | 8.66 | 5.29 | 5.29 | 0.02 | 0.02 |
| gene10334 | https://www.ncbi.nlm.nih.gov/gene/?term=LOC105283973 | 1536 | 30 | 195.31 | 1 | 6.51 | 3.33 | 61.25 | 55.99 | 5.26 | 5.26 | 0.1 | 0.1 |
| gene5924 | https://www.ncbi.nlm.nih.gov/gene/?term=LOC105278853 | 1035 | 6 | 57.97 | 1 | 9.66 | 16.67 | 22.54 | 17.31 | 5.23 | 5.23 | 0.03 | 0.03 |
| gene6814 | https://www.ncbi.nlm.nih.gov/gene/?term=LOC105279898 | 1343 | 9 | 67.01 | 1 | 7.45 | 11.11 | 79.86 | 74.67 | 5.19 | 5.19 | 0.03 | 0.03 |
| gene7962 | https://www.ncbi.nlm.nih.gov/gene/?term=LOC105281226 | 1280 | 7 | 54.69 | 1 | 7.81 | 14.29 | 69.47 | 64.31 | 5.16 | 5.16 | 0.03 | 0.03 |
| gene3048 | https://www.ncbi.nlm.nih.gov/gene/?term=LOC105275585 | 2468 | 25 | 101.3 | 1 | 4.05 | 4 | 31.83 | 26.81 | 5.02 | 5.02 | 0.05 | 0.05 |
| gene13210 | https://www.ncbi.nlm.nih.gov/gene/?term=LOC105287253 | 757 | 8 | 105.68 | 2 | 26.42 | 25 | 52.09 | 47.22 | 4.87 | 4.87 | 0.05 | 0.05 |
| gene10192 | https://www.ncbi.nlm.nih.gov/gene/?term=LOC105283803 | 1636 | 7 | 42.79 | 1 | 6.11 | 14.29 | 37.31 | 32.5 | 4.82 | 4.82 | 0.02 | 0.02 |

...
... only first 25 lines shown ...

Directory: OGL File: ogl-Cb_LI_BCphaseA2.txt

| gene_ID | gene_link | gwidth | #Sites | #per10Kb | %perSite | %pNucl |
|---|---|---|---|---|---|---|
| gene12538 | https://www.ncbi.nlm.nih.gov/gene/?term=LOC105286531 | 372 | 42 | 1129.03 | 38.35 | 4.33 |
| gene5917 | https://www.ncbi.nlm.nih.gov/gene/?term=LOC105278849 | 823 | 136 | 1652.49 | 25.47 | 4.21 |
| gene13384 | https://www.ncbi.nlm.nih.gov/gene/?term=LOC105287447 | 1877 | 240 | 1278.64 | 32.67 | 4.18 |
| gene5911 | https://www.ncbi.nlm.nih.gov/gene/?term=LOC105278844 | 943 | 97 | 1028.63 | 37.69 | 3.88 |
| gene5309 | https://www.ncbi.nlm.nih.gov/gene/?term=LOC105278160 | 1704 | 161 | 944.84 | 37.8 | 3.57 |
| gene211 | https://www.ncbi.nlm.nih.gov/gene/?term=LOC105274534 | 1280 | 91 | 710.94 | 49.01 | 3.48 |
| gene12364 | https://www.ncbi.nlm.nih.gov/gene/?term=LOC105286269 | 1403 | 91 | 648.61 | 51.53 | 3.34 |
| gene1654 | https://www.ncbi.nlm.nih.gov/gene/?term=LOC105287811 | 2285 | 165 | 722.1 | 45.4 | 3.28 |
| gene12330 | https://www.ncbi.nlm.nih.gov/gene/?term=LOC105286235 | 1501 | 110 | 732.84 | 44.1 | 3.23 |
| gene10742 | https://www.ncbi.nlm.nih.gov/gene/?term=LOC109611309 | 776 | 58 | 747.42 | 43.13 | 3.22 |
| gene12746 | https://www.ncbi.nlm.nih.gov/gene/?term=LOC105286737 | 2688 | 368 | 1369.05 | 23.56 | 3.22 |
| gene3731 | https://www.ncbi.nlm.nih.gov/gene/?term=LOC105276371 | 865 | 56 | 647.4 | 49.67 | 3.22 |
| gene6078 | https://www.ncbi.nlm.nih.gov/gene/?term=LOC105279052 | 1130 | 139 | 1230.09 | 26.01 | 3.2 |
| gene12805 | https://www.ncbi.nlm.nih.gov/gene/?term=LOC105286801 | 1379 | 115 | 833.94 | 37.24 | 3.11 |
| gene5301 | https://www.ncbi.nlm.nih.gov/gene/?term=LOC105278148 | 1881 | 208 | 1105.79 | 27.74 | 3.07 |
| gene3719 | https://www.ncbi.nlm.nih.gov/gene/?term=LOC105276360 | 1471 | 153 | 1040.11 | 29.45 | 3.06 |
| gene2797 | https://www.ncbi.nlm.nih.gov/gene/?term=LOC105275314 | 1489 | 93 | 624.58 | 47.6 | 2.97 |
| gene7527 | https://www.ncbi.nlm.nih.gov/gene/?term=LOC105280739 | 2686 | 145 | 539.84 | 54.6 | 2.95 |
| gene1171 | https://www.ncbi.nlm.nih.gov/gene/?term=LOC105285174 | 844 | 58 | 687.2 | 41.33 | 2.84 |
| gene1562 | https://www.ncbi.nlm.nih.gov/gene/?term=LOC105287692 | 1491 | 129 | 865.19 | 32.49 | 2.81 |
| gene4635 | https://www.ncbi.nlm.nih.gov/gene/?term=LOC105277390 | 2067 | 126 | 609.58 | 46.08 | 2.81 |
| gene570 | https://www.ncbi.nlm.nih.gov/gene/?term=LOC105278549 | 1673 | 112 | 669.46 | 41.85 | 2.8 |
| gene10743 | https://www.ncbi.nlm.nih.gov/gene/?term=LOC105284456 | 1028 | 65 | 632.3 | 43.96 | 2.78 |
| gene8453 | https://www.ncbi.nlm.nih.gov/gene/?term=LOC105281826 | 1750 | 160 | 914.29 | 30.44 | 2.78 |

...

... only first 25 lines shown ...

Directory: OGL File: ogl-Cb_LI_BCphaseA2.vs.BCphaseB1.txt

| gene_ID | gene_link | gwidth | #Sites | #per10Kb | #dmSites | #dmsp10kb | %dmSites | %pSite1 | %pSite2 | DMpSite | ADMpSite | DMpNucl | ADMpNucl |
|---|---|---|---|---|---|---|---|---|---|---|---|---|---|
| gene7199 | https://www.ncbi.nlm.nih.gov/gene/?term=LOC105280366 | 1414 | 30 | 212.16 | 1 | 7.07 | 3.33 | 43.5 | 41.71 | 1.79 | 1.79 | 0.04 | 0.04 |
| gene10606 | https://www.ncbi.nlm.nih.gov/gene/?term=LOC105284273 | 6119 | 258 | 421.64 | 1 | 1.63 | 0.39 | 19.65 | 18.03 | 1.63 | 1.63 | 0.07 | 0.07 |
| gene2856 | https://www.ncbi.nlm.nih.gov/gene/?term=LOC105275384 | 1987 | 91 | 457.98 | 1 | 5.03 | 1.1 | 33.74 | 32.36 | 1.37 | 1.37 | 0.06 | 0.06 |
| gene3793 | https://www.ncbi.nlm.nih.gov/gene/?term=LOC105276429 | 2901 | 119 | 410.2 | 1 | 3.45 | 0.84 | 23.8 | 22.52 | 1.28 | 1.28 | 0.05 | 0.05 |
| gene9740 | https://www.ncbi.nlm.nih.gov/gene/?term=LOC105283277 | 2887 | 143 | 495.32 | 1 | 3.46 | 0.7 | 19.74 | 18.46 | 1.28 | 1.28 | 0.06 | 0.06 |
| gene564 | https://www.ncbi.nlm.nih.gov/gene/?term=LOC105278478 | 2802 | 17 | 60.67 | 1 | 3.57 | 5.88 | 14.91 | 14.04 | 0.87 | 0.87 | 0.01 | 0.01 |
| gene10844 | https://www.ncbi.nlm.nih.gov/gene/?term=LOC105284535 | 9086 | 246 | 270.75 | 1 | 1.1 | 0.41 | 16.16 | 15.32 | 0.84 | 0.84 | 0.02 | 0.02 |
| gene570 | https://www.ncbi.nlm.nih.gov/gene/?term=LOC105278549 | 1673 | 70 | 418.41 | 1 | 5.98 | 1.43 | 44.03 | 43.19 | 0.83 | 0.83 | 0.03 | 0.03 |
| gene10326 | https://www.ncbi.nlm.nih.gov/gene/?term=LOC105283962 | 3042 | 130 | 427.35 | 1 | 3.29 | 0.77 | 8.82 | 8.26 | 0.56 | 0.56 | 0.02 | 0.02 |
| gene1855 | https://www.ncbi.nlm.nih.gov/gene/?term=LOC105288036 | 2387 | 89 | 372.85 | 1 | 4.19 | 1.12 | 30.86 | 30.35 | 0.51 | 0.51 | 0.02 | 0.02 |
| gene7166 | https://www.ncbi.nlm.nih.gov/gene/?term=LOC105280326 | 6117 | 235 | 384.18 | 1 | 1.63 | 0.43 | 15.36 | 14.92 | 0.43 | 0.43 | 0.02 | 0.02 |
| gene12203 | https://www.ncbi.nlm.nih.gov/gene/?term=LOC105286099 | 5967 | 84 | 140.77 | 1 | 1.68 | 1.19 | 11.21 | 10.88 | 0.34 | 0.34 | 0 | 0 |
| gene5101 | https://www.ncbi.nlm.nih.gov/gene/?term=LOC105277903 | 1813 | 34 | 187.53 | 1 | 5.52 | 2.94 | 48.71 | 48.5 | 0.21 | 0.21 | 0 | 0 |
| gene7183 | https://www.ncbi.nlm.nih.gov/gene/?term=LOC105280348 | 3925 | 72 | 183.44 | 1 | 2.55 | 1.39 | 20.87 | 20.69 | 0.19 | 0.19 | 0 | 0 |
| gene7245 | https://www.ncbi.nlm.nih.gov/gene/?term=LOC105280410 | 12478 | 277 | 221.99 | 1 | 0.8 | 0.36 | 19.64 | 19.47 | 0.17 | 0.17 | 0 | 0 |
| gene9755 | https://www.ncbi.nlm.nih.gov/gene/?term=LOC105283296 | 2177 | 92 | 422.6 | 1 | 4.59 | 1.09 | 18.09 | 17.95 | 0.14 | 0.14 | 0.01 | 0.01 |
| gene3744 | https://www.ncbi.nlm.nih.gov/gene/?term=LOC105276385 | 9376 | 459 | 489.55 | 1 | 1.07 | 0.22 | 5.46 | 5.7 | -0.24 | 0.24 | -0.01 | 0.01 |
| gene4892 | https://www.ncbi.nlm.nih.gov/gene/?term=LOC105277667 | 3898 | 126 | 323.24 | 1 | 2.57 | 0.79 | 27.95 | 28.19 | -0.24 | 0.24 | -0.01 | 0.01 |
| gene8814 | https://www.ncbi.nlm.nih.gov/gene/?term=LOC105282240 | 6910 | 175 | 253.26 | 1 | 1.45 | 0.57 | 9.54 | 9.87 | -0.32 | 0.32 | -0.01 | 0.01 |
| gene10819 | https://www.ncbi.nlm.nih.gov/gene/?term=LOC105284507 | 5767 | 94 | 163 | 1 | 1.73 | 1.06 | 20.43 | 21.18 | -0.75 | 0.75 | -0.01 | 0.01 |
| gene11447 | https://www.ncbi.nlm.nih.gov/gene/?term=LOC105285252 | 7437 | 131 | 176.15 | 1 | 1.34 | 0.76 | 12.71 | 13.67 | -0.95 | 0.95 | -0.02 | 0.02 |
| gene9722 | https://www.ncbi.nlm.nih.gov/gene/?term=LOC105283264 | 3951 | 88 | 222.73 | 1 | 2.53 | 1.14 | 18.85 | 20.39 | -1.54 | 1.54 | -0.03 | 0.03 |

Directory: OGL File: ogl-Cb_LI_BCphaseA2.vs.BCphaseB2.txt

| gene_ID | gene_link | gwidth | #Sites | #per10Kb | #dmSites | #dmsp10kb | %dmSites | %pSite1 | %pSite2 | DMpSite | ADMpSite | DMpNucl | ADMpNucl |
|---|---|---|---|---|---|---|---|---|---|---|---|---|---|
| gene2593 | https://www.ncbi.nlm.nih.gov/gene/?term=LOC105275070 | 1774 | 5 | 28.18 | 5 | 28.18 | 100 | 80.28 | 10.96 | 69.32 | 69.32 | 0.2 | 0.2 |
| gene3165 | https://www.ncbi.nlm.nih.gov/gene/?term=LOC105275706 | 2374 | 67 | 282.22 | 5 | 21.06 | 7.46 | 36.03 | 23.6 | 12.43 | 12.43 | 0.35 | 0.35 |
| gene1957 | https://www.ncbi.nlm.nih.gov/gene/?term=LOC105288153 | 1044 | 16 | 153.26 | 1 | 9.58 | 6.25 | 15.68 | 4.26 | 11.42 | 11.42 | 0.18 | 0.18 |
| gene2824 | https://www.ncbi.nlm.nih.gov/gene/?term=LOC105275337 | 1772 | 64 | 361.17 | 2 | 11.29 | 3.12 | 37.53 | 29.09 | 8.44 | 8.44 | 0.3 | 0.3 |
| gene9468 | https://www.ncbi.nlm.nih.gov/gene/?term=LOC105282976 | 1636 | 53 | 323.96 | 1 | 6.11 | 1.89 | 28.76 | 22.1 | 6.66 | 6.66 | 0.22 | 0.22 |
| gene12395 | https://www.ncbi.nlm.nih.gov/gene/?term=LOC105286302 | 4093 | 17 | 41.53 | 1 | 2.44 | 5.88 | 21.99 | 16.58 | 5.41 | 5.41 | 0.02 | 0.02 |
| gene5714 | https://www.ncbi.nlm.nih.gov/gene/?term=LOC105278729 | 710 | 24 | 338.03 | 1 | 14.08 | 4.17 | 61.56 | 56.79 | 4.77 | 4.77 | 0.16 | 0.16 |
| gene1518 | https://www.ncbi.nlm.nih.gov/gene/?term=LOC105287648 | 2195 | 27 | 123.01 | 1 | 4.56 | 3.7 | 41.41 | 38.92 | 2.49 | 2.49 | 0.03 | 0.03 |
| gene3385 | https://www.ncbi.nlm.nih.gov/gene/?term=LOC105275966 | 3305 | 61 | 184.57 | 1 | 3.03 | 1.64 | 49.58 | 47.17 | 2.41 | 2.41 | 0.04 | 0.04 |
| gene5877 | https://www.ncbi.nlm.nih.gov/gene/?term=LOC105278807 | 1719 | 49 | 285.05 | 1 | 5.82 | 2.04 | 23.71 | 21.48 | 2.23 | 2.23 | 0.06 | 0.06 |
| gene5052 | https://www.ncbi.nlm.nih.gov/gene/?term=LOC105277856 | 3167 | 34 | 107.36 | 1 | 3.16 | 2.94 | 38.79 | 36.66 | 2.13 | 2.13 | 0.02 | 0.02 |
| gene3817 | https://www.ncbi.nlm.nih.gov/gene/?term=LOC105276449 | 1230 | 42 | 341.46 | 1 | 8.13 | 2.38 | 35.93 | 33.84 | 2.09 | 2.09 | 0.07 | 0.07 |
| gene8115 | https://www.ncbi.nlm.nih.gov/gene/?term=LOC105281420 | 3623 | 51 | 140.77 | 1 | 2.76 | 1.96 | 12.6 | 10.51 | 2.09 | 2.09 | 0.03 | 0.03 |
| gene2970 | https://www.ncbi.nlm.nih.gov/gene/?term=LOC105275518 | 2340 | 19 | 81.2 | 1 | 4.27 | 5.26 | 46.39 | 44.36 | 2.03 | 2.03 | 0.02 | 0.02 |
| gene8742 | https://www.ncbi.nlm.nih.gov/gene/?term=LOC105282151 | 2563 | 41 | 159.97 | 2 | 7.8 | 4.88 | 36.1 | 34.31 | 1.79 | 1.79 | 0.03 | 0.03 |
| gene7556 | https://www.ncbi.nlm.nih.gov/gene/?term=LOC105280774 | 2792 | 102 | 365.33 | 1 | 3.58 | 0.98 | 32.11 | 30.45 | 1.66 | 1.66 | 0.06 | 0.06 |
| gene7398 | https://www.ncbi.nlm.nih.gov/gene/?term=LOC105280589 | 3452 | 39 | 112.98 | 1 | 2.9 | 2.56 | 23.26 | 21.67 | 1.58 | 1.58 | 0.02 | 0.02 |
| gene9462 | https://www.ncbi.nlm.nih.gov/gene/?term=LOC105282970 | 1469 | 39 | 265.49 | 1 | 6.81 | 2.56 | 53.94 | 52.44 | 1.5 | 1.5 | 0.04 | 0.04 |
| gene12572 | https://www.ncbi.nlm.nih.gov/gene/?term=LOC105286514 | 2374 | 42 | 176.92 | 1 | 4.21 | 2.38 | 34.39 | 32.92 | 1.47 | 1.47 | 0.03 | 0.03 |
| gene927 | https://www.ncbi.nlm.nih.gov/gene/?term=LOC105282600 | 3727 | 53 | 142.21 | 1 | 2.68 | 1.89 | 32.11 | 30.67 | 1.44 | 1.44 | 0.02 | 0.02 |
| gene6528 | https://www.ncbi.nlm.nih.gov/gene/?term=LOC105279591 | 7222 | 55 | 76.16 | 1 | 1.38 | 1.82 | 29.5 | 28.12 | 1.38 | 1.38 | 0.01 | 0.01 |
| gene7568 | https://www.ncbi.nlm.nih.gov/gene/?term=LOC105280786 | 2525 | 111 | 439.6 | 1 | 3.96 | 0.9 | 33.28 | 31.99 | 1.29 | 1.29 | 0.06 | 0.06 |
| gene4892 | https://www.ncbi.nlm.nih.gov/gene/?term=LOC105277667 | 3898 | 125 | 320.68 | 1 | 2.57 | 0.8 | 27.74 | 26.47 | 1.27 | 1.27 | 0.04 | 0.04 |
| gene10606 | https://www.ncbi.nlm.nih.gov/gene/?term=LOC105284273 | 6119 | 251 | 410.2 | 1 | 1.63 | 0.4 | 20.03 | 18.79 | 1.24 | 1.24 | 0.05 | 0.05 |
| gene12290 | https://www.ncbi.nlm.nih.gov/gene/?term=LOC105286190 | 6212 | 304 | 489.38 | 1 | 1.61 | 0.33 | 8.69 | 7.5 | 1.18 | 1.18 | 0.06 | 0.06 |
| gene1884 | https://www.ncbi.nlm.nih.gov/gene/?term=LOC105288072 | 4169 | 71 | 170.3 | 1 | 2.4 | 1.41 | 25.5 | 24.39 | 1.1 | 1.1 | 0.02 | 0.02 |
| gene3470 | https://www.ncbi.nlm.nih.gov/gene/?term=LOC105276059 | 8716 | 145 | 166.36 | 1 | 1.15 | 0.69 | 10.01 | 8.94 | 1.08 | 1.08 | 0.02 | 0.02 |
| gene5967 | https://www.ncbi.nlm.nih.gov/gene/?term=LOC105278918 | 4393 | 121 | 275.44 | 1 | 2.28 | 0.83 | 15.97 | 14.91 | 1.05 | 1.05 | 0.03 | 0.03 |
| gene940 | https://www.ncbi.nlm.nih.gov/gene/?term=LOC105282794 | 3578 | 85 | 237.56 | 1 | 2.79 | 1.18 | 16.1 | 15.1 | 1 | 1 | 0.02 | 0.02 |
| gene10678 | https://www.ncbi.nlm.nih.gov/gene/?term=LOC105284355 | 4031 | 167 | 414.29 | 1 | 2.48 | 0.6 | 19.59 | 18.65 | 0.94 | 0.94 | 0.04 | 0.04 |
| gene4034 | https://www.ncbi.nlm.nih.gov/gene/?term=LOC105276704 | 2065 | 101 | 489.1 | 1 | 4.84 | 0.99 | 13.4 | 12.57 | 0.83 | 0.83 | 0.04 | 0.04 |
| gene8729 | https://www.ncbi.nlm.nih.gov/gene/?term=LOC105282137 | 5005 | 197 | 393.61 | 1 | 2 | 0.51 | 17.44 | 16.65 | 0.8 | 0.8 | 0.03 | 0.03 |
| gene10520 | https://www.ncbi.nlm.nih.gov/gene/?term=LOC105284184 | 2451 | 85 | 346.8 | 1 | 4.08 | 1.18 | 34.37 | 33.59 | 0.78 | 0.78 | 0.03 | 0.03 |
| gene13033 | https://www.ncbi.nlm.nih.gov/gene/?term=LOC105287063 | 1877 | 49 | 261.05 | 1 | 5.33 | 2.04 | 31.12 | 30.35 | 0.77 | 0.77 | 0.02 | 0.02 |
| gene2825 | https://www.ncbi.nlm.nih.gov/gene/?term=LOC105275343 | 5545 | 96 | 173.13 | 2 | 3.61 | 2.08 | 11.79 | 11.04 | 0.75 | 0.75 | 0.01 | 0.01 |
| gene9336 | https://www.ncbi.nlm.nih.gov/gene/?term=LOC105282834 | 5993 | 124 | 206.91 | 1 | 1.67 | 0.81 | 16.06 | 15.3 | 0.75 | 0.75 | 0.02 | 0.02 |
| gene12516 | https://www.ncbi.nlm.nih.gov/gene/?term=LOC105286452 | 1529 | 38 | 248.53 | 1 | 6.54 | 2.63 | 52.34 | 51.61 | 0.73 | 0.73 | 0.02 | 0.02 |
| gene6620 | https://www.ncbi.nlm.nih.gov/gene/?term=LOC105279687 | 17998 | 336 | 186.69 | 1 | 0.56 | 0.3 | 5.15 | 4.42 | 0.73 | 0.73 | 0.01 | 0.01 |
| gene4376 | https://www.ncbi.nlm.nih.gov/gene/?term=LOC105277083 | 4307 | 127 | 294.87 | 1 | 2.32 | 0.79 | 19.77 | 19.11 | 0.65 | 0.65 | 0.02 | 0.02 |
| gene12307 | https://www.ncbi.nlm.nih.gov/gene/?term=LOC105286216 | 10313 | 166 | 160.96 | 1 | 0.97 | 0.6 | 16.86 | 16.32 | 0.54 | 0.54 | 0.01 | 0.01 |
| gene7368 | https://www.ncbi.nlm.nih.gov/gene/?term=LOC105280556 | 4105 | 50 | 121.8 | 1 | 2.44 | 2 | 18.71 | 18.17 | 0.54 | 0.54 | 0.01 | 0.01 |
| gene7514 | https://www.ncbi.nlm.nih.gov/gene/?term=LOC105280723 | 4471 | 200 | 447.33 | 1 | 2.24 | 0.5 | 14.49 | 14.01 | 0.49 | 0.49 | 0.02 | 0.02 |
| gene2056 | https://www.ncbi.nlm.nih.gov/gene/?term=LOC105274452 | 6584 | 263 | 399.45 | 2 | 3.04 | 0.76 | 13.53 | 13.06 | 0.47 | 0.47 | 0.02 | 0.02 |
| gene3286 | https://www.ncbi.nlm.nih.gov/gene/?term=LOC105275848 | 3744 | 140 | 373.93 | 1 | 2.67 | 0.71 | 23 | 22.56 | 0.45 | 0.45 | 0.02 | 0.02 |
| gene292 | https://www.ncbi.nlm.nih.gov/gene/?term=LOC105275398 | 2825 | 86 | 304.42 | 1 | 3.54 | 1.16 | 21.94 | 21.53 | 0.4 | 0.4 | 0.01 | 0.01 |
| gene4962 | https://www.ncbi.nlm.nih.gov/gene/?term=LOC105277761 | 4645 | 150 | 322.93 | 1 | 2.15 | 0.67 | 15.92 | 15.52 | 0.4 | 0.4 | 0.01 | 0.01 |
| gene7245 | https://www.ncbi.nlm.nih.gov/gene/?term=LOC105280410 | 12478 | 237 | 189.93 | 1 | 0.8 | 0.42 | 21.04 | 20.64 | 0.4 | 0.4 | 0.01 | 0.01 |
| gene10654 | https://www.ncbi.nlm.nih.gov/gene/?term=LOC105284329 | 4027 | 50 | 124.16 | 1 | 2.48 | 2 | 19.81 | 19.43 | 0.39 | 0.39 | 0 | 0 |
| gene2173 | https://www.ncbi.nlm.nih.gov/gene/?term=LOC105274590 | 4387 | 64 | 145.89 | 2 | 4.56 | 3.12 | 34.09 | 33.7 | 0.39 | 0.39 | 0.01 | 0.01 |
| gene6031 | https://www.ncbi.nlm.nih.gov/gene/?term=LOC105279000 | 5691 | 195 | 342.65 | 1 | 1.76 | 0.51 | 14.39 | 14 | 0.39 | 0.39 | 0.01 | 0.01 |
| gene5746 | https://www.ncbi.nlm.nih.gov/gene/?term=LOC105278666 | 10343 | 461 | 445.71 | 1 | 0.97 | 0.22 | 7.31 | 6.93 | 0.37 | 0.37 | 0.02 | 0.02 |
| gene1430 | https://www.ncbi.nlm.nih.gov/gene/?term=LOC105287551 | 6433 | 231 | 359.09 | 1 | 1.55 | 0.43 | 9.26 | 8.92 | 0.34 | 0.34 | 0.01 | 0.01 |
| gene10531 | https://www.ncbi.nlm.nih.gov/gene/?term=LOC105284190 | 6881 | 173 | 251.42 | 1 | 1.45 | 0.58 | 12.82 | 12.5 | 0.32 | 0.32 | 0.01 | 0.01 |
| gene1310 | https://www.ncbi.nlm.nih.gov/gene/?term=LOC105286661 | 2582 | 66 | 255.62 | 1 | 3.87 | 1.52 | 24.45 | 24.13 | 0.32 | 0.32 | 0.01 | 0.01 |
| gene4246 | https://www.ncbi.nlm.nih.gov/gene/?term=LOC105276941 | 16412 | 552 | 336.34 | 1 | 0.61 | 0.18 | 5.12 | 4.8 | 0.32 | 0.32 | 0.01 | 0.01 |
| gene11568 | https://www.ncbi.nlm.nih.gov/gene/?term=LOC105285395 | 8086 | 192 | 237.45 | 1 | 1.24 | 0.52 | 15.07 | 14.77 | 0.31 | 0.31 | 0.01 | 0.01 |
| gene1470 | https://www.ncbi.nlm.nih.gov/gene/?term=LOC105287597 | 2974 | 204 | 685.94 | 1 | 3.36 | 0.49 | 17.73 | 17.44 | 0.3 | 0.3 | 0.02 | 0.02 |
| gene3856 | https://www.ncbi.nlm.nih.gov/gene/?term=LOC105276499 | 5236 | 190 | 362.87 | 1 | 1.91 | 0.53 | 9.53 | 9.24 | 0.29 | 0.29 | 0.01 | 0.01 |
| gene3599 | https://www.ncbi.nlm.nih.gov/gene/?term=LOC105276196 | 2955 | 48 | 162.44 | 1 | 3.38 | 2.08 | 45.56 | 45.28 | 0.28 | 0.28 | 0 | 0 |
| gene7249 | https://www.ncbi.nlm.nih.gov/gene/?term=LOC105280422 | 1775 | 63 | 354.93 | 1 | 5.63 | 1.59 | 17.96 | 17.74 | 0.23 | 0.23 | 0.01 | 0.01 |

| | | | | | | | | | | | | | |
|---|---|---|---|---|---|---|---|---|---|---|---|---|---|
| gene8600 | https://www.ncbi.nlm.nih.gov/gene/?term=LOC105282001 | 8513 | 156 | 183.25 | 1 | 1.17 | 0.64 | 12.04 | 11.82 | 0.22 | 0.22 | 0 | 0 |
| gene2806 | https://www.ncbi.nlm.nih.gov/gene/?term=LOC105275327 | 14257 | 494 | 346.5 1 | 0.7 | 0.2 | 6.88 | 6.68 | 0.2 | 0.2 | 0.01 | 0.01 | |
| gene11275 | https://www.ncbi.nlm.nih.gov/gene/?term=LOC105285055 | 4119 | 255 | 619.08 | 1 | 2.43 | 0.39 | 11.36 | 11.16 | 0.19 | 0.19 | 0.01 | 0.01 |
| gene2846 | https://www.ncbi.nlm.nih.gov/gene/?term=LOC105275374 | 11246 | 502 | 446.38 | 1 | 0.89 | 0.2 | 11.4 | 11.21 | 0.19 | 0.19 | 0.01 | 0.01 |
| gene7166 | https://www.ncbi.nlm.nih.gov/gene/?term=LOC105280326 | 6117 | 233 | 380.91 | 1 | 1.63 | 0.43 | 15.64 | 15.46 | 0.19 | 0.19 | 0.01 | 0.01 |
| gene12439 | https://www.ncbi.nlm.nih.gov/gene/?term=LOC105286357 | 4761 | 81 | 170.13 | 1 | 2.1 | 1.23 | 32.58 | 32.4 | 0.18 | 0.18 | 0 | 0 |
| gene11579 | https://www.ncbi.nlm.nih.gov/gene/?term=LOC105285406 | 13571 | 267 | 196.74 | 1 | 0.74 | 0.37 | 6.68 | 6.51 | 0.17 | 0.17 | 0 | 0 |
| gene8343 | https://www.ncbi.nlm.nih.gov/gene/?term=LOC105281700 | 2172 | 37 | 170.35 | 1 | 4.6 | 2.7 | 39.71 | 39.54 | 0.16 | 0.16 | 0 | 0 |
| gene11401 | https://www.ncbi.nlm.nih.gov/gene/?term=LOC105285202 | 1841 | 82 | 445.41 | 1 | 5.43 | 1.22 | 25.44 | 25.28 | 0.15 | 0.15 | 0.01 | 0.01 |
| gene12203 | https://www.ncbi.nlm.nih.gov/gene/?term=LOC105286099 | 5967 | 72 | 120.66 | 1 | 1.68 | 1.39 | 10.74 | 10.59 | 0.15 | 0.15 | 0 | 0 |
| gene12806 | https://www.ncbi.nlm.nih.gov/gene/?term=LOC105286803 | 4196 | 143 | 340.8 1 | 2.38 | 0.7 | 11.07 | 10.92 | 0.15 | 0.15 | 0.01 | 0.01 | |
| gene4190 | https://www.ncbi.nlm.nih.gov/gene/?term=LOC105276875 | 3150 | 107 | 339.68 | 1 | 3.17 | 0.93 | 23.8 | 23.66 | 0.15 | 0.15 | 0 | 0 |
| gene1198 | https://www.ncbi.nlm.nih.gov/gene/?term=LOC105285507 | 2738 | 88 | 321.4 1 | 3.65 | 1.14 | 23.48 | 23.34 | 0.14 | 0.14 | 0 | 0 | |
| gene5524 | https://www.ncbi.nlm.nih.gov/gene/?term=LOC105278397 | 5177 | 203 | 392.12 | 1 | 1.93 | 0.49 | 25.25 | 25.12 | 0.13 | 0.13 | 0.01 | 0.01 |
| gene12802 | https://www.ncbi.nlm.nih.gov/gene/?term=LOC105286799 | 3183 | 185 | 581.21 | 2 | 6.28 | 1.08 | 16.06 | 15.94 | 0.12 | 0.12 | 0.01 | 0.01 |
| gene10392 | https://www.ncbi.nlm.nih.gov/gene/?term=LOC105284036 | 3216 | 94 | 292.29 | 2 | 6.22 | 2.13 | 22.55 | 22.51 | 0.04 | 0.04 | 0 | 0 |
| gene53 | https://www.ncbi.nlm.nih.gov/gene/?term=LOC105277148 | 19283 | 287 | 148.84 | 1 | 0.52 | 0.35 | 5.29 | 5.27 | 0.03 | 0.03 | 0 | 0 |
| gene6286 | https://www.ncbi.nlm.nih.gov/gene/?term=LOC105279298 | 2861 | 125 | 436.91 | 2 | 6.99 | 1.6 | 25.22 | 25.19 | 0.03 | 0.03 | 0 | 0 |
| gene13044 | https://www.ncbi.nlm.nih.gov/gene/?term=LOC105287070 | 5094 | 241 | 473.11 | 1 | 1.96 | 0.41 | 5.28 | 5.27 | 0.02 | 0.02 | 0 | 0 |
| gene2844 | https://www.ncbi.nlm.nih.gov/gene/?term=LOC105275363 | 4330 | 192 | 443.42 | 1 | 2.31 | 0.52 | 14.25 | 14.23 | 0.02 | 0.02 | 0 | 0 |
| gene5525 | https://www.ncbi.nlm.nih.gov/gene/?term=LOC105278396 | 3090 | 188 | 608.41 | 1 | 3.24 | 0.53 | 24.8 | 24.79 | 0.01 | 0.01 | 0 | 0 |
| gene9330 | https://www.ncbi.nlm.nih.gov/gene/?term=LOC105282825 | 3936 | 80 | 203.25 | 1 | 2.54 | 1.25 | 12.57 | 12.58 | -0.01 | 0.01 | 0 | 0 |
| gene9755 | https://www.ncbi.nlm.nih.gov/gene/?term=LOC105283296 | 2177 | 85 | 390.45 | 1 | 4.59 | 1.18 | 21.44 | 21.45 | -0.01 | 0.01 | 0 | 0 |
| gene298 | https://www.ncbi.nlm.nih.gov/gene/?term=LOC105275460 | 2473 | 114 | 460.98 | 1 | 4.04 | 0.88 | 13.89 | 13.91 | -0.02 | 0.02 | 0 | 0 |
| gene10549 | https://www.ncbi.nlm.nih.gov/gene/?term=LOC105284210 | 7145 | 325 | 454.86 | 2 | 2.8 | 0.62 | 10.71 | 10.76 | -0.05 | 0.05 | 0 | 0 |
| gene3848 | https://www.ncbi.nlm.nih.gov/gene/?term=LOC105276486 | 5899 | 133 | 225.46 | 1 | 1.7 | 0.75 | 24.81 | 24.86 | -0.05 | 0.05 | 0 | 0 |
| gene9177 | https://www.ncbi.nlm.nih.gov/gene/?term=LOC105282646 | 3320 | 98 | 295.18 | 1 | 3.01 | 1.02 | 22.82 | 22.88 | -0.05 | 0.05 | 0 | 0 |
| gene4641 | https://www.ncbi.nlm.nih.gov/gene/?term=LOC105277398 | 1937 | 77 | 397.52 | 1 | 5.16 | 1.3 | 34.01 | 34.07 | -0.06 | 0.06 | 0 | 0 |
| gene11688 | https://www.ncbi.nlm.nih.gov/gene/?term=LOC105285533 | 1637 | 60 | 366.52 | 1 | 6.11 | 1.67 | 26.05 | 26.13 | -0.07 | 0.07 | 0 | 0 |
| gene3107 | https://www.ncbi.nlm.nih.gov/gene/?term=LOC105275650 | 5103 | 181 | 354.69 | 1 | 1.96 | 0.55 | 15.45 | 15.53 | -0.08 | 0.08 | 0 | 0 |
| gene5831 | https://www.ncbi.nlm.nih.gov/gene/?term=LOC105278761 | 2506 | 119 | 474.86 | 2 | 7.98 | 1.68 | 21.61 | 21.68 | -0.08 | 0.08 | 0 | 0 |
| gene9445 | https://www.ncbi.nlm.nih.gov/gene/?term=LOC105282951 | 2285 | 115 | 503.28 | 1 | 4.38 | 0.87 | 20.19 | 20.28 | -0.09 | 0.09 | 0 | 0 |
| gene12674 | https://www.ncbi.nlm.nih.gov/gene/?term=LOC105286649 | 18283 | 811 | 443.58 | 1 | 0.55 | 0.12 | 0.93 | 1.06 | -0.13 | 0.13 | -0.01 | 0.01 |
| gene13114 | https://www.ncbi.nlm.nih.gov/gene/?term=LOC105287147 | 7609 | 409 | 537.52 | 1 | 1.31 | 0.24 | 9.57 | 9.7 | -0.13 | 0.13 | -0.01 | 0.01 |
| gene6652 | https://www.ncbi.nlm.nih.gov/gene/?term=LOC105279719 | 2831 | 79 | 279.05 | 1 | 3.53 | 1.27 | 24.78 | 24.91 | -0.14 | 0.14 | 0 | 0 |
| gene1632 | https://www.ncbi.nlm.nih.gov/gene/?term=LOC105287764 | 18061 | 393 | 217.6 2 | 1.11 | 0.51 | 2.97 | 3.13 | -0.17 | 0.17 | 0 | 0 | |
| gene8661 | https://www.ncbi.nlm.nih.gov/gene/?term=LOC105282063 | 1926 | 32 | 166.15 | 1 | 5.19 | 3.12 | 29.26 | 29.43 | -0.17 | 0.17 | 0 | 0 |
| gene3885 | https://www.ncbi.nlm.nih.gov/gene/?term=LOC105276527 | 7928 | 317 | 399.85 | 1 | 1.26 | 0.32 | 9.68 | 9.86 | -0.18 | 0.18 | -0.01 | 0.01 |
| gene5777 | https://www.ncbi.nlm.nih.gov/gene/?term=LOC105278691 | 3883 | 148 | 381.15 | 1 | 2.58 | 0.68 | 14.2 | 14.38 | -0.18 | 0.18 | -0.01 | 0.01 |
| gene6583 | https://www.ncbi.nlm.nih.gov/gene/?term=LOC105279649 | 7444 | 332 | 446 1 | 1.34 | 0.3 | 10.76 | 10.98 | -0.22 | 0.22 | -0.01 | 0.01 | |
| gene11454 | https://www.ncbi.nlm.nih.gov/gene/?term=LOC105285256 | 1906 | 47 | 246.59 | 1 | 5.25 | 2.13 | 45.38 | 45.61 | -0.23 | 0.23 | -0.01 | 0.01 |
| gene5375 | https://www.ncbi.nlm.nih.gov/gene/?term=LOC105278228 | 8640 | 163 | 188.66 | 1 | 1.16 | 0.61 | 21.78 | 22.01 | -0.23 | 0.23 | 0 | 0 |
| gene13036 | https://www.ncbi.nlm.nih.gov/gene/?term=LOC105287059 | 4959 | 163 | 328.7 1 | 2.02 | 0.61 | 16.39 | 16.65 | -0.26 | 0.26 | -0.01 | 0.01 | |
| gene3727 | https://www.ncbi.nlm.nih.gov/gene/?term=LOC105276367 | 2484 | 69 | 277.78 | 1 | 4.03 | 1.45 | 46.73 | 46.99 | -0.27 | 0.27 | -0.01 | 0.01 |
| gene6237 | https://www.ncbi.nlm.nih.gov/gene/?term=LOC105279244 | 1484 | 24 | 161.73 | 1 | 6.74 | 4.17 | 52.96 | 53.25 | -0.29 | 0.29 | 0 | 0 |
| gene2683 | https://www.ncbi.nlm.nih.gov/gene/?term=LOC105275187 | 5492 | 70 | 127.46 | 1 | 1.82 | 1.43 | 14.08 | 14.4 | -0.31 | 0.31 | 0 | 0 |
| gene10155 | https://www.ncbi.nlm.nih.gov/gene/?term=LOC105283756 | 7030 | 93 | 132.29 | 1 | 1.42 | 1.08 | 5.63 | 5.96 | -0.34 | 0.34 | 0 | 0 |
| gene2257 | https://www.ncbi.nlm.nih.gov/gene/?term=LOC105274689 | 3658 | 71 | 194.1 1 | 2.73 | 1.41 | 14.85 | 15.18 | -0.34 | 0.34 | -0.01 | 0.01 | |
| gene7600 | https://www.ncbi.nlm.nih.gov/gene/?term=LOC105280819 | 1615 | 39 | 241.49 | 1 | 6.19 | 2.56 | 35.76 | 36.1 | -0.34 | 0.34 | -0.01 | 0.01 |
| gene11082 | https://www.ncbi.nlm.nih.gov/gene/?term=LOC105284824 | 7436 | 200 | 268.96 | 1 | 1.34 | 0.5 | 5.46 | 5.81 | -0.35 | 0.35 | -0.01 | 0.01 |
| gene12573 | https://www.ncbi.nlm.nih.gov/gene/?term=LOC105286515 | 3647 | 113 | 309.84 | 1 | 2.74 | 0.88 | 10.7 | 11.07 | -0.37 | 0.37 | -0.01 | 0.01 |
| gene10600 | https://www.ncbi.nlm.nih.gov/gene/?term=LOC105284270 | 4837 | 124 | 256.36 | 1 | 2.07 | 0.81 | 10.47 | 10.85 | -0.38 | 0.38 | -0.01 | 0.01 |
| gene3046 | https://www.ncbi.nlm.nih.gov/gene/?term=LOC105275587 | 2394 | 56 | 233.92 | 1 | 4.18 | 1.79 | 33.82 | 34.2 | -0.38 | 0.38 | -0.01 | 0.01 |
| gene12981 | https://www.ncbi.nlm.nih.gov/gene/?term=LOC105286999 | 2063 | 40 | 193.89 | 1 | 4.85 | 2.5 | 35.8 | 36.25 | -0.45 | 0.45 | -0.01 | 0.01 |
| gene10107 | https://www.ncbi.nlm.nih.gov/gene/?term=LOC105283712 | 1883 | 32 | 169.94 | 1 | 5.31 | 3.12 | 29.98 | 30.44 | -0.46 | 0.46 | -0.01 | 0.01 |
| gene11561 | https://www.ncbi.nlm.nih.gov/gene/?term=LOC105285387 | 11528 | 194 | 168.29 | 1 | 0.87 | 0.52 | 8.65 | 9.11 | -0.47 | 0.47 | -0.01 | 0.01 |
| gene8015 | https://www.ncbi.nlm.nih.gov/gene/?term=LOC105281312 | 2998 | 72 | 240.16 | 1 | 3.34 | 1.39 | 29.67 | 30.18 | -0.5 | 0.5 | -0.01 | 0.01 |
| gene8814 | https://www.ncbi.nlm.nih.gov/gene/?term=LOC105282240 | 6910 | 170 | 246.02 | 1 | 1.45 | 0.59 | 9.16 | 9.68 | -0.52 | 0.52 | -0.01 | 0.01 |
| gene4720 | https://www.ncbi.nlm.nih.gov/gene/?term=LOC105277479 | 1697 | 47 | 276.96 | 1 | 5.89 | 2.13 | 37.56 | 38.12 | -0.55 | 0.55 | -0.02 | 0.02 |
| gene10391 | https://www.ncbi.nlm.nih.gov/gene/?term=LOC109611283 | 1994 | 44 | 220.66 | 1 | 5.02 | 2.27 | 32.96 | 33.52 | -0.56 | 0.56 | -0.01 | 0.01 |
| gene3671 | https://www.ncbi.nlm.nih.gov/gene/?term=LOC105276272 | 1966 | 36 | 183.11 | 1 | 5.09 | 2.78 | 32.08 | 32.66 | -0.58 | 0.58 | -0.01 | 0.01 |
| gene9826 | https://www.ncbi.nlm.nih.gov/gene/?term=LOC105283385 | 9219 | 305 | 330.84 | 1 | 1.08 | 0.33 | 12.6 | 13.19 | -0.59 | 0.59 | -0.02 | 0.02 |
| gene6966 | https://www.ncbi.nlm.nih.gov/gene/?term=LOC105280104 | 3747 | 122 | 325.59 | 1 | 2.67 | 0.82 | 15.27 | 15.88 | -0.62 | 0.62 | -0.02 | 0.02 |
| gene8274 | https://www.ncbi.nlm.nih.gov/gene/?term=LOC105281624 | 5425 | 230 | 423.96 | 1 | 1.84 | 0.43 | 9.61 | 10.33 | -0.73 | 0.73 | -0.03 | 0.03 |

| | | | | | | | | | | | | | |
|---|---|---|---|---|---|---|---|---|---|---|---|---|---|
| gene10018 | https://www.ncbi.nlm.nih.gov/gene/?term=LOC105283607 | 4414 | 191 | 432.71 | 1 | 2.27 | 0.52 | 15.56 | 16.32 | -0.76 | 0.76 | -0.03 | 0.03 |
| gene5298 | https://www.ncbi.nlm.nih.gov/gene/?term=LOC105278145 | 4060 | 244 | 600.99 | 2 | 4.93 | 0.82 | 10.55 | 11.35 | -0.8 | 0.8 | -0.05 | 0.05 |
| gene10213 | https://www.ncbi.nlm.nih.gov/gene/?term=LOC105283826 | 1503 | 64 | 425.82 | 1 | 6.65 | 1.56 | 14.04 | 14.85 | -0.81 | 0.81 | -0.03 | 0.03 |
| gene1147 | https://www.ncbi.nlm.nih.gov/gene/?term=LOC105284954 | 4022 | 95 | 236.2 | 1 | 2.49 | 1.05 | 15.81 | 16.65 | -0.84 | 0.84 | -0.02 | 0.02 |
| gene7915 | https://www.ncbi.nlm.nih.gov/gene/?term=LOC105281173 | 1597 | 38 | 237.95 | 1 | 6.26 | 2.63 | 59.94 | 60.79 | -0.85 | 0.85 | -0.02 | 0.02 |
| gene2413 | https://www.ncbi.nlm.nih.gov/gene/?term=LOC105274863 | 2006 | 44 | 219.34 | 1 | 4.99 | 2.27 | 63.14 | 64.02 | -0.87 | 0.87 | -0.02 | 0.02 |
| gene4498 | https://www.ncbi.nlm.nih.gov/gene/?term=LOC105277239 | 2831 | 129 | 455.67 | 1 | 3.53 | 0.78 | 14.44 | 15.34 | -0.89 | 0.89 | -0.04 | 0.04 |
| gene6639 | https://www.ncbi.nlm.nih.gov/gene/?term=LOC105279705 | 7656 | 62 | 80.98 | 1 | 1.31 | 1.61 | 22.96 | 23.88 | -0.92 | 0.92 | -0.01 | 0.01 |
| gene1993 | https://www.ncbi.nlm.nih.gov/gene/?term=LOC105288193 | 1449 | 75 | 517.6 | 1 | 6.9 | 1.33 | 35.76 | 36.74 | -0.98 | 0.98 | -0.05 | 0.05 |
| gene11340 | https://www.ncbi.nlm.nih.gov/gene/?term=LOC105285131 | 2918 | 101 | 346.13 | 1 | 3.43 | 0.99 | 10.9 | 11.93 | -1.02 | 1.02 | -0.04 | 0.04 |
| gene12406 | https://www.ncbi.nlm.nih.gov/gene/?term=LOC105286316 | 1386 | 28 | 202.02 | 1 | 7.22 | 3.57 | 47.22 | 48.43 | -1.22 | 1.22 | -0.02 | 0.02 |
| gene2271 | https://www.ncbi.nlm.nih.gov/gene/?term=LOC105274706 | 1448 | 56 | 386.74 | 1 | 6.91 | 1.79 | 36.89 | 38.13 | -1.24 | 1.24 | -0.05 | 0.05 |
| gene10345 | https://www.ncbi.nlm.nih.gov/gene/?term=LOC105283989 | 4444 | 23 | 51.76 | 1 | 2.25 | 4.35 | 43.82 | 45.07 | -1.26 | 1.26 | -0.01 | 0.01 |
| gene3168 | https://www.ncbi.nlm.nih.gov/gene/?term=LOC105275711 | 6150 | 70 | 113.82 | 1 | 1.63 | 1.43 | 12.53 | 13.84 | -1.31 | 1.31 | -0.01 | 0.01 |
| gene5646 | https://www.ncbi.nlm.nih.gov/gene/?term=LOC105278534 | 7475 | 181 | 242.14 | 1 | 1.34 | 0.55 | 17.19 | 18.52 | -1.32 | 1.32 | -0.03 | 0.03 |
| gene322 | https://www.ncbi.nlm.nih.gov/gene/?term=LOC105275777 | 1606 | 24 | 149.44 | 1 | 6.23 | 4.17 | 45 | 46.35 | -1.35 | 1.35 | -0.02 | 0.02 |
| gene388 | https://www.ncbi.nlm.nih.gov/gene/?term=LOC105276538 | 1558 | 20 | 128.37 | 2 | 12.84 | 10 | 64.73 | 66.14 | -1.41 | 1.41 | -0.02 | 0.02 |
| gene11447 | https://www.ncbi.nlm.nih.gov/gene/?term=LOC105285252 | 7437 | 120 | 161.36 | 1 | 1.34 | 0.83 | 13.34 | 14.97 | -1.63 | 1.63 | -0.03 | 0.03 |
| gene6422 | https://www.ncbi.nlm.nih.gov/gene/?term=LOC105279458 | 2419 | 34 | 140.55 | 1 | 4.13 | 2.94 | 51.48 | 53.11 | -1.63 | 1.63 | -0.02 | 0.02 |
| gene10819 | https://www.ncbi.nlm.nih.gov/gene/?term=LOC105284507 | 5767 | 77 | 133.52 | 1 | 1.73 | 1.3 | 20.73 | 22.56 | -1.84 | 1.84 | -0.02 | 0.02 |
| gene6455 | https://www.ncbi.nlm.nih.gov/gene/?term=LOC105279500 | 2612 | 33 | 126.34 | 1 | 3.83 | 3.03 | 45.07 | 47.06 | -1.99 | 1.99 | -0.03 | 0.03 |
| gene1220 | https://www.ncbi.nlm.nih.gov/gene/?term=LOC105285742 | 3453 | 108 | 312.77 | 1 | 2.9 | 0.93 | 21.23 | 23.26 | -2.03 | 2.03 | -0.06 | 0.06 |
| gene2748 | https://www.ncbi.nlm.nih.gov/gene/?term=LOC105275261 | 3845 | 52 | 135.24 | 1 | 2.6 | 1.92 | 36.08 | 38.33 | -2.25 | 2.25 | -0.03 | 0.03 |
| gene920 | https://www.ncbi.nlm.nih.gov/gene/?term=LOC105282518 | 1197 | 40 | 334.17 | 1 | 8.35 | 2.5 | 37.69 | 40.21 | -2.52 | 2.52 | -0.08 | 0.08 |
| gene2404 | https://www.ncbi.nlm.nih.gov/gene/?term=LOC105274853 | 1767 | 36 | 203.74 | 1 | 5.66 | 2.78 | 27.62 | 31.08 | -3.47 | 3.47 | -0.07 | 0.07 |
| gene1199 | https://www.ncbi.nlm.nih.gov/gene/?term=LOC105285498 | 2220 | 26 | 117.12 | 1 | 4.5 | 3.85 | 33.05 | 37.03 | -3.99 | 3.99 | -0.05 | 0.05 |
| gene12021 | https://www.ncbi.nlm.nih.gov/gene/?term=LOC105285905 | 1399 | 20 | 142.96 | 1 | 7.15 | 5 | 32.7 | 38.03 | -5.34 | 5.34 | -0.08 | 0.08 |
| gene10816 | https://www.ncbi.nlm.nih.gov/gene/?term=LOC105284502 | 1974 | 18 | 91.19 | 1 | 5.07 | 5.56 | 38.87 | 46.78 | -7.91 | 7.91 | -0.07 | 0.07 |
| gene10963 | https://www.ncbi.nlm.nih.gov/gene/?term=LOC105284677 | 1576 | 28 | 177.66 | 2 | 12.69 | 7.14 | 19.34 | 35.66 | -16.32 | 16.32 | -0.29 | 0.29 |

Directory: OGL File: ogl-Cb_LI_BCphaseA2.vs.RphaseA1.txt

| gene_ID | gene_link | gwidth | #Sites | #per10Kb | #dmSites | #dmsp10kb | %dmSites | %pSite1 | %pSite2 | DMpSite | ADMpSite | DMpNucl | ADMpNucl |
|---|---|---|---|---|---|---|---|---|---|---|---|---|---|
| gene7414 | https://www.ncbi.nlm.nih.gov/gene/?term=LOC105280604 | 746 | 1 | 13.4 | 1 | 13.4 | 100 | 75 | 42.31 | 32.69 | 32.69 | 0.04 | 0.04 |
| gene9265 | https://www.ncbi.nlm.nih.gov/gene/?term=LOC105282752 | 1389 | 1 | 7.2 | 1 | 7.2 | 100 | 46.67 | 22.86 | 23.81 | 23.81 | 0.02 | 0.02 |
| gene11618 | https://www.ncbi.nlm.nih.gov/gene/?term=LOC105285452 | 1046 | 1 | 9.56 | 1 | 9.56 | 100 | 68.42 | 47.83 | 20.59 | 20.59 | 0.02 | 0.02 |
| gene12624 | https://www.ncbi.nlm.nih.gov/gene/?term=LOC105286586 | 981 | 1 | 10.19 | 1 | 10.19 | 100 | 35.71 | 15.56 | 20.15 | 20.15 | 0.02 | 0.02 |
| gene1087 | https://www.ncbi.nlm.nih.gov/gene/?term=LOC105284346 | 2052 | 2 | 9.75 | 1 | 4.87 | 50 | 28.11 | 14.29 | 13.83 | 13.83 | 0.01 | 0.01 |
| gene1024 | https://www.ncbi.nlm.nih.gov/gene/?term=LOC105283693 | 1307 | 12 | 91.81 | 4 | 30.6 | 33.33 | 71.36 | 58.1 | 13.26 | 13.26 | 0.12 | 0.12 |
| gene5029 | https://www.ncbi.nlm.nih.gov/gene/?term=LOC105277834 | 1242 | 2 | 16.1 | 1 | 8.05 | 50 | 24.28 | 12.73 | 11.56 | 11.56 | 0.02 | 0.02 |
| gene7864 | https://www.ncbi.nlm.nih.gov/gene/?term=LOC105281110 | 946 | 4 | 42.28 | 1 | 10.57 | 25 | 40.14 | 29.24 | 10.91 | 10.91 | 0.05 | 0.05 |
| gene11932 | https://www.ncbi.nlm.nih.gov/gene/?term=LOC109611391 | 917 | 8 | 87.24 | 1 | 10.91 | 12.5 | 32.02 | 22.62 | 9.4 | 9.4 | 0.08 | 0.08 |
| gene11023 | https://www.ncbi.nlm.nih.gov/gene/?term=LOC105284759 | 1222 | 9 | 73.65 | 1 | 8.18 | 11.11 | 38.59 | 30.71 | 7.87 | 7.87 | 0.06 | 0.06 |
| gene10833 | https://www.ncbi.nlm.nih.gov/gene/?term=LOC105284521 | 2093 | 14 | 66.89 | 1 | 4.78 | 7.14 | 62.65 | 54.81 | 7.84 | 7.84 | 0.05 | 0.05 |
| gene764 | https://www.ncbi.nlm.nih.gov/gene/?term=LOC105280683 | 2178 | 21 | 96.42 | 4 | 18.37 | 19.05 | 54.64 | 46.81 | 7.83 | 7.83 | 0.08 | 0.08 |
| gene11073 | https://www.ncbi.nlm.nih.gov/gene/?term=LOC105284817 | 1692 | 5 | 29.55 | 1 | 5.91 | 20 | 33.92 | 26.99 | 6.92 | 6.92 | 0.02 | 0.02 |
| gene3015 | https://www.ncbi.nlm.nih.gov/gene/?term=LOC105275557 | 811 | 11 | 135.64 | 1 | 12.33 | 9.09 | 74.6 | 67.82 | 6.78 | 6.78 | 0.09 | 0.09 |
| gene2380 | https://www.ncbi.nlm.nih.gov/gene/?term=LOC105274834 | 1214 | 9 | 74.14 | 3 | 24.71 | 33.33 | 53.69 | 46.94 | 6.75 | 6.75 | 0.05 | 0.05 |
| gene5188 | https://www.ncbi.nlm.nih.gov/gene/?term=LOC105277995 | 2486 | 13 | 52.29 | 2 | 8.05 | 15.38 | 18.76 | 12.14 | 6.63 | 6.63 | 0.03 | 0.03 |
| gene2855 | https://www.ncbi.nlm.nih.gov/gene/?term=LOC105275378 | 1189 | 20 | 168.21 | 1 | 8.41 | 5 | 32.85 | 26.65 | 6.2 | 6.2 | 0.1 | 0.1 |
| gene12357 | https://www.ncbi.nlm.nih.gov/gene/?term=LOC105286263 | 1877 | 23 | 122.54 | 4 | 21.31 | 17.39 | 38.1 | 31.96 | 6.14 | 6.14 | 0.08 | 0.08 |
| gene1780 | https://www.ncbi.nlm.nih.gov/gene/?term=LOC105287955 | 2042 | 12 | 58.77 | 1 | 4.9 | 8.33 | 91.37 | 85.34 | 6.03 | 6.03 | 0.04 | 0.04 |
| gene7611 | https://www.ncbi.nlm.nih.gov/gene/?term=LOC105280835 | 1890 | 6 | 31.75 | 1 | 5.29 | 16.67 | 6.39 | 0.36 | 6.03 | 6.03 | 0.02 | 0.02 |
| gene9903 | https://www.ncbi.nlm.nih.gov/gene/?term=LOC105283475 | 1725 | 7 | 40.58 | 1 | 5.8 | 14.29 | 61.36 | 55.71 | 5.65 | 5.65 | 0.02 | 0.02 |
| gene7657 | https://www.ncbi.nlm.nih.gov/gene/?term=LOC105280890 | 1109 | 15 | 135.26 | 2 | 18.03 | 13.33 | 31.21 | 25.71 | 5.5 | 5.5 | 0.07 | 0.07 |
| gene5066 | https://www.ncbi.nlm.nih.gov/gene/?term=LOC105277868 | 2120 | 17 | 80.19 | 1 | 4.72 | 5.88 | 54.78 | 49.31 | 5.48 | 5.48 | 0.04 | 0.04 |
| gene4708 | https://www.ncbi.nlm.nih.gov/gene/?term=LOC105277468 | 1712 | 31 | 181.07 | 3 | 17.52 | 9.68 | 40.08 | 34.7 | 5.39 | 5.39 | 0.1 | 0.1 |

...

... only first 25 lines shown ...

Directory: OGL File: ogl-Cb_LI_BCphaseA2.vs.RphaseA2.txt

| gene_ID | gene_link | gwidth | #Sites | #per10Kb | #dmSites | #dmsp10kb | %dmSites | %pSite1 | %pSite2 | DMpSite | ADMpSite | DMpNucl | ADMpNucl |
|---|---|---|---|---|---|---|---|---|---|---|---|---|---|
| gene7414 | https://www.ncbi.nlm.nih.gov/gene/?term=LOC105280604 | 746 | 1 | 13.4 | 1 | 13.4 | 100 | 75 | 50 | 25 | 25 | 0.03 | 0.03 |
| gene8330 | https://www.ncbi.nlm.nih.gov/gene/?term=LOC105281688 | 1179 | 4 | 33.93 | 1 | 8.48 | 25 | 83.9 | 74.11 | 9.79 | 9.79 | 0.03 | 0.03 |
| gene1517 | https://www.ncbi.nlm.nih.gov/gene/?term=LOC105287649 | 1566 | 7 | 44.7 | 1 | 6.39 | 14.29 | 47.32 | 40.38 | 6.93 | 6.93 | 0.03 | 0.03 |
| gene11269 | https://www.ncbi.nlm.nih.gov/gene/?term=LOC105285051 | 1363 | 15 | 110.05 | 2 | 14.67 | 13.33 | 37.1 | 31.4 | 5.69 | 5.69 | 0.06 | 0.06 |
| gene6755 | https://www.ncbi.nlm.nih.gov/gene/?term=LOC105279839 | 2112 | 11 | 52.08 | 2 | 9.47 | 18.18 | 28.82 | 23.92 | 4.9 | 4.9 | 0.03 | 0.03 |
| gene10237 | https://www.ncbi.nlm.nih.gov/gene/?term=LOC105283856 | 1259 | 26 | 206.51 | 1 | 7.94 | 3.85 | 64.95 | 60.12 | 4.82 | 4.82 | 0.1 | 0.1 |
| gene3297 | https://www.ncbi.nlm.nih.gov/gene/?term=LOC105275861 | 2795 | 16 | 57.25 | 1 | 3.58 | 6.25 | 17.96 | 13.14 | 4.82 | 4.82 | 0.03 | 0.03 |
| gene7727 | https://www.ncbi.nlm.nih.gov/gene/?term=LOC105280969 | 1006 | 9 | 89.46 | 1 | 9.94 | 11.11 | 58.85 | 54.45 | 4.4 | 4.4 | 0.04 | 0.04 |
| gene7300 | https://www.ncbi.nlm.nih.gov/gene/?term=LOC105280472 | 1589 | 6 | 37.76 | 1 | 6.29 | 16.67 | 78.84 | 74.62 | 4.22 | 4.22 | 0.02 | 0.02 |
| gene7398 | https://www.ncbi.nlm.nih.gov/gene/?term=LOC105280589 | 3452 | 45 | 130.36 | 2 | 5.79 | 4.44 | 28.17 | 23.98 | 4.19 | 4.19 | 0.05 | 0.05 |
| gene8608 | https://www.ncbi.nlm.nih.gov/gene/?term=LOC105282002 | 1467 | 26 | 177.23 | 1 | 6.82 | 3.85 | 67.13 | 63.11 | 4.02 | 4.02 | 0.07 | 0.07 |
| gene9464 | https://www.ncbi.nlm.nih.gov/gene/?term=LOC105282975 | 1450 | 22 | 151.72 | 2 | 13.79 | 9.09 | 49.9 | 46.07 | 3.84 | 3.84 | 0.06 | 0.06 |
| gene1024 | https://www.ncbi.nlm.nih.gov/gene/?term=LOC105283693 | 1307 | 12 | 91.81 | 1 | 7.65 | 8.33 | 71.36 | 67.66 | 3.7 | 3.7 | 0.03 | 0.03 |
| gene2959 | https://www.ncbi.nlm.nih.gov/gene/?term=LOC105275501 | 1057 | 7 | 66.23 | 1 | 9.46 | 14.29 | 25.2 | 21.53 | 3.67 | 3.67 | 0.02 | 0.02 |
| gene7190 | https://www.ncbi.nlm.nih.gov/gene/?term=LOC105280355 | 1026 | 14 | 136.45 | 1 | 9.75 | 7.14 | 18.6 | 15.18 | 3.42 | 3.42 | 0.05 | 0.05 |
| gene8198 | https://www.ncbi.nlm.nih.gov/gene/?term=LOC105281539 | 4803 | 18 | 37.48 | 1 | 2.08 | 5.56 | 62.51 | 59.26 | 3.25 | 3.25 | 0.01 | 0.01 |
| gene4156 | https://www.ncbi.nlm.nih.gov/gene/?term=LOC105276840 | 3307 | 11 | 33.26 | 1 | 3.02 | 9.09 | 6.96 | 3.82 | 3.14 | 3.14 | 0.01 | 0.01 |
| gene416 | https://www.ncbi.nlm.nih.gov/gene/?term=LOC105276798 | 2864 | 8 | 27.93 | 1 | 3.49 | 12.5 | 6.95 | 3.82 | 3.14 | 3.14 | 0.01 | 0.01 |
| gene11816 | https://www.ncbi.nlm.nih.gov/gene/?term=LOC105285668 | 1912 | 22 | 115.06 | 3 | 15.69 | 13.64 | 41.69 | 38.58 | 3.11 | 3.11 | 0.04 | 0.04 |
| gene10743 | https://www.ncbi.nlm.nih.gov/gene/?term=LOC105284456 | 1028 | 43 | 418.29 | 3 | 29.18 | 6.98 | 44.49 | 41.39 | 3.1 | 3.1 | 0.13 | 0.13 |
| gene12357 | https://www.ncbi.nlm.nih.gov/gene/?term=LOC105286263 | 1877 | 29 | 154.5 | 1 | 5.33 | 3.45 | 33.04 | 29.95 | 3.09 | 3.09 | 0.05 | 0.05 |
| gene2009 | https://www.ncbi.nlm.nih.gov/gene/?term=LOC105288212 | 1653 | 24 | 145.19 | 1 | 6.05 | 4.17 | 26.6 | 23.57 | 3.03 | 3.03 | 0.04 | 0.04 |
| gene8891 | https://www.ncbi.nlm.nih.gov/gene/?term=LOC105282325 | 2254 | 21 | 93.17 | 1 | 4.44 | 4.76 | 36.49 | 33.5 | 2.99 | 2.99 | 0.03 | 0.03 |
| gene7266 | https://www.ncbi.nlm.nih.gov/gene/?term=LOC105280436 | 4823 | 41 | 85.01 | 3 | 6.22 | 7.32 | 18.51 | 15.53 | 2.98 | 2.98 | 0.03 | 0.03 |

...
... only first 25 lines shown ...

Directory: OGL File: ogl-Cb_LI_BCphaseA2.vs.RphaseB1.txt

| gene_ID | gene_link | gwidth | #Sites | #per10Kb | #dmSites | #dmsp10kb | %dmSites | %pSite1 | %pSite2 | DMpSite | ADMpSite | DMpNucl | ADMpNucl |
|---|---|---|---|---|---|---|---|---|---|---|---|---|---|
| gene2593 | https://www.ncbi.nlm.nih.gov/gene/?term=LOC105275070 | 1774 | 9 | 50.73 | 4 | 22.55 | 44.44 | 77.93 | 29.94 | 47.99 | 47.99 | 0.24 | 0.24 |
| gene3165 | https://www.ncbi.nlm.nih.gov/gene/?term=LOC105275706 | 2374 | 67 | 282.22 | 6 | 25.27 | 8.96 | 35.39 | 24.21 | 11.18 | 11.18 | 0.32 | 0.32 |
| gene2824 | https://www.ncbi.nlm.nih.gov/gene/?term=LOC105275337 | 1772 | 62 | 349.89 | 2 | 11.29 | 3.23 | 38.77 | 30.45 | 8.31 | 8.31 | 0.29 | 0.29 |
| gene5714 | https://www.ncbi.nlm.nih.gov/gene/?term=LOC105278729 | 710 | 16 | 225.35 | 1 | 14.08 | 6.25 | 57.91 | 51.9 | 6.01 | 6.01 | 0.14 | 0.14 |
| gene1993 | https://www.ncbi.nlm.nih.gov/gene/?term=LOC105288193 | 1449 | 75 | 517.6 | 1 | 6.9 | 1.33 | 35.76 | 32.82 | 2.94 | 2.94 | 0.15 | 0.15 |
| gene13033 | https://www.ncbi.nlm.nih.gov/gene/?term=LOC105287063 | 1877 | 49 | 261.05 | 1 | 5.33 | 2.04 | 32.47 | 29.74 | 2.73 | 2.73 | 0.07 | 0.07 |
| gene10984 | https://www.ncbi.nlm.nih.gov/gene/?term=LOC105284704 | 1987 | 62 | 312.03 | 1 | 5.03 | 1.61 | 26.65 | 24.35 | 2.3 | 2.3 | 0.07 | 0.07 |
| gene3671 | https://www.ncbi.nlm.nih.gov/gene/?term=LOC105276272 | 1966 | 38 | 193.29 | 1 | 5.09 | 2.63 | 30.65 | 28.5 | 2.15 | 2.15 | 0.04 | 0.04 |
| gene7556 | https://www.ncbi.nlm.nih.gov/gene/?term=LOC105280774 | 2792 | 101 | 361.75 | 1 | 3.58 | 0.99 | 32.38 | 30.39 | 1.99 | 1.99 | 0.07 | 0.07 |
| gene7398 | https://www.ncbi.nlm.nih.gov/gene/?term=LOC105280589 | 3452 | 42 | 121.67 | 1 | 2.9 | 2.38 | 27.89 | 25.91 | 1.98 | 1.98 | 0.02 | 0.02 |
| gene4034 | https://www.ncbi.nlm.nih.gov/gene/?term=LOC105276704 | 2065 | 86 | 416.46 | 1 | 4.84 | 1.16 | 15.38 | 13.53 | 1.85 | 1.85 | 0.08 | 0.08 |
| gene6652 | https://www.ncbi.nlm.nih.gov/gene/?term=LOC105279719 | 2831 | 78 | 275.52 | 1 | 3.53 | 1.28 | 25.09 | 23.41 | 1.68 | 1.68 | 0.05 | 0.05 |
| gene10520 | https://www.ncbi.nlm.nih.gov/gene/?term=LOC105284184 | 2451 | 87 | 354.96 | 1 | 4.08 | 1.15 | 34.29 | 32.67 | 1.61 | 1.61 | 0.06 | 0.06 |
| gene10091 | https://www.ncbi.nlm.nih.gov/gene/?term=LOC105283697 | 4988 | 137 | 274.66 | 1 | 2 | 0.73 | 17.98 | 16.41 | 1.57 | 1.57 | 0.04 | 0.04 |
| gene4502 | https://www.ncbi.nlm.nih.gov/gene/?term=LOC105277243 | 2289 | 32 | 139.8 | 1 | 4.37 | 3.12 | 44.88 | 43.35 | 1.53 | 1.53 | 0.02 | 0.02 |
| gene5877 | https://www.ncbi.nlm.nih.gov/gene/?term=LOC105278807 | 1719 | 49 | 285.05 | 1 | 5.82 | 2.04 | 25.27 | 23.74 | 1.53 | 1.53 | 0.04 | 0.04 |
| gene9336 | https://www.ncbi.nlm.nih.gov/gene/?term=LOC105282834 | 5993 | 124 | 206.91 | 1 | 1.67 | 0.81 | 15.53 | 14.01 | 1.52 | 1.52 | 0.03 | 0.03 |
| gene6455 | https://www.ncbi.nlm.nih.gov/gene/?term=LOC105279500 | 2612 | 33 | 126.34 | 1 | 3.83 | 3.03 | 51.99 | 50.58 | 1.41 | 1.41 | 0.02 | 0.02 |
| gene7734 | https://www.ncbi.nlm.nih.gov/gene/?term=LOC105280972 | 1528 | 32 | 209.42 | 1 | 6.54 | 3.12 | 19.4 | 18.05 | 1.35 | 1.35 | 0.03 | 0.03 |
| gene2833 | https://www.ncbi.nlm.nih.gov/gene/?term=LOC105275352 | 5528 | 38 | 68.74 | 1 | 1.81 | 2.63 | 25.27 | 24 | 1.27 | 1.27 | 0.01 | 0.01 |
| gene7888 | https://www.ncbi.nlm.nih.gov/gene/?term=LOC105281141 | 4357 | 157 | 360.34 | 1 | 2.3 | 0.64 | 31.42 | 30.25 | 1.17 | 1.17 | 0.04 | 0.04 |
| gene10606 | https://www.ncbi.nlm.nih.gov/gene/?term=LOC105284273 | 6119 | 260 | 424.91 | 1 | 1.63 | 0.38 | 19.55 | 18.4 | 1.15 | 1.15 | 0.05 | 0.05 |
| gene2970 | https://www.ncbi.nlm.nih.gov/gene/?term=LOC105275518 | 2340 | 19 | 81.2 | 1 | 4.27 | 5.26 | 51.65 | 50.52 | 1.13 | 1.13 | 0.01 | 0.01 |
| gene10107 | https://www.ncbi.nlm.nih.gov/gene/?term=LOC105283712 | 1883 | 30 | 159.32 | 1 | 5.31 | 3.33 | 31.98 | 30.87 | 1.11 | 1.11 | 0.02 | 0.02 |
| gene3817 | https://www.ncbi.nlm.nih.gov/gene/?term=LOC105276449 | 1230 | 43 | 349.59 | 1 | 8.13 | 2.33 | 30.78 | 29.68 | 1.1 | 1.1 | 0.04 | 0.04 |
| gene7568 | https://www.ncbi.nlm.nih.gov/gene/?term=LOC105280786 | 2525 | 122 | 483.17 | 1 | 3.96 | 0.82 | 30.88 | 29.81 | 1.07 | 1.07 | 0.05 | 0.05 |
| gene940 | https://www.ncbi.nlm.nih.gov/gene/?term=LOC105282794 | 3578 | 86 | 240.36 | 1 | 2.79 | 1.16 | 16.54 | 15.48 | 1.06 | 1.06 | 0.03 | 0.03 |
| gene7249 | https://www.ncbi.nlm.nih.gov/gene/?term=LOC105280422 | 1775 | 62 | 349.3 | 1 | 5.63 | 1.61 | 17.51 | 16.57 | 0.94 | 0.94 | 0.03 | 0.03 |
| gene5831 | https://www.ncbi.nlm.nih.gov/gene/?term=LOC105278761 | 2506 | 112 | 446.93 | 2 | 7.98 | 1.79 | 21.67 | 20.75 | 0.92 | 0.92 | 0.04 | 0.04 |
| gene10222 | https://www.ncbi.nlm.nih.gov/gene/?term=LOC105283838 | 1748 | 21 | 120.14 | 1 | 5.72 | 4.76 | 46.37 | 45.47 | 0.9 | 0.9 | 0.01 | 0.01 |
| gene322 | https://www.ncbi.nlm.nih.gov/gene/?term=LOC105275777 | 1606 | 23 | 143.21 | 1 | 6.23 | 4.35 | 44.64 | 43.87 | 0.76 | 0.76 | 0.01 | 0.01 |
| gene6286 | https://www.ncbi.nlm.nih.gov/gene/?term=LOC105279298 | 2861 | 121 | 422.93 | 2 | 6.99 | 1.65 | 24.64 | 23.9 | 0.74 | 0.74 | 0.03 | 0.03 |
| gene3912 | https://www.ncbi.nlm.nih.gov/gene/?term=LOC105276560 | 17900 | 309 | 172.63 | 1 | 0.56 | 0.32 | 2.52 | 1.81 | 0.7 | 0.7 | 0.01 | 0.01 |
| gene1147 | https://www.ncbi.nlm.nih.gov/gene/?term=LOC105284954 | 4022 | 96 | 238.69 | 1 | 2.49 | 1.04 | 15.76 | 15.07 | 0.69 | 0.69 | 0.02 | 0.02 |
| gene6031 | https://www.ncbi.nlm.nih.gov/gene/?term=LOC105279000 | 5691 | 208 | 365.49 | 1 | 1.76 | 0.48 | 12.77 | 12.08 | 0.69 | 0.69 | 0.03 | 0.03 |
| gene8015 | https://www.ncbi.nlm.nih.gov/gene/?term=LOC105281312 | 2998 | 68 | 226.82 | 1 | 3.34 | 1.47 | 31.47 | 30.8 | 0.67 | 0.67 | 0.02 | 0.02 |
| gene292 | https://www.ncbi.nlm.nih.gov/gene/?term=LOC105275398 | 2825 | 92 | 325.66 | 1 | 3.54 | 1.09 | 23.29 | 22.65 | 0.64 | 0.64 | 0.02 | 0.02 |
| gene7657 | https://www.ncbi.nlm.nih.gov/gene/?term=LOC105280890 | 1109 | 21 | 189.36 | 1 | 9.02 | 4.76 | 27.2 | 26.57 | 0.64 | 0.64 | 0.01 | 0.01 |
| gene1310 | https://www.ncbi.nlm.nih.gov/gene/?term=LOC105286661 | 2582 | 71 | 274.98 | 1 | 3.87 | 1.41 | 21.34 | 20.74 | 0.61 | 0.61 | 0.02 | 0.02 |
| gene10678 | https://www.ncbi.nlm.nih.gov/gene/?term=LOC105284355 | 4031 | 161 | 399.4 | 1 | 2.48 | 0.62 | 20.49 | 19.95 | 0.54 | 0.54 | 0.02 | 0.02 |
| gene12572 | https://www.ncbi.nlm.nih.gov/gene/?term=LOC105286514 | 2374 | 44 | 185.34 | 1 | 4.21 | 2.27 | 33.89 | 33.38 | 0.52 | 0.52 | 0.01 | 0.01 |
| gene4892 | https://www.ncbi.nlm.nih.gov/gene/?term=LOC105277667 | 3898 | 123 | 315.55 | 1 | 2.57 | 0.81 | 28.28 | 27.77 | 0.51 | 0.51 | 0.02 | 0.02 |
| gene298 | https://www.ncbi.nlm.nih.gov/gene/?term=LOC105275460 | 2473 | 118 | 477.15 | 1 | 4.04 | 0.85 | 13.46 | 12.97 | 0.49 | 0.49 | 0.02 | 0.02 |
| gene289 | https://www.ncbi.nlm.nih.gov/gene/?term=LOC105275347 | 2533 | 72 | 284.25 | 1 | 3.95 | 1.39 | 24.33 | 23.84 | 0.48 | 0.48 | 0.01 | 0.01 |
| gene11568 | https://www.ncbi.nlm.nih.gov/gene/?term=LOC105285395 | 8086 | 202 | 249.81 | 1 | 1.24 | 0.5 | 14.4 | 13.94 | 0.46 | 0.46 | 0.01 | 0.01 |
| gene2056 | https://www.ncbi.nlm.nih.gov/gene/?term=LOC105274452 | 6584 | 236 | 358.44 | 1 | 1.52 | 0.42 | 14.57 | 14.16 | 0.42 | 0.42 | 0.01 | 0.01 |
| gene3470 | https://www.ncbi.nlm.nih.gov/gene/?term=LOC105276059 | 8716 | 151 | 173.24 | 1 | 1.15 | 0.66 | 10.19 | 9.77 | 0.42 | 0.42 | 0.01 | 0.01 |
| gene8729 | https://www.ncbi.nlm.nih.gov/gene/?term=LOC105282137 | 5005 | 211 | 421.58 | 1 | 2 | 0.47 | 16 | 15.6 | 0.4 | 0.4 | 0.02 | 0.02 |
| gene7245 | https://www.ncbi.nlm.nih.gov/gene/?term=LOC105280410 | 12478 | 264 | 211.57 | 1 | 0.8 | 0.38 | 19.99 | 19.61 | 0.38 | 0.38 | 0.01 | 0.01 |
| gene8600 | https://www.ncbi.nlm.nih.gov/gene/?term=LOC105282001 | 8513 | 152 | 178.55 | 1 | 1.17 | 0.66 | 12.95 | 12.59 | 0.36 | 0.36 | 0.01 | 0.01 |
| gene3747 | https://www.ncbi.nlm.nih.gov/gene/?term=LOC105276387 | 3736 | 188 | 503.21 | 1 | 2.68 | 0.53 | 14.43 | 14.09 | 0.35 | 0.35 | 0.02 | 0.02 |
| gene8178 | https://www.ncbi.nlm.nih.gov/gene/?term=LOC105281515 | 8647 | 52 | 60.14 | 1 | 1.16 | 1.92 | 27.71 | 27.4 | 0.32 | 0.32 | 0 | 0 |
| gene2683 | https://www.ncbi.nlm.nih.gov/gene/?term=LOC105275187 | 5492 | 69 | 125.64 | 1 | 1.82 | 1.45 | 14.14 | 13.85 | 0.29 | 0.29 | 0 | 0 |
| gene6237 | https://www.ncbi.nlm.nih.gov/gene/?term=LOC105279244 | 1484 | 27 | 181.94 | 1 | 6.74 | 3.7 | 50.23 | 49.93 | 0.29 | 0.29 | 0.01 | 0.01 |
| gene2846 | https://www.ncbi.nlm.nih.gov/gene/?term=LOC105275374 | 11246 | 500 | 444.6 | 1 | 0.89 | 0.2 | 10.21 | 9.93 | 0.28 | 0.28 | 0.01 | 0.01 |
| gene3727 | https://www.ncbi.nlm.nih.gov/gene/?term=LOC105276367 | 2484 | 68 | 273.75 | 1 | 4.03 | 1.47 | 46.01 | 45.74 | 0.27 | 0.27 | 0.01 | 0.01 |
| gene8343 | https://www.ncbi.nlm.nih.gov/gene/?term=LOC105281700 | 2172 | 38 | 174.95 | 1 | 4.6 | 2.63 | 39.23 | 38.97 | 0.26 | 0.26 | 0 | 0 |
| gene5746 | https://www.ncbi.nlm.nih.gov/gene/?term=LOC105278666 | 10343 | 455 | 439.91 | 1 | 0.97 | 0.22 | 7.39 | 7.14 | 0.25 | 0.25 | 0.01 | 0.01 |
| gene11819 | https://www.ncbi.nlm.nih.gov/gene/?term=LOC105285671 | 2366 | 98 | 414.2 | 1 | 4.23 | 1.02 | 30.44 | 30.22 | 0.23 | 0.23 | 0.01 | 0.01 |
| gene9755 | https://www.ncbi.nlm.nih.gov/gene/?term=LOC105283296 | 2177 | 87 | 399.63 | 1 | 4.59 | 1.15 | 15.89 | 15.67 | 0.23 | 0.23 | 0.01 | 0.01 |

| | | | | | | | | | | | | | |
|---|---|---|---|---|---|---|---|---|---|---|---|---|---|
| gene474 | https://www.ncbi.nlm.nih.gov/gene/?term=LOC105277435 | 3625 | 173 | 477.24 | 1 | 2.76 | 0.58 | 10.83 | 10.64 | 0.2 | 0.2 | 0.01 | 0.01 |
| gene6620 | https://www.ncbi.nlm.nih.gov/gene/?term=LOC105279687 | 17998 | 349 | 193.91 | 1 | 0.56 | 0.29 | 4.97 | 4.84 | 0.13 | 0.13 | 0 | 0 |
| gene11275 | https://www.ncbi.nlm.nih.gov/gene/?term=LOC105285055 | 4119 | 239 | 580.24 | 1 | 2.43 | 0.42 | 11.8 | 11.69 | 0.12 | 0.12 | 0.01 | 0.01 |
| gene2271 | https://www.ncbi.nlm.nih.gov/gene/?term=LOC105274706 | 1448 | 51 | 352.21 | 1 | 6.91 | 1.96 | 38.8 | 38.68 | 0.12 | 0.12 | 0 | 0 |
| gene12307 | https://www.ncbi.nlm.nih.gov/gene/?term=LOC105286216 | 10313 | 178 | 172.6 1 | 0.97 | 0.56 | 15.99 | 15.89 | 0.1 | 0.1 | 0 | 0 | |
| gene3856 | https://www.ncbi.nlm.nih.gov/gene/?term=LOC105276499 | 5236 | 177 | 338.04 | 1 | 1.91 | 0.56 | 10.08 | 9.99 | 0.09 | 0.09 | 0 | 0 |
| gene7166 | https://www.ncbi.nlm.nih.gov/gene/?term=LOC105280326 | 6117 | 222 | 362.92 | 1 | 1.63 | 0.45 | 16.24 | 16.15 | 0.09 | 0.09 | 0 | 0 |
| gene9330 | https://www.ncbi.nlm.nih.gov/gene/?term=LOC105282825 | 3936 | 78 | 198.17 | 1 | 2.54 | 1.28 | 12.99 | 12.9 | 0.09 | 0.09 | 0 | 0 |
| gene53 | https://www.ncbi.nlm.nih.gov/gene/?term=LOC105277148 | 19283 | 283 | 146.76 | 1 | 0.52 | 0.35 | 5.45 | 5.37 | 0.08 | 0.08 | 0 | 0 |
| gene10344 | https://www.ncbi.nlm.nih.gov/gene/?term=LOC105283987 | 4841 | 71 | 146.66 | 1 | 2.07 | 1.41 | 32.31 | 32.24 | 0.07 | 0.07 | 0 | 0 |
| gene10654 | https://www.ncbi.nlm.nih.gov/gene/?term=LOC105284329 | 4027 | 51 | 126.65 | 1 | 2.48 | 1.96 | 19.38 | 19.32 | 0.06 | 0.06 | 0 | 0 |
| gene10600 | https://www.ncbi.nlm.nih.gov/gene/?term=LOC105284270 | 4837 | 100 | 206.74 | 1 | 2.07 | 1 | 12.53 | 12.48 | 0.05 | 0.05 | 0 | 0 |
| gene12981 | https://www.ncbi.nlm.nih.gov/gene/?term=LOC105286999 | 2063 | 44 | 213.28 | 1 | 4.85 | 2.27 | 35.25 | 35.21 | 0.04 | 0.04 | 0 | 0 |
| gene3885 | https://www.ncbi.nlm.nih.gov/gene/?term=LOC105276527 | 7928 | 333 | 420.03 | 1 | 1.26 | 0.3 | 8.6 | 8.57 | 0.03 | 0.03 | 0 | 0 |
| gene10392 | https://www.ncbi.nlm.nih.gov/gene/?term=LOC105284036 | 3216 | 87 | 270.52 | 2 | 6.22 | 2.3 | 24.3 | 24.3 | 0.01 | 0.01 | 0 | 0 |
| gene10549 | https://www.ncbi.nlm.nih.gov/gene/?term=LOC105284210 | 7145 | 327 | 457.66 | 2 | 2.8 | 0.61 | 10.82 | 10.83 | -0.01 | 0.01 | 0 | 0 |
| gene7600 | https://www.ncbi.nlm.nih.gov/gene/?term=LOC105280819 | 1615 | 38 | 235.29 | 1 | 6.19 | 2.63 | 35.56 | 35.58 | -0.02 | 0.02 | 0 | 0 |
| gene2806 | https://www.ncbi.nlm.nih.gov/gene/?term=LOC105275327 | 14257 | 472 | 331.07 | 1 | 0.7 | 0.21 | 7.12 | 7.15 | -0.03 | 0.03 | 0 | 0 |
| gene4190 | https://www.ncbi.nlm.nih.gov/gene/?term=LOC105276875 | 3150 | 101 | 320.63 | 1 | 3.17 | 0.99 | 24.58 | 24.61 | -0.03 | 0.03 | 0 | 0 |
| gene1632 | https://www.ncbi.nlm.nih.gov/gene/?term=LOC105287764 | 18061 | 391 | 216.49 | 1 | 0.55 | 0.26 | 3.3 | 3.35 | -0.05 | 0.05 | 0 | 0 |
| gene5524 | https://www.ncbi.nlm.nih.gov/gene/?term=LOC105278397 | 5177 | 200 | 386.32 | 1 | 1.93 | 0.5 | 24.93 | 25 | -0.07 | 0.07 | 0 | 0 |
| gene12516 | https://www.ncbi.nlm.nih.gov/gene/?term=LOC105286452 | 1529 | 42 | 274.69 | 1 | 6.54 | 2.38 | 46.48 | 46.6 | -0.11 | 0.11 | 0 | 0 |
| gene5525 | https://www.ncbi.nlm.nih.gov/gene/?term=LOC105278396 | 3090 | 186 | 601.94 | 1 | 3.24 | 0.54 | 24.32 | 24.44 | -0.12 | 0.12 | -0.01 | 0.01 |
| gene12802 | https://www.ncbi.nlm.nih.gov/gene/?term=LOC105286799 | 3183 | 173 | 543.51 | 1 | 3.14 | 0.58 | 17.3 | 17.43 | -0.13 | 0.13 | -0.01 | 0.01 |
| gene9029 | https://www.ncbi.nlm.nih.gov/gene/?term=LOC105282491 | 4762 | 169 | 354.89 | 1 | 2.1 | 0.59 | 7.49 | 7.61 | -0.13 | 0.13 | 0 | 0 |
| gene4641 | https://www.ncbi.nlm.nih.gov/gene/?term=LOC105277398 | 1937 | 67 | 345.9 1 | 5.16 | 1.49 | 35.24 | 35.39 | -0.14 | 0.14 | 0 | 0 | |
| gene1884 | https://www.ncbi.nlm.nih.gov/gene/?term=LOC105288072 | 4169 | 77 | 184.7 1 | 2.4 | 1.3 | 24.07 | 24.23 | -0.16 | 0.16 | 0 | 0 | |
| gene10018 | https://www.ncbi.nlm.nih.gov/gene/?term=LOC105283607 | 4414 | 192 | 434.98 | 1 | 2.27 | 0.52 | 15.73 | 15.92 | -0.19 | 0.19 | -0.01 | 0.01 |
| gene13036 | https://www.ncbi.nlm.nih.gov/gene/?term=LOC105287059 | 4959 | 172 | 346.84 | 1 | 2.02 | 0.58 | 16.67 | 16.87 | -0.2 | 0.2 | -0.01 | 0.01 |
| gene2173 | https://www.ncbi.nlm.nih.gov/gene/?term=LOC105274590 | 4387 | 67 | 152.72 | 2 | 4.56 | 2.99 | 31.19 | 31.42 | -0.23 | 0.23 | 0 | 0 |
| gene1470 | https://www.ncbi.nlm.nih.gov/gene/?term=LOC105287597 | 2974 | 192 | 645.6 1 | 3.36 | 0.52 | 18.97 | 19.22 | -0.25 | 0.25 | -0.02 | 0.02 | |
| gene4376 | https://www.ncbi.nlm.nih.gov/gene/?term=LOC105277083 | 4307 | 123 | 285.58 | 1 | 2.32 | 0.81 | 19.63 | 19.89 | -0.26 | 0.26 | -0.01 | 0.01 |
| gene2748 | https://www.ncbi.nlm.nih.gov/gene/?term=LOC105275261 | 3845 | 50 | 130.04 | 1 | 2.6 | 2 | 37.52 | 37.82 | -0.3 | 0.3 | 0 | 0 |
| gene3599 | https://www.ncbi.nlm.nih.gov/gene/?term=LOC105276196 | 2955 | 45 | 152.28 | 1 | 3.38 | 2.22 | 48.59 | 48.91 | -0.31 | 0.31 | 0 | 0 |
| gene10490 | https://www.ncbi.nlm.nih.gov/gene/?term=LOC105284146 | 13960 | 149 | 106.73 | 1 | 0.72 | 0.67 | 10.54 | 10.87 | -0.33 | 0.33 | 0 | 0 |
| gene5646 | https://www.ncbi.nlm.nih.gov/gene/?term=LOC105278534 | 7475 | 180 | 240.8 1 | 1.34 | 0.56 | 17.08 | 17.41 | -0.33 | 0.33 | -0.01 | 0.01 | |
| gene13044 | https://www.ncbi.nlm.nih.gov/gene/?term=LOC105287070 | 5094 | 224 | 439.73 | 1 | 1.96 | 0.45 | 5.36 | 5.7 | -0.34 | 0.34 | -0.01 | 0.01 |
| gene6639 | https://www.ncbi.nlm.nih.gov/gene/?term=LOC105279705 | 7656 | 66 | 86.21 1 | 1.31 | 1.52 | 21.87 | 22.21 | -0.34 | 0.34 | 0 | 0 | |
| gene12674 | https://www.ncbi.nlm.nih.gov/gene/?term=LOC105286649 | 18283 | 764 | 417.87 | 1 | 0.55 | 0.13 | 0.96 | 1.31 | -0.35 | 0.35 | -0.01 | 0.01 |
| gene1571 | https://www.ncbi.nlm.nih.gov/gene/?term=LOC105287700 | 3392 | 160 | 471.7 1 | 2.95 | 0.62 | 15.1 | 15.46 | -0.36 | 0.36 | -0.02 | 0.02 | |
| gene12406 | https://www.ncbi.nlm.nih.gov/gene/?term=LOC105286316 | 1386 | 29 | 209.24 | 1 | 7.22 | 3.45 | 45.59 | 46 | -0.41 | 0.41 | -0.01 | 0.01 |
| gene5298 | https://www.ncbi.nlm.nih.gov/gene/?term=LOC105278145 | 4060 | 251 | 618.23 | 2 | 4.93 | 0.8 | 10.52 | 10.93 | -0.42 | 0.42 | -0.03 | 0.03 |
| gene4962 | https://www.ncbi.nlm.nih.gov/gene/?term=LOC105277761 | 4645 | 161 | 346.61 | 1 | 2.15 | 0.62 | 14.28 | 14.77 | -0.48 | 0.48 | -0.02 | 0.02 |
| gene13114 | https://www.ncbi.nlm.nih.gov/gene/?term=LOC105287147 | 7609 | 398 | 523.06 | 1 | 1.31 | 0.25 | 9.41 | 9.93 | -0.52 | 0.52 | -0.03 | 0.03 |
| gene8274 | https://www.ncbi.nlm.nih.gov/gene/?term=LOC105281624 | 5425 | 221 | 407.37 | 1 | 1.84 | 0.45 | 9.57 | 10.1 | -0.54 | 0.54 | -0.02 | 0.02 |
| gene9326 | https://www.ncbi.nlm.nih.gov/gene/?term=LOC105282818 | 16612 | 217 | 130.63 | 1 | 0.6 | 0.46 | 13.52 | 14.06 | -0.54 | 0.54 | -0.01 | 0.01 |
| gene3107 | https://www.ncbi.nlm.nih.gov/gene/?term=LOC105275650 | 5103 | 172 | 337.06 | 1 | 1.96 | 0.58 | 16.37 | 16.93 | -0.56 | 0.56 | -0.02 | 0.02 |
| gene5375 | https://www.ncbi.nlm.nih.gov/gene/?term=LOC105278228 | 8640 | 167 | 193.29 | 1 | 1.16 | 0.6 | 22.05 | 22.68 | -0.63 | 0.63 | -0.01 | 0.01 |
| gene11561 | https://www.ncbi.nlm.nih.gov/gene/?term=LOC105285387 | 11528 | 200 | 173.49 | 1 | 0.87 | 0.5 | 9.19 | 9.83 | -0.64 | 0.64 | -0.01 | 0.01 |
| gene12573 | https://www.ncbi.nlm.nih.gov/gene/?term=LOC105286515 | 3647 | 108 | 296.13 | 1 | 2.74 | 0.93 | 11.17 | 11.84 | -0.67 | 0.67 | -0.02 | 0.02 |
| gene9327 | https://www.ncbi.nlm.nih.gov/gene/?term=LOC105282821 | 7793 | 161 | 206.6 1 | 1.28 | 0.62 | 12.72 | 13.4 | -0.68 | 0.68 | -0.01 | 0.01 | |
| gene1198 | https://www.ncbi.nlm.nih.gov/gene/?term=LOC105285507 | 2738 | 92 | 336.01 | 1 | 3.65 | 1.09 | 20.74 | 21.44 | -0.7 | 0.7 | -0.02 | 0.02 |
| gene11454 | https://www.ncbi.nlm.nih.gov/gene/?term=LOC105285256 | 1906 | 48 | 251.84 | 1 | 5.25 | 2.08 | 44.93 | 45.73 | -0.8 | 0.8 | -0.02 | 0.02 |
| gene5581 | https://www.ncbi.nlm.nih.gov/gene/?term=LOC105278459 | 2935 | 97 | 330.49 | 1 | 3.41 | 1.03 | 20.44 | 21.26 | -0.83 | 0.83 | -0.03 | 0.03 |
| gene7368 | https://www.ncbi.nlm.nih.gov/gene/?term=LOC105280556 | 4105 | 58 | 141.29 | 1 | 2.44 | 1.72 | 18.1 | 19.01 | -0.91 | 0.91 | -0.01 | 0.01 |
| gene6966 | https://www.ncbi.nlm.nih.gov/gene/?term=LOC105280104 | 3747 | 124 | 330.93 | 1 | 2.67 | 0.81 | 16.15 | 17.09 | -0.94 | 0.94 | -0.03 | 0.03 |
| gene8814 | https://www.ncbi.nlm.nih.gov/gene/?term=LOC105282240 | 6910 | 166 | 240.23 | 1 | 1.45 | 0.6 | 9.27 | 10.23 | -0.96 | 0.96 | -0.02 | 0.02 |
| gene12203 | https://www.ncbi.nlm.nih.gov/gene/?term=LOC105286099 | 5967 | 79 | 132.39 | 1 | 1.68 | 1.27 | 11.92 | 12.93 | -1.01 | 1.01 | -0.01 | 0.01 |
| gene2835 | https://www.ncbi.nlm.nih.gov/gene/?term=LOC105275357 | 2492 | 114 | 457.46 | 1 | 4.01 | 0.88 | 13.76 | 14.82 | -1.05 | 1.05 | -0.05 | 0.05 |
| gene7915 | https://www.ncbi.nlm.nih.gov/gene/?term=LOC105281173 | 1597 | 39 | 244.21 | 1 | 6.26 | 2.56 | 60.85 | 62.01 | -1.16 | 1.16 | -0.03 | 0.03 |
| gene11447 | https://www.ncbi.nlm.nih.gov/gene/?term=LOC105285252 | 7437 | 126 | 169.42 | 2 | 2.69 | 1.59 | 12.66 | 13.85 | -1.19 | 1.19 | -0.02 | 0.02 |
| gene6583 | https://www.ncbi.nlm.nih.gov/gene/?term=LOC105279649 | 7444 | 343 | 460.77 | 1 | 1.34 | 0.29 | 10.51 | 11.78 | -1.26 | 1.26 | -0.06 | 0.06 |
| gene11340 | https://www.ncbi.nlm.nih.gov/gene/?term=LOC105285131 | 2918 | 89 | 305 1 | 3.43 | 1.12 | 11.73 | 13.01 | -1.28 | 1.28 | -0.04 | 0.04 | |
| gene3335 | https://www.ncbi.nlm.nih.gov/gene/?term=LOC105275898 | 4665 | 147 | 315.11 | 1 | 2.14 | 0.68 | 11.48 | 12.76 | -1.29 | 1.29 | -0.04 | 0.04 |

| | | | | | | | | | | | | | |
|---|---|---|---|---|---|---|---|---|---|---|---|---|---|
| gene3848 | https://www.ncbi.nlm.nih.gov/gene/?term=LOC105276486 | 5899 | 145 | 245.8 | 1 | 1.7 | 0.69 | 23.24 | 24.56 | -1.32 | 1.32 | -0.03 | 0.03 |
| gene10345 | https://www.ncbi.nlm.nih.gov/gene/?term=LOC105283989 | 4444 | 21 | 47.25 | 1 | 2.25 | 4.76 | 48.56 | 49.94 | -1.37 | 1.37 | -0.01 | 0.01 |
| gene7207 | https://www.ncbi.nlm.nih.gov/gene/?term=LOC105280376 | 4659 | 112 | 240.39 | 1 | 2.15 | 0.89 | 28.49 | 29.89 | -1.4 | 1.4 | -0.03 | 0.03 |
| gene10155 | https://www.ncbi.nlm.nih.gov/gene/?term=LOC105283756 | 7030 | 98 | 139.4 | 1 | 1.42 | 1.02 | 6.09 | 7.87 | -1.78 | 1.78 | -0.02 | 0.02 |
| gene920 | https://www.ncbi.nlm.nih.gov/gene/?term=LOC105282518 | 1197 | 39 | 325.81 | 1 | 8.35 | 2.56 | 36.44 | 38.28 | -1.85 | 1.85 | -0.06 | 0.06 |
| gene3168 | https://www.ncbi.nlm.nih.gov/gene/?term=LOC105275711 | 6150 | 80 | 130.08 | 1 | 1.63 | 1.25 | 11.3 | 13.17 | -1.86 | 1.86 | -0.02 | 0.02 |
| gene10819 | https://www.ncbi.nlm.nih.gov/gene/?term=LOC105284507 | 5767 | 86 | 149.12 | 1 | 1.73 | 1.16 | 21.64 | 23.53 | -1.89 | 1.89 | -0.03 | 0.03 |
| gene1220 | https://www.ncbi.nlm.nih.gov/gene/?term=LOC105285742 | 3453 | 98 | 283.81 | 1 | 2.9 | 1.02 | 19.04 | 20.94 | -1.9 | 1.9 | -0.05 | 0.05 |
| gene388 | https://www.ncbi.nlm.nih.gov/gene/?term=LOC105276538 | 1558 | 19 | 121.95 | 2 | 12.84 | 10.53 | 66.07 | 68.07 | -1.99 | 1.99 | -0.02 | 0.02 |
| gene10879 | https://www.ncbi.nlm.nih.gov/gene/?term=LOC105284574 | 2423 | 45 | 185.72 | 1 | 4.13 | 2.22 | 57.53 | 59.68 | -2.15 | 2.15 | -0.04 | 0.04 |
| gene10552 | https://www.ncbi.nlm.nih.gov/gene/?term=LOC105284213 | 878 | 15 | 170.84 | 1 | 11.39 | 6.67 | 15.38 | 18.37 | -2.99 | 2.99 | -0.05 | 0.05 |
| gene12021 | https://www.ncbi.nlm.nih.gov/gene/?term=LOC105285905 | 1399 | 27 | 192.99 | 1 | 7.15 | 3.7 | 31.19 | 34.36 | -3.17 | 3.17 | -0.06 | 0.06 |
| gene1199 | https://www.ncbi.nlm.nih.gov/gene/?term=LOC105285498 | 2220 | 25 | 112.61 | 1 | 4.5 | 4 | 32.49 | 35.74 | -3.26 | 3.26 | -0.04 | 0.04 |
| gene2413 | https://www.ncbi.nlm.nih.gov/gene/?term=LOC105274863 | 2006 | 46 | 229.31 | 1 | 4.99 | 2.17 | 60.76 | 64.28 | -3.53 | 3.53 | -0.08 | 0.08 |
| gene2404 | https://www.ncbi.nlm.nih.gov/gene/?term=LOC105274853 | 1767 | 34 | 192.42 | 1 | 5.66 | 2.94 | 29.24 | 33.79 | -4.55 | 4.55 | -0.09 | 0.09 |
| gene3166 | https://www.ncbi.nlm.nih.gov/gene/?term=LOC105275707 | 2393 | 11 | 45.97 | 1 | 4.18 | 9.09 | 62.4 | 71.24 | -8.83 | 8.83 | -0.04 | 0.04 |
| gene9109 | https://www.ncbi.nlm.nih.gov/gene/?term=LOC105282561 | 2400 | 11 | 45.83 | 1 | 4.17 | 9.09 | 17.27 | 26.23 | -8.97 | 8.97 | -0.04 | 0.04 |
| gene11584 | https://www.ncbi.nlm.nih.gov/gene/?term=LOC105285416 | 3709 | 6 | 16.18 | 2 | 5.39 | 33.33 | 18.09 | 32.8 | -14.71 | 14.71 | -0.02 | 0.02 |
| gene393 | https://www.ncbi.nlm.nih.gov/gene/?term=LOC105276566 | 1123 | 2 | 17.81 | 1 | 8.9 | 50 | 48.08 | 74.3 | -26.22 | 26.22 | -0.05 | 0.05 |

Directory: OGL File: ogl-Cb_LI_BCphaseA2.vs.RphaseB2.txt

| gene_ID | gene_link | gwidth | #Sites | #per10Kb | #dmSites | #dmsp10kb | %dmSites | %pSite1 | %pSite2 | DMpSite | ADMpSite | DMpNucl | ADMpNucl |
|---|---|---|---|---|---|---|---|---|---|---|---|---|---|
| gene2593 | https://www.ncbi.nlm.nih.gov/gene/?term=LOC105275070 | 1774 | 7 | 39.46 | 5 | 28.18 | 71.43 | 66.52 | 4.09 | 62.43 | 62.43 | 0.25 | 0.25 |
| gene3165 | https://www.ncbi.nlm.nih.gov/gene/?term=LOC105275706 | 2374 | 72 | 303.29 | 6 | 25.27 | 8.33 | 33.88 | 22.12 | 11.76 | 11.76 | 0.36 | 0.36 |
| gene2824 | https://www.ncbi.nlm.nih.gov/gene/?term=LOC105275337 | 1772 | 60 | 338.6 | 2 | 11.29 | 3.33 | 38.89 | 29.82 | 9.07 | 9.07 | 0.31 | 0.31 |
| gene5714 | https://www.ncbi.nlm.nih.gov/gene/?term=LOC105278729 | 710 | 17 | 239.44 | 1 | 14.08 | 5.88 | 60.22 | 53.54 | 6.67 | 6.67 | 0.16 | 0.16 |
| gene7398 | https://www.ncbi.nlm.nih.gov/gene/?term=LOC105280589 | 3452 | 34 | 98.49 | 1 | 2.9 | 2.94 | 23.83 | 19.97 | 3.86 | 3.86 | 0.04 | 0.04 |
| gene2970 | https://www.ncbi.nlm.nih.gov/gene/?term=LOC105275518 | 2340 | 18 | 76.92 | 1 | 4.27 | 5.56 | 54.32 | 51.43 | 2.89 | 2.89 | 0.02 | 0.02 |
| gene940 | https://www.ncbi.nlm.nih.gov/gene/?term=LOC105282794 | 3578 | 87 | 243.15 | 1 | 2.79 | 1.15 | 16.03 | 14.13 | 1.9 | 1.9 | 0.05 | 0.05 |
| gene6455 | https://www.ncbi.nlm.nih.gov/gene/?term=LOC105279500 | 2612 | 33 | 126.34 | 1 | 3.83 | 3.03 | 51.77 | 49.92 | 1.86 | 1.86 | 0.02 | 0.02 |
| gene7568 | https://www.ncbi.nlm.nih.gov/gene/?term=LOC105280786 | 2525 | 112 | 443.56 | 1 | 3.96 | 0.89 | 32.64 | 30.88 | 1.76 | 1.76 | 0.08 | 0.08 |
| gene10520 | https://www.ncbi.nlm.nih.gov/gene/?term=LOC105284184 | 2451 | 78 | 318.24 | 1 | 4.08 | 1.28 | 37.44 | 35.71 | 1.72 | 1.72 | 0.05 | 0.05 |
| gene9462 | https://www.ncbi.nlm.nih.gov/gene/?term=LOC105282970 | 1469 | 39 | 265.49 | 1 | 6.81 | 2.56 | 51.12 | 49.47 | 1.65 | 1.65 | 0.04 | 0.04 |
| gene3385 | https://www.ncbi.nlm.nih.gov/gene/?term=LOC105275966 | 3305 | 53 | 160.36 | 1 | 3.03 | 1.89 | 54.5 | 52.88 | 1.62 | 1.62 | 0.03 | 0.03 |
| gene3594 | https://www.ncbi.nlm.nih.gov/gene/?term=LOC105276190 | 4910 | 83 | 169.04 | 1 | 2.04 | 1.2 | 11.44 | 9.83 | 1.62 | 1.62 | 0.03 | 0.03 |
| gene11819 | https://www.ncbi.nlm.nih.gov/gene/?term=LOC105285671 | 2366 | 100 | 422.65 | 1 | 4.23 | 1 | 27.82 | 26.43 | 1.39 | 1.39 | 0.06 | 0.06 |
| gene5877 | https://www.ncbi.nlm.nih.gov/gene/?term=LOC105278807 | 1719 | 46 | 267.6 | 1 | 5.82 | 2.17 | 26.78 | 25.52 | 1.25 | 1.25 | 0.03 | 0.03 |
| gene35 | https://www.ncbi.nlm.nih.gov/gene/?term=LOC105275197 | 2643 | 43 | 162.69 | 1 | 3.78 | 2.33 | 44.25 | 43.02 | 1.22 | 1.22 | 0.02 | 0.02 |
| gene13033 | https://www.ncbi.nlm.nih.gov/gene/?term=LOC105287063 | 1877 | 48 | 255.73 | 1 | 5.33 | 2.08 | 31.55 | 30.37 | 1.19 | 1.19 | 0.03 | 0.03 |
| gene3470 | https://www.ncbi.nlm.nih.gov/gene/?term=LOC105276059 | 8716 | 145 | 166.36 | 1 | 1.15 | 0.69 | 9.2 | 8.01 | 1.19 | 1.19 | 0.02 | 0.02 |
| gene7915 | https://www.ncbi.nlm.nih.gov/gene/?term=LOC105281173 | 1597 | 38 | 237.95 | 1 | 6.26 | 2.63 | 59.94 | 58.74 | 1.19 | 1.19 | 0.03 | 0.03 |
| gene12806 | https://www.ncbi.nlm.nih.gov/gene/?term=LOC105286803 | 4196 | 137 | 326.5 | 1 | 2.38 | 0.73 | 10.84 | 9.75 | 1.09 | 1.09 | 0.04 | 0.04 |
| gene6652 | https://www.ncbi.nlm.nih.gov/gene/?term=LOC105279719 | 2831 | 79 | 279.05 | 1 | 3.53 | 1.27 | 24.35 | 23.32 | 1.03 | 1.03 | 0.03 | 0.03 |
| gene7556 | https://www.ncbi.nlm.nih.gov/gene/?term=LOC105280774 | 2792 | 89 | 318.77 | 1 | 3.58 | 1.12 | 36.33 | 35.33 | 1 | 1 | 0.03 | 0.03 |
| gene3671 | https://www.ncbi.nlm.nih.gov/gene/?term=LOC105276272 | 1966 | 34 | 172.94 | 1 | 5.09 | 2.94 | 33.29 | 32.32 | 0.98 | 0.98 | 0.02 | 0.02 |
| gene10678 | https://www.ncbi.nlm.nih.gov/gene/?term=LOC105284355 | 4031 | 158 | 391.96 | 1 | 2.48 | 0.63 | 20.75 | 19.82 | 0.93 | 0.93 | 0.04 | 0.04 |
| gene3817 | https://www.ncbi.nlm.nih.gov/gene/?term=LOC105276449 | 1230 | 42 | 341.46 | 1 | 8.13 | 2.38 | 35.93 | 35.04 | 0.88 | 0.88 | 0.03 | 0.03 |
| gene10344 | https://www.ncbi.nlm.nih.gov/gene/?term=LOC105283987 | 4841 | 64 | 132.2 | 1 | 2.07 | 1.56 | 30.28 | 29.42 | 0.86 | 0.86 | 0.01 | 0.01 |
| gene10606 | https://www.ncbi.nlm.nih.gov/gene/?term=LOC105284273 | 6119 | 249 | 406.93 | 1 | 1.63 | 0.4 | 18.46 | 17.61 | 0.86 | 0.86 | 0.03 | 0.03 |
| gene1470 | https://www.ncbi.nlm.nih.gov/gene/?term=LOC105287597 | 2974 | 195 | 655.68 | 1 | 3.36 | 0.51 | 18.31 | 17.52 | 0.8 | 0.8 | 0.05 | 0.05 |
| gene4962 | https://www.ncbi.nlm.nih.gov/gene/?term=LOC105277761 | 4645 | 142 | 305.71 | 1 | 2.15 | 0.7 | 14.73 | 13.94 | 0.79 | 0.79 | 0.02 | 0.02 |
| gene4892 | https://www.ncbi.nlm.nih.gov/gene/?term=LOC105277667 | 3898 | 124 | 318.11 | 1 | 2.57 | 0.81 | 27.97 | 27.19 | 0.78 | 0.78 | 0.02 | 0.02 |
| gene11401 | https://www.ncbi.nlm.nih.gov/gene/?term=LOC105285202 | 1841 | 82 | 445.41 | 1 | 5.43 | 1.22 | 26.05 | 25.33 | 0.73 | 0.73 | 0.03 | 0.03 |
| gene9336 | https://www.ncbi.nlm.nih.gov/gene/?term=LOC105282834 | 5993 | 121 | 201.9 | 1 | 1.67 | 0.83 | 16.02 | 15.31 | 0.72 | 0.72 | 0.01 | 0.01 |
| gene10600 | https://www.ncbi.nlm.nih.gov/gene/?term=LOC105284270 | 4837 | 119 | 246.02 | 1 | 2.07 | 0.84 | 10.7 | 10.02 | 0.68 | 0.68 | 0.02 | 0.02 |
| gene2846 | https://www.ncbi.nlm.nih.gov/gene/?term=LOC105275374 | 11246 | 464 | 412.59 | 1 | 0.89 | 0.22 | 12.19 | 11.56 | 0.63 | 0.63 | 0.03 | 0.03 |
| gene2173 | https://www.ncbi.nlm.nih.gov/gene/?term=LOC105274590 | 4387 | 57 | 129.93 | 2 | 4.56 | 3.51 | 33.32 | 32.7 | 0.62 | 0.62 | 0.01 | 0.01 |
| gene3727 | https://www.ncbi.nlm.nih.gov/gene/?term=LOC105276367 | 2484 | 66 | 265.7 | 1 | 4.03 | 1.52 | 47.44 | 46.84 | 0.61 | 0.61 | 0.02 | 0.02 |
| gene11568 | https://www.ncbi.nlm.nih.gov/gene/?term=LOC105285395 | 8086 | 180 | 222.61 | 1 | 1.24 | 0.56 | 17.19 | 16.59 | 0.6 | 0.6 | 0.01 | 0.01 |
| gene10984 | https://www.ncbi.nlm.nih.gov/gene/?term=LOC105284704 | 1987 | 56 | 281.83 | 1 | 5.03 | 1.79 | 25.7 | 25.1 | 0.59 | 0.59 | 0.02 | 0.02 |
| gene2844 | https://www.ncbi.nlm.nih.gov/gene/?term=LOC105275363 | 4330 | 181 | 418.01 | 1 | 2.31 | 0.55 | 15.09 | 14.51 | 0.58 | 0.58 | 0.02 | 0.02 |
| gene8274 | https://www.ncbi.nlm.nih.gov/gene/?term=LOC105281624 | 5425 | 204 | 376.04 | 1 | 1.84 | 0.49 | 10.02 | 9.47 | 0.56 | 0.56 | 0.02 | 0.02 |
| gene7245 | https://www.ncbi.nlm.nih.gov/gene/?term=LOC105280410 | 12478 | 234 | 187.53 | 1 | 0.8 | 0.43 | 21.44 | 20.93 | 0.52 | 0.52 | 0.01 | 0.01 |
| gene5573 | https://www.ncbi.nlm.nih.gov/gene/?term=LOC105278454 | 2688 | 35 | 130.21 | 1 | 3.72 | 2.86 | 35.99 | 35.53 | 0.47 | 0.47 | 0.01 | 0.01 |
| gene12572 | https://www.ncbi.nlm.nih.gov/gene/?term=LOC105286514 | 2374 | 45 | 189.55 | 1 | 4.21 | 2.22 | 33.13 | 32.67 | 0.45 | 0.45 | 0.01 | 0.01 |
| gene34 | https://www.ncbi.nlm.nih.gov/gene/?term=LOC105278416 | 7120 | 124 | 174.16 | 1 | 1.4 | 0.81 | 15.53 | 15.09 | 0.43 | 0.43 | 0.01 | 0.01 |
| gene2056 | https://www.ncbi.nlm.nih.gov/gene/?term=LOC105274452 | 6584 | 251 | 381.23 | 2 | 3.04 | 0.8 | 13.77 | 13.35 | 0.42 | 0.42 | 0.02 | 0.02 |
| gene6286 | https://www.ncbi.nlm.nih.gov/gene/?term=LOC105279298 | 2861 | 128 | 447.4 | 2 | 6.99 | 1.56 | 24.57 | 24.17 | 0.4 | 0.4 | 0.02 | 0.02 |
| gene7166 | https://www.ncbi.nlm.nih.gov/gene/?term=LOC105280326 | 6117 | 214 | 349.84 | 1 | 1.63 | 0.47 | 16.62 | 16.23 | 0.4 | 0.4 | 0.01 | 0.01 |
| gene4246 | https://www.ncbi.nlm.nih.gov/gene/?term=LOC105276941 | 16412 | 493 | 300.39 | 1 | 0.61 | 0.2 | 5.46 | 5.07 | 0.38 | 0.38 | 0.01 | 0.01 |
| gene5746 | https://www.ncbi.nlm.nih.gov/gene/?term=LOC105278666 | 10343 | 425 | 410.91 | 1 | 0.97 | 0.24 | 7.62 | 7.27 | 0.35 | 0.35 | 0.01 | 0.01 |
| gene10392 | https://www.ncbi.nlm.nih.gov/gene/?term=LOC105284036 | 3216 | 91 | 282.96 | 2 | 6.22 | 2.2 | 22.63 | 22.31 | 0.32 | 0.32 | 0.01 | 0.01 |
| gene473 | https://www.ncbi.nlm.nih.gov/gene/?term=LOC105277423 | 2342 | 108 | 461.14 | 1 | 4.27 | 0.93 | 30.31 | 29.99 | 0.32 | 0.32 | 0.01 | 0.01 |
| gene11579 | https://www.ncbi.nlm.nih.gov/gene/?term=LOC105285406 | 13571 | 228 | 168.01 | 1 | 0.74 | 0.44 | 7.6 | 7.29 | 0.3 | 0.3 | 0.01 | 0.01 |
| gene2271 | https://www.ncbi.nlm.nih.gov/gene/?term=LOC105274706 | 1448 | 56 | 386.74 | 1 | 6.91 | 1.79 | 36.69 | 36.4 | 0.3 | 0.3 | 0.01 | 0.01 |
| gene10107 | https://www.ncbi.nlm.nih.gov/gene/?term=LOC105283712 | 1883 | 32 | 169.94 | 1 | 5.31 | 3.12 | 29.98 | 29.7 | 0.28 | 0.28 | 0 | 0 |
| gene10531 | https://www.ncbi.nlm.nih.gov/gene/?term=LOC105284190 | 6881 | 168 | 244.15 | 1 | 1.45 | 0.6 | 13.32 | 13.04 | 0.27 | 0.27 | 0.01 | 0.01 |
| gene7211 | https://www.ncbi.nlm.nih.gov/gene/?term=LOC105280377 | 2024 | 57 | 281.62 | 1 | 4.94 | 1.75 | 50.41 | 50.16 | 0.25 | 0.25 | 0.01 | 0.01 |
| gene298 | https://www.ncbi.nlm.nih.gov/gene/?term=LOC105275460 | 2473 | 116 | 469.07 | 1 | 4.04 | 0.86 | 13.65 | 13.43 | 0.23 | 0.23 | 0.01 | 0.01 |
| gene5525 | https://www.ncbi.nlm.nih.gov/gene/?term=LOC105278396 | 3090 | 181 | 585.76 | 1 | 3.24 | 0.55 | 24.85 | 24.63 | 0.22 | 0.22 | 0.01 | 0.01 |
| gene5524 | https://www.ncbi.nlm.nih.gov/gene/?term=LOC105278397 | 5177 | 196 | 378.6 | 1 | 1.93 | 0.51 | 24.88 | 24.69 | 0.18 | 0.18 | 0.01 | 0.01 |
| gene8600 | https://www.ncbi.nlm.nih.gov/gene/?term=LOC105282001 | 8513 | 139 | 163.28 | 1 | 1.17 | 0.72 | 11.37 | 11.2 | 0.17 | 0.17 | 0 | 0 |

| | | | | | | | | | | | | | |
|---|---|---|---|---|---|---|---|---|---|---|---|---|---|
| gene9330 | https://www.ncbi.nlm.nih.gov/gene/?term=LOC105282825 | 3936 | 69 | 175.3 1 | 2.54 | 1.45 | 13.52 | 13.35 | 0.17 | 0.17 | 0 | 0 | |
| gene12307 | https://www.ncbi.nlm.nih.gov/gene/?term=LOC105286216 | 10313 | 167 | 161.93 | 1 | 0.97 | 0.6 | 17.01 | 16.86 | 0.15 | 0.15 | 0 | 0 |
| gene6620 | https://www.ncbi.nlm.nih.gov/gene/?term=LOC105279687 | 17998 | 312 | 173.35 | 1 | 0.56 | 0.32 | 5.46 | 5.32 | 0.14 | 0.14 | 0 | 0 |
| gene12265 | https://www.ncbi.nlm.nih.gov/gene/?term=LOC105286166 | 3050 | 104 | 340.98 | 1 | 3.28 | 0.96 | 25.25 | 25.13 | 0.13 | 0.13 | 0 | 0 |
| gene292 | https://www.ncbi.nlm.nih.gov/gene/?term=LOC105275398 | 2825 | 81 | 286.73 | 1 | 3.54 | 1.23 | 26.31 | 26.19 | 0.11 | 0.11 | 0 | 0 |
| gene11275 | https://www.ncbi.nlm.nih.gov/gene/?term=LOC105285055 | 4119 | 252 | 611.8 1 | 2.43 | 0.4 | 11.37 | 11.3 | 0.07 | 0.07 | 0 | 0 | |
| gene2806 | https://www.ncbi.nlm.nih.gov/gene/?term=LOC105275327 | 14257 | 460 | 322.65 | 1 | 0.7 | 0.22 | 7.25 | 7.19 | 0.06 | 0.06 | 0 | 0 |
| gene12981 | https://www.ncbi.nlm.nih.gov/gene/?term=LOC105286999 | 2063 | 42 | 203.59 | 1 | 4.85 | 2.38 | 36.12 | 36.07 | 0.05 | 0.05 | 0 | 0 |
| gene13114 | https://www.ncbi.nlm.nih.gov/gene/?term=LOC105287147 | 7609 | 383 | 503.35 | 1 | 1.31 | 0.26 | 9.68 | 9.63 | 0.04 | 0.04 | 0 | 0 |
| gene53 | https://www.ncbi.nlm.nih.gov/gene/?term=LOC105277148 | 19283 | 258 | 133.8 1 | 0.52 | 0.39 | 4.46 | 4.42 | 0.04 | 0.04 | 0 | 0 | |
| gene11082 | https://www.ncbi.nlm.nih.gov/gene/?term=LOC105284824 | 7436 | 185 | 248.79 | 1 | 1.34 | 0.54 | 6.28 | 6.27 | 0.01 | 0.01 | 0 | 0 |
| gene1632 | https://www.ncbi.nlm.nih.gov/gene/?term=LOC105287764 | 18061 | 371 | 205.41 | 2 | 1.11 | 0.54 | 3.28 | 3.27 | 0.01 | 0.01 | 0 | 0 |
| gene6966 | https://www.ncbi.nlm.nih.gov/gene/?term=LOC105280104 | 3747 | 123 | 328.26 | 1 | 2.67 | 0.81 | 16.59 | 16.58 | 0.01 | 0.01 | 0 | 0 |
| gene4190 | https://www.ncbi.nlm.nih.gov/gene/?term=LOC105276875 | 3150 | 109 | 346.03 | 1 | 3.17 | 0.92 | 24.02 | 24.04 | -0.02 | 0.02 | 0 | 0 |
| gene1993 | https://www.ncbi.nlm.nih.gov/gene/?term=LOC105288193 | 1449 | 74 | 510.7 1 | 6.9 | 1.35 | 34.89 | 34.92 | -0.03 | 0.03 | 0 | 0 | |
| gene8729 | https://www.ncbi.nlm.nih.gov/gene/?term=LOC105282137 | 5005 | 198 | 395.6 1 | 2 | 0.51 | 16.91 | 16.95 | -0.04 | 0.04 | 0 | 0 | |
| gene5831 | https://www.ncbi.nlm.nih.gov/gene/?term=LOC105278761 | 2506 | 109 | 434.96 | 1 | 3.99 | 0.92 | 24.44 | 24.49 | -0.06 | 0.06 | 0 | 0 |
| gene9177 | https://www.ncbi.nlm.nih.gov/gene/?term=LOC105282646 | 3320 | 99 | 298.19 | 1 | 3.01 | 1.01 | 25.05 | 25.11 | -0.06 | 0.06 | 0 | 0 |
| gene7249 | https://www.ncbi.nlm.nih.gov/gene/?term=LOC105280422 | 1775 | 60 | 338.03 | 1 | 5.63 | 1.67 | 16.97 | 17.06 | -0.09 | 0.09 | 0 | 0 |
| gene3286 | https://www.ncbi.nlm.nih.gov/gene/?term=LOC105275848 | 3744 | 134 | 357.91 | 1 | 2.67 | 0.75 | 22.27 | 22.41 | -0.14 | 0.14 | -0.01 | 0.01 |
| gene5967 | https://www.ncbi.nlm.nih.gov/gene/?term=LOC105278918 | 4393 | 116 | 264.06 | 1 | 2.28 | 0.86 | 14.39 | 14.54 | -0.15 | 0.15 | 0 | 0 |
| gene10549 | https://www.ncbi.nlm.nih.gov/gene/?term=LOC105284210 | 7145 | 286 | 400.28 | 2 | 2.8 | 0.7 | 11.97 | 12.18 | -0.21 | 0.21 | -0.01 | 0.01 |
| gene12802 | https://www.ncbi.nlm.nih.gov/gene/?term=LOC105286799 | 3183 | 160 | 502.67 | 1 | 3.14 | 0.62 | 17.6 | 17.81 | -0.21 | 0.21 | -0.01 | 0.01 |
| gene3107 | https://www.ncbi.nlm.nih.gov/gene/?term=LOC105275650 | 5103 | 166 | 325.3 1 | 1.96 | 0.6 | 14.75 | 14.95 | -0.21 | 0.21 | -0.01 | 0.01 | |
| gene9755 | https://www.ncbi.nlm.nih.gov/gene/?term=LOC105283296 | 2177 | 85 | 390.45 | 1 | 4.59 | 1.18 | 19.16 | 19.37 | -0.21 | 0.21 | -0.01 | 0.01 |
| gene8661 | https://www.ncbi.nlm.nih.gov/gene/?term=LOC105282063 | 1926 | 35 | 181.72 | 1 | 5.19 | 2.86 | 30.68 | 30.89 | -0.22 | 0.22 | 0 | 0 |
| gene1199 | https://www.ncbi.nlm.nih.gov/gene/?term=LOC105285498 | 2220 | 26 | 117.12 | 1 | 4.5 | 3.85 | 31.35 | 31.59 | -0.24 | 0.24 | 0 | 0 |
| gene5733 | https://www.ncbi.nlm.nih.gov/gene/?term=LOC105278649 | 5221 | 114 | 218.35 | 1 | 1.92 | 0.88 | 16.23 | 16.48 | -0.25 | 0.25 | -0.01 | 0.01 |
| gene1310 | https://www.ncbi.nlm.nih.gov/gene/?term=LOC105286661 | 2582 | 72 | 278.85 | 1 | 3.87 | 1.39 | 21.23 | 21.48 | -0.26 | 0.26 | -0.01 | 0.01 |
| gene9326 | https://www.ncbi.nlm.nih.gov/gene/?term=LOC105282818 | 16612 | 173 | 104.14 | 1 | 0.6 | 0.58 | 12.61 | 12.88 | -0.27 | 0.27 | 0 | 0 |
| gene12674 | https://www.ncbi.nlm.nih.gov/gene/?term=LOC105286649 | 18283 | 747 | 408.58 | 1 | 0.55 | 0.13 | 0.89 | 1.18 | -0.29 | 0.29 | -0.01 | 0.01 |
| gene13036 | https://www.ncbi.nlm.nih.gov/gene/?term=LOC105287059 | 4959 | 146 | 294.41 | 1 | 2.02 | 0.68 | 18.9 | 19.21 | -0.31 | 0.31 | -0.01 | 0.01 |
| gene3885 | https://www.ncbi.nlm.nih.gov/gene/?term=LOC105276527 | 7928 | 305 | 384.71 | 1 | 1.26 | 0.33 | 9.75 | 10.06 | -0.32 | 0.32 | -0.01 | 0.01 |
| gene6583 | https://www.ncbi.nlm.nih.gov/gene/?term=LOC105279649 | 7444 | 303 | 407.04 | 1 | 1.34 | 0.33 | 11.36 | 11.69 | -0.33 | 0.33 | -0.01 | 0.01 |
| gene10018 | https://www.ncbi.nlm.nih.gov/gene/?term=LOC105283607 | 4414 | 187 | 423.65 | 1 | 2.27 | 0.53 | 16.13 | 16.48 | -0.35 | 0.35 | -0.01 | 0.01 |
| gene2122 | https://www.ncbi.nlm.nih.gov/gene/?term=LOC105274531 | 8153 | 289 | 354.47 | 1 | 1.23 | 0.35 | 10.34 | 10.69 | -0.35 | 0.35 | -0.01 | 0.01 |
| gene6031 | https://www.ncbi.nlm.nih.gov/gene/?term=LOC105279000 | 5691 | 187 | 328.59 | 1 | 1.76 | 0.53 | 16.08 | 16.46 | -0.38 | 0.38 | -0.01 | 0.01 |
| gene12573 | https://www.ncbi.nlm.nih.gov/gene/?term=LOC105286515 | 3647 | 98 | 268.71 | 1 | 2.74 | 1.02 | 13.27 | 13.66 | -0.39 | 0.39 | -0.01 | 0.01 |
| gene12021 | https://www.ncbi.nlm.nih.gov/gene/?term=LOC105285905 | 1399 | 16 | 114.37 | 1 | 7.15 | 6.25 | 47.33 | 47.75 | -0.42 | 0.42 | 0 | 0 |
| gene6639 | https://www.ncbi.nlm.nih.gov/gene/?term=LOC105279705 | 7656 | 61 | 79.68 1 | 1.31 | 1.64 | 23.52 | 23.98 | -0.46 | 0.46 | 0 | 0 | |
| gene5298 | https://www.ncbi.nlm.nih.gov/gene/?term=LOC105278145 | 4060 | 245 | 603.45 | 2 | 4.93 | 0.82 | 10.68 | 11.19 | -0.51 | 0.51 | -0.03 | 0.03 |
| gene4376 | https://www.ncbi.nlm.nih.gov/gene/?term=LOC105277083 | 4307 | 102 | 236.82 | 1 | 2.32 | 0.98 | 24.57 | 25.16 | -0.58 | 0.58 | -0.01 | 0.01 |
| gene11340 | https://www.ncbi.nlm.nih.gov/gene/?term=LOC105285131 | 2918 | 86 | 294.72 | 1 | 3.43 | 1.16 | 11.93 | 12.54 | -0.61 | 0.61 | -0.02 | 0.02 |
| gene5273 | https://www.ncbi.nlm.nih.gov/gene/?term=LOC105278108 | 4067 | 44 | 108.19 | 1 | 2.46 | 2.27 | 24.76 | 25.36 | -0.61 | 0.61 | -0.01 | 0.01 |
| gene4498 | https://www.ncbi.nlm.nih.gov/gene/?term=LOC105277239 | 2831 | 122 | 430.94 | 1 | 3.53 | 0.82 | 12.06 | 12.7 | -0.64 | 0.64 | -0.03 | 0.03 |
| gene12203 | https://www.ncbi.nlm.nih.gov/gene/?term=LOC105286099 | 5967 | 73 | 122.34 | 1 | 1.68 | 1.37 | 10.19 | 10.87 | -0.68 | 0.68 | -0.01 | 0.01 |
| gene5646 | https://www.ncbi.nlm.nih.gov/gene/?term=LOC105278534 | 7475 | 175 | 234.11 | 1 | 1.34 | 0.57 | 17.8 | 18.5 | -0.7 | 0.7 | -0.02 | 0.02 |
| gene12406 | https://www.ncbi.nlm.nih.gov/gene/?term=LOC105286316 | 1386 | 28 | 202.02 | 1 | 7.22 | 3.57 | 44.18 | 44.91 | -0.73 | 0.73 | -0.01 | 0.01 |
| gene2748 | https://www.ncbi.nlm.nih.gov/gene/?term=LOC105275261 | 3845 | 51 | 132.64 | 1 | 2.6 | 1.96 | 36.93 | 37.66 | -0.74 | 0.74 | -0.01 | 0.01 |
| gene2683 | https://www.ncbi.nlm.nih.gov/gene/?term=LOC105275187 | 5492 | 66 | 120.17 | 1 | 1.82 | 1.52 | 11.92 | 12.72 | -0.8 | 0.8 | -0.01 | 0.01 |
| gene7368 | https://www.ncbi.nlm.nih.gov/gene/?term=LOC105280556 | 4105 | 35 | 85.26 1 | 2.44 | 2.86 | 17.18 | 18.02 | -0.84 | 0.84 | -0.01 | 0.01 | |
| gene9101 | https://www.ncbi.nlm.nih.gov/gene/?term=LOC105282553 | 3068 | 108 | 352.02 | 1 | 3.26 | 0.93 | 22.57 | 23.41 | -0.84 | 0.84 | -0.03 | 0.03 |
| gene9327 | https://www.ncbi.nlm.nih.gov/gene/?term=LOC105282821 | 7793 | 132 | 169.38 | 1 | 1.28 | 0.76 | 11.87 | 12.7 | -0.84 | 0.84 | -0.01 | 0.01 |
| gene8343 | https://www.ncbi.nlm.nih.gov/gene/?term=LOC105281700 | 2172 | 34 | 156.54 | 1 | 4.6 | 2.94 | 35.91 | 36.82 | -0.9 | 0.9 | -0.01 | 0.01 |
| gene11447 | https://www.ncbi.nlm.nih.gov/gene/?term=LOC105285252 | 7437 | 114 | 153.29 | 2 | 2.69 | 1.75 | 13.96 | 14.93 | -0.97 | 0.97 | -0.01 | 0.01 |
| gene11976 | https://www.ncbi.nlm.nih.gov/gene/?term=LOC105285860 | 3112 | 52 | 167.1 1 | 3.21 | 1.92 | 21.02 | 22 | -0.98 | 0.98 | -0.02 | 0.02 | |
| gene12516 | https://www.ncbi.nlm.nih.gov/gene/?term=LOC105286452 | 1529 | 40 | 261.61 | 1 | 6.54 | 2.5 | 50.21 | 51.25 | -1.04 | 1.04 | -0.03 | 0.03 |
| gene1147 | https://www.ncbi.nlm.nih.gov/gene/?term=LOC105284954 | 4022 | 84 | 208.85 | 1 | 2.49 | 1.19 | 17.29 | 18.34 | -1.06 | 1.06 | -0.02 | 0.02 |
| gene10391 | https://www.ncbi.nlm.nih.gov/gene/?term=LOC109611283 | 1994 | 42 | 210.63 | 1 | 5.02 | 2.38 | 32.6 | 33.67 | -1.07 | 1.07 | -0.02 | 0.02 |
| gene10155 | https://www.ncbi.nlm.nih.gov/gene/?term=LOC105283756 | 7030 | 100 | 142.25 | 1 | 1.42 | 1 | 5.99 | 7.12 | -1.12 | 1.12 | -0.02 | 0.02 |
| gene1220 | https://www.ncbi.nlm.nih.gov/gene/?term=LOC105285742 | 3453 | 107 | 309.88 | 1 | 2.9 | 0.93 | 22.27 | 23.44 | -1.17 | 1.17 | -0.04 | 0.04 |
| gene10819 | https://www.ncbi.nlm.nih.gov/gene/?term=LOC105284507 | 5767 | 76 | 131.78 | 1 | 1.73 | 1.32 | 23.17 | 24.52 | -1.35 | 1.35 | -0.02 | 0.02 |
| gene7207 | https://www.ncbi.nlm.nih.gov/gene/?term=LOC105280376 | 4659 | 109 | 233.96 | 1 | 2.15 | 0.92 | 29.99 | 31.37 | -1.38 | 1.38 | -0.03 | 0.03 |
| gene7600 | https://www.ncbi.nlm.nih.gov/gene/?term=LOC105280819 | 1615 | 37 | 229.1 1 | 6.19 | 2.7 | 36.56 | 38.16 | -1.6 | 1.6 | -0.04 | 0.04 | |

| | | | | | | | | | | | | | |
|---|---|---|---|---|---|---|---|---|---|---|---|---|---|
| gene7850 | https://www.ncbi.nlm.nih.gov/gene/?term=LOC105281095 | 3050 | 63 | 206.56 | 1 | 3.28 | 1.59 | 33.9 | 35.57 | -1.67 | 1.67 | -0.03 | 0.03 |
| gene388 | https://www.ncbi.nlm.nih.gov/gene/?term=LOC105276538 | 1558 | 19 | 121.95 | 2 | 12.84 | 10.53 | 63.26 | 65.13 | -1.87 | 1.87 | -0.02 | 0.02 |
| gene2413 | https://www.ncbi.nlm.nih.gov/gene/?term=LOC105274863 | 2006 | 40 | 199.4 | 1 | 4.99 | 2.5 | 64.23 | 66.19 | -1.96 | 1.96 | -0.04 | 0.04 |
| gene920 | https://www.ncbi.nlm.nih.gov/gene/?term=LOC105282518 | 1197 | 39 | 325.81 | 1 | 8.35 | 2.56 | 38.65 | 40.64 | -1.98 | 1.98 | -0.06 | 0.06 |
| gene10213 | https://www.ncbi.nlm.nih.gov/gene/?term=LOC105283826 | 1503 | 54 | 359.28 | 1 | 6.65 | 1.85 | 13.66 | 15.73 | -2.07 | 2.07 | -0.07 | 0.07 |
| gene322 | https://www.ncbi.nlm.nih.gov/gene/?term=LOC105275777 | 1606 | 23 | 143.21 | 1 | 6.23 | 4.35 | 46.95 | 49.13 | -2.17 | 2.17 | -0.03 | 0.03 |
| gene3168 | https://www.ncbi.nlm.nih.gov/gene/?term=LOC105275711 | 6150 | 67 | 108.94 | 1 | 1.63 | 1.49 | 13.09 | 15.36 | -2.27 | 2.27 | -0.02 | 0.02 |
| gene3599 | https://www.ncbi.nlm.nih.gov/gene/?term=LOC105276196 | 2955 | 47 | 159.05 | 1 | 3.38 | 2.13 | 46.53 | 48.84 | -2.31 | 2.31 | -0.04 | 0.04 |
| gene10879 | https://www.ncbi.nlm.nih.gov/gene/?term=LOC105284574 | 2423 | 46 | 189.85 | 1 | 4.13 | 2.17 | 56.45 | 59.11 | -2.65 | 2.65 | -0.05 | 0.05 |
| gene2404 | https://www.ncbi.nlm.nih.gov/gene/?term=LOC105274853 | 1767 | 34 | 192.42 | 1 | 5.66 | 2.94 | 29.24 | 32.2 | -2.96 | 2.96 | -0.06 | 0.06 |
| gene4641 | https://www.ncbi.nlm.nih.gov/gene/?term=LOC105277398 | 1937 | 68 | 351.06 | 1 | 5.16 | 1.47 | 37.35 | 40.44 | -3.09 | 3.09 | -0.11 | 0.11 |
| gene10345 | https://www.ncbi.nlm.nih.gov/gene/?term=LOC105283989 | 4444 | 22 | 49.5 | 1 | 2.25 | 4.55 | 37.81 | 42.87 | -5.06 | 5.06 | -0.03 | 0.03 |
| gene10963 | https://www.ncbi.nlm.nih.gov/gene/?term=LOC105284677 | 1576 | 27 | 171.32 | 3 | 19.04 | 11.11 | 20.06 | 37.04 | -16.98 | 16.98 | -0.29 | 0.29 |
| gene393 | https://www.ncbi.nlm.nih.gov/gene/?term=LOC105276566 | 1123 | 2 | 17.81 | 1 | 8.9 | 50 | 48.08 | 76.78 | -28.71 | 28.71 | -0.05 | 0.05 |

Directory: OGL File: ogl-Cb_LI_BCphaseB1.txt

| gene_ID | gene_link | gwidth | #Sites | #per10Kb | %perSite | %pNucl |
|---|---|---|---|---|---|---|
| gene12538 | https://www.ncbi.nlm.nih.gov/gene/?term=LOC105286531 | 372 | 43 | 1155.91 | 37.6 | 4.35 |
| gene5911 | https://www.ncbi.nlm.nih.gov/gene/?term=LOC105278844 | 943 | 109 | 1155.89 | 33.35 | 3.86 |
| gene13384 | https://www.ncbi.nlm.nih.gov/gene/?term=LOC105287447 | 1877 | 240 | 1278.64 | 29.63 | 3.79 |
| gene5309 | https://www.ncbi.nlm.nih.gov/gene/?term=LOC105278160 | 1704 | 163 | 956.57 | 39.24 | 3.75 |
| gene5917 | https://www.ncbi.nlm.nih.gov/gene/?term=LOC105278849 | 823 | 115 | 1397.33 | 25.77 | 3.6 |
| gene211 | https://www.ncbi.nlm.nih.gov/gene/?term=LOC105274534 | 1280 | 103 | 804.69 | 41.41 | 3.33 |
| gene4878 | https://www.ncbi.nlm.nih.gov/gene/?term=LOC105277648 | 675 | 40 | 592.59 | 55.76 | 3.3 |
| gene12330 | https://www.ncbi.nlm.nih.gov/gene/?term=LOC105286235 | 1501 | 111 | 739.51 | 44.42 | 3.28 |
| gene1562 | https://www.ncbi.nlm.nih.gov/gene/?term=LOC105287692 | 1491 | 142 | 952.38 | 34.46 | 3.28 |
| gene6078 | https://www.ncbi.nlm.nih.gov/gene/?term=LOC105279052 | 1130 | 139 | 1230.09 | 26.68 | 3.28 |
| gene12746 | https://www.ncbi.nlm.nih.gov/gene/?term=LOC105286737 | 2688 | 393 | 1462.05 | 22.36 | 3.27 |
| gene1654 | https://www.ncbi.nlm.nih.gov/gene/?term=LOC105287811 | 2285 | 165 | 722.1 | 43.88 | 3.17 |
| gene12364 | https://www.ncbi.nlm.nih.gov/gene/?term=LOC105286269 | 1403 | 85 | 605.84 | 51.17 | 3.1 |
| gene12805 | https://www.ncbi.nlm.nih.gov/gene/?term=LOC105286801 | 1379 | 121 | 877.45 | 35.03 | 3.07 |
| gene9471 | https://www.ncbi.nlm.nih.gov/gene/?term=LOC105282986 | 1605 | 135 | 841.12 | 36.19 | 3.04 |
| gene570 | https://www.ncbi.nlm.nih.gov/gene/?term=LOC105278549 | 1673 | 124 | 741.18 | 40.89 | 3.03 |
| gene12652 | https://www.ncbi.nlm.nih.gov/gene/?term=LOC109610640 | 1122 | 63 | 561.5 | 53.52 | 3.01 |
| gene3731 | https://www.ncbi.nlm.nih.gov/gene/?term=LOC105276371 | 865 | 55 | 635.84 | 47.18 | 3 |
| gene3719 | https://www.ncbi.nlm.nih.gov/gene/?term=LOC105276360 | 1471 | 163 | 1108.09 | 26.81 | 2.97 |
| gene7405 | https://www.ncbi.nlm.nih.gov/gene/?term=LOC105280594 | 1263 | 87 | 688.84 | 42.66 | 2.94 |
| gene7527 | https://www.ncbi.nlm.nih.gov/gene/?term=LOC105280739 | 2686 | 159 | 591.96 | 49.69 | 2.94 |
| gene5301 | https://www.ncbi.nlm.nih.gov/gene/?term=LOC105278148 | 1881 | 201 | 1068.58 | 27.41 | 2.93 |
| gene8453 | https://www.ncbi.nlm.nih.gov/gene/?term=LOC105281826 | 1750 | 169 | 965.71 | 30.13 | 2.91 |
| gene4633 | https://www.ncbi.nlm.nih.gov/gene/?term=LOC105277389 | 2426 | 183 | 754.33 | 38.21 | 2.88 |

...

... only first 25 lines shown ...

Directory: OGL File: ogl-Cb_LI_BCphaseB1.vs.BCphaseB2.txt

| gene_ID | gene_link | gwidth | #Sites | #per10Kb | #dmSites | #dmsp10kb | %dmSites | %pSite1 | %pSite2 | DMpSite | ADMpSite | DMpNucl | ADMpNucl |
|---|---|---|---|---|---|---|---|---|---|---|---|---|---|
| gene2593 | https://www.ncbi.nlm.nih.gov/gene/?term=LOC105275070 | 1774 | 5 | 28.18 | 5 | 28.18 | 100 | 72.92 | 10.96 | 61.96 | 61.96 | 0.17 | 0.17 |
| gene3165 | https://www.ncbi.nlm.nih.gov/gene/?term=LOC105275706 | 2374 | 66 | 278.01 | 5 | 21.06 | 7.58 | 34.05 | 24.01 | 10.05 | 10.05 | 0.28 | 0.28 |
| gene1962 | https://www.ncbi.nlm.nih.gov/gene/?term=LOC105288157 | 1246 | 17 | 136.44 | 1 | 8.03 | 5.88 | 18.51 | 13.14 | 5.37 | 5.37 | 0.07 | 0.07 |
| gene2907 | https://www.ncbi.nlm.nih.gov/gene/?term=LOC105275447 | 2539 | 40 | 157.54 | 1 | 3.94 | 2.5 | 7.22 | 3.04 | 4.18 | 4.18 | 0.07 | 0.07 |
| gene2970 | https://www.ncbi.nlm.nih.gov/gene/?term=LOC105275518 | 2340 | 25 | 106.84 | 1 | 4.27 | 4 | 43.33 | 39.71 | 3.62 | 3.62 | 0.04 | 0.04 |
| gene5714 | https://www.ncbi.nlm.nih.gov/gene/?term=LOC105278729 | 710 | 24 | 338.03 | 1 | 14.08 | 4.17 | 59.89 | 56.79 | 3.1 | 3.1 | 0.1 | 0.1 |
| gene7525 | https://www.ncbi.nlm.nih.gov/gene/?term=LOC105280738 | 4237 | 41 | 96.77 | 1 | 2.36 | 2.44 | 26.13 | 23.59 | 2.54 | 2.54 | 0.02 | 0.02 |
| gene9478 | https://www.ncbi.nlm.nih.gov/gene/?term=LOC105282991 | 3726 | 57 | 152.98 | 1 | 2.68 | 1.75 | 35.88 | 33.61 | 2.27 | 2.27 | 0.03 | 0.03 |
| gene3817 | https://www.ncbi.nlm.nih.gov/gene/?term=LOC105276449 | 1230 | 51 | 414.63 | 1 | 8.13 | 1.96 | 27.91 | 26.06 | 1.85 | 1.85 | 0.08 | 0.08 |
| gene8178 | https://www.ncbi.nlm.nih.gov/gene/?term=LOC105281515 | 8647 | 61 | 70.54 | 1 | 1.16 | 1.64 | 27.1 | 25.25 | 1.85 | 1.85 | 0.01 | 0.01 |
| gene1518 | https://www.ncbi.nlm.nih.gov/gene/?term=LOC105287648 | 2195 | 30 | 136.67 | 1 | 4.56 | 3.33 | 43.53 | 41.69 | 1.83 | 1.83 | 0.03 | 0.03 |
| gene940 | https://www.ncbi.nlm.nih.gov/gene/?term=LOC105282794 | 3578 | 88 | 245.95 | 1 | 2.79 | 1.14 | 16.46 | 14.64 | 1.82 | 1.82 | 0.04 | 0.04 |
| gene7368 | https://www.ncbi.nlm.nih.gov/gene/?term=LOC105280556 | 4105 | 54 | 131.55 | 1 | 2.44 | 1.85 | 18.52 | 16.73 | 1.79 | 1.79 | 0.02 | 0.02 |
| gene1884 | https://www.ncbi.nlm.nih.gov/gene/?term=LOC105288072 | 4169 | 76 | 182.3 | 1 | 2.4 | 1.32 | 24.99 | 23.26 | 1.72 | 1.72 | 0.03 | 0.03 |
| gene5877 | https://www.ncbi.nlm.nih.gov/gene/?term=LOC105278807 | 1719 | 53 | 308.32 | 1 | 5.82 | 1.89 | 23.93 | 22.25 | 1.68 | 1.68 | 0.05 | 0.05 |
| gene5101 | https://www.ncbi.nlm.nih.gov/gene/?term=LOC105277903 | 1813 | 37 | 204.08 | 1 | 5.52 | 2.7 | 54.27 | 52.77 | 1.5 | 1.5 | 0.03 | 0.03 |
| gene4892 | https://www.ncbi.nlm.nih.gov/gene/?term=LOC105277667 | 3898 | 127 | 325.81 | 1 | 2.57 | 0.79 | 27.62 | 26.17 | 1.45 | 1.45 | 0.05 | 0.05 |
| gene12572 | https://www.ncbi.nlm.nih.gov/gene/?term=LOC105286514 | 2374 | 46 | 193.77 | 1 | 4.21 | 2.17 | 36.88 | 35.48 | 1.4 | 1.4 | 0.03 | 0.03 |
| gene35 | https://www.ncbi.nlm.nih.gov/gene/?term=LOC105275197 | 2643 | 50 | 189.18 | 1 | 3.78 | 2 | 38.68 | 37.33 | 1.34 | 1.34 | 0.03 | 0.03 |
| gene10345 | https://www.ncbi.nlm.nih.gov/gene/?term=LOC105283989 | 4444 | 29 | 65.26 | 1 | 2.25 | 3.45 | 49.14 | 47.94 | 1.2 | 1.2 | 0.01 | 0.01 |
| gene10678 | https://www.ncbi.nlm.nih.gov/gene/?term=LOC105284355 | 4031 | 174 | 431.65 | 1 | 2.48 | 0.57 | 19.29 | 18.18 | 1.11 | 1.11 | 0.05 | 0.05 |
| gene5967 | https://www.ncbi.nlm.nih.gov/gene/?term=LOC105278918 | 4393 | 146 | 332.35 | 1 | 2.28 | 0.68 | 13.82 | 12.77 | 1.06 | 1.06 | 0.04 | 0.04 |
| gene7568 | https://www.ncbi.nlm.nih.gov/gene/?term=LOC105280786 | 2525 | 119 | 471.29 | 1 | 3.96 | 0.84 | 30.86 | 29.84 | 1.03 | 1.03 | 0.05 | 0.05 |
| gene7888 | https://www.ncbi.nlm.nih.gov/gene/?term=LOC105281141 | 4357 | 164 | 376.41 | 1 | 2.3 | 0.61 | 29.56 | 28.67 | 0.9 | 0.9 | 0.03 | 0.03 |
| gene2856 | https://www.ncbi.nlm.nih.gov/gene/?term=LOC105275384 | 1987 | 92 | 463.01 | 1 | 5.03 | 1.09 | 32.79 | 31.9 | 0.89 | 0.89 | 0.04 | 0.04 |
| gene927 | https://www.ncbi.nlm.nih.gov/gene/?term=LOC105282600 | 3727 | 50 | 134.16 | 1 | 2.68 | 2 | 33.71 | 32.83 | 0.88 | 0.88 | 0.01 | 0.01 |
| gene13033 | https://www.ncbi.nlm.nih.gov/gene/?term=LOC105287063 | 1877 | 53 | 282.37 | 1 | 5.33 | 1.89 | 30.73 | 29.86 | 0.87 | 0.87 | 0.02 | 0.02 |
| gene4720 | https://www.ncbi.nlm.nih.gov/gene/?term=LOC105277479 | 1697 | 40 | 235.71 | 1 | 5.89 | 2.5 | 43.85 | 42.99 | 0.86 | 0.86 | 0.02 | 0.02 |
| gene12516 | https://www.ncbi.nlm.nih.gov/gene/?term=LOC105286452 | 1529 | 37 | 241.99 | 1 | 6.54 | 2.7 | 49.81 | 49.09 | 0.73 | 0.73 | 0.02 | 0.02 |
| gene292 | https://www.ncbi.nlm.nih.gov/gene/?term=LOC105275398 | 2825 | 85 | 300.88 | 1 | 3.54 | 1.18 | 21.01 | 20.28 | 0.73 | 0.73 | 0.02 | 0.02 |
| gene8742 | https://www.ncbi.nlm.nih.gov/gene/?term=LOC105282151 | 2563 | 44 | 171.67 | 2 | 7.8 | 4.55 | 36.03 | 35.31 | 0.72 | 0.72 | 0.01 | 0.01 |
| gene9336 | https://www.ncbi.nlm.nih.gov/gene/?term=LOC105282834 | 5993 | 129 | 215.25 | 1 | 1.67 | 0.78 | 15.82 | 15.16 | 0.66 | 0.66 | 0.01 | 0.01 |
| gene10540 | https://www.ncbi.nlm.nih.gov/gene/?term=LOC105284200 | 5267 | 204 | 387.32 | 1 | 1.9 | 0.49 | 12.98 | 12.34 | 0.64 | 0.64 | 0.02 | 0.02 |
| gene11401 | https://www.ncbi.nlm.nih.gov/gene/?term=LOC105285202 | 1841 | 77 | 418.25 | 1 | 5.43 | 1.3 | 24.18 | 23.56 | 0.62 | 0.62 | 0.03 | 0.03 |
| gene12439 | https://www.ncbi.nlm.nih.gov/gene/?term=LOC105286357 | 4761 | 86 | 180.63 | 1 | 2.1 | 1.16 | 26.67 | 26.07 | 0.6 | 0.6 | 0.01 | 0.01 |
| gene34 | https://www.ncbi.nlm.nih.gov/gene/?term=LOC105278416 | 7120 | 146 | 205.06 | 1 | 1.4 | 0.68 | 13.63 | 13.03 | 0.6 | 0.6 | 0.01 | 0.01 |
| gene5654 | https://www.ncbi.nlm.nih.gov/gene/?term=LOC105278543 | 4019 | 160 | 398.11 | 1 | 2.49 | 0.62 | 15.95 | 15.36 | 0.6 | 0.6 | 0.02 | 0.02 |
| gene6583 | https://www.ncbi.nlm.nih.gov/gene/?term=LOC105279649 | 7444 | 350 | 470.18 | 1 | 1.34 | 0.29 | 11.44 | 10.85 | 0.59 | 0.59 | 0.03 | 0.03 |
| gene2683 | https://www.ncbi.nlm.nih.gov/gene/?term=LOC105275187 | 5492 | 75 | 136.56 | 1 | 1.82 | 1.33 | 15.98 | 15.4 | 0.58 | 0.58 | 0.01 | 0.01 |
| gene4023 | https://www.ncbi.nlm.nih.gov/gene/?term=LOC105276692 | 1759 | 37 | 210.35 | 1 | 5.69 | 2.7 | 20.35 | 19.78 | 0.57 | 0.57 | 0.01 | 0.01 |
| gene7884 | https://www.ncbi.nlm.nih.gov/gene/?term=LOC105281266 | 3390 | 79 | 233.04 | 1 | 2.95 | 1.27 | 16.01 | 15.44 | 0.57 | 0.57 | 0.01 | 0.01 |
| gene4376 | https://www.ncbi.nlm.nih.gov/gene/?term=LOC105277083 | 4307 | 129 | 299.51 | 1 | 2.32 | 0.78 | 19.34 | 18.78 | 0.56 | 0.56 | 0.02 | 0.02 |
| gene1855 | https://www.ncbi.nlm.nih.gov/gene/?term=LOC105288036 | 2387 | 86 | 360.28 | 1 | 4.19 | 1.16 | 29.99 | 29.44 | 0.55 | 0.55 | 0.02 | 0.02 |
| gene3107 | https://www.ncbi.nlm.nih.gov/gene/?term=LOC105275650 | 5103 | 178 | 348.81 | 1 | 1.96 | 0.56 | 16.67 | 16.15 | 0.52 | 0.52 | 0.02 | 0.02 |
| gene3206 | https://www.ncbi.nlm.nih.gov/gene/?term=LOC105275752 | 3474 | 129 | 371.33 | 1 | 2.88 | 0.78 | 32.13 | 31.62 | 0.52 | 0.52 | 0.02 | 0.02 |
| gene1332 | https://www.ncbi.nlm.nih.gov/gene/?term=LOC105286872 | 2460 | 158 | 642.28 | 1 | 4.07 | 0.63 | 20.79 | 20.28 | 0.51 | 0.51 | 0.03 | 0.03 |
| gene2710 | https://www.ncbi.nlm.nih.gov/gene/?term=LOC105275218 | 3222 | 98 | 304.16 | 1 | 3.1 | 1.02 | 22.4 | 21.9 | 0.5 | 0.5 | 0.02 | 0.02 |
| gene2668 | https://www.ncbi.nlm.nih.gov/gene/?term=LOC105275170 | 12496 | 301 | 240.88 | 1 | 0.8 | 0.33 | 2.32 | 1.82 | 0.49 | 0.49 | 0.01 | 0.01 |
| gene6966 | https://www.ncbi.nlm.nih.gov/gene/?term=LOC105280104 | 3747 | 133 | 354.95 | 1 | 2.67 | 0.75 | 15.81 | 15.32 | 0.49 | 0.49 | 0.02 | 0.02 |
| gene9445 | https://www.ncbi.nlm.nih.gov/gene/?term=LOC105282951 | 2285 | 115 | 503.28 | 1 | 4.38 | 0.87 | 20.4 | 19.9 | 0.49 | 0.49 | 0.02 | 0.02 |
| gene738 | https://www.ncbi.nlm.nih.gov/gene/?term=LOC105280421 | 4927 | 222 | 450.58 | 1 | 2.03 | 0.45 | 14.83 | 14.35 | 0.48 | 0.48 | 0.02 | 0.02 |
| gene8729 | https://www.ncbi.nlm.nih.gov/gene/?term=LOC105282137 | 5005 | 202 | 403.6 | 1 | 2 | 0.5 | 16.8 | 16.33 | 0.46 | 0.46 | 0.02 | 0.02 |
| gene3470 | https://www.ncbi.nlm.nih.gov/gene/?term=LOC105276059 | 8716 | 164 | 188.16 | 1 | 1.15 | 0.61 | 7.87 | 7.47 | 0.4 | 0.4 | 0.01 | 0.01 |
| gene11454 | https://www.ncbi.nlm.nih.gov/gene/?term=LOC105285256 | 1906 | 51 | 267.58 | 1 | 5.25 | 1.96 | 42.4 | 42.03 | 0.37 | 0.37 | 0.01 | 0.01 |
| gene5831 | https://www.ncbi.nlm.nih.gov/gene/?term=LOC105278761 | 2506 | 126 | 502.79 | 1 | 3.99 | 0.79 | 21.41 | 21.05 | 0.36 | 0.36 | 0.02 | 0.02 |
| gene6620 | https://www.ncbi.nlm.nih.gov/gene/?term=LOC105279687 | 17998 | 384 | 213.36 | 1 | 0.56 | 0.26 | 4.83 | 4.47 | 0.36 | 0.36 | 0.01 | 0.01 |
| gene5375 | https://www.ncbi.nlm.nih.gov/gene/?term=LOC105278228 | 8640 | 173 | 200.23 | 1 | 1.16 | 0.58 | 23.79 | 23.44 | 0.35 | 0.35 | 0.01 | 0.01 |
| gene12802 | https://www.ncbi.nlm.nih.gov/gene/?term=LOC105286799 | 3183 | 189 | 593.78 | 1 | 3.14 | 0.53 | 15.7 | 15.39 | 0.32 | 0.32 | 0.02 | 0.02 |
| gene2844 | https://www.ncbi.nlm.nih.gov/gene/?term=LOC105275363 | 4330 | 206 | 475.75 | 1 | 2.31 | 0.49 | 14.15 | 13.85 | 0.3 | 0.3 | 0.01 | 0.01 |
| gene11568 | https://www.ncbi.nlm.nih.gov/gene/?term=LOC105285395 | 8086 | 204 | 252.29 | 1 | 1.24 | 0.49 | 14.79 | 14.52 | 0.27 | 0.27 | 0.01 | 0.01 |

| | | | | | | | | | | | | | |
|---|---|---|---|---|---|---|---|---|---|---|---|---|---|
| gene1310 | https://www.ncbi.nlm.nih.gov/gene/?term=LOC105286661 | 2582 | 65 | 251.74 | 1 | 3.87 | 1.54 | 23.22 | 22.96 | 0.26 | 0.26 | 0.01 | 0.01 |
| gene10531 | https://www.ncbi.nlm.nih.gov/gene/?term=LOC105284190 | 6881 | 195 | 283.39 | 1 | 1.45 | 0.51 | 11.74 | 11.5 | 0.24 | 0.24 | 0.01 | 0.01 |
| gene11579 | https://www.ncbi.nlm.nih.gov/gene/?term=LOC105285406 | 13571 | 291 | 214.43 | 1 | 0.74 | 0.34 | 6.38 | 6.14 | 0.24 | 0.24 | 0.01 | 0.01 |
| gene10844 | https://www.ncbi.nlm.nih.gov/gene/?term=LOC105284535 | 9086 | 256 | 281.75 | 1 | 1.1 | 0.39 | 14.59 | 14.36 | 0.23 | 0.23 | 0.01 | 0.01 |
| gene2806 | https://www.ncbi.nlm.nih.gov/gene/?term=LOC105275327 | 14257 | 542 | 380.16 | 1 | 0.7 | 0.18 | 6.63 | 6.42 | 0.21 | 0.21 | 0.01 | 0.01 |
| gene3599 | https://www.ncbi.nlm.nih.gov/gene/?term=LOC105276196 | 2955 | 59 | 199.66 | 1 | 3.38 | 1.69 | 37.86 | 37.72 | 0.15 | 0.15 | 0 | 0 |
| gene53 | https://www.ncbi.nlm.nih.gov/gene/?term=LOC105277148 | 19283 | 324 | 168.02 | 1 | 0.52 | 0.31 | 4.91 | 4.75 | 0.15 | 0.15 | 0 | 0 |
| gene9462 | https://www.ncbi.nlm.nih.gov/gene/?term=LOC105282970 | 1469 | 42 | 285.91 | 1 | 6.81 | 2.38 | 51.11 | 50.96 | 0.15 | 0.15 | 0 | 0 |
| gene3885 | https://www.ncbi.nlm.nih.gov/gene/?term=LOC105276527 | 7928 | 334 | 421.29 | 1 | 1.26 | 0.3 | 9.88 | 9.74 | 0.14 | 0.14 | 0.01 | 0.01 |
| gene7183 | https://www.ncbi.nlm.nih.gov/gene/?term=LOC105280348 | 3925 | 76 | 193.63 | 1 | 2.55 | 1.32 | 19.42 | 19.29 | 0.13 | 0.13 | 0 | 0 |
| gene12307 | https://www.ncbi.nlm.nih.gov/gene/?term=LOC105286216 | 10313 | 174 | 168.72 | 1 | 0.97 | 0.57 | 15.92 | 15.81 | 0.12 | 0.12 | 0 | 0 |
| gene5184 | https://www.ncbi.nlm.nih.gov/gene/?term=LOC105277999 | 5528 | 146 | 264.11 | 1 | 1.81 | 0.68 | 11.67 | 11.57 | 0.1 | 0.1 | 0 | 0 |
| gene5746 | https://www.ncbi.nlm.nih.gov/gene/?term=LOC105278666 | 10343 | 482 | 466.02 | 1 | 0.97 | 0.21 | 6.71 | 6.61 | 0.1 | 0.1 | 0 | 0 |
| gene12573 | https://www.ncbi.nlm.nih.gov/gene/?term=LOC105286515 | 3647 | 103 | 282.42 | 1 | 2.74 | 0.97 | 10.56 | 10.47 | 0.09 | 0.09 | 0 | 0 |
| gene8343 | https://www.ncbi.nlm.nih.gov/gene/?term=LOC105281700 | 2172 | 37 | 170.35 | 1 | 4.6 | 2.7 | 39.73 | 39.63 | 0.09 | 0.09 | 0 | 0 |
| gene298 | https://www.ncbi.nlm.nih.gov/gene/?term=LOC105275460 | 2473 | 122 | 493.33 | 1 | 4.04 | 0.82 | 13.15 | 13.07 | 0.08 | 0.08 | 0 | 0 |
| gene10155 | https://www.ncbi.nlm.nih.gov/gene/?term=LOC105283756 | 7030 | 109 | 155.05 | 1 | 1.42 | 0.92 | 5.36 | 5.29 | 0.07 | 0.07 | 0 | 0 |
| gene1632 | https://www.ncbi.nlm.nih.gov/gene/?term=LOC105287764 | 18061 | 406 | 224.79 | 2 | 1.11 | 0.49 | 3.27 | 3.2 | 0.07 | 0.07 | 0 | 0 |
| gene9326 | https://www.ncbi.nlm.nih.gov/gene/?term=LOC105282818 | 16612 | 217 | 130.63 | 1 | 0.6 | 0.46 | 11.57 | 11.5 | 0.07 | 0.07 | 0 | 0 |
| gene10596 | https://www.ncbi.nlm.nih.gov/gene/?term=LOC105284262 | 6520 | 234 | 358.9 1 | 1.53 | 0.43 | 3.38 | 3.34 | 0.04 | 0.04 | 0 | 0 | |
| gene2056 | https://www.ncbi.nlm.nih.gov/gene/?term=LOC105274452 | 6584 | 263 | 399.45 | 2 | 3.04 | 0.76 | 12.97 | 12.95 | 0.02 | 0.02 | 0 | 0 |
| gene7398 | https://www.ncbi.nlm.nih.gov/gene/?term=LOC105280589 | 3452 | 37 | 107.18 | 1 | 2.9 | 2.7 | 19.84 | 19.83 | 0.01 | 0.01 | 0 | 0 |
| gene10018 | https://www.ncbi.nlm.nih.gov/gene/?term=LOC105283607 | 4414 | 208 | 471.23 | 1 | 2.27 | 0.48 | 15.08 | 15.07 | 0 | 0 | 0 | 0 |
| gene1430 | https://www.ncbi.nlm.nih.gov/gene/?term=LOC105287551 | 6433 | 232 | 360.64 | 1 | 1.55 | 0.43 | 8.43 | 8.44 | 0 | 0 | 0 | 0 |
| gene11275 | https://www.ncbi.nlm.nih.gov/gene/?term=LOC105285055 | 4119 | 248 | 602.09 | 1 | 2.43 | 0.4 | 12.13 | 12.15 | -0.02 | 0.02 | 0 | 0 |
| gene13114 | https://www.ncbi.nlm.nih.gov/gene/?term=LOC105287147 | 7609 | 430 | 565.12 | 1 | 1.31 | 0.23 | 9.08 | 9.1 | -0.02 | 0.02 | 0 | 0 |
| gene3856 | https://www.ncbi.nlm.nih.gov/gene/?term=LOC105276499 | 5236 | 194 | 370.51 | 1 | 1.91 | 0.52 | 10.61 | 10.65 | -0.04 | 0.04 | 0 | 0 |
| gene9327 | https://www.ncbi.nlm.nih.gov/gene/?term=LOC105282821 | 7793 | 173 | 221.99 | 1 | 1.28 | 0.58 | 10.79 | 10.87 | -0.08 | 0.08 | 0 | 0 |
| gene7166 | https://www.ncbi.nlm.nih.gov/gene/?term=LOC105280326 | 6117 | 253 | 413.6 1 | 1.63 | 0.4 | 14.52 | 14.62 | -0.1 | 0.1 | 0 | 0 | |
| gene10886 | https://www.ncbi.nlm.nih.gov/gene/?term=LOC105284582 | 1475 | 47 | 318.64 | 1 | 6.78 | 2.13 | 22.72 | 22.83 | -0.11 | 0.11 | 0 | 0 |
| gene6528 | https://www.ncbi.nlm.nih.gov/gene/?term=LOC105279591 | 7222 | 59 | 81.69 1 | 1.38 | 1.69 | 26.07 | 26.22 | -0.14 | 0.14 | 0 | 0 | |
| gene2633 | https://www.ncbi.nlm.nih.gov/gene/?term=LOC105275127 | 7457 | 270 | 362.08 | 1 | 1.34 | 0.37 | 7.9 | 8.07 | -0.16 | 0.16 | -0.01 | 0.01 |
| gene2173 | https://www.ncbi.nlm.nih.gov/gene/?term=LOC105274590 | 4387 | 84 | 191.47 | 2 | 4.56 | 2.38 | 27.83 | 27.99 | -0.17 | 0.17 | 0 | 0 |
| gene11561 | https://www.ncbi.nlm.nih.gov/gene/?term=LOC105285387 | 11528 | 202 | 175.23 | 1 | 0.87 | 0.5 | 8.85 | 9.03 | -0.18 | 0.18 | 0 | 0 |
| gene7514 | https://www.ncbi.nlm.nih.gov/gene/?term=LOC105280723 | 4471 | 210 | 469.69 | 1 | 2.24 | 0.48 | 13.2 | 13.38 | -0.18 | 0.18 | -0.01 | 0.01 |
| gene10549 | https://www.ncbi.nlm.nih.gov/gene/?term=LOC105284210 | 7145 | 340 | 475.86 | 2 | 2.8 | 0.59 | 9.99 | 10.19 | -0.2 | 0.2 | -0.01 | 0.01 |
| gene8814 | https://www.ncbi.nlm.nih.gov/gene/?term=LOC105282240 | 6910 | 198 | 286.54 | 1 | 1.45 | 0.51 | 8.79 | 9 | -0.21 | 0.21 | -0.01 | 0.01 |
| gene2846 | https://www.ncbi.nlm.nih.gov/gene/?term=LOC105275374 | 11246 | 512 | 455.27 | 1 | 0.89 | 0.2 | 9.81 | 10.04 | -0.23 | 0.23 | -0.01 | 0.01 |
| gene10213 | https://www.ncbi.nlm.nih.gov/gene/?term=LOC105283826 | 1503 | 65 | 432.47 | 1 | 6.65 | 1.54 | 15.16 | 15.4 | -0.24 | 0.24 | -0.01 | 0.01 |
| gene13036 | https://www.ncbi.nlm.nih.gov/gene/?term=LOC105287059 | 4959 | 176 | 354.91 | 1 | 2.02 | 0.57 | 16.13 | 16.41 | -0.28 | 0.28 | -0.01 | 0.01 |
| gene1470 | https://www.ncbi.nlm.nih.gov/gene/?term=LOC105287597 | 2974 | 204 | 685.94 | 1 | 3.36 | 0.49 | 17.06 | 17.34 | -0.28 | 0.28 | -0.02 | 0.02 |
| gene10392 | https://www.ncbi.nlm.nih.gov/gene/?term=LOC105284036 | 3216 | 106 | 329.6 2 | 6.22 | 1.89 | 20.71 | 21 | -0.29 | 0.29 | -0.01 | 0.01 | |
| gene10520 | https://www.ncbi.nlm.nih.gov/gene/?term=LOC105284184 | 2451 | 101 | 412.08 | 1 | 4.08 | 0.99 | 29.13 | 29.43 | -0.3 | 0.3 | -0.01 | 0.01 |
| gene11082 | https://www.ncbi.nlm.nih.gov/gene/?term=LOC105284824 | 7436 | 214 | 287.79 | 1 | 1.34 | 0.47 | 6.56 | 6.86 | -0.31 | 0.31 | -0.01 | 0.01 |
| gene3046 | https://www.ncbi.nlm.nih.gov/gene/?term=LOC105275587 | 2394 | 55 | 229.74 | 1 | 4.18 | 1.82 | 36.59 | 36.9 | -0.31 | 0.31 | -0.01 | 0.01 |
| gene1907 | https://www.ncbi.nlm.nih.gov/gene/?term=LOC105288099 | 1238 | 36 | 290.79 | 1 | 8.08 | 2.78 | 33.72 | 34.04 | -0.32 | 0.32 | -0.01 | 0.01 |
| gene5777 | https://www.ncbi.nlm.nih.gov/gene/?term=LOC105278691 | 3883 | 168 | 432.66 | 1 | 2.58 | 0.6 | 14.66 | 14.98 | -0.33 | 0.33 | -0.01 | 0.01 |
| gene10600 | https://www.ncbi.nlm.nih.gov/gene/?term=LOC105284270 | 4837 | 124 | 256.36 | 1 | 2.07 | 0.81 | 11.8 | 12.15 | -0.35 | 0.35 | -0.01 | 0.01 |
| gene12203 | https://www.ncbi.nlm.nih.gov/gene/?term=LOC105286099 | 5967 | 75 | 125.69 | 1 | 1.68 | 1.33 | 9.95 | 10.3 | -0.35 | 0.35 | 0 | 0 |
| gene6031 | https://www.ncbi.nlm.nih.gov/gene/?term=LOC105279000 | 5691 | 197 | 346.16 | 1 | 1.76 | 0.51 | 12.37 | 12.73 | -0.35 | 0.35 | -0.01 | 0.01 |
| gene5524 | https://www.ncbi.nlm.nih.gov/gene/?term=LOC105278397 | 5177 | 205 | 395.98 | 1 | 1.93 | 0.49 | 24.95 | 25.35 | -0.39 | 0.39 | -0.02 | 0.02 |
| gene6286 | https://www.ncbi.nlm.nih.gov/gene/?term=LOC105279298 | 2861 | 133 | 464.87 | 2 | 6.99 | 1.5 | 23.95 | 24.34 | -0.39 | 0.39 | -0.02 | 0.02 |
| gene4190 | https://www.ncbi.nlm.nih.gov/gene/?term=LOC105276875 | 3150 | 123 | 390.48 | 1 | 3.17 | 0.81 | 20.99 | 21.41 | -0.42 | 0.42 | -0.02 | 0.02 |
| gene570 | https://www.ncbi.nlm.nih.gov/gene/?term=LOC105278549 | 1673 | 71 | 424.39 | 1 | 5.98 | 1.41 | 43.51 | 43.92 | -0.42 | 0.42 | -0.02 | 0.02 |
| gene12806 | https://www.ncbi.nlm.nih.gov/gene/?term=LOC105286803 | 4196 | 160 | 381.32 | 1 | 2.38 | 0.62 | 9.57 | 10.02 | -0.45 | 0.45 | -0.02 | 0.02 |
| gene8274 | https://www.ncbi.nlm.nih.gov/gene/?term=LOC105281624 | 5425 | 236 | 435.02 | 1 | 1.84 | 0.42 | 9.29 | 9.74 | -0.45 | 0.45 | -0.02 | 0.02 |
| gene10326 | https://www.ncbi.nlm.nih.gov/gene/?term=LOC105283962 | 3042 | 132 | 433.93 | 1 | 3.29 | 0.76 | 8.33 | 8.79 | -0.46 | 0.46 | -0.02 | 0.02 |
| gene2413 | https://www.ncbi.nlm.nih.gov/gene/?term=LOC105274863 | 2006 | 54 | 269.19 | 1 | 4.99 | 1.85 | 52.24 | 52.76 | -0.51 | 0.51 | -0.01 | 0.01 |
| gene4962 | https://www.ncbi.nlm.nih.gov/gene/?term=LOC105277761 | 4645 | 151 | 325.08 | 1 | 2.15 | 0.66 | 14.59 | 15.1 | -0.51 | 0.51 | -0.02 | 0.02 |
| gene388 | https://www.ncbi.nlm.nih.gov/gene/?term=LOC105276538 | 1558 | 19 | 121.95 | 2 | 12.84 | 10.53 | 66.25 | 66.78 | -0.53 | 0.53 | -0.01 | 0.01 |
| gene2748 | https://www.ncbi.nlm.nih.gov/gene/?term=LOC105275261 | 3845 | 55 | 143.04 | 1 | 2.6 | 1.82 | 38.33 | 38.87 | -0.54 | 0.54 | -0.01 | 0.01 |
| gene10606 | https://www.ncbi.nlm.nih.gov/gene/?term=LOC105284273 | 6119 | 264 | 431.44 | 1 | 1.63 | 0.38 | 18.32 | 18.9 | -0.57 | 0.57 | -0.02 | 0.02 |
| gene5525 | https://www.ncbi.nlm.nih.gov/gene/?term=LOC105278396 | 3090 | 189 | 611.65 | 1 | 3.24 | 0.53 | 24.57 | 25.15 | -0.58 | 0.58 | -0.04 | 0.04 |
| gene11583 | https://www.ncbi.nlm.nih.gov/gene/?term=LOC105285414 | 8192 | 184 | 224.61 | 1 | 1.22 | 0.54 | 8.68 | 9.27 | -0.59 | 0.59 | -0.01 | 0.01 |

| | | | | | | | | | | | | | |
|---|---|---|---|---|---|---|---|---|---|---|---|---|---|
| gene9101 | https://www.ncbi.nlm.nih.gov/gene/?term=LOC105282553 | 3068 | 108 | 352.02 | 1 | 3.26 | 0.93 | 22.97 | 23.62 | -0.64 | 0.64 | -0.02 | 0.02 |
| gene4034 | https://www.ncbi.nlm.nih.gov/gene/?term=LOC105276704 | 2065 | 103 | 498.79 | 2 | 9.69 | 1.94 | 12.96 | 13.6 | -0.65 | 0.65 | -0.03 | 0.03 |
| gene7249 | https://www.ncbi.nlm.nih.gov/gene/?term=LOC105280422 | 1775 | 62 | 349.3 | 1 | 5.63 | 1.61 | 17.31 | 17.96 | -0.66 | 0.66 | -0.02 | 0.02 |
| gene6639 | https://www.ncbi.nlm.nih.gov/gene/?term=LOC105279705 | 7656 | 70 | 91.43 | 1 | 1.31 | 1.43 | 24.79 | 25.46 | -0.67 | 0.67 | -0.01 | 0.01 |
| gene5646 | https://www.ncbi.nlm.nih.gov/gene/?term=LOC105278534 | 7475 | 195 | 260.87 | 1 | 1.34 | 0.51 | 17.04 | 17.72 | -0.68 | 0.68 | -0.02 | 0.02 |
| gene322 | https://www.ncbi.nlm.nih.gov/gene/?term=LOC105275777 | 1606 | 27 | 168.12 | 1 | 6.23 | 3.7 | 40.46 | 41.2 | -0.74 | 0.74 | -0.01 | 0.01 |
| gene2404 | https://www.ncbi.nlm.nih.gov/gene/?term=LOC105274853 | 1767 | 38 | 215.05 | 1 | 5.66 | 2.63 | 34.09 | 34.89 | -0.8 | 0.8 | -0.02 | 0.02 |
| gene11688 | https://www.ncbi.nlm.nih.gov/gene/?term=LOC105285533 | 1637 | 70 | 427.61 | 1 | 6.11 | 1.43 | 22.69 | 23.5 | -0.81 | 0.81 | -0.03 | 0.03 |
| gene10107 | https://www.ncbi.nlm.nih.gov/gene/?term=LOC105283712 | 1883 | 36 | 191.18 | 1 | 5.31 | 2.78 | 28.27 | 29.18 | -0.91 | 0.91 | -0.02 | 0.02 |
| gene3168 | https://www.ncbi.nlm.nih.gov/gene/?term=LOC105275711 | 6150 | 75 | 121.95 | 1 | 1.63 | 1.33 | 12.84 | 13.77 | -0.93 | 0.93 | -0.01 | 0.01 |
| gene3793 | https://www.ncbi.nlm.nih.gov/gene/?term=LOC105276429 | 2901 | 110 | 379.18 | 1 | 3.45 | 0.91 | 25.42 | 26.35 | -0.93 | 0.93 | -0.04 | 0.04 |
| gene7247 | https://www.ncbi.nlm.nih.gov/gene/?term=LOC105280407 | 10486 | 212 | 202.17 | 1 | 0.95 | 0.47 | 15.9 | 16.83 | -0.93 | 0.93 | -0.02 | 0.02 |
| gene7199 | https://www.ncbi.nlm.nih.gov/gene/?term=LOC105280366 | 1414 | 32 | 226.31 | 1 | 7.07 | 3.12 | 42.1 | 43.06 | -0.96 | 0.96 | -0.02 | 0.02 |
| gene6652 | https://www.ncbi.nlm.nih.gov/gene/?term=LOC105279719 | 2831 | 85 | 300.25 | 1 | 3.53 | 1.18 | 22.31 | 23.3 | -0.99 | 0.99 | -0.03 | 0.03 |
| gene11340 | https://www.ncbi.nlm.nih.gov/gene/?term=LOC105285131 | 2918 | 105 | 359.84 | 1 | 3.43 | 0.95 | 13.12 | 14.12 | -1 | 1 | -0.04 | 0.04 |
| gene1147 | https://www.ncbi.nlm.nih.gov/gene/?term=LOC105284954 | 4022 | 114 | 283.44 | 1 | 2.49 | 0.88 | 13.79 | 14.79 | -1 | 1 | -0.03 | 0.03 |
| gene5298 | https://www.ncbi.nlm.nih.gov/gene/?term=LOC105278145 | 4060 | 236 | 581.28 | 2 | 4.93 | 0.85 | 10.36 | 11.46 | -1.1 | 1.1 | -0.06 | 0.06 |
| gene9740 | https://www.ncbi.nlm.nih.gov/gene/?term=LOC105283277 | 2887 | 140 | 484.93 | 1 | 3.46 | 0.71 | 19.86 | 21.12 | -1.26 | 1.26 | -0.06 | 0.06 |
| gene920 | https://www.ncbi.nlm.nih.gov/gene/?term=LOC105282518 | 1197 | 42 | 350.88 | 1 | 8.35 | 2.38 | 37.02 | 38.3 | -1.28 | 1.28 | -0.04 | 0.04 |
| gene8015 | https://www.ncbi.nlm.nih.gov/gene/?term=LOC105281312 | 2998 | 65 | 216.81 | 1 | 3.34 | 1.54 | 31.99 | 33.36 | -1.36 | 1.36 | -0.03 | 0.03 |
| gene12406 | https://www.ncbi.nlm.nih.gov/gene/?term=LOC105286316 | 1386 | 32 | 230.88 | 1 | 7.22 | 3.12 | 45.77 | 47.17 | -1.4 | 1.4 | -0.03 | 0.03 |
| gene1220 | https://www.ncbi.nlm.nih.gov/gene/?term=LOC105285742 | 3453 | 125 | 362 | 1 | 2.9 | 0.8 | 19.11 | 20.58 | -1.47 | 1.47 | -0.05 | 0.05 |
| gene7915 | https://www.ncbi.nlm.nih.gov/gene/?term=LOC105281173 | 1597 | 38 | 237.95 | 1 | 6.26 | 2.63 | 59.22 | 60.79 | -1.57 | 1.57 | -0.04 | 0.04 |
| gene13235 | https://www.ncbi.nlm.nih.gov/gene/?term=LOC105287278 | 2679 | 53 | 197.84 | 1 | 3.73 | 1.89 | 24.79 | 26.36 | -1.58 | 1.58 | -0.03 | 0.03 |
| gene2271 | https://www.ncbi.nlm.nih.gov/gene/?term=LOC105274706 | 1448 | 51 | 352.21 | 1 | 6.91 | 1.96 | 40.05 | 41.71 | -1.67 | 1.67 | -0.06 | 0.06 |
| gene4498 | https://www.ncbi.nlm.nih.gov/gene/?term=LOC105277239 | 2831 | 140 | 494.52 | 1 | 3.53 | 0.71 | 13.31 | 15.32 | -2.01 | 2.01 | -0.1 | 0.1 |
| gene7248 | https://www.ncbi.nlm.nih.gov/gene/?term=LOC105280408 | 5175 | 57 | 110.14 | 1 | 1.93 | 1.75 | 18.23 | 20.49 | -2.26 | 2.26 | -0.02 | 0.02 |
| gene3727 | https://www.ncbi.nlm.nih.gov/gene/?term=LOC105276367 | 2484 | 66 | 265.7 | 1 | 4.03 | 1.52 | 44.56 | 46.97 | -2.41 | 2.41 | -0.06 | 0.06 |
| gene6422 | https://www.ncbi.nlm.nih.gov/gene/?term=LOC105279458 | 2419 | 37 | 152.96 | 1 | 4.13 | 2.7 | 49.83 | 52.5 | -2.66 | 2.66 | -0.04 | 0.04 |
| gene1505 | https://www.ncbi.nlm.nih.gov/gene/?term=LOC105287633 | 1693 | 27 | 159.48 | 1 | 5.91 | 3.7 | 42.55 | 45.31 | -2.76 | 2.76 | -0.04 | 0.04 |
| gene9411 | https://www.ncbi.nlm.nih.gov/gene/?term=LOC105282914 | 5534 | 41 | 74.09 | 1 | 1.81 | 2.44 | 49.41 | 52.21 | -2.8 | 2.8 | -0.02 | 0.02 |
| gene6237 | https://www.ncbi.nlm.nih.gov/gene/?term=LOC105279244 | 1484 | 25 | 168.46 | 1 | 6.74 | 4 | 50.3 | 53.48 | -3.18 | 3.18 | -0.05 | 0.05 |
| gene6455 | https://www.ncbi.nlm.nih.gov/gene/?term=LOC105279500 | 2612 | 33 | 126.34 | 1 | 3.83 | 3.03 | 43.71 | 47.06 | -3.35 | 3.35 | -0.04 | 0.04 |
| gene564 | https://www.ncbi.nlm.nih.gov/gene/?term=LOC105278478 | 2802 | 17 | 60.67 | 1 | 3.57 | 5.88 | 14.04 | 17.66 | -3.63 | 3.63 | -0.02 | 0.02 |
| gene10602 | https://www.ncbi.nlm.nih.gov/gene/?term=LOC105284268 | 2122 | 18 | 84.83 | 1 | 4.71 | 5.56 | 26.04 | 29.95 | -3.91 | 3.91 | -0.03 | 0.03 |
| gene1199 | https://www.ncbi.nlm.nih.gov/gene/?term=LOC105285498 | 2220 | 26 | 117.12 | 1 | 4.5 | 3.85 | 33.11 | 37.03 | -3.92 | 3.92 | -0.05 | 0.05 |
| gene12021 | https://www.ncbi.nlm.nih.gov/gene/?term=LOC105285905 | 1399 | 20 | 142.96 | 1 | 7.15 | 5 | 33.2 | 38.03 | -4.83 | 4.83 | -0.07 | 0.07 |
| gene12430 | https://www.ncbi.nlm.nih.gov/gene/?term=LOC105286348 | 3389 | 19 | 56.06 | 1 | 2.95 | 5.26 | 6.26 | 11.43 | -5.17 | 5.17 | -0.03 | 0.03 |
| gene10963 | https://www.ncbi.nlm.nih.gov/gene/?term=LOC105284677 | 1576 | 29 | 184.01 | 1 | 6.35 | 3.45 | 18.87 | 36.99 | -18.12 | 18.12 | -0.33 | 0.33 |
| gene393 | https://www.ncbi.nlm.nih.gov/gene/?term=LOC105276566 | 1123 | 2 | 17.81 | 1 | 8.9 | 50 | 47.22 | 70.57 | -23.35 | 23.35 | -0.04 | 0.04 |

Directory: OGL File: ogl-Cb_LI_BCphaseB1.vs.RphaseA1.txt

| gene_ID | gene_link | gwidth | #Sites | #per10Kb | #dmSites | #dmsp10kb | %dmSites | %pSite1 | %pSite2 | DMpSite | ADMpSite | DMpNucl | ADMpNucl |
|---|---|---|---|---|---|---|---|---|---|---|---|---|---|
| gene12518 | https://www.ncbi.nlm.nih.gov/gene/?term=LOC105286453 | 1109 | 6 | 54.1 | 1 | 9.02 | 16.67 | 54.64 | 39.73 | 14.91 | 14.91 | 0.08 | 0.08 |
| gene9354 | https://www.ncbi.nlm.nih.gov/gene/?term=LOC105282854 | 1836 | 6 | 32.68 | 1 | 5.45 | 16.67 | 52.41 | 41.89 | 10.52 | 10.52 | 0.03 | 0.03 |
| gene12173 | https://www.ncbi.nlm.nih.gov/gene/?term=LOC105286067 | 518 | 3 | 57.92 | 1 | 19.31 | 33.33 | 13.69 | 3.17 | 10.51 | 10.51 | 0.06 | 0.06 |
| gene5950 | https://www.ncbi.nlm.nih.gov/gene/?term=LOC105278893 | 2674 | 6 | 22.44 | 1 | 3.74 | 16.67 | 34.71 | 25.34 | 9.37 | 9.37 | 0.02 | 0.02 |
| gene2855 | https://www.ncbi.nlm.nih.gov/gene/?term=LOC105275378 | 1189 | 21 | 176.62 | 3 | 25.23 | 14.29 | 38.9 | 30.15 | 8.75 | 8.75 | 0.15 | 0.15 |
| gene9690 | https://www.ncbi.nlm.nih.gov/gene/?term=LOC105283241 | 887 | 7 | 78.92 | 1 | 11.27 | 14.29 | 55.79 | 47.4 | 8.39 | 8.39 | 0.07 | 0.07 |
| gene5966 | https://www.ncbi.nlm.nih.gov/gene/?term=LOC105278917 | 2178 | 3 | 13.77 | 1 | 4.59 | 33.33 | 33.22 | 25.32 | 7.9 | 7.9 | 0.01 | 0.01 |
| gene13301 | https://www.ncbi.nlm.nih.gov/gene/?term=LOC105287352 | 656 | 7 | 106.71 | 1 | 15.24 | 14.29 | 24.74 | 17.08 | 7.67 | 7.67 | 0.08 | 0.08 |
| gene11073 | https://www.ncbi.nlm.nih.gov/gene/?term=LOC105284817 | 1692 | 9 | 53.19 | 1 | 5.91 | 11.11 | 40.23 | 32.79 | 7.44 | 7.44 | 0.04 | 0.04 |
| gene3128 | https://www.ncbi.nlm.nih.gov/gene/?term=LOC105275669 | 1566 | 8 | 51.09 | 1 | 6.39 | 12.5 | 54.98 | 48.31 | 6.67 | 6.67 | 0.03 | 0.03 |
| gene7488 | https://www.ncbi.nlm.nih.gov/gene/?term=LOC105280699 | 1563 | 5 | 31.99 | 1 | 6.4 | 20 | 31.65 | 25.04 | 6.61 | 6.61 | 0.02 | 0.02 |
| gene4732 | https://www.ncbi.nlm.nih.gov/gene/?term=LOC105277491 | 1647 | 11 | 66.79 | 1 | 6.07 | 9.09 | 19.89 | 14.07 | 5.82 | 5.82 | 0.04 | 0.04 |
| gene12610 | https://www.ncbi.nlm.nih.gov/gene/?term=LOC105286572 | 2000 | 6 | 30 | 1 | 5 | 16.67 | 16.76 | 11.15 | 5.61 | 5.61 | 0.02 | 0.02 |
| gene9255 | https://www.ncbi.nlm.nih.gov/gene/?term=LOC105282733 | 3562 | 21 | 58.96 | 2 | 5.61 | 9.52 | 36 | 30.71 | 5.29 | 5.29 | 0.03 | 0.03 |
| gene12818 | https://www.ncbi.nlm.nih.gov/gene/?term=LOC105286815 | 1095 | 11 | 100.46 | 1 | 9.13 | 9.09 | 63.21 | 58.15 | 5.07 | 5.07 | 0.05 | 0.05 |
| gene7300 | https://www.ncbi.nlm.nih.gov/gene/?term=LOC105280472 | 1589 | 12 | 75.52 | 1 | 6.29 | 8.33 | 42.17 | 37.11 | 5.06 | 5.06 | 0.04 | 0.04 |
| gene1656 | https://www.ncbi.nlm.nih.gov/gene/?term=LOC105287814 | 812 | 8 | 98.52 | 1 | 12.32 | 12.5 | 35.83 | 30.99 | 4.84 | 4.84 | 0.05 | 0.05 |
| gene5306 | https://www.ncbi.nlm.nih.gov/gene/?term=LOC105278153 | 481 | 17 | 353.43 | 2 | 41.58 | 11.76 | 21.3 | 16.46 | 4.84 | 4.84 | 0.17 | 0.17 |
| gene10176 | https://www.ncbi.nlm.nih.gov/gene/?term=LOC105283785 | 1637 | 35 | 213.81 | 1 | 6.11 | 2.86 | 18.49 | 13.7 | 4.79 | 4.79 | 0.1 | 0.1 |
| gene9684 | https://www.ncbi.nlm.nih.gov/gene/?term=LOC105283227 | 2228 | 15 | 67.32 | 2 | 8.98 | 13.33 | 34.73 | 29.95 | 4.79 | 4.79 | 0.03 | 0.03 |
| gene1742 | https://www.ncbi.nlm.nih.gov/gene/?term=LOC105287978 | 3019 | 7 | 23.19 | 1 | 3.31 | 14.29 | 17.41 | 12.85 | 4.56 | 4.56 | 0.01 | 0.01 |
| gene2970 | https://www.ncbi.nlm.nih.gov/gene/?term=LOC105275518 | 2340 | 26 | 111.11 | 2 | 8.55 | 7.69 | 47.9 | 43.34 | 4.56 | 4.56 | 0.05 | 0.05 |
| gene9852 | https://www.ncbi.nlm.nih.gov/gene/?term=LOC105283419 | 1060 | 16 | 150.94 | 2 | 18.87 | 12.5 | 6.02 | 1.53 | 4.48 | 4.48 | 0.07 | 0.07 |
| gene4032 | https://www.ncbi.nlm.nih.gov/gene/?term=LOC105276707 | 1993 | 16 | 80.28 | 2 | 10.04 | 12.5 | 37.03 | 32.6 | 4.43 | 4.43 | 0.04 | 0.04 |

...

... only first 25 lines shown ...

Directory: OGL File: ogl-Cb_LI_BCphaseB1.vs.RphaseA2.txt

| gene_ID | gene_link | gwidth | #Sites | #per10Kb | #dmSites | #dmsp10kb | %dmSites | %pSite1 | %pSite2 | DMpSite | ADMpSite | DMpNucl | ADMpNucl |
|---|---|---|---|---|---|---|---|---|---|---|---|---|---|
| gene7300 | https://www.ncbi.nlm.nih.gov/gene/?term=LOC105280472 | 1589 | 12 | 75.52 | 2 | 12.59 | 16.67 | 42.17 | 34.37 | 7.8 | 7.8 | 0.06 | 0.06 |
| gene5695 | https://www.ncbi.nlm.nih.gov/gene/?term=LOC105278607 | 4140 | 22 | 53.14 | 2 | 4.83 | 9.09 | 13.44 | 6.4 | 7.04 | 7.04 | 0.04 | 0.04 |
| gene10343 | https://www.ncbi.nlm.nih.gov/gene/?term=LOC105283986 | 1763 | 7 | 39.71 | 1 | 5.67 | 14.29 | 29.98 | 22.98 | 7 | 7 | 0.03 | 0.03 |
| gene12518 | https://www.ncbi.nlm.nih.gov/gene/?term=LOC105286453 | 1109 | 13 | 117.22 | 2 | 18.03 | 15.38 | 34.82 | 28.73 | 6.08 | 6.08 | 0.07 | 0.07 |
| gene1742 | https://www.ncbi.nlm.nih.gov/gene/?term=LOC105287978 | 3019 | 7 | 23.19 | 2 | 6.62 | 28.57 | 17.41 | 11.62 | 5.79 | 5.79 | 0.01 | 0.01 |
| gene11185 | https://www.ncbi.nlm.nih.gov/gene/?term=LOC105284939 | 1723 | 17 | 98.67 | 1 | 5.8 | 5.88 | 37.14 | 31.9 | 5.24 | 5.24 | 0.05 | 0.05 |
| gene1894 | https://www.ncbi.nlm.nih.gov/gene/?term=LOC105288083 | 445 | 17 | 382.02 | 2 | 44.94 | 11.76 | 56.3 | 51.18 | 5.11 | 5.11 | 0.2 | 0.2 |
| gene4621 | https://www.ncbi.nlm.nih.gov/gene/?term=LOC105277369 | 1157 | 8 | 69.14 | 1 | 8.64 | 12.5 | 67.07 | 62.16 | 4.9 | 4.9 | 0.03 | 0.03 |
| gene11850 | https://www.ncbi.nlm.nih.gov/gene/?term=LOC105285707 | 2053 | 31 | 151 | 2 | 9.74 | 6.45 | 60.03 | 55.77 | 4.26 | 4.26 | 0.06 | 0.06 |
| gene12173 | https://www.ncbi.nlm.nih.gov/gene/?term=LOC105286067 | 518 | 6 | 115.83 | 1 | 19.31 | 16.67 | 11.97 | 7.8 | 4.17 | 4.17 | 0.05 | 0.05 |
| gene7727 | https://www.ncbi.nlm.nih.gov/gene/?term=LOC105280969 | 1006 | 15 | 149.11 | 1 | 9.94 | 6.67 | 36.57 | 32.67 | 3.89 | 3.89 | 0.06 | 0.06 |
| gene2837 | https://www.ncbi.nlm.nih.gov/gene/?term=LOC105275355 | 2872 | 160 | 557.1 | 10 | 34.82 | 6.25 | 29.18 | 25.36 | 3.82 | 3.82 | 0.21 | 0.21 |
| gene2855 | https://www.ncbi.nlm.nih.gov/gene/?term=LOC105275378 | 1189 | 28 | 235.49 | 2 | 16.82 | 7.14 | 43.5 | 39.96 | 3.54 | 3.54 | 0.08 | 0.08 |
| gene9854 | https://www.ncbi.nlm.nih.gov/gene/?term=LOC105283423 | 2931 | 16 | 54.59 | 1 | 3.41 | 6.25 | 37.51 | 33.96 | 3.54 | 3.54 | 0.02 | 0.02 |
| gene3282 | https://www.ncbi.nlm.nih.gov/gene/?term=LOC105275847 | 1942 | 25 | 128.73 | 1 | 5.15 | 4 | 47.32 | 43.85 | 3.47 | 3.47 | 0.04 | 0.04 |
| gene6879 | https://www.ncbi.nlm.nih.gov/gene/?term=LOC105280001 | 2322 | 22 | 94.75 | 1 | 4.31 | 4.55 | 25.57 | 22.33 | 3.24 | 3.24 | 0.03 | 0.03 |
| gene7840 | https://www.ncbi.nlm.nih.gov/gene/?term=LOC105281081 | 2338 | 15 | 64.16 | 1 | 4.28 | 6.67 | 19.85 | 16.65 | 3.2 | 3.2 | 0.02 | 0.02 |
| gene7549 | https://www.ncbi.nlm.nih.gov/gene/?term=LOC105280764 | 2668 | 23 | 86.21 | 2 | 7.5 | 8.7 | 13.43 | 10.25 | 3.17 | 3.17 | 0.03 | 0.03 |
| gene3000 | https://www.ncbi.nlm.nih.gov/gene/?term=LOC105275544 | 1577 | 23 | 145.85 | 3 | 19.02 | 13.04 | 37.23 | 34.08 | 3.15 | 3.15 | 0.05 | 0.05 |
| gene2220 | https://www.ncbi.nlm.nih.gov/gene/?term=LOC105274640 | 2263 | 9 | 39.77 | 1 | 4.42 | 11.11 | 71.96 | 68.85 | 3.11 | 3.11 | 0.01 | 0.01 |
| gene108 | https://www.ncbi.nlm.nih.gov/gene/?term=LOC105284604 | 1903 | 17 | 89.33 | 2 | 10.51 | 11.76 | 3.09 | 0 | 3.09 | 3.09 | 0.03 | 0.03 |
| gene1962 | https://www.ncbi.nlm.nih.gov/gene/?term=LOC105288157 | 1246 | 19 | 152.49 | 1 | 8.03 | 5.26 | 16.56 | 13.6 | 2.96 | 2.96 | 0.05 | 0.05 |
| gene4999 | https://www.ncbi.nlm.nih.gov/gene/?term=LOC105277805 | 6695 | 14 | 20.91 | 1 | 1.49 | 7.14 | 45.95 | 43.05 | 2.9 | 2.9 | 0.01 | 0.01 |
| gene10237 | https://www.ncbi.nlm.nih.gov/gene/?term=LOC105283856 | 1259 | 23 | 182.68 | 1 | 7.94 | 4.35 | 64.51 | 61.63 | 2.88 | 2.88 | 0.05 | 0.05 |

...
... only first 25 lines shown ...

Directory: OGL File: ogl-Cb_LI_BCphaseB1.vs.RphaseB1.txt

| gene_ID | gene_link | gwidth | #Sites | #per10Kb | #dmSites | #dmsp10kb | %dmSites | %pSite1 | %pSite2 | DMpSite | ADMpSite | DMpNucl | ADMpNucl | |
|---|---|---|---|---|---|---|---|---|---|---|---|---|---|---|
| gene2593 | https://www.ncbi.nlm.nih.gov/gene/?term=LOC105275070 | 1774 | 11 | 62.01 | 1 | 5.64 | 9.09 | 61.58 | 25.28 | 36.29 | 36.29 | 0.23 | 0.23 | |
| gene3165 | https://www.ncbi.nlm.nih.gov/gene/?term=LOC105275706 | 2374 | 65 | 273.8 | 4 | 16.85 | 6.15 | 33.82 | 25.03 | 8.79 | 8.79 | 0.24 | 0.24 | |
| gene2837 | https://www.ncbi.nlm.nih.gov/gene/?term=LOC105275355 | 2872 | 165 | 574.51 | | 1 | 3.48 | 0.61 | 28.35 | 22.92 | 5.43 | 5.43 | 0.31 | 0.31 |
| gene2970 | https://www.ncbi.nlm.nih.gov/gene/?term=LOC105275518 | 2340 | 25 | 106.84 | | 1 | 4.27 | 4 | 44.93 | 41.23 | 3.69 | 3.69 | 0.04 | 0.04 |
| gene5714 | https://www.ncbi.nlm.nih.gov/gene/?term=LOC105278729 | 710 | 16 | 225.35 | | 1 | 14.08 | 6.25 | 55.52 | 51.9 | 3.62 | 3.62 | 0.08 | 0.08 |
| gene2856 | https://www.ncbi.nlm.nih.gov/gene/?term=LOC105275384 | 1987 | 92 | 463.01 | | 1 | 5.03 | 1.09 | 34.19 | 31.19 | 3 | 3 | 0.14 | 0.14 |
| gene3671 | https://www.ncbi.nlm.nih.gov/gene/?term=LOC105276272 | 1966 | 40 | 203.46 | | 1 | 5.09 | 2.5 | 30.25 | 27.31 | 2.94 | 2.94 | 0.06 | 0.06 |
| gene396 | https://www.ncbi.nlm.nih.gov/gene/?term=LOC105276611 | 1722 | 36 | 209.06 | | 1 | 5.81 | 2.78 | 30.31 | 27.55 | 2.76 | 2.76 | 0.06 | 0.06 |
| gene13033 | https://www.ncbi.nlm.nih.gov/gene/?term=LOC105287063 | 1877 | 50 | 266.38 | | 1 | 5.33 | 2 | 31.92 | 29.55 | 2.37 | 2.37 | 0.06 | 0.06 |
| gene1855 | https://www.ncbi.nlm.nih.gov/gene/?term=LOC105288036 | 2387 | 91 | 381.23 | | 1 | 4.19 | 1.1 | 29.43 | 27.13 | 2.3 | 2.3 | 0.09 | 0.09 |
| gene2241 | https://www.ncbi.nlm.nih.gov/gene/?term=LOC105274666 | 2598 | 61 | 234.8 | 1 | 3.85 | 1.64 | 30.46 | 28.61 | 1.85 | 1.85 | 0.04 | 0.04 | |
| gene10345 | https://www.ncbi.nlm.nih.gov/gene/?term=LOC105283989 | 4444 | 41 | 92.26 | 1 | 2.25 | 2.44 | 38.24 | 36.42 | 1.83 | 1.83 | 0.02 | 0.02 | |
| gene7493 | https://www.ncbi.nlm.nih.gov/gene/?term=LOC105280704 | 4225 | 147 | 347.93 | | 1 | 2.37 | 0.68 | 18.36 | 16.63 | 1.72 | 1.72 | 0.06 | 0.06 |
| gene5831 | https://www.ncbi.nlm.nih.gov/gene/?term=LOC105278761 | 2506 | 118 | 470.87 | | 2 | 7.98 | 1.69 | 22.67 | 21.1 | 1.58 | 1.58 | 0.07 | 0.07 |
| gene940 | https://www.ncbi.nlm.nih.gov/gene/?term=LOC105282794 | 3578 | 94 | 262.72 | | 1 | 2.79 | 1.06 | 15.53 | 14.17 | 1.36 | 1.36 | 0.04 | 0.04 |
| gene4023 | https://www.ncbi.nlm.nih.gov/gene/?term=LOC105276692 | 1759 | 37 | 210.35 | | 1 | 5.69 | 2.7 | 23.22 | 21.93 | 1.3 | 1.3 | 0.03 | 0.03 |
| gene9336 | https://www.ncbi.nlm.nih.gov/gene/?term=LOC105282834 | 5993 | 134 | 223.59 | | 1 | 1.67 | 0.75 | 14.88 | 13.64 | 1.24 | 1.24 | 0.03 | 0.03 |
| gene2748 | https://www.ncbi.nlm.nih.gov/gene/?term=LOC105275261 | 3845 | 53 | 137.84 | | 1 | 2.6 | 1.89 | 39.61 | 38.42 | 1.19 | 1.19 | 0.02 | 0.02 |
| gene10107 | https://www.ncbi.nlm.nih.gov/gene/?term=LOC105283712 | 1883 | 36 | 191.18 | | 1 | 5.31 | 2.78 | 30.43 | 29.26 | 1.17 | 1.17 | 0.02 | 0.02 |
| gene1884 | https://www.ncbi.nlm.nih.gov/gene/?term=LOC105288072 | 4169 | 84 | 201.49 | | 1 | 2.4 | 1.19 | 24.86 | 23.72 | 1.14 | 1.14 | 0.02 | 0.02 |
| gene8178 | https://www.ncbi.nlm.nih.gov/gene/?term=LOC105281515 | 8647 | 60 | 69.39 | 1 | 1.16 | 1.67 | 30.54 | 29.47 | 1.06 | 1.06 | 0.01 | 0.01 | |
| gene7734 | https://www.ncbi.nlm.nih.gov/gene/?term=LOC105280972 | 1528 | 35 | 229.06 | | 1 | 6.54 | 2.86 | 17.63 | 16.59 | 1.04 | 1.04 | 0.02 | 0.02 |
| gene7884 | https://www.ncbi.nlm.nih.gov/gene/?term=LOC105281266 | 3390 | 87 | 256.64 | | 1 | 2.95 | 1.15 | 15.85 | 14.89 | 0.96 | 0.96 | 0.02 | 0.02 |
| gene4641 | https://www.ncbi.nlm.nih.gov/gene/?term=LOC105277398 | 1937 | 72 | 371.71 | | 1 | 5.16 | 1.39 | 36.3 | 35.35 | 0.95 | 0.95 | 0.04 | 0.04 |
| gene5877 | https://www.ncbi.nlm.nih.gov/gene/?term=LOC105278807 | 1719 | 51 | 296.68 | | 1 | 5.82 | 1.96 | 26.14 | 25.26 | 0.89 | 0.89 | 0.03 | 0.03 |
| gene1147 | https://www.ncbi.nlm.nih.gov/gene/?term=LOC105284954 | 4022 | 118 | 293.39 | | 1 | 2.49 | 0.85 | 15.13 | 14.31 | 0.81 | 0.81 | 0.02 | 0.02 |
| gene3817 | https://www.ncbi.nlm.nih.gov/gene/?term=LOC105276449 | 1230 | 55 | 447.15 | | 1 | 8.13 | 1.82 | 22.28 | 21.47 | 0.8 | 0.8 | 0.04 | 0.04 |
| gene474 | https://www.ncbi.nlm.nih.gov/gene/?term=LOC105277435 | 3625 | 176 | 485.52 | | 1 | 2.76 | 0.57 | 10.76 | 9.98 | 0.78 | 0.78 | 0.04 | 0.04 |
| gene292 | https://www.ncbi.nlm.nih.gov/gene/?term=LOC105275398 | 2825 | 88 | 311.5 | 1 | 3.54 | 1.14 | 22.66 | 21.93 | 0.73 | 0.73 | 0.02 | 0.02 | |
| gene7183 | https://www.ncbi.nlm.nih.gov/gene/?term=LOC105280348 | 3925 | 80 | 203.82 | | 1 | 2.55 | 1.25 | 17.35 | 16.62 | 0.73 | 0.73 | 0.01 | 0.01 |
| gene322 | https://www.ncbi.nlm.nih.gov/gene/?term=LOC105275777 | 1606 | 28 | 174.35 | | 1 | 6.23 | 3.57 | 36.76 | 36.04 | 0.72 | 0.72 | 0.01 | 0.01 |
| gene298 | https://www.ncbi.nlm.nih.gov/gene/?term=LOC105275460 | 2473 | 126 | 509.5 | 1 | 4.04 | 0.79 | 13.09 | 12.43 | 0.67 | 0.67 | 0.03 | 0.03 | |
| gene10678 | https://www.ncbi.nlm.nih.gov/gene/?term=LOC105284355 | 4031 | 169 | 419.25 | | 1 | 2.48 | 0.59 | 20.31 | 19.66 | 0.65 | 0.65 | 0.03 | 0.03 |
| gene4034 | https://www.ncbi.nlm.nih.gov/gene/?term=LOC105276704 | 2065 | 87 | 421.31 | | 2 | 9.69 | 2.3 | 14.01 | 13.38 | 0.63 | 0.63 | 0.03 | 0.03 |
| gene7888 | https://www.ncbi.nlm.nih.gov/gene/?term=LOC105281141 | 4357 | 170 | 390.18 | | 1 | 2.3 | 0.59 | 28.6 | 28 | 0.6 | 0.6 | 0.02 | 0.02 |
| gene6652 | https://www.ncbi.nlm.nih.gov/gene/?term=LOC105279719 | 2831 | 88 | 310.84 | | 1 | 3.53 | 1.14 | 21.65 | 21.08 | 0.57 | 0.57 | 0.02 | 0.02 |
| gene4892 | https://www.ncbi.nlm.nih.gov/gene/?term=LOC105277667 | 3898 | 127 | 325.81 | | 1 | 2.57 | 0.79 | 27.72 | 27.21 | 0.52 | 0.52 | 0.02 | 0.02 |
| gene12978 | https://www.ncbi.nlm.nih.gov/gene/?term=LOC105286997 | 2858 | 150 | 524.84 | | 1 | 3.5 | 0.67 | 11.28 | 10.84 | 0.44 | 0.44 | 0.02 | 0.02 |
| gene11819 | https://www.ncbi.nlm.nih.gov/gene/?term=LOC105285671 | 2366 | 104 | 439.56 | | 1 | 4.23 | 0.96 | 30.06 | 29.63 | 0.43 | 0.43 | 0.02 | 0.02 |
| gene2683 | https://www.ncbi.nlm.nih.gov/gene/?term=LOC105275187 | 5492 | 78 | 142.02 | | 1 | 1.82 | 1.28 | 16.44 | 16.02 | 0.43 | 0.43 | 0.01 | 0.01 |
| gene3107 | https://www.ncbi.nlm.nih.gov/gene/?term=LOC105275650 | 5103 | 182 | 356.65 | | 1 | 1.96 | 0.55 | 16.89 | 16.49 | 0.4 | 0.4 | 0.01 | 0.01 |
| gene6455 | https://www.ncbi.nlm.nih.gov/gene/?term=LOC105279500 | 2612 | 35 | 134 | 1 | 3.83 | 2.86 | 48.92 | 48.53 | 0.39 | 0.39 | 0.01 | 0.01 | |
| gene3885 | https://www.ncbi.nlm.nih.gov/gene/?term=LOC105276527 | 7928 | 369 | 465.44 | | 1 | 1.26 | 0.27 | 8.26 | 7.93 | 0.33 | 0.33 | 0.02 | 0.02 |
| gene13036 | https://www.ncbi.nlm.nih.gov/gene/?term=LOC105287059 | 4959 | 193 | 389.19 | | 1 | 2.02 | 0.52 | 16.75 | 16.43 | 0.32 | 0.32 | 0.01 | 0.01 |
| gene8729 | https://www.ncbi.nlm.nih.gov/gene/?term=LOC105282137 | 5005 | 226 | 451.55 | | 1 | 2 | 0.44 | 14.98 | 14.68 | 0.29 | 0.29 | 0.01 | 0.01 |
| gene12572 | https://www.ncbi.nlm.nih.gov/gene/?term=LOC105286514 | 2374 | 52 | 219.04 | | 1 | 4.21 | 1.92 | 35.13 | 34.85 | 0.28 | 0.28 | 0.01 | 0.01 |
| gene9101 | https://www.ncbi.nlm.nih.gov/gene/?term=LOC105282553 | 3068 | 106 | 345.5 | 1 | 3.26 | 0.94 | 22.9 | 22.63 | 0.27 | 0.27 | 0.01 | 0.01 | |
| gene10018 | https://www.ncbi.nlm.nih.gov/gene/?term=LOC105283607 | 4414 | 228 | 516.54 | | 1 | 2.27 | 0.44 | 14.1 | 13.84 | 0.26 | 0.26 | 0.01 | 0.01 |
| gene2668 | https://www.ncbi.nlm.nih.gov/gene/?term=LOC105275170 | 12496 | 356 | 284.89 | | 1 | 0.8 | 0.28 | 2.04 | 1.8 | 0.24 | 0.24 | 0.01 | 0.01 |
| gene10540 | https://www.ncbi.nlm.nih.gov/gene/?term=LOC105284200 | 5267 | 214 | 406.3 | 1 | 1.9 | 0.47 | 13.24 | 13.01 | 0.23 | 0.23 | 0.01 | 0.01 | |
| gene10596 | https://www.ncbi.nlm.nih.gov/gene/?term=LOC105284262 | 6520 | 239 | 366.56 | | 1 | 1.53 | 0.42 | 4.23 | 4.05 | 0.19 | 0.19 | 0.01 | 0.01 |
| gene6286 | https://www.ncbi.nlm.nih.gov/gene/?term=LOC105279298 | 2861 | 128 | 447.4 | 1 | 3.5 | 0.78 | 23.43 | 23.27 | 0.16 | 0.16 | 0.01 | 0.01 | |
| gene7368 | https://www.ncbi.nlm.nih.gov/gene/?term=LOC105280556 | 4105 | 71 | 172.96 | | 1 | 2.44 | 1.41 | 15.89 | 15.73 | 0.16 | 0.16 | 0 | 0 |
| gene12406 | https://www.ncbi.nlm.nih.gov/gene/?term=LOC105286316 | 1386 | 44 | 317.46 | | 1 | 7.22 | 2.27 | 32.85 | 32.71 | 0.14 | 0.14 | 0 | 0 |
| gene10600 | https://www.ncbi.nlm.nih.gov/gene/?term=LOC105284270 | 4837 | 111 | 229.48 | | 1 | 2.07 | 0.9 | 11.96 | 11.83 | 0.13 | 0.13 | 0 | 0 |
| gene927 | https://www.ncbi.nlm.nih.gov/gene/?term=LOC105282600 | 3727 | 60 | 160.99 | | 1 | 2.68 | 1.67 | 28.88 | 28.75 | 0.13 | 0.13 | 0 | 0 |
| gene9327 | https://www.ncbi.nlm.nih.gov/gene/?term=LOC105282821 | 7793 | 183 | 234.83 | | 1 | 1.28 | 0.55 | 12.55 | 12.42 | 0.13 | 0.13 | 0 | 0 |
| gene11275 | https://www.ncbi.nlm.nih.gov/gene/?term=LOC105285055 | 4119 | 246 | 597.23 | | 1 | 2.43 | 0.41 | 12.22 | 12.12 | 0.1 | 0.1 | 0.01 | 0.01 |
| gene5375 | https://www.ncbi.nlm.nih.gov/gene/?term=LOC105278228 | 8640 | 191 | 221.06 | | 1 | 1.16 | 0.52 | 22.09 | 21.99 | 0.1 | 0.1 | 0 | 0 |
| gene10222 | https://www.ncbi.nlm.nih.gov/gene/?term=LOC105283838 | 1748 | 30 | 171.62 | | 1 | 5.72 | 3.33 | 35.37 | 35.28 | 0.09 | 0.09 | 0 | 0 |

| | | | | | | | | | | | | | |
|---|---|---|---|---|---|---|---|---|---|---|---|---|---|
| gene10520 | https://www.ncbi.nlm.nih.gov/gene/?term=LOC105284184 | 2451 | 102 | 416.16 | 1 | 4.08 | 0.98 | 30 | 29.93 | 0.07 | 0.07 | 0 | 0 |
| gene9326 | https://www.ncbi.nlm.nih.gov/gene/?term=LOC105282818 | 16612 | 236 | 142.07 | 1 | 0.6 | 0.42 | 13.17 | 13.1 | 0.07 | 0.07 | 0 | 0 |
| gene2443 | https://www.ncbi.nlm.nih.gov/gene/?term=LOC105274895 | 10433 | 424 | 406.4 1 | 0.96 | 0.24 | 5.17 | 5.13 | 0.03 | 0.03 | 0 | 0 | |
| gene53 | https://www.ncbi.nlm.nih.gov/gene/?term=LOC105277148 | 19283 | 333 | 172.69 | 1 | 0.52 | 0.3 | 4.68 | 4.68 | 0 | 0 | 0 | 0 |
| gene11454 | https://www.ncbi.nlm.nih.gov/gene/?term=LOC105285256 | 1906 | 50 | 262.33 | 1 | 5.25 | 2 | 44.07 | 44.08 | -0.01 | 0.01 | 0 | 0 |
| gene13114 | https://www.ncbi.nlm.nih.gov/gene/?term=LOC105287147 | 7609 | 432 | 567.75 | 1 | 1.31 | 0.23 | 9.01 | 9.02 | -0.01 | 0.01 | 0 | 0 |
| gene2271 | https://www.ncbi.nlm.nih.gov/gene/?term=LOC105274706 | 1448 | 47 | 324.59 | 1 | 6.91 | 2.13 | 41.96 | 41.97 | -0.01 | 0.01 | 0 | 0 |
| gene8600 | https://www.ncbi.nlm.nih.gov/gene/?term=LOC105282001 | 8513 | 206 | 241.98 | 1 | 1.17 | 0.49 | 9.71 | 9.75 | -0.04 | 0.04 | 0 | 0 |
| gene5746 | https://www.ncbi.nlm.nih.gov/gene/?term=LOC105278666 | 10343 | 483 | 466.98 | 1 | 0.97 | 0.21 | 6.37 | 6.41 | -0.05 | 0.05 | 0 | 0 |
| gene6620 | https://www.ncbi.nlm.nih.gov/gene/?term=LOC105279687 | 17998 | 448 | 248.92 | 1 | 0.56 | 0.22 | 4.43 | 4.47 | -0.05 | 0.05 | 0 | 0 |
| gene1632 | https://www.ncbi.nlm.nih.gov/gene/?term=LOC105287764 | 18061 | 431 | 238.64 | 1 | 0.55 | 0.23 | 3.3 | 3.37 | -0.07 | 0.07 | 0 | 0 |
| gene2806 | https://www.ncbi.nlm.nih.gov/gene/?term=LOC105275327 | 14257 | 552 | 387.18 | 1 | 0.7 | 0.18 | 6.34 | 6.41 | -0.07 | 0.07 | 0 | 0 |
| gene3470 | https://www.ncbi.nlm.nih.gov/gene/?term=LOC105276059 | 8716 | 182 | 208.81 | 1 | 1.15 | 0.55 | 9.23 | 9.32 | -0.09 | 0.09 | 0 | 0 |
| gene5646 | https://www.ncbi.nlm.nih.gov/gene/?term=LOC105278534 | 7475 | 202 | 270.23 | 1 | 1.34 | 0.5 | 16.69 | 16.79 | -0.1 | 0.1 | 0 | 0 |
| gene12427 | https://www.ncbi.nlm.nih.gov/gene/?term=LOC105286339 | 5220 | 187 | 358.24 | 1 | 1.92 | 0.53 | 15.91 | 16.02 | -0.12 | 0.12 | 0 | 0 |
| gene12802 | https://www.ncbi.nlm.nih.gov/gene/?term=LOC105286799 | 3183 | 180 | 565.5 1 | 3.14 | 0.56 | 16.81 | 16.93 | -0.12 | 0.12 | -0.01 | 0.01 | |
| gene5654 | https://www.ncbi.nlm.nih.gov/gene/?term=LOC105278543 | 4019 | 181 | 450.36 | 1 | 2.49 | 0.55 | 15.7 | 15.83 | -0.13 | 0.13 | -0.01 | 0.01 |
| gene10844 | https://www.ncbi.nlm.nih.gov/gene/?term=LOC105284535 | 9086 | 259 | 285.05 | 1 | 1.1 | 0.39 | 14.4 | 14.56 | -0.16 | 0.16 | 0 | 0 |
| gene8274 | https://www.ncbi.nlm.nih.gov/gene/?term=LOC105281624 | 5425 | 227 | 418.43 | 1 | 1.84 | 0.44 | 9.53 | 9.7 | -0.17 | 0.17 | -0.01 | 0.01 |
| gene13039 | https://www.ncbi.nlm.nih.gov/gene/?term=LOC105287065 | 4902 | 253 | 516.12 | 1 | 2.04 | 0.4 | 12.12 | 12.3 | -0.18 | 0.18 | -0.01 | 0.01 |
| gene11579 | https://www.ncbi.nlm.nih.gov/gene/?term=LOC105285406 | 13571 | 316 | 232.85 | 1 | 0.74 | 0.32 | 6.01 | 6.26 | -0.25 | 0.25 | -0.01 | 0.01 |
| gene7249 | https://www.ncbi.nlm.nih.gov/gene/?term=LOC105280422 | 1775 | 66 | 371.83 | 1 | 5.63 | 1.52 | 15.78 | 16.04 | -0.26 | 0.26 | -0.01 | 0.01 |
| gene6639 | https://www.ncbi.nlm.nih.gov/gene/?term=LOC105279705 | 7656 | 83 | 108.41 | 1 | 1.31 | 1.2 | 19.9 | 20.17 | -0.28 | 0.28 | 0 | 0 |
| gene7199 | https://www.ncbi.nlm.nih.gov/gene/?term=LOC105280366 | 1414 | 34 | 240.45 | 1 | 7.07 | 2.94 | 44.65 | 44.98 | -0.33 | 0.33 | -0.01 | 0.01 |
| gene10549 | https://www.ncbi.nlm.nih.gov/gene/?term=LOC105284210 | 7145 | 365 | 510.85 | 2 | 2.8 | 0.55 | 9.55 | 9.9 | -0.35 | 0.35 | -0.02 | 0.02 |
| gene5524 | https://www.ncbi.nlm.nih.gov/gene/?term=LOC105278397 | 5177 | 209 | 403.71 | 1 | 1.93 | 0.48 | 23.99 | 24.35 | -0.36 | 0.36 | -0.01 | 0.01 |
| gene10326 | https://www.ncbi.nlm.nih.gov/gene/?term=LOC105283962 | 3042 | 137 | 450.36 | 1 | 3.29 | 0.73 | 7.31 | 7.68 | -0.37 | 0.37 | -0.02 | 0.02 |
| gene4376 | https://www.ncbi.nlm.nih.gov/gene/?term=LOC105277083 | 4307 | 129 | 299.51 | 1 | 2.32 | 0.78 | 18.69 | 19.06 | -0.38 | 0.38 | -0.01 | 0.01 |
| gene11583 | https://www.ncbi.nlm.nih.gov/gene/?term=LOC105285414 | 8192 | 192 | 234.38 | 1 | 1.22 | 0.52 | 8.5 | 8.89 | -0.39 | 0.39 | -0.01 | 0.01 |
| gene7915 | https://www.ncbi.nlm.nih.gov/gene/?term=LOC105281173 | 1597 | 42 | 262.99 | 1 | 6.26 | 2.38 | 62.33 | 62.75 | -0.42 | 0.42 | -0.01 | 0.01 |
| gene3793 | https://www.ncbi.nlm.nih.gov/gene/?term=LOC105276429 | 2901 | 131 | 451.57 | 1 | 3.45 | 0.76 | 22.23 | 22.66 | -0.43 | 0.43 | -0.02 | 0.02 |
| gene5525 | https://www.ncbi.nlm.nih.gov/gene/?term=LOC105278396 | 3090 | 189 | 611.65 | 1 | 3.24 | 0.53 | 24.08 | 24.52 | -0.44 | 0.44 | -0.03 | 0.03 |
| gene9330 | https://www.ncbi.nlm.nih.gov/gene/?term=LOC105282825 | 3936 | 98 | 248.98 | 1 | 2.54 | 1.02 | 13.47 | 13.91 | -0.44 | 0.44 | -0.01 | 0.01 |
| gene10606 | https://www.ncbi.nlm.nih.gov/gene/?term=LOC105284273 | 6119 | 272 | 444.52 | 1 | 1.63 | 0.37 | 17.87 | 18.32 | -0.45 | 0.45 | -0.02 | 0.02 |
| gene6583 | https://www.ncbi.nlm.nih.gov/gene/?term=LOC105279649 | 7444 | 372 | 499.73 | 1 | 1.34 | 0.27 | 10.85 | 11.31 | -0.45 | 0.45 | -0.02 | 0.02 |
| gene4190 | https://www.ncbi.nlm.nih.gov/gene/?term=LOC105276875 | 3150 | 134 | 425.4 1 | 3.17 | 0.75 | 20.21 | 20.67 | -0.46 | 0.46 | -0.02 | 0.02 | |
| gene7247 | https://www.ncbi.nlm.nih.gov/gene/?term=LOC105280407 | 10486 | 227 | 216.48 | 1 | 0.95 | 0.44 | 14.79 | 15.25 | -0.46 | 0.46 | -0.01 | 0.01 |
| gene7166 | https://www.ncbi.nlm.nih.gov/gene/?term=LOC105280326 | 6117 | 259 | 423.41 | 1 | 1.63 | 0.39 | 14.46 | 14.93 | -0.47 | 0.47 | -0.02 | 0.02 |
| gene2173 | https://www.ncbi.nlm.nih.gov/gene/?term=LOC105274590 | 4387 | 87 | 198.31 | 2 | 4.56 | 2.3 | 23.56 | 24.1 | -0.54 | 0.54 | -0.01 | 0.01 |
| gene4033 | https://www.ncbi.nlm.nih.gov/gene/?term=LOC105276705 | 10546 | 155 | 146.98 | 1 | 0.95 | 0.65 | 5.56 | 6.1 | -0.55 | 0.55 | -0.01 | 0.01 |
| gene6966 | https://www.ncbi.nlm.nih.gov/gene/?term=LOC105280104 | 3747 | 132 | 352.28 | 1 | 2.67 | 0.76 | 16.91 | 17.46 | -0.55 | 0.55 | -0.02 | 0.02 |
| gene11561 | https://www.ncbi.nlm.nih.gov/gene/?term=LOC105285387 | 11528 | 218 | 189.1 1 | 0.87 | 0.46 | 8.62 | 9.18 | -0.56 | 0.56 | -0.01 | 0.01 | |
| gene8814 | https://www.ncbi.nlm.nih.gov/gene/?term=LOC105282240 | 6910 | 204 | 295.22 | 1 | 1.45 | 0.49 | 8.01 | 8.64 | -0.63 | 0.63 | -0.02 | 0.02 |
| gene5184 | https://www.ncbi.nlm.nih.gov/gene/?term=LOC105277999 | 5528 | 165 | 298.48 | 1 | 1.81 | 0.61 | 11.31 | 11.98 | -0.67 | 0.67 | -0.02 | 0.02 |
| gene11235 | https://www.ncbi.nlm.nih.gov/gene/?term=LOC105285004 | 1339 | 36 | 268.86 | 1 | 7.47 | 2.78 | 19.18 | 19.88 | -0.7 | 0.7 | -0.02 | 0.02 |
| gene1310 | https://www.ncbi.nlm.nih.gov/gene/?term=LOC105286661 | 2582 | 68 | 263.36 | 1 | 3.87 | 1.47 | 18.99 | 19.74 | -0.75 | 0.75 | -0.02 | 0.02 |
| gene12573 | https://www.ncbi.nlm.nih.gov/gene/?term=LOC105286515 | 3647 | 107 | 293.39 | 1 | 2.74 | 0.93 | 10.59 | 11.4 | -0.82 | 0.82 | -0.02 | 0.02 |
| gene1470 | https://www.ncbi.nlm.nih.gov/gene/?term=LOC105287597 | 2974 | 192 | 645.6 1 | 3.36 | 0.52 | 18.41 | 19.24 | -0.83 | 0.83 | -0.05 | 0.05 | |
| gene5298 | https://www.ncbi.nlm.nih.gov/gene/?term=LOC105278145 | 4060 | 262 | 645.32 | 2 | 4.93 | 0.76 | 9.65 | 10.52 | -0.86 | 0.86 | -0.06 | 0.06 |
| gene6698 | https://www.ncbi.nlm.nih.gov/gene/?term=LOC105279778 | 4099 | 161 | 392.78 | 1 | 2.44 | 0.62 | 8.69 | 9.57 | -0.88 | 0.88 | -0.03 | 0.03 |
| gene920 | https://www.ncbi.nlm.nih.gov/gene/?term=LOC105282518 | 1197 | 44 | 367.59 | 1 | 8.35 | 2.27 | 33.07 | 33.97 | -0.89 | 0.89 | -0.03 | 0.03 |
| gene5101 | https://www.ncbi.nlm.nih.gov/gene/?term=LOC105277903 | 1813 | 37 | 204.08 | 1 | 5.52 | 2.7 | 54.12 | 55.06 | -0.94 | 0.94 | -0.02 | 0.02 |
| gene10155 | https://www.ncbi.nlm.nih.gov/gene/?term=LOC105283756 | 7030 | 116 | 165.01 | 1 | 1.42 | 0.86 | 7.15 | 8.11 | -0.96 | 0.96 | -0.02 | 0.02 |
| gene10392 | https://www.ncbi.nlm.nih.gov/gene/?term=LOC105284036 | 3216 | 103 | 320.27 | 2 | 6.22 | 1.94 | 23.68 | 24.65 | -0.97 | 0.97 | -0.03 | 0.03 |
| gene570 | https://www.ncbi.nlm.nih.gov/gene/?term=LOC105278549 | 1673 | 72 | 430.36 | 1 | 5.98 | 1.39 | 42.9 | 43.9 | -0.99 | 0.99 | -0.04 | 0.04 |
| gene1220 | https://www.ncbi.nlm.nih.gov/gene/?term=LOC105285742 | 3453 | 119 | 344.63 | 1 | 2.9 | 0.84 | 16.38 | 17.43 | -1.05 | 1.05 | -0.04 | 0.04 |
| gene9598 | https://www.ncbi.nlm.nih.gov/gene/?term=LOC105283132 | 1769 | 56 | 316.56 | 1 | 5.65 | 1.79 | 39.06 | 40.18 | -1.12 | 1.12 | -0.04 | 0.04 |
| gene4962 | https://www.ncbi.nlm.nih.gov/gene/?term=LOC105277761 | 4645 | 156 | 335.84 | 1 | 2.15 | 0.64 | 14.16 | 15.31 | -1.15 | 1.15 | -0.04 | 0.04 |
| gene3168 | https://www.ncbi.nlm.nih.gov/gene/?term=LOC105275711 | 6150 | 97 | 157.72 | 1 | 1.63 | 1.03 | 10.51 | 11.68 | -1.17 | 1.17 | -0.02 | 0.02 |
| gene388 | https://www.ncbi.nlm.nih.gov/gene/?term=LOC105276538 | 1558 | 21 | 134.79 | 1 | 6.42 | 4.76 | 69.07 | 70.25 | -1.18 | 1.18 | -0.02 | 0.02 |
| gene3599 | https://www.ncbi.nlm.nih.gov/gene/?term=LOC105276196 | 2955 | 60 | 203.05 | 1 | 3.38 | 1.67 | 37.66 | 38.86 | -1.2 | 1.2 | -0.02 | 0.02 |
| gene12203 | https://www.ncbi.nlm.nih.gov/gene/?term=LOC105286099 | 5967 | 87 | 145.8 1 | 1.68 | 1.15 | 10.81 | 12.06 | -1.25 | 1.25 | -0.02 | 0.02 | |
| gene7398 | https://www.ncbi.nlm.nih.gov/gene/?term=LOC105280589 | 3452 | 42 | 121.67 | 1 | 2.9 | 2.38 | 24.95 | 26.26 | -1.31 | 1.31 | -0.02 | 0.02 |
| gene1198 | https://www.ncbi.nlm.nih.gov/gene/?term=LOC105285507 | 2738 | 97 | 354.27 | 1 | 3.65 | 1.03 | 20.27 | 21.73 | -1.46 | 1.46 | -0.05 | 0.05 |

| | | | | | | | | | | | | | |
|---|---|---|---|---|---|---|---|---|---|---|---|---|---|
| gene12021 | https://www.ncbi.nlm.nih.gov/gene/?term=LOC105285905 | 1399 | 29 | 207.29 | 1 | 7.15 | 3.45 | 32.77 | 34.25 | -1.48 | 1.48 | -0.03 | 0.03 |
| gene6422 | https://www.ncbi.nlm.nih.gov/gene/?term=LOC105279458 | 2419 | 42 | 173.63 | 1 | 4.13 | 2.38 | 44.07 | 45.58 | -1.51 | 1.51 | -0.03 | 0.03 |
| gene11340 | https://www.ncbi.nlm.nih.gov/gene/?term=LOC105285131 | 2918 | 98 | 335.85 | 1 | 3.43 | 1.02 | 13.78 | 15.33 | -1.55 | 1.55 | -0.05 | 0.05 |
| gene3727 | https://www.ncbi.nlm.nih.gov/gene/?term=LOC105276367 | 2484 | 65 | 261.67 | 1 | 4.03 | 1.54 | 43.95 | 45.54 | -1.59 | 1.59 | -0.04 | 0.04 |
| gene9411 | https://www.ncbi.nlm.nih.gov/gene/?term=LOC105282914 | 5534 | 42 | 75.89 | 1 | 1.81 | 2.38 | 49.22 | 50.92 | -1.7 | 1.7 | -0.01 | 0.01 |
| gene2404 | https://www.ncbi.nlm.nih.gov/gene/?term=LOC105274853 | 1767 | 36 | 203.74 | 1 | 5.66 | 2.78 | 33.17 | 35.01 | -1.84 | 1.84 | -0.04 | 0.04 |
| gene7248 | https://www.ncbi.nlm.nih.gov/gene/?term=LOC105280408 | 5175 | 59 | 114.01 | 1 | 1.93 | 1.69 | 15.82 | 17.71 | -1.89 | 1.89 | -0.02 | 0.02 |
| gene9740 | https://www.ncbi.nlm.nih.gov/gene/?term=LOC105283277 | 2887 | 139 | 481.47 | 1 | 3.46 | 0.72 | 18.49 | 20.55 | -2.05 | 2.05 | -0.1 | 0.1 |
| gene652 | https://www.ncbi.nlm.nih.gov/gene/?term=LOC105279528 | 5806 | 39 | 67.17 | 1 | 1.72 | 2.56 | 17.73 | 19.93 | -2.2 | 2.2 | -0.01 | 0.01 |
| gene9464 | https://www.ncbi.nlm.nih.gov/gene/?term=LOC105282975 | 1450 | 24 | 165.52 | 1 | 6.9 | 4.17 | 39.26 | 42.21 | -2.95 | 2.95 | -0.05 | 0.05 |
| gene1199 | https://www.ncbi.nlm.nih.gov/gene/?term=LOC105285498 | 2220 | 30 | 135.14 | 1 | 4.5 | 3.33 | 33.52 | 36.79 | -3.27 | 3.27 | -0.04 | 0.04 |
| gene6237 | https://www.ncbi.nlm.nih.gov/gene/?term=LOC105279244 | 1484 | 33 | 222.37 | 1 | 6.74 | 3.03 | 44.64 | 48.04 | -3.4 | 3.4 | -0.08 | 0.08 |
| gene1505 | https://www.ncbi.nlm.nih.gov/gene/?term=LOC105287633 | 1693 | 22 | 129.95 | 1 | 5.91 | 4.55 | 51.59 | 55.25 | -3.65 | 3.65 | -0.05 | 0.05 |
| gene2413 | https://www.ncbi.nlm.nih.gov/gene/?term=LOC105274863 | 2006 | 62 | 309.07 | 1 | 4.99 | 1.61 | 48.7 | 53.36 | -4.66 | 4.66 | -0.14 | 0.14 |
| gene564 | https://www.ncbi.nlm.nih.gov/gene/?term=LOC105278478 | 2802 | 16 | 57.1 | 1 | 3.57 | 6.25 | 14.91 | 19.95 | -5.03 | 5.03 | -0.03 | 0.03 |
| gene11584 | https://www.ncbi.nlm.nih.gov/gene/?term=LOC105285416 | 3709 | 16 | 43.14 | 2 | 5.39 | 12.5 | 21.86 | 29.47 | -7.61 | 7.61 | -0.03 | 0.03 |
| gene9109 | https://www.ncbi.nlm.nih.gov/gene/?term=LOC105282561 | 2400 | 14 | 58.33 | 1 | 4.17 | 7.14 | 13.59 | 21.35 | -7.76 | 7.76 | -0.05 | 0.05 |
| gene3166 | https://www.ncbi.nlm.nih.gov/gene/?term=LOC105275707 | 2393 | 10 | 41.79 | 1 | 4.18 | 10 | 65.97 | 76.22 | -10.25 | 10.25 | -0.04 | 0.04 |
| gene10963 | https://www.ncbi.nlm.nih.gov/gene/?term=LOC105284677 | 1576 | 30 | 190.36 | 1 | 6.35 | 3.33 | 19.35 | 34.92 | -15.57 | 15.57 | -0.3 | 0.3 |
| gene393 | https://www.ncbi.nlm.nih.gov/gene/?term=LOC105276566 | 1123 | 2 | 17.81 | 1 | 8.9 | 50 | 47.22 | 74.3 | -27.08 | 27.08 | -0.05 | 0.05 |

Directory: OGL File: ogl-Cb_LI_BCphaseB1.vs.RphaseB2.txt

| gene_ID | gene_link | gwidth | #Sites | #per10Kb | #dmSites | #dmsp10kb | %dmSites | %pSite1 | %pSite2 | DMpSite | ADMpSite | DMpNucl | ADMpNucl |
|---|---|---|---|---|---|---|---|---|---|---|---|---|---|
| gene2593 | https://www.ncbi.nlm.nih.gov/gene/?term=LOC105275070 | 1774 | 7 | 39.46 | 5 | 28.18 | 71.43 | 58.65 | 4.09 | 54.56 | 54.56 | 0.22 | 0.22 |
| gene3165 | https://www.ncbi.nlm.nih.gov/gene/?term=LOC105275706 | 2374 | 68 | 286.44 | 5 | 21.06 | 7.35 | 33.32 | 23.13 | 10.19 | 10.19 | 0.29 | 0.29 |
| gene2970 | https://www.ncbi.nlm.nih.gov/gene/?term=LOC105275518 | 2340 | 20 | 85.47 | 1 | 4.27 | 5 | 58.11 | 52.81 | 5.3 | 5.3 | 0.05 | 0.05 |
| gene5714 | https://www.ncbi.nlm.nih.gov/gene/?term=LOC105278729 | 710 | 17 | 239.44 | 1 | 14.08 | 5.88 | 58.14 | 53.54 | 4.59 | 4.59 | 0.11 | 0.11 |
| gene1518 | https://www.ncbi.nlm.nih.gov/gene/?term=LOC105287648 | 2195 | 25 | 113.9 | 1 | 4.56 | 4 | 44.43 | 41.18 | 3.25 | 3.25 | 0.04 | 0.04 |
| gene940 | https://www.ncbi.nlm.nih.gov/gene/?term=LOC105282794 | 3578 | 83 | 231.97 | 1 | 2.79 | 1.2 | 17.02 | 13.98 | 3.04 | 3.04 | 0.07 | 0.07 |
| gene5647 | https://www.ncbi.nlm.nih.gov/gene/?term=LOC105278567 | 3118 | 36 | 115.46 | 1 | 3.21 | 2.78 | 16.65 | 14.27 | 2.38 | 2.38 | 0.03 | 0.03 |
| gene5273 | https://www.ncbi.nlm.nih.gov/gene/?term=LOC105278108 | 4067 | 46 | 113.11 | 1 | 2.46 | 2.17 | 27.18 | 24.92 | 2.26 | 2.26 | 0.03 | 0.03 |
| gene6334 | https://www.ncbi.nlm.nih.gov/gene/?term=LOC105279356 | 1471 | 48 | 326.31 | 1 | 6.8 | 2.08 | 28.86 | 26.95 | 1.91 | 1.91 | 0.06 | 0.06 |
| gene8178 | https://www.ncbi.nlm.nih.gov/gene/?term=LOC105281515 | 8647 | 44 | 50.88 | 1 | 1.16 | 2.27 | 25.92 | 24.04 | 1.88 | 1.88 | 0.01 | 0.01 |
| gene7734 | https://www.ncbi.nlm.nih.gov/gene/?term=LOC105280972 | 1528 | 30 | 196.34 | 1 | 6.54 | 3.33 | 20.49 | 18.79 | 1.71 | 1.71 | 0.03 | 0.03 |
| gene1855 | https://www.ncbi.nlm.nih.gov/gene/?term=LOC105288036 | 2387 | 78 | 326.77 | 1 | 4.19 | 1.28 | 33.45 | 31.79 | 1.66 | 1.66 | 0.05 | 0.05 |
| gene8661 | https://www.ncbi.nlm.nih.gov/gene/?term=LOC105282063 | 1926 | 34 | 176.53 | 1 | 5.19 | 2.94 | 30.6 | 28.98 | 1.62 | 1.62 | 0.03 | 0.03 |
| gene7199 | https://www.ncbi.nlm.nih.gov/gene/?term=LOC105280366 | 1414 | 22 | 155.59 | 1 | 7.07 | 4.55 | 43.24 | 41.76 | 1.47 | 1.47 | 0.02 | 0.02 |
| gene11819 | https://www.ncbi.nlm.nih.gov/gene/?term=LOC105285671 | 2366 | 102 | 431.11 | 1 | 4.23 | 0.98 | 27.3 | 25.91 | 1.39 | 1.39 | 0.06 | 0.06 |
| gene7568 | https://www.ncbi.nlm.nih.gov/gene/?term=LOC105280786 | 2525 | 122 | 483.17 | 1 | 3.96 | 0.82 | 29.81 | 28.44 | 1.38 | 1.38 | 0.07 | 0.07 |
| gene6966 | https://www.ncbi.nlm.nih.gov/gene/?term=LOC105280104 | 3747 | 135 | 360.29 | 1 | 2.67 | 0.74 | 17.67 | 16.38 | 1.29 | 1.29 | 0.05 | 0.05 |
| gene2748 | https://www.ncbi.nlm.nih.gov/gene/?term=LOC105275261 | 3845 | 54 | 140.44 | 1 | 2.6 | 1.85 | 40.26 | 39.17 | 1.09 | 1.09 | 0.02 | 0.02 |
| gene4213 | https://www.ncbi.nlm.nih.gov/gene/?term=LOC105276901 | 3257 | 81 | 248.7 | 1 | 3.07 | 1.23 | 28.04 | 26.96 | 1.09 | 1.09 | 0.03 | 0.03 |
| gene1332 | https://www.ncbi.nlm.nih.gov/gene/?term=LOC105286872 | 2460 | 154 | 626.02 | 1 | 4.07 | 0.65 | 21.26 | 20.18 | 1.08 | 1.08 | 0.07 | 0.07 |
| gene6455 | https://www.ncbi.nlm.nih.gov/gene/?term=LOC105279500 | 2612 | 35 | 134 | 1 | 3.83 | 2.86 | 49.33 | 48.29 | 1.04 | 1.04 | 0.01 | 0.01 |
| gene2844 | https://www.ncbi.nlm.nih.gov/gene/?term=LOC105275363 | 4330 | 190 | 438.8 | 1 | 2.31 | 0.53 | 14.79 | 13.86 | 0.93 | 0.93 | 0.04 | 0.04 |
| gene10678 | https://www.ncbi.nlm.nih.gov/gene/?term=LOC105284355 | 4031 | 166 | 411.81 | 1 | 2.48 | 0.6 | 19.71 | 18.79 | 0.92 | 0.92 | 0.04 | 0.04 |
| gene10600 | https://www.ncbi.nlm.nih.gov/gene/?term=LOC105284270 | 4837 | 119 | 246.02 | 1 | 2.07 | 0.84 | 10.98 | 10.08 | 0.9 | 0.9 | 0.02 | 0.02 |
| gene7398 | https://www.ncbi.nlm.nih.gov/gene/?term=LOC105280589 | 3452 | 33 | 95.6 | 1 | 2.9 | 3.03 | 21.34 | 20.45 | 0.9 | 0.9 | 0.01 | 0.01 |
| gene4034 | https://www.ncbi.nlm.nih.gov/gene/?term=LOC105276704 | 2065 | 87 | 421.31 | 1 | 4.84 | 1.15 | 14.97 | 14.08 | 0.89 | 0.89 | 0.04 | 0.04 |
| gene5831 | https://www.ncbi.nlm.nih.gov/gene/?term=LOC105278761 | 2506 | 110 | 438.95 | 1 | 3.99 | 0.91 | 23.47 | 22.6 | 0.88 | 0.88 | 0.04 | 0.04 |
| gene11401 | https://www.ncbi.nlm.nih.gov/gene/?term=LOC105285202 | 1841 | 79 | 429.11 | 1 | 5.43 | 1.27 | 24.33 | 23.48 | 0.85 | 0.85 | 0.04 | 0.04 |
| gene9336 | https://www.ncbi.nlm.nih.gov/gene/?term=LOC105282834 | 5993 | 124 | 206.91 | 1 | 1.67 | 0.81 | 16.5 | 15.65 | 0.85 | 0.85 | 0.02 | 0.02 |
| gene8274 | https://www.ncbi.nlm.nih.gov/gene/?term=LOC105281624 | 5425 | 211 | 388.94 | 1 | 1.84 | 0.47 | 9.95 | 9.18 | 0.77 | 0.77 | 0.03 | 0.03 |
| gene3470 | https://www.ncbi.nlm.nih.gov/gene/?term=LOC105276059 | 8716 | 157 | 180.13 | 1 | 1.15 | 0.64 | 8.72 | 7.97 | 0.76 | 0.76 | 0.01 | 0.01 |
| gene303 | https://www.ncbi.nlm.nih.gov/gene/?term=LOC105275584 | 3081 | 86 | 279.13 | 1 | 3.25 | 1.16 | 20.08 | 19.37 | 0.71 | 0.71 | 0.02 | 0.02 |
| gene12516 | https://www.ncbi.nlm.nih.gov/gene/?term=LOC105286452 | 1529 | 38 | 248.53 | 1 | 6.54 | 2.63 | 50.05 | 49.37 | 0.68 | 0.68 | 0.02 | 0.02 |
| gene11568 | https://www.ncbi.nlm.nih.gov/gene/?term=LOC105285395 | 8086 | 190 | 234.97 | 1 | 1.24 | 0.53 | 16.67 | 16 | 0.67 | 0.67 | 0.02 | 0.02 |
| gene4892 | https://www.ncbi.nlm.nih.gov/gene/?term=LOC105277667 | 3898 | 126 | 323.24 | 1 | 2.57 | 0.79 | 27.81 | 27.16 | 0.65 | 0.65 | 0.02 | 0.02 |
| gene10520 | https://www.ncbi.nlm.nih.gov/gene/?term=LOC105284184 | 2451 | 84 | 342.72 | 1 | 4.08 | 1.19 | 33.8 | 33.16 | 0.64 | 0.64 | 0.02 | 0.02 |
| gene8252 | https://www.ncbi.nlm.nih.gov/gene/?term=LOC105281599 | 3813 | 49 | 128.51 | 1 | 2.62 | 2.04 | 4.72 | 4.12 | 0.6 | 0.6 | 0.01 | 0.01 |
| gene473 | https://www.ncbi.nlm.nih.gov/gene/?term=LOC105277423 | 2342 | 106 | 452.6 | 1 | 4.27 | 0.94 | 28.94 | 28.4 | 0.54 | 0.54 | 0.02 | 0.02 |
| gene7884 | https://www.ncbi.nlm.nih.gov/gene/?term=LOC105281266 | 3390 | 76 | 224.19 | 1 | 2.95 | 1.32 | 16.63 | 16.11 | 0.52 | 0.52 | 0.01 | 0.01 |
| gene11579 | https://www.ncbi.nlm.nih.gov/gene/?term=LOC105285406 | 13571 | 226 | 166.53 | 1 | 0.74 | 0.44 | 7.69 | 7.18 | 0.51 | 0.51 | 0.01 | 0.01 |
| gene5877 | https://www.ncbi.nlm.nih.gov/gene/?term=LOC105278807 | 1719 | 49 | 285.05 | 1 | 5.82 | 2.04 | 27.11 | 26.65 | 0.46 | 0.46 | 0.01 | 0.01 |
| gene10540 | https://www.ncbi.nlm.nih.gov/gene/?term=LOC105284200 | 5267 | 220 | 417.7 | 1 | 1.9 | 0.45 | 12.54 | 12.08 | 0.45 | 0.45 | 0.02 | 0.02 |
| gene11082 | https://www.ncbi.nlm.nih.gov/gene/?term=LOC105284824 | 7436 | 200 | 268.96 | 1 | 1.34 | 0.5 | 7.89 | 7.46 | 0.43 | 0.43 | 0.01 | 0.01 |
| gene7552 | https://www.ncbi.nlm.nih.gov/gene/?term=LOC105280769 | 8566 | 287 | 335.05 | 1 | 1.17 | 0.35 | 2.06 | 1.63 | 0.43 | 0.43 | 0.01 | 0.01 |
| gene6583 | https://www.ncbi.nlm.nih.gov/gene/?term=LOC105279649 | 7444 | 320 | 429.88 | 1 | 1.34 | 0.31 | 11.91 | 11.5 | 0.41 | 0.41 | 0.02 | 0.02 |
| gene10531 | https://www.ncbi.nlm.nih.gov/gene/?term=LOC105284190 | 6881 | 186 | 270.31 | 1 | 1.45 | 0.54 | 12.46 | 12.07 | 0.39 | 0.39 | 0.01 | 0.01 |
| gene298 | https://www.ncbi.nlm.nih.gov/gene/?term=LOC105275460 | 2473 | 117 | 473.11 | 1 | 4.04 | 0.85 | 13.71 | 13.31 | 0.39 | 0.39 | 0.02 | 0.02 |
| gene7556 | https://www.ncbi.nlm.nih.gov/gene/?term=LOC105280774 | 2792 | 97 | 347.42 | 1 | 3.58 | 1.03 | 33.33 | 32.94 | 0.39 | 0.39 | 0.01 | 0.01 |
| gene12806 | https://www.ncbi.nlm.nih.gov/gene/?term=LOC105286803 | 4196 | 141 | 336.03 | 1 | 2.38 | 0.71 | 9.78 | 9.4 | 0.38 | 0.38 | 0.01 | 0.01 |
| gene10844 | https://www.ncbi.nlm.nih.gov/gene/?term=LOC105284535 | 9086 | 231 | 254.24 | 1 | 1.1 | 0.43 | 15.03 | 14.66 | 0.37 | 0.37 | 0.01 | 0.01 |
| gene13114 | https://www.ncbi.nlm.nih.gov/gene/?term=LOC105287147 | 7609 | 398 | 523.06 | 1 | 1.31 | 0.25 | 9.72 | 9.4 | 0.33 | 0.33 | 0.02 | 0.02 |
| gene292 | https://www.ncbi.nlm.nih.gov/gene/?term=LOC105275398 | 2825 | 81 | 286.73 | 1 | 3.54 | 1.23 | 24.98 | 24.69 | 0.3 | 0.3 | 0.01 | 0.01 |
| gene5524 | https://www.ncbi.nlm.nih.gov/gene/?term=LOC105278397 | 5177 | 198 | 382.46 | 1 | 1.93 | 0.51 | 24.98 | 24.69 | 0.29 | 0.29 | 0.01 | 0.01 |
| gene2173 | https://www.ncbi.nlm.nih.gov/gene/?term=LOC105274590 | 4387 | 70 | 159.56 | 2 | 4.56 | 2.86 | 26.31 | 26.05 | 0.27 | 0.27 | 0 | 0 |
| gene2271 | https://www.ncbi.nlm.nih.gov/gene/?term=LOC105274706 | 1448 | 51 | 352.21 | 1 | 6.91 | 1.96 | 40.12 | 39.86 | 0.26 | 0.26 | 0.01 | 0.01 |
| gene4216 | https://www.ncbi.nlm.nih.gov/gene/?term=LOC105276906 | 2127 | 90 | 423.13 | 1 | 4.7 | 1.11 | 21.35 | 21.12 | 0.23 | 0.23 | 0.01 | 0.01 |
| gene10018 | https://www.ncbi.nlm.nih.gov/gene/?term=LOC105283607 | 4414 | 207 | 468.96 | 1 | 2.27 | 0.48 | 15.29 | 15.09 | 0.2 | 0.2 | 0.01 | 0.01 |
| gene6639 | https://www.ncbi.nlm.nih.gov/gene/?term=LOC105279705 | 7656 | 67 | 87.51 | 1 | 1.31 | 1.49 | 22.85 | 22.67 | 0.17 | 0.17 | 0 | 0 |
| gene10596 | https://www.ncbi.nlm.nih.gov/gene/?term=LOC105284262 | 6520 | 208 | 319.02 | 1 | 1.53 | 0.48 | 4.3 | 4.14 | 0.16 | 0.16 | 0.01 | 0.01 |
| gene1632 | https://www.ncbi.nlm.nih.gov/gene/?term=LOC105287764 | 18061 | 391 | 216.49 | 2 | 1.11 | 0.51 | 3.3 | 3.14 | 0.16 | 0.16 | 0 | 0 |

| | | | | | | | | | | | | | |
|---|---|---|---|---|---|---|---|---|---|---|---|---|---|
| gene3206 | https://www.ncbi.nlm.nih.gov/gene/?term=LOC105275752 | 3474 | 118 | 339.67 | 1 | 2.88 | 0.85 | 32.15 | 31.99 | 0.16 | 0.16 | 0.01 | 0.01 |
| gene570 | https://www.ncbi.nlm.nih.gov/gene/?term=LOC105278549 | 1673 | 63 | 376.57 | 1 | 5.98 | 1.59 | 43.38 | 43.23 | 0.15 | 0.15 | 0.01 | 0.01 |
| gene7888 | https://www.ncbi.nlm.nih.gov/gene/?term=LOC105281141 | 4357 | 153 | 351.16 | 1 | 2.3 | 0.65 | 32.3 | 32.17 | 0.13 | 0.13 | 0 | 0 |
| gene5525 | https://www.ncbi.nlm.nih.gov/gene/?term=LOC105278396 | 3090 | 182 | 589 | 1 | 3.24 | 0.55 | 24.88 | 24.76 | 0.12 | 0.12 | 0.01 | 0.01 |
| gene5746 | https://www.ncbi.nlm.nih.gov/gene/?term=LOC105278666 | 10343 | 441 | 426.38 | 1 | 0.97 | 0.23 | 6.88 | 6.76 | 0.12 | 0.12 | 0.01 | 0.01 |
| gene10392 | https://www.ncbi.nlm.nih.gov/gene/?term=LOC105284036 | 3216 | 99 | 307.84 | 2 | 6.22 | 2.02 | 22.54 | 22.43 | 0.11 | 0.11 | 0 | 0 |
| gene7368 | https://www.ncbi.nlm.nih.gov/gene/?term=LOC105280556 | 4105 | 41 | 99.88 | 1 | 2.44 | 2.44 | 16.06 | 15.95 | 0.11 | 0.11 | 0 | 0 |
| gene2806 | https://www.ncbi.nlm.nih.gov/gene/?term=LOC105275327 | 14257 | 473 | 331.77 | 1 | 0.7 | 0.21 | 7.14 | 7.04 | 0.1 | 0.1 | 0 | 0 |
| gene1470 | https://www.ncbi.nlm.nih.gov/gene/?term=LOC105287597 | 2974 | 186 | 625.42 | 1 | 3.36 | 0.54 | 18.48 | 18.39 | 0.09 | 0.09 | 0.01 | 0.01 |
| gene3885 | https://www.ncbi.nlm.nih.gov/gene/?term=LOC105276527 | 7928 | 318 | 401.11 | 1 | 1.26 | 0.31 | 9.68 | 9.6 | 0.09 | 0.09 | 0 | 0 |
| gene9101 | https://www.ncbi.nlm.nih.gov/gene/?term=LOC105282553 | 3068 | 104 | 338.98 | 1 | 3.26 | 0.96 | 23.44 | 23.39 | 0.06 | 0.06 | 0 | 0 |
| gene1408 | https://www.ncbi.nlm.nih.gov/gene/?term=LOC109611520 | 8645 | 306 | 353.96 | 1 | 1.16 | 0.33 | 6.02 | 5.98 | 0.05 | 0.05 | 0 | 0 |
| gene2633 | https://www.ncbi.nlm.nih.gov/gene/?term=LOC105275127 | 7457 | 252 | 337.94 | 1 | 1.34 | 0.4 | 8.36 | 8.33 | 0.03 | 0.03 | 0 | 0 |
| gene10326 | https://www.ncbi.nlm.nih.gov/gene/?term=LOC105283962 | 3042 | 124 | 407.63 | 1 | 3.29 | 0.81 | 8.49 | 8.5 | -0.01 | 0.01 | 0 | 0 |
| gene11275 | https://www.ncbi.nlm.nih.gov/gene/?term=LOC105285055 | 4119 | 252 | 611.8 | 1 | 2.43 | 0.4 | 11.84 | 11.86 | -0.03 | 0.03 | 0 | 0 |
| gene6620 | https://www.ncbi.nlm.nih.gov/gene/?term=LOC105279687 | 17998 | 355 | 197.24 | 1 | 0.56 | 0.28 | 5.15 | 5.19 | -0.04 | 0.04 | 0 | 0 |
| gene7915 | https://www.ncbi.nlm.nih.gov/gene/?term=LOC105281173 | 1597 | 39 | 244.21 | 1 | 6.26 | 2.56 | 59.75 | 59.8 | -0.05 | 0.05 | 0 | 0 |
| gene12572 | https://www.ncbi.nlm.nih.gov/gene/?term=LOC105286514 | 2374 | 49 | 206.4 | 1 | 4.21 | 2.04 | 35.44 | 35.51 | -0.07 | 0.07 | 0 | 0 |
| gene2683 | https://www.ncbi.nlm.nih.gov/gene/?term=LOC105275187 | 5492 | 70 | 127.46 | 1 | 1.82 | 1.43 | 12.3 | 12.38 | -0.07 | 0.07 | 0 | 0 |
| gene5387 | https://www.ncbi.nlm.nih.gov/gene/?term=LOC105278245 | 6451 | 112 | 173.62 | 1 | 1.55 | 0.89 | 22.85 | 22.92 | -0.07 | 0.07 | 0 | 0 |
| gene8814 | https://www.ncbi.nlm.nih.gov/gene/?term=LOC105282240 | 6910 | 176 | 254.7 | 1 | 1.45 | 0.57 | 9.18 | 9.27 | -0.08 | 0.08 | 0 | 0 |
| gene8600 | https://www.ncbi.nlm.nih.gov/gene/?term=LOC105282001 | 8513 | 151 | 177.38 | 1 | 1.17 | 0.66 | 10.34 | 10.43 | -0.09 | 0.09 | 0 | 0 |
| gene8729 | https://www.ncbi.nlm.nih.gov/gene/?term=LOC105282137 | 5005 | 211 | 421.58 | 1 | 2 | 0.47 | 15.81 | 15.91 | -0.09 | 0.09 | 0 | 0 |
| gene2404 | https://www.ncbi.nlm.nih.gov/gene/?term=LOC105274853 | 1767 | 35 | 198.08 | 1 | 5.66 | 2.86 | 33.43 | 33.53 | -0.1 | 0.1 | 0 | 0 |
| gene5101 | https://www.ncbi.nlm.nih.gov/gene/?term=LOC105277903 | 1813 | 35 | 193.05 | 1 | 5.52 | 2.86 | 51.16 | 51.27 | -0.1 | 0.1 | 0 | 0 |
| gene53 | https://www.ncbi.nlm.nih.gov/gene/?term=LOC105277148 | 19283 | 295 | 152.98 | 1 | 0.52 | 0.34 | 3.8 | 3.9 | -0.1 | 0.1 | 0 | 0 |
| gene5646 | https://www.ncbi.nlm.nih.gov/gene/?term=LOC105278534 | 7475 | 187 | 250.17 | 1 | 1.34 | 0.53 | 18.3 | 18.4 | -0.1 | 0.1 | 0 | 0 |
| gene2846 | https://www.ncbi.nlm.nih.gov/gene/?term=LOC105275374 | 11246 | 477 | 424.15 | 1 | 0.89 | 0.21 | 10.56 | 10.68 | -0.11 | 0.11 | 0 | 0 |
| gene34 | https://www.ncbi.nlm.nih.gov/gene/?term=LOC105278416 | 7120 | 140 | 196.63 | 1 | 1.4 | 0.71 | 13.56 | 13.67 | -0.11 | 0.11 | 0 | 0 |
| gene7166 | https://www.ncbi.nlm.nih.gov/gene/?term=LOC105280326 | 6117 | 243 | 397.25 | 1 | 1.63 | 0.41 | 14.54 | 14.65 | -0.11 | 0.11 | 0 | 0 |
| gene4190 | https://www.ncbi.nlm.nih.gov/gene/?term=LOC105276875 | 3150 | 117 | 371.43 | 1 | 3.17 | 0.85 | 21.8 | 21.95 | -0.15 | 0.15 | -0.01 | 0.01 |
| gene12802 | https://www.ncbi.nlm.nih.gov/gene/?term=LOC105286799 | 3183 | 164 | 515.24 | 1 | 3.14 | 0.61 | 17.26 | 17.43 | -0.17 | 0.17 | -0.01 | 0.01 |
| gene1430 | https://www.ncbi.nlm.nih.gov/gene/?term=LOC105287551 | 6433 | 223 | 346.65 | 1 | 1.55 | 0.45 | 8.23 | 8.41 | -0.18 | 0.18 | -0.01 | 0.01 |
| gene9326 | https://www.ncbi.nlm.nih.gov/gene/?term=LOC105282818 | 16612 | 178 | 107.15 | 1 | 0.6 | 0.56 | 13.41 | 13.58 | -0.18 | 0.18 | 0 | 0 |
| gene6286 | https://www.ncbi.nlm.nih.gov/gene/?term=LOC105279298 | 2861 | 129 | 450.89 | 2 | 6.99 | 1.55 | 23.63 | 23.86 | -0.23 | 0.23 | -0.01 | 0.01 |
| gene4962 | https://www.ncbi.nlm.nih.gov/gene/?term=LOC105277761 | 4645 | 144 | 310.01 | 1 | 2.15 | 0.69 | 13.44 | 13.75 | -0.31 | 0.31 | -0.01 | 0.01 |
| gene9327 | https://www.ncbi.nlm.nih.gov/gene/?term=LOC105282821 | 7793 | 145 | 186.06 | 1 | 1.28 | 0.69 | 12.57 | 12.89 | -0.32 | 0.32 | -0.01 | 0.01 |
| gene2957 | https://www.ncbi.nlm.nih.gov/gene/?term=LOC105275500 | 10926 | 240 | 219.66 | 1 | 0.92 | 0.42 | 6.99 | 7.33 | -0.34 | 0.34 | -0.01 | 0.01 |
| gene12307 | https://www.ncbi.nlm.nih.gov/gene/?term=LOC105286216 | 10313 | 187 | 181.32 | 1 | 0.97 | 0.53 | 14.69 | 15.04 | -0.35 | 0.35 | -0.01 | 0.01 |
| gene10155 | https://www.ncbi.nlm.nih.gov/gene/?term=LOC105283756 | 7030 | 112 | 159.32 | 1 | 1.42 | 0.89 | 5.92 | 6.29 | -0.37 | 0.37 | -0.01 | 0.01 |
| gene4023 | https://www.ncbi.nlm.nih.gov/gene/?term=LOC105276692 | 1759 | 31 | 176.24 | 1 | 5.69 | 3.23 | 23.25 | 23.63 | -0.38 | 0.38 | -0.01 | 0.01 |
| gene1310 | https://www.ncbi.nlm.nih.gov/gene/?term=LOC105286661 | 2582 | 71 | 274.98 | 1 | 3.87 | 1.41 | 21.29 | 21.68 | -0.39 | 0.39 | -0.01 | 0.01 |
| gene12573 | https://www.ncbi.nlm.nih.gov/gene/?term=LOC105286515 | 3647 | 100 | 274.2 | 1 | 2.74 | 1 | 13.03 | 13.44 | -0.42 | 0.42 | -0.01 | 0.01 |
| gene13036 | https://www.ncbi.nlm.nih.gov/gene/?term=LOC105287059 | 4959 | 160 | 322.65 | 1 | 2.02 | 0.62 | 19.69 | 20.15 | -0.46 | 0.46 | -0.01 | 0.01 |
| gene920 | https://www.ncbi.nlm.nih.gov/gene/?term=LOC105282518 | 1197 | 52 | 434.42 | 1 | 8.35 | 1.92 | 30.27 | 30.77 | -0.49 | 0.49 | -0.02 | 0.02 |
| gene10549 | https://www.ncbi.nlm.nih.gov/gene/?term=LOC105284210 | 7145 | 290 | 405.88 | 2 | 2.8 | 0.69 | 11.49 | 12 | -0.51 | 0.51 | -0.02 | 0.02 |
| gene1199 | https://www.ncbi.nlm.nih.gov/gene/?term=LOC105285498 | 2220 | 28 | 126.13 | 1 | 4.5 | 3.57 | 27.44 | 27.99 | -0.55 | 0.55 | -0.01 | 0.01 |
| gene657 | https://www.ncbi.nlm.nih.gov/gene/?term=LOC105279667 | 6421 | 81 | 126.15 | 1 | 1.56 | 1.23 | 2.49 | 3.12 | -0.63 | 0.63 | -0.01 | 0.01 |
| gene7183 | https://www.ncbi.nlm.nih.gov/gene/?term=LOC105280348 | 3925 | 70 | 178.34 | 1 | 2.55 | 1.43 | 20.85 | 21.48 | -0.63 | 0.63 | -0.01 | 0.01 |
| gene7249 | https://www.ncbi.nlm.nih.gov/gene/?term=LOC105280422 | 1775 | 59 | 332.39 | 1 | 5.63 | 1.69 | 16.45 | 17.08 | -0.63 | 0.63 | -0.02 | 0.02 |
| gene12021 | https://www.ncbi.nlm.nih.gov/gene/?term=LOC105285905 | 1399 | 15 | 107.22 | 1 | 7.15 | 6.67 | 50 | 50.68 | -0.68 | 0.68 | -0.01 | 0.01 |
| gene7247 | https://www.ncbi.nlm.nih.gov/gene/?term=LOC105280407 | 10486 | 206 | 196.45 | 1 | 0.95 | 0.49 | 16.14 | 16.82 | -0.68 | 0.68 | -0.01 | 0.01 |
| gene11561 | https://www.ncbi.nlm.nih.gov/gene/?term=LOC105285387 | 11528 | 178 | 154.41 | 1 | 0.87 | 0.56 | 9.89 | 10.61 | -0.72 | 0.72 | -0.01 | 0.01 |
| gene9740 | https://www.ncbi.nlm.nih.gov/gene/?term=LOC105283277 | 2887 | 137 | 474.54 | 1 | 3.46 | 0.73 | 19.95 | 20.71 | -0.76 | 0.76 | -0.04 | 0.04 |
| gene12265 | https://www.ncbi.nlm.nih.gov/gene/?term=LOC105286166 | 3050 | 110 | 360.66 | 1 | 3.28 | 0.91 | 22.97 | 23.8 | -0.82 | 0.82 | -0.03 | 0.03 |
| gene10606 | https://www.ncbi.nlm.nih.gov/gene/?term=LOC105284273 | 6119 | 248 | 405.29 | 1 | 1.63 | 0.4 | 17.15 | 17.98 | -0.83 | 0.83 | -0.03 | 0.03 |
| gene35 | https://www.ncbi.nlm.nih.gov/gene/?term=LOC105275197 | 2643 | 50 | 189.18 | 1 | 3.78 | 2 | 37.01 | 37.84 | -0.83 | 0.83 | -0.02 | 0.02 |
| gene5298 | https://www.ncbi.nlm.nih.gov/gene/?term=LOC105278145 | 4060 | 235 | 578.82 | 2 | 4.93 | 0.85 | 10.4 | 11.25 | -0.85 | 0.85 | -0.05 | 0.05 |
| gene4376 | https://www.ncbi.nlm.nih.gov/gene/?term=LOC105277083 | 4307 | 102 | 236.82 | 1 | 2.32 | 0.98 | 24.25 | 25.14 | -0.89 | 0.89 | -0.02 | 0.02 |
| gene1220 | https://www.ncbi.nlm.nih.gov/gene/?term=LOC105285742 | 3453 | 111 | 321.46 | 1 | 2.9 | 0.9 | 21.24 | 22.19 | -0.96 | 0.96 | -0.03 | 0.03 |
| gene12203 | https://www.ncbi.nlm.nih.gov/gene/?term=LOC105286099 | 5967 | 75 | 125.69 | 1 | 1.68 | 1.33 | 9.56 | 10.52 | -0.96 | 0.96 | -0.01 | 0.01 |
| gene6031 | https://www.ncbi.nlm.nih.gov/gene/?term=LOC105279000 | 5691 | 179 | 314.53 | 1 | 1.76 | 0.56 | 13.44 | 14.41 | -0.97 | 0.97 | -0.03 | 0.03 |
| gene7211 | https://www.ncbi.nlm.nih.gov/gene/?term=LOC105280377 | 2024 | 67 | 331.03 | 1 | 4.94 | 1.49 | 44.45 | 45.48 | -1.03 | 1.03 | -0.03 | 0.03 |
| gene4033 | https://www.ncbi.nlm.nih.gov/gene/?term=LOC105276705 | 10546 | 142 | 134.65 | 1 | 0.95 | 0.7 | 7.83 | 8.89 | -1.06 | 1.06 | -0.01 | 0.01 |

| | | | | | | | | | | | | | |
|---|---|---|---|---|---|---|---|---|---|---|---|---|---|
| gene3385 | https://www.ncbi.nlm.nih.gov/gene/?term=LOC105275966 | 3305 | 58 | 175.49 | 1 | 3.03 | 1.72 | 49.09 | 50.19 | -1.1 | 1.1 | -0.02 | 0.02 |
| gene12981 | https://www.ncbi.nlm.nih.gov/gene/?term=LOC105286999 | 2063 | 46 | 222.98 | 1 | 4.85 | 2.17 | 31.8 | 32.93 | -1.13 | 1.13 | -0.03 | 0.03 |
| gene12406 | https://www.ncbi.nlm.nih.gov/gene/?term=LOC105286316 | 1386 | 33 | 238.1 1 | 7.22 | 3.03 | 40.83 | 41.98 | -1.15 | 1.15 | -0.03 | 0.03 | |
| gene10213 | https://www.ncbi.nlm.nih.gov/gene/?term=LOC105283826 | 1503 | 54 | 359.28 | 1 | 6.65 | 1.85 | 14.52 | 15.73 | -1.21 | 1.21 | -0.04 | 0.04 |
| gene3793 | https://www.ncbi.nlm.nih.gov/gene/?term=LOC105276429 | 2901 | 112 | 386.07 | 1 | 3.45 | 0.89 | 25.16 | 26.39 | -1.23 | 1.23 | -0.05 | 0.05 |
| gene11340 | https://www.ncbi.nlm.nih.gov/gene/?term=LOC105285131 | 2918 | 88 | 301.58 | 1 | 3.43 | 1.14 | 10.13 | 11.45 | -1.32 | 1.32 | -0.04 | 0.04 |
| gene322 | https://www.ncbi.nlm.nih.gov/gene/?term=LOC105275777 | 1606 | 28 | 174.35 | 1 | 6.23 | 3.57 | 42.75 | 44.07 | -1.32 | 1.32 | -0.02 | 0.02 |
| gene7248 | https://www.ncbi.nlm.nih.gov/gene/?term=LOC105280408 | 5175 | 52 | 100.48 | 1 | 1.93 | 1.92 | 16.62 | 18.28 | -1.65 | 1.65 | -0.02 | 0.02 |
| gene8343 | https://www.ncbi.nlm.nih.gov/gene/?term=LOC105281700 | 2172 | 32 | 147.33 | 1 | 4.6 | 3.12 | 37.31 | 38.97 | -1.66 | 1.66 | -0.02 | 0.02 |
| gene1147 | https://www.ncbi.nlm.nih.gov/gene/?term=LOC105284954 | 4022 | 90 | 223.77 | 1 | 2.49 | 1.11 | 16.68 | 18.39 | -1.71 | 1.71 | -0.04 | 0.04 |
| gene2413 | https://www.ncbi.nlm.nih.gov/gene/?term=LOC105274863 | 2006 | 49 | 244.27 | 1 | 4.99 | 2.04 | 53.75 | 55.47 | -1.72 | 1.72 | -0.04 | 0.04 |
| gene13235 | https://www.ncbi.nlm.nih.gov/gene/?term=LOC105287278 | 2679 | 34 | 126.91 | 1 | 3.73 | 2.94 | 22.53 | 24.28 | -1.75 | 1.75 | -0.02 | 0.02 |
| gene3727 | https://www.ncbi.nlm.nih.gov/gene/?term=LOC105276367 | 2484 | 63 | 253.62 | 1 | 4.03 | 1.59 | 45.38 | 47.24 | -1.87 | 1.87 | -0.05 | 0.05 |
| gene4641 | https://www.ncbi.nlm.nih.gov/gene/?term=LOC105277398 | 1937 | 71 | 366.55 | 1 | 5.16 | 1.41 | 37.78 | 39.69 | -1.91 | 1.91 | -0.07 | 0.07 |
| gene10107 | https://www.ncbi.nlm.nih.gov/gene/?term=LOC105283712 | 1883 | 35 | 185.87 | 1 | 5.31 | 2.86 | 26.3 | 28.27 | -1.97 | 1.97 | -0.04 | 0.04 |
| gene4498 | https://www.ncbi.nlm.nih.gov/gene/?term=LOC105277239 | 2831 | 121 | 427.41 | 1 | 3.53 | 0.83 | 11.26 | 13.23 | -1.97 | 1.97 | -0.08 | 0.08 |
| gene10345 | https://www.ncbi.nlm.nih.gov/gene/?term=LOC105283989 | 4444 | 27 | 60.76 1 | 2.25 | 3.7 | 36.46 | 38.69 | -2.24 | 2.24 | -0.01 | 0.01 | |
| gene388 | https://www.ncbi.nlm.nih.gov/gene/?term=LOC105276538 | 1558 | 18 | 115.53 | 2 | 12.84 | 11.11 | 66.09 | 68.41 | -2.32 | 2.32 | -0.03 | 0.03 |
| gene3168 | https://www.ncbi.nlm.nih.gov/gene/?term=LOC105275711 | 6150 | 69 | 112.2 1 | 1.63 | 1.45 | 12.54 | 15.09 | -2.54 | 2.54 | -0.03 | 0.03 | |
| gene3599 | https://www.ncbi.nlm.nih.gov/gene/?term=LOC105276196 | 2955 | 59 | 199.66 | 1 | 3.38 | 1.69 | 37.76 | 40.32 | -2.56 | 2.56 | -0.05 | 0.05 |
| gene1505 | https://www.ncbi.nlm.nih.gov/gene/?term=LOC105287633 | 1693 | 27 | 159.48 | 1 | 5.91 | 3.7 | 42.55 | 45.35 | -2.8 | 2.8 | -0.04 | 0.04 |
| gene6237 | https://www.ncbi.nlm.nih.gov/gene/?term=LOC105279244 | 1484 | 25 | 168.46 | 1 | 6.74 | 4 | 41.13 | 44.31 | -3.18 | 3.18 | -0.05 | 0.05 |
| gene564 | https://www.ncbi.nlm.nih.gov/gene/?term=LOC105278478 | 2802 | 17 | 60.67 1 | 3.57 | 5.88 | 14.04 | 17.34 | -3.31 | 3.31 | -0.02 | 0.02 | |
| gene6422 | https://www.ncbi.nlm.nih.gov/gene/?term=LOC105279458 | 2419 | 38 | 157.09 | 1 | 4.13 | 2.63 | 46.62 | 50.37 | -3.75 | 3.75 | -0.06 | 0.06 |
| gene10602 | https://www.ncbi.nlm.nih.gov/gene/?term=LOC105284268 | 2122 | 19 | 89.54 1 | 4.71 | 5.26 | 26.09 | 32.99 | -6.9 | 6.9 | -0.06 | 0.06 | |
| gene1008 | https://www.ncbi.nlm.nih.gov/gene/?term=LOC105283492 | 1479 | 10 | 67.61 1 | 6.76 | 10 | 34.7 | 42 | -7.3 | 7.3 | -0.05 | 0.05 | |
| gene393 | https://www.ncbi.nlm.nih.gov/gene/?term=LOC105276566 | 1123 | 2 | 17.81 1 | 8.9 | 50 | 47.22 | 76.78 | -29.56 | | 29.56 | -0.05 | 0.05 |

Directory: OGL File: ogl-Cb_LI_BCphaseB2.txt

| gene_ID | gene_link | gwidth | #Sites | #per10Kb | %perSite | %pNucl |
|---|---|---|---|---|---|---|
| gene12538 | https://www.ncbi.nlm.nih.gov/gene/?term=LOC105286531 | 372 | 43 | 1155.91 | 35.54 | 4.11 |
| gene13384 | https://www.ncbi.nlm.nih.gov/gene/?term=LOC105287447 | 1877 | 245 | 1305.27 | 28.47 | 3.72 |
| gene5917 | https://www.ncbi.nlm.nih.gov/gene/?term=LOC105278849 | 823 | 126 | 1530.98 | 23.79 | 3.64 |
| gene5911 | https://www.ncbi.nlm.nih.gov/gene/?term=LOC105278844 | 943 | 98 | 1039.24 | 34.92 | 3.63 |
| gene211 | https://www.ncbi.nlm.nih.gov/gene/?term=LOC105274534 | 1280 | 101 | 789.06 | 45.75 | 3.61 |
| gene11569 | https://www.ncbi.nlm.nih.gov/gene/?term=LOC105285394 | 1311 | 61 | 465.29 | 72.04 | 3.35 |
| gene12364 | https://www.ncbi.nlm.nih.gov/gene/?term=LOC105286269 | 1403 | 90 | 641.48 | 50.55 | 3.24 |
| gene5309 | https://www.ncbi.nlm.nih.gov/gene/?term=LOC105278160 | 1704 | 155 | 909.62 | 35.44 | 3.22 |
| gene6078 | https://www.ncbi.nlm.nih.gov/gene/?term=LOC105279052 | 1130 | 138 | 1221.24 | 26.05 | 3.18 |
| gene3719 | https://www.ncbi.nlm.nih.gov/gene/?term=LOC105276360 | 1471 | 160 | 1087.7 | 29.08 | 3.16 |
| gene12330 | https://www.ncbi.nlm.nih.gov/gene/?term=LOC105286235 | 1501 | 112 | 746.17 | 41.99 | 3.13 |
| gene1562 | https://www.ncbi.nlm.nih.gov/gene/?term=LOC105287692 | 1491 | 129 | 865.19 | 35.56 | 3.08 |
| gene12746 | https://www.ncbi.nlm.nih.gov/gene/?term=LOC105286737 | 2688 | 363 | 1350.45 | 22.13 | 2.99 |
| gene190 | https://www.ncbi.nlm.nih.gov/gene/?term=LOC105288088 | 1797 | 127 | 706.73 | 41.77 | 2.95 |
| gene3731 | https://www.ncbi.nlm.nih.gov/gene/?term=LOC105276371 | 865 | 57 | 658.96 | 43.94 | 2.9 |
| gene1654 | https://www.ncbi.nlm.nih.gov/gene/?term=LOC105287811 | 2285 | 150 | 656.46 | 43.71 | 2.87 |
| gene2797 | https://www.ncbi.nlm.nih.gov/gene/?term=LOC105275314 | 1489 | 93 | 624.58 | 45.82 | 2.86 |
| gene9471 | https://www.ncbi.nlm.nih.gov/gene/?term=LOC105282986 | 1605 | 140 | 872.27 | 32.66 | 2.85 |
| gene570 | https://www.ncbi.nlm.nih.gov/gene/?term=LOC105278549 | 1673 | 107 | 639.57 | 44.01 | 2.81 |
| gene5301 | https://www.ncbi.nlm.nih.gov/gene/?term=LOC105278148 | 1881 | 192 | 1020.73 | 27.32 | 2.79 |
| gene6978 | https://www.ncbi.nlm.nih.gov/gene/?term=LOC105280117 | 1816 | 96 | 528.63 | 52.73 | 2.79 |
| gene12403 | https://www.ncbi.nlm.nih.gov/gene/?term=LOC105286311 | 1413 | 64 | 452.94 | 61.18 | 2.77 |
| gene4878 | https://www.ncbi.nlm.nih.gov/gene/?term=LOC105277648 | 675 | 36 | 533.33 | 51.21 | 2.73 |
| gene4635 | https://www.ncbi.nlm.nih.gov/gene/?term=LOC105277390 | 2067 | 125 | 604.74 | 44.98 | 2.72 |

...
... only first 25 lines shown ...

Directory: OGL File: ogl-Cb_LI_BCphaseB2.vs.RphaseA1.txt

| gene_ID | gene_link | gwidth | #Sites | #per10Kb | #dmSites | #dmsp10kb | %dmSites | %pSite1 | %pSite2 | DMpSite | ADMpSite | DMpNucl | ADMpNucl |
|---|---|---|---|---|---|---|---|---|---|---|---|---|---|
| gene393 | https://www.ncbi.nlm.nih.gov/gene/?term=LOC105276566 | 1123 | 2 | 17.81 | 1 | 8.9 | 50 | 70.57 | 50 | 20.57 | 20.57 | 0.04 | 0.04 |
| gene12021 | https://www.ncbi.nlm.nih.gov/gene/?term=LOC105285905 | 1399 | 21 | 150.11 | 1 | 7.15 | 4.76 | 36.22 | 31.01 | 5.22 | 5.22 | 0.08 | 0.08 |
| gene9411 | https://www.ncbi.nlm.nih.gov/gene/?term=LOC105282914 | 5534 | 41 | 74.09 | 1 | 1.81 | 2.44 | 52.21 | 48.58 | 3.63 | 3.63 | 0.03 | 0.03 |
| gene1199 | https://www.ncbi.nlm.nih.gov/gene/?term=LOC105285498 | 2220 | 25 | 112.61 | 1 | 4.5 | 4 | 37.37 | 33.76 | 3.61 | 3.61 | 0.04 | 0.04 |
| gene388 | https://www.ncbi.nlm.nih.gov/gene/?term=LOC105276538 | 1558 | 20 | 128.37 | 2 | 12.84 | 10 | 66.14 | 62.93 | 3.2 | 3.2 | 0.04 | 0.04 |
| gene10714 | https://www.ncbi.nlm.nih.gov/gene/?term=LOC105284396 | 1966 | 62 | 315.36 | 1 | 5.09 | 1.61 | 36.35 | 33.21 | 3.14 | 3.14 | 0.1 | 0.1 |
| gene322 | https://www.ncbi.nlm.nih.gov/gene/?term=LOC105275777 | 1606 | 25 | 155.67 | 1 | 6.23 | 4 | 44.37 | 41.49 | 2.89 | 2.89 | 0.04 | 0.04 |
| gene12406 | https://www.ncbi.nlm.nih.gov/gene/?term=LOC105286316 | 1386 | 31 | 223.67 | 1 | 7.22 | 3.23 | 46.61 | 43.84 | 2.77 | 2.77 | 0.06 | 0.06 |
| gene6652 | https://www.ncbi.nlm.nih.gov/gene/?term=LOC105279719 | 2831 | 86 | 303.78 | 1 | 3.53 | 1.16 | 23.08 | 20.39 | 2.69 | 2.69 | 0.08 | 0.08 |
| gene3168 | https://www.ncbi.nlm.nih.gov/gene/?term=LOC105275711 | 6150 | 73 | 118.7 | 1 | 1.63 | 1.37 | 14.69 | 12.05 | 2.64 | 2.64 | 0.03 | 0.03 |
| gene6422 | https://www.ncbi.nlm.nih.gov/gene/?term=LOC105279458 | 2419 | 37 | 152.96 | 1 | 4.13 | 2.7 | 52.5 | 49.97 | 2.53 | 2.53 | 0.04 | 0.04 |
| gene6455 | https://www.ncbi.nlm.nih.gov/gene/?term=LOC105279500 | 2612 | 30 | 114.85 | 1 | 3.83 | 3.33 | 48.51 | 46 | 2.5 | 2.5 | 0.03 | 0.03 |
| gene564 | https://www.ncbi.nlm.nih.gov/gene/?term=LOC105278478 | 2802 | 18 | 64.24 | 1 | 3.57 | 5.56 | 17.32 | 14.97 | 2.35 | 2.35 | 0.02 | 0.02 |
| gene6806 | https://www.ncbi.nlm.nih.gov/gene/?term=LOC105279894 | 3649 | 35 | 95.92 | 1 | 2.74 | 2.86 | 40.58 | 38.26 | 2.32 | 2.32 | 0.02 | 0.02 |
| gene9101 | https://www.ncbi.nlm.nih.gov/gene/?term=LOC105282553 | 3068 | 106 | 345.5 | 1 | 3.26 | 0.94 | 23.35 | 21.05 | 2.29 | 2.29 | 0.08 | 0.08 |
| gene7199 | https://www.ncbi.nlm.nih.gov/gene/?term=LOC105280366 | 1414 | 33 | 233.38 | 1 | 7.07 | 3.03 | 43.27 | 41 | 2.28 | 2.28 | 0.05 | 0.05 |
| gene2404 | https://www.ncbi.nlm.nih.gov/gene/?term=LOC105274853 | 1767 | 43 | 243.35 | 1 | 5.66 | 2.33 | 31.85 | 29.62 | 2.23 | 2.23 | 0.05 | 0.05 |
| gene6237 | https://www.ncbi.nlm.nih.gov/gene/?term=LOC105279244 | 1484 | 23 | 154.99 | 1 | 6.74 | 4.35 | 50.08 | 47.92 | 2.16 | 2.16 | 0.03 | 0.03 |
| gene2271 | https://www.ncbi.nlm.nih.gov/gene/?term=LOC105274706 | 1448 | 51 | 352.21 | 1 | 6.91 | 1.96 | 39.73 | 37.8 | 1.94 | 1.94 | 0.07 | 0.07 |
| gene2748 | https://www.ncbi.nlm.nih.gov/gene/?term=LOC105275261 | 3845 | 54 | 140.44 | 1 | 2.6 | 1.85 | 39.59 | 37.65 | 1.94 | 1.94 | 0.03 | 0.03 |
| gene10107 | https://www.ncbi.nlm.nih.gov/gene/?term=LOC105283712 | 1883 | 35 | 185.87 | 1 | 5.31 | 2.86 | 27.16 | 25.31 | 1.85 | 1.85 | 0.03 | 0.03 |
| gene6639 | https://www.ncbi.nlm.nih.gov/gene/?term=LOC105279705 | 7656 | 68 | 88.82 | 1 | 1.31 | 1.47 | 23.58 | 21.76 | 1.82 | 1.82 | 0.02 | 0.02 |
| gene4498 | https://www.ncbi.nlm.nih.gov/gene/?term=LOC105277239 | 2831 | 130 | 459.2 | 1 | 3.53 | 0.77 | 12.47 | 10.69 | 1.78 | 1.78 | 0.08 | 0.08 |
| gene1907 | https://www.ncbi.nlm.nih.gov/gene/?term=LOC105288099 | 1238 | 35 | 282.71 | 1 | 8.08 | 2.86 | 34.89 | 33.16 | 1.73 | 1.73 | 0.05 | 0.05 |
| gene920 | https://www.ncbi.nlm.nih.gov/gene/?term=LOC105282518 | 1197 | 41 | 342.52 | 1 | 8.35 | 2.44 | 39.23 | 37.74 | 1.49 | 1.49 | 0.05 | 0.05 |
| gene7550 | https://www.ncbi.nlm.nih.gov/gene/?term=LOC105280763 | 2173 | 98 | 450.99 | 1 | 4.6 | 1.02 | 27.13 | 25.7 | 1.43 | 1.43 | 0.06 | 0.06 |
| gene10602 | https://www.ncbi.nlm.nih.gov/gene/?term=LOC105284268 | 2122 | 21 | 98.96 | 1 | 4.71 | 4.76 | 25.67 | 24.45 | 1.22 | 1.22 | 0.01 | 0.01 |
| gene3671 | https://www.ncbi.nlm.nih.gov/gene/?term=LOC105276272 | 1966 | 46 | 233.98 | 1 | 5.09 | 2.17 | 27.89 | 26.72 | 1.16 | 1.16 | 0.03 | 0.03 |
| gene5646 | https://www.ncbi.nlm.nih.gov/gene/?term=LOC105278534 | 7475 | 185 | 247.49 | 1 | 1.34 | 0.54 | 18.24 | 17.21 | 1.03 | 1.03 | 0.03 | 0.03 |
| gene3046 | https://www.ncbi.nlm.nih.gov/gene/?term=LOC105275587 | 2394 | 58 | 242.27 | 1 | 4.18 | 1.72 | 36.72 | 35.74 | 0.98 | 0.98 | 0.02 | 0.02 |
| gene7915 | https://www.ncbi.nlm.nih.gov/gene/?term=LOC105281173 | 1597 | 38 | 237.95 | 1 | 6.26 | 2.63 | 60.79 | 59.81 | 0.98 | 0.98 | 0.02 | 0.02 |
| gene8015 | https://www.ncbi.nlm.nih.gov/gene/?term=LOC105281312 | 2998 | 65 | 216.81 | 1 | 3.34 | 1.54 | 33.24 | 32.3 | 0.93 | 0.93 | 0.02 | 0.02 |
| gene7398 | https://www.ncbi.nlm.nih.gov/gene/?term=LOC105280589 | 3452 | 36 | 104.29 | 1 | 2.9 | 2.78 | 20.14 | 19.24 | 0.9 | 0.9 | 0.01 | 0.01 |
| gene10392 | https://www.ncbi.nlm.nih.gov/gene/?term=LOC105284036 | 3216 | 106 | 329.6 | 2 | 6.22 | 1.89 | 21.2 | 20.33 | 0.87 | 0.87 | 0.03 | 0.03 |
| gene1310 | https://www.ncbi.nlm.nih.gov/gene/?term=LOC105286661 | 2582 | 62 | 240.12 | 1 | 3.87 | 1.61 | 22.65 | 21.78 | 0.87 | 0.87 | 0.02 | 0.02 |
| gene9462 | https://www.ncbi.nlm.nih.gov/gene/?term=LOC105282970 | 1469 | 41 | 279.1 | 1 | 6.81 | 2.44 | 50 | 49.15 | 0.85 | 0.85 | 0.02 | 0.02 |
| gene11340 | https://www.ncbi.nlm.nih.gov/gene/?term=LOC105285131 | 2918 | 105 | 359.84 | 1 | 3.43 | 0.95 | 14.38 | 13.54 | 0.84 | 0.84 | 0.03 | 0.03 |
| gene6286 | https://www.ncbi.nlm.nih.gov/gene/?term=LOC105279298 | 2861 | 131 | 457.88 | 2 | 6.99 | 1.53 | 24.99 | 24.15 | 0.84 | 0.84 | 0.04 | 0.04 |
| gene1220 | https://www.ncbi.nlm.nih.gov/gene/?term=LOC105285742 | 3453 | 106 | 306.98 | 1 | 2.9 | 0.94 | 18.53 | 17.74 | 0.79 | 0.79 | 0.02 | 0.02 |
| gene8729 | https://www.ncbi.nlm.nih.gov/gene/?term=LOC105282137 | 5005 | 200 | 399.6 | 1 | 2 | 0.5 | 16.14 | 15.41 | 0.72 | 0.72 | 0.03 | 0.03 |
| gene10018 | https://www.ncbi.nlm.nih.gov/gene/?term=LOC105283607 | 4414 | 201 | 455.37 | 1 | 2.27 | 0.5 | 15.61 | 14.98 | 0.63 | 0.63 | 0.03 | 0.03 |
| gene4190 | https://www.ncbi.nlm.nih.gov/gene/?term=LOC105276875 | 3150 | 111 | 352.38 | 1 | 3.17 | 0.9 | 22.75 | 22.14 | 0.61 | 0.61 | 0.02 | 0.02 |
| gene12203 | https://www.ncbi.nlm.nih.gov/gene/?term=LOC105286099 | 5967 | 72 | 120.66 | 1 | 1.68 | 1.39 | 10.57 | 10 | 0.57 | 0.57 | 0.01 | 0.01 |
| gene5298 | https://www.ncbi.nlm.nih.gov/gene/?term=LOC105278145 | 4060 | 237 | 583.74 | 3 | 7.39 | 1.27 | 11.65 | 11.08 | 0.57 | 0.57 | 0.03 | 0.03 |
| gene4962 | https://www.ncbi.nlm.nih.gov/gene/?term=LOC105277761 | 4645 | 147 | 316.47 | 1 | 2.15 | 0.68 | 14.43 | 13.91 | 0.52 | 0.52 | 0.02 | 0.02 |
| gene10520 | https://www.ncbi.nlm.nih.gov/gene/?term=LOC105284184 | 2451 | 93 | 379.44 | 1 | 4.08 | 1.08 | 31.28 | 30.8 | 0.49 | 0.49 | 0.02 | 0.02 |
| gene10844 | https://www.ncbi.nlm.nih.gov/gene/?term=LOC105284535 | 9086 | 238 | 261.94 | 1 | 1.1 | 0.42 | 14.5 | 14.01 | 0.49 | 0.49 | 0.01 | 0.01 |
| gene2173 | https://www.ncbi.nlm.nih.gov/gene/?term=LOC105274590 | 4387 | 80 | 182.36 | 1 | 2.28 | 1.25 | 27.39 | 26.91 | 0.48 | 0.48 | 0.01 | 0.01 |
| gene10326 | https://www.ncbi.nlm.nih.gov/gene/?term=LOC105283962 | 3042 | 126 | 414.2 | 1 | 3.29 | 0.79 | 9.42 | 8.95 | 0.47 | 0.47 | 0.02 | 0.02 |
| gene6031 | https://www.ncbi.nlm.nih.gov/gene/?term=LOC105279000 | 5691 | 195 | 342.65 | 1 | 1.76 | 0.51 | 13.76 | 13.29 | 0.47 | 0.47 | 0.02 | 0.02 |
| gene10600 | https://www.ncbi.nlm.nih.gov/gene/?term=LOC105284270 | 4837 | 115 | 237.75 | 1 | 2.07 | 0.87 | 11.71 | 11.26 | 0.46 | 0.46 | 0.01 | 0.01 |
| gene2257 | https://www.ncbi.nlm.nih.gov/gene/?term=LOC105274689 | 3658 | 73 | 199.56 | 1 | 2.73 | 1.37 | 16.2 | 15.75 | 0.46 | 0.46 | 0.01 | 0.01 |
| gene3885 | https://www.ncbi.nlm.nih.gov/gene/?term=LOC105276527 | 7928 | 318 | 401.11 | 1 | 1.26 | 0.31 | 10.14 | 9.68 | 0.46 | 0.46 | 0.02 | 0.02 |
| gene12572 | https://www.ncbi.nlm.nih.gov/gene/?term=LOC105286514 | 2374 | 44 | 185.34 | 1 | 4.21 | 2.27 | 36.23 | 35.78 | 0.45 | 0.45 | 0.01 | 0.01 |
| gene7600 | https://www.ncbi.nlm.nih.gov/gene/?term=LOC105280819 | 1615 | 45 | 278.64 | 1 | 6.19 | 2.22 | 37.23 | 36.8 | 0.43 | 0.43 | 0.01 | 0.01 |
| gene8274 | https://www.ncbi.nlm.nih.gov/gene/?term=LOC105281624 | 5425 | 219 | 403.69 | 1 | 1.84 | 0.46 | 10.56 | 10.14 | 0.42 | 0.42 | 0.02 | 0.02 |
| gene9740 | https://www.ncbi.nlm.nih.gov/gene/?term=LOC105283277 | 2887 | 136 | 471.08 | 1 | 3.46 | 0.74 | 19.63 | 19.22 | 0.41 | 0.41 | 0.02 | 0.02 |
| gene10213 | https://www.ncbi.nlm.nih.gov/gene/?term=LOC105283826 | 1503 | 64 | 425.82 | 1 | 6.65 | 1.56 | 16.48 | 16.1 | 0.38 | 0.38 | 0.02 | 0.02 |
| gene3599 | https://www.ncbi.nlm.nih.gov/gene/?term=LOC105276196 | 2955 | 57 | 192.89 | 1 | 3.38 | 1.75 | 38.84 | 38.45 | 0.38 | 0.38 | 0.01 | 0.01 |
| gene3107 | https://www.ncbi.nlm.nih.gov/gene/?term=LOC105275650 | 5103 | 174 | 340.98 | 1 | 1.96 | 0.57 | 16.52 | 16.19 | 0.33 | 0.33 | 0.01 | 0.01 |

| | | | | | | | | | | | | | |
|---|---|---|---|---|---|---|---|---|---|---|---|---|---|
| gene10155 | https://www.ncbi.nlm.nih.gov/gene/?term=LOC105283756 | 7030 | 104 | 147.94 | 1 | 1.42 | 0.96 | 5.54 | 5.23 | 0.31 | 0.31 | 0 | 0 |
| gene6966 | https://www.ncbi.nlm.nih.gov/gene/?term=LOC105280104 | 3747 | 121 | 322.93 | 1 | 2.67 | 0.83 | 16.24 | 15.96 | 0.28 | 0.28 | 0.01 | 0.01 |
| gene9445 | https://www.ncbi.nlm.nih.gov/gene/?term=LOC105282951 | 2285 | 114 | 498.91 | 1 | 4.38 | 0.88 | 20.31 | 20.03 | 0.28 | 0.28 | 0.01 | 0.01 |
| gene1470 | https://www.ncbi.nlm.nih.gov/gene/?term=LOC105287597 | 2974 | 195 | 655.68 | 1 | 3.36 | 0.51 | 18.15 | 17.88 | 0.27 | 0.27 | 0.02 | 0.02 |
| gene11454 | https://www.ncbi.nlm.nih.gov/gene/?term=LOC105285256 | 1906 | 51 | 267.58 | 1 | 5.25 | 1.96 | 42.03 | 41.77 | 0.26 | 0.26 | 0.01 | 0.01 |
| gene12307 | https://www.ncbi.nlm.nih.gov/gene/?term=LOC105286216 | 10313 | 160 | 155.14 | 1 | 0.97 | 0.62 | 16.94 | 16.73 | 0.22 | 0.22 | 0 | 0 |
| gene5524 | https://www.ncbi.nlm.nih.gov/gene/?term=LOC105278397 | 5177 | 202 | 390.19 | 1 | 1.93 | 0.5 | 25.27 | 25.04 | 0.22 | 0.22 | 0.01 | 0.01 |
| gene570 | https://www.ncbi.nlm.nih.gov/gene/?term=LOC105278549 | 1673 | 69 | 412.43 | 1 | 5.98 | 1.45 | 43.78 | 43.57 | 0.21 | 0.21 | 0.01 | 0.01 |
| gene9177 | https://www.ncbi.nlm.nih.gov/gene/?term=LOC105282646 | 3320 | 101 | 304.22 | 1 | 3.01 | 0.99 | 21.24 | 21.05 | 0.2 | 0.2 | 0.01 | 0.01 |
| gene10886 | https://www.ncbi.nlm.nih.gov/gene/?term=LOC105284582 | 1475 | 49 | 332.2 1 | 6.78 | 2.04 | 22 | 21.8 | 0.19 | 0.19 | 0.01 | 0.01 | |
| gene5525 | https://www.ncbi.nlm.nih.gov/gene/?term=LOC105278396 | 3090 | 186 | 601.94 | 1 | 3.24 | 0.54 | 25.06 | 24.87 | 0.18 | 0.18 | 0.01 | 0.01 |
| gene7183 | https://www.ncbi.nlm.nih.gov/gene/?term=LOC105280348 | 3925 | 70 | 178.34 | 1 | 2.55 | 1.43 | 18.6 | 18.42 | 0.18 | 0.18 | 0 | 0 |
| gene7249 | https://www.ncbi.nlm.nih.gov/gene/?term=LOC105280422 | 1775 | 63 | 354.93 | 1 | 5.63 | 1.59 | 17.74 | 17.55 | 0.18 | 0.18 | 0.01 | 0.01 |
| gene13036 | https://www.ncbi.nlm.nih.gov/gene/?term=LOC105287059 | 4959 | 165 | 332.73 | 1 | 2.02 | 0.61 | 17.15 | 16.98 | 0.17 | 0.17 | 0.01 | 0.01 |
| gene10531 | https://www.ncbi.nlm.nih.gov/gene/?term=LOC105284190 | 6881 | 175 | 254.32 | 1 | 1.45 | 0.57 | 12.25 | 12.09 | 0.16 | 0.16 | 0 | 0 |
| gene10606 | https://www.ncbi.nlm.nih.gov/gene/?term=LOC105284273 | 6119 | 255 | 416.73 | 1 | 1.63 | 0.39 | 19.24 | 19.09 | 0.16 | 0.16 | 0.01 | 0.01 |
| gene5777 | https://www.ncbi.nlm.nih.gov/gene/?term=LOC105278691 | 3883 | 152 | 391.45 | 1 | 2.58 | 0.66 | 14.89 | 14.73 | 0.16 | 0.16 | 0.01 | 0.01 |
| gene11082 | https://www.ncbi.nlm.nih.gov/gene/?term=LOC105284824 | 7436 | 212 | 285.1 1 | 1.34 | 0.47 | 6.95 | 6.8 | 0.15 | 0.15 | 0 | 0 | |
| gene7166 | https://www.ncbi.nlm.nih.gov/gene/?term=LOC105280326 | 6117 | 238 | 389.08 | 1 | 1.63 | 0.42 | 15.04 | 14.89 | 0.15 | 0.15 | 0.01 | 0.01 |
| gene8343 | https://www.ncbi.nlm.nih.gov/gene/?term=LOC105281700 | 2172 | 37 | 170.35 | 1 | 4.6 | 2.7 | 39.63 | 39.48 | 0.15 | 0.15 | 0 | 0 |
| gene13114 | https://www.ncbi.nlm.nih.gov/gene/?term=LOC105287147 | 7609 | 419 | 550.66 | 1 | 1.31 | 0.24 | 8.81 | 8.67 | 0.14 | 0.14 | 0.01 | 0.01 |
| gene53 | https://www.ncbi.nlm.nih.gov/gene/?term=LOC105277148 | 19283 | 291 | 150.91 | 1 | 0.52 | 0.34 | 4.8 | 4.67 | 0.13 | 0.13 | 0 | 0 |
| gene1632 | https://www.ncbi.nlm.nih.gov/gene/?term=LOC105287764 | 18061 | 379 | 209.84 | 1 | 0.55 | 0.26 | 3.62 | 3.51 | 0.11 | 0.11 | 0 | 0 |
| gene10549 | https://www.ncbi.nlm.nih.gov/gene/?term=LOC105284210 | 7145 | 335 | 468.86 | 2 | 2.8 | 0.6 | 10.35 | 10.25 | 0.09 | 0.09 | 0 | 0 |
| gene2413 | https://www.ncbi.nlm.nih.gov/gene/?term=LOC105274863 | 2006 | 53 | 264.21 | 1 | 4.99 | 1.89 | 52.76 | 52.69 | 0.07 | 0.07 | 0 | 0 |
| gene11275 | https://www.ncbi.nlm.nih.gov/gene/?term=LOC105285055 | 4119 | 252 | 611.8 1 | 2.43 | 0.4 | 10.71 | 10.67 | 0.05 | 0.05 | 0 | 0 | |
| gene1332 | https://www.ncbi.nlm.nih.gov/gene/?term=LOC105286872 | 2460 | 158 | 642.28 | 1 | 4.07 | 0.63 | 20.39 | 20.35 | 0.04 | 0.04 | 0 | 0 |
| gene3856 | https://www.ncbi.nlm.nih.gov/gene/?term=LOC105276499 | 5236 | 196 | 374.33 | 1 | 1.91 | 0.51 | 10.18 | 10.15 | 0.03 | 0.03 | 0 | 0 |
| gene1147 | https://www.ncbi.nlm.nih.gov/gene/?term=LOC105284954 | 4022 | 111 | 275.98 | 1 | 2.49 | 0.9 | 11.93 | 11.9 | 0.02 | 0.02 | 0 | 0 |
| gene11583 | https://www.ncbi.nlm.nih.gov/gene/?term=LOC105285414 | 8192 | 178 | 217.29 | 1 | 1.22 | 0.56 | 9.44 | 9.48 | -0.04 | 0.04 | 0 | 0 |
| gene5375 | https://www.ncbi.nlm.nih.gov/gene/?term=LOC105278228 | 8640 | 154 | 178.24 | 1 | 1.16 | 0.65 | 24.99 | 25.04 | -0.05 | 0.05 | 0 | 0 |
| gene5746 | https://www.ncbi.nlm.nih.gov/gene/?term=LOC105278666 | 10343 | 477 | 461.18 | 1 | 0.97 | 0.21 | 6.71 | 6.77 | -0.05 | 0.05 | 0 | 0 |
| gene2844 | https://www.ncbi.nlm.nih.gov/gene/?term=LOC105275363 | 4330 | 197 | 454.97 | 1 | 2.31 | 0.51 | 14.26 | 14.32 | -0.06 | 0.06 | 0 | 0 |
| gene10345 | https://www.ncbi.nlm.nih.gov/gene/?term=LOC105283989 | 4444 | 26 | 58.51 1 | 2.25 | 3.85 | 49.48 | 49.58 | -0.09 | 0.09 | 0 | 0 | |
| gene2846 | https://www.ncbi.nlm.nih.gov/gene/?term=LOC105275374 | 11246 | 465 | 413.48 | 1 | 0.89 | 0.22 | 10.6 | 10.73 | -0.13 | 0.13 | -0.01 | 0.01 |
| gene12802 | https://www.ncbi.nlm.nih.gov/gene/?term=LOC105286799 | 3183 | 190 | 596.92 | 1 | 3.14 | 0.53 | 15.49 | 15.63 | -0.14 | 0.14 | -0.01 | 0.01 |
| gene11579 | https://www.ncbi.nlm.nih.gov/gene/?term=LOC105285406 | 13571 | 280 | 206.32 | 1 | 0.74 | 0.36 | 6.48 | 6.65 | -0.17 | 0.17 | 0 | 0 |
| gene8814 | https://www.ncbi.nlm.nih.gov/gene/?term=LOC105282240 | 6910 | 185 | 267.73 | 1 | 1.45 | 0.54 | 9.48 | 9.66 | -0.18 | 0.18 | 0 | 0 |
| gene11561 | https://www.ncbi.nlm.nih.gov/gene/?term=LOC105285387 | 11528 | 207 | 179.56 | 1 | 0.87 | 0.48 | 8.9 | 9.12 | -0.22 | 0.22 | 0 | 0 |
| gene10678 | https://www.ncbi.nlm.nih.gov/gene/?term=LOC105284355 | 4031 | 182 | 451.5 1 | 2.48 | 0.55 | 17.39 | 17.62 | -0.23 | 0.23 | -0.01 | 0.01 | |
| gene34 | https://www.ncbi.nlm.nih.gov/gene/?term=LOC105278416 | 7120 | 132 | 185.39 | 1 | 1.4 | 0.76 | 14.08 | 14.34 | -0.26 | 0.26 | 0 | 0 |
| gene7556 | https://www.ncbi.nlm.nih.gov/gene/?term=LOC105280774 | 2792 | 118 | 422.64 | 1 | 3.58 | 0.85 | 27.01 | 27.27 | -0.26 | 0.26 | -0.01 | 0.01 |
| gene8600 | https://www.ncbi.nlm.nih.gov/gene/?term=LOC105282001 | 8513 | 186 | 218.49 | 1 | 1.17 | 0.54 | 10 | 10.27 | -0.27 | 0.27 | -0.01 | 0.01 |
| gene5052 | https://www.ncbi.nlm.nih.gov/gene/?term=LOC105277856 | 3167 | 33 | 104.2 1 | 3.16 | 3.03 | 39.08 | 39.37 | -0.29 | 0.29 | 0 | 0 | |
| gene2806 | https://www.ncbi.nlm.nih.gov/gene/?term=LOC105275327 | 14257 | 511 | 358.42 | 1 | 0.7 | 0.2 | 6.7 | 7.03 | -0.33 | 0.33 | -0.01 | 0.01 |
| gene35 | https://www.ncbi.nlm.nih.gov/gene/?term=LOC105275197 | 2643 | 48 | 181.61 | 1 | 3.78 | 2.08 | 38.22 | 38.55 | -0.33 | 0.33 | -0.01 | 0.01 |
| gene2668 | https://www.ncbi.nlm.nih.gov/gene/?term=LOC105275170 | 12496 | 279 | 223.27 | 1 | 0.8 | 0.36 | 1.97 | 2.32 | -0.35 | 0.35 | -0.01 | 0.01 |
| gene738 | https://www.ncbi.nlm.nih.gov/gene/?term=LOC105280421 | 4927 | 216 | 438.4 1 | 2.03 | 0.46 | 14.22 | 14.61 | -0.39 | 0.39 | -0.02 | 0.02 | |
| gene1855 | https://www.ncbi.nlm.nih.gov/gene/?term=LOC105288036 | 2387 | 84 | 351.91 | 1 | 4.19 | 1.19 | 29.18 | 29.59 | -0.41 | 0.41 | -0.01 | 0.01 |
| gene7248 | https://www.ncbi.nlm.nih.gov/gene/?term=LOC105280408 | 5175 | 61 | 117.87 | 1 | 1.93 | 1.64 | 20.52 | 20.93 | -0.41 | 0.41 | 0 | 0 |
| gene6620 | https://www.ncbi.nlm.nih.gov/gene/?term=LOC105279687 | 17998 | 368 | 204.47 | 1 | 0.56 | 0.27 | 4.64 | 5.1 | -0.46 | 0.46 | -0.01 | 0.01 |
| gene7247 | https://www.ncbi.nlm.nih.gov/gene/?term=LOC105280407 | 10486 | 218 | 207.9 1 | 0.95 | 0.46 | 16.54 | 17.03 | -0.49 | 0.49 | -0.01 | 0.01 | |
| gene9336 | https://www.ncbi.nlm.nih.gov/gene/?term=LOC105282834 | 5993 | 129 | 215.25 | 1 | 1.67 | 0.78 | 14.53 | 15.02 | -0.49 | 0.49 | -0.01 | 0.01 |
| gene3206 | https://www.ncbi.nlm.nih.gov/gene/?term=LOC105275752 | 3474 | 129 | 371.33 | 1 | 2.88 | 0.78 | 29.67 | 30.17 | -0.5 | 0.5 | -0.02 | 0.02 |
| gene940 | https://www.ncbi.nlm.nih.gov/gene/?term=LOC105282794 | 3578 | 85 | 237.56 | 1 | 2.79 | 1.18 | 13.96 | 14.47 | -0.51 | 0.51 | -0.01 | 0.01 |
| gene298 | https://www.ncbi.nlm.nih.gov/gene/?term=LOC105275460 | 2473 | 120 | 485.24 | 1 | 4.04 | 0.83 | 13.22 | 13.75 | -0.53 | 0.53 | -0.03 | 0.03 |
| gene3470 | https://www.ncbi.nlm.nih.gov/gene/?term=LOC105276059 | 8716 | 152 | 174.39 | 1 | 1.15 | 0.66 | 7.9 | 8.44 | -0.54 | 0.54 | -0.01 | 0.01 |
| gene6583 | https://www.ncbi.nlm.nih.gov/gene/?term=LOC105279649 | 7444 | 340 | 456.74 | 1 | 1.34 | 0.29 | 10.48 | 11.08 | -0.59 | 0.59 | -0.03 | 0.03 |
| gene7514 | https://www.ncbi.nlm.nih.gov/gene/?term=LOC105280723 | 4471 | 208 | 465.22 | 1 | 2.24 | 0.48 | 13.49 | 14.1 | -0.6 | 0.6 | -0.03 | 0.03 |
| gene4376 | https://www.ncbi.nlm.nih.gov/gene/?term=LOC105277083 | 4307 | 114 | 264.69 | 1 | 2.32 | 0.88 | 20.15 | 20.76 | -0.61 | 0.61 | -0.02 | 0.02 |
| gene12516 | https://www.ncbi.nlm.nih.gov/gene/?term=LOC105286452 | 1529 | 35 | 228.91 | 1 | 6.54 | 2.86 | 51.89 | 52.54 | -0.65 | 0.65 | -0.01 | 0.01 |
| gene13033 | https://www.ncbi.nlm.nih.gov/gene/?term=LOC105287063 | 1877 | 51 | 271.71 | 1 | 5.33 | 1.96 | 30.35 | 31.05 | -0.7 | 0.7 | -0.02 | 0.02 |
| gene2056 | https://www.ncbi.nlm.nih.gov/gene/?term=LOC105274452 | 6584 | 243 | 369.08 | 1 | 1.52 | 0.41 | 13.17 | 13.88 | -0.71 | 0.71 | -0.03 | 0.03 |
| gene5733 | https://www.ncbi.nlm.nih.gov/gene/?term=LOC105278649 | 5221 | 91 | 174.3 1 | 1.92 | 1.1 | 19.36 | 20.1 | -0.74 | 0.74 | -0.01 | 0.01 | |

| | | | | | | | | | | | | | | | |
|---|---|---|---|---|---|---|---|---|---|---|---|---|---|---|---|
| gene8742 | https://www.ncbi.nlm.nih.gov/gene/?term=LOC105282151 | 2563 | 42 | 163.87 | | 1 | 3.9 | 2.38 | 34.83 | 35.57 | -0.75 | 0.75 | -0.01 | 0.01 | |
| gene1430 | https://www.ncbi.nlm.nih.gov/gene/?term=LOC105287551 | 6433 | 235 | 365.3 | 1 | 1.55 | 0.43 | 8.56 | 9.58 | -1.02 | 1.02 | -0.04 | 0.04 | | |
| gene11401 | https://www.ncbi.nlm.nih.gov/gene/?term=LOC105285202 | 1841 | 75 | 407.39 | | 1 | 5.43 | 1.33 | 24.08 | 25.12 | -1.03 | 1.03 | -0.04 | 0.04 | |
| gene12439 | https://www.ncbi.nlm.nih.gov/gene/?term=LOC105286357 | 4761 | 81 | 170.13 | | 1 | 2.1 | 1.23 | 29.49 | 30.52 | -1.03 | 1.03 | -0.02 | 0.02 | |
| gene2683 | https://www.ncbi.nlm.nih.gov/gene/?term=LOC105275187 | 5492 | 70 | 127.46 | | 1 | 1.82 | 1.43 | 13.71 | 14.76 | -1.05 | 1.05 | -0.01 | 0.01 | |
| gene6528 | https://www.ncbi.nlm.nih.gov/gene/?term=LOC105279591 | 7222 | 59 | 81.69 | 1 | 1.38 | 1.69 | 26.22 | 27.32 | -1.1 | 1.1 | -0.01 | 0.01 | | |
| gene7884 | https://www.ncbi.nlm.nih.gov/gene/?term=LOC105281266 | 3390 | 81 | 238.94 | | 1 | 2.95 | 1.23 | 16.21 | 17.33 | -1.12 | 1.12 | -0.03 | 0.03 | |
| gene2071 | https://www.ncbi.nlm.nih.gov/gene/?term=LOC105274471 | 2446 | 121 | 494.69 | | 1 | 4.09 | 0.83 | 19.71 | 20.9 | -1.19 | 1.19 | -0.06 | 0.06 | |
| gene4023 | https://www.ncbi.nlm.nih.gov/gene/?term=LOC105276692 | 1759 | 36 | 204.66 | | 1 | 5.69 | 2.78 | 20.19 | 21.39 | -1.2 | 1.2 | -0.02 | 0.02 | |
| gene7368 | https://www.ncbi.nlm.nih.gov/gene/?term=LOC105280556 | 4105 | 50 | 121.8 | 1 | 2.44 | 2 | 14.06 | 15.36 | -1.3 | 1.3 | -0.02 | 0.02 | | |
| gene4720 | https://www.ncbi.nlm.nih.gov/gene/?term=LOC105277479 | 1697 | 50 | 294.64 | | 1 | 5.89 | 2 | 35.87 | 37.27 | -1.4 | 1.4 | -0.04 | 0.04 | |
| gene10654 | https://www.ncbi.nlm.nih.gov/gene/?term=LOC105284329 | 4027 | 51 | 126.65 | | 1 | 2.48 | 1.96 | 17.3 | 18.75 | -1.45 | 1.45 | -0.02 | 0.02 | |
| gene2856 | https://www.ncbi.nlm.nih.gov/gene/?term=LOC105275384 | 1987 | 87 | 437.85 | | 1 | 5.03 | 1.15 | 34.24 | 35.77 | -1.53 | 1.53 | -0.07 | 0.07 | |
| gene7568 | https://www.ncbi.nlm.nih.gov/gene/?term=LOC105280786 | 2525 | 109 | 431.68 | | 1 | 3.96 | 0.92 | 28.7 | 30.24 | -1.54 | 1.54 | -0.07 | 0.07 | |
| gene4892 | https://www.ncbi.nlm.nih.gov/gene/?term=LOC105277667 | 3898 | 120 | 307.85 | | 1 | 2.57 | 0.83 | 27.15 | 28.74 | -1.59 | 1.59 | -0.05 | 0.05 | |
| gene1518 | https://www.ncbi.nlm.nih.gov/gene/?term=LOC105287648 | 2195 | 35 | 159.45 | | 1 | 4.56 | 2.86 | 35.74 | 37.41 | -1.67 | 1.67 | -0.03 | 0.03 | |
| gene9478 | https://www.ncbi.nlm.nih.gov/gene/?term=LOC105282991 | 3726 | 58 | 155.66 | | 1 | 2.68 | 1.72 | 33.94 | 35.62 | -1.68 | 1.68 | -0.03 | 0.03 | |
| gene927 | https://www.ncbi.nlm.nih.gov/gene/?term=LOC105282600 | 3727 | 52 | 139.52 | | 1 | 2.68 | 1.92 | 31.26 | 33.17 | -1.91 | 1.91 | -0.03 | 0.03 | |
| gene1884 | https://www.ncbi.nlm.nih.gov/gene/?term=LOC105288072 | 4169 | 78 | 187.1 | 1 | 2.4 | 1.28 | 22.55 | 24.51 | -1.96 | 1.96 | -0.04 | 0.04 | | |
| gene3817 | https://www.ncbi.nlm.nih.gov/gene/?term=LOC105276449 | 1230 | 45 | 365.85 | | 1 | 8.13 | 2.22 | 27.32 | 29.29 | -1.97 | 1.97 | -0.07 | 0.07 | |
| gene9647 | https://www.ncbi.nlm.nih.gov/gene/?term=LOC109611233 | 3094 | 35 | 113.12 | | 1 | 3.23 | 2.86 | 38.21 | 40.18 | -1.97 | 1.97 | -0.02 | 0.02 | |
| gene9648 | https://www.ncbi.nlm.nih.gov/gene/?term=LOC105283183 | 2602 | 33 | 126.83 | | 1 | 3.84 | 3.03 | 40.53 | 42.61 | -2.08 | 2.08 | -0.03 | 0.03 | |
| gene5877 | https://www.ncbi.nlm.nih.gov/gene/?term=LOC105278807 | 1719 | 53 | 308.32 | | 1 | 5.82 | 1.89 | 22.63 | 24.74 | -2.11 | 2.11 | -0.06 | 0.06 | |
| gene8661 | https://www.ncbi.nlm.nih.gov/gene/?term=LOC105282063 | 1926 | 30 | 155.76 | | 1 | 5.19 | 3.33 | 28.06 | 30.33 | -2.28 | 2.28 | -0.04 | 0.04 | |
| gene8178 | https://www.ncbi.nlm.nih.gov/gene/?term=LOC105281515 | 8647 | 63 | 72.86 | 1 | 1.16 | 1.59 | 24.53 | 26.83 | -2.3 | 2.3 | -0.02 | 0.02 | | |
| gene5101 | https://www.ncbi.nlm.nih.gov/gene/?term=LOC105277903 | 1813 | 36 | 198.57 | | 1 | 5.52 | 2.78 | 53.23 | 56.32 | -3.09 | 3.09 | -0.06 | 0.06 | |
| gene5714 | https://www.ncbi.nlm.nih.gov/gene/?term=LOC105278729 | 710 | 21 | 295.77 | | 1 | 14.08 | 4.76 | 56.35 | 60.77 | -4.42 | 4.42 | -0.13 | 0.13 | |
| gene5647 | https://www.ncbi.nlm.nih.gov/gene/?term=LOC105278567 | 3118 | 29 | 93.01 | 1 | 3.21 | 3.45 | 13.21 | 19.43 | -6.21 | 6.21 | -0.06 | 0.06 | | |
| gene3165 | https://www.ncbi.nlm.nih.gov/gene/?term=LOC105275706 | 2374 | 60 | 252.74 | | 3 | 12.64 | 5 | 22.21 | 32.53 | -10.32 | | 10.32 | -0.26 | 0.26 |
| gene2593 | https://www.ncbi.nlm.nih.gov/gene/?term=LOC105275070 | 1774 | 5 | 28.18 | 5 | 28.18 | 100 | 10.96 | 79.22 | -68.26 | | 68.26 | -0.19 | 0.19 | |

Directory: OGL File: ogl-Cb_LI_BCphaseB2.vs.RphaseA2.txt

| gene_ID | gene_link | gwidth | #Sites | #per10Kb | #dmSites | #dmsp10kb | %dmSites | %pSite1 | %pSite2 | DMpSite | ADMpSite | DMpNucl | ADMpNucl |
|---|---|---|---|---|---|---|---|---|---|---|---|---|---|
| gene393 | https://www.ncbi.nlm.nih.gov/gene/?term=LOC105276566 | 1123 | 2 | 17.81 | 1 | 8.9 | 50 | 70.57 | 47.62 | 22.95 | 22.95 | 0.04 | 0.04 |
| gene10963 | https://www.ncbi.nlm.nih.gov/gene/?term=LOC105284677 | 1576 | 30 | 190.36 | 2 | 12.69 | 6.67 | 38.74 | 23.8 | 14.95 | 14.95 | 0.28 | 0.28 |
| gene9109 | https://www.ncbi.nlm.nih.gov/gene/?term=LOC105282561 | 2400 | 7 | 29.17 | 1 | 4.17 | 14.29 | 63.87 | 57.68 | 6.19 | 6.19 | 0.02 | 0.02 |
| gene10816 | https://www.ncbi.nlm.nih.gov/gene/?term=LOC105284502 | 1974 | 21 | 106.38 | 1 | 5.07 | 4.76 | 45.81 | 40.14 | 5.67 | 5.67 | 0.06 | 0.06 |
| gene1199 | https://www.ncbi.nlm.nih.gov/gene/?term=LOC105285498 | 2220 | 26 | 117.12 | 1 | 4.5 | 3.85 | 37.03 | 32.91 | 4.12 | 4.12 | 0.05 | 0.05 |
| gene9411 | https://www.ncbi.nlm.nih.gov/gene/?term=LOC105282914 | 5534 | 43 | 77.7 | 1 | 1.81 | 2.33 | 49.78 | 47.15 | 2.63 | 2.63 | 0.02 | 0.02 |
| gene920 | https://www.ncbi.nlm.nih.gov/gene/?term=LOC105282518 | 1197 | 42 | 350.88 | 1 | 8.35 | 2.38 | 38.3 | 35.67 | 2.62 | 2.62 | 0.09 | 0.09 |
| gene6237 | https://www.ncbi.nlm.nih.gov/gene/?term=LOC105279244 | 1484 | 25 | 168.46 | 1 | 6.74 | 4 | 53.48 | 51.47 | 2.01 | 2.01 | 0.03 | 0.03 |
| gene6652 | https://www.ncbi.nlm.nih.gov/gene/?term=LOC105279719 | 2831 | 82 | 289.65 | 1 | 3.53 | 1.22 | 24.05 | 22.34 | 1.72 | 1.72 | 0.05 | 0.05 |
| gene12021 | https://www.ncbi.nlm.nih.gov/gene/?term=LOC105285905 | 1399 | 22 | 157.26 | 1 | 7.15 | 4.55 | 34.58 | 32.98 | 1.6 | 1.6 | 0.03 | 0.03 |
| gene3200 | https://www.ncbi.nlm.nih.gov/gene/?term=LOC105275743 | 1786 | 76 | 425.53 | 1 | 5.6 | 1.32 | 23.84 | 22.24 | 1.6 | 1.6 | 0.07 | 0.07 |
| gene9101 | https://www.ncbi.nlm.nih.gov/gene/?term=LOC105282553 | 3068 | 115 | 374.84 | 1 | 3.26 | 0.87 | 23.82 | 22.25 | 1.57 | 1.57 | 0.06 | 0.06 |
| gene6455 | https://www.ncbi.nlm.nih.gov/gene/?term=LOC105279500 | 2612 | 34 | 130.17 | 1 | 3.83 | 2.94 | 45.97 | 44.48 | 1.5 | 1.5 | 0.02 | 0.02 |
| gene10714 | https://www.ncbi.nlm.nih.gov/gene/?term=LOC105284396 | 1966 | 66 | 335.71 | 1 | 5.09 | 1.52 | 34.65 | 33.2 | 1.45 | 1.45 | 0.05 | 0.05 |
| gene7398 | https://www.ncbi.nlm.nih.gov/gene/?term=LOC105280589 | 3452 | 39 | 112.98 | 1 | 2.9 | 2.56 | 21.67 | 20.29 | 1.38 | 1.38 | 0.02 | 0.02 |
| gene1505 | https://www.ncbi.nlm.nih.gov/gene/?term=LOC105287633 | 1693 | 31 | 183.11 | 1 | 5.91 | 3.23 | 40.09 | 38.72 | 1.37 | 1.37 | 0.03 | 0.03 |
| gene4498 | https://www.ncbi.nlm.nih.gov/gene/?term=LOC105277239 | 2831 | 141 | 498.06 | 1 | 3.53 | 0.71 | 15.21 | 13.89 | 1.32 | 1.32 | 0.07 | 0.07 |
| gene2404 | https://www.ncbi.nlm.nih.gov/gene/?term=LOC105274853 | 1767 | 47 | 265.99 | 1 | 5.66 | 2.13 | 31.76 | 30.45 | 1.31 | 1.31 | 0.03 | 0.03 |
| gene11340 | https://www.ncbi.nlm.nih.gov/gene/?term=LOC105285131 | 2918 | 110 | 376.97 | 1 | 3.43 | 0.91 | 14.31 | 13.13 | 1.19 | 1.19 | 0.04 | 0.04 |
| gene564 | https://www.ncbi.nlm.nih.gov/gene/?term=LOC105278478 | 2802 | 24 | 85.65 | 1 | 3.57 | 4.17 | 23.58 | 22.43 | 1.15 | 1.15 | 0.01 | 0.01 |
| gene1768 | https://www.ncbi.nlm.nih.gov/gene/?term=LOC105287938 | 2792 | 79 | 282.95 | 1 | 3.58 | 1.27 | 29.01 | 27.92 | 1.09 | 1.09 | 0.03 | 0.03 |
| gene11447 | https://www.ncbi.nlm.nih.gov/gene/?term=LOC105285252 | 7437 | 128 | 172.11 | 1 | 1.34 | 0.78 | 14.37 | 13.39 | 0.98 | 0.98 | 0.02 | 0.02 |
| gene814 | https://www.ncbi.nlm.nih.gov/gene/?term=LOC105281296 | 2722 | 52 | 191.04 | 1 | 3.67 | 1.92 | 24.92 | 24.03 | 0.89 | 0.89 | 0.02 | 0.02 |
| gene3671 | https://www.ncbi.nlm.nih.gov/gene/?term=LOC105276272 | 1966 | 51 | 259.41 | 1 | 5.09 | 1.96 | 27.37 | 26.55 | 0.82 | 0.82 | 0.02 | 0.02 |
| gene5646 | https://www.ncbi.nlm.nih.gov/gene/?term=LOC105278534 | 7475 | 200 | 267.56 | 1 | 1.34 | 0.5 | 17.47 | 16.68 | 0.79 | 0.79 | 0.02 | 0.02 |
| gene5298 | https://www.ncbi.nlm.nih.gov/gene/?term=LOC105278145 | 4060 | 254 | 625.62 | 3 | 7.39 | 1.18 | 11.08 | 10.34 | 0.75 | 0.75 | 0.05 | 0.05 |
| gene7600 | https://www.ncbi.nlm.nih.gov/gene/?term=LOC105280819 | 1615 | 43 | 266.25 | 1 | 6.19 | 2.33 | 38.96 | 38.22 | 0.75 | 0.75 | 0.02 | 0.02 |
| gene1220 | https://www.ncbi.nlm.nih.gov/gene/?term=LOC105285742 | 3453 | 133 | 385.17 | 1 | 2.9 | 0.75 | 19.92 | 19.19 | 0.74 | 0.74 | 0.03 | 0.03 |
| gene3046 | https://www.ncbi.nlm.nih.gov/gene/?term=LOC105275587 | 2394 | 58 | 242.27 | 1 | 4.18 | 1.72 | 36.02 | 35.31 | 0.71 | 0.71 | 0.02 | 0.02 |
| gene7199 | https://www.ncbi.nlm.nih.gov/gene/?term=LOC105280366 | 1414 | 33 | 233.38 | 1 | 7.07 | 3.03 | 43.27 | 42.56 | 0.71 | 0.71 | 0.02 | 0.02 |
| gene7915 | https://www.ncbi.nlm.nih.gov/gene/?term=LOC105281173 | 1597 | 38 | 237.95 | 1 | 6.26 | 2.63 | 60.79 | 60.11 | 0.68 | 0.68 | 0.02 | 0.02 |
| gene10379 | https://www.ncbi.nlm.nih.gov/gene/?term=LOC105284028 | 6974 | 95 | 136.22 | 1 | 1.43 | 1.05 | 15.2 | 14.55 | 0.66 | 0.66 | 0.01 | 0.01 |
| gene657 | https://www.ncbi.nlm.nih.gov/gene/?term=LOC105279667 | 6421 | 104 | 161.97 | 1 | 1.56 | 0.96 | 3.28 | 2.76 | 0.52 | 0.52 | 0.01 | 0.01 |
| gene3727 | https://www.ncbi.nlm.nih.gov/gene/?term=LOC105276367 | 2484 | 77 | 309.98 | 1 | 4.03 | 1.3 | 42.16 | 41.65 | 0.51 | 0.51 | 0.02 | 0.02 |
| gene9327 | https://www.ncbi.nlm.nih.gov/gene/?term=LOC105282821 | 7793 | 174 | 223.28 | 1 | 1.28 | 0.57 | 10.81 | 10.29 | 0.51 | 0.51 | 0.01 | 0.01 |
| gene6639 | https://www.ncbi.nlm.nih.gov/gene/?term=LOC105279705 | 7656 | 73 | 95.35 | 1 | 1.31 | 1.37 | 24.42 | 23.96 | 0.46 | 0.46 | 0 | 0 |
| gene11583 | https://www.ncbi.nlm.nih.gov/gene/?term=LOC105285414 | 8192 | 188 | 229.49 | 1 | 1.22 | 0.53 | 8.66 | 8.2 | 0.45 | 0.45 | 0.01 | 0.01 |
| gene10213 | https://www.ncbi.nlm.nih.gov/gene/?term=LOC105283826 | 1503 | 66 | 439.12 | 1 | 6.65 | 1.52 | 15.98 | 15.58 | 0.4 | 0.4 | 0.02 | 0.02 |
| gene10819 | https://www.ncbi.nlm.nih.gov/gene/?term=LOC105284507 | 5767 | 94 | 163 | 1 | 1.73 | 1.06 | 19.47 | 19.06 | 0.4 | 0.4 | 0.01 | 0.01 |
| gene12406 | https://www.ncbi.nlm.nih.gov/gene/?term=LOC105286316 | 1386 | 32 | 230.88 | 1 | 7.22 | 3.12 | 47.17 | 46.8 | 0.37 | 0.37 | 0.01 | 0.01 |
| gene2748 | https://www.ncbi.nlm.nih.gov/gene/?term=LOC105275261 | 3845 | 56 | 145.64 | 1 | 2.6 | 1.79 | 39.96 | 39.6 | 0.36 | 0.36 | 0.01 | 0.01 |
| gene7148 | https://www.ncbi.nlm.nih.gov/gene/?term=LOC105280304 | 8920 | 123 | 137.89 | 1 | 1.12 | 0.81 | 7.45 | 7.09 | 0.36 | 0.36 | 0 | 0 |
| gene3937 | https://www.ncbi.nlm.nih.gov/gene/?term=LOC105276592 | 9576 | 86 | 89.81 | 1 | 1.04 | 1.16 | 7.94 | 7.61 | 0.34 | 0.34 | 0 | 0 |
| gene11454 | https://www.ncbi.nlm.nih.gov/gene/?term=LOC105285256 | 1906 | 50 | 262.33 | 1 | 5.25 | 2 | 42.87 | 42.54 | 0.33 | 0.33 | 0.01 | 0.01 |
| gene3168 | https://www.ncbi.nlm.nih.gov/gene/?term=LOC105275711 | 6150 | 79 | 128.46 | 1 | 1.63 | 1.27 | 13.78 | 13.47 | 0.32 | 0.32 | 0 | 0 |
| gene8274 | https://www.ncbi.nlm.nih.gov/gene/?term=LOC105281624 | 5425 | 255 | 470.05 | 1 | 1.84 | 0.39 | 9.6 | 9.29 | 0.31 | 0.31 | 0.01 | 0.01 |
| gene9755 | https://www.ncbi.nlm.nih.gov/gene/?term=LOC105283296 | 2177 | 97 | 445.57 | 1 | 4.59 | 1.03 | 20.64 | 20.32 | 0.31 | 0.31 | 0.01 | 0.01 |
| gene2413 | https://www.ncbi.nlm.nih.gov/gene/?term=LOC105274863 | 2006 | 55 | 274.18 | 1 | 4.99 | 1.82 | 52.75 | 52.45 | 0.3 | 0.3 | 0.01 | 0.01 |
| gene322 | https://www.ncbi.nlm.nih.gov/gene/?term=LOC105275777 | 1606 | 28 | 174.35 | 1 | 6.23 | 3.57 | 39.73 | 39.43 | 0.3 | 0.3 | 0.01 | 0.01 |
| gene8015 | https://www.ncbi.nlm.nih.gov/gene/?term=LOC105281312 | 2998 | 74 | 246.83 | 1 | 3.34 | 1.35 | 29.49 | 29.21 | 0.28 | 0.28 | 0.01 | 0.01 |
| gene2835 | https://www.ncbi.nlm.nih.gov/gene/?term=LOC105275357 | 2492 | 129 | 517.66 | 1 | 4.01 | 0.78 | 13.74 | 13.5 | 0.25 | 0.25 | 0.01 | 0.01 |
| gene7249 | https://www.ncbi.nlm.nih.gov/gene/?term=LOC105280422 | 1775 | 63 | 354.93 | 1 | 5.63 | 1.59 | 17.74 | 17.51 | 0.23 | 0.23 | 0.01 | 0.01 |
| gene10018 | https://www.ncbi.nlm.nih.gov/gene/?term=LOC105283607 | 4414 | 208 | 471.23 | 1 | 2.27 | 0.48 | 15.07 | 14.94 | 0.13 | 0.13 | 0.01 | 0.01 |
| gene4190 | https://www.ncbi.nlm.nih.gov/gene/?term=LOC105276875 | 3150 | 124 | 393.65 | 1 | 3.17 | 0.81 | 21.27 | 21.17 | 0.1 | 0.1 | 0 | 0 |
| gene1632 | https://www.ncbi.nlm.nih.gov/gene/?term=LOC105287764 | 18061 | 434 | 240.3 | 1 | 0.55 | 0.23 | 3.44 | 3.34 | 0.09 | 0.09 | 0 | 0 |
| gene10549 | https://www.ncbi.nlm.nih.gov/gene/?term=LOC105284210 | 7145 | 358 | 501.05 | 2 | 2.8 | 0.56 | 10 | 9.93 | 0.07 | 0.07 | 0 | 0 |
| gene9326 | https://www.ncbi.nlm.nih.gov/gene/?term=LOC105282818 | 16612 | 222 | 133.64 | 1 | 0.6 | 0.45 | 11.24 | 11.17 | 0.06 | 0.06 | 0 | 0 |
| gene10520 | https://www.ncbi.nlm.nih.gov/gene/?term=LOC105284184 | 2451 | 104 | 424.32 | 1 | 4.08 | 0.96 | 28.9 | 28.86 | 0.04 | 0.04 | 0 | 0 |
| gene10155 | https://www.ncbi.nlm.nih.gov/gene/?term=LOC105283756 | 7030 | 110 | 156.47 | 1 | 1.42 | 0.91 | 5.47 | 5.44 | 0.03 | 0.03 | 0 | 0 |
| gene1147 | https://www.ncbi.nlm.nih.gov/gene/?term=LOC105284954 | 4022 | 118 | 293.39 | 1 | 2.49 | 0.85 | 14.22 | 14.19 | 0.03 | 0.03 | 0 | 0 |

| | | | | | | | | | | | | | |
|---|---|---|---|---|---|---|---|---|---|---|---|---|---|
| gene53 | https://www.ncbi.nlm.nih.gov/gene/?term=LOC105277148 | 19283 | 331 | 171.65 | 1 | 0.52 | 0.3 | 4.76 | 4.72 | 0.03 | 0.03 | 0 | 0 |
| gene11688 | https://www.ncbi.nlm.nih.gov/gene/?term=LOC105285533 | 1637 | 71 | 433.72 | 1 | 6.11 | 1.41 | 23.23 | 23.21 | 0.02 | 0.02 | 0 | 0 |
| gene7550 | https://www.ncbi.nlm.nih.gov/gene/?term=LOC105280763 | 2173 | 109 | 501.61 | 1 | 4.6 | 0.92 | 25.93 | 25.92 | 0.01 | 0.01 | 0 | 0 |
| gene9445 | https://www.ncbi.nlm.nih.gov/gene/?term=LOC105282951 | 2285 | 122 | 533.92 | 1 | 4.38 | 0.82 | 19.11 | 19.11 | 0.01 | 0.01 | 0 | 0 |
| gene4527 | https://www.ncbi.nlm.nih.gov/gene/?term=LOC105277273 | 3686 | 170 | 461.2 1 | 2.71 | 0.59 | 12.09 | 12.09 | 0 | 0 | 0 | 0 | |
| gene3885 | https://www.ncbi.nlm.nih.gov/gene/?term=LOC105276527 | 7928 | 341 | 430.12 | 1 | 1.26 | 0.29 | 9.68 | 9.69 | -0.01 | 0.01 | 0 | 0 |
| gene1310 | https://www.ncbi.nlm.nih.gov/gene/?term=LOC105286661 | 2582 | 67 | 259.49 | 1 | 3.87 | 1.49 | 24.89 | 24.92 | -0.03 | 0.03 | 0 | 0 |
| gene13114 | https://www.ncbi.nlm.nih.gov/gene/?term=LOC105287147 | 7609 | 446 | 586.15 | 1 | 1.31 | 0.22 | 9.09 | 9.12 | -0.03 | 0.03 | 0 | 0 |
| gene12674 | https://www.ncbi.nlm.nih.gov/gene/?term=LOC105286649 | 18283 | 886 | 484.6 1 | 0.55 | 0.11 | 1.03 | 1.06 | -0.04 | 0.04 | 0 | 0 | |
| gene1470 | https://www.ncbi.nlm.nih.gov/gene/?term=LOC105287597 | 2974 | 221 | 743.11 | 1 | 3.36 | 0.45 | 16.16 | 16.21 | -0.04 | 0.04 | 0 | 0 |
| gene2122 | https://www.ncbi.nlm.nih.gov/gene/?term=LOC105274531 | 8153 | 308 | 377.78 | 1 | 1.23 | 0.32 | 10.22 | 10.26 | -0.04 | 0.04 | 0 | 0 |
| gene6649 | https://www.ncbi.nlm.nih.gov/gene/?term=LOC105279717 | 6017 | 191 | 317.43 | 1 | 1.66 | 0.52 | 12.72 | 12.75 | -0.04 | 0.04 | 0 | 0 |
| gene3856 | https://www.ncbi.nlm.nih.gov/gene/?term=LOC105276499 | 5236 | 217 | 414.44 | 1 | 1.91 | 0.46 | 9.77 | 9.82 | -0.05 | 0.05 | 0 | 0 |
| gene5777 | https://www.ncbi.nlm.nih.gov/gene/?term=LOC105278691 | 3883 | 175 | 450.68 | 1 | 2.58 | 0.57 | 14.25 | 14.3 | -0.05 | 0.05 | 0 | 0 |
| gene10596 | https://www.ncbi.nlm.nih.gov/gene/?term=LOC105284262 | 6520 | 245 | 375.77 | 1 | 1.53 | 0.41 | 3.24 | 3.34 | -0.09 | 0.09 | 0 | 0 |
| gene3286 | https://www.ncbi.nlm.nih.gov/gene/?term=LOC105275848 | 3744 | 151 | 403.31 | 1 | 2.67 | 0.66 | 21.78 | 21.89 | -0.11 | 0.11 | 0 | 0 |
| gene11082 | https://www.ncbi.nlm.nih.gov/gene/?term=LOC105284824 | 7436 | 224 | 301.24 | 1 | 1.34 | 0.45 | 7.03 | 7.19 | -0.15 | 0.15 | 0 | 0 |
| gene12573 | https://www.ncbi.nlm.nih.gov/gene/?term=LOC105286515 | 3647 | 121 | 331.78 | 1 | 2.74 | 0.83 | 10.48 | 10.65 | -0.17 | 0.17 | -0.01 | 0.01 |
| gene8343 | https://www.ncbi.nlm.nih.gov/gene/?term=LOC105281700 | 2172 | 39 | 179.56 | 1 | 4.6 | 2.56 | 37.71 | 37.89 | -0.17 | 0.17 | 0 | 0 |
| gene10345 | https://www.ncbi.nlm.nih.gov/gene/?term=LOC105283989 | 4444 | 29 | 65.26 1 | 2.25 | 3.45 | 47.94 | 48.12 | -0.18 | 0.18 | 0 | 0 | |
| gene11561 | https://www.ncbi.nlm.nih.gov/gene/?term=LOC105285387 | 11528 | 218 | 189.1 1 | 0.87 | 0.46 | 9.03 | 9.22 | -0.19 | 0.19 | 0 | 0 | |
| gene7245 | https://www.ncbi.nlm.nih.gov/gene/?term=LOC105280410 | 12478 | 265 | 212.37 | 1 | 0.8 | 0.38 | 19.68 | 19.87 | -0.19 | 0.19 | 0 | 0 |
| gene10392 | https://www.ncbi.nlm.nih.gov/gene/?term=LOC105284036 | 3216 | 111 | 345.15 | 2 | 6.22 | 1.8 | 22.59 | 22.79 | -0.2 | 0.2 | -0.01 | 0.01 |
| gene11401 | https://www.ncbi.nlm.nih.gov/gene/?term=LOC105285202 | 1841 | 82 | 445.41 | 1 | 5.43 | 1.22 | 25.28 | 25.48 | -0.2 | 0.2 | -0.01 | 0.01 |
| gene13036 | https://www.ncbi.nlm.nih.gov/gene/?term=LOC105287059 | 4959 | 179 | 360.96 | 1 | 2.02 | 0.56 | 16.67 | 16.87 | -0.2 | 0.2 | -0.01 | 0.01 |
| gene11579 | https://www.ncbi.nlm.nih.gov/gene/?term=LOC105285406 | 13571 | 312 | 229.9 1 | 0.74 | 0.32 | 5.9 | 6.13 | -0.23 | 0.23 | -0.01 | 0.01 | |
| gene2844 | https://www.ncbi.nlm.nih.gov/gene/?term=LOC105275363 | 4330 | 215 | 496.54 | 1 | 2.31 | 0.47 | 13.31 | 13.55 | -0.24 | 0.24 | -0.01 | 0.01 |
| gene6528 | https://www.ncbi.nlm.nih.gov/gene/?term=LOC105279591 | 7222 | 60 | 83.08 1 | 1.38 | 1.67 | 26.03 | 26.28 | -0.25 | 0.25 | 0 | 0 | |
| gene6583 | https://www.ncbi.nlm.nih.gov/gene/?term=LOC105279649 | 7444 | 372 | 499.73 | 1 | 1.34 | 0.27 | 10.77 | 11.02 | -0.25 | 0.25 | -0.01 | 0.01 |
| gene12203 | https://www.ncbi.nlm.nih.gov/gene/?term=LOC105286099 | 5967 | 74 | 124.02 | 1 | 1.68 | 1.35 | 10.44 | 10.7 | -0.27 | 0.27 | 0 | 0 |
| gene5613 | https://www.ncbi.nlm.nih.gov/gene/?term=LOC105278504 | 8524 | 313 | 367.2 1 | 1.17 | 0.32 | 4.41 | 4.68 | -0.27 | 0.27 | -0.01 | 0.01 | |
| gene7166 | https://www.ncbi.nlm.nih.gov/gene/?term=LOC105280326 | 6117 | 260 | 425.04 | 1 | 1.63 | 0.38 | 14.15 | 14.42 | -0.27 | 0.27 | -0.01 | 0.01 |
| gene2846 | https://www.ncbi.nlm.nih.gov/gene/?term=LOC105275374 | 11246 | 548 | 487.28 | 1 | 0.89 | 0.18 | 10.73 | 11.01 | -0.28 | 0.28 | -0.01 | 0.01 |
| gene8600 | https://www.ncbi.nlm.nih.gov/gene/?term=LOC105282001 | 8513 | 197 | 231.41 | 1 | 1.17 | 0.51 | 9.89 | 10.16 | -0.28 | 0.28 | -0.01 | 0.01 |
| gene9177 | https://www.ncbi.nlm.nih.gov/gene/?term=LOC105282646 | 3320 | 113 | 340.36 | 1 | 3.01 | 0.88 | 20.04 | 20.32 | -0.28 | 0.28 | -0.01 | 0.01 |
| gene11275 | https://www.ncbi.nlm.nih.gov/gene/?term=LOC105285055 | 4119 | 268 | 650.64 | 1 | 2.43 | 0.37 | 11.27 | 11.56 | -0.29 | 0.29 | -0.02 | 0.02 |
| gene6286 | https://www.ncbi.nlm.nih.gov/gene/?term=LOC105279298 | 2861 | 138 | 482.35 | 2 | 6.99 | 1.45 | 23.81 | 24.1 | -0.29 | 0.29 | -0.01 | 0.01 |
| gene8814 | https://www.ncbi.nlm.nih.gov/gene/?term=LOC105282240 | 6910 | 204 | 295.22 | 1 | 1.45 | 0.49 | 8.85 | 9.15 | -0.3 | 0.3 | -0.01 | 0.01 |
| gene1198 | https://www.ncbi.nlm.nih.gov/gene/?term=LOC105285507 | 2738 | 90 | 328.71 | 1 | 3.65 | 1.11 | 23.4 | 23.71 | -0.31 | 0.31 | -0.01 | 0.01 |
| gene5746 | https://www.ncbi.nlm.nih.gov/gene/?term=LOC105278666 | 10343 | 497 | 480.52 | 1 | 0.97 | 0.2 | 6.84 | 7.15 | -0.31 | 0.31 | -0.01 | 0.01 |
| gene3107 | https://www.ncbi.nlm.nih.gov/gene/?term=LOC105275650 | 5103 | 197 | 386.05 | 1 | 1.96 | 0.51 | 15.38 | 15.71 | -0.33 | 0.33 | -0.01 | 0.01 |
| gene7707 | https://www.ncbi.nlm.nih.gov/gene/?term=LOC105280953 | 4925 | 133 | 270.05 | 1 | 2.03 | 0.75 | 26.19 | 26.53 | -0.33 | 0.33 | -0.01 | 0.01 |
| gene2266 | https://www.ncbi.nlm.nih.gov/gene/?term=LOC105274699 | 2391 | 65 | 271.85 | 1 | 4.18 | 1.54 | 22.79 | 23.14 | -0.35 | 0.35 | -0.01 | 0.01 |
| gene34 | https://www.ncbi.nlm.nih.gov/gene/?term=LOC105278416 | 7120 | 153 | 214.89 | 1 | 1.4 | 0.65 | 12.92 | 13.27 | -0.35 | 0.35 | -0.01 | 0.01 |
| gene3848 | https://www.ncbi.nlm.nih.gov/gene/?term=LOC105276486 | 5899 | 170 | 288.18 | 1 | 1.7 | 0.59 | 21.62 | 21.98 | -0.37 | 0.37 | -0.01 | 0.01 |
| gene5831 | https://www.ncbi.nlm.nih.gov/gene/?term=LOC105278761 | 2506 | 128 | 510.77 | 1 | 3.99 | 0.78 | 21.72 | 22.12 | -0.39 | 0.39 | -0.02 | 0.02 |
| gene2806 | https://www.ncbi.nlm.nih.gov/gene/?term=LOC105275327 | 14257 | 571 | 400.51 | 1 | 0.7 | 0.18 | 6.14 | 6.54 | -0.4 | 0.4 | -0.02 | 0.02 |
| gene4962 | https://www.ncbi.nlm.nih.gov/gene/?term=LOC105277761 | 4645 | 164 | 353.07 | 1 | 2.15 | 0.61 | 14.72 | 15.12 | -0.4 | 0.4 | -0.01 | 0.01 |
| gene738 | https://www.ncbi.nlm.nih.gov/gene/?term=LOC105280421 | 4927 | 229 | 464.79 | 1 | 2.03 | 0.44 | 14.03 | 14.44 | -0.4 | 0.4 | -0.02 | 0.02 |
| gene2056 | https://www.ncbi.nlm.nih.gov/gene/?term=LOC105274452 | 6584 | 282 | 428.31 | 2 | 3.04 | 0.71 | 12.46 | 12.87 | -0.41 | 0.41 | -0.02 | 0.02 |
| gene10600 | https://www.ncbi.nlm.nih.gov/gene/?term=LOC105284270 | 4837 | 134 | 277.03 | 1 | 2.07 | 0.75 | 11.69 | 12.12 | -0.43 | 0.43 | -0.01 | 0.01 |
| gene7207 | https://www.ncbi.nlm.nih.gov/gene/?term=LOC105280376 | 4659 | 110 | 236.1 1 | 2.15 | 0.91 | 30.27 | 30.71 | -0.45 | 0.45 | -0.01 | 0.01 | |
| gene10531 | https://www.ncbi.nlm.nih.gov/gene/?term=LOC105284190 | 6881 | 203 | 295.02 | 1 | 1.45 | 0.49 | 11.33 | 11.8 | -0.47 | 0.47 | -0.01 | 0.01 |
| gene2271 | https://www.ncbi.nlm.nih.gov/gene/?term=LOC105274706 | 1448 | 58 | 400.55 | 1 | 6.91 | 1.72 | 36.82 | 37.28 | -0.47 | 0.47 | -0.02 | 0.02 |
| gene3470 | https://www.ncbi.nlm.nih.gov/gene/?term=LOC105276059 | 8716 | 175 | 200.78 | 1 | 1.15 | 0.57 | 7.55 | 8.01 | -0.47 | 0.47 | -0.01 | 0.01 |
| gene10606 | https://www.ncbi.nlm.nih.gov/gene/?term=LOC105284273 | 6119 | 278 | 454.32 | 1 | 1.63 | 0.36 | 18.22 | 18.7 | -0.48 | 0.48 | -0.02 | 0.02 |
| gene5877 | https://www.ncbi.nlm.nih.gov/gene/?term=LOC105278807 | 1719 | 54 | 314.14 | 1 | 5.82 | 1.85 | 24.15 | 24.64 | -0.49 | 0.49 | -0.02 | 0.02 |
| gene298 | https://www.ncbi.nlm.nih.gov/gene/?term=LOC105275460 | 2473 | 121 | 489.28 | 1 | 4.04 | 0.83 | 13.07 | 13.58 | -0.51 | 0.51 | -0.03 | 0.03 |
| gene4996 | https://www.ncbi.nlm.nih.gov/gene/?term=LOC105277798 | 5617 | 58 | 103.26 | 1 | 1.78 | 1.72 | 13.2 | 13.71 | -0.51 | 0.51 | -0.01 | 0.01 |
| gene6031 | https://www.ncbi.nlm.nih.gov/gene/?term=LOC105279000 | 5691 | 209 | 367.25 | 1 | 1.76 | 0.48 | 13.23 | 13.74 | -0.51 | 0.51 | -0.02 | 0.02 |
| gene7183 | https://www.ncbi.nlm.nih.gov/gene/?term=LOC105280348 | 3925 | 79 | 201.27 | 1 | 2.55 | 1.27 | 19.74 | 20.25 | -0.51 | 0.51 | -0.01 | 0.01 |
| gene1430 | https://www.ncbi.nlm.nih.gov/gene/?term=LOC105287551 | 6433 | 256 | 397.95 | 1 | 1.55 | 0.39 | 8.96 | 9.5 | -0.53 | 0.53 | -0.02 | 0.02 |
| gene5375 | https://www.ncbi.nlm.nih.gov/gene/?term=LOC105278228 | 8640 | 177 | 204.86 | 1 | 1.16 | 0.56 | 22.94 | 23.49 | -0.55 | 0.55 | -0.01 | 0.01 |
| gene7514 | https://www.ncbi.nlm.nih.gov/gene/?term=LOC105280723 | 4471 | 216 | 483.11 | 1 | 2.24 | 0.46 | 13.02 | 13.57 | -0.55 | 0.55 | -0.03 | 0.03 |

| | | | | | | | | | | | | | |
|---|---|---|---|---|---|---|---|---|---|---|---|---|---|
| gene8729 | https://www.ncbi.nlm.nih.gov/gene/?term=LOC105282137 | 5005 | 209 | 417.58 | 1 | 2 | 0.48 | 15.97 | 16.54 | -0.56 | 0.56 | -0.02 | 0.02 |
| gene12307 | https://www.ncbi.nlm.nih.gov/gene/?term=LOC105286216 | 10313 | 173 | 167.75 | 1 | 0.97 | 0.58 | 16 | 16.58 | -0.58 | 0.58 | -0.01 | 0.01 |
| gene6966 | https://www.ncbi.nlm.nih.gov/gene/?term=LOC105280104 | 3747 | 140 | 373.63 | 1 | 2.67 | 0.71 | 14.7 | 15.28 | -0.58 | 0.58 | -0.02 | 0.02 |
| gene6620 | https://www.ncbi.nlm.nih.gov/gene/?term=LOC105279687 | 17998 | 402 | 223.36 | 1 | 0.56 | 0.25 | 4.4 | 5 | -0.6 | 0.6 | -0.01 | 0.01 |
| gene10107 | https://www.ncbi.nlm.nih.gov/gene/?term=LOC105283712 | 1883 | 36 | 191.18 | 1 | 5.31 | 2.78 | 29.18 | 29.79 | -0.61 | 0.61 | -0.01 | 0.01 |
| gene12439 | https://www.ncbi.nlm.nih.gov/gene/?term=LOC105286357 | 4761 | 94 | 197.44 | 1 | 2.1 | 1.06 | 30.53 | 31.14 | -0.61 | 0.61 | -0.01 | 0.01 |
| gene35 | https://www.ncbi.nlm.nih.gov/gene/?term=LOC105275197 | 2643 | 53 | 200.53 | 1 | 3.78 | 1.89 | 36.78 | 37.42 | -0.64 | 0.64 | -0.01 | 0.01 |
| gene2173 | https://www.ncbi.nlm.nih.gov/gene/?term=LOC105274590 | 4387 | 89 | 202.87 | 2 | 4.56 | 2.25 | 28.89 | 29.55 | -0.66 | 0.66 | -0.01 | 0.01 |
| gene1518 | https://www.ncbi.nlm.nih.gov/gene/?term=LOC105287648 | 2195 | 35 | 159.45 | 1 | 4.56 | 2.86 | 35.74 | 36.45 | -0.71 | 0.71 | -0.01 | 0.01 |
| gene3206 | https://www.ncbi.nlm.nih.gov/gene/?term=LOC105275752 | 3474 | 142 | 408.75 | 1 | 2.88 | 0.7 | 30.85 | 31.58 | -0.73 | 0.73 | -0.03 | 0.03 |
| gene2825 | https://www.ncbi.nlm.nih.gov/gene/?term=LOC105275343 | 5545 | 105 | 189.36 | 1 | 1.8 | 0.95 | 10.15 | 10.91 | -0.77 | 0.77 | -0.01 | 0.01 |
| gene4376 | https://www.ncbi.nlm.nih.gov/gene/?term=LOC105277083 | 4307 | 136 | 315.77 | 1 | 2.32 | 0.74 | 17.99 | 18.79 | -0.8 | 0.8 | -0.03 | 0.03 |
| gene935 | https://www.ncbi.nlm.nih.gov/gene/?term=LOC105282682 | 4368 | 241 | 551.74 | 1 | 2.29 | 0.41 | 12.96 | 13.79 | -0.83 | 0.83 | -0.05 | 0.05 |
| gene11568 | https://www.ncbi.nlm.nih.gov/gene/?term=LOC105285395 | 8086 | 226 | 279.5 1 | 1.24 | 0.44 | 14.03 | 14.87 | -0.84 | 0.84 | -0.02 | 0.02 | |
| gene9330 | https://www.ncbi.nlm.nih.gov/gene/?term=LOC105282825 | 3936 | 98 | 248.98 | 1 | 2.54 | 1.02 | 14.19 | 15.06 | -0.86 | 0.86 | -0.02 | 0.02 |
| gene11801 | https://www.ncbi.nlm.nih.gov/gene/?term=LOC105285656 | 2497 | 137 | 548.66 | 1 | 4 | 0.73 | 8 | 8.86 | -0.87 | 0.87 | -0.05 | 0.05 |
| gene7556 | https://www.ncbi.nlm.nih.gov/gene/?term=LOC105280774 | 2792 | 123 | 440.54 | 1 | 3.58 | 0.81 | 26.02 | 26.91 | -0.89 | 0.89 | -0.04 | 0.04 |
| gene4720 | https://www.ncbi.nlm.nih.gov/gene/?term=LOC105277479 | 1697 | 45 | 265.17 | 1 | 5.89 | 2.22 | 39.86 | 40.77 | -0.91 | 0.91 | -0.02 | 0.02 |
| gene2683 | https://www.ncbi.nlm.nih.gov/gene/?term=LOC105275187 | 5492 | 78 | 142.02 | 1 | 1.82 | 1.28 | 15.12 | 16.04 | -0.92 | 0.92 | -0.01 | 0.01 |
| gene12802 | https://www.ncbi.nlm.nih.gov/gene/?term=LOC105286799 | 3183 | 193 | 606.35 | 2 | 6.28 | 1.04 | 15.46 | 16.4 | -0.94 | 0.94 | -0.06 | 0.06 |
| gene3599 | https://www.ncbi.nlm.nih.gov/gene/?term=LOC105276196 | 2955 | 60 | 203.05 | 1 | 3.38 | 1.67 | 37.09 | 38.05 | -0.97 | 0.97 | -0.02 | 0.02 |
| gene811 | https://www.ncbi.nlm.nih.gov/gene/?term=LOC105281261 | 5614 | 170 | 302.81 | 1 | 1.78 | 0.59 | 13.19 | 14.19 | -1 | 1 | -0.03 | 0.03 |
| gene4225 | https://www.ncbi.nlm.nih.gov/gene/?term=LOC105276917 | 2443 | 30 | 122.8 1 | 4.09 | 3.33 | 32.94 | 33.99 | -1.05 | 1.05 | -0.01 | 0.01 | |
| gene940 | https://www.ncbi.nlm.nih.gov/gene/?term=LOC105282794 | 3578 | 92 | 257.13 | 1 | 2.79 | 1.09 | 14.15 | 15.24 | -1.1 | 1.1 | -0.03 | 0.03 |
| gene4048 | https://www.ncbi.nlm.nih.gov/gene/?term=LOC105276724 | 2955 | 42 | 142.13 | 1 | 3.38 | 2.38 | 10.45 | 11.57 | -1.12 | 1.12 | -0.02 | 0.02 |
| gene10678 | https://www.ncbi.nlm.nih.gov/gene/?term=LOC105284355 | 4031 | 183 | 453.98 | 1 | 2.48 | 0.55 | 17.56 | 18.71 | -1.15 | 1.15 | -0.05 | 0.05 |
| gene10540 | https://www.ncbi.nlm.nih.gov/gene/?term=LOC105284200 | 5267 | 213 | 404.4 1 | 1.9 | 0.47 | 12.38 | 13.55 | -1.16 | 1.16 | -0.05 | 0.05 | |
| gene2833 | https://www.ncbi.nlm.nih.gov/gene/?term=LOC105275352 | 5528 | 41 | 74.17 1 | 1.81 | 2.44 | 22.41 | 23.6 | -1.2 | 1.2 | -0.01 | 0.01 | |
| gene5524 | https://www.ncbi.nlm.nih.gov/gene/?term=LOC105278397 | 5177 | 206 | 397.91 | 1 | 1.93 | 0.49 | 25.22 | 26.43 | -1.21 | 1.21 | -0.05 | 0.05 |
| gene9462 | https://www.ncbi.nlm.nih.gov/gene/?term=LOC105282970 | 1469 | 42 | 285.91 | 1 | 6.81 | 2.38 | 50.96 | 52.17 | -1.21 | 1.21 | -0.03 | 0.03 |
| gene3859 | https://www.ncbi.nlm.nih.gov/gene/?term=LOC105276501 | 5788 | 106 | 183.14 | 1 | 1.73 | 0.94 | 14.91 | 16.17 | -1.26 | 1.26 | -0.02 | 0.02 |
| gene4034 | https://www.ncbi.nlm.nih.gov/gene/?term=LOC105276704 | 2065 | 106 | 513.32 | 2 | 9.69 | 1.89 | 12.65 | 13.91 | -1.26 | 1.26 | -0.06 | 0.06 |
| gene2257 | https://www.ncbi.nlm.nih.gov/gene/?term=LOC105274689 | 3658 | 86 | 235.1 1 | 2.73 | 1.16 | 16.99 | 18.26 | -1.27 | 1.27 | -0.03 | 0.03 | |
| gene10391 | https://www.ncbi.nlm.nih.gov/gene/?term=LOC109611283 | 1994 | 46 | 230.69 | 1 | 5.02 | 2.17 | 34.01 | 35.36 | -1.35 | 1.35 | -0.03 | 0.03 |
| gene3032 | https://www.ncbi.nlm.nih.gov/gene/?term=LOC105275575 | 1713 | 29 | 169.29 | 1 | 5.84 | 3.45 | 12.34 | 13.69 | -1.35 | 1.35 | -0.02 | 0.02 |
| gene5525 | https://www.ncbi.nlm.nih.gov/gene/?term=LOC105278396 | 3090 | 190 | 614.89 | 1 | 3.24 | 0.53 | 25.02 | 26.4 | -1.38 | 1.38 | -0.08 | 0.08 |
| gene4892 | https://www.ncbi.nlm.nih.gov/gene/?term=LOC105277667 | 3898 | 131 | 336.07 | 1 | 2.57 | 0.76 | 26.19 | 27.61 | -1.42 | 1.42 | -0.05 | 0.05 |
| gene12516 | https://www.ncbi.nlm.nih.gov/gene/?term=LOC105286452 | 1529 | 38 | 248.53 | 1 | 6.54 | 2.63 | 51.61 | 53.08 | -1.47 | 1.47 | -0.04 | 0.04 |
| gene2970 | https://www.ncbi.nlm.nih.gov/gene/?term=LOC105275518 | 2340 | 28 | 119.66 | 1 | 4.27 | 3.57 | 37.12 | 38.59 | -1.47 | 1.47 | -0.02 | 0.02 |
| gene7368 | https://www.ncbi.nlm.nih.gov/gene/?term=LOC105280556 | 4105 | 55 | 133.98 | 1 | 2.44 | 1.82 | 17.06 | 18.54 | -1.48 | 1.48 | -0.02 | 0.02 |
| gene8661 | https://www.ncbi.nlm.nih.gov/gene/?term=LOC105282063 | 1926 | 32 | 166.15 | 1 | 5.19 | 3.12 | 32.4 | 33.92 | -1.52 | 1.52 | -0.03 | 0.03 |
| gene10864 | https://www.ncbi.nlm.nih.gov/gene/?term=LOC105284557 | 1666 | 84 | 504.2 1 | 6 | 1.19 | 28.16 | 29.73 | -1.56 | 1.56 | -0.08 | 0.08 | |
| gene5967 | https://www.ncbi.nlm.nih.gov/gene/?term=LOC105278918 | 4393 | 148 | 336.9 1 | 2.28 | 0.68 | 12.61 | 14.19 | -1.58 | 1.58 | -0.05 | 0.05 | |
| gene1884 | https://www.ncbi.nlm.nih.gov/gene/?term=LOC105288072 | 4169 | 83 | 199.09 | 1 | 2.4 | 1.2 | 21.36 | 22.96 | -1.6 | 1.6 | -0.03 | 0.03 |
| gene9336 | https://www.ncbi.nlm.nih.gov/gene/?term=LOC105282834 | 5993 | 141 | 235.27 | 1 | 1.67 | 0.71 | 14.11 | 15.72 | -1.61 | 1.61 | -0.04 | 0.04 |
| gene13033 | https://www.ncbi.nlm.nih.gov/gene/?term=LOC105287063 | 1877 | 56 | 298.35 | 1 | 5.33 | 1.79 | 28.26 | 30.08 | -1.82 | 1.82 | -0.05 | 0.05 |
| gene927 | https://www.ncbi.nlm.nih.gov/gene/?term=LOC105282600 | 3727 | 56 | 150.25 | 1 | 2.68 | 1.79 | 29.88 | 31.8 | -1.92 | 1.92 | -0.03 | 0.03 |
| gene1962 | https://www.ncbi.nlm.nih.gov/gene/?term=LOC105288157 | 1246 | 18 | 144.46 | 1 | 8.03 | 5.56 | 12.41 | 14.36 | -1.95 | 1.95 | -0.03 | 0.03 |
| gene7525 | https://www.ncbi.nlm.nih.gov/gene/?term=LOC105280738 | 4237 | 43 | 101.49 | 1 | 2.36 | 2.33 | 24.72 | 26.71 | -1.99 | 1.99 | -0.02 | 0.02 |
| gene3385 | https://www.ncbi.nlm.nih.gov/gene/?term=LOC105275966 | 3305 | 73 | 220.88 | 1 | 3.03 | 1.37 | 43.5 | 45.58 | -2.08 | 2.08 | -0.05 | 0.05 |
| gene8742 | https://www.ncbi.nlm.nih.gov/gene/?term=LOC105282151 | 2563 | 45 | 175.58 | 2 | 7.8 | 4.44 | 34.53 | 36.64 | -2.11 | 2.11 | -0.04 | 0.04 |
| gene8178 | https://www.ncbi.nlm.nih.gov/gene/?term=LOC105281515 | 8647 | 58 | 67.08 1 | 1.16 | 1.72 | 25.41 | 27.56 | -2.15 | 2.15 | -0.01 | 0.01 | |
| gene388 | https://www.ncbi.nlm.nih.gov/gene/?term=LOC105276538 | 1558 | 21 | 134.79 | 2 | 12.84 | 9.52 | 67.26 | 69.44 | -2.18 | 2.18 | -0.03 | 0.03 |
| gene3817 | https://www.ncbi.nlm.nih.gov/gene/?term=LOC105276449 | 1230 | 57 | 463.41 | 1 | 8.13 | 1.75 | 31.59 | 33.84 | -2.25 | 2.25 | -0.1 | 0.1 |
| gene5573 | https://www.ncbi.nlm.nih.gov/gene/?term=LOC105278454 | 2688 | 42 | 156.25 | 1 | 3.72 | 2.38 | 29.78 | 32.15 | -2.37 | 2.37 | -0.04 | 0.04 |
| gene9647 | https://www.ncbi.nlm.nih.gov/gene/?term=LOC109611233 | 3094 | 42 | 135.75 | 1 | 3.23 | 2.38 | 33.15 | 35.66 | -2.51 | 2.51 | -0.03 | 0.03 |
| gene5052 | https://www.ncbi.nlm.nih.gov/gene/?term=LOC105277856 | 3167 | 39 | 123.14 | 1 | 3.16 | 2.56 | 38.42 | 40.95 | -2.53 | 2.53 | -0.03 | 0.03 |
| gene7568 | https://www.ncbi.nlm.nih.gov/gene/?term=LOC105280786 | 2525 | 119 | 471.29 | 1 | 3.96 | 0.84 | 29.84 | 32.5 | -2.66 | 2.66 | -0.13 | 0.13 |
| gene9478 | https://www.ncbi.nlm.nih.gov/gene/?term=LOC105282991 | 3726 | 61 | 163.71 | 1 | 2.68 | 1.64 | 35.67 | 38.35 | -2.68 | 2.68 | -0.04 | 0.04 |
| gene9648 | https://www.ncbi.nlm.nih.gov/gene/?term=LOC105283183 | 2602 | 36 | 138.36 | 1 | 3.84 | 2.78 | 38.67 | 41.6 | -2.92 | 2.92 | -0.04 | 0.04 |
| gene7165 | https://www.ncbi.nlm.nih.gov/gene/?term=LOC105280327 | 2688 | 109 | 405.51 | 1 | 3.72 | 0.92 | 26.1 | 29.06 | -2.96 | 2.96 | -0.12 | 0.12 |
| gene12572 | https://www.ncbi.nlm.nih.gov/gene/?term=LOC105286514 | 2374 | 47 | 197.98 | 1 | 4.21 | 2.13 | 34.72 | 38.06 | -3.34 | 3.34 | -0.07 | 0.07 |
| gene2907 | https://www.ncbi.nlm.nih.gov/gene/?term=LOC105275447 | 2539 | 43 | 169.36 | 1 | 3.94 | 2.33 | 3.37 | 7.11 | -3.74 | 3.74 | -0.06 | 0.06 |
| gene11619 | https://www.ncbi.nlm.nih.gov/gene/?term=LOC105285454 | 1649 | 29 | 175.86 | 1 | 6.06 | 3.45 | 26.22 | 30.26 | -4.03 | 4.03 | -0.07 | 0.07 |

| | | | | | | | | | | | | | | | |
|---|---|---|---|---|---|---|---|---|---|---|---|---|---|---|---|
| gene5647 | https://www.ncbi.nlm.nih.gov/gene/?term=LOC105278567 | 3118 | 32 | 102.63 | | 1 | 3.21 | 3.12 | 11.48 | 15.59 | -4.11 | 4.11 | -0.04 | 0.04 | |
| gene5714 | https://www.ncbi.nlm.nih.gov/gene/?term=LOC105278729 | 710 | 24 | 338.03 | | 1 | 14.08 | 4.17 | 56.79 | 61.08 | -4.29 | 4.29 | -0.14 | 0.14 | |
| gene12961 | https://www.ncbi.nlm.nih.gov/gene/?term=LOC105286983 | 3094 | 69 | 223.01 | | 2 | 6.46 | 2.9 | 0.97 | 5.84 | -4.87 | 4.87 | -0.11 | 0.11 | |
| gene2824 | https://www.ncbi.nlm.nih.gov/gene/?term=LOC105275337 | 1772 | 64 | 361.17 | | 2 | 11.29 | 3.12 | 29.09 | 37.7 | -8.61 | 8.61 | -0.31 | 0.31 | |
| gene3165 | https://www.ncbi.nlm.nih.gov/gene/?term=LOC105275706 | 2374 | 69 | 290.65 | | 5 | 21.06 | 7.25 | 23.81 | 34.06 | -10.25 | | 10.25 | -0.3 | 0.3 |
| gene7988 | https://www.ncbi.nlm.nih.gov/gene/?term=LOC105281282 | 3237 | 3 | 9.27 | 1 | 3.09 | 33.33 | 23.25 | 65.16 | -41.92 | | 41.92 | -0.04 | 0.04 | |
| gene2593 | https://www.ncbi.nlm.nih.gov/gene/?term=LOC105275070 | 1774 | 5 | 28.18 | 5 | 28.18 | 100 | 10.96 | 81.03 | -70.07 | | 70.07 | -0.2 | 0.2 | |

Directory: OGL File: ogl-Cb_LI_BCphaseB2.vs.RphaseB1.txt

| gene_ID | gene_link | gwidth | #Sites | #per10Kb | #dmSites | #dmsp10kb | %dmSites | %pSite1 | %pSite2 | DMpSite | ADMpSite | DMpNucl | ADMpNucl |
|---|---|---|---|---|---|---|---|---|---|---|---|---|---|
| gene5877 | https://www.ncbi.nlm.nih.gov/gene/?term=LOC105278807 | 1719 | 50 | 290.87 | 1 | 5.82 | 2 | 23.49 | 23.85 | -0.36 | 0.36 | -0.01 | 0.01 |
| gene1470 | https://www.ncbi.nlm.nih.gov/gene/?term=LOC105287597 | 2974 | 190 | 638.87 | 1 | 3.36 | 0.53 | 18.6 | 19.2 | -0.6 | 0.6 | -0.04 | 0.04 |

Directory: OGL File: ogl-Cb_LI_BCphaseB2.vs.RphaseB2.txt

| gene_ID | gene_link | gwidth | #Sites | #per10Kb | #dmSites | #dmsp10kb | %dmSites | %pSite1 | %pSite2 | DMpSite | ADMpSite | DMpNucl | ADMpNucl |
|---|---|---|---|---|---|---|---|---|---|---|---|---|---|
| gene13225 | https://www.ncbi.nlm.nih.gov/gene/?term=LOC105287265 | 1631 | 2 | 12.26 | 1 | 6.13 | 50 | 88.97 | 71.1 | 17.88 | 17.88 | 0.02 | 0.02 |
| gene6576 | https://www.ncbi.nlm.nih.gov/gene/?term=LOC105279639 | 2616 | 3 | 11.47 | 2 | 7.65 | 66.67 | 79.09 | 65.15 | 13.93 | 13.93 | 0.02 | 0.02 |
| gene12853 | https://www.ncbi.nlm.nih.gov/gene/?term=LOC105286857 | 1589 | 2 | 12.59 | 1 | 6.29 | 50 | 69.58 | 57.77 | 11.81 | 11.81 | 0.01 | 0.01 |
| gene13119 | https://www.ncbi.nlm.nih.gov/gene/?term=LOC105287152 | 1525 | 2 | 13.11 | 1 | 6.56 | 50 | 30.95 | 20.38 | 10.57 | 10.57 | 0.01 | 0.01 |
| gene10278 | https://www.ncbi.nlm.nih.gov/gene/?term=LOC105283908 | 902 | 14 | 155.21 | 4 | 44.35 | 28.57 | 48.14 | 38.93 | 9.21 | 9.21 | 0.14 | 0.14 |
| gene4648 | https://www.ncbi.nlm.nih.gov/gene/?term=LOC105277406 | 2653 | 8 | 30.15 | 1 | 3.77 | 12.5 | 50.8 | 42.14 | 8.66 | 8.66 | 0.03 | 0.03 |
| gene6530 | https://www.ncbi.nlm.nih.gov/gene/?term=LOC105279592 | 1123 | 7 | 62.33 | 1 | 8.9 | 14.29 | 23.02 | 14.43 | 8.59 | 8.59 | 0.05 | 0.05 |
| gene12355 | https://www.ncbi.nlm.nih.gov/gene/?term=LOC105286261 | 1301 | 8 | 61.49 | 1 | 7.69 | 12.5 | 76.73 | 68.31 | 8.42 | 8.42 | 0.05 | 0.05 |
| gene12481 | https://www.ncbi.nlm.nih.gov/gene/?term=LOC105286408 | 356 | 3 | 84.27 | 1 | 28.09 | 33.33 | 17.04 | 8.98 | 8.06 | 8.06 | 0.07 | 0.07 |
| gene13178 | https://www.ncbi.nlm.nih.gov/gene/?term=LOC105287216 | 2535 | 6 | 23.67 | 1 | 3.94 | 16.67 | 27.44 | 19.82 | 7.62 | 7.62 | 0.02 | 0.02 |
| gene13131 | https://www.ncbi.nlm.nih.gov/gene/?term=LOC105287168 | 1392 | 8 | 57.47 | 2 | 14.37 | 25 | 60.44 | 53.29 | 7.15 | 7.15 | 0.04 | 0.04 |
| gene8174 | https://www.ncbi.nlm.nih.gov/gene/?term=LOC105281512 | 2154 | 12 | 55.71 | 2 | 9.29 | 16.67 | 33.85 | 26.91 | 6.94 | 6.94 | 0.04 | 0.04 |
| gene9483 | https://www.ncbi.nlm.nih.gov/gene/?term=LOC105282996 | 3739 | 6 | 16.05 | 1 | 2.67 | 16.67 | 23.81 | 17.02 | 6.8 | 6.8 | 0.01 | 0.01 |
| gene2636 | https://www.ncbi.nlm.nih.gov/gene/?term=LOC105275141 | 947 | 10 | 105.6 | 2 | 21.12 | 20 | 26.02 | 19.35 | 6.68 | 6.68 | 0.07 | 0.07 |
| gene11537 | https://www.ncbi.nlm.nih.gov/gene/?term=LOC105285354 | 1747 | 2 | 11.45 | 1 | 5.72 | 50 | 65.6 | 58.97 | 6.63 | 6.63 | 0.01 | 0.01 |
| gene3483 | https://www.ncbi.nlm.nih.gov/gene/?term=LOC105276075 | 1444 | 10 | 69.25 | 1 | 6.93 | 10 | 39.38 | 33.04 | 6.34 | 6.34 | 0.04 | 0.04 |
| gene9845 | https://www.ncbi.nlm.nih.gov/gene/?term=LOC105283410 | 1401 | 9 | 64.24 | 2 | 14.28 | 22.22 | 48.37 | 42.07 | 6.3 | 6.3 | 0.04 | 0.04 |
| gene3916 | https://www.ncbi.nlm.nih.gov/gene/?term=LOC105276569 | 1967 | 15 | 76.26 | 1 | 5.08 | 6.67 | 26.08 | 19.99 | 6.08 | 6.08 | 0.05 | 0.05 |
| gene5162 | https://www.ncbi.nlm.nih.gov/gene/?term=LOC105277980 | 2407 | 29 | 120.48 | 2 | 8.31 | 6.9 | 50.46 | 44.38 | 6.08 | 6.08 | 0.07 | 0.07 |
| gene11426 | https://www.ncbi.nlm.nih.gov/gene/?term=LOC105285229 | 1413 | 5 | 35.39 | 1 | 7.08 | 20 | 23.57 | 17.61 | 5.95 | 5.95 | 0.02 | 0.02 |
| gene11399 | https://www.ncbi.nlm.nih.gov/gene/?term=LOC105285203 | 1029 | 28 | 272.11 | 2 | 19.44 | 7.14 | 30.21 | 24.31 | 5.9 | 5.9 | 0.16 | 0.16 |
| gene11818 | https://www.ncbi.nlm.nih.gov/gene/?term=LOC105285669 | 2185 | 16 | 73.23 | 1 | 4.58 | 6.25 | 26.14 | 20.26 | 5.88 | 5.88 | 0.04 | 0.04 |
| gene1350 | https://www.ncbi.nlm.nih.gov/gene/?term=LOC105287003 | 2212 | 12 | 54.25 | 3 | 13.56 | 25 | 39.02 | 33.3 | 5.71 | 5.71 | 0.03 | 0.03 |
| gene11024 | https://www.ncbi.nlm.nih.gov/gene/?term=LOC105284763 | 4090 | 12 | 29.34 | 1 | 2.44 | 8.33 | 34.19 | 28.79 | 5.4 | 5.4 | 0.02 | 0.02 |

...
... only first 25 lines shown ...

Directory: OGL File: ogl-Cb_LI_RphaseA1.txt

| gene_ID | gene_link | gwidth | #Sites | #per10Kb | %perSite | %pNucl |
|---|---|---|---|---|---|---|
| gene12538 | https://www.ncbi.nlm.nih.gov/gene/?term=LOC105286531 | 372 | 43 | 1155.91 | 36.48 | 4.22 |
| gene211 | https://www.ncbi.nlm.nih.gov/gene/?term=LOC105274534 | 1280 | 100 | 781.25 | 47.61 | 3.72 |
| gene5911 | https://www.ncbi.nlm.nih.gov/gene/?term=LOC105278844 | 943 | 95 | 1007.42 | 36.67 | 3.69 |
| gene13384 | https://www.ncbi.nlm.nih.gov/gene/?term=LOC105287447 | 1877 | 228 | 1214.7 | 29.65 | 3.6 |
| gene12330 | https://www.ncbi.nlm.nih.gov/gene/?term=LOC105286235 | 1501 | 118 | 786.14 | 42.69 | 3.36 |
| gene5309 | https://www.ncbi.nlm.nih.gov/gene/?term=LOC105278160 | 1704 | 158 | 927.23 | 35.33 | 3.28 |
| gene5917 | https://www.ncbi.nlm.nih.gov/gene/?term=LOC105278849 | 823 | 113 | 1373.03 | 23.86 | 3.28 |
| gene12364 | https://www.ncbi.nlm.nih.gov/gene/?term=LOC105286269 | 1403 | 87 | 620.1 | 52.62 | 3.26 |
| gene12746 | https://www.ncbi.nlm.nih.gov/gene/?term=LOC105286737 | 2688 | 366 | 1361.61 | 23.78 | 3.24 |
| gene4878 | https://www.ncbi.nlm.nih.gov/gene/?term=LOC105277648 | 675 | 38 | 562.96 | 57.09 | 3.21 |
| gene1654 | https://www.ncbi.nlm.nih.gov/gene/?term=LOC105287811 | 2285 | 165 | 722.1 | 44.27 | 3.2 |
| gene2797 | https://www.ncbi.nlm.nih.gov/gene/?term=LOC105275314 | 1489 | 97 | 651.44 | 47.77 | 3.11 |
| gene3731 | https://www.ncbi.nlm.nih.gov/gene/?term=LOC105276371 | 865 | 55 | 635.84 | 48.54 | 3.09 |
| gene6078 | https://www.ncbi.nlm.nih.gov/gene/?term=LOC105279052 | 1130 | 122 | 1079.65 | 28.5 | 3.08 |
| gene3719 | https://www.ncbi.nlm.nih.gov/gene/?term=LOC105276360 | 1471 | 155 | 1053.7 | 27.58 | 2.91 |
| gene458 | https://www.ncbi.nlm.nih.gov/gene/?term=LOC105277254 | 1378 | 113 | 820.03 | 35.06 | 2.87 |
| gene5301 | https://www.ncbi.nlm.nih.gov/gene/?term=LOC105278148 | 1881 | 199 | 1057.95 | 27.12 | 2.87 |
| gene1562 | https://www.ncbi.nlm.nih.gov/gene/?term=LOC105287692 | 1491 | 117 | 784.71 | 35.76 | 2.81 |
| gene7527 | https://www.ncbi.nlm.nih.gov/gene/?term=LOC105280739 | 2686 | 145 | 539.84 | 52.04 | 2.81 |
| gene11566 | https://www.ncbi.nlm.nih.gov/gene/?term=LOC105285392 | 1603 | 138 | 860.89 | 32.41 | 2.79 |
| gene4633 | https://www.ncbi.nlm.nih.gov/gene/?term=LOC105277389 | 2426 | 167 | 688.38 | 40.41 | 2.78 |
| gene8208 | https://www.ncbi.nlm.nih.gov/gene/?term=LOC105281548 | 1240 | 97 | 782.26 | 34.99 | 2.74 |
| gene9471 | https://www.ncbi.nlm.nih.gov/gene/?term=LOC105282986 | 1605 | 121 | 753.89 | 36.41 | 2.74 |
| gene11569 | https://www.ncbi.nlm.nih.gov/gene/?term=LOC105285394 | 1311 | 53 | 404.27 | 67.05 | 2.71 |

...
... only first 25 lines shown ...

Directory: OGL File: ogl-Cb_LI_RphaseA1.vs.RphaseA2.txt

| gene_ID | gene_link | gwidth | #Sites | #per10Kb | #dmSites | #dmsp10kb | %dmSites | %pSite1 | %pSite2 | DMpSite | ADMpSite | DMpNucl | ADMpNucl |
|---|---|---|---|---|---|---|---|---|---|---|---|---|---|
| gene4630 | https://www.ncbi.nlm.nih.gov/gene/?term=LOC105277380 | 1893 | 5 | 26.41 | 1 | 5.28 | 20 | 48.18 | 35.31 | 12.87 | 12.87 | 0.03 | 0.03 |
| gene4631 | https://www.ncbi.nlm.nih.gov/gene/?term=LOC105277379 | 1348 | 5 | 37.09 | 1 | 7.42 | 20 | 51.95 | 41.8 | 10.16 | 10.16 | 0.04 | 0.04 |
| gene7684 | https://www.ncbi.nlm.nih.gov/gene/?term=LOC105280926 | 1727 | 9 | 52.11 | 1 | 5.79 | 11.11 | 57.47 | 49.81 | 7.66 | 7.66 | 0.04 | 0.04 |
| gene5695 | https://www.ncbi.nlm.nih.gov/gene/?term=LOC105278607 | 4140 | 22 | 53.14 | 2 | 4.83 | 9.09 | 16.79 | 9.48 | 7.31 | 7.31 | 0.04 | 0.04 |
| gene266 | https://www.ncbi.nlm.nih.gov/gene/?term=LOC105275133 | 1445 | 3 | 20.76 | 1 | 6.92 | 33.33 | 50.67 | 43.59 | 7.08 | 7.08 | 0.01 | 0.01 |
| gene6423 | https://www.ncbi.nlm.nih.gov/gene/?term=LOC105279461 | 1638 | 21 | 128.21 | 1 | 6.11 | 4.76 | 39.13 | 33.51 | 5.62 | 5.62 | 0.07 | 0.07 |
| gene5374 | https://www.ncbi.nlm.nih.gov/gene/?term=LOC105278227 | 2529 | 6 | 23.72 | 1 | 3.95 | 16.67 | 77.84 | 72.62 | 5.22 | 5.22 | 0.01 | 0.01 |
| gene477 | https://www.ncbi.nlm.nih.gov/gene/?term=LOC105277498 | 1683 | 7 | 41.59 | 1 | 5.94 | 14.29 | 28.69 | 23.62 | 5.07 | 5.07 | 0.02 | 0.02 |
| gene11035 | https://www.ncbi.nlm.nih.gov/gene/?term=LOC105284773 | 3419 | 5 | 14.62 | 1 | 2.92 | 20 | 8.77 | 3.9 | 4.87 | 4.87 | 0.01 | 0.01 |
| gene1924 | https://www.ncbi.nlm.nih.gov/gene/?term=LOC105288116 | 1820 | 10 | 54.95 | 1 | 5.49 | 10 | 7.47 | 3 | 4.48 | 4.48 | 0.02 | 0.02 |
| gene11800 | https://www.ncbi.nlm.nih.gov/gene/?term=LOC105285653 | 2693 | 13 | 48.27 | 1 | 3.71 | 7.69 | 56.71 | 52.38 | 4.32 | 4.32 | 0.02 | 0.02 |
| gene224 | https://www.ncbi.nlm.nih.gov/gene/?term=LOC109611562 | 1010 | 5 | 49.5 | 1 | 9.9 | 20 | 5.71 | 1.55 | 4.17 | 4.17 | 0.02 | 0.02 |
| gene9283 | https://www.ncbi.nlm.nih.gov/gene/?term=LOC105282769 | 2782 | 43 | 154.57 | 3 | 10.78 | 6.98 | 51.3 | 47.2 | 4.11 | 4.11 | 0.06 | 0.06 |
| gene7190 | https://www.ncbi.nlm.nih.gov/gene/?term=LOC105280355 | 1026 | 14 | 136.45 | 1 | 9.75 | 7.14 | 6.85 | 2.9 | 3.95 | 3.95 | 0.05 | 0.05 |
| gene4999 | https://www.ncbi.nlm.nih.gov/gene/?term=LOC105277805 | 6695 | 15 | 22.4 | 1 | 1.49 | 6.67 | 55.88 | 52 | 3.88 | 3.88 | 0.01 | 0.01 |
| gene9545 | https://www.ncbi.nlm.nih.gov/gene/?term=LOC105283071 | 1986 | 9 | 45.32 | 1 | 5.04 | 11.11 | 22.29 | 18.66 | 3.63 | 3.63 | 0.02 | 0.02 |
| gene4119 | https://www.ncbi.nlm.nih.gov/gene/?term=LOC105276806 | 1608 | 27 | 167.91 | 3 | 18.66 | 11.11 | 40.05 | 36.43 | 3.62 | 3.62 | 0.06 | 0.06 |
| gene4409 | https://www.ncbi.nlm.nih.gov/gene/?term=LOC105277140 | 1157 | 24 | 207.43 | 1 | 8.64 | 4.17 | 8.39 | 5.09 | 3.3 | 3.3 | 0.07 | 0.07 |
| gene7949 | https://www.ncbi.nlm.nih.gov/gene/?term=LOC105281206 | 1405 | 36 | 256.23 | 2 | 14.23 | 5.56 | 48.07 | 44.81 | 3.26 | 3.26 | 0.08 | 0.08 |
| gene11185 | https://www.ncbi.nlm.nih.gov/gene/?term=LOC105284939 | 1723 | 16 | 92.86 | 1 | 5.8 | 6.25 | 20.86 | 17.65 | 3.22 | 3.22 | 0.03 | 0.03 |
| gene11671 | https://www.ncbi.nlm.nih.gov/gene/?term=LOC105285514 | 1273 | 20 | 157.11 | 1 | 7.86 | 5 | 44.44 | 41.28 | 3.16 | 3.16 | 0.05 | 0.05 |
| gene8267 | https://www.ncbi.nlm.nih.gov/gene/?term=LOC105281612 | 1014 | 12 | 118.34 | 1 | 9.86 | 8.33 | 22.91 | 19.75 | 3.16 | 3.16 | 0.04 | 0.04 |
| gene265 | https://www.ncbi.nlm.nih.gov/gene/?term=LOC105275144 | 2128 | 42 | 197.37 | 4 | 18.8 | 9.52 | 32.47 | 29.32 | 3.15 | 3.15 | 0.06 | 0.06 |
| gene8113 | https://www.ncbi.nlm.nih.gov/gene/?term=LOC105281419 | 2170 | 25 | 115.21 | 2 | 9.22 | 8 | 7.95 | 4.92 | 3.03 | 3.03 | 0.03 | 0.03 |

...
... only first 25 lines shown ...

Directory: OGL File: ogl-Cb_LI_RphaseA1.vs.RphaseB1.txt

| gene_ID | gene_link | gwidth | #Sites | #per10Kb | #dmSites | #dmsp10kb | %dmSites | %pSite1 | %pSite2 | DMpSite | ADMpSite | DMpNucl | ADMpNucl |
|---|---|---|---|---|---|---|---|---|---|---|---|---|---|
| gene2593 | https://www.ncbi.nlm.nih.gov/gene/?term=LOC105275070 | 1774 | 11 | 62.01 | 3 | 16.91 | 27.27 | 69.15 | 25.28 | 43.86 | 43.86 | 0.27 | 0.27 |
| gene3165 | https://www.ncbi.nlm.nih.gov/gene/?term=LOC105275706 | 2374 | 63 | 265.37 | 3 | 12.64 | 4.76 | 31.1 | 21.89 | 9.2 | 9.2 | 0.24 | 0.24 |
| gene2837 | https://www.ncbi.nlm.nih.gov/gene/?term=LOC105275355 | 2872 | 164 | 571.03 | 1 | 3.48 | 0.61 | 27.39 | 23.06 | 4.33 | 4.33 | 0.25 | 0.25 |
| gene5714 | https://www.ncbi.nlm.nih.gov/gene/?term=LOC105278729 | 710 | 16 | 225.35 | 1 | 14.08 | 6.25 | 55.81 | 51.9 | 3.91 | 3.91 | 0.09 | 0.09 |
| gene9478 | https://www.ncbi.nlm.nih.gov/gene/?term=LOC105282991 | 3726 | 58 | 155.66 | 1 | 2.68 | 1.72 | 31.92 | 28.27 | 3.65 | 3.65 | 0.06 | 0.06 |
| gene2856 | https://www.ncbi.nlm.nih.gov/gene/?term=LOC105275384 | 1987 | 88 | 442.88 | 1 | 5.03 | 1.14 | 36.35 | 32.79 | 3.56 | 3.56 | 0.16 | 0.16 |
| gene13033 | https://www.ncbi.nlm.nih.gov/gene/?term=LOC105287063 | 1877 | 49 | 261.05 | 1 | 5.33 | 2.04 | 32.74 | 30.05 | 2.68 | 2.68 | 0.07 | 0.07 |
| gene5877 | https://www.ncbi.nlm.nih.gov/gene/?term=LOC105278807 | 1719 | 51 | 296.68 | 1 | 5.82 | 1.96 | 27.19 | 24.91 | 2.27 | 2.27 | 0.07 | 0.07 |
| gene3671 | https://www.ncbi.nlm.nih.gov/gene/?term=LOC105276272 | 1966 | 44 | 223.8 | 1 | 5.09 | 2.27 | 27.22 | 24.96 | 2.26 | 2.26 | 0.05 | 0.05 |
| gene1855 | https://www.ncbi.nlm.nih.gov/gene/?term=LOC105288036 | 2387 | 90 | 377.04 | 1 | 4.19 | 1.11 | 28.76 | 26.75 | 2.02 | 2.02 | 0.08 | 0.08 |
| gene9189 | https://www.ncbi.nlm.nih.gov/gene/?term=LOC105282652 | 4699 | 84 | 178.76 | 1 | 2.13 | 1.19 | 16.38 | 14.61 | 1.78 | 1.78 | 0.03 | 0.03 |
| gene12516 | https://www.ncbi.nlm.nih.gov/gene/?term=LOC105286452 | 1529 | 38 | 248.53 | 1 | 6.54 | 2.63 | 49.76 | 48.05 | 1.71 | 1.71 | 0.04 | 0.04 |
| gene8178 | https://www.ncbi.nlm.nih.gov/gene/?term=LOC105281515 | 8647 | 60 | 69.39 | 1 | 1.16 | 1.67 | 29.73 | 28.04 | 1.69 | 1.69 | 0.01 | 0.01 |
| gene4023 | https://www.ncbi.nlm.nih.gov/gene/?term=LOC105276692 | 1759 | 36 | 204.66 | 1 | 5.69 | 2.78 | 23.9 | 22.4 | 1.5 | 1.5 | 0.03 | 0.03 |
| gene927 | https://www.ncbi.nlm.nih.gov/gene/?term=LOC105282600 | 3727 | 60 | 160.99 | 1 | 2.68 | 1.67 | 27.81 | 26.32 | 1.49 | 1.49 | 0.02 | 0.02 |
| gene1884 | https://www.ncbi.nlm.nih.gov/gene/?term=LOC105288072 | 4169 | 82 | 196.69 | 1 | 2.4 | 1.22 | 24.39 | 23.08 | 1.31 | 1.31 | 0.03 | 0.03 |
| gene11819 | https://www.ncbi.nlm.nih.gov/gene/?term=LOC105285671 | 2366 | 100 | 422.65 | 1 | 4.23 | 1 | 31.41 | 30.12 | 1.29 | 1.29 | 0.05 | 0.05 |
| gene4641 | https://www.ncbi.nlm.nih.gov/gene/?term=LOC105277398 | 1937 | 73 | 376.87 | 1 | 5.16 | 1.37 | 36.14 | 34.87 | 1.27 | 1.27 | 0.05 | 0.05 |
| gene10091 | https://www.ncbi.nlm.nih.gov/gene/?term=LOC105283697 | 4988 | 154 | 308.74 | 1 | 2 | 0.65 | 16.48 | 15.36 | 1.12 | 1.12 | 0.03 | 0.03 |
| gene9336 | https://www.ncbi.nlm.nih.gov/gene/?term=LOC105282834 | 5993 | 134 | 223.59 | 1 | 1.67 | 0.75 | 14.48 | 13.37 | 1.1 | 1.1 | 0.02 | 0.02 |
| gene2683 | https://www.ncbi.nlm.nih.gov/gene/?term=LOC105275187 | 5492 | 73 | 132.92 | 1 | 1.82 | 1.37 | 14.7 | 13.62 | 1.08 | 1.08 | 0.01 | 0.01 |
| gene3817 | https://www.ncbi.nlm.nih.gov/gene/?term=LOC105276449 | 1230 | 48 | 390.24 | 1 | 8.13 | 2.08 | 25.64 | 24.6 | 1.04 | 1.04 | 0.04 | 0.04 |
| gene298 | https://www.ncbi.nlm.nih.gov/gene/?term=LOC105275460 | 2473 | 126 | 509.5 | 1 | 4.04 | 0.79 | 13.41 | 12.43 | 0.98 | 0.98 | 0.05 | 0.05 |
| gene652 | https://www.ncbi.nlm.nih.gov/gene/?term=LOC105279528 | 5806 | 39 | 67.17 | 1 | 1.72 | 2.56 | 22.38 | 21.58 | 0.8 | 0.8 | 0.01 | 0.01 |
| gene10107 | https://www.ncbi.nlm.nih.gov/gene/?term=LOC105283712 | 1883 | 36 | 191.18 | 1 | 5.31 | 2.78 | 27.63 | 26.89 | 0.74 | 0.74 | 0.01 | 0.01 |
| gene5831 | https://www.ncbi.nlm.nih.gov/gene/?term=LOC105278761 | 2506 | 115 | 458.9 | 2 | 7.98 | 1.74 | 22.22 | 21.51 | 0.71 | 0.71 | 0.03 | 0.03 |
| gene940 | https://www.ncbi.nlm.nih.gov/gene/?term=LOC105282794 | 3578 | 93 | 259.92 | 1 | 2.79 | 1.08 | 14.3 | 13.63 | 0.67 | 0.67 | 0.02 | 0.02 |
| gene10540 | https://www.ncbi.nlm.nih.gov/gene/?term=LOC105284200 | 5267 | 209 | 396.81 | 1 | 1.9 | 0.48 | 13.12 | 12.47 | 0.65 | 0.65 | 0.03 | 0.03 |
| gene4892 | https://www.ncbi.nlm.nih.gov/gene/?term=LOC105277667 | 3898 | 121 | 310.42 | 1 | 2.57 | 0.83 | 29.02 | 28.4 | 0.62 | 0.62 | 0.02 | 0.02 |
| gene7600 | https://www.ncbi.nlm.nih.gov/gene/?term=LOC105280819 | 1615 | 41 | 253.87 | 1 | 6.19 | 2.44 | 39.36 | 38.88 | 0.48 | 0.48 | 0.01 | 0.01 |
| gene1147 | https://www.ncbi.nlm.nih.gov/gene/?term=LOC105284954 | 4022 | 118 | 293.39 | 1 | 2.49 | 0.85 | 11.22 | 10.78 | 0.44 | 0.44 | 0.01 | 0.01 |
| gene13036 | https://www.ncbi.nlm.nih.gov/gene/?term=LOC105287059 | 4959 | 180 | 362.98 | 1 | 2.02 | 0.56 | 17.65 | 17.23 | 0.42 | 0.42 | 0.02 | 0.02 |
| gene12441 | https://www.ncbi.nlm.nih.gov/gene/?term=LOC105286355 | 3252 | 149 | 458.18 | 1 | 3.08 | 0.67 | 21.45 | 21.08 | 0.38 | 0.38 | 0.02 | 0.02 |
| gene10222 | https://www.ncbi.nlm.nih.gov/gene/?term=LOC105283838 | 1748 | 30 | 171.62 | 1 | 5.72 | 3.33 | 33.06 | 32.69 | 0.37 | 0.37 | 0.01 | 0.01 |
| gene10243 | https://www.ncbi.nlm.nih.gov/gene/?term=LOC105283862 | 3239 | 43 | 132.76 | 1 | 3.09 | 2.33 | 29.28 | 28.91 | 0.37 | 0.37 | 0 | 0 |
| gene5101 | https://www.ncbi.nlm.nih.gov/gene/?term=LOC105277903 | 1813 | 36 | 198.57 | 1 | 5.52 | 2.78 | 56.09 | 55.74 | 0.35 | 0.35 | 0.01 | 0.01 |
| gene2844 | https://www.ncbi.nlm.nih.gov/gene/?term=LOC105275363 | 4330 | 218 | 503.46 | 1 | 2.31 | 0.46 | 13.23 | 12.92 | 0.3 | 0.3 | 0.02 | 0.02 |
| gene7247 | https://www.ncbi.nlm.nih.gov/gene/?term=LOC105280407 | 10486 | 226 | 215.53 | 1 | 0.95 | 0.44 | 15.61 | 15.31 | 0.3 | 0.3 | 0.01 | 0.01 |
| gene10654 | https://www.ncbi.nlm.nih.gov/gene/?term=LOC105284329 | 4027 | 71 | 176.31 | 1 | 2.48 | 1.41 | 19.25 | 18.99 | 0.26 | 0.26 | 0 | 0 |
| gene6455 | https://www.ncbi.nlm.nih.gov/gene/?term=LOC105279500 | 2612 | 31 | 118.68 | 1 | 3.83 | 3.23 | 47.76 | 47.54 | 0.22 | 0.22 | 0 | 0 |
| gene10345 | https://www.ncbi.nlm.nih.gov/gene/?term=LOC105283989 | 4444 | 29 | 65.26 | 1 | 2.25 | 3.45 | 49.14 | 48.92 | 0.21 | 0.21 | 0 | 0 |
| gene10520 | https://www.ncbi.nlm.nih.gov/gene/?term=LOC105284184 | 2451 | 92 | 375.36 | 1 | 4.08 | 1.09 | 31.61 | 31.4 | 0.21 | 0.21 | 0.01 | 0.01 |
| gene7183 | https://www.ncbi.nlm.nih.gov/gene/?term=LOC105280348 | 3925 | 79 | 201.27 | 1 | 2.55 | 1.27 | 17.02 | 16.83 | 0.19 | 0.19 | 0 | 0 |
| gene2668 | https://www.ncbi.nlm.nih.gov/gene/?term=LOC105275170 | 12496 | 348 | 278.49 | 1 | 0.8 | 0.29 | 1.95 | 1.78 | 0.18 | 0.18 | 0 | 0 |
| gene2846 | https://www.ncbi.nlm.nih.gov/gene/?term=LOC105275374 | 11246 | 493 | 438.38 | 1 | 0.89 | 0.2 | 9.9 | 9.75 | 0.16 | 0.16 | 0.01 | 0.01 |
| gene9327 | https://www.ncbi.nlm.nih.gov/gene/?term=LOC105282821 | 7793 | 162 | 207.88 | 1 | 1.28 | 0.62 | 13.28 | 13.13 | 0.16 | 0.16 | 0 | 0 |
| gene12978 | https://www.ncbi.nlm.nih.gov/gene/?term=LOC105286997 | 2858 | 142 | 496.85 | 1 | 3.5 | 0.7 | 11.15 | 11.02 | 0.14 | 0.14 | 0.01 | 0.01 |
| gene7556 | https://www.ncbi.nlm.nih.gov/gene/?term=LOC105280774 | 2792 | 132 | 472.78 | 1 | 3.58 | 0.76 | 25.2 | 25.1 | 0.1 | 0.1 | 0 | 0 |
| gene10606 | https://www.ncbi.nlm.nih.gov/gene/?term=LOC105284273 | 6119 | 261 | 426.54 | 1 | 1.63 | 0.38 | 19.09 | 19 | 0.08 | 0.08 | 0 | 0 |
| gene8600 | https://www.ncbi.nlm.nih.gov/gene/?term=LOC105282001 | 8513 | 189 | 222.01 | 1 | 1.17 | 0.53 | 10.4 | 10.32 | 0.08 | 0.08 | 0 | 0 |
| gene2806 | https://www.ncbi.nlm.nih.gov/gene/?term=LOC105275327 | 14257 | 529 | 371.05 | 1 | 0.7 | 0.19 | 6.66 | 6.63 | 0.03 | 0.03 | 0 | 0 |
| gene2748 | https://www.ncbi.nlm.nih.gov/gene/?term=LOC105275261 | 3845 | 55 | 143.04 | 1 | 2.6 | 1.82 | 38.35 | 38.32 | 0.02 | 0.02 | 0 | 0 |
| gene7368 | https://www.ncbi.nlm.nih.gov/gene/?term=LOC105280556 | 4105 | 65 | 158.34 | 1 | 2.44 | 1.54 | 14.12 | 14.11 | 0.01 | 0.01 | 0 | 0 |
| gene11583 | https://www.ncbi.nlm.nih.gov/gene/?term=LOC105285414 | 8192 | 189 | 230.71 | 1 | 1.22 | 0.53 | 8.96 | 8.96 | 0 | 0 | 0 | 0 |
| gene10879 | https://www.ncbi.nlm.nih.gov/gene/?term=LOC105284574 | 2423 | 50 | 206.36 | 1 | 4.13 | 2 | 52.18 | 52.2 | -0.01 | 0.01 | 0 | 0 |
| gene3470 | https://www.ncbi.nlm.nih.gov/gene/?term=LOC105276059 | 8716 | 161 | 184.72 | 1 | 1.15 | 0.62 | 9.23 | 9.23 | -0.01 | 0.01 | 0 | 0 |
| gene5746 | https://www.ncbi.nlm.nih.gov/gene/?term=LOC105278666 | 10343 | 473 | 457.31 | 1 | 0.97 | 0.21 | 6.89 | 6.9 | -0.01 | 0.01 | 0 | 0 |
| gene5375 | https://www.ncbi.nlm.nih.gov/gene/?term=LOC105278228 | 8640 | 170 | 196.76 | 1 | 1.16 | 0.59 | 23.53 | 23.55 | -0.02 | 0.02 | 0 | 0 |
| gene9326 | https://www.ncbi.nlm.nih.gov/gene/?term=LOC105282818 | 16612 | 210 | 126.41 | 1 | 0.6 | 0.48 | 13.96 | 13.99 | -0.03 | 0.03 | 0 | 0 |
| gene12573 | https://www.ncbi.nlm.nih.gov/gene/?term=LOC105286515 | 3647 | 112 | 307.1 | 1 | 2.74 | 0.89 | 11.12 | 11.16 | -0.04 | 0.04 | 0 | 0 |

| | | | | | | | | | | | | | |
|---|---|---|---|---|---|---|---|---|---|---|---|---|---|
| gene2443 | https://www.ncbi.nlm.nih.gov/gene/?term=LOC105274895 | 10433 | 409 | 392.03 | 1 | 0.96 | 0.24 | 4.52 | 4.57 | -0.04 | 0.04 | 0 | 0 |
| gene7249 | https://www.ncbi.nlm.nih.gov/gene/?term=LOC105280422 | 1775 | 66 | 371.83 | 1 | 5.63 | 1.52 | 15.99 | 16.04 | -0.05 | 0.05 | 0 | 0 |
| gene11275 | https://www.ncbi.nlm.nih.gov/gene/?term=LOC105285055 | 4119 | 251 | 609.37 | 1 | 2.43 | 0.4 | 10.55 | 10.62 | -0.07 | 0.07 | 0 | 0 |
| gene1220 | https://www.ncbi.nlm.nih.gov/gene/?term=LOC105285742 | 3453 | 109 | 315.67 | 1 | 2.9 | 0.92 | 16.97 | 17.05 | -0.09 | 0.09 | 0 | 0 |
| gene6620 | https://www.ncbi.nlm.nih.gov/gene/?term=LOC105279687 | 17998 | 433 | 240.58 | 1 | 0.56 | 0.23 | 4.48 | 4.58 | -0.1 | 0.1 | 0 | 0 |
| gene53 | https://www.ncbi.nlm.nih.gov/gene/?term=LOC105277148 | 19283 | 305 | 158.17 | 1 | 0.52 | 0.33 | 4.34 | 4.47 | -0.13 | 0.13 | 0 | 0 |
| gene5052 | https://www.ncbi.nlm.nih.gov/gene/?term=LOC105277856 | 3167 | 33 | 104.2 | 1 | 3.16 | 3.03 | 39.38 | 39.53 | -0.15 | 0.15 | 0 | 0 |
| gene11561 | https://www.ncbi.nlm.nih.gov/gene/?term=LOC105285387 | 11528 | 219 | 189.97 | 1 | 0.87 | 0.46 | 9.39 | 9.54 | -0.16 | 0.16 | 0 | 0 |
| gene292 | https://www.ncbi.nlm.nih.gov/gene/?term=LOC105275398 | 2825 | 83 | 293.81 | 1 | 3.54 | 1.2 | 20.15 | 20.32 | -0.16 | 0.16 | 0 | 0 |
| gene2271 | https://www.ncbi.nlm.nih.gov/gene/?term=LOC105274706 | 1448 | 51 | 352.21 | 1 | 6.91 | 1.96 | 37.35 | 37.52 | -0.17 | 0.17 | -0.01 | 0.01 |
| gene3885 | https://www.ncbi.nlm.nih.gov/gene/?term=LOC105276527 | 7928 | 336 | 423.81 | 1 | 1.26 | 0.3 | 8.31 | 8.48 | -0.17 | 0.17 | -0.01 | 0.01 |
| gene8274 | https://www.ncbi.nlm.nih.gov/gene/?term=LOC105281624 | 5425 | 216 | 398.16 | 1 | 1.84 | 0.46 | 9.81 | 9.99 | -0.18 | 0.18 | -0.01 | 0.01 |
| gene1632 | https://www.ncbi.nlm.nih.gov/gene/?term=LOC105287764 | 18061 | 398 | 220.36 | 1 | 0.55 | 0.25 | 3.49 | 3.69 | -0.19 | 0.19 | 0 | 0 |
| gene8814 | https://www.ncbi.nlm.nih.gov/gene/?term=LOC105282240 | 6910 | 188 | 272.07 | 1 | 1.45 | 0.53 | 8.71 | 8.93 | -0.22 | 0.22 | -0.01 | 0.01 |
| gene10018 | https://www.ncbi.nlm.nih.gov/gene/?term=LOC105283607 | 4414 | 219 | 496.15 | 1 | 2.27 | 0.46 | 13.97 | 14.21 | -0.23 | 0.23 | -0.01 | 0.01 |
| gene10600 | https://www.ncbi.nlm.nih.gov/gene/?term=LOC105284270 | 4837 | 103 | 212.94 | 1 | 2.07 | 0.97 | 12.43 | 12.66 | -0.23 | 0.23 | 0 | 0 |
| gene13114 | https://www.ncbi.nlm.nih.gov/gene/?term=LOC105287147 | 7609 | 415 | 545.41 | 1 | 1.31 | 0.24 | 8.59 | 8.85 | -0.26 | 0.26 | -0.01 | 0.01 |
| gene12802 | https://www.ncbi.nlm.nih.gov/gene/?term=LOC105286799 | 3183 | 178 | 559.22 | 1 | 3.14 | 0.56 | 16.8 | 17.1 | -0.29 | 0.29 | -0.02 | 0.02 |
| gene6286 | https://www.ncbi.nlm.nih.gov/gene/?term=LOC105279298 | 2861 | 125 | 436.91 | 2 | 6.99 | 1.6 | 23.63 | 23.91 | -0.29 | 0.29 | -0.01 | 0.01 |
| gene10549 | https://www.ncbi.nlm.nih.gov/gene/?term=LOC105284210 | 7145 | 348 | 487.05 | 2 | 2.8 | 0.57 | 9.98 | 10.29 | -0.31 | 0.31 | -0.01 | 0.01 |
| gene5525 | https://www.ncbi.nlm.nih.gov/gene/?term=LOC105278396 | 3090 | 186 | 601.94 | 1 | 3.24 | 0.54 | 24.12 | 24.45 | -0.33 | 0.33 | -0.02 | 0.02 |
| gene9330 | https://www.ncbi.nlm.nih.gov/gene/?term=LOC105282825 | 3936 | 96 | 243.9 | 1 | 2.54 | 1.04 | 14.55 | 14.91 | -0.36 | 0.36 | -0.01 | 0.01 |
| gene4376 | https://www.ncbi.nlm.nih.gov/gene/?term=LOC105277083 | 4307 | 120 | 278.62 | 1 | 2.32 | 0.83 | 19.12 | 19.51 | -0.39 | 0.39 | -0.01 | 0.01 |
| gene5524 | https://www.ncbi.nlm.nih.gov/gene/?term=LOC105278397 | 5177 | 205 | 395.98 | 1 | 1.93 | 0.49 | 24.01 | 24.4 | -0.39 | 0.39 | -0.02 | 0.02 |
| gene6031 | https://www.ncbi.nlm.nih.gov/gene/?term=LOC105279000 | 5691 | 210 | 369 | 1 | 1.76 | 0.48 | 11.92 | 12.31 | -0.4 | 0.4 | -0.01 | 0.01 |
| gene6583 | https://www.ncbi.nlm.nih.gov/gene/?term=LOC105279649 | 7444 | 364 | 488.98 | 1 | 1.34 | 0.27 | 10.46 | 10.89 | -0.43 | 0.43 | -0.02 | 0.02 |
| gene10678 | https://www.ncbi.nlm.nih.gov/gene/?term=LOC105284355 | 4031 | 173 | 429.17 | 1 | 2.48 | 0.58 | 18.28 | 18.72 | -0.44 | 0.44 | -0.02 | 0.02 |
| gene12307 | https://www.ncbi.nlm.nih.gov/gene/?term=LOC105286216 | 10313 | 186 | 180.35 | 1 | 0.97 | 0.54 | 14.88 | 15.35 | -0.47 | 0.47 | -0.01 | 0.01 |
| gene5298 | https://www.ncbi.nlm.nih.gov/gene/?term=LOC105278145 | 4060 | 253 | 623.15 | 3 | 7.39 | 1.19 | 10.61 | 11.09 | -0.49 | 0.49 | -0.03 | 0.03 |
| gene9647 | https://www.ncbi.nlm.nih.gov/gene/?term=LOC109611233 | 3094 | 33 | 106.66 | 1 | 3.23 | 3.03 | 41.64 | 42.16 | -0.52 | 0.52 | -0.01 | 0.01 |
| gene9648 | https://www.ncbi.nlm.nih.gov/gene/?term=LOC105283183 | 2602 | 31 | 119.14 | 1 | 3.84 | 3.23 | 44.33 | 44.88 | -0.55 | 0.55 | -0.01 | 0.01 |
| gene10326 | https://www.ncbi.nlm.nih.gov/gene/?term=LOC105283962 | 3042 | 132 | 433.93 | 1 | 3.29 | 0.76 | 8.06 | 8.65 | -0.59 | 0.59 | -0.03 | 0.03 |
| gene3107 | https://www.ncbi.nlm.nih.gov/gene/?term=LOC105275650 | 5103 | 180 | 352.73 | 1 | 1.96 | 0.56 | 15.71 | 16.3 | -0.59 | 0.59 | -0.02 | 0.02 |
| gene5646 | https://www.ncbi.nlm.nih.gov/gene/?term=LOC105278534 | 7475 | 198 | 264.88 | 1 | 1.34 | 0.51 | 17.1 | 17.7 | -0.6 | 0.6 | -0.02 | 0.02 |
| gene7166 | https://www.ncbi.nlm.nih.gov/gene/?term=LOC105280326 | 6117 | 249 | 407.06 | 1 | 1.63 | 0.4 | 14.56 | 15.24 | -0.68 | 0.68 | -0.03 | 0.03 |
| gene6237 | https://www.ncbi.nlm.nih.gov/gene/?term=LOC105279244 | 1484 | 25 | 168.46 | 1 | 6.74 | 4 | 45.17 | 45.86 | -0.69 | 0.69 | -0.01 | 0.01 |
| gene10844 | https://www.ncbi.nlm.nih.gov/gene/?term=LOC105284535 | 9086 | 258 | 283.95 | 1 | 1.1 | 0.39 | 13.99 | 14.72 | -0.73 | 0.73 | -0.02 | 0.02 |
| gene9740 | https://www.ncbi.nlm.nih.gov/gene/?term=LOC105283277 | 2887 | 138 | 478 | 1 | 3.46 | 0.72 | 17.69 | 18.47 | -0.78 | 0.78 | -0.04 | 0.04 |
| gene4190 | https://www.ncbi.nlm.nih.gov/gene/?term=LOC105276875 | 3150 | 125 | 396.83 | 1 | 3.17 | 0.8 | 19.56 | 20.35 | -0.79 | 0.79 | -0.03 | 0.03 |
| gene1470 | https://www.ncbi.nlm.nih.gov/gene/?term=LOC105287597 | 2974 | 182 | 611.97 | 1 | 3.36 | 0.55 | 19.45 | 20.26 | -0.81 | 0.81 | -0.05 | 0.05 |
| gene6652 | https://www.ncbi.nlm.nih.gov/gene/?term=LOC105279719 | 2831 | 87 | 307.31 | 1 | 3.53 | 1.15 | 20.49 | 21.33 | -0.83 | 0.83 | -0.03 | 0.03 |
| gene8729 | https://www.ncbi.nlm.nih.gov/gene/?term=LOC105282137 | 5005 | 225 | 449.55 | 1 | 2 | 0.44 | 13.97 | 14.84 | -0.86 | 0.86 | -0.04 | 0.04 |
| gene11454 | https://www.ncbi.nlm.nih.gov/gene/?term=LOC105285256 | 1906 | 50 | 262.33 | 1 | 5.25 | 2 | 43.22 | 44.08 | -0.87 | 0.87 | -0.02 | 0.02 |
| gene10155 | https://www.ncbi.nlm.nih.gov/gene/?term=LOC105283756 | 7030 | 112 | 159.32 | 1 | 1.42 | 0.89 | 7.35 | 8.24 | -0.89 | 0.89 | -0.01 | 0.01 |
| gene570 | https://www.ncbi.nlm.nih.gov/gene/?term=LOC105278549 | 1673 | 70 | 418.41 | 1 | 5.98 | 1.43 | 42.95 | 43.99 | -1.04 | 1.04 | -0.04 | 0.04 |
| gene6422 | https://www.ncbi.nlm.nih.gov/gene/?term=LOC105279458 | 2419 | 43 | 177.76 | 1 | 4.13 | 2.33 | 44.62 | 45.68 | -1.05 | 1.05 | -0.02 | 0.02 |
| gene920 | https://www.ncbi.nlm.nih.gov/gene/?term=LOC105282518 | 1197 | 44 | 367.59 | 1 | 8.35 | 2.27 | 32.92 | 33.98 | -1.06 | 1.06 | -0.04 | 0.04 |
| gene2173 | https://www.ncbi.nlm.nih.gov/gene/?term=LOC105274590 | 4387 | 84 | 191.47 | 1 | 2.28 | 1.19 | 23.89 | 24.96 | -1.07 | 1.07 | -0.02 | 0.02 |
| gene4962 | https://www.ncbi.nlm.nih.gov/gene/?term=LOC105277761 | 4645 | 160 | 344.46 | 1 | 2.15 | 0.62 | 12.8 | 13.96 | -1.16 | 1.16 | -0.04 | 0.04 |
| gene7915 | https://www.ncbi.nlm.nih.gov/gene/?term=LOC105281173 | 1597 | 38 | 237.95 | 1 | 6.26 | 2.63 | 59.81 | 61.01 | -1.2 | 1.2 | -0.03 | 0.03 |
| gene3335 | https://www.ncbi.nlm.nih.gov/gene/?term=LOC105275898 | 4665 | 157 | 336.55 | 1 | 2.14 | 0.64 | 12.01 | 13.22 | -1.21 | 1.21 | -0.04 | 0.04 |
| gene10392 | https://www.ncbi.nlm.nih.gov/gene/?term=LOC105284036 | 3216 | 98 | 304.73 | 1 | 3.11 | 1.02 | 22.15 | 23.39 | -1.25 | 1.25 | -0.04 | 0.04 |
| gene6639 | https://www.ncbi.nlm.nih.gov/gene/?term=LOC105279705 | 7656 | 81 | 105.8 | 1 | 1.31 | 1.23 | 19.09 | 20.33 | -1.25 | 1.25 | -0.01 | 0.01 |
| gene3599 | https://www.ncbi.nlm.nih.gov/gene/?term=LOC105276196 | 2955 | 57 | 192.89 | 1 | 3.38 | 1.75 | 39.57 | 40.89 | -1.32 | 1.32 | -0.03 | 0.03 |
| gene9101 | https://www.ncbi.nlm.nih.gov/gene/?term=LOC105282553 | 3068 | 106 | 345.5 | 1 | 3.26 | 0.94 | 21.02 | 22.34 | -1.33 | 1.33 | -0.05 | 0.05 |
| gene1310 | https://www.ncbi.nlm.nih.gov/gene/?term=LOC105286661 | 2582 | 67 | 259.49 | 1 | 3.87 | 1.49 | 17.48 | 18.89 | -1.41 | 1.41 | -0.04 | 0.04 |
| gene7248 | https://www.ncbi.nlm.nih.gov/gene/?term=LOC105280408 | 5175 | 60 | 115.94 | 1 | 1.93 | 1.67 | 15.92 | 17.41 | -1.49 | 1.49 | -0.02 | 0.02 |
| gene11340 | https://www.ncbi.nlm.nih.gov/gene/?term=LOC105285131 | 2918 | 98 | 335.85 | 1 | 3.43 | 1.02 | 13.77 | 15.33 | -1.56 | 1.56 | -0.05 | 0.05 |
| gene2413 | https://www.ncbi.nlm.nih.gov/gene/?term=LOC105274863 | 2006 | 60 | 299.1 | 1 | 4.99 | 1.67 | 50.89 | 52.54 | -1.65 | 1.65 | -0.05 | 0.05 |
| gene1198 | https://www.ncbi.nlm.nih.gov/gene/?term=LOC105285507 | 2738 | 98 | 357.93 | 1 | 3.65 | 1.02 | 19.77 | 21.55 | -1.78 | 1.78 | -0.06 | 0.06 |
| gene2404 | https://www.ncbi.nlm.nih.gov/gene/?term=LOC105274853 | 1767 | 42 | 237.69 | 1 | 5.66 | 2.38 | 30.33 | 32.12 | -1.8 | 1.8 | -0.04 | 0.04 |
| gene1575 | https://www.ncbi.nlm.nih.gov/gene/?term=LOC105287706 | 3783 | 44 | 116.31 | 1 | 2.64 | 2.27 | 12.72 | 14.57 | -1.85 | 1.85 | -0.02 | 0.02 |
| gene7550 | https://www.ncbi.nlm.nih.gov/gene/?term=LOC105280763 | 2173 | 101 | 464.8 | 1 | 4.6 | 0.99 | 26.35 | 28.2 | -1.85 | 1.85 | -0.09 | 0.09 |
| gene7398 | https://www.ncbi.nlm.nih.gov/gene/?term=LOC105280589 | 3452 | 41 | 118.77 | 1 | 2.9 | 2.44 | 24.33 | 26.31 | -1.98 | 1.98 | -0.02 | 0.02 |

| | | | | | | | | | | | | | |
|---|---|---|---|---|---|---|---|---|---|---|---|---|---|
| gene12203 | https://www.ncbi.nlm.nih.gov/gene/?term=LOC105286099 | 5967 | 80 | 134.07 | 1 | 1.68 | 1.25 | 10.4 | 12.39 | -2 | 2 | -0.03 | 0.03 |
| gene7199 | https://www.ncbi.nlm.nih.gov/gene/?term=LOC105280366 | 1414 | 37 | 261.67 | 1 | 7.07 | 2.7 | 44.35 | 46.38 | -2.03 | 2.03 | -0.05 | 0.05 |
| gene12572 | https://www.ncbi.nlm.nih.gov/gene/?term=LOC105286514 | 2374 | 51 | 214.83 | 1 | 4.21 | 1.96 | 34.55 | 36.64 | -2.09 | 2.09 | -0.04 | 0.04 |
| gene3028 | https://www.ncbi.nlm.nih.gov/gene/?term=LOC105275571 | 1924 | 30 | 155.93 | 1 | 5.2 | 3.33 | 42.66 | 44.75 | -2.09 | 2.09 | -0.03 | 0.03 |
| gene3168 | https://www.ncbi.nlm.nih.gov/gene/?term=LOC105275711 | 6150 | 87 | 141.46 | 1 | 1.63 | 1.15 | 10.39 | 12.71 | -2.33 | 2.33 | -0.03 | 0.03 |
| gene12406 | https://www.ncbi.nlm.nih.gov/gene/?term=LOC105286316 | 1386 | 35 | 252.53 | 1 | 7.22 | 2.86 | 40.6 | 43.17 | -2.57 | 2.57 | -0.06 | 0.06 |
| gene11235 | https://www.ncbi.nlm.nih.gov/gene/?term=LOC105285004 | 1339 | 38 | 283.79 | 1 | 7.47 | 2.63 | 16.34 | 18.94 | -2.6 | 2.6 | -0.07 | 0.07 |
| gene12021 | https://www.ncbi.nlm.nih.gov/gene/?term=LOC105285905 | 1399 | 30 | 214.44 | 1 | 7.15 | 3.33 | 30.6 | 33.22 | -2.62 | 2.62 | -0.06 | 0.06 |
| gene9411 | https://www.ncbi.nlm.nih.gov/gene/?term=LOC105282914 | 5534 | 42 | 75.89 | 1 | 1.81 | 2.38 | 47.45 | 50.92 | -3.47 | 3.47 | -0.03 | 0.03 |
| gene564 | https://www.ncbi.nlm.nih.gov/gene/?term=LOC105278478 | 2802 | 17 | 60.67 | 1 | 3.57 | 5.88 | 15.85 | 19.36 | -3.51 | 3.51 | -0.02 | 0.02 |
| gene388 | https://www.ncbi.nlm.nih.gov/gene/?term=LOC105276538 | 1558 | 22 | 141.21 | 2 | 12.84 | 9.09 | 54.75 | 58.78 | -4.03 | 4.03 | -0.06 | 0.06 |
| gene11584 | https://www.ncbi.nlm.nih.gov/gene/?term=LOC105285416 | 3709 | 13 | 35.05 | 2 | 5.39 | 15.38 | 23.21 | 33.23 | -10.02 | 10.02 | -0.04 | 0.04 |
| gene393 | https://www.ncbi.nlm.nih.gov/gene/?term=LOC105276566 | 1123 | 2 | 17.81 | 1 | 8.9 | 50 | 50 | 74.3 | -24.3 | 24.3 | -0.04 | 0.04 |

Directory: OGL File: ogl-Cb_LI_RphaseA1.vs.RphaseB2.txt

| gene_ID | gene_link | gwidth | #Sites | #per10Kb | #dmSites | #dmsp10kb | %dmSites | %pSite1 | %pSite2 | DMpSite | ADMpSite | DMpNucl | ADMpNucl | |
|---|---|---|---|---|---|---|---|---|---|---|---|---|---|---|
| gene2593 | https://www.ncbi.nlm.nih.gov/gene/?term=LOC105275070 | 1774 | 7 | 39.46 | 5 | 28.18 | 71.43 | 68.01 | 4.09 | 63.92 | 63.92 | 0.25 | 0.25 | |
| gene3165 | https://www.ncbi.nlm.nih.gov/gene/?term=LOC105275706 | 2374 | 65 | 273.8 | 5 | 21.06 | 7.69 | 30.28 | 21 | 9.28 | 9.28 | 0.25 | 0.25 | |
| gene2824 | https://www.ncbi.nlm.nih.gov/gene/?term=LOC105275337 | 1772 | 58 | 327.31 | | 1 | 5.64 | 1.72 | 38.96 | 30.77 | 8.18 | 8.18 | 0.27 | 0.27 |
| gene6334 | https://www.ncbi.nlm.nih.gov/gene/?term=LOC105279356 | 1471 | 45 | 305.91 | | 1 | 6.8 | 2.22 | 31.17 | 28.75 | 2.42 | 2.42 | 0.07 | 0.07 |
| gene1884 | https://www.ncbi.nlm.nih.gov/gene/?term=LOC105288072 | 4169 | 70 | 167.91 | | 1 | 2.4 | 1.43 | 26.72 | 24.36 | 2.36 | 2.36 | 0.04 | 0.04 |
| gene8178 | https://www.ncbi.nlm.nih.gov/gene/?term=LOC105281515 | 8647 | 43 | 49.73 | 1 | 1.16 | 2.33 | 27.02 | 24.75 | 2.27 | 2.27 | 0.01 | 0.01 | |
| gene11819 | https://www.ncbi.nlm.nih.gov/gene/?term=LOC105285671 | 2366 | 98 | 414.2 | 1 | 4.23 | 1.02 | 29.1 | 26.97 | 2.13 | 2.13 | 0.09 | 0.09 | |
| gene5877 | https://www.ncbi.nlm.nih.gov/gene/?term=LOC105278807 | 1719 | 48 | 279.23 | | 1 | 5.82 | 2.08 | 26.96 | 25.13 | 1.83 | 1.83 | 0.05 | 0.05 |
| gene7884 | https://www.ncbi.nlm.nih.gov/gene/?term=LOC105281266 | 3390 | 76 | 224.19 | | 1 | 2.95 | 1.32 | 18.91 | 17.21 | 1.7 | 1.7 | 0.04 | 0.04 |
| gene11401 | https://www.ncbi.nlm.nih.gov/gene/?term=LOC105285202 | 1841 | 78 | 423.68 | | 1 | 5.43 | 1.28 | 26.41 | 24.78 | 1.62 | 1.62 | 0.07 | 0.07 |
| gene1855 | https://www.ncbi.nlm.nih.gov/gene/?term=LOC105288036 | 2387 | 77 | 322.58 | | 1 | 4.19 | 1.3 | 32.99 | 31.48 | 1.51 | 1.51 | 0.05 | 0.05 |
| gene5101 | https://www.ncbi.nlm.nih.gov/gene/?term=LOC105277903 | 1813 | 35 | 193.05 | | 1 | 5.52 | 2.86 | 52.77 | 51.27 | 1.51 | 1.51 | 0.03 | 0.03 |
| gene940 | https://www.ncbi.nlm.nih.gov/gene/?term=LOC105282794 | 3578 | 85 | 237.56 | | 1 | 2.79 | 1.18 | 14.76 | 13.28 | 1.48 | 1.48 | 0.04 | 0.04 |
| gene10654 | https://www.ncbi.nlm.nih.gov/gene/?term=LOC105284329 | 4027 | 51 | 126.65 | | 1 | 2.48 | 1.96 | 20.19 | 18.78 | 1.41 | 1.41 | 0.02 | 0.02 |
| gene7568 | https://www.ncbi.nlm.nih.gov/gene/?term=LOC105280786 | 2525 | 114 | 451.49 | | 1 | 3.96 | 0.88 | 28.64 | 27.23 | 1.41 | 1.41 | 0.06 | 0.06 |
| gene7915 | https://www.ncbi.nlm.nih.gov/gene/?term=LOC105281173 | 1597 | 38 | 237.95 | | 1 | 6.26 | 2.63 | 59.81 | 58.74 | 1.06 | 1.06 | 0.03 | 0.03 |
| gene12516 | https://www.ncbi.nlm.nih.gov/gene/?term=LOC105286452 | 1529 | 38 | 248.53 | | 1 | 6.54 | 2.63 | 50.31 | 49.37 | 0.94 | 0.94 | 0.02 | 0.02 |
| gene8661 | https://www.ncbi.nlm.nih.gov/gene/?term=LOC105282063 | 1926 | 26 | 134.99 | | 1 | 5.19 | 3.85 | 38.79 | 37.9 | 0.89 | 0.89 | 0.01 | 0.01 |
| gene2844 | https://www.ncbi.nlm.nih.gov/gene/?term=LOC105275363 | 4330 | 186 | 429.56 | | 1 | 2.31 | 0.54 | 15.2 | 14.36 | 0.85 | 0.85 | 0.04 | 0.04 |
| gene11454 | https://www.ncbi.nlm.nih.gov/gene/?term=LOC105285256 | 1906 | 45 | 236.1 | 1 | 5.25 | 2.22 | 47.83 | 47.02 | 0.81 | 0.81 | 0.02 | 0.02 | |
| gene4023 | https://www.ncbi.nlm.nih.gov/gene/?term=LOC105276692 | 1759 | 31 | 176.24 | | 1 | 5.69 | 3.23 | 24.44 | 23.63 | 0.81 | 0.81 | 0.01 | 0.01 |
| gene4892 | https://www.ncbi.nlm.nih.gov/gene/?term=LOC105277667 | 3898 | 119 | 305.28 | | 1 | 2.57 | 0.84 | 28.98 | 28.19 | 0.8 | 0.8 | 0.02 | 0.02 |
| gene3470 | https://www.ncbi.nlm.nih.gov/gene/?term=LOC105276059 | 8716 | 146 | 167.51 | | 1 | 1.15 | 0.68 | 9.33 | 8.54 | 0.79 | 0.79 | 0.01 | 0.01 |
| gene10540 | https://www.ncbi.nlm.nih.gov/gene/?term=LOC105284200 | 5267 | 215 | 408.2 | 1 | 1.9 | 0.47 | 12.8 | 12.05 | 0.75 | 0.75 | 0.03 | 0.03 | |
| gene4213 | https://www.ncbi.nlm.nih.gov/gene/?term=LOC105276901 | 3257 | 83 | 254.84 | | 1 | 3.07 | 1.2 | 27.23 | 26.49 | 0.74 | 0.74 | 0.02 | 0.02 |
| gene10520 | https://www.ncbi.nlm.nih.gov/gene/?term=LOC105284184 | 2451 | 79 | 322.32 | | 1 | 4.08 | 1.27 | 35.68 | 34.99 | 0.69 | 0.69 | 0.02 | 0.02 |
| gene298 | https://www.ncbi.nlm.nih.gov/gene/?term=LOC105275460 | 2473 | 118 | 477.15 | | 1 | 4.04 | 0.85 | 13.95 | 13.28 | 0.67 | 0.67 | 0.03 | 0.03 |
| gene9336 | https://www.ncbi.nlm.nih.gov/gene/?term=LOC105282834 | 5993 | 123 | 205.24 | | 1 | 1.67 | 0.81 | 15.27 | 14.61 | 0.66 | 0.66 | 0.01 | 0.01 |
| gene10600 | https://www.ncbi.nlm.nih.gov/gene/?term=LOC105284270 | 4837 | 115 | 237.75 | | 1 | 2.07 | 0.87 | 11.06 | 10.43 | 0.63 | 0.63 | 0.02 | 0.02 |
| gene6966 | https://www.ncbi.nlm.nih.gov/gene/?term=LOC105280104 | 3747 | 124 | 330.93 | | 1 | 2.67 | 0.81 | 17.23 | 16.6 | 0.63 | 0.63 | 0.02 | 0.02 |
| gene1430 | https://www.ncbi.nlm.nih.gov/gene/?term=LOC105287551 | 6433 | 216 | 335.77 | | 1 | 1.55 | 0.46 | 9.29 | 8.67 | 0.62 | 0.62 | 0.02 | 0.02 |
| gene570 | https://www.ncbi.nlm.nih.gov/gene/?term=LOC105278549 | 1673 | 62 | 370.59 | | 1 | 5.98 | 1.61 | 43.95 | 43.37 | 0.58 | 0.58 | 0.02 | 0.02 |
| gene5273 | https://www.ncbi.nlm.nih.gov/gene/?term=LOC105278108 | 4067 | 43 | 105.73 | | 1 | 2.46 | 2.33 | 26.66 | 26.12 | 0.55 | 0.55 | 0.01 | 0.01 |
| gene6583 | https://www.ncbi.nlm.nih.gov/gene/?term=LOC105279649 | 7444 | 315 | 423.16 | | 1 | 1.34 | 0.32 | 11.55 | 11.04 | 0.51 | 0.51 | 0.02 | 0.02 |
| gene12978 | https://www.ncbi.nlm.nih.gov/gene/?term=LOC105286997 | 2858 | 106 | 370.89 | | 1 | 3.5 | 0.94 | 12 | 11.5 | 0.5 | 0.5 | 0.02 | 0.02 |
| gene5831 | https://www.ncbi.nlm.nih.gov/gene/?term=LOC105278761 | 2506 | 105 | 418.99 | | 1 | 3.99 | 0.95 | 24.55 | 24.07 | 0.48 | 0.48 | 0.02 | 0.02 |
| gene8274 | https://www.ncbi.nlm.nih.gov/gene/?term=LOC105281624 | 5425 | 198 | 364.98 | | 1 | 1.84 | 0.51 | 10.15 | 9.67 | 0.48 | 0.48 | 0.02 | 0.02 |
| gene1332 | https://www.ncbi.nlm.nih.gov/gene/?term=LOC105286872 | 2460 | 156 | 634.15 | | 1 | 4.07 | 0.64 | 20.52 | 20.12 | 0.41 | 0.41 | 0.03 | 0.03 |
| gene2846 | https://www.ncbi.nlm.nih.gov/gene/?term=LOC105275374 | 11246 | 432 | 384.14 | | 1 | 0.89 | 0.23 | 11.15 | 10.76 | 0.39 | 0.39 | 0.02 | 0.02 |
| gene11579 | https://www.ncbi.nlm.nih.gov/gene/?term=LOC105285406 | 13571 | 229 | 168.74 | | 1 | 0.74 | 0.44 | 7.6 | 7.22 | 0.38 | 0.38 | 0.01 | 0.01 |
| gene12439 | https://www.ncbi.nlm.nih.gov/gene/?term=LOC105286357 | 4761 | 87 | 182.73 | | 1 | 2.1 | 1.15 | 25.3 | 24.92 | 0.38 | 0.38 | 0.01 | 0.01 |
| gene7247 | https://www.ncbi.nlm.nih.gov/gene/?term=LOC105280407 | 10486 | 208 | 198.36 | | 1 | 0.95 | 0.48 | 16.58 | 16.25 | 0.33 | 0.33 | 0.01 | 0.01 |
| gene2266 | https://www.ncbi.nlm.nih.gov/gene/?term=LOC105274699 | 2391 | 49 | 204.94 | | 1 | 4.18 | 2.04 | 23.32 | 23 | 0.32 | 0.32 | 0.01 | 0.01 |
| gene13114 | https://www.ncbi.nlm.nih.gov/gene/?term=LOC105287147 | 7609 | 393 | 516.49 | | 1 | 1.31 | 0.25 | 9.24 | 9.02 | 0.23 | 0.23 | 0.01 | 0.01 |
| gene5525 | https://www.ncbi.nlm.nih.gov/gene/?term=LOC105278396 | 3090 | 180 | 582.52 | | 1 | 3.24 | 0.56 | 24.99 | 24.77 | 0.22 | 0.22 | 0.01 | 0.01 |
| gene2683 | https://www.ncbi.nlm.nih.gov/gene/?term=LOC105275187 | 5492 | 65 | 118.35 | | 1 | 1.82 | 1.54 | 11.46 | 11.25 | 0.21 | 0.21 | 0 | 0 |
| gene5052 | https://www.ncbi.nlm.nih.gov/gene/?term=LOC105277856 | 3167 | 28 | 88.41 | 1 | 3.16 | 3.57 | 33.66 | 33.45 | 0.21 | 0.21 | 0 | 0 | |
| gene11082 | https://www.ncbi.nlm.nih.gov/gene/?term=LOC105284824 | 7436 | 189 | 254.17 | | 1 | 1.34 | 0.53 | 8.04 | 7.84 | 0.2 | 0.2 | 0.01 | 0.01 |
| gene11055 | https://www.ncbi.nlm.nih.gov/gene/?term=LOC105284796 | 2283 | 121 | 530 | 1 | 4.38 | 0.83 | 34.82 | 34.65 | 0.17 | 0.17 | 0.01 | 0.01 | |
| gene1470 | https://www.ncbi.nlm.nih.gov/gene/?term=LOC105287597 | 2974 | 181 | 608.61 | | 1 | 3.36 | 0.55 | 19.06 | 18.9 | 0.16 | 0.16 | 0.01 | 0.01 |
| gene2806 | https://www.ncbi.nlm.nih.gov/gene/?term=LOC105275327 | 14257 | 456 | 319.84 | | 1 | 0.7 | 0.22 | 7.38 | 7.24 | 0.14 | 0.14 | 0 | 0 |
| gene1220 | https://www.ncbi.nlm.nih.gov/gene/?term=LOC105285742 | 3453 | 100 | 289.6 | 1 | 2.9 | 1 | 18.13 | 18.02 | 0.11 | 0.11 | 0 | 0 | |
| gene5733 | https://www.ncbi.nlm.nih.gov/gene/?term=LOC105278649 | 5221 | 92 | 176.21 | | 1 | 1.92 | 1.09 | 19.86 | 19.75 | 0.11 | 0.11 | 0 | 0 |
| gene5746 | https://www.ncbi.nlm.nih.gov/gene/?term=LOC105278666 | 10343 | 436 | 421.54 | | 1 | 0.97 | 0.23 | 6.97 | 6.87 | 0.11 | 0.11 | 0 | 0 |
| gene8600 | https://www.ncbi.nlm.nih.gov/gene/?term=LOC105282001 | 8513 | 151 | 177.38 | | 1 | 1.17 | 0.66 | 10.5 | 10.38 | 0.11 | 0.11 | 0 | 0 |
| gene10844 | https://www.ncbi.nlm.nih.gov/gene/?term=LOC105284535 | 9086 | 222 | 244.33 | | 1 | 1.1 | 0.45 | 15.15 | 15.05 | 0.1 | 0.1 | 0 | 0 |
| gene1632 | https://www.ncbi.nlm.nih.gov/gene/?term=LOC105287764 | 18061 | 367 | 203.2 | 1 | 0.55 | 0.27 | 3.44 | 3.36 | 0.08 | 0.08 | 0 | 0 | |
| gene5524 | https://www.ncbi.nlm.nih.gov/gene/?term=LOC105278397 | 5177 | 196 | 378.6 | 1 | 1.93 | 0.51 | 24.77 | 24.69 | 0.07 | 0.07 | 0 | 0 | |
| gene1408 | https://www.ncbi.nlm.nih.gov/gene/?term=LOC109611520 | 8645 | 289 | 334.3 | 1 | 1.16 | 0.35 | 6.21 | 6.16 | 0.05 | 0.05 | 0 | 0 | |
| gene7368 | https://www.ncbi.nlm.nih.gov/gene/?term=LOC105280556 | 4105 | 38 | 92.57 | 1 | 2.44 | 2.63 | 16.79 | 16.73 | 0.05 | 0.05 | 0 | 0 | |

| | | | | | | | | | | | | | |
|---|---|---|---|---|---|---|---|---|---|---|---|---|---|
| gene9740 | https://www.ncbi.nlm.nih.gov/gene/?term=LOC105283277 | 2887 | 131 | 453.76 | 1 | 3.46 | 0.76 | 19.31 | 19.27 | 0.04 | 0.04 | 0 | 0 |
| gene8814 | https://www.ncbi.nlm.nih.gov/gene/?term=LOC105282240 | 6910 | 163 | 235.89 | 1 | 1.45 | 0.61 | 9.99 | 9.96 | 0.03 | 0.03 | 0 | 0 |
| gene2271 | https://www.ncbi.nlm.nih.gov/gene/?term=LOC105274706 | 1448 | 50 | 345.3 | 1 | 6.91 | 2 | 38.1 | 38.09 | 0.01 | 0.01 | 0 | 0 |
| gene2173 | https://www.ncbi.nlm.nih.gov/gene/?term=LOC105274590 | 4387 | 68 | 155 | 1 | 2.28 | 1.47 | 26.72 | 26.74 | -0.02 | 0.02 | 0 | 0 |
| gene10678 | https://www.ncbi.nlm.nih.gov/gene/?term=LOC105284355 | 4031 | 168 | 416.77 | 1 | 2.48 | 0.6 | 18.14 | 18.16 | -0.03 | 0.03 | 0 | 0 |
| gene10984 | https://www.ncbi.nlm.nih.gov/gene/?term=LOC105284704 | 1987 | 58 | 291.9 | 1 | 5.03 | 1.72 | 23.02 | 23.06 | -0.04 | 0.04 | 0 | 0 |
| gene10326 | https://www.ncbi.nlm.nih.gov/gene/?term=LOC105283962 | 3042 | 119 | 391.19 | 1 | 3.29 | 0.84 | 8.91 | 8.98 | -0.07 | 0.07 | 0 | 0 |
| gene11275 | https://www.ncbi.nlm.nih.gov/gene/?term=LOC105285055 | 4119 | 257 | 623.94 | 1 | 2.43 | 0.39 | 10.45 | 10.51 | -0.07 | 0.07 | 0 | 0 |
| gene4962 | https://www.ncbi.nlm.nih.gov/gene/?term=LOC105277761 | 4645 | 142 | 305.71 | 1 | 2.15 | 0.7 | 13.6 | 13.69 | -0.09 | 0.09 | 0 | 0 |
| gene7398 | https://www.ncbi.nlm.nih.gov/gene/?term=LOC105280589 | 3452 | 33 | 95.6 | 1 | 2.9 | 3.03 | 20.49 | 20.58 | -0.09 | 0.09 | 0 | 0 |
| gene6620 | https://www.ncbi.nlm.nih.gov/gene/?term=LOC105279687 | 17998 | 344 | 191.13 | 1 | 0.56 | 0.29 | 5.2 | 5.3 | -0.1 | 0.1 | 0 | 0 |
| gene7556 | https://www.ncbi.nlm.nih.gov/gene/?term=LOC105280774 | 2792 | 98 | 351 | 1 | 3.58 | 1.02 | 32.45 | 32.61 | -0.17 | 0.17 | -0.01 | 0.01 |
| gene53 | https://www.ncbi.nlm.nih.gov/gene/?term=LOC105277148 | 19283 | 268 | 138.98 | 1 | 0.52 | 0.37 | 3.71 | 3.95 | -0.23 | 0.23 | 0 | 0 |
| gene10018 | https://www.ncbi.nlm.nih.gov/gene/?term=LOC105283607 | 4414 | 202 | 457.63 | 1 | 2.27 | 0.5 | 14.99 | 15.23 | -0.24 | 0.24 | -0.01 | 0.01 |
| gene6286 | https://www.ncbi.nlm.nih.gov/gene/?term=LOC105279298 | 2861 | 130 | 454.39 | 2 | 6.99 | 1.54 | 23.66 | 23.9 | -0.24 | 0.24 | -0.01 | 0.01 |
| gene9326 | https://www.ncbi.nlm.nih.gov/gene/?term=LOC105282818 | 16612 | 169 | 101.73 | 1 | 0.6 | 0.59 | 13.49 | 13.76 | -0.26 | 0.26 | 0 | 0 |
| gene10606 | https://www.ncbi.nlm.nih.gov/gene/?term=LOC105284273 | 6119 | 244 | 398.76 | 1 | 1.63 | 0.41 | 18.11 | 18.39 | -0.29 | 0.29 | -0.01 | 0.01 |
| gene34 | https://www.ncbi.nlm.nih.gov/gene/?term=LOC105278416 | 7120 | 124 | 174.16 | 1 | 1.4 | 0.81 | 14.41 | 14.72 | -0.3 | 0.3 | -0.01 | 0.01 |
| gene12573 | https://www.ncbi.nlm.nih.gov/gene/?term=LOC105286515 | 3647 | 105 | 287.91 | 1 | 2.74 | 0.95 | 12.53 | 12.84 | -0.31 | 0.31 | -0.01 | 0.01 |
| gene3885 | https://www.ncbi.nlm.nih.gov/gene/?term=LOC105276527 | 7928 | 299 | 377.14 | 1 | 1.26 | 0.33 | 9.83 | 10.14 | -0.31 | 0.31 | -0.01 | 0.01 |
| gene7166 | https://www.ncbi.nlm.nih.gov/gene/?term=LOC105280326 | 6117 | 226 | 369.46 | 1 | 1.63 | 0.44 | 14.91 | 15.21 | -0.31 | 0.31 | -0.01 | 0.01 |
| gene10549 | https://www.ncbi.nlm.nih.gov/gene/?term=LOC105284210 | 7145 | 284 | 397.48 | 2 | 2.8 | 0.7 | 11.85 | 12.19 | -0.33 | 0.33 | -0.01 | 0.01 |
| gene2748 | https://www.ncbi.nlm.nih.gov/gene/?term=LOC105275261 | 3845 | 53 | 137.84 | 1 | 2.6 | 1.89 | 38.36 | 38.68 | -0.33 | 0.33 | 0 | 0 |
| gene5646 | https://www.ncbi.nlm.nih.gov/gene/?term=LOC105278534 | 7475 | 176 | 235.45 | 1 | 1.34 | 0.57 | 18.67 | 19 | -0.33 | 0.33 | -0.01 | 0.01 |
| gene2404 | https://www.ncbi.nlm.nih.gov/gene/?term=LOC105274853 | 1767 | 41 | 232.03 | 1 | 5.66 | 2.44 | 30.51 | 30.87 | -0.36 | 0.36 | -0.01 | 0.01 |
| gene9327 | https://www.ncbi.nlm.nih.gov/gene/?term=LOC105282821 | 7793 | 140 | 179.65 | 1 | 1.28 | 0.71 | 12.3 | 12.71 | -0.41 | 0.41 | -0.01 | 0.01 |
| gene5375 | https://www.ncbi.nlm.nih.gov/gene/?term=LOC105278228 | 8640 | 152 | 175.93 | 1 | 1.16 | 0.66 | 22.96 | 23.39 | -0.42 | 0.42 | -0.01 | 0.01 |
| gene10879 | https://www.ncbi.nlm.nih.gov/gene/?term=LOC105284574 | 2423 | 50 | 206.36 | 1 | 4.13 | 2 | 52.82 | 53.31 | -0.49 | 0.49 | -0.01 | 0.01 |
| gene7211 | https://www.ncbi.nlm.nih.gov/gene/?term=LOC105280377 | 2024 | 63 | 311.26 | 1 | 4.94 | 1.59 | 48.25 | 48.74 | -0.49 | 0.49 | -0.02 | 0.02 |
| gene7550 | https://www.ncbi.nlm.nih.gov/gene/?term=LOC105280763 | 2173 | 76 | 349.75 | 1 | 4.6 | 1.32 | 32.2 | 32.7 | -0.49 | 0.49 | -0.02 | 0.02 |
| gene12307 | https://www.ncbi.nlm.nih.gov/gene/?term=LOC105286216 | 10313 | 164 | 159.02 | 1 | 0.97 | 0.61 | 16.27 | 16.77 | -0.5 | 0.5 | -0.01 | 0.01 |
| gene12802 | https://www.ncbi.nlm.nih.gov/gene/?term=LOC105286799 | 3183 | 159 | 499.53 | 1 | 3.14 | 0.63 | 17.45 | 17.99 | -0.54 | 0.54 | -0.03 | 0.03 |
| gene292 | https://www.ncbi.nlm.nih.gov/gene/?term=LOC105275398 | 2825 | 78 | 276.11 | 1 | 3.54 | 1.28 | 22.03 | 22.57 | -0.55 | 0.55 | -0.02 | 0.02 |
| gene4216 | https://www.ncbi.nlm.nih.gov/gene/?term=LOC105276906 | 2127 | 86 | 404.33 | 1 | 4.7 | 1.16 | 19.06 | 19.63 | -0.57 | 0.57 | -0.02 | 0.02 |
| gene5298 | https://www.ncbi.nlm.nih.gov/gene/?term=LOC105278145 | 4060 | 235 | 578.82 | 2 | 4.93 | 0.85 | 10.98 | 11.56 | -0.58 | 0.58 | -0.03 | 0.03 |
| gene13036 | https://www.ncbi.nlm.nih.gov/gene/?term=LOC105287059 | 4959 | 148 | 298.45 | 1 | 2.02 | 0.68 | 20.35 | 20.94 | -0.59 | 0.59 | -0.02 | 0.02 |
| gene3107 | https://www.ncbi.nlm.nih.gov/gene/?term=LOC105275650 | 5103 | 152 | 297.86 | 1 | 1.96 | 0.66 | 16.19 | 16.78 | -0.59 | 0.59 | -0.02 | 0.02 |
| gene4190 | https://www.ncbi.nlm.nih.gov/gene/?term=LOC105276875 | 3150 | 105 | 333.33 | 1 | 3.17 | 0.95 | 22.87 | 23.46 | -0.6 | 0.6 | -0.02 | 0.02 |
| gene7249 | https://www.ncbi.nlm.nih.gov/gene/?term=LOC105280422 | 1775 | 60 | 338.03 | 1 | 5.63 | 1.67 | 16.46 | 17.06 | -0.6 | 0.6 | -0.02 | 0.02 |
| gene1147 | https://www.ncbi.nlm.nih.gov/gene/?term=LOC105284954 | 4022 | 91 | 226.26 | 1 | 2.49 | 1.1 | 14.14 | 14.77 | -0.63 | 0.63 | -0.01 | 0.01 |
| gene9647 | https://www.ncbi.nlm.nih.gov/gene/?term=LOC109611233 | 3094 | 32 | 103.43 | 1 | 3.23 | 3.12 | 37.53 | 38.18 | -0.65 | 0.65 | -0.01 | 0.01 |
| gene9648 | https://www.ncbi.nlm.nih.gov/gene/?term=LOC105283183 | 2602 | 30 | 115.3 | 1 | 3.84 | 3.33 | 40.03 | 40.73 | -0.7 | 0.7 | -0.01 | 0.01 |
| gene4376 | https://www.ncbi.nlm.nih.gov/gene/?term=LOC105277083 | 4307 | 97 | 225.21 | 1 | 2.32 | 1.03 | 24.28 | 25.05 | -0.77 | 0.77 | -0.02 | 0.02 |
| gene920 | https://www.ncbi.nlm.nih.gov/gene/?term=LOC105282518 | 1197 | 48 | 401 | 1 | 8.35 | 2.08 | 32.24 | 33.02 | -0.78 | 0.78 | -0.03 | 0.03 |
| gene10392 | https://www.ncbi.nlm.nih.gov/gene/?term=LOC105284036 | 3216 | 94 | 292.29 | 2 | 6.22 | 2.13 | 22.41 | 23.26 | -0.84 | 0.84 | -0.02 | 0.02 |
| gene12203 | https://www.ncbi.nlm.nih.gov/gene/?term=LOC105286099 | 5967 | 70 | 117.31 | 1 | 1.68 | 1.43 | 9.91 | 10.79 | -0.88 | 0.88 | -0.01 | 0.01 |
| gene11340 | https://www.ncbi.nlm.nih.gov/gene/?term=LOC105285131 | 2918 | 87 | 298.15 | 1 | 3.43 | 1.15 | 10.66 | 11.58 | -0.92 | 0.92 | -0.03 | 0.03 |
| gene7248 | https://www.ncbi.nlm.nih.gov/gene/?term=LOC105280408 | 5175 | 52 | 100.48 | 1 | 1.93 | 1.92 | 17.59 | 18.54 | -0.95 | 0.95 | -0.01 | 0.01 |
| gene10155 | https://www.ncbi.nlm.nih.gov/gene/?term=LOC105283756 | 7030 | 108 | 153.63 | 1 | 1.42 | 0.93 | 5.58 | 6.54 | -0.96 | 0.96 | -0.01 | 0.01 |
| gene12021 | https://www.ncbi.nlm.nih.gov/gene/?term=LOC105285905 | 1399 | 16 | 114.37 | 1 | 7.15 | 6.25 | 46.76 | 47.75 | -0.98 | 0.98 | -0.01 | 0.01 |
| gene1310 | https://www.ncbi.nlm.nih.gov/gene/?term=LOC105286661 | 2582 | 68 | 263.36 | 1 | 3.87 | 1.47 | 19.86 | 20.85 | -1 | 1 | -0.03 | 0.03 |
| gene8729 | https://www.ncbi.nlm.nih.gov/gene/?term=LOC105282137 | 5005 | 213 | 425.57 | 1 | 2 | 0.47 | 14.62 | 15.67 | -1.05 | 1.05 | -0.04 | 0.04 |
| gene7600 | https://www.ncbi.nlm.nih.gov/gene/?term=LOC105280819 | 1615 | 37 | 229.1 | 1 | 6.19 | 2.7 | 36.99 | 38.16 | -1.16 | 1.16 | -0.03 | 0.03 |
| gene10213 | https://www.ncbi.nlm.nih.gov/gene/?term=LOC105283826 | 1503 | 53 | 352.63 | 1 | 6.65 | 1.89 | 14.78 | 15.95 | -1.17 | 1.17 | -0.04 | 0.04 |
| gene6639 | https://www.ncbi.nlm.nih.gov/gene/?term=LOC105279705 | 7656 | 69 | 90.13 | 1 | 1.31 | 1.45 | 20.74 | 21.96 | -1.21 | 1.21 | -0.01 | 0.01 |
| gene35 | https://www.ncbi.nlm.nih.gov/gene/?term=LOC105275197 | 2643 | 45 | 170.26 | 1 | 3.78 | 2.22 | 38.91 | 40.15 | -1.23 | 1.23 | -0.02 | 0.02 |
| gene6031 | https://www.ncbi.nlm.nih.gov/gene/?term=LOC105279000 | 5691 | 180 | 316.29 | 1 | 1.76 | 0.56 | 14.61 | 15.92 | -1.31 | 1.31 | -0.04 | 0.04 |
| gene564 | https://www.ncbi.nlm.nih.gov/gene/?term=LOC105278478 | 2802 | 18 | 64.24 | 1 | 3.57 | 5.56 | 14.97 | 16.38 | -1.41 | 1.41 | -0.01 | 0.01 |
| gene4641 | https://www.ncbi.nlm.nih.gov/gene/?term=LOC105277398 | 1937 | 71 | 366.55 | 1 | 5.16 | 1.41 | 38.26 | 39.69 | -1.43 | 1.43 | -0.05 | 0.05 |
| gene7183 | https://www.ncbi.nlm.nih.gov/gene/?term=LOC105280348 | 3925 | 70 | 178.34 | 1 | 2.55 | 1.43 | 19.96 | 21.48 | -1.52 | 1.52 | -0.03 | 0.03 |
| gene8343 | https://www.ncbi.nlm.nih.gov/gene/?term=LOC105281700 | 2172 | 32 | 147.33 | 1 | 4.6 | 3.12 | 37.43 | 38.97 | -1.55 | 1.55 | -0.02 | 0.02 |
| gene7199 | https://www.ncbi.nlm.nih.gov/gene/?term=LOC105280366 | 1414 | 24 | 169.73 | 1 | 7.07 | 4.17 | 43.55 | 45.17 | -1.62 | 1.62 | -0.03 | 0.03 |
| gene12572 | https://www.ncbi.nlm.nih.gov/gene/?term=LOC105286514 | 2374 | 47 | 197.98 | 1 | 4.21 | 2.13 | 35.17 | 36.86 | -1.69 | 1.69 | -0.03 | 0.03 |
| gene9101 | https://www.ncbi.nlm.nih.gov/gene/?term=LOC105282553 | 3068 | 102 | 332.46 | 1 | 3.26 | 0.98 | 21.53 | 23.25 | -1.72 | 1.72 | -0.06 | 0.06 |

| | | | | | | | | | | | | | |
|---|---|---|---|---|---|---|---|---|---|---|---|---|---|
| gene6237 | https://www.ncbi.nlm.nih.gov/gene/?term=LOC105279244 | 1484 | 22 | 148.25 | 1 | 6.74 | 4.55 | 40.19 | 41.99 | -1.8 | 1.8 | -0.03 | 0.03 |
| gene2413 | https://www.ncbi.nlm.nih.gov/gene/?term=LOC105274863 | 2006 | 49 | 244.27 | 1 | 4.99 | 2.04 | 53.65 | 55.47 | -1.81 | 1.81 | -0.04 | 0.04 |
| gene7406 | https://www.ncbi.nlm.nih.gov/gene/?term=LOC105280596 | 3432 | 106 | 308.86 | 1 | 2.91 | 0.94 | 17.02 | 19.17 | -2.15 | 2.15 | -0.07 | 0.07 |
| gene12981 | https://www.ncbi.nlm.nih.gov/gene/?term=LOC105286999 | 2063 | 42 | 203.59 | 1 | 4.85 | 2.38 | 33.61 | 35.96 | -2.34 | 2.34 | -0.05 | 0.05 |
| gene12406 | https://www.ncbi.nlm.nih.gov/gene/?term=LOC105286316 | 1386 | 31 | 223.67 | 1 | 7.22 | 3.23 | 42.03 | 44.54 | -2.51 | 2.51 | -0.06 | 0.06 |
| gene10107 | https://www.ncbi.nlm.nih.gov/gene/?term=LOC105283712 | 1883 | 35 | 185.87 | 1 | 5.31 | 2.86 | 23.29 | 25.95 | -2.66 | 2.66 | -0.05 | 0.05 |
| gene3599 | https://www.ncbi.nlm.nih.gov/gene/?term=LOC105276196 | 2955 | 58 | 196.28 | 1 | 3.38 | 1.72 | 37.85 | 40.55 | -2.7 | 2.7 | -0.05 | 0.05 |
| gene6422 | https://www.ncbi.nlm.nih.gov/gene/?term=LOC105279458 | 2419 | 38 | 157.09 | 1 | 4.13 | 2.63 | 47.43 | 50.37 | -2.94 | 2.94 | -0.05 | 0.05 |
| gene3168 | https://www.ncbi.nlm.nih.gov/gene/?term=LOC105275711 | 6150 | 69 | 112.2 | 1 | 1.63 | 1.45 | 12.17 | 15.18 | -3.01 | 3.01 | -0.03 | 0.03 |
| gene322 | https://www.ncbi.nlm.nih.gov/gene/?term=LOC105275777 | 1606 | 27 | 168.12 | 1 | 6.23 | 3.7 | 38.42 | 42 | -3.59 | 3.59 | -0.06 | 0.06 |
| gene388 | https://www.ncbi.nlm.nih.gov/gene/?term=LOC105276538 | 1558 | 19 | 121.95 | 2 | 12.84 | 10.53 | 61.29 | 65.13 | -3.85 | 3.85 | -0.05 | 0.05 |
| gene10345 | https://www.ncbi.nlm.nih.gov/gene/?term=LOC105283989 | 4444 | 22 | 49.5 | 1 | 2.25 | 4.55 | 37.68 | 41.66 | -3.99 | 3.99 | -0.02 | 0.02 |
| gene10602 | https://www.ncbi.nlm.nih.gov/gene/?term=LOC105284268 | 2122 | 19 | 89.54 | 1 | 4.71 | 5.26 | 27.9 | 32.99 | -5.08 | 5.08 | -0.05 | 0.05 |
| gene1008 | https://www.ncbi.nlm.nih.gov/gene/?term=LOC105283492 | 1479 | 9 | 60.85 | 1 | 6.76 | 11.11 | 30.57 | 36.08 | -5.51 | 5.51 | -0.03 | 0.03 |
| gene393 | https://www.ncbi.nlm.nih.gov/gene/?term=LOC105276566 | 1123 | 2 | 17.81 | 1 | 8.9 | 50 | 50 | 76.78 | -26.78 | 26.78 | -0.05 | 0.05 |

Directory: OGL File: ogl-Cb_LI_RphaseA2.txt

| gene_ID | gene_link | gwidth | #Sites | #per10Kb | %perSite | %pNucl |
|---|---|---|---|---|---|---|
| gene13384 | https://www.ncbi.nlm.nih.gov/gene/?term=LOC105287447 | 1877 | 296 | 1576.98 | 27.63 | 4.36 |
| gene5917 | https://www.ncbi.nlm.nih.gov/gene/?term=LOC105278849 | 823 | 139 | 1688.94 | 25.03 | 4.23 |
| gene10219 | https://www.ncbi.nlm.nih.gov/gene/?term=LOC105283830 | 235 | 22 | 936.17 | 44.27 | 4.14 |
| gene12364 | https://www.ncbi.nlm.nih.gov/gene/?term=LOC105286269 | 1403 | 110 | 784.03 | 52.64 | 4.13 |
| gene12538 | https://www.ncbi.nlm.nih.gov/gene/?term=LOC105286531 | 372 | 43 | 1155.91 | 35.68 | 4.12 |
| gene5911 | https://www.ncbi.nlm.nih.gov/gene/?term=LOC105278844 | 943 | 122 | 1293.74 | 31.64 | 4.09 |
| gene5309 | https://www.ncbi.nlm.nih.gov/gene/?term=LOC105278160 | 1704 | 183 | 1073.94 | 37.02 | 3.98 |
| gene3719 | https://www.ncbi.nlm.nih.gov/gene/?term=LOC105276360 | 1471 | 205 | 1393.61 | 28.45 | 3.97 |
| gene11569 | https://www.ncbi.nlm.nih.gov/gene/?term=LOC105285394 | 1311 | 75 | 572.08 | 68.68 | 3.93 |
| gene4878 | https://www.ncbi.nlm.nih.gov/gene/?term=LOC105277648 | 675 | 45 | 666.67 | 57.93 | 3.86 |
| gene1654 | https://www.ncbi.nlm.nih.gov/gene/?term=LOC105287811 | 2285 | 205 | 897.16 | 42.64 | 3.83 |
| gene211 | https://www.ncbi.nlm.nih.gov/gene/?term=LOC105274534 | 1280 | 111 | 867.19 | 44.2 | 3.83 |
| gene12652 | https://www.ncbi.nlm.nih.gov/gene/?term=LOC109610640 | 1122 | 71 | 632.8 | 57.98 | 3.67 |
| gene2797 | https://www.ncbi.nlm.nih.gov/gene/?term=LOC105275314 | 1489 | 132 | 886.5 | 41.45 | 3.67 |
| gene570 | https://www.ncbi.nlm.nih.gov/gene/?term=LOC105278549 | 1673 | 145 | 866.71 | 41.46 | 3.59 |
| gene6078 | https://www.ncbi.nlm.nih.gov/gene/?term=LOC105279052 | 1130 | 163 | 1442.48 | 24.83 | 3.58 |
| gene10943 | https://www.ncbi.nlm.nih.gov/gene/?term=LOC105284659 | 1759 | 129 | 733.37 | 48.52 | 3.56 |
| gene12330 | https://www.ncbi.nlm.nih.gov/gene/?term=LOC105286235 | 1501 | 122 | 812.79 | 42.08 | 3.42 |
| gene12746 | https://www.ncbi.nlm.nih.gov/gene/?term=LOC105286737 | 2688 | 406 | 1510.42 | 22.57 | 3.41 |
| gene1562 | https://www.ncbi.nlm.nih.gov/gene/?term=LOC105287692 | 1491 | 153 | 1026.16 | 33.21 | 3.41 |
| gene3731 | https://www.ncbi.nlm.nih.gov/gene/?term=LOC105276371 | 865 | 65 | 751.45 | 44.86 | 3.37 |
| gene5301 | https://www.ncbi.nlm.nih.gov/gene/?term=LOC105278148 | 1881 | 240 | 1275.92 | 26.28 | 3.35 |
| gene11566 | https://www.ncbi.nlm.nih.gov/gene/?term=LOC105285392 | 1603 | 157 | 979.41 | 33.58 | 3.29 |
| gene4635 | https://www.ncbi.nlm.nih.gov/gene/?term=LOC105277390 | 2067 | 168 | 812.77 | 40.39 | 3.28 |

...

... only first 25 lines shown ...

Directory: OGL File: ogl-Cb_LI_RphaseA2.vs.RphaseB1.txt

| gene_ID | gene_link | gwidth | #Sites | #per10Kb | #dmSites | #dmsp10kb | %dmSites | %pSite1 | %pSite2 | DMpSite | ADMpSite | DMpNucl | ADMpNucl |
|---|---|---|---|---|---|---|---|---|---|---|---|---|---|
| gene2593 | https://www.ncbi.nlm.nih.gov/gene/?term=LOC105275070 | 1774 | 11 | 62.01 | 6 | 33.82 | 54.55 | 67.77 | 25.28 | 42.49 | 42.49 | 0.26 | 0.26 |
| gene3165 | https://www.ncbi.nlm.nih.gov/gene/?term=LOC105275706 | 2374 | 70 | 294.86 | 5 | 21.06 | 7.14 | 33.14 | 24.38 | 8.76 | 8.76 | 0.26 | 0.26 |
| gene2824 | https://www.ncbi.nlm.nih.gov/gene/?term=LOC105275337 | 1772 | 63 | 355.53 | 2 | 11.29 | 3.17 | 38.53 | 29.97 | 8.56 | 8.56 | 0.3 | 0.3 |
| gene12395 | https://www.ncbi.nlm.nih.gov/gene/?term=LOC105286302 | 4093 | 24 | 58.64 | 1 | 2.44 | 4.17 | 20.4 | 15.15 | 5.24 | 5.24 | 0.03 | 0.03 |
| gene12961 | https://www.ncbi.nlm.nih.gov/gene/?term=LOC105286983 | 3094 | 71 | 229.48 | 1 | 3.23 | 1.41 | 6.3 | 1.17 | 5.12 | 5.12 | 0.12 | 0.12 |
| gene5714 | https://www.ncbi.nlm.nih.gov/gene/?term=LOC105278729 | 710 | 16 | 225.35 | 1 | 14.08 | 6.25 | 56.86 | 51.9 | 4.96 | 4.96 | 0.11 | 0.11 |
| gene13033 | https://www.ncbi.nlm.nih.gov/gene/?term=LOC105287063 | 1877 | 55 | 293.02 | 1 | 5.33 | 1.82 | 30.5 | 27.04 | 3.46 | 3.46 | 0.1 | 0.1 |
| gene1007 | https://www.ncbi.nlm.nih.gov/gene/?term=LOC105283457 | 1878 | 49 | 260.92 | 1 | 5.32 | 2.04 | 35.2 | 31.9 | 3.3 | 3.3 | 0.09 | 0.09 |
| gene8178 | https://www.ncbi.nlm.nih.gov/gene/?term=LOC105281515 | 8647 | 59 | 68.23 | 1 | 1.16 | 1.69 | 30.95 | 28.43 | 2.52 | 2.52 | 0.02 | 0.02 |
| gene10107 | https://www.ncbi.nlm.nih.gov/gene/?term=LOC105283712 | 1883 | 34 | 180.56 | 1 | 5.31 | 2.94 | 31.9 | 29.48 | 2.42 | 2.42 | 0.04 | 0.04 |
| gene10984 | https://www.ncbi.nlm.nih.gov/gene/?term=LOC105284704 | 1987 | 66 | 332.16 | 1 | 5.03 | 1.52 | 25.73 | 23.31 | 2.42 | 2.42 | 0.08 | 0.08 |
| gene274 | https://www.ncbi.nlm.nih.gov/gene/?term=LOC105275190 | 3488 | 74 | 212.16 | 1 | 2.87 | 1.35 | 22.52 | 20.11 | 2.4 | 2.4 | 0.05 | 0.05 |
| gene2970 | https://www.ncbi.nlm.nih.gov/gene/?term=LOC105275518 | 2340 | 28 | 119.66 | 1 | 4.27 | 3.57 | 41.75 | 39.36 | 2.39 | 2.39 | 0.03 | 0.03 |
| gene2271 | https://www.ncbi.nlm.nih.gov/gene/?term=LOC105274706 | 1448 | 57 | 393.65 | 1 | 6.91 | 1.75 | 39.53 | 37.37 | 2.16 | 2.16 | 0.09 | 0.09 |
| gene4034 | https://www.ncbi.nlm.nih.gov/gene/?term=LOC105276704 | 2065 | 92 | 445.52 | 5 | 24.21 | 5.43 | 14.65 | 12.65 | 2 | 2 | 0.09 | 0.09 |
| gene7568 | https://www.ncbi.nlm.nih.gov/gene/?term=LOC105280786 | 2525 | 143 | 566.34 | 1 | 3.96 | 0.7 | 27.63 | 25.63 | 2 | 2 | 0.11 | 0.11 |
| gene3671 | https://www.ncbi.nlm.nih.gov/gene/?term=LOC105276272 | 1966 | 47 | 239.06 | 1 | 5.09 | 2.13 | 25.43 | 23.44 | 1.98 | 1.98 | 0.05 | 0.05 |
| gene9336 | https://www.ncbi.nlm.nih.gov/gene/?term=LOC105282834 | 5993 | 137 | 228.6 | 1 | 1.67 | 0.73 | 15.64 | 13.69 | 1.95 | 1.95 | 0.04 | 0.04 |
| gene3774 | https://www.ncbi.nlm.nih.gov/gene/?term=LOC105276553 | 2716 | 84 | 309.28 | 1 | 3.68 | 1.19 | 43.69 | 41.82 | 1.87 | 1.87 | 0.06 | 0.06 |
| gene388 | https://www.ncbi.nlm.nih.gov/gene/?term=LOC105276538 | 1558 | 24 | 154.04 | 2 | 12.84 | 8.33 | 66.6 | 64.8 | 1.8 | 1.8 | 0.03 | 0.03 |
| gene4996 | https://www.ncbi.nlm.nih.gov/gene/?term=LOC105277798 | 5617 | 74 | 131.74 | 1 | 1.78 | 1.35 | 11.47 | 9.83 | 1.64 | 1.64 | 0.02 | 0.02 |
| gene10879 | https://www.ncbi.nlm.nih.gov/gene/?term=LOC105284574 | 2423 | 52 | 214.61 | 1 | 4.13 | 1.92 | 56.31 | 54.67 | 1.63 | 1.63 | 0.04 | 0.04 |
| gene8742 | https://www.ncbi.nlm.nih.gov/gene/?term=LOC105282151 | 2563 | 44 | 171.67 | 1 | 3.9 | 2.27 | 43.82 | 42.21 | 1.61 | 1.61 | 0.03 | 0.03 |
| gene940 | https://www.ncbi.nlm.nih.gov/gene/?term=LOC105282794 | 3578 | 95 | 265.51 | 1 | 2.79 | 1.05 | 15.95 | 14.34 | 1.61 | 1.61 | 0.04 | 0.04 |
| gene7493 | https://www.ncbi.nlm.nih.gov/gene/?term=LOC105280704 | 4225 | 155 | 366.86 | 1 | 2.37 | 0.65 | 17.59 | 16.02 | 1.58 | 1.58 | 0.06 | 0.06 |
| gene3817 | https://www.ncbi.nlm.nih.gov/gene/?term=LOC105276449 | 1230 | 55 | 447.15 | 1 | 8.13 | 1.82 | 24.78 | 23.21 | 1.57 | 1.57 | 0.07 | 0.07 |
| gene2241 | https://www.ncbi.nlm.nih.gov/gene/?term=LOC105274666 | 2598 | 64 | 246.34 | 1 | 3.85 | 1.56 | 28.81 | 27.27 | 1.54 | 1.54 | 0.04 | 0.04 |
| gene2748 | https://www.ncbi.nlm.nih.gov/gene/?term=LOC105275261 | 3845 | 55 | 143.04 | 1 | 2.6 | 1.82 | 41.25 | 39.71 | 1.54 | 1.54 | 0.02 | 0.02 |
| gene5273 | https://www.ncbi.nlm.nih.gov/gene/?term=LOC105278108 | 4067 | 57 | 140.15 | 1 | 2.46 | 1.75 | 25.93 | 24.39 | 1.54 | 1.54 | 0.02 | 0.02 |
| gene2825 | https://www.ncbi.nlm.nih.gov/gene/?term=LOC105275343 | 5545 | 98 | 176.74 | 1 | 1.8 | 1.02 | 9.51 | 8.1 | 1.42 | 1.42 | 0.03 | 0.03 |
| gene1147 | https://www.ncbi.nlm.nih.gov/gene/?term=LOC105284954 | 4022 | 114 | 283.44 | 1 | 2.49 | 0.88 | 14.88 | 13.52 | 1.36 | 1.36 | 0.04 | 0.04 |
| gene322 | https://www.ncbi.nlm.nih.gov/gene/?term=LOC105275777 | 1606 | 29 | 180.57 | 1 | 6.23 | 3.45 | 36.14 | 34.8 | 1.35 | 1.35 | 0.02 | 0.02 |
| gene5525 | https://www.ncbi.nlm.nih.gov/gene/?term=LOC105278396 | 3090 | 189 | 611.65 | 1 | 3.24 | 0.53 | 25.87 | 24.52 | 1.35 | 1.35 | 0.08 | 0.08 |
| gene12106 | https://www.ncbi.nlm.nih.gov/gene/?term=LOC105286002 | 8556 | 83 | 97.01 | 1 | 1.17 | 1.2 | 3.39 | 2.1 | 1.29 | 1.29 | 0.01 | 0.01 |
| gene927 | https://www.ncbi.nlm.nih.gov/gene/?term=LOC105282600 | 3727 | 67 | 179.77 | 1 | 2.68 | 1.49 | 27.6 | 26.32 | 1.28 | 1.28 | 0.02 | 0.02 |
| gene12572 | https://www.ncbi.nlm.nih.gov/gene/?term=LOC105286514 | 2374 | 56 | 235.89 | 1 | 4.21 | 1.79 | 36.49 | 35.22 | 1.27 | 1.27 | 0.03 | 0.03 |
| gene6455 | https://www.ncbi.nlm.nih.gov/gene/?term=LOC105279500 | 2612 | 36 | 137.83 | 1 | 3.83 | 2.78 | 48.83 | 47.56 | 1.26 | 1.26 | 0.02 | 0.02 |
| gene4048 | https://www.ncbi.nlm.nih.gov/gene/?term=LOC105276724 | 2955 | 47 | 159.05 | 1 | 3.38 | 2.13 | 12.29 | 11.14 | 1.14 | 1.14 | 0.02 | 0.02 |
| gene5052 | https://www.ncbi.nlm.nih.gov/gene/?term=LOC105277856 | 3167 | 36 | 113.67 | 1 | 3.16 | 2.78 | 37.89 | 36.75 | 1.14 | 1.14 | 0.01 | 0.01 |
| gene935 | https://www.ncbi.nlm.nih.gov/gene/?term=LOC105282682 | 4368 | 240 | 549.45 | 1 | 2.29 | 0.42 | 13.64 | 12.5 | 1.14 | 1.14 | 0.06 | 0.06 |
| gene5524 | https://www.ncbi.nlm.nih.gov/gene/?term=LOC105278397 | 5177 | 206 | 397.91 | 1 | 1.93 | 0.49 | 25.82 | 24.7 | 1.12 | 1.12 | 0.04 | 0.04 |
| gene5831 | https://www.ncbi.nlm.nih.gov/gene/?term=LOC105278761 | 2506 | 119 | 474.86 | 1 | 3.99 | 0.84 | 22.74 | 21.63 | 1.11 | 1.11 | 0.05 | 0.05 |
| gene10222 | https://www.ncbi.nlm.nih.gov/gene/?term=LOC105283838 | 1748 | 33 | 188.79 | 1 | 5.72 | 3.03 | 33.58 | 32.49 | 1.1 | 1.1 | 0.02 | 0.02 |
| gene12406 | https://www.ncbi.nlm.nih.gov/gene/?term=LOC105286316 | 1386 | 52 | 375.18 | 1 | 7.22 | 1.92 | 33.87 | 32.77 | 1.1 | 1.1 | 0.04 | 0.04 |
| gene5877 | https://www.ncbi.nlm.nih.gov/gene/?term=LOC105278807 | 1719 | 53 | 308.32 | 1 | 5.82 | 1.89 | 27.62 | 26.53 | 1.08 | 1.08 | 0.03 | 0.03 |
| gene12021 | https://www.ncbi.nlm.nih.gov/gene/?term=LOC105285905 | 1399 | 27 | 192.99 | 1 | 7.15 | 3.7 | 37.97 | 36.9 | 1.06 | 1.06 | 0.02 | 0.02 |
| gene10243 | https://www.ncbi.nlm.nih.gov/gene/?term=LOC105283862 | 3239 | 45 | 138.93 | 1 | 3.09 | 2.22 | 29.81 | 28.76 | 1.05 | 1.05 | 0.01 | 0.01 |
| gene7183 | https://www.ncbi.nlm.nih.gov/gene/?term=LOC105280348 | 3925 | 79 | 201.27 | 1 | 2.55 | 1.27 | 18.1 | 17.08 | 1.01 | 1.01 | 0.02 | 0.02 |
| gene2833 | https://www.ncbi.nlm.nih.gov/gene/?term=LOC105275352 | 5528 | 43 | 77.79 | 1 | 1.81 | 2.33 | 22.51 | 21.53 | 0.98 | 0.98 | 0.01 | 0.01 |
| gene3773 | https://www.ncbi.nlm.nih.gov/gene/?term=LOC105276408 | 5522 | 200 | 362.19 | 1 | 1.81 | 0.5 | 21.6 | 20.62 | 0.98 | 0.98 | 0.04 | 0.04 |
| gene298 | https://www.ncbi.nlm.nih.gov/gene/?term=LOC105275460 | 2473 | 123 | 497.37 | 1 | 4.04 | 0.81 | 13.29 | 12.33 | 0.96 | 0.96 | 0.05 | 0.05 |
| gene7556 | https://www.ncbi.nlm.nih.gov/gene/?term=LOC105280774 | 2792 | 135 | 483.52 | 1 | 3.58 | 0.74 | 25.37 | 24.41 | 0.96 | 0.96 | 0.05 | 0.05 |
| gene11819 | https://www.ncbi.nlm.nih.gov/gene/?term=LOC105285671 | 2366 | 103 | 435.33 | 1 | 4.23 | 0.97 | 30.19 | 29.25 | 0.95 | 0.95 | 0.04 | 0.04 |
| gene1884 | https://www.ncbi.nlm.nih.gov/gene/?term=LOC105288072 | 4169 | 88 | 211.08 | 1 | 2.4 | 1.14 | 22.65 | 21.71 | 0.94 | 0.94 | 0.02 | 0.02 |
| gene10540 | https://www.ncbi.nlm.nih.gov/gene/?term=LOC105284200 | 5267 | 219 | 415.8 | 1 | 1.9 | 0.46 | 13.98 | 13.1 | 0.88 | 0.88 | 0.04 | 0.04 |
| gene12516 | https://www.ncbi.nlm.nih.gov/gene/?term=LOC105286452 | 1529 | 42 | 274.69 | 1 | 6.54 | 2.38 | 47.45 | 46.6 | 0.86 | 0.86 | 0.02 | 0.02 |
| gene6286 | https://www.ncbi.nlm.nih.gov/gene/?term=LOC105279298 | 2861 | 133 | 464.87 | 2 | 6.99 | 1.5 | 23.83 | 22.97 | 0.86 | 0.86 | 0.04 | 0.04 |
| gene10678 | https://www.ncbi.nlm.nih.gov/gene/?term=LOC105284355 | 4031 | 172 | 426.69 | 1 | 2.48 | 0.58 | 20.12 | 19.3 | 0.83 | 0.83 | 0.04 | 0.04 |
| gene4892 | https://www.ncbi.nlm.nih.gov/gene/?term=LOC105277667 | 3898 | 127 | 325.81 | 1 | 2.57 | 0.79 | 28.13 | 27.36 | 0.77 | 0.77 | 0.03 | 0.03 |
| gene10600 | https://www.ncbi.nlm.nih.gov/gene/?term=LOC105284270 | 4837 | 114 | 235.68 | 1 | 2.07 | 0.88 | 12.4 | 11.65 | 0.75 | 0.75 | 0.02 | 0.02 |

| | | | | | | | | | | | | | |
|---|---|---|---|---|---|---|---|---|---|---|---|---|---|
| gene10606 | https://www.ncbi.nlm.nih.gov/gene/?term=LOC105284273 | 6119 | 286 | 467.4 1 | 1.63 | 0.35 | 18.29 | 17.53 | 0.75 | 0.75 | 0.04 | 0.04 | |
| gene8015 | https://www.ncbi.nlm.nih.gov/gene/?term=LOC105281312 | 2998 | 69 | 230.15 | 1 | 3.34 | 1.45 | 31.17 | 30.43 | 0.74 | 0.74 | 0.02 | 0.02 |
| gene289 | https://www.ncbi.nlm.nih.gov/gene/?term=LOC105275347 | 2533 | 90 | 355.31 | 1 | 3.95 | 1.11 | 24.42 | 23.69 | 0.73 | 0.73 | 0.03 | 0.03 |
| gene10864 | https://www.ncbi.nlm.nih.gov/gene/?term=LOC105284557 | 1666 | 91 | 546.22 | 1 | 6 | 1.1 | 27.88 | 27.16 | 0.72 | 0.72 | 0.04 | 0.04 |
| gene13036 | https://www.ncbi.nlm.nih.gov/gene/?term=LOC105287059 | 4959 | 200 | 403.31 | 1 | 2.02 | 0.5 | 17.56 | 16.85 | 0.71 | 0.71 | 0.03 | 0.03 |
| gene12802 | https://www.ncbi.nlm.nih.gov/gene/?term=LOC105286799 | 3183 | 184 | 578.07 | 1 | 3.14 | 0.54 | 17.39 | 16.7 | 0.69 | 0.69 | 0.04 | 0.04 |
| gene9330 | https://www.ncbi.nlm.nih.gov/gene/?term=LOC105282825 | 3936 | 102 | 259.15 | 1 | 2.54 | 0.98 | 14.91 | 14.23 | 0.68 | 0.68 | 0.02 | 0.02 |
| gene3912 | https://www.ncbi.nlm.nih.gov/gene/?term=LOC105276560 | 17900 | 373 | 208.38 | 3 | 1.68 | 0.8 | 2.56 | 1.89 | 0.67 | 0.67 | 0.01 | 0.01 |
| gene814 | https://www.ncbi.nlm.nih.gov/gene/?term=LOC105281296 | 2722 | 57 | 209.4 1 | 3.67 | 1.75 | 23.86 | 23.2 | 0.65 | 0.65 | 0.01 | 0.01 | |
| gene8343 | https://www.ncbi.nlm.nih.gov/gene/?term=LOC105281700 | 2172 | 43 | 197.97 | 1 | 4.6 | 2.33 | 36.96 | 36.39 | 0.57 | 0.57 | 0.01 | 0.01 |
| gene1573 | https://www.ncbi.nlm.nih.gov/gene/?term=LOC105287704 | 3030 | 143 | 471.95 | 1 | 3.3 | 0.7 | 19.92 | 19.38 | 0.54 | 0.54 | 0.03 | 0.03 |
| gene2257 | https://www.ncbi.nlm.nih.gov/gene/?term=LOC105274689 | 3658 | 90 | 246.04 | 1 | 2.73 | 1.11 | 15.54 | 15.05 | 0.49 | 0.49 | 0.01 | 0.01 |
| gene6031 | https://www.ncbi.nlm.nih.gov/gene/?term=LOC105279000 | 5691 | 226 | 397.12 | 1 | 1.76 | 0.44 | 12.2 | 11.72 | 0.48 | 0.48 | 0.02 | 0.02 |
| gene3206 | https://www.ncbi.nlm.nih.gov/gene/?term=LOC105275752 | 3474 | 138 | 397.24 | 1 | 2.88 | 0.72 | 31.47 | 31.01 | 0.46 | 0.46 | 0.02 | 0.02 |
| gene7888 | https://www.ncbi.nlm.nih.gov/gene/?term=LOC105281141 | 4357 | 180 | 413.13 | 1 | 2.3 | 0.56 | 27.03 | 26.57 | 0.45 | 0.45 | 0.02 | 0.02 |
| gene4641 | https://www.ncbi.nlm.nih.gov/gene/?term=LOC105277398 | 1937 | 74 | 382.03 | 1 | 5.16 | 1.35 | 35.22 | 34.78 | 0.44 | 0.44 | 0.02 | 0.02 |
| gene7249 | https://www.ncbi.nlm.nih.gov/gene/?term=LOC105280422 | 1775 | 66 | 371.83 | 1 | 5.63 | 1.52 | 16.48 | 16.04 | 0.44 | 0.44 | 0.02 | 0.02 |
| gene9648 | https://www.ncbi.nlm.nih.gov/gene/?term=LOC105283183 | 2602 | 33 | 126.83 | 1 | 3.84 | 3.03 | 45.18 | 44.75 | 0.44 | 0.44 | 0.01 | 0.01 |
| gene2330 | https://www.ncbi.nlm.nih.gov/gene/?term=LOC105274773 | 13104 | 208 | 158.73 | 1 | 0.76 | 0.48 | 15.28 | 14.85 | 0.43 | 0.43 | 0.01 | 0.01 |
| gene9647 | https://www.ncbi.nlm.nih.gov/gene/?term=LOC109611233 | 3094 | 35 | 113.12 | 1 | 3.23 | 2.86 | 42.6 | 42.19 | 0.41 | 0.41 | 0 | 0 |
| gene10520 | https://www.ncbi.nlm.nih.gov/gene/?term=LOC105284184 | 2451 | 105 | 428.4 1 | 4.08 | 0.95 | 28.89 | 28.49 | 0.4 | 0.4 | 0.02 | 0.02 | |
| gene2846 | https://www.ncbi.nlm.nih.gov/gene/?term=LOC105275374 | 11246 | 566 | 503.29 | 1 | 0.89 | 0.18 | 9.56 | 9.16 | 0.4 | 0.4 | 0.02 | 0.02 |
| gene8814 | https://www.ncbi.nlm.nih.gov/gene/?term=LOC105282240 | 6910 | 209 | 302.46 | 1 | 1.45 | 0.48 | 8.7 | 8.29 | 0.4 | 0.4 | 0.01 | 0.01 |
| gene570 | https://www.ncbi.nlm.nih.gov/gene/?term=LOC105278549 | 1673 | 74 | 442.32 | 1 | 5.98 | 1.35 | 44.93 | 44.55 | 0.38 | 0.38 | 0.02 | 0.02 |
| gene3885 | https://www.ncbi.nlm.nih.gov/gene/?term=LOC105276527 | 7928 | 380 | 479.31 | 1 | 1.26 | 0.26 | 8.1 | 7.75 | 0.34 | 0.34 | 0.02 | 0.02 |
| gene9029 | https://www.ncbi.nlm.nih.gov/gene/?term=LOC105282491 | 4762 | 190 | 398.99 | 1 | 2.1 | 0.53 | 8.59 | 8.25 | 0.34 | 0.34 | 0.01 | 0.01 |
| gene2056 | https://www.ncbi.nlm.nih.gov/gene/?term=LOC105274452 | 6584 | 268 | 407.05 | 1 | 1.52 | 0.37 | 13.37 | 13.03 | 0.33 | 0.33 | 0.01 | 0.01 |
| gene2844 | https://www.ncbi.nlm.nih.gov/gene/?term=LOC105275363 | 4330 | 258 | 595.84 | 1 | 2.31 | 0.39 | 11.65 | 11.32 | 0.33 | 0.33 | 0.02 | 0.02 |
| gene11568 | https://www.ncbi.nlm.nih.gov/gene/?term=LOC105285395 | 8086 | 257 | 317.83 | 1 | 1.24 | 0.39 | 13.19 | 12.93 | 0.27 | 0.27 | 0.01 | 0.01 |
| gene10018 | https://www.ncbi.nlm.nih.gov/gene/?term=LOC105283607 | 4414 | 222 | 502.95 | 1 | 2.27 | 0.45 | 14.44 | 14.18 | 0.26 | 0.26 | 0.01 | 0.01 |
| gene6620 | https://www.ncbi.nlm.nih.gov/gene/?term=LOC105279687 | 17998 | 456 | 253.36 | 1 | 0.56 | 0.22 | 4.79 | 4.53 | 0.26 | 0.26 | 0.01 | 0.01 |
| gene2683 | https://www.ncbi.nlm.nih.gov/gene/?term=LOC105275187 | 5492 | 87 | 158.41 | 1 | 1.82 | 1.15 | 15.13 | 14.88 | 0.25 | 0.25 | 0 | 0 |
| gene8600 | https://www.ncbi.nlm.nih.gov/gene/?term=LOC105282001 | 8513 | 211 | 247.86 | 1 | 1.17 | 0.47 | 9.85 | 9.6 | 0.25 | 0.25 | 0.01 | 0.01 |
| gene1310 | https://www.ncbi.nlm.nih.gov/gene/?term=LOC105286661 | 2582 | 72 | 278.85 | 1 | 3.87 | 1.39 | 20.69 | 20.45 | 0.24 | 0.24 | 0.01 | 0.01 |
| gene292 | https://www.ncbi.nlm.nih.gov/gene/?term=LOC105275398 | 2825 | 92 | 325.66 | 1 | 3.54 | 1.09 | 21.85 | 21.62 | 0.23 | 0.23 | 0.01 | 0.01 |
| gene11275 | https://www.ncbi.nlm.nih.gov/gene/?term=LOC105285055 | 4119 | 262 | 636.08 | 1 | 2.43 | 0.38 | 11.65 | 11.43 | 0.22 | 0.22 | 0.01 | 0.01 |
| gene8729 | https://www.ncbi.nlm.nih.gov/gene/?term=LOC105282137 | 5005 | 236 | 471.53 | 1 | 2 | 0.42 | 14.47 | 14.26 | 0.21 | 0.21 | 0.01 | 0.01 |
| gene2806 | https://www.ncbi.nlm.nih.gov/gene/?term=LOC105275327 | 14257 | 550 | 385.78 | 1 | 0.7 | 0.18 | 6.58 | 6.38 | 0.2 | 0.2 | 0.01 | 0.01 |
| gene3747 | https://www.ncbi.nlm.nih.gov/gene/?term=LOC105276387 | 3736 | 198 | 529.98 | 1 | 2.68 | 0.51 | 14.52 | 14.33 | 0.19 | 0.19 | 0.01 | 0.01 |
| gene5746 | https://www.ncbi.nlm.nih.gov/gene/?term=LOC105278666 | 10343 | 491 | 474.72 | 1 | 0.97 | 0.2 | 6.89 | 6.71 | 0.19 | 0.19 | 0.01 | 0.01 |
| gene10596 | https://www.ncbi.nlm.nih.gov/gene/?term=LOC105284262 | 6520 | 247 | 378.83 | 1 | 1.53 | 0.4 | 4.51 | 4.34 | 0.18 | 0.18 | 0.01 | 0.01 |
| gene7245 | https://www.ncbi.nlm.nih.gov/gene/?term=LOC105280410 | 12478 | 319 | 255.65 | 1 | 0.8 | 0.31 | 17.86 | 17.71 | 0.15 | 0.15 | 0 | 0 |
| gene2173 | https://www.ncbi.nlm.nih.gov/gene/?term=LOC105274590 | 4387 | 91 | 207.43 | 2 | 4.56 | 2.2 | 24.1 | 23.96 | 0.14 | 0.14 | 0 | 0 |
| gene4225 | https://www.ncbi.nlm.nih.gov/gene/?term=LOC105276917 | 2443 | 40 | 163.73 | 1 | 4.09 | 2.5 | 29.99 | 29.86 | 0.14 | 0.14 | 0 | 0 |
| gene6966 | https://www.ncbi.nlm.nih.gov/gene/?term=LOC105280104 | 3747 | 133 | 354.95 | 1 | 2.67 | 0.75 | 17.09 | 16.96 | 0.13 | 0.13 | 0 | 0 |
| gene6652 | https://www.ncbi.nlm.nih.gov/gene/?term=LOC105279719 | 2831 | 83 | 293.18 | 1 | 3.53 | 1.2 | 23.57 | 23.44 | 0.12 | 0.12 | 0 | 0 |
| gene9722 | https://www.ncbi.nlm.nih.gov/gene/?term=LOC105283264 | 3951 | 94 | 237.91 | 1 | 2.53 | 1.06 | 22.96 | 22.87 | 0.1 | 0.1 | 0 | 0 |
| gene4033 | https://www.ncbi.nlm.nih.gov/gene/?term=LOC105276705 | 10546 | 157 | 148.87 | 1 | 0.95 | 0.64 | 6.45 | 6.38 | 0.06 | 0.06 | 0 | 0 |
| gene3727 | https://www.ncbi.nlm.nih.gov/gene/?term=LOC105276367 | 2484 | 76 | 305.96 | 1 | 4.03 | 1.32 | 41.02 | 40.98 | 0.04 | 0.04 | 0 | 0 |
| gene12307 | https://www.ncbi.nlm.nih.gov/gene/?term=LOC105286216 | 10313 | 209 | 202.66 | 1 | 0.97 | 0.48 | 14.07 | 14.04 | 0.02 | 0.02 | 0 | 0 |
| gene3470 | https://www.ncbi.nlm.nih.gov/gene/?term=LOC105276059 | 8716 | 186 | 213.4 1 | 1.15 | 0.54 | 9.64 | 9.62 | 0.02 | 0.02 | 0 | 0 | |
| gene3107 | https://www.ncbi.nlm.nih.gov/gene/?term=LOC105275650 | 5103 | 187 | 366.45 | 1 | 1.96 | 0.53 | 16.49 | 16.48 | 0.01 | 0.01 | 0 | 0 |
| gene3335 | https://www.ncbi.nlm.nih.gov/gene/?term=LOC105275898 | 4665 | 172 | 368.7 1 | 2.14 | 0.58 | 12.53 | 12.51 | 0.01 | 0.01 | 0 | 0 | |
| gene9326 | https://www.ncbi.nlm.nih.gov/gene/?term=LOC105282818 | 16612 | 251 | 151.1 1 | 0.6 | 0.4 | 12.99 | 12.99 | 0 | 0 | 0 | 0 | |
| gene11561 | https://www.ncbi.nlm.nih.gov/gene/?term=LOC105285387 | 11528 | 226 | 196.04 | 1 | 0.87 | 0.44 | 9.46 | 9.47 | -0.01 | 0.01 | 0 | 0 |
| gene13039 | https://www.ncbi.nlm.nih.gov/gene/?term=LOC105287065 | 4902 | 263 | 536.52 | 1 | 2.04 | 0.38 | 11.98 | 12.01 | -0.02 | 0.02 | 0 | 0 |
| gene7166 | https://www.ncbi.nlm.nih.gov/gene/?term=LOC105280326 | 6117 | 258 | 421.78 | 1 | 1.63 | 0.39 | 14.79 | 14.83 | -0.04 | 0.04 | 0 | 0 |
| gene6649 | https://www.ncbi.nlm.nih.gov/gene/?term=LOC105279717 | 6017 | 181 | 300.81 | 1 | 1.66 | 0.55 | 11.29 | 11.35 | -0.06 | 0.06 | 0 | 0 |
| gene3856 | https://www.ncbi.nlm.nih.gov/gene/?term=LOC105276499 | 5236 | 206 | 393.43 | 1 | 1.91 | 0.49 | 10.22 | 10.29 | -0.07 | 0.07 | 0 | 0 |
| gene10326 | https://www.ncbi.nlm.nih.gov/gene/?term=LOC105283962 | 3042 | 145 | 476.66 | 1 | 3.29 | 0.69 | 9.18 | 9.26 | -0.08 | 0.08 | 0 | 0 |
| gene10549 | https://www.ncbi.nlm.nih.gov/gene/?term=LOC105284210 | 7145 | 374 | 523.44 | 2 | 2.8 | 0.53 | 9.66 | 9.75 | -0.09 | 0.09 | 0 | 0 |
| gene2835 | https://www.ncbi.nlm.nih.gov/gene/?term=LOC105275357 | 2492 | 122 | 489.57 | 1 | 4.01 | 0.82 | 14.14 | 14.23 | -0.09 | 0.09 | 0 | 0 |
| gene4190 | https://www.ncbi.nlm.nih.gov/gene/?term=LOC105276875 | 3150 | 135 | 428.57 | 1 | 3.17 | 0.74 | 20.47 | 20.57 | -0.09 | 0.09 | 0 | 0 |
| gene5613 | https://www.ncbi.nlm.nih.gov/gene/?term=LOC105278504 | 8524 | 274 | 321.45 | 1 | 1.17 | 0.36 | 5.89 | 5.98 | -0.09 | 0.09 | 0 | 0 |

| | | | | | | | | | | | | | |
|---|---|---|---|---|---|---|---|---|---|---|---|---|---|
| gene9755 | https://www.ncbi.nlm.nih.gov/gene/?term=LOC105283296 | 2177 | 97 | 445.57 | 1 | 4.59 | 1.03 | 14.1 | 14.18 | -0.09 | 0.09 | 0 | 0 |
| gene13114 | https://www.ncbi.nlm.nih.gov/gene/?term=LOC105287147 | 7609 | 447 | 587.46 | 1 | 1.31 | 0.22 | 9.07 | 9.17 | -0.1 | 0.1 | -0.01 | 0.01 |
| gene4376 | https://www.ncbi.nlm.nih.gov/gene/?term=LOC105277083 | 4307 | 140 | 325.05 | 1 | 2.32 | 0.71 | 17.67 | 17.76 | -0.1 | 0.1 | 0 | 0 |
| gene53 | https://www.ncbi.nlm.nih.gov/gene/?term=LOC105277148 | 19283 | 339 | 175.8 1 | 0.52 | 0.29 | 4.53 | 4.65 | -0.11 | 0.11 | 0 | 0 | |
| gene1632 | https://www.ncbi.nlm.nih.gov/gene/?term=LOC105287764 | 18061 | 462 | 255.8 1 | 0.55 | 0.22 | 3.51 | 3.63 | -0.12 | 0.12 | 0 | 0 | |
| gene8274 | https://www.ncbi.nlm.nih.gov/gene/?term=LOC105281624 | 5425 | 244 | 449.77 | 1 | 1.84 | 0.41 | 9.34 | 9.46 | -0.12 | 0.12 | -0.01 | 0.01 |
| gene3599 | https://www.ncbi.nlm.nih.gov/gene/?term=LOC105276196 | 2955 | 61 | 206.43 | 1 | 3.38 | 1.64 | 38.99 | 39.11 | -0.13 | 0.13 | 0 | 0 |
| gene9327 | https://www.ncbi.nlm.nih.gov/gene/?term=LOC105282821 | 7793 | 184 | 236.11 | 1 | 1.28 | 0.54 | 12.73 | 12.87 | -0.14 | 0.14 | 0 | 0 |
| gene11579 | https://www.ncbi.nlm.nih.gov/gene/?term=LOC105285406 | 13571 | 341 | 251.27 | 1 | 0.74 | 0.29 | 6.04 | 6.2 | -0.15 | 0.15 | 0 | 0 |
| gene12674 | https://www.ncbi.nlm.nih.gov/gene/?term=LOC105286649 | 18283 | 836 | 457.26 | 1 | 0.55 | 0.12 | 1.34 | 1.5 | -0.16 | 0.16 | -0.01 | 0.01 |
| gene11082 | https://www.ncbi.nlm.nih.gov/gene/?term=LOC105284824 | 7436 | 245 | 329.48 | 1 | 1.34 | 0.41 | 7.32 | 7.57 | -0.25 | 0.25 | -0.01 | 0.01 |
| gene1198 | https://www.ncbi.nlm.nih.gov/gene/?term=LOC105285507 | 2738 | 98 | 357.93 | 1 | 3.65 | 1.02 | 21.45 | 21.7 | -0.25 | 0.25 | -0.01 | 0.01 |
| gene7368 | https://www.ncbi.nlm.nih.gov/gene/?term=LOC105280556 | 4105 | 76 | 185.14 | 1 | 2.44 | 1.32 | 15.19 | 15.44 | -0.25 | 0.25 | 0 | 0 |
| gene9464 | https://www.ncbi.nlm.nih.gov/gene/?term=LOC105282975 | 1450 | 25 | 172.41 | 1 | 6.9 | 4 | 40.46 | 40.7 | -0.25 | 0.25 | 0 | 0 |
| gene12407 | https://www.ncbi.nlm.nih.gov/gene/?term=LOC105286315 | 8593 | 227 | 264.17 | 1 | 1.16 | 0.44 | 13.44 | 13.72 | -0.28 | 0.28 | -0.01 | 0.01 |
| gene11583 | https://www.ncbi.nlm.nih.gov/gene/?term=LOC105285414 | 8192 | 204 | 249.02 | 1 | 1.22 | 0.49 | 7.78 | 8.09 | -0.31 | 0.31 | -0.01 | 0.01 |
| gene12573 | https://www.ncbi.nlm.nih.gov/gene/?term=LOC105286515 | 3647 | 128 | 350.97 | 1 | 2.74 | 0.78 | 10.64 | 10.97 | -0.33 | 0.33 | -0.01 | 0.01 |
| gene9598 | https://www.ncbi.nlm.nih.gov/gene/?term=LOC105283132 | 1769 | 56 | 316.56 | 1 | 5.65 | 1.79 | 39.81 | 40.18 | -0.38 | 0.38 | -0.01 | 0.01 |
| gene1220 | https://www.ncbi.nlm.nih.gov/gene/?term=LOC105285742 | 3453 | 123 | 356.21 | 1 | 2.9 | 0.81 | 16.77 | 17.17 | -0.4 | 0.4 | -0.01 | 0.01 |
| gene657 | https://www.ncbi.nlm.nih.gov/gene/?term=LOC105279667 | 6421 | 106 | 165.08 | 1 | 1.56 | 0.94 | 2.17 | 2.57 | -0.4 | 0.4 | -0.01 | 0.01 |
| gene5646 | https://www.ncbi.nlm.nih.gov/gene/?term=LOC105278534 | 7475 | 214 | 286.29 | 1 | 1.34 | 0.47 | 16.15 | 16.57 | -0.41 | 0.41 | -0.01 | 0.01 |
| gene7148 | https://www.ncbi.nlm.nih.gov/gene/?term=LOC105280304 | 8920 | 143 | 160.31 | 1 | 1.12 | 0.7 | 6.95 | 7.37 | -0.42 | 0.42 | -0.01 | 0.01 |
| gene7600 | https://www.ncbi.nlm.nih.gov/gene/?term=LOC105280819 | 1615 | 43 | 266.25 | 1 | 6.19 | 2.33 | 37.46 | 37.88 | -0.42 | 0.42 | -0.01 | 0.01 |
| gene1470 | https://www.ncbi.nlm.nih.gov/gene/?term=LOC105287597 | 2974 | 200 | 672.49 | 1 | 3.36 | 0.5 | 18.11 | 18.56 | -0.46 | 0.46 | -0.03 | 0.03 |
| gene6583 | https://www.ncbi.nlm.nih.gov/gene/?term=LOC105279649 | 7444 | 393 | 527.94 | 1 | 1.34 | 0.25 | 10.67 | 11.14 | -0.47 | 0.47 | -0.02 | 0.02 |
| gene10345 | https://www.ncbi.nlm.nih.gov/gene/?term=LOC105283989 | 4444 | 40 | 90.01 1 | 2.25 | 2.5 | 37.64 | 38.15 | -0.5 | 0.5 | 0 | 0 | |
| gene5298 | https://www.ncbi.nlm.nih.gov/gene/?term=LOC105278145 | 4060 | 275 | 677.34 | 2 | 4.93 | 0.73 | 9.89 | 10.42 | -0.53 | 0.53 | -0.04 | 0.04 |
| gene3848 | https://www.ncbi.nlm.nih.gov/gene/?term=LOC105276486 | 5899 | 176 | 298.36 | 1 | 1.7 | 0.57 | 23.6 | 24.17 | -0.57 | 0.57 | -0.02 | 0.02 |
| gene4962 | https://www.ncbi.nlm.nih.gov/gene/?term=LOC105277761 | 4645 | 175 | 376.75 | 1 | 2.15 | 0.57 | 13.58 | 14.2 | -0.62 | 0.62 | -0.02 | 0.02 |
| gene7207 | https://www.ncbi.nlm.nih.gov/gene/?term=LOC105280376 | 4659 | 124 | 266.15 | 1 | 2.15 | 0.81 | 26.54 | 27.17 | -0.63 | 0.63 | -0.02 | 0.02 |
| gene12377 | https://www.ncbi.nlm.nih.gov/gene/?term=LOC105286279 | 5846 | 278 | 475.54 | 1 | 1.71 | 0.36 | 2.15 | 2.82 | -0.67 | 0.67 | -0.03 | 0.03 |
| gene7550 | https://www.ncbi.nlm.nih.gov/gene/?term=LOC105280763 | 2173 | 109 | 501.61 | 1 | 4.6 | 0.92 | 27.44 | 28.14 | -0.7 | 0.7 | -0.04 | 0.04 |
| gene10819 | https://www.ncbi.nlm.nih.gov/gene/?term=LOC105284507 | 5767 | 122 | 211.55 | 1 | 1.73 | 0.82 | 18.54 | 19.25 | -0.71 | 0.71 | -0.01 | 0.01 |
| gene3385 | https://www.ncbi.nlm.nih.gov/gene/?term=LOC105275966 | 3305 | 75 | 226.93 | 1 | 3.03 | 1.33 | 43.86 | 44.6 | -0.75 | 0.75 | -0.02 | 0.02 |
| gene10213 | https://www.ncbi.nlm.nih.gov/gene/?term=LOC105283826 | 1503 | 62 | 412.51 | 1 | 6.65 | 1.61 | 14.74 | 15.54 | -0.8 | 0.8 | -0.03 | 0.03 |
| gene11454 | https://www.ncbi.nlm.nih.gov/gene/?term=LOC105285256 | 1906 | 50 | 262.33 | 1 | 5.25 | 2 | 43.28 | 44.08 | -0.8 | 0.8 | -0.02 | 0.02 |
| gene10392 | https://www.ncbi.nlm.nih.gov/gene/?term=LOC105284036 | 3216 | 103 | 320.27 | 2 | 6.22 | 1.94 | 24.04 | 24.86 | -0.81 | 0.81 | -0.03 | 0.03 |
| gene7915 | https://www.ncbi.nlm.nih.gov/gene/?term=LOC105281173 | 1597 | 39 | 244.21 | 1 | 6.26 | 2.56 | 61.13 | 62.01 | -0.88 | 0.88 | -0.02 | 0.02 |
| gene11401 | https://www.ncbi.nlm.nih.gov/gene/?term=LOC105285202 | 1841 | 89 | 483.43 | 1 | 5.43 | 1.12 | 23.71 | 24.59 | -0.89 | 0.89 | -0.04 | 0.04 |
| gene11447 | https://www.ncbi.nlm.nih.gov/gene/?term=LOC105285252 | 7437 | 150 | 201.69 | 1 | 1.34 | 0.67 | 11.78 | 12.78 | -0.99 | 0.99 | -0.02 | 0.02 |
| gene3168 | https://www.ncbi.nlm.nih.gov/gene/?term=LOC105275711 | 6150 | 95 | 154.47 | 1 | 1.63 | 1.05 | 10.97 | 11.98 | -1 | 1 | -0.02 | 0.02 |
| gene12203 | https://www.ncbi.nlm.nih.gov/gene/?term=LOC105286099 | 5967 | 87 | 145.8 1 | 1.68 | 1.15 | 11.05 | 12.06 | -1.01 | 1.01 | -0.01 | 0.01 | |
| gene7657 | https://www.ncbi.nlm.nih.gov/gene/?term=LOC105280890 | 1109 | 23 | 207.39 | 1 | 9.02 | 4.35 | 23.17 | 24.26 | -1.09 | 1.09 | -0.02 | 0.02 |
| gene652 | https://www.ncbi.nlm.nih.gov/gene/?term=LOC105279528 | 5806 | 39 | 67.17 1 | 1.72 | 2.56 | 19.31 | 20.42 | -1.11 | 1.11 | -0.01 | 0.01 | |
| gene6639 | https://www.ncbi.nlm.nih.gov/gene/?term=LOC105279705 | 7656 | 88 | 114.94 | 1 | 1.31 | 1.14 | 19.96 | 21.09 | -1.13 | 1.13 | -0.01 | 0.01 |
| gene9101 | https://www.ncbi.nlm.nih.gov/gene/?term=LOC105282553 | 3068 | 110 | 358.54 | 1 | 3.26 | 0.91 | 21.48 | 22.61 | -1.13 | 1.13 | -0.04 | 0.04 |
| gene10155 | https://www.ncbi.nlm.nih.gov/gene/?term=LOC105283756 | 7030 | 119 | 169.27 | 1 | 1.42 | 0.84 | 6.89 | 8.19 | -1.3 | 1.3 | -0.02 | 0.02 |
| gene2404 | https://www.ncbi.nlm.nih.gov/gene/?term=LOC105274853 | 1767 | 45 | 254.67 | 1 | 5.66 | 2.22 | 30.61 | 31.94 | -1.33 | 1.33 | -0.03 | 0.03 |
| gene10654 | https://www.ncbi.nlm.nih.gov/gene/?term=LOC105284329 | 4027 | 72 | 178.79 | 1 | 2.48 | 1.39 | 18.86 | 20.42 | -1.56 | 1.56 | -0.03 | 0.03 |
| gene564 | https://www.ncbi.nlm.nih.gov/gene/?term=LOC105278478 | 2802 | 22 | 78.52 1 | 3.57 | 4.55 | 23.15 | 24.78 | -1.63 | 1.63 | -0.01 | 0.01 | |
| gene11340 | https://www.ncbi.nlm.nih.gov/gene/?term=LOC105285131 | 2918 | 100 | 342.7 1 | 3.43 | 1 | 13.44 | 15.08 | -1.64 | 1.64 | -0.06 | 0.06 | |
| gene6237 | https://www.ncbi.nlm.nih.gov/gene/?term=LOC105279244 | 1484 | 30 | 202.16 | 1 | 6.74 | 3.33 | 49.36 | 51.05 | -1.69 | 1.69 | -0.03 | 0.03 |
| gene920 | https://www.ncbi.nlm.nih.gov/gene/?term=LOC105282518 | 1197 | 46 | 384.29 | 1 | 8.35 | 2.17 | 30.45 | 32.64 | -2.19 | 2.19 | -0.08 | 0.08 |
| gene9411 | https://www.ncbi.nlm.nih.gov/gene/?term=LOC105282914 | 5534 | 42 | 75.89 1 | 1.81 | 2.38 | 48.54 | 50.92 | -2.38 | 2.38 | -0.02 | 0.02 | |
| gene1768 | https://www.ncbi.nlm.nih.gov/gene/?term=LOC105287938 | 2792 | 89 | 318.77 | 1 | 3.58 | 1.12 | 25.55 | 28.06 | -2.51 | 2.51 | -0.08 | 0.08 |
| gene10552 | https://www.ncbi.nlm.nih.gov/gene/?term=LOC105284213 | 878 | 23 | 261.96 | 1 | 11.39 | 4.35 | 9.72 | 12.3 | -2.58 | 2.58 | -0.07 | 0.07 |
| gene7398 | https://www.ncbi.nlm.nih.gov/gene/?term=LOC105280589 | 3452 | 42 | 121.67 | 1 | 2.9 | 2.38 | 23.2 | 25.91 | -2.71 | 2.71 | -0.03 | 0.03 |
| gene2413 | https://www.ncbi.nlm.nih.gov/gene/?term=LOC105274863 | 2006 | 65 | 324.03 | 1 | 4.99 | 1.54 | 49.74 | 52.89 | -3.14 | 3.14 | -0.1 | 0.1 |
| gene1199 | https://www.ncbi.nlm.nih.gov/gene/?term=LOC105285498 | 2220 | 29 | 130.63 | 1 | 4.5 | 3.45 | 32.78 | 35.99 | -3.21 | 3.21 | -0.04 | 0.04 |
| gene1505 | https://www.ncbi.nlm.nih.gov/gene/?term=LOC105287633 | 1693 | 24 | 141.76 | 1 | 5.91 | 4.17 | 48.26 | 51.48 | -3.22 | 3.22 | -0.05 | 0.05 |
| gene9109 | https://www.ncbi.nlm.nih.gov/gene/?term=LOC105282561 | 2400 | 17 | 70.83 1 | 4.17 | 5.88 | 28.52 | 31.86 | -3.34 | 3.34 | -0.02 | 0.02 | |
| gene11584 | https://www.ncbi.nlm.nih.gov/gene/?term=LOC105285416 | 3709 | 15 | 40.44 2 | 5.39 | 13.33 | 24.1 | 29.33 | -5.23 | 5.23 | -0.02 | 0.02 | |
| gene3166 | https://www.ncbi.nlm.nih.gov/gene/?term=LOC105275707 | 2393 | 12 | 50.15 1 | 4.18 | 8.33 | 62.45 | 73.29 | -10.83 | | 10.83 | -0.05 | 0.05 |
| gene8691 | https://www.ncbi.nlm.nih.gov/gene/?term=LOC105282094 | 2028 | 6 | 29.59 1 | 4.93 | 16.67 | 86.17 | 97.58 | -11.41 | | 11.41 | -0.03 | 0.03 |

| | | | | | | | | | | | | | |
|---|---|---|---|---|---|---|---|---|---|---|---|---|---|
| gene393 | https://www.ncbi.nlm.nih.gov/gene/?term=LOC105276566 | 1123 | 2 | 17.81 | 1 | 8.9 | 50 | 47.62 | 74.3 | -26.68 | 26.68 | -0.05 | 0.05 |

Directory: OGL File: ogl-Cb_LI_RphaseA2.vs.RphaseB2.txt

| gene_ID | gene_link | gwidth | #Sites | #per10Kb | #dmSites | #dmsp10kb | %dmSites | %pSite1 | %pSite2 | DMpSite | ADMpSite | DMpNucl | ADMpNucl |
|---|---|---|---|---|---|---|---|---|---|---|---|---|---|
| gene2593 | https://www.ncbi.nlm.nih.gov/gene/?term=LOC105275070 | 1774 | 7 | 39.46 | 5 | 28.18 | 71.43 | 65.68 | 4.09 | 61.59 | 61.59 | 0.24 | 0.24 |
| gene3165 | https://www.ncbi.nlm.nih.gov/gene/?term=LOC105275706 | 2374 | 74 | 311.71 | 6 | 25.27 | 8.11 | 32.18 | 22.24 | 9.94 | 9.94 | 0.31 | 0.31 |
| gene2824 | https://www.ncbi.nlm.nih.gov/gene/?term=LOC105275337 | 1772 | 60 | 338.6 | 2 | 11.29 | 3.33 | 38.84 | 29.82 | 9.02 | 9.02 | 0.31 | 0.31 |
| gene2907 | https://www.ncbi.nlm.nih.gov/gene/?term=LOC105275447 | 2539 | 37 | 145.73 | 1 | 3.94 | 2.7 | 8.22 | 2.41 | 5.8 | 5.8 | 0.08 | 0.08 |
| gene5714 | https://www.ncbi.nlm.nih.gov/gene/?term=LOC105278729 | 710 | 17 | 239.44 | 1 | 14.08 | 5.88 | 59.17 | 53.54 | 5.62 | 5.62 | 0.13 | 0.13 |
| gene8943 | https://www.ncbi.nlm.nih.gov/gene/?term=LOC105282388 | 1220 | 33 | 270.49 | 1 | 8.2 | 3.03 | 49.82 | 45.11 | 4.71 | 4.71 | 0.13 | 0.13 |
| gene2335 | https://www.ncbi.nlm.nih.gov/gene/?term=LOC105274778 | 3945 | 38 | 96.32 | 1 | 2.53 | 2.63 | 48.07 | 44.68 | 3.39 | 3.39 | 0.03 | 0.03 |
| gene12021 | https://www.ncbi.nlm.nih.gov/gene/?term=LOC105285905 | 1399 | 16 | 114.37 | 1 | 7.15 | 6.25 | 51.11 | 47.75 | 3.36 | 3.36 | 0.04 | 0.04 |
| gene2970 | https://www.ncbi.nlm.nih.gov/gene/?term=LOC105275518 | 2340 | 24 | 102.56 | 1 | 4.27 | 4.17 | 49.7 | 46.61 | 3.09 | 3.09 | 0.03 | 0.03 |
| gene7568 | https://www.ncbi.nlm.nih.gov/gene/?term=LOC105280786 | 2525 | 126 | 499.01 | 1 | 3.96 | 0.79 | 30.35 | 27.57 | 2.78 | 2.78 | 0.14 | 0.14 |
| gene2271 | https://www.ncbi.nlm.nih.gov/gene/?term=LOC105274706 | 1448 | 56 | 386.74 | 1 | 6.91 | 1.79 | 38.93 | 36.4 | 2.53 | 2.53 | 0.1 | 0.1 |
| gene4225 | https://www.ncbi.nlm.nih.gov/gene/?term=LOC105276917 | 2443 | 36 | 147.36 | 1 | 4.09 | 2.78 | 30.16 | 27.69 | 2.47 | 2.47 | 0.04 | 0.04 |
| gene940 | https://www.ncbi.nlm.nih.gov/gene/?term=LOC105282794 | 3578 | 88 | 245.95 | 1 | 2.79 | 1.14 | 16.61 | 14.17 | 2.44 | 2.44 | 0.06 | 0.06 |
| gene13033 | https://www.ncbi.nlm.nih.gov/gene/?term=LOC105287063 | 1877 | 48 | 255.73 | 1 | 5.33 | 2.08 | 32.8 | 30.37 | 2.43 | 2.43 | 0.06 | 0.06 |
| gene4034 | https://www.ncbi.nlm.nih.gov/gene/?term=LOC105276704 | 2065 | 88 | 426.15 | 1 | 4.84 | 1.14 | 16.14 | 13.92 | 2.22 | 2.22 | 0.09 | 0.09 |
| gene5052 | https://www.ncbi.nlm.nih.gov/gene/?term=LOC105277856 | 3167 | 34 | 107.36 | 1 | 3.16 | 2.94 | 35.71 | 33.56 | 2.15 | 2.15 | 0.02 | 0.02 |
| gene5273 | https://www.ncbi.nlm.nih.gov/gene/?term=LOC105278108 | 4067 | 51 | 125.4 | 1 | 2.46 | 1.96 | 25.16 | 23.04 | 2.12 | 2.12 | 0.03 | 0.03 |
| gene9598 | https://www.ncbi.nlm.nih.gov/gene/?term=LOC105283132 | 1769 | 57 | 322.22 | 1 | 5.65 | 1.75 | 39.92 | 37.87 | 2.05 | 2.05 | 0.07 | 0.07 |
| gene12106 | https://www.ncbi.nlm.nih.gov/gene/?term=LOC105286002 | 8556 | 68 | 79.48 | 1 | 1.17 | 1.47 | 4.38 | 2.38 | 2 | 2 | 0.02 | 0.02 |
| gene11819 | https://www.ncbi.nlm.nih.gov/gene/?term=LOC105285671 | 2366 | 103 | 435.33 | 1 | 4.23 | 0.97 | 27.63 | 25.66 | 1.97 | 1.97 | 0.09 | 0.09 |
| gene7211 | https://www.ncbi.nlm.nih.gov/gene/?term=LOC105280377 | 2024 | 69 | 340.91 | 1 | 4.94 | 1.45 | 48.01 | 46.12 | 1.89 | 1.89 | 0.06 | 0.06 |
| gene927 | https://www.ncbi.nlm.nih.gov/gene/?term=LOC105282600 | 3727 | 54 | 144.89 | 1 | 2.68 | 1.85 | 28.21 | 26.36 | 1.85 | 1.85 | 0.03 | 0.03 |
| gene388 | https://www.ncbi.nlm.nih.gov/gene/?term=LOC105276538 | 1558 | 21 | 134.79 | 2 | 12.84 | 9.52 | 69.8 | 67.98 | 1.82 | 1.82 | 0.02 | 0.02 |
| gene5525 | https://www.ncbi.nlm.nih.gov/gene/?term=LOC105278396 | 3090 | 184 | 595.47 | 1 | 3.24 | 0.54 | 26.64 | 24.83 | 1.81 | 1.81 | 0.11 | 0.11 |
| gene8178 | https://www.ncbi.nlm.nih.gov/gene/?term=LOC105281515 | 8647 | 45 | 52.04 | 1 | 1.16 | 2.22 | 25.4 | 23.65 | 1.75 | 1.75 | 0.01 | 0.01 |
| gene3594 | https://www.ncbi.nlm.nih.gov/gene/?term=LOC105276190 | 4910 | 95 | 193.48 | 1 | 2.04 | 1.05 | 10.96 | 9.28 | 1.68 | 1.68 | 0.03 | 0.03 |
| gene5524 | https://www.ncbi.nlm.nih.gov/gene/?term=LOC105278397 | 5177 | 201 | 388.26 | 1 | 1.93 | 0.5 | 26.23 | 24.63 | 1.61 | 1.61 | 0.06 | 0.06 |
| gene9646 | https://www.ncbi.nlm.nih.gov/gene/?term=LOC105283182 | 3588 | 133 | 370.68 | 1 | 2.79 | 0.75 | 13.97 | 12.43 | 1.54 | 1.54 | 0.06 | 0.06 |
| gene1518 | https://www.ncbi.nlm.nih.gov/gene/?term=LOC105287648 | 2195 | 26 | 118.45 | 1 | 4.56 | 3.85 | 41.13 | 39.6 | 1.53 | 1.53 | 0.02 | 0.02 |
| gene564 | https://www.ncbi.nlm.nih.gov/gene/?term=LOC105278478 | 2802 | 28 | 99.93 | 1 | 3.57 | 3.57 | 18.19 | 16.66 | 1.53 | 1.53 | 0.02 | 0.02 |
| gene5647 | https://www.ncbi.nlm.nih.gov/gene/?term=LOC105278567 | 3118 | 38 | 121.87 | 1 | 3.21 | 2.63 | 15.22 | 13.72 | 1.5 | 1.5 | 0.02 | 0.02 |
| gene3817 | https://www.ncbi.nlm.nih.gov/gene/?term=LOC105276449 | 1230 | 52 | 422.76 | 1 | 8.13 | 1.92 | 31.62 | 30.14 | 1.48 | 1.48 | 0.06 | 0.06 |
| gene6455 | https://www.ncbi.nlm.nih.gov/gene/?term=LOC105279500 | 2612 | 35 | 134 | 1 | 3.83 | 2.86 | 49.73 | 48.29 | 1.45 | 1.45 | 0.02 | 0.02 |
| gene10879 | https://www.ncbi.nlm.nih.gov/gene/?term=LOC105284574 | 2423 | 53 | 218.74 | 1 | 4.13 | 1.89 | 56.3 | 54.86 | 1.44 | 1.44 | 0.03 | 0.03 |
| gene10984 | https://www.ncbi.nlm.nih.gov/gene/?term=LOC105284704 | 1987 | 61 | 307 | 1 | 5.03 | 1.64 | 24.7 | 23.27 | 1.43 | 1.43 | 0.04 | 0.04 |
| gene8742 | https://www.ncbi.nlm.nih.gov/gene/?term=LOC105282151 | 2563 | 39 | 152.17 | 2 | 7.8 | 5.13 | 42.27 | 40.86 | 1.41 | 1.41 | 0.02 | 0.02 |
| gene9462 | https://www.ncbi.nlm.nih.gov/gene/?term=LOC105282970 | 1469 | 40 | 272.29 | 1 | 6.81 | 2.5 | 49.62 | 48.23 | 1.39 | 1.39 | 0.04 | 0.04 |
| gene4996 | https://www.ncbi.nlm.nih.gov/gene/?term=LOC105277798 | 5617 | 56 | 99.7 | 1 | 1.78 | 1.79 | 13.35 | 11.98 | 1.38 | 1.38 | 0.01 | 0.01 |
| gene8661 | https://www.ncbi.nlm.nih.gov/gene/?term=LOC105282063 | 1926 | 36 | 186.92 | 1 | 5.19 | 2.78 | 33.96 | 32.69 | 1.28 | 1.28 | 0.02 | 0.02 |
| gene9336 | https://www.ncbi.nlm.nih.gov/gene/?term=LOC105282834 | 5993 | 129 | 215.25 | 1 | 1.67 | 0.78 | 16.18 | 14.94 | 1.24 | 1.24 | 0.03 | 0.03 |
| gene10600 | https://www.ncbi.nlm.nih.gov/gene/?term=LOC105284270 | 4837 | 127 | 262.56 | 1 | 2.07 | 0.79 | 10.81 | 9.59 | 1.22 | 1.22 | 0.03 | 0.03 |
| gene2748 | https://www.ncbi.nlm.nih.gov/gene/?term=LOC105275261 | 3845 | 58 | 150.85 | 1 | 2.6 | 1.72 | 39.72 | 38.52 | 1.2 | 1.2 | 0.02 | 0.02 |
| gene10678 | https://www.ncbi.nlm.nih.gov/gene/?term=LOC105284355 | 4031 | 169 | 419.25 | 1 | 2.48 | 0.59 | 19.82 | 18.64 | 1.18 | 1.18 | 0.05 | 0.05 |
| gene1588 | https://www.ncbi.nlm.nih.gov/gene/?term=LOC105287720 | 4942 | 99 | 200.32 | 1 | 2.02 | 1.01 | 16.41 | 15.25 | 1.16 | 1.16 | 0.02 | 0.02 |
| gene4023 | https://www.ncbi.nlm.nih.gov/gene/?term=LOC105276692 | 1759 | 37 | 210.35 | 1 | 5.69 | 2.7 | 20.96 | 19.79 | 1.16 | 1.16 | 0.02 | 0.02 |
| gene6966 | https://www.ncbi.nlm.nih.gov/gene/?term=LOC105280104 | 3747 | 139 | 370.96 | 1 | 2.67 | 0.72 | 16.72 | 15.56 | 1.16 | 1.16 | 0.04 | 0.04 |
| gene7915 | https://www.ncbi.nlm.nih.gov/gene/?term=LOC105281173 | 1597 | 39 | 244.21 | 1 | 6.26 | 2.56 | 60.96 | 59.8 | 1.15 | 1.15 | 0.03 | 0.03 |
| gene12572 | https://www.ncbi.nlm.nih.gov/gene/?term=LOC105286514 | 2374 | 50 | 210.61 | 1 | 4.21 | 2 | 36.01 | 34.88 | 1.14 | 1.14 | 0.02 | 0.02 |
| gene9648 | https://www.ncbi.nlm.nih.gov/gene/?term=LOC105283183 | 2602 | 31 | 119.14 | 1 | 3.84 | 3.23 | 40.75 | 39.71 | 1.04 | 1.04 | 0.01 | 0.01 |
| gene1993 | https://www.ncbi.nlm.nih.gov/gene/?term=LOC105288193 | 1449 | 77 | 531.4 | 1 | 6.9 | 1.3 | 34.64 | 33.62 | 1.02 | 1.02 | 0.05 | 0.05 |
| gene4500 | https://www.ncbi.nlm.nih.gov/gene/?term=LOC105277238 | 2708 | 85 | 313.88 | 1 | 3.69 | 1.18 | 26.93 | 25.91 | 1.02 | 1.02 | 0.03 | 0.03 |
| gene11454 | https://www.ncbi.nlm.nih.gov/gene/?term=LOC105285256 | 1906 | 45 | 236.1 | 1 | 5.25 | 2.22 | 48.03 | 47.02 | 1 | 1 | 0.02 | 0.02 |
| gene10540 | https://www.ncbi.nlm.nih.gov/gene/?term=LOC105284200 | 5267 | 230 | 436.68 | 1 | 1.9 | 0.43 | 12.62 | 11.63 | 0.99 | 0.99 | 0.04 | 0.04 |
| gene935 | https://www.ncbi.nlm.nih.gov/gene/?term=LOC105282682 | 4368 | 222 | 508.24 | 1 | 2.29 | 0.45 | 13.71 | 12.73 | 0.98 | 0.98 | 0.05 | 0.05 |
| gene2173 | https://www.ncbi.nlm.nih.gov/gene/?term=LOC105274590 | 4387 | 71 | 161.84 | 2 | 4.56 | 2.82 | 27.99 | 27.02 | 0.97 | 0.97 | 0.02 | 0.02 |
| gene10520 | https://www.ncbi.nlm.nih.gov/gene/?term=LOC105284184 | 2451 | 84 | 342.72 | 1 | 4.08 | 1.19 | 34.09 | 33.16 | 0.93 | 0.93 | 0.03 | 0.03 |
| gene11568 | https://www.ncbi.nlm.nih.gov/gene/?term=LOC105285395 | 8086 | 215 | 265.89 | 1 | 1.24 | 0.47 | 16.71 | 15.82 | 0.89 | 0.89 | 0.02 | 0.02 |
| gene2844 | https://www.ncbi.nlm.nih.gov/gene/?term=LOC105275363 | 4330 | 196 | 452.66 | 1 | 2.31 | 0.51 | 14.69 | 13.8 | 0.89 | 0.89 | 0.04 | 0.04 |
| gene9647 | https://www.ncbi.nlm.nih.gov/gene/?term=LOC109611233 | 3094 | 37 | 119.59 | 1 | 3.23 | 2.7 | 34.14 | 33.27 | 0.87 | 0.87 | 0.01 | 0.01 |
| gene5831 | https://www.ncbi.nlm.nih.gov/gene/?term=LOC105278761 | 2506 | 117 | 466.88 | 2 | 7.98 | 1.71 | 24.49 | 23.66 | 0.83 | 0.83 | 0.04 | 0.04 |

| | | | | | | | | | | | | | |
|---|---|---|---|---|---|---|---|---|---|---|---|---|---|
| gene3385 | https://www.ncbi.nlm.nih.gov/gene/?term=LOC105275966 | 3305 | 62 | 187.59 | 1 | 3.03 | 1.61 | 48.06 | 47.24 | 0.82 | 0.82 | 0.02 | 0.02 |
| gene6528 | https://www.ncbi.nlm.nih.gov/gene/?term=LOC105279591 | 7222 | 55 | 76.16 1 | 1.38 | 1.82 | 30.56 | 29.74 | 0.82 | 0.82 | 0.01 | 0.01 | |
| gene11801 | https://www.ncbi.nlm.nih.gov/gene/?term=LOC105285656 | 2497 | 140 | 560.67 | 1 | 4 | 0.71 | 9.18 | 8.36 | 0.81 | 0.81 | 0.05 | 0.05 |
| gene10344 | https://www.ncbi.nlm.nih.gov/gene/?term=LOC105283987 | 4841 | 67 | 138.4 1 | 2.07 | 1.49 | 29.63 | 28.82 | 0.8 | 0.8 | 0.01 | 0.01 | |
| gene7556 | https://www.ncbi.nlm.nih.gov/gene/?term=LOC105280774 | 2792 | 98 | 351 1 | 3.58 | 1.02 | 33.4 | 32.6 | 0.8 | 0.8 | 0.03 | 0.03 | |
| gene298 | https://www.ncbi.nlm.nih.gov/gene/?term=LOC105275460 | 2473 | 117 | 473.11 | 1 | 4.04 | 0.85 | 14.05 | 13.27 | 0.78 | 0.78 | 0.04 | 0.04 |
| gene10391 | https://www.ncbi.nlm.nih.gov/gene/?term=LOC109611283 | 1994 | 42 | 210.63 | 1 | 5.02 | 2.38 | 34.44 | 33.67 | 0.76 | 0.76 | 0.02 | 0.02 |
| gene3470 | https://www.ncbi.nlm.nih.gov/gene/?term=LOC105276059 | 8716 | 160 | 183.57 | 1 | 1.15 | 0.62 | 8.58 | 7.82 | 0.76 | 0.76 | 0.01 | 0.01 |
| gene4892 | https://www.ncbi.nlm.nih.gov/gene/?term=LOC105277667 | 3898 | 129 | 330.94 | 1 | 2.57 | 0.78 | 28.03 | 27.27 | 0.76 | 0.76 | 0.03 | 0.03 |
| gene8274 | https://www.ncbi.nlm.nih.gov/gene/?term=LOC105281624 | 5425 | 219 | 403.69 | 1 | 1.84 | 0.46 | 9.87 | 9.12 | 0.75 | 0.75 | 0.03 | 0.03 |
| gene2257 | https://www.ncbi.nlm.nih.gov/gene/?term=LOC105274689 | 3658 | 70 | 191.36 | 1 | 2.73 | 1.43 | 15.64 | 14.92 | 0.72 | 0.72 | 0.01 | 0.01 |
| gene2266 | https://www.ncbi.nlm.nih.gov/gene/?term=LOC105274699 | 2391 | 53 | 221.66 | 1 | 4.18 | 1.89 | 24.7 | 23.99 | 0.72 | 0.72 | 0.02 | 0.02 |
| gene3107 | https://www.ncbi.nlm.nih.gov/gene/?term=LOC105275650 | 5103 | 167 | 327.26 | 1 | 1.96 | 0.6 | 16.25 | 15.53 | 0.72 | 0.72 | 0.02 | 0.02 |
| gene2633 | https://www.ncbi.nlm.nih.gov/gene/?term=LOC105275127 | 7457 | 259 | 347.32 | 1 | 1.34 | 0.39 | 8.88 | 8.23 | 0.66 | 0.66 | 0.02 | 0.02 |
| gene8814 | https://www.ncbi.nlm.nih.gov/gene/?term=LOC105282240 | 6910 | 181 | 261.94 | 1 | 1.45 | 0.55 | 9.72 | 9.08 | 0.64 | 0.64 | 0.02 | 0.02 |
| gene2846 | https://www.ncbi.nlm.nih.gov/gene/?term=LOC105275374 | 11246 | 498 | 442.82 | 1 | 0.89 | 0.2 | 12.01 | 11.42 | 0.6 | 0.6 | 0.03 | 0.03 |
| gene10531 | https://www.ncbi.nlm.nih.gov/gene/?term=LOC105284190 | 6881 | 191 | 277.58 | 1 | 1.45 | 0.52 | 12.39 | 11.8 | 0.58 | 0.58 | 0.02 | 0.02 |
| gene10864 | https://www.ncbi.nlm.nih.gov/gene/?term=LOC105284557 | 1666 | 83 | 498.2 1 | 6 | 1.2 | 29.52 | 28.94 | 0.58 | 0.58 | 0.03 | 0.03 | |
| gene9330 | https://www.ncbi.nlm.nih.gov/gene/?term=LOC105282825 | 3936 | 85 | 215.96 | 1 | 2.54 | 1.18 | 14.44 | 13.87 | 0.57 | 0.57 | 0.01 | 0.01 |
| gene1470 | https://www.ncbi.nlm.nih.gov/gene/?term=LOC105287597 | 2974 | 197 | 662.41 | 1 | 3.36 | 0.51 | 17.93 | 17.38 | 0.55 | 0.55 | 0.04 | 0.04 |
| gene3727 | https://www.ncbi.nlm.nih.gov/gene/?term=LOC105276367 | 2484 | 74 | 297.91 | 1 | 4.03 | 1.35 | 42.37 | 41.83 | 0.54 | 0.54 | 0.02 | 0.02 |
| gene11579 | https://www.ncbi.nlm.nih.gov/gene/?term=LOC105285406 | 13571 | 245 | 180.53 | 1 | 0.74 | 0.41 | 7.35 | 6.84 | 0.5 | 0.5 | 0.01 | 0.01 |
| gene12802 | https://www.ncbi.nlm.nih.gov/gene/?term=LOC105286799 | 3183 | 166 | 521.52 | 1 | 3.14 | 0.6 | 17.88 | 17.37 | 0.5 | 0.5 | 0.03 | 0.03 |
| gene7245 | https://www.ncbi.nlm.nih.gov/gene/?term=LOC105280410 | 12478 | 260 | 208.37 | 1 | 0.8 | 0.38 | 20.02 | 19.52 | 0.5 | 0.5 | 0.01 | 0.01 |
| gene4962 | https://www.ncbi.nlm.nih.gov/gene/?term=LOC105277761 | 4645 | 149 | 320.78 | 1 | 2.15 | 0.67 | 13.8 | 13.32 | 0.49 | 0.49 | 0.02 | 0.02 |
| gene6583 | https://www.ncbi.nlm.nih.gov/gene/?term=LOC105279649 | 7444 | 333 | 447.34 | 1 | 1.34 | 0.3 | 12.03 | 11.53 | 0.49 | 0.49 | 0.02 | 0.02 |
| gene8252 | https://www.ncbi.nlm.nih.gov/gene/?term=LOC105281599 | 3813 | 49 | 128.51 | 1 | 2.62 | 2.04 | 4.61 | 4.12 | 0.49 | 0.49 | 0.01 | 0.01 |
| gene12516 | https://www.ncbi.nlm.nih.gov/gene/?term=LOC105286452 | 1529 | 40 | 261.61 | 1 | 6.54 | 2.5 | 51.71 | 51.25 | 0.47 | 0.47 | 0.01 | 0.01 |
| gene10606 | https://www.ncbi.nlm.nih.gov/gene/?term=LOC105284273 | 6119 | 260 | 424.91 | 1 | 1.63 | 0.38 | 17.76 | 17.3 | 0.45 | 0.45 | 0.02 | 0.02 |
| gene11401 | https://www.ncbi.nlm.nih.gov/gene/?term=LOC105285202 | 1841 | 87 | 472.57 | 1 | 5.43 | 1.15 | 25.64 | 25.2 | 0.44 | 0.44 | 0.02 | 0.02 |
| gene11976 | https://www.ncbi.nlm.nih.gov/gene/?term=LOC105285860 | 3112 | 55 | 176.74 | 1 | 3.21 | 1.82 | 24.17 | 23.73 | 0.43 | 0.43 | 0.01 | 0.01 |
| gene6286 | https://www.ncbi.nlm.nih.gov/gene/?term=LOC105279298 | 2861 | 132 | 461.38 | 2 | 6.99 | 1.52 | 23.96 | 23.54 | 0.42 | 0.42 | 0.02 | 0.02 |
| gene5967 | https://www.ncbi.nlm.nih.gov/gene/?term=LOC105278918 | 4393 | 137 | 311.86 | 1 | 2.28 | 0.73 | 13.27 | 12.86 | 0.41 | 0.41 | 0.01 | 0.01 |
| gene13114 | https://www.ncbi.nlm.nih.gov/gene/?term=LOC105287147 | 7609 | 413 | 542.78 | 1 | 1.31 | 0.24 | 9.55 | 9.19 | 0.36 | 0.36 | 0.02 | 0.02 |
| gene7550 | https://www.ncbi.nlm.nih.gov/gene/?term=LOC105280763 | 2173 | 84 | 386.56 | 1 | 4.6 | 1.19 | 32.91 | 32.56 | 0.35 | 0.35 | 0.01 | 0.01 |
| gene2806 | https://www.ncbi.nlm.nih.gov/gene/?term=LOC105275327 | 14257 | 490 | 343.69 | 1 | 0.7 | 0.2 | 7.15 | 6.82 | 0.34 | 0.34 | 0.01 | 0.01 |
| gene1408 | https://www.ncbi.nlm.nih.gov/gene/?term=LOC109611520 | 8645 | 325 | 375.94 | 1 | 1.16 | 0.31 | 6.16 | 5.83 | 0.33 | 0.33 | 0.01 | 0.01 |
| gene1430 | https://www.ncbi.nlm.nih.gov/gene/?term=LOC105287551 | 6433 | 237 | 368.41 | 1 | 1.55 | 0.42 | 8.71 | 8.38 | 0.33 | 0.33 | 0.01 | 0.01 |
| gene10107 | https://www.ncbi.nlm.nih.gov/gene/?term=LOC105283712 | 1883 | 36 | 191.18 | 1 | 5.31 | 2.78 | 27.71 | 27.4 | 0.31 | 0.31 | 0.01 | 0.01 |
| gene5613 | https://www.ncbi.nlm.nih.gov/gene/?term=LOC105278504 | 8524 | 250 | 293.29 | 1 | 1.17 | 0.4 | 4.85 | 4.54 | 0.31 | 0.31 | 0.01 | 0.01 |
| gene8600 | https://www.ncbi.nlm.nih.gov/gene/?term=LOC105282001 | 8513 | 158 | 185.6 1 | 1.17 | 0.63 | 10.54 | 10.23 | 0.31 | 0.31 | 0.01 | 0.01 | |
| gene10392 | https://www.ncbi.nlm.nih.gov/gene/?term=LOC105284036 | 3216 | 104 | 323.38 | 2 | 6.22 | 1.92 | 22.48 | 22.18 | 0.3 | 0.3 | 0.01 | 0.01 |
| gene2056 | https://www.ncbi.nlm.nih.gov/gene/?term=LOC105274452 | 6584 | 257 | 390.34 | 1 | 1.52 | 0.39 | 13.38 | 13.08 | 0.3 | 0.3 | 0.01 | 0.01 |
| gene5746 | https://www.ncbi.nlm.nih.gov/gene/?term=LOC105278666 | 10343 | 447 | 432.18 | 1 | 0.97 | 0.22 | 7.23 | 6.93 | 0.3 | 0.3 | 0.01 | 0.01 |
| gene11082 | https://www.ncbi.nlm.nih.gov/gene/?term=LOC105284824 | 7436 | 205 | 275.69 | 1 | 1.34 | 0.49 | 7.69 | 7.4 | 0.29 | 0.29 | 0.01 | 0.01 |
| gene12265 | https://www.ncbi.nlm.nih.gov/gene/?term=LOC105286166 | 3050 | 110 | 360.66 | 1 | 3.28 | 0.91 | 24.09 | 23.8 | 0.29 | 0.29 | 0.01 | 0.01 |
| gene12307 | https://www.ncbi.nlm.nih.gov/gene/?term=LOC105286216 | 10313 | 190 | 184.23 | 1 | 0.97 | 0.53 | 15.65 | 15.38 | 0.26 | 0.26 | 0 | 0 |
| gene4190 | https://www.ncbi.nlm.nih.gov/gene/?term=LOC105276875 | 3150 | 118 | 374.6 1 | 3.17 | 0.85 | 22.65 | 22.39 | 0.26 | 0.26 | 0.01 | 0.01 | |
| gene10596 | https://www.ncbi.nlm.nih.gov/gene/?term=LOC105284262 | 6520 | 215 | 329.75 | 1 | 1.53 | 0.47 | 4.27 | 4.03 | 0.24 | 0.24 | 0.01 | 0.01 |
| gene7166 | https://www.ncbi.nlm.nih.gov/gene/?term=LOC105280326 | 6117 | 245 | 400.52 | 1 | 1.63 | 0.41 | 14.73 | 14.53 | 0.2 | 0.2 | 0.01 | 0.01 |
| gene3856 | https://www.ncbi.nlm.nih.gov/gene/?term=LOC105276499 | 5236 | 184 | 351.41 | 1 | 1.91 | 0.54 | 10.07 | 9.88 | 0.18 | 0.18 | 0.01 | 0.01 |
| gene6620 | https://www.ncbi.nlm.nih.gov/gene/?term=LOC105279687 | 17998 | 368 | 204.47 | 1 | 0.56 | 0.27 | 5.31 | 5.13 | 0.18 | 0.18 | 0 | 0 |
| gene5387 | https://www.ncbi.nlm.nih.gov/gene/?term=LOC105278245 | 6451 | 113 | 175.17 | 1 | 1.55 | 0.88 | 22.95 | 22.78 | 0.17 | 0.17 | 0 | 0 |
| gene3671 | https://www.ncbi.nlm.nih.gov/gene/?term=LOC105276272 | 1966 | 48 | 244.15 | 1 | 5.09 | 2.08 | 27.74 | 27.59 | 0.16 | 0.16 | 0 | 0 |
| gene11275 | https://www.ncbi.nlm.nih.gov/gene/?term=LOC105285055 | 4119 | 272 | 660.35 | 1 | 2.43 | 0.37 | 11.12 | 11.01 | 0.12 | 0.12 | 0.01 | 0.01 |
| gene3206 | https://www.ncbi.nlm.nih.gov/gene/?term=LOC105275752 | 3474 | 132 | 379.97 | 1 | 2.88 | 0.76 | 32.74 | 32.62 | 0.12 | 0.12 | 0 | 0 |
| gene1632 | https://www.ncbi.nlm.nih.gov/gene/?term=LOC105287764 | 18061 | 416 | 230.33 | 1 | 0.55 | 0.24 | 3.35 | 3.24 | 0.11 | 0.11 | 0 | 0 |
| gene5877 | https://www.ncbi.nlm.nih.gov/gene/?term=LOC105278807 | 1719 | 50 | 290.87 | 1 | 5.82 | 2 | 27.13 | 27.02 | 0.11 | 0.11 | 0 | 0 |
| gene10018 | https://www.ncbi.nlm.nih.gov/gene/?term=LOC105283607 | 4414 | 207 | 468.96 | 1 | 2.27 | 0.48 | 15.18 | 15.09 | 0.09 | 0.09 | 0 | 0 |
| gene3885 | https://www.ncbi.nlm.nih.gov/gene/?term=LOC105276527 | 7928 | 322 | 406.16 | 1 | 1.26 | 0.31 | 9.68 | 9.59 | 0.09 | 0.09 | 0 | 0 |
| gene9029 | https://www.ncbi.nlm.nih.gov/gene/?term=LOC105282491 | 4762 | 167 | 350.69 | 1 | 2.1 | 0.6 | 9.69 | 9.6 | 0.09 | 0.09 | 0 | 0 |
| gene2122 | https://www.ncbi.nlm.nih.gov/gene/?term=LOC105274531 | 8153 | 304 | 372.87 | 1 | 1.23 | 0.33 | 10.7 | 10.62 | 0.08 | 0.08 | 0 | 0 |
| gene13036 | https://www.ncbi.nlm.nih.gov/gene/?term=LOC105287059 | 4959 | 163 | 328.7 1 | 2.02 | 0.61 | 20.67 | 20.67 | 0.01 | 0.01 | 0 | 0 | |
| gene7850 | https://www.ncbi.nlm.nih.gov/gene/?term=LOC105281095 | 3050 | 67 | 219.67 | 1 | 3.28 | 1.49 | 33.74 | 33.73 | 0.01 | 0.01 | 0 | 0 |

| | | | | | | | | | | | | | |
|---|---|---|---|---|---|---|---|---|---|---|---|---|---|
| gene10549 | https://www.ncbi.nlm.nih.gov/gene/?term=LOC105284210 | 7145 | 298 | 417.07 | 2 | 2.8 | 0.67 | 11.73 | 11.76 | -0.04 | 0.04 | 0 | 0 |
| gene4048 | https://www.ncbi.nlm.nih.gov/gene/?term=LOC105276724 | 2955 | 32 | 108.29 | 1 | 3.38 | 3.12 | 11.95 | 11.99 | -0.04 | 0.04 | 0 | 0 |
| gene7249 | https://www.ncbi.nlm.nih.gov/gene/?term=LOC105280422 | 1775 | 60 | 338.03 | 1 | 5.63 | 1.67 | 17.01 | 17.06 | -0.05 | 0.05 | 0 | 0 |
| gene1220 | https://www.ncbi.nlm.nih.gov/gene/?term=LOC105285742 | 3453 | 123 | 356.21 | 1 | 2.9 | 0.81 | 20.51 | 20.57 | -0.06 | 0.06 | 0 | 0 |
| gene34 | https://www.ncbi.nlm.nih.gov/gene/?term=LOC105278416 | 7120 | 141 | 198.03 | 1 | 1.4 | 0.71 | 14.13 | 14.19 | -0.06 | 0.06 | 0 | 0 |
| gene12573 | https://www.ncbi.nlm.nih.gov/gene/?term=LOC105286515 | 3647 | 112 | 307.1 1 | 2.74 | 0.89 | 12.05 | 12.13 | -0.07 | 0.07 | 0 | 0 | |
| gene7368 | https://www.ncbi.nlm.nih.gov/gene/?term=LOC105280556 | 4105 | 40 | 97.44 1 | 2.44 | 2.5 | 15.83 | 15.9 | -0.07 | 0.07 | 0 | 0 | |
| gene9177 | https://www.ncbi.nlm.nih.gov/gene/?term=LOC105282646 | 3320 | 116 | 349.4 1 | 3.01 | 0.86 | 24.81 | 24.88 | -0.07 | 0.07 | 0 | 0 | |
| gene2683 | https://www.ncbi.nlm.nih.gov/gene/?term=LOC105275187 | 5492 | 71 | 129.28 | 1 | 1.82 | 1.41 | 12.33 | 12.41 | -0.08 | 0.08 | 0 | 0 |
| gene1147 | https://www.ncbi.nlm.nih.gov/gene/?term=LOC105284954 | 4022 | 101 | 251.12 | 1 | 2.49 | 0.99 | 15.4 | 15.49 | -0.09 | 0.09 | 0 | 0 |
| gene3286 | https://www.ncbi.nlm.nih.gov/gene/?term=LOC105275848 | 3744 | 144 | 384.62 | 1 | 2.67 | 0.69 | 21.45 | 21.54 | -0.09 | 0.09 | 0 | 0 |
| gene12439 | https://www.ncbi.nlm.nih.gov/gene/?term=LOC105286357 | 4761 | 97 | 203.74 | 1 | 2.1 | 1.03 | 28.47 | 28.59 | -0.12 | 0.12 | 0 | 0 |
| gene12674 | https://www.ncbi.nlm.nih.gov/gene/?term=LOC105286649 | 18283 | 788 | 431 1 | 0.55 | 0.13 | 1.01 | 1.14 | -0.13 | 0.13 | -0.01 | 0.01 | |
| gene6649 | https://www.ncbi.nlm.nih.gov/gene/?term=LOC105279717 | 6017 | 171 | 284.19 | 1 | 1.66 | 0.58 | 12.42 | 12.54 | -0.13 | 0.13 | 0 | 0 |
| gene5573 | https://www.ncbi.nlm.nih.gov/gene/?term=LOC105278454 | 2688 | 37 | 137.65 | 1 | 3.72 | 2.7 | 33.59 | 33.73 | -0.14 | 0.14 | 0 | 0 |
| gene8729 | https://www.ncbi.nlm.nih.gov/gene/?term=LOC105282137 | 5005 | 220 | 439.56 | 1 | 2 | 0.45 | 15.19 | 15.33 | -0.14 | 0.14 | -0.01 | 0.01 |
| gene9755 | https://www.ncbi.nlm.nih.gov/gene/?term=LOC105283296 | 2177 | 92 | 422.6 1 | 4.59 | 1.09 | 18.56 | 18.74 | -0.18 | 0.18 | -0.01 | 0.01 | |
| gene7493 | https://www.ncbi.nlm.nih.gov/gene/?term=LOC105280704 | 4225 | 132 | 312.43 | 1 | 2.37 | 0.76 | 20.13 | 20.33 | -0.21 | 0.21 | -0.01 | 0.01 |
| gene10195 | https://www.ncbi.nlm.nih.gov/gene/?term=LOC105283806 | 10532 | 75 | 71.21 1 | 0.95 | 1.33 | 14.09 | 14.33 | -0.24 | 0.24 | 0 | 0 | |
| gene1310 | https://www.ncbi.nlm.nih.gov/gene/?term=LOC105286661 | 2582 | 72 | 278.85 | 1 | 3.87 | 1.39 | 21.24 | 21.48 | -0.25 | 0.25 | -0.01 | 0.01 |
| gene2404 | https://www.ncbi.nlm.nih.gov/gene/?term=LOC105274853 | 1767 | 43 | 243.35 | 1 | 5.66 | 2.33 | 29.37 | 29.63 | -0.26 | 0.26 | -0.01 | 0.01 |
| gene289 | https://www.ncbi.nlm.nih.gov/gene/?term=LOC105275347 | 2533 | 69 | 272.4 1 | 3.95 | 1.45 | 31.43 | 31.68 | -0.26 | 0.26 | -0.01 | 0.01 | |
| gene6031 | https://www.ncbi.nlm.nih.gov/gene/?term=LOC105279000 | 5691 | 190 | 333.86 | 1 | 1.76 | 0.53 | 15.94 | 16.2 | -0.26 | 0.26 | -0.01 | 0.01 |
| gene5646 | https://www.ncbi.nlm.nih.gov/gene/?term=LOC105278534 | 7475 | 197 | 263.55 | 1 | 1.34 | 0.51 | 17.35 | 17.62 | -0.27 | 0.27 | -0.01 | 0.01 |
| gene10819 | https://www.ncbi.nlm.nih.gov/gene/?term=LOC105284507 | 5767 | 84 | 145.66 | 1 | 1.73 | 1.19 | 22.04 | 22.33 | -0.3 | 0.3 | 0 | 0 |
| gene9326 | https://www.ncbi.nlm.nih.gov/gene/?term=LOC105282818 | 16612 | 192 | 115.58 | 1 | 0.6 | 0.52 | 12.9 | 13.21 | -0.3 | 0.3 | 0 | 0 |
| gene53 | https://www.ncbi.nlm.nih.gov/gene/?term=LOC105277148 | 19283 | 303 | 157.13 | 1 | 0.52 | 0.33 | 3.79 | 4.11 | -0.32 | 0.32 | -0.01 | 0.01 |
| gene11583 | https://www.ncbi.nlm.nih.gov/gene/?term=LOC105285414 | 8192 | 182 | 222.17 | 1 | 1.22 | 0.55 | 8.25 | 8.59 | -0.33 | 0.33 | -0.01 | 0.01 |
| gene8015 | https://www.ncbi.nlm.nih.gov/gene/?term=LOC105281312 | 2998 | 62 | 206.8 1 | 3.34 | 1.61 | 34.79 | 35.12 | -0.33 | 0.33 | -0.01 | 0.01 | |
| gene2835 | https://www.ncbi.nlm.nih.gov/gene/?term=LOC105275357 | 2492 | 128 | 513.64 | 1 | 4.01 | 0.78 | 14.7 | 15.03 | -0.34 | 0.34 | -0.02 | 0.02 |
| gene11561 | https://www.ncbi.nlm.nih.gov/gene/?term=LOC105285387 | 11528 | 184 | 159.61 | 1 | 0.87 | 0.54 | 10.15 | 10.52 | -0.37 | 0.37 | -0.01 | 0.01 |
| gene12981 | https://www.ncbi.nlm.nih.gov/gene/?term=LOC105286999 | 2063 | 44 | 213.28 | 1 | 4.85 | 2.27 | 34 | 34.43 | -0.43 | 0.43 | -0.01 | 0.01 |
| gene6652 | https://www.ncbi.nlm.nih.gov/gene/?term=LOC105279719 | 2831 | 80 | 282.59 | 1 | 3.53 | 1.25 | 22.97 | 23.4 | -0.43 | 0.43 | -0.01 | 0.01 |
| gene1199 | https://www.ncbi.nlm.nih.gov/gene/?term=LOC105285498 | 2220 | 30 | 135.14 | 1 | 4.5 | 3.33 | 26.9 | 27.37 | -0.47 | 0.47 | -0.01 | 0.01 |
| gene4376 | https://www.ncbi.nlm.nih.gov/gene/?term=LOC105277083 | 4307 | 111 | 257.72 | 1 | 2.32 | 0.9 | 22.97 | 23.44 | -0.47 | 0.47 | -0.01 | 0.01 |
| gene657 | https://www.ncbi.nlm.nih.gov/gene/?term=LOC105279667 | 6421 | 85 | 132.38 | 1 | 1.56 | 1.18 | 2.59 | 3.08 | -0.49 | 0.49 | -0.01 | 0.01 |
| gene12203 | https://www.ncbi.nlm.nih.gov/gene/?term=LOC105286099 | 5967 | 78 | 130.72 | 1 | 1.68 | 1.28 | 9.64 | 10.17 | -0.53 | 0.53 | -0.01 | 0.01 |
| gene814 | https://www.ncbi.nlm.nih.gov/gene/?term=LOC105281296 | 2722 | 44 | 161.65 | 1 | 3.67 | 2.27 | 26.1 | 26.63 | -0.53 | 0.53 | -0.01 | 0.01 |
| gene3747 | https://www.ncbi.nlm.nih.gov/gene/?term=LOC105276387 | 3736 | 168 | 449.68 | 1 | 2.68 | 0.6 | 14.1 | 14.65 | -0.54 | 0.54 | -0.02 | 0.02 |
| gene12406 | https://www.ncbi.nlm.nih.gov/gene/?term=LOC105286316 | 1386 | 36 | 259.74 | 1 | 7.22 | 2.78 | 38.06 | 38.61 | -0.55 | 0.55 | -0.01 | 0.01 |
| gene35 | https://www.ncbi.nlm.nih.gov/gene/?term=LOC105275197 | 2643 | 51 | 192.96 | 1 | 3.78 | 1.96 | 38.21 | 38.8 | -0.59 | 0.59 | -0.01 | 0.01 |
| gene6639 | https://www.ncbi.nlm.nih.gov/gene/?term=LOC105279705 | 7656 | 73 | 95.35 1 | 1.31 | 1.37 | 23.16 | 23.78 | -0.62 | 0.62 | -0.01 | 0.01 | |
| gene11447 | https://www.ncbi.nlm.nih.gov/gene/?term=LOC105285252 | 7437 | 119 | 160.01 | 1 | 1.34 | 0.84 | 14.25 | 14.93 | -0.68 | 0.68 | -0.01 | 0.01 |
| gene5298 | https://www.ncbi.nlm.nih.gov/gene/?term=LOC105278145 | 4060 | 250 | 615.76 | 2 | 4.93 | 0.8 | 10.45 | 11.12 | -0.68 | 0.68 | -0.04 | 0.04 |
| gene4033 | https://www.ncbi.nlm.nih.gov/gene/?term=LOC105276705 | 10546 | 136 | 128.96 | 1 | 0.95 | 0.74 | 8.69 | 9.42 | -0.73 | 0.73 | -0.01 | 0.01 |
| gene9327 | https://www.ncbi.nlm.nih.gov/gene/?term=LOC105282821 | 7793 | 149 | 191.2 1 | 1.28 | 0.67 | 12.54 | 13.29 | -0.74 | 0.74 | -0.01 | 0.01 | |
| gene7207 | https://www.ncbi.nlm.nih.gov/gene/?term=LOC105280376 | 4659 | 111 | 238.25 | 1 | 2.15 | 0.9 | 30.17 | 30.93 | -0.76 | 0.76 | -0.02 | 0.02 |
| gene6237 | https://www.ncbi.nlm.nih.gov/gene/?term=LOC105279244 | 1484 | 26 | 175.2 1 | 6.74 | 3.85 | 41.81 | 42.6 | -0.8 | 0.8 | -0.01 | 0.01 | |
| gene10155 | https://www.ncbi.nlm.nih.gov/gene/?term=LOC105283756 | 7030 | 114 | 162.16 | 1 | 1.42 | 0.88 | 5.48 | 6.39 | -0.91 | 0.91 | -0.01 | 0.01 |
| gene4498 | https://www.ncbi.nlm.nih.gov/gene/?term=LOC105277239 | 2831 | 120 | 423.88 | 1 | 3.53 | 0.83 | 12.33 | 13.34 | -1.01 | 1.01 | -0.04 | 0.04 |
| gene322 | https://www.ncbi.nlm.nih.gov/gene/?term=LOC105275777 | 1606 | 28 | 174.35 | 1 | 6.23 | 3.57 | 39.43 | 40.5 | -1.07 | 1.07 | -0.02 | 0.02 |
| gene7398 | https://www.ncbi.nlm.nih.gov/gene/?term=LOC105280589 | 3452 | 34 | 98.49 1 | 2.9 | 2.94 | 18.86 | 19.97 | -1.12 | 1.12 | -0.01 | 0.01 | |
| gene1768 | https://www.ncbi.nlm.nih.gov/gene/?term=LOC105287938 | 2792 | 87 | 311.6 1 | 3.58 | 1.15 | 25.07 | 26.23 | -1.17 | 1.17 | -0.04 | 0.04 | |
| gene3168 | https://www.ncbi.nlm.nih.gov/gene/?term=LOC105275711 | 6150 | 75 | 121.95 | 1 | 1.63 | 1.33 | 13.14 | 14.33 | -1.18 | 1.18 | -0.01 | 0.01 |
| gene7406 | https://www.ncbi.nlm.nih.gov/gene/?term=LOC105280596 | 3432 | 115 | 335.08 | 2 | 5.83 | 1.74 | 17.75 | 19.02 | -1.26 | 1.26 | -0.04 | 0.04 |
| gene11340 | https://www.ncbi.nlm.nih.gov/gene/?term=LOC105285131 | 2918 | 91 | 311.86 | 1 | 3.43 | 1.1 | 11.55 | 12.83 | -1.28 | 1.28 | -0.04 | 0.04 |
| gene9101 | https://www.ncbi.nlm.nih.gov/gene/?term=LOC105282553 | 3068 | 108 | 352.02 | 1 | 3.26 | 0.93 | 22.02 | 23.41 | -1.4 | 1.4 | -0.05 | 0.05 |
| gene8343 | https://www.ncbi.nlm.nih.gov/gene/?term=LOC105281700 | 2172 | 34 | 156.54 | 1 | 4.6 | 2.94 | 35.34 | 36.82 | -1.48 | 1.48 | -0.02 | 0.02 |
| gene10213 | https://www.ncbi.nlm.nih.gov/gene/?term=LOC105283826 | 1503 | 54 | 359.28 | 1 | 6.65 | 1.85 | 14.21 | 15.73 | -1.51 | 1.51 | -0.05 | 0.05 |
| gene7600 | https://www.ncbi.nlm.nih.gov/gene/?term=LOC105280819 | 1615 | 38 | 235.29 | 1 | 6.19 | 2.63 | 35.64 | 37.15 | -1.52 | 1.52 | -0.04 | 0.04 |
| gene3827 | https://www.ncbi.nlm.nih.gov/gene/?term=LOC105276463 | 2749 | 82 | 298.29 | 1 | 3.64 | 1.22 | 27.66 | 29.29 | -1.63 | 1.63 | -0.05 | 0.05 |
| gene3599 | https://www.ncbi.nlm.nih.gov/gene/?term=LOC105276196 | 2955 | 62 | 209.81 | 1 | 3.38 | 1.61 | 36.96 | 38.6 | -1.64 | 1.64 | -0.03 | 0.03 |
| gene1505 | https://www.ncbi.nlm.nih.gov/gene/?term=LOC105287633 | 1693 | 29 | 171.29 | 1 | 5.91 | 3.45 | 41.25 | 42.91 | -1.66 | 1.66 | -0.03 | 0.03 |
| gene920 | https://www.ncbi.nlm.nih.gov/gene/?term=LOC105282518 | 1197 | 52 | 434.42 | 1 | 8.35 | 1.92 | 29.01 | 30.77 | -1.76 | 1.76 | -0.08 | 0.08 |

| | | | | | | | | | | | | | |
|---|---|---|---|---|---|---|---|---|---|---|---|---|---|
| gene2413 | https://www.ncbi.nlm.nih.gov/gene/?term=LOC105274863 | 2006 | 50 | 249.25 | 1 | 4.99 | 2 | 53.71 | 55.57 | -1.87 | 1.87 | -0.05 | 0.05 |
| gene10396 | https://www.ncbi.nlm.nih.gov/gene/?term=LOC105284047 | 1227 | 23 | 187.45 | 1 | 8.15 | 4.35 | 26.88 | 28.88 | -2 | 2 | -0.04 | 0.04 |
| gene4641 | https://www.ncbi.nlm.nih.gov/gene/?term=LOC105277398 | 1937 | 72 | 371.71 | 1 | 5.16 | 1.39 | 36.58 | 39.26 | -2.68 | 2.68 | -0.1 | 0.1 |
| gene10345 | https://www.ncbi.nlm.nih.gov/gene/?term=LOC105283989 | 4444 | 27 | 60.76 | 1 | 2.25 | 3.7 | 34.86 | 38.69 | -3.83 | 3.83 | -0.02 | 0.02 |
| gene1008 | https://www.ncbi.nlm.nih.gov/gene/?term=LOC105283492 | 1479 | 10 | 67.61 | 1 | 6.76 | 10 | 37.74 | 42 | -4.25 | 4.25 | -0.03 | 0.03 |
| gene9702 | https://www.ncbi.nlm.nih.gov/gene/?term=LOC105283234 | 2318 | 7 | 30.2 | 1 | 4.31 | 14.29 | 42.66 | 51.92 | -9.26 | 9.26 | -0.03 | 0.03 |
| gene10963 | https://www.ncbi.nlm.nih.gov/gene/?term=LOC105284677 | 1576 | 30 | 190.36 | 1 | 6.35 | 3.33 | 25.25 | 41.23 | -15.97 | 15.97 | -0.3 | 0.3 |
| gene393 | https://www.ncbi.nlm.nih.gov/gene/?term=LOC105276566 | 1123 | 2 | 17.81 | 1 | 8.9 | 50 | 47.62 | 76.78 | -29.16 | 29.16 | -0.05 | 0.05 |

Directory: OGL File: ogl-Cb_LI_RphaseB1.txt

| gene_ID | gene_link | gwidth | #Sites | #per10Kb | %perSite | %pNucl |
|---|---|---|---|---|---|---|
| gene12538 | https://www.ncbi.nlm.nih.gov/gene/?term=LOC105286531 | 372 | 40 | 1075.27 | 35.13 | 3.78 |
| gene5911 | https://www.ncbi.nlm.nih.gov/gene/?term=LOC105278844 | 943 | 102 | 1081.65 | 34.66 | 3.75 |
| gene13384 | https://www.ncbi.nlm.nih.gov/gene/?term=LOC105287447 | 1877 | 230 | 1225.36 | 29.95 | 3.67 |
| gene211 | https://www.ncbi.nlm.nih.gov/gene/?term=LOC105274534 | 1280 | 99 | 773.44 | 46.14 | 3.57 |
| gene12364 | https://www.ncbi.nlm.nih.gov/gene/?term=LOC105286269 | 1403 | 95 | 677.12 | 50.19 | 3.4 |
| gene5917 | https://www.ncbi.nlm.nih.gov/gene/?term=LOC105278849 | 823 | 125 | 1518.83 | 22.36 | 3.4 |
| gene3719 | https://www.ncbi.nlm.nih.gov/gene/?term=LOC105276360 | 1471 | 169 | 1148.88 | 29.3 | 3.37 |
| gene5309 | https://www.ncbi.nlm.nih.gov/gene/?term=LOC105278160 | 1704 | 167 | 980.05 | 34.27 | 3.36 |
| gene570 | https://www.ncbi.nlm.nih.gov/gene/?term=LOC105278549 | 1673 | 120 | 717.27 | 46.56 | 3.34 |
| gene4878 | https://www.ncbi.nlm.nih.gov/gene/?term=LOC105277648 | 675 | 40 | 592.59 | 54.59 | 3.23 |
| gene1654 | https://www.ncbi.nlm.nih.gov/gene/?term=LOC105287811 | 2285 | 147 | 643.33 | 49.46 | 3.18 |
| gene11566 | https://www.ncbi.nlm.nih.gov/gene/?term=LOC105285392 | 1603 | 134 | 835.93 | 36.82 | 3.08 |
| gene12330 | https://www.ncbi.nlm.nih.gov/gene/?term=LOC105286235 | 1501 | 105 | 699.53 | 44.05 | 3.08 |
| gene12746 | https://www.ncbi.nlm.nih.gov/gene/?term=LOC105286737 | 2688 | 370 | 1376.49 | 22.28 | 3.07 |
| gene6078 | https://www.ncbi.nlm.nih.gov/gene/?term=LOC105279052 | 1130 | 126 | 1115.04 | 27.5 | 3.07 |
| gene12652 | https://www.ncbi.nlm.nih.gov/gene/?term=LOC109610640 | 1122 | 60 | 534.76 | 56.29 | 3.01 |
| gene1562 | https://www.ncbi.nlm.nih.gov/gene/?term=LOC105287692 | 1491 | 134 | 898.73 | 33.27 | 2.99 |
| gene2797 | https://www.ncbi.nlm.nih.gov/gene/?term=LOC105275314 | 1489 | 104 | 698.46 | 41.67 | 2.91 |
| gene190 | https://www.ncbi.nlm.nih.gov/gene/?term=LOC105288088 | 1797 | 123 | 684.47 | 42.11 | 2.88 |
| gene3731 | https://www.ncbi.nlm.nih.gov/gene/?term=LOC105276371 | 865 | 58 | 670.52 | 42.64 | 2.86 |
| gene7527 | https://www.ncbi.nlm.nih.gov/gene/?term=LOC105280739 | 2686 | 148 | 551.01 | 51.09 | 2.81 |
| gene6978 | https://www.ncbi.nlm.nih.gov/gene/?term=LOC105280117 | 1816 | 92 | 506.61 | 55.33 | 2.8 |
| gene12805 | https://www.ncbi.nlm.nih.gov/gene/?term=LOC105286801 | 1379 | 110 | 797.68 | 34.81 | 2.78 |
| gene12403 | https://www.ncbi.nlm.nih.gov/gene/?term=LOC105286311 | 1413 | 65 | 460.01 | 60.28 | 2.77 |

...

... only first 25 lines shown ...

Directory: OGL File: ogl-Cb_LI_RphaseB1.vs.RphaseB2.txt

| gene_ID | gene_link | gwidth | #Sites | #per10Kb | #dmSites | #dmsp10kb | %dmSites | %pSite1 | %pSite2 | DMpSite | ADMpSite | DMpNucl | ADMpNucl |
|---|---|---|---|---|---|---|---|---|---|---|---|---|---|
| gene2593 | https://www.ncbi.nlm.nih.gov/gene/?term=LOC105275070 | 1774 | 6 | 33.82 | 2 | 11.27 | 33.33 | 23.92 | 4.77 | 19.15 | 19.15 | 0.06 | 0.06 |
| gene11599 | https://www.ncbi.nlm.nih.gov/gene/?term=LOC105285431 | 1550 | 6 | 38.71 | 3 | 19.35 | 50 | 46.72 | 30.29 | 16.43 | 16.43 | 0.06 | 0.06 |
| gene11124 | https://www.ncbi.nlm.nih.gov/gene/?term=LOC105284871 | 1889 | 3 | 15.88 | 1 | 5.29 | 33.33 | 46.42 | 31.48 | 14.94 | 14.94 | 0.02 | 0.02 |
| gene6576 | https://www.ncbi.nlm.nih.gov/gene/?term=LOC105279639 | 2616 | 3 | 11.47 | 1 | 3.82 | 33.33 | 77.78 | 65.15 | 12.62 | 12.62 | 0.01 | 0.01 |
| gene2636 | https://www.ncbi.nlm.nih.gov/gene/?term=LOC105275141 | 947 | 11 | 116.16 | 4 | 42.24 | 36.36 | 32.51 | 19.93 | 12.58 | 12.58 | 0.15 | 0.15 |
| gene10194 | https://www.ncbi.nlm.nih.gov/gene/?term=LOC105283805 | 1309 | 6 | 45.84 | 3 | 22.92 | 50 | 87.61 | 75.59 | 12.01 | 12.01 | 0.06 | 0.06 |
| gene4732 | https://www.ncbi.nlm.nih.gov/gene/?term=LOC105277491 | 1647 | 12 | 72.86 | 2 | 12.14 | 16.67 | 22.08 | 12.22 | 9.85 | 9.85 | 0.07 | 0.07 |
| gene9354 | https://www.ncbi.nlm.nih.gov/gene/?term=LOC105282854 | 1836 | 2 | 10.89 | 1 | 5.45 | 50 | 38.24 | 29.01 | 9.22 | 9.22 | 0.01 | 0.01 |
| gene9109 | https://www.ncbi.nlm.nih.gov/gene/?term=LOC105282561 | 2400 | 5 | 20.83 | 1 | 4.17 | 20 | 36.89 | 27.8 | 9.09 | 9.09 | 0.02 | 0.02 |
| gene7955 | https://www.ncbi.nlm.nih.gov/gene/?term=LOC105281216 | 3744 | 12 | 32.05 | 2 | 5.34 | 16.67 | 39.32 | 30.29 | 9.02 | 9.02 | 0.03 | 0.03 |
| gene9192 | https://www.ncbi.nlm.nih.gov/gene/?term=LOC105282657 | 1404 | 6 | 42.74 | 1 | 7.12 | 16.67 | 58.95 | 50.62 | 8.33 | 8.33 | 0.04 | 0.04 |
| gene9845 | https://www.ncbi.nlm.nih.gov/gene/?term=LOC105283410 | 1401 | 9 | 64.24 | 2 | 14.28 | 22.22 | 50.4 | 42.07 | 8.33 | 8.33 | 0.05 | 0.05 |
| gene4621 | https://www.ncbi.nlm.nih.gov/gene/?term=LOC105277369 | 1157 | 4 | 34.57 | 1 | 8.64 | 25 | 86.56 | 78.3 | 8.25 | 8.25 | 0.03 | 0.03 |
| gene4515 | https://www.ncbi.nlm.nih.gov/gene/?term=LOC109610839 | 335 | 7 | 208.96 | 1 | 29.85 | 14.29 | 66.57 | 58.9 | 7.67 | 7.67 | 0.16 | 0.16 |
| gene5060 | https://www.ncbi.nlm.nih.gov/gene/?term=LOC105277862 | 1458 | 7 | 48.01 | 1 | 6.86 | 14.29 | 63.47 | 55.84 | 7.64 | 7.64 | 0.04 | 0.04 |
| gene240 | https://www.ncbi.nlm.nih.gov/gene/?term=LOC105274857 | 1382 | 4 | 28.94 | 1 | 7.24 | 25 | 52.94 | 45.79 | 7.15 | 7.15 | 0.02 | 0.02 |
| gene10278 | https://www.ncbi.nlm.nih.gov/gene/?term=LOC105283908 | 902 | 12 | 133.04 | 4 | 44.35 | 33.33 | 41.95 | 34.88 | 7.07 | 7.07 | 0.09 | 0.09 |
| gene12853 | https://www.ncbi.nlm.nih.gov/gene/?term=LOC105286857 | 1589 | 4 | 25.17 | 1 | 6.29 | 25 | 74.57 | 67.52 | 7.05 | 7.05 | 0.02 | 0.02 |
| gene12355 | https://www.ncbi.nlm.nih.gov/gene/?term=LOC105286261 | 1301 | 8 | 61.49 | 1 | 7.69 | 12.5 | 74.9 | 68.31 | 6.59 | 6.59 | 0.04 | 0.04 |
| gene4044 | https://www.ncbi.nlm.nih.gov/gene/?term=LOC105276718 | 1729 | 10 | 57.84 | 1 | 5.78 | 10 | 28.19 | 21.81 | 6.38 | 6.38 | 0.04 | 0.04 |
| gene10189 | https://www.ncbi.nlm.nih.gov/gene/?term=LOC105283800 | 871 | 13 | 149.25 | 1 | 11.48 | 7.69 | 18.91 | 13.21 | 5.7 | 5.7 | 0.09 | 0.09 |
| gene3250 | https://www.ncbi.nlm.nih.gov/gene/?term=LOC105275812 | 2029 | 13 | 64.07 | 1 | 4.93 | 7.69 | 22.84 | 17.14 | 5.7 | 5.7 | 0.04 | 0.04 |
| gene7967 | https://www.ncbi.nlm.nih.gov/gene/?term=LOC105281233 | 4322 | 13 | 30.08 | 1 | 2.31 | 7.69 | 21.1 | 15.42 | 5.68 | 5.68 | 0.02 | 0.02 |
| gene10199 | https://www.ncbi.nlm.nih.gov/gene/?term=LOC105283809 | 2716 | 13 | 47.86 | 1 | 3.68 | 7.69 | 69.81 | 64.23 | 5.58 | 5.58 | 0.03 | 0.03 |

...
... only first 25 lines shown ...

Directory: OGL File: ogl-Cb_LI_RphaseB2.txt

| gene_ID | gene_link | gwidth | #Sites | #per10Kb | %perSite | %pNucl |
|---|---|---|---|---|---|---|
| gene12538 | https://www.ncbi.nlm.nih.gov/gene/?term=LOC105286531 | 372 | 41 | 1102.15 | 36.19 | 3.99 |
| gene13384 | https://www.ncbi.nlm.nih.gov/gene/?term=LOC105287447 | 1877 | 236 | 1257.33 | 28.98 | 3.64 |
| gene5917 | https://www.ncbi.nlm.nih.gov/gene/?term=LOC105278849 | 823 | 124 | 1506.68 | 23.86 | 3.6 |
| gene5911 | https://www.ncbi.nlm.nih.gov/gene/?term=LOC105278844 | 943 | 95 | 1007.42 | 35.49 | 3.58 |
| gene3719 | https://www.ncbi.nlm.nih.gov/gene/?term=LOC105276360 | 1471 | 141 | 958.53 | 35.05 | 3.36 |
| gene5309 | https://www.ncbi.nlm.nih.gov/gene/?term=LOC105278160 | 1704 | 158 | 927.23 | 36.29 | 3.36 |
| gene211 | https://www.ncbi.nlm.nih.gov/gene/?term=LOC105274534 | 1280 | 86 | 671.88 | 49.53 | 3.33 |
| gene3731 | https://www.ncbi.nlm.nih.gov/gene/?term=LOC105276371 | 865 | 57 | 658.96 | 46.13 | 3.04 |
| gene1562 | https://www.ncbi.nlm.nih.gov/gene/?term=LOC105287692 | 1491 | 129 | 865.19 | 34.84 | 3.01 |
| gene1654 | https://www.ncbi.nlm.nih.gov/gene/?term=LOC105287811 | 2285 | 161 | 704.6 | 42.7 | 3.01 |
| gene12805 | https://www.ncbi.nlm.nih.gov/gene/?term=LOC105286801 | 1379 | 108 | 783.18 | 37.84 | 2.96 |
| gene6078 | https://www.ncbi.nlm.nih.gov/gene/?term=LOC105279052 | 1130 | 111 | 982.3 | 29.64 | 2.91 |
| gene12330 | https://www.ncbi.nlm.nih.gov/gene/?term=LOC105286235 | 1501 | 95 | 632.91 | 45.44 | 2.88 |
| gene12746 | https://www.ncbi.nlm.nih.gov/gene/?term=LOC105286737 | 2688 | 337 | 1253.72 | 22.91 | 2.87 |
| gene2797 | https://www.ncbi.nlm.nih.gov/gene/?term=LOC105275314 | 1489 | 89 | 597.72 | 47.65 | 2.85 |
| gene12403 | https://www.ncbi.nlm.nih.gov/gene/?term=LOC105286311 | 1413 | 63 | 445.86 | 62.76 | 2.8 |
| gene5301 | https://www.ncbi.nlm.nih.gov/gene/?term=LOC105278148 | 1881 | 182 | 967.57 | 27.83 | 2.69 |
| gene343 | https://www.ncbi.nlm.nih.gov/gene/?term=LOC105275996 | 1620 | 64 | 395.06 | 67.56 | 2.67 |
| gene12364 | https://www.ncbi.nlm.nih.gov/gene/?term=LOC105286269 | 1403 | 67 | 477.55 | 54.92 | 2.62 |
| gene3334 | https://www.ncbi.nlm.nih.gov/gene/?term=LOC105275897 | 1689 | 116 | 686.8 | 38.03 | 2.61 |
| gene7527 | https://www.ncbi.nlm.nih.gov/gene/?term=LOC105280739 | 2686 | 128 | 476.55 | 54.67 | 2.61 |
| gene6978 | https://www.ncbi.nlm.nih.gov/gene/?term=LOC105280117 | 1816 | 82 | 451.54 | 57.37 | 2.59 |
| gene5497 | https://www.ncbi.nlm.nih.gov/gene/?term=LOC105278368 | 1690 | 64 | 378.7 | 67.97 | 2.57 |
| gene2279 | https://www.ncbi.nlm.nih.gov/gene/?term=LOC105274711 | 2037 | 116 | 569.46 | 44.99 | 2.56 |

...

... only first 25 lines shown ...

Directory: OGL File: rnk-dmg-Cb_LI_BCphaseA1.vs.BCphaseA2.txt

| gene_ID | gene_link | gwidth | #Sites | #per10Kb | #dmSites | #dmsp10kb | %dmSites | %pSite1 | %pSite2 | DMpSite | ADMpSite | DMpNucl | ADMpNucl |
|---|---|---|---|---|---|---|---|---|---|---|---|---|---|
| gene3165 | https://www.ncbi.nlm.nih.gov/gene/?term=LOC105275706 | 2374 | 65 | 273.8 | 6 | 25.27 | 9.23 | 24.65 | 38.08 | -13.43 | 13.43 | -0.37 | 0.37 |
| gene2824 | https://www.ncbi.nlm.nih.gov/gene/?term=LOC105275337 | 1772 | 58 | 327.31 | 1 | 5.64 | 1.72 | 31.86 | 38.08 | -6.22 | 6.22 | -0.2 | 0.2 |
| gene2593 | https://www.ncbi.nlm.nih.gov/gene/?term=LOC105275070 | 1774 | 5 | 28.18 | 2 | 11.27 | 40 | 19.71 | 71.28 | -51.57 | 51.57 | -0.15 | 0.15 |
| gene5714 | https://www.ncbi.nlm.nih.gov/gene/?term=LOC105278729 | 710 | 14 | 197.18 | 1 | 14.08 | 7.14 | 51.67 | 57.98 | -6.31 | 6.31 | -0.12 | 0.12 |
| gene10213 | https://www.ncbi.nlm.nih.gov/gene/?term=LOC105283826 | 1503 | 53 | 352.63 | 1 | 6.65 | 1.89 | 16.45 | 13.62 | 2.82 | 2.82 | 0.1 | 0.1 |
| gene920 | https://www.ncbi.nlm.nih.gov/gene/?term=LOC105282518 | 1197 | 40 | 334.17 | 1 | 8.35 | 2.5 | 39.94 | 37.69 | 2.25 | 2.25 | 0.08 | 0.08 |
| gene6334 | https://www.ncbi.nlm.nih.gov/gene/?term=LOC105279356 | 1471 | 40 | 271.92 | 1 | 6.8 | 2.5 | 29.12 | 32.21 | -3.09 | 3.09 | -0.08 | 0.08 |
| gene7734 | https://www.ncbi.nlm.nih.gov/gene/?term=LOC105280972 | 1528 | 26 | 170.16 | 1 | 6.54 | 3.85 | 20.04 | 23.88 | -3.84 | 3.84 | -0.07 | 0.07 |
| gene2676 | https://www.ncbi.nlm.nih.gov/gene/?term=LOC105275181 | 1353 | 15 | 110.86 | 1 | 7.39 | 6.67 | 67.39 | 74.05 | -6.66 | 6.66 | -0.07 | 0.07 |
| gene7556 | https://www.ncbi.nlm.nih.gov/gene/?term=LOC105280774 | 2792 | 98 | 351 | 1 | 3.58 | 1.02 | 31.47 | 33.13 | -1.66 | 1.66 | -0.06 | 0.06 |
| gene5877 | https://www.ncbi.nlm.nih.gov/gene/?term=LOC105278807 | 1719 | 50 | 290.87 | 1 | 5.82 | 2 | 21.66 | 23.62 | -1.95 | 1.95 | -0.06 | 0.06 |
| gene12406 | https://www.ncbi.nlm.nih.gov/gene/?term=LOC105286316 | 1386 | 29 | 209.24 | 1 | 7.22 | 3.45 | 41.65 | 44.38 | -2.73 | 2.73 | -0.06 | 0.06 |
| gene393 | https://www.ncbi.nlm.nih.gov/gene/?term=LOC105276566 | 1123 | 2 | 17.81 | 1 | 8.9 | 50 | 77.65 | 48.08 | 29.58 | 29.58 | 0.05 | 0.05 |
| gene7915 | https://www.ncbi.nlm.nih.gov/gene/?term=LOC105281173 | 1597 | 25 | 156.54 | 1 | 6.26 | 4 | 64.87 | 61.62 | 3.25 | 3.25 | 0.05 | 0.05 |
| gene1220 | https://www.ncbi.nlm.nih.gov/gene/?term=LOC105285742 | 3453 | 90 | 260.64 | 1 | 2.9 | 1.11 | 27.79 | 25.88 | 1.92 | 1.92 | 0.05 | 0.05 |
| gene11401 | https://www.ncbi.nlm.nih.gov/gene/?term=LOC105285202 | 1841 | 47 | 255.3 | 1 | 5.43 | 2.13 | 22.36 | 20.6 | 1.77 | 1.77 | 0.05 | 0.05 |
| gene6966 | https://www.ncbi.nlm.nih.gov/gene/?term=LOC105280104 | 3747 | 116 | 309.58 | 1 | 2.67 | 0.86 | 15.51 | 13.86 | 1.65 | 1.65 | 0.05 | 0.05 |
| gene5298 | https://www.ncbi.nlm.nih.gov/gene/?term=LOC105278145 | 4060 | 245 | 603.45 | 2 | 4.93 | 0.82 | 11.36 | 10.5 | 0.86 | 0.86 | 0.05 | 0.05 |
| gene10984 | https://www.ncbi.nlm.nih.gov/gene/?term=LOC105284704 | 1987 | 61 | 307 | 1 | 5.03 | 1.64 | 25.94 | 27.43 | -1.49 | 1.49 | -0.05 | 0.05 |
| gene10520 | https://www.ncbi.nlm.nih.gov/gene/?term=LOC105284184 | 2451 | 67 | 273.36 | 1 | 4.08 | 1.49 | 41.63 | 43.45 | -1.81 | 1.81 | -0.05 | 0.05 |
| gene12516 | https://www.ncbi.nlm.nih.gov/gene/?term=LOC105286452 | 1529 | 35 | 228.91 | 1 | 6.54 | 2.86 | 51.01 | 53.03 | -2.03 | 2.03 | -0.05 | 0.05 |
| gene9109 | https://www.ncbi.nlm.nih.gov/gene/?term=LOC105282561 | 2400 | 10 | 41.67 | 1 | 4.17 | 10 | 26.72 | 17.84 | 8.89 | 8.89 | 0.04 | 0.04 |
| gene7952 | https://www.ncbi.nlm.nih.gov/gene/?term=LOC105281213 | 2963 | 31 | 104.62 | 1 | 3.37 | 3.23 | 42.96 | 39.57 | 3.39 | 3.39 | 0.04 | 0.04 |
| gene12981 | https://www.ncbi.nlm.nih.gov/gene/?term=LOC105286999 | 2063 | 39 | 189.05 | 1 | 4.85 | 2.56 | 33.12 | 30.76 | 2.36 | 2.36 | 0.04 | 0.04 |
| gene2404 | https://www.ncbi.nlm.nih.gov/gene/?term=LOC105274853 | 1767 | 33 | 186.76 | 1 | 5.66 | 3.03 | 31.53 | 29.22 | 2.31 | 2.31 | 0.04 | 0.04 |
| gene10391 | https://www.ncbi.nlm.nih.gov/gene/?term=LOC109611283 | 1994 | 41 | 205.62 | 1 | 5.02 | 2.44 | 35.16 | 33.4 | 1.76 | 1.76 | 0.04 | 0.04 |
| gene11454 | https://www.ncbi.nlm.nih.gov/gene/?term=LOC105285256 | 1906 | 48 | 251.84 | 1 | 5.25 | 2.08 | 46.56 | 44.93 | 1.63 | 1.63 | 0.04 | 0.04 |
| gene11275 | https://www.ncbi.nlm.nih.gov/gene/?term=LOC105285055 | 4119 | 196 | 475.84 | 1 | 2.43 | 0.51 | 13.91 | 14.65 | -0.74 | 0.74 | -0.04 | 0.04 |
| gene298 | https://www.ncbi.nlm.nih.gov/gene/?term=LOC105275460 | 2473 | 107 | 432.67 | 1 | 4.04 | 0.93 | 13.52 | 14.43 | -0.91 | 0.91 | -0.04 | 0.04 |
| gene11819 | https://www.ncbi.nlm.nih.gov/gene/?term=LOC105285671 | 2366 | 91 | 384.62 | 1 | 4.23 | 1.1 | 28.12 | 29.18 | -1.06 | 1.06 | -0.04 | 0.04 |
| gene1310 | https://www.ncbi.nlm.nih.gov/gene/?term=LOC105286661 | 2582 | 65 | 251.74 | 1 | 3.87 | 1.54 | 19.96 | 21.58 | -1.62 | 1.62 | -0.04 | 0.04 |
| gene274 | https://www.ncbi.nlm.nih.gov/gene/?term=LOC105275190 | 3488 | 66 | 189.22 | 1 | 2.87 | 1.52 | 20.22 | 22.09 | -1.87 | 1.87 | -0.04 | 0.04 |
| gene3727 | https://www.ncbi.nlm.nih.gov/gene/?term=LOC105276367 | 2484 | 54 | 217.39 | 1 | 4.03 | 1.85 | 48.38 | 50.33 | -1.95 | 1.95 | -0.04 | 0.04 |
| gene7398 | https://www.ncbi.nlm.nih.gov/gene/?term=LOC105280589 | 3452 | 41 | 118.77 | 1 | 2.9 | 2.44 | 25.07 | 28.12 | -3.05 | 3.05 | -0.04 | 0.04 |
| gene7368 | https://www.ncbi.nlm.nih.gov/gene/?term=LOC105280556 | 4105 | 40 | 97.44 | 1 | 2.44 | 2.5 | 14.82 | 11.87 | 2.94 | 2.94 | 0.03 | 0.03 |
| gene3168 | https://www.ncbi.nlm.nih.gov/gene/?term=LOC105275711 | 6150 | 71 | 115.45 | 1 | 1.63 | 1.41 | 14.71 | 11.97 | 2.74 | 2.74 | 0.03 | 0.03 |
| gene5490 | https://www.ncbi.nlm.nih.gov/gene/?term=LOC105278358 | 2095 | 25 | 119.33 | 1 | 4.77 | 4 | 36.86 | 34.23 | 2.63 | 2.63 | 0.03 | 0.03 |
| gene388 | https://www.ncbi.nlm.nih.gov/gene/?term=LOC105276538 | 1558 | 19 | 121.95 | 2 | 12.84 | 10.53 | 69.41 | 66.83 | 2.58 | 2.58 | 0.03 | 0.03 |
| gene6422 | https://www.ncbi.nlm.nih.gov/gene/?term=LOC105279458 | 2419 | 33 | 136.42 | 1 | 4.13 | 3.03 | 55.1 | 53.04 | 2.06 | 2.06 | 0.03 | 0.03 |
| gene3107 | https://www.ncbi.nlm.nih.gov/gene/?term=LOC105275650 | 5103 | 143 | 280.23 | 1 | 1.96 | 0.7 | 16.9 | 15.68 | 1.21 | 1.21 | 0.03 | 0.03 |
| gene4190 | https://www.ncbi.nlm.nih.gov/gene/?term=LOC105276875 | 3150 | 104 | 330.16 | 1 | 3.17 | 0.96 | 25.99 | 25.04 | 0.95 | 0.95 | 0.03 | 0.03 |
| gene3286 | https://www.ncbi.nlm.nih.gov/gene/?term=LOC105275848 | 3744 | 121 | 323.18 | 1 | 2.67 | 0.83 | 22.9 | 22 | 0.9 | 0.9 | 0.03 | 0.03 |
| gene1993 | https://www.ncbi.nlm.nih.gov/gene/?term=LOC105288193 | 1449 | 62 | 427.88 | 1 | 6.9 | 1.61 | 35.55 | 34.76 | 0.78 | 0.78 | 0.03 | 0.03 |
| gene474 | https://www.ncbi.nlm.nih.gov/gene/?term=LOC105277435 | 3625 | 150 | 413.79 | 1 | 2.76 | 0.67 | 12.65 | 11.92 | 0.73 | 0.73 | 0.03 | 0.03 |
| gene2844 | https://www.ncbi.nlm.nih.gov/gene/?term=LOC105275363 | 4330 | 179 | 413.39 | 1 | 2.31 | 0.56 | 14.53 | 15.15 | -0.61 | 0.61 | -0.03 | 0.03 |
| gene8729 | https://www.ncbi.nlm.nih.gov/gene/?term=LOC105282137 | 5005 | 195 | 389.61 | 1 | 2 | 0.51 | 16.47 | 17.35 | -0.89 | 0.89 | -0.03 | 0.03 |
| gene7166 | https://www.ncbi.nlm.nih.gov/gene/?term=LOC105280326 | 6117 | 195 | 318.78 | 1 | 1.63 | 0.51 | 16.86 | 17.78 | -0.92 | 0.92 | -0.03 | 0.03 |
| gene9101 | https://www.ncbi.nlm.nih.gov/gene/?term=LOC105282553 | 3068 | 104 | 338.98 | 1 | 3.26 | 0.96 | 20.36 | 21.29 | -0.93 | 0.93 | -0.03 | 0.03 |
| gene4023 | https://www.ncbi.nlm.nih.gov/gene/?term=LOC105276692 | 1759 | 35 | 198.98 | 1 | 5.69 | 2.86 | 23.93 | 25.35 | -1.42 | 1.42 | -0.03 | 0.03 |
| gene7183 | https://www.ncbi.nlm.nih.gov/gene/?term=LOC105280348 | 3925 | 51 | 129.94 | 1 | 2.55 | 1.96 | 25.44 | 27.74 | -2.3 | 2.3 | -0.03 | 0.03 |
| gene10107 | https://www.ncbi.nlm.nih.gov/gene/?term=LOC105283712 | 1883 | 23 | 122.15 | 1 | 5.31 | 4.35 | 27.23 | 29.73 | -2.49 | 2.49 | -0.03 | 0.03 |
| gene1199 | https://www.ncbi.nlm.nih.gov/gene/?term=LOC105285498 | 2220 | 23 | 103.6 | 1 | 4.5 | 4.35 | 35.45 | 33.87 | 1.58 | 1.58 | 0.02 | 0.02 |
| gene12021 | https://www.ncbi.nlm.nih.gov/gene/?term=LOC105285905 | 1399 | 21 | 150.11 | 1 | 7.15 | 4.76 | 41.18 | 39.62 | 1.55 | 1.55 | 0.02 | 0.02 |
| gene2748 | https://www.ncbi.nlm.nih.gov/gene/?term=LOC105275261 | 3845 | 54 | 140.44 | 1 | 2.6 | 1.85 | 38.15 | 36.73 | 1.42 | 1.42 | 0.02 | 0.02 |
| gene11447 | https://www.ncbi.nlm.nih.gov/gene/?term=LOC105285252 | 7437 | 119 | 160.01 | 1 | 1.34 | 0.84 | 14.91 | 13.69 | 1.22 | 1.22 | 0.02 | 0.02 |
| gene5375 | https://www.ncbi.nlm.nih.gov/gene/?term=LOC105278228 | 8640 | 143 | 165.51 | 1 | 1.16 | 0.7 | 24.34 | 23.23 | 1.11 | 1.11 | 0.02 | 0.02 |
| gene5646 | https://www.ncbi.nlm.nih.gov/gene/?term=LOC105278534 | 7475 | 166 | 222.07 | 1 | 1.34 | 0.6 | 18.54 | 17.53 | 1.02 | 1.02 | 0.02 | 0.02 |
| gene4376 | https://www.ncbi.nlm.nih.gov/gene/?term=LOC105277083 | 4307 | 92 | 213.61 | 1 | 2.32 | 1.09 | 26.82 | 26.06 | 0.76 | 0.76 | 0.02 | 0.02 |
| gene11340 | https://www.ncbi.nlm.nih.gov/gene/?term=LOC105285131 | 2918 | 93 | 318.71 | 1 | 3.43 | 1.08 | 12.16 | 11.4 | 0.75 | 0.75 | 0.02 | 0.02 |
| gene2271 | https://www.ncbi.nlm.nih.gov/gene/?term=LOC105274706 | 1448 | 52 | 359.12 | 1 | 6.91 | 1.92 | 37.3 | 36.65 | 0.64 | 0.64 | 0.02 | 0.02 |

| | | | | | | | | | | | | | |
|---|---|---|---|---|---|---|---|---|---|---|---|---|---|
| gene12573 | https://www.ncbi.nlm.nih.gov/gene/?term=LOC105286515 | 3647 | 106 | 290.65 | 1 | 2.74 | 0.94 | 10.61 | 10.01 | 0.6 | 0.6 | 0.02 | 0.02 |
| gene9645 | https://www.ncbi.nlm.nih.gov/gene/?term=LOC105283181 | 3875 | 118 | 304.52 | 1 | 2.58 | 0.85 | 12.99 | 12.4 | 0.59 | 0.59 | 0.02 | 0.02 |
| gene13036 | https://www.ncbi.nlm.nih.gov/gene/?term=LOC105287059 | 4959 | 150 | 302.48 | 1 | 2.02 | 0.67 | 17.57 | 17 | 0.57 | 0.57 | 0.02 | 0.02 |
| gene5831 | https://www.ncbi.nlm.nih.gov/gene/?term=LOC105278761 | 2506 | 100 | 399.04 | 2 | 7.98 | 2 | 24.32 | 23.77 | 0.54 | 0.54 | 0.02 | 0.02 |
| gene13114 | https://www.ncbi.nlm.nih.gov/gene/?term=LOC105287147 | 7609 | 360 | 473.12 | 1 | 1.31 | 0.28 | 10.34 | 9.95 | 0.39 | 0.39 | 0.02 | 0.02 |
| gene7568 | https://www.ncbi.nlm.nih.gov/gene/?term=LOC105280786 | 2525 | 107 | 423.76 | 1 | 3.96 | 0.93 | 33.06 | 32.68 | 0.38 | 0.38 | 0.02 | 0.02 |
| gene10606 | https://www.ncbi.nlm.nih.gov/gene/?term=LOC105284273 | 6119 | 227 | 370.98 | 1 | 1.63 | 0.44 | 21.11 | 21.54 | -0.43 | 0.43 | -0.02 | 0.02 |
| gene5649 | https://www.ncbi.nlm.nih.gov/gene/?term=LOC105278538 | 2140 | 77 | 359.81 | 1 | 4.67 | 1.3 | 17.91 | 18.36 | -0.46 | 0.46 | -0.02 | 0.02 |
| gene9755 | https://www.ncbi.nlm.nih.gov/gene/?term=LOC105283296 | 2177 | 79 | 362.88 | 1 | 4.59 | 1.27 | 20.77 | 21.25 | -0.48 | 0.48 | -0.02 | 0.02 |
| gene10678 | https://www.ncbi.nlm.nih.gov/gene/?term=LOC105284355 | 4031 | 152 | 377.08 | 1 | 2.48 | 0.66 | 20.4 | 20.91 | -0.51 | 0.51 | -0.02 | 0.02 |
| gene7888 | https://www.ncbi.nlm.nih.gov/gene/?term=LOC105281141 | 4357 | 153 | 351.16 | 1 | 2.3 | 0.65 | 30.21 | 30.79 | -0.57 | 0.57 | -0.02 | 0.02 |
| gene292 | https://www.ncbi.nlm.nih.gov/gene/?term=LOC105275398 | 2825 | 77 | 272.57 | 1 | 3.54 | 1.3 | 19.76 | 20.59 | -0.82 | 0.82 | -0.02 | 0.02 |
| gene10531 | https://www.ncbi.nlm.nih.gov/gene/?term=LOC105284190 | 6881 | 164 | 238.34 | 1 | 1.45 | 0.61 | 10.48 | 11.52 | -1.03 | 1.03 | -0.02 | 0.02 |
| gene2173 | https://www.ncbi.nlm.nih.gov/gene/?term=LOC105274590 | 4387 | 64 | 145.89 | 1 | 2.28 | 1.56 | 26 | 27.13 | -1.13 | 1.13 | -0.02 | 0.02 |
| gene6455 | https://www.ncbi.nlm.nih.gov/gene/?term=LOC105279500 | 2612 | 31 | 118.68 | 1 | 3.83 | 3.23 | 51.05 | 52.76 | -1.71 | 1.71 | -0.02 | 0.02 |
| gene10819 | https://www.ncbi.nlm.nih.gov/gene/?term=LOC105284507 | 5767 | 69 | 119.65 | 1 | 1.73 | 1.45 | 22.66 | 21.47 | 1.19 | 1.19 | 0.01 | 0.01 |
| gene322 | https://www.ncbi.nlm.nih.gov/gene/?term=LOC105275777 | 1606 | 21 | 130.76 | 1 | 6.23 | 4.76 | 49.88 | 48.85 | 1.04 | 1.04 | 0.01 | 0.01 |
| gene10155 | https://www.ncbi.nlm.nih.gov/gene/?term=LOC105283756 | 7030 | 86 | 122.33 | 1 | 1.42 | 1.16 | 7.44 | 6.49 | 0.95 | 0.95 | 0.01 | 0.01 |
| gene3671 | https://www.ncbi.nlm.nih.gov/gene/?term=LOC105276272 | 1966 | 25 | 127.16 | 1 | 5.09 | 4 | 43.89 | 43.09 | 0.8 | 0.8 | 0.01 | 0.01 |
| gene3848 | https://www.ncbi.nlm.nih.gov/gene/?term=LOC105276486 | 5899 | 107 | 181.39 | 1 | 1.7 | 0.93 | 25.75 | 24.98 | 0.77 | 0.77 | 0.01 | 0.01 |
| gene2683 | https://www.ncbi.nlm.nih.gov/gene/?term=LOC105275187 | 5492 | 66 | 120.17 | 1 | 1.82 | 1.52 | 14.03 | 13.27 | 0.76 | 0.76 | 0.01 | 0.01 |
| gene1147 | https://www.ncbi.nlm.nih.gov/gene/?term=LOC105284954 | 4022 | 75 | 186.47 | 1 | 2.49 | 1.33 | 14.93 | 14.22 | 0.71 | 0.71 | 0.01 | 0.01 |
| gene7207 | https://www.ncbi.nlm.nih.gov/gene/?term=LOC105280376 | 4659 | 98 | 210.35 | 1 | 2.15 | 1.02 | 30.66 | 30.04 | 0.62 | 0.62 | 0.01 | 0.01 |
| gene657 | https://www.ncbi.nlm.nih.gov/gene/?term=LOC105279667 | 6421 | 64 | 99.67 1 | 1.56 | 1.56 | 3.41 | 2.8 | 0.61 | 0.61 | 0.01 | 0.01 | |
| gene3599 | https://www.ncbi.nlm.nih.gov/gene/?term=LOC105276196 | 2955 | 40 | 135.36 | 1 | 3.38 | 2.5 | 52.66 | 52.1 | 0.56 | 0.56 | 0.01 | 0.01 |
| gene12307 | https://www.ncbi.nlm.nih.gov/gene/?term=LOC105286216 | 10313 | 146 | 141.57 | 1 | 0.97 | 0.68 | 17.27 | 16.76 | 0.51 | 0.51 | 0.01 | 0.01 |
| gene8661 | https://www.ncbi.nlm.nih.gov/gene/?term=LOC105282063 | 1926 | 39 | 202.49 | 1 | 5.19 | 2.56 | 23.72 | 23.24 | 0.48 | 0.48 | 0.01 | 0.01 |
| gene10326 | https://www.ncbi.nlm.nih.gov/gene/?term=LOC105283962 | 3042 | 92 | 302.43 | 1 | 3.29 | 1.09 | 15.29 | 14.85 | 0.44 | 0.44 | 0.01 | 0.01 |
| gene13044 | https://www.ncbi.nlm.nih.gov/gene/?term=LOC105287070 | 5094 | 159 | 312.13 | 1 | 1.96 | 0.63 | 5.69 | 5.29 | 0.4 | 0.4 | 0.01 | 0.01 |
| gene3817 | https://www.ncbi.nlm.nih.gov/gene/?term=LOC105276449 | 1230 | 39 | 317.07 | 1 | 8.13 | 2.56 | 32.77 | 32.39 | 0.38 | 0.38 | 0.01 | 0.01 |
| gene3206 | https://www.ncbi.nlm.nih.gov/gene/?term=LOC105275752 | 3474 | 117 | 336.79 | 1 | 2.88 | 0.85 | 29.75 | 29.38 | 0.37 | 0.37 | 0.01 | 0.01 |
| gene6583 | https://www.ncbi.nlm.nih.gov/gene/?term=LOC105279649 | 7444 | 308 | 413.76 | 1 | 1.34 | 0.32 | 11.75 | 11.41 | 0.34 | 0.34 | 0.01 | 0.01 |
| gene10392 | https://www.ncbi.nlm.nih.gov/gene/?term=LOC105284036 | 3216 | 76 | 236.32 | 1 | 3.11 | 1.32 | 25.16 | 24.85 | 0.31 | 0.31 | 0.01 | 0.01 |
| gene12674 | https://www.ncbi.nlm.nih.gov/gene/?term=LOC105286649 | 18283 | 638 | 348.96 | 1 | 0.55 | 0.16 | 1.31 | 1.05 | 0.26 | 0.26 | 0.01 | 0.01 |
| gene10549 | https://www.ncbi.nlm.nih.gov/gene/?term=LOC105284210 | 7145 | 298 | 417.07 | 1 | 1.4 | 0.34 | 11.27 | 11.01 | 0.25 | 0.25 | 0.01 | 0.01 |
| gene7249 | https://www.ncbi.nlm.nih.gov/gene/?term=LOC105280422 | 1775 | 58 | 326.76 | 1 | 5.63 | 1.72 | 15.27 | 15.08 | 0.19 | 0.19 | 0.01 | 0.01 |
| gene10018 | https://www.ncbi.nlm.nih.gov/gene/?term=LOC105283607 | 4414 | 181 | 410.06 | 1 | 2.27 | 0.55 | 16.23 | 16.36 | -0.13 | 0.13 | -0.01 | 0.01 |
| gene1470 | https://www.ncbi.nlm.nih.gov/gene/?term=LOC105287597 | 2974 | 188 | 632.15 | 1 | 3.36 | 0.53 | 17.15 | 17.35 | -0.19 | 0.19 | -0.01 | 0.01 |
| gene3885 | https://www.ncbi.nlm.nih.gov/gene/?term=LOC105276527 | 7928 | 323 | 407.42 | 1 | 1.26 | 0.31 | 8.78 | 9 | -0.22 | 0.22 | -0.01 | 0.01 |
| gene10540 | https://www.ncbi.nlm.nih.gov/gene/?term=LOC105284200 | 5267 | 183 | 347.45 | 1 | 1.9 | 0.55 | 14.63 | 14.93 | -0.3 | 0.3 | -0.01 | 0.01 |
| gene6286 | https://www.ncbi.nlm.nih.gov/gene/?term=LOC105279298 | 2861 | 128 | 447.4 2 | 6.99 | 1.56 | 23.41 | 23.73 | -0.32 | 0.32 | -0.01 | 0.01 | |
| gene3856 | https://www.ncbi.nlm.nih.gov/gene/?term=LOC105276499 | 5236 | 178 | 339.95 | 1 | 1.91 | 0.56 | 9.73 | 10.06 | -0.33 | 0.33 | -0.01 | 0.01 |
| gene4892 | https://www.ncbi.nlm.nih.gov/gene/?term=LOC105277667 | 3898 | 119 | 305.28 | 1 | 2.57 | 0.84 | 27.91 | 28.29 | -0.38 | 0.38 | -0.01 | 0.01 |
| gene12317 | https://www.ncbi.nlm.nih.gov/gene/?term=LOC105286223 | 2235 | 79 | 353.47 | 1 | 4.47 | 1.27 | 30.56 | 30.96 | -0.4 | 0.4 | -0.01 | 0.01 |
| gene1884 | https://www.ncbi.nlm.nih.gov/gene/?term=LOC105288072 | 4169 | 76 | 182.3 1 | 2.4 | 1.32 | 23.55 | 23.95 | -0.4 | 0.4 | -0.01 | 0.01 | |
| gene2056 | https://www.ncbi.nlm.nih.gov/gene/?term=LOC105274452 | 6584 | 206 | 312.88 | 1 | 1.52 | 0.49 | 14.33 | 14.74 | -0.41 | 0.41 | -0.01 | 0.01 |
| gene4641 | https://www.ncbi.nlm.nih.gov/gene/?term=LOC105277398 | 1937 | 68 | 351.06 | 1 | 5.16 | 1.47 | 34.8 | 35.21 | -0.41 | 0.41 | -0.01 | 0.01 |
| gene303 | https://www.ncbi.nlm.nih.gov/gene/?term=LOC105275584 | 3081 | 71 | 230.44 | 1 | 3.25 | 1.41 | 26.68 | 27.17 | -0.49 | 0.49 | -0.01 | 0.01 |
| gene13033 | https://www.ncbi.nlm.nih.gov/gene/?term=LOC105287063 | 1877 | 45 | 239.74 | 1 | 5.33 | 2.22 | 34.91 | 35.54 | -0.62 | 0.62 | -0.01 | 0.01 |
| gene8600 | https://www.ncbi.nlm.nih.gov/gene/?term=LOC105282001 | 8513 | 126 | 148.01 | 1 | 1.17 | 0.79 | 11.25 | 11.9 | -0.65 | 0.65 | -0.01 | 0.01 |
| gene7245 | https://www.ncbi.nlm.nih.gov/gene/?term=LOC105280410 | 12478 | 234 | 187.53 | 1 | 0.8 | 0.43 | 20.56 | 21.23 | -0.67 | 0.67 | -0.01 | 0.01 |
| gene8343 | https://www.ncbi.nlm.nih.gov/gene/?term=LOC105281700 | 2172 | 40 | 184.16 | 1 | 4.6 | 2.5 | 41.18 | 41.86 | -0.68 | 0.68 | -0.01 | 0.01 |
| gene9336 | https://www.ncbi.nlm.nih.gov/gene/?term=LOC105282834 | 5993 | 106 | 176.87 | 1 | 1.67 | 0.94 | 15.44 | 16.15 | -0.71 | 0.71 | -0.01 | 0.01 |
| gene10344 | https://www.ncbi.nlm.nih.gov/gene/?term=LOC105283987 | 4841 | 64 | 132.2 1 | 2.07 | 1.56 | 32.11 | 32.92 | -0.81 | 0.81 | -0.01 | 0.01 | |
| gene6528 | https://www.ncbi.nlm.nih.gov/gene/?term=LOC105279591 | 7222 | 67 | 92.77 1 | 1.38 | 1.49 | 22.84 | 24.15 | -1.32 | 1.32 | -0.01 | 0.01 | |
| gene2970 | https://www.ncbi.nlm.nih.gov/gene/?term=LOC105275518 | 2340 | 18 | 76.92 1 | 4.27 | 5.56 | 47.45 | 48.96 | -1.51 | 1.51 | -0.01 | 0.01 | |
| gene10345 | https://www.ncbi.nlm.nih.gov/gene/?term=LOC105283989 | 4444 | 21 | 47.25 1 | 2.25 | 4.76 | 48.99 | 48.08 | 0.91 | 0.91 | 0 | 0 | |
| gene10600 | https://www.ncbi.nlm.nih.gov/gene/?term=LOC105284270 | 4837 | 97 | 200.54 | 1 | 2.07 | 1.03 | 13.04 | 12.82 | 0.22 | 0.22 | 0 | 0 |
| gene4962 | https://www.ncbi.nlm.nih.gov/gene/?term=LOC105277761 | 4645 | 140 | 301.4 1 | 2.15 | 0.71 | 14.56 | 14.4 | 0.16 | 0.16 | 0 | 0 | |
| gene5967 | https://www.ncbi.nlm.nih.gov/gene/?term=LOC105278918 | 4393 | 121 | 275.44 | 1 | 2.28 | 0.83 | 16.38 | 16.32 | 0.06 | 0.06 | 0 | 0 |
| gene6031 | https://www.ncbi.nlm.nih.gov/gene/?term=LOC105279000 | 5691 | 174 | 305.75 | 1 | 1.76 | 0.57 | 14.03 | 13.97 | 0.06 | 0.06 | 0 | 0 |
| gene11583 | https://www.ncbi.nlm.nih.gov/gene/?term=LOC105285414 | 8192 | 168 | 205.08 | 1 | 1.22 | 0.6 | 8.3 | 8.25 | 0.05 | 0.05 | 0 | 0 |
| gene940 | https://www.ncbi.nlm.nih.gov/gene/?term=LOC105282794 | 3578 | 82 | 229.18 | 1 | 2.79 | 1.22 | 16.49 | 16.45 | 0.04 | 0.04 | 0 | 0 |
| gene5524 | https://www.ncbi.nlm.nih.gov/gene/?term=LOC105278397 | 5177 | 200 | 386.32 | 1 | 1.93 | 0.5 | 24.39 | 24.36 | 0.03 | 0.03 | 0 | 0 |

| | | | | | | | | | | | | | | |
|---|---|---|---|---|---|---|---|---|---|---|---|---|---|---|
| gene5746 | https://www.ncbi.nlm.nih.gov/gene/?term=LOC105278666 | 10343 | 387 | 374.17 | | 1 | 0.97 | 0.26 | 7.75 | 7.72 | 0.03 | 0.03 | 0 | 0 |
| gene6639 | https://www.ncbi.nlm.nih.gov/gene/?term=LOC105279705 | 7656 | 58 | 75.76 | 1 | 1.31 | 1.72 | 24.54 | 24.54 | 0 | 0 | 0 | 0 | |
| gene12802 | https://www.ncbi.nlm.nih.gov/gene/?term=LOC105286799 | 3183 | 165 | 518.38 | | 1 | 3.14 | 0.61 | 16.21 | 16.23 | -0.02 | 0.02 | 0 | 0 |
| gene2806 | https://www.ncbi.nlm.nih.gov/gene/?term=LOC105275327 | 14257 | 414 | 290.38 | | 1 | 0.7 | 0.24 | 7.7 | 7.73 | -0.02 | 0.02 | 0 | 0 |
| gene11579 | https://www.ncbi.nlm.nih.gov/gene/?term=LOC105285406 | 13571 | 248 | 182.74 | | 1 | 0.74 | 0.4 | 5.98 | 6.04 | -0.05 | 0.05 | 0 | 0 |
| gene7148 | https://www.ncbi.nlm.nih.gov/gene/?term=LOC105280304 | 8920 | 90 | 100.9 | 1 | 1.12 | 1.11 | 7.28 | 7.34 | -0.05 | 0.05 | 0 | 0 | |
| gene53 | https://www.ncbi.nlm.nih.gov/gene/?term=LOC105277148 | 19283 | 236 | 122.39 | | 1 | 0.52 | 0.42 | 4.4 | 4.47 | -0.07 | 0.07 | 0 | 0 |
| gene5525 | https://www.ncbi.nlm.nih.gov/gene/?term=LOC105278396 | 3090 | 186 | 601.94 | | 1 | 3.24 | 0.54 | 24.51 | 24.57 | -0.07 | 0.07 | 0 | 0 |
| gene6620 | https://www.ncbi.nlm.nih.gov/gene/?term=LOC105279687 | 17998 | 294 | 163.35 | | 1 | 0.56 | 0.34 | 5.33 | 5.42 | -0.1 | 0.1 | 0 | 0 |
| gene4246 | https://www.ncbi.nlm.nih.gov/gene/?term=LOC105276941 | 16412 | 446 | 271.75 | | 1 | 0.61 | 0.22 | 5.36 | 5.47 | -0.11 | 0.11 | 0 | 0 |
| gene9330 | https://www.ncbi.nlm.nih.gov/gene/?term=LOC105282825 | 3936 | 80 | 203.25 | | 1 | 2.54 | 1.25 | 13.05 | 13.17 | -0.11 | 0.11 | 0 | 0 |
| gene12203 | https://www.ncbi.nlm.nih.gov/gene/?term=LOC105286099 | 5967 | 70 | 117.31 | | 1 | 1.68 | 1.43 | 11.64 | 11.77 | -0.13 | 0.13 | 0 | 0 |
| gene11568 | https://www.ncbi.nlm.nih.gov/gene/?term=LOC105285395 | 8086 | 160 | 197.87 | | 1 | 1.24 | 0.62 | 17.75 | 17.97 | -0.23 | 0.23 | 0 | 0 |

Directory: OGL File: rnk-dmg-Cb_LI_BCphaseA1.vs.BCphaseB1.txt

| gene_ID | gene_link | gwidth | #Sites | #per10Kb | #dmSites | #dmsp10kb | %dmSites | %pSite1 | %pSite2 | DMpSite | ADMpSite | DMpNucl | ADMpNucl |
|---|---|---|---|---|---|---|---|---|---|---|---|---|---|
| gene3165 | https://www.ncbi.nlm.nih.gov/gene/?term=LOC105275706 | 2374 | 66 | 278.01 | 4 | 16.85 | 6.06 | 24.58 | 35.31 | -10.73 | 10.73 | -0.3 | 0.3 |
| gene2593 | https://www.ncbi.nlm.nih.gov/gene/?term=LOC105275070 | 1774 | 6 | 33.82 | 1 | 5.64 | 16.67 | 16.42 | 54.02 | -37.6 | 37.6 | -0.13 | 0.13 |
| gene9740 | https://www.ncbi.nlm.nih.gov/gene/?term=LOC105283277 | 2887 | 138 | 478 | 1 | 3.46 | 0.72 | 23.22 | 20.78 | 2.44 | 2.44 | 0.12 | 0.12 |
| gene7199 | https://www.ncbi.nlm.nih.gov/gene/?term=LOC105280366 | 1414 | 24 | 169.73 | 1 | 7.07 | 4.17 | 45.18 | 38.6 | 6.59 | 6.59 | 0.11 | 0.11 |
| gene8929 | https://www.ncbi.nlm.nih.gov/gene/?term=LOC105282371 | 1849 | 8 | 43.27 | 1 | 5.41 | 12.5 | 22.25 | 44.14 | -21.89 | 21.89 | -0.09 | 0.09 |
| gene10602 | https://www.ncbi.nlm.nih.gov/gene/?term=LOC105284268 | 2122 | 20 | 94.25 | 1 | 4.71 | 5 | 32.63 | 25.61 | 7.02 | 7.02 | 0.07 | 0.07 |
| gene5298 | https://www.ncbi.nlm.nih.gov/gene/?term=LOC105278145 | 4060 | 250 | 615.76 | 2 | 4.93 | 0.8 | 10.79 | 9.61 | 1.17 | 1.17 | 0.07 | 0.07 |
| gene5714 | https://www.ncbi.nlm.nih.gov/gene/?term=LOC105278729 | 710 | 14 | 197.18 | 1 | 14.08 | 7.14 | 51.67 | 55.07 | -3.41 | 3.41 | -0.07 | 0.07 |
| gene9464 | https://www.ncbi.nlm.nih.gov/gene/?term=LOC105282975 | 1450 | 24 | 165.52 | 1 | 6.9 | 4.17 | 42.63 | 39.26 | 3.37 | 3.37 | 0.06 | 0.06 |
| gene10213 | https://www.ncbi.nlm.nih.gov/gene/?term=LOC105283826 | 1503 | 54 | 359.28 | 1 | 6.65 | 1.85 | 16.14 | 14.48 | 1.66 | 1.66 | 0.06 | 0.06 |
| gene298 | https://www.ncbi.nlm.nih.gov/gene/?term=LOC105275460 | 2473 | 115 | 465.02 | 1 | 4.04 | 0.87 | 12.68 | 13.89 | -1.21 | 1.21 | -0.06 | 0.06 |
| gene1855 | https://www.ncbi.nlm.nih.gov/gene/?term=LOC105288036 | 2387 | 87 | 364.47 | 1 | 4.19 | 1.15 | 28.59 | 30.36 | -1.76 | 1.76 | -0.06 | 0.06 |
| gene6334 | https://www.ncbi.nlm.nih.gov/gene/?term=LOC105279356 | 1471 | 43 | 292.32 | 1 | 6.8 | 2.33 | 28.25 | 30.28 | -2.03 | 2.03 | -0.06 | 0.06 |
| gene12516 | https://www.ncbi.nlm.nih.gov/gene/?term=LOC105286452 | 1529 | 35 | 228.91 | 1 | 6.54 | 2.86 | 51.01 | 53.47 | -2.46 | 2.46 | -0.06 | 0.06 |
| gene1962 | https://www.ncbi.nlm.nih.gov/gene/?term=LOC105288157 | 1246 | 16 | 128.41 | 1 | 8.03 | 6.25 | 13.95 | 18.55 | -4.6 | 4.6 | -0.06 | 0.06 |
| gene393 | https://www.ncbi.nlm.nih.gov/gene/?term=LOC105276566 | 1123 | 2 | 17.81 | 1 | 8.9 | 50 | 77.65 | 47.22 | 30.43 | 30.43 | 0.05 | 0.05 |
| gene11401 | https://www.ncbi.nlm.nih.gov/gene/?term=LOC105285202 | 1841 | 50 | 271.59 | 1 | 5.43 | 2 | 21.11 | 19.1 | 2.01 | 2.01 | 0.05 | 0.05 |
| gene10326 | https://www.ncbi.nlm.nih.gov/gene/?term=LOC105283962 | 3042 | 93 | 305.72 | 1 | 3.29 | 1.08 | 11.63 | 10.03 | 1.6 | 1.6 | 0.05 | 0.05 |
| gene11340 | https://www.ncbi.nlm.nih.gov/gene/?term=LOC105285131 | 2918 | 97 | 332.42 | 1 | 3.43 | 1.03 | 14.75 | 13.34 | 1.41 | 1.41 | 0.05 | 0.05 |
| gene4190 | https://www.ncbi.nlm.nih.gov/gene/?term=LOC105276875 | 3150 | 130 | 412.7 | 1 | 3.17 | 0.77 | 21.07 | 19.84 | 1.23 | 1.23 | 0.05 | 0.05 |
| gene10606 | https://www.ncbi.nlm.nih.gov/gene/?term=LOC105284273 | 6119 | 249 | 406.93 | 2 | 3.27 | 0.8 | 20.32 | 19.15 | 1.16 | 1.16 | 0.05 | 0.05 |
| gene4034 | https://www.ncbi.nlm.nih.gov/gene/?term=LOC105276704 | 2065 | 84 | 406.78 | 1 | 4.84 | 1.19 | 12.86 | 14.01 | -1.15 | 1.15 | -0.05 | 0.05 |
| gene570 | https://www.ncbi.nlm.nih.gov/gene/?term=LOC105278549 | 1673 | 63 | 376.57 | 1 | 5.98 | 1.59 | 40.52 | 41.79 | -1.27 | 1.27 | -0.05 | 0.05 |
| gene2856 | https://www.ncbi.nlm.nih.gov/gene/?term=LOC105275384 | 1987 | 76 | 382.49 | 1 | 5.03 | 1.32 | 32.95 | 34.23 | -1.28 | 1.28 | -0.05 | 0.05 |
| gene4641 | https://www.ncbi.nlm.nih.gov/gene/?term=LOC105277398 | 1937 | 70 | 361.38 | 1 | 5.16 | 1.43 | 35.36 | 36.76 | -1.4 | 1.4 | -0.05 | 0.05 |
| gene9109 | https://www.ncbi.nlm.nih.gov/gene/?term=LOC105282561 | 2400 | 12 | 50 | 1 | 4.17 | 8.33 | 20.55 | 12.93 | 7.62 | 7.62 | 0.04 | 0.04 |
| gene3168 | https://www.ncbi.nlm.nih.gov/gene/?term=LOC105275711 | 6150 | 79 | 128.46 | 1 | 1.63 | 1.27 | 14.18 | 11.09 | 3.09 | 3.09 | 0.04 | 0.04 |
| gene6422 | https://www.ncbi.nlm.nih.gov/gene/?term=LOC105279458 | 2419 | 36 | 148.82 | 1 | 4.13 | 2.78 | 53.83 | 51.06 | 2.78 | 2.78 | 0.04 | 0.04 |
| gene7915 | https://www.ncbi.nlm.nih.gov/gene/?term=LOC105281173 | 1597 | 25 | 156.54 | 1 | 6.26 | 4 | 64.87 | 62.27 | 2.61 | 2.61 | 0.04 | 0.04 |
| gene920 | https://www.ncbi.nlm.nih.gov/gene/?term=LOC105282518 | 1197 | 43 | 359.23 | 1 | 8.35 | 2.33 | 37.5 | 36.41 | 1.09 | 1.09 | 0.04 | 0.04 |
| gene1470 | https://www.ncbi.nlm.nih.gov/gene/?term=LOC105287597 | 2974 | 180 | 605.25 | 1 | 3.36 | 0.56 | 17.88 | 17.25 | 0.63 | 0.63 | 0.04 | 0.04 |
| gene1332 | https://www.ncbi.nlm.nih.gov/gene/?term=LOC105286872 | 2460 | 141 | 573.17 | 1 | 4.07 | 0.71 | 21.92 | 22.68 | -0.76 | 0.76 | -0.04 | 0.04 |
| gene2844 | https://www.ncbi.nlm.nih.gov/gene/?term=LOC105275363 | 4330 | 187 | 431.87 | 1 | 2.31 | 0.53 | 14.36 | 15.35 | -0.99 | 0.99 | -0.04 | 0.04 |
| gene9101 | https://www.ncbi.nlm.nih.gov/gene/?term=LOC105282553 | 3068 | 103 | 335.72 | 1 | 3.26 | 0.97 | 20.56 | 21.9 | -1.34 | 1.34 | -0.04 | 0.04 |
| gene5877 | https://www.ncbi.nlm.nih.gov/gene/?term=LOC105278807 | 1719 | 51 | 296.68 | 1 | 5.82 | 1.96 | 22.99 | 24.43 | -1.44 | 1.44 | -0.04 | 0.04 |
| gene5647 | https://www.ncbi.nlm.nih.gov/gene/?term=LOC105278567 | 3118 | 38 | 121.87 | 1 | 3.21 | 2.63 | 12.56 | 16.06 | -3.5 | 3.5 | -0.04 | 0.04 |
| gene7368 | https://www.ncbi.nlm.nih.gov/gene/?term=LOC105280556 | 4105 | 49 | 119.37 | 1 | 2.44 | 2.04 | 12.67 | 10.47 | 2.2 | 2.2 | 0.03 | 0.03 |
| gene3727 | https://www.ncbi.nlm.nih.gov/gene/?term=LOC105276367 | 2484 | 50 | 201.29 | 1 | 4.03 | 2 | 50.02 | 48.5 | 1.52 | 1.52 | 0.03 | 0.03 |
| gene7345 | https://www.ncbi.nlm.nih.gov/gene/?term=LOC105280529 | 3833 | 87 | 226.98 | 1 | 2.61 | 1.15 | 18.24 | 16.79 | 1.44 | 1.44 | 0.03 | 0.03 |
| gene3599 | https://www.ncbi.nlm.nih.gov/gene/?term=LOC105276196 | 2955 | 54 | 182.74 | 1 | 3.38 | 1.85 | 40.6 | 39.18 | 1.42 | 1.42 | 0.03 | 0.03 |
| gene2271 | https://www.ncbi.nlm.nih.gov/gene/?term=LOC105274706 | 1448 | 47 | 324.59 | 1 | 6.91 | 2.13 | 41.16 | 40.14 | 1.03 | 1.03 | 0.03 | 0.03 |
| gene1220 | https://www.ncbi.nlm.nih.gov/gene/?term=LOC105285742 | 3453 | 94 | 272.23 | 1 | 2.9 | 1.06 | 26.72 | 25.74 | 0.97 | 0.97 | 0.03 | 0.03 |
| gene6031 | https://www.ncbi.nlm.nih.gov/gene/?term=LOC105279000 | 5691 | 180 | 316.29 | 1 | 1.76 | 0.56 | 14.05 | 13.11 | 0.94 | 0.94 | 0.03 | 0.03 |
| gene4962 | https://www.ncbi.nlm.nih.gov/gene/?term=LOC105277761 | 4645 | 140 | 301.4 | 1 | 2.15 | 0.71 | 15.27 | 14.38 | 0.89 | 0.89 | 0.03 | 0.03 |
| gene7249 | https://www.ncbi.nlm.nih.gov/gene/?term=LOC105280422 | 1775 | 61 | 343.66 | 1 | 5.63 | 1.64 | 15.16 | 14.36 | 0.81 | 0.81 | 0.03 | 0.03 |
| gene3793 | https://www.ncbi.nlm.nih.gov/gene/?term=LOC105276429 | 2901 | 112 | 386.07 | 1 | 3.45 | 0.89 | 26.42 | 25.76 | 0.65 | 0.65 | 0.03 | 0.03 |
| gene5525 | https://www.ncbi.nlm.nih.gov/gene/?term=LOC105278396 | 3090 | 187 | 605.18 | 1 | 3.24 | 0.53 | 24.38 | 23.89 | 0.49 | 0.49 | 0.03 | 0.03 |
| gene3885 | https://www.ncbi.nlm.nih.gov/gene/?term=LOC105276527 | 7928 | 346 | 436.43 | 1 | 1.26 | 0.29 | 8.3 | 8.89 | -0.6 | 0.6 | -0.03 | 0.03 |
| gene10018 | https://www.ncbi.nlm.nih.gov/gene/?term=LOC105283607 | 4414 | 204 | 462.17 | 1 | 2.27 | 0.49 | 14.8 | 15.47 | -0.67 | 0.67 | -0.03 | 0.03 |
| gene8729 | https://www.ncbi.nlm.nih.gov/gene/?term=LOC105282137 | 5005 | 199 | 397.6 | 1 | 2 | 0.5 | 16.19 | 16.99 | -0.79 | 0.79 | -0.03 | 0.03 |
| gene11819 | https://www.ncbi.nlm.nih.gov/gene/?term=LOC105285671 | 2366 | 96 | 405.75 | 1 | 4.23 | 1.04 | 27.73 | 28.57 | -0.84 | 0.84 | -0.03 | 0.03 |
| gene1310 | https://www.ncbi.nlm.nih.gov/gene/?term=LOC105286661 | 2582 | 65 | 251.74 | 1 | 3.87 | 1.54 | 19.96 | 20.95 | -0.99 | 0.99 | -0.03 | 0.03 |
| gene292 | https://www.ncbi.nlm.nih.gov/gene/?term=LOC105275398 | 2825 | 76 | 269.03 | 1 | 3.54 | 1.32 | 20.33 | 21.34 | -1.01 | 1.01 | -0.03 | 0.03 |
| gene12406 | https://www.ncbi.nlm.nih.gov/gene/?term=LOC105286316 | 1386 | 30 | 216.45 | 1 | 7.22 | 3.33 | 39.2 | 40.74 | -1.55 | 1.55 | -0.03 | 0.03 |
| gene1884 | https://www.ncbi.nlm.nih.gov/gene/?term=LOC105288072 | 4169 | 80 | 191.89 | 1 | 2.4 | 1.25 | 22.74 | 24.44 | -1.7 | 1.7 | -0.03 | 0.03 |
| gene2676 | https://www.ncbi.nlm.nih.gov/gene/?term=LOC105275181 | 1353 | 14 | 103.47 | 1 | 7.39 | 7.14 | 65.95 | 68.91 | -2.96 | 2.96 | -0.03 | 0.03 |
| gene2970 | https://www.ncbi.nlm.nih.gov/gene/?term=LOC105275518 | 2340 | 20 | 85.47 | 1 | 4.27 | 5 | 47.36 | 50.74 | -3.38 | 3.38 | -0.03 | 0.03 |
| gene652 | https://www.ncbi.nlm.nih.gov/gene/?term=LOC105279528 | 5806 | 42 | 72.34 | 1 | 1.72 | 2.38 | 16.48 | 14.36 | 2.12 | 2.12 | 0.02 | 0.02 |
| gene1199 | https://www.ncbi.nlm.nih.gov/gene/?term=LOC105285498 | 2220 | 26 | 117.12 | 1 | 4.5 | 3.85 | 33.11 | 31.11 | 2 | 2 | 0.02 | 0.02 |
| gene388 | https://www.ncbi.nlm.nih.gov/gene/?term=LOC105276538 | 1558 | 19 | 121.95 | 1 | 6.42 | 5.26 | 69.93 | 68.17 | 1.77 | 1.77 | 0.02 | 0.02 |

| | | | | | | | | | | | | | |
|---|---|---|---|---|---|---|---|---|---|---|---|---|---|
| gene322 | https://www.ncbi.nlm.nih.gov/gene/?term=LOC105275777 | 1606 | 22 | 136.99 | 1 | 6.23 | 4.55 | 47.62 | 46.19 | 1.43 | 1.43 | 0.02 | 0.02 |
| gene7247 | https://www.ncbi.nlm.nih.gov/gene/?term=LOC105280407 | 10486 | 205 | 195.5 1 | 0.95 | 0.49 | 17 | 15.99 | 1.01 | 1.01 | 0.02 | 0.02 | |
| gene4376 | https://www.ncbi.nlm.nih.gov/gene/?term=LOC105277083 | 4307 | 92 | 213.61 | 1 | 2.32 | 1.09 | 26.68 | 25.69 | 0.99 | 0.99 | 0.02 | 0.02 |
| gene10392 | https://www.ncbi.nlm.nih.gov/gene/?term=LOC105284036 | 3216 | 84 | 261.19 | 2 | 6.22 | 2.38 | 23.38 | 22.62 | 0.76 | 0.76 | 0.02 | 0.02 |
| gene10844 | https://www.ncbi.nlm.nih.gov/gene/?term=LOC105284535 | 9086 | 220 | 242.13 | 1 | 1.1 | 0.45 | 16.85 | 16.12 | 0.73 | 0.73 | 0.02 | 0.02 |
| gene6966 | https://www.ncbi.nlm.nih.gov/gene/?term=LOC105280104 | 3747 | 124 | 330.93 | 1 | 2.67 | 0.81 | 15.19 | 14.51 | 0.68 | 0.68 | 0.02 | 0.02 |
| gene11454 | https://www.ncbi.nlm.nih.gov/gene/?term=LOC105285256 | 1906 | 52 | 272.82 | 1 | 5.25 | 1.92 | 42.98 | 42.36 | 0.62 | 0.62 | 0.02 | 0.02 |
| gene7568 | https://www.ncbi.nlm.nih.gov/gene/?term=LOC105280786 | 2525 | 121 | 479.21 | 1 | 3.96 | 0.83 | 29.4 | 28.89 | 0.52 | 0.52 | 0.02 | 0.02 |
| gene13036 | https://www.ncbi.nlm.nih.gov/gene/?term=LOC105287059 | 4959 | 166 | 334.74 | 1 | 2.02 | 0.6 | 17.36 | 16.86 | 0.5 | 0.5 | 0.02 | 0.02 |
| gene474 | https://www.ncbi.nlm.nih.gov/gene/?term=LOC105277435 | 3625 | 155 | 427.59 | 1 | 2.76 | 0.65 | 12.2 | 11.71 | 0.49 | 0.49 | 0.02 | 0.02 |
| gene10549 | https://www.ncbi.nlm.nih.gov/gene/?term=LOC105284210 | 7145 | 319 | 446.47 | 1 | 1.4 | 0.31 | 10.59 | 10.18 | 0.4 | 0.4 | 0.02 | 0.02 |
| gene6286 | https://www.ncbi.nlm.nih.gov/gene/?term=LOC105279298 | 2861 | 134 | 468.37 | 1 | 3.5 | 0.75 | 23.06 | 22.69 | 0.37 | 0.37 | 0.02 | 0.02 |
| gene3856 | https://www.ncbi.nlm.nih.gov/gene/?term=LOC105276499 | 5236 | 197 | 376.24 | 1 | 1.91 | 0.51 | 9.94 | 10.34 | -0.41 | 0.41 | -0.02 | 0.02 |
| gene6583 | https://www.ncbi.nlm.nih.gov/gene/?term=LOC105279649 | 7444 | 317 | 425.85 | 1 | 1.34 | 0.32 | 11.9 | 12.34 | -0.44 | 0.44 | -0.02 | 0.02 |
| gene11275 | https://www.ncbi.nlm.nih.gov/gene/?term=LOC105285055 | 4119 | 213 | 517.12 | 1 | 2.43 | 0.47 | 13.26 | 13.73 | -0.47 | 0.47 | -0.02 | 0.02 |
| gene10678 | https://www.ncbi.nlm.nih.gov/gene/?term=LOC105284355 | 4031 | 170 | 421.73 | 1 | 2.48 | 0.59 | 18.52 | 19.03 | -0.5 | 0.5 | -0.02 | 0.02 |
| gene9336 | https://www.ncbi.nlm.nih.gov/gene/?term=LOC105282834 | 5993 | 115 | 191.89 | 1 | 1.67 | 0.87 | 14.71 | 15.58 | -0.87 | 0.87 | -0.02 | 0.02 |
| gene8343 | https://www.ncbi.nlm.nih.gov/gene/?term=LOC105281700 | 2172 | 44 | 202.58 | 1 | 4.6 | 2.27 | 37.43 | 38.56 | -1.13 | 1.13 | -0.02 | 0.02 |
| gene5101 | https://www.ncbi.nlm.nih.gov/gene/?term=LOC105277903 | 1813 | 34 | 187.53 | 1 | 5.52 | 2.94 | 52.57 | 53.85 | -1.27 | 1.27 | -0.02 | 0.02 |
| gene5402 | https://www.ncbi.nlm.nih.gov/gene/?term=LOC105278260 | 3068 | 42 | 136.9 1 | 3.26 | 2.38 | 12.8 | 14.3 | -1.5 | 1.5 | -0.02 | 0.02 | |
| gene7248 | https://www.ncbi.nlm.nih.gov/gene/?term=LOC105280408 | 5175 | 44 | 85.02 1 | 1.93 | 2.27 | 21.35 | 20.08 | 1.27 | 1.27 | 0.01 | 0.01 | |
| gene657 | https://www.ncbi.nlm.nih.gov/gene/?term=LOC105279667 | 6421 | 78 | 121.48 | 1 | 1.56 | 1.28 | 4.46 | 3.72 | 0.74 | 0.74 | 0.01 | 0.01 |
| gene12203 | https://www.ncbi.nlm.nih.gov/gene/?term=LOC105286099 | 5967 | 78 | 130.72 | 1 | 1.68 | 1.28 | 11.28 | 10.61 | 0.67 | 0.67 | 0.01 | 0.01 |
| gene1147 | https://www.ncbi.nlm.nih.gov/gene/?term=LOC105284954 | 4022 | 87 | 216.31 | 1 | 2.49 | 1.15 | 15.12 | 14.51 | 0.61 | 0.61 | 0.01 | 0.01 |
| gene12021 | https://www.ncbi.nlm.nih.gov/gene/?term=LOC105285905 | 1399 | 21 | 150.11 | 1 | 7.15 | 4.76 | 44.76 | 44.21 | 0.55 | 0.55 | 0.01 | 0.01 |
| gene2683 | https://www.ncbi.nlm.nih.gov/gene/?term=LOC105275187 | 5492 | 75 | 136.56 | 1 | 1.82 | 1.33 | 14.15 | 13.62 | 0.53 | 0.53 | 0.01 | 0.01 |
| gene11583 | https://www.ncbi.nlm.nih.gov/gene/?term=LOC105285414 | 8192 | 179 | 218.51 | 1 | 1.22 | 0.56 | 9.12 | 8.6 | 0.52 | 0.52 | 0.01 | 0.01 |
| gene2748 | https://www.ncbi.nlm.nih.gov/gene/?term=LOC105275261 | 3845 | 54 | 140.44 | 1 | 2.6 | 1.85 | 38.38 | 37.86 | 0.52 | 0.52 | 0.01 | 0.01 |
| gene7801 | https://www.ncbi.nlm.nih.gov/gene/?term=LOC105281047 | 5137 | 113 | 219.97 | 1 | 1.95 | 0.88 | 10.88 | 10.38 | 0.5 | 0.5 | 0.01 | 0.01 |
| gene5524 | https://www.ncbi.nlm.nih.gov/gene/?term=LOC105278397 | 5177 | 201 | 388.26 | 1 | 1.93 | 0.5 | 24.26 | 23.91 | 0.35 | 0.35 | 0.01 | 0.01 |
| gene927 | https://www.ncbi.nlm.nih.gov/gene/?term=LOC105282600 | 3727 | 57 | 152.94 | 1 | 2.68 | 1.75 | 29.17 | 28.82 | 0.35 | 0.35 | 0.01 | 0.01 |
| gene12573 | https://www.ncbi.nlm.nih.gov/gene/?term=LOC105286515 | 3647 | 102 | 279.68 | 1 | 2.74 | 0.98 | 10.87 | 10.55 | 0.32 | 0.32 | 0.01 | 0.01 |
| gene5746 | https://www.ncbi.nlm.nih.gov/gene/?term=LOC105278666 | 10343 | 405 | 391.57 | 1 | 0.97 | 0.25 | 7.03 | 6.73 | 0.3 | 0.3 | 0.01 | 0.01 |
| gene4213 | https://www.ncbi.nlm.nih.gov/gene/?term=LOC105276901 | 3257 | 95 | 291.68 | 1 | 3.07 | 1.05 | 23.09 | 22.87 | 0.22 | 0.22 | 0.01 | 0.01 |
| gene5831 | https://www.ncbi.nlm.nih.gov/gene/?term=LOC105278761 | 2506 | 103 | 411.01 | 2 | 7.98 | 1.94 | 24.82 | 24.61 | 0.21 | 0.21 | 0.01 | 0.01 |
| gene8274 | https://www.ncbi.nlm.nih.gov/gene/?term=LOC105281624 | 5425 | 203 | 374.19 | 1 | 1.84 | 0.49 | 10.66 | 10.47 | 0.2 | 0.2 | 0.01 | 0.01 |
| gene3206 | https://www.ncbi.nlm.nih.gov/gene/?term=LOC105275752 | 3474 | 123 | 354.06 | 1 | 2.88 | 0.81 | 28.92 | 28.73 | 0.19 | 0.19 | 0.01 | 0.01 |
| gene10596 | https://www.ncbi.nlm.nih.gov/gene/?term=LOC105284262 | 6520 | 228 | 349.69 | 1 | 1.53 | 0.44 | 3.34 | 3.5 | -0.16 | 0.16 | -0.01 | 0.01 |
| gene10540 | https://www.ncbi.nlm.nih.gov/gene/?term=LOC105284200 | 5267 | 193 | 366.43 | 1 | 1.9 | 0.52 | 13.99 | 14.21 | -0.22 | 0.22 | -0.01 | 0.01 |
| gene303 | https://www.ncbi.nlm.nih.gov/gene/?term=LOC105275584 | 3081 | 86 | 279.13 | 1 | 3.25 | 1.16 | 23.78 | 24 | -0.22 | 0.22 | -0.01 | 0.01 |
| gene5184 | https://www.ncbi.nlm.nih.gov/gene/?term=LOC105277999 | 5528 | 147 | 265.92 | 1 | 1.81 | 0.68 | 12.4 | 12.64 | -0.24 | 0.24 | -0.01 | 0.01 |
| gene7888 | https://www.ncbi.nlm.nih.gov/gene/?term=LOC105281141 | 4357 | 155 | 355.75 | 1 | 2.3 | 0.65 | 29.83 | 30.06 | -0.24 | 0.24 | -0.01 | 0.01 |
| gene7166 | https://www.ncbi.nlm.nih.gov/gene/?term=LOC105280326 | 6117 | 208 | 340.04 | 1 | 1.63 | 0.48 | 15.97 | 16.24 | -0.27 | 0.27 | -0.01 | 0.01 |
| gene4023 | https://www.ncbi.nlm.nih.gov/gene/?term=LOC105276692 | 1759 | 38 | 216.03 | 1 | 5.69 | 2.63 | 22.43 | 22.73 | -0.3 | 0.3 | -0.01 | 0.01 |
| gene2443 | https://www.ncbi.nlm.nih.gov/gene/?term=LOC105274895 | 10433 | 345 | 330.68 | 1 | 0.96 | 0.29 | 6.76 | 7.09 | -0.33 | 0.33 | -0.01 | 0.01 |
| gene10520 | https://www.ncbi.nlm.nih.gov/gene/?term=LOC105284184 | 2451 | 82 | 334.56 | 1 | 4.08 | 1.22 | 34.55 | 34.9 | -0.35 | 0.35 | -0.01 | 0.01 |
| gene2404 | https://www.ncbi.nlm.nih.gov/gene/?term=LOC105274853 | 1767 | 37 | 209.39 | 1 | 5.66 | 2.7 | 31.99 | 32.37 | -0.37 | 0.37 | -0.01 | 0.01 |
| gene10531 | https://www.ncbi.nlm.nih.gov/gene/?term=LOC105284190 | 6881 | 192 | 279.03 | 1 | 1.45 | 0.52 | 9.5 | 9.91 | -0.4 | 0.4 | -0.01 | 0.01 |
| gene4892 | https://www.ncbi.nlm.nih.gov/gene/?term=LOC105277667 | 3898 | 120 | 307.85 | 1 | 2.57 | 0.83 | 28.07 | 28.49 | -0.42 | 0.42 | -0.01 | 0.01 |
| gene8600 | https://www.ncbi.nlm.nih.gov/gene/?term=LOC105282001 | 8513 | 153 | 179.73 | 1 | 1.17 | 0.65 | 9.37 | 9.87 | -0.5 | 0.5 | -0.01 | 0.01 |
| gene274 | https://www.ncbi.nlm.nih.gov/gene/?term=LOC105275190 | 3488 | 70 | 200.69 | 1 | 2.87 | 1.43 | 19.35 | 20.02 | -0.67 | 0.67 | -0.01 | 0.01 |
| gene10107 | https://www.ncbi.nlm.nih.gov/gene/?term=LOC105283712 | 1883 | 28 | 148.7 1 | 5.31 | 3.57 | 28.01 | 28.9 | -0.88 | 0.88 | -0.01 | 0.01 | |
| gene7183 | https://www.ncbi.nlm.nih.gov/gene/?term=LOC105280348 | 3925 | 52 | 132.48 | 2 | 5.1 | 3.85 | 25.66 | 26.67 | -1.01 | 1.01 | -0.01 | 0.01 |
| gene10345 | https://www.ncbi.nlm.nih.gov/gene/?term=LOC105283989 | 4444 | 29 | 65.26 1 | 2.25 | 3.45 | 48.19 | 50.2 | -2.01 | 2.01 | -0.01 | 0.01 | |
| gene564 | https://www.ncbi.nlm.nih.gov/gene/?term=LOC105278478 | 2802 | 13 | 46.4 1 | 3.57 | 7.69 | 25.47 | 24.48 | 0.99 | 0.99 | 0 | 0 | |
| gene10155 | https://www.ncbi.nlm.nih.gov/gene/?term=LOC105283756 | 7030 | 97 | 137.98 | 1 | 1.42 | 1.03 | 6.83 | 6.6 | 0.23 | 0.23 | 0 | 0 |
| gene6620 | https://www.ncbi.nlm.nih.gov/gene/?term=LOC105279687 | 17998 | 363 | 201.69 | 1 | 0.56 | 0.28 | 5.06 | 4.84 | 0.22 | 0.22 | 0 | 0 |
| gene10600 | https://www.ncbi.nlm.nih.gov/gene/?term=LOC105284270 | 4837 | 103 | 212.94 | 1 | 2.07 | 0.97 | 12.92 | 12.79 | 0.13 | 0.13 | 0 | 0 |
| gene5646 | https://www.ncbi.nlm.nih.gov/gene/?term=LOC105278534 | 7475 | 185 | 247.49 | 1 | 1.34 | 0.54 | 17.64 | 17.56 | 0.08 | 0.08 | 0 | 0 |
| gene5967 | https://www.ncbi.nlm.nih.gov/gene/?term=LOC105278918 | 4393 | 132 | 300.48 | 1 | 2.28 | 0.76 | 15.7 | 15.62 | 0.08 | 0.08 | 0 | 0 |
| gene12802 | https://www.ncbi.nlm.nih.gov/gene/?term=LOC105286799 | 3183 | 170 | 534.09 | 1 | 3.14 | 0.59 | 15.91 | 15.85 | 0.06 | 0.06 | 0 | 0 |
| gene3470 | https://www.ncbi.nlm.nih.gov/gene/?term=LOC105276059 | 8716 | 159 | 182.42 | 1 | 1.15 | 0.63 | 9.24 | 9.2 | 0.04 | 0.04 | 0 | 0 |
| gene2806 | https://www.ncbi.nlm.nih.gov/gene/?term=LOC105275327 | 14257 | 479 | 335.98 | 1 | 0.7 | 0.21 | 7.13 | 7.12 | 0.01 | 0.01 | 0 | 0 |
| gene5375 | https://www.ncbi.nlm.nih.gov/gene/?term=LOC105278228 | 8640 | 161 | 186.34 | 1 | 1.16 | 0.62 | 23.31 | 23.3 | 0.01 | 0.01 | 0 | 0 |

| | | | | | | | | | | | | | | |
|---|---|---|---|---|---|---|---|---|---|---|---|---|---|---|
| gene2056 | https://www.ncbi.nlm.nih.gov/gene/?term=LOC105274452 | 6584 | 206 | 312.88 | | 1 | 1.52 | 0.49 | 14.18 | 14.18 | 0 | 0 | 0 | 0 |
| gene11579 | https://www.ncbi.nlm.nih.gov/gene/?term=LOC105285406 | 13571 | 282 | 207.8 | 1 | 0.74 | 0.35 | 5.68 | 5.69 | -0.01 | 0.01 | 0 | 0 | |
| gene3107 | https://www.ncbi.nlm.nih.gov/gene/?term=LOC105275650 | 5103 | 150 | 293.94 | | 1 | 1.96 | 0.67 | 17.09 | 17.11 | -0.02 | 0.02 | 0 | 0 |
| gene2173 | https://www.ncbi.nlm.nih.gov/gene/?term=LOC105274590 | 4387 | 79 | 180.08 | | 1 | 2.28 | 1.27 | 22.08 | 22.11 | -0.03 | 0.03 | 0 | 0 |
| gene13114 | https://www.ncbi.nlm.nih.gov/gene/?term=LOC105287147 | 7609 | 381 | 500.72 | | 1 | 1.31 | 0.26 | 9.67 | 9.7 | -0.04 | 0.04 | 0 | 0 |
| gene2668 | https://www.ncbi.nlm.nih.gov/gene/?term=LOC105275170 | 12496 | 336 | 268.89 | | 1 | 0.8 | 0.3 | 2.05 | 2.12 | -0.07 | 0.07 | 0 | 0 |
| gene6698 | https://www.ncbi.nlm.nih.gov/gene/?term=LOC105279778 | 4099 | 138 | 336.67 | | 1 | 2.44 | 0.72 | 9.41 | 9.48 | -0.07 | 0.07 | 0 | 0 |
| gene53 | https://www.ncbi.nlm.nih.gov/gene/?term=LOC105277148 | 19283 | 270 | 140.02 | | 1 | 0.52 | 0.37 | 3.92 | 4.02 | -0.1 | 0.1 | 0 | 0 |
| gene11568 | https://www.ncbi.nlm.nih.gov/gene/?term=LOC105285395 | 8086 | 176 | 217.66 | | 1 | 1.24 | 0.57 | 15.78 | 15.89 | -0.11 | 0.11 | 0 | 0 |
| gene7398 | https://www.ncbi.nlm.nih.gov/gene/?term=LOC105280589 | 3452 | 41 | 118.77 | | 1 | 2.9 | 2.44 | 25.07 | 25.33 | -0.26 | 0.26 | 0 | 0 |
| gene6528 | https://www.ncbi.nlm.nih.gov/gene/?term=LOC105279591 | 7222 | 67 | 92.77 | 1 | 1.38 | 1.49 | 23.03 | 23.32 | -0.28 | 0.28 | 0 | 0 | |
| gene6455 | https://www.ncbi.nlm.nih.gov/gene/?term=LOC105279500 | 2612 | 32 | 122.51 | | 1 | 3.83 | 3.12 | 49.77 | 50.07 | -0.3 | 0.3 | 0 | 0 |
| gene9411 | https://www.ncbi.nlm.nih.gov/gene/?term=LOC105282914 | 5534 | 41 | 74.09 | 1 | 1.81 | 2.44 | 49.68 | 50.18 | -0.49 | 0.49 | 0 | 0 | |
| gene8178 | https://www.ncbi.nlm.nih.gov/gene/?term=LOC105281515 | 8647 | 44 | 50.88 | 1 | 1.16 | 2.27 | 29.36 | 29.86 | -0.5 | 0.5 | 0 | 0 | |
| gene6639 | https://www.ncbi.nlm.nih.gov/gene/?term=LOC105279705 | 7656 | 68 | 88.82 | 1 | 1.31 | 1.47 | 22.19 | 22.7 | -0.51 | 0.51 | 0 | 0 | |

Directory: OGL File: rnk-dmg-Cb_LI_BCphaseA1.vs.BCphaseB2.txt

| gene_ID | gene_link | gwidth | #Sites | #per10Kb | #dmSites | #dmsp10kb | %dmSites | %pSite1 | %pSite2 | DMpSite | ADMpSite | DMpNucl | ADMpNucl |
|---|---|---|---|---|---|---|---|---|---|---|---|---|---|
| gene10219 | https://www.ncbi.nlm.nih.gov/gene/?term=LOC105283830 | 235 | 12 | 510.64 | 1 | 42.55 | 8.33 | 42.96 | 33.59 | 9.36 | 9.36 | 0.48 | 0.48 |
| gene5917 | https://www.ncbi.nlm.nih.gov/gene/?term=LOC105278849 | 823 | 81 | 984.2 | 2 | 24.3 | 2.47 | 24.78 | 23.07 | 1.71 | 1.71 | 0.17 | 0.17 |
| gene5809 | https://www.ncbi.nlm.nih.gov/gene/?term=LOC105278737 | 1159 | 83 | 716.13 | 1 | 8.63 | 1.2 | 16.21 | 18.32 | -2.11 | 2.11 | -0.15 | 0.15 |
| gene558 | https://www.ncbi.nlm.nih.gov/gene/?term=LOC105278458 | 216 | 3 | 138.89 | 1 | 46.3 | 33.33 | 20.87 | 10.44 | 10.43 | 10.43 | 0.14 | 0.14 |
| gene8965 | https://www.ncbi.nlm.nih.gov/gene/?term=LOC105282405 | 1310 | 31 | 236.64 | 3 | 22.9 | 9.68 | 51.28 | 45.18 | 6.1 | 6.1 | 0.14 | 0.14 |
| gene1696 | https://www.ncbi.nlm.nih.gov/gene/?term=LOC105287856 | 1917 | 118 | 615.55 | 2 | 10.43 | 1.69 | 12.7 | 14.93 | -2.23 | 2.23 | -0.14 | 0.14 |
| gene10509 | https://www.ncbi.nlm.nih.gov/gene/?term=LOC105284168 | 1350 | 71 | 525.93 | 1 | 7.41 | 1.41 | 22.24 | 24.86 | -2.62 | 2.62 | -0.14 | 0.14 |
| gene1534 | https://www.ncbi.nlm.nih.gov/gene/?term=LOC105287662 | 1711 | 32 | 187.03 | 3 | 17.53 | 9.38 | 32.94 | 39.9 | -6.95 | 6.95 | -0.13 | 0.13 |
| gene9471 | https://www.ncbi.nlm.nih.gov/gene/?term=LOC105282986 | 1605 | 79 | 492.21 | 3 | 18.69 | 3.8 | 31.65 | 29.21 | 2.44 | 2.44 | 0.12 | 0.12 |
| gene4013 | https://www.ncbi.nlm.nih.gov/gene/?term=LOC105276679 | 2007 | 65 | 323.87 | 3 | 14.95 | 4.62 | 39.73 | 36.29 | 3.44 | 3.44 | 0.11 | 0.11 |
| gene952 | https://www.ncbi.nlm.nih.gov/gene/?term=LOC105283016 | 2375 | 111 | 467.37 | 3 | 12.63 | 2.7 | 23.65 | 21.4 | 2.25 | 2.25 | 0.11 | 0.11 |
| gene12842 | https://www.ncbi.nlm.nih.gov/gene/?term=LOC105286845 | 1157 | 60 | 518.58 | 3 | 25.93 | 5 | 17.37 | 19.47 | -2.1 | 2.1 | -0.11 | 0.11 |
| gene12374 | https://www.ncbi.nlm.nih.gov/gene/?term=LOC105286277 | 852 | 27 | 316.9 | 1 | 11.74 | 3.7 | 37.34 | 40.66 | -3.32 | 3.32 | -0.11 | 0.11 |
| gene168 | https://www.ncbi.nlm.nih.gov/gene/?term=LOC105287895 | 2303 | 50 | 217.11 | 1 | 4.34 | 2 | 28.58 | 33.64 | -5.06 | 5.06 | -0.11 | 0.11 |
| gene10721 | https://www.ncbi.nlm.nih.gov/gene/?term=LOC105284403 | 1537 | 24 | 156.15 | 3 | 19.52 | 12.5 | 43.65 | 50.5 | -6.85 | 6.85 | -0.11 | 0.11 |
| gene2824 | https://www.ncbi.nlm.nih.gov/gene/?term=LOC105275337 | 1772 | 57 | 321.67 | 5 | 28.22 | 8.77 | 32.13 | 29.09 | 3.04 | 3.04 | 0.1 | 0.1 |
| gene1802 | https://www.ncbi.nlm.nih.gov/gene/?term=LOC105287981 | 3173 | 135 | 425.46 | 2 | 6.3 | 1.48 | 18.91 | 16.6 | 2.31 | 2.31 | 0.1 | 0.1 |
| gene7568 | https://www.ncbi.nlm.nih.gov/gene/?term=LOC105280786 | 2525 | 110 | 435.64 | 3 | 11.88 | 2.73 | 32.15 | 29.95 | 2.21 | 2.21 | 0.1 | 0.1 |
| gene7401 | https://www.ncbi.nlm.nih.gov/gene/?term=LOC105280591 | 2018 | 91 | 450.94 | 1 | 4.96 | 1.1 | 14.79 | 12.63 | 2.16 | 2.16 | 0.1 | 0.1 |
| gene9781 | https://www.ncbi.nlm.nih.gov/gene/?term=LOC105283351 | 1245 | 60 | 481.93 | 1 | 8.03 | 1.67 | 24.95 | 22.93 | 2.02 | 2.02 | 0.1 | 0.1 |
| gene6057 | https://www.ncbi.nlm.nih.gov/gene/?term=LOC105279035 | 2105 | 146 | 693.59 | 6 | 28.5 | 4.11 | 19.02 | 17.55 | 1.48 | 1.48 | 0.1 | 0.1 |
| gene570 | https://www.ncbi.nlm.nih.gov/gene/?term=LOC105278549 | 1673 | 65 | 388.52 | 3 | 17.93 | 4.62 | 41.78 | 44.3 | -2.52 | 2.52 | -0.1 | 0.1 |
| gene4643 | https://www.ncbi.nlm.nih.gov/gene/?term=LOC109610847 | 429 | 5 | 116.55 | 1 | 23.31 | 20 | 29.86 | 22.36 | 7.5 | 7.5 | 0.09 | 0.09 |
| gene1914 | https://www.ncbi.nlm.nih.gov/gene/?term=LOC105288105 | 1289 | 23 | 178.43 | 2 | 15.52 | 8.7 | 60.89 | 56.04 | 4.85 | 4.85 | 0.09 | 0.09 |

...

... only first 25 lines shown ...

Directory: OGL File: rnk-dmg-Cb_LI_BCphaseA1.vs.RphaseA1.txt

| gene_ID | gene_link | gwidth | #Sites | #per10Kb | #dmSites | #dmsp10kb | %dmSites | %pSite1 | %pSite2 | DMpSite | ADMpSite | DMpNucl | ADMpNucl |
|---|---|---|---|---|---|---|---|---|---|---|---|---|---|
| gene3165 | https://www.ncbi.nlm.nih.gov/gene/?term=LOC105275706 | 2374 | 59 | 248.53 | 4 | 16.85 | 6.78 | 23.46 | 34.48 | -11.02 | 11.02 | -0.27 | 0.27 |
| gene7199 | https://www.ncbi.nlm.nih.gov/gene/?term=LOC105280366 | 1414 | 26 | 183.88 | 1 | 7.07 | 3.85 | 47.43 | 38.76 | 8.67 | 8.67 | 0.16 | 0.16 |
| gene2593 | https://www.ncbi.nlm.nih.gov/gene/?term=LOC105275070 | 1774 | 6 | 33.82 | 2 | 11.27 | 33.33 | 16.42 | 60.02 | -43.6 | 43.6 | -0.15 | 0.15 |
| gene11819 | https://www.ncbi.nlm.nih.gov/gene/?term=LOC105285671 | 2366 | 90 | 380.39 | 1 | 4.23 | 1.11 | 28.49 | 30.76 | -2.27 | 2.27 | -0.09 | 0.09 |
| gene298 | https://www.ncbi.nlm.nih.gov/gene/?term=LOC105275460 | 2473 | 115 | 465.02 | 1 | 4.04 | 0.87 | 12.68 | 14.4 | -1.73 | 1.73 | -0.08 | 0.08 |
| gene5877 | https://www.ncbi.nlm.nih.gov/gene/?term=LOC105278807 | 1719 | 51 | 296.68 | 1 | 5.82 | 1.96 | 21.68 | 24.24 | -2.55 | 2.55 | -0.08 | 0.08 |
| gene5101 | https://www.ncbi.nlm.nih.gov/gene/?term=LOC105277903 | 1813 | 34 | 187.53 | 1 | 5.52 | 2.94 | 52.57 | 56.73 | -4.16 | 4.16 | -0.08 | 0.08 |
| gene10714 | https://www.ncbi.nlm.nih.gov/gene/?term=LOC105284396 | 1966 | 53 | 269.58 | 1 | 5.09 | 1.89 | 40.95 | 38.4 | 2.55 | 2.55 | 0.07 | 0.07 |
| gene10213 | https://www.ncbi.nlm.nih.gov/gene/?term=LOC105283826 | 1503 | 52 | 345.97 | 1 | 6.65 | 1.92 | 16.76 | 14.79 | 1.97 | 1.97 | 0.07 | 0.07 |
| gene4034 | https://www.ncbi.nlm.nih.gov/gene/?term=LOC105276704 | 2065 | 80 | 387.41 | 1 | 4.84 | 1.25 | 13.38 | 15.07 | -1.69 | 1.69 | -0.07 | 0.07 |
| gene4641 | https://www.ncbi.nlm.nih.gov/gene/?term=LOC105277398 | 1937 | 70 | 361.38 | 1 | 5.16 | 1.43 | 35.36 | 37.39 | -2.03 | 2.03 | -0.07 | 0.07 |
| gene388 | https://www.ncbi.nlm.nih.gov/gene/?term=LOC105276538 | 1558 | 19 | 121.95 | 1 | 6.42 | 5.26 | 69.41 | 64.89 | 4.52 | 4.52 | 0.06 | 0.06 |
| gene7915 | https://www.ncbi.nlm.nih.gov/gene/?term=LOC105281173 | 1597 | 25 | 156.54 | 1 | 6.26 | 4 | 64.87 | 60.99 | 3.88 | 3.88 | 0.06 | 0.06 |
| gene11401 | https://www.ncbi.nlm.nih.gov/gene/?term=LOC105285202 | 1841 | 49 | 266.16 | 1 | 5.43 | 2.04 | 21.54 | 19.38 | 2.16 | 2.16 | 0.06 | 0.06 |
| gene2271 | https://www.ncbi.nlm.nih.gov/gene/?term=LOC105274706 | 1448 | 49 | 338.4 | 1 | 6.91 | 2.04 | 38.87 | 37.01 | 1.86 | 1.86 | 0.06 | 0.06 |
| gene4190 | https://www.ncbi.nlm.nih.gov/gene/?term=LOC105276875 | 3150 | 124 | 393.65 | 1 | 3.17 | 0.81 | 21.46 | 19.89 | 1.58 | 1.58 | 0.06 | 0.06 |
| gene9740 | https://www.ncbi.nlm.nih.gov/gene/?term=LOC105283277 | 2887 | 135 | 467.61 | 1 | 3.46 | 0.74 | 21.88 | 20.52 | 1.37 | 1.37 | 0.06 | 0.06 |
| gene5298 | https://www.ncbi.nlm.nih.gov/gene/?term=LOC105278145 | 4060 | 240 | 591.13 | 2 | 4.93 | 0.83 | 11.5 | 10.44 | 1.06 | 1.06 | 0.06 | 0.06 |
| gene570 | https://www.ncbi.nlm.nih.gov/gene/?term=LOC105278549 | 1673 | 63 | 376.57 | 1 | 5.98 | 1.59 | 41.36 | 42.83 | -1.47 | 1.47 | -0.06 | 0.06 |
| gene6334 | https://www.ncbi.nlm.nih.gov/gene/?term=LOC105279356 | 1471 | 43 | 292.32 | 1 | 6.8 | 2.33 | 28.25 | 30.45 | -2.2 | 2.2 | -0.06 | 0.06 |
| gene12516 | https://www.ncbi.nlm.nih.gov/gene/?term=LOC105286452 | 1529 | 35 | 228.91 | 1 | 6.54 | 2.86 | 51.01 | 53.56 | -2.56 | 2.56 | -0.06 | 0.06 |
| gene393 | https://www.ncbi.nlm.nih.gov/gene/?term=LOC105276566 | 1123 | 2 | 17.81 | 1 | 8.9 | 50 | 77.65 | 50 | 27.65 | 27.65 | 0.05 | 0.05 |
| gene12517 | https://www.ncbi.nlm.nih.gov/gene/?term=LOC105286451 | 1626 | 10 | 61.5 | 1 | 6.15 | 10 | 52.19 | 44.01 | 8.18 | 8.18 | 0.05 | 0.05 |
| gene920 | https://www.ncbi.nlm.nih.gov/gene/?term=LOC105282518 | 1197 | 42 | 350.88 | 1 | 8.35 | 2.38 | 38.27 | 36.85 | 1.43 | 1.43 | 0.05 | 0.05 |
| gene5649 | https://www.ncbi.nlm.nih.gov/gene/?term=LOC105278538 | 2140 | 76 | 355.14 | 1 | 4.67 | 1.32 | 18.24 | 16.87 | 1.37 | 1.37 | 0.05 | 0.05 |
| gene8873 | https://www.ncbi.nlm.nih.gov/gene/?term=LOC105282297 | 2055 | 78 | 379.56 | 1 | 4.87 | 1.28 | 33.62 | 34.98 | -1.37 | 1.37 | -0.05 | 0.05 |
| gene5647 | https://www.ncbi.nlm.nih.gov/gene/?term=LOC105278567 | 3118 | 32 | 102.63 | 1 | 3.21 | 3.12 | 13.07 | 18.25 | -5.18 | 5.18 | -0.05 | 0.05 |
| gene3168 | https://www.ncbi.nlm.nih.gov/gene/?term=LOC105275711 | 6150 | 77 | 125.2 | 1 | 1.63 | 1.3 | 14.51 | 10.99 | 3.51 | 3.51 | 0.04 | 0.04 |
| gene6422 | https://www.ncbi.nlm.nih.gov/gene/?term=LOC105279458 | 2419 | 37 | 152.96 | 1 | 4.13 | 2.7 | 54.1 | 51.26 | 2.83 | 2.83 | 0.04 | 0.04 |
| gene322 | https://www.ncbi.nlm.nih.gov/gene/?term=LOC105275777 | 1606 | 23 | 143.21 | 1 | 6.23 | 4.35 | 45.55 | 43.01 | 2.54 | 2.54 | 0.04 | 0.04 |
| gene11454 | https://www.ncbi.nlm.nih.gov/gene/?term=LOC105285256 | 1906 | 52 | 272.82 | 1 | 5.25 | 1.92 | 42.98 | 41.47 | 1.51 | 1.51 | 0.04 | 0.04 |
| gene10326 | https://www.ncbi.nlm.nih.gov/gene/?term=LOC105283962 | 3042 | 94 | 309.01 | 1 | 3.29 | 1.06 | 12.22 | 10.91 | 1.31 | 1.31 | 0.04 | 0.04 |
| gene6031 | https://www.ncbi.nlm.nih.gov/gene/?term=LOC105279000 | 5691 | 175 | 307.5 | 1 | 1.76 | 0.57 | 14.52 | 13.23 | 1.29 | 1.29 | 0.04 | 0.04 |
| gene7568 | https://www.ncbi.nlm.nih.gov/gene/?term=LOC105280786 | 2525 | 116 | 459.41 | 1 | 3.96 | 0.86 | 28.86 | 28.06 | 0.8 | 0.8 | 0.04 | 0.04 |
| gene2844 | https://www.ncbi.nlm.nih.gov/gene/?term=LOC105275363 | 4330 | 185 | 427.25 | 1 | 2.31 | 0.54 | 14.56 | 15.46 | -0.9 | 0.9 | -0.04 | 0.04 |
| gene1855 | https://www.ncbi.nlm.nih.gov/gene/?term=LOC105288036 | 2387 | 86 | 360.28 | 1 | 4.19 | 1.16 | 28.48 | 29.73 | -1.24 | 1.24 | -0.04 | 0.04 |
| gene12021 | https://www.ncbi.nlm.nih.gov/gene/?term=LOC105285905 | 1399 | 23 | 164.4 | 1 | 7.15 | 4.35 | 41.24 | 39.35 | 1.89 | 1.89 | 0.03 | 0.03 |
| gene2752 | https://www.ncbi.nlm.nih.gov/gene/?term=LOC105275266 | 2111 | 49 | 232.12 | 1 | 4.74 | 2.04 | 23.87 | 22.43 | 1.44 | 1.44 | 0.03 | 0.03 |
| gene10844 | https://www.ncbi.nlm.nih.gov/gene/?term=LOC105284535 | 9086 | 218 | 239.93 | 1 | 1.1 | 0.46 | 16.62 | 15.4 | 1.22 | 1.22 | 0.03 | 0.03 |
| gene4376 | https://www.ncbi.nlm.nih.gov/gene/?term=LOC105277083 | 4307 | 90 | 208.96 | 1 | 2.32 | 1.11 | 26.1 | 24.89 | 1.21 | 1.21 | 0.03 | 0.03 |
| gene4962 | https://www.ncbi.nlm.nih.gov/gene/?term=LOC105277761 | 4645 | 147 | 316.47 | 1 | 2.15 | 0.68 | 13.9 | 13.02 | 0.88 | 0.88 | 0.03 | 0.03 |
| gene6966 | https://www.ncbi.nlm.nih.gov/gene/?term=LOC105280104 | 3747 | 119 | 317.59 | 1 | 2.67 | 0.84 | 15.37 | 14.52 | 0.85 | 0.85 | 0.03 | 0.03 |
| gene7550 | https://www.ncbi.nlm.nih.gov/gene/?term=LOC105280763 | 2173 | 93 | 427.98 | 1 | 4.6 | 1.08 | 27 | 26.23 | 0.77 | 0.77 | 0.03 | 0.03 |
| gene7249 | https://www.ncbi.nlm.nih.gov/gene/?term=LOC105280422 | 1775 | 61 | 343.66 | 1 | 5.63 | 1.64 | 15.16 | 14.42 | 0.75 | 0.75 | 0.03 | 0.03 |
| gene10606 | https://www.ncbi.nlm.nih.gov/gene/?term=LOC105284273 | 6119 | 243 | 397.12 | 2 | 3.27 | 0.82 | 20.74 | 20.03 | 0.71 | 0.71 | 0.03 | 0.03 |
| gene6286 | https://www.ncbi.nlm.nih.gov/gene/?term=LOC105279298 | 2861 | 136 | 475.36 | 1 | 3.5 | 0.74 | 22.78 | 22.14 | 0.64 | 0.64 | 0.03 | 0.03 |
| gene1470 | https://www.ncbi.nlm.nih.gov/gene/?term=LOC105287597 | 2974 | 181 | 608.61 | 1 | 3.36 | 0.55 | 17.88 | 17.42 | 0.47 | 0.47 | 0.03 | 0.03 |
| gene274 | https://www.ncbi.nlm.nih.gov/gene/?term=LOC105275190 | 3488 | 65 | 186.35 | 1 | 2.87 | 1.54 | 21.06 | 22.74 | -1.69 | 1.69 | -0.03 | 0.03 |
| gene1884 | https://www.ncbi.nlm.nih.gov/gene/?term=LOC105288072 | 4169 | 79 | 189.49 | 1 | 2.4 | 1.27 | 22.97 | 24.67 | -1.7 | 1.7 | -0.03 | 0.03 |
| gene2676 | https://www.ncbi.nlm.nih.gov/gene/?term=LOC105275181 | 1353 | 14 | 103.47 | 1 | 7.39 | 7.14 | 65.95 | 69.25 | -3.3 | 3.3 | -0.03 | 0.03 |
| gene7368 | https://www.ncbi.nlm.nih.gov/gene/?term=LOC105280556 | 4105 | 46 | 112.06 | 1 | 2.44 | 2.17 | 13.39 | 11.38 | 2 | 2 | 0.02 | 0.02 |
| gene2748 | https://www.ncbi.nlm.nih.gov/gene/?term=LOC105275261 | 3845 | 55 | 143.04 | 1 | 2.6 | 1.82 | 38.68 | 37.31 | 1.36 | 1.36 | 0.02 | 0.02 |
| gene3599 | https://www.ncbi.nlm.nih.gov/gene/?term=LOC105276196 | 2955 | 55 | 186.13 | 1 | 3.38 | 1.82 | 40.49 | 39.41 | 1.07 | 1.07 | 0.02 | 0.02 |
| gene10392 | https://www.ncbi.nlm.nih.gov/gene/?term=LOC105284036 | 3216 | 83 | 258.08 | 1 | 3.11 | 1.2 | 24.77 | 23.81 | 0.96 | 0.96 | 0.02 | 0.02 |
| gene3107 | https://www.ncbi.nlm.nih.gov/gene/?term=LOC105275650 | 5103 | 145 | 284.15 | 1 | 1.96 | 0.69 | 17.36 | 16.74 | 0.62 | 0.62 | 0.02 | 0.02 |
| gene11340 | https://www.ncbi.nlm.nih.gov/gene/?term=LOC105285131 | 2918 | 97 | 332.42 | 1 | 3.43 | 1.03 | 14.71 | 14.15 | 0.56 | 0.56 | 0.02 | 0.02 |
| gene3206 | https://www.ncbi.nlm.nih.gov/gene/?term=LOC105275752 | 3474 | 126 | 362.69 | 1 | 2.88 | 0.79 | 28.38 | 27.85 | 0.53 | 0.53 | 0.02 | 0.02 |
| gene5746 | https://www.ncbi.nlm.nih.gov/gene/?term=LOC105278666 | 10343 | 393 | 379.97 | 1 | 0.97 | 0.25 | 7.69 | 7.22 | 0.47 | 0.47 | 0.02 | 0.02 |
| gene8729 | https://www.ncbi.nlm.nih.gov/gene/?term=LOC105282137 | 5005 | 200 | 399.6 | 1 | 2 | 0.5 | 15.61 | 15.22 | 0.39 | 0.39 | 0.02 | 0.02 |
| gene13114 | https://www.ncbi.nlm.nih.gov/gene/?term=LOC105287147 | 7609 | 370 | 486.27 | 1 | 1.31 | 0.27 | 9.54 | 9.22 | 0.32 | 0.32 | 0.02 | 0.02 |

| | | | | | | | | | | | | | |
|---|---|---|---|---|---|---|---|---|---|---|---|---|---|
| gene1332 | https://www.ncbi.nlm.nih.gov/gene/?term=LOC105286872 | 2460 | 142 | 577.24 | 1 | 4.07 | 0.7 | 21.72 | 21.98 | -0.26 | 0.26 | -0.02 | 0.02 |
| gene6583 | https://www.ncbi.nlm.nih.gov/gene/?term=LOC105279649 | 7444 | 308 | 413.76 | 1 | 1.34 | 0.32 | 11.09 | 11.46 | -0.37 | 0.37 | -0.02 | 0.02 |
| gene10018 | https://www.ncbi.nlm.nih.gov/gene/?term=LOC105283607 | 4414 | 194 | 439.51 | 1 | 2.27 | 0.52 | 15.27 | 15.67 | -0.4 | 0.4 | -0.02 | 0.02 |
| gene11275 | https://www.ncbi.nlm.nih.gov/gene/?term=LOC105285055 | 4119 | 202 | 490.41 | 1 | 2.43 | 0.5 | 12.81 | 13.23 | -0.42 | 0.42 | -0.02 | 0.02 |
| gene12978 | https://www.ncbi.nlm.nih.gov/gene/?term=LOC105286997 | 2858 | 134 | 468.86 | 1 | 3.5 | 0.75 | 11.8 | 12.24 | -0.44 | 0.44 | -0.02 | 0.02 |
| gene10540 | https://www.ncbi.nlm.nih.gov/gene/?term=LOC105284200 | 5267 | 195 | 370.23 | 1 | 1.9 | 0.51 | 13.62 | 14.09 | -0.47 | 0.47 | -0.02 | 0.02 |
| gene10520 | https://www.ncbi.nlm.nih.gov/gene/?term=LOC105284184 | 2451 | 80 | 326.4 1 | 4.08 | 1.25 | 34.85 | 35.33 | -0.48 | 0.48 | -0.02 | 0.02 | |
| gene9478 | https://www.ncbi.nlm.nih.gov/gene/?term=LOC105282991 | 3726 | 48 | 128.82 | 1 | 2.68 | 2.08 | 31.5 | 33.03 | -1.53 | 1.53 | -0.02 | 0.02 |
| gene7248 | https://www.ncbi.nlm.nih.gov/gene/?term=LOC105280408 | 5175 | 44 | 85.02 1 | 1.93 | 2.27 | 22.41 | 21.41 | 0.99 | 0.99 | 0.01 | 0.01 | |
| gene6806 | https://www.ncbi.nlm.nih.gov/gene/?term=LOC105279894 | 3649 | 39 | 106.88 | 1 | 2.74 | 2.56 | 35.25 | 34.29 | 0.97 | 0.97 | 0.01 | 0.01 |
| gene12203 | https://www.ncbi.nlm.nih.gov/gene/?term=LOC105286099 | 5967 | 69 | 115.64 | 1 | 1.68 | 1.45 | 11.63 | 10.68 | 0.96 | 0.96 | 0.01 | 0.01 |
| gene12307 | https://www.ncbi.nlm.nih.gov/gene/?term=LOC105286216 | 10313 | 145 | 140.6 1 | 0.97 | 0.69 | 17.77 | 16.9 | 0.87 | 0.87 | 0.01 | 0.01 | |
| gene6639 | https://www.ncbi.nlm.nih.gov/gene/?term=LOC105279705 | 7656 | 70 | 91.43 1 | 1.31 | 1.43 | 22.25 | 21.42 | 0.84 | 0.84 | 0.01 | 0.01 | |
| gene10600 | https://www.ncbi.nlm.nih.gov/gene/?term=LOC105284270 | 4837 | 95 | 196.4 1 | 2.07 | 1.05 | 13.92 | 13.23 | 0.69 | 0.69 | 0.01 | 0.01 | |
| gene2404 | https://www.ncbi.nlm.nih.gov/gene/?term=LOC105274853 | 1767 | 39 | 220.71 | 1 | 5.66 | 2.56 | 31.07 | 30.51 | 0.56 | 0.56 | 0.01 | 0.01 |
| gene10155 | https://www.ncbi.nlm.nih.gov/gene/?term=LOC105283756 | 7030 | 93 | 132.29 | 1 | 1.42 | 1.08 | 7.03 | 6.49 | 0.54 | 0.54 | 0.01 | 0.01 |
| gene657 | https://www.ncbi.nlm.nih.gov/gene/?term=LOC105279667 | 6421 | 77 | 119.92 | 1 | 1.56 | 1.3 | 5.63 | 5.1 | 0.53 | 0.53 | 0.01 | 0.01 |
| gene7801 | https://www.ncbi.nlm.nih.gov/gene/?term=LOC105281047 | 5137 | 109 | 212.19 | 1 | 1.95 | 0.92 | 11.18 | 10.67 | 0.5 | 0.5 | 0.01 | 0.01 |
| gene7398 | https://www.ncbi.nlm.nih.gov/gene/?term=LOC105280589 | 3452 | 40 | 115.87 | 1 | 2.9 | 2.5 | 25.2 | 24.72 | 0.48 | 0.48 | 0.01 | 0.01 |
| gene940 | https://www.ncbi.nlm.nih.gov/gene/?term=LOC105282794 | 3578 | 84 | 234.77 | 1 | 2.79 | 1.19 | 14.53 | 14.06 | 0.47 | 0.47 | 0.01 | 0.01 |
| gene5524 | https://www.ncbi.nlm.nih.gov/gene/?term=LOC105278397 | 5177 | 199 | 384.39 | 1 | 1.93 | 0.5 | 24.51 | 24.16 | 0.35 | 0.35 | 0.01 | 0.01 |
| gene5831 | https://www.ncbi.nlm.nih.gov/gene/?term=LOC105278761 | 2506 | 99 | 395.05 | 1 | 3.99 | 1.01 | 25.53 | 25.18 | 0.35 | 0.35 | 0.01 | 0.01 |
| gene7600 | https://www.ncbi.nlm.nih.gov/gene/?term=LOC105280819 | 1615 | 41 | 253.87 | 1 | 6.19 | 2.44 | 40.74 | 40.39 | 0.35 | 0.35 | 0.01 | 0.01 |
| gene1147 | https://www.ncbi.nlm.nih.gov/gene/?term=LOC105284954 | 4022 | 87 | 216.31 | 1 | 2.49 | 1.15 | 14.67 | 14.36 | 0.3 | 0.3 | 0.01 | 0.01 |
| gene2173 | https://www.ncbi.nlm.nih.gov/gene/?term=LOC105274590 | 4387 | 78 | 177.8 1 | 2.28 | 1.28 | 22.31 | 22.01 | 0.3 | 0.3 | 0.01 | 0.01 | |
| gene9101 | https://www.ncbi.nlm.nih.gov/gene/?term=LOC105282553 | 3068 | 103 | 335.72 | 1 | 3.26 | 0.97 | 20.28 | 19.99 | 0.29 | 0.29 | 0.01 | 0.01 |
| gene13036 | https://www.ncbi.nlm.nih.gov/gene/?term=LOC105287059 | 4959 | 157 | 316.6 1 | 2.02 | 0.64 | 17.83 | 17.59 | 0.25 | 0.25 | 0.01 | 0.01 | |
| gene10549 | https://www.ncbi.nlm.nih.gov/gene/?term=LOC105284210 | 7145 | 313 | 438.07 | 1 | 1.4 | 0.32 | 10.77 | 10.55 | 0.22 | 0.22 | 0.01 | 0.01 |
| gene8274 | https://www.ncbi.nlm.nih.gov/gene/?term=LOC105281624 | 5425 | 189 | 348.39 | 1 | 1.84 | 0.53 | 11.18 | 10.97 | 0.21 | 0.21 | 0.01 | 0.01 |
| gene12802 | https://www.ncbi.nlm.nih.gov/gene/?term=LOC105286799 | 3183 | 170 | 534.09 | 1 | 3.14 | 0.59 | 15.96 | 15.78 | 0.18 | 0.18 | 0.01 | 0.01 |
| gene5525 | https://www.ncbi.nlm.nih.gov/gene/?term=LOC105278396 | 3090 | 185 | 598.71 | 1 | 3.24 | 0.54 | 24.64 | 24.46 | 0.18 | 0.18 | 0.01 | 0.01 |
| gene2443 | https://www.ncbi.nlm.nih.gov/gene/?term=LOC105274895 | 10433 | 330 | 316.3 1 | 0.96 | 0.3 | 5.47 | 5.65 | -0.18 | 0.18 | -0.01 | 0.01 | |
| gene4892 | https://www.ncbi.nlm.nih.gov/gene/?term=LOC105277667 | 3898 | 116 | 297.59 | 1 | 2.57 | 0.86 | 28.81 | 28.99 | -0.19 | 0.19 | -0.01 | 0.01 |
| gene3335 | https://www.ncbi.nlm.nih.gov/gene/?term=LOC105275898 | 4665 | 128 | 274.38 | 1 | 2.14 | 0.78 | 13.41 | 13.77 | -0.36 | 0.36 | -0.01 | 0.01 |
| gene3856 | https://www.ncbi.nlm.nih.gov/gene/?term=LOC105276499 | 5236 | 188 | 359.05 | 1 | 1.91 | 0.53 | 10.14 | 10.5 | -0.36 | 0.36 | -0.01 | 0.01 |
| gene10243 | https://www.ncbi.nlm.nih.gov/gene/?term=LOC105283862 | 3239 | 38 | 117.32 | 1 | 3.09 | 2.63 | 30.66 | 31.1 | -0.44 | 0.44 | -0.01 | 0.01 |
| gene8600 | https://www.ncbi.nlm.nih.gov/gene/?term=LOC105282001 | 8513 | 148 | 173.85 | 1 | 1.17 | 0.68 | 9.45 | 9.9 | -0.45 | 0.45 | -0.01 | 0.01 |
| gene13033 | https://www.ncbi.nlm.nih.gov/gene/?term=LOC105287063 | 1877 | 45 | 239.74 | 1 | 5.33 | 2.22 | 34.91 | 35.5 | -0.59 | 0.59 | -0.01 | 0.01 |
| gene4023 | https://www.ncbi.nlm.nih.gov/gene/?term=LOC105276692 | 1759 | 36 | 204.66 | 1 | 5.69 | 2.78 | 23.26 | 23.9 | -0.64 | 0.64 | -0.01 | 0.01 |
| gene8343 | https://www.ncbi.nlm.nih.gov/gene/?term=LOC105281700 | 2172 | 45 | 207.18 | 1 | 4.6 | 2.22 | 38.72 | 39.37 | -0.65 | 0.65 | -0.01 | 0.01 |
| gene6528 | https://www.ncbi.nlm.nih.gov/gene/?term=LOC105279591 | 7222 | 68 | 94.16 1 | 1.38 | 1.47 | 22.7 | 23.89 | -1.19 | 1.19 | -0.01 | 0.01 | |
| gene5052 | https://www.ncbi.nlm.nih.gov/gene/?term=LOC105277856 | 3167 | 27 | 85.25 1 | 3.16 | 3.7 | 36.57 | 38.1 | -1.52 | 1.52 | -0.01 | 0.01 | |
| gene10345 | https://www.ncbi.nlm.nih.gov/gene/?term=LOC105283989 | 4444 | 27 | 60.76 1 | 2.25 | 3.7 | 48.21 | 50.36 | -2.14 | 2.14 | -0.01 | 0.01 | |
| gene1220 | https://www.ncbi.nlm.nih.gov/gene/?term=LOC105285742 | 3453 | 86 | 249.06 | 1 | 2.9 | 1.16 | 21.68 | 21.51 | 0.17 | 0.17 | 0 | 0 |
| gene7247 | https://www.ncbi.nlm.nih.gov/gene/?term=LOC105280407 | 10486 | 206 | 196.45 | 1 | 0.95 | 0.49 | 17.14 | 16.97 | 0.17 | 0.17 | 0 | 0 |
| gene4213 | https://www.ncbi.nlm.nih.gov/gene/?term=LOC105276901 | 3257 | 96 | 294.75 | 1 | 3.07 | 1.04 | 22.87 | 22.72 | 0.14 | 0.14 | 0 | 0 |
| gene10107 | https://www.ncbi.nlm.nih.gov/gene/?term=LOC105283712 | 1883 | 29 | 154.01 | 1 | 5.31 | 3.45 | 27.22 | 27.09 | 0.13 | 0.13 | 0 | 0 |
| gene9645 | https://www.ncbi.nlm.nih.gov/gene/?term=LOC105283181 | 3875 | 134 | 345.81 | 1 | 2.58 | 0.75 | 11.45 | 11.37 | 0.08 | 0.08 | 0 | 0 |
| gene6620 | https://www.ncbi.nlm.nih.gov/gene/?term=LOC105279687 | 17998 | 354 | 196.69 | 1 | 0.56 | 0.28 | 5.08 | 5.05 | 0.03 | 0.03 | 0 | 0 |
| gene11583 | https://www.ncbi.nlm.nih.gov/gene/?term=LOC105285414 | 8192 | 175 | 213.62 | 1 | 1.22 | 0.57 | 9.21 | 9.24 | -0.03 | 0.03 | 0 | 0 |
| gene2806 | https://www.ncbi.nlm.nih.gov/gene/?term=LOC105275327 | 14257 | 462 | 324.05 | 1 | 0.7 | 0.22 | 7.37 | 7.4 | -0.03 | 0.03 | 0 | 0 |
| gene11579 | https://www.ncbi.nlm.nih.gov/gene/?term=LOC105285406 | 13571 | 260 | 191.58 | 1 | 0.74 | 0.38 | 5.75 | 5.79 | -0.04 | 0.04 | 0 | 0 |
| gene7166 | https://www.ncbi.nlm.nih.gov/gene/?term=LOC105280326 | 6117 | 208 | 340.04 | 1 | 1.63 | 0.48 | 16.25 | 16.28 | -0.04 | 0.04 | 0 | 0 |
| gene12406 | https://www.ncbi.nlm.nih.gov/gene/?term=LOC105286316 | 1386 | 30 | 216.45 | 1 | 7.22 | 3.33 | 39.2 | 39.29 | -0.09 | 0.09 | 0 | 0 |
| gene5375 | https://www.ncbi.nlm.nih.gov/gene/?term=LOC105278228 | 8640 | 148 | 171.3 1 | 1.16 | 0.68 | 24.78 | 24.89 | -0.11 | 0.11 | 0 | 0 | |
| gene12573 | https://www.ncbi.nlm.nih.gov/gene/?term=LOC105286515 | 3647 | 105 | 287.91 | 1 | 2.74 | 0.95 | 10.51 | 10.65 | -0.14 | 0.14 | 0 | 0 |
| gene1310 | https://www.ncbi.nlm.nih.gov/gene/?term=LOC105286661 | 2582 | 63 | 244 1 | 3.87 | 1.59 | 19.4 | 19.54 | -0.14 | 0.14 | 0 | 0 | |
| gene10531 | https://www.ncbi.nlm.nih.gov/gene/?term=LOC105284190 | 6881 | 180 | 261.59 | 1 | 1.45 | 0.56 | 10.06 | 10.22 | -0.16 | 0.16 | 0 | 0 |
| gene1632 | https://www.ncbi.nlm.nih.gov/gene/?term=LOC105287764 | 18061 | 334 | 184.93 | 1 | 0.55 | 0.3 | 2.71 | 2.98 | -0.27 | 0.27 | 0 | 0 |
| gene7183 | https://www.ncbi.nlm.nih.gov/gene/?term=LOC105280348 | 3925 | 53 | 135.03 | 1 | 2.55 | 1.89 | 25.26 | 25.53 | -0.27 | 0.27 | 0 | 0 |
| gene6455 | https://www.ncbi.nlm.nih.gov/gene/?term=LOC105279500 | 2612 | 29 | 111.03 | 1 | 3.83 | 3.45 | 48.01 | 48.34 | -0.33 | 0.33 | 0 | 0 |
| gene564 | https://www.ncbi.nlm.nih.gov/gene/?term=LOC105278478 | 2802 | 12 | 42.83 1 | 3.57 | 8.33 | 20.51 | 21.62 | -1.11 | 1.11 | 0 | 0 | |

Directory: OGL File: rnk-dmg-Cb_LI_BCphaseA1.vs.RphaseA2.txt

| gene_ID | gene_link | gwidth | #Sites | #per10Kb | #dmSites | #dmsp10kb | %dmSites | %pSite1 | %pSite2 | DMpSite | ADMpSite | DMpNucl | ADMpNucl |
|---|---|---|---|---|---|---|---|---|---|---|---|---|---|
| gene3165 | https://www.ncbi.nlm.nih.gov/gene/?term=LOC105275706 | 2374 | 67 | 282.22 | 4 | 16.85 | 5.97 | 25.11 | 35.96 | -10.84 | 10.84 | -0.31 | 0.31 |
| gene2593 | https://www.ncbi.nlm.nih.gov/gene/?term=LOC105275070 | 1774 | 6 | 33.82 | 3 | 16.91 | 50 | 16.42 | 57.88 | -41.46 | 41.46 | -0.14 | 0.14 |
| gene3200 | https://www.ncbi.nlm.nih.gov/gene/?term=LOC105275743 | 1786 | 77 | 431.13 | 1 | 5.6 | 1.3 | 22.67 | 19.88 | 2.79 | 2.79 | 0.12 | 0.12 |
| gene4034 | https://www.ncbi.nlm.nih.gov/gene/?term=LOC105276704 | 2065 | 87 | 421.31 | 2 | 9.69 | 2.3 | 12.36 | 15.2 | -2.84 | 2.84 | -0.12 | 0.12 |
| gene12961 | https://www.ncbi.nlm.nih.gov/gene/?term=LOC105286983 | 3094 | 70 | 226.24 | 1 | 3.23 | 1.43 | 0.98 | 5.77 | -4.79 | 4.79 | -0.11 | 0.11 |
| gene5714 | https://www.ncbi.nlm.nih.gov/gene/?term=LOC105278729 | 710 | 14 | 197.18 | 1 | 14.08 | 7.14 | 51.67 | 57.2 | -5.53 | 5.53 | -0.11 | 0.11 |
| gene12558 | https://www.ncbi.nlm.nih.gov/gene/?term=LOC105286502 | 1285 | 43 | 334.63 | 1 | 7.78 | 2.33 | 24.55 | 27.41 | -2.86 | 2.86 | -0.1 | 0.1 |
| gene920 | https://www.ncbi.nlm.nih.gov/gene/?term=LOC105282518 | 1197 | 43 | 359.23 | 1 | 8.35 | 2.33 | 37.5 | 34.97 | 2.52 | 2.52 | 0.09 | 0.09 |
| gene5525 | https://www.ncbi.nlm.nih.gov/gene/?term=LOC105278396 | 3090 | 186 | 601.94 | 1 | 3.24 | 0.54 | 24.51 | 25.99 | -1.48 | 1.48 | -0.09 | 0.09 |
| gene274 | https://www.ncbi.nlm.nih.gov/gene/?term=LOC105275190 | 3488 | 73 | 209.29 | 1 | 2.87 | 1.37 | 18.68 | 22.58 | -3.9 | 3.9 | -0.08 | 0.08 |
| gene6814 | https://www.ncbi.nlm.nih.gov/gene/?term=LOC105279898 | 1343 | 15 | 111.69 | 1 | 7.45 | 6.67 | 59.84 | 67.04 | -7.2 | 7.2 | -0.08 | 0.08 |
| gene10213 | https://www.ncbi.nlm.nih.gov/gene/?term=LOC105283826 | 1503 | 54 | 359.28 | 1 | 6.65 | 1.85 | 16.14 | 14.22 | 1.92 | 1.92 | 0.07 | 0.07 |
| gene11819 | https://www.ncbi.nlm.nih.gov/gene/?term=LOC105285671 | 2366 | 95 | 401.52 | 1 | 4.23 | 1.05 | 27.1 | 28.85 | -1.76 | 1.76 | -0.07 | 0.07 |
| gene12317 | https://www.ncbi.nlm.nih.gov/gene/?term=LOC105286223 | 2235 | 84 | 375.84 | 1 | 4.47 | 1.19 | 29.87 | 31.79 | -1.93 | 1.93 | -0.07 | 0.07 |
| gene12406 | https://www.ncbi.nlm.nih.gov/gene/?term=LOC105286316 | 1386 | 34 | 245.31 | 1 | 7.22 | 2.94 | 39.04 | 41.82 | -2.78 | 2.78 | -0.07 | 0.07 |
| gene12516 | https://www.ncbi.nlm.nih.gov/gene/?term=LOC105286452 | 1529 | 35 | 228.91 | 1 | 6.54 | 2.86 | 51.01 | 54.04 | -3.03 | 3.03 | -0.07 | 0.07 |
| gene5298 | https://www.ncbi.nlm.nih.gov/gene/?term=LOC105278145 | 4060 | 257 | 633 | 2 | 4.93 | 0.78 | 10.97 | 10.08 | 0.88 | 0.88 | 0.06 | 0.06 |
| gene298 | https://www.ncbi.nlm.nih.gov/gene/?term=LOC105275460 | 2473 | 112 | 452.89 | 1 | 4.04 | 0.89 | 12.88 | 14.18 | -1.3 | 1.3 | -0.06 | 0.06 |
| gene10984 | https://www.ncbi.nlm.nih.gov/gene/?term=LOC105284704 | 1987 | 64 | 322.09 | 1 | 5.03 | 1.56 | 25.47 | 27.27 | -1.8 | 1.8 | -0.06 | 0.06 |
| gene393 | https://www.ncbi.nlm.nih.gov/gene/?term=LOC105276566 | 1123 | 2 | 17.81 | 1 | 8.9 | 50 | 77.65 | 47.62 | 30.03 | 30.03 | 0.05 | 0.05 |
| gene7952 | https://www.ncbi.nlm.nih.gov/gene/?term=LOC105281213 | 2963 | 43 | 145.12 | 1 | 3.37 | 2.33 | 42.11 | 38.78 | 3.33 | 3.33 | 0.05 | 0.05 |
| gene4190 | https://www.ncbi.nlm.nih.gov/gene/?term=LOC105276875 | 3150 | 132 | 419.05 | 1 | 3.17 | 0.76 | 21.24 | 20.15 | 1.09 | 1.09 | 0.05 | 0.05 |
| gene5524 | https://www.ncbi.nlm.nih.gov/gene/?term=LOC105278397 | 5177 | 200 | 386.32 | 1 | 1.93 | 0.5 | 24.39 | 25.66 | -1.27 | 1.27 | -0.05 | 0.05 |
| gene6334 | https://www.ncbi.nlm.nih.gov/gene/?term=LOC105279356 | 1471 | 43 | 292.32 | 1 | 6.8 | 2.33 | 28.25 | 29.87 | -1.62 | 1.62 | -0.05 | 0.05 |
| gene10107 | https://www.ncbi.nlm.nih.gov/gene/?term=LOC105283712 | 1883 | 27 | 143.39 | 1 | 5.31 | 3.7 | 26.21 | 29.43 | -3.23 | 3.23 | -0.05 | 0.05 |
| gene5052 | https://www.ncbi.nlm.nih.gov/gene/?term=LOC105277856 | 3167 | 28 | 88.41 | 1 | 3.16 | 3.57 | 35.73 | 41.1 | -5.37 | 5.37 | -0.05 | 0.05 |
| gene4630 | https://www.ncbi.nlm.nih.gov/gene/?term=LOC105277380 | 1893 | 5 | 26.41 | 1 | 5.28 | 20 | 69.42 | 53.65 | 15.77 | 15.77 | 0.04 | 0.04 |
| gene7915 | https://www.ncbi.nlm.nih.gov/gene/?term=LOC105281173 | 1597 | 25 | 156.54 | 1 | 6.26 | 4 | 64.87 | 62.52 | 2.35 | 2.35 | 0.04 | 0.04 |
| gene11454 | https://www.ncbi.nlm.nih.gov/gene/?term=LOC105285256 | 1906 | 51 | 267.58 | 1 | 5.25 | 1.96 | 43.83 | 42.42 | 1.4 | 1.4 | 0.04 | 0.04 |
| gene10326 | https://www.ncbi.nlm.nih.gov/gene/?term=LOC105283962 | 3042 | 102 | 335.31 | 1 | 3.29 | 0.98 | 14.13 | 13.04 | 1.09 | 1.09 | 0.04 | 0.04 |
| gene10540 | https://www.ncbi.nlm.nih.gov/gene/?term=LOC105284200 | 5267 | 199 | 377.82 | 1 | 1.9 | 0.5 | 13.8 | 14.75 | -0.95 | 0.95 | -0.04 | 0.04 |
| gene5877 | https://www.ncbi.nlm.nih.gov/gene/?term=LOC105278807 | 1719 | 52 | 302.5 | 1 | 5.82 | 1.92 | 22.91 | 24.37 | -1.46 | 1.46 | -0.04 | 0.04 |
| gene1310 | https://www.ncbi.nlm.nih.gov/gene/?term=LOC105286661 | 2582 | 65 | 251.74 | 1 | 3.87 | 1.54 | 19.96 | 21.59 | -1.63 | 1.63 | -0.04 | 0.04 |
| gene4023 | https://www.ncbi.nlm.nih.gov/gene/?term=LOC105276692 | 1759 | 39 | 221.72 | 1 | 5.69 | 2.56 | 19.85 | 21.5 | -1.65 | 1.65 | -0.04 | 0.04 |
| gene3727 | https://www.ncbi.nlm.nih.gov/gene/?term=LOC105276367 | 2484 | 54 | 217.39 | 1 | 4.03 | 1.85 | 48.38 | 50.37 | -1.99 | 1.99 | -0.04 | 0.04 |
| gene12021 | https://www.ncbi.nlm.nih.gov/gene/?term=LOC105285905 | 1399 | 23 | 164.4 | 1 | 7.15 | 4.35 | 41.24 | 43.7 | -2.46 | 2.46 | -0.04 | 0.04 |
| gene2335 | https://www.ncbi.nlm.nih.gov/gene/?term=LOC105274778 | 3945 | 38 | 96.32 | 1 | 2.53 | 2.63 | 45.06 | 48.85 | -3.79 | 3.79 | -0.04 | 0.04 |
| gene9109 | https://www.ncbi.nlm.nih.gov/gene/?term=LOC105282561 | 2400 | 15 | 62.5 | 1 | 4.17 | 6.67 | 34.05 | 29.4 | 4.65 | 4.65 | 0.03 | 0.03 |
| gene7368 | https://www.ncbi.nlm.nih.gov/gene/?term=LOC105280556 | 4105 | 49 | 119.37 | 1 | 2.44 | 2.04 | 12.57 | 9.91 | 2.66 | 2.66 | 0.03 | 0.03 |
| gene3168 | https://www.ncbi.nlm.nih.gov/gene/?term=LOC105275711 | 6150 | 80 | 130.08 | 1 | 1.63 | 1.25 | 14.39 | 12.36 | 2.03 | 2.03 | 0.03 | 0.03 |
| gene7345 | https://www.ncbi.nlm.nih.gov/gene/?term=LOC105280529 | 3833 | 92 | 240.02 | 1 | 2.61 | 1.09 | 17.99 | 16.93 | 1.05 | 1.05 | 0.03 | 0.03 |
| gene3286 | https://www.ncbi.nlm.nih.gov/gene/?term=LOC105275848 | 3744 | 132 | 352.56 | 1 | 2.67 | 0.76 | 21.71 | 20.73 | 0.98 | 0.98 | 0.03 | 0.03 |
| gene13429 | https://www.ncbi.nlm.nih.gov/gene/?term=LOC105287501 | 3714 | 102 | 274.64 | 1 | 2.69 | 0.98 | 22.78 | 21.81 | 0.97 | 0.97 | 0.03 | 0.03 |
| gene11275 | https://www.ncbi.nlm.nih.gov/gene/?term=LOC105285055 | 4119 | 216 | 524.4 | 1 | 2.43 | 0.46 | 13.08 | 13.7 | -0.62 | 0.62 | -0.03 | 0.03 |
| gene12802 | https://www.ncbi.nlm.nih.gov/gene/?term=LOC105286799 | 3183 | 172 | 540.37 | 1 | 3.14 | 0.58 | 15.78 | 16.4 | -0.62 | 0.62 | -0.03 | 0.03 |
| gene3817 | https://www.ncbi.nlm.nih.gov/gene/?term=LOC105276449 | 1230 | 48 | 390.24 | 1 | 8.13 | 2.08 | 26.73 | 27.42 | -0.69 | 0.69 | -0.03 | 0.03 |
| gene10018 | https://www.ncbi.nlm.nih.gov/gene/?term=LOC105283607 | 4414 | 198 | 448.57 | 1 | 2.27 | 0.51 | 15.25 | 15.95 | -0.7 | 0.7 | -0.03 | 0.03 |
| gene2271 | https://www.ncbi.nlm.nih.gov/gene/?term=LOC105274706 | 1448 | 53 | 366.02 | 1 | 6.91 | 1.89 | 36.91 | 37.62 | -0.71 | 0.71 | -0.03 | 0.03 |
| gene3206 | https://www.ncbi.nlm.nih.gov/gene/?term=LOC105275752 | 3474 | 125 | 359.82 | 1 | 2.88 | 0.8 | 28.34 | 29.06 | -0.72 | 0.72 | -0.03 | 0.03 |
| gene2844 | https://www.ncbi.nlm.nih.gov/gene/?term=LOC105275363 | 4330 | 192 | 443.42 | 1 | 2.31 | 0.52 | 14.13 | 14.88 | -0.75 | 0.75 | -0.03 | 0.03 |
| gene7556 | https://www.ncbi.nlm.nih.gov/gene/?term=LOC105280774 | 2792 | 117 | 419.05 | 1 | 3.58 | 0.85 | 27.44 | 28.23 | -0.79 | 0.79 | -0.03 | 0.03 |
| gene8729 | https://www.ncbi.nlm.nih.gov/gene/?term=LOC105282137 | 5005 | 205 | 409.59 | 1 | 2 | 0.49 | 15.76 | 16.6 | -0.84 | 0.84 | -0.03 | 0.03 |
| gene4641 | https://www.ncbi.nlm.nih.gov/gene/?term=LOC105277398 | 1937 | 70 | 361.38 | 1 | 5.16 | 1.43 | 35.36 | 36.26 | -0.9 | 0.9 | -0.03 | 0.03 |
| gene10531 | https://www.ncbi.nlm.nih.gov/gene/?term=LOC105284190 | 6881 | 199 | 289.2 | 1 | 1.45 | 0.5 | 9.56 | 10.66 | -1.1 | 1.1 | -0.03 | 0.03 |
| gene13033 | https://www.ncbi.nlm.nih.gov/gene/?term=LOC105287063 | 1877 | 45 | 239.74 | 1 | 5.33 | 2.22 | 34.91 | 36.28 | -1.37 | 1.37 | -0.03 | 0.03 |
| gene1884 | https://www.ncbi.nlm.nih.gov/gene/?term=LOC105288072 | 4169 | 82 | 196.69 | 1 | 2.4 | 1.22 | 22.32 | 23.72 | -1.41 | 1.41 | -0.03 | 0.03 |
| gene2257 | https://www.ncbi.nlm.nih.gov/gene/?term=LOC105274689 | 3658 | 75 | 205.03 | 1 | 2.73 | 1.33 | 12.35 | 13.76 | -1.41 | 1.41 | -0.03 | 0.03 |
| gene8343 | https://www.ncbi.nlm.nih.gov/gene/?term=LOC105281700 | 2172 | 46 | 211.79 | 1 | 4.6 | 2.17 | 39.86 | 41.32 | -1.46 | 1.46 | -0.03 | 0.03 |
| gene5647 | https://www.ncbi.nlm.nih.gov/gene/?term=LOC105278567 | 3118 | 37 | 118.67 | 1 | 3.21 | 2.7 | 11.3 | 14.2 | -2.9 | 2.9 | -0.03 | 0.03 |
| gene1199 | https://www.ncbi.nlm.nih.gov/gene/?term=LOC105285498 | 2220 | 26 | 117.12 | 1 | 4.5 | 3.85 | 33.11 | 31.3 | 1.81 | 1.81 | 0.02 | 0.02 |

| | | | | | | | | | | | | | |
|---|---|---|---|---|---|---|---|---|---|---|---|---|---|
| gene7398 | https://www.ncbi.nlm.nih.gov/gene/?term=LOC105280589 | 3452 | 41 | 118.77 | 1 | 2.9 | 2.44 | 25.07 | 23.32 | 1.75 | 1.75 | 0.02 | 0.02 |
| gene3671 | https://www.ncbi.nlm.nih.gov/gene/?term=LOC105276272 | 1966 | 27 | 137.33 | 1 | 5.09 | 3.7 | 41.96 | 40.71 | 1.25 | 1.25 | 0.02 | 0.02 |
| gene322 | https://www.ncbi.nlm.nih.gov/gene/?term=LOC105275777 | 1606 | 23 | 143.21 | 1 | 6.23 | 4.35 | 45.55 | 44.33 | 1.21 | 1.21 | 0.02 | 0.02 |
| gene6237 | https://www.ncbi.nlm.nih.gov/gene/?term=LOC105279244 | 1484 | 26 | 175.2 | 1 | 6.74 | 3.85 | 46.7 | 45.51 | 1.19 | 1.19 | 0.02 | 0.02 |
| gene10819 | https://www.ncbi.nlm.nih.gov/gene/?term=LOC105284507 | 5767 | 83 | 143.92 | 1 | 1.73 | 1.2 | 21.4 | 20.36 | 1.04 | 1.04 | 0.02 | 0.02 |
| gene6966 | https://www.ncbi.nlm.nih.gov/gene/?term=LOC105280104 | 3747 | 125 | 333.6 | 1 | 2.67 | 0.8 | 14.94 | 14.21 | 0.73 | 0.73 | 0.02 | 0.02 |
| gene9722 | https://www.ncbi.nlm.nih.gov/gene/?term=LOC105283264 | 3951 | 88 | 222.73 | 1 | 2.53 | 1.14 | 24.81 | 24.09 | 0.72 | 0.72 | 0.02 | 0.02 |
| gene11340 | https://www.ncbi.nlm.nih.gov/gene/?term=LOC105285131 | 2918 | 102 | 349.55 | 1 | 3.43 | 0.98 | 14.35 | 13.65 | 0.7 | 0.7 | 0.02 | 0.02 |
| gene12573 | https://www.ncbi.nlm.nih.gov/gene/?term=LOC105286515 | 3647 | 115 | 315.33 | 1 | 2.74 | 0.87 | 11.99 | 11.33 | 0.66 | 0.66 | 0.02 | 0.02 |
| gene11401 | https://www.ncbi.nlm.nih.gov/gene/?term=LOC105285202 | 1841 | 50 | 271.59 | 1 | 5.43 | 2 | 21.11 | 20.46 | 0.65 | 0.65 | 0.02 | 0.02 |
| gene7600 | https://www.ncbi.nlm.nih.gov/gene/?term=LOC105280819 | 1615 | 41 | 253.87 | 1 | 6.19 | 2.44 | 40.74 | 40.08 | 0.65 | 0.65 | 0.02 | 0.02 |
| gene10714 | https://www.ncbi.nlm.nih.gov/gene/?term=LOC105284396 | 1966 | 57 | 289.93 | 1 | 5.09 | 1.75 | 38.23 | 37.64 | 0.59 | 0.59 | 0.02 | 0.02 |
| gene4962 | https://www.ncbi.nlm.nih.gov/gene/?term=LOC105277761 | 4645 | 150 | 322.93 | 1 | 2.15 | 0.67 | 14.77 | 14.2 | 0.57 | 0.57 | 0.02 | 0.02 |
| gene6286 | https://www.ncbi.nlm.nih.gov/gene/?term=LOC105279298 | 2861 | 139 | 485.84 | 2 | 6.99 | 1.44 | 22.43 | 22.79 | -0.36 | 0.36 | -0.02 | 0.02 |
| gene7568 | https://www.ncbi.nlm.nih.gov/gene/?term=LOC105280786 | 2525 | 123 | 487.13 | 1 | 3.96 | 0.81 | 28.92 | 29.32 | -0.39 | 0.39 | -0.02 | 0.02 |
| gene7888 | https://www.ncbi.nlm.nih.gov/gene/?term=LOC105281141 | 4357 | 165 | 378.7 | 1 | 2.3 | 0.61 | 28.04 | 28.47 | -0.43 | 0.43 | -0.02 | 0.02 |
| gene4892 | https://www.ncbi.nlm.nih.gov/gene/?term=LOC105277667 | 3898 | 122 | 312.98 | 1 | 2.57 | 0.82 | 27.68 | 28.18 | -0.5 | 0.5 | -0.02 | 0.02 |
| gene7550 | https://www.ncbi.nlm.nih.gov/gene/?term=LOC105280763 | 2173 | 98 | 450.99 | 1 | 4.6 | 1.02 | 26.91 | 27.42 | -0.51 | 0.51 | -0.02 | 0.02 |
| gene3885 | https://www.ncbi.nlm.nih.gov/gene/?term=LOC105276527 | 7928 | 352 | 444 | 1 | 1.26 | 0.28 | 8.21 | 8.74 | -0.52 | 0.52 | -0.02 | 0.02 |
| gene5967 | https://www.ncbi.nlm.nih.gov/gene/?term=LOC105278918 | 4393 | 135 | 307.31 | 1 | 2.28 | 0.74 | 16.02 | 16.55 | -0.53 | 0.53 | -0.02 | 0.02 |
| gene7166 | https://www.ncbi.nlm.nih.gov/gene/?term=LOC105280326 | 6117 | 213 | 348.21 | 1 | 1.63 | 0.47 | 15.89 | 16.43 | -0.54 | 0.54 | -0.02 | 0.02 |
| gene474 | https://www.ncbi.nlm.nih.gov/gene/?term=LOC105277435 | 3625 | 158 | 435.86 | 1 | 2.76 | 0.63 | 12.73 | 13.28 | -0.55 | 0.55 | -0.02 | 0.02 |
| gene10678 | https://www.ncbi.nlm.nih.gov/gene/?term=LOC105284355 | 4031 | 164 | 406.85 | 1 | 2.48 | 0.61 | 19.18 | 19.78 | -0.6 | 0.6 | -0.02 | 0.02 |
| gene8661 | https://www.ncbi.nlm.nih.gov/gene/?term=LOC105282063 | 1926 | 43 | 223.26 | 1 | 5.19 | 2.33 | 23.84 | 24.54 | -0.71 | 0.71 | -0.02 | 0.02 |
| gene2173 | https://www.ncbi.nlm.nih.gov/gene/?term=LOC105274590 | 4387 | 84 | 191.47 | 1 | 2.28 | 1.19 | 20.89 | 21.7 | -0.81 | 0.81 | -0.02 | 0.02 |
| gene9598 | https://www.ncbi.nlm.nih.gov/gene/?term=LOC105283132 | 1769 | 54 | 305.26 | 1 | 5.65 | 1.85 | 40.78 | 41.59 | -0.81 | 0.81 | -0.02 | 0.02 |
| gene4070 | https://www.ncbi.nlm.nih.gov/gene/?term=LOC105276745 | 3957 | 72 | 181.96 | 1 | 2.53 | 1.39 | 20.98 | 22.28 | -1.3 | 1.3 | -0.02 | 0.02 |
| gene9336 | https://www.ncbi.nlm.nih.gov/gene/?term=LOC105282834 | 5993 | 109 | 181.88 | 1 | 1.67 | 0.92 | 15.48 | 16.78 | -1.31 | 1.31 | -0.02 | 0.02 |
| gene1962 | https://www.ncbi.nlm.nih.gov/gene/?term=LOC105288157 | 1246 | 16 | 128.41 | 1 | 8.03 | 6.25 | 13.95 | 15.31 | -1.36 | 1.36 | -0.02 | 0.02 |
| gene7183 | https://www.ncbi.nlm.nih.gov/gene/?term=LOC105280348 | 3925 | 53 | 135.03 | 1 | 2.55 | 1.89 | 25.26 | 26.63 | -1.37 | 1.37 | -0.02 | 0.02 |
| gene6455 | https://www.ncbi.nlm.nih.gov/gene/?term=LOC105279500 | 2612 | 32 | 122.51 | 1 | 3.83 | 3.12 | 49.77 | 51.44 | -1.68 | 1.68 | -0.02 | 0.02 |
| gene5402 | https://www.ncbi.nlm.nih.gov/gene/?term=LOC105278260 | 3068 | 41 | 133.64 | 1 | 3.26 | 2.44 | 13.01 | 14.77 | -1.77 | 1.77 | -0.02 | 0.02 |
| gene10243 | https://www.ncbi.nlm.nih.gov/gene/?term=LOC105283862 | 3239 | 37 | 114.23 | 1 | 3.09 | 2.7 | 31.49 | 33.45 | -1.96 | 1.96 | -0.02 | 0.02 |
| gene2970 | https://www.ncbi.nlm.nih.gov/gene/?term=LOC105275518 | 2340 | 21 | 89.74 | 1 | 4.27 | 4.76 | 45.1 | 47.46 | -2.36 | 2.36 | -0.02 | 0.02 |
| gene814 | https://www.ncbi.nlm.nih.gov/gene/?term=LOC105281296 | 2722 | 43 | 157.97 | 1 | 3.67 | 2.33 | 29.73 | 28.81 | 0.92 | 0.92 | 0.01 | 0.01 |
| gene11447 | https://www.ncbi.nlm.nih.gov/gene/?term=LOC105285252 | 7437 | 135 | 181.52 | 1 | 1.34 | 0.74 | 13.54 | 12.74 | 0.8 | 0.8 | 0.01 | 0.01 |
| gene1159 | https://www.ncbi.nlm.nih.gov/gene/?term=LOC105285084 | 3282 | 27 | 82.27 | 1 | 3.05 | 3.7 | 22 | 21.23 | 0.77 | 0.77 | 0.01 | 0.01 |
| gene652 | https://www.ncbi.nlm.nih.gov/gene/?term=LOC105279528 | 5806 | 44 | 75.78 | 1 | 1.72 | 2.27 | 16.35 | 15.59 | 0.76 | 0.76 | 0.01 | 0.01 |
| gene9464 | https://www.ncbi.nlm.nih.gov/gene/?term=LOC105282975 | 1450 | 25 | 172.41 | 1 | 6.9 | 4 | 41.13 | 40.46 | 0.67 | 0.67 | 0.01 | 0.01 |
| gene4376 | https://www.ncbi.nlm.nih.gov/gene/?term=LOC105277083 | 4307 | 96 | 222.89 | 1 | 2.32 | 1.04 | 25.73 | 25.08 | 0.66 | 0.66 | 0.01 | 0.01 |
| gene10155 | https://www.ncbi.nlm.nih.gov/gene/?term=LOC105283756 | 7030 | 97 | 137.98 | 1 | 1.42 | 1.03 | 6.83 | 6.25 | 0.59 | 0.59 | 0.01 | 0.01 |
| gene12203 | https://www.ncbi.nlm.nih.gov/gene/?term=LOC105286099 | 5967 | 78 | 130.72 | 1 | 1.68 | 1.28 | 11.28 | 10.77 | 0.51 | 0.51 | 0.01 | 0.01 |
| gene3107 | https://www.ncbi.nlm.nih.gov/gene/?term=LOC105275650 | 5103 | 151 | 295.9 | 1 | 1.96 | 0.66 | 17.08 | 16.7 | 0.38 | 0.38 | 0.01 | 0.01 |
| gene5646 | https://www.ncbi.nlm.nih.gov/gene/?term=LOC105278534 | 7475 | 186 | 248.83 | 1 | 1.34 | 0.54 | 18.03 | 17.67 | 0.36 | 0.36 | 0.01 | 0.01 |
| gene10392 | https://www.ncbi.nlm.nih.gov/gene/?term=LOC105284036 | 3216 | 83 | 258.08 | 2 | 6.22 | 2.41 | 23.78 | 23.42 | 0.35 | 0.35 | 0.01 | 0.01 |
| gene1220 | https://www.ncbi.nlm.nih.gov/gene/?term=LOC105285742 | 3453 | 99 | 286.71 | 1 | 2.9 | 1.01 | 25.65 | 25.32 | 0.33 | 0.33 | 0.01 | 0.01 |
| gene3848 | https://www.ncbi.nlm.nih.gov/gene/?term=LOC105276486 | 5899 | 121 | 205.12 | 1 | 1.7 | 0.83 | 23.22 | 22.92 | 0.3 | 0.3 | 0.01 | 0.01 |
| gene11583 | https://www.ncbi.nlm.nih.gov/gene/?term=LOC105285414 | 8192 | 177 | 216.06 | 1 | 1.22 | 0.56 | 8.82 | 8.53 | 0.29 | 0.29 | 0.01 | 0.01 |
| gene5831 | https://www.ncbi.nlm.nih.gov/gene/?term=LOC105278761 | 2506 | 104 | 415 | 1 | 3.99 | 0.96 | 25.24 | 24.96 | 0.28 | 0.28 | 0.01 | 0.01 |
| gene13114 | https://www.ncbi.nlm.nih.gov/gene/?term=LOC105287147 | 7609 | 389 | 511.24 | 1 | 1.31 | 0.26 | 9.77 | 9.58 | 0.19 | 0.19 | 0.01 | 0.01 |
| gene5746 | https://www.ncbi.nlm.nih.gov/gene/?term=LOC105278666 | 10343 | 409 | 395.44 | 1 | 0.97 | 0.24 | 7.43 | 7.26 | 0.17 | 0.17 | 0.01 | 0.01 |
| gene12674 | https://www.ncbi.nlm.nih.gov/gene/?term=LOC105286649 | 18283 | 680 | 371.93 | 1 | 0.55 | 0.15 | 1.31 | 1.16 | 0.15 | 0.15 | 0.01 | 0.01 |
| gene10864 | https://www.ncbi.nlm.nih.gov/gene/?term=LOC105284557 | 1666 | 78 | 468.19 | 1 | 6 | 1.28 | 30.45 | 30.59 | -0.13 | 0.13 | -0.01 | 0.01 |
| gene5649 | https://www.ncbi.nlm.nih.gov/gene/?term=LOC105278538 | 2140 | 84 | 392.52 | 1 | 4.67 | 1.19 | 18.7 | 18.83 | -0.13 | 0.13 | -0.01 | 0.01 |
| gene6583 | https://www.ncbi.nlm.nih.gov/gene/?term=LOC105279649 | 7444 | 330 | 443.31 | 1 | 1.34 | 0.3 | 11.47 | 11.66 | -0.2 | 0.2 | -0.01 | 0.01 |
| gene2806 | https://www.ncbi.nlm.nih.gov/gene/?term=LOC105275327 | 14257 | 480 | 336.68 | 1 | 0.7 | 0.21 | 7.1 | 7.32 | -0.22 | 0.22 | -0.01 | 0.01 |
| gene7249 | https://www.ncbi.nlm.nih.gov/gene/?term=LOC105280422 | 1775 | 61 | 343.66 | 1 | 5.63 | 1.64 | 15.16 | 15.4 | -0.24 | 0.24 | -0.01 | 0.01 |
| gene9101 | https://www.ncbi.nlm.nih.gov/gene/?term=LOC105282553 | 3068 | 104 | 338.98 | 1 | 3.26 | 0.96 | 20.36 | 20.64 | -0.27 | 0.27 | -0.01 | 0.01 |
| gene5613 | https://www.ncbi.nlm.nih.gov/gene/?term=LOC105278504 | 8524 | 211 | 247.54 | 1 | 1.17 | 0.47 | 5.46 | 5.74 | -0.28 | 0.28 | -0.01 | 0.01 |
| gene6620 | https://www.ncbi.nlm.nih.gov/gene/?term=LOC105279687 | 17998 | 360 | 200.02 | 1 | 0.56 | 0.28 | 5.06 | 5.34 | -0.28 | 0.28 | -0.01 | 0.01 |
| gene1573 | https://www.ncbi.nlm.nih.gov/gene/?term=LOC105287704 | 3030 | 131 | 432.34 | 1 | 3.3 | 0.76 | 20.77 | 21.09 | -0.32 | 0.32 | -0.01 | 0.01 |
| gene2056 | https://www.ncbi.nlm.nih.gov/gene/?term=LOC105274452 | 6584 | 212 | 321.99 | 1 | 1.52 | 0.47 | 13.94 | 14.26 | -0.33 | 0.33 | -0.01 | 0.01 |
| gene1147 | https://www.ncbi.nlm.nih.gov/gene/?term=LOC105284954 | 4022 | 82 | 203.88 | 1 | 2.49 | 1.22 | 15.94 | 16.29 | -0.34 | 0.34 | -0.01 | 0.01 |
| gene10600 | https://www.ncbi.nlm.nih.gov/gene/?term=LOC105284270 | 4837 | 105 | 217.08 | 1 | 2.07 | 0.95 | 12.85 | 13.22 | -0.37 | 0.37 | -0.01 | 0.01 |

| | | | | | | | | | | | | | |
|---|---|---|---|---|---|---|---|---|---|---|---|---|---|
| gene3856 | https://www.ncbi.nlm.nih.gov/gene/?term=LOC105276499 | 5236 | 201 | 383.88 | 1 | 1.91 | 0.5 | 10.34 | 10.72 | -0.39 | 0.39 | -0.01 | 0.01 |
| gene10520 | https://www.ncbi.nlm.nih.gov/gene/?term=LOC105284184 | 2451 | 82 | 334.56 | 1 | 4.08 | 1.22 | 34.55 | 34.95 | -0.4 | 0.4 | -0.01 | 0.01 |
| gene5375 | https://www.ncbi.nlm.nih.gov/gene/?term=LOC105278228 | 8640 | 159 | 184.03 | 1 | 1.16 | 0.63 | 24.17 | 24.62 | -0.45 | 0.45 | -0.01 | 0.01 |
| gene4225 | https://www.ncbi.nlm.nih.gov/gene/?term=LOC105276917 | 2443 | 26 | 106.43 | 1 | 4.09 | 3.85 | 36.73 | 37.26 | -0.53 | 0.53 | -0.01 | 0.01 |
| gene9330 | https://www.ncbi.nlm.nih.gov/gene/?term=LOC105282825 | 3936 | 90 | 228.66 | 1 | 2.54 | 1.11 | 15.33 | 15.86 | -0.53 | 0.53 | -0.01 | 0.01 |
| gene8600 | https://www.ncbi.nlm.nih.gov/gene/?term=LOC105282001 | 8513 | 158 | 185.6 | 1 | 1.17 | 0.63 | 9.54 | 10.09 | -0.55 | 0.55 | -0.01 | 0.01 |
| gene7245 | https://www.ncbi.nlm.nih.gov/gene/?term=LOC105280410 | 12478 | 260 | 208.37 | 1 | 0.8 | 0.38 | 19.03 | 19.62 | -0.59 | 0.59 | -0.01 | 0.01 |
| gene3385 | https://www.ncbi.nlm.nih.gov/gene/?term=LOC105275966 | 3305 | 59 | 178.52 | 1 | 3.03 | 1.69 | 50.29 | 50.9 | -0.61 | 0.61 | -0.01 | 0.01 |
| gene10379 | https://www.ncbi.nlm.nih.gov/gene/?term=LOC105284028 | 6974 | 74 | 106.11 | 1 | 1.43 | 1.35 | 8.99 | 9.61 | -0.62 | 0.62 | -0.01 | 0.01 |
| gene388 | https://www.ncbi.nlm.nih.gov/gene/?term=LOC105276538 | 1558 | 20 | 128.37 | 2 | 12.84 | 10 | 70.94 | 71.62 | -0.68 | 0.68 | -0.01 | 0.01 |
| gene927 | https://www.ncbi.nlm.nih.gov/gene/?term=LOC105282600 | 3727 | 65 | 174.4 | 1 | 2.68 | 1.54 | 26.3 | 27.02 | -0.73 | 0.73 | -0.01 | 0.01 |
| gene2330 | https://www.ncbi.nlm.nih.gov/gene/?term=LOC105274773 | 13104 | 172 | 131.26 | 1 | 0.76 | 0.58 | 15.95 | 16.78 | -0.83 | 0.83 | -0.01 | 0.01 |
| gene6528 | https://www.ncbi.nlm.nih.gov/gene/?term=LOC105279591 | 7222 | 75 | 103.85 | 1 | 1.38 | 1.33 | 20.71 | 21.6 | -0.89 | 0.89 | -0.01 | 0.01 |
| gene10345 | https://www.ncbi.nlm.nih.gov/gene/?term=LOC105283989 | 4444 | 30 | 67.51 | 1 | 2.25 | 3.33 | 46.58 | 47.61 | -1.03 | 1.03 | -0.01 | 0.01 |
| gene8178 | https://www.ncbi.nlm.nih.gov/gene/?term=LOC105281515 | 8647 | 46 | 53.2 | 1 | 1.16 | 2.17 | 28.18 | 29.56 | -1.38 | 1.38 | -0.01 | 0.01 |
| gene1809 | https://www.ncbi.nlm.nih.gov/gene/?term=LOC105287989 | 7010 | 73 | 104.14 | 1 | 1.43 | 1.37 | 23.7 | 23.28 | 0.42 | 0.42 | 0 | 0 |
| gene12307 | https://www.ncbi.nlm.nih.gov/gene/?term=LOC105286216 | 10313 | 167 | 161.93 | 1 | 0.97 | 0.6 | 15.75 | 15.52 | 0.23 | 0.23 | 0 | 0 |
| gene657 | https://www.ncbi.nlm.nih.gov/gene/?term=LOC105279667 | 6421 | 80 | 124.59 | 1 | 1.56 | 1.25 | 3.15 | 2.93 | 0.21 | 0.21 | 0 | 0 |
| gene940 | https://www.ncbi.nlm.nih.gov/gene/?term=LOC105282794 | 3578 | 88 | 245.95 | 1 | 2.79 | 1.14 | 15.65 | 15.49 | 0.16 | 0.16 | 0 | 0 |
| gene6031 | https://www.ncbi.nlm.nih.gov/gene/?term=LOC105279000 | 5691 | 185 | 325.07 | 1 | 1.76 | 0.54 | 13.85 | 13.69 | 0.15 | 0.15 | 0 | 0 |
| gene3599 | https://www.ncbi.nlm.nih.gov/gene/?term=LOC105276196 | 2955 | 56 | 189.51 | 1 | 3.38 | 1.79 | 40.12 | 39.99 | 0.13 | 0.13 | 0 | 0 |
| gene2404 | https://www.ncbi.nlm.nih.gov/gene/?term=LOC105274853 | 1767 | 42 | 237.69 | 1 | 5.66 | 2.38 | 28.85 | 28.79 | 0.07 | 0.07 | 0 | 0 |
| gene2991 | https://www.ncbi.nlm.nih.gov/gene/?term=LOC105275536 | 11224 | 214 | 190.66 | 1 | 0.89 | 0.47 | 5.64 | 5.58 | 0.05 | 0.05 | 0 | 0 |
| gene4527 | https://www.ncbi.nlm.nih.gov/gene/?term=LOC105277273 | 3686 | 151 | 409.66 | 1 | 2.71 | 0.66 | 12.71 | 12.65 | 0.05 | 0.05 | 0 | 0 |
| gene4996 | https://www.ncbi.nlm.nih.gov/gene/?term=LOC105277798 | 5617 | 64 | 113.94 | 1 | 1.78 | 1.56 | 10.86 | 10.81 | 0.05 | 0.05 | 0 | 0 |
| gene2266 | https://www.ncbi.nlm.nih.gov/gene/?term=LOC105274699 | 2391 | 63 | 263.49 | 1 | 4.18 | 1.59 | 23.04 | 22.99 | 0.04 | 0.04 | 0 | 0 |
| gene7148 | https://www.ncbi.nlm.nih.gov/gene/?term=LOC105280304 | 8920 | 123 | 137.89 | 1 | 1.12 | 0.81 | 5.79 | 5.75 | 0.04 | 0.04 | 0 | 0 |
| gene53 | https://www.ncbi.nlm.nih.gov/gene/?term=LOC105277148 | 19283 | 269 | 139.5 | 1 | 0.52 | 0.37 | 3.9 | 3.86 | 0.03 | 0.03 | 0 | 0 |
| gene9645 | https://www.ncbi.nlm.nih.gov/gene/?term=LOC105283181 | 3875 | 138 | 356.13 | 1 | 2.58 | 0.72 | 11.99 | 11.98 | 0.01 | 0.01 | 0 | 0 |
| gene10549 | https://www.ncbi.nlm.nih.gov/gene/?term=LOC105284210 | 7145 | 323 | 452.06 | 1 | 1.4 | 0.31 | 10.47 | 10.47 | 0 | 0 | 0 | 0 |
| gene1470 | https://www.ncbi.nlm.nih.gov/gene/?term=LOC105287597 | 2974 | 192 | 645.6 | 1 | 3.36 | 0.52 | 16.86 | 16.86 | 0 | 0 | 0 | 0 |
| gene9755 | https://www.ncbi.nlm.nih.gov/gene/?term=LOC105283296 | 2177 | 90 | 413.41 | 1 | 4.59 | 1.11 | 19.36 | 19.36 | 0 | 0 | 0 | 0 |
| gene9411 | https://www.ncbi.nlm.nih.gov/gene/?term=LOC105282914 | 5534 | 41 | 74.09 | 1 | 1.81 | 2.44 | 49.68 | 49.72 | -0.04 | 0.04 | 0 | 0 |
| gene13036 | https://www.ncbi.nlm.nih.gov/gene/?term=LOC105287059 | 4959 | 165 | 332.73 | 1 | 2.02 | 0.61 | 17.35 | 17.4 | -0.05 | 0.05 | 0 | 0 |
| gene7207 | https://www.ncbi.nlm.nih.gov/gene/?term=LOC105280376 | 4659 | 105 | 225.37 | 1 | 2.15 | 0.95 | 28.77 | 28.82 | -0.05 | 0.05 | 0 | 0 |
| gene10606 | https://www.ncbi.nlm.nih.gov/gene/?term=LOC105284273 | 6119 | 252 | 411.83 | 2 | 3.27 | 0.79 | 19.91 | 19.99 | -0.08 | 0.08 | 0 | 0 |
| gene11568 | https://www.ncbi.nlm.nih.gov/gene/?term=LOC105285395 | 8086 | 184 | 227.55 | 1 | 1.24 | 0.54 | 16.42 | 16.51 | -0.08 | 0.08 | 0 | 0 |
| gene3470 | https://www.ncbi.nlm.nih.gov/gene/?term=LOC105276059 | 8716 | 161 | 184.72 | 1 | 1.15 | 0.62 | 9.12 | 9.21 | -0.08 | 0.08 | 0 | 0 |
| gene8274 | https://www.ncbi.nlm.nih.gov/gene/?term=LOC105281624 | 5425 | 208 | 383.41 | 1 | 1.84 | 0.48 | 10.51 | 10.6 | -0.09 | 0.09 | 0 | 0 |
| gene2748 | https://www.ncbi.nlm.nih.gov/gene/?term=LOC105275261 | 3845 | 58 | 150.85 | 1 | 2.6 | 1.72 | 39.98 | 40.08 | -0.1 | 0.1 | 0 | 0 |
| gene1632 | https://www.ncbi.nlm.nih.gov/gene/?term=LOC105287764 | 18061 | 354 | 196 | 1 | 0.55 | 0.28 | 2.69 | 2.8 | -0.11 | 0.11 | 0 | 0 |
| gene10596 | https://www.ncbi.nlm.nih.gov/gene/?term=LOC105284262 | 6520 | 235 | 360.43 | 1 | 1.53 | 0.43 | 3.69 | 3.81 | -0.12 | 0.12 | 0 | 0 |
| gene2683 | https://www.ncbi.nlm.nih.gov/gene/?term=LOC105275187 | 5492 | 75 | 136.56 | 1 | 1.82 | 1.33 | 14.14 | 14.34 | -0.2 | 0.2 | 0 | 0 |
| gene11579 | https://www.ncbi.nlm.nih.gov/gene/?term=LOC105285406 | 13571 | 293 | 215.9 | 1 | 0.74 | 0.34 | 5.86 | 6.07 | -0.21 | 0.21 | 0 | 0 |
| gene6639 | https://www.ncbi.nlm.nih.gov/gene/?term=LOC105279705 | 7656 | 69 | 90.13 | 1 | 1.31 | 1.45 | 21.96 | 22.32 | -0.36 | 0.36 | 0 | 0 |

Directory: OGL File: rnk-dmg-Cb_LI_BCphaseA1.vs.RphaseB1.txt

| gene_ID | gene_link | gwidth | #Sites | #per10Kb | #dmSites | #dmsp10kb | %dmSites | %pSite1 | %pSite2 | DMpSite | ADMpSite | DMpNucl | ADMpNucl |
|---|---|---|---|---|---|---|---|---|---|---|---|---|---|
| gene10237 | https://www.ncbi.nlm.nih.gov/gene/?term=LOC105283856 | 1259 | 25 | 198.57 | 5 | 39.71 | 20 | 64.65 | 55.66 | 9 | 9 | 0.18 | 0.18 |
| gene8965 | https://www.ncbi.nlm.nih.gov/gene/?term=LOC105282405 | 1310 | 31 | 236.64 | 2 | 15.27 | 6.45 | 51.28 | 44.69 | 6.59 | 6.59 | 0.16 | 0.16 |
| gene1993 | https://www.ncbi.nlm.nih.gov/gene/?term=LOC105288193 | 1449 | 64 | 441.68 | 4 | 27.61 | 6.25 | 34.43 | 31.28 | 3.16 | 3.16 | 0.14 | 0.14 |
| gene13384 | https://www.ncbi.nlm.nih.gov/gene/?term=LOC105287447 | 1877 | 145 | 772.51 | 7 | 37.29 | 4.83 | 29.16 | 30.96 | -1.8 | 1.8 | -0.14 | 0.14 |
| gene5809 | https://www.ncbi.nlm.nih.gov/gene/?term=LOC105278737 | 1159 | 83 | 716.13 | 1 | 8.63 | 1.2 | 16.21 | 18.18 | -1.97 | 1.97 | -0.14 | 0.14 |
| gene6314 | https://www.ncbi.nlm.nih.gov/gene/?term=LOC105279335 | 718 | 34 | 473.54 | 2 | 27.86 | 5.88 | 18.89 | 16.04 | 2.85 | 2.85 | 0.13 | 0.13 |
| gene5309 | https://www.ncbi.nlm.nih.gov/gene/?term=LOC105278160 | 1704 | 85 | 498.83 | 2 | 11.74 | 2.35 | 38.16 | 35.69 | 2.46 | 2.46 | 0.12 | 0.12 |
| gene560 | https://www.ncbi.nlm.nih.gov/gene/?term=LOC109610897 | 1584 | 112 | 707.07 | 3 | 18.94 | 2.68 | 15.45 | 13.95 | 1.5 | 1.5 | 0.11 | 0.11 |
| gene5412 | https://www.ncbi.nlm.nih.gov/gene/?term=LOC105278266 | 1868 | 111 | 594.22 | 2 | 10.71 | 1.8 | 25.18 | 27.02 | -1.84 | 1.84 | -0.11 | 0.11 |
| gene5650 | https://www.ncbi.nlm.nih.gov/gene/?term=LOC105278537 | 1672 | 61 | 364.83 | 5 | 29.9 | 8.2 | 28.88 | 31.76 | -2.89 | 2.89 | -0.11 | 0.11 |
| gene11413 | https://www.ncbi.nlm.nih.gov/gene/?term=LOC105285213 | 1472 | 37 | 251.36 | 4 | 27.17 | 10.81 | 13.65 | 18.03 | -4.37 | 4.37 | -0.11 | 0.11 |
| gene305 | https://www.ncbi.nlm.nih.gov/gene/?term=LOC105275539 | 1277 | 23 | 180.11 | 2 | 15.66 | 8.7 | 20.8 | 15.48 | 5.32 | 5.32 | 0.1 | 0.1 |
| gene10334 | https://www.ncbi.nlm.nih.gov/gene/?term=LOC105283973 | 1536 | 33 | 214.84 | 4 | 26.04 | 12.12 | 58.47 | 53.86 | 4.61 | 4.61 | 0.1 | 0.1 |
| gene3213 | https://www.ncbi.nlm.nih.gov/gene/?term=LOC105275761 | 2635 | 101 | 383.3 | 6 | 22.77 | 5.94 | 29.66 | 26.96 | 2.7 | 2.7 | 0.1 | 0.1 |
| gene2140 | https://www.ncbi.nlm.nih.gov/gene/?term=LOC105274555 | 1379 | 56 | 406.09 | 2 | 14.5 | 3.57 | 36.51 | 34.17 | 2.35 | 2.35 | 0.1 | 0.1 |
| gene460 | https://www.ncbi.nlm.nih.gov/gene/?term=LOC105277371 | 1819 | 76 | 417.81 | 1 | 5.5 | 1.32 | 19.38 | 17.09 | 2.29 | 2.29 | 0.1 | 0.1 |
| gene1562 | https://www.ncbi.nlm.nih.gov/gene/?term=LOC105287692 | 1491 | 97 | 650.57 | 3 | 20.12 | 3.09 | 30.3 | 28.78 | 1.52 | 1.52 | 0.1 | 0.1 |
| gene5207 | https://www.ncbi.nlm.nih.gov/gene/?term=LOC105278043 | 1205 | 73 | 605.81 | 1 | 8.3 | 1.37 | 20.73 | 22.42 | -1.69 | 1.69 | -0.1 | 0.1 |
| gene762 | https://www.ncbi.nlm.nih.gov/gene/?term=LOC105280710 | 1173 | 66 | 562.66 | 5 | 42.63 | 7.58 | 21.68 | 23.45 | -1.77 | 1.77 | -0.1 | 0.1 |
| gene12330 | https://www.ncbi.nlm.nih.gov/gene/?term=LOC105286235 | 1501 | 76 | 506.33 | 3 | 19.99 | 3.95 | 37.88 | 39.89 | -2.01 | 2.01 | -0.1 | 0.1 |
| gene6110 | https://www.ncbi.nlm.nih.gov/gene/?term=LOC105279087 | 1077 | 28 | 259.98 | 1 | 9.29 | 3.57 | 52.85 | 56.81 | -3.96 | 3.96 | -0.1 | 0.1 |
| gene3520 | https://www.ncbi.nlm.nih.gov/gene/?term=LOC105276115 | 1525 | 39 | 255.74 | 3 | 19.67 | 7.69 | 23.35 | 27.32 | -3.97 | 3.97 | -0.1 | 0.1 |
| gene4643 | https://www.ncbi.nlm.nih.gov/gene/?term=LOC109610847 | 429 | 5 | 116.55 | 1 | 23.31 | 20 | 29.86 | 22.57 | 7.29 | 7.29 | 0.09 | 0.09 |
| gene13026 | https://www.ncbi.nlm.nih.gov/gene/?term=LOC105287046 | 1786 | 55 | 307.95 | 1 | 5.6 | 1.82 | 21.15 | 18.07 | 3.07 | 3.07 | 0.09 | 0.09 |

...
... only first 25 lines shown ...

Directory: OGL File: rnk-dmg-Cb_LI_BCphaseA1.vs.RphaseB2.txt

| gene_ID | gene_link | gwidth | #Sites | #per10Kb | #dmSites | #dmsp10kb | %dmSites | %pSite1 | %pSite2 | DMpSite | ADMpSite | DMpNucl | ADMpNucl |
|---|---|---|---|---|---|---|---|---|---|---|---|---|---|
| gene5412 | https://www.ncbi.nlm.nih.gov/gene/?term=LOC105278266 | 1868 | 105 | 562.1 | 4 | 21.41 | 3.81 | 26.49 | 29.78 | -3.29 | 3.29 | -0.18 | 0.18 |
| gene8965 | https://www.ncbi.nlm.nih.gov/gene/?term=LOC105282405 | 1310 | 31 | 236.64 | 5 | 38.17 | 16.13 | 51.28 | 44.76 | 6.52 | 6.52 | 0.15 | 0.15 |
| gene5917 | https://www.ncbi.nlm.nih.gov/gene/?term=LOC105278849 | 823 | 80 | 972.05 | 4 | 48.6 | 5 | 24.88 | 23.29 | 1.59 | 1.59 | 0.15 | 0.15 |
| gene3929 | https://www.ncbi.nlm.nih.gov/gene/?term=LOC105276585 | 1525 | 115 | 754.1 | 1 | 6.56 | 0.87 | 7.17 | 5.29 | 1.88 | 1.88 | 0.14 | 0.14 |
| gene11399 | https://www.ncbi.nlm.nih.gov/gene/?term=LOC105285203 | 1029 | 27 | 262.39 | 1 | 9.72 | 3.7 | 30.13 | 25.46 | 4.67 | 4.67 | 0.12 | 0.12 |
| gene1696 | https://www.ncbi.nlm.nih.gov/gene/?term=LOC105287856 | 1917 | 114 | 594.68 | 6 | 31.3 | 5.26 | 12.21 | 14.28 | -2.07 | 2.07 | -0.12 | 0.12 |
| gene6776 | https://www.ncbi.nlm.nih.gov/gene/?term=LOC105279863 | 1193 | 17 | 142.5 | 4 | 33.53 | 23.53 | 11.89 | 20.13 | -8.24 | 8.24 | -0.12 | 0.12 |
| gene558 | https://www.ncbi.nlm.nih.gov/gene/?term=LOC105278458 | 216 | 3 | 138.89 | 1 | 46.3 | 33.33 | 20.87 | 29.54 | -8.67 | 8.67 | -0.12 | 0.12 |
| gene10363 | https://www.ncbi.nlm.nih.gov/gene/?term=LOC105284009 | 1503 | 26 | 172.99 | 3 | 19.96 | 11.54 | 22.01 | 15.79 | 6.22 | 6.22 | 0.11 | 0.11 |
| gene1802 | https://www.ncbi.nlm.nih.gov/gene/?term=LOC105287981 | 3173 | 131 | 412.86 | 2 | 6.3 | 1.53 | 19.33 | 16.77 | 2.56 | 2.56 | 0.11 | 0.11 |
| gene5379 | https://www.ncbi.nlm.nih.gov/gene/?term=LOC105278234 | 2307 | 119 | 515.82 | 1 | 4.33 | 0.84 | 9.63 | 7.57 | 2.06 | 2.06 | 0.11 | 0.11 |
| gene13384 | https://www.ncbi.nlm.nih.gov/gene/?term=LOC105287447 | 1877 | 144 | 767.18 | 1 | 5.33 | 0.69 | 29.36 | 30.8 | -1.44 | 1.44 | -0.11 | 0.11 |
| gene762 | https://www.ncbi.nlm.nih.gov/gene/?term=LOC105280710 | 1173 | 61 | 520.03 | 1 | 8.53 | 1.64 | 23.4 | 25.56 | -2.17 | 2.17 | -0.11 | 0.11 |
| gene10334 | https://www.ncbi.nlm.nih.gov/gene/?term=LOC105283973 | 1536 | 30 | 195.31 | 1 | 6.51 | 3.33 | 61.25 | 55.99 | 5.26 | 5.26 | 0.1 | 0.1 |
| gene5937 | https://www.ncbi.nlm.nih.gov/gene/?term=LOC105278873 | 1902 | 141 | 741.32 | 4 | 21.03 | 2.84 | 20.24 | 21.57 | -1.32 | 1.32 | -0.1 | 0.1 |
| gene4641 | https://www.ncbi.nlm.nih.gov/gene/?term=LOC105277398 | 1937 | 61 | 314.92 | 5 | 25.81 | 8.2 | 38.08 | 41.19 | -3.11 | 3.11 | -0.1 | 0.1 |
| gene10584 | https://www.ncbi.nlm.nih.gov/gene/?term=LOC105284253 | 1255 | 36 | 286.85 | 2 | 15.94 | 5.56 | 27.9 | 31.27 | -3.36 | 3.36 | -0.1 | 0.1 |
| gene1222 | https://www.ncbi.nlm.nih.gov/gene/?term=LOC105285716 | 1097 | 25 | 227.89 | 2 | 18.23 | 8 | 39.46 | 43.8 | -4.34 | 4.34 | -0.1 | 0.1 |
| gene1179 | https://www.ncbi.nlm.nih.gov/gene/?term=LOC105285207 | 2994 | 60 | 200.4 | 5 | 16.7 | 8.33 | 21.02 | 25.85 | -4.83 | 4.83 | -0.1 | 0.1 |
| gene1534 | https://www.ncbi.nlm.nih.gov/gene/?term=LOC105287662 | 1711 | 31 | 181.18 | 1 | 5.84 | 3.23 | 32.16 | 37.64 | -5.47 | 5.47 | -0.1 | 0.1 |
| gene5006 | https://www.ncbi.nlm.nih.gov/gene/?term=LOC105277810 | 1382 | 21 | 151.95 | 1 | 7.24 | 4.76 | 45.33 | 39.66 | 5.66 | 5.66 | 0.09 | 0.09 |
| gene9280 | https://www.ncbi.nlm.nih.gov/gene/?term=LOC105282764 | 1635 | 35 | 214.07 | 3 | 18.35 | 8.57 | 40.89 | 36.66 | 4.23 | 4.23 | 0.09 | 0.09 |
| gene1589 | https://www.ncbi.nlm.nih.gov/gene/?term=LOC105287721 | 1407 | 39 | 277.19 | 1 | 7.11 | 2.56 | 25.67 | 22.55 | 3.13 | 3.13 | 0.09 | 0.09 |
| gene2824 | https://www.ncbi.nlm.nih.gov/gene/?term=LOC105275337 | 1772 | 54 | 304.74 | 3 | 16.93 | 5.56 | 32.91 | 29.82 | 3.09 | 3.09 | 0.09 | 0.09 |

...

... only first 25 lines shown ...

Directory: OGL File: rnk-dmg-Cb_LI_BCphaseA2.vs.BCphaseB1.txt

| gene_ID | gene_link | gwidth | #Sites | #per10Kb | #dmSites | #dmsp10kb | %dmSites | %pSite1 | %pSite2 | DMpSite | ADMpSite | DMpNucl | ADMpNucl |
|---|---|---|---|---|---|---|---|---|---|---|---|---|---|
| gene10606 | https://www.ncbi.nlm.nih.gov/gene/?term=LOC105284273 | 6119 | 258 | 421.64 | 1 | 1.63 | 0.39 | 19.65 | 18.03 | 1.63 | 1.63 | 0.07 | 0.07 |
| gene2856 | https://www.ncbi.nlm.nih.gov/gene/?term=LOC105275384 | 1987 | 91 | 457.98 | 1 | 5.03 | 1.1 | 33.74 | 32.36 | 1.37 | 1.37 | 0.06 | 0.06 |
| gene9740 | https://www.ncbi.nlm.nih.gov/gene/?term=LOC105283277 | 2887 | 143 | 495.32 | 1 | 3.46 | 0.7 | 19.74 | 18.46 | 1.28 | 1.28 | 0.06 | 0.06 |
| gene3793 | https://www.ncbi.nlm.nih.gov/gene/?term=LOC105276429 | 2901 | 119 | 410.2 | 1 | 3.45 | 0.84 | 23.8 | 22.52 | 1.28 | 1.28 | 0.05 | 0.05 |
| gene7199 | https://www.ncbi.nlm.nih.gov/gene/?term=LOC105280366 | 1414 | 30 | 212.16 | 1 | 7.07 | 3.33 | 43.5 | 41.71 | 1.79 | 1.79 | 0.04 | 0.04 |
| gene570 | https://www.ncbi.nlm.nih.gov/gene/?term=LOC105278549 | 1673 | 70 | 418.41 | 1 | 5.98 | 1.43 | 44.03 | 43.19 | 0.83 | 0.83 | 0.03 | 0.03 |
| gene9722 | https://www.ncbi.nlm.nih.gov/gene/?term=LOC105283264 | 3951 | 88 | 222.73 | 1 | 2.53 | 1.14 | 18.85 | 20.39 | -1.54 | 1.54 | -0.03 | 0.03 |
| gene10844 | https://www.ncbi.nlm.nih.gov/gene/?term=LOC105284535 | 9086 | 246 | 270.75 | 1 | 1.1 | 0.41 | 16.16 | 15.32 | 0.84 | 0.84 | 0.02 | 0.02 |
| gene10326 | https://www.ncbi.nlm.nih.gov/gene/?term=LOC105283962 | 3042 | 130 | 427.35 | 1 | 3.29 | 0.77 | 8.82 | 8.26 | 0.56 | 0.56 | 0.02 | 0.02 |
| gene1855 | https://www.ncbi.nlm.nih.gov/gene/?term=LOC105288036 | 2387 | 89 | 372.85 | 1 | 4.19 | 1.12 | 30.86 | 30.35 | 0.51 | 0.51 | 0.02 | 0.02 |
| gene7166 | https://www.ncbi.nlm.nih.gov/gene/?term=LOC105280326 | 6117 | 235 | 384.18 | 1 | 1.63 | 0.43 | 15.36 | 14.92 | 0.43 | 0.43 | 0.02 | 0.02 |
| gene11447 | https://www.ncbi.nlm.nih.gov/gene/?term=LOC105285252 | 7437 | 131 | 176.15 | 1 | 1.34 | 0.76 | 12.71 | 13.67 | -0.95 | 0.95 | -0.02 | 0.02 |
| gene564 | https://www.ncbi.nlm.nih.gov/gene/?term=LOC105278478 | 2802 | 17 | 60.67 | 1 | 3.57 | 5.88 | 14.91 | 14.04 | 0.87 | 0.87 | 0.01 | 0.01 |
| gene9755 | https://www.ncbi.nlm.nih.gov/gene/?term=LOC105283296 | 2177 | 92 | 422.6 | 1 | 4.59 | 1.09 | 18.09 | 17.95 | 0.14 | 0.14 | 0.01 | 0.01 |
| gene3744 | https://www.ncbi.nlm.nih.gov/gene/?term=LOC105276385 | 9376 | 459 | 489.55 | 1 | 1.07 | 0.22 | 5.46 | 5.7 | -0.24 | 0.24 | -0.01 | 0.01 |
| gene4892 | https://www.ncbi.nlm.nih.gov/gene/?term=LOC105277667 | 3898 | 126 | 323.24 | 1 | 2.57 | 0.79 | 27.95 | 28.19 | -0.24 | 0.24 | -0.01 | 0.01 |
| gene8814 | https://www.ncbi.nlm.nih.gov/gene/?term=LOC105282240 | 6910 | 175 | 253.26 | 1 | 1.45 | 0.57 | 9.54 | 9.87 | -0.32 | 0.32 | -0.01 | 0.01 |
| gene10819 | https://www.ncbi.nlm.nih.gov/gene/?term=LOC105284507 | 5767 | 94 | 163 | 1 | 1.73 | 1.06 | 20.43 | 21.18 | -0.75 | 0.75 | -0.01 | 0.01 |
| gene12203 | https://www.ncbi.nlm.nih.gov/gene/?term=LOC105286099 | 5967 | 84 | 140.77 | 1 | 1.68 | 1.19 | 11.21 | 10.88 | 0.34 | 0.34 | 0 | 0 |
| gene5101 | https://www.ncbi.nlm.nih.gov/gene/?term=LOC105277903 | 1813 | 34 | 187.53 | 1 | 5.52 | 2.94 | 48.71 | 48.5 | 0.21 | 0.21 | 0 | 0 |
| gene7183 | https://www.ncbi.nlm.nih.gov/gene/?term=LOC105280348 | 3925 | 72 | 183.44 | 1 | 2.55 | 1.39 | 20.87 | 20.69 | 0.19 | 0.19 | 0 | 0 |
| gene7245 | https://www.ncbi.nlm.nih.gov/gene/?term=LOC105280410 | 12478 | 277 | 221.99 | 1 | 0.8 | 0.36 | 19.64 | 19.47 | 0.17 | 0.17 | 0 | 0 |

Directory: OGL File: rnk-dmg-Cb_LI_BCphaseA2.vs.BCphaseB2.txt

| gene_ID | gene_link | gwidth | #Sites | #per10Kb | #dmSites | #dmsp10kb | %dmSites | %pSite1 | %pSite2 | DMpSite | ADMpSite | DMpNucl | ADMpNucl |
|---|---|---|---|---|---|---|---|---|---|---|---|---|---|
| gene3165 | https://www.ncbi.nlm.nih.gov/gene/?term=LOC105275706 | 2374 | 67 | 282.22 | 5 | 21.06 | 7.46 | 36.03 | 23.6 | 12.43 | 12.43 | 0.35 | 0.35 |
| gene2824 | https://www.ncbi.nlm.nih.gov/gene/?term=LOC105275337 | 1772 | 64 | 361.17 | 2 | 11.29 | 3.12 | 37.53 | 29.09 | 8.44 | 8.44 | 0.3 | 0.3 |
| gene10963 | https://www.ncbi.nlm.nih.gov/gene/?term=LOC105284677 | 1576 | 28 | 177.66 | 2 | 12.69 | 7.14 | 19.34 | 35.66 | -16.32 | 16.32 | -0.29 | 0.29 |
| gene9468 | https://www.ncbi.nlm.nih.gov/gene/?term=LOC105282976 | 1636 | 53 | 323.96 | 1 | 6.11 | 1.89 | 28.76 | 22.1 | 6.66 | 6.66 | 0.22 | 0.22 |
| gene2593 | https://www.ncbi.nlm.nih.gov/gene/?term=LOC105275070 | 1774 | 5 | 28.18 | 5 | 28.18 | 100 | 80.28 | 10.96 | 69.32 | 69.32 | 0.2 | 0.2 |
| gene1957 | https://www.ncbi.nlm.nih.gov/gene/?term=LOC105288153 | 1044 | 16 | 153.26 | 1 | 9.58 | 6.25 | 15.68 | 4.26 | 11.42 | 11.42 | 0.18 | 0.18 |
| gene5714 | https://www.ncbi.nlm.nih.gov/gene/?term=LOC105278729 | 710 | 24 | 338.03 | 1 | 14.08 | 4.17 | 61.56 | 56.79 | 4.77 | 4.77 | 0.16 | 0.16 |
| gene920 | https://www.ncbi.nlm.nih.gov/gene/?term=LOC105282518 | 1197 | 40 | 334.17 | 1 | 8.35 | 2.5 | 37.69 | 40.21 | -2.52 | 2.52 | -0.08 | 0.08 |
| gene12021 | https://www.ncbi.nlm.nih.gov/gene/?term=LOC105285905 | 1399 | 20 | 142.96 | 1 | 7.15 | 5 | 32.7 | 38.03 | -5.34 | 5.34 | -0.08 | 0.08 |
| gene3817 | https://www.ncbi.nlm.nih.gov/gene/?term=LOC105276449 | 1230 | 42 | 341.46 | 1 | 8.13 | 2.38 | 35.93 | 33.84 | 2.09 | 2.09 | 0.07 | 0.07 |
| gene2404 | https://www.ncbi.nlm.nih.gov/gene/?term=LOC105274853 | 1767 | 36 | 203.74 | 1 | 5.66 | 2.78 | 27.62 | 31.08 | -3.47 | 3.47 | -0.07 | 0.07 |
| gene10816 | https://www.ncbi.nlm.nih.gov/gene/?term=LOC105284502 | 1974 | 18 | 91.19 | 1 | 5.07 | 5.56 | 38.87 | 46.78 | -7.91 | 7.91 | -0.07 | 0.07 |
| gene5877 | https://www.ncbi.nlm.nih.gov/gene/?term=LOC105278807 | 1719 | 49 | 285.05 | 1 | 5.82 | 2.04 | 23.71 | 21.48 | 2.23 | 2.23 | 0.06 | 0.06 |
| gene7556 | https://www.ncbi.nlm.nih.gov/gene/?term=LOC105280774 | 2792 | 102 | 365.33 | 1 | 3.58 | 0.98 | 32.11 | 30.45 | 1.66 | 1.66 | 0.06 | 0.06 |
| gene7568 | https://www.ncbi.nlm.nih.gov/gene/?term=LOC105280786 | 2525 | 111 | 439.6 | 1 | 3.96 | 0.9 | 33.28 | 31.99 | 1.29 | 1.29 | 0.06 | 0.06 |
| gene12290 | https://www.ncbi.nlm.nih.gov/gene/?term=LOC105286190 | 6212 | 304 | 489.38 | 1 | 1.61 | 0.33 | 8.69 | 7.5 | 1.18 | 1.18 | 0.06 | 0.06 |
| gene1220 | https://www.ncbi.nlm.nih.gov/gene/?term=LOC105285742 | 3453 | 108 | 312.77 | 1 | 2.9 | 0.93 | 21.23 | 23.26 | -2.03 | 2.03 | -0.06 | 0.06 |
| gene10606 | https://www.ncbi.nlm.nih.gov/gene/?term=LOC105284273 | 6119 | 251 | 410.2 | 1 | 1.63 | 0.4 | 20.03 | 18.79 | 1.24 | 1.24 | 0.05 | 0.05 |
| gene5298 | https://www.ncbi.nlm.nih.gov/gene/?term=LOC105278145 | 4060 | 244 | 600.99 | 2 | 4.93 | 0.82 | 10.55 | 11.35 | -0.8 | 0.8 | -0.05 | 0.05 |
| gene1993 | https://www.ncbi.nlm.nih.gov/gene/?term=LOC105288193 | 1449 | 75 | 517.6 | 1 | 6.9 | 1.33 | 35.76 | 36.74 | -0.98 | 0.98 | -0.05 | 0.05 |
| gene2271 | https://www.ncbi.nlm.nih.gov/gene/?term=LOC105274706 | 1448 | 56 | 386.74 | 1 | 6.91 | 1.79 | 36.89 | 38.13 | -1.24 | 1.24 | -0.05 | 0.05 |
| gene1199 | https://www.ncbi.nlm.nih.gov/gene/?term=LOC105285498 | 2220 | 26 | 117.12 | 1 | 4.5 | 3.85 | 33.05 | 37.03 | -3.99 | 3.99 | -0.05 | 0.05 |
| gene3385 | https://www.ncbi.nlm.nih.gov/gene/?term=LOC105275966 | 3305 | 61 | 184.57 | 1 | 3.03 | 1.64 | 49.58 | 47.17 | 2.41 | 2.41 | 0.04 | 0.04 |
| gene9462 | https://www.ncbi.nlm.nih.gov/gene/?term=LOC105282970 | 1469 | 39 | 265.49 | 1 | 6.81 | 2.56 | 53.94 | 52.44 | 1.5 | 1.5 | 0.04 | 0.04 |
| gene4892 | https://www.ncbi.nlm.nih.gov/gene/?term=LOC105277667 | 3898 | 125 | 320.68 | 1 | 2.57 | 0.8 | 27.74 | 26.47 | 1.27 | 1.27 | 0.04 | 0.04 |
| gene10678 | https://www.ncbi.nlm.nih.gov/gene/?term=LOC105284355 | 4031 | 167 | 414.29 | 1 | 2.48 | 0.6 | 19.59 | 18.65 | 0.94 | 0.94 | 0.04 | 0.04 |
| gene4034 | https://www.ncbi.nlm.nih.gov/gene/?term=LOC105276704 | 2065 | 101 | 489.1 | 1 | 4.84 | 0.99 | 13.4 | 12.57 | 0.83 | 0.83 | 0.04 | 0.04 |
| gene4498 | https://www.ncbi.nlm.nih.gov/gene/?term=LOC105277239 | 2831 | 129 | 455.67 | 1 | 3.53 | 0.78 | 14.44 | 15.34 | -0.89 | 0.89 | -0.04 | 0.04 |
| gene11340 | https://www.ncbi.nlm.nih.gov/gene/?term=LOC105285131 | 2918 | 101 | 346.13 | 1 | 3.43 | 0.99 | 10.9 | 11.93 | -1.02 | 1.02 | -0.04 | 0.04 |
| gene1518 | https://www.ncbi.nlm.nih.gov/gene/?term=LOC105287648 | 2195 | 27 | 123.01 | 1 | 4.56 | 3.7 | 41.41 | 38.92 | 2.49 | 2.49 | 0.03 | 0.03 |
| gene8115 | https://www.ncbi.nlm.nih.gov/gene/?term=LOC105281420 | 3623 | 51 | 140.77 | 1 | 2.76 | 1.96 | 12.6 | 10.51 | 2.09 | 2.09 | 0.03 | 0.03 |
| gene8742 | https://www.ncbi.nlm.nih.gov/gene/?term=LOC105282151 | 2563 | 41 | 159.97 | 2 | 7.8 | 4.88 | 36.1 | 34.31 | 1.79 | 1.79 | 0.03 | 0.03 |
| gene12572 | https://www.ncbi.nlm.nih.gov/gene/?term=LOC105286514 | 2374 | 42 | 176.92 | 1 | 4.21 | 2.38 | 34.39 | 32.92 | 1.47 | 1.47 | 0.03 | 0.03 |
| gene5967 | https://www.ncbi.nlm.nih.gov/gene/?term=LOC105278918 | 4393 | 121 | 275.44 | 1 | 2.28 | 0.83 | 15.97 | 14.91 | 1.05 | 1.05 | 0.03 | 0.03 |
| gene8729 | https://www.ncbi.nlm.nih.gov/gene/?term=LOC105282137 | 5005 | 197 | 393.61 | 1 | 2 | 0.51 | 17.44 | 16.65 | 0.8 | 0.8 | 0.03 | 0.03 |
| gene10520 | https://www.ncbi.nlm.nih.gov/gene/?term=LOC105284184 | 2451 | 85 | 346.8 | 1 | 4.08 | 1.18 | 34.37 | 33.59 | 0.78 | 0.78 | 0.03 | 0.03 |
| gene8274 | https://www.ncbi.nlm.nih.gov/gene/?term=LOC105281624 | 5425 | 230 | 423.96 | 1 | 1.84 | 0.43 | 9.61 | 10.33 | -0.73 | 0.73 | -0.03 | 0.03 |
| gene10018 | https://www.ncbi.nlm.nih.gov/gene/?term=LOC105283607 | 4414 | 191 | 432.71 | 1 | 2.27 | 0.52 | 15.56 | 16.32 | -0.76 | 0.76 | -0.03 | 0.03 |
| gene10213 | https://www.ncbi.nlm.nih.gov/gene/?term=LOC105283826 | 1503 | 64 | 425.82 | 1 | 6.65 | 1.56 | 14.04 | 14.85 | -0.81 | 0.81 | -0.03 | 0.03 |
| gene5646 | https://www.ncbi.nlm.nih.gov/gene/?term=LOC105278534 | 7475 | 181 | 242.14 | 1 | 1.34 | 0.55 | 17.19 | 18.52 | -1.32 | 1.32 | -0.03 | 0.03 |
| gene11447 | https://www.ncbi.nlm.nih.gov/gene/?term=LOC105285252 | 7437 | 120 | 161.36 | 1 | 1.34 | 0.83 | 13.34 | 14.97 | -1.63 | 1.63 | -0.03 | 0.03 |
| gene6455 | https://www.ncbi.nlm.nih.gov/gene/?term=LOC105279500 | 2612 | 33 | 126.34 | 1 | 3.83 | 3.03 | 45.07 | 47.06 | -1.99 | 1.99 | -0.03 | 0.03 |
| gene2748 | https://www.ncbi.nlm.nih.gov/gene/?term=LOC105275261 | 3845 | 52 | 135.24 | 1 | 2.6 | 1.92 | 36.08 | 38.33 | -2.25 | 2.25 | -0.03 | 0.03 |
| gene12395 | https://www.ncbi.nlm.nih.gov/gene/?term=LOC105286302 | 4093 | 17 | 41.53 | 1 | 2.44 | 5.88 | 21.99 | 16.58 | 5.41 | 5.41 | 0.02 | 0.02 |
| gene5052 | https://www.ncbi.nlm.nih.gov/gene/?term=LOC105277856 | 3167 | 34 | 107.36 | 1 | 3.16 | 2.94 | 38.79 | 36.66 | 2.13 | 2.13 | 0.02 | 0.02 |
| gene2970 | https://www.ncbi.nlm.nih.gov/gene/?term=LOC105275518 | 2340 | 19 | 81.2 | 1 | 4.27 | 5.26 | 46.39 | 44.36 | 2.03 | 2.03 | 0.02 | 0.02 |
| gene7398 | https://www.ncbi.nlm.nih.gov/gene/?term=LOC105280589 | 3452 | 39 | 112.98 | 1 | 2.9 | 2.56 | 23.26 | 21.67 | 1.58 | 1.58 | 0.02 | 0.02 |
| gene927 | https://www.ncbi.nlm.nih.gov/gene/?term=LOC105282600 | 3727 | 53 | 142.21 | 1 | 2.68 | 1.89 | 32.11 | 30.67 | 1.44 | 1.44 | 0.02 | 0.02 |
| gene1884 | https://www.ncbi.nlm.nih.gov/gene/?term=LOC105288072 | 4169 | 71 | 170.3 | 1 | 2.4 | 1.41 | 25.5 | 24.39 | 1.1 | 1.1 | 0.02 | 0.02 |
| gene3470 | https://www.ncbi.nlm.nih.gov/gene/?term=LOC105276059 | 8716 | 145 | 166.36 | 1 | 1.15 | 0.69 | 10.01 | 8.94 | 1.08 | 1.08 | 0.02 | 0.02 |
| gene940 | https://www.ncbi.nlm.nih.gov/gene/?term=LOC105282794 | 3578 | 85 | 237.56 | 1 | 2.79 | 1.18 | 16.1 | 15.1 | 1 | 1 | 0.02 | 0.02 |
| gene13033 | https://www.ncbi.nlm.nih.gov/gene/?term=LOC105287063 | 1877 | 49 | 261.05 | 1 | 5.33 | 2.04 | 31.12 | 30.35 | 0.77 | 0.77 | 0.02 | 0.02 |
| gene9336 | https://www.ncbi.nlm.nih.gov/gene/?term=LOC105282834 | 5993 | 124 | 206.91 | 1 | 1.67 | 0.81 | 16.06 | 15.3 | 0.75 | 0.75 | 0.02 | 0.02 |
| gene12516 | https://www.ncbi.nlm.nih.gov/gene/?term=LOC105286452 | 1529 | 38 | 248.53 | 1 | 6.54 | 2.63 | 52.34 | 51.61 | 0.73 | 0.73 | 0.02 | 0.02 |
| gene4376 | https://www.ncbi.nlm.nih.gov/gene/?term=LOC105277083 | 4307 | 127 | 294.87 | 1 | 2.32 | 0.79 | 19.77 | 19.11 | 0.65 | 0.65 | 0.02 | 0.02 |
| gene7514 | https://www.ncbi.nlm.nih.gov/gene/?term=LOC105280723 | 4471 | 200 | 447.33 | 1 | 2.24 | 0.5 | 14.49 | 14.01 | 0.49 | 0.49 | 0.02 | 0.02 |
| gene2056 | https://www.ncbi.nlm.nih.gov/gene/?term=LOC105274452 | 6584 | 263 | 399.45 | 2 | 3.04 | 0.76 | 13.53 | 13.06 | 0.47 | 0.47 | 0.02 | 0.02 |
| gene3286 | https://www.ncbi.nlm.nih.gov/gene/?term=LOC105275848 | 3744 | 140 | 373.93 | 1 | 2.67 | 0.71 | 23 | 22.56 | 0.45 | 0.45 | 0.02 | 0.02 |
| gene5746 | https://www.ncbi.nlm.nih.gov/gene/?term=LOC105278666 | 10343 | 461 | 445.71 | 1 | 0.97 | 0.22 | 7.31 | 6.93 | 0.37 | 0.37 | 0.02 | 0.02 |
| gene1470 | https://www.ncbi.nlm.nih.gov/gene/?term=LOC105287597 | 2974 | 204 | 685.94 | 1 | 3.36 | 0.49 | 17.73 | 17.44 | 0.3 | 0.3 | 0.02 | 0.02 |

| | | | | | | | | | | | | | |
|---|---|---|---|---|---|---|---|---|---|---|---|---|---|
| gene4720 | https://www.ncbi.nlm.nih.gov/gene/?term=LOC105277479 | 1697 | 47 | 276.96 | 1 | 5.89 | 2.13 | 37.56 | 38.12 | -0.55 | 0.55 | -0.02 | 0.02 |
| gene9826 | https://www.ncbi.nlm.nih.gov/gene/?term=LOC105283385 | 9219 | 305 | 330.84 | 1 | 1.08 | 0.33 | 12.6 | 13.19 | -0.59 | 0.59 | -0.02 | 0.02 |
| gene6966 | https://www.ncbi.nlm.nih.gov/gene/?term=LOC105280104 | 3747 | 122 | 325.59 | 1 | 2.67 | 0.82 | 15.27 | 15.88 | -0.62 | 0.62 | -0.02 | 0.02 |
| gene1147 | https://www.ncbi.nlm.nih.gov/gene/?term=LOC105284954 | 4022 | 95 | 236.2 1 | 2.49 | 1.05 | 15.81 | 16.65 | -0.84 | 0.84 | -0.02 | 0.02 | |
| gene7915 | https://www.ncbi.nlm.nih.gov/gene/?term=LOC105281173 | 1597 | 38 | 237.95 | 1 | 6.26 | 2.63 | 59.94 | 60.79 | -0.85 | 0.85 | -0.02 | 0.02 |
| gene2413 | https://www.ncbi.nlm.nih.gov/gene/?term=LOC105274863 | 2006 | 44 | 219.34 | 1 | 4.99 | 2.27 | 63.14 | 64.02 | -0.87 | 0.87 | -0.02 | 0.02 |
| gene12406 | https://www.ncbi.nlm.nih.gov/gene/?term=LOC105286316 | 1386 | 28 | 202.02 | 1 | 7.22 | 3.57 | 47.22 | 48.43 | -1.22 | 1.22 | -0.02 | 0.02 |
| gene322 | https://www.ncbi.nlm.nih.gov/gene/?term=LOC105275777 | 1606 | 24 | 149.44 | 1 | 6.23 | 4.17 | 45 | 46.35 | -1.35 | 1.35 | -0.02 | 0.02 |
| gene388 | https://www.ncbi.nlm.nih.gov/gene/?term=LOC105276538 | 1558 | 20 | 128.37 | 2 | 12.84 | 10 | 64.73 | 66.14 | -1.41 | 1.41 | -0.02 | 0.02 |
| gene6422 | https://www.ncbi.nlm.nih.gov/gene/?term=LOC105279458 | 2419 | 34 | 140.55 | 1 | 4.13 | 2.94 | 51.48 | 53.11 | -1.63 | 1.63 | -0.02 | 0.02 |
| gene10819 | https://www.ncbi.nlm.nih.gov/gene/?term=LOC105284507 | 5767 | 77 | 133.52 | 1 | 1.73 | 1.3 | 20.73 | 22.56 | -1.84 | 1.84 | -0.02 | 0.02 |
| gene6528 | https://www.ncbi.nlm.nih.gov/gene/?term=LOC105279591 | 7222 | 55 | 76.16 1 | 1.38 | 1.82 | 29.5 | 28.12 | 1.38 | 1.38 | 0.01 | 0.01 | |
| gene2825 | https://www.ncbi.nlm.nih.gov/gene/?term=LOC105275343 | 5545 | 96 | 173.13 | 2 | 3.61 | 2.08 | 11.79 | 11.04 | 0.75 | 0.75 | 0.01 | 0.01 |
| gene6620 | https://www.ncbi.nlm.nih.gov/gene/?term=LOC105279687 | 17998 | 336 | 186.69 | 1 | 0.56 | 0.3 | 5.15 | 4.42 | 0.73 | 0.73 | 0.01 | 0.01 |
| gene12307 | https://www.ncbi.nlm.nih.gov/gene/?term=LOC105286216 | 10313 | 166 | 160.96 | 1 | 0.97 | 0.6 | 16.86 | 16.32 | 0.54 | 0.54 | 0.01 | 0.01 |
| gene7368 | https://www.ncbi.nlm.nih.gov/gene/?term=LOC105280556 | 4105 | 50 | 121.8 1 | 2.44 | 2 | 18.71 | 18.17 | 0.54 | 0.54 | 0.01 | 0.01 | |
| gene292 | https://www.ncbi.nlm.nih.gov/gene/?term=LOC105275398 | 2825 | 86 | 304.42 | 1 | 3.54 | 1.16 | 21.94 | 21.53 | 0.4 | 0.4 | 0.01 | 0.01 |
| gene4962 | https://www.ncbi.nlm.nih.gov/gene/?term=LOC105277761 | 4645 | 150 | 322.93 | 1 | 2.15 | 0.67 | 15.92 | 15.52 | 0.4 | 0.4 | 0.01 | 0.01 |
| gene7245 | https://www.ncbi.nlm.nih.gov/gene/?term=LOC105280410 | 12478 | 237 | 189.93 | 1 | 0.8 | 0.42 | 21.04 | 20.64 | 0.4 | 0.4 | 0.01 | 0.01 |
| gene2173 | https://www.ncbi.nlm.nih.gov/gene/?term=LOC105274590 | 4387 | 64 | 145.89 | 2 | 4.56 | 3.12 | 34.09 | 33.7 | 0.39 | 0.39 | 0.01 | 0.01 |
| gene6031 | https://www.ncbi.nlm.nih.gov/gene/?term=LOC105279000 | 5691 | 195 | 342.65 | 1 | 1.76 | 0.51 | 14.39 | 14 | 0.39 | 0.39 | 0.01 | 0.01 |
| gene1430 | https://www.ncbi.nlm.nih.gov/gene/?term=LOC105287551 | 6433 | 231 | 359.09 | 1 | 1.55 | 0.43 | 9.26 | 8.92 | 0.34 | 0.34 | 0.01 | 0.01 |
| gene10531 | https://www.ncbi.nlm.nih.gov/gene/?term=LOC105284190 | 6881 | 173 | 251.42 | 1 | 1.45 | 0.58 | 12.82 | 12.5 | 0.32 | 0.32 | 0.01 | 0.01 |
| gene1310 | https://www.ncbi.nlm.nih.gov/gene/?term=LOC105286661 | 2582 | 66 | 255.62 | 1 | 3.87 | 1.52 | 24.45 | 24.13 | 0.32 | 0.32 | 0.01 | 0.01 |
| gene4246 | https://www.ncbi.nlm.nih.gov/gene/?term=LOC105276941 | 16412 | 552 | 336.34 | 1 | 0.61 | 0.18 | 5.12 | 4.8 | 0.32 | 0.32 | 0.01 | 0.01 |
| gene11568 | https://www.ncbi.nlm.nih.gov/gene/?term=LOC105285395 | 8086 | 192 | 237.45 | 1 | 1.24 | 0.52 | 15.07 | 14.77 | 0.31 | 0.31 | 0.01 | 0.01 |
| gene3856 | https://www.ncbi.nlm.nih.gov/gene/?term=LOC105276499 | 5236 | 190 | 362.87 | 1 | 1.91 | 0.53 | 9.53 | 9.24 | 0.29 | 0.29 | 0.01 | 0.01 |
| gene7249 | https://www.ncbi.nlm.nih.gov/gene/?term=LOC105280422 | 1775 | 63 | 354.93 | 1 | 5.63 | 1.59 | 17.96 | 17.74 | 0.23 | 0.23 | 0.01 | 0.01 |
| gene2806 | https://www.ncbi.nlm.nih.gov/gene/?term=LOC105275327 | 14257 | 494 | 346.5 1 | 0.7 | 0.2 | 6.88 | 6.68 | 0.2 | 0.2 | 0.01 | 0.01 | |
| gene11275 | https://www.ncbi.nlm.nih.gov/gene/?term=LOC105285055 | 4119 | 255 | 619.08 | 1 | 2.43 | 0.39 | 11.36 | 11.16 | 0.19 | 0.19 | 0.01 | 0.01 |
| gene2846 | https://www.ncbi.nlm.nih.gov/gene/?term=LOC105275374 | 11246 | 502 | 446.38 | 1 | 0.89 | 0.2 | 11.4 | 11.21 | 0.19 | 0.19 | 0.01 | 0.01 |
| gene7166 | https://www.ncbi.nlm.nih.gov/gene/?term=LOC105280326 | 6117 | 233 | 380.91 | 1 | 1.63 | 0.43 | 15.64 | 15.46 | 0.19 | 0.19 | 0.01 | 0.01 |
| gene11401 | https://www.ncbi.nlm.nih.gov/gene/?term=LOC105285202 | 1841 | 82 | 445.41 | 1 | 5.43 | 1.22 | 25.44 | 25.28 | 0.15 | 0.15 | 0.01 | 0.01 |
| gene12806 | https://www.ncbi.nlm.nih.gov/gene/?term=LOC105286803 | 4196 | 143 | 340.8 1 | 2.38 | 0.7 | 11.07 | 10.92 | 0.15 | 0.15 | 0.01 | 0.01 | |
| gene5524 | https://www.ncbi.nlm.nih.gov/gene/?term=LOC105278397 | 5177 | 203 | 392.12 | 1 | 1.93 | 0.49 | 25.25 | 25.12 | 0.13 | 0.13 | 0.01 | 0.01 |
| gene12802 | https://www.ncbi.nlm.nih.gov/gene/?term=LOC105286799 | 3183 | 185 | 581.21 | 2 | 6.28 | 1.08 | 16.06 | 15.94 | 0.12 | 0.12 | 0.01 | 0.01 |
| gene12674 | https://www.ncbi.nlm.nih.gov/gene/?term=LOC105286649 | 18283 | 811 | 443.58 | 1 | 0.55 | 0.12 | 0.93 | 1.06 | -0.13 | 0.13 | -0.01 | 0.01 |
| gene13114 | https://www.ncbi.nlm.nih.gov/gene/?term=LOC105287147 | 7609 | 409 | 537.52 | 1 | 1.31 | 0.24 | 9.57 | 9.7 | -0.13 | 0.13 | -0.01 | 0.01 |
| gene3885 | https://www.ncbi.nlm.nih.gov/gene/?term=LOC105276527 | 7928 | 317 | 399.85 | 1 | 1.26 | 0.32 | 9.68 | 9.86 | -0.18 | 0.18 | -0.01 | 0.01 |
| gene5777 | https://www.ncbi.nlm.nih.gov/gene/?term=LOC105278691 | 3883 | 148 | 381.15 | 1 | 2.58 | 0.68 | 14.2 | 14.38 | -0.18 | 0.18 | -0.01 | 0.01 |
| gene6583 | https://www.ncbi.nlm.nih.gov/gene/?term=LOC105279649 | 7444 | 332 | 446 1 | 1.34 | 0.3 | 10.76 | 10.98 | -0.22 | 0.22 | -0.01 | 0.01 | |
| gene11454 | https://www.ncbi.nlm.nih.gov/gene/?term=LOC105285256 | 1906 | 47 | 246.59 | 1 | 5.25 | 2.13 | 45.38 | 45.61 | -0.23 | 0.23 | -0.01 | 0.01 |
| gene13036 | https://www.ncbi.nlm.nih.gov/gene/?term=LOC105287059 | 4959 | 163 | 328.7 1 | 2.02 | 0.61 | 16.39 | 16.65 | -0.26 | 0.26 | -0.01 | 0.01 | |
| gene3727 | https://www.ncbi.nlm.nih.gov/gene/?term=LOC105276367 | 2484 | 69 | 277.78 | 1 | 4.03 | 1.45 | 46.73 | 46.99 | -0.27 | 0.27 | -0.01 | 0.01 |
| gene2257 | https://www.ncbi.nlm.nih.gov/gene/?term=LOC105274689 | 3658 | 71 | 194.1 1 | 2.73 | 1.41 | 14.85 | 15.18 | -0.34 | 0.34 | -0.01 | 0.01 | |
| gene7600 | https://www.ncbi.nlm.nih.gov/gene/?term=LOC105280819 | 1615 | 39 | 241.49 | 1 | 6.19 | 2.56 | 35.76 | 36.1 | -0.34 | 0.34 | -0.01 | 0.01 |
| gene11082 | https://www.ncbi.nlm.nih.gov/gene/?term=LOC105284824 | 7436 | 200 | 268.96 | 1 | 1.34 | 0.5 | 5.46 | 5.81 | -0.35 | 0.35 | -0.01 | 0.01 |
| gene12573 | https://www.ncbi.nlm.nih.gov/gene/?term=LOC105286515 | 3647 | 113 | 309.84 | 1 | 2.74 | 0.88 | 10.7 | 11.07 | -0.37 | 0.37 | -0.01 | 0.01 |
| gene10600 | https://www.ncbi.nlm.nih.gov/gene/?term=LOC105284270 | 4837 | 124 | 256.36 | 1 | 2.07 | 0.81 | 10.47 | 10.85 | -0.38 | 0.38 | -0.01 | 0.01 |
| gene3046 | https://www.ncbi.nlm.nih.gov/gene/?term=LOC105275587 | 2394 | 56 | 233.92 | 1 | 4.18 | 1.79 | 33.82 | 34.2 | -0.38 | 0.38 | -0.01 | 0.01 |
| gene12981 | https://www.ncbi.nlm.nih.gov/gene/?term=LOC105286999 | 2063 | 40 | 193.89 | 1 | 4.85 | 2.5 | 35.8 | 36.25 | -0.45 | 0.45 | -0.01 | 0.01 |
| gene10107 | https://www.ncbi.nlm.nih.gov/gene/?term=LOC105283712 | 1883 | 32 | 169.94 | 1 | 5.31 | 3.12 | 29.98 | 30.44 | -0.46 | 0.46 | -0.01 | 0.01 |
| gene11561 | https://www.ncbi.nlm.nih.gov/gene/?term=LOC105285387 | 11528 | 194 | 168.29 | 1 | 0.87 | 0.52 | 8.65 | 9.11 | -0.47 | 0.47 | -0.01 | 0.01 |
| gene8015 | https://www.ncbi.nlm.nih.gov/gene/?term=LOC105281312 | 2998 | 72 | 240.16 | 1 | 3.34 | 1.39 | 29.67 | 30.18 | -0.5 | 0.5 | -0.01 | 0.01 |
| gene8814 | https://www.ncbi.nlm.nih.gov/gene/?term=LOC105282240 | 6910 | 170 | 246.02 | 1 | 1.45 | 0.59 | 9.16 | 9.68 | -0.52 | 0.52 | -0.01 | 0.01 |
| gene10391 | https://www.ncbi.nlm.nih.gov/gene/?term=LOC109611283 | 1994 | 44 | 220.66 | 1 | 5.02 | 2.27 | 32.96 | 33.52 | -0.56 | 0.56 | -0.01 | 0.01 |
| gene3671 | https://www.ncbi.nlm.nih.gov/gene/?term=LOC105276272 | 1966 | 36 | 183.11 | 1 | 5.09 | 2.78 | 32.08 | 32.66 | -0.58 | 0.58 | -0.01 | 0.01 |
| gene6639 | https://www.ncbi.nlm.nih.gov/gene/?term=LOC105279705 | 7656 | 62 | 80.98 1 | 1.31 | 1.61 | 22.96 | 23.88 | -0.92 | 0.92 | -0.01 | 0.01 | |
| gene10345 | https://www.ncbi.nlm.nih.gov/gene/?term=LOC105283989 | 4444 | 23 | 51.76 1 | 2.25 | 4.35 | 43.82 | 45.07 | -1.26 | 1.26 | -0.01 | 0.01 | |
| gene3168 | https://www.ncbi.nlm.nih.gov/gene/?term=LOC105275711 | 6150 | 70 | 113.82 | 1 | 1.63 | 1.43 | 12.53 | 13.84 | -1.31 | 1.31 | -0.01 | 0.01 |
| gene10654 | https://www.ncbi.nlm.nih.gov/gene/?term=LOC105284329 | 4027 | 50 | 124.16 | 1 | 2.48 | 2 | 19.81 | 19.43 | 0.39 | 0.39 | 0 | 0 |
| gene3599 | https://www.ncbi.nlm.nih.gov/gene/?term=LOC105276196 | 2955 | 48 | 162.44 | 1 | 3.38 | 2.08 | 45.56 | 45.28 | 0.28 | 0.28 | 0 | 0 |
| gene8600 | https://www.ncbi.nlm.nih.gov/gene/?term=LOC105282001 | 8513 | 156 | 183.25 | 1 | 1.17 | 0.64 | 12.04 | 11.82 | 0.22 | 0.22 | 0 | 0 |
| gene12439 | https://www.ncbi.nlm.nih.gov/gene/?term=LOC105286357 | 4761 | 81 | 170.13 | 1 | 2.1 | 1.23 | 32.58 | 32.4 | 0.18 | 0.18 | 0 | 0 |

| | | | | | | | | | | | | | |
|---|---|---|---|---|---|---|---|---|---|---|---|---|---|
| gene11579 | https://www.ncbi.nlm.nih.gov/gene/?term=LOC105285406 | 13571 | 267 | 196.74 | 1 | 0.74 | 0.37 | 6.68 | 6.51 | 0.17 | 0.17 | 0 | 0 |
| gene8343 | https://www.ncbi.nlm.nih.gov/gene/?term=LOC105281700 | 2172 | 37 | 170.35 | 1 | 4.6 | 2.7 | 39.71 | 39.54 | 0.16 | 0.16 | 0 | 0 |
| gene12203 | https://www.ncbi.nlm.nih.gov/gene/?term=LOC105286099 | 5967 | 72 | 120.66 | 1 | 1.68 | 1.39 | 10.74 | 10.59 | 0.15 | 0.15 | 0 | 0 |
| gene4190 | https://www.ncbi.nlm.nih.gov/gene/?term=LOC105276875 | 3150 | 107 | 339.68 | 1 | 3.17 | 0.93 | 23.8 | 23.66 | 0.15 | 0.15 | 0 | 0 |
| gene1198 | https://www.ncbi.nlm.nih.gov/gene/?term=LOC105285507 | 2738 | 88 | 321.4 1 | 3.65 | 1.14 | 23.48 | 23.34 | 0.14 | 0.14 | 0 | 0 | |
| gene10392 | https://www.ncbi.nlm.nih.gov/gene/?term=LOC105284036 | 3216 | 94 | 292.29 | 2 | 6.22 | 2.13 | 22.55 | 22.51 | 0.04 | 0.04 | 0 | 0 |
| gene53 | https://www.ncbi.nlm.nih.gov/gene/?term=LOC105277148 | 19283 | 287 | 148.84 | 1 | 0.52 | 0.35 | 5.29 | 5.27 | 0.03 | 0.03 | 0 | 0 |
| gene6286 | https://www.ncbi.nlm.nih.gov/gene/?term=LOC105279298 | 2861 | 125 | 436.91 | 2 | 6.99 | 1.6 | 25.22 | 25.19 | 0.03 | 0.03 | 0 | 0 |
| gene13044 | https://www.ncbi.nlm.nih.gov/gene/?term=LOC105287070 | 5094 | 241 | 473.11 | 1 | 1.96 | 0.41 | 5.28 | 5.27 | 0.02 | 0.02 | 0 | 0 |
| gene2844 | https://www.ncbi.nlm.nih.gov/gene/?term=LOC105275363 | 4330 | 192 | 443.42 | 1 | 2.31 | 0.52 | 14.25 | 14.23 | 0.02 | 0.02 | 0 | 0 |
| gene5525 | https://www.ncbi.nlm.nih.gov/gene/?term=LOC105278396 | 3090 | 188 | 608.41 | 1 | 3.24 | 0.53 | 24.8 | 24.79 | 0.01 | 0.01 | 0 | 0 |
| gene9330 | https://www.ncbi.nlm.nih.gov/gene/?term=LOC105282825 | 3936 | 80 | 203.25 | 1 | 2.54 | 1.25 | 12.57 | 12.58 | -0.01 | 0.01 | 0 | 0 |
| gene9755 | https://www.ncbi.nlm.nih.gov/gene/?term=LOC105283296 | 2177 | 85 | 390.45 | 1 | 4.59 | 1.18 | 21.44 | 21.45 | -0.01 | 0.01 | 0 | 0 |
| gene298 | https://www.ncbi.nlm.nih.gov/gene/?term=LOC105275460 | 2473 | 114 | 460.98 | 1 | 4.04 | 0.88 | 13.89 | 13.91 | -0.02 | 0.02 | 0 | 0 |
| gene10549 | https://www.ncbi.nlm.nih.gov/gene/?term=LOC105284210 | 7145 | 325 | 454.86 | 2 | 2.8 | 0.62 | 10.71 | 10.76 | -0.05 | 0.05 | 0 | 0 |
| gene3848 | https://www.ncbi.nlm.nih.gov/gene/?term=LOC105276486 | 5899 | 133 | 225.46 | 1 | 1.7 | 0.75 | 24.81 | 24.86 | -0.05 | 0.05 | 0 | 0 |
| gene9177 | https://www.ncbi.nlm.nih.gov/gene/?term=LOC105282646 | 3320 | 98 | 295.18 | 1 | 3.01 | 1.02 | 22.82 | 22.88 | -0.05 | 0.05 | 0 | 0 |
| gene4641 | https://www.ncbi.nlm.nih.gov/gene/?term=LOC105277398 | 1937 | 77 | 397.52 | 1 | 5.16 | 1.3 | 34.01 | 34.07 | -0.06 | 0.06 | 0 | 0 |
| gene11688 | https://www.ncbi.nlm.nih.gov/gene/?term=LOC105285533 | 1637 | 60 | 366.52 | 1 | 6.11 | 1.67 | 26.05 | 26.13 | -0.07 | 0.07 | 0 | 0 |
| gene3107 | https://www.ncbi.nlm.nih.gov/gene/?term=LOC105275650 | 5103 | 181 | 354.69 | 1 | 1.96 | 0.55 | 15.45 | 15.53 | -0.08 | 0.08 | 0 | 0 |
| gene5831 | https://www.ncbi.nlm.nih.gov/gene/?term=LOC105278761 | 2506 | 119 | 474.86 | 2 | 7.98 | 1.68 | 21.61 | 21.68 | -0.08 | 0.08 | 0 | 0 |
| gene9445 | https://www.ncbi.nlm.nih.gov/gene/?term=LOC105282951 | 2285 | 115 | 503.28 | 1 | 4.38 | 0.87 | 20.19 | 20.28 | -0.09 | 0.09 | 0 | 0 |
| gene6652 | https://www.ncbi.nlm.nih.gov/gene/?term=LOC105279719 | 2831 | 79 | 279.05 | 1 | 3.53 | 1.27 | 24.78 | 24.91 | -0.14 | 0.14 | 0 | 0 |
| gene1632 | https://www.ncbi.nlm.nih.gov/gene/?term=LOC105287764 | 18061 | 393 | 217.6 2 | 1.11 | 0.51 | 2.97 | 3.13 | -0.17 | 0.17 | 0 | 0 | |
| gene8661 | https://www.ncbi.nlm.nih.gov/gene/?term=LOC105282063 | 1926 | 32 | 166.15 | 1 | 5.19 | 3.12 | 29.26 | 29.43 | -0.17 | 0.17 | 0 | 0 |
| gene5375 | https://www.ncbi.nlm.nih.gov/gene/?term=LOC105278228 | 8640 | 163 | 188.66 | 1 | 1.16 | 0.61 | 21.78 | 22.01 | -0.23 | 0.23 | 0 | 0 |
| gene6237 | https://www.ncbi.nlm.nih.gov/gene/?term=LOC105279244 | 1484 | 24 | 161.73 | 1 | 6.74 | 4.17 | 52.96 | 53.25 | -0.29 | 0.29 | 0 | 0 |
| gene2683 | https://www.ncbi.nlm.nih.gov/gene/?term=LOC105275187 | 5492 | 70 | 127.46 | 1 | 1.82 | 1.43 | 14.08 | 14.4 | -0.31 | 0.31 | 0 | 0 |
| gene10155 | https://www.ncbi.nlm.nih.gov/gene/?term=LOC105283756 | 7030 | 93 | 132.29 | 1 | 1.42 | 1.08 | 5.63 | 5.96 | -0.34 | 0.34 | 0 | 0 |

Directory: OGL File: rnk-dmg-Cb_LI_BCphaseA2.vs.RphaseA1.txt

| gene_ID | gene_link | gwidth | #Sites | #per10Kb | #dmSites | #dmsp10kb | %dmSites | %pSite1 | %pSite2 | DMpSite | ADMpSite | DMpNucl | ADMpNucl |
|---|---|---|---|---|---|---|---|---|---|---|---|---|---|
| gene558 | https://www.ncbi.nlm.nih.gov/gene/?term=LOC105278458 | 216 | 3 | 138.89 | 1 | 46.3 | 33.33 | 10.57 | 24.69 | -14.12 | 14.12 | -0.2 | 0.2 |
| gene12538 | https://www.ncbi.nlm.nih.gov/gene/?term=LOC105286531 | 372 | 23 | 618.28 | 1 | 26.88 | 4.35 | 39.65 | 36.5 | 3.15 | 3.15 | 0.19 | 0.19 |
| gene11566 | https://www.ncbi.nlm.nih.gov/gene/?term=LOC105285392 | 1603 | 82 | 511.54 | 3 | 18.71 | 3.66 | 38.11 | 34.95 | 3.16 | 3.16 | 0.16 | 0.16 |
| gene2837 | https://www.ncbi.nlm.nih.gov/gene/?term=LOC105275355 | 2872 | 154 | 536.21 | 7 | 24.37 | 4.55 | 26.21 | 29.04 | -2.82 | 2.82 | -0.15 | 0.15 |
| gene10731 | https://www.ncbi.nlm.nih.gov/gene/?term=LOC105284411 | 1794 | 63 | 351.17 | 4 | 22.3 | 6.35 | 36.85 | 33 | 3.85 | 3.85 | 0.14 | 0.14 |
| gene12411 | https://www.ncbi.nlm.nih.gov/gene/?term=LOC105286320 | 1660 | 60 | 361.45 | 4 | 24.1 | 6.67 | 41.91 | 38.27 | 3.64 | 3.64 | 0.13 | 0.13 |
| gene4635 | https://www.ncbi.nlm.nih.gov/gene/?term=LOC105277390 | 2067 | 81 | 391.87 | 3 | 14.51 | 3.7 | 45.5 | 42.07 | 3.43 | 3.43 | 0.13 | 0.13 |
| gene3268 | https://www.ncbi.nlm.nih.gov/gene/?term=LOC105275833 | 1040 | 41 | 394.23 | 2 | 19.23 | 4.88 | 38.46 | 35.12 | 3.35 | 3.35 | 0.13 | 0.13 |
| gene12925 | https://www.ncbi.nlm.nih.gov/gene/?term=LOC109611487 | 391 | 27 | 690.54 | 1 | 25.58 | 3.7 | 17.14 | 15.18 | 1.95 | 1.95 | 0.13 | 0.13 |
| gene1024 | https://www.ncbi.nlm.nih.gov/gene/?term=LOC105283693 | 1307 | 12 | 91.81 | 4 | 30.6 | 33.33 | 71.36 | 58.1 | 13.26 | 13.26 | 0.12 | 0.12 |
| gene12927 | https://www.ncbi.nlm.nih.gov/gene/?term=LOC105286946 | 1231 | 49 | 398.05 | 3 | 24.37 | 6.12 | 34.73 | 31.65 | 3.08 | 3.08 | 0.12 | 0.12 |
| gene5447 | https://www.ncbi.nlm.nih.gov/gene/?term=LOC105278314 | 1482 | 85 | 573.55 | 2 | 13.5 | 2.35 | 25.47 | 23.3 | 2.17 | 2.17 | 0.12 | 0.12 |
| gene3910 | https://www.ncbi.nlm.nih.gov/gene/?term=LOC105276563 | 1957 | 115 | 587.63 | 6 | 30.66 | 5.22 | 18.73 | 16.62 | 2.11 | 2.11 | 0.12 | 0.12 |
| gene5937 | https://www.ncbi.nlm.nih.gov/gene/?term=LOC105278873 | 1902 | 146 | 767.61 | 6 | 31.55 | 4.11 | 21.31 | 19.68 | 1.63 | 1.63 | 0.12 | 0.12 |
| gene2855 | https://www.ncbi.nlm.nih.gov/gene/?term=LOC105275378 | 1189 | 20 | 168.21 | 1 | 8.41 | 5 | 32.85 | 26.65 | 6.2 | 6.2 | 0.1 | 0.1 |
| gene4708 | https://www.ncbi.nlm.nih.gov/gene/?term=LOC105277468 | 1712 | 31 | 181.07 | 3 | 17.52 | 9.68 | 40.08 | 34.7 | 5.39 | 5.39 | 0.1 | 0.1 |
| gene4013 | https://www.ncbi.nlm.nih.gov/gene/?term=LOC105276679 | 2007 | 64 | 318.88 | 1 | 4.98 | 1.56 | 39.86 | 36.65 | 3.22 | 3.22 | 0.1 | 0.1 |
| gene10112 | https://www.ncbi.nlm.nih.gov/gene/?term=LOC105283717 | 1483 | 80 | 539.45 | 5 | 33.72 | 6.25 | 29.96 | 28.17 | 1.79 | 1.79 | 0.1 | 0.1 |
| gene13157 | https://www.ncbi.nlm.nih.gov/gene/?term=LOC105287195 | 837 | 21 | 250.9 | 1 | 11.95 | 4.76 | 20.59 | 24.57 | -3.98 | 3.98 | -0.1 | 0.1 |
| gene3015 | https://www.ncbi.nlm.nih.gov/gene/?term=LOC105275557 | 811 | 11 | 135.64 | 1 | 12.33 | 9.09 | 74.6 | 67.82 | 6.78 | 6.78 | 0.09 | 0.09 |
| gene6974 | https://www.ncbi.nlm.nih.gov/gene/?term=LOC105280111 | 1003 | 21 | 209.37 | 1 | 9.97 | 4.76 | 44.89 | 40.66 | 4.23 | 4.23 | 0.09 | 0.09 |
| gene11109 | https://www.ncbi.nlm.nih.gov/gene/?term=LOC105284855 | 1786 | 42 | 235.16 | 2 | 11.2 | 4.76 | 43.39 | 39.73 | 3.66 | 3.66 | 0.09 | 0.09 |
| gene7588 | https://www.ncbi.nlm.nih.gov/gene/?term=LOC105280808 | 1612 | 43 | 266.75 | 1 | 6.2 | 2.33 | 30.22 | 26.92 | 3.3 | 3.3 | 0.09 | 0.09 |
| gene11934 | https://www.ncbi.nlm.nih.gov/gene/?term=LOC105285812 | 1594 | 46 | 288.58 | 1 | 6.27 | 2.17 | 22.97 | 19.95 | 3.02 | 3.02 | 0.09 | 0.09 |

...

... only first 25 lines shown ...

Directory: OGL File: rnk-dmg-Cb_LI_BCphaseA2.vs.RphaseA2.txt

| gene_ID | gene_link | gwidth | #Sites | #per10Kb | #dmSites | #dmsp10kb | %dmSites | %pSite1 | %pSite2 | DMpSite | ADMpSite | DMpNucl | ADMpNucl |
|---|---|---|---|---|---|---|---|---|---|---|---|---|---|
| gene12733 | https://www.ncbi.nlm.nih.gov/gene/?term=LOC109611472 | 1351 | 27 | 199.85 | 2 | 14.8 | 7.41 | 32.9 | 39.85 | -6.95 | 6.95 | -0.14 | 0.14 |
| gene10743 | https://www.ncbi.nlm.nih.gov/gene/?term=LOC105284456 | 1028 | 43 | 418.29 | 3 | 29.18 | 6.98 | 44.49 | 41.39 | 3.1 | 3.1 | 0.13 | 0.13 |
| gene9121 | https://www.ncbi.nlm.nih.gov/gene/?term=LOC105282572 | 771 | 35 | 453.96 | 1 | 12.97 | 2.86 | 29.82 | 32.76 | -2.94 | 2.94 | -0.13 | 0.13 |
| gene12311 | https://www.ncbi.nlm.nih.gov/gene/?term=LOC105286212 | 1005 | 36 | 358.21 | 1 | 9.95 | 2.78 | 37.91 | 41.67 | -3.76 | 3.76 | -0.13 | 0.13 |
| gene1171 | https://www.ncbi.nlm.nih.gov/gene/?term=LOC105285174 | 844 | 39 | 462.09 | 1 | 11.85 | 2.56 | 36.65 | 39.21 | -2.56 | 2.56 | -0.12 | 0.12 |
| gene1562 | https://www.ncbi.nlm.nih.gov/gene/?term=LOC105287692 | 1491 | 98 | 657.28 | 3 | 20.12 | 3.06 | 28.39 | 30.05 | -1.66 | 1.66 | -0.11 | 0.11 |
| gene4227 | https://www.ncbi.nlm.nih.gov/gene/?term=LOC105276920 | 2587 | 95 | 367.22 | 1 | 3.87 | 1.05 | 19.9 | 22.92 | -3.01 | 3.01 | -0.11 | 0.11 |
| gene25 | https://www.ncbi.nlm.nih.gov/gene/?term=LOC105274524 | 1118 | 38 | 339.89 | 1 | 8.94 | 2.63 | 28.99 | 32.2 | -3.2 | 3.2 | -0.11 | 0.11 |
| gene11980 | https://www.ncbi.nlm.nih.gov/gene/?term=LOC109611395 | 571 | 5 | 87.57 | 1 | 17.51 | 20 | 33.3 | 45.97 | -12.67 | 12.67 | -0.11 | 0.11 |
| gene10237 | https://www.ncbi.nlm.nih.gov/gene/?term=LOC105283856 | 1259 | 26 | 206.51 | 1 | 7.94 | 3.85 | 64.95 | 60.12 | 4.82 | 4.82 | 0.1 | 0.1 |
| gene10731 | https://www.ncbi.nlm.nih.gov/gene/?term=LOC105284411 | 1794 | 63 | 351.17 | 1 | 5.57 | 1.59 | 35.73 | 32.92 | 2.81 | 2.81 | 0.1 | 0.1 |
| gene10742 | https://www.ncbi.nlm.nih.gov/gene/?term=LOC109611309 | 776 | 37 | 476.8 | 2 | 25.77 | 5.41 | 42.09 | 40.01 | 2.09 | 2.09 | 0.1 | 0.1 |
| gene3719 | https://www.ncbi.nlm.nih.gov/gene/?term=LOC105276360 | 1471 | 108 | 734.19 | 1 | 6.8 | 0.93 | 29.44 | 30.83 | -1.39 | 1.39 | -0.1 | 0.1 |
| gene5912 | https://www.ncbi.nlm.nih.gov/gene/?term=LOC105278847 | 2089 | 125 | 598.37 | 1 | 4.79 | 0.8 | 8.99 | 10.64 | -1.65 | 1.65 | -0.1 | 0.1 |
| gene3787 | https://www.ncbi.nlm.nih.gov/gene/?term=LOC105276423 | 1679 | 77 | 458.61 | 1 | 5.96 | 1.3 | 16.64 | 18.77 | -2.13 | 2.13 | -0.1 | 0.1 |
| gene7767 | https://www.ncbi.nlm.nih.gov/gene/?term=LOC105281009 | 1509 | 40 | 265.08 | 3 | 19.88 | 7.5 | 53.17 | 56.81 | -3.65 | 3.65 | -0.1 | 0.1 |
| gene3165 | https://www.ncbi.nlm.nih.gov/gene/?term=LOC105275706 | 2374 | 82 | 345.41 | 3 | 12.64 | 3.66 | 33.31 | 30.79 | 2.52 | 2.52 | 0.09 | 0.09 |
| gene9183 | https://www.ncbi.nlm.nih.gov/gene/?term=LOC105282640 | 2752 | 211 | 766.72 | 2 | 7.27 | 0.95 | 21.27 | 22.49 | -1.22 | 1.22 | -0.09 | 0.09 |
| gene2790 | https://www.ncbi.nlm.nih.gov/gene/?term=LOC105275307 | 1429 | 54 | 377.89 | 2 | 14 | 3.7 | 39.26 | 41.56 | -2.3 | 2.3 | -0.09 | 0.09 |
| gene4313 | https://www.ncbi.nlm.nih.gov/gene/?term=LOC105277012 | 1491 | 52 | 348.76 | 3 | 20.12 | 5.77 | 24.71 | 27.22 | -2.51 | 2.51 | -0.09 | 0.09 |
| gene10720 | https://www.ncbi.nlm.nih.gov/gene/?term=LOC105284400 | 1914 | 44 | 229.89 | 3 | 15.67 | 6.82 | 27.76 | 31.46 | -3.7 | 3.7 | -0.09 | 0.09 |
| gene10721 | https://www.ncbi.nlm.nih.gov/gene/?term=LOC105284403 | 1537 | 29 | 188.68 | 2 | 13.01 | 6.9 | 50.07 | 54.88 | -4.81 | 4.81 | -0.09 | 0.09 |
| gene11521 | https://www.ncbi.nlm.nih.gov/gene/?term=LOC105285336 | 476 | 7 | 147.06 | 1 | 21.01 | 14.29 | 45.58 | 51.56 | -5.98 | 5.98 | -0.09 | 0.09 |
| gene7810 | https://www.ncbi.nlm.nih.gov/gene/?term=LOC105281055 | 1333 | 50 | 375.09 | 1 | 7.5 | 2 | 26.8 | 24.6 | 2.2 | 2.2 | 0.08 | 0.08 |

...
... only first 25 lines shown ...

Directory: OGL File: rnk-dmg-Cb_LI_BCphaseA2.vs.RphaseB1.txt

| gene_ID | gene_link | gwidth | #Sites | #per10Kb | #dmSites | #dmsp10kb | %dmSites | %pSite1 | %pSite2 | DMpSite | ADMpSite | DMpNucl | ADMpNucl |
|---|---|---|---|---|---|---|---|---|---|---|---|---|---|
| gene3165 | https://www.ncbi.nlm.nih.gov/gene/?term=LOC105275706 | 2374 | 67 | 282.22 | 6 | 25.27 | 8.96 | 35.39 | 24.21 | 11.18 | 11.18 | 0.32 | 0.32 |
| gene2824 | https://www.ncbi.nlm.nih.gov/gene/?term=LOC105275337 | 1772 | 62 | 349.89 | 2 | 11.29 | 3.23 | 38.77 | 30.45 | 8.31 | 8.31 | 0.29 | 0.29 |
| gene2593 | https://www.ncbi.nlm.nih.gov/gene/?term=LOC105275070 | 1774 | 9 | 50.73 | 4 | 22.55 | 44.44 | 77.93 | 29.94 | 47.99 | 47.99 | 0.24 | 0.24 |
| gene1993 | https://www.ncbi.nlm.nih.gov/gene/?term=LOC105288193 | 1449 | 75 | 517.6 | 1 | 6.9 | 1.33 | 35.76 | 32.82 | 2.94 | 2.94 | 0.15 | 0.15 |
| gene5714 | https://www.ncbi.nlm.nih.gov/gene/?term=LOC105278729 | 710 | 16 | 225.35 | 1 | 14.08 | 6.25 | 57.91 | 51.9 | 6.01 | 6.01 | 0.14 | 0.14 |
| gene2404 | https://www.ncbi.nlm.nih.gov/gene/?term=LOC105274853 | 1767 | 34 | 192.42 | 1 | 5.66 | 2.94 | 29.24 | 33.79 | -4.55 | 4.55 | -0.09 | 0.09 |
| gene4034 | https://www.ncbi.nlm.nih.gov/gene/?term=LOC105276704 | 2065 | 86 | 416.46 | 1 | 4.84 | 1.16 | 15.38 | 13.53 | 1.85 | 1.85 | 0.08 | 0.08 |
| gene2413 | https://www.ncbi.nlm.nih.gov/gene/?term=LOC105274863 | 2006 | 46 | 229.31 | 1 | 4.99 | 2.17 | 60.76 | 64.28 | -3.53 | 3.53 | -0.08 | 0.08 |
| gene13033 | https://www.ncbi.nlm.nih.gov/gene/?term=LOC105287063 | 1877 | 49 | 261.05 | 1 | 5.33 | 2.04 | 32.47 | 29.74 | 2.73 | 2.73 | 0.07 | 0.07 |
| gene10984 | https://www.ncbi.nlm.nih.gov/gene/?term=LOC105284704 | 1987 | 62 | 312.03 | 1 | 5.03 | 1.61 | 26.65 | 24.35 | 2.3 | 2.3 | 0.07 | 0.07 |
| gene7556 | https://www.ncbi.nlm.nih.gov/gene/?term=LOC105280774 | 2792 | 101 | 361.75 | 1 | 3.58 | 0.99 | 32.38 | 30.39 | 1.99 | 1.99 | 0.07 | 0.07 |
| gene10520 | https://www.ncbi.nlm.nih.gov/gene/?term=LOC105284184 | 2451 | 87 | 354.96 | 1 | 4.08 | 1.15 | 34.29 | 32.67 | 1.61 | 1.61 | 0.06 | 0.06 |
| gene6583 | https://www.ncbi.nlm.nih.gov/gene/?term=LOC105279649 | 7444 | 343 | 460.77 | 1 | 1.34 | 0.29 | 10.51 | 11.78 | -1.26 | 1.26 | -0.06 | 0.06 |
| gene920 | https://www.ncbi.nlm.nih.gov/gene/?term=LOC105282518 | 1197 | 39 | 325.81 | 1 | 8.35 | 2.56 | 36.44 | 38.28 | -1.85 | 1.85 | -0.06 | 0.06 |
| gene12021 | https://www.ncbi.nlm.nih.gov/gene/?term=LOC105285905 | 1399 | 27 | 192.99 | 1 | 7.15 | 3.7 | 31.19 | 34.36 | -3.17 | 3.17 | -0.06 | 0.06 |
| gene6652 | https://www.ncbi.nlm.nih.gov/gene/?term=LOC105279719 | 2831 | 78 | 275.52 | 1 | 3.53 | 1.28 | 25.09 | 23.41 | 1.68 | 1.68 | 0.05 | 0.05 |
| gene10606 | https://www.ncbi.nlm.nih.gov/gene/?term=LOC105284273 | 6119 | 260 | 424.91 | 1 | 1.63 | 0.38 | 19.55 | 18.4 | 1.15 | 1.15 | 0.05 | 0.05 |
| gene7568 | https://www.ncbi.nlm.nih.gov/gene/?term=LOC105280786 | 2525 | 122 | 483.17 | 1 | 3.96 | 0.82 | 30.88 | 29.81 | 1.07 | 1.07 | 0.05 | 0.05 |
| gene2835 | https://www.ncbi.nlm.nih.gov/gene/?term=LOC105275357 | 2492 | 114 | 457.46 | 1 | 4.01 | 0.88 | 13.76 | 14.82 | -1.05 | 1.05 | -0.05 | 0.05 |
| gene1220 | https://www.ncbi.nlm.nih.gov/gene/?term=LOC105285742 | 3453 | 98 | 283.81 | 1 | 2.9 | 1.02 | 19.04 | 20.94 | -1.9 | 1.9 | -0.05 | 0.05 |
| gene10552 | https://www.ncbi.nlm.nih.gov/gene/?term=LOC105284213 | 878 | 15 | 170.84 | 1 | 11.39 | 6.67 | 15.38 | 18.37 | -2.99 | 2.99 | -0.05 | 0.05 |
| gene393 | https://www.ncbi.nlm.nih.gov/gene/?term=LOC105276566 | 1123 | 2 | 17.81 | 1 | 8.9 | 50 | 48.08 | 74.3 | -26.22 | 26.22 | -0.05 | 0.05 |
| gene3671 | https://www.ncbi.nlm.nih.gov/gene/?term=LOC105276272 | 1966 | 38 | 193.29 | 1 | 5.09 | 2.63 | 30.65 | 28.5 | 2.15 | 2.15 | 0.04 | 0.04 |
| gene10091 | https://www.ncbi.nlm.nih.gov/gene/?term=LOC105283697 | 4988 | 137 | 274.66 | 1 | 2 | 0.73 | 17.98 | 16.41 | 1.57 | 1.57 | 0.04 | 0.04 |
| gene5877 | https://www.ncbi.nlm.nih.gov/gene/?term=LOC105278807 | 1719 | 49 | 285.05 | 1 | 5.82 | 2.04 | 25.27 | 23.74 | 1.53 | 1.53 | 0.04 | 0.04 |
| gene7888 | https://www.ncbi.nlm.nih.gov/gene/?term=LOC105281141 | 4357 | 157 | 360.34 | 1 | 2.3 | 0.64 | 31.42 | 30.25 | 1.17 | 1.17 | 0.04 | 0.04 |
| gene3817 | https://www.ncbi.nlm.nih.gov/gene/?term=LOC105276449 | 1230 | 43 | 349.59 | 1 | 8.13 | 2.33 | 30.78 | 29.68 | 1.1 | 1.1 | 0.04 | 0.04 |
| gene5831 | https://www.ncbi.nlm.nih.gov/gene/?term=LOC105278761 | 2506 | 112 | 446.93 | 2 | 7.98 | 1.79 | 21.67 | 20.75 | 0.92 | 0.92 | 0.04 | 0.04 |
| gene11340 | https://www.ncbi.nlm.nih.gov/gene/?term=LOC105285131 | 2918 | 89 | 305 | 1 | 3.43 | 1.12 | 11.73 | 13.01 | -1.28 | 1.28 | -0.04 | 0.04 |
| gene3335 | https://www.ncbi.nlm.nih.gov/gene/?term=LOC105275898 | 4665 | 147 | 315.11 | 1 | 2.14 | 0.68 | 11.48 | 12.76 | -1.29 | 1.29 | -0.04 | 0.04 |
| gene10879 | https://www.ncbi.nlm.nih.gov/gene/?term=LOC105284574 | 2423 | 45 | 185.72 | 1 | 4.13 | 2.22 | 57.53 | 59.68 | -2.15 | 2.15 | -0.04 | 0.04 |
| gene1199 | https://www.ncbi.nlm.nih.gov/gene/?term=LOC105285498 | 2220 | 25 | 112.61 | 1 | 4.5 | 4 | 32.49 | 35.74 | -3.26 | 3.26 | -0.04 | 0.04 |
| gene3166 | https://www.ncbi.nlm.nih.gov/gene/?term=LOC105275707 | 2393 | 11 | 45.97 | 1 | 4.18 | 9.09 | 62.4 | 71.24 | -8.83 | 8.83 | -0.04 | 0.04 |
| gene9109 | https://www.ncbi.nlm.nih.gov/gene/?term=LOC105282561 | 2400 | 11 | 45.83 | 1 | 4.17 | 9.09 | 17.27 | 26.23 | -8.97 | 8.97 | -0.04 | 0.04 |
| gene9336 | https://www.ncbi.nlm.nih.gov/gene/?term=LOC105282834 | 5993 | 124 | 206.91 | 1 | 1.67 | 0.81 | 15.53 | 14.01 | 1.52 | 1.52 | 0.03 | 0.03 |
| gene7734 | https://www.ncbi.nlm.nih.gov/gene/?term=LOC105280972 | 1528 | 32 | 209.42 | 1 | 6.54 | 3.12 | 19.4 | 18.05 | 1.35 | 1.35 | 0.03 | 0.03 |
| gene940 | https://www.ncbi.nlm.nih.gov/gene/?term=LOC105282794 | 3578 | 86 | 240.36 | 1 | 2.79 | 1.16 | 16.54 | 15.48 | 1.06 | 1.06 | 0.03 | 0.03 |
| gene7249 | https://www.ncbi.nlm.nih.gov/gene/?term=LOC105280422 | 1775 | 62 | 349.3 | 1 | 5.63 | 1.61 | 17.51 | 16.57 | 0.94 | 0.94 | 0.03 | 0.03 |
| gene6286 | https://www.ncbi.nlm.nih.gov/gene/?term=LOC105279298 | 2861 | 121 | 422.93 | 2 | 6.99 | 1.65 | 24.64 | 23.9 | 0.74 | 0.74 | 0.03 | 0.03 |
| gene6031 | https://www.ncbi.nlm.nih.gov/gene/?term=LOC105279000 | 5691 | 208 | 365.49 | 1 | 1.76 | 0.48 | 12.77 | 12.08 | 0.69 | 0.69 | 0.03 | 0.03 |
| gene5298 | https://www.ncbi.nlm.nih.gov/gene/?term=LOC105278145 | 4060 | 251 | 618.23 | 2 | 4.93 | 0.8 | 10.52 | 10.93 | -0.42 | 0.42 | -0.03 | 0.03 |
| gene13114 | https://www.ncbi.nlm.nih.gov/gene/?term=LOC105287147 | 7609 | 398 | 523.06 | 1 | 1.31 | 0.25 | 9.41 | 9.93 | -0.52 | 0.52 | -0.03 | 0.03 |
| gene5581 | https://www.ncbi.nlm.nih.gov/gene/?term=LOC105278459 | 2935 | 97 | 330.49 | 1 | 3.41 | 1.03 | 20.44 | 21.26 | -0.83 | 0.83 | -0.03 | 0.03 |
| gene6966 | https://www.ncbi.nlm.nih.gov/gene/?term=LOC105280104 | 3747 | 124 | 330.93 | 1 | 2.67 | 0.81 | 16.15 | 17.09 | -0.94 | 0.94 | -0.03 | 0.03 |
| gene7915 | https://www.ncbi.nlm.nih.gov/gene/?term=LOC105281173 | 1597 | 39 | 244.21 | 1 | 6.26 | 2.56 | 60.85 | 62.01 | -1.16 | 1.16 | -0.03 | 0.03 |
| gene3848 | https://www.ncbi.nlm.nih.gov/gene/?term=LOC105276486 | 5899 | 145 | 245.8 | 1 | 1.7 | 0.69 | 23.24 | 24.56 | -1.32 | 1.32 | -0.03 | 0.03 |
| gene7207 | https://www.ncbi.nlm.nih.gov/gene/?term=LOC105280376 | 4659 | 112 | 240.39 | 1 | 2.15 | 0.89 | 28.49 | 29.89 | -1.4 | 1.4 | -0.03 | 0.03 |
| gene10819 | https://www.ncbi.nlm.nih.gov/gene/?term=LOC105284507 | 5767 | 86 | 149.12 | 1 | 1.73 | 1.16 | 21.64 | 23.53 | -1.89 | 1.89 | -0.03 | 0.03 |
| gene7398 | https://www.ncbi.nlm.nih.gov/gene/?term=LOC105280589 | 3452 | 42 | 121.67 | 1 | 2.9 | 2.38 | 27.89 | 25.91 | 1.98 | 1.98 | 0.02 | 0.02 |
| gene4502 | https://www.ncbi.nlm.nih.gov/gene/?term=LOC105277243 | 2289 | 32 | 139.8 | 1 | 4.37 | 3.12 | 44.88 | 43.35 | 1.53 | 1.53 | 0.02 | 0.02 |
| gene6455 | https://www.ncbi.nlm.nih.gov/gene/?term=LOC105279500 | 2612 | 33 | 126.34 | 1 | 3.83 | 3.03 | 51.99 | 50.58 | 1.41 | 1.41 | 0.02 | 0.02 |
| gene10107 | https://www.ncbi.nlm.nih.gov/gene/?term=LOC105283712 | 1883 | 30 | 159.32 | 1 | 5.31 | 3.33 | 31.98 | 30.87 | 1.11 | 1.11 | 0.02 | 0.02 |
| gene1147 | https://www.ncbi.nlm.nih.gov/gene/?term=LOC105284954 | 4022 | 96 | 238.69 | 1 | 2.49 | 1.04 | 15.76 | 15.07 | 0.69 | 0.69 | 0.02 | 0.02 |
| gene8015 | https://www.ncbi.nlm.nih.gov/gene/?term=LOC105281312 | 2998 | 68 | 226.82 | 1 | 3.34 | 1.47 | 31.47 | 30.8 | 0.67 | 0.67 | 0.02 | 0.02 |
| gene292 | https://www.ncbi.nlm.nih.gov/gene/?term=LOC105275398 | 2825 | 92 | 325.66 | 1 | 3.54 | 1.09 | 23.29 | 22.65 | 0.64 | 0.64 | 0.02 | 0.02 |
| gene1310 | https://www.ncbi.nlm.nih.gov/gene/?term=LOC105286661 | 2582 | 71 | 274.98 | 1 | 3.87 | 1.41 | 21.34 | 20.74 | 0.61 | 0.61 | 0.02 | 0.02 |
| gene10678 | https://www.ncbi.nlm.nih.gov/gene/?term=LOC105284355 | 4031 | 161 | 399.4 | 1 | 2.48 | 0.62 | 20.49 | 19.95 | 0.54 | 0.54 | 0.02 | 0.02 |
| gene4892 | https://www.ncbi.nlm.nih.gov/gene/?term=LOC105277667 | 3898 | 123 | 315.55 | 1 | 2.57 | 0.81 | 28.28 | 27.77 | 0.51 | 0.51 | 0.02 | 0.02 |
| gene298 | https://www.ncbi.nlm.nih.gov/gene/?term=LOC105275460 | 2473 | 118 | 477.15 | 1 | 4.04 | 0.85 | 13.46 | 12.97 | 0.49 | 0.49 | 0.02 | 0.02 |
| gene8729 | https://www.ncbi.nlm.nih.gov/gene/?term=LOC105282137 | 5005 | 211 | 421.58 | 1 | 2 | 0.47 | 16 | 15.6 | 0.4 | 0.4 | 0.02 | 0.02 |

| | | | | | | | | | | | | | |
|---|---|---|---|---|---|---|---|---|---|---|---|---|---|
| gene3747 | https://www.ncbi.nlm.nih.gov/gene/?term=LOC105276387 | 3736 | 188 | 503.21 | 1 | 2.68 | 0.53 | 14.43 | 14.09 | 0.35 | 0.35 | 0.02 | 0.02 |
| gene1470 | https://www.ncbi.nlm.nih.gov/gene/?term=LOC105287597 | 2974 | 192 | 645.6 1 | 3.36 | 0.52 | 18.97 | 19.22 | -0.25 | 0.25 | -0.02 | 0.02 | |
| gene1571 | https://www.ncbi.nlm.nih.gov/gene/?term=LOC105287700 | 3392 | 160 | 471.7 1 | 2.95 | 0.62 | 15.1 | 15.46 | -0.36 | 0.36 | -0.02 | 0.02 | |
| gene4962 | https://www.ncbi.nlm.nih.gov/gene/?term=LOC105277761 | 4645 | 161 | 346.61 | 1 | 2.15 | 0.62 | 14.28 | 14.77 | -0.48 | 0.48 | -0.02 | 0.02 |
| gene8274 | https://www.ncbi.nlm.nih.gov/gene/?term=LOC105281624 | 5425 | 221 | 407.37 | 1 | 1.84 | 0.45 | 9.57 | 10.1 | -0.54 | 0.54 | -0.02 | 0.02 |
| gene3107 | https://www.ncbi.nlm.nih.gov/gene/?term=LOC105275650 | 5103 | 172 | 337.06 | 1 | 1.96 | 0.58 | 16.37 | 16.93 | -0.56 | 0.56 | -0.02 | 0.02 |
| gene12573 | https://www.ncbi.nlm.nih.gov/gene/?term=LOC105286515 | 3647 | 108 | 296.13 | 1 | 2.74 | 0.93 | 11.17 | 11.84 | -0.67 | 0.67 | -0.02 | 0.02 |
| gene1198 | https://www.ncbi.nlm.nih.gov/gene/?term=LOC105285507 | 2738 | 92 | 336.01 | 1 | 3.65 | 1.09 | 20.74 | 21.44 | -0.7 | 0.7 | -0.02 | 0.02 |
| gene11454 | https://www.ncbi.nlm.nih.gov/gene/?term=LOC105285256 | 1906 | 48 | 251.84 | 1 | 5.25 | 2.08 | 44.93 | 45.73 | -0.8 | 0.8 | -0.02 | 0.02 |
| gene8814 | https://www.ncbi.nlm.nih.gov/gene/?term=LOC105282240 | 6910 | 166 | 240.23 | 1 | 1.45 | 0.6 | 9.27 | 10.23 | -0.96 | 0.96 | -0.02 | 0.02 |
| gene11447 | https://www.ncbi.nlm.nih.gov/gene/?term=LOC105285252 | 7437 | 126 | 169.42 | 2 | 2.69 | 1.59 | 12.66 | 13.85 | -1.19 | 1.19 | -0.02 | 0.02 |
| gene10155 | https://www.ncbi.nlm.nih.gov/gene/?term=LOC105283756 | 7030 | 98 | 139.4 1 | 1.42 | 1.02 | 6.09 | 7.87 | -1.78 | 1.78 | -0.02 | 0.02 | |
| gene3168 | https://www.ncbi.nlm.nih.gov/gene/?term=LOC105275711 | 6150 | 80 | 130.08 | 1 | 1.63 | 1.25 | 11.3 | 13.17 | -1.86 | 1.86 | -0.02 | 0.02 |
| gene388 | https://www.ncbi.nlm.nih.gov/gene/?term=LOC105276538 | 1558 | 19 | 121.95 | 2 | 12.84 | 10.53 | 66.07 | 68.07 | -1.99 | 1.99 | -0.02 | 0.02 |
| gene11584 | https://www.ncbi.nlm.nih.gov/gene/?term=LOC105285416 | 3709 | 6 | 16.18 2 | 5.39 | 33.33 | 18.09 | 32.8 | -14.71 | | 14.71 | -0.02 | 0.02 |
| gene2833 | https://www.ncbi.nlm.nih.gov/gene/?term=LOC105275352 | 5528 | 38 | 68.74 1 | 1.81 | 2.63 | 25.27 | 24 | 1.27 | 1.27 | 0.01 | 0.01 | |
| gene2970 | https://www.ncbi.nlm.nih.gov/gene/?term=LOC105275518 | 2340 | 19 | 81.2 1 | 4.27 | 5.26 | 51.65 | 50.52 | 1.13 | 1.13 | 0.01 | 0.01 | |
| gene10222 | https://www.ncbi.nlm.nih.gov/gene/?term=LOC105283838 | 1748 | 21 | 120.14 | 1 | 5.72 | 4.76 | 46.37 | 45.47 | 0.9 | 0.9 | 0.01 | 0.01 |
| gene322 | https://www.ncbi.nlm.nih.gov/gene/?term=LOC105275777 | 1606 | 23 | 143.21 | 1 | 6.23 | 4.35 | 44.64 | 43.87 | 0.76 | 0.76 | 0.01 | 0.01 |
| gene3912 | https://www.ncbi.nlm.nih.gov/gene/?term=LOC105276560 | 17900 | 309 | 172.63 | 1 | 0.56 | 0.32 | 2.52 | 1.81 | 0.7 | 0.7 | 0.01 | 0.01 |
| gene7657 | https://www.ncbi.nlm.nih.gov/gene/?term=LOC105280890 | 1109 | 21 | 189.36 | 1 | 9.02 | 4.76 | 27.2 | 26.57 | 0.64 | 0.64 | 0.01 | 0.01 |
| gene12572 | https://www.ncbi.nlm.nih.gov/gene/?term=LOC105286514 | 2374 | 44 | 185.34 | 1 | 4.21 | 2.27 | 33.89 | 33.38 | 0.52 | 0.52 | 0.01 | 0.01 |
| gene289 | https://www.ncbi.nlm.nih.gov/gene/?term=LOC105275347 | 2533 | 72 | 284.25 | 1 | 3.95 | 1.39 | 24.33 | 23.84 | 0.48 | 0.48 | 0.01 | 0.01 |
| gene11568 | https://www.ncbi.nlm.nih.gov/gene/?term=LOC105285395 | 8086 | 202 | 249.81 | 1 | 1.24 | 0.5 | 14.4 | 13.94 | 0.46 | 0.46 | 0.01 | 0.01 |
| gene2056 | https://www.ncbi.nlm.nih.gov/gene/?term=LOC105274452 | 6584 | 236 | 358.44 | 1 | 1.52 | 0.42 | 14.57 | 14.16 | 0.42 | 0.42 | 0.01 | 0.01 |
| gene3470 | https://www.ncbi.nlm.nih.gov/gene/?term=LOC105276059 | 8716 | 151 | 173.24 | 1 | 1.15 | 0.66 | 10.19 | 9.77 | 0.42 | 0.42 | 0.01 | 0.01 |
| gene7245 | https://www.ncbi.nlm.nih.gov/gene/?term=LOC105280410 | 12478 | 264 | 211.57 | 1 | 0.8 | 0.38 | 19.99 | 19.61 | 0.38 | 0.38 | 0.01 | 0.01 |
| gene8600 | https://www.ncbi.nlm.nih.gov/gene/?term=LOC105282001 | 8513 | 152 | 178.55 | 1 | 1.17 | 0.66 | 12.95 | 12.59 | 0.36 | 0.36 | 0.01 | 0.01 |
| gene6237 | https://www.ncbi.nlm.nih.gov/gene/?term=LOC105279244 | 1484 | 27 | 181.94 | 1 | 6.74 | 3.7 | 50.23 | 49.93 | 0.29 | 0.29 | 0.01 | 0.01 |
| gene2846 | https://www.ncbi.nlm.nih.gov/gene/?term=LOC105275374 | 11246 | 500 | 444.6 1 | 0.89 | 0.2 | 10.21 | 9.93 | 0.28 | 0.28 | 0.01 | 0.01 | |
| gene3727 | https://www.ncbi.nlm.nih.gov/gene/?term=LOC105276367 | 2484 | 68 | 273.75 | 1 | 4.03 | 1.47 | 46.01 | 45.74 | 0.27 | 0.27 | 0.01 | 0.01 |
| gene5746 | https://www.ncbi.nlm.nih.gov/gene/?term=LOC105278666 | 10343 | 455 | 439.91 | 1 | 0.97 | 0.22 | 7.39 | 7.14 | 0.25 | 0.25 | 0.01 | 0.01 |
| gene11819 | https://www.ncbi.nlm.nih.gov/gene/?term=LOC105285671 | 2366 | 98 | 414.2 1 | 4.23 | 1.02 | 30.44 | 30.22 | 0.23 | 0.23 | 0.01 | 0.01 | |
| gene9755 | https://www.ncbi.nlm.nih.gov/gene/?term=LOC105283296 | 2177 | 87 | 399.63 | 1 | 4.59 | 1.15 | 15.89 | 15.67 | 0.23 | 0.23 | 0.01 | 0.01 |
| gene474 | https://www.ncbi.nlm.nih.gov/gene/?term=LOC105277435 | 3625 | 173 | 477.24 | 1 | 2.76 | 0.58 | 10.83 | 10.64 | 0.2 | 0.2 | 0.01 | 0.01 |
| gene11275 | https://www.ncbi.nlm.nih.gov/gene/?term=LOC105285055 | 4119 | 239 | 580.24 | 1 | 2.43 | 0.42 | 11.8 | 11.69 | 0.12 | 0.12 | 0.01 | 0.01 |
| gene5525 | https://www.ncbi.nlm.nih.gov/gene/?term=LOC105278396 | 3090 | 186 | 601.94 | 1 | 3.24 | 0.54 | 24.32 | 24.44 | -0.12 | 0.12 | -0.01 | 0.01 |
| gene12802 | https://www.ncbi.nlm.nih.gov/gene/?term=LOC105286799 | 3183 | 173 | 543.51 | 1 | 3.14 | 0.58 | 17.3 | 17.43 | -0.13 | 0.13 | -0.01 | 0.01 |
| gene10018 | https://www.ncbi.nlm.nih.gov/gene/?term=LOC105283607 | 4414 | 192 | 434.98 | 1 | 2.27 | 0.52 | 15.73 | 15.92 | -0.19 | 0.19 | -0.01 | 0.01 |
| gene13036 | https://www.ncbi.nlm.nih.gov/gene/?term=LOC105287059 | 4959 | 172 | 346.84 | 1 | 2.02 | 0.58 | 16.67 | 16.87 | -0.2 | 0.2 | -0.01 | 0.01 |
| gene4376 | https://www.ncbi.nlm.nih.gov/gene/?term=LOC105277083 | 4307 | 123 | 285.58 | 1 | 2.32 | 0.81 | 19.63 | 19.89 | -0.26 | 0.26 | -0.01 | 0.01 |
| gene5646 | https://www.ncbi.nlm.nih.gov/gene/?term=LOC105278534 | 7475 | 180 | 240.8 1 | 1.34 | 0.56 | 17.08 | 17.41 | -0.33 | 0.33 | -0.01 | 0.01 | |
| gene13044 | https://www.ncbi.nlm.nih.gov/gene/?term=LOC105287070 | 5094 | 224 | 439.73 | 1 | 1.96 | 0.45 | 5.36 | 5.7 | -0.34 | 0.34 | -0.01 | 0.01 |
| gene12674 | https://www.ncbi.nlm.nih.gov/gene/?term=LOC105286649 | 18283 | 764 | 417.87 | 1 | 0.55 | 0.13 | 0.96 | 1.31 | -0.35 | 0.35 | -0.01 | 0.01 |
| gene12406 | https://www.ncbi.nlm.nih.gov/gene/?term=LOC105286316 | 1386 | 29 | 209.24 | 1 | 7.22 | 3.45 | 45.59 | 46 | -0.41 | 0.41 | -0.01 | 0.01 |
| gene9326 | https://www.ncbi.nlm.nih.gov/gene/?term=LOC105282818 | 16612 | 217 | 130.63 | 1 | 0.6 | 0.46 | 13.52 | 14.06 | -0.54 | 0.54 | -0.01 | 0.01 |
| gene5375 | https://www.ncbi.nlm.nih.gov/gene/?term=LOC105278228 | 8640 | 167 | 193.29 | 1 | 1.16 | 0.6 | 22.05 | 22.68 | -0.63 | 0.63 | -0.01 | 0.01 |
| gene11561 | https://www.ncbi.nlm.nih.gov/gene/?term=LOC105285387 | 11528 | 200 | 173.49 | 1 | 0.87 | 0.5 | 9.19 | 9.83 | -0.64 | 0.64 | -0.01 | 0.01 |
| gene9327 | https://www.ncbi.nlm.nih.gov/gene/?term=LOC105282821 | 7793 | 161 | 206.6 1 | 1.28 | 0.62 | 12.72 | 13.4 | -0.68 | 0.68 | -0.01 | 0.01 | |
| gene7368 | https://www.ncbi.nlm.nih.gov/gene/?term=LOC105280556 | 4105 | 58 | 141.29 | 1 | 2.44 | 1.72 | 18.1 | 19.01 | -0.91 | 0.91 | -0.01 | 0.01 |
| gene12203 | https://www.ncbi.nlm.nih.gov/gene/?term=LOC105286099 | 5967 | 79 | 132.39 | 1 | 1.68 | 1.27 | 11.92 | 12.93 | -1.01 | 1.01 | -0.01 | 0.01 |
| gene10345 | https://www.ncbi.nlm.nih.gov/gene/?term=LOC105283989 | 4444 | 21 | 47.25 1 | 2.25 | 4.76 | 48.56 | 49.94 | -1.37 | 1.37 | -0.01 | 0.01 | |
| gene8178 | https://www.ncbi.nlm.nih.gov/gene/?term=LOC105281515 | 8647 | 52 | 60.14 1 | 1.16 | 1.92 | 27.71 | 27.4 | 0.32 | 0.32 | 0 | 0 | |
| gene2683 | https://www.ncbi.nlm.nih.gov/gene/?term=LOC105275187 | 5492 | 69 | 125.64 | 1 | 1.82 | 1.45 | 14.14 | 13.85 | 0.29 | 0.29 | 0 | 0 |
| gene8343 | https://www.ncbi.nlm.nih.gov/gene/?term=LOC105281700 | 2172 | 38 | 174.95 | 1 | 4.6 | 2.63 | 39.23 | 38.97 | 0.26 | 0.26 | 0 | 0 |
| gene6620 | https://www.ncbi.nlm.nih.gov/gene/?term=LOC105279687 | 17998 | 349 | 193.91 | 1 | 0.56 | 0.29 | 4.97 | 4.84 | 0.13 | 0.13 | 0 | 0 |
| gene2271 | https://www.ncbi.nlm.nih.gov/gene/?term=LOC105274706 | 1448 | 51 | 352.21 | 1 | 6.91 | 1.96 | 38.8 | 38.68 | 0.12 | 0.12 | 0 | 0 |
| gene12307 | https://www.ncbi.nlm.nih.gov/gene/?term=LOC105286216 | 10313 | 178 | 172.6 1 | 0.97 | 0.56 | 15.99 | 15.89 | 0.1 | 0.1 | 0 | 0 | |
| gene3856 | https://www.ncbi.nlm.nih.gov/gene/?term=LOC105276499 | 5236 | 177 | 338.04 | 1 | 1.91 | 0.56 | 10.08 | 9.99 | 0.09 | 0.09 | 0 | 0 |
| gene7166 | https://www.ncbi.nlm.nih.gov/gene/?term=LOC105280326 | 6117 | 222 | 362.92 | 1 | 1.63 | 0.45 | 16.24 | 16.15 | 0.09 | 0.09 | 0 | 0 |
| gene9330 | https://www.ncbi.nlm.nih.gov/gene/?term=LOC105282825 | 3936 | 78 | 198.17 | 1 | 2.54 | 1.28 | 12.99 | 12.9 | 0.09 | 0.09 | 0 | 0 |
| gene53 | https://www.ncbi.nlm.nih.gov/gene/?term=LOC105277148 | 19283 | 283 | 146.76 | 1 | 0.52 | 0.35 | 5.45 | 5.37 | 0.08 | 0.08 | 0 | 0 |
| gene10344 | https://www.ncbi.nlm.nih.gov/gene/?term=LOC105283987 | 4841 | 71 | 146.66 | 1 | 2.07 | 1.41 | 32.31 | 32.24 | 0.07 | 0.07 | 0 | 0 |
| gene10654 | https://www.ncbi.nlm.nih.gov/gene/?term=LOC105284329 | 4027 | 51 | 126.65 | 1 | 2.48 | 1.96 | 19.38 | 19.32 | 0.06 | 0.06 | 0 | 0 |

| | | | | | | | | | | | | | |
|---|---|---|---|---|---|---|---|---|---|---|---|---|---|
| gene10600 | https://www.ncbi.nlm.nih.gov/gene/?term=LOC105284270 | 4837 | 100 | 206.74 | 1 | 2.07 | 1 | 12.53 | 12.48 | 0.05 | 0.05 | 0 | 0 |
| gene12981 | https://www.ncbi.nlm.nih.gov/gene/?term=LOC105286999 | 2063 | 44 | 213.28 | 1 | 4.85 | 2.27 | 35.25 | 35.21 | 0.04 | 0.04 | 0 | 0 |
| gene3885 | https://www.ncbi.nlm.nih.gov/gene/?term=LOC105276527 | 7928 | 333 | 420.03 | 1 | 1.26 | 0.3 | 8.6 | 8.57 | 0.03 | 0.03 | 0 | 0 |
| gene10392 | https://www.ncbi.nlm.nih.gov/gene/?term=LOC105284036 | 3216 | 87 | 270.52 | 2 | 6.22 | 2.3 | 24.3 | 24.3 | 0.01 | 0.01 | 0 | 0 |
| gene10549 | https://www.ncbi.nlm.nih.gov/gene/?term=LOC105284210 | 7145 | 327 | 457.66 | 2 | 2.8 | 0.61 | 10.82 | 10.83 | -0.01 | 0.01 | 0 | 0 |
| gene7600 | https://www.ncbi.nlm.nih.gov/gene/?term=LOC105280819 | 1615 | 38 | 235.29 | 1 | 6.19 | 2.63 | 35.56 | 35.58 | -0.02 | 0.02 | 0 | 0 |
| gene2806 | https://www.ncbi.nlm.nih.gov/gene/?term=LOC105275327 | 14257 | 472 | 331.07 | 1 | 0.7 | 0.21 | 7.12 | 7.15 | -0.03 | 0.03 | 0 | 0 |
| gene4190 | https://www.ncbi.nlm.nih.gov/gene/?term=LOC105276875 | 3150 | 101 | 320.63 | 1 | 3.17 | 0.99 | 24.58 | 24.61 | -0.03 | 0.03 | 0 | 0 |
| gene1632 | https://www.ncbi.nlm.nih.gov/gene/?term=LOC105287764 | 18061 | 391 | 216.49 | 1 | 0.55 | 0.26 | 3.3 | 3.35 | -0.05 | 0.05 | 0 | 0 |
| gene5524 | https://www.ncbi.nlm.nih.gov/gene/?term=LOC105278397 | 5177 | 200 | 386.32 | 1 | 1.93 | 0.5 | 24.93 | 25 | -0.07 | 0.07 | 0 | 0 |
| gene12516 | https://www.ncbi.nlm.nih.gov/gene/?term=LOC105286452 | 1529 | 42 | 274.69 | 1 | 6.54 | 2.38 | 46.48 | 46.6 | -0.11 | 0.11 | 0 | 0 |
| gene9029 | https://www.ncbi.nlm.nih.gov/gene/?term=LOC105282491 | 4762 | 169 | 354.89 | 1 | 2.1 | 0.59 | 7.49 | 7.61 | -0.13 | 0.13 | 0 | 0 |
| gene4641 | https://www.ncbi.nlm.nih.gov/gene/?term=LOC105277398 | 1937 | 67 | 345.9 | 1 | 5.16 | 1.49 | 35.24 | 35.39 | -0.14 | 0.14 | 0 | 0 |
| gene1884 | https://www.ncbi.nlm.nih.gov/gene/?term=LOC105288072 | 4169 | 77 | 184.7 | 1 | 2.4 | 1.3 | 24.07 | 24.23 | -0.16 | 0.16 | 0 | 0 |
| gene2173 | https://www.ncbi.nlm.nih.gov/gene/?term=LOC105274590 | 4387 | 67 | 152.72 | 2 | 4.56 | 2.99 | 31.19 | 31.42 | -0.23 | 0.23 | 0 | 0 |
| gene2748 | https://www.ncbi.nlm.nih.gov/gene/?term=LOC105275261 | 3845 | 50 | 130.04 | 1 | 2.6 | 2 | 37.52 | 37.82 | -0.3 | 0.3 | 0 | 0 |
| gene3599 | https://www.ncbi.nlm.nih.gov/gene/?term=LOC105276196 | 2955 | 45 | 152.28 | 1 | 3.38 | 2.22 | 48.59 | 48.91 | -0.31 | 0.31 | 0 | 0 |
| gene10490 | https://www.ncbi.nlm.nih.gov/gene/?term=LOC105284146 | 13960 | 149 | 106.73 | 1 | 0.72 | 0.67 | 10.54 | 10.87 | -0.33 | 0.33 | 0 | 0 |
| gene6639 | https://www.ncbi.nlm.nih.gov/gene/?term=LOC105279705 | 7656 | 66 | 86.21 | 1 | 1.31 | 1.52 | 21.87 | 22.21 | -0.34 | 0.34 | 0 | 0 |

Directory: OGL File: rnk-dmg-Cb_LI_BCphaseA2.vs.RphaseB2.txt

| gene_ID | gene_link | gwidth | #Sites | #per10Kb | #dmSites | #dmsp10kb | %dmSites | %pSite1 | %pSite2 | DMpSite | ADMpSite | DMpNucl | ADMpNucl |
|---|---|---|---|---|---|---|---|---|---|---|---|---|---|
| gene3165 | https://www.ncbi.nlm.nih.gov/gene/?term=LOC105275706 | 2374 | 72 | 303.29 | 6 | 25.27 | 8.33 | 33.88 | 22.12 | 11.76 | 11.76 | 0.36 | 0.36 |
| gene2824 | https://www.ncbi.nlm.nih.gov/gene/?term=LOC105275337 | 1772 | 60 | 338.6 | 2 | 11.29 | 3.33 | 38.89 | 29.82 | 9.07 | 9.07 | 0.31 | 0.31 |
| gene10963 | https://www.ncbi.nlm.nih.gov/gene/?term=LOC105284677 | 1576 | 27 | 171.32 | 3 | 19.04 | 11.11 | 20.06 | 37.04 | -16.98 | 16.98 | -0.29 | 0.29 |
| gene2593 | https://www.ncbi.nlm.nih.gov/gene/?term=LOC105275070 | 1774 | 7 | 39.46 | 5 | 28.18 | 71.43 | 66.52 | 4.09 | 62.43 | 62.43 | 0.25 | 0.25 |
| gene5714 | https://www.ncbi.nlm.nih.gov/gene/?term=LOC105278729 | 710 | 17 | 239.44 | 1 | 14.08 | 5.88 | 60.22 | 53.54 | 6.67 | 6.67 | 0.16 | 0.16 |
| gene4641 | https://www.ncbi.nlm.nih.gov/gene/?term=LOC105277398 | 1937 | 68 | 351.06 | 1 | 5.16 | 1.47 | 37.35 | 40.44 | -3.09 | 3.09 | -0.11 | 0.11 |
| gene7568 | https://www.ncbi.nlm.nih.gov/gene/?term=LOC105280786 | 2525 | 112 | 443.56 | 1 | 3.96 | 0.89 | 32.64 | 30.88 | 1.76 | 1.76 | 0.08 | 0.08 |
| gene10213 | https://www.ncbi.nlm.nih.gov/gene/?term=LOC105283826 | 1503 | 54 | 359.28 | 1 | 6.65 | 1.85 | 13.66 | 15.73 | -2.07 | 2.07 | -0.07 | 0.07 |
| gene11819 | https://www.ncbi.nlm.nih.gov/gene/?term=LOC105285671 | 2366 | 100 | 422.65 | 1 | 4.23 | 1 | 27.82 | 26.43 | 1.39 | 1.39 | 0.06 | 0.06 |
| gene920 | https://www.ncbi.nlm.nih.gov/gene/?term=LOC105282518 | 1197 | 39 | 325.81 | 1 | 8.35 | 2.56 | 38.65 | 40.64 | -1.98 | 1.98 | -0.06 | 0.06 |
| gene2404 | https://www.ncbi.nlm.nih.gov/gene/?term=LOC105274853 | 1767 | 34 | 192.42 | 1 | 5.66 | 2.94 | 29.24 | 32.2 | -2.96 | 2.96 | -0.06 | 0.06 |
| gene940 | https://www.ncbi.nlm.nih.gov/gene/?term=LOC105282794 | 3578 | 87 | 243.15 | 1 | 2.79 | 1.15 | 16.03 | 14.13 | 1.9 | 1.9 | 0.05 | 0.05 |
| gene10520 | https://www.ncbi.nlm.nih.gov/gene/?term=LOC105284184 | 2451 | 78 | 318.24 | 1 | 4.08 | 1.28 | 37.44 | 35.71 | 1.72 | 1.72 | 0.05 | 0.05 |
| gene1470 | https://www.ncbi.nlm.nih.gov/gene/?term=LOC105287597 | 2974 | 195 | 655.68 | 1 | 3.36 | 0.51 | 18.31 | 17.52 | 0.8 | 0.8 | 0.05 | 0.05 |
| gene10879 | https://www.ncbi.nlm.nih.gov/gene/?term=LOC105284574 | 2423 | 46 | 189.85 | 1 | 4.13 | 2.17 | 56.45 | 59.11 | -2.65 | 2.65 | -0.05 | 0.05 |
| gene393 | https://www.ncbi.nlm.nih.gov/gene/?term=LOC105276566 | 1123 | 2 | 17.81 | 1 | 8.9 | 50 | 48.08 | 76.78 | -28.71 | 28.71 | -0.05 | 0.05 |
| gene7398 | https://www.ncbi.nlm.nih.gov/gene/?term=LOC105280589 | 3452 | 34 | 98.49 | 1 | 2.9 | 2.94 | 23.83 | 19.97 | 3.86 | 3.86 | 0.04 | 0.04 |
| gene9462 | https://www.ncbi.nlm.nih.gov/gene/?term=LOC105282970 | 1469 | 39 | 265.49 | 1 | 6.81 | 2.56 | 51.12 | 49.47 | 1.65 | 1.65 | 0.04 | 0.04 |
| gene12806 | https://www.ncbi.nlm.nih.gov/gene/?term=LOC105286803 | 4196 | 137 | 326.5 | 1 | 2.38 | 0.73 | 10.84 | 9.75 | 1.09 | 1.09 | 0.04 | 0.04 |
| gene10678 | https://www.ncbi.nlm.nih.gov/gene/?term=LOC105284355 | 4031 | 158 | 391.96 | 1 | 2.48 | 0.63 | 20.75 | 19.82 | 0.93 | 0.93 | 0.04 | 0.04 |
| gene1220 | https://www.ncbi.nlm.nih.gov/gene/?term=LOC105285742 | 3453 | 107 | 309.88 | 1 | 2.9 | 0.93 | 22.27 | 23.44 | -1.17 | 1.17 | -0.04 | 0.04 |
| gene7600 | https://www.ncbi.nlm.nih.gov/gene/?term=LOC105280819 | 1615 | 37 | 229.1 | 1 | 6.19 | 2.7 | 36.56 | 38.16 | -1.6 | 1.6 | -0.04 | 0.04 |
| gene2413 | https://www.ncbi.nlm.nih.gov/gene/?term=LOC105274863 | 2006 | 40 | 199.4 | 1 | 4.99 | 2.5 | 64.23 | 66.19 | -1.96 | 1.96 | -0.04 | 0.04 |
| gene3599 | https://www.ncbi.nlm.nih.gov/gene/?term=LOC105276196 | 2955 | 47 | 159.05 | 1 | 3.38 | 2.13 | 46.53 | 48.84 | -2.31 | 2.31 | -0.04 | 0.04 |
| gene3385 | https://www.ncbi.nlm.nih.gov/gene/?term=LOC105275966 | 3305 | 53 | 160.36 | 1 | 3.03 | 1.89 | 54.5 | 52.88 | 1.62 | 1.62 | 0.03 | 0.03 |
| gene3594 | https://www.ncbi.nlm.nih.gov/gene/?term=LOC105276190 | 4910 | 83 | 169.04 | 1 | 2.04 | 1.2 | 11.44 | 9.83 | 1.62 | 1.62 | 0.03 | 0.03 |
| gene5877 | https://www.ncbi.nlm.nih.gov/gene/?term=LOC105278807 | 1719 | 46 | 267.6 | 1 | 5.82 | 2.17 | 26.78 | 25.52 | 1.25 | 1.25 | 0.03 | 0.03 |
| gene13033 | https://www.ncbi.nlm.nih.gov/gene/?term=LOC105287063 | 1877 | 48 | 255.73 | 1 | 5.33 | 2.08 | 31.55 | 30.37 | 1.19 | 1.19 | 0.03 | 0.03 |
| gene7915 | https://www.ncbi.nlm.nih.gov/gene/?term=LOC105281173 | 1597 | 38 | 237.95 | 1 | 6.26 | 2.63 | 59.94 | 58.74 | 1.19 | 1.19 | 0.03 | 0.03 |
| gene6652 | https://www.ncbi.nlm.nih.gov/gene/?term=LOC105279719 | 2831 | 79 | 279.05 | 1 | 3.53 | 1.27 | 24.35 | 23.32 | 1.03 | 1.03 | 0.03 | 0.03 |
| gene7556 | https://www.ncbi.nlm.nih.gov/gene/?term=LOC105280774 | 2792 | 89 | 318.77 | 1 | 3.58 | 1.12 | 36.33 | 35.33 | 1 | 1 | 0.03 | 0.03 |
| gene3817 | https://www.ncbi.nlm.nih.gov/gene/?term=LOC105276449 | 1230 | 42 | 341.46 | 1 | 8.13 | 2.38 | 35.93 | 35.04 | 0.88 | 0.88 | 0.03 | 0.03 |
| gene10606 | https://www.ncbi.nlm.nih.gov/gene/?term=LOC105284273 | 6119 | 249 | 406.93 | 1 | 1.63 | 0.4 | 18.46 | 17.61 | 0.86 | 0.86 | 0.03 | 0.03 |
| gene11401 | https://www.ncbi.nlm.nih.gov/gene/?term=LOC105285202 | 1841 | 82 | 445.41 | 1 | 5.43 | 1.22 | 26.05 | 25.33 | 0.73 | 0.73 | 0.03 | 0.03 |
| gene2846 | https://www.ncbi.nlm.nih.gov/gene/?term=LOC105275374 | 11246 | 464 | 412.59 | 1 | 0.89 | 0.22 | 12.19 | 11.56 | 0.63 | 0.63 | 0.03 | 0.03 |
| gene5298 | https://www.ncbi.nlm.nih.gov/gene/?term=LOC105278145 | 4060 | 245 | 603.45 | 2 | 4.93 | 0.82 | 10.68 | 11.19 | -0.51 | 0.51 | -0.03 | 0.03 |
| gene4498 | https://www.ncbi.nlm.nih.gov/gene/?term=LOC105277239 | 2831 | 122 | 430.94 | 1 | 3.53 | 0.82 | 12.06 | 12.7 | -0.64 | 0.64 | -0.03 | 0.03 |
| gene9101 | https://www.ncbi.nlm.nih.gov/gene/?term=LOC105282553 | 3068 | 108 | 352.02 | 1 | 3.26 | 0.93 | 22.57 | 23.41 | -0.84 | 0.84 | -0.03 | 0.03 |
| gene12516 | https://www.ncbi.nlm.nih.gov/gene/?term=LOC105286452 | 1529 | 40 | 261.61 | 1 | 6.54 | 2.5 | 50.21 | 51.25 | -1.04 | 1.04 | -0.03 | 0.03 |
| gene7207 | https://www.ncbi.nlm.nih.gov/gene/?term=LOC105280376 | 4659 | 109 | 233.96 | 1 | 2.15 | 0.92 | 29.99 | 31.37 | -1.38 | 1.38 | -0.03 | 0.03 |
| gene7850 | https://www.ncbi.nlm.nih.gov/gene/?term=LOC105281095 | 3050 | 63 | 206.56 | 1 | 3.28 | 1.59 | 33.9 | 35.57 | -1.67 | 1.67 | -0.03 | 0.03 |
| gene322 | https://www.ncbi.nlm.nih.gov/gene/?term=LOC105275777 | 1606 | 23 | 143.21 | 1 | 6.23 | 4.35 | 46.95 | 49.13 | -2.17 | 2.17 | -0.03 | 0.03 |
| gene10345 | https://www.ncbi.nlm.nih.gov/gene/?term=LOC105283989 | 4444 | 22 | 49.5 | 1 | 2.25 | 4.55 | 37.81 | 42.87 | -5.06 | 5.06 | -0.03 | 0.03 |
| gene2970 | https://www.ncbi.nlm.nih.gov/gene/?term=LOC105275518 | 2340 | 18 | 76.92 | 1 | 4.27 | 5.56 | 54.32 | 51.43 | 2.89 | 2.89 | 0.02 | 0.02 |
| gene6455 | https://www.ncbi.nlm.nih.gov/gene/?term=LOC105279500 | 2612 | 33 | 126.34 | 1 | 3.83 | 3.03 | 51.77 | 49.92 | 1.86 | 1.86 | 0.02 | 0.02 |
| gene35 | https://www.ncbi.nlm.nih.gov/gene/?term=LOC105275197 | 2643 | 43 | 162.69 | 1 | 3.78 | 2.33 | 44.25 | 43.02 | 1.22 | 1.22 | 0.02 | 0.02 |
| gene3470 | https://www.ncbi.nlm.nih.gov/gene/?term=LOC105276059 | 8716 | 145 | 166.36 | 1 | 1.15 | 0.69 | 9.2 | 8.01 | 1.19 | 1.19 | 0.02 | 0.02 |
| gene3671 | https://www.ncbi.nlm.nih.gov/gene/?term=LOC105276272 | 1966 | 34 | 172.94 | 1 | 5.09 | 2.94 | 33.29 | 32.32 | 0.98 | 0.98 | 0.02 | 0.02 |
| gene4962 | https://www.ncbi.nlm.nih.gov/gene/?term=LOC105277761 | 4645 | 142 | 305.71 | 1 | 2.15 | 0.7 | 14.73 | 13.94 | 0.79 | 0.79 | 0.02 | 0.02 |
| gene4892 | https://www.ncbi.nlm.nih.gov/gene/?term=LOC105277667 | 3898 | 124 | 318.11 | 1 | 2.57 | 0.81 | 27.97 | 27.19 | 0.78 | 0.78 | 0.02 | 0.02 |
| gene10600 | https://www.ncbi.nlm.nih.gov/gene/?term=LOC105284270 | 4837 | 119 | 246.02 | 1 | 2.07 | 0.84 | 10.7 | 10.02 | 0.68 | 0.68 | 0.02 | 0.02 |
| gene3727 | https://www.ncbi.nlm.nih.gov/gene/?term=LOC105276367 | 2484 | 66 | 265.7 | 1 | 4.03 | 1.52 | 47.44 | 46.84 | 0.61 | 0.61 | 0.02 | 0.02 |
| gene10984 | https://www.ncbi.nlm.nih.gov/gene/?term=LOC105284704 | 1987 | 56 | 281.83 | 1 | 5.03 | 1.79 | 25.7 | 25.1 | 0.59 | 0.59 | 0.02 | 0.02 |
| gene2844 | https://www.ncbi.nlm.nih.gov/gene/?term=LOC105275363 | 4330 | 181 | 418.01 | 1 | 2.31 | 0.55 | 15.09 | 14.51 | 0.58 | 0.58 | 0.02 | 0.02 |
| gene8274 | https://www.ncbi.nlm.nih.gov/gene/?term=LOC105281624 | 5425 | 204 | 376.04 | 1 | 1.84 | 0.49 | 10.02 | 9.47 | 0.56 | 0.56 | 0.02 | 0.02 |
| gene2056 | https://www.ncbi.nlm.nih.gov/gene/?term=LOC105274452 | 6584 | 251 | 381.23 | 2 | 3.04 | 0.8 | 13.77 | 13.35 | 0.42 | 0.42 | 0.02 | 0.02 |
| gene6286 | https://www.ncbi.nlm.nih.gov/gene/?term=LOC105279298 | 2861 | 128 | 447.4 | 2 | 6.99 | 1.56 | 24.57 | 24.17 | 0.4 | 0.4 | 0.02 | 0.02 |
| gene11340 | https://www.ncbi.nlm.nih.gov/gene/?term=LOC105285131 | 2918 | 86 | 294.72 | 1 | 3.43 | 1.16 | 11.93 | 12.54 | -0.61 | 0.61 | -0.02 | 0.02 |
| gene5646 | https://www.ncbi.nlm.nih.gov/gene/?term=LOC105278534 | 7475 | 175 | 234.11 | 1 | 1.34 | 0.57 | 17.8 | 18.5 | -0.7 | 0.7 | -0.02 | 0.02 |
| gene11976 | https://www.ncbi.nlm.nih.gov/gene/?term=LOC105285860 | 3112 | 52 | 167.1 | 1 | 3.21 | 1.92 | 21.02 | 22 | -0.98 | 0.98 | -0.02 | 0.02 |

| | | | | | | | | | | | | | |
|---|---|---|---|---|---|---|---|---|---|---|---|---|---|
| gene1147 | https://www.ncbi.nlm.nih.gov/gene/?term=LOC105284954 | 4022 | 84 | 208.85 | 1 | 2.49 | 1.19 | 17.29 | 18.34 | -1.06 | 1.06 | -0.02 | 0.02 |
| gene10391 | https://www.ncbi.nlm.nih.gov/gene/?term=LOC109611283 | 1994 | 42 | 210.63 | 1 | 5.02 | 2.38 | 32.6 | 33.67 | -1.07 | 1.07 | -0.02 | 0.02 |
| gene10155 | https://www.ncbi.nlm.nih.gov/gene/?term=LOC105283756 | 7030 | 100 | 142.25 | 1 | 1.42 | 1 | 5.99 | 7.12 | -1.12 | 1.12 | -0.02 | 0.02 |
| gene10819 | https://www.ncbi.nlm.nih.gov/gene/?term=LOC105284507 | 5767 | 76 | 131.78 | 1 | 1.73 | 1.32 | 23.17 | 24.52 | -1.35 | 1.35 | -0.02 | 0.02 |
| gene388 | https://www.ncbi.nlm.nih.gov/gene/?term=LOC105276538 | 1558 | 19 | 121.95 | 2 | 12.84 | 10.53 | 63.26 | 65.13 | -1.87 | 1.87 | -0.02 | 0.02 |
| gene3168 | https://www.ncbi.nlm.nih.gov/gene/?term=LOC105275711 | 6150 | 67 | 108.94 | 1 | 1.63 | 1.49 | 13.09 | 15.36 | -2.27 | 2.27 | -0.02 | 0.02 |
| gene10344 | https://www.ncbi.nlm.nih.gov/gene/?term=LOC105283987 | 4841 | 64 | 132.2 1 | 2.07 | 1.56 | 30.28 | 29.42 | 0.86 | 0.86 | 0.01 | 0.01 | |
| gene9336 | https://www.ncbi.nlm.nih.gov/gene/?term=LOC105282834 | 5993 | 121 | 201.9 1 | 1.67 | 0.83 | 16.02 | 15.31 | 0.72 | 0.72 | 0.01 | 0.01 | |
| gene2173 | https://www.ncbi.nlm.nih.gov/gene/?term=LOC105274590 | 4387 | 57 | 129.93 | 2 | 4.56 | 3.51 | 33.32 | 32.7 | 0.62 | 0.62 | 0.01 | 0.01 |
| gene11568 | https://www.ncbi.nlm.nih.gov/gene/?term=LOC105285395 | 8086 | 180 | 222.61 | 1 | 1.24 | 0.56 | 17.19 | 16.59 | 0.6 | 0.6 | 0.01 | 0.01 |
| gene7245 | https://www.ncbi.nlm.nih.gov/gene/?term=LOC105280410 | 12478 | 234 | 187.53 | 1 | 0.8 | 0.43 | 21.44 | 20.93 | 0.52 | 0.52 | 0.01 | 0.01 |
| gene5573 | https://www.ncbi.nlm.nih.gov/gene/?term=LOC105278454 | 2688 | 35 | 130.21 | 1 | 3.72 | 2.86 | 35.99 | 35.53 | 0.47 | 0.47 | 0.01 | 0.01 |
| gene12572 | https://www.ncbi.nlm.nih.gov/gene/?term=LOC105286514 | 2374 | 45 | 189.55 | 1 | 4.21 | 2.22 | 33.13 | 32.67 | 0.45 | 0.45 | 0.01 | 0.01 |
| gene34 | https://www.ncbi.nlm.nih.gov/gene/?term=LOC105278416 | 7120 | 124 | 174.16 | 1 | 1.4 | 0.81 | 15.53 | 15.09 | 0.43 | 0.43 | 0.01 | 0.01 |
| gene7166 | https://www.ncbi.nlm.nih.gov/gene/?term=LOC105280326 | 6117 | 214 | 349.84 | 1 | 1.63 | 0.47 | 16.62 | 16.23 | 0.4 | 0.4 | 0.01 | 0.01 |
| gene4246 | https://www.ncbi.nlm.nih.gov/gene/?term=LOC105276941 | 16412 | 493 | 300.39 | 1 | 0.61 | 0.2 | 5.46 | 5.07 | 0.38 | 0.38 | 0.01 | 0.01 |
| gene5746 | https://www.ncbi.nlm.nih.gov/gene/?term=LOC105278666 | 10343 | 425 | 410.91 | 1 | 0.97 | 0.24 | 7.62 | 7.27 | 0.35 | 0.35 | 0.01 | 0.01 |
| gene10392 | https://www.ncbi.nlm.nih.gov/gene/?term=LOC105284036 | 3216 | 91 | 282.96 | 2 | 6.22 | 2.2 | 22.63 | 22.31 | 0.32 | 0.32 | 0.01 | 0.01 |
| gene473 | https://www.ncbi.nlm.nih.gov/gene/?term=LOC105277423 | 2342 | 108 | 461.14 | 1 | 4.27 | 0.93 | 30.31 | 29.99 | 0.32 | 0.32 | 0.01 | 0.01 |
| gene11579 | https://www.ncbi.nlm.nih.gov/gene/?term=LOC105285406 | 13571 | 228 | 168.01 | 1 | 0.74 | 0.44 | 7.6 | 7.29 | 0.3 | 0.3 | 0.01 | 0.01 |
| gene2271 | https://www.ncbi.nlm.nih.gov/gene/?term=LOC105274706 | 1448 | 56 | 386.74 | 1 | 6.91 | 1.79 | 36.69 | 36.4 | 0.3 | 0.3 | 0.01 | 0.01 |
| gene10531 | https://www.ncbi.nlm.nih.gov/gene/?term=LOC105284190 | 6881 | 168 | 244.15 | 1 | 1.45 | 0.6 | 13.32 | 13.04 | 0.27 | 0.27 | 0.01 | 0.01 |
| gene7211 | https://www.ncbi.nlm.nih.gov/gene/?term=LOC105280377 | 2024 | 57 | 281.62 | 1 | 4.94 | 1.75 | 50.41 | 50.16 | 0.25 | 0.25 | 0.01 | 0.01 |
| gene298 | https://www.ncbi.nlm.nih.gov/gene/?term=LOC105275460 | 2473 | 116 | 469.07 | 1 | 4.04 | 0.86 | 13.65 | 13.43 | 0.23 | 0.23 | 0.01 | 0.01 |
| gene5525 | https://www.ncbi.nlm.nih.gov/gene/?term=LOC105278396 | 3090 | 181 | 585.76 | 1 | 3.24 | 0.55 | 24.85 | 24.63 | 0.22 | 0.22 | 0.01 | 0.01 |
| gene5524 | https://www.ncbi.nlm.nih.gov/gene/?term=LOC105278397 | 5177 | 196 | 378.6 1 | 1.93 | 0.51 | 24.88 | 24.69 | 0.18 | 0.18 | 0.01 | 0.01 | |
| gene3286 | https://www.ncbi.nlm.nih.gov/gene/?term=LOC105275848 | 3744 | 134 | 357.91 | 1 | 2.67 | 0.75 | 22.27 | 22.41 | -0.14 | 0.14 | -0.01 | 0.01 |
| gene10549 | https://www.ncbi.nlm.nih.gov/gene/?term=LOC105284210 | 7145 | 286 | 400.28 | 2 | 2.8 | 0.7 | 11.97 | 12.18 | -0.21 | 0.21 | -0.01 | 0.01 |
| gene12802 | https://www.ncbi.nlm.nih.gov/gene/?term=LOC105286799 | 3183 | 160 | 502.67 | 1 | 3.14 | 0.62 | 17.6 | 17.81 | -0.21 | 0.21 | -0.01 | 0.01 |
| gene3107 | https://www.ncbi.nlm.nih.gov/gene/?term=LOC105275650 | 5103 | 166 | 325.3 1 | 1.96 | 0.6 | 14.75 | 14.95 | -0.21 | 0.21 | -0.01 | 0.01 | |
| gene9755 | https://www.ncbi.nlm.nih.gov/gene/?term=LOC105283296 | 2177 | 85 | 390.45 | 1 | 4.59 | 1.18 | 19.16 | 19.37 | -0.21 | 0.21 | -0.01 | 0.01 |
| gene5733 | https://www.ncbi.nlm.nih.gov/gene/?term=LOC105278649 | 5221 | 114 | 218.35 | 1 | 1.92 | 0.88 | 16.23 | 16.48 | -0.25 | 0.25 | -0.01 | 0.01 |
| gene1310 | https://www.ncbi.nlm.nih.gov/gene/?term=LOC105286661 | 2582 | 72 | 278.85 | 1 | 3.87 | 1.39 | 21.23 | 21.48 | -0.26 | 0.26 | -0.01 | 0.01 |
| gene12674 | https://www.ncbi.nlm.nih.gov/gene/?term=LOC105286649 | 18283 | 747 | 408.58 | 1 | 0.55 | 0.13 | 0.89 | 1.18 | -0.29 | 0.29 | -0.01 | 0.01 |
| gene13036 | https://www.ncbi.nlm.nih.gov/gene/?term=LOC105287059 | 4959 | 146 | 294.41 | 1 | 2.02 | 0.68 | 18.9 | 19.21 | -0.31 | 0.31 | -0.01 | 0.01 |
| gene3885 | https://www.ncbi.nlm.nih.gov/gene/?term=LOC105276527 | 7928 | 305 | 384.71 | 1 | 1.26 | 0.33 | 9.75 | 10.06 | -0.32 | 0.32 | -0.01 | 0.01 |
| gene6583 | https://www.ncbi.nlm.nih.gov/gene/?term=LOC105279649 | 7444 | 303 | 407.04 | 1 | 1.34 | 0.33 | 11.36 | 11.69 | -0.33 | 0.33 | -0.01 | 0.01 |
| gene10018 | https://www.ncbi.nlm.nih.gov/gene/?term=LOC105283607 | 4414 | 187 | 423.65 | 1 | 2.27 | 0.53 | 16.13 | 16.48 | -0.35 | 0.35 | -0.01 | 0.01 |
| gene2122 | https://www.ncbi.nlm.nih.gov/gene/?term=LOC105274531 | 8153 | 289 | 354.47 | 1 | 1.23 | 0.35 | 10.34 | 10.69 | -0.35 | 0.35 | -0.01 | 0.01 |
| gene6031 | https://www.ncbi.nlm.nih.gov/gene/?term=LOC105279000 | 5691 | 187 | 328.59 | 1 | 1.76 | 0.53 | 16.08 | 16.46 | -0.38 | 0.38 | -0.01 | 0.01 |
| gene12573 | https://www.ncbi.nlm.nih.gov/gene/?term=LOC105286515 | 3647 | 98 | 268.71 | 1 | 2.74 | 1.02 | 13.27 | 13.66 | -0.39 | 0.39 | -0.01 | 0.01 |
| gene4376 | https://www.ncbi.nlm.nih.gov/gene/?term=LOC105277083 | 4307 | 102 | 236.82 | 1 | 2.32 | 0.98 | 24.57 | 25.16 | -0.58 | 0.58 | -0.01 | 0.01 |
| gene5273 | https://www.ncbi.nlm.nih.gov/gene/?term=LOC105278108 | 4067 | 44 | 108.19 | 1 | 2.46 | 2.27 | 24.76 | 25.36 | -0.61 | 0.61 | -0.01 | 0.01 |
| gene12203 | https://www.ncbi.nlm.nih.gov/gene/?term=LOC105286099 | 5967 | 73 | 122.34 | 1 | 1.68 | 1.37 | 10.19 | 10.87 | -0.68 | 0.68 | -0.01 | 0.01 |
| gene12406 | https://www.ncbi.nlm.nih.gov/gene/?term=LOC105286316 | 1386 | 28 | 202.02 | 1 | 7.22 | 3.57 | 44.18 | 44.91 | -0.73 | 0.73 | -0.01 | 0.01 |
| gene2748 | https://www.ncbi.nlm.nih.gov/gene/?term=LOC105275261 | 3845 | 51 | 132.64 | 1 | 2.6 | 1.96 | 36.93 | 37.66 | -0.74 | 0.74 | -0.01 | 0.01 |
| gene2683 | https://www.ncbi.nlm.nih.gov/gene/?term=LOC105275187 | 5492 | 66 | 120.17 | 1 | 1.82 | 1.52 | 11.92 | 12.72 | -0.8 | 0.8 | -0.01 | 0.01 |
| gene7368 | https://www.ncbi.nlm.nih.gov/gene/?term=LOC105280556 | 4105 | 35 | 85.26 1 | 2.44 | 2.86 | 17.18 | 18.02 | -0.84 | 0.84 | -0.01 | 0.01 | |
| gene9327 | https://www.ncbi.nlm.nih.gov/gene/?term=LOC105282821 | 7793 | 132 | 169.38 | 1 | 1.28 | 0.76 | 11.87 | 12.7 | -0.84 | 0.84 | -0.01 | 0.01 |
| gene8343 | https://www.ncbi.nlm.nih.gov/gene/?term=LOC105281700 | 2172 | 34 | 156.54 | 1 | 4.6 | 2.94 | 35.91 | 36.82 | -0.9 | 0.9 | -0.01 | 0.01 |
| gene11447 | https://www.ncbi.nlm.nih.gov/gene/?term=LOC105285252 | 7437 | 114 | 153.29 | 2 | 2.69 | 1.75 | 13.96 | 14.93 | -0.97 | 0.97 | -0.01 | 0.01 |
| gene10107 | https://www.ncbi.nlm.nih.gov/gene/?term=LOC105283712 | 1883 | 32 | 169.94 | 1 | 5.31 | 3.12 | 29.98 | 29.7 | 0.28 | 0.28 | 0 | 0 |
| gene8600 | https://www.ncbi.nlm.nih.gov/gene/?term=LOC105282001 | 8513 | 139 | 163.28 | 1 | 1.17 | 0.72 | 11.37 | 11.2 | 0.17 | 0.17 | 0 | 0 |
| gene9330 | https://www.ncbi.nlm.nih.gov/gene/?term=LOC105282825 | 3936 | 69 | 175.3 1 | 2.54 | 1.45 | 13.52 | 13.35 | 0.17 | 0.17 | 0 | 0 | |
| gene12307 | https://www.ncbi.nlm.nih.gov/gene/?term=LOC105286216 | 10313 | 167 | 161.93 | 1 | 0.97 | 0.6 | 17.01 | 16.86 | 0.15 | 0.15 | 0 | 0 |
| gene6620 | https://www.ncbi.nlm.nih.gov/gene/?term=LOC105279687 | 17998 | 312 | 173.35 | 1 | 0.56 | 0.32 | 5.46 | 5.32 | 0.14 | 0.14 | 0 | 0 |
| gene12265 | https://www.ncbi.nlm.nih.gov/gene/?term=LOC105286166 | 3050 | 104 | 340.98 | 1 | 3.28 | 0.96 | 25.25 | 25.13 | 0.13 | 0.13 | 0 | 0 |
| gene292 | https://www.ncbi.nlm.nih.gov/gene/?term=LOC105275398 | 2825 | 81 | 286.73 | 1 | 3.54 | 1.23 | 26.31 | 26.19 | 0.11 | 0.11 | 0 | 0 |
| gene11275 | https://www.ncbi.nlm.nih.gov/gene/?term=LOC105285055 | 4119 | 252 | 611.8 1 | 2.43 | 0.4 | 11.37 | 11.3 | 0.07 | 0.07 | 0 | 0 | |
| gene2806 | https://www.ncbi.nlm.nih.gov/gene/?term=LOC105275327 | 14257 | 460 | 322.65 | 1 | 0.7 | 0.22 | 7.25 | 7.19 | 0.06 | 0.06 | 0 | 0 |
| gene12981 | https://www.ncbi.nlm.nih.gov/gene/?term=LOC105286999 | 2063 | 42 | 203.59 | 1 | 4.85 | 2.38 | 36.12 | 36.07 | 0.05 | 0.05 | 0 | 0 |
| gene13114 | https://www.ncbi.nlm.nih.gov/gene/?term=LOC105287147 | 7609 | 383 | 503.35 | 1 | 1.31 | 0.26 | 9.68 | 9.63 | 0.04 | 0.04 | 0 | 0 |
| gene53 | https://www.ncbi.nlm.nih.gov/gene/?term=LOC105277148 | 19283 | 258 | 133.8 1 | 0.52 | 0.39 | 4.46 | 4.42 | 0.04 | 0.04 | 0 | 0 | |
| gene11082 | https://www.ncbi.nlm.nih.gov/gene/?term=LOC105284824 | 7436 | 185 | 248.79 | 1 | 1.34 | 0.54 | 6.28 | 6.27 | 0.01 | 0.01 | 0 | 0 |

| | | | | | | | | | | | | | | |
|---|---|---|---|---|---|---|---|---|---|---|---|---|---|---|
| gene1632 | https://www.ncbi.nlm.nih.gov/gene/?term=LOC105287764 | 18061 | 371 | 205.41 | | 2 | 1.11 | 0.54 | 3.28 | 3.27 | 0.01 | 0.01 | 0 | 0 |
| gene6966 | https://www.ncbi.nlm.nih.gov/gene/?term=LOC105280104 | 3747 | 123 | 328.26 | | 1 | 2.67 | 0.81 | 16.59 | 16.58 | 0.01 | 0.01 | 0 | 0 |
| gene4190 | https://www.ncbi.nlm.nih.gov/gene/?term=LOC105276875 | 3150 | 109 | 346.03 | | 1 | 3.17 | 0.92 | 24.02 | 24.04 | -0.02 | 0.02 | 0 | 0 |
| gene1993 | https://www.ncbi.nlm.nih.gov/gene/?term=LOC105288193 | 1449 | 74 | 510.7 | 1 | 6.9 | 1.35 | 34.89 | 34.92 | -0.03 | 0.03 | 0 | 0 | |
| gene8729 | https://www.ncbi.nlm.nih.gov/gene/?term=LOC105282137 | 5005 | 198 | 395.6 | 1 | 2 | 0.51 | 16.91 | 16.95 | -0.04 | 0.04 | 0 | 0 | |
| gene5831 | https://www.ncbi.nlm.nih.gov/gene/?term=LOC105278761 | 2506 | 109 | 434.96 | | 1 | 3.99 | 0.92 | 24.44 | 24.49 | -0.06 | 0.06 | 0 | 0 |
| gene9177 | https://www.ncbi.nlm.nih.gov/gene/?term=LOC105282646 | 3320 | 99 | 298.19 | | 1 | 3.01 | 1.01 | 25.05 | 25.11 | -0.06 | 0.06 | 0 | 0 |
| gene7249 | https://www.ncbi.nlm.nih.gov/gene/?term=LOC105280422 | 1775 | 60 | 338.03 | | 1 | 5.63 | 1.67 | 16.97 | 17.06 | -0.09 | 0.09 | 0 | 0 |
| gene5967 | https://www.ncbi.nlm.nih.gov/gene/?term=LOC105278918 | 4393 | 116 | 264.06 | | 1 | 2.28 | 0.86 | 14.39 | 14.54 | -0.15 | 0.15 | 0 | 0 |
| gene8661 | https://www.ncbi.nlm.nih.gov/gene/?term=LOC105282063 | 1926 | 35 | 181.72 | | 1 | 5.19 | 2.86 | 30.68 | 30.89 | -0.22 | 0.22 | 0 | 0 |
| gene1199 | https://www.ncbi.nlm.nih.gov/gene/?term=LOC105285498 | 2220 | 26 | 117.12 | | 1 | 4.5 | 3.85 | 31.35 | 31.59 | -0.24 | 0.24 | 0 | 0 |
| gene9326 | https://www.ncbi.nlm.nih.gov/gene/?term=LOC105282818 | 16612 | 173 | 104.14 | | 1 | 0.6 | 0.58 | 12.61 | 12.88 | -0.27 | 0.27 | 0 | 0 |
| gene12021 | https://www.ncbi.nlm.nih.gov/gene/?term=LOC105285905 | 1399 | 16 | 114.37 | | 1 | 7.15 | 6.25 | 47.33 | 47.75 | -0.42 | 0.42 | 0 | 0 |
| gene6639 | https://www.ncbi.nlm.nih.gov/gene/?term=LOC105279705 | 7656 | 61 | 79.68 | 1 | 1.31 | 1.64 | 23.52 | 23.98 | -0.46 | 0.46 | 0 | 0 | |

Directory: OGL File: rnk-dmg-Cb_LI_BCphaseB1.vs.BCphaseB2.txt

| gene_ID | gene_link | gwidth | #Sites | #per10Kb | #dmSites | #dmsp10kb | %dmSites | %pSite1 | %pSite2 | DMpSite | ADMpSite | DMpNucl | ADMpNucl |
|---|---|---|---|---|---|---|---|---|---|---|---|---|---|
| gene10963 | https://www.ncbi.nlm.nih.gov/gene/?term=LOC105284677 | 1576 | 29 | 184.01 | 1 | 6.35 | 3.45 | 18.87 | 36.99 | -18.12 | 18.12 | -0.33 | 0.33 |
| gene3165 | https://www.ncbi.nlm.nih.gov/gene/?term=LOC105275706 | 2374 | 66 | 278.01 | 5 | 21.06 | 7.58 | 34.05 | 24.01 | 10.05 | 10.05 | 0.28 | 0.28 |
| gene2593 | https://www.ncbi.nlm.nih.gov/gene/?term=LOC105275070 | 1774 | 5 | 28.18 | 5 | 28.18 | 100 | 72.92 | 10.96 | 61.96 | 61.96 | 0.17 | 0.17 |
| gene5714 | https://www.ncbi.nlm.nih.gov/gene/?term=LOC105278729 | 710 | 24 | 338.03 | 1 | 14.08 | 4.17 | 59.89 | 56.79 | 3.1 | 3.1 | 0.1 | 0.1 |
| gene4498 | https://www.ncbi.nlm.nih.gov/gene/?term=LOC105277239 | 2831 | 140 | 494.52 | 1 | 3.53 | 0.71 | 13.31 | 15.32 | -2.01 | 2.01 | -0.1 | 0.1 |
| gene3817 | https://www.ncbi.nlm.nih.gov/gene/?term=LOC105276449 | 1230 | 51 | 414.63 | 1 | 8.13 | 1.96 | 27.91 | 26.06 | 1.85 | 1.85 | 0.08 | 0.08 |
| gene1962 | https://www.ncbi.nlm.nih.gov/gene/?term=LOC105288157 | 1246 | 17 | 136.44 | 1 | 8.03 | 5.88 | 18.51 | 13.14 | 5.37 | 5.37 | 0.07 | 0.07 |
| gene2907 | https://www.ncbi.nlm.nih.gov/gene/?term=LOC105275447 | 2539 | 40 | 157.54 | 1 | 3.94 | 2.5 | 7.22 | 3.04 | 4.18 | 4.18 | 0.07 | 0.07 |
| gene12021 | https://www.ncbi.nlm.nih.gov/gene/?term=LOC105285905 | 1399 | 20 | 142.96 | 1 | 7.15 | 5 | 33.2 | 38.03 | -4.83 | 4.83 | -0.07 | 0.07 |
| gene5298 | https://www.ncbi.nlm.nih.gov/gene/?term=LOC105278145 | 4060 | 236 | 581.28 | 2 | 4.93 | 0.85 | 10.36 | 11.46 | -1.1 | 1.1 | -0.06 | 0.06 |
| gene9740 | https://www.ncbi.nlm.nih.gov/gene/?term=LOC105283277 | 2887 | 140 | 484.93 | 1 | 3.46 | 0.71 | 19.86 | 21.12 | -1.26 | 1.26 | -0.06 | 0.06 |
| gene2271 | https://www.ncbi.nlm.nih.gov/gene/?term=LOC105274706 | 1448 | 51 | 352.21 | 1 | 6.91 | 1.96 | 40.05 | 41.71 | -1.67 | 1.67 | -0.06 | 0.06 |
| gene3727 | https://www.ncbi.nlm.nih.gov/gene/?term=LOC105276367 | 2484 | 66 | 265.7 | 1 | 4.03 | 1.52 | 44.56 | 46.97 | -2.41 | 2.41 | -0.06 | 0.06 |
| gene5877 | https://www.ncbi.nlm.nih.gov/gene/?term=LOC105278807 | 1719 | 53 | 308.32 | 1 | 5.82 | 1.89 | 23.93 | 22.25 | 1.68 | 1.68 | 0.05 | 0.05 |
| gene4892 | https://www.ncbi.nlm.nih.gov/gene/?term=LOC105277667 | 3898 | 127 | 325.81 | 1 | 2.57 | 0.79 | 27.62 | 26.17 | 1.45 | 1.45 | 0.05 | 0.05 |
| gene10678 | https://www.ncbi.nlm.nih.gov/gene/?term=LOC105284355 | 4031 | 174 | 431.65 | 1 | 2.48 | 0.57 | 19.29 | 18.18 | 1.11 | 1.11 | 0.05 | 0.05 |
| gene7568 | https://www.ncbi.nlm.nih.gov/gene/?term=LOC105280786 | 2525 | 119 | 471.29 | 1 | 3.96 | 0.84 | 30.86 | 29.84 | 1.03 | 1.03 | 0.05 | 0.05 |
| gene1220 | https://www.ncbi.nlm.nih.gov/gene/?term=LOC105285742 | 3453 | 125 | 362 | 1 | 2.9 | 0.8 | 19.11 | 20.58 | -1.47 | 1.47 | -0.05 | 0.05 |
| gene6237 | https://www.ncbi.nlm.nih.gov/gene/?term=LOC105279244 | 1484 | 25 | 168.46 | 1 | 6.74 | 4 | 50.3 | 53.48 | -3.18 | 3.18 | -0.05 | 0.05 |
| gene1199 | https://www.ncbi.nlm.nih.gov/gene/?term=LOC105285498 | 2220 | 26 | 117.12 | 1 | 4.5 | 3.85 | 33.11 | 37.03 | -3.92 | 3.92 | -0.05 | 0.05 |
| gene2970 | https://www.ncbi.nlm.nih.gov/gene/?term=LOC105275518 | 2340 | 25 | 106.84 | 1 | 4.27 | 4 | 43.33 | 39.71 | 3.62 | 3.62 | 0.04 | 0.04 |
| gene940 | https://www.ncbi.nlm.nih.gov/gene/?term=LOC105282794 | 3578 | 88 | 245.95 | 1 | 2.79 | 1.14 | 16.46 | 14.64 | 1.82 | 1.82 | 0.04 | 0.04 |
| gene5967 | https://www.ncbi.nlm.nih.gov/gene/?term=LOC105278918 | 4393 | 146 | 332.35 | 1 | 2.28 | 0.68 | 13.82 | 12.77 | 1.06 | 1.06 | 0.04 | 0.04 |
| gene2856 | https://www.ncbi.nlm.nih.gov/gene/?term=LOC105275384 | 1987 | 92 | 463.01 | 1 | 5.03 | 1.09 | 32.79 | 31.9 | 0.89 | 0.89 | 0.04 | 0.04 |
| gene5525 | https://www.ncbi.nlm.nih.gov/gene/?term=LOC105278396 | 3090 | 189 | 611.65 | 1 | 3.24 | 0.53 | 24.57 | 25.15 | -0.58 | 0.58 | -0.04 | 0.04 |
| gene3793 | https://www.ncbi.nlm.nih.gov/gene/?term=LOC105276429 | 2901 | 110 | 379.18 | 1 | 3.45 | 0.91 | 25.42 | 26.35 | -0.93 | 0.93 | -0.04 | 0.04 |
| gene11340 | https://www.ncbi.nlm.nih.gov/gene/?term=LOC105285131 | 2918 | 105 | 359.84 | 1 | 3.43 | 0.95 | 13.12 | 14.12 | -1 | 1 | -0.04 | 0.04 |
| gene920 | https://www.ncbi.nlm.nih.gov/gene/?term=LOC105282518 | 1197 | 42 | 350.88 | 1 | 8.35 | 2.38 | 37.02 | 38.3 | -1.28 | 1.28 | -0.04 | 0.04 |
| gene7915 | https://www.ncbi.nlm.nih.gov/gene/?term=LOC105281173 | 1597 | 38 | 237.95 | 1 | 6.26 | 2.63 | 59.22 | 60.79 | -1.57 | 1.57 | -0.04 | 0.04 |
| gene6422 | https://www.ncbi.nlm.nih.gov/gene/?term=LOC105279458 | 2419 | 37 | 152.96 | 1 | 4.13 | 2.7 | 49.83 | 52.5 | -2.66 | 2.66 | -0.04 | 0.04 |
| gene1505 | https://www.ncbi.nlm.nih.gov/gene/?term=LOC105287633 | 1693 | 27 | 159.48 | 1 | 5.91 | 3.7 | 42.55 | 45.31 | -2.76 | 2.76 | -0.04 | 0.04 |
| gene6455 | https://www.ncbi.nlm.nih.gov/gene/?term=LOC105279500 | 2612 | 33 | 126.34 | 1 | 3.83 | 3.03 | 43.71 | 47.06 | -3.35 | 3.35 | -0.04 | 0.04 |
| gene393 | https://www.ncbi.nlm.nih.gov/gene/?term=LOC105276566 | 1123 | 2 | 17.81 | 1 | 8.9 | 50 | 47.22 | 70.57 | -23.35 | 23.35 | -0.04 | 0.04 |
| gene9478 | https://www.ncbi.nlm.nih.gov/gene/?term=LOC105282991 | 3726 | 57 | 152.98 | 1 | 2.68 | 1.75 | 35.88 | 33.61 | 2.27 | 2.27 | 0.03 | 0.03 |
| gene1518 | https://www.ncbi.nlm.nih.gov/gene/?term=LOC105287648 | 2195 | 30 | 136.67 | 1 | 4.56 | 3.33 | 43.53 | 41.69 | 1.83 | 1.83 | 0.03 | 0.03 |
| gene1884 | https://www.ncbi.nlm.nih.gov/gene/?term=LOC105288072 | 4169 | 76 | 182.3 | 1 | 2.4 | 1.32 | 24.99 | 23.26 | 1.72 | 1.72 | 0.03 | 0.03 |
| gene5101 | https://www.ncbi.nlm.nih.gov/gene/?term=LOC105277903 | 1813 | 37 | 204.08 | 1 | 5.52 | 2.7 | 54.27 | 52.77 | 1.5 | 1.5 | 0.03 | 0.03 |
| gene12572 | https://www.ncbi.nlm.nih.gov/gene/?term=LOC105286514 | 2374 | 46 | 193.77 | 1 | 4.21 | 2.17 | 36.88 | 35.48 | 1.4 | 1.4 | 0.03 | 0.03 |
| gene35 | https://www.ncbi.nlm.nih.gov/gene/?term=LOC105275197 | 2643 | 50 | 189.18 | 1 | 3.78 | 2 | 38.68 | 37.33 | 1.34 | 1.34 | 0.03 | 0.03 |
| gene7888 | https://www.ncbi.nlm.nih.gov/gene/?term=LOC105281141 | 4357 | 164 | 376.41 | 1 | 2.3 | 0.61 | 29.56 | 28.67 | 0.9 | 0.9 | 0.03 | 0.03 |
| gene11401 | https://www.ncbi.nlm.nih.gov/gene/?term=LOC105285202 | 1841 | 77 | 418.25 | 1 | 5.43 | 1.3 | 24.18 | 23.56 | 0.62 | 0.62 | 0.03 | 0.03 |
| gene6583 | https://www.ncbi.nlm.nih.gov/gene/?term=LOC105279649 | 7444 | 350 | 470.18 | 1 | 1.34 | 0.29 | 11.44 | 10.85 | 0.59 | 0.59 | 0.03 | 0.03 |
| gene1332 | https://www.ncbi.nlm.nih.gov/gene/?term=LOC105286872 | 2460 | 158 | 642.28 | 1 | 4.07 | 0.63 | 20.79 | 20.28 | 0.51 | 0.51 | 0.03 | 0.03 |
| gene4034 | https://www.ncbi.nlm.nih.gov/gene/?term=LOC105276704 | 2065 | 103 | 498.79 | 2 | 9.69 | 1.94 | 12.96 | 13.6 | -0.65 | 0.65 | -0.03 | 0.03 |
| gene11688 | https://www.ncbi.nlm.nih.gov/gene/?term=LOC105285533 | 1637 | 70 | 427.61 | 1 | 6.11 | 1.43 | 22.69 | 23.5 | -0.81 | 0.81 | -0.03 | 0.03 |
| gene6652 | https://www.ncbi.nlm.nih.gov/gene/?term=LOC105279719 | 2831 | 85 | 300.25 | 1 | 3.53 | 1.18 | 22.31 | 23.3 | -0.99 | 0.99 | -0.03 | 0.03 |
| gene1147 | https://www.ncbi.nlm.nih.gov/gene/?term=LOC105284954 | 4022 | 114 | 283.44 | 1 | 2.49 | 0.88 | 13.79 | 14.79 | -1 | 1 | -0.03 | 0.03 |
| gene8015 | https://www.ncbi.nlm.nih.gov/gene/?term=LOC105281312 | 2998 | 65 | 216.81 | 1 | 3.34 | 1.54 | 31.99 | 33.36 | -1.36 | 1.36 | -0.03 | 0.03 |
| gene12406 | https://www.ncbi.nlm.nih.gov/gene/?term=LOC105286316 | 1386 | 32 | 230.88 | 1 | 7.22 | 3.12 | 45.77 | 47.17 | -1.4 | 1.4 | -0.03 | 0.03 |
| gene13235 | https://www.ncbi.nlm.nih.gov/gene/?term=LOC105287278 | 2679 | 53 | 197.84 | 1 | 3.73 | 1.89 | 24.79 | 26.36 | -1.58 | 1.58 | -0.03 | 0.03 |
| gene10602 | https://www.ncbi.nlm.nih.gov/gene/?term=LOC105284268 | 2122 | 18 | 84.83 | 1 | 4.71 | 5.56 | 26.04 | 29.95 | -3.91 | 3.91 | -0.03 | 0.03 |
| gene12430 | https://www.ncbi.nlm.nih.gov/gene/?term=LOC105286348 | 3389 | 19 | 56.06 | 1 | 2.95 | 5.26 | 6.26 | 11.43 | -5.17 | 5.17 | -0.03 | 0.03 |
| gene7525 | https://www.ncbi.nlm.nih.gov/gene/?term=LOC105280738 | 4237 | 41 | 96.77 | 1 | 2.36 | 2.44 | 26.13 | 23.59 | 2.54 | 2.54 | 0.02 | 0.02 |
| gene7368 | https://www.ncbi.nlm.nih.gov/gene/?term=LOC105280556 | 4105 | 54 | 131.55 | 1 | 2.44 | 1.85 | 18.52 | 16.73 | 1.79 | 1.79 | 0.02 | 0.02 |
| gene13033 | https://www.ncbi.nlm.nih.gov/gene/?term=LOC105287063 | 1877 | 53 | 282.37 | 1 | 5.33 | 1.89 | 30.73 | 29.86 | 0.87 | 0.87 | 0.02 | 0.02 |
| gene4720 | https://www.ncbi.nlm.nih.gov/gene/?term=LOC105277479 | 1697 | 40 | 235.71 | 1 | 5.89 | 2.5 | 43.85 | 42.99 | 0.86 | 0.86 | 0.02 | 0.02 |
| gene12516 | https://www.ncbi.nlm.nih.gov/gene/?term=LOC105286452 | 1529 | 37 | 241.99 | 1 | 6.54 | 2.7 | 49.81 | 49.09 | 0.73 | 0.73 | 0.02 | 0.02 |
| gene292 | https://www.ncbi.nlm.nih.gov/gene/?term=LOC105275398 | 2825 | 85 | 300.88 | 1 | 3.54 | 1.18 | 21.01 | 20.28 | 0.73 | 0.73 | 0.02 | 0.02 |
| gene10540 | https://www.ncbi.nlm.nih.gov/gene/?term=LOC105284200 | 5267 | 204 | 387.32 | 1 | 1.9 | 0.49 | 12.98 | 12.34 | 0.64 | 0.64 | 0.02 | 0.02 |
| gene5654 | https://www.ncbi.nlm.nih.gov/gene/?term=LOC105278543 | 4019 | 160 | 398.11 | 1 | 2.49 | 0.62 | 15.95 | 15.36 | 0.6 | 0.6 | 0.02 | 0.02 |

| | | | | | | | | | | | | | |
|---|---|---|---|---|---|---|---|---|---|---|---|---|---|
| gene4376 | https://www.ncbi.nlm.nih.gov/gene/?term=LOC105277083 | 4307 | 129 | 299.51 | 1 | 2.32 | 0.78 | 19.34 | 18.78 | 0.56 | 0.56 | 0.02 | 0.02 |
| gene1855 | https://www.ncbi.nlm.nih.gov/gene/?term=LOC105288036 | 2387 | 86 | 360.28 | 1 | 4.19 | 1.16 | 29.99 | 29.44 | 0.55 | 0.55 | 0.02 | 0.02 |
| gene3107 | https://www.ncbi.nlm.nih.gov/gene/?term=LOC105275650 | 5103 | 178 | 348.81 | 1 | 1.96 | 0.56 | 16.67 | 16.15 | 0.52 | 0.52 | 0.02 | 0.02 |
| gene3206 | https://www.ncbi.nlm.nih.gov/gene/?term=LOC105275752 | 3474 | 129 | 371.33 | 1 | 2.88 | 0.78 | 32.13 | 31.62 | 0.52 | 0.52 | 0.02 | 0.02 |
| gene2710 | https://www.ncbi.nlm.nih.gov/gene/?term=LOC105275218 | 3222 | 98 | 304.16 | 1 | 3.1 | 1.02 | 22.4 | 21.9 | 0.5 | 0.5 | 0.02 | 0.02 |
| gene6966 | https://www.ncbi.nlm.nih.gov/gene/?term=LOC105280104 | 3747 | 133 | 354.95 | 1 | 2.67 | 0.75 | 15.81 | 15.32 | 0.49 | 0.49 | 0.02 | 0.02 |
| gene9445 | https://www.ncbi.nlm.nih.gov/gene/?term=LOC105282951 | 2285 | 115 | 503.28 | 1 | 4.38 | 0.87 | 20.4 | 19.9 | 0.49 | 0.49 | 0.02 | 0.02 |
| gene738 | https://www.ncbi.nlm.nih.gov/gene/?term=LOC105280421 | 4927 | 222 | 450.58 | 1 | 2.03 | 0.45 | 14.83 | 14.35 | 0.48 | 0.48 | 0.02 | 0.02 |
| gene8729 | https://www.ncbi.nlm.nih.gov/gene/?term=LOC105282137 | 5005 | 202 | 403.6 1 | 2 | 0.5 | 16.8 | 16.33 | 0.46 | 0.46 | 0.02 | 0.02 | |
| gene5831 | https://www.ncbi.nlm.nih.gov/gene/?term=LOC105278761 | 2506 | 126 | 502.79 | 1 | 3.99 | 0.79 | 21.41 | 21.05 | 0.36 | 0.36 | 0.02 | 0.02 |
| gene12802 | https://www.ncbi.nlm.nih.gov/gene/?term=LOC105286799 | 3183 | 189 | 593.78 | 1 | 3.14 | 0.53 | 15.7 | 15.39 | 0.32 | 0.32 | 0.02 | 0.02 |
| gene1470 | https://www.ncbi.nlm.nih.gov/gene/?term=LOC105287597 | 2974 | 204 | 685.94 | 1 | 3.36 | 0.49 | 17.06 | 17.34 | -0.28 | 0.28 | -0.02 | 0.02 |
| gene5524 | https://www.ncbi.nlm.nih.gov/gene/?term=LOC105278397 | 5177 | 205 | 395.98 | 1 | 1.93 | 0.49 | 24.95 | 25.35 | -0.39 | 0.39 | -0.02 | 0.02 |
| gene6286 | https://www.ncbi.nlm.nih.gov/gene/?term=LOC105279298 | 2861 | 133 | 464.87 | 2 | 6.99 | 1.5 | 23.95 | 24.34 | -0.39 | 0.39 | -0.02 | 0.02 |
| gene4190 | https://www.ncbi.nlm.nih.gov/gene/?term=LOC105276875 | 3150 | 123 | 390.48 | 1 | 3.17 | 0.81 | 20.99 | 21.41 | -0.42 | 0.42 | -0.02 | 0.02 |
| gene570 | https://www.ncbi.nlm.nih.gov/gene/?term=LOC105278549 | 1673 | 71 | 424.39 | 1 | 5.98 | 1.41 | 43.51 | 43.92 | -0.42 | 0.42 | -0.02 | 0.02 |
| gene12806 | https://www.ncbi.nlm.nih.gov/gene/?term=LOC105286803 | 4196 | 160 | 381.32 | 1 | 2.38 | 0.62 | 9.57 | 10.02 | -0.45 | 0.45 | -0.02 | 0.02 |
| gene8274 | https://www.ncbi.nlm.nih.gov/gene/?term=LOC105281624 | 5425 | 236 | 435.02 | 1 | 1.84 | 0.42 | 9.29 | 9.74 | -0.45 | 0.45 | -0.02 | 0.02 |
| gene10326 | https://www.ncbi.nlm.nih.gov/gene/?term=LOC105283962 | 3042 | 132 | 433.93 | 1 | 3.29 | 0.76 | 8.33 | 8.79 | -0.46 | 0.46 | -0.02 | 0.02 |
| gene4962 | https://www.ncbi.nlm.nih.gov/gene/?term=LOC105277761 | 4645 | 151 | 325.08 | 1 | 2.15 | 0.66 | 14.59 | 15.1 | -0.51 | 0.51 | -0.02 | 0.02 |
| gene10606 | https://www.ncbi.nlm.nih.gov/gene/?term=LOC105284273 | 6119 | 264 | 431.44 | 1 | 1.63 | 0.38 | 18.32 | 18.9 | -0.57 | 0.57 | -0.02 | 0.02 |
| gene9101 | https://www.ncbi.nlm.nih.gov/gene/?term=LOC105282553 | 3068 | 108 | 352.02 | 1 | 3.26 | 0.93 | 22.97 | 23.62 | -0.64 | 0.64 | -0.02 | 0.02 |
| gene7249 | https://www.ncbi.nlm.nih.gov/gene/?term=LOC105280422 | 1775 | 62 | 349.3 1 | 5.63 | 1.61 | 17.31 | 17.96 | -0.66 | 0.66 | -0.02 | 0.02 | |
| gene5646 | https://www.ncbi.nlm.nih.gov/gene/?term=LOC105278534 | 7475 | 195 | 260.87 | 1 | 1.34 | 0.51 | 17.04 | 17.72 | -0.68 | 0.68 | -0.02 | 0.02 |
| gene2404 | https://www.ncbi.nlm.nih.gov/gene/?term=LOC105274853 | 1767 | 38 | 215.05 | 1 | 5.66 | 2.63 | 34.09 | 34.89 | -0.8 | 0.8 | -0.02 | 0.02 |
| gene10107 | https://www.ncbi.nlm.nih.gov/gene/?term=LOC105283712 | 1883 | 36 | 191.18 | 1 | 5.31 | 2.78 | 28.27 | 29.18 | -0.91 | 0.91 | -0.02 | 0.02 |
| gene7247 | https://www.ncbi.nlm.nih.gov/gene/?term=LOC105280407 | 10486 | 212 | 202.17 | 1 | 0.95 | 0.47 | 15.9 | 16.83 | -0.93 | 0.93 | -0.02 | 0.02 |
| gene7199 | https://www.ncbi.nlm.nih.gov/gene/?term=LOC105280366 | 1414 | 32 | 226.31 | 1 | 7.07 | 3.12 | 42.1 | 43.06 | -0.96 | 0.96 | -0.02 | 0.02 |
| gene7248 | https://www.ncbi.nlm.nih.gov/gene/?term=LOC105280408 | 5175 | 57 | 110.14 | 1 | 1.93 | 1.75 | 18.23 | 20.49 | -2.26 | 2.26 | -0.02 | 0.02 |
| gene9411 | https://www.ncbi.nlm.nih.gov/gene/?term=LOC105282914 | 5534 | 41 | 74.09 1 | 1.81 | 2.44 | 49.41 | 52.21 | -2.8 | 2.8 | -0.02 | 0.02 | |
| gene564 | https://www.ncbi.nlm.nih.gov/gene/?term=LOC105278478 | 2802 | 17 | 60.67 1 | 3.57 | 5.88 | 14.04 | 17.66 | -3.63 | 3.63 | -0.02 | 0.02 | |
| gene8178 | https://www.ncbi.nlm.nih.gov/gene/?term=LOC105281515 | 8647 | 61 | 70.54 1 | 1.16 | 1.64 | 27.1 | 25.25 | 1.85 | 1.85 | 0.01 | 0.01 | |
| gene10345 | https://www.ncbi.nlm.nih.gov/gene/?term=LOC105283989 | 4444 | 29 | 65.26 1 | 2.25 | 3.45 | 49.14 | 47.94 | 1.2 | 1.2 | 0.01 | 0.01 | |
| gene927 | https://www.ncbi.nlm.nih.gov/gene/?term=LOC105282600 | 3727 | 50 | 134.16 | 1 | 2.68 | 2 | 33.71 | 32.83 | 0.88 | 0.88 | 0.01 | 0.01 |
| gene8742 | https://www.ncbi.nlm.nih.gov/gene/?term=LOC105282151 | 2563 | 44 | 171.67 | 2 | 7.8 | 4.55 | 36.03 | 35.31 | 0.72 | 0.72 | 0.01 | 0.01 |
| gene9336 | https://www.ncbi.nlm.nih.gov/gene/?term=LOC105282834 | 5993 | 129 | 215.25 | 1 | 1.67 | 0.78 | 15.82 | 15.16 | 0.66 | 0.66 | 0.01 | 0.01 |
| gene12439 | https://www.ncbi.nlm.nih.gov/gene/?term=LOC105286357 | 4761 | 86 | 180.63 | 1 | 2.1 | 1.16 | 26.67 | 26.07 | 0.6 | 0.6 | 0.01 | 0.01 |
| gene34 | https://www.ncbi.nlm.nih.gov/gene/?term=LOC105278416 | 7120 | 146 | 205.06 | 1 | 1.4 | 0.68 | 13.63 | 13.03 | 0.6 | 0.6 | 0.01 | 0.01 |
| gene2683 | https://www.ncbi.nlm.nih.gov/gene/?term=LOC105275187 | 5492 | 75 | 136.56 | 1 | 1.82 | 1.33 | 15.98 | 15.4 | 0.58 | 0.58 | 0.01 | 0.01 |
| gene4023 | https://www.ncbi.nlm.nih.gov/gene/?term=LOC105276692 | 1759 | 37 | 210.35 | 1 | 5.69 | 2.7 | 20.35 | 19.78 | 0.57 | 0.57 | 0.01 | 0.01 |
| gene7884 | https://www.ncbi.nlm.nih.gov/gene/?term=LOC105281266 | 3390 | 79 | 233.04 | 1 | 2.95 | 1.27 | 16.01 | 15.44 | 0.57 | 0.57 | 0.01 | 0.01 |
| gene2668 | https://www.ncbi.nlm.nih.gov/gene/?term=LOC105275170 | 12496 | 301 | 240.88 | 1 | 0.8 | 0.33 | 2.32 | 1.82 | 0.49 | 0.49 | 0.01 | 0.01 |
| gene3470 | https://www.ncbi.nlm.nih.gov/gene/?term=LOC105276059 | 8716 | 164 | 188.16 | 1 | 1.15 | 0.61 | 7.87 | 7.47 | 0.4 | 0.4 | 0.01 | 0.01 |
| gene11454 | https://www.ncbi.nlm.nih.gov/gene/?term=LOC105285256 | 1906 | 51 | 267.58 | 1 | 5.25 | 1.96 | 42.4 | 42.03 | 0.37 | 0.37 | 0.01 | 0.01 |
| gene6620 | https://www.ncbi.nlm.nih.gov/gene/?term=LOC105279687 | 17998 | 384 | 213.36 | 1 | 0.56 | 0.26 | 4.83 | 4.47 | 0.36 | 0.36 | 0.01 | 0.01 |
| gene5375 | https://www.ncbi.nlm.nih.gov/gene/?term=LOC105278228 | 8640 | 173 | 200.23 | 1 | 1.16 | 0.58 | 23.79 | 23.44 | 0.35 | 0.35 | 0.01 | 0.01 |
| gene2844 | https://www.ncbi.nlm.nih.gov/gene/?term=LOC105275363 | 4330 | 206 | 475.75 | 1 | 2.31 | 0.49 | 14.15 | 13.85 | 0.3 | 0.3 | 0.01 | 0.01 |
| gene11568 | https://www.ncbi.nlm.nih.gov/gene/?term=LOC105285395 | 8086 | 204 | 252.29 | 1 | 1.24 | 0.49 | 14.79 | 14.52 | 0.27 | 0.27 | 0.01 | 0.01 |
| gene1310 | https://www.ncbi.nlm.nih.gov/gene/?term=LOC105286661 | 2582 | 65 | 251.74 | 1 | 3.87 | 1.54 | 23.22 | 22.96 | 0.26 | 0.26 | 0.01 | 0.01 |
| gene10531 | https://www.ncbi.nlm.nih.gov/gene/?term=LOC105284190 | 6881 | 195 | 283.39 | 1 | 1.45 | 0.51 | 11.74 | 11.5 | 0.24 | 0.24 | 0.01 | 0.01 |
| gene11579 | https://www.ncbi.nlm.nih.gov/gene/?term=LOC105285406 | 13571 | 291 | 214.43 | 1 | 0.74 | 0.34 | 6.38 | 6.14 | 0.24 | 0.24 | 0.01 | 0.01 |
| gene10844 | https://www.ncbi.nlm.nih.gov/gene/?term=LOC105284535 | 9086 | 256 | 281.75 | 1 | 1.1 | 0.39 | 14.59 | 14.36 | 0.23 | 0.23 | 0.01 | 0.01 |
| gene2806 | https://www.ncbi.nlm.nih.gov/gene/?term=LOC105275327 | 14257 | 542 | 380.16 | 1 | 0.7 | 0.18 | 6.63 | 6.42 | 0.21 | 0.21 | 0.01 | 0.01 |
| gene3885 | https://www.ncbi.nlm.nih.gov/gene/?term=LOC105276527 | 7928 | 334 | 421.29 | 1 | 1.26 | 0.3 | 9.88 | 9.74 | 0.14 | 0.14 | 0.01 | 0.01 |
| gene2633 | https://www.ncbi.nlm.nih.gov/gene/?term=LOC105275127 | 7457 | 270 | 362.08 | 1 | 1.34 | 0.37 | 7.9 | 8.07 | -0.16 | 0.16 | -0.01 | 0.01 |
| gene7514 | https://www.ncbi.nlm.nih.gov/gene/?term=LOC105280723 | 4471 | 210 | 469.69 | 1 | 2.24 | 0.48 | 13.2 | 13.38 | -0.18 | 0.18 | -0.01 | 0.01 |
| gene10549 | https://www.ncbi.nlm.nih.gov/gene/?term=LOC105284210 | 7145 | 340 | 475.86 | 2 | 2.8 | 0.59 | 9.99 | 10.19 | -0.2 | 0.2 | -0.01 | 0.01 |
| gene8814 | https://www.ncbi.nlm.nih.gov/gene/?term=LOC105282240 | 6910 | 198 | 286.54 | 1 | 1.45 | 0.51 | 8.79 | 9 | -0.21 | 0.21 | -0.01 | 0.01 |
| gene2846 | https://www.ncbi.nlm.nih.gov/gene/?term=LOC105275374 | 11246 | 512 | 455.27 | 1 | 0.89 | 0.2 | 9.81 | 10.04 | -0.23 | 0.23 | -0.01 | 0.01 |
| gene10213 | https://www.ncbi.nlm.nih.gov/gene/?term=LOC105283826 | 1503 | 65 | 432.47 | 1 | 6.65 | 1.54 | 15.16 | 15.4 | -0.24 | 0.24 | -0.01 | 0.01 |
| gene13036 | https://www.ncbi.nlm.nih.gov/gene/?term=LOC105287059 | 4959 | 176 | 354.91 | 1 | 2.02 | 0.57 | 16.13 | 16.41 | -0.28 | 0.28 | -0.01 | 0.01 |
| gene10392 | https://www.ncbi.nlm.nih.gov/gene/?term=LOC105284036 | 3216 | 106 | 329.6 2 | 6.22 | 1.89 | 20.71 | 21 | -0.29 | 0.29 | -0.01 | 0.01 | |
| gene10520 | https://www.ncbi.nlm.nih.gov/gene/?term=LOC105284184 | 2451 | 101 | 412.08 | 1 | 4.08 | 0.99 | 29.13 | 29.43 | -0.3 | 0.3 | -0.01 | 0.01 |
| gene11082 | https://www.ncbi.nlm.nih.gov/gene/?term=LOC105284824 | 7436 | 214 | 287.79 | 1 | 1.34 | 0.47 | 6.56 | 6.86 | -0.31 | 0.31 | -0.01 | 0.01 |

| | | | | | | | | | | | | | |
|---|---|---|---|---|---|---|---|---|---|---|---|---|---|
| gene3046 | https://www.ncbi.nlm.nih.gov/gene/?term=LOC105275587 | 2394 | 55 | 229.74 | 1 | 4.18 | 1.82 | 36.59 | 36.9 | -0.31 | 0.31 | -0.01 | 0.01 |
| gene1907 | https://www.ncbi.nlm.nih.gov/gene/?term=LOC105288099 | 1238 | 36 | 290.79 | 1 | 8.08 | 2.78 | 33.72 | 34.04 | -0.32 | 0.32 | -0.01 | 0.01 |
| gene5777 | https://www.ncbi.nlm.nih.gov/gene/?term=LOC105278691 | 3883 | 168 | 432.66 | 1 | 2.58 | 0.6 | 14.66 | 14.98 | -0.33 | 0.33 | -0.01 | 0.01 |
| gene10600 | https://www.ncbi.nlm.nih.gov/gene/?term=LOC105284270 | 4837 | 124 | 256.36 | 1 | 2.07 | 0.81 | 11.8 | 12.15 | -0.35 | 0.35 | -0.01 | 0.01 |
| gene6031 | https://www.ncbi.nlm.nih.gov/gene/?term=LOC105279000 | 5691 | 197 | 346.16 | 1 | 1.76 | 0.51 | 12.37 | 12.73 | -0.35 | 0.35 | -0.01 | 0.01 |
| gene2413 | https://www.ncbi.nlm.nih.gov/gene/?term=LOC105274863 | 2006 | 54 | 269.19 | 1 | 4.99 | 1.85 | 52.24 | 52.76 | -0.51 | 0.51 | -0.01 | 0.01 |
| gene388 | https://www.ncbi.nlm.nih.gov/gene/?term=LOC105276538 | 1558 | 19 | 121.95 | 2 | 12.84 | 10.53 | 66.25 | 66.78 | -0.53 | 0.53 | -0.01 | 0.01 |
| gene2748 | https://www.ncbi.nlm.nih.gov/gene/?term=LOC105275261 | 3845 | 55 | 143.04 | 1 | 2.6 | 1.82 | 38.33 | 38.87 | -0.54 | 0.54 | -0.01 | 0.01 |
| gene11583 | https://www.ncbi.nlm.nih.gov/gene/?term=LOC105285414 | 8192 | 184 | 224.61 | 1 | 1.22 | 0.54 | 8.68 | 9.27 | -0.59 | 0.59 | -0.01 | 0.01 |
| gene6639 | https://www.ncbi.nlm.nih.gov/gene/?term=LOC105279705 | 7656 | 70 | 91.43 1 | 1.31 | 1.43 | 24.79 | 25.46 | -0.67 | 0.67 | -0.01 | 0.01 | |
| gene322 | https://www.ncbi.nlm.nih.gov/gene/?term=LOC105275777 | 1606 | 27 | 168.12 | 1 | 6.23 | 3.7 | 40.46 | 41.2 | -0.74 | 0.74 | -0.01 | 0.01 |
| gene3168 | https://www.ncbi.nlm.nih.gov/gene/?term=LOC105275711 | 6150 | 75 | 121.95 | 1 | 1.63 | 1.33 | 12.84 | 13.77 | -0.93 | 0.93 | -0.01 | 0.01 |
| gene3599 | https://www.ncbi.nlm.nih.gov/gene/?term=LOC105276196 | 2955 | 59 | 199.66 | 1 | 3.38 | 1.69 | 37.86 | 37.72 | 0.15 | 0.15 | 0 | 0 |
| gene53 | https://www.ncbi.nlm.nih.gov/gene/?term=LOC105277148 | 19283 | 324 | 168.02 | 1 | 0.52 | 0.31 | 4.91 | 4.75 | 0.15 | 0.15 | 0 | 0 |
| gene9462 | https://www.ncbi.nlm.nih.gov/gene/?term=LOC105282970 | 1469 | 42 | 285.91 | 1 | 6.81 | 2.38 | 51.11 | 50.96 | 0.15 | 0.15 | 0 | 0 |
| gene7183 | https://www.ncbi.nlm.nih.gov/gene/?term=LOC105280348 | 3925 | 76 | 193.63 | 1 | 2.55 | 1.32 | 19.42 | 19.29 | 0.13 | 0.13 | 0 | 0 |
| gene12307 | https://www.ncbi.nlm.nih.gov/gene/?term=LOC105286216 | 10313 | 174 | 168.72 | 1 | 0.97 | 0.57 | 15.92 | 15.81 | 0.12 | 0.12 | 0 | 0 |
| gene5184 | https://www.ncbi.nlm.nih.gov/gene/?term=LOC105277999 | 5528 | 146 | 264.11 | 1 | 1.81 | 0.68 | 11.67 | 11.57 | 0.1 | 0.1 | 0 | 0 |
| gene5746 | https://www.ncbi.nlm.nih.gov/gene/?term=LOC105278666 | 10343 | 482 | 466.02 | 1 | 0.97 | 0.21 | 6.71 | 6.61 | 0.1 | 0.1 | 0 | 0 |
| gene12573 | https://www.ncbi.nlm.nih.gov/gene/?term=LOC105286515 | 3647 | 103 | 282.42 | 1 | 2.74 | 0.97 | 10.56 | 10.47 | 0.09 | 0.09 | 0 | 0 |
| gene8343 | https://www.ncbi.nlm.nih.gov/gene/?term=LOC105281700 | 2172 | 37 | 170.35 | 1 | 4.6 | 2.7 | 39.73 | 39.63 | 0.09 | 0.09 | 0 | 0 |
| gene298 | https://www.ncbi.nlm.nih.gov/gene/?term=LOC105275460 | 2473 | 122 | 493.33 | 1 | 4.04 | 0.82 | 13.15 | 13.07 | 0.08 | 0.08 | 0 | 0 |
| gene10155 | https://www.ncbi.nlm.nih.gov/gene/?term=LOC105283756 | 7030 | 109 | 155.05 | 1 | 1.42 | 0.92 | 5.36 | 5.29 | 0.07 | 0.07 | 0 | 0 |
| gene1632 | https://www.ncbi.nlm.nih.gov/gene/?term=LOC105287764 | 18061 | 406 | 224.79 | 2 | 1.11 | 0.49 | 3.27 | 3.2 | 0.07 | 0.07 | 0 | 0 |
| gene9326 | https://www.ncbi.nlm.nih.gov/gene/?term=LOC105282818 | 16612 | 217 | 130.63 | 1 | 0.6 | 0.46 | 11.57 | 11.5 | 0.07 | 0.07 | 0 | 0 |
| gene10596 | https://www.ncbi.nlm.nih.gov/gene/?term=LOC105284262 | 6520 | 234 | 358.9 1 | 1.53 | 0.43 | 3.38 | 3.34 | 0.04 | 0.04 | 0 | 0 | |
| gene2056 | https://www.ncbi.nlm.nih.gov/gene/?term=LOC105274452 | 6584 | 263 | 399.45 | 2 | 3.04 | 0.76 | 12.97 | 12.95 | 0.02 | 0.02 | 0 | 0 |
| gene7398 | https://www.ncbi.nlm.nih.gov/gene/?term=LOC105280589 | 3452 | 37 | 107.18 | 1 | 2.9 | 2.7 | 19.84 | 19.83 | 0.01 | 0.01 | 0 | 0 |
| gene10018 | https://www.ncbi.nlm.nih.gov/gene/?term=LOC105283607 | 4414 | 208 | 471.23 | 1 | 2.27 | 0.48 | 15.08 | 15.07 | 0 | 0 | 0 | 0 |
| gene1430 | https://www.ncbi.nlm.nih.gov/gene/?term=LOC105287551 | 6433 | 232 | 360.64 | 1 | 1.55 | 0.43 | 8.43 | 8.44 | 0 | 0 | 0 | 0 |
| gene11275 | https://www.ncbi.nlm.nih.gov/gene/?term=LOC105285055 | 4119 | 248 | 602.09 | 1 | 2.43 | 0.4 | 12.13 | 12.15 | -0.02 | 0.02 | 0 | 0 |
| gene13114 | https://www.ncbi.nlm.nih.gov/gene/?term=LOC105287147 | 7609 | 430 | 565.12 | 1 | 1.31 | 0.23 | 9.08 | 9.1 | -0.02 | 0.02 | 0 | 0 |
| gene3856 | https://www.ncbi.nlm.nih.gov/gene/?term=LOC105276499 | 5236 | 194 | 370.51 | 1 | 1.91 | 0.52 | 10.61 | 10.65 | -0.04 | 0.04 | 0 | 0 |
| gene9327 | https://www.ncbi.nlm.nih.gov/gene/?term=LOC105282821 | 7793 | 173 | 221.99 | 1 | 1.28 | 0.58 | 10.79 | 10.87 | -0.08 | 0.08 | 0 | 0 |
| gene7166 | https://www.ncbi.nlm.nih.gov/gene/?term=LOC105280326 | 6117 | 253 | 413.6 1 | 1.63 | 0.4 | 14.52 | 14.62 | -0.1 | 0.1 | 0 | 0 | |
| gene10886 | https://www.ncbi.nlm.nih.gov/gene/?term=LOC105284582 | 1475 | 47 | 318.64 | 1 | 6.78 | 2.13 | 22.72 | 22.83 | -0.11 | 0.11 | 0 | 0 |
| gene6528 | https://www.ncbi.nlm.nih.gov/gene/?term=LOC105279591 | 7222 | 59 | 81.69 1 | 1.38 | 1.69 | 26.07 | 26.22 | -0.14 | 0.14 | 0 | 0 | |
| gene2173 | https://www.ncbi.nlm.nih.gov/gene/?term=LOC105274590 | 4387 | 84 | 191.47 | 2 | 4.56 | 2.38 | 27.83 | 27.99 | -0.17 | 0.17 | 0 | 0 |
| gene11561 | https://www.ncbi.nlm.nih.gov/gene/?term=LOC105285387 | 11528 | 202 | 175.23 | 1 | 0.87 | 0.5 | 8.85 | 9.03 | -0.18 | 0.18 | 0 | 0 |
| gene12203 | https://www.ncbi.nlm.nih.gov/gene/?term=LOC105286099 | 5967 | 75 | 125.69 | 1 | 1.68 | 1.33 | 9.95 | 10.3 | -0.35 | 0.35 | 0 | 0 |

Directory: OGL File: rnk-dmg-Cb_LI_BCphaseB1.vs.RphaseA1.txt

| gene_ID | gene_link | gwidth | #Sites | #per10Kb | #dmSites | #dmsp10kb | %dmSites | %pSite1 | %pSite2 | DMpSite | ADMpSite | DMpNucl | ADMpNucl |
|---|---|---|---|---|---|---|---|---|---|---|---|---|---|
| gene11566 | https://www.ncbi.nlm.nih.gov/gene/?term=LOC105285392 | 1603 | 91 | 567.69 | 6 | 37.43 | 6.59 | 35.74 | 31.54 | 4.2 | 4.2 | 0.24 | 0.24 |
| gene5306 | https://www.ncbi.nlm.nih.gov/gene/?term=LOC105278153 | 481 | 17 | 353.43 | 2 | 41.58 | 11.76 | 21.3 | 16.46 | 4.84 | 4.84 | 0.17 | 0.17 |
| gene2855 | https://www.ncbi.nlm.nih.gov/gene/?term=LOC105275378 | 1189 | 21 | 176.62 | 3 | 25.23 | 14.29 | 38.9 | 30.15 | 8.75 | 8.75 | 0.15 | 0.15 |
| gene2111 | https://www.ncbi.nlm.nih.gov/gene/?term=LOC105274528 | 2028 | 144 | 710.06 | 3 | 14.79 | 2.08 | 19.76 | 17.65 | 2.1 | 2.1 | 0.15 | 0.15 |
| gene11402 | https://www.ncbi.nlm.nih.gov/gene/?term=LOC105285201 | 335 | 16 | 477.61 | 1 | 29.85 | 6.25 | 21.09 | 18.26 | 2.83 | 2.83 | 0.14 | 0.14 |
| gene10112 | https://www.ncbi.nlm.nih.gov/gene/?term=LOC105283717 | 1483 | 81 | 546.19 | 4 | 26.97 | 4.94 | 29.9 | 27.32 | 2.58 | 2.58 | 0.14 | 0.14 |
| gene560 | https://www.ncbi.nlm.nih.gov/gene/?term=LOC109610897 | 1584 | 125 | 789.14 | 2 | 12.63 | 1.6 | 14.21 | 16.02 | -1.81 | 1.81 | -0.14 | 0.14 |
| gene12670 | https://www.ncbi.nlm.nih.gov/gene/?term=LOC105286645 | 1506 | 55 | 365.21 | 1 | 6.64 | 1.82 | 37.73 | 41.37 | -3.64 | 3.64 | -0.13 | 0.13 |
| gene10513 | https://www.ncbi.nlm.nih.gov/gene/?term=LOC105284174 | 1606 | 77 | 479.45 | 1 | 6.23 | 1.3 | 21.46 | 23.96 | -2.5 | 2.5 | -0.12 | 0.12 |
| gene12364 | https://www.ncbi.nlm.nih.gov/gene/?term=LOC105286269 | 1403 | 51 | 363.51 | 4 | 28.51 | 7.84 | 50.68 | 53.95 | -3.27 | 3.27 | -0.12 | 0.12 |
| gene9468 | https://www.ncbi.nlm.nih.gov/gene/?term=LOC105282976 | 1636 | 63 | 385.09 | 2 | 12.22 | 3.17 | 23.75 | 20.8 | 2.95 | 2.95 | 0.11 | 0.11 |
| gene5447 | https://www.ncbi.nlm.nih.gov/gene/?term=LOC105278314 | 1482 | 93 | 627.53 | 1 | 6.75 | 1.08 | 23.89 | 22.18 | 1.72 | 1.72 | 0.11 | 0.11 |
| gene4380 | https://www.ncbi.nlm.nih.gov/gene/?term=LOC105277087 | 334 | 10 | 299.4 | 1 | 29.94 | 10 | 6.29 | 9.99 | -3.7 | 3.7 | -0.11 | 0.11 |
| gene2036 | https://www.ncbi.nlm.nih.gov/gene/?term=LOC105288242 | 1260 | 36 | 285.71 | 1 | 7.94 | 2.78 | 42.3 | 46.17 | -3.87 | 3.87 | -0.11 | 0.11 |
| gene10176 | https://www.ncbi.nlm.nih.gov/gene/?term=LOC105283785 | 1637 | 35 | 213.81 | 1 | 6.11 | 2.86 | 18.49 | 13.7 | 4.79 | 4.79 | 0.1 | 0.1 |
| gene11109 | https://www.ncbi.nlm.nih.gov/gene/?term=LOC105284855 | 1786 | 56 | 313.55 | 2 | 11.2 | 3.57 | 33.74 | 30.6 | 3.13 | 3.13 | 0.1 | 0.1 |
| gene13266 | https://www.ncbi.nlm.nih.gov/gene/?term=LOC105287313 | 1615 | 87 | 538.7 | 3 | 18.58 | 3.45 | 28.84 | 27.02 | 1.82 | 1.82 | 0.1 | 0.1 |
| gene3910 | https://www.ncbi.nlm.nih.gov/gene/?term=LOC105276563 | 1957 | 116 | 592.74 | 5 | 25.55 | 4.31 | 17.71 | 16.03 | 1.69 | 1.69 | 0.1 | 0.1 |
| gene276 | https://www.ncbi.nlm.nih.gov/gene/?term=LOC105275214 | 1299 | 44 | 338.72 | 1 | 7.7 | 2.27 | 33.69 | 30.91 | 2.78 | 2.78 | 0.09 | 0.09 |
| gene7165 | https://www.ncbi.nlm.nih.gov/gene/?term=LOC105280327 | 2688 | 111 | 412.95 | 6 | 22.32 | 5.41 | 28.5 | 26.42 | 2.08 | 2.08 | 0.09 | 0.09 |
| gene1564 | https://www.ncbi.nlm.nih.gov/gene/?term=LOC105287690 | 1386 | 109 | 786.44 | 3 | 21.65 | 2.75 | 18.55 | 17.35 | 1.21 | 1.21 | 0.09 | 0.09 |
| gene5937 | https://www.ncbi.nlm.nih.gov/gene/?term=LOC105278873 | 1902 | 149 | 783.39 | 6 | 31.55 | 4.03 | 20.65 | 19.44 | 1.21 | 1.21 | 0.09 | 0.09 |
| gene10743 | https://www.ncbi.nlm.nih.gov/gene/?term=LOC105284456 | 1028 | 42 | 408.56 | 3 | 29.18 | 7.14 | 38.04 | 40.31 | -2.27 | 2.27 | -0.09 | 0.09 |
| gene1769 | https://www.ncbi.nlm.nih.gov/gene/?term=LOC105287942 | 2154 | 78 | 362.12 | 5 | 23.21 | 6.41 | 31.01 | 33.47 | -2.46 | 2.46 | -0.09 | 0.09 |

...

... only first 25 lines shown ...

Directory: OGL File: rnk-dmg-Cb_LI_BCphaseB1.vs.RphaseA2.txt

| gene_ID | gene_link | gwidth | #Sites | #per10Kb | #dmSites | #dmsp10kb | %dmSites | %pSite1 | %pSite2 | DMpSite | ADMpSite | DMpNucl | ADMpNucl |
|---|---|---|---|---|---|---|---|---|---|---|---|---|---|
| gene2837 | https://www.ncbi.nlm.nih.gov/gene/?term=LOC105275355 | 2872 | 160 | 557.1 | 10 | 34.82 | 6.25 | 29.18 | 25.36 | 3.82 | 3.82 | 0.21 | 0.21 |
| gene1894 | https://www.ncbi.nlm.nih.gov/gene/?term=LOC105288083 | 445 | 17 | 382.02 | 2 | 44.94 | 11.76 | 56.3 | 51.18 | 5.11 | 5.11 | 0.2 | 0.2 |
| gene12925 | https://www.ncbi.nlm.nih.gov/gene/?term=LOC109611487 | 391 | 27 | 690.54 | 1 | 25.58 | 3.7 | 14.67 | 17.34 | -2.67 | 2.67 | -0.18 | 0.18 |
| gene12311 | https://www.ncbi.nlm.nih.gov/gene/?term=LOC105286212 | 1005 | 39 | 388.06 | 3 | 29.85 | 7.69 | 33.27 | 37.99 | -4.72 | 4.72 | -0.18 | 0.18 |
| gene3268 | https://www.ncbi.nlm.nih.gov/gene/?term=LOC105275833 | 1040 | 43 | 413.46 | 1 | 9.62 | 2.33 | 35.22 | 39.24 | -4.03 | 4.03 | -0.17 | 0.17 |
| gene6075 | https://www.ncbi.nlm.nih.gov/gene/?term=LOC105279049 | 1285 | 99 | 770.43 | 1 | 7.78 | 1.01 | 16.76 | 18.86 | -2.11 | 2.11 | -0.16 | 0.16 |
| gene3815 | https://www.ncbi.nlm.nih.gov/gene/?term=LOC105276451 | 1324 | 39 | 294.56 | 3 | 22.66 | 7.69 | 36.8 | 42.26 | -5.46 | 5.46 | -0.16 | 0.16 |
| gene9740 | https://www.ncbi.nlm.nih.gov/gene/?term=LOC105283277 | 2887 | 145 | 502.25 | 3 | 10.39 | 2.07 | 19.78 | 22.8 | -3.02 | 3.02 | -0.15 | 0.15 |
| gene8798 | https://www.ncbi.nlm.nih.gov/gene/?term=LOC105282220 | 2300 | 235 | 1021.74 | 3 | 13.04 | 1.28 | 7.15 | 8.56 | -1.4 | 1.4 | -0.14 | 0.14 |
| gene2790 | https://www.ncbi.nlm.nih.gov/gene/?term=LOC105275307 | 1429 | 55 | 384.88 | 2 | 14 | 3.64 | 37.41 | 41.04 | -3.64 | 3.64 | -0.14 | 0.14 |
| gene12403 | https://www.ncbi.nlm.nih.gov/gene/?term=LOC105286311 | 1413 | 52 | 368.01 | 3 | 21.23 | 5.77 | 55.61 | 59.38 | -3.77 | 3.77 | -0.14 | 0.14 |
| gene5917 | https://www.ncbi.nlm.nih.gov/gene/?term=LOC105278849 | 823 | 82 | 996.35 | 1 | 12.15 | 1.22 | 25.33 | 24.02 | 1.31 | 1.31 | 0.13 | 0.13 |
| gene7917 | https://www.ncbi.nlm.nih.gov/gene/?term=LOC105281172 | 1304 | 30 | 230.06 | 3 | 23.01 | 10 | 42.68 | 48.2 | -5.52 | 5.52 | -0.13 | 0.13 |
| gene11779 | https://www.ncbi.nlm.nih.gov/gene/?term=LOC105285628 | 1037 | 22 | 212.15 | 3 | 28.93 | 13.64 | 32.81 | 38.84 | -6.03 | 6.03 | -0.13 | 0.13 |
| gene2111 | https://www.ncbi.nlm.nih.gov/gene/?term=LOC105274528 | 2028 | 151 | 744.58 | 2 | 9.86 | 1.32 | 19 | 17.45 | 1.55 | 1.55 | 0.12 | 0.12 |
| gene560 | https://www.ncbi.nlm.nih.gov/gene/?term=LOC109610897 | 1584 | 148 | 934.34 | 2 | 12.63 | 1.35 | 13.32 | 14.65 | -1.33 | 1.33 | -0.12 | 0.12 |
| gene6057 | https://www.ncbi.nlm.nih.gov/gene/?term=LOC105279035 | 2105 | 166 | 788.6 | 4 | 19 | 2.41 | 18.44 | 19.91 | -1.47 | 1.47 | -0.12 | 0.12 |
| gene5525 | https://www.ncbi.nlm.nih.gov/gene/?term=LOC105278396 | 3090 | 191 | 618.12 | 4 | 12.94 | 2.09 | 24.34 | 26.26 | -1.92 | 1.92 | -0.12 | 0.12 |
| gene5423 | https://www.ncbi.nlm.nih.gov/gene/?term=LOC105278282 | 1776 | 67 | 377.25 | 3 | 16.89 | 4.48 | 31.84 | 34.9 | -3.06 | 3.06 | -0.12 | 0.12 |
| gene8333 | https://www.ncbi.nlm.nih.gov/gene/?term=LOC105281691 | 1264 | 34 | 268.99 | 4 | 31.65 | 11.76 | 48.64 | 53.25 | -4.6 | 4.6 | -0.12 | 0.12 |
| gene3269 | https://www.ncbi.nlm.nih.gov/gene/?term=LOC105275832 | 728 | 55 | 755.49 | 2 | 27.47 | 3.64 | 8.05 | 9.52 | -1.47 | 1.47 | -0.11 | 0.11 |
| gene9183 | https://www.ncbi.nlm.nih.gov/gene/?term=LOC105282640 | 2752 | 209 | 759.45 | 2 | 7.27 | 0.96 | 21 | 22.49 | -1.49 | 1.49 | -0.11 | 0.11 |
| gene8453 | https://www.ncbi.nlm.nih.gov/gene/?term=LOC105281826 | 1750 | 99 | 565.71 | 1 | 5.71 | 1.01 | 28.31 | 30.17 | -1.86 | 1.86 | -0.11 | 0.11 |
| gene10513 | https://www.ncbi.nlm.nih.gov/gene/?term=LOC105284174 | 1606 | 89 | 554.17 | 1 | 6.23 | 1.12 | 19.86 | 21.82 | -1.96 | 1.96 | -0.11 | 0.11 |

...

... only first 25 lines shown ...

Directory: OGL File: rnk-dmg-Cb_LI_BCphaseB1.vs.RphaseB1.txt

| gene_ID | gene_link | gwidth | #Sites | #per10Kb | #dmSites | #dmsp10kb | %dmSites | %pSite1 | %pSite2 | DMpSite | ADMpSite | DMpNucl | ADMpNucl |
|---|---|---|---|---|---|---|---|---|---|---|---|---|---|
| gene2837 | https://www.ncbi.nlm.nih.gov/gene/?term=LOC105275355 | 2872 | 165 | 574.51 | 1 | 3.48 | 0.61 | 28.35 | 22.92 | 5.43 | 5.43 | 0.31 | 0.31 |
| gene10963 | https://www.ncbi.nlm.nih.gov/gene/?term=LOC105284677 | 1576 | 30 | 190.36 | 1 | 6.35 | 3.33 | 19.35 | 34.92 | -15.57 | 15.57 | -0.3 | 0.3 |
| gene3165 | https://www.ncbi.nlm.nih.gov/gene/?term=LOC105275706 | 2374 | 65 | 273.8 | 4 | 16.85 | 6.15 | 33.82 | 25.03 | 8.79 | 8.79 | 0.24 | 0.24 |
| gene2593 | https://www.ncbi.nlm.nih.gov/gene/?term=LOC105275070 | 1774 | 11 | 62.01 | 1 | 5.64 | 9.09 | 61.58 | 25.28 | 36.29 | 36.29 | 0.23 | 0.23 |
| gene2856 | https://www.ncbi.nlm.nih.gov/gene/?term=LOC105275384 | 1987 | 92 | 463.01 | 1 | 5.03 | 1.09 | 34.19 | 31.19 | 3 | 3 | 0.14 | 0.14 |
| gene2413 | https://www.ncbi.nlm.nih.gov/gene/?term=LOC105274863 | 2006 | 62 | 309.07 | 1 | 4.99 | 1.61 | 48.7 | 53.36 | -4.66 | 4.66 | -0.14 | 0.14 |
| gene9740 | https://www.ncbi.nlm.nih.gov/gene/?term=LOC105283277 | 2887 | 139 | 481.47 | 1 | 3.46 | 0.72 | 18.49 | 20.55 | -2.05 | 2.05 | -0.1 | 0.1 |
| gene1855 | https://www.ncbi.nlm.nih.gov/gene/?term=LOC105288036 | 2387 | 91 | 381.23 | 1 | 4.19 | 1.1 | 29.43 | 27.13 | 2.3 | 2.3 | 0.09 | 0.09 |
| gene5714 | https://www.ncbi.nlm.nih.gov/gene/?term=LOC105278729 | 710 | 16 | 225.35 | 1 | 14.08 | 6.25 | 55.52 | 51.9 | 3.62 | 3.62 | 0.08 | 0.08 |
| gene6237 | https://www.ncbi.nlm.nih.gov/gene/?term=LOC105279244 | 1484 | 33 | 222.37 | 1 | 6.74 | 3.03 | 44.64 | 48.04 | -3.4 | 3.4 | -0.08 | 0.08 |
| gene5831 | https://www.ncbi.nlm.nih.gov/gene/?term=LOC105278761 | 2506 | 118 | 470.87 | 2 | 7.98 | 1.69 | 22.67 | 21.1 | 1.58 | 1.58 | 0.07 | 0.07 |
| gene3671 | https://www.ncbi.nlm.nih.gov/gene/?term=LOC105276272 | 1966 | 40 | 203.46 | 1 | 5.09 | 2.5 | 30.25 | 27.31 | 2.94 | 2.94 | 0.06 | 0.06 |
| gene396 | https://www.ncbi.nlm.nih.gov/gene/?term=LOC105276611 | 1722 | 36 | 209.06 | 1 | 5.81 | 2.78 | 30.31 | 27.55 | 2.76 | 2.76 | 0.06 | 0.06 |
| gene13033 | https://www.ncbi.nlm.nih.gov/gene/?term=LOC105287063 | 1877 | 50 | 266.38 | 1 | 5.33 | 2 | 31.92 | 29.55 | 2.37 | 2.37 | 0.06 | 0.06 |
| gene7493 | https://www.ncbi.nlm.nih.gov/gene/?term=LOC105280704 | 4225 | 147 | 347.93 | 1 | 2.37 | 0.68 | 18.36 | 16.63 | 1.72 | 1.72 | 0.06 | 0.06 |
| gene5298 | https://www.ncbi.nlm.nih.gov/gene/?term=LOC105278145 | 4060 | 262 | 645.32 | 2 | 4.93 | 0.76 | 9.65 | 10.52 | -0.86 | 0.86 | -0.06 | 0.06 |
| gene1470 | https://www.ncbi.nlm.nih.gov/gene/?term=LOC105287597 | 2974 | 192 | 645.6 | 1 | 3.36 | 0.52 | 18.41 | 19.24 | -0.83 | 0.83 | -0.05 | 0.05 |
| gene1198 | https://www.ncbi.nlm.nih.gov/gene/?term=LOC105285507 | 2738 | 97 | 354.27 | 1 | 3.65 | 1.03 | 20.27 | 21.73 | -1.46 | 1.46 | -0.05 | 0.05 |
| gene11340 | https://www.ncbi.nlm.nih.gov/gene/?term=LOC105285131 | 2918 | 98 | 335.85 | 1 | 3.43 | 1.02 | 13.78 | 15.33 | -1.55 | 1.55 | -0.05 | 0.05 |
| gene9464 | https://www.ncbi.nlm.nih.gov/gene/?term=LOC105282975 | 1450 | 24 | 165.52 | 1 | 6.9 | 4.17 | 39.26 | 42.21 | -2.95 | 2.95 | -0.05 | 0.05 |
| gene1505 | https://www.ncbi.nlm.nih.gov/gene/?term=LOC105287633 | 1693 | 22 | 129.95 | 1 | 5.91 | 4.55 | 51.59 | 55.25 | -3.65 | 3.65 | -0.05 | 0.05 |
| gene9109 | https://www.ncbi.nlm.nih.gov/gene/?term=LOC105282561 | 2400 | 14 | 58.33 | 1 | 4.17 | 7.14 | 13.59 | 21.35 | -7.76 | 7.76 | -0.05 | 0.05 |
| gene393 | https://www.ncbi.nlm.nih.gov/gene/?term=LOC105276566 | 1123 | 2 | 17.81 | 1 | 8.9 | 50 | 47.22 | 74.3 | -27.08 | 27.08 | -0.05 | 0.05 |
| gene2970 | https://www.ncbi.nlm.nih.gov/gene/?term=LOC105275518 | 2340 | 25 | 106.84 | 1 | 4.27 | 4 | 44.93 | 41.23 | 3.69 | 3.69 | 0.04 | 0.04 |
| gene2241 | https://www.ncbi.nlm.nih.gov/gene/?term=LOC105274666 | 2598 | 61 | 234.8 | 1 | 3.85 | 1.64 | 30.46 | 28.61 | 1.85 | 1.85 | 0.04 | 0.04 |
| gene940 | https://www.ncbi.nlm.nih.gov/gene/?term=LOC105282794 | 3578 | 94 | 262.72 | 1 | 2.79 | 1.06 | 15.53 | 14.17 | 1.36 | 1.36 | 0.04 | 0.04 |
| gene4641 | https://www.ncbi.nlm.nih.gov/gene/?term=LOC105277398 | 1937 | 72 | 371.71 | 1 | 5.16 | 1.39 | 36.3 | 35.35 | 0.95 | 0.95 | 0.04 | 0.04 |
| gene3817 | https://www.ncbi.nlm.nih.gov/gene/?term=LOC105276449 | 1230 | 55 | 447.15 | 1 | 8.13 | 1.82 | 22.28 | 21.47 | 0.8 | 0.8 | 0.04 | 0.04 |
| gene474 | https://www.ncbi.nlm.nih.gov/gene/?term=LOC105277435 | 3625 | 176 | 485.52 | 1 | 2.76 | 0.57 | 10.76 | 9.98 | 0.78 | 0.78 | 0.04 | 0.04 |
| gene570 | https://www.ncbi.nlm.nih.gov/gene/?term=LOC105278549 | 1673 | 72 | 430.36 | 1 | 5.98 | 1.39 | 42.9 | 43.9 | -0.99 | 0.99 | -0.04 | 0.04 |
| gene1220 | https://www.ncbi.nlm.nih.gov/gene/?term=LOC105285742 | 3453 | 119 | 344.63 | 1 | 2.9 | 0.84 | 16.38 | 17.43 | -1.05 | 1.05 | -0.04 | 0.04 |
| gene9598 | https://www.ncbi.nlm.nih.gov/gene/?term=LOC105283132 | 1769 | 56 | 316.56 | 1 | 5.65 | 1.79 | 39.06 | 40.18 | -1.12 | 1.12 | -0.04 | 0.04 |
| gene4962 | https://www.ncbi.nlm.nih.gov/gene/?term=LOC105277761 | 4645 | 156 | 335.84 | 1 | 2.15 | 0.64 | 14.16 | 15.31 | -1.15 | 1.15 | -0.04 | 0.04 |
| gene3727 | https://www.ncbi.nlm.nih.gov/gene/?term=LOC105276367 | 2484 | 65 | 261.67 | 1 | 4.03 | 1.54 | 43.95 | 45.54 | -1.59 | 1.59 | -0.04 | 0.04 |
| gene2404 | https://www.ncbi.nlm.nih.gov/gene/?term=LOC105274853 | 1767 | 36 | 203.74 | 1 | 5.66 | 2.78 | 33.17 | 35.01 | -1.84 | 1.84 | -0.04 | 0.04 |
| gene1199 | https://www.ncbi.nlm.nih.gov/gene/?term=LOC105285498 | 2220 | 30 | 135.14 | 1 | 4.5 | 3.33 | 33.52 | 36.79 | -3.27 | 3.27 | -0.04 | 0.04 |
| gene3166 | https://www.ncbi.nlm.nih.gov/gene/?term=LOC105275707 | 2393 | 10 | 41.79 | 1 | 4.18 | 10 | 65.97 | 76.22 | -10.25 | 10.25 | -0.04 | 0.04 |
| gene4023 | https://www.ncbi.nlm.nih.gov/gene/?term=LOC105276692 | 1759 | 37 | 210.35 | 1 | 5.69 | 2.7 | 23.22 | 21.93 | 1.3 | 1.3 | 0.03 | 0.03 |
| gene9336 | https://www.ncbi.nlm.nih.gov/gene/?term=LOC105282834 | 5993 | 134 | 223.59 | 1 | 1.67 | 0.75 | 14.88 | 13.64 | 1.24 | 1.24 | 0.03 | 0.03 |
| gene5877 | https://www.ncbi.nlm.nih.gov/gene/?term=LOC105278807 | 1719 | 51 | 296.68 | 1 | 5.82 | 1.96 | 26.14 | 25.26 | 0.89 | 0.89 | 0.03 | 0.03 |
| gene298 | https://www.ncbi.nlm.nih.gov/gene/?term=LOC105275460 | 2473 | 126 | 509.5 | 1 | 4.04 | 0.79 | 13.09 | 12.43 | 0.67 | 0.67 | 0.03 | 0.03 |
| gene10678 | https://www.ncbi.nlm.nih.gov/gene/?term=LOC105284355 | 4031 | 169 | 419.25 | 1 | 2.48 | 0.59 | 20.31 | 19.66 | 0.65 | 0.65 | 0.03 | 0.03 |
| gene4034 | https://www.ncbi.nlm.nih.gov/gene/?term=LOC105276704 | 2065 | 87 | 421.31 | 2 | 9.69 | 2.3 | 14.01 | 13.38 | 0.63 | 0.63 | 0.03 | 0.03 |
| gene5525 | https://www.ncbi.nlm.nih.gov/gene/?term=LOC105278396 | 3090 | 189 | 611.65 | 1 | 3.24 | 0.53 | 24.08 | 24.52 | -0.44 | 0.44 | -0.03 | 0.03 |
| gene6698 | https://www.ncbi.nlm.nih.gov/gene/?term=LOC105279778 | 4099 | 161 | 392.78 | 1 | 2.44 | 0.62 | 8.69 | 9.57 | -0.88 | 0.88 | -0.03 | 0.03 |
| gene920 | https://www.ncbi.nlm.nih.gov/gene/?term=LOC105282518 | 1197 | 44 | 367.59 | 1 | 8.35 | 2.27 | 33.07 | 33.97 | -0.89 | 0.89 | -0.03 | 0.03 |
| gene10392 | https://www.ncbi.nlm.nih.gov/gene/?term=LOC105284036 | 3216 | 103 | 320.27 | 2 | 6.22 | 1.94 | 23.68 | 24.65 | -0.97 | 0.97 | -0.03 | 0.03 |
| gene12021 | https://www.ncbi.nlm.nih.gov/gene/?term=LOC105285905 | 1399 | 29 | 207.29 | 1 | 7.15 | 3.45 | 32.77 | 34.25 | -1.48 | 1.48 | -0.03 | 0.03 |
| gene6422 | https://www.ncbi.nlm.nih.gov/gene/?term=LOC105279458 | 2419 | 42 | 173.63 | 1 | 4.13 | 2.38 | 44.07 | 45.58 | -1.51 | 1.51 | -0.03 | 0.03 |
| gene564 | https://www.ncbi.nlm.nih.gov/gene/?term=LOC105278478 | 2802 | 16 | 57.1 | 1 | 3.57 | 6.25 | 14.91 | 19.95 | -5.03 | 5.03 | -0.03 | 0.03 |
| gene11584 | https://www.ncbi.nlm.nih.gov/gene/?term=LOC105285416 | 3709 | 16 | 43.14 | 2 | 5.39 | 12.5 | 21.86 | 29.47 | -7.61 | 7.61 | -0.03 | 0.03 |
| gene10345 | https://www.ncbi.nlm.nih.gov/gene/?term=LOC105283989 | 4444 | 41 | 92.26 | 1 | 2.25 | 2.44 | 38.24 | 36.42 | 1.83 | 1.83 | 0.02 | 0.02 |
| gene2748 | https://www.ncbi.nlm.nih.gov/gene/?term=LOC105275261 | 3845 | 53 | 137.84 | 1 | 2.6 | 1.89 | 39.61 | 38.42 | 1.19 | 1.19 | 0.02 | 0.02 |
| gene10107 | https://www.ncbi.nlm.nih.gov/gene/?term=LOC105283712 | 1883 | 36 | 191.18 | 1 | 5.31 | 2.78 | 30.43 | 29.26 | 1.17 | 1.17 | 0.02 | 0.02 |
| gene1884 | https://www.ncbi.nlm.nih.gov/gene/?term=LOC105288072 | 4169 | 84 | 201.49 | 1 | 2.4 | 1.19 | 24.86 | 23.72 | 1.14 | 1.14 | 0.02 | 0.02 |
| gene7734 | https://www.ncbi.nlm.nih.gov/gene/?term=LOC105280972 | 1528 | 35 | 229.06 | 1 | 6.54 | 2.86 | 17.63 | 16.59 | 1.04 | 1.04 | 0.02 | 0.02 |
| gene7884 | https://www.ncbi.nlm.nih.gov/gene/?term=LOC105281266 | 3390 | 87 | 256.64 | 1 | 2.95 | 1.15 | 15.85 | 14.89 | 0.96 | 0.96 | 0.02 | 0.02 |
| gene1147 | https://www.ncbi.nlm.nih.gov/gene/?term=LOC105284954 | 4022 | 118 | 293.39 | 1 | 2.49 | 0.85 | 15.13 | 14.31 | 0.81 | 0.81 | 0.02 | 0.02 |
| gene292 | https://www.ncbi.nlm.nih.gov/gene/?term=LOC105275398 | 2825 | 88 | 311.5 | 1 | 3.54 | 1.14 | 22.66 | 21.93 | 0.73 | 0.73 | 0.02 | 0.02 |
| gene7888 | https://www.ncbi.nlm.nih.gov/gene/?term=LOC105281141 | 4357 | 170 | 390.18 | 1 | 2.3 | 0.59 | 28.6 | 28 | 0.6 | 0.6 | 0.02 | 0.02 |

| | | | | | | | | | | | | | |
|---|---|---|---|---|---|---|---|---|---|---|---|---|---|
| gene6652 | https://www.ncbi.nlm.nih.gov/gene/?term=LOC105279719 | 2831 | 88 | 310.84 | 1 | 3.53 | 1.14 | 21.65 | 21.08 | 0.57 | 0.57 | 0.02 | 0.02 |
| gene4892 | https://www.ncbi.nlm.nih.gov/gene/?term=LOC105277667 | 3898 | 127 | 325.81 | 1 | 2.57 | 0.79 | 27.72 | 27.21 | 0.52 | 0.52 | 0.02 | 0.02 |
| gene12978 | https://www.ncbi.nlm.nih.gov/gene/?term=LOC105286997 | 2858 | 150 | 524.84 | 1 | 3.5 | 0.67 | 11.28 | 10.84 | 0.44 | 0.44 | 0.02 | 0.02 |
| gene11819 | https://www.ncbi.nlm.nih.gov/gene/?term=LOC105285671 | 2366 | 104 | 439.56 | 1 | 4.23 | 0.96 | 30.06 | 29.63 | 0.43 | 0.43 | 0.02 | 0.02 |
| gene3885 | https://www.ncbi.nlm.nih.gov/gene/?term=LOC105276527 | 7928 | 369 | 465.44 | 1 | 1.26 | 0.27 | 8.26 | 7.93 | 0.33 | 0.33 | 0.02 | 0.02 |
| gene10549 | https://www.ncbi.nlm.nih.gov/gene/?term=LOC105284210 | 7145 | 365 | 510.85 | 2 | 2.8 | 0.55 | 9.55 | 9.9 | -0.35 | 0.35 | -0.02 | 0.02 |
| gene10326 | https://www.ncbi.nlm.nih.gov/gene/?term=LOC105283962 | 3042 | 137 | 450.36 | 1 | 3.29 | 0.73 | 7.31 | 7.68 | -0.37 | 0.37 | -0.02 | 0.02 |
| gene3793 | https://www.ncbi.nlm.nih.gov/gene/?term=LOC105276429 | 2901 | 131 | 451.57 | 1 | 3.45 | 0.76 | 22.23 | 22.66 | -0.43 | 0.43 | -0.02 | 0.02 |
| gene10606 | https://www.ncbi.nlm.nih.gov/gene/?term=LOC105284273 | 6119 | 272 | 444.52 | 1 | 1.63 | 0.37 | 17.87 | 18.32 | -0.45 | 0.45 | -0.02 | 0.02 |
| gene6583 | https://www.ncbi.nlm.nih.gov/gene/?term=LOC105279649 | 7444 | 372 | 499.73 | 1 | 1.34 | 0.27 | 10.85 | 11.31 | -0.45 | 0.45 | -0.02 | 0.02 |
| gene4190 | https://www.ncbi.nlm.nih.gov/gene/?term=LOC105276875 | 3150 | 134 | 425.4 | 1 | 3.17 | 0.75 | 20.21 | 20.67 | -0.46 | 0.46 | -0.02 | 0.02 |
| gene7166 | https://www.ncbi.nlm.nih.gov/gene/?term=LOC105280326 | 6117 | 259 | 423.41 | 1 | 1.63 | 0.39 | 14.46 | 14.93 | -0.47 | 0.47 | -0.02 | 0.02 |
| gene6966 | https://www.ncbi.nlm.nih.gov/gene/?term=LOC105280104 | 3747 | 132 | 352.28 | 1 | 2.67 | 0.76 | 16.91 | 17.46 | -0.55 | 0.55 | -0.02 | 0.02 |
| gene8814 | https://www.ncbi.nlm.nih.gov/gene/?term=LOC105282240 | 6910 | 204 | 295.22 | 1 | 1.45 | 0.49 | 8.01 | 8.64 | -0.63 | 0.63 | -0.02 | 0.02 |
| gene5184 | https://www.ncbi.nlm.nih.gov/gene/?term=LOC105277999 | 5528 | 165 | 298.48 | 1 | 1.81 | 0.61 | 11.31 | 11.98 | -0.67 | 0.67 | -0.02 | 0.02 |
| gene11235 | https://www.ncbi.nlm.nih.gov/gene/?term=LOC105285004 | 1339 | 36 | 268.86 | 1 | 7.47 | 2.78 | 19.18 | 19.88 | -0.7 | 0.7 | -0.02 | 0.02 |
| gene1310 | https://www.ncbi.nlm.nih.gov/gene/?term=LOC105286661 | 2582 | 68 | 263.36 | 1 | 3.87 | 1.47 | 18.99 | 19.74 | -0.75 | 0.75 | -0.02 | 0.02 |
| gene12573 | https://www.ncbi.nlm.nih.gov/gene/?term=LOC105286515 | 3647 | 107 | 293.39 | 1 | 2.74 | 0.93 | 10.59 | 11.4 | -0.82 | 0.82 | -0.02 | 0.02 |
| gene5101 | https://www.ncbi.nlm.nih.gov/gene/?term=LOC105277903 | 1813 | 37 | 204.08 | 1 | 5.52 | 2.7 | 54.12 | 55.06 | -0.94 | 0.94 | -0.02 | 0.02 |
| gene10155 | https://www.ncbi.nlm.nih.gov/gene/?term=LOC105283756 | 7030 | 116 | 165.01 | 1 | 1.42 | 0.86 | 7.15 | 8.11 | -0.96 | 0.96 | -0.02 | 0.02 |
| gene3168 | https://www.ncbi.nlm.nih.gov/gene/?term=LOC105275711 | 6150 | 97 | 157.72 | 1 | 1.63 | 1.03 | 10.51 | 11.68 | -1.17 | 1.17 | -0.02 | 0.02 |
| gene388 | https://www.ncbi.nlm.nih.gov/gene/?term=LOC105276538 | 1558 | 21 | 134.79 | 1 | 6.42 | 4.76 | 69.07 | 70.25 | -1.18 | 1.18 | -0.02 | 0.02 |
| gene3599 | https://www.ncbi.nlm.nih.gov/gene/?term=LOC105276196 | 2955 | 60 | 203.05 | 1 | 3.38 | 1.67 | 37.66 | 38.86 | -1.2 | 1.2 | -0.02 | 0.02 |
| gene12203 | https://www.ncbi.nlm.nih.gov/gene/?term=LOC105286099 | 5967 | 87 | 145.8 | 1 | 1.68 | 1.15 | 10.81 | 12.06 | -1.25 | 1.25 | -0.02 | 0.02 |
| gene7398 | https://www.ncbi.nlm.nih.gov/gene/?term=LOC105280589 | 3452 | 42 | 121.67 | 1 | 2.9 | 2.38 | 24.95 | 26.26 | -1.31 | 1.31 | -0.02 | 0.02 |
| gene7248 | https://www.ncbi.nlm.nih.gov/gene/?term=LOC105280408 | 5175 | 59 | 114.01 | 1 | 1.93 | 1.69 | 15.82 | 17.71 | -1.89 | 1.89 | -0.02 | 0.02 |
| gene8178 | https://www.ncbi.nlm.nih.gov/gene/?term=LOC105281515 | 8647 | 60 | 69.39 | 1 | 1.16 | 1.67 | 30.54 | 29.47 | 1.06 | 1.06 | 0.01 | 0.01 |
| gene7183 | https://www.ncbi.nlm.nih.gov/gene/?term=LOC105280348 | 3925 | 80 | 203.82 | 1 | 2.55 | 1.25 | 17.35 | 16.62 | 0.73 | 0.73 | 0.01 | 0.01 |
| gene322 | https://www.ncbi.nlm.nih.gov/gene/?term=LOC105275777 | 1606 | 28 | 174.35 | 1 | 6.23 | 3.57 | 36.76 | 36.04 | 0.72 | 0.72 | 0.01 | 0.01 |
| gene2683 | https://www.ncbi.nlm.nih.gov/gene/?term=LOC105275187 | 5492 | 78 | 142.02 | 1 | 1.82 | 1.28 | 16.44 | 16.02 | 0.43 | 0.43 | 0.01 | 0.01 |
| gene3107 | https://www.ncbi.nlm.nih.gov/gene/?term=LOC105275650 | 5103 | 182 | 356.65 | 1 | 1.96 | 0.55 | 16.89 | 16.49 | 0.4 | 0.4 | 0.01 | 0.01 |
| gene6455 | https://www.ncbi.nlm.nih.gov/gene/?term=LOC105279500 | 2612 | 35 | 134 | 1 | 3.83 | 2.86 | 48.92 | 48.53 | 0.39 | 0.39 | 0.01 | 0.01 |
| gene13036 | https://www.ncbi.nlm.nih.gov/gene/?term=LOC105287059 | 4959 | 193 | 389.19 | 1 | 2.02 | 0.52 | 16.75 | 16.43 | 0.32 | 0.32 | 0.01 | 0.01 |
| gene8729 | https://www.ncbi.nlm.nih.gov/gene/?term=LOC105282137 | 5005 | 226 | 451.55 | 1 | 2 | 0.44 | 14.98 | 14.68 | 0.29 | 0.29 | 0.01 | 0.01 |
| gene12572 | https://www.ncbi.nlm.nih.gov/gene/?term=LOC105286514 | 2374 | 52 | 219.04 | 1 | 4.21 | 1.92 | 35.13 | 34.85 | 0.28 | 0.28 | 0.01 | 0.01 |
| gene9101 | https://www.ncbi.nlm.nih.gov/gene/?term=LOC105282553 | 3068 | 106 | 345.5 | 1 | 3.26 | 0.94 | 22.9 | 22.63 | 0.27 | 0.27 | 0.01 | 0.01 |
| gene10018 | https://www.ncbi.nlm.nih.gov/gene/?term=LOC105283607 | 4414 | 228 | 516.54 | 1 | 2.27 | 0.44 | 14.1 | 13.84 | 0.26 | 0.26 | 0.01 | 0.01 |
| gene2668 | https://www.ncbi.nlm.nih.gov/gene/?term=LOC105275170 | 12496 | 356 | 284.89 | 1 | 0.8 | 0.28 | 2.04 | 1.8 | 0.24 | 0.24 | 0.01 | 0.01 |
| gene10540 | https://www.ncbi.nlm.nih.gov/gene/?term=LOC105284200 | 5267 | 214 | 406.3 | 1 | 1.9 | 0.47 | 13.24 | 13.01 | 0.23 | 0.23 | 0.01 | 0.01 |
| gene10596 | https://www.ncbi.nlm.nih.gov/gene/?term=LOC105284262 | 6520 | 239 | 366.56 | 1 | 1.53 | 0.42 | 4.23 | 4.05 | 0.19 | 0.19 | 0.01 | 0.01 |
| gene6286 | https://www.ncbi.nlm.nih.gov/gene/?term=LOC105279298 | 2861 | 128 | 447.4 | 1 | 3.5 | 0.78 | 23.43 | 23.27 | 0.16 | 0.16 | 0.01 | 0.01 |
| gene11275 | https://www.ncbi.nlm.nih.gov/gene/?term=LOC105285055 | 4119 | 246 | 597.23 | 1 | 2.43 | 0.41 | 12.22 | 12.12 | 0.1 | 0.1 | 0.01 | 0.01 |
| gene12802 | https://www.ncbi.nlm.nih.gov/gene/?term=LOC105286799 | 3183 | 180 | 565.5 | 1 | 3.14 | 0.56 | 16.81 | 16.93 | -0.12 | 0.12 | -0.01 | 0.01 |
| gene5654 | https://www.ncbi.nlm.nih.gov/gene/?term=LOC105278543 | 4019 | 181 | 450.36 | 1 | 2.49 | 0.55 | 15.7 | 15.83 | -0.13 | 0.13 | -0.01 | 0.01 |
| gene8274 | https://www.ncbi.nlm.nih.gov/gene/?term=LOC105281624 | 5425 | 227 | 418.43 | 1 | 1.84 | 0.44 | 9.53 | 9.7 | -0.17 | 0.17 | -0.01 | 0.01 |
| gene13039 | https://www.ncbi.nlm.nih.gov/gene/?term=LOC105287065 | 4902 | 253 | 516.12 | 1 | 2.04 | 0.4 | 12.12 | 12.3 | -0.18 | 0.18 | -0.01 | 0.01 |
| gene11579 | https://www.ncbi.nlm.nih.gov/gene/?term=LOC105285406 | 13571 | 316 | 232.85 | 1 | 0.74 | 0.32 | 6.01 | 6.26 | -0.25 | 0.25 | -0.01 | 0.01 |
| gene7249 | https://www.ncbi.nlm.nih.gov/gene/?term=LOC105280422 | 1775 | 66 | 371.83 | 1 | 5.63 | 1.52 | 15.78 | 16.04 | -0.26 | 0.26 | -0.01 | 0.01 |
| gene7199 | https://www.ncbi.nlm.nih.gov/gene/?term=LOC105280366 | 1414 | 34 | 240.45 | 1 | 7.07 | 2.94 | 44.65 | 44.98 | -0.33 | 0.33 | -0.01 | 0.01 |
| gene5524 | https://www.ncbi.nlm.nih.gov/gene/?term=LOC105278397 | 5177 | 209 | 403.71 | 1 | 1.93 | 0.48 | 23.99 | 24.35 | -0.36 | 0.36 | -0.01 | 0.01 |
| gene4376 | https://www.ncbi.nlm.nih.gov/gene/?term=LOC105277083 | 4307 | 129 | 299.51 | 1 | 2.32 | 0.78 | 18.69 | 19.06 | -0.38 | 0.38 | -0.01 | 0.01 |
| gene11583 | https://www.ncbi.nlm.nih.gov/gene/?term=LOC105285414 | 8192 | 192 | 234.38 | 1 | 1.22 | 0.52 | 8.5 | 8.89 | -0.39 | 0.39 | -0.01 | 0.01 |
| gene7915 | https://www.ncbi.nlm.nih.gov/gene/?term=LOC105281173 | 1597 | 42 | 262.99 | 1 | 6.26 | 2.38 | 62.33 | 62.75 | -0.42 | 0.42 | -0.01 | 0.01 |
| gene9330 | https://www.ncbi.nlm.nih.gov/gene/?term=LOC105282825 | 3936 | 98 | 248.98 | 1 | 2.54 | 1.02 | 13.47 | 13.91 | -0.44 | 0.44 | -0.01 | 0.01 |
| gene7247 | https://www.ncbi.nlm.nih.gov/gene/?term=LOC105280407 | 10486 | 227 | 216.48 | 1 | 0.95 | 0.44 | 14.79 | 15.25 | -0.46 | 0.46 | -0.01 | 0.01 |
| gene2173 | https://www.ncbi.nlm.nih.gov/gene/?term=LOC105274590 | 4387 | 87 | 198.31 | 2 | 4.56 | 2.3 | 23.56 | 24.1 | -0.54 | 0.54 | -0.01 | 0.01 |
| gene4033 | https://www.ncbi.nlm.nih.gov/gene/?term=LOC105276705 | 10546 | 155 | 146.98 | 1 | 0.95 | 0.65 | 5.56 | 6.1 | -0.55 | 0.55 | -0.01 | 0.01 |
| gene11561 | https://www.ncbi.nlm.nih.gov/gene/?term=LOC105285387 | 11528 | 218 | 189.1 | 1 | 0.87 | 0.46 | 8.62 | 9.18 | -0.56 | 0.56 | -0.01 | 0.01 |
| gene9411 | https://www.ncbi.nlm.nih.gov/gene/?term=LOC105282914 | 5534 | 42 | 75.89 | 1 | 1.81 | 2.38 | 49.22 | 50.92 | -1.7 | 1.7 | -0.01 | 0.01 |
| gene652 | https://www.ncbi.nlm.nih.gov/gene/?term=LOC105279528 | 5806 | 39 | 67.17 | 1 | 1.72 | 2.56 | 17.73 | 19.93 | -2.2 | 2.2 | -0.01 | 0.01 |
| gene7368 | https://www.ncbi.nlm.nih.gov/gene/?term=LOC105280556 | 4105 | 71 | 172.96 | 1 | 2.44 | 1.41 | 15.89 | 15.73 | 0.16 | 0.16 | 0 | 0 |
| gene12406 | https://www.ncbi.nlm.nih.gov/gene/?term=LOC105286316 | 1386 | 44 | 317.46 | 1 | 7.22 | 2.27 | 32.85 | 32.71 | 0.14 | 0.14 | 0 | 0 |
| gene10600 | https://www.ncbi.nlm.nih.gov/gene/?term=LOC105284270 | 4837 | 111 | 229.48 | 1 | 2.07 | 0.9 | 11.96 | 11.83 | 0.13 | 0.13 | 0 | 0 |
| gene927 | https://www.ncbi.nlm.nih.gov/gene/?term=LOC105282600 | 3727 | 60 | 160.99 | 1 | 2.68 | 1.67 | 28.88 | 28.75 | 0.13 | 0.13 | 0 | 0 |

| | | | | | | | | | | | | | |
|---|---|---|---|---|---|---|---|---|---|---|---|---|---|
| gene9327 | https://www.ncbi.nlm.nih.gov/gene/?term=LOC105282821 | 7793 | 183 | 234.83 | 1 | 1.28 | 0.55 | 12.55 | 12.42 | 0.13 | 0.13 | 0 | 0 |
| gene5375 | https://www.ncbi.nlm.nih.gov/gene/?term=LOC105278228 | 8640 | 191 | 221.06 | 1 | 1.16 | 0.52 | 22.09 | 21.99 | 0.1 | 0.1 | 0 | 0 |
| gene10222 | https://www.ncbi.nlm.nih.gov/gene/?term=LOC105283838 | 1748 | 30 | 171.62 | 1 | 5.72 | 3.33 | 35.37 | 35.28 | 0.09 | 0.09 | 0 | 0 |
| gene10520 | https://www.ncbi.nlm.nih.gov/gene/?term=LOC105284184 | 2451 | 102 | 416.16 | 1 | 4.08 | 0.98 | 30 | 29.93 | 0.07 | 0.07 | 0 | 0 |
| gene9326 | https://www.ncbi.nlm.nih.gov/gene/?term=LOC105282818 | 16612 | 236 | 142.07 | 1 | 0.6 | 0.42 | 13.17 | 13.1 | 0.07 | 0.07 | 0 | 0 |
| gene2443 | https://www.ncbi.nlm.nih.gov/gene/?term=LOC105274895 | 10433 | 424 | 406.4 1 | 0.96 | 0.24 | 5.17 | 5.13 | 0.03 | 0.03 | 0 | 0 | |
| gene53 | https://www.ncbi.nlm.nih.gov/gene/?term=LOC105277148 | 19283 | 333 | 172.69 | 1 | 0.52 | 0.3 | 4.68 | 4.68 | 0 | 0 | 0 | 0 |
| gene11454 | https://www.ncbi.nlm.nih.gov/gene/?term=LOC105285256 | 1906 | 50 | 262.33 | 1 | 5.25 | 2 | 44.07 | 44.08 | -0.01 | 0.01 | 0 | 0 |
| gene13114 | https://www.ncbi.nlm.nih.gov/gene/?term=LOC105287147 | 7609 | 432 | 567.75 | 1 | 1.31 | 0.23 | 9.01 | 9.02 | -0.01 | 0.01 | 0 | 0 |
| gene2271 | https://www.ncbi.nlm.nih.gov/gene/?term=LOC105274706 | 1448 | 47 | 324.59 | 1 | 6.91 | 2.13 | 41.96 | 41.97 | -0.01 | 0.01 | 0 | 0 |
| gene8600 | https://www.ncbi.nlm.nih.gov/gene/?term=LOC105282001 | 8513 | 206 | 241.98 | 1 | 1.17 | 0.49 | 9.71 | 9.75 | -0.04 | 0.04 | 0 | 0 |
| gene5746 | https://www.ncbi.nlm.nih.gov/gene/?term=LOC105278666 | 10343 | 483 | 466.98 | 1 | 0.97 | 0.21 | 6.37 | 6.41 | -0.05 | 0.05 | 0 | 0 |
| gene6620 | https://www.ncbi.nlm.nih.gov/gene/?term=LOC105279687 | 17998 | 448 | 248.92 | 1 | 0.56 | 0.22 | 4.43 | 4.47 | -0.05 | 0.05 | 0 | 0 |
| gene1632 | https://www.ncbi.nlm.nih.gov/gene/?term=LOC105287764 | 18061 | 431 | 238.64 | 1 | 0.55 | 0.23 | 3.3 | 3.37 | -0.07 | 0.07 | 0 | 0 |
| gene2806 | https://www.ncbi.nlm.nih.gov/gene/?term=LOC105275327 | 14257 | 552 | 387.18 | 1 | 0.7 | 0.18 | 6.34 | 6.41 | -0.07 | 0.07 | 0 | 0 |
| gene3470 | https://www.ncbi.nlm.nih.gov/gene/?term=LOC105276059 | 8716 | 182 | 208.81 | 1 | 1.15 | 0.55 | 9.23 | 9.32 | -0.09 | 0.09 | 0 | 0 |
| gene5646 | https://www.ncbi.nlm.nih.gov/gene/?term=LOC105278534 | 7475 | 202 | 270.23 | 1 | 1.34 | 0.5 | 16.69 | 16.79 | -0.1 | 0.1 | 0 | 0 |
| gene12427 | https://www.ncbi.nlm.nih.gov/gene/?term=LOC105286339 | 5220 | 187 | 358.24 | 1 | 1.92 | 0.53 | 15.91 | 16.02 | -0.12 | 0.12 | 0 | 0 |
| gene10844 | https://www.ncbi.nlm.nih.gov/gene/?term=LOC105284535 | 9086 | 259 | 285.05 | 1 | 1.1 | 0.39 | 14.4 | 14.56 | -0.16 | 0.16 | 0 | 0 |
| gene6639 | https://www.ncbi.nlm.nih.gov/gene/?term=LOC105279705 | 7656 | 83 | 108.41 | 1 | 1.31 | 1.2 | 19.9 | 20.17 | -0.28 | 0.28 | 0 | 0 |

Directory: OGL File: rnk-dmg-Cb_LI_BCphaseB1.vs.RphaseB2.txt

| gene_ID | gene_link | gwidth | #Sites | #per10Kb | #dmSites | #dmsp10kb | %dmSites | %pSite1 | %pSite2 | DMpSite | ADMpSite | DMpNucl | ADMpNucl |
|---|---|---|---|---|---|---|---|---|---|---|---|---|---|
| gene3165 | https://www.ncbi.nlm.nih.gov/gene/?term=LOC105275706 | 2374 | 68 | 286.44 | 5 | 21.06 | 7.35 | 33.32 | 23.13 | 10.19 | 10.19 | 0.29 | 0.29 |
| gene2593 | https://www.ncbi.nlm.nih.gov/gene/?term=LOC105275070 | 1774 | 7 | 39.46 | 5 | 28.18 | 71.43 | 58.65 | 4.09 | 54.56 | 54.56 | 0.22 | 0.22 |
| gene5714 | https://www.ncbi.nlm.nih.gov/gene/?term=LOC105278729 | 710 | 17 | 239.44 | 1 | 14.08 | 5.88 | 58.14 | 53.54 | 4.59 | 4.59 | 0.11 | 0.11 |
| gene4498 | https://www.ncbi.nlm.nih.gov/gene/?term=LOC105277239 | 2831 | 121 | 427.41 | 1 | 3.53 | 0.83 | 11.26 | 13.23 | -1.97 | 1.97 | -0.08 | 0.08 |
| gene940 | https://www.ncbi.nlm.nih.gov/gene/?term=LOC105282794 | 3578 | 83 | 231.97 | 1 | 2.79 | 1.2 | 17.02 | 13.98 | 3.04 | 3.04 | 0.07 | 0.07 |
| gene7568 | https://www.ncbi.nlm.nih.gov/gene/?term=LOC105280786 | 2525 | 122 | 483.17 | 1 | 3.96 | 0.82 | 29.81 | 28.44 | 1.38 | 1.38 | 0.07 | 0.07 |
| gene1332 | https://www.ncbi.nlm.nih.gov/gene/?term=LOC105286872 | 2460 | 154 | 626.02 | 1 | 4.07 | 0.65 | 21.26 | 20.18 | 1.08 | 1.08 | 0.07 | 0.07 |
| gene4641 | https://www.ncbi.nlm.nih.gov/gene/?term=LOC105277398 | 1937 | 71 | 366.55 | 1 | 5.16 | 1.41 | 37.78 | 39.69 | -1.91 | 1.91 | -0.07 | 0.07 |
| gene6334 | https://www.ncbi.nlm.nih.gov/gene/?term=LOC105279356 | 1471 | 48 | 326.31 | 1 | 6.8 | 2.08 | 28.86 | 26.95 | 1.91 | 1.91 | 0.06 | 0.06 |
| gene11819 | https://www.ncbi.nlm.nih.gov/gene/?term=LOC105285671 | 2366 | 102 | 431.11 | 1 | 4.23 | 0.98 | 27.3 | 25.91 | 1.39 | 1.39 | 0.06 | 0.06 |
| gene6422 | https://www.ncbi.nlm.nih.gov/gene/?term=LOC105279458 | 2419 | 38 | 157.09 | 1 | 4.13 | 2.63 | 46.62 | 50.37 | -3.75 | 3.75 | -0.06 | 0.06 |
| gene10602 | https://www.ncbi.nlm.nih.gov/gene/?term=LOC105284268 | 2122 | 19 | 89.54 | 1 | 4.71 | 5.26 | 26.09 | 32.99 | -6.9 | 6.9 | -0.06 | 0.06 |
| gene2970 | https://www.ncbi.nlm.nih.gov/gene/?term=LOC105275518 | 2340 | 20 | 85.47 | 1 | 4.27 | 5 | 58.11 | 52.81 | 5.3 | 5.3 | 0.05 | 0.05 |
| gene1855 | https://www.ncbi.nlm.nih.gov/gene/?term=LOC105288036 | 2387 | 78 | 326.77 | 1 | 4.19 | 1.28 | 33.45 | 31.79 | 1.66 | 1.66 | 0.05 | 0.05 |
| gene6966 | https://www.ncbi.nlm.nih.gov/gene/?term=LOC105280104 | 3747 | 135 | 360.29 | 1 | 2.67 | 0.74 | 17.67 | 16.38 | 1.29 | 1.29 | 0.05 | 0.05 |
| gene5298 | https://www.ncbi.nlm.nih.gov/gene/?term=LOC105278145 | 4060 | 235 | 578.82 | 2 | 4.93 | 0.85 | 10.4 | 11.25 | -0.85 | 0.85 | -0.05 | 0.05 |
| gene3793 | https://www.ncbi.nlm.nih.gov/gene/?term=LOC105276429 | 2901 | 112 | 386.07 | 1 | 3.45 | 0.89 | 25.16 | 26.39 | -1.23 | 1.23 | -0.05 | 0.05 |
| gene3727 | https://www.ncbi.nlm.nih.gov/gene/?term=LOC105276367 | 2484 | 63 | 253.62 | 1 | 4.03 | 1.59 | 45.38 | 47.24 | -1.87 | 1.87 | -0.05 | 0.05 |
| gene3599 | https://www.ncbi.nlm.nih.gov/gene/?term=LOC105276196 | 2955 | 59 | 199.66 | 1 | 3.38 | 1.69 | 37.76 | 40.32 | -2.56 | 2.56 | -0.05 | 0.05 |
| gene6237 | https://www.ncbi.nlm.nih.gov/gene/?term=LOC105279244 | 1484 | 25 | 168.46 | 1 | 6.74 | 4 | 41.13 | 44.31 | -3.18 | 3.18 | -0.05 | 0.05 |
| gene1008 | https://www.ncbi.nlm.nih.gov/gene/?term=LOC105283492 | 1479 | 10 | 67.61 | 1 | 6.76 | 10 | 34.7 | 42 | -7.3 | 7.3 | -0.05 | 0.05 |
| gene393 | https://www.ncbi.nlm.nih.gov/gene/?term=LOC105276566 | 1123 | 2 | 17.81 | 1 | 8.9 | 50 | 47.22 | 76.78 | -29.56 | 29.56 | -0.05 | 0.05 |
| gene1518 | https://www.ncbi.nlm.nih.gov/gene/?term=LOC105287648 | 2195 | 25 | 113.9 | 1 | 4.56 | 4 | 44.43 | 41.18 | 3.25 | 3.25 | 0.04 | 0.04 |
| gene2844 | https://www.ncbi.nlm.nih.gov/gene/?term=LOC105275363 | 4330 | 190 | 438.8 | 1 | 2.31 | 0.53 | 14.79 | 13.86 | 0.93 | 0.93 | 0.04 | 0.04 |
| gene10678 | https://www.ncbi.nlm.nih.gov/gene/?term=LOC105284355 | 4031 | 166 | 411.81 | 1 | 2.48 | 0.6 | 19.71 | 18.79 | 0.92 | 0.92 | 0.04 | 0.04 |
| gene4034 | https://www.ncbi.nlm.nih.gov/gene/?term=LOC105276704 | 2065 | 87 | 421.31 | 1 | 4.84 | 1.15 | 14.97 | 14.08 | 0.89 | 0.89 | 0.04 | 0.04 |
| gene5831 | https://www.ncbi.nlm.nih.gov/gene/?term=LOC105278761 | 2506 | 110 | 438.95 | 1 | 3.99 | 0.91 | 23.47 | 22.6 | 0.88 | 0.88 | 0.04 | 0.04 |
| gene11401 | https://www.ncbi.nlm.nih.gov/gene/?term=LOC105285202 | 1841 | 79 | 429.11 | 1 | 5.43 | 1.27 | 24.33 | 23.48 | 0.85 | 0.85 | 0.04 | 0.04 |
| gene9740 | https://www.ncbi.nlm.nih.gov/gene/?term=LOC105283277 | 2887 | 137 | 474.54 | 1 | 3.46 | 0.73 | 19.95 | 20.71 | -0.76 | 0.76 | -0.04 | 0.04 |
| gene10213 | https://www.ncbi.nlm.nih.gov/gene/?term=LOC105283826 | 1503 | 54 | 359.28 | 1 | 6.65 | 1.85 | 14.52 | 15.73 | -1.21 | 1.21 | -0.04 | 0.04 |
| gene11340 | https://www.ncbi.nlm.nih.gov/gene/?term=LOC105285131 | 2918 | 88 | 301.58 | 1 | 3.43 | 1.14 | 10.13 | 11.45 | -1.32 | 1.32 | -0.04 | 0.04 |
| gene1147 | https://www.ncbi.nlm.nih.gov/gene/?term=LOC105284954 | 4022 | 90 | 223.77 | 1 | 2.49 | 1.11 | 16.68 | 18.39 | -1.71 | 1.71 | -0.04 | 0.04 |
| gene2413 | https://www.ncbi.nlm.nih.gov/gene/?term=LOC105274863 | 2006 | 49 | 244.27 | 1 | 4.99 | 2.04 | 53.75 | 55.47 | -1.72 | 1.72 | -0.04 | 0.04 |
| gene10107 | https://www.ncbi.nlm.nih.gov/gene/?term=LOC105283712 | 1883 | 35 | 185.87 | 1 | 5.31 | 2.86 | 26.3 | 28.27 | -1.97 | 1.97 | -0.04 | 0.04 |
| gene1505 | https://www.ncbi.nlm.nih.gov/gene/?term=LOC105287633 | 1693 | 27 | 159.48 | 1 | 5.91 | 3.7 | 42.55 | 45.35 | -2.8 | 2.8 | -0.04 | 0.04 |
| gene5647 | https://www.ncbi.nlm.nih.gov/gene/?term=LOC105278567 | 3118 | 36 | 115.46 | 1 | 3.21 | 2.78 | 16.65 | 14.27 | 2.38 | 2.38 | 0.03 | 0.03 |
| gene5273 | https://www.ncbi.nlm.nih.gov/gene/?term=LOC105278108 | 4067 | 46 | 113.11 | 1 | 2.46 | 2.17 | 27.18 | 24.92 | 2.26 | 2.26 | 0.03 | 0.03 |
| gene7734 | https://www.ncbi.nlm.nih.gov/gene/?term=LOC105280972 | 1528 | 30 | 196.34 | 1 | 6.54 | 3.33 | 20.49 | 18.79 | 1.71 | 1.71 | 0.03 | 0.03 |
| gene8661 | https://www.ncbi.nlm.nih.gov/gene/?term=LOC105282063 | 1926 | 34 | 176.53 | 1 | 5.19 | 2.94 | 30.6 | 28.98 | 1.62 | 1.62 | 0.03 | 0.03 |
| gene4213 | https://www.ncbi.nlm.nih.gov/gene/?term=LOC105276901 | 3257 | 81 | 248.7 | 1 | 3.07 | 1.23 | 28.04 | 26.96 | 1.09 | 1.09 | 0.03 | 0.03 |
| gene8274 | https://www.ncbi.nlm.nih.gov/gene/?term=LOC105281624 | 5425 | 211 | 388.94 | 1 | 1.84 | 0.47 | 9.95 | 9.18 | 0.77 | 0.77 | 0.03 | 0.03 |
| gene12265 | https://www.ncbi.nlm.nih.gov/gene/?term=LOC105286166 | 3050 | 110 | 360.66 | 1 | 3.28 | 0.91 | 22.97 | 23.8 | -0.82 | 0.82 | -0.03 | 0.03 |
| gene10606 | https://www.ncbi.nlm.nih.gov/gene/?term=LOC105284273 | 6119 | 248 | 405.29 | 1 | 1.63 | 0.4 | 17.15 | 17.98 | -0.83 | 0.83 | -0.03 | 0.03 |
| gene1220 | https://www.ncbi.nlm.nih.gov/gene/?term=LOC105285742 | 3453 | 111 | 321.46 | 1 | 2.9 | 0.9 | 21.24 | 22.19 | -0.96 | 0.96 | -0.03 | 0.03 |
| gene6031 | https://www.ncbi.nlm.nih.gov/gene/?term=LOC105279000 | 5691 | 179 | 314.53 | 1 | 1.76 | 0.56 | 13.44 | 14.41 | -0.97 | 0.97 | -0.03 | 0.03 |
| gene7211 | https://www.ncbi.nlm.nih.gov/gene/?term=LOC105280377 | 2024 | 67 | 331.03 | 1 | 4.94 | 1.49 | 44.45 | 45.48 | -1.03 | 1.03 | -0.03 | 0.03 |
| gene12981 | https://www.ncbi.nlm.nih.gov/gene/?term=LOC105286999 | 2063 | 46 | 222.98 | 1 | 4.85 | 2.17 | 31.8 | 32.93 | -1.13 | 1.13 | -0.03 | 0.03 |
| gene12406 | https://www.ncbi.nlm.nih.gov/gene/?term=LOC105286316 | 1386 | 33 | 238.1 | 1 | 7.22 | 3.03 | 40.83 | 41.98 | -1.15 | 1.15 | -0.03 | 0.03 |
| gene388 | https://www.ncbi.nlm.nih.gov/gene/?term=LOC105276538 | 1558 | 18 | 115.53 | 2 | 12.84 | 11.11 | 66.09 | 68.41 | -2.32 | 2.32 | -0.03 | 0.03 |
| gene3168 | https://www.ncbi.nlm.nih.gov/gene/?term=LOC105275711 | 6150 | 69 | 112.2 | 1 | 1.63 | 1.45 | 12.54 | 15.09 | -2.54 | 2.54 | -0.03 | 0.03 |
| gene7199 | https://www.ncbi.nlm.nih.gov/gene/?term=LOC105280366 | 1414 | 22 | 155.59 | 1 | 7.07 | 4.55 | 43.24 | 41.76 | 1.47 | 1.47 | 0.02 | 0.02 |
| gene2748 | https://www.ncbi.nlm.nih.gov/gene/?term=LOC105275261 | 3845 | 54 | 140.44 | 1 | 2.6 | 1.85 | 40.26 | 39.17 | 1.09 | 1.09 | 0.02 | 0.02 |
| gene10600 | https://www.ncbi.nlm.nih.gov/gene/?term=LOC105284270 | 4837 | 119 | 246.02 | 1 | 2.07 | 0.84 | 10.98 | 10.08 | 0.9 | 0.9 | 0.02 | 0.02 |
| gene9336 | https://www.ncbi.nlm.nih.gov/gene/?term=LOC105282834 | 5993 | 124 | 206.91 | 1 | 1.67 | 0.81 | 16.5 | 15.65 | 0.85 | 0.85 | 0.02 | 0.02 |
| gene303 | https://www.ncbi.nlm.nih.gov/gene/?term=LOC105275584 | 3081 | 86 | 279.13 | 1 | 3.25 | 1.16 | 20.08 | 19.37 | 0.71 | 0.71 | 0.02 | 0.02 |
| gene12516 | https://www.ncbi.nlm.nih.gov/gene/?term=LOC105286452 | 1529 | 38 | 248.53 | 1 | 6.54 | 2.63 | 50.05 | 49.37 | 0.68 | 0.68 | 0.02 | 0.02 |
| gene11568 | https://www.ncbi.nlm.nih.gov/gene/?term=LOC105285395 | 8086 | 190 | 234.97 | 1 | 1.24 | 0.53 | 16.67 | 16 | 0.67 | 0.67 | 0.02 | 0.02 |
| gene4892 | https://www.ncbi.nlm.nih.gov/gene/?term=LOC105277667 | 3898 | 126 | 323.24 | 1 | 2.57 | 0.79 | 27.81 | 27.16 | 0.65 | 0.65 | 0.02 | 0.02 |
| gene10520 | https://www.ncbi.nlm.nih.gov/gene/?term=LOC105284184 | 2451 | 84 | 342.72 | 1 | 4.08 | 1.19 | 33.8 | 33.16 | 0.64 | 0.64 | 0.02 | 0.02 |
| gene473 | https://www.ncbi.nlm.nih.gov/gene/?term=LOC105277423 | 2342 | 106 | 452.6 | 1 | 4.27 | 0.94 | 28.94 | 28.4 | 0.54 | 0.54 | 0.02 | 0.02 |

| | | | | | | | | | | | | | |
|---|---|---|---|---|---|---|---|---|---|---|---|---|---|
| gene10540 | https://www.ncbi.nlm.nih.gov/gene/?term=LOC105284200 | 5267 | 220 | 417.7 | 1 | 1.9 | 0.45 | 12.54 | 12.08 | 0.45 | 0.45 | 0.02 | 0.02 | |
| gene6583 | https://www.ncbi.nlm.nih.gov/gene/?term=LOC105279649 | 7444 | 320 | 429.88 | | 1 | 1.34 | 0.31 | 11.91 | 11.5 | 0.41 | 0.41 | 0.02 | 0.02 |
| gene298 | https://www.ncbi.nlm.nih.gov/gene/?term=LOC105275460 | 2473 | 117 | 473.11 | | 1 | 4.04 | 0.85 | 13.71 | 13.31 | 0.39 | 0.39 | 0.02 | 0.02 |
| gene13114 | https://www.ncbi.nlm.nih.gov/gene/?term=LOC105287147 | 7609 | 398 | 523.06 | | 1 | 1.31 | 0.25 | 9.72 | 9.4 | 0.33 | 0.33 | 0.02 | 0.02 |
| gene920 | https://www.ncbi.nlm.nih.gov/gene/?term=LOC105282518 | 1197 | 52 | 434.42 | | 1 | 8.35 | 1.92 | 30.27 | 30.77 | -0.49 | 0.49 | -0.02 | 0.02 |
| gene10549 | https://www.ncbi.nlm.nih.gov/gene/?term=LOC105284210 | 7145 | 290 | 405.88 | | 2 | 2.8 | 0.69 | 11.49 | 12 | -0.51 | 0.51 | -0.02 | 0.02 |
| gene7249 | https://www.ncbi.nlm.nih.gov/gene/?term=LOC105280422 | 1775 | 59 | 332.39 | | 1 | 5.63 | 1.69 | 16.45 | 17.08 | -0.63 | 0.63 | -0.02 | 0.02 |
| gene35 | https://www.ncbi.nlm.nih.gov/gene/?term=LOC105275197 | 2643 | 50 | 189.18 | | 1 | 3.78 | 2 | 37.01 | 37.84 | -0.83 | 0.83 | -0.02 | 0.02 |
| gene4376 | https://www.ncbi.nlm.nih.gov/gene/?term=LOC105277083 | 4307 | 102 | 236.82 | | 1 | 2.32 | 0.98 | 24.25 | 25.14 | -0.89 | 0.89 | -0.02 | 0.02 |
| gene3385 | https://www.ncbi.nlm.nih.gov/gene/?term=LOC105275966 | 3305 | 58 | 175.49 | | 1 | 3.03 | 1.72 | 49.09 | 50.19 | -1.1 | 1.1 | -0.02 | 0.02 |
| gene322 | https://www.ncbi.nlm.nih.gov/gene/?term=LOC105275777 | 1606 | 28 | 174.35 | | 1 | 6.23 | 3.57 | 42.75 | 44.07 | -1.32 | 1.32 | -0.02 | 0.02 |
| gene7248 | https://www.ncbi.nlm.nih.gov/gene/?term=LOC105280408 | 5175 | 52 | 100.48 | | 1 | 1.93 | 1.92 | 16.62 | 18.28 | -1.65 | 1.65 | -0.02 | 0.02 |
| gene8343 | https://www.ncbi.nlm.nih.gov/gene/?term=LOC105281700 | 2172 | 32 | 147.33 | | 1 | 4.6 | 3.12 | 37.31 | 38.97 | -1.66 | 1.66 | -0.02 | 0.02 |
| gene13235 | https://www.ncbi.nlm.nih.gov/gene/?term=LOC105287278 | 2679 | 34 | 126.91 | | 1 | 3.73 | 2.94 | 22.53 | 24.28 | -1.75 | 1.75 | -0.02 | 0.02 |
| gene564 | https://www.ncbi.nlm.nih.gov/gene/?term=LOC105278478 | 2802 | 17 | 60.67 | 1 | 3.57 | 5.88 | 14.04 | 17.34 | -3.31 | 3.31 | -0.02 | 0.02 | |
| gene8178 | https://www.ncbi.nlm.nih.gov/gene/?term=LOC105281515 | 8647 | 44 | 50.88 | 1 | 1.16 | 2.27 | 25.92 | 24.04 | 1.88 | 1.88 | 0.01 | 0.01 | |
| gene6455 | https://www.ncbi.nlm.nih.gov/gene/?term=LOC105279500 | 2612 | 35 | 134 | 1 | 3.83 | 2.86 | 49.33 | 48.29 | 1.04 | 1.04 | 0.01 | 0.01 | |
| gene7398 | https://www.ncbi.nlm.nih.gov/gene/?term=LOC105280589 | 3452 | 33 | 95.6 | 1 | 2.9 | 3.03 | 21.34 | 20.45 | 0.9 | 0.9 | 0.01 | 0.01 | |
| gene3470 | https://www.ncbi.nlm.nih.gov/gene/?term=LOC105276059 | 8716 | 157 | 180.13 | | 1 | 1.15 | 0.64 | 8.72 | 7.97 | 0.76 | 0.76 | 0.01 | 0.01 |
| gene8252 | https://www.ncbi.nlm.nih.gov/gene/?term=LOC105281599 | 3813 | 49 | 128.51 | | 1 | 2.62 | 2.04 | 4.72 | 4.12 | 0.6 | 0.6 | 0.01 | 0.01 |
| gene7884 | https://www.ncbi.nlm.nih.gov/gene/?term=LOC105281266 | 3390 | 76 | 224.19 | | 1 | 2.95 | 1.32 | 16.63 | 16.11 | 0.52 | 0.52 | 0.01 | 0.01 |
| gene11579 | https://www.ncbi.nlm.nih.gov/gene/?term=LOC105285406 | 13571 | 226 | 166.53 | | 1 | 0.74 | 0.44 | 7.69 | 7.18 | 0.51 | 0.51 | 0.01 | 0.01 |
| gene5877 | https://www.ncbi.nlm.nih.gov/gene/?term=LOC105278807 | 1719 | 49 | 285.05 | | 1 | 5.82 | 2.04 | 27.11 | 26.65 | 0.46 | 0.46 | 0.01 | 0.01 |
| gene11082 | https://www.ncbi.nlm.nih.gov/gene/?term=LOC105284824 | 7436 | 200 | 268.96 | | 1 | 1.34 | 0.5 | 7.89 | 7.46 | 0.43 | 0.43 | 0.01 | 0.01 |
| gene7552 | https://www.ncbi.nlm.nih.gov/gene/?term=LOC105280769 | 8566 | 287 | 335.05 | | 1 | 1.17 | 0.35 | 2.06 | 1.63 | 0.43 | 0.43 | 0.01 | 0.01 |
| gene10531 | https://www.ncbi.nlm.nih.gov/gene/?term=LOC105284190 | 6881 | 186 | 270.31 | | 1 | 1.45 | 0.54 | 12.46 | 12.07 | 0.39 | 0.39 | 0.01 | 0.01 |
| gene7556 | https://www.ncbi.nlm.nih.gov/gene/?term=LOC105280774 | 2792 | 97 | 347.42 | | 1 | 3.58 | 1.03 | 33.33 | 32.94 | 0.39 | 0.39 | 0.01 | 0.01 |
| gene12806 | https://www.ncbi.nlm.nih.gov/gene/?term=LOC105286803 | 4196 | 141 | 336.03 | | 1 | 2.38 | 0.71 | 9.78 | 9.4 | 0.38 | 0.38 | 0.01 | 0.01 |
| gene10844 | https://www.ncbi.nlm.nih.gov/gene/?term=LOC105284535 | 9086 | 231 | 254.24 | | 1 | 1.1 | 0.43 | 15.03 | 14.66 | 0.37 | 0.37 | 0.01 | 0.01 |
| gene292 | https://www.ncbi.nlm.nih.gov/gene/?term=LOC105275398 | 2825 | 81 | 286.73 | | 1 | 3.54 | 1.23 | 24.98 | 24.69 | 0.3 | 0.3 | 0.01 | 0.01 |
| gene5524 | https://www.ncbi.nlm.nih.gov/gene/?term=LOC105278397 | 5177 | 198 | 382.46 | | 1 | 1.93 | 0.51 | 24.98 | 24.69 | 0.29 | 0.29 | 0.01 | 0.01 |
| gene2271 | https://www.ncbi.nlm.nih.gov/gene/?term=LOC105274706 | 1448 | 51 | 352.21 | | 1 | 6.91 | 1.96 | 40.12 | 39.86 | 0.26 | 0.26 | 0.01 | 0.01 |
| gene4216 | https://www.ncbi.nlm.nih.gov/gene/?term=LOC105276906 | 2127 | 90 | 423.13 | | 1 | 4.7 | 1.11 | 21.35 | 21.12 | 0.23 | 0.23 | 0.01 | 0.01 |
| gene10018 | https://www.ncbi.nlm.nih.gov/gene/?term=LOC105283607 | 4414 | 207 | 468.96 | | 1 | 2.27 | 0.48 | 15.29 | 15.09 | 0.2 | 0.2 | 0.01 | 0.01 |
| gene10596 | https://www.ncbi.nlm.nih.gov/gene/?term=LOC105284262 | 6520 | 208 | 319.02 | | 1 | 1.53 | 0.48 | 4.3 | 4.14 | 0.16 | 0.16 | 0.01 | 0.01 |
| gene3206 | https://www.ncbi.nlm.nih.gov/gene/?term=LOC105275752 | 3474 | 118 | 339.67 | | 1 | 2.88 | 0.85 | 32.15 | 31.99 | 0.16 | 0.16 | 0.01 | 0.01 |
| gene570 | https://www.ncbi.nlm.nih.gov/gene/?term=LOC105278549 | 1673 | 63 | 376.57 | | 1 | 5.98 | 1.59 | 43.38 | 43.23 | 0.15 | 0.15 | 0.01 | 0.01 |
| gene5525 | https://www.ncbi.nlm.nih.gov/gene/?term=LOC105278396 | 3090 | 182 | 589 | 1 | 3.24 | 0.55 | 24.88 | 24.76 | 0.12 | 0.12 | 0.01 | 0.01 | |
| gene5746 | https://www.ncbi.nlm.nih.gov/gene/?term=LOC105278666 | 10343 | 441 | 426.38 | | 1 | 0.97 | 0.23 | 6.88 | 6.76 | 0.12 | 0.12 | 0.01 | 0.01 |
| gene1470 | https://www.ncbi.nlm.nih.gov/gene/?term=LOC105287597 | 2974 | 186 | 625.42 | | 1 | 3.36 | 0.54 | 18.48 | 18.39 | 0.09 | 0.09 | 0.01 | 0.01 |
| gene4190 | https://www.ncbi.nlm.nih.gov/gene/?term=LOC105276875 | 3150 | 117 | 371.43 | | 1 | 3.17 | 0.85 | 21.8 | 21.95 | -0.15 | 0.15 | -0.01 | 0.01 |
| gene12802 | https://www.ncbi.nlm.nih.gov/gene/?term=LOC105286799 | 3183 | 164 | 515.24 | | 1 | 3.14 | 0.61 | 17.26 | 17.43 | -0.17 | 0.17 | -0.01 | 0.01 |
| gene1430 | https://www.ncbi.nlm.nih.gov/gene/?term=LOC105287551 | 6433 | 223 | 346.65 | | 1 | 1.55 | 0.45 | 8.23 | 8.41 | -0.18 | 0.18 | -0.01 | 0.01 |
| gene6286 | https://www.ncbi.nlm.nih.gov/gene/?term=LOC105279298 | 2861 | 129 | 450.89 | | 2 | 6.99 | 1.55 | 23.63 | 23.86 | -0.23 | 0.23 | -0.01 | 0.01 |
| gene4962 | https://www.ncbi.nlm.nih.gov/gene/?term=LOC105277761 | 4645 | 144 | 310.01 | | 1 | 2.15 | 0.69 | 13.44 | 13.75 | -0.31 | 0.31 | -0.01 | 0.01 |
| gene9327 | https://www.ncbi.nlm.nih.gov/gene/?term=LOC105282821 | 7793 | 145 | 186.06 | | 1 | 1.28 | 0.69 | 12.57 | 12.89 | -0.32 | 0.32 | -0.01 | 0.01 |
| gene2957 | https://www.ncbi.nlm.nih.gov/gene/?term=LOC105275500 | 10926 | 240 | 219.66 | | 1 | 0.92 | 0.42 | 6.99 | 7.33 | -0.34 | 0.34 | -0.01 | 0.01 |
| gene12307 | https://www.ncbi.nlm.nih.gov/gene/?term=LOC105286216 | 10313 | 187 | 181.32 | | 1 | 0.97 | 0.53 | 14.69 | 15.04 | -0.35 | 0.35 | -0.01 | 0.01 |
| gene10155 | https://www.ncbi.nlm.nih.gov/gene/?term=LOC105283756 | 7030 | 112 | 159.32 | | 1 | 1.42 | 0.89 | 5.92 | 6.29 | -0.37 | 0.37 | -0.01 | 0.01 |
| gene4023 | https://www.ncbi.nlm.nih.gov/gene/?term=LOC105276692 | 1759 | 31 | 176.24 | | 1 | 5.69 | 3.23 | 23.25 | 23.63 | -0.38 | 0.38 | -0.01 | 0.01 |
| gene1310 | https://www.ncbi.nlm.nih.gov/gene/?term=LOC105286661 | 2582 | 71 | 274.98 | | 1 | 3.87 | 1.41 | 21.29 | 21.68 | -0.39 | 0.39 | -0.01 | 0.01 |
| gene12573 | https://www.ncbi.nlm.nih.gov/gene/?term=LOC105286515 | 3647 | 100 | 274.2 | 1 | 2.74 | 1 | 13.03 | 13.44 | -0.42 | 0.42 | -0.01 | 0.01 | |
| gene13036 | https://www.ncbi.nlm.nih.gov/gene/?term=LOC105287059 | 4959 | 160 | 322.65 | | 1 | 2.02 | 0.62 | 19.69 | 20.15 | -0.46 | 0.46 | -0.01 | 0.01 |
| gene1199 | https://www.ncbi.nlm.nih.gov/gene/?term=LOC105285498 | 2220 | 28 | 126.13 | | 1 | 4.5 | 3.57 | 27.44 | 27.99 | -0.55 | 0.55 | -0.01 | 0.01 |
| gene657 | https://www.ncbi.nlm.nih.gov/gene/?term=LOC105279667 | 6421 | 81 | 126.15 | | 1 | 1.56 | 1.23 | 2.49 | 3.12 | -0.63 | 0.63 | -0.01 | 0.01 |
| gene7183 | https://www.ncbi.nlm.nih.gov/gene/?term=LOC105280348 | 3925 | 70 | 178.34 | | 1 | 2.55 | 1.43 | 20.85 | 21.48 | -0.63 | 0.63 | -0.01 | 0.01 |
| gene12021 | https://www.ncbi.nlm.nih.gov/gene/?term=LOC105285905 | 1399 | 15 | 107.22 | | 1 | 7.15 | 6.67 | 50 | 50.68 | -0.68 | 0.68 | -0.01 | 0.01 |
| gene7247 | https://www.ncbi.nlm.nih.gov/gene/?term=LOC105280407 | 10486 | 206 | 196.45 | | 1 | 0.95 | 0.49 | 16.14 | 16.82 | -0.68 | 0.68 | -0.01 | 0.01 |
| gene11561 | https://www.ncbi.nlm.nih.gov/gene/?term=LOC105285387 | 11528 | 178 | 154.41 | | 1 | 0.87 | 0.56 | 9.89 | 10.61 | -0.72 | 0.72 | -0.01 | 0.01 |
| gene12203 | https://www.ncbi.nlm.nih.gov/gene/?term=LOC105286099 | 5967 | 75 | 125.69 | | 1 | 1.68 | 1.33 | 9.56 | 10.52 | -0.96 | 0.96 | -0.01 | 0.01 |
| gene4033 | https://www.ncbi.nlm.nih.gov/gene/?term=LOC105276705 | 10546 | 142 | 134.65 | | 1 | 0.95 | 0.7 | 7.83 | 8.89 | -1.06 | 1.06 | -0.01 | 0.01 |
| gene10345 | https://www.ncbi.nlm.nih.gov/gene/?term=LOC105283989 | 4444 | 27 | 60.76 | 1 | 2.25 | 3.7 | 36.46 | 38.69 | -2.24 | 2.24 | -0.01 | 0.01 | |
| gene2173 | https://www.ncbi.nlm.nih.gov/gene/?term=LOC105274590 | 4387 | 70 | 159.56 | | 2 | 4.56 | 2.86 | 26.31 | 26.05 | 0.27 | 0.27 | 0 | 0 |
| gene6639 | https://www.ncbi.nlm.nih.gov/gene/?term=LOC105279705 | 7656 | 67 | 87.51 | 1 | 1.31 | 1.49 | 22.85 | 22.67 | 0.17 | 0.17 | 0 | 0 | |

| | | | | | | | | | | | | | |
|---|---|---|---|---|---|---|---|---|---|---|---|---|---|
| gene1632 | https://www.ncbi.nlm.nih.gov/gene/?term=LOC105287764 | 18061 | 391 | 216.49 | 2 | 1.11 | 0.51 | 3.3 | 3.14 | 0.16 | 0.16 | 0 | 0 |
| gene7888 | https://www.ncbi.nlm.nih.gov/gene/?term=LOC105281141 | 4357 | 153 | 351.16 | 1 | 2.3 | 0.65 | 32.3 | 32.17 | 0.13 | 0.13 | 0 | 0 |
| gene10392 | https://www.ncbi.nlm.nih.gov/gene/?term=LOC105284036 | 3216 | 99 | 307.84 | 2 | 6.22 | 2.02 | 22.54 | 22.43 | 0.11 | 0.11 | 0 | 0 |
| gene7368 | https://www.ncbi.nlm.nih.gov/gene/?term=LOC105280556 | 4105 | 41 | 99.88 1 | 2.44 | 2.44 | 16.06 | 15.95 | 0.11 | 0.11 | 0 | 0 | |
| gene2806 | https://www.ncbi.nlm.nih.gov/gene/?term=LOC105275327 | 14257 | 473 | 331.77 | 1 | 0.7 | 0.21 | 7.14 | 7.04 | 0.1 | 0.1 | 0 | 0 |
| gene3885 | https://www.ncbi.nlm.nih.gov/gene/?term=LOC105276527 | 7928 | 318 | 401.11 | 1 | 1.26 | 0.31 | 9.68 | 9.6 | 0.09 | 0.09 | 0 | 0 |
| gene9101 | https://www.ncbi.nlm.nih.gov/gene/?term=LOC105282553 | 3068 | 104 | 338.98 | 1 | 3.26 | 0.96 | 23.44 | 23.39 | 0.06 | 0.06 | 0 | 0 |
| gene1408 | https://www.ncbi.nlm.nih.gov/gene/?term=LOC109611520 | 8645 | 306 | 353.96 | 1 | 1.16 | 0.33 | 6.02 | 5.98 | 0.05 | 0.05 | 0 | 0 |
| gene2633 | https://www.ncbi.nlm.nih.gov/gene/?term=LOC105275127 | 7457 | 252 | 337.94 | 1 | 1.34 | 0.4 | 8.36 | 8.33 | 0.03 | 0.03 | 0 | 0 |
| gene10326 | https://www.ncbi.nlm.nih.gov/gene/?term=LOC105283962 | 3042 | 124 | 407.63 | 1 | 3.29 | 0.81 | 8.49 | 8.5 | -0.01 | 0.01 | 0 | 0 |
| gene11275 | https://www.ncbi.nlm.nih.gov/gene/?term=LOC105285055 | 4119 | 252 | 611.8 1 | 2.43 | 0.4 | 11.84 | 11.86 | -0.03 | 0.03 | 0 | 0 | |
| gene6620 | https://www.ncbi.nlm.nih.gov/gene/?term=LOC105279687 | 17998 | 355 | 197.24 | 1 | 0.56 | 0.28 | 5.15 | 5.19 | -0.04 | 0.04 | 0 | 0 |
| gene7915 | https://www.ncbi.nlm.nih.gov/gene/?term=LOC105281173 | 1597 | 39 | 244.21 | 1 | 6.26 | 2.56 | 59.75 | 59.8 | -0.05 | 0.05 | 0 | 0 |
| gene12572 | https://www.ncbi.nlm.nih.gov/gene/?term=LOC105286514 | 2374 | 49 | 206.4 1 | 4.21 | 2.04 | 35.44 | 35.51 | -0.07 | 0.07 | 0 | 0 | |
| gene2683 | https://www.ncbi.nlm.nih.gov/gene/?term=LOC105275187 | 5492 | 70 | 127.46 | 1 | 1.82 | 1.43 | 12.3 | 12.38 | -0.07 | 0.07 | 0 | 0 |
| gene5387 | https://www.ncbi.nlm.nih.gov/gene/?term=LOC105278245 | 6451 | 112 | 173.62 | 1 | 1.55 | 0.89 | 22.85 | 22.92 | -0.07 | 0.07 | 0 | 0 |
| gene8814 | https://www.ncbi.nlm.nih.gov/gene/?term=LOC105282240 | 6910 | 176 | 254.7 1 | 1.45 | 0.57 | 9.18 | 9.27 | -0.08 | 0.08 | 0 | 0 | |
| gene8600 | https://www.ncbi.nlm.nih.gov/gene/?term=LOC105282001 | 8513 | 151 | 177.38 | 1 | 1.17 | 0.66 | 10.34 | 10.43 | -0.09 | 0.09 | 0 | 0 |
| gene8729 | https://www.ncbi.nlm.nih.gov/gene/?term=LOC105282137 | 5005 | 211 | 421.58 | 1 | 2 | 0.47 | 15.81 | 15.91 | -0.09 | 0.09 | 0 | 0 |
| gene2404 | https://www.ncbi.nlm.nih.gov/gene/?term=LOC105274853 | 1767 | 35 | 198.08 | 1 | 5.66 | 2.86 | 33.43 | 33.53 | -0.1 | 0.1 | 0 | 0 |
| gene5101 | https://www.ncbi.nlm.nih.gov/gene/?term=LOC105277903 | 1813 | 35 | 193.05 | 1 | 5.52 | 2.86 | 51.16 | 51.27 | -0.1 | 0.1 | 0 | 0 |
| gene53 | https://www.ncbi.nlm.nih.gov/gene/?term=LOC105277148 | 19283 | 295 | 152.98 | 1 | 0.52 | 0.34 | 3.8 | 3.9 | -0.1 | 0.1 | 0 | 0 |
| gene5646 | https://www.ncbi.nlm.nih.gov/gene/?term=LOC105278534 | 7475 | 187 | 250.17 | 1 | 1.34 | 0.53 | 18.3 | 18.4 | -0.1 | 0.1 | 0 | 0 |
| gene2846 | https://www.ncbi.nlm.nih.gov/gene/?term=LOC105275374 | 11246 | 477 | 424.15 | 1 | 0.89 | 0.21 | 10.56 | 10.68 | -0.11 | 0.11 | 0 | 0 |
| gene34 | https://www.ncbi.nlm.nih.gov/gene/?term=LOC105278416 | 7120 | 140 | 196.63 | 1 | 1.4 | 0.71 | 13.56 | 13.67 | -0.11 | 0.11 | 0 | 0 |
| gene7166 | https://www.ncbi.nlm.nih.gov/gene/?term=LOC105280326 | 6117 | 243 | 397.25 | 1 | 1.63 | 0.41 | 14.54 | 14.65 | -0.11 | 0.11 | 0 | 0 |
| gene9326 | https://www.ncbi.nlm.nih.gov/gene/?term=LOC105282818 | 16612 | 178 | 107.15 | 1 | 0.6 | 0.56 | 13.41 | 13.58 | -0.18 | 0.18 | 0 | 0 |

Directory: OGL File: rnk-dmg-Cb_LI_BCphaseB2.vs.RphaseA1.txt

| gene_ID | gene_link | gwidth | #Sites | #per10Kb | #dmSites | #dmsp10kb | %dmSites | %pSite1 | %pSite2 | DMpSite | ADMpSite | DMpNucl | ADMpNucl |
|---|---|---|---|---|---|---|---|---|---|---|---|---|---|
| gene3165 | https://www.ncbi.nlm.nih.gov/gene/?term=LOC105275706 | 2374 | 60 | 252.74 | 3 | 12.64 | 5 | 22.21 | 32.53 | -10.32 | 10.32 | -0.26 | 0.26 |
| gene2593 | https://www.ncbi.nlm.nih.gov/gene/?term=LOC105275070 | 1774 | 5 | 28.18 | 5 | 28.18 | 100 | 10.96 | 79.22 | -68.26 | 68.26 | -0.19 | 0.19 |
| gene5714 | https://www.ncbi.nlm.nih.gov/gene/?term=LOC105278729 | 710 | 21 | 295.77 | 1 | 14.08 | 4.76 | 56.35 | 60.77 | -4.42 | 4.42 | -0.13 | 0.13 |
| gene10714 | https://www.ncbi.nlm.nih.gov/gene/?term=LOC105284396 | 1966 | 62 | 315.36 | 1 | 5.09 | 1.61 | 36.35 | 33.21 | 3.14 | 3.14 | 0.1 | 0.1 |
| gene12021 | https://www.ncbi.nlm.nih.gov/gene/?term=LOC105285905 | 1399 | 21 | 150.11 | 1 | 7.15 | 4.76 | 36.22 | 31.01 | 5.22 | 5.22 | 0.08 | 0.08 |
| gene6652 | https://www.ncbi.nlm.nih.gov/gene/?term=LOC105279719 | 2831 | 86 | 303.78 | 1 | 3.53 | 1.16 | 23.08 | 20.39 | 2.69 | 2.69 | 0.08 | 0.08 |
| gene9101 | https://www.ncbi.nlm.nih.gov/gene/?term=LOC105282553 | 3068 | 106 | 345.5 | 1 | 3.26 | 0.94 | 23.35 | 21.05 | 2.29 | 2.29 | 0.08 | 0.08 |
| gene4498 | https://www.ncbi.nlm.nih.gov/gene/?term=LOC105277239 | 2831 | 130 | 459.2 | 1 | 3.53 | 0.77 | 12.47 | 10.69 | 1.78 | 1.78 | 0.08 | 0.08 |
| gene2271 | https://www.ncbi.nlm.nih.gov/gene/?term=LOC105274706 | 1448 | 51 | 352.21 | 1 | 6.91 | 1.96 | 39.73 | 37.8 | 1.94 | 1.94 | 0.07 | 0.07 |
| gene2856 | https://www.ncbi.nlm.nih.gov/gene/?term=LOC105275384 | 1987 | 87 | 437.85 | 1 | 5.03 | 1.15 | 34.24 | 35.77 | -1.53 | 1.53 | -0.07 | 0.07 |
| gene7568 | https://www.ncbi.nlm.nih.gov/gene/?term=LOC105280786 | 2525 | 109 | 431.68 | 1 | 3.96 | 0.92 | 28.7 | 30.24 | -1.54 | 1.54 | -0.07 | 0.07 |
| gene3817 | https://www.ncbi.nlm.nih.gov/gene/?term=LOC105276449 | 1230 | 45 | 365.85 | 1 | 8.13 | 2.22 | 27.32 | 29.29 | -1.97 | 1.97 | -0.07 | 0.07 |
| gene12406 | https://www.ncbi.nlm.nih.gov/gene/?term=LOC105286316 | 1386 | 31 | 223.67 | 1 | 7.22 | 3.23 | 46.61 | 43.84 | 2.77 | 2.77 | 0.06 | 0.06 |
| gene7550 | https://www.ncbi.nlm.nih.gov/gene/?term=LOC105280763 | 2173 | 98 | 450.99 | 1 | 4.6 | 1.02 | 27.13 | 25.7 | 1.43 | 1.43 | 0.06 | 0.06 |
| gene2071 | https://www.ncbi.nlm.nih.gov/gene/?term=LOC105274471 | 2446 | 121 | 494.69 | 1 | 4.09 | 0.83 | 19.71 | 20.9 | -1.19 | 1.19 | -0.06 | 0.06 |
| gene5877 | https://www.ncbi.nlm.nih.gov/gene/?term=LOC105278807 | 1719 | 53 | 308.32 | 1 | 5.82 | 1.89 | 22.63 | 24.74 | -2.11 | 2.11 | -0.06 | 0.06 |
| gene5101 | https://www.ncbi.nlm.nih.gov/gene/?term=LOC105277903 | 1813 | 36 | 198.57 | 1 | 5.52 | 2.78 | 53.23 | 56.32 | -3.09 | 3.09 | -0.06 | 0.06 |
| gene5647 | https://www.ncbi.nlm.nih.gov/gene/?term=LOC105278567 | 3118 | 29 | 93.01 | 1 | 3.21 | 3.45 | 13.21 | 19.43 | -6.21 | 6.21 | -0.06 | 0.06 |
| gene7199 | https://www.ncbi.nlm.nih.gov/gene/?term=LOC105280366 | 1414 | 33 | 233.38 | 1 | 7.07 | 3.03 | 43.27 | 41 | 2.28 | 2.28 | 0.05 | 0.05 |
| gene2404 | https://www.ncbi.nlm.nih.gov/gene/?term=LOC105274853 | 1767 | 43 | 243.35 | 1 | 5.66 | 2.33 | 31.85 | 29.62 | 2.23 | 2.23 | 0.05 | 0.05 |
| gene1907 | https://www.ncbi.nlm.nih.gov/gene/?term=LOC105288099 | 1238 | 35 | 282.71 | 1 | 8.08 | 2.86 | 34.89 | 33.16 | 1.73 | 1.73 | 0.05 | 0.05 |
| gene920 | https://www.ncbi.nlm.nih.gov/gene/?term=LOC105282518 | 1197 | 41 | 342.52 | 1 | 8.35 | 2.44 | 39.23 | 37.74 | 1.49 | 1.49 | 0.05 | 0.05 |
| gene4892 | https://www.ncbi.nlm.nih.gov/gene/?term=LOC105277667 | 3898 | 120 | 307.85 | 1 | 2.57 | 0.83 | 27.15 | 28.74 | -1.59 | 1.59 | -0.05 | 0.05 |
| gene393 | https://www.ncbi.nlm.nih.gov/gene/?term=LOC105276566 | 1123 | 2 | 17.81 | 1 | 8.9 | 50 | 70.57 | 50 | 20.57 | 20.57 | 0.04 | 0.04 |
| gene1199 | https://www.ncbi.nlm.nih.gov/gene/?term=LOC105285498 | 2220 | 25 | 112.61 | 1 | 4.5 | 4 | 37.37 | 33.76 | 3.61 | 3.61 | 0.04 | 0.04 |
| gene388 | https://www.ncbi.nlm.nih.gov/gene/?term=LOC105276538 | 1558 | 20 | 128.37 | 2 | 12.84 | 10 | 66.14 | 62.93 | 3.2 | 3.2 | 0.04 | 0.04 |
| gene322 | https://www.ncbi.nlm.nih.gov/gene/?term=LOC105275777 | 1606 | 25 | 155.67 | 1 | 6.23 | 4 | 44.37 | 41.49 | 2.89 | 2.89 | 0.04 | 0.04 |
| gene6422 | https://www.ncbi.nlm.nih.gov/gene/?term=LOC105279458 | 2419 | 37 | 152.96 | 1 | 4.13 | 2.7 | 52.5 | 49.97 | 2.53 | 2.53 | 0.04 | 0.04 |
| gene6286 | https://www.ncbi.nlm.nih.gov/gene/?term=LOC105279298 | 2861 | 131 | 457.88 | 2 | 6.99 | 1.53 | 24.99 | 24.15 | 0.84 | 0.84 | 0.04 | 0.04 |
| gene1430 | https://www.ncbi.nlm.nih.gov/gene/?term=LOC105287551 | 6433 | 235 | 365.3 | 1 | 1.55 | 0.43 | 8.56 | 9.58 | -1.02 | 1.02 | -0.04 | 0.04 |
| gene11401 | https://www.ncbi.nlm.nih.gov/gene/?term=LOC105285202 | 1841 | 75 | 407.39 | 1 | 5.43 | 1.33 | 24.08 | 25.12 | -1.03 | 1.03 | -0.04 | 0.04 |
| gene4720 | https://www.ncbi.nlm.nih.gov/gene/?term=LOC105277479 | 1697 | 50 | 294.64 | 1 | 5.89 | 2 | 35.87 | 37.27 | -1.4 | 1.4 | -0.04 | 0.04 |
| gene1884 | https://www.ncbi.nlm.nih.gov/gene/?term=LOC105288072 | 4169 | 78 | 187.1 | 1 | 2.4 | 1.28 | 22.55 | 24.51 | -1.96 | 1.96 | -0.04 | 0.04 |
| gene8661 | https://www.ncbi.nlm.nih.gov/gene/?term=LOC105282063 | 1926 | 30 | 155.76 | 1 | 5.19 | 3.33 | 28.06 | 30.33 | -2.28 | 2.28 | -0.04 | 0.04 |
| gene9411 | https://www.ncbi.nlm.nih.gov/gene/?term=LOC105282914 | 5534 | 41 | 74.09 | 1 | 1.81 | 2.44 | 52.21 | 48.58 | 3.63 | 3.63 | 0.03 | 0.03 |
| gene3168 | https://www.ncbi.nlm.nih.gov/gene/?term=LOC105275711 | 6150 | 73 | 118.7 | 1 | 1.63 | 1.37 | 14.69 | 12.05 | 2.64 | 2.64 | 0.03 | 0.03 |
| gene6455 | https://www.ncbi.nlm.nih.gov/gene/?term=LOC105279500 | 2612 | 30 | 114.85 | 1 | 3.83 | 3.33 | 48.51 | 46 | 2.5 | 2.5 | 0.03 | 0.03 |
| gene6237 | https://www.ncbi.nlm.nih.gov/gene/?term=LOC105279244 | 1484 | 23 | 154.99 | 1 | 6.74 | 4.35 | 50.08 | 47.92 | 2.16 | 2.16 | 0.03 | 0.03 |
| gene2748 | https://www.ncbi.nlm.nih.gov/gene/?term=LOC105275261 | 3845 | 54 | 140.44 | 1 | 2.6 | 1.85 | 39.59 | 37.65 | 1.94 | 1.94 | 0.03 | 0.03 |
| gene10107 | https://www.ncbi.nlm.nih.gov/gene/?term=LOC105283712 | 1883 | 35 | 185.87 | 1 | 5.31 | 2.86 | 27.16 | 25.31 | 1.85 | 1.85 | 0.03 | 0.03 |
| gene3671 | https://www.ncbi.nlm.nih.gov/gene/?term=LOC105276272 | 1966 | 46 | 233.98 | 1 | 5.09 | 2.17 | 27.89 | 26.72 | 1.16 | 1.16 | 0.03 | 0.03 |
| gene5646 | https://www.ncbi.nlm.nih.gov/gene/?term=LOC105278534 | 7475 | 185 | 247.49 | 1 | 1.34 | 0.54 | 18.24 | 17.21 | 1.03 | 1.03 | 0.03 | 0.03 |
| gene10392 | https://www.ncbi.nlm.nih.gov/gene/?term=LOC105284036 | 3216 | 106 | 329.6 | 2 | 6.22 | 1.89 | 21.2 | 20.33 | 0.87 | 0.87 | 0.03 | 0.03 |
| gene11340 | https://www.ncbi.nlm.nih.gov/gene/?term=LOC105285131 | 2918 | 105 | 359.84 | 1 | 3.43 | 0.95 | 14.38 | 13.54 | 0.84 | 0.84 | 0.03 | 0.03 |
| gene8729 | https://www.ncbi.nlm.nih.gov/gene/?term=LOC105282137 | 5005 | 200 | 399.6 | 1 | 2 | 0.5 | 16.14 | 15.41 | 0.72 | 0.72 | 0.03 | 0.03 |
| gene10018 | https://www.ncbi.nlm.nih.gov/gene/?term=LOC105283607 | 4414 | 201 | 455.37 | 1 | 2.27 | 0.5 | 15.61 | 14.98 | 0.63 | 0.63 | 0.03 | 0.03 |
| gene5298 | https://www.ncbi.nlm.nih.gov/gene/?term=LOC105278145 | 4060 | 237 | 583.74 | 3 | 7.39 | 1.27 | 11.65 | 11.08 | 0.57 | 0.57 | 0.03 | 0.03 |
| gene298 | https://www.ncbi.nlm.nih.gov/gene/?term=LOC105275460 | 2473 | 120 | 485.24 | 1 | 4.04 | 0.83 | 13.22 | 13.75 | -0.53 | 0.53 | -0.03 | 0.03 |
| gene6583 | https://www.ncbi.nlm.nih.gov/gene/?term=LOC105279649 | 7444 | 340 | 456.74 | 1 | 1.34 | 0.29 | 10.48 | 11.08 | -0.59 | 0.59 | -0.03 | 0.03 |
| gene7514 | https://www.ncbi.nlm.nih.gov/gene/?term=LOC105280723 | 4471 | 208 | 465.22 | 1 | 2.24 | 0.48 | 13.49 | 14.1 | -0.6 | 0.6 | -0.03 | 0.03 |
| gene2056 | https://www.ncbi.nlm.nih.gov/gene/?term=LOC105274452 | 6584 | 243 | 369.08 | 1 | 1.52 | 0.41 | 13.17 | 13.88 | -0.71 | 0.71 | -0.03 | 0.03 |
| gene7884 | https://www.ncbi.nlm.nih.gov/gene/?term=LOC105281266 | 3390 | 81 | 238.94 | 1 | 2.95 | 1.23 | 16.21 | 17.33 | -1.12 | 1.12 | -0.03 | 0.03 |
| gene1518 | https://www.ncbi.nlm.nih.gov/gene/?term=LOC105287648 | 2195 | 35 | 159.45 | 1 | 4.56 | 2.86 | 35.74 | 37.41 | -1.67 | 1.67 | -0.03 | 0.03 |
| gene9478 | https://www.ncbi.nlm.nih.gov/gene/?term=LOC105282991 | 3726 | 58 | 155.66 | 1 | 2.68 | 1.72 | 33.94 | 35.62 | -1.68 | 1.68 | -0.03 | 0.03 |
| gene927 | https://www.ncbi.nlm.nih.gov/gene/?term=LOC105282600 | 3727 | 52 | 139.52 | 1 | 2.68 | 1.92 | 31.26 | 33.17 | -1.91 | 1.91 | -0.03 | 0.03 |
| gene9648 | https://www.ncbi.nlm.nih.gov/gene/?term=LOC105283183 | 2602 | 33 | 126.83 | 1 | 3.84 | 3.03 | 40.53 | 42.61 | -2.08 | 2.08 | -0.03 | 0.03 |
| gene564 | https://www.ncbi.nlm.nih.gov/gene/?term=LOC105278478 | 2802 | 18 | 64.24 | 1 | 3.57 | 5.56 | 17.32 | 14.97 | 2.35 | 2.35 | 0.02 | 0.02 |
| gene6806 | https://www.ncbi.nlm.nih.gov/gene/?term=LOC105279894 | 3649 | 35 | 95.92 | 1 | 2.74 | 2.86 | 40.58 | 38.26 | 2.32 | 2.32 | 0.02 | 0.02 |
| gene6639 | https://www.ncbi.nlm.nih.gov/gene/?term=LOC105279705 | 7656 | 68 | 88.82 | 1 | 1.31 | 1.47 | 23.58 | 21.76 | 1.82 | 1.82 | 0.02 | 0.02 |
| gene3046 | https://www.ncbi.nlm.nih.gov/gene/?term=LOC105275587 | 2394 | 58 | 242.27 | 1 | 4.18 | 1.72 | 36.72 | 35.74 | 0.98 | 0.98 | 0.02 | 0.02 |

| | | | | | | | | | | | | | |
|---|---|---|---|---|---|---|---|---|---|---|---|---|---|
| gene7915 | https://www.ncbi.nlm.nih.gov/gene/?term=LOC105281173 | 1597 | 38 | 237.95 | 1 | 6.26 | 2.63 | 60.79 | 59.81 | 0.98 | 0.98 | 0.02 | 0.02 |
| gene8015 | https://www.ncbi.nlm.nih.gov/gene/?term=LOC105281312 | 2998 | 65 | 216.81 | 1 | 3.34 | 1.54 | 33.24 | 32.3 | 0.93 | 0.93 | 0.02 | 0.02 |
| gene1310 | https://www.ncbi.nlm.nih.gov/gene/?term=LOC105286661 | 2582 | 62 | 240.12 | 1 | 3.87 | 1.61 | 22.65 | 21.78 | 0.87 | 0.87 | 0.02 | 0.02 |
| gene9462 | https://www.ncbi.nlm.nih.gov/gene/?term=LOC105282970 | 1469 | 41 | 279.1 1 | 6.81 | 2.44 | 50 | 49.15 | 0.85 | 0.85 | 0.02 | 0.02 | |
| gene1220 | https://www.ncbi.nlm.nih.gov/gene/?term=LOC105285742 | 3453 | 106 | 306.98 | 1 | 2.9 | 0.94 | 18.53 | 17.74 | 0.79 | 0.79 | 0.02 | 0.02 |
| gene4190 | https://www.ncbi.nlm.nih.gov/gene/?term=LOC105276875 | 3150 | 111 | 352.38 | 1 | 3.17 | 0.9 | 22.75 | 22.14 | 0.61 | 0.61 | 0.02 | 0.02 |
| gene4962 | https://www.ncbi.nlm.nih.gov/gene/?term=LOC105277761 | 4645 | 147 | 316.47 | 1 | 2.15 | 0.68 | 14.43 | 13.91 | 0.52 | 0.52 | 0.02 | 0.02 |
| gene10520 | https://www.ncbi.nlm.nih.gov/gene/?term=LOC105284184 | 2451 | 93 | 379.44 | 1 | 4.08 | 1.08 | 31.28 | 30.8 | 0.49 | 0.49 | 0.02 | 0.02 |
| gene10326 | https://www.ncbi.nlm.nih.gov/gene/?term=LOC105283962 | 3042 | 126 | 414.2 1 | 3.29 | 0.79 | 9.42 | 8.95 | 0.47 | 0.47 | 0.02 | 0.02 | |
| gene6031 | https://www.ncbi.nlm.nih.gov/gene/?term=LOC105279000 | 5691 | 195 | 342.65 | 1 | 1.76 | 0.51 | 13.76 | 13.29 | 0.47 | 0.47 | 0.02 | 0.02 |
| gene3885 | https://www.ncbi.nlm.nih.gov/gene/?term=LOC105276527 | 7928 | 318 | 401.11 | 1 | 1.26 | 0.31 | 10.14 | 9.68 | 0.46 | 0.46 | 0.02 | 0.02 |
| gene8274 | https://www.ncbi.nlm.nih.gov/gene/?term=LOC105281624 | 5425 | 219 | 403.69 | 1 | 1.84 | 0.46 | 10.56 | 10.14 | 0.42 | 0.42 | 0.02 | 0.02 |
| gene9740 | https://www.ncbi.nlm.nih.gov/gene/?term=LOC105283277 | 2887 | 136 | 471.08 | 1 | 3.46 | 0.74 | 19.63 | 19.22 | 0.41 | 0.41 | 0.02 | 0.02 |
| gene10213 | https://www.ncbi.nlm.nih.gov/gene/?term=LOC105283826 | 1503 | 64 | 425.82 | 1 | 6.65 | 1.56 | 16.48 | 16.1 | 0.38 | 0.38 | 0.02 | 0.02 |
| gene1470 | https://www.ncbi.nlm.nih.gov/gene/?term=LOC105287597 | 2974 | 195 | 655.68 | 1 | 3.36 | 0.51 | 18.15 | 17.88 | 0.27 | 0.27 | 0.02 | 0.02 |
| gene738 | https://www.ncbi.nlm.nih.gov/gene/?term=LOC105280421 | 4927 | 216 | 438.4 1 | 2.03 | 0.46 | 14.22 | 14.61 | -0.39 | 0.39 | -0.02 | 0.02 | |
| gene3206 | https://www.ncbi.nlm.nih.gov/gene/?term=LOC105275752 | 3474 | 129 | 371.33 | 1 | 2.88 | 0.78 | 29.67 | 30.17 | -0.5 | 0.5 | -0.02 | 0.02 |
| gene4376 | https://www.ncbi.nlm.nih.gov/gene/?term=LOC105277083 | 4307 | 114 | 264.69 | 1 | 2.32 | 0.88 | 20.15 | 20.76 | -0.61 | 0.61 | -0.02 | 0.02 |
| gene13033 | https://www.ncbi.nlm.nih.gov/gene/?term=LOC105287063 | 1877 | 51 | 271.71 | 1 | 5.33 | 1.96 | 30.35 | 31.05 | -0.7 | 0.7 | -0.02 | 0.02 |
| gene12439 | https://www.ncbi.nlm.nih.gov/gene/?term=LOC105286357 | 4761 | 81 | 170.13 | 1 | 2.1 | 1.23 | 29.49 | 30.52 | -1.03 | 1.03 | -0.02 | 0.02 |
| gene4023 | https://www.ncbi.nlm.nih.gov/gene/?term=LOC105276692 | 1759 | 36 | 204.66 | 1 | 5.69 | 2.78 | 20.19 | 21.39 | -1.2 | 1.2 | -0.02 | 0.02 |
| gene7368 | https://www.ncbi.nlm.nih.gov/gene/?term=LOC105280556 | 4105 | 50 | 121.8 1 | 2.44 | 2 | 14.06 | 15.36 | -1.3 | 1.3 | -0.02 | 0.02 | |
| gene10654 | https://www.ncbi.nlm.nih.gov/gene/?term=LOC105284329 | 4027 | 51 | 126.65 | 1 | 2.48 | 1.96 | 17.3 | 18.75 | -1.45 | 1.45 | -0.02 | 0.02 |
| gene9647 | https://www.ncbi.nlm.nih.gov/gene/?term=LOC109611233 | 3094 | 35 | 113.12 | 1 | 3.23 | 2.86 | 38.21 | 40.18 | -1.97 | 1.97 | -0.02 | 0.02 |
| gene8178 | https://www.ncbi.nlm.nih.gov/gene/?term=LOC105281515 | 8647 | 63 | 72.86 1 | 1.16 | 1.59 | 24.53 | 26.83 | -2.3 | 2.3 | -0.02 | 0.02 | |
| gene10602 | https://www.ncbi.nlm.nih.gov/gene/?term=LOC105284268 | 2122 | 21 | 98.96 1 | 4.71 | 4.76 | 25.67 | 24.45 | 1.22 | 1.22 | 0.01 | 0.01 | |
| gene7398 | https://www.ncbi.nlm.nih.gov/gene/?term=LOC105280589 | 3452 | 36 | 104.29 | 1 | 2.9 | 2.78 | 20.14 | 19.24 | 0.9 | 0.9 | 0.01 | 0.01 |
| gene12203 | https://www.ncbi.nlm.nih.gov/gene/?term=LOC105286099 | 5967 | 72 | 120.66 | 1 | 1.68 | 1.39 | 10.57 | 10 | 0.57 | 0.57 | 0.01 | 0.01 |
| gene10844 | https://www.ncbi.nlm.nih.gov/gene/?term=LOC105284535 | 9086 | 238 | 261.94 | 1 | 1.1 | 0.42 | 14.5 | 14.01 | 0.49 | 0.49 | 0.01 | 0.01 |
| gene2173 | https://www.ncbi.nlm.nih.gov/gene/?term=LOC105274590 | 4387 | 80 | 182.36 | 1 | 2.28 | 1.25 | 27.39 | 26.91 | 0.48 | 0.48 | 0.01 | 0.01 |
| gene10600 | https://www.ncbi.nlm.nih.gov/gene/?term=LOC105284270 | 4837 | 115 | 237.75 | 1 | 2.07 | 0.87 | 11.71 | 11.26 | 0.46 | 0.46 | 0.01 | 0.01 |
| gene2257 | https://www.ncbi.nlm.nih.gov/gene/?term=LOC105274689 | 3658 | 73 | 199.56 | 1 | 2.73 | 1.37 | 16.2 | 15.75 | 0.46 | 0.46 | 0.01 | 0.01 |
| gene12572 | https://www.ncbi.nlm.nih.gov/gene/?term=LOC105286514 | 2374 | 44 | 185.34 | 1 | 4.21 | 2.27 | 36.23 | 35.78 | 0.45 | 0.45 | 0.01 | 0.01 |
| gene7600 | https://www.ncbi.nlm.nih.gov/gene/?term=LOC105280819 | 1615 | 45 | 278.64 | 1 | 6.19 | 2.22 | 37.23 | 36.8 | 0.43 | 0.43 | 0.01 | 0.01 |
| gene3599 | https://www.ncbi.nlm.nih.gov/gene/?term=LOC105276196 | 2955 | 57 | 192.89 | 1 | 3.38 | 1.75 | 38.84 | 38.45 | 0.38 | 0.38 | 0.01 | 0.01 |
| gene3107 | https://www.ncbi.nlm.nih.gov/gene/?term=LOC105275650 | 5103 | 174 | 340.98 | 1 | 1.96 | 0.57 | 16.52 | 16.19 | 0.33 | 0.33 | 0.01 | 0.01 |
| gene6966 | https://www.ncbi.nlm.nih.gov/gene/?term=LOC105280104 | 3747 | 121 | 322.93 | 1 | 2.67 | 0.83 | 16.24 | 15.96 | 0.28 | 0.28 | 0.01 | 0.01 |
| gene9445 | https://www.ncbi.nlm.nih.gov/gene/?term=LOC105282951 | 2285 | 114 | 498.91 | 1 | 4.38 | 0.88 | 20.31 | 20.03 | 0.28 | 0.28 | 0.01 | 0.01 |
| gene11454 | https://www.ncbi.nlm.nih.gov/gene/?term=LOC105285256 | 1906 | 51 | 267.58 | 1 | 5.25 | 1.96 | 42.03 | 41.77 | 0.26 | 0.26 | 0.01 | 0.01 |
| gene5524 | https://www.ncbi.nlm.nih.gov/gene/?term=LOC105278397 | 5177 | 202 | 390.19 | 1 | 1.93 | 0.5 | 25.27 | 25.04 | 0.22 | 0.22 | 0.01 | 0.01 |
| gene570 | https://www.ncbi.nlm.nih.gov/gene/?term=LOC105278549 | 1673 | 69 | 412.43 | 1 | 5.98 | 1.45 | 43.78 | 43.57 | 0.21 | 0.21 | 0.01 | 0.01 |
| gene9177 | https://www.ncbi.nlm.nih.gov/gene/?term=LOC105282646 | 3320 | 101 | 304.22 | 1 | 3.01 | 0.99 | 21.24 | 21.05 | 0.2 | 0.2 | 0.01 | 0.01 |
| gene10886 | https://www.ncbi.nlm.nih.gov/gene/?term=LOC105284582 | 1475 | 49 | 332.2 1 | 6.78 | 2.04 | 22 | 21.8 | 0.19 | 0.19 | 0.01 | 0.01 | |
| gene5525 | https://www.ncbi.nlm.nih.gov/gene/?term=LOC105278396 | 3090 | 186 | 601.94 | 1 | 3.24 | 0.54 | 25.06 | 24.87 | 0.18 | 0.18 | 0.01 | 0.01 |
| gene7249 | https://www.ncbi.nlm.nih.gov/gene/?term=LOC105280422 | 1775 | 63 | 354.93 | 1 | 5.63 | 1.59 | 17.74 | 17.55 | 0.18 | 0.18 | 0.01 | 0.01 |
| gene13036 | https://www.ncbi.nlm.nih.gov/gene/?term=LOC105287059 | 4959 | 165 | 332.73 | 1 | 2.02 | 0.61 | 17.15 | 16.98 | 0.17 | 0.17 | 0.01 | 0.01 |
| gene10606 | https://www.ncbi.nlm.nih.gov/gene/?term=LOC105284273 | 6119 | 255 | 416.73 | 1 | 1.63 | 0.39 | 19.24 | 19.09 | 0.16 | 0.16 | 0.01 | 0.01 |
| gene5777 | https://www.ncbi.nlm.nih.gov/gene/?term=LOC105278691 | 3883 | 152 | 391.45 | 1 | 2.58 | 0.66 | 14.89 | 14.73 | 0.16 | 0.16 | 0.01 | 0.01 |
| gene7166 | https://www.ncbi.nlm.nih.gov/gene/?term=LOC105280326 | 6117 | 238 | 389.08 | 1 | 1.63 | 0.42 | 15.04 | 14.89 | 0.15 | 0.15 | 0.01 | 0.01 |
| gene13114 | https://www.ncbi.nlm.nih.gov/gene/?term=LOC105287147 | 7609 | 419 | 550.66 | 1 | 1.31 | 0.24 | 8.81 | 8.67 | 0.14 | 0.14 | 0.01 | 0.01 |
| gene2846 | https://www.ncbi.nlm.nih.gov/gene/?term=LOC105275374 | 11246 | 465 | 413.48 | 1 | 0.89 | 0.22 | 10.6 | 10.73 | -0.13 | 0.13 | -0.01 | 0.01 |
| gene12802 | https://www.ncbi.nlm.nih.gov/gene/?term=LOC105286799 | 3183 | 190 | 596.92 | 1 | 3.14 | 0.53 | 15.49 | 15.63 | -0.14 | 0.14 | -0.01 | 0.01 |
| gene10678 | https://www.ncbi.nlm.nih.gov/gene/?term=LOC105284355 | 4031 | 182 | 451.5 1 | 2.48 | 0.55 | 17.39 | 17.62 | -0.23 | 0.23 | -0.01 | 0.01 | |
| gene7556 | https://www.ncbi.nlm.nih.gov/gene/?term=LOC105280774 | 2792 | 118 | 422.64 | 1 | 3.58 | 0.85 | 27.01 | 27.27 | -0.26 | 0.26 | -0.01 | 0.01 |
| gene8600 | https://www.ncbi.nlm.nih.gov/gene/?term=LOC105282001 | 8513 | 186 | 218.49 | 1 | 1.17 | 0.54 | 10 | 10.27 | -0.27 | 0.27 | -0.01 | 0.01 |
| gene2806 | https://www.ncbi.nlm.nih.gov/gene/?term=LOC105275327 | 14257 | 511 | 358.42 | 1 | 0.7 | 0.2 | 6.7 | 7.03 | -0.33 | 0.33 | -0.01 | 0.01 |
| gene35 | https://www.ncbi.nlm.nih.gov/gene/?term=LOC105275197 | 2643 | 48 | 181.61 | 1 | 3.78 | 2.08 | 38.22 | 38.55 | -0.33 | 0.33 | -0.01 | 0.01 |
| gene2668 | https://www.ncbi.nlm.nih.gov/gene/?term=LOC105275170 | 12496 | 279 | 223.27 | 1 | 0.8 | 0.36 | 1.97 | 2.32 | -0.35 | 0.35 | -0.01 | 0.01 |
| gene1855 | https://www.ncbi.nlm.nih.gov/gene/?term=LOC105288036 | 2387 | 84 | 351.91 | 1 | 4.19 | 1.19 | 29.18 | 29.59 | -0.41 | 0.41 | -0.01 | 0.01 |
| gene6620 | https://www.ncbi.nlm.nih.gov/gene/?term=LOC105279687 | 17998 | 368 | 204.47 | 1 | 0.56 | 0.27 | 4.64 | 5.1 | -0.46 | 0.46 | -0.01 | 0.01 |
| gene7247 | https://www.ncbi.nlm.nih.gov/gene/?term=LOC105280407 | 10486 | 218 | 207.9 1 | 0.95 | 0.46 | 16.54 | 17.03 | -0.49 | 0.49 | -0.01 | 0.01 | |
| gene9336 | https://www.ncbi.nlm.nih.gov/gene/?term=LOC105282834 | 5993 | 129 | 215.25 | 1 | 1.67 | 0.78 | 14.53 | 15.02 | -0.49 | 0.49 | -0.01 | 0.01 |
| gene940 | https://www.ncbi.nlm.nih.gov/gene/?term=LOC105282794 | 3578 | 85 | 237.56 | 1 | 2.79 | 1.18 | 13.96 | 14.47 | -0.51 | 0.51 | -0.01 | 0.01 |
| gene3470 | https://www.ncbi.nlm.nih.gov/gene/?term=LOC105276059 | 8716 | 152 | 174.39 | 1 | 1.15 | 0.66 | 7.9 | 8.44 | -0.54 | 0.54 | -0.01 | 0.01 |

| | | | | | | | | | | | | | | |
|---|---|---|---|---|---|---|---|---|---|---|---|---|---|---|
| gene12516 | https://www.ncbi.nlm.nih.gov/gene/?term=LOC105286452 | 1529 | 35 | 228.91 | | 1 | 6.54 | 2.86 | 51.89 | 52.54 | -0.65 | 0.65 | -0.01 | 0.01 |
| gene5733 | https://www.ncbi.nlm.nih.gov/gene/?term=LOC105278649 | 5221 | 91 | 174.3 | 1 | 1.92 | 1.1 | 19.36 | 20.1 | -0.74 | 0.74 | -0.01 | 0.01 | |
| gene8742 | https://www.ncbi.nlm.nih.gov/gene/?term=LOC105282151 | 2563 | 42 | 163.87 | | 1 | 3.9 | 2.38 | 34.83 | 35.57 | -0.75 | 0.75 | -0.01 | 0.01 |
| gene2683 | https://www.ncbi.nlm.nih.gov/gene/?term=LOC105275187 | 5492 | 70 | 127.46 | | 1 | 1.82 | 1.43 | 13.71 | 14.76 | -1.05 | 1.05 | -0.01 | 0.01 |
| gene6528 | https://www.ncbi.nlm.nih.gov/gene/?term=LOC105279591 | 7222 | 59 | 81.69 | 1 | 1.38 | 1.69 | 26.22 | 27.32 | -1.1 | 1.1 | -0.01 | 0.01 | |
| gene10155 | https://www.ncbi.nlm.nih.gov/gene/?term=LOC105283756 | 7030 | 104 | 147.94 | | 1 | 1.42 | 0.96 | 5.54 | 5.23 | 0.31 | 0.31 | 0 | 0 |
| gene12307 | https://www.ncbi.nlm.nih.gov/gene/?term=LOC105286216 | 10313 | 160 | 155.14 | | 1 | 0.97 | 0.62 | 16.94 | 16.73 | 0.22 | 0.22 | 0 | 0 |
| gene7183 | https://www.ncbi.nlm.nih.gov/gene/?term=LOC105280348 | 3925 | 70 | 178.34 | | 1 | 2.55 | 1.43 | 18.6 | 18.42 | 0.18 | 0.18 | 0 | 0 |
| gene10531 | https://www.ncbi.nlm.nih.gov/gene/?term=LOC105284190 | 6881 | 175 | 254.32 | | 1 | 1.45 | 0.57 | 12.25 | 12.09 | 0.16 | 0.16 | 0 | 0 |
| gene11082 | https://www.ncbi.nlm.nih.gov/gene/?term=LOC105284824 | 7436 | 212 | 285.1 | 1 | 1.34 | 0.47 | 6.95 | 6.8 | 0.15 | 0.15 | 0 | 0 | |
| gene8343 | https://www.ncbi.nlm.nih.gov/gene/?term=LOC105281700 | 2172 | 37 | 170.35 | | 1 | 4.6 | 2.7 | 39.63 | 39.48 | 0.15 | 0.15 | 0 | 0 |
| gene53 | https://www.ncbi.nlm.nih.gov/gene/?term=LOC105277148 | 19283 | 291 | 150.91 | | 1 | 0.52 | 0.34 | 4.8 | 4.67 | 0.13 | 0.13 | 0 | 0 |
| gene1632 | https://www.ncbi.nlm.nih.gov/gene/?term=LOC105287764 | 18061 | 379 | 209.84 | | 1 | 0.55 | 0.26 | 3.62 | 3.51 | 0.11 | 0.11 | 0 | 0 |
| gene10549 | https://www.ncbi.nlm.nih.gov/gene/?term=LOC105284210 | 7145 | 335 | 468.86 | | 2 | 2.8 | 0.6 | 10.35 | 10.25 | 0.09 | 0.09 | 0 | 0 |
| gene2413 | https://www.ncbi.nlm.nih.gov/gene/?term=LOC105274863 | 2006 | 53 | 264.21 | | 1 | 4.99 | 1.89 | 52.76 | 52.69 | 0.07 | 0.07 | 0 | 0 |
| gene11275 | https://www.ncbi.nlm.nih.gov/gene/?term=LOC105285055 | 4119 | 252 | 611.8 | 1 | 2.43 | 0.4 | 10.71 | 10.67 | 0.05 | 0.05 | 0 | 0 | |
| gene1332 | https://www.ncbi.nlm.nih.gov/gene/?term=LOC105286872 | 2460 | 158 | 642.28 | | 1 | 4.07 | 0.63 | 20.39 | 20.35 | 0.04 | 0.04 | 0 | 0 |
| gene3856 | https://www.ncbi.nlm.nih.gov/gene/?term=LOC105276499 | 5236 | 196 | 374.33 | | 1 | 1.91 | 0.51 | 10.18 | 10.15 | 0.03 | 0.03 | 0 | 0 |
| gene1147 | https://www.ncbi.nlm.nih.gov/gene/?term=LOC105284954 | 4022 | 111 | 275.98 | | 1 | 2.49 | 0.9 | 11.93 | 11.9 | 0.02 | 0.02 | 0 | 0 |
| gene11583 | https://www.ncbi.nlm.nih.gov/gene/?term=LOC105285414 | 8192 | 178 | 217.29 | | 1 | 1.22 | 0.56 | 9.44 | 9.48 | -0.04 | 0.04 | 0 | 0 |
| gene5375 | https://www.ncbi.nlm.nih.gov/gene/?term=LOC105278228 | 8640 | 154 | 178.24 | | 1 | 1.16 | 0.65 | 24.99 | 25.04 | -0.05 | 0.05 | 0 | 0 |
| gene5746 | https://www.ncbi.nlm.nih.gov/gene/?term=LOC105278666 | 10343 | 477 | 461.18 | | 1 | 0.97 | 0.21 | 6.71 | 6.77 | -0.05 | 0.05 | 0 | 0 |
| gene2844 | https://www.ncbi.nlm.nih.gov/gene/?term=LOC105275363 | 4330 | 197 | 454.97 | | 1 | 2.31 | 0.51 | 14.26 | 14.32 | -0.06 | 0.06 | 0 | 0 |
| gene10345 | https://www.ncbi.nlm.nih.gov/gene/?term=LOC105283989 | 4444 | 26 | 58.51 | 1 | 2.25 | 3.85 | 49.48 | 49.58 | -0.09 | 0.09 | 0 | 0 | |
| gene11579 | https://www.ncbi.nlm.nih.gov/gene/?term=LOC105285406 | 13571 | 280 | 206.32 | | 1 | 0.74 | 0.36 | 6.48 | 6.65 | -0.17 | 0.17 | 0 | 0 |
| gene8814 | https://www.ncbi.nlm.nih.gov/gene/?term=LOC105282240 | 6910 | 185 | 267.73 | | 1 | 1.45 | 0.54 | 9.48 | 9.66 | -0.18 | 0.18 | 0 | 0 |
| gene11561 | https://www.ncbi.nlm.nih.gov/gene/?term=LOC105285387 | 11528 | 207 | 179.56 | | 1 | 0.87 | 0.48 | 8.9 | 9.12 | -0.22 | 0.22 | 0 | 0 |
| gene34 | https://www.ncbi.nlm.nih.gov/gene/?term=LOC105278416 | 7120 | 132 | 185.39 | | 1 | 1.4 | 0.76 | 14.08 | 14.34 | -0.26 | 0.26 | 0 | 0 |
| gene5052 | https://www.ncbi.nlm.nih.gov/gene/?term=LOC105277856 | 3167 | 33 | 104.2 | 1 | 3.16 | 3.03 | 39.08 | 39.37 | -0.29 | 0.29 | 0 | 0 | |
| gene7248 | https://www.ncbi.nlm.nih.gov/gene/?term=LOC105280408 | 5175 | 61 | 117.87 | | 1 | 1.93 | 1.64 | 20.52 | 20.93 | -0.41 | 0.41 | 0 | 0 |

Directory: OGL File: rnk-dmg-Cb_LI_BCphaseB2.vs.RphaseA2.txt

| gene_ID | gene_link | gwidth | #Sites | #per10Kb | #dmSites | #dmsp10kb | %dmSites | %pSite1 | %pSite2 | DMpSite | ADMpSite | DMpNucl | ADMpNucl |
|---|---|---|---|---|---|---|---|---|---|---|---|---|---|
| gene2824 | https://www.ncbi.nlm.nih.gov/gene/?term=LOC105275337 | 1772 | 64 | 361.17 | 2 | 11.29 | 3.12 | 29.09 | 37.7 | -8.61 | 8.61 | -0.31 | 0.31 |
| gene3165 | https://www.ncbi.nlm.nih.gov/gene/?term=LOC105275706 | 2374 | 69 | 290.65 | 5 | 21.06 | 7.25 | 23.81 | 34.06 | -10.25 | 10.25 | -0.3 | 0.3 |
| gene10963 | https://www.ncbi.nlm.nih.gov/gene/?term=LOC105284677 | 1576 | 30 | 190.36 | 2 | 12.69 | 6.67 | 38.74 | 23.8 | 14.95 | 14.95 | 0.28 | 0.28 |
| gene2593 | https://www.ncbi.nlm.nih.gov/gene/?term=LOC105275070 | 1774 | 5 | 28.18 | 5 | 28.18 | 100 | 10.96 | 81.03 | -70.07 | 70.07 | -0.2 | 0.2 |
| gene5714 | https://www.ncbi.nlm.nih.gov/gene/?term=LOC105278729 | 710 | 24 | 338.03 | 1 | 14.08 | 4.17 | 56.79 | 61.08 | -4.29 | 4.29 | -0.14 | 0.14 |
| gene7568 | https://www.ncbi.nlm.nih.gov/gene/?term=LOC105280786 | 2525 | 119 | 471.29 | 1 | 3.96 | 0.84 | 29.84 | 32.5 | -2.66 | 2.66 | -0.13 | 0.13 |
| gene7165 | https://www.ncbi.nlm.nih.gov/gene/?term=LOC105280327 | 2688 | 109 | 405.51 | 1 | 3.72 | 0.92 | 26.1 | 29.06 | -2.96 | 2.96 | -0.12 | 0.12 |
| gene12961 | https://www.ncbi.nlm.nih.gov/gene/?term=LOC105286983 | 3094 | 69 | 223.01 | 2 | 6.46 | 2.9 | 0.97 | 5.84 | -4.87 | 4.87 | -0.11 | 0.11 |
| gene3817 | https://www.ncbi.nlm.nih.gov/gene/?term=LOC105276449 | 1230 | 57 | 463.41 | 1 | 8.13 | 1.75 | 31.59 | 33.84 | -2.25 | 2.25 | -0.1 | 0.1 |
| gene920 | https://www.ncbi.nlm.nih.gov/gene/?term=LOC105282518 | 1197 | 42 | 350.88 | 1 | 8.35 | 2.38 | 38.3 | 35.67 | 2.62 | 2.62 | 0.09 | 0.09 |
| gene5525 | https://www.ncbi.nlm.nih.gov/gene/?term=LOC105278396 | 3090 | 190 | 614.89 | 1 | 3.24 | 0.53 | 25.02 | 26.4 | -1.38 | 1.38 | -0.08 | 0.08 |
| gene10864 | https://www.ncbi.nlm.nih.gov/gene/?term=LOC105284557 | 1666 | 84 | 504.2 | 1 | 6 | 1.19 | 28.16 | 29.73 | -1.56 | 1.56 | -0.08 | 0.08 |
| gene3200 | https://www.ncbi.nlm.nih.gov/gene/?term=LOC105275743 | 1786 | 76 | 425.53 | 1 | 5.6 | 1.32 | 23.84 | 22.24 | 1.6 | 1.6 | 0.07 | 0.07 |
| gene4498 | https://www.ncbi.nlm.nih.gov/gene/?term=LOC105277239 | 2831 | 141 | 498.06 | 1 | 3.53 | 0.71 | 15.21 | 13.89 | 1.32 | 1.32 | 0.07 | 0.07 |
| gene12572 | https://www.ncbi.nlm.nih.gov/gene/?term=LOC105286514 | 2374 | 47 | 197.98 | 1 | 4.21 | 2.13 | 34.72 | 38.06 | -3.34 | 3.34 | -0.07 | 0.07 |
| gene11619 | https://www.ncbi.nlm.nih.gov/gene/?term=LOC105285454 | 1649 | 29 | 175.86 | 1 | 6.06 | 3.45 | 26.22 | 30.26 | -4.03 | 4.03 | -0.07 | 0.07 |
| gene10816 | https://www.ncbi.nlm.nih.gov/gene/?term=LOC105284502 | 1974 | 21 | 106.38 | 1 | 5.07 | 4.76 | 45.81 | 40.14 | 5.67 | 5.67 | 0.06 | 0.06 |
| gene9101 | https://www.ncbi.nlm.nih.gov/gene/?term=LOC105282553 | 3068 | 115 | 374.84 | 1 | 3.26 | 0.87 | 23.82 | 22.25 | 1.57 | 1.57 | 0.06 | 0.06 |
| gene12802 | https://www.ncbi.nlm.nih.gov/gene/?term=LOC105286799 | 3183 | 193 | 606.35 | 2 | 6.28 | 1.04 | 15.46 | 16.4 | -0.94 | 0.94 | -0.06 | 0.06 |
| gene4034 | https://www.ncbi.nlm.nih.gov/gene/?term=LOC105276704 | 2065 | 106 | 513.32 | 2 | 9.69 | 1.89 | 12.65 | 13.91 | -1.26 | 1.26 | -0.06 | 0.06 |
| gene2907 | https://www.ncbi.nlm.nih.gov/gene/?term=LOC105275447 | 2539 | 43 | 169.36 | 1 | 3.94 | 2.33 | 3.37 | 7.11 | -3.74 | 3.74 | -0.06 | 0.06 |
| gene1199 | https://www.ncbi.nlm.nih.gov/gene/?term=LOC105285498 | 2220 | 26 | 117.12 | 1 | 4.5 | 3.85 | 37.03 | 32.91 | 4.12 | 4.12 | 0.05 | 0.05 |
| gene6652 | https://www.ncbi.nlm.nih.gov/gene/?term=LOC105279719 | 2831 | 82 | 289.65 | 1 | 3.53 | 1.22 | 24.05 | 22.34 | 1.72 | 1.72 | 0.05 | 0.05 |
| gene10714 | https://www.ncbi.nlm.nih.gov/gene/?term=LOC105284396 | 1966 | 66 | 335.71 | 1 | 5.09 | 1.52 | 34.65 | 33.2 | 1.45 | 1.45 | 0.05 | 0.05 |
| gene5298 | https://www.ncbi.nlm.nih.gov/gene/?term=LOC105278145 | 4060 | 254 | 625.62 | 3 | 7.39 | 1.18 | 11.08 | 10.34 | 0.75 | 0.75 | 0.05 | 0.05 |
| gene935 | https://www.ncbi.nlm.nih.gov/gene/?term=LOC105282682 | 4368 | 241 | 551.74 | 1 | 2.29 | 0.41 | 12.96 | 13.79 | -0.83 | 0.83 | -0.05 | 0.05 |
| gene11801 | https://www.ncbi.nlm.nih.gov/gene/?term=LOC105285656 | 2497 | 137 | 548.66 | 1 | 4 | 0.73 | 8 | 8.86 | -0.87 | 0.87 | -0.05 | 0.05 |
| gene10678 | https://www.ncbi.nlm.nih.gov/gene/?term=LOC105284355 | 4031 | 183 | 453.98 | 1 | 2.48 | 0.55 | 17.56 | 18.71 | -1.15 | 1.15 | -0.05 | 0.05 |
| gene10540 | https://www.ncbi.nlm.nih.gov/gene/?term=LOC105284200 | 5267 | 213 | 404.4 | 1 | 1.9 | 0.47 | 12.38 | 13.55 | -1.16 | 1.16 | -0.05 | 0.05 |
| gene5524 | https://www.ncbi.nlm.nih.gov/gene/?term=LOC105278397 | 5177 | 206 | 397.91 | 1 | 1.93 | 0.49 | 25.22 | 26.43 | -1.21 | 1.21 | -0.05 | 0.05 |
| gene4892 | https://www.ncbi.nlm.nih.gov/gene/?term=LOC105277667 | 3898 | 131 | 336.07 | 1 | 2.57 | 0.76 | 26.19 | 27.61 | -1.42 | 1.42 | -0.05 | 0.05 |
| gene5967 | https://www.ncbi.nlm.nih.gov/gene/?term=LOC105278918 | 4393 | 148 | 336.9 | 1 | 2.28 | 0.68 | 12.61 | 14.19 | -1.58 | 1.58 | -0.05 | 0.05 |
| gene13033 | https://www.ncbi.nlm.nih.gov/gene/?term=LOC105287063 | 1877 | 56 | 298.35 | 1 | 5.33 | 1.79 | 28.26 | 30.08 | -1.82 | 1.82 | -0.05 | 0.05 |
| gene3385 | https://www.ncbi.nlm.nih.gov/gene/?term=LOC105275966 | 3305 | 73 | 220.88 | 1 | 3.03 | 1.37 | 43.5 | 45.58 | -2.08 | 2.08 | -0.05 | 0.05 |
| gene393 | https://www.ncbi.nlm.nih.gov/gene/?term=LOC105276566 | 1123 | 2 | 17.81 | 1 | 8.9 | 50 | 70.57 | 47.62 | 22.95 | 22.95 | 0.04 | 0.04 |
| gene11340 | https://www.ncbi.nlm.nih.gov/gene/?term=LOC105285131 | 2918 | 110 | 376.97 | 1 | 3.43 | 0.91 | 14.31 | 13.13 | 1.19 | 1.19 | 0.04 | 0.04 |
| gene7556 | https://www.ncbi.nlm.nih.gov/gene/?term=LOC105280774 | 2792 | 123 | 440.54 | 1 | 3.58 | 0.81 | 26.02 | 26.91 | -0.89 | 0.89 | -0.04 | 0.04 |
| gene12516 | https://www.ncbi.nlm.nih.gov/gene/?term=LOC105286452 | 1529 | 38 | 248.53 | 1 | 6.54 | 2.63 | 51.61 | 53.08 | -1.47 | 1.47 | -0.04 | 0.04 |
| gene9336 | https://www.ncbi.nlm.nih.gov/gene/?term=LOC105282834 | 5993 | 141 | 235.27 | 1 | 1.67 | 0.71 | 14.11 | 15.72 | -1.61 | 1.61 | -0.04 | 0.04 |
| gene8742 | https://www.ncbi.nlm.nih.gov/gene/?term=LOC105282151 | 2563 | 45 | 175.58 | 2 | 7.8 | 4.44 | 34.53 | 36.64 | -2.11 | 2.11 | -0.04 | 0.04 |
| gene5573 | https://www.ncbi.nlm.nih.gov/gene/?term=LOC105278454 | 2688 | 42 | 156.25 | 1 | 3.72 | 2.38 | 29.78 | 32.15 | -2.37 | 2.37 | -0.04 | 0.04 |
| gene9478 | https://www.ncbi.nlm.nih.gov/gene/?term=LOC105282991 | 3726 | 61 | 163.71 | 1 | 2.68 | 1.64 | 35.67 | 38.35 | -2.68 | 2.68 | -0.04 | 0.04 |
| gene9648 | https://www.ncbi.nlm.nih.gov/gene/?term=LOC105283183 | 2602 | 36 | 138.36 | 1 | 3.84 | 2.78 | 38.67 | 41.6 | -2.92 | 2.92 | -0.04 | 0.04 |
| gene5647 | https://www.ncbi.nlm.nih.gov/gene/?term=LOC105278567 | 3118 | 32 | 102.63 | 1 | 3.21 | 3.12 | 11.48 | 15.59 | -4.11 | 4.11 | -0.04 | 0.04 |
| gene7988 | https://www.ncbi.nlm.nih.gov/gene/?term=LOC105281282 | 3237 | 3 | 9.27 | 1 | 3.09 | 33.33 | 23.25 | 65.16 | -41.92 | 41.92 | -0.04 | 0.04 |
| gene6237 | https://www.ncbi.nlm.nih.gov/gene/?term=LOC105279244 | 1484 | 25 | 168.46 | 1 | 6.74 | 4 | 53.48 | 51.47 | 2.01 | 2.01 | 0.03 | 0.03 |
| gene12021 | https://www.ncbi.nlm.nih.gov/gene/?term=LOC105285905 | 1399 | 22 | 157.26 | 1 | 7.15 | 4.55 | 34.58 | 32.98 | 1.6 | 1.6 | 0.03 | 0.03 |
| gene1505 | https://www.ncbi.nlm.nih.gov/gene/?term=LOC105287633 | 1693 | 31 | 183.11 | 1 | 5.91 | 3.23 | 40.09 | 38.72 | 1.37 | 1.37 | 0.03 | 0.03 |
| gene2404 | https://www.ncbi.nlm.nih.gov/gene/?term=LOC105274853 | 1767 | 47 | 265.99 | 1 | 5.66 | 2.13 | 31.76 | 30.45 | 1.31 | 1.31 | 0.03 | 0.03 |
| gene1768 | https://www.ncbi.nlm.nih.gov/gene/?term=LOC105287938 | 2792 | 79 | 282.95 | 1 | 3.58 | 1.27 | 29.01 | 27.92 | 1.09 | 1.09 | 0.03 | 0.03 |
| gene1220 | https://www.ncbi.nlm.nih.gov/gene/?term=LOC105285742 | 3453 | 133 | 385.17 | 1 | 2.9 | 0.75 | 19.92 | 19.19 | 0.74 | 0.74 | 0.03 | 0.03 |
| gene298 | https://www.ncbi.nlm.nih.gov/gene/?term=LOC105275460 | 2473 | 121 | 489.28 | 1 | 4.04 | 0.83 | 13.07 | 13.58 | -0.51 | 0.51 | -0.03 | 0.03 |
| gene7514 | https://www.ncbi.nlm.nih.gov/gene/?term=LOC105280723 | 4471 | 216 | 483.11 | 1 | 2.24 | 0.46 | 13.02 | 13.57 | -0.55 | 0.55 | -0.03 | 0.03 |
| gene3206 | https://www.ncbi.nlm.nih.gov/gene/?term=LOC105275752 | 3474 | 142 | 408.75 | 1 | 2.88 | 0.7 | 30.85 | 31.58 | -0.73 | 0.73 | -0.03 | 0.03 |
| gene4376 | https://www.ncbi.nlm.nih.gov/gene/?term=LOC105277083 | 4307 | 136 | 315.77 | 1 | 2.32 | 0.74 | 17.99 | 18.79 | -0.8 | 0.8 | -0.03 | 0.03 |
| gene811 | https://www.ncbi.nlm.nih.gov/gene/?term=LOC105281261 | 5614 | 170 | 302.81 | 1 | 1.78 | 0.59 | 13.19 | 14.19 | -1 | 1 | -0.03 | 0.03 |
| gene940 | https://www.ncbi.nlm.nih.gov/gene/?term=LOC105282794 | 3578 | 92 | 257.13 | 1 | 2.79 | 1.09 | 14.15 | 15.24 | -1.1 | 1.1 | -0.03 | 0.03 |
| gene9462 | https://www.ncbi.nlm.nih.gov/gene/?term=LOC105282970 | 1469 | 42 | 285.91 | 1 | 6.81 | 2.38 | 50.96 | 52.17 | -1.21 | 1.21 | -0.03 | 0.03 |
| gene2257 | https://www.ncbi.nlm.nih.gov/gene/?term=LOC105274689 | 3658 | 86 | 235.1 | 1 | 2.73 | 1.16 | 16.99 | 18.26 | -1.27 | 1.27 | -0.03 | 0.03 |
| gene10391 | https://www.ncbi.nlm.nih.gov/gene/?term=LOC109611283 | 1994 | 46 | 230.69 | 1 | 5.02 | 2.17 | 34.01 | 35.36 | -1.35 | 1.35 | -0.03 | 0.03 |

| | | | | | | | | | | | | | |
|---|---|---|---|---|---|---|---|---|---|---|---|---|---|
| gene8661 | https://www.ncbi.nlm.nih.gov/gene/?term=LOC105282063 | 1926 | 32 | 166.15 | 1 | 5.19 | 3.12 | 32.4 | 33.92 | -1.52 | 1.52 | -0.03 | 0.03 |
| gene1884 | https://www.ncbi.nlm.nih.gov/gene/?term=LOC105288072 | 4169 | 83 | 199.09 | 1 | 2.4 | 1.2 | 21.36 | 22.96 | -1.6 | 1.6 | -0.03 | 0.03 |
| gene927 | https://www.ncbi.nlm.nih.gov/gene/?term=LOC105282600 | 3727 | 56 | 150.25 | 1 | 2.68 | 1.79 | 29.88 | 31.8 | -1.92 | 1.92 | -0.03 | 0.03 |
| gene1962 | https://www.ncbi.nlm.nih.gov/gene/?term=LOC105288157 | 1246 | 18 | 144.46 | 1 | 8.03 | 5.56 | 12.41 | 14.36 | -1.95 | 1.95 | -0.03 | 0.03 |
| gene388 | https://www.ncbi.nlm.nih.gov/gene/?term=LOC105276538 | 1558 | 21 | 134.79 | 2 | 12.84 | 9.52 | 67.26 | 69.44 | -2.18 | 2.18 | -0.03 | 0.03 |
| gene9647 | https://www.ncbi.nlm.nih.gov/gene/?term=LOC109611233 | 3094 | 42 | 135.75 | 1 | 3.23 | 2.38 | 33.15 | 35.66 | -2.51 | 2.51 | -0.03 | 0.03 |
| gene5052 | https://www.ncbi.nlm.nih.gov/gene/?term=LOC105277856 | 3167 | 39 | 123.14 | 1 | 3.16 | 2.56 | 38.42 | 40.95 | -2.53 | 2.53 | -0.03 | 0.03 |
| gene9109 | https://www.ncbi.nlm.nih.gov/gene/?term=LOC105282561 | 2400 | 7 | 29.17 | 1 | 4.17 | 14.29 | 63.87 | 57.68 | 6.19 | 6.19 | 0.02 | 0.02 |
| gene9411 | https://www.ncbi.nlm.nih.gov/gene/?term=LOC105282914 | 5534 | 43 | 77.7 | 1 | 1.81 | 2.33 | 49.78 | 47.15 | 2.63 | 2.63 | 0.02 | 0.02 |
| gene6455 | https://www.ncbi.nlm.nih.gov/gene/?term=LOC105279500 | 2612 | 34 | 130.17 | 1 | 3.83 | 2.94 | 45.97 | 44.48 | 1.5 | 1.5 | 0.02 | 0.02 |
| gene7398 | https://www.ncbi.nlm.nih.gov/gene/?term=LOC105280589 | 3452 | 39 | 112.98 | 1 | 2.9 | 2.56 | 21.67 | 20.29 | 1.38 | 1.38 | 0.02 | 0.02 |
| gene11447 | https://www.ncbi.nlm.nih.gov/gene/?term=LOC105285252 | 7437 | 128 | 172.11 | 1 | 1.34 | 0.78 | 14.37 | 13.39 | 0.98 | 0.98 | 0.02 | 0.02 |
| gene814 | https://www.ncbi.nlm.nih.gov/gene/?term=LOC105281296 | 2722 | 52 | 191.04 | 1 | 3.67 | 1.92 | 24.92 | 24.03 | 0.89 | 0.89 | 0.02 | 0.02 |
| gene3671 | https://www.ncbi.nlm.nih.gov/gene/?term=LOC105276272 | 1966 | 51 | 259.41 | 1 | 5.09 | 1.96 | 27.37 | 26.55 | 0.82 | 0.82 | 0.02 | 0.02 |
| gene5646 | https://www.ncbi.nlm.nih.gov/gene/?term=LOC105278534 | 7475 | 200 | 267.56 | 1 | 1.34 | 0.5 | 17.47 | 16.68 | 0.79 | 0.79 | 0.02 | 0.02 |
| gene7600 | https://www.ncbi.nlm.nih.gov/gene/?term=LOC105280819 | 1615 | 43 | 266.25 | 1 | 6.19 | 2.33 | 38.96 | 38.22 | 0.75 | 0.75 | 0.02 | 0.02 |
| gene3046 | https://www.ncbi.nlm.nih.gov/gene/?term=LOC105275587 | 2394 | 58 | 242.27 | 1 | 4.18 | 1.72 | 36.02 | 35.31 | 0.71 | 0.71 | 0.02 | 0.02 |
| gene7199 | https://www.ncbi.nlm.nih.gov/gene/?term=LOC105280366 | 1414 | 33 | 233.38 | 1 | 7.07 | 3.03 | 43.27 | 42.56 | 0.71 | 0.71 | 0.02 | 0.02 |
| gene7915 | https://www.ncbi.nlm.nih.gov/gene/?term=LOC105281173 | 1597 | 38 | 237.95 | 1 | 6.26 | 2.63 | 60.79 | 60.11 | 0.68 | 0.68 | 0.02 | 0.02 |
| gene3727 | https://www.ncbi.nlm.nih.gov/gene/?term=LOC105276367 | 2484 | 77 | 309.98 | 1 | 4.03 | 1.3 | 42.16 | 41.65 | 0.51 | 0.51 | 0.02 | 0.02 |
| gene10213 | https://www.ncbi.nlm.nih.gov/gene/?term=LOC105283826 | 1503 | 66 | 439.12 | 1 | 6.65 | 1.52 | 15.98 | 15.58 | 0.4 | 0.4 | 0.02 | 0.02 |
| gene11275 | https://www.ncbi.nlm.nih.gov/gene/?term=LOC105285055 | 4119 | 268 | 650.64 | 1 | 2.43 | 0.37 | 11.27 | 11.56 | -0.29 | 0.29 | -0.02 | 0.02 |
| gene5831 | https://www.ncbi.nlm.nih.gov/gene/?term=LOC105278761 | 2506 | 128 | 510.77 | 1 | 3.99 | 0.78 | 21.72 | 22.12 | -0.39 | 0.39 | -0.02 | 0.02 |
| gene2806 | https://www.ncbi.nlm.nih.gov/gene/?term=LOC105275327 | 14257 | 571 | 400.51 | 1 | 0.7 | 0.18 | 6.14 | 6.54 | -0.4 | 0.4 | -0.02 | 0.02 |
| gene738 | https://www.ncbi.nlm.nih.gov/gene/?term=LOC105280421 | 4927 | 229 | 464.79 | 1 | 2.03 | 0.44 | 14.03 | 14.44 | -0.4 | 0.4 | -0.02 | 0.02 |
| gene2056 | https://www.ncbi.nlm.nih.gov/gene/?term=LOC105274452 | 6584 | 282 | 428.31 | 2 | 3.04 | 0.71 | 12.46 | 12.87 | -0.41 | 0.41 | -0.02 | 0.02 |
| gene2271 | https://www.ncbi.nlm.nih.gov/gene/?term=LOC105274706 | 1448 | 58 | 400.55 | 1 | 6.91 | 1.72 | 36.82 | 37.28 | -0.47 | 0.47 | -0.02 | 0.02 |
| gene10606 | https://www.ncbi.nlm.nih.gov/gene/?term=LOC105284273 | 6119 | 278 | 454.32 | 1 | 1.63 | 0.36 | 18.22 | 18.7 | -0.48 | 0.48 | -0.02 | 0.02 |
| gene5877 | https://www.ncbi.nlm.nih.gov/gene/?term=LOC105278807 | 1719 | 54 | 314.14 | 1 | 5.82 | 1.85 | 24.15 | 24.64 | -0.49 | 0.49 | -0.02 | 0.02 |
| gene6031 | https://www.ncbi.nlm.nih.gov/gene/?term=LOC105279000 | 5691 | 209 | 367.25 | 1 | 1.76 | 0.48 | 13.23 | 13.74 | -0.51 | 0.51 | -0.02 | 0.02 |
| gene1430 | https://www.ncbi.nlm.nih.gov/gene/?term=LOC105287551 | 6433 | 256 | 397.95 | 1 | 1.55 | 0.39 | 8.96 | 9.5 | -0.53 | 0.53 | -0.02 | 0.02 |
| gene8729 | https://www.ncbi.nlm.nih.gov/gene/?term=LOC105282137 | 5005 | 209 | 417.58 | 1 | 2 | 0.48 | 15.97 | 16.54 | -0.56 | 0.56 | -0.02 | 0.02 |
| gene6966 | https://www.ncbi.nlm.nih.gov/gene/?term=LOC105280104 | 3747 | 140 | 373.63 | 1 | 2.67 | 0.71 | 14.7 | 15.28 | -0.58 | 0.58 | -0.02 | 0.02 |
| gene11568 | https://www.ncbi.nlm.nih.gov/gene/?term=LOC105285395 | 8086 | 226 | 279.5 | 1 | 1.24 | 0.44 | 14.03 | 14.87 | -0.84 | 0.84 | -0.02 | 0.02 |
| gene9330 | https://www.ncbi.nlm.nih.gov/gene/?term=LOC105282825 | 3936 | 98 | 248.98 | 1 | 2.54 | 1.02 | 14.19 | 15.06 | -0.86 | 0.86 | -0.02 | 0.02 |
| gene4720 | https://www.ncbi.nlm.nih.gov/gene/?term=LOC105277479 | 1697 | 45 | 265.17 | 1 | 5.89 | 2.22 | 39.86 | 40.77 | -0.91 | 0.91 | -0.02 | 0.02 |
| gene3599 | https://www.ncbi.nlm.nih.gov/gene/?term=LOC105276196 | 2955 | 60 | 203.05 | 1 | 3.38 | 1.67 | 37.09 | 38.05 | -0.97 | 0.97 | -0.02 | 0.02 |
| gene4048 | https://www.ncbi.nlm.nih.gov/gene/?term=LOC105276724 | 2955 | 42 | 142.13 | 1 | 3.38 | 2.38 | 10.45 | 11.57 | -1.12 | 1.12 | -0.02 | 0.02 |
| gene3859 | https://www.ncbi.nlm.nih.gov/gene/?term=LOC105276501 | 5788 | 106 | 183.14 | 1 | 1.73 | 0.94 | 14.91 | 16.17 | -1.26 | 1.26 | -0.02 | 0.02 |
| gene3032 | https://www.ncbi.nlm.nih.gov/gene/?term=LOC105275575 | 1713 | 29 | 169.29 | 1 | 5.84 | 3.45 | 12.34 | 13.69 | -1.35 | 1.35 | -0.02 | 0.02 |
| gene2970 | https://www.ncbi.nlm.nih.gov/gene/?term=LOC105275518 | 2340 | 28 | 119.66 | 1 | 4.27 | 3.57 | 37.12 | 38.59 | -1.47 | 1.47 | -0.02 | 0.02 |
| gene7368 | https://www.ncbi.nlm.nih.gov/gene/?term=LOC105280556 | 4105 | 55 | 133.98 | 1 | 2.44 | 1.82 | 17.06 | 18.54 | -1.48 | 1.48 | -0.02 | 0.02 |
| gene7525 | https://www.ncbi.nlm.nih.gov/gene/?term=LOC105280738 | 4237 | 43 | 101.49 | 1 | 2.36 | 2.33 | 24.72 | 26.71 | -1.99 | 1.99 | -0.02 | 0.02 |
| gene564 | https://www.ncbi.nlm.nih.gov/gene/?term=LOC105278478 | 2802 | 24 | 85.65 | 1 | 3.57 | 4.17 | 23.58 | 22.43 | 1.15 | 1.15 | 0.01 | 0.01 |
| gene10379 | https://www.ncbi.nlm.nih.gov/gene/?term=LOC105284028 | 6974 | 95 | 136.22 | 1 | 1.43 | 1.05 | 15.2 | 14.55 | 0.66 | 0.66 | 0.01 | 0.01 |
| gene657 | https://www.ncbi.nlm.nih.gov/gene/?term=LOC105279667 | 6421 | 104 | 161.97 | 1 | 1.56 | 0.96 | 3.28 | 2.76 | 0.52 | 0.52 | 0.01 | 0.01 |
| gene9327 | https://www.ncbi.nlm.nih.gov/gene/?term=LOC105282821 | 7793 | 174 | 223.28 | 1 | 1.28 | 0.57 | 10.81 | 10.29 | 0.51 | 0.51 | 0.01 | 0.01 |
| gene11583 | https://www.ncbi.nlm.nih.gov/gene/?term=LOC105285414 | 8192 | 188 | 229.49 | 1 | 1.22 | 0.53 | 8.66 | 8.2 | 0.45 | 0.45 | 0.01 | 0.01 |
| gene10819 | https://www.ncbi.nlm.nih.gov/gene/?term=LOC105284507 | 5767 | 94 | 163 | 1 | 1.73 | 1.06 | 19.47 | 19.06 | 0.4 | 0.4 | 0.01 | 0.01 |
| gene12406 | https://www.ncbi.nlm.nih.gov/gene/?term=LOC105286316 | 1386 | 32 | 230.88 | 1 | 7.22 | 3.12 | 47.17 | 46.8 | 0.37 | 0.37 | 0.01 | 0.01 |
| gene2748 | https://www.ncbi.nlm.nih.gov/gene/?term=LOC105275261 | 3845 | 56 | 145.64 | 1 | 2.6 | 1.79 | 39.96 | 39.6 | 0.36 | 0.36 | 0.01 | 0.01 |
| gene11454 | https://www.ncbi.nlm.nih.gov/gene/?term=LOC105285256 | 1906 | 50 | 262.33 | 1 | 5.25 | 2 | 42.87 | 42.54 | 0.33 | 0.33 | 0.01 | 0.01 |
| gene8274 | https://www.ncbi.nlm.nih.gov/gene/?term=LOC105281624 | 5425 | 255 | 470.05 | 1 | 1.84 | 0.39 | 9.6 | 9.29 | 0.31 | 0.31 | 0.01 | 0.01 |
| gene9755 | https://www.ncbi.nlm.nih.gov/gene/?term=LOC105283296 | 2177 | 97 | 445.57 | 1 | 4.59 | 1.03 | 20.64 | 20.32 | 0.31 | 0.31 | 0.01 | 0.01 |
| gene2413 | https://www.ncbi.nlm.nih.gov/gene/?term=LOC105274863 | 2006 | 55 | 274.18 | 1 | 4.99 | 1.82 | 52.75 | 52.45 | 0.3 | 0.3 | 0.01 | 0.01 |
| gene322 | https://www.ncbi.nlm.nih.gov/gene/?term=LOC105275777 | 1606 | 28 | 174.35 | 1 | 6.23 | 3.57 | 39.73 | 39.43 | 0.3 | 0.3 | 0.01 | 0.01 |
| gene8015 | https://www.ncbi.nlm.nih.gov/gene/?term=LOC105281312 | 2998 | 74 | 246.83 | 1 | 3.34 | 1.35 | 29.49 | 29.21 | 0.28 | 0.28 | 0.01 | 0.01 |
| gene2835 | https://www.ncbi.nlm.nih.gov/gene/?term=LOC105275357 | 2492 | 129 | 517.66 | 1 | 4.01 | 0.78 | 13.74 | 13.5 | 0.25 | 0.25 | 0.01 | 0.01 |
| gene7249 | https://www.ncbi.nlm.nih.gov/gene/?term=LOC105280422 | 1775 | 63 | 354.93 | 1 | 5.63 | 1.59 | 17.74 | 17.51 | 0.23 | 0.23 | 0.01 | 0.01 |
| gene10018 | https://www.ncbi.nlm.nih.gov/gene/?term=LOC105283607 | 4414 | 208 | 471.23 | 1 | 2.27 | 0.48 | 15.07 | 14.94 | 0.13 | 0.13 | 0.01 | 0.01 |
| gene12573 | https://www.ncbi.nlm.nih.gov/gene/?term=LOC105286515 | 3647 | 121 | 331.78 | 1 | 2.74 | 0.83 | 10.48 | 10.65 | -0.17 | 0.17 | -0.01 | 0.01 |
| gene10392 | https://www.ncbi.nlm.nih.gov/gene/?term=LOC105284036 | 3216 | 111 | 345.15 | 2 | 6.22 | 1.8 | 22.59 | 22.79 | -0.2 | 0.2 | -0.01 | 0.01 |
| gene11401 | https://www.ncbi.nlm.nih.gov/gene/?term=LOC105285202 | 1841 | 82 | 445.41 | 1 | 5.43 | 1.22 | 25.28 | 25.48 | -0.2 | 0.2 | -0.01 | 0.01 |
| gene13036 | https://www.ncbi.nlm.nih.gov/gene/?term=LOC105287059 | 4959 | 179 | 360.96 | 1 | 2.02 | 0.56 | 16.67 | 16.87 | -0.2 | 0.2 | -0.01 | 0.01 |

| | | | | | | | | | | | | | |
|---|---|---|---|---|---|---|---|---|---|---|---|---|---|
| gene11579 | https://www.ncbi.nlm.nih.gov/gene/?term=LOC105285406 | 13571 | 312 | 229.9 | 1 | 0.74 | 0.32 | 5.9 | 6.13 | -0.23 | 0.23 | -0.01 | 0.01 |
| gene2844 | https://www.ncbi.nlm.nih.gov/gene/?term=LOC105275363 | 4330 | 215 | 496.54 | 1 | 2.31 | 0.47 | 13.31 | 13.55 | -0.24 | 0.24 | -0.01 | 0.01 |
| gene6583 | https://www.ncbi.nlm.nih.gov/gene/?term=LOC105279649 | 7444 | 372 | 499.73 | 1 | 1.34 | 0.27 | 10.77 | 11.02 | -0.25 | 0.25 | -0.01 | 0.01 |
| gene5613 | https://www.ncbi.nlm.nih.gov/gene/?term=LOC105278504 | 8524 | 313 | 367.2 | 1 | 1.17 | 0.32 | 4.41 | 4.68 | -0.27 | 0.27 | -0.01 | 0.01 |
| gene7166 | https://www.ncbi.nlm.nih.gov/gene/?term=LOC105280326 | 6117 | 260 | 425.04 | 1 | 1.63 | 0.38 | 14.15 | 14.42 | -0.27 | 0.27 | -0.01 | 0.01 |
| gene2846 | https://www.ncbi.nlm.nih.gov/gene/?term=LOC105275374 | 11246 | 548 | 487.28 | 1 | 0.89 | 0.18 | 10.73 | 11.01 | -0.28 | 0.28 | -0.01 | 0.01 |
| gene8600 | https://www.ncbi.nlm.nih.gov/gene/?term=LOC105282001 | 8513 | 197 | 231.41 | 1 | 1.17 | 0.51 | 9.89 | 10.16 | -0.28 | 0.28 | -0.01 | 0.01 |
| gene9177 | https://www.ncbi.nlm.nih.gov/gene/?term=LOC105282646 | 3320 | 113 | 340.36 | 1 | 3.01 | 0.88 | 20.04 | 20.32 | -0.28 | 0.28 | -0.01 | 0.01 |
| gene6286 | https://www.ncbi.nlm.nih.gov/gene/?term=LOC105279298 | 2861 | 138 | 482.35 | 2 | 6.99 | 1.45 | 23.81 | 24.1 | -0.29 | 0.29 | -0.01 | 0.01 |
| gene8814 | https://www.ncbi.nlm.nih.gov/gene/?term=LOC105282240 | 6910 | 204 | 295.22 | 1 | 1.45 | 0.49 | 8.85 | 9.15 | -0.3 | 0.3 | -0.01 | 0.01 |
| gene1198 | https://www.ncbi.nlm.nih.gov/gene/?term=LOC105285507 | 2738 | 90 | 328.71 | 1 | 3.65 | 1.11 | 23.4 | 23.71 | -0.31 | 0.31 | -0.01 | 0.01 |
| gene5746 | https://www.ncbi.nlm.nih.gov/gene/?term=LOC105278666 | 10343 | 497 | 480.52 | 1 | 0.97 | 0.2 | 6.84 | 7.15 | -0.31 | 0.31 | -0.01 | 0.01 |
| gene3107 | https://www.ncbi.nlm.nih.gov/gene/?term=LOC105275650 | 5103 | 197 | 386.05 | 1 | 1.96 | 0.51 | 15.38 | 15.71 | -0.33 | 0.33 | -0.01 | 0.01 |
| gene7707 | https://www.ncbi.nlm.nih.gov/gene/?term=LOC105280953 | 4925 | 133 | 270.05 | 1 | 2.03 | 0.75 | 26.19 | 26.53 | -0.33 | 0.33 | -0.01 | 0.01 |
| gene2266 | https://www.ncbi.nlm.nih.gov/gene/?term=LOC105274699 | 2391 | 65 | 271.85 | 1 | 4.18 | 1.54 | 22.79 | 23.14 | -0.35 | 0.35 | -0.01 | 0.01 |
| gene34 | https://www.ncbi.nlm.nih.gov/gene/?term=LOC105278416 | 7120 | 153 | 214.89 | 1 | 1.4 | 0.65 | 12.92 | 13.27 | -0.35 | 0.35 | -0.01 | 0.01 |
| gene3848 | https://www.ncbi.nlm.nih.gov/gene/?term=LOC105276486 | 5899 | 170 | 288.18 | 1 | 1.7 | 0.59 | 21.62 | 21.98 | -0.37 | 0.37 | -0.01 | 0.01 |
| gene4962 | https://www.ncbi.nlm.nih.gov/gene/?term=LOC105277761 | 4645 | 164 | 353.07 | 1 | 2.15 | 0.61 | 14.72 | 15.12 | -0.4 | 0.4 | -0.01 | 0.01 |
| gene10600 | https://www.ncbi.nlm.nih.gov/gene/?term=LOC105284270 | 4837 | 134 | 277.03 | 1 | 2.07 | 0.75 | 11.69 | 12.12 | -0.43 | 0.43 | -0.01 | 0.01 |
| gene7207 | https://www.ncbi.nlm.nih.gov/gene/?term=LOC105280376 | 4659 | 110 | 236.1 | 1 | 2.15 | 0.91 | 30.27 | 30.71 | -0.45 | 0.45 | -0.01 | 0.01 |
| gene10531 | https://www.ncbi.nlm.nih.gov/gene/?term=LOC105284190 | 6881 | 203 | 295.02 | 1 | 1.45 | 0.49 | 11.33 | 11.8 | -0.47 | 0.47 | -0.01 | 0.01 |
| gene3470 | https://www.ncbi.nlm.nih.gov/gene/?term=LOC105276059 | 8716 | 175 | 200.78 | 1 | 1.15 | 0.57 | 7.55 | 8.01 | -0.47 | 0.47 | -0.01 | 0.01 |
| gene4996 | https://www.ncbi.nlm.nih.gov/gene/?term=LOC105277798 | 5617 | 58 | 103.26 | 1 | 1.78 | 1.72 | 13.2 | 13.71 | -0.51 | 0.51 | -0.01 | 0.01 |
| gene7183 | https://www.ncbi.nlm.nih.gov/gene/?term=LOC105280348 | 3925 | 79 | 201.27 | 1 | 2.55 | 1.27 | 19.74 | 20.25 | -0.51 | 0.51 | -0.01 | 0.01 |
| gene5375 | https://www.ncbi.nlm.nih.gov/gene/?term=LOC105278228 | 8640 | 177 | 204.86 | 1 | 1.16 | 0.56 | 22.94 | 23.49 | -0.55 | 0.55 | -0.01 | 0.01 |
| gene12307 | https://www.ncbi.nlm.nih.gov/gene/?term=LOC105286216 | 10313 | 173 | 167.75 | 1 | 0.97 | 0.58 | 16 | 16.58 | -0.58 | 0.58 | -0.01 | 0.01 |
| gene6620 | https://www.ncbi.nlm.nih.gov/gene/?term=LOC105279687 | 17998 | 402 | 223.36 | 1 | 0.56 | 0.25 | 4.4 | 5 | -0.6 | 0.6 | -0.01 | 0.01 |
| gene10107 | https://www.ncbi.nlm.nih.gov/gene/?term=LOC105283712 | 1883 | 36 | 191.18 | 1 | 5.31 | 2.78 | 29.18 | 29.79 | -0.61 | 0.61 | -0.01 | 0.01 |
| gene12439 | https://www.ncbi.nlm.nih.gov/gene/?term=LOC105286357 | 4761 | 94 | 197.44 | 1 | 2.1 | 1.06 | 30.53 | 31.14 | -0.61 | 0.61 | -0.01 | 0.01 |
| gene35 | https://www.ncbi.nlm.nih.gov/gene/?term=LOC105275197 | 2643 | 53 | 200.53 | 1 | 3.78 | 1.89 | 36.78 | 37.42 | -0.64 | 0.64 | -0.01 | 0.01 |
| gene2173 | https://www.ncbi.nlm.nih.gov/gene/?term=LOC105274590 | 4387 | 89 | 202.87 | 2 | 4.56 | 2.25 | 28.89 | 29.55 | -0.66 | 0.66 | -0.01 | 0.01 |
| gene1518 | https://www.ncbi.nlm.nih.gov/gene/?term=LOC105287648 | 2195 | 35 | 159.45 | 1 | 4.56 | 2.86 | 35.74 | 36.45 | -0.71 | 0.71 | -0.01 | 0.01 |
| gene2825 | https://www.ncbi.nlm.nih.gov/gene/?term=LOC105275343 | 5545 | 105 | 189.36 | 1 | 1.8 | 0.95 | 10.15 | 10.91 | -0.77 | 0.77 | -0.01 | 0.01 |
| gene2683 | https://www.ncbi.nlm.nih.gov/gene/?term=LOC105275187 | 5492 | 78 | 142.02 | 1 | 1.82 | 1.28 | 15.12 | 16.04 | -0.92 | 0.92 | -0.01 | 0.01 |
| gene4225 | https://www.ncbi.nlm.nih.gov/gene/?term=LOC105276917 | 2443 | 30 | 122.8 | 1 | 4.09 | 3.33 | 32.94 | 33.99 | -1.05 | 1.05 | -0.01 | 0.01 |
| gene2833 | https://www.ncbi.nlm.nih.gov/gene/?term=LOC105275352 | 5528 | 41 | 74.17 | 1 | 1.81 | 2.44 | 22.41 | 23.6 | -1.2 | 1.2 | -0.01 | 0.01 |
| gene8178 | https://www.ncbi.nlm.nih.gov/gene/?term=LOC105281515 | 8647 | 58 | 67.08 | 1 | 1.16 | 1.72 | 25.41 | 27.56 | -2.15 | 2.15 | -0.01 | 0.01 |
| gene6639 | https://www.ncbi.nlm.nih.gov/gene/?term=LOC105279705 | 7656 | 73 | 95.35 | 1 | 1.31 | 1.37 | 24.42 | 23.96 | 0.46 | 0.46 | 0 | 0 |
| gene7148 | https://www.ncbi.nlm.nih.gov/gene/?term=LOC105280304 | 8920 | 123 | 137.89 | 1 | 1.12 | 0.81 | 7.45 | 7.09 | 0.36 | 0.36 | 0 | 0 |
| gene3937 | https://www.ncbi.nlm.nih.gov/gene/?term=LOC105276592 | 9576 | 86 | 89.81 | 1 | 1.04 | 1.16 | 7.94 | 7.61 | 0.34 | 0.34 | 0 | 0 |
| gene3168 | https://www.ncbi.nlm.nih.gov/gene/?term=LOC105275711 | 6150 | 79 | 128.46 | 1 | 1.63 | 1.27 | 13.78 | 13.47 | 0.32 | 0.32 | 0 | 0 |
| gene4190 | https://www.ncbi.nlm.nih.gov/gene/?term=LOC105276875 | 3150 | 124 | 393.65 | 1 | 3.17 | 0.81 | 21.27 | 21.17 | 0.1 | 0.1 | 0 | 0 |
| gene1632 | https://www.ncbi.nlm.nih.gov/gene/?term=LOC105287764 | 18061 | 434 | 240.3 | 1 | 0.55 | 0.23 | 3.44 | 3.34 | 0.09 | 0.09 | 0 | 0 |
| gene10549 | https://www.ncbi.nlm.nih.gov/gene/?term=LOC105284210 | 7145 | 358 | 501.05 | 2 | 2.8 | 0.56 | 10 | 9.93 | 0.07 | 0.07 | 0 | 0 |
| gene9326 | https://www.ncbi.nlm.nih.gov/gene/?term=LOC105282818 | 16612 | 222 | 133.64 | 1 | 0.6 | 0.45 | 11.24 | 11.17 | 0.06 | 0.06 | 0 | 0 |
| gene10520 | https://www.ncbi.nlm.nih.gov/gene/?term=LOC105284184 | 2451 | 104 | 424.32 | 1 | 4.08 | 0.96 | 28.9 | 28.86 | 0.04 | 0.04 | 0 | 0 |
| gene10155 | https://www.ncbi.nlm.nih.gov/gene/?term=LOC105283756 | 7030 | 110 | 156.47 | 1 | 1.42 | 0.91 | 5.47 | 5.44 | 0.03 | 0.03 | 0 | 0 |
| gene1147 | https://www.ncbi.nlm.nih.gov/gene/?term=LOC105284954 | 4022 | 118 | 293.39 | 1 | 2.49 | 0.85 | 14.22 | 14.19 | 0.03 | 0.03 | 0 | 0 |
| gene53 | https://www.ncbi.nlm.nih.gov/gene/?term=LOC105277148 | 19283 | 331 | 171.65 | 1 | 0.52 | 0.3 | 4.76 | 4.72 | 0.03 | 0.03 | 0 | 0 |
| gene11688 | https://www.ncbi.nlm.nih.gov/gene/?term=LOC105285533 | 1637 | 71 | 433.72 | 1 | 6.11 | 1.41 | 23.23 | 23.21 | 0.02 | 0.02 | 0 | 0 |
| gene7550 | https://www.ncbi.nlm.nih.gov/gene/?term=LOC105280763 | 2173 | 109 | 501.61 | 1 | 4.6 | 0.92 | 25.93 | 25.92 | 0.01 | 0.01 | 0 | 0 |
| gene9445 | https://www.ncbi.nlm.nih.gov/gene/?term=LOC105282951 | 2285 | 122 | 533.92 | 1 | 4.38 | 0.82 | 19.11 | 19.11 | 0.01 | 0.01 | 0 | 0 |
| gene4527 | https://www.ncbi.nlm.nih.gov/gene/?term=LOC105277273 | 3686 | 170 | 461.2 | 1 | 2.71 | 0.59 | 12.09 | 12.09 | 0 | 0 | 0 | 0 |
| gene3885 | https://www.ncbi.nlm.nih.gov/gene/?term=LOC105276527 | 7928 | 341 | 430.12 | 1 | 1.26 | 0.29 | 9.68 | 9.69 | -0.01 | 0.01 | 0 | 0 |
| gene1310 | https://www.ncbi.nlm.nih.gov/gene/?term=LOC105286661 | 2582 | 67 | 259.49 | 1 | 3.87 | 1.49 | 24.89 | 24.92 | -0.03 | 0.03 | 0 | 0 |
| gene13114 | https://www.ncbi.nlm.nih.gov/gene/?term=LOC105287147 | 7609 | 446 | 586.15 | 1 | 1.31 | 0.22 | 9.09 | 9.12 | -0.03 | 0.03 | 0 | 0 |
| gene12674 | https://www.ncbi.nlm.nih.gov/gene/?term=LOC105286649 | 18283 | 886 | 484.6 | 1 | 0.55 | 0.11 | 1.03 | 1.06 | -0.04 | 0.04 | 0 | 0 |
| gene1470 | https://www.ncbi.nlm.nih.gov/gene/?term=LOC105287597 | 2974 | 221 | 743.11 | 1 | 3.36 | 0.45 | 16.16 | 16.21 | -0.04 | 0.04 | 0 | 0 |
| gene2122 | https://www.ncbi.nlm.nih.gov/gene/?term=LOC105274531 | 8153 | 308 | 377.78 | 1 | 1.23 | 0.32 | 10.22 | 10.26 | -0.04 | 0.04 | 0 | 0 |
| gene6649 | https://www.ncbi.nlm.nih.gov/gene/?term=LOC105279717 | 6017 | 191 | 317.43 | 1 | 1.66 | 0.52 | 12.72 | 12.75 | -0.04 | 0.04 | 0 | 0 |
| gene3856 | https://www.ncbi.nlm.nih.gov/gene/?term=LOC105276499 | 5236 | 217 | 414.44 | 1 | 1.91 | 0.46 | 9.77 | 9.82 | -0.05 | 0.05 | 0 | 0 |
| gene5777 | https://www.ncbi.nlm.nih.gov/gene/?term=LOC105278691 | 3883 | 175 | 450.68 | 1 | 2.58 | 0.57 | 14.25 | 14.3 | -0.05 | 0.05 | 0 | 0 |
| gene10596 | https://www.ncbi.nlm.nih.gov/gene/?term=LOC105284262 | 6520 | 245 | 375.77 | 1 | 1.53 | 0.41 | 3.24 | 3.34 | -0.09 | 0.09 | 0 | 0 |
| gene3286 | https://www.ncbi.nlm.nih.gov/gene/?term=LOC105275848 | 3744 | 151 | 403.31 | 1 | 2.67 | 0.66 | 21.78 | 21.89 | -0.11 | 0.11 | 0 | 0 |

| | | | | | | | | | | | | | |
|---|---|---|---|---|---|---|---|---|---|---|---|---|---|
| gene11082 | https://www.ncbi.nlm.nih.gov/gene/?term=LOC105284824 | 7436 | 224 | 301.24 | 1 | 1.34 | 0.45 | 7.03 | 7.19 | -0.15 | 0.15 | 0 | 0 |
| gene8343 | https://www.ncbi.nlm.nih.gov/gene/?term=LOC105281700 | 2172 | 39 | 179.56 | 1 | 4.6 | 2.56 | 37.71 | 37.89 | -0.17 | 0.17 | 0 | 0 |
| gene10345 | https://www.ncbi.nlm.nih.gov/gene/?term=LOC105283989 | 4444 | 29 | 65.26 | 1 | 2.25 | 3.45 | 47.94 | 48.12 | -0.18 | 0.18 | 0 | 0 |
| gene11561 | https://www.ncbi.nlm.nih.gov/gene/?term=LOC105285387 | 11528 | 218 | 189.1 | 1 | 0.87 | 0.46 | 9.03 | 9.22 | -0.19 | 0.19 | 0 | 0 |
| gene7245 | https://www.ncbi.nlm.nih.gov/gene/?term=LOC105280410 | 12478 | 265 | 212.37 | 1 | 0.8 | 0.38 | 19.68 | 19.87 | -0.19 | 0.19 | 0 | 0 |
| gene6528 | https://www.ncbi.nlm.nih.gov/gene/?term=LOC105279591 | 7222 | 60 | 83.08 | 1 | 1.38 | 1.67 | 26.03 | 26.28 | -0.25 | 0.25 | 0 | 0 |
| gene12203 | https://www.ncbi.nlm.nih.gov/gene/?term=LOC105286099 | 5967 | 74 | 124.02 | 1 | 1.68 | 1.35 | 10.44 | 10.7 | -0.27 | 0.27 | 0 | 0 |

Directory: OGL File: rnk-dmg-Cb_LI_BCphaseB2.vs.RphaseB1.txt

| gene_ID | gene_link | gwidth | #Sites | #per10Kb | #dmSites | #dmsp10kb | %dmSites | %pSite1 | %pSite2 | DMpSite | ADMpSite | DMpNucl | ADMpNucl |
|---|---|---|---|---|---|---|---|---|---|---|---|---|---|
| gene1470 | https://www.ncbi.nlm.nih.gov/gene/?term=LOC105287597 | 2974 | 190 | 638.87 | 1 | 3.36 | 0.53 | 18.6 | 19.2 | -0.6 | 0.6 | -0.04 | 0.04 |
| gene5877 | https://www.ncbi.nlm.nih.gov/gene/?term=LOC105278807 | 1719 | 50 | 290.87 | 1 | 5.82 | 2 | 23.49 | 23.85 | -0.36 | 0.36 | -0.01 | 0.01 |

Directory: OGL File: rnk-dmg-Cb_LI_BCphaseB2.vs.RphaseB2.txt

| gene_ID | gene_link | gwidth | #Sites | #per10Kb | #dmSites | #dmsp10kb | %dmSites | %pSite1 | %pSite2 | DMpSite | ADMpSite | DMpNucl | ADMpNucl |
|---|---|---|---|---|---|---|---|---|---|---|---|---|---|
| gene558 | https://www.ncbi.nlm.nih.gov/gene/?term=LOC105278458 | 216 | 3 | 138.89 | 1 | 46.3 | 33.33 | 10.44 | 29.54 | -19.1 | 19.1 | -0.27 | 0.27 |
| gene10219 | https://www.ncbi.nlm.nih.gov/gene/?term=LOC105283830 | 235 | 4 | 170.21 | 1 | 42.55 | 25 | 29.84 | 43.16 | -13.31 | 13.31 | -0.23 | 0.23 |
| gene11399 | https://www.ncbi.nlm.nih.gov/gene/?term=LOC105285203 | 1029 | 28 | 272.11 | 2 | 19.44 | 7.14 | 30.21 | 24.31 | 5.9 | 5.9 | 0.16 | 0.16 |
| gene3334 | https://www.ncbi.nlm.nih.gov/gene/?term=LOC105275897 | 1689 | 72 | 426.29 | 2 | 11.84 | 2.78 | 36.11 | 39.55 | -3.44 | 3.44 | -0.15 | 0.15 |
| gene10278 | https://www.ncbi.nlm.nih.gov/gene/?term=LOC105283908 | 902 | 14 | 155.21 | 4 | 44.35 | 28.57 | 48.14 | 38.93 | 9.21 | 9.21 | 0.14 | 0.14 |
| gene5987 | https://www.ncbi.nlm.nih.gov/gene/?term=LOC105278938 | 1882 | 110 | 584.48 | 1 | 5.31 | 0.91 | 21.13 | 23.32 | -2.19 | 2.19 | -0.13 | 0.13 |
| gene1562 | https://www.ncbi.nlm.nih.gov/gene/?term=LOC105287692 | 1491 | 89 | 596.91 | 4 | 26.83 | 4.49 | 31.38 | 29.32 | 2.06 | 2.06 | 0.12 | 0.12 |
| gene3910 | https://www.ncbi.nlm.nih.gov/gene/?term=LOC105276563 | 1957 | 112 | 572.3 | 6 | 30.66 | 5.36 | 19.61 | 17.59 | 2.02 | 2.02 | 0.12 | 0.12 |
| gene762 | https://www.ncbi.nlm.nih.gov/gene/?term=LOC105280710 | 1173 | 60 | 511.51 | 2 | 17.05 | 3.33 | 23.52 | 25.93 | -2.41 | 2.41 | -0.12 | 0.12 |
| gene5191 | https://www.ncbi.nlm.nih.gov/gene/?term=LOC105278012 | 411 | 6 | 145.99 | 1 | 24.33 | 16.67 | 44.15 | 52.1 | -7.95 | 7.95 | -0.12 | 0.12 |
| gene9965 | https://www.ncbi.nlm.nih.gov/gene/?term=LOC105283552 | 1563 | 37 | 236.72 | 1 | 6.4 | 2.7 | 55.71 | 50.95 | 4.76 | 4.76 | 0.11 | 0.11 |
| gene5809 | https://www.ncbi.nlm.nih.gov/gene/?term=LOC105278737 | 1159 | 91 | 785.16 | 1 | 8.63 | 1.1 | 17.55 | 16.13 | 1.42 | 1.42 | 0.11 | 0.11 |
| gene12660 | https://www.ncbi.nlm.nih.gov/gene/?term=LOC105286633 | 1293 | 47 | 363.5 | 2 | 15.47 | 4.26 | 13.29 | 16.18 | -2.89 | 2.89 | -0.11 | 0.11 |
| gene9019 | https://www.ncbi.nlm.nih.gov/gene/?term=LOC105282481 | 1743 | 57 | 327.02 | 1 | 5.74 | 1.75 | 34.58 | 37.81 | -3.22 | 3.22 | -0.11 | 0.11 |
| gene10743 | https://www.ncbi.nlm.nih.gov/gene/?term=LOC105284456 | 1028 | 30 | 291.83 | 2 | 19.46 | 6.67 | 47.46 | 51.1 | -3.64 | 3.64 | -0.11 | 0.11 |
| gene10644 | https://www.ncbi.nlm.nih.gov/gene/?term=LOC105284320 | 358 | 9 | 251.4 | 1 | 27.93 | 11.11 | 24.22 | 28.77 | -4.55 | 4.55 | -0.11 | 0.11 |
| gene12033 | https://www.ncbi.nlm.nih.gov/gene/?term=LOC105285914 | 844 | 28 | 331.75 | 2 | 23.7 | 7.14 | 16.23 | 13.21 | 3.02 | 3.02 | 0.1 | 0.1 |
| gene8463 | https://www.ncbi.nlm.nih.gov/gene/?term=LOC105281837 | 2474 | 106 | 428.46 | 1 | 4.04 | 0.94 | 22.3 | 19.98 | 2.31 | 2.31 | 0.1 | 0.1 |
| gene6075 | https://www.ncbi.nlm.nih.gov/gene/?term=LOC105279049 | 1285 | 90 | 700.39 | 2 | 15.56 | 2.22 | 17.35 | 18.81 | -1.46 | 1.46 | -0.1 | 0.1 |
| gene5412 | https://www.ncbi.nlm.nih.gov/gene/?term=LOC105278266 | 1868 | 115 | 615.63 | 1 | 5.35 | 0.87 | 29.91 | 31.54 | -1.63 | 1.63 | -0.1 | 0.1 |
| gene3268 | https://www.ncbi.nlm.nih.gov/gene/?term=LOC105275833 | 1040 | 42 | 403.85 | 2 | 19.23 | 4.76 | 32.43 | 34.9 | -2.47 | 2.47 | -0.1 | 0.1 |
| gene10584 | https://www.ncbi.nlm.nih.gov/gene/?term=LOC105284253 | 1255 | 41 | 326.69 | 1 | 7.97 | 2.44 | 28.64 | 31.83 | -3.18 | 3.18 | -0.1 | 0.1 |
| gene2481 | https://www.ncbi.nlm.nih.gov/gene/?term=LOC105274938 | 1937 | 54 | 278.78 | 5 | 25.81 | 9.26 | 26.94 | 30.66 | -3.73 | 3.73 | -0.1 | 0.1 |
| gene2709 | https://www.ncbi.nlm.nih.gov/gene/?term=LOC105275219 | 999 | 18 | 180.18 | 1 | 10.01 | 5.56 | 37.14 | 42.54 | -5.4 | 5.4 | -0.1 | 0.1 |

...

... only first 25 lines shown ...

Directory: OGL File: rnk-dmg-Cb_LI_RphaseA1.vs.RphaseA2.txt

| gene_ID | gene_link | gwidth | #Sites | #per10Kb | #dmSites | #dmsp10kb | %dmSites | %pSite1 | %pSite2 | DMpSite | ADMpSite | DMpNucl | ADMpNucl |
|---|---|---|---|---|---|---|---|---|---|---|---|---|---|
| gene10232 | https://www.ncbi.nlm.nih.gov/gene/?term=LOC105283847 | 1591 | 72 | 452.55 | 2 | 12.57 | 2.78 | 32.03 | 35.7 | -3.67 | 3.67 | -0.17 | 0.17 |
| gene12411 | https://www.ncbi.nlm.nih.gov/gene/?term=LOC105286320 | 1660 | 64 | 385.54 | 4 | 24.1 | 6.25 | 38.65 | 42.95 | -4.3 | 4.3 | -0.17 | 0.17 |
| gene3268 | https://www.ncbi.nlm.nih.gov/gene/?term=LOC105275833 | 1040 | 41 | 394.23 | 1 | 9.62 | 2.44 | 35.12 | 39.5 | -4.38 | 4.38 | -0.17 | 0.17 |
| gene3719 | https://www.ncbi.nlm.nih.gov/gene/?term=LOC105276360 | 1471 | 107 | 727.4 | 1 | 6.8 | 0.93 | 30.24 | 32.33 | -2.08 | 2.08 | -0.15 | 0.15 |
| gene12925 | https://www.ncbi.nlm.nih.gov/gene/?term=LOC109611487 | 391 | 27 | 690.54 | 1 | 25.58 | 3.7 | 15.18 | 17.34 | -2.15 | 2.15 | -0.15 | 0.15 |
| gene3803 | https://www.ncbi.nlm.nih.gov/gene/?term=LOC105276438 | 2613 | 162 | 619.98 | 2 | 7.65 | 1.23 | 19.62 | 21.97 | -2.35 | 2.35 | -0.15 | 0.15 |
| gene3910 | https://www.ncbi.nlm.nih.gov/gene/?term=LOC105276563 | 1957 | 118 | 602.96 | 7 | 35.77 | 5.93 | 16.25 | 18.68 | -2.43 | 2.43 | -0.15 | 0.15 |
| gene11566 | https://www.ncbi.nlm.nih.gov/gene/?term=LOC105285392 | 1603 | 91 | 567.69 | 4 | 24.95 | 4.4 | 31.49 | 34.19 | -2.7 | 2.7 | -0.15 | 0.15 |
| gene2837 | https://www.ncbi.nlm.nih.gov/gene/?term=LOC105275355 | 2872 | 161 | 560.58 | 8 | 27.86 | 4.97 | 27.89 | 25.3 | 2.58 | 2.58 | 0.14 | 0.14 |
| gene6781 | https://www.ncbi.nlm.nih.gov/gene/?term=LOC105279857 | 241 | 12 | 497.93 | 1 | 41.49 | 8.33 | 7.9 | 10.68 | -2.78 | 2.78 | -0.14 | 0.14 |
| gene5027 | https://www.ncbi.nlm.nih.gov/gene/?term=LOC105277833 | 905 | 61 | 674.03 | 1 | 11.05 | 1.64 | 2.47 | 4.36 | -1.89 | 1.89 | -0.13 | 0.13 |
| gene10176 | https://www.ncbi.nlm.nih.gov/gene/?term=LOC105283785 | 1637 | 42 | 256.57 | 3 | 18.33 | 7.14 | 12.82 | 18.05 | -5.23 | 5.23 | -0.13 | 0.13 |
| gene5828 | https://www.ncbi.nlm.nih.gov/gene/?term=LOC105278756 | 4586 | 413 | 900.57 | 3 | 6.54 | 0.73 | 9.45 | 10.77 | -1.33 | 1.33 | -0.12 | 0.12 |
| gene9183 | https://www.ncbi.nlm.nih.gov/gene/?term=LOC105282640 | 2752 | 206 | 748.55 | 3 | 10.9 | 1.46 | 21.54 | 23.12 | -1.58 | 1.58 | -0.12 | 0.12 |
| gene12652 | https://www.ncbi.nlm.nih.gov/gene/?term=LOC109610640 | 1122 | 45 | 401.07 | 4 | 35.65 | 8.89 | 47.28 | 50.3 | -3.02 | 3.02 | -0.12 | 0.12 |
| gene2274 | https://www.ncbi.nlm.nih.gov/gene/?term=LOC109610680 | 1467 | 31 | 211.32 | 2 | 13.63 | 6.45 | 65.47 | 71.37 | -5.89 | 5.89 | -0.12 | 0.12 |
| gene8798 | https://www.ncbi.nlm.nih.gov/gene/?term=LOC105282220 | 2300 | 232 | 1008.7 | 2 | 8.7 | 0.86 | 7.56 | 8.63 | -1.07 | 1.07 | -0.11 | 0.11 |
| gene12416 | https://www.ncbi.nlm.nih.gov/gene/?term=LOC105286327 | 3689 | 219 | 593.66 | 2 | 5.42 | 0.91 | 9 | 10.78 | -1.78 | 1.78 | -0.11 | 0.11 |
| gene5447 | https://www.ncbi.nlm.nih.gov/gene/?term=LOC105278314 | 1482 | 95 | 641.03 | 2 | 13.5 | 2.11 | 21.76 | 23.54 | -1.78 | 1.78 | -0.11 | 0.11 |
| gene3314 | https://www.ncbi.nlm.nih.gov/gene/?term=LOC105275878 | 2314 | 133 | 574.76 | 2 | 8.64 | 1.5 | 29.12 | 30.97 | -1.84 | 1.84 | -0.11 | 0.11 |
| gene10084 | https://www.ncbi.nlm.nih.gov/gene/?term=LOC105283683 | 1448 | 36 | 248.62 | 4 | 27.62 | 11.11 | 29.43 | 33.99 | -4.56 | 4.56 | -0.11 | 0.11 |
| gene8333 | https://www.ncbi.nlm.nih.gov/gene/?term=LOC105281691 | 1264 | 28 | 221.52 | 3 | 23.73 | 10.71 | 52.62 | 57.6 | -4.98 | 4.98 | -0.11 | 0.11 |
| gene1024 | https://www.ncbi.nlm.nih.gov/gene/?term=LOC105283693 | 1307 | 16 | 122.42 | 4 | 30.6 | 25 | 60.96 | 69.71 | -8.75 | 8.75 | -0.11 | 0.11 |
| gene13341 | https://www.ncbi.nlm.nih.gov/gene/?term=LOC105287399 | 534 | 6 | 112.36 | 1 | 18.73 | 16.67 | 23.22 | 33.07 | -9.85 | 9.85 | -0.11 | 0.11 |

...

... only first 25 lines shown ...

Directory: OGL File: rnk-dmg-Cb_LI_RphaseA1.vs.RphaseB1.txt

| gene_ID | gene_link | gwidth | #Sites | #per10Kb | #dmSites | #dmsp10kb | %dmSites | %pSite1 | %pSite2 | DMpSite | ADMpSite | DMpNucl | ADMpNucl |
|---|---|---|---|---|---|---|---|---|---|---|---|---|---|
| gene2593 | https://www.ncbi.nlm.nih.gov/gene/?term=LOC105275070 | 1774 | 11 | 62.01 | 3 | 16.91 | 27.27 | 69.15 | 25.28 | 43.86 | 43.86 | 0.27 | 0.27 |
| gene2837 | https://www.ncbi.nlm.nih.gov/gene/?term=LOC105275355 | 2872 | 164 | 571.03 | 1 | 3.48 | 0.61 | 27.39 | 23.06 | 4.33 | 4.33 | 0.25 | 0.25 |
| gene3165 | https://www.ncbi.nlm.nih.gov/gene/?term=LOC105275706 | 2374 | 63 | 265.37 | 3 | 12.64 | 4.76 | 31.1 | 21.89 | 9.2 | 9.2 | 0.24 | 0.24 |
| gene2856 | https://www.ncbi.nlm.nih.gov/gene/?term=LOC105275384 | 1987 | 88 | 442.88 | 1 | 5.03 | 1.14 | 36.35 | 32.79 | 3.56 | 3.56 | 0.16 | 0.16 |
| gene5714 | https://www.ncbi.nlm.nih.gov/gene/?term=LOC105278729 | 710 | 16 | 225.35 | 1 | 14.08 | 6.25 | 55.81 | 51.9 | 3.91 | 3.91 | 0.09 | 0.09 |
| gene7550 | https://www.ncbi.nlm.nih.gov/gene/?term=LOC105280763 | 2173 | 101 | 464.8 | 1 | 4.6 | 0.99 | 26.35 | 28.2 | -1.85 | 1.85 | -0.09 | 0.09 |
| gene1855 | https://www.ncbi.nlm.nih.gov/gene/?term=LOC105288036 | 2387 | 90 | 377.04 | 1 | 4.19 | 1.11 | 28.76 | 26.75 | 2.02 | 2.02 | 0.08 | 0.08 |
| gene13033 | https://www.ncbi.nlm.nih.gov/gene/?term=LOC105287063 | 1877 | 49 | 261.05 | 1 | 5.33 | 2.04 | 32.74 | 30.05 | 2.68 | 2.68 | 0.07 | 0.07 |
| gene5877 | https://www.ncbi.nlm.nih.gov/gene/?term=LOC105278807 | 1719 | 51 | 296.68 | 1 | 5.82 | 1.96 | 27.19 | 24.91 | 2.27 | 2.27 | 0.07 | 0.07 |
| gene11235 | https://www.ncbi.nlm.nih.gov/gene/?term=LOC105285004 | 1339 | 38 | 283.79 | 1 | 7.47 | 2.63 | 16.34 | 18.94 | -2.6 | 2.6 | -0.07 | 0.07 |
| gene9478 | https://www.ncbi.nlm.nih.gov/gene/?term=LOC105282991 | 3726 | 58 | 155.66 | 1 | 2.68 | 1.72 | 31.92 | 28.27 | 3.65 | 3.65 | 0.06 | 0.06 |
| gene1198 | https://www.ncbi.nlm.nih.gov/gene/?term=LOC105285507 | 2738 | 98 | 357.93 | 1 | 3.65 | 1.02 | 19.77 | 21.55 | -1.78 | 1.78 | -0.06 | 0.06 |
| gene12406 | https://www.ncbi.nlm.nih.gov/gene/?term=LOC105286316 | 1386 | 35 | 252.53 | 1 | 7.22 | 2.86 | 40.6 | 43.17 | -2.57 | 2.57 | -0.06 | 0.06 |
| gene12021 | https://www.ncbi.nlm.nih.gov/gene/?term=LOC105285905 | 1399 | 30 | 214.44 | 1 | 7.15 | 3.33 | 30.6 | 33.22 | -2.62 | 2.62 | -0.06 | 0.06 |
| gene388 | https://www.ncbi.nlm.nih.gov/gene/?term=LOC105276538 | 1558 | 22 | 141.21 | 2 | 12.84 | 9.09 | 54.75 | 58.78 | -4.03 | 4.03 | -0.06 | 0.06 |
| gene3671 | https://www.ncbi.nlm.nih.gov/gene/?term=LOC105276272 | 1966 | 44 | 223.8 | 1 | 5.09 | 2.27 | 27.22 | 24.96 | 2.26 | 2.26 | 0.05 | 0.05 |
| gene11819 | https://www.ncbi.nlm.nih.gov/gene/?term=LOC105285671 | 2366 | 100 | 422.65 | 1 | 4.23 | 1 | 31.41 | 30.12 | 1.29 | 1.29 | 0.05 | 0.05 |
| gene4641 | https://www.ncbi.nlm.nih.gov/gene/?term=LOC105277398 | 1937 | 73 | 376.87 | 1 | 5.16 | 1.37 | 36.14 | 34.87 | 1.27 | 1.27 | 0.05 | 0.05 |
| gene298 | https://www.ncbi.nlm.nih.gov/gene/?term=LOC105275460 | 2473 | 126 | 509.5 | 1 | 4.04 | 0.79 | 13.41 | 12.43 | 0.98 | 0.98 | 0.05 | 0.05 |
| gene1470 | https://www.ncbi.nlm.nih.gov/gene/?term=LOC105287597 | 2974 | 182 | 611.97 | 1 | 3.36 | 0.55 | 19.45 | 20.26 | -0.81 | 0.81 | -0.05 | 0.05 |
| gene9101 | https://www.ncbi.nlm.nih.gov/gene/?term=LOC105282553 | 3068 | 106 | 345.5 | 1 | 3.26 | 0.94 | 21.02 | 22.34 | -1.33 | 1.33 | -0.05 | 0.05 |
| gene11340 | https://www.ncbi.nlm.nih.gov/gene/?term=LOC105285131 | 2918 | 98 | 335.85 | 1 | 3.43 | 1.02 | 13.77 | 15.33 | -1.56 | 1.56 | -0.05 | 0.05 |
| gene2413 | https://www.ncbi.nlm.nih.gov/gene/?term=LOC105274863 | 2006 | 60 | 299.1 | 1 | 4.99 | 1.67 | 50.89 | 52.54 | -1.65 | 1.65 | -0.05 | 0.05 |
| gene7199 | https://www.ncbi.nlm.nih.gov/gene/?term=LOC105280366 | 1414 | 37 | 261.67 | 1 | 7.07 | 2.7 | 44.35 | 46.38 | -2.03 | 2.03 | -0.05 | 0.05 |
| gene12516 | https://www.ncbi.nlm.nih.gov/gene/?term=LOC105286452 | 1529 | 38 | 248.53 | 1 | 6.54 | 2.63 | 49.76 | 48.05 | 1.71 | 1.71 | 0.04 | 0.04 |
| gene3817 | https://www.ncbi.nlm.nih.gov/gene/?term=LOC105276449 | 1230 | 48 | 390.24 | 1 | 8.13 | 2.08 | 25.64 | 24.6 | 1.04 | 1.04 | 0.04 | 0.04 |
| gene9740 | https://www.ncbi.nlm.nih.gov/gene/?term=LOC105283277 | 2887 | 138 | 478 | 1 | 3.46 | 0.72 | 17.69 | 18.47 | -0.78 | 0.78 | -0.04 | 0.04 |
| gene8729 | https://www.ncbi.nlm.nih.gov/gene/?term=LOC105282137 | 5005 | 225 | 449.55 | 1 | 2 | 0.44 | 13.97 | 14.84 | -0.86 | 0.86 | -0.04 | 0.04 |
| gene570 | https://www.ncbi.nlm.nih.gov/gene/?term=LOC105278549 | 1673 | 70 | 418.41 | 1 | 5.98 | 1.43 | 42.95 | 43.99 | -1.04 | 1.04 | -0.04 | 0.04 |
| gene920 | https://www.ncbi.nlm.nih.gov/gene/?term=LOC105282518 | 1197 | 44 | 367.59 | 1 | 8.35 | 2.27 | 32.92 | 33.98 | -1.06 | 1.06 | -0.04 | 0.04 |
| gene4962 | https://www.ncbi.nlm.nih.gov/gene/?term=LOC105277761 | 4645 | 160 | 344.46 | 1 | 2.15 | 0.62 | 12.8 | 13.96 | -1.16 | 1.16 | -0.04 | 0.04 |
| gene3335 | https://www.ncbi.nlm.nih.gov/gene/?term=LOC105275898 | 4665 | 157 | 336.55 | 1 | 2.14 | 0.64 | 12.01 | 13.22 | -1.21 | 1.21 | -0.04 | 0.04 |
| gene10392 | https://www.ncbi.nlm.nih.gov/gene/?term=LOC105284036 | 3216 | 98 | 304.73 | 1 | 3.11 | 1.02 | 22.15 | 23.39 | -1.25 | 1.25 | -0.04 | 0.04 |
| gene1310 | https://www.ncbi.nlm.nih.gov/gene/?term=LOC105286661 | 2582 | 67 | 259.49 | 1 | 3.87 | 1.49 | 17.48 | 18.89 | -1.41 | 1.41 | -0.04 | 0.04 |
| gene2404 | https://www.ncbi.nlm.nih.gov/gene/?term=LOC105274853 | 1767 | 42 | 237.69 | 1 | 5.66 | 2.38 | 30.33 | 32.12 | -1.8 | 1.8 | -0.04 | 0.04 |
| gene12572 | https://www.ncbi.nlm.nih.gov/gene/?term=LOC105286514 | 2374 | 51 | 214.83 | 1 | 4.21 | 1.96 | 34.55 | 36.64 | -2.09 | 2.09 | -0.04 | 0.04 |
| gene11584 | https://www.ncbi.nlm.nih.gov/gene/?term=LOC105285416 | 3709 | 13 | 35.05 | 2 | 5.39 | 15.38 | 23.21 | 33.23 | -10.02 | 10.02 | -0.04 | 0.04 |
| gene393 | https://www.ncbi.nlm.nih.gov/gene/?term=LOC105276566 | 1123 | 2 | 17.81 | 1 | 8.9 | 50 | 50 | 74.3 | -24.3 | 24.3 | -0.04 | 0.04 |
| gene9189 | https://www.ncbi.nlm.nih.gov/gene/?term=LOC105282652 | 4699 | 84 | 178.76 | 1 | 2.13 | 1.19 | 16.38 | 14.61 | 1.78 | 1.78 | 0.03 | 0.03 |
| gene4023 | https://www.ncbi.nlm.nih.gov/gene/?term=LOC105276692 | 1759 | 36 | 204.66 | 1 | 5.69 | 2.78 | 23.9 | 22.4 | 1.5 | 1.5 | 0.03 | 0.03 |
| gene1884 | https://www.ncbi.nlm.nih.gov/gene/?term=LOC105288072 | 4169 | 82 | 196.69 | 1 | 2.4 | 1.22 | 24.39 | 23.08 | 1.31 | 1.31 | 0.03 | 0.03 |
| gene10091 | https://www.ncbi.nlm.nih.gov/gene/?term=LOC105283697 | 4988 | 154 | 308.74 | 1 | 2 | 0.65 | 16.48 | 15.36 | 1.12 | 1.12 | 0.03 | 0.03 |
| gene5831 | https://www.ncbi.nlm.nih.gov/gene/?term=LOC105278761 | 2506 | 115 | 458.9 | 2 | 7.98 | 1.74 | 22.22 | 21.51 | 0.71 | 0.71 | 0.03 | 0.03 |
| gene10540 | https://www.ncbi.nlm.nih.gov/gene/?term=LOC105284200 | 5267 | 209 | 396.81 | 1 | 1.9 | 0.48 | 13.12 | 12.47 | 0.65 | 0.65 | 0.03 | 0.03 |
| gene5298 | https://www.ncbi.nlm.nih.gov/gene/?term=LOC105278145 | 4060 | 253 | 623.15 | 3 | 7.39 | 1.19 | 10.61 | 11.09 | -0.49 | 0.49 | -0.03 | 0.03 |
| gene10326 | https://www.ncbi.nlm.nih.gov/gene/?term=LOC105283962 | 3042 | 132 | 433.93 | 1 | 3.29 | 0.76 | 8.06 | 8.65 | -0.59 | 0.59 | -0.03 | 0.03 |
| gene7166 | https://www.ncbi.nlm.nih.gov/gene/?term=LOC105280326 | 6117 | 249 | 407.06 | 1 | 1.63 | 0.4 | 14.56 | 15.24 | -0.68 | 0.68 | -0.03 | 0.03 |
| gene4190 | https://www.ncbi.nlm.nih.gov/gene/?term=LOC105276875 | 3150 | 125 | 396.83 | 1 | 3.17 | 0.8 | 19.56 | 20.35 | -0.79 | 0.79 | -0.03 | 0.03 |
| gene6652 | https://www.ncbi.nlm.nih.gov/gene/?term=LOC105279719 | 2831 | 87 | 307.31 | 1 | 3.53 | 1.15 | 20.49 | 21.33 | -0.83 | 0.83 | -0.03 | 0.03 |
| gene7915 | https://www.ncbi.nlm.nih.gov/gene/?term=LOC105281173 | 1597 | 38 | 237.95 | 1 | 6.26 | 2.63 | 59.81 | 61.01 | -1.2 | 1.2 | -0.03 | 0.03 |
| gene3599 | https://www.ncbi.nlm.nih.gov/gene/?term=LOC105276196 | 2955 | 57 | 192.89 | 1 | 3.38 | 1.75 | 39.57 | 40.89 | -1.32 | 1.32 | -0.03 | 0.03 |
| gene12203 | https://www.ncbi.nlm.nih.gov/gene/?term=LOC105286099 | 5967 | 80 | 134.07 | 1 | 1.68 | 1.25 | 10.4 | 12.39 | -2 | 2 | -0.03 | 0.03 |
| gene3028 | https://www.ncbi.nlm.nih.gov/gene/?term=LOC105275571 | 1924 | 30 | 155.93 | 1 | 5.2 | 3.33 | 42.66 | 44.75 | -2.09 | 2.09 | -0.03 | 0.03 |
| gene3168 | https://www.ncbi.nlm.nih.gov/gene/?term=LOC105275711 | 6150 | 87 | 141.46 | 1 | 1.63 | 1.15 | 10.39 | 12.71 | -2.33 | 2.33 | -0.03 | 0.03 |
| gene9411 | https://www.ncbi.nlm.nih.gov/gene/?term=LOC105282914 | 5534 | 42 | 75.89 | 1 | 1.81 | 2.38 | 47.45 | 50.92 | -3.47 | 3.47 | -0.03 | 0.03 |
| gene927 | https://www.ncbi.nlm.nih.gov/gene/?term=LOC105282600 | 3727 | 60 | 160.99 | 1 | 2.68 | 1.67 | 27.81 | 26.32 | 1.49 | 1.49 | 0.02 | 0.02 |
| gene9336 | https://www.ncbi.nlm.nih.gov/gene/?term=LOC105282834 | 5993 | 134 | 223.59 | 1 | 1.67 | 0.75 | 14.48 | 13.37 | 1.1 | 1.1 | 0.02 | 0.02 |
| gene940 | https://www.ncbi.nlm.nih.gov/gene/?term=LOC105282794 | 3578 | 93 | 259.92 | 1 | 2.79 | 1.08 | 14.3 | 13.63 | 0.67 | 0.67 | 0.02 | 0.02 |
| gene4892 | https://www.ncbi.nlm.nih.gov/gene/?term=LOC105277667 | 3898 | 121 | 310.42 | 1 | 2.57 | 0.83 | 29.02 | 28.4 | 0.62 | 0.62 | 0.02 | 0.02 |
| gene13036 | https://www.ncbi.nlm.nih.gov/gene/?term=LOC105287059 | 4959 | 180 | 362.98 | 1 | 2.02 | 0.56 | 17.65 | 17.23 | 0.42 | 0.42 | 0.02 | 0.02 |

| | | | | | | | | | | | | | |
|---|---|---|---|---|---|---|---|---|---|---|---|---|---|
| gene12441 | https://www.ncbi.nlm.nih.gov/gene/?term=LOC105286355 | 3252 | 149 | 458.18 | 1 | 3.08 | 0.67 | 21.45 | 21.08 | 0.38 | 0.38 | 0.02 | 0.02 |
| gene2844 | https://www.ncbi.nlm.nih.gov/gene/?term=LOC105275363 | 4330 | 218 | 503.46 | 1 | 2.31 | 0.46 | 13.23 | 12.92 | 0.3 | 0.3 | 0.02 | 0.02 |
| gene12802 | https://www.ncbi.nlm.nih.gov/gene/?term=LOC105286799 | 3183 | 178 | 559.22 | 1 | 3.14 | 0.56 | 16.8 | 17.1 | -0.29 | 0.29 | -0.02 | 0.02 |
| gene5525 | https://www.ncbi.nlm.nih.gov/gene/?term=LOC105278396 | 3090 | 186 | 601.94 | 1 | 3.24 | 0.54 | 24.12 | 24.45 | -0.33 | 0.33 | -0.02 | 0.02 |
| gene5524 | https://www.ncbi.nlm.nih.gov/gene/?term=LOC105278397 | 5177 | 205 | 395.98 | 1 | 1.93 | 0.49 | 24.01 | 24.4 | -0.39 | 0.39 | -0.02 | 0.02 |
| gene6583 | https://www.ncbi.nlm.nih.gov/gene/?term=LOC105279649 | 7444 | 364 | 488.98 | 1 | 1.34 | 0.27 | 10.46 | 10.89 | -0.43 | 0.43 | -0.02 | 0.02 |
| gene10678 | https://www.ncbi.nlm.nih.gov/gene/?term=LOC105284355 | 4031 | 173 | 429.17 | 1 | 2.48 | 0.58 | 18.28 | 18.72 | -0.44 | 0.44 | -0.02 | 0.02 |
| gene3107 | https://www.ncbi.nlm.nih.gov/gene/?term=LOC105275650 | 5103 | 180 | 352.73 | 1 | 1.96 | 0.56 | 15.71 | 16.3 | -0.59 | 0.59 | -0.02 | 0.02 |
| gene5646 | https://www.ncbi.nlm.nih.gov/gene/?term=LOC105278534 | 7475 | 198 | 264.88 | 1 | 1.34 | 0.51 | 17.1 | 17.7 | -0.6 | 0.6 | -0.02 | 0.02 |
| gene10844 | https://www.ncbi.nlm.nih.gov/gene/?term=LOC105284535 | 9086 | 258 | 283.95 | 1 | 1.1 | 0.39 | 13.99 | 14.72 | -0.73 | 0.73 | -0.02 | 0.02 |
| gene11454 | https://www.ncbi.nlm.nih.gov/gene/?term=LOC105285256 | 1906 | 50 | 262.33 | 1 | 5.25 | 2 | 43.22 | 44.08 | -0.87 | 0.87 | -0.02 | 0.02 |
| gene6422 | https://www.ncbi.nlm.nih.gov/gene/?term=LOC105279458 | 2419 | 43 | 177.76 | 1 | 4.13 | 2.33 | 44.62 | 45.68 | -1.05 | 1.05 | -0.02 | 0.02 |
| gene2173 | https://www.ncbi.nlm.nih.gov/gene/?term=LOC105274590 | 4387 | 84 | 191.47 | 1 | 2.28 | 1.19 | 23.89 | 24.96 | -1.07 | 1.07 | -0.02 | 0.02 |
| gene7248 | https://www.ncbi.nlm.nih.gov/gene/?term=LOC105280408 | 5175 | 60 | 115.94 | 1 | 1.93 | 1.67 | 15.92 | 17.41 | -1.49 | 1.49 | -0.02 | 0.02 |
| gene1575 | https://www.ncbi.nlm.nih.gov/gene/?term=LOC105287706 | 3783 | 44 | 116.31 | 1 | 2.64 | 2.27 | 12.72 | 14.57 | -1.85 | 1.85 | -0.02 | 0.02 |
| gene7398 | https://www.ncbi.nlm.nih.gov/gene/?term=LOC105280589 | 3452 | 41 | 118.77 | 1 | 2.9 | 2.44 | 24.33 | 26.31 | -1.98 | 1.98 | -0.02 | 0.02 |
| gene564 | https://www.ncbi.nlm.nih.gov/gene/?term=LOC105278478 | 2802 | 17 | 60.67 1 | 3.57 | 5.88 | 15.85 | 19.36 | -3.51 | 3.51 | -0.02 | 0.02 | |
| gene8178 | https://www.ncbi.nlm.nih.gov/gene/?term=LOC105281515 | 8647 | 60 | 69.39 1 | 1.16 | 1.67 | 29.73 | 28.04 | 1.69 | 1.69 | 0.01 | 0.01 | |
| gene2683 | https://www.ncbi.nlm.nih.gov/gene/?term=LOC105275187 | 5492 | 73 | 132.92 | 1 | 1.82 | 1.37 | 14.7 | 13.62 | 1.08 | 1.08 | 0.01 | 0.01 |
| gene652 | https://www.ncbi.nlm.nih.gov/gene/?term=LOC105279528 | 5806 | 39 | 67.17 1 | 1.72 | 2.56 | 22.38 | 21.58 | 0.8 | 0.8 | 0.01 | 0.01 | |
| gene10107 | https://www.ncbi.nlm.nih.gov/gene/?term=LOC105283712 | 1883 | 36 | 191.18 | 1 | 5.31 | 2.78 | 27.63 | 26.89 | 0.74 | 0.74 | 0.01 | 0.01 |
| gene7600 | https://www.ncbi.nlm.nih.gov/gene/?term=LOC105280819 | 1615 | 41 | 253.87 | 1 | 6.19 | 2.44 | 39.36 | 38.88 | 0.48 | 0.48 | 0.01 | 0.01 |
| gene1147 | https://www.ncbi.nlm.nih.gov/gene/?term=LOC105284954 | 4022 | 118 | 293.39 | 1 | 2.49 | 0.85 | 11.22 | 10.78 | 0.44 | 0.44 | 0.01 | 0.01 |
| gene10222 | https://www.ncbi.nlm.nih.gov/gene/?term=LOC105283838 | 1748 | 30 | 171.62 | 1 | 5.72 | 3.33 | 33.06 | 32.69 | 0.37 | 0.37 | 0.01 | 0.01 |
| gene5101 | https://www.ncbi.nlm.nih.gov/gene/?term=LOC105277903 | 1813 | 36 | 198.57 | 1 | 5.52 | 2.78 | 56.09 | 55.74 | 0.35 | 0.35 | 0.01 | 0.01 |
| gene7247 | https://www.ncbi.nlm.nih.gov/gene/?term=LOC105280407 | 10486 | 226 | 215.53 | 1 | 0.95 | 0.44 | 15.61 | 15.31 | 0.3 | 0.3 | 0.01 | 0.01 |
| gene10520 | https://www.ncbi.nlm.nih.gov/gene/?term=LOC105284184 | 2451 | 92 | 375.36 | 1 | 4.08 | 1.09 | 31.61 | 31.4 | 0.21 | 0.21 | 0.01 | 0.01 |
| gene2846 | https://www.ncbi.nlm.nih.gov/gene/?term=LOC105275374 | 11246 | 493 | 438.38 | 1 | 0.89 | 0.2 | 9.9 | 9.75 | 0.16 | 0.16 | 0.01 | 0.01 |
| gene12978 | https://www.ncbi.nlm.nih.gov/gene/?term=LOC105286997 | 2858 | 142 | 496.85 | 1 | 3.5 | 0.7 | 11.15 | 11.02 | 0.14 | 0.14 | 0.01 | 0.01 |
| gene2271 | https://www.ncbi.nlm.nih.gov/gene/?term=LOC105274706 | 1448 | 51 | 352.21 | 1 | 6.91 | 1.96 | 37.35 | 37.52 | -0.17 | 0.17 | -0.01 | 0.01 |
| gene3885 | https://www.ncbi.nlm.nih.gov/gene/?term=LOC105276527 | 7928 | 336 | 423.81 | 1 | 1.26 | 0.3 | 8.31 | 8.48 | -0.17 | 0.17 | -0.01 | 0.01 |
| gene8274 | https://www.ncbi.nlm.nih.gov/gene/?term=LOC105281624 | 5425 | 216 | 398.16 | 1 | 1.84 | 0.46 | 9.81 | 9.99 | -0.18 | 0.18 | -0.01 | 0.01 |
| gene8814 | https://www.ncbi.nlm.nih.gov/gene/?term=LOC105282240 | 6910 | 188 | 272.07 | 1 | 1.45 | 0.53 | 8.71 | 8.93 | -0.22 | 0.22 | -0.01 | 0.01 |
| gene10018 | https://www.ncbi.nlm.nih.gov/gene/?term=LOC105283607 | 4414 | 219 | 496.15 | 1 | 2.27 | 0.46 | 13.97 | 14.21 | -0.23 | 0.23 | -0.01 | 0.01 |
| gene13114 | https://www.ncbi.nlm.nih.gov/gene/?term=LOC105287147 | 7609 | 415 | 545.41 | 1 | 1.31 | 0.24 | 8.59 | 8.85 | -0.26 | 0.26 | -0.01 | 0.01 |
| gene6286 | https://www.ncbi.nlm.nih.gov/gene/?term=LOC105279298 | 2861 | 125 | 436.91 | 2 | 6.99 | 1.6 | 23.63 | 23.91 | -0.29 | 0.29 | -0.01 | 0.01 |
| gene10549 | https://www.ncbi.nlm.nih.gov/gene/?term=LOC105284210 | 7145 | 348 | 487.05 | 2 | 2.8 | 0.57 | 9.98 | 10.29 | -0.31 | 0.31 | -0.01 | 0.01 |
| gene9330 | https://www.ncbi.nlm.nih.gov/gene/?term=LOC105282825 | 3936 | 96 | 243.9 1 | 2.54 | 1.04 | 14.55 | 14.91 | -0.36 | 0.36 | -0.01 | 0.01 | |
| gene4376 | https://www.ncbi.nlm.nih.gov/gene/?term=LOC105277083 | 4307 | 120 | 278.62 | 1 | 2.32 | 0.83 | 19.12 | 19.51 | -0.39 | 0.39 | -0.01 | 0.01 |
| gene6031 | https://www.ncbi.nlm.nih.gov/gene/?term=LOC105279000 | 5691 | 210 | 369 1 | 1.76 | 0.48 | 11.92 | 12.31 | -0.4 | 0.4 | -0.01 | 0.01 | |
| gene12307 | https://www.ncbi.nlm.nih.gov/gene/?term=LOC105286216 | 10313 | 186 | 180.35 | 1 | 0.97 | 0.54 | 14.88 | 15.35 | -0.47 | 0.47 | -0.01 | 0.01 |
| gene9647 | https://www.ncbi.nlm.nih.gov/gene/?term=LOC109611233 | 3094 | 33 | 106.66 | 1 | 3.23 | 3.03 | 41.64 | 42.16 | -0.52 | 0.52 | -0.01 | 0.01 |
| gene9648 | https://www.ncbi.nlm.nih.gov/gene/?term=LOC105283183 | 2602 | 31 | 119.14 | 1 | 3.84 | 3.23 | 44.33 | 44.88 | -0.55 | 0.55 | -0.01 | 0.01 |
| gene6237 | https://www.ncbi.nlm.nih.gov/gene/?term=LOC105279244 | 1484 | 25 | 168.46 | 1 | 6.74 | 4 | 45.17 | 45.86 | -0.69 | 0.69 | -0.01 | 0.01 |
| gene10155 | https://www.ncbi.nlm.nih.gov/gene/?term=LOC105283756 | 7030 | 112 | 159.32 | 1 | 1.42 | 0.89 | 7.35 | 8.24 | -0.89 | 0.89 | -0.01 | 0.01 |
| gene6639 | https://www.ncbi.nlm.nih.gov/gene/?term=LOC105279705 | 7656 | 81 | 105.8 1 | 1.31 | 1.23 | 19.09 | 20.33 | -1.25 | 1.25 | -0.01 | 0.01 | |
| gene10243 | https://www.ncbi.nlm.nih.gov/gene/?term=LOC105283862 | 3239 | 43 | 132.76 | 1 | 3.09 | 2.33 | 29.28 | 28.91 | 0.37 | 0.37 | 0 | 0 |
| gene10654 | https://www.ncbi.nlm.nih.gov/gene/?term=LOC105284329 | 4027 | 71 | 176.31 | 1 | 2.48 | 1.41 | 19.25 | 18.99 | 0.26 | 0.26 | 0 | 0 |
| gene6455 | https://www.ncbi.nlm.nih.gov/gene/?term=LOC105279500 | 2612 | 31 | 118.68 | 1 | 3.83 | 3.23 | 47.76 | 47.54 | 0.22 | 0.22 | 0 | 0 |
| gene10345 | https://www.ncbi.nlm.nih.gov/gene/?term=LOC105283989 | 4444 | 29 | 65.26 1 | 2.25 | 3.45 | 49.14 | 48.92 | 0.21 | 0.21 | 0 | 0 | |
| gene7183 | https://www.ncbi.nlm.nih.gov/gene/?term=LOC105280348 | 3925 | 79 | 201.27 | 1 | 2.55 | 1.27 | 17.02 | 16.83 | 0.19 | 0.19 | 0 | 0 |
| gene2668 | https://www.ncbi.nlm.nih.gov/gene/?term=LOC105275170 | 12496 | 348 | 278.49 | 1 | 0.8 | 0.29 | 1.95 | 1.78 | 0.18 | 0.18 | 0 | 0 |
| gene9327 | https://www.ncbi.nlm.nih.gov/gene/?term=LOC105282821 | 7793 | 162 | 207.88 | 1 | 1.28 | 0.62 | 13.28 | 13.13 | 0.16 | 0.16 | 0 | 0 |
| gene7556 | https://www.ncbi.nlm.nih.gov/gene/?term=LOC105280774 | 2792 | 132 | 472.78 | 1 | 3.58 | 0.76 | 25.2 | 25.1 | 0.1 | 0.1 | 0 | 0 |
| gene10606 | https://www.ncbi.nlm.nih.gov/gene/?term=LOC105284273 | 6119 | 261 | 426.54 | 1 | 1.63 | 0.38 | 19.09 | 19 | 0.08 | 0.08 | 0 | 0 |
| gene8600 | https://www.ncbi.nlm.nih.gov/gene/?term=LOC105282001 | 8513 | 189 | 222.01 | 1 | 1.17 | 0.53 | 10.4 | 10.32 | 0.08 | 0.08 | 0 | 0 |
| gene2806 | https://www.ncbi.nlm.nih.gov/gene/?term=LOC105275327 | 14257 | 529 | 371.05 | 1 | 0.7 | 0.19 | 6.66 | 6.63 | 0.03 | 0.03 | 0 | 0 |
| gene2748 | https://www.ncbi.nlm.nih.gov/gene/?term=LOC105275261 | 3845 | 55 | 143.04 | 1 | 2.6 | 1.82 | 38.35 | 38.32 | 0.02 | 0.02 | 0 | 0 |
| gene7368 | https://www.ncbi.nlm.nih.gov/gene/?term=LOC105280556 | 4105 | 65 | 158.34 | 1 | 2.44 | 1.54 | 14.12 | 14.11 | 0.01 | 0.01 | 0 | 0 |
| gene11583 | https://www.ncbi.nlm.nih.gov/gene/?term=LOC105285414 | 8192 | 189 | 230.71 | 1 | 1.22 | 0.53 | 8.96 | 8.96 | 0 | 0 | 0 | 0 |
| gene10879 | https://www.ncbi.nlm.nih.gov/gene/?term=LOC105284574 | 2423 | 50 | 206.36 | 1 | 4.13 | 2 | 52.18 | 52.2 | -0.01 | 0.01 | 0 | 0 |
| gene3470 | https://www.ncbi.nlm.nih.gov/gene/?term=LOC105276059 | 8716 | 161 | 184.72 | 1 | 1.15 | 0.62 | 9.23 | 9.23 | -0.01 | 0.01 | 0 | 0 |
| gene5746 | https://www.ncbi.nlm.nih.gov/gene/?term=LOC105278666 | 10343 | 473 | 457.31 | 1 | 0.97 | 0.21 | 6.89 | 6.9 | -0.01 | 0.01 | 0 | 0 |
| gene5375 | https://www.ncbi.nlm.nih.gov/gene/?term=LOC105278228 | 8640 | 170 | 196.76 | 1 | 1.16 | 0.59 | 23.53 | 23.55 | -0.02 | 0.02 | 0 | 0 |

| | | | | | | | | | | | | | |
|---|---|---|---|---|---|---|---|---|---|---|---|---|---|
| gene9326 | https://www.ncbi.nlm.nih.gov/gene/?term=LOC105282818 | 16612 | 210 | 126.41 | 1 | 0.6 | 0.48 | 13.96 | 13.99 | -0.03 | 0.03 | 0 | 0 |
| gene12573 | https://www.ncbi.nlm.nih.gov/gene/?term=LOC105286515 | 3647 | 112 | 307.1 1 | 2.74 | 0.89 | 11.12 | 11.16 | -0.04 | 0.04 | 0 | 0 | |
| gene2443 | https://www.ncbi.nlm.nih.gov/gene/?term=LOC105274895 | 10433 | 409 | 392.03 | 1 | 0.96 | 0.24 | 4.52 | 4.57 | -0.04 | 0.04 | 0 | 0 |
| gene7249 | https://www.ncbi.nlm.nih.gov/gene/?term=LOC105280422 | 1775 | 66 | 371.83 | 1 | 5.63 | 1.52 | 15.99 | 16.04 | -0.05 | 0.05 | 0 | 0 |
| gene11275 | https://www.ncbi.nlm.nih.gov/gene/?term=LOC105285055 | 4119 | 251 | 609.37 | 1 | 2.43 | 0.4 | 10.55 | 10.62 | -0.07 | 0.07 | 0 | 0 |
| gene1220 | https://www.ncbi.nlm.nih.gov/gene/?term=LOC105285742 | 3453 | 109 | 315.67 | 1 | 2.9 | 0.92 | 16.97 | 17.05 | -0.09 | 0.09 | 0 | 0 |
| gene6620 | https://www.ncbi.nlm.nih.gov/gene/?term=LOC105279687 | 17998 | 433 | 240.58 | 1 | 0.56 | 0.23 | 4.48 | 4.58 | -0.1 | 0.1 | 0 | 0 |
| gene53 | https://www.ncbi.nlm.nih.gov/gene/?term=LOC105277148 | 19283 | 305 | 158.17 | 1 | 0.52 | 0.33 | 4.34 | 4.47 | -0.13 | 0.13 | 0 | 0 |
| gene5052 | https://www.ncbi.nlm.nih.gov/gene/?term=LOC105277856 | 3167 | 33 | 104.2 1 | 3.16 | 3.03 | 39.38 | 39.53 | -0.15 | 0.15 | 0 | 0 | |
| gene11561 | https://www.ncbi.nlm.nih.gov/gene/?term=LOC105285387 | 11528 | 219 | 189.97 | 1 | 0.87 | 0.46 | 9.39 | 9.54 | -0.16 | 0.16 | 0 | 0 |
| gene292 | https://www.ncbi.nlm.nih.gov/gene/?term=LOC105275398 | 2825 | 83 | 293.81 | 1 | 3.54 | 1.2 | 20.15 | 20.32 | -0.16 | 0.16 | 0 | 0 |
| gene1632 | https://www.ncbi.nlm.nih.gov/gene/?term=LOC105287764 | 18061 | 398 | 220.36 | 1 | 0.55 | 0.25 | 3.49 | 3.69 | -0.19 | 0.19 | 0 | 0 |
| gene10600 | https://www.ncbi.nlm.nih.gov/gene/?term=LOC105284270 | 4837 | 103 | 212.94 | 1 | 2.07 | 0.97 | 12.43 | 12.66 | -0.23 | 0.23 | 0 | 0 |

Directory: OGL File: rnk-dmg-Cb_LI_RphaseA1.vs.RphaseB2.txt

| gene_ID | gene_link | gwidth | #Sites | #per10Kb | #dmSites | #dmsp10kb | %dmSites | %pSite1 | %pSite2 | DMpSite | ADMpSite | DMpNucl | ADMpNucl |
|---|---|---|---|---|---|---|---|---|---|---|---|---|---|
| gene2824 | https://www.ncbi.nlm.nih.gov/gene/?term=LOC105275337 | 1772 | 58 | 327.31 | 1 | 5.64 | 1.72 | 38.96 | 30.77 | 8.18 | 8.18 | 0.27 | 0.27 |
| gene2593 | https://www.ncbi.nlm.nih.gov/gene/?term=LOC105275070 | 1774 | 7 | 39.46 | 5 | 28.18 | 71.43 | 68.01 | 4.09 | 63.92 | 63.92 | 0.25 | 0.25 |
| gene3165 | https://www.ncbi.nlm.nih.gov/gene/?term=LOC105275706 | 2374 | 65 | 273.8 | 5 | 21.06 | 7.69 | 30.28 | 21 | 9.28 | 9.28 | 0.25 | 0.25 |
| gene11819 | https://www.ncbi.nlm.nih.gov/gene/?term=LOC105285671 | 2366 | 98 | 414.2 | 1 | 4.23 | 1.02 | 29.1 | 26.97 | 2.13 | 2.13 | 0.09 | 0.09 |
| gene6334 | https://www.ncbi.nlm.nih.gov/gene/?term=LOC105279356 | 1471 | 45 | 305.91 | 1 | 6.8 | 2.22 | 31.17 | 28.75 | 2.42 | 2.42 | 0.07 | 0.07 |
| gene11401 | https://www.ncbi.nlm.nih.gov/gene/?term=LOC105285202 | 1841 | 78 | 423.68 | 1 | 5.43 | 1.28 | 26.41 | 24.78 | 1.62 | 1.62 | 0.07 | 0.07 |
| gene7406 | https://www.ncbi.nlm.nih.gov/gene/?term=LOC105280596 | 3432 | 106 | 308.86 | 1 | 2.91 | 0.94 | 17.02 | 19.17 | -2.15 | 2.15 | -0.07 | 0.07 |
| gene7568 | https://www.ncbi.nlm.nih.gov/gene/?term=LOC105280786 | 2525 | 114 | 451.49 | 1 | 3.96 | 0.88 | 28.64 | 27.23 | 1.41 | 1.41 | 0.06 | 0.06 |
| gene9101 | https://www.ncbi.nlm.nih.gov/gene/?term=LOC105282553 | 3068 | 102 | 332.46 | 1 | 3.26 | 0.98 | 21.53 | 23.25 | -1.72 | 1.72 | -0.06 | 0.06 |
| gene12406 | https://www.ncbi.nlm.nih.gov/gene/?term=LOC105286316 | 1386 | 31 | 223.67 | 1 | 7.22 | 3.23 | 42.03 | 44.54 | -2.51 | 2.51 | -0.06 | 0.06 |
| gene322 | https://www.ncbi.nlm.nih.gov/gene/?term=LOC105275777 | 1606 | 27 | 168.12 | 1 | 6.23 | 3.7 | 38.42 | 42 | -3.59 | 3.59 | -0.06 | 0.06 |
| gene5877 | https://www.ncbi.nlm.nih.gov/gene/?term=LOC105278807 | 1719 | 48 | 279.23 | 1 | 5.82 | 2.08 | 26.96 | 25.13 | 1.83 | 1.83 | 0.05 | 0.05 |
| gene1855 | https://www.ncbi.nlm.nih.gov/gene/?term=LOC105288036 | 2387 | 77 | 322.58 | 1 | 4.19 | 1.3 | 32.99 | 31.48 | 1.51 | 1.51 | 0.05 | 0.05 |
| gene4641 | https://www.ncbi.nlm.nih.gov/gene/?term=LOC105277398 | 1937 | 71 | 366.55 | 1 | 5.16 | 1.41 | 38.26 | 39.69 | -1.43 | 1.43 | -0.05 | 0.05 |
| gene12981 | https://www.ncbi.nlm.nih.gov/gene/?term=LOC105286999 | 2063 | 42 | 203.59 | 1 | 4.85 | 2.38 | 33.61 | 35.96 | -2.34 | 2.34 | -0.05 | 0.05 |
| gene10107 | https://www.ncbi.nlm.nih.gov/gene/?term=LOC105283712 | 1883 | 35 | 185.87 | 1 | 5.31 | 2.86 | 23.29 | 25.95 | -2.66 | 2.66 | -0.05 | 0.05 |
| gene3599 | https://www.ncbi.nlm.nih.gov/gene/?term=LOC105276196 | 2955 | 58 | 196.28 | 1 | 3.38 | 1.72 | 37.85 | 40.55 | -2.7 | 2.7 | -0.05 | 0.05 |
| gene6422 | https://www.ncbi.nlm.nih.gov/gene/?term=LOC105279458 | 2419 | 38 | 157.09 | 1 | 4.13 | 2.63 | 47.43 | 50.37 | -2.94 | 2.94 | -0.05 | 0.05 |
| gene388 | https://www.ncbi.nlm.nih.gov/gene/?term=LOC105276538 | 1558 | 19 | 121.95 | 2 | 12.84 | 10.53 | 61.29 | 65.13 | -3.85 | 3.85 | -0.05 | 0.05 |
| gene10602 | https://www.ncbi.nlm.nih.gov/gene/?term=LOC105284268 | 2122 | 19 | 89.54 | 1 | 4.71 | 5.26 | 27.9 | 32.99 | -5.08 | 5.08 | -0.05 | 0.05 |
| gene393 | https://www.ncbi.nlm.nih.gov/gene/?term=LOC105276566 | 1123 | 2 | 17.81 | 1 | 8.9 | 50 | 50 | 76.78 | -26.78 | 26.78 | -0.05 | 0.05 |
| gene1884 | https://www.ncbi.nlm.nih.gov/gene/?term=LOC105288072 | 4169 | 70 | 167.91 | 1 | 2.4 | 1.43 | 26.72 | 24.36 | 2.36 | 2.36 | 0.04 | 0.04 |
| gene7884 | https://www.ncbi.nlm.nih.gov/gene/?term=LOC105281266 | 3390 | 76 | 224.19 | 1 | 2.95 | 1.32 | 18.91 | 17.21 | 1.7 | 1.7 | 0.04 | 0.04 |
| gene940 | https://www.ncbi.nlm.nih.gov/gene/?term=LOC105282794 | 3578 | 85 | 237.56 | 1 | 2.79 | 1.18 | 14.76 | 13.28 | 1.48 | 1.48 | 0.04 | 0.04 |
| gene2844 | https://www.ncbi.nlm.nih.gov/gene/?term=LOC105275363 | 4330 | 186 | 429.56 | 1 | 2.31 | 0.54 | 15.2 | 14.36 | 0.85 | 0.85 | 0.04 | 0.04 |
| gene8729 | https://www.ncbi.nlm.nih.gov/gene/?term=LOC105282137 | 5005 | 213 | 425.57 | 1 | 2 | 0.47 | 14.62 | 15.67 | -1.05 | 1.05 | -0.04 | 0.04 |
| gene10213 | https://www.ncbi.nlm.nih.gov/gene/?term=LOC105283826 | 1503 | 53 | 352.63 | 1 | 6.65 | 1.89 | 14.78 | 15.95 | -1.17 | 1.17 | -0.04 | 0.04 |
| gene6031 | https://www.ncbi.nlm.nih.gov/gene/?term=LOC105279000 | 5691 | 180 | 316.29 | 1 | 1.76 | 0.56 | 14.61 | 15.92 | -1.31 | 1.31 | -0.04 | 0.04 |
| gene2413 | https://www.ncbi.nlm.nih.gov/gene/?term=LOC105274863 | 2006 | 49 | 244.27 | 1 | 4.99 | 2.04 | 53.65 | 55.47 | -1.81 | 1.81 | -0.04 | 0.04 |
| gene5101 | https://www.ncbi.nlm.nih.gov/gene/?term=LOC105277903 | 1813 | 35 | 193.05 | 1 | 5.52 | 2.86 | 52.77 | 51.27 | 1.51 | 1.51 | 0.03 | 0.03 |
| gene7915 | https://www.ncbi.nlm.nih.gov/gene/?term=LOC105281173 | 1597 | 38 | 237.95 | 1 | 6.26 | 2.63 | 59.81 | 58.74 | 1.06 | 1.06 | 0.03 | 0.03 |
| gene10540 | https://www.ncbi.nlm.nih.gov/gene/?term=LOC105284200 | 5267 | 215 | 408.2 | 1 | 1.9 | 0.47 | 12.8 | 12.05 | 0.75 | 0.75 | 0.03 | 0.03 |
| gene298 | https://www.ncbi.nlm.nih.gov/gene/?term=LOC105275460 | 2473 | 118 | 477.15 | 1 | 4.04 | 0.85 | 13.95 | 13.28 | 0.67 | 0.67 | 0.03 | 0.03 |
| gene1332 | https://www.ncbi.nlm.nih.gov/gene/?term=LOC105286872 | 2460 | 156 | 634.15 | 1 | 4.07 | 0.64 | 20.52 | 20.12 | 0.41 | 0.41 | 0.03 | 0.03 |
| gene12802 | https://www.ncbi.nlm.nih.gov/gene/?term=LOC105286799 | 3183 | 159 | 499.53 | 1 | 3.14 | 0.63 | 17.45 | 17.99 | -0.54 | 0.54 | -0.03 | 0.03 |
| gene5298 | https://www.ncbi.nlm.nih.gov/gene/?term=LOC105278145 | 4060 | 235 | 578.82 | 2 | 4.93 | 0.85 | 10.98 | 11.56 | -0.58 | 0.58 | -0.03 | 0.03 |
| gene920 | https://www.ncbi.nlm.nih.gov/gene/?term=LOC105282518 | 1197 | 48 | 401 | 1 | 8.35 | 2.08 | 32.24 | 33.02 | -0.78 | 0.78 | -0.03 | 0.03 |
| gene11340 | https://www.ncbi.nlm.nih.gov/gene/?term=LOC105285131 | 2918 | 87 | 298.15 | 1 | 3.43 | 1.15 | 10.66 | 11.58 | -0.92 | 0.92 | -0.03 | 0.03 |
| gene1310 | https://www.ncbi.nlm.nih.gov/gene/?term=LOC105286661 | 2582 | 68 | 263.36 | 1 | 3.87 | 1.47 | 19.86 | 20.85 | -1 | 1 | -0.03 | 0.03 |
| gene7600 | https://www.ncbi.nlm.nih.gov/gene/?term=LOC105280819 | 1615 | 37 | 229.1 | 1 | 6.19 | 2.7 | 36.99 | 38.16 | -1.16 | 1.16 | -0.03 | 0.03 |
| gene7183 | https://www.ncbi.nlm.nih.gov/gene/?term=LOC105280348 | 3925 | 70 | 178.34 | 1 | 2.55 | 1.43 | 19.96 | 21.48 | -1.52 | 1.52 | -0.03 | 0.03 |
| gene7199 | https://www.ncbi.nlm.nih.gov/gene/?term=LOC105280366 | 1414 | 24 | 169.73 | 1 | 7.07 | 4.17 | 43.55 | 45.17 | -1.62 | 1.62 | -0.03 | 0.03 |
| gene12572 | https://www.ncbi.nlm.nih.gov/gene/?term=LOC105286514 | 2374 | 47 | 197.98 | 1 | 4.21 | 2.13 | 35.17 | 36.86 | -1.69 | 1.69 | -0.03 | 0.03 |
| gene6237 | https://www.ncbi.nlm.nih.gov/gene/?term=LOC105279244 | 1484 | 22 | 148.25 | 1 | 6.74 | 4.55 | 40.19 | 41.99 | -1.8 | 1.8 | -0.03 | 0.03 |
| gene3168 | https://www.ncbi.nlm.nih.gov/gene/?term=LOC105275711 | 6150 | 69 | 112.2 | 1 | 1.63 | 1.45 | 12.17 | 15.18 | -3.01 | 3.01 | -0.03 | 0.03 |
| gene1008 | https://www.ncbi.nlm.nih.gov/gene/?term=LOC105283492 | 1479 | 9 | 60.85 | 1 | 6.76 | 11.11 | 30.57 | 36.08 | -5.51 | 5.51 | -0.03 | 0.03 |
| gene10654 | https://www.ncbi.nlm.nih.gov/gene/?term=LOC105284329 | 4027 | 51 | 126.65 | 1 | 2.48 | 1.96 | 20.19 | 18.78 | 1.41 | 1.41 | 0.02 | 0.02 |
| gene12516 | https://www.ncbi.nlm.nih.gov/gene/?term=LOC105286452 | 1529 | 38 | 248.53 | 1 | 6.54 | 2.63 | 50.31 | 49.37 | 0.94 | 0.94 | 0.02 | 0.02 |
| gene11454 | https://www.ncbi.nlm.nih.gov/gene/?term=LOC105285256 | 1906 | 45 | 236.1 | 1 | 5.25 | 2.22 | 47.83 | 47.02 | 0.81 | 0.81 | 0.02 | 0.02 |
| gene4892 | https://www.ncbi.nlm.nih.gov/gene/?term=LOC105277667 | 3898 | 119 | 305.28 | 1 | 2.57 | 0.84 | 28.98 | 28.19 | 0.8 | 0.8 | 0.02 | 0.02 |
| gene4213 | https://www.ncbi.nlm.nih.gov/gene/?term=LOC105276901 | 3257 | 83 | 254.84 | 1 | 3.07 | 1.2 | 27.23 | 26.49 | 0.74 | 0.74 | 0.02 | 0.02 |
| gene10520 | https://www.ncbi.nlm.nih.gov/gene/?term=LOC105284184 | 2451 | 79 | 322.32 | 1 | 4.08 | 1.27 | 35.68 | 34.99 | 0.69 | 0.69 | 0.02 | 0.02 |
| gene10600 | https://www.ncbi.nlm.nih.gov/gene/?term=LOC105284270 | 4837 | 115 | 237.75 | 1 | 2.07 | 0.87 | 11.06 | 10.43 | 0.63 | 0.63 | 0.02 | 0.02 |
| gene6966 | https://www.ncbi.nlm.nih.gov/gene/?term=LOC105280104 | 3747 | 124 | 330.93 | 1 | 2.67 | 0.81 | 17.23 | 16.6 | 0.63 | 0.63 | 0.02 | 0.02 |
| gene1430 | https://www.ncbi.nlm.nih.gov/gene/?term=LOC105287551 | 6433 | 216 | 335.77 | 1 | 1.55 | 0.46 | 9.29 | 8.67 | 0.62 | 0.62 | 0.02 | 0.02 |
| gene570 | https://www.ncbi.nlm.nih.gov/gene/?term=LOC105278549 | 1673 | 62 | 370.59 | 1 | 5.98 | 1.61 | 43.95 | 43.37 | 0.58 | 0.58 | 0.02 | 0.02 |
| gene6583 | https://www.ncbi.nlm.nih.gov/gene/?term=LOC105279649 | 7444 | 315 | 423.16 | 1 | 1.34 | 0.32 | 11.55 | 11.04 | 0.51 | 0.51 | 0.02 | 0.02 |
| gene12978 | https://www.ncbi.nlm.nih.gov/gene/?term=LOC105286997 | 2858 | 106 | 370.89 | 1 | 3.5 | 0.94 | 12 | 11.5 | 0.5 | 0.5 | 0.02 | 0.02 |
| gene5831 | https://www.ncbi.nlm.nih.gov/gene/?term=LOC105278761 | 2506 | 105 | 418.99 | 1 | 3.99 | 0.95 | 24.55 | 24.07 | 0.48 | 0.48 | 0.02 | 0.02 |
| gene8274 | https://www.ncbi.nlm.nih.gov/gene/?term=LOC105281624 | 5425 | 198 | 364.98 | 1 | 1.84 | 0.51 | 10.15 | 9.67 | 0.48 | 0.48 | 0.02 | 0.02 |

| | | | | | | | | | | | | | |
|---|---|---|---|---|---|---|---|---|---|---|---|---|---|
| gene2846 | https://www.ncbi.nlm.nih.gov/gene/?term=LOC105275374 | 11246 | 432 | 384.14 | 1 | 0.89 | 0.23 | 11.15 | 10.76 | 0.39 | 0.39 | 0.02 | 0.02 |
| gene7211 | https://www.ncbi.nlm.nih.gov/gene/?term=LOC105280377 | 2024 | 63 | 311.26 | 1 | 4.94 | 1.59 | 48.25 | 48.74 | -0.49 | 0.49 | -0.02 | 0.02 |
| gene7550 | https://www.ncbi.nlm.nih.gov/gene/?term=LOC105280763 | 2173 | 76 | 349.75 | 1 | 4.6 | 1.32 | 32.2 | 32.7 | -0.49 | 0.49 | -0.02 | 0.02 |
| gene292 | https://www.ncbi.nlm.nih.gov/gene/?term=LOC105275398 | 2825 | 78 | 276.11 | 1 | 3.54 | 1.28 | 22.03 | 22.57 | -0.55 | 0.55 | -0.02 | 0.02 |
| gene4216 | https://www.ncbi.nlm.nih.gov/gene/?term=LOC105276906 | 2127 | 86 | 404.33 | 1 | 4.7 | 1.16 | 19.06 | 19.63 | -0.57 | 0.57 | -0.02 | 0.02 |
| gene13036 | https://www.ncbi.nlm.nih.gov/gene/?term=LOC105287059 | 4959 | 148 | 298.45 | 1 | 2.02 | 0.68 | 20.35 | 20.94 | -0.59 | 0.59 | -0.02 | 0.02 |
| gene3107 | https://www.ncbi.nlm.nih.gov/gene/?term=LOC105275650 | 5103 | 152 | 297.86 | 1 | 1.96 | 0.66 | 16.19 | 16.78 | -0.59 | 0.59 | -0.02 | 0.02 |
| gene4190 | https://www.ncbi.nlm.nih.gov/gene/?term=LOC105276875 | 3150 | 105 | 333.33 | 1 | 3.17 | 0.95 | 22.87 | 23.46 | -0.6 | 0.6 | -0.02 | 0.02 |
| gene7249 | https://www.ncbi.nlm.nih.gov/gene/?term=LOC105280422 | 1775 | 60 | 338.03 | 1 | 5.63 | 1.67 | 16.46 | 17.06 | -0.6 | 0.6 | -0.02 | 0.02 |
| gene4376 | https://www.ncbi.nlm.nih.gov/gene/?term=LOC105277083 | 4307 | 97 | 225.21 | 1 | 2.32 | 1.03 | 24.28 | 25.05 | -0.77 | 0.77 | -0.02 | 0.02 |
| gene10392 | https://www.ncbi.nlm.nih.gov/gene/?term=LOC105284036 | 3216 | 94 | 292.29 | 2 | 6.22 | 2.13 | 22.41 | 23.26 | -0.84 | 0.84 | -0.02 | 0.02 |
| gene35 | https://www.ncbi.nlm.nih.gov/gene/?term=LOC105275197 | 2643 | 45 | 170.26 | 1 | 3.78 | 2.22 | 38.91 | 40.15 | -1.23 | 1.23 | -0.02 | 0.02 |
| gene8343 | https://www.ncbi.nlm.nih.gov/gene/?term=LOC105281700 | 2172 | 32 | 147.33 | 1 | 4.6 | 3.12 | 37.43 | 38.97 | -1.55 | 1.55 | -0.02 | 0.02 |
| gene10345 | https://www.ncbi.nlm.nih.gov/gene/?term=LOC105283989 | 4444 | 22 | 49.5 | 1 | 2.25 | 4.55 | 37.68 | 41.66 | -3.99 | 3.99 | -0.02 | 0.02 |
| gene8178 | https://www.ncbi.nlm.nih.gov/gene/?term=LOC105281515 | 8647 | 43 | 49.73 | 1 | 1.16 | 2.33 | 27.02 | 24.75 | 2.27 | 2.27 | 0.01 | 0.01 |
| gene8661 | https://www.ncbi.nlm.nih.gov/gene/?term=LOC105282063 | 1926 | 26 | 134.99 | 1 | 5.19 | 3.85 | 38.79 | 37.9 | 0.89 | 0.89 | 0.01 | 0.01 |
| gene4023 | https://www.ncbi.nlm.nih.gov/gene/?term=LOC105276692 | 1759 | 31 | 176.24 | 1 | 5.69 | 3.23 | 24.44 | 23.63 | 0.81 | 0.81 | 0.01 | 0.01 |
| gene3470 | https://www.ncbi.nlm.nih.gov/gene/?term=LOC105276059 | 8716 | 146 | 167.51 | 1 | 1.15 | 0.68 | 9.33 | 8.54 | 0.79 | 0.79 | 0.01 | 0.01 |
| gene9336 | https://www.ncbi.nlm.nih.gov/gene/?term=LOC105282834 | 5993 | 123 | 205.24 | 1 | 1.67 | 0.81 | 15.27 | 14.61 | 0.66 | 0.66 | 0.01 | 0.01 |
| gene5273 | https://www.ncbi.nlm.nih.gov/gene/?term=LOC105278108 | 4067 | 43 | 105.73 | 1 | 2.46 | 2.33 | 26.66 | 26.12 | 0.55 | 0.55 | 0.01 | 0.01 |
| gene11579 | https://www.ncbi.nlm.nih.gov/gene/?term=LOC105285406 | 13571 | 229 | 168.74 | 1 | 0.74 | 0.44 | 7.6 | 7.22 | 0.38 | 0.38 | 0.01 | 0.01 |
| gene12439 | https://www.ncbi.nlm.nih.gov/gene/?term=LOC105286357 | 4761 | 87 | 182.73 | 1 | 2.1 | 1.15 | 25.3 | 24.92 | 0.38 | 0.38 | 0.01 | 0.01 |
| gene7247 | https://www.ncbi.nlm.nih.gov/gene/?term=LOC105280407 | 10486 | 208 | 198.36 | 1 | 0.95 | 0.48 | 16.58 | 16.25 | 0.33 | 0.33 | 0.01 | 0.01 |
| gene2266 | https://www.ncbi.nlm.nih.gov/gene/?term=LOC105274699 | 2391 | 49 | 204.94 | 1 | 4.18 | 2.04 | 23.32 | 23 | 0.32 | 0.32 | 0.01 | 0.01 |
| gene13114 | https://www.ncbi.nlm.nih.gov/gene/?term=LOC105287147 | 7609 | 393 | 516.49 | 1 | 1.31 | 0.25 | 9.24 | 9.02 | 0.23 | 0.23 | 0.01 | 0.01 |
| gene5525 | https://www.ncbi.nlm.nih.gov/gene/?term=LOC105278396 | 3090 | 180 | 582.52 | 1 | 3.24 | 0.56 | 24.99 | 24.77 | 0.22 | 0.22 | 0.01 | 0.01 |
| gene11082 | https://www.ncbi.nlm.nih.gov/gene/?term=LOC105284824 | 7436 | 189 | 254.17 | 1 | 1.34 | 0.53 | 8.04 | 7.84 | 0.2 | 0.2 | 0.01 | 0.01 |
| gene11055 | https://www.ncbi.nlm.nih.gov/gene/?term=LOC105284796 | 2283 | 121 | 530 | 1 | 4.38 | 0.83 | 34.82 | 34.65 | 0.17 | 0.17 | 0.01 | 0.01 |
| gene1470 | https://www.ncbi.nlm.nih.gov/gene/?term=LOC105287597 | 2974 | 181 | 608.61 | 1 | 3.36 | 0.55 | 19.06 | 18.9 | 0.16 | 0.16 | 0.01 | 0.01 |
| gene7556 | https://www.ncbi.nlm.nih.gov/gene/?term=LOC105280774 | 2792 | 98 | 351 | 1 | 3.58 | 1.02 | 32.45 | 32.61 | -0.17 | 0.17 | -0.01 | 0.01 |
| gene10018 | https://www.ncbi.nlm.nih.gov/gene/?term=LOC105283607 | 4414 | 202 | 457.63 | 1 | 2.27 | 0.5 | 14.99 | 15.23 | -0.24 | 0.24 | -0.01 | 0.01 |
| gene6286 | https://www.ncbi.nlm.nih.gov/gene/?term=LOC105279298 | 2861 | 130 | 454.39 | 2 | 6.99 | 1.54 | 23.66 | 23.9 | -0.24 | 0.24 | -0.01 | 0.01 |
| gene10606 | https://www.ncbi.nlm.nih.gov/gene/?term=LOC105284273 | 6119 | 244 | 398.76 | 1 | 1.63 | 0.41 | 18.11 | 18.39 | -0.29 | 0.29 | -0.01 | 0.01 |
| gene34 | https://www.ncbi.nlm.nih.gov/gene/?term=LOC105278416 | 7120 | 124 | 174.16 | 1 | 1.4 | 0.81 | 14.41 | 14.72 | -0.3 | 0.3 | -0.01 | 0.01 |
| gene12573 | https://www.ncbi.nlm.nih.gov/gene/?term=LOC105286515 | 3647 | 105 | 287.91 | 1 | 2.74 | 0.95 | 12.53 | 12.84 | -0.31 | 0.31 | -0.01 | 0.01 |
| gene3885 | https://www.ncbi.nlm.nih.gov/gene/?term=LOC105276527 | 7928 | 299 | 377.14 | 1 | 1.26 | 0.33 | 9.83 | 10.14 | -0.31 | 0.31 | -0.01 | 0.01 |
| gene7166 | https://www.ncbi.nlm.nih.gov/gene/?term=LOC105280326 | 6117 | 226 | 369.46 | 1 | 1.63 | 0.44 | 14.91 | 15.21 | -0.31 | 0.31 | -0.01 | 0.01 |
| gene10549 | https://www.ncbi.nlm.nih.gov/gene/?term=LOC105284210 | 7145 | 284 | 397.48 | 2 | 2.8 | 0.7 | 11.85 | 12.19 | -0.33 | 0.33 | -0.01 | 0.01 |
| gene5646 | https://www.ncbi.nlm.nih.gov/gene/?term=LOC105278534 | 7475 | 176 | 235.45 | 1 | 1.34 | 0.57 | 18.67 | 19 | -0.33 | 0.33 | -0.01 | 0.01 |
| gene2404 | https://www.ncbi.nlm.nih.gov/gene/?term=LOC105274853 | 1767 | 41 | 232.03 | 1 | 5.66 | 2.44 | 30.51 | 30.87 | -0.36 | 0.36 | -0.01 | 0.01 |
| gene9327 | https://www.ncbi.nlm.nih.gov/gene/?term=LOC105282821 | 7793 | 140 | 179.65 | 1 | 1.28 | 0.71 | 12.3 | 12.71 | -0.41 | 0.41 | -0.01 | 0.01 |
| gene5375 | https://www.ncbi.nlm.nih.gov/gene/?term=LOC105278228 | 8640 | 152 | 175.93 | 1 | 1.16 | 0.66 | 22.96 | 23.39 | -0.42 | 0.42 | -0.01 | 0.01 |
| gene10879 | https://www.ncbi.nlm.nih.gov/gene/?term=LOC105284574 | 2423 | 50 | 206.36 | 1 | 4.13 | 2 | 52.82 | 53.31 | -0.49 | 0.49 | -0.01 | 0.01 |
| gene12307 | https://www.ncbi.nlm.nih.gov/gene/?term=LOC105286216 | 10313 | 164 | 159.02 | 1 | 0.97 | 0.61 | 16.27 | 16.77 | -0.5 | 0.5 | -0.01 | 0.01 |
| gene1147 | https://www.ncbi.nlm.nih.gov/gene/?term=LOC105284954 | 4022 | 91 | 226.26 | 1 | 2.49 | 1.1 | 14.14 | 14.77 | -0.63 | 0.63 | -0.01 | 0.01 |
| gene9647 | https://www.ncbi.nlm.nih.gov/gene/?term=LOC109611233 | 3094 | 32 | 103.43 | 1 | 3.23 | 3.12 | 37.53 | 38.18 | -0.65 | 0.65 | -0.01 | 0.01 |
| gene9648 | https://www.ncbi.nlm.nih.gov/gene/?term=LOC105283183 | 2602 | 30 | 115.3 | 1 | 3.84 | 3.33 | 40.03 | 40.73 | -0.7 | 0.7 | -0.01 | 0.01 |
| gene12203 | https://www.ncbi.nlm.nih.gov/gene/?term=LOC105286099 | 5967 | 70 | 117.31 | 1 | 1.68 | 1.43 | 9.91 | 10.79 | -0.88 | 0.88 | -0.01 | 0.01 |
| gene7248 | https://www.ncbi.nlm.nih.gov/gene/?term=LOC105280408 | 5175 | 52 | 100.48 | 1 | 1.93 | 1.92 | 17.59 | 18.54 | -0.95 | 0.95 | -0.01 | 0.01 |
| gene10155 | https://www.ncbi.nlm.nih.gov/gene/?term=LOC105283756 | 7030 | 108 | 153.63 | 1 | 1.42 | 0.93 | 5.58 | 6.54 | -0.96 | 0.96 | -0.01 | 0.01 |
| gene12021 | https://www.ncbi.nlm.nih.gov/gene/?term=LOC105285905 | 1399 | 16 | 114.37 | 1 | 7.15 | 6.25 | 46.76 | 47.75 | -0.98 | 0.98 | -0.01 | 0.01 |
| gene6639 | https://www.ncbi.nlm.nih.gov/gene/?term=LOC105279705 | 7656 | 69 | 90.13 | 1 | 1.31 | 1.45 | 20.74 | 21.96 | -1.21 | 1.21 | -0.01 | 0.01 |
| gene564 | https://www.ncbi.nlm.nih.gov/gene/?term=LOC105278478 | 2802 | 18 | 64.24 | 1 | 3.57 | 5.56 | 14.97 | 16.38 | -1.41 | 1.41 | -0.01 | 0.01 |
| gene2683 | https://www.ncbi.nlm.nih.gov/gene/?term=LOC105275187 | 5492 | 65 | 118.35 | 1 | 1.82 | 1.54 | 11.46 | 11.25 | 0.21 | 0.21 | 0 | 0 |
| gene5052 | https://www.ncbi.nlm.nih.gov/gene/?term=LOC105277856 | 3167 | 28 | 88.41 | 1 | 3.16 | 3.57 | 33.66 | 33.45 | 0.21 | 0.21 | 0 | 0 |
| gene2806 | https://www.ncbi.nlm.nih.gov/gene/?term=LOC105275327 | 14257 | 456 | 319.84 | 1 | 0.7 | 0.22 | 7.38 | 7.24 | 0.14 | 0.14 | 0 | 0 |
| gene1220 | https://www.ncbi.nlm.nih.gov/gene/?term=LOC105285742 | 3453 | 100 | 289.6 | 1 | 2.9 | 1 | 18.13 | 18.02 | 0.11 | 0.11 | 0 | 0 |
| gene5733 | https://www.ncbi.nlm.nih.gov/gene/?term=LOC105278649 | 5221 | 92 | 176.21 | 1 | 1.92 | 1.09 | 19.86 | 19.75 | 0.11 | 0.11 | 0 | 0 |
| gene5746 | https://www.ncbi.nlm.nih.gov/gene/?term=LOC105278666 | 10343 | 436 | 421.54 | 1 | 0.97 | 0.23 | 6.97 | 6.87 | 0.11 | 0.11 | 0 | 0 |
| gene8600 | https://www.ncbi.nlm.nih.gov/gene/?term=LOC105282001 | 8513 | 151 | 177.38 | 1 | 1.17 | 0.66 | 10.5 | 10.38 | 0.11 | 0.11 | 0 | 0 |
| gene10844 | https://www.ncbi.nlm.nih.gov/gene/?term=LOC105284535 | 9086 | 222 | 244.33 | 1 | 1.1 | 0.45 | 15.15 | 15.05 | 0.1 | 0.1 | 0 | 0 |
| gene1632 | https://www.ncbi.nlm.nih.gov/gene/?term=LOC105287764 | 18061 | 367 | 203.2 | 1 | 0.55 | 0.27 | 3.44 | 3.36 | 0.08 | 0.08 | 0 | 0 |
| gene5524 | https://www.ncbi.nlm.nih.gov/gene/?term=LOC105278397 | 5177 | 196 | 378.6 | 1 | 1.93 | 0.51 | 24.77 | 24.69 | 0.07 | 0.07 | 0 | 0 |
| gene1408 | https://www.ncbi.nlm.nih.gov/gene/?term=LOC109611520 | 8645 | 289 | 334.3 | 1 | 1.16 | 0.35 | 6.21 | 6.16 | 0.05 | 0.05 | 0 | 0 |

| | | | | | | | | | | | | | |
|---|---|---|---|---|---|---|---|---|---|---|---|---|---|
| gene7368 | https://www.ncbi.nlm.nih.gov/gene/?term=LOC105280556 | 4105 | 38 | 92.57 | 1 | 2.44 | 2.63 | 16.79 | 16.73 | 0.05 | 0.05 | 0 | 0 |
| gene9740 | https://www.ncbi.nlm.nih.gov/gene/?term=LOC105283277 | 2887 | 131 | 453.76 | 1 | 3.46 | 0.76 | 19.31 | 19.27 | 0.04 | 0.04 | 0 | 0 |
| gene8814 | https://www.ncbi.nlm.nih.gov/gene/?term=LOC105282240 | 6910 | 163 | 235.89 | 1 | 1.45 | 0.61 | 9.99 | 9.96 | 0.03 | 0.03 | 0 | 0 |
| gene2271 | https://www.ncbi.nlm.nih.gov/gene/?term=LOC105274706 | 1448 | 50 | 345.3 | 1 | 6.91 | 2 | 38.1 | 38.09 | 0.01 | 0.01 | 0 | 0 |
| gene2173 | https://www.ncbi.nlm.nih.gov/gene/?term=LOC105274590 | 4387 | 68 | 155 | 1 | 2.28 | 1.47 | 26.72 | 26.74 | -0.02 | 0.02 | 0 | 0 |
| gene10678 | https://www.ncbi.nlm.nih.gov/gene/?term=LOC105284355 | 4031 | 168 | 416.77 | 1 | 2.48 | 0.6 | 18.14 | 18.16 | -0.03 | 0.03 | 0 | 0 |
| gene10984 | https://www.ncbi.nlm.nih.gov/gene/?term=LOC105284704 | 1987 | 58 | 291.9 | 1 | 5.03 | 1.72 | 23.02 | 23.06 | -0.04 | 0.04 | 0 | 0 |
| gene10326 | https://www.ncbi.nlm.nih.gov/gene/?term=LOC105283962 | 3042 | 119 | 391.19 | 1 | 3.29 | 0.84 | 8.91 | 8.98 | -0.07 | 0.07 | 0 | 0 |
| gene11275 | https://www.ncbi.nlm.nih.gov/gene/?term=LOC105285055 | 4119 | 257 | 623.94 | 1 | 2.43 | 0.39 | 10.45 | 10.51 | -0.07 | 0.07 | 0 | 0 |
| gene4962 | https://www.ncbi.nlm.nih.gov/gene/?term=LOC105277761 | 4645 | 142 | 305.71 | 1 | 2.15 | 0.7 | 13.6 | 13.69 | -0.09 | 0.09 | 0 | 0 |
| gene7398 | https://www.ncbi.nlm.nih.gov/gene/?term=LOC105280589 | 3452 | 33 | 95.6 | 1 | 2.9 | 3.03 | 20.49 | 20.58 | -0.09 | 0.09 | 0 | 0 |
| gene6620 | https://www.ncbi.nlm.nih.gov/gene/?term=LOC105279687 | 17998 | 344 | 191.13 | 1 | 0.56 | 0.29 | 5.2 | 5.3 | -0.1 | 0.1 | 0 | 0 |
| gene53 | https://www.ncbi.nlm.nih.gov/gene/?term=LOC105277148 | 19283 | 268 | 138.98 | 1 | 0.52 | 0.37 | 3.71 | 3.95 | -0.23 | 0.23 | 0 | 0 |
| gene9326 | https://www.ncbi.nlm.nih.gov/gene/?term=LOC105282818 | 16612 | 169 | 101.73 | 1 | 0.6 | 0.59 | 13.49 | 13.76 | -0.26 | 0.26 | 0 | 0 |
| gene2748 | https://www.ncbi.nlm.nih.gov/gene/?term=LOC105275261 | 3845 | 53 | 137.84 | 1 | 2.6 | 1.89 | 38.36 | 38.68 | -0.33 | 0.33 | 0 | 0 |

Directory: OGL File: rnk-dmg-Cb_LI_RphaseA2.vs.RphaseB1.txt

| gene_ID | gene_link | gwidth | #Sites | #per10Kb | #dmSites | #dmsp10kb | %dmSites | %pSite1 | %pSite2 | DMpSite | ADMpSite | DMpNucl | ADMpNucl |
|---|---|---|---|---|---|---|---|---|---|---|---|---|---|
| gene2824 | https://www.ncbi.nlm.nih.gov/gene/?term=LOC105275337 | 1772 | 63 | 355.53 | 2 | 11.29 | 3.17 | 38.53 | 29.97 | 8.56 | 8.56 | 0.3 | 0.3 |
| gene2593 | https://www.ncbi.nlm.nih.gov/gene/?term=LOC105275070 | 1774 | 11 | 62.01 | 6 | 33.82 | 54.55 | 67.77 | 25.28 | 42.49 | 42.49 | 0.26 | 0.26 |
| gene3165 | https://www.ncbi.nlm.nih.gov/gene/?term=LOC105275706 | 2374 | 70 | 294.86 | 5 | 21.06 | 7.14 | 33.14 | 24.38 | 8.76 | 8.76 | 0.26 | 0.26 |
| gene12961 | https://www.ncbi.nlm.nih.gov/gene/?term=LOC105286983 | 3094 | 71 | 229.48 | 1 | 3.23 | 1.41 | 6.3 | 1.17 | 5.12 | 5.12 | 0.12 | 0.12 |
| gene5714 | https://www.ncbi.nlm.nih.gov/gene/?term=LOC105278729 | 710 | 16 | 225.35 | 1 | 14.08 | 6.25 | 56.86 | 51.9 | 4.96 | 4.96 | 0.11 | 0.11 |
| gene7568 | https://www.ncbi.nlm.nih.gov/gene/?term=LOC105280786 | 2525 | 143 | 566.34 | 1 | 3.96 | 0.7 | 27.63 | 25.63 | 2 | 2 | 0.11 | 0.11 |
| gene13033 | https://www.ncbi.nlm.nih.gov/gene/?term=LOC105287063 | 1877 | 55 | 293.02 | 1 | 5.33 | 1.82 | 30.5 | 27.04 | 3.46 | 3.46 | 0.1 | 0.1 |
| gene2413 | https://www.ncbi.nlm.nih.gov/gene/?term=LOC105274863 | 2006 | 65 | 324.03 | 1 | 4.99 | 1.54 | 49.74 | 52.89 | -3.14 | 3.14 | -0.1 | 0.1 |
| gene1007 | https://www.ncbi.nlm.nih.gov/gene/?term=LOC105283457 | 1878 | 49 | 260.92 | 1 | 5.32 | 2.04 | 35.2 | 31.9 | 3.3 | 3.3 | 0.09 | 0.09 |
| gene2271 | https://www.ncbi.nlm.nih.gov/gene/?term=LOC105274706 | 1448 | 57 | 393.65 | 1 | 6.91 | 1.75 | 39.53 | 37.37 | 2.16 | 2.16 | 0.09 | 0.09 |
| gene4034 | https://www.ncbi.nlm.nih.gov/gene/?term=LOC105276704 | 2065 | 92 | 445.52 | 5 | 24.21 | 5.43 | 14.65 | 12.65 | 2 | 2 | 0.09 | 0.09 |
| gene10984 | https://www.ncbi.nlm.nih.gov/gene/?term=LOC105284704 | 1987 | 66 | 332.16 | 1 | 5.03 | 1.52 | 25.73 | 23.31 | 2.42 | 2.42 | 0.08 | 0.08 |
| gene5525 | https://www.ncbi.nlm.nih.gov/gene/?term=LOC105278396 | 3090 | 189 | 611.65 | 1 | 3.24 | 0.53 | 25.87 | 24.52 | 1.35 | 1.35 | 0.08 | 0.08 |
| gene920 | https://www.ncbi.nlm.nih.gov/gene/?term=LOC105282518 | 1197 | 46 | 384.29 | 1 | 8.35 | 2.17 | 30.45 | 32.64 | -2.19 | 2.19 | -0.08 | 0.08 |
| gene1768 | https://www.ncbi.nlm.nih.gov/gene/?term=LOC105287938 | 2792 | 89 | 318.77 | 1 | 3.58 | 1.12 | 25.55 | 28.06 | -2.51 | 2.51 | -0.08 | 0.08 |
| gene3817 | https://www.ncbi.nlm.nih.gov/gene/?term=LOC105276449 | 1230 | 55 | 447.15 | 1 | 8.13 | 1.82 | 24.78 | 23.21 | 1.57 | 1.57 | 0.07 | 0.07 |
| gene10552 | https://www.ncbi.nlm.nih.gov/gene/?term=LOC105284213 | 878 | 23 | 261.96 | 1 | 11.39 | 4.35 | 9.72 | 12.3 | -2.58 | 2.58 | -0.07 | 0.07 |
| gene3774 | https://www.ncbi.nlm.nih.gov/gene/?term=LOC105276553 | 2716 | 84 | 309.28 | 1 | 3.68 | 1.19 | 43.69 | 41.82 | 1.87 | 1.87 | 0.06 | 0.06 |
| gene7493 | https://www.ncbi.nlm.nih.gov/gene/?term=LOC105280704 | 4225 | 155 | 366.86 | 1 | 2.37 | 0.65 | 17.59 | 16.02 | 1.58 | 1.58 | 0.06 | 0.06 |
| gene935 | https://www.ncbi.nlm.nih.gov/gene/?term=LOC105282682 | 4368 | 240 | 549.45 | 1 | 2.29 | 0.42 | 13.64 | 12.5 | 1.14 | 1.14 | 0.06 | 0.06 |
| gene11340 | https://www.ncbi.nlm.nih.gov/gene/?term=LOC105285131 | 2918 | 100 | 342.7 | 1 | 3.43 | 1 | 13.44 | 15.08 | -1.64 | 1.64 | -0.06 | 0.06 |
| gene274 | https://www.ncbi.nlm.nih.gov/gene/?term=LOC105275190 | 3488 | 74 | 212.16 | 1 | 2.87 | 1.35 | 22.52 | 20.11 | 2.4 | 2.4 | 0.05 | 0.05 |
| gene3671 | https://www.ncbi.nlm.nih.gov/gene/?term=LOC105276272 | 1966 | 47 | 239.06 | 1 | 5.09 | 2.13 | 25.43 | 23.44 | 1.98 | 1.98 | 0.05 | 0.05 |
| gene5831 | https://www.ncbi.nlm.nih.gov/gene/?term=LOC105278761 | 2506 | 119 | 474.86 | 1 | 3.99 | 0.84 | 22.74 | 21.63 | 1.11 | 1.11 | 0.05 | 0.05 |
| gene298 | https://www.ncbi.nlm.nih.gov/gene/?term=LOC105275460 | 2473 | 123 | 497.37 | 1 | 4.04 | 0.81 | 13.29 | 12.33 | 0.96 | 0.96 | 0.05 | 0.05 |
| gene7556 | https://www.ncbi.nlm.nih.gov/gene/?term=LOC105280774 | 2792 | 135 | 483.52 | 1 | 3.58 | 0.74 | 25.37 | 24.41 | 0.96 | 0.96 | 0.05 | 0.05 |
| gene1505 | https://www.ncbi.nlm.nih.gov/gene/?term=LOC105287633 | 1693 | 24 | 141.76 | 1 | 5.91 | 4.17 | 48.26 | 51.48 | -3.22 | 3.22 | -0.05 | 0.05 |
| gene3166 | https://www.ncbi.nlm.nih.gov/gene/?term=LOC105275707 | 2393 | 12 | 50.15 | 1 | 4.18 | 8.33 | 62.45 | 73.29 | -10.83 | 10.83 | -0.05 | 0.05 |
| gene393 | https://www.ncbi.nlm.nih.gov/gene/?term=LOC105276566 | 1123 | 2 | 17.81 | 1 | 8.9 | 50 | 47.62 | 74.3 | -26.68 | 26.68 | -0.05 | 0.05 |
| gene10107 | https://www.ncbi.nlm.nih.gov/gene/?term=LOC105283712 | 1883 | 34 | 180.56 | 1 | 5.31 | 2.94 | 31.9 | 29.48 | 2.42 | 2.42 | 0.04 | 0.04 |
| gene9336 | https://www.ncbi.nlm.nih.gov/gene/?term=LOC105282834 | 5993 | 137 | 228.6 | 1 | 1.67 | 0.73 | 15.64 | 13.69 | 1.95 | 1.95 | 0.04 | 0.04 |
| gene10879 | https://www.ncbi.nlm.nih.gov/gene/?term=LOC105284574 | 2423 | 52 | 214.61 | 1 | 4.13 | 1.92 | 56.31 | 54.67 | 1.63 | 1.63 | 0.04 | 0.04 |
| gene940 | https://www.ncbi.nlm.nih.gov/gene/?term=LOC105282794 | 3578 | 95 | 265.51 | 1 | 2.79 | 1.05 | 15.95 | 14.34 | 1.61 | 1.61 | 0.04 | 0.04 |
| gene2241 | https://www.ncbi.nlm.nih.gov/gene/?term=LOC105274666 | 2598 | 64 | 246.34 | 1 | 3.85 | 1.56 | 28.81 | 27.27 | 1.54 | 1.54 | 0.04 | 0.04 |
| gene1147 | https://www.ncbi.nlm.nih.gov/gene/?term=LOC105284954 | 4022 | 114 | 283.44 | 1 | 2.49 | 0.88 | 14.88 | 13.52 | 1.36 | 1.36 | 0.04 | 0.04 |
| gene5524 | https://www.ncbi.nlm.nih.gov/gene/?term=LOC105278397 | 5177 | 206 | 397.91 | 1 | 1.93 | 0.49 | 25.82 | 24.7 | 1.12 | 1.12 | 0.04 | 0.04 |
| gene12406 | https://www.ncbi.nlm.nih.gov/gene/?term=LOC105286316 | 1386 | 52 | 375.18 | 1 | 7.22 | 1.92 | 33.87 | 32.77 | 1.1 | 1.1 | 0.04 | 0.04 |
| gene3773 | https://www.ncbi.nlm.nih.gov/gene/?term=LOC105276408 | 5522 | 200 | 362.19 | 1 | 1.81 | 0.5 | 21.6 | 20.62 | 0.98 | 0.98 | 0.04 | 0.04 |
| gene11819 | https://www.ncbi.nlm.nih.gov/gene/?term=LOC105285671 | 2366 | 103 | 435.33 | 1 | 4.23 | 0.97 | 30.19 | 29.25 | 0.95 | 0.95 | 0.04 | 0.04 |
| gene10540 | https://www.ncbi.nlm.nih.gov/gene/?term=LOC105284200 | 5267 | 219 | 415.8 | 1 | 1.9 | 0.46 | 13.98 | 13.1 | 0.88 | 0.88 | 0.04 | 0.04 |
| gene6286 | https://www.ncbi.nlm.nih.gov/gene/?term=LOC105279298 | 2861 | 133 | 464.87 | 2 | 6.99 | 1.5 | 23.83 | 22.97 | 0.86 | 0.86 | 0.04 | 0.04 |
| gene10678 | https://www.ncbi.nlm.nih.gov/gene/?term=LOC105284355 | 4031 | 172 | 426.69 | 1 | 2.48 | 0.58 | 20.12 | 19.3 | 0.83 | 0.83 | 0.04 | 0.04 |
| gene10606 | https://www.ncbi.nlm.nih.gov/gene/?term=LOC105284273 | 6119 | 286 | 467.4 | 1 | 1.63 | 0.35 | 18.29 | 17.53 | 0.75 | 0.75 | 0.04 | 0.04 |
| gene10864 | https://www.ncbi.nlm.nih.gov/gene/?term=LOC105284557 | 1666 | 91 | 546.22 | 1 | 6 | 1.1 | 27.88 | 27.16 | 0.72 | 0.72 | 0.04 | 0.04 |
| gene12802 | https://www.ncbi.nlm.nih.gov/gene/?term=LOC105286799 | 3183 | 184 | 578.07 | 1 | 3.14 | 0.54 | 17.39 | 16.7 | 0.69 | 0.69 | 0.04 | 0.04 |
| gene5298 | https://www.ncbi.nlm.nih.gov/gene/?term=LOC105278145 | 4060 | 275 | 677.34 | 2 | 4.93 | 0.73 | 9.89 | 10.42 | -0.53 | 0.53 | -0.04 | 0.04 |
| gene7550 | https://www.ncbi.nlm.nih.gov/gene/?term=LOC105280763 | 2173 | 109 | 501.61 | 1 | 4.6 | 0.92 | 27.44 | 28.14 | -0.7 | 0.7 | -0.04 | 0.04 |
| gene11401 | https://www.ncbi.nlm.nih.gov/gene/?term=LOC105285202 | 1841 | 89 | 483.43 | 1 | 5.43 | 1.12 | 23.71 | 24.59 | -0.89 | 0.89 | -0.04 | 0.04 |
| gene9101 | https://www.ncbi.nlm.nih.gov/gene/?term=LOC105282553 | 3068 | 110 | 358.54 | 1 | 3.26 | 0.91 | 21.48 | 22.61 | -1.13 | 1.13 | -0.04 | 0.04 |
| gene1199 | https://www.ncbi.nlm.nih.gov/gene/?term=LOC105285498 | 2220 | 29 | 130.63 | 1 | 4.5 | 3.45 | 32.78 | 35.99 | -3.21 | 3.21 | -0.04 | 0.04 |
| gene12395 | https://www.ncbi.nlm.nih.gov/gene/?term=LOC105286302 | 4093 | 24 | 58.64 | 1 | 2.44 | 4.17 | 20.4 | 15.15 | 5.24 | 5.24 | 0.03 | 0.03 |
| gene2970 | https://www.ncbi.nlm.nih.gov/gene/?term=LOC105275518 | 2340 | 28 | 119.66 | 1 | 4.27 | 3.57 | 41.75 | 39.36 | 2.39 | 2.39 | 0.03 | 0.03 |
| gene388 | https://www.ncbi.nlm.nih.gov/gene/?term=LOC105276538 | 1558 | 24 | 154.04 | 2 | 12.84 | 8.33 | 66.6 | 64.8 | 1.8 | 1.8 | 0.03 | 0.03 |
| gene8742 | https://www.ncbi.nlm.nih.gov/gene/?term=LOC105282151 | 2563 | 44 | 171.67 | 1 | 3.9 | 2.27 | 43.82 | 42.21 | 1.61 | 1.61 | 0.03 | 0.03 |
| gene2825 | https://www.ncbi.nlm.nih.gov/gene/?term=LOC105275343 | 5545 | 98 | 176.74 | 1 | 1.8 | 1.02 | 9.51 | 8.1 | 1.42 | 1.42 | 0.03 | 0.03 |
| gene12572 | https://www.ncbi.nlm.nih.gov/gene/?term=LOC105286514 | 2374 | 56 | 235.89 | 1 | 4.21 | 1.79 | 36.49 | 35.22 | 1.27 | 1.27 | 0.03 | 0.03 |
| gene5877 | https://www.ncbi.nlm.nih.gov/gene/?term=LOC105278807 | 1719 | 53 | 308.32 | 1 | 5.82 | 1.89 | 27.62 | 26.53 | 1.08 | 1.08 | 0.03 | 0.03 |
| gene4892 | https://www.ncbi.nlm.nih.gov/gene/?term=LOC105277667 | 3898 | 127 | 325.81 | 1 | 2.57 | 0.79 | 28.13 | 27.36 | 0.77 | 0.77 | 0.03 | 0.03 |
| gene289 | https://www.ncbi.nlm.nih.gov/gene/?term=LOC105275347 | 2533 | 90 | 355.31 | 1 | 3.95 | 1.11 | 24.42 | 23.69 | 0.73 | 0.73 | 0.03 | 0.03 |
| gene13036 | https://www.ncbi.nlm.nih.gov/gene/?term=LOC105287059 | 4959 | 200 | 403.31 | 1 | 2.02 | 0.5 | 17.56 | 16.85 | 0.71 | 0.71 | 0.03 | 0.03 |

| | | | | | | | | | | | | | |
|---|---|---|---|---|---|---|---|---|---|---|---|---|---|
| gene1573 | https://www.ncbi.nlm.nih.gov/gene/?term=LOC105287704 | 3030 | 143 | 471.95 | 1 | 3.3 | 0.7 | 19.92 | 19.38 | 0.54 | 0.54 | 0.03 | 0.03 |
| gene1470 | https://www.ncbi.nlm.nih.gov/gene/?term=LOC105287597 | 2974 | 200 | 672.49 | 1 | 3.36 | 0.5 | 18.11 | 18.56 | -0.46 | 0.46 | -0.03 | 0.03 |
| gene12377 | https://www.ncbi.nlm.nih.gov/gene/?term=LOC105286279 | 5846 | 278 | 475.54 | 1 | 1.71 | 0.36 | 2.15 | 2.82 | -0.67 | 0.67 | -0.03 | 0.03 |
| gene10213 | https://www.ncbi.nlm.nih.gov/gene/?term=LOC105283826 | 1503 | 62 | 412.51 | 1 | 6.65 | 1.61 | 14.74 | 15.54 | -0.8 | 0.8 | -0.03 | 0.03 |
| gene10392 | https://www.ncbi.nlm.nih.gov/gene/?term=LOC105284036 | 3216 | 103 | 320.27 | 2 | 6.22 | 1.94 | 24.04 | 24.86 | -0.81 | 0.81 | -0.03 | 0.03 |
| gene2404 | https://www.ncbi.nlm.nih.gov/gene/?term=LOC105274853 | 1767 | 45 | 254.67 | 1 | 5.66 | 2.22 | 30.61 | 31.94 | -1.33 | 1.33 | -0.03 | 0.03 |
| gene10654 | https://www.ncbi.nlm.nih.gov/gene/?term=LOC105284329 | 4027 | 72 | 178.79 | 1 | 2.48 | 1.39 | 18.86 | 20.42 | -1.56 | 1.56 | -0.03 | 0.03 |
| gene6237 | https://www.ncbi.nlm.nih.gov/gene/?term=LOC105279244 | 1484 | 30 | 202.16 | 1 | 6.74 | 3.33 | 49.36 | 51.05 | -1.69 | 1.69 | -0.03 | 0.03 |
| gene7398 | https://www.ncbi.nlm.nih.gov/gene/?term=LOC105280589 | 3452 | 42 | 121.67 | 1 | 2.9 | 2.38 | 23.2 | 25.91 | -2.71 | 2.71 | -0.03 | 0.03 |
| gene8691 | https://www.ncbi.nlm.nih.gov/gene/?term=LOC105282094 | 2028 | 6 | 29.59 1 | 4.93 | 16.67 | 86.17 | 97.58 | -11.41 | | 11.41 | -0.03 | 0.03 |
| gene8178 | https://www.ncbi.nlm.nih.gov/gene/?term=LOC105281515 | 8647 | 59 | 68.23 1 | 1.16 | 1.69 | 30.95 | 28.43 | 2.52 | 2.52 | 0.02 | 0.02 | |
| gene4996 | https://www.ncbi.nlm.nih.gov/gene/?term=LOC105277798 | 5617 | 74 | 131.74 | 1 | 1.78 | 1.35 | 11.47 | 9.83 | 1.64 | 1.64 | 0.02 | 0.02 |
| gene2748 | https://www.ncbi.nlm.nih.gov/gene/?term=LOC105275261 | 3845 | 55 | 143.04 | 1 | 2.6 | 1.82 | 41.25 | 39.71 | 1.54 | 1.54 | 0.02 | 0.02 |
| gene5273 | https://www.ncbi.nlm.nih.gov/gene/?term=LOC105278108 | 4067 | 57 | 140.15 | 1 | 2.46 | 1.75 | 25.93 | 24.39 | 1.54 | 1.54 | 0.02 | 0.02 |
| gene322 | https://www.ncbi.nlm.nih.gov/gene/?term=LOC105275777 | 1606 | 29 | 180.57 | 1 | 6.23 | 3.45 | 36.14 | 34.8 | 1.35 | 1.35 | 0.02 | 0.02 |
| gene927 | https://www.ncbi.nlm.nih.gov/gene/?term=LOC105282600 | 3727 | 67 | 179.77 | 1 | 2.68 | 1.49 | 27.6 | 26.32 | 1.28 | 1.28 | 0.02 | 0.02 |
| gene6455 | https://www.ncbi.nlm.nih.gov/gene/?term=LOC105279500 | 2612 | 36 | 137.83 | 1 | 3.83 | 2.78 | 48.83 | 47.56 | 1.26 | 1.26 | 0.02 | 0.02 |
| gene4048 | https://www.ncbi.nlm.nih.gov/gene/?term=LOC105276724 | 2955 | 47 | 159.05 | 1 | 3.38 | 2.13 | 12.29 | 11.14 | 1.14 | 1.14 | 0.02 | 0.02 |
| gene10222 | https://www.ncbi.nlm.nih.gov/gene/?term=LOC105283838 | 1748 | 33 | 188.79 | 1 | 5.72 | 3.03 | 33.58 | 32.49 | 1.1 | 1.1 | 0.02 | 0.02 |
| gene12021 | https://www.ncbi.nlm.nih.gov/gene/?term=LOC105285905 | 1399 | 27 | 192.99 | 1 | 7.15 | 3.7 | 37.97 | 36.9 | 1.06 | 1.06 | 0.02 | 0.02 |
| gene7183 | https://www.ncbi.nlm.nih.gov/gene/?term=LOC105280348 | 3925 | 79 | 201.27 | 1 | 2.55 | 1.27 | 18.1 | 17.08 | 1.01 | 1.01 | 0.02 | 0.02 |
| gene1884 | https://www.ncbi.nlm.nih.gov/gene/?term=LOC105288072 | 4169 | 88 | 211.08 | 1 | 2.4 | 1.14 | 22.65 | 21.71 | 0.94 | 0.94 | 0.02 | 0.02 |
| gene12516 | https://www.ncbi.nlm.nih.gov/gene/?term=LOC105286452 | 1529 | 42 | 274.69 | 1 | 6.54 | 2.38 | 47.45 | 46.6 | 0.86 | 0.86 | 0.02 | 0.02 |
| gene10600 | https://www.ncbi.nlm.nih.gov/gene/?term=LOC105284270 | 4837 | 114 | 235.68 | 1 | 2.07 | 0.88 | 12.4 | 11.65 | 0.75 | 0.75 | 0.02 | 0.02 |
| gene8015 | https://www.ncbi.nlm.nih.gov/gene/?term=LOC105281312 | 2998 | 69 | 230.15 | 1 | 3.34 | 1.45 | 31.17 | 30.43 | 0.74 | 0.74 | 0.02 | 0.02 |
| gene9330 | https://www.ncbi.nlm.nih.gov/gene/?term=LOC105282825 | 3936 | 102 | 259.15 | 1 | 2.54 | 0.98 | 14.91 | 14.23 | 0.68 | 0.68 | 0.02 | 0.02 |
| gene6031 | https://www.ncbi.nlm.nih.gov/gene/?term=LOC105279000 | 5691 | 226 | 397.12 | 1 | 1.76 | 0.44 | 12.2 | 11.72 | 0.48 | 0.48 | 0.02 | 0.02 |
| gene3206 | https://www.ncbi.nlm.nih.gov/gene/?term=LOC105275752 | 3474 | 138 | 397.24 | 1 | 2.88 | 0.72 | 31.47 | 31.01 | 0.46 | 0.46 | 0.02 | 0.02 |
| gene7888 | https://www.ncbi.nlm.nih.gov/gene/?term=LOC105281141 | 4357 | 180 | 413.13 | 1 | 2.3 | 0.56 | 27.03 | 26.57 | 0.45 | 0.45 | 0.02 | 0.02 |
| gene4641 | https://www.ncbi.nlm.nih.gov/gene/?term=LOC105277398 | 1937 | 74 | 382.03 | 1 | 5.16 | 1.35 | 35.22 | 34.78 | 0.44 | 0.44 | 0.02 | 0.02 |
| gene7249 | https://www.ncbi.nlm.nih.gov/gene/?term=LOC105280422 | 1775 | 66 | 371.83 | 1 | 5.63 | 1.52 | 16.48 | 16.04 | 0.44 | 0.44 | 0.02 | 0.02 |
| gene10520 | https://www.ncbi.nlm.nih.gov/gene/?term=LOC105284184 | 2451 | 105 | 428.4 1 | 4.08 | 0.95 | 28.89 | 28.49 | 0.4 | 0.4 | 0.02 | 0.02 | |
| gene2846 | https://www.ncbi.nlm.nih.gov/gene/?term=LOC105275374 | 11246 | 566 | 503.29 | 1 | 0.89 | 0.18 | 9.56 | 9.16 | 0.4 | 0.4 | 0.02 | 0.02 |
| gene570 | https://www.ncbi.nlm.nih.gov/gene/?term=LOC105278549 | 1673 | 74 | 442.32 | 1 | 5.98 | 1.35 | 44.93 | 44.55 | 0.38 | 0.38 | 0.02 | 0.02 |
| gene3885 | https://www.ncbi.nlm.nih.gov/gene/?term=LOC105276527 | 7928 | 380 | 479.31 | 1 | 1.26 | 0.26 | 8.1 | 7.75 | 0.34 | 0.34 | 0.02 | 0.02 |
| gene2844 | https://www.ncbi.nlm.nih.gov/gene/?term=LOC105275363 | 4330 | 258 | 595.84 | 1 | 2.31 | 0.39 | 11.65 | 11.32 | 0.33 | 0.33 | 0.02 | 0.02 |
| gene6583 | https://www.ncbi.nlm.nih.gov/gene/?term=LOC105279649 | 7444 | 393 | 527.94 | 1 | 1.34 | 0.25 | 10.67 | 11.14 | -0.47 | 0.47 | -0.02 | 0.02 |
| gene3848 | https://www.ncbi.nlm.nih.gov/gene/?term=LOC105276486 | 5899 | 176 | 298.36 | 1 | 1.7 | 0.57 | 23.6 | 24.17 | -0.57 | 0.57 | -0.02 | 0.02 |
| gene4962 | https://www.ncbi.nlm.nih.gov/gene/?term=LOC105277761 | 4645 | 175 | 376.75 | 1 | 2.15 | 0.57 | 13.58 | 14.2 | -0.62 | 0.62 | -0.02 | 0.02 |
| gene7207 | https://www.ncbi.nlm.nih.gov/gene/?term=LOC105280376 | 4659 | 124 | 266.15 | 1 | 2.15 | 0.81 | 26.54 | 27.17 | -0.63 | 0.63 | -0.02 | 0.02 |
| gene3385 | https://www.ncbi.nlm.nih.gov/gene/?term=LOC105275966 | 3305 | 75 | 226.93 | 1 | 3.03 | 1.33 | 43.86 | 44.6 | -0.75 | 0.75 | -0.02 | 0.02 |
| gene11454 | https://www.ncbi.nlm.nih.gov/gene/?term=LOC105285256 | 1906 | 50 | 262.33 | 1 | 5.25 | 2 | 43.28 | 44.08 | -0.8 | 0.8 | -0.02 | 0.02 |
| gene7915 | https://www.ncbi.nlm.nih.gov/gene/?term=LOC105281173 | 1597 | 39 | 244.21 | 1 | 6.26 | 2.56 | 61.13 | 62.01 | -0.88 | 0.88 | -0.02 | 0.02 |
| gene11447 | https://www.ncbi.nlm.nih.gov/gene/?term=LOC105285252 | 7437 | 150 | 201.69 | 1 | 1.34 | 0.67 | 11.78 | 12.78 | -0.99 | 0.99 | -0.02 | 0.02 |
| gene3168 | https://www.ncbi.nlm.nih.gov/gene/?term=LOC105275711 | 6150 | 95 | 154.47 | 1 | 1.63 | 1.05 | 10.97 | 11.98 | -1 | 1 | -0.02 | 0.02 |
| gene7657 | https://www.ncbi.nlm.nih.gov/gene/?term=LOC105280890 | 1109 | 23 | 207.39 | 1 | 9.02 | 4.35 | 23.17 | 24.26 | -1.09 | 1.09 | -0.02 | 0.02 |
| gene10155 | https://www.ncbi.nlm.nih.gov/gene/?term=LOC105283756 | 7030 | 119 | 169.27 | 1 | 1.42 | 0.84 | 6.89 | 8.19 | -1.3 | 1.3 | -0.02 | 0.02 |
| gene9411 | https://www.ncbi.nlm.nih.gov/gene/?term=LOC105282914 | 5534 | 42 | 75.89 1 | 1.81 | 2.38 | 48.54 | 50.92 | -2.38 | 2.38 | -0.02 | 0.02 | |
| gene9109 | https://www.ncbi.nlm.nih.gov/gene/?term=LOC105282561 | 2400 | 17 | 70.83 1 | 4.17 | 5.88 | 28.52 | 31.86 | -3.34 | 3.34 | -0.02 | 0.02 | |
| gene11584 | https://www.ncbi.nlm.nih.gov/gene/?term=LOC105285416 | 3709 | 15 | 40.44 2 | 5.39 | 13.33 | 24.1 | 29.33 | -5.23 | 5.23 | -0.02 | 0.02 | |
| gene12106 | https://www.ncbi.nlm.nih.gov/gene/?term=LOC105286002 | 8556 | 83 | 97.01 1 | 1.17 | 1.2 | 3.39 | 2.1 | 1.29 | 1.29 | 0.01 | 0.01 | |
| gene5052 | https://www.ncbi.nlm.nih.gov/gene/?term=LOC105277856 | 3167 | 36 | 113.67 | 1 | 3.16 | 2.78 | 37.89 | 36.75 | 1.14 | 1.14 | 0.01 | 0.01 |
| gene10243 | https://www.ncbi.nlm.nih.gov/gene/?term=LOC105283862 | 3239 | 45 | 138.93 | 1 | 3.09 | 2.22 | 29.81 | 28.76 | 1.05 | 1.05 | 0.01 | 0.01 |
| gene2833 | https://www.ncbi.nlm.nih.gov/gene/?term=LOC105275352 | 5528 | 43 | 77.79 1 | 1.81 | 2.33 | 22.51 | 21.53 | 0.98 | 0.98 | 0.01 | 0.01 | |
| gene3912 | https://www.ncbi.nlm.nih.gov/gene/?term=LOC105276560 | 17900 | 373 | 208.38 | 3 | 1.68 | 0.8 | 2.56 | 1.89 | 0.67 | 0.67 | 0.01 | 0.01 |
| gene814 | https://www.ncbi.nlm.nih.gov/gene/?term=LOC105281296 | 2722 | 57 | 209.4 1 | 3.67 | 1.75 | 23.86 | 23.2 | 0.65 | 0.65 | 0.01 | 0.01 | |
| gene8343 | https://www.ncbi.nlm.nih.gov/gene/?term=LOC105281700 | 2172 | 43 | 197.97 | 1 | 4.6 | 2.33 | 36.96 | 36.39 | 0.57 | 0.57 | 0.01 | 0.01 |
| gene2257 | https://www.ncbi.nlm.nih.gov/gene/?term=LOC105274689 | 3658 | 90 | 246.04 | 1 | 2.73 | 1.11 | 15.54 | 15.05 | 0.49 | 0.49 | 0.01 | 0.01 |
| gene9648 | https://www.ncbi.nlm.nih.gov/gene/?term=LOC105283183 | 2602 | 33 | 126.83 | 1 | 3.84 | 3.03 | 45.18 | 44.75 | 0.44 | 0.44 | 0.01 | 0.01 |
| gene2330 | https://www.ncbi.nlm.nih.gov/gene/?term=LOC105274773 | 13104 | 208 | 158.73 | 1 | 0.76 | 0.48 | 15.28 | 14.85 | 0.43 | 0.43 | 0.01 | 0.01 |
| gene8814 | https://www.ncbi.nlm.nih.gov/gene/?term=LOC105282240 | 6910 | 209 | 302.46 | 1 | 1.45 | 0.48 | 8.7 | 8.29 | 0.4 | 0.4 | 0.01 | 0.01 |
| gene9029 | https://www.ncbi.nlm.nih.gov/gene/?term=LOC105282491 | 4762 | 190 | 398.99 | 1 | 2.1 | 0.53 | 8.59 | 8.25 | 0.34 | 0.34 | 0.01 | 0.01 |
| gene2056 | https://www.ncbi.nlm.nih.gov/gene/?term=LOC105274452 | 6584 | 268 | 407.05 | 1 | 1.52 | 0.37 | 13.37 | 13.03 | 0.33 | 0.33 | 0.01 | 0.01 |
| gene11568 | https://www.ncbi.nlm.nih.gov/gene/?term=LOC105285395 | 8086 | 257 | 317.83 | 1 | 1.24 | 0.39 | 13.19 | 12.93 | 0.27 | 0.27 | 0.01 | 0.01 |

| | | | | | | | | | | | | | |
|---|---|---|---|---|---|---|---|---|---|---|---|---|---|
| gene10018 | https://www.ncbi.nlm.nih.gov/gene/?term=LOC105283607 | 4414 | 222 | 502.95 | 1 | 2.27 | 0.45 | 14.44 | 14.18 | 0.26 | 0.26 | 0.01 | 0.01 |
| gene6620 | https://www.ncbi.nlm.nih.gov/gene/?term=LOC105279687 | 17998 | 456 | 253.36 | 1 | 0.56 | 0.22 | 4.79 | 4.53 | 0.26 | 0.26 | 0.01 | 0.01 |
| gene8600 | https://www.ncbi.nlm.nih.gov/gene/?term=LOC105282001 | 8513 | 211 | 247.86 | 1 | 1.17 | 0.47 | 9.85 | 9.6 | 0.25 | 0.25 | 0.01 | 0.01 |
| gene1310 | https://www.ncbi.nlm.nih.gov/gene/?term=LOC105286661 | 2582 | 72 | 278.85 | 1 | 3.87 | 1.39 | 20.69 | 20.45 | 0.24 | 0.24 | 0.01 | 0.01 |
| gene292 | https://www.ncbi.nlm.nih.gov/gene/?term=LOC105275398 | 2825 | 92 | 325.66 | 1 | 3.54 | 1.09 | 21.85 | 21.62 | 0.23 | 0.23 | 0.01 | 0.01 |
| gene11275 | https://www.ncbi.nlm.nih.gov/gene/?term=LOC105285055 | 4119 | 262 | 636.08 | 1 | 2.43 | 0.38 | 11.65 | 11.43 | 0.22 | 0.22 | 0.01 | 0.01 |
| gene8729 | https://www.ncbi.nlm.nih.gov/gene/?term=LOC105282137 | 5005 | 236 | 471.53 | 1 | 2 | 0.42 | 14.47 | 14.26 | 0.21 | 0.21 | 0.01 | 0.01 |
| gene2806 | https://www.ncbi.nlm.nih.gov/gene/?term=LOC105275327 | 14257 | 550 | 385.78 | 1 | 0.7 | 0.18 | 6.58 | 6.38 | 0.2 | 0.2 | 0.01 | 0.01 |
| gene3747 | https://www.ncbi.nlm.nih.gov/gene/?term=LOC105276387 | 3736 | 198 | 529.98 | 1 | 2.68 | 0.51 | 14.52 | 14.33 | 0.19 | 0.19 | 0.01 | 0.01 |
| gene5746 | https://www.ncbi.nlm.nih.gov/gene/?term=LOC105278666 | 10343 | 491 | 474.72 | 1 | 0.97 | 0.2 | 6.89 | 6.71 | 0.19 | 0.19 | 0.01 | 0.01 |
| gene10596 | https://www.ncbi.nlm.nih.gov/gene/?term=LOC105284262 | 6520 | 247 | 378.83 | 1 | 1.53 | 0.4 | 4.51 | 4.34 | 0.18 | 0.18 | 0.01 | 0.01 |
| gene13114 | https://www.ncbi.nlm.nih.gov/gene/?term=LOC105287147 | 7609 | 447 | 587.46 | 1 | 1.31 | 0.22 | 9.07 | 9.17 | -0.1 | 0.1 | -0.01 | 0.01 |
| gene8274 | https://www.ncbi.nlm.nih.gov/gene/?term=LOC105281624 | 5425 | 244 | 449.77 | 1 | 1.84 | 0.41 | 9.34 | 9.46 | -0.12 | 0.12 | -0.01 | 0.01 |
| gene12674 | https://www.ncbi.nlm.nih.gov/gene/?term=LOC105286649 | 18283 | 836 | 457.26 | 1 | 0.55 | 0.12 | 1.34 | 1.5 | -0.16 | 0.16 | -0.01 | 0.01 |
| gene11082 | https://www.ncbi.nlm.nih.gov/gene/?term=LOC105284824 | 7436 | 245 | 329.48 | 1 | 1.34 | 0.41 | 7.32 | 7.57 | -0.25 | 0.25 | -0.01 | 0.01 |
| gene1198 | https://www.ncbi.nlm.nih.gov/gene/?term=LOC105285507 | 2738 | 98 | 357.93 | 1 | 3.65 | 1.02 | 21.45 | 21.7 | -0.25 | 0.25 | -0.01 | 0.01 |
| gene12407 | https://www.ncbi.nlm.nih.gov/gene/?term=LOC105286315 | 8593 | 227 | 264.17 | 1 | 1.16 | 0.44 | 13.44 | 13.72 | -0.28 | 0.28 | -0.01 | 0.01 |
| gene11583 | https://www.ncbi.nlm.nih.gov/gene/?term=LOC105285414 | 8192 | 204 | 249.02 | 1 | 1.22 | 0.49 | 7.78 | 8.09 | -0.31 | 0.31 | -0.01 | 0.01 |
| gene12573 | https://www.ncbi.nlm.nih.gov/gene/?term=LOC105286515 | 3647 | 128 | 350.97 | 1 | 2.74 | 0.78 | 10.64 | 10.97 | -0.33 | 0.33 | -0.01 | 0.01 |
| gene9598 | https://www.ncbi.nlm.nih.gov/gene/?term=LOC105283132 | 1769 | 56 | 316.56 | 1 | 5.65 | 1.79 | 39.81 | 40.18 | -0.38 | 0.38 | -0.01 | 0.01 |
| gene1220 | https://www.ncbi.nlm.nih.gov/gene/?term=LOC105285742 | 3453 | 123 | 356.21 | 1 | 2.9 | 0.81 | 16.77 | 17.17 | -0.4 | 0.4 | -0.01 | 0.01 |
| gene657 | https://www.ncbi.nlm.nih.gov/gene/?term=LOC105279667 | 6421 | 106 | 165.08 | 1 | 1.56 | 0.94 | 2.17 | 2.57 | -0.4 | 0.4 | -0.01 | 0.01 |
| gene5646 | https://www.ncbi.nlm.nih.gov/gene/?term=LOC105278534 | 7475 | 214 | 286.29 | 1 | 1.34 | 0.47 | 16.15 | 16.57 | -0.41 | 0.41 | -0.01 | 0.01 |
| gene7148 | https://www.ncbi.nlm.nih.gov/gene/?term=LOC105280304 | 8920 | 143 | 160.31 | 1 | 1.12 | 0.7 | 6.95 | 7.37 | -0.42 | 0.42 | -0.01 | 0.01 |
| gene7600 | https://www.ncbi.nlm.nih.gov/gene/?term=LOC105280819 | 1615 | 43 | 266.25 | 1 | 6.19 | 2.33 | 37.46 | 37.88 | -0.42 | 0.42 | -0.01 | 0.01 |
| gene10819 | https://www.ncbi.nlm.nih.gov/gene/?term=LOC105284507 | 5767 | 122 | 211.55 | 1 | 1.73 | 0.82 | 18.54 | 19.25 | -0.71 | 0.71 | -0.01 | 0.01 |
| gene12203 | https://www.ncbi.nlm.nih.gov/gene/?term=LOC105286099 | 5967 | 87 | 145.8 1 | 1.68 | 1.15 | 11.05 | 12.06 | -1.01 | 1.01 | -0.01 | 0.01 | |
| gene652 | https://www.ncbi.nlm.nih.gov/gene/?term=LOC105279528 | 5806 | 39 | 67.17 1 | 1.72 | 2.56 | 19.31 | 20.42 | -1.11 | 1.11 | -0.01 | 0.01 | |
| gene6639 | https://www.ncbi.nlm.nih.gov/gene/?term=LOC105279705 | 7656 | 88 | 114.94 | 1 | 1.31 | 1.14 | 19.96 | 21.09 | -1.13 | 1.13 | -0.01 | 0.01 |
| gene564 | https://www.ncbi.nlm.nih.gov/gene/?term=LOC105278478 | 2802 | 22 | 78.52 1 | 3.57 | 4.55 | 23.15 | 24.78 | -1.63 | 1.63 | -0.01 | 0.01 | |
| gene9647 | https://www.ncbi.nlm.nih.gov/gene/?term=LOC109611233 | 3094 | 35 | 113.12 | 1 | 3.23 | 2.86 | 42.6 | 42.19 | 0.41 | 0.41 | 0 | 0 |
| gene2683 | https://www.ncbi.nlm.nih.gov/gene/?term=LOC105275187 | 5492 | 87 | 158.41 | 1 | 1.82 | 1.15 | 15.13 | 14.88 | 0.25 | 0.25 | 0 | 0 |
| gene7245 | https://www.ncbi.nlm.nih.gov/gene/?term=LOC105280410 | 12478 | 319 | 255.65 | 1 | 0.8 | 0.31 | 17.86 | 17.71 | 0.15 | 0.15 | 0 | 0 |
| gene2173 | https://www.ncbi.nlm.nih.gov/gene/?term=LOC105274590 | 4387 | 91 | 207.43 | 2 | 4.56 | 2.2 | 24.1 | 23.96 | 0.14 | 0.14 | 0 | 0 |
| gene4225 | https://www.ncbi.nlm.nih.gov/gene/?term=LOC105276917 | 2443 | 40 | 163.73 | 1 | 4.09 | 2.5 | 29.99 | 29.86 | 0.14 | 0.14 | 0 | 0 |
| gene6966 | https://www.ncbi.nlm.nih.gov/gene/?term=LOC105280104 | 3747 | 133 | 354.95 | 1 | 2.67 | 0.75 | 17.09 | 16.96 | 0.13 | 0.13 | 0 | 0 |
| gene6652 | https://www.ncbi.nlm.nih.gov/gene/?term=LOC105279719 | 2831 | 83 | 293.18 | 1 | 3.53 | 1.2 | 23.57 | 23.44 | 0.12 | 0.12 | 0 | 0 |
| gene9722 | https://www.ncbi.nlm.nih.gov/gene/?term=LOC105283264 | 3951 | 94 | 237.91 | 1 | 2.53 | 1.06 | 22.96 | 22.87 | 0.1 | 0.1 | 0 | 0 |
| gene4033 | https://www.ncbi.nlm.nih.gov/gene/?term=LOC105276705 | 10546 | 157 | 148.87 | 1 | 0.95 | 0.64 | 6.45 | 6.38 | 0.06 | 0.06 | 0 | 0 |
| gene3727 | https://www.ncbi.nlm.nih.gov/gene/?term=LOC105276367 | 2484 | 76 | 305.96 | 1 | 4.03 | 1.32 | 41.02 | 40.98 | 0.04 | 0.04 | 0 | 0 |
| gene12307 | https://www.ncbi.nlm.nih.gov/gene/?term=LOC105286216 | 10313 | 209 | 202.66 | 1 | 0.97 | 0.48 | 14.07 | 14.04 | 0.02 | 0.02 | 0 | 0 |
| gene3470 | https://www.ncbi.nlm.nih.gov/gene/?term=LOC105276059 | 8716 | 186 | 213.4 1 | 1.15 | 0.54 | 9.64 | 9.62 | 0.02 | 0.02 | 0 | 0 | |
| gene3107 | https://www.ncbi.nlm.nih.gov/gene/?term=LOC105275650 | 5103 | 187 | 366.45 | 1 | 1.96 | 0.53 | 16.49 | 16.48 | 0.01 | 0.01 | 0 | 0 |
| gene3335 | https://www.ncbi.nlm.nih.gov/gene/?term=LOC105275898 | 4665 | 172 | 368.7 1 | 2.14 | 0.58 | 12.53 | 12.51 | 0.01 | 0.01 | 0 | 0 | |
| gene9326 | https://www.ncbi.nlm.nih.gov/gene/?term=LOC105282818 | 16612 | 251 | 151.1 1 | 0.6 | 0.4 | 12.99 | 12.99 | 0 | 0 | 0 | 0 | |
| gene11561 | https://www.ncbi.nlm.nih.gov/gene/?term=LOC105285387 | 11528 | 226 | 196.04 | 1 | 0.87 | 0.44 | 9.46 | 9.47 | -0.01 | 0.01 | 0 | 0 |
| gene13039 | https://www.ncbi.nlm.nih.gov/gene/?term=LOC105287065 | 4902 | 263 | 536.52 | 1 | 2.04 | 0.38 | 11.98 | 12.01 | -0.02 | 0.02 | 0 | 0 |
| gene7166 | https://www.ncbi.nlm.nih.gov/gene/?term=LOC105280326 | 6117 | 258 | 421.78 | 1 | 1.63 | 0.39 | 14.79 | 14.83 | -0.04 | 0.04 | 0 | 0 |
| gene6649 | https://www.ncbi.nlm.nih.gov/gene/?term=LOC105279717 | 6017 | 181 | 300.81 | 1 | 1.66 | 0.55 | 11.29 | 11.35 | -0.06 | 0.06 | 0 | 0 |
| gene3856 | https://www.ncbi.nlm.nih.gov/gene/?term=LOC105276499 | 5236 | 206 | 393.43 | 1 | 1.91 | 0.49 | 10.22 | 10.29 | -0.07 | 0.07 | 0 | 0 |
| gene10326 | https://www.ncbi.nlm.nih.gov/gene/?term=LOC105283962 | 3042 | 145 | 476.66 | 1 | 3.29 | 0.69 | 9.18 | 9.26 | -0.08 | 0.08 | 0 | 0 |
| gene10549 | https://www.ncbi.nlm.nih.gov/gene/?term=LOC105284210 | 7145 | 374 | 523.44 | 2 | 2.8 | 0.53 | 9.66 | 9.75 | -0.09 | 0.09 | 0 | 0 |
| gene2835 | https://www.ncbi.nlm.nih.gov/gene/?term=LOC105275357 | 2492 | 122 | 489.57 | 1 | 4.01 | 0.82 | 14.14 | 14.23 | -0.09 | 0.09 | 0 | 0 |
| gene4190 | https://www.ncbi.nlm.nih.gov/gene/?term=LOC105276875 | 3150 | 135 | 428.57 | 1 | 3.17 | 0.74 | 20.47 | 20.57 | -0.09 | 0.09 | 0 | 0 |
| gene5613 | https://www.ncbi.nlm.nih.gov/gene/?term=LOC105278504 | 8524 | 274 | 321.45 | 1 | 1.17 | 0.36 | 5.89 | 5.98 | -0.09 | 0.09 | 0 | 0 |
| gene9755 | https://www.ncbi.nlm.nih.gov/gene/?term=LOC105283296 | 2177 | 97 | 445.57 | 1 | 4.59 | 1.03 | 14.1 | 14.18 | -0.09 | 0.09 | 0 | 0 |
| gene4376 | https://www.ncbi.nlm.nih.gov/gene/?term=LOC105277083 | 4307 | 140 | 325.05 | 1 | 2.32 | 0.71 | 17.67 | 17.76 | -0.1 | 0.1 | 0 | 0 |
| gene53 | https://www.ncbi.nlm.nih.gov/gene/?term=LOC105277148 | 19283 | 339 | 175.8 1 | 0.52 | 0.29 | 4.53 | 4.65 | -0.11 | 0.11 | 0 | 0 | |
| gene1632 | https://www.ncbi.nlm.nih.gov/gene/?term=LOC105287764 | 18061 | 462 | 255.8 1 | 0.55 | 0.22 | 3.51 | 3.63 | -0.12 | 0.12 | 0 | 0 | |
| gene3599 | https://www.ncbi.nlm.nih.gov/gene/?term=LOC105276196 | 2955 | 61 | 206.43 | 1 | 3.38 | 1.64 | 38.99 | 39.11 | -0.13 | 0.13 | 0 | 0 |
| gene9327 | https://www.ncbi.nlm.nih.gov/gene/?term=LOC105282821 | 7793 | 184 | 236.11 | 1 | 1.28 | 0.54 | 12.73 | 12.87 | -0.14 | 0.14 | 0 | 0 |
| gene11579 | https://www.ncbi.nlm.nih.gov/gene/?term=LOC105285406 | 13571 | 341 | 251.27 | 1 | 0.74 | 0.29 | 6.04 | 6.2 | -0.15 | 0.15 | 0 | 0 |
| gene7368 | https://www.ncbi.nlm.nih.gov/gene/?term=LOC105280556 | 4105 | 76 | 185.14 | 1 | 2.44 | 1.32 | 15.19 | 15.44 | -0.25 | 0.25 | 0 | 0 |
| gene9464 | https://www.ncbi.nlm.nih.gov/gene/?term=LOC105282975 | 1450 | 25 | 172.41 | 1 | 6.9 | 4 | 40.46 | 40.7 | -0.25 | 0.25 | 0 | 0 |

| | | | | | | | | | | | | | |
|---|---|---|---|---|---|---|---|---|---|---|---|---|---|
| gene10345 | https://www.ncbi.nlm.nih.gov/gene/?term=LOC105283989 | 4444 | 40 | 90.01 | 1 | 2.25 | 2.5 | 37.64 | 38.15 | -0.5 | 0.5 | 0 | 0 |

Directory: OGL File: rnk-dmg-Cb_LI_RphaseA2.vs.RphaseB2.txt

| gene_ID | gene_link | gwidth | #Sites | #per10Kb | #dmSites | #dmsp10kb | %dmSites | %pSite1 | %pSite2 | DMpSite | ADMpSite | DMpNucl | ADMpNucl |
|---|---|---|---|---|---|---|---|---|---|---|---|---|---|
| gene3165 | https://www.ncbi.nlm.nih.gov/gene/?term=LOC105275706 | 2374 | 74 | 311.71 | 6 | 25.27 | 8.11 | 32.18 | 22.24 | 9.94 | 9.94 | 0.31 | 0.31 |
| gene2824 | https://www.ncbi.nlm.nih.gov/gene/?term=LOC105275337 | 1772 | 60 | 338.6 | 2 | 11.29 | 3.33 | 38.84 | 29.82 | 9.02 | 9.02 | 0.31 | 0.31 |
| gene10963 | https://www.ncbi.nlm.nih.gov/gene/?term=LOC105284677 | 1576 | 30 | 190.36 | 1 | 6.35 | 3.33 | 25.25 | 41.23 | -15.97 | 15.97 | -0.3 | 0.3 |
| gene2593 | https://www.ncbi.nlm.nih.gov/gene/?term=LOC105275070 | 1774 | 7 | 39.46 | 5 | 28.18 | 71.43 | 65.68 | 4.09 | 61.59 | 61.59 | 0.24 | 0.24 |
| gene7568 | https://www.ncbi.nlm.nih.gov/gene/?term=LOC105280786 | 2525 | 126 | 499.01 | 1 | 3.96 | 0.79 | 30.35 | 27.57 | 2.78 | 2.78 | 0.14 | 0.14 |
| gene5714 | https://www.ncbi.nlm.nih.gov/gene/?term=LOC105278729 | 710 | 17 | 239.44 | 1 | 14.08 | 5.88 | 59.17 | 53.54 | 5.62 | 5.62 | 0.13 | 0.13 |
| gene8943 | https://www.ncbi.nlm.nih.gov/gene/?term=LOC105282388 | 1220 | 33 | 270.49 | 1 | 8.2 | 3.03 | 49.82 | 45.11 | 4.71 | 4.71 | 0.13 | 0.13 |
| gene5525 | https://www.ncbi.nlm.nih.gov/gene/?term=LOC105278396 | 3090 | 184 | 595.47 | 1 | 3.24 | 0.54 | 26.64 | 24.83 | 1.81 | 1.81 | 0.11 | 0.11 |
| gene2271 | https://www.ncbi.nlm.nih.gov/gene/?term=LOC105274706 | 1448 | 56 | 386.74 | 1 | 6.91 | 1.79 | 38.93 | 36.4 | 2.53 | 2.53 | 0.1 | 0.1 |
| gene4641 | https://www.ncbi.nlm.nih.gov/gene/?term=LOC105277398 | 1937 | 72 | 371.71 | 1 | 5.16 | 1.39 | 36.58 | 39.26 | -2.68 | 2.68 | -0.1 | 0.1 |
| gene4034 | https://www.ncbi.nlm.nih.gov/gene/?term=LOC105276704 | 2065 | 88 | 426.15 | 1 | 4.84 | 1.14 | 16.14 | 13.92 | 2.22 | 2.22 | 0.09 | 0.09 |
| gene11819 | https://www.ncbi.nlm.nih.gov/gene/?term=LOC105285671 | 2366 | 103 | 435.33 | 1 | 4.23 | 0.97 | 27.63 | 25.66 | 1.97 | 1.97 | 0.09 | 0.09 |
| gene2907 | https://www.ncbi.nlm.nih.gov/gene/?term=LOC105275447 | 2539 | 37 | 145.73 | 1 | 3.94 | 2.7 | 8.22 | 2.41 | 5.8 | 5.8 | 0.08 | 0.08 |
| gene920 | https://www.ncbi.nlm.nih.gov/gene/?term=LOC105282518 | 1197 | 52 | 434.42 | 1 | 8.35 | 1.92 | 29.01 | 30.77 | -1.76 | 1.76 | -0.08 | 0.08 |
| gene9598 | https://www.ncbi.nlm.nih.gov/gene/?term=LOC105283132 | 1769 | 57 | 322.22 | 1 | 5.65 | 1.75 | 39.92 | 37.87 | 2.05 | 2.05 | 0.07 | 0.07 |
| gene940 | https://www.ncbi.nlm.nih.gov/gene/?term=LOC105282794 | 3578 | 88 | 245.95 | 1 | 2.79 | 1.14 | 16.61 | 14.17 | 2.44 | 2.44 | 0.06 | 0.06 |
| gene13033 | https://www.ncbi.nlm.nih.gov/gene/?term=LOC105287063 | 1877 | 48 | 255.73 | 1 | 5.33 | 2.08 | 32.8 | 30.37 | 2.43 | 2.43 | 0.06 | 0.06 |
| gene7211 | https://www.ncbi.nlm.nih.gov/gene/?term=LOC105280377 | 2024 | 69 | 340.91 | 1 | 4.94 | 1.45 | 48.01 | 46.12 | 1.89 | 1.89 | 0.06 | 0.06 |
| gene5524 | https://www.ncbi.nlm.nih.gov/gene/?term=LOC105278397 | 5177 | 201 | 388.26 | 1 | 1.93 | 0.5 | 26.23 | 24.63 | 1.61 | 1.61 | 0.06 | 0.06 |
| gene9646 | https://www.ncbi.nlm.nih.gov/gene/?term=LOC105283182 | 3588 | 133 | 370.68 | 1 | 2.79 | 0.75 | 13.97 | 12.43 | 1.54 | 1.54 | 0.06 | 0.06 |
| gene3817 | https://www.ncbi.nlm.nih.gov/gene/?term=LOC105276449 | 1230 | 52 | 422.76 | 1 | 8.13 | 1.92 | 31.62 | 30.14 | 1.48 | 1.48 | 0.06 | 0.06 |
| gene10678 | https://www.ncbi.nlm.nih.gov/gene/?term=LOC105284355 | 4031 | 169 | 419.25 | 1 | 2.48 | 0.59 | 19.82 | 18.64 | 1.18 | 1.18 | 0.05 | 0.05 |
| gene1993 | https://www.ncbi.nlm.nih.gov/gene/?term=LOC105288193 | 1449 | 77 | 531.4 | 1 | 6.9 | 1.3 | 34.64 | 33.62 | 1.02 | 1.02 | 0.05 | 0.05 |
| gene935 | https://www.ncbi.nlm.nih.gov/gene/?term=LOC105282682 | 4368 | 222 | 508.24 | 1 | 2.29 | 0.45 | 13.71 | 12.73 | 0.98 | 0.98 | 0.05 | 0.05 |
| gene11801 | https://www.ncbi.nlm.nih.gov/gene/?term=LOC105285656 | 2497 | 140 | 560.67 | 1 | 4 | 0.71 | 9.18 | 8.36 | 0.81 | 0.81 | 0.05 | 0.05 |
| gene9101 | https://www.ncbi.nlm.nih.gov/gene/?term=LOC105282553 | 3068 | 108 | 352.02 | 1 | 3.26 | 0.93 | 22.02 | 23.41 | -1.4 | 1.4 | -0.05 | 0.05 |
| gene10213 | https://www.ncbi.nlm.nih.gov/gene/?term=LOC105283826 | 1503 | 54 | 359.28 | 1 | 6.65 | 1.85 | 14.21 | 15.73 | -1.51 | 1.51 | -0.05 | 0.05 |
| gene3827 | https://www.ncbi.nlm.nih.gov/gene/?term=LOC105276463 | 2749 | 82 | 298.29 | 1 | 3.64 | 1.22 | 27.66 | 29.29 | -1.63 | 1.63 | -0.05 | 0.05 |
| gene2413 | https://www.ncbi.nlm.nih.gov/gene/?term=LOC105274863 | 2006 | 50 | 249.25 | 1 | 4.99 | 2 | 53.71 | 55.57 | -1.87 | 1.87 | -0.05 | 0.05 |
| gene393 | https://www.ncbi.nlm.nih.gov/gene/?term=LOC105276566 | 1123 | 2 | 17.81 | 1 | 8.9 | 50 | 47.62 | 76.78 | -29.16 | 29.16 | -0.05 | 0.05 |
| gene12021 | https://www.ncbi.nlm.nih.gov/gene/?term=LOC105285905 | 1399 | 16 | 114.37 | 1 | 7.15 | 6.25 | 51.11 | 47.75 | 3.36 | 3.36 | 0.04 | 0.04 |
| gene4225 | https://www.ncbi.nlm.nih.gov/gene/?term=LOC105276917 | 2443 | 36 | 147.36 | 1 | 4.09 | 2.78 | 30.16 | 27.69 | 2.47 | 2.47 | 0.04 | 0.04 |
| gene10984 | https://www.ncbi.nlm.nih.gov/gene/?term=LOC105284704 | 1987 | 61 | 307 | 1 | 5.03 | 1.64 | 24.7 | 23.27 | 1.43 | 1.43 | 0.04 | 0.04 |
| gene9462 | https://www.ncbi.nlm.nih.gov/gene/?term=LOC105282970 | 1469 | 40 | 272.29 | 1 | 6.81 | 2.5 | 49.62 | 48.23 | 1.39 | 1.39 | 0.04 | 0.04 |
| gene6966 | https://www.ncbi.nlm.nih.gov/gene/?term=LOC105280104 | 3747 | 139 | 370.96 | 1 | 2.67 | 0.72 | 16.72 | 15.56 | 1.16 | 1.16 | 0.04 | 0.04 |
| gene10540 | https://www.ncbi.nlm.nih.gov/gene/?term=LOC105284200 | 5267 | 230 | 436.68 | 1 | 1.9 | 0.43 | 12.62 | 11.63 | 0.99 | 0.99 | 0.04 | 0.04 |
| gene2844 | https://www.ncbi.nlm.nih.gov/gene/?term=LOC105275363 | 4330 | 196 | 452.66 | 1 | 2.31 | 0.51 | 14.69 | 13.8 | 0.89 | 0.89 | 0.04 | 0.04 |
| gene5831 | https://www.ncbi.nlm.nih.gov/gene/?term=LOC105278761 | 2506 | 117 | 466.88 | 2 | 7.98 | 1.71 | 24.49 | 23.66 | 0.83 | 0.83 | 0.04 | 0.04 |
| gene298 | https://www.ncbi.nlm.nih.gov/gene/?term=LOC105275460 | 2473 | 117 | 473.11 | 1 | 4.04 | 0.85 | 14.05 | 13.27 | 0.78 | 0.78 | 0.04 | 0.04 |
| gene1470 | https://www.ncbi.nlm.nih.gov/gene/?term=LOC105287597 | 2974 | 197 | 662.41 | 1 | 3.36 | 0.51 | 17.93 | 17.38 | 0.55 | 0.55 | 0.04 | 0.04 |
| gene5298 | https://www.ncbi.nlm.nih.gov/gene/?term=LOC105278145 | 4060 | 250 | 615.76 | 2 | 4.93 | 0.8 | 10.45 | 11.12 | -0.68 | 0.68 | -0.04 | 0.04 |
| gene4498 | https://www.ncbi.nlm.nih.gov/gene/?term=LOC105277239 | 2831 | 120 | 423.88 | 1 | 3.53 | 0.83 | 12.33 | 13.34 | -1.01 | 1.01 | -0.04 | 0.04 |
| gene1768 | https://www.ncbi.nlm.nih.gov/gene/?term=LOC105287938 | 2792 | 87 | 311.6 | 1 | 3.58 | 1.15 | 25.07 | 26.23 | -1.17 | 1.17 | -0.04 | 0.04 |
| gene7406 | https://www.ncbi.nlm.nih.gov/gene/?term=LOC105280596 | 3432 | 115 | 335.08 | 2 | 5.83 | 1.74 | 17.75 | 19.02 | -1.26 | 1.26 | -0.04 | 0.04 |
| gene11340 | https://www.ncbi.nlm.nih.gov/gene/?term=LOC105285131 | 2918 | 91 | 311.86 | 1 | 3.43 | 1.1 | 11.55 | 12.83 | -1.28 | 1.28 | -0.04 | 0.04 |
| gene7600 | https://www.ncbi.nlm.nih.gov/gene/?term=LOC105280819 | 1615 | 38 | 235.29 | 1 | 6.19 | 2.63 | 35.64 | 37.15 | -1.52 | 1.52 | -0.04 | 0.04 |
| gene10396 | https://www.ncbi.nlm.nih.gov/gene/?term=LOC105284047 | 1227 | 23 | 187.45 | 1 | 8.15 | 4.35 | 26.88 | 28.88 | -2 | 2 | -0.04 | 0.04 |
| gene2335 | https://www.ncbi.nlm.nih.gov/gene/?term=LOC105274778 | 3945 | 38 | 96.32 | 1 | 2.53 | 2.63 | 48.07 | 44.68 | 3.39 | 3.39 | 0.03 | 0.03 |
| gene2970 | https://www.ncbi.nlm.nih.gov/gene/?term=LOC105275518 | 2340 | 24 | 102.56 | 1 | 4.27 | 4.17 | 49.7 | 46.61 | 3.09 | 3.09 | 0.03 | 0.03 |
| gene5273 | https://www.ncbi.nlm.nih.gov/gene/?term=LOC105278108 | 4067 | 51 | 125.4 | 1 | 2.46 | 1.96 | 25.16 | 23.04 | 2.12 | 2.12 | 0.03 | 0.03 |
| gene927 | https://www.ncbi.nlm.nih.gov/gene/?term=LOC105282600 | 3727 | 54 | 144.89 | 1 | 2.68 | 1.85 | 28.21 | 26.36 | 1.85 | 1.85 | 0.03 | 0.03 |
| gene3594 | https://www.ncbi.nlm.nih.gov/gene/?term=LOC105276190 | 4910 | 95 | 193.48 | 1 | 2.04 | 1.05 | 10.96 | 9.28 | 1.68 | 1.68 | 0.03 | 0.03 |
| gene10879 | https://www.ncbi.nlm.nih.gov/gene/?term=LOC105284574 | 2423 | 53 | 218.74 | 1 | 4.13 | 1.89 | 56.3 | 54.86 | 1.44 | 1.44 | 0.03 | 0.03 |
| gene9336 | https://www.ncbi.nlm.nih.gov/gene/?term=LOC105282834 | 5993 | 129 | 215.25 | 1 | 1.67 | 0.78 | 16.18 | 14.94 | 1.24 | 1.24 | 0.03 | 0.03 |
| gene10600 | https://www.ncbi.nlm.nih.gov/gene/?term=LOC105284270 | 4837 | 127 | 262.56 | 1 | 2.07 | 0.79 | 10.81 | 9.59 | 1.22 | 1.22 | 0.03 | 0.03 |
| gene7915 | https://www.ncbi.nlm.nih.gov/gene/?term=LOC105281173 | 1597 | 39 | 244.21 | 1 | 6.26 | 2.56 | 60.96 | 59.8 | 1.15 | 1.15 | 0.03 | 0.03 |
| gene4500 | https://www.ncbi.nlm.nih.gov/gene/?term=LOC105277238 | 2708 | 85 | 313.88 | 1 | 3.69 | 1.18 | 26.93 | 25.91 | 1.02 | 1.02 | 0.03 | 0.03 |
| gene10520 | https://www.ncbi.nlm.nih.gov/gene/?term=LOC105284184 | 2451 | 84 | 342.72 | 1 | 4.08 | 1.19 | 34.09 | 33.16 | 0.93 | 0.93 | 0.03 | 0.03 |
| gene7556 | https://www.ncbi.nlm.nih.gov/gene/?term=LOC105280774 | 2792 | 98 | 351 | 1 | 3.58 | 1.02 | 33.4 | 32.6 | 0.8 | 0.8 | 0.03 | 0.03 |
| gene4892 | https://www.ncbi.nlm.nih.gov/gene/?term=LOC105277667 | 3898 | 129 | 330.94 | 1 | 2.57 | 0.78 | 28.03 | 27.27 | 0.76 | 0.76 | 0.03 | 0.03 |

| | | | | | | | | | | | | | |
|---|---|---|---|---|---|---|---|---|---|---|---|---|---|
| gene8274 | https://www.ncbi.nlm.nih.gov/gene/?term=LOC105281624 | 5425 | 219 | 403.69 | 1 | 1.84 | 0.46 | 9.87 | 9.12 | 0.75 | 0.75 | 0.03 | 0.03 |
| gene2846 | https://www.ncbi.nlm.nih.gov/gene/?term=LOC105275374 | 11246 | 498 | 442.82 | 1 | 0.89 | 0.2 | 12.01 | 11.42 | 0.6 | 0.6 | 0.03 | 0.03 |
| gene10864 | https://www.ncbi.nlm.nih.gov/gene/?term=LOC105284557 | 1666 | 83 | 498.2 1 | 6 | 1.2 | 29.52 | 28.94 | 0.58 | 0.58 | 0.03 | 0.03 | |
| gene12802 | https://www.ncbi.nlm.nih.gov/gene/?term=LOC105286799 | 3183 | 166 | 521.52 | 1 | 3.14 | 0.6 | 17.88 | 17.37 | 0.5 | 0.5 | 0.03 | 0.03 |
| gene3599 | https://www.ncbi.nlm.nih.gov/gene/?term=LOC105276196 | 2955 | 62 | 209.81 | 1 | 3.38 | 1.61 | 36.96 | 38.6 | -1.64 | 1.64 | -0.03 | 0.03 |
| gene1505 | https://www.ncbi.nlm.nih.gov/gene/?term=LOC105287633 | 1693 | 29 | 171.29 | 1 | 5.91 | 3.45 | 41.25 | 42.91 | -1.66 | 1.66 | -0.03 | 0.03 |
| gene1008 | https://www.ncbi.nlm.nih.gov/gene/?term=LOC105283492 | 1479 | 10 | 67.61 1 | 6.76 | 10 | 37.74 | 42 | -4.25 | 4.25 | -0.03 | 0.03 | |
| gene9702 | https://www.ncbi.nlm.nih.gov/gene/?term=LOC105283234 | 2318 | 7 | 30.2 1 | 4.31 | 14.29 | 42.66 | 51.92 | -9.26 | 9.26 | -0.03 | 0.03 | |
| gene5052 | https://www.ncbi.nlm.nih.gov/gene/?term=LOC105277856 | 3167 | 34 | 107.36 | 1 | 3.16 | 2.94 | 35.71 | 33.56 | 2.15 | 2.15 | 0.02 | 0.02 |
| gene12106 | https://www.ncbi.nlm.nih.gov/gene/?term=LOC105286002 | 8556 | 68 | 79.48 1 | 1.17 | 1.47 | 4.38 | 2.38 | 2 | 2 | 0.02 | 0.02 | |
| gene388 | https://www.ncbi.nlm.nih.gov/gene/?term=LOC105276538 | 1558 | 21 | 134.79 | 2 | 12.84 | 9.52 | 69.8 | 67.98 | 1.82 | 1.82 | 0.02 | 0.02 |
| gene1518 | https://www.ncbi.nlm.nih.gov/gene/?term=LOC105287648 | 2195 | 26 | 118.45 | 1 | 4.56 | 3.85 | 41.13 | 39.6 | 1.53 | 1.53 | 0.02 | 0.02 |
| gene564 | https://www.ncbi.nlm.nih.gov/gene/?term=LOC105278478 | 2802 | 28 | 99.93 1 | 3.57 | 3.57 | 18.19 | 16.66 | 1.53 | 1.53 | 0.02 | 0.02 | |
| gene5647 | https://www.ncbi.nlm.nih.gov/gene/?term=LOC105278567 | 3118 | 38 | 121.87 | 1 | 3.21 | 2.63 | 15.22 | 13.72 | 1.5 | 1.5 | 0.02 | 0.02 |
| gene6455 | https://www.ncbi.nlm.nih.gov/gene/?term=LOC105279500 | 2612 | 35 | 134 1 | 3.83 | 2.86 | 49.73 | 48.29 | 1.45 | 1.45 | 0.02 | 0.02 | |
| gene8742 | https://www.ncbi.nlm.nih.gov/gene/?term=LOC105282151 | 2563 | 39 | 152.17 | 2 | 7.8 | 5.13 | 42.27 | 40.86 | 1.41 | 1.41 | 0.02 | 0.02 |
| gene8661 | https://www.ncbi.nlm.nih.gov/gene/?term=LOC105282063 | 1926 | 36 | 186.92 | 1 | 5.19 | 2.78 | 33.96 | 32.69 | 1.28 | 1.28 | 0.02 | 0.02 |
| gene2748 | https://www.ncbi.nlm.nih.gov/gene/?term=LOC105275261 | 3845 | 58 | 150.85 | 1 | 2.6 | 1.72 | 39.72 | 38.52 | 1.2 | 1.2 | 0.02 | 0.02 |
| gene1588 | https://www.ncbi.nlm.nih.gov/gene/?term=LOC105287720 | 4942 | 99 | 200.32 | 1 | 2.02 | 1.01 | 16.41 | 15.25 | 1.16 | 1.16 | 0.02 | 0.02 |
| gene4023 | https://www.ncbi.nlm.nih.gov/gene/?term=LOC105276692 | 1759 | 37 | 210.35 | 1 | 5.69 | 2.7 | 20.96 | 19.79 | 1.16 | 1.16 | 0.02 | 0.02 |
| gene12572 | https://www.ncbi.nlm.nih.gov/gene/?term=LOC105286514 | 2374 | 50 | 210.61 | 1 | 4.21 | 2 | 36.01 | 34.88 | 1.14 | 1.14 | 0.02 | 0.02 |
| gene11454 | https://www.ncbi.nlm.nih.gov/gene/?term=LOC105285256 | 1906 | 45 | 236.1 1 | 5.25 | 2.22 | 48.03 | 47.02 | 1 | 1 | 0.02 | 0.02 | |
| gene2173 | https://www.ncbi.nlm.nih.gov/gene/?term=LOC105274590 | 4387 | 71 | 161.84 | 2 | 4.56 | 2.82 | 27.99 | 27.02 | 0.97 | 0.97 | 0.02 | 0.02 |
| gene11568 | https://www.ncbi.nlm.nih.gov/gene/?term=LOC105285395 | 8086 | 215 | 265.89 | 1 | 1.24 | 0.47 | 16.71 | 15.82 | 0.89 | 0.89 | 0.02 | 0.02 |
| gene3385 | https://www.ncbi.nlm.nih.gov/gene/?term=LOC105275966 | 3305 | 62 | 187.59 | 1 | 3.03 | 1.61 | 48.06 | 47.24 | 0.82 | 0.82 | 0.02 | 0.02 |
| gene10391 | https://www.ncbi.nlm.nih.gov/gene/?term=LOC109611283 | 1994 | 42 | 210.63 | 1 | 5.02 | 2.38 | 34.44 | 33.67 | 0.76 | 0.76 | 0.02 | 0.02 |
| gene2266 | https://www.ncbi.nlm.nih.gov/gene/?term=LOC105274699 | 2391 | 53 | 221.66 | 1 | 4.18 | 1.89 | 24.7 | 23.99 | 0.72 | 0.72 | 0.02 | 0.02 |
| gene3107 | https://www.ncbi.nlm.nih.gov/gene/?term=LOC105275650 | 5103 | 167 | 327.26 | 1 | 1.96 | 0.6 | 16.25 | 15.53 | 0.72 | 0.72 | 0.02 | 0.02 |
| gene2633 | https://www.ncbi.nlm.nih.gov/gene/?term=LOC105275127 | 7457 | 259 | 347.32 | 1 | 1.34 | 0.39 | 8.88 | 8.23 | 0.66 | 0.66 | 0.02 | 0.02 |
| gene8814 | https://www.ncbi.nlm.nih.gov/gene/?term=LOC105282240 | 6910 | 181 | 261.94 | 1 | 1.45 | 0.55 | 9.72 | 9.08 | 0.64 | 0.64 | 0.02 | 0.02 |
| gene10531 | https://www.ncbi.nlm.nih.gov/gene/?term=LOC105284190 | 6881 | 191 | 277.58 | 1 | 1.45 | 0.52 | 12.39 | 11.8 | 0.58 | 0.58 | 0.02 | 0.02 |
| gene3727 | https://www.ncbi.nlm.nih.gov/gene/?term=LOC105276367 | 2484 | 74 | 297.91 | 1 | 4.03 | 1.35 | 42.37 | 41.83 | 0.54 | 0.54 | 0.02 | 0.02 |
| gene4962 | https://www.ncbi.nlm.nih.gov/gene/?term=LOC105277761 | 4645 | 149 | 320.78 | 1 | 2.15 | 0.67 | 13.8 | 13.32 | 0.49 | 0.49 | 0.02 | 0.02 |
| gene6583 | https://www.ncbi.nlm.nih.gov/gene/?term=LOC105279649 | 7444 | 333 | 447.34 | 1 | 1.34 | 0.3 | 12.03 | 11.53 | 0.49 | 0.49 | 0.02 | 0.02 |
| gene10606 | https://www.ncbi.nlm.nih.gov/gene/?term=LOC105284273 | 6119 | 260 | 424.91 | 1 | 1.63 | 0.38 | 17.76 | 17.3 | 0.45 | 0.45 | 0.02 | 0.02 |
| gene11401 | https://www.ncbi.nlm.nih.gov/gene/?term=LOC105285202 | 1841 | 87 | 472.57 | 1 | 5.43 | 1.15 | 25.64 | 25.2 | 0.44 | 0.44 | 0.02 | 0.02 |
| gene6286 | https://www.ncbi.nlm.nih.gov/gene/?term=LOC105279298 | 2861 | 132 | 461.38 | 2 | 6.99 | 1.52 | 23.96 | 23.54 | 0.42 | 0.42 | 0.02 | 0.02 |
| gene13114 | https://www.ncbi.nlm.nih.gov/gene/?term=LOC105287147 | 7609 | 413 | 542.78 | 1 | 1.31 | 0.24 | 9.55 | 9.19 | 0.36 | 0.36 | 0.02 | 0.02 |
| gene2835 | https://www.ncbi.nlm.nih.gov/gene/?term=LOC105275357 | 2492 | 128 | 513.64 | 1 | 4.01 | 0.78 | 14.7 | 15.03 | -0.34 | 0.34 | -0.02 | 0.02 |
| gene3747 | https://www.ncbi.nlm.nih.gov/gene/?term=LOC105276387 | 3736 | 168 | 449.68 | 1 | 2.68 | 0.6 | 14.1 | 14.65 | -0.54 | 0.54 | -0.02 | 0.02 |
| gene7207 | https://www.ncbi.nlm.nih.gov/gene/?term=LOC105280376 | 4659 | 111 | 238.25 | 1 | 2.15 | 0.9 | 30.17 | 30.93 | -0.76 | 0.76 | -0.02 | 0.02 |
| gene322 | https://www.ncbi.nlm.nih.gov/gene/?term=LOC105275777 | 1606 | 28 | 174.35 | 1 | 6.23 | 3.57 | 39.43 | 40.5 | -1.07 | 1.07 | -0.02 | 0.02 |
| gene8343 | https://www.ncbi.nlm.nih.gov/gene/?term=LOC105281700 | 2172 | 34 | 156.54 | 1 | 4.6 | 2.94 | 35.34 | 36.82 | -1.48 | 1.48 | -0.02 | 0.02 |
| gene10345 | https://www.ncbi.nlm.nih.gov/gene/?term=LOC105283989 | 4444 | 27 | 60.76 1 | 2.25 | 3.7 | 34.86 | 38.69 | -3.83 | 3.83 | -0.02 | 0.02 | |
| gene8178 | https://www.ncbi.nlm.nih.gov/gene/?term=LOC105281515 | 8647 | 45 | 52.04 1 | 1.16 | 2.22 | 25.4 | 23.65 | 1.75 | 1.75 | 0.01 | 0.01 | |
| gene4996 | https://www.ncbi.nlm.nih.gov/gene/?term=LOC105277798 | 5617 | 56 | 99.7 1 | 1.78 | 1.79 | 13.35 | 11.98 | 1.38 | 1.38 | 0.01 | 0.01 | |
| gene9648 | https://www.ncbi.nlm.nih.gov/gene/?term=LOC105283183 | 2602 | 31 | 119.14 | 1 | 3.84 | 3.23 | 40.75 | 39.71 | 1.04 | 1.04 | 0.01 | 0.01 |
| gene9647 | https://www.ncbi.nlm.nih.gov/gene/?term=LOC109611233 | 3094 | 37 | 119.59 | 1 | 3.23 | 2.7 | 34.14 | 33.27 | 0.87 | 0.87 | 0.01 | 0.01 |
| gene6528 | https://www.ncbi.nlm.nih.gov/gene/?term=LOC105279591 | 7222 | 55 | 76.16 1 | 1.38 | 1.82 | 30.56 | 29.74 | 0.82 | 0.82 | 0.01 | 0.01 | |
| gene10344 | https://www.ncbi.nlm.nih.gov/gene/?term=LOC105283987 | 4841 | 67 | 138.4 1 | 2.07 | 1.49 | 29.63 | 28.82 | 0.8 | 0.8 | 0.01 | 0.01 | |
| gene3470 | https://www.ncbi.nlm.nih.gov/gene/?term=LOC105276059 | 8716 | 160 | 183.57 | 1 | 1.15 | 0.62 | 8.58 | 7.82 | 0.76 | 0.76 | 0.01 | 0.01 |
| gene2257 | https://www.ncbi.nlm.nih.gov/gene/?term=LOC105274689 | 3658 | 70 | 191.36 | 1 | 2.73 | 1.43 | 15.64 | 14.92 | 0.72 | 0.72 | 0.01 | 0.01 |
| gene9330 | https://www.ncbi.nlm.nih.gov/gene/?term=LOC105282825 | 3936 | 85 | 215.96 | 1 | 2.54 | 1.18 | 14.44 | 13.87 | 0.57 | 0.57 | 0.01 | 0.01 |
| gene11579 | https://www.ncbi.nlm.nih.gov/gene/?term=LOC105285406 | 13571 | 245 | 180.53 | 1 | 0.74 | 0.41 | 7.35 | 6.84 | 0.5 | 0.5 | 0.01 | 0.01 |
| gene7245 | https://www.ncbi.nlm.nih.gov/gene/?term=LOC105280410 | 12478 | 260 | 208.37 | 1 | 0.8 | 0.38 | 20.02 | 19.52 | 0.5 | 0.5 | 0.01 | 0.01 |
| gene8252 | https://www.ncbi.nlm.nih.gov/gene/?term=LOC105281599 | 3813 | 49 | 128.51 | 1 | 2.62 | 2.04 | 4.61 | 4.12 | 0.49 | 0.49 | 0.01 | 0.01 |
| gene12516 | https://www.ncbi.nlm.nih.gov/gene/?term=LOC105286452 | 1529 | 40 | 261.61 | 1 | 6.54 | 2.5 | 51.71 | 51.25 | 0.47 | 0.47 | 0.01 | 0.01 |
| gene11976 | https://www.ncbi.nlm.nih.gov/gene/?term=LOC105285860 | 3112 | 55 | 176.74 | 1 | 3.21 | 1.82 | 24.17 | 23.73 | 0.43 | 0.43 | 0.01 | 0.01 |
| gene5967 | https://www.ncbi.nlm.nih.gov/gene/?term=LOC105278918 | 4393 | 137 | 311.86 | 1 | 2.28 | 0.73 | 13.27 | 12.86 | 0.41 | 0.41 | 0.01 | 0.01 |
| gene7550 | https://www.ncbi.nlm.nih.gov/gene/?term=LOC105280763 | 2173 | 84 | 386.56 | 1 | 4.6 | 1.19 | 32.91 | 32.56 | 0.35 | 0.35 | 0.01 | 0.01 |
| gene2806 | https://www.ncbi.nlm.nih.gov/gene/?term=LOC105275327 | 14257 | 490 | 343.69 | 1 | 0.7 | 0.2 | 7.15 | 6.82 | 0.34 | 0.34 | 0.01 | 0.01 |
| gene1408 | https://www.ncbi.nlm.nih.gov/gene/?term=LOC109611520 | 8645 | 325 | 375.94 | 1 | 1.16 | 0.31 | 6.16 | 5.83 | 0.33 | 0.33 | 0.01 | 0.01 |
| gene1430 | https://www.ncbi.nlm.nih.gov/gene/?term=LOC105287551 | 6433 | 237 | 368.41 | 1 | 1.55 | 0.42 | 8.71 | 8.38 | 0.33 | 0.33 | 0.01 | 0.01 |
| gene10107 | https://www.ncbi.nlm.nih.gov/gene/?term=LOC105283712 | 1883 | 36 | 191.18 | 1 | 5.31 | 2.78 | 27.71 | 27.4 | 0.31 | 0.31 | 0.01 | 0.01 |

| | | | | | | | | | | | | | |
|---|---|---|---|---|---|---|---|---|---|---|---|---|---|
| gene5613 | https://www.ncbi.nlm.nih.gov/gene/?term=LOC105278504 | 8524 | 250 | 293.29 | 1 | 1.17 | 0.4 | 4.85 | 4.54 | 0.31 | 0.31 | 0.01 | 0.01 |
| gene8600 | https://www.ncbi.nlm.nih.gov/gene/?term=LOC105282001 | 8513 | 158 | 185.6 | 1 | 1.17 | 0.63 | 10.54 | 10.23 | 0.31 | 0.31 | 0.01 | 0.01 |
| gene10392 | https://www.ncbi.nlm.nih.gov/gene/?term=LOC105284036 | 3216 | 104 | 323.38 | 2 | 6.22 | 1.92 | 22.48 | 22.18 | 0.3 | 0.3 | 0.01 | 0.01 |
| gene2056 | https://www.ncbi.nlm.nih.gov/gene/?term=LOC105274452 | 6584 | 257 | 390.34 | 1 | 1.52 | 0.39 | 13.38 | 13.08 | 0.3 | 0.3 | 0.01 | 0.01 |
| gene5746 | https://www.ncbi.nlm.nih.gov/gene/?term=LOC105278666 | 10343 | 447 | 432.18 | 1 | 0.97 | 0.22 | 7.23 | 6.93 | 0.3 | 0.3 | 0.01 | 0.01 |
| gene11082 | https://www.ncbi.nlm.nih.gov/gene/?term=LOC105284824 | 7436 | 205 | 275.69 | 1 | 1.34 | 0.49 | 7.69 | 7.4 | 0.29 | 0.29 | 0.01 | 0.01 |
| gene12265 | https://www.ncbi.nlm.nih.gov/gene/?term=LOC105286166 | 3050 | 110 | 360.66 | 1 | 3.28 | 0.91 | 24.09 | 23.8 | 0.29 | 0.29 | 0.01 | 0.01 |
| gene4190 | https://www.ncbi.nlm.nih.gov/gene/?term=LOC105276875 | 3150 | 118 | 374.6 | 1 | 3.17 | 0.85 | 22.65 | 22.39 | 0.26 | 0.26 | 0.01 | 0.01 |
| gene10596 | https://www.ncbi.nlm.nih.gov/gene/?term=LOC105284262 | 6520 | 215 | 329.75 | 1 | 1.53 | 0.47 | 4.27 | 4.03 | 0.24 | 0.24 | 0.01 | 0.01 |
| gene7166 | https://www.ncbi.nlm.nih.gov/gene/?term=LOC105280326 | 6117 | 245 | 400.52 | 1 | 1.63 | 0.41 | 14.73 | 14.53 | 0.2 | 0.2 | 0.01 | 0.01 |
| gene3856 | https://www.ncbi.nlm.nih.gov/gene/?term=LOC105276499 | 5236 | 184 | 351.41 | 1 | 1.91 | 0.54 | 10.07 | 9.88 | 0.18 | 0.18 | 0.01 | 0.01 |
| gene11275 | https://www.ncbi.nlm.nih.gov/gene/?term=LOC105285055 | 4119 | 272 | 660.35 | 1 | 2.43 | 0.37 | 11.12 | 11.01 | 0.12 | 0.12 | 0.01 | 0.01 |
| gene12674 | https://www.ncbi.nlm.nih.gov/gene/?term=LOC105286649 | 18283 | 788 | 431 | 1 | 0.55 | 0.13 | 1.01 | 1.14 | -0.13 | 0.13 | -0.01 | 0.01 |
| gene8729 | https://www.ncbi.nlm.nih.gov/gene/?term=LOC105282137 | 5005 | 220 | 439.56 | 1 | 2 | 0.45 | 15.19 | 15.33 | -0.14 | 0.14 | -0.01 | 0.01 |
| gene9755 | https://www.ncbi.nlm.nih.gov/gene/?term=LOC105283296 | 2177 | 92 | 422.6 | 1 | 4.59 | 1.09 | 18.56 | 18.74 | -0.18 | 0.18 | -0.01 | 0.01 |
| gene7493 | https://www.ncbi.nlm.nih.gov/gene/?term=LOC105280704 | 4225 | 132 | 312.43 | 1 | 2.37 | 0.76 | 20.13 | 20.33 | -0.21 | 0.21 | -0.01 | 0.01 |
| gene1310 | https://www.ncbi.nlm.nih.gov/gene/?term=LOC105286661 | 2582 | 72 | 278.85 | 1 | 3.87 | 1.39 | 21.24 | 21.48 | -0.25 | 0.25 | -0.01 | 0.01 |
| gene2404 | https://www.ncbi.nlm.nih.gov/gene/?term=LOC105274853 | 1767 | 43 | 243.35 | 1 | 5.66 | 2.33 | 29.37 | 29.63 | -0.26 | 0.26 | -0.01 | 0.01 |
| gene289 | https://www.ncbi.nlm.nih.gov/gene/?term=LOC105275347 | 2533 | 69 | 272.4 | 1 | 3.95 | 1.45 | 31.43 | 31.68 | -0.26 | 0.26 | -0.01 | 0.01 |
| gene6031 | https://www.ncbi.nlm.nih.gov/gene/?term=LOC105279000 | 5691 | 190 | 333.86 | 1 | 1.76 | 0.53 | 15.94 | 16.2 | -0.26 | 0.26 | -0.01 | 0.01 |
| gene5646 | https://www.ncbi.nlm.nih.gov/gene/?term=LOC105278534 | 7475 | 197 | 263.55 | 1 | 1.34 | 0.51 | 17.35 | 17.62 | -0.27 | 0.27 | -0.01 | 0.01 |
| gene53 | https://www.ncbi.nlm.nih.gov/gene/?term=LOC105277148 | 19283 | 303 | 157.13 | 1 | 0.52 | 0.33 | 3.79 | 4.11 | -0.32 | 0.32 | -0.01 | 0.01 |
| gene11583 | https://www.ncbi.nlm.nih.gov/gene/?term=LOC105285414 | 8192 | 182 | 222.17 | 1 | 1.22 | 0.55 | 8.25 | 8.59 | -0.33 | 0.33 | -0.01 | 0.01 |
| gene8015 | https://www.ncbi.nlm.nih.gov/gene/?term=LOC105281312 | 2998 | 62 | 206.8 | 1 | 3.34 | 1.61 | 34.79 | 35.12 | -0.33 | 0.33 | -0.01 | 0.01 |
| gene11561 | https://www.ncbi.nlm.nih.gov/gene/?term=LOC105285387 | 11528 | 184 | 159.61 | 1 | 0.87 | 0.54 | 10.15 | 10.52 | -0.37 | 0.37 | -0.01 | 0.01 |
| gene12981 | https://www.ncbi.nlm.nih.gov/gene/?term=LOC105286999 | 2063 | 44 | 213.28 | 1 | 4.85 | 2.27 | 34 | 34.43 | -0.43 | 0.43 | -0.01 | 0.01 |
| gene6652 | https://www.ncbi.nlm.nih.gov/gene/?term=LOC105279719 | 2831 | 80 | 282.59 | 1 | 3.53 | 1.25 | 22.97 | 23.4 | -0.43 | 0.43 | -0.01 | 0.01 |
| gene1199 | https://www.ncbi.nlm.nih.gov/gene/?term=LOC105285498 | 2220 | 30 | 135.14 | 1 | 4.5 | 3.33 | 26.9 | 27.37 | -0.47 | 0.47 | -0.01 | 0.01 |
| gene4376 | https://www.ncbi.nlm.nih.gov/gene/?term=LOC105277083 | 4307 | 111 | 257.72 | 1 | 2.32 | 0.9 | 22.97 | 23.44 | -0.47 | 0.47 | -0.01 | 0.01 |
| gene657 | https://www.ncbi.nlm.nih.gov/gene/?term=LOC105279667 | 6421 | 85 | 132.38 | 1 | 1.56 | 1.18 | 2.59 | 3.08 | -0.49 | 0.49 | -0.01 | 0.01 |
| gene12203 | https://www.ncbi.nlm.nih.gov/gene/?term=LOC105286099 | 5967 | 78 | 130.72 | 1 | 1.68 | 1.28 | 9.64 | 10.17 | -0.53 | 0.53 | -0.01 | 0.01 |
| gene814 | https://www.ncbi.nlm.nih.gov/gene/?term=LOC105281296 | 2722 | 44 | 161.65 | 1 | 3.67 | 2.27 | 26.1 | 26.63 | -0.53 | 0.53 | -0.01 | 0.01 |
| gene12406 | https://www.ncbi.nlm.nih.gov/gene/?term=LOC105286316 | 1386 | 36 | 259.74 | 1 | 7.22 | 2.78 | 38.06 | 38.61 | -0.55 | 0.55 | -0.01 | 0.01 |
| gene35 | https://www.ncbi.nlm.nih.gov/gene/?term=LOC105275197 | 2643 | 51 | 192.96 | 1 | 3.78 | 1.96 | 38.21 | 38.8 | -0.59 | 0.59 | -0.01 | 0.01 |
| gene6639 | https://www.ncbi.nlm.nih.gov/gene/?term=LOC105279705 | 7656 | 73 | 95.35 | 1 | 1.31 | 1.37 | 23.16 | 23.78 | -0.62 | 0.62 | -0.01 | 0.01 |
| gene11447 | https://www.ncbi.nlm.nih.gov/gene/?term=LOC105285252 | 7437 | 119 | 160.01 | 1 | 1.34 | 0.84 | 14.25 | 14.93 | -0.68 | 0.68 | -0.01 | 0.01 |
| gene4033 | https://www.ncbi.nlm.nih.gov/gene/?term=LOC105276705 | 10546 | 136 | 128.96 | 1 | 0.95 | 0.74 | 8.69 | 9.42 | -0.73 | 0.73 | -0.01 | 0.01 |
| gene9327 | https://www.ncbi.nlm.nih.gov/gene/?term=LOC105282821 | 7793 | 149 | 191.2 | 1 | 1.28 | 0.67 | 12.54 | 13.29 | -0.74 | 0.74 | -0.01 | 0.01 |
| gene6237 | https://www.ncbi.nlm.nih.gov/gene/?term=LOC105279244 | 1484 | 26 | 175.2 | 1 | 6.74 | 3.85 | 41.81 | 42.6 | -0.8 | 0.8 | -0.01 | 0.01 |
| gene10155 | https://www.ncbi.nlm.nih.gov/gene/?term=LOC105283756 | 7030 | 114 | 162.16 | 1 | 1.42 | 0.88 | 5.48 | 6.39 | -0.91 | 0.91 | -0.01 | 0.01 |
| gene7398 | https://www.ncbi.nlm.nih.gov/gene/?term=LOC105280589 | 3452 | 34 | 98.49 | 1 | 2.9 | 2.94 | 18.86 | 19.97 | -1.12 | 1.12 | -0.01 | 0.01 |
| gene3168 | https://www.ncbi.nlm.nih.gov/gene/?term=LOC105275711 | 6150 | 75 | 121.95 | 1 | 1.63 | 1.33 | 13.14 | 14.33 | -1.18 | 1.18 | -0.01 | 0.01 |
| gene12307 | https://www.ncbi.nlm.nih.gov/gene/?term=LOC105286216 | 10313 | 190 | 184.23 | 1 | 0.97 | 0.53 | 15.65 | 15.38 | 0.26 | 0.26 | 0 | 0 |
| gene6620 | https://www.ncbi.nlm.nih.gov/gene/?term=LOC105279687 | 17998 | 368 | 204.47 | 1 | 0.56 | 0.27 | 5.31 | 5.13 | 0.18 | 0.18 | 0 | 0 |
| gene5387 | https://www.ncbi.nlm.nih.gov/gene/?term=LOC105278245 | 6451 | 113 | 175.17 | 1 | 1.55 | 0.88 | 22.95 | 22.78 | 0.17 | 0.17 | 0 | 0 |
| gene3671 | https://www.ncbi.nlm.nih.gov/gene/?term=LOC105276272 | 1966 | 48 | 244.15 | 1 | 5.09 | 2.08 | 27.74 | 27.59 | 0.16 | 0.16 | 0 | 0 |
| gene3206 | https://www.ncbi.nlm.nih.gov/gene/?term=LOC105275752 | 3474 | 132 | 379.97 | 1 | 2.88 | 0.76 | 32.74 | 32.62 | 0.12 | 0.12 | 0 | 0 |
| gene1632 | https://www.ncbi.nlm.nih.gov/gene/?term=LOC105287764 | 18061 | 416 | 230.33 | 1 | 0.55 | 0.24 | 3.35 | 3.24 | 0.11 | 0.11 | 0 | 0 |
| gene5877 | https://www.ncbi.nlm.nih.gov/gene/?term=LOC105278807 | 1719 | 50 | 290.87 | 1 | 5.82 | 2 | 27.13 | 27.02 | 0.11 | 0.11 | 0 | 0 |
| gene10018 | https://www.ncbi.nlm.nih.gov/gene/?term=LOC105283607 | 4414 | 207 | 468.96 | 1 | 2.27 | 0.48 | 15.18 | 15.09 | 0.09 | 0.09 | 0 | 0 |
| gene3885 | https://www.ncbi.nlm.nih.gov/gene/?term=LOC105276527 | 7928 | 322 | 406.16 | 1 | 1.26 | 0.31 | 9.68 | 9.59 | 0.09 | 0.09 | 0 | 0 |
| gene9029 | https://www.ncbi.nlm.nih.gov/gene/?term=LOC105282491 | 4762 | 167 | 350.69 | 1 | 2.1 | 0.6 | 9.69 | 9.6 | 0.09 | 0.09 | 0 | 0 |
| gene2122 | https://www.ncbi.nlm.nih.gov/gene/?term=LOC105274531 | 8153 | 304 | 372.87 | 1 | 1.23 | 0.33 | 10.7 | 10.62 | 0.08 | 0.08 | 0 | 0 |
| gene13036 | https://www.ncbi.nlm.nih.gov/gene/?term=LOC105287059 | 4959 | 163 | 328.7 | 1 | 2.02 | 0.61 | 20.67 | 20.67 | 0.01 | 0.01 | 0 | 0 |
| gene7850 | https://www.ncbi.nlm.nih.gov/gene/?term=LOC105281095 | 3050 | 67 | 219.67 | 1 | 3.28 | 1.49 | 33.74 | 33.73 | 0.01 | 0.01 | 0 | 0 |
| gene10549 | https://www.ncbi.nlm.nih.gov/gene/?term=LOC105284210 | 7145 | 298 | 417.07 | 2 | 2.8 | 0.67 | 11.73 | 11.76 | -0.04 | 0.04 | 0 | 0 |
| gene4048 | https://www.ncbi.nlm.nih.gov/gene/?term=LOC105276724 | 2955 | 32 | 108.29 | 1 | 3.38 | 3.12 | 11.95 | 11.99 | -0.04 | 0.04 | 0 | 0 |
| gene7249 | https://www.ncbi.nlm.nih.gov/gene/?term=LOC105280422 | 1775 | 60 | 338.03 | 1 | 5.63 | 1.67 | 17.01 | 17.06 | -0.05 | 0.05 | 0 | 0 |
| gene1220 | https://www.ncbi.nlm.nih.gov/gene/?term=LOC105285742 | 3453 | 123 | 356.21 | 1 | 2.9 | 0.81 | 20.51 | 20.57 | -0.06 | 0.06 | 0 | 0 |
| gene34 | https://www.ncbi.nlm.nih.gov/gene/?term=LOC105278416 | 7120 | 141 | 198.03 | 1 | 1.4 | 0.71 | 14.13 | 14.19 | -0.06 | 0.06 | 0 | 0 |
| gene12573 | https://www.ncbi.nlm.nih.gov/gene/?term=LOC105286515 | 3647 | 112 | 307.1 | 1 | 2.74 | 0.89 | 12.05 | 12.13 | -0.07 | 0.07 | 0 | 0 |
| gene7368 | https://www.ncbi.nlm.nih.gov/gene/?term=LOC105280556 | 4105 | 40 | 97.44 | 1 | 2.44 | 2.5 | 15.83 | 15.9 | -0.07 | 0.07 | 0 | 0 |
| gene9177 | https://www.ncbi.nlm.nih.gov/gene/?term=LOC105282646 | 3320 | 116 | 349.4 | 1 | 3.01 | 0.86 | 24.81 | 24.88 | -0.07 | 0.07 | 0 | 0 |
| gene2683 | https://www.ncbi.nlm.nih.gov/gene/?term=LOC105275187 | 5492 | 71 | 129.28 | 1 | 1.82 | 1.41 | 12.33 | 12.41 | -0.08 | 0.08 | 0 | 0 |

| | | | | | | | | | | | | | |
|---|---|---|---|---|---|---|---|---|---|---|---|---|---|
| gene1147 | https://www.ncbi.nlm.nih.gov/gene/?term=LOC105284954 | 4022 | 101 | 251.12 | 1 | 2.49 | 0.99 | 15.4 | 15.49 | -0.09 | 0.09 | 0 | 0 |
| gene3286 | https://www.ncbi.nlm.nih.gov/gene/?term=LOC105275848 | 3744 | 144 | 384.62 | 1 | 2.67 | 0.69 | 21.45 | 21.54 | -0.09 | 0.09 | 0 | 0 |
| gene12439 | https://www.ncbi.nlm.nih.gov/gene/?term=LOC105286357 | 4761 | 97 | 203.74 | 1 | 2.1 | 1.03 | 28.47 | 28.59 | -0.12 | 0.12 | 0 | 0 |
| gene6649 | https://www.ncbi.nlm.nih.gov/gene/?term=LOC105279717 | 6017 | 171 | 284.19 | 1 | 1.66 | 0.58 | 12.42 | 12.54 | -0.13 | 0.13 | 0 | 0 |
| gene5573 | https://www.ncbi.nlm.nih.gov/gene/?term=LOC105278454 | 2688 | 37 | 137.65 | 1 | 3.72 | 2.7 | 33.59 | 33.73 | -0.14 | 0.14 | 0 | 0 |
| gene10195 | https://www.ncbi.nlm.nih.gov/gene/?term=LOC105283806 | 10532 | 75 | 71.21 | 1 | 0.95 | 1.33 | 14.09 | 14.33 | -0.24 | 0.24 | 0 | 0 |
| gene10819 | https://www.ncbi.nlm.nih.gov/gene/?term=LOC105284507 | 5767 | 84 | 145.66 | 1 | 1.73 | 1.19 | 22.04 | 22.33 | -0.3 | 0.3 | 0 | 0 |
| gene9326 | https://www.ncbi.nlm.nih.gov/gene/?term=LOC105282818 | 16612 | 192 | 115.58 | 1 | 0.6 | 0.52 | 12.9 | 13.21 | -0.3 | 0.3 | 0 | 0 |

Directory: OGL File: rnk-dmg-Cb_LI_RphaseB1.vs.RphaseB2.txt

| gene_ID | gene_link | gwidth | #Sites | #per10Kb | #dmSites | #dmsp10kb | %dmSites | %pSite1 | %pSite2 | DMpSite | ADMpSite | DMpNucl | ADMpNucl |
|---|---|---|---|---|---|---|---|---|---|---|---|---|---|
| gene10442 | https://www.ncbi.nlm.nih.gov/gene/?term=LOC105284093 | 809 | 61 | 754.02 | 2 | 24.72 | 3.28 | 10.04 | 7.71 | 2.34 | 2.34 | 0.18 | 0.18 |
| gene10584 | https://www.ncbi.nlm.nih.gov/gene/?term=LOC105284253 | 1255 | 41 | 326.69 | 5 | 39.84 | 12.2 | 26.29 | 31.83 | -5.54 | 5.54 | -0.18 | 0.18 |
| gene5917 | https://www.ncbi.nlm.nih.gov/gene/?term=LOC105278849 | 823 | 80 | 972.05 | 2 | 24.3 | 2.5 | 21.53 | 23.29 | -1.76 | 1.76 | -0.17 | 0.17 |
| gene10644 | https://www.ncbi.nlm.nih.gov/gene/?term=LOC105284320 | 358 | 10 | 279.33 | 2 | 55.87 | 20 | 24.37 | 30.4 | -6.02 | 6.02 | -0.17 | 0.17 |
| gene4515 | https://www.ncbi.nlm.nih.gov/gene/?term=LOC109610839 | 335 | 7 | 208.96 | 1 | 29.85 | 14.29 | 66.57 | 58.9 | 7.67 | 7.67 | 0.16 | 0.16 |
| gene1993 | https://www.ncbi.nlm.nih.gov/gene/?term=LOC105288193 | 1449 | 77 | 531.4 | 5 | 34.51 | 6.49 | 30.7 | 33.62 | -2.92 | 2.92 | -0.16 | 0.16 |
| gene9019 | https://www.ncbi.nlm.nih.gov/gene/?term=LOC105282481 | 1743 | 58 | 332.76 | 4 | 22.95 | 6.9 | 32.56 | 37.33 | -4.77 | 4.77 | -0.16 | 0.16 |
| gene2636 | https://www.ncbi.nlm.nih.gov/gene/?term=LOC105275141 | 947 | 11 | 116.16 | 4 | 42.24 | 36.36 | 32.51 | 19.93 | 12.58 | 12.58 | 0.15 | 0.15 |
| gene9291 | https://www.ncbi.nlm.nih.gov/gene/?term=LOC105282787 | 1045 | 42 | 401.91 | 3 | 28.71 | 7.14 | 37.93 | 34.07 | 3.85 | 3.85 | 0.15 | 0.15 |
| gene5309 | https://www.ncbi.nlm.nih.gov/gene/?term=LOC105278160 | 1704 | 91 | 534.04 | 1 | 5.87 | 1.1 | 34.43 | 37.28 | -2.85 | 2.85 | -0.15 | 0.15 |
| gene10529 | https://www.ncbi.nlm.nih.gov/gene/?term=LOC105284189 | 2602 | 133 | 511.15 | 3 | 11.53 | 2.26 | 17.44 | 14.96 | 2.49 | 2.49 | 0.13 | 0.13 |
| gene1589 | https://www.ncbi.nlm.nih.gov/gene/?term=LOC105287721 | 1407 | 41 | 291.4 | 2 | 14.21 | 4.88 | 25.63 | 21.54 | 4.09 | 4.09 | 0.12 | 0.12 |
| gene4597 | https://www.ncbi.nlm.nih.gov/gene/?term=LOC105277345 | 2173 | 78 | 358.95 | 3 | 13.81 | 3.85 | 18.24 | 14.85 | 3.38 | 3.38 | 0.12 | 0.12 |
| gene6324 | https://www.ncbi.nlm.nih.gov/gene/?term=LOC105279346 | 4650 | 253 | 544.09 | 7 | 15.05 | 2.77 | 14.36 | 16.48 | -2.12 | 2.12 | -0.12 | 0.12 |
| gene558 | https://www.ncbi.nlm.nih.gov/gene/?term=LOC105278458 | 216 | 3 | 138.89 | 1 | 46.3 | 33.33 | 20.77 | 29.54 | -8.77 | 8.77 | -0.12 | 0.12 |
| gene11399 | https://www.ncbi.nlm.nih.gov/gene/?term=LOC105285203 | 1029 | 30 | 291.55 | 2 | 19.44 | 6.67 | 30.87 | 26.95 | 3.93 | 3.93 | 0.11 | 0.11 |
| gene2559 | https://www.ncbi.nlm.nih.gov/gene/?term=LOC105275030 | 1897 | 119 | 627.31 | 6 | 31.63 | 5.04 | 19.12 | 20.91 | -1.79 | 1.79 | -0.11 | 0.11 |
| gene1223 | https://www.ncbi.nlm.nih.gov/gene/?term=LOC105285713 | 1048 | 38 | 362.6 | 2 | 19.08 | 5.26 | 46.12 | 43.33 | 2.79 | 2.79 | 0.1 | 0.1 |
| gene2130 | https://www.ncbi.nlm.nih.gov/gene/?term=LOC105274542 | 1254 | 52 | 414.67 | 3 | 23.92 | 5.77 | 30.14 | 27.83 | 2.32 | 2.32 | 0.1 | 0.1 |
| gene6441 | https://www.ncbi.nlm.nih.gov/gene/?term=LOC105279487 | 2117 | 100 | 472.37 | 2 | 9.45 | 2 | 15.08 | 13.01 | 2.07 | 2.07 | 0.1 | 0.1 |
| gene3910 | https://www.ncbi.nlm.nih.gov/gene/?term=LOC105276563 | 1957 | 112 | 572.3 | 4 | 20.44 | 3.57 | 18.76 | 17 | 1.77 | 1.77 | 0.1 | 0.1 |
| gene12842 | https://www.ncbi.nlm.nih.gov/gene/?term=LOC105286845 | 1157 | 78 | 674.16 | 2 | 17.29 | 2.56 | 11.01 | 12.53 | -1.53 | 1.53 | -0.1 | 0.1 |
| gene5987 | https://www.ncbi.nlm.nih.gov/gene/?term=LOC105278938 | 1882 | 109 | 579.17 | 4 | 21.25 | 3.67 | 19.75 | 21.46 | -1.71 | 1.71 | -0.1 | 0.1 |
| gene6781 | https://www.ncbi.nlm.nih.gov/gene/?term=LOC105279857 | 241 | 10 | 414.94 | 1 | 41.49 | 10 | 8.85 | 11.17 | -2.32 | 2.32 | -0.1 | 0.1 |

...
... only first 25 lines shown ...

```
Directory: OGL File: wrt-Cb_LI.txt

... comparing  BCphaseA1.vs.BCphaseA2 ...

Number of genes <=  20000  and with #dmsites >=  1 :  137
sample1     sum of percent methylation:  3379.84     average:    24.67
sample2     sum of percent methylation:  3413.89     average:    24.92

sum difference:   -34.02      average difference:  -0.2483

      Wilcoxon signed rank test with continuity correction

data:  pw_summary$`%pSite1` and pw_summary$`%pSite2`
V = 4806, p-value = 0.7
alternative hypothesis: true location shift is not equal to 0

... comparing  BCphaseA1.vs.BCphaseB1 ...

Number of genes <=  20000  and with #dmsites >=  1 :  139
sample1     sum of percent methylation:  3279.30     average:    23.59
sample2     sum of percent methylation:  3296.07     average:    23.71

sum difference:   -16.72      average difference:  -0.1203

      Wilcoxon signed rank test with continuity correction
...
... only first 25 lines shown ...
```

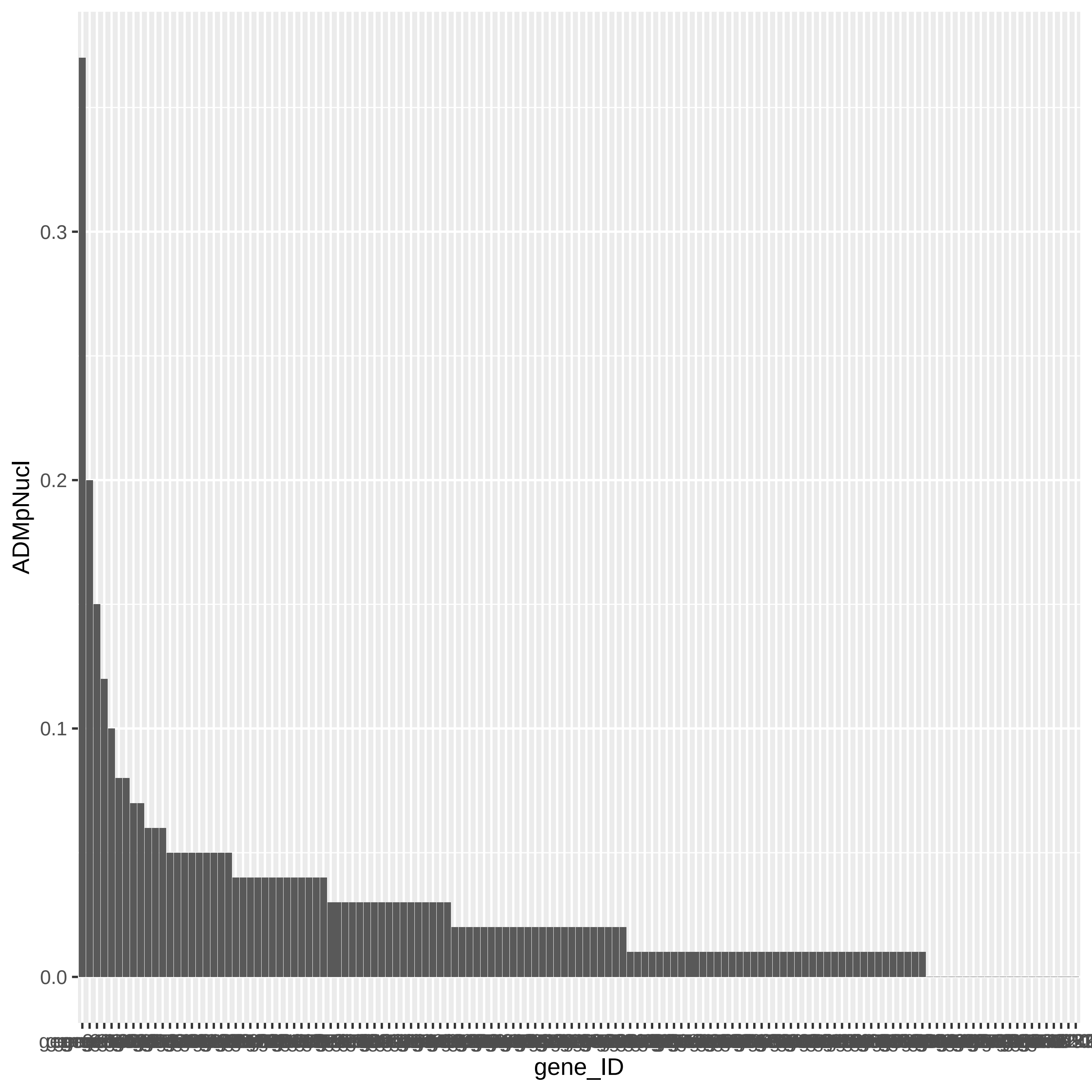

ADMpNucl
0.3
0.2
0.1
0.0
gene_ID

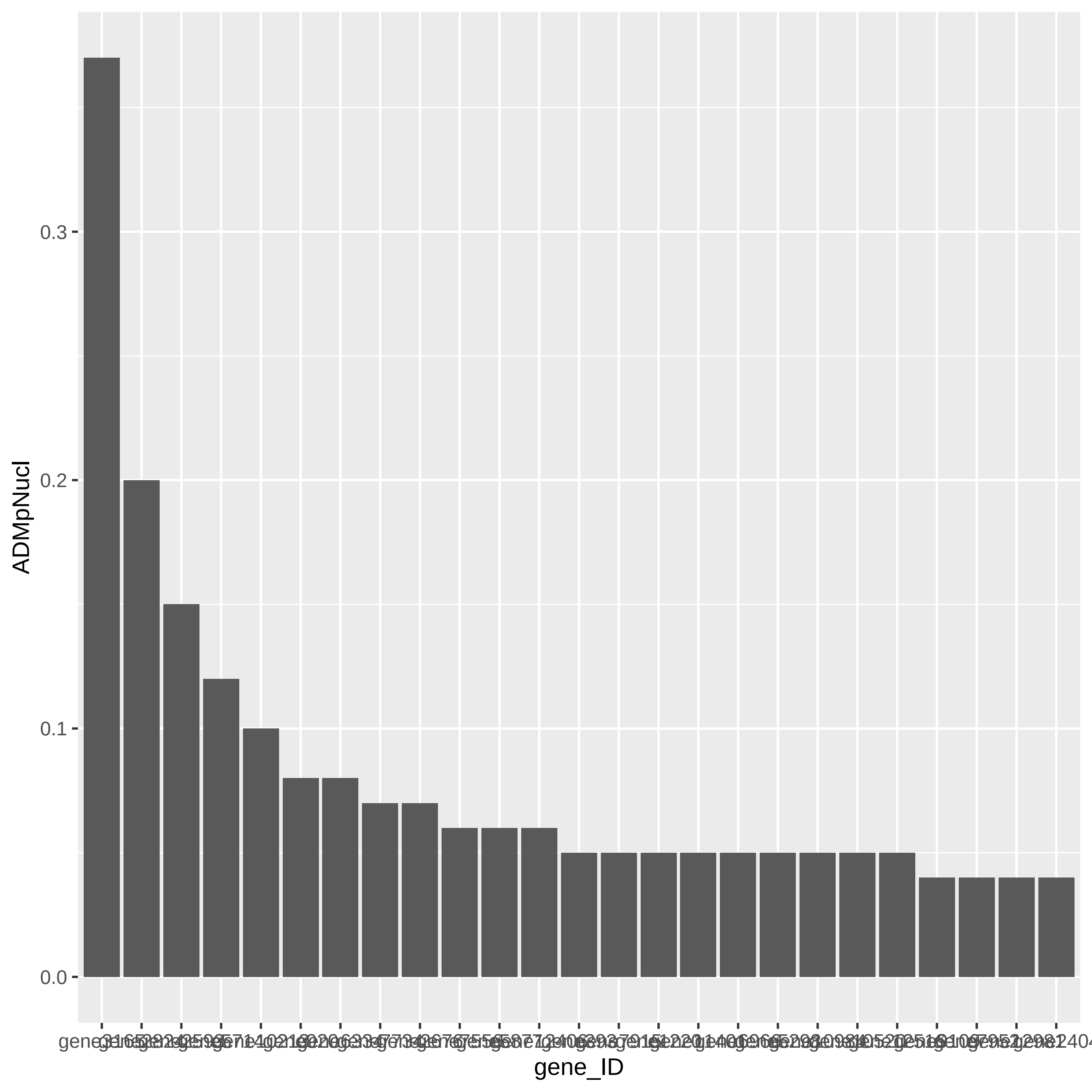

0.3
0.2
0.1
0.0
ADMpNucl
gene_ID

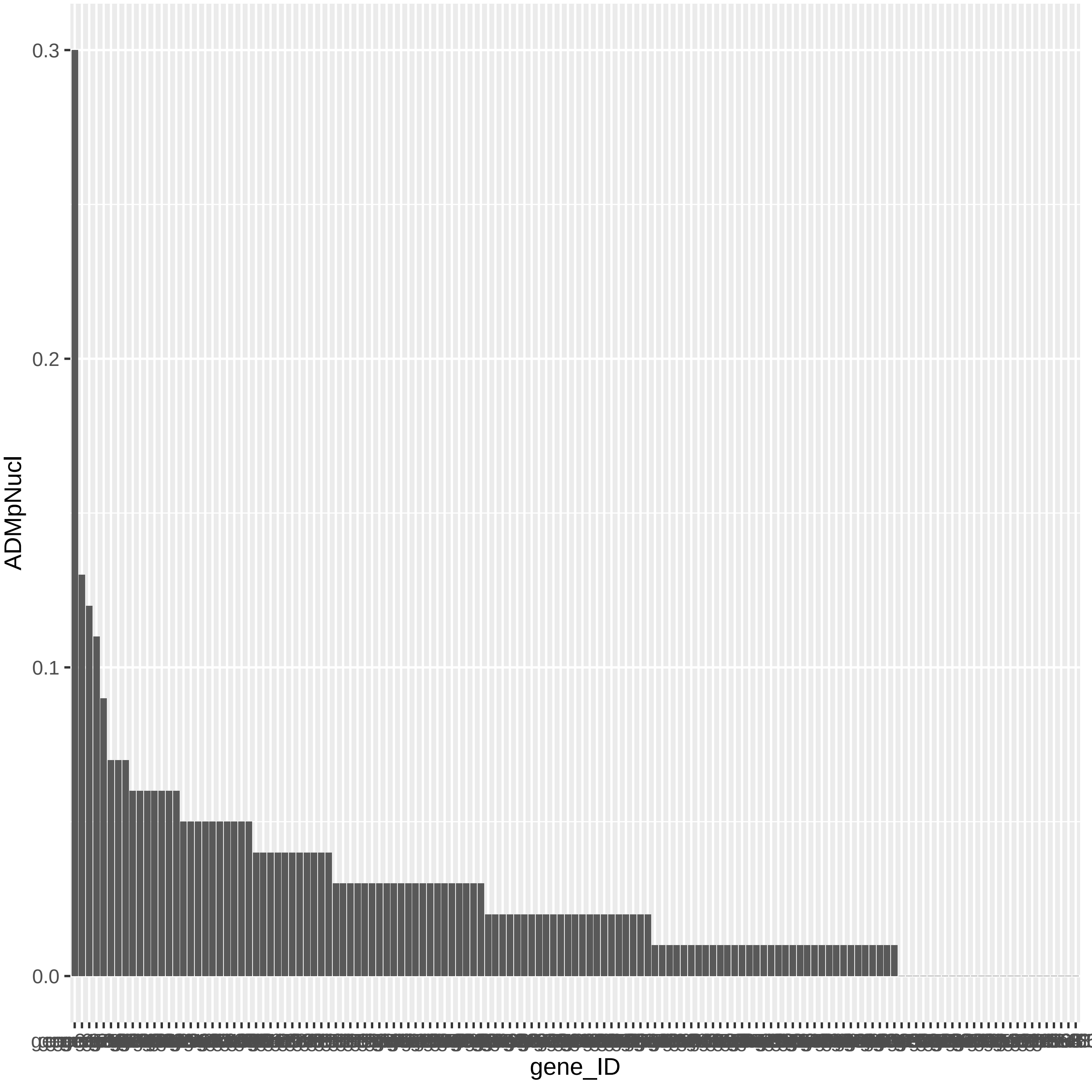

ADMpNucl
0.3
0.2
0.1
0.0
gene_ID

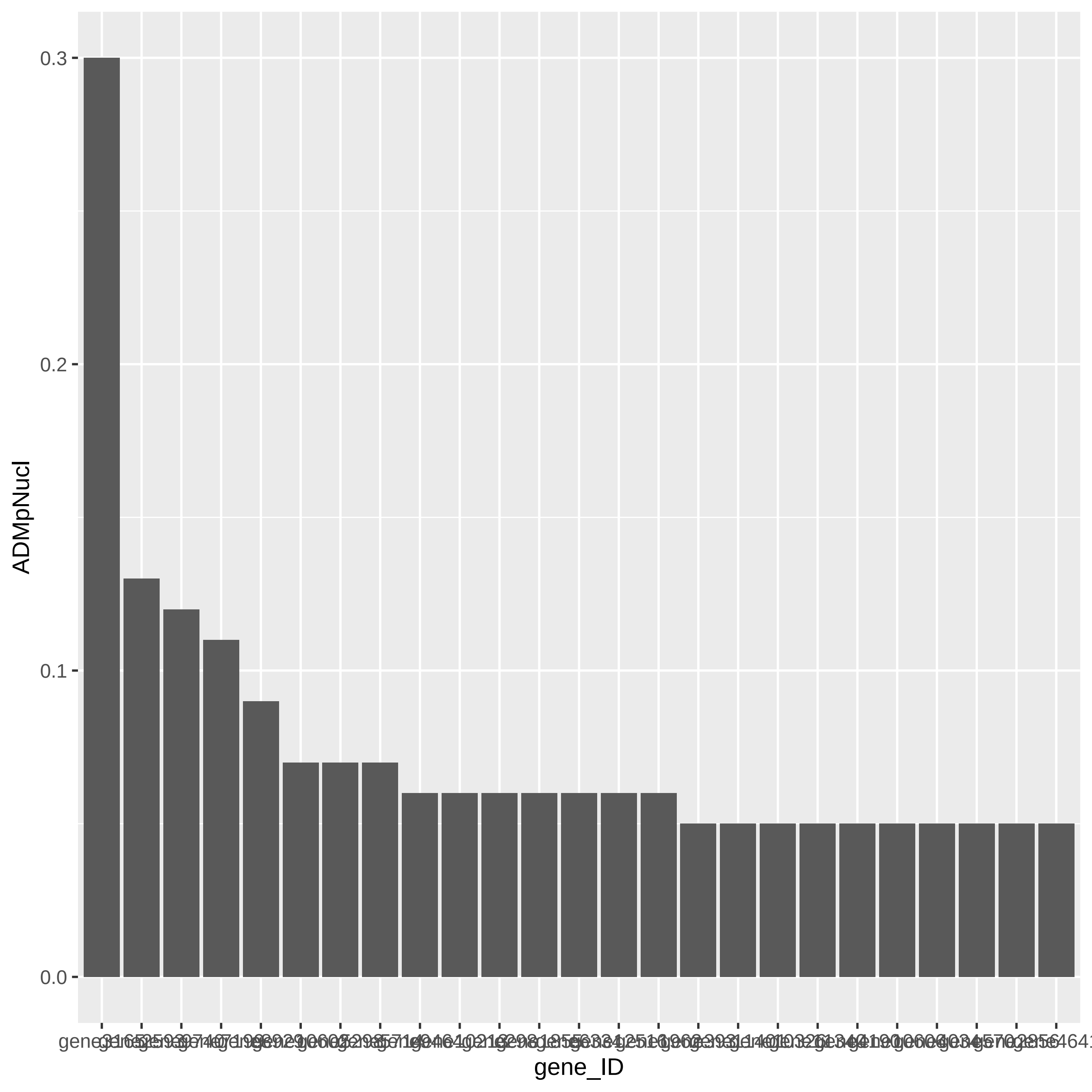

0.3
0.2
0.1
0.0
ADMpNucl
gene_ID

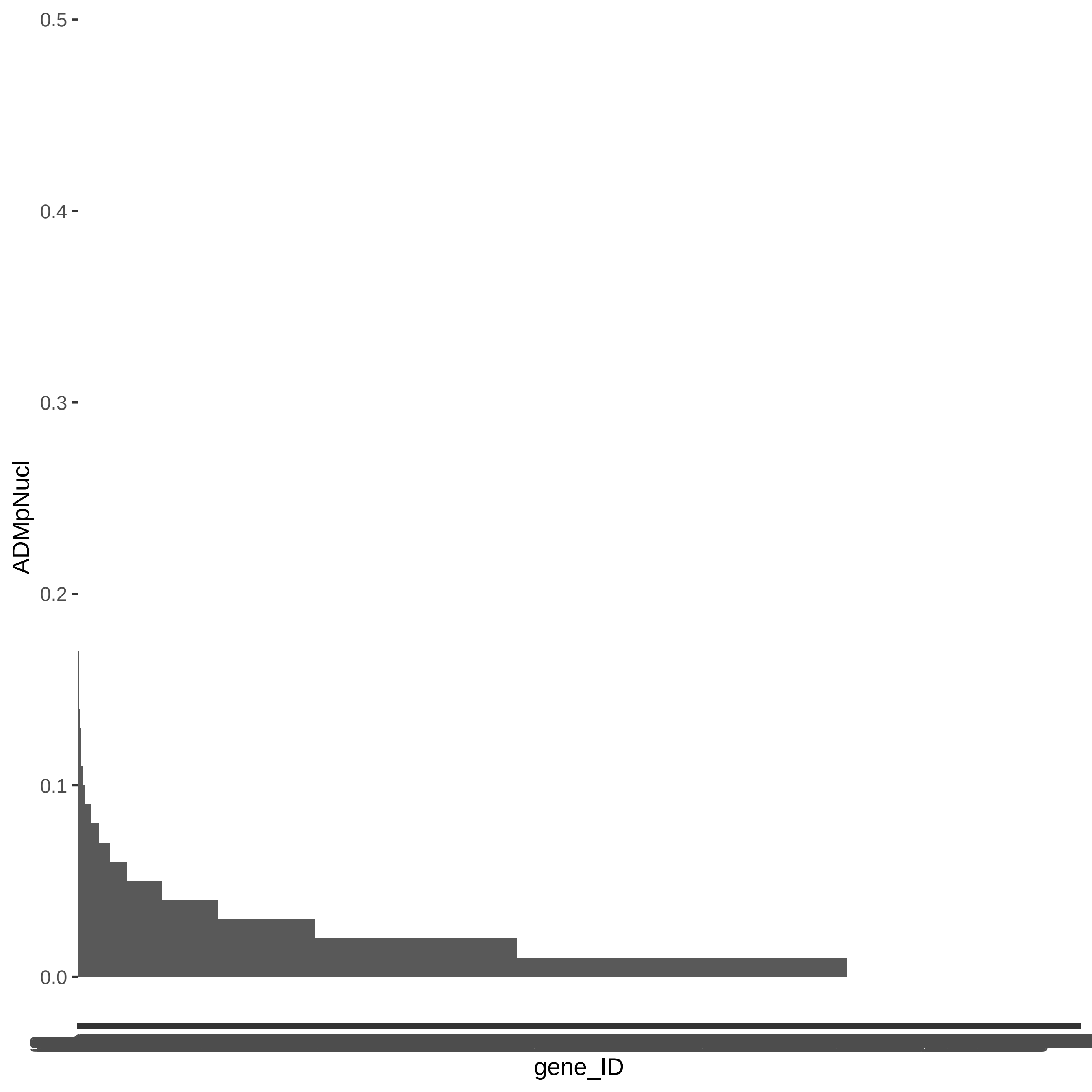
0.5
0.4
0.3
0.2
0.1
0.0
ADMpNucl
gene_ID

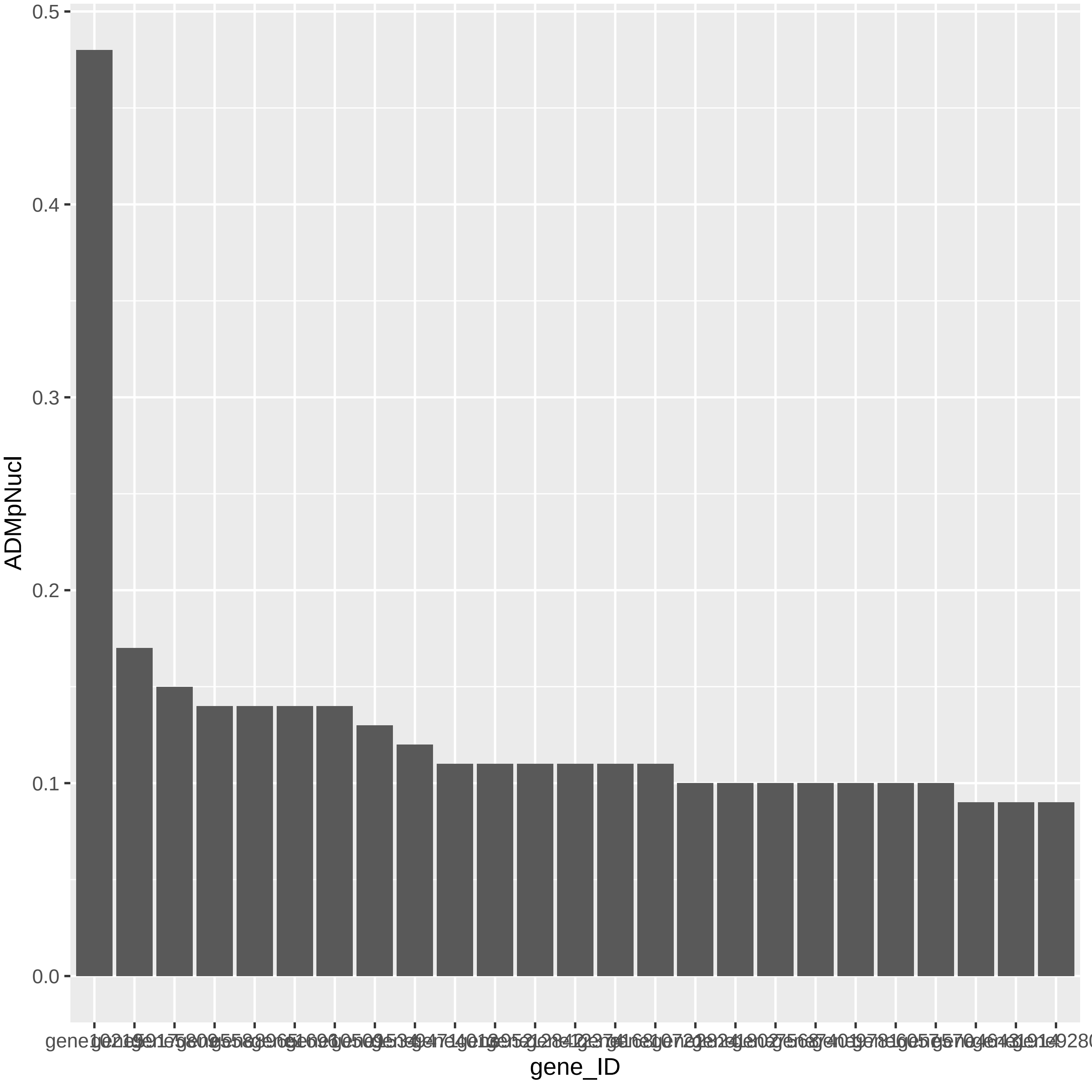
0.5
0.4
0.3
0.2
0.1
0.0
ADMpNucl
gene_ID

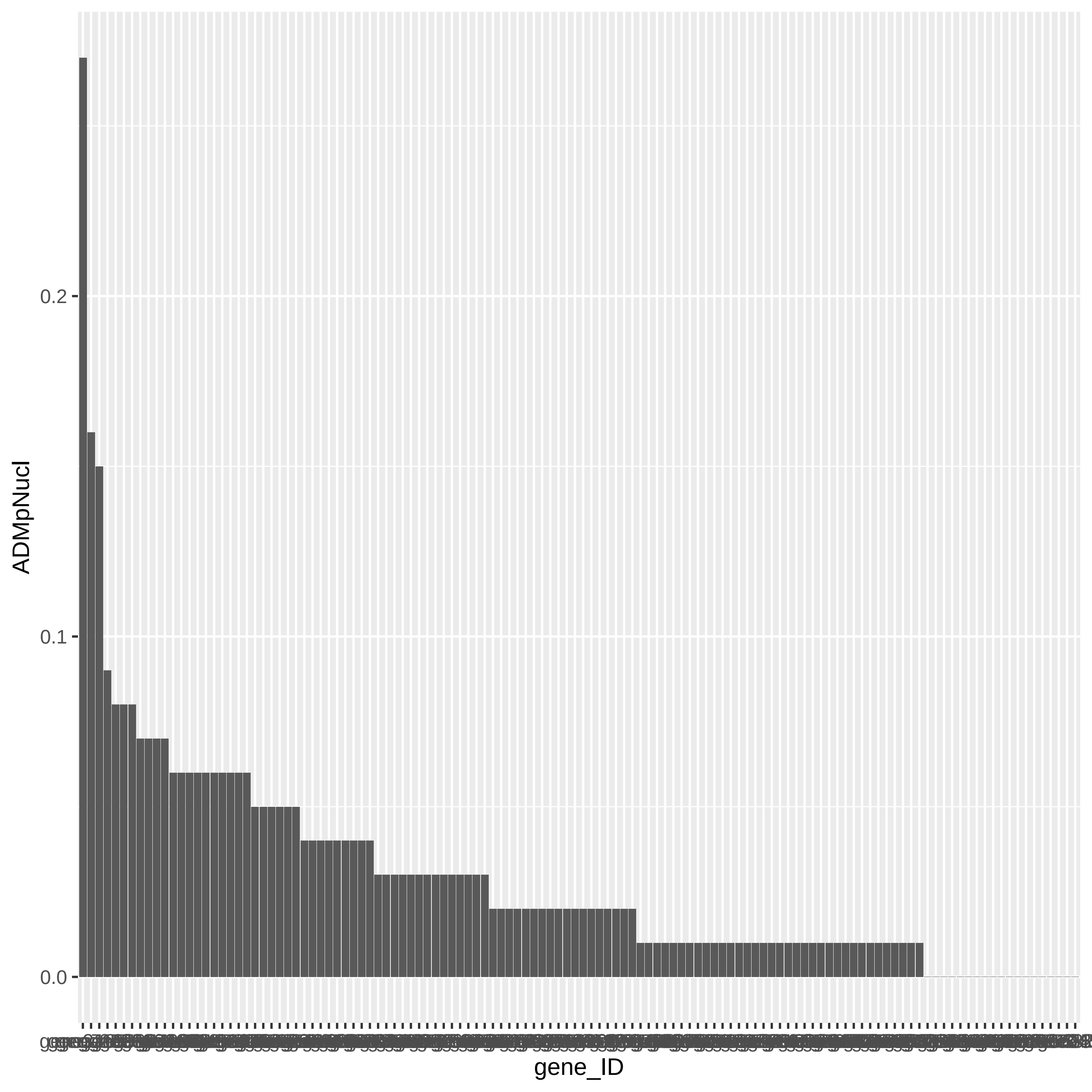
ADMpNucl
0.2
0.1
0.0
gene_ID

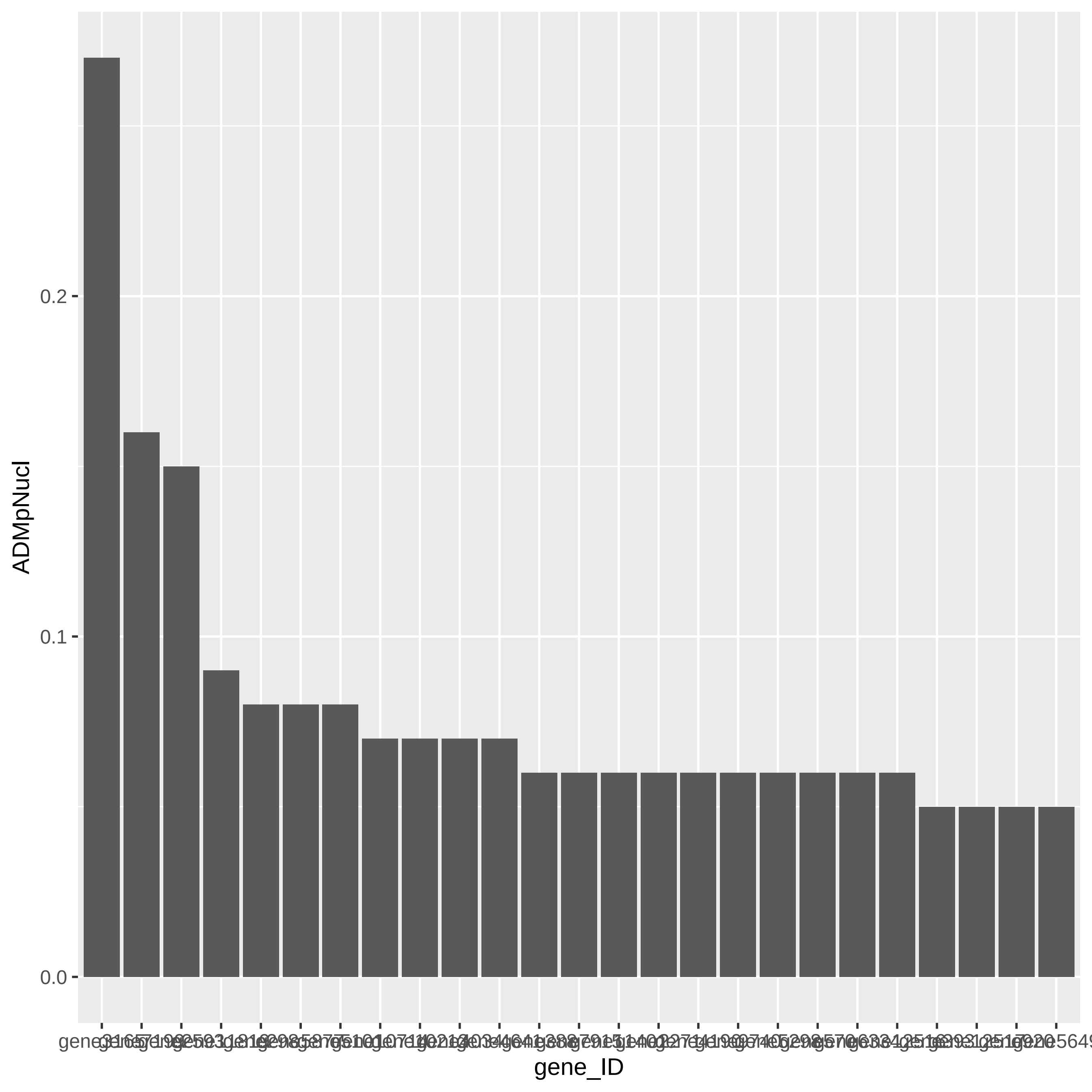
ADMpNucl
0.2
0.1
0.0
gene_ID

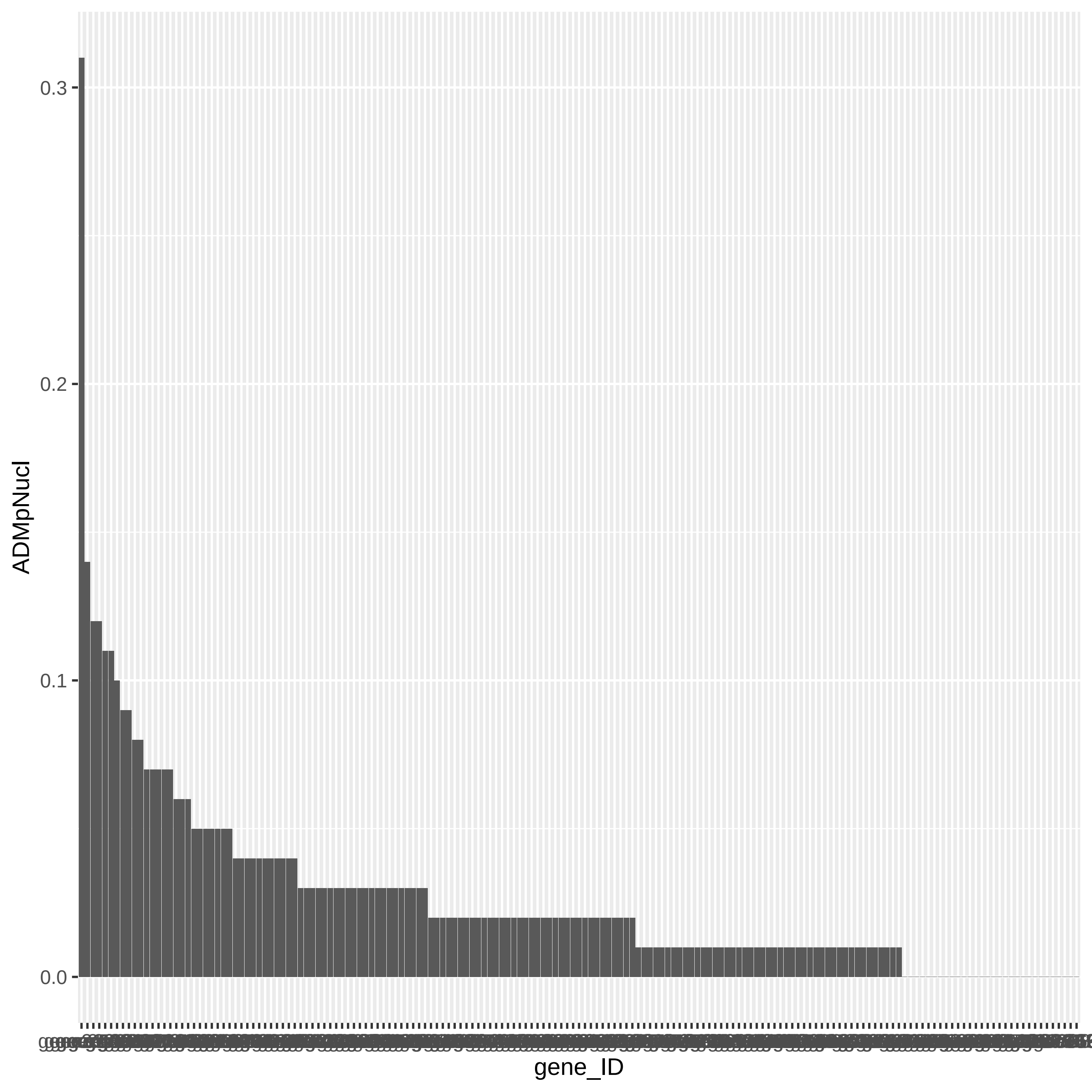
0.3
0.2
0.1
0.0
ADMpNucl
gene_ID

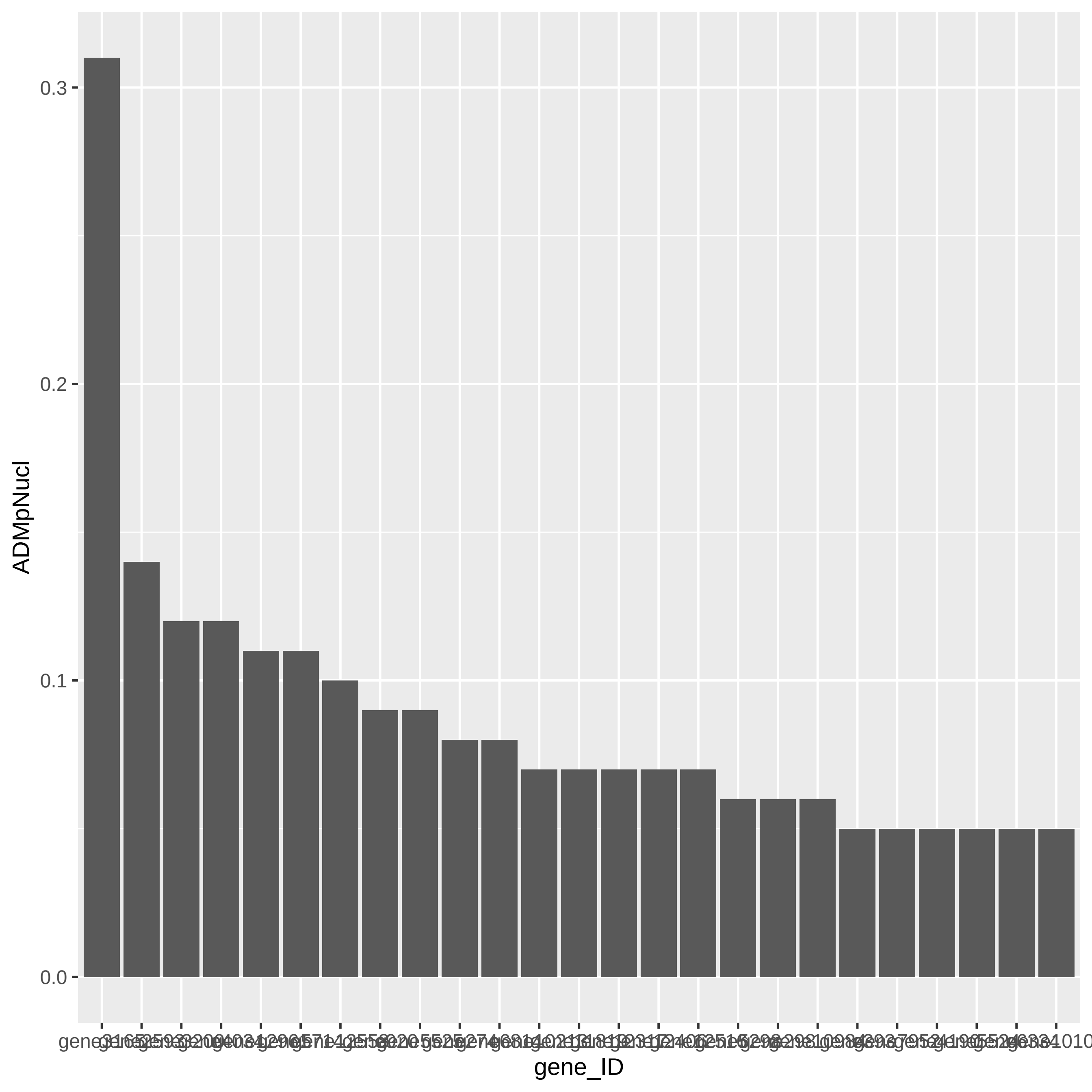
ADMpNucl
0.3
0.2
0.1
0.0
gene_ID

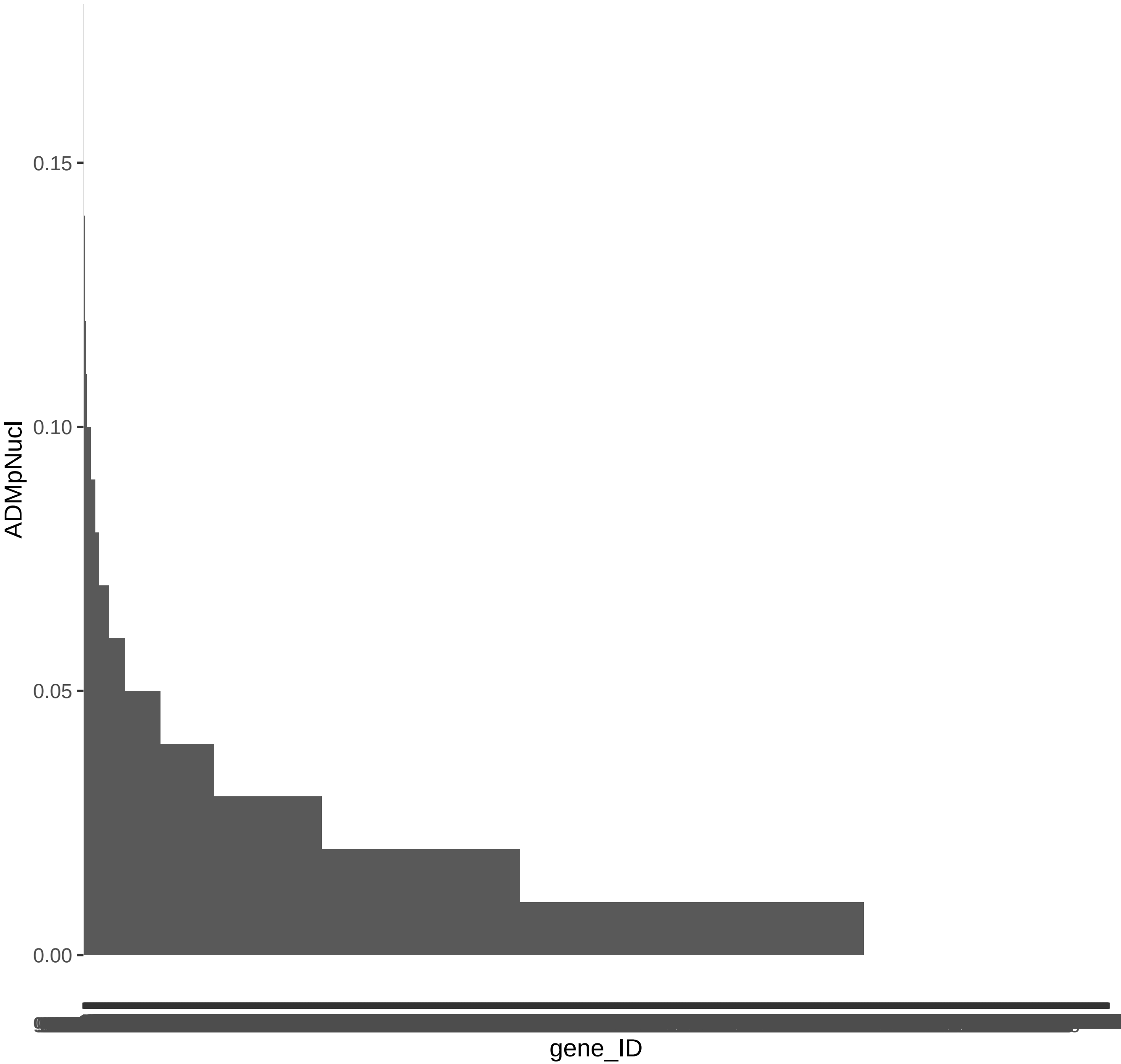

0.15
0.10
0.05
0.00
ADMpNucl
gene_ID

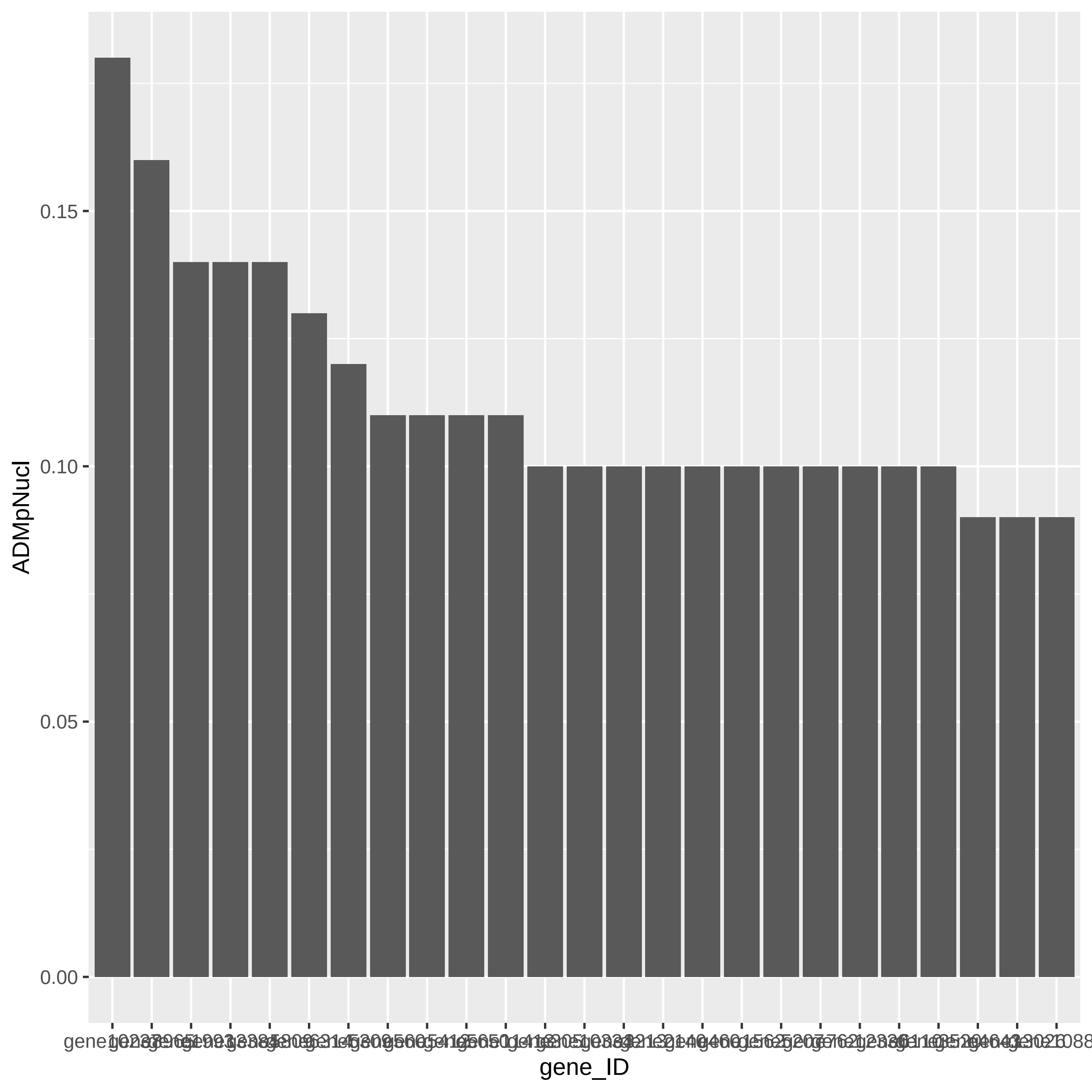

0.15
0.10
0.05
0.00
ADMpNucl
gene_ID

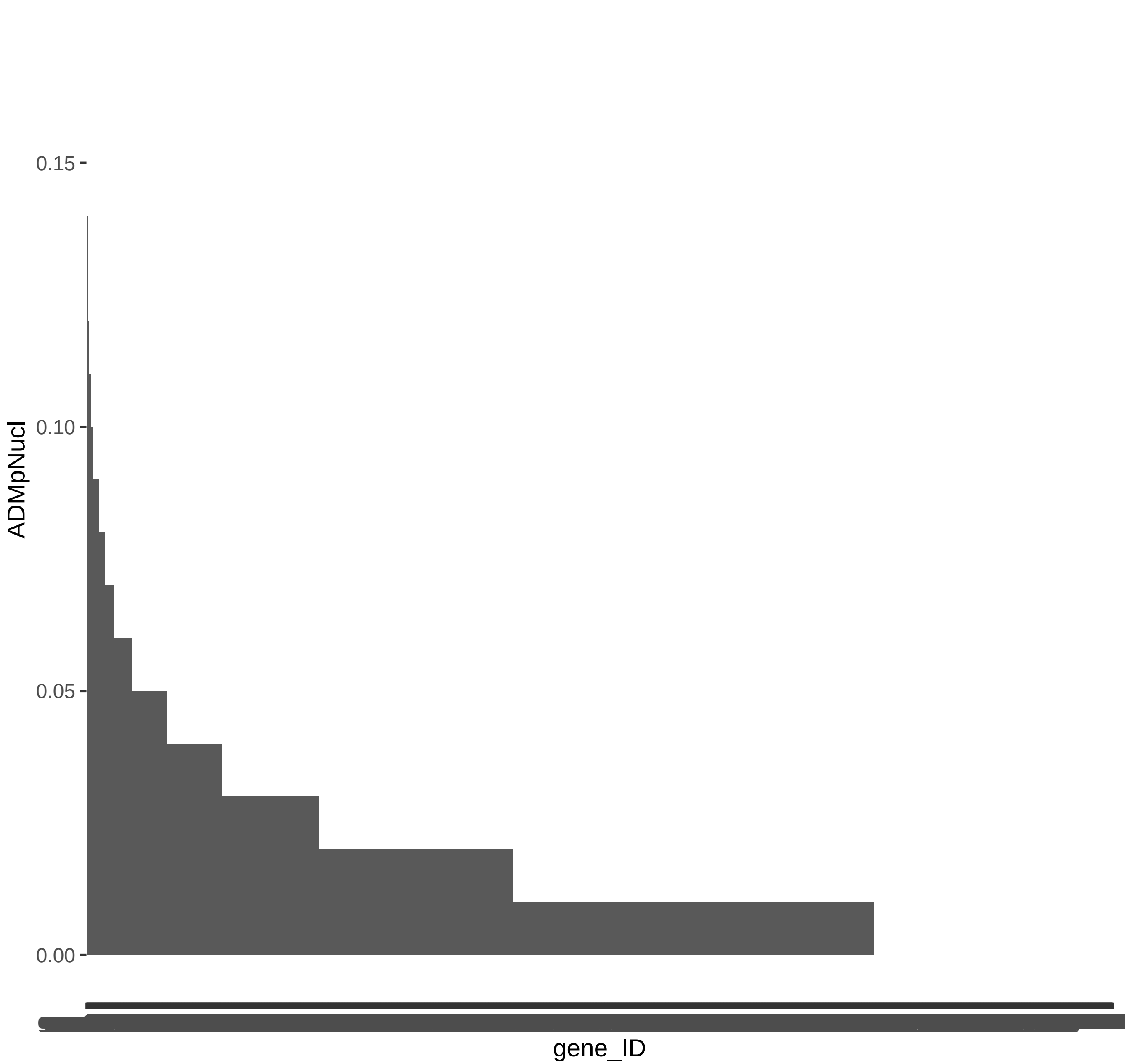
ADMpNucl
0.15
0.10
0.05
0.00
gene_ID

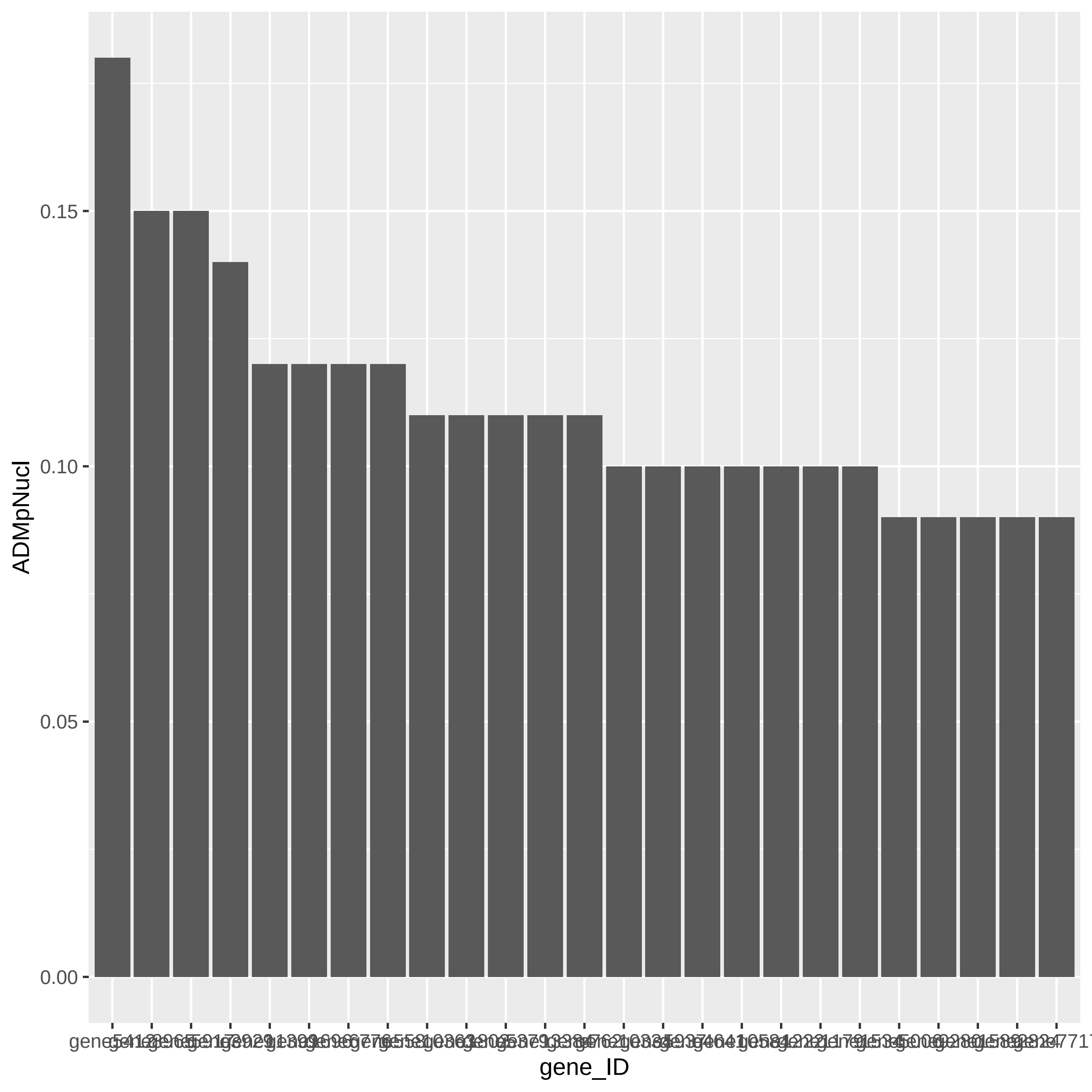
ADMpNucl
0.00
0.05
0.10
0.15
gene_ID

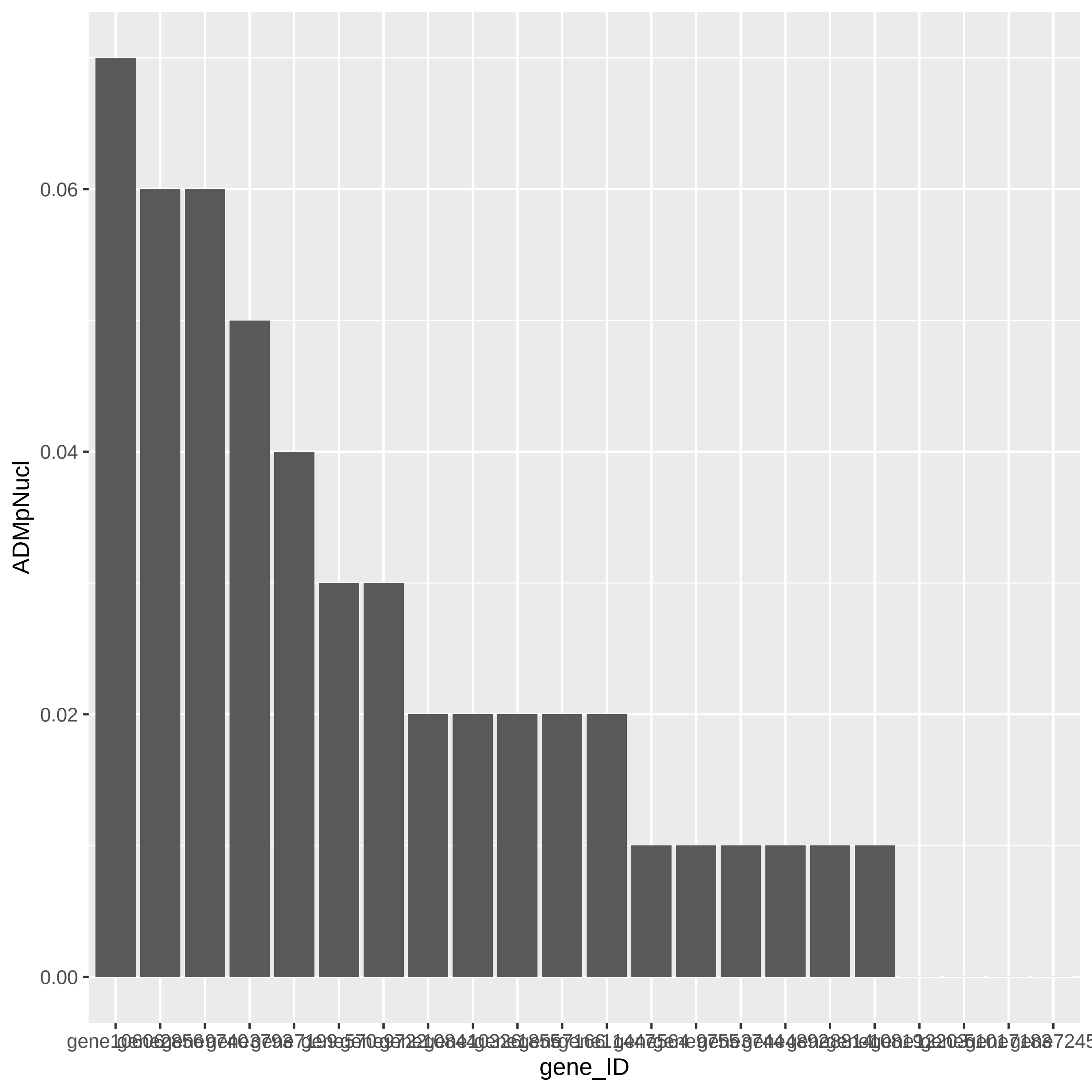
ADMpNucl
0.06
0.04
0.02
0.00
gene_ID

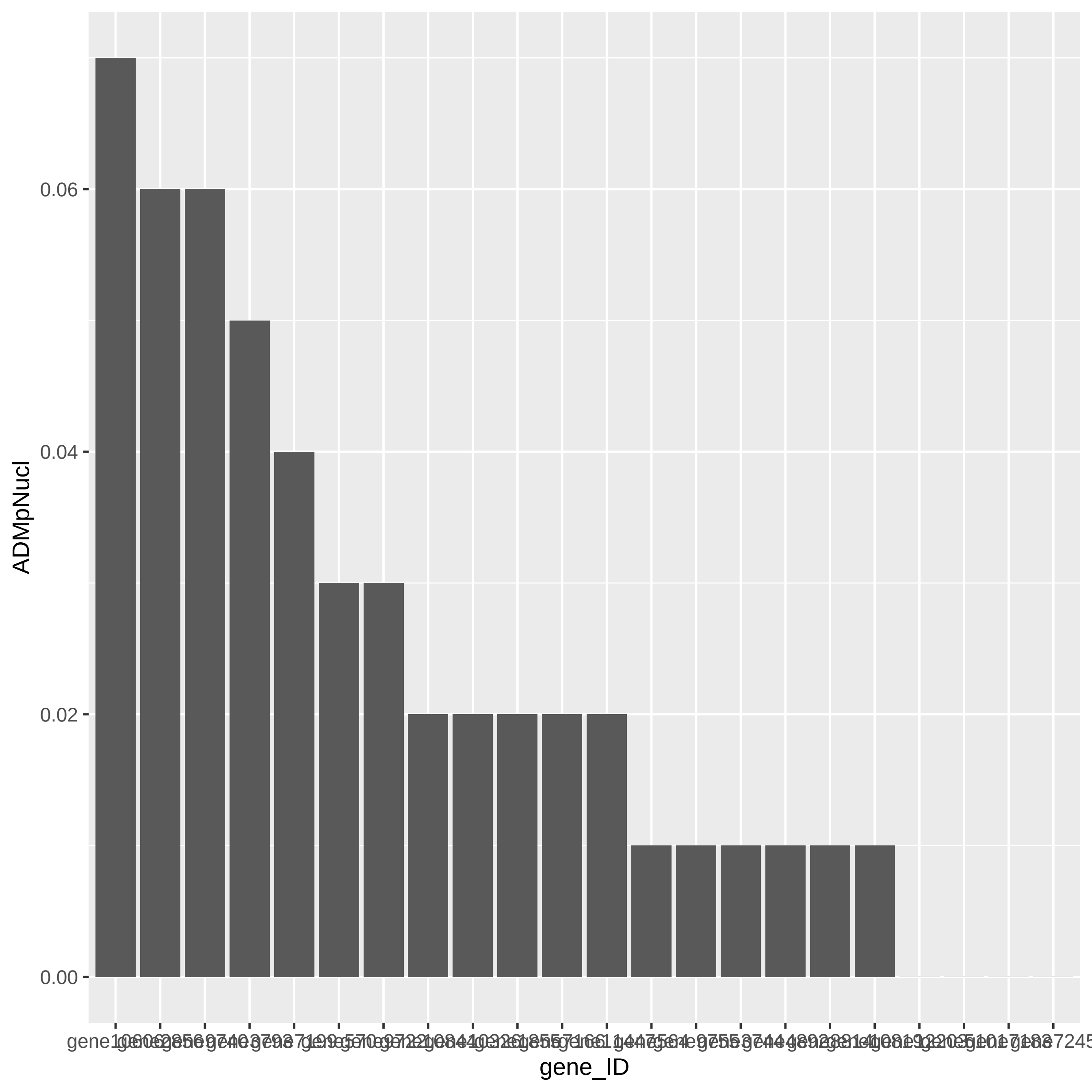

0.06
0.04
0.02
0.00
ADMpNucl
gene_ID

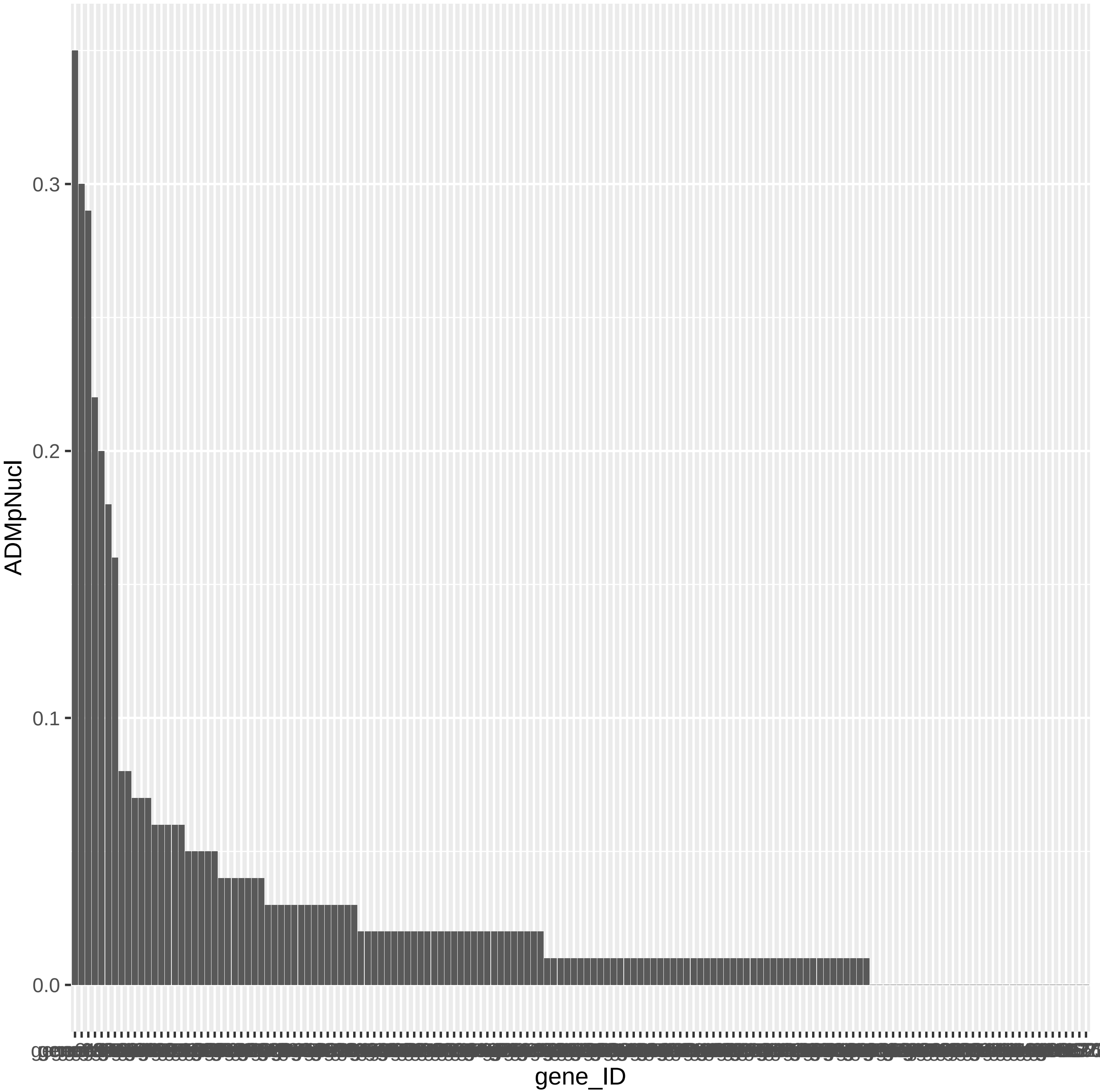

0.3
0.2
0.1
0.0
ADMpNucl
gene_ID

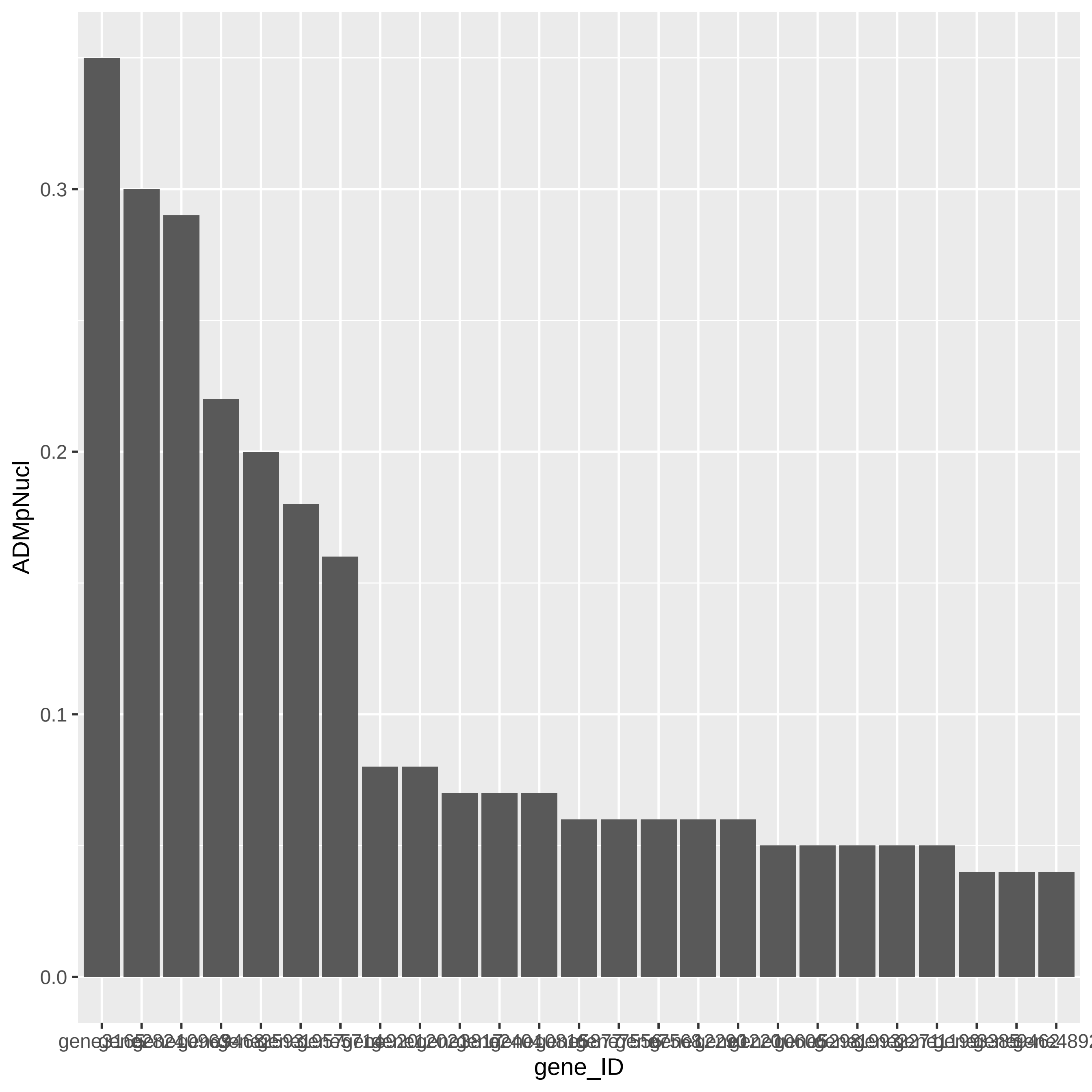
ADMpNucl
0.0
0.1
0.2
0.3
gene_ID

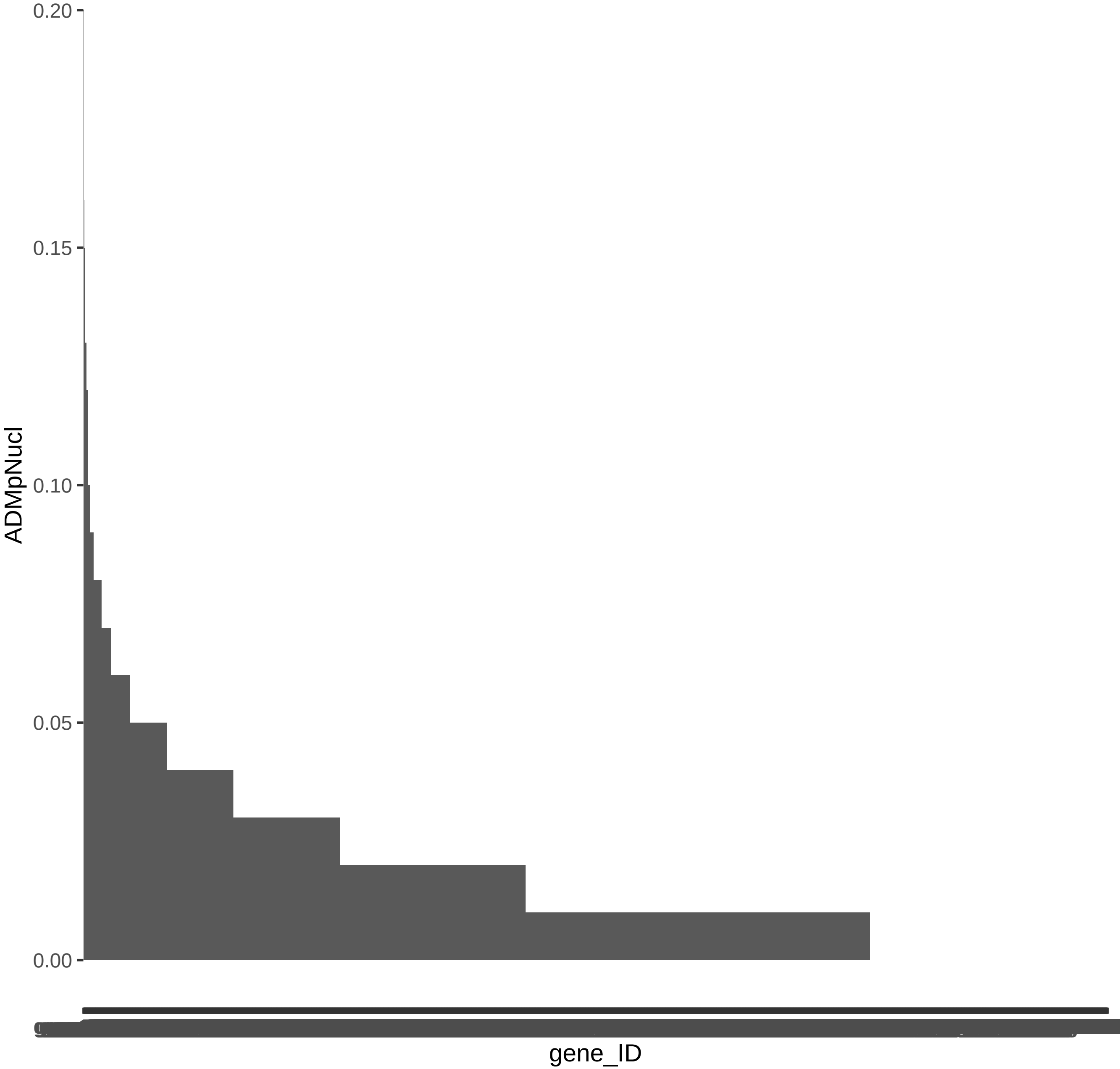

0.20
0.15
0.10
0.05
0.00
ADMpNucl
gene_ID

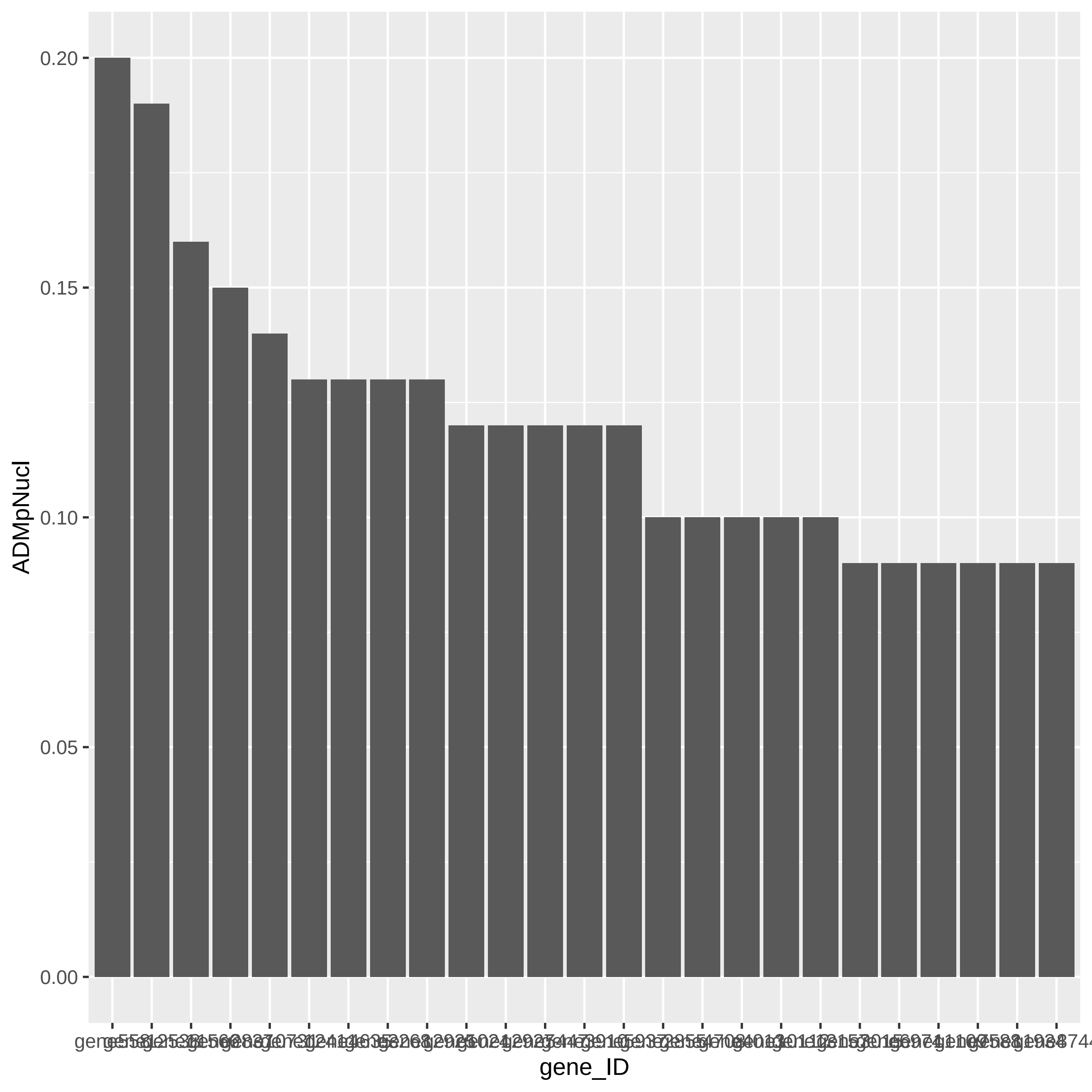
0.20
0.15
0.10
0.05
0.00
ADMpNucl
gene_ID

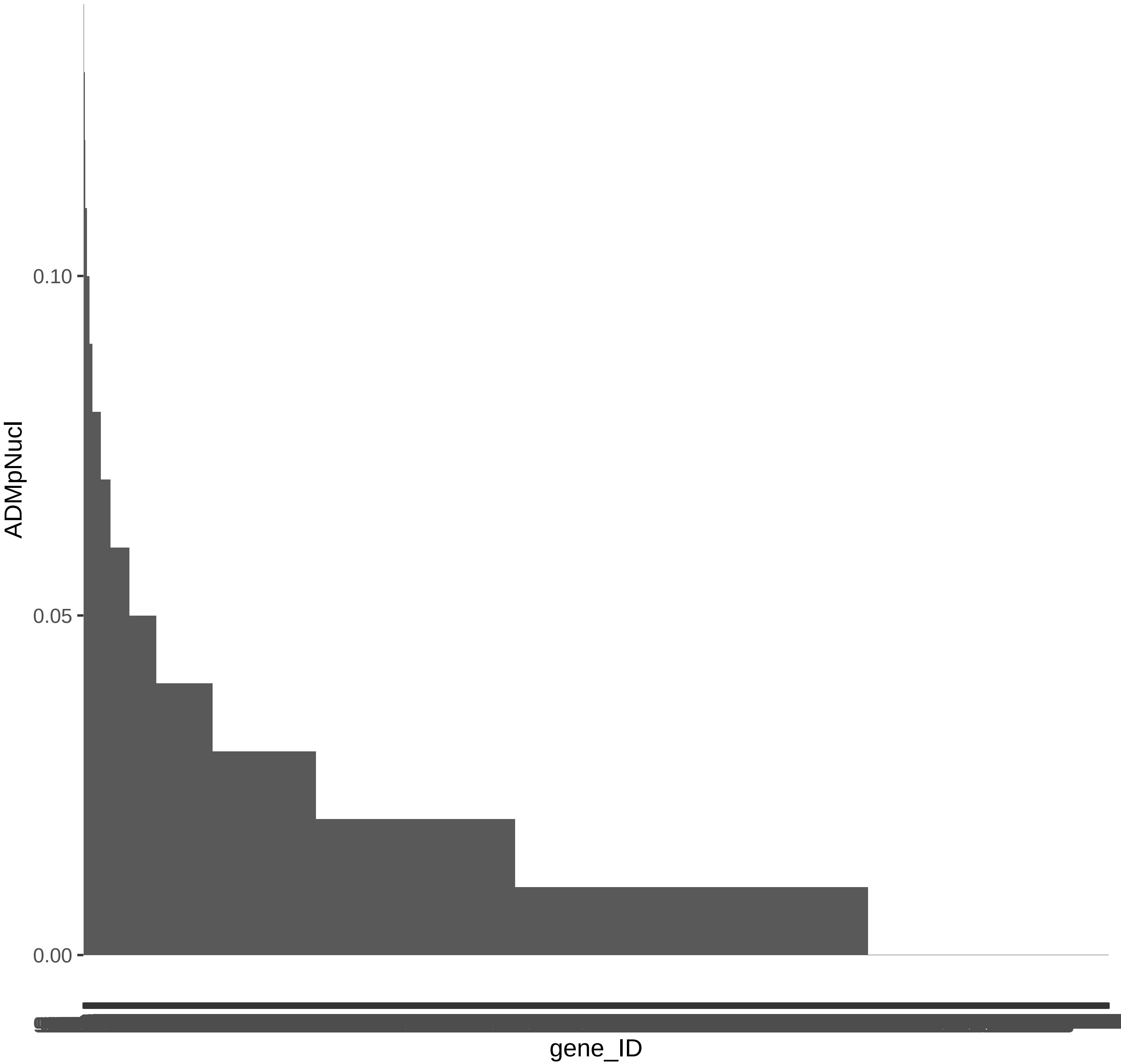

ADMpNucl
0.10
0.05
0.00
gene_ID

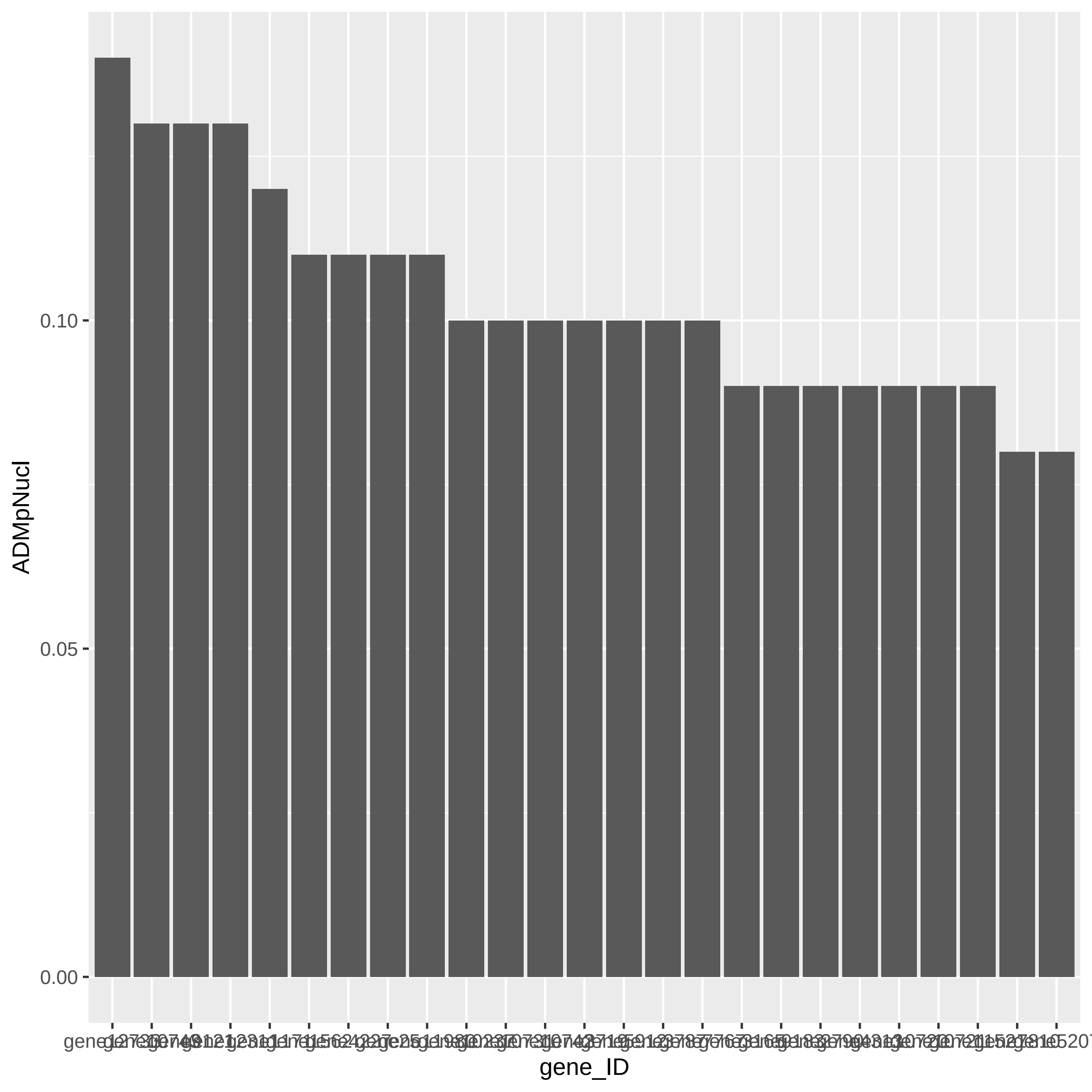
ADMpNucl
0.00
0.05
0.10
gene_ID

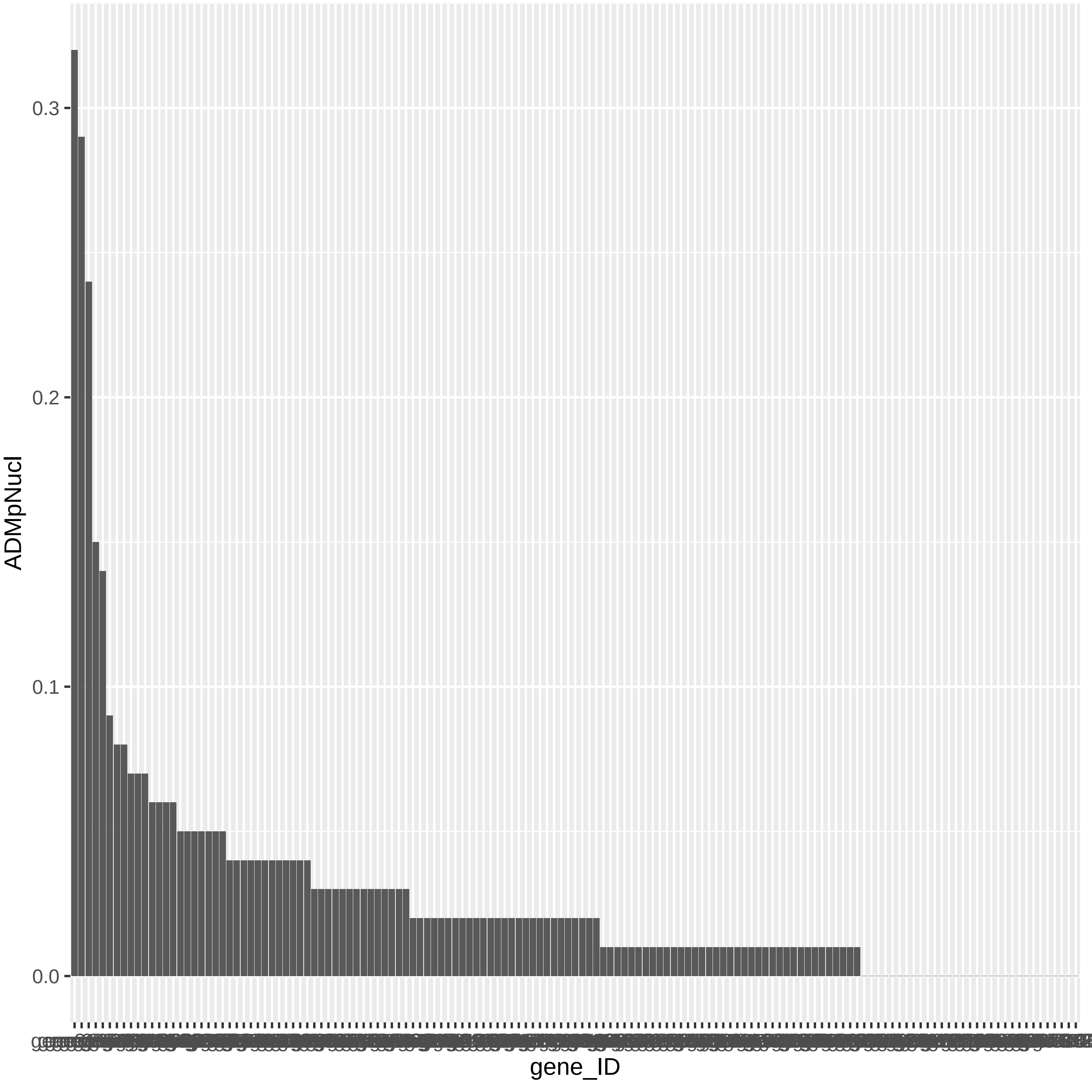
0.3
0.2
0.1
0.0
ADMpNucl
gene_ID

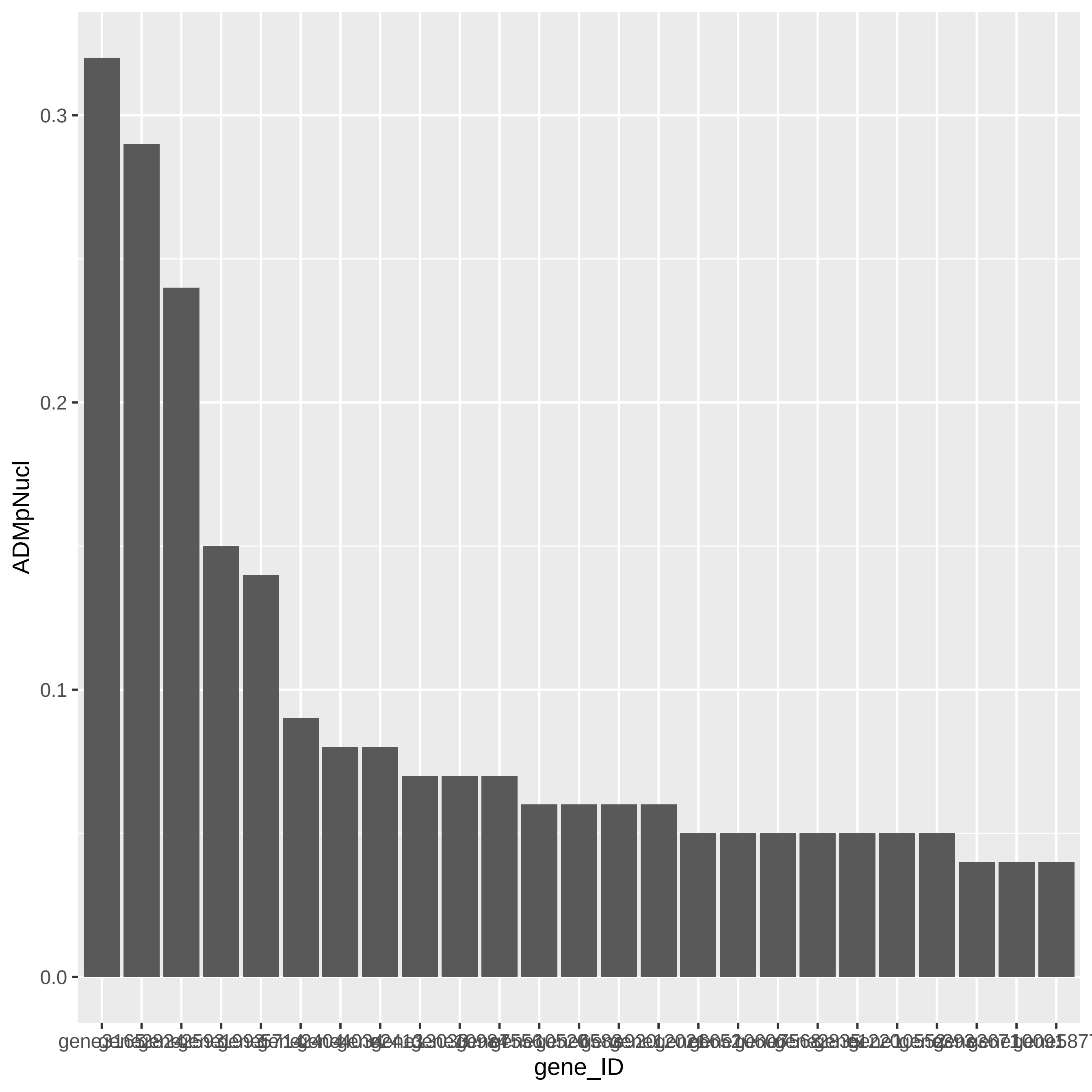

0.3
0.2
0.1
0.0
ADMpNucl
gene_ID

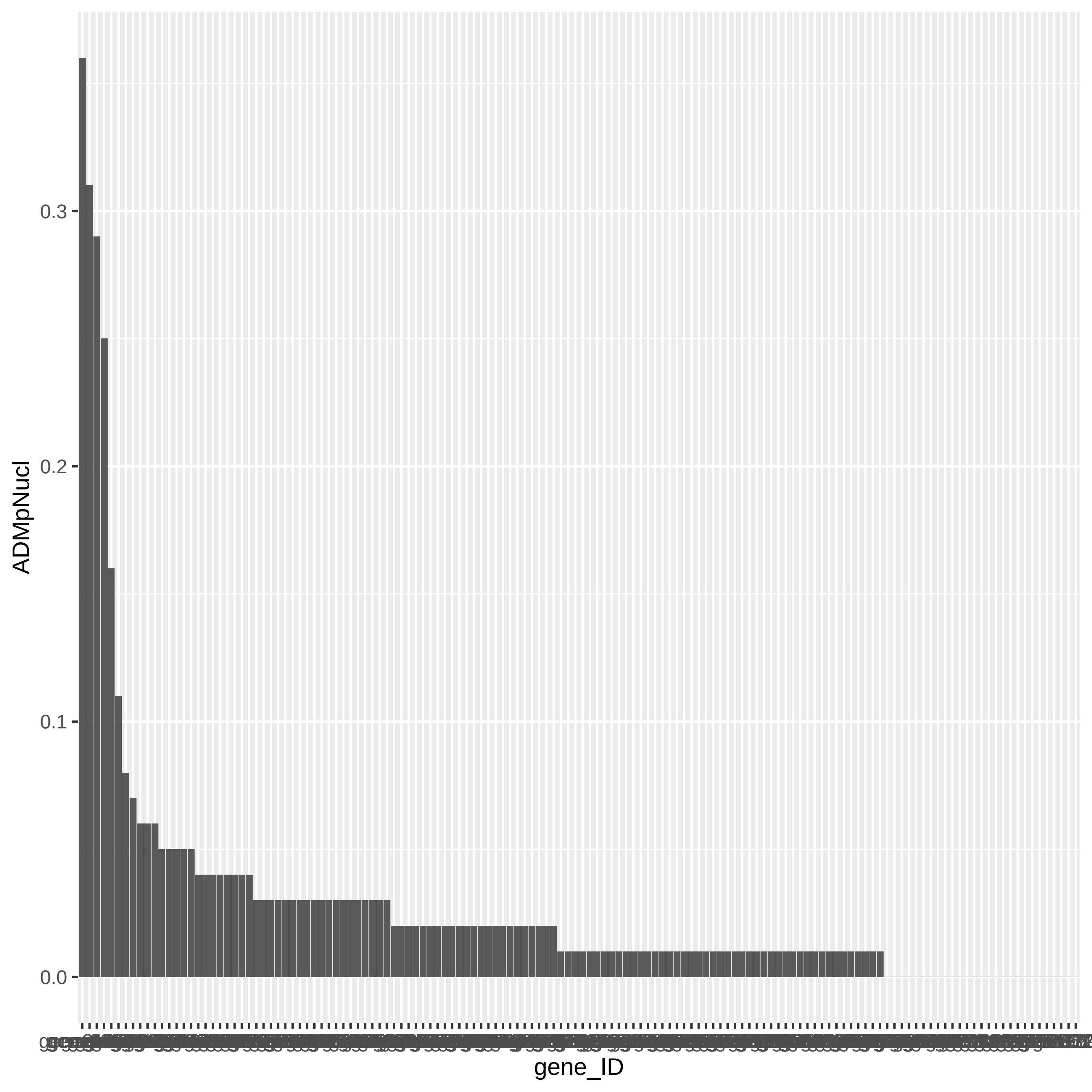

ADMpNucl
0.3
0.2
0.1
0.0
gene_ID

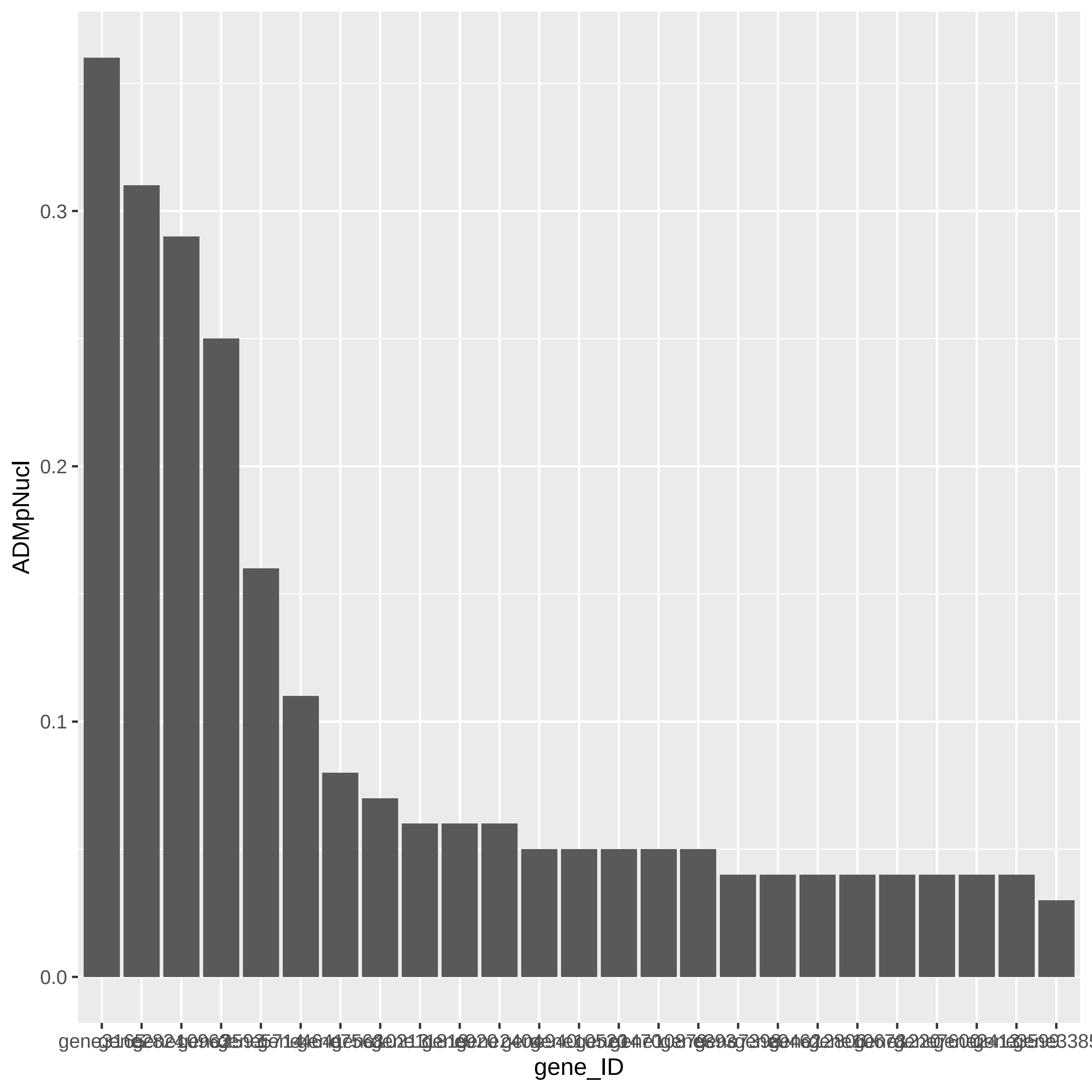
ADMpNucl
0.3
0.2
0.1
0.0
gene_ID

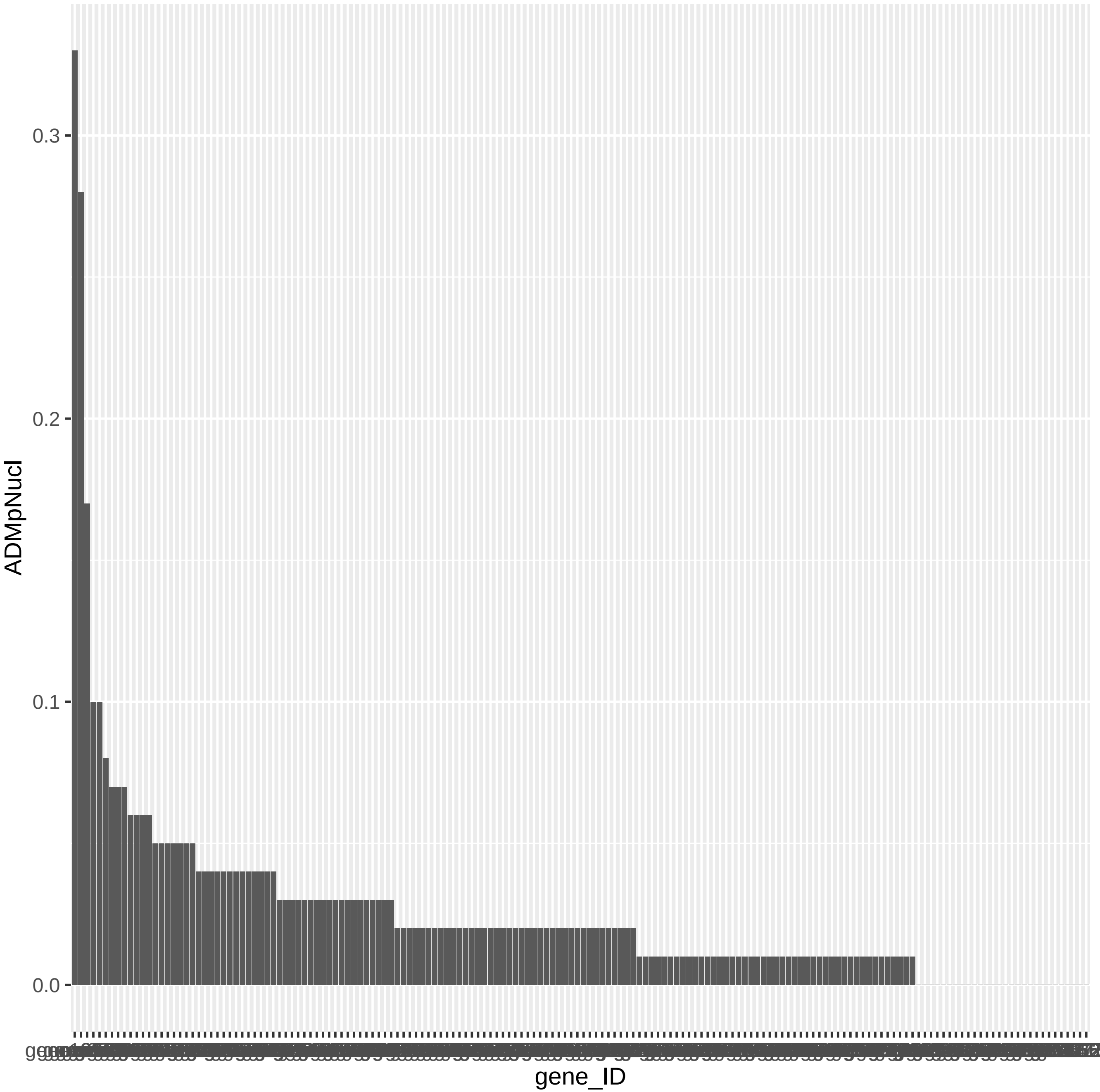
ADMpNucl
0.3
0.2
0.1
0.0
gene_ID

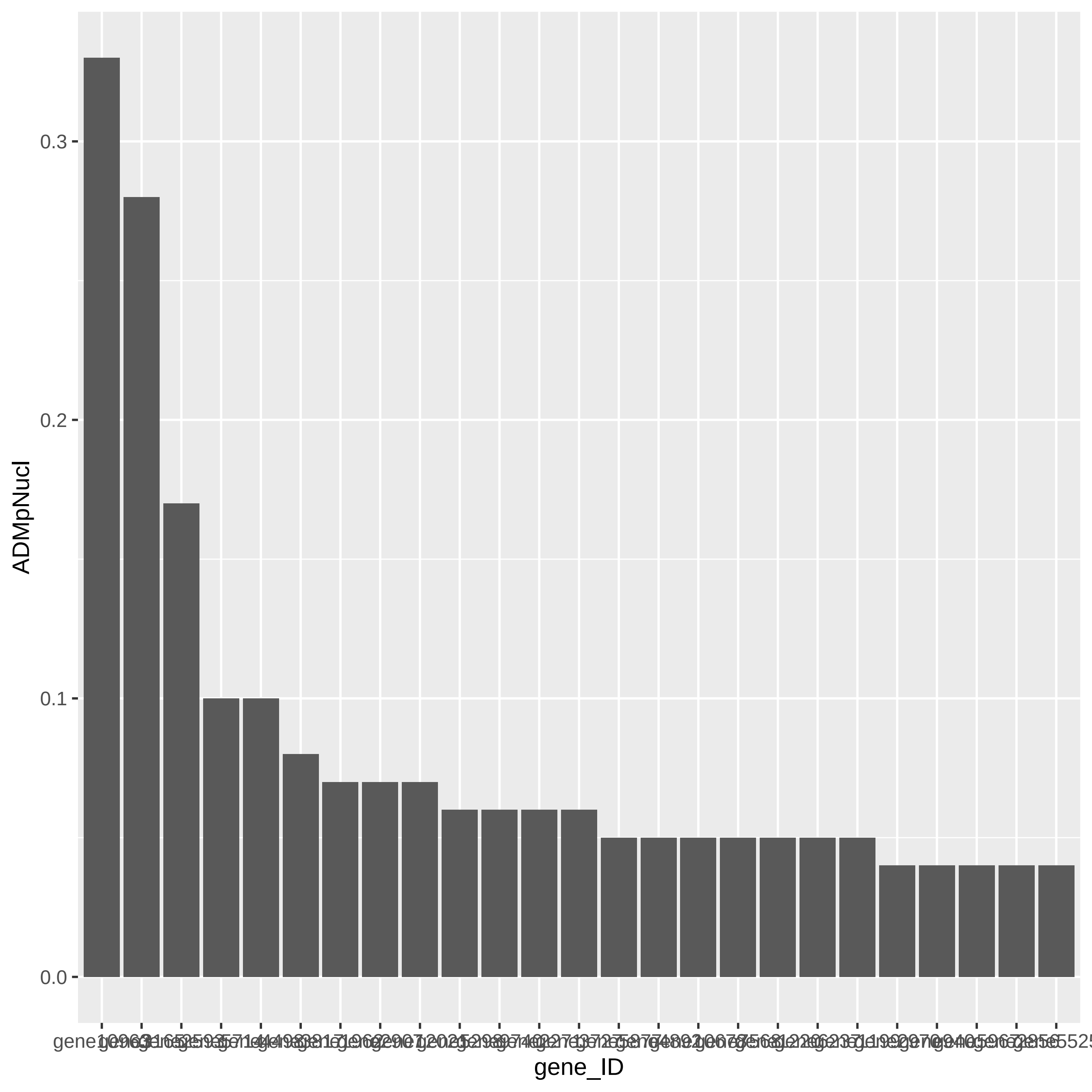

ADMpNucl
0.0
0.1
0.2
0.3
gene_ID

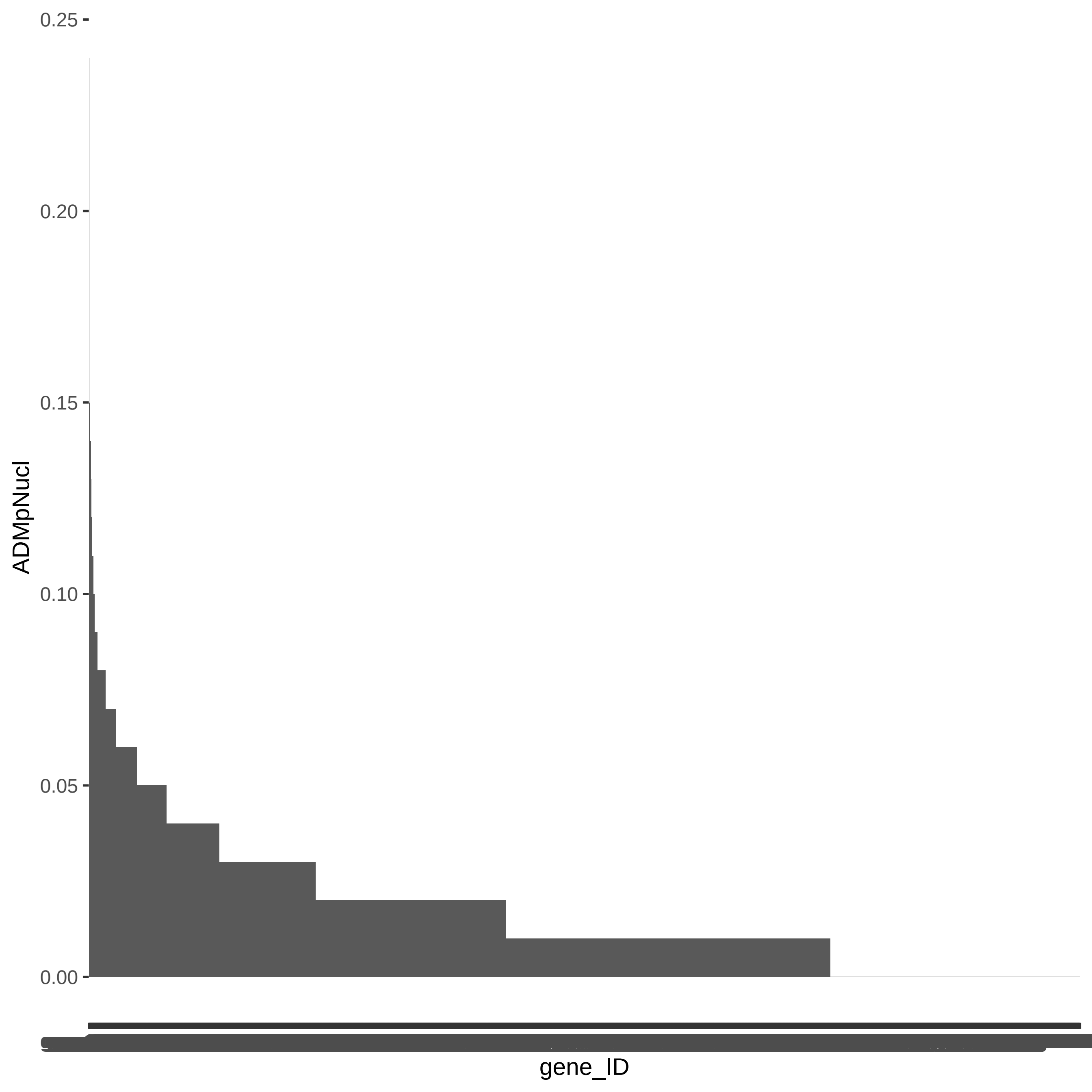

0.25
0.20
0.15
0.10
0.05
0.00
ADMpNucl
gene_ID

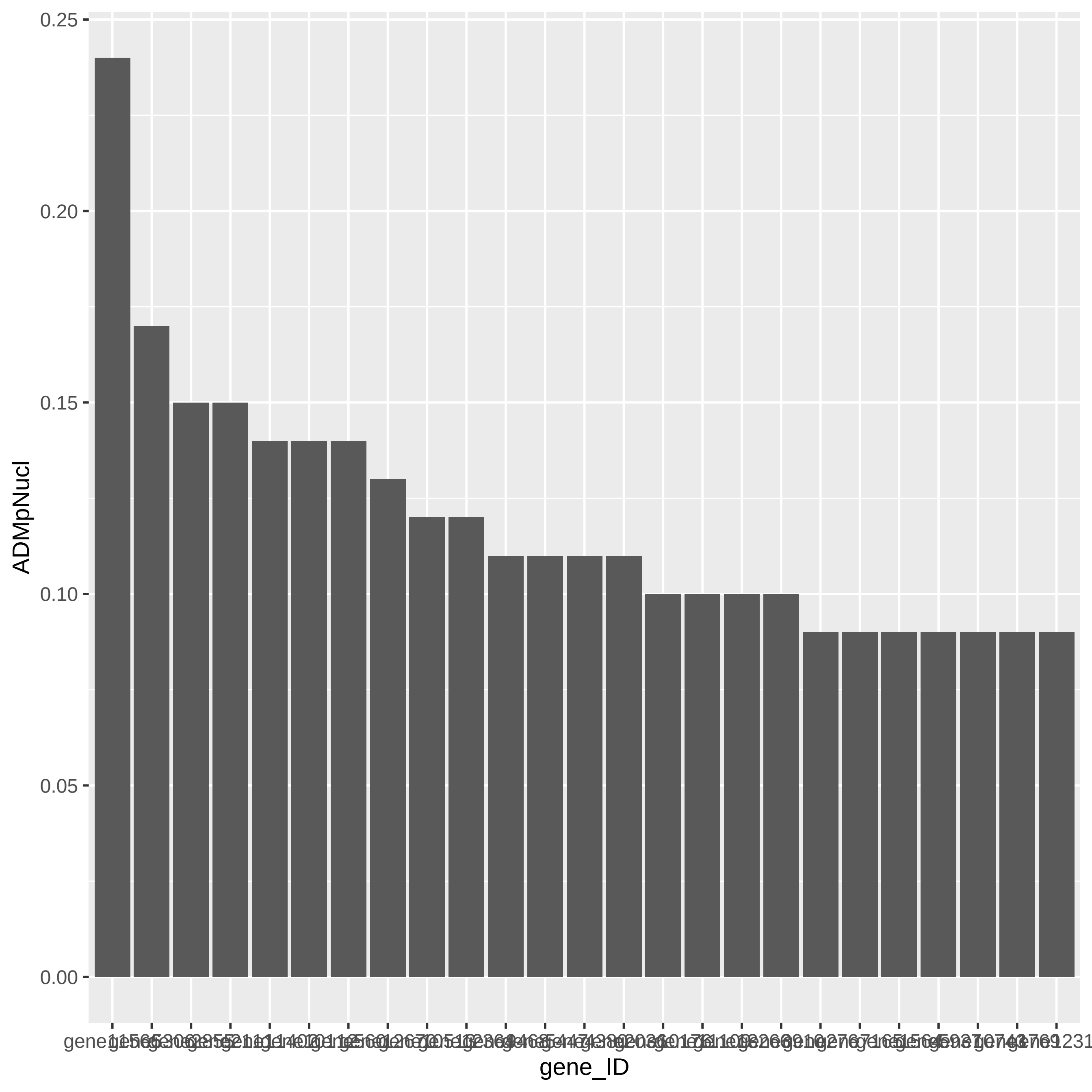

0.25
0.20
0.15
0.10
0.05
0.00
ADMpNucl
gene_ID

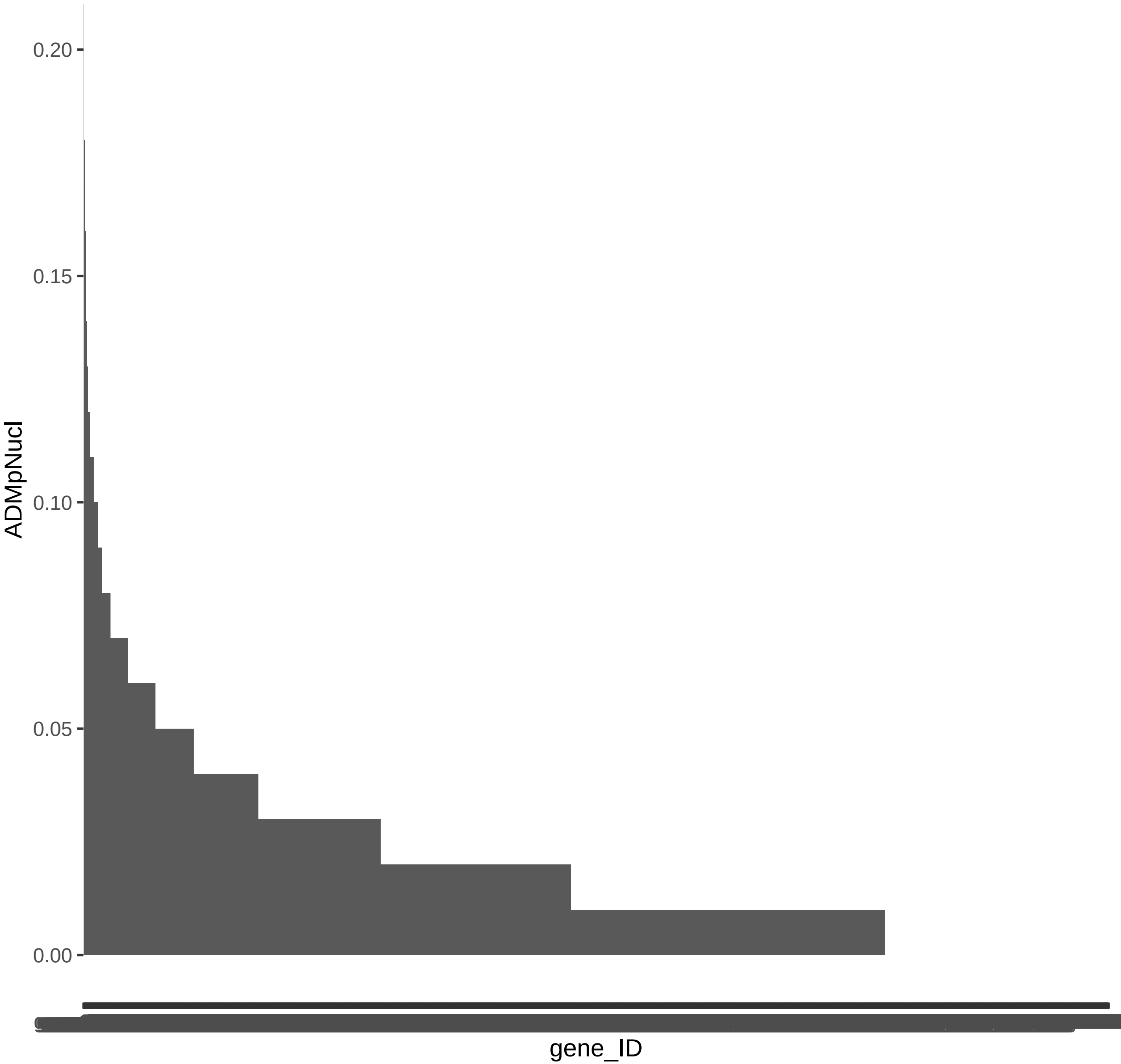

ADMpNucl
0.20
0.15
0.10
0.05
0.00
gene_ID

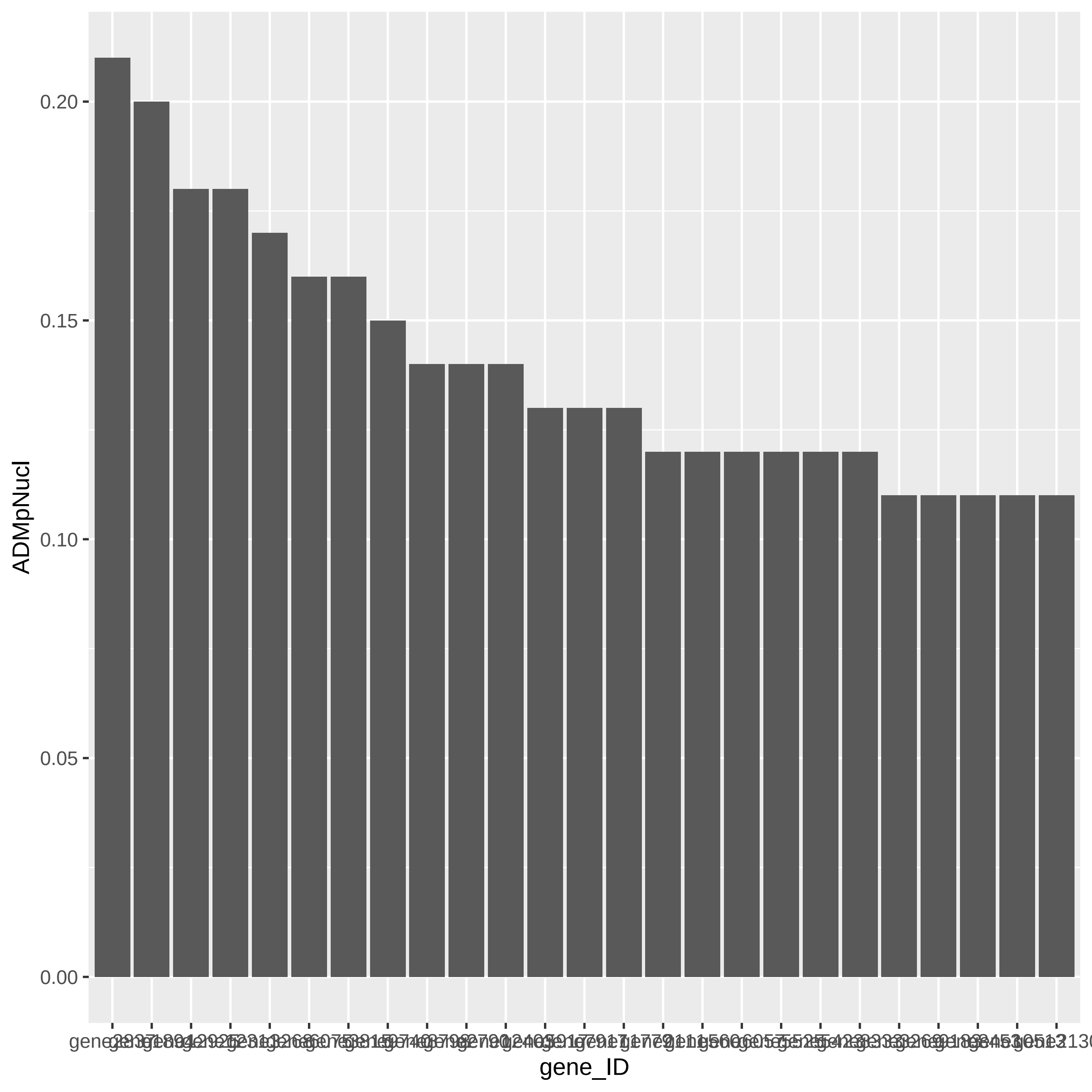
0.20
0.15
0.10
0.05
0.00
ADMpNucl
gene_ID

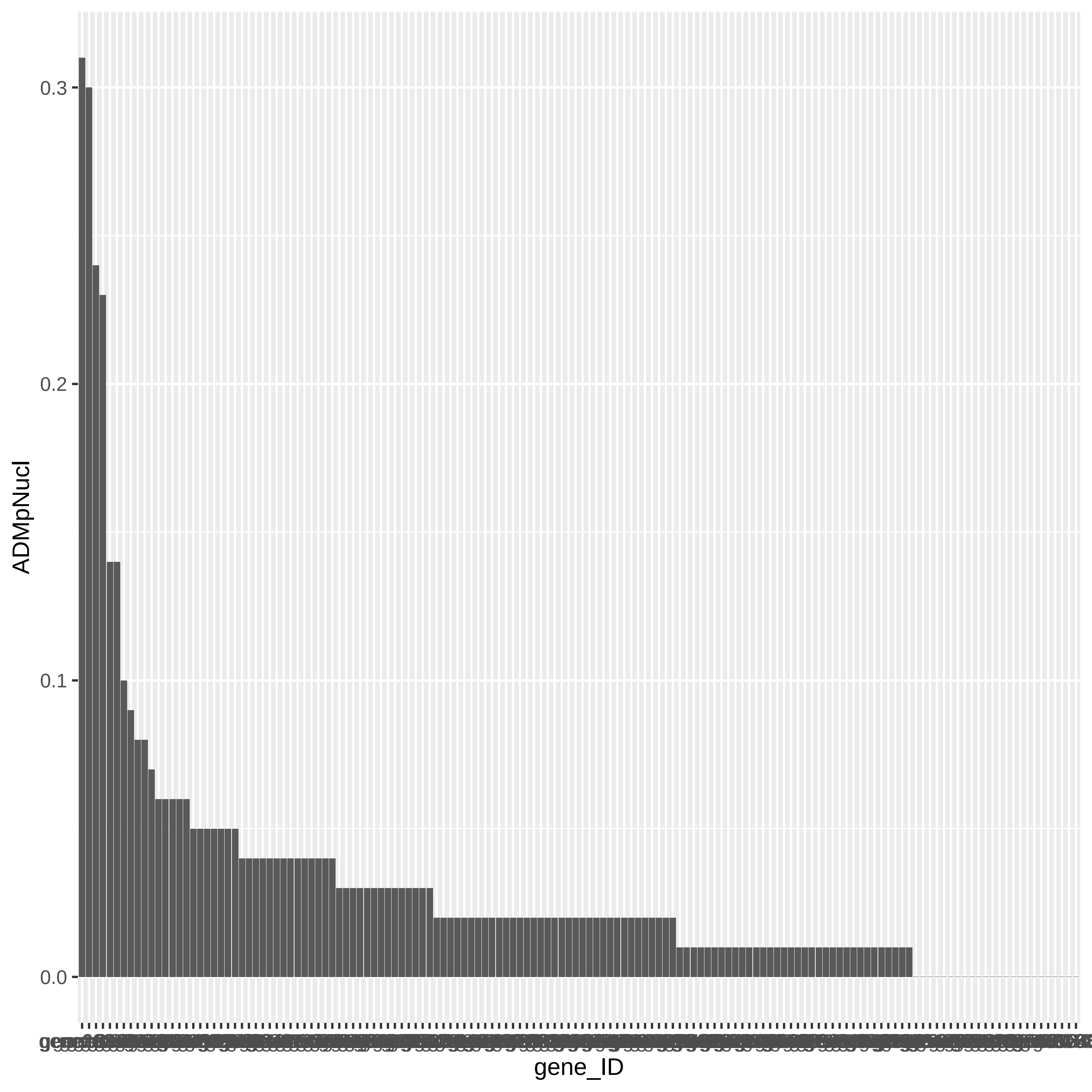

ADMpNucl
0.3
0.2
0.1
0.0
gene_ID

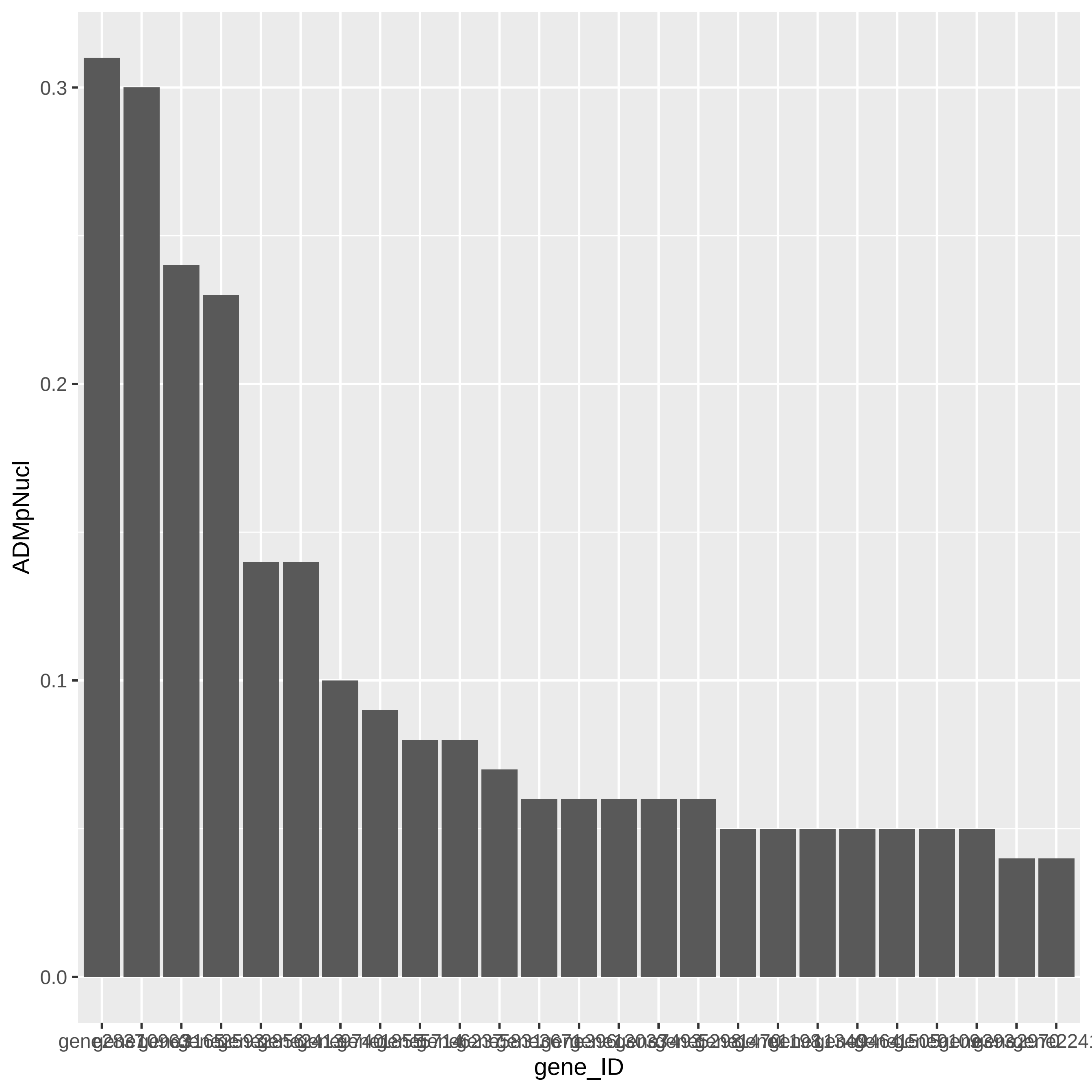
0.3
0.2
0.1
0.0
ADMpNucl
gene_ID

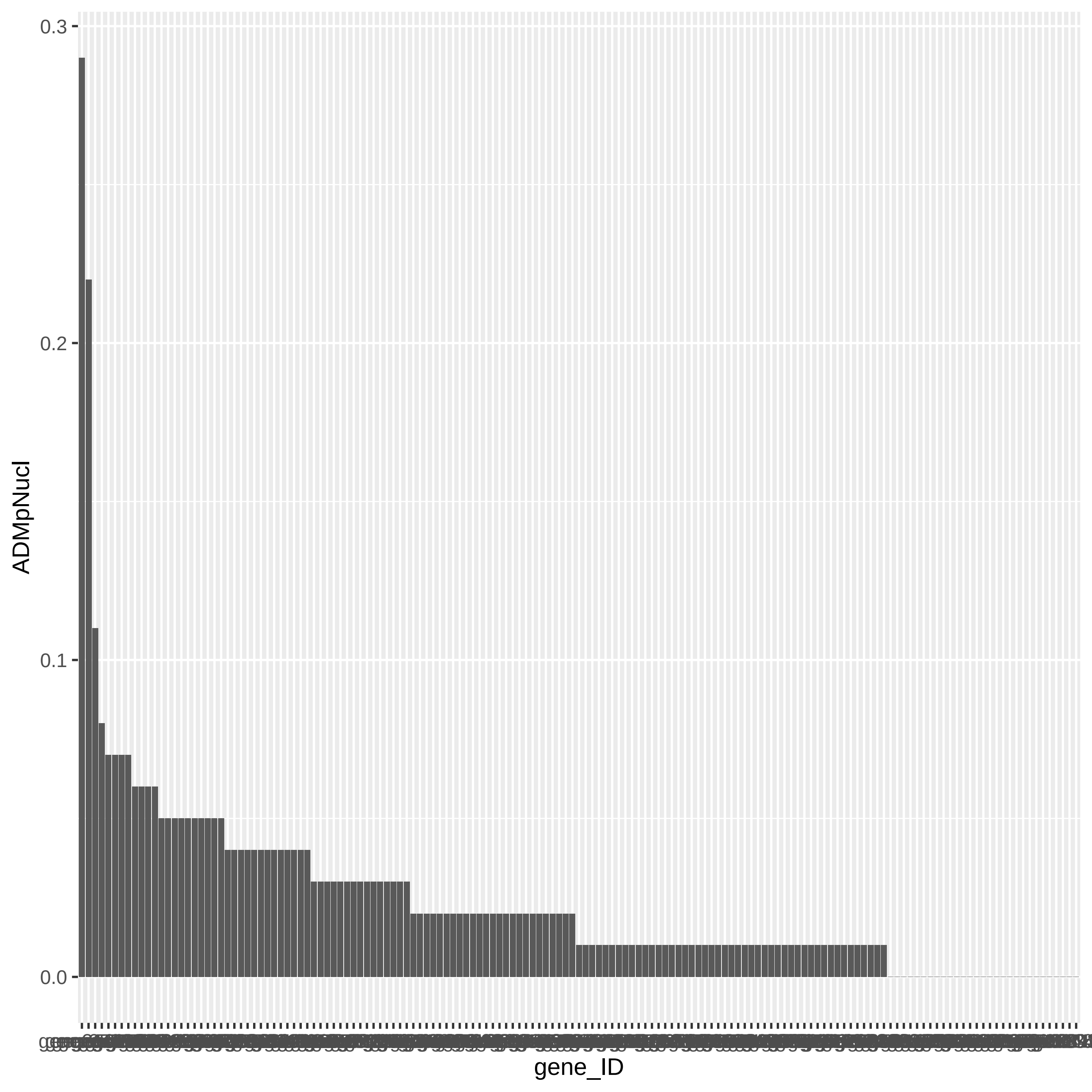
0.3
0.2
0.1
0.0
ADMpNucl
gene_ID

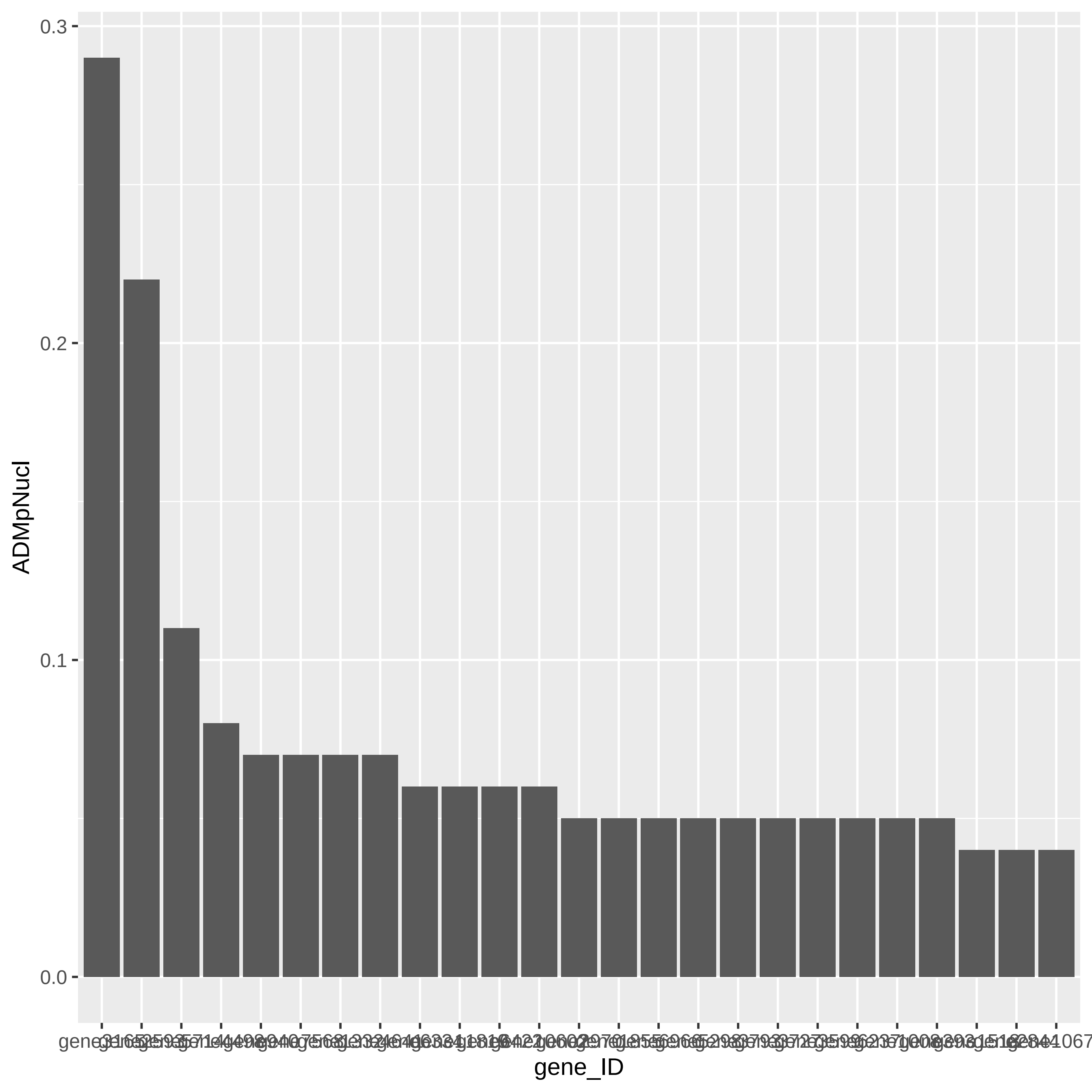

ADMpNucl
0.3
0.2
0.1
0.0
gene_ID

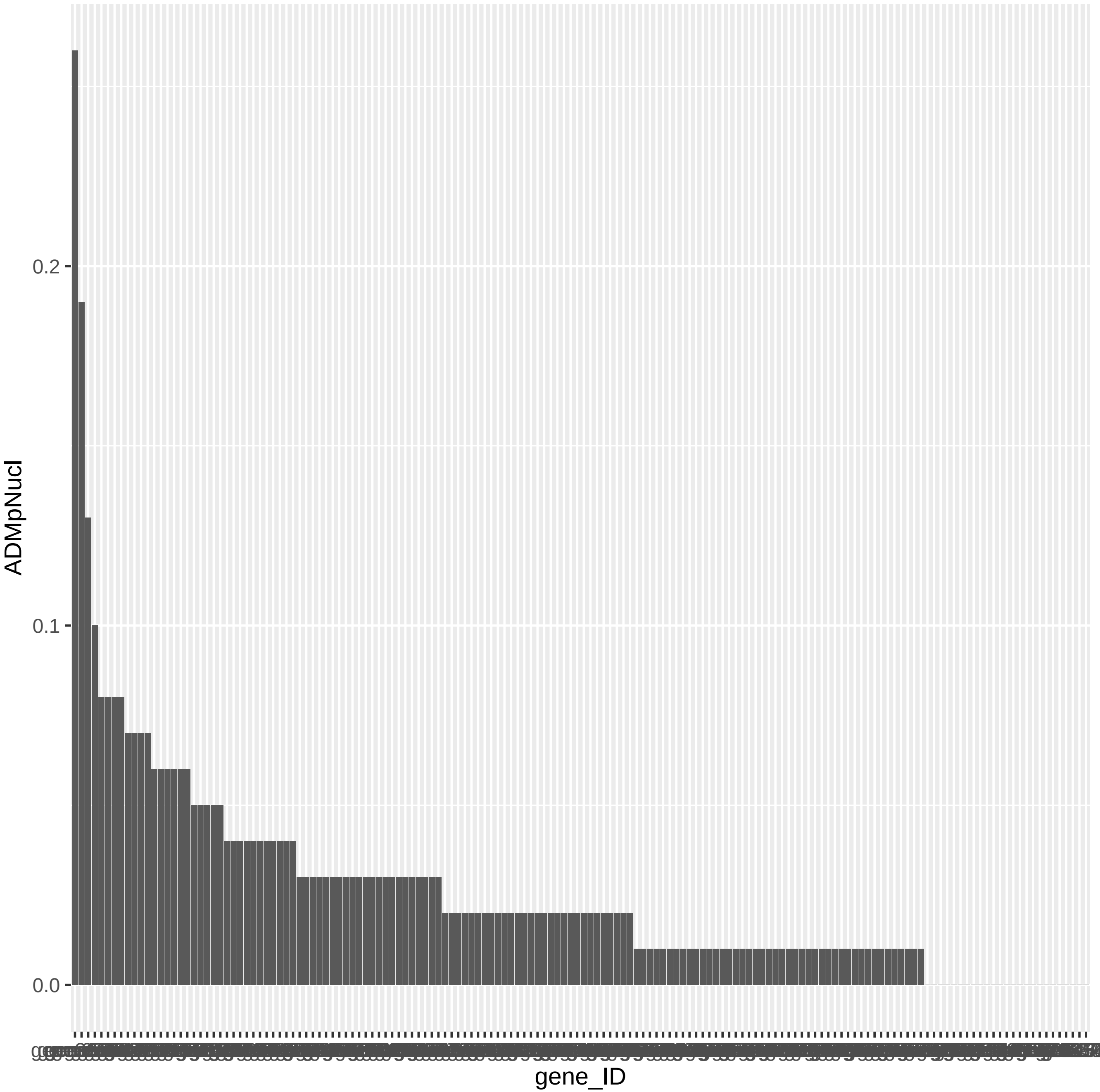

0.2
0.1
0.0
ADMpNucl
gene_ID

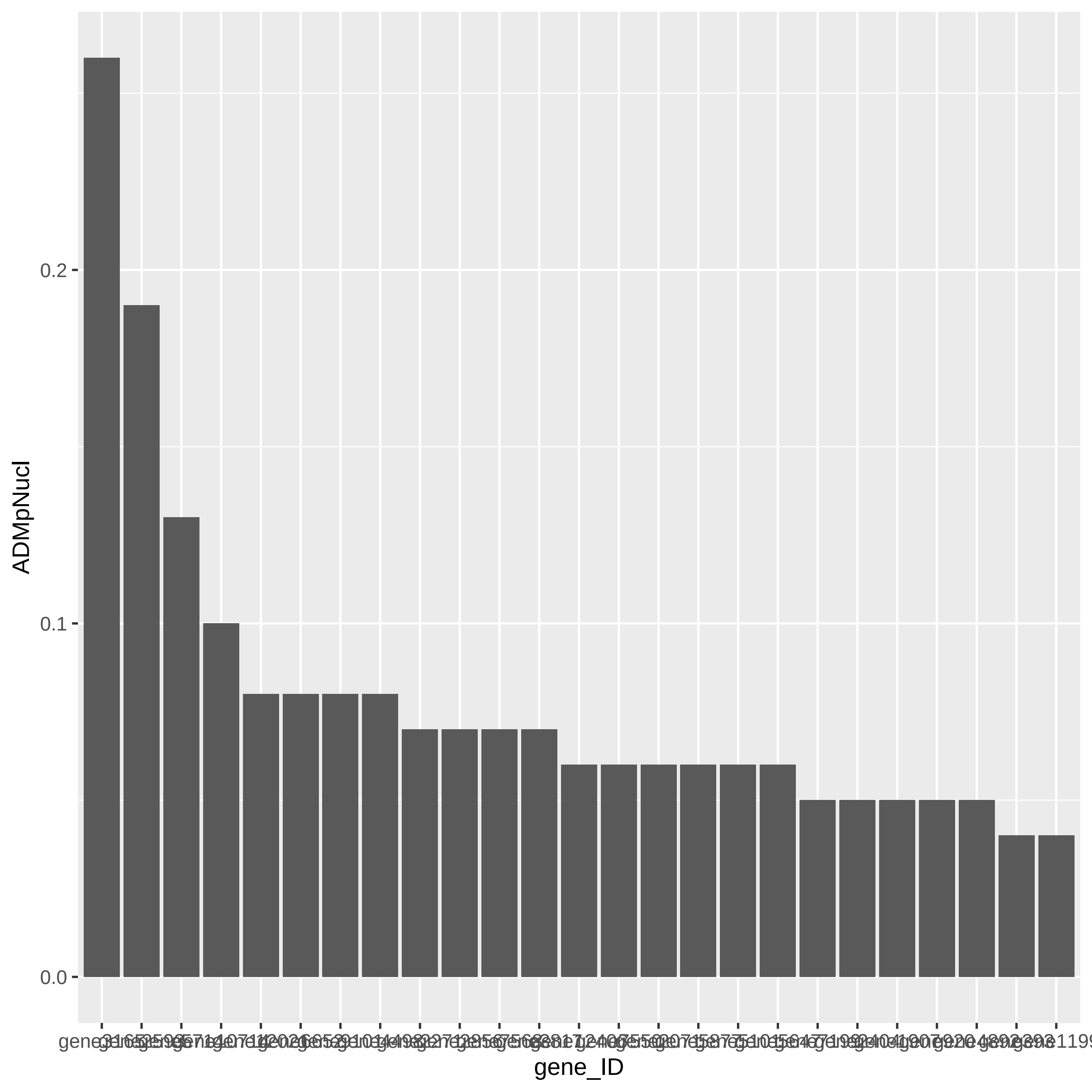
ADMpNucl
0.2
0.1
0.0
gene_ID

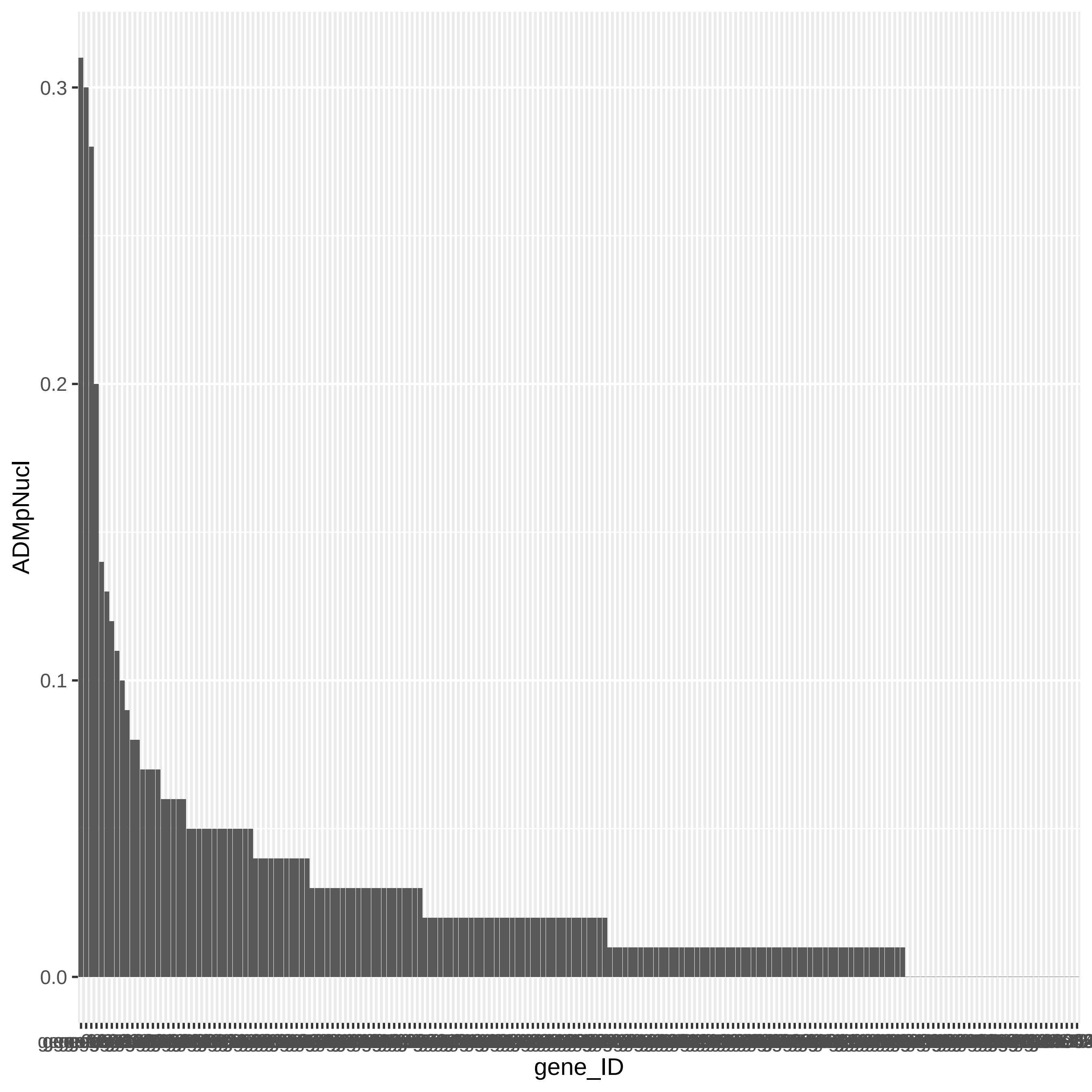

ADMpNucl
0.3
0.2
0.1
0.0
gene_ID

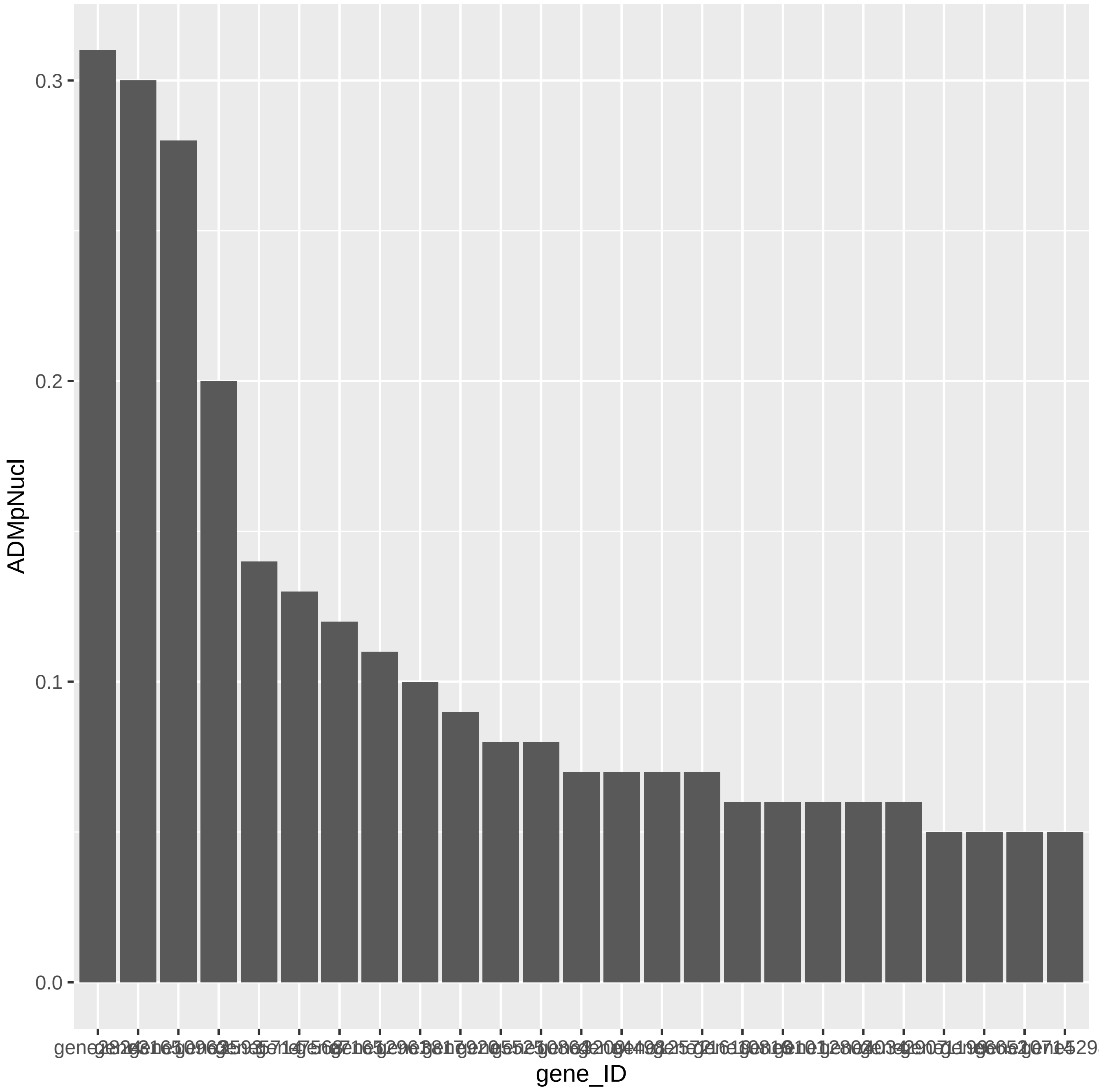

ADMpNucl
0.3
0.2
0.1
0.0
gene_ID

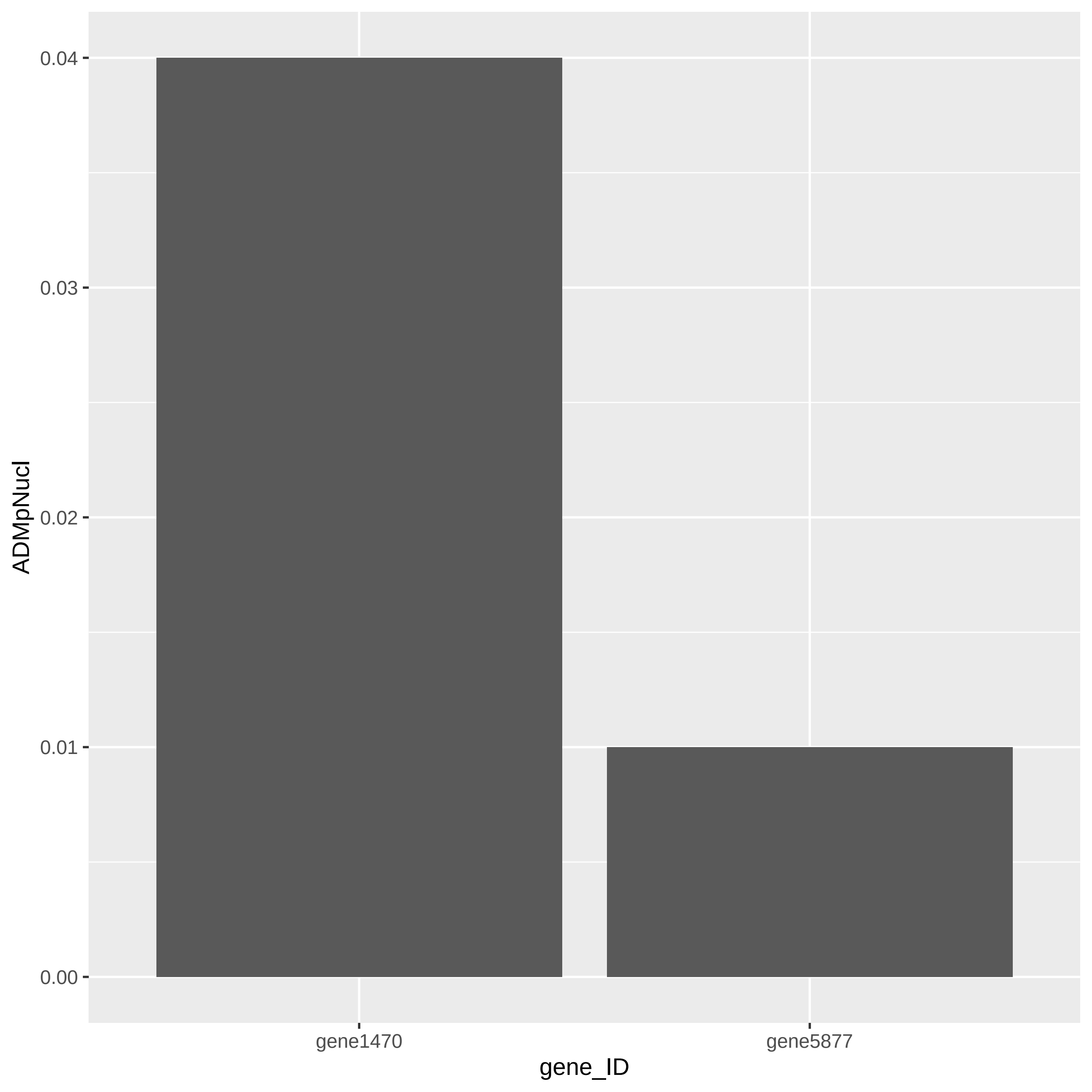
0.04
0.03
0.02
0.01
0.00
ADMpNucl
gene1470
gene5877
gene_ID

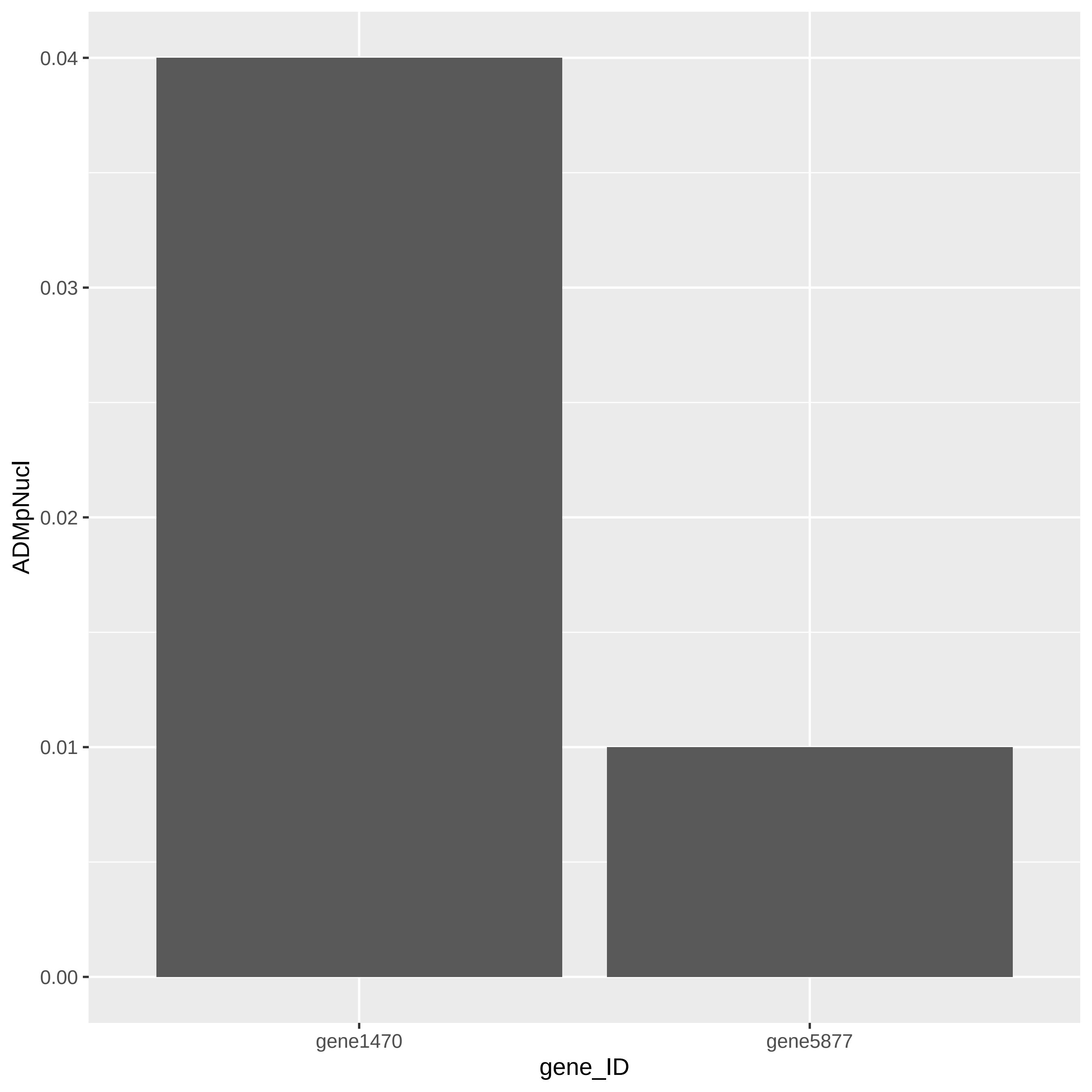

ADMpNucl
0.04
0.03
0.02
0.01
0.00
gene1470
gene5877
gene_ID

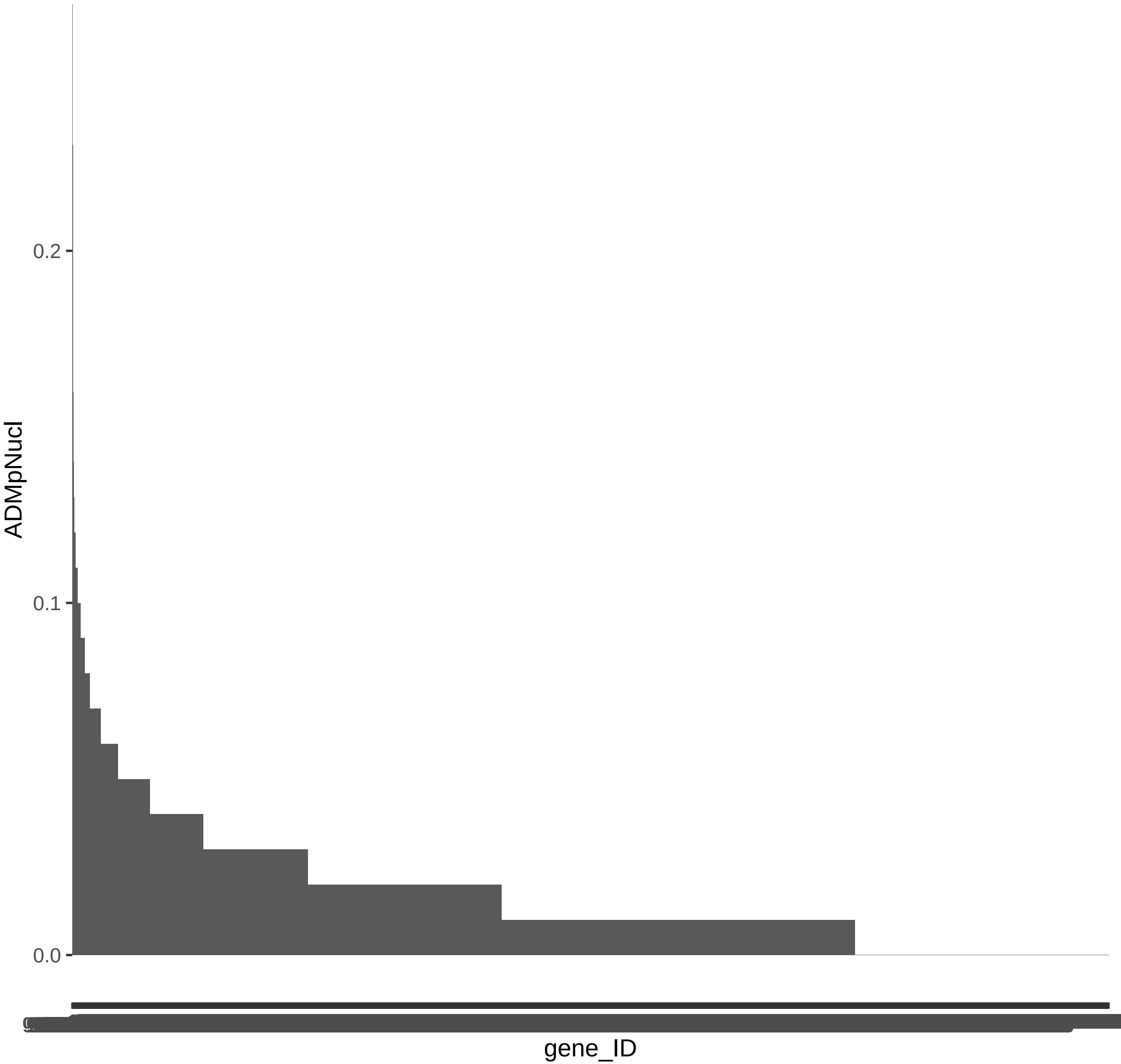

ADMpNucl
0.2
0.1
0.0
gene_ID

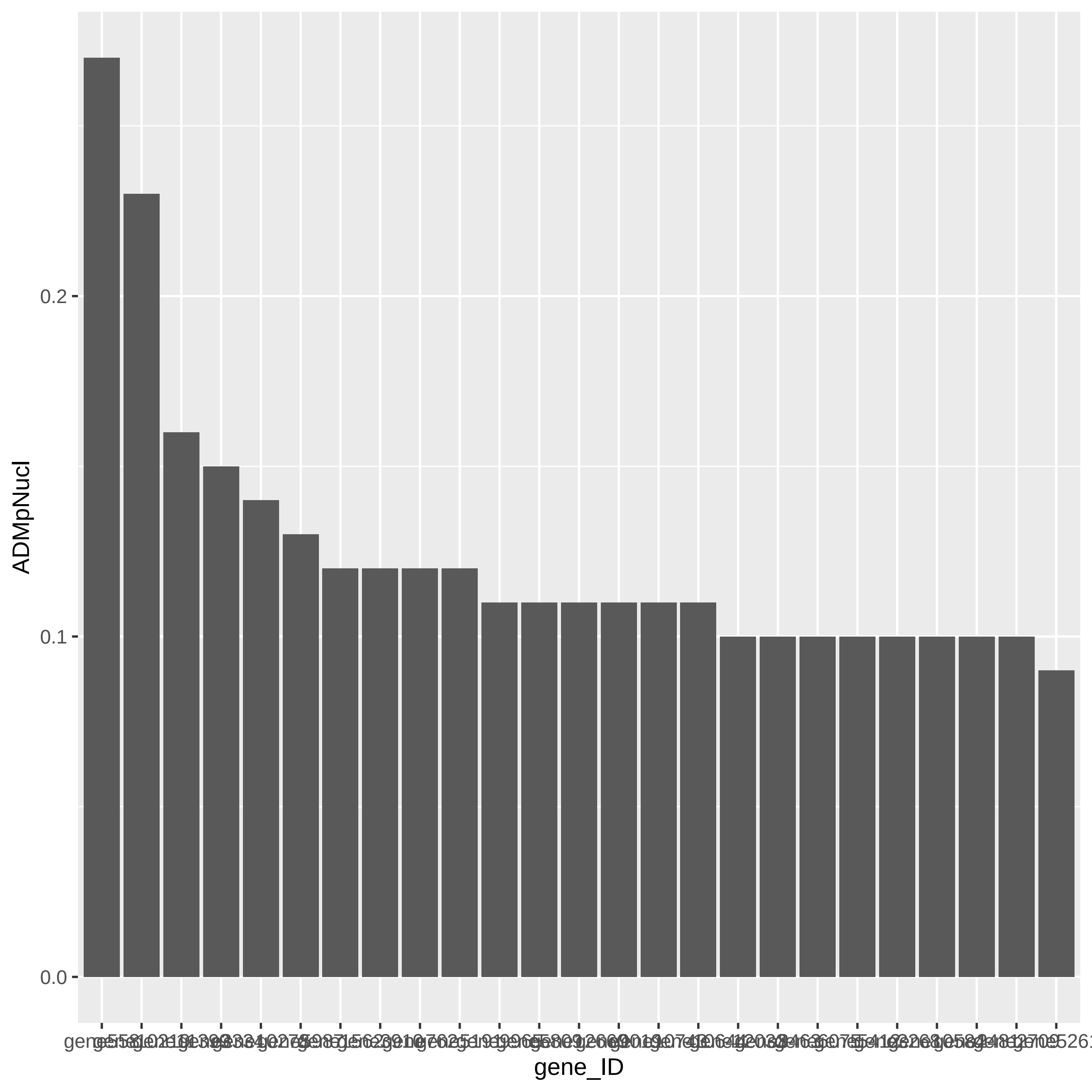
ADMpNucl
0.0
0.1
0.2
gene_ID

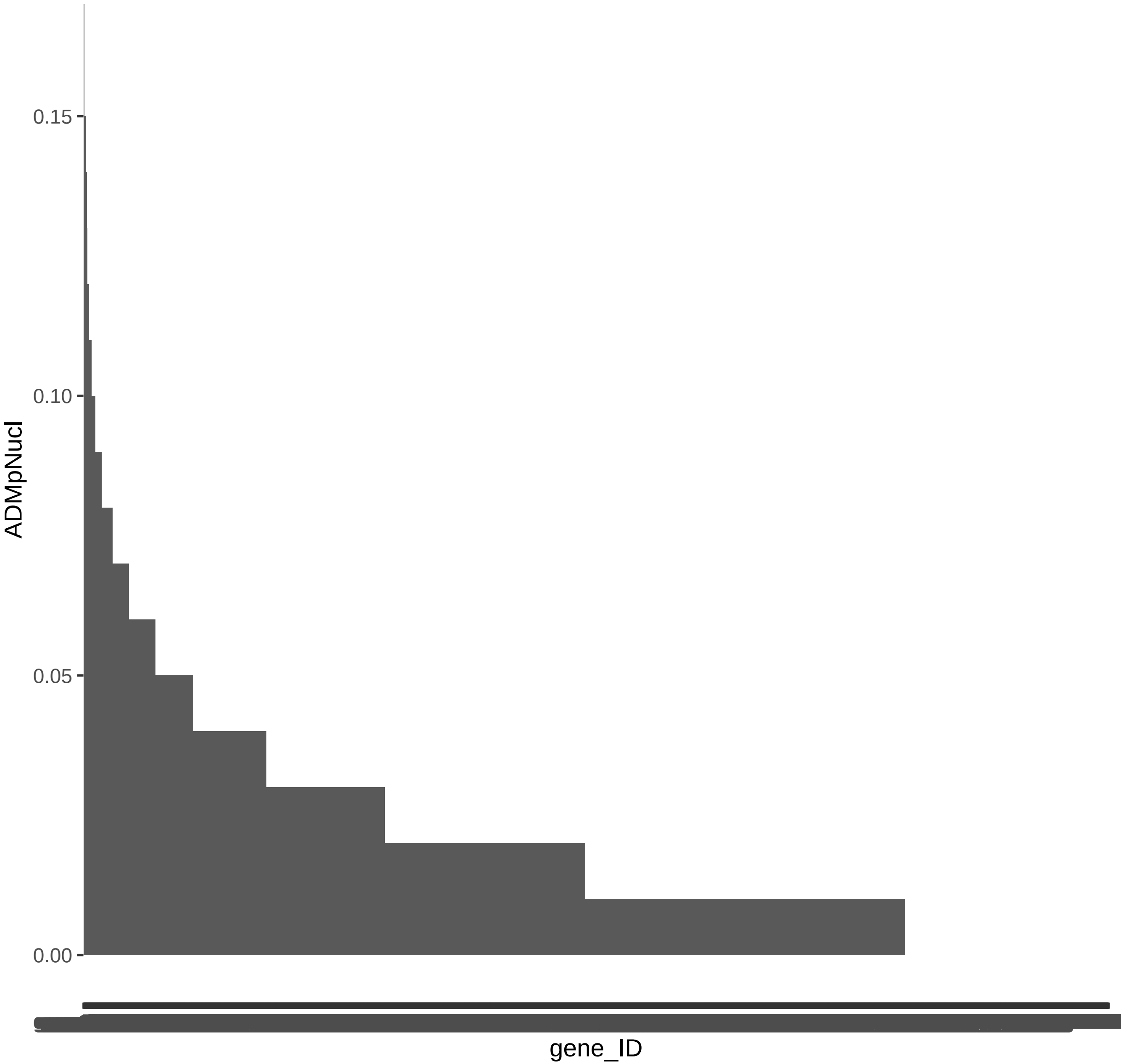

0.15
0.10
0.05
0.00
ADMpNucl
gene_ID

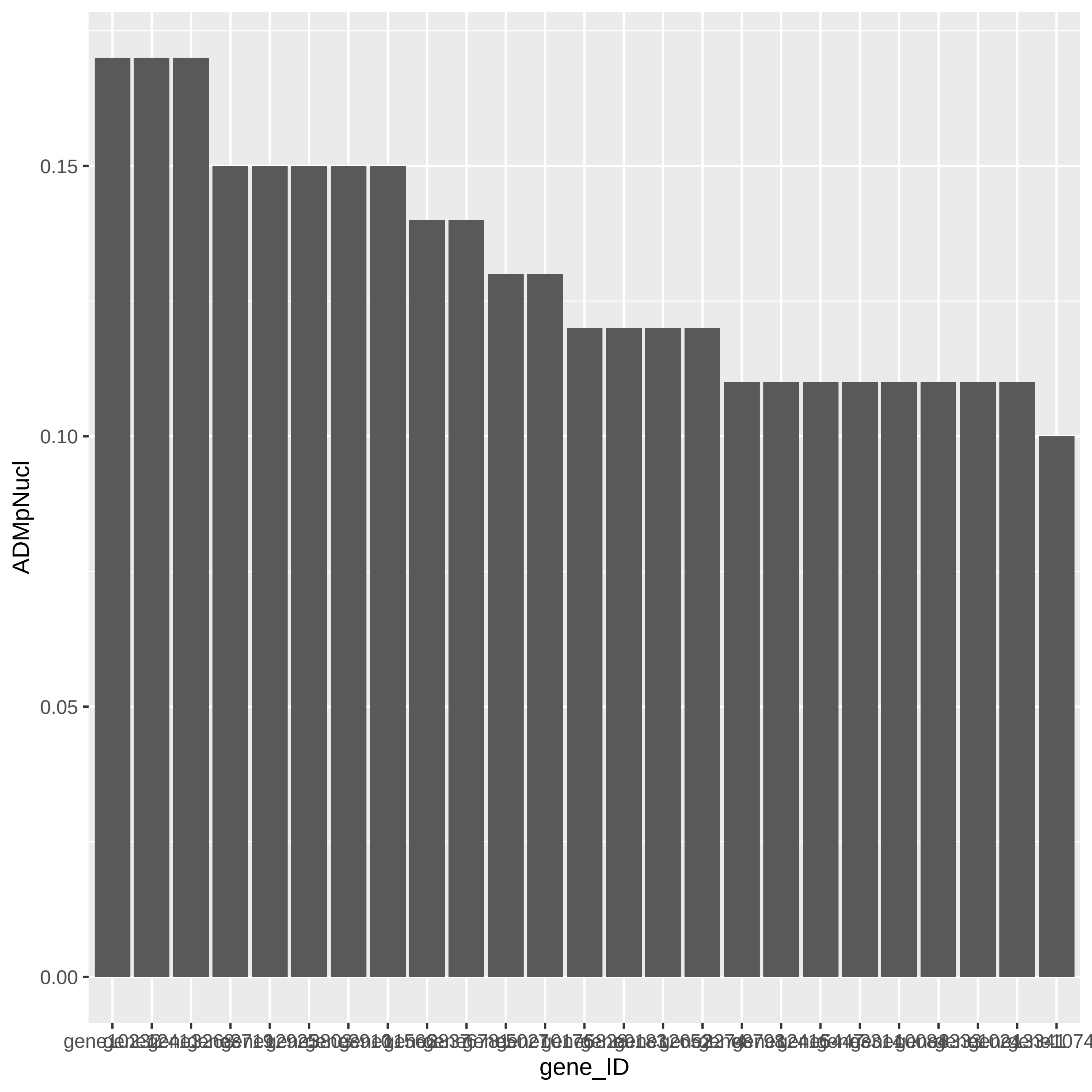

ADMpNucl
0.15
0.10
0.05
0.00
gene_ID

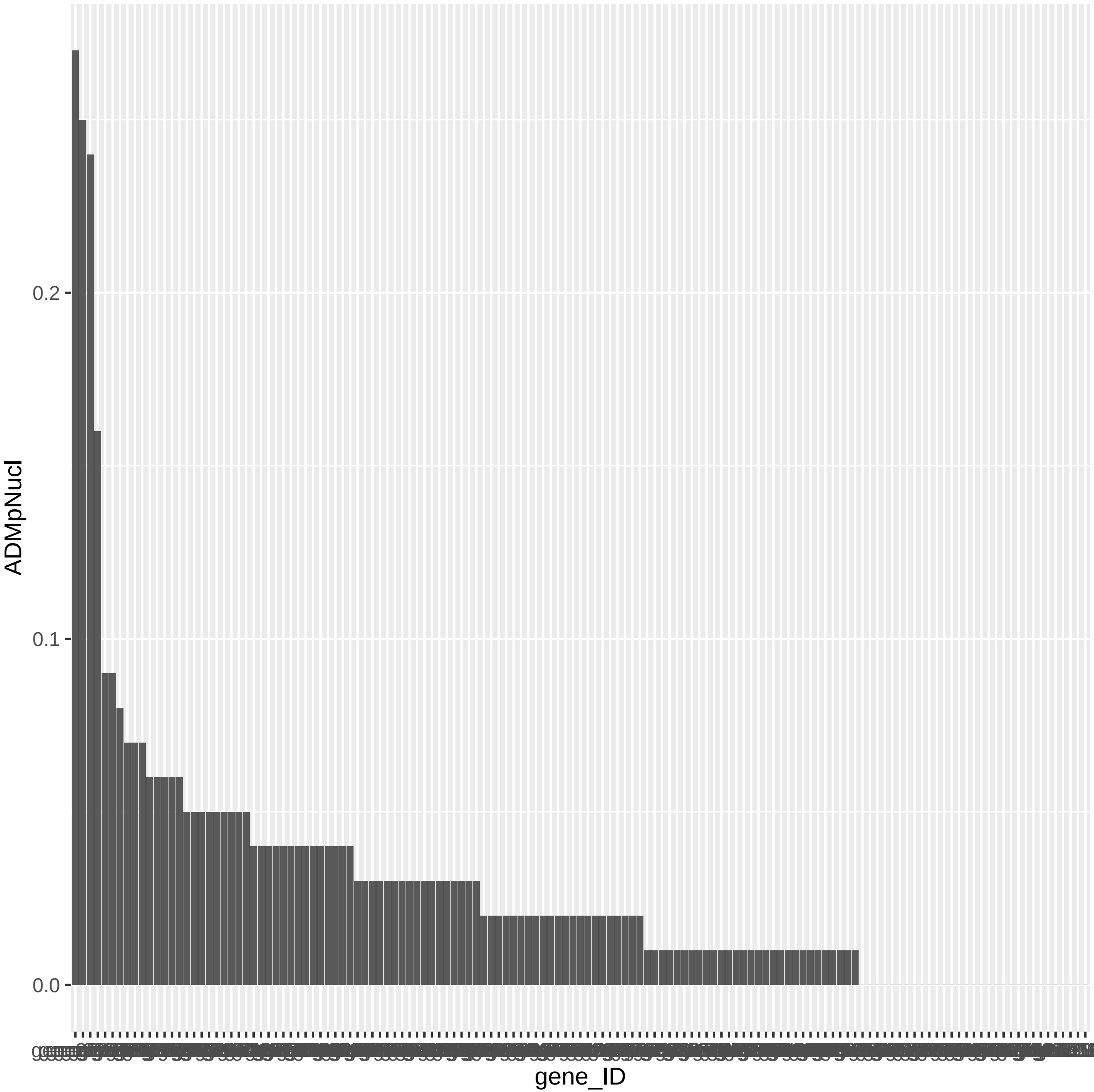
ADMpNucl
0.2
0.1
0.0
gene_ID

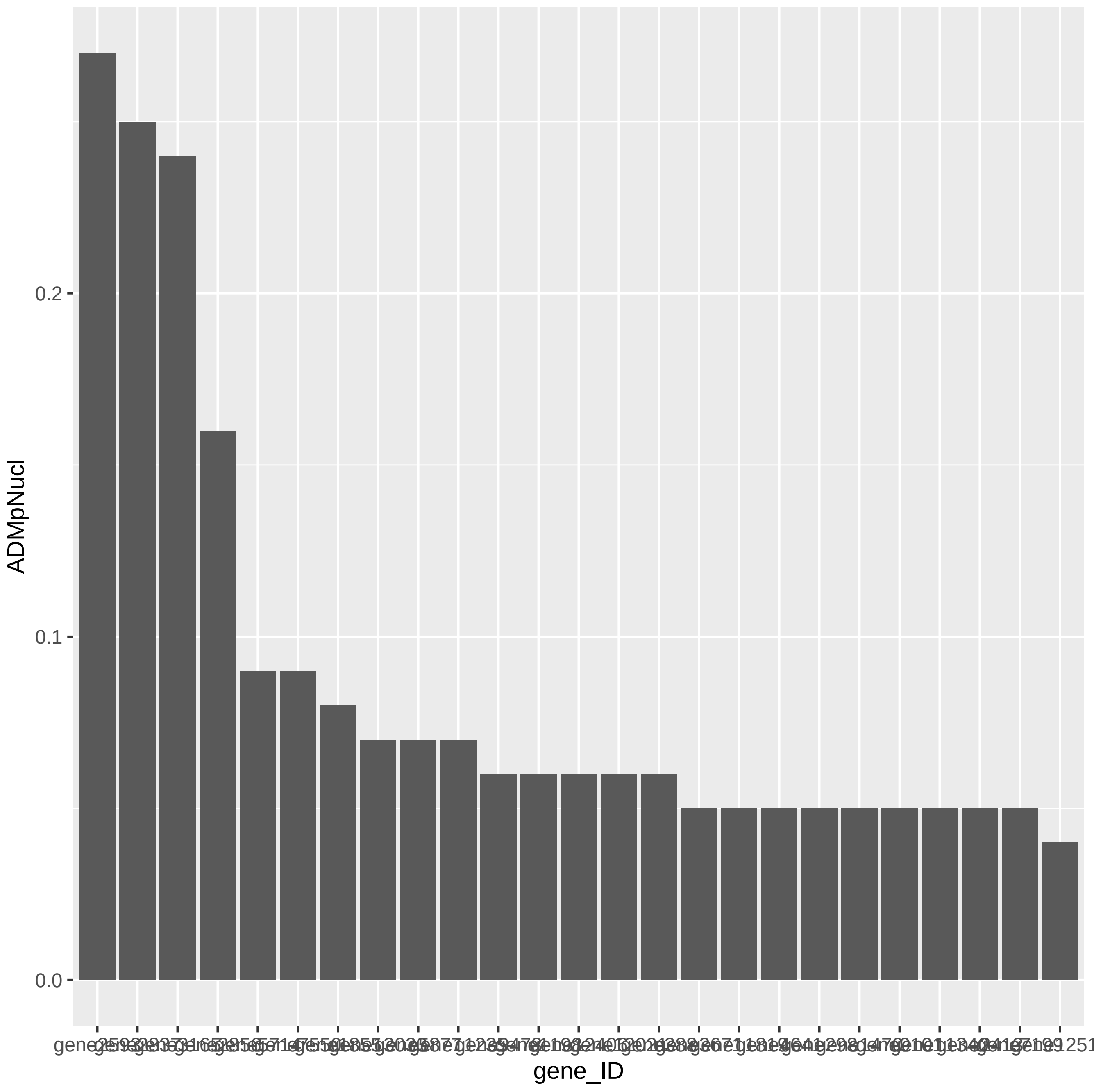
ADMpNucl
0.2
0.1
0.0
gene_ID

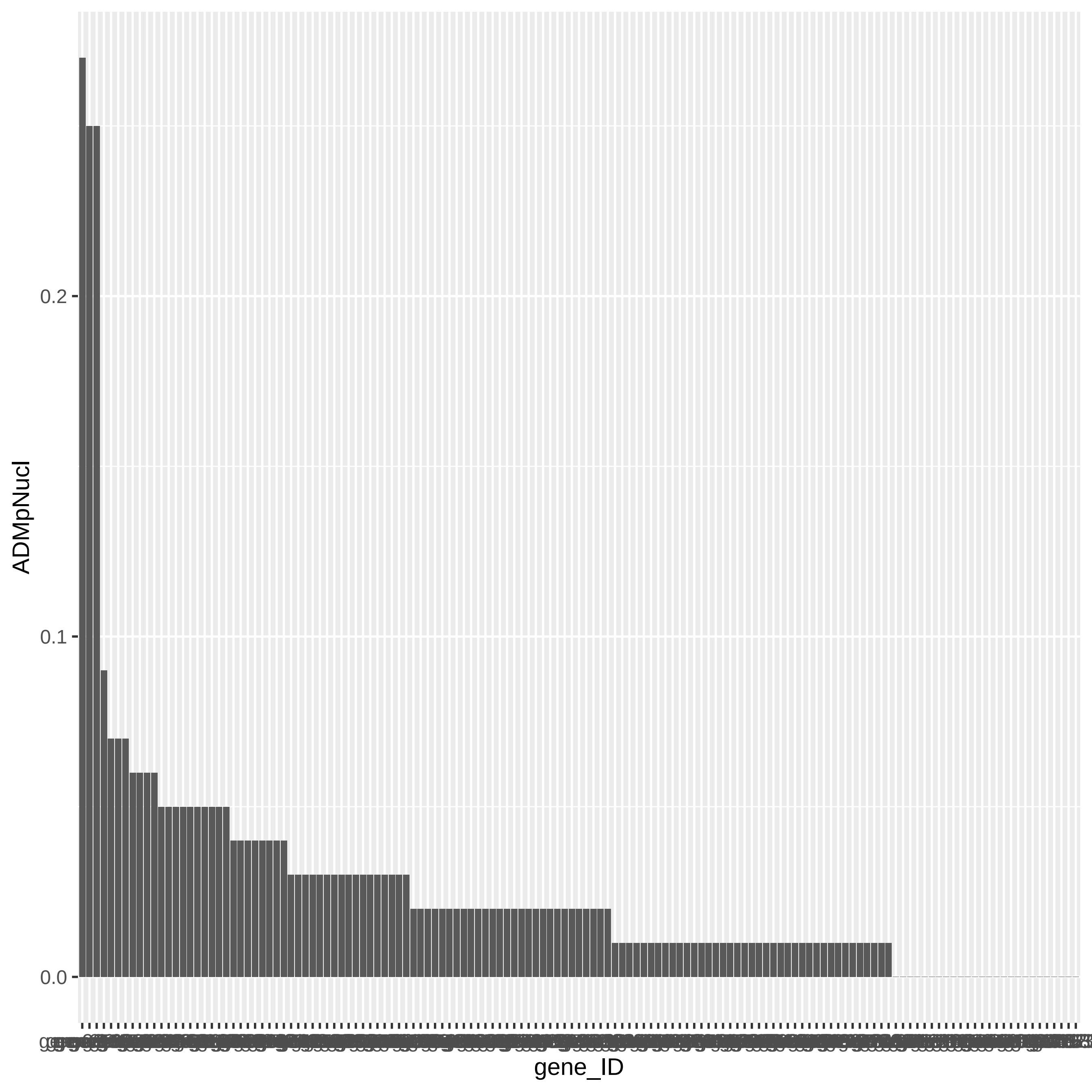

ADMpNucl
0.2
0.1
0.0
gene_ID

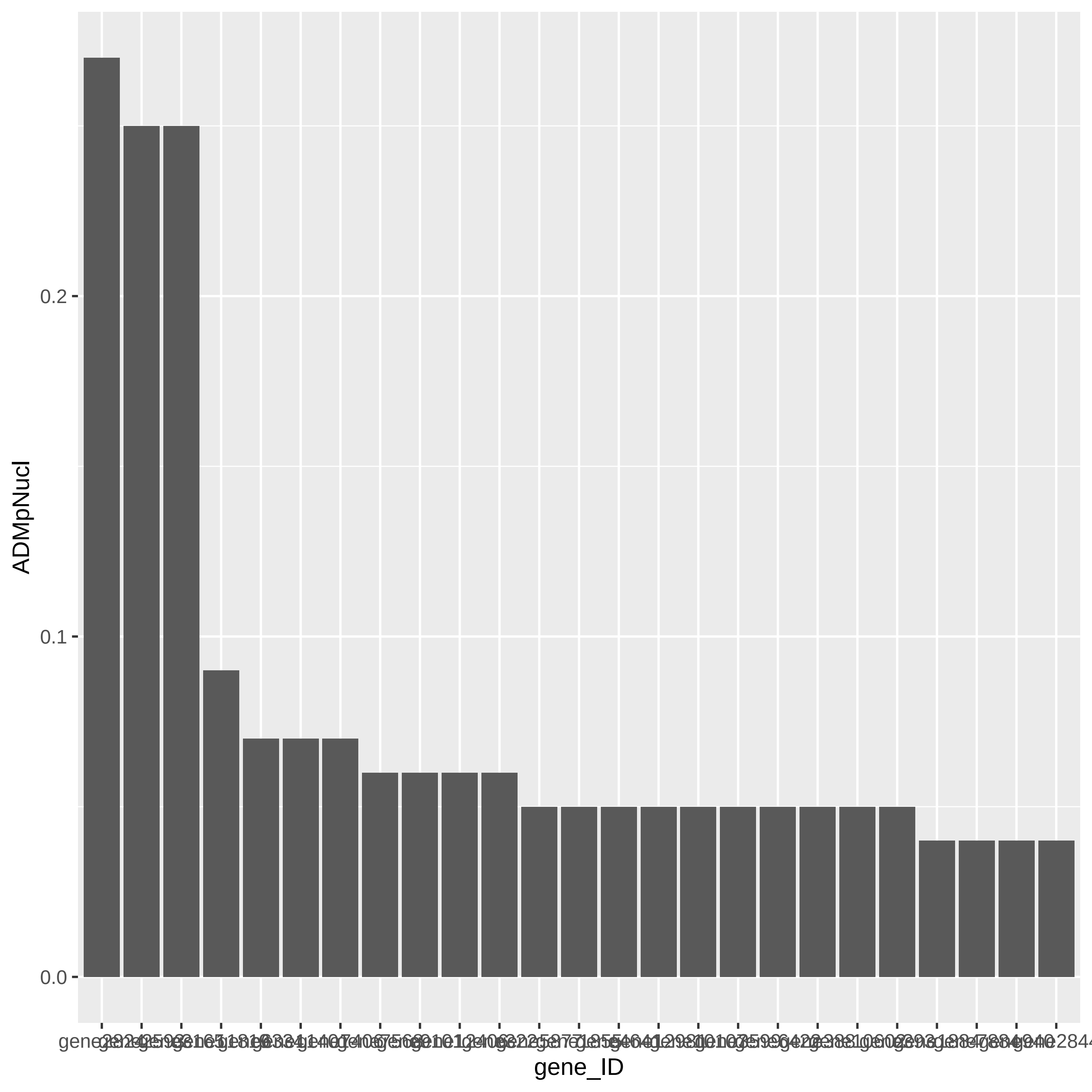

ADMpNucl
0.0
0.1
0.2
gene_ID

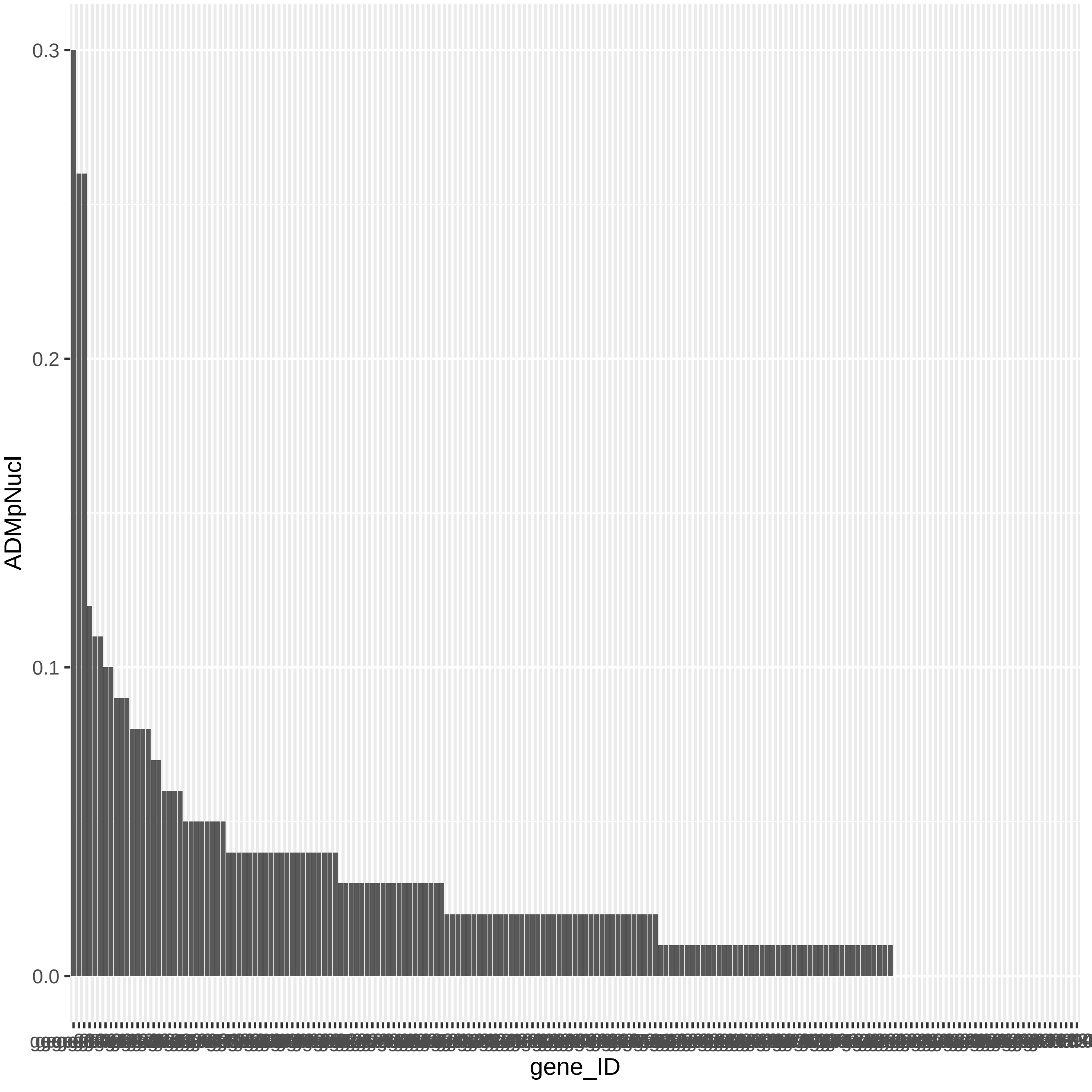

0.3
0.2
0.1
0.0
ADMpNucl
gene_ID

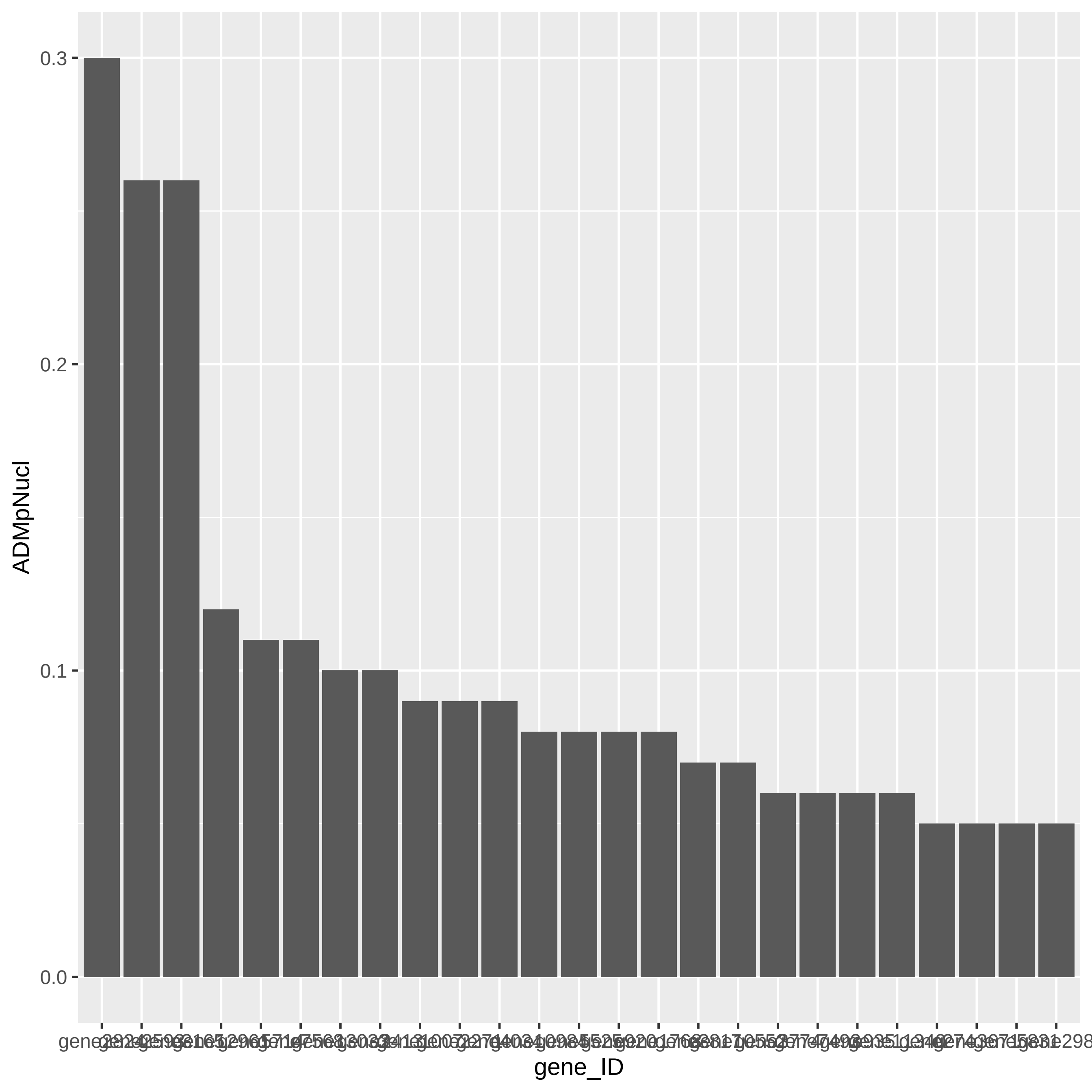
ADMpNucl
0.3
0.2
0.1
0.0
gene_ID

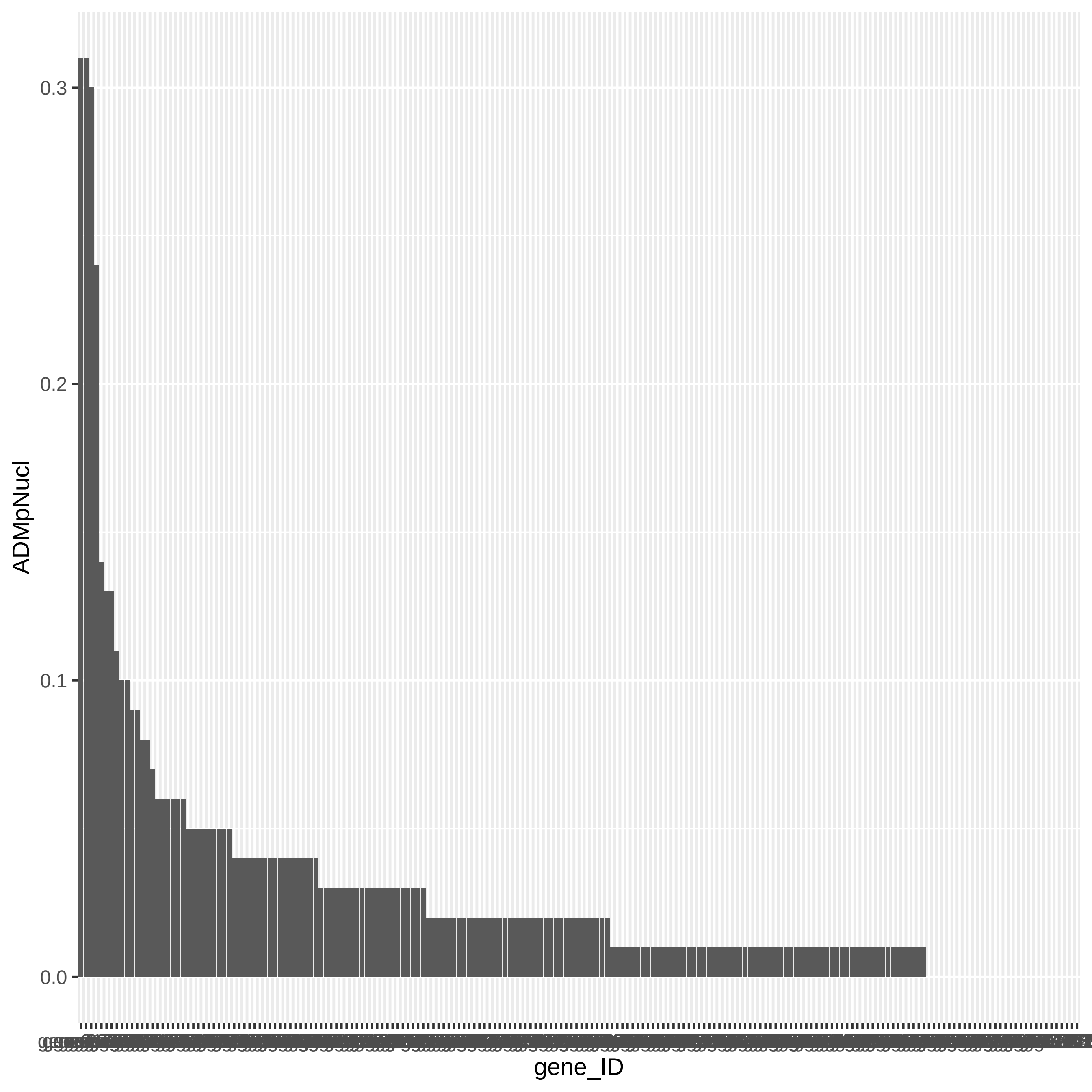
0.3
0.2
0.1
0.0
ADMpNucl
gene_ID

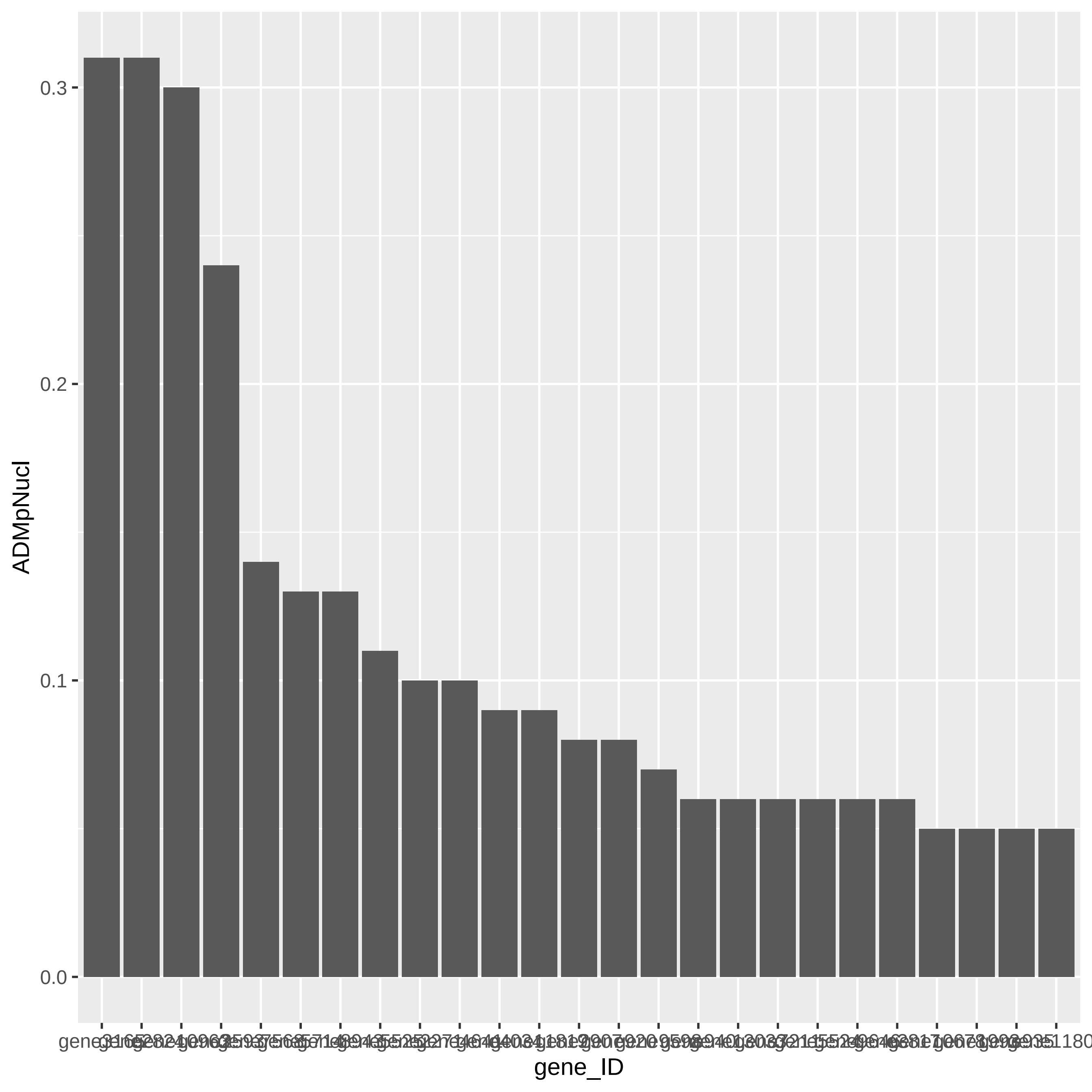

0.3
0.2
0.1
0.0
ADMpNucl
gene_ID

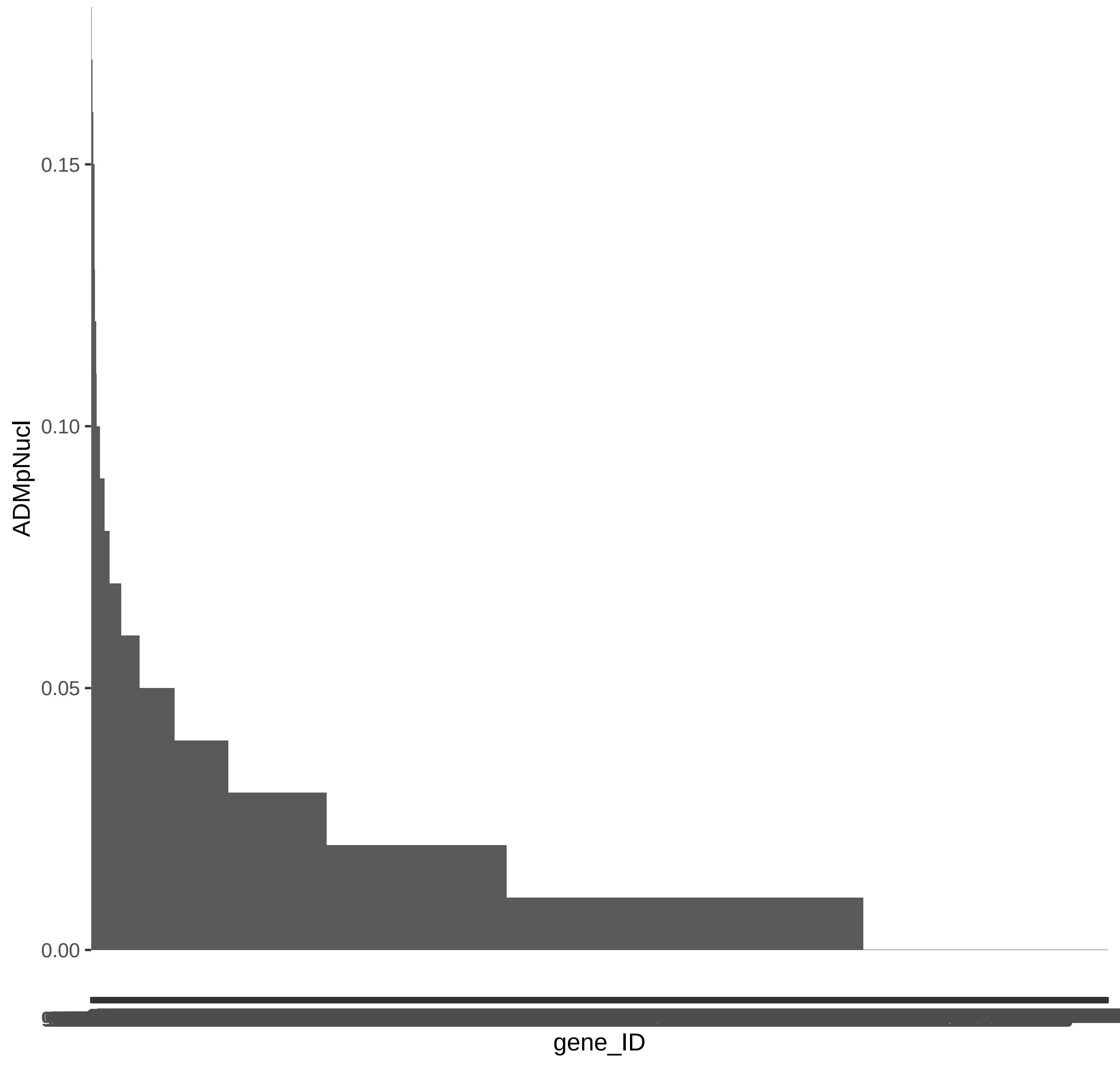
0.15
0.10
0.05
0.00
ADMpNucl
gene_ID

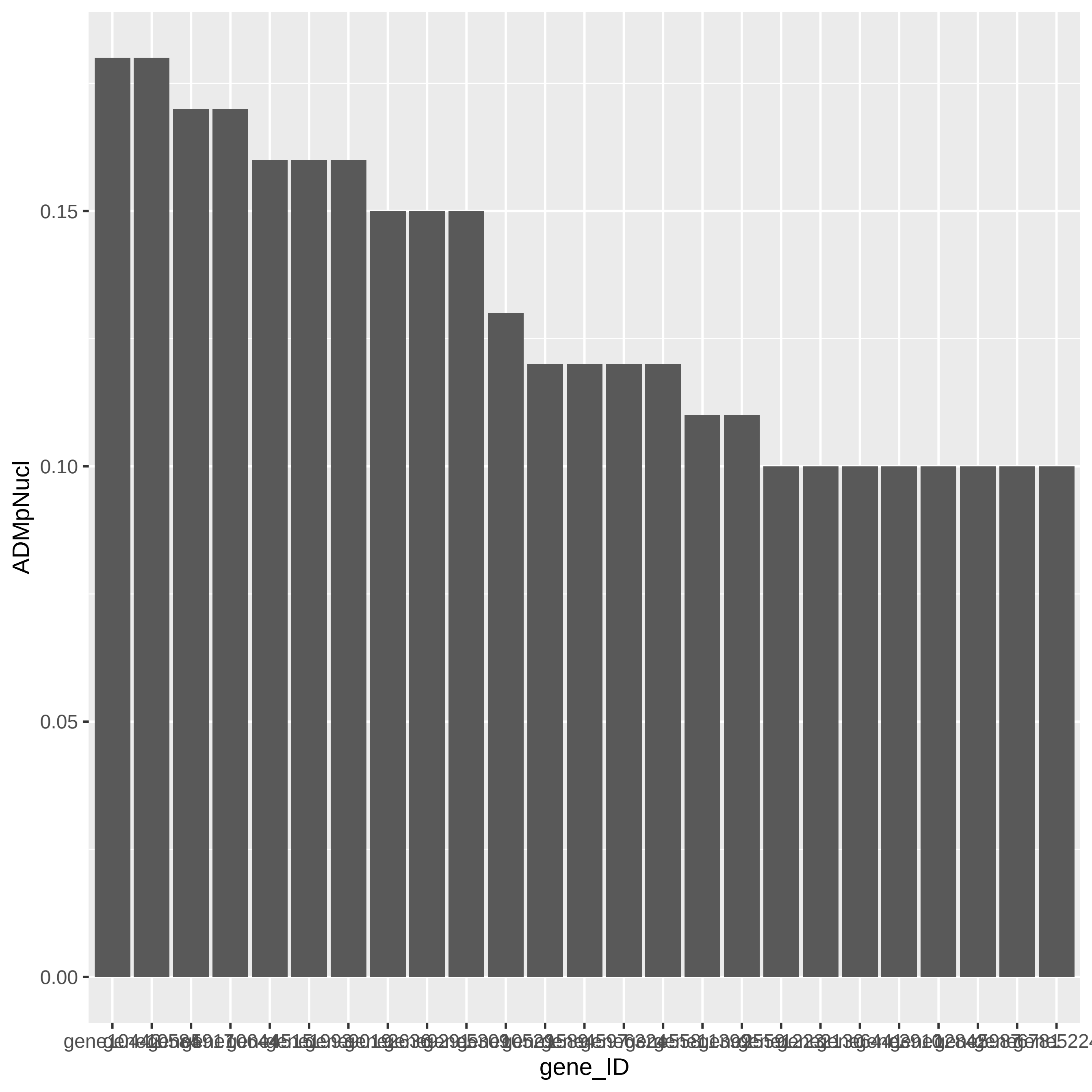
ADMpNucl
0.15
0.10
0.05
0.00
gene_ID

```
Directory: OGLall File: 0READMEogl

OGL - Ordered gene lists.

Input: studyhc annotation   maxgwidth minnbrdmsites
       dmgprp (= output of BWASPR:show_dmsg())

Output: files ogl-*.txt rnk-dmg-*.txt
         ogl-<study>_<sample>.txt ogl-<study>_<sample1>.vs.<sample2>.txt
         rnk-dmg--<study>_<sample1>.vs.<sample2>.txt rnk-dmg--<study>_<sample1>.vs.<sample2>.pdf
         wrt-<study>.txt

Notes: Output files ogl-<study>_<sample>.txt give tables for each sample with columns

        gene_ID gwidth #Sites #per10Kb %perSite %pNucl

       ordered by %pNucl.  If available, a link to an NCBI entry of gene_ID is inserted as second column.
       Abbreviations used: %perSite, percent methylation per site
                           %perNucl, percent methylation per nucleotide of the gene

       Output files ogl-<study>_<sample1>.vs.<sample2>.txt give tables for each comparison with columns

        gene_ID gwidth #Sites #per10Kb #dmSites #dmsp10kb %dmSites %pSite1 %pSite2 DMpSite ADMpSite DMpNucl ADMpNucl

       ordered by DMpSite.  If available, a link to an NCBI entry of gene_ID is inserted as second column.
       Abbreviations used: %dmSites, percent sites that are differentially methylated
                           %pSite1, average per site % methylation for sample1
                           %pSite2, average per site % methylation for sample2
                           DMpSite, average per site difference in % methylation between sample1 and sample2
                           ADMpSite, absolute value of DMpSite
                           DMpNucl, average per nucleotide difference in % methylation between sample1 and sample2
                           ADMpNucl, absolute value of DMpNucl

       Output files rnk-dmg-<study>_<sample1>.vs.<sample2>.txt are equivalent to files
        ogl-<study>_<sample1>.vs.<sample2>.txt but ordered by ADMpNucl.
       Output files rnk-dmg-<study>_<sample1>.vs.<sample2>.pdf provide visualization of the distribution
        of ADMpNucl values.

       Output file wrt-<study>.txt gives results of the Wilcoxon signed rank test comparing
        the %pSite1 and %pSite2 vectors

       In output directory OGL, tables are restricted to genes with gwidth <= maxgwidth and, for
        pairwise comparisons, #dmSites >= minnbrdmsites

       If either maxgwidth or minnbrdmsites is set, then output directory OGLall will show the full tables
        with all genes (for reference).
```

Directory: OGLall File: ogl-Cb_LI_BCphaseA1.txt

| gene_ID | gene_link | gwidth | #Sites | #per10Kb | %perSite | %pNucl |
|---|---|---|---|---|---|---|
| gene5911 | https://www.ncbi.nlm.nih.gov/gene/?term=LOC105278844 | 943 | 105 | 1113.47 | 31.24 | 3.48 |
| gene10219 | https://www.ncbi.nlm.nih.gov/gene/?term=LOC105283830 | 235 | 21 | 893.62 | 38.08 | 3.4 |
| gene13384 | https://www.ncbi.nlm.nih.gov/gene/?term=LOC105287447 | 1877 | 216 | 1150.77 | 28.75 | 3.31 |
| gene12538 | https://www.ncbi.nlm.nih.gov/gene/?term=LOC105286531 | 372 | 35 | 940.86 | 34.82 | 3.28 |
| gene5309 | https://www.ncbi.nlm.nih.gov/gene/?term=LOC105278160 | 1704 | 148 | 868.54 | 36.82 | 3.2 |
| gene211 | https://www.ncbi.nlm.nih.gov/gene/?term=LOC105274534 | 1280 | 86 | 671.88 | 45.78 | 3.08 |
| gene12746 | https://www.ncbi.nlm.nih.gov/gene/?term=LOC105286737 | 2688 | 310 | 1153.27 | 25.77 | 2.97 |
| gene3719 | https://www.ncbi.nlm.nih.gov/gene/?term=LOC105276360 | 1471 | 132 | 897.35 | 33.01 | 2.96 |
| gene5917 | https://www.ncbi.nlm.nih.gov/gene/?term=LOC105278849 | 823 | 99 | 1202.92 | 24.3 | 2.92 |
| gene1654 | https://www.ncbi.nlm.nih.gov/gene/?term=LOC105287811 | 2285 | 134 | 586.43 | 48.38 | 2.84 |
| gene3731 | https://www.ncbi.nlm.nih.gov/gene/?term=LOC105276371 | 865 | 55 | 635.84 | 44.05 | 2.8 |
| gene12330 | https://www.ncbi.nlm.nih.gov/gene/?term=LOC105286235 | 1501 | 97 | 646.24 | 43.04 | 2.78 |
| gene1562 | https://www.ncbi.nlm.nih.gov/gene/?term=LOC105287692 | 1491 | 120 | 804.83 | 33.36 | 2.68 |
| gene2797 | https://www.ncbi.nlm.nih.gov/gene/?term=LOC105275314 | 1489 | 79 | 530.56 | 50.1 | 2.66 |
| gene12364 | https://www.ncbi.nlm.nih.gov/gene/?term=LOC105286269 | 1403 | 68 | 484.68 | 54.55 | 2.64 |
| gene458 | https://www.ncbi.nlm.nih.gov/gene/?term=LOC105277254 | 1378 | 109 | 791 | 33.34 | 2.64 |
| gene5301 | https://www.ncbi.nlm.nih.gov/gene/?term=LOC105278148 | 1881 | 182 | 967.57 | 26.39 | 2.55 |
| gene9471 | https://www.ncbi.nlm.nih.gov/gene/?term=LOC105282986 | 1605 | 124 | 772.59 | 32.83 | 2.54 |
| gene7527 | https://www.ncbi.nlm.nih.gov/gene/?term=LOC105280739 | 2686 | 131 | 487.71 | 51.92 | 2.53 |
| gene12652 | https://www.ncbi.nlm.nih.gov/gene/?term=LOC109610640 | 1122 | 55 | 490.2 | 51.51 | 2.52 |
| gene3806 | https://www.ncbi.nlm.nih.gov/gene/?term=LOC105276439 | 2385 | 143 | 599.58 | 42.03 | 2.52 |
| gene2129 | https://www.ncbi.nlm.nih.gov/gene/?term=LOC105274543 | 1631 | 104 | 637.65 | 38.35 | 2.45 |
| gene11569 | https://www.ncbi.nlm.nih.gov/gene/?term=LOC105285394 | 1311 | 46 | 350.88 | 69.57 | 2.44 |
| gene5825 | https://www.ncbi.nlm.nih.gov/gene/?term=LOC105278753 | 1754 | 100 | 570.13 | 42.54 | 2.43 |

...

... only first 25 lines shown ...

Directory: OGLall File: ogl-Cb_LI_BCphaseA1.vs.BCphaseA2.txt

| gene_ID | gene_link | gwidth | #Sites | #per10Kb | #dmSites | #dmsp10kb | %dmSites | %pSite1 | %pSite2 | DMpSite | ADMpSite | DMpNucl | ADMpNucl |
|---|---|---|---|---|---|---|---|---|---|---|---|---|---|
| gene393 | https://www.ncbi.nlm.nih.gov/gene/?term=LOC105276566 | 1123 | 2 | 17.81 | 1 | 8.9 | 50 | 77.65 | 48.08 | 29.58 | 29.58 | 0.05 | 0.05 |
| gene9109 | https://www.ncbi.nlm.nih.gov/gene/?term=LOC105282561 | 2400 | 10 | 41.67 | 1 | 4.17 | 10 | 26.72 | 17.84 | 8.89 | 8.89 | 0.04 | 0.04 |
| gene7952 | https://www.ncbi.nlm.nih.gov/gene/?term=LOC105281213 | 2963 | 31 | 104.62 | 1 | 3.37 | 3.23 | 42.96 | 39.57 | 3.39 | 3.39 | 0.04 | 0.04 |
| gene7915 | https://www.ncbi.nlm.nih.gov/gene/?term=LOC105281173 | 1597 | 25 | 156.54 | 1 | 6.26 | 4 | 64.87 | 61.62 | 3.25 | 3.25 | 0.05 | 0.05 |
| gene7368 | https://www.ncbi.nlm.nih.gov/gene/?term=LOC105280556 | 4105 | 40 | 97.44 | 1 | 2.44 | 2.5 | 14.82 | 11.87 | 2.94 | 2.94 | 0.03 | 0.03 |
| gene10213 | https://www.ncbi.nlm.nih.gov/gene/?term=LOC105283826 | 1503 | 53 | 352.63 | 1 | 6.65 | 1.89 | 16.45 | 13.62 | 2.82 | 2.82 | 0.1 | 0.1 |
| gene3168 | https://www.ncbi.nlm.nih.gov/gene/?term=LOC105275711 | 6150 | 71 | 115.45 | 1 | 1.63 | 1.41 | 14.71 | 11.97 | 2.74 | 2.74 | 0.03 | 0.03 |
| gene5490 | https://www.ncbi.nlm.nih.gov/gene/?term=LOC105278358 | 2095 | 25 | 119.33 | 1 | 4.77 | 4 | 36.86 | 34.23 | 2.63 | 2.63 | 0.03 | 0.03 |
| gene388 | https://www.ncbi.nlm.nih.gov/gene/?term=LOC105276538 | 1558 | 19 | 121.95 | 2 | 12.84 | 10.53 | 69.41 | 66.83 | 2.58 | 2.58 | 0.03 | 0.03 |
| gene12981 | https://www.ncbi.nlm.nih.gov/gene/?term=LOC105286999 | 2063 | 39 | 189.05 | 1 | 4.85 | 2.56 | 33.12 | 30.76 | 2.36 | 2.36 | 0.04 | 0.04 |
| gene2404 | https://www.ncbi.nlm.nih.gov/gene/?term=LOC105274853 | 1767 | 33 | 186.76 | 1 | 5.66 | 3.03 | 31.53 | 29.22 | 2.31 | 2.31 | 0.04 | 0.04 |
| gene920 | https://www.ncbi.nlm.nih.gov/gene/?term=LOC105282518 | 1197 | 40 | 334.17 | 1 | 8.35 | 2.5 | 39.94 | 37.69 | 2.25 | 2.25 | 0.08 | 0.08 |
| gene6422 | https://www.ncbi.nlm.nih.gov/gene/?term=LOC105279458 | 2419 | 33 | 136.42 | 1 | 4.13 | 3.03 | 55.1 | 53.04 | 2.06 | 2.06 | 0.03 | 0.03 |
| gene1220 | https://www.ncbi.nlm.nih.gov/gene/?term=LOC105285742 | 3453 | 90 | 260.64 | 1 | 2.9 | 1.11 | 27.79 | 25.88 | 1.92 | 1.92 | 0.05 | 0.05 |
| gene11401 | https://www.ncbi.nlm.nih.gov/gene/?term=LOC105285202 | 1841 | 47 | 255.3 | 1 | 5.43 | 2.13 | 22.36 | 20.6 | 1.77 | 1.77 | 0.05 | 0.05 |
| gene10391 | https://www.ncbi.nlm.nih.gov/gene/?term=LOC109611283 | 1994 | 41 | 205.62 | 1 | 5.02 | 2.44 | 35.16 | 33.4 | 1.76 | 1.76 | 0.04 | 0.04 |
| gene6966 | https://www.ncbi.nlm.nih.gov/gene/?term=LOC105280104 | 3747 | 116 | 309.58 | 1 | 2.67 | 0.86 | 15.51 | 13.86 | 1.65 | 1.65 | 0.05 | 0.05 |
| gene11454 | https://www.ncbi.nlm.nih.gov/gene/?term=LOC105285256 | 1906 | 48 | 251.84 | 1 | 5.25 | 2.08 | 46.56 | 44.93 | 1.63 | 1.63 | 0.04 | 0.04 |
| gene1199 | https://www.ncbi.nlm.nih.gov/gene/?term=LOC105285498 | 2220 | 23 | 103.6 | 1 | 4.5 | 4.35 | 35.45 | 33.87 | 1.58 | 1.58 | 0.02 | 0.02 |
| gene12021 | https://www.ncbi.nlm.nih.gov/gene/?term=LOC105285905 | 1399 | 21 | 150.11 | 1 | 7.15 | 4.76 | 41.18 | 39.62 | 1.55 | 1.55 | 0.02 | 0.02 |
| gene2748 | https://www.ncbi.nlm.nih.gov/gene/?term=LOC105275261 | 3845 | 54 | 140.44 | 1 | 2.6 | 1.85 | 38.15 | 36.73 | 1.42 | 1.42 | 0.02 | 0.02 |
| gene11447 | https://www.ncbi.nlm.nih.gov/gene/?term=LOC105285252 | 7437 | 119 | 160.01 | 1 | 1.34 | 0.84 | 14.91 | 13.69 | 1.22 | 1.22 | 0.02 | 0.02 |
| gene3107 | https://www.ncbi.nlm.nih.gov/gene/?term=LOC105275650 | 5103 | 143 | 280.23 | 1 | 1.96 | 0.7 | 16.9 | 15.68 | 1.21 | 1.21 | 0.03 | 0.03 |
| gene10819 | https://www.ncbi.nlm.nih.gov/gene/?term=LOC105284507 | 5767 | 69 | 119.65 | 1 | 1.73 | 1.45 | 22.66 | 21.47 | 1.19 | 1.19 | 0.01 | 0.01 |
| gene5375 | https://www.ncbi.nlm.nih.gov/gene/?term=LOC105278228 | 8640 | 143 | 165.51 | 1 | 1.16 | 0.7 | 24.34 | 23.23 | 1.11 | 1.11 | 0.02 | 0.02 |
| gene322 | https://www.ncbi.nlm.nih.gov/gene/?term=LOC105275777 | 1606 | 21 | 130.76 | 1 | 6.23 | 4.76 | 49.88 | 48.85 | 1.04 | 1.04 | 0.01 | 0.01 |
| gene5646 | https://www.ncbi.nlm.nih.gov/gene/?term=LOC105278534 | 7475 | 166 | 222.07 | 1 | 1.34 | 0.6 | 18.54 | 17.53 | 1.02 | 1.02 | 0.02 | 0.02 |
| gene10155 | https://www.ncbi.nlm.nih.gov/gene/?term=LOC105283756 | 7030 | 86 | 122.33 | 1 | 1.42 | 1.16 | 7.44 | 6.49 | 0.95 | 0.95 | 0.01 | 0.01 |
| gene4190 | https://www.ncbi.nlm.nih.gov/gene/?term=LOC105276875 | 3150 | 104 | 330.16 | 1 | 3.17 | 0.96 | 25.99 | 25.04 | 0.95 | 0.95 | 0.03 | 0.03 |
| gene10345 | https://www.ncbi.nlm.nih.gov/gene/?term=LOC105283989 | 4444 | 21 | 47.25 | 1 | 2.25 | 4.76 | 48.99 | 48.08 | 0.91 | 0.91 | 0 | 0 |
| gene3286 | https://www.ncbi.nlm.nih.gov/gene/?term=LOC105275848 | 3744 | 121 | 323.18 | 1 | 2.67 | 0.83 | 22.9 | 22 | 0.9 | 0.9 | 0.03 | 0.03 |
| gene5298 | https://www.ncbi.nlm.nih.gov/gene/?term=LOC105278145 | 4060 | 245 | 603.45 | 2 | 4.93 | 0.82 | 11.36 | 10.5 | 0.86 | 0.86 | 0.05 | 0.05 |
| gene3671 | https://www.ncbi.nlm.nih.gov/gene/?term=LOC105276272 | 1966 | 25 | 127.16 | 1 | 5.09 | 4 | 43.89 | 43.09 | 0.8 | 0.8 | 0.01 | 0.01 |
| gene1993 | https://www.ncbi.nlm.nih.gov/gene/?term=LOC105288193 | 1449 | 62 | 427.88 | 1 | 6.9 | 1.61 | 35.55 | 34.76 | 0.78 | 0.78 | 0.03 | 0.03 |
| gene3848 | https://www.ncbi.nlm.nih.gov/gene/?term=LOC105276486 | 5899 | 107 | 181.39 | 1 | 1.7 | 0.93 | 25.75 | 24.98 | 0.77 | 0.77 | 0.01 | 0.01 |
| gene2683 | https://www.ncbi.nlm.nih.gov/gene/?term=LOC105275187 | 5492 | 66 | 120.17 | 1 | 1.82 | 1.52 | 14.03 | 13.27 | 0.76 | 0.76 | 0.01 | 0.01 |
| gene4376 | https://www.ncbi.nlm.nih.gov/gene/?term=LOC105277083 | 4307 | 92 | 213.61 | 1 | 2.32 | 1.09 | 26.82 | 26.06 | 0.76 | 0.76 | 0.02 | 0.02 |
| gene11340 | https://www.ncbi.nlm.nih.gov/gene/?term=LOC105285131 | 2918 | 93 | 318.71 | 1 | 3.43 | 1.08 | 12.16 | 11.4 | 0.75 | 0.75 | 0.02 | 0.02 |
| gene474 | https://www.ncbi.nlm.nih.gov/gene/?term=LOC105277435 | 3625 | 150 | 413.79 | 1 | 2.76 | 0.67 | 12.65 | 11.92 | 0.73 | 0.73 | 0.03 | 0.03 |
| gene1147 | https://www.ncbi.nlm.nih.gov/gene/?term=LOC105284954 | 4022 | 75 | 186.47 | 1 | 2.49 | 1.33 | 14.93 | 14.22 | 0.71 | 0.71 | 0.01 | 0.01 |
| gene2271 | https://www.ncbi.nlm.nih.gov/gene/?term=LOC105274706 | 1448 | 52 | 359.12 | 1 | 6.91 | 1.92 | 37.3 | 36.65 | 0.64 | 0.64 | 0.02 | 0.02 |
| gene7207 | https://www.ncbi.nlm.nih.gov/gene/?term=LOC105280376 | 4659 | 98 | 210.35 | 1 | 2.15 | 1.02 | 30.66 | 30.04 | 0.62 | 0.62 | 0.01 | 0.01 |
| gene657 | https://www.ncbi.nlm.nih.gov/gene/?term=LOC105279667 | 6421 | 64 | 99.67 | 1 | 1.56 | 1.56 | 3.41 | 2.8 | 0.61 | 0.61 | 0.01 | 0.01 |
| gene12573 | https://www.ncbi.nlm.nih.gov/gene/?term=LOC105286515 | 3647 | 106 | 290.65 | 1 | 2.74 | 0.94 | 10.61 | 10.01 | 0.6 | 0.6 | 0.02 | 0.02 |
| gene9645 | https://www.ncbi.nlm.nih.gov/gene/?term=LOC105283181 | 3875 | 118 | 304.52 | 1 | 2.58 | 0.85 | 12.99 | 12.4 | 0.59 | 0.59 | 0.02 | 0.02 |
| gene13036 | https://www.ncbi.nlm.nih.gov/gene/?term=LOC105287059 | 4959 | 150 | 302.48 | 1 | 2.02 | 0.67 | 17.57 | 17 | 0.57 | 0.57 | 0.02 | 0.02 |
| gene3599 | https://www.ncbi.nlm.nih.gov/gene/?term=LOC105276196 | 2955 | 40 | 135.36 | 1 | 3.38 | 2.5 | 52.66 | 52.1 | 0.56 | 0.56 | 0.01 | 0.01 |
| gene5831 | https://www.ncbi.nlm.nih.gov/gene/?term=LOC105278761 | 2506 | 100 | 399.04 | 2 | 7.98 | 2 | 24.32 | 23.77 | 0.54 | 0.54 | 0.02 | 0.02 |
| gene12307 | https://www.ncbi.nlm.nih.gov/gene/?term=LOC105286216 | 10313 | 146 | 141.57 | 1 | 0.97 | 0.68 | 17.27 | 16.76 | 0.51 | 0.51 | 0.01 | 0.01 |
| gene8661 | https://www.ncbi.nlm.nih.gov/gene/?term=LOC105282063 | 1926 | 39 | 202.49 | 1 | 5.19 | 2.56 | 23.72 | 23.24 | 0.48 | 0.48 | 0.01 | 0.01 |
| gene10326 | https://www.ncbi.nlm.nih.gov/gene/?term=LOC105283962 | 3042 | 92 | 302.43 | 1 | 3.29 | 1.09 | 15.29 | 14.85 | 0.44 | 0.44 | 0.01 | 0.01 |
| gene13044 | https://www.ncbi.nlm.nih.gov/gene/?term=LOC105287070 | 5094 | 159 | 312.13 | 1 | 1.96 | 0.63 | 5.69 | 5.29 | 0.4 | 0.4 | 0.01 | 0.01 |
| gene13114 | https://www.ncbi.nlm.nih.gov/gene/?term=LOC105287147 | 7609 | 360 | 473.12 | 1 | 1.31 | 0.28 | 10.34 | 9.95 | 0.39 | 0.39 | 0.02 | 0.02 |
| gene3817 | https://www.ncbi.nlm.nih.gov/gene/?term=LOC105276449 | 1230 | 39 | 317.07 | 1 | 8.13 | 2.56 | 32.77 | 32.39 | 0.38 | 0.38 | 0.01 | 0.01 |
| gene7568 | https://www.ncbi.nlm.nih.gov/gene/?term=LOC105280786 | 2525 | 107 | 423.76 | 1 | 3.96 | 0.93 | 33.06 | 32.68 | 0.38 | 0.38 | 0.02 | 0.02 |
| gene3206 | https://www.ncbi.nlm.nih.gov/gene/?term=LOC105275752 | 3474 | 117 | 336.79 | 1 | 2.88 | 0.85 | 29.75 | 29.38 | 0.37 | 0.37 | 0.01 | 0.01 |
| gene6583 | https://www.ncbi.nlm.nih.gov/gene/?term=LOC105279649 | 7444 | 308 | 413.76 | 1 | 1.34 | 0.32 | 11.75 | 11.41 | 0.34 | 0.34 | 0.01 | 0.01 |
| gene10392 | https://www.ncbi.nlm.nih.gov/gene/?term=LOC105284036 | 3216 | 76 | 236.32 | 1 | 3.11 | 1.32 | 25.16 | 24.85 | 0.31 | 0.31 | 0.01 | 0.01 |
| gene5539 | https://www.ncbi.nlm.nih.gov/gene/?term=LOC105278415 | 25024 | 542 | 216.59 | 1 | 0.4 | 0.18 | 4.22 | 3.95 | 0.27 | 0.27 | 0.01 | 0.01 |
| gene12674 | https://www.ncbi.nlm.nih.gov/gene/?term=LOC105286649 | 18283 | 638 | 348.96 | 1 | 0.55 | 0.16 | 1.31 | 1.05 | 0.26 | 0.26 | 0.01 | 0.01 |

| | | | | | | | | | | | | | |
|---|---|---|---|---|---|---|---|---|---|---|---|---|---|
| gene10549 | https://www.ncbi.nlm.nih.gov/gene/?term=LOC105284210 | 7145 | 298 | 417.07 | 1 | 1.4 | 0.34 | 11.27 | 11.01 | 0.25 | 0.25 | 0.01 | 0.01 |
| gene10600 | https://www.ncbi.nlm.nih.gov/gene/?term=LOC105284270 | 4837 | 97 | 200.54 | 1 | 2.07 | 1.03 | 13.04 | 12.82 | 0.22 | 0.22 | 0 | 0 |
| gene7249 | https://www.ncbi.nlm.nih.gov/gene/?term=LOC105280422 | 1775 | 58 | 326.76 | 1 | 5.63 | 1.72 | 15.27 | 15.08 | 0.19 | 0.19 | 0.01 | 0.01 |
| gene4962 | https://www.ncbi.nlm.nih.gov/gene/?term=LOC105277761 | 4645 | 140 | 301.4 | 1 | 2.15 | 0.71 | 14.56 | 14.4 | 0.16 | 0.16 | 0 | 0 |
| gene5967 | https://www.ncbi.nlm.nih.gov/gene/?term=LOC105278918 | 4393 | 121 | 275.44 | 1 | 2.28 | 0.83 | 16.38 | 16.32 | 0.06 | 0.06 | 0 | 0 |
| gene6031 | https://www.ncbi.nlm.nih.gov/gene/?term=LOC105279000 | 5691 | 174 | 305.75 | 1 | 1.76 | 0.57 | 14.03 | 13.97 | 0.06 | 0.06 | 0 | 0 |
| gene11583 | https://www.ncbi.nlm.nih.gov/gene/?term=LOC105285414 | 8192 | 168 | 205.08 | 1 | 1.22 | 0.6 | 8.3 | 8.25 | 0.05 | 0.05 | 0 | 0 |
| gene940 | https://www.ncbi.nlm.nih.gov/gene/?term=LOC105282794 | 3578 | 82 | 229.18 | 1 | 2.79 | 1.22 | 16.49 | 16.45 | 0.04 | 0.04 | 0 | 0 |
| gene5524 | https://www.ncbi.nlm.nih.gov/gene/?term=LOC105278397 | 5177 | 200 | 386.32 | 1 | 1.93 | 0.5 | 24.39 | 24.36 | 0.03 | 0.03 | 0 | 0 |
| gene5746 | https://www.ncbi.nlm.nih.gov/gene/?term=LOC105278666 | 10343 | 387 | 374.17 | 1 | 0.97 | 0.26 | 7.75 | 7.72 | 0.03 | 0.03 | 0 | 0 |
| gene4209 | https://www.ncbi.nlm.nih.gov/gene/?term=LOC105277125 | 50550 | 1576 | 311.77 | 1 | 0.2 | 0.06 | 1.66 | 1.64 | 0.02 | 0.02 | 0 | 0 |
| gene6639 | https://www.ncbi.nlm.nih.gov/gene/?term=LOC105279705 | 7656 | 58 | 75.76 | 1 | 1.31 | 1.72 | 24.54 | 24.54 | 0 | 0 | 0 | 0 |
| gene12802 | https://www.ncbi.nlm.nih.gov/gene/?term=LOC105286799 | 3183 | 165 | 518.38 | 1 | 3.14 | 0.61 | 16.21 | 16.23 | -0.02 | 0.02 | 0 | 0 |
| gene2806 | https://www.ncbi.nlm.nih.gov/gene/?term=LOC105275327 | 14257 | 414 | 290.38 | 1 | 0.7 | 0.24 | 7.7 | 7.73 | -0.02 | 0.02 | 0 | 0 |
| gene11579 | https://www.ncbi.nlm.nih.gov/gene/?term=LOC105285406 | 13571 | 248 | 182.74 | 1 | 0.74 | 0.4 | 5.98 | 6.04 | -0.05 | 0.05 | 0 | 0 |
| gene7148 | https://www.ncbi.nlm.nih.gov/gene/?term=LOC105280304 | 8920 | 90 | 100.9 | 1 | 1.12 | 1.11 | 7.28 | 7.34 | -0.05 | 0.05 | 0 | 0 |
| gene53 | https://www.ncbi.nlm.nih.gov/gene/?term=LOC105277148 | 19283 | 236 | 122.39 | 1 | 0.52 | 0.42 | 4.4 | 4.47 | -0.07 | 0.07 | 0 | 0 |
| gene5525 | https://www.ncbi.nlm.nih.gov/gene/?term=LOC105278396 | 3090 | 186 | 601.94 | 1 | 3.24 | 0.54 | 24.51 | 24.57 | -0.07 | 0.07 | 0 | 0 |
| gene6620 | https://www.ncbi.nlm.nih.gov/gene/?term=LOC105279687 | 17998 | 294 | 163.35 | 1 | 0.56 | 0.34 | 5.33 | 5.42 | -0.1 | 0.1 | 0 | 0 |
| gene4246 | https://www.ncbi.nlm.nih.gov/gene/?term=LOC105276941 | 16412 | 446 | 271.75 | 1 | 0.61 | 0.22 | 5.36 | 5.47 | -0.11 | 0.11 | 0 | 0 |
| gene9330 | https://www.ncbi.nlm.nih.gov/gene/?term=LOC105282825 | 3936 | 80 | 203.25 | 1 | 2.54 | 1.25 | 13.05 | 13.17 | -0.11 | 0.11 | 0 | 0 |
| gene10018 | https://www.ncbi.nlm.nih.gov/gene/?term=LOC105283607 | 4414 | 181 | 410.06 | 1 | 2.27 | 0.55 | 16.23 | 16.36 | -0.13 | 0.13 | -0.01 | 0.01 |
| gene12203 | https://www.ncbi.nlm.nih.gov/gene/?term=LOC105286099 | 5967 | 70 | 117.31 | 1 | 1.68 | 1.43 | 11.64 | 11.77 | -0.13 | 0.13 | 0 | 0 |
| gene1470 | https://www.ncbi.nlm.nih.gov/gene/?term=LOC105287597 | 2974 | 188 | 632.15 | 1 | 3.36 | 0.53 | 17.15 | 17.35 | -0.19 | 0.19 | -0.01 | 0.01 |
| gene3885 | https://www.ncbi.nlm.nih.gov/gene/?term=LOC105276527 | 7928 | 323 | 407.42 | 1 | 1.26 | 0.31 | 8.78 | 9 | -0.22 | 0.22 | -0.01 | 0.01 |
| gene11568 | https://www.ncbi.nlm.nih.gov/gene/?term=LOC105285395 | 8086 | 160 | 197.87 | 1 | 1.24 | 0.62 | 17.75 | 17.97 | -0.23 | 0.23 | 0 | 0 |
| gene10540 | https://www.ncbi.nlm.nih.gov/gene/?term=LOC105284200 | 5267 | 183 | 347.45 | 1 | 1.9 | 0.55 | 14.63 | 14.93 | -0.3 | 0.3 | -0.01 | 0.01 |
| gene6286 | https://www.ncbi.nlm.nih.gov/gene/?term=LOC105279298 | 2861 | 128 | 447.4 | 2 | 6.99 | 1.56 | 23.41 | 23.73 | -0.32 | 0.32 | -0.01 | 0.01 |
| gene3856 | https://www.ncbi.nlm.nih.gov/gene/?term=LOC105276499 | 5236 | 178 | 339.95 | 1 | 1.91 | 0.56 | 9.73 | 10.06 | -0.33 | 0.33 | -0.01 | 0.01 |
| gene4892 | https://www.ncbi.nlm.nih.gov/gene/?term=LOC105277667 | 3898 | 119 | 305.28 | 1 | 2.57 | 0.84 | 27.91 | 28.29 | -0.38 | 0.38 | -0.01 | 0.01 |
| gene12317 | https://www.ncbi.nlm.nih.gov/gene/?term=LOC105286223 | 2235 | 79 | 353.47 | 1 | 4.47 | 1.27 | 30.56 | 30.96 | -0.4 | 0.4 | -0.01 | 0.01 |
| gene1884 | https://www.ncbi.nlm.nih.gov/gene/?term=LOC105288072 | 4169 | 76 | 182.3 | 1 | 2.4 | 1.32 | 23.55 | 23.95 | -0.4 | 0.4 | -0.01 | 0.01 |
| gene2056 | https://www.ncbi.nlm.nih.gov/gene/?term=LOC105274452 | 6584 | 206 | 312.88 | 1 | 1.52 | 0.49 | 14.33 | 14.74 | -0.41 | 0.41 | -0.01 | 0.01 |
| gene4641 | https://www.ncbi.nlm.nih.gov/gene/?term=LOC105277398 | 1937 | 68 | 351.06 | 1 | 5.16 | 1.47 | 34.8 | 35.21 | -0.41 | 0.41 | -0.01 | 0.01 |
| gene10606 | https://www.ncbi.nlm.nih.gov/gene/?term=LOC105284273 | 6119 | 227 | 370.98 | 1 | 1.63 | 0.44 | 21.11 | 21.54 | -0.43 | 0.43 | -0.02 | 0.02 |
| gene5649 | https://www.ncbi.nlm.nih.gov/gene/?term=LOC105278538 | 2140 | 77 | 359.81 | 1 | 4.67 | 1.3 | 17.91 | 18.36 | -0.46 | 0.46 | -0.02 | 0.02 |
| gene9755 | https://www.ncbi.nlm.nih.gov/gene/?term=LOC105283296 | 2177 | 79 | 362.88 | 1 | 4.59 | 1.27 | 20.77 | 21.25 | -0.48 | 0.48 | -0.02 | 0.02 |
| gene303 | https://www.ncbi.nlm.nih.gov/gene/?term=LOC105275584 | 3081 | 71 | 230.45 | 1 | 3.25 | 1.41 | 26.68 | 27.17 | -0.49 | 0.49 | -0.01 | 0.01 |
| gene10678 | https://www.ncbi.nlm.nih.gov/gene/?term=LOC105284355 | 4031 | 152 | 377.08 | 1 | 2.48 | 0.66 | 20.4 | 20.91 | -0.51 | 0.51 | -0.02 | 0.02 |
| gene7888 | https://www.ncbi.nlm.nih.gov/gene/?term=LOC105281141 | 4357 | 153 | 351.16 | 1 | 2.3 | 0.65 | 30.21 | 30.79 | -0.57 | 0.57 | -0.02 | 0.02 |
| gene2844 | https://www.ncbi.nlm.nih.gov/gene/?term=LOC105275363 | 4330 | 179 | 413.39 | 1 | 2.31 | 0.56 | 14.53 | 15.15 | -0.61 | 0.61 | -0.03 | 0.03 |
| gene13033 | https://www.ncbi.nlm.nih.gov/gene/?term=LOC105287063 | 1877 | 45 | 239.74 | 1 | 5.33 | 2.22 | 34.91 | 35.54 | -0.62 | 0.62 | -0.01 | 0.01 |
| gene8600 | https://www.ncbi.nlm.nih.gov/gene/?term=LOC105282001 | 8513 | 126 | 148.01 | 1 | 1.17 | 0.79 | 11.25 | 11.9 | -0.65 | 0.65 | -0.01 | 0.01 |
| gene7245 | https://www.ncbi.nlm.nih.gov/gene/?term=LOC105280410 | 12478 | 234 | 187.53 | 1 | 0.8 | 0.43 | 20.56 | 21.23 | -0.67 | 0.67 | -0.01 | 0.01 |
| gene8343 | https://www.ncbi.nlm.nih.gov/gene/?term=LOC105281700 | 2172 | 40 | 184.16 | 1 | 4.6 | 2.5 | 41.18 | 41.86 | -0.68 | 0.68 | -0.01 | 0.01 |
| gene9336 | https://www.ncbi.nlm.nih.gov/gene/?term=LOC105282834 | 5993 | 106 | 176.87 | 1 | 1.67 | 0.94 | 15.44 | 16.15 | -0.71 | 0.71 | -0.01 | 0.01 |
| gene11275 | https://www.ncbi.nlm.nih.gov/gene/?term=LOC105285055 | 4119 | 196 | 475.84 | 1 | 2.43 | 0.51 | 13.91 | 14.65 | -0.74 | 0.74 | -0.04 | 0.04 |
| gene10344 | https://www.ncbi.nlm.nih.gov/gene/?term=LOC105283987 | 4841 | 64 | 132.2 | 1 | 2.07 | 1.56 | 32.11 | 32.92 | -0.81 | 0.81 | -0.01 | 0.01 |
| gene292 | https://www.ncbi.nlm.nih.gov/gene/?term=LOC105275398 | 2825 | 77 | 272.57 | 1 | 3.54 | 1.3 | 19.76 | 20.59 | -0.82 | 0.82 | -0.02 | 0.02 |
| gene8729 | https://www.ncbi.nlm.nih.gov/gene/?term=LOC105282137 | 5005 | 195 | 389.61 | 1 | 2 | 0.51 | 16.47 | 17.35 | -0.89 | 0.89 | -0.03 | 0.03 |
| gene298 | https://www.ncbi.nlm.nih.gov/gene/?term=LOC105275460 | 2473 | 107 | 432.67 | 1 | 4.04 | 0.93 | 13.52 | 14.43 | -0.91 | 0.91 | -0.04 | 0.04 |
| gene7166 | https://www.ncbi.nlm.nih.gov/gene/?term=LOC105280326 | 6117 | 195 | 318.78 | 1 | 1.63 | 0.51 | 16.86 | 17.78 | -0.92 | 0.92 | -0.03 | 0.03 |
| gene9101 | https://www.ncbi.nlm.nih.gov/gene/?term=LOC105282553 | 3068 | 104 | 338.98 | 1 | 3.26 | 0.96 | 20.36 | 21.29 | -0.93 | 0.93 | -0.03 | 0.03 |
| gene10531 | https://www.ncbi.nlm.nih.gov/gene/?term=LOC105284190 | 6881 | 164 | 238.34 | 1 | 1.45 | 0.61 | 10.48 | 11.52 | -1.03 | 1.03 | -0.02 | 0.02 |
| gene11819 | https://www.ncbi.nlm.nih.gov/gene/?term=LOC105285671 | 2366 | 91 | 384.62 | 1 | 4.23 | 1.1 | 28.12 | 29.18 | -1.06 | 1.06 | -0.04 | 0.04 |
| gene2173 | https://www.ncbi.nlm.nih.gov/gene/?term=LOC105274590 | 4387 | 64 | 145.89 | 1 | 2.28 | 1.56 | 26 | 27.13 | -1.13 | 1.13 | -0.02 | 0.02 |
| gene6528 | https://www.ncbi.nlm.nih.gov/gene/?term=LOC105279591 | 7222 | 67 | 92.77 | 1 | 1.38 | 1.49 | 22.84 | 24.15 | -1.32 | 1.32 | -0.01 | 0.01 |
| gene4023 | https://www.ncbi.nlm.nih.gov/gene/?term=LOC105276692 | 1759 | 35 | 198.98 | 1 | 5.69 | 2.86 | 23.93 | 25.35 | -1.42 | 1.42 | -0.03 | 0.03 |
| gene10984 | https://www.ncbi.nlm.nih.gov/gene/?term=LOC105284704 | 1987 | 61 | 307 | 1 | 5.03 | 1.64 | 25.94 | 27.43 | -1.49 | 1.49 | -0.05 | 0.05 |
| gene2970 | https://www.ncbi.nlm.nih.gov/gene/?term=LOC105275518 | 2340 | 18 | 76.92 | 1 | 4.27 | 5.56 | 47.45 | 48.96 | -1.51 | 1.51 | -0.01 | 0.01 |
| gene1310 | https://www.ncbi.nlm.nih.gov/gene/?term=LOC105286661 | 2582 | 65 | 251.74 | 1 | 3.87 | 1.54 | 19.96 | 21.58 | -1.62 | 1.62 | -0.04 | 0.04 |
| gene7556 | https://www.ncbi.nlm.nih.gov/gene/?term=LOC105280774 | 2792 | 98 | 351 | 1 | 3.58 | 1.02 | 31.47 | 33.13 | -1.66 | 1.66 | -0.06 | 0.06 |
| gene6455 | https://www.ncbi.nlm.nih.gov/gene/?term=LOC105279500 | 2612 | 31 | 118.68 | 1 | 3.83 | 3.23 | 51.05 | 52.76 | -1.71 | 1.71 | -0.02 | 0.02 |
| gene10520 | https://www.ncbi.nlm.nih.gov/gene/?term=LOC105284184 | 2451 | 67 | 273.36 | 1 | 4.08 | 1.49 | 41.63 | 43.45 | -1.81 | 1.81 | -0.05 | 0.05 |

| | | | | | | | | | | | | | | |
|---|---|---|---|---|---|---|---|---|---|---|---|---|---|---|
| gene274 | https://www.ncbi.nlm.nih.gov/gene/?term=LOC105275190 | 3488 | 66 | 189.22 | | 1 | 2.87 | 1.52 | 20.22 | 22.09 | -1.87 | 1.87 | -0.04 | 0.04 |
| gene3727 | https://www.ncbi.nlm.nih.gov/gene/?term=LOC105276367 | 2484 | 54 | 217.39 | | 1 | 4.03 | 1.85 | 48.38 | 50.33 | -1.95 | 1.95 | -0.04 | 0.04 |
| gene5877 | https://www.ncbi.nlm.nih.gov/gene/?term=LOC105278807 | 1719 | 50 | 290.87 | | 1 | 5.82 | 2 | 21.66 | 23.62 | -1.95 | 1.95 | -0.06 | 0.06 |
| gene12516 | https://www.ncbi.nlm.nih.gov/gene/?term=LOC105286452 | 1529 | 35 | 228.91 | | 1 | 6.54 | 2.86 | 51.01 | 53.03 | -2.03 | 2.03 | -0.05 | 0.05 |
| gene7183 | https://www.ncbi.nlm.nih.gov/gene/?term=LOC105280348 | 3925 | 51 | 129.94 | | 1 | 2.55 | 1.96 | 25.44 | 27.74 | -2.3 | 2.3 | -0.03 | 0.03 |
| gene10107 | https://www.ncbi.nlm.nih.gov/gene/?term=LOC105283712 | 1883 | 23 | 122.15 | | 1 | 5.31 | 4.35 | 27.23 | 29.73 | -2.49 | 2.49 | -0.03 | 0.03 |
| gene12406 | https://www.ncbi.nlm.nih.gov/gene/?term=LOC105286316 | 1386 | 29 | 209.24 | | 1 | 7.22 | 3.45 | 41.65 | 44.38 | -2.73 | 2.73 | -0.06 | 0.06 |
| gene7398 | https://www.ncbi.nlm.nih.gov/gene/?term=LOC105280589 | 3452 | 41 | 118.77 | | 1 | 2.9 | 2.44 | 25.07 | 28.12 | -3.05 | 3.05 | -0.04 | 0.04 |
| gene6334 | https://www.ncbi.nlm.nih.gov/gene/?term=LOC105279356 | 1471 | 40 | 271.92 | | 1 | 6.8 | 2.5 | 29.12 | 32.21 | -3.09 | 3.09 | -0.08 | 0.08 |
| gene7734 | https://www.ncbi.nlm.nih.gov/gene/?term=LOC105280972 | 1528 | 26 | 170.16 | | 1 | 6.54 | 3.85 | 20.04 | 23.88 | -3.84 | 3.84 | -0.07 | 0.07 |
| gene2824 | https://www.ncbi.nlm.nih.gov/gene/?term=LOC105275337 | 1772 | 58 | 327.31 | | 1 | 5.64 | 1.72 | 31.86 | 38.08 | -6.22 | 6.22 | -0.2 | 0.2 |
| gene5714 | https://www.ncbi.nlm.nih.gov/gene/?term=LOC105278729 | 710 | 14 | 197.18 | | 1 | 14.08 | 7.14 | 51.67 | 57.98 | -6.31 | 6.31 | -0.12 | 0.12 |
| gene2676 | https://www.ncbi.nlm.nih.gov/gene/?term=LOC105275181 | 1353 | 15 | 110.86 | | 1 | 7.39 | 6.67 | 67.39 | 74.05 | -6.66 | 6.66 | -0.07 | 0.07 |
| gene3165 | https://www.ncbi.nlm.nih.gov/gene/?term=LOC105275706 | 2374 | 65 | 273.8 | 6 | 25.27 | 9.23 | 24.65 | 38.08 | -13.43 | | 13.43 | -0.37 | 0.37 |
| gene2593 | https://www.ncbi.nlm.nih.gov/gene/?term=LOC105275070 | 1774 | 5 | 28.18 | 2 | 11.27 | 40 | 19.71 | 71.28 | -51.57 | | 51.57 | -0.15 | 0.15 |

Directory: OGLall File: ogl-Cb_LI_BCphaseA1.vs.BCphaseB1.txt

| gene_ID | gene_link | gwidth | #Sites | #per10Kb | #dmSites | #dmsp10kb | %dmSites | %pSite1 | %pSite2 | DMpSite | ADMpSite | DMpNucl | ADMpNucl |
|---|---|---|---|---|---|---|---|---|---|---|---|---|---|
| gene393 | https://www.ncbi.nlm.nih.gov/gene/?term=LOC105276566 | 1123 | 2 | 17.81 | 1 | 8.9 | 50 | 77.65 | 47.22 | 30.43 | 30.43 | 0.05 | 0.05 |
| gene9109 | https://www.ncbi.nlm.nih.gov/gene/?term=LOC105282561 | 2400 | 12 | 50 | 1 | 4.17 | 8.33 | 20.55 | 12.93 | 7.62 | 7.62 | 0.04 | 0.04 |
| gene10602 | https://www.ncbi.nlm.nih.gov/gene/?term=LOC105284268 | 2122 | 20 | 94.25 | 1 | 4.71 | 5 | 32.63 | 25.61 | 7.02 | 7.02 | 0.07 | 0.07 |
| gene7199 | https://www.ncbi.nlm.nih.gov/gene/?term=LOC105280366 | 1414 | 24 | 169.73 | 1 | 7.07 | 4.17 | 45.18 | 38.6 | 6.59 | 6.59 | 0.11 | 0.11 |
| gene9464 | https://www.ncbi.nlm.nih.gov/gene/?term=LOC105282975 | 1450 | 24 | 165.52 | 1 | 6.9 | 4.17 | 42.63 | 39.26 | 3.37 | 3.37 | 0.06 | 0.06 |
| gene3168 | https://www.ncbi.nlm.nih.gov/gene/?term=LOC105275711 | 6150 | 79 | 128.46 | 1 | 1.63 | 1.27 | 14.18 | 11.09 | 3.09 | 3.09 | 0.04 | 0.04 |
| gene6422 | https://www.ncbi.nlm.nih.gov/gene/?term=LOC105279458 | 2419 | 36 | 148.82 | 1 | 4.13 | 2.78 | 53.83 | 51.06 | 2.78 | 2.78 | 0.04 | 0.04 |
| gene7915 | https://www.ncbi.nlm.nih.gov/gene/?term=LOC105281173 | 1597 | 25 | 156.54 | 1 | 6.26 | 4 | 64.87 | 62.27 | 2.61 | 2.61 | 0.04 | 0.04 |
| gene9740 | https://www.ncbi.nlm.nih.gov/gene/?term=LOC105283277 | 2887 | 138 | 478 | 1 | 3.46 | 0.72 | 23.22 | 20.78 | 2.44 | 2.44 | 0.12 | 0.12 |
| gene7368 | https://www.ncbi.nlm.nih.gov/gene/?term=LOC105280556 | 4105 | 49 | 119.37 | 1 | 2.44 | 2.04 | 12.67 | 10.47 | 2.2 | 2.2 | 0.03 | 0.03 |
| gene652 | https://www.ncbi.nlm.nih.gov/gene/?term=LOC105279528 | 5806 | 42 | 72.34 | 1 | 1.72 | 2.38 | 16.48 | 14.36 | 2.12 | 2.12 | 0.02 | 0.02 |
| gene11401 | https://www.ncbi.nlm.nih.gov/gene/?term=LOC105285202 | 1841 | 50 | 271.59 | 1 | 5.43 | 2 | 21.11 | 19.1 | 2.01 | 2.01 | 0.05 | 0.05 |
| gene1199 | https://www.ncbi.nlm.nih.gov/gene/?term=LOC105285498 | 2220 | 26 | 117.12 | 1 | 4.5 | 3.85 | 33.11 | 31.11 | 2 | 2 | 0.02 | 0.02 |
| gene388 | https://www.ncbi.nlm.nih.gov/gene/?term=LOC105276538 | 1558 | 19 | 121.95 | 1 | 6.42 | 5.26 | 69.93 | 68.17 | 1.77 | 1.77 | 0.02 | 0.02 |
| gene10213 | https://www.ncbi.nlm.nih.gov/gene/?term=LOC105283826 | 1503 | 54 | 359.28 | 1 | 6.65 | 1.85 | 16.14 | 14.48 | 1.66 | 1.66 | 0.06 | 0.06 |
| gene10326 | https://www.ncbi.nlm.nih.gov/gene/?term=LOC105283962 | 3042 | 93 | 305.72 | 1 | 3.29 | 1.08 | 11.63 | 10.03 | 1.6 | 1.6 | 0.05 | 0.05 |
| gene3727 | https://www.ncbi.nlm.nih.gov/gene/?term=LOC105276367 | 2484 | 50 | 201.29 | 1 | 4.03 | 2 | 50.02 | 48.5 | 1.52 | 1.52 | 0.03 | 0.03 |
| gene7345 | https://www.ncbi.nlm.nih.gov/gene/?term=LOC105280529 | 3833 | 87 | 226.98 | 1 | 2.61 | 1.15 | 18.24 | 16.79 | 1.44 | 1.44 | 0.03 | 0.03 |
| gene322 | https://www.ncbi.nlm.nih.gov/gene/?term=LOC105275777 | 1606 | 22 | 136.99 | 1 | 6.23 | 4.55 | 47.62 | 46.19 | 1.43 | 1.43 | 0.02 | 0.02 |
| gene3599 | https://www.ncbi.nlm.nih.gov/gene/?term=LOC105276196 | 2955 | 54 | 182.74 | 1 | 3.38 | 1.85 | 40.6 | 39.18 | 1.42 | 1.42 | 0.03 | 0.03 |
| gene11340 | https://www.ncbi.nlm.nih.gov/gene/?term=LOC105285131 | 2918 | 97 | 332.42 | 1 | 3.43 | 1.03 | 14.75 | 13.34 | 1.41 | 1.41 | 0.05 | 0.05 |
| gene7248 | https://www.ncbi.nlm.nih.gov/gene/?term=LOC105280408 | 5175 | 44 | 85.02 | 1 | 1.93 | 2.27 | 21.35 | 20.08 | 1.27 | 1.27 | 0.01 | 0.01 |
| gene4190 | https://www.ncbi.nlm.nih.gov/gene/?term=LOC105276875 | 3150 | 130 | 412.7 | 1 | 3.17 | 0.77 | 21.07 | 19.84 | 1.23 | 1.23 | 0.05 | 0.05 |
| gene5298 | https://www.ncbi.nlm.nih.gov/gene/?term=LOC105278145 | 4060 | 250 | 615.76 | 2 | 4.93 | 0.8 | 10.79 | 9.61 | 1.17 | 1.17 | 0.07 | 0.07 |
| gene10606 | https://www.ncbi.nlm.nih.gov/gene/?term=LOC105284273 | 6119 | 249 | 406.93 | 2 | 3.27 | 0.8 | 20.32 | 19.15 | 1.16 | 1.16 | 0.05 | 0.05 |
| gene920 | https://www.ncbi.nlm.nih.gov/gene/?term=LOC105282518 | 1197 | 43 | 359.23 | 1 | 8.35 | 2.33 | 37.5 | 36.41 | 1.09 | 1.09 | 0.04 | 0.04 |
| gene2271 | https://www.ncbi.nlm.nih.gov/gene/?term=LOC105274706 | 1448 | 47 | 324.59 | 1 | 6.91 | 2.13 | 41.16 | 40.14 | 1.03 | 1.03 | 0.03 | 0.03 |
| gene7247 | https://www.ncbi.nlm.nih.gov/gene/?term=LOC105280407 | 10486 | 205 | 195.5 | 1 | 0.95 | 0.49 | 17 | 15.99 | 1.01 | 1.01 | 0.02 | 0.02 |
| gene4376 | https://www.ncbi.nlm.nih.gov/gene/?term=LOC105277083 | 4307 | 92 | 213.61 | 1 | 2.32 | 1.09 | 26.68 | 25.69 | 0.99 | 0.99 | 0.02 | 0.02 |
| gene564 | https://www.ncbi.nlm.nih.gov/gene/?term=LOC105278478 | 2802 | 13 | 46.4 | 1 | 3.57 | 7.69 | 25.47 | 24.48 | 0.99 | 0.99 | 0 | 0 |
| gene1220 | https://www.ncbi.nlm.nih.gov/gene/?term=LOC105285742 | 3453 | 94 | 272.23 | 1 | 2.9 | 1.06 | 26.72 | 25.74 | 0.97 | 0.97 | 0.03 | 0.03 |
| gene6031 | https://www.ncbi.nlm.nih.gov/gene/?term=LOC105279000 | 5691 | 180 | 316.29 | 1 | 1.76 | 0.56 | 14.05 | 13.11 | 0.94 | 0.94 | 0.03 | 0.03 |
| gene4962 | https://www.ncbi.nlm.nih.gov/gene/?term=LOC105277761 | 4645 | 140 | 301.4 | 1 | 2.15 | 0.71 | 15.27 | 14.38 | 0.89 | 0.89 | 0.03 | 0.03 |
| gene7249 | https://www.ncbi.nlm.nih.gov/gene/?term=LOC105280422 | 1775 | 61 | 343.66 | 1 | 5.63 | 1.64 | 15.16 | 14.36 | 0.81 | 0.81 | 0.03 | 0.03 |
| gene10392 | https://www.ncbi.nlm.nih.gov/gene/?term=LOC105284036 | 3216 | 84 | 261.19 | 2 | 6.22 | 2.38 | 23.38 | 22.62 | 0.76 | 0.76 | 0.02 | 0.02 |
| gene657 | https://www.ncbi.nlm.nih.gov/gene/?term=LOC105279667 | 6421 | 78 | 121.48 | 1 | 1.56 | 1.28 | 4.46 | 3.72 | 0.74 | 0.74 | 0.01 | 0.01 |
| gene10844 | https://www.ncbi.nlm.nih.gov/gene/?term=LOC105284535 | 9086 | 220 | 242.13 | 1 | 1.1 | 0.45 | 16.85 | 16.12 | 0.73 | 0.73 | 0.02 | 0.02 |
| gene6966 | https://www.ncbi.nlm.nih.gov/gene/?term=LOC105280104 | 3747 | 124 | 330.93 | 1 | 2.67 | 0.81 | 15.19 | 14.51 | 0.68 | 0.68 | 0.02 | 0.02 |
| gene12203 | https://www.ncbi.nlm.nih.gov/gene/?term=LOC105286099 | 5967 | 78 | 130.72 | 1 | 1.68 | 1.28 | 11.28 | 10.61 | 0.67 | 0.67 | 0.01 | 0.01 |
| gene3793 | https://www.ncbi.nlm.nih.gov/gene/?term=LOC105276429 | 2901 | 112 | 386.07 | 1 | 3.45 | 0.89 | 26.42 | 25.76 | 0.65 | 0.65 | 0.03 | 0.03 |
| gene1470 | https://www.ncbi.nlm.nih.gov/gene/?term=LOC105287597 | 2974 | 180 | 605.25 | 1 | 3.36 | 0.56 | 17.88 | 17.25 | 0.63 | 0.63 | 0.04 | 0.04 |
| gene11454 | https://www.ncbi.nlm.nih.gov/gene/?term=LOC105285256 | 1906 | 52 | 272.82 | 1 | 5.25 | 1.92 | 42.98 | 42.36 | 0.62 | 0.62 | 0.02 | 0.02 |
| gene1147 | https://www.ncbi.nlm.nih.gov/gene/?term=LOC105284954 | 4022 | 87 | 216.31 | 1 | 2.49 | 1.15 | 15.12 | 14.51 | 0.61 | 0.61 | 0.01 | 0.01 |
| gene12021 | https://www.ncbi.nlm.nih.gov/gene/?term=LOC105285905 | 1399 | 21 | 150.11 | 1 | 7.15 | 4.76 | 44.76 | 44.21 | 0.55 | 0.55 | 0.01 | 0.01 |
| gene2683 | https://www.ncbi.nlm.nih.gov/gene/?term=LOC105275187 | 5492 | 75 | 136.56 | 1 | 1.82 | 1.33 | 14.15 | 13.62 | 0.53 | 0.53 | 0.01 | 0.01 |
| gene11583 | https://www.ncbi.nlm.nih.gov/gene/?term=LOC105285414 | 8192 | 179 | 218.51 | 1 | 1.22 | 0.56 | 9.12 | 8.6 | 0.52 | 0.52 | 0.01 | 0.01 |
| gene2748 | https://www.ncbi.nlm.nih.gov/gene/?term=LOC105275261 | 3845 | 54 | 140.44 | 1 | 2.6 | 1.85 | 38.38 | 37.86 | 0.52 | 0.52 | 0.01 | 0.01 |
| gene7568 | https://www.ncbi.nlm.nih.gov/gene/?term=LOC105280786 | 2525 | 121 | 479.21 | 1 | 3.96 | 0.83 | 29.4 | 28.89 | 0.52 | 0.52 | 0.02 | 0.02 |
| gene13036 | https://www.ncbi.nlm.nih.gov/gene/?term=LOC105287059 | 4959 | 166 | 334.74 | 1 | 2.02 | 0.6 | 17.36 | 16.86 | 0.5 | 0.5 | 0.02 | 0.02 |
| gene7801 | https://www.ncbi.nlm.nih.gov/gene/?term=LOC105281047 | 5137 | 113 | 219.97 | 1 | 1.95 | 0.88 | 10.88 | 10.38 | 0.5 | 0.5 | 0.01 | 0.01 |
| gene474 | https://www.ncbi.nlm.nih.gov/gene/?term=LOC105277435 | 3625 | 155 | 427.59 | 1 | 2.76 | 0.65 | 12.2 | 11.71 | 0.49 | 0.49 | 0.02 | 0.02 |
| gene5525 | https://www.ncbi.nlm.nih.gov/gene/?term=LOC105278396 | 3090 | 187 | 605.18 | 1 | 3.24 | 0.53 | 24.38 | 23.89 | 0.49 | 0.49 | 0.03 | 0.03 |
| gene10549 | https://www.ncbi.nlm.nih.gov/gene/?term=LOC105284210 | 7145 | 319 | 446.47 | 1 | 1.4 | 0.31 | 10.59 | 10.18 | 0.4 | 0.4 | 0.02 | 0.02 |
| gene6286 | https://www.ncbi.nlm.nih.gov/gene/?term=LOC105279298 | 2861 | 134 | 468.37 | 1 | 3.5 | 0.75 | 23.06 | 22.69 | 0.37 | 0.37 | 0.02 | 0.02 |
| gene5524 | https://www.ncbi.nlm.nih.gov/gene/?term=LOC105278397 | 5177 | 201 | 388.26 | 1 | 1.93 | 0.5 | 24.26 | 23.91 | 0.35 | 0.35 | 0.01 | 0.01 |
| gene927 | https://www.ncbi.nlm.nih.gov/gene/?term=LOC105282600 | 3727 | 57 | 152.94 | 1 | 2.68 | 1.75 | 29.17 | 28.82 | 0.35 | 0.35 | 0.01 | 0.01 |
| gene12573 | https://www.ncbi.nlm.nih.gov/gene/?term=LOC105286515 | 3647 | 102 | 279.68 | 1 | 2.74 | 0.98 | 10.87 | 10.55 | 0.32 | 0.32 | 0.01 | 0.01 |
| gene5746 | https://www.ncbi.nlm.nih.gov/gene/?term=LOC105278666 | 10343 | 405 | 391.57 | 1 | 0.97 | 0.25 | 7.03 | 6.73 | 0.3 | 0.3 | 0.01 | 0.01 |
| gene10155 | https://www.ncbi.nlm.nih.gov/gene/?term=LOC105283756 | 7030 | 97 | 137.98 | 1 | 1.42 | 1.03 | 6.83 | 6.6 | 0.23 | 0.23 | 0 | 0 |
| gene5539 | https://www.ncbi.nlm.nih.gov/gene/?term=LOC105278415 | 25024 | 582 | 232.58 | 1 | 0.4 | 0.17 | 3.85 | 3.62 | 0.23 | 0.23 | 0.01 | 0.01 |

| | | | | | | | | | | | | | |
|---|---|---|---|---|---|---|---|---|---|---|---|---|---|
| gene4213 | https://www.ncbi.nlm.nih.gov/gene/?term=LOC105276901 | 3257 | 95 | 291.68 | 1 | 3.07 | 1.05 | 23.09 | 22.87 | 0.22 | 0.22 | 0.01 | 0.01 |
| gene6620 | https://www.ncbi.nlm.nih.gov/gene/?term=LOC105279687 | 17998 | 363 | 201.69 | 1 | 0.56 | 0.28 | 5.06 | 4.84 | 0.22 | 0.22 | 0 | 0 |
| gene5831 | https://www.ncbi.nlm.nih.gov/gene/?term=LOC105278761 | 2506 | 103 | 411.01 | 2 | 7.98 | 1.94 | 24.82 | 24.61 | 0.21 | 0.21 | 0.01 | 0.01 |
| gene8274 | https://www.ncbi.nlm.nih.gov/gene/?term=LOC105281624 | 5425 | 203 | 374.19 | 1 | 1.84 | 0.49 | 10.66 | 10.47 | 0.2 | 0.2 | 0.01 | 0.01 |
| gene3206 | https://www.ncbi.nlm.nih.gov/gene/?term=LOC105275752 | 3474 | 123 | 354.06 | 1 | 2.88 | 0.81 | 28.92 | 28.73 | 0.19 | 0.19 | 0.01 | 0.01 |
| gene10600 | https://www.ncbi.nlm.nih.gov/gene/?term=LOC105284270 | 4837 | 103 | 212.94 | 1 | 2.07 | 0.97 | 12.92 | 12.79 | 0.13 | 0.13 | 0 | 0 |
| gene5646 | https://www.ncbi.nlm.nih.gov/gene/?term=LOC105278534 | 7475 | 185 | 247.49 | 1 | 1.34 | 0.54 | 17.64 | 17.56 | 0.08 | 0.08 | 0 | 0 |
| gene5967 | https://www.ncbi.nlm.nih.gov/gene/?term=LOC105278918 | 4393 | 132 | 300.48 | 1 | 2.28 | 0.76 | 15.7 | 15.62 | 0.08 | 0.08 | 0 | 0 |
| gene12802 | https://www.ncbi.nlm.nih.gov/gene/?term=LOC105286799 | 3183 | 170 | 534.09 | 1 | 3.14 | 0.59 | 15.91 | 15.85 | 0.06 | 0.06 | 0 | 0 |
| gene3470 | https://www.ncbi.nlm.nih.gov/gene/?term=LOC105276059 | 8716 | 159 | 182.42 | 1 | 1.15 | 0.63 | 9.24 | 9.2 | 0.04 | 0.04 | 0 | 0 |
| gene2806 | https://www.ncbi.nlm.nih.gov/gene/?term=LOC105275327 | 14257 | 479 | 335.98 | 1 | 0.7 | 0.21 | 7.13 | 7.12 | 0.01 | 0.01 | 0 | 0 |
| gene5375 | https://www.ncbi.nlm.nih.gov/gene/?term=LOC105278228 | 8640 | 161 | 186.34 | 1 | 1.16 | 0.62 | 23.31 | 23.3 | 0.01 | 0.01 | 0 | 0 |
| gene2056 | https://www.ncbi.nlm.nih.gov/gene/?term=LOC105274452 | 6584 | 206 | 312.88 | 1 | 1.52 | 0.49 | 14.18 | 14.18 | 0 | 0 | 0 | 0 |
| gene11579 | https://www.ncbi.nlm.nih.gov/gene/?term=LOC105285406 | 13571 | 282 | 207.8 1 | 0.74 | 0.35 | 5.68 | 5.69 | -0.01 | 0.01 | 0 | 0 | |
| gene3107 | https://www.ncbi.nlm.nih.gov/gene/?term=LOC105275650 | 5103 | 150 | 293.94 | 1 | 1.96 | 0.67 | 17.09 | 17.11 | -0.02 | 0.02 | 0 | 0 |
| gene2173 | https://www.ncbi.nlm.nih.gov/gene/?term=LOC105274590 | 4387 | 79 | 180.08 | 1 | 2.28 | 1.27 | 22.08 | 22.11 | -0.03 | 0.03 | 0 | 0 |
| gene13114 | https://www.ncbi.nlm.nih.gov/gene/?term=LOC105287147 | 7609 | 381 | 500.72 | 1 | 1.31 | 0.26 | 9.67 | 9.7 | -0.04 | 0.04 | 0 | 0 |
| gene2668 | https://www.ncbi.nlm.nih.gov/gene/?term=LOC105275170 | 12496 | 336 | 268.89 | 1 | 0.8 | 0.3 | 2.05 | 2.12 | -0.07 | 0.07 | 0 | 0 |
| gene6698 | https://www.ncbi.nlm.nih.gov/gene/?term=LOC105279778 | 4099 | 138 | 336.67 | 1 | 2.44 | 0.72 | 9.41 | 9.48 | -0.07 | 0.07 | 0 | 0 |
| gene6783 | https://www.ncbi.nlm.nih.gov/gene/?term=LOC105279865 | 62267 | 1495 | 240.1 1 | 0.16 | 0.07 | 1.69 | 1.78 | -0.09 | 0.09 | 0 | 0 | |
| gene53 | https://www.ncbi.nlm.nih.gov/gene/?term=LOC105277148 | 19283 | 270 | 140.02 | 1 | 0.52 | 0.37 | 3.92 | 4.02 | -0.1 | 0.1 | 0 | 0 |
| gene11568 | https://www.ncbi.nlm.nih.gov/gene/?term=LOC105285395 | 8086 | 176 | 217.66 | 1 | 1.24 | 0.57 | 15.78 | 15.89 | -0.11 | 0.11 | 0 | 0 |
| gene10596 | https://www.ncbi.nlm.nih.gov/gene/?term=LOC105284262 | 6520 | 228 | 349.69 | 1 | 1.53 | 0.44 | 3.34 | 3.5 | -0.16 | 0.16 | -0.01 | 0.01 |
| gene10540 | https://www.ncbi.nlm.nih.gov/gene/?term=LOC105284200 | 5267 | 193 | 366.43 | 1 | 1.9 | 0.52 | 13.99 | 14.21 | -0.22 | 0.22 | -0.01 | 0.01 |
| gene303 | https://www.ncbi.nlm.nih.gov/gene/?term=LOC105275584 | 3081 | 86 | 279.13 | 1 | 3.25 | 1.16 | 23.78 | 24 | -0.22 | 0.22 | -0.01 | 0.01 |
| gene5184 | https://www.ncbi.nlm.nih.gov/gene/?term=LOC105277999 | 5528 | 147 | 265.92 | 1 | 1.81 | 0.68 | 12.4 | 12.64 | -0.24 | 0.24 | -0.01 | 0.01 |
| gene7888 | https://www.ncbi.nlm.nih.gov/gene/?term=LOC105281141 | 4357 | 155 | 355.75 | 1 | 2.3 | 0.65 | 29.83 | 30.06 | -0.24 | 0.24 | -0.01 | 0.01 |
| gene7398 | https://www.ncbi.nlm.nih.gov/gene/?term=LOC105280589 | 3452 | 41 | 118.77 | 1 | 2.9 | 2.44 | 25.07 | 25.33 | -0.26 | 0.26 | 0 | 0 |
| gene7166 | https://www.ncbi.nlm.nih.gov/gene/?term=LOC105280326 | 6117 | 208 | 340.04 | 1 | 1.63 | 0.48 | 15.97 | 16.24 | -0.27 | 0.27 | -0.01 | 0.01 |
| gene6528 | https://www.ncbi.nlm.nih.gov/gene/?term=LOC105279591 | 7222 | 67 | 92.77 1 | 1.38 | 1.49 | 23.03 | 23.32 | -0.28 | 0.28 | 0 | 0 | |
| gene4023 | https://www.ncbi.nlm.nih.gov/gene/?term=LOC105276692 | 1759 | 38 | 216.03 | 1 | 5.69 | 2.63 | 22.43 | 22.73 | -0.3 | 0.3 | -0.01 | 0.01 |
| gene6455 | https://www.ncbi.nlm.nih.gov/gene/?term=LOC105279500 | 2612 | 32 | 122.51 | 1 | 3.83 | 3.12 | 49.77 | 50.07 | -0.3 | 0.3 | 0 | 0 |
| gene2443 | https://www.ncbi.nlm.nih.gov/gene/?term=LOC105274895 | 10433 | 345 | 330.68 | 1 | 0.96 | 0.29 | 6.76 | 7.09 | -0.33 | 0.33 | -0.01 | 0.01 |
| gene10520 | https://www.ncbi.nlm.nih.gov/gene/?term=LOC105284184 | 2451 | 82 | 334.56 | 1 | 4.08 | 1.22 | 34.55 | 34.9 | -0.35 | 0.35 | -0.01 | 0.01 |
| gene2404 | https://www.ncbi.nlm.nih.gov/gene/?term=LOC105274853 | 1767 | 37 | 209.39 | 1 | 5.66 | 2.7 | 31.99 | 32.37 | -0.37 | 0.37 | -0.01 | 0.01 |
| gene10531 | https://www.ncbi.nlm.nih.gov/gene/?term=LOC105284190 | 6881 | 192 | 279.03 | 1 | 1.45 | 0.52 | 9.5 | 9.91 | -0.4 | 0.4 | -0.01 | 0.01 |
| gene3856 | https://www.ncbi.nlm.nih.gov/gene/?term=LOC105276499 | 5236 | 197 | 376.24 | 1 | 1.91 | 0.51 | 9.94 | 10.34 | -0.41 | 0.41 | -0.02 | 0.02 |
| gene4892 | https://www.ncbi.nlm.nih.gov/gene/?term=LOC105277667 | 3898 | 120 | 307.85 | 1 | 2.57 | 0.83 | 28.07 | 28.49 | -0.42 | 0.42 | -0.01 | 0.01 |
| gene6583 | https://www.ncbi.nlm.nih.gov/gene/?term=LOC105279649 | 7444 | 317 | 425.85 | 1 | 1.34 | 0.32 | 11.9 | 12.34 | -0.44 | 0.44 | -0.02 | 0.02 |
| gene11275 | https://www.ncbi.nlm.nih.gov/gene/?term=LOC105285055 | 4119 | 213 | 517.12 | 1 | 2.43 | 0.47 | 13.26 | 13.73 | -0.47 | 0.47 | -0.02 | 0.02 |
| gene9411 | https://www.ncbi.nlm.nih.gov/gene/?term=LOC105282914 | 5534 | 41 | 74.09 1 | 1.81 | 2.44 | 49.68 | 50.18 | -0.49 | 0.49 | 0 | 0 | |
| gene10678 | https://www.ncbi.nlm.nih.gov/gene/?term=LOC105284355 | 4031 | 170 | 421.73 | 1 | 2.48 | 0.59 | 18.52 | 19.03 | -0.5 | 0.5 | -0.02 | 0.02 |
| gene8178 | https://www.ncbi.nlm.nih.gov/gene/?term=LOC105281515 | 8647 | 44 | 50.88 1 | 1.16 | 2.27 | 29.36 | 29.86 | -0.5 | 0.5 | 0 | 0 | |
| gene8600 | https://www.ncbi.nlm.nih.gov/gene/?term=LOC105282001 | 8513 | 153 | 179.73 | 1 | 1.17 | 0.65 | 9.37 | 9.87 | -0.5 | 0.5 | -0.01 | 0.01 |
| gene6639 | https://www.ncbi.nlm.nih.gov/gene/?term=LOC105279705 | 7656 | 68 | 88.82 1 | 1.31 | 1.47 | 22.19 | 22.7 | -0.51 | 0.51 | 0 | 0 | |
| gene3885 | https://www.ncbi.nlm.nih.gov/gene/?term=LOC105276527 | 7928 | 346 | 436.43 | 1 | 1.26 | 0.29 | 8.3 | 8.89 | -0.6 | 0.6 | -0.03 | 0.03 |
| gene10018 | https://www.ncbi.nlm.nih.gov/gene/?term=LOC105283607 | 4414 | 204 | 462.17 | 1 | 2.27 | 0.49 | 14.8 | 15.47 | -0.67 | 0.67 | -0.03 | 0.03 |
| gene274 | https://www.ncbi.nlm.nih.gov/gene/?term=LOC105275190 | 3488 | 70 | 200.69 | 1 | 2.87 | 1.43 | 19.35 | 20.02 | -0.67 | 0.67 | -0.01 | 0.01 |
| gene1332 | https://www.ncbi.nlm.nih.gov/gene/?term=LOC105286872 | 2460 | 141 | 573.17 | 1 | 4.07 | 0.71 | 21.92 | 22.68 | -0.76 | 0.76 | -0.04 | 0.04 |
| gene8729 | https://www.ncbi.nlm.nih.gov/gene/?term=LOC105282137 | 5005 | 199 | 397.6 1 | 2 | 0.5 | 16.19 | 16.99 | -0.79 | 0.79 | -0.03 | 0.03 | |
| gene11819 | https://www.ncbi.nlm.nih.gov/gene/?term=LOC105285671 | 2366 | 96 | 405.75 | 1 | 4.23 | 1.04 | 27.73 | 28.57 | -0.84 | 0.84 | -0.03 | 0.03 |
| gene9336 | https://www.ncbi.nlm.nih.gov/gene/?term=LOC105282834 | 5993 | 115 | 191.89 | 1 | 1.67 | 0.87 | 14.71 | 15.58 | -0.87 | 0.87 | -0.02 | 0.02 |
| gene10107 | https://www.ncbi.nlm.nih.gov/gene/?term=LOC105283712 | 1883 | 28 | 148.7 1 | 5.31 | 3.57 | 28.01 | 28.9 | -0.88 | 0.88 | -0.01 | 0.01 | |
| gene1310 | https://www.ncbi.nlm.nih.gov/gene/?term=LOC105286661 | 2582 | 65 | 251.74 | 1 | 3.87 | 1.54 | 19.96 | 20.95 | -0.99 | 0.99 | -0.03 | 0.03 |
| gene2844 | https://www.ncbi.nlm.nih.gov/gene/?term=LOC105275363 | 4330 | 187 | 431.87 | 1 | 2.31 | 0.53 | 14.36 | 15.35 | -0.99 | 0.99 | -0.04 | 0.04 |
| gene292 | https://www.ncbi.nlm.nih.gov/gene/?term=LOC105275398 | 2825 | 76 | 269.03 | 1 | 3.54 | 1.32 | 20.33 | 21.34 | -1.01 | 1.01 | -0.03 | 0.03 |
| gene7183 | https://www.ncbi.nlm.nih.gov/gene/?term=LOC105280348 | 3925 | 52 | 132.48 | 2 | 5.1 | 3.85 | 25.66 | 26.67 | -1.01 | 1.01 | -0.01 | 0.01 |
| gene8343 | https://www.ncbi.nlm.nih.gov/gene/?term=LOC105281700 | 2172 | 44 | 202.58 | 1 | 4.6 | 2.27 | 37.43 | 38.56 | -1.13 | 1.13 | -0.02 | 0.02 |
| gene4034 | https://www.ncbi.nlm.nih.gov/gene/?term=LOC105276704 | 2065 | 84 | 406.78 | 1 | 4.84 | 1.19 | 12.86 | 14.01 | -1.15 | 1.15 | -0.05 | 0.05 |
| gene298 | https://www.ncbi.nlm.nih.gov/gene/?term=LOC105275460 | 2473 | 115 | 465.02 | 1 | 4.04 | 0.87 | 12.68 | 13.89 | -1.21 | 1.21 | -0.06 | 0.06 |
| gene5101 | https://www.ncbi.nlm.nih.gov/gene/?term=LOC105277903 | 1813 | 34 | 187.53 | 1 | 5.52 | 2.94 | 52.57 | 53.85 | -1.27 | 1.27 | -0.02 | 0.02 |
| gene570 | https://www.ncbi.nlm.nih.gov/gene/?term=LOC105278549 | 1673 | 63 | 376.57 | 1 | 5.98 | 1.59 | 40.52 | 41.79 | -1.27 | 1.27 | -0.05 | 0.05 |
| gene2856 | https://www.ncbi.nlm.nih.gov/gene/?term=LOC105275384 | 1987 | 76 | 382.49 | 1 | 5.03 | 1.32 | 32.95 | 34.23 | -1.28 | 1.28 | -0.05 | 0.05 |
| gene9101 | https://www.ncbi.nlm.nih.gov/gene/?term=LOC105282553 | 3068 | 103 | 335.72 | 1 | 3.26 | 0.97 | 20.56 | 21.9 | -1.34 | 1.34 | -0.04 | 0.04 |

| | | | | | | | | | | | | | |
|---|---|---|---|---|---|---|---|---|---|---|---|---|---|
| gene4641 | https://www.ncbi.nlm.nih.gov/gene/?term=LOC105277398 | 1937 | 70 | 361.38 | 1 | 5.16 | 1.43 | 35.36 | 36.76 | -1.4 | 1.4 | -0.05 | 0.05 |
| gene5877 | https://www.ncbi.nlm.nih.gov/gene/?term=LOC105278807 | 1719 | 51 | 296.68 | 1 | 5.82 | 1.96 | 22.99 | 24.43 | -1.44 | 1.44 | -0.04 | 0.04 |
| gene5402 | https://www.ncbi.nlm.nih.gov/gene/?term=LOC105278260 | 3068 | 42 | 136.9 | 1 | 3.26 | 2.38 | 12.8 | 14.3 | -1.5 | 1.5 | -0.02 | 0.02 |
| gene12406 | https://www.ncbi.nlm.nih.gov/gene/?term=LOC105286316 | 1386 | 30 | 216.45 | 1 | 7.22 | 3.33 | 39.2 | 40.74 | -1.55 | 1.55 | -0.03 | 0.03 |
| gene1884 | https://www.ncbi.nlm.nih.gov/gene/?term=LOC105288072 | 4169 | 80 | 191.89 | 1 | 2.4 | 1.25 | 22.74 | 24.44 | -1.7 | 1.7 | -0.03 | 0.03 |
| gene1855 | https://www.ncbi.nlm.nih.gov/gene/?term=LOC105288036 | 2387 | 87 | 364.47 | 1 | 4.19 | 1.15 | 28.59 | 30.36 | -1.76 | 1.76 | -0.06 | 0.06 |
| gene10345 | https://www.ncbi.nlm.nih.gov/gene/?term=LOC105283989 | 4444 | 29 | 65.26 | 1 | 2.25 | 3.45 | 48.19 | 50.2 | -2.01 | 2.01 | -0.01 | 0.01 |
| gene6334 | https://www.ncbi.nlm.nih.gov/gene/?term=LOC105279356 | 1471 | 43 | 292.32 | 1 | 6.8 | 2.33 | 28.25 | 30.28 | -2.03 | 2.03 | -0.06 | 0.06 |
| gene12516 | https://www.ncbi.nlm.nih.gov/gene/?term=LOC105286452 | 1529 | 35 | 228.91 | 1 | 6.54 | 2.86 | 51.01 | 53.47 | -2.46 | 2.46 | -0.06 | 0.06 |
| gene2676 | https://www.ncbi.nlm.nih.gov/gene/?term=LOC105275181 | 1353 | 14 | 103.47 | 1 | 7.39 | 7.14 | 65.95 | 68.91 | -2.96 | 2.96 | -0.03 | 0.03 |
| gene2970 | https://www.ncbi.nlm.nih.gov/gene/?term=LOC105275518 | 2340 | 20 | 85.47 | 1 | 4.27 | 5 | 47.36 | 50.74 | -3.38 | 3.38 | -0.03 | 0.03 |
| gene5714 | https://www.ncbi.nlm.nih.gov/gene/?term=LOC105278729 | 710 | 14 | 197.18 | 1 | 14.08 | 7.14 | 51.67 | 55.07 | -3.41 | 3.41 | -0.07 | 0.07 |
| gene5647 | https://www.ncbi.nlm.nih.gov/gene/?term=LOC105278567 | 3118 | 38 | 121.87 | 1 | 3.21 | 2.63 | 12.56 | 16.06 | -3.5 | 3.5 | -0.04 | 0.04 |
| gene1962 | https://www.ncbi.nlm.nih.gov/gene/?term=LOC105288157 | 1246 | 16 | 128.41 | 1 | 8.03 | 6.25 | 13.95 | 18.55 | -4.6 | 4.6 | -0.06 | 0.06 |
| gene3165 | https://www.ncbi.nlm.nih.gov/gene/?term=LOC105275706 | 2374 | 66 | 278.01 | 4 | 16.85 | 6.06 | 24.58 | 35.31 | -10.73 | 10.73 | -0.3 | 0.3 |
| gene8929 | https://www.ncbi.nlm.nih.gov/gene/?term=LOC105282371 | 1849 | 8 | 43.27 | 1 | 5.41 | 12.5 | 22.25 | 44.14 | -21.89 | 21.89 | -0.09 | 0.09 |
| gene2593 | https://www.ncbi.nlm.nih.gov/gene/?term=LOC105275070 | 1774 | 6 | 33.82 | 1 | 5.64 | 16.67 | 16.42 | 54.02 | -37.6 | 37.6 | -0.13 | 0.13 |

Directory: OGLall File: ogl-Cb_LI_BCphaseA1.vs.BCphaseB2.txt

| gene_ID | gene_link | gwidth | #Sites | #per10Kb | #dmSites | #dmsp10kb | %dmSites | %pSite1 | %pSite2 | DMpSite | ADMpSite | DMpNucl | ADMpNucl |
|---|---|---|---|---|---|---|---|---|---|---|---|---|---|
| gene7106 | https://www.ncbi.nlm.nih.gov/gene/?term=LOC105280261 | 970 | 4 | 41.24 | 1 | 10.31 | 25 | 82.16 | 67.27 | 14.89 | 14.89 | 0.06 | 0.06 |
| gene11980 | https://www.ncbi.nlm.nih.gov/gene/?term=LOC109611395 | 571 | 3 | 52.54 | 1 | 17.51 | 33.33 | 22.75 | 11.76 | 10.99 | 10.99 | 0.06 | 0.06 |
| gene558 | https://www.ncbi.nlm.nih.gov/gene/?term=LOC105278458 | 216 | 3 | 138.89 | 1 | 46.3 | 33.33 | 20.87 | 10.44 | 10.43 | 10.43 | 0.14 | 0.14 |
| gene7684 | https://www.ncbi.nlm.nih.gov/gene/?term=LOC105280926 | 1727 | 6 | 34.74 | 2 | 11.58 | 33.33 | 47.53 | 37.53 | 10 | 10 | 0.03 | 0.03 |
| gene10219 | https://www.ncbi.nlm.nih.gov/gene/?term=LOC105283830 | 235 | 12 | 510.64 | 1 | 42.55 | 8.33 | 42.96 | 33.59 | 9.36 | 9.36 | 0.48 | 0.48 |
| gene3395 | https://www.ncbi.nlm.nih.gov/gene/?term=LOC105275978 | 2468 | 5 | 20.26 | 1 | 4.05 | 20 | 19.06 | 10.64 | 8.42 | 8.42 | 0.02 | 0.02 |
| gene12003 | https://www.ncbi.nlm.nih.gov/gene/?term=LOC105285885 | 1336 | 11 | 82.34 | 2 | 14.97 | 18.18 | 63.58 | 55.36 | 8.22 | 8.22 | 0.07 | 0.07 |
| gene1656 | https://www.ncbi.nlm.nih.gov/gene/?term=LOC105287814 | 812 | 7 | 86.21 | 1 | 12.32 | 14.29 | 37.14 | 29.51 | 7.63 | 7.63 | 0.07 | 0.07 |
| gene4643 | https://www.ncbi.nlm.nih.gov/gene/?term=LOC109610847 | 429 | 5 | 116.55 | 1 | 23.31 | 20 | 29.86 | 22.36 | 7.5 | 7.5 | 0.09 | 0.09 |
| gene10325 | https://www.ncbi.nlm.nih.gov/gene/?term=LOC105283963 | 1286 | 9 | 69.98 | 1 | 7.78 | 11.11 | 57.17 | 49.99 | 7.19 | 7.19 | 0.05 | 0.05 |
| gene8172 | https://www.ncbi.nlm.nih.gov/gene/?term=LOC105281507 | 946 | 5 | 52.85 | 1 | 10.57 | 20 | 11.82 | 4.83 | 6.99 | 6.99 | 0.04 | 0.04 |
| gene405 | https://www.ncbi.nlm.nih.gov/gene/?term=LOC105276706 | 1442 | 16 | 110.96 | 1 | 6.93 | 6.25 | 82.68 | 76.42 | 6.27 | 6.27 | 0.07 | 0.07 |
| gene8965 | https://www.ncbi.nlm.nih.gov/gene/?term=LOC105282405 | 1310 | 31 | 236.64 | 3 | 22.9 | 9.68 | 51.28 | 45.18 | 6.1 | 6.1 | 0.14 | 0.14 |
| gene539 | https://www.ncbi.nlm.nih.gov/gene/?term=LOC105278243 | 1952 | 18 | 92.21 | 1 | 5.12 | 5.56 | 20.77 | 14.99 | 5.77 | 5.77 | 0.05 | 0.05 |
| gene10310 | https://www.ncbi.nlm.nih.gov/gene/?term=LOC109611277 | 1730 | 8 | 46.24 | 1 | 5.78 | 12.5 | 12.29 | 6.59 | 5.7 | 5.7 | 0.03 | 0.03 |
| gene11269 | https://www.ncbi.nlm.nih.gov/gene/?term=LOC105285051 | 1363 | 11 | 80.7 | 3 | 22.01 | 27.27 | 46.02 | 40.39 | 5.64 | 5.64 | 0.05 | 0.05 |
| gene8458 | https://www.ncbi.nlm.nih.gov/gene/?term=LOC105281831 | 1053 | 8 | 75.97 | 1 | 9.5 | 12.5 | 87.04 | 81.5 | 5.54 | 5.54 | 0.04 | 0.04 |
| gene7962 | https://www.ncbi.nlm.nih.gov/gene/?term=LOC105281226 | 1280 | 6 | 46.88 | 1 | 7.81 | 16.67 | 62.79 | 57.27 | 5.53 | 5.53 | 0.03 | 0.03 |
| gene10204 | https://www.ncbi.nlm.nih.gov/gene/?term=LOC105283813 | 1312 | 8 | 60.98 | 1 | 7.62 | 12.5 | 29.71 | 24.26 | 5.45 | 5.45 | 0.03 | 0.03 |
| gene10792 | https://www.ncbi.nlm.nih.gov/gene/?term=LOC105284483 | 2910 | 24 | 82.47 | 1 | 3.44 | 4.17 | 22.77 | 17.6 | 5.17 | 5.17 | 0.04 | 0.04 |
| gene10357 | https://www.ncbi.nlm.nih.gov/gene/?term=LOC105284002 | 1605 | 12 | 74.77 | 1 | 6.23 | 8.33 | 44.27 | 39.3 | 4.98 | 4.98 | 0.04 | 0.04 |
| gene1914 | https://www.ncbi.nlm.nih.gov/gene/?term=LOC105288105 | 1289 | 23 | 178.43 | 2 | 15.52 | 8.7 | 60.89 | 56.04 | 4.85 | 4.85 | 0.09 | 0.09 |
| gene3296 | https://www.ncbi.nlm.nih.gov/gene/?term=LOC105275859 | 2340 | 21 | 89.74 | 2 | 8.55 | 9.52 | 53.4 | 48.62 | 4.78 | 4.78 | 0.04 | 0.04 |
| gene939 | https://www.ncbi.nlm.nih.gov/gene/?term=LOC105282729 | 1450 | 11 | 75.86 | 1 | 6.9 | 9.09 | 28.64 | 23.91 | 4.73 | 4.73 | 0.04 | 0.04 |

...
... only first 25 lines shown ...

Directory: OGLall File: ogl-Cb_LI_BCphaseA1.vs.RphaseA1.txt

| gene_ID | gene_link | gwidth | #Sites | #per10Kb | #dmSites | #dmsp10kb | %dmSites | %pSite1 | %pSite2 | DMpSite | ADMpSite | DMpNucl | ADMpNucl |
|---|---|---|---|---|---|---|---|---|---|---|---|---|---|
| gene393 | https://www.ncbi.nlm.nih.gov/gene/?term=LOC105276566 | 1123 | 2 | 17.81 | 1 | 8.9 | 50 | 77.65 | 50 | 27.65 | 27.65 | 0.05 | 0.05 |
| gene7199 | https://www.ncbi.nlm.nih.gov/gene/?term=LOC105280366 | 1414 | 26 | 183.88 | 1 | 7.07 | 3.85 | 47.43 | 38.76 | 8.67 | 8.67 | 0.16 | 0.16 |
| gene12517 | https://www.ncbi.nlm.nih.gov/gene/?term=LOC105286451 | 1626 | 10 | 61.5 | 1 | 6.15 | 10 | 52.19 | 44.01 | 8.18 | 8.18 | 0.05 | 0.05 |
| gene388 | https://www.ncbi.nlm.nih.gov/gene/?term=LOC105276538 | 1558 | 19 | 121.95 | 1 | 6.42 | 5.26 | 69.41 | 64.89 | 4.52 | 4.52 | 0.06 | 0.06 |
| gene7915 | https://www.ncbi.nlm.nih.gov/gene/?term=LOC105281173 | 1597 | 25 | 156.54 | 1 | 6.26 | 4 | 64.87 | 60.99 | 3.88 | 3.88 | 0.06 | 0.06 |
| gene3168 | https://www.ncbi.nlm.nih.gov/gene/?term=LOC105275711 | 6150 | 77 | 125.2 | 1 | 1.63 | 1.3 | 14.51 | 10.99 | 3.51 | 3.51 | 0.04 | 0.04 |
| gene6422 | https://www.ncbi.nlm.nih.gov/gene/?term=LOC105279458 | 2419 | 37 | 152.96 | 1 | 4.13 | 2.7 | 54.1 | 51.26 | 2.83 | 2.83 | 0.04 | 0.04 |
| gene10714 | https://www.ncbi.nlm.nih.gov/gene/?term=LOC105284396 | 1966 | 53 | 269.58 | 1 | 5.09 | 1.89 | 40.95 | 38.4 | 2.55 | 2.55 | 0.07 | 0.07 |
| gene322 | https://www.ncbi.nlm.nih.gov/gene/?term=LOC105275777 | 1606 | 23 | 143.21 | 1 | 6.23 | 4.35 | 45.55 | 43.01 | 2.54 | 2.54 | 0.04 | 0.04 |
| gene11401 | https://www.ncbi.nlm.nih.gov/gene/?term=LOC105285202 | 1841 | 49 | 266.16 | 1 | 5.43 | 2.04 | 21.54 | 19.38 | 2.16 | 2.16 | 0.06 | 0.06 |
| gene7368 | https://www.ncbi.nlm.nih.gov/gene/?term=LOC105280556 | 4105 | 46 | 112.06 | 1 | 2.44 | 2.17 | 13.39 | 11.38 | 2 | 2 | 0.02 | 0.02 |
| gene10213 | https://www.ncbi.nlm.nih.gov/gene/?term=LOC105283826 | 1503 | 52 | 345.97 | 1 | 6.65 | 1.92 | 16.76 | 14.79 | 1.97 | 1.97 | 0.07 | 0.07 |
| gene12021 | https://www.ncbi.nlm.nih.gov/gene/?term=LOC105285905 | 1399 | 23 | 164.4 | 1 | 7.15 | 4.35 | 41.24 | 39.35 | 1.89 | 1.89 | 0.03 | 0.03 |
| gene2271 | https://www.ncbi.nlm.nih.gov/gene/?term=LOC105274706 | 1448 | 49 | 338.4 | 1 | 6.91 | 2.04 | 38.87 | 37.01 | 1.86 | 1.86 | 0.06 | 0.06 |
| gene4190 | https://www.ncbi.nlm.nih.gov/gene/?term=LOC105276875 | 3150 | 124 | 393.65 | 1 | 3.17 | 0.81 | 21.46 | 19.89 | 1.58 | 1.58 | 0.06 | 0.06 |
| gene11454 | https://www.ncbi.nlm.nih.gov/gene/?term=LOC105285256 | 1906 | 52 | 272.82 | 1 | 5.25 | 1.92 | 42.98 | 41.47 | 1.51 | 1.51 | 0.04 | 0.04 |
| gene2752 | https://www.ncbi.nlm.nih.gov/gene/?term=LOC105275266 | 2111 | 49 | 232.12 | 1 | 4.74 | 2.04 | 23.87 | 22.43 | 1.44 | 1.44 | 0.03 | 0.03 |
| gene920 | https://www.ncbi.nlm.nih.gov/gene/?term=LOC105282518 | 1197 | 42 | 350.88 | 1 | 8.35 | 2.38 | 38.27 | 36.85 | 1.43 | 1.43 | 0.05 | 0.05 |
| gene5649 | https://www.ncbi.nlm.nih.gov/gene/?term=LOC105278538 | 2140 | 76 | 355.14 | 1 | 4.67 | 1.32 | 18.24 | 16.87 | 1.37 | 1.37 | 0.05 | 0.05 |
| gene9740 | https://www.ncbi.nlm.nih.gov/gene/?term=LOC105283277 | 2887 | 135 | 467.61 | 1 | 3.46 | 0.74 | 21.88 | 20.52 | 1.37 | 1.37 | 0.06 | 0.06 |
| gene2748 | https://www.ncbi.nlm.nih.gov/gene/?term=LOC105275261 | 3845 | 55 | 143.04 | 1 | 2.6 | 1.82 | 38.68 | 37.31 | 1.36 | 1.36 | 0.02 | 0.02 |
| gene10326 | https://www.ncbi.nlm.nih.gov/gene/?term=LOC105283962 | 3042 | 94 | 309.01 | 1 | 3.29 | 1.06 | 12.22 | 10.91 | 1.31 | 1.31 | 0.04 | 0.04 |
| gene6031 | https://www.ncbi.nlm.nih.gov/gene/?term=LOC105279000 | 5691 | 175 | 307.5 | 1 | 1.76 | 0.57 | 14.52 | 13.23 | 1.29 | 1.29 | 0.04 | 0.04 |
| gene10844 | https://www.ncbi.nlm.nih.gov/gene/?term=LOC105284535 | 9086 | 218 | 239.93 | 1 | 1.1 | 0.46 | 16.62 | 15.4 | 1.22 | 1.22 | 0.03 | 0.03 |
| gene4376 | https://www.ncbi.nlm.nih.gov/gene/?term=LOC105277083 | 4307 | 90 | 208.96 | 1 | 2.32 | 1.11 | 26.1 | 24.89 | 1.21 | 1.21 | 0.03 | 0.03 |
| gene3599 | https://www.ncbi.nlm.nih.gov/gene/?term=LOC105276196 | 2955 | 55 | 186.13 | 1 | 3.38 | 1.82 | 40.49 | 39.41 | 1.07 | 1.07 | 0.02 | 0.02 |
| gene5298 | https://www.ncbi.nlm.nih.gov/gene/?term=LOC105278145 | 4060 | 240 | 591.13 | 2 | 4.93 | 0.83 | 11.5 | 10.44 | 1.06 | 1.06 | 0.06 | 0.06 |
| gene7248 | https://www.ncbi.nlm.nih.gov/gene/?term=LOC105280408 | 5175 | 44 | 85.02 | 1 | 1.93 | 2.27 | 22.41 | 21.41 | 0.99 | 0.99 | 0.01 | 0.01 |
| gene6806 | https://www.ncbi.nlm.nih.gov/gene/?term=LOC105279894 | 3649 | 39 | 106.88 | 1 | 2.74 | 2.56 | 35.25 | 34.29 | 0.97 | 0.97 | 0.01 | 0.01 |
| gene10392 | https://www.ncbi.nlm.nih.gov/gene/?term=LOC105284036 | 3216 | 83 | 258.08 | 1 | 3.11 | 1.2 | 24.77 | 23.81 | 0.96 | 0.96 | 0.02 | 0.02 |
| gene12203 | https://www.ncbi.nlm.nih.gov/gene/?term=LOC105286099 | 5967 | 69 | 115.64 | 1 | 1.68 | 1.45 | 11.63 | 10.68 | 0.96 | 0.96 | 0.01 | 0.01 |
| gene4962 | https://www.ncbi.nlm.nih.gov/gene/?term=LOC105277761 | 4645 | 147 | 316.47 | 1 | 2.15 | 0.68 | 13.9 | 13.02 | 0.88 | 0.88 | 0.03 | 0.03 |
| gene12307 | https://www.ncbi.nlm.nih.gov/gene/?term=LOC105286216 | 10313 | 145 | 140.6 | 1 | 0.97 | 0.69 | 17.77 | 16.9 | 0.87 | 0.87 | 0.01 | 0.01 |
| gene6966 | https://www.ncbi.nlm.nih.gov/gene/?term=LOC105280104 | 3747 | 119 | 317.59 | 1 | 2.67 | 0.84 | 15.37 | 14.52 | 0.85 | 0.85 | 0.03 | 0.03 |
| gene6639 | https://www.ncbi.nlm.nih.gov/gene/?term=LOC105279705 | 7656 | 70 | 91.43 | 1 | 1.31 | 1.43 | 22.25 | 21.42 | 0.84 | 0.84 | 0.01 | 0.01 |
| gene7568 | https://www.ncbi.nlm.nih.gov/gene/?term=LOC105280786 | 2525 | 116 | 459.41 | 1 | 3.96 | 0.86 | 28.86 | 28.06 | 0.8 | 0.8 | 0.04 | 0.04 |
| gene7550 | https://www.ncbi.nlm.nih.gov/gene/?term=LOC105280763 | 2173 | 93 | 427.98 | 1 | 4.6 | 1.08 | 27 | 26.23 | 0.77 | 0.77 | 0.03 | 0.03 |
| gene7249 | https://www.ncbi.nlm.nih.gov/gene/?term=LOC105280422 | 1775 | 61 | 343.66 | 1 | 5.63 | 1.64 | 15.16 | 14.42 | 0.75 | 0.75 | 0.03 | 0.03 |
| gene10606 | https://www.ncbi.nlm.nih.gov/gene/?term=LOC105284273 | 6119 | 243 | 397.12 | 2 | 3.27 | 0.82 | 20.74 | 20.03 | 0.71 | 0.71 | 0.03 | 0.03 |
| gene10600 | https://www.ncbi.nlm.nih.gov/gene/?term=LOC105284270 | 4837 | 95 | 196.4 | 1 | 2.07 | 1.05 | 13.92 | 13.23 | 0.69 | 0.69 | 0.01 | 0.01 |
| gene6286 | https://www.ncbi.nlm.nih.gov/gene/?term=LOC105279298 | 2861 | 136 | 475.36 | 1 | 3.5 | 0.74 | 22.78 | 22.14 | 0.64 | 0.64 | 0.03 | 0.03 |
| gene3107 | https://www.ncbi.nlm.nih.gov/gene/?term=LOC105275650 | 5103 | 145 | 284.15 | 1 | 1.96 | 0.69 | 17.36 | 16.74 | 0.62 | 0.62 | 0.02 | 0.02 |
| gene11340 | https://www.ncbi.nlm.nih.gov/gene/?term=LOC105285131 | 2918 | 97 | 332.42 | 1 | 3.43 | 1.03 | 14.71 | 14.15 | 0.56 | 0.56 | 0.02 | 0.02 |
| gene2404 | https://www.ncbi.nlm.nih.gov/gene/?term=LOC105274853 | 1767 | 39 | 220.71 | 1 | 5.66 | 2.56 | 31.07 | 30.51 | 0.56 | 0.56 | 0.01 | 0.01 |
| gene10155 | https://www.ncbi.nlm.nih.gov/gene/?term=LOC105283756 | 7030 | 93 | 132.29 | 1 | 1.42 | 1.08 | 7.03 | 6.49 | 0.54 | 0.54 | 0.01 | 0.01 |
| gene3206 | https://www.ncbi.nlm.nih.gov/gene/?term=LOC105275752 | 3474 | 126 | 362.69 | 1 | 2.88 | 0.79 | 28.38 | 27.85 | 0.53 | 0.53 | 0.02 | 0.02 |
| gene657 | https://www.ncbi.nlm.nih.gov/gene/?term=LOC105279667 | 6421 | 77 | 119.92 | 1 | 1.56 | 1.3 | 5.63 | 5.1 | 0.53 | 0.53 | 0.01 | 0.01 |
| gene7801 | https://www.ncbi.nlm.nih.gov/gene/?term=LOC105281047 | 5137 | 109 | 212.19 | 1 | 1.95 | 0.92 | 11.18 | 10.67 | 0.5 | 0.5 | 0.01 | 0.01 |
| gene7398 | https://www.ncbi.nlm.nih.gov/gene/?term=LOC105280589 | 3452 | 40 | 115.87 | 1 | 2.9 | 2.5 | 25.2 | 24.72 | 0.48 | 0.48 | 0.01 | 0.01 |
| gene1470 | https://www.ncbi.nlm.nih.gov/gene/?term=LOC105287597 | 2974 | 181 | 608.61 | 1 | 3.36 | 0.55 | 17.88 | 17.42 | 0.47 | 0.47 | 0.03 | 0.03 |
| gene5746 | https://www.ncbi.nlm.nih.gov/gene/?term=LOC105278666 | 10343 | 393 | 379.97 | 1 | 0.97 | 0.25 | 7.69 | 7.22 | 0.47 | 0.47 | 0.02 | 0.02 |
| gene940 | https://www.ncbi.nlm.nih.gov/gene/?term=LOC105282794 | 3578 | 84 | 234.77 | 1 | 2.79 | 1.19 | 14.53 | 14.06 | 0.47 | 0.47 | 0.01 | 0.01 |
| gene8729 | https://www.ncbi.nlm.nih.gov/gene/?term=LOC105282137 | 5005 | 200 | 399.6 | 1 | 2 | 0.5 | 15.61 | 15.22 | 0.39 | 0.39 | 0.02 | 0.02 |
| gene5524 | https://www.ncbi.nlm.nih.gov/gene/?term=LOC105278397 | 5177 | 199 | 384.39 | 1 | 1.93 | 0.5 | 24.51 | 24.16 | 0.35 | 0.35 | 0.01 | 0.01 |
| gene5831 | https://www.ncbi.nlm.nih.gov/gene/?term=LOC105278761 | 2506 | 99 | 395.05 | 1 | 3.99 | 1.01 | 25.53 | 25.18 | 0.35 | 0.35 | 0.01 | 0.01 |
| gene7600 | https://www.ncbi.nlm.nih.gov/gene/?term=LOC105280819 | 1615 | 41 | 253.87 | 1 | 6.19 | 2.44 | 40.74 | 40.39 | 0.35 | 0.35 | 0.01 | 0.01 |
| gene13114 | https://www.ncbi.nlm.nih.gov/gene/?term=LOC105287147 | 7609 | 370 | 486.27 | 1 | 1.31 | 0.27 | 9.54 | 9.22 | 0.32 | 0.32 | 0.02 | 0.02 |
| gene5539 | https://www.ncbi.nlm.nih.gov/gene/?term=LOC105278415 | 25024 | 562 | 224.58 | 1 | 0.4 | 0.18 | 4.11 | 3.79 | 0.31 | 0.31 | 0.01 | 0.01 |
| gene1147 | https://www.ncbi.nlm.nih.gov/gene/?term=LOC105284954 | 4022 | 87 | 216.31 | 1 | 2.49 | 1.15 | 14.67 | 14.36 | 0.3 | 0.3 | 0.01 | 0.01 |
| gene2173 | https://www.ncbi.nlm.nih.gov/gene/?term=LOC105274590 | 4387 | 78 | 177.8 | 1 | 2.28 | 1.28 | 22.31 | 22.01 | 0.3 | 0.3 | 0.01 | 0.01 |

| | | | | | | | | | | | | | |
|---|---|---|---|---|---|---|---|---|---|---|---|---|---|
| gene9101 | https://www.ncbi.nlm.nih.gov/gene/?term=LOC105282553 | 3068 | 103 | 335.72 | 1 | 3.26 | 0.97 | 20.28 | 19.99 | 0.29 | 0.29 | 0.01 | 0.01 |
| gene13036 | https://www.ncbi.nlm.nih.gov/gene/?term=LOC105287059 | 4959 | 157 | 316.6 | 1 | 2.02 | 0.64 | 17.83 | 17.59 | 0.25 | 0.25 | 0.01 | 0.01 |
| gene10549 | https://www.ncbi.nlm.nih.gov/gene/?term=LOC105284210 | 7145 | 313 | 438.07 | 1 | 1.4 | 0.32 | 10.77 | 10.55 | 0.22 | 0.22 | 0.01 | 0.01 |
| gene8274 | https://www.ncbi.nlm.nih.gov/gene/?term=LOC105281624 | 5425 | 189 | 348.39 | 1 | 1.84 | 0.53 | 11.18 | 10.97 | 0.21 | 0.21 | 0.01 | 0.01 |
| gene7547 | https://www.ncbi.nlm.nih.gov/gene/?term=LOC105280767 | 24576 | 683 | 277.91 | 1 | 0.41 | 0.15 | 4.82 | 4.61 | 0.2 | 0.2 | 0.01 | 0.01 |
| gene12802 | https://www.ncbi.nlm.nih.gov/gene/?term=LOC105286799 | 3183 | 170 | 534.09 | 1 | 3.14 | 0.59 | 15.96 | 15.78 | 0.18 | 0.18 | 0.01 | 0.01 |
| gene5525 | https://www.ncbi.nlm.nih.gov/gene/?term=LOC105278396 | 3090 | 185 | 598.71 | 1 | 3.24 | 0.54 | 24.64 | 24.46 | 0.18 | 0.18 | 0.01 | 0.01 |
| gene1220 | https://www.ncbi.nlm.nih.gov/gene/?term=LOC105285742 | 3453 | 86 | 249.06 | 1 | 2.9 | 1.16 | 21.68 | 21.51 | 0.17 | 0.17 | 0 | 0 |
| gene7247 | https://www.ncbi.nlm.nih.gov/gene/?term=LOC105280407 | 10486 | 206 | 196.45 | 1 | 0.95 | 0.49 | 17.14 | 16.97 | 0.17 | 0.17 | 0 | 0 |
| gene4213 | https://www.ncbi.nlm.nih.gov/gene/?term=LOC105276901 | 3257 | 96 | 294.75 | 1 | 3.07 | 1.04 | 22.87 | 22.72 | 0.14 | 0.14 | 0 | 0 |
| gene6805 | https://www.ncbi.nlm.nih.gov/gene/?term=LOC105279895 | 87888 | 975 | 110.94 | 1 | 0.11 | 0.1 | 1.92 | 1.78 | 0.14 | 0.14 | 0 | 0 |
| gene10107 | https://www.ncbi.nlm.nih.gov/gene/?term=LOC105283712 | 1883 | 29 | 154.01 | 1 | 5.31 | 3.45 | 27.22 | 27.09 | 0.13 | 0.13 | 0 | 0 |
| gene9645 | https://www.ncbi.nlm.nih.gov/gene/?term=LOC105283181 | 3875 | 134 | 345.81 | 1 | 2.58 | 0.75 | 11.45 | 11.37 | 0.08 | 0.08 | 0 | 0 |
| gene6620 | https://www.ncbi.nlm.nih.gov/gene/?term=LOC105279687 | 17998 | 354 | 196.69 | 1 | 0.56 | 0.28 | 5.08 | 5.05 | 0.03 | 0.03 | 0 | 0 |
| gene11583 | https://www.ncbi.nlm.nih.gov/gene/?term=LOC105285414 | 8192 | 175 | 213.62 | 1 | 1.22 | 0.57 | 9.21 | 9.24 | -0.03 | 0.03 | 0 | 0 |
| gene2806 | https://www.ncbi.nlm.nih.gov/gene/?term=LOC105275327 | 14257 | 462 | 324.05 | 1 | 0.7 | 0.22 | 7.37 | 7.4 | -0.03 | 0.03 | 0 | 0 |
| gene11579 | https://www.ncbi.nlm.nih.gov/gene/?term=LOC105285406 | 13571 | 260 | 191.58 | 1 | 0.74 | 0.38 | 5.75 | 5.79 | -0.04 | 0.04 | 0 | 0 |
| gene7166 | https://www.ncbi.nlm.nih.gov/gene/?term=LOC105280326 | 6117 | 208 | 340.04 | 1 | 1.63 | 0.48 | 16.25 | 16.28 | -0.04 | 0.04 | 0 | 0 |
| gene12406 | https://www.ncbi.nlm.nih.gov/gene/?term=LOC105286316 | 1386 | 30 | 216.45 | 1 | 7.22 | 3.33 | 39.2 | 39.29 | -0.09 | 0.09 | 0 | 0 |
| gene5375 | https://www.ncbi.nlm.nih.gov/gene/?term=LOC105278228 | 8640 | 148 | 171.3 | 1 | 1.16 | 0.68 | 24.78 | 24.89 | -0.11 | 0.11 | 0 | 0 |
| gene12573 | https://www.ncbi.nlm.nih.gov/gene/?term=LOC105286515 | 3647 | 105 | 287.91 | 1 | 2.74 | 0.95 | 10.51 | 10.65 | -0.14 | 0.14 | 0 | 0 |
| gene1310 | https://www.ncbi.nlm.nih.gov/gene/?term=LOC105286661 | 2582 | 63 | 244 | 1 | 3.87 | 1.59 | 19.4 | 19.54 | -0.14 | 0.14 | 0 | 0 |
| gene10531 | https://www.ncbi.nlm.nih.gov/gene/?term=LOC105284190 | 6881 | 180 | 261.59 | 1 | 1.45 | 0.56 | 10.06 | 10.22 | -0.16 | 0.16 | 0 | 0 |
| gene2443 | https://www.ncbi.nlm.nih.gov/gene/?term=LOC105274895 | 10433 | 330 | 316.3 | 1 | 0.96 | 0.3 | 5.47 | 5.65 | -0.18 | 0.18 | -0.01 | 0.01 |
| gene4892 | https://www.ncbi.nlm.nih.gov/gene/?term=LOC105277667 | 3898 | 116 | 297.59 | 1 | 2.57 | 0.86 | 28.81 | 28.99 | -0.19 | 0.19 | -0.01 | 0.01 |
| gene1332 | https://www.ncbi.nlm.nih.gov/gene/?term=LOC105286872 | 2460 | 142 | 577.24 | 1 | 4.07 | 0.7 | 21.72 | 21.98 | -0.26 | 0.26 | -0.02 | 0.02 |
| gene1632 | https://www.ncbi.nlm.nih.gov/gene/?term=LOC105287764 | 18061 | 334 | 184.93 | 1 | 0.55 | 0.3 | 2.71 | 2.98 | -0.27 | 0.27 | 0 | 0 |
| gene7183 | https://www.ncbi.nlm.nih.gov/gene/?term=LOC105280348 | 3925 | 53 | 135.03 | 1 | 2.55 | 1.89 | 25.26 | 25.53 | -0.27 | 0.27 | 0 | 0 |
| gene6455 | https://www.ncbi.nlm.nih.gov/gene/?term=LOC105279500 | 2612 | 29 | 111.03 | 1 | 3.83 | 3.45 | 48.01 | 48.34 | -0.33 | 0.33 | 0 | 0 |
| gene3335 | https://www.ncbi.nlm.nih.gov/gene/?term=LOC105275898 | 4665 | 128 | 274.38 | 1 | 2.14 | 0.78 | 13.41 | 13.77 | -0.36 | 0.36 | -0.01 | 0.01 |
| gene3856 | https://www.ncbi.nlm.nih.gov/gene/?term=LOC105276499 | 5236 | 188 | 359.05 | 1 | 1.91 | 0.53 | 10.14 | 10.5 | -0.36 | 0.36 | -0.01 | 0.01 |
| gene6583 | https://www.ncbi.nlm.nih.gov/gene/?term=LOC105279649 | 7444 | 308 | 413.76 | 1 | 1.34 | 0.32 | 11.09 | 11.46 | -0.37 | 0.37 | -0.02 | 0.02 |
| gene10018 | https://www.ncbi.nlm.nih.gov/gene/?term=LOC105283607 | 4414 | 194 | 439.51 | 1 | 2.27 | 0.52 | 15.27 | 15.67 | -0.4 | 0.4 | -0.02 | 0.02 |
| gene11275 | https://www.ncbi.nlm.nih.gov/gene/?term=LOC105285055 | 4119 | 202 | 490.41 | 1 | 2.43 | 0.5 | 12.81 | 13.23 | -0.42 | 0.42 | -0.02 | 0.02 |
| gene10243 | https://www.ncbi.nlm.nih.gov/gene/?term=LOC105283862 | 3239 | 38 | 117.32 | 1 | 3.09 | 2.63 | 30.66 | 31.1 | -0.44 | 0.44 | -0.01 | 0.01 |
| gene12978 | https://www.ncbi.nlm.nih.gov/gene/?term=LOC105286997 | 2858 | 134 | 468.86 | 1 | 3.5 | 0.75 | 11.8 | 12.24 | -0.44 | 0.44 | -0.02 | 0.02 |
| gene8600 | https://www.ncbi.nlm.nih.gov/gene/?term=LOC105282001 | 8513 | 148 | 173.85 | 1 | 1.17 | 0.68 | 9.45 | 9.9 | -0.45 | 0.45 | -0.01 | 0.01 |
| gene10540 | https://www.ncbi.nlm.nih.gov/gene/?term=LOC105284200 | 5267 | 195 | 370.23 | 1 | 1.9 | 0.51 | 13.62 | 14.09 | -0.47 | 0.47 | -0.02 | 0.02 |
| gene10520 | https://www.ncbi.nlm.nih.gov/gene/?term=LOC105284184 | 2451 | 80 | 326.4 | 1 | 4.08 | 1.25 | 34.85 | 35.33 | -0.48 | 0.48 | -0.02 | 0.02 |
| gene13033 | https://www.ncbi.nlm.nih.gov/gene/?term=LOC105287063 | 1877 | 45 | 239.74 | 1 | 5.33 | 2.22 | 34.91 | 35.5 | -0.59 | 0.59 | -0.01 | 0.01 |
| gene4023 | https://www.ncbi.nlm.nih.gov/gene/?term=LOC105276692 | 1759 | 36 | 204.66 | 1 | 5.69 | 2.78 | 23.26 | 23.9 | -0.64 | 0.64 | -0.01 | 0.01 |
| gene8343 | https://www.ncbi.nlm.nih.gov/gene/?term=LOC105281700 | 2172 | 45 | 207.18 | 1 | 4.6 | 2.22 | 38.72 | 39.37 | -0.65 | 0.65 | -0.01 | 0.01 |
| gene2844 | https://www.ncbi.nlm.nih.gov/gene/?term=LOC105275363 | 4330 | 185 | 427.25 | 1 | 2.31 | 0.54 | 14.56 | 15.46 | -0.9 | 0.9 | -0.04 | 0.04 |
| gene564 | https://www.ncbi.nlm.nih.gov/gene/?term=LOC105278478 | 2802 | 12 | 42.83 | 1 | 3.57 | 8.33 | 20.51 | 21.62 | -1.11 | 1.11 | 0 | 0 |
| gene6528 | https://www.ncbi.nlm.nih.gov/gene/?term=LOC105279591 | 7222 | 68 | 94.16 | 1 | 1.38 | 1.47 | 22.7 | 23.89 | -1.19 | 1.19 | -0.01 | 0.01 |
| gene1855 | https://www.ncbi.nlm.nih.gov/gene/?term=LOC105288036 | 2387 | 86 | 360.28 | 1 | 4.19 | 1.16 | 28.48 | 29.73 | -1.24 | 1.24 | -0.04 | 0.04 |
| gene8873 | https://www.ncbi.nlm.nih.gov/gene/?term=LOC105282297 | 2055 | 78 | 379.56 | 1 | 4.87 | 1.28 | 33.62 | 34.98 | -1.37 | 1.37 | -0.05 | 0.05 |
| gene570 | https://www.ncbi.nlm.nih.gov/gene/?term=LOC105278549 | 1673 | 63 | 376.57 | 1 | 5.98 | 1.59 | 41.36 | 42.83 | -1.47 | 1.47 | -0.06 | 0.06 |
| gene5052 | https://www.ncbi.nlm.nih.gov/gene/?term=LOC105277856 | 3167 | 27 | 85.25 | 1 | 3.16 | 3.7 | 36.57 | 38.1 | -1.52 | 1.52 | -0.01 | 0.01 |
| gene9478 | https://www.ncbi.nlm.nih.gov/gene/?term=LOC105282991 | 3726 | 48 | 128.82 | 1 | 2.68 | 2.08 | 31.5 | 33.03 | -1.53 | 1.53 | -0.02 | 0.02 |
| gene274 | https://www.ncbi.nlm.nih.gov/gene/?term=LOC105275190 | 3488 | 65 | 186.35 | 1 | 2.87 | 1.54 | 21.06 | 22.74 | -1.69 | 1.69 | -0.03 | 0.03 |
| gene4034 | https://www.ncbi.nlm.nih.gov/gene/?term=LOC105276704 | 2065 | 80 | 387.41 | 1 | 4.84 | 1.25 | 13.38 | 15.07 | -1.69 | 1.69 | -0.07 | 0.07 |
| gene1884 | https://www.ncbi.nlm.nih.gov/gene/?term=LOC105288072 | 4169 | 79 | 189.49 | 1 | 2.4 | 1.27 | 22.97 | 24.67 | -1.7 | 1.7 | -0.03 | 0.03 |
| gene298 | https://www.ncbi.nlm.nih.gov/gene/?term=LOC105275460 | 2473 | 115 | 465.02 | 1 | 4.04 | 0.87 | 12.68 | 14.4 | -1.73 | 1.73 | -0.08 | 0.08 |
| gene4641 | https://www.ncbi.nlm.nih.gov/gene/?term=LOC105277398 | 1937 | 70 | 361.38 | 1 | 5.16 | 1.43 | 35.36 | 37.39 | -2.03 | 2.03 | -0.07 | 0.07 |
| gene10345 | https://www.ncbi.nlm.nih.gov/gene/?term=LOC105283989 | 4444 | 27 | 60.76 | 1 | 2.25 | 3.7 | 48.21 | 50.36 | -2.14 | 2.14 | -0.01 | 0.01 |
| gene6334 | https://www.ncbi.nlm.nih.gov/gene/?term=LOC105279356 | 1471 | 43 | 292.32 | 1 | 6.8 | 2.33 | 28.25 | 30.45 | -2.2 | 2.2 | -0.06 | 0.06 |
| gene11819 | https://www.ncbi.nlm.nih.gov/gene/?term=LOC105285671 | 2366 | 90 | 380.39 | 1 | 4.23 | 1.11 | 28.49 | 30.76 | -2.27 | 2.27 | -0.09 | 0.09 |
| gene5877 | https://www.ncbi.nlm.nih.gov/gene/?term=LOC105278807 | 1719 | 51 | 296.68 | 1 | 5.82 | 1.96 | 21.68 | 24.24 | -2.55 | 2.55 | -0.08 | 0.08 |
| gene12516 | https://www.ncbi.nlm.nih.gov/gene/?term=LOC105286452 | 1529 | 35 | 228.91 | 1 | 6.54 | 2.86 | 51.01 | 53.56 | -2.56 | 2.56 | -0.06 | 0.06 |
| gene2676 | https://www.ncbi.nlm.nih.gov/gene/?term=LOC105275181 | 1353 | 14 | 103.47 | 1 | 7.39 | 7.14 | 65.95 | 69.25 | -3.3 | 3.3 | -0.03 | 0.03 |
| gene5101 | https://www.ncbi.nlm.nih.gov/gene/?term=LOC105277903 | 1813 | 34 | 187.53 | 1 | 5.52 | 2.94 | 52.57 | 56.73 | -4.16 | 4.16 | -0.08 | 0.08 |
| gene5647 | https://www.ncbi.nlm.nih.gov/gene/?term=LOC105278567 | 3118 | 32 | 102.63 | 1 | 3.21 | 3.12 | 13.07 | 18.25 | -5.18 | 5.18 | -0.05 | 0.05 |
| gene3165 | https://www.ncbi.nlm.nih.gov/gene/?term=LOC105275706 | 2374 | 59 | 248.53 | 4 | 16.85 | 6.78 | 23.46 | 34.48 | -11.02 | 11.02 | -0.27 | 0.27 |

| | | | | | | | | | | | | | |
|---|---|---|---|---|---|---|---|---|---|---|---|---|---|
| gene2593 | https://www.ncbi.nlm.nih.gov/gene/?term=LOC105275070 | 1774 | 6 | 33.82 | 2 | 11.27 | 33.33 | 16.42 | 60.02 | -43.6 | 43.6 | -0.15 | 0.15 |

Directory: OGLall File: ogl-Cb_LI_BCphaseA1.vs.RphaseA2.txt

| gene_ID | gene_link | gwidth | #Sites | #per10Kb | #dmSites | #dmsp10kb | %dmSites | %pSite1 | %pSite2 | DMpSite | ADMpSite | DMpNucl | ADMpNucl |
|---|---|---|---|---|---|---|---|---|---|---|---|---|---|
| gene393 | https://www.ncbi.nlm.nih.gov/gene/?term=LOC105276566 | 1123 | 2 | 17.81 | 1 | 8.9 | 50 | 77.65 | 47.62 | 30.03 | 30.03 | 0.05 | 0.05 |
| gene4630 | https://www.ncbi.nlm.nih.gov/gene/?term=LOC105277380 | 1893 | 5 | 26.41 | 1 | 5.28 | 20 | 69.42 | 53.65 | 15.77 | 15.77 | 0.04 | 0.04 |
| gene9109 | https://www.ncbi.nlm.nih.gov/gene/?term=LOC105282561 | 2400 | 15 | 62.5 | 1 | 4.17 | 6.67 | 34.05 | 29.4 | 4.65 | 4.65 | 0.03 | 0.03 |
| gene7952 | https://www.ncbi.nlm.nih.gov/gene/?term=LOC105281213 | 2963 | 43 | 145.12 | 1 | 3.37 | 2.33 | 42.11 | 38.78 | 3.33 | 3.33 | 0.05 | 0.05 |
| gene3200 | https://www.ncbi.nlm.nih.gov/gene/?term=LOC105275743 | 1786 | 77 | 431.13 | 1 | 5.6 | 1.3 | 22.67 | 19.88 | 2.79 | 2.79 | 0.12 | 0.12 |
| gene7368 | https://www.ncbi.nlm.nih.gov/gene/?term=LOC105280556 | 4105 | 49 | 119.37 | 1 | 2.44 | 2.04 | 12.57 | 9.91 | 2.66 | 2.66 | 0.03 | 0.03 |
| gene920 | https://www.ncbi.nlm.nih.gov/gene/?term=LOC105282518 | 1197 | 43 | 359.23 | 1 | 8.35 | 2.33 | 37.5 | 34.97 | 2.52 | 2.52 | 0.09 | 0.09 |
| gene7915 | https://www.ncbi.nlm.nih.gov/gene/?term=LOC105281173 | 1597 | 25 | 156.54 | 1 | 6.26 | 4 | 64.87 | 62.52 | 2.35 | 2.35 | 0.04 | 0.04 |
| gene3168 | https://www.ncbi.nlm.nih.gov/gene/?term=LOC105275711 | 6150 | 80 | 130.08 | 1 | 1.63 | 1.25 | 14.39 | 12.36 | 2.03 | 2.03 | 0.03 | 0.03 |
| gene10213 | https://www.ncbi.nlm.nih.gov/gene/?term=LOC105283826 | 1503 | 54 | 359.28 | 1 | 6.65 | 1.85 | 16.14 | 14.22 | 1.92 | 1.92 | 0.07 | 0.07 |
| gene1199 | https://www.ncbi.nlm.nih.gov/gene/?term=LOC105285498 | 2220 | 26 | 117.12 | 1 | 4.5 | 3.85 | 33.11 | 31.3 | 1.81 | 1.81 | 0.02 | 0.02 |
| gene7398 | https://www.ncbi.nlm.nih.gov/gene/?term=LOC105280589 | 3452 | 41 | 118.77 | 1 | 2.9 | 2.44 | 25.07 | 23.32 | 1.75 | 1.75 | 0.02 | 0.02 |
| gene11454 | https://www.ncbi.nlm.nih.gov/gene/?term=LOC105285256 | 1906 | 51 | 267.58 | 1 | 5.25 | 1.96 | 43.83 | 42.42 | 1.4 | 1.4 | 0.04 | 0.04 |
| gene3671 | https://www.ncbi.nlm.nih.gov/gene/?term=LOC105276272 | 1966 | 27 | 137.33 | 1 | 5.09 | 3.7 | 41.96 | 40.71 | 1.25 | 1.25 | 0.02 | 0.02 |
| gene322 | https://www.ncbi.nlm.nih.gov/gene/?term=LOC105275777 | 1606 | 23 | 143.21 | 1 | 6.23 | 4.35 | 45.55 | 44.33 | 1.21 | 1.21 | 0.02 | 0.02 |
| gene6237 | https://www.ncbi.nlm.nih.gov/gene/?term=LOC105279244 | 1484 | 26 | 175.2 | 1 | 6.74 | 3.85 | 46.7 | 45.51 | 1.19 | 1.19 | 0.02 | 0.02 |
| gene10326 | https://www.ncbi.nlm.nih.gov/gene/?term=LOC105283962 | 3042 | 102 | 335.31 | 1 | 3.29 | 0.98 | 14.13 | 13.04 | 1.09 | 1.09 | 0.04 | 0.04 |
| gene4190 | https://www.ncbi.nlm.nih.gov/gene/?term=LOC105276875 | 3150 | 132 | 419.05 | 1 | 3.17 | 0.76 | 21.24 | 20.15 | 1.09 | 1.09 | 0.05 | 0.05 |
| gene7345 | https://www.ncbi.nlm.nih.gov/gene/?term=LOC105280529 | 3833 | 92 | 240.02 | 1 | 2.61 | 1.09 | 17.99 | 16.93 | 1.05 | 1.05 | 0.03 | 0.03 |
| gene10819 | https://www.ncbi.nlm.nih.gov/gene/?term=LOC105284507 | 5767 | 83 | 143.92 | 1 | 1.73 | 1.2 | 21.4 | 20.36 | 1.04 | 1.04 | 0.02 | 0.02 |
| gene3286 | https://www.ncbi.nlm.nih.gov/gene/?term=LOC105275848 | 3744 | 132 | 352.56 | 1 | 2.67 | 0.76 | 21.71 | 20.73 | 0.98 | 0.98 | 0.03 | 0.03 |
| gene13429 | https://www.ncbi.nlm.nih.gov/gene/?term=LOC105287501 | 3714 | 102 | 274.64 | 1 | 2.69 | 0.98 | 22.78 | 21.81 | 0.97 | 0.97 | 0.03 | 0.03 |
| gene814 | https://www.ncbi.nlm.nih.gov/gene/?term=LOC105281296 | 2722 | 43 | 157.97 | 1 | 3.67 | 2.33 | 29.73 | 28.81 | 0.92 | 0.92 | 0.01 | 0.01 |
| gene5298 | https://www.ncbi.nlm.nih.gov/gene/?term=LOC105278145 | 4060 | 257 | 633 | 2 | 4.93 | 0.78 | 10.97 | 10.08 | 0.88 | 0.88 | 0.06 | 0.06 |
| gene11447 | https://www.ncbi.nlm.nih.gov/gene/?term=LOC105285252 | 7437 | 135 | 181.52 | 1 | 1.34 | 0.74 | 13.54 | 12.74 | 0.8 | 0.8 | 0.01 | 0.01 |
| gene1159 | https://www.ncbi.nlm.nih.gov/gene/?term=LOC105285084 | 3282 | 27 | 82.27 | 1 | 3.05 | 3.7 | 22 | 21.23 | 0.77 | 0.77 | 0.01 | 0.01 |
| gene652 | https://www.ncbi.nlm.nih.gov/gene/?term=LOC105279528 | 5806 | 44 | 75.78 | 1 | 1.72 | 2.27 | 16.35 | 15.59 | 0.76 | 0.76 | 0.01 | 0.01 |
| gene6966 | https://www.ncbi.nlm.nih.gov/gene/?term=LOC105280104 | 3747 | 125 | 333.6 | 1 | 2.67 | 0.8 | 14.94 | 14.21 | 0.73 | 0.73 | 0.02 | 0.02 |
| gene9722 | https://www.ncbi.nlm.nih.gov/gene/?term=LOC105283264 | 3951 | 88 | 222.73 | 1 | 2.53 | 1.14 | 24.81 | 24.09 | 0.72 | 0.72 | 0.02 | 0.02 |
| gene11340 | https://www.ncbi.nlm.nih.gov/gene/?term=LOC105285131 | 2918 | 102 | 349.55 | 1 | 3.43 | 0.98 | 14.35 | 13.65 | 0.7 | 0.7 | 0.02 | 0.02 |
| gene9464 | https://www.ncbi.nlm.nih.gov/gene/?term=LOC105282975 | 1450 | 25 | 172.41 | 1 | 6.9 | 4 | 41.13 | 40.46 | 0.67 | 0.67 | 0.01 | 0.01 |
| gene12573 | https://www.ncbi.nlm.nih.gov/gene/?term=LOC105286515 | 3647 | 115 | 315.33 | 1 | 2.74 | 0.87 | 11.99 | 11.33 | 0.66 | 0.66 | 0.02 | 0.02 |
| gene4376 | https://www.ncbi.nlm.nih.gov/gene/?term=LOC105277083 | 4307 | 96 | 222.89 | 1 | 2.32 | 1.04 | 25.73 | 25.08 | 0.66 | 0.66 | 0.01 | 0.01 |
| gene11401 | https://www.ncbi.nlm.nih.gov/gene/?term=LOC105285202 | 1841 | 50 | 271.59 | 1 | 5.43 | 2 | 21.11 | 20.46 | 0.65 | 0.65 | 0.02 | 0.02 |
| gene7600 | https://www.ncbi.nlm.nih.gov/gene/?term=LOC105280819 | 1615 | 41 | 253.87 | 1 | 6.19 | 2.44 | 40.74 | 40.08 | 0.65 | 0.65 | 0.02 | 0.02 |
| gene10155 | https://www.ncbi.nlm.nih.gov/gene/?term=LOC105283756 | 7030 | 97 | 137.98 | 1 | 1.42 | 1.03 | 6.83 | 6.25 | 0.59 | 0.59 | 0.01 | 0.01 |
| gene10714 | https://www.ncbi.nlm.nih.gov/gene/?term=LOC105284396 | 1966 | 57 | 289.93 | 1 | 5.09 | 1.75 | 38.23 | 37.64 | 0.59 | 0.59 | 0.02 | 0.02 |
| gene4962 | https://www.ncbi.nlm.nih.gov/gene/?term=LOC105277761 | 4645 | 150 | 322.93 | 1 | 2.15 | 0.67 | 14.77 | 14.2 | 0.57 | 0.57 | 0.02 | 0.02 |
| gene12203 | https://www.ncbi.nlm.nih.gov/gene/?term=LOC105286099 | 5967 | 78 | 130.72 | 1 | 1.68 | 1.28 | 11.28 | 10.77 | 0.51 | 0.51 | 0.01 | 0.01 |
| gene1809 | https://www.ncbi.nlm.nih.gov/gene/?term=LOC105287989 | 7010 | 73 | 104.14 | 1 | 1.43 | 1.37 | 23.7 | 23.28 | 0.42 | 0.42 | 0 | 0 |
| gene3107 | https://www.ncbi.nlm.nih.gov/gene/?term=LOC105275650 | 5103 | 151 | 295.9 | 1 | 1.96 | 0.66 | 17.08 | 16.7 | 0.38 | 0.38 | 0.01 | 0.01 |
| gene5646 | https://www.ncbi.nlm.nih.gov/gene/?term=LOC105278534 | 7475 | 186 | 248.83 | 1 | 1.34 | 0.54 | 18.03 | 17.67 | 0.36 | 0.36 | 0.01 | 0.01 |
| gene10392 | https://www.ncbi.nlm.nih.gov/gene/?term=LOC105284036 | 3216 | 83 | 258.08 | 2 | 6.22 | 2.41 | 23.78 | 23.42 | 0.35 | 0.35 | 0.01 | 0.01 |
| gene1220 | https://www.ncbi.nlm.nih.gov/gene/?term=LOC105285742 | 3453 | 99 | 286.71 | 1 | 2.9 | 1.01 | 25.65 | 25.32 | 0.33 | 0.33 | 0.01 | 0.01 |
| gene3848 | https://www.ncbi.nlm.nih.gov/gene/?term=LOC105276486 | 5899 | 121 | 205.12 | 1 | 1.7 | 0.83 | 23.22 | 22.92 | 0.3 | 0.3 | 0.01 | 0.01 |
| gene11583 | https://www.ncbi.nlm.nih.gov/gene/?term=LOC105285414 | 8192 | 177 | 216.06 | 1 | 1.22 | 0.56 | 8.82 | 8.53 | 0.29 | 0.29 | 0.01 | 0.01 |
| gene5539 | https://www.ncbi.nlm.nih.gov/gene/?term=LOC105278415 | 25024 | 604 | 241.37 | 1 | 0.4 | 0.17 | 3.91 | 3.62 | 0.29 | 0.29 | 0.01 | 0.01 |
| gene5831 | https://www.ncbi.nlm.nih.gov/gene/?term=LOC105278761 | 2506 | 104 | 415 | 1 | 3.99 | 0.96 | 25.24 | 24.96 | 0.28 | 0.28 | 0.01 | 0.01 |
| gene12307 | https://www.ncbi.nlm.nih.gov/gene/?term=LOC105286216 | 10313 | 167 | 161.93 | 1 | 0.97 | 0.6 | 15.75 | 15.52 | 0.23 | 0.23 | 0 | 0 |
| gene657 | https://www.ncbi.nlm.nih.gov/gene/?term=LOC105279667 | 6421 | 80 | 124.59 | 1 | 1.56 | 1.25 | 3.15 | 2.93 | 0.21 | 0.21 | 0 | 0 |
| gene13114 | https://www.ncbi.nlm.nih.gov/gene/?term=LOC105287147 | 7609 | 389 | 511.24 | 1 | 1.31 | 0.26 | 9.77 | 9.58 | 0.19 | 0.19 | 0.01 | 0.01 |
| gene5746 | https://www.ncbi.nlm.nih.gov/gene/?term=LOC105278666 | 10343 | 409 | 395.44 | 1 | 0.97 | 0.24 | 7.43 | 7.26 | 0.17 | 0.17 | 0.01 | 0.01 |
| gene940 | https://www.ncbi.nlm.nih.gov/gene/?term=LOC105282794 | 3578 | 88 | 245.95 | 1 | 2.79 | 1.14 | 15.65 | 15.49 | 0.16 | 0.16 | 0 | 0 |
| gene12674 | https://www.ncbi.nlm.nih.gov/gene/?term=LOC105286649 | 18283 | 680 | 371.93 | 1 | 0.55 | 0.15 | 1.31 | 1.16 | 0.15 | 0.15 | 0.01 | 0.01 |
| gene6031 | https://www.ncbi.nlm.nih.gov/gene/?term=LOC105279000 | 5691 | 185 | 325.07 | 1 | 1.76 | 0.54 | 13.85 | 13.69 | 0.15 | 0.15 | 0 | 0 |
| gene3599 | https://www.ncbi.nlm.nih.gov/gene/?term=LOC105276196 | 2955 | 56 | 189.51 | 1 | 3.38 | 1.79 | 40.12 | 39.99 | 0.13 | 0.13 | 0 | 0 |
| gene7547 | https://www.ncbi.nlm.nih.gov/gene/?term=LOC105280767 | 24576 | 753 | 306.4 | 1 | 0.41 | 0.13 | 4.75 | 4.63 | 0.12 | 0.12 | 0 | 0 |
| gene2404 | https://www.ncbi.nlm.nih.gov/gene/?term=LOC105274853 | 1767 | 42 | 237.69 | 1 | 5.66 | 2.38 | 28.85 | 28.79 | 0.07 | 0.07 | 0 | 0 |
| gene2991 | https://www.ncbi.nlm.nih.gov/gene/?term=LOC105275536 | 11224 | 214 | 190.66 | 1 | 0.89 | 0.47 | 5.64 | 5.58 | 0.05 | 0.05 | 0 | 0 |
| gene4527 | https://www.ncbi.nlm.nih.gov/gene/?term=LOC105277273 | 3686 | 151 | 409.66 | 1 | 2.71 | 0.66 | 12.71 | 12.65 | 0.05 | 0.05 | 0 | 0 |

| | | | | | | | | | | | | | |
|---|---|---|---|---|---|---|---|---|---|---|---|---|---|
| gene4996 | https://www.ncbi.nlm.nih.gov/gene/?term=LOC105277798 | 5617 | 64 | 113.94 | 1 | 1.78 | 1.56 | 10.86 | 10.81 | 0.05 | 0.05 | 0 | 0 |
| gene2266 | https://www.ncbi.nlm.nih.gov/gene/?term=LOC105274699 | 2391 | 63 | 263.49 | 1 | 4.18 | 1.59 | 23.04 | 22.99 | 0.04 | 0.04 | 0 | 0 |
| gene7148 | https://www.ncbi.nlm.nih.gov/gene/?term=LOC105280304 | 8920 | 123 | 137.89 | 1 | 1.12 | 0.81 | 5.79 | 5.75 | 0.04 | 0.04 | 0 | 0 |
| gene53 | https://www.ncbi.nlm.nih.gov/gene/?term=LOC105277148 | 19283 | 269 | 139.5 | 1 | 0.52 | 0.37 | 3.9 | 3.86 | 0.03 | 0.03 | 0 | 0 |
| gene9645 | https://www.ncbi.nlm.nih.gov/gene/?term=LOC105283181 | 3875 | 138 | 356.13 | 1 | 2.58 | 0.72 | 11.99 | 11.98 | 0.01 | 0.01 | 0 | 0 |
| gene10549 | https://www.ncbi.nlm.nih.gov/gene/?term=LOC105284210 | 7145 | 323 | 452.06 | 1 | 1.4 | 0.31 | 10.47 | 10.47 | 0 | 0 | 0 | 0 |
| gene1470 | https://www.ncbi.nlm.nih.gov/gene/?term=LOC105287597 | 2974 | 192 | 645.6 | 1 | 3.36 | 0.52 | 16.86 | 16.86 | 0 | 0 | 0 | 0 |
| gene9755 | https://www.ncbi.nlm.nih.gov/gene/?term=LOC105283296 | 2177 | 90 | 413.41 | 1 | 4.59 | 1.11 | 19.36 | 19.36 | 0 | 0 | 0 | 0 |
| gene4209 | https://www.ncbi.nlm.nih.gov/gene/?term=LOC105277125 | 50550 | 1735 | 343.22 | 1 | 0.2 | 0.06 | 1.59 | 1.62 | -0.03 | 0.03 | 0 | 0 |
| gene9411 | https://www.ncbi.nlm.nih.gov/gene/?term=LOC105282914 | 5534 | 41 | 74.09 | 1 | 1.81 | 2.44 | 49.68 | 49.72 | -0.04 | 0.04 | 0 | 0 |
| gene13036 | https://www.ncbi.nlm.nih.gov/gene/?term=LOC105287059 | 4959 | 165 | 332.73 | 1 | 2.02 | 0.61 | 17.35 | 17.4 | -0.05 | 0.05 | 0 | 0 |
| gene7207 | https://www.ncbi.nlm.nih.gov/gene/?term=LOC105280376 | 4659 | 105 | 225.37 | 1 | 2.15 | 0.95 | 28.77 | 28.82 | -0.05 | 0.05 | 0 | 0 |
| gene10606 | https://www.ncbi.nlm.nih.gov/gene/?term=LOC105284273 | 6119 | 252 | 411.83 | 2 | 3.27 | 0.79 | 19.91 | 19.99 | -0.08 | 0.08 | 0 | 0 |
| gene11568 | https://www.ncbi.nlm.nih.gov/gene/?term=LOC105285395 | 8086 | 184 | 227.55 | 1 | 1.24 | 0.54 | 16.42 | 16.51 | -0.08 | 0.08 | 0 | 0 |
| gene3470 | https://www.ncbi.nlm.nih.gov/gene/?term=LOC105276059 | 8716 | 161 | 184.72 | 1 | 1.15 | 0.62 | 9.12 | 9.21 | -0.08 | 0.08 | 0 | 0 |
| gene8274 | https://www.ncbi.nlm.nih.gov/gene/?term=LOC105281624 | 5425 | 208 | 383.41 | 1 | 1.84 | 0.48 | 10.51 | 10.6 | -0.09 | 0.09 | 0 | 0 |
| gene2748 | https://www.ncbi.nlm.nih.gov/gene/?term=LOC105275261 | 3845 | 58 | 150.85 | 1 | 2.6 | 1.72 | 39.98 | 40.08 | -0.1 | 0.1 | 0 | 0 |
| gene1632 | https://www.ncbi.nlm.nih.gov/gene/?term=LOC105287764 | 18061 | 354 | 196 | 1 | 0.55 | 0.28 | 2.69 | 2.8 | -0.11 | 0.11 | 0 | 0 |
| gene10596 | https://www.ncbi.nlm.nih.gov/gene/?term=LOC105284262 | 6520 | 235 | 360.43 | 1 | 1.53 | 0.43 | 3.69 | 3.81 | -0.12 | 0.12 | 0 | 0 |
| gene10864 | https://www.ncbi.nlm.nih.gov/gene/?term=LOC105284557 | 1666 | 78 | 468.19 | 1 | 6 | 1.28 | 30.45 | 30.59 | -0.13 | 0.13 | -0.01 | 0.01 |
| gene5649 | https://www.ncbi.nlm.nih.gov/gene/?term=LOC105278538 | 2140 | 84 | 392.52 | 1 | 4.67 | 1.19 | 18.7 | 18.83 | -0.13 | 0.13 | -0.01 | 0.01 |
| gene3195 | https://www.ncbi.nlm.nih.gov/gene/?term=LOC105275750 | 28072 | 713 | 253.99 | 1 | 0.36 | 0.14 | 11.56 | 11.74 | -0.18 | 0.18 | 0 | 0 |
| gene2683 | https://www.ncbi.nlm.nih.gov/gene/?term=LOC105275187 | 5492 | 75 | 136.56 | 1 | 1.82 | 1.33 | 14.14 | 14.34 | -0.2 | 0.2 | 0 | 0 |
| gene6583 | https://www.ncbi.nlm.nih.gov/gene/?term=LOC105279649 | 7444 | 330 | 443.31 | 1 | 1.34 | 0.3 | 11.47 | 11.66 | -0.2 | 0.2 | -0.01 | 0.01 |
| gene11579 | https://www.ncbi.nlm.nih.gov/gene/?term=LOC105285406 | 13571 | 293 | 215.9 | 1 | 0.74 | 0.34 | 5.86 | 6.07 | -0.21 | 0.21 | 0 | 0 |
| gene2806 | https://www.ncbi.nlm.nih.gov/gene/?term=LOC105275327 | 14257 | 480 | 336.68 | 1 | 0.7 | 0.21 | 7.1 | 7.32 | -0.22 | 0.22 | -0.01 | 0.01 |
| gene7249 | https://www.ncbi.nlm.nih.gov/gene/?term=LOC105280422 | 1775 | 61 | 343.66 | 1 | 5.63 | 1.64 | 15.16 | 15.4 | -0.24 | 0.24 | -0.01 | 0.01 |
| gene9101 | https://www.ncbi.nlm.nih.gov/gene/?term=LOC105282553 | 3068 | 104 | 338.98 | 1 | 3.26 | 0.96 | 20.36 | 20.64 | -0.27 | 0.27 | -0.01 | 0.01 |
| gene5613 | https://www.ncbi.nlm.nih.gov/gene/?term=LOC105278504 | 8524 | 211 | 247.54 | 1 | 1.17 | 0.47 | 5.46 | 5.74 | -0.28 | 0.28 | -0.01 | 0.01 |
| gene6620 | https://www.ncbi.nlm.nih.gov/gene/?term=LOC105279687 | 17998 | 360 | 200.02 | 1 | 0.56 | 0.28 | 5.06 | 5.34 | -0.28 | 0.28 | -0.01 | 0.01 |
| gene1573 | https://www.ncbi.nlm.nih.gov/gene/?term=LOC105287704 | 3030 | 131 | 432.34 | 1 | 3.3 | 0.76 | 20.77 | 21.09 | -0.32 | 0.32 | -0.01 | 0.01 |
| gene2056 | https://www.ncbi.nlm.nih.gov/gene/?term=LOC105274452 | 6584 | 212 | 321.99 | 1 | 1.52 | 0.47 | 13.94 | 14.26 | -0.33 | 0.33 | -0.01 | 0.01 |
| gene1147 | https://www.ncbi.nlm.nih.gov/gene/?term=LOC105284954 | 4022 | 82 | 203.88 | 1 | 2.49 | 1.22 | 15.94 | 16.29 | -0.34 | 0.34 | -0.01 | 0.01 |
| gene6286 | https://www.ncbi.nlm.nih.gov/gene/?term=LOC105279298 | 2861 | 139 | 485.84 | 2 | 6.99 | 1.44 | 22.43 | 22.79 | -0.36 | 0.36 | -0.02 | 0.02 |
| gene6639 | https://www.ncbi.nlm.nih.gov/gene/?term=LOC105279705 | 7656 | 69 | 90.13 | 1 | 1.31 | 1.45 | 21.96 | 22.32 | -0.36 | 0.36 | 0 | 0 |
| gene10600 | https://www.ncbi.nlm.nih.gov/gene/?term=LOC105284270 | 4837 | 105 | 217.08 | 1 | 2.07 | 0.95 | 12.85 | 13.22 | -0.37 | 0.37 | -0.01 | 0.01 |
| gene3856 | https://www.ncbi.nlm.nih.gov/gene/?term=LOC105276499 | 5236 | 201 | 383.88 | 1 | 1.91 | 0.5 | 10.34 | 10.72 | -0.39 | 0.39 | -0.01 | 0.01 |
| gene7568 | https://www.ncbi.nlm.nih.gov/gene/?term=LOC105280786 | 2525 | 123 | 487.13 | 1 | 3.96 | 0.81 | 28.92 | 29.32 | -0.39 | 0.39 | -0.02 | 0.02 |
| gene10520 | https://www.ncbi.nlm.nih.gov/gene/?term=LOC105284184 | 2451 | 82 | 334.56 | 1 | 4.08 | 1.22 | 34.55 | 34.95 | -0.4 | 0.4 | -0.01 | 0.01 |
| gene7888 | https://www.ncbi.nlm.nih.gov/gene/?term=LOC105281141 | 4357 | 165 | 378.7 | 1 | 2.3 | 0.61 | 28.04 | 28.47 | -0.43 | 0.43 | -0.02 | 0.02 |
| gene5375 | https://www.ncbi.nlm.nih.gov/gene/?term=LOC105278228 | 8640 | 159 | 184.03 | 1 | 1.16 | 0.63 | 24.17 | 24.62 | -0.45 | 0.45 | -0.01 | 0.01 |
| gene4892 | https://www.ncbi.nlm.nih.gov/gene/?term=LOC105277667 | 3898 | 122 | 312.98 | 1 | 2.57 | 0.82 | 27.68 | 28.18 | -0.5 | 0.5 | -0.02 | 0.02 |
| gene7550 | https://www.ncbi.nlm.nih.gov/gene/?term=LOC105280763 | 2173 | 98 | 450.99 | 1 | 4.6 | 1.02 | 26.91 | 27.42 | -0.51 | 0.51 | -0.02 | 0.02 |
| gene3885 | https://www.ncbi.nlm.nih.gov/gene/?term=LOC105276527 | 7928 | 352 | 444 | 1 | 1.26 | 0.28 | 8.21 | 8.74 | -0.52 | 0.52 | -0.02 | 0.02 |
| gene4225 | https://www.ncbi.nlm.nih.gov/gene/?term=LOC105276917 | 2443 | 26 | 106.43 | 1 | 4.09 | 3.85 | 36.73 | 37.26 | -0.53 | 0.53 | -0.01 | 0.01 |
| gene5967 | https://www.ncbi.nlm.nih.gov/gene/?term=LOC105278918 | 4393 | 135 | 307.31 | 1 | 2.28 | 0.74 | 16.02 | 16.55 | -0.53 | 0.53 | -0.02 | 0.02 |
| gene9330 | https://www.ncbi.nlm.nih.gov/gene/?term=LOC105282825 | 3936 | 90 | 228.66 | 1 | 2.54 | 1.11 | 15.33 | 15.86 | -0.53 | 0.53 | -0.01 | 0.01 |
| gene7166 | https://www.ncbi.nlm.nih.gov/gene/?term=LOC105280326 | 6117 | 213 | 348.21 | 1 | 1.63 | 0.47 | 15.89 | 16.43 | -0.54 | 0.54 | -0.02 | 0.02 |
| gene474 | https://www.ncbi.nlm.nih.gov/gene/?term=LOC105277435 | 3625 | 158 | 435.86 | 1 | 2.76 | 0.63 | 12.73 | 13.28 | -0.55 | 0.55 | -0.02 | 0.02 |
| gene8600 | https://www.ncbi.nlm.nih.gov/gene/?term=LOC105282001 | 8513 | 158 | 185.6 | 1 | 1.17 | 0.63 | 9.54 | 10.09 | -0.55 | 0.55 | -0.01 | 0.01 |
| gene7245 | https://www.ncbi.nlm.nih.gov/gene/?term=LOC105280410 | 12478 | 260 | 208.37 | 1 | 0.8 | 0.38 | 19.03 | 19.62 | -0.59 | 0.59 | -0.01 | 0.01 |
| gene10678 | https://www.ncbi.nlm.nih.gov/gene/?term=LOC105284355 | 4031 | 164 | 406.85 | 1 | 2.48 | 0.61 | 19.18 | 19.78 | -0.6 | 0.6 | -0.02 | 0.02 |
| gene3385 | https://www.ncbi.nlm.nih.gov/gene/?term=LOC105275966 | 3305 | 59 | 178.52 | 1 | 3.03 | 1.69 | 50.29 | 50.9 | -0.61 | 0.61 | -0.01 | 0.01 |
| gene10379 | https://www.ncbi.nlm.nih.gov/gene/?term=LOC105284028 | 6974 | 74 | 106.11 | 1 | 1.43 | 1.35 | 8.99 | 9.61 | -0.62 | 0.62 | -0.01 | 0.01 |
| gene11275 | https://www.ncbi.nlm.nih.gov/gene/?term=LOC105285055 | 4119 | 216 | 524.4 | 1 | 2.43 | 0.46 | 13.08 | 13.7 | -0.62 | 0.62 | -0.03 | 0.03 |
| gene12802 | https://www.ncbi.nlm.nih.gov/gene/?term=LOC105286799 | 3183 | 172 | 540.37 | 1 | 3.14 | 0.58 | 15.78 | 16.4 | -0.62 | 0.62 | -0.03 | 0.03 |
| gene388 | https://www.ncbi.nlm.nih.gov/gene/?term=LOC105276538 | 1558 | 20 | 128.37 | 2 | 12.84 | 10 | 70.94 | 71.62 | -0.68 | 0.68 | -0.01 | 0.01 |
| gene3817 | https://www.ncbi.nlm.nih.gov/gene/?term=LOC105276449 | 1230 | 48 | 390.24 | 1 | 8.13 | 2.08 | 26.73 | 27.42 | -0.69 | 0.69 | -0.03 | 0.03 |
| gene10018 | https://www.ncbi.nlm.nih.gov/gene/?term=LOC105283607 | 4414 | 198 | 448.57 | 1 | 2.27 | 0.51 | 15.25 | 15.95 | -0.7 | 0.7 | -0.03 | 0.03 |
| gene2271 | https://www.ncbi.nlm.nih.gov/gene/?term=LOC105274706 | 1448 | 53 | 366.02 | 1 | 6.91 | 1.89 | 36.91 | 37.62 | -0.71 | 0.71 | -0.03 | 0.03 |
| gene8661 | https://www.ncbi.nlm.nih.gov/gene/?term=LOC105282063 | 1926 | 43 | 223.26 | 1 | 5.19 | 2.33 | 23.84 | 24.54 | -0.71 | 0.71 | -0.02 | 0.02 |
| gene3206 | https://www.ncbi.nlm.nih.gov/gene/?term=LOC105275752 | 3474 | 125 | 359.82 | 1 | 2.88 | 0.8 | 28.34 | 29.06 | -0.72 | 0.72 | -0.03 | 0.03 |
| gene927 | https://www.ncbi.nlm.nih.gov/gene/?term=LOC105282600 | 3727 | 65 | 174.4 | 1 | 2.68 | 1.54 | 26.3 | 27.02 | -0.73 | 0.73 | -0.01 | 0.01 |
| gene2844 | https://www.ncbi.nlm.nih.gov/gene/?term=LOC105275363 | 4330 | 192 | 443.42 | 1 | 2.31 | 0.52 | 14.13 | 14.88 | -0.75 | 0.75 | -0.03 | 0.03 |

| | | | | | | | | | | | | | |
|---|---|---|---|---|---|---|---|---|---|---|---|---|---|
| gene7556 | https://www.ncbi.nlm.nih.gov/gene/?term=LOC105280774 | 2792 | 117 | 419.05 | 1 | 3.58 | 0.85 | 27.44 | 28.23 | -0.79 | 0.79 | -0.03 | 0.03 |
| gene2173 | https://www.ncbi.nlm.nih.gov/gene/?term=LOC105274590 | 4387 | 84 | 191.47 | 1 | 2.28 | 1.19 | 20.89 | 21.7 | -0.81 | 0.81 | -0.02 | 0.02 |
| gene9598 | https://www.ncbi.nlm.nih.gov/gene/?term=LOC105283132 | 1769 | 54 | 305.26 | 1 | 5.65 | 1.85 | 40.78 | 41.59 | -0.81 | 0.81 | -0.02 | 0.02 |
| gene2330 | https://www.ncbi.nlm.nih.gov/gene/?term=LOC105274773 | 13104 | 172 | 131.26 | 1 | 0.76 | 0.58 | 15.95 | 16.78 | -0.83 | 0.83 | -0.01 | 0.01 |
| gene8729 | https://www.ncbi.nlm.nih.gov/gene/?term=LOC105282137 | 5005 | 205 | 409.59 | 1 | 2 | 0.49 | 15.76 | 16.6 | -0.84 | 0.84 | -0.03 | 0.03 |
| gene6528 | https://www.ncbi.nlm.nih.gov/gene/?term=LOC105279591 | 7222 | 75 | 103.85 | 1 | 1.38 | 1.33 | 20.71 | 21.6 | -0.89 | 0.89 | -0.01 | 0.01 |
| gene4641 | https://www.ncbi.nlm.nih.gov/gene/?term=LOC105277398 | 1937 | 70 | 361.38 | 1 | 5.16 | 1.43 | 35.36 | 36.26 | -0.9 | 0.9 | -0.03 | 0.03 |
| gene10540 | https://www.ncbi.nlm.nih.gov/gene/?term=LOC105284200 | 5267 | 199 | 377.82 | 1 | 1.9 | 0.5 | 13.8 | 14.75 | -0.95 | 0.95 | -0.04 | 0.04 |
| gene10345 | https://www.ncbi.nlm.nih.gov/gene/?term=LOC105283989 | 4444 | 30 | 67.51 | 1 | 2.25 | 3.33 | 46.58 | 47.61 | -1.03 | 1.03 | -0.01 | 0.01 |
| gene10531 | https://www.ncbi.nlm.nih.gov/gene/?term=LOC105284190 | 6881 | 199 | 289.2 | 1 | 1.45 | 0.5 | 9.56 | 10.66 | -1.1 | 1.1 | -0.03 | 0.03 |
| gene5524 | https://www.ncbi.nlm.nih.gov/gene/?term=LOC105278397 | 5177 | 200 | 386.32 | 1 | 1.93 | 0.5 | 24.39 | 25.66 | -1.27 | 1.27 | -0.05 | 0.05 |
| gene298 | https://www.ncbi.nlm.nih.gov/gene/?term=LOC105275460 | 2473 | 112 | 452.89 | 1 | 4.04 | 0.89 | 12.88 | 14.18 | -1.3 | 1.3 | -0.06 | 0.06 |
| gene4070 | https://www.ncbi.nlm.nih.gov/gene/?term=LOC105276745 | 3957 | 72 | 181.96 | 1 | 2.53 | 1.39 | 20.98 | 22.28 | -1.3 | 1.3 | -0.02 | 0.02 |
| gene9336 | https://www.ncbi.nlm.nih.gov/gene/?term=LOC105282834 | 5993 | 109 | 181.88 | 1 | 1.67 | 0.92 | 15.48 | 16.78 | -1.31 | 1.31 | -0.02 | 0.02 |
| gene1962 | https://www.ncbi.nlm.nih.gov/gene/?term=LOC105288157 | 1246 | 16 | 128.41 | 1 | 8.03 | 6.25 | 13.95 | 15.31 | -1.36 | 1.36 | -0.02 | 0.02 |
| gene13033 | https://www.ncbi.nlm.nih.gov/gene/?term=LOC105287063 | 1877 | 45 | 239.74 | 1 | 5.33 | 2.22 | 34.91 | 36.28 | -1.37 | 1.37 | -0.03 | 0.03 |
| gene7183 | https://www.ncbi.nlm.nih.gov/gene/?term=LOC105280348 | 3925 | 53 | 135.03 | 1 | 2.55 | 1.89 | 25.26 | 26.63 | -1.37 | 1.37 | -0.02 | 0.02 |
| gene8178 | https://www.ncbi.nlm.nih.gov/gene/?term=LOC105281515 | 8647 | 46 | 53.2 | 1 | 1.16 | 2.17 | 28.18 | 29.56 | -1.38 | 1.38 | -0.01 | 0.01 |
| gene1884 | https://www.ncbi.nlm.nih.gov/gene/?term=LOC105288072 | 4169 | 82 | 196.69 | 1 | 2.4 | 1.22 | 22.32 | 23.72 | -1.41 | 1.41 | -0.03 | 0.03 |
| gene2257 | https://www.ncbi.nlm.nih.gov/gene/?term=LOC105274689 | 3658 | 75 | 205.03 | 1 | 2.73 | 1.33 | 12.35 | 13.76 | -1.41 | 1.41 | -0.03 | 0.03 |
| gene5877 | https://www.ncbi.nlm.nih.gov/gene/?term=LOC105278807 | 1719 | 52 | 302.5 | 1 | 5.82 | 1.92 | 22.91 | 24.37 | -1.46 | 1.46 | -0.04 | 0.04 |
| gene8343 | https://www.ncbi.nlm.nih.gov/gene/?term=LOC105281700 | 2172 | 46 | 211.79 | 1 | 4.6 | 2.17 | 39.86 | 41.32 | -1.46 | 1.46 | -0.03 | 0.03 |
| gene5525 | https://www.ncbi.nlm.nih.gov/gene/?term=LOC105278396 | 3090 | 186 | 601.94 | 1 | 3.24 | 0.54 | 24.51 | 25.99 | -1.48 | 1.48 | -0.09 | 0.09 |
| gene6334 | https://www.ncbi.nlm.nih.gov/gene/?term=LOC105279356 | 1471 | 43 | 292.32 | 1 | 6.8 | 2.33 | 28.25 | 29.87 | -1.62 | 1.62 | -0.05 | 0.05 |
| gene1310 | https://www.ncbi.nlm.nih.gov/gene/?term=LOC105286661 | 2582 | 65 | 251.74 | 1 | 3.87 | 1.54 | 19.96 | 21.59 | -1.63 | 1.63 | -0.04 | 0.04 |
| gene4023 | https://www.ncbi.nlm.nih.gov/gene/?term=LOC105276692 | 1759 | 39 | 221.72 | 1 | 5.69 | 2.56 | 19.85 | 21.5 | -1.65 | 1.65 | -0.04 | 0.04 |
| gene6455 | https://www.ncbi.nlm.nih.gov/gene/?term=LOC105279500 | 2612 | 32 | 122.51 | 1 | 3.83 | 3.12 | 49.77 | 51.44 | -1.68 | 1.68 | -0.02 | 0.02 |
| gene11819 | https://www.ncbi.nlm.nih.gov/gene/?term=LOC105285671 | 2366 | 95 | 401.52 | 1 | 4.23 | 1.05 | 27.1 | 28.85 | -1.76 | 1.76 | -0.07 | 0.07 |
| gene5402 | https://www.ncbi.nlm.nih.gov/gene/?term=LOC105278260 | 3068 | 41 | 133.64 | 1 | 3.26 | 2.44 | 13.01 | 14.77 | -1.77 | 1.77 | -0.02 | 0.02 |
| gene10984 | https://www.ncbi.nlm.nih.gov/gene/?term=LOC105284704 | 1987 | 64 | 322.09 | 1 | 5.03 | 1.56 | 25.47 | 27.27 | -1.8 | 1.8 | -0.06 | 0.06 |
| gene12317 | https://www.ncbi.nlm.nih.gov/gene/?term=LOC105286223 | 2235 | 84 | 375.84 | 1 | 4.47 | 1.19 | 29.87 | 31.79 | -1.93 | 1.93 | -0.07 | 0.07 |
| gene10243 | https://www.ncbi.nlm.nih.gov/gene/?term=LOC105283862 | 3239 | 37 | 114.23 | 1 | 3.09 | 2.7 | 31.49 | 33.45 | -1.96 | 1.96 | -0.02 | 0.02 |
| gene3727 | https://www.ncbi.nlm.nih.gov/gene/?term=LOC105276367 | 2484 | 54 | 217.39 | 1 | 4.03 | 1.85 | 48.38 | 50.37 | -1.99 | 1.99 | -0.04 | 0.04 |
| gene2970 | https://www.ncbi.nlm.nih.gov/gene/?term=LOC105275518 | 2340 | 21 | 89.74 | 1 | 4.27 | 4.76 | 45.1 | 47.46 | -2.36 | 2.36 | -0.02 | 0.02 |
| gene12021 | https://www.ncbi.nlm.nih.gov/gene/?term=LOC105285905 | 1399 | 23 | 164.4 | 1 | 7.15 | 4.35 | 41.24 | 43.7 | -2.46 | 2.46 | -0.04 | 0.04 |
| gene12406 | https://www.ncbi.nlm.nih.gov/gene/?term=LOC105286316 | 1386 | 34 | 245.31 | 1 | 7.22 | 2.94 | 39.04 | 41.82 | -2.78 | 2.78 | -0.07 | 0.07 |
| gene4034 | https://www.ncbi.nlm.nih.gov/gene/?term=LOC105276704 | 2065 | 87 | 421.31 | 2 | 9.69 | 2.3 | 12.36 | 15.2 | -2.84 | 2.84 | -0.12 | 0.12 |
| gene12558 | https://www.ncbi.nlm.nih.gov/gene/?term=LOC105286502 | 1285 | 43 | 334.63 | 1 | 7.78 | 2.33 | 24.55 | 27.41 | -2.86 | 2.86 | -0.1 | 0.1 |
| gene5647 | https://www.ncbi.nlm.nih.gov/gene/?term=LOC105278567 | 3118 | 37 | 118.67 | 1 | 3.21 | 2.7 | 11.3 | 14.2 | -2.9 | 2.9 | -0.03 | 0.03 |
| gene12516 | https://www.ncbi.nlm.nih.gov/gene/?term=LOC105286452 | 1529 | 35 | 228.91 | 1 | 6.54 | 2.86 | 51.01 | 54.04 | -3.03 | 3.03 | -0.07 | 0.07 |
| gene10107 | https://www.ncbi.nlm.nih.gov/gene/?term=LOC105283712 | 1883 | 27 | 143.39 | 1 | 5.31 | 3.7 | 26.21 | 29.43 | -3.23 | 3.23 | -0.05 | 0.05 |
| gene2335 | https://www.ncbi.nlm.nih.gov/gene/?term=LOC105274778 | 3945 | 38 | 96.32 | 1 | 2.53 | 2.63 | 45.06 | 48.85 | -3.79 | 3.79 | -0.04 | 0.04 |
| gene274 | https://www.ncbi.nlm.nih.gov/gene/?term=LOC105275190 | 3488 | 73 | 209.29 | 1 | 2.87 | 1.37 | 18.68 | 22.58 | -3.9 | 3.9 | -0.08 | 0.08 |
| gene12961 | https://www.ncbi.nlm.nih.gov/gene/?term=LOC105286983 | 3094 | 70 | 226.24 | 1 | 3.23 | 1.43 | 0.98 | 5.77 | -4.79 | 4.79 | -0.11 | 0.11 |
| gene5052 | https://www.ncbi.nlm.nih.gov/gene/?term=LOC105277856 | 3167 | 28 | 88.41 | 1 | 3.16 | 3.57 | 35.73 | 41.1 | -5.37 | 5.37 | -0.05 | 0.05 |
| gene5714 | https://www.ncbi.nlm.nih.gov/gene/?term=LOC105278729 | 710 | 14 | 197.18 | 1 | 14.08 | 7.14 | 51.67 | 57.2 | -5.53 | 5.53 | -0.11 | 0.11 |
| gene6814 | https://www.ncbi.nlm.nih.gov/gene/?term=LOC105279898 | 1343 | 15 | 111.69 | 1 | 7.45 | 6.67 | 59.84 | 67.04 | -7.2 | 7.2 | -0.08 | 0.08 |
| gene3165 | https://www.ncbi.nlm.nih.gov/gene/?term=LOC105275706 | 2374 | 67 | 282.22 | 4 | 16.85 | 5.97 | 25.11 | 35.96 | -10.84 | 10.84 | -0.31 | 0.31 |
| gene2593 | https://www.ncbi.nlm.nih.gov/gene/?term=LOC105275070 | 1774 | 6 | 33.82 | 3 | 16.91 | 50 | 16.42 | 57.88 | -41.46 | 41.46 | -0.14 | 0.14 |

Directory: OGLall File: ogl-Cb_LI_BCphaseA1.vs.RphaseB1.txt

| gene_ID | gene_link | gwidth | #Sites | #per10Kb | #dmSites | #dmsp10kb | %dmSites | %pSite1 | %pSite2 | DMpSite | ADMpSite | DMpNucl | ADMpNucl |
|---|---|---|---|---|---|---|---|---|---|---|---|---|---|
| gene10784 | https://www.ncbi.nlm.nih.gov/gene/?term=LOC105284474 | 481 | 1 | 20.79 | 1 | 20.79 | 100 | 66.67 | 38.1 | 28.57 | 28.57 | 0.06 | 0.06 |
| gene2022 | https://www.ncbi.nlm.nih.gov/gene/?term=LOC105288227 | 1245 | 3 | 24.1 | 1 | 8.03 | 33.33 | 86.21 | 76.94 | 9.26 | 9.26 | 0.02 | 0.02 |
| gene10237 | https://www.ncbi.nlm.nih.gov/gene/?term=LOC105283856 | 1259 | 25 | 198.57 | 5 | 39.71 | 20 | 64.65 | 55.66 | 9 | 9 | 0.18 | 0.18 |
| gene12003 | https://www.ncbi.nlm.nih.gov/gene/?term=LOC105285885 | 1336 | 10 | 74.85 | 1 | 7.49 | 10 | 64.94 | 57.31 | 7.63 | 7.63 | 0.06 | 0.06 |
| gene4643 | https://www.ncbi.nlm.nih.gov/gene/?term=LOC109610847 | 429 | 5 | 116.55 | 1 | 23.31 | 20 | 29.86 | 22.57 | 7.29 | 7.29 | 0.09 | 0.09 |
| gene8965 | https://www.ncbi.nlm.nih.gov/gene/?term=LOC105282405 | 1310 | 31 | 236.64 | 2 | 15.27 | 6.45 | 51.28 | 44.69 | 6.59 | 6.59 | 0.16 | 0.16 |
| gene11436 | https://www.ncbi.nlm.nih.gov/gene/?term=LOC105285241 | 2207 | 14 | 63.43 | 2 | 9.06 | 14.29 | 48.57 | 42.07 | 6.5 | 6.5 | 0.04 | 0.04 |
| gene7962 | https://www.ncbi.nlm.nih.gov/gene/?term=LOC105281226 | 1280 | 10 | 78.12 | 1 | 7.81 | 10 | 77.35 | 71.1 | 6.25 | 6.25 | 0.05 | 0.05 |
| gene12819 | https://www.ncbi.nlm.nih.gov/gene/?term=LOC105286816 | 4989 | 10 | 20.04 | 1 | 2 | 10 | 37.75 | 31.69 | 6.06 | 6.06 | 0.01 | 0.01 |
| gene571 | https://www.ncbi.nlm.nih.gov/gene/?term=LOC105278541 | 1569 | 16 | 101.98 | 1 | 6.37 | 6.25 | 28.82 | 23.23 | 5.58 | 5.58 | 0.06 | 0.06 |
| gene305 | https://www.ncbi.nlm.nih.gov/gene/?term=LOC105275539 | 1277 | 23 | 180.11 | 2 | 15.66 | 8.7 | 20.8 | 15.48 | 5.32 | 5.32 | 0.1 | 0.1 |
| gene8172 | https://www.ncbi.nlm.nih.gov/gene/?term=LOC105281507 | 946 | 5 | 52.85 | 1 | 10.57 | 20 | 11.82 | 6.67 | 5.15 | 5.15 | 0.03 | 0.03 |
| gene10187 | https://www.ncbi.nlm.nih.gov/gene/?term=LOC105283798 | 2511 | 20 | 79.65 | 1 | 3.98 | 5 | 33.53 | 28.81 | 4.72 | 4.72 | 0.04 | 0.04 |
| gene7684 | https://www.ncbi.nlm.nih.gov/gene/?term=LOC105280926 | 1727 | 7 | 40.53 | 1 | 5.79 | 14.29 | 49.96 | 45.26 | 4.69 | 4.69 | 0.02 | 0.02 |
| gene9483 | https://www.ncbi.nlm.nih.gov/gene/?term=LOC105282996 | 3739 | 9 | 24.07 | 2 | 5.35 | 22.22 | 28.24 | 23.6 | 4.64 | 4.64 | 0.01 | 0.01 |
| gene10334 | https://www.ncbi.nlm.nih.gov/gene/?term=LOC105283973 | 1536 | 33 | 214.84 | 4 | 26.04 | 12.12 | 58.47 | 53.86 | 4.61 | 4.61 | 0.1 | 0.1 |
| gene4999 | https://www.ncbi.nlm.nih.gov/gene/?term=LOC105277805 | 6695 | 13 | 19.42 | 1 | 1.49 | 7.69 | 45.92 | 41.43 | 4.49 | 4.49 | 0.01 | 0.01 |
| gene5066 | https://www.ncbi.nlm.nih.gov/gene/?term=LOC105277868 | 2120 | 15 | 70.75 | 2 | 9.43 | 13.33 | 54.5 | 50.07 | 4.44 | 4.44 | 0.03 | 0.03 |
| gene11269 | https://www.ncbi.nlm.nih.gov/gene/?term=LOC105285051 | 1363 | 10 | 73.37 | 1 | 7.34 | 10 | 46.13 | 41.71 | 4.41 | 4.41 | 0.03 | 0.03 |
| gene470 | https://www.ncbi.nlm.nih.gov/gene/?term=LOC105277393 | 799 | 14 | 175.22 | 1 | 12.52 | 7.14 | 32.95 | 28.75 | 4.2 | 4.2 | 0.07 | 0.07 |
| gene4986 | https://www.ncbi.nlm.nih.gov/gene/?term=LOC105277787 | 1212 | 7 | 57.76 | 1 | 8.25 | 14.29 | 24.61 | 20.42 | 4.18 | 4.18 | 0.02 | 0.02 |
| gene10346 | https://www.ncbi.nlm.nih.gov/gene/?term=LOC105283988 | 2518 | 11 | 43.69 | 1 | 3.97 | 9.09 | 45.73 | 41.61 | 4.12 | 4.12 | 0.02 | 0.02 |
| gene9905 | https://www.ncbi.nlm.nih.gov/gene/?term=LOC105283473 | 937 | 9 | 96.05 | 2 | 21.34 | 22.22 | 34.37 | 30.37 | 4 | 4 | 0.04 | 0.04 |
| gene11135 | https://www.ncbi.nlm.nih.gov/gene/?term=LOC105284881 | 2024 | 21 | 103.75 | 2 | 9.88 | 9.52 | 26.62 | 22.75 | 3.88 | 3.88 | 0.04 | 0.04 |

...
... only first 25 lines shown ...

Directory: OGLall File: ogl-Cb_LI_BCphaseA1.vs.RphaseB2.txt

| gene_ID | gene_link | gwidth | #Sites | #per10Kb | #dmSites | #dmsp10kb | %dmSites | %pSite1 | %pSite2 | DMpSite | ADMpSite | DMpNucl | ADMpNucl |
|---|---|---|---|---|---|---|---|---|---|---|---|---|---|
| gene2022 | https://www.ncbi.nlm.nih.gov/gene/?term=LOC105288227 | 1245 | 2 | 16.06 | 1 | 8.03 | 50 | 79.31 | 55.55 | 23.76 | 23.76 | 0.04 | 0.04 |
| gene2593 | https://www.ncbi.nlm.nih.gov/gene/?term=LOC105275070 | 1774 | 5 | 28.18 | 1 | 5.64 | 20 | 19.71 | 4.7 | 15.01 | 15.01 | 0.04 | 0.04 |
| gene11599 | https://www.ncbi.nlm.nih.gov/gene/?term=LOC105285431 | 1550 | 5 | 32.26 | 2 | 12.9 | 40 | 49.86 | 35.35 | 14.51 | 14.51 | 0.05 | 0.05 |
| gene1656 | https://www.ncbi.nlm.nih.gov/gene/?term=LOC105287814 | 812 | 5 | 61.58 | 2 | 24.63 | 40 | 35.51 | 23.39 | 12.12 | 12.12 | 0.07 | 0.07 |
| gene9109 | https://www.ncbi.nlm.nih.gov/gene/?term=LOC105282561 | 2400 | 4 | 16.67 | 1 | 4.17 | 25 | 43.84 | 31.75 | 12.09 | 12.09 | 0.02 | 0.02 |
| gene9845 | https://www.ncbi.nlm.nih.gov/gene/?term=LOC105283410 | 1401 | 9 | 64.24 | 2 | 14.28 | 22.22 | 50.57 | 42.07 | 8.5 | 8.5 | 0.05 | 0.05 |
| gene4044 | https://www.ncbi.nlm.nih.gov/gene/?term=LOC105276718 | 1729 | 9 | 52.05 | 3 | 17.35 | 33.33 | 30.94 | 22.72 | 8.22 | 8.22 | 0.04 | 0.04 |
| gene4620 | https://www.ncbi.nlm.nih.gov/gene/?term=LOC105277368 | 1739 | 6 | 34.5 | 1 | 5.75 | 16.67 | 22.3 | 14.79 | 7.5 | 7.5 | 0.03 | 0.03 |
| gene8965 | https://www.ncbi.nlm.nih.gov/gene/?term=LOC105282405 | 1310 | 31 | 236.64 | 5 | 38.17 | 16.13 | 51.28 | 44.76 | 6.52 | 6.52 | 0.15 | 0.15 |
| gene11145 | https://www.ncbi.nlm.nih.gov/gene/?term=LOC105284899 | 1781 | 14 | 78.61 | 1 | 5.61 | 7.14 | 40.04 | 33.58 | 6.46 | 6.46 | 0.05 | 0.05 |
| gene5060 | https://www.ncbi.nlm.nih.gov/gene/?term=LOC105277862 | 1458 | 7 | 48.01 | 1 | 6.86 | 14.29 | 62.13 | 55.84 | 6.3 | 6.3 | 0.03 | 0.03 |
| gene10363 | https://www.ncbi.nlm.nih.gov/gene/?term=LOC105284009 | 1503 | 26 | 172.99 | 3 | 19.96 | 11.54 | 22.01 | 15.79 | 6.22 | 6.22 | 0.11 | 0.11 |
| gene12853 | https://www.ncbi.nlm.nih.gov/gene/?term=LOC105286857 | 1589 | 4 | 25.17 | 1 | 6.29 | 25 | 73.58 | 67.52 | 6.05 | 6.05 | 0.02 | 0.02 |
| gene11598 | https://www.ncbi.nlm.nih.gov/gene/?term=LOC105285429 | 1324 | 8 | 60.42 | 1 | 7.55 | 12.5 | 53.59 | 47.67 | 5.92 | 5.92 | 0.04 | 0.04 |
| gene5006 | https://www.ncbi.nlm.nih.gov/gene/?term=LOC105277810 | 1382 | 21 | 151.95 | 1 | 7.24 | 4.76 | 45.33 | 39.66 | 5.66 | 5.66 | 0.09 | 0.09 |
| gene4944 | https://www.ncbi.nlm.nih.gov/gene/?term=LOC105277745 | 1344 | 8 | 59.52 | 1 | 7.44 | 12.5 | 67.79 | 62.31 | 5.48 | 5.48 | 0.03 | 0.03 |
| gene2226 | https://www.ncbi.nlm.nih.gov/gene/?term=LOC109611583 | 3276 | 11 | 33.58 | 1 | 3.05 | 9.09 | 13.95 | 8.66 | 5.29 | 5.29 | 0.02 | 0.02 |
| gene10334 | https://www.ncbi.nlm.nih.gov/gene/?term=LOC105283973 | 1536 | 30 | 195.31 | 1 | 6.51 | 3.33 | 61.25 | 55.99 | 5.26 | 5.26 | 0.1 | 0.1 |
| gene5924 | https://www.ncbi.nlm.nih.gov/gene/?term=LOC105278853 | 1035 | 6 | 57.97 | 1 | 9.66 | 16.67 | 22.54 | 17.31 | 5.23 | 5.23 | 0.03 | 0.03 |
| gene6814 | https://www.ncbi.nlm.nih.gov/gene/?term=LOC105279898 | 1343 | 9 | 67.01 | 1 | 7.45 | 11.11 | 79.86 | 74.67 | 5.19 | 5.19 | 0.03 | 0.03 |
| gene7962 | https://www.ncbi.nlm.nih.gov/gene/?term=LOC105281226 | 1280 | 7 | 54.69 | 1 | 7.81 | 14.29 | 69.47 | 64.31 | 5.16 | 5.16 | 0.03 | 0.03 |
| gene3048 | https://www.ncbi.nlm.nih.gov/gene/?term=LOC105275585 | 2468 | 25 | 101.3 | 1 | 4.05 | 4 | 31.83 | 26.81 | 5.02 | 5.02 | 0.05 | 0.05 |
| gene13210 | https://www.ncbi.nlm.nih.gov/gene/?term=LOC105287253 | 757 | 8 | 105.68 | 2 | 26.42 | 25 | 52.09 | 47.22 | 4.87 | 4.87 | 0.05 | 0.05 |
| gene10192 | https://www.ncbi.nlm.nih.gov/gene/?term=LOC105283803 | 1636 | 7 | 42.79 | 1 | 6.11 | 14.29 | 37.31 | 32.5 | 4.82 | 4.82 | 0.02 | 0.02 |

...
... only first 25 lines shown ...

Directory: OGLall File: ogl-Cb_LI_BCphaseA2.txt

| gene_ID | gene_link | gwidth | #Sites | #per10Kb | %perSite | %pNucl |
|---|---|---|---|---|---|---|
| gene12538 | https://www.ncbi.nlm.nih.gov/gene/?term=LOC105286531 | 372 | 42 | 1129.03 | 38.35 | 4.33 |
| gene5917 | https://www.ncbi.nlm.nih.gov/gene/?term=LOC105278849 | 823 | 136 | 1652.49 | 25.47 | 4.21 |
| gene13384 | https://www.ncbi.nlm.nih.gov/gene/?term=LOC105287447 | 1877 | 240 | 1278.64 | 32.67 | 4.18 |
| gene5911 | https://www.ncbi.nlm.nih.gov/gene/?term=LOC105278844 | 943 | 97 | 1028.63 | 37.69 | 3.88 |
| gene5309 | https://www.ncbi.nlm.nih.gov/gene/?term=LOC105278160 | 1704 | 161 | 944.84 | 37.8 | 3.57 |
| gene211 | https://www.ncbi.nlm.nih.gov/gene/?term=LOC105274534 | 1280 | 91 | 710.94 | 49.01 | 3.48 |
| gene12364 | https://www.ncbi.nlm.nih.gov/gene/?term=LOC105286269 | 1403 | 91 | 648.61 | 51.53 | 3.34 |
| gene1654 | https://www.ncbi.nlm.nih.gov/gene/?term=LOC105287811 | 2285 | 165 | 722.1 | 45.4 | 3.28 |
| gene12330 | https://www.ncbi.nlm.nih.gov/gene/?term=LOC105286235 | 1501 | 110 | 732.84 | 44.1 | 3.23 |
| gene10742 | https://www.ncbi.nlm.nih.gov/gene/?term=LOC109611309 | 776 | 58 | 747.42 | 43.13 | 3.22 |
| gene12746 | https://www.ncbi.nlm.nih.gov/gene/?term=LOC105286737 | 2688 | 368 | 1369.05 | 23.56 | 3.22 |
| gene3731 | https://www.ncbi.nlm.nih.gov/gene/?term=LOC105276371 | 865 | 56 | 647.4 | 49.67 | 3.22 |
| gene6078 | https://www.ncbi.nlm.nih.gov/gene/?term=LOC105279052 | 1130 | 139 | 1230.09 | 26.01 | 3.2 |
| gene12805 | https://www.ncbi.nlm.nih.gov/gene/?term=LOC105286801 | 1379 | 115 | 833.94 | 37.24 | 3.11 |
| gene5301 | https://www.ncbi.nlm.nih.gov/gene/?term=LOC105278148 | 1881 | 208 | 1105.79 | 27.74 | 3.07 |
| gene3719 | https://www.ncbi.nlm.nih.gov/gene/?term=LOC105276360 | 1471 | 153 | 1040.11 | 29.45 | 3.06 |
| gene2797 | https://www.ncbi.nlm.nih.gov/gene/?term=LOC105275314 | 1489 | 93 | 624.58 | 47.6 | 2.97 |
| gene7527 | https://www.ncbi.nlm.nih.gov/gene/?term=LOC105280739 | 2686 | 145 | 539.84 | 54.6 | 2.95 |
| gene1171 | https://www.ncbi.nlm.nih.gov/gene/?term=LOC105285174 | 844 | 58 | 687.2 | 41.33 | 2.84 |
| gene1562 | https://www.ncbi.nlm.nih.gov/gene/?term=LOC105287692 | 1491 | 129 | 865.19 | 32.49 | 2.81 |
| gene4635 | https://www.ncbi.nlm.nih.gov/gene/?term=LOC105277390 | 2067 | 126 | 609.58 | 46.08 | 2.81 |
| gene570 | https://www.ncbi.nlm.nih.gov/gene/?term=LOC105278549 | 1673 | 112 | 669.46 | 41.85 | 2.8 |
| gene10743 | https://www.ncbi.nlm.nih.gov/gene/?term=LOC105284456 | 1028 | 65 | 632.3 | 43.96 | 2.78 |
| gene8453 | https://www.ncbi.nlm.nih.gov/gene/?term=LOC105281826 | 1750 | 160 | 914.29 | 30.44 | 2.78 |

...
... only first 25 lines shown ...

Directory: OGLall File: ogl-Cb_LI_BCphaseA2.vs.BCphaseB1.txt

| gene_ID | gene_link | gwidth | #Sites | #per10Kb | #dmSites | #dmsp10kb | %dmSites | %pSite1 | %pSite2 | DMpSite | ADMpSite | DMpNucl | ADMpNucl |
|---|---|---|---|---|---|---|---|---|---|---|---|---|---|
| gene7199 | https://www.ncbi.nlm.nih.gov/gene/?term=LOC105280366 | 1414 | 30 | 212.16 | 1 | 7.07 | 3.33 | 43.5 | 41.71 | 1.79 | 1.79 | 0.04 | 0.04 |
| gene10606 | https://www.ncbi.nlm.nih.gov/gene/?term=LOC105284273 | 6119 | 258 | 421.64 | 1 | 1.63 | 0.39 | 19.65 | 18.03 | 1.63 | 1.63 | 0.07 | 0.07 |
| gene2856 | https://www.ncbi.nlm.nih.gov/gene/?term=LOC105275384 | 1987 | 91 | 457.98 | 1 | 5.03 | 1.1 | 33.74 | 32.36 | 1.37 | 1.37 | 0.06 | 0.06 |
| gene3793 | https://www.ncbi.nlm.nih.gov/gene/?term=LOC105276429 | 2901 | 119 | 410.2 | 1 | 3.45 | 0.84 | 23.8 | 22.52 | 1.28 | 1.28 | 0.05 | 0.05 |
| gene9740 | https://www.ncbi.nlm.nih.gov/gene/?term=LOC105283277 | 2887 | 143 | 495.32 | 1 | 3.46 | 0.7 | 19.74 | 18.46 | 1.28 | 1.28 | 0.06 | 0.06 |
| gene564 | https://www.ncbi.nlm.nih.gov/gene/?term=LOC105278478 | 2802 | 17 | 60.67 | 1 | 3.57 | 5.88 | 14.91 | 14.04 | 0.87 | 0.87 | 0.01 | 0.01 |
| gene10844 | https://www.ncbi.nlm.nih.gov/gene/?term=LOC105284535 | 9086 | 246 | 270.75 | 1 | 1.1 | 0.41 | 16.16 | 15.32 | 0.84 | 0.84 | 0.02 | 0.02 |
| gene570 | https://www.ncbi.nlm.nih.gov/gene/?term=LOC105278549 | 1673 | 70 | 418.41 | 1 | 5.98 | 1.43 | 44.03 | 43.19 | 0.83 | 0.83 | 0.03 | 0.03 |
| gene10326 | https://www.ncbi.nlm.nih.gov/gene/?term=LOC105283962 | 3042 | 130 | 427.35 | 1 | 3.29 | 0.77 | 8.82 | 8.26 | 0.56 | 0.56 | 0.02 | 0.02 |
| gene1855 | https://www.ncbi.nlm.nih.gov/gene/?term=LOC105288036 | 2387 | 89 | 372.85 | 1 | 4.19 | 1.12 | 30.86 | 30.35 | 0.51 | 0.51 | 0.02 | 0.02 |
| gene7166 | https://www.ncbi.nlm.nih.gov/gene/?term=LOC105280326 | 6117 | 235 | 384.18 | 1 | 1.63 | 0.43 | 15.36 | 14.92 | 0.43 | 0.43 | 0.02 | 0.02 |
| gene12203 | https://www.ncbi.nlm.nih.gov/gene/?term=LOC105286099 | 5967 | 84 | 140.77 | 1 | 1.68 | 1.19 | 11.21 | 10.88 | 0.34 | 0.34 | 0 | 0 |
| gene5101 | https://www.ncbi.nlm.nih.gov/gene/?term=LOC105277903 | 1813 | 34 | 187.53 | 1 | 5.52 | 2.94 | 48.71 | 48.5 | 0.21 | 0.21 | 0 | 0 |
| gene7183 | https://www.ncbi.nlm.nih.gov/gene/?term=LOC105280348 | 3925 | 72 | 183.44 | 1 | 2.55 | 1.39 | 20.87 | 20.69 | 0.19 | 0.19 | 0 | 0 |
| gene7245 | https://www.ncbi.nlm.nih.gov/gene/?term=LOC105280410 | 12478 | 277 | 221.99 | 1 | 0.8 | 0.36 | 19.64 | 19.47 | 0.17 | 0.17 | 0 | 0 |
| gene9755 | https://www.ncbi.nlm.nih.gov/gene/?term=LOC105283296 | 2177 | 92 | 422.6 | 1 | 4.59 | 1.09 | 18.09 | 17.95 | 0.14 | 0.14 | 0.01 | 0.01 |
| gene4209 | https://www.ncbi.nlm.nih.gov/gene/?term=LOC105277125 | 50550 | 1862 | 368.35 | 1 | 0.2 | 0.05 | 1.6 | 1.65 | -0.05 | 0.05 | 0 | 0 |
| gene3744 | https://www.ncbi.nlm.nih.gov/gene/?term=LOC105276385 | 9376 | 459 | 489.55 | 1 | 1.07 | 0.22 | 5.46 | 5.7 | -0.24 | 0.24 | -0.01 | 0.01 |
| gene4892 | https://www.ncbi.nlm.nih.gov/gene/?term=LOC105277667 | 3898 | 126 | 323.24 | 1 | 2.57 | 0.79 | 27.95 | 28.19 | -0.24 | 0.24 | -0.01 | 0.01 |
| gene8814 | https://www.ncbi.nlm.nih.gov/gene/?term=LOC105282240 | 6910 | 175 | 253.26 | 1 | 1.45 | 0.57 | 9.54 | 9.87 | -0.32 | 0.32 | -0.01 | 0.01 |
| gene10819 | https://www.ncbi.nlm.nih.gov/gene/?term=LOC105284507 | 5767 | 94 | 163 | 1 | 1.73 | 1.06 | 20.43 | 21.18 | -0.75 | 0.75 | -0.01 | 0.01 |
| gene11447 | https://www.ncbi.nlm.nih.gov/gene/?term=LOC105285252 | 7437 | 131 | 176.15 | 1 | 1.34 | 0.76 | 12.71 | 13.67 | -0.95 | 0.95 | -0.02 | 0.02 |
| gene9722 | https://www.ncbi.nlm.nih.gov/gene/?term=LOC105283264 | 3951 | 88 | 222.73 | 1 | 2.53 | 1.14 | 18.85 | 20.39 | -1.54 | 1.54 | -0.03 | 0.03 |

Directory: OGLall File: ogl-Cb_LI_BCphaseA2.vs.BCphaseB2.txt

| gene_ID | gene_link | gwidth | #Sites | #per10Kb | #dmSites | #dmsp10kb | %dmSites | %pSite1 | %pSite2 | DMpSite | ADMpSite | DMpNucl | ADMpNucl |
|---|---|---|---|---|---|---|---|---|---|---|---|---|---|
| gene2593 | https://www.ncbi.nlm.nih.gov/gene/?term=LOC105275070 | 1774 | 5 | 28.18 | 5 | 28.18 | 100 | 80.28 | 10.96 | 69.32 | 69.32 | 0.2 | 0.2 |
| gene3165 | https://www.ncbi.nlm.nih.gov/gene/?term=LOC105275706 | 2374 | 67 | 282.22 | 5 | 21.06 | 7.46 | 36.03 | 23.6 | 12.43 | 12.43 | 0.35 | 0.35 |
| gene1957 | https://www.ncbi.nlm.nih.gov/gene/?term=LOC105288153 | 1044 | 16 | 153.26 | 1 | 9.58 | 6.25 | 15.68 | 4.26 | 11.42 | 11.42 | 0.18 | 0.18 |
| gene2824 | https://www.ncbi.nlm.nih.gov/gene/?term=LOC105275337 | 1772 | 64 | 361.17 | 2 | 11.29 | 3.12 | 37.53 | 29.09 | 8.44 | 8.44 | 0.3 | 0.3 |
| gene9468 | https://www.ncbi.nlm.nih.gov/gene/?term=LOC105282976 | 1636 | 53 | 323.96 | 1 | 6.11 | 1.89 | 28.76 | 22.1 | 6.66 | 6.66 | 0.22 | 0.22 |
| gene12395 | https://www.ncbi.nlm.nih.gov/gene/?term=LOC105286302 | 4093 | 17 | 41.53 | 1 | 2.44 | 5.88 | 21.99 | 16.58 | 5.41 | 5.41 | 0.02 | 0.02 |
| gene5714 | https://www.ncbi.nlm.nih.gov/gene/?term=LOC105278729 | 710 | 24 | 338.03 | 1 | 14.08 | 4.17 | 61.56 | 56.79 | 4.77 | 4.77 | 0.16 | 0.16 |
| gene1518 | https://www.ncbi.nlm.nih.gov/gene/?term=LOC105287648 | 2195 | 27 | 123.01 | 1 | 4.56 | 3.7 | 41.41 | 38.92 | 2.49 | 2.49 | 0.03 | 0.03 |
| gene3385 | https://www.ncbi.nlm.nih.gov/gene/?term=LOC105275966 | 3305 | 61 | 184.57 | 1 | 3.03 | 1.64 | 49.58 | 47.17 | 2.41 | 2.41 | 0.04 | 0.04 |
| gene5877 | https://www.ncbi.nlm.nih.gov/gene/?term=LOC105278807 | 1719 | 49 | 285.05 | 1 | 5.82 | 2.04 | 23.71 | 21.48 | 2.23 | 2.23 | 0.06 | 0.06 |
| gene5052 | https://www.ncbi.nlm.nih.gov/gene/?term=LOC105277856 | 3167 | 34 | 107.36 | 1 | 3.16 | 2.94 | 38.79 | 36.66 | 2.13 | 2.13 | 0.02 | 0.02 |
| gene3817 | https://www.ncbi.nlm.nih.gov/gene/?term=LOC105276449 | 1230 | 42 | 341.46 | 1 | 8.13 | 2.38 | 35.93 | 33.84 | 2.09 | 2.09 | 0.07 | 0.07 |
| gene8115 | https://www.ncbi.nlm.nih.gov/gene/?term=LOC105281420 | 3623 | 51 | 140.77 | 1 | 2.76 | 1.96 | 12.6 | 10.51 | 2.09 | 2.09 | 0.03 | 0.03 |
| gene2970 | https://www.ncbi.nlm.nih.gov/gene/?term=LOC105275518 | 2340 | 19 | 81.2 | 1 | 4.27 | 5.26 | 46.39 | 44.36 | 2.03 | 2.03 | 0.02 | 0.02 |
| gene8742 | https://www.ncbi.nlm.nih.gov/gene/?term=LOC105282151 | 2563 | 41 | 159.97 | 2 | 7.8 | 4.88 | 36.1 | 34.31 | 1.79 | 1.79 | 0.03 | 0.03 |
| gene7556 | https://www.ncbi.nlm.nih.gov/gene/?term=LOC105280774 | 2792 | 102 | 365.33 | 1 | 3.58 | 0.98 | 32.11 | 30.45 | 1.66 | 1.66 | 0.06 | 0.06 |
| gene7398 | https://www.ncbi.nlm.nih.gov/gene/?term=LOC105280589 | 3452 | 39 | 112.98 | 1 | 2.9 | 2.56 | 23.26 | 21.67 | 1.58 | 1.58 | 0.02 | 0.02 |
| gene9462 | https://www.ncbi.nlm.nih.gov/gene/?term=LOC105282970 | 1469 | 39 | 265.49 | 1 | 6.81 | 2.56 | 53.94 | 52.44 | 1.5 | 1.5 | 0.04 | 0.04 |
| gene12572 | https://www.ncbi.nlm.nih.gov/gene/?term=LOC105286514 | 2374 | 42 | 176.92 | 1 | 4.21 | 2.38 | 34.39 | 32.92 | 1.47 | 1.47 | 0.03 | 0.03 |
| gene927 | https://www.ncbi.nlm.nih.gov/gene/?term=LOC105282600 | 3727 | 53 | 142.21 | 1 | 2.68 | 1.89 | 32.11 | 30.67 | 1.44 | 1.44 | 0.02 | 0.02 |
| gene6528 | https://www.ncbi.nlm.nih.gov/gene/?term=LOC105279591 | 7222 | 55 | 76.16 | 1 | 1.38 | 1.82 | 29.5 | 28.12 | 1.38 | 1.38 | 0.01 | 0.01 |
| gene7568 | https://www.ncbi.nlm.nih.gov/gene/?term=LOC105280786 | 2525 | 111 | 439.6 | 1 | 3.96 | 0.9 | 33.28 | 31.99 | 1.29 | 1.29 | 0.06 | 0.06 |
| gene4892 | https://www.ncbi.nlm.nih.gov/gene/?term=LOC105277667 | 3898 | 125 | 320.68 | 1 | 2.57 | 0.8 | 27.74 | 26.47 | 1.27 | 1.27 | 0.04 | 0.04 |
| gene10606 | https://www.ncbi.nlm.nih.gov/gene/?term=LOC105284273 | 6119 | 251 | 410.2 | 1 | 1.63 | 0.4 | 20.03 | 18.79 | 1.24 | 1.24 | 0.05 | 0.05 |
| gene12290 | https://www.ncbi.nlm.nih.gov/gene/?term=LOC105286190 | 6212 | 304 | 489.38 | 1 | 1.61 | 0.33 | 8.69 | 7.5 | 1.18 | 1.18 | 0.06 | 0.06 |
| gene1884 | https://www.ncbi.nlm.nih.gov/gene/?term=LOC105288072 | 4169 | 71 | 170.3 | 1 | 2.4 | 1.41 | 25.5 | 24.39 | 1.1 | 1.1 | 0.02 | 0.02 |
| gene3470 | https://www.ncbi.nlm.nih.gov/gene/?term=LOC105276059 | 8716 | 145 | 166.36 | 1 | 1.15 | 0.69 | 10.01 | 8.94 | 1.08 | 1.08 | 0.02 | 0.02 |
| gene5967 | https://www.ncbi.nlm.nih.gov/gene/?term=LOC105278918 | 4393 | 121 | 275.44 | 1 | 2.28 | 0.83 | 15.97 | 14.91 | 1.05 | 1.05 | 0.03 | 0.03 |
| gene940 | https://www.ncbi.nlm.nih.gov/gene/?term=LOC105282794 | 3578 | 85 | 237.56 | 1 | 2.79 | 1.18 | 16.1 | 15.1 | 1 | 1 | 0.02 | 0.02 |
| gene10678 | https://www.ncbi.nlm.nih.gov/gene/?term=LOC105284355 | 4031 | 167 | 414.29 | 1 | 2.48 | 0.6 | 19.59 | 18.65 | 0.94 | 0.94 | 0.04 | 0.04 |
| gene4034 | https://www.ncbi.nlm.nih.gov/gene/?term=LOC105276704 | 2065 | 101 | 489.1 | 1 | 4.84 | 0.99 | 13.4 | 12.57 | 0.83 | 0.83 | 0.04 | 0.04 |
| gene8729 | https://www.ncbi.nlm.nih.gov/gene/?term=LOC105282137 | 5005 | 197 | 393.61 | 1 | 2 | 0.51 | 17.44 | 16.65 | 0.8 | 0.8 | 0.03 | 0.03 |
| gene10520 | https://www.ncbi.nlm.nih.gov/gene/?term=LOC105284184 | 2451 | 85 | 346.8 | 1 | 4.08 | 1.18 | 34.37 | 33.59 | 0.78 | 0.78 | 0.03 | 0.03 |
| gene13033 | https://www.ncbi.nlm.nih.gov/gene/?term=LOC105287063 | 1877 | 49 | 261.05 | 1 | 5.33 | 2.04 | 31.12 | 30.35 | 0.77 | 0.77 | 0.02 | 0.02 |
| gene2825 | https://www.ncbi.nlm.nih.gov/gene/?term=LOC105275343 | 5545 | 96 | 173.13 | 2 | 3.61 | 2.08 | 11.79 | 11.04 | 0.75 | 0.75 | 0.01 | 0.01 |
| gene9336 | https://www.ncbi.nlm.nih.gov/gene/?term=LOC105282834 | 5993 | 124 | 206.91 | 1 | 1.67 | 0.81 | 16.06 | 15.3 | 0.75 | 0.75 | 0.02 | 0.02 |
| gene12516 | https://www.ncbi.nlm.nih.gov/gene/?term=LOC105286452 | 1529 | 38 | 248.53 | 1 | 6.54 | 2.63 | 52.34 | 51.61 | 0.73 | 0.73 | 0.02 | 0.02 |
| gene6620 | https://www.ncbi.nlm.nih.gov/gene/?term=LOC105279687 | 17998 | 336 | 186.69 | 1 | 0.56 | 0.3 | 5.15 | 4.42 | 0.73 | 0.73 | 0.01 | 0.01 |
| gene4376 | https://www.ncbi.nlm.nih.gov/gene/?term=LOC105277083 | 4307 | 127 | 294.87 | 1 | 2.32 | 0.79 | 19.77 | 19.11 | 0.65 | 0.65 | 0.02 | 0.02 |
| gene12307 | https://www.ncbi.nlm.nih.gov/gene/?term=LOC105286216 | 10313 | 166 | 160.96 | 1 | 0.97 | 0.6 | 16.86 | 16.32 | 0.54 | 0.54 | 0.01 | 0.01 |
| gene7368 | https://www.ncbi.nlm.nih.gov/gene/?term=LOC105280556 | 4105 | 50 | 121.8 | 1 | 2.44 | 2 | 18.71 | 18.17 | 0.54 | 0.54 | 0.01 | 0.01 |
| gene7514 | https://www.ncbi.nlm.nih.gov/gene/?term=LOC105280723 | 4471 | 200 | 447.33 | 1 | 2.24 | 0.5 | 14.49 | 14.01 | 0.49 | 0.49 | 0.02 | 0.02 |
| gene2056 | https://www.ncbi.nlm.nih.gov/gene/?term=LOC105274452 | 6584 | 263 | 399.45 | 2 | 3.04 | 0.76 | 13.53 | 13.06 | 0.47 | 0.47 | 0.02 | 0.02 |
| gene3286 | https://www.ncbi.nlm.nih.gov/gene/?term=LOC105275848 | 3744 | 140 | 373.93 | 1 | 2.67 | 0.71 | 23 | 22.56 | 0.45 | 0.45 | 0.02 | 0.02 |
| gene292 | https://www.ncbi.nlm.nih.gov/gene/?term=LOC105275398 | 2825 | 86 | 304.42 | 1 | 3.54 | 1.16 | 21.94 | 21.53 | 0.4 | 0.4 | 0.01 | 0.01 |
| gene4962 | https://www.ncbi.nlm.nih.gov/gene/?term=LOC105277761 | 4645 | 150 | 322.93 | 1 | 2.15 | 0.67 | 15.92 | 15.52 | 0.4 | 0.4 | 0.01 | 0.01 |
| gene7245 | https://www.ncbi.nlm.nih.gov/gene/?term=LOC105280410 | 12478 | 237 | 189.93 | 1 | 0.8 | 0.42 | 21.04 | 20.64 | 0.4 | 0.4 | 0.01 | 0.01 |
| gene10654 | https://www.ncbi.nlm.nih.gov/gene/?term=LOC105284329 | 4027 | 50 | 124.16 | 1 | 2.48 | 2 | 19.81 | 19.43 | 0.39 | 0.39 | 0 | 0 |
| gene2173 | https://www.ncbi.nlm.nih.gov/gene/?term=LOC105274590 | 4387 | 64 | 145.89 | 2 | 4.56 | 3.12 | 34.09 | 33.7 | 0.39 | 0.39 | 0.01 | 0.01 |
| gene6031 | https://www.ncbi.nlm.nih.gov/gene/?term=LOC105279000 | 5691 | 195 | 342.65 | 1 | 1.76 | 0.51 | 14.39 | 14 | 0.39 | 0.39 | 0.01 | 0.01 |
| gene5746 | https://www.ncbi.nlm.nih.gov/gene/?term=LOC105278666 | 10343 | 461 | 445.71 | 1 | 0.97 | 0.22 | 7.31 | 6.93 | 0.37 | 0.37 | 0.02 | 0.02 |
| gene1430 | https://www.ncbi.nlm.nih.gov/gene/?term=LOC105287551 | 6433 | 231 | 359.09 | 1 | 1.55 | 0.43 | 9.26 | 8.92 | 0.34 | 0.34 | 0.01 | 0.01 |
| gene10531 | https://www.ncbi.nlm.nih.gov/gene/?term=LOC105284190 | 6881 | 173 | 251.42 | 1 | 1.45 | 0.58 | 12.82 | 12.5 | 0.32 | 0.32 | 0.01 | 0.01 |
| gene1310 | https://www.ncbi.nlm.nih.gov/gene/?term=LOC105286661 | 2582 | 66 | 255.62 | 1 | 3.87 | 1.52 | 24.45 | 24.13 | 0.32 | 0.32 | 0.01 | 0.01 |
| gene4246 | https://www.ncbi.nlm.nih.gov/gene/?term=LOC105276941 | 16412 | 552 | 336.34 | 1 | 0.61 | 0.18 | 5.12 | 4.8 | 0.32 | 0.32 | 0.01 | 0.01 |
| gene11568 | https://www.ncbi.nlm.nih.gov/gene/?term=LOC105285395 | 8086 | 192 | 237.45 | 1 | 1.24 | 0.52 | 15.07 | 14.77 | 0.31 | 0.31 | 0.01 | 0.01 |
| gene1470 | https://www.ncbi.nlm.nih.gov/gene/?term=LOC105287597 | 2974 | 204 | 685.94 | 1 | 3.36 | 0.49 | 17.73 | 17.44 | 0.3 | 0.3 | 0.02 | 0.02 |
| gene3856 | https://www.ncbi.nlm.nih.gov/gene/?term=LOC105276499 | 5236 | 190 | 362.87 | 1 | 1.91 | 0.53 | 9.53 | 9.24 | 0.29 | 0.29 | 0.01 | 0.01 |
| gene3599 | https://www.ncbi.nlm.nih.gov/gene/?term=LOC105276196 | 2955 | 48 | 162.44 | 1 | 3.38 | 2.08 | 45.56 | 45.28 | 0.28 | 0.28 | 0 | 0 |
| gene7249 | https://www.ncbi.nlm.nih.gov/gene/?term=LOC105280422 | 1775 | 63 | 354.93 | 1 | 5.63 | 1.59 | 17.96 | 17.74 | 0.23 | 0.23 | 0.01 | 0.01 |

| | | | | | | | | | | | | | |
|---|---|---|---|---|---|---|---|---|---|---|---|---|---|
| gene8600 | https://www.ncbi.nlm.nih.gov/gene/?term=LOC105282001 | 8513 | 156 | 183.25 | 1 | 1.17 | 0.64 | 12.04 | 11.82 | 0.22 | 0.22 | 0 | 0 |
| gene2806 | https://www.ncbi.nlm.nih.gov/gene/?term=LOC105275327 | 14257 | 494 | 346.5 1 | 0.7 | 0.2 | 6.88 | 6.68 | 0.2 | 0.2 | 0.01 | 0.01 | |
| gene11275 | https://www.ncbi.nlm.nih.gov/gene/?term=LOC105285055 | 4119 | 255 | 619.08 | 1 | 2.43 | 0.39 | 11.36 | 11.16 | 0.19 | 0.19 | 0.01 | 0.01 |
| gene2846 | https://www.ncbi.nlm.nih.gov/gene/?term=LOC105275374 | 11246 | 502 | 446.38 | 1 | 0.89 | 0.2 | 11.4 | 11.21 | 0.19 | 0.19 | 0.01 | 0.01 |
| gene7166 | https://www.ncbi.nlm.nih.gov/gene/?term=LOC105280326 | 6117 | 233 | 380.91 | 1 | 1.63 | 0.43 | 15.64 | 15.46 | 0.19 | 0.19 | 0.01 | 0.01 |
| gene12439 | https://www.ncbi.nlm.nih.gov/gene/?term=LOC105286357 | 4761 | 81 | 170.13 | 1 | 2.1 | 1.23 | 32.58 | 32.4 | 0.18 | 0.18 | 0 | 0 |
| gene11579 | https://www.ncbi.nlm.nih.gov/gene/?term=LOC105285406 | 13571 | 267 | 196.74 | 1 | 0.74 | 0.37 | 6.68 | 6.51 | 0.17 | 0.17 | 0 | 0 |
| gene8343 | https://www.ncbi.nlm.nih.gov/gene/?term=LOC105281700 | 2172 | 37 | 170.35 | 1 | 4.6 | 2.7 | 39.71 | 39.54 | 0.16 | 0.16 | 0 | 0 |
| gene11401 | https://www.ncbi.nlm.nih.gov/gene/?term=LOC105285202 | 1841 | 82 | 445.41 | 1 | 5.43 | 1.22 | 25.44 | 25.28 | 0.15 | 0.15 | 0.01 | 0.01 |
| gene12203 | https://www.ncbi.nlm.nih.gov/gene/?term=LOC105286099 | 5967 | 72 | 120.66 | 1 | 1.68 | 1.39 | 10.74 | 10.59 | 0.15 | 0.15 | 0 | 0 |
| gene12806 | https://www.ncbi.nlm.nih.gov/gene/?term=LOC105286803 | 4196 | 143 | 340.8 1 | 2.38 | 0.7 | 11.07 | 10.92 | 0.15 | 0.15 | 0.01 | 0.01 | |
| gene4190 | https://www.ncbi.nlm.nih.gov/gene/?term=LOC105276875 | 3150 | 107 | 339.68 | 1 | 3.17 | 0.93 | 23.8 | 23.66 | 0.15 | 0.15 | 0 | 0 |
| gene1198 | https://www.ncbi.nlm.nih.gov/gene/?term=LOC105285507 | 2738 | 88 | 321.4 1 | 3.65 | 1.14 | 23.48 | 23.34 | 0.14 | 0.14 | 0 | 0 | |
| gene5524 | https://www.ncbi.nlm.nih.gov/gene/?term=LOC105278397 | 5177 | 203 | 392.12 | 1 | 1.93 | 0.49 | 25.25 | 25.12 | 0.13 | 0.13 | 0.01 | 0.01 |
| gene12802 | https://www.ncbi.nlm.nih.gov/gene/?term=LOC105286799 | 3183 | 185 | 581.21 | 2 | 6.28 | 1.08 | 16.06 | 15.94 | 0.12 | 0.12 | 0.01 | 0.01 |
| gene10392 | https://www.ncbi.nlm.nih.gov/gene/?term=LOC105284036 | 3216 | 94 | 292.29 | 2 | 6.22 | 2.13 | 22.55 | 22.51 | 0.04 | 0.04 | 0 | 0 |
| gene53 | https://www.ncbi.nlm.nih.gov/gene/?term=LOC105277148 | 19283 | 287 | 148.84 | 1 | 0.52 | 0.35 | 5.29 | 5.27 | 0.03 | 0.03 | 0 | 0 |
| gene6286 | https://www.ncbi.nlm.nih.gov/gene/?term=LOC105279298 | 2861 | 125 | 436.91 | 2 | 6.99 | 1.6 | 25.22 | 25.19 | 0.03 | 0.03 | 0 | 0 |
| gene13044 | https://www.ncbi.nlm.nih.gov/gene/?term=LOC105287070 | 5094 | 241 | 473.11 | 1 | 1.96 | 0.41 | 5.28 | 5.27 | 0.02 | 0.02 | 0 | 0 |
| gene2844 | https://www.ncbi.nlm.nih.gov/gene/?term=LOC105275363 | 4330 | 192 | 443.42 | 1 | 2.31 | 0.52 | 14.25 | 14.23 | 0.02 | 0.02 | 0 | 0 |
| gene5525 | https://www.ncbi.nlm.nih.gov/gene/?term=LOC105278396 | 3090 | 188 | 608.41 | 1 | 3.24 | 0.53 | 24.8 | 24.79 | 0.01 | 0.01 | 0 | 0 |
| gene9330 | https://www.ncbi.nlm.nih.gov/gene/?term=LOC105282825 | 3936 | 80 | 203.25 | 1 | 2.54 | 1.25 | 12.57 | 12.58 | -0.01 | 0.01 | 0 | 0 |
| gene9755 | https://www.ncbi.nlm.nih.gov/gene/?term=LOC105283296 | 2177 | 85 | 390.45 | 1 | 4.59 | 1.18 | 21.44 | 21.45 | -0.01 | 0.01 | 0 | 0 |
| gene298 | https://www.ncbi.nlm.nih.gov/gene/?term=LOC105275460 | 2473 | 114 | 460.98 | 1 | 4.04 | 0.88 | 13.89 | 13.91 | -0.02 | 0.02 | 0 | 0 |
| gene7319 | https://www.ncbi.nlm.nih.gov/gene/?term=LOC105280513 | 68997 | 1241 | 179.86 | 1 | 0.14 | 0.08 | 0.56 | 0.6 | -0.04 | 0.04 | 0 | 0 |
| gene10549 | https://www.ncbi.nlm.nih.gov/gene/?term=LOC105284210 | 7145 | 325 | 454.86 | 2 | 2.8 | 0.62 | 10.71 | 10.76 | -0.05 | 0.05 | 0 | 0 |
| gene3848 | https://www.ncbi.nlm.nih.gov/gene/?term=LOC105276486 | 5899 | 133 | 225.46 | 1 | 1.7 | 0.75 | 24.81 | 24.86 | -0.05 | 0.05 | 0 | 0 |
| gene9177 | https://www.ncbi.nlm.nih.gov/gene/?term=LOC105282646 | 3320 | 98 | 295.18 | 1 | 3.01 | 1.02 | 22.82 | 22.88 | -0.05 | 0.05 | 0 | 0 |
| gene4209 | https://www.ncbi.nlm.nih.gov/gene/?term=LOC105277125 | 50550 | 1852 | 366.37 | 1 | 0.2 | 0.05 | 1.57 | 1.63 | -0.06 | 0.06 | 0 | 0 |
| gene4641 | https://www.ncbi.nlm.nih.gov/gene/?term=LOC105277398 | 1937 | 77 | 397.52 | 1 | 5.16 | 1.3 | 34.01 | 34.07 | -0.06 | 0.06 | 0 | 0 |
| gene11688 | https://www.ncbi.nlm.nih.gov/gene/?term=LOC105285533 | 1637 | 60 | 366.52 | 1 | 6.11 | 1.67 | 26.05 | 26.13 | -0.07 | 0.07 | 0 | 0 |
| gene3107 | https://www.ncbi.nlm.nih.gov/gene/?term=LOC105275650 | 5103 | 181 | 354.69 | 1 | 1.96 | 0.55 | 15.45 | 15.53 | -0.08 | 0.08 | 0 | 0 |
| gene5831 | https://www.ncbi.nlm.nih.gov/gene/?term=LOC105278761 | 2506 | 119 | 474.86 | 2 | 7.98 | 1.68 | 21.61 | 21.68 | -0.08 | 0.08 | 0 | 0 |
| gene9445 | https://www.ncbi.nlm.nih.gov/gene/?term=LOC105282951 | 2285 | 115 | 503.28 | 1 | 4.38 | 0.87 | 20.19 | 20.28 | -0.09 | 0.09 | 0 | 0 |
| gene12674 | https://www.ncbi.nlm.nih.gov/gene/?term=LOC105286649 | 18283 | 811 | 443.58 | 1 | 0.55 | 0.12 | 0.93 | 1.06 | -0.13 | 0.13 | -0.01 | 0.01 |
| gene13114 | https://www.ncbi.nlm.nih.gov/gene/?term=LOC105287147 | 7609 | 409 | 537.52 | 1 | 1.31 | 0.24 | 9.57 | 9.7 | -0.13 | 0.13 | -0.01 | 0.01 |
| gene6652 | https://www.ncbi.nlm.nih.gov/gene/?term=LOC105279719 | 2831 | 79 | 279.05 | 1 | 3.53 | 1.27 | 24.78 | 24.91 | -0.14 | 0.14 | 0 | 0 |
| gene1632 | https://www.ncbi.nlm.nih.gov/gene/?term=LOC105287764 | 18061 | 393 | 217.6 2 | 1.11 | 0.51 | 2.97 | 3.13 | -0.17 | 0.17 | 0 | 0 | |
| gene8661 | https://www.ncbi.nlm.nih.gov/gene/?term=LOC105282063 | 1926 | 32 | 166.15 | 1 | 5.19 | 3.12 | 29.26 | 29.43 | -0.17 | 0.17 | 0 | 0 |
| gene3885 | https://www.ncbi.nlm.nih.gov/gene/?term=LOC105276527 | 7928 | 317 | 399.85 | 1 | 1.26 | 0.32 | 9.68 | 9.86 | -0.18 | 0.18 | -0.01 | 0.01 |
| gene5777 | https://www.ncbi.nlm.nih.gov/gene/?term=LOC105278691 | 3883 | 148 | 381.15 | 1 | 2.58 | 0.68 | 14.2 | 14.38 | -0.18 | 0.18 | -0.01 | 0.01 |
| gene6583 | https://www.ncbi.nlm.nih.gov/gene/?term=LOC105279649 | 7444 | 332 | 446 1 | 1.34 | 0.3 | 10.76 | 10.98 | -0.22 | 0.22 | -0.01 | 0.01 | |
| gene11454 | https://www.ncbi.nlm.nih.gov/gene/?term=LOC105285256 | 1906 | 47 | 246.59 | 1 | 5.25 | 2.13 | 45.38 | 45.61 | -0.23 | 0.23 | -0.01 | 0.01 |
| gene5375 | https://www.ncbi.nlm.nih.gov/gene/?term=LOC105278228 | 8640 | 163 | 188.66 | 1 | 1.16 | 0.61 | 21.78 | 22.01 | -0.23 | 0.23 | 0 | 0 |
| gene13036 | https://www.ncbi.nlm.nih.gov/gene/?term=LOC105287059 | 4959 | 163 | 328.7 1 | 2.02 | 0.61 | 16.39 | 16.65 | -0.26 | 0.26 | -0.01 | 0.01 | |
| gene3727 | https://www.ncbi.nlm.nih.gov/gene/?term=LOC105276367 | 2484 | 69 | 277.78 | 1 | 4.03 | 1.45 | 46.73 | 46.99 | -0.27 | 0.27 | -0.01 | 0.01 |
| gene6237 | https://www.ncbi.nlm.nih.gov/gene/?term=LOC105279244 | 1484 | 24 | 161.73 | 1 | 6.74 | 4.17 | 52.96 | 53.25 | -0.29 | 0.29 | 0 | 0 |
| gene2683 | https://www.ncbi.nlm.nih.gov/gene/?term=LOC105275187 | 5492 | 70 | 127.46 | 1 | 1.82 | 1.43 | 14.08 | 14.4 | -0.31 | 0.31 | 0 | 0 |
| gene10155 | https://www.ncbi.nlm.nih.gov/gene/?term=LOC105283756 | 7030 | 93 | 132.29 | 1 | 1.42 | 1.08 | 5.63 | 5.96 | -0.34 | 0.34 | 0 | 0 |
| gene2257 | https://www.ncbi.nlm.nih.gov/gene/?term=LOC105274689 | 3658 | 71 | 194.1 1 | 2.73 | 1.41 | 14.85 | 15.18 | -0.34 | 0.34 | -0.01 | 0.01 | |
| gene7600 | https://www.ncbi.nlm.nih.gov/gene/?term=LOC105280819 | 1615 | 39 | 241.49 | 1 | 6.19 | 2.56 | 35.76 | 36.1 | -0.34 | 0.34 | -0.01 | 0.01 |
| gene11082 | https://www.ncbi.nlm.nih.gov/gene/?term=LOC105284824 | 7436 | 200 | 268.96 | 1 | 1.34 | 0.5 | 5.46 | 5.81 | -0.35 | 0.35 | -0.01 | 0.01 |
| gene12573 | https://www.ncbi.nlm.nih.gov/gene/?term=LOC105286515 | 3647 | 113 | 309.84 | 1 | 2.74 | 0.88 | 10.7 | 11.07 | -0.37 | 0.37 | -0.01 | 0.01 |
| gene10600 | https://www.ncbi.nlm.nih.gov/gene/?term=LOC105284270 | 4837 | 124 | 256.36 | 1 | 2.07 | 0.81 | 10.47 | 10.85 | -0.38 | 0.38 | -0.01 | 0.01 |
| gene3046 | https://www.ncbi.nlm.nih.gov/gene/?term=LOC105275587 | 2394 | 56 | 233.92 | 1 | 4.18 | 1.79 | 33.82 | 34.2 | -0.38 | 0.38 | -0.01 | 0.01 |
| gene12981 | https://www.ncbi.nlm.nih.gov/gene/?term=LOC105286999 | 2063 | 40 | 193.89 | 1 | 4.85 | 2.5 | 35.8 | 36.25 | -0.45 | 0.45 | -0.01 | 0.01 |
| gene10107 | https://www.ncbi.nlm.nih.gov/gene/?term=LOC105283712 | 1883 | 32 | 169.94 | 1 | 5.31 | 3.12 | 29.98 | 30.44 | -0.46 | 0.46 | -0.01 | 0.01 |
| gene11561 | https://www.ncbi.nlm.nih.gov/gene/?term=LOC105285387 | 11528 | 194 | 168.29 | 1 | 0.87 | 0.52 | 8.65 | 9.11 | -0.47 | 0.47 | -0.01 | 0.01 |
| gene8015 | https://www.ncbi.nlm.nih.gov/gene/?term=LOC105281312 | 2998 | 72 | 240.16 | 1 | 3.34 | 1.39 | 29.67 | 30.18 | -0.5 | 0.5 | -0.01 | 0.01 |
| gene8814 | https://www.ncbi.nlm.nih.gov/gene/?term=LOC105282240 | 6910 | 170 | 246.02 | 1 | 1.45 | 0.59 | 9.16 | 9.68 | -0.52 | 0.52 | -0.01 | 0.01 |
| gene4720 | https://www.ncbi.nlm.nih.gov/gene/?term=LOC105277479 | 1697 | 47 | 276.96 | 1 | 5.89 | 2.13 | 37.56 | 38.12 | -0.55 | 0.55 | -0.02 | 0.02 |
| gene10391 | https://www.ncbi.nlm.nih.gov/gene/?term=LOC109611283 | 1994 | 44 | 220.66 | 1 | 5.02 | 2.27 | 32.96 | 33.52 | -0.56 | 0.56 | -0.01 | 0.01 |
| gene3671 | https://www.ncbi.nlm.nih.gov/gene/?term=LOC105276272 | 1966 | 36 | 183.11 | 1 | 5.09 | 2.78 | 32.08 | 32.66 | -0.58 | 0.58 | -0.01 | 0.01 |
| gene9826 | https://www.ncbi.nlm.nih.gov/gene/?term=LOC105283385 | 9219 | 305 | 330.84 | 1 | 1.08 | 0.33 | 12.6 | 13.19 | -0.59 | 0.59 | -0.02 | 0.02 |

| | | | | | | | | | | | | | | |
|---|---|---|---|---|---|---|---|---|---|---|---|---|---|---|
| gene6966 | https://www.ncbi.nlm.nih.gov/gene/?term=LOC105280104 | 3747 | 122 | 325.59 | 1 | 2.67 | 0.82 | 15.27 | 15.88 | -0.62 | 0.62 | -0.02 | 0.02 | |
| gene8274 | https://www.ncbi.nlm.nih.gov/gene/?term=LOC105281624 | 5425 | 230 | 423.96 | 1 | 1.84 | 0.43 | 9.61 | 10.33 | -0.73 | 0.73 | -0.03 | 0.03 | |
| gene10018 | https://www.ncbi.nlm.nih.gov/gene/?term=LOC105283607 | 4414 | 191 | 432.71 | 1 | 2.27 | 0.52 | 15.56 | 16.32 | -0.76 | 0.76 | -0.03 | 0.03 | |
| gene5298 | https://www.ncbi.nlm.nih.gov/gene/?term=LOC105278145 | 4060 | 244 | 600.99 | 2 | 4.93 | 0.82 | 10.55 | 11.35 | -0.8 | 0.8 | -0.05 | 0.05 | |
| gene10213 | https://www.ncbi.nlm.nih.gov/gene/?term=LOC105283826 | 1503 | 64 | 425.82 | 1 | 6.65 | 1.56 | 14.04 | 14.85 | -0.81 | 0.81 | -0.03 | 0.03 | |
| gene1147 | https://www.ncbi.nlm.nih.gov/gene/?term=LOC105284954 | 4022 | 95 | 236.2 1 | 2.49 | 1.05 | 15.81 | 16.65 | -0.84 | 0.84 | -0.02 | 0.02 | | |
| gene7915 | https://www.ncbi.nlm.nih.gov/gene/?term=LOC105281173 | 1597 | 38 | 237.95 | 1 | 6.26 | 2.63 | 59.94 | 60.79 | -0.85 | 0.85 | -0.02 | 0.02 | |
| gene2413 | https://www.ncbi.nlm.nih.gov/gene/?term=LOC105274863 | 2006 | 44 | 219.34 | 1 | 4.99 | 2.27 | 63.14 | 64.02 | -0.87 | 0.87 | -0.02 | 0.02 | |
| gene4498 | https://www.ncbi.nlm.nih.gov/gene/?term=LOC105277239 | 2831 | 129 | 455.67 | 1 | 3.53 | 0.78 | 14.44 | 15.34 | -0.89 | 0.89 | -0.04 | 0.04 | |
| gene6639 | https://www.ncbi.nlm.nih.gov/gene/?term=LOC105279705 | 7656 | 62 | 80.98 1 | 1.31 | 1.61 | 22.96 | 23.88 | -0.92 | 0.92 | -0.01 | 0.01 | | |
| gene1993 | https://www.ncbi.nlm.nih.gov/gene/?term=LOC105288193 | 1449 | 75 | 517.6 1 | 6.9 | 1.33 | 35.76 | 36.74 | -0.98 | 0.98 | -0.05 | 0.05 | | |
| gene11340 | https://www.ncbi.nlm.nih.gov/gene/?term=LOC105285131 | 2918 | 101 | 346.13 | 1 | 3.43 | 0.99 | 10.9 | 11.93 | -1.02 | 1.02 | -0.04 | 0.04 | |
| gene12406 | https://www.ncbi.nlm.nih.gov/gene/?term=LOC105286316 | 1386 | 28 | 202.02 | 1 | 7.22 | 3.57 | 47.22 | 48.43 | -1.22 | 1.22 | -0.02 | 0.02 | |
| gene2271 | https://www.ncbi.nlm.nih.gov/gene/?term=LOC105274706 | 1448 | 56 | 386.74 | 1 | 6.91 | 1.79 | 36.89 | 38.13 | -1.24 | 1.24 | -0.05 | 0.05 | |
| gene10345 | https://www.ncbi.nlm.nih.gov/gene/?term=LOC105283989 | 4444 | 23 | 51.76 1 | 2.25 | 4.35 | 43.82 | 45.07 | -1.26 | 1.26 | -0.01 | 0.01 | | |
| gene3168 | https://www.ncbi.nlm.nih.gov/gene/?term=LOC105275711 | 6150 | 70 | 113.82 | 1 | 1.63 | 1.43 | 12.53 | 13.84 | -1.31 | 1.31 | -0.01 | 0.01 | |
| gene5646 | https://www.ncbi.nlm.nih.gov/gene/?term=LOC105278534 | 7475 | 181 | 242.14 | 1 | 1.34 | 0.55 | 17.19 | 18.52 | -1.32 | 1.32 | -0.03 | 0.03 | |
| gene322 | https://www.ncbi.nlm.nih.gov/gene/?term=LOC105275777 | 1606 | 24 | 149.44 | 1 | 6.23 | 4.17 | 45 | 46.35 | -1.35 | 1.35 | -0.02 | 0.02 | |
| gene388 | https://www.ncbi.nlm.nih.gov/gene/?term=LOC105276538 | 1558 | 20 | 128.37 | 2 | 12.84 | 10 | 64.73 | 66.14 | -1.41 | 1.41 | -0.02 | 0.02 | |
| gene11447 | https://www.ncbi.nlm.nih.gov/gene/?term=LOC105285252 | 7437 | 120 | 161.36 | 1 | 1.34 | 0.83 | 13.34 | 14.97 | -1.63 | 1.63 | -0.03 | 0.03 | |
| gene6422 | https://www.ncbi.nlm.nih.gov/gene/?term=LOC105279458 | 2419 | 34 | 140.55 | 1 | 4.13 | 2.94 | 51.48 | 53.11 | -1.63 | 1.63 | -0.02 | 0.02 | |
| gene10819 | https://www.ncbi.nlm.nih.gov/gene/?term=LOC105284507 | 5767 | 77 | 133.52 | 1 | 1.73 | 1.3 | 20.73 | 22.56 | -1.84 | 1.84 | -0.02 | 0.02 | |
| gene6455 | https://www.ncbi.nlm.nih.gov/gene/?term=LOC105279500 | 2612 | 33 | 126.34 | 1 | 3.83 | 3.03 | 45.07 | 47.06 | -1.99 | 1.99 | -0.03 | 0.03 | |
| gene1220 | https://www.ncbi.nlm.nih.gov/gene/?term=LOC105285742 | 3453 | 108 | 312.77 | 1 | 2.9 | 0.93 | 21.23 | 23.26 | -2.03 | 2.03 | -0.06 | 0.06 | |
| gene2748 | https://www.ncbi.nlm.nih.gov/gene/?term=LOC105275261 | 3845 | 52 | 135.24 | 1 | 2.6 | 1.92 | 36.08 | 38.33 | -2.25 | 2.25 | -0.03 | 0.03 | |
| gene920 | https://www.ncbi.nlm.nih.gov/gene/?term=LOC105282518 | 1197 | 40 | 334.17 | 1 | 8.35 | 2.5 | 37.69 | 40.21 | -2.52 | 2.52 | -0.08 | 0.08 | |
| gene2404 | https://www.ncbi.nlm.nih.gov/gene/?term=LOC105274853 | 1767 | 36 | 203.74 | 1 | 5.66 | 2.78 | 27.62 | 31.08 | -3.47 | 3.47 | -0.07 | 0.07 | |
| gene1199 | https://www.ncbi.nlm.nih.gov/gene/?term=LOC105285498 | 2220 | 26 | 117.12 | 1 | 4.5 | 3.85 | 33.05 | 37.03 | -3.99 | 3.99 | -0.05 | 0.05 | |
| gene12021 | https://www.ncbi.nlm.nih.gov/gene/?term=LOC105285905 | 1399 | 20 | 142.96 | 1 | 7.15 | 5 | 32.7 | 38.03 | -5.34 | 5.34 | -0.08 | 0.08 | |
| gene10816 | https://www.ncbi.nlm.nih.gov/gene/?term=LOC105284502 | 1974 | 18 | 91.19 1 | 5.07 | 5.56 | 38.87 | 46.78 | -7.91 | 7.91 | -0.07 | 0.07 | | |
| gene10963 | https://www.ncbi.nlm.nih.gov/gene/?term=LOC105284677 | 1576 | 28 | 177.66 | 2 | 12.69 | 7.14 | 19.34 | 35.66 | -16.32 | | 16.32 | -0.29 | 0.29 |

Directory: OGLall File: ogl-Cb_LI_BCphaseA2.vs.RphaseA1.txt

| gene_ID | gene_link | gwidth | #Sites | #per10Kb | #dmSites | #dmsp10kb | %dmSites | %pSite1 | %pSite2 | DMpSite | ADMpSite | DMpNucl | ADMpNucl |
|---|---|---|---|---|---|---|---|---|---|---|---|---|---|
| gene7414 | https://www.ncbi.nlm.nih.gov/gene/?term=LOC105280604 | 746 | 1 | 13.4 | 1 | 13.4 | 100 | 75 | 42.31 | 32.69 | 32.69 | 0.04 | 0.04 |
| gene9265 | https://www.ncbi.nlm.nih.gov/gene/?term=LOC105282752 | 1389 | 1 | 7.2 | 1 | 7.2 | 100 | 46.67 | 22.86 | 23.81 | 23.81 | 0.02 | 0.02 |
| gene11618 | https://www.ncbi.nlm.nih.gov/gene/?term=LOC105285452 | 1046 | 1 | 9.56 | 1 | 9.56 | 100 | 68.42 | 47.83 | 20.59 | 20.59 | 0.02 | 0.02 |
| gene12624 | https://www.ncbi.nlm.nih.gov/gene/?term=LOC105286586 | 981 | 1 | 10.19 | 1 | 10.19 | 100 | 35.71 | 15.56 | 20.15 | 20.15 | 0.02 | 0.02 |
| gene1087 | https://www.ncbi.nlm.nih.gov/gene/?term=LOC105284346 | 2052 | 2 | 9.75 | 1 | 4.87 | 50 | 28.11 | 14.29 | 13.83 | 13.83 | 0.01 | 0.01 |
| gene1024 | https://www.ncbi.nlm.nih.gov/gene/?term=LOC105283693 | 1307 | 12 | 91.81 | 4 | 30.6 | 33.33 | 71.36 | 58.1 | 13.26 | 13.26 | 0.12 | 0.12 |
| gene5029 | https://www.ncbi.nlm.nih.gov/gene/?term=LOC105277834 | 1242 | 2 | 16.1 | 1 | 8.05 | 50 | 24.28 | 12.73 | 11.56 | 11.56 | 0.02 | 0.02 |
| gene7864 | https://www.ncbi.nlm.nih.gov/gene/?term=LOC105281110 | 946 | 4 | 42.28 | 1 | 10.57 | 25 | 40.14 | 29.24 | 10.91 | 10.91 | 0.05 | 0.05 |
| gene11932 | https://www.ncbi.nlm.nih.gov/gene/?term=LOC109611391 | 917 | 8 | 87.24 | 1 | 10.91 | 12.5 | 32.02 | 22.62 | 9.4 | 9.4 | 0.08 | 0.08 |
| gene11023 | https://www.ncbi.nlm.nih.gov/gene/?term=LOC105284759 | 1222 | 9 | 73.65 | 1 | 8.18 | 11.11 | 38.59 | 30.71 | 7.87 | 7.87 | 0.06 | 0.06 |
| gene10833 | https://www.ncbi.nlm.nih.gov/gene/?term=LOC105284521 | 2093 | 14 | 66.89 | 1 | 4.78 | 7.14 | 62.65 | 54.81 | 7.84 | 7.84 | 0.05 | 0.05 |
| gene764 | https://www.ncbi.nlm.nih.gov/gene/?term=LOC105280683 | 2178 | 21 | 96.42 | 4 | 18.37 | 19.05 | 54.64 | 46.81 | 7.83 | 7.83 | 0.08 | 0.08 |
| gene11073 | https://www.ncbi.nlm.nih.gov/gene/?term=LOC105284817 | 1692 | 5 | 29.55 | 1 | 5.91 | 20 | 33.92 | 26.99 | 6.92 | 6.92 | 0.02 | 0.02 |
| gene3015 | https://www.ncbi.nlm.nih.gov/gene/?term=LOC105275557 | 811 | 11 | 135.64 | 1 | 12.33 | 9.09 | 74.6 | 67.82 | 6.78 | 6.78 | 0.09 | 0.09 |
| gene2380 | https://www.ncbi.nlm.nih.gov/gene/?term=LOC105274834 | 1214 | 9 | 74.14 | 3 | 24.71 | 33.33 | 53.69 | 46.94 | 6.75 | 6.75 | 0.05 | 0.05 |
| gene5188 | https://www.ncbi.nlm.nih.gov/gene/?term=LOC105277995 | 2486 | 13 | 52.29 | 2 | 8.05 | 15.38 | 18.76 | 12.14 | 6.63 | 6.63 | 0.03 | 0.03 |
| gene2855 | https://www.ncbi.nlm.nih.gov/gene/?term=LOC105275378 | 1189 | 20 | 168.21 | 1 | 8.41 | 5 | 32.85 | 26.65 | 6.2 | 6.2 | 0.1 | 0.1 |
| gene12357 | https://www.ncbi.nlm.nih.gov/gene/?term=LOC105286263 | 1877 | 23 | 122.54 | 4 | 21.31 | 17.39 | 38.1 | 31.96 | 6.14 | 6.14 | 0.08 | 0.08 |
| gene1780 | https://www.ncbi.nlm.nih.gov/gene/?term=LOC105287955 | 2042 | 12 | 58.77 | 1 | 4.9 | 8.33 | 91.37 | 85.34 | 6.03 | 6.03 | 0.04 | 0.04 |
| gene7611 | https://www.ncbi.nlm.nih.gov/gene/?term=LOC105280835 | 1890 | 6 | 31.75 | 1 | 5.29 | 16.67 | 6.39 | 0.36 | 6.03 | 6.03 | 0.02 | 0.02 |
| gene9903 | https://www.ncbi.nlm.nih.gov/gene/?term=LOC105283475 | 1725 | 7 | 40.58 | 1 | 5.8 | 14.29 | 61.36 | 55.71 | 5.65 | 5.65 | 0.02 | 0.02 |
| gene7657 | https://www.ncbi.nlm.nih.gov/gene/?term=LOC105280890 | 1109 | 15 | 135.26 | 2 | 18.03 | 13.33 | 31.21 | 25.71 | 5.5 | 5.5 | 0.07 | 0.07 |
| gene5066 | https://www.ncbi.nlm.nih.gov/gene/?term=LOC105277868 | 2120 | 17 | 80.19 | 1 | 4.72 | 5.88 | 54.78 | 49.31 | 5.48 | 5.48 | 0.04 | 0.04 |
| gene4708 | https://www.ncbi.nlm.nih.gov/gene/?term=LOC105277468 | 1712 | 31 | 181.07 | 3 | 17.52 | 9.68 | 40.08 | 34.7 | 5.39 | 5.39 | 0.1 | 0.1 |

...

... only first 25 lines shown ...

Directory: OGLall File: ogl-Cb_LI_BCphaseA2.vs.RphaseA2.txt

| gene_ID | gene_link | gwidth | #Sites | #per10Kb | #dmSites | #dmsp10kb | %dmSites | %pSite1 | %pSite2 | DMpSite | ADMpSite | DMpNucl | ADMpNucl |
|---|---|---|---|---|---|---|---|---|---|---|---|---|---|
| gene7414 | https://www.ncbi.nlm.nih.gov/gene/?term=LOC105280604 | 746 | 1 | 13.4 | 1 | 13.4 | 100 | 75 | 50 | 25 | 25 | 0.03 | 0.03 |
| gene8330 | https://www.ncbi.nlm.nih.gov/gene/?term=LOC105281688 | 1179 | 4 | 33.93 | 1 | 8.48 | 25 | 83.9 | 74.11 | 9.79 | 9.79 | 0.03 | 0.03 |
| gene1517 | https://www.ncbi.nlm.nih.gov/gene/?term=LOC105287649 | 1566 | 7 | 44.7 | 1 | 6.39 | 14.29 | 47.32 | 40.38 | 6.93 | 6.93 | 0.03 | 0.03 |
| gene11269 | https://www.ncbi.nlm.nih.gov/gene/?term=LOC105285051 | 1363 | 15 | 110.05 | 2 | 14.67 | 13.33 | 37.1 | 31.4 | 5.69 | 5.69 | 0.06 | 0.06 |
| gene6755 | https://www.ncbi.nlm.nih.gov/gene/?term=LOC105279839 | 2112 | 11 | 52.08 | 2 | 9.47 | 18.18 | 28.82 | 23.92 | 4.9 | 4.9 | 0.03 | 0.03 |
| gene10237 | https://www.ncbi.nlm.nih.gov/gene/?term=LOC105283856 | 1259 | 26 | 206.51 | 1 | 7.94 | 3.85 | 64.95 | 60.12 | 4.82 | 4.82 | 0.1 | 0.1 |
| gene3297 | https://www.ncbi.nlm.nih.gov/gene/?term=LOC105275861 | 2795 | 16 | 57.25 | 1 | 3.58 | 6.25 | 17.96 | 13.14 | 4.82 | 4.82 | 0.03 | 0.03 |
| gene7727 | https://www.ncbi.nlm.nih.gov/gene/?term=LOC105280969 | 1006 | 9 | 89.46 | 1 | 9.94 | 11.11 | 58.85 | 54.45 | 4.4 | 4.4 | 0.04 | 0.04 |
| gene7300 | https://www.ncbi.nlm.nih.gov/gene/?term=LOC105280472 | 1589 | 6 | 37.76 | 1 | 6.29 | 16.67 | 78.84 | 74.62 | 4.22 | 4.22 | 0.02 | 0.02 |
| gene7398 | https://www.ncbi.nlm.nih.gov/gene/?term=LOC105280589 | 3452 | 45 | 130.36 | 2 | 5.79 | 4.44 | 28.17 | 23.98 | 4.19 | 4.19 | 0.05 | 0.05 |
| gene8608 | https://www.ncbi.nlm.nih.gov/gene/?term=LOC105282002 | 1467 | 26 | 177.23 | 1 | 6.82 | 3.85 | 67.13 | 63.11 | 4.02 | 4.02 | 0.07 | 0.07 |
| gene9464 | https://www.ncbi.nlm.nih.gov/gene/?term=LOC105282975 | 1450 | 22 | 151.72 | 2 | 13.79 | 9.09 | 49.9 | 46.07 | 3.84 | 3.84 | 0.06 | 0.06 |
| gene1024 | https://www.ncbi.nlm.nih.gov/gene/?term=LOC105283693 | 1307 | 12 | 91.81 | 1 | 7.65 | 8.33 | 71.36 | 67.66 | 3.7 | 3.7 | 0.03 | 0.03 |
| gene2959 | https://www.ncbi.nlm.nih.gov/gene/?term=LOC105275501 | 1057 | 7 | 66.23 | 1 | 9.46 | 14.29 | 25.2 | 21.53 | 3.67 | 3.67 | 0.02 | 0.02 |
| gene7190 | https://www.ncbi.nlm.nih.gov/gene/?term=LOC105280355 | 1026 | 14 | 136.45 | 1 | 9.75 | 7.14 | 18.6 | 15.18 | 3.42 | 3.42 | 0.05 | 0.05 |
| gene8198 | https://www.ncbi.nlm.nih.gov/gene/?term=LOC105281539 | 4803 | 18 | 37.48 | 1 | 2.08 | 5.56 | 62.51 | 59.26 | 3.25 | 3.25 | 0.01 | 0.01 |
| gene4156 | https://www.ncbi.nlm.nih.gov/gene/?term=LOC105276840 | 3307 | 11 | 33.26 | 1 | 3.02 | 9.09 | 6.96 | 3.82 | 3.14 | 3.14 | 0.01 | 0.01 |
| gene416 | https://www.ncbi.nlm.nih.gov/gene/?term=LOC105276798 | 2864 | 8 | 27.93 | 1 | 3.49 | 12.5 | 6.95 | 3.82 | 3.14 | 3.14 | 0.01 | 0.01 |
| gene11816 | https://www.ncbi.nlm.nih.gov/gene/?term=LOC105285668 | 1912 | 22 | 115.06 | 3 | 15.69 | 13.64 | 41.69 | 38.58 | 3.11 | 3.11 | 0.04 | 0.04 |
| gene10743 | https://www.ncbi.nlm.nih.gov/gene/?term=LOC105284456 | 1028 | 43 | 418.29 | 3 | 29.18 | 6.98 | 44.49 | 41.39 | 3.1 | 3.1 | 0.13 | 0.13 |
| gene12357 | https://www.ncbi.nlm.nih.gov/gene/?term=LOC105286263 | 1877 | 29 | 154.5 | 1 | 5.33 | 3.45 | 33.04 | 29.95 | 3.09 | 3.09 | 0.05 | 0.05 |
| gene2009 | https://www.ncbi.nlm.nih.gov/gene/?term=LOC105288212 | 1653 | 24 | 145.19 | 1 | 6.05 | 4.17 | 26.6 | 23.57 | 3.03 | 3.03 | 0.04 | 0.04 |
| gene8891 | https://www.ncbi.nlm.nih.gov/gene/?term=LOC105282325 | 2254 | 21 | 93.17 | 1 | 4.44 | 4.76 | 36.49 | 33.5 | 2.99 | 2.99 | 0.03 | 0.03 |
| gene7266 | https://www.ncbi.nlm.nih.gov/gene/?term=LOC105280436 | 4823 | 41 | 85.01 | 3 | 6.22 | 7.32 | 18.51 | 15.53 | 2.98 | 2.98 | 0.03 | 0.03 |

...

... only first 25 lines shown ...

Directory: OGLall File: ogl-Cb_LI_BCphaseA2.vs.RphaseB1.txt

| gene_ID | gene_link | gwidth | #Sites | #per10Kb | #dmSites | #dmsp10kb | %dmSites | %pSite1 | %pSite2 | DMpSite | ADMpSite | DMpNucl | ADMpNucl |
|---|---|---|---|---|---|---|---|---|---|---|---|---|---|
| gene2593 | https://www.ncbi.nlm.nih.gov/gene/?term=LOC105275070 | 1774 | 9 | 50.73 | 4 | 22.55 | 44.44 | 77.93 | 29.94 | 47.99 | 47.99 | 0.24 | 0.24 |
| gene3165 | https://www.ncbi.nlm.nih.gov/gene/?term=LOC105275706 | 2374 | 67 | 282.22 | 6 | 25.27 | 8.96 | 35.39 | 24.21 | 11.18 | 11.18 | 0.32 | 0.32 |
| gene2824 | https://www.ncbi.nlm.nih.gov/gene/?term=LOC105275337 | 1772 | 62 | 349.89 | 2 | 11.29 | 3.23 | 38.77 | 30.45 | 8.31 | 8.31 | 0.29 | 0.29 |
| gene5714 | https://www.ncbi.nlm.nih.gov/gene/?term=LOC105278729 | 710 | 16 | 225.35 | 1 | 14.08 | 6.25 | 57.91 | 51.9 | 6.01 | 6.01 | 0.14 | 0.14 |
| gene1993 | https://www.ncbi.nlm.nih.gov/gene/?term=LOC105288193 | 1449 | 75 | 517.6 | 1 | 6.9 | 1.33 | 35.76 | 32.82 | 2.94 | 2.94 | 0.15 | 0.15 |
| gene13033 | https://www.ncbi.nlm.nih.gov/gene/?term=LOC105287063 | 1877 | 49 | 261.05 | 1 | 5.33 | 2.04 | 32.47 | 29.74 | 2.73 | 2.73 | 0.07 | 0.07 |
| gene10984 | https://www.ncbi.nlm.nih.gov/gene/?term=LOC105284704 | 1987 | 62 | 312.03 | 1 | 5.03 | 1.61 | 26.65 | 24.35 | 2.3 | 2.3 | 0.07 | 0.07 |
| gene3671 | https://www.ncbi.nlm.nih.gov/gene/?term=LOC105276272 | 1966 | 38 | 193.29 | 1 | 5.09 | 2.63 | 30.65 | 28.5 | 2.15 | 2.15 | 0.04 | 0.04 |
| gene7556 | https://www.ncbi.nlm.nih.gov/gene/?term=LOC105280774 | 2792 | 101 | 361.75 | 1 | 3.58 | 0.99 | 32.38 | 30.39 | 1.99 | 1.99 | 0.07 | 0.07 |
| gene7398 | https://www.ncbi.nlm.nih.gov/gene/?term=LOC105280589 | 3452 | 42 | 121.67 | 1 | 2.9 | 2.38 | 27.89 | 25.91 | 1.98 | 1.98 | 0.02 | 0.02 |
| gene4034 | https://www.ncbi.nlm.nih.gov/gene/?term=LOC105276704 | 2065 | 86 | 416.46 | 1 | 4.84 | 1.16 | 15.38 | 13.53 | 1.85 | 1.85 | 0.08 | 0.08 |
| gene6652 | https://www.ncbi.nlm.nih.gov/gene/?term=LOC105279719 | 2831 | 78 | 275.52 | 1 | 3.53 | 1.28 | 25.09 | 23.41 | 1.68 | 1.68 | 0.05 | 0.05 |
| gene10520 | https://www.ncbi.nlm.nih.gov/gene/?term=LOC105284184 | 2451 | 87 | 354.96 | 1 | 4.08 | 1.15 | 34.29 | 32.67 | 1.61 | 1.61 | 0.06 | 0.06 |
| gene10091 | https://www.ncbi.nlm.nih.gov/gene/?term=LOC105283697 | 4988 | 137 | 274.66 | 1 | 2 | 0.73 | 17.98 | 16.41 | 1.57 | 1.57 | 0.04 | 0.04 |
| gene4502 | https://www.ncbi.nlm.nih.gov/gene/?term=LOC105277243 | 2289 | 32 | 139.8 | 1 | 4.37 | 3.12 | 44.88 | 43.35 | 1.53 | 1.53 | 0.02 | 0.02 |
| gene5877 | https://www.ncbi.nlm.nih.gov/gene/?term=LOC105278807 | 1719 | 49 | 285.05 | 1 | 5.82 | 2.04 | 25.27 | 23.74 | 1.53 | 1.53 | 0.04 | 0.04 |
| gene9336 | https://www.ncbi.nlm.nih.gov/gene/?term=LOC105282834 | 5993 | 124 | 206.91 | 1 | 1.67 | 0.81 | 15.53 | 14.01 | 1.52 | 1.52 | 0.03 | 0.03 |
| gene6455 | https://www.ncbi.nlm.nih.gov/gene/?term=LOC105279500 | 2612 | 33 | 126.34 | 1 | 3.83 | 3.03 | 51.99 | 50.58 | 1.41 | 1.41 | 0.02 | 0.02 |
| gene7734 | https://www.ncbi.nlm.nih.gov/gene/?term=LOC105280972 | 1528 | 32 | 209.42 | 1 | 6.54 | 3.12 | 19.4 | 18.05 | 1.35 | 1.35 | 0.03 | 0.03 |
| gene2833 | https://www.ncbi.nlm.nih.gov/gene/?term=LOC105275352 | 5528 | 38 | 68.74 | 1 | 1.81 | 2.63 | 25.27 | 24 | 1.27 | 1.27 | 0.01 | 0.01 |
| gene7888 | https://www.ncbi.nlm.nih.gov/gene/?term=LOC105281141 | 4357 | 157 | 360.34 | 1 | 2.3 | 0.64 | 31.42 | 30.25 | 1.17 | 1.17 | 0.04 | 0.04 |
| gene10606 | https://www.ncbi.nlm.nih.gov/gene/?term=LOC105284273 | 6119 | 260 | 424.91 | 1 | 1.63 | 0.38 | 19.55 | 18.4 | 1.15 | 1.15 | 0.05 | 0.05 |
| gene2970 | https://www.ncbi.nlm.nih.gov/gene/?term=LOC105275518 | 2340 | 19 | 81.2 | 1 | 4.27 | 5.26 | 51.65 | 50.52 | 1.13 | 1.13 | 0.01 | 0.01 |
| gene10107 | https://www.ncbi.nlm.nih.gov/gene/?term=LOC105283712 | 1883 | 30 | 159.32 | 1 | 5.31 | 3.33 | 31.98 | 30.87 | 1.11 | 1.11 | 0.02 | 0.02 |
| gene3817 | https://www.ncbi.nlm.nih.gov/gene/?term=LOC105276449 | 1230 | 43 | 349.59 | 1 | 8.13 | 2.33 | 30.78 | 29.68 | 1.1 | 1.1 | 0.04 | 0.04 |
| gene7568 | https://www.ncbi.nlm.nih.gov/gene/?term=LOC105280786 | 2525 | 122 | 483.17 | 1 | 3.96 | 0.82 | 30.88 | 29.81 | 1.07 | 1.07 | 0.05 | 0.05 |
| gene940 | https://www.ncbi.nlm.nih.gov/gene/?term=LOC105282794 | 3578 | 86 | 240.36 | 1 | 2.79 | 1.16 | 16.54 | 15.48 | 1.06 | 1.06 | 0.03 | 0.03 |
| gene7249 | https://www.ncbi.nlm.nih.gov/gene/?term=LOC105280422 | 1775 | 62 | 349.3 | 1 | 5.63 | 1.61 | 17.51 | 16.57 | 0.94 | 0.94 | 0.03 | 0.03 |
| gene5831 | https://www.ncbi.nlm.nih.gov/gene/?term=LOC105278761 | 2506 | 112 | 446.93 | 2 | 7.98 | 1.79 | 21.67 | 20.75 | 0.92 | 0.92 | 0.04 | 0.04 |
| gene10222 | https://www.ncbi.nlm.nih.gov/gene/?term=LOC105283838 | 1748 | 21 | 120.14 | 1 | 5.72 | 4.76 | 46.37 | 45.47 | 0.9 | 0.9 | 0.01 | 0.01 |
| gene322 | https://www.ncbi.nlm.nih.gov/gene/?term=LOC105275777 | 1606 | 23 | 143.21 | 1 | 6.23 | 4.35 | 44.64 | 43.87 | 0.76 | 0.76 | 0.01 | 0.01 |
| gene6286 | https://www.ncbi.nlm.nih.gov/gene/?term=LOC105279298 | 2861 | 121 | 422.93 | 2 | 6.99 | 1.65 | 24.64 | 23.9 | 0.74 | 0.74 | 0.03 | 0.03 |
| gene3912 | https://www.ncbi.nlm.nih.gov/gene/?term=LOC105276560 | 17900 | 309 | 172.63 | 1 | 0.56 | 0.32 | 2.52 | 1.81 | 0.7 | 0.7 | 0.01 | 0.01 |
| gene1147 | https://www.ncbi.nlm.nih.gov/gene/?term=LOC105284954 | 4022 | 96 | 238.69 | 1 | 2.49 | 1.04 | 15.76 | 15.07 | 0.69 | 0.69 | 0.02 | 0.02 |
| gene6031 | https://www.ncbi.nlm.nih.gov/gene/?term=LOC105279000 | 5691 | 208 | 365.49 | 1 | 1.76 | 0.48 | 12.77 | 12.08 | 0.69 | 0.69 | 0.03 | 0.03 |
| gene8015 | https://www.ncbi.nlm.nih.gov/gene/?term=LOC105281312 | 2998 | 68 | 226.82 | 1 | 3.34 | 1.47 | 31.47 | 30.8 | 0.67 | 0.67 | 0.02 | 0.02 |
| gene292 | https://www.ncbi.nlm.nih.gov/gene/?term=LOC105275398 | 2825 | 92 | 325.66 | 1 | 3.54 | 1.09 | 23.29 | 22.65 | 0.64 | 0.64 | 0.02 | 0.02 |
| gene7657 | https://www.ncbi.nlm.nih.gov/gene/?term=LOC105280890 | 1109 | 21 | 189.36 | 1 | 9.02 | 4.76 | 27.2 | 26.57 | 0.64 | 0.64 | 0.01 | 0.01 |
| gene1310 | https://www.ncbi.nlm.nih.gov/gene/?term=LOC105286661 | 2582 | 71 | 274.98 | 1 | 3.87 | 1.41 | 21.34 | 20.74 | 0.61 | 0.61 | 0.02 | 0.02 |
| gene10678 | https://www.ncbi.nlm.nih.gov/gene/?term=LOC105284355 | 4031 | 161 | 399.4 | 1 | 2.48 | 0.62 | 20.49 | 19.95 | 0.54 | 0.54 | 0.02 | 0.02 |
| gene12572 | https://www.ncbi.nlm.nih.gov/gene/?term=LOC105286514 | 2374 | 44 | 185.34 | 1 | 4.21 | 2.27 | 33.89 | 33.38 | 0.52 | 0.52 | 0.01 | 0.01 |
| gene4892 | https://www.ncbi.nlm.nih.gov/gene/?term=LOC105277667 | 3898 | 123 | 315.55 | 1 | 2.57 | 0.81 | 28.28 | 27.77 | 0.51 | 0.51 | 0.02 | 0.02 |
| gene298 | https://www.ncbi.nlm.nih.gov/gene/?term=LOC105275460 | 2473 | 118 | 477.15 | 1 | 4.04 | 0.85 | 13.46 | 12.97 | 0.49 | 0.49 | 0.02 | 0.02 |
| gene289 | https://www.ncbi.nlm.nih.gov/gene/?term=LOC105275347 | 2533 | 72 | 284.25 | 1 | 3.95 | 1.39 | 24.33 | 23.84 | 0.48 | 0.48 | 0.01 | 0.01 |
| gene11568 | https://www.ncbi.nlm.nih.gov/gene/?term=LOC105285395 | 8086 | 202 | 249.81 | 1 | 1.24 | 0.5 | 14.4 | 13.94 | 0.46 | 0.46 | 0.01 | 0.01 |
| gene2056 | https://www.ncbi.nlm.nih.gov/gene/?term=LOC105274452 | 6584 | 236 | 358.44 | 1 | 1.52 | 0.42 | 14.57 | 14.16 | 0.42 | 0.42 | 0.01 | 0.01 |
| gene3470 | https://www.ncbi.nlm.nih.gov/gene/?term=LOC105276059 | 8716 | 151 | 173.24 | 1 | 1.15 | 0.66 | 10.19 | 9.77 | 0.42 | 0.42 | 0.01 | 0.01 |
| gene8729 | https://www.ncbi.nlm.nih.gov/gene/?term=LOC105282137 | 5005 | 211 | 421.58 | 1 | 2 | 0.47 | 16 | 15.6 | 0.4 | 0.4 | 0.02 | 0.02 |
| gene7245 | https://www.ncbi.nlm.nih.gov/gene/?term=LOC105280410 | 12478 | 264 | 211.57 | 1 | 0.8 | 0.38 | 19.99 | 19.61 | 0.38 | 0.38 | 0.01 | 0.01 |
| gene8600 | https://www.ncbi.nlm.nih.gov/gene/?term=LOC105282001 | 8513 | 152 | 178.55 | 1 | 1.17 | 0.66 | 12.95 | 12.59 | 0.36 | 0.36 | 0.01 | 0.01 |
| gene3747 | https://www.ncbi.nlm.nih.gov/gene/?term=LOC105276387 | 3736 | 188 | 503.21 | 1 | 2.68 | 0.53 | 14.43 | 14.09 | 0.35 | 0.35 | 0.02 | 0.02 |
| gene8178 | https://www.ncbi.nlm.nih.gov/gene/?term=LOC105281515 | 8647 | 52 | 60.14 | 1 | 1.16 | 1.92 | 27.71 | 27.4 | 0.32 | 0.32 | 0 | 0 |
| gene2683 | https://www.ncbi.nlm.nih.gov/gene/?term=LOC105275187 | 5492 | 69 | 125.64 | 1 | 1.82 | 1.45 | 14.14 | 13.85 | 0.29 | 0.29 | 0 | 0 |
| gene6237 | https://www.ncbi.nlm.nih.gov/gene/?term=LOC105279244 | 1484 | 27 | 181.94 | 1 | 6.74 | 3.7 | 50.23 | 49.93 | 0.29 | 0.29 | 0.01 | 0.01 |
| gene2846 | https://www.ncbi.nlm.nih.gov/gene/?term=LOC105275374 | 11246 | 500 | 444.6 | 1 | 0.89 | 0.2 | 10.21 | 9.93 | 0.28 | 0.28 | 0.01 | 0.01 |
| gene3727 | https://www.ncbi.nlm.nih.gov/gene/?term=LOC105276367 | 2484 | 68 | 273.75 | 1 | 4.03 | 1.47 | 46.01 | 45.74 | 0.27 | 0.27 | 0.01 | 0.01 |
| gene8343 | https://www.ncbi.nlm.nih.gov/gene/?term=LOC105281700 | 2172 | 38 | 174.95 | 1 | 4.6 | 2.63 | 39.23 | 38.97 | 0.26 | 0.26 | 0 | 0 |
| gene5746 | https://www.ncbi.nlm.nih.gov/gene/?term=LOC105278666 | 10343 | 455 | 439.91 | 1 | 0.97 | 0.22 | 7.39 | 7.14 | 0.25 | 0.25 | 0.01 | 0.01 |
| gene11819 | https://www.ncbi.nlm.nih.gov/gene/?term=LOC105285671 | 2366 | 98 | 414.2 | 1 | 4.23 | 1.02 | 30.44 | 30.22 | 0.23 | 0.23 | 0.01 | 0.01 |
| gene9755 | https://www.ncbi.nlm.nih.gov/gene/?term=LOC105283296 | 2177 | 87 | 399.63 | 1 | 4.59 | 1.15 | 15.89 | 15.67 | 0.23 | 0.23 | 0.01 | 0.01 |

| | | | | | | | | | | | | | |
|---|---|---|---|---|---|---|---|---|---|---|---|---|---|
| gene474 | https://www.ncbi.nlm.nih.gov/gene/?term=LOC105277435 | 3625 | 173 | 477.24 | 1 | 2.76 | 0.58 | 10.83 | 10.64 | 0.2 | 0.2 | 0.01 | 0.01 |
| gene6620 | https://www.ncbi.nlm.nih.gov/gene/?term=LOC105279687 | 17998 | 349 | 193.91 | 1 | 0.56 | 0.29 | 4.97 | 4.84 | 0.13 | 0.13 | 0 | 0 |
| gene11275 | https://www.ncbi.nlm.nih.gov/gene/?term=LOC105285055 | 4119 | 239 | 580.24 | 1 | 2.43 | 0.42 | 11.8 | 11.69 | 0.12 | 0.12 | 0.01 | 0.01 |
| gene2271 | https://www.ncbi.nlm.nih.gov/gene/?term=LOC105274706 | 1448 | 51 | 352.21 | 1 | 6.91 | 1.96 | 38.8 | 38.68 | 0.12 | 0.12 | 0 | 0 |
| gene12307 | https://www.ncbi.nlm.nih.gov/gene/?term=LOC105286216 | 10313 | 178 | 172.6 | 1 | 0.97 | 0.56 | 15.99 | 15.89 | 0.1 | 0.1 | 0 | 0 |
| gene3856 | https://www.ncbi.nlm.nih.gov/gene/?term=LOC105276499 | 5236 | 177 | 338.04 | 1 | 1.91 | 0.56 | 10.08 | 9.99 | 0.09 | 0.09 | 0 | 0 |
| gene7166 | https://www.ncbi.nlm.nih.gov/gene/?term=LOC105280326 | 6117 | 222 | 362.92 | 1 | 1.63 | 0.45 | 16.24 | 16.15 | 0.09 | 0.09 | 0 | 0 |
| gene9330 | https://www.ncbi.nlm.nih.gov/gene/?term=LOC105282825 | 3936 | 78 | 198.17 | 1 | 2.54 | 1.28 | 12.99 | 12.9 | 0.09 | 0.09 | 0 | 0 |
| gene53 | https://www.ncbi.nlm.nih.gov/gene/?term=LOC105277148 | 19283 | 283 | 146.76 | 1 | 0.52 | 0.35 | 5.45 | 5.37 | 0.08 | 0.08 | 0 | 0 |
| gene10344 | https://www.ncbi.nlm.nih.gov/gene/?term=LOC105283987 | 4841 | 71 | 146.66 | 1 | 2.07 | 1.41 | 32.31 | 32.24 | 0.07 | 0.07 | 0 | 0 |
| gene10654 | https://www.ncbi.nlm.nih.gov/gene/?term=LOC105284329 | 4027 | 51 | 126.65 | 1 | 2.48 | 1.96 | 19.38 | 19.32 | 0.06 | 0.06 | 0 | 0 |
| gene10600 | https://www.ncbi.nlm.nih.gov/gene/?term=LOC105284270 | 4837 | 100 | 206.74 | 1 | 2.07 | 1 | 12.53 | 12.48 | 0.05 | 0.05 | 0 | 0 |
| gene12981 | https://www.ncbi.nlm.nih.gov/gene/?term=LOC105286999 | 2063 | 44 | 213.28 | 1 | 4.85 | 2.27 | 35.25 | 35.21 | 0.04 | 0.04 | 0 | 0 |
| gene3885 | https://www.ncbi.nlm.nih.gov/gene/?term=LOC105276527 | 7928 | 333 | 420.03 | 1 | 1.26 | 0.3 | 8.6 | 8.57 | 0.03 | 0.03 | 0 | 0 |
| gene10392 | https://www.ncbi.nlm.nih.gov/gene/?term=LOC105284036 | 3216 | 87 | 270.52 | 2 | 6.22 | 2.3 | 24.3 | 24.3 | 0.01 | 0.01 | 0 | 0 |
| gene10549 | https://www.ncbi.nlm.nih.gov/gene/?term=LOC105284210 | 7145 | 327 | 457.66 | 2 | 2.8 | 0.61 | 10.82 | 10.83 | -0.01 | 0.01 | 0 | 0 |
| gene7600 | https://www.ncbi.nlm.nih.gov/gene/?term=LOC105280819 | 1615 | 38 | 235.29 | 1 | 6.19 | 2.63 | 35.56 | 35.58 | -0.02 | 0.02 | 0 | 0 |
| gene2806 | https://www.ncbi.nlm.nih.gov/gene/?term=LOC105275327 | 14257 | 472 | 331.07 | 1 | 0.7 | 0.21 | 7.12 | 7.15 | -0.03 | 0.03 | 0 | 0 |
| gene4190 | https://www.ncbi.nlm.nih.gov/gene/?term=LOC105276875 | 3150 | 101 | 320.63 | 1 | 3.17 | 0.99 | 24.58 | 24.61 | -0.03 | 0.03 | 0 | 0 |
| gene1632 | https://www.ncbi.nlm.nih.gov/gene/?term=LOC105287764 | 18061 | 391 | 216.49 | 1 | 0.55 | 0.26 | 3.3 | 3.35 | -0.05 | 0.05 | 0 | 0 |
| gene5524 | https://www.ncbi.nlm.nih.gov/gene/?term=LOC105278397 | 5177 | 200 | 386.32 | 1 | 1.93 | 0.5 | 24.93 | 25 | -0.07 | 0.07 | 0 | 0 |
| gene4209 | https://www.ncbi.nlm.nih.gov/gene/?term=LOC105277125 | 50550 | 1783 | 352.72 | 1 | 0.2 | 0.06 | 1.63 | 1.71 | -0.08 | 0.08 | 0 | 0 |
| gene12516 | https://www.ncbi.nlm.nih.gov/gene/?term=LOC105286452 | 1529 | 42 | 274.69 | 1 | 6.54 | 2.38 | 46.48 | 46.6 | -0.11 | 0.11 | 0 | 0 |
| gene5525 | https://www.ncbi.nlm.nih.gov/gene/?term=LOC105278396 | 3090 | 186 | 601.94 | 1 | 3.24 | 0.54 | 24.32 | 24.44 | -0.12 | 0.12 | -0.01 | 0.01 |
| gene12802 | https://www.ncbi.nlm.nih.gov/gene/?term=LOC105286799 | 3183 | 173 | 543.51 | 1 | 3.14 | 0.58 | 17.3 | 17.43 | -0.13 | 0.13 | -0.01 | 0.01 |
| gene9029 | https://www.ncbi.nlm.nih.gov/gene/?term=LOC105282491 | 4762 | 169 | 354.89 | 1 | 2.1 | 0.59 | 7.49 | 7.61 | -0.13 | 0.13 | 0 | 0 |
| gene4641 | https://www.ncbi.nlm.nih.gov/gene/?term=LOC105277398 | 1937 | 67 | 345.9 | 1 | 5.16 | 1.49 | 35.24 | 35.39 | -0.14 | 0.14 | 0 | 0 |
| gene1884 | https://www.ncbi.nlm.nih.gov/gene/?term=LOC105288072 | 4169 | 77 | 184.7 | 1 | 2.4 | 1.3 | 24.07 | 24.23 | -0.16 | 0.16 | 0 | 0 |
| gene5539 | https://www.ncbi.nlm.nih.gov/gene/?term=LOC105278415 | 25024 | 661 | 264.15 | 1 | 0.4 | 0.15 | 3.65 | 3.82 | -0.17 | 0.17 | 0 | 0 |
| gene10018 | https://www.ncbi.nlm.nih.gov/gene/?term=LOC105283607 | 4414 | 192 | 434.98 | 1 | 2.27 | 0.52 | 15.73 | 15.92 | -0.19 | 0.19 | -0.01 | 0.01 |
| gene10489 | https://www.ncbi.nlm.nih.gov/gene/?term=LOC105284147 | 36992 | 478 | 129.22 | 1 | 0.27 | 0.21 | 10.06 | 10.25 | -0.19 | 0.19 | 0 | 0 |
| gene13036 | https://www.ncbi.nlm.nih.gov/gene/?term=LOC105287059 | 4959 | 172 | 346.84 | 1 | 2.02 | 0.58 | 16.67 | 16.87 | -0.2 | 0.2 | -0.01 | 0.01 |
| gene2173 | https://www.ncbi.nlm.nih.gov/gene/?term=LOC105274590 | 4387 | 67 | 152.72 | 2 | 4.56 | 2.99 | 31.19 | 31.42 | -0.23 | 0.23 | 0 | 0 |
| gene1470 | https://www.ncbi.nlm.nih.gov/gene/?term=LOC105287597 | 2974 | 192 | 645.6 | 1 | 3.36 | 0.52 | 18.97 | 19.22 | -0.25 | 0.25 | -0.02 | 0.02 |
| gene4376 | https://www.ncbi.nlm.nih.gov/gene/?term=LOC105277083 | 4307 | 123 | 285.58 | 1 | 2.32 | 0.81 | 19.63 | 19.89 | -0.26 | 0.26 | -0.01 | 0.01 |
| gene2748 | https://www.ncbi.nlm.nih.gov/gene/?term=LOC105275261 | 3845 | 50 | 130.04 | 1 | 2.6 | 2 | 37.52 | 37.82 | -0.3 | 0.3 | 0 | 0 |
| gene3599 | https://www.ncbi.nlm.nih.gov/gene/?term=LOC105276196 | 2955 | 45 | 152.28 | 1 | 3.38 | 2.22 | 48.59 | 48.91 | -0.31 | 0.31 | 0 | 0 |
| gene10490 | https://www.ncbi.nlm.nih.gov/gene/?term=LOC105284146 | 13960 | 149 | 106.73 | 1 | 0.72 | 0.67 | 10.54 | 10.87 | -0.33 | 0.33 | 0 | 0 |
| gene5646 | https://www.ncbi.nlm.nih.gov/gene/?term=LOC105278534 | 7475 | 180 | 240.8 | 1 | 1.34 | 0.56 | 17.08 | 17.41 | -0.33 | 0.33 | -0.01 | 0.01 |
| gene13044 | https://www.ncbi.nlm.nih.gov/gene/?term=LOC105287070 | 5094 | 224 | 439.73 | 1 | 1.96 | 0.45 | 5.36 | 5.7 | -0.34 | 0.34 | -0.01 | 0.01 |
| gene6639 | https://www.ncbi.nlm.nih.gov/gene/?term=LOC105279705 | 7656 | 66 | 86.21 | 1 | 1.31 | 1.52 | 21.87 | 22.21 | -0.34 | 0.34 | 0 | 0 |
| gene12674 | https://www.ncbi.nlm.nih.gov/gene/?term=LOC105286649 | 18283 | 764 | 417.87 | 1 | 0.55 | 0.13 | 0.96 | 1.31 | -0.35 | 0.35 | -0.01 | 0.01 |
| gene1571 | https://www.ncbi.nlm.nih.gov/gene/?term=LOC105287700 | 3392 | 160 | 471.7 | 1 | 2.95 | 0.62 | 15.1 | 15.46 | -0.36 | 0.36 | -0.02 | 0.02 |
| gene12406 | https://www.ncbi.nlm.nih.gov/gene/?term=LOC105286316 | 1386 | 29 | 209.24 | 1 | 7.22 | 3.45 | 45.59 | 46 | -0.41 | 0.41 | -0.01 | 0.01 |
| gene5298 | https://www.ncbi.nlm.nih.gov/gene/?term=LOC105278145 | 4060 | 251 | 618.23 | 2 | 4.93 | 0.8 | 10.52 | 10.93 | -0.42 | 0.42 | -0.03 | 0.03 |
| gene4962 | https://www.ncbi.nlm.nih.gov/gene/?term=LOC105277761 | 4645 | 161 | 346.61 | 1 | 2.15 | 0.62 | 14.28 | 14.77 | -0.48 | 0.48 | -0.02 | 0.02 |
| gene13114 | https://www.ncbi.nlm.nih.gov/gene/?term=LOC105287147 | 7609 | 398 | 523.06 | 1 | 1.31 | 0.25 | 9.41 | 9.93 | -0.52 | 0.52 | -0.03 | 0.03 |
| gene8274 | https://www.ncbi.nlm.nih.gov/gene/?term=LOC105281624 | 5425 | 221 | 407.37 | 1 | 1.84 | 0.45 | 9.57 | 10.1 | -0.54 | 0.54 | -0.02 | 0.02 |
| gene9326 | https://www.ncbi.nlm.nih.gov/gene/?term=LOC105282818 | 16612 | 217 | 130.63 | 1 | 0.6 | 0.46 | 13.52 | 14.06 | -0.54 | 0.54 | -0.01 | 0.01 |
| gene3107 | https://www.ncbi.nlm.nih.gov/gene/?term=LOC105275650 | 5103 | 172 | 337.06 | 1 | 1.96 | 0.58 | 16.37 | 16.93 | -0.56 | 0.56 | -0.02 | 0.02 |
| gene5375 | https://www.ncbi.nlm.nih.gov/gene/?term=LOC105278228 | 8640 | 167 | 193.29 | 1 | 1.16 | 0.6 | 22.05 | 22.68 | -0.63 | 0.63 | -0.01 | 0.01 |
| gene11561 | https://www.ncbi.nlm.nih.gov/gene/?term=LOC105285387 | 11528 | 200 | 173.49 | 1 | 0.87 | 0.5 | 9.19 | 9.83 | -0.64 | 0.64 | -0.01 | 0.01 |
| gene12573 | https://www.ncbi.nlm.nih.gov/gene/?term=LOC105286515 | 3647 | 108 | 296.13 | 1 | 2.74 | 0.93 | 11.17 | 11.84 | -0.67 | 0.67 | -0.02 | 0.02 |
| gene9327 | https://www.ncbi.nlm.nih.gov/gene/?term=LOC105282821 | 7793 | 161 | 206.6 | 1 | 1.28 | 0.62 | 12.72 | 13.4 | -0.68 | 0.68 | -0.01 | 0.01 |
| gene1198 | https://www.ncbi.nlm.nih.gov/gene/?term=LOC105285507 | 2738 | 92 | 336.01 | 1 | 3.65 | 1.09 | 20.74 | 21.44 | -0.7 | 0.7 | -0.02 | 0.02 |
| gene11454 | https://www.ncbi.nlm.nih.gov/gene/?term=LOC105285256 | 1906 | 48 | 251.84 | 1 | 5.25 | 2.08 | 44.93 | 45.73 | -0.8 | 0.8 | -0.02 | 0.02 |
| gene5581 | https://www.ncbi.nlm.nih.gov/gene/?term=LOC105278459 | 2935 | 97 | 330.49 | 1 | 3.41 | 1.03 | 20.44 | 21.26 | -0.83 | 0.83 | -0.03 | 0.03 |
| gene7368 | https://www.ncbi.nlm.nih.gov/gene/?term=LOC105280556 | 4105 | 58 | 141.29 | 1 | 2.44 | 1.72 | 18.1 | 19.01 | -0.91 | 0.91 | -0.01 | 0.01 |
| gene6966 | https://www.ncbi.nlm.nih.gov/gene/?term=LOC105280104 | 3747 | 124 | 330.93 | 1 | 2.67 | 0.81 | 16.15 | 17.09 | -0.94 | 0.94 | -0.03 | 0.03 |
| gene8814 | https://www.ncbi.nlm.nih.gov/gene/?term=LOC105282240 | 6910 | 166 | 240.23 | 1 | 1.45 | 0.6 | 9.27 | 10.23 | -0.96 | 0.96 | -0.02 | 0.02 |
| gene12203 | https://www.ncbi.nlm.nih.gov/gene/?term=LOC105286099 | 5967 | 79 | 132.39 | 1 | 1.68 | 1.27 | 11.92 | 12.93 | -1.01 | 1.01 | -0.01 | 0.01 |
| gene2835 | https://www.ncbi.nlm.nih.gov/gene/?term=LOC105275357 | 2492 | 114 | 457.46 | 1 | 4.01 | 0.88 | 13.76 | 14.82 | -1.05 | 1.05 | -0.05 | 0.05 |
| gene7915 | https://www.ncbi.nlm.nih.gov/gene/?term=LOC105281173 | 1597 | 39 | 244.21 | 1 | 6.26 | 2.56 | 60.85 | 62.01 | -1.16 | 1.16 | -0.03 | 0.03 |
| gene11447 | https://www.ncbi.nlm.nih.gov/gene/?term=LOC105285252 | 7437 | 126 | 169.42 | 2 | 2.69 | 1.59 | 12.66 | 13.85 | -1.19 | 1.19 | -0.02 | 0.02 |

| | | | | | | | | | | | | | |
|---|---|---|---|---|---|---|---|---|---|---|---|---|---|
| gene6583 | https://www.ncbi.nlm.nih.gov/gene/?term=LOC105279649 | 7444 | 343 | 460.77 | 1 | 1.34 | 0.29 | 10.51 | 11.78 | -1.26 | 1.26 | -0.06 | 0.06 |
| gene11340 | https://www.ncbi.nlm.nih.gov/gene/?term=LOC105285131 | 2918 | 89 | 305 | 1 | 3.43 | 1.12 | 11.73 | 13.01 | -1.28 | 1.28 | -0.04 | 0.04 |
| gene3335 | https://www.ncbi.nlm.nih.gov/gene/?term=LOC105275898 | 4665 | 147 | 315.11 | 1 | 2.14 | 0.68 | 11.48 | 12.76 | -1.29 | 1.29 | -0.04 | 0.04 |
| gene3848 | https://www.ncbi.nlm.nih.gov/gene/?term=LOC105276486 | 5899 | 145 | 245.8 | 1 | 1.7 | 0.69 | 23.24 | 24.56 | -1.32 | 1.32 | -0.03 | 0.03 |
| gene10345 | https://www.ncbi.nlm.nih.gov/gene/?term=LOC105283989 | 4444 | 21 | 47.25 | 1 | 2.25 | 4.76 | 48.56 | 49.94 | -1.37 | 1.37 | -0.01 | 0.01 |
| gene7207 | https://www.ncbi.nlm.nih.gov/gene/?term=LOC105280376 | 4659 | 112 | 240.39 | 1 | 2.15 | 0.89 | 28.49 | 29.89 | -1.4 | 1.4 | -0.03 | 0.03 |
| gene10155 | https://www.ncbi.nlm.nih.gov/gene/?term=LOC105283756 | 7030 | 98 | 139.4 | 1 | 1.42 | 1.02 | 6.09 | 7.87 | -1.78 | 1.78 | -0.02 | 0.02 |
| gene920 | https://www.ncbi.nlm.nih.gov/gene/?term=LOC105282518 | 1197 | 39 | 325.81 | 1 | 8.35 | 2.56 | 36.44 | 38.28 | -1.85 | 1.85 | -0.06 | 0.06 |
| gene3168 | https://www.ncbi.nlm.nih.gov/gene/?term=LOC105275711 | 6150 | 80 | 130.08 | 1 | 1.63 | 1.25 | 11.3 | 13.17 | -1.86 | 1.86 | -0.02 | 0.02 |
| gene10819 | https://www.ncbi.nlm.nih.gov/gene/?term=LOC105284507 | 5767 | 86 | 149.12 | 1 | 1.73 | 1.16 | 21.64 | 23.53 | -1.89 | 1.89 | -0.03 | 0.03 |
| gene1220 | https://www.ncbi.nlm.nih.gov/gene/?term=LOC105285742 | 3453 | 98 | 283.81 | 1 | 2.9 | 1.02 | 19.04 | 20.94 | -1.9 | 1.9 | -0.05 | 0.05 |
| gene388 | https://www.ncbi.nlm.nih.gov/gene/?term=LOC105276538 | 1558 | 19 | 121.95 | 2 | 12.84 | 10.53 | 66.07 | 68.07 | -1.99 | 1.99 | -0.02 | 0.02 |
| gene10879 | https://www.ncbi.nlm.nih.gov/gene/?term=LOC105284574 | 2423 | 45 | 185.72 | 1 | 4.13 | 2.22 | 57.53 | 59.68 | -2.15 | 2.15 | -0.04 | 0.04 |
| gene10552 | https://www.ncbi.nlm.nih.gov/gene/?term=LOC105284213 | 878 | 15 | 170.84 | 1 | 11.39 | 6.67 | 15.38 | 18.37 | -2.99 | 2.99 | -0.05 | 0.05 |
| gene12021 | https://www.ncbi.nlm.nih.gov/gene/?term=LOC105285905 | 1399 | 27 | 192.99 | 1 | 7.15 | 3.7 | 31.19 | 34.36 | -3.17 | 3.17 | -0.06 | 0.06 |
| gene1199 | https://www.ncbi.nlm.nih.gov/gene/?term=LOC105285498 | 2220 | 25 | 112.61 | 1 | 4.5 | 4 | 32.49 | 35.74 | -3.26 | 3.26 | -0.04 | 0.04 |
| gene2413 | https://www.ncbi.nlm.nih.gov/gene/?term=LOC105274863 | 2006 | 46 | 229.31 | 1 | 4.99 | 2.17 | 60.76 | 64.28 | -3.53 | 3.53 | -0.08 | 0.08 |
| gene2404 | https://www.ncbi.nlm.nih.gov/gene/?term=LOC105274853 | 1767 | 34 | 192.42 | 1 | 5.66 | 2.94 | 29.24 | 33.79 | -4.55 | 4.55 | -0.09 | 0.09 |
| gene3166 | https://www.ncbi.nlm.nih.gov/gene/?term=LOC105275707 | 2393 | 11 | 45.97 | 1 | 4.18 | 9.09 | 62.4 | 71.24 | -8.83 | 8.83 | -0.04 | 0.04 |
| gene9109 | https://www.ncbi.nlm.nih.gov/gene/?term=LOC105282561 | 2400 | 11 | 45.83 | 1 | 4.17 | 9.09 | 17.27 | 26.23 | -8.97 | 8.97 | -0.04 | 0.04 |
| gene11584 | https://www.ncbi.nlm.nih.gov/gene/?term=LOC105285416 | 3709 | 6 | 16.18 | 2 | 5.39 | 33.33 | 18.09 | 32.8 | -14.71 | 14.71 | -0.02 | 0.02 |
| gene393 | https://www.ncbi.nlm.nih.gov/gene/?term=LOC105276566 | 1123 | 2 | 17.81 | 1 | 8.9 | 50 | 48.08 | 74.3 | -26.22 | 26.22 | -0.05 | 0.05 |

Directory: OGLall File: ogl-Cb_LI_BCphaseA2.vs.RphaseB2.txt

| gene_ID | gene_link | gwidth | #Sites | #per10Kb | #dmSites | #dmsp10kb | %dmSites | %pSite1 | %pSite2 | DMpSite | ADMpSite | DMpNucl | ADMpNucl | |
|---|---|---|---|---|---|---|---|---|---|---|---|---|---|---|
| gene2593 | https://www.ncbi.nlm.nih.gov/gene/?term=LOC105275070 | 1774 | 7 | 39.46 | 5 | 28.18 | 71.43 | 66.52 | 4.09 | 62.43 | 62.43 | 0.25 | 0.25 | |
| gene3165 | https://www.ncbi.nlm.nih.gov/gene/?term=LOC105275706 | 2374 | 72 | 303.29 | | 6 | 25.27 | 8.33 | 33.88 | 22.12 | 11.76 | 11.76 | 0.36 | 0.36 |
| gene2824 | https://www.ncbi.nlm.nih.gov/gene/?term=LOC105275337 | 1772 | 60 | 338.6 | 2 | 11.29 | 3.33 | 38.89 | 29.82 | 9.07 | 9.07 | 0.31 | 0.31 | |
| gene5714 | https://www.ncbi.nlm.nih.gov/gene/?term=LOC105278729 | 710 | 17 | 239.44 | | 1 | 14.08 | 5.88 | 60.22 | 53.54 | 6.67 | 6.67 | 0.16 | 0.16 |
| gene7398 | https://www.ncbi.nlm.nih.gov/gene/?term=LOC105280589 | 3452 | 34 | 98.49 | 1 | 2.9 | 2.94 | 23.83 | 19.97 | 3.86 | 3.86 | 0.04 | 0.04 | |
| gene2970 | https://www.ncbi.nlm.nih.gov/gene/?term=LOC105275518 | 2340 | 18 | 76.92 | 1 | 4.27 | 5.56 | 54.32 | 51.43 | 2.89 | 2.89 | 0.02 | 0.02 | |
| gene940 | https://www.ncbi.nlm.nih.gov/gene/?term=LOC105282794 | 3578 | 87 | 243.15 | | 1 | 2.79 | 1.15 | 16.03 | 14.13 | 1.9 | 1.9 | 0.05 | 0.05 |
| gene6455 | https://www.ncbi.nlm.nih.gov/gene/?term=LOC105279500 | 2612 | 33 | 126.34 | | 1 | 3.83 | 3.03 | 51.77 | 49.92 | 1.86 | 1.86 | 0.02 | 0.02 |
| gene7568 | https://www.ncbi.nlm.nih.gov/gene/?term=LOC105280786 | 2525 | 112 | 443.56 | | 1 | 3.96 | 0.89 | 32.64 | 30.88 | 1.76 | 1.76 | 0.08 | 0.08 |
| gene10520 | https://www.ncbi.nlm.nih.gov/gene/?term=LOC105284184 | 2451 | 78 | 318.24 | | 1 | 4.08 | 1.28 | 37.44 | 35.71 | 1.72 | 1.72 | 0.05 | 0.05 |
| gene9462 | https://www.ncbi.nlm.nih.gov/gene/?term=LOC105282970 | 1469 | 39 | 265.49 | | 1 | 6.81 | 2.56 | 51.12 | 49.47 | 1.65 | 1.65 | 0.04 | 0.04 |
| gene3385 | https://www.ncbi.nlm.nih.gov/gene/?term=LOC105275966 | 3305 | 53 | 160.36 | | 1 | 3.03 | 1.89 | 54.5 | 52.88 | 1.62 | 1.62 | 0.03 | 0.03 |
| gene3594 | https://www.ncbi.nlm.nih.gov/gene/?term=LOC105276190 | 4910 | 83 | 169.04 | | 1 | 2.04 | 1.2 | 11.44 | 9.83 | 1.62 | 1.62 | 0.03 | 0.03 |
| gene11819 | https://www.ncbi.nlm.nih.gov/gene/?term=LOC105285671 | 2366 | 100 | 422.65 | | 1 | 4.23 | 1 | 27.82 | 26.43 | 1.39 | 1.39 | 0.06 | 0.06 |
| gene5877 | https://www.ncbi.nlm.nih.gov/gene/?term=LOC105278807 | 1719 | 46 | 267.6 | 1 | 5.82 | 2.17 | 26.78 | 25.52 | 1.25 | 1.25 | 0.03 | 0.03 | |
| gene35 | https://www.ncbi.nlm.nih.gov/gene/?term=LOC105275197 | 2643 | 43 | 162.69 | | 1 | 3.78 | 2.33 | 44.25 | 43.02 | 1.22 | 1.22 | 0.02 | 0.02 |
| gene13033 | https://www.ncbi.nlm.nih.gov/gene/?term=LOC105287063 | 1877 | 48 | 255.73 | | 1 | 5.33 | 2.08 | 31.55 | 30.37 | 1.19 | 1.19 | 0.03 | 0.03 |
| gene3470 | https://www.ncbi.nlm.nih.gov/gene/?term=LOC105276059 | 8716 | 145 | 166.36 | | 1 | 1.15 | 0.69 | 9.2 | 8.01 | 1.19 | 1.19 | 0.02 | 0.02 |
| gene7915 | https://www.ncbi.nlm.nih.gov/gene/?term=LOC105281173 | 1597 | 38 | 237.95 | | 1 | 6.26 | 2.63 | 59.94 | 58.74 | 1.19 | 1.19 | 0.03 | 0.03 |
| gene12806 | https://www.ncbi.nlm.nih.gov/gene/?term=LOC105286803 | 4196 | 137 | 326.5 | 1 | 2.38 | 0.73 | 10.84 | 9.75 | 1.09 | 1.09 | 0.04 | 0.04 | |
| gene6652 | https://www.ncbi.nlm.nih.gov/gene/?term=LOC105279719 | 2831 | 79 | 279.05 | | 1 | 3.53 | 1.27 | 24.35 | 23.32 | 1.03 | 1.03 | 0.03 | 0.03 |
| gene7556 | https://www.ncbi.nlm.nih.gov/gene/?term=LOC105280774 | 2792 | 89 | 318.77 | | 1 | 3.58 | 1.12 | 36.33 | 35.33 | 1 | 1 | 0.03 | 0.03 |
| gene3671 | https://www.ncbi.nlm.nih.gov/gene/?term=LOC105276272 | 1966 | 34 | 172.94 | | 1 | 5.09 | 2.94 | 33.29 | 32.32 | 0.98 | 0.98 | 0.02 | 0.02 |
| gene10678 | https://www.ncbi.nlm.nih.gov/gene/?term=LOC105284355 | 4031 | 158 | 391.96 | | 1 | 2.48 | 0.63 | 20.75 | 19.82 | 0.93 | 0.93 | 0.04 | 0.04 |
| gene3817 | https://www.ncbi.nlm.nih.gov/gene/?term=LOC105276449 | 1230 | 42 | 341.46 | | 1 | 8.13 | 2.38 | 35.93 | 35.04 | 0.88 | 0.88 | 0.03 | 0.03 |
| gene10344 | https://www.ncbi.nlm.nih.gov/gene/?term=LOC105283987 | 4841 | 64 | 132.2 | 1 | 2.07 | 1.56 | 30.28 | 29.42 | 0.86 | 0.86 | 0.01 | 0.01 | |
| gene10606 | https://www.ncbi.nlm.nih.gov/gene/?term=LOC105284273 | 6119 | 249 | 406.93 | | 1 | 1.63 | 0.4 | 18.46 | 17.61 | 0.86 | 0.86 | 0.03 | 0.03 |
| gene1470 | https://www.ncbi.nlm.nih.gov/gene/?term=LOC105287597 | 2974 | 195 | 655.68 | | 1 | 3.36 | 0.51 | 18.31 | 17.52 | 0.8 | 0.8 | 0.05 | 0.05 |
| gene4962 | https://www.ncbi.nlm.nih.gov/gene/?term=LOC105277761 | 4645 | 142 | 305.71 | | 1 | 2.15 | 0.7 | 14.73 | 13.94 | 0.79 | 0.79 | 0.02 | 0.02 |
| gene4892 | https://www.ncbi.nlm.nih.gov/gene/?term=LOC105277667 | 3898 | 124 | 318.11 | | 1 | 2.57 | 0.81 | 27.97 | 27.19 | 0.78 | 0.78 | 0.02 | 0.02 |
| gene11401 | https://www.ncbi.nlm.nih.gov/gene/?term=LOC105285202 | 1841 | 82 | 445.41 | | 1 | 5.43 | 1.22 | 26.05 | 25.33 | 0.73 | 0.73 | 0.03 | 0.03 |
| gene9336 | https://www.ncbi.nlm.nih.gov/gene/?term=LOC105282834 | 5993 | 121 | 201.9 | 1 | 1.67 | 0.83 | 16.02 | 15.31 | 0.72 | 0.72 | 0.01 | 0.01 | |
| gene10600 | https://www.ncbi.nlm.nih.gov/gene/?term=LOC105284270 | 4837 | 119 | 246.02 | | 1 | 2.07 | 0.84 | 10.7 | 10.02 | 0.68 | 0.68 | 0.02 | 0.02 |
| gene2846 | https://www.ncbi.nlm.nih.gov/gene/?term=LOC105275374 | 11246 | 464 | 412.59 | | 1 | 0.89 | 0.22 | 12.19 | 11.56 | 0.63 | 0.63 | 0.03 | 0.03 |
| gene2173 | https://www.ncbi.nlm.nih.gov/gene/?term=LOC105274590 | 4387 | 57 | 129.93 | | 2 | 4.56 | 3.51 | 33.32 | 32.7 | 0.62 | 0.62 | 0.01 | 0.01 |
| gene3727 | https://www.ncbi.nlm.nih.gov/gene/?term=LOC105276367 | 2484 | 66 | 265.7 | 1 | 4.03 | 1.52 | 47.44 | 46.84 | 0.61 | 0.61 | 0.02 | 0.02 | |
| gene11568 | https://www.ncbi.nlm.nih.gov/gene/?term=LOC105285395 | 8086 | 180 | 222.61 | | 1 | 1.24 | 0.56 | 17.19 | 16.59 | 0.6 | 0.6 | 0.01 | 0.01 |
| gene10984 | https://www.ncbi.nlm.nih.gov/gene/?term=LOC105284704 | 1987 | 56 | 281.83 | | 1 | 5.03 | 1.79 | 25.7 | 25.1 | 0.59 | 0.59 | 0.02 | 0.02 |
| gene2844 | https://www.ncbi.nlm.nih.gov/gene/?term=LOC105275363 | 4330 | 181 | 418.01 | | 1 | 2.31 | 0.55 | 15.09 | 14.51 | 0.58 | 0.58 | 0.02 | 0.02 |
| gene8274 | https://www.ncbi.nlm.nih.gov/gene/?term=LOC105281624 | 5425 | 204 | 376.04 | | 1 | 1.84 | 0.49 | 10.02 | 9.47 | 0.56 | 0.56 | 0.02 | 0.02 |
| gene7245 | https://www.ncbi.nlm.nih.gov/gene/?term=LOC105280410 | 12478 | 234 | 187.53 | | 1 | 0.8 | 0.43 | 21.44 | 20.93 | 0.52 | 0.52 | 0.01 | 0.01 |
| gene5573 | https://www.ncbi.nlm.nih.gov/gene/?term=LOC105278454 | 2688 | 35 | 130.21 | | 1 | 3.72 | 2.86 | 35.99 | 35.53 | 0.47 | 0.47 | 0.01 | 0.01 |
| gene12572 | https://www.ncbi.nlm.nih.gov/gene/?term=LOC105286514 | 2374 | 45 | 189.55 | | 1 | 4.21 | 2.22 | 33.13 | 32.67 | 0.45 | 0.45 | 0.01 | 0.01 |
| gene34 | https://www.ncbi.nlm.nih.gov/gene/?term=LOC105278416 | 7120 | 124 | 174.16 | | 1 | 1.4 | 0.81 | 15.53 | 15.09 | 0.43 | 0.43 | 0.01 | 0.01 |
| gene2056 | https://www.ncbi.nlm.nih.gov/gene/?term=LOC105274452 | 6584 | 251 | 381.23 | | 2 | 3.04 | 0.8 | 13.77 | 13.35 | 0.42 | 0.42 | 0.02 | 0.02 |
| gene6286 | https://www.ncbi.nlm.nih.gov/gene/?term=LOC105279298 | 2861 | 128 | 447.4 | 2 | 6.99 | 1.56 | 24.57 | 24.17 | 0.4 | 0.4 | 0.02 | 0.02 | |
| gene7166 | https://www.ncbi.nlm.nih.gov/gene/?term=LOC105280326 | 6117 | 214 | 349.84 | | 1 | 1.63 | 0.47 | 16.62 | 16.23 | 0.4 | 0.4 | 0.01 | 0.01 |
| gene4246 | https://www.ncbi.nlm.nih.gov/gene/?term=LOC105276941 | 16412 | 493 | 300.39 | | 1 | 0.61 | 0.2 | 5.46 | 5.07 | 0.38 | 0.38 | 0.01 | 0.01 |
| gene5746 | https://www.ncbi.nlm.nih.gov/gene/?term=LOC105278666 | 10343 | 425 | 410.91 | | 1 | 0.97 | 0.24 | 7.62 | 7.27 | 0.35 | 0.35 | 0.01 | 0.01 |
| gene10392 | https://www.ncbi.nlm.nih.gov/gene/?term=LOC105284036 | 3216 | 91 | 282.96 | | 2 | 6.22 | 2.2 | 22.63 | 22.31 | 0.32 | 0.32 | 0.01 | 0.01 |
| gene473 | https://www.ncbi.nlm.nih.gov/gene/?term=LOC105277423 | 2342 | 108 | 461.14 | | 1 | 4.27 | 0.93 | 30.31 | 29.99 | 0.32 | 0.32 | 0.01 | 0.01 |
| gene11579 | https://www.ncbi.nlm.nih.gov/gene/?term=LOC105285406 | 13571 | 228 | 168.01 | | 1 | 0.74 | 0.44 | 7.6 | 7.29 | 0.3 | 0.3 | 0.01 | 0.01 |
| gene2271 | https://www.ncbi.nlm.nih.gov/gene/?term=LOC105274706 | 1448 | 56 | 386.74 | | 1 | 6.91 | 1.79 | 36.69 | 36.4 | 0.3 | 0.3 | 0.01 | 0.01 |
| gene10107 | https://www.ncbi.nlm.nih.gov/gene/?term=LOC105283712 | 1883 | 32 | 169.94 | | 1 | 5.31 | 3.12 | 29.98 | 29.7 | 0.28 | 0.28 | 0 | 0 |
| gene10531 | https://www.ncbi.nlm.nih.gov/gene/?term=LOC105284190 | 6881 | 168 | 244.15 | | 1 | 1.45 | 0.6 | 13.32 | 13.04 | 0.27 | 0.27 | 0.01 | 0.01 |
| gene7211 | https://www.ncbi.nlm.nih.gov/gene/?term=LOC105280377 | 2024 | 57 | 281.62 | | 1 | 4.94 | 1.75 | 50.41 | 50.16 | 0.25 | 0.25 | 0.01 | 0.01 |
| gene298 | https://www.ncbi.nlm.nih.gov/gene/?term=LOC105275460 | 2473 | 116 | 469.07 | | 1 | 4.04 | 0.86 | 13.65 | 13.43 | 0.23 | 0.23 | 0.01 | 0.01 |
| gene5525 | https://www.ncbi.nlm.nih.gov/gene/?term=LOC105278396 | 3090 | 181 | 585.76 | | 1 | 3.24 | 0.55 | 24.85 | 24.63 | 0.22 | 0.22 | 0.01 | 0.01 |
| gene5524 | https://www.ncbi.nlm.nih.gov/gene/?term=LOC105278397 | 5177 | 196 | 378.6 | 1 | 1.93 | 0.51 | 24.88 | 24.69 | 0.18 | 0.18 | 0.01 | 0.01 | |
| gene8600 | https://www.ncbi.nlm.nih.gov/gene/?term=LOC105282001 | 8513 | 139 | 163.28 | | 1 | 1.17 | 0.72 | 11.37 | 11.2 | 0.17 | 0.17 | 0 | 0 |

| | | | | | | | | | | | | | |
|---|---|---|---|---|---|---|---|---|---|---|---|---|---|
| gene9330 | https://www.ncbi.nlm.nih.gov/gene/?term=LOC105282825 | 3936 | 69 | 175.3 | 1 | 2.54 | 1.45 | 13.52 | 13.35 | 0.17 | 0.17 | 0 | 0 |
| gene12307 | https://www.ncbi.nlm.nih.gov/gene/?term=LOC105286216 | 10313 | 167 | 161.93 | 1 | 0.97 | 0.6 | 17.01 | 16.86 | 0.15 | 0.15 | 0 | 0 |
| gene6620 | https://www.ncbi.nlm.nih.gov/gene/?term=LOC105279687 | 17998 | 312 | 173.35 | 1 | 0.56 | 0.32 | 5.46 | 5.32 | 0.14 | 0.14 | 0 | 0 |
| gene12265 | https://www.ncbi.nlm.nih.gov/gene/?term=LOC105286166 | 3050 | 104 | 340.98 | 1 | 3.28 | 0.96 | 25.25 | 25.13 | 0.13 | 0.13 | 0 | 0 |
| gene292 | https://www.ncbi.nlm.nih.gov/gene/?term=LOC105275398 | 2825 | 81 | 286.73 | 1 | 3.54 | 1.23 | 26.31 | 26.19 | 0.11 | 0.11 | 0 | 0 |
| gene11275 | https://www.ncbi.nlm.nih.gov/gene/?term=LOC105285055 | 4119 | 252 | 611.8 | 1 | 2.43 | 0.4 | 11.37 | 11.3 | 0.07 | 0.07 | 0 | 0 |
| gene2806 | https://www.ncbi.nlm.nih.gov/gene/?term=LOC105275327 | 14257 | 460 | 322.65 | 1 | 0.7 | 0.22 | 7.25 | 7.19 | 0.06 | 0.06 | 0 | 0 |
| gene12981 | https://www.ncbi.nlm.nih.gov/gene/?term=LOC105286999 | 2063 | 42 | 203.59 | 1 | 4.85 | 2.38 | 36.12 | 36.07 | 0.05 | 0.05 | 0 | 0 |
| gene13114 | https://www.ncbi.nlm.nih.gov/gene/?term=LOC105287147 | 7609 | 383 | 503.35 | 1 | 1.31 | 0.26 | 9.68 | 9.63 | 0.04 | 0.04 | 0 | 0 |
| gene53 | https://www.ncbi.nlm.nih.gov/gene/?term=LOC105277148 | 19283 | 258 | 133.8 | 1 | 0.52 | 0.39 | 4.46 | 4.42 | 0.04 | 0.04 | 0 | 0 |
| gene5572 | https://www.ncbi.nlm.nih.gov/gene/?term=LOC105278455 | 27887 | 864 | 309.82 | 1 | 0.36 | 0.12 | 6.83 | 6.79 | 0.04 | 0.04 | 0 | 0 |
| gene11082 | https://www.ncbi.nlm.nih.gov/gene/?term=LOC105284824 | 7436 | 185 | 248.79 | 1 | 1.34 | 0.54 | 6.28 | 6.27 | 0.01 | 0.01 | 0 | 0 |
| gene1632 | https://www.ncbi.nlm.nih.gov/gene/?term=LOC105287764 | 18061 | 371 | 205.41 | 2 | 1.11 | 0.54 | 3.28 | 3.27 | 0.01 | 0.01 | 0 | 0 |
| gene4209 | https://www.ncbi.nlm.nih.gov/gene/?term=LOC105277125 | 50550 | 1672 | 330.76 | 1 | 0.2 | 0.06 | 1.66 | 1.65 | 0.01 | 0.01 | 0 | 0 |
| gene6966 | https://www.ncbi.nlm.nih.gov/gene/?term=LOC105280104 | 3747 | 123 | 328.26 | 1 | 2.67 | 0.81 | 16.59 | 16.58 | 0.01 | 0.01 | 0 | 0 |
| gene4190 | https://www.ncbi.nlm.nih.gov/gene/?term=LOC105276875 | 3150 | 109 | 346.03 | 1 | 3.17 | 0.92 | 24.02 | 24.04 | -0.02 | 0.02 | 0 | 0 |
| gene1993 | https://www.ncbi.nlm.nih.gov/gene/?term=LOC105288193 | 1449 | 74 | 510.7 | 1 | 6.9 | 1.35 | 34.89 | 34.92 | -0.03 | 0.03 | 0 | 0 |
| gene8729 | https://www.ncbi.nlm.nih.gov/gene/?term=LOC105282137 | 5005 | 198 | 395.6 | 1 | 2 | 0.51 | 16.91 | 16.95 | -0.04 | 0.04 | 0 | 0 |
| gene5831 | https://www.ncbi.nlm.nih.gov/gene/?term=LOC105278761 | 2506 | 109 | 434.96 | 1 | 3.99 | 0.92 | 24.44 | 24.49 | -0.06 | 0.06 | 0 | 0 |
| gene9177 | https://www.ncbi.nlm.nih.gov/gene/?term=LOC105282646 | 3320 | 99 | 298.19 | 1 | 3.01 | 1.01 | 25.05 | 25.11 | -0.06 | 0.06 | 0 | 0 |
| gene7249 | https://www.ncbi.nlm.nih.gov/gene/?term=LOC105280422 | 1775 | 60 | 338.03 | 1 | 5.63 | 1.67 | 16.97 | 17.06 | -0.09 | 0.09 | 0 | 0 |
| gene3286 | https://www.ncbi.nlm.nih.gov/gene/?term=LOC105275848 | 3744 | 134 | 357.91 | 1 | 2.67 | 0.75 | 22.27 | 22.41 | -0.14 | 0.14 | -0.01 | 0.01 |
| gene5967 | https://www.ncbi.nlm.nih.gov/gene/?term=LOC105278918 | 4393 | 116 | 264.06 | 1 | 2.28 | 0.86 | 14.39 | 14.54 | -0.15 | 0.15 | 0 | 0 |
| gene10549 | https://www.ncbi.nlm.nih.gov/gene/?term=LOC105284210 | 7145 | 286 | 400.28 | 2 | 2.8 | 0.7 | 11.97 | 12.18 | -0.21 | 0.21 | -0.01 | 0.01 |
| gene12802 | https://www.ncbi.nlm.nih.gov/gene/?term=LOC105286799 | 3183 | 160 | 502.67 | 1 | 3.14 | 0.62 | 17.6 | 17.81 | -0.21 | 0.21 | -0.01 | 0.01 |
| gene3107 | https://www.ncbi.nlm.nih.gov/gene/?term=LOC105275650 | 5103 | 166 | 325.3 | 1 | 1.96 | 0.6 | 14.75 | 14.95 | -0.21 | 0.21 | -0.01 | 0.01 |
| gene9755 | https://www.ncbi.nlm.nih.gov/gene/?term=LOC105283296 | 2177 | 85 | 390.45 | 1 | 4.59 | 1.18 | 19.16 | 19.37 | -0.21 | 0.21 | -0.01 | 0.01 |
| gene8661 | https://www.ncbi.nlm.nih.gov/gene/?term=LOC105282063 | 1926 | 35 | 181.72 | 1 | 5.19 | 2.86 | 30.68 | 30.89 | -0.22 | 0.22 | 0 | 0 |
| gene1199 | https://www.ncbi.nlm.nih.gov/gene/?term=LOC105285498 | 2220 | 26 | 117.12 | 1 | 4.5 | 3.85 | 31.35 | 31.59 | -0.24 | 0.24 | 0 | 0 |
| gene5733 | https://www.ncbi.nlm.nih.gov/gene/?term=LOC105278649 | 5221 | 114 | 218.35 | 1 | 1.92 | 0.88 | 16.23 | 16.48 | -0.25 | 0.25 | -0.01 | 0.01 |
| gene1310 | https://www.ncbi.nlm.nih.gov/gene/?term=LOC105286661 | 2582 | 72 | 278.85 | 1 | 3.87 | 1.39 | 21.23 | 21.48 | -0.26 | 0.26 | -0.01 | 0.01 |
| gene9326 | https://www.ncbi.nlm.nih.gov/gene/?term=LOC105282818 | 16612 | 173 | 104.14 | 1 | 0.6 | 0.58 | 12.61 | 12.88 | -0.27 | 0.27 | 0 | 0 |
| gene12674 | https://www.ncbi.nlm.nih.gov/gene/?term=LOC105286649 | 18283 | 747 | 408.58 | 1 | 0.55 | 0.13 | 0.89 | 1.18 | -0.29 | 0.29 | -0.01 | 0.01 |
| gene13036 | https://www.ncbi.nlm.nih.gov/gene/?term=LOC105287059 | 4959 | 146 | 294.41 | 1 | 2.02 | 0.68 | 18.9 | 19.21 | -0.31 | 0.31 | -0.01 | 0.01 |
| gene3885 | https://www.ncbi.nlm.nih.gov/gene/?term=LOC105276527 | 7928 | 305 | 384.71 | 1 | 1.26 | 0.33 | 9.75 | 10.06 | -0.32 | 0.32 | -0.01 | 0.01 |
| gene6583 | https://www.ncbi.nlm.nih.gov/gene/?term=LOC105279649 | 7444 | 303 | 407.04 | 1 | 1.34 | 0.33 | 11.36 | 11.69 | -0.33 | 0.33 | -0.01 | 0.01 |
| gene10018 | https://www.ncbi.nlm.nih.gov/gene/?term=LOC105283607 | 4414 | 187 | 423.65 | 1 | 2.27 | 0.53 | 16.13 | 16.48 | -0.35 | 0.35 | -0.01 | 0.01 |
| gene2122 | https://www.ncbi.nlm.nih.gov/gene/?term=LOC105274531 | 8153 | 289 | 354.47 | 1 | 1.23 | 0.35 | 10.34 | 10.69 | -0.35 | 0.35 | -0.01 | 0.01 |
| gene6031 | https://www.ncbi.nlm.nih.gov/gene/?term=LOC105279000 | 5691 | 187 | 328.59 | 1 | 1.76 | 0.53 | 16.08 | 16.46 | -0.38 | 0.38 | -0.01 | 0.01 |
| gene12573 | https://www.ncbi.nlm.nih.gov/gene/?term=LOC105286515 | 3647 | 98 | 268.71 | 1 | 2.74 | 1.02 | 13.27 | 13.66 | -0.39 | 0.39 | -0.01 | 0.01 |
| gene12021 | https://www.ncbi.nlm.nih.gov/gene/?term=LOC105285905 | 1399 | 16 | 114.37 | 1 | 7.15 | 6.25 | 47.33 | 47.75 | -0.42 | 0.42 | 0 | 0 |
| gene6639 | https://www.ncbi.nlm.nih.gov/gene/?term=LOC105279705 | 7656 | 61 | 79.68 | 1 | 1.31 | 1.64 | 23.52 | 23.98 | -0.46 | 0.46 | 0 | 0 |
| gene5298 | https://www.ncbi.nlm.nih.gov/gene/?term=LOC105278145 | 4060 | 245 | 603.45 | 2 | 4.93 | 0.82 | 10.68 | 11.19 | -0.51 | 0.51 | -0.03 | 0.03 |
| gene4376 | https://www.ncbi.nlm.nih.gov/gene/?term=LOC105277083 | 4307 | 102 | 236.82 | 1 | 2.32 | 0.98 | 24.57 | 25.16 | -0.58 | 0.58 | -0.01 | 0.01 |
| gene11340 | https://www.ncbi.nlm.nih.gov/gene/?term=LOC105285131 | 2918 | 86 | 294.72 | 1 | 3.43 | 1.16 | 11.93 | 12.54 | -0.61 | 0.61 | -0.02 | 0.02 |
| gene5273 | https://www.ncbi.nlm.nih.gov/gene/?term=LOC105278108 | 4067 | 44 | 108.19 | 1 | 2.46 | 2.27 | 24.76 | 25.36 | -0.61 | 0.61 | -0.01 | 0.01 |
| gene4498 | https://www.ncbi.nlm.nih.gov/gene/?term=LOC105277239 | 2831 | 122 | 430.94 | 1 | 3.53 | 0.82 | 12.06 | 12.7 | -0.64 | 0.64 | -0.03 | 0.03 |
| gene12203 | https://www.ncbi.nlm.nih.gov/gene/?term=LOC105286099 | 5967 | 73 | 122.34 | 1 | 1.68 | 1.37 | 10.19 | 10.87 | -0.68 | 0.68 | -0.01 | 0.01 |
| gene5646 | https://www.ncbi.nlm.nih.gov/gene/?term=LOC105278534 | 7475 | 175 | 234.11 | 1 | 1.34 | 0.57 | 17.8 | 18.5 | -0.7 | 0.7 | -0.02 | 0.02 |
| gene12406 | https://www.ncbi.nlm.nih.gov/gene/?term=LOC105286316 | 1386 | 28 | 202.02 | 1 | 7.22 | 3.57 | 44.18 | 44.91 | -0.73 | 0.73 | -0.01 | 0.01 |
| gene2748 | https://www.ncbi.nlm.nih.gov/gene/?term=LOC105275261 | 3845 | 51 | 132.64 | 1 | 2.6 | 1.96 | 36.93 | 37.66 | -0.74 | 0.74 | -0.01 | 0.01 |
| gene2683 | https://www.ncbi.nlm.nih.gov/gene/?term=LOC105275187 | 5492 | 66 | 120.17 | 1 | 1.82 | 1.52 | 11.92 | 12.72 | -0.8 | 0.8 | -0.01 | 0.01 |
| gene7368 | https://www.ncbi.nlm.nih.gov/gene/?term=LOC105280556 | 4105 | 35 | 85.26 | 1 | 2.44 | 2.86 | 17.18 | 18.02 | -0.84 | 0.84 | -0.01 | 0.01 |
| gene9101 | https://www.ncbi.nlm.nih.gov/gene/?term=LOC105282553 | 3068 | 108 | 352.02 | 1 | 3.26 | 0.93 | 22.57 | 23.41 | -0.84 | 0.84 | -0.03 | 0.03 |
| gene9327 | https://www.ncbi.nlm.nih.gov/gene/?term=LOC105282821 | 7793 | 132 | 169.38 | 1 | 1.28 | 0.76 | 11.87 | 12.7 | -0.84 | 0.84 | -0.01 | 0.01 |
| gene8343 | https://www.ncbi.nlm.nih.gov/gene/?term=LOC105281700 | 2172 | 34 | 156.54 | 1 | 4.6 | 2.94 | 35.91 | 36.82 | -0.9 | 0.9 | -0.01 | 0.01 |
| gene11447 | https://www.ncbi.nlm.nih.gov/gene/?term=LOC105285252 | 7437 | 114 | 153.29 | 2 | 2.69 | 1.75 | 13.96 | 14.93 | -0.97 | 0.97 | -0.01 | 0.01 |
| gene11976 | https://www.ncbi.nlm.nih.gov/gene/?term=LOC105285860 | 3112 | 52 | 167.1 | 1 | 3.21 | 1.92 | 21.02 | 22 | -0.98 | 0.98 | -0.02 | 0.02 |
| gene12516 | https://www.ncbi.nlm.nih.gov/gene/?term=LOC105286452 | 1529 | 40 | 261.61 | 1 | 6.54 | 2.5 | 50.21 | 51.25 | -1.04 | 1.04 | -0.03 | 0.03 |
| gene1147 | https://www.ncbi.nlm.nih.gov/gene/?term=LOC105284954 | 4022 | 84 | 208.85 | 1 | 2.49 | 1.19 | 17.29 | 18.34 | -1.06 | 1.06 | -0.02 | 0.02 |
| gene10391 | https://www.ncbi.nlm.nih.gov/gene/?term=LOC109611283 | 1994 | 42 | 210.63 | 1 | 5.02 | 2.38 | 32.6 | 33.67 | -1.07 | 1.07 | -0.02 | 0.02 |
| gene10155 | https://www.ncbi.nlm.nih.gov/gene/?term=LOC105283756 | 7030 | 100 | 142.25 | 1 | 1.42 | 1 | 5.99 | 7.12 | -1.12 | 1.12 | -0.02 | 0.02 |
| gene1220 | https://www.ncbi.nlm.nih.gov/gene/?term=LOC105285742 | 3453 | 107 | 309.88 | 1 | 2.9 | 0.93 | 22.27 | 23.44 | -1.17 | 1.17 | -0.04 | 0.04 |
| gene10819 | https://www.ncbi.nlm.nih.gov/gene/?term=LOC105284507 | 5767 | 76 | 131.78 | 1 | 1.73 | 1.32 | 23.17 | 24.52 | -1.35 | 1.35 | -0.02 | 0.02 |

| | | | | | | | | | | | | | |
|---|---|---|---|---|---|---|---|---|---|---|---|---|---|
| gene7207 | https://www.ncbi.nlm.nih.gov/gene/?term=LOC105280376 | 4659 | 109 | 233.96 | 1 | 2.15 | 0.92 | 29.99 | 31.37 | -1.38 | 1.38 | -0.03 | 0.03 |
| gene7600 | https://www.ncbi.nlm.nih.gov/gene/?term=LOC105280819 | 1615 | 37 | 229.1 | 1 | 6.19 | 2.7 | 36.56 | 38.16 | -1.6 | 1.6 | -0.04 | 0.04 |
| gene7850 | https://www.ncbi.nlm.nih.gov/gene/?term=LOC105281095 | 3050 | 63 | 206.56 | 1 | 3.28 | 1.59 | 33.9 | 35.57 | -1.67 | 1.67 | -0.03 | 0.03 |
| gene388 | https://www.ncbi.nlm.nih.gov/gene/?term=LOC105276538 | 1558 | 19 | 121.95 | 2 | 12.84 | 10.53 | 63.26 | 65.13 | -1.87 | 1.87 | -0.02 | 0.02 |
| gene2413 | https://www.ncbi.nlm.nih.gov/gene/?term=LOC105274863 | 2006 | 40 | 199.4 | 1 | 4.99 | 2.5 | 64.23 | 66.19 | -1.96 | 1.96 | -0.04 | 0.04 |
| gene920 | https://www.ncbi.nlm.nih.gov/gene/?term=LOC105282518 | 1197 | 39 | 325.81 | 1 | 8.35 | 2.56 | 38.65 | 40.64 | -1.98 | 1.98 | -0.06 | 0.06 |
| gene10213 | https://www.ncbi.nlm.nih.gov/gene/?term=LOC105283826 | 1503 | 54 | 359.28 | 1 | 6.65 | 1.85 | 13.66 | 15.73 | -2.07 | 2.07 | -0.07 | 0.07 |
| gene322 | https://www.ncbi.nlm.nih.gov/gene/?term=LOC105275777 | 1606 | 23 | 143.21 | 1 | 6.23 | 4.35 | 46.95 | 49.13 | -2.17 | 2.17 | -0.03 | 0.03 |
| gene3168 | https://www.ncbi.nlm.nih.gov/gene/?term=LOC105275711 | 6150 | 67 | 108.94 | 1 | 1.63 | 1.49 | 13.09 | 15.36 | -2.27 | 2.27 | -0.02 | 0.02 |
| gene3599 | https://www.ncbi.nlm.nih.gov/gene/?term=LOC105276196 | 2955 | 47 | 159.05 | 1 | 3.38 | 2.13 | 46.53 | 48.84 | -2.31 | 2.31 | -0.04 | 0.04 |
| gene10879 | https://www.ncbi.nlm.nih.gov/gene/?term=LOC105284574 | 2423 | 46 | 189.85 | 1 | 4.13 | 2.17 | 56.45 | 59.11 | -2.65 | 2.65 | -0.05 | 0.05 |
| gene2404 | https://www.ncbi.nlm.nih.gov/gene/?term=LOC105274853 | 1767 | 34 | 192.42 | 1 | 5.66 | 2.94 | 29.24 | 32.2 | -2.96 | 2.96 | -0.06 | 0.06 |
| gene4641 | https://www.ncbi.nlm.nih.gov/gene/?term=LOC105277398 | 1937 | 68 | 351.06 | 1 | 5.16 | 1.47 | 37.35 | 40.44 | -3.09 | 3.09 | -0.11 | 0.11 |
| gene10345 | https://www.ncbi.nlm.nih.gov/gene/?term=LOC105283989 | 4444 | 22 | 49.5 | 1 | 2.25 | 4.55 | 37.81 | 42.87 | -5.06 | 5.06 | -0.03 | 0.03 |
| gene10963 | https://www.ncbi.nlm.nih.gov/gene/?term=LOC105284677 | 1576 | 27 | 171.32 | 3 | 19.04 | 11.11 | 20.06 | 37.04 | -16.98 | 16.98 | -0.29 | 0.29 |
| gene393 | https://www.ncbi.nlm.nih.gov/gene/?term=LOC105276566 | 1123 | 2 | 17.81 | 1 | 8.9 | 50 | 48.08 | 76.78 | -28.71 | 28.71 | -0.05 | 0.05 |

Directory: OGLall File: ogl-Cb_LI_BCphaseB1.txt

| gene_ID | gene_link | gwidth | #Sites | #per10Kb | %perSite | %pNucl |
|---|---|---|---|---|---|---|
| gene12538 | https://www.ncbi.nlm.nih.gov/gene/?term=LOC105286531 | 372 | 43 | 1155.91 | 37.6 | 4.35 |
| gene5911 | https://www.ncbi.nlm.nih.gov/gene/?term=LOC105278844 | 943 | 109 | 1155.89 | 33.35 | 3.86 |
| gene13384 | https://www.ncbi.nlm.nih.gov/gene/?term=LOC105287447 | 1877 | 240 | 1278.64 | 29.63 | 3.79 |
| gene5309 | https://www.ncbi.nlm.nih.gov/gene/?term=LOC105278160 | 1704 | 163 | 956.57 | 39.24 | 3.75 |
| gene5917 | https://www.ncbi.nlm.nih.gov/gene/?term=LOC105278849 | 823 | 115 | 1397.33 | 25.77 | 3.6 |
| gene211 | https://www.ncbi.nlm.nih.gov/gene/?term=LOC105274534 | 1280 | 103 | 804.69 | 41.41 | 3.33 |
| gene4878 | https://www.ncbi.nlm.nih.gov/gene/?term=LOC105277648 | 675 | 40 | 592.59 | 55.76 | 3.3 |
| gene12330 | https://www.ncbi.nlm.nih.gov/gene/?term=LOC105286235 | 1501 | 111 | 739.51 | 44.42 | 3.28 |
| gene1562 | https://www.ncbi.nlm.nih.gov/gene/?term=LOC105287692 | 1491 | 142 | 952.38 | 34.46 | 3.28 |
| gene6078 | https://www.ncbi.nlm.nih.gov/gene/?term=LOC105279052 | 1130 | 139 | 1230.09 | 26.68 | 3.28 |
| gene12746 | https://www.ncbi.nlm.nih.gov/gene/?term=LOC105286737 | 2688 | 393 | 1462.05 | 22.36 | 3.27 |
| gene1654 | https://www.ncbi.nlm.nih.gov/gene/?term=LOC105287811 | 2285 | 165 | 722.1 | 43.88 | 3.17 |
| gene12364 | https://www.ncbi.nlm.nih.gov/gene/?term=LOC105286269 | 1403 | 85 | 605.84 | 51.17 | 3.1 |
| gene12805 | https://www.ncbi.nlm.nih.gov/gene/?term=LOC105286801 | 1379 | 121 | 877.45 | 35.03 | 3.07 |
| gene9471 | https://www.ncbi.nlm.nih.gov/gene/?term=LOC105282986 | 1605 | 135 | 841.12 | 36.19 | 3.04 |
| gene570 | https://www.ncbi.nlm.nih.gov/gene/?term=LOC105278549 | 1673 | 124 | 741.18 | 40.89 | 3.03 |
| gene12652 | https://www.ncbi.nlm.nih.gov/gene/?term=LOC109610640 | 1122 | 63 | 561.5 | 53.52 | 3.01 |
| gene3731 | https://www.ncbi.nlm.nih.gov/gene/?term=LOC105276371 | 865 | 55 | 635.84 | 47.18 | 3 |
| gene3719 | https://www.ncbi.nlm.nih.gov/gene/?term=LOC105276360 | 1471 | 163 | 1108.09 | 26.81 | 2.97 |
| gene7405 | https://www.ncbi.nlm.nih.gov/gene/?term=LOC105280594 | 1263 | 87 | 688.84 | 42.66 | 2.94 |
| gene7527 | https://www.ncbi.nlm.nih.gov/gene/?term=LOC105280739 | 2686 | 159 | 591.96 | 49.69 | 2.94 |
| gene5301 | https://www.ncbi.nlm.nih.gov/gene/?term=LOC105278148 | 1881 | 201 | 1068.58 | 27.41 | 2.93 |
| gene8453 | https://www.ncbi.nlm.nih.gov/gene/?term=LOC105281826 | 1750 | 169 | 965.71 | 30.13 | 2.91 |
| gene4633 | https://www.ncbi.nlm.nih.gov/gene/?term=LOC105277389 | 2426 | 183 | 754.33 | 38.21 | 2.88 |

...
... only first 25 lines shown ...

Directory: OGLall File: ogl-Cb_LI_BCphaseB1.vs.BCphaseB2.txt

| gene_ID | gene_link | gwidth | #Sites | #per10Kb | #dmSites | #dmsp10kb | %dmSites | %pSite1 | %pSite2 | DMpSite | ADMpSite | DMpNucl | ADMpNucl |
|---|---|---|---|---|---|---|---|---|---|---|---|---|---|
| gene2593 | https://www.ncbi.nlm.nih.gov/gene/?term=LOC105275070 | 1774 | 5 | 28.18 | 5 | 28.18 | 100 | 72.92 | 10.96 | 61.96 | 61.96 | 0.17 | 0.17 |
| gene3165 | https://www.ncbi.nlm.nih.gov/gene/?term=LOC105275706 | 2374 | 66 | 278.01 | 5 | 21.06 | 7.58 | 34.05 | 24.01 | 10.05 | 10.05 | 0.28 | 0.28 |
| gene1962 | https://www.ncbi.nlm.nih.gov/gene/?term=LOC105288157 | 1246 | 17 | 136.44 | 1 | 8.03 | 5.88 | 18.51 | 13.14 | 5.37 | 5.37 | 0.07 | 0.07 |
| gene2907 | https://www.ncbi.nlm.nih.gov/gene/?term=LOC105275447 | 2539 | 40 | 157.54 | 1 | 3.94 | 2.5 | 7.22 | 3.04 | 4.18 | 4.18 | 0.07 | 0.07 |
| gene2970 | https://www.ncbi.nlm.nih.gov/gene/?term=LOC105275518 | 2340 | 25 | 106.84 | 1 | 4.27 | 4 | 43.33 | 39.71 | 3.62 | 3.62 | 0.04 | 0.04 |
| gene5714 | https://www.ncbi.nlm.nih.gov/gene/?term=LOC105278729 | 710 | 24 | 338.03 | 1 | 14.08 | 4.17 | 59.89 | 56.79 | 3.1 | 3.1 | 0.1 | 0.1 |
| gene7525 | https://www.ncbi.nlm.nih.gov/gene/?term=LOC105280738 | 4237 | 41 | 96.77 | 1 | 2.36 | 2.44 | 26.13 | 23.59 | 2.54 | 2.54 | 0.02 | 0.02 |
| gene9478 | https://www.ncbi.nlm.nih.gov/gene/?term=LOC105282991 | 3726 | 57 | 152.98 | 1 | 2.68 | 1.75 | 35.88 | 33.61 | 2.27 | 2.27 | 0.03 | 0.03 |
| gene3817 | https://www.ncbi.nlm.nih.gov/gene/?term=LOC105276449 | 1230 | 51 | 414.63 | 1 | 8.13 | 1.96 | 27.91 | 26.06 | 1.85 | 1.85 | 0.08 | 0.08 |
| gene8178 | https://www.ncbi.nlm.nih.gov/gene/?term=LOC105281515 | 8647 | 61 | 70.54 | 1 | 1.16 | 1.64 | 27.1 | 25.25 | 1.85 | 1.85 | 0.01 | 0.01 |
| gene1518 | https://www.ncbi.nlm.nih.gov/gene/?term=LOC105287648 | 2195 | 30 | 136.67 | 1 | 4.56 | 3.33 | 43.53 | 41.69 | 1.83 | 1.83 | 0.03 | 0.03 |
| gene940 | https://www.ncbi.nlm.nih.gov/gene/?term=LOC105282794 | 3578 | 88 | 245.95 | 1 | 2.79 | 1.14 | 16.46 | 14.64 | 1.82 | 1.82 | 0.04 | 0.04 |
| gene7368 | https://www.ncbi.nlm.nih.gov/gene/?term=LOC105280556 | 4105 | 54 | 131.55 | 1 | 2.44 | 1.85 | 18.52 | 16.73 | 1.79 | 1.79 | 0.02 | 0.02 |
| gene1884 | https://www.ncbi.nlm.nih.gov/gene/?term=LOC105288072 | 4169 | 76 | 182.3 | 1 | 2.4 | 1.32 | 24.99 | 23.26 | 1.72 | 1.72 | 0.03 | 0.03 |
| gene5877 | https://www.ncbi.nlm.nih.gov/gene/?term=LOC105278807 | 1719 | 53 | 308.32 | 1 | 5.82 | 1.89 | 23.93 | 22.25 | 1.68 | 1.68 | 0.05 | 0.05 |
| gene5101 | https://www.ncbi.nlm.nih.gov/gene/?term=LOC105277903 | 1813 | 37 | 204.08 | 1 | 5.52 | 2.7 | 54.27 | 52.77 | 1.5 | 1.5 | 0.03 | 0.03 |
| gene4892 | https://www.ncbi.nlm.nih.gov/gene/?term=LOC105277667 | 3898 | 127 | 325.81 | 1 | 2.57 | 0.79 | 27.62 | 26.17 | 1.45 | 1.45 | 0.05 | 0.05 |
| gene12572 | https://www.ncbi.nlm.nih.gov/gene/?term=LOC105286514 | 2374 | 46 | 193.77 | 1 | 4.21 | 2.17 | 36.88 | 35.48 | 1.4 | 1.4 | 0.03 | 0.03 |
| gene35 | https://www.ncbi.nlm.nih.gov/gene/?term=LOC105275197 | 2643 | 50 | 189.18 | 1 | 3.78 | 2 | 38.68 | 37.33 | 1.34 | 1.34 | 0.03 | 0.03 |
| gene10345 | https://www.ncbi.nlm.nih.gov/gene/?term=LOC105283989 | 4444 | 29 | 65.26 | 1 | 2.25 | 3.45 | 49.14 | 47.94 | 1.2 | 1.2 | 0.01 | 0.01 |
| gene10678 | https://www.ncbi.nlm.nih.gov/gene/?term=LOC105284355 | 4031 | 174 | 431.65 | 1 | 2.48 | 0.57 | 19.29 | 18.18 | 1.11 | 1.11 | 0.05 | 0.05 |
| gene5967 | https://www.ncbi.nlm.nih.gov/gene/?term=LOC105278918 | 4393 | 146 | 332.35 | 1 | 2.28 | 0.68 | 13.82 | 12.77 | 1.06 | 1.06 | 0.04 | 0.04 |
| gene7568 | https://www.ncbi.nlm.nih.gov/gene/?term=LOC105280786 | 2525 | 119 | 471.29 | 1 | 3.96 | 0.84 | 30.86 | 29.84 | 1.03 | 1.03 | 0.05 | 0.05 |
| gene7888 | https://www.ncbi.nlm.nih.gov/gene/?term=LOC105281141 | 4357 | 164 | 376.41 | 1 | 2.3 | 0.61 | 29.56 | 28.67 | 0.9 | 0.9 | 0.03 | 0.03 |
| gene2856 | https://www.ncbi.nlm.nih.gov/gene/?term=LOC105275384 | 1987 | 92 | 463.01 | 1 | 5.03 | 1.09 | 32.79 | 31.9 | 0.89 | 0.89 | 0.04 | 0.04 |
| gene927 | https://www.ncbi.nlm.nih.gov/gene/?term=LOC105282600 | 3727 | 50 | 134.16 | 1 | 2.68 | 2 | 33.71 | 32.83 | 0.88 | 0.88 | 0.01 | 0.01 |
| gene13033 | https://www.ncbi.nlm.nih.gov/gene/?term=LOC105287063 | 1877 | 53 | 282.37 | 1 | 5.33 | 1.89 | 30.73 | 29.86 | 0.87 | 0.87 | 0.02 | 0.02 |
| gene4720 | https://www.ncbi.nlm.nih.gov/gene/?term=LOC105277479 | 1697 | 40 | 235.71 | 1 | 5.89 | 2.5 | 43.85 | 42.99 | 0.86 | 0.86 | 0.02 | 0.02 |
| gene12516 | https://www.ncbi.nlm.nih.gov/gene/?term=LOC105286452 | 1529 | 37 | 241.99 | 1 | 6.54 | 2.7 | 49.81 | 49.09 | 0.73 | 0.73 | 0.02 | 0.02 |
| gene292 | https://www.ncbi.nlm.nih.gov/gene/?term=LOC105275398 | 2825 | 85 | 300.88 | 1 | 3.54 | 1.18 | 21.01 | 20.28 | 0.73 | 0.73 | 0.02 | 0.02 |
| gene8742 | https://www.ncbi.nlm.nih.gov/gene/?term=LOC105282151 | 2563 | 44 | 171.67 | 2 | 7.8 | 4.55 | 36.03 | 35.31 | 0.72 | 0.72 | 0.01 | 0.01 |
| gene9336 | https://www.ncbi.nlm.nih.gov/gene/?term=LOC105282834 | 5993 | 129 | 215.25 | 1 | 1.67 | 0.78 | 15.82 | 15.16 | 0.66 | 0.66 | 0.01 | 0.01 |
| gene10540 | https://www.ncbi.nlm.nih.gov/gene/?term=LOC105284200 | 5267 | 204 | 387.32 | 1 | 1.9 | 0.49 | 12.98 | 12.34 | 0.64 | 0.64 | 0.02 | 0.02 |
| gene11401 | https://www.ncbi.nlm.nih.gov/gene/?term=LOC105285202 | 1841 | 77 | 418.25 | 1 | 5.43 | 1.3 | 24.18 | 23.56 | 0.62 | 0.62 | 0.03 | 0.03 |
| gene12439 | https://www.ncbi.nlm.nih.gov/gene/?term=LOC105286357 | 4761 | 86 | 180.63 | 1 | 2.1 | 1.16 | 26.67 | 26.07 | 0.6 | 0.6 | 0.01 | 0.01 |
| gene34 | https://www.ncbi.nlm.nih.gov/gene/?term=LOC105278416 | 7120 | 146 | 205.06 | 1 | 1.4 | 0.68 | 13.63 | 13.03 | 0.6 | 0.6 | 0.01 | 0.01 |
| gene5654 | https://www.ncbi.nlm.nih.gov/gene/?term=LOC105278543 | 4019 | 160 | 398.11 | 1 | 2.49 | 0.62 | 15.95 | 15.36 | 0.6 | 0.6 | 0.02 | 0.02 |
| gene6583 | https://www.ncbi.nlm.nih.gov/gene/?term=LOC105279649 | 7444 | 350 | 470.18 | 1 | 1.34 | 0.29 | 11.44 | 10.85 | 0.59 | 0.59 | 0.03 | 0.03 |
| gene2683 | https://www.ncbi.nlm.nih.gov/gene/?term=LOC105275187 | 5492 | 75 | 136.56 | 1 | 1.82 | 1.33 | 15.98 | 15.4 | 0.58 | 0.58 | 0.01 | 0.01 |
| gene4023 | https://www.ncbi.nlm.nih.gov/gene/?term=LOC105276692 | 1759 | 37 | 210.35 | 1 | 5.69 | 2.7 | 20.35 | 19.78 | 0.57 | 0.57 | 0.01 | 0.01 |
| gene7884 | https://www.ncbi.nlm.nih.gov/gene/?term=LOC105281266 | 3390 | 79 | 233.04 | 1 | 2.95 | 1.27 | 16.01 | 15.44 | 0.57 | 0.57 | 0.01 | 0.01 |
| gene4376 | https://www.ncbi.nlm.nih.gov/gene/?term=LOC105277083 | 4307 | 129 | 299.51 | 1 | 2.32 | 0.78 | 19.34 | 18.78 | 0.56 | 0.56 | 0.02 | 0.02 |
| gene1855 | https://www.ncbi.nlm.nih.gov/gene/?term=LOC105288036 | 2387 | 86 | 360.28 | 1 | 4.19 | 1.16 | 29.99 | 29.44 | 0.55 | 0.55 | 0.02 | 0.02 |
| gene3107 | https://www.ncbi.nlm.nih.gov/gene/?term=LOC105275650 | 5103 | 178 | 348.81 | 1 | 1.96 | 0.56 | 16.67 | 16.15 | 0.52 | 0.52 | 0.02 | 0.02 |
| gene3206 | https://www.ncbi.nlm.nih.gov/gene/?term=LOC105275752 | 3474 | 129 | 371.33 | 1 | 2.88 | 0.78 | 32.13 | 31.62 | 0.52 | 0.52 | 0.02 | 0.02 |
| gene1332 | https://www.ncbi.nlm.nih.gov/gene/?term=LOC105286872 | 2460 | 158 | 642.28 | 1 | 4.07 | 0.63 | 20.79 | 20.28 | 0.51 | 0.51 | 0.03 | 0.03 |
| gene2710 | https://www.ncbi.nlm.nih.gov/gene/?term=LOC105275218 | 3222 | 98 | 304.16 | 1 | 3.1 | 1.02 | 22.4 | 21.9 | 0.5 | 0.5 | 0.02 | 0.02 |
| gene2668 | https://www.ncbi.nlm.nih.gov/gene/?term=LOC105275170 | 12496 | 301 | 240.88 | 1 | 0.8 | 0.33 | 2.32 | 1.82 | 0.49 | 0.49 | 0.01 | 0.01 |
| gene6966 | https://www.ncbi.nlm.nih.gov/gene/?term=LOC105280104 | 3747 | 133 | 354.95 | 1 | 2.67 | 0.75 | 15.81 | 15.32 | 0.49 | 0.49 | 0.02 | 0.02 |
| gene9445 | https://www.ncbi.nlm.nih.gov/gene/?term=LOC105282951 | 2285 | 115 | 503.28 | 1 | 4.38 | 0.87 | 20.4 | 19.9 | 0.49 | 0.49 | 0.02 | 0.02 |
| gene738 | https://www.ncbi.nlm.nih.gov/gene/?term=LOC105280421 | 4927 | 222 | 450.58 | 1 | 2.03 | 0.45 | 14.83 | 14.35 | 0.48 | 0.48 | 0.02 | 0.02 |
| gene8729 | https://www.ncbi.nlm.nih.gov/gene/?term=LOC105282137 | 5005 | 202 | 403.6 | 1 | 2 | 0.5 | 16.8 | 16.33 | 0.46 | 0.46 | 0.02 | 0.02 |
| gene3470 | https://www.ncbi.nlm.nih.gov/gene/?term=LOC105276059 | 8716 | 164 | 188.16 | 1 | 1.15 | 0.61 | 7.87 | 7.47 | 0.4 | 0.4 | 0.01 | 0.01 |
| gene11454 | https://www.ncbi.nlm.nih.gov/gene/?term=LOC105285256 | 1906 | 51 | 267.58 | 1 | 5.25 | 1.96 | 42.4 | 42.03 | 0.37 | 0.37 | 0.01 | 0.01 |
| gene5831 | https://www.ncbi.nlm.nih.gov/gene/?term=LOC105278761 | 2506 | 126 | 502.79 | 1 | 3.99 | 0.79 | 21.41 | 21.05 | 0.36 | 0.36 | 0.02 | 0.02 |
| gene6620 | https://www.ncbi.nlm.nih.gov/gene/?term=LOC105279687 | 17998 | 384 | 213.36 | 1 | 0.56 | 0.26 | 4.83 | 4.47 | 0.36 | 0.36 | 0.01 | 0.01 |
| gene5375 | https://www.ncbi.nlm.nih.gov/gene/?term=LOC105278228 | 8640 | 173 | 200.23 | 1 | 1.16 | 0.58 | 23.79 | 23.44 | 0.35 | 0.35 | 0.01 | 0.01 |
| gene12802 | https://www.ncbi.nlm.nih.gov/gene/?term=LOC105286799 | 3183 | 189 | 593.78 | 1 | 3.14 | 0.53 | 15.7 | 15.39 | 0.32 | 0.32 | 0.02 | 0.02 |
| gene2844 | https://www.ncbi.nlm.nih.gov/gene/?term=LOC105275363 | 4330 | 206 | 475.75 | 1 | 2.31 | 0.49 | 14.15 | 13.85 | 0.3 | 0.3 | 0.01 | 0.01 |
| gene11568 | https://www.ncbi.nlm.nih.gov/gene/?term=LOC105285395 | 8086 | 204 | 252.29 | 1 | 1.24 | 0.49 | 14.79 | 14.52 | 0.27 | 0.27 | 0.01 | 0.01 |

| | | | | | | | | | | | | | |
|---|---|---|---|---|---|---|---|---|---|---|---|---|---|
| gene1310 | https://www.ncbi.nlm.nih.gov/gene/?term=LOC105286661 | 2582 | 65 | 251.74 | 1 | 3.87 | 1.54 | 23.22 | 22.96 | 0.26 | 0.26 | 0.01 | 0.01 |
| gene10531 | https://www.ncbi.nlm.nih.gov/gene/?term=LOC105284190 | 6881 | 195 | 283.39 | 1 | 1.45 | 0.51 | 11.74 | 11.5 | 0.24 | 0.24 | 0.01 | 0.01 |
| gene11579 | https://www.ncbi.nlm.nih.gov/gene/?term=LOC105285406 | 13571 | 291 | 214.43 | 1 | 0.74 | 0.34 | 6.38 | 6.14 | 0.24 | 0.24 | 0.01 | 0.01 |
| gene10844 | https://www.ncbi.nlm.nih.gov/gene/?term=LOC105284535 | 9086 | 256 | 281.75 | 1 | 1.1 | 0.39 | 14.59 | 14.36 | 0.23 | 0.23 | 0.01 | 0.01 |
| gene2806 | https://www.ncbi.nlm.nih.gov/gene/?term=LOC105275327 | 14257 | 542 | 380.16 | 1 | 0.7 | 0.18 | 6.63 | 6.42 | 0.21 | 0.21 | 0.01 | 0.01 |
| gene3599 | https://www.ncbi.nlm.nih.gov/gene/?term=LOC105276196 | 2955 | 59 | 199.66 | 1 | 3.38 | 1.69 | 37.86 | 37.72 | 0.15 | 0.15 | 0 | 0 |
| gene53 | https://www.ncbi.nlm.nih.gov/gene/?term=LOC105277148 | 19283 | 324 | 168.02 | 1 | 0.52 | 0.31 | 4.91 | 4.75 | 0.15 | 0.15 | 0 | 0 |
| gene9462 | https://www.ncbi.nlm.nih.gov/gene/?term=LOC105282970 | 1469 | 42 | 285.91 | 1 | 6.81 | 2.38 | 51.11 | 50.96 | 0.15 | 0.15 | 0 | 0 |
| gene3885 | https://www.ncbi.nlm.nih.gov/gene/?term=LOC105276527 | 7928 | 334 | 421.29 | 1 | 1.26 | 0.3 | 9.88 | 9.74 | 0.14 | 0.14 | 0.01 | 0.01 |
| gene5539 | https://www.ncbi.nlm.nih.gov/gene/?term=LOC105278415 | 25024 | 720 | 287.72 | 1 | 0.4 | 0.14 | 3.22 | 3.07 | 0.14 | 0.14 | 0 | 0 |
| gene7183 | https://www.ncbi.nlm.nih.gov/gene/?term=LOC105280348 | 3925 | 76 | 193.63 | 1 | 2.55 | 1.32 | 19.42 | 19.29 | 0.13 | 0.13 | 0 | 0 |
| gene862 | https://www.ncbi.nlm.nih.gov/gene/?term=LOC105281802 | 73922 | 2171 | 293.69 | 1 | 0.14 | 0.05 | 2.2 | 2.07 | 0.13 | 0.13 | 0 | 0 |
| gene12307 | https://www.ncbi.nlm.nih.gov/gene/?term=LOC105286216 | 10313 | 174 | 168.72 | 1 | 0.97 | 0.57 | 15.92 | 15.81 | 0.12 | 0.12 | 0 | 0 |
| gene5184 | https://www.ncbi.nlm.nih.gov/gene/?term=LOC105277999 | 5528 | 146 | 264.11 | 1 | 1.81 | 0.68 | 11.67 | 11.57 | 0.1 | 0.1 | 0 | 0 |
| gene5746 | https://www.ncbi.nlm.nih.gov/gene/?term=LOC105278666 | 10343 | 482 | 466.02 | 1 | 0.97 | 0.21 | 6.71 | 6.61 | 0.1 | 0.1 | 0 | 0 |
| gene12573 | https://www.ncbi.nlm.nih.gov/gene/?term=LOC105286515 | 3647 | 103 | 282.42 | 1 | 2.74 | 0.97 | 10.56 | 10.47 | 0.09 | 0.09 | 0 | 0 |
| gene8343 | https://www.ncbi.nlm.nih.gov/gene/?term=LOC105281700 | 2172 | 37 | 170.35 | 1 | 4.6 | 2.7 | 39.73 | 39.63 | 0.09 | 0.09 | 0 | 0 |
| gene298 | https://www.ncbi.nlm.nih.gov/gene/?term=LOC105275460 | 2473 | 122 | 493.33 | 1 | 4.04 | 0.82 | 13.15 | 13.07 | 0.08 | 0.08 | 0 | 0 |
| gene10155 | https://www.ncbi.nlm.nih.gov/gene/?term=LOC105283756 | 7030 | 109 | 155.05 | 1 | 1.42 | 0.92 | 5.36 | 5.29 | 0.07 | 0.07 | 0 | 0 |
| gene1632 | https://www.ncbi.nlm.nih.gov/gene/?term=LOC105287764 | 18061 | 406 | 224.79 | 2 | 1.11 | 0.49 | 3.27 | 3.2 | 0.07 | 0.07 | 0 | 0 |
| gene7319 | https://www.ncbi.nlm.nih.gov/gene/?term=LOC105280513 | 68997 | 1376 | 199.43 | 1 | 0.14 | 0.07 | 0.74 | 0.67 | 0.07 | 0.07 | 0 | 0 |
| gene9326 | https://www.ncbi.nlm.nih.gov/gene/?term=LOC105282818 | 16612 | 217 | 130.63 | 1 | 0.6 | 0.46 | 11.57 | 11.5 | 0.07 | 0.07 | 0 | 0 |
| gene10596 | https://www.ncbi.nlm.nih.gov/gene/?term=LOC105284262 | 6520 | 234 | 358.9 1 | 1.53 | 0.43 | 3.38 | 3.34 | 0.04 | 0.04 | 0 | 0 | |
| gene2056 | https://www.ncbi.nlm.nih.gov/gene/?term=LOC105274452 | 6584 | 263 | 399.45 | 2 | 3.04 | 0.76 | 12.97 | 12.95 | 0.02 | 0.02 | 0 | 0 |
| gene5197 | https://www.ncbi.nlm.nih.gov/gene/?term=LOC105278018 | 89233 | 3838 | 430.11 | 1 | 0.11 | 0.03 | 0.67 | 0.66 | 0.01 | 0.01 | 0 | 0 |
| gene7398 | https://www.ncbi.nlm.nih.gov/gene/?term=LOC105280589 | 3452 | 37 | 107.18 | 1 | 2.9 | 2.7 | 19.84 | 19.83 | 0.01 | 0.01 | 0 | 0 |
| gene10018 | https://www.ncbi.nlm.nih.gov/gene/?term=LOC105283607 | 4414 | 208 | 471.23 | 1 | 2.27 | 0.48 | 15.08 | 15.07 | 0 | 0 | 0 | 0 |
| gene1430 | https://www.ncbi.nlm.nih.gov/gene/?term=LOC105287551 | 6433 | 232 | 360.64 | 1 | 1.55 | 0.43 | 8.43 | 8.44 | 0 | 0 | 0 | 0 |
| gene11275 | https://www.ncbi.nlm.nih.gov/gene/?term=LOC105285055 | 4119 | 248 | 602.09 | 1 | 2.43 | 0.4 | 12.13 | 12.15 | -0.02 | 0.02 | 0 | 0 |
| gene13114 | https://www.ncbi.nlm.nih.gov/gene/?term=LOC105287147 | 7609 | 430 | 565.12 | 1 | 1.31 | 0.23 | 9.08 | 9.1 | -0.02 | 0.02 | 0 | 0 |
| gene3856 | https://www.ncbi.nlm.nih.gov/gene/?term=LOC105276499 | 5236 | 194 | 370.51 | 1 | 1.91 | 0.52 | 10.61 | 10.65 | -0.04 | 0.04 | 0 | 0 |
| gene9327 | https://www.ncbi.nlm.nih.gov/gene/?term=LOC105282821 | 7793 | 173 | 221.99 | 1 | 1.28 | 0.58 | 10.79 | 10.87 | -0.08 | 0.08 | 0 | 0 |
| gene7166 | https://www.ncbi.nlm.nih.gov/gene/?term=LOC105280326 | 6117 | 253 | 413.6 1 | 1.63 | 0.4 | 14.52 | 14.62 | -0.1 | 0.1 | 0 | 0 | |
| gene10886 | https://www.ncbi.nlm.nih.gov/gene/?term=LOC105284582 | 1475 | 47 | 318.64 | 1 | 6.78 | 2.13 | 22.72 | 22.83 | -0.11 | 0.11 | 0 | 0 |
| gene6528 | https://www.ncbi.nlm.nih.gov/gene/?term=LOC105279591 | 7222 | 59 | 81.69 1 | 1.38 | 1.69 | 26.07 | 26.22 | -0.14 | 0.14 | 0 | 0 | |
| gene2633 | https://www.ncbi.nlm.nih.gov/gene/?term=LOC105275127 | 7457 | 270 | 362.08 | 1 | 1.34 | 0.37 | 7.9 | 8.07 | -0.16 | 0.16 | -0.01 | 0.01 |
| gene2173 | https://www.ncbi.nlm.nih.gov/gene/?term=LOC105274590 | 4387 | 84 | 191.47 | 2 | 4.56 | 2.38 | 27.83 | 27.99 | -0.17 | 0.17 | 0 | 0 |
| gene11561 | https://www.ncbi.nlm.nih.gov/gene/?term=LOC105285387 | 11528 | 202 | 175.23 | 1 | 0.87 | 0.5 | 8.85 | 9.03 | -0.18 | 0.18 | 0 | 0 |
| gene7514 | https://www.ncbi.nlm.nih.gov/gene/?term=LOC105280723 | 4471 | 210 | 469.69 | 1 | 2.24 | 0.48 | 13.2 | 13.38 | -0.18 | 0.18 | -0.01 | 0.01 |
| gene10549 | https://www.ncbi.nlm.nih.gov/gene/?term=LOC105284210 | 7145 | 340 | 475.86 | 2 | 2.8 | 0.59 | 9.99 | 10.19 | -0.2 | 0.2 | -0.01 | 0.01 |
| gene8814 | https://www.ncbi.nlm.nih.gov/gene/?term=LOC105282240 | 6910 | 198 | 286.54 | 1 | 1.45 | 0.51 | 8.79 | 9 | -0.21 | 0.21 | -0.01 | 0.01 |
| gene2846 | https://www.ncbi.nlm.nih.gov/gene/?term=LOC105275374 | 11246 | 512 | 455.27 | 1 | 0.89 | 0.2 | 9.81 | 10.04 | -0.23 | 0.23 | -0.01 | 0.01 |
| gene10213 | https://www.ncbi.nlm.nih.gov/gene/?term=LOC105283826 | 1503 | 65 | 432.47 | 1 | 6.65 | 1.54 | 15.16 | 15.4 | -0.24 | 0.24 | -0.01 | 0.01 |
| gene13036 | https://www.ncbi.nlm.nih.gov/gene/?term=LOC105287059 | 4959 | 176 | 354.91 | 1 | 2.02 | 0.57 | 16.13 | 16.41 | -0.28 | 0.28 | -0.01 | 0.01 |
| gene1470 | https://www.ncbi.nlm.nih.gov/gene/?term=LOC105287597 | 2974 | 204 | 685.94 | 1 | 3.36 | 0.49 | 17.06 | 17.34 | -0.28 | 0.28 | -0.02 | 0.02 |
| gene10392 | https://www.ncbi.nlm.nih.gov/gene/?term=LOC105284036 | 3216 | 106 | 329.6 2 | 6.22 | 1.89 | 20.71 | 21 | -0.29 | 0.29 | -0.01 | 0.01 | |
| gene10520 | https://www.ncbi.nlm.nih.gov/gene/?term=LOC105284184 | 2451 | 101 | 412.08 | 1 | 4.08 | 0.99 | 29.13 | 29.43 | -0.3 | 0.3 | -0.01 | 0.01 |
| gene11082 | https://www.ncbi.nlm.nih.gov/gene/?term=LOC105284824 | 7436 | 214 | 287.79 | 1 | 1.34 | 0.47 | 6.56 | 6.86 | -0.31 | 0.31 | -0.01 | 0.01 |
| gene3046 | https://www.ncbi.nlm.nih.gov/gene/?term=LOC105275587 | 2394 | 55 | 229.74 | 1 | 4.18 | 1.82 | 36.59 | 36.9 | -0.31 | 0.31 | -0.01 | 0.01 |
| gene1907 | https://www.ncbi.nlm.nih.gov/gene/?term=LOC105288099 | 1238 | 36 | 290.79 | 1 | 8.08 | 2.78 | 33.72 | 34.04 | -0.32 | 0.32 | -0.01 | 0.01 |
| gene5777 | https://www.ncbi.nlm.nih.gov/gene/?term=LOC105278691 | 3883 | 168 | 432.66 | 1 | 2.58 | 0.6 | 14.66 | 14.98 | -0.33 | 0.33 | -0.01 | 0.01 |
| gene10600 | https://www.ncbi.nlm.nih.gov/gene/?term=LOC105284270 | 4837 | 124 | 256.36 | 1 | 2.07 | 0.81 | 11.8 | 12.15 | -0.35 | 0.35 | -0.01 | 0.01 |
| gene12203 | https://www.ncbi.nlm.nih.gov/gene/?term=LOC105286099 | 5967 | 75 | 125.69 | 1 | 1.68 | 1.33 | 9.95 | 10.3 | -0.35 | 0.35 | 0 | 0 |
| gene6031 | https://www.ncbi.nlm.nih.gov/gene/?term=LOC105279000 | 5691 | 197 | 346.16 | 1 | 1.76 | 0.51 | 12.37 | 12.73 | -0.35 | 0.35 | -0.01 | 0.01 |
| gene5524 | https://www.ncbi.nlm.nih.gov/gene/?term=LOC105278397 | 5177 | 205 | 395.98 | 1 | 1.93 | 0.49 | 24.95 | 25.35 | -0.39 | 0.39 | -0.02 | 0.02 |
| gene6286 | https://www.ncbi.nlm.nih.gov/gene/?term=LOC105279298 | 2861 | 133 | 464.87 | 2 | 6.99 | 1.5 | 23.95 | 24.34 | -0.39 | 0.39 | -0.02 | 0.02 |
| gene4190 | https://www.ncbi.nlm.nih.gov/gene/?term=LOC105276875 | 3150 | 123 | 390.48 | 1 | 3.17 | 0.81 | 20.99 | 21.41 | -0.42 | 0.42 | -0.02 | 0.02 |
| gene570 | https://www.ncbi.nlm.nih.gov/gene/?term=LOC105278549 | 1673 | 71 | 424.39 | 1 | 5.98 | 1.41 | 43.51 | 43.92 | -0.42 | 0.42 | -0.02 | 0.02 |
| gene12806 | https://www.ncbi.nlm.nih.gov/gene/?term=LOC105286803 | 4196 | 160 | 381.32 | 1 | 2.38 | 0.62 | 9.57 | 10.02 | -0.45 | 0.45 | -0.02 | 0.02 |
| gene8274 | https://www.ncbi.nlm.nih.gov/gene/?term=LOC105281624 | 5425 | 236 | 435.02 | 1 | 1.84 | 0.42 | 9.29 | 9.74 | -0.45 | 0.45 | -0.02 | 0.02 |
| gene10326 | https://www.ncbi.nlm.nih.gov/gene/?term=LOC105283962 | 3042 | 132 | 433.93 | 1 | 3.29 | 0.76 | 8.33 | 8.79 | -0.46 | 0.46 | -0.02 | 0.02 |
| gene2413 | https://www.ncbi.nlm.nih.gov/gene/?term=LOC105274863 | 2006 | 54 | 269.19 | 1 | 4.99 | 1.85 | 52.24 | 52.76 | -0.51 | 0.51 | -0.01 | 0.01 |
| gene4962 | https://www.ncbi.nlm.nih.gov/gene/?term=LOC105277761 | 4645 | 151 | 325.08 | 1 | 2.15 | 0.66 | 14.59 | 15.1 | -0.51 | 0.51 | -0.02 | 0.02 |
| gene388 | https://www.ncbi.nlm.nih.gov/gene/?term=LOC105276538 | 1558 | 19 | 121.95 | 2 | 12.84 | 10.53 | 66.25 | 66.78 | -0.53 | 0.53 | -0.01 | 0.01 |

| | | | | | | | | | | | | | | |
|---|---|---|---|---|---|---|---|---|---|---|---|---|---|---|
| gene2748 | https://www.ncbi.nlm.nih.gov/gene/?term=LOC105275261 | 3845 | 55 | 143.04 | 1 | 2.6 | 1.82 | 38.33 | 38.87 | -0.54 | 0.54 | -0.01 | 0.01 | |
| gene10606 | https://www.ncbi.nlm.nih.gov/gene/?term=LOC105284273 | 6119 | 264 | 431.44 | 1 | 1.63 | 0.38 | 18.32 | 18.9 | -0.57 | 0.57 | -0.02 | 0.02 | |
| gene5525 | https://www.ncbi.nlm.nih.gov/gene/?term=LOC105278396 | 3090 | 189 | 611.65 | 1 | 3.24 | 0.53 | 24.57 | 25.15 | -0.58 | 0.58 | -0.04 | 0.04 | |
| gene11583 | https://www.ncbi.nlm.nih.gov/gene/?term=LOC105285414 | 8192 | 184 | 224.61 | 1 | 1.22 | 0.54 | 8.68 | 9.27 | -0.59 | 0.59 | -0.01 | 0.01 | |
| gene9101 | https://www.ncbi.nlm.nih.gov/gene/?term=LOC105282553 | 3068 | 108 | 352.02 | 1 | 3.26 | 0.93 | 22.97 | 23.62 | -0.64 | 0.64 | -0.02 | 0.02 | |
| gene4034 | https://www.ncbi.nlm.nih.gov/gene/?term=LOC105276704 | 2065 | 103 | 498.79 | 2 | 9.69 | 1.94 | 12.96 | 13.6 | -0.65 | 0.65 | -0.03 | 0.03 | |
| gene7249 | https://www.ncbi.nlm.nih.gov/gene/?term=LOC105280422 | 1775 | 62 | 349.3 1 | 5.63 | 1.61 | 17.31 | 17.96 | -0.66 | 0.66 | -0.02 | 0.02 | | |
| gene6639 | https://www.ncbi.nlm.nih.gov/gene/?term=LOC105279705 | 7656 | 70 | 91.43 1 | 1.31 | 1.43 | 24.79 | 25.46 | -0.67 | 0.67 | -0.01 | 0.01 | | |
| gene5646 | https://www.ncbi.nlm.nih.gov/gene/?term=LOC105278534 | 7475 | 195 | 260.87 | 1 | 1.34 | 0.51 | 17.04 | 17.72 | -0.68 | 0.68 | -0.02 | 0.02 | |
| gene322 | https://www.ncbi.nlm.nih.gov/gene/?term=LOC105275777 | 1606 | 27 | 168.12 | 1 | 6.23 | 3.7 | 40.46 | 41.2 | -0.74 | 0.74 | -0.01 | 0.01 | |
| gene2404 | https://www.ncbi.nlm.nih.gov/gene/?term=LOC105274853 | 1767 | 38 | 215.05 | 1 | 5.66 | 2.63 | 34.09 | 34.89 | -0.8 | 0.8 | -0.02 | 0.02 | |
| gene11688 | https://www.ncbi.nlm.nih.gov/gene/?term=LOC105285533 | 1637 | 70 | 427.61 | 1 | 6.11 | 1.43 | 22.69 | 23.5 | -0.81 | 0.81 | -0.03 | 0.03 | |
| gene10107 | https://www.ncbi.nlm.nih.gov/gene/?term=LOC105283712 | 1883 | 36 | 191.18 | 1 | 5.31 | 2.78 | 28.27 | 29.18 | -0.91 | 0.91 | -0.02 | 0.02 | |
| gene3168 | https://www.ncbi.nlm.nih.gov/gene/?term=LOC105275711 | 6150 | 75 | 121.95 | 1 | 1.63 | 1.33 | 12.84 | 13.77 | -0.93 | 0.93 | -0.01 | 0.01 | |
| gene3793 | https://www.ncbi.nlm.nih.gov/gene/?term=LOC105276429 | 2901 | 110 | 379.18 | 1 | 3.45 | 0.91 | 25.42 | 26.35 | -0.93 | 0.93 | -0.04 | 0.04 | |
| gene7247 | https://www.ncbi.nlm.nih.gov/gene/?term=LOC105280407 | 10486 | 212 | 202.17 | 1 | 0.95 | 0.47 | 15.9 | 16.83 | -0.93 | 0.93 | -0.02 | 0.02 | |
| gene7199 | https://www.ncbi.nlm.nih.gov/gene/?term=LOC105280366 | 1414 | 32 | 226.31 | 1 | 7.07 | 3.12 | 42.1 | 43.06 | -0.96 | 0.96 | -0.02 | 0.02 | |
| gene6652 | https://www.ncbi.nlm.nih.gov/gene/?term=LOC105279719 | 2831 | 85 | 300.25 | 1 | 3.53 | 1.18 | 22.31 | 23.3 | -0.99 | 0.99 | -0.03 | 0.03 | |
| gene11340 | https://www.ncbi.nlm.nih.gov/gene/?term=LOC105285131 | 2918 | 105 | 359.84 | 1 | 3.43 | 0.95 | 13.12 | 14.12 | -1 | 1 | -0.04 | 0.04 | |
| gene1147 | https://www.ncbi.nlm.nih.gov/gene/?term=LOC105284954 | 4022 | 114 | 283.44 | 1 | 2.49 | 0.88 | 13.79 | 14.79 | -1 | 1 | -0.03 | 0.03 | |
| gene5298 | https://www.ncbi.nlm.nih.gov/gene/?term=LOC105278145 | 4060 | 236 | 581.28 | 2 | 4.93 | 0.85 | 10.36 | 11.46 | -1.1 | 1.1 | -0.06 | 0.06 | |
| gene9740 | https://www.ncbi.nlm.nih.gov/gene/?term=LOC105283277 | 2887 | 140 | 484.93 | 1 | 3.46 | 0.71 | 19.86 | 21.12 | -1.26 | 1.26 | -0.06 | 0.06 | |
| gene920 | https://www.ncbi.nlm.nih.gov/gene/?term=LOC105282518 | 1197 | 42 | 350.88 | 1 | 8.35 | 2.38 | 37.02 | 38.3 | -1.28 | 1.28 | -0.04 | 0.04 | |
| gene8015 | https://www.ncbi.nlm.nih.gov/gene/?term=LOC105281312 | 2998 | 65 | 216.81 | 1 | 3.34 | 1.54 | 31.99 | 33.36 | -1.36 | 1.36 | -0.03 | 0.03 | |
| gene12406 | https://www.ncbi.nlm.nih.gov/gene/?term=LOC105286316 | 1386 | 32 | 230.88 | 1 | 7.22 | 3.12 | 45.77 | 47.17 | -1.4 | 1.4 | -0.03 | 0.03 | |
| gene1220 | https://www.ncbi.nlm.nih.gov/gene/?term=LOC105285742 | 3453 | 125 | 362 1 | 2.9 | 0.8 | 19.11 | 20.58 | -1.47 | 1.47 | -0.05 | 0.05 | | |
| gene7915 | https://www.ncbi.nlm.nih.gov/gene/?term=LOC105281173 | 1597 | 38 | 237.95 | 1 | 6.26 | 2.63 | 59.22 | 60.79 | -1.57 | 1.57 | -0.04 | 0.04 | |
| gene13235 | https://www.ncbi.nlm.nih.gov/gene/?term=LOC105287278 | 2679 | 53 | 197.84 | 1 | 3.73 | 1.89 | 24.79 | 26.36 | -1.58 | 1.58 | -0.03 | 0.03 | |
| gene2271 | https://www.ncbi.nlm.nih.gov/gene/?term=LOC105274706 | 1448 | 51 | 352.21 | 1 | 6.91 | 1.96 | 40.05 | 41.71 | -1.67 | 1.67 | -0.06 | 0.06 | |
| gene4498 | https://www.ncbi.nlm.nih.gov/gene/?term=LOC105277239 | 2831 | 140 | 494.52 | 1 | 3.53 | 0.71 | 13.31 | 15.32 | -2.01 | 2.01 | -0.1 | 0.1 | |
| gene7248 | https://www.ncbi.nlm.nih.gov/gene/?term=LOC105280408 | 5175 | 57 | 110.14 | 1 | 1.93 | 1.75 | 18.23 | 20.49 | -2.26 | 2.26 | -0.02 | 0.02 | |
| gene3727 | https://www.ncbi.nlm.nih.gov/gene/?term=LOC105276367 | 2484 | 66 | 265.7 1 | 4.03 | 1.52 | 44.56 | 46.97 | -2.41 | 2.41 | -0.06 | 0.06 | | |
| gene6422 | https://www.ncbi.nlm.nih.gov/gene/?term=LOC105279458 | 2419 | 37 | 152.96 | 1 | 4.13 | 2.7 | 49.83 | 52.5 | -2.66 | 2.66 | -0.04 | 0.04 | |
| gene1505 | https://www.ncbi.nlm.nih.gov/gene/?term=LOC105287633 | 1693 | 27 | 159.48 | 1 | 5.91 | 3.7 | 42.55 | 45.31 | -2.76 | 2.76 | -0.04 | 0.04 | |
| gene9411 | https://www.ncbi.nlm.nih.gov/gene/?term=LOC105282914 | 5534 | 41 | 74.09 1 | 1.81 | 2.44 | 49.41 | 52.21 | -2.8 | 2.8 | -0.02 | 0.02 | | |
| gene6237 | https://www.ncbi.nlm.nih.gov/gene/?term=LOC105279244 | 1484 | 25 | 168.46 | 1 | 6.74 | 4 | 50.3 | 53.48 | -3.18 | 3.18 | -0.05 | 0.05 | |
| gene6455 | https://www.ncbi.nlm.nih.gov/gene/?term=LOC105279500 | 2612 | 33 | 126.34 | 1 | 3.83 | 3.03 | 43.71 | 47.06 | -3.35 | 3.35 | -0.04 | 0.04 | |
| gene564 | https://www.ncbi.nlm.nih.gov/gene/?term=LOC105278478 | 2802 | 17 | 60.67 1 | 3.57 | 5.88 | 14.04 | 17.66 | -3.63 | 3.63 | -0.02 | 0.02 | | |
| gene10602 | https://www.ncbi.nlm.nih.gov/gene/?term=LOC105284268 | 2122 | 18 | 84.83 1 | 4.71 | 5.56 | 26.04 | 29.95 | -3.91 | 3.91 | -0.03 | 0.03 | | |
| gene1199 | https://www.ncbi.nlm.nih.gov/gene/?term=LOC105285498 | 2220 | 26 | 117.12 | 1 | 4.5 | 3.85 | 33.11 | 37.03 | -3.92 | 3.92 | -0.05 | 0.05 | |
| gene12021 | https://www.ncbi.nlm.nih.gov/gene/?term=LOC105285905 | 1399 | 20 | 142.96 | 1 | 7.15 | 5 | 33.2 | 38.03 | -4.83 | 4.83 | -0.07 | 0.07 | |
| gene12430 | https://www.ncbi.nlm.nih.gov/gene/?term=LOC105286348 | 3389 | 19 | 56.06 1 | 2.95 | 5.26 | 6.26 | 11.43 | -5.17 | 5.17 | -0.03 | 0.03 | | |
| gene10963 | https://www.ncbi.nlm.nih.gov/gene/?term=LOC105284677 | 1576 | 29 | 184.01 | 1 | 6.35 | 3.45 | 18.87 | 36.99 | -18.12 | | 18.12 | -0.33 | 0.33 |
| gene393 | https://www.ncbi.nlm.nih.gov/gene/?term=LOC105276566 | 1123 | 2 | 17.81 1 | 8.9 | 50 | 47.22 | 70.57 | -23.35 | | 23.35 | -0.04 | 0.04 | |

Directory: OGLall File: ogl-Cb_LI_BCphaseB1.vs.RphaseA1.txt

| gene_ID | gene_link | gwidth | #Sites | #per10Kb | #dmSites | #dmsp10kb | %dmSites | %pSite1 | %pSite2 | DMpSite | ADMpSite | DMpNucl | ADMpNucl |
|---|---|---|---|---|---|---|---|---|---|---|---|---|---|
| gene12518 | https://www.ncbi.nlm.nih.gov/gene/?term=LOC105286453 | 1109 | 6 | 54.1 | 1 | 9.02 | 16.67 | 54.64 | 39.73 | 14.91 | 14.91 | 0.08 | 0.08 |
| gene9354 | https://www.ncbi.nlm.nih.gov/gene/?term=LOC105282854 | 1836 | 6 | 32.68 | 1 | 5.45 | 16.67 | 52.41 | 41.89 | 10.52 | 10.52 | 0.03 | 0.03 |
| gene12173 | https://www.ncbi.nlm.nih.gov/gene/?term=LOC105286067 | 518 | 3 | 57.92 | 1 | 19.31 | 33.33 | 13.69 | 3.17 | 10.51 | 10.51 | 0.06 | 0.06 |
| gene5950 | https://www.ncbi.nlm.nih.gov/gene/?term=LOC105278893 | 2674 | 6 | 22.44 | 1 | 3.74 | 16.67 | 34.71 | 25.34 | 9.37 | 9.37 | 0.02 | 0.02 |
| gene2855 | https://www.ncbi.nlm.nih.gov/gene/?term=LOC105275378 | 1189 | 21 | 176.62 | 3 | 25.23 | 14.29 | 38.9 | 30.15 | 8.75 | 8.75 | 0.15 | 0.15 |
| gene9690 | https://www.ncbi.nlm.nih.gov/gene/?term=LOC105283241 | 887 | 7 | 78.92 | 1 | 11.27 | 14.29 | 55.79 | 47.4 | 8.39 | 8.39 | 0.07 | 0.07 |
| gene5966 | https://www.ncbi.nlm.nih.gov/gene/?term=LOC105278917 | 2178 | 3 | 13.77 | 1 | 4.59 | 33.33 | 33.22 | 25.32 | 7.9 | 7.9 | 0.01 | 0.01 |
| gene13301 | https://www.ncbi.nlm.nih.gov/gene/?term=LOC105287352 | 656 | 7 | 106.71 | 1 | 15.24 | 14.29 | 24.74 | 17.08 | 7.67 | 7.67 | 0.08 | 0.08 |
| gene11073 | https://www.ncbi.nlm.nih.gov/gene/?term=LOC105284817 | 1692 | 9 | 53.19 | 1 | 5.91 | 11.11 | 40.23 | 32.79 | 7.44 | 7.44 | 0.04 | 0.04 |
| gene3128 | https://www.ncbi.nlm.nih.gov/gene/?term=LOC105275669 | 1566 | 8 | 51.09 | 1 | 6.39 | 12.5 | 54.98 | 48.31 | 6.67 | 6.67 | 0.03 | 0.03 |
| gene7488 | https://www.ncbi.nlm.nih.gov/gene/?term=LOC105280699 | 1563 | 5 | 31.99 | 1 | 6.4 | 20 | 31.65 | 25.04 | 6.61 | 6.61 | 0.02 | 0.02 |
| gene4732 | https://www.ncbi.nlm.nih.gov/gene/?term=LOC105277491 | 1647 | 11 | 66.79 | 1 | 6.07 | 9.09 | 19.89 | 14.07 | 5.82 | 5.82 | 0.04 | 0.04 |
| gene12610 | https://www.ncbi.nlm.nih.gov/gene/?term=LOC105286572 | 2000 | 6 | 30 | 1 | 5 | 16.67 | 16.76 | 11.15 | 5.61 | 5.61 | 0.02 | 0.02 |
| gene9255 | https://www.ncbi.nlm.nih.gov/gene/?term=LOC105282733 | 3562 | 21 | 58.96 | 2 | 5.61 | 9.52 | 36 | 30.71 | 5.29 | 5.29 | 0.03 | 0.03 |
| gene12818 | https://www.ncbi.nlm.nih.gov/gene/?term=LOC105286815 | 1095 | 11 | 100.46 | 1 | 9.13 | 9.09 | 63.21 | 58.15 | 5.07 | 5.07 | 0.05 | 0.05 |
| gene7300 | https://www.ncbi.nlm.nih.gov/gene/?term=LOC105280472 | 1589 | 12 | 75.52 | 1 | 6.29 | 8.33 | 42.17 | 37.11 | 5.06 | 5.06 | 0.04 | 0.04 |
| gene1656 | https://www.ncbi.nlm.nih.gov/gene/?term=LOC105287814 | 812 | 8 | 98.52 | 1 | 12.32 | 12.5 | 35.83 | 30.99 | 4.84 | 4.84 | 0.05 | 0.05 |
| gene5306 | https://www.ncbi.nlm.nih.gov/gene/?term=LOC105278153 | 481 | 17 | 353.43 | 2 | 41.58 | 11.76 | 21.3 | 16.46 | 4.84 | 4.84 | 0.17 | 0.17 |
| gene10176 | https://www.ncbi.nlm.nih.gov/gene/?term=LOC105283785 | 1637 | 35 | 213.81 | 1 | 6.11 | 2.86 | 18.49 | 13.7 | 4.79 | 4.79 | 0.1 | 0.1 |
| gene9684 | https://www.ncbi.nlm.nih.gov/gene/?term=LOC105283227 | 2228 | 15 | 67.32 | 2 | 8.98 | 13.33 | 34.73 | 29.95 | 4.79 | 4.79 | 0.03 | 0.03 |
| gene1742 | https://www.ncbi.nlm.nih.gov/gene/?term=LOC105287978 | 3019 | 7 | 23.19 | 1 | 3.31 | 14.29 | 17.41 | 12.85 | 4.56 | 4.56 | 0.01 | 0.01 |
| gene2970 | https://www.ncbi.nlm.nih.gov/gene/?term=LOC105275518 | 2340 | 26 | 111.11 | 2 | 8.55 | 7.69 | 47.9 | 43.34 | 4.56 | 4.56 | 0.05 | 0.05 |
| gene9852 | https://www.ncbi.nlm.nih.gov/gene/?term=LOC105283419 | 1060 | 16 | 150.94 | 2 | 18.87 | 12.5 | 6.02 | 1.53 | 4.48 | 4.48 | 0.07 | 0.07 |
| gene4032 | https://www.ncbi.nlm.nih.gov/gene/?term=LOC105276707 | 1993 | 16 | 80.28 | 2 | 10.04 | 12.5 | 37.03 | 32.6 | 4.43 | 4.43 | 0.04 | 0.04 |

...
... only first 25 lines shown ...

Directory: OGLall File: ogl-Cb_LI_BCphaseB1.vs.RphaseA2.txt

| gene_ID | gene_link | gwidth | #Sites | #per10Kb | #dmSites | #dmsp10kb | %dmSites | %pSite1 | %pSite2 | DMpSite | ADMpSite | DMpNucl | ADMpNucl |
|---|---|---|---|---|---|---|---|---|---|---|---|---|---|
| gene7300 | https://www.ncbi.nlm.nih.gov/gene/?term=LOC105280472 | 1589 | 12 | 75.52 | 2 | 12.59 | 16.67 | 42.17 | 34.37 | 7.8 | 7.8 | 0.06 | 0.06 |
| gene5695 | https://www.ncbi.nlm.nih.gov/gene/?term=LOC105278607 | 4140 | 22 | 53.14 | 2 | 4.83 | 9.09 | 13.44 | 6.4 | 7.04 | 7.04 | 0.04 | 0.04 |
| gene10343 | https://www.ncbi.nlm.nih.gov/gene/?term=LOC105283986 | 1763 | 7 | 39.71 | 1 | 5.67 | 14.29 | 29.98 | 22.98 | 7 | 7 | 0.03 | 0.03 |
| gene12518 | https://www.ncbi.nlm.nih.gov/gene/?term=LOC105286453 | 1109 | 13 | 117.22 | 2 | 18.03 | 15.38 | 34.82 | 28.73 | 6.08 | 6.08 | 0.07 | 0.07 |
| gene1742 | https://www.ncbi.nlm.nih.gov/gene/?term=LOC105287978 | 3019 | 7 | 23.19 | 2 | 6.62 | 28.57 | 17.41 | 11.62 | 5.79 | 5.79 | 0.01 | 0.01 |
| gene11185 | https://www.ncbi.nlm.nih.gov/gene/?term=LOC105284939 | 1723 | 17 | 98.67 | 1 | 5.8 | 5.88 | 37.14 | 31.9 | 5.24 | 5.24 | 0.05 | 0.05 |
| gene1894 | https://www.ncbi.nlm.nih.gov/gene/?term=LOC105288083 | 445 | 17 | 382.02 | 2 | 44.94 | 11.76 | 56.3 | 51.18 | 5.11 | 5.11 | 0.2 | 0.2 |
| gene4621 | https://www.ncbi.nlm.nih.gov/gene/?term=LOC105277369 | 1157 | 8 | 69.14 | 1 | 8.64 | 12.5 | 67.07 | 62.16 | 4.9 | 4.9 | 0.03 | 0.03 |
| gene11850 | https://www.ncbi.nlm.nih.gov/gene/?term=LOC105285707 | 2053 | 31 | 151 | 2 | 9.74 | 6.45 | 60.03 | 55.77 | 4.26 | 4.26 | 0.06 | 0.06 |
| gene12173 | https://www.ncbi.nlm.nih.gov/gene/?term=LOC105286067 | 518 | 6 | 115.83 | 1 | 19.31 | 16.67 | 11.97 | 7.8 | 4.17 | 4.17 | 0.05 | 0.05 |
| gene7727 | https://www.ncbi.nlm.nih.gov/gene/?term=LOC105280969 | 1006 | 15 | 149.11 | 1 | 9.94 | 6.67 | 36.57 | 32.67 | 3.89 | 3.89 | 0.06 | 0.06 |
| gene2837 | https://www.ncbi.nlm.nih.gov/gene/?term=LOC105275355 | 2872 | 160 | 557.1 | 10 | 34.82 | 6.25 | 29.18 | 25.36 | 3.82 | 3.82 | 0.21 | 0.21 |
| gene2855 | https://www.ncbi.nlm.nih.gov/gene/?term=LOC105275378 | 1189 | 28 | 235.49 | 2 | 16.82 | 7.14 | 43.5 | 39.96 | 3.54 | 3.54 | 0.08 | 0.08 |
| gene9854 | https://www.ncbi.nlm.nih.gov/gene/?term=LOC105283423 | 2931 | 16 | 54.59 | 1 | 3.41 | 6.25 | 37.51 | 33.96 | 3.54 | 3.54 | 0.02 | 0.02 |
| gene3282 | https://www.ncbi.nlm.nih.gov/gene/?term=LOC105275847 | 1942 | 25 | 128.73 | 1 | 5.15 | 4 | 47.32 | 43.85 | 3.47 | 3.47 | 0.04 | 0.04 |
| gene6879 | https://www.ncbi.nlm.nih.gov/gene/?term=LOC105280001 | 2322 | 22 | 94.75 | 1 | 4.31 | 4.55 | 25.57 | 22.33 | 3.24 | 3.24 | 0.03 | 0.03 |
| gene7840 | https://www.ncbi.nlm.nih.gov/gene/?term=LOC105281081 | 2338 | 15 | 64.16 | 1 | 4.28 | 6.67 | 19.85 | 16.65 | 3.2 | 3.2 | 0.02 | 0.02 |
| gene7549 | https://www.ncbi.nlm.nih.gov/gene/?term=LOC105280764 | 2668 | 23 | 86.21 | 2 | 7.5 | 8.7 | 13.43 | 10.25 | 3.17 | 3.17 | 0.03 | 0.03 |
| gene3000 | https://www.ncbi.nlm.nih.gov/gene/?term=LOC105275544 | 1577 | 23 | 145.85 | 3 | 19.02 | 13.04 | 37.23 | 34.08 | 3.15 | 3.15 | 0.05 | 0.05 |
| gene2220 | https://www.ncbi.nlm.nih.gov/gene/?term=LOC105274640 | 2263 | 9 | 39.77 | 1 | 4.42 | 11.11 | 71.96 | 68.85 | 3.11 | 3.11 | 0.01 | 0.01 |
| gene108 | https://www.ncbi.nlm.nih.gov/gene/?term=LOC105284604 | 1903 | 17 | 89.33 | 2 | 10.51 | 11.76 | 3.09 | 0 | 3.09 | 3.09 | 0.03 | 0.03 |
| gene1962 | https://www.ncbi.nlm.nih.gov/gene/?term=LOC105288157 | 1246 | 19 | 152.49 | 1 | 8.03 | 5.26 | 16.56 | 13.6 | 2.96 | 2.96 | 0.05 | 0.05 |
| gene4999 | https://www.ncbi.nlm.nih.gov/gene/?term=LOC105277805 | 6695 | 14 | 20.91 | 1 | 1.49 | 7.14 | 45.95 | 43.05 | 2.9 | 2.9 | 0.01 | 0.01 |
| gene10237 | https://www.ncbi.nlm.nih.gov/gene/?term=LOC105283856 | 1259 | 23 | 182.68 | 1 | 7.94 | 4.35 | 64.51 | 61.63 | 2.88 | 2.88 | 0.05 | 0.05 |

...
... only first 25 lines shown ...

Directory: OGLall File: ogl-Cb_LI_BCphaseB1.vs.RphaseB1.txt

| gene_ID | gene_link | gwidth | #Sites | #per10Kb | #dmSites | #dmsp10kb | %dmSites | %pSite1 | %pSite2 | DMpSite | ADMpSite | DMpNucl | ADMpNucl |
|---|---|---|---|---|---|---|---|---|---|---|---|---|---|
| gene2593 | https://www.ncbi.nlm.nih.gov/gene/?term=LOC105275070 | 1774 | 11 | 62.01 | 1 | 5.64 | 9.09 | 61.58 | 25.28 | 36.29 | 36.29 | 0.23 | 0.23 |
| gene3165 | https://www.ncbi.nlm.nih.gov/gene/?term=LOC105275706 | 2374 | 65 | 273.8 | 4 | 16.85 | 6.15 | 33.82 | 25.03 | 8.79 | 8.79 | 0.24 | 0.24 |
| gene2837 | https://www.ncbi.nlm.nih.gov/gene/?term=LOC105275355 | 2872 | 165 | 574.51 | 1 | 3.48 | 0.61 | 28.35 | 22.92 | 5.43 | 5.43 | 0.31 | 0.31 |
| gene2970 | https://www.ncbi.nlm.nih.gov/gene/?term=LOC105275518 | 2340 | 25 | 106.84 | 1 | 4.27 | 4 | 44.93 | 41.23 | 3.69 | 3.69 | 0.04 | 0.04 |
| gene5714 | https://www.ncbi.nlm.nih.gov/gene/?term=LOC105278729 | 710 | 16 | 225.35 | 1 | 14.08 | 6.25 | 55.52 | 51.9 | 3.62 | 3.62 | 0.08 | 0.08 |
| gene2856 | https://www.ncbi.nlm.nih.gov/gene/?term=LOC105275384 | 1987 | 92 | 463.01 | 1 | 5.03 | 1.09 | 34.19 | 31.19 | 3 | 3 | 0.14 | 0.14 |
| gene3671 | https://www.ncbi.nlm.nih.gov/gene/?term=LOC105276272 | 1966 | 40 | 203.46 | 1 | 5.09 | 2.5 | 30.25 | 27.31 | 2.94 | 2.94 | 0.06 | 0.06 |
| gene396 | https://www.ncbi.nlm.nih.gov/gene/?term=LOC105276611 | 1722 | 36 | 209.06 | 1 | 5.81 | 2.78 | 30.31 | 27.55 | 2.76 | 2.76 | 0.06 | 0.06 |
| gene13033 | https://www.ncbi.nlm.nih.gov/gene/?term=LOC105287063 | 1877 | 50 | 266.38 | 1 | 5.33 | 2 | 31.92 | 29.55 | 2.37 | 2.37 | 0.06 | 0.06 |
| gene1855 | https://www.ncbi.nlm.nih.gov/gene/?term=LOC105288036 | 2387 | 91 | 381.23 | 1 | 4.19 | 1.1 | 29.43 | 27.13 | 2.3 | 2.3 | 0.09 | 0.09 |
| gene2241 | https://www.ncbi.nlm.nih.gov/gene/?term=LOC105274666 | 2598 | 61 | 234.8 | 1 | 3.85 | 1.64 | 30.46 | 28.61 | 1.85 | 1.85 | 0.04 | 0.04 |
| gene10345 | https://www.ncbi.nlm.nih.gov/gene/?term=LOC105283989 | 4444 | 41 | 92.26 | 1 | 2.25 | 2.44 | 38.24 | 36.42 | 1.83 | 1.83 | 0.02 | 0.02 |
| gene7493 | https://www.ncbi.nlm.nih.gov/gene/?term=LOC105280704 | 4225 | 147 | 347.93 | 1 | 2.37 | 0.68 | 18.36 | 16.63 | 1.72 | 1.72 | 0.06 | 0.06 |
| gene5831 | https://www.ncbi.nlm.nih.gov/gene/?term=LOC105278761 | 2506 | 118 | 470.87 | 2 | 7.98 | 1.69 | 22.67 | 21.1 | 1.58 | 1.58 | 0.07 | 0.07 |
| gene940 | https://www.ncbi.nlm.nih.gov/gene/?term=LOC105282794 | 3578 | 94 | 262.72 | 1 | 2.79 | 1.06 | 15.53 | 14.17 | 1.36 | 1.36 | 0.04 | 0.04 |
| gene4023 | https://www.ncbi.nlm.nih.gov/gene/?term=LOC105276692 | 1759 | 37 | 210.35 | 1 | 5.69 | 2.7 | 23.22 | 21.93 | 1.3 | 1.3 | 0.03 | 0.03 |
| gene9336 | https://www.ncbi.nlm.nih.gov/gene/?term=LOC105282834 | 5993 | 134 | 223.59 | 1 | 1.67 | 0.75 | 14.88 | 13.64 | 1.24 | 1.24 | 0.03 | 0.03 |
| gene2748 | https://www.ncbi.nlm.nih.gov/gene/?term=LOC105275261 | 3845 | 53 | 137.84 | 1 | 2.6 | 1.89 | 39.61 | 38.42 | 1.19 | 1.19 | 0.02 | 0.02 |
| gene10107 | https://www.ncbi.nlm.nih.gov/gene/?term=LOC105283712 | 1883 | 36 | 191.18 | 1 | 5.31 | 2.78 | 30.43 | 29.26 | 1.17 | 1.17 | 0.02 | 0.02 |
| gene1884 | https://www.ncbi.nlm.nih.gov/gene/?term=LOC105288072 | 4169 | 84 | 201.49 | 1 | 2.4 | 1.19 | 24.86 | 23.72 | 1.14 | 1.14 | 0.02 | 0.02 |
| gene8178 | https://www.ncbi.nlm.nih.gov/gene/?term=LOC105281515 | 8647 | 60 | 69.39 | 1 | 1.16 | 1.67 | 30.54 | 29.47 | 1.06 | 1.06 | 0.01 | 0.01 |
| gene7734 | https://www.ncbi.nlm.nih.gov/gene/?term=LOC105280972 | 1528 | 35 | 229.06 | 1 | 6.54 | 2.86 | 17.63 | 16.59 | 1.04 | 1.04 | 0.02 | 0.02 |
| gene7884 | https://www.ncbi.nlm.nih.gov/gene/?term=LOC105281266 | 3390 | 87 | 256.64 | 1 | 2.95 | 1.15 | 15.85 | 14.89 | 0.96 | 0.96 | 0.02 | 0.02 |
| gene4641 | https://www.ncbi.nlm.nih.gov/gene/?term=LOC105277398 | 1937 | 72 | 371.71 | 1 | 5.16 | 1.39 | 36.3 | 35.35 | 0.95 | 0.95 | 0.04 | 0.04 |
| gene5877 | https://www.ncbi.nlm.nih.gov/gene/?term=LOC105278807 | 1719 | 51 | 296.68 | 1 | 5.82 | 1.96 | 26.14 | 25.26 | 0.89 | 0.89 | 0.03 | 0.03 |
| gene1147 | https://www.ncbi.nlm.nih.gov/gene/?term=LOC105284954 | 4022 | 118 | 293.39 | 1 | 2.49 | 0.85 | 15.13 | 14.31 | 0.81 | 0.81 | 0.02 | 0.02 |
| gene3817 | https://www.ncbi.nlm.nih.gov/gene/?term=LOC105276449 | 1230 | 55 | 447.15 | 1 | 8.13 | 1.82 | 22.28 | 21.47 | 0.8 | 0.8 | 0.04 | 0.04 |
| gene474 | https://www.ncbi.nlm.nih.gov/gene/?term=LOC105277435 | 3625 | 176 | 485.52 | 1 | 2.76 | 0.57 | 10.76 | 9.98 | 0.78 | 0.78 | 0.04 | 0.04 |
| gene292 | https://www.ncbi.nlm.nih.gov/gene/?term=LOC105275398 | 2825 | 88 | 311.5 | 1 | 3.54 | 1.14 | 22.66 | 21.93 | 0.73 | 0.73 | 0.02 | 0.02 |
| gene7183 | https://www.ncbi.nlm.nih.gov/gene/?term=LOC105280348 | 3925 | 80 | 203.82 | 1 | 2.55 | 1.25 | 17.35 | 16.62 | 0.73 | 0.73 | 0.01 | 0.01 |
| gene322 | https://www.ncbi.nlm.nih.gov/gene/?term=LOC105275777 | 1606 | 28 | 174.35 | 1 | 6.23 | 3.57 | 36.76 | 36.04 | 0.72 | 0.72 | 0.01 | 0.01 |
| gene298 | https://www.ncbi.nlm.nih.gov/gene/?term=LOC105275460 | 2473 | 126 | 509.5 | 1 | 4.04 | 0.79 | 13.09 | 12.43 | 0.67 | 0.67 | 0.03 | 0.03 |
| gene10678 | https://www.ncbi.nlm.nih.gov/gene/?term=LOC105284355 | 4031 | 169 | 419.25 | 1 | 2.48 | 0.59 | 20.31 | 19.66 | 0.65 | 0.65 | 0.03 | 0.03 |
| gene4034 | https://www.ncbi.nlm.nih.gov/gene/?term=LOC105276704 | 2065 | 87 | 421.31 | 2 | 9.69 | 2.3 | 14.01 | 13.38 | 0.63 | 0.63 | 0.03 | 0.03 |
| gene7888 | https://www.ncbi.nlm.nih.gov/gene/?term=LOC105281141 | 4357 | 170 | 390.18 | 1 | 2.3 | 0.59 | 28.6 | 28 | 0.6 | 0.6 | 0.02 | 0.02 |
| gene6652 | https://www.ncbi.nlm.nih.gov/gene/?term=LOC105279719 | 2831 | 88 | 310.84 | 1 | 3.53 | 1.14 | 21.65 | 21.08 | 0.57 | 0.57 | 0.02 | 0.02 |
| gene4892 | https://www.ncbi.nlm.nih.gov/gene/?term=LOC105277667 | 3898 | 127 | 325.81 | 1 | 2.57 | 0.79 | 27.72 | 27.21 | 0.52 | 0.52 | 0.02 | 0.02 |
| gene12978 | https://www.ncbi.nlm.nih.gov/gene/?term=LOC105286997 | 2858 | 150 | 524.84 | 1 | 3.5 | 0.67 | 11.28 | 10.84 | 0.44 | 0.44 | 0.02 | 0.02 |
| gene11819 | https://www.ncbi.nlm.nih.gov/gene/?term=LOC105285671 | 2366 | 104 | 439.56 | 1 | 4.23 | 0.96 | 30.06 | 29.63 | 0.43 | 0.43 | 0.02 | 0.02 |
| gene2683 | https://www.ncbi.nlm.nih.gov/gene/?term=LOC105275187 | 5492 | 78 | 142.02 | 1 | 1.82 | 1.28 | 16.44 | 16.02 | 0.43 | 0.43 | 0.01 | 0.01 |
| gene3107 | https://www.ncbi.nlm.nih.gov/gene/?term=LOC105275650 | 5103 | 182 | 356.65 | 1 | 1.96 | 0.55 | 16.89 | 16.49 | 0.4 | 0.4 | 0.01 | 0.01 |
| gene6455 | https://www.ncbi.nlm.nih.gov/gene/?term=LOC105279500 | 2612 | 35 | 134 | 1 | 3.83 | 2.86 | 48.92 | 48.53 | 0.39 | 0.39 | 0.01 | 0.01 |
| gene3885 | https://www.ncbi.nlm.nih.gov/gene/?term=LOC105276527 | 7928 | 369 | 465.44 | 1 | 1.26 | 0.27 | 8.26 | 7.93 | 0.33 | 0.33 | 0.02 | 0.02 |
| gene13036 | https://www.ncbi.nlm.nih.gov/gene/?term=LOC105287059 | 4959 | 193 | 389.19 | 1 | 2.02 | 0.52 | 16.75 | 16.43 | 0.32 | 0.32 | 0.01 | 0.01 |
| gene8729 | https://www.ncbi.nlm.nih.gov/gene/?term=LOC105282137 | 5005 | 226 | 451.55 | 1 | 2 | 0.44 | 14.98 | 14.68 | 0.29 | 0.29 | 0.01 | 0.01 |
| gene12572 | https://www.ncbi.nlm.nih.gov/gene/?term=LOC105286514 | 2374 | 52 | 219.04 | 1 | 4.21 | 1.92 | 35.13 | 34.85 | 0.28 | 0.28 | 0.01 | 0.01 |
| gene9101 | https://www.ncbi.nlm.nih.gov/gene/?term=LOC105282553 | 3068 | 106 | 345.5 | 1 | 3.26 | 0.94 | 22.9 | 22.63 | 0.27 | 0.27 | 0.01 | 0.01 |
| gene10018 | https://www.ncbi.nlm.nih.gov/gene/?term=LOC105283607 | 4414 | 228 | 516.54 | 1 | 2.27 | 0.44 | 14.1 | 13.84 | 0.26 | 0.26 | 0.01 | 0.01 |
| gene2668 | https://www.ncbi.nlm.nih.gov/gene/?term=LOC105275170 | 12496 | 356 | 284.89 | 1 | 0.8 | 0.28 | 2.04 | 1.8 | 0.24 | 0.24 | 0.01 | 0.01 |
| gene10540 | https://www.ncbi.nlm.nih.gov/gene/?term=LOC105284200 | 5267 | 214 | 406.3 | 1 | 1.9 | 0.47 | 13.24 | 13.01 | 0.23 | 0.23 | 0.01 | 0.01 |
| gene10596 | https://www.ncbi.nlm.nih.gov/gene/?term=LOC105284262 | 6520 | 239 | 366.56 | 1 | 1.53 | 0.42 | 4.23 | 4.05 | 0.19 | 0.19 | 0.01 | 0.01 |
| gene6286 | https://www.ncbi.nlm.nih.gov/gene/?term=LOC105279298 | 2861 | 128 | 447.4 | 1 | 3.5 | 0.78 | 23.43 | 23.27 | 0.16 | 0.16 | 0.01 | 0.01 |
| gene7368 | https://www.ncbi.nlm.nih.gov/gene/?term=LOC105280556 | 4105 | 71 | 172.96 | 1 | 2.44 | 1.41 | 15.89 | 15.73 | 0.16 | 0.16 | 0 | 0 |
| gene12406 | https://www.ncbi.nlm.nih.gov/gene/?term=LOC105286316 | 1386 | 44 | 317.46 | 1 | 7.22 | 2.27 | 32.85 | 32.71 | 0.14 | 0.14 | 0 | 0 |
| gene10600 | https://www.ncbi.nlm.nih.gov/gene/?term=LOC105284270 | 4837 | 111 | 229.48 | 1 | 2.07 | 0.9 | 11.96 | 11.83 | 0.13 | 0.13 | 0 | 0 |
| gene927 | https://www.ncbi.nlm.nih.gov/gene/?term=LOC105282600 | 3727 | 60 | 160.99 | 1 | 2.68 | 1.67 | 28.88 | 28.75 | 0.13 | 0.13 | 0 | 0 |
| gene9327 | https://www.ncbi.nlm.nih.gov/gene/?term=LOC105282821 | 7793 | 183 | 234.83 | 1 | 1.28 | 0.55 | 12.55 | 12.42 | 0.13 | 0.13 | 0 | 0 |
| gene11275 | https://www.ncbi.nlm.nih.gov/gene/?term=LOC105285055 | 4119 | 246 | 597.23 | 1 | 2.43 | 0.41 | 12.22 | 12.12 | 0.1 | 0.1 | 0.01 | 0.01 |
| gene5375 | https://www.ncbi.nlm.nih.gov/gene/?term=LOC105278228 | 8640 | 191 | 221.06 | 1 | 1.16 | 0.52 | 22.09 | 21.99 | 0.1 | 0.1 | 0 | 0 |
| gene10222 | https://www.ncbi.nlm.nih.gov/gene/?term=LOC105283838 | 1748 | 30 | 171.62 | 1 | 5.72 | 3.33 | 35.37 | 35.28 | 0.09 | 0.09 | 0 | 0 |

| | | | | | | | | | | | | | |
|---|---|---|---|---|---|---|---|---|---|---|---|---|---|
| gene10520 | https://www.ncbi.nlm.nih.gov/gene/?term=LOC105284184 | 2451 | 102 | 416.16 | 1 | 4.08 | 0.98 | 30 | 29.93 | 0.07 | 0.07 | 0 | 0 |
| gene9326 | https://www.ncbi.nlm.nih.gov/gene/?term=LOC105282818 | 16612 | 236 | 142.07 | 1 | 0.6 | 0.42 | 13.17 | 13.1 | 0.07 | 0.07 | 0 | 0 |
| gene2443 | https://www.ncbi.nlm.nih.gov/gene/?term=LOC105274895 | 10433 | 424 | 406.4 1 | 0.96 | 0.24 | 5.17 | 5.13 | 0.03 | 0.03 | 0 | 0 | |
| gene53 | https://www.ncbi.nlm.nih.gov/gene/?term=LOC105277148 | 19283 | 333 | 172.69 | 1 | 0.52 | 0.3 | 4.68 | 4.68 | 0 | 0 | 0 | 0 |
| gene11454 | https://www.ncbi.nlm.nih.gov/gene/?term=LOC105285256 | 1906 | 50 | 262.33 | 1 | 5.25 | 2 | 44.07 | 44.08 | -0.01 | 0.01 | 0 | 0 |
| gene13114 | https://www.ncbi.nlm.nih.gov/gene/?term=LOC105287147 | 7609 | 432 | 567.75 | 1 | 1.31 | 0.23 | 9.01 | 9.02 | -0.01 | 0.01 | 0 | 0 |
| gene2271 | https://www.ncbi.nlm.nih.gov/gene/?term=LOC105274706 | 1448 | 47 | 324.59 | 1 | 6.91 | 2.13 | 41.96 | 41.97 | -0.01 | 0.01 | 0 | 0 |
| gene862 | https://www.ncbi.nlm.nih.gov/gene/?term=LOC105281802 | 73922 | 2318 | 313.57 | 1 | 0.14 | 0.04 | 2.22 | 2.23 | -0.01 | 0.01 | 0 | 0 |
| gene8600 | https://www.ncbi.nlm.nih.gov/gene/?term=LOC105282001 | 8513 | 206 | 241.98 | 1 | 1.17 | 0.49 | 9.71 | 9.75 | -0.04 | 0.04 | 0 | 0 |
| gene5746 | https://www.ncbi.nlm.nih.gov/gene/?term=LOC105278666 | 10343 | 483 | 466.98 | 1 | 0.97 | 0.21 | 6.37 | 6.41 | -0.05 | 0.05 | 0 | 0 |
| gene6620 | https://www.ncbi.nlm.nih.gov/gene/?term=LOC105279687 | 17998 | 448 | 248.92 | 1 | 0.56 | 0.22 | 4.43 | 4.47 | -0.05 | 0.05 | 0 | 0 |
| gene5539 | https://www.ncbi.nlm.nih.gov/gene/?term=LOC105278415 | 25024 | 790 | 315.7 1 | 0.4 | 0.13 | 3.02 | 3.08 | -0.06 | 0.06 | 0 | 0 | |
| gene1632 | https://www.ncbi.nlm.nih.gov/gene/?term=LOC105287764 | 18061 | 431 | 238.64 | 1 | 0.55 | 0.23 | 3.3 | 3.37 | -0.07 | 0.07 | 0 | 0 |
| gene2806 | https://www.ncbi.nlm.nih.gov/gene/?term=LOC105275327 | 14257 | 552 | 387.18 | 1 | 0.7 | 0.18 | 6.34 | 6.41 | -0.07 | 0.07 | 0 | 0 |
| gene5197 | https://www.ncbi.nlm.nih.gov/gene/?term=LOC105278018 | 89233 | 3791 | 424.84 | 1 | 0.11 | 0.03 | 0.65 | 0.72 | -0.07 | 0.07 | 0 | 0 |
| gene3470 | https://www.ncbi.nlm.nih.gov/gene/?term=LOC105276059 | 8716 | 182 | 208.81 | 1 | 1.15 | 0.55 | 9.23 | 9.32 | -0.09 | 0.09 | 0 | 0 |
| gene5646 | https://www.ncbi.nlm.nih.gov/gene/?term=LOC105278534 | 7475 | 202 | 270.23 | 1 | 1.34 | 0.5 | 16.69 | 16.79 | -0.1 | 0.1 | 0 | 0 |
| gene12427 | https://www.ncbi.nlm.nih.gov/gene/?term=LOC105286339 | 5220 | 187 | 358.24 | 1 | 1.92 | 0.53 | 15.91 | 16.02 | -0.12 | 0.12 | 0 | 0 |
| gene12802 | https://www.ncbi.nlm.nih.gov/gene/?term=LOC105286799 | 3183 | 180 | 565.5 1 | 3.14 | 0.56 | 16.81 | 16.93 | -0.12 | 0.12 | -0.01 | 0.01 | |
| gene5654 | https://www.ncbi.nlm.nih.gov/gene/?term=LOC105278543 | 4019 | 181 | 450.36 | 1 | 2.49 | 0.55 | 15.7 | 15.83 | -0.13 | 0.13 | -0.01 | 0.01 |
| gene10844 | https://www.ncbi.nlm.nih.gov/gene/?term=LOC105284535 | 9086 | 259 | 285.05 | 1 | 1.1 | 0.39 | 14.4 | 14.56 | -0.16 | 0.16 | 0 | 0 |
| gene8274 | https://www.ncbi.nlm.nih.gov/gene/?term=LOC105281624 | 5425 | 227 | 418.43 | 1 | 1.84 | 0.44 | 9.53 | 9.7 | -0.17 | 0.17 | -0.01 | 0.01 |
| gene13039 | https://www.ncbi.nlm.nih.gov/gene/?term=LOC105287065 | 4902 | 253 | 516.12 | 1 | 2.04 | 0.4 | 12.12 | 12.3 | -0.18 | 0.18 | -0.01 | 0.01 |
| gene11579 | https://www.ncbi.nlm.nih.gov/gene/?term=LOC105285406 | 13571 | 316 | 232.85 | 1 | 0.74 | 0.32 | 6.01 | 6.26 | -0.25 | 0.25 | -0.01 | 0.01 |
| gene7249 | https://www.ncbi.nlm.nih.gov/gene/?term=LOC105280422 | 1775 | 66 | 371.83 | 1 | 5.63 | 1.52 | 15.78 | 16.04 | -0.26 | 0.26 | -0.01 | 0.01 |
| gene6639 | https://www.ncbi.nlm.nih.gov/gene/?term=LOC105279705 | 7656 | 83 | 108.41 | 1 | 1.31 | 1.2 | 19.9 | 20.17 | -0.28 | 0.28 | 0 | 0 |
| gene7199 | https://www.ncbi.nlm.nih.gov/gene/?term=LOC105280366 | 1414 | 34 | 240.45 | 1 | 7.07 | 2.94 | 44.65 | 44.98 | -0.33 | 0.33 | -0.01 | 0.01 |
| gene10549 | https://www.ncbi.nlm.nih.gov/gene/?term=LOC105284210 | 7145 | 365 | 510.85 | 2 | 2.8 | 0.55 | 9.55 | 9.9 | -0.35 | 0.35 | -0.02 | 0.02 |
| gene5524 | https://www.ncbi.nlm.nih.gov/gene/?term=LOC105278397 | 5177 | 209 | 403.71 | 1 | 1.93 | 0.48 | 23.99 | 24.35 | -0.36 | 0.36 | -0.01 | 0.01 |
| gene10326 | https://www.ncbi.nlm.nih.gov/gene/?term=LOC105283962 | 3042 | 137 | 450.36 | 1 | 3.29 | 0.73 | 7.31 | 7.68 | -0.37 | 0.37 | -0.02 | 0.02 |
| gene4376 | https://www.ncbi.nlm.nih.gov/gene/?term=LOC105277083 | 4307 | 129 | 299.51 | 1 | 2.32 | 0.78 | 18.69 | 19.06 | -0.38 | 0.38 | -0.01 | 0.01 |
| gene11583 | https://www.ncbi.nlm.nih.gov/gene/?term=LOC105285414 | 8192 | 192 | 234.38 | 1 | 1.22 | 0.52 | 8.5 | 8.89 | -0.39 | 0.39 | -0.01 | 0.01 |
| gene7915 | https://www.ncbi.nlm.nih.gov/gene/?term=LOC105281173 | 1597 | 42 | 262.99 | 1 | 6.26 | 2.38 | 62.33 | 62.75 | -0.42 | 0.42 | -0.01 | 0.01 |
| gene3793 | https://www.ncbi.nlm.nih.gov/gene/?term=LOC105276429 | 2901 | 131 | 451.57 | 1 | 3.45 | 0.76 | 22.23 | 22.66 | -0.43 | 0.43 | -0.02 | 0.02 |
| gene5525 | https://www.ncbi.nlm.nih.gov/gene/?term=LOC105278396 | 3090 | 189 | 611.65 | 1 | 3.24 | 0.53 | 24.08 | 24.52 | -0.44 | 0.44 | -0.03 | 0.03 |
| gene9330 | https://www.ncbi.nlm.nih.gov/gene/?term=LOC105282825 | 3936 | 98 | 248.98 | 1 | 2.54 | 1.02 | 13.47 | 13.91 | -0.44 | 0.44 | -0.01 | 0.01 |
| gene10606 | https://www.ncbi.nlm.nih.gov/gene/?term=LOC105284273 | 6119 | 272 | 444.52 | 1 | 1.63 | 0.37 | 17.87 | 18.32 | -0.45 | 0.45 | -0.02 | 0.02 |
| gene6583 | https://www.ncbi.nlm.nih.gov/gene/?term=LOC105279649 | 7444 | 372 | 499.73 | 1 | 1.34 | 0.27 | 10.85 | 11.31 | -0.45 | 0.45 | -0.02 | 0.02 |
| gene4190 | https://www.ncbi.nlm.nih.gov/gene/?term=LOC105276875 | 3150 | 134 | 425.4 1 | 3.17 | 0.75 | 20.21 | 20.67 | -0.46 | 0.46 | -0.02 | 0.02 | |
| gene7247 | https://www.ncbi.nlm.nih.gov/gene/?term=LOC105280407 | 10486 | 227 | 216.48 | 1 | 0.95 | 0.44 | 14.79 | 15.25 | -0.46 | 0.46 | -0.01 | 0.01 |
| gene7166 | https://www.ncbi.nlm.nih.gov/gene/?term=LOC105280326 | 6117 | 259 | 423.41 | 1 | 1.63 | 0.39 | 14.46 | 14.93 | -0.47 | 0.47 | -0.02 | 0.02 |
| gene2173 | https://www.ncbi.nlm.nih.gov/gene/?term=LOC105274590 | 4387 | 87 | 198.31 | 2 | 4.56 | 2.3 | 23.56 | 24.1 | -0.54 | 0.54 | -0.01 | 0.01 |
| gene4033 | https://www.ncbi.nlm.nih.gov/gene/?term=LOC105276705 | 10546 | 155 | 146.98 | 1 | 0.95 | 0.65 | 5.56 | 6.1 | -0.55 | 0.55 | -0.01 | 0.01 |
| gene6966 | https://www.ncbi.nlm.nih.gov/gene/?term=LOC105280104 | 3747 | 132 | 352.28 | 1 | 2.67 | 0.76 | 16.91 | 17.46 | -0.55 | 0.55 | -0.02 | 0.02 |
| gene11561 | https://www.ncbi.nlm.nih.gov/gene/?term=LOC105285387 | 11528 | 218 | 189.1 1 | 0.87 | 0.46 | 8.62 | 9.18 | -0.56 | 0.56 | -0.01 | 0.01 | |
| gene8814 | https://www.ncbi.nlm.nih.gov/gene/?term=LOC105282240 | 6910 | 204 | 295.22 | 1 | 1.45 | 0.49 | 8.01 | 8.64 | -0.63 | 0.63 | -0.02 | 0.02 |
| gene5184 | https://www.ncbi.nlm.nih.gov/gene/?term=LOC105277999 | 5528 | 165 | 298.48 | 1 | 1.81 | 0.61 | 11.31 | 11.98 | -0.67 | 0.67 | -0.02 | 0.02 |
| gene11235 | https://www.ncbi.nlm.nih.gov/gene/?term=LOC105285004 | 1339 | 36 | 268.86 | 1 | 7.47 | 2.78 | 19.18 | 19.88 | -0.7 | 0.7 | -0.02 | 0.02 |
| gene1310 | https://www.ncbi.nlm.nih.gov/gene/?term=LOC105286661 | 2582 | 68 | 263.36 | 1 | 3.87 | 1.47 | 18.99 | 19.74 | -0.75 | 0.75 | -0.02 | 0.02 |
| gene12573 | https://www.ncbi.nlm.nih.gov/gene/?term=LOC105286515 | 3647 | 107 | 293.39 | 1 | 2.74 | 0.93 | 10.59 | 11.4 | -0.82 | 0.82 | -0.02 | 0.02 |
| gene1470 | https://www.ncbi.nlm.nih.gov/gene/?term=LOC105287597 | 2974 | 192 | 645.6 1 | 3.36 | 0.52 | 18.41 | 19.24 | -0.83 | 0.83 | -0.05 | 0.05 | |
| gene5298 | https://www.ncbi.nlm.nih.gov/gene/?term=LOC105278145 | 4060 | 262 | 645.32 | 2 | 4.93 | 0.76 | 9.65 | 10.52 | -0.86 | 0.86 | -0.06 | 0.06 |
| gene6698 | https://www.ncbi.nlm.nih.gov/gene/?term=LOC105279778 | 4099 | 161 | 392.78 | 1 | 2.44 | 0.62 | 8.69 | 9.57 | -0.88 | 0.88 | -0.03 | 0.03 |
| gene920 | https://www.ncbi.nlm.nih.gov/gene/?term=LOC105282518 | 1197 | 44 | 367.59 | 1 | 8.35 | 2.27 | 33.07 | 33.97 | -0.89 | 0.89 | -0.03 | 0.03 |
| gene5101 | https://www.ncbi.nlm.nih.gov/gene/?term=LOC105277903 | 1813 | 37 | 204.08 | 1 | 5.52 | 2.7 | 54.12 | 55.06 | -0.94 | 0.94 | -0.02 | 0.02 |
| gene10155 | https://www.ncbi.nlm.nih.gov/gene/?term=LOC105283756 | 7030 | 116 | 165.01 | 1 | 1.42 | 0.86 | 7.15 | 8.11 | -0.96 | 0.96 | -0.02 | 0.02 |
| gene10392 | https://www.ncbi.nlm.nih.gov/gene/?term=LOC105284036 | 3216 | 103 | 320.27 | 2 | 6.22 | 1.94 | 23.68 | 24.65 | -0.97 | 0.97 | -0.03 | 0.03 |
| gene570 | https://www.ncbi.nlm.nih.gov/gene/?term=LOC105278549 | 1673 | 72 | 430.36 | 1 | 5.98 | 1.39 | 42.9 | 43.9 | -0.99 | 0.99 | -0.04 | 0.04 |
| gene1220 | https://www.ncbi.nlm.nih.gov/gene/?term=LOC105285742 | 3453 | 119 | 344.63 | 1 | 2.9 | 0.84 | 16.38 | 17.43 | -1.05 | 1.05 | -0.04 | 0.04 |
| gene9598 | https://www.ncbi.nlm.nih.gov/gene/?term=LOC105283132 | 1769 | 56 | 316.56 | 1 | 5.65 | 1.79 | 39.06 | 40.18 | -1.12 | 1.12 | -0.04 | 0.04 |
| gene4962 | https://www.ncbi.nlm.nih.gov/gene/?term=LOC105277761 | 4645 | 156 | 335.84 | 1 | 2.15 | 0.64 | 14.16 | 15.31 | -1.15 | 1.15 | -0.04 | 0.04 |
| gene3168 | https://www.ncbi.nlm.nih.gov/gene/?term=LOC105275711 | 6150 | 97 | 157.72 | 1 | 1.63 | 1.03 | 10.51 | 11.68 | -1.17 | 1.17 | -0.02 | 0.02 |
| gene388 | https://www.ncbi.nlm.nih.gov/gene/?term=LOC105276538 | 1558 | 21 | 134.79 | 1 | 6.42 | 4.76 | 69.07 | 70.25 | -1.18 | 1.18 | -0.02 | 0.02 |
| gene3599 | https://www.ncbi.nlm.nih.gov/gene/?term=LOC105276196 | 2955 | 60 | 203.05 | 1 | 3.38 | 1.67 | 37.66 | 38.86 | -1.2 | 1.2 | -0.02 | 0.02 |

| | | | | | | | | | | | | | |
|---|---|---|---|---|---|---|---|---|---|---|---|---|---|
| gene12203 | https://www.ncbi.nlm.nih.gov/gene/?term=LOC105286099 | 5967 | 87 | 145.8 | 1 | 1.68 | 1.15 | 10.81 | 12.06 | -1.25 | 1.25 | -0.02 | 0.02 |
| gene7398 | https://www.ncbi.nlm.nih.gov/gene/?term=LOC105280589 | 3452 | 42 | 121.67 | 1 | 2.9 | 2.38 | 24.95 | 26.26 | -1.31 | 1.31 | -0.02 | 0.02 |
| gene1198 | https://www.ncbi.nlm.nih.gov/gene/?term=LOC105285507 | 2738 | 97 | 354.27 | 1 | 3.65 | 1.03 | 20.27 | 21.73 | -1.46 | 1.46 | -0.05 | 0.05 |
| gene12021 | https://www.ncbi.nlm.nih.gov/gene/?term=LOC105285905 | 1399 | 29 | 207.29 | 1 | 7.15 | 3.45 | 32.77 | 34.25 | -1.48 | 1.48 | -0.03 | 0.03 |
| gene6422 | https://www.ncbi.nlm.nih.gov/gene/?term=LOC105279458 | 2419 | 42 | 173.63 | 1 | 4.13 | 2.38 | 44.07 | 45.58 | -1.51 | 1.51 | -0.03 | 0.03 |
| gene11340 | https://www.ncbi.nlm.nih.gov/gene/?term=LOC105285131 | 2918 | 98 | 335.85 | 1 | 3.43 | 1.02 | 13.78 | 15.33 | -1.55 | 1.55 | -0.05 | 0.05 |
| gene3727 | https://www.ncbi.nlm.nih.gov/gene/?term=LOC105276367 | 2484 | 65 | 261.67 | 1 | 4.03 | 1.54 | 43.95 | 45.54 | -1.59 | 1.59 | -0.04 | 0.04 |
| gene9411 | https://www.ncbi.nlm.nih.gov/gene/?term=LOC105282914 | 5534 | 42 | 75.89 | 1 | 1.81 | 2.38 | 49.22 | 50.92 | -1.7 | 1.7 | -0.01 | 0.01 |
| gene2404 | https://www.ncbi.nlm.nih.gov/gene/?term=LOC105274853 | 1767 | 36 | 203.74 | 1 | 5.66 | 2.78 | 33.17 | 35.01 | -1.84 | 1.84 | -0.04 | 0.04 |
| gene7248 | https://www.ncbi.nlm.nih.gov/gene/?term=LOC105280408 | 5175 | 59 | 114.01 | 1 | 1.93 | 1.69 | 15.82 | 17.71 | -1.89 | 1.89 | -0.02 | 0.02 |
| gene9740 | https://www.ncbi.nlm.nih.gov/gene/?term=LOC105283277 | 2887 | 139 | 481.47 | 1 | 3.46 | 0.72 | 18.49 | 20.55 | -2.05 | 2.05 | -0.1 | 0.1 |
| gene652 | https://www.ncbi.nlm.nih.gov/gene/?term=LOC105279528 | 5806 | 39 | 67.17 | 1 | 1.72 | 2.56 | 17.73 | 19.93 | -2.2 | 2.2 | -0.01 | 0.01 |
| gene9464 | https://www.ncbi.nlm.nih.gov/gene/?term=LOC105282975 | 1450 | 24 | 165.52 | 1 | 6.9 | 4.17 | 39.26 | 42.21 | -2.95 | 2.95 | -0.05 | 0.05 |
| gene1199 | https://www.ncbi.nlm.nih.gov/gene/?term=LOC105285498 | 2220 | 30 | 135.14 | 1 | 4.5 | 3.33 | 33.52 | 36.79 | -3.27 | 3.27 | -0.04 | 0.04 |
| gene6237 | https://www.ncbi.nlm.nih.gov/gene/?term=LOC105279244 | 1484 | 33 | 222.37 | 1 | 6.74 | 3.03 | 44.64 | 48.04 | -3.4 | 3.4 | -0.08 | 0.08 |
| gene1505 | https://www.ncbi.nlm.nih.gov/gene/?term=LOC105287633 | 1693 | 22 | 129.95 | 1 | 5.91 | 4.55 | 51.59 | 55.25 | -3.65 | 3.65 | -0.05 | 0.05 |
| gene2413 | https://www.ncbi.nlm.nih.gov/gene/?term=LOC105274863 | 2006 | 62 | 309.07 | 1 | 4.99 | 1.61 | 48.7 | 53.36 | -4.66 | 4.66 | -0.14 | 0.14 |
| gene564 | https://www.ncbi.nlm.nih.gov/gene/?term=LOC105278478 | 2802 | 16 | 57.1 | 1 | 3.57 | 6.25 | 14.91 | 19.95 | -5.03 | 5.03 | -0.03 | 0.03 |
| gene11584 | https://www.ncbi.nlm.nih.gov/gene/?term=LOC105285416 | 3709 | 16 | 43.14 | 2 | 5.39 | 12.5 | 21.86 | 29.47 | -7.61 | 7.61 | -0.03 | 0.03 |
| gene9109 | https://www.ncbi.nlm.nih.gov/gene/?term=LOC105282561 | 2400 | 14 | 58.33 | 1 | 4.17 | 7.14 | 13.59 | 21.35 | -7.76 | 7.76 | -0.05 | 0.05 |
| gene3166 | https://www.ncbi.nlm.nih.gov/gene/?term=LOC105275707 | 2393 | 10 | 41.79 | 1 | 4.18 | 10 | 65.97 | 76.22 | -10.25 | 10.25 | -0.04 | 0.04 |
| gene10963 | https://www.ncbi.nlm.nih.gov/gene/?term=LOC105284677 | 1576 | 30 | 190.36 | 1 | 6.35 | 3.33 | 19.35 | 34.92 | -15.57 | 15.57 | -0.3 | 0.3 |
| gene393 | https://www.ncbi.nlm.nih.gov/gene/?term=LOC105276566 | 1123 | 2 | 17.81 | 1 | 8.9 | 50 | 47.22 | 74.3 | -27.08 | 27.08 | -0.05 | 0.05 |

Directory: OGLall File: ogl-Cb_LI_BCphaseB1.vs.RphaseB2.txt

| gene_ID | gene_link | gwidth | #Sites | #per10Kb | #dmSites | #dmsp10kb | %dmSites | %pSite1 | %pSite2 | DMpSite | ADMpSite | DMpNucl | ADMpNucl |
|---|---|---|---|---|---|---|---|---|---|---|---|---|---|
| gene2593 | https://www.ncbi.nlm.nih.gov/gene/?term=LOC105275070 | 1774 | 7 | 39.46 | 5 | 28.18 | 71.43 | 58.65 | 4.09 | 54.56 | 54.56 | 0.22 | 0.22 |
| gene3165 | https://www.ncbi.nlm.nih.gov/gene/?term=LOC105275706 | 2374 | 68 | 286.44 | 5 | 21.06 | 7.35 | 33.32 | 23.13 | 10.19 | 10.19 | 0.29 | 0.29 |
| gene2970 | https://www.ncbi.nlm.nih.gov/gene/?term=LOC105275518 | 2340 | 20 | 85.47 | 1 | 4.27 | 5 | 58.11 | 52.81 | 5.3 | 5.3 | 0.05 | 0.05 |
| gene5714 | https://www.ncbi.nlm.nih.gov/gene/?term=LOC105278729 | 710 | 17 | 239.44 | 1 | 14.08 | 5.88 | 58.14 | 53.54 | 4.59 | 4.59 | 0.11 | 0.11 |
| gene1518 | https://www.ncbi.nlm.nih.gov/gene/?term=LOC105287648 | 2195 | 25 | 113.9 | 1 | 4.56 | 4 | 44.43 | 41.18 | 3.25 | 3.25 | 0.04 | 0.04 |
| gene940 | https://www.ncbi.nlm.nih.gov/gene/?term=LOC105282794 | 3578 | 83 | 231.97 | 1 | 2.79 | 1.2 | 17.02 | 13.98 | 3.04 | 3.04 | 0.07 | 0.07 |
| gene5647 | https://www.ncbi.nlm.nih.gov/gene/?term=LOC105278567 | 3118 | 36 | 115.46 | 1 | 3.21 | 2.78 | 16.65 | 14.27 | 2.38 | 2.38 | 0.03 | 0.03 |
| gene5273 | https://www.ncbi.nlm.nih.gov/gene/?term=LOC105278108 | 4067 | 46 | 113.11 | 1 | 2.46 | 2.17 | 27.18 | 24.92 | 2.26 | 2.26 | 0.03 | 0.03 |
| gene6334 | https://www.ncbi.nlm.nih.gov/gene/?term=LOC105279356 | 1471 | 48 | 326.31 | 1 | 6.8 | 2.08 | 28.86 | 26.95 | 1.91 | 1.91 | 0.06 | 0.06 |
| gene8178 | https://www.ncbi.nlm.nih.gov/gene/?term=LOC105281515 | 8647 | 44 | 50.88 | 1 | 1.16 | 2.27 | 25.92 | 24.04 | 1.88 | 1.88 | 0.01 | 0.01 |
| gene7734 | https://www.ncbi.nlm.nih.gov/gene/?term=LOC105280972 | 1528 | 30 | 196.34 | 1 | 6.54 | 3.33 | 20.49 | 18.79 | 1.71 | 1.71 | 0.03 | 0.03 |
| gene1855 | https://www.ncbi.nlm.nih.gov/gene/?term=LOC105288036 | 2387 | 78 | 326.77 | 1 | 4.19 | 1.28 | 33.45 | 31.79 | 1.66 | 1.66 | 0.05 | 0.05 |
| gene8661 | https://www.ncbi.nlm.nih.gov/gene/?term=LOC105282063 | 1926 | 34 | 176.53 | 1 | 5.19 | 2.94 | 30.6 | 28.98 | 1.62 | 1.62 | 0.03 | 0.03 |
| gene7199 | https://www.ncbi.nlm.nih.gov/gene/?term=LOC105280366 | 1414 | 22 | 155.59 | 1 | 7.07 | 4.55 | 43.24 | 41.76 | 1.47 | 1.47 | 0.02 | 0.02 |
| gene11819 | https://www.ncbi.nlm.nih.gov/gene/?term=LOC105285671 | 2366 | 102 | 431.11 | 1 | 4.23 | 0.98 | 27.3 | 25.91 | 1.39 | 1.39 | 0.06 | 0.06 |
| gene7568 | https://www.ncbi.nlm.nih.gov/gene/?term=LOC105280786 | 2525 | 122 | 483.17 | 1 | 3.96 | 0.82 | 29.81 | 28.44 | 1.38 | 1.38 | 0.07 | 0.07 |
| gene6966 | https://www.ncbi.nlm.nih.gov/gene/?term=LOC105280104 | 3747 | 135 | 360.29 | 1 | 2.67 | 0.74 | 17.67 | 16.38 | 1.29 | 1.29 | 0.05 | 0.05 |
| gene2748 | https://www.ncbi.nlm.nih.gov/gene/?term=LOC105275261 | 3845 | 54 | 140.44 | 1 | 2.6 | 1.85 | 40.26 | 39.17 | 1.09 | 1.09 | 0.02 | 0.02 |
| gene4213 | https://www.ncbi.nlm.nih.gov/gene/?term=LOC105276901 | 3257 | 81 | 248.7 | 1 | 3.07 | 1.23 | 28.04 | 26.96 | 1.09 | 1.09 | 0.03 | 0.03 |
| gene1332 | https://www.ncbi.nlm.nih.gov/gene/?term=LOC105286872 | 2460 | 154 | 626.02 | 1 | 4.07 | 0.65 | 21.26 | 20.18 | 1.08 | 1.08 | 0.07 | 0.07 |
| gene6455 | https://www.ncbi.nlm.nih.gov/gene/?term=LOC105279500 | 2612 | 35 | 134 | 1 | 3.83 | 2.86 | 49.33 | 48.29 | 1.04 | 1.04 | 0.01 | 0.01 |
| gene2844 | https://www.ncbi.nlm.nih.gov/gene/?term=LOC105275363 | 4330 | 190 | 438.8 | 1 | 2.31 | 0.53 | 14.79 | 13.86 | 0.93 | 0.93 | 0.04 | 0.04 |
| gene10678 | https://www.ncbi.nlm.nih.gov/gene/?term=LOC105284355 | 4031 | 166 | 411.81 | 1 | 2.48 | 0.6 | 19.71 | 18.79 | 0.92 | 0.92 | 0.04 | 0.04 |
| gene10600 | https://www.ncbi.nlm.nih.gov/gene/?term=LOC105284270 | 4837 | 119 | 246.02 | 1 | 2.07 | 0.84 | 10.98 | 10.08 | 0.9 | 0.9 | 0.02 | 0.02 |
| gene7398 | https://www.ncbi.nlm.nih.gov/gene/?term=LOC105280589 | 3452 | 33 | 95.6 | 1 | 2.9 | 3.03 | 21.34 | 20.45 | 0.9 | 0.9 | 0.01 | 0.01 |
| gene4034 | https://www.ncbi.nlm.nih.gov/gene/?term=LOC105276704 | 2065 | 87 | 421.31 | 1 | 4.84 | 1.15 | 14.97 | 14.08 | 0.89 | 0.89 | 0.04 | 0.04 |
| gene5831 | https://www.ncbi.nlm.nih.gov/gene/?term=LOC105278761 | 2506 | 110 | 438.95 | 1 | 3.99 | 0.91 | 23.47 | 22.6 | 0.88 | 0.88 | 0.04 | 0.04 |
| gene11401 | https://www.ncbi.nlm.nih.gov/gene/?term=LOC105285202 | 1841 | 79 | 429.11 | 1 | 5.43 | 1.27 | 24.33 | 23.48 | 0.85 | 0.85 | 0.04 | 0.04 |
| gene9336 | https://www.ncbi.nlm.nih.gov/gene/?term=LOC105282834 | 5993 | 124 | 206.91 | 1 | 1.67 | 0.81 | 16.5 | 15.65 | 0.85 | 0.85 | 0.02 | 0.02 |
| gene8274 | https://www.ncbi.nlm.nih.gov/gene/?term=LOC105281624 | 5425 | 211 | 388.94 | 1 | 1.84 | 0.47 | 9.95 | 9.18 | 0.77 | 0.77 | 0.03 | 0.03 |
| gene3470 | https://www.ncbi.nlm.nih.gov/gene/?term=LOC105276059 | 8716 | 157 | 180.13 | 1 | 1.15 | 0.64 | 8.72 | 7.97 | 0.76 | 0.76 | 0.01 | 0.01 |
| gene303 | https://www.ncbi.nlm.nih.gov/gene/?term=LOC105275584 | 3081 | 86 | 279.13 | 1 | 3.25 | 1.16 | 20.08 | 19.37 | 0.71 | 0.71 | 0.02 | 0.02 |
| gene12516 | https://www.ncbi.nlm.nih.gov/gene/?term=LOC105286452 | 1529 | 38 | 248.53 | 1 | 6.54 | 2.63 | 50.05 | 49.37 | 0.68 | 0.68 | 0.02 | 0.02 |
| gene11568 | https://www.ncbi.nlm.nih.gov/gene/?term=LOC105285395 | 8086 | 190 | 234.97 | 1 | 1.24 | 0.53 | 16.67 | 16 | 0.67 | 0.67 | 0.02 | 0.02 |
| gene4892 | https://www.ncbi.nlm.nih.gov/gene/?term=LOC105277667 | 3898 | 126 | 323.24 | 1 | 2.57 | 0.79 | 27.81 | 27.16 | 0.65 | 0.65 | 0.02 | 0.02 |
| gene10520 | https://www.ncbi.nlm.nih.gov/gene/?term=LOC105284184 | 2451 | 84 | 342.72 | 1 | 4.08 | 1.19 | 33.8 | 33.16 | 0.64 | 0.64 | 0.02 | 0.02 |
| gene8252 | https://www.ncbi.nlm.nih.gov/gene/?term=LOC105281599 | 3813 | 49 | 128.51 | 1 | 2.62 | 2.04 | 4.72 | 4.12 | 0.6 | 0.6 | 0.01 | 0.01 |
| gene473 | https://www.ncbi.nlm.nih.gov/gene/?term=LOC105277423 | 2342 | 106 | 452.6 | 1 | 4.27 | 0.94 | 28.94 | 28.4 | 0.54 | 0.54 | 0.02 | 0.02 |
| gene7884 | https://www.ncbi.nlm.nih.gov/gene/?term=LOC105281266 | 3390 | 76 | 224.19 | 1 | 2.95 | 1.32 | 16.63 | 16.11 | 0.52 | 0.52 | 0.01 | 0.01 |
| gene11579 | https://www.ncbi.nlm.nih.gov/gene/?term=LOC105285406 | 13571 | 226 | 166.53 | 1 | 0.74 | 0.44 | 7.69 | 7.18 | 0.51 | 0.51 | 0.01 | 0.01 |
| gene5877 | https://www.ncbi.nlm.nih.gov/gene/?term=LOC105278807 | 1719 | 49 | 285.05 | 1 | 5.82 | 2.04 | 27.11 | 26.65 | 0.46 | 0.46 | 0.01 | 0.01 |
| gene10540 | https://www.ncbi.nlm.nih.gov/gene/?term=LOC105284200 | 5267 | 220 | 417.7 | 1 | 1.9 | 0.45 | 12.54 | 12.08 | 0.45 | 0.45 | 0.02 | 0.02 |
| gene11082 | https://www.ncbi.nlm.nih.gov/gene/?term=LOC105284824 | 7436 | 200 | 268.96 | 1 | 1.34 | 0.5 | 7.89 | 7.46 | 0.43 | 0.43 | 0.01 | 0.01 |
| gene7552 | https://www.ncbi.nlm.nih.gov/gene/?term=LOC105280769 | 8566 | 287 | 335.05 | 1 | 1.17 | 0.35 | 2.06 | 1.63 | 0.43 | 0.43 | 0.01 | 0.01 |
| gene6583 | https://www.ncbi.nlm.nih.gov/gene/?term=LOC105279649 | 7444 | 320 | 429.88 | 1 | 1.34 | 0.31 | 11.91 | 11.5 | 0.41 | 0.41 | 0.02 | 0.02 |
| gene10531 | https://www.ncbi.nlm.nih.gov/gene/?term=LOC105284190 | 6881 | 186 | 270.31 | 1 | 1.45 | 0.54 | 12.46 | 12.07 | 0.39 | 0.39 | 0.01 | 0.01 |
| gene298 | https://www.ncbi.nlm.nih.gov/gene/?term=LOC105275460 | 2473 | 117 | 473.11 | 1 | 4.04 | 0.85 | 13.71 | 13.31 | 0.39 | 0.39 | 0.02 | 0.02 |
| gene7556 | https://www.ncbi.nlm.nih.gov/gene/?term=LOC105280774 | 2792 | 97 | 347.42 | 1 | 3.58 | 1.03 | 33.33 | 32.94 | 0.39 | 0.39 | 0.01 | 0.01 |
| gene12806 | https://www.ncbi.nlm.nih.gov/gene/?term=LOC105286803 | 4196 | 141 | 336.03 | 1 | 2.38 | 0.71 | 9.78 | 9.4 | 0.38 | 0.38 | 0.01 | 0.01 |
| gene10844 | https://www.ncbi.nlm.nih.gov/gene/?term=LOC105284535 | 9086 | 231 | 254.24 | 1 | 1.1 | 0.43 | 15.03 | 14.66 | 0.37 | 0.37 | 0.01 | 0.01 |
| gene13114 | https://www.ncbi.nlm.nih.gov/gene/?term=LOC105287147 | 7609 | 398 | 523.06 | 1 | 1.31 | 0.25 | 9.72 | 9.4 | 0.33 | 0.33 | 0.02 | 0.02 |
| gene292 | https://www.ncbi.nlm.nih.gov/gene/?term=LOC105275398 | 2825 | 81 | 286.73 | 1 | 3.54 | 1.23 | 24.98 | 24.69 | 0.3 | 0.3 | 0.01 | 0.01 |
| gene5524 | https://www.ncbi.nlm.nih.gov/gene/?term=LOC105278397 | 5177 | 198 | 382.46 | 1 | 1.93 | 0.51 | 24.98 | 24.69 | 0.29 | 0.29 | 0.01 | 0.01 |
| gene2173 | https://www.ncbi.nlm.nih.gov/gene/?term=LOC105274590 | 4387 | 70 | 159.56 | 2 | 4.56 | 2.86 | 26.31 | 26.05 | 0.27 | 0.27 | 0 | 0 |
| gene2271 | https://www.ncbi.nlm.nih.gov/gene/?term=LOC105274706 | 1448 | 51 | 352.21 | 1 | 6.91 | 1.96 | 40.12 | 39.86 | 0.26 | 0.26 | 0.01 | 0.01 |
| gene4216 | https://www.ncbi.nlm.nih.gov/gene/?term=LOC105276906 | 2127 | 90 | 423.13 | 1 | 4.7 | 1.11 | 21.35 | 21.12 | 0.23 | 0.23 | 0.01 | 0.01 |
| gene10018 | https://www.ncbi.nlm.nih.gov/gene/?term=LOC105283607 | 4414 | 207 | 468.96 | 1 | 2.27 | 0.48 | 15.29 | 15.09 | 0.2 | 0.2 | 0.01 | 0.01 |
| gene6639 | https://www.ncbi.nlm.nih.gov/gene/?term=LOC105279705 | 7656 | 67 | 87.51 | 1 | 1.31 | 1.49 | 22.85 | 22.67 | 0.17 | 0.17 | 0 | 0 |
| gene10596 | https://www.ncbi.nlm.nih.gov/gene/?term=LOC105284262 | 6520 | 208 | 319.02 | 1 | 1.53 | 0.48 | 4.3 | 4.14 | 0.16 | 0.16 | 0.01 | 0.01 |
| gene1632 | https://www.ncbi.nlm.nih.gov/gene/?term=LOC105287764 | 18061 | 391 | 216.49 | 2 | 1.11 | 0.51 | 3.3 | 3.14 | 0.16 | 0.16 | 0 | 0 |

| | | | | | | | | | | | | | |
|---|---|---|---|---|---|---|---|---|---|---|---|---|---|
| gene3206 | https://www.ncbi.nlm.nih.gov/gene/?term=LOC105275752 | 3474 | 118 | 339.67 | 1 | 2.88 | 0.85 | 32.15 | 31.99 | 0.16 | 0.16 | 0.01 | 0.01 |
| gene570 | https://www.ncbi.nlm.nih.gov/gene/?term=LOC105278549 | 1673 | 63 | 376.57 | 1 | 5.98 | 1.59 | 43.38 | 43.23 | 0.15 | 0.15 | 0.01 | 0.01 |
| gene862 | https://www.ncbi.nlm.nih.gov/gene/?term=LOC105281802 | 73922 | 1907 | 257.97 | 1 | 0.14 | 0.05 | 2.33 | 2.19 | 0.14 | 0.14 | 0 | 0 |
| gene7888 | https://www.ncbi.nlm.nih.gov/gene/?term=LOC105281141 | 4357 | 153 | 351.16 | 1 | 2.3 | 0.65 | 32.3 | 32.17 | 0.13 | 0.13 | 0 | 0 |
| gene5525 | https://www.ncbi.nlm.nih.gov/gene/?term=LOC105278396 | 3090 | 182 | 589 | 1 | 3.24 | 0.55 | 24.88 | 24.76 | 0.12 | 0.12 | 0.01 | 0.01 |
| gene5746 | https://www.ncbi.nlm.nih.gov/gene/?term=LOC105278666 | 10343 | 441 | 426.38 | 1 | 0.97 | 0.23 | 6.88 | 6.76 | 0.12 | 0.12 | 0.01 | 0.01 |
| gene10392 | https://www.ncbi.nlm.nih.gov/gene/?term=LOC105284036 | 3216 | 99 | 307.84 | 2 | 6.22 | 2.02 | 22.54 | 22.43 | 0.11 | 0.11 | 0 | 0 |
| gene7368 | https://www.ncbi.nlm.nih.gov/gene/?term=LOC105280556 | 4105 | 41 | 99.88 | 1 | 2.44 | 2.44 | 16.06 | 15.95 | 0.11 | 0.11 | 0 | 0 |
| gene2806 | https://www.ncbi.nlm.nih.gov/gene/?term=LOC105275327 | 14257 | 473 | 331.77 | 1 | 0.7 | 0.21 | 7.14 | 7.04 | 0.1 | 0.1 | 0 | 0 |
| gene1470 | https://www.ncbi.nlm.nih.gov/gene/?term=LOC105287597 | 2974 | 186 | 625.42 | 1 | 3.36 | 0.54 | 18.48 | 18.39 | 0.09 | 0.09 | 0.01 | 0.01 |
| gene3885 | https://www.ncbi.nlm.nih.gov/gene/?term=LOC105276527 | 7928 | 318 | 401.11 | 1 | 1.26 | 0.31 | 9.68 | 9.6 | 0.09 | 0.09 | 0 | 0 |
| gene9101 | https://www.ncbi.nlm.nih.gov/gene/?term=LOC105282553 | 3068 | 104 | 338.98 | 1 | 3.26 | 0.96 | 23.44 | 23.39 | 0.06 | 0.06 | 0 | 0 |
| gene1408 | https://www.ncbi.nlm.nih.gov/gene/?term=LOC109611520 | 8645 | 306 | 353.96 | 1 | 1.16 | 0.33 | 6.02 | 5.98 | 0.05 | 0.05 | 0 | 0 |
| gene2633 | https://www.ncbi.nlm.nih.gov/gene/?term=LOC105275127 | 7457 | 252 | 337.94 | 1 | 1.34 | 0.4 | 8.36 | 8.33 | 0.03 | 0.03 | 0 | 0 |
| gene10326 | https://www.ncbi.nlm.nih.gov/gene/?term=LOC105283962 | 3042 | 124 | 407.63 | 1 | 3.29 | 0.81 | 8.49 | 8.5 | -0.01 | 0.01 | 0 | 0 |
| gene11275 | https://www.ncbi.nlm.nih.gov/gene/?term=LOC105285055 | 4119 | 252 | 611.8 | 1 | 2.43 | 0.4 | 11.84 | 11.86 | -0.03 | 0.03 | 0 | 0 |
| gene6620 | https://www.ncbi.nlm.nih.gov/gene/?term=LOC105279687 | 17998 | 355 | 197.24 | 1 | 0.56 | 0.28 | 5.15 | 5.19 | -0.04 | 0.04 | 0 | 0 |
| gene7915 | https://www.ncbi.nlm.nih.gov/gene/?term=LOC105281173 | 1597 | 39 | 244.21 | 1 | 6.26 | 2.56 | 59.75 | 59.8 | -0.05 | 0.05 | 0 | 0 |
| gene12572 | https://www.ncbi.nlm.nih.gov/gene/?term=LOC105286514 | 2374 | 49 | 206.4 | 1 | 4.21 | 2.04 | 35.44 | 35.51 | -0.07 | 0.07 | 0 | 0 |
| gene2683 | https://www.ncbi.nlm.nih.gov/gene/?term=LOC105275187 | 5492 | 70 | 127.46 | 1 | 1.82 | 1.43 | 12.3 | 12.38 | -0.07 | 0.07 | 0 | 0 |
| gene5387 | https://www.ncbi.nlm.nih.gov/gene/?term=LOC105278245 | 6451 | 112 | 173.62 | 1 | 1.55 | 0.89 | 22.85 | 22.92 | -0.07 | 0.07 | 0 | 0 |
| gene8814 | https://www.ncbi.nlm.nih.gov/gene/?term=LOC105282240 | 6910 | 176 | 254.7 | 1 | 1.45 | 0.57 | 9.18 | 9.27 | -0.08 | 0.08 | 0 | 0 |
| gene8600 | https://www.ncbi.nlm.nih.gov/gene/?term=LOC105282001 | 8513 | 151 | 177.38 | 1 | 1.17 | 0.66 | 10.34 | 10.43 | -0.09 | 0.09 | 0 | 0 |
| gene8729 | https://www.ncbi.nlm.nih.gov/gene/?term=LOC105282137 | 5005 | 211 | 421.58 | 1 | 2 | 0.47 | 15.81 | 15.91 | -0.09 | 0.09 | 0 | 0 |
| gene2404 | https://www.ncbi.nlm.nih.gov/gene/?term=LOC105274853 | 1767 | 35 | 198.08 | 1 | 5.66 | 2.86 | 33.43 | 33.53 | -0.1 | 0.1 | 0 | 0 |
| gene5101 | https://www.ncbi.nlm.nih.gov/gene/?term=LOC105277903 | 1813 | 35 | 193.05 | 1 | 5.52 | 2.86 | 51.16 | 51.27 | -0.1 | 0.1 | 0 | 0 |
| gene53 | https://www.ncbi.nlm.nih.gov/gene/?term=LOC105277148 | 19283 | 295 | 152.98 | 1 | 0.52 | 0.34 | 3.8 | 3.9 | -0.1 | 0.1 | 0 | 0 |
| gene5646 | https://www.ncbi.nlm.nih.gov/gene/?term=LOC105278534 | 7475 | 187 | 250.17 | 1 | 1.34 | 0.53 | 18.3 | 18.4 | -0.1 | 0.1 | 0 | 0 |
| gene2846 | https://www.ncbi.nlm.nih.gov/gene/?term=LOC105275374 | 11246 | 477 | 424.15 | 1 | 0.89 | 0.21 | 10.56 | 10.68 | -0.11 | 0.11 | 0 | 0 |
| gene34 | https://www.ncbi.nlm.nih.gov/gene/?term=LOC105278416 | 7120 | 140 | 196.63 | 1 | 1.4 | 0.71 | 13.56 | 13.67 | -0.11 | 0.11 | 0 | 0 |
| gene7166 | https://www.ncbi.nlm.nih.gov/gene/?term=LOC105280326 | 6117 | 243 | 397.25 | 1 | 1.63 | 0.41 | 14.54 | 14.65 | -0.11 | 0.11 | 0 | 0 |
| gene4190 | https://www.ncbi.nlm.nih.gov/gene/?term=LOC105276875 | 3150 | 117 | 371.43 | 1 | 3.17 | 0.85 | 21.8 | 21.95 | -0.15 | 0.15 | -0.01 | 0.01 |
| gene12802 | https://www.ncbi.nlm.nih.gov/gene/?term=LOC105286799 | 3183 | 164 | 515.24 | 1 | 3.14 | 0.61 | 17.26 | 17.43 | -0.17 | 0.17 | -0.01 | 0.01 |
| gene1430 | https://www.ncbi.nlm.nih.gov/gene/?term=LOC105287551 | 6433 | 223 | 346.65 | 1 | 1.55 | 0.45 | 8.23 | 8.41 | -0.18 | 0.18 | -0.01 | 0.01 |
| gene9326 | https://www.ncbi.nlm.nih.gov/gene/?term=LOC105282818 | 16612 | 178 | 107.15 | 1 | 0.6 | 0.56 | 13.41 | 13.58 | -0.18 | 0.18 | 0 | 0 |
| gene6286 | https://www.ncbi.nlm.nih.gov/gene/?term=LOC105279298 | 2861 | 129 | 450.89 | 2 | 6.99 | 1.55 | 23.63 | 23.86 | -0.23 | 0.23 | -0.01 | 0.01 |
| gene4962 | https://www.ncbi.nlm.nih.gov/gene/?term=LOC105277761 | 4645 | 144 | 310.01 | 1 | 2.15 | 0.69 | 13.44 | 13.75 | -0.31 | 0.31 | -0.01 | 0.01 |
| gene9327 | https://www.ncbi.nlm.nih.gov/gene/?term=LOC105282821 | 7793 | 145 | 186.06 | 1 | 1.28 | 0.69 | 12.57 | 12.89 | -0.32 | 0.32 | -0.01 | 0.01 |
| gene2957 | https://www.ncbi.nlm.nih.gov/gene/?term=LOC105275500 | 10926 | 240 | 219.66 | 1 | 0.92 | 0.42 | 6.99 | 7.33 | -0.34 | 0.34 | -0.01 | 0.01 |
| gene12307 | https://www.ncbi.nlm.nih.gov/gene/?term=LOC105286216 | 10313 | 187 | 181.32 | 1 | 0.97 | 0.53 | 14.69 | 15.04 | -0.35 | 0.35 | -0.01 | 0.01 |
| gene10155 | https://www.ncbi.nlm.nih.gov/gene/?term=LOC105283756 | 7030 | 112 | 159.32 | 1 | 1.42 | 0.89 | 5.92 | 6.29 | -0.37 | 0.37 | -0.01 | 0.01 |
| gene4023 | https://www.ncbi.nlm.nih.gov/gene/?term=LOC105276692 | 1759 | 31 | 176.24 | 1 | 5.69 | 3.23 | 23.25 | 23.63 | -0.38 | 0.38 | -0.01 | 0.01 |
| gene1310 | https://www.ncbi.nlm.nih.gov/gene/?term=LOC105286661 | 2582 | 71 | 274.98 | 1 | 3.87 | 1.41 | 21.29 | 21.68 | -0.39 | 0.39 | -0.01 | 0.01 |
| gene12573 | https://www.ncbi.nlm.nih.gov/gene/?term=LOC105286515 | 3647 | 100 | 274.2 | 1 | 2.74 | 1 | 13.03 | 13.44 | -0.42 | 0.42 | -0.01 | 0.01 |
| gene13036 | https://www.ncbi.nlm.nih.gov/gene/?term=LOC105287059 | 4959 | 160 | 322.65 | 1 | 2.02 | 0.62 | 19.69 | 20.15 | -0.46 | 0.46 | -0.01 | 0.01 |
| gene920 | https://www.ncbi.nlm.nih.gov/gene/?term=LOC105282518 | 1197 | 52 | 434.42 | 1 | 8.35 | 1.92 | 30.27 | 30.77 | -0.49 | 0.49 | -0.02 | 0.02 |
| gene10549 | https://www.ncbi.nlm.nih.gov/gene/?term=LOC105284210 | 7145 | 290 | 405.88 | 2 | 2.8 | 0.69 | 11.49 | 12 | -0.51 | 0.51 | -0.02 | 0.02 |
| gene1199 | https://www.ncbi.nlm.nih.gov/gene/?term=LOC105285498 | 2220 | 28 | 126.13 | 1 | 4.5 | 3.57 | 27.44 | 27.99 | -0.55 | 0.55 | -0.01 | 0.01 |
| gene657 | https://www.ncbi.nlm.nih.gov/gene/?term=LOC105279667 | 6421 | 81 | 126.15 | 1 | 1.56 | 1.23 | 2.49 | 3.12 | -0.63 | 0.63 | -0.01 | 0.01 |
| gene7183 | https://www.ncbi.nlm.nih.gov/gene/?term=LOC105280348 | 3925 | 70 | 178.34 | 1 | 2.55 | 1.43 | 20.85 | 21.48 | -0.63 | 0.63 | -0.01 | 0.01 |
| gene7249 | https://www.ncbi.nlm.nih.gov/gene/?term=LOC105280422 | 1775 | 59 | 332.39 | 1 | 5.63 | 1.69 | 16.45 | 17.08 | -0.63 | 0.63 | -0.02 | 0.02 |
| gene12021 | https://www.ncbi.nlm.nih.gov/gene/?term=LOC105285905 | 1399 | 15 | 107.22 | 1 | 7.15 | 6.67 | 50 | 50.68 | -0.68 | 0.68 | -0.01 | 0.01 |
| gene7247 | https://www.ncbi.nlm.nih.gov/gene/?term=LOC105280407 | 10486 | 206 | 196.45 | 1 | 0.95 | 0.49 | 16.14 | 16.82 | -0.68 | 0.68 | -0.01 | 0.01 |
| gene11561 | https://www.ncbi.nlm.nih.gov/gene/?term=LOC105285387 | 11528 | 178 | 154.41 | 1 | 0.87 | 0.56 | 9.89 | 10.61 | -0.72 | 0.72 | -0.01 | 0.01 |
| gene9740 | https://www.ncbi.nlm.nih.gov/gene/?term=LOC105283277 | 2887 | 137 | 474.54 | 1 | 3.46 | 0.73 | 19.95 | 20.71 | -0.76 | 0.76 | -0.04 | 0.04 |
| gene12265 | https://www.ncbi.nlm.nih.gov/gene/?term=LOC105286166 | 3050 | 110 | 360.66 | 1 | 3.28 | 0.91 | 22.97 | 23.8 | -0.82 | 0.82 | -0.03 | 0.03 |
| gene10606 | https://www.ncbi.nlm.nih.gov/gene/?term=LOC105284273 | 6119 | 248 | 405.29 | 1 | 1.63 | 0.4 | 17.15 | 17.98 | -0.83 | 0.83 | -0.03 | 0.03 |
| gene35 | https://www.ncbi.nlm.nih.gov/gene/?term=LOC105275197 | 2643 | 50 | 189.18 | 1 | 3.78 | 2 | 37.01 | 37.84 | -0.83 | 0.83 | -0.02 | 0.02 |
| gene5298 | https://www.ncbi.nlm.nih.gov/gene/?term=LOC105278145 | 4060 | 235 | 578.82 | 2 | 4.93 | 0.85 | 10.4 | 11.25 | -0.85 | 0.85 | -0.05 | 0.05 |
| gene4376 | https://www.ncbi.nlm.nih.gov/gene/?term=LOC105277083 | 4307 | 102 | 236.82 | 1 | 2.32 | 0.98 | 24.25 | 25.14 | -0.89 | 0.89 | -0.02 | 0.02 |
| gene1220 | https://www.ncbi.nlm.nih.gov/gene/?term=LOC105285742 | 3453 | 111 | 321.46 | 1 | 2.9 | 0.9 | 21.24 | 22.19 | -0.96 | 0.96 | -0.03 | 0.03 |
| gene12203 | https://www.ncbi.nlm.nih.gov/gene/?term=LOC105286099 | 5967 | 75 | 125.69 | 1 | 1.68 | 1.33 | 9.56 | 10.52 | -0.96 | 0.96 | -0.01 | 0.01 |
| gene6031 | https://www.ncbi.nlm.nih.gov/gene/?term=LOC105279000 | 5691 | 179 | 314.53 | 1 | 1.76 | 0.56 | 13.44 | 14.41 | -0.97 | 0.97 | -0.03 | 0.03 |
| gene7211 | https://www.ncbi.nlm.nih.gov/gene/?term=LOC105280377 | 2024 | 67 | 331.03 | 1 | 4.94 | 1.49 | 44.45 | 45.48 | -1.03 | 1.03 | -0.03 | 0.03 |

| | | | | | | | | | | | | | |
|---|---|---|---|---|---|---|---|---|---|---|---|---|---|
| gene4033 | https://www.ncbi.nlm.nih.gov/gene/?term=LOC105276705 | 10546 | 142 | 134.65 | 1 | 0.95 | 0.7 | 7.83 | 8.89 | -1.06 | 1.06 | -0.01 | 0.01 |
| gene3385 | https://www.ncbi.nlm.nih.gov/gene/?term=LOC105275966 | 3305 | 58 | 175.49 | 1 | 3.03 | 1.72 | 49.09 | 50.19 | -1.1 | 1.1 | -0.02 | 0.02 |
| gene12981 | https://www.ncbi.nlm.nih.gov/gene/?term=LOC105286999 | 2063 | 46 | 222.98 | 1 | 4.85 | 2.17 | 31.8 | 32.93 | -1.13 | 1.13 | -0.03 | 0.03 |
| gene12406 | https://www.ncbi.nlm.nih.gov/gene/?term=LOC105286316 | 1386 | 33 | 238.1 | 1 | 7.22 | 3.03 | 40.83 | 41.98 | -1.15 | 1.15 | -0.03 | 0.03 |
| gene10213 | https://www.ncbi.nlm.nih.gov/gene/?term=LOC105283826 | 1503 | 54 | 359.28 | 1 | 6.65 | 1.85 | 14.52 | 15.73 | -1.21 | 1.21 | -0.04 | 0.04 |
| gene3793 | https://www.ncbi.nlm.nih.gov/gene/?term=LOC105276429 | 2901 | 112 | 386.07 | 1 | 3.45 | 0.89 | 25.16 | 26.39 | -1.23 | 1.23 | -0.05 | 0.05 |
| gene11340 | https://www.ncbi.nlm.nih.gov/gene/?term=LOC105285131 | 2918 | 88 | 301.58 | 1 | 3.43 | 1.14 | 10.13 | 11.45 | -1.32 | 1.32 | -0.04 | 0.04 |
| gene322 | https://www.ncbi.nlm.nih.gov/gene/?term=LOC105275777 | 1606 | 28 | 174.35 | 1 | 6.23 | 3.57 | 42.75 | 44.07 | -1.32 | 1.32 | -0.02 | 0.02 |
| gene7248 | https://www.ncbi.nlm.nih.gov/gene/?term=LOC105280408 | 5175 | 52 | 100.48 | 1 | 1.93 | 1.92 | 16.62 | 18.28 | -1.65 | 1.65 | -0.02 | 0.02 |
| gene8343 | https://www.ncbi.nlm.nih.gov/gene/?term=LOC105281700 | 2172 | 32 | 147.33 | 1 | 4.6 | 3.12 | 37.31 | 38.97 | -1.66 | 1.66 | -0.02 | 0.02 |
| gene1147 | https://www.ncbi.nlm.nih.gov/gene/?term=LOC105284954 | 4022 | 90 | 223.77 | 1 | 2.49 | 1.11 | 16.68 | 18.39 | -1.71 | 1.71 | -0.04 | 0.04 |
| gene2413 | https://www.ncbi.nlm.nih.gov/gene/?term=LOC105274863 | 2006 | 49 | 244.27 | 1 | 4.99 | 2.04 | 53.75 | 55.47 | -1.72 | 1.72 | -0.04 | 0.04 |
| gene13235 | https://www.ncbi.nlm.nih.gov/gene/?term=LOC105287278 | 2679 | 34 | 126.91 | 1 | 3.73 | 2.94 | 22.53 | 24.28 | -1.75 | 1.75 | -0.02 | 0.02 |
| gene3727 | https://www.ncbi.nlm.nih.gov/gene/?term=LOC105276367 | 2484 | 63 | 253.62 | 1 | 4.03 | 1.59 | 45.38 | 47.24 | -1.87 | 1.87 | -0.05 | 0.05 |
| gene4641 | https://www.ncbi.nlm.nih.gov/gene/?term=LOC105277398 | 1937 | 71 | 366.55 | 1 | 5.16 | 1.41 | 37.78 | 39.69 | -1.91 | 1.91 | -0.07 | 0.07 |
| gene10107 | https://www.ncbi.nlm.nih.gov/gene/?term=LOC105283712 | 1883 | 35 | 185.87 | 1 | 5.31 | 2.86 | 26.3 | 28.27 | -1.97 | 1.97 | -0.04 | 0.04 |
| gene4498 | https://www.ncbi.nlm.nih.gov/gene/?term=LOC105277239 | 2831 | 121 | 427.41 | 1 | 3.53 | 0.83 | 11.26 | 13.23 | -1.97 | 1.97 | -0.08 | 0.08 |
| gene10345 | https://www.ncbi.nlm.nih.gov/gene/?term=LOC105283989 | 4444 | 27 | 60.76 | 1 | 2.25 | 3.7 | 36.46 | 38.69 | -2.24 | 2.24 | -0.01 | 0.01 |
| gene388 | https://www.ncbi.nlm.nih.gov/gene/?term=LOC105276538 | 1558 | 18 | 115.53 | 2 | 12.84 | 11.11 | 66.09 | 68.41 | -2.32 | 2.32 | -0.03 | 0.03 |
| gene3168 | https://www.ncbi.nlm.nih.gov/gene/?term=LOC105275711 | 6150 | 69 | 112.2 | 1 | 1.63 | 1.45 | 12.54 | 15.09 | -2.54 | 2.54 | -0.03 | 0.03 |
| gene3599 | https://www.ncbi.nlm.nih.gov/gene/?term=LOC105276196 | 2955 | 59 | 199.66 | 1 | 3.38 | 1.69 | 37.76 | 40.32 | -2.56 | 2.56 | -0.05 | 0.05 |
| gene1505 | https://www.ncbi.nlm.nih.gov/gene/?term=LOC105287633 | 1693 | 27 | 159.48 | 1 | 5.91 | 3.7 | 42.55 | 45.35 | -2.8 | 2.8 | -0.04 | 0.04 |
| gene6237 | https://www.ncbi.nlm.nih.gov/gene/?term=LOC105279244 | 1484 | 25 | 168.46 | 1 | 6.74 | 4 | 41.13 | 44.31 | -3.18 | 3.18 | -0.05 | 0.05 |
| gene564 | https://www.ncbi.nlm.nih.gov/gene/?term=LOC105278478 | 2802 | 17 | 60.67 | 1 | 3.57 | 5.88 | 14.04 | 17.34 | -3.31 | 3.31 | -0.02 | 0.02 |
| gene6422 | https://www.ncbi.nlm.nih.gov/gene/?term=LOC105279458 | 2419 | 38 | 157.09 | 1 | 4.13 | 2.63 | 46.62 | 50.37 | -3.75 | 3.75 | -0.06 | 0.06 |
| gene10602 | https://www.ncbi.nlm.nih.gov/gene/?term=LOC105284268 | 2122 | 19 | 89.54 | 1 | 4.71 | 5.26 | 26.09 | 32.99 | -6.9 | 6.9 | -0.06 | 0.06 |
| gene1008 | https://www.ncbi.nlm.nih.gov/gene/?term=LOC105283492 | 1479 | 10 | 67.61 | 1 | 6.76 | 10 | 34.7 | 42 | -7.3 | 7.3 | -0.05 | 0.05 |
| gene393 | https://www.ncbi.nlm.nih.gov/gene/?term=LOC105276566 | 1123 | 2 | 17.81 | 1 | 8.9 | 50 | 47.22 | 76.78 | -29.56 | 29.56 | -0.05 | 0.05 |

Directory: OGLall File: ogl-Cb_LI_BCphaseB2.txt

| gene_ID | gene_link | gwidth | #Sites | #per10Kb | %perSite | %pNucl |
|---|---|---|---|---|---|---|
| gene12538 | https://www.ncbi.nlm.nih.gov/gene/?term=LOC105286531 | 372 | 43 | 1155.91 | 35.54 | 4.11 |
| gene13384 | https://www.ncbi.nlm.nih.gov/gene/?term=LOC105287447 | 1877 | 245 | 1305.27 | 28.47 | 3.72 |
| gene5917 | https://www.ncbi.nlm.nih.gov/gene/?term=LOC105278849 | 823 | 126 | 1530.98 | 23.79 | 3.64 |
| gene5911 | https://www.ncbi.nlm.nih.gov/gene/?term=LOC105278844 | 943 | 98 | 1039.24 | 34.92 | 3.63 |
| gene211 | https://www.ncbi.nlm.nih.gov/gene/?term=LOC105274534 | 1280 | 101 | 789.06 | 45.75 | 3.61 |
| gene11569 | https://www.ncbi.nlm.nih.gov/gene/?term=LOC105285394 | 1311 | 61 | 465.29 | 72.04 | 3.35 |
| gene12364 | https://www.ncbi.nlm.nih.gov/gene/?term=LOC105286269 | 1403 | 90 | 641.48 | 50.55 | 3.24 |
| gene5309 | https://www.ncbi.nlm.nih.gov/gene/?term=LOC105278160 | 1704 | 155 | 909.62 | 35.44 | 3.22 |
| gene6078 | https://www.ncbi.nlm.nih.gov/gene/?term=LOC105279052 | 1130 | 138 | 1221.24 | 26.05 | 3.18 |
| gene3719 | https://www.ncbi.nlm.nih.gov/gene/?term=LOC105276360 | 1471 | 160 | 1087.7 | 29.08 | 3.16 |
| gene12330 | https://www.ncbi.nlm.nih.gov/gene/?term=LOC105286235 | 1501 | 112 | 746.17 | 41.99 | 3.13 |
| gene1562 | https://www.ncbi.nlm.nih.gov/gene/?term=LOC105287692 | 1491 | 129 | 865.19 | 35.56 | 3.08 |
| gene12746 | https://www.ncbi.nlm.nih.gov/gene/?term=LOC105286737 | 2688 | 363 | 1350.45 | 22.13 | 2.99 |
| gene190 | https://www.ncbi.nlm.nih.gov/gene/?term=LOC105288088 | 1797 | 127 | 706.73 | 41.77 | 2.95 |
| gene3731 | https://www.ncbi.nlm.nih.gov/gene/?term=LOC105276371 | 865 | 57 | 658.96 | 43.94 | 2.9 |
| gene1654 | https://www.ncbi.nlm.nih.gov/gene/?term=LOC105287811 | 2285 | 150 | 656.46 | 43.71 | 2.87 |
| gene2797 | https://www.ncbi.nlm.nih.gov/gene/?term=LOC105275314 | 1489 | 93 | 624.58 | 45.82 | 2.86 |
| gene9471 | https://www.ncbi.nlm.nih.gov/gene/?term=LOC105282986 | 1605 | 140 | 872.27 | 32.66 | 2.85 |
| gene570 | https://www.ncbi.nlm.nih.gov/gene/?term=LOC105278549 | 1673 | 107 | 639.57 | 44.01 | 2.81 |
| gene5301 | https://www.ncbi.nlm.nih.gov/gene/?term=LOC105278148 | 1881 | 192 | 1020.73 | 27.32 | 2.79 |
| gene6978 | https://www.ncbi.nlm.nih.gov/gene/?term=LOC105280117 | 1816 | 96 | 528.63 | 52.73 | 2.79 |
| gene12403 | https://www.ncbi.nlm.nih.gov/gene/?term=LOC105286311 | 1413 | 64 | 452.94 | 61.18 | 2.77 |
| gene4878 | https://www.ncbi.nlm.nih.gov/gene/?term=LOC105277648 | 675 | 36 | 533.33 | 51.21 | 2.73 |
| gene4635 | https://www.ncbi.nlm.nih.gov/gene/?term=LOC105277390 | 2067 | 125 | 604.74 | 44.98 | 2.72 |

...
... only first 25 lines shown ...

Directory: OGLall File: ogl-Cb_LI_BCphaseB2.vs.RphaseA1.txt

| gene_ID | gene_link | gwidth | #Sites | #per10Kb | #dmSites | #dmsp10kb | %dmSites | %pSite1 | %pSite2 | DMpSite | ADMpSite | DMpNucl | ADMpNucl |
|---|---|---|---|---|---|---|---|---|---|---|---|---|---|
| gene393 | https://www.ncbi.nlm.nih.gov/gene/?term=LOC105276566 | 1123 | 2 | 17.81 | 1 | 8.9 | 50 | 70.57 | 50 | 20.57 | 20.57 | 0.04 | 0.04 |
| gene12021 | https://www.ncbi.nlm.nih.gov/gene/?term=LOC105285905 | 1399 | 21 | 150.11 | 1 | 7.15 | 4.76 | 36.22 | 31.01 | 5.22 | 5.22 | 0.08 | 0.08 |
| gene9411 | https://www.ncbi.nlm.nih.gov/gene/?term=LOC105282914 | 5534 | 41 | 74.09 | 1 | 1.81 | 2.44 | 52.21 | 48.58 | 3.63 | 3.63 | 0.03 | 0.03 |
| gene1199 | https://www.ncbi.nlm.nih.gov/gene/?term=LOC105285498 | 2220 | 25 | 112.61 | 1 | 4.5 | 4 | 37.37 | 33.76 | 3.61 | 3.61 | 0.04 | 0.04 |
| gene388 | https://www.ncbi.nlm.nih.gov/gene/?term=LOC105276538 | 1558 | 20 | 128.37 | 2 | 12.84 | 10 | 66.14 | 62.93 | 3.2 | 3.2 | 0.04 | 0.04 |
| gene10714 | https://www.ncbi.nlm.nih.gov/gene/?term=LOC105284396 | 1966 | 62 | 315.36 | 1 | 5.09 | 1.61 | 36.35 | 33.21 | 3.14 | 3.14 | 0.1 | 0.1 |
| gene322 | https://www.ncbi.nlm.nih.gov/gene/?term=LOC105275777 | 1606 | 25 | 155.67 | 1 | 6.23 | 4 | 44.37 | 41.49 | 2.89 | 2.89 | 0.04 | 0.04 |
| gene12406 | https://www.ncbi.nlm.nih.gov/gene/?term=LOC105286316 | 1386 | 31 | 223.67 | 1 | 7.22 | 3.23 | 46.61 | 43.84 | 2.77 | 2.77 | 0.06 | 0.06 |
| gene6652 | https://www.ncbi.nlm.nih.gov/gene/?term=LOC105279719 | 2831 | 86 | 303.78 | 1 | 3.53 | 1.16 | 23.08 | 20.39 | 2.69 | 2.69 | 0.08 | 0.08 |
| gene3168 | https://www.ncbi.nlm.nih.gov/gene/?term=LOC105275711 | 6150 | 73 | 118.7 | 1 | 1.63 | 1.37 | 14.69 | 12.05 | 2.64 | 2.64 | 0.03 | 0.03 |
| gene6422 | https://www.ncbi.nlm.nih.gov/gene/?term=LOC105279458 | 2419 | 37 | 152.96 | 1 | 4.13 | 2.7 | 52.5 | 49.97 | 2.53 | 2.53 | 0.04 | 0.04 |
| gene6455 | https://www.ncbi.nlm.nih.gov/gene/?term=LOC105279500 | 2612 | 30 | 114.85 | 1 | 3.83 | 3.33 | 48.51 | 46 | 2.5 | 2.5 | 0.03 | 0.03 |
| gene564 | https://www.ncbi.nlm.nih.gov/gene/?term=LOC105278478 | 2802 | 18 | 64.24 | 1 | 3.57 | 5.56 | 17.32 | 14.97 | 2.35 | 2.35 | 0.02 | 0.02 |
| gene6806 | https://www.ncbi.nlm.nih.gov/gene/?term=LOC105279894 | 3649 | 35 | 95.92 | 1 | 2.74 | 2.86 | 40.58 | 38.26 | 2.32 | 2.32 | 0.02 | 0.02 |
| gene9101 | https://www.ncbi.nlm.nih.gov/gene/?term=LOC105282553 | 3068 | 106 | 345.5 | 1 | 3.26 | 0.94 | 23.35 | 21.05 | 2.29 | 2.29 | 0.08 | 0.08 |
| gene7199 | https://www.ncbi.nlm.nih.gov/gene/?term=LOC105280366 | 1414 | 33 | 233.38 | 1 | 7.07 | 3.03 | 43.27 | 41 | 2.28 | 2.28 | 0.05 | 0.05 |
| gene2404 | https://www.ncbi.nlm.nih.gov/gene/?term=LOC105274853 | 1767 | 43 | 243.35 | 1 | 5.66 | 2.33 | 31.85 | 29.62 | 2.23 | 2.23 | 0.05 | 0.05 |
| gene6237 | https://www.ncbi.nlm.nih.gov/gene/?term=LOC105279244 | 1484 | 23 | 154.99 | 1 | 6.74 | 4.35 | 50.08 | 47.92 | 2.16 | 2.16 | 0.03 | 0.03 |
| gene2271 | https://www.ncbi.nlm.nih.gov/gene/?term=LOC105274706 | 1448 | 51 | 352.21 | 1 | 6.91 | 1.96 | 39.73 | 37.8 | 1.94 | 1.94 | 0.07 | 0.07 |
| gene2748 | https://www.ncbi.nlm.nih.gov/gene/?term=LOC105275261 | 3845 | 54 | 140.44 | 1 | 2.6 | 1.85 | 39.59 | 37.65 | 1.94 | 1.94 | 0.03 | 0.03 |
| gene10107 | https://www.ncbi.nlm.nih.gov/gene/?term=LOC105283712 | 1883 | 35 | 185.87 | 1 | 5.31 | 2.86 | 27.16 | 25.31 | 1.85 | 1.85 | 0.03 | 0.03 |
| gene6639 | https://www.ncbi.nlm.nih.gov/gene/?term=LOC105279705 | 7656 | 68 | 88.82 | 1 | 1.31 | 1.47 | 23.58 | 21.76 | 1.82 | 1.82 | 0.02 | 0.02 |
| gene4498 | https://www.ncbi.nlm.nih.gov/gene/?term=LOC105277239 | 2831 | 130 | 459.2 | 1 | 3.53 | 0.77 | 12.47 | 10.69 | 1.78 | 1.78 | 0.08 | 0.08 |
| gene1907 | https://www.ncbi.nlm.nih.gov/gene/?term=LOC105288099 | 1238 | 35 | 282.71 | 1 | 8.08 | 2.86 | 34.89 | 33.16 | 1.73 | 1.73 | 0.05 | 0.05 |
| gene920 | https://www.ncbi.nlm.nih.gov/gene/?term=LOC105282518 | 1197 | 41 | 342.52 | 1 | 8.35 | 2.44 | 39.23 | 37.74 | 1.49 | 1.49 | 0.05 | 0.05 |
| gene7550 | https://www.ncbi.nlm.nih.gov/gene/?term=LOC105280763 | 2173 | 98 | 450.99 | 1 | 4.6 | 1.02 | 27.13 | 25.7 | 1.43 | 1.43 | 0.06 | 0.06 |
| gene10602 | https://www.ncbi.nlm.nih.gov/gene/?term=LOC105284268 | 2122 | 21 | 98.96 | 1 | 4.71 | 4.76 | 25.67 | 24.45 | 1.22 | 1.22 | 0.01 | 0.01 |
| gene3671 | https://www.ncbi.nlm.nih.gov/gene/?term=LOC105276272 | 1966 | 46 | 233.98 | 1 | 5.09 | 2.17 | 27.89 | 26.72 | 1.16 | 1.16 | 0.03 | 0.03 |
| gene5646 | https://www.ncbi.nlm.nih.gov/gene/?term=LOC105278534 | 7475 | 185 | 247.49 | 1 | 1.34 | 0.54 | 18.24 | 17.21 | 1.03 | 1.03 | 0.03 | 0.03 |
| gene3046 | https://www.ncbi.nlm.nih.gov/gene/?term=LOC105275587 | 2394 | 58 | 242.27 | 1 | 4.18 | 1.72 | 36.72 | 35.74 | 0.98 | 0.98 | 0.02 | 0.02 |
| gene7915 | https://www.ncbi.nlm.nih.gov/gene/?term=LOC105281173 | 1597 | 38 | 237.95 | 1 | 6.26 | 2.63 | 60.79 | 59.81 | 0.98 | 0.98 | 0.02 | 0.02 |
| gene8015 | https://www.ncbi.nlm.nih.gov/gene/?term=LOC105281312 | 2998 | 65 | 216.81 | 1 | 3.34 | 1.54 | 33.24 | 32.3 | 0.93 | 0.93 | 0.02 | 0.02 |
| gene7398 | https://www.ncbi.nlm.nih.gov/gene/?term=LOC105280589 | 3452 | 36 | 104.29 | 1 | 2.9 | 2.78 | 20.14 | 19.24 | 0.9 | 0.9 | 0.01 | 0.01 |
| gene10392 | https://www.ncbi.nlm.nih.gov/gene/?term=LOC105284036 | 3216 | 106 | 329.6 | 2 | 6.22 | 1.89 | 21.2 | 20.33 | 0.87 | 0.87 | 0.03 | 0.03 |
| gene1310 | https://www.ncbi.nlm.nih.gov/gene/?term=LOC105286661 | 2582 | 62 | 240.12 | 1 | 3.87 | 1.61 | 22.65 | 21.78 | 0.87 | 0.87 | 0.02 | 0.02 |
| gene9462 | https://www.ncbi.nlm.nih.gov/gene/?term=LOC105282970 | 1469 | 41 | 279.1 | 1 | 6.81 | 2.44 | 50 | 49.15 | 0.85 | 0.85 | 0.02 | 0.02 |
| gene11340 | https://www.ncbi.nlm.nih.gov/gene/?term=LOC105285131 | 2918 | 105 | 359.84 | 1 | 3.43 | 0.95 | 14.38 | 13.54 | 0.84 | 0.84 | 0.03 | 0.03 |
| gene6286 | https://www.ncbi.nlm.nih.gov/gene/?term=LOC105279298 | 2861 | 131 | 457.88 | 2 | 6.99 | 1.53 | 24.99 | 24.15 | 0.84 | 0.84 | 0.04 | 0.04 |
| gene1220 | https://www.ncbi.nlm.nih.gov/gene/?term=LOC105285742 | 3453 | 106 | 306.98 | 1 | 2.9 | 0.94 | 18.53 | 17.74 | 0.79 | 0.79 | 0.02 | 0.02 |
| gene8729 | https://www.ncbi.nlm.nih.gov/gene/?term=LOC105282137 | 5005 | 200 | 399.6 | 1 | 2 | 0.5 | 16.14 | 15.41 | 0.72 | 0.72 | 0.03 | 0.03 |
| gene10018 | https://www.ncbi.nlm.nih.gov/gene/?term=LOC105283607 | 4414 | 201 | 455.37 | 1 | 2.27 | 0.5 | 15.61 | 14.98 | 0.63 | 0.63 | 0.03 | 0.03 |
| gene4190 | https://www.ncbi.nlm.nih.gov/gene/?term=LOC105276875 | 3150 | 111 | 352.38 | 1 | 3.17 | 0.9 | 22.75 | 22.14 | 0.61 | 0.61 | 0.02 | 0.02 |
| gene12203 | https://www.ncbi.nlm.nih.gov/gene/?term=LOC105286099 | 5967 | 72 | 120.66 | 1 | 1.68 | 1.39 | 10.57 | 10 | 0.57 | 0.57 | 0.01 | 0.01 |
| gene5298 | https://www.ncbi.nlm.nih.gov/gene/?term=LOC105278145 | 4060 | 237 | 583.74 | 3 | 7.39 | 1.27 | 11.65 | 11.08 | 0.57 | 0.57 | 0.03 | 0.03 |
| gene4962 | https://www.ncbi.nlm.nih.gov/gene/?term=LOC105277761 | 4645 | 147 | 316.47 | 1 | 2.15 | 0.68 | 14.43 | 13.91 | 0.52 | 0.52 | 0.02 | 0.02 |
| gene10520 | https://www.ncbi.nlm.nih.gov/gene/?term=LOC105284184 | 2451 | 93 | 379.44 | 1 | 4.08 | 1.08 | 31.28 | 30.8 | 0.49 | 0.49 | 0.02 | 0.02 |
| gene10844 | https://www.ncbi.nlm.nih.gov/gene/?term=LOC105284535 | 9086 | 238 | 261.94 | 1 | 1.1 | 0.42 | 14.5 | 14.01 | 0.49 | 0.49 | 0.01 | 0.01 |
| gene2173 | https://www.ncbi.nlm.nih.gov/gene/?term=LOC105274590 | 4387 | 80 | 182.36 | 1 | 2.28 | 1.25 | 27.39 | 26.91 | 0.48 | 0.48 | 0.01 | 0.01 |
| gene10326 | https://www.ncbi.nlm.nih.gov/gene/?term=LOC105283962 | 3042 | 126 | 414.2 | 1 | 3.29 | 0.79 | 9.42 | 8.95 | 0.47 | 0.47 | 0.02 | 0.02 |
| gene6031 | https://www.ncbi.nlm.nih.gov/gene/?term=LOC105279000 | 5691 | 195 | 342.65 | 1 | 1.76 | 0.51 | 13.76 | 13.29 | 0.47 | 0.47 | 0.02 | 0.02 |
| gene10600 | https://www.ncbi.nlm.nih.gov/gene/?term=LOC105284270 | 4837 | 115 | 237.75 | 1 | 2.07 | 0.87 | 11.71 | 11.26 | 0.46 | 0.46 | 0.01 | 0.01 |
| gene2257 | https://www.ncbi.nlm.nih.gov/gene/?term=LOC105274689 | 3658 | 73 | 199.56 | 1 | 2.73 | 1.37 | 16.2 | 15.75 | 0.46 | 0.46 | 0.01 | 0.01 |
| gene3885 | https://www.ncbi.nlm.nih.gov/gene/?term=LOC105276527 | 7928 | 318 | 401.11 | 1 | 1.26 | 0.31 | 10.14 | 9.68 | 0.46 | 0.46 | 0.02 | 0.02 |
| gene12572 | https://www.ncbi.nlm.nih.gov/gene/?term=LOC105286514 | 2374 | 44 | 185.34 | 1 | 4.21 | 2.27 | 36.23 | 35.78 | 0.45 | 0.45 | 0.01 | 0.01 |
| gene7600 | https://www.ncbi.nlm.nih.gov/gene/?term=LOC105280819 | 1615 | 45 | 278.64 | 1 | 6.19 | 2.22 | 37.23 | 36.8 | 0.43 | 0.43 | 0.01 | 0.01 |
| gene8274 | https://www.ncbi.nlm.nih.gov/gene/?term=LOC105281624 | 5425 | 219 | 403.69 | 1 | 1.84 | 0.46 | 10.56 | 10.14 | 0.42 | 0.42 | 0.02 | 0.02 |
| gene9740 | https://www.ncbi.nlm.nih.gov/gene/?term=LOC105283277 | 2887 | 136 | 471.08 | 1 | 3.46 | 0.74 | 19.63 | 19.22 | 0.41 | 0.41 | 0.02 | 0.02 |
| gene10213 | https://www.ncbi.nlm.nih.gov/gene/?term=LOC105283826 | 1503 | 64 | 425.82 | 1 | 6.65 | 1.56 | 16.48 | 16.1 | 0.38 | 0.38 | 0.02 | 0.02 |
| gene3599 | https://www.ncbi.nlm.nih.gov/gene/?term=LOC105276196 | 2955 | 57 | 192.89 | 1 | 3.38 | 1.75 | 38.84 | 38.45 | 0.38 | 0.38 | 0.01 | 0.01 |
| gene3107 | https://www.ncbi.nlm.nih.gov/gene/?term=LOC105275650 | 5103 | 174 | 340.98 | 1 | 1.96 | 0.57 | 16.52 | 16.19 | 0.33 | 0.33 | 0.01 | 0.01 |

| | | | | | | | | | | | | | |
|---|---|---|---|---|---|---|---|---|---|---|---|---|---|
| gene10155 | https://www.ncbi.nlm.nih.gov/gene/?term=LOC105283756 | 7030 | 104 | 147.94 | 1 | 1.42 | 0.96 | 5.54 | 5.23 | 0.31 | 0.31 | 0 | 0 |
| gene6966 | https://www.ncbi.nlm.nih.gov/gene/?term=LOC105280104 | 3747 | 121 | 322.93 | 1 | 2.67 | 0.83 | 16.24 | 15.96 | 0.28 | 0.28 | 0.01 | 0.01 |
| gene9445 | https://www.ncbi.nlm.nih.gov/gene/?term=LOC105282951 | 2285 | 114 | 498.91 | 1 | 4.38 | 0.88 | 20.31 | 20.03 | 0.28 | 0.28 | 0.01 | 0.01 |
| gene1470 | https://www.ncbi.nlm.nih.gov/gene/?term=LOC105287597 | 2974 | 195 | 655.68 | 1 | 3.36 | 0.51 | 18.15 | 17.88 | 0.27 | 0.27 | 0.02 | 0.02 |
| gene11454 | https://www.ncbi.nlm.nih.gov/gene/?term=LOC105285256 | 1906 | 51 | 267.58 | 1 | 5.25 | 1.96 | 42.03 | 41.77 | 0.26 | 0.26 | 0.01 | 0.01 |
| gene12307 | https://www.ncbi.nlm.nih.gov/gene/?term=LOC105286216 | 10313 | 160 | 155.14 | 1 | 0.97 | 0.62 | 16.94 | 16.73 | 0.22 | 0.22 | 0 | 0 |
| gene5524 | https://www.ncbi.nlm.nih.gov/gene/?term=LOC105278397 | 5177 | 202 | 390.19 | 1 | 1.93 | 0.5 | 25.27 | 25.04 | 0.22 | 0.22 | 0.01 | 0.01 |
| gene570 | https://www.ncbi.nlm.nih.gov/gene/?term=LOC105278549 | 1673 | 69 | 412.43 | 1 | 5.98 | 1.45 | 43.78 | 43.57 | 0.21 | 0.21 | 0.01 | 0.01 |
| gene7547 | https://www.ncbi.nlm.nih.gov/gene/?term=LOC105280767 | 24576 | 785 | 319.42 | 1 | 0.41 | 0.13 | 4.48 | 4.27 | 0.21 | 0.21 | 0.01 | 0.01 |
| gene9177 | https://www.ncbi.nlm.nih.gov/gene/?term=LOC105282646 | 3320 | 101 | 304.22 | 1 | 3.01 | 0.99 | 21.24 | 21.05 | 0.2 | 0.2 | 0.01 | 0.01 |
| gene10886 | https://www.ncbi.nlm.nih.gov/gene/?term=LOC105284582 | 1475 | 49 | 332.2 1 | 6.78 | 2.04 | 22 | 21.8 | 0.19 | 0.19 | 0.01 | 0.01 | |
| gene5525 | https://www.ncbi.nlm.nih.gov/gene/?term=LOC105278396 | 3090 | 186 | 601.94 | 1 | 3.24 | 0.54 | 25.06 | 24.87 | 0.18 | 0.18 | 0.01 | 0.01 |
| gene7183 | https://www.ncbi.nlm.nih.gov/gene/?term=LOC105280348 | 3925 | 70 | 178.34 | 1 | 2.55 | 1.43 | 18.6 | 18.42 | 0.18 | 0.18 | 0 | 0 |
| gene7249 | https://www.ncbi.nlm.nih.gov/gene/?term=LOC105280422 | 1775 | 63 | 354.93 | 1 | 5.63 | 1.59 | 17.74 | 17.55 | 0.18 | 0.18 | 0.01 | 0.01 |
| gene13036 | https://www.ncbi.nlm.nih.gov/gene/?term=LOC105287059 | 4959 | 165 | 332.73 | 1 | 2.02 | 0.61 | 17.15 | 16.98 | 0.17 | 0.17 | 0.01 | 0.01 |
| gene10531 | https://www.ncbi.nlm.nih.gov/gene/?term=LOC105284190 | 6881 | 175 | 254.32 | 1 | 1.45 | 0.57 | 12.25 | 12.09 | 0.16 | 0.16 | 0 | 0 |
| gene10606 | https://www.ncbi.nlm.nih.gov/gene/?term=LOC105284273 | 6119 | 255 | 416.73 | 1 | 1.63 | 0.39 | 19.24 | 19.09 | 0.16 | 0.16 | 0.01 | 0.01 |
| gene5777 | https://www.ncbi.nlm.nih.gov/gene/?term=LOC105278691 | 3883 | 152 | 391.45 | 1 | 2.58 | 0.66 | 14.89 | 14.73 | 0.16 | 0.16 | 0.01 | 0.01 |
| gene11082 | https://www.ncbi.nlm.nih.gov/gene/?term=LOC105284824 | 7436 | 212 | 285.1 1 | 1.34 | 0.47 | 6.95 | 6.8 | 0.15 | 0.15 | 0 | 0 | |
| gene7166 | https://www.ncbi.nlm.nih.gov/gene/?term=LOC105280326 | 6117 | 238 | 389.08 | 1 | 1.63 | 0.42 | 15.04 | 14.89 | 0.15 | 0.15 | 0.01 | 0.01 |
| gene8343 | https://www.ncbi.nlm.nih.gov/gene/?term=LOC105281700 | 2172 | 37 | 170.35 | 1 | 4.6 | 2.7 | 39.63 | 39.48 | 0.15 | 0.15 | 0 | 0 |
| gene13114 | https://www.ncbi.nlm.nih.gov/gene/?term=LOC105287147 | 7609 | 419 | 550.66 | 1 | 1.31 | 0.24 | 8.81 | 8.67 | 0.14 | 0.14 | 0.01 | 0.01 |
| gene53 | https://www.ncbi.nlm.nih.gov/gene/?term=LOC105277148 | 19283 | 291 | 150.91 | 1 | 0.52 | 0.34 | 4.8 | 4.67 | 0.13 | 0.13 | 0 | 0 |
| gene1632 | https://www.ncbi.nlm.nih.gov/gene/?term=LOC105287764 | 18061 | 379 | 209.84 | 1 | 0.55 | 0.26 | 3.62 | 3.51 | 0.11 | 0.11 | 0 | 0 |
| gene10549 | https://www.ncbi.nlm.nih.gov/gene/?term=LOC105284210 | 7145 | 335 | 468.86 | 2 | 2.8 | 0.6 | 10.35 | 10.25 | 0.09 | 0.09 | 0 | 0 |
| gene2413 | https://www.ncbi.nlm.nih.gov/gene/?term=LOC105274863 | 2006 | 53 | 264.21 | 1 | 4.99 | 1.89 | 52.76 | 52.69 | 0.07 | 0.07 | 0 | 0 |
| gene6805 | https://www.ncbi.nlm.nih.gov/gene/?term=LOC105279895 | 87888 | 973 | 110.71 | 1 | 0.11 | 0.1 | 1.87 | 1.81 | 0.06 | 0.06 | 0 | 0 |
| gene11275 | https://www.ncbi.nlm.nih.gov/gene/?term=LOC105285055 | 4119 | 252 | 611.8 1 | 2.43 | 0.4 | 10.71 | 10.67 | 0.05 | 0.05 | 0 | 0 | |
| gene862 | https://www.ncbi.nlm.nih.gov/gene/?term=LOC105281802 | 73922 | 1969 | 266.36 | 1 | 0.14 | 0.05 | 2.17 | 2.12 | 0.05 | 0.05 | 0 | 0 |
| gene1332 | https://www.ncbi.nlm.nih.gov/gene/?term=LOC105286872 | 2460 | 158 | 642.28 | 1 | 4.07 | 0.63 | 20.39 | 20.35 | 0.04 | 0.04 | 0 | 0 |
| gene3856 | https://www.ncbi.nlm.nih.gov/gene/?term=LOC105276499 | 5236 | 196 | 374.33 | 1 | 1.91 | 0.51 | 10.18 | 10.15 | 0.03 | 0.03 | 0 | 0 |
| gene1147 | https://www.ncbi.nlm.nih.gov/gene/?term=LOC105284954 | 4022 | 111 | 275.98 | 1 | 2.49 | 0.9 | 11.93 | 11.9 | 0.02 | 0.02 | 0 | 0 |
| gene5197 | https://www.ncbi.nlm.nih.gov/gene/?term=LOC105278018 | 89233 | 3377 | 378.45 | 1 | 0.11 | 0.03 | 0.69 | 0.68 | 0 | 0 | 0 | 0 |
| gene11583 | https://www.ncbi.nlm.nih.gov/gene/?term=LOC105285414 | 8192 | 178 | 217.29 | 1 | 1.22 | 0.56 | 9.44 | 9.48 | -0.04 | 0.04 | 0 | 0 |
| gene5375 | https://www.ncbi.nlm.nih.gov/gene/?term=LOC105278228 | 8640 | 154 | 178.24 | 1 | 1.16 | 0.65 | 24.99 | 25.04 | -0.05 | 0.05 | 0 | 0 |
| gene5746 | https://www.ncbi.nlm.nih.gov/gene/?term=LOC105278666 | 10343 | 477 | 461.18 | 1 | 0.97 | 0.21 | 6.71 | 6.77 | -0.05 | 0.05 | 0 | 0 |
| gene2844 | https://www.ncbi.nlm.nih.gov/gene/?term=LOC105275363 | 4330 | 197 | 454.97 | 1 | 2.31 | 0.51 | 14.26 | 14.32 | -0.06 | 0.06 | 0 | 0 |
| gene7319 | https://www.ncbi.nlm.nih.gov/gene/?term=LOC105280513 | 68997 | 1336 | 193.63 | 1 | 0.14 | 0.07 | 0.68 | 0.75 | -0.06 | 0.06 | 0 | 0 |
| gene10345 | https://www.ncbi.nlm.nih.gov/gene/?term=LOC105283989 | 4444 | 26 | 58.51 1 | 2.25 | 3.85 | 49.48 | 49.58 | -0.09 | 0.09 | 0 | 0 | |
| gene2846 | https://www.ncbi.nlm.nih.gov/gene/?term=LOC105275374 | 11246 | 465 | 413.48 | 1 | 0.89 | 0.22 | 10.6 | 10.73 | -0.13 | 0.13 | -0.01 | 0.01 |
| gene12802 | https://www.ncbi.nlm.nih.gov/gene/?term=LOC105286799 | 3183 | 190 | 596.92 | 1 | 3.14 | 0.53 | 15.49 | 15.63 | -0.14 | 0.14 | -0.01 | 0.01 |
| gene11579 | https://www.ncbi.nlm.nih.gov/gene/?term=LOC105285406 | 13571 | 280 | 206.32 | 1 | 0.74 | 0.36 | 6.48 | 6.65 | -0.17 | 0.17 | 0 | 0 |
| gene8814 | https://www.ncbi.nlm.nih.gov/gene/?term=LOC105282240 | 6910 | 185 | 267.73 | 1 | 1.45 | 0.54 | 9.48 | 9.66 | -0.18 | 0.18 | 0 | 0 |
| gene11561 | https://www.ncbi.nlm.nih.gov/gene/?term=LOC105285387 | 11528 | 207 | 179.56 | 1 | 0.87 | 0.48 | 8.9 | 9.12 | -0.22 | 0.22 | 0 | 0 |
| gene10678 | https://www.ncbi.nlm.nih.gov/gene/?term=LOC105284355 | 4031 | 182 | 451.5 1 | 2.48 | 0.55 | 17.39 | 17.62 | -0.23 | 0.23 | -0.01 | 0.01 | |
| gene34 | https://www.ncbi.nlm.nih.gov/gene/?term=LOC105278416 | 7120 | 132 | 185.39 | 1 | 1.4 | 0.76 | 14.08 | 14.34 | -0.26 | 0.26 | 0 | 0 |
| gene7556 | https://www.ncbi.nlm.nih.gov/gene/?term=LOC105280774 | 2792 | 118 | 422.64 | 1 | 3.58 | 0.85 | 27.01 | 27.27 | -0.26 | 0.26 | -0.01 | 0.01 |
| gene8600 | https://www.ncbi.nlm.nih.gov/gene/?term=LOC105282001 | 8513 | 186 | 218.49 | 1 | 1.17 | 0.54 | 10 | 10.27 | -0.27 | 0.27 | -0.01 | 0.01 |
| gene5052 | https://www.ncbi.nlm.nih.gov/gene/?term=LOC105277856 | 3167 | 33 | 104.2 1 | 3.16 | 3.03 | 39.08 | 39.37 | -0.29 | 0.29 | 0 | 0 | |
| gene2806 | https://www.ncbi.nlm.nih.gov/gene/?term=LOC105275327 | 14257 | 511 | 358.42 | 1 | 0.7 | 0.2 | 6.7 | 7.03 | -0.33 | 0.33 | -0.01 | 0.01 |
| gene35 | https://www.ncbi.nlm.nih.gov/gene/?term=LOC105275197 | 2643 | 48 | 181.61 | 1 | 3.78 | 2.08 | 38.22 | 38.55 | -0.33 | 0.33 | -0.01 | 0.01 |
| gene2668 | https://www.ncbi.nlm.nih.gov/gene/?term=LOC105275170 | 12496 | 279 | 223.27 | 1 | 0.8 | 0.36 | 1.97 | 2.32 | -0.35 | 0.35 | -0.01 | 0.01 |
| gene738 | https://www.ncbi.nlm.nih.gov/gene/?term=LOC105280421 | 4927 | 216 | 438.4 1 | 2.03 | 0.46 | 14.22 | 14.61 | -0.39 | 0.39 | -0.02 | 0.02 | |
| gene1855 | https://www.ncbi.nlm.nih.gov/gene/?term=LOC105288036 | 2387 | 84 | 351.91 | 1 | 4.19 | 1.19 | 29.18 | 29.59 | -0.41 | 0.41 | -0.01 | 0.01 |
| gene7248 | https://www.ncbi.nlm.nih.gov/gene/?term=LOC105280408 | 5175 | 61 | 117.87 | 1 | 1.93 | 1.64 | 20.52 | 20.93 | -0.41 | 0.41 | 0 | 0 |
| gene6620 | https://www.ncbi.nlm.nih.gov/gene/?term=LOC105279687 | 17998 | 368 | 204.47 | 1 | 0.56 | 0.27 | 4.64 | 5.1 | -0.46 | 0.46 | -0.01 | 0.01 |
| gene7247 | https://www.ncbi.nlm.nih.gov/gene/?term=LOC105280407 | 10486 | 218 | 207.9 1 | 0.95 | 0.46 | 16.54 | 17.03 | -0.49 | 0.49 | -0.01 | 0.01 | |
| gene9336 | https://www.ncbi.nlm.nih.gov/gene/?term=LOC105282834 | 5993 | 129 | 215.25 | 1 | 1.67 | 0.78 | 14.53 | 15.02 | -0.49 | 0.49 | -0.01 | 0.01 |
| gene3206 | https://www.ncbi.nlm.nih.gov/gene/?term=LOC105275752 | 3474 | 129 | 371.33 | 1 | 2.88 | 0.78 | 29.67 | 30.17 | -0.5 | 0.5 | -0.02 | 0.02 |
| gene940 | https://www.ncbi.nlm.nih.gov/gene/?term=LOC105282794 | 3578 | 85 | 237.56 | 1 | 2.79 | 1.18 | 13.96 | 14.47 | -0.51 | 0.51 | -0.01 | 0.01 |
| gene298 | https://www.ncbi.nlm.nih.gov/gene/?term=LOC105275460 | 2473 | 120 | 485.24 | 1 | 4.04 | 0.83 | 13.22 | 13.75 | -0.53 | 0.53 | -0.03 | 0.03 |
| gene3470 | https://www.ncbi.nlm.nih.gov/gene/?term=LOC105276059 | 8716 | 152 | 174.39 | 1 | 1.15 | 0.66 | 7.9 | 8.44 | -0.54 | 0.54 | -0.01 | 0.01 |
| gene6583 | https://www.ncbi.nlm.nih.gov/gene/?term=LOC105279649 | 7444 | 340 | 456.74 | 1 | 1.34 | 0.29 | 10.48 | 11.08 | -0.59 | 0.59 | -0.03 | 0.03 |
| gene7514 | https://www.ncbi.nlm.nih.gov/gene/?term=LOC105280723 | 4471 | 208 | 465.22 | 1 | 2.24 | 0.48 | 13.49 | 14.1 | -0.6 | 0.6 | -0.03 | 0.03 |

| | | | | | | | | | | | | | | | |
|---|---|---|---|---|---|---|---|---|---|---|---|---|---|---|---|
| gene4376 | https://www.ncbi.nlm.nih.gov/gene/?term=LOC105277083 | 4307 | 114 | 264.69 | | 1 | 2.32 | 0.88 | 20.15 | 20.76 | -0.61 | 0.61 | -0.02 | 0.02 | |
| gene12516 | https://www.ncbi.nlm.nih.gov/gene/?term=LOC105286452 | 1529 | 35 | 228.91 | | 1 | 6.54 | 2.86 | 51.89 | 52.54 | -0.65 | 0.65 | -0.01 | 0.01 | |
| gene13033 | https://www.ncbi.nlm.nih.gov/gene/?term=LOC105287063 | 1877 | 51 | 271.71 | | 1 | 5.33 | 1.96 | 30.35 | 31.05 | -0.7 | 0.7 | -0.02 | 0.02 | |
| gene2056 | https://www.ncbi.nlm.nih.gov/gene/?term=LOC105274452 | 6584 | 243 | 369.08 | | 1 | 1.52 | 0.41 | 13.17 | 13.88 | -0.71 | 0.71 | -0.03 | 0.03 | |
| gene5733 | https://www.ncbi.nlm.nih.gov/gene/?term=LOC105278649 | 5221 | 91 | 174.3 | 1 | 1.92 | 1.1 | 19.36 | 20.1 | -0.74 | 0.74 | -0.01 | 0.01 | | |
| gene8742 | https://www.ncbi.nlm.nih.gov/gene/?term=LOC105282151 | 2563 | 42 | 163.87 | | 1 | 3.9 | 2.38 | 34.83 | 35.57 | -0.75 | 0.75 | -0.01 | 0.01 | |
| gene1430 | https://www.ncbi.nlm.nih.gov/gene/?term=LOC105287551 | 6433 | 235 | 365.3 | 1 | 1.55 | 0.43 | 8.56 | 9.58 | -1.02 | 1.02 | -0.04 | 0.04 | | |
| gene11401 | https://www.ncbi.nlm.nih.gov/gene/?term=LOC105285202 | 1841 | 75 | 407.39 | | 1 | 5.43 | 1.33 | 24.08 | 25.12 | -1.03 | 1.03 | -0.04 | 0.04 | |
| gene12439 | https://www.ncbi.nlm.nih.gov/gene/?term=LOC105286357 | 4761 | 81 | 170.13 | | 1 | 2.1 | 1.23 | 29.49 | 30.52 | -1.03 | 1.03 | -0.02 | 0.02 | |
| gene2683 | https://www.ncbi.nlm.nih.gov/gene/?term=LOC105275187 | 5492 | 70 | 127.46 | | 1 | 1.82 | 1.43 | 13.71 | 14.76 | -1.05 | 1.05 | -0.01 | 0.01 | |
| gene6528 | https://www.ncbi.nlm.nih.gov/gene/?term=LOC105279591 | 7222 | 59 | 81.69 | 1 | 1.38 | 1.69 | 26.22 | 27.32 | -1.1 | 1.1 | -0.01 | 0.01 | | |
| gene7884 | https://www.ncbi.nlm.nih.gov/gene/?term=LOC105281266 | 3390 | 81 | 238.94 | | 1 | 2.95 | 1.23 | 16.21 | 17.33 | -1.12 | 1.12 | -0.03 | 0.03 | |
| gene2071 | https://www.ncbi.nlm.nih.gov/gene/?term=LOC105274471 | 2446 | 121 | 494.69 | | 1 | 4.09 | 0.83 | 19.71 | 20.9 | -1.19 | 1.19 | -0.06 | 0.06 | |
| gene4023 | https://www.ncbi.nlm.nih.gov/gene/?term=LOC105276692 | 1759 | 36 | 204.66 | | 1 | 5.69 | 2.78 | 20.19 | 21.39 | -1.2 | 1.2 | -0.02 | 0.02 | |
| gene7368 | https://www.ncbi.nlm.nih.gov/gene/?term=LOC105280556 | 4105 | 50 | 121.8 | 1 | 2.44 | 2 | 14.06 | 15.36 | -1.3 | 1.3 | -0.02 | 0.02 | | |
| gene4720 | https://www.ncbi.nlm.nih.gov/gene/?term=LOC105277479 | 1697 | 50 | 294.64 | | 1 | 5.89 | 2 | 35.87 | 37.27 | -1.4 | 1.4 | -0.04 | 0.04 | |
| gene10654 | https://www.ncbi.nlm.nih.gov/gene/?term=LOC105284329 | 4027 | 51 | 126.65 | | 1 | 2.48 | 1.96 | 17.3 | 18.75 | -1.45 | 1.45 | -0.02 | 0.02 | |
| gene2856 | https://www.ncbi.nlm.nih.gov/gene/?term=LOC105275384 | 1987 | 87 | 437.85 | | 1 | 5.03 | 1.15 | 34.24 | 35.77 | -1.53 | 1.53 | -0.07 | 0.07 | |
| gene7568 | https://www.ncbi.nlm.nih.gov/gene/?term=LOC105280786 | 2525 | 109 | 431.68 | | 1 | 3.96 | 0.92 | 28.7 | 30.24 | -1.54 | 1.54 | -0.07 | 0.07 | |
| gene4892 | https://www.ncbi.nlm.nih.gov/gene/?term=LOC105277667 | 3898 | 120 | 307.85 | | 1 | 2.57 | 0.83 | 27.15 | 28.74 | -1.59 | 1.59 | -0.05 | 0.05 | |
| gene1518 | https://www.ncbi.nlm.nih.gov/gene/?term=LOC105287648 | 2195 | 35 | 159.45 | | 1 | 4.56 | 2.86 | 35.74 | 37.41 | -1.67 | 1.67 | -0.03 | 0.03 | |
| gene9478 | https://www.ncbi.nlm.nih.gov/gene/?term=LOC105282991 | 3726 | 58 | 155.66 | | 1 | 2.68 | 1.72 | 33.94 | 35.62 | -1.68 | 1.68 | -0.03 | 0.03 | |
| gene927 | https://www.ncbi.nlm.nih.gov/gene/?term=LOC105282600 | 3727 | 52 | 139.52 | | 1 | 2.68 | 1.92 | 31.26 | 33.17 | -1.91 | 1.91 | -0.03 | 0.03 | |
| gene1884 | https://www.ncbi.nlm.nih.gov/gene/?term=LOC105288072 | 4169 | 78 | 187.1 | 1 | 2.4 | 1.28 | 22.55 | 24.51 | -1.96 | 1.96 | -0.04 | 0.04 | | |
| gene3817 | https://www.ncbi.nlm.nih.gov/gene/?term=LOC105276449 | 1230 | 45 | 365.85 | | 1 | 8.13 | 2.22 | 27.32 | 29.29 | -1.97 | 1.97 | -0.07 | 0.07 | |
| gene9647 | https://www.ncbi.nlm.nih.gov/gene/?term=LOC109611233 | 3094 | 35 | 113.12 | | 1 | 3.23 | 2.86 | 38.21 | 40.18 | -1.97 | 1.97 | -0.02 | 0.02 | |
| gene9648 | https://www.ncbi.nlm.nih.gov/gene/?term=LOC105283183 | 2602 | 33 | 126.83 | | 1 | 3.84 | 3.03 | 40.53 | 42.61 | -2.08 | 2.08 | -0.03 | 0.03 | |
| gene5877 | https://www.ncbi.nlm.nih.gov/gene/?term=LOC105278807 | 1719 | 53 | 308.32 | | 1 | 5.82 | 1.89 | 22.63 | 24.74 | -2.11 | 2.11 | -0.06 | 0.06 | |
| gene8661 | https://www.ncbi.nlm.nih.gov/gene/?term=LOC105282063 | 1926 | 30 | 155.76 | | 1 | 5.19 | 3.33 | 28.06 | 30.33 | -2.28 | 2.28 | -0.04 | 0.04 | |
| gene8178 | https://www.ncbi.nlm.nih.gov/gene/?term=LOC105281515 | 8647 | 63 | 72.86 | 1 | 1.16 | 1.59 | 24.53 | 26.83 | -2.3 | 2.3 | -0.02 | 0.02 | | |
| gene5101 | https://www.ncbi.nlm.nih.gov/gene/?term=LOC105277903 | 1813 | 36 | 198.57 | | 1 | 5.52 | 2.78 | 53.23 | 56.32 | -3.09 | 3.09 | -0.06 | 0.06 | |
| gene5714 | https://www.ncbi.nlm.nih.gov/gene/?term=LOC105278729 | 710 | 21 | 295.77 | | 1 | 14.08 | 4.76 | 56.35 | 60.77 | -4.42 | 4.42 | -0.13 | 0.13 | |
| gene5647 | https://www.ncbi.nlm.nih.gov/gene/?term=LOC105278567 | 3118 | 29 | 93.01 | 1 | 3.21 | 3.45 | 13.21 | 19.43 | -6.21 | 6.21 | -0.06 | 0.06 | | |
| gene3165 | https://www.ncbi.nlm.nih.gov/gene/?term=LOC105275706 | 2374 | 60 | 252.74 | | 3 | 12.64 | 5 | 22.21 | 32.53 | -10.32 | | 10.32 | -0.26 | 0.26 |
| gene2593 | https://www.ncbi.nlm.nih.gov/gene/?term=LOC105275070 | 1774 | 5 | 28.18 | 5 | 28.18 | 100 | 10.96 | 79.22 | -68.26 | | 68.26 | -0.19 | 0.19 | |

Directory: OGLall File: ogl-Cb_LI_BCphaseB2.vs.RphaseA2.txt

| gene_ID | gene_link | gwidth | #Sites | #per10Kb | #dmSites | #dmsp10kb | %dmSites | %pSite1 | %pSite2 | DMpSite | ADMpSite | DMpNucl | ADMpNucl |
|---|---|---|---|---|---|---|---|---|---|---|---|---|---|
| gene393 | https://www.ncbi.nlm.nih.gov/gene/?term=LOC105276566 | 1123 | 2 | 17.81 | 1 | 8.9 | 50 | 70.57 | 47.62 | 22.95 | 22.95 | 0.04 | 0.04 |
| gene10963 | https://www.ncbi.nlm.nih.gov/gene/?term=LOC105284677 | 1576 | 30 | 190.36 | 2 | 12.69 | 6.67 | 38.74 | 23.8 | 14.95 | 14.95 | 0.28 | 0.28 |
| gene9109 | https://www.ncbi.nlm.nih.gov/gene/?term=LOC105282561 | 2400 | 7 | 29.17 | 1 | 4.17 | 14.29 | 63.87 | 57.68 | 6.19 | 6.19 | 0.02 | 0.02 |
| gene10816 | https://www.ncbi.nlm.nih.gov/gene/?term=LOC105284502 | 1974 | 21 | 106.38 | 1 | 5.07 | 4.76 | 45.81 | 40.14 | 5.67 | 5.67 | 0.06 | 0.06 |
| gene1199 | https://www.ncbi.nlm.nih.gov/gene/?term=LOC105285498 | 2220 | 26 | 117.12 | 1 | 4.5 | 3.85 | 37.03 | 32.91 | 4.12 | 4.12 | 0.05 | 0.05 |
| gene9411 | https://www.ncbi.nlm.nih.gov/gene/?term=LOC105282914 | 5534 | 43 | 77.7 | 1 | 1.81 | 2.33 | 49.78 | 47.15 | 2.63 | 2.63 | 0.02 | 0.02 |
| gene920 | https://www.ncbi.nlm.nih.gov/gene/?term=LOC105282518 | 1197 | 42 | 350.88 | 1 | 8.35 | 2.38 | 38.3 | 35.67 | 2.62 | 2.62 | 0.09 | 0.09 |
| gene6237 | https://www.ncbi.nlm.nih.gov/gene/?term=LOC105279244 | 1484 | 25 | 168.46 | 1 | 6.74 | 4 | 53.48 | 51.47 | 2.01 | 2.01 | 0.03 | 0.03 |
| gene6652 | https://www.ncbi.nlm.nih.gov/gene/?term=LOC105279719 | 2831 | 82 | 289.65 | 1 | 3.53 | 1.22 | 24.05 | 22.34 | 1.72 | 1.72 | 0.05 | 0.05 |
| gene12021 | https://www.ncbi.nlm.nih.gov/gene/?term=LOC105285905 | 1399 | 22 | 157.26 | 1 | 7.15 | 4.55 | 34.58 | 32.98 | 1.6 | 1.6 | 0.03 | 0.03 |
| gene3200 | https://www.ncbi.nlm.nih.gov/gene/?term=LOC105275743 | 1786 | 76 | 425.53 | 1 | 5.6 | 1.32 | 23.84 | 22.24 | 1.6 | 1.6 | 0.07 | 0.07 |
| gene9101 | https://www.ncbi.nlm.nih.gov/gene/?term=LOC105282553 | 3068 | 115 | 374.84 | 1 | 3.26 | 0.87 | 23.82 | 22.25 | 1.57 | 1.57 | 0.06 | 0.06 |
| gene6455 | https://www.ncbi.nlm.nih.gov/gene/?term=LOC105279500 | 2612 | 34 | 130.17 | 1 | 3.83 | 2.94 | 45.97 | 44.48 | 1.5 | 1.5 | 0.02 | 0.02 |
| gene10714 | https://www.ncbi.nlm.nih.gov/gene/?term=LOC105284396 | 1966 | 66 | 335.71 | 1 | 5.09 | 1.52 | 34.65 | 33.2 | 1.45 | 1.45 | 0.05 | 0.05 |
| gene7398 | https://www.ncbi.nlm.nih.gov/gene/?term=LOC105280589 | 3452 | 39 | 112.98 | 1 | 2.9 | 2.56 | 21.67 | 20.29 | 1.38 | 1.38 | 0.02 | 0.02 |
| gene1505 | https://www.ncbi.nlm.nih.gov/gene/?term=LOC105287633 | 1693 | 31 | 183.11 | 1 | 5.91 | 3.23 | 40.09 | 38.72 | 1.37 | 1.37 | 0.03 | 0.03 |
| gene4498 | https://www.ncbi.nlm.nih.gov/gene/?term=LOC105277239 | 2831 | 141 | 498.06 | 1 | 3.53 | 0.71 | 15.21 | 13.89 | 1.32 | 1.32 | 0.07 | 0.07 |
| gene2404 | https://www.ncbi.nlm.nih.gov/gene/?term=LOC105274853 | 1767 | 47 | 265.99 | 1 | 5.66 | 2.13 | 31.76 | 30.45 | 1.31 | 1.31 | 0.03 | 0.03 |
| gene11340 | https://www.ncbi.nlm.nih.gov/gene/?term=LOC105285131 | 2918 | 110 | 376.97 | 1 | 3.43 | 0.91 | 14.31 | 13.13 | 1.19 | 1.19 | 0.04 | 0.04 |
| gene564 | https://www.ncbi.nlm.nih.gov/gene/?term=LOC105278478 | 2802 | 24 | 85.65 | 1 | 3.57 | 4.17 | 23.58 | 22.43 | 1.15 | 1.15 | 0.01 | 0.01 |
| gene1768 | https://www.ncbi.nlm.nih.gov/gene/?term=LOC105287938 | 2792 | 79 | 282.95 | 1 | 3.58 | 1.27 | 29.01 | 27.92 | 1.09 | 1.09 | 0.03 | 0.03 |
| gene11447 | https://www.ncbi.nlm.nih.gov/gene/?term=LOC105285252 | 7437 | 128 | 172.11 | 1 | 1.34 | 0.78 | 14.37 | 13.39 | 0.98 | 0.98 | 0.02 | 0.02 |
| gene814 | https://www.ncbi.nlm.nih.gov/gene/?term=LOC105281296 | 2722 | 52 | 191.04 | 1 | 3.67 | 1.92 | 24.92 | 24.03 | 0.89 | 0.89 | 0.02 | 0.02 |
| gene3671 | https://www.ncbi.nlm.nih.gov/gene/?term=LOC105276272 | 1966 | 51 | 259.41 | 1 | 5.09 | 1.96 | 27.37 | 26.55 | 0.82 | 0.82 | 0.02 | 0.02 |

...

... only first 25 lines shown ...

Directory: OGLall File: ogl-Cb_LI_BCphaseB2.vs.RphaseB1.txt

| gene_ID | gene_link | gwidth | #Sites | #per10Kb | #dmSites | #dmsp10kb | %dmSites | %pSite1 | %pSite2 | DMpSite | ADMpSite | DMpNucl | ADMpNucl |
|---|---|---|---|---|---|---|---|---|---|---|---|---|---|
| gene5877 | https://www.ncbi.nlm.nih.gov/gene/?term=LOC105278807 | 1719 | 50 | 290.87 | 1 | 5.82 | 2 | 23.49 | 23.85 | -0.36 | 0.36 | -0.01 | 0.01 |
| gene1470 | https://www.ncbi.nlm.nih.gov/gene/?term=LOC105287597 | 2974 | 190 | 638.87 | 1 | 3.36 | 0.53 | 18.6 | 19.2 | -0.6 | 0.6 | -0.04 | 0.04 |

Directory: OGLall File: ogl-Cb_LI_BCphaseB2.vs.RphaseB2.txt

| gene_ID | gene_link | gwidth | #Sites | #per10Kb | #dmSites | #dmsp10kb | %dmSites | %pSite1 | %pSite2 | DMpSite | ADMpSite | DMpNucl | ADMpNucl |
|---|---|---|---|---|---|---|---|---|---|---|---|---|---|
| gene13225 | https://www.ncbi.nlm.nih.gov/gene/?term=LOC105287265 | 1631 | 2 | 12.26 | 1 | 6.13 | 50 | 88.97 | 71.1 | 17.88 | 17.88 | 0.02 | 0.02 |
| gene6576 | https://www.ncbi.nlm.nih.gov/gene/?term=LOC105279639 | 2616 | 3 | 11.47 | 2 | 7.65 | 66.67 | 79.09 | 65.15 | 13.93 | 13.93 | 0.02 | 0.02 |
| gene12853 | https://www.ncbi.nlm.nih.gov/gene/?term=LOC105286857 | 1589 | 2 | 12.59 | 1 | 6.29 | 50 | 69.58 | 57.77 | 11.81 | 11.81 | 0.01 | 0.01 |
| gene13119 | https://www.ncbi.nlm.nih.gov/gene/?term=LOC105287152 | 1525 | 2 | 13.11 | 1 | 6.56 | 50 | 30.95 | 20.38 | 10.57 | 10.57 | 0.01 | 0.01 |
| gene10278 | https://www.ncbi.nlm.nih.gov/gene/?term=LOC105283908 | 902 | 14 | 155.21 | 4 | 44.35 | 28.57 | 48.14 | 38.93 | 9.21 | 9.21 | 0.14 | 0.14 |
| gene4648 | https://www.ncbi.nlm.nih.gov/gene/?term=LOC105277406 | 2653 | 8 | 30.15 | 1 | 3.77 | 12.5 | 50.8 | 42.14 | 8.66 | 8.66 | 0.03 | 0.03 |
| gene6530 | https://www.ncbi.nlm.nih.gov/gene/?term=LOC105279592 | 1123 | 7 | 62.33 | 1 | 8.9 | 14.29 | 23.02 | 14.43 | 8.59 | 8.59 | 0.05 | 0.05 |
| gene12355 | https://www.ncbi.nlm.nih.gov/gene/?term=LOC105286261 | 1301 | 8 | 61.49 | 1 | 7.69 | 12.5 | 76.73 | 68.31 | 8.42 | 8.42 | 0.05 | 0.05 |
| gene12481 | https://www.ncbi.nlm.nih.gov/gene/?term=LOC105286408 | 356 | 3 | 84.27 | 1 | 28.09 | 33.33 | 17.04 | 8.98 | 8.06 | 8.06 | 0.07 | 0.07 |
| gene13178 | https://www.ncbi.nlm.nih.gov/gene/?term=LOC105287216 | 2535 | 6 | 23.67 | 1 | 3.94 | 16.67 | 27.44 | 19.82 | 7.62 | 7.62 | 0.02 | 0.02 |
| gene13131 | https://www.ncbi.nlm.nih.gov/gene/?term=LOC105287168 | 1392 | 8 | 57.47 | 2 | 14.37 | 25 | 60.44 | 53.29 | 7.15 | 7.15 | 0.04 | 0.04 |
| gene8174 | https://www.ncbi.nlm.nih.gov/gene/?term=LOC105281512 | 2154 | 12 | 55.71 | 2 | 9.29 | 16.67 | 33.85 | 26.91 | 6.94 | 6.94 | 0.04 | 0.04 |
| gene9483 | https://www.ncbi.nlm.nih.gov/gene/?term=LOC105282996 | 3739 | 6 | 16.05 | 1 | 2.67 | 16.67 | 23.81 | 17.02 | 6.8 | 6.8 | 0.01 | 0.01 |
| gene2636 | https://www.ncbi.nlm.nih.gov/gene/?term=LOC105275141 | 947 | 10 | 105.6 | 2 | 21.12 | 20 | 26.02 | 19.35 | 6.68 | 6.68 | 0.07 | 0.07 |
| gene11537 | https://www.ncbi.nlm.nih.gov/gene/?term=LOC105285354 | 1747 | 2 | 11.45 | 1 | 5.72 | 50 | 65.6 | 58.97 | 6.63 | 6.63 | 0.01 | 0.01 |
| gene3483 | https://www.ncbi.nlm.nih.gov/gene/?term=LOC105276075 | 1444 | 10 | 69.25 | 1 | 6.93 | 10 | 39.38 | 33.04 | 6.34 | 6.34 | 0.04 | 0.04 |
| gene9845 | https://www.ncbi.nlm.nih.gov/gene/?term=LOC105283410 | 1401 | 9 | 64.24 | 2 | 14.28 | 22.22 | 48.37 | 42.07 | 6.3 | 6.3 | 0.04 | 0.04 |
| gene3916 | https://www.ncbi.nlm.nih.gov/gene/?term=LOC105276569 | 1967 | 15 | 76.26 | 1 | 5.08 | 6.67 | 26.08 | 19.99 | 6.08 | 6.08 | 0.05 | 0.05 |
| gene5162 | https://www.ncbi.nlm.nih.gov/gene/?term=LOC105277980 | 2407 | 29 | 120.48 | 2 | 8.31 | 6.9 | 50.46 | 44.38 | 6.08 | 6.08 | 0.07 | 0.07 |
| gene11426 | https://www.ncbi.nlm.nih.gov/gene/?term=LOC105285229 | 1413 | 5 | 35.39 | 1 | 7.08 | 20 | 23.57 | 17.61 | 5.95 | 5.95 | 0.02 | 0.02 |
| gene11399 | https://www.ncbi.nlm.nih.gov/gene/?term=LOC105285203 | 1029 | 28 | 272.11 | 2 | 19.44 | 7.14 | 30.21 | 24.31 | 5.9 | 5.9 | 0.16 | 0.16 |
| gene11818 | https://www.ncbi.nlm.nih.gov/gene/?term=LOC105285669 | 2185 | 16 | 73.23 | 1 | 4.58 | 6.25 | 26.14 | 20.26 | 5.88 | 5.88 | 0.04 | 0.04 |
| gene1350 | https://www.ncbi.nlm.nih.gov/gene/?term=LOC105287003 | 2212 | 12 | 54.25 | 3 | 13.56 | 25 | 39.02 | 33.3 | 5.71 | 5.71 | 0.03 | 0.03 |
| gene11024 | https://www.ncbi.nlm.nih.gov/gene/?term=LOC105284763 | 4090 | 12 | 29.34 | 1 | 2.44 | 8.33 | 34.19 | 28.79 | 5.4 | 5.4 | 0.02 | 0.02 |

...
... only first 25 lines shown ...

Directory: OGLall File: ogl-Cb_LI_RphaseA1.txt

| gene_ID | gene_link | gwidth | #Sites | #per10Kb | %perSite | %pNucl |
|---|---|---|---|---|---|---|
| gene12538 | https://www.ncbi.nlm.nih.gov/gene/?term=LOC105286531 | 372 | 43 | 1155.91 | 36.48 | 4.22 |
| gene211 | https://www.ncbi.nlm.nih.gov/gene/?term=LOC105274534 | 1280 | 100 | 781.25 | 47.61 | 3.72 |
| gene5911 | https://www.ncbi.nlm.nih.gov/gene/?term=LOC105278844 | 943 | 95 | 1007.42 | 36.67 | 3.69 |
| gene13384 | https://www.ncbi.nlm.nih.gov/gene/?term=LOC105287447 | 1877 | 228 | 1214.7 | 29.65 | 3.6 |
| gene12330 | https://www.ncbi.nlm.nih.gov/gene/?term=LOC105286235 | 1501 | 118 | 786.14 | 42.69 | 3.36 |
| gene5309 | https://www.ncbi.nlm.nih.gov/gene/?term=LOC105278160 | 1704 | 158 | 927.23 | 35.33 | 3.28 |
| gene5917 | https://www.ncbi.nlm.nih.gov/gene/?term=LOC105278849 | 823 | 113 | 1373.03 | 23.86 | 3.28 |
| gene12364 | https://www.ncbi.nlm.nih.gov/gene/?term=LOC105286269 | 1403 | 87 | 620.1 | 52.62 | 3.26 |
| gene12746 | https://www.ncbi.nlm.nih.gov/gene/?term=LOC105286737 | 2688 | 366 | 1361.61 | 23.78 | 3.24 |
| gene4878 | https://www.ncbi.nlm.nih.gov/gene/?term=LOC105277648 | 675 | 38 | 562.96 | 57.09 | 3.21 |
| gene1654 | https://www.ncbi.nlm.nih.gov/gene/?term=LOC105287811 | 2285 | 165 | 722.1 | 44.27 | 3.2 |
| gene2797 | https://www.ncbi.nlm.nih.gov/gene/?term=LOC105275314 | 1489 | 97 | 651.44 | 47.77 | 3.11 |
| gene3731 | https://www.ncbi.nlm.nih.gov/gene/?term=LOC105276371 | 865 | 55 | 635.84 | 48.54 | 3.09 |
| gene6078 | https://www.ncbi.nlm.nih.gov/gene/?term=LOC105279052 | 1130 | 122 | 1079.65 | 28.5 | 3.08 |
| gene3719 | https://www.ncbi.nlm.nih.gov/gene/?term=LOC105276360 | 1471 | 155 | 1053.7 | 27.58 | 2.91 |
| gene458 | https://www.ncbi.nlm.nih.gov/gene/?term=LOC105277254 | 1378 | 113 | 820.03 | 35.06 | 2.87 |
| gene5301 | https://www.ncbi.nlm.nih.gov/gene/?term=LOC105278148 | 1881 | 199 | 1057.95 | 27.12 | 2.87 |
| gene1562 | https://www.ncbi.nlm.nih.gov/gene/?term=LOC105287692 | 1491 | 117 | 784.71 | 35.76 | 2.81 |
| gene7527 | https://www.ncbi.nlm.nih.gov/gene/?term=LOC105280739 | 2686 | 145 | 539.84 | 52.04 | 2.81 |
| gene11566 | https://www.ncbi.nlm.nih.gov/gene/?term=LOC105285392 | 1603 | 138 | 860.89 | 32.41 | 2.79 |
| gene4633 | https://www.ncbi.nlm.nih.gov/gene/?term=LOC105277389 | 2426 | 167 | 688.38 | 40.41 | 2.78 |
| gene8208 | https://www.ncbi.nlm.nih.gov/gene/?term=LOC105281548 | 1240 | 97 | 782.26 | 34.99 | 2.74 |
| gene9471 | https://www.ncbi.nlm.nih.gov/gene/?term=LOC105282986 | 1605 | 121 | 753.89 | 36.41 | 2.74 |
| gene11569 | https://www.ncbi.nlm.nih.gov/gene/?term=LOC105285394 | 1311 | 53 | 404.27 | 67.05 | 2.71 |

...

... only first 25 lines shown ...

Directory: OGLall File: ogl-Cb_LI_RphaseA1.vs.RphaseA2.txt

| gene_ID | gene_link | gwidth | #Sites | #per10Kb | #dmSites | #dmsp10kb | %dmSites | %pSite1 | %pSite2 | DMpSite | ADMpSite | DMpNucl | ADMpNucl |
|---|---|---|---|---|---|---|---|---|---|---|---|---|---|
| gene4630 | https://www.ncbi.nlm.nih.gov/gene/?term=LOC105277380 | 1893 | 5 | 26.41 | 1 | 5.28 | 20 | 48.18 | 35.31 | 12.87 | 12.87 | 0.03 | 0.03 |
| gene4631 | https://www.ncbi.nlm.nih.gov/gene/?term=LOC105277379 | 1348 | 5 | 37.09 | 1 | 7.42 | 20 | 51.95 | 41.8 | 10.16 | 10.16 | 0.04 | 0.04 |
| gene7684 | https://www.ncbi.nlm.nih.gov/gene/?term=LOC105280926 | 1727 | 9 | 52.11 | 1 | 5.79 | 11.11 | 57.47 | 49.81 | 7.66 | 7.66 | 0.04 | 0.04 |
| gene5695 | https://www.ncbi.nlm.nih.gov/gene/?term=LOC105278607 | 4140 | 22 | 53.14 | 2 | 4.83 | 9.09 | 16.79 | 9.48 | 7.31 | 7.31 | 0.04 | 0.04 |
| gene266 | https://www.ncbi.nlm.nih.gov/gene/?term=LOC105275133 | 1445 | 3 | 20.76 | 1 | 6.92 | 33.33 | 50.67 | 43.59 | 7.08 | 7.08 | 0.01 | 0.01 |
| gene6423 | https://www.ncbi.nlm.nih.gov/gene/?term=LOC105279461 | 1638 | 21 | 128.21 | 1 | 6.11 | 4.76 | 39.13 | 33.51 | 5.62 | 5.62 | 0.07 | 0.07 |
| gene5374 | https://www.ncbi.nlm.nih.gov/gene/?term=LOC105278227 | 2529 | 6 | 23.72 | 1 | 3.95 | 16.67 | 77.84 | 72.62 | 5.22 | 5.22 | 0.01 | 0.01 |
| gene477 | https://www.ncbi.nlm.nih.gov/gene/?term=LOC105277498 | 1683 | 7 | 41.59 | 1 | 5.94 | 14.29 | 28.69 | 23.62 | 5.07 | 5.07 | 0.02 | 0.02 |
| gene11035 | https://www.ncbi.nlm.nih.gov/gene/?term=LOC105284773 | 3419 | 5 | 14.62 | 1 | 2.92 | 20 | 8.77 | 3.9 | 4.87 | 4.87 | 0.01 | 0.01 |
| gene1924 | https://www.ncbi.nlm.nih.gov/gene/?term=LOC105288116 | 1820 | 10 | 54.95 | 1 | 5.49 | 10 | 7.47 | 3 | 4.48 | 4.48 | 0.02 | 0.02 |
| gene11800 | https://www.ncbi.nlm.nih.gov/gene/?term=LOC105285653 | 2693 | 13 | 48.27 | 1 | 3.71 | 7.69 | 56.71 | 52.38 | 4.32 | 4.32 | 0.02 | 0.02 |
| gene224 | https://www.ncbi.nlm.nih.gov/gene/?term=LOC109611562 | 1010 | 5 | 49.5 | 1 | 9.9 | 20 | 5.71 | 1.55 | 4.17 | 4.17 | 0.02 | 0.02 |
| gene9283 | https://www.ncbi.nlm.nih.gov/gene/?term=LOC105282769 | 2782 | 43 | 154.57 | 3 | 10.78 | 6.98 | 51.3 | 47.2 | 4.11 | 4.11 | 0.06 | 0.06 |
| gene7190 | https://www.ncbi.nlm.nih.gov/gene/?term=LOC105280355 | 1026 | 14 | 136.45 | 1 | 9.75 | 7.14 | 6.85 | 2.9 | 3.95 | 3.95 | 0.05 | 0.05 |
| gene4999 | https://www.ncbi.nlm.nih.gov/gene/?term=LOC105277805 | 6695 | 15 | 22.4 | 1 | 1.49 | 6.67 | 55.88 | 52 | 3.88 | 3.88 | 0.01 | 0.01 |
| gene9545 | https://www.ncbi.nlm.nih.gov/gene/?term=LOC105283071 | 1986 | 9 | 45.32 | 1 | 5.04 | 11.11 | 22.29 | 18.66 | 3.63 | 3.63 | 0.02 | 0.02 |
| gene4119 | https://www.ncbi.nlm.nih.gov/gene/?term=LOC105276806 | 1608 | 27 | 167.91 | 3 | 18.66 | 11.11 | 40.05 | 36.43 | 3.62 | 3.62 | 0.06 | 0.06 |
| gene4409 | https://www.ncbi.nlm.nih.gov/gene/?term=LOC105277140 | 1157 | 24 | 207.43 | 1 | 8.64 | 4.17 | 8.39 | 5.09 | 3.3 | 3.3 | 0.07 | 0.07 |
| gene7949 | https://www.ncbi.nlm.nih.gov/gene/?term=LOC105281206 | 1405 | 36 | 256.23 | 2 | 14.23 | 5.56 | 48.07 | 44.81 | 3.26 | 3.26 | 0.08 | 0.08 |
| gene11185 | https://www.ncbi.nlm.nih.gov/gene/?term=LOC105284939 | 1723 | 16 | 92.86 | 1 | 5.8 | 6.25 | 20.86 | 17.65 | 3.22 | 3.22 | 0.03 | 0.03 |
| gene11671 | https://www.ncbi.nlm.nih.gov/gene/?term=LOC105285514 | 1273 | 20 | 157.11 | 1 | 7.86 | 5 | 44.44 | 41.28 | 3.16 | 3.16 | 0.05 | 0.05 |
| gene8267 | https://www.ncbi.nlm.nih.gov/gene/?term=LOC105281612 | 1014 | 12 | 118.34 | 1 | 9.86 | 8.33 | 22.91 | 19.75 | 3.16 | 3.16 | 0.04 | 0.04 |
| gene265 | https://www.ncbi.nlm.nih.gov/gene/?term=LOC105275144 | 2128 | 42 | 197.37 | 4 | 18.8 | 9.52 | 32.47 | 29.32 | 3.15 | 3.15 | 0.06 | 0.06 |
| gene8113 | https://www.ncbi.nlm.nih.gov/gene/?term=LOC105281419 | 2170 | 25 | 115.21 | 2 | 9.22 | 8 | 7.95 | 4.92 | 3.03 | 3.03 | 0.03 | 0.03 |

...
... only first 25 lines shown ...

Directory: OGLall File: ogl-Cb_LI_RphaseA1.vs.RphaseB1.txt

| gene_ID | gene_link | gwidth | #Sites | #per10Kb | #dmSites | #dmsp10kb | %dmSites | %pSite1 | %pSite2 | DMpSite | ADMpSite | DMpNucl | ADMpNucl |
|---|---|---|---|---|---|---|---|---|---|---|---|---|---|
| gene2593 | https://www.ncbi.nlm.nih.gov/gene/?term=LOC105275070 | 1774 | 11 | 62.01 | 3 | 16.91 | 27.27 | 69.15 | 25.28 | 43.86 | 43.86 | 0.27 | 0.27 |
| gene3165 | https://www.ncbi.nlm.nih.gov/gene/?term=LOC105275706 | 2374 | 63 | 265.37 | 3 | 12.64 | 4.76 | 31.1 | 21.89 | 9.2 | 9.2 | 0.24 | 0.24 |
| gene2837 | https://www.ncbi.nlm.nih.gov/gene/?term=LOC105275355 | 2872 | 164 | 571.03 | 1 | 3.48 | 0.61 | 27.39 | 23.06 | 4.33 | 4.33 | 0.25 | 0.25 |
| gene5714 | https://www.ncbi.nlm.nih.gov/gene/?term=LOC105278729 | 710 | 16 | 225.35 | 1 | 14.08 | 6.25 | 55.81 | 51.9 | 3.91 | 3.91 | 0.09 | 0.09 |
| gene9478 | https://www.ncbi.nlm.nih.gov/gene/?term=LOC105282991 | 3726 | 58 | 155.66 | 1 | 2.68 | 1.72 | 31.92 | 28.27 | 3.65 | 3.65 | 0.06 | 0.06 |
| gene2856 | https://www.ncbi.nlm.nih.gov/gene/?term=LOC105275384 | 1987 | 88 | 442.88 | 1 | 5.03 | 1.14 | 36.35 | 32.79 | 3.56 | 3.56 | 0.16 | 0.16 |
| gene13033 | https://www.ncbi.nlm.nih.gov/gene/?term=LOC105287063 | 1877 | 49 | 261.05 | 1 | 5.33 | 2.04 | 32.74 | 30.05 | 2.68 | 2.68 | 0.07 | 0.07 |
| gene5877 | https://www.ncbi.nlm.nih.gov/gene/?term=LOC105278807 | 1719 | 51 | 296.68 | 1 | 5.82 | 1.96 | 27.19 | 24.91 | 2.27 | 2.27 | 0.07 | 0.07 |
| gene3671 | https://www.ncbi.nlm.nih.gov/gene/?term=LOC105276272 | 1966 | 44 | 223.8 | 1 | 5.09 | 2.27 | 27.22 | 24.96 | 2.26 | 2.26 | 0.05 | 0.05 |
| gene1855 | https://www.ncbi.nlm.nih.gov/gene/?term=LOC105288036 | 2387 | 90 | 377.04 | 1 | 4.19 | 1.11 | 28.76 | 26.75 | 2.02 | 2.02 | 0.08 | 0.08 |
| gene9189 | https://www.ncbi.nlm.nih.gov/gene/?term=LOC105282652 | 4699 | 84 | 178.76 | 1 | 2.13 | 1.19 | 16.38 | 14.61 | 1.78 | 1.78 | 0.03 | 0.03 |
| gene12516 | https://www.ncbi.nlm.nih.gov/gene/?term=LOC105286452 | 1529 | 38 | 248.53 | 1 | 6.54 | 2.63 | 49.76 | 48.05 | 1.71 | 1.71 | 0.04 | 0.04 |
| gene8178 | https://www.ncbi.nlm.nih.gov/gene/?term=LOC105281515 | 8647 | 60 | 69.39 | 1 | 1.16 | 1.67 | 29.73 | 28.04 | 1.69 | 1.69 | 0.01 | 0.01 |
| gene4023 | https://www.ncbi.nlm.nih.gov/gene/?term=LOC105276692 | 1759 | 36 | 204.66 | 1 | 5.69 | 2.78 | 23.9 | 22.4 | 1.5 | 1.5 | 0.03 | 0.03 |
| gene927 | https://www.ncbi.nlm.nih.gov/gene/?term=LOC105282600 | 3727 | 60 | 160.99 | 1 | 2.68 | 1.67 | 27.81 | 26.32 | 1.49 | 1.49 | 0.02 | 0.02 |
| gene1884 | https://www.ncbi.nlm.nih.gov/gene/?term=LOC105288072 | 4169 | 82 | 196.69 | 1 | 2.4 | 1.22 | 24.39 | 23.08 | 1.31 | 1.31 | 0.03 | 0.03 |
| gene11819 | https://www.ncbi.nlm.nih.gov/gene/?term=LOC105285671 | 2366 | 100 | 422.65 | 1 | 4.23 | 1 | 31.41 | 30.12 | 1.29 | 1.29 | 0.05 | 0.05 |
| gene4641 | https://www.ncbi.nlm.nih.gov/gene/?term=LOC105277398 | 1937 | 73 | 376.87 | 1 | 5.16 | 1.37 | 36.14 | 34.87 | 1.27 | 1.27 | 0.05 | 0.05 |
| gene10091 | https://www.ncbi.nlm.nih.gov/gene/?term=LOC105283697 | 4988 | 154 | 308.74 | 1 | 2 | 0.65 | 16.48 | 15.36 | 1.12 | 1.12 | 0.03 | 0.03 |
| gene9336 | https://www.ncbi.nlm.nih.gov/gene/?term=LOC105282834 | 5993 | 134 | 223.59 | 1 | 1.67 | 0.75 | 14.48 | 13.37 | 1.1 | 1.1 | 0.02 | 0.02 |
| gene2683 | https://www.ncbi.nlm.nih.gov/gene/?term=LOC105275187 | 5492 | 73 | 132.92 | 1 | 1.82 | 1.37 | 14.7 | 13.62 | 1.08 | 1.08 | 0.01 | 0.01 |
| gene3817 | https://www.ncbi.nlm.nih.gov/gene/?term=LOC105276449 | 1230 | 48 | 390.24 | 1 | 8.13 | 2.08 | 25.64 | 24.6 | 1.04 | 1.04 | 0.04 | 0.04 |
| gene298 | https://www.ncbi.nlm.nih.gov/gene/?term=LOC105275460 | 2473 | 126 | 509.5 | 1 | 4.04 | 0.79 | 13.41 | 12.43 | 0.98 | 0.98 | 0.05 | 0.05 |
| gene652 | https://www.ncbi.nlm.nih.gov/gene/?term=LOC105279528 | 5806 | 39 | 67.17 | 1 | 1.72 | 2.56 | 22.38 | 21.58 | 0.8 | 0.8 | 0.01 | 0.01 |
| gene10107 | https://www.ncbi.nlm.nih.gov/gene/?term=LOC105283712 | 1883 | 36 | 191.18 | 1 | 5.31 | 2.78 | 27.63 | 26.89 | 0.74 | 0.74 | 0.01 | 0.01 |
| gene5831 | https://www.ncbi.nlm.nih.gov/gene/?term=LOC105278761 | 2506 | 115 | 458.9 | 2 | 7.98 | 1.74 | 22.22 | 21.51 | 0.71 | 0.71 | 0.03 | 0.03 |
| gene940 | https://www.ncbi.nlm.nih.gov/gene/?term=LOC105282794 | 3578 | 93 | 259.92 | 1 | 2.79 | 1.08 | 14.3 | 13.63 | 0.67 | 0.67 | 0.02 | 0.02 |
| gene10540 | https://www.ncbi.nlm.nih.gov/gene/?term=LOC105284200 | 5267 | 209 | 396.81 | 1 | 1.9 | 0.48 | 13.12 | 12.47 | 0.65 | 0.65 | 0.03 | 0.03 |
| gene4892 | https://www.ncbi.nlm.nih.gov/gene/?term=LOC105277667 | 3898 | 121 | 310.42 | 1 | 2.57 | 0.83 | 29.02 | 28.4 | 0.62 | 0.62 | 0.02 | 0.02 |
| gene7600 | https://www.ncbi.nlm.nih.gov/gene/?term=LOC105280819 | 1615 | 41 | 253.87 | 1 | 6.19 | 2.44 | 39.36 | 38.88 | 0.48 | 0.48 | 0.01 | 0.01 |
| gene1147 | https://www.ncbi.nlm.nih.gov/gene/?term=LOC105284954 | 4022 | 118 | 293.39 | 1 | 2.49 | 0.85 | 11.22 | 10.78 | 0.44 | 0.44 | 0.01 | 0.01 |
| gene13036 | https://www.ncbi.nlm.nih.gov/gene/?term=LOC105287059 | 4959 | 180 | 362.98 | 1 | 2.02 | 0.56 | 17.65 | 17.23 | 0.42 | 0.42 | 0.02 | 0.02 |
| gene12441 | https://www.ncbi.nlm.nih.gov/gene/?term=LOC105286355 | 3252 | 149 | 458.18 | 1 | 3.08 | 0.67 | 21.45 | 21.08 | 0.38 | 0.38 | 0.02 | 0.02 |
| gene10222 | https://www.ncbi.nlm.nih.gov/gene/?term=LOC105283838 | 1748 | 30 | 171.62 | 1 | 5.72 | 3.33 | 33.06 | 32.69 | 0.37 | 0.37 | 0.01 | 0.01 |
| gene10243 | https://www.ncbi.nlm.nih.gov/gene/?term=LOC105283862 | 3239 | 43 | 132.76 | 1 | 3.09 | 2.33 | 29.28 | 28.91 | 0.37 | 0.37 | 0 | 0 |
| gene5101 | https://www.ncbi.nlm.nih.gov/gene/?term=LOC105277903 | 1813 | 36 | 198.57 | 1 | 5.52 | 2.78 | 56.09 | 55.74 | 0.35 | 0.35 | 0.01 | 0.01 |
| gene2844 | https://www.ncbi.nlm.nih.gov/gene/?term=LOC105275363 | 4330 | 218 | 503.46 | 1 | 2.31 | 0.46 | 13.23 | 12.92 | 0.3 | 0.3 | 0.02 | 0.02 |
| gene7247 | https://www.ncbi.nlm.nih.gov/gene/?term=LOC105280407 | 10486 | 226 | 215.53 | 1 | 0.95 | 0.44 | 15.61 | 15.31 | 0.3 | 0.3 | 0.01 | 0.01 |
| gene10654 | https://www.ncbi.nlm.nih.gov/gene/?term=LOC105284329 | 4027 | 71 | 176.31 | 1 | 2.48 | 1.41 | 19.25 | 18.99 | 0.26 | 0.26 | 0 | 0 |
| gene6455 | https://www.ncbi.nlm.nih.gov/gene/?term=LOC105279500 | 2612 | 31 | 118.68 | 1 | 3.83 | 3.23 | 47.76 | 47.54 | 0.22 | 0.22 | 0 | 0 |
| gene10345 | https://www.ncbi.nlm.nih.gov/gene/?term=LOC105283989 | 4444 | 29 | 65.26 | 1 | 2.25 | 3.45 | 49.14 | 48.92 | 0.21 | 0.21 | 0 | 0 |
| gene10520 | https://www.ncbi.nlm.nih.gov/gene/?term=LOC105284184 | 2451 | 92 | 375.36 | 1 | 4.08 | 1.09 | 31.61 | 31.4 | 0.21 | 0.21 | 0.01 | 0.01 |
| gene7183 | https://www.ncbi.nlm.nih.gov/gene/?term=LOC105280348 | 3925 | 79 | 201.27 | 1 | 2.55 | 1.27 | 17.02 | 16.83 | 0.19 | 0.19 | 0 | 0 |
| gene2668 | https://www.ncbi.nlm.nih.gov/gene/?term=LOC105275170 | 12496 | 348 | 278.49 | 1 | 0.8 | 0.29 | 1.95 | 1.78 | 0.18 | 0.18 | 0 | 0 |
| gene2846 | https://www.ncbi.nlm.nih.gov/gene/?term=LOC105275374 | 11246 | 493 | 438.38 | 1 | 0.89 | 0.2 | 9.9 | 9.75 | 0.16 | 0.16 | 0.01 | 0.01 |
| gene9327 | https://www.ncbi.nlm.nih.gov/gene/?term=LOC105282821 | 7793 | 162 | 207.88 | 1 | 1.28 | 0.62 | 13.28 | 13.13 | 0.16 | 0.16 | 0 | 0 |
| gene12978 | https://www.ncbi.nlm.nih.gov/gene/?term=LOC105286997 | 2858 | 142 | 496.85 | 1 | 3.5 | 0.7 | 11.15 | 11.02 | 0.14 | 0.14 | 0.01 | 0.01 |
| gene7556 | https://www.ncbi.nlm.nih.gov/gene/?term=LOC105280774 | 2792 | 132 | 472.78 | 1 | 3.58 | 0.76 | 25.2 | 25.1 | 0.1 | 0.1 | 0 | 0 |
| gene10606 | https://www.ncbi.nlm.nih.gov/gene/?term=LOC105284273 | 6119 | 261 | 426.54 | 1 | 1.63 | 0.38 | 19.09 | 19 | 0.08 | 0.08 | 0 | 0 |
| gene8600 | https://www.ncbi.nlm.nih.gov/gene/?term=LOC105282001 | 8513 | 189 | 222.01 | 1 | 1.17 | 0.53 | 10.4 | 10.32 | 0.08 | 0.08 | 0 | 0 |
| gene2806 | https://www.ncbi.nlm.nih.gov/gene/?term=LOC105275327 | 14257 | 529 | 371.05 | 1 | 0.7 | 0.19 | 6.66 | 6.63 | 0.03 | 0.03 | 0 | 0 |
| gene2748 | https://www.ncbi.nlm.nih.gov/gene/?term=LOC105275261 | 3845 | 55 | 143.04 | 1 | 2.6 | 1.82 | 38.35 | 38.32 | 0.02 | 0.02 | 0 | 0 |
| gene7368 | https://www.ncbi.nlm.nih.gov/gene/?term=LOC105280556 | 4105 | 65 | 158.34 | 1 | 2.44 | 1.54 | 14.12 | 14.11 | 0.01 | 0.01 | 0 | 0 |
| gene11583 | https://www.ncbi.nlm.nih.gov/gene/?term=LOC105285414 | 8192 | 189 | 230.71 | 1 | 1.22 | 0.53 | 8.96 | 8.96 | 0 | 0 | 0 | 0 |
| gene10879 | https://www.ncbi.nlm.nih.gov/gene/?term=LOC105284574 | 2423 | 50 | 206.36 | 1 | 4.13 | 2 | 52.18 | 52.2 | -0.01 | 0.01 | 0 | 0 |
| gene3470 | https://www.ncbi.nlm.nih.gov/gene/?term=LOC105276059 | 8716 | 161 | 184.72 | 1 | 1.15 | 0.62 | 9.23 | 9.23 | -0.01 | 0.01 | 0 | 0 |
| gene5746 | https://www.ncbi.nlm.nih.gov/gene/?term=LOC105278666 | 10343 | 473 | 457.31 | 1 | 0.97 | 0.21 | 6.89 | 6.9 | -0.01 | 0.01 | 0 | 0 |
| gene5375 | https://www.ncbi.nlm.nih.gov/gene/?term=LOC105278228 | 8640 | 170 | 196.76 | 1 | 1.16 | 0.59 | 23.53 | 23.55 | -0.02 | 0.02 | 0 | 0 |
| gene9326 | https://www.ncbi.nlm.nih.gov/gene/?term=LOC105282818 | 16612 | 210 | 126.41 | 1 | 0.6 | 0.48 | 13.96 | 13.99 | -0.03 | 0.03 | 0 | 0 |
| gene12573 | https://www.ncbi.nlm.nih.gov/gene/?term=LOC105286515 | 3647 | 112 | 307.1 | 1 | 2.74 | 0.89 | 11.12 | 11.16 | -0.04 | 0.04 | 0 | 0 |

| | | | | | | | | | | | | | |
|---|---|---|---|---|---|---|---|---|---|---|---|---|---|
| gene2443 | https://www.ncbi.nlm.nih.gov/gene/?term=LOC105274895 | 10433 | 409 | 392.03 | 1 | 0.96 | 0.24 | 4.52 | 4.57 | -0.04 | 0.04 | 0 | 0 |
| gene7249 | https://www.ncbi.nlm.nih.gov/gene/?term=LOC105280422 | 1775 | 66 | 371.83 | 1 | 5.63 | 1.52 | 15.99 | 16.04 | -0.05 | 0.05 | 0 | 0 |
| gene11275 | https://www.ncbi.nlm.nih.gov/gene/?term=LOC105285055 | 4119 | 251 | 609.37 | 1 | 2.43 | 0.4 | 10.55 | 10.62 | -0.07 | 0.07 | 0 | 0 |
| gene5539 | https://www.ncbi.nlm.nih.gov/gene/?term=LOC105278415 | 25024 | 724 | 289.32 | 1 | 0.4 | 0.14 | 3.42 | 3.48 | -0.07 | 0.07 | 0 | 0 |
| gene5197 | https://www.ncbi.nlm.nih.gov/gene/?term=LOC105278018 | 89233 | 3396 | 380.58 | 1 | 0.11 | 0.03 | 0.66 | 0.73 | -0.08 | 0.08 | 0 | 0 |
| gene1220 | https://www.ncbi.nlm.nih.gov/gene/?term=LOC105285742 | 3453 | 109 | 315.67 | 1 | 2.9 | 0.92 | 16.97 | 17.05 | -0.09 | 0.09 | 0 | 0 |
| gene6620 | https://www.ncbi.nlm.nih.gov/gene/?term=LOC105279687 | 17998 | 433 | 240.58 | 1 | 0.56 | 0.23 | 4.48 | 4.58 | -0.1 | 0.1 | 0 | 0 |
| gene53 | https://www.ncbi.nlm.nih.gov/gene/?term=LOC105277148 | 19283 | 305 | 158.17 | 1 | 0.52 | 0.33 | 4.34 | 4.47 | -0.13 | 0.13 | 0 | 0 |
| gene5052 | https://www.ncbi.nlm.nih.gov/gene/?term=LOC105277856 | 3167 | 33 | 104.2 1 | 3.16 | 3.03 | 39.38 | 39.53 | -0.15 | 0.15 | 0 | 0 | |
| gene862 | https://www.ncbi.nlm.nih.gov/gene/?term=LOC105281802 | 73922 | 2087 | 282.32 | 1 | 0.14 | 0.05 | 2.14 | 2.3 | -0.15 | 0.15 | 0 | 0 |
| gene11561 | https://www.ncbi.nlm.nih.gov/gene/?term=LOC105285387 | 11528 | 219 | 189.97 | 1 | 0.87 | 0.46 | 9.39 | 9.54 | -0.16 | 0.16 | 0 | 0 |
| gene292 | https://www.ncbi.nlm.nih.gov/gene/?term=LOC105275398 | 2825 | 83 | 293.81 | 1 | 3.54 | 1.2 | 20.15 | 20.32 | -0.16 | 0.16 | 0 | 0 |
| gene2271 | https://www.ncbi.nlm.nih.gov/gene/?term=LOC105274706 | 1448 | 51 | 352.21 | 1 | 6.91 | 1.96 | 37.35 | 37.52 | -0.17 | 0.17 | -0.01 | 0.01 |
| gene3885 | https://www.ncbi.nlm.nih.gov/gene/?term=LOC105276527 | 7928 | 336 | 423.81 | 1 | 1.26 | 0.3 | 8.31 | 8.48 | -0.17 | 0.17 | -0.01 | 0.01 |
| gene8274 | https://www.ncbi.nlm.nih.gov/gene/?term=LOC105281624 | 5425 | 216 | 398.16 | 1 | 1.84 | 0.46 | 9.81 | 9.99 | -0.18 | 0.18 | -0.01 | 0.01 |
| gene1632 | https://www.ncbi.nlm.nih.gov/gene/?term=LOC105287764 | 18061 | 398 | 220.36 | 1 | 0.55 | 0.25 | 3.49 | 3.69 | -0.19 | 0.19 | 0 | 0 |
| gene7547 | https://www.ncbi.nlm.nih.gov/gene/?term=LOC105280767 | 24576 | 808 | 328.78 | 1 | 0.41 | 0.12 | 4.37 | 4.58 | -0.21 | 0.21 | -0.01 | 0.01 |
| gene8814 | https://www.ncbi.nlm.nih.gov/gene/?term=LOC105282240 | 6910 | 188 | 272.07 | 1 | 1.45 | 0.53 | 8.71 | 8.93 | -0.22 | 0.22 | -0.01 | 0.01 |
| gene10018 | https://www.ncbi.nlm.nih.gov/gene/?term=LOC105283607 | 4414 | 219 | 496.15 | 1 | 2.27 | 0.46 | 13.97 | 14.21 | -0.23 | 0.23 | -0.01 | 0.01 |
| gene10600 | https://www.ncbi.nlm.nih.gov/gene/?term=LOC105284270 | 4837 | 103 | 212.94 | 1 | 2.07 | 0.97 | 12.43 | 12.66 | -0.23 | 0.23 | 0 | 0 |
| gene13114 | https://www.ncbi.nlm.nih.gov/gene/?term=LOC105287147 | 7609 | 415 | 545.41 | 1 | 1.31 | 0.24 | 8.59 | 8.85 | -0.26 | 0.26 | -0.01 | 0.01 |
| gene12802 | https://www.ncbi.nlm.nih.gov/gene/?term=LOC105286799 | 3183 | 178 | 559.22 | 1 | 3.14 | 0.56 | 16.8 | 17.1 | -0.29 | 0.29 | -0.02 | 0.02 |
| gene6286 | https://www.ncbi.nlm.nih.gov/gene/?term=LOC105279298 | 2861 | 125 | 436.91 | 2 | 6.99 | 1.6 | 23.63 | 23.91 | -0.29 | 0.29 | -0.01 | 0.01 |
| gene10549 | https://www.ncbi.nlm.nih.gov/gene/?term=LOC105284210 | 7145 | 348 | 487.05 | 2 | 2.8 | 0.57 | 9.98 | 10.29 | -0.31 | 0.31 | -0.01 | 0.01 |
| gene5525 | https://www.ncbi.nlm.nih.gov/gene/?term=LOC105278396 | 3090 | 186 | 601.94 | 1 | 3.24 | 0.54 | 24.12 | 24.45 | -0.33 | 0.33 | -0.02 | 0.02 |
| gene9330 | https://www.ncbi.nlm.nih.gov/gene/?term=LOC105282825 | 3936 | 96 | 243.9 1 | 2.54 | 1.04 | 14.55 | 14.91 | -0.36 | 0.36 | -0.01 | 0.01 | |
| gene4376 | https://www.ncbi.nlm.nih.gov/gene/?term=LOC105277083 | 4307 | 120 | 278.62 | 1 | 2.32 | 0.83 | 19.12 | 19.51 | -0.39 | 0.39 | -0.01 | 0.01 |
| gene5524 | https://www.ncbi.nlm.nih.gov/gene/?term=LOC105278397 | 5177 | 205 | 395.98 | 1 | 1.93 | 0.49 | 24.01 | 24.4 | -0.39 | 0.39 | -0.02 | 0.02 |
| gene6031 | https://www.ncbi.nlm.nih.gov/gene/?term=LOC105279000 | 5691 | 210 | 369 1 | 1.76 | 0.48 | 11.92 | 12.31 | -0.4 | 0.4 | -0.01 | 0.01 | |
| gene6583 | https://www.ncbi.nlm.nih.gov/gene/?term=LOC105279649 | 7444 | 364 | 488.98 | 1 | 1.34 | 0.27 | 10.46 | 10.89 | -0.43 | 0.43 | -0.02 | 0.02 |
| gene10678 | https://www.ncbi.nlm.nih.gov/gene/?term=LOC105284355 | 4031 | 173 | 429.17 | 1 | 2.48 | 0.58 | 18.28 | 18.72 | -0.44 | 0.44 | -0.02 | 0.02 |
| gene12307 | https://www.ncbi.nlm.nih.gov/gene/?term=LOC105286216 | 10313 | 186 | 180.35 | 1 | 0.97 | 0.54 | 14.88 | 15.35 | -0.47 | 0.47 | -0.01 | 0.01 |
| gene5298 | https://www.ncbi.nlm.nih.gov/gene/?term=LOC105278145 | 4060 | 253 | 623.15 | 3 | 7.39 | 1.19 | 10.61 | 11.09 | -0.49 | 0.49 | -0.03 | 0.03 |
| gene9647 | https://www.ncbi.nlm.nih.gov/gene/?term=LOC109611233 | 3094 | 33 | 106.66 | 1 | 3.23 | 3.03 | 41.64 | 42.16 | -0.52 | 0.52 | -0.01 | 0.01 |
| gene9648 | https://www.ncbi.nlm.nih.gov/gene/?term=LOC105283183 | 2602 | 31 | 119.14 | 1 | 3.84 | 3.23 | 44.33 | 44.88 | -0.55 | 0.55 | -0.01 | 0.01 |
| gene10326 | https://www.ncbi.nlm.nih.gov/gene/?term=LOC105283962 | 3042 | 132 | 433.93 | 1 | 3.29 | 0.76 | 8.06 | 8.65 | -0.59 | 0.59 | -0.03 | 0.03 |
| gene3107 | https://www.ncbi.nlm.nih.gov/gene/?term=LOC105275650 | 5103 | 180 | 352.73 | 1 | 1.96 | 0.56 | 15.71 | 16.3 | -0.59 | 0.59 | -0.02 | 0.02 |
| gene5646 | https://www.ncbi.nlm.nih.gov/gene/?term=LOC105278534 | 7475 | 198 | 264.88 | 1 | 1.34 | 0.51 | 17.1 | 17.7 | -0.6 | 0.6 | -0.02 | 0.02 |
| gene7166 | https://www.ncbi.nlm.nih.gov/gene/?term=LOC105280326 | 6117 | 249 | 407.06 | 1 | 1.63 | 0.4 | 14.56 | 15.24 | -0.68 | 0.68 | -0.03 | 0.03 |
| gene6237 | https://www.ncbi.nlm.nih.gov/gene/?term=LOC105279244 | 1484 | 25 | 168.46 | 1 | 6.74 | 4 | 45.17 | 45.86 | -0.69 | 0.69 | -0.01 | 0.01 |
| gene10844 | https://www.ncbi.nlm.nih.gov/gene/?term=LOC105284535 | 9086 | 258 | 283.95 | 1 | 1.1 | 0.39 | 13.99 | 14.72 | -0.73 | 0.73 | -0.02 | 0.02 |
| gene9740 | https://www.ncbi.nlm.nih.gov/gene/?term=LOC105283277 | 2887 | 138 | 478 1 | 3.46 | 0.72 | 17.69 | 18.47 | -0.78 | 0.78 | -0.04 | 0.04 | |
| gene4190 | https://www.ncbi.nlm.nih.gov/gene/?term=LOC105276875 | 3150 | 125 | 396.83 | 1 | 3.17 | 0.8 | 19.56 | 20.35 | -0.79 | 0.79 | -0.03 | 0.03 |
| gene1470 | https://www.ncbi.nlm.nih.gov/gene/?term=LOC105287597 | 2974 | 182 | 611.97 | 1 | 3.36 | 0.55 | 19.45 | 20.26 | -0.81 | 0.81 | -0.05 | 0.05 |
| gene6652 | https://www.ncbi.nlm.nih.gov/gene/?term=LOC105279719 | 2831 | 87 | 307.31 | 1 | 3.53 | 1.15 | 20.49 | 21.33 | -0.83 | 0.83 | -0.03 | 0.03 |
| gene8729 | https://www.ncbi.nlm.nih.gov/gene/?term=LOC105282137 | 5005 | 225 | 449.55 | 1 | 2 | 0.44 | 13.97 | 14.84 | -0.86 | 0.86 | -0.04 | 0.04 |
| gene11454 | https://www.ncbi.nlm.nih.gov/gene/?term=LOC105285256 | 1906 | 50 | 262.33 | 1 | 5.25 | 2 | 43.22 | 44.08 | -0.87 | 0.87 | -0.02 | 0.02 |
| gene10155 | https://www.ncbi.nlm.nih.gov/gene/?term=LOC105283756 | 7030 | 112 | 159.32 | 1 | 1.42 | 0.89 | 7.35 | 8.24 | -0.89 | 0.89 | -0.01 | 0.01 |
| gene570 | https://www.ncbi.nlm.nih.gov/gene/?term=LOC105278549 | 1673 | 70 | 418.41 | 1 | 5.98 | 1.43 | 42.95 | 43.99 | -1.04 | 1.04 | -0.04 | 0.04 |
| gene6422 | https://www.ncbi.nlm.nih.gov/gene/?term=LOC105279458 | 2419 | 43 | 177.76 | 1 | 4.13 | 2.33 | 44.62 | 45.68 | -1.05 | 1.05 | -0.02 | 0.02 |
| gene920 | https://www.ncbi.nlm.nih.gov/gene/?term=LOC105282518 | 1197 | 44 | 367.59 | 1 | 8.35 | 2.27 | 32.92 | 33.98 | -1.06 | 1.06 | -0.04 | 0.04 |
| gene2173 | https://www.ncbi.nlm.nih.gov/gene/?term=LOC105274590 | 4387 | 84 | 191.47 | 1 | 2.28 | 1.19 | 23.89 | 24.96 | -1.07 | 1.07 | -0.02 | 0.02 |
| gene4962 | https://www.ncbi.nlm.nih.gov/gene/?term=LOC105277761 | 4645 | 160 | 344.46 | 1 | 2.15 | 0.62 | 12.8 | 13.96 | -1.16 | 1.16 | -0.04 | 0.04 |
| gene7915 | https://www.ncbi.nlm.nih.gov/gene/?term=LOC105281173 | 1597 | 38 | 237.95 | 1 | 6.26 | 2.63 | 59.81 | 61.01 | -1.2 | 1.2 | -0.03 | 0.03 |
| gene3335 | https://www.ncbi.nlm.nih.gov/gene/?term=LOC105275898 | 4665 | 157 | 336.55 | 1 | 2.14 | 0.64 | 12.01 | 13.22 | -1.21 | 1.21 | -0.04 | 0.04 |
| gene10392 | https://www.ncbi.nlm.nih.gov/gene/?term=LOC105284036 | 3216 | 98 | 304.73 | 1 | 3.11 | 1.02 | 22.15 | 23.39 | -1.25 | 1.25 | -0.04 | 0.04 |
| gene6639 | https://www.ncbi.nlm.nih.gov/gene/?term=LOC105279705 | 7656 | 81 | 105.8 1 | 1.31 | 1.23 | 19.09 | 20.33 | -1.25 | 1.25 | -0.01 | 0.01 | |
| gene3599 | https://www.ncbi.nlm.nih.gov/gene/?term=LOC105276196 | 2955 | 57 | 192.89 | 1 | 3.38 | 1.75 | 39.57 | 40.89 | -1.32 | 1.32 | -0.03 | 0.03 |
| gene9101 | https://www.ncbi.nlm.nih.gov/gene/?term=LOC105282553 | 3068 | 106 | 345.5 1 | 3.26 | 0.94 | 21.02 | 22.34 | -1.33 | 1.33 | -0.05 | 0.05 | |
| gene1310 | https://www.ncbi.nlm.nih.gov/gene/?term=LOC105286661 | 2582 | 67 | 259.49 | 1 | 3.87 | 1.49 | 17.48 | 18.89 | -1.41 | 1.41 | -0.04 | 0.04 |
| gene7248 | https://www.ncbi.nlm.nih.gov/gene/?term=LOC105280408 | 5175 | 60 | 115.94 | 1 | 1.93 | 1.67 | 15.92 | 17.41 | -1.49 | 1.49 | -0.02 | 0.02 |
| gene11340 | https://www.ncbi.nlm.nih.gov/gene/?term=LOC105285131 | 2918 | 98 | 335.85 | 1 | 3.43 | 1.02 | 13.77 | 15.33 | -1.56 | 1.56 | -0.05 | 0.05 |
| gene2413 | https://www.ncbi.nlm.nih.gov/gene/?term=LOC105274863 | 2006 | 60 | 299.1 1 | 4.99 | 1.67 | 50.89 | 52.54 | -1.65 | 1.65 | -0.05 | 0.05 | |
| gene1198 | https://www.ncbi.nlm.nih.gov/gene/?term=LOC105285507 | 2738 | 98 | 357.93 | 1 | 3.65 | 1.02 | 19.77 | 21.55 | -1.78 | 1.78 | -0.06 | 0.06 |

| | | | | | | | | | | | | | |
|---|---|---|---|---|---|---|---|---|---|---|---|---|---|
| gene2404 | https://www.ncbi.nlm.nih.gov/gene/?term=LOC105274853 | 1767 | 42 | 237.69 | 1 | 5.66 | 2.38 | 30.33 | 32.12 | -1.8 | 1.8 | -0.04 | 0.04 |
| gene1575 | https://www.ncbi.nlm.nih.gov/gene/?term=LOC105287706 | 3783 | 44 | 116.31 | 1 | 2.64 | 2.27 | 12.72 | 14.57 | -1.85 | 1.85 | -0.02 | 0.02 |
| gene7550 | https://www.ncbi.nlm.nih.gov/gene/?term=LOC105280763 | 2173 | 101 | 464.8 | 1 | 4.6 | 0.99 | 26.35 | 28.2 | -1.85 | 1.85 | -0.09 | 0.09 |
| gene7398 | https://www.ncbi.nlm.nih.gov/gene/?term=LOC105280589 | 3452 | 41 | 118.77 | 1 | 2.9 | 2.44 | 24.33 | 26.31 | -1.98 | 1.98 | -0.02 | 0.02 |
| gene12203 | https://www.ncbi.nlm.nih.gov/gene/?term=LOC105286099 | 5967 | 80 | 134.07 | 1 | 1.68 | 1.25 | 10.4 | 12.39 | -2 | 2 | -0.03 | 0.03 |
| gene7199 | https://www.ncbi.nlm.nih.gov/gene/?term=LOC105280366 | 1414 | 37 | 261.67 | 1 | 7.07 | 2.7 | 44.35 | 46.38 | -2.03 | 2.03 | -0.05 | 0.05 |
| gene12572 | https://www.ncbi.nlm.nih.gov/gene/?term=LOC105286514 | 2374 | 51 | 214.83 | 1 | 4.21 | 1.96 | 34.55 | 36.64 | -2.09 | 2.09 | -0.04 | 0.04 |
| gene3028 | https://www.ncbi.nlm.nih.gov/gene/?term=LOC105275571 | 1924 | 30 | 155.93 | 1 | 5.2 | 3.33 | 42.66 | 44.75 | -2.09 | 2.09 | -0.03 | 0.03 |
| gene3168 | https://www.ncbi.nlm.nih.gov/gene/?term=LOC105275711 | 6150 | 87 | 141.46 | 1 | 1.63 | 1.15 | 10.39 | 12.71 | -2.33 | 2.33 | -0.03 | 0.03 |
| gene12406 | https://www.ncbi.nlm.nih.gov/gene/?term=LOC105286316 | 1386 | 35 | 252.53 | 1 | 7.22 | 2.86 | 40.6 | 43.17 | -2.57 | 2.57 | -0.06 | 0.06 |
| gene11235 | https://www.ncbi.nlm.nih.gov/gene/?term=LOC105285004 | 1339 | 38 | 283.79 | 1 | 7.47 | 2.63 | 16.34 | 18.94 | -2.6 | 2.6 | -0.07 | 0.07 |
| gene12021 | https://www.ncbi.nlm.nih.gov/gene/?term=LOC105285905 | 1399 | 30 | 214.44 | 1 | 7.15 | 3.33 | 30.6 | 33.22 | -2.62 | 2.62 | -0.06 | 0.06 |
| gene9411 | https://www.ncbi.nlm.nih.gov/gene/?term=LOC105282914 | 5534 | 42 | 75.89 | 1 | 1.81 | 2.38 | 47.45 | 50.92 | -3.47 | 3.47 | -0.03 | 0.03 |
| gene564 | https://www.ncbi.nlm.nih.gov/gene/?term=LOC105278478 | 2802 | 17 | 60.67 | 1 | 3.57 | 5.88 | 15.85 | 19.36 | -3.51 | 3.51 | -0.02 | 0.02 |
| gene388 | https://www.ncbi.nlm.nih.gov/gene/?term=LOC105276538 | 1558 | 22 | 141.21 | 2 | 12.84 | 9.09 | 54.75 | 58.78 | -4.03 | 4.03 | -0.06 | 0.06 |
| gene11584 | https://www.ncbi.nlm.nih.gov/gene/?term=LOC105285416 | 3709 | 13 | 35.05 | 2 | 5.39 | 15.38 | 23.21 | 33.23 | -10.02 | 10.02 | -0.04 | 0.04 |
| gene393 | https://www.ncbi.nlm.nih.gov/gene/?term=LOC105276566 | 1123 | 2 | 17.81 | 1 | 8.9 | 50 | 50 | 74.3 | -24.3 | 24.3 | -0.04 | 0.04 |

Directory: OGLall File: ogl-Cb_LI_RphaseA1.vs.RphaseB2.txt

| gene_ID | gene_link | gwidth | #Sites | #per10Kb | #dmSites | #dmsp10kb | %dmSites | %pSite1 | %pSite2 | DMpSite | ADMpSite | DMpNucl | ADMpNucl | |
|---|---|---|---|---|---|---|---|---|---|---|---|---|---|---|
| gene2593 | https://www.ncbi.nlm.nih.gov/gene/?term=LOC105275070 | 1774 | 7 | 39.46 | 5 | 28.18 | 71.43 | 68.01 | 4.09 | 63.92 | 63.92 | 0.25 | 0.25 | |
| gene3165 | https://www.ncbi.nlm.nih.gov/gene/?term=LOC105275706 | 2374 | 65 | 273.8 | 5 | 21.06 | 7.69 | 30.28 | 21 | 9.28 | 9.28 | 0.25 | 0.25 | |
| gene2824 | https://www.ncbi.nlm.nih.gov/gene/?term=LOC105275337 | 1772 | 58 | 327.31 | | 1 | 5.64 | 1.72 | 38.96 | 30.77 | 8.18 | 8.18 | 0.27 | 0.27 |
| gene6334 | https://www.ncbi.nlm.nih.gov/gene/?term=LOC105279356 | 1471 | 45 | 305.91 | | 1 | 6.8 | 2.22 | 31.17 | 28.75 | 2.42 | 2.42 | 0.07 | 0.07 |
| gene1884 | https://www.ncbi.nlm.nih.gov/gene/?term=LOC105288072 | 4169 | 70 | 167.91 | | 1 | 2.4 | 1.43 | 26.72 | 24.36 | 2.36 | 2.36 | 0.04 | 0.04 |
| gene8178 | https://www.ncbi.nlm.nih.gov/gene/?term=LOC105281515 | 8647 | 43 | 49.73 | 1 | 1.16 | 2.33 | 27.02 | 24.75 | 2.27 | 2.27 | 0.01 | 0.01 | |
| gene11819 | https://www.ncbi.nlm.nih.gov/gene/?term=LOC105285671 | 2366 | 98 | 414.2 | 1 | 4.23 | 1.02 | 29.1 | 26.97 | 2.13 | 2.13 | 0.09 | 0.09 | |
| gene5877 | https://www.ncbi.nlm.nih.gov/gene/?term=LOC105278807 | 1719 | 48 | 279.23 | | 1 | 5.82 | 2.08 | 26.96 | 25.13 | 1.83 | 1.83 | 0.05 | 0.05 |
| gene7884 | https://www.ncbi.nlm.nih.gov/gene/?term=LOC105281266 | 3390 | 76 | 224.19 | | 1 | 2.95 | 1.32 | 18.91 | 17.21 | 1.7 | 1.7 | 0.04 | 0.04 |
| gene11401 | https://www.ncbi.nlm.nih.gov/gene/?term=LOC105285202 | 1841 | 78 | 423.68 | | 1 | 5.43 | 1.28 | 26.41 | 24.78 | 1.62 | 1.62 | 0.07 | 0.07 |
| gene1855 | https://www.ncbi.nlm.nih.gov/gene/?term=LOC105288036 | 2387 | 77 | 322.58 | | 1 | 4.19 | 1.3 | 32.99 | 31.48 | 1.51 | 1.51 | 0.05 | 0.05 |
| gene5101 | https://www.ncbi.nlm.nih.gov/gene/?term=LOC105277903 | 1813 | 35 | 193.05 | | 1 | 5.52 | 2.86 | 52.77 | 51.27 | 1.51 | 1.51 | 0.03 | 0.03 |
| gene940 | https://www.ncbi.nlm.nih.gov/gene/?term=LOC105282794 | 3578 | 85 | 237.56 | | 1 | 2.79 | 1.18 | 14.76 | 13.28 | 1.48 | 1.48 | 0.04 | 0.04 |
| gene10654 | https://www.ncbi.nlm.nih.gov/gene/?term=LOC105284329 | 4027 | 51 | 126.65 | | 1 | 2.48 | 1.96 | 20.19 | 18.78 | 1.41 | 1.41 | 0.02 | 0.02 |
| gene7568 | https://www.ncbi.nlm.nih.gov/gene/?term=LOC105280786 | 2525 | 114 | 451.49 | | 1 | 3.96 | 0.88 | 28.64 | 27.23 | 1.41 | 1.41 | 0.06 | 0.06 |
| gene7915 | https://www.ncbi.nlm.nih.gov/gene/?term=LOC105281173 | 1597 | 38 | 237.95 | | 1 | 6.26 | 2.63 | 59.81 | 58.74 | 1.06 | 1.06 | 0.03 | 0.03 |
| gene12516 | https://www.ncbi.nlm.nih.gov/gene/?term=LOC105286452 | 1529 | 38 | 248.53 | | 1 | 6.54 | 2.63 | 50.31 | 49.37 | 0.94 | 0.94 | 0.02 | 0.02 |
| gene8661 | https://www.ncbi.nlm.nih.gov/gene/?term=LOC105282063 | 1926 | 26 | 134.99 | | 1 | 5.19 | 3.85 | 38.79 | 37.9 | 0.89 | 0.89 | 0.01 | 0.01 |
| gene2844 | https://www.ncbi.nlm.nih.gov/gene/?term=LOC105275363 | 4330 | 186 | 429.56 | | 1 | 2.31 | 0.54 | 15.2 | 14.36 | 0.85 | 0.85 | 0.04 | 0.04 |
| gene11454 | https://www.ncbi.nlm.nih.gov/gene/?term=LOC105285256 | 1906 | 45 | 236.1 | 1 | 5.25 | 2.22 | 47.83 | 47.02 | 0.81 | 0.81 | 0.02 | 0.02 | |
| gene4023 | https://www.ncbi.nlm.nih.gov/gene/?term=LOC105276692 | 1759 | 31 | 176.24 | | 1 | 5.69 | 3.23 | 24.44 | 23.63 | 0.81 | 0.81 | 0.01 | 0.01 |
| gene4892 | https://www.ncbi.nlm.nih.gov/gene/?term=LOC105277667 | 3898 | 119 | 305.28 | | 1 | 2.57 | 0.84 | 28.98 | 28.19 | 0.8 | 0.8 | 0.02 | 0.02 |
| gene3470 | https://www.ncbi.nlm.nih.gov/gene/?term=LOC105276059 | 8716 | 146 | 167.51 | | 1 | 1.15 | 0.68 | 9.33 | 8.54 | 0.79 | 0.79 | 0.01 | 0.01 |
| gene10540 | https://www.ncbi.nlm.nih.gov/gene/?term=LOC105284200 | 5267 | 215 | 408.2 | 1 | 1.9 | 0.47 | 12.8 | 12.05 | 0.75 | 0.75 | 0.03 | 0.03 | |
| gene4213 | https://www.ncbi.nlm.nih.gov/gene/?term=LOC105276901 | 3257 | 83 | 254.84 | | 1 | 3.07 | 1.2 | 27.23 | 26.49 | 0.74 | 0.74 | 0.02 | 0.02 |
| gene10520 | https://www.ncbi.nlm.nih.gov/gene/?term=LOC105284184 | 2451 | 79 | 322.32 | | 1 | 4.08 | 1.27 | 35.68 | 34.99 | 0.69 | 0.69 | 0.02 | 0.02 |
| gene298 | https://www.ncbi.nlm.nih.gov/gene/?term=LOC105275460 | 2473 | 118 | 477.15 | | 1 | 4.04 | 0.85 | 13.95 | 13.28 | 0.67 | 0.67 | 0.03 | 0.03 |
| gene9336 | https://www.ncbi.nlm.nih.gov/gene/?term=LOC105282834 | 5993 | 123 | 205.24 | | 1 | 1.67 | 0.81 | 15.27 | 14.61 | 0.66 | 0.66 | 0.01 | 0.01 |
| gene10600 | https://www.ncbi.nlm.nih.gov/gene/?term=LOC105284270 | 4837 | 115 | 237.75 | | 1 | 2.07 | 0.87 | 11.06 | 10.43 | 0.63 | 0.63 | 0.02 | 0.02 |
| gene6966 | https://www.ncbi.nlm.nih.gov/gene/?term=LOC105280104 | 3747 | 124 | 330.93 | | 1 | 2.67 | 0.81 | 17.23 | 16.6 | 0.63 | 0.63 | 0.02 | 0.02 |
| gene1430 | https://www.ncbi.nlm.nih.gov/gene/?term=LOC105287551 | 6433 | 216 | 335.77 | | 1 | 1.55 | 0.46 | 9.29 | 8.67 | 0.62 | 0.62 | 0.02 | 0.02 |
| gene570 | https://www.ncbi.nlm.nih.gov/gene/?term=LOC105278549 | 1673 | 62 | 370.59 | | 1 | 5.98 | 1.61 | 43.95 | 43.37 | 0.58 | 0.58 | 0.02 | 0.02 |
| gene5273 | https://www.ncbi.nlm.nih.gov/gene/?term=LOC105278108 | 4067 | 43 | 105.73 | | 1 | 2.46 | 2.33 | 26.66 | 26.12 | 0.55 | 0.55 | 0.01 | 0.01 |
| gene6583 | https://www.ncbi.nlm.nih.gov/gene/?term=LOC105279649 | 7444 | 315 | 423.16 | | 1 | 1.34 | 0.32 | 11.55 | 11.04 | 0.51 | 0.51 | 0.02 | 0.02 |
| gene12978 | https://www.ncbi.nlm.nih.gov/gene/?term=LOC105286997 | 2858 | 106 | 370.89 | | 1 | 3.5 | 0.94 | 12 | 11.5 | 0.5 | 0.5 | 0.02 | 0.02 |
| gene5831 | https://www.ncbi.nlm.nih.gov/gene/?term=LOC105278761 | 2506 | 105 | 418.99 | | 1 | 3.99 | 0.95 | 24.55 | 24.07 | 0.48 | 0.48 | 0.02 | 0.02 |
| gene8274 | https://www.ncbi.nlm.nih.gov/gene/?term=LOC105281624 | 5425 | 198 | 364.98 | | 1 | 1.84 | 0.51 | 10.15 | 9.67 | 0.48 | 0.48 | 0.02 | 0.02 |
| gene1332 | https://www.ncbi.nlm.nih.gov/gene/?term=LOC105286872 | 2460 | 156 | 634.15 | | 1 | 4.07 | 0.64 | 20.52 | 20.12 | 0.41 | 0.41 | 0.03 | 0.03 |
| gene2846 | https://www.ncbi.nlm.nih.gov/gene/?term=LOC105275374 | 11246 | 432 | 384.14 | | 1 | 0.89 | 0.23 | 11.15 | 10.76 | 0.39 | 0.39 | 0.02 | 0.02 |
| gene11579 | https://www.ncbi.nlm.nih.gov/gene/?term=LOC105285406 | 13571 | 229 | 168.74 | | 1 | 0.74 | 0.44 | 7.6 | 7.22 | 0.38 | 0.38 | 0.01 | 0.01 |
| gene12439 | https://www.ncbi.nlm.nih.gov/gene/?term=LOC105286357 | 4761 | 87 | 182.73 | | 1 | 2.1 | 1.15 | 25.3 | 24.92 | 0.38 | 0.38 | 0.01 | 0.01 |
| gene7247 | https://www.ncbi.nlm.nih.gov/gene/?term=LOC105280407 | 10486 | 208 | 198.36 | | 1 | 0.95 | 0.48 | 16.58 | 16.25 | 0.33 | 0.33 | 0.01 | 0.01 |
| gene2266 | https://www.ncbi.nlm.nih.gov/gene/?term=LOC105274699 | 2391 | 49 | 204.94 | | 1 | 4.18 | 2.04 | 23.32 | 23 | 0.32 | 0.32 | 0.01 | 0.01 |
| gene13114 | https://www.ncbi.nlm.nih.gov/gene/?term=LOC105287147 | 7609 | 393 | 516.49 | | 1 | 1.31 | 0.25 | 9.24 | 9.02 | 0.23 | 0.23 | 0.01 | 0.01 |
| gene5525 | https://www.ncbi.nlm.nih.gov/gene/?term=LOC105278396 | 3090 | 180 | 582.52 | | 1 | 3.24 | 0.56 | 24.99 | 24.77 | 0.22 | 0.22 | 0.01 | 0.01 |
| gene2683 | https://www.ncbi.nlm.nih.gov/gene/?term=LOC105275187 | 5492 | 65 | 118.35 | | 1 | 1.82 | 1.54 | 11.46 | 11.25 | 0.21 | 0.21 | 0 | 0 |
| gene5052 | https://www.ncbi.nlm.nih.gov/gene/?term=LOC105277856 | 3167 | 28 | 88.41 | 1 | 3.16 | 3.57 | 33.66 | 33.45 | 0.21 | 0.21 | 0 | 0 | |
| gene11082 | https://www.ncbi.nlm.nih.gov/gene/?term=LOC105284824 | 7436 | 189 | 254.17 | | 1 | 1.34 | 0.53 | 8.04 | 7.84 | 0.2 | 0.2 | 0.01 | 0.01 |
| gene11055 | https://www.ncbi.nlm.nih.gov/gene/?term=LOC105284796 | 2283 | 121 | 530 | 1 | 4.38 | 0.83 | 34.82 | 34.65 | 0.17 | 0.17 | 0.01 | 0.01 | |
| gene1470 | https://www.ncbi.nlm.nih.gov/gene/?term=LOC105287597 | 2974 | 181 | 608.61 | | 1 | 3.36 | 0.55 | 19.06 | 18.9 | 0.16 | 0.16 | 0.01 | 0.01 |
| gene2806 | https://www.ncbi.nlm.nih.gov/gene/?term=LOC105275327 | 14257 | 456 | 319.84 | | 1 | 0.7 | 0.22 | 7.38 | 7.24 | 0.14 | 0.14 | 0 | 0 |
| gene1220 | https://www.ncbi.nlm.nih.gov/gene/?term=LOC105285742 | 3453 | 100 | 289.6 | 1 | 2.9 | 1 | 18.13 | 18.02 | 0.11 | 0.11 | 0 | 0 | |
| gene5733 | https://www.ncbi.nlm.nih.gov/gene/?term=LOC105278649 | 5221 | 92 | 176.21 | | 1 | 1.92 | 1.09 | 19.86 | 19.75 | 0.11 | 0.11 | 0 | 0 |
| gene5746 | https://www.ncbi.nlm.nih.gov/gene/?term=LOC105278666 | 10343 | 436 | 421.54 | | 1 | 0.97 | 0.23 | 6.97 | 6.87 | 0.11 | 0.11 | 0 | 0 |
| gene8600 | https://www.ncbi.nlm.nih.gov/gene/?term=LOC105282001 | 8513 | 151 | 177.38 | | 1 | 1.17 | 0.66 | 10.5 | 10.38 | 0.11 | 0.11 | 0 | 0 |
| gene10844 | https://www.ncbi.nlm.nih.gov/gene/?term=LOC105284535 | 9086 | 222 | 244.33 | | 1 | 1.1 | 0.45 | 15.15 | 15.05 | 0.1 | 0.1 | 0 | 0 |
| gene1632 | https://www.ncbi.nlm.nih.gov/gene/?term=LOC105287764 | 18061 | 367 | 203.2 | 1 | 0.55 | 0.27 | 3.44 | 3.36 | 0.08 | 0.08 | 0 | 0 | |
| gene5524 | https://www.ncbi.nlm.nih.gov/gene/?term=LOC105278397 | 5177 | 196 | 378.6 | 1 | 1.93 | 0.51 | 24.77 | 24.69 | 0.07 | 0.07 | 0 | 0 | |
| gene1408 | https://www.ncbi.nlm.nih.gov/gene/?term=LOC109611520 | 8645 | 289 | 334.3 | 1 | 1.16 | 0.35 | 6.21 | 6.16 | 0.05 | 0.05 | 0 | 0 | |
| gene7368 | https://www.ncbi.nlm.nih.gov/gene/?term=LOC105280556 | 4105 | 38 | 92.57 | 1 | 2.44 | 2.63 | 16.79 | 16.73 | 0.05 | 0.05 | 0 | 0 | |

| | | | | | | | | | | | | | |
|---|---|---|---|---|---|---|---|---|---|---|---|---|---|
| gene9740 | https://www.ncbi.nlm.nih.gov/gene/?term=LOC105283277 | 2887 | 131 | 453.76 | 1 | 3.46 | 0.76 | 19.31 | 19.27 | 0.04 | 0.04 | 0 | 0 |
| gene8814 | https://www.ncbi.nlm.nih.gov/gene/?term=LOC105282240 | 6910 | 163 | 235.89 | 1 | 1.45 | 0.61 | 9.99 | 9.96 | 0.03 | 0.03 | 0 | 0 |
| gene2271 | https://www.ncbi.nlm.nih.gov/gene/?term=LOC105274706 | 1448 | 50 | 345.3 1 | 6.91 | 2 | 38.1 | 38.09 | 0.01 | 0.01 | 0 | 0 | |
| gene2173 | https://www.ncbi.nlm.nih.gov/gene/?term=LOC105274590 | 4387 | 68 | 155 1 | 2.28 | 1.47 | 26.72 | 26.74 | -0.02 | 0.02 | 0 | 0 | |
| gene7547 | https://www.ncbi.nlm.nih.gov/gene/?term=LOC105280767 | 24576 | 671 | 273.03 | 1 | 0.41 | 0.15 | 4.75 | 4.77 | -0.02 | 0.02 | 0 | 0 |
| gene10678 | https://www.ncbi.nlm.nih.gov/gene/?term=LOC105284355 | 4031 | 168 | 416.77 | 1 | 2.48 | 0.6 | 18.14 | 18.16 | -0.03 | 0.03 | 0 | 0 |
| gene10984 | https://www.ncbi.nlm.nih.gov/gene/?term=LOC105284704 | 1987 | 58 | 291.9 1 | 5.03 | 1.72 | 23.02 | 23.06 | -0.04 | 0.04 | 0 | 0 | |
| gene862 | https://www.ncbi.nlm.nih.gov/gene/?term=LOC105281802 | 73922 | 1763 | 238.49 | 1 | 0.14 | 0.06 | 2.22 | 2.27 | -0.05 | 0.05 | 0 | 0 |
| gene10326 | https://www.ncbi.nlm.nih.gov/gene/?term=LOC105283962 | 3042 | 119 | 391.19 | 1 | 3.29 | 0.84 | 8.91 | 8.98 | -0.07 | 0.07 | 0 | 0 |
| gene11275 | https://www.ncbi.nlm.nih.gov/gene/?term=LOC105285055 | 4119 | 257 | 623.94 | 1 | 2.43 | 0.39 | 10.45 | 10.51 | -0.07 | 0.07 | 0 | 0 |
| gene4962 | https://www.ncbi.nlm.nih.gov/gene/?term=LOC105277761 | 4645 | 142 | 305.71 | 1 | 2.15 | 0.7 | 13.6 | 13.69 | -0.09 | 0.09 | 0 | 0 |
| gene7398 | https://www.ncbi.nlm.nih.gov/gene/?term=LOC105280589 | 3452 | 33 | 95.6 1 | 2.9 | 3.03 | 20.49 | 20.58 | -0.09 | 0.09 | 0 | 0 | |
| gene6620 | https://www.ncbi.nlm.nih.gov/gene/?term=LOC105279687 | 17998 | 344 | 191.13 | 1 | 0.56 | 0.29 | 5.2 | 5.3 | -0.1 | 0.1 | 0 | 0 |
| gene7556 | https://www.ncbi.nlm.nih.gov/gene/?term=LOC105280774 | 2792 | 98 | 351 1 | 3.58 | 1.02 | 32.45 | 32.61 | -0.17 | 0.17 | -0.01 | 0.01 | |
| gene53 | https://www.ncbi.nlm.nih.gov/gene/?term=LOC105277148 | 19283 | 268 | 138.98 | 1 | 0.52 | 0.37 | 3.71 | 3.95 | -0.23 | 0.23 | 0 | 0 |
| gene10018 | https://www.ncbi.nlm.nih.gov/gene/?term=LOC105283607 | 4414 | 202 | 457.63 | 1 | 2.27 | 0.5 | 14.99 | 15.23 | -0.24 | 0.24 | -0.01 | 0.01 |
| gene6286 | https://www.ncbi.nlm.nih.gov/gene/?term=LOC105279298 | 2861 | 130 | 454.39 | 2 | 6.99 | 1.54 | 23.66 | 23.9 | -0.24 | 0.24 | -0.01 | 0.01 |
| gene9326 | https://www.ncbi.nlm.nih.gov/gene/?term=LOC105282818 | 16612 | 169 | 101.73 | 1 | 0.6 | 0.59 | 13.49 | 13.76 | -0.26 | 0.26 | 0 | 0 |
| gene10606 | https://www.ncbi.nlm.nih.gov/gene/?term=LOC105284273 | 6119 | 244 | 398.76 | 1 | 1.63 | 0.41 | 18.11 | 18.39 | -0.29 | 0.29 | -0.01 | 0.01 |
| gene34 | https://www.ncbi.nlm.nih.gov/gene/?term=LOC105278416 | 7120 | 124 | 174.16 | 1 | 1.4 | 0.81 | 14.41 | 14.72 | -0.3 | 0.3 | -0.01 | 0.01 |
| gene12573 | https://www.ncbi.nlm.nih.gov/gene/?term=LOC105286515 | 3647 | 105 | 287.91 | 1 | 2.74 | 0.95 | 12.53 | 12.84 | -0.31 | 0.31 | -0.01 | 0.01 |
| gene3885 | https://www.ncbi.nlm.nih.gov/gene/?term=LOC105276527 | 7928 | 299 | 377.14 | 1 | 1.26 | 0.33 | 9.83 | 10.14 | -0.31 | 0.31 | -0.01 | 0.01 |
| gene7166 | https://www.ncbi.nlm.nih.gov/gene/?term=LOC105280326 | 6117 | 226 | 369.46 | 1 | 1.63 | 0.44 | 14.91 | 15.21 | -0.31 | 0.31 | -0.01 | 0.01 |
| gene10549 | https://www.ncbi.nlm.nih.gov/gene/?term=LOC105284210 | 7145 | 284 | 397.48 | 2 | 2.8 | 0.7 | 11.85 | 12.19 | -0.33 | 0.33 | -0.01 | 0.01 |
| gene2748 | https://www.ncbi.nlm.nih.gov/gene/?term=LOC105275261 | 3845 | 53 | 137.84 | 1 | 2.6 | 1.89 | 38.36 | 38.68 | -0.33 | 0.33 | 0 | 0 |
| gene5646 | https://www.ncbi.nlm.nih.gov/gene/?term=LOC105278534 | 7475 | 176 | 235.45 | 1 | 1.34 | 0.57 | 18.67 | 19 | -0.33 | 0.33 | -0.01 | 0.01 |
| gene2404 | https://www.ncbi.nlm.nih.gov/gene/?term=LOC105274853 | 1767 | 41 | 232.03 | 1 | 5.66 | 2.44 | 30.51 | 30.87 | -0.36 | 0.36 | -0.01 | 0.01 |
| gene9327 | https://www.ncbi.nlm.nih.gov/gene/?term=LOC105282821 | 7793 | 140 | 179.65 | 1 | 1.28 | 0.71 | 12.3 | 12.71 | -0.41 | 0.41 | -0.01 | 0.01 |
| gene5375 | https://www.ncbi.nlm.nih.gov/gene/?term=LOC105278228 | 8640 | 152 | 175.93 | 1 | 1.16 | 0.66 | 22.96 | 23.39 | -0.42 | 0.42 | -0.01 | 0.01 |
| gene10879 | https://www.ncbi.nlm.nih.gov/gene/?term=LOC105284574 | 2423 | 50 | 206.36 | 1 | 4.13 | 2 | 52.82 | 53.31 | -0.49 | 0.49 | -0.01 | 0.01 |
| gene7211 | https://www.ncbi.nlm.nih.gov/gene/?term=LOC105280377 | 2024 | 63 | 311.26 | 1 | 4.94 | 1.59 | 48.25 | 48.74 | -0.49 | 0.49 | -0.02 | 0.02 |
| gene7550 | https://www.ncbi.nlm.nih.gov/gene/?term=LOC105280763 | 2173 | 76 | 349.75 | 1 | 4.6 | 1.32 | 32.2 | 32.7 | -0.49 | 0.49 | -0.02 | 0.02 |
| gene12307 | https://www.ncbi.nlm.nih.gov/gene/?term=LOC105286216 | 10313 | 164 | 159.02 | 1 | 0.97 | 0.61 | 16.27 | 16.77 | -0.5 | 0.5 | -0.01 | 0.01 |
| gene12802 | https://www.ncbi.nlm.nih.gov/gene/?term=LOC105286799 | 3183 | 159 | 499.53 | 1 | 3.14 | 0.63 | 17.45 | 17.99 | -0.54 | 0.54 | -0.03 | 0.03 |
| gene292 | https://www.ncbi.nlm.nih.gov/gene/?term=LOC105275398 | 2825 | 78 | 276.11 | 1 | 3.54 | 1.28 | 22.03 | 22.57 | -0.55 | 0.55 | -0.02 | 0.02 |
| gene4216 | https://www.ncbi.nlm.nih.gov/gene/?term=LOC105276906 | 2127 | 86 | 404.33 | 1 | 4.7 | 1.16 | 19.06 | 19.63 | -0.57 | 0.57 | -0.02 | 0.02 |
| gene5298 | https://www.ncbi.nlm.nih.gov/gene/?term=LOC105278145 | 4060 | 235 | 578.82 | 2 | 4.93 | 0.85 | 10.98 | 11.56 | -0.58 | 0.58 | -0.03 | 0.03 |
| gene13036 | https://www.ncbi.nlm.nih.gov/gene/?term=LOC105287059 | 4959 | 148 | 298.45 | 1 | 2.02 | 0.68 | 20.35 | 20.94 | -0.59 | 0.59 | -0.02 | 0.02 |
| gene3107 | https://www.ncbi.nlm.nih.gov/gene/?term=LOC105275650 | 5103 | 152 | 297.86 | 1 | 1.96 | 0.66 | 16.19 | 16.78 | -0.59 | 0.59 | -0.02 | 0.02 |
| gene4190 | https://www.ncbi.nlm.nih.gov/gene/?term=LOC105276875 | 3150 | 105 | 333.33 | 1 | 3.17 | 0.95 | 22.87 | 23.46 | -0.6 | 0.6 | -0.02 | 0.02 |
| gene7249 | https://www.ncbi.nlm.nih.gov/gene/?term=LOC105280422 | 1775 | 60 | 338.03 | 1 | 5.63 | 1.67 | 16.46 | 17.06 | -0.6 | 0.6 | -0.02 | 0.02 |
| gene1147 | https://www.ncbi.nlm.nih.gov/gene/?term=LOC105284954 | 4022 | 91 | 226.26 | 1 | 2.49 | 1.1 | 14.14 | 14.77 | -0.63 | 0.63 | -0.01 | 0.01 |
| gene9647 | https://www.ncbi.nlm.nih.gov/gene/?term=LOC109611233 | 3094 | 32 | 103.43 | 1 | 3.23 | 3.12 | 37.53 | 38.18 | -0.65 | 0.65 | -0.01 | 0.01 |
| gene9648 | https://www.ncbi.nlm.nih.gov/gene/?term=LOC105283183 | 2602 | 30 | 115.3 1 | 3.84 | 3.33 | 40.03 | 40.73 | -0.7 | 0.7 | -0.01 | 0.01 | |
| gene4376 | https://www.ncbi.nlm.nih.gov/gene/?term=LOC105277083 | 4307 | 97 | 225.21 | 1 | 2.32 | 1.03 | 24.28 | 25.05 | -0.77 | 0.77 | -0.02 | 0.02 |
| gene920 | https://www.ncbi.nlm.nih.gov/gene/?term=LOC105282518 | 1197 | 48 | 401 1 | 8.35 | 2.08 | 32.24 | 33.02 | -0.78 | 0.78 | -0.03 | 0.03 | |
| gene10392 | https://www.ncbi.nlm.nih.gov/gene/?term=LOC105284036 | 3216 | 94 | 292.29 | 2 | 6.22 | 2.13 | 22.41 | 23.26 | -0.84 | 0.84 | -0.02 | 0.02 |
| gene12203 | https://www.ncbi.nlm.nih.gov/gene/?term=LOC105286099 | 5967 | 70 | 117.31 | 1 | 1.68 | 1.43 | 9.91 | 10.79 | -0.88 | 0.88 | -0.01 | 0.01 |
| gene11340 | https://www.ncbi.nlm.nih.gov/gene/?term=LOC105285131 | 2918 | 87 | 298.15 | 1 | 3.43 | 1.15 | 10.66 | 11.58 | -0.92 | 0.92 | -0.03 | 0.03 |
| gene7248 | https://www.ncbi.nlm.nih.gov/gene/?term=LOC105280408 | 5175 | 52 | 100.48 | 1 | 1.93 | 1.92 | 17.59 | 18.54 | -0.95 | 0.95 | -0.01 | 0.01 |
| gene10155 | https://www.ncbi.nlm.nih.gov/gene/?term=LOC105283756 | 7030 | 108 | 153.63 | 1 | 1.42 | 0.93 | 5.58 | 6.54 | -0.96 | 0.96 | -0.01 | 0.01 |
| gene12021 | https://www.ncbi.nlm.nih.gov/gene/?term=LOC105285905 | 1399 | 16 | 114.37 | 1 | 7.15 | 6.25 | 46.76 | 47.75 | -0.98 | 0.98 | -0.01 | 0.01 |
| gene1310 | https://www.ncbi.nlm.nih.gov/gene/?term=LOC105286661 | 2582 | 68 | 263.36 | 1 | 3.87 | 1.47 | 19.86 | 20.85 | -1 | 1 | -0.03 | 0.03 |
| gene8729 | https://www.ncbi.nlm.nih.gov/gene/?term=LOC105282137 | 5005 | 213 | 425.57 | 1 | 2 | 0.47 | 14.62 | 15.67 | -1.05 | 1.05 | -0.04 | 0.04 |
| gene7600 | https://www.ncbi.nlm.nih.gov/gene/?term=LOC105280819 | 1615 | 37 | 229.1 1 | 6.19 | 2.7 | 36.99 | 38.16 | -1.16 | 1.16 | -0.03 | 0.03 | |
| gene10213 | https://www.ncbi.nlm.nih.gov/gene/?term=LOC105283826 | 1503 | 53 | 352.63 | 1 | 6.65 | 1.89 | 14.78 | 15.95 | -1.17 | 1.17 | -0.04 | 0.04 |
| gene6639 | https://www.ncbi.nlm.nih.gov/gene/?term=LOC105279705 | 7656 | 69 | 90.13 1 | 1.31 | 1.45 | 20.74 | 21.96 | -1.21 | 1.21 | -0.01 | 0.01 | |
| gene35 | https://www.ncbi.nlm.nih.gov/gene/?term=LOC105275197 | 2643 | 45 | 170.26 | 1 | 3.78 | 2.22 | 38.91 | 40.15 | -1.23 | 1.23 | -0.02 | 0.02 |
| gene6031 | https://www.ncbi.nlm.nih.gov/gene/?term=LOC105279000 | 5691 | 180 | 316.29 | 1 | 1.76 | 0.56 | 14.61 | 15.92 | -1.31 | 1.31 | -0.04 | 0.04 |
| gene564 | https://www.ncbi.nlm.nih.gov/gene/?term=LOC105278478 | 2802 | 18 | 64.24 1 | 3.57 | 5.56 | 14.97 | 16.38 | -1.41 | 1.41 | -0.01 | 0.01 | |
| gene4641 | https://www.ncbi.nlm.nih.gov/gene/?term=LOC105277398 | 1937 | 71 | 366.55 | 1 | 5.16 | 1.41 | 38.26 | 39.69 | -1.43 | 1.43 | -0.05 | 0.05 |
| gene7183 | https://www.ncbi.nlm.nih.gov/gene/?term=LOC105280348 | 3925 | 70 | 178.34 | 1 | 2.55 | 1.43 | 19.96 | 21.48 | -1.52 | 1.52 | -0.03 | 0.03 |
| gene8343 | https://www.ncbi.nlm.nih.gov/gene/?term=LOC105281700 | 2172 | 32 | 147.33 | 1 | 4.6 | 3.12 | 37.43 | 38.97 | -1.55 | 1.55 | -0.02 | 0.02 |
| gene7199 | https://www.ncbi.nlm.nih.gov/gene/?term=LOC105280366 | 1414 | 24 | 169.73 | 1 | 7.07 | 4.17 | 43.55 | 45.17 | -1.62 | 1.62 | -0.03 | 0.03 |

| | | | | | | | | | | | | | | |
|---|---|---|---|---|---|---|---|---|---|---|---|---|---|---|
| gene12572 | https://www.ncbi.nlm.nih.gov/gene/?term=LOC105286514 | 2374 | 47 | 197.98 | | 1 | 4.21 | 2.13 | 35.17 | 36.86 | -1.69 | 1.69 | -0.03 | 0.03 |
| gene9101 | https://www.ncbi.nlm.nih.gov/gene/?term=LOC105282553 | 3068 | 102 | 332.46 | | 1 | 3.26 | 0.98 | 21.53 | 23.25 | -1.72 | 1.72 | -0.06 | 0.06 |
| gene6237 | https://www.ncbi.nlm.nih.gov/gene/?term=LOC105279244 | 1484 | 22 | 148.25 | | 1 | 6.74 | 4.55 | 40.19 | 41.99 | -1.8 | 1.8 | -0.03 | 0.03 |
| gene2413 | https://www.ncbi.nlm.nih.gov/gene/?term=LOC105274863 | 2006 | 49 | 244.27 | | 1 | 4.99 | 2.04 | 53.65 | 55.47 | -1.81 | 1.81 | -0.04 | 0.04 |
| gene7406 | https://www.ncbi.nlm.nih.gov/gene/?term=LOC105280596 | 3432 | 106 | 308.86 | | 1 | 2.91 | 0.94 | 17.02 | 19.17 | -2.15 | 2.15 | -0.07 | 0.07 |
| gene12981 | https://www.ncbi.nlm.nih.gov/gene/?term=LOC105286999 | 2063 | 42 | 203.59 | | 1 | 4.85 | 2.38 | 33.61 | 35.96 | -2.34 | 2.34 | -0.05 | 0.05 |
| gene12406 | https://www.ncbi.nlm.nih.gov/gene/?term=LOC105286316 | 1386 | 31 | 223.67 | | 1 | 7.22 | 3.23 | 42.03 | 44.54 | -2.51 | 2.51 | -0.06 | 0.06 |
| gene10107 | https://www.ncbi.nlm.nih.gov/gene/?term=LOC105283712 | 1883 | 35 | 185.87 | | 1 | 5.31 | 2.86 | 23.29 | 25.95 | -2.66 | 2.66 | -0.05 | 0.05 |
| gene3599 | https://www.ncbi.nlm.nih.gov/gene/?term=LOC105276196 | 2955 | 58 | 196.28 | | 1 | 3.38 | 1.72 | 37.85 | 40.55 | -2.7 | 2.7 | -0.05 | 0.05 |
| gene6422 | https://www.ncbi.nlm.nih.gov/gene/?term=LOC105279458 | 2419 | 38 | 157.09 | | 1 | 4.13 | 2.63 | 47.43 | 50.37 | -2.94 | 2.94 | -0.05 | 0.05 |
| gene3168 | https://www.ncbi.nlm.nih.gov/gene/?term=LOC105275711 | 6150 | 69 | 112.2 | 1 | 1.63 | 1.45 | 12.17 | 15.18 | -3.01 | 3.01 | -0.03 | 0.03 | |
| gene322 | https://www.ncbi.nlm.nih.gov/gene/?term=LOC105275777 | 1606 | 27 | 168.12 | | 1 | 6.23 | 3.7 | 38.42 | 42 | -3.59 | 3.59 | -0.06 | 0.06 |
| gene388 | https://www.ncbi.nlm.nih.gov/gene/?term=LOC105276538 | 1558 | 19 | 121.95 | | 2 | 12.84 | 10.53 | 61.29 | 65.13 | -3.85 | 3.85 | -0.05 | 0.05 |
| gene10345 | https://www.ncbi.nlm.nih.gov/gene/?term=LOC105283989 | 4444 | 22 | 49.5 | 1 | 2.25 | 4.55 | 37.68 | 41.66 | -3.99 | 3.99 | -0.02 | 0.02 | |
| gene10602 | https://www.ncbi.nlm.nih.gov/gene/?term=LOC105284268 | 2122 | 19 | 89.54 | 1 | 4.71 | 5.26 | 27.9 | 32.99 | -5.08 | 5.08 | -0.05 | 0.05 | |
| gene1008 | https://www.ncbi.nlm.nih.gov/gene/?term=LOC105283492 | 1479 | 9 | 60.85 | 1 | 6.76 | 11.11 | 30.57 | 36.08 | -5.51 | 5.51 | -0.03 | 0.03 | |
| gene393 | https://www.ncbi.nlm.nih.gov/gene/?term=LOC105276566 | 1123 | 2 | 17.81 | 1 | 8.9 | 50 | 50 | 76.78 | -26.78 | | 26.78 | -0.05 | 0.05 |

Directory: OGLall File: ogl-Cb_LI_RphaseA2.txt

| gene_ID | gene_link | gwidth | #Sites | #per10Kb | %perSite | %pNucl |
|---|---|---|---|---|---|---|
| gene13384 | https://www.ncbi.nlm.nih.gov/gene/?term=LOC105287447 | 1877 | 296 | 1576.98 | 27.63 | 4.36 |
| gene5917 | https://www.ncbi.nlm.nih.gov/gene/?term=LOC105278849 | 823 | 139 | 1688.94 | 25.03 | 4.23 |
| gene10219 | https://www.ncbi.nlm.nih.gov/gene/?term=LOC105283830 | 235 | 22 | 936.17 | 44.27 | 4.14 |
| gene12364 | https://www.ncbi.nlm.nih.gov/gene/?term=LOC105286269 | 1403 | 110 | 784.03 | 52.64 | 4.13 |
| gene12538 | https://www.ncbi.nlm.nih.gov/gene/?term=LOC105286531 | 372 | 43 | 1155.91 | 35.68 | 4.12 |
| gene5911 | https://www.ncbi.nlm.nih.gov/gene/?term=LOC105278844 | 943 | 122 | 1293.74 | 31.64 | 4.09 |
| gene5309 | https://www.ncbi.nlm.nih.gov/gene/?term=LOC105278160 | 1704 | 183 | 1073.94 | 37.02 | 3.98 |
| gene3719 | https://www.ncbi.nlm.nih.gov/gene/?term=LOC105276360 | 1471 | 205 | 1393.61 | 28.45 | 3.97 |
| gene11569 | https://www.ncbi.nlm.nih.gov/gene/?term=LOC105285394 | 1311 | 75 | 572.08 | 68.68 | 3.93 |
| gene4878 | https://www.ncbi.nlm.nih.gov/gene/?term=LOC105277648 | 675 | 45 | 666.67 | 57.93 | 3.86 |
| gene1654 | https://www.ncbi.nlm.nih.gov/gene/?term=LOC105287811 | 2285 | 205 | 897.16 | 42.64 | 3.83 |
| gene211 | https://www.ncbi.nlm.nih.gov/gene/?term=LOC105274534 | 1280 | 111 | 867.19 | 44.2 | 3.83 |
| gene12652 | https://www.ncbi.nlm.nih.gov/gene/?term=LOC109610640 | 1122 | 71 | 632.8 | 57.98 | 3.67 |
| gene2797 | https://www.ncbi.nlm.nih.gov/gene/?term=LOC105275314 | 1489 | 132 | 886.5 | 41.45 | 3.67 |
| gene570 | https://www.ncbi.nlm.nih.gov/gene/?term=LOC105278549 | 1673 | 145 | 866.71 | 41.46 | 3.59 |
| gene6078 | https://www.ncbi.nlm.nih.gov/gene/?term=LOC105279052 | 1130 | 163 | 1442.48 | 24.83 | 3.58 |
| gene10943 | https://www.ncbi.nlm.nih.gov/gene/?term=LOC105284659 | 1759 | 129 | 733.37 | 48.52 | 3.56 |
| gene12330 | https://www.ncbi.nlm.nih.gov/gene/?term=LOC105286235 | 1501 | 122 | 812.79 | 42.08 | 3.42 |
| gene12746 | https://www.ncbi.nlm.nih.gov/gene/?term=LOC105286737 | 2688 | 406 | 1510.42 | 22.57 | 3.41 |
| gene1562 | https://www.ncbi.nlm.nih.gov/gene/?term=LOC105287692 | 1491 | 153 | 1026.16 | 33.21 | 3.41 |
| gene3731 | https://www.ncbi.nlm.nih.gov/gene/?term=LOC105276371 | 865 | 65 | 751.45 | 44.86 | 3.37 |
| gene5301 | https://www.ncbi.nlm.nih.gov/gene/?term=LOC105278148 | 1881 | 240 | 1275.92 | 26.28 | 3.35 |
| gene11566 | https://www.ncbi.nlm.nih.gov/gene/?term=LOC105285392 | 1603 | 157 | 979.41 | 33.58 | 3.29 |
| gene4635 | https://www.ncbi.nlm.nih.gov/gene/?term=LOC105277390 | 2067 | 168 | 812.77 | 40.39 | 3.28 |

...

... only first 25 lines shown ...

Directory: OGLall File: ogl-Cb_LI_RphaseA2.vs.RphaseB1.txt

| gene_ID | gene_link | gwidth | #Sites | #per10Kb | #dmSites | #dmsp10kb | %dmSites | %pSite1 | %pSite2 | DMpSite | ADMpSite | DMpNucl | ADMpNucl |
|---|---|---|---|---|---|---|---|---|---|---|---|---|---|
| gene2593 | https://www.ncbi.nlm.nih.gov/gene/?term=LOC105275070 | 1774 | 11 | 62.01 | 6 | 33.82 | 54.55 | 67.77 | 25.28 | 42.49 | 42.49 | 0.26 | 0.26 |
| gene3165 | https://www.ncbi.nlm.nih.gov/gene/?term=LOC105275706 | 2374 | 70 | 294.86 | 5 | 21.06 | 7.14 | 33.14 | 24.38 | 8.76 | 8.76 | 0.26 | 0.26 |
| gene2824 | https://www.ncbi.nlm.nih.gov/gene/?term=LOC105275337 | 1772 | 63 | 355.53 | 2 | 11.29 | 3.17 | 38.53 | 29.97 | 8.56 | 8.56 | 0.3 | 0.3 |
| gene12395 | https://www.ncbi.nlm.nih.gov/gene/?term=LOC105286302 | 4093 | 24 | 58.64 | 1 | 2.44 | 4.17 | 20.4 | 15.15 | 5.24 | 5.24 | 0.03 | 0.03 |
| gene12961 | https://www.ncbi.nlm.nih.gov/gene/?term=LOC105286983 | 3094 | 71 | 229.48 | 1 | 3.23 | 1.41 | 6.3 | 1.17 | 5.12 | 5.12 | 0.12 | 0.12 |
| gene5714 | https://www.ncbi.nlm.nih.gov/gene/?term=LOC105278729 | 710 | 16 | 225.35 | 1 | 14.08 | 6.25 | 56.86 | 51.9 | 4.96 | 4.96 | 0.11 | 0.11 |
| gene13033 | https://www.ncbi.nlm.nih.gov/gene/?term=LOC105287063 | 1877 | 55 | 293.02 | 1 | 5.33 | 1.82 | 30.5 | 27.04 | 3.46 | 3.46 | 0.1 | 0.1 |
| gene1007 | https://www.ncbi.nlm.nih.gov/gene/?term=LOC105283457 | 1878 | 49 | 260.92 | 1 | 5.32 | 2.04 | 35.2 | 31.9 | 3.3 | 3.3 | 0.09 | 0.09 |
| gene8178 | https://www.ncbi.nlm.nih.gov/gene/?term=LOC105281515 | 8647 | 59 | 68.23 | 1 | 1.16 | 1.69 | 30.95 | 28.43 | 2.52 | 2.52 | 0.02 | 0.02 |
| gene10107 | https://www.ncbi.nlm.nih.gov/gene/?term=LOC105283712 | 1883 | 34 | 180.56 | 1 | 5.31 | 2.94 | 31.9 | 29.48 | 2.42 | 2.42 | 0.04 | 0.04 |
| gene10984 | https://www.ncbi.nlm.nih.gov/gene/?term=LOC105284704 | 1987 | 66 | 332.16 | 1 | 5.03 | 1.52 | 25.73 | 23.31 | 2.42 | 2.42 | 0.08 | 0.08 |
| gene274 | https://www.ncbi.nlm.nih.gov/gene/?term=LOC105275190 | 3488 | 74 | 212.16 | 1 | 2.87 | 1.35 | 22.52 | 20.11 | 2.4 | 2.4 | 0.05 | 0.05 |
| gene2970 | https://www.ncbi.nlm.nih.gov/gene/?term=LOC105275518 | 2340 | 28 | 119.66 | 1 | 4.27 | 3.57 | 41.75 | 39.36 | 2.39 | 2.39 | 0.03 | 0.03 |
| gene2271 | https://www.ncbi.nlm.nih.gov/gene/?term=LOC105274706 | 1448 | 57 | 393.65 | 1 | 6.91 | 1.75 | 39.53 | 37.37 | 2.16 | 2.16 | 0.09 | 0.09 |
| gene4034 | https://www.ncbi.nlm.nih.gov/gene/?term=LOC105276704 | 2065 | 92 | 445.52 | 5 | 24.21 | 5.43 | 14.65 | 12.65 | 2 | 2 | 0.09 | 0.09 |
| gene7568 | https://www.ncbi.nlm.nih.gov/gene/?term=LOC105280786 | 2525 | 143 | 566.34 | 1 | 3.96 | 0.7 | 27.63 | 25.63 | 2 | 2 | 0.11 | 0.11 |
| gene3671 | https://www.ncbi.nlm.nih.gov/gene/?term=LOC105276272 | 1966 | 47 | 239.06 | 1 | 5.09 | 2.13 | 25.43 | 23.44 | 1.98 | 1.98 | 0.05 | 0.05 |
| gene9336 | https://www.ncbi.nlm.nih.gov/gene/?term=LOC105282834 | 5993 | 137 | 228.6 | 1 | 1.67 | 0.73 | 15.64 | 13.69 | 1.95 | 1.95 | 0.04 | 0.04 |
| gene3774 | https://www.ncbi.nlm.nih.gov/gene/?term=LOC105276553 | 2716 | 84 | 309.28 | 1 | 3.68 | 1.19 | 43.69 | 41.82 | 1.87 | 1.87 | 0.06 | 0.06 |
| gene388 | https://www.ncbi.nlm.nih.gov/gene/?term=LOC105276538 | 1558 | 24 | 154.04 | 2 | 12.84 | 8.33 | 66.6 | 64.8 | 1.8 | 1.8 | 0.03 | 0.03 |
| gene4996 | https://www.ncbi.nlm.nih.gov/gene/?term=LOC105277798 | 5617 | 74 | 131.74 | 1 | 1.78 | 1.35 | 11.47 | 9.83 | 1.64 | 1.64 | 0.02 | 0.02 |
| gene10879 | https://www.ncbi.nlm.nih.gov/gene/?term=LOC105284574 | 2423 | 52 | 214.61 | 1 | 4.13 | 1.92 | 56.31 | 54.67 | 1.63 | 1.63 | 0.04 | 0.04 |
| gene8742 | https://www.ncbi.nlm.nih.gov/gene/?term=LOC105282151 | 2563 | 44 | 171.67 | 1 | 3.9 | 2.27 | 43.82 | 42.21 | 1.61 | 1.61 | 0.03 | 0.03 |
| gene940 | https://www.ncbi.nlm.nih.gov/gene/?term=LOC105282794 | 3578 | 95 | 265.51 | 1 | 2.79 | 1.05 | 15.95 | 14.34 | 1.61 | 1.61 | 0.04 | 0.04 |
| gene7493 | https://www.ncbi.nlm.nih.gov/gene/?term=LOC105280704 | 4225 | 155 | 366.86 | 1 | 2.37 | 0.65 | 17.59 | 16.02 | 1.58 | 1.58 | 0.06 | 0.06 |
| gene3817 | https://www.ncbi.nlm.nih.gov/gene/?term=LOC105276449 | 1230 | 55 | 447.15 | 1 | 8.13 | 1.82 | 24.78 | 23.21 | 1.57 | 1.57 | 0.07 | 0.07 |
| gene2241 | https://www.ncbi.nlm.nih.gov/gene/?term=LOC105274666 | 2598 | 64 | 246.34 | 1 | 3.85 | 1.56 | 28.81 | 27.27 | 1.54 | 1.54 | 0.04 | 0.04 |
| gene2748 | https://www.ncbi.nlm.nih.gov/gene/?term=LOC105275261 | 3845 | 55 | 143.04 | 1 | 2.6 | 1.82 | 41.25 | 39.71 | 1.54 | 1.54 | 0.02 | 0.02 |
| gene5273 | https://www.ncbi.nlm.nih.gov/gene/?term=LOC105278108 | 4067 | 57 | 140.15 | 1 | 2.46 | 1.75 | 25.93 | 24.39 | 1.54 | 1.54 | 0.02 | 0.02 |
| gene2825 | https://www.ncbi.nlm.nih.gov/gene/?term=LOC105275343 | 5545 | 98 | 176.74 | 1 | 1.8 | 1.02 | 9.51 | 8.1 | 1.42 | 1.42 | 0.03 | 0.03 |
| gene1147 | https://www.ncbi.nlm.nih.gov/gene/?term=LOC105284954 | 4022 | 114 | 283.44 | 1 | 2.49 | 0.88 | 14.88 | 13.52 | 1.36 | 1.36 | 0.04 | 0.04 |
| gene322 | https://www.ncbi.nlm.nih.gov/gene/?term=LOC105275777 | 1606 | 29 | 180.57 | 1 | 6.23 | 3.45 | 36.14 | 34.8 | 1.35 | 1.35 | 0.02 | 0.02 |
| gene5525 | https://www.ncbi.nlm.nih.gov/gene/?term=LOC105278396 | 3090 | 189 | 611.65 | 1 | 3.24 | 0.53 | 25.87 | 24.52 | 1.35 | 1.35 | 0.08 | 0.08 |
| gene12106 | https://www.ncbi.nlm.nih.gov/gene/?term=LOC105286002 | 8556 | 83 | 97.01 | 1 | 1.17 | 1.2 | 3.39 | 2.1 | 1.29 | 1.29 | 0.01 | 0.01 |
| gene927 | https://www.ncbi.nlm.nih.gov/gene/?term=LOC105282600 | 3727 | 67 | 179.77 | 1 | 2.68 | 1.49 | 27.6 | 26.32 | 1.28 | 1.28 | 0.02 | 0.02 |
| gene12572 | https://www.ncbi.nlm.nih.gov/gene/?term=LOC105286514 | 2374 | 56 | 235.89 | 1 | 4.21 | 1.79 | 36.49 | 35.22 | 1.27 | 1.27 | 0.03 | 0.03 |
| gene6455 | https://www.ncbi.nlm.nih.gov/gene/?term=LOC105279500 | 2612 | 36 | 137.83 | 1 | 3.83 | 2.78 | 48.83 | 47.56 | 1.26 | 1.26 | 0.02 | 0.02 |
| gene4048 | https://www.ncbi.nlm.nih.gov/gene/?term=LOC105276724 | 2955 | 47 | 159.05 | 1 | 3.38 | 2.13 | 12.29 | 11.14 | 1.14 | 1.14 | 0.02 | 0.02 |
| gene5052 | https://www.ncbi.nlm.nih.gov/gene/?term=LOC105277856 | 3167 | 36 | 113.67 | 1 | 3.16 | 2.78 | 37.89 | 36.75 | 1.14 | 1.14 | 0.01 | 0.01 |
| gene935 | https://www.ncbi.nlm.nih.gov/gene/?term=LOC105282682 | 4368 | 240 | 549.45 | 1 | 2.29 | 0.42 | 13.64 | 12.5 | 1.14 | 1.14 | 0.06 | 0.06 |
| gene5524 | https://www.ncbi.nlm.nih.gov/gene/?term=LOC105278397 | 5177 | 206 | 397.91 | 1 | 1.93 | 0.49 | 25.82 | 24.7 | 1.12 | 1.12 | 0.04 | 0.04 |
| gene5831 | https://www.ncbi.nlm.nih.gov/gene/?term=LOC105278761 | 2506 | 119 | 474.86 | 1 | 3.99 | 0.84 | 22.74 | 21.63 | 1.11 | 1.11 | 0.05 | 0.05 |
| gene10222 | https://www.ncbi.nlm.nih.gov/gene/?term=LOC105283838 | 1748 | 33 | 188.79 | 1 | 5.72 | 3.03 | 33.58 | 32.49 | 1.1 | 1.1 | 0.02 | 0.02 |
| gene12406 | https://www.ncbi.nlm.nih.gov/gene/?term=LOC105286316 | 1386 | 52 | 375.18 | 1 | 7.22 | 1.92 | 33.87 | 32.77 | 1.1 | 1.1 | 0.04 | 0.04 |
| gene5877 | https://www.ncbi.nlm.nih.gov/gene/?term=LOC105278807 | 1719 | 53 | 308.32 | 1 | 5.82 | 1.89 | 27.62 | 26.53 | 1.08 | 1.08 | 0.03 | 0.03 |
| gene12021 | https://www.ncbi.nlm.nih.gov/gene/?term=LOC105285905 | 1399 | 27 | 192.99 | 1 | 7.15 | 3.7 | 37.97 | 36.9 | 1.06 | 1.06 | 0.02 | 0.02 |
| gene10243 | https://www.ncbi.nlm.nih.gov/gene/?term=LOC105283862 | 3239 | 45 | 138.93 | 1 | 3.09 | 2.22 | 29.81 | 28.76 | 1.05 | 1.05 | 0.01 | 0.01 |
| gene7183 | https://www.ncbi.nlm.nih.gov/gene/?term=LOC105280348 | 3925 | 79 | 201.27 | 1 | 2.55 | 1.27 | 18.1 | 17.08 | 1.01 | 1.01 | 0.02 | 0.02 |
| gene2833 | https://www.ncbi.nlm.nih.gov/gene/?term=LOC105275352 | 5528 | 43 | 77.79 | 1 | 1.81 | 2.33 | 22.51 | 21.53 | 0.98 | 0.98 | 0.01 | 0.01 |
| gene3773 | https://www.ncbi.nlm.nih.gov/gene/?term=LOC105276408 | 5522 | 200 | 362.19 | 1 | 1.81 | 0.5 | 21.6 | 20.62 | 0.98 | 0.98 | 0.04 | 0.04 |
| gene298 | https://www.ncbi.nlm.nih.gov/gene/?term=LOC105275460 | 2473 | 123 | 497.37 | 1 | 4.04 | 0.81 | 13.29 | 12.33 | 0.96 | 0.96 | 0.05 | 0.05 |
| gene7556 | https://www.ncbi.nlm.nih.gov/gene/?term=LOC105280774 | 2792 | 135 | 483.52 | 1 | 3.58 | 0.74 | 25.37 | 24.41 | 0.96 | 0.96 | 0.05 | 0.05 |
| gene11819 | https://www.ncbi.nlm.nih.gov/gene/?term=LOC105285671 | 2366 | 103 | 435.33 | 1 | 4.23 | 0.97 | 30.19 | 29.25 | 0.95 | 0.95 | 0.04 | 0.04 |
| gene1884 | https://www.ncbi.nlm.nih.gov/gene/?term=LOC105288072 | 4169 | 88 | 211.08 | 1 | 2.4 | 1.14 | 22.65 | 21.71 | 0.94 | 0.94 | 0.02 | 0.02 |
| gene10540 | https://www.ncbi.nlm.nih.gov/gene/?term=LOC105284200 | 5267 | 219 | 415.8 | 1 | 1.9 | 0.46 | 13.98 | 13.1 | 0.88 | 0.88 | 0.04 | 0.04 |
| gene12516 | https://www.ncbi.nlm.nih.gov/gene/?term=LOC105286452 | 1529 | 42 | 274.69 | 1 | 6.54 | 2.38 | 47.45 | 46.6 | 0.86 | 0.86 | 0.02 | 0.02 |
| gene6286 | https://www.ncbi.nlm.nih.gov/gene/?term=LOC105279298 | 2861 | 133 | 464.87 | 2 | 6.99 | 1.5 | 23.83 | 22.97 | 0.86 | 0.86 | 0.04 | 0.04 |
| gene10678 | https://www.ncbi.nlm.nih.gov/gene/?term=LOC105284355 | 4031 | 172 | 426.69 | 1 | 2.48 | 0.58 | 20.12 | 19.3 | 0.83 | 0.83 | 0.04 | 0.04 |
| gene4892 | https://www.ncbi.nlm.nih.gov/gene/?term=LOC105277667 | 3898 | 127 | 325.81 | 1 | 2.57 | 0.79 | 28.13 | 27.36 | 0.77 | 0.77 | 0.03 | 0.03 |
| gene10600 | https://www.ncbi.nlm.nih.gov/gene/?term=LOC105284270 | 4837 | 114 | 235.68 | 1 | 2.07 | 0.88 | 12.4 | 11.65 | 0.75 | 0.75 | 0.02 | 0.02 |

| | | | | | | | | | | | | | |
|---|---|---|---|---|---|---|---|---|---|---|---|---|---|
| gene10606 | https://www.ncbi.nlm.nih.gov/gene/?term=LOC105284273 | 6119 | 286 | 467.4 | 1 | 1.63 | 0.35 | 18.29 | 17.53 | 0.75 | 0.75 | 0.04 | 0.04 |
| gene8015 | https://www.ncbi.nlm.nih.gov/gene/?term=LOC105281312 | 2998 | 69 | 230.15 | 1 | 3.34 | 1.45 | 31.17 | 30.43 | 0.74 | 0.74 | 0.02 | 0.02 |
| gene289 | https://www.ncbi.nlm.nih.gov/gene/?term=LOC105275347 | 2533 | 90 | 355.31 | 1 | 3.95 | 1.11 | 24.42 | 23.69 | 0.73 | 0.73 | 0.03 | 0.03 |
| gene10864 | https://www.ncbi.nlm.nih.gov/gene/?term=LOC105284557 | 1666 | 91 | 546.22 | 1 | 6 | 1.1 | 27.88 | 27.16 | 0.72 | 0.72 | 0.04 | 0.04 |
| gene13036 | https://www.ncbi.nlm.nih.gov/gene/?term=LOC105287059 | 4959 | 200 | 403.31 | 1 | 2.02 | 0.5 | 17.56 | 16.85 | 0.71 | 0.71 | 0.03 | 0.03 |
| gene12802 | https://www.ncbi.nlm.nih.gov/gene/?term=LOC105286799 | 3183 | 184 | 578.07 | 1 | 3.14 | 0.54 | 17.39 | 16.7 | 0.69 | 0.69 | 0.04 | 0.04 |
| gene9330 | https://www.ncbi.nlm.nih.gov/gene/?term=LOC105282825 | 3936 | 102 | 259.15 | 1 | 2.54 | 0.98 | 14.91 | 14.23 | 0.68 | 0.68 | 0.02 | 0.02 |
| gene3912 | https://www.ncbi.nlm.nih.gov/gene/?term=LOC105276560 | 17900 | 373 | 208.38 | 3 | 1.68 | 0.8 | 2.56 | 1.89 | 0.67 | 0.67 | 0.01 | 0.01 |
| gene814 | https://www.ncbi.nlm.nih.gov/gene/?term=LOC105281296 | 2722 | 57 | 209.4 | 1 | 3.67 | 1.75 | 23.86 | 23.2 | 0.65 | 0.65 | 0.01 | 0.01 |
| gene8343 | https://www.ncbi.nlm.nih.gov/gene/?term=LOC105281700 | 2172 | 43 | 197.97 | 1 | 4.6 | 2.33 | 36.96 | 36.39 | 0.57 | 0.57 | 0.01 | 0.01 |
| gene1573 | https://www.ncbi.nlm.nih.gov/gene/?term=LOC105287704 | 3030 | 143 | 471.95 | 1 | 3.3 | 0.7 | 19.92 | 19.38 | 0.54 | 0.54 | 0.03 | 0.03 |
| gene2257 | https://www.ncbi.nlm.nih.gov/gene/?term=LOC105274689 | 3658 | 90 | 246.04 | 1 | 2.73 | 1.11 | 15.54 | 15.05 | 0.49 | 0.49 | 0.01 | 0.01 |
| gene6031 | https://www.ncbi.nlm.nih.gov/gene/?term=LOC105279000 | 5691 | 226 | 397.12 | 1 | 1.76 | 0.44 | 12.2 | 11.72 | 0.48 | 0.48 | 0.02 | 0.02 |
| gene3206 | https://www.ncbi.nlm.nih.gov/gene/?term=LOC105275752 | 3474 | 138 | 397.24 | 1 | 2.88 | 0.72 | 31.47 | 31.01 | 0.46 | 0.46 | 0.02 | 0.02 |
| gene7888 | https://www.ncbi.nlm.nih.gov/gene/?term=LOC105281141 | 4357 | 180 | 413.13 | 1 | 2.3 | 0.56 | 27.03 | 26.57 | 0.45 | 0.45 | 0.02 | 0.02 |
| gene4641 | https://www.ncbi.nlm.nih.gov/gene/?term=LOC105277398 | 1937 | 74 | 382.03 | 1 | 5.16 | 1.35 | 35.22 | 34.78 | 0.44 | 0.44 | 0.02 | 0.02 |
| gene7249 | https://www.ncbi.nlm.nih.gov/gene/?term=LOC105280422 | 1775 | 66 | 371.83 | 1 | 5.63 | 1.52 | 16.48 | 16.04 | 0.44 | 0.44 | 0.02 | 0.02 |
| gene9648 | https://www.ncbi.nlm.nih.gov/gene/?term=LOC105283183 | 2602 | 33 | 126.83 | 1 | 3.84 | 3.03 | 45.18 | 44.75 | 0.44 | 0.44 | 0.01 | 0.01 |
| gene2330 | https://www.ncbi.nlm.nih.gov/gene/?term=LOC105274773 | 13104 | 208 | 158.73 | 1 | 0.76 | 0.48 | 15.28 | 14.85 | 0.43 | 0.43 | 0.01 | 0.01 |
| gene9647 | https://www.ncbi.nlm.nih.gov/gene/?term=LOC109611233 | 3094 | 35 | 113.12 | 1 | 3.23 | 2.86 | 42.6 | 42.19 | 0.41 | 0.41 | 0 | 0 |
| gene10520 | https://www.ncbi.nlm.nih.gov/gene/?term=LOC105284184 | 2451 | 105 | 428.4 | 1 | 4.08 | 0.95 | 28.89 | 28.49 | 0.4 | 0.4 | 0.02 | 0.02 |
| gene2846 | https://www.ncbi.nlm.nih.gov/gene/?term=LOC105275374 | 11246 | 566 | 503.29 | 1 | 0.89 | 0.18 | 9.56 | 9.16 | 0.4 | 0.4 | 0.02 | 0.02 |
| gene8814 | https://www.ncbi.nlm.nih.gov/gene/?term=LOC105282240 | 6910 | 209 | 302.46 | 1 | 1.45 | 0.48 | 8.7 | 8.29 | 0.4 | 0.4 | 0.01 | 0.01 |
| gene570 | https://www.ncbi.nlm.nih.gov/gene/?term=LOC105278549 | 1673 | 74 | 442.32 | 1 | 5.98 | 1.35 | 44.93 | 44.55 | 0.38 | 0.38 | 0.02 | 0.02 |
| gene3885 | https://www.ncbi.nlm.nih.gov/gene/?term=LOC105276527 | 7928 | 380 | 479.31 | 1 | 1.26 | 0.26 | 8.1 | 7.75 | 0.34 | 0.34 | 0.02 | 0.02 |
| gene9029 | https://www.ncbi.nlm.nih.gov/gene/?term=LOC105282491 | 4762 | 190 | 398.99 | 1 | 2.1 | 0.53 | 8.59 | 8.25 | 0.34 | 0.34 | 0.01 | 0.01 |
| gene2056 | https://www.ncbi.nlm.nih.gov/gene/?term=LOC105274452 | 6584 | 268 | 407.05 | 1 | 1.52 | 0.37 | 13.37 | 13.03 | 0.33 | 0.33 | 0.01 | 0.01 |
| gene2844 | https://www.ncbi.nlm.nih.gov/gene/?term=LOC105275363 | 4330 | 258 | 595.84 | 1 | 2.31 | 0.39 | 11.65 | 11.32 | 0.33 | 0.33 | 0.02 | 0.02 |
| gene11568 | https://www.ncbi.nlm.nih.gov/gene/?term=LOC105285395 | 8086 | 257 | 317.83 | 1 | 1.24 | 0.39 | 13.19 | 12.93 | 0.27 | 0.27 | 0.01 | 0.01 |
| gene10018 | https://www.ncbi.nlm.nih.gov/gene/?term=LOC105283607 | 4414 | 222 | 502.95 | 1 | 2.27 | 0.45 | 14.44 | 14.18 | 0.26 | 0.26 | 0.01 | 0.01 |
| gene6620 | https://www.ncbi.nlm.nih.gov/gene/?term=LOC105279687 | 17998 | 456 | 253.36 | 1 | 0.56 | 0.22 | 4.79 | 4.53 | 0.26 | 0.26 | 0.01 | 0.01 |
| gene2683 | https://www.ncbi.nlm.nih.gov/gene/?term=LOC105275187 | 5492 | 87 | 158.41 | 1 | 1.82 | 1.15 | 15.13 | 14.88 | 0.25 | 0.25 | 0 | 0 |
| gene8600 | https://www.ncbi.nlm.nih.gov/gene/?term=LOC105282001 | 8513 | 211 | 247.86 | 1 | 1.17 | 0.47 | 9.85 | 9.6 | 0.25 | 0.25 | 0.01 | 0.01 |
| gene1310 | https://www.ncbi.nlm.nih.gov/gene/?term=LOC105286661 | 2582 | 72 | 278.85 | 1 | 3.87 | 1.39 | 20.69 | 20.45 | 0.24 | 0.24 | 0.01 | 0.01 |
| gene292 | https://www.ncbi.nlm.nih.gov/gene/?term=LOC105275398 | 2825 | 92 | 325.66 | 1 | 3.54 | 1.09 | 21.85 | 21.62 | 0.23 | 0.23 | 0.01 | 0.01 |
| gene11275 | https://www.ncbi.nlm.nih.gov/gene/?term=LOC105285055 | 4119 | 262 | 636.08 | 1 | 2.43 | 0.38 | 11.65 | 11.43 | 0.22 | 0.22 | 0.01 | 0.01 |
| gene8729 | https://www.ncbi.nlm.nih.gov/gene/?term=LOC105282137 | 5005 | 236 | 471.53 | 1 | 2 | 0.42 | 14.47 | 14.26 | 0.21 | 0.21 | 0.01 | 0.01 |
| gene2806 | https://www.ncbi.nlm.nih.gov/gene/?term=LOC105275327 | 14257 | 550 | 385.78 | 1 | 0.7 | 0.18 | 6.58 | 6.38 | 0.2 | 0.2 | 0.01 | 0.01 |
| gene3747 | https://www.ncbi.nlm.nih.gov/gene/?term=LOC105276387 | 3736 | 198 | 529.98 | 1 | 2.68 | 0.51 | 14.52 | 14.33 | 0.19 | 0.19 | 0.01 | 0.01 |
| gene5746 | https://www.ncbi.nlm.nih.gov/gene/?term=LOC105278666 | 10343 | 491 | 474.72 | 1 | 0.97 | 0.2 | 6.89 | 6.71 | 0.19 | 0.19 | 0.01 | 0.01 |
| gene10596 | https://www.ncbi.nlm.nih.gov/gene/?term=LOC105284262 | 6520 | 247 | 378.83 | 1 | 1.53 | 0.4 | 4.51 | 4.34 | 0.18 | 0.18 | 0.01 | 0.01 |
| gene7245 | https://www.ncbi.nlm.nih.gov/gene/?term=LOC105280410 | 12478 | 319 | 255.65 | 1 | 0.8 | 0.31 | 17.86 | 17.71 | 0.15 | 0.15 | 0 | 0 |
| gene2173 | https://www.ncbi.nlm.nih.gov/gene/?term=LOC105274590 | 4387 | 91 | 207.43 | 2 | 4.56 | 2.2 | 24.1 | 23.96 | 0.14 | 0.14 | 0 | 0 |
| gene4225 | https://www.ncbi.nlm.nih.gov/gene/?term=LOC105276917 | 2443 | 40 | 163.73 | 1 | 4.09 | 2.5 | 29.99 | 29.86 | 0.14 | 0.14 | 0 | 0 |
| gene6966 | https://www.ncbi.nlm.nih.gov/gene/?term=LOC105280104 | 3747 | 133 | 354.95 | 1 | 2.67 | 0.75 | 17.09 | 16.96 | 0.13 | 0.13 | 0 | 0 |
| gene6652 | https://www.ncbi.nlm.nih.gov/gene/?term=LOC105279719 | 2831 | 83 | 293.18 | 1 | 3.53 | 1.2 | 23.57 | 23.44 | 0.12 | 0.12 | 0 | 0 |
| gene9722 | https://www.ncbi.nlm.nih.gov/gene/?term=LOC105283264 | 3951 | 94 | 237.91 | 1 | 2.53 | 1.06 | 22.96 | 22.87 | 0.1 | 0.1 | 0 | 0 |
| gene4033 | https://www.ncbi.nlm.nih.gov/gene/?term=LOC105276705 | 10546 | 157 | 148.87 | 1 | 0.95 | 0.64 | 6.45 | 6.38 | 0.06 | 0.06 | 0 | 0 |
| gene3727 | https://www.ncbi.nlm.nih.gov/gene/?term=LOC105276367 | 2484 | 76 | 305.96 | 1 | 4.03 | 1.32 | 41.02 | 40.98 | 0.04 | 0.04 | 0 | 0 |
| gene7319 | https://www.ncbi.nlm.nih.gov/gene/?term=LOC105280513 | 68997 | 1592 | 230.73 | 1 | 0.14 | 0.06 | 0.75 | 0.72 | 0.03 | 0.03 | 0 | 0 |
| gene12307 | https://www.ncbi.nlm.nih.gov/gene/?term=LOC105286216 | 10313 | 209 | 202.66 | 1 | 0.97 | 0.48 | 14.07 | 14.04 | 0.02 | 0.02 | 0 | 0 |
| gene3470 | https://www.ncbi.nlm.nih.gov/gene/?term=LOC105276059 | 8716 | 186 | 213.4 | 1 | 1.15 | 0.54 | 9.64 | 9.62 | 0.02 | 0.02 | 0 | 0 |
| gene3107 | https://www.ncbi.nlm.nih.gov/gene/?term=LOC105275650 | 5103 | 187 | 366.45 | 1 | 1.96 | 0.53 | 16.49 | 16.48 | 0.01 | 0.01 | 0 | 0 |
| gene3335 | https://www.ncbi.nlm.nih.gov/gene/?term=LOC105275898 | 4665 | 172 | 368.7 | 1 | 2.14 | 0.58 | 12.53 | 12.51 | 0.01 | 0.01 | 0 | 0 |
| gene9326 | https://www.ncbi.nlm.nih.gov/gene/?term=LOC105282818 | 16612 | 251 | 151.1 | 1 | 0.6 | 0.4 | 12.99 | 12.99 | 0 | 0 | 0 | 0 |
| gene11561 | https://www.ncbi.nlm.nih.gov/gene/?term=LOC105285387 | 11528 | 226 | 196.04 | 1 | 0.87 | 0.44 | 9.46 | 9.47 | -0.01 | 0.01 | 0 | 0 |
| gene13039 | https://www.ncbi.nlm.nih.gov/gene/?term=LOC105287065 | 4902 | 263 | 536.52 | 1 | 2.04 | 0.38 | 11.98 | 12.01 | -0.02 | 0.02 | 0 | 0 |
| gene4209 | https://www.ncbi.nlm.nih.gov/gene/?term=LOC105277125 | 50550 | 2067 | 408.9 | 1 | 0.2 | 0.05 | 1.64 | 1.66 | -0.02 | 0.02 | 0 | 0 |
| gene7166 | https://www.ncbi.nlm.nih.gov/gene/?term=LOC105280326 | 6117 | 258 | 421.78 | 1 | 1.63 | 0.39 | 14.79 | 14.83 | -0.04 | 0.04 | 0 | 0 |
| gene6649 | https://www.ncbi.nlm.nih.gov/gene/?term=LOC105279717 | 6017 | 181 | 300.81 | 1 | 1.66 | 0.55 | 11.29 | 11.35 | -0.06 | 0.06 | 0 | 0 |
| gene3856 | https://www.ncbi.nlm.nih.gov/gene/?term=LOC105276499 | 5236 | 206 | 393.43 | 1 | 1.91 | 0.49 | 10.22 | 10.29 | -0.07 | 0.07 | 0 | 0 |
| gene10326 | https://www.ncbi.nlm.nih.gov/gene/?term=LOC105283962 | 3042 | 145 | 476.66 | 1 | 3.29 | 0.69 | 9.18 | 9.26 | -0.08 | 0.08 | 0 | 0 |
| gene10549 | https://www.ncbi.nlm.nih.gov/gene/?term=LOC105284210 | 7145 | 374 | 523.44 | 2 | 2.8 | 0.53 | 9.66 | 9.75 | -0.09 | 0.09 | 0 | 0 |
| gene2835 | https://www.ncbi.nlm.nih.gov/gene/?term=LOC105275357 | 2492 | 122 | 489.57 | 1 | 4.01 | 0.82 | 14.14 | 14.23 | -0.09 | 0.09 | 0 | 0 |

| | | | | | | | | | | | | | |
|---|---|---|---|---|---|---|---|---|---|---|---|---|---|
| gene4190 | https://www.ncbi.nlm.nih.gov/gene/?term=LOC105276875 | 3150 | 135 | 428.57 | 1 | 3.17 | 0.74 | 20.47 | 20.57 | -0.09 | 0.09 | 0 | 0 |
| gene5613 | https://www.ncbi.nlm.nih.gov/gene/?term=LOC105278504 | 8524 | 274 | 321.45 | 1 | 1.17 | 0.36 | 5.89 | 5.98 | -0.09 | 0.09 | 0 | 0 |
| gene9755 | https://www.ncbi.nlm.nih.gov/gene/?term=LOC105283296 | 2177 | 97 | 445.57 | 1 | 4.59 | 1.03 | 14.1 | 14.18 | -0.09 | 0.09 | 0 | 0 |
| gene13114 | https://www.ncbi.nlm.nih.gov/gene/?term=LOC105287147 | 7609 | 447 | 587.46 | 1 | 1.31 | 0.22 | 9.07 | 9.17 | -0.1 | 0.1 | -0.01 | 0.01 |
| gene4376 | https://www.ncbi.nlm.nih.gov/gene/?term=LOC105277083 | 4307 | 140 | 325.05 | 1 | 2.32 | 0.71 | 17.67 | 17.76 | -0.1 | 0.1 | 0 | 0 |
| gene53 | https://www.ncbi.nlm.nih.gov/gene/?term=LOC105277148 | 19283 | 339 | 175.8 1 | 0.52 | 0.29 | 4.53 | 4.65 | -0.11 | 0.11 | 0 | 0 | |
| gene1632 | https://www.ncbi.nlm.nih.gov/gene/?term=LOC105287764 | 18061 | 462 | 255.8 1 | 0.55 | 0.22 | 3.51 | 3.63 | -0.12 | 0.12 | 0 | 0 | |
| gene7547 | https://www.ncbi.nlm.nih.gov/gene/?term=LOC105280767 | 24576 | 950 | 386.56 | 1 | 0.41 | 0.11 | 4.24 | 4.36 | -0.12 | 0.12 | 0 | 0 |
| gene8274 | https://www.ncbi.nlm.nih.gov/gene/?term=LOC105281624 | 5425 | 244 | 449.77 | 1 | 1.84 | 0.41 | 9.34 | 9.46 | -0.12 | 0.12 | -0.01 | 0.01 |
| gene3599 | https://www.ncbi.nlm.nih.gov/gene/?term=LOC105276196 | 2955 | 61 | 206.43 | 1 | 3.38 | 1.64 | 38.99 | 39.11 | -0.13 | 0.13 | 0 | 0 |
| gene9327 | https://www.ncbi.nlm.nih.gov/gene/?term=LOC105282821 | 7793 | 184 | 236.11 | 1 | 1.28 | 0.54 | 12.73 | 12.87 | -0.14 | 0.14 | 0 | 0 |
| gene11579 | https://www.ncbi.nlm.nih.gov/gene/?term=LOC105285406 | 13571 | 341 | 251.27 | 1 | 0.74 | 0.29 | 6.04 | 6.2 | -0.15 | 0.15 | 0 | 0 |
| gene12674 | https://www.ncbi.nlm.nih.gov/gene/?term=LOC105286649 | 18283 | 836 | 457.26 | 1 | 0.55 | 0.12 | 1.34 | 1.5 | -0.16 | 0.16 | -0.01 | 0.01 |
| gene5539 | https://www.ncbi.nlm.nih.gov/gene/?term=LOC105278415 | 25024 | 835 | 333.68 | 1 | 0.4 | 0.12 | 3.01 | 3.18 | -0.17 | 0.17 | -0.01 | 0.01 |
| gene11082 | https://www.ncbi.nlm.nih.gov/gene/?term=LOC105284824 | 7436 | 245 | 329.48 | 1 | 1.34 | 0.41 | 7.32 | 7.57 | -0.25 | 0.25 | -0.01 | 0.01 |
| gene1198 | https://www.ncbi.nlm.nih.gov/gene/?term=LOC105285507 | 2738 | 98 | 357.93 | 1 | 3.65 | 1.02 | 21.45 | 21.7 | -0.25 | 0.25 | -0.01 | 0.01 |
| gene7368 | https://www.ncbi.nlm.nih.gov/gene/?term=LOC105280556 | 4105 | 76 | 185.14 | 1 | 2.44 | 1.32 | 15.19 | 15.44 | -0.25 | 0.25 | 0 | 0 |
| gene9464 | https://www.ncbi.nlm.nih.gov/gene/?term=LOC105282975 | 1450 | 25 | 172.41 | 1 | 6.9 | 4 | 40.46 | 40.7 | -0.25 | 0.25 | 0 | 0 |
| gene12407 | https://www.ncbi.nlm.nih.gov/gene/?term=LOC105286315 | 8593 | 227 | 264.17 | 1 | 1.16 | 0.44 | 13.44 | 13.72 | -0.28 | 0.28 | -0.01 | 0.01 |
| gene11583 | https://www.ncbi.nlm.nih.gov/gene/?term=LOC105285414 | 8192 | 204 | 249.02 | 1 | 1.22 | 0.49 | 7.78 | 8.09 | -0.31 | 0.31 | -0.01 | 0.01 |
| gene12573 | https://www.ncbi.nlm.nih.gov/gene/?term=LOC105286515 | 3647 | 128 | 350.97 | 1 | 2.74 | 0.78 | 10.64 | 10.97 | -0.33 | 0.33 | -0.01 | 0.01 |
| gene9598 | https://www.ncbi.nlm.nih.gov/gene/?term=LOC105283132 | 1769 | 56 | 316.56 | 1 | 5.65 | 1.79 | 39.81 | 40.18 | -0.38 | 0.38 | -0.01 | 0.01 |
| gene1220 | https://www.ncbi.nlm.nih.gov/gene/?term=LOC105285742 | 3453 | 123 | 356.21 | 1 | 2.9 | 0.81 | 16.77 | 17.17 | -0.4 | 0.4 | -0.01 | 0.01 |
| gene657 | https://www.ncbi.nlm.nih.gov/gene/?term=LOC105279667 | 6421 | 106 | 165.08 | 1 | 1.56 | 0.94 | 2.17 | 2.57 | -0.4 | 0.4 | -0.01 | 0.01 |
| gene5646 | https://www.ncbi.nlm.nih.gov/gene/?term=LOC105278534 | 7475 | 214 | 286.29 | 1 | 1.34 | 0.47 | 16.15 | 16.57 | -0.41 | 0.41 | -0.01 | 0.01 |
| gene7148 | https://www.ncbi.nlm.nih.gov/gene/?term=LOC105280304 | 8920 | 143 | 160.31 | 1 | 1.12 | 0.7 | 6.95 | 7.37 | -0.42 | 0.42 | -0.01 | 0.01 |
| gene7600 | https://www.ncbi.nlm.nih.gov/gene/?term=LOC105280819 | 1615 | 43 | 266.25 | 1 | 6.19 | 2.33 | 37.46 | 37.88 | -0.42 | 0.42 | -0.01 | 0.01 |
| gene1470 | https://www.ncbi.nlm.nih.gov/gene/?term=LOC105287597 | 2974 | 200 | 672.49 | 1 | 3.36 | 0.5 | 18.11 | 18.56 | -0.46 | 0.46 | -0.03 | 0.03 |
| gene6583 | https://www.ncbi.nlm.nih.gov/gene/?term=LOC105279649 | 7444 | 393 | 527.94 | 1 | 1.34 | 0.25 | 10.67 | 11.14 | -0.47 | 0.47 | -0.02 | 0.02 |
| gene10345 | https://www.ncbi.nlm.nih.gov/gene/?term=LOC105283989 | 4444 | 40 | 90.01 1 | 2.25 | 2.5 | 37.64 | 38.15 | -0.5 | 0.5 | 0 | 0 | |
| gene5298 | https://www.ncbi.nlm.nih.gov/gene/?term=LOC105278145 | 4060 | 275 | 677.34 | 2 | 4.93 | 0.73 | 9.89 | 10.42 | -0.53 | 0.53 | -0.04 | 0.04 |
| gene3848 | https://www.ncbi.nlm.nih.gov/gene/?term=LOC105276486 | 5899 | 176 | 298.36 | 1 | 1.7 | 0.57 | 23.6 | 24.17 | -0.57 | 0.57 | -0.02 | 0.02 |
| gene4962 | https://www.ncbi.nlm.nih.gov/gene/?term=LOC105277761 | 4645 | 175 | 376.75 | 1 | 2.15 | 0.57 | 13.58 | 14.2 | -0.62 | 0.62 | -0.02 | 0.02 |
| gene7207 | https://www.ncbi.nlm.nih.gov/gene/?term=LOC105280376 | 4659 | 124 | 266.15 | 1 | 2.15 | 0.81 | 26.54 | 27.17 | -0.63 | 0.63 | -0.02 | 0.02 |
| gene12377 | https://www.ncbi.nlm.nih.gov/gene/?term=LOC105286279 | 5846 | 278 | 475.54 | 1 | 1.71 | 0.36 | 2.15 | 2.82 | -0.67 | 0.67 | -0.03 | 0.03 |
| gene7550 | https://www.ncbi.nlm.nih.gov/gene/?term=LOC105280763 | 2173 | 109 | 501.61 | 1 | 4.6 | 0.92 | 27.44 | 28.14 | -0.7 | 0.7 | -0.04 | 0.04 |
| gene10819 | https://www.ncbi.nlm.nih.gov/gene/?term=LOC105284507 | 5767 | 122 | 211.55 | 1 | 1.73 | 0.82 | 18.54 | 19.25 | -0.71 | 0.71 | -0.01 | 0.01 |
| gene3385 | https://www.ncbi.nlm.nih.gov/gene/?term=LOC105275966 | 3305 | 75 | 226.93 | 1 | 3.03 | 1.33 | 43.86 | 44.6 | -0.75 | 0.75 | -0.02 | 0.02 |
| gene10213 | https://www.ncbi.nlm.nih.gov/gene/?term=LOC105283826 | 1503 | 62 | 412.51 | 1 | 6.65 | 1.61 | 14.74 | 15.54 | -0.8 | 0.8 | -0.03 | 0.03 |
| gene11454 | https://www.ncbi.nlm.nih.gov/gene/?term=LOC105285256 | 1906 | 50 | 262.33 | 1 | 5.25 | 2 | 43.28 | 44.08 | -0.8 | 0.8 | -0.02 | 0.02 |
| gene10392 | https://www.ncbi.nlm.nih.gov/gene/?term=LOC105284036 | 3216 | 103 | 320.27 | 2 | 6.22 | 1.94 | 24.04 | 24.86 | -0.81 | 0.81 | -0.03 | 0.03 |
| gene7915 | https://www.ncbi.nlm.nih.gov/gene/?term=LOC105281173 | 1597 | 39 | 244.21 | 1 | 6.26 | 2.56 | 61.13 | 62.01 | -0.88 | 0.88 | -0.02 | 0.02 |
| gene11401 | https://www.ncbi.nlm.nih.gov/gene/?term=LOC105285202 | 1841 | 89 | 483.43 | 1 | 5.43 | 1.12 | 23.71 | 24.59 | -0.89 | 0.89 | -0.04 | 0.04 |
| gene11447 | https://www.ncbi.nlm.nih.gov/gene/?term=LOC105285252 | 7437 | 150 | 201.69 | 1 | 1.34 | 0.67 | 11.78 | 12.78 | -0.99 | 0.99 | -0.02 | 0.02 |
| gene3168 | https://www.ncbi.nlm.nih.gov/gene/?term=LOC105275711 | 6150 | 95 | 154.47 | 1 | 1.63 | 1.05 | 10.97 | 11.98 | -1 | 1 | -0.02 | 0.02 |
| gene12203 | https://www.ncbi.nlm.nih.gov/gene/?term=LOC105286099 | 5967 | 87 | 145.8 1 | 1.68 | 1.15 | 11.05 | 12.06 | -1.01 | 1.01 | -0.01 | 0.01 | |
| gene7657 | https://www.ncbi.nlm.nih.gov/gene/?term=LOC105280890 | 1109 | 23 | 207.39 | 1 | 9.02 | 4.35 | 23.17 | 24.26 | -1.09 | 1.09 | -0.02 | 0.02 |
| gene652 | https://www.ncbi.nlm.nih.gov/gene/?term=LOC105279528 | 5806 | 39 | 67.17 1 | 1.72 | 2.56 | 19.31 | 20.42 | -1.11 | 1.11 | -0.01 | 0.01 | |
| gene6639 | https://www.ncbi.nlm.nih.gov/gene/?term=LOC105279705 | 7656 | 88 | 114.94 | 1 | 1.31 | 1.14 | 19.96 | 21.09 | -1.13 | 1.13 | -0.01 | 0.01 |
| gene9101 | https://www.ncbi.nlm.nih.gov/gene/?term=LOC105282553 | 3068 | 110 | 358.54 | 1 | 3.26 | 0.91 | 21.48 | 22.61 | -1.13 | 1.13 | -0.04 | 0.04 |
| gene10155 | https://www.ncbi.nlm.nih.gov/gene/?term=LOC105283756 | 7030 | 119 | 169.27 | 1 | 1.42 | 0.84 | 6.89 | 8.19 | -1.3 | 1.3 | -0.02 | 0.02 |
| gene2404 | https://www.ncbi.nlm.nih.gov/gene/?term=LOC105274853 | 1767 | 45 | 254.67 | 1 | 5.66 | 2.22 | 30.61 | 31.94 | -1.33 | 1.33 | -0.03 | 0.03 |
| gene10654 | https://www.ncbi.nlm.nih.gov/gene/?term=LOC105284329 | 4027 | 72 | 178.79 | 1 | 2.48 | 1.39 | 18.86 | 20.42 | -1.56 | 1.56 | -0.03 | 0.03 |
| gene564 | https://www.ncbi.nlm.nih.gov/gene/?term=LOC105278478 | 2802 | 22 | 78.52 1 | 3.57 | 4.55 | 23.15 | 24.78 | -1.63 | 1.63 | -0.01 | 0.01 | |
| gene11340 | https://www.ncbi.nlm.nih.gov/gene/?term=LOC105285131 | 2918 | 100 | 342.7 1 | 3.43 | 1 | 13.44 | 15.08 | -1.64 | 1.64 | -0.06 | 0.06 | |
| gene6237 | https://www.ncbi.nlm.nih.gov/gene/?term=LOC105279244 | 1484 | 30 | 202.16 | 1 | 6.74 | 3.33 | 49.36 | 51.05 | -1.69 | 1.69 | -0.03 | 0.03 |
| gene920 | https://www.ncbi.nlm.nih.gov/gene/?term=LOC105282518 | 1197 | 46 | 384.29 | 1 | 8.35 | 2.17 | 30.45 | 32.64 | -2.19 | 2.19 | -0.08 | 0.08 |
| gene9411 | https://www.ncbi.nlm.nih.gov/gene/?term=LOC105282914 | 5534 | 42 | 75.89 1 | 1.81 | 2.38 | 48.54 | 50.92 | -2.38 | 2.38 | -0.02 | 0.02 | |
| gene1768 | https://www.ncbi.nlm.nih.gov/gene/?term=LOC105287938 | 2792 | 89 | 318.77 | 1 | 3.58 | 1.12 | 25.55 | 28.06 | -2.51 | 2.51 | -0.08 | 0.08 |
| gene10552 | https://www.ncbi.nlm.nih.gov/gene/?term=LOC105284213 | 878 | 23 | 261.96 | 1 | 11.39 | 4.35 | 9.72 | 12.3 | -2.58 | 2.58 | -0.07 | 0.07 |
| gene7398 | https://www.ncbi.nlm.nih.gov/gene/?term=LOC105280589 | 3452 | 42 | 121.67 | 1 | 2.9 | 2.38 | 23.2 | 25.91 | -2.71 | 2.71 | -0.03 | 0.03 |
| gene2413 | https://www.ncbi.nlm.nih.gov/gene/?term=LOC105274863 | 2006 | 65 | 324.03 | 1 | 4.99 | 1.54 | 49.74 | 52.89 | -3.14 | 3.14 | -0.1 | 0.1 |
| gene1199 | https://www.ncbi.nlm.nih.gov/gene/?term=LOC105285498 | 2220 | 29 | 130.63 | 1 | 4.5 | 3.45 | 32.78 | 35.99 | -3.21 | 3.21 | -0.04 | 0.04 |
| gene1505 | https://www.ncbi.nlm.nih.gov/gene/?term=LOC105287633 | 1693 | 24 | 141.76 | 1 | 5.91 | 4.17 | 48.26 | 51.48 | -3.22 | 3.22 | -0.05 | 0.05 |

| | | | | | | | | | | | | | | |
|---|---|---|---|---|---|---|---|---|---|---|---|---|---|---|
| gene9109 | https://www.ncbi.nlm.nih.gov/gene/?term=LOC105282561 | 2400 | 17 | 70.83 | 1 | 4.17 | 5.88 | 28.52 | 31.86 | -3.34 | 3.34 | -0.02 | 0.02 | |
| gene11584 | https://www.ncbi.nlm.nih.gov/gene/?term=LOC105285416 | 3709 | 15 | 40.44 | 2 | 5.39 | 13.33 | 24.1 | 29.33 | -5.23 | 5.23 | -0.02 | 0.02 | |
| gene3166 | https://www.ncbi.nlm.nih.gov/gene/?term=LOC105275707 | 2393 | 12 | 50.15 | 1 | 4.18 | 8.33 | 62.45 | 73.29 | -10.83 | | 10.83 | -0.05 | 0.05 |
| gene8691 | https://www.ncbi.nlm.nih.gov/gene/?term=LOC105282094 | 2028 | 6 | 29.59 | 1 | 4.93 | 16.67 | 86.17 | 97.58 | -11.41 | | 11.41 | -0.03 | 0.03 |
| gene393 | https://www.ncbi.nlm.nih.gov/gene/?term=LOC105276566 | 1123 | 2 | 17.81 | 1 | 8.9 | 50 | 47.62 | 74.3 | -26.68 | | 26.68 | -0.05 | 0.05 |

Directory: OGLall File: ogl-Cb_LI_RphaseA2.vs.RphaseB2.txt

| gene_ID | gene_link | gwidth | #Sites | #per10Kb | #dmSites | #dmsp10kb | %dmSites | %pSite1 | %pSite2 | DMpSite | ADMpSite | DMpNucl | ADMpNucl |
|---|---|---|---|---|---|---|---|---|---|---|---|---|---|
| gene2593 | https://www.ncbi.nlm.nih.gov/gene/?term=LOC105275070 | 1774 | 7 | 39.46 | 5 | 28.18 | 71.43 | 65.68 | 4.09 | 61.59 | 61.59 | 0.24 | 0.24 |
| gene3165 | https://www.ncbi.nlm.nih.gov/gene/?term=LOC105275706 | 2374 | 74 | 311.71 | 6 | 25.27 | 8.11 | 32.18 | 22.24 | 9.94 | 9.94 | 0.31 | 0.31 |
| gene2824 | https://www.ncbi.nlm.nih.gov/gene/?term=LOC105275337 | 1772 | 60 | 338.6 | 2 | 11.29 | 3.33 | 38.84 | 29.82 | 9.02 | 9.02 | 0.31 | 0.31 |
| gene2907 | https://www.ncbi.nlm.nih.gov/gene/?term=LOC105275447 | 2539 | 37 | 145.73 | 1 | 3.94 | 2.7 | 8.22 | 2.41 | 5.8 | 5.8 | 0.08 | 0.08 |
| gene5714 | https://www.ncbi.nlm.nih.gov/gene/?term=LOC105278729 | 710 | 17 | 239.44 | 1 | 14.08 | 5.88 | 59.17 | 53.54 | 5.62 | 5.62 | 0.13 | 0.13 |
| gene8943 | https://www.ncbi.nlm.nih.gov/gene/?term=LOC105282388 | 1220 | 33 | 270.49 | 1 | 8.2 | 3.03 | 49.82 | 45.11 | 4.71 | 4.71 | 0.13 | 0.13 |
| gene2335 | https://www.ncbi.nlm.nih.gov/gene/?term=LOC105274778 | 3945 | 38 | 96.32 | 1 | 2.53 | 2.63 | 48.07 | 44.68 | 3.39 | 3.39 | 0.03 | 0.03 |
| gene12021 | https://www.ncbi.nlm.nih.gov/gene/?term=LOC105285905 | 1399 | 16 | 114.37 | 1 | 7.15 | 6.25 | 51.11 | 47.75 | 3.36 | 3.36 | 0.04 | 0.04 |
| gene2970 | https://www.ncbi.nlm.nih.gov/gene/?term=LOC105275518 | 2340 | 24 | 102.56 | 1 | 4.27 | 4.17 | 49.7 | 46.61 | 3.09 | 3.09 | 0.03 | 0.03 |
| gene7568 | https://www.ncbi.nlm.nih.gov/gene/?term=LOC105280786 | 2525 | 126 | 499.01 | 1 | 3.96 | 0.79 | 30.35 | 27.57 | 2.78 | 2.78 | 0.14 | 0.14 |
| gene2271 | https://www.ncbi.nlm.nih.gov/gene/?term=LOC105274706 | 1448 | 56 | 386.74 | 1 | 6.91 | 1.79 | 38.93 | 36.4 | 2.53 | 2.53 | 0.1 | 0.1 |
| gene4225 | https://www.ncbi.nlm.nih.gov/gene/?term=LOC105276917 | 2443 | 36 | 147.36 | 1 | 4.09 | 2.78 | 30.16 | 27.69 | 2.47 | 2.47 | 0.04 | 0.04 |
| gene940 | https://www.ncbi.nlm.nih.gov/gene/?term=LOC105282794 | 3578 | 88 | 245.95 | 1 | 2.79 | 1.14 | 16.61 | 14.17 | 2.44 | 2.44 | 0.06 | 0.06 |
| gene13033 | https://www.ncbi.nlm.nih.gov/gene/?term=LOC105287063 | 1877 | 48 | 255.73 | 1 | 5.33 | 2.08 | 32.8 | 30.37 | 2.43 | 2.43 | 0.06 | 0.06 |
| gene4034 | https://www.ncbi.nlm.nih.gov/gene/?term=LOC105276704 | 2065 | 88 | 426.15 | 1 | 4.84 | 1.14 | 16.14 | 13.92 | 2.22 | 2.22 | 0.09 | 0.09 |
| gene5052 | https://www.ncbi.nlm.nih.gov/gene/?term=LOC105277856 | 3167 | 34 | 107.36 | 1 | 3.16 | 2.94 | 35.71 | 33.56 | 2.15 | 2.15 | 0.02 | 0.02 |
| gene5273 | https://www.ncbi.nlm.nih.gov/gene/?term=LOC105278108 | 4067 | 51 | 125.4 | 1 | 2.46 | 1.96 | 25.16 | 23.04 | 2.12 | 2.12 | 0.03 | 0.03 |
| gene9598 | https://www.ncbi.nlm.nih.gov/gene/?term=LOC105283132 | 1769 | 57 | 322.22 | 1 | 5.65 | 1.75 | 39.92 | 37.87 | 2.05 | 2.05 | 0.07 | 0.07 |
| gene12106 | https://www.ncbi.nlm.nih.gov/gene/?term=LOC105286002 | 8556 | 68 | 79.48 | 1 | 1.17 | 1.47 | 4.38 | 2.38 | 2 | 2 | 0.02 | 0.02 |
| gene11819 | https://www.ncbi.nlm.nih.gov/gene/?term=LOC105285671 | 2366 | 103 | 435.33 | 1 | 4.23 | 0.97 | 27.63 | 25.66 | 1.97 | 1.97 | 0.09 | 0.09 |
| gene7211 | https://www.ncbi.nlm.nih.gov/gene/?term=LOC105280377 | 2024 | 69 | 340.91 | 1 | 4.94 | 1.45 | 48.01 | 46.12 | 1.89 | 1.89 | 0.06 | 0.06 |
| gene927 | https://www.ncbi.nlm.nih.gov/gene/?term=LOC105282600 | 3727 | 54 | 144.89 | 1 | 2.68 | 1.85 | 28.21 | 26.36 | 1.85 | 1.85 | 0.03 | 0.03 |
| gene388 | https://www.ncbi.nlm.nih.gov/gene/?term=LOC105276538 | 1558 | 21 | 134.79 | 2 | 12.84 | 9.52 | 69.8 | 67.98 | 1.82 | 1.82 | 0.02 | 0.02 |
| gene5525 | https://www.ncbi.nlm.nih.gov/gene/?term=LOC105278396 | 3090 | 184 | 595.47 | 1 | 3.24 | 0.54 | 26.64 | 24.83 | 1.81 | 1.81 | 0.11 | 0.11 |

...
... only first 25 lines shown ...

Directory: OGLall File: ogl-Cb_LI_RphaseB1.txt

| gene_ID | gene_link | gwidth | #Sites | #per10Kb | %perSite | %pNucl |
|---|---|---|---|---|---|---|
| gene12538 | https://www.ncbi.nlm.nih.gov/gene/?term=LOC105286531 | 372 | 40 | 1075.27 | 35.13 | 3.78 |
| gene5911 | https://www.ncbi.nlm.nih.gov/gene/?term=LOC105278844 | 943 | 102 | 1081.65 | 34.66 | 3.75 |
| gene13384 | https://www.ncbi.nlm.nih.gov/gene/?term=LOC105287447 | 1877 | 230 | 1225.36 | 29.95 | 3.67 |
| gene211 | https://www.ncbi.nlm.nih.gov/gene/?term=LOC105274534 | 1280 | 99 | 773.44 | 46.14 | 3.57 |
| gene12364 | https://www.ncbi.nlm.nih.gov/gene/?term=LOC105286269 | 1403 | 95 | 677.12 | 50.19 | 3.4 |
| gene5917 | https://www.ncbi.nlm.nih.gov/gene/?term=LOC105278849 | 823 | 125 | 1518.83 | 22.36 | 3.4 |
| gene3719 | https://www.ncbi.nlm.nih.gov/gene/?term=LOC105276360 | 1471 | 169 | 1148.88 | 29.3 | 3.37 |
| gene5309 | https://www.ncbi.nlm.nih.gov/gene/?term=LOC105278160 | 1704 | 167 | 980.05 | 34.27 | 3.36 |
| gene570 | https://www.ncbi.nlm.nih.gov/gene/?term=LOC105278549 | 1673 | 120 | 717.27 | 46.56 | 3.34 |
| gene4878 | https://www.ncbi.nlm.nih.gov/gene/?term=LOC105277648 | 675 | 40 | 592.59 | 54.59 | 3.23 |
| gene1654 | https://www.ncbi.nlm.nih.gov/gene/?term=LOC105287811 | 2285 | 147 | 643.33 | 49.46 | 3.18 |
| gene11566 | https://www.ncbi.nlm.nih.gov/gene/?term=LOC105285392 | 1603 | 134 | 835.93 | 36.82 | 3.08 |
| gene12330 | https://www.ncbi.nlm.nih.gov/gene/?term=LOC105286235 | 1501 | 105 | 699.53 | 44.05 | 3.08 |
| gene12746 | https://www.ncbi.nlm.nih.gov/gene/?term=LOC105286737 | 2688 | 370 | 1376.49 | 22.28 | 3.07 |
| gene6078 | https://www.ncbi.nlm.nih.gov/gene/?term=LOC105279052 | 1130 | 126 | 1115.04 | 27.5 | 3.07 |
| gene12652 | https://www.ncbi.nlm.nih.gov/gene/?term=LOC109610640 | 1122 | 60 | 534.76 | 56.29 | 3.01 |
| gene1562 | https://www.ncbi.nlm.nih.gov/gene/?term=LOC105287692 | 1491 | 134 | 898.73 | 33.27 | 2.99 |
| gene2797 | https://www.ncbi.nlm.nih.gov/gene/?term=LOC105275314 | 1489 | 104 | 698.46 | 41.67 | 2.91 |
| gene190 | https://www.ncbi.nlm.nih.gov/gene/?term=LOC105288088 | 1797 | 123 | 684.47 | 42.11 | 2.88 |
| gene3731 | https://www.ncbi.nlm.nih.gov/gene/?term=LOC105276371 | 865 | 58 | 670.52 | 42.64 | 2.86 |
| gene7527 | https://www.ncbi.nlm.nih.gov/gene/?term=LOC105280739 | 2686 | 148 | 551.01 | 51.09 | 2.81 |
| gene6978 | https://www.ncbi.nlm.nih.gov/gene/?term=LOC105280117 | 1816 | 92 | 506.61 | 55.33 | 2.8 |
| gene12805 | https://www.ncbi.nlm.nih.gov/gene/?term=LOC105286801 | 1379 | 110 | 797.68 | 34.81 | 2.78 |
| gene12403 | https://www.ncbi.nlm.nih.gov/gene/?term=LOC105286311 | 1413 | 65 | 460.01 | 60.28 | 2.77 |

...
... only first 25 lines shown ...

Directory: OGLall File: ogl-Cb_LI_RphaseB1.vs.RphaseB2.txt

| gene_ID | gene_link | gwidth | #Sites | #per10Kb | #dmSites | #dmsp10kb | %dmSites | %pSite1 | %pSite2 | DMpSite | ADMpSite | DMpNucl | ADMpNucl |
|---|---|---|---|---|---|---|---|---|---|---|---|---|---|
| gene2593 | https://www.ncbi.nlm.nih.gov/gene/?term=LOC105275070 | 1774 | 6 | 33.82 | 2 | 11.27 | 33.33 | 23.92 | 4.77 | 19.15 | 19.15 | 0.06 | 0.06 |
| gene11599 | https://www.ncbi.nlm.nih.gov/gene/?term=LOC105285431 | 1550 | 6 | 38.71 | 3 | 19.35 | 50 | 46.72 | 30.29 | 16.43 | 16.43 | 0.06 | 0.06 |
| gene11124 | https://www.ncbi.nlm.nih.gov/gene/?term=LOC105284871 | 1889 | 3 | 15.88 | 1 | 5.29 | 33.33 | 46.42 | 31.48 | 14.94 | 14.94 | 0.02 | 0.02 |
| gene6576 | https://www.ncbi.nlm.nih.gov/gene/?term=LOC105279639 | 2616 | 3 | 11.47 | 1 | 3.82 | 33.33 | 77.78 | 65.15 | 12.62 | 12.62 | 0.01 | 0.01 |
| gene2636 | https://www.ncbi.nlm.nih.gov/gene/?term=LOC105275141 | 947 | 11 | 116.16 | 4 | 42.24 | 36.36 | 32.51 | 19.93 | 12.58 | 12.58 | 0.15 | 0.15 |
| gene10194 | https://www.ncbi.nlm.nih.gov/gene/?term=LOC105283805 | 1309 | 6 | 45.84 | 3 | 22.92 | 50 | 87.61 | 75.59 | 12.01 | 12.01 | 0.06 | 0.06 |
| gene4732 | https://www.ncbi.nlm.nih.gov/gene/?term=LOC105277491 | 1647 | 12 | 72.86 | 2 | 12.14 | 16.67 | 22.08 | 12.22 | 9.85 | 9.85 | 0.07 | 0.07 |
| gene9354 | https://www.ncbi.nlm.nih.gov/gene/?term=LOC105282854 | 1836 | 2 | 10.89 | 1 | 5.45 | 50 | 38.24 | 29.01 | 9.22 | 9.22 | 0.01 | 0.01 |
| gene9109 | https://www.ncbi.nlm.nih.gov/gene/?term=LOC105282561 | 2400 | 5 | 20.83 | 1 | 4.17 | 20 | 36.89 | 27.8 | 9.09 | 9.09 | 0.02 | 0.02 |
| gene7955 | https://www.ncbi.nlm.nih.gov/gene/?term=LOC105281216 | 3744 | 12 | 32.05 | 2 | 5.34 | 16.67 | 39.32 | 30.29 | 9.02 | 9.02 | 0.03 | 0.03 |
| gene9192 | https://www.ncbi.nlm.nih.gov/gene/?term=LOC105282657 | 1404 | 6 | 42.74 | 1 | 7.12 | 16.67 | 58.95 | 50.62 | 8.33 | 8.33 | 0.04 | 0.04 |
| gene9845 | https://www.ncbi.nlm.nih.gov/gene/?term=LOC105283410 | 1401 | 9 | 64.24 | 2 | 14.28 | 22.22 | 50.4 | 42.07 | 8.33 | 8.33 | 0.05 | 0.05 |
| gene4621 | https://www.ncbi.nlm.nih.gov/gene/?term=LOC105277369 | 1157 | 4 | 34.57 | 1 | 8.64 | 25 | 86.56 | 78.3 | 8.25 | 8.25 | 0.03 | 0.03 |
| gene4515 | https://www.ncbi.nlm.nih.gov/gene/?term=LOC109610839 | 335 | 7 | 208.96 | 1 | 29.85 | 14.29 | 66.57 | 58.9 | 7.67 | 7.67 | 0.16 | 0.16 |
| gene5060 | https://www.ncbi.nlm.nih.gov/gene/?term=LOC105277862 | 1458 | 7 | 48.01 | 1 | 6.86 | 14.29 | 63.47 | 55.84 | 7.64 | 7.64 | 0.04 | 0.04 |
| gene240 | https://www.ncbi.nlm.nih.gov/gene/?term=LOC105274857 | 1382 | 4 | 28.94 | 1 | 7.24 | 25 | 52.94 | 45.79 | 7.15 | 7.15 | 0.02 | 0.02 |
| gene10278 | https://www.ncbi.nlm.nih.gov/gene/?term=LOC105283908 | 902 | 12 | 133.04 | 4 | 44.35 | 33.33 | 41.95 | 34.88 | 7.07 | 7.07 | 0.09 | 0.09 |
| gene12853 | https://www.ncbi.nlm.nih.gov/gene/?term=LOC105286857 | 1589 | 4 | 25.17 | 1 | 6.29 | 25 | 74.57 | 67.52 | 7.05 | 7.05 | 0.02 | 0.02 |
| gene12355 | https://www.ncbi.nlm.nih.gov/gene/?term=LOC105286261 | 1301 | 8 | 61.49 | 1 | 7.69 | 12.5 | 74.9 | 68.31 | 6.59 | 6.59 | 0.04 | 0.04 |
| gene4044 | https://www.ncbi.nlm.nih.gov/gene/?term=LOC105276718 | 1729 | 10 | 57.84 | 1 | 5.78 | 10 | 28.19 | 21.81 | 6.38 | 6.38 | 0.04 | 0.04 |
| gene10189 | https://www.ncbi.nlm.nih.gov/gene/?term=LOC105283800 | 871 | 13 | 149.25 | 1 | 11.48 | 7.69 | 18.91 | 13.21 | 5.7 | 5.7 | 0.09 | 0.09 |
| gene3250 | https://www.ncbi.nlm.nih.gov/gene/?term=LOC105275812 | 2029 | 13 | 64.07 | 1 | 4.93 | 7.69 | 22.84 | 17.14 | 5.7 | 5.7 | 0.04 | 0.04 |
| gene7967 | https://www.ncbi.nlm.nih.gov/gene/?term=LOC105281233 | 4322 | 13 | 30.08 | 1 | 2.31 | 7.69 | 21.1 | 15.42 | 5.68 | 5.68 | 0.02 | 0.02 |
| gene10199 | https://www.ncbi.nlm.nih.gov/gene/?term=LOC105283809 | 2716 | 13 | 47.86 | 1 | 3.68 | 7.69 | 69.81 | 64.23 | 5.58 | 5.58 | 0.03 | 0.03 |

...
... only first 25 lines shown ...

Directory: OGLall File: ogl-Cb_LI_RphaseB2.txt

| gene_ID | gene_link | gwidth | #Sites | #per10Kb | %perSite | %pNucl |
|---|---|---|---|---|---|---|
| gene12538 | https://www.ncbi.nlm.nih.gov/gene/?term=LOC105286531 | 372 | 41 | 1102.15 | 36.19 | 3.99 |
| gene13384 | https://www.ncbi.nlm.nih.gov/gene/?term=LOC105287447 | 1877 | 236 | 1257.33 | 28.98 | 3.64 |
| gene5917 | https://www.ncbi.nlm.nih.gov/gene/?term=LOC105278849 | 823 | 124 | 1506.68 | 23.86 | 3.6 |
| gene5911 | https://www.ncbi.nlm.nih.gov/gene/?term=LOC105278844 | 943 | 95 | 1007.42 | 35.49 | 3.58 |
| gene3719 | https://www.ncbi.nlm.nih.gov/gene/?term=LOC105276360 | 1471 | 141 | 958.53 | 35.05 | 3.36 |
| gene5309 | https://www.ncbi.nlm.nih.gov/gene/?term=LOC105278160 | 1704 | 158 | 927.23 | 36.29 | 3.36 |
| gene211 | https://www.ncbi.nlm.nih.gov/gene/?term=LOC105274534 | 1280 | 86 | 671.88 | 49.53 | 3.33 |
| gene3731 | https://www.ncbi.nlm.nih.gov/gene/?term=LOC105276371 | 865 | 57 | 658.96 | 46.13 | 3.04 |
| gene1562 | https://www.ncbi.nlm.nih.gov/gene/?term=LOC105287692 | 1491 | 129 | 865.19 | 34.84 | 3.01 |
| gene1654 | https://www.ncbi.nlm.nih.gov/gene/?term=LOC105287811 | 2285 | 161 | 704.6 | 42.7 | 3.01 |
| gene12805 | https://www.ncbi.nlm.nih.gov/gene/?term=LOC105286801 | 1379 | 108 | 783.18 | 37.84 | 2.96 |
| gene6078 | https://www.ncbi.nlm.nih.gov/gene/?term=LOC105279052 | 1130 | 111 | 982.3 | 29.64 | 2.91 |
| gene12330 | https://www.ncbi.nlm.nih.gov/gene/?term=LOC105286235 | 1501 | 95 | 632.91 | 45.44 | 2.88 |
| gene12746 | https://www.ncbi.nlm.nih.gov/gene/?term=LOC105286737 | 2688 | 337 | 1253.72 | 22.91 | 2.87 |
| gene2797 | https://www.ncbi.nlm.nih.gov/gene/?term=LOC105275314 | 1489 | 89 | 597.72 | 47.65 | 2.85 |
| gene12403 | https://www.ncbi.nlm.nih.gov/gene/?term=LOC105286311 | 1413 | 63 | 445.86 | 62.76 | 2.8 |
| gene5301 | https://www.ncbi.nlm.nih.gov/gene/?term=LOC105278148 | 1881 | 182 | 967.57 | 27.83 | 2.69 |
| gene343 | https://www.ncbi.nlm.nih.gov/gene/?term=LOC105275996 | 1620 | 64 | 395.06 | 67.56 | 2.67 |
| gene12364 | https://www.ncbi.nlm.nih.gov/gene/?term=LOC105286269 | 1403 | 67 | 477.55 | 54.92 | 2.62 |
| gene3334 | https://www.ncbi.nlm.nih.gov/gene/?term=LOC105275897 | 1689 | 116 | 686.8 | 38.03 | 2.61 |
| gene7527 | https://www.ncbi.nlm.nih.gov/gene/?term=LOC105280739 | 2686 | 128 | 476.55 | 54.67 | 2.61 |
| gene6978 | https://www.ncbi.nlm.nih.gov/gene/?term=LOC105280117 | 1816 | 82 | 451.54 | 57.37 | 2.59 |
| gene5497 | https://www.ncbi.nlm.nih.gov/gene/?term=LOC105278368 | 1690 | 64 | 378.7 | 67.97 | 2.57 |
| gene2279 | https://www.ncbi.nlm.nih.gov/gene/?term=LOC105274711 | 2037 | 116 | 569.46 | 44.99 | 2.56 |

...
... only first 25 lines shown ...

Directory: OGLall File: rnk-dmg-Cb_LI_BCphaseA1.vs.BCphaseA2.txt

| gene_ID | gene_link | gwidth | #Sites | #per10Kb | #dmSites | #dmsp10kb | %dmSites | %pSite1 | %pSite2 | DMpSite | ADMpSite | DMpNucl | ADMpNucl |
|---|---|---|---|---|---|---|---|---|---|---|---|---|---|
| gene3165 | https://www.ncbi.nlm.nih.gov/gene/?term=LOC105275706 | 2374 | 65 | 273.8 | 6 | 25.27 | 9.23 | 24.65 | 38.08 | -13.43 | 13.43 | -0.37 | 0.37 |
| gene2824 | https://www.ncbi.nlm.nih.gov/gene/?term=LOC105275337 | 1772 | 58 | 327.31 | 1 | 5.64 | 1.72 | 31.86 | 38.08 | -6.22 | 6.22 | -0.2 | 0.2 |
| gene2593 | https://www.ncbi.nlm.nih.gov/gene/?term=LOC105275070 | 1774 | 5 | 28.18 | 2 | 11.27 | 40 | 19.71 | 71.28 | -51.57 | 51.57 | -0.15 | 0.15 |
| gene5714 | https://www.ncbi.nlm.nih.gov/gene/?term=LOC105278729 | 710 | 14 | 197.18 | 1 | 14.08 | 7.14 | 51.67 | 57.98 | -6.31 | 6.31 | -0.12 | 0.12 |
| gene10213 | https://www.ncbi.nlm.nih.gov/gene/?term=LOC105283826 | 1503 | 53 | 352.63 | 1 | 6.65 | 1.89 | 16.45 | 13.62 | 2.82 | 2.82 | 0.1 | 0.1 |
| gene920 | https://www.ncbi.nlm.nih.gov/gene/?term=LOC105282518 | 1197 | 40 | 334.17 | 1 | 8.35 | 2.5 | 39.94 | 37.69 | 2.25 | 2.25 | 0.08 | 0.08 |
| gene6334 | https://www.ncbi.nlm.nih.gov/gene/?term=LOC105279356 | 1471 | 40 | 271.92 | 1 | 6.8 | 2.5 | 29.12 | 32.21 | -3.09 | 3.09 | -0.08 | 0.08 |
| gene7734 | https://www.ncbi.nlm.nih.gov/gene/?term=LOC105280972 | 1528 | 26 | 170.16 | 1 | 6.54 | 3.85 | 20.04 | 23.88 | -3.84 | 3.84 | -0.07 | 0.07 |
| gene2676 | https://www.ncbi.nlm.nih.gov/gene/?term=LOC105275181 | 1353 | 15 | 110.86 | 1 | 7.39 | 6.67 | 67.39 | 74.05 | -6.66 | 6.66 | -0.07 | 0.07 |
| gene7556 | https://www.ncbi.nlm.nih.gov/gene/?term=LOC105280774 | 2792 | 98 | 351 | 1 | 3.58 | 1.02 | 31.47 | 33.13 | -1.66 | 1.66 | -0.06 | 0.06 |
| gene5877 | https://www.ncbi.nlm.nih.gov/gene/?term=LOC105278807 | 1719 | 50 | 290.87 | 1 | 5.82 | 2 | 21.66 | 23.62 | -1.95 | 1.95 | -0.06 | 0.06 |
| gene12406 | https://www.ncbi.nlm.nih.gov/gene/?term=LOC105286316 | 1386 | 29 | 209.24 | 1 | 7.22 | 3.45 | 41.65 | 44.38 | -2.73 | 2.73 | -0.06 | 0.06 |
| gene393 | https://www.ncbi.nlm.nih.gov/gene/?term=LOC105276566 | 1123 | 2 | 17.81 | 1 | 8.9 | 50 | 77.65 | 48.08 | 29.58 | 29.58 | 0.05 | 0.05 |
| gene7915 | https://www.ncbi.nlm.nih.gov/gene/?term=LOC105281173 | 1597 | 25 | 156.54 | 1 | 6.26 | 4 | 64.87 | 61.62 | 3.25 | 3.25 | 0.05 | 0.05 |
| gene1220 | https://www.ncbi.nlm.nih.gov/gene/?term=LOC105285742 | 3453 | 90 | 260.64 | 1 | 2.9 | 1.11 | 27.79 | 25.88 | 1.92 | 1.92 | 0.05 | 0.05 |
| gene11401 | https://www.ncbi.nlm.nih.gov/gene/?term=LOC105285202 | 1841 | 47 | 255.3 | 1 | 5.43 | 2.13 | 22.36 | 20.6 | 1.77 | 1.77 | 0.05 | 0.05 |
| gene6966 | https://www.ncbi.nlm.nih.gov/gene/?term=LOC105280104 | 3747 | 116 | 309.58 | 1 | 2.67 | 0.86 | 15.51 | 13.86 | 1.65 | 1.65 | 0.05 | 0.05 |
| gene5298 | https://www.ncbi.nlm.nih.gov/gene/?term=LOC105278145 | 4060 | 245 | 603.45 | 2 | 4.93 | 0.82 | 11.36 | 10.5 | 0.86 | 0.86 | 0.05 | 0.05 |
| gene10984 | https://www.ncbi.nlm.nih.gov/gene/?term=LOC105284704 | 1987 | 61 | 307 | 1 | 5.03 | 1.64 | 25.94 | 27.43 | -1.49 | 1.49 | -0.05 | 0.05 |
| gene10520 | https://www.ncbi.nlm.nih.gov/gene/?term=LOC105284184 | 2451 | 67 | 273.36 | 1 | 4.08 | 1.49 | 41.63 | 43.45 | -1.81 | 1.81 | -0.05 | 0.05 |
| gene12516 | https://www.ncbi.nlm.nih.gov/gene/?term=LOC105286452 | 1529 | 35 | 228.91 | 1 | 6.54 | 2.86 | 51.01 | 53.03 | -2.03 | 2.03 | -0.05 | 0.05 |
| gene9109 | https://www.ncbi.nlm.nih.gov/gene/?term=LOC105282561 | 2400 | 10 | 41.67 | 1 | 4.17 | 10 | 26.72 | 17.84 | 8.89 | 8.89 | 0.04 | 0.04 |
| gene7952 | https://www.ncbi.nlm.nih.gov/gene/?term=LOC105281213 | 2963 | 31 | 104.62 | 1 | 3.37 | 3.23 | 42.96 | 39.57 | 3.39 | 3.39 | 0.04 | 0.04 |
| gene12981 | https://www.ncbi.nlm.nih.gov/gene/?term=LOC105286999 | 2063 | 39 | 189.05 | 1 | 4.85 | 2.56 | 33.12 | 30.76 | 2.36 | 2.36 | 0.04 | 0.04 |
| gene2404 | https://www.ncbi.nlm.nih.gov/gene/?term=LOC105274853 | 1767 | 33 | 186.76 | 1 | 5.66 | 3.03 | 31.53 | 29.22 | 2.31 | 2.31 | 0.04 | 0.04 |
| gene10391 | https://www.ncbi.nlm.nih.gov/gene/?term=LOC109611283 | 1994 | 41 | 205.62 | 1 | 5.02 | 2.44 | 35.16 | 33.4 | 1.76 | 1.76 | 0.04 | 0.04 |
| gene11454 | https://www.ncbi.nlm.nih.gov/gene/?term=LOC105285256 | 1906 | 48 | 251.84 | 1 | 5.25 | 2.08 | 46.56 | 44.93 | 1.63 | 1.63 | 0.04 | 0.04 |
| gene11275 | https://www.ncbi.nlm.nih.gov/gene/?term=LOC105285055 | 4119 | 196 | 475.84 | 1 | 2.43 | 0.51 | 13.91 | 14.65 | -0.74 | 0.74 | -0.04 | 0.04 |
| gene298 | https://www.ncbi.nlm.nih.gov/gene/?term=LOC105275460 | 2473 | 107 | 432.67 | 1 | 4.04 | 0.93 | 13.52 | 14.43 | -0.91 | 0.91 | -0.04 | 0.04 |
| gene11819 | https://www.ncbi.nlm.nih.gov/gene/?term=LOC105285671 | 2366 | 91 | 384.62 | 1 | 4.23 | 1.1 | 28.12 | 29.18 | -1.06 | 1.06 | -0.04 | 0.04 |
| gene1310 | https://www.ncbi.nlm.nih.gov/gene/?term=LOC105286661 | 2582 | 65 | 251.74 | 1 | 3.87 | 1.54 | 19.96 | 21.58 | -1.62 | 1.62 | -0.04 | 0.04 |
| gene274 | https://www.ncbi.nlm.nih.gov/gene/?term=LOC105275190 | 3488 | 66 | 189.22 | 1 | 2.87 | 1.52 | 20.22 | 22.09 | -1.87 | 1.87 | -0.04 | 0.04 |
| gene3727 | https://www.ncbi.nlm.nih.gov/gene/?term=LOC105276367 | 2484 | 54 | 217.39 | 1 | 4.03 | 1.85 | 48.38 | 50.33 | -1.95 | 1.95 | -0.04 | 0.04 |
| gene7398 | https://www.ncbi.nlm.nih.gov/gene/?term=LOC105280589 | 3452 | 41 | 118.77 | 1 | 2.9 | 2.44 | 25.07 | 28.12 | -3.05 | 3.05 | -0.04 | 0.04 |
| gene7368 | https://www.ncbi.nlm.nih.gov/gene/?term=LOC105280556 | 4105 | 40 | 97.44 | 1 | 2.44 | 2.5 | 14.82 | 11.87 | 2.94 | 2.94 | 0.03 | 0.03 |
| gene3168 | https://www.ncbi.nlm.nih.gov/gene/?term=LOC105275711 | 6150 | 71 | 115.45 | 1 | 1.63 | 1.41 | 14.71 | 11.97 | 2.74 | 2.74 | 0.03 | 0.03 |
| gene5490 | https://www.ncbi.nlm.nih.gov/gene/?term=LOC105278358 | 2095 | 25 | 119.33 | 1 | 4.77 | 4 | 36.86 | 34.23 | 2.63 | 2.63 | 0.03 | 0.03 |
| gene388 | https://www.ncbi.nlm.nih.gov/gene/?term=LOC105276538 | 1558 | 19 | 121.95 | 2 | 12.84 | 10.53 | 69.41 | 66.83 | 2.58 | 2.58 | 0.03 | 0.03 |
| gene6422 | https://www.ncbi.nlm.nih.gov/gene/?term=LOC105279458 | 2419 | 33 | 136.42 | 1 | 4.13 | 3.03 | 55.1 | 53.04 | 2.06 | 2.06 | 0.03 | 0.03 |
| gene3107 | https://www.ncbi.nlm.nih.gov/gene/?term=LOC105275650 | 5103 | 143 | 280.23 | 1 | 1.96 | 0.7 | 16.9 | 15.68 | 1.21 | 1.21 | 0.03 | 0.03 |
| gene4190 | https://www.ncbi.nlm.nih.gov/gene/?term=LOC105276875 | 3150 | 104 | 330.16 | 1 | 3.17 | 0.96 | 25.99 | 25.04 | 0.95 | 0.95 | 0.03 | 0.03 |
| gene3286 | https://www.ncbi.nlm.nih.gov/gene/?term=LOC105275848 | 3744 | 121 | 323.18 | 1 | 2.67 | 0.83 | 22.9 | 22 | 0.9 | 0.9 | 0.03 | 0.03 |
| gene1993 | https://www.ncbi.nlm.nih.gov/gene/?term=LOC105288193 | 1449 | 62 | 427.88 | 1 | 6.9 | 1.61 | 35.55 | 34.76 | 0.78 | 0.78 | 0.03 | 0.03 |
| gene474 | https://www.ncbi.nlm.nih.gov/gene/?term=LOC105277435 | 3625 | 150 | 413.79 | 1 | 2.76 | 0.67 | 12.65 | 11.92 | 0.73 | 0.73 | 0.03 | 0.03 |
| gene2844 | https://www.ncbi.nlm.nih.gov/gene/?term=LOC105275363 | 4330 | 179 | 413.39 | 1 | 2.31 | 0.56 | 14.53 | 15.15 | -0.61 | 0.61 | -0.03 | 0.03 |
| gene8729 | https://www.ncbi.nlm.nih.gov/gene/?term=LOC105282137 | 5005 | 195 | 389.61 | 1 | 2 | 0.51 | 16.47 | 17.35 | -0.89 | 0.89 | -0.03 | 0.03 |
| gene7166 | https://www.ncbi.nlm.nih.gov/gene/?term=LOC105280326 | 6117 | 195 | 318.78 | 1 | 1.63 | 0.51 | 16.86 | 17.78 | -0.92 | 0.92 | -0.03 | 0.03 |
| gene9101 | https://www.ncbi.nlm.nih.gov/gene/?term=LOC105282553 | 3068 | 104 | 338.98 | 1 | 3.26 | 0.96 | 20.36 | 21.29 | -0.93 | 0.93 | -0.03 | 0.03 |
| gene4023 | https://www.ncbi.nlm.nih.gov/gene/?term=LOC105276692 | 1759 | 35 | 198.98 | 1 | 5.69 | 2.86 | 23.93 | 25.35 | -1.42 | 1.42 | -0.03 | 0.03 |
| gene7183 | https://www.ncbi.nlm.nih.gov/gene/?term=LOC105280348 | 3925 | 51 | 129.94 | 1 | 2.55 | 1.96 | 25.44 | 27.74 | -2.3 | 2.3 | -0.03 | 0.03 |
| gene10107 | https://www.ncbi.nlm.nih.gov/gene/?term=LOC105283712 | 1883 | 23 | 122.15 | 1 | 5.31 | 4.35 | 27.23 | 29.73 | -2.49 | 2.49 | -0.03 | 0.03 |
| gene1199 | https://www.ncbi.nlm.nih.gov/gene/?term=LOC105285498 | 2220 | 23 | 103.6 | 1 | 4.5 | 4.35 | 35.45 | 33.87 | 1.58 | 1.58 | 0.02 | 0.02 |
| gene12021 | https://www.ncbi.nlm.nih.gov/gene/?term=LOC105285905 | 1399 | 21 | 150.11 | 1 | 7.15 | 4.76 | 41.18 | 39.62 | 1.55 | 1.55 | 0.02 | 0.02 |
| gene2748 | https://www.ncbi.nlm.nih.gov/gene/?term=LOC105275261 | 3845 | 54 | 140.44 | 1 | 2.6 | 1.85 | 38.15 | 36.73 | 1.42 | 1.42 | 0.02 | 0.02 |
| gene11447 | https://www.ncbi.nlm.nih.gov/gene/?term=LOC105285252 | 7437 | 119 | 160.01 | 1 | 1.34 | 0.84 | 14.91 | 13.69 | 1.22 | 1.22 | 0.02 | 0.02 |
| gene5375 | https://www.ncbi.nlm.nih.gov/gene/?term=LOC105278228 | 8640 | 143 | 165.51 | 1 | 1.16 | 0.7 | 24.34 | 23.23 | 1.11 | 1.11 | 0.02 | 0.02 |
| gene5646 | https://www.ncbi.nlm.nih.gov/gene/?term=LOC105278534 | 7475 | 166 | 222.07 | 1 | 1.34 | 0.6 | 18.54 | 17.53 | 1.02 | 1.02 | 0.02 | 0.02 |
| gene4376 | https://www.ncbi.nlm.nih.gov/gene/?term=LOC105277083 | 4307 | 92 | 213.61 | 1 | 2.32 | 1.09 | 26.82 | 26.06 | 0.76 | 0.76 | 0.02 | 0.02 |
| gene11340 | https://www.ncbi.nlm.nih.gov/gene/?term=LOC105285131 | 2918 | 93 | 318.71 | 1 | 3.43 | 1.08 | 12.16 | 11.4 | 0.75 | 0.75 | 0.02 | 0.02 |
| gene2271 | https://www.ncbi.nlm.nih.gov/gene/?term=LOC105274706 | 1448 | 52 | 359.12 | 1 | 6.91 | 1.92 | 37.3 | 36.65 | 0.64 | 0.64 | 0.02 | 0.02 |

| | | | | | | | | | | | | | |
|---|---|---|---|---|---|---|---|---|---|---|---|---|---|
| gene12573 | https://www.ncbi.nlm.nih.gov/gene/?term=LOC105286515 | 3647 | 106 | 290.65 | 1 | 2.74 | 0.94 | 10.61 | 10.01 | 0.6 | 0.6 | 0.02 | 0.02 |
| gene9645 | https://www.ncbi.nlm.nih.gov/gene/?term=LOC105283181 | 3875 | 118 | 304.52 | 1 | 2.58 | 0.85 | 12.99 | 12.4 | 0.59 | 0.59 | 0.02 | 0.02 |
| gene13036 | https://www.ncbi.nlm.nih.gov/gene/?term=LOC105287059 | 4959 | 150 | 302.48 | 1 | 2.02 | 0.67 | 17.57 | 17 | 0.57 | 0.57 | 0.02 | 0.02 |
| gene5831 | https://www.ncbi.nlm.nih.gov/gene/?term=LOC105278761 | 2506 | 100 | 399.04 | 2 | 7.98 | 2 | 24.32 | 23.77 | 0.54 | 0.54 | 0.02 | 0.02 |
| gene13114 | https://www.ncbi.nlm.nih.gov/gene/?term=LOC105287147 | 7609 | 360 | 473.12 | 1 | 1.31 | 0.28 | 10.34 | 9.95 | 0.39 | 0.39 | 0.02 | 0.02 |
| gene7568 | https://www.ncbi.nlm.nih.gov/gene/?term=LOC105280786 | 2525 | 107 | 423.76 | 1 | 3.96 | 0.93 | 33.06 | 32.68 | 0.38 | 0.38 | 0.02 | 0.02 |
| gene10606 | https://www.ncbi.nlm.nih.gov/gene/?term=LOC105284273 | 6119 | 227 | 370.98 | 1 | 1.63 | 0.44 | 21.11 | 21.54 | -0.43 | 0.43 | -0.02 | 0.02 |
| gene5649 | https://www.ncbi.nlm.nih.gov/gene/?term=LOC105278538 | 2140 | 77 | 359.81 | 1 | 4.67 | 1.3 | 17.91 | 18.36 | -0.46 | 0.46 | -0.02 | 0.02 |
| gene9755 | https://www.ncbi.nlm.nih.gov/gene/?term=LOC105283296 | 2177 | 79 | 362.88 | 1 | 4.59 | 1.27 | 20.77 | 21.25 | -0.48 | 0.48 | -0.02 | 0.02 |
| gene10678 | https://www.ncbi.nlm.nih.gov/gene/?term=LOC105284355 | 4031 | 152 | 377.08 | 1 | 2.48 | 0.66 | 20.4 | 20.91 | -0.51 | 0.51 | -0.02 | 0.02 |
| gene7888 | https://www.ncbi.nlm.nih.gov/gene/?term=LOC105281141 | 4357 | 153 | 351.16 | 1 | 2.3 | 0.65 | 30.21 | 30.79 | -0.57 | 0.57 | -0.02 | 0.02 |
| gene292 | https://www.ncbi.nlm.nih.gov/gene/?term=LOC105275398 | 2825 | 77 | 272.57 | 1 | 3.54 | 1.3 | 19.76 | 20.59 | -0.82 | 0.82 | -0.02 | 0.02 |
| gene10531 | https://www.ncbi.nlm.nih.gov/gene/?term=LOC105284190 | 6881 | 164 | 238.34 | 1 | 1.45 | 0.61 | 10.48 | 11.52 | -1.03 | 1.03 | -0.02 | 0.02 |
| gene2173 | https://www.ncbi.nlm.nih.gov/gene/?term=LOC105274590 | 4387 | 64 | 145.89 | 1 | 2.28 | 1.56 | 26 | 27.13 | -1.13 | 1.13 | -0.02 | 0.02 |
| gene6455 | https://www.ncbi.nlm.nih.gov/gene/?term=LOC105279500 | 2612 | 31 | 118.68 | 1 | 3.83 | 3.23 | 51.05 | 52.76 | -1.71 | 1.71 | -0.02 | 0.02 |
| gene10819 | https://www.ncbi.nlm.nih.gov/gene/?term=LOC105284507 | 5767 | 69 | 119.65 | 1 | 1.73 | 1.45 | 22.66 | 21.47 | 1.19 | 1.19 | 0.01 | 0.01 |
| gene322 | https://www.ncbi.nlm.nih.gov/gene/?term=LOC105275777 | 1606 | 21 | 130.76 | 1 | 6.23 | 4.76 | 49.88 | 48.85 | 1.04 | 1.04 | 0.01 | 0.01 |
| gene10155 | https://www.ncbi.nlm.nih.gov/gene/?term=LOC105283756 | 7030 | 86 | 122.33 | 1 | 1.42 | 1.16 | 7.44 | 6.49 | 0.95 | 0.95 | 0.01 | 0.01 |
| gene3671 | https://www.ncbi.nlm.nih.gov/gene/?term=LOC105276272 | 1966 | 25 | 127.16 | 1 | 5.09 | 4 | 43.89 | 43.09 | 0.8 | 0.8 | 0.01 | 0.01 |
| gene3848 | https://www.ncbi.nlm.nih.gov/gene/?term=LOC105276486 | 5899 | 107 | 181.39 | 1 | 1.7 | 0.93 | 25.75 | 24.98 | 0.77 | 0.77 | 0.01 | 0.01 |
| gene2683 | https://www.ncbi.nlm.nih.gov/gene/?term=LOC105275187 | 5492 | 66 | 120.17 | 1 | 1.82 | 1.52 | 14.03 | 13.27 | 0.76 | 0.76 | 0.01 | 0.01 |
| gene1147 | https://www.ncbi.nlm.nih.gov/gene/?term=LOC105284954 | 4022 | 75 | 186.47 | 1 | 2.49 | 1.33 | 14.93 | 14.22 | 0.71 | 0.71 | 0.01 | 0.01 |
| gene7207 | https://www.ncbi.nlm.nih.gov/gene/?term=LOC105280376 | 4659 | 98 | 210.35 | 1 | 2.15 | 1.02 | 30.66 | 30.04 | 0.62 | 0.62 | 0.01 | 0.01 |
| gene657 | https://www.ncbi.nlm.nih.gov/gene/?term=LOC105279667 | 6421 | 64 | 99.67 1 | 1.56 | 1.56 | 3.41 | 2.8 | 0.61 | 0.61 | 0.01 | 0.01 | |
| gene3599 | https://www.ncbi.nlm.nih.gov/gene/?term=LOC105276196 | 2955 | 40 | 135.36 | 1 | 3.38 | 2.5 | 52.66 | 52.1 | 0.56 | 0.56 | 0.01 | 0.01 |
| gene12307 | https://www.ncbi.nlm.nih.gov/gene/?term=LOC105286216 | 10313 | 146 | 141.57 | 1 | 0.97 | 0.68 | 17.27 | 16.76 | 0.51 | 0.51 | 0.01 | 0.01 |
| gene8661 | https://www.ncbi.nlm.nih.gov/gene/?term=LOC105282063 | 1926 | 39 | 202.49 | 1 | 5.19 | 2.56 | 23.72 | 23.24 | 0.48 | 0.48 | 0.01 | 0.01 |
| gene10326 | https://www.ncbi.nlm.nih.gov/gene/?term=LOC105283962 | 3042 | 92 | 302.43 | 1 | 3.29 | 1.09 | 15.29 | 14.85 | 0.44 | 0.44 | 0.01 | 0.01 |
| gene13044 | https://www.ncbi.nlm.nih.gov/gene/?term=LOC105287070 | 5094 | 159 | 312.13 | 1 | 1.96 | 0.63 | 5.69 | 5.29 | 0.4 | 0.4 | 0.01 | 0.01 |
| gene3817 | https://www.ncbi.nlm.nih.gov/gene/?term=LOC105276449 | 1230 | 39 | 317.07 | 1 | 8.13 | 2.56 | 32.77 | 32.39 | 0.38 | 0.38 | 0.01 | 0.01 |
| gene3206 | https://www.ncbi.nlm.nih.gov/gene/?term=LOC105275752 | 3474 | 117 | 336.79 | 1 | 2.88 | 0.85 | 29.75 | 29.38 | 0.37 | 0.37 | 0.01 | 0.01 |
| gene6583 | https://www.ncbi.nlm.nih.gov/gene/?term=LOC105279649 | 7444 | 308 | 413.76 | 1 | 1.34 | 0.32 | 11.75 | 11.41 | 0.34 | 0.34 | 0.01 | 0.01 |
| gene10392 | https://www.ncbi.nlm.nih.gov/gene/?term=LOC105284036 | 3216 | 76 | 236.32 | 1 | 3.11 | 1.32 | 25.16 | 24.85 | 0.31 | 0.31 | 0.01 | 0.01 |
| gene5539 | https://www.ncbi.nlm.nih.gov/gene/?term=LOC105278415 | 25024 | 542 | 216.59 | 1 | 0.4 | 0.18 | 4.22 | 3.95 | 0.27 | 0.27 | 0.01 | 0.01 |
| gene12674 | https://www.ncbi.nlm.nih.gov/gene/?term=LOC105286649 | 18283 | 638 | 348.96 | 1 | 0.55 | 0.16 | 1.31 | 1.05 | 0.26 | 0.26 | 0.01 | 0.01 |
| gene10549 | https://www.ncbi.nlm.nih.gov/gene/?term=LOC105284210 | 7145 | 298 | 417.07 | 1 | 1.4 | 0.34 | 11.27 | 11.01 | 0.25 | 0.25 | 0.01 | 0.01 |
| gene7249 | https://www.ncbi.nlm.nih.gov/gene/?term=LOC105280422 | 1775 | 58 | 326.76 | 1 | 5.63 | 1.72 | 15.27 | 15.08 | 0.19 | 0.19 | 0.01 | 0.01 |
| gene10018 | https://www.ncbi.nlm.nih.gov/gene/?term=LOC105283607 | 4414 | 181 | 410.06 | 1 | 2.27 | 0.55 | 16.23 | 16.36 | -0.13 | 0.13 | -0.01 | 0.01 |
| gene1470 | https://www.ncbi.nlm.nih.gov/gene/?term=LOC105287597 | 2974 | 188 | 632.15 | 1 | 3.36 | 0.53 | 17.15 | 17.35 | -0.19 | 0.19 | -0.01 | 0.01 |
| gene3885 | https://www.ncbi.nlm.nih.gov/gene/?term=LOC105276527 | 7928 | 323 | 407.42 | 1 | 1.26 | 0.31 | 8.78 | 9 | -0.22 | 0.22 | -0.01 | 0.01 |
| gene10540 | https://www.ncbi.nlm.nih.gov/gene/?term=LOC105284200 | 5267 | 183 | 347.45 | 1 | 1.9 | 0.55 | 14.63 | 14.93 | -0.3 | 0.3 | -0.01 | 0.01 |
| gene6286 | https://www.ncbi.nlm.nih.gov/gene/?term=LOC105279298 | 2861 | 128 | 447.4 2 | 6.99 | 1.56 | 23.41 | 23.73 | -0.32 | 0.32 | -0.01 | 0.01 | |
| gene3856 | https://www.ncbi.nlm.nih.gov/gene/?term=LOC105276499 | 5236 | 178 | 339.95 | 1 | 1.91 | 0.56 | 9.73 | 10.06 | -0.33 | 0.33 | -0.01 | 0.01 |
| gene4892 | https://www.ncbi.nlm.nih.gov/gene/?term=LOC105277667 | 3898 | 119 | 305.28 | 1 | 2.57 | 0.84 | 27.91 | 28.29 | -0.38 | 0.38 | -0.01 | 0.01 |
| gene12317 | https://www.ncbi.nlm.nih.gov/gene/?term=LOC105286223 | 2235 | 79 | 353.47 | 1 | 4.47 | 1.27 | 30.56 | 30.96 | -0.4 | 0.4 | -0.01 | 0.01 |
| gene1884 | https://www.ncbi.nlm.nih.gov/gene/?term=LOC105288072 | 4169 | 76 | 182.3 1 | 2.4 | 1.32 | 23.55 | 23.95 | -0.4 | 0.4 | -0.01 | 0.01 | |
| gene2056 | https://www.ncbi.nlm.nih.gov/gene/?term=LOC105274452 | 6584 | 206 | 312.88 | 1 | 1.52 | 0.49 | 14.33 | 14.74 | -0.41 | 0.41 | -0.01 | 0.01 |
| gene4641 | https://www.ncbi.nlm.nih.gov/gene/?term=LOC105277398 | 1937 | 68 | 351.06 | 1 | 5.16 | 1.47 | 34.8 | 35.21 | -0.41 | 0.41 | -0.01 | 0.01 |
| gene303 | https://www.ncbi.nlm.nih.gov/gene/?term=LOC105275584 | 3081 | 71 | 230.44 | 1 | 3.25 | 1.41 | 26.68 | 27.17 | -0.49 | 0.49 | -0.01 | 0.01 |
| gene13033 | https://www.ncbi.nlm.nih.gov/gene/?term=LOC105287063 | 1877 | 45 | 239.74 | 1 | 5.33 | 2.22 | 34.91 | 35.54 | -0.62 | 0.62 | -0.01 | 0.01 |
| gene8600 | https://www.ncbi.nlm.nih.gov/gene/?term=LOC105282001 | 8513 | 126 | 148.01 | 1 | 1.17 | 0.79 | 11.25 | 11.9 | -0.65 | 0.65 | -0.01 | 0.01 |
| gene7245 | https://www.ncbi.nlm.nih.gov/gene/?term=LOC105280410 | 12478 | 234 | 187.53 | 1 | 0.8 | 0.43 | 20.56 | 21.23 | -0.67 | 0.67 | -0.01 | 0.01 |
| gene8343 | https://www.ncbi.nlm.nih.gov/gene/?term=LOC105281700 | 2172 | 40 | 184.16 | 1 | 4.6 | 2.5 | 41.18 | 41.86 | -0.68 | 0.68 | -0.01 | 0.01 |
| gene9336 | https://www.ncbi.nlm.nih.gov/gene/?term=LOC105282834 | 5993 | 106 | 176.87 | 1 | 1.67 | 0.94 | 15.44 | 16.15 | -0.71 | 0.71 | -0.01 | 0.01 |
| gene10344 | https://www.ncbi.nlm.nih.gov/gene/?term=LOC105283987 | 4841 | 64 | 132.2 1 | 2.07 | 1.56 | 32.11 | 32.92 | -0.81 | 0.81 | -0.01 | 0.01 | |
| gene6528 | https://www.ncbi.nlm.nih.gov/gene/?term=LOC105279591 | 7222 | 67 | 92.77 1 | 1.38 | 1.49 | 22.84 | 24.15 | -1.32 | 1.32 | -0.01 | 0.01 | |
| gene2970 | https://www.ncbi.nlm.nih.gov/gene/?term=LOC105275518 | 2340 | 18 | 76.92 1 | 4.27 | 5.56 | 47.45 | 48.96 | -1.51 | 1.51 | -0.01 | 0.01 | |
| gene10345 | https://www.ncbi.nlm.nih.gov/gene/?term=LOC105283989 | 4444 | 21 | 47.25 1 | 2.25 | 4.76 | 48.99 | 48.08 | 0.91 | 0.91 | 0 | 0 | |
| gene10600 | https://www.ncbi.nlm.nih.gov/gene/?term=LOC105284270 | 4837 | 97 | 200.54 | 1 | 2.07 | 1.03 | 13.04 | 12.82 | 0.22 | 0.22 | 0 | 0 |
| gene4962 | https://www.ncbi.nlm.nih.gov/gene/?term=LOC105277761 | 4645 | 140 | 301.4 1 | 2.15 | 0.71 | 14.56 | 14.4 | 0.16 | 0.16 | 0 | 0 | |
| gene5967 | https://www.ncbi.nlm.nih.gov/gene/?term=LOC105278918 | 4393 | 121 | 275.44 | 1 | 2.28 | 0.83 | 16.38 | 16.32 | 0.06 | 0.06 | 0 | 0 |
| gene6031 | https://www.ncbi.nlm.nih.gov/gene/?term=LOC105279000 | 5691 | 174 | 305.75 | 1 | 1.76 | 0.57 | 14.03 | 13.97 | 0.06 | 0.06 | 0 | 0 |
| gene11583 | https://www.ncbi.nlm.nih.gov/gene/?term=LOC105285414 | 8192 | 168 | 205.08 | 1 | 1.22 | 0.6 | 8.3 | 8.25 | 0.05 | 0.05 | 0 | 0 |
| gene940 | https://www.ncbi.nlm.nih.gov/gene/?term=LOC105282794 | 3578 | 82 | 229.18 | 1 | 2.79 | 1.22 | 16.49 | 16.45 | 0.04 | 0.04 | 0 | 0 |

| | | | | | | | | | | | | | |
|---|---|---|---|---|---|---|---|---|---|---|---|---|---|
| gene5524 | https://www.ncbi.nlm.nih.gov/gene/?term=LOC105278397 | 5177 | 200 | 386.32 | 1 | 1.93 | 0.5 | 24.39 | 24.36 | 0.03 | 0.03 | 0 | 0 |
| gene5746 | https://www.ncbi.nlm.nih.gov/gene/?term=LOC105278666 | 10343 | 387 | 374.17 | 1 | 0.97 | 0.26 | 7.75 | 7.72 | 0.03 | 0.03 | 0 | 0 |
| gene4209 | https://www.ncbi.nlm.nih.gov/gene/?term=LOC105277125 | 50550 | 1576 | 311.77 | 1 | 0.2 | 0.06 | 1.66 | 1.64 | 0.02 | 0.02 | 0 | 0 |
| gene6639 | https://www.ncbi.nlm.nih.gov/gene/?term=LOC105279705 | 7656 | 58 | 75.76 1 | 1.31 | 1.72 | 24.54 | 24.54 | 0 | 0 | 0 | 0 | |
| gene12802 | https://www.ncbi.nlm.nih.gov/gene/?term=LOC105286799 | 3183 | 165 | 518.38 | 1 | 3.14 | 0.61 | 16.21 | 16.23 | -0.02 | 0.02 | 0 | 0 |
| gene2806 | https://www.ncbi.nlm.nih.gov/gene/?term=LOC105275327 | 14257 | 414 | 290.38 | 1 | 0.7 | 0.24 | 7.7 | 7.73 | -0.02 | 0.02 | 0 | 0 |
| gene11579 | https://www.ncbi.nlm.nih.gov/gene/?term=LOC105285406 | 13571 | 248 | 182.74 | 1 | 0.74 | 0.4 | 5.98 | 6.04 | -0.05 | 0.05 | 0 | 0 |
| gene7148 | https://www.ncbi.nlm.nih.gov/gene/?term=LOC105280304 | 8920 | 90 | 100.9 1 | 1.12 | 1.11 | 7.28 | 7.34 | -0.05 | 0.05 | 0 | 0 | |
| gene53 | https://www.ncbi.nlm.nih.gov/gene/?term=LOC105277148 | 19283 | 236 | 122.39 | 1 | 0.52 | 0.42 | 4.4 | 4.47 | -0.07 | 0.07 | 0 | 0 |
| gene5525 | https://www.ncbi.nlm.nih.gov/gene/?term=LOC105278396 | 3090 | 186 | 601.94 | 1 | 3.24 | 0.54 | 24.51 | 24.57 | -0.07 | 0.07 | 0 | 0 |
| gene6620 | https://www.ncbi.nlm.nih.gov/gene/?term=LOC105279687 | 17998 | 294 | 163.35 | 1 | 0.56 | 0.34 | 5.33 | 5.42 | -0.1 | 0.1 | 0 | 0 |
| gene4246 | https://www.ncbi.nlm.nih.gov/gene/?term=LOC105276941 | 16412 | 446 | 271.75 | 1 | 0.61 | 0.22 | 5.36 | 5.47 | -0.11 | 0.11 | 0 | 0 |
| gene9330 | https://www.ncbi.nlm.nih.gov/gene/?term=LOC105282825 | 3936 | 80 | 203.25 | 1 | 2.54 | 1.25 | 13.05 | 13.17 | -0.11 | 0.11 | 0 | 0 |
| gene12203 | https://www.ncbi.nlm.nih.gov/gene/?term=LOC105286099 | 5967 | 70 | 117.31 | 1 | 1.68 | 1.43 | 11.64 | 11.77 | -0.13 | 0.13 | 0 | 0 |
| gene11568 | https://www.ncbi.nlm.nih.gov/gene/?term=LOC105285395 | 8086 | 160 | 197.87 | 1 | 1.24 | 0.62 | 17.75 | 17.97 | -0.23 | 0.23 | 0 | 0 |

Directory: OGLall File: rnk-dmg-Cb_LI_BCphaseA1.vs.BCphaseB1.txt

| gene_ID | gene_link | gwidth | #Sites | #per10Kb | #dmSites | #dmsp10kb | %dmSites | %pSite1 | %pSite2 | DMpSite | ADMpSite | DMpNucl | ADMpNucl |
|---|---|---|---|---|---|---|---|---|---|---|---|---|---|
| gene3165 | https://www.ncbi.nlm.nih.gov/gene/?term=LOC105275706 | 2374 | 66 | 278.01 | 4 | 16.85 | 6.06 | 24.58 | 35.31 | -10.73 | 10.73 | -0.3 | 0.3 |
| gene2593 | https://www.ncbi.nlm.nih.gov/gene/?term=LOC105275070 | 1774 | 6 | 33.82 | 1 | 5.64 | 16.67 | 16.42 | 54.02 | -37.6 | 37.6 | -0.13 | 0.13 |
| gene9740 | https://www.ncbi.nlm.nih.gov/gene/?term=LOC105283277 | 2887 | 138 | 478 | 1 | 3.46 | 0.72 | 23.22 | 20.78 | 2.44 | 2.44 | 0.12 | 0.12 |
| gene7199 | https://www.ncbi.nlm.nih.gov/gene/?term=LOC105280366 | 1414 | 24 | 169.73 | 1 | 7.07 | 4.17 | 45.18 | 38.6 | 6.59 | 6.59 | 0.11 | 0.11 |
| gene8929 | https://www.ncbi.nlm.nih.gov/gene/?term=LOC105282371 | 1849 | 8 | 43.27 | 1 | 5.41 | 12.5 | 22.25 | 44.14 | -21.89 | 21.89 | -0.09 | 0.09 |
| gene10602 | https://www.ncbi.nlm.nih.gov/gene/?term=LOC105284268 | 2122 | 20 | 94.25 | 1 | 4.71 | 5 | 32.63 | 25.61 | 7.02 | 7.02 | 0.07 | 0.07 |
| gene5298 | https://www.ncbi.nlm.nih.gov/gene/?term=LOC105278145 | 4060 | 250 | 615.76 | 2 | 4.93 | 0.8 | 10.79 | 9.61 | 1.17 | 1.17 | 0.07 | 0.07 |
| gene5714 | https://www.ncbi.nlm.nih.gov/gene/?term=LOC105278729 | 710 | 14 | 197.18 | 1 | 14.08 | 7.14 | 51.67 | 55.07 | -3.41 | 3.41 | -0.07 | 0.07 |
| gene9464 | https://www.ncbi.nlm.nih.gov/gene/?term=LOC105282975 | 1450 | 24 | 165.52 | 1 | 6.9 | 4.17 | 42.63 | 39.26 | 3.37 | 3.37 | 0.06 | 0.06 |
| gene10213 | https://www.ncbi.nlm.nih.gov/gene/?term=LOC105283826 | 1503 | 54 | 359.28 | 1 | 6.65 | 1.85 | 16.14 | 14.48 | 1.66 | 1.66 | 0.06 | 0.06 |
| gene298 | https://www.ncbi.nlm.nih.gov/gene/?term=LOC105275460 | 2473 | 115 | 465.02 | 1 | 4.04 | 0.87 | 12.68 | 13.89 | -1.21 | 1.21 | -0.06 | 0.06 |
| gene1855 | https://www.ncbi.nlm.nih.gov/gene/?term=LOC105288036 | 2387 | 87 | 364.47 | 1 | 4.19 | 1.15 | 28.59 | 30.36 | -1.76 | 1.76 | -0.06 | 0.06 |
| gene6334 | https://www.ncbi.nlm.nih.gov/gene/?term=LOC105279356 | 1471 | 43 | 292.32 | 1 | 6.8 | 2.33 | 28.25 | 30.28 | -2.03 | 2.03 | -0.06 | 0.06 |
| gene12516 | https://www.ncbi.nlm.nih.gov/gene/?term=LOC105286452 | 1529 | 35 | 228.91 | 1 | 6.54 | 2.86 | 51.01 | 53.47 | -2.46 | 2.46 | -0.06 | 0.06 |
| gene1962 | https://www.ncbi.nlm.nih.gov/gene/?term=LOC105288157 | 1246 | 16 | 128.41 | 1 | 8.03 | 6.25 | 13.95 | 18.55 | -4.6 | 4.6 | -0.06 | 0.06 |
| gene393 | https://www.ncbi.nlm.nih.gov/gene/?term=LOC105276566 | 1123 | 2 | 17.81 | 1 | 8.9 | 50 | 77.65 | 47.22 | 30.43 | 30.43 | 0.05 | 0.05 |
| gene11401 | https://www.ncbi.nlm.nih.gov/gene/?term=LOC105285202 | 1841 | 50 | 271.59 | 1 | 5.43 | 2 | 21.11 | 19.1 | 2.01 | 2.01 | 0.05 | 0.05 |
| gene10326 | https://www.ncbi.nlm.nih.gov/gene/?term=LOC105283962 | 3042 | 93 | 305.72 | 1 | 3.29 | 1.08 | 11.63 | 10.03 | 1.6 | 1.6 | 0.05 | 0.05 |
| gene11340 | https://www.ncbi.nlm.nih.gov/gene/?term=LOC105285131 | 2918 | 97 | 332.42 | 1 | 3.43 | 1.03 | 14.75 | 13.34 | 1.41 | 1.41 | 0.05 | 0.05 |
| gene4190 | https://www.ncbi.nlm.nih.gov/gene/?term=LOC105276875 | 3150 | 130 | 412.7 | 1 | 3.17 | 0.77 | 21.07 | 19.84 | 1.23 | 1.23 | 0.05 | 0.05 |
| gene10606 | https://www.ncbi.nlm.nih.gov/gene/?term=LOC105284273 | 6119 | 249 | 406.93 | 2 | 3.27 | 0.8 | 20.32 | 19.15 | 1.16 | 1.16 | 0.05 | 0.05 |
| gene4034 | https://www.ncbi.nlm.nih.gov/gene/?term=LOC105276704 | 2065 | 84 | 406.78 | 1 | 4.84 | 1.19 | 12.86 | 14.01 | -1.15 | 1.15 | -0.05 | 0.05 |
| gene570 | https://www.ncbi.nlm.nih.gov/gene/?term=LOC105278549 | 1673 | 63 | 376.57 | 1 | 5.98 | 1.59 | 40.52 | 41.79 | -1.27 | 1.27 | -0.05 | 0.05 |
| gene2856 | https://www.ncbi.nlm.nih.gov/gene/?term=LOC105275384 | 1987 | 76 | 382.49 | 1 | 5.03 | 1.32 | 32.95 | 34.23 | -1.28 | 1.28 | -0.05 | 0.05 |
| gene4641 | https://www.ncbi.nlm.nih.gov/gene/?term=LOC105277398 | 1937 | 70 | 361.38 | 1 | 5.16 | 1.43 | 35.36 | 36.76 | -1.4 | 1.4 | -0.05 | 0.05 |
| gene9109 | https://www.ncbi.nlm.nih.gov/gene/?term=LOC105282561 | 2400 | 12 | 50 | 1 | 4.17 | 8.33 | 20.55 | 12.93 | 7.62 | 7.62 | 0.04 | 0.04 |
| gene3168 | https://www.ncbi.nlm.nih.gov/gene/?term=LOC105275711 | 6150 | 79 | 128.46 | 1 | 1.63 | 1.27 | 14.18 | 11.09 | 3.09 | 3.09 | 0.04 | 0.04 |
| gene6422 | https://www.ncbi.nlm.nih.gov/gene/?term=LOC105279458 | 2419 | 36 | 148.82 | 1 | 4.13 | 2.78 | 53.83 | 51.06 | 2.78 | 2.78 | 0.04 | 0.04 |
| gene7915 | https://www.ncbi.nlm.nih.gov/gene/?term=LOC105281173 | 1597 | 25 | 156.54 | 1 | 6.26 | 4 | 64.87 | 62.27 | 2.61 | 2.61 | 0.04 | 0.04 |
| gene920 | https://www.ncbi.nlm.nih.gov/gene/?term=LOC105282518 | 1197 | 43 | 359.23 | 1 | 8.35 | 2.33 | 37.5 | 36.41 | 1.09 | 1.09 | 0.04 | 0.04 |
| gene1470 | https://www.ncbi.nlm.nih.gov/gene/?term=LOC105287597 | 2974 | 180 | 605.25 | 1 | 3.36 | 0.56 | 17.88 | 17.25 | 0.63 | 0.63 | 0.04 | 0.04 |
| gene1332 | https://www.ncbi.nlm.nih.gov/gene/?term=LOC105286872 | 2460 | 141 | 573.17 | 1 | 4.07 | 0.71 | 21.92 | 22.68 | -0.76 | 0.76 | -0.04 | 0.04 |
| gene2844 | https://www.ncbi.nlm.nih.gov/gene/?term=LOC105275363 | 4330 | 187 | 431.87 | 1 | 2.31 | 0.53 | 14.36 | 15.35 | -0.99 | 0.99 | -0.04 | 0.04 |
| gene9101 | https://www.ncbi.nlm.nih.gov/gene/?term=LOC105282553 | 3068 | 103 | 335.72 | 1 | 3.26 | 0.97 | 20.56 | 21.9 | -1.34 | 1.34 | -0.04 | 0.04 |
| gene5877 | https://www.ncbi.nlm.nih.gov/gene/?term=LOC105278807 | 1719 | 51 | 296.68 | 1 | 5.82 | 1.96 | 22.99 | 24.43 | -1.44 | 1.44 | -0.04 | 0.04 |
| gene5647 | https://www.ncbi.nlm.nih.gov/gene/?term=LOC105278567 | 3118 | 38 | 121.87 | 1 | 3.21 | 2.63 | 12.56 | 16.06 | -3.5 | 3.5 | -0.04 | 0.04 |
| gene7368 | https://www.ncbi.nlm.nih.gov/gene/?term=LOC105280556 | 4105 | 49 | 119.37 | 1 | 2.44 | 2.04 | 12.67 | 10.47 | 2.2 | 2.2 | 0.03 | 0.03 |
| gene3727 | https://www.ncbi.nlm.nih.gov/gene/?term=LOC105276367 | 2484 | 50 | 201.29 | 1 | 4.03 | 2 | 50.02 | 48.5 | 1.52 | 1.52 | 0.03 | 0.03 |
| gene7345 | https://www.ncbi.nlm.nih.gov/gene/?term=LOC105280529 | 3833 | 87 | 226.98 | 1 | 2.61 | 1.15 | 18.24 | 16.79 | 1.44 | 1.44 | 0.03 | 0.03 |
| gene3599 | https://www.ncbi.nlm.nih.gov/gene/?term=LOC105276196 | 2955 | 54 | 182.74 | 1 | 3.38 | 1.85 | 40.6 | 39.18 | 1.42 | 1.42 | 0.03 | 0.03 |
| gene2271 | https://www.ncbi.nlm.nih.gov/gene/?term=LOC105274706 | 1448 | 47 | 324.59 | 1 | 6.91 | 2.13 | 41.16 | 40.14 | 1.03 | 1.03 | 0.03 | 0.03 |
| gene1220 | https://www.ncbi.nlm.nih.gov/gene/?term=LOC105285742 | 3453 | 94 | 272.23 | 1 | 2.9 | 1.06 | 26.72 | 25.74 | 0.97 | 0.97 | 0.03 | 0.03 |
| gene6031 | https://www.ncbi.nlm.nih.gov/gene/?term=LOC105279000 | 5691 | 180 | 316.29 | 1 | 1.76 | 0.56 | 14.05 | 13.11 | 0.94 | 0.94 | 0.03 | 0.03 |
| gene4962 | https://www.ncbi.nlm.nih.gov/gene/?term=LOC105277761 | 4645 | 140 | 301.4 | 1 | 2.15 | 0.71 | 15.27 | 14.38 | 0.89 | 0.89 | 0.03 | 0.03 |
| gene7249 | https://www.ncbi.nlm.nih.gov/gene/?term=LOC105280422 | 1775 | 61 | 343.66 | 1 | 5.63 | 1.64 | 15.16 | 14.36 | 0.81 | 0.81 | 0.03 | 0.03 |
| gene3793 | https://www.ncbi.nlm.nih.gov/gene/?term=LOC105276429 | 2901 | 112 | 386.07 | 1 | 3.45 | 0.89 | 26.42 | 25.76 | 0.65 | 0.65 | 0.03 | 0.03 |
| gene5525 | https://www.ncbi.nlm.nih.gov/gene/?term=LOC105278396 | 3090 | 187 | 605.18 | 1 | 3.24 | 0.53 | 24.38 | 23.89 | 0.49 | 0.49 | 0.03 | 0.03 |
| gene3885 | https://www.ncbi.nlm.nih.gov/gene/?term=LOC105276527 | 7928 | 346 | 436.43 | 1 | 1.26 | 0.29 | 8.3 | 8.89 | -0.6 | 0.6 | -0.03 | 0.03 |
| gene10018 | https://www.ncbi.nlm.nih.gov/gene/?term=LOC105283607 | 4414 | 204 | 462.17 | 1 | 2.27 | 0.49 | 14.8 | 15.47 | -0.67 | 0.67 | -0.03 | 0.03 |
| gene8729 | https://www.ncbi.nlm.nih.gov/gene/?term=LOC105282137 | 5005 | 199 | 397.6 | 1 | 2 | 0.5 | 16.19 | 16.99 | -0.79 | 0.79 | -0.03 | 0.03 |
| gene11819 | https://www.ncbi.nlm.nih.gov/gene/?term=LOC105285671 | 2366 | 96 | 405.75 | 1 | 4.23 | 1.04 | 27.73 | 28.57 | -0.84 | 0.84 | -0.03 | 0.03 |
| gene1310 | https://www.ncbi.nlm.nih.gov/gene/?term=LOC105286661 | 2582 | 65 | 251.74 | 1 | 3.87 | 1.54 | 19.96 | 20.95 | -0.99 | 0.99 | -0.03 | 0.03 |
| gene292 | https://www.ncbi.nlm.nih.gov/gene/?term=LOC105275398 | 2825 | 76 | 269.03 | 1 | 3.54 | 1.32 | 20.33 | 21.34 | -1.01 | 1.01 | -0.03 | 0.03 |
| gene12406 | https://www.ncbi.nlm.nih.gov/gene/?term=LOC105286316 | 1386 | 30 | 216.45 | 1 | 7.22 | 3.33 | 39.2 | 40.74 | -1.55 | 1.55 | -0.03 | 0.03 |
| gene1884 | https://www.ncbi.nlm.nih.gov/gene/?term=LOC105288072 | 4169 | 80 | 191.89 | 1 | 2.4 | 1.25 | 22.74 | 24.44 | -1.7 | 1.7 | -0.03 | 0.03 |
| gene2676 | https://www.ncbi.nlm.nih.gov/gene/?term=LOC105275181 | 1353 | 14 | 103.47 | 1 | 7.39 | 7.14 | 65.95 | 68.91 | -2.96 | 2.96 | -0.03 | 0.03 |
| gene2970 | https://www.ncbi.nlm.nih.gov/gene/?term=LOC105275518 | 2340 | 20 | 85.47 | 1 | 4.27 | 5 | 47.36 | 50.74 | -3.38 | 3.38 | -0.03 | 0.03 |
| gene652 | https://www.ncbi.nlm.nih.gov/gene/?term=LOC105279528 | 5806 | 42 | 72.34 | 1 | 1.72 | 2.38 | 16.48 | 14.36 | 2.12 | 2.12 | 0.02 | 0.02 |
| gene1199 | https://www.ncbi.nlm.nih.gov/gene/?term=LOC105285498 | 2220 | 26 | 117.12 | 1 | 4.5 | 3.85 | 33.11 | 31.11 | 2 | 2 | 0.02 | 0.02 |
| gene388 | https://www.ncbi.nlm.nih.gov/gene/?term=LOC105276538 | 1558 | 19 | 121.95 | 1 | 6.42 | 5.26 | 69.93 | 68.17 | 1.77 | 1.77 | 0.02 | 0.02 |

| | | | | | | | | | | | | | |
|---|---|---|---|---|---|---|---|---|---|---|---|---|---|
| gene322 | https://www.ncbi.nlm.nih.gov/gene/?term=LOC105275777 | 1606 | 22 | 136.99 | 1 | 6.23 | 4.55 | 47.62 | 46.19 | 1.43 | 1.43 | 0.02 | 0.02 |
| gene7247 | https://www.ncbi.nlm.nih.gov/gene/?term=LOC105280407 | 10486 | 205 | 195.5 | 1 | 0.95 | 0.49 | 17 | 15.99 | 1.01 | 1.01 | 0.02 | 0.02 |
| gene4376 | https://www.ncbi.nlm.nih.gov/gene/?term=LOC105277083 | 4307 | 92 | 213.61 | 1 | 2.32 | 1.09 | 26.68 | 25.69 | 0.99 | 0.99 | 0.02 | 0.02 |
| gene10392 | https://www.ncbi.nlm.nih.gov/gene/?term=LOC105284036 | 3216 | 84 | 261.19 | 2 | 6.22 | 2.38 | 23.38 | 22.62 | 0.76 | 0.76 | 0.02 | 0.02 |
| gene10844 | https://www.ncbi.nlm.nih.gov/gene/?term=LOC105284535 | 9086 | 220 | 242.13 | 1 | 1.1 | 0.45 | 16.85 | 16.12 | 0.73 | 0.73 | 0.02 | 0.02 |
| gene6966 | https://www.ncbi.nlm.nih.gov/gene/?term=LOC105280104 | 3747 | 124 | 330.93 | 1 | 2.67 | 0.81 | 15.19 | 14.51 | 0.68 | 0.68 | 0.02 | 0.02 |
| gene11454 | https://www.ncbi.nlm.nih.gov/gene/?term=LOC105285256 | 1906 | 52 | 272.82 | 1 | 5.25 | 1.92 | 42.98 | 42.36 | 0.62 | 0.62 | 0.02 | 0.02 |
| gene7568 | https://www.ncbi.nlm.nih.gov/gene/?term=LOC105280786 | 2525 | 121 | 479.21 | 1 | 3.96 | 0.83 | 29.4 | 28.89 | 0.52 | 0.52 | 0.02 | 0.02 |
| gene13036 | https://www.ncbi.nlm.nih.gov/gene/?term=LOC105287059 | 4959 | 166 | 334.74 | 1 | 2.02 | 0.6 | 17.36 | 16.86 | 0.5 | 0.5 | 0.02 | 0.02 |
| gene474 | https://www.ncbi.nlm.nih.gov/gene/?term=LOC105277435 | 3625 | 155 | 427.59 | 1 | 2.76 | 0.65 | 12.2 | 11.71 | 0.49 | 0.49 | 0.02 | 0.02 |
| gene10549 | https://www.ncbi.nlm.nih.gov/gene/?term=LOC105284210 | 7145 | 319 | 446.47 | 1 | 1.4 | 0.31 | 10.59 | 10.18 | 0.4 | 0.4 | 0.02 | 0.02 |
| gene6286 | https://www.ncbi.nlm.nih.gov/gene/?term=LOC105279298 | 2861 | 134 | 468.37 | 1 | 3.5 | 0.75 | 23.06 | 22.69 | 0.37 | 0.37 | 0.02 | 0.02 |
| gene3856 | https://www.ncbi.nlm.nih.gov/gene/?term=LOC105276499 | 5236 | 197 | 376.24 | 1 | 1.91 | 0.51 | 9.94 | 10.34 | -0.41 | 0.41 | -0.02 | 0.02 |
| gene6583 | https://www.ncbi.nlm.nih.gov/gene/?term=LOC105279649 | 7444 | 317 | 425.85 | 1 | 1.34 | 0.32 | 11.9 | 12.34 | -0.44 | 0.44 | -0.02 | 0.02 |
| gene11275 | https://www.ncbi.nlm.nih.gov/gene/?term=LOC105285055 | 4119 | 213 | 517.12 | 1 | 2.43 | 0.47 | 13.26 | 13.73 | -0.47 | 0.47 | -0.02 | 0.02 |
| gene10678 | https://www.ncbi.nlm.nih.gov/gene/?term=LOC105284355 | 4031 | 170 | 421.73 | 1 | 2.48 | 0.59 | 18.52 | 19.03 | -0.5 | 0.5 | -0.02 | 0.02 |
| gene9336 | https://www.ncbi.nlm.nih.gov/gene/?term=LOC105282834 | 5993 | 115 | 191.89 | 1 | 1.67 | 0.87 | 14.71 | 15.58 | -0.87 | 0.87 | -0.02 | 0.02 |
| gene8343 | https://www.ncbi.nlm.nih.gov/gene/?term=LOC105281700 | 2172 | 44 | 202.58 | 1 | 4.6 | 2.27 | 37.43 | 38.56 | -1.13 | 1.13 | -0.02 | 0.02 |
| gene5101 | https://www.ncbi.nlm.nih.gov/gene/?term=LOC105277903 | 1813 | 34 | 187.53 | 1 | 5.52 | 2.94 | 52.57 | 53.85 | -1.27 | 1.27 | -0.02 | 0.02 |
| gene5402 | https://www.ncbi.nlm.nih.gov/gene/?term=LOC105278260 | 3068 | 42 | 136.9 | 1 | 3.26 | 2.38 | 12.8 | 14.3 | -1.5 | 1.5 | -0.02 | 0.02 |
| gene7248 | https://www.ncbi.nlm.nih.gov/gene/?term=LOC105280408 | 5175 | 44 | 85.02 | 1 | 1.93 | 2.27 | 21.35 | 20.08 | 1.27 | 1.27 | 0.01 | 0.01 |
| gene657 | https://www.ncbi.nlm.nih.gov/gene/?term=LOC105279667 | 6421 | 78 | 121.48 | 1 | 1.56 | 1.28 | 4.46 | 3.72 | 0.74 | 0.74 | 0.01 | 0.01 |
| gene12203 | https://www.ncbi.nlm.nih.gov/gene/?term=LOC105286099 | 5967 | 78 | 130.72 | 1 | 1.68 | 1.28 | 11.28 | 10.61 | 0.67 | 0.67 | 0.01 | 0.01 |
| gene1147 | https://www.ncbi.nlm.nih.gov/gene/?term=LOC105284954 | 4022 | 87 | 216.31 | 1 | 2.49 | 1.15 | 15.12 | 14.51 | 0.61 | 0.61 | 0.01 | 0.01 |
| gene12021 | https://www.ncbi.nlm.nih.gov/gene/?term=LOC105285905 | 1399 | 21 | 150.11 | 1 | 7.15 | 4.76 | 44.76 | 44.21 | 0.55 | 0.55 | 0.01 | 0.01 |
| gene2683 | https://www.ncbi.nlm.nih.gov/gene/?term=LOC105275187 | 5492 | 75 | 136.56 | 1 | 1.82 | 1.33 | 14.15 | 13.62 | 0.53 | 0.53 | 0.01 | 0.01 |
| gene11583 | https://www.ncbi.nlm.nih.gov/gene/?term=LOC105285414 | 8192 | 179 | 218.51 | 1 | 1.22 | 0.56 | 9.12 | 8.6 | 0.52 | 0.52 | 0.01 | 0.01 |
| gene2748 | https://www.ncbi.nlm.nih.gov/gene/?term=LOC105275261 | 3845 | 54 | 140.44 | 1 | 2.6 | 1.85 | 38.38 | 37.86 | 0.52 | 0.52 | 0.01 | 0.01 |
| gene7801 | https://www.ncbi.nlm.nih.gov/gene/?term=LOC105281047 | 5137 | 113 | 219.97 | 1 | 1.95 | 0.88 | 10.88 | 10.38 | 0.5 | 0.5 | 0.01 | 0.01 |
| gene5524 | https://www.ncbi.nlm.nih.gov/gene/?term=LOC105278397 | 5177 | 201 | 388.26 | 1 | 1.93 | 0.5 | 24.26 | 23.91 | 0.35 | 0.35 | 0.01 | 0.01 |
| gene927 | https://www.ncbi.nlm.nih.gov/gene/?term=LOC105282600 | 3727 | 57 | 152.94 | 1 | 2.68 | 1.75 | 29.17 | 28.82 | 0.35 | 0.35 | 0.01 | 0.01 |
| gene12573 | https://www.ncbi.nlm.nih.gov/gene/?term=LOC105286515 | 3647 | 102 | 279.68 | 1 | 2.74 | 0.98 | 10.87 | 10.55 | 0.32 | 0.32 | 0.01 | 0.01 |
| gene5746 | https://www.ncbi.nlm.nih.gov/gene/?term=LOC105278666 | 10343 | 405 | 391.57 | 1 | 0.97 | 0.25 | 7.03 | 6.73 | 0.3 | 0.3 | 0.01 | 0.01 |
| gene5539 | https://www.ncbi.nlm.nih.gov/gene/?term=LOC105278415 | 25024 | 582 | 232.58 | 1 | 0.4 | 0.17 | 3.85 | 3.62 | 0.23 | 0.23 | 0.01 | 0.01 |
| gene4213 | https://www.ncbi.nlm.nih.gov/gene/?term=LOC105276901 | 3257 | 95 | 291.68 | 1 | 3.07 | 1.05 | 23.09 | 22.87 | 0.22 | 0.22 | 0.01 | 0.01 |
| gene5831 | https://www.ncbi.nlm.nih.gov/gene/?term=LOC105278761 | 2506 | 103 | 411.01 | 2 | 7.98 | 1.94 | 24.82 | 24.61 | 0.21 | 0.21 | 0.01 | 0.01 |
| gene8274 | https://www.ncbi.nlm.nih.gov/gene/?term=LOC105281624 | 5425 | 203 | 374.19 | 1 | 1.84 | 0.49 | 10.66 | 10.47 | 0.2 | 0.2 | 0.01 | 0.01 |
| gene3206 | https://www.ncbi.nlm.nih.gov/gene/?term=LOC105275752 | 3474 | 123 | 354.06 | 1 | 2.88 | 0.81 | 28.92 | 28.73 | 0.19 | 0.19 | 0.01 | 0.01 |
| gene10596 | https://www.ncbi.nlm.nih.gov/gene/?term=LOC105284262 | 6520 | 228 | 349.69 | 1 | 1.53 | 0.44 | 3.34 | 3.5 | -0.16 | 0.16 | -0.01 | 0.01 |
| gene10540 | https://www.ncbi.nlm.nih.gov/gene/?term=LOC105284200 | 5267 | 193 | 366.43 | 1 | 1.9 | 0.52 | 13.99 | 14.21 | -0.22 | 0.22 | -0.01 | 0.01 |
| gene303 | https://www.ncbi.nlm.nih.gov/gene/?term=LOC105275584 | 3081 | 86 | 279.13 | 1 | 3.25 | 1.16 | 23.78 | 24 | -0.22 | 0.22 | -0.01 | 0.01 |
| gene5184 | https://www.ncbi.nlm.nih.gov/gene/?term=LOC105277999 | 5528 | 147 | 265.92 | 1 | 1.81 | 0.68 | 12.4 | 12.64 | -0.24 | 0.24 | -0.01 | 0.01 |
| gene7888 | https://www.ncbi.nlm.nih.gov/gene/?term=LOC105281141 | 4357 | 155 | 355.75 | 1 | 2.3 | 0.65 | 29.83 | 30.06 | -0.24 | 0.24 | -0.01 | 0.01 |
| gene7166 | https://www.ncbi.nlm.nih.gov/gene/?term=LOC105280326 | 6117 | 208 | 340.04 | 1 | 1.63 | 0.48 | 15.97 | 16.24 | -0.27 | 0.27 | -0.01 | 0.01 |
| gene4023 | https://www.ncbi.nlm.nih.gov/gene/?term=LOC105276692 | 1759 | 38 | 216.03 | 1 | 5.69 | 2.63 | 22.43 | 22.73 | -0.3 | 0.3 | -0.01 | 0.01 |
| gene2443 | https://www.ncbi.nlm.nih.gov/gene/?term=LOC105274895 | 10433 | 345 | 330.68 | 1 | 0.96 | 0.29 | 6.76 | 7.09 | -0.33 | 0.33 | -0.01 | 0.01 |
| gene10520 | https://www.ncbi.nlm.nih.gov/gene/?term=LOC105284184 | 2451 | 82 | 334.56 | 1 | 4.08 | 1.22 | 34.55 | 34.9 | -0.35 | 0.35 | -0.01 | 0.01 |
| gene2404 | https://www.ncbi.nlm.nih.gov/gene/?term=LOC105274853 | 1767 | 37 | 209.39 | 1 | 5.66 | 2.7 | 31.99 | 32.37 | -0.37 | 0.37 | -0.01 | 0.01 |
| gene10531 | https://www.ncbi.nlm.nih.gov/gene/?term=LOC105284190 | 6881 | 192 | 279.03 | 1 | 1.45 | 0.52 | 9.5 | 9.91 | -0.4 | 0.4 | -0.01 | 0.01 |
| gene4892 | https://www.ncbi.nlm.nih.gov/gene/?term=LOC105277667 | 3898 | 120 | 307.85 | 1 | 2.57 | 0.83 | 28.07 | 28.49 | -0.42 | 0.42 | -0.01 | 0.01 |
| gene8600 | https://www.ncbi.nlm.nih.gov/gene/?term=LOC105282001 | 8513 | 153 | 179.73 | 1 | 1.17 | 0.65 | 9.37 | 9.87 | -0.5 | 0.5 | -0.01 | 0.01 |
| gene274 | https://www.ncbi.nlm.nih.gov/gene/?term=LOC105275190 | 3488 | 70 | 200.69 | 1 | 2.87 | 1.43 | 19.35 | 20.02 | -0.67 | 0.67 | -0.01 | 0.01 |
| gene10107 | https://www.ncbi.nlm.nih.gov/gene/?term=LOC105283712 | 1883 | 28 | 148.7 | 1 | 5.31 | 3.57 | 28.01 | 28.9 | -0.88 | 0.88 | -0.01 | 0.01 |
| gene7183 | https://www.ncbi.nlm.nih.gov/gene/?term=LOC105280348 | 3925 | 52 | 132.48 | 2 | 5.1 | 3.85 | 25.66 | 26.67 | -1.01 | 1.01 | -0.01 | 0.01 |
| gene10345 | https://www.ncbi.nlm.nih.gov/gene/?term=LOC105283989 | 4444 | 29 | 65.26 | 1 | 2.25 | 3.45 | 48.19 | 50.2 | -2.01 | 2.01 | -0.01 | 0.01 |
| gene564 | https://www.ncbi.nlm.nih.gov/gene/?term=LOC105278478 | 2802 | 13 | 46.4 | 1 | 3.57 | 7.69 | 25.47 | 24.48 | 0.99 | 0.99 | 0 | 0 |
| gene10155 | https://www.ncbi.nlm.nih.gov/gene/?term=LOC105283756 | 7030 | 97 | 137.98 | 1 | 1.42 | 1.03 | 6.83 | 6.6 | 0.23 | 0.23 | 0 | 0 |
| gene6620 | https://www.ncbi.nlm.nih.gov/gene/?term=LOC105279687 | 17998 | 363 | 201.69 | 1 | 0.56 | 0.28 | 5.06 | 4.84 | 0.22 | 0.22 | 0 | 0 |
| gene10600 | https://www.ncbi.nlm.nih.gov/gene/?term=LOC105284270 | 4837 | 103 | 212.94 | 1 | 2.07 | 0.97 | 12.92 | 12.79 | 0.13 | 0.13 | 0 | 0 |
| gene5646 | https://www.ncbi.nlm.nih.gov/gene/?term=LOC105278534 | 7475 | 185 | 247.49 | 1 | 1.34 | 0.54 | 17.64 | 17.56 | 0.08 | 0.08 | 0 | 0 |
| gene5967 | https://www.ncbi.nlm.nih.gov/gene/?term=LOC105278918 | 4393 | 132 | 300.48 | 1 | 2.28 | 0.76 | 15.7 | 15.62 | 0.08 | 0.08 | 0 | 0 |
| gene12802 | https://www.ncbi.nlm.nih.gov/gene/?term=LOC105286799 | 3183 | 170 | 534.09 | 1 | 3.14 | 0.59 | 15.91 | 15.85 | 0.06 | 0.06 | 0 | 0 |
| gene3470 | https://www.ncbi.nlm.nih.gov/gene/?term=LOC105276059 | 8716 | 159 | 182.42 | 1 | 1.15 | 0.63 | 9.24 | 9.2 | 0.04 | 0.04 | 0 | 0 |
| gene2806 | https://www.ncbi.nlm.nih.gov/gene/?term=LOC105275327 | 14257 | 479 | 335.98 | 1 | 0.7 | 0.21 | 7.13 | 7.12 | 0.01 | 0.01 | 0 | 0 |

| | | | | | | | | | | | | | | |
|---|---|---|---|---|---|---|---|---|---|---|---|---|---|---|
| gene5375 | https://www.ncbi.nlm.nih.gov/gene/?term=LOC105278228 | 8640 | 161 | 186.34 | | 1 | 1.16 | 0.62 | 23.31 | 23.3 | 0.01 | 0.01 | 0 | 0 |
| gene2056 | https://www.ncbi.nlm.nih.gov/gene/?term=LOC105274452 | 6584 | 206 | 312.88 | | 1 | 1.52 | 0.49 | 14.18 | 14.18 | 0 | 0 | 0 | 0 |
| gene11579 | https://www.ncbi.nlm.nih.gov/gene/?term=LOC105285406 | 13571 | 282 | 207.8 | 1 | 0.74 | 0.35 | 5.68 | 5.69 | -0.01 | 0.01 | 0 | 0 | |
| gene3107 | https://www.ncbi.nlm.nih.gov/gene/?term=LOC105275650 | 5103 | 150 | 293.94 | | 1 | 1.96 | 0.67 | 17.09 | 17.11 | -0.02 | 0.02 | 0 | 0 |
| gene2173 | https://www.ncbi.nlm.nih.gov/gene/?term=LOC105274590 | 4387 | 79 | 180.08 | | 1 | 2.28 | 1.27 | 22.08 | 22.11 | -0.03 | 0.03 | 0 | 0 |
| gene13114 | https://www.ncbi.nlm.nih.gov/gene/?term=LOC105287147 | 7609 | 381 | 500.72 | | 1 | 1.31 | 0.26 | 9.67 | 9.7 | -0.04 | 0.04 | 0 | 0 |
| gene2668 | https://www.ncbi.nlm.nih.gov/gene/?term=LOC105275170 | 12496 | 336 | 268.89 | | 1 | 0.8 | 0.3 | 2.05 | 2.12 | -0.07 | 0.07 | 0 | 0 |
| gene6698 | https://www.ncbi.nlm.nih.gov/gene/?term=LOC105279778 | 4099 | 138 | 336.67 | | 1 | 2.44 | 0.72 | 9.41 | 9.48 | -0.07 | 0.07 | 0 | 0 |
| gene6783 | https://www.ncbi.nlm.nih.gov/gene/?term=LOC105279865 | 62267 | 1495 | 240.1 | 1 | 0.16 | 0.07 | 1.69 | 1.78 | -0.09 | 0.09 | 0 | 0 | |
| gene53 | https://www.ncbi.nlm.nih.gov/gene/?term=LOC105277148 | 19283 | 270 | 140.02 | | 1 | 0.52 | 0.37 | 3.92 | 4.02 | -0.1 | 0.1 | 0 | 0 |
| gene11568 | https://www.ncbi.nlm.nih.gov/gene/?term=LOC105285395 | 8086 | 176 | 217.66 | | 1 | 1.24 | 0.57 | 15.78 | 15.89 | -0.11 | 0.11 | 0 | 0 |
| gene7398 | https://www.ncbi.nlm.nih.gov/gene/?term=LOC105280589 | 3452 | 41 | 118.77 | | 1 | 2.9 | 2.44 | 25.07 | 25.33 | -0.26 | 0.26 | 0 | 0 |
| gene6528 | https://www.ncbi.nlm.nih.gov/gene/?term=LOC105279591 | 7222 | 67 | 92.77 | 1 | 1.38 | 1.49 | 23.03 | 23.32 | -0.28 | 0.28 | 0 | 0 | |
| gene6455 | https://www.ncbi.nlm.nih.gov/gene/?term=LOC105279500 | 2612 | 32 | 122.51 | | 1 | 3.83 | 3.12 | 49.77 | 50.07 | -0.3 | 0.3 | 0 | 0 |
| gene9411 | https://www.ncbi.nlm.nih.gov/gene/?term=LOC105282914 | 5534 | 41 | 74.09 | 1 | 1.81 | 2.44 | 49.68 | 50.18 | -0.49 | 0.49 | 0 | 0 | |
| gene8178 | https://www.ncbi.nlm.nih.gov/gene/?term=LOC105281515 | 8647 | 44 | 50.88 | 1 | 1.16 | 2.27 | 29.36 | 29.86 | -0.5 | 0.5 | 0 | 0 | |
| gene6639 | https://www.ncbi.nlm.nih.gov/gene/?term=LOC105279705 | 7656 | 68 | 88.82 | 1 | 1.31 | 1.47 | 22.19 | 22.7 | -0.51 | 0.51 | 0 | 0 | |

Directory: OGLall File: rnk-dmg-Cb_LI_BCphaseA1.vs.BCphaseB2.txt

| gene_ID | gene_link | gwidth | #Sites | #per10Kb | #dmSites | #dmsp10kb | %dmSites | %pSite1 | %pSite2 | DMpSite | ADMpSite | DMpNucl | ADMpNucl |
|---|---|---|---|---|---|---|---|---|---|---|---|---|---|
| gene10219 | https://www.ncbi.nlm.nih.gov/gene/?term=LOC105283830 | 235 | 12 | 510.64 | 1 | 42.55 | 8.33 | 42.96 | 33.59 | 9.36 | 9.36 | 0.48 | 0.48 |
| gene5917 | https://www.ncbi.nlm.nih.gov/gene/?term=LOC105278849 | 823 | 81 | 984.2 | 2 | 24.3 | 2.47 | 24.78 | 23.07 | 1.71 | 1.71 | 0.17 | 0.17 |
| gene5809 | https://www.ncbi.nlm.nih.gov/gene/?term=LOC105278737 | 1159 | 83 | 716.13 | 1 | 8.63 | 1.2 | 16.21 | 18.32 | -2.11 | 2.11 | -0.15 | 0.15 |
| gene558 | https://www.ncbi.nlm.nih.gov/gene/?term=LOC105278458 | 216 | 3 | 138.89 | 1 | 46.3 | 33.33 | 20.87 | 10.44 | 10.43 | 10.43 | 0.14 | 0.14 |
| gene8965 | https://www.ncbi.nlm.nih.gov/gene/?term=LOC105282405 | 1310 | 31 | 236.64 | 3 | 22.9 | 9.68 | 51.28 | 45.18 | 6.1 | 6.1 | 0.14 | 0.14 |
| gene1696 | https://www.ncbi.nlm.nih.gov/gene/?term=LOC105287856 | 1917 | 118 | 615.55 | 2 | 10.43 | 1.69 | 12.7 | 14.93 | -2.23 | 2.23 | -0.14 | 0.14 |
| gene10509 | https://www.ncbi.nlm.nih.gov/gene/?term=LOC105284168 | 1350 | 71 | 525.93 | 1 | 7.41 | 1.41 | 22.24 | 24.86 | -2.62 | 2.62 | -0.14 | 0.14 |
| gene1534 | https://www.ncbi.nlm.nih.gov/gene/?term=LOC105287662 | 1711 | 32 | 187.03 | 3 | 17.53 | 9.38 | 32.94 | 39.9 | -6.95 | 6.95 | -0.13 | 0.13 |
| gene9471 | https://www.ncbi.nlm.nih.gov/gene/?term=LOC105282986 | 1605 | 79 | 492.21 | 3 | 18.69 | 3.8 | 31.65 | 29.21 | 2.44 | 2.44 | 0.12 | 0.12 |
| gene4013 | https://www.ncbi.nlm.nih.gov/gene/?term=LOC105276679 | 2007 | 65 | 323.87 | 3 | 14.95 | 4.62 | 39.73 | 36.29 | 3.44 | 3.44 | 0.11 | 0.11 |
| gene952 | https://www.ncbi.nlm.nih.gov/gene/?term=LOC105283016 | 2375 | 111 | 467.37 | 3 | 12.63 | 2.7 | 23.65 | 21.4 | 2.25 | 2.25 | 0.11 | 0.11 |
| gene12842 | https://www.ncbi.nlm.nih.gov/gene/?term=LOC105286845 | 1157 | 60 | 518.58 | 3 | 25.93 | 5 | 17.37 | 19.47 | -2.1 | 2.1 | -0.11 | 0.11 |
| gene12374 | https://www.ncbi.nlm.nih.gov/gene/?term=LOC105286277 | 852 | 27 | 316.9 | 1 | 11.74 | 3.7 | 37.34 | 40.66 | -3.32 | 3.32 | -0.11 | 0.11 |
| gene168 | https://www.ncbi.nlm.nih.gov/gene/?term=LOC105287895 | 2303 | 50 | 217.11 | 1 | 4.34 | 2 | 28.58 | 33.64 | -5.06 | 5.06 | -0.11 | 0.11 |
| gene10721 | https://www.ncbi.nlm.nih.gov/gene/?term=LOC105284403 | 1537 | 24 | 156.15 | 3 | 19.52 | 12.5 | 43.65 | 50.5 | -6.85 | 6.85 | -0.11 | 0.11 |
| gene2824 | https://www.ncbi.nlm.nih.gov/gene/?term=LOC105275337 | 1772 | 57 | 321.67 | 5 | 28.22 | 8.77 | 32.13 | 29.09 | 3.04 | 3.04 | 0.1 | 0.1 |
| gene1802 | https://www.ncbi.nlm.nih.gov/gene/?term=LOC105287981 | 3173 | 135 | 425.46 | 2 | 6.3 | 1.48 | 18.91 | 16.6 | 2.31 | 2.31 | 0.1 | 0.1 |
| gene7568 | https://www.ncbi.nlm.nih.gov/gene/?term=LOC105280786 | 2525 | 110 | 435.64 | 3 | 11.88 | 2.73 | 32.15 | 29.95 | 2.21 | 2.21 | 0.1 | 0.1 |
| gene7401 | https://www.ncbi.nlm.nih.gov/gene/?term=LOC105280591 | 2018 | 91 | 450.94 | 1 | 4.96 | 1.1 | 14.79 | 12.63 | 2.16 | 2.16 | 0.1 | 0.1 |
| gene9781 | https://www.ncbi.nlm.nih.gov/gene/?term=LOC105283351 | 1245 | 60 | 481.93 | 1 | 8.03 | 1.67 | 24.95 | 22.93 | 2.02 | 2.02 | 0.1 | 0.1 |
| gene6057 | https://www.ncbi.nlm.nih.gov/gene/?term=LOC105279035 | 2105 | 146 | 693.59 | 6 | 28.5 | 4.11 | 19.02 | 17.55 | 1.48 | 1.48 | 0.1 | 0.1 |
| gene570 | https://www.ncbi.nlm.nih.gov/gene/?term=LOC105278549 | 1673 | 65 | 388.52 | 3 | 17.93 | 4.62 | 41.78 | 44.3 | -2.52 | 2.52 | -0.1 | 0.1 |
| gene4643 | https://www.ncbi.nlm.nih.gov/gene/?term=LOC109610847 | 429 | 5 | 116.55 | 1 | 23.31 | 20 | 29.86 | 22.36 | 7.5 | 7.5 | 0.09 | 0.09 |
| gene1914 | https://www.ncbi.nlm.nih.gov/gene/?term=LOC105288105 | 1289 | 23 | 178.43 | 2 | 15.52 | 8.7 | 60.89 | 56.04 | 4.85 | 4.85 | 0.09 | 0.09 |

...
... only first 25 lines shown ...

Directory: OGLall File: rnk-dmg-Cb_LI_BCphaseA1.vs.RphaseA1.txt

| gene_ID | gene_link | gwidth | #Sites | #per10Kb | #dmSites | #dmsp10kb | %dmSites | %pSite1 | %pSite2 | DMpSite | ADMpSite | DMpNucl | ADMpNucl |
|---|---|---|---|---|---|---|---|---|---|---|---|---|---|
| gene3165 | https://www.ncbi.nlm.nih.gov/gene/?term=LOC105275706 | 2374 | 59 | 248.53 | 4 | 16.85 | 6.78 | 23.46 | 34.48 | -11.02 | 11.02 | -0.27 | 0.27 |
| gene7199 | https://www.ncbi.nlm.nih.gov/gene/?term=LOC105280366 | 1414 | 26 | 183.88 | 1 | 7.07 | 3.85 | 47.43 | 38.76 | 8.67 | 8.67 | 0.16 | 0.16 |
| gene2593 | https://www.ncbi.nlm.nih.gov/gene/?term=LOC105275070 | 1774 | 6 | 33.82 | 2 | 11.27 | 33.33 | 16.42 | 60.02 | -43.6 | 43.6 | -0.15 | 0.15 |
| gene11819 | https://www.ncbi.nlm.nih.gov/gene/?term=LOC105285671 | 2366 | 90 | 380.39 | 1 | 4.23 | 1.11 | 28.49 | 30.76 | -2.27 | 2.27 | -0.09 | 0.09 |
| gene298 | https://www.ncbi.nlm.nih.gov/gene/?term=LOC105275460 | 2473 | 115 | 465.02 | 1 | 4.04 | 0.87 | 12.68 | 14.4 | -1.73 | 1.73 | -0.08 | 0.08 |
| gene5877 | https://www.ncbi.nlm.nih.gov/gene/?term=LOC105278807 | 1719 | 51 | 296.68 | 1 | 5.82 | 1.96 | 21.68 | 24.24 | -2.55 | 2.55 | -0.08 | 0.08 |
| gene5101 | https://www.ncbi.nlm.nih.gov/gene/?term=LOC105277903 | 1813 | 34 | 187.53 | 1 | 5.52 | 2.94 | 52.57 | 56.73 | -4.16 | 4.16 | -0.08 | 0.08 |
| gene10714 | https://www.ncbi.nlm.nih.gov/gene/?term=LOC105284396 | 1966 | 53 | 269.58 | 1 | 5.09 | 1.89 | 40.95 | 38.4 | 2.55 | 2.55 | 0.07 | 0.07 |
| gene10213 | https://www.ncbi.nlm.nih.gov/gene/?term=LOC105283826 | 1503 | 52 | 345.97 | 1 | 6.65 | 1.92 | 16.76 | 14.79 | 1.97 | 1.97 | 0.07 | 0.07 |
| gene4034 | https://www.ncbi.nlm.nih.gov/gene/?term=LOC105276704 | 2065 | 80 | 387.41 | 1 | 4.84 | 1.25 | 13.38 | 15.07 | -1.69 | 1.69 | -0.07 | 0.07 |
| gene4641 | https://www.ncbi.nlm.nih.gov/gene/?term=LOC105277398 | 1937 | 70 | 361.38 | 1 | 5.16 | 1.43 | 35.36 | 37.39 | -2.03 | 2.03 | -0.07 | 0.07 |
| gene388 | https://www.ncbi.nlm.nih.gov/gene/?term=LOC105276538 | 1558 | 19 | 121.95 | 1 | 6.42 | 5.26 | 69.41 | 64.89 | 4.52 | 4.52 | 0.06 | 0.06 |
| gene7915 | https://www.ncbi.nlm.nih.gov/gene/?term=LOC105281173 | 1597 | 25 | 156.54 | 1 | 6.26 | 4 | 64.87 | 60.99 | 3.88 | 3.88 | 0.06 | 0.06 |
| gene11401 | https://www.ncbi.nlm.nih.gov/gene/?term=LOC105285202 | 1841 | 49 | 266.16 | 1 | 5.43 | 2.04 | 21.54 | 19.38 | 2.16 | 2.16 | 0.06 | 0.06 |
| gene2271 | https://www.ncbi.nlm.nih.gov/gene/?term=LOC105274706 | 1448 | 49 | 338.4 | 1 | 6.91 | 2.04 | 38.87 | 37.01 | 1.86 | 1.86 | 0.06 | 0.06 |
| gene4190 | https://www.ncbi.nlm.nih.gov/gene/?term=LOC105276875 | 3150 | 124 | 393.65 | 1 | 3.17 | 0.81 | 21.46 | 19.89 | 1.58 | 1.58 | 0.06 | 0.06 |
| gene9740 | https://www.ncbi.nlm.nih.gov/gene/?term=LOC105283277 | 2887 | 135 | 467.61 | 1 | 3.46 | 0.74 | 21.88 | 20.52 | 1.37 | 1.37 | 0.06 | 0.06 |
| gene5298 | https://www.ncbi.nlm.nih.gov/gene/?term=LOC105278145 | 4060 | 240 | 591.13 | 2 | 4.93 | 0.83 | 11.5 | 10.44 | 1.06 | 1.06 | 0.06 | 0.06 |
| gene570 | https://www.ncbi.nlm.nih.gov/gene/?term=LOC105278549 | 1673 | 63 | 376.57 | 1 | 5.98 | 1.59 | 41.36 | 42.83 | -1.47 | 1.47 | -0.06 | 0.06 |
| gene6334 | https://www.ncbi.nlm.nih.gov/gene/?term=LOC105279356 | 1471 | 43 | 292.32 | 1 | 6.8 | 2.33 | 28.25 | 30.45 | -2.2 | 2.2 | -0.06 | 0.06 |
| gene12516 | https://www.ncbi.nlm.nih.gov/gene/?term=LOC105286452 | 1529 | 35 | 228.91 | 1 | 6.54 | 2.86 | 51.01 | 53.56 | -2.56 | 2.56 | -0.06 | 0.06 |
| gene393 | https://www.ncbi.nlm.nih.gov/gene/?term=LOC105276566 | 1123 | 2 | 17.81 | 1 | 8.9 | 50 | 77.65 | 50 | 27.65 | 27.65 | 0.05 | 0.05 |
| gene12517 | https://www.ncbi.nlm.nih.gov/gene/?term=LOC105286451 | 1626 | 10 | 61.5 | 1 | 6.15 | 10 | 52.19 | 44.01 | 8.18 | 8.18 | 0.05 | 0.05 |
| gene920 | https://www.ncbi.nlm.nih.gov/gene/?term=LOC105282518 | 1197 | 42 | 350.88 | 1 | 8.35 | 2.38 | 38.27 | 36.85 | 1.43 | 1.43 | 0.05 | 0.05 |
| gene5649 | https://www.ncbi.nlm.nih.gov/gene/?term=LOC105278538 | 2140 | 76 | 355.14 | 1 | 4.67 | 1.32 | 18.24 | 16.87 | 1.37 | 1.37 | 0.05 | 0.05 |
| gene8873 | https://www.ncbi.nlm.nih.gov/gene/?term=LOC105282297 | 2055 | 78 | 379.56 | 1 | 4.87 | 1.28 | 33.62 | 34.98 | -1.37 | 1.37 | -0.05 | 0.05 |
| gene5647 | https://www.ncbi.nlm.nih.gov/gene/?term=LOC105278567 | 3118 | 32 | 102.63 | 1 | 3.21 | 3.12 | 13.07 | 18.25 | -5.18 | 5.18 | -0.05 | 0.05 |
| gene3168 | https://www.ncbi.nlm.nih.gov/gene/?term=LOC105275711 | 6150 | 77 | 125.2 | 1 | 1.63 | 1.3 | 14.51 | 10.99 | 3.51 | 3.51 | 0.04 | 0.04 |
| gene6422 | https://www.ncbi.nlm.nih.gov/gene/?term=LOC105279458 | 2419 | 37 | 152.96 | 1 | 4.13 | 2.7 | 54.1 | 51.26 | 2.83 | 2.83 | 0.04 | 0.04 |
| gene322 | https://www.ncbi.nlm.nih.gov/gene/?term=LOC105275777 | 1606 | 23 | 143.21 | 1 | 6.23 | 4.35 | 45.55 | 43.01 | 2.54 | 2.54 | 0.04 | 0.04 |
| gene11454 | https://www.ncbi.nlm.nih.gov/gene/?term=LOC105285256 | 1906 | 52 | 272.82 | 1 | 5.25 | 1.92 | 42.98 | 41.47 | 1.51 | 1.51 | 0.04 | 0.04 |
| gene10326 | https://www.ncbi.nlm.nih.gov/gene/?term=LOC105283962 | 3042 | 94 | 309.01 | 1 | 3.29 | 1.06 | 12.22 | 10.91 | 1.31 | 1.31 | 0.04 | 0.04 |
| gene6031 | https://www.ncbi.nlm.nih.gov/gene/?term=LOC105279000 | 5691 | 175 | 307.5 | 1 | 1.76 | 0.57 | 14.52 | 13.23 | 1.29 | 1.29 | 0.04 | 0.04 |
| gene7568 | https://www.ncbi.nlm.nih.gov/gene/?term=LOC105280786 | 2525 | 116 | 459.41 | 1 | 3.96 | 0.86 | 28.86 | 28.06 | 0.8 | 0.8 | 0.04 | 0.04 |
| gene2844 | https://www.ncbi.nlm.nih.gov/gene/?term=LOC105275363 | 4330 | 185 | 427.25 | 1 | 2.31 | 0.54 | 14.56 | 15.46 | -0.9 | 0.9 | -0.04 | 0.04 |
| gene1855 | https://www.ncbi.nlm.nih.gov/gene/?term=LOC105288036 | 2387 | 86 | 360.28 | 1 | 4.19 | 1.16 | 28.48 | 29.73 | -1.24 | 1.24 | -0.04 | 0.04 |
| gene12021 | https://www.ncbi.nlm.nih.gov/gene/?term=LOC105285905 | 1399 | 23 | 164.4 | 1 | 7.15 | 4.35 | 41.24 | 39.35 | 1.89 | 1.89 | 0.03 | 0.03 |
| gene2752 | https://www.ncbi.nlm.nih.gov/gene/?term=LOC105275266 | 2111 | 49 | 232.12 | 1 | 4.74 | 2.04 | 23.87 | 22.43 | 1.44 | 1.44 | 0.03 | 0.03 |
| gene10844 | https://www.ncbi.nlm.nih.gov/gene/?term=LOC105284535 | 9086 | 218 | 239.93 | 1 | 1.1 | 0.46 | 16.62 | 15.4 | 1.22 | 1.22 | 0.03 | 0.03 |
| gene4376 | https://www.ncbi.nlm.nih.gov/gene/?term=LOC105277083 | 4307 | 90 | 208.96 | 1 | 2.32 | 1.11 | 26.1 | 24.89 | 1.21 | 1.21 | 0.03 | 0.03 |
| gene4962 | https://www.ncbi.nlm.nih.gov/gene/?term=LOC105277761 | 4645 | 147 | 316.47 | 1 | 2.15 | 0.68 | 13.9 | 13.02 | 0.88 | 0.88 | 0.03 | 0.03 |
| gene6966 | https://www.ncbi.nlm.nih.gov/gene/?term=LOC105280104 | 3747 | 119 | 317.59 | 1 | 2.67 | 0.84 | 15.37 | 14.52 | 0.85 | 0.85 | 0.03 | 0.03 |
| gene7550 | https://www.ncbi.nlm.nih.gov/gene/?term=LOC105280763 | 2173 | 93 | 427.98 | 1 | 4.6 | 1.08 | 27 | 26.23 | 0.77 | 0.77 | 0.03 | 0.03 |
| gene7249 | https://www.ncbi.nlm.nih.gov/gene/?term=LOC105280422 | 1775 | 61 | 343.66 | 1 | 5.63 | 1.64 | 15.16 | 14.42 | 0.75 | 0.75 | 0.03 | 0.03 |
| gene10606 | https://www.ncbi.nlm.nih.gov/gene/?term=LOC105284273 | 6119 | 243 | 397.12 | 2 | 3.27 | 0.82 | 20.74 | 20.03 | 0.71 | 0.71 | 0.03 | 0.03 |
| gene6286 | https://www.ncbi.nlm.nih.gov/gene/?term=LOC105279298 | 2861 | 136 | 475.36 | 1 | 3.5 | 0.74 | 22.78 | 22.14 | 0.64 | 0.64 | 0.03 | 0.03 |
| gene1470 | https://www.ncbi.nlm.nih.gov/gene/?term=LOC105287597 | 2974 | 181 | 608.61 | 1 | 3.36 | 0.55 | 17.88 | 17.42 | 0.47 | 0.47 | 0.03 | 0.03 |
| gene274 | https://www.ncbi.nlm.nih.gov/gene/?term=LOC105275190 | 3488 | 65 | 186.35 | 1 | 2.87 | 1.54 | 21.06 | 22.74 | -1.69 | 1.69 | -0.03 | 0.03 |
| gene1884 | https://www.ncbi.nlm.nih.gov/gene/?term=LOC105288072 | 4169 | 79 | 189.49 | 1 | 2.4 | 1.27 | 22.97 | 24.67 | -1.7 | 1.7 | -0.03 | 0.03 |
| gene2676 | https://www.ncbi.nlm.nih.gov/gene/?term=LOC105275181 | 1353 | 14 | 103.47 | 1 | 7.39 | 7.14 | 65.95 | 69.25 | -3.3 | 3.3 | -0.03 | 0.03 |
| gene7368 | https://www.ncbi.nlm.nih.gov/gene/?term=LOC105280556 | 4105 | 46 | 112.06 | 1 | 2.44 | 2.17 | 13.39 | 11.38 | 2 | 2 | 0.02 | 0.02 |
| gene2748 | https://www.ncbi.nlm.nih.gov/gene/?term=LOC105275261 | 3845 | 55 | 143.04 | 1 | 2.6 | 1.82 | 38.68 | 37.31 | 1.36 | 1.36 | 0.02 | 0.02 |
| gene3599 | https://www.ncbi.nlm.nih.gov/gene/?term=LOC105276196 | 2955 | 55 | 186.13 | 1 | 3.38 | 1.82 | 40.49 | 39.41 | 1.07 | 1.07 | 0.02 | 0.02 |
| gene10392 | https://www.ncbi.nlm.nih.gov/gene/?term=LOC105284036 | 3216 | 83 | 258.08 | 1 | 3.11 | 1.2 | 24.77 | 23.81 | 0.96 | 0.96 | 0.02 | 0.02 |
| gene3107 | https://www.ncbi.nlm.nih.gov/gene/?term=LOC105275650 | 5103 | 145 | 284.15 | 1 | 1.96 | 0.69 | 17.36 | 16.74 | 0.62 | 0.62 | 0.02 | 0.02 |
| gene11340 | https://www.ncbi.nlm.nih.gov/gene/?term=LOC105285131 | 2918 | 97 | 332.42 | 1 | 3.43 | 1.03 | 14.71 | 14.15 | 0.56 | 0.56 | 0.02 | 0.02 |
| gene3206 | https://www.ncbi.nlm.nih.gov/gene/?term=LOC105275752 | 3474 | 126 | 362.69 | 1 | 2.88 | 0.79 | 28.38 | 27.85 | 0.53 | 0.53 | 0.02 | 0.02 |
| gene5746 | https://www.ncbi.nlm.nih.gov/gene/?term=LOC105278666 | 10343 | 393 | 379.97 | 1 | 0.97 | 0.25 | 7.69 | 7.22 | 0.47 | 0.47 | 0.02 | 0.02 |
| gene8729 | https://www.ncbi.nlm.nih.gov/gene/?term=LOC105282137 | 5005 | 200 | 399.6 | 1 | 2 | 0.5 | 15.61 | 15.22 | 0.39 | 0.39 | 0.02 | 0.02 |
| gene13114 | https://www.ncbi.nlm.nih.gov/gene/?term=LOC105287147 | 7609 | 370 | 486.27 | 1 | 1.31 | 0.27 | 9.54 | 9.22 | 0.32 | 0.32 | 0.02 | 0.02 |

| | | | | | | | | | | | | | |
|---|---|---|---|---|---|---|---|---|---|---|---|---|---|
| gene1332 | https://www.ncbi.nlm.nih.gov/gene/?term=LOC105286872 | 2460 | 142 | 577.24 | 1 | 4.07 | 0.7 | 21.72 | 21.98 | -0.26 | 0.26 | -0.02 | 0.02 |
| gene6583 | https://www.ncbi.nlm.nih.gov/gene/?term=LOC105279649 | 7444 | 308 | 413.76 | 1 | 1.34 | 0.32 | 11.09 | 11.46 | -0.37 | 0.37 | -0.02 | 0.02 |
| gene10018 | https://www.ncbi.nlm.nih.gov/gene/?term=LOC105283607 | 4414 | 194 | 439.51 | 1 | 2.27 | 0.52 | 15.27 | 15.67 | -0.4 | 0.4 | -0.02 | 0.02 |
| gene11275 | https://www.ncbi.nlm.nih.gov/gene/?term=LOC105285055 | 4119 | 202 | 490.41 | 1 | 2.43 | 0.5 | 12.81 | 13.23 | -0.42 | 0.42 | -0.02 | 0.02 |
| gene12978 | https://www.ncbi.nlm.nih.gov/gene/?term=LOC105286997 | 2858 | 134 | 468.86 | 1 | 3.5 | 0.75 | 11.8 | 12.24 | -0.44 | 0.44 | -0.02 | 0.02 |
| gene10540 | https://www.ncbi.nlm.nih.gov/gene/?term=LOC105284200 | 5267 | 195 | 370.23 | 1 | 1.9 | 0.51 | 13.62 | 14.09 | -0.47 | 0.47 | -0.02 | 0.02 |
| gene10520 | https://www.ncbi.nlm.nih.gov/gene/?term=LOC105284184 | 2451 | 80 | 326.4 | 1 | 4.08 | 1.25 | 34.85 | 35.33 | -0.48 | 0.48 | -0.02 | 0.02 |
| gene9478 | https://www.ncbi.nlm.nih.gov/gene/?term=LOC105282991 | 3726 | 48 | 128.82 | 1 | 2.68 | 2.08 | 31.5 | 33.03 | -1.53 | 1.53 | -0.02 | 0.02 |
| gene7248 | https://www.ncbi.nlm.nih.gov/gene/?term=LOC105280408 | 5175 | 44 | 85.02 | 1 | 1.93 | 2.27 | 22.41 | 21.41 | 0.99 | 0.99 | 0.01 | 0.01 |
| gene6806 | https://www.ncbi.nlm.nih.gov/gene/?term=LOC105279894 | 3649 | 39 | 106.88 | 1 | 2.74 | 2.56 | 35.25 | 34.29 | 0.97 | 0.97 | 0.01 | 0.01 |
| gene12203 | https://www.ncbi.nlm.nih.gov/gene/?term=LOC105286099 | 5967 | 69 | 115.64 | 1 | 1.68 | 1.45 | 11.63 | 10.68 | 0.96 | 0.96 | 0.01 | 0.01 |
| gene12307 | https://www.ncbi.nlm.nih.gov/gene/?term=LOC105286216 | 10313 | 145 | 140.6 | 1 | 0.97 | 0.69 | 17.77 | 16.9 | 0.87 | 0.87 | 0.01 | 0.01 |
| gene6639 | https://www.ncbi.nlm.nih.gov/gene/?term=LOC105279705 | 7656 | 70 | 91.43 | 1 | 1.31 | 1.43 | 22.25 | 21.42 | 0.84 | 0.84 | 0.01 | 0.01 |
| gene10600 | https://www.ncbi.nlm.nih.gov/gene/?term=LOC105284270 | 4837 | 95 | 196.4 | 1 | 2.07 | 1.05 | 13.92 | 13.23 | 0.69 | 0.69 | 0.01 | 0.01 |
| gene2404 | https://www.ncbi.nlm.nih.gov/gene/?term=LOC105274853 | 1767 | 39 | 220.71 | 1 | 5.66 | 2.56 | 31.07 | 30.51 | 0.56 | 0.56 | 0.01 | 0.01 |
| gene10155 | https://www.ncbi.nlm.nih.gov/gene/?term=LOC105283756 | 7030 | 93 | 132.29 | 1 | 1.42 | 1.08 | 7.03 | 6.49 | 0.54 | 0.54 | 0.01 | 0.01 |
| gene657 | https://www.ncbi.nlm.nih.gov/gene/?term=LOC105279667 | 6421 | 77 | 119.92 | 1 | 1.56 | 1.3 | 5.63 | 5.1 | 0.53 | 0.53 | 0.01 | 0.01 |
| gene7801 | https://www.ncbi.nlm.nih.gov/gene/?term=LOC105281047 | 5137 | 109 | 212.19 | 1 | 1.95 | 0.92 | 11.18 | 10.67 | 0.5 | 0.5 | 0.01 | 0.01 |
| gene7398 | https://www.ncbi.nlm.nih.gov/gene/?term=LOC105280589 | 3452 | 40 | 115.87 | 1 | 2.9 | 2.5 | 25.2 | 24.72 | 0.48 | 0.48 | 0.01 | 0.01 |
| gene940 | https://www.ncbi.nlm.nih.gov/gene/?term=LOC105282794 | 3578 | 84 | 234.77 | 1 | 2.79 | 1.19 | 14.53 | 14.06 | 0.47 | 0.47 | 0.01 | 0.01 |
| gene5524 | https://www.ncbi.nlm.nih.gov/gene/?term=LOC105278397 | 5177 | 199 | 384.39 | 1 | 1.93 | 0.5 | 24.51 | 24.16 | 0.35 | 0.35 | 0.01 | 0.01 |
| gene5831 | https://www.ncbi.nlm.nih.gov/gene/?term=LOC105278761 | 2506 | 99 | 395.05 | 1 | 3.99 | 1.01 | 25.53 | 25.18 | 0.35 | 0.35 | 0.01 | 0.01 |
| gene7600 | https://www.ncbi.nlm.nih.gov/gene/?term=LOC105280819 | 1615 | 41 | 253.87 | 1 | 6.19 | 2.44 | 40.74 | 40.39 | 0.35 | 0.35 | 0.01 | 0.01 |
| gene5539 | https://www.ncbi.nlm.nih.gov/gene/?term=LOC105278415 | 25024 | 562 | 224.58 | 1 | 0.4 | 0.18 | 4.11 | 3.79 | 0.31 | 0.31 | 0.01 | 0.01 |
| gene1147 | https://www.ncbi.nlm.nih.gov/gene/?term=LOC105284954 | 4022 | 87 | 216.31 | 1 | 2.49 | 1.15 | 14.67 | 14.36 | 0.3 | 0.3 | 0.01 | 0.01 |
| gene2173 | https://www.ncbi.nlm.nih.gov/gene/?term=LOC105274590 | 4387 | 78 | 177.8 | 1 | 2.28 | 1.28 | 22.31 | 22.01 | 0.3 | 0.3 | 0.01 | 0.01 |
| gene9101 | https://www.ncbi.nlm.nih.gov/gene/?term=LOC105282553 | 3068 | 103 | 335.72 | 1 | 3.26 | 0.97 | 20.28 | 19.99 | 0.29 | 0.29 | 0.01 | 0.01 |
| gene13036 | https://www.ncbi.nlm.nih.gov/gene/?term=LOC105287059 | 4959 | 157 | 316.6 | 1 | 2.02 | 0.64 | 17.83 | 17.59 | 0.25 | 0.25 | 0.01 | 0.01 |
| gene10549 | https://www.ncbi.nlm.nih.gov/gene/?term=LOC105284210 | 7145 | 313 | 438.07 | 1 | 1.4 | 0.32 | 10.77 | 10.55 | 0.22 | 0.22 | 0.01 | 0.01 |
| gene8274 | https://www.ncbi.nlm.nih.gov/gene/?term=LOC105281624 | 5425 | 189 | 348.39 | 1 | 1.84 | 0.53 | 11.18 | 10.97 | 0.21 | 0.21 | 0.01 | 0.01 |
| gene7547 | https://www.ncbi.nlm.nih.gov/gene/?term=LOC105280767 | 24576 | 683 | 277.91 | 1 | 0.41 | 0.15 | 4.82 | 4.61 | 0.2 | 0.2 | 0.01 | 0.01 |
| gene12802 | https://www.ncbi.nlm.nih.gov/gene/?term=LOC105286799 | 3183 | 170 | 534.09 | 1 | 3.14 | 0.59 | 15.96 | 15.78 | 0.18 | 0.18 | 0.01 | 0.01 |
| gene5525 | https://www.ncbi.nlm.nih.gov/gene/?term=LOC105278396 | 3090 | 185 | 598.71 | 1 | 3.24 | 0.54 | 24.64 | 24.46 | 0.18 | 0.18 | 0.01 | 0.01 |
| gene2443 | https://www.ncbi.nlm.nih.gov/gene/?term=LOC105274895 | 10433 | 330 | 316.3 | 1 | 0.96 | 0.3 | 5.47 | 5.65 | -0.18 | 0.18 | -0.01 | 0.01 |
| gene4892 | https://www.ncbi.nlm.nih.gov/gene/?term=LOC105277667 | 3898 | 116 | 297.59 | 1 | 2.57 | 0.86 | 28.81 | 28.99 | -0.19 | 0.19 | -0.01 | 0.01 |
| gene3335 | https://www.ncbi.nlm.nih.gov/gene/?term=LOC105275898 | 4665 | 128 | 274.38 | 1 | 2.14 | 0.78 | 13.41 | 13.77 | -0.36 | 0.36 | -0.01 | 0.01 |
| gene3856 | https://www.ncbi.nlm.nih.gov/gene/?term=LOC105276499 | 5236 | 188 | 359.05 | 1 | 1.91 | 0.53 | 10.14 | 10.5 | -0.36 | 0.36 | -0.01 | 0.01 |
| gene10243 | https://www.ncbi.nlm.nih.gov/gene/?term=LOC105283862 | 3239 | 38 | 117.32 | 1 | 3.09 | 2.63 | 30.66 | 31.1 | -0.44 | 0.44 | -0.01 | 0.01 |
| gene8600 | https://www.ncbi.nlm.nih.gov/gene/?term=LOC105282001 | 8513 | 148 | 173.85 | 1 | 1.17 | 0.68 | 9.45 | 9.9 | -0.45 | 0.45 | -0.01 | 0.01 |
| gene13033 | https://www.ncbi.nlm.nih.gov/gene/?term=LOC105287063 | 1877 | 45 | 239.74 | 1 | 5.33 | 2.22 | 34.91 | 35.5 | -0.59 | 0.59 | -0.01 | 0.01 |
| gene4023 | https://www.ncbi.nlm.nih.gov/gene/?term=LOC105276692 | 1759 | 36 | 204.66 | 1 | 5.69 | 2.78 | 23.26 | 23.9 | -0.64 | 0.64 | -0.01 | 0.01 |
| gene8343 | https://www.ncbi.nlm.nih.gov/gene/?term=LOC105281700 | 2172 | 45 | 207.18 | 1 | 4.6 | 2.22 | 38.72 | 39.37 | -0.65 | 0.65 | -0.01 | 0.01 |
| gene6528 | https://www.ncbi.nlm.nih.gov/gene/?term=LOC105279591 | 7222 | 68 | 94.16 | 1 | 1.38 | 1.47 | 22.7 | 23.89 | -1.19 | 1.19 | -0.01 | 0.01 |
| gene5052 | https://www.ncbi.nlm.nih.gov/gene/?term=LOC105277856 | 3167 | 27 | 85.25 | 1 | 3.16 | 3.7 | 36.57 | 38.1 | -1.52 | 1.52 | -0.01 | 0.01 |
| gene10345 | https://www.ncbi.nlm.nih.gov/gene/?term=LOC105283989 | 4444 | 27 | 60.76 | 1 | 2.25 | 3.7 | 48.21 | 50.36 | -2.14 | 2.14 | -0.01 | 0.01 |
| gene1220 | https://www.ncbi.nlm.nih.gov/gene/?term=LOC105285742 | 3453 | 86 | 249.06 | 1 | 2.9 | 1.16 | 21.68 | 21.51 | 0.17 | 0.17 | 0 | 0 |
| gene7247 | https://www.ncbi.nlm.nih.gov/gene/?term=LOC105280407 | 10486 | 206 | 196.45 | 1 | 0.95 | 0.49 | 17.14 | 16.97 | 0.17 | 0.17 | 0 | 0 |
| gene4213 | https://www.ncbi.nlm.nih.gov/gene/?term=LOC105276901 | 3257 | 96 | 294.75 | 1 | 3.07 | 1.04 | 22.87 | 22.72 | 0.14 | 0.14 | 0 | 0 |
| gene6805 | https://www.ncbi.nlm.nih.gov/gene/?term=LOC105279895 | 87888 | 975 | 110.94 | 1 | 0.11 | 0.1 | 1.92 | 1.78 | 0.14 | 0.14 | 0 | 0 |
| gene10107 | https://www.ncbi.nlm.nih.gov/gene/?term=LOC105283712 | 1883 | 29 | 154.01 | 1 | 5.31 | 3.45 | 27.22 | 27.09 | 0.13 | 0.13 | 0 | 0 |
| gene9645 | https://www.ncbi.nlm.nih.gov/gene/?term=LOC105283181 | 3875 | 134 | 345.81 | 1 | 2.58 | 0.75 | 11.45 | 11.37 | 0.08 | 0.08 | 0 | 0 |
| gene6620 | https://www.ncbi.nlm.nih.gov/gene/?term=LOC105279687 | 17998 | 354 | 196.69 | 1 | 0.56 | 0.28 | 5.08 | 5.05 | 0.03 | 0.03 | 0 | 0 |
| gene11583 | https://www.ncbi.nlm.nih.gov/gene/?term=LOC105285414 | 8192 | 175 | 213.62 | 1 | 1.22 | 0.57 | 9.21 | 9.24 | -0.03 | 0.03 | 0 | 0 |
| gene2806 | https://www.ncbi.nlm.nih.gov/gene/?term=LOC105275327 | 14257 | 462 | 324.05 | 1 | 0.7 | 0.22 | 7.37 | 7.4 | -0.03 | 0.03 | 0 | 0 |
| gene11579 | https://www.ncbi.nlm.nih.gov/gene/?term=LOC105285406 | 13571 | 260 | 191.58 | 1 | 0.74 | 0.38 | 5.75 | 5.79 | -0.04 | 0.04 | 0 | 0 |
| gene7166 | https://www.ncbi.nlm.nih.gov/gene/?term=LOC105280326 | 6117 | 208 | 340.04 | 1 | 1.63 | 0.48 | 16.25 | 16.28 | -0.04 | 0.04 | 0 | 0 |
| gene12406 | https://www.ncbi.nlm.nih.gov/gene/?term=LOC105286316 | 1386 | 30 | 216.45 | 1 | 7.22 | 3.33 | 39.2 | 39.29 | -0.09 | 0.09 | 0 | 0 |
| gene5375 | https://www.ncbi.nlm.nih.gov/gene/?term=LOC105278228 | 8640 | 148 | 171.3 | 1 | 1.16 | 0.68 | 24.78 | 24.89 | -0.11 | 0.11 | 0 | 0 |
| gene12573 | https://www.ncbi.nlm.nih.gov/gene/?term=LOC105286515 | 3647 | 105 | 287.91 | 1 | 2.74 | 0.95 | 10.51 | 10.65 | -0.14 | 0.14 | 0 | 0 |
| gene1310 | https://www.ncbi.nlm.nih.gov/gene/?term=LOC105286661 | 2582 | 63 | 244 | 1 | 3.87 | 1.59 | 19.4 | 19.54 | -0.14 | 0.14 | 0 | 0 |
| gene10531 | https://www.ncbi.nlm.nih.gov/gene/?term=LOC105284190 | 6881 | 180 | 261.59 | 1 | 1.45 | 0.56 | 10.06 | 10.22 | -0.16 | 0.16 | 0 | 0 |
| gene1632 | https://www.ncbi.nlm.nih.gov/gene/?term=LOC105287764 | 18061 | 334 | 184.93 | 1 | 0.55 | 0.3 | 2.71 | 2.98 | -0.27 | 0.27 | 0 | 0 |
| gene7183 | https://www.ncbi.nlm.nih.gov/gene/?term=LOC105280348 | 3925 | 53 | 135.03 | 1 | 2.55 | 1.89 | 25.26 | 25.53 | -0.27 | 0.27 | 0 | 0 |
| gene6455 | https://www.ncbi.nlm.nih.gov/gene/?term=LOC105279500 | 2612 | 29 | 111.03 | 1 | 3.83 | 3.45 | 48.01 | 48.34 | -0.33 | 0.33 | 0 | 0 |

| | | | | | | | | | | | | | |
|---|---|---|---|---|---|---|---|---|---|---|---|---|---|
| gene564 | https://www.ncbi.nlm.nih.gov/gene/?term=LOC105278478 | 2802 | 12 | 42.83 | 1 | 3.57 | 8.33 | 20.51 | 21.62 | -1.11 | 1.11 | 0 | 0 |

Directory: OGLall File: rnk-dmg-Cb_LI_BCphaseA1.vs.RphaseA2.txt

| gene_ID | gene_link | gwidth | #Sites | #per10Kb | #dmSites | #dmsp10kb | %dmSites | %pSite1 | %pSite2 | DMpSite | ADMpSite | DMpNucl | ADMpNucl |
|---|---|---|---|---|---|---|---|---|---|---|---|---|---|
| gene3165 | https://www.ncbi.nlm.nih.gov/gene/?term=LOC105275706 | 2374 | 67 | 282.22 | 4 | 16.85 | 5.97 | 25.11 | 35.96 | -10.84 | 10.84 | -0.31 | 0.31 |
| gene2593 | https://www.ncbi.nlm.nih.gov/gene/?term=LOC105275070 | 1774 | 6 | 33.82 | 3 | 16.91 | 50 | 16.42 | 57.88 | -41.46 | 41.46 | -0.14 | 0.14 |
| gene3200 | https://www.ncbi.nlm.nih.gov/gene/?term=LOC105275743 | 1786 | 77 | 431.13 | 1 | 5.6 | 1.3 | 22.67 | 19.88 | 2.79 | 2.79 | 0.12 | 0.12 |
| gene4034 | https://www.ncbi.nlm.nih.gov/gene/?term=LOC105276704 | 2065 | 87 | 421.31 | 2 | 9.69 | 2.3 | 12.36 | 15.2 | -2.84 | 2.84 | -0.12 | 0.12 |
| gene12961 | https://www.ncbi.nlm.nih.gov/gene/?term=LOC105286983 | 3094 | 70 | 226.24 | 1 | 3.23 | 1.43 | 0.98 | 5.77 | -4.79 | 4.79 | -0.11 | 0.11 |
| gene5714 | https://www.ncbi.nlm.nih.gov/gene/?term=LOC105278729 | 710 | 14 | 197.18 | 1 | 14.08 | 7.14 | 51.67 | 57.2 | -5.53 | 5.53 | -0.11 | 0.11 |
| gene12558 | https://www.ncbi.nlm.nih.gov/gene/?term=LOC105286502 | 1285 | 43 | 334.63 | 1 | 7.78 | 2.33 | 24.55 | 27.41 | -2.86 | 2.86 | -0.1 | 0.1 |
| gene920 | https://www.ncbi.nlm.nih.gov/gene/?term=LOC105282518 | 1197 | 43 | 359.23 | 1 | 8.35 | 2.33 | 37.5 | 34.97 | 2.52 | 2.52 | 0.09 | 0.09 |
| gene5525 | https://www.ncbi.nlm.nih.gov/gene/?term=LOC105278396 | 3090 | 186 | 601.94 | 1 | 3.24 | 0.54 | 24.51 | 25.99 | -1.48 | 1.48 | -0.09 | 0.09 |
| gene274 | https://www.ncbi.nlm.nih.gov/gene/?term=LOC105275190 | 3488 | 73 | 209.29 | 1 | 2.87 | 1.37 | 18.68 | 22.58 | -3.9 | 3.9 | -0.08 | 0.08 |
| gene6814 | https://www.ncbi.nlm.nih.gov/gene/?term=LOC105279898 | 1343 | 15 | 111.69 | 1 | 7.45 | 6.67 | 59.84 | 67.04 | -7.2 | 7.2 | -0.08 | 0.08 |
| gene10213 | https://www.ncbi.nlm.nih.gov/gene/?term=LOC105283826 | 1503 | 54 | 359.28 | 1 | 6.65 | 1.85 | 16.14 | 14.22 | 1.92 | 1.92 | 0.07 | 0.07 |
| gene11819 | https://www.ncbi.nlm.nih.gov/gene/?term=LOC105285671 | 2366 | 95 | 401.52 | 1 | 4.23 | 1.05 | 27.1 | 28.85 | -1.76 | 1.76 | -0.07 | 0.07 |
| gene12317 | https://www.ncbi.nlm.nih.gov/gene/?term=LOC105286223 | 2235 | 84 | 375.84 | 1 | 4.47 | 1.19 | 29.87 | 31.79 | -1.93 | 1.93 | -0.07 | 0.07 |
| gene12406 | https://www.ncbi.nlm.nih.gov/gene/?term=LOC105286316 | 1386 | 34 | 245.31 | 1 | 7.22 | 2.94 | 39.04 | 41.82 | -2.78 | 2.78 | -0.07 | 0.07 |
| gene12516 | https://www.ncbi.nlm.nih.gov/gene/?term=LOC105286452 | 1529 | 35 | 228.91 | 1 | 6.54 | 2.86 | 51.01 | 54.04 | -3.03 | 3.03 | -0.07 | 0.07 |
| gene5298 | https://www.ncbi.nlm.nih.gov/gene/?term=LOC105278145 | 4060 | 257 | 633 | 2 | 4.93 | 0.78 | 10.97 | 10.08 | 0.88 | 0.88 | 0.06 | 0.06 |
| gene298 | https://www.ncbi.nlm.nih.gov/gene/?term=LOC105275460 | 2473 | 112 | 452.89 | 1 | 4.04 | 0.89 | 12.88 | 14.18 | -1.3 | 1.3 | -0.06 | 0.06 |
| gene10984 | https://www.ncbi.nlm.nih.gov/gene/?term=LOC105284704 | 1987 | 64 | 322.09 | 1 | 5.03 | 1.56 | 25.47 | 27.27 | -1.8 | 1.8 | -0.06 | 0.06 |
| gene393 | https://www.ncbi.nlm.nih.gov/gene/?term=LOC105276566 | 1123 | 2 | 17.81 | 1 | 8.9 | 50 | 77.65 | 47.62 | 30.03 | 30.03 | 0.05 | 0.05 |
| gene7952 | https://www.ncbi.nlm.nih.gov/gene/?term=LOC105281213 | 2963 | 43 | 145.12 | 1 | 3.37 | 2.33 | 42.11 | 38.78 | 3.33 | 3.33 | 0.05 | 0.05 |
| gene4190 | https://www.ncbi.nlm.nih.gov/gene/?term=LOC105276875 | 3150 | 132 | 419.05 | 1 | 3.17 | 0.76 | 21.24 | 20.15 | 1.09 | 1.09 | 0.05 | 0.05 |
| gene5524 | https://www.ncbi.nlm.nih.gov/gene/?term=LOC105278397 | 5177 | 200 | 386.32 | 1 | 1.93 | 0.5 | 24.39 | 25.66 | -1.27 | 1.27 | -0.05 | 0.05 |
| gene6334 | https://www.ncbi.nlm.nih.gov/gene/?term=LOC105279356 | 1471 | 43 | 292.32 | 1 | 6.8 | 2.33 | 28.25 | 29.87 | -1.62 | 1.62 | -0.05 | 0.05 |
| gene10107 | https://www.ncbi.nlm.nih.gov/gene/?term=LOC105283712 | 1883 | 27 | 143.39 | 1 | 5.31 | 3.7 | 26.21 | 29.43 | -3.23 | 3.23 | -0.05 | 0.05 |
| gene5052 | https://www.ncbi.nlm.nih.gov/gene/?term=LOC105277856 | 3167 | 28 | 88.41 | 1 | 3.16 | 3.57 | 35.73 | 41.1 | -5.37 | 5.37 | -0.05 | 0.05 |
| gene4630 | https://www.ncbi.nlm.nih.gov/gene/?term=LOC105277380 | 1893 | 5 | 26.41 | 1 | 5.28 | 20 | 69.42 | 53.65 | 15.77 | 15.77 | 0.04 | 0.04 |
| gene7915 | https://www.ncbi.nlm.nih.gov/gene/?term=LOC105281173 | 1597 | 25 | 156.54 | 1 | 6.26 | 4 | 64.87 | 62.52 | 2.35 | 2.35 | 0.04 | 0.04 |
| gene11454 | https://www.ncbi.nlm.nih.gov/gene/?term=LOC105285256 | 1906 | 51 | 267.58 | 1 | 5.25 | 1.96 | 43.83 | 42.42 | 1.4 | 1.4 | 0.04 | 0.04 |
| gene10326 | https://www.ncbi.nlm.nih.gov/gene/?term=LOC105283962 | 3042 | 102 | 335.31 | 1 | 3.29 | 0.98 | 14.13 | 13.04 | 1.09 | 1.09 | 0.04 | 0.04 |
| gene10540 | https://www.ncbi.nlm.nih.gov/gene/?term=LOC105284200 | 5267 | 199 | 377.82 | 1 | 1.9 | 0.5 | 13.8 | 14.75 | -0.95 | 0.95 | -0.04 | 0.04 |
| gene5877 | https://www.ncbi.nlm.nih.gov/gene/?term=LOC105278807 | 1719 | 52 | 302.5 | 1 | 5.82 | 1.92 | 22.91 | 24.37 | -1.46 | 1.46 | -0.04 | 0.04 |
| gene1310 | https://www.ncbi.nlm.nih.gov/gene/?term=LOC105286661 | 2582 | 65 | 251.74 | 1 | 3.87 | 1.54 | 19.96 | 21.59 | -1.63 | 1.63 | -0.04 | 0.04 |
| gene4023 | https://www.ncbi.nlm.nih.gov/gene/?term=LOC105276692 | 1759 | 39 | 221.72 | 1 | 5.69 | 2.56 | 19.85 | 21.5 | -1.65 | 1.65 | -0.04 | 0.04 |
| gene3727 | https://www.ncbi.nlm.nih.gov/gene/?term=LOC105276367 | 2484 | 54 | 217.39 | 1 | 4.03 | 1.85 | 48.38 | 50.37 | -1.99 | 1.99 | -0.04 | 0.04 |
| gene12021 | https://www.ncbi.nlm.nih.gov/gene/?term=LOC105285905 | 1399 | 23 | 164.4 | 1 | 7.15 | 4.35 | 41.24 | 43.7 | -2.46 | 2.46 | -0.04 | 0.04 |
| gene2335 | https://www.ncbi.nlm.nih.gov/gene/?term=LOC105274778 | 3945 | 38 | 96.32 | 1 | 2.53 | 2.63 | 45.06 | 48.85 | -3.79 | 3.79 | -0.04 | 0.04 |
| gene9109 | https://www.ncbi.nlm.nih.gov/gene/?term=LOC105282561 | 2400 | 15 | 62.5 | 1 | 4.17 | 6.67 | 34.05 | 29.4 | 4.65 | 4.65 | 0.03 | 0.03 |
| gene7368 | https://www.ncbi.nlm.nih.gov/gene/?term=LOC105280556 | 4105 | 49 | 119.37 | 1 | 2.44 | 2.04 | 12.57 | 9.91 | 2.66 | 2.66 | 0.03 | 0.03 |
| gene3168 | https://www.ncbi.nlm.nih.gov/gene/?term=LOC105275711 | 6150 | 80 | 130.08 | 1 | 1.63 | 1.25 | 14.39 | 12.36 | 2.03 | 2.03 | 0.03 | 0.03 |
| gene7345 | https://www.ncbi.nlm.nih.gov/gene/?term=LOC105280529 | 3833 | 92 | 240.02 | 1 | 2.61 | 1.09 | 17.99 | 16.93 | 1.05 | 1.05 | 0.03 | 0.03 |
| gene3286 | https://www.ncbi.nlm.nih.gov/gene/?term=LOC105275848 | 3744 | 132 | 352.56 | 1 | 2.67 | 0.76 | 21.71 | 20.73 | 0.98 | 0.98 | 0.03 | 0.03 |
| gene13429 | https://www.ncbi.nlm.nih.gov/gene/?term=LOC105287501 | 3714 | 102 | 274.64 | 1 | 2.69 | 0.98 | 22.78 | 21.81 | 0.97 | 0.97 | 0.03 | 0.03 |
| gene11275 | https://www.ncbi.nlm.nih.gov/gene/?term=LOC105285055 | 4119 | 216 | 524.4 | 1 | 2.43 | 0.46 | 13.08 | 13.7 | -0.62 | 0.62 | -0.03 | 0.03 |
| gene12802 | https://www.ncbi.nlm.nih.gov/gene/?term=LOC105286799 | 3183 | 172 | 540.37 | 1 | 3.14 | 0.58 | 15.78 | 16.4 | -0.62 | 0.62 | -0.03 | 0.03 |
| gene3817 | https://www.ncbi.nlm.nih.gov/gene/?term=LOC105276449 | 1230 | 48 | 390.24 | 1 | 8.13 | 2.08 | 26.73 | 27.42 | -0.69 | 0.69 | -0.03 | 0.03 |
| gene10018 | https://www.ncbi.nlm.nih.gov/gene/?term=LOC105283607 | 4414 | 198 | 448.57 | 1 | 2.27 | 0.51 | 15.25 | 15.95 | -0.7 | 0.7 | -0.03 | 0.03 |
| gene2271 | https://www.ncbi.nlm.nih.gov/gene/?term=LOC105274706 | 1448 | 53 | 366.02 | 1 | 6.91 | 1.89 | 36.91 | 37.62 | -0.71 | 0.71 | -0.03 | 0.03 |
| gene3206 | https://www.ncbi.nlm.nih.gov/gene/?term=LOC105275752 | 3474 | 125 | 359.82 | 1 | 2.88 | 0.8 | 28.34 | 29.06 | -0.72 | 0.72 | -0.03 | 0.03 |
| gene2844 | https://www.ncbi.nlm.nih.gov/gene/?term=LOC105275363 | 4330 | 192 | 443.42 | 1 | 2.31 | 0.52 | 14.13 | 14.88 | -0.75 | 0.75 | -0.03 | 0.03 |
| gene7556 | https://www.ncbi.nlm.nih.gov/gene/?term=LOC105280774 | 2792 | 117 | 419.05 | 1 | 3.58 | 0.85 | 27.44 | 28.23 | -0.79 | 0.79 | -0.03 | 0.03 |
| gene8729 | https://www.ncbi.nlm.nih.gov/gene/?term=LOC105282137 | 5005 | 205 | 409.59 | 1 | 2 | 0.49 | 15.76 | 16.6 | -0.84 | 0.84 | -0.03 | 0.03 |
| gene4641 | https://www.ncbi.nlm.nih.gov/gene/?term=LOC105277398 | 1937 | 70 | 361.38 | 1 | 5.16 | 1.43 | 35.36 | 36.26 | -0.9 | 0.9 | -0.03 | 0.03 |
| gene10531 | https://www.ncbi.nlm.nih.gov/gene/?term=LOC105284190 | 6881 | 199 | 289.2 | 1 | 1.45 | 0.5 | 9.56 | 10.66 | -1.1 | 1.1 | -0.03 | 0.03 |
| gene13033 | https://www.ncbi.nlm.nih.gov/gene/?term=LOC105287063 | 1877 | 45 | 239.74 | 1 | 5.33 | 2.22 | 34.91 | 36.28 | -1.37 | 1.37 | -0.03 | 0.03 |
| gene1884 | https://www.ncbi.nlm.nih.gov/gene/?term=LOC105288072 | 4169 | 82 | 196.69 | 1 | 2.4 | 1.22 | 22.32 | 23.72 | -1.41 | 1.41 | -0.03 | 0.03 |
| gene2257 | https://www.ncbi.nlm.nih.gov/gene/?term=LOC105274689 | 3658 | 75 | 205.03 | 1 | 2.73 | 1.33 | 12.35 | 13.76 | -1.41 | 1.41 | -0.03 | 0.03 |
| gene8343 | https://www.ncbi.nlm.nih.gov/gene/?term=LOC105281700 | 2172 | 46 | 211.79 | 1 | 4.6 | 2.17 | 39.86 | 41.32 | -1.46 | 1.46 | -0.03 | 0.03 |
| gene5647 | https://www.ncbi.nlm.nih.gov/gene/?term=LOC105278567 | 3118 | 37 | 118.67 | 1 | 3.21 | 2.7 | 11.3 | 14.2 | -2.9 | 2.9 | -0.03 | 0.03 |
| gene1199 | https://www.ncbi.nlm.nih.gov/gene/?term=LOC105285498 | 2220 | 26 | 117.12 | 1 | 4.5 | 3.85 | 33.11 | 31.3 | 1.81 | 1.81 | 0.02 | 0.02 |

| | | | | | | | | | | | | | |
|---|---|---|---|---|---|---|---|---|---|---|---|---|---|
| gene7398 | https://www.ncbi.nlm.nih.gov/gene/?term=LOC105280589 | 3452 | 41 | 118.77 | 1 | 2.9 | 2.44 | 25.07 | 23.32 | 1.75 | 1.75 | 0.02 | 0.02 |
| gene3671 | https://www.ncbi.nlm.nih.gov/gene/?term=LOC105276272 | 1966 | 27 | 137.33 | 1 | 5.09 | 3.7 | 41.96 | 40.71 | 1.25 | 1.25 | 0.02 | 0.02 |
| gene322 | https://www.ncbi.nlm.nih.gov/gene/?term=LOC105275777 | 1606 | 23 | 143.21 | 1 | 6.23 | 4.35 | 45.55 | 44.33 | 1.21 | 1.21 | 0.02 | 0.02 |
| gene6237 | https://www.ncbi.nlm.nih.gov/gene/?term=LOC105279244 | 1484 | 26 | 175.2 1 | 6.74 | 3.85 | 46.7 | 45.51 | 1.19 | 1.19 | 0.02 | 0.02 | |
| gene10819 | https://www.ncbi.nlm.nih.gov/gene/?term=LOC105284507 | 5767 | 83 | 143.92 | 1 | 1.73 | 1.2 | 21.4 | 20.36 | 1.04 | 1.04 | 0.02 | 0.02 |
| gene6966 | https://www.ncbi.nlm.nih.gov/gene/?term=LOC105280104 | 3747 | 125 | 333.6 1 | 2.67 | 0.8 | 14.94 | 14.21 | 0.73 | 0.73 | 0.02 | 0.02 | |
| gene9722 | https://www.ncbi.nlm.nih.gov/gene/?term=LOC105283264 | 3951 | 88 | 222.73 | 1 | 2.53 | 1.14 | 24.81 | 24.09 | 0.72 | 0.72 | 0.02 | 0.02 |
| gene11340 | https://www.ncbi.nlm.nih.gov/gene/?term=LOC105285131 | 2918 | 102 | 349.55 | 1 | 3.43 | 0.98 | 14.35 | 13.65 | 0.7 | 0.7 | 0.02 | 0.02 |
| gene12573 | https://www.ncbi.nlm.nih.gov/gene/?term=LOC105286515 | 3647 | 115 | 315.33 | 1 | 2.74 | 0.87 | 11.99 | 11.33 | 0.66 | 0.66 | 0.02 | 0.02 |
| gene11401 | https://www.ncbi.nlm.nih.gov/gene/?term=LOC105285202 | 1841 | 50 | 271.59 | 1 | 5.43 | 2 | 21.11 | 20.46 | 0.65 | 0.65 | 0.02 | 0.02 |
| gene7600 | https://www.ncbi.nlm.nih.gov/gene/?term=LOC105280819 | 1615 | 41 | 253.87 | 1 | 6.19 | 2.44 | 40.74 | 40.08 | 0.65 | 0.65 | 0.02 | 0.02 |
| gene10714 | https://www.ncbi.nlm.nih.gov/gene/?term=LOC105284396 | 1966 | 57 | 289.93 | 1 | 5.09 | 1.75 | 38.23 | 37.64 | 0.59 | 0.59 | 0.02 | 0.02 |
| gene4962 | https://www.ncbi.nlm.nih.gov/gene/?term=LOC105277761 | 4645 | 150 | 322.93 | 1 | 2.15 | 0.67 | 14.77 | 14.2 | 0.57 | 0.57 | 0.02 | 0.02 |
| gene6286 | https://www.ncbi.nlm.nih.gov/gene/?term=LOC105279298 | 2861 | 139 | 485.84 | 2 | 6.99 | 1.44 | 22.43 | 22.79 | -0.36 | 0.36 | -0.02 | 0.02 |
| gene7568 | https://www.ncbi.nlm.nih.gov/gene/?term=LOC105280786 | 2525 | 123 | 487.13 | 1 | 3.96 | 0.81 | 28.92 | 29.32 | -0.39 | 0.39 | -0.02 | 0.02 |
| gene7888 | https://www.ncbi.nlm.nih.gov/gene/?term=LOC105281141 | 4357 | 165 | 378.7 1 | 2.3 | 0.61 | 28.04 | 28.47 | -0.43 | 0.43 | -0.02 | 0.02 | |
| gene4892 | https://www.ncbi.nlm.nih.gov/gene/?term=LOC105277667 | 3898 | 122 | 312.98 | 1 | 2.57 | 0.82 | 27.68 | 28.18 | -0.5 | 0.5 | -0.02 | 0.02 |
| gene7550 | https://www.ncbi.nlm.nih.gov/gene/?term=LOC105280763 | 2173 | 98 | 450.99 | 1 | 4.6 | 1.02 | 26.91 | 27.42 | -0.51 | 0.51 | -0.02 | 0.02 |
| gene3885 | https://www.ncbi.nlm.nih.gov/gene/?term=LOC105276527 | 7928 | 352 | 444 1 | 1.26 | 0.28 | 8.21 | 8.74 | -0.52 | 0.52 | -0.02 | 0.02 | |
| gene5967 | https://www.ncbi.nlm.nih.gov/gene/?term=LOC105278918 | 4393 | 135 | 307.31 | 1 | 2.28 | 0.74 | 16.02 | 16.55 | -0.53 | 0.53 | -0.02 | 0.02 |
| gene7166 | https://www.ncbi.nlm.nih.gov/gene/?term=LOC105280326 | 6117 | 213 | 348.21 | 1 | 1.63 | 0.47 | 15.89 | 16.43 | -0.54 | 0.54 | -0.02 | 0.02 |
| gene474 | https://www.ncbi.nlm.nih.gov/gene/?term=LOC105277435 | 3625 | 158 | 435.86 | 1 | 2.76 | 0.63 | 12.73 | 13.28 | -0.55 | 0.55 | -0.02 | 0.02 |
| gene10678 | https://www.ncbi.nlm.nih.gov/gene/?term=LOC105284355 | 4031 | 164 | 406.85 | 1 | 2.48 | 0.61 | 19.18 | 19.78 | -0.6 | 0.6 | -0.02 | 0.02 |
| gene8661 | https://www.ncbi.nlm.nih.gov/gene/?term=LOC105282063 | 1926 | 43 | 223.26 | 1 | 5.19 | 2.33 | 23.84 | 24.54 | -0.71 | 0.71 | -0.02 | 0.02 |
| gene2173 | https://www.ncbi.nlm.nih.gov/gene/?term=LOC105274590 | 4387 | 84 | 191.47 | 1 | 2.28 | 1.19 | 20.89 | 21.7 | -0.81 | 0.81 | -0.02 | 0.02 |
| gene9598 | https://www.ncbi.nlm.nih.gov/gene/?term=LOC105283132 | 1769 | 54 | 305.26 | 1 | 5.65 | 1.85 | 40.78 | 41.59 | -0.81 | 0.81 | -0.02 | 0.02 |
| gene4070 | https://www.ncbi.nlm.nih.gov/gene/?term=LOC105276745 | 3957 | 72 | 181.96 | 1 | 2.53 | 1.39 | 20.98 | 22.28 | -1.3 | 1.3 | -0.02 | 0.02 |
| gene9336 | https://www.ncbi.nlm.nih.gov/gene/?term=LOC105282834 | 5993 | 109 | 181.88 | 1 | 1.67 | 0.92 | 15.48 | 16.78 | -1.31 | 1.31 | -0.02 | 0.02 |
| gene1962 | https://www.ncbi.nlm.nih.gov/gene/?term=LOC105288157 | 1246 | 16 | 128.41 | 1 | 8.03 | 6.25 | 13.95 | 15.31 | -1.36 | 1.36 | -0.02 | 0.02 |
| gene7183 | https://www.ncbi.nlm.nih.gov/gene/?term=LOC105280348 | 3925 | 53 | 135.03 | 1 | 2.55 | 1.89 | 25.26 | 26.63 | -1.37 | 1.37 | -0.02 | 0.02 |
| gene6455 | https://www.ncbi.nlm.nih.gov/gene/?term=LOC105279500 | 2612 | 32 | 122.51 | 1 | 3.83 | 3.12 | 49.77 | 51.44 | -1.68 | 1.68 | -0.02 | 0.02 |
| gene5402 | https://www.ncbi.nlm.nih.gov/gene/?term=LOC105278260 | 3068 | 41 | 133.64 | 1 | 3.26 | 2.44 | 13.01 | 14.77 | -1.77 | 1.77 | -0.02 | 0.02 |
| gene10243 | https://www.ncbi.nlm.nih.gov/gene/?term=LOC105283862 | 3239 | 37 | 114.23 | 1 | 3.09 | 2.7 | 31.49 | 33.45 | -1.96 | 1.96 | -0.02 | 0.02 |
| gene2970 | https://www.ncbi.nlm.nih.gov/gene/?term=LOC105275518 | 2340 | 21 | 89.74 1 | 4.27 | 4.76 | 45.1 | 47.46 | -2.36 | 2.36 | -0.02 | 0.02 | |
| gene814 | https://www.ncbi.nlm.nih.gov/gene/?term=LOC105281296 | 2722 | 43 | 157.97 | 1 | 3.67 | 2.33 | 29.73 | 28.81 | 0.92 | 0.92 | 0.01 | 0.01 |
| gene11447 | https://www.ncbi.nlm.nih.gov/gene/?term=LOC105285252 | 7437 | 135 | 181.52 | 1 | 1.34 | 0.74 | 13.54 | 12.74 | 0.8 | 0.8 | 0.01 | 0.01 |
| gene1159 | https://www.ncbi.nlm.nih.gov/gene/?term=LOC105285084 | 3282 | 27 | 82.27 1 | 3.05 | 3.7 | 22 | 21.23 | 0.77 | 0.77 | 0.01 | 0.01 | |
| gene652 | https://www.ncbi.nlm.nih.gov/gene/?term=LOC105279528 | 5806 | 44 | 75.78 1 | 1.72 | 2.27 | 16.35 | 15.59 | 0.76 | 0.76 | 0.01 | 0.01 | |
| gene9464 | https://www.ncbi.nlm.nih.gov/gene/?term=LOC105282975 | 1450 | 25 | 172.41 | 1 | 6.9 | 4 | 41.13 | 40.46 | 0.67 | 0.67 | 0.01 | 0.01 |
| gene4376 | https://www.ncbi.nlm.nih.gov/gene/?term=LOC105277083 | 4307 | 96 | 222.89 | 1 | 2.32 | 1.04 | 25.73 | 25.08 | 0.66 | 0.66 | 0.01 | 0.01 |
| gene10155 | https://www.ncbi.nlm.nih.gov/gene/?term=LOC105283756 | 7030 | 97 | 137.98 | 1 | 1.42 | 1.03 | 6.83 | 6.25 | 0.59 | 0.59 | 0.01 | 0.01 |
| gene12203 | https://www.ncbi.nlm.nih.gov/gene/?term=LOC105286099 | 5967 | 78 | 130.72 | 1 | 1.68 | 1.28 | 11.28 | 10.77 | 0.51 | 0.51 | 0.01 | 0.01 |
| gene3107 | https://www.ncbi.nlm.nih.gov/gene/?term=LOC105275650 | 5103 | 151 | 295.9 1 | 1.96 | 0.66 | 17.08 | 16.7 | 0.38 | 0.38 | 0.01 | 0.01 | |
| gene5646 | https://www.ncbi.nlm.nih.gov/gene/?term=LOC105278534 | 7475 | 186 | 248.83 | 1 | 1.34 | 0.54 | 18.03 | 17.67 | 0.36 | 0.36 | 0.01 | 0.01 |
| gene10392 | https://www.ncbi.nlm.nih.gov/gene/?term=LOC105284036 | 3216 | 83 | 258.08 | 2 | 6.22 | 2.41 | 23.78 | 23.42 | 0.35 | 0.35 | 0.01 | 0.01 |
| gene1220 | https://www.ncbi.nlm.nih.gov/gene/?term=LOC105285742 | 3453 | 99 | 286.71 | 1 | 2.9 | 1.01 | 25.65 | 25.32 | 0.33 | 0.33 | 0.01 | 0.01 |
| gene3848 | https://www.ncbi.nlm.nih.gov/gene/?term=LOC105276486 | 5899 | 121 | 205.12 | 1 | 1.7 | 0.83 | 23.22 | 22.92 | 0.3 | 0.3 | 0.01 | 0.01 |
| gene11583 | https://www.ncbi.nlm.nih.gov/gene/?term=LOC105285414 | 8192 | 177 | 216.06 | 1 | 1.22 | 0.56 | 8.82 | 8.53 | 0.29 | 0.29 | 0.01 | 0.01 |
| gene5539 | https://www.ncbi.nlm.nih.gov/gene/?term=LOC105278415 | 25024 | 604 | 241.37 | 1 | 0.4 | 0.17 | 3.91 | 3.62 | 0.29 | 0.29 | 0.01 | 0.01 |
| gene5831 | https://www.ncbi.nlm.nih.gov/gene/?term=LOC105278761 | 2506 | 104 | 415 1 | 3.99 | 0.96 | 25.24 | 24.96 | 0.28 | 0.28 | 0.01 | 0.01 | |
| gene13114 | https://www.ncbi.nlm.nih.gov/gene/?term=LOC105287147 | 7609 | 389 | 511.24 | 1 | 1.31 | 0.26 | 9.77 | 9.58 | 0.19 | 0.19 | 0.01 | 0.01 |
| gene5746 | https://www.ncbi.nlm.nih.gov/gene/?term=LOC105278666 | 10343 | 409 | 395.44 | 1 | 0.97 | 0.24 | 7.43 | 7.26 | 0.17 | 0.17 | 0.01 | 0.01 |
| gene12674 | https://www.ncbi.nlm.nih.gov/gene/?term=LOC105286649 | 18283 | 680 | 371.93 | 1 | 0.55 | 0.15 | 1.31 | 1.16 | 0.15 | 0.15 | 0.01 | 0.01 |
| gene10864 | https://www.ncbi.nlm.nih.gov/gene/?term=LOC105284557 | 1666 | 78 | 468.19 | 1 | 6 | 1.28 | 30.45 | 30.59 | -0.13 | 0.13 | -0.01 | 0.01 |
| gene5649 | https://www.ncbi.nlm.nih.gov/gene/?term=LOC105278538 | 2140 | 84 | 392.52 | 1 | 4.67 | 1.19 | 18.7 | 18.83 | -0.13 | 0.13 | -0.01 | 0.01 |
| gene6583 | https://www.ncbi.nlm.nih.gov/gene/?term=LOC105279649 | 7444 | 330 | 443.31 | 1 | 1.34 | 0.3 | 11.47 | 11.66 | -0.2 | 0.2 | -0.01 | 0.01 |
| gene2806 | https://www.ncbi.nlm.nih.gov/gene/?term=LOC105275327 | 14257 | 480 | 336.68 | 1 | 0.7 | 0.21 | 7.1 | 7.32 | -0.22 | 0.22 | -0.01 | 0.01 |
| gene7249 | https://www.ncbi.nlm.nih.gov/gene/?term=LOC105280422 | 1775 | 61 | 343.66 | 1 | 5.63 | 1.64 | 15.16 | 15.4 | -0.24 | 0.24 | -0.01 | 0.01 |
| gene9101 | https://www.ncbi.nlm.nih.gov/gene/?term=LOC105282553 | 3068 | 104 | 338.98 | 1 | 3.26 | 0.96 | 20.36 | 20.64 | -0.27 | 0.27 | -0.01 | 0.01 |
| gene5613 | https://www.ncbi.nlm.nih.gov/gene/?term=LOC105278504 | 8524 | 211 | 247.54 | 1 | 1.17 | 0.47 | 5.46 | 5.74 | -0.28 | 0.28 | -0.01 | 0.01 |
| gene6620 | https://www.ncbi.nlm.nih.gov/gene/?term=LOC105279687 | 17998 | 360 | 200.02 | 1 | 0.56 | 0.28 | 5.06 | 5.34 | -0.28 | 0.28 | -0.01 | 0.01 |
| gene1573 | https://www.ncbi.nlm.nih.gov/gene/?term=LOC105287704 | 3030 | 131 | 432.34 | 1 | 3.3 | 0.76 | 20.77 | 21.09 | -0.32 | 0.32 | -0.01 | 0.01 |
| gene2056 | https://www.ncbi.nlm.nih.gov/gene/?term=LOC105274452 | 6584 | 212 | 321.99 | 1 | 1.52 | 0.47 | 13.94 | 14.26 | -0.33 | 0.33 | -0.01 | 0.01 |
| gene1147 | https://www.ncbi.nlm.nih.gov/gene/?term=LOC105284954 | 4022 | 82 | 203.88 | 1 | 2.49 | 1.22 | 15.94 | 16.29 | -0.34 | 0.34 | -0.01 | 0.01 |

| | | | | | | | | | | | | | | |
|---|---|---|---|---|---|---|---|---|---|---|---|---|---|---|
| gene10600 | https://www.ncbi.nlm.nih.gov/gene/?term=LOC105284270 | 4837 | 105 | 217.08 | | 1 | 2.07 | 0.95 | 12.85 | 13.22 | -0.37 | 0.37 | -0.01 | 0.01 |
| gene3856 | https://www.ncbi.nlm.nih.gov/gene/?term=LOC105276499 | 5236 | 201 | 383.88 | | 1 | 1.91 | 0.5 | 10.34 | 10.72 | -0.39 | 0.39 | -0.01 | 0.01 |
| gene10520 | https://www.ncbi.nlm.nih.gov/gene/?term=LOC105284184 | 2451 | 82 | 334.56 | | 1 | 4.08 | 1.22 | 34.55 | 34.95 | -0.4 | 0.4 | -0.01 | 0.01 |
| gene5375 | https://www.ncbi.nlm.nih.gov/gene/?term=LOC105278228 | 8640 | 159 | 184.03 | | 1 | 1.16 | 0.63 | 24.17 | 24.62 | -0.45 | 0.45 | -0.01 | 0.01 |
| gene4225 | https://www.ncbi.nlm.nih.gov/gene/?term=LOC105276917 | 2443 | 26 | 106.43 | | 1 | 4.09 | 3.85 | 36.73 | 37.26 | -0.53 | 0.53 | -0.01 | 0.01 |
| gene9330 | https://www.ncbi.nlm.nih.gov/gene/?term=LOC105282825 | 3936 | 90 | 228.66 | | 1 | 2.54 | 1.11 | 15.33 | 15.86 | -0.53 | 0.53 | -0.01 | 0.01 |
| gene8600 | https://www.ncbi.nlm.nih.gov/gene/?term=LOC105282001 | 8513 | 158 | 185.6 | 1 | 1.17 | 0.63 | 9.54 | 10.09 | -0.55 | 0.55 | -0.01 | 0.01 | |
| gene7245 | https://www.ncbi.nlm.nih.gov/gene/?term=LOC105280410 | 12478 | 260 | 208.37 | | 1 | 0.8 | 0.38 | 19.03 | 19.62 | -0.59 | 0.59 | -0.01 | 0.01 |
| gene3385 | https://www.ncbi.nlm.nih.gov/gene/?term=LOC105275966 | 3305 | 59 | 178.52 | | 1 | 3.03 | 1.69 | 50.29 | 50.9 | -0.61 | 0.61 | -0.01 | 0.01 |
| gene10379 | https://www.ncbi.nlm.nih.gov/gene/?term=LOC105284028 | 6974 | 74 | 106.11 | | 1 | 1.43 | 1.35 | 8.99 | 9.61 | -0.62 | 0.62 | -0.01 | 0.01 |
| gene388 | https://www.ncbi.nlm.nih.gov/gene/?term=LOC105276538 | 1558 | 20 | 128.37 | | 2 | 12.84 | 10 | 70.94 | 71.62 | -0.68 | 0.68 | -0.01 | 0.01 |
| gene927 | https://www.ncbi.nlm.nih.gov/gene/?term=LOC105282600 | 3727 | 65 | 174.4 | 1 | 2.68 | 1.54 | 26.3 | 27.02 | -0.73 | 0.73 | -0.01 | 0.01 | |
| gene2330 | https://www.ncbi.nlm.nih.gov/gene/?term=LOC105274773 | 13104 | 172 | 131.26 | | 1 | 0.76 | 0.58 | 15.95 | 16.78 | -0.83 | 0.83 | -0.01 | 0.01 |
| gene6528 | https://www.ncbi.nlm.nih.gov/gene/?term=LOC105279591 | 7222 | 75 | 103.85 | | 1 | 1.38 | 1.33 | 20.71 | 21.6 | -0.89 | 0.89 | -0.01 | 0.01 |
| gene10345 | https://www.ncbi.nlm.nih.gov/gene/?term=LOC105283989 | 4444 | 30 | 67.51 | 1 | 2.25 | 3.33 | 46.58 | 47.61 | -1.03 | 1.03 | -0.01 | 0.01 | |
| gene8178 | https://www.ncbi.nlm.nih.gov/gene/?term=LOC105281515 | 8647 | 46 | 53.2 | 1 | 1.16 | 2.17 | 28.18 | 29.56 | -1.38 | 1.38 | -0.01 | 0.01 | |
| gene1809 | https://www.ncbi.nlm.nih.gov/gene/?term=LOC105287989 | 7010 | 73 | 104.14 | | 1 | 1.43 | 1.37 | 23.7 | 23.28 | 0.42 | 0.42 | 0 | 0 |
| gene12307 | https://www.ncbi.nlm.nih.gov/gene/?term=LOC105286216 | 10313 | 167 | 161.93 | | 1 | 0.97 | 0.6 | 15.75 | 15.52 | 0.23 | 0.23 | 0 | 0 |
| gene657 | https://www.ncbi.nlm.nih.gov/gene/?term=LOC105279667 | 6421 | 80 | 124.59 | | 1 | 1.56 | 1.25 | 3.15 | 2.93 | 0.21 | 0.21 | 0 | 0 |
| gene940 | https://www.ncbi.nlm.nih.gov/gene/?term=LOC105282794 | 3578 | 88 | 245.95 | | 1 | 2.79 | 1.14 | 15.65 | 15.49 | 0.16 | 0.16 | 0 | 0 |
| gene6031 | https://www.ncbi.nlm.nih.gov/gene/?term=LOC105279000 | 5691 | 185 | 325.07 | | 1 | 1.76 | 0.54 | 13.85 | 13.69 | 0.15 | 0.15 | 0 | 0 |
| gene3599 | https://www.ncbi.nlm.nih.gov/gene/?term=LOC105276196 | 2955 | 56 | 189.51 | | 1 | 3.38 | 1.79 | 40.12 | 39.99 | 0.13 | 0.13 | 0 | 0 |
| gene7547 | https://www.ncbi.nlm.nih.gov/gene/?term=LOC105280767 | 24576 | 753 | 306.4 | 1 | 0.41 | 0.13 | 4.75 | 4.63 | 0.12 | 0.12 | 0 | 0 | |
| gene2404 | https://www.ncbi.nlm.nih.gov/gene/?term=LOC105274853 | 1767 | 42 | 237.69 | | 1 | 5.66 | 2.38 | 28.85 | 28.79 | 0.07 | 0.07 | 0 | 0 |
| gene2991 | https://www.ncbi.nlm.nih.gov/gene/?term=LOC105275536 | 11224 | 214 | 190.66 | | 1 | 0.89 | 0.47 | 5.64 | 5.58 | 0.05 | 0.05 | 0 | 0 |
| gene4527 | https://www.ncbi.nlm.nih.gov/gene/?term=LOC105277273 | 3686 | 151 | 409.66 | | 1 | 2.71 | 0.66 | 12.71 | 12.65 | 0.05 | 0.05 | 0 | 0 |
| gene4996 | https://www.ncbi.nlm.nih.gov/gene/?term=LOC105277798 | 5617 | 64 | 113.94 | | 1 | 1.78 | 1.56 | 10.86 | 10.81 | 0.05 | 0.05 | 0 | 0 |
| gene2266 | https://www.ncbi.nlm.nih.gov/gene/?term=LOC105274699 | 2391 | 63 | 263.49 | | 1 | 4.18 | 1.59 | 23.04 | 22.99 | 0.04 | 0.04 | 0 | 0 |
| gene7148 | https://www.ncbi.nlm.nih.gov/gene/?term=LOC105280304 | 8920 | 123 | 137.89 | | 1 | 1.12 | 0.81 | 5.79 | 5.75 | 0.04 | 0.04 | 0 | 0 |
| gene53 | https://www.ncbi.nlm.nih.gov/gene/?term=LOC105277148 | 19283 | 269 | 139.5 | 1 | 0.52 | 0.37 | 3.9 | 3.86 | 0.03 | 0.03 | 0 | 0 | |
| gene9645 | https://www.ncbi.nlm.nih.gov/gene/?term=LOC105283181 | 3875 | 138 | 356.13 | | 1 | 2.58 | 0.72 | 11.99 | 11.98 | 0.01 | 0.01 | 0 | 0 |
| gene10549 | https://www.ncbi.nlm.nih.gov/gene/?term=LOC105284210 | 7145 | 323 | 452.06 | | 1 | 1.4 | 0.31 | 10.47 | 10.47 | 0 | 0 | 0 | 0 |
| gene1470 | https://www.ncbi.nlm.nih.gov/gene/?term=LOC105287597 | 2974 | 192 | 645.6 | 1 | 3.36 | 0.52 | 16.86 | 16.86 | 0 | 0 | 0 | 0 | |
| gene9755 | https://www.ncbi.nlm.nih.gov/gene/?term=LOC105283296 | 2177 | 90 | 413.41 | | 1 | 4.59 | 1.11 | 19.36 | 19.36 | 0 | 0 | 0 | 0 |
| gene4209 | https://www.ncbi.nlm.nih.gov/gene/?term=LOC105277125 | 50550 | 1735 | 343.22 | | 1 | 0.2 | 0.06 | 1.59 | 1.62 | -0.03 | 0.03 | 0 | 0 |
| gene9411 | https://www.ncbi.nlm.nih.gov/gene/?term=LOC105282914 | 5534 | 41 | 74.09 | 1 | 1.81 | 2.44 | 49.68 | 49.72 | -0.04 | 0.04 | 0 | 0 | |
| gene13036 | https://www.ncbi.nlm.nih.gov/gene/?term=LOC105287059 | 4959 | 165 | 332.73 | | 1 | 2.02 | 0.61 | 17.35 | 17.4 | -0.05 | 0.05 | 0 | 0 |
| gene7207 | https://www.ncbi.nlm.nih.gov/gene/?term=LOC105280376 | 4659 | 105 | 225.37 | | 1 | 2.15 | 0.95 | 28.77 | 28.82 | -0.05 | 0.05 | 0 | 0 |
| gene10606 | https://www.ncbi.nlm.nih.gov/gene/?term=LOC105284273 | 6119 | 252 | 411.83 | | 2 | 3.27 | 0.79 | 19.91 | 19.99 | -0.08 | 0.08 | 0 | 0 |
| gene11568 | https://www.ncbi.nlm.nih.gov/gene/?term=LOC105285395 | 8086 | 184 | 227.55 | | 1 | 1.24 | 0.54 | 16.42 | 16.51 | -0.08 | 0.08 | 0 | 0 |
| gene3470 | https://www.ncbi.nlm.nih.gov/gene/?term=LOC105276059 | 8716 | 161 | 184.72 | | 1 | 1.15 | 0.62 | 9.12 | 9.21 | -0.08 | 0.08 | 0 | 0 |
| gene8274 | https://www.ncbi.nlm.nih.gov/gene/?term=LOC105281624 | 5425 | 208 | 383.41 | | 1 | 1.84 | 0.48 | 10.51 | 10.6 | -0.09 | 0.09 | 0 | 0 |
| gene2748 | https://www.ncbi.nlm.nih.gov/gene/?term=LOC105275261 | 3845 | 58 | 150.85 | | 1 | 2.6 | 1.72 | 39.98 | 40.08 | -0.1 | 0.1 | 0 | 0 |
| gene1632 | https://www.ncbi.nlm.nih.gov/gene/?term=LOC105287764 | 18061 | 354 | 196 | 1 | 0.55 | 0.28 | 2.69 | 2.8 | -0.11 | 0.11 | 0 | 0 | |
| gene10596 | https://www.ncbi.nlm.nih.gov/gene/?term=LOC105284262 | 6520 | 235 | 360.43 | | 1 | 1.53 | 0.43 | 3.69 | 3.81 | -0.12 | 0.12 | 0 | 0 |
| gene3195 | https://www.ncbi.nlm.nih.gov/gene/?term=LOC105275750 | 28072 | 713 | 253.99 | | 1 | 0.36 | 0.14 | 11.56 | 11.74 | -0.18 | 0.18 | 0 | 0 |
| gene2683 | https://www.ncbi.nlm.nih.gov/gene/?term=LOC105275187 | 5492 | 75 | 136.56 | | 1 | 1.82 | 1.33 | 14.14 | 14.34 | -0.2 | 0.2 | 0 | 0 |
| gene11579 | https://www.ncbi.nlm.nih.gov/gene/?term=LOC105285406 | 13571 | 293 | 215.9 | 1 | 0.74 | 0.34 | 5.86 | 6.07 | -0.21 | 0.21 | 0 | 0 | |
| gene6639 | https://www.ncbi.nlm.nih.gov/gene/?term=LOC105279705 | 7656 | 69 | 90.13 | 1 | 1.31 | 1.45 | 21.96 | 22.32 | -0.36 | 0.36 | 0 | 0 | |

Directory: OGLall File: rnk-dmg-Cb_LI_BCphaseA1.vs.RphaseB1.txt

| gene_ID | gene_link | gwidth | #Sites | #per10Kb | #dmSites | #dmsp10kb | %dmSites | %pSite1 | %pSite2 | DMpSite | ADMpSite | DMpNucl | ADMpNucl |
|---|---|---|---|---|---|---|---|---|---|---|---|---|---|
| gene10237 | https://www.ncbi.nlm.nih.gov/gene/?term=LOC105283856 | 1259 | 25 | 198.57 | 5 | 39.71 | 20 | 64.65 | 55.66 | 9 | 9 | 0.18 | 0.18 |
| gene8965 | https://www.ncbi.nlm.nih.gov/gene/?term=LOC105282405 | 1310 | 31 | 236.64 | 2 | 15.27 | 6.45 | 51.28 | 44.69 | 6.59 | 6.59 | 0.16 | 0.16 |
| gene1993 | https://www.ncbi.nlm.nih.gov/gene/?term=LOC105288193 | 1449 | 64 | 441.68 | 4 | 27.61 | 6.25 | 34.43 | 31.28 | 3.16 | 3.16 | 0.14 | 0.14 |
| gene13384 | https://www.ncbi.nlm.nih.gov/gene/?term=LOC105287447 | 1877 | 145 | 772.51 | 7 | 37.29 | 4.83 | 29.16 | 30.96 | -1.8 | 1.8 | -0.14 | 0.14 |
| gene5809 | https://www.ncbi.nlm.nih.gov/gene/?term=LOC105278737 | 1159 | 83 | 716.13 | 1 | 8.63 | 1.2 | 16.21 | 18.18 | -1.97 | 1.97 | -0.14 | 0.14 |
| gene6314 | https://www.ncbi.nlm.nih.gov/gene/?term=LOC105279335 | 718 | 34 | 473.54 | 2 | 27.86 | 5.88 | 18.89 | 16.04 | 2.85 | 2.85 | 0.13 | 0.13 |
| gene5309 | https://www.ncbi.nlm.nih.gov/gene/?term=LOC105278160 | 1704 | 85 | 498.83 | 2 | 11.74 | 2.35 | 38.16 | 35.69 | 2.46 | 2.46 | 0.12 | 0.12 |
| gene560 | https://www.ncbi.nlm.nih.gov/gene/?term=LOC109610897 | 1584 | 112 | 707.07 | 3 | 18.94 | 2.68 | 15.45 | 13.95 | 1.5 | 1.5 | 0.11 | 0.11 |
| gene5412 | https://www.ncbi.nlm.nih.gov/gene/?term=LOC105278266 | 1868 | 111 | 594.22 | 2 | 10.71 | 1.8 | 25.18 | 27.02 | -1.84 | 1.84 | -0.11 | 0.11 |
| gene5650 | https://www.ncbi.nlm.nih.gov/gene/?term=LOC105278537 | 1672 | 61 | 364.83 | 5 | 29.9 | 8.2 | 28.88 | 31.76 | -2.89 | 2.89 | -0.11 | 0.11 |
| gene11413 | https://www.ncbi.nlm.nih.gov/gene/?term=LOC105285213 | 1472 | 37 | 251.36 | 4 | 27.17 | 10.81 | 13.65 | 18.03 | -4.37 | 4.37 | -0.11 | 0.11 |
| gene305 | https://www.ncbi.nlm.nih.gov/gene/?term=LOC105275539 | 1277 | 23 | 180.11 | 2 | 15.66 | 8.7 | 20.8 | 15.48 | 5.32 | 5.32 | 0.1 | 0.1 |
| gene10334 | https://www.ncbi.nlm.nih.gov/gene/?term=LOC105283973 | 1536 | 33 | 214.84 | 4 | 26.04 | 12.12 | 58.47 | 53.86 | 4.61 | 4.61 | 0.1 | 0.1 |
| gene3213 | https://www.ncbi.nlm.nih.gov/gene/?term=LOC105275761 | 2635 | 101 | 383.3 | 6 | 22.77 | 5.94 | 29.66 | 26.96 | 2.7 | 2.7 | 0.1 | 0.1 |
| gene2140 | https://www.ncbi.nlm.nih.gov/gene/?term=LOC105274555 | 1379 | 56 | 406.09 | 2 | 14.5 | 3.57 | 36.51 | 34.17 | 2.35 | 2.35 | 0.1 | 0.1 |
| gene460 | https://www.ncbi.nlm.nih.gov/gene/?term=LOC105277371 | 1819 | 76 | 417.81 | 1 | 5.5 | 1.32 | 19.38 | 17.09 | 2.29 | 2.29 | 0.1 | 0.1 |
| gene1562 | https://www.ncbi.nlm.nih.gov/gene/?term=LOC105287692 | 1491 | 97 | 650.57 | 3 | 20.12 | 3.09 | 30.3 | 28.78 | 1.52 | 1.52 | 0.1 | 0.1 |
| gene5207 | https://www.ncbi.nlm.nih.gov/gene/?term=LOC105278043 | 1205 | 73 | 605.81 | 1 | 8.3 | 1.37 | 20.73 | 22.42 | -1.69 | 1.69 | -0.1 | 0.1 |
| gene762 | https://www.ncbi.nlm.nih.gov/gene/?term=LOC105280710 | 1173 | 66 | 562.66 | 5 | 42.63 | 7.58 | 21.68 | 23.45 | -1.77 | 1.77 | -0.1 | 0.1 |
| gene12330 | https://www.ncbi.nlm.nih.gov/gene/?term=LOC105286235 | 1501 | 76 | 506.33 | 3 | 19.99 | 3.95 | 37.88 | 39.89 | -2.01 | 2.01 | -0.1 | 0.1 |
| gene6110 | https://www.ncbi.nlm.nih.gov/gene/?term=LOC105279087 | 1077 | 28 | 259.98 | 1 | 9.29 | 3.57 | 52.85 | 56.81 | -3.96 | 3.96 | -0.1 | 0.1 |
| gene3520 | https://www.ncbi.nlm.nih.gov/gene/?term=LOC105276115 | 1525 | 39 | 255.74 | 3 | 19.67 | 7.69 | 23.35 | 27.32 | -3.97 | 3.97 | -0.1 | 0.1 |
| gene4643 | https://www.ncbi.nlm.nih.gov/gene/?term=LOC109610847 | 429 | 5 | 116.55 | 1 | 23.31 | 20 | 29.86 | 22.57 | 7.29 | 7.29 | 0.09 | 0.09 |
| gene13026 | https://www.ncbi.nlm.nih.gov/gene/?term=LOC105287046 | 1786 | 55 | 307.95 | 1 | 5.6 | 1.82 | 21.15 | 18.07 | 3.07 | 3.07 | 0.09 | 0.09 |

...
... only first 25 lines shown ...

Directory: OGLall File: rnk-dmg-Cb_LI_BCphaseA1.vs.RphaseB2.txt

| gene_ID | gene_link | gwidth | #Sites | #per10Kb | #dmSites | #dmsp10kb | %dmSites | %pSite1 | %pSite2 | DMpSite | ADMpSite | DMpNucl | ADMpNucl |
|---|---|---|---|---|---|---|---|---|---|---|---|---|---|
| gene5412 | https://www.ncbi.nlm.nih.gov/gene/?term=LOC105278266 | 1868 | 105 | 562.1 | 4 | 21.41 | 3.81 | 26.49 | 29.78 | -3.29 | 3.29 | -0.18 | 0.18 |
| gene8965 | https://www.ncbi.nlm.nih.gov/gene/?term=LOC105282405 | 1310 | 31 | 236.64 | 5 | 38.17 | 16.13 | 51.28 | 44.76 | 6.52 | 6.52 | 0.15 | 0.15 |
| gene5917 | https://www.ncbi.nlm.nih.gov/gene/?term=LOC105278849 | 823 | 80 | 972.05 | 4 | 48.6 | 5 | 24.88 | 23.29 | 1.59 | 1.59 | 0.15 | 0.15 |
| gene3929 | https://www.ncbi.nlm.nih.gov/gene/?term=LOC105276585 | 1525 | 115 | 754.1 | 1 | 6.56 | 0.87 | 7.17 | 5.29 | 1.88 | 1.88 | 0.14 | 0.14 |
| gene11399 | https://www.ncbi.nlm.nih.gov/gene/?term=LOC105285203 | 1029 | 27 | 262.39 | 1 | 9.72 | 3.7 | 30.13 | 25.46 | 4.67 | 4.67 | 0.12 | 0.12 |
| gene1696 | https://www.ncbi.nlm.nih.gov/gene/?term=LOC105287856 | 1917 | 114 | 594.68 | 6 | 31.3 | 5.26 | 12.21 | 14.28 | -2.07 | 2.07 | -0.12 | 0.12 |
| gene6776 | https://www.ncbi.nlm.nih.gov/gene/?term=LOC105279863 | 1193 | 17 | 142.5 | 4 | 33.53 | 23.53 | 11.89 | 20.13 | -8.24 | 8.24 | -0.12 | 0.12 |
| gene558 | https://www.ncbi.nlm.nih.gov/gene/?term=LOC105278458 | 216 | 3 | 138.89 | 1 | 46.3 | 33.33 | 20.87 | 29.54 | -8.67 | 8.67 | -0.12 | 0.12 |
| gene10363 | https://www.ncbi.nlm.nih.gov/gene/?term=LOC105284009 | 1503 | 26 | 172.99 | 3 | 19.96 | 11.54 | 22.01 | 15.79 | 6.22 | 6.22 | 0.11 | 0.11 |
| gene1802 | https://www.ncbi.nlm.nih.gov/gene/?term=LOC105287981 | 3173 | 131 | 412.86 | 2 | 6.3 | 1.53 | 19.33 | 16.77 | 2.56 | 2.56 | 0.11 | 0.11 |
| gene5379 | https://www.ncbi.nlm.nih.gov/gene/?term=LOC105278234 | 2307 | 119 | 515.82 | 1 | 4.33 | 0.84 | 9.63 | 7.57 | 2.06 | 2.06 | 0.11 | 0.11 |
| gene13384 | https://www.ncbi.nlm.nih.gov/gene/?term=LOC105287447 | 1877 | 144 | 767.18 | 1 | 5.33 | 0.69 | 29.36 | 30.8 | -1.44 | 1.44 | -0.11 | 0.11 |
| gene762 | https://www.ncbi.nlm.nih.gov/gene/?term=LOC105280710 | 1173 | 61 | 520.03 | 1 | 8.53 | 1.64 | 23.4 | 25.56 | -2.17 | 2.17 | -0.11 | 0.11 |
| gene10334 | https://www.ncbi.nlm.nih.gov/gene/?term=LOC105283973 | 1536 | 30 | 195.31 | 1 | 6.51 | 3.33 | 61.25 | 55.99 | 5.26 | 5.26 | 0.1 | 0.1 |
| gene5937 | https://www.ncbi.nlm.nih.gov/gene/?term=LOC105278873 | 1902 | 141 | 741.32 | 4 | 21.03 | 2.84 | 20.24 | 21.57 | -1.32 | 1.32 | -0.1 | 0.1 |
| gene4641 | https://www.ncbi.nlm.nih.gov/gene/?term=LOC105277398 | 1937 | 61 | 314.92 | 5 | 25.81 | 8.2 | 38.08 | 41.19 | -3.11 | 3.11 | -0.1 | 0.1 |
| gene10584 | https://www.ncbi.nlm.nih.gov/gene/?term=LOC105284253 | 1255 | 36 | 286.85 | 2 | 15.94 | 5.56 | 27.9 | 31.27 | -3.36 | 3.36 | -0.1 | 0.1 |
| gene1222 | https://www.ncbi.nlm.nih.gov/gene/?term=LOC105285716 | 1097 | 25 | 227.89 | 2 | 18.23 | 8 | 39.46 | 43.8 | -4.34 | 4.34 | -0.1 | 0.1 |
| gene1179 | https://www.ncbi.nlm.nih.gov/gene/?term=LOC105285207 | 2994 | 60 | 200.4 | 5 | 16.7 | 8.33 | 21.02 | 25.85 | -4.83 | 4.83 | -0.1 | 0.1 |
| gene1534 | https://www.ncbi.nlm.nih.gov/gene/?term=LOC105287662 | 1711 | 31 | 181.18 | 1 | 5.84 | 3.23 | 32.16 | 37.64 | -5.47 | 5.47 | -0.1 | 0.1 |
| gene5006 | https://www.ncbi.nlm.nih.gov/gene/?term=LOC105277810 | 1382 | 21 | 151.95 | 1 | 7.24 | 4.76 | 45.33 | 39.66 | 5.66 | 5.66 | 0.09 | 0.09 |
| gene9280 | https://www.ncbi.nlm.nih.gov/gene/?term=LOC105282764 | 1635 | 35 | 214.07 | 3 | 18.35 | 8.57 | 40.89 | 36.66 | 4.23 | 4.23 | 0.09 | 0.09 |
| gene1589 | https://www.ncbi.nlm.nih.gov/gene/?term=LOC105287721 | 1407 | 39 | 277.19 | 1 | 7.11 | 2.56 | 25.67 | 22.55 | 3.13 | 3.13 | 0.09 | 0.09 |
| gene2824 | https://www.ncbi.nlm.nih.gov/gene/?term=LOC105275337 | 1772 | 54 | 304.74 | 3 | 16.93 | 5.56 | 32.91 | 29.82 | 3.09 | 3.09 | 0.09 | 0.09 |

...

... only first 25 lines shown ...

Directory: OGLall File: rnk-dmg-Cb_LI_BCphaseA2.vs.BCphaseB1.txt

| gene_ID | gene_link | gwidth | #Sites | #per10Kb | #dmSites | #dmsp10kb | %dmSites | %pSite1 | %pSite2 | DMpSite | ADMpSite | DMpNucl | ADMpNucl |
|---|---|---|---|---|---|---|---|---|---|---|---|---|---|
| gene10606 | https://www.ncbi.nlm.nih.gov/gene/?term=LOC105284273 | 6119 | 258 | 421.64 | 1 | 1.63 | 0.39 | 19.65 | 18.03 | 1.63 | 1.63 | 0.07 | 0.07 |
| gene2856 | https://www.ncbi.nlm.nih.gov/gene/?term=LOC105275384 | 1987 | 91 | 457.98 | 1 | 5.03 | 1.1 | 33.74 | 32.36 | 1.37 | 1.37 | 0.06 | 0.06 |
| gene9740 | https://www.ncbi.nlm.nih.gov/gene/?term=LOC105283277 | 2887 | 143 | 495.32 | 1 | 3.46 | 0.7 | 19.74 | 18.46 | 1.28 | 1.28 | 0.06 | 0.06 |
| gene3793 | https://www.ncbi.nlm.nih.gov/gene/?term=LOC105276429 | 2901 | 119 | 410.2 | 1 | 3.45 | 0.84 | 23.8 | 22.52 | 1.28 | 1.28 | 0.05 | 0.05 |
| gene7199 | https://www.ncbi.nlm.nih.gov/gene/?term=LOC105280366 | 1414 | 30 | 212.16 | 1 | 7.07 | 3.33 | 43.5 | 41.71 | 1.79 | 1.79 | 0.04 | 0.04 |
| gene570 | https://www.ncbi.nlm.nih.gov/gene/?term=LOC105278549 | 1673 | 70 | 418.41 | 1 | 5.98 | 1.43 | 44.03 | 43.19 | 0.83 | 0.83 | 0.03 | 0.03 |
| gene9722 | https://www.ncbi.nlm.nih.gov/gene/?term=LOC105283264 | 3951 | 88 | 222.73 | 1 | 2.53 | 1.14 | 18.85 | 20.39 | -1.54 | 1.54 | -0.03 | 0.03 |
| gene10844 | https://www.ncbi.nlm.nih.gov/gene/?term=LOC105284535 | 9086 | 246 | 270.75 | 1 | 1.1 | 0.41 | 16.16 | 15.32 | 0.84 | 0.84 | 0.02 | 0.02 |
| gene10326 | https://www.ncbi.nlm.nih.gov/gene/?term=LOC105283962 | 3042 | 130 | 427.35 | 1 | 3.29 | 0.77 | 8.82 | 8.26 | 0.56 | 0.56 | 0.02 | 0.02 |
| gene1855 | https://www.ncbi.nlm.nih.gov/gene/?term=LOC105288036 | 2387 | 89 | 372.85 | 1 | 4.19 | 1.12 | 30.86 | 30.35 | 0.51 | 0.51 | 0.02 | 0.02 |
| gene7166 | https://www.ncbi.nlm.nih.gov/gene/?term=LOC105280326 | 6117 | 235 | 384.18 | 1 | 1.63 | 0.43 | 15.36 | 14.92 | 0.43 | 0.43 | 0.02 | 0.02 |
| gene11447 | https://www.ncbi.nlm.nih.gov/gene/?term=LOC105285252 | 7437 | 131 | 176.15 | 1 | 1.34 | 0.76 | 12.71 | 13.67 | -0.95 | 0.95 | -0.02 | 0.02 |
| gene564 | https://www.ncbi.nlm.nih.gov/gene/?term=LOC105278478 | 2802 | 17 | 60.67 | 1 | 3.57 | 5.88 | 14.91 | 14.04 | 0.87 | 0.87 | 0.01 | 0.01 |
| gene9755 | https://www.ncbi.nlm.nih.gov/gene/?term=LOC105283296 | 2177 | 92 | 422.6 | 1 | 4.59 | 1.09 | 18.09 | 17.95 | 0.14 | 0.14 | 0.01 | 0.01 |
| gene3744 | https://www.ncbi.nlm.nih.gov/gene/?term=LOC105276385 | 9376 | 459 | 489.55 | 1 | 1.07 | 0.22 | 5.46 | 5.7 | -0.24 | 0.24 | -0.01 | 0.01 |
| gene4892 | https://www.ncbi.nlm.nih.gov/gene/?term=LOC105277667 | 3898 | 126 | 323.24 | 1 | 2.57 | 0.79 | 27.95 | 28.19 | -0.24 | 0.24 | -0.01 | 0.01 |
| gene8814 | https://www.ncbi.nlm.nih.gov/gene/?term=LOC105282240 | 6910 | 175 | 253.26 | 1 | 1.45 | 0.57 | 9.54 | 9.87 | -0.32 | 0.32 | -0.01 | 0.01 |
| gene10819 | https://www.ncbi.nlm.nih.gov/gene/?term=LOC105284507 | 5767 | 94 | 163 | 1 | 1.73 | 1.06 | 20.43 | 21.18 | -0.75 | 0.75 | -0.01 | 0.01 |
| gene12203 | https://www.ncbi.nlm.nih.gov/gene/?term=LOC105286099 | 5967 | 84 | 140.77 | 1 | 1.68 | 1.19 | 11.21 | 10.88 | 0.34 | 0.34 | 0 | 0 |
| gene5101 | https://www.ncbi.nlm.nih.gov/gene/?term=LOC105277903 | 1813 | 34 | 187.53 | 1 | 5.52 | 2.94 | 48.71 | 48.5 | 0.21 | 0.21 | 0 | 0 |
| gene7183 | https://www.ncbi.nlm.nih.gov/gene/?term=LOC105280348 | 3925 | 72 | 183.44 | 1 | 2.55 | 1.39 | 20.87 | 20.69 | 0.19 | 0.19 | 0 | 0 |
| gene7245 | https://www.ncbi.nlm.nih.gov/gene/?term=LOC105280410 | 12478 | 277 | 221.99 | 1 | 0.8 | 0.36 | 19.64 | 19.47 | 0.17 | 0.17 | 0 | 0 |
| gene4209 | https://www.ncbi.nlm.nih.gov/gene/?term=LOC105277125 | 50550 | 1862 | 368.35 | 1 | 0.2 | 0.05 | 1.6 | 1.65 | -0.05 | 0.05 | 0 | 0 |

Directory: OGLall File: rnk-dmg-Cb_LI_BCphaseA2.vs.BCphaseB2.txt

| gene_ID | gene_link | gwidth | #Sites | #per10Kb | #dmSites | #dmsp10kb | %dmSites | %pSite1 | %pSite2 | DMpSite | ADMpSite | DMpNucl | ADMpNucl |
|---|---|---|---|---|---|---|---|---|---|---|---|---|---|
| gene3165 | https://www.ncbi.nlm.nih.gov/gene/?term=LOC105275706 | 2374 | 67 | 282.22 | 5 | 21.06 | 7.46 | 36.03 | 23.6 | 12.43 | 12.43 | 0.35 | 0.35 |
| gene2824 | https://www.ncbi.nlm.nih.gov/gene/?term=LOC105275337 | 1772 | 64 | 361.17 | 2 | 11.29 | 3.12 | 37.53 | 29.09 | 8.44 | 8.44 | 0.3 | 0.3 |
| gene10963 | https://www.ncbi.nlm.nih.gov/gene/?term=LOC105284677 | 1576 | 28 | 177.66 | 2 | 12.69 | 7.14 | 19.34 | 35.66 | -16.32 | 16.32 | -0.29 | 0.29 |
| gene9468 | https://www.ncbi.nlm.nih.gov/gene/?term=LOC105282976 | 1636 | 53 | 323.96 | 1 | 6.11 | 1.89 | 28.76 | 22.1 | 6.66 | 6.66 | 0.22 | 0.22 |
| gene2593 | https://www.ncbi.nlm.nih.gov/gene/?term=LOC105275070 | 1774 | 5 | 28.18 | 5 | 28.18 | 100 | 80.28 | 10.96 | 69.32 | 69.32 | 0.2 | 0.2 |
| gene1957 | https://www.ncbi.nlm.nih.gov/gene/?term=LOC105288153 | 1044 | 16 | 153.26 | 1 | 9.58 | 6.25 | 15.68 | 4.26 | 11.42 | 11.42 | 0.18 | 0.18 |
| gene5714 | https://www.ncbi.nlm.nih.gov/gene/?term=LOC105278729 | 710 | 24 | 338.03 | 1 | 14.08 | 4.17 | 61.56 | 56.79 | 4.77 | 4.77 | 0.16 | 0.16 |
| gene920 | https://www.ncbi.nlm.nih.gov/gene/?term=LOC105282518 | 1197 | 40 | 334.17 | 1 | 8.35 | 2.5 | 37.69 | 40.21 | -2.52 | 2.52 | -0.08 | 0.08 |
| gene12021 | https://www.ncbi.nlm.nih.gov/gene/?term=LOC105285905 | 1399 | 20 | 142.96 | 1 | 7.15 | 5 | 32.7 | 38.03 | -5.34 | 5.34 | -0.08 | 0.08 |
| gene3817 | https://www.ncbi.nlm.nih.gov/gene/?term=LOC105276449 | 1230 | 42 | 341.46 | 1 | 8.13 | 2.38 | 35.93 | 33.84 | 2.09 | 2.09 | 0.07 | 0.07 |
| gene2404 | https://www.ncbi.nlm.nih.gov/gene/?term=LOC105274853 | 1767 | 36 | 203.74 | 1 | 5.66 | 2.78 | 27.62 | 31.08 | -3.47 | 3.47 | -0.07 | 0.07 |
| gene10816 | https://www.ncbi.nlm.nih.gov/gene/?term=LOC105284502 | 1974 | 18 | 91.19 | 1 | 5.07 | 5.56 | 38.87 | 46.78 | -7.91 | 7.91 | -0.07 | 0.07 |
| gene5877 | https://www.ncbi.nlm.nih.gov/gene/?term=LOC105278807 | 1719 | 49 | 285.05 | 1 | 5.82 | 2.04 | 23.71 | 21.48 | 2.23 | 2.23 | 0.06 | 0.06 |
| gene7556 | https://www.ncbi.nlm.nih.gov/gene/?term=LOC105280774 | 2792 | 102 | 365.33 | 1 | 3.58 | 0.98 | 32.11 | 30.45 | 1.66 | 1.66 | 0.06 | 0.06 |
| gene7568 | https://www.ncbi.nlm.nih.gov/gene/?term=LOC105280786 | 2525 | 111 | 439.6 | 1 | 3.96 | 0.9 | 33.28 | 31.99 | 1.29 | 1.29 | 0.06 | 0.06 |
| gene12290 | https://www.ncbi.nlm.nih.gov/gene/?term=LOC105286190 | 6212 | 304 | 489.38 | 1 | 1.61 | 0.33 | 8.69 | 7.5 | 1.18 | 1.18 | 0.06 | 0.06 |
| gene1220 | https://www.ncbi.nlm.nih.gov/gene/?term=LOC105285742 | 3453 | 108 | 312.77 | 1 | 2.9 | 0.93 | 21.23 | 23.26 | -2.03 | 2.03 | -0.06 | 0.06 |
| gene10606 | https://www.ncbi.nlm.nih.gov/gene/?term=LOC105284273 | 6119 | 251 | 410.2 | 1 | 1.63 | 0.4 | 20.03 | 18.79 | 1.24 | 1.24 | 0.05 | 0.05 |
| gene5298 | https://www.ncbi.nlm.nih.gov/gene/?term=LOC105278145 | 4060 | 244 | 600.99 | 2 | 4.93 | 0.82 | 10.55 | 11.35 | -0.8 | 0.8 | -0.05 | 0.05 |
| gene1993 | https://www.ncbi.nlm.nih.gov/gene/?term=LOC105288193 | 1449 | 75 | 517.6 | 1 | 6.9 | 1.33 | 35.76 | 36.74 | -0.98 | 0.98 | -0.05 | 0.05 |
| gene2271 | https://www.ncbi.nlm.nih.gov/gene/?term=LOC105274706 | 1448 | 56 | 386.74 | 1 | 6.91 | 1.79 | 36.89 | 38.13 | -1.24 | 1.24 | -0.05 | 0.05 |
| gene1199 | https://www.ncbi.nlm.nih.gov/gene/?term=LOC105285498 | 2220 | 26 | 117.12 | 1 | 4.5 | 3.85 | 33.05 | 37.03 | -3.99 | 3.99 | -0.05 | 0.05 |
| gene3385 | https://www.ncbi.nlm.nih.gov/gene/?term=LOC105275966 | 3305 | 61 | 184.57 | 1 | 3.03 | 1.64 | 49.58 | 47.17 | 2.41 | 2.41 | 0.04 | 0.04 |
| gene9462 | https://www.ncbi.nlm.nih.gov/gene/?term=LOC105282970 | 1469 | 39 | 265.49 | 1 | 6.81 | 2.56 | 53.94 | 52.44 | 1.5 | 1.5 | 0.04 | 0.04 |
| gene4892 | https://www.ncbi.nlm.nih.gov/gene/?term=LOC105277667 | 3898 | 125 | 320.68 | 1 | 2.57 | 0.8 | 27.74 | 26.47 | 1.27 | 1.27 | 0.04 | 0.04 |
| gene10678 | https://www.ncbi.nlm.nih.gov/gene/?term=LOC105284355 | 4031 | 167 | 414.29 | 1 | 2.48 | 0.6 | 19.59 | 18.65 | 0.94 | 0.94 | 0.04 | 0.04 |
| gene4034 | https://www.ncbi.nlm.nih.gov/gene/?term=LOC105276704 | 2065 | 101 | 489.1 | 1 | 4.84 | 0.99 | 13.4 | 12.57 | 0.83 | 0.83 | 0.04 | 0.04 |
| gene4498 | https://www.ncbi.nlm.nih.gov/gene/?term=LOC105277239 | 2831 | 129 | 455.67 | 1 | 3.53 | 0.78 | 14.44 | 15.34 | -0.89 | 0.89 | -0.04 | 0.04 |
| gene11340 | https://www.ncbi.nlm.nih.gov/gene/?term=LOC105285131 | 2918 | 101 | 346.13 | 1 | 3.43 | 0.99 | 10.9 | 11.93 | -1.02 | 1.02 | -0.04 | 0.04 |
| gene1518 | https://www.ncbi.nlm.nih.gov/gene/?term=LOC105287648 | 2195 | 27 | 123.01 | 1 | 4.56 | 3.7 | 41.41 | 38.92 | 2.49 | 2.49 | 0.03 | 0.03 |
| gene8115 | https://www.ncbi.nlm.nih.gov/gene/?term=LOC105281420 | 3623 | 51 | 140.77 | 1 | 2.76 | 1.96 | 12.6 | 10.51 | 2.09 | 2.09 | 0.03 | 0.03 |
| gene8742 | https://www.ncbi.nlm.nih.gov/gene/?term=LOC105282151 | 2563 | 41 | 159.97 | 2 | 7.8 | 4.88 | 36.1 | 34.31 | 1.79 | 1.79 | 0.03 | 0.03 |
| gene12572 | https://www.ncbi.nlm.nih.gov/gene/?term=LOC105286514 | 2374 | 42 | 176.92 | 1 | 4.21 | 2.38 | 34.39 | 32.92 | 1.47 | 1.47 | 0.03 | 0.03 |
| gene5967 | https://www.ncbi.nlm.nih.gov/gene/?term=LOC105278918 | 4393 | 121 | 275.44 | 1 | 2.28 | 0.83 | 15.97 | 14.91 | 1.05 | 1.05 | 0.03 | 0.03 |
| gene8729 | https://www.ncbi.nlm.nih.gov/gene/?term=LOC105282137 | 5005 | 197 | 393.61 | 1 | 2 | 0.51 | 17.44 | 16.65 | 0.8 | 0.8 | 0.03 | 0.03 |
| gene10520 | https://www.ncbi.nlm.nih.gov/gene/?term=LOC105284184 | 2451 | 85 | 346.8 | 1 | 4.08 | 1.18 | 34.37 | 33.59 | 0.78 | 0.78 | 0.03 | 0.03 |
| gene8274 | https://www.ncbi.nlm.nih.gov/gene/?term=LOC105281624 | 5425 | 230 | 423.96 | 1 | 1.84 | 0.43 | 9.61 | 10.33 | -0.73 | 0.73 | -0.03 | 0.03 |
| gene10018 | https://www.ncbi.nlm.nih.gov/gene/?term=LOC105283607 | 4414 | 191 | 432.71 | 1 | 2.27 | 0.52 | 15.56 | 16.32 | -0.76 | 0.76 | -0.03 | 0.03 |
| gene10213 | https://www.ncbi.nlm.nih.gov/gene/?term=LOC105283826 | 1503 | 64 | 425.82 | 1 | 6.65 | 1.56 | 14.04 | 14.85 | -0.81 | 0.81 | -0.03 | 0.03 |
| gene5646 | https://www.ncbi.nlm.nih.gov/gene/?term=LOC105278534 | 7475 | 181 | 242.14 | 1 | 1.34 | 0.55 | 17.19 | 18.52 | -1.32 | 1.32 | -0.03 | 0.03 |
| gene11447 | https://www.ncbi.nlm.nih.gov/gene/?term=LOC105285252 | 7437 | 120 | 161.36 | 1 | 1.34 | 0.83 | 13.34 | 14.97 | -1.63 | 1.63 | -0.03 | 0.03 |
| gene6455 | https://www.ncbi.nlm.nih.gov/gene/?term=LOC105279500 | 2612 | 33 | 126.34 | 1 | 3.83 | 3.03 | 45.07 | 47.06 | -1.99 | 1.99 | -0.03 | 0.03 |
| gene2748 | https://www.ncbi.nlm.nih.gov/gene/?term=LOC105275261 | 3845 | 52 | 135.24 | 1 | 2.6 | 1.92 | 36.08 | 38.33 | -2.25 | 2.25 | -0.03 | 0.03 |
| gene12395 | https://www.ncbi.nlm.nih.gov/gene/?term=LOC105286302 | 4093 | 17 | 41.53 | 1 | 2.44 | 5.88 | 21.99 | 16.58 | 5.41 | 5.41 | 0.02 | 0.02 |
| gene5052 | https://www.ncbi.nlm.nih.gov/gene/?term=LOC105277856 | 3167 | 34 | 107.36 | 1 | 3.16 | 2.94 | 38.79 | 36.66 | 2.13 | 2.13 | 0.02 | 0.02 |
| gene2970 | https://www.ncbi.nlm.nih.gov/gene/?term=LOC105275518 | 2340 | 19 | 81.2 | 1 | 4.27 | 5.26 | 46.39 | 44.36 | 2.03 | 2.03 | 0.02 | 0.02 |
| gene7398 | https://www.ncbi.nlm.nih.gov/gene/?term=LOC105280589 | 3452 | 39 | 112.98 | 1 | 2.9 | 2.56 | 23.26 | 21.67 | 1.58 | 1.58 | 0.02 | 0.02 |
| gene927 | https://www.ncbi.nlm.nih.gov/gene/?term=LOC105282600 | 3727 | 53 | 142.21 | 1 | 2.68 | 1.89 | 32.11 | 30.67 | 1.44 | 1.44 | 0.02 | 0.02 |
| gene1884 | https://www.ncbi.nlm.nih.gov/gene/?term=LOC105288072 | 4169 | 71 | 170.3 | 1 | 2.4 | 1.41 | 25.5 | 24.39 | 1.1 | 1.1 | 0.02 | 0.02 |
| gene3470 | https://www.ncbi.nlm.nih.gov/gene/?term=LOC105276059 | 8716 | 145 | 166.36 | 1 | 1.15 | 0.69 | 10.01 | 8.94 | 1.08 | 1.08 | 0.02 | 0.02 |
| gene940 | https://www.ncbi.nlm.nih.gov/gene/?term=LOC105282794 | 3578 | 85 | 237.56 | 1 | 2.79 | 1.18 | 16.1 | 15.1 | 1 | 1 | 0.02 | 0.02 |
| gene13033 | https://www.ncbi.nlm.nih.gov/gene/?term=LOC105287063 | 1877 | 49 | 261.05 | 1 | 5.33 | 2.04 | 31.12 | 30.35 | 0.77 | 0.77 | 0.02 | 0.02 |
| gene9336 | https://www.ncbi.nlm.nih.gov/gene/?term=LOC105282834 | 5993 | 124 | 206.91 | 1 | 1.67 | 0.81 | 16.06 | 15.3 | 0.75 | 0.75 | 0.02 | 0.02 |
| gene12516 | https://www.ncbi.nlm.nih.gov/gene/?term=LOC105286452 | 1529 | 38 | 248.53 | 1 | 6.54 | 2.63 | 52.34 | 51.61 | 0.73 | 0.73 | 0.02 | 0.02 |
| gene4376 | https://www.ncbi.nlm.nih.gov/gene/?term=LOC105277083 | 4307 | 127 | 294.87 | 1 | 2.32 | 0.79 | 19.77 | 19.11 | 0.65 | 0.65 | 0.02 | 0.02 |
| gene7514 | https://www.ncbi.nlm.nih.gov/gene/?term=LOC105280723 | 4471 | 200 | 447.33 | 1 | 2.24 | 0.5 | 14.49 | 14.01 | 0.49 | 0.49 | 0.02 | 0.02 |
| gene2056 | https://www.ncbi.nlm.nih.gov/gene/?term=LOC105274452 | 6584 | 263 | 399.45 | 2 | 3.04 | 0.76 | 13.53 | 13.06 | 0.47 | 0.47 | 0.02 | 0.02 |
| gene3286 | https://www.ncbi.nlm.nih.gov/gene/?term=LOC105275848 | 3744 | 140 | 373.93 | 1 | 2.67 | 0.71 | 23 | 22.56 | 0.45 | 0.45 | 0.02 | 0.02 |
| gene5746 | https://www.ncbi.nlm.nih.gov/gene/?term=LOC105278666 | 10343 | 461 | 445.71 | 1 | 0.97 | 0.22 | 7.31 | 6.93 | 0.37 | 0.37 | 0.02 | 0.02 |
| gene1470 | https://www.ncbi.nlm.nih.gov/gene/?term=LOC105287597 | 2974 | 204 | 685.94 | 1 | 3.36 | 0.49 | 17.73 | 17.44 | 0.3 | 0.3 | 0.02 | 0.02 |

| | | | | | | | | | | | | | |
|---|---|---|---|---|---|---|---|---|---|---|---|---|---|
| gene4720 | https://www.ncbi.nlm.nih.gov/gene/?term=LOC105277479 | 1697 | 47 | 276.96 | 1 | 5.89 | 2.13 | 37.56 | 38.12 | -0.55 | 0.55 | -0.02 | 0.02 |
| gene9826 | https://www.ncbi.nlm.nih.gov/gene/?term=LOC105283385 | 9219 | 305 | 330.84 | 1 | 1.08 | 0.33 | 12.6 | 13.19 | -0.59 | 0.59 | -0.02 | 0.02 |
| gene6966 | https://www.ncbi.nlm.nih.gov/gene/?term=LOC105280104 | 3747 | 122 | 325.59 | 1 | 2.67 | 0.82 | 15.27 | 15.88 | -0.62 | 0.62 | -0.02 | 0.02 |
| gene1147 | https://www.ncbi.nlm.nih.gov/gene/?term=LOC105284954 | 4022 | 95 | 236.2 1 | 2.49 | 1.05 | 15.81 | 16.65 | -0.84 | 0.84 | -0.02 | 0.02 | |
| gene7915 | https://www.ncbi.nlm.nih.gov/gene/?term=LOC105281173 | 1597 | 38 | 237.95 | 1 | 6.26 | 2.63 | 59.94 | 60.79 | -0.85 | 0.85 | -0.02 | 0.02 |
| gene2413 | https://www.ncbi.nlm.nih.gov/gene/?term=LOC105274863 | 2006 | 44 | 219.34 | 1 | 4.99 | 2.27 | 63.14 | 64.02 | -0.87 | 0.87 | -0.02 | 0.02 |
| gene12406 | https://www.ncbi.nlm.nih.gov/gene/?term=LOC105286316 | 1386 | 28 | 202.02 | 1 | 7.22 | 3.57 | 47.22 | 48.43 | -1.22 | 1.22 | -0.02 | 0.02 |
| gene322 | https://www.ncbi.nlm.nih.gov/gene/?term=LOC105275777 | 1606 | 24 | 149.44 | 1 | 6.23 | 4.17 | 45 | 46.35 | -1.35 | 1.35 | -0.02 | 0.02 |
| gene388 | https://www.ncbi.nlm.nih.gov/gene/?term=LOC105276538 | 1558 | 20 | 128.37 | 2 | 12.84 | 10 | 64.73 | 66.14 | -1.41 | 1.41 | -0.02 | 0.02 |
| gene6422 | https://www.ncbi.nlm.nih.gov/gene/?term=LOC105279458 | 2419 | 34 | 140.55 | 1 | 4.13 | 2.94 | 51.48 | 53.11 | -1.63 | 1.63 | -0.02 | 0.02 |
| gene10819 | https://www.ncbi.nlm.nih.gov/gene/?term=LOC105284507 | 5767 | 77 | 133.52 | 1 | 1.73 | 1.3 | 20.73 | 22.56 | -1.84 | 1.84 | -0.02 | 0.02 |
| gene6528 | https://www.ncbi.nlm.nih.gov/gene/?term=LOC105279591 | 7222 | 55 | 76.16 1 | 1.38 | 1.82 | 29.5 | 28.12 | 1.38 | 1.38 | 0.01 | 0.01 | |
| gene2825 | https://www.ncbi.nlm.nih.gov/gene/?term=LOC105275343 | 5545 | 96 | 173.13 | 2 | 3.61 | 2.08 | 11.79 | 11.04 | 0.75 | 0.75 | 0.01 | 0.01 |
| gene6620 | https://www.ncbi.nlm.nih.gov/gene/?term=LOC105279687 | 17998 | 336 | 186.69 | 1 | 0.56 | 0.3 | 5.15 | 4.42 | 0.73 | 0.73 | 0.01 | 0.01 |
| gene12307 | https://www.ncbi.nlm.nih.gov/gene/?term=LOC105286216 | 10313 | 166 | 160.96 | 1 | 0.97 | 0.6 | 16.86 | 16.32 | 0.54 | 0.54 | 0.01 | 0.01 |
| gene7368 | https://www.ncbi.nlm.nih.gov/gene/?term=LOC105280556 | 4105 | 50 | 121.8 1 | 2.44 | 2 | 18.71 | 18.17 | 0.54 | 0.54 | 0.01 | 0.01 | |
| gene292 | https://www.ncbi.nlm.nih.gov/gene/?term=LOC105275398 | 2825 | 86 | 304.42 | 1 | 3.54 | 1.16 | 21.94 | 21.53 | 0.4 | 0.4 | 0.01 | 0.01 |
| gene4962 | https://www.ncbi.nlm.nih.gov/gene/?term=LOC105277761 | 4645 | 150 | 322.93 | 1 | 2.15 | 0.67 | 15.92 | 15.52 | 0.4 | 0.4 | 0.01 | 0.01 |
| gene7245 | https://www.ncbi.nlm.nih.gov/gene/?term=LOC105280410 | 12478 | 237 | 189.93 | 1 | 0.8 | 0.42 | 21.04 | 20.64 | 0.4 | 0.4 | 0.01 | 0.01 |
| gene2173 | https://www.ncbi.nlm.nih.gov/gene/?term=LOC105274590 | 4387 | 64 | 145.89 | 2 | 4.56 | 3.12 | 34.09 | 33.7 | 0.39 | 0.39 | 0.01 | 0.01 |
| gene6031 | https://www.ncbi.nlm.nih.gov/gene/?term=LOC105279000 | 5691 | 195 | 342.65 | 1 | 1.76 | 0.51 | 14.39 | 14 | 0.39 | 0.39 | 0.01 | 0.01 |
| gene1430 | https://www.ncbi.nlm.nih.gov/gene/?term=LOC105287551 | 6433 | 231 | 359.09 | 1 | 1.55 | 0.43 | 9.26 | 8.92 | 0.34 | 0.34 | 0.01 | 0.01 |
| gene10531 | https://www.ncbi.nlm.nih.gov/gene/?term=LOC105284190 | 6881 | 173 | 251.42 | 1 | 1.45 | 0.58 | 12.82 | 12.5 | 0.32 | 0.32 | 0.01 | 0.01 |
| gene1310 | https://www.ncbi.nlm.nih.gov/gene/?term=LOC105286661 | 2582 | 66 | 255.62 | 1 | 3.87 | 1.52 | 24.45 | 24.13 | 0.32 | 0.32 | 0.01 | 0.01 |
| gene4246 | https://www.ncbi.nlm.nih.gov/gene/?term=LOC105276941 | 16412 | 552 | 336.34 | 1 | 0.61 | 0.18 | 5.12 | 4.8 | 0.32 | 0.32 | 0.01 | 0.01 |
| gene11568 | https://www.ncbi.nlm.nih.gov/gene/?term=LOC105285395 | 8086 | 192 | 237.45 | 1 | 1.24 | 0.52 | 15.07 | 14.77 | 0.31 | 0.31 | 0.01 | 0.01 |
| gene3856 | https://www.ncbi.nlm.nih.gov/gene/?term=LOC105276499 | 5236 | 190 | 362.87 | 1 | 1.91 | 0.53 | 9.53 | 9.24 | 0.29 | 0.29 | 0.01 | 0.01 |
| gene7249 | https://www.ncbi.nlm.nih.gov/gene/?term=LOC105280422 | 1775 | 63 | 354.93 | 1 | 5.63 | 1.59 | 17.96 | 17.74 | 0.23 | 0.23 | 0.01 | 0.01 |
| gene2806 | https://www.ncbi.nlm.nih.gov/gene/?term=LOC105275327 | 14257 | 494 | 346.5 1 | 0.7 | 0.2 | 6.88 | 6.68 | 0.2 | 0.2 | 0.01 | 0.01 | |
| gene11275 | https://www.ncbi.nlm.nih.gov/gene/?term=LOC105285055 | 4119 | 255 | 619.08 | 1 | 2.43 | 0.39 | 11.36 | 11.16 | 0.19 | 0.19 | 0.01 | 0.01 |
| gene2846 | https://www.ncbi.nlm.nih.gov/gene/?term=LOC105275374 | 11246 | 502 | 446.38 | 1 | 0.89 | 0.2 | 11.4 | 11.21 | 0.19 | 0.19 | 0.01 | 0.01 |
| gene7166 | https://www.ncbi.nlm.nih.gov/gene/?term=LOC105280326 | 6117 | 233 | 380.91 | 1 | 1.63 | 0.43 | 15.64 | 15.46 | 0.19 | 0.19 | 0.01 | 0.01 |
| gene11401 | https://www.ncbi.nlm.nih.gov/gene/?term=LOC105285202 | 1841 | 82 | 445.41 | 1 | 5.43 | 1.22 | 25.44 | 25.28 | 0.15 | 0.15 | 0.01 | 0.01 |
| gene12806 | https://www.ncbi.nlm.nih.gov/gene/?term=LOC105286803 | 4196 | 143 | 340.8 1 | 2.38 | 0.7 | 11.07 | 10.92 | 0.15 | 0.15 | 0.01 | 0.01 | |
| gene5524 | https://www.ncbi.nlm.nih.gov/gene/?term=LOC105278397 | 5177 | 203 | 392.12 | 1 | 1.93 | 0.49 | 25.25 | 25.12 | 0.13 | 0.13 | 0.01 | 0.01 |
| gene12802 | https://www.ncbi.nlm.nih.gov/gene/?term=LOC105286799 | 3183 | 185 | 581.21 | 2 | 6.28 | 1.08 | 16.06 | 15.94 | 0.12 | 0.12 | 0.01 | 0.01 |
| gene12674 | https://www.ncbi.nlm.nih.gov/gene/?term=LOC105286649 | 18283 | 811 | 443.58 | 1 | 0.55 | 0.12 | 0.93 | 1.06 | -0.13 | 0.13 | -0.01 | 0.01 |
| gene13114 | https://www.ncbi.nlm.nih.gov/gene/?term=LOC105287147 | 7609 | 409 | 537.52 | 1 | 1.31 | 0.24 | 9.57 | 9.7 | -0.13 | 0.13 | -0.01 | 0.01 |
| gene3885 | https://www.ncbi.nlm.nih.gov/gene/?term=LOC105276527 | 7928 | 317 | 399.85 | 1 | 1.26 | 0.32 | 9.68 | 9.86 | -0.18 | 0.18 | -0.01 | 0.01 |
| gene5777 | https://www.ncbi.nlm.nih.gov/gene/?term=LOC105278691 | 3883 | 148 | 381.15 | 1 | 2.58 | 0.68 | 14.2 | 14.38 | -0.18 | 0.18 | -0.01 | 0.01 |
| gene6583 | https://www.ncbi.nlm.nih.gov/gene/?term=LOC105279649 | 7444 | 332 | 446 1 | 1.34 | 0.3 | 10.76 | 10.98 | -0.22 | 0.22 | -0.01 | 0.01 | |
| gene11454 | https://www.ncbi.nlm.nih.gov/gene/?term=LOC105285256 | 1906 | 47 | 246.59 | 1 | 5.25 | 2.13 | 45.38 | 45.61 | -0.23 | 0.23 | -0.01 | 0.01 |
| gene13036 | https://www.ncbi.nlm.nih.gov/gene/?term=LOC105287059 | 4959 | 163 | 328.7 1 | 2.02 | 0.61 | 16.39 | 16.65 | -0.26 | 0.26 | -0.01 | 0.01 | |
| gene3727 | https://www.ncbi.nlm.nih.gov/gene/?term=LOC105276367 | 2484 | 69 | 277.78 | 1 | 4.03 | 1.45 | 46.73 | 46.99 | -0.27 | 0.27 | -0.01 | 0.01 |
| gene2257 | https://www.ncbi.nlm.nih.gov/gene/?term=LOC105274689 | 3658 | 71 | 194.1 1 | 2.73 | 1.41 | 14.85 | 15.18 | -0.34 | 0.34 | -0.01 | 0.01 | |
| gene7600 | https://www.ncbi.nlm.nih.gov/gene/?term=LOC105280819 | 1615 | 39 | 241.49 | 1 | 6.19 | 2.56 | 35.76 | 36.1 | -0.34 | 0.34 | -0.01 | 0.01 |
| gene11082 | https://www.ncbi.nlm.nih.gov/gene/?term=LOC105284824 | 7436 | 200 | 268.96 | 1 | 1.34 | 0.5 | 5.46 | 5.81 | -0.35 | 0.35 | -0.01 | 0.01 |
| gene12573 | https://www.ncbi.nlm.nih.gov/gene/?term=LOC105286515 | 3647 | 113 | 309.84 | 1 | 2.74 | 0.88 | 10.7 | 11.07 | -0.37 | 0.37 | -0.01 | 0.01 |
| gene10600 | https://www.ncbi.nlm.nih.gov/gene/?term=LOC105284270 | 4837 | 124 | 256.36 | 1 | 2.07 | 0.81 | 10.47 | 10.85 | -0.38 | 0.38 | -0.01 | 0.01 |
| gene3046 | https://www.ncbi.nlm.nih.gov/gene/?term=LOC105275587 | 2394 | 56 | 233.92 | 1 | 4.18 | 1.79 | 33.82 | 34.2 | -0.38 | 0.38 | -0.01 | 0.01 |
| gene12981 | https://www.ncbi.nlm.nih.gov/gene/?term=LOC105286999 | 2063 | 40 | 193.89 | 1 | 4.85 | 2.5 | 35.8 | 36.25 | -0.45 | 0.45 | -0.01 | 0.01 |
| gene10107 | https://www.ncbi.nlm.nih.gov/gene/?term=LOC105283712 | 1883 | 32 | 169.94 | 1 | 5.31 | 3.12 | 29.98 | 30.44 | -0.46 | 0.46 | -0.01 | 0.01 |
| gene11561 | https://www.ncbi.nlm.nih.gov/gene/?term=LOC105285387 | 11528 | 194 | 168.29 | 1 | 0.87 | 0.52 | 8.65 | 9.11 | -0.47 | 0.47 | -0.01 | 0.01 |
| gene8015 | https://www.ncbi.nlm.nih.gov/gene/?term=LOC105281312 | 2998 | 72 | 240.16 | 1 | 3.34 | 1.39 | 29.67 | 30.18 | -0.5 | 0.5 | -0.01 | 0.01 |
| gene8814 | https://www.ncbi.nlm.nih.gov/gene/?term=LOC105282240 | 6910 | 170 | 246.02 | 1 | 1.45 | 0.59 | 9.16 | 9.68 | -0.52 | 0.52 | -0.01 | 0.01 |
| gene10391 | https://www.ncbi.nlm.nih.gov/gene/?term=LOC109611283 | 1994 | 44 | 220.66 | 1 | 5.02 | 2.27 | 32.96 | 33.52 | -0.56 | 0.56 | -0.01 | 0.01 |
| gene3671 | https://www.ncbi.nlm.nih.gov/gene/?term=LOC105276272 | 1966 | 36 | 183.11 | 1 | 5.09 | 2.78 | 32.08 | 32.66 | -0.58 | 0.58 | -0.01 | 0.01 |
| gene6639 | https://www.ncbi.nlm.nih.gov/gene/?term=LOC105279705 | 7656 | 62 | 80.98 1 | 1.31 | 1.61 | 22.96 | 23.88 | -0.92 | 0.92 | -0.01 | 0.01 | |
| gene10345 | https://www.ncbi.nlm.nih.gov/gene/?term=LOC105283989 | 4444 | 23 | 51.76 1 | 2.25 | 4.35 | 43.82 | 45.07 | -1.26 | 1.26 | -0.01 | 0.01 | |
| gene3168 | https://www.ncbi.nlm.nih.gov/gene/?term=LOC105275711 | 6150 | 70 | 113.82 | 1 | 1.63 | 1.43 | 12.53 | 13.84 | -1.31 | 1.31 | -0.01 | 0.01 |
| gene10654 | https://www.ncbi.nlm.nih.gov/gene/?term=LOC105284329 | 4027 | 50 | 124.16 | 1 | 2.48 | 2 | 19.81 | 19.43 | 0.39 | 0.39 | 0 | 0 |
| gene3599 | https://www.ncbi.nlm.nih.gov/gene/?term=LOC105276196 | 2955 | 48 | 162.44 | 1 | 3.38 | 2.08 | 45.56 | 45.28 | 0.28 | 0.28 | 0 | 0 |
| gene8600 | https://www.ncbi.nlm.nih.gov/gene/?term=LOC105282001 | 8513 | 156 | 183.25 | 1 | 1.17 | 0.64 | 12.04 | 11.82 | 0.22 | 0.22 | 0 | 0 |
| gene12439 | https://www.ncbi.nlm.nih.gov/gene/?term=LOC105286357 | 4761 | 81 | 170.13 | 1 | 2.1 | 1.23 | 32.58 | 32.4 | 0.18 | 0.18 | 0 | 0 |

| | | | | | | | | | | | | | |
|---|---|---|---|---|---|---|---|---|---|---|---|---|---|
| gene11579 | https://www.ncbi.nlm.nih.gov/gene/?term=LOC105285406 | 13571 | 267 | 196.74 | 1 | 0.74 | 0.37 | 6.68 | 6.51 | 0.17 | 0.17 | 0 | 0 |
| gene8343 | https://www.ncbi.nlm.nih.gov/gene/?term=LOC105281700 | 2172 | 37 | 170.35 | 1 | 4.6 | 2.7 | 39.71 | 39.54 | 0.16 | 0.16 | 0 | 0 |
| gene12203 | https://www.ncbi.nlm.nih.gov/gene/?term=LOC105286099 | 5967 | 72 | 120.66 | 1 | 1.68 | 1.39 | 10.74 | 10.59 | 0.15 | 0.15 | 0 | 0 |
| gene4190 | https://www.ncbi.nlm.nih.gov/gene/?term=LOC105276875 | 3150 | 107 | 339.68 | 1 | 3.17 | 0.93 | 23.8 | 23.66 | 0.15 | 0.15 | 0 | 0 |
| gene1198 | https://www.ncbi.nlm.nih.gov/gene/?term=LOC105285507 | 2738 | 88 | 321.4 1 | 3.65 | 1.14 | 23.48 | 23.34 | 0.14 | 0.14 | 0 | 0 | |
| gene10392 | https://www.ncbi.nlm.nih.gov/gene/?term=LOC105284036 | 3216 | 94 | 292.29 | 2 | 6.22 | 2.13 | 22.55 | 22.51 | 0.04 | 0.04 | 0 | 0 |
| gene53 | https://www.ncbi.nlm.nih.gov/gene/?term=LOC105277148 | 19283 | 287 | 148.84 | 1 | 0.52 | 0.35 | 5.29 | 5.27 | 0.03 | 0.03 | 0 | 0 |
| gene6286 | https://www.ncbi.nlm.nih.gov/gene/?term=LOC105279298 | 2861 | 125 | 436.91 | 2 | 6.99 | 1.6 | 25.22 | 25.19 | 0.03 | 0.03 | 0 | 0 |
| gene13044 | https://www.ncbi.nlm.nih.gov/gene/?term=LOC105287070 | 5094 | 241 | 473.11 | 1 | 1.96 | 0.41 | 5.28 | 5.27 | 0.02 | 0.02 | 0 | 0 |
| gene2844 | https://www.ncbi.nlm.nih.gov/gene/?term=LOC105275363 | 4330 | 192 | 443.42 | 1 | 2.31 | 0.52 | 14.25 | 14.23 | 0.02 | 0.02 | 0 | 0 |
| gene5525 | https://www.ncbi.nlm.nih.gov/gene/?term=LOC105278396 | 3090 | 188 | 608.41 | 1 | 3.24 | 0.53 | 24.8 | 24.79 | 0.01 | 0.01 | 0 | 0 |
| gene9330 | https://www.ncbi.nlm.nih.gov/gene/?term=LOC105282825 | 3936 | 80 | 203.25 | 1 | 2.54 | 1.25 | 12.57 | 12.58 | -0.01 | 0.01 | 0 | 0 |
| gene9755 | https://www.ncbi.nlm.nih.gov/gene/?term=LOC105283296 | 2177 | 85 | 390.45 | 1 | 4.59 | 1.18 | 21.44 | 21.45 | -0.01 | 0.01 | 0 | 0 |
| gene298 | https://www.ncbi.nlm.nih.gov/gene/?term=LOC105275460 | 2473 | 114 | 460.98 | 1 | 4.04 | 0.88 | 13.89 | 13.91 | -0.02 | 0.02 | 0 | 0 |
| gene7319 | https://www.ncbi.nlm.nih.gov/gene/?term=LOC105280513 | 68997 | 1241 | 179.86 | 1 | 0.14 | 0.08 | 0.56 | 0.6 | -0.04 | 0.04 | 0 | 0 |
| gene10549 | https://www.ncbi.nlm.nih.gov/gene/?term=LOC105284210 | 7145 | 325 | 454.86 | 2 | 2.8 | 0.62 | 10.71 | 10.76 | -0.05 | 0.05 | 0 | 0 |
| gene3848 | https://www.ncbi.nlm.nih.gov/gene/?term=LOC105276486 | 5899 | 133 | 225.46 | 1 | 1.7 | 0.75 | 24.81 | 24.86 | -0.05 | 0.05 | 0 | 0 |
| gene9177 | https://www.ncbi.nlm.nih.gov/gene/?term=LOC105282646 | 3320 | 98 | 295.18 | 1 | 3.01 | 1.02 | 22.82 | 22.88 | -0.05 | 0.05 | 0 | 0 |
| gene4209 | https://www.ncbi.nlm.nih.gov/gene/?term=LOC105277125 | 50550 | 1852 | 366.37 | 1 | 0.2 | 0.05 | 1.57 | 1.63 | -0.06 | 0.06 | 0 | 0 |
| gene4641 | https://www.ncbi.nlm.nih.gov/gene/?term=LOC105277398 | 1937 | 77 | 397.52 | 1 | 5.16 | 1.3 | 34.01 | 34.07 | -0.06 | 0.06 | 0 | 0 |
| gene11688 | https://www.ncbi.nlm.nih.gov/gene/?term=LOC105285533 | 1637 | 60 | 366.52 | 1 | 6.11 | 1.67 | 26.05 | 26.13 | -0.07 | 0.07 | 0 | 0 |
| gene3107 | https://www.ncbi.nlm.nih.gov/gene/?term=LOC105275650 | 5103 | 181 | 354.69 | 1 | 1.96 | 0.55 | 15.45 | 15.53 | -0.08 | 0.08 | 0 | 0 |
| gene5831 | https://www.ncbi.nlm.nih.gov/gene/?term=LOC105278761 | 2506 | 119 | 474.86 | 2 | 7.98 | 1.68 | 21.61 | 21.68 | -0.08 | 0.08 | 0 | 0 |
| gene9445 | https://www.ncbi.nlm.nih.gov/gene/?term=LOC105282951 | 2285 | 115 | 503.28 | 1 | 4.38 | 0.87 | 20.19 | 20.28 | -0.09 | 0.09 | 0 | 0 |
| gene6652 | https://www.ncbi.nlm.nih.gov/gene/?term=LOC105279719 | 2831 | 79 | 279.05 | 1 | 3.53 | 1.27 | 24.78 | 24.91 | -0.14 | 0.14 | 0 | 0 |
| gene1632 | https://www.ncbi.nlm.nih.gov/gene/?term=LOC105287764 | 18061 | 393 | 217.6 2 | 1.11 | 0.51 | 2.97 | 3.13 | -0.17 | 0.17 | 0 | 0 | |
| gene8661 | https://www.ncbi.nlm.nih.gov/gene/?term=LOC105282063 | 1926 | 32 | 166.15 | 1 | 5.19 | 3.12 | 29.26 | 29.43 | -0.17 | 0.17 | 0 | 0 |
| gene5375 | https://www.ncbi.nlm.nih.gov/gene/?term=LOC105278228 | 8640 | 163 | 188.66 | 1 | 1.16 | 0.61 | 21.78 | 22.01 | -0.23 | 0.23 | 0 | 0 |
| gene6237 | https://www.ncbi.nlm.nih.gov/gene/?term=LOC105279244 | 1484 | 24 | 161.73 | 1 | 6.74 | 4.17 | 52.96 | 53.25 | -0.29 | 0.29 | 0 | 0 |
| gene2683 | https://www.ncbi.nlm.nih.gov/gene/?term=LOC105275187 | 5492 | 70 | 127.46 | 1 | 1.82 | 1.43 | 14.08 | 14.4 | -0.31 | 0.31 | 0 | 0 |
| gene10155 | https://www.ncbi.nlm.nih.gov/gene/?term=LOC105283756 | 7030 | 93 | 132.29 | 1 | 1.42 | 1.08 | 5.63 | 5.96 | -0.34 | 0.34 | 0 | 0 |

Directory: OGLall File: rnk-dmg-Cb_LI_BCphaseA2.vs.RphaseA1.txt

| gene_ID | gene_link | gwidth | #Sites | #per10Kb | #dmSites | #dmsp10kb | %dmSites | %pSite1 | %pSite2 | DMpSite | ADMpSite | DMpNucl | ADMpNucl |
|---|---|---|---|---|---|---|---|---|---|---|---|---|---|
| gene558 | https://www.ncbi.nlm.nih.gov/gene/?term=LOC105278458 | 216 | 3 | 138.89 | 1 | 46.3 | 33.33 | 10.57 | 24.69 | -14.12 | 14.12 | -0.2 | 0.2 |
| gene12538 | https://www.ncbi.nlm.nih.gov/gene/?term=LOC105286531 | 372 | 23 | 618.28 | 1 | 26.88 | 4.35 | 39.65 | 36.5 | 3.15 | 3.15 | 0.19 | 0.19 |
| gene11566 | https://www.ncbi.nlm.nih.gov/gene/?term=LOC105285392 | 1603 | 82 | 511.54 | 3 | 18.71 | 3.66 | 38.11 | 34.95 | 3.16 | 3.16 | 0.16 | 0.16 |
| gene2837 | https://www.ncbi.nlm.nih.gov/gene/?term=LOC105275355 | 2872 | 154 | 536.21 | 7 | 24.37 | 4.55 | 26.21 | 29.04 | -2.82 | 2.82 | -0.15 | 0.15 |
| gene10731 | https://www.ncbi.nlm.nih.gov/gene/?term=LOC105284411 | 1794 | 63 | 351.17 | 4 | 22.3 | 6.35 | 36.85 | 33 | 3.85 | 3.85 | 0.14 | 0.14 |
| gene12411 | https://www.ncbi.nlm.nih.gov/gene/?term=LOC105286320 | 1660 | 60 | 361.45 | 4 | 24.1 | 6.67 | 41.91 | 38.27 | 3.64 | 3.64 | 0.13 | 0.13 |
| gene4635 | https://www.ncbi.nlm.nih.gov/gene/?term=LOC105277390 | 2067 | 81 | 391.87 | 3 | 14.51 | 3.7 | 45.5 | 42.07 | 3.43 | 3.43 | 0.13 | 0.13 |
| gene3268 | https://www.ncbi.nlm.nih.gov/gene/?term=LOC105275833 | 1040 | 41 | 394.23 | 2 | 19.23 | 4.88 | 38.46 | 35.12 | 3.35 | 3.35 | 0.13 | 0.13 |
| gene12925 | https://www.ncbi.nlm.nih.gov/gene/?term=LOC109611487 | 391 | 27 | 690.54 | 1 | 25.58 | 3.7 | 17.14 | 15.18 | 1.95 | 1.95 | 0.13 | 0.13 |
| gene1024 | https://www.ncbi.nlm.nih.gov/gene/?term=LOC105283693 | 1307 | 12 | 91.81 | 4 | 30.6 | 33.33 | 71.36 | 58.1 | 13.26 | 13.26 | 0.12 | 0.12 |
| gene12927 | https://www.ncbi.nlm.nih.gov/gene/?term=LOC105286946 | 1231 | 49 | 398.05 | 3 | 24.37 | 6.12 | 34.73 | 31.65 | 3.08 | 3.08 | 0.12 | 0.12 |
| gene5447 | https://www.ncbi.nlm.nih.gov/gene/?term=LOC105278314 | 1482 | 85 | 573.55 | 2 | 13.5 | 2.35 | 25.47 | 23.3 | 2.17 | 2.17 | 0.12 | 0.12 |
| gene3910 | https://www.ncbi.nlm.nih.gov/gene/?term=LOC105276563 | 1957 | 115 | 587.63 | 6 | 30.66 | 5.22 | 18.73 | 16.62 | 2.11 | 2.11 | 0.12 | 0.12 |
| gene5937 | https://www.ncbi.nlm.nih.gov/gene/?term=LOC105278873 | 1902 | 146 | 767.61 | 6 | 31.55 | 4.11 | 21.31 | 19.68 | 1.63 | 1.63 | 0.12 | 0.12 |
| gene2855 | https://www.ncbi.nlm.nih.gov/gene/?term=LOC105275378 | 1189 | 20 | 168.21 | 1 | 8.41 | 5 | 32.85 | 26.65 | 6.2 | 6.2 | 0.1 | 0.1 |
| gene4708 | https://www.ncbi.nlm.nih.gov/gene/?term=LOC105277468 | 1712 | 31 | 181.07 | 3 | 17.52 | 9.68 | 40.08 | 34.7 | 5.39 | 5.39 | 0.1 | 0.1 |
| gene4013 | https://www.ncbi.nlm.nih.gov/gene/?term=LOC105276679 | 2007 | 64 | 318.88 | 1 | 4.98 | 1.56 | 39.86 | 36.65 | 3.22 | 3.22 | 0.1 | 0.1 |
| gene10112 | https://www.ncbi.nlm.nih.gov/gene/?term=LOC105283717 | 1483 | 80 | 539.45 | 5 | 33.72 | 6.25 | 29.96 | 28.17 | 1.79 | 1.79 | 0.1 | 0.1 |
| gene13157 | https://www.ncbi.nlm.nih.gov/gene/?term=LOC105287195 | 837 | 21 | 250.9 | 1 | 11.95 | 4.76 | 20.59 | 24.57 | -3.98 | 3.98 | -0.1 | 0.1 |
| gene3015 | https://www.ncbi.nlm.nih.gov/gene/?term=LOC105275557 | 811 | 11 | 135.64 | 1 | 12.33 | 9.09 | 74.6 | 67.82 | 6.78 | 6.78 | 0.09 | 0.09 |
| gene6974 | https://www.ncbi.nlm.nih.gov/gene/?term=LOC105280111 | 1003 | 21 | 209.37 | 1 | 9.97 | 4.76 | 44.89 | 40.66 | 4.23 | 4.23 | 0.09 | 0.09 |
| gene11109 | https://www.ncbi.nlm.nih.gov/gene/?term=LOC105284855 | 1786 | 42 | 235.16 | 2 | 11.2 | 4.76 | 43.39 | 39.73 | 3.66 | 3.66 | 0.09 | 0.09 |
| gene7588 | https://www.ncbi.nlm.nih.gov/gene/?term=LOC105280808 | 1612 | 43 | 266.75 | 1 | 6.2 | 2.33 | 30.22 | 26.92 | 3.3 | 3.3 | 0.09 | 0.09 |
| gene11934 | https://www.ncbi.nlm.nih.gov/gene/?term=LOC105285812 | 1594 | 46 | 288.58 | 1 | 6.27 | 2.17 | 22.97 | 19.95 | 3.02 | 3.02 | 0.09 | 0.09 |

...

... only first 25 lines shown ...

Directory: OGLall File: rnk-dmg-Cb_LI_BCphaseA2.vs.RphaseA2.txt

| gene_ID | gene_link | gwidth | #Sites | #per10Kb | #dmSites | #dmsp10kb | %dmSites | %pSite1 | %pSite2 | DMpSite | ADMpSite | DMpNucl | ADMpNucl |
|---|---|---|---|---|---|---|---|---|---|---|---|---|---|
| gene12733 | https://www.ncbi.nlm.nih.gov/gene/?term=LOC109611472 | 1351 | 27 | 199.85 | 2 | 14.8 | 7.41 | 32.9 | 39.85 | -6.95 | 6.95 | -0.14 | 0.14 |
| gene10743 | https://www.ncbi.nlm.nih.gov/gene/?term=LOC105284456 | 1028 | 43 | 418.29 | 3 | 29.18 | 6.98 | 44.49 | 41.39 | 3.1 | 3.1 | 0.13 | 0.13 |
| gene9121 | https://www.ncbi.nlm.nih.gov/gene/?term=LOC105282572 | 771 | 35 | 453.96 | 1 | 12.97 | 2.86 | 29.82 | 32.76 | -2.94 | 2.94 | -0.13 | 0.13 |
| gene12311 | https://www.ncbi.nlm.nih.gov/gene/?term=LOC105286212 | 1005 | 36 | 358.21 | 1 | 9.95 | 2.78 | 37.91 | 41.67 | -3.76 | 3.76 | -0.13 | 0.13 |
| gene1171 | https://www.ncbi.nlm.nih.gov/gene/?term=LOC105285174 | 844 | 39 | 462.09 | 1 | 11.85 | 2.56 | 36.65 | 39.21 | -2.56 | 2.56 | -0.12 | 0.12 |
| gene1562 | https://www.ncbi.nlm.nih.gov/gene/?term=LOC105287692 | 1491 | 98 | 657.28 | 3 | 20.12 | 3.06 | 28.39 | 30.05 | -1.66 | 1.66 | -0.11 | 0.11 |
| gene4227 | https://www.ncbi.nlm.nih.gov/gene/?term=LOC105276920 | 2587 | 95 | 367.22 | 1 | 3.87 | 1.05 | 19.9 | 22.92 | -3.01 | 3.01 | -0.11 | 0.11 |
| gene25 | https://www.ncbi.nlm.nih.gov/gene/?term=LOC105274524 | 1118 | 38 | 339.89 | 1 | 8.94 | 2.63 | 28.99 | 32.2 | -3.2 | 3.2 | -0.11 | 0.11 |
| gene11980 | https://www.ncbi.nlm.nih.gov/gene/?term=LOC109611395 | 571 | 5 | 87.57 | 1 | 17.51 | 20 | 33.3 | 45.97 | -12.67 | 12.67 | -0.11 | 0.11 |
| gene10237 | https://www.ncbi.nlm.nih.gov/gene/?term=LOC105283856 | 1259 | 26 | 206.51 | 1 | 7.94 | 3.85 | 64.95 | 60.12 | 4.82 | 4.82 | 0.1 | 0.1 |
| gene10731 | https://www.ncbi.nlm.nih.gov/gene/?term=LOC105284411 | 1794 | 63 | 351.17 | 1 | 5.57 | 1.59 | 35.73 | 32.92 | 2.81 | 2.81 | 0.1 | 0.1 |
| gene10742 | https://www.ncbi.nlm.nih.gov/gene/?term=LOC109611309 | 776 | 37 | 476.8 | 2 | 25.77 | 5.41 | 42.09 | 40.01 | 2.09 | 2.09 | 0.1 | 0.1 |
| gene3719 | https://www.ncbi.nlm.nih.gov/gene/?term=LOC105276360 | 1471 | 108 | 734.19 | 1 | 6.8 | 0.93 | 29.44 | 30.83 | -1.39 | 1.39 | -0.1 | 0.1 |
| gene5912 | https://www.ncbi.nlm.nih.gov/gene/?term=LOC105278847 | 2089 | 125 | 598.37 | 1 | 4.79 | 0.8 | 8.99 | 10.64 | -1.65 | 1.65 | -0.1 | 0.1 |
| gene3787 | https://www.ncbi.nlm.nih.gov/gene/?term=LOC105276423 | 1679 | 77 | 458.61 | 1 | 5.96 | 1.3 | 16.64 | 18.77 | -2.13 | 2.13 | -0.1 | 0.1 |
| gene7767 | https://www.ncbi.nlm.nih.gov/gene/?term=LOC105281009 | 1509 | 40 | 265.08 | 3 | 19.88 | 7.5 | 53.17 | 56.81 | -3.65 | 3.65 | -0.1 | 0.1 |
| gene3165 | https://www.ncbi.nlm.nih.gov/gene/?term=LOC105275706 | 2374 | 82 | 345.41 | 3 | 12.64 | 3.66 | 33.31 | 30.79 | 2.52 | 2.52 | 0.09 | 0.09 |
| gene9183 | https://www.ncbi.nlm.nih.gov/gene/?term=LOC105282640 | 2752 | 211 | 766.72 | 2 | 7.27 | 0.95 | 21.27 | 22.49 | -1.22 | 1.22 | -0.09 | 0.09 |
| gene2790 | https://www.ncbi.nlm.nih.gov/gene/?term=LOC105275307 | 1429 | 54 | 377.89 | 2 | 14 | 3.7 | 39.26 | 41.56 | -2.3 | 2.3 | -0.09 | 0.09 |
| gene4313 | https://www.ncbi.nlm.nih.gov/gene/?term=LOC105277012 | 1491 | 52 | 348.76 | 3 | 20.12 | 5.77 | 24.71 | 27.22 | -2.51 | 2.51 | -0.09 | 0.09 |
| gene10720 | https://www.ncbi.nlm.nih.gov/gene/?term=LOC105284400 | 1914 | 44 | 229.89 | 3 | 15.67 | 6.82 | 27.76 | 31.46 | -3.7 | 3.7 | -0.09 | 0.09 |
| gene10721 | https://www.ncbi.nlm.nih.gov/gene/?term=LOC105284403 | 1537 | 29 | 188.68 | 2 | 13.01 | 6.9 | 50.07 | 54.88 | -4.81 | 4.81 | -0.09 | 0.09 |
| gene11521 | https://www.ncbi.nlm.nih.gov/gene/?term=LOC105285336 | 476 | 7 | 147.06 | 1 | 21.01 | 14.29 | 45.58 | 51.56 | -5.98 | 5.98 | -0.09 | 0.09 |
| gene7810 | https://www.ncbi.nlm.nih.gov/gene/?term=LOC105281055 | 1333 | 50 | 375.09 | 1 | 7.5 | 2 | 26.8 | 24.6 | 2.2 | 2.2 | 0.08 | 0.08 |

...

... only first 25 lines shown ...

Directory: OGLall File: rnk-dmg-Cb_LI_BCphaseA2.vs.RphaseB1.txt

| gene_ID | gene_link | gwidth | #Sites | #per10Kb | #dmSites | #dmsp10kb | %dmSites | %pSite1 | %pSite2 | DMpSite | ADMpSite | DMpNucl | ADMpNucl |
|---|---|---|---|---|---|---|---|---|---|---|---|---|---|
| gene3165 | https://www.ncbi.nlm.nih.gov/gene/?term=LOC105275706 | 2374 | 67 | 282.22 | 6 | 25.27 | 8.96 | 35.39 | 24.21 | 11.18 | 11.18 | 0.32 | 0.32 |
| gene2824 | https://www.ncbi.nlm.nih.gov/gene/?term=LOC105275337 | 1772 | 62 | 349.89 | 2 | 11.29 | 3.23 | 38.77 | 30.45 | 8.31 | 8.31 | 0.29 | 0.29 |
| gene2593 | https://www.ncbi.nlm.nih.gov/gene/?term=LOC105275070 | 1774 | 9 | 50.73 | 4 | 22.55 | 44.44 | 77.93 | 29.94 | 47.99 | 47.99 | 0.24 | 0.24 |
| gene1993 | https://www.ncbi.nlm.nih.gov/gene/?term=LOC105288193 | 1449 | 75 | 517.6 | 1 | 6.9 | 1.33 | 35.76 | 32.82 | 2.94 | 2.94 | 0.15 | 0.15 |
| gene5714 | https://www.ncbi.nlm.nih.gov/gene/?term=LOC105278729 | 710 | 16 | 225.35 | 1 | 14.08 | 6.25 | 57.91 | 51.9 | 6.01 | 6.01 | 0.14 | 0.14 |
| gene2404 | https://www.ncbi.nlm.nih.gov/gene/?term=LOC105274853 | 1767 | 34 | 192.42 | 1 | 5.66 | 2.94 | 29.24 | 33.79 | -4.55 | 4.55 | -0.09 | 0.09 |
| gene4034 | https://www.ncbi.nlm.nih.gov/gene/?term=LOC105276704 | 2065 | 86 | 416.46 | 1 | 4.84 | 1.16 | 15.38 | 13.53 | 1.85 | 1.85 | 0.08 | 0.08 |
| gene2413 | https://www.ncbi.nlm.nih.gov/gene/?term=LOC105274863 | 2006 | 46 | 229.31 | 1 | 4.99 | 2.17 | 60.76 | 64.28 | -3.53 | 3.53 | -0.08 | 0.08 |
| gene13033 | https://www.ncbi.nlm.nih.gov/gene/?term=LOC105287063 | 1877 | 49 | 261.05 | 1 | 5.33 | 2.04 | 32.47 | 29.74 | 2.73 | 2.73 | 0.07 | 0.07 |
| gene10984 | https://www.ncbi.nlm.nih.gov/gene/?term=LOC105284704 | 1987 | 62 | 312.03 | 1 | 5.03 | 1.61 | 26.65 | 24.35 | 2.3 | 2.3 | 0.07 | 0.07 |
| gene7556 | https://www.ncbi.nlm.nih.gov/gene/?term=LOC105280774 | 2792 | 101 | 361.75 | 1 | 3.58 | 0.99 | 32.38 | 30.39 | 1.99 | 1.99 | 0.07 | 0.07 |
| gene10520 | https://www.ncbi.nlm.nih.gov/gene/?term=LOC105284184 | 2451 | 87 | 354.96 | 1 | 4.08 | 1.15 | 34.29 | 32.67 | 1.61 | 1.61 | 0.06 | 0.06 |
| gene6583 | https://www.ncbi.nlm.nih.gov/gene/?term=LOC105279649 | 7444 | 343 | 460.77 | 1 | 1.34 | 0.29 | 10.51 | 11.78 | -1.26 | 1.26 | -0.06 | 0.06 |
| gene920 | https://www.ncbi.nlm.nih.gov/gene/?term=LOC105282518 | 1197 | 39 | 325.81 | 1 | 8.35 | 2.56 | 36.44 | 38.28 | -1.85 | 1.85 | -0.06 | 0.06 |
| gene12021 | https://www.ncbi.nlm.nih.gov/gene/?term=LOC105285905 | 1399 | 27 | 192.99 | 1 | 7.15 | 3.7 | 31.19 | 34.36 | -3.17 | 3.17 | -0.06 | 0.06 |
| gene6652 | https://www.ncbi.nlm.nih.gov/gene/?term=LOC105279719 | 2831 | 78 | 275.52 | 1 | 3.53 | 1.28 | 25.09 | 23.41 | 1.68 | 1.68 | 0.05 | 0.05 |
| gene10606 | https://www.ncbi.nlm.nih.gov/gene/?term=LOC105284273 | 6119 | 260 | 424.91 | 1 | 1.63 | 0.38 | 19.55 | 18.4 | 1.15 | 1.15 | 0.05 | 0.05 |
| gene7568 | https://www.ncbi.nlm.nih.gov/gene/?term=LOC105280786 | 2525 | 122 | 483.17 | 1 | 3.96 | 0.82 | 30.88 | 29.81 | 1.07 | 1.07 | 0.05 | 0.05 |
| gene2835 | https://www.ncbi.nlm.nih.gov/gene/?term=LOC105275357 | 2492 | 114 | 457.46 | 1 | 4.01 | 0.88 | 13.76 | 14.82 | -1.05 | 1.05 | -0.05 | 0.05 |
| gene1220 | https://www.ncbi.nlm.nih.gov/gene/?term=LOC105285742 | 3453 | 98 | 283.81 | 1 | 2.9 | 1.02 | 19.04 | 20.94 | -1.9 | 1.9 | -0.05 | 0.05 |
| gene10552 | https://www.ncbi.nlm.nih.gov/gene/?term=LOC105284213 | 878 | 15 | 170.84 | 1 | 11.39 | 6.67 | 15.38 | 18.37 | -2.99 | 2.99 | -0.05 | 0.05 |
| gene393 | https://www.ncbi.nlm.nih.gov/gene/?term=LOC105276566 | 1123 | 2 | 17.81 | 1 | 8.9 | 50 | 48.08 | 74.3 | -26.22 | 26.22 | -0.05 | 0.05 |
| gene3671 | https://www.ncbi.nlm.nih.gov/gene/?term=LOC105276272 | 1966 | 38 | 193.29 | 1 | 5.09 | 2.63 | 30.65 | 28.5 | 2.15 | 2.15 | 0.04 | 0.04 |
| gene10091 | https://www.ncbi.nlm.nih.gov/gene/?term=LOC105283697 | 4988 | 137 | 274.66 | 1 | 2 | 0.73 | 17.98 | 16.41 | 1.57 | 1.57 | 0.04 | 0.04 |
| gene5877 | https://www.ncbi.nlm.nih.gov/gene/?term=LOC105278807 | 1719 | 49 | 285.05 | 1 | 5.82 | 2.04 | 25.27 | 23.74 | 1.53 | 1.53 | 0.04 | 0.04 |
| gene7888 | https://www.ncbi.nlm.nih.gov/gene/?term=LOC105281141 | 4357 | 157 | 360.34 | 1 | 2.3 | 0.64 | 31.42 | 30.25 | 1.17 | 1.17 | 0.04 | 0.04 |
| gene3817 | https://www.ncbi.nlm.nih.gov/gene/?term=LOC105276449 | 1230 | 43 | 349.59 | 1 | 8.13 | 2.33 | 30.78 | 29.68 | 1.1 | 1.1 | 0.04 | 0.04 |
| gene5831 | https://www.ncbi.nlm.nih.gov/gene/?term=LOC105278761 | 2506 | 112 | 446.93 | 2 | 7.98 | 1.79 | 21.67 | 20.75 | 0.92 | 0.92 | 0.04 | 0.04 |
| gene11340 | https://www.ncbi.nlm.nih.gov/gene/?term=LOC105285131 | 2918 | 89 | 305 | 1 | 3.43 | 1.12 | 11.73 | 13.01 | -1.28 | 1.28 | -0.04 | 0.04 |
| gene3335 | https://www.ncbi.nlm.nih.gov/gene/?term=LOC105275898 | 4665 | 147 | 315.11 | 1 | 2.14 | 0.68 | 11.48 | 12.76 | -1.29 | 1.29 | -0.04 | 0.04 |
| gene10879 | https://www.ncbi.nlm.nih.gov/gene/?term=LOC105284574 | 2423 | 45 | 185.72 | 1 | 4.13 | 2.22 | 57.53 | 59.68 | -2.15 | 2.15 | -0.04 | 0.04 |
| gene1199 | https://www.ncbi.nlm.nih.gov/gene/?term=LOC105285498 | 2220 | 25 | 112.61 | 1 | 4.5 | 4 | 32.49 | 35.74 | -3.26 | 3.26 | -0.04 | 0.04 |
| gene3166 | https://www.ncbi.nlm.nih.gov/gene/?term=LOC105275707 | 2393 | 11 | 45.97 | 1 | 4.18 | 9.09 | 62.4 | 71.24 | -8.83 | 8.83 | -0.04 | 0.04 |
| gene9109 | https://www.ncbi.nlm.nih.gov/gene/?term=LOC105282561 | 2400 | 11 | 45.83 | 1 | 4.17 | 9.09 | 17.27 | 26.23 | -8.97 | 8.97 | -0.04 | 0.04 |
| gene9336 | https://www.ncbi.nlm.nih.gov/gene/?term=LOC105282834 | 5993 | 124 | 206.91 | 1 | 1.67 | 0.81 | 15.53 | 14.01 | 1.52 | 1.52 | 0.03 | 0.03 |
| gene7734 | https://www.ncbi.nlm.nih.gov/gene/?term=LOC105280972 | 1528 | 32 | 209.42 | 1 | 6.54 | 3.12 | 19.4 | 18.05 | 1.35 | 1.35 | 0.03 | 0.03 |
| gene940 | https://www.ncbi.nlm.nih.gov/gene/?term=LOC105282794 | 3578 | 86 | 240.36 | 1 | 2.79 | 1.16 | 16.54 | 15.48 | 1.06 | 1.06 | 0.03 | 0.03 |
| gene7249 | https://www.ncbi.nlm.nih.gov/gene/?term=LOC105280422 | 1775 | 62 | 349.3 | 1 | 5.63 | 1.61 | 17.51 | 16.57 | 0.94 | 0.94 | 0.03 | 0.03 |
| gene6286 | https://www.ncbi.nlm.nih.gov/gene/?term=LOC105279298 | 2861 | 121 | 422.93 | 2 | 6.99 | 1.65 | 24.64 | 23.9 | 0.74 | 0.74 | 0.03 | 0.03 |
| gene6031 | https://www.ncbi.nlm.nih.gov/gene/?term=LOC105279000 | 5691 | 208 | 365.49 | 1 | 1.76 | 0.48 | 12.77 | 12.08 | 0.69 | 0.69 | 0.03 | 0.03 |
| gene5298 | https://www.ncbi.nlm.nih.gov/gene/?term=LOC105278145 | 4060 | 251 | 618.23 | 2 | 4.93 | 0.8 | 10.52 | 10.93 | -0.42 | 0.42 | -0.03 | 0.03 |
| gene13114 | https://www.ncbi.nlm.nih.gov/gene/?term=LOC105287147 | 7609 | 398 | 523.06 | 1 | 1.31 | 0.25 | 9.41 | 9.93 | -0.52 | 0.52 | -0.03 | 0.03 |
| gene5581 | https://www.ncbi.nlm.nih.gov/gene/?term=LOC105278459 | 2935 | 97 | 330.49 | 1 | 3.41 | 1.03 | 20.44 | 21.26 | -0.83 | 0.83 | -0.03 | 0.03 |
| gene6966 | https://www.ncbi.nlm.nih.gov/gene/?term=LOC105280104 | 3747 | 124 | 330.93 | 1 | 2.67 | 0.81 | 16.15 | 17.09 | -0.94 | 0.94 | -0.03 | 0.03 |
| gene7915 | https://www.ncbi.nlm.nih.gov/gene/?term=LOC105281173 | 1597 | 39 | 244.21 | 1 | 6.26 | 2.56 | 60.85 | 62.01 | -1.16 | 1.16 | -0.03 | 0.03 |
| gene3848 | https://www.ncbi.nlm.nih.gov/gene/?term=LOC105276486 | 5899 | 145 | 245.8 | 1 | 1.7 | 0.69 | 23.24 | 24.56 | -1.32 | 1.32 | -0.03 | 0.03 |
| gene7207 | https://www.ncbi.nlm.nih.gov/gene/?term=LOC105280376 | 4659 | 112 | 240.39 | 1 | 2.15 | 0.89 | 28.49 | 29.89 | -1.4 | 1.4 | -0.03 | 0.03 |
| gene10819 | https://www.ncbi.nlm.nih.gov/gene/?term=LOC105284507 | 5767 | 86 | 149.12 | 1 | 1.73 | 1.16 | 21.64 | 23.53 | -1.89 | 1.89 | -0.03 | 0.03 |
| gene7398 | https://www.ncbi.nlm.nih.gov/gene/?term=LOC105280589 | 3452 | 42 | 121.67 | 1 | 2.9 | 2.38 | 27.89 | 25.91 | 1.98 | 1.98 | 0.02 | 0.02 |
| gene4502 | https://www.ncbi.nlm.nih.gov/gene/?term=LOC105277243 | 2289 | 32 | 139.8 | 1 | 4.37 | 3.12 | 44.88 | 43.35 | 1.53 | 1.53 | 0.02 | 0.02 |
| gene6455 | https://www.ncbi.nlm.nih.gov/gene/?term=LOC105279500 | 2612 | 33 | 126.34 | 1 | 3.83 | 3.03 | 51.99 | 50.58 | 1.41 | 1.41 | 0.02 | 0.02 |
| gene10107 | https://www.ncbi.nlm.nih.gov/gene/?term=LOC105283712 | 1883 | 30 | 159.32 | 1 | 5.31 | 3.33 | 31.98 | 30.87 | 1.11 | 1.11 | 0.02 | 0.02 |
| gene1147 | https://www.ncbi.nlm.nih.gov/gene/?term=LOC105284954 | 4022 | 96 | 238.69 | 1 | 2.49 | 1.04 | 15.76 | 15.07 | 0.69 | 0.69 | 0.02 | 0.02 |
| gene8015 | https://www.ncbi.nlm.nih.gov/gene/?term=LOC105281312 | 2998 | 68 | 226.82 | 1 | 3.34 | 1.47 | 31.47 | 30.8 | 0.67 | 0.67 | 0.02 | 0.02 |
| gene292 | https://www.ncbi.nlm.nih.gov/gene/?term=LOC105275398 | 2825 | 92 | 325.66 | 1 | 3.54 | 1.09 | 23.29 | 22.65 | 0.64 | 0.64 | 0.02 | 0.02 |
| gene1310 | https://www.ncbi.nlm.nih.gov/gene/?term=LOC105286661 | 2582 | 71 | 274.98 | 1 | 3.87 | 1.41 | 21.34 | 20.74 | 0.61 | 0.61 | 0.02 | 0.02 |
| gene10678 | https://www.ncbi.nlm.nih.gov/gene/?term=LOC105284355 | 4031 | 161 | 399.4 | 1 | 2.48 | 0.62 | 20.49 | 19.95 | 0.54 | 0.54 | 0.02 | 0.02 |
| gene4892 | https://www.ncbi.nlm.nih.gov/gene/?term=LOC105277667 | 3898 | 123 | 315.55 | 1 | 2.57 | 0.81 | 28.28 | 27.77 | 0.51 | 0.51 | 0.02 | 0.02 |
| gene298 | https://www.ncbi.nlm.nih.gov/gene/?term=LOC105275460 | 2473 | 118 | 477.15 | 1 | 4.04 | 0.85 | 13.46 | 12.97 | 0.49 | 0.49 | 0.02 | 0.02 |
| gene8729 | https://www.ncbi.nlm.nih.gov/gene/?term=LOC105282137 | 5005 | 211 | 421.58 | 1 | 2 | 0.47 | 16 | 15.6 | 0.4 | 0.4 | 0.02 | 0.02 |

| | | | | | | | | | | | | | |
|---|---|---|---|---|---|---|---|---|---|---|---|---|---|
| gene3747 | https://www.ncbi.nlm.nih.gov/gene/?term=LOC105276387 | 3736 | 188 | 503.21 | 1 | 2.68 | 0.53 | 14.43 | 14.09 | 0.35 | 0.35 | 0.02 | 0.02 |
| gene1470 | https://www.ncbi.nlm.nih.gov/gene/?term=LOC105287597 | 2974 | 192 | 645.6 1 | 3.36 | 0.52 | 18.97 | 19.22 | -0.25 | 0.25 | -0.02 | 0.02 | |
| gene1571 | https://www.ncbi.nlm.nih.gov/gene/?term=LOC105287700 | 3392 | 160 | 471.7 1 | 2.95 | 0.62 | 15.1 | 15.46 | -0.36 | 0.36 | -0.02 | 0.02 | |
| gene4962 | https://www.ncbi.nlm.nih.gov/gene/?term=LOC105277761 | 4645 | 161 | 346.61 | 1 | 2.15 | 0.62 | 14.28 | 14.77 | -0.48 | 0.48 | -0.02 | 0.02 |
| gene8274 | https://www.ncbi.nlm.nih.gov/gene/?term=LOC105281624 | 5425 | 221 | 407.37 | 1 | 1.84 | 0.45 | 9.57 | 10.1 | -0.54 | 0.54 | -0.02 | 0.02 |
| gene3107 | https://www.ncbi.nlm.nih.gov/gene/?term=LOC105275650 | 5103 | 172 | 337.06 | 1 | 1.96 | 0.58 | 16.37 | 16.93 | -0.56 | 0.56 | -0.02 | 0.02 |
| gene12573 | https://www.ncbi.nlm.nih.gov/gene/?term=LOC105286515 | 3647 | 108 | 296.13 | 1 | 2.74 | 0.93 | 11.17 | 11.84 | -0.67 | 0.67 | -0.02 | 0.02 |
| gene1198 | https://www.ncbi.nlm.nih.gov/gene/?term=LOC105285507 | 2738 | 92 | 336.01 | 1 | 3.65 | 1.09 | 20.74 | 21.44 | -0.7 | 0.7 | -0.02 | 0.02 |
| gene11454 | https://www.ncbi.nlm.nih.gov/gene/?term=LOC105285256 | 1906 | 48 | 251.84 | 1 | 5.25 | 2.08 | 44.93 | 45.73 | -0.8 | 0.8 | -0.02 | 0.02 |
| gene8814 | https://www.ncbi.nlm.nih.gov/gene/?term=LOC105282240 | 6910 | 166 | 240.23 | 1 | 1.45 | 0.6 | 9.27 | 10.23 | -0.96 | 0.96 | -0.02 | 0.02 |
| gene11447 | https://www.ncbi.nlm.nih.gov/gene/?term=LOC105285252 | 7437 | 126 | 169.42 | 2 | 2.69 | 1.59 | 12.66 | 13.85 | -1.19 | 1.19 | -0.02 | 0.02 |
| gene10155 | https://www.ncbi.nlm.nih.gov/gene/?term=LOC105283756 | 7030 | 98 | 139.4 1 | 1.42 | 1.02 | 6.09 | 7.87 | -1.78 | 1.78 | -0.02 | 0.02 | |
| gene3168 | https://www.ncbi.nlm.nih.gov/gene/?term=LOC105275711 | 6150 | 80 | 130.08 | 1 | 1.63 | 1.25 | 11.3 | 13.17 | -1.86 | 1.86 | -0.02 | 0.02 |
| gene388 | https://www.ncbi.nlm.nih.gov/gene/?term=LOC105276538 | 1558 | 19 | 121.95 | 2 | 12.84 | 10.53 | 66.07 | 68.07 | -1.99 | 1.99 | -0.02 | 0.02 |
| gene11584 | https://www.ncbi.nlm.nih.gov/gene/?term=LOC105285416 | 3709 | 6 | 16.18 2 | 5.39 | 33.33 | 18.09 | 32.8 | -14.71 | | 14.71 | -0.02 | 0.02 |
| gene2833 | https://www.ncbi.nlm.nih.gov/gene/?term=LOC105275352 | 5528 | 38 | 68.74 1 | 1.81 | 2.63 | 25.27 | 24 | 1.27 | 1.27 | 0.01 | 0.01 | |
| gene2970 | https://www.ncbi.nlm.nih.gov/gene/?term=LOC105275518 | 2340 | 19 | 81.2 1 | 4.27 | 5.26 | 51.65 | 50.52 | 1.13 | 1.13 | 0.01 | 0.01 | |
| gene10222 | https://www.ncbi.nlm.nih.gov/gene/?term=LOC105283838 | 1748 | 21 | 120.14 | 1 | 5.72 | 4.76 | 46.37 | 45.47 | 0.9 | 0.9 | 0.01 | 0.01 |
| gene322 | https://www.ncbi.nlm.nih.gov/gene/?term=LOC105275777 | 1606 | 23 | 143.21 | 1 | 6.23 | 4.35 | 44.64 | 43.87 | 0.76 | 0.76 | 0.01 | 0.01 |
| gene3912 | https://www.ncbi.nlm.nih.gov/gene/?term=LOC105276560 | 17900 | 309 | 172.63 | 1 | 0.56 | 0.32 | 2.52 | 1.81 | 0.7 | 0.7 | 0.01 | 0.01 |
| gene7657 | https://www.ncbi.nlm.nih.gov/gene/?term=LOC105280890 | 1109 | 21 | 189.36 | 1 | 9.02 | 4.76 | 27.2 | 26.57 | 0.64 | 0.64 | 0.01 | 0.01 |
| gene12572 | https://www.ncbi.nlm.nih.gov/gene/?term=LOC105286514 | 2374 | 44 | 185.34 | 1 | 4.21 | 2.27 | 33.89 | 33.38 | 0.52 | 0.52 | 0.01 | 0.01 |
| gene289 | https://www.ncbi.nlm.nih.gov/gene/?term=LOC105275347 | 2533 | 72 | 284.25 | 1 | 3.95 | 1.39 | 24.33 | 23.84 | 0.48 | 0.48 | 0.01 | 0.01 |
| gene11568 | https://www.ncbi.nlm.nih.gov/gene/?term=LOC105285395 | 8086 | 202 | 249.81 | 1 | 1.24 | 0.5 | 14.4 | 13.94 | 0.46 | 0.46 | 0.01 | 0.01 |
| gene2056 | https://www.ncbi.nlm.nih.gov/gene/?term=LOC105274452 | 6584 | 236 | 358.44 | 1 | 1.52 | 0.42 | 14.57 | 14.16 | 0.42 | 0.42 | 0.01 | 0.01 |
| gene3470 | https://www.ncbi.nlm.nih.gov/gene/?term=LOC105276059 | 8716 | 151 | 173.24 | 1 | 1.15 | 0.66 | 10.19 | 9.77 | 0.42 | 0.42 | 0.01 | 0.01 |
| gene7245 | https://www.ncbi.nlm.nih.gov/gene/?term=LOC105280410 | 12478 | 264 | 211.57 | 1 | 0.8 | 0.38 | 19.99 | 19.61 | 0.38 | 0.38 | 0.01 | 0.01 |
| gene8600 | https://www.ncbi.nlm.nih.gov/gene/?term=LOC105282001 | 8513 | 152 | 178.55 | 1 | 1.17 | 0.66 | 12.95 | 12.59 | 0.36 | 0.36 | 0.01 | 0.01 |
| gene6237 | https://www.ncbi.nlm.nih.gov/gene/?term=LOC105279244 | 1484 | 27 | 181.94 | 1 | 6.74 | 3.7 | 50.23 | 49.93 | 0.29 | 0.29 | 0.01 | 0.01 |
| gene2846 | https://www.ncbi.nlm.nih.gov/gene/?term=LOC105275374 | 11246 | 500 | 444.6 1 | 0.89 | 0.2 | 10.21 | 9.93 | 0.28 | 0.28 | 0.01 | 0.01 | |
| gene3727 | https://www.ncbi.nlm.nih.gov/gene/?term=LOC105276367 | 2484 | 68 | 273.75 | 1 | 4.03 | 1.47 | 46.01 | 45.74 | 0.27 | 0.27 | 0.01 | 0.01 |
| gene5746 | https://www.ncbi.nlm.nih.gov/gene/?term=LOC105278666 | 10343 | 455 | 439.91 | 1 | 0.97 | 0.22 | 7.39 | 7.14 | 0.25 | 0.25 | 0.01 | 0.01 |
| gene11819 | https://www.ncbi.nlm.nih.gov/gene/?term=LOC105285671 | 2366 | 98 | 414.2 1 | 4.23 | 1.02 | 30.44 | 30.22 | 0.23 | 0.23 | 0.01 | 0.01 | |
| gene9755 | https://www.ncbi.nlm.nih.gov/gene/?term=LOC105283296 | 2177 | 87 | 399.63 | 1 | 4.59 | 1.15 | 15.89 | 15.67 | 0.23 | 0.23 | 0.01 | 0.01 |
| gene474 | https://www.ncbi.nlm.nih.gov/gene/?term=LOC105277435 | 3625 | 173 | 477.24 | 1 | 2.76 | 0.58 | 10.83 | 10.64 | 0.2 | 0.2 | 0.01 | 0.01 |
| gene11275 | https://www.ncbi.nlm.nih.gov/gene/?term=LOC105285055 | 4119 | 239 | 580.24 | 1 | 2.43 | 0.42 | 11.8 | 11.69 | 0.12 | 0.12 | 0.01 | 0.01 |
| gene5525 | https://www.ncbi.nlm.nih.gov/gene/?term=LOC105278396 | 3090 | 186 | 601.94 | 1 | 3.24 | 0.54 | 24.32 | 24.44 | -0.12 | 0.12 | -0.01 | 0.01 |
| gene12802 | https://www.ncbi.nlm.nih.gov/gene/?term=LOC105286799 | 3183 | 173 | 543.51 | 1 | 3.14 | 0.58 | 17.3 | 17.43 | -0.13 | 0.13 | -0.01 | 0.01 |
| gene10018 | https://www.ncbi.nlm.nih.gov/gene/?term=LOC105283607 | 4414 | 192 | 434.98 | 1 | 2.27 | 0.52 | 15.73 | 15.92 | -0.19 | 0.19 | -0.01 | 0.01 |
| gene13036 | https://www.ncbi.nlm.nih.gov/gene/?term=LOC105287059 | 4959 | 172 | 346.84 | 1 | 2.02 | 0.58 | 16.67 | 16.87 | -0.2 | 0.2 | -0.01 | 0.01 |
| gene4376 | https://www.ncbi.nlm.nih.gov/gene/?term=LOC105277083 | 4307 | 123 | 285.58 | 1 | 2.32 | 0.81 | 19.63 | 19.89 | -0.26 | 0.26 | -0.01 | 0.01 |
| gene5646 | https://www.ncbi.nlm.nih.gov/gene/?term=LOC105278534 | 7475 | 180 | 240.8 1 | 1.34 | 0.56 | 17.08 | 17.41 | -0.33 | 0.33 | -0.01 | 0.01 | |
| gene13044 | https://www.ncbi.nlm.nih.gov/gene/?term=LOC105287070 | 5094 | 224 | 439.73 | 1 | 1.96 | 0.45 | 5.36 | 5.7 | -0.34 | 0.34 | -0.01 | 0.01 |
| gene12674 | https://www.ncbi.nlm.nih.gov/gene/?term=LOC105286649 | 18283 | 764 | 417.87 | 1 | 0.55 | 0.13 | 0.96 | 1.31 | -0.35 | 0.35 | -0.01 | 0.01 |
| gene12406 | https://www.ncbi.nlm.nih.gov/gene/?term=LOC105286316 | 1386 | 29 | 209.24 | 1 | 7.22 | 3.45 | 45.59 | 46 | -0.41 | 0.41 | -0.01 | 0.01 |
| gene9326 | https://www.ncbi.nlm.nih.gov/gene/?term=LOC105282818 | 16612 | 217 | 130.63 | 1 | 0.6 | 0.46 | 13.52 | 14.06 | -0.54 | 0.54 | -0.01 | 0.01 |
| gene5375 | https://www.ncbi.nlm.nih.gov/gene/?term=LOC105278228 | 8640 | 167 | 193.29 | 1 | 1.16 | 0.6 | 22.05 | 22.68 | -0.63 | 0.63 | -0.01 | 0.01 |
| gene11561 | https://www.ncbi.nlm.nih.gov/gene/?term=LOC105285387 | 11528 | 200 | 173.49 | 1 | 0.87 | 0.5 | 9.19 | 9.83 | -0.64 | 0.64 | -0.01 | 0.01 |
| gene9327 | https://www.ncbi.nlm.nih.gov/gene/?term=LOC105282821 | 7793 | 161 | 206.6 1 | 1.28 | 0.62 | 12.72 | 13.4 | -0.68 | 0.68 | -0.01 | 0.01 | |
| gene7368 | https://www.ncbi.nlm.nih.gov/gene/?term=LOC105280556 | 4105 | 58 | 141.29 | 1 | 2.44 | 1.72 | 18.1 | 19.01 | -0.91 | 0.91 | -0.01 | 0.01 |
| gene12203 | https://www.ncbi.nlm.nih.gov/gene/?term=LOC105286099 | 5967 | 79 | 132.39 | 1 | 1.68 | 1.27 | 11.92 | 12.93 | -1.01 | 1.01 | -0.01 | 0.01 |
| gene10345 | https://www.ncbi.nlm.nih.gov/gene/?term=LOC105283989 | 4444 | 21 | 47.25 1 | 2.25 | 4.76 | 48.56 | 49.94 | -1.37 | 1.37 | -0.01 | 0.01 | |
| gene8178 | https://www.ncbi.nlm.nih.gov/gene/?term=LOC105281515 | 8647 | 52 | 60.14 1 | 1.16 | 1.92 | 27.71 | 27.4 | 0.32 | 0.32 | 0 | 0 | |
| gene2683 | https://www.ncbi.nlm.nih.gov/gene/?term=LOC105275187 | 5492 | 69 | 125.64 | 1 | 1.82 | 1.45 | 14.14 | 13.85 | 0.29 | 0.29 | 0 | 0 |
| gene8343 | https://www.ncbi.nlm.nih.gov/gene/?term=LOC105281700 | 2172 | 38 | 174.95 | 1 | 4.6 | 2.63 | 39.23 | 38.97 | 0.26 | 0.26 | 0 | 0 |
| gene6620 | https://www.ncbi.nlm.nih.gov/gene/?term=LOC105279687 | 17998 | 349 | 193.91 | 1 | 0.56 | 0.29 | 4.97 | 4.84 | 0.13 | 0.13 | 0 | 0 |
| gene2271 | https://www.ncbi.nlm.nih.gov/gene/?term=LOC105274706 | 1448 | 51 | 352.21 | 1 | 6.91 | 1.96 | 38.8 | 38.68 | 0.12 | 0.12 | 0 | 0 |
| gene12307 | https://www.ncbi.nlm.nih.gov/gene/?term=LOC105286216 | 10313 | 178 | 172.6 1 | 0.97 | 0.56 | 15.99 | 15.89 | 0.1 | 0.1 | 0 | 0 | |
| gene3856 | https://www.ncbi.nlm.nih.gov/gene/?term=LOC105276499 | 5236 | 177 | 338.04 | 1 | 1.91 | 0.56 | 10.08 | 9.99 | 0.09 | 0.09 | 0 | 0 |
| gene7166 | https://www.ncbi.nlm.nih.gov/gene/?term=LOC105280326 | 6117 | 222 | 362.92 | 1 | 1.63 | 0.45 | 16.24 | 16.15 | 0.09 | 0.09 | 0 | 0 |
| gene9330 | https://www.ncbi.nlm.nih.gov/gene/?term=LOC105282825 | 3936 | 78 | 198.17 | 1 | 2.54 | 1.28 | 12.99 | 12.9 | 0.09 | 0.09 | 0 | 0 |
| gene53 | https://www.ncbi.nlm.nih.gov/gene/?term=LOC105277148 | 19283 | 283 | 146.76 | 1 | 0.52 | 0.35 | 5.45 | 5.37 | 0.08 | 0.08 | 0 | 0 |
| gene10344 | https://www.ncbi.nlm.nih.gov/gene/?term=LOC105283987 | 4841 | 71 | 146.66 | 1 | 2.07 | 1.41 | 32.31 | 32.24 | 0.07 | 0.07 | 0 | 0 |
| gene10654 | https://www.ncbi.nlm.nih.gov/gene/?term=LOC105284329 | 4027 | 51 | 126.65 | 1 | 2.48 | 1.96 | 19.38 | 19.32 | 0.06 | 0.06 | 0 | 0 |

| | | | | | | | | | | | | | |
|---|---|---|---|---|---|---|---|---|---|---|---|---|---|
| gene10600 | https://www.ncbi.nlm.nih.gov/gene/?term=LOC105284270 | 4837 | 100 | 206.74 | 1 | 2.07 | 1 | 12.53 | 12.48 | 0.05 | 0.05 | 0 | 0 |
| gene12981 | https://www.ncbi.nlm.nih.gov/gene/?term=LOC105286999 | 2063 | 44 | 213.28 | 1 | 4.85 | 2.27 | 35.25 | 35.21 | 0.04 | 0.04 | 0 | 0 |
| gene3885 | https://www.ncbi.nlm.nih.gov/gene/?term=LOC105276527 | 7928 | 333 | 420.03 | 1 | 1.26 | 0.3 | 8.6 | 8.57 | 0.03 | 0.03 | 0 | 0 |
| gene10392 | https://www.ncbi.nlm.nih.gov/gene/?term=LOC105284036 | 3216 | 87 | 270.52 | 2 | 6.22 | 2.3 | 24.3 | 24.3 | 0.01 | 0.01 | 0 | 0 |
| gene10549 | https://www.ncbi.nlm.nih.gov/gene/?term=LOC105284210 | 7145 | 327 | 457.66 | 2 | 2.8 | 0.61 | 10.82 | 10.83 | -0.01 | 0.01 | 0 | 0 |
| gene7600 | https://www.ncbi.nlm.nih.gov/gene/?term=LOC105280819 | 1615 | 38 | 235.29 | 1 | 6.19 | 2.63 | 35.56 | 35.58 | -0.02 | 0.02 | 0 | 0 |
| gene2806 | https://www.ncbi.nlm.nih.gov/gene/?term=LOC105275327 | 14257 | 472 | 331.07 | 1 | 0.7 | 0.21 | 7.12 | 7.15 | -0.03 | 0.03 | 0 | 0 |
| gene4190 | https://www.ncbi.nlm.nih.gov/gene/?term=LOC105276875 | 3150 | 101 | 320.63 | 1 | 3.17 | 0.99 | 24.58 | 24.61 | -0.03 | 0.03 | 0 | 0 |
| gene1632 | https://www.ncbi.nlm.nih.gov/gene/?term=LOC105287764 | 18061 | 391 | 216.49 | 1 | 0.55 | 0.26 | 3.3 | 3.35 | -0.05 | 0.05 | 0 | 0 |
| gene5524 | https://www.ncbi.nlm.nih.gov/gene/?term=LOC105278397 | 5177 | 200 | 386.32 | 1 | 1.93 | 0.5 | 24.93 | 25 | -0.07 | 0.07 | 0 | 0 |
| gene4209 | https://www.ncbi.nlm.nih.gov/gene/?term=LOC105277125 | 50550 | 1783 | 352.72 | 1 | 0.2 | 0.06 | 1.63 | 1.71 | -0.08 | 0.08 | 0 | 0 |
| gene12516 | https://www.ncbi.nlm.nih.gov/gene/?term=LOC105286452 | 1529 | 42 | 274.69 | 1 | 6.54 | 2.38 | 46.48 | 46.6 | -0.11 | 0.11 | 0 | 0 |
| gene9029 | https://www.ncbi.nlm.nih.gov/gene/?term=LOC105282491 | 4762 | 169 | 354.89 | 1 | 2.1 | 0.59 | 7.49 | 7.61 | -0.13 | 0.13 | 0 | 0 |
| gene4641 | https://www.ncbi.nlm.nih.gov/gene/?term=LOC105277398 | 1937 | 67 | 345.9 | 1 | 5.16 | 1.49 | 35.24 | 35.39 | -0.14 | 0.14 | 0 | 0 |
| gene1884 | https://www.ncbi.nlm.nih.gov/gene/?term=LOC105288072 | 4169 | 77 | 184.7 | 1 | 2.4 | 1.3 | 24.07 | 24.23 | -0.16 | 0.16 | 0 | 0 |
| gene5539 | https://www.ncbi.nlm.nih.gov/gene/?term=LOC105278415 | 25024 | 661 | 264.15 | 1 | 0.4 | 0.15 | 3.65 | 3.82 | -0.17 | 0.17 | 0 | 0 |
| gene10489 | https://www.ncbi.nlm.nih.gov/gene/?term=LOC105284147 | 36992 | 478 | 129.22 | 1 | 0.27 | 0.21 | 10.06 | 10.25 | -0.19 | 0.19 | 0 | 0 |
| gene2173 | https://www.ncbi.nlm.nih.gov/gene/?term=LOC105274590 | 4387 | 67 | 152.72 | 2 | 4.56 | 2.99 | 31.19 | 31.42 | -0.23 | 0.23 | 0 | 0 |
| gene2748 | https://www.ncbi.nlm.nih.gov/gene/?term=LOC105275261 | 3845 | 50 | 130.04 | 1 | 2.6 | 2 | 37.52 | 37.82 | -0.3 | 0.3 | 0 | 0 |
| gene3599 | https://www.ncbi.nlm.nih.gov/gene/?term=LOC105276196 | 2955 | 45 | 152.28 | 1 | 3.38 | 2.22 | 48.59 | 48.91 | -0.31 | 0.31 | 0 | 0 |
| gene10490 | https://www.ncbi.nlm.nih.gov/gene/?term=LOC105284146 | 13960 | 149 | 106.73 | 1 | 0.72 | 0.67 | 10.54 | 10.87 | -0.33 | 0.33 | 0 | 0 |
| gene6639 | https://www.ncbi.nlm.nih.gov/gene/?term=LOC105279705 | 7656 | 66 | 86.21 | 1 | 1.31 | 1.52 | 21.87 | 22.21 | -0.34 | 0.34 | 0 | 0 |

Directory: OGLall File: rnk-dmg-Cb_LI_BCphaseA2.vs.RphaseB2.txt

| gene_ID | gene_link | gwidth | #Sites | #per10Kb | #dmSites | #dmsp10kb | %dmSites | %pSite1 | %pSite2 | DMpSite | ADMpSite | DMpNucl | ADMpNucl |
|---|---|---|---|---|---|---|---|---|---|---|---|---|---|
| gene3165 | https://www.ncbi.nlm.nih.gov/gene/?term=LOC105275706 | 2374 | 72 | 303.29 | 6 | 25.27 | 8.33 | 33.88 | 22.12 | 11.76 | 11.76 | 0.36 | 0.36 |
| gene2824 | https://www.ncbi.nlm.nih.gov/gene/?term=LOC105275337 | 1772 | 60 | 338.6 | 2 | 11.29 | 3.33 | 38.89 | 29.82 | 9.07 | 9.07 | 0.31 | 0.31 |
| gene10963 | https://www.ncbi.nlm.nih.gov/gene/?term=LOC105284677 | 1576 | 27 | 171.32 | 3 | 19.04 | 11.11 | 20.06 | 37.04 | -16.98 | 16.98 | -0.29 | 0.29 |
| gene2593 | https://www.ncbi.nlm.nih.gov/gene/?term=LOC105275070 | 1774 | 7 | 39.46 | 5 | 28.18 | 71.43 | 66.52 | 4.09 | 62.43 | 62.43 | 0.25 | 0.25 |
| gene5714 | https://www.ncbi.nlm.nih.gov/gene/?term=LOC105278729 | 710 | 17 | 239.44 | 1 | 14.08 | 5.88 | 60.22 | 53.54 | 6.67 | 6.67 | 0.16 | 0.16 |
| gene4641 | https://www.ncbi.nlm.nih.gov/gene/?term=LOC105277398 | 1937 | 68 | 351.06 | 1 | 5.16 | 1.47 | 37.35 | 40.44 | -3.09 | 3.09 | -0.11 | 0.11 |
| gene7568 | https://www.ncbi.nlm.nih.gov/gene/?term=LOC105280786 | 2525 | 112 | 443.56 | 1 | 3.96 | 0.89 | 32.64 | 30.88 | 1.76 | 1.76 | 0.08 | 0.08 |
| gene10213 | https://www.ncbi.nlm.nih.gov/gene/?term=LOC105283826 | 1503 | 54 | 359.28 | 1 | 6.65 | 1.85 | 13.66 | 15.73 | -2.07 | 2.07 | -0.07 | 0.07 |
| gene11819 | https://www.ncbi.nlm.nih.gov/gene/?term=LOC105285671 | 2366 | 100 | 422.65 | 1 | 4.23 | 1 | 27.82 | 26.43 | 1.39 | 1.39 | 0.06 | 0.06 |
| gene920 | https://www.ncbi.nlm.nih.gov/gene/?term=LOC105282518 | 1197 | 39 | 325.81 | 1 | 8.35 | 2.56 | 38.65 | 40.64 | -1.98 | 1.98 | -0.06 | 0.06 |
| gene2404 | https://www.ncbi.nlm.nih.gov/gene/?term=LOC105274853 | 1767 | 34 | 192.42 | 1 | 5.66 | 2.94 | 29.24 | 32.2 | -2.96 | 2.96 | -0.06 | 0.06 |
| gene940 | https://www.ncbi.nlm.nih.gov/gene/?term=LOC105282794 | 3578 | 87 | 243.15 | 1 | 2.79 | 1.15 | 16.03 | 14.13 | 1.9 | 1.9 | 0.05 | 0.05 |
| gene10520 | https://www.ncbi.nlm.nih.gov/gene/?term=LOC105284184 | 2451 | 78 | 318.24 | 1 | 4.08 | 1.28 | 37.44 | 35.71 | 1.72 | 1.72 | 0.05 | 0.05 |
| gene1470 | https://www.ncbi.nlm.nih.gov/gene/?term=LOC105287597 | 2974 | 195 | 655.68 | 1 | 3.36 | 0.51 | 18.31 | 17.52 | 0.8 | 0.8 | 0.05 | 0.05 |
| gene10879 | https://www.ncbi.nlm.nih.gov/gene/?term=LOC105284574 | 2423 | 46 | 189.85 | 1 | 4.13 | 2.17 | 56.45 | 59.11 | -2.65 | 2.65 | -0.05 | 0.05 |
| gene393 | https://www.ncbi.nlm.nih.gov/gene/?term=LOC105276566 | 1123 | 2 | 17.81 | 1 | 8.9 | 50 | 48.08 | 76.78 | -28.71 | 28.71 | -0.05 | 0.05 |
| gene7398 | https://www.ncbi.nlm.nih.gov/gene/?term=LOC105280589 | 3452 | 34 | 98.49 | 1 | 2.9 | 2.94 | 23.83 | 19.97 | 3.86 | 3.86 | 0.04 | 0.04 |
| gene9462 | https://www.ncbi.nlm.nih.gov/gene/?term=LOC105282970 | 1469 | 39 | 265.49 | 1 | 6.81 | 2.56 | 51.12 | 49.47 | 1.65 | 1.65 | 0.04 | 0.04 |
| gene12806 | https://www.ncbi.nlm.nih.gov/gene/?term=LOC105286803 | 4196 | 137 | 326.5 | 1 | 2.38 | 0.73 | 10.84 | 9.75 | 1.09 | 1.09 | 0.04 | 0.04 |
| gene10678 | https://www.ncbi.nlm.nih.gov/gene/?term=LOC105284355 | 4031 | 158 | 391.96 | 1 | 2.48 | 0.63 | 20.75 | 19.82 | 0.93 | 0.93 | 0.04 | 0.04 |
| gene1220 | https://www.ncbi.nlm.nih.gov/gene/?term=LOC105285742 | 3453 | 107 | 309.88 | 1 | 2.9 | 0.93 | 22.27 | 23.44 | -1.17 | 1.17 | -0.04 | 0.04 |
| gene7600 | https://www.ncbi.nlm.nih.gov/gene/?term=LOC105280819 | 1615 | 37 | 229.1 | 1 | 6.19 | 2.7 | 36.56 | 38.16 | -1.6 | 1.6 | -0.04 | 0.04 |
| gene2413 | https://www.ncbi.nlm.nih.gov/gene/?term=LOC105274863 | 2006 | 40 | 199.4 | 1 | 4.99 | 2.5 | 64.23 | 66.19 | -1.96 | 1.96 | -0.04 | 0.04 |
| gene3599 | https://www.ncbi.nlm.nih.gov/gene/?term=LOC105276196 | 2955 | 47 | 159.05 | 1 | 3.38 | 2.13 | 46.53 | 48.84 | -2.31 | 2.31 | -0.04 | 0.04 |
| gene3385 | https://www.ncbi.nlm.nih.gov/gene/?term=LOC105275966 | 3305 | 53 | 160.36 | 1 | 3.03 | 1.89 | 54.5 | 52.88 | 1.62 | 1.62 | 0.03 | 0.03 |
| gene3594 | https://www.ncbi.nlm.nih.gov/gene/?term=LOC105276190 | 4910 | 83 | 169.04 | 1 | 2.04 | 1.2 | 11.44 | 9.83 | 1.62 | 1.62 | 0.03 | 0.03 |
| gene5877 | https://www.ncbi.nlm.nih.gov/gene/?term=LOC105278807 | 1719 | 46 | 267.6 | 1 | 5.82 | 2.17 | 26.78 | 25.52 | 1.25 | 1.25 | 0.03 | 0.03 |
| gene13033 | https://www.ncbi.nlm.nih.gov/gene/?term=LOC105287063 | 1877 | 48 | 255.73 | 1 | 5.33 | 2.08 | 31.55 | 30.37 | 1.19 | 1.19 | 0.03 | 0.03 |
| gene7915 | https://www.ncbi.nlm.nih.gov/gene/?term=LOC105281173 | 1597 | 38 | 237.95 | 1 | 6.26 | 2.63 | 59.94 | 58.74 | 1.19 | 1.19 | 0.03 | 0.03 |
| gene6652 | https://www.ncbi.nlm.nih.gov/gene/?term=LOC105279719 | 2831 | 79 | 279.05 | 1 | 3.53 | 1.27 | 24.35 | 23.32 | 1.03 | 1.03 | 0.03 | 0.03 |
| gene7556 | https://www.ncbi.nlm.nih.gov/gene/?term=LOC105280774 | 2792 | 89 | 318.77 | 1 | 3.58 | 1.12 | 36.33 | 35.33 | 1 | 1 | 0.03 | 0.03 |
| gene3817 | https://www.ncbi.nlm.nih.gov/gene/?term=LOC105276449 | 1230 | 42 | 341.46 | 1 | 8.13 | 2.38 | 35.93 | 35.04 | 0.88 | 0.88 | 0.03 | 0.03 |
| gene10606 | https://www.ncbi.nlm.nih.gov/gene/?term=LOC105284273 | 6119 | 249 | 406.93 | 1 | 1.63 | 0.4 | 18.46 | 17.61 | 0.86 | 0.86 | 0.03 | 0.03 |
| gene11401 | https://www.ncbi.nlm.nih.gov/gene/?term=LOC105285202 | 1841 | 82 | 445.41 | 1 | 5.43 | 1.22 | 26.05 | 25.33 | 0.73 | 0.73 | 0.03 | 0.03 |
| gene2846 | https://www.ncbi.nlm.nih.gov/gene/?term=LOC105275374 | 11246 | 464 | 412.59 | 1 | 0.89 | 0.22 | 12.19 | 11.56 | 0.63 | 0.63 | 0.03 | 0.03 |
| gene5298 | https://www.ncbi.nlm.nih.gov/gene/?term=LOC105278145 | 4060 | 245 | 603.45 | 2 | 4.93 | 0.82 | 10.68 | 11.19 | -0.51 | 0.51 | -0.03 | 0.03 |
| gene4498 | https://www.ncbi.nlm.nih.gov/gene/?term=LOC105277239 | 2831 | 122 | 430.94 | 1 | 3.53 | 0.82 | 12.06 | 12.7 | -0.64 | 0.64 | -0.03 | 0.03 |
| gene9101 | https://www.ncbi.nlm.nih.gov/gene/?term=LOC105282553 | 3068 | 108 | 352.02 | 1 | 3.26 | 0.93 | 22.57 | 23.41 | -0.84 | 0.84 | -0.03 | 0.03 |
| gene12516 | https://www.ncbi.nlm.nih.gov/gene/?term=LOC105286452 | 1529 | 40 | 261.61 | 1 | 6.54 | 2.5 | 50.21 | 51.25 | -1.04 | 1.04 | -0.03 | 0.03 |
| gene7207 | https://www.ncbi.nlm.nih.gov/gene/?term=LOC105280376 | 4659 | 109 | 233.96 | 1 | 2.15 | 0.92 | 29.99 | 31.37 | -1.38 | 1.38 | -0.03 | 0.03 |
| gene7850 | https://www.ncbi.nlm.nih.gov/gene/?term=LOC105281095 | 3050 | 63 | 206.56 | 1 | 3.28 | 1.59 | 33.9 | 35.57 | -1.67 | 1.67 | -0.03 | 0.03 |
| gene322 | https://www.ncbi.nlm.nih.gov/gene/?term=LOC105275777 | 1606 | 23 | 143.21 | 1 | 6.23 | 4.35 | 46.95 | 49.13 | -2.17 | 2.17 | -0.03 | 0.03 |
| gene10345 | https://www.ncbi.nlm.nih.gov/gene/?term=LOC105283989 | 4444 | 22 | 49.5 | 1 | 2.25 | 4.55 | 37.81 | 42.87 | -5.06 | 5.06 | -0.03 | 0.03 |
| gene2970 | https://www.ncbi.nlm.nih.gov/gene/?term=LOC105275518 | 2340 | 18 | 76.92 | 1 | 4.27 | 5.56 | 54.32 | 51.43 | 2.89 | 2.89 | 0.02 | 0.02 |
| gene6455 | https://www.ncbi.nlm.nih.gov/gene/?term=LOC105279500 | 2612 | 33 | 126.34 | 1 | 3.83 | 3.03 | 51.77 | 49.92 | 1.86 | 1.86 | 0.02 | 0.02 |
| gene35 | https://www.ncbi.nlm.nih.gov/gene/?term=LOC105275197 | 2643 | 43 | 162.69 | 1 | 3.78 | 2.33 | 44.25 | 43.02 | 1.22 | 1.22 | 0.02 | 0.02 |
| gene3470 | https://www.ncbi.nlm.nih.gov/gene/?term=LOC105276059 | 8716 | 145 | 166.36 | 1 | 1.15 | 0.69 | 9.2 | 8.01 | 1.19 | 1.19 | 0.02 | 0.02 |
| gene3671 | https://www.ncbi.nlm.nih.gov/gene/?term=LOC105276272 | 1966 | 34 | 172.94 | 1 | 5.09 | 2.94 | 33.29 | 32.32 | 0.98 | 0.98 | 0.02 | 0.02 |
| gene4962 | https://www.ncbi.nlm.nih.gov/gene/?term=LOC105277761 | 4645 | 142 | 305.71 | 1 | 2.15 | 0.7 | 14.73 | 13.94 | 0.79 | 0.79 | 0.02 | 0.02 |
| gene4892 | https://www.ncbi.nlm.nih.gov/gene/?term=LOC105277667 | 3898 | 124 | 318.11 | 1 | 2.57 | 0.81 | 27.97 | 27.19 | 0.78 | 0.78 | 0.02 | 0.02 |
| gene10600 | https://www.ncbi.nlm.nih.gov/gene/?term=LOC105284270 | 4837 | 119 | 246.02 | 1 | 2.07 | 0.84 | 10.7 | 10.02 | 0.68 | 0.68 | 0.02 | 0.02 |
| gene3727 | https://www.ncbi.nlm.nih.gov/gene/?term=LOC105276367 | 2484 | 66 | 265.7 | 1 | 4.03 | 1.52 | 47.44 | 46.84 | 0.61 | 0.61 | 0.02 | 0.02 |
| gene10984 | https://www.ncbi.nlm.nih.gov/gene/?term=LOC105284704 | 1987 | 56 | 281.83 | 1 | 5.03 | 1.79 | 25.7 | 25.1 | 0.59 | 0.59 | 0.02 | 0.02 |
| gene2844 | https://www.ncbi.nlm.nih.gov/gene/?term=LOC105275363 | 4330 | 181 | 418.01 | 1 | 2.31 | 0.55 | 15.09 | 14.51 | 0.58 | 0.58 | 0.02 | 0.02 |
| gene8274 | https://www.ncbi.nlm.nih.gov/gene/?term=LOC105281624 | 5425 | 204 | 376.04 | 1 | 1.84 | 0.49 | 10.02 | 9.47 | 0.56 | 0.56 | 0.02 | 0.02 |
| gene2056 | https://www.ncbi.nlm.nih.gov/gene/?term=LOC105274452 | 6584 | 251 | 381.23 | 2 | 3.04 | 0.8 | 13.77 | 13.35 | 0.42 | 0.42 | 0.02 | 0.02 |
| gene6286 | https://www.ncbi.nlm.nih.gov/gene/?term=LOC105279298 | 2861 | 128 | 447.4 | 2 | 6.99 | 1.56 | 24.57 | 24.17 | 0.4 | 0.4 | 0.02 | 0.02 |
| gene11340 | https://www.ncbi.nlm.nih.gov/gene/?term=LOC105285131 | 2918 | 86 | 294.72 | 1 | 3.43 | 1.16 | 11.93 | 12.54 | -0.61 | 0.61 | -0.02 | 0.02 |
| gene5646 | https://www.ncbi.nlm.nih.gov/gene/?term=LOC105278534 | 7475 | 175 | 234.11 | 1 | 1.34 | 0.57 | 17.8 | 18.5 | -0.7 | 0.7 | -0.02 | 0.02 |
| gene11976 | https://www.ncbi.nlm.nih.gov/gene/?term=LOC105285860 | 3112 | 52 | 167.1 | 1 | 3.21 | 1.92 | 21.02 | 22 | -0.98 | 0.98 | -0.02 | 0.02 |

| | | | | | | | | | | | | | |
|---|---|---|---|---|---|---|---|---|---|---|---|---|---|
| gene1147 | https://www.ncbi.nlm.nih.gov/gene/?term=LOC105284954 | 4022 | 84 | 208.85 | 1 | 2.49 | 1.19 | 17.29 | 18.34 | -1.06 | 1.06 | -0.02 | 0.02 |
| gene10391 | https://www.ncbi.nlm.nih.gov/gene/?term=LOC109611283 | 1994 | 42 | 210.63 | 1 | 5.02 | 2.38 | 32.6 | 33.67 | -1.07 | 1.07 | -0.02 | 0.02 |
| gene10155 | https://www.ncbi.nlm.nih.gov/gene/?term=LOC105283756 | 7030 | 100 | 142.25 | 1 | 1.42 | 1 | 5.99 | 7.12 | -1.12 | 1.12 | -0.02 | 0.02 |
| gene10819 | https://www.ncbi.nlm.nih.gov/gene/?term=LOC105284507 | 5767 | 76 | 131.78 | 1 | 1.73 | 1.32 | 23.17 | 24.52 | -1.35 | 1.35 | -0.02 | 0.02 |
| gene388 | https://www.ncbi.nlm.nih.gov/gene/?term=LOC105276538 | 1558 | 19 | 121.95 | 2 | 12.84 | 10.53 | 63.26 | 65.13 | -1.87 | 1.87 | -0.02 | 0.02 |
| gene3168 | https://www.ncbi.nlm.nih.gov/gene/?term=LOC105275711 | 6150 | 67 | 108.94 | 1 | 1.63 | 1.49 | 13.09 | 15.36 | -2.27 | 2.27 | -0.02 | 0.02 |
| gene10344 | https://www.ncbi.nlm.nih.gov/gene/?term=LOC105283987 | 4841 | 64 | 132.2 1 | 2.07 | 1.56 | 30.28 | 29.42 | 0.86 | 0.86 | 0.01 | 0.01 | |
| gene9336 | https://www.ncbi.nlm.nih.gov/gene/?term=LOC105282834 | 5993 | 121 | 201.9 1 | 1.67 | 0.83 | 16.02 | 15.31 | 0.72 | 0.72 | 0.01 | 0.01 | |
| gene2173 | https://www.ncbi.nlm.nih.gov/gene/?term=LOC105274590 | 4387 | 57 | 129.93 | 2 | 4.56 | 3.51 | 33.32 | 32.7 | 0.62 | 0.62 | 0.01 | 0.01 |
| gene11568 | https://www.ncbi.nlm.nih.gov/gene/?term=LOC105285395 | 8086 | 180 | 222.61 | 1 | 1.24 | 0.56 | 17.19 | 16.59 | 0.6 | 0.6 | 0.01 | 0.01 |
| gene7245 | https://www.ncbi.nlm.nih.gov/gene/?term=LOC105280410 | 12478 | 234 | 187.53 | 1 | 0.8 | 0.43 | 21.44 | 20.93 | 0.52 | 0.52 | 0.01 | 0.01 |
| gene5573 | https://www.ncbi.nlm.nih.gov/gene/?term=LOC105278454 | 2688 | 35 | 130.21 | 1 | 3.72 | 2.86 | 35.99 | 35.53 | 0.47 | 0.47 | 0.01 | 0.01 |
| gene12572 | https://www.ncbi.nlm.nih.gov/gene/?term=LOC105286514 | 2374 | 45 | 189.55 | 1 | 4.21 | 2.22 | 33.13 | 32.67 | 0.45 | 0.45 | 0.01 | 0.01 |
| gene34 | https://www.ncbi.nlm.nih.gov/gene/?term=LOC105278416 | 7120 | 124 | 174.16 | 1 | 1.4 | 0.81 | 15.53 | 15.09 | 0.43 | 0.43 | 0.01 | 0.01 |
| gene7166 | https://www.ncbi.nlm.nih.gov/gene/?term=LOC105280326 | 6117 | 214 | 349.84 | 1 | 1.63 | 0.47 | 16.62 | 16.23 | 0.4 | 0.4 | 0.01 | 0.01 |
| gene4246 | https://www.ncbi.nlm.nih.gov/gene/?term=LOC105276941 | 16412 | 493 | 300.39 | 1 | 0.61 | 0.2 | 5.46 | 5.07 | 0.38 | 0.38 | 0.01 | 0.01 |
| gene5746 | https://www.ncbi.nlm.nih.gov/gene/?term=LOC105278666 | 10343 | 425 | 410.91 | 1 | 0.97 | 0.24 | 7.62 | 7.27 | 0.35 | 0.35 | 0.01 | 0.01 |
| gene10392 | https://www.ncbi.nlm.nih.gov/gene/?term=LOC105284036 | 3216 | 91 | 282.96 | 2 | 6.22 | 2.2 | 22.63 | 22.31 | 0.32 | 0.32 | 0.01 | 0.01 |
| gene473 | https://www.ncbi.nlm.nih.gov/gene/?term=LOC105277423 | 2342 | 108 | 461.14 | 1 | 4.27 | 0.93 | 30.31 | 29.99 | 0.32 | 0.32 | 0.01 | 0.01 |
| gene11579 | https://www.ncbi.nlm.nih.gov/gene/?term=LOC105285406 | 13571 | 228 | 168.01 | 1 | 0.74 | 0.44 | 7.6 | 7.29 | 0.3 | 0.3 | 0.01 | 0.01 |
| gene2271 | https://www.ncbi.nlm.nih.gov/gene/?term=LOC105274706 | 1448 | 56 | 386.74 | 1 | 6.91 | 1.79 | 36.69 | 36.4 | 0.3 | 0.3 | 0.01 | 0.01 |
| gene10531 | https://www.ncbi.nlm.nih.gov/gene/?term=LOC105284190 | 6881 | 168 | 244.15 | 1 | 1.45 | 0.6 | 13.32 | 13.04 | 0.27 | 0.27 | 0.01 | 0.01 |
| gene7211 | https://www.ncbi.nlm.nih.gov/gene/?term=LOC105280377 | 2024 | 57 | 281.62 | 1 | 4.94 | 1.75 | 50.41 | 50.16 | 0.25 | 0.25 | 0.01 | 0.01 |
| gene298 | https://www.ncbi.nlm.nih.gov/gene/?term=LOC105275460 | 2473 | 116 | 469.07 | 1 | 4.04 | 0.86 | 13.65 | 13.43 | 0.23 | 0.23 | 0.01 | 0.01 |
| gene5525 | https://www.ncbi.nlm.nih.gov/gene/?term=LOC105278396 | 3090 | 181 | 585.76 | 1 | 3.24 | 0.55 | 24.85 | 24.63 | 0.22 | 0.22 | 0.01 | 0.01 |
| gene5524 | https://www.ncbi.nlm.nih.gov/gene/?term=LOC105278397 | 5177 | 196 | 378.6 1 | 1.93 | 0.51 | 24.88 | 24.69 | 0.18 | 0.18 | 0.01 | 0.01 | |
| gene3286 | https://www.ncbi.nlm.nih.gov/gene/?term=LOC105275848 | 3744 | 134 | 357.91 | 1 | 2.67 | 0.75 | 22.27 | 22.41 | -0.14 | 0.14 | -0.01 | 0.01 |
| gene10549 | https://www.ncbi.nlm.nih.gov/gene/?term=LOC105284210 | 7145 | 286 | 400.28 | 2 | 2.8 | 0.7 | 11.97 | 12.18 | -0.21 | 0.21 | -0.01 | 0.01 |
| gene12802 | https://www.ncbi.nlm.nih.gov/gene/?term=LOC105286799 | 3183 | 160 | 502.67 | 1 | 3.14 | 0.62 | 17.6 | 17.81 | -0.21 | 0.21 | -0.01 | 0.01 |
| gene3107 | https://www.ncbi.nlm.nih.gov/gene/?term=LOC105275650 | 5103 | 166 | 325.3 1 | 1.96 | 0.6 | 14.75 | 14.95 | -0.21 | 0.21 | -0.01 | 0.01 | |
| gene9755 | https://www.ncbi.nlm.nih.gov/gene/?term=LOC105283296 | 2177 | 85 | 390.45 | 1 | 4.59 | 1.18 | 19.16 | 19.37 | -0.21 | 0.21 | -0.01 | 0.01 |
| gene5733 | https://www.ncbi.nlm.nih.gov/gene/?term=LOC105278649 | 5221 | 114 | 218.35 | 1 | 1.92 | 0.88 | 16.23 | 16.48 | -0.25 | 0.25 | -0.01 | 0.01 |
| gene1310 | https://www.ncbi.nlm.nih.gov/gene/?term=LOC105286661 | 2582 | 72 | 278.85 | 1 | 3.87 | 1.39 | 21.23 | 21.48 | -0.26 | 0.26 | -0.01 | 0.01 |
| gene12674 | https://www.ncbi.nlm.nih.gov/gene/?term=LOC105286649 | 18283 | 747 | 408.58 | 1 | 0.55 | 0.13 | 0.89 | 1.18 | -0.29 | 0.29 | -0.01 | 0.01 |
| gene13036 | https://www.ncbi.nlm.nih.gov/gene/?term=LOC105287059 | 4959 | 146 | 294.41 | 1 | 2.02 | 0.68 | 18.9 | 19.21 | -0.31 | 0.31 | -0.01 | 0.01 |
| gene3885 | https://www.ncbi.nlm.nih.gov/gene/?term=LOC105276527 | 7928 | 305 | 384.71 | 1 | 1.26 | 0.33 | 9.75 | 10.06 | -0.32 | 0.32 | -0.01 | 0.01 |
| gene6583 | https://www.ncbi.nlm.nih.gov/gene/?term=LOC105279649 | 7444 | 303 | 407.04 | 1 | 1.34 | 0.33 | 11.36 | 11.69 | -0.33 | 0.33 | -0.01 | 0.01 |
| gene10018 | https://www.ncbi.nlm.nih.gov/gene/?term=LOC105283607 | 4414 | 187 | 423.65 | 1 | 2.27 | 0.53 | 16.13 | 16.48 | -0.35 | 0.35 | -0.01 | 0.01 |
| gene2122 | https://www.ncbi.nlm.nih.gov/gene/?term=LOC105274531 | 8153 | 289 | 354.47 | 1 | 1.23 | 0.35 | 10.34 | 10.69 | -0.35 | 0.35 | -0.01 | 0.01 |
| gene6031 | https://www.ncbi.nlm.nih.gov/gene/?term=LOC105279000 | 5691 | 187 | 328.59 | 1 | 1.76 | 0.53 | 16.08 | 16.46 | -0.38 | 0.38 | -0.01 | 0.01 |
| gene12573 | https://www.ncbi.nlm.nih.gov/gene/?term=LOC105286515 | 3647 | 98 | 268.71 | 1 | 2.74 | 1.02 | 13.27 | 13.66 | -0.39 | 0.39 | -0.01 | 0.01 |
| gene4376 | https://www.ncbi.nlm.nih.gov/gene/?term=LOC105277083 | 4307 | 102 | 236.82 | 1 | 2.32 | 0.98 | 24.57 | 25.16 | -0.58 | 0.58 | -0.01 | 0.01 |
| gene5273 | https://www.ncbi.nlm.nih.gov/gene/?term=LOC105278108 | 4067 | 44 | 108.19 | 1 | 2.46 | 2.27 | 24.76 | 25.36 | -0.61 | 0.61 | -0.01 | 0.01 |
| gene12203 | https://www.ncbi.nlm.nih.gov/gene/?term=LOC105286099 | 5967 | 73 | 122.34 | 1 | 1.68 | 1.37 | 10.19 | 10.87 | -0.68 | 0.68 | -0.01 | 0.01 |
| gene12406 | https://www.ncbi.nlm.nih.gov/gene/?term=LOC105286316 | 1386 | 28 | 202.02 | 1 | 7.22 | 3.57 | 44.18 | 44.91 | -0.73 | 0.73 | -0.01 | 0.01 |
| gene2748 | https://www.ncbi.nlm.nih.gov/gene/?term=LOC105275261 | 3845 | 51 | 132.64 | 1 | 2.6 | 1.96 | 36.93 | 37.66 | -0.74 | 0.74 | -0.01 | 0.01 |
| gene2683 | https://www.ncbi.nlm.nih.gov/gene/?term=LOC105275187 | 5492 | 66 | 120.17 | 1 | 1.82 | 1.52 | 11.92 | 12.72 | -0.8 | 0.8 | -0.01 | 0.01 |
| gene7368 | https://www.ncbi.nlm.nih.gov/gene/?term=LOC105280556 | 4105 | 35 | 85.26 1 | 2.44 | 2.86 | 17.18 | 18.02 | -0.84 | 0.84 | -0.01 | 0.01 | |
| gene9327 | https://www.ncbi.nlm.nih.gov/gene/?term=LOC105282821 | 7793 | 132 | 169.38 | 1 | 1.28 | 0.76 | 11.87 | 12.7 | -0.84 | 0.84 | -0.01 | 0.01 |
| gene8343 | https://www.ncbi.nlm.nih.gov/gene/?term=LOC105281700 | 2172 | 34 | 156.54 | 1 | 4.6 | 2.94 | 35.91 | 36.82 | -0.9 | 0.9 | -0.01 | 0.01 |
| gene11447 | https://www.ncbi.nlm.nih.gov/gene/?term=LOC105285252 | 7437 | 114 | 153.29 | 2 | 2.69 | 1.75 | 13.96 | 14.93 | -0.97 | 0.97 | -0.01 | 0.01 |
| gene10107 | https://www.ncbi.nlm.nih.gov/gene/?term=LOC105283712 | 1883 | 32 | 169.94 | 1 | 5.31 | 3.12 | 29.98 | 29.7 | 0.28 | 0.28 | 0 | 0 |
| gene8600 | https://www.ncbi.nlm.nih.gov/gene/?term=LOC105282001 | 8513 | 139 | 163.28 | 1 | 1.17 | 0.72 | 11.37 | 11.2 | 0.17 | 0.17 | 0 | 0 |
| gene9330 | https://www.ncbi.nlm.nih.gov/gene/?term=LOC105282825 | 3936 | 69 | 175.3 1 | 2.54 | 1.45 | 13.52 | 13.35 | 0.17 | 0.17 | 0 | 0 | |
| gene12307 | https://www.ncbi.nlm.nih.gov/gene/?term=LOC105286216 | 10313 | 167 | 161.93 | 1 | 0.97 | 0.6 | 17.01 | 16.86 | 0.15 | 0.15 | 0 | 0 |
| gene6620 | https://www.ncbi.nlm.nih.gov/gene/?term=LOC105279687 | 17998 | 312 | 173.35 | 1 | 0.56 | 0.32 | 5.46 | 5.32 | 0.14 | 0.14 | 0 | 0 |
| gene12265 | https://www.ncbi.nlm.nih.gov/gene/?term=LOC105286166 | 3050 | 104 | 340.98 | 1 | 3.28 | 0.96 | 25.25 | 25.13 | 0.13 | 0.13 | 0 | 0 |
| gene292 | https://www.ncbi.nlm.nih.gov/gene/?term=LOC105275398 | 2825 | 81 | 286.73 | 1 | 3.54 | 1.23 | 26.31 | 26.19 | 0.11 | 0.11 | 0 | 0 |
| gene11275 | https://www.ncbi.nlm.nih.gov/gene/?term=LOC105285055 | 4119 | 252 | 611.8 1 | 2.43 | 0.4 | 11.37 | 11.3 | 0.07 | 0.07 | 0 | 0 | |
| gene2806 | https://www.ncbi.nlm.nih.gov/gene/?term=LOC105275327 | 14257 | 460 | 322.65 | 1 | 0.7 | 0.22 | 7.25 | 7.19 | 0.06 | 0.06 | 0 | 0 |
| gene12981 | https://www.ncbi.nlm.nih.gov/gene/?term=LOC105286999 | 2063 | 42 | 203.59 | 1 | 4.85 | 2.38 | 36.12 | 36.07 | 0.05 | 0.05 | 0 | 0 |
| gene13114 | https://www.ncbi.nlm.nih.gov/gene/?term=LOC105287147 | 7609 | 383 | 503.35 | 1 | 1.31 | 0.26 | 9.68 | 9.63 | 0.04 | 0.04 | 0 | 0 |
| gene53 | https://www.ncbi.nlm.nih.gov/gene/?term=LOC105277148 | 19283 | 258 | 133.8 1 | 0.52 | 0.39 | 4.46 | 4.42 | 0.04 | 0.04 | 0 | 0 | |
| gene5572 | https://www.ncbi.nlm.nih.gov/gene/?term=LOC105278455 | 27887 | 864 | 309.82 | 1 | 0.36 | 0.12 | 6.83 | 6.79 | 0.04 | 0.04 | 0 | 0 |

| | | | | | | | | | | | | | | |
|---|---|---|---|---|---|---|---|---|---|---|---|---|---|---|
| gene11082 | https://www.ncbi.nlm.nih.gov/gene/?term=LOC105284824 | 7436 | 185 | 248.79 | | 1 | 1.34 | 0.54 | 6.28 | 6.27 | 0.01 | 0.01 | 0 | 0 |
| gene1632 | https://www.ncbi.nlm.nih.gov/gene/?term=LOC105287764 | 18061 | 371 | 205.41 | | 2 | 1.11 | 0.54 | 3.28 | 3.27 | 0.01 | 0.01 | 0 | 0 |
| gene4209 | https://www.ncbi.nlm.nih.gov/gene/?term=LOC105277125 | 50550 | 1672 | 330.76 | | 1 | 0.2 | 0.06 | 1.66 | 1.65 | 0.01 | 0.01 | 0 | 0 |
| gene6966 | https://www.ncbi.nlm.nih.gov/gene/?term=LOC105280104 | 3747 | 123 | 328.26 | | 1 | 2.67 | 0.81 | 16.59 | 16.58 | 0.01 | 0.01 | 0 | 0 |
| gene4190 | https://www.ncbi.nlm.nih.gov/gene/?term=LOC105276875 | 3150 | 109 | 346.03 | | 1 | 3.17 | 0.92 | 24.02 | 24.04 | -0.02 | 0.02 | 0 | 0 |
| gene1993 | https://www.ncbi.nlm.nih.gov/gene/?term=LOC105288193 | 1449 | 74 | 510.7 | 1 | 6.9 | 1.35 | 34.89 | 34.92 | -0.03 | 0.03 | 0 | 0 | |
| gene8729 | https://www.ncbi.nlm.nih.gov/gene/?term=LOC105282137 | 5005 | 198 | 395.6 | 1 | 2 | 0.51 | 16.91 | 16.95 | -0.04 | 0.04 | 0 | 0 | |
| gene5831 | https://www.ncbi.nlm.nih.gov/gene/?term=LOC105278761 | 2506 | 109 | 434.96 | | 1 | 3.99 | 0.92 | 24.44 | 24.49 | -0.06 | 0.06 | 0 | 0 |
| gene9177 | https://www.ncbi.nlm.nih.gov/gene/?term=LOC105282646 | 3320 | 99 | 298.19 | | 1 | 3.01 | 1.01 | 25.05 | 25.11 | -0.06 | 0.06 | 0 | 0 |
| gene7249 | https://www.ncbi.nlm.nih.gov/gene/?term=LOC105280422 | 1775 | 60 | 338.03 | | 1 | 5.63 | 1.67 | 16.97 | 17.06 | -0.09 | 0.09 | 0 | 0 |
| gene5967 | https://www.ncbi.nlm.nih.gov/gene/?term=LOC105278918 | 4393 | 116 | 264.06 | | 1 | 2.28 | 0.86 | 14.39 | 14.54 | -0.15 | 0.15 | 0 | 0 |
| gene8661 | https://www.ncbi.nlm.nih.gov/gene/?term=LOC105282063 | 1926 | 35 | 181.72 | | 1 | 5.19 | 2.86 | 30.68 | 30.89 | -0.22 | 0.22 | 0 | 0 |
| gene1199 | https://www.ncbi.nlm.nih.gov/gene/?term=LOC105285498 | 2220 | 26 | 117.12 | | 1 | 4.5 | 3.85 | 31.35 | 31.59 | -0.24 | 0.24 | 0 | 0 |
| gene9326 | https://www.ncbi.nlm.nih.gov/gene/?term=LOC105282818 | 16612 | 173 | 104.14 | | 1 | 0.6 | 0.58 | 12.61 | 12.88 | -0.27 | 0.27 | 0 | 0 |
| gene12021 | https://www.ncbi.nlm.nih.gov/gene/?term=LOC105285905 | 1399 | 16 | 114.37 | | 1 | 7.15 | 6.25 | 47.33 | 47.75 | -0.42 | 0.42 | 0 | 0 |
| gene6639 | https://www.ncbi.nlm.nih.gov/gene/?term=LOC105279705 | 7656 | 61 | 79.68 | 1 | 1.31 | 1.64 | 23.52 | 23.98 | -0.46 | 0.46 | 0 | 0 | |

Directory: OGLall File: rnk-dmg-Cb_LI_BCphaseB1.vs.BCphaseB2.txt

| gene_ID | gene_link | gwidth | #Sites | #per10Kb | #dmSites | #dmsp10kb | %dmSites | %pSite1 | %pSite2 | DMpSite | ADMpSite | DMpNucl | ADMpNucl |
|---|---|---|---|---|---|---|---|---|---|---|---|---|---|
| gene10963 | https://www.ncbi.nlm.nih.gov/gene/?term=LOC105284677 | 1576 | 29 | 184.01 | 1 | 6.35 | 3.45 | 18.87 | 36.99 | -18.12 | 18.12 | -0.33 | 0.33 |
| gene3165 | https://www.ncbi.nlm.nih.gov/gene/?term=LOC105275706 | 2374 | 66 | 278.01 | 5 | 21.06 | 7.58 | 34.05 | 24.01 | 10.05 | 10.05 | 0.28 | 0.28 |
| gene2593 | https://www.ncbi.nlm.nih.gov/gene/?term=LOC105275070 | 1774 | 5 | 28.18 | 5 | 28.18 | 100 | 72.92 | 10.96 | 61.96 | 61.96 | 0.17 | 0.17 |
| gene5714 | https://www.ncbi.nlm.nih.gov/gene/?term=LOC105278729 | 710 | 24 | 338.03 | 1 | 14.08 | 4.17 | 59.89 | 56.79 | 3.1 | 3.1 | 0.1 | 0.1 |
| gene4498 | https://www.ncbi.nlm.nih.gov/gene/?term=LOC105277239 | 2831 | 140 | 494.52 | 1 | 3.53 | 0.71 | 13.31 | 15.32 | -2.01 | 2.01 | -0.1 | 0.1 |
| gene3817 | https://www.ncbi.nlm.nih.gov/gene/?term=LOC105276449 | 1230 | 51 | 414.63 | 1 | 8.13 | 1.96 | 27.91 | 26.06 | 1.85 | 1.85 | 0.08 | 0.08 |
| gene1962 | https://www.ncbi.nlm.nih.gov/gene/?term=LOC105288157 | 1246 | 17 | 136.44 | 1 | 8.03 | 5.88 | 18.51 | 13.14 | 5.37 | 5.37 | 0.07 | 0.07 |
| gene2907 | https://www.ncbi.nlm.nih.gov/gene/?term=LOC105275447 | 2539 | 40 | 157.54 | 1 | 3.94 | 2.5 | 7.22 | 3.04 | 4.18 | 4.18 | 0.07 | 0.07 |
| gene12021 | https://www.ncbi.nlm.nih.gov/gene/?term=LOC105285905 | 1399 | 20 | 142.96 | 1 | 7.15 | 5 | 33.2 | 38.03 | -4.83 | 4.83 | -0.07 | 0.07 |
| gene5298 | https://www.ncbi.nlm.nih.gov/gene/?term=LOC105278145 | 4060 | 236 | 581.28 | 2 | 4.93 | 0.85 | 10.36 | 11.46 | -1.1 | 1.1 | -0.06 | 0.06 |
| gene9740 | https://www.ncbi.nlm.nih.gov/gene/?term=LOC105283277 | 2887 | 140 | 484.93 | 1 | 3.46 | 0.71 | 19.86 | 21.12 | -1.26 | 1.26 | -0.06 | 0.06 |
| gene2271 | https://www.ncbi.nlm.nih.gov/gene/?term=LOC105274706 | 1448 | 51 | 352.21 | 1 | 6.91 | 1.96 | 40.05 | 41.71 | -1.67 | 1.67 | -0.06 | 0.06 |
| gene3727 | https://www.ncbi.nlm.nih.gov/gene/?term=LOC105276367 | 2484 | 66 | 265.7 | 1 | 4.03 | 1.52 | 44.56 | 46.97 | -2.41 | 2.41 | -0.06 | 0.06 |
| gene5877 | https://www.ncbi.nlm.nih.gov/gene/?term=LOC105278807 | 1719 | 53 | 308.32 | 1 | 5.82 | 1.89 | 23.93 | 22.25 | 1.68 | 1.68 | 0.05 | 0.05 |
| gene4892 | https://www.ncbi.nlm.nih.gov/gene/?term=LOC105277667 | 3898 | 127 | 325.81 | 1 | 2.57 | 0.79 | 27.62 | 26.17 | 1.45 | 1.45 | 0.05 | 0.05 |
| gene10678 | https://www.ncbi.nlm.nih.gov/gene/?term=LOC105284355 | 4031 | 174 | 431.65 | 1 | 2.48 | 0.57 | 19.29 | 18.18 | 1.11 | 1.11 | 0.05 | 0.05 |
| gene7568 | https://www.ncbi.nlm.nih.gov/gene/?term=LOC105280786 | 2525 | 119 | 471.29 | 1 | 3.96 | 0.84 | 30.86 | 29.84 | 1.03 | 1.03 | 0.05 | 0.05 |
| gene1220 | https://www.ncbi.nlm.nih.gov/gene/?term=LOC105285742 | 3453 | 125 | 362 | 1 | 2.9 | 0.8 | 19.11 | 20.58 | -1.47 | 1.47 | -0.05 | 0.05 |
| gene6237 | https://www.ncbi.nlm.nih.gov/gene/?term=LOC105279244 | 1484 | 25 | 168.46 | 1 | 6.74 | 4 | 50.3 | 53.48 | -3.18 | 3.18 | -0.05 | 0.05 |
| gene1199 | https://www.ncbi.nlm.nih.gov/gene/?term=LOC105285498 | 2220 | 26 | 117.12 | 1 | 4.5 | 3.85 | 33.11 | 37.03 | -3.92 | 3.92 | -0.05 | 0.05 |
| gene2970 | https://www.ncbi.nlm.nih.gov/gene/?term=LOC105275518 | 2340 | 25 | 106.84 | 1 | 4.27 | 4 | 43.33 | 39.71 | 3.62 | 3.62 | 0.04 | 0.04 |
| gene940 | https://www.ncbi.nlm.nih.gov/gene/?term=LOC105282794 | 3578 | 88 | 245.95 | 1 | 2.79 | 1.14 | 16.46 | 14.64 | 1.82 | 1.82 | 0.04 | 0.04 |
| gene5967 | https://www.ncbi.nlm.nih.gov/gene/?term=LOC105278918 | 4393 | 146 | 332.35 | 1 | 2.28 | 0.68 | 13.82 | 12.77 | 1.06 | 1.06 | 0.04 | 0.04 |
| gene2856 | https://www.ncbi.nlm.nih.gov/gene/?term=LOC105275384 | 1987 | 92 | 463.01 | 1 | 5.03 | 1.09 | 32.79 | 31.9 | 0.89 | 0.89 | 0.04 | 0.04 |
| gene5525 | https://www.ncbi.nlm.nih.gov/gene/?term=LOC105278396 | 3090 | 189 | 611.65 | 1 | 3.24 | 0.53 | 24.57 | 25.15 | -0.58 | 0.58 | -0.04 | 0.04 |
| gene3793 | https://www.ncbi.nlm.nih.gov/gene/?term=LOC105276429 | 2901 | 110 | 379.18 | 1 | 3.45 | 0.91 | 25.42 | 26.35 | -0.93 | 0.93 | -0.04 | 0.04 |
| gene11340 | https://www.ncbi.nlm.nih.gov/gene/?term=LOC105285131 | 2918 | 105 | 359.84 | 1 | 3.43 | 0.95 | 13.12 | 14.12 | -1 | 1 | -0.04 | 0.04 |
| gene920 | https://www.ncbi.nlm.nih.gov/gene/?term=LOC105282518 | 1197 | 42 | 350.88 | 1 | 8.35 | 2.38 | 37.02 | 38.3 | -1.28 | 1.28 | -0.04 | 0.04 |
| gene7915 | https://www.ncbi.nlm.nih.gov/gene/?term=LOC105281173 | 1597 | 38 | 237.95 | 1 | 6.26 | 2.63 | 59.22 | 60.79 | -1.57 | 1.57 | -0.04 | 0.04 |
| gene6422 | https://www.ncbi.nlm.nih.gov/gene/?term=LOC105279458 | 2419 | 37 | 152.96 | 1 | 4.13 | 2.7 | 49.83 | 52.5 | -2.66 | 2.66 | -0.04 | 0.04 |
| gene1505 | https://www.ncbi.nlm.nih.gov/gene/?term=LOC105287633 | 1693 | 27 | 159.48 | 1 | 5.91 | 3.7 | 42.55 | 45.31 | -2.76 | 2.76 | -0.04 | 0.04 |
| gene6455 | https://www.ncbi.nlm.nih.gov/gene/?term=LOC105279500 | 2612 | 33 | 126.34 | 1 | 3.83 | 3.03 | 43.71 | 47.06 | -3.35 | 3.35 | -0.04 | 0.04 |
| gene393 | https://www.ncbi.nlm.nih.gov/gene/?term=LOC105276566 | 1123 | 2 | 17.81 | 1 | 8.9 | 50 | 47.22 | 70.57 | -23.35 | 23.35 | -0.04 | 0.04 |
| gene9478 | https://www.ncbi.nlm.nih.gov/gene/?term=LOC105282991 | 3726 | 57 | 152.98 | 1 | 2.68 | 1.75 | 35.88 | 33.61 | 2.27 | 2.27 | 0.03 | 0.03 |
| gene1518 | https://www.ncbi.nlm.nih.gov/gene/?term=LOC105287648 | 2195 | 30 | 136.67 | 1 | 4.56 | 3.33 | 43.53 | 41.69 | 1.83 | 1.83 | 0.03 | 0.03 |
| gene1884 | https://www.ncbi.nlm.nih.gov/gene/?term=LOC105288072 | 4169 | 76 | 182.3 | 1 | 2.4 | 1.32 | 24.99 | 23.26 | 1.72 | 1.72 | 0.03 | 0.03 |
| gene5101 | https://www.ncbi.nlm.nih.gov/gene/?term=LOC105277903 | 1813 | 37 | 204.08 | 1 | 5.52 | 2.7 | 54.27 | 52.77 | 1.5 | 1.5 | 0.03 | 0.03 |
| gene12572 | https://www.ncbi.nlm.nih.gov/gene/?term=LOC105286514 | 2374 | 46 | 193.77 | 1 | 4.21 | 2.17 | 36.88 | 35.48 | 1.4 | 1.4 | 0.03 | 0.03 |
| gene35 | https://www.ncbi.nlm.nih.gov/gene/?term=LOC105275197 | 2643 | 50 | 189.18 | 1 | 3.78 | 2 | 38.68 | 37.33 | 1.34 | 1.34 | 0.03 | 0.03 |
| gene7888 | https://www.ncbi.nlm.nih.gov/gene/?term=LOC105281141 | 4357 | 164 | 376.41 | 1 | 2.3 | 0.61 | 29.56 | 28.67 | 0.9 | 0.9 | 0.03 | 0.03 |
| gene11401 | https://www.ncbi.nlm.nih.gov/gene/?term=LOC105285202 | 1841 | 77 | 418.25 | 1 | 5.43 | 1.3 | 24.18 | 23.56 | 0.62 | 0.62 | 0.03 | 0.03 |
| gene6583 | https://www.ncbi.nlm.nih.gov/gene/?term=LOC105279649 | 7444 | 350 | 470.18 | 1 | 1.34 | 0.29 | 11.44 | 10.85 | 0.59 | 0.59 | 0.03 | 0.03 |
| gene1332 | https://www.ncbi.nlm.nih.gov/gene/?term=LOC105286872 | 2460 | 158 | 642.28 | 1 | 4.07 | 0.63 | 20.79 | 20.28 | 0.51 | 0.51 | 0.03 | 0.03 |
| gene4034 | https://www.ncbi.nlm.nih.gov/gene/?term=LOC105276704 | 2065 | 103 | 498.79 | 2 | 9.69 | 1.94 | 12.96 | 13.6 | -0.65 | 0.65 | -0.03 | 0.03 |
| gene11688 | https://www.ncbi.nlm.nih.gov/gene/?term=LOC105285533 | 1637 | 70 | 427.61 | 1 | 6.11 | 1.43 | 22.69 | 23.5 | -0.81 | 0.81 | -0.03 | 0.03 |
| gene6652 | https://www.ncbi.nlm.nih.gov/gene/?term=LOC105279719 | 2831 | 85 | 300.25 | 1 | 3.53 | 1.18 | 22.31 | 23.3 | -0.99 | 0.99 | -0.03 | 0.03 |
| gene1147 | https://www.ncbi.nlm.nih.gov/gene/?term=LOC105284954 | 4022 | 114 | 283.44 | 1 | 2.49 | 0.88 | 13.79 | 14.79 | -1 | 1 | -0.03 | 0.03 |
| gene8015 | https://www.ncbi.nlm.nih.gov/gene/?term=LOC105281312 | 2998 | 65 | 216.81 | 1 | 3.34 | 1.54 | 31.99 | 33.36 | -1.36 | 1.36 | -0.03 | 0.03 |
| gene12406 | https://www.ncbi.nlm.nih.gov/gene/?term=LOC105286316 | 1386 | 32 | 230.88 | 1 | 7.22 | 3.12 | 45.77 | 47.17 | -1.4 | 1.4 | -0.03 | 0.03 |
| gene13235 | https://www.ncbi.nlm.nih.gov/gene/?term=LOC105287278 | 2679 | 53 | 197.84 | 1 | 3.73 | 1.89 | 24.79 | 26.36 | -1.58 | 1.58 | -0.03 | 0.03 |
| gene10602 | https://www.ncbi.nlm.nih.gov/gene/?term=LOC105284268 | 2122 | 18 | 84.83 | 1 | 4.71 | 5.56 | 26.04 | 29.95 | -3.91 | 3.91 | -0.03 | 0.03 |
| gene12430 | https://www.ncbi.nlm.nih.gov/gene/?term=LOC105286348 | 3389 | 19 | 56.06 | 1 | 2.95 | 5.26 | 6.26 | 11.43 | -5.17 | 5.17 | -0.03 | 0.03 |
| gene7525 | https://www.ncbi.nlm.nih.gov/gene/?term=LOC105280738 | 4237 | 41 | 96.77 | 1 | 2.36 | 2.44 | 26.13 | 23.59 | 2.54 | 2.54 | 0.02 | 0.02 |
| gene7368 | https://www.ncbi.nlm.nih.gov/gene/?term=LOC105280556 | 4105 | 54 | 131.55 | 1 | 2.44 | 1.85 | 18.52 | 16.73 | 1.79 | 1.79 | 0.02 | 0.02 |
| gene13033 | https://www.ncbi.nlm.nih.gov/gene/?term=LOC105287063 | 1877 | 53 | 282.37 | 1 | 5.33 | 1.89 | 30.73 | 29.86 | 0.87 | 0.87 | 0.02 | 0.02 |
| gene4720 | https://www.ncbi.nlm.nih.gov/gene/?term=LOC105277479 | 1697 | 40 | 235.71 | 1 | 5.89 | 2.5 | 43.85 | 42.99 | 0.86 | 0.86 | 0.02 | 0.02 |
| gene12516 | https://www.ncbi.nlm.nih.gov/gene/?term=LOC105286452 | 1529 | 37 | 241.99 | 1 | 6.54 | 2.7 | 49.81 | 49.09 | 0.73 | 0.73 | 0.02 | 0.02 |
| gene292 | https://www.ncbi.nlm.nih.gov/gene/?term=LOC105275398 | 2825 | 85 | 300.88 | 1 | 3.54 | 1.18 | 21.01 | 20.28 | 0.73 | 0.73 | 0.02 | 0.02 |
| gene10540 | https://www.ncbi.nlm.nih.gov/gene/?term=LOC105284200 | 5267 | 204 | 387.32 | 1 | 1.9 | 0.49 | 12.98 | 12.34 | 0.64 | 0.64 | 0.02 | 0.02 |
| gene5654 | https://www.ncbi.nlm.nih.gov/gene/?term=LOC105278543 | 4019 | 160 | 398.11 | 1 | 2.49 | 0.62 | 15.95 | 15.36 | 0.6 | 0.6 | 0.02 | 0.02 |

| | | | | | | | | | | | | | |
|---|---|---|---|---|---|---|---|---|---|---|---|---|---|
| gene4376 | https://www.ncbi.nlm.nih.gov/gene/?term=LOC105277083 | 4307 | 129 | 299.51 | 1 | 2.32 | 0.78 | 19.34 | 18.78 | 0.56 | 0.56 | 0.02 | 0.02 |
| gene1855 | https://www.ncbi.nlm.nih.gov/gene/?term=LOC105288036 | 2387 | 86 | 360.28 | 1 | 4.19 | 1.16 | 29.99 | 29.44 | 0.55 | 0.55 | 0.02 | 0.02 |
| gene3107 | https://www.ncbi.nlm.nih.gov/gene/?term=LOC105275650 | 5103 | 178 | 348.81 | 1 | 1.96 | 0.56 | 16.67 | 16.15 | 0.52 | 0.52 | 0.02 | 0.02 |
| gene3206 | https://www.ncbi.nlm.nih.gov/gene/?term=LOC105275752 | 3474 | 129 | 371.33 | 1 | 2.88 | 0.78 | 32.13 | 31.62 | 0.52 | 0.52 | 0.02 | 0.02 |
| gene2710 | https://www.ncbi.nlm.nih.gov/gene/?term=LOC105275218 | 3222 | 98 | 304.16 | 1 | 3.1 | 1.02 | 22.4 | 21.9 | 0.5 | 0.5 | 0.02 | 0.02 |
| gene6966 | https://www.ncbi.nlm.nih.gov/gene/?term=LOC105280104 | 3747 | 133 | 354.95 | 1 | 2.67 | 0.75 | 15.81 | 15.32 | 0.49 | 0.49 | 0.02 | 0.02 |
| gene9445 | https://www.ncbi.nlm.nih.gov/gene/?term=LOC105282951 | 2285 | 115 | 503.28 | 1 | 4.38 | 0.87 | 20.4 | 19.9 | 0.49 | 0.49 | 0.02 | 0.02 |
| gene738 | https://www.ncbi.nlm.nih.gov/gene/?term=LOC105280421 | 4927 | 222 | 450.58 | 1 | 2.03 | 0.45 | 14.83 | 14.35 | 0.48 | 0.48 | 0.02 | 0.02 |
| gene8729 | https://www.ncbi.nlm.nih.gov/gene/?term=LOC105282137 | 5005 | 202 | 403.6 1 | 2 | 0.5 | 16.8 | 16.33 | 0.46 | 0.46 | 0.02 | 0.02 | |
| gene5831 | https://www.ncbi.nlm.nih.gov/gene/?term=LOC105278761 | 2506 | 126 | 502.79 | 1 | 3.99 | 0.79 | 21.41 | 21.05 | 0.36 | 0.36 | 0.02 | 0.02 |
| gene12802 | https://www.ncbi.nlm.nih.gov/gene/?term=LOC105286799 | 3183 | 189 | 593.78 | 1 | 3.14 | 0.53 | 15.7 | 15.39 | 0.32 | 0.32 | 0.02 | 0.02 |
| gene1470 | https://www.ncbi.nlm.nih.gov/gene/?term=LOC105287597 | 2974 | 204 | 685.94 | 1 | 3.36 | 0.49 | 17.06 | 17.34 | -0.28 | 0.28 | -0.02 | 0.02 |
| gene5524 | https://www.ncbi.nlm.nih.gov/gene/?term=LOC105278397 | 5177 | 205 | 395.98 | 1 | 1.93 | 0.49 | 24.95 | 25.35 | -0.39 | 0.39 | -0.02 | 0.02 |
| gene6286 | https://www.ncbi.nlm.nih.gov/gene/?term=LOC105279298 | 2861 | 133 | 464.87 | 2 | 6.99 | 1.5 | 23.95 | 24.34 | -0.39 | 0.39 | -0.02 | 0.02 |
| gene4190 | https://www.ncbi.nlm.nih.gov/gene/?term=LOC105276875 | 3150 | 123 | 390.48 | 1 | 3.17 | 0.81 | 20.99 | 21.41 | -0.42 | 0.42 | -0.02 | 0.02 |
| gene570 | https://www.ncbi.nlm.nih.gov/gene/?term=LOC105278549 | 1673 | 71 | 424.39 | 1 | 5.98 | 1.41 | 43.51 | 43.92 | -0.42 | 0.42 | -0.02 | 0.02 |
| gene12806 | https://www.ncbi.nlm.nih.gov/gene/?term=LOC105286803 | 4196 | 160 | 381.32 | 1 | 2.38 | 0.62 | 9.57 | 10.02 | -0.45 | 0.45 | -0.02 | 0.02 |
| gene8274 | https://www.ncbi.nlm.nih.gov/gene/?term=LOC105281624 | 5425 | 236 | 435.02 | 1 | 1.84 | 0.42 | 9.29 | 9.74 | -0.45 | 0.45 | -0.02 | 0.02 |
| gene10326 | https://www.ncbi.nlm.nih.gov/gene/?term=LOC105283962 | 3042 | 132 | 433.93 | 1 | 3.29 | 0.76 | 8.33 | 8.79 | -0.46 | 0.46 | -0.02 | 0.02 |
| gene4962 | https://www.ncbi.nlm.nih.gov/gene/?term=LOC105277761 | 4645 | 151 | 325.08 | 1 | 2.15 | 0.66 | 14.59 | 15.1 | -0.51 | 0.51 | -0.02 | 0.02 |
| gene10606 | https://www.ncbi.nlm.nih.gov/gene/?term=LOC105284273 | 6119 | 264 | 431.44 | 1 | 1.63 | 0.38 | 18.32 | 18.9 | -0.57 | 0.57 | -0.02 | 0.02 |
| gene9101 | https://www.ncbi.nlm.nih.gov/gene/?term=LOC105282553 | 3068 | 108 | 352.02 | 1 | 3.26 | 0.93 | 22.97 | 23.62 | -0.64 | 0.64 | -0.02 | 0.02 |
| gene7249 | https://www.ncbi.nlm.nih.gov/gene/?term=LOC105280422 | 1775 | 62 | 349.3 1 | 5.63 | 1.61 | 17.31 | 17.96 | -0.66 | 0.66 | -0.02 | 0.02 | |
| gene5646 | https://www.ncbi.nlm.nih.gov/gene/?term=LOC105278534 | 7475 | 195 | 260.87 | 1 | 1.34 | 0.51 | 17.04 | 17.72 | -0.68 | 0.68 | -0.02 | 0.02 |
| gene2404 | https://www.ncbi.nlm.nih.gov/gene/?term=LOC105274853 | 1767 | 38 | 215.05 | 1 | 5.66 | 2.63 | 34.09 | 34.89 | -0.8 | 0.8 | -0.02 | 0.02 |
| gene10107 | https://www.ncbi.nlm.nih.gov/gene/?term=LOC105283712 | 1883 | 36 | 191.18 | 1 | 5.31 | 2.78 | 28.27 | 29.18 | -0.91 | 0.91 | -0.02 | 0.02 |
| gene7247 | https://www.ncbi.nlm.nih.gov/gene/?term=LOC105280407 | 10486 | 212 | 202.17 | 1 | 0.95 | 0.47 | 15.9 | 16.83 | -0.93 | 0.93 | -0.02 | 0.02 |
| gene7199 | https://www.ncbi.nlm.nih.gov/gene/?term=LOC105280366 | 1414 | 32 | 226.31 | 1 | 7.07 | 3.12 | 42.1 | 43.06 | -0.96 | 0.96 | -0.02 | 0.02 |
| gene7248 | https://www.ncbi.nlm.nih.gov/gene/?term=LOC105280408 | 5175 | 57 | 110.14 | 1 | 1.93 | 1.75 | 18.23 | 20.49 | -2.26 | 2.26 | -0.02 | 0.02 |
| gene9411 | https://www.ncbi.nlm.nih.gov/gene/?term=LOC105282914 | 5534 | 41 | 74.09 1 | 1.81 | 2.44 | 49.41 | 52.21 | -2.8 | 2.8 | -0.02 | 0.02 | |
| gene564 | https://www.ncbi.nlm.nih.gov/gene/?term=LOC105278478 | 2802 | 17 | 60.67 1 | 3.57 | 5.88 | 14.04 | 17.66 | -3.63 | 3.63 | -0.02 | 0.02 | |
| gene8178 | https://www.ncbi.nlm.nih.gov/gene/?term=LOC105281515 | 8647 | 61 | 70.54 1 | 1.16 | 1.64 | 27.1 | 25.25 | 1.85 | 1.85 | 0.01 | 0.01 | |
| gene10345 | https://www.ncbi.nlm.nih.gov/gene/?term=LOC105283989 | 4444 | 29 | 65.26 1 | 2.25 | 3.45 | 49.14 | 47.94 | 1.2 | 1.2 | 0.01 | 0.01 | |
| gene927 | https://www.ncbi.nlm.nih.gov/gene/?term=LOC105282600 | 3727 | 50 | 134.16 | 1 | 2.68 | 2 | 33.71 | 32.83 | 0.88 | 0.88 | 0.01 | 0.01 |
| gene8742 | https://www.ncbi.nlm.nih.gov/gene/?term=LOC105282151 | 2563 | 44 | 171.67 | 2 | 7.8 | 4.55 | 36.03 | 35.31 | 0.72 | 0.72 | 0.01 | 0.01 |
| gene9336 | https://www.ncbi.nlm.nih.gov/gene/?term=LOC105282834 | 5993 | 129 | 215.25 | 1 | 1.67 | 0.78 | 15.82 | 15.16 | 0.66 | 0.66 | 0.01 | 0.01 |
| gene12439 | https://www.ncbi.nlm.nih.gov/gene/?term=LOC105286357 | 4761 | 86 | 180.63 | 1 | 2.1 | 1.16 | 26.67 | 26.07 | 0.6 | 0.6 | 0.01 | 0.01 |
| gene34 | https://www.ncbi.nlm.nih.gov/gene/?term=LOC105278416 | 7120 | 146 | 205.06 | 1 | 1.4 | 0.68 | 13.63 | 13.03 | 0.6 | 0.6 | 0.01 | 0.01 |
| gene2683 | https://www.ncbi.nlm.nih.gov/gene/?term=LOC105275187 | 5492 | 75 | 136.56 | 1 | 1.82 | 1.33 | 15.98 | 15.4 | 0.58 | 0.58 | 0.01 | 0.01 |
| gene4023 | https://www.ncbi.nlm.nih.gov/gene/?term=LOC105276692 | 1759 | 37 | 210.35 | 1 | 5.69 | 2.7 | 20.35 | 19.78 | 0.57 | 0.57 | 0.01 | 0.01 |
| gene7884 | https://www.ncbi.nlm.nih.gov/gene/?term=LOC105281266 | 3390 | 79 | 233.04 | 1 | 2.95 | 1.27 | 16.01 | 15.44 | 0.57 | 0.57 | 0.01 | 0.01 |
| gene2668 | https://www.ncbi.nlm.nih.gov/gene/?term=LOC105275170 | 12496 | 301 | 240.88 | 1 | 0.8 | 0.33 | 2.32 | 1.82 | 0.49 | 0.49 | 0.01 | 0.01 |
| gene3470 | https://www.ncbi.nlm.nih.gov/gene/?term=LOC105276059 | 8716 | 164 | 188.16 | 1 | 1.15 | 0.61 | 7.87 | 7.47 | 0.4 | 0.4 | 0.01 | 0.01 |
| gene11454 | https://www.ncbi.nlm.nih.gov/gene/?term=LOC105285256 | 1906 | 51 | 267.58 | 1 | 5.25 | 1.96 | 42.4 | 42.03 | 0.37 | 0.37 | 0.01 | 0.01 |
| gene6620 | https://www.ncbi.nlm.nih.gov/gene/?term=LOC105279687 | 17998 | 384 | 213.36 | 1 | 0.56 | 0.26 | 4.83 | 4.47 | 0.36 | 0.36 | 0.01 | 0.01 |
| gene5375 | https://www.ncbi.nlm.nih.gov/gene/?term=LOC105278228 | 8640 | 173 | 200.23 | 1 | 1.16 | 0.58 | 23.79 | 23.44 | 0.35 | 0.35 | 0.01 | 0.01 |
| gene2844 | https://www.ncbi.nlm.nih.gov/gene/?term=LOC105275363 | 4330 | 206 | 475.75 | 1 | 2.31 | 0.49 | 14.15 | 13.85 | 0.3 | 0.3 | 0.01 | 0.01 |
| gene11568 | https://www.ncbi.nlm.nih.gov/gene/?term=LOC105285395 | 8086 | 204 | 252.29 | 1 | 1.24 | 0.49 | 14.79 | 14.52 | 0.27 | 0.27 | 0.01 | 0.01 |
| gene1310 | https://www.ncbi.nlm.nih.gov/gene/?term=LOC105286661 | 2582 | 65 | 251.74 | 1 | 3.87 | 1.54 | 23.22 | 22.96 | 0.26 | 0.26 | 0.01 | 0.01 |
| gene10531 | https://www.ncbi.nlm.nih.gov/gene/?term=LOC105284190 | 6881 | 195 | 283.39 | 1 | 1.45 | 0.51 | 11.74 | 11.5 | 0.24 | 0.24 | 0.01 | 0.01 |
| gene11579 | https://www.ncbi.nlm.nih.gov/gene/?term=LOC105285406 | 13571 | 291 | 214.43 | 1 | 0.74 | 0.34 | 6.38 | 6.14 | 0.24 | 0.24 | 0.01 | 0.01 |
| gene10844 | https://www.ncbi.nlm.nih.gov/gene/?term=LOC105284535 | 9086 | 256 | 281.75 | 1 | 1.1 | 0.39 | 14.59 | 14.36 | 0.23 | 0.23 | 0.01 | 0.01 |
| gene2806 | https://www.ncbi.nlm.nih.gov/gene/?term=LOC105275327 | 14257 | 542 | 380.16 | 1 | 0.7 | 0.18 | 6.63 | 6.42 | 0.21 | 0.21 | 0.01 | 0.01 |
| gene3885 | https://www.ncbi.nlm.nih.gov/gene/?term=LOC105276527 | 7928 | 334 | 421.29 | 1 | 1.26 | 0.3 | 9.88 | 9.74 | 0.14 | 0.14 | 0.01 | 0.01 |
| gene2633 | https://www.ncbi.nlm.nih.gov/gene/?term=LOC105275127 | 7457 | 270 | 362.08 | 1 | 1.34 | 0.37 | 7.9 | 8.07 | -0.16 | 0.16 | -0.01 | 0.01 |
| gene7514 | https://www.ncbi.nlm.nih.gov/gene/?term=LOC105280723 | 4471 | 210 | 469.69 | 1 | 2.24 | 0.48 | 13.2 | 13.38 | -0.18 | 0.18 | -0.01 | 0.01 |
| gene10549 | https://www.ncbi.nlm.nih.gov/gene/?term=LOC105284210 | 7145 | 340 | 475.86 | 2 | 2.8 | 0.59 | 9.99 | 10.19 | -0.2 | 0.2 | -0.01 | 0.01 |
| gene8814 | https://www.ncbi.nlm.nih.gov/gene/?term=LOC105282240 | 6910 | 198 | 286.54 | 1 | 1.45 | 0.51 | 8.79 | 9 | -0.21 | 0.21 | -0.01 | 0.01 |
| gene2846 | https://www.ncbi.nlm.nih.gov/gene/?term=LOC105275374 | 11246 | 512 | 455.27 | 1 | 0.89 | 0.2 | 9.81 | 10.04 | -0.23 | 0.23 | -0.01 | 0.01 |
| gene10213 | https://www.ncbi.nlm.nih.gov/gene/?term=LOC105283826 | 1503 | 65 | 432.47 | 1 | 6.65 | 1.54 | 15.16 | 15.4 | -0.24 | 0.24 | -0.01 | 0.01 |
| gene13036 | https://www.ncbi.nlm.nih.gov/gene/?term=LOC105287059 | 4959 | 176 | 354.91 | 1 | 2.02 | 0.57 | 16.13 | 16.41 | -0.28 | 0.28 | -0.01 | 0.01 |
| gene10392 | https://www.ncbi.nlm.nih.gov/gene/?term=LOC105284036 | 3216 | 106 | 329.6 2 | 6.22 | 1.89 | 20.71 | 21 | -0.29 | 0.29 | -0.01 | 0.01 | |
| gene10520 | https://www.ncbi.nlm.nih.gov/gene/?term=LOC105284184 | 2451 | 101 | 412.08 | 1 | 4.08 | 0.99 | 29.13 | 29.43 | -0.3 | 0.3 | -0.01 | 0.01 |
| gene11082 | https://www.ncbi.nlm.nih.gov/gene/?term=LOC105284824 | 7436 | 214 | 287.79 | 1 | 1.34 | 0.47 | 6.56 | 6.86 | -0.31 | 0.31 | -0.01 | 0.01 |

| | | | | | | | | | | | | | |
|---|---|---|---|---|---|---|---|---|---|---|---|---|---|
| gene3046 | https://www.ncbi.nlm.nih.gov/gene/?term=LOC105275587 | 2394 | 55 | 229.74 | 1 | 4.18 | 1.82 | 36.59 | 36.9 | -0.31 | 0.31 | -0.01 | 0.01 |
| gene1907 | https://www.ncbi.nlm.nih.gov/gene/?term=LOC105288099 | 1238 | 36 | 290.79 | 1 | 8.08 | 2.78 | 33.72 | 34.04 | -0.32 | 0.32 | -0.01 | 0.01 |
| gene5777 | https://www.ncbi.nlm.nih.gov/gene/?term=LOC105278691 | 3883 | 168 | 432.66 | 1 | 2.58 | 0.6 | 14.66 | 14.98 | -0.33 | 0.33 | -0.01 | 0.01 |
| gene10600 | https://www.ncbi.nlm.nih.gov/gene/?term=LOC105284270 | 4837 | 124 | 256.36 | 1 | 2.07 | 0.81 | 11.8 | 12.15 | -0.35 | 0.35 | -0.01 | 0.01 |
| gene6031 | https://www.ncbi.nlm.nih.gov/gene/?term=LOC105279000 | 5691 | 197 | 346.16 | 1 | 1.76 | 0.51 | 12.37 | 12.73 | -0.35 | 0.35 | -0.01 | 0.01 |
| gene2413 | https://www.ncbi.nlm.nih.gov/gene/?term=LOC105274863 | 2006 | 54 | 269.19 | 1 | 4.99 | 1.85 | 52.24 | 52.76 | -0.51 | 0.51 | -0.01 | 0.01 |
| gene388 | https://www.ncbi.nlm.nih.gov/gene/?term=LOC105276538 | 1558 | 19 | 121.95 | 2 | 12.84 | 10.53 | 66.25 | 66.78 | -0.53 | 0.53 | -0.01 | 0.01 |
| gene2748 | https://www.ncbi.nlm.nih.gov/gene/?term=LOC105275261 | 3845 | 55 | 143.04 | 1 | 2.6 | 1.82 | 38.33 | 38.87 | -0.54 | 0.54 | -0.01 | 0.01 |
| gene11583 | https://www.ncbi.nlm.nih.gov/gene/?term=LOC105285414 | 8192 | 184 | 224.61 | 1 | 1.22 | 0.54 | 8.68 | 9.27 | -0.59 | 0.59 | -0.01 | 0.01 |
| gene6639 | https://www.ncbi.nlm.nih.gov/gene/?term=LOC105279705 | 7656 | 70 | 91.43 1 | 1.31 | 1.43 | 24.79 | 25.46 | -0.67 | 0.67 | -0.01 | 0.01 | |
| gene322 | https://www.ncbi.nlm.nih.gov/gene/?term=LOC105275777 | 1606 | 27 | 168.12 | 1 | 6.23 | 3.7 | 40.46 | 41.2 | -0.74 | 0.74 | -0.01 | 0.01 |
| gene3168 | https://www.ncbi.nlm.nih.gov/gene/?term=LOC105275711 | 6150 | 75 | 121.95 | 1 | 1.63 | 1.33 | 12.84 | 13.77 | -0.93 | 0.93 | -0.01 | 0.01 |
| gene3599 | https://www.ncbi.nlm.nih.gov/gene/?term=LOC105276196 | 2955 | 59 | 199.66 | 1 | 3.38 | 1.69 | 37.86 | 37.72 | 0.15 | 0.15 | 0 | 0 |
| gene53 | https://www.ncbi.nlm.nih.gov/gene/?term=LOC105277148 | 19283 | 324 | 168.02 | 1 | 0.52 | 0.31 | 4.91 | 4.75 | 0.15 | 0.15 | 0 | 0 |
| gene9462 | https://www.ncbi.nlm.nih.gov/gene/?term=LOC105282970 | 1469 | 42 | 285.91 | 1 | 6.81 | 2.38 | 51.11 | 50.96 | 0.15 | 0.15 | 0 | 0 |
| gene5539 | https://www.ncbi.nlm.nih.gov/gene/?term=LOC105278415 | 25024 | 720 | 287.72 | 1 | 0.4 | 0.14 | 3.22 | 3.07 | 0.14 | 0.14 | 0 | 0 |
| gene7183 | https://www.ncbi.nlm.nih.gov/gene/?term=LOC105280348 | 3925 | 76 | 193.63 | 1 | 2.55 | 1.32 | 19.42 | 19.29 | 0.13 | 0.13 | 0 | 0 |
| gene862 | https://www.ncbi.nlm.nih.gov/gene/?term=LOC105281802 | 73922 | 2171 | 293.69 | 1 | 0.14 | 0.05 | 2.2 | 2.07 | 0.13 | 0.13 | 0 | 0 |
| gene12307 | https://www.ncbi.nlm.nih.gov/gene/?term=LOC105286216 | 10313 | 174 | 168.72 | 1 | 0.97 | 0.57 | 15.92 | 15.81 | 0.12 | 0.12 | 0 | 0 |
| gene5184 | https://www.ncbi.nlm.nih.gov/gene/?term=LOC105277999 | 5528 | 146 | 264.11 | 1 | 1.81 | 0.68 | 11.67 | 11.57 | 0.1 | 0.1 | 0 | 0 |
| gene5746 | https://www.ncbi.nlm.nih.gov/gene/?term=LOC105278666 | 10343 | 482 | 466.02 | 1 | 0.97 | 0.21 | 6.71 | 6.61 | 0.1 | 0.1 | 0 | 0 |
| gene12573 | https://www.ncbi.nlm.nih.gov/gene/?term=LOC105286515 | 3647 | 103 | 282.42 | 1 | 2.74 | 0.97 | 10.56 | 10.47 | 0.09 | 0.09 | 0 | 0 |
| gene8343 | https://www.ncbi.nlm.nih.gov/gene/?term=LOC105281700 | 2172 | 37 | 170.35 | 1 | 4.6 | 2.7 | 39.73 | 39.63 | 0.09 | 0.09 | 0 | 0 |
| gene298 | https://www.ncbi.nlm.nih.gov/gene/?term=LOC105275460 | 2473 | 122 | 493.33 | 1 | 4.04 | 0.82 | 13.15 | 13.07 | 0.08 | 0.08 | 0 | 0 |
| gene10155 | https://www.ncbi.nlm.nih.gov/gene/?term=LOC105283756 | 7030 | 109 | 155.05 | 1 | 1.42 | 0.92 | 5.36 | 5.29 | 0.07 | 0.07 | 0 | 0 |
| gene1632 | https://www.ncbi.nlm.nih.gov/gene/?term=LOC105287764 | 18061 | 406 | 224.79 | 2 | 1.11 | 0.49 | 3.27 | 3.2 | 0.07 | 0.07 | 0 | 0 |
| gene7319 | https://www.ncbi.nlm.nih.gov/gene/?term=LOC105280513 | 68997 | 1376 | 199.43 | 1 | 0.14 | 0.07 | 0.74 | 0.67 | 0.07 | 0.07 | 0 | 0 |
| gene9326 | https://www.ncbi.nlm.nih.gov/gene/?term=LOC105282818 | 16612 | 217 | 130.63 | 1 | 0.6 | 0.46 | 11.57 | 11.5 | 0.07 | 0.07 | 0 | 0 |
| gene10596 | https://www.ncbi.nlm.nih.gov/gene/?term=LOC105284262 | 6520 | 234 | 358.9 1 | 1.53 | 0.43 | 3.38 | 3.34 | 0.04 | 0.04 | 0 | 0 | |
| gene2056 | https://www.ncbi.nlm.nih.gov/gene/?term=LOC105274452 | 6584 | 263 | 399.45 | 2 | 3.04 | 0.76 | 12.97 | 12.95 | 0.02 | 0.02 | 0 | 0 |
| gene5197 | https://www.ncbi.nlm.nih.gov/gene/?term=LOC105278018 | 89233 | 3838 | 430.11 | 1 | 0.11 | 0.03 | 0.67 | 0.66 | 0.01 | 0.01 | 0 | 0 |
| gene7398 | https://www.ncbi.nlm.nih.gov/gene/?term=LOC105280589 | 3452 | 37 | 107.18 | 1 | 2.9 | 2.7 | 19.84 | 19.83 | 0.01 | 0.01 | 0 | 0 |
| gene10018 | https://www.ncbi.nlm.nih.gov/gene/?term=LOC105283607 | 4414 | 208 | 471.23 | 1 | 2.27 | 0.48 | 15.08 | 15.07 | 0 | 0 | 0 | 0 |
| gene1430 | https://www.ncbi.nlm.nih.gov/gene/?term=LOC105287551 | 6433 | 232 | 360.64 | 1 | 1.55 | 0.43 | 8.43 | 8.44 | 0 | 0 | 0 | 0 |
| gene11275 | https://www.ncbi.nlm.nih.gov/gene/?term=LOC105285055 | 4119 | 248 | 602.09 | 1 | 2.43 | 0.4 | 12.13 | 12.15 | -0.02 | 0.02 | 0 | 0 |
| gene13114 | https://www.ncbi.nlm.nih.gov/gene/?term=LOC105287147 | 7609 | 430 | 565.12 | 1 | 1.31 | 0.23 | 9.08 | 9.1 | -0.02 | 0.02 | 0 | 0 |
| gene3856 | https://www.ncbi.nlm.nih.gov/gene/?term=LOC105276499 | 5236 | 194 | 370.51 | 1 | 1.91 | 0.52 | 10.61 | 10.65 | -0.04 | 0.04 | 0 | 0 |
| gene9327 | https://www.ncbi.nlm.nih.gov/gene/?term=LOC105282821 | 7793 | 173 | 221.99 | 1 | 1.28 | 0.58 | 10.79 | 10.87 | -0.08 | 0.08 | 0 | 0 |
| gene7166 | https://www.ncbi.nlm.nih.gov/gene/?term=LOC105280326 | 6117 | 253 | 413.6 1 | 1.63 | 0.4 | 14.52 | 14.62 | -0.1 | 0.1 | 0 | 0 | |
| gene10886 | https://www.ncbi.nlm.nih.gov/gene/?term=LOC105284582 | 1475 | 47 | 318.64 | 1 | 6.78 | 2.13 | 22.72 | 22.83 | -0.11 | 0.11 | 0 | 0 |
| gene6528 | https://www.ncbi.nlm.nih.gov/gene/?term=LOC105279591 | 7222 | 59 | 81.69 1 | 1.38 | 1.69 | 26.07 | 26.22 | -0.14 | 0.14 | 0 | 0 | |
| gene2173 | https://www.ncbi.nlm.nih.gov/gene/?term=LOC105274590 | 4387 | 84 | 191.47 | 2 | 4.56 | 2.38 | 27.83 | 27.99 | -0.17 | 0.17 | 0 | 0 |
| gene11561 | https://www.ncbi.nlm.nih.gov/gene/?term=LOC105285387 | 11528 | 202 | 175.23 | 1 | 0.87 | 0.5 | 8.85 | 9.03 | -0.18 | 0.18 | 0 | 0 |
| gene12203 | https://www.ncbi.nlm.nih.gov/gene/?term=LOC105286099 | 5967 | 75 | 125.69 | 1 | 1.68 | 1.33 | 9.95 | 10.3 | -0.35 | 0.35 | 0 | 0 |

Directory: OGLall File: rnk-dmg-Cb_LI_BCphaseB1.vs.RphaseA1.txt

| gene_ID | gene_link | gwidth | #Sites | #per10Kb | #dmSites | #dmsp10kb | %dmSites | %pSite1 | %pSite2 | DMpSite | ADMpSite | DMpNucl | ADMpNucl |
|---|---|---|---|---|---|---|---|---|---|---|---|---|---|
| gene11566 | https://www.ncbi.nlm.nih.gov/gene/?term=LOC105285392 | 1603 | 91 | 567.69 | 6 | 37.43 | 6.59 | 35.74 | 31.54 | 4.2 | 4.2 | 0.24 | 0.24 |
| gene5306 | https://www.ncbi.nlm.nih.gov/gene/?term=LOC105278153 | 481 | 17 | 353.43 | 2 | 41.58 | 11.76 | 21.3 | 16.46 | 4.84 | 4.84 | 0.17 | 0.17 |
| gene2855 | https://www.ncbi.nlm.nih.gov/gene/?term=LOC105275378 | 1189 | 21 | 176.62 | 3 | 25.23 | 14.29 | 38.9 | 30.15 | 8.75 | 8.75 | 0.15 | 0.15 |
| gene2111 | https://www.ncbi.nlm.nih.gov/gene/?term=LOC105274528 | 2028 | 144 | 710.06 | 3 | 14.79 | 2.08 | 19.76 | 17.65 | 2.1 | 2.1 | 0.15 | 0.15 |
| gene11402 | https://www.ncbi.nlm.nih.gov/gene/?term=LOC105285201 | 335 | 16 | 477.61 | 1 | 29.85 | 6.25 | 21.09 | 18.26 | 2.83 | 2.83 | 0.14 | 0.14 |
| gene10112 | https://www.ncbi.nlm.nih.gov/gene/?term=LOC105283717 | 1483 | 81 | 546.19 | 4 | 26.97 | 4.94 | 29.9 | 27.32 | 2.58 | 2.58 | 0.14 | 0.14 |
| gene560 | https://www.ncbi.nlm.nih.gov/gene/?term=LOC109610897 | 1584 | 125 | 789.14 | 2 | 12.63 | 1.6 | 14.21 | 16.02 | -1.81 | 1.81 | -0.14 | 0.14 |
| gene12670 | https://www.ncbi.nlm.nih.gov/gene/?term=LOC105286645 | 1506 | 55 | 365.21 | 1 | 6.64 | 1.82 | 37.73 | 41.37 | -3.64 | 3.64 | -0.13 | 0.13 |
| gene10513 | https://www.ncbi.nlm.nih.gov/gene/?term=LOC105284174 | 1606 | 77 | 479.45 | 1 | 6.23 | 1.3 | 21.46 | 23.96 | -2.5 | 2.5 | -0.12 | 0.12 |
| gene12364 | https://www.ncbi.nlm.nih.gov/gene/?term=LOC105286269 | 1403 | 51 | 363.51 | 4 | 28.51 | 7.84 | 50.68 | 53.95 | -3.27 | 3.27 | -0.12 | 0.12 |
| gene9468 | https://www.ncbi.nlm.nih.gov/gene/?term=LOC105282976 | 1636 | 63 | 385.09 | 2 | 12.22 | 3.17 | 23.75 | 20.8 | 2.95 | 2.95 | 0.11 | 0.11 |
| gene5447 | https://www.ncbi.nlm.nih.gov/gene/?term=LOC105278314 | 1482 | 93 | 627.53 | 1 | 6.75 | 1.08 | 23.89 | 22.18 | 1.72 | 1.72 | 0.11 | 0.11 |
| gene4380 | https://www.ncbi.nlm.nih.gov/gene/?term=LOC105277087 | 334 | 10 | 299.4 | 1 | 29.94 | 10 | 6.29 | 9.99 | -3.7 | 3.7 | -0.11 | 0.11 |
| gene2036 | https://www.ncbi.nlm.nih.gov/gene/?term=LOC105288242 | 1260 | 36 | 285.71 | 1 | 7.94 | 2.78 | 42.3 | 46.17 | -3.87 | 3.87 | -0.11 | 0.11 |
| gene10176 | https://www.ncbi.nlm.nih.gov/gene/?term=LOC105283785 | 1637 | 35 | 213.81 | 1 | 6.11 | 2.86 | 18.49 | 13.7 | 4.79 | 4.79 | 0.1 | 0.1 |
| gene11109 | https://www.ncbi.nlm.nih.gov/gene/?term=LOC105284855 | 1786 | 56 | 313.55 | 2 | 11.2 | 3.57 | 33.74 | 30.6 | 3.13 | 3.13 | 0.1 | 0.1 |
| gene13266 | https://www.ncbi.nlm.nih.gov/gene/?term=LOC105287313 | 1615 | 87 | 538.7 | 3 | 18.58 | 3.45 | 28.84 | 27.02 | 1.82 | 1.82 | 0.1 | 0.1 |
| gene3910 | https://www.ncbi.nlm.nih.gov/gene/?term=LOC105276563 | 1957 | 116 | 592.74 | 5 | 25.55 | 4.31 | 17.71 | 16.03 | 1.69 | 1.69 | 0.1 | 0.1 |
| gene276 | https://www.ncbi.nlm.nih.gov/gene/?term=LOC105275214 | 1299 | 44 | 338.72 | 1 | 7.7 | 2.27 | 33.69 | 30.91 | 2.78 | 2.78 | 0.09 | 0.09 |
| gene7165 | https://www.ncbi.nlm.nih.gov/gene/?term=LOC105280327 | 2688 | 111 | 412.95 | 6 | 22.32 | 5.41 | 28.5 | 26.42 | 2.08 | 2.08 | 0.09 | 0.09 |
| gene1564 | https://www.ncbi.nlm.nih.gov/gene/?term=LOC105287690 | 1386 | 109 | 786.44 | 3 | 21.65 | 2.75 | 18.55 | 17.35 | 1.21 | 1.21 | 0.09 | 0.09 |
| gene5937 | https://www.ncbi.nlm.nih.gov/gene/?term=LOC105278873 | 1902 | 149 | 783.39 | 6 | 31.55 | 4.03 | 20.65 | 19.44 | 1.21 | 1.21 | 0.09 | 0.09 |
| gene10743 | https://www.ncbi.nlm.nih.gov/gene/?term=LOC105284456 | 1028 | 42 | 408.56 | 3 | 29.18 | 7.14 | 38.04 | 40.31 | -2.27 | 2.27 | -0.09 | 0.09 |
| gene1769 | https://www.ncbi.nlm.nih.gov/gene/?term=LOC105287942 | 2154 | 78 | 362.12 | 5 | 23.21 | 6.41 | 31.01 | 33.47 | -2.46 | 2.46 | -0.09 | 0.09 |

...

... only first 25 lines shown ...

Directory: OGLall File: rnk-dmg-Cb_LI_BCphaseB1.vs.RphaseA2.txt

| gene_ID | gene_link | gwidth | #Sites | #per10Kb | #dmSites | #dmsp10kb | %dmSites | %pSite1 | %pSite2 | DMpSite | ADMpSite | DMpNucl | ADMpNucl |
|---|---|---|---|---|---|---|---|---|---|---|---|---|---|
| gene2837 | https://www.ncbi.nlm.nih.gov/gene/?term=LOC105275355 | 2872 | 160 | 557.1 | 10 | 34.82 | 6.25 | 29.18 | 25.36 | 3.82 | 3.82 | 0.21 | 0.21 |
| gene1894 | https://www.ncbi.nlm.nih.gov/gene/?term=LOC105288083 | 445 | 17 | 382.02 | 2 | 44.94 | 11.76 | 56.3 | 51.18 | 5.11 | 5.11 | 0.2 | 0.2 |
| gene12925 | https://www.ncbi.nlm.nih.gov/gene/?term=LOC109611487 | 391 | 27 | 690.54 | 1 | 25.58 | 3.7 | 14.67 | 17.34 | -2.67 | 2.67 | -0.18 | 0.18 |
| gene12311 | https://www.ncbi.nlm.nih.gov/gene/?term=LOC105286212 | 1005 | 39 | 388.06 | 3 | 29.85 | 7.69 | 33.27 | 37.99 | -4.72 | 4.72 | -0.18 | 0.18 |
| gene3268 | https://www.ncbi.nlm.nih.gov/gene/?term=LOC105275833 | 1040 | 43 | 413.46 | 1 | 9.62 | 2.33 | 35.22 | 39.24 | -4.03 | 4.03 | -0.17 | 0.17 |
| gene6075 | https://www.ncbi.nlm.nih.gov/gene/?term=LOC105279049 | 1285 | 99 | 770.43 | 1 | 7.78 | 1.01 | 16.76 | 18.86 | -2.11 | 2.11 | -0.16 | 0.16 |
| gene3815 | https://www.ncbi.nlm.nih.gov/gene/?term=LOC105276451 | 1324 | 39 | 294.56 | 3 | 22.66 | 7.69 | 36.8 | 42.26 | -5.46 | 5.46 | -0.16 | 0.16 |
| gene9740 | https://www.ncbi.nlm.nih.gov/gene/?term=LOC105283277 | 2887 | 145 | 502.25 | 3 | 10.39 | 2.07 | 19.78 | 22.8 | -3.02 | 3.02 | -0.15 | 0.15 |
| gene8798 | https://www.ncbi.nlm.nih.gov/gene/?term=LOC105282220 | 2300 | 235 | 1021.74 | 3 | 13.04 | 1.28 | 7.15 | 8.56 | -1.4 | 1.4 | -0.14 | 0.14 |
| gene2790 | https://www.ncbi.nlm.nih.gov/gene/?term=LOC105275307 | 1429 | 55 | 384.88 | 2 | 14 | 3.64 | 37.41 | 41.04 | -3.64 | 3.64 | -0.14 | 0.14 |
| gene12403 | https://www.ncbi.nlm.nih.gov/gene/?term=LOC105286311 | 1413 | 52 | 368.01 | 3 | 21.23 | 5.77 | 55.61 | 59.38 | -3.77 | 3.77 | -0.14 | 0.14 |
| gene5917 | https://www.ncbi.nlm.nih.gov/gene/?term=LOC105278849 | 823 | 82 | 996.35 | 1 | 12.15 | 1.22 | 25.33 | 24.02 | 1.31 | 1.31 | 0.13 | 0.13 |
| gene7917 | https://www.ncbi.nlm.nih.gov/gene/?term=LOC105281172 | 1304 | 30 | 230.06 | 3 | 23.01 | 10 | 42.68 | 48.2 | -5.52 | 5.52 | -0.13 | 0.13 |
| gene11779 | https://www.ncbi.nlm.nih.gov/gene/?term=LOC105285628 | 1037 | 22 | 212.15 | 3 | 28.93 | 13.64 | 32.81 | 38.84 | -6.03 | 6.03 | -0.13 | 0.13 |
| gene2111 | https://www.ncbi.nlm.nih.gov/gene/?term=LOC105274528 | 2028 | 151 | 744.58 | 2 | 9.86 | 1.32 | 19 | 17.45 | 1.55 | 1.55 | 0.12 | 0.12 |
| gene560 | https://www.ncbi.nlm.nih.gov/gene/?term=LOC109610897 | 1584 | 148 | 934.34 | 2 | 12.63 | 1.35 | 13.32 | 14.65 | -1.33 | 1.33 | -0.12 | 0.12 |
| gene6057 | https://www.ncbi.nlm.nih.gov/gene/?term=LOC105279035 | 2105 | 166 | 788.6 | 4 | 19 | 2.41 | 18.44 | 19.91 | -1.47 | 1.47 | -0.12 | 0.12 |
| gene5525 | https://www.ncbi.nlm.nih.gov/gene/?term=LOC105278396 | 3090 | 191 | 618.12 | 4 | 12.94 | 2.09 | 24.34 | 26.26 | -1.92 | 1.92 | -0.12 | 0.12 |
| gene5423 | https://www.ncbi.nlm.nih.gov/gene/?term=LOC105278282 | 1776 | 67 | 377.25 | 3 | 16.89 | 4.48 | 31.84 | 34.9 | -3.06 | 3.06 | -0.12 | 0.12 |
| gene8333 | https://www.ncbi.nlm.nih.gov/gene/?term=LOC105281691 | 1264 | 34 | 268.99 | 4 | 31.65 | 11.76 | 48.64 | 53.25 | -4.6 | 4.6 | -0.12 | 0.12 |
| gene3269 | https://www.ncbi.nlm.nih.gov/gene/?term=LOC105275832 | 728 | 55 | 755.49 | 2 | 27.47 | 3.64 | 8.05 | 9.52 | -1.47 | 1.47 | -0.11 | 0.11 |
| gene9183 | https://www.ncbi.nlm.nih.gov/gene/?term=LOC105282640 | 2752 | 209 | 759.45 | 2 | 7.27 | 0.96 | 21 | 22.49 | -1.49 | 1.49 | -0.11 | 0.11 |
| gene8453 | https://www.ncbi.nlm.nih.gov/gene/?term=LOC105281826 | 1750 | 99 | 565.71 | 1 | 5.71 | 1.01 | 28.31 | 30.17 | -1.86 | 1.86 | -0.11 | 0.11 |
| gene10513 | https://www.ncbi.nlm.nih.gov/gene/?term=LOC105284174 | 1606 | 89 | 554.17 | 1 | 6.23 | 1.12 | 19.86 | 21.82 | -1.96 | 1.96 | -0.11 | 0.11 |

...

... only first 25 lines shown ...

Directory: OGLall File: rnk-dmg-Cb_LI_BCphaseB1.vs.RphaseB1.txt

| gene_ID | gene_link | gwidth | #Sites | #per10Kb | #dmSites | #dmsp10kb | %dmSites | %pSite1 | %pSite2 | DMpSite | ADMpSite | DMpNucl | ADMpNucl |
|---|---|---|---|---|---|---|---|---|---|---|---|---|---|
| gene2837 | https://www.ncbi.nlm.nih.gov/gene/?term=LOC105275355 | 2872 | 165 | 574.51 | 1 | 3.48 | 0.61 | 28.35 | 22.92 | 5.43 | 5.43 | 0.31 | 0.31 |
| gene10963 | https://www.ncbi.nlm.nih.gov/gene/?term=LOC105284677 | 1576 | 30 | 190.36 | 1 | 6.35 | 3.33 | 19.35 | 34.92 | -15.57 | 15.57 | -0.3 | 0.3 |
| gene3165 | https://www.ncbi.nlm.nih.gov/gene/?term=LOC105275706 | 2374 | 65 | 273.8 | 4 | 16.85 | 6.15 | 33.82 | 25.03 | 8.79 | 8.79 | 0.24 | 0.24 |
| gene2593 | https://www.ncbi.nlm.nih.gov/gene/?term=LOC105275070 | 1774 | 11 | 62.01 | 1 | 5.64 | 9.09 | 61.58 | 25.28 | 36.29 | 36.29 | 0.23 | 0.23 |
| gene2856 | https://www.ncbi.nlm.nih.gov/gene/?term=LOC105275384 | 1987 | 92 | 463.01 | 1 | 5.03 | 1.09 | 34.19 | 31.19 | 3 | 3 | 0.14 | 0.14 |
| gene2413 | https://www.ncbi.nlm.nih.gov/gene/?term=LOC105274863 | 2006 | 62 | 309.07 | 1 | 4.99 | 1.61 | 48.7 | 53.36 | -4.66 | 4.66 | -0.14 | 0.14 |
| gene9740 | https://www.ncbi.nlm.nih.gov/gene/?term=LOC105283277 | 2887 | 139 | 481.47 | 1 | 3.46 | 0.72 | 18.49 | 20.55 | -2.05 | 2.05 | -0.1 | 0.1 |
| gene1855 | https://www.ncbi.nlm.nih.gov/gene/?term=LOC105288036 | 2387 | 91 | 381.23 | 1 | 4.19 | 1.1 | 29.43 | 27.13 | 2.3 | 2.3 | 0.09 | 0.09 |
| gene5714 | https://www.ncbi.nlm.nih.gov/gene/?term=LOC105278729 | 710 | 16 | 225.35 | 1 | 14.08 | 6.25 | 55.52 | 51.9 | 3.62 | 3.62 | 0.08 | 0.08 |
| gene6237 | https://www.ncbi.nlm.nih.gov/gene/?term=LOC105279244 | 1484 | 33 | 222.37 | 1 | 6.74 | 3.03 | 44.64 | 48.04 | -3.4 | 3.4 | -0.08 | 0.08 |
| gene5831 | https://www.ncbi.nlm.nih.gov/gene/?term=LOC105278761 | 2506 | 118 | 470.87 | 2 | 7.98 | 1.69 | 22.67 | 21.1 | 1.58 | 1.58 | 0.07 | 0.07 |
| gene3671 | https://www.ncbi.nlm.nih.gov/gene/?term=LOC105276272 | 1966 | 40 | 203.46 | 1 | 5.09 | 2.5 | 30.25 | 27.31 | 2.94 | 2.94 | 0.06 | 0.06 |
| gene396 | https://www.ncbi.nlm.nih.gov/gene/?term=LOC105276611 | 1722 | 36 | 209.06 | 1 | 5.81 | 2.78 | 30.31 | 27.55 | 2.76 | 2.76 | 0.06 | 0.06 |
| gene13033 | https://www.ncbi.nlm.nih.gov/gene/?term=LOC105287063 | 1877 | 50 | 266.38 | 1 | 5.33 | 2 | 31.92 | 29.55 | 2.37 | 2.37 | 0.06 | 0.06 |
| gene7493 | https://www.ncbi.nlm.nih.gov/gene/?term=LOC105280704 | 4225 | 147 | 347.93 | 1 | 2.37 | 0.68 | 18.36 | 16.63 | 1.72 | 1.72 | 0.06 | 0.06 |
| gene5298 | https://www.ncbi.nlm.nih.gov/gene/?term=LOC105278145 | 4060 | 262 | 645.32 | 2 | 4.93 | 0.76 | 9.65 | 10.52 | -0.86 | 0.86 | -0.06 | 0.06 |
| gene1470 | https://www.ncbi.nlm.nih.gov/gene/?term=LOC105287597 | 2974 | 192 | 645.6 | 1 | 3.36 | 0.52 | 18.41 | 19.24 | -0.83 | 0.83 | -0.05 | 0.05 |
| gene1198 | https://www.ncbi.nlm.nih.gov/gene/?term=LOC105285507 | 2738 | 97 | 354.27 | 1 | 3.65 | 1.03 | 20.27 | 21.73 | -1.46 | 1.46 | -0.05 | 0.05 |
| gene11340 | https://www.ncbi.nlm.nih.gov/gene/?term=LOC105285131 | 2918 | 98 | 335.85 | 1 | 3.43 | 1.02 | 13.78 | 15.33 | -1.55 | 1.55 | -0.05 | 0.05 |
| gene9464 | https://www.ncbi.nlm.nih.gov/gene/?term=LOC105282975 | 1450 | 24 | 165.52 | 1 | 6.9 | 4.17 | 39.26 | 42.21 | -2.95 | 2.95 | -0.05 | 0.05 |
| gene1505 | https://www.ncbi.nlm.nih.gov/gene/?term=LOC105287633 | 1693 | 22 | 129.95 | 1 | 5.91 | 4.55 | 51.59 | 55.25 | -3.65 | 3.65 | -0.05 | 0.05 |
| gene9109 | https://www.ncbi.nlm.nih.gov/gene/?term=LOC105282561 | 2400 | 14 | 58.33 | 1 | 4.17 | 7.14 | 13.59 | 21.35 | -7.76 | 7.76 | -0.05 | 0.05 |
| gene393 | https://www.ncbi.nlm.nih.gov/gene/?term=LOC105276566 | 1123 | 2 | 17.81 | 1 | 8.9 | 50 | 47.22 | 74.3 | -27.08 | 27.08 | -0.05 | 0.05 |
| gene2970 | https://www.ncbi.nlm.nih.gov/gene/?term=LOC105275518 | 2340 | 25 | 106.84 | 1 | 4.27 | 4 | 44.93 | 41.23 | 3.69 | 3.69 | 0.04 | 0.04 |
| gene2241 | https://www.ncbi.nlm.nih.gov/gene/?term=LOC105274666 | 2598 | 61 | 234.8 | 1 | 3.85 | 1.64 | 30.46 | 28.61 | 1.85 | 1.85 | 0.04 | 0.04 |
| gene940 | https://www.ncbi.nlm.nih.gov/gene/?term=LOC105282794 | 3578 | 94 | 262.72 | 1 | 2.79 | 1.06 | 15.53 | 14.17 | 1.36 | 1.36 | 0.04 | 0.04 |
| gene4641 | https://www.ncbi.nlm.nih.gov/gene/?term=LOC105277398 | 1937 | 72 | 371.71 | 1 | 5.16 | 1.39 | 36.3 | 35.35 | 0.95 | 0.95 | 0.04 | 0.04 |
| gene3817 | https://www.ncbi.nlm.nih.gov/gene/?term=LOC105276449 | 1230 | 55 | 447.15 | 1 | 8.13 | 1.82 | 22.28 | 21.47 | 0.8 | 0.8 | 0.04 | 0.04 |
| gene474 | https://www.ncbi.nlm.nih.gov/gene/?term=LOC105277435 | 3625 | 176 | 485.52 | 1 | 2.76 | 0.57 | 10.76 | 9.98 | 0.78 | 0.78 | 0.04 | 0.04 |
| gene570 | https://www.ncbi.nlm.nih.gov/gene/?term=LOC105278549 | 1673 | 72 | 430.36 | 1 | 5.98 | 1.39 | 42.9 | 43.9 | -0.99 | 0.99 | -0.04 | 0.04 |
| gene1220 | https://www.ncbi.nlm.nih.gov/gene/?term=LOC105285742 | 3453 | 119 | 344.63 | 1 | 2.9 | 0.84 | 16.38 | 17.43 | -1.05 | 1.05 | -0.04 | 0.04 |
| gene9598 | https://www.ncbi.nlm.nih.gov/gene/?term=LOC105283132 | 1769 | 56 | 316.56 | 1 | 5.65 | 1.79 | 39.06 | 40.18 | -1.12 | 1.12 | -0.04 | 0.04 |
| gene4962 | https://www.ncbi.nlm.nih.gov/gene/?term=LOC105277761 | 4645 | 156 | 335.84 | 1 | 2.15 | 0.64 | 14.16 | 15.31 | -1.15 | 1.15 | -0.04 | 0.04 |
| gene3727 | https://www.ncbi.nlm.nih.gov/gene/?term=LOC105276367 | 2484 | 65 | 261.67 | 1 | 4.03 | 1.54 | 43.95 | 45.54 | -1.59 | 1.59 | -0.04 | 0.04 |
| gene2404 | https://www.ncbi.nlm.nih.gov/gene/?term=LOC105274853 | 1767 | 36 | 203.74 | 1 | 5.66 | 2.78 | 33.17 | 35.01 | -1.84 | 1.84 | -0.04 | 0.04 |
| gene1199 | https://www.ncbi.nlm.nih.gov/gene/?term=LOC105285498 | 2220 | 30 | 135.14 | 1 | 4.5 | 3.33 | 33.52 | 36.79 | -3.27 | 3.27 | -0.04 | 0.04 |
| gene3166 | https://www.ncbi.nlm.nih.gov/gene/?term=LOC105275707 | 2393 | 10 | 41.79 | 1 | 4.18 | 10 | 65.97 | 76.22 | -10.25 | 10.25 | -0.04 | 0.04 |
| gene4023 | https://www.ncbi.nlm.nih.gov/gene/?term=LOC105276692 | 1759 | 37 | 210.35 | 1 | 5.69 | 2.7 | 23.22 | 21.93 | 1.3 | 1.3 | 0.03 | 0.03 |
| gene9336 | https://www.ncbi.nlm.nih.gov/gene/?term=LOC105282834 | 5993 | 134 | 223.59 | 1 | 1.67 | 0.75 | 14.88 | 13.64 | 1.24 | 1.24 | 0.03 | 0.03 |
| gene5877 | https://www.ncbi.nlm.nih.gov/gene/?term=LOC105278807 | 1719 | 51 | 296.68 | 1 | 5.82 | 1.96 | 26.14 | 25.26 | 0.89 | 0.89 | 0.03 | 0.03 |
| gene298 | https://www.ncbi.nlm.nih.gov/gene/?term=LOC105275460 | 2473 | 126 | 509.5 | 1 | 4.04 | 0.79 | 13.09 | 12.43 | 0.67 | 0.67 | 0.03 | 0.03 |
| gene10678 | https://www.ncbi.nlm.nih.gov/gene/?term=LOC105284355 | 4031 | 169 | 419.25 | 1 | 2.48 | 0.59 | 20.31 | 19.66 | 0.65 | 0.65 | 0.03 | 0.03 |
| gene4034 | https://www.ncbi.nlm.nih.gov/gene/?term=LOC105276704 | 2065 | 87 | 421.31 | 2 | 9.69 | 2.3 | 14.01 | 13.38 | 0.63 | 0.63 | 0.03 | 0.03 |
| gene5525 | https://www.ncbi.nlm.nih.gov/gene/?term=LOC105278396 | 3090 | 189 | 611.65 | 1 | 3.24 | 0.53 | 24.08 | 24.52 | -0.44 | 0.44 | -0.03 | 0.03 |
| gene6698 | https://www.ncbi.nlm.nih.gov/gene/?term=LOC105279778 | 4099 | 161 | 392.78 | 1 | 2.44 | 0.62 | 8.69 | 9.57 | -0.88 | 0.88 | -0.03 | 0.03 |
| gene920 | https://www.ncbi.nlm.nih.gov/gene/?term=LOC105282518 | 1197 | 44 | 367.59 | 1 | 8.35 | 2.27 | 33.07 | 33.97 | -0.89 | 0.89 | -0.03 | 0.03 |
| gene10392 | https://www.ncbi.nlm.nih.gov/gene/?term=LOC105284036 | 3216 | 103 | 320.27 | 2 | 6.22 | 1.94 | 23.68 | 24.65 | -0.97 | 0.97 | -0.03 | 0.03 |
| gene12021 | https://www.ncbi.nlm.nih.gov/gene/?term=LOC105285905 | 1399 | 29 | 207.29 | 1 | 7.15 | 3.45 | 32.77 | 34.25 | -1.48 | 1.48 | -0.03 | 0.03 |
| gene6422 | https://www.ncbi.nlm.nih.gov/gene/?term=LOC105279458 | 2419 | 42 | 173.63 | 1 | 4.13 | 2.38 | 44.07 | 45.58 | -1.51 | 1.51 | -0.03 | 0.03 |
| gene564 | https://www.ncbi.nlm.nih.gov/gene/?term=LOC105278478 | 2802 | 16 | 57.1 | 1 | 3.57 | 6.25 | 14.91 | 19.95 | -5.03 | 5.03 | -0.03 | 0.03 |
| gene11584 | https://www.ncbi.nlm.nih.gov/gene/?term=LOC105285416 | 3709 | 16 | 43.14 | 2 | 5.39 | 12.5 | 21.86 | 29.47 | -7.61 | 7.61 | -0.03 | 0.03 |
| gene10345 | https://www.ncbi.nlm.nih.gov/gene/?term=LOC105283989 | 4444 | 41 | 92.26 | 1 | 2.25 | 2.44 | 38.24 | 36.42 | 1.83 | 1.83 | 0.02 | 0.02 |
| gene2748 | https://www.ncbi.nlm.nih.gov/gene/?term=LOC105275261 | 3845 | 53 | 137.84 | 1 | 2.6 | 1.89 | 39.61 | 38.42 | 1.19 | 1.19 | 0.02 | 0.02 |
| gene10107 | https://www.ncbi.nlm.nih.gov/gene/?term=LOC105283712 | 1883 | 36 | 191.18 | 1 | 5.31 | 2.78 | 30.43 | 29.26 | 1.17 | 1.17 | 0.02 | 0.02 |
| gene1884 | https://www.ncbi.nlm.nih.gov/gene/?term=LOC105288072 | 4169 | 84 | 201.49 | 1 | 2.4 | 1.19 | 24.86 | 23.72 | 1.14 | 1.14 | 0.02 | 0.02 |
| gene7734 | https://www.ncbi.nlm.nih.gov/gene/?term=LOC105280972 | 1528 | 35 | 229.06 | 1 | 6.54 | 2.86 | 17.63 | 16.59 | 1.04 | 1.04 | 0.02 | 0.02 |
| gene7884 | https://www.ncbi.nlm.nih.gov/gene/?term=LOC105281266 | 3390 | 87 | 256.64 | 1 | 2.95 | 1.15 | 15.85 | 14.89 | 0.96 | 0.96 | 0.02 | 0.02 |
| gene1147 | https://www.ncbi.nlm.nih.gov/gene/?term=LOC105284954 | 4022 | 118 | 293.39 | 1 | 2.49 | 0.85 | 15.13 | 14.31 | 0.81 | 0.81 | 0.02 | 0.02 |
| gene292 | https://www.ncbi.nlm.nih.gov/gene/?term=LOC105275398 | 2825 | 88 | 311.5 | 1 | 3.54 | 1.14 | 22.66 | 21.93 | 0.73 | 0.73 | 0.02 | 0.02 |
| gene7888 | https://www.ncbi.nlm.nih.gov/gene/?term=LOC105281141 | 4357 | 170 | 390.18 | 1 | 2.3 | 0.59 | 28.6 | 28 | 0.6 | 0.6 | 0.02 | 0.02 |

| | | | | | | | | | | | | | |
|---|---|---|---|---|---|---|---|---|---|---|---|---|---|
| gene6652 | https://www.ncbi.nlm.nih.gov/gene/?term=LOC105279719 | 2831 | 88 | 310.84 | 1 | 3.53 | 1.14 | 21.65 | 21.08 | 0.57 | 0.57 | 0.02 | 0.02 |
| gene4892 | https://www.ncbi.nlm.nih.gov/gene/?term=LOC105277667 | 3898 | 127 | 325.81 | 1 | 2.57 | 0.79 | 27.72 | 27.21 | 0.52 | 0.52 | 0.02 | 0.02 |
| gene12978 | https://www.ncbi.nlm.nih.gov/gene/?term=LOC105286997 | 2858 | 150 | 524.84 | 1 | 3.5 | 0.67 | 11.28 | 10.84 | 0.44 | 0.44 | 0.02 | 0.02 |
| gene11819 | https://www.ncbi.nlm.nih.gov/gene/?term=LOC105285671 | 2366 | 104 | 439.56 | 1 | 4.23 | 0.96 | 30.06 | 29.63 | 0.43 | 0.43 | 0.02 | 0.02 |
| gene3885 | https://www.ncbi.nlm.nih.gov/gene/?term=LOC105276527 | 7928 | 369 | 465.44 | 1 | 1.26 | 0.27 | 8.26 | 7.93 | 0.33 | 0.33 | 0.02 | 0.02 |
| gene10549 | https://www.ncbi.nlm.nih.gov/gene/?term=LOC105284210 | 7145 | 365 | 510.85 | 2 | 2.8 | 0.55 | 9.55 | 9.9 | -0.35 | 0.35 | -0.02 | 0.02 |
| gene10326 | https://www.ncbi.nlm.nih.gov/gene/?term=LOC105283962 | 3042 | 137 | 450.36 | 1 | 3.29 | 0.73 | 7.31 | 7.68 | -0.37 | 0.37 | -0.02 | 0.02 |
| gene3793 | https://www.ncbi.nlm.nih.gov/gene/?term=LOC105276429 | 2901 | 131 | 451.57 | 1 | 3.45 | 0.76 | 22.23 | 22.66 | -0.43 | 0.43 | -0.02 | 0.02 |
| gene10606 | https://www.ncbi.nlm.nih.gov/gene/?term=LOC105284273 | 6119 | 272 | 444.52 | 1 | 1.63 | 0.37 | 17.87 | 18.32 | -0.45 | 0.45 | -0.02 | 0.02 |
| gene6583 | https://www.ncbi.nlm.nih.gov/gene/?term=LOC105279649 | 7444 | 372 | 499.73 | 1 | 1.34 | 0.27 | 10.85 | 11.31 | -0.45 | 0.45 | -0.02 | 0.02 |
| gene4190 | https://www.ncbi.nlm.nih.gov/gene/?term=LOC105276875 | 3150 | 134 | 425.4 1 | 3.17 | 0.75 | 20.21 | 20.67 | -0.46 | 0.46 | -0.02 | 0.02 | |
| gene7166 | https://www.ncbi.nlm.nih.gov/gene/?term=LOC105280326 | 6117 | 259 | 423.41 | 1 | 1.63 | 0.39 | 14.46 | 14.93 | -0.47 | 0.47 | -0.02 | 0.02 |
| gene6966 | https://www.ncbi.nlm.nih.gov/gene/?term=LOC105280104 | 3747 | 132 | 352.28 | 1 | 2.67 | 0.76 | 16.91 | 17.46 | -0.55 | 0.55 | -0.02 | 0.02 |
| gene8814 | https://www.ncbi.nlm.nih.gov/gene/?term=LOC105282240 | 6910 | 204 | 295.22 | 1 | 1.45 | 0.49 | 8.01 | 8.64 | -0.63 | 0.63 | -0.02 | 0.02 |
| gene5184 | https://www.ncbi.nlm.nih.gov/gene/?term=LOC105277999 | 5528 | 165 | 298.48 | 1 | 1.81 | 0.61 | 11.31 | 11.98 | -0.67 | 0.67 | -0.02 | 0.02 |
| gene11235 | https://www.ncbi.nlm.nih.gov/gene/?term=LOC105285004 | 1339 | 36 | 268.86 | 1 | 7.47 | 2.78 | 19.18 | 19.88 | -0.7 | 0.7 | -0.02 | 0.02 |
| gene1310 | https://www.ncbi.nlm.nih.gov/gene/?term=LOC105286661 | 2582 | 68 | 263.36 | 1 | 3.87 | 1.47 | 18.99 | 19.74 | -0.75 | 0.75 | -0.02 | 0.02 |
| gene12573 | https://www.ncbi.nlm.nih.gov/gene/?term=LOC105286515 | 3647 | 107 | 293.39 | 1 | 2.74 | 0.93 | 10.59 | 11.4 | -0.82 | 0.82 | -0.02 | 0.02 |
| gene5101 | https://www.ncbi.nlm.nih.gov/gene/?term=LOC105277903 | 1813 | 37 | 204.08 | 1 | 5.52 | 2.7 | 54.12 | 55.06 | -0.94 | 0.94 | -0.02 | 0.02 |
| gene10155 | https://www.ncbi.nlm.nih.gov/gene/?term=LOC105283756 | 7030 | 116 | 165.01 | 1 | 1.42 | 0.86 | 7.15 | 8.11 | -0.96 | 0.96 | -0.02 | 0.02 |
| gene3168 | https://www.ncbi.nlm.nih.gov/gene/?term=LOC105275711 | 6150 | 97 | 157.72 | 1 | 1.63 | 1.03 | 10.51 | 11.68 | -1.17 | 1.17 | -0.02 | 0.02 |
| gene388 | https://www.ncbi.nlm.nih.gov/gene/?term=LOC105276538 | 1558 | 21 | 134.79 | 1 | 6.42 | 4.76 | 69.07 | 70.25 | -1.18 | 1.18 | -0.02 | 0.02 |
| gene3599 | https://www.ncbi.nlm.nih.gov/gene/?term=LOC105276196 | 2955 | 60 | 203.05 | 1 | 3.38 | 1.67 | 37.66 | 38.86 | -1.2 | 1.2 | -0.02 | 0.02 |
| gene12203 | https://www.ncbi.nlm.nih.gov/gene/?term=LOC105286099 | 5967 | 87 | 145.8 1 | 1.68 | 1.15 | 10.81 | 12.06 | -1.25 | 1.25 | -0.02 | 0.02 | |
| gene7398 | https://www.ncbi.nlm.nih.gov/gene/?term=LOC105280589 | 3452 | 42 | 121.67 | 1 | 2.9 | 2.38 | 24.95 | 26.26 | -1.31 | 1.31 | -0.02 | 0.02 |
| gene7248 | https://www.ncbi.nlm.nih.gov/gene/?term=LOC105280408 | 5175 | 59 | 114.01 | 1 | 1.93 | 1.69 | 15.82 | 17.71 | -1.89 | 1.89 | -0.02 | 0.02 |
| gene8178 | https://www.ncbi.nlm.nih.gov/gene/?term=LOC105281515 | 8647 | 60 | 69.39 1 | 1.16 | 1.67 | 30.54 | 29.47 | 1.06 | 1.06 | 0.01 | 0.01 | |
| gene7183 | https://www.ncbi.nlm.nih.gov/gene/?term=LOC105280348 | 3925 | 80 | 203.82 | 1 | 2.55 | 1.25 | 17.35 | 16.62 | 0.73 | 0.73 | 0.01 | 0.01 |
| gene322 | https://www.ncbi.nlm.nih.gov/gene/?term=LOC105275777 | 1606 | 28 | 174.35 | 1 | 6.23 | 3.57 | 36.76 | 36.04 | 0.72 | 0.72 | 0.01 | 0.01 |
| gene2683 | https://www.ncbi.nlm.nih.gov/gene/?term=LOC105275187 | 5492 | 78 | 142.02 | 1 | 1.82 | 1.28 | 16.44 | 16.02 | 0.43 | 0.43 | 0.01 | 0.01 |
| gene3107 | https://www.ncbi.nlm.nih.gov/gene/?term=LOC105275650 | 5103 | 182 | 356.65 | 1 | 1.96 | 0.55 | 16.89 | 16.49 | 0.4 | 0.4 | 0.01 | 0.01 |
| gene6455 | https://www.ncbi.nlm.nih.gov/gene/?term=LOC105279500 | 2612 | 35 | 134 1 | 3.83 | 2.86 | 48.92 | 48.53 | 0.39 | 0.39 | 0.01 | 0.01 | |
| gene13036 | https://www.ncbi.nlm.nih.gov/gene/?term=LOC105287059 | 4959 | 193 | 389.19 | 1 | 2.02 | 0.52 | 16.75 | 16.43 | 0.32 | 0.32 | 0.01 | 0.01 |
| gene8729 | https://www.ncbi.nlm.nih.gov/gene/?term=LOC105282137 | 5005 | 226 | 451.55 | 1 | 2 | 0.44 | 14.98 | 14.68 | 0.29 | 0.29 | 0.01 | 0.01 |
| gene12572 | https://www.ncbi.nlm.nih.gov/gene/?term=LOC105286514 | 2374 | 52 | 219.04 | 1 | 4.21 | 1.92 | 35.13 | 34.85 | 0.28 | 0.28 | 0.01 | 0.01 |
| gene9101 | https://www.ncbi.nlm.nih.gov/gene/?term=LOC105282553 | 3068 | 106 | 345.5 1 | 3.26 | 0.94 | 22.9 | 22.63 | 0.27 | 0.27 | 0.01 | 0.01 | |
| gene10018 | https://www.ncbi.nlm.nih.gov/gene/?term=LOC105283607 | 4414 | 228 | 516.54 | 1 | 2.27 | 0.44 | 14.1 | 13.84 | 0.26 | 0.26 | 0.01 | 0.01 |
| gene2668 | https://www.ncbi.nlm.nih.gov/gene/?term=LOC105275170 | 12496 | 356 | 284.89 | 1 | 0.8 | 0.28 | 2.04 | 1.8 | 0.24 | 0.24 | 0.01 | 0.01 |
| gene10540 | https://www.ncbi.nlm.nih.gov/gene/?term=LOC105284200 | 5267 | 214 | 406.3 1 | 1.9 | 0.47 | 13.24 | 13.01 | 0.23 | 0.23 | 0.01 | 0.01 | |
| gene10596 | https://www.ncbi.nlm.nih.gov/gene/?term=LOC105284262 | 6520 | 239 | 366.56 | 1 | 1.53 | 0.42 | 4.23 | 4.05 | 0.19 | 0.19 | 0.01 | 0.01 |
| gene6286 | https://www.ncbi.nlm.nih.gov/gene/?term=LOC105279298 | 2861 | 128 | 447.4 1 | 3.5 | 0.78 | 23.43 | 23.27 | 0.16 | 0.16 | 0.01 | 0.01 | |
| gene11275 | https://www.ncbi.nlm.nih.gov/gene/?term=LOC105285055 | 4119 | 246 | 597.23 | 1 | 2.43 | 0.41 | 12.22 | 12.12 | 0.1 | 0.1 | 0.01 | 0.01 |
| gene12802 | https://www.ncbi.nlm.nih.gov/gene/?term=LOC105286799 | 3183 | 180 | 565.5 1 | 3.14 | 0.56 | 16.81 | 16.93 | -0.12 | 0.12 | -0.01 | 0.01 | |
| gene5654 | https://www.ncbi.nlm.nih.gov/gene/?term=LOC105278543 | 4019 | 181 | 450.36 | 1 | 2.49 | 0.55 | 15.7 | 15.83 | -0.13 | 0.13 | -0.01 | 0.01 |
| gene8274 | https://www.ncbi.nlm.nih.gov/gene/?term=LOC105281624 | 5425 | 227 | 418.43 | 1 | 1.84 | 0.44 | 9.53 | 9.7 | -0.17 | 0.17 | -0.01 | 0.01 |
| gene13039 | https://www.ncbi.nlm.nih.gov/gene/?term=LOC105287065 | 4902 | 253 | 516.12 | 1 | 2.04 | 0.4 | 12.12 | 12.3 | -0.18 | 0.18 | -0.01 | 0.01 |
| gene11579 | https://www.ncbi.nlm.nih.gov/gene/?term=LOC105285406 | 13571 | 316 | 232.85 | 1 | 0.74 | 0.32 | 6.01 | 6.26 | -0.25 | 0.25 | -0.01 | 0.01 |
| gene7249 | https://www.ncbi.nlm.nih.gov/gene/?term=LOC105280422 | 1775 | 66 | 371.83 | 1 | 5.63 | 1.52 | 15.78 | 16.04 | -0.26 | 0.26 | -0.01 | 0.01 |
| gene7199 | https://www.ncbi.nlm.nih.gov/gene/?term=LOC105280366 | 1414 | 34 | 240.45 | 1 | 7.07 | 2.94 | 44.65 | 44.98 | -0.33 | 0.33 | -0.01 | 0.01 |
| gene5524 | https://www.ncbi.nlm.nih.gov/gene/?term=LOC105278397 | 5177 | 209 | 403.71 | 1 | 1.93 | 0.48 | 23.99 | 24.35 | -0.36 | 0.36 | -0.01 | 0.01 |
| gene4376 | https://www.ncbi.nlm.nih.gov/gene/?term=LOC105277083 | 4307 | 129 | 299.51 | 1 | 2.32 | 0.78 | 18.69 | 19.06 | -0.38 | 0.38 | -0.01 | 0.01 |
| gene11583 | https://www.ncbi.nlm.nih.gov/gene/?term=LOC105285414 | 8192 | 192 | 234.38 | 1 | 1.22 | 0.52 | 8.5 | 8.89 | -0.39 | 0.39 | -0.01 | 0.01 |
| gene7915 | https://www.ncbi.nlm.nih.gov/gene/?term=LOC105281173 | 1597 | 42 | 262.99 | 1 | 6.26 | 2.38 | 62.33 | 62.75 | -0.42 | 0.42 | -0.01 | 0.01 |
| gene9330 | https://www.ncbi.nlm.nih.gov/gene/?term=LOC105282825 | 3936 | 98 | 248.98 | 1 | 2.54 | 1.02 | 13.47 | 13.91 | -0.44 | 0.44 | -0.01 | 0.01 |
| gene7247 | https://www.ncbi.nlm.nih.gov/gene/?term=LOC105280407 | 10486 | 227 | 216.48 | 1 | 0.95 | 0.44 | 14.79 | 15.25 | -0.46 | 0.46 | -0.01 | 0.01 |
| gene2173 | https://www.ncbi.nlm.nih.gov/gene/?term=LOC105274590 | 4387 | 87 | 198.31 | 2 | 4.56 | 2.3 | 23.56 | 24.1 | -0.54 | 0.54 | -0.01 | 0.01 |
| gene4033 | https://www.ncbi.nlm.nih.gov/gene/?term=LOC105276705 | 10546 | 155 | 146.98 | 1 | 0.95 | 0.65 | 5.56 | 6.1 | -0.55 | 0.55 | -0.01 | 0.01 |
| gene11561 | https://www.ncbi.nlm.nih.gov/gene/?term=LOC105285387 | 11528 | 218 | 189.1 1 | 0.87 | 0.46 | 8.62 | 9.18 | -0.56 | 0.56 | -0.01 | 0.01 | |
| gene9411 | https://www.ncbi.nlm.nih.gov/gene/?term=LOC105282914 | 5534 | 42 | 75.89 1 | 1.81 | 2.38 | 49.22 | 50.92 | -1.7 | 1.7 | -0.01 | 0.01 | |
| gene652 | https://www.ncbi.nlm.nih.gov/gene/?term=LOC105279528 | 5806 | 39 | 67.17 1 | 1.72 | 2.56 | 17.73 | 19.93 | -2.2 | 2.2 | -0.01 | 0.01 | |
| gene7368 | https://www.ncbi.nlm.nih.gov/gene/?term=LOC105280556 | 4105 | 71 | 172.96 | 1 | 2.44 | 1.41 | 15.89 | 15.73 | 0.16 | 0.16 | 0 | 0 |
| gene12406 | https://www.ncbi.nlm.nih.gov/gene/?term=LOC105286316 | 1386 | 44 | 317.46 | 1 | 7.22 | 2.27 | 32.85 | 32.71 | 0.14 | 0.14 | 0 | 0 |
| gene10600 | https://www.ncbi.nlm.nih.gov/gene/?term=LOC105284270 | 4837 | 111 | 229.48 | 1 | 2.07 | 0.9 | 11.96 | 11.83 | 0.13 | 0.13 | 0 | 0 |
| gene927 | https://www.ncbi.nlm.nih.gov/gene/?term=LOC105282600 | 3727 | 60 | 160.99 | 1 | 2.68 | 1.67 | 28.88 | 28.75 | 0.13 | 0.13 | 0 | 0 |

| | | | | | | | | | | | | | | |
|---|---|---|---|---|---|---|---|---|---|---|---|---|---|---|
| gene9327 | https://www.ncbi.nlm.nih.gov/gene/?term=LOC105282821 | 7793 | 183 | 234.83 | | 1 | 1.28 | 0.55 | 12.55 | 12.42 | 0.13 | 0.13 | 0 | 0 |
| gene5375 | https://www.ncbi.nlm.nih.gov/gene/?term=LOC105278228 | 8640 | 191 | 221.06 | | 1 | 1.16 | 0.52 | 22.09 | 21.99 | 0.1 | 0.1 | 0 | 0 |
| gene10222 | https://www.ncbi.nlm.nih.gov/gene/?term=LOC105283838 | 1748 | 30 | 171.62 | | 1 | 5.72 | 3.33 | 35.37 | 35.28 | 0.09 | 0.09 | 0 | 0 |
| gene10520 | https://www.ncbi.nlm.nih.gov/gene/?term=LOC105284184 | 2451 | 102 | 416.16 | | 1 | 4.08 | 0.98 | 30 | 29.93 | 0.07 | 0.07 | 0 | 0 |
| gene9326 | https://www.ncbi.nlm.nih.gov/gene/?term=LOC105282818 | 16612 | 236 | 142.07 | | 1 | 0.6 | 0.42 | 13.17 | 13.1 | 0.07 | 0.07 | 0 | 0 |
| gene2443 | https://www.ncbi.nlm.nih.gov/gene/?term=LOC105274895 | 10433 | 424 | 406.4 | 1 | 0.96 | 0.24 | 5.17 | 5.13 | 0.03 | 0.03 | 0 | 0 | |
| gene53 | https://www.ncbi.nlm.nih.gov/gene/?term=LOC105277148 | 19283 | 333 | 172.69 | | 1 | 0.52 | 0.3 | 4.68 | 4.68 | 0 | 0 | 0 | 0 |
| gene11454 | https://www.ncbi.nlm.nih.gov/gene/?term=LOC105285256 | 1906 | 50 | 262.33 | | 1 | 5.25 | 2 | 44.07 | 44.08 | -0.01 | 0.01 | 0 | 0 |
| gene13114 | https://www.ncbi.nlm.nih.gov/gene/?term=LOC105287147 | 7609 | 432 | 567.75 | | 1 | 1.31 | 0.23 | 9.01 | 9.02 | -0.01 | 0.01 | 0 | 0 |
| gene2271 | https://www.ncbi.nlm.nih.gov/gene/?term=LOC105274706 | 1448 | 47 | 324.59 | | 1 | 6.91 | 2.13 | 41.96 | 41.97 | -0.01 | 0.01 | 0 | 0 |
| gene862 | https://www.ncbi.nlm.nih.gov/gene/?term=LOC105281802 | 73922 | 2318 | 313.57 | | 1 | 0.14 | 0.04 | 2.22 | 2.23 | -0.01 | 0.01 | 0 | 0 |
| gene8600 | https://www.ncbi.nlm.nih.gov/gene/?term=LOC105282001 | 8513 | 206 | 241.98 | | 1 | 1.17 | 0.49 | 9.71 | 9.75 | -0.04 | 0.04 | 0 | 0 |
| gene5746 | https://www.ncbi.nlm.nih.gov/gene/?term=LOC105278666 | 10343 | 483 | 466.98 | | 1 | 0.97 | 0.21 | 6.37 | 6.41 | -0.05 | 0.05 | 0 | 0 |
| gene6620 | https://www.ncbi.nlm.nih.gov/gene/?term=LOC105279687 | 17998 | 448 | 248.92 | | 1 | 0.56 | 0.22 | 4.43 | 4.47 | -0.05 | 0.05 | 0 | 0 |
| gene5539 | https://www.ncbi.nlm.nih.gov/gene/?term=LOC105278415 | 25024 | 790 | 315.7 | 1 | 0.4 | 0.13 | 3.02 | 3.08 | -0.06 | 0.06 | 0 | 0 | |
| gene1632 | https://www.ncbi.nlm.nih.gov/gene/?term=LOC105287764 | 18061 | 431 | 238.64 | | 1 | 0.55 | 0.23 | 3.3 | 3.37 | -0.07 | 0.07 | 0 | 0 |
| gene2806 | https://www.ncbi.nlm.nih.gov/gene/?term=LOC105275327 | 14257 | 552 | 387.18 | | 1 | 0.7 | 0.18 | 6.34 | 6.41 | -0.07 | 0.07 | 0 | 0 |
| gene5197 | https://www.ncbi.nlm.nih.gov/gene/?term=LOC105278018 | 89233 | 3791 | 424.84 | | 1 | 0.11 | 0.03 | 0.65 | 0.72 | -0.07 | 0.07 | 0 | 0 |
| gene3470 | https://www.ncbi.nlm.nih.gov/gene/?term=LOC105276059 | 8716 | 182 | 208.81 | | 1 | 1.15 | 0.55 | 9.23 | 9.32 | -0.09 | 0.09 | 0 | 0 |
| gene5646 | https://www.ncbi.nlm.nih.gov/gene/?term=LOC105278534 | 7475 | 202 | 270.23 | | 1 | 1.34 | 0.5 | 16.69 | 16.79 | -0.1 | 0.1 | 0 | 0 |
| gene12427 | https://www.ncbi.nlm.nih.gov/gene/?term=LOC105286339 | 5220 | 187 | 358.24 | | 1 | 1.92 | 0.53 | 15.91 | 16.02 | -0.12 | 0.12 | 0 | 0 |
| gene10844 | https://www.ncbi.nlm.nih.gov/gene/?term=LOC105284535 | 9086 | 259 | 285.05 | | 1 | 1.1 | 0.39 | 14.4 | 14.56 | -0.16 | 0.16 | 0 | 0 |
| gene6639 | https://www.ncbi.nlm.nih.gov/gene/?term=LOC105279705 | 7656 | 83 | 108.41 | | 1 | 1.31 | 1.2 | 19.9 | 20.17 | -0.28 | 0.28 | 0 | 0 |

Directory: OGLall File: rnk-dmg-Cb_LI_BCphaseB1.vs.RphaseB2.txt

| gene_ID | gene_link | gwidth | #Sites | #per10Kb | #dmSites | #dmsp10kb | %dmSites | %pSite1 | %pSite2 | DMpSite | ADMpSite | DMpNucl | ADMpNucl |
|---|---|---|---|---|---|---|---|---|---|---|---|---|---|
| gene3165 | https://www.ncbi.nlm.nih.gov/gene/?term=LOC105275706 | 2374 | 68 | 286.44 | 5 | 21.06 | 7.35 | 33.32 | 23.13 | 10.19 | 10.19 | 0.29 | 0.29 |
| gene2593 | https://www.ncbi.nlm.nih.gov/gene/?term=LOC105275070 | 1774 | 7 | 39.46 | 5 | 28.18 | 71.43 | 58.65 | 4.09 | 54.56 | 54.56 | 0.22 | 0.22 |
| gene5714 | https://www.ncbi.nlm.nih.gov/gene/?term=LOC105278729 | 710 | 17 | 239.44 | 1 | 14.08 | 5.88 | 58.14 | 53.54 | 4.59 | 4.59 | 0.11 | 0.11 |
| gene4498 | https://www.ncbi.nlm.nih.gov/gene/?term=LOC105277239 | 2831 | 121 | 427.41 | 1 | 3.53 | 0.83 | 11.26 | 13.23 | -1.97 | 1.97 | -0.08 | 0.08 |
| gene940 | https://www.ncbi.nlm.nih.gov/gene/?term=LOC105282794 | 3578 | 83 | 231.97 | 1 | 2.79 | 1.2 | 17.02 | 13.98 | 3.04 | 3.04 | 0.07 | 0.07 |
| gene7568 | https://www.ncbi.nlm.nih.gov/gene/?term=LOC105280786 | 2525 | 122 | 483.17 | 1 | 3.96 | 0.82 | 29.81 | 28.44 | 1.38 | 1.38 | 0.07 | 0.07 |
| gene1332 | https://www.ncbi.nlm.nih.gov/gene/?term=LOC105286872 | 2460 | 154 | 626.02 | 1 | 4.07 | 0.65 | 21.26 | 20.18 | 1.08 | 1.08 | 0.07 | 0.07 |
| gene4641 | https://www.ncbi.nlm.nih.gov/gene/?term=LOC105277398 | 1937 | 71 | 366.55 | 1 | 5.16 | 1.41 | 37.78 | 39.69 | -1.91 | 1.91 | -0.07 | 0.07 |
| gene6334 | https://www.ncbi.nlm.nih.gov/gene/?term=LOC105279356 | 1471 | 48 | 326.31 | 1 | 6.8 | 2.08 | 28.86 | 26.95 | 1.91 | 1.91 | 0.06 | 0.06 |
| gene11819 | https://www.ncbi.nlm.nih.gov/gene/?term=LOC105285671 | 2366 | 102 | 431.11 | 1 | 4.23 | 0.98 | 27.3 | 25.91 | 1.39 | 1.39 | 0.06 | 0.06 |
| gene6422 | https://www.ncbi.nlm.nih.gov/gene/?term=LOC105279458 | 2419 | 38 | 157.09 | 1 | 4.13 | 2.63 | 46.62 | 50.37 | -3.75 | 3.75 | -0.06 | 0.06 |
| gene10602 | https://www.ncbi.nlm.nih.gov/gene/?term=LOC105284268 | 2122 | 19 | 89.54 | 1 | 4.71 | 5.26 | 26.09 | 32.99 | -6.9 | 6.9 | -0.06 | 0.06 |
| gene2970 | https://www.ncbi.nlm.nih.gov/gene/?term=LOC105275518 | 2340 | 20 | 85.47 | 1 | 4.27 | 5 | 58.11 | 52.81 | 5.3 | 5.3 | 0.05 | 0.05 |
| gene1855 | https://www.ncbi.nlm.nih.gov/gene/?term=LOC105288036 | 2387 | 78 | 326.77 | 1 | 4.19 | 1.28 | 33.45 | 31.79 | 1.66 | 1.66 | 0.05 | 0.05 |
| gene6966 | https://www.ncbi.nlm.nih.gov/gene/?term=LOC105280104 | 3747 | 135 | 360.29 | 1 | 2.67 | 0.74 | 17.67 | 16.38 | 1.29 | 1.29 | 0.05 | 0.05 |
| gene5298 | https://www.ncbi.nlm.nih.gov/gene/?term=LOC105278145 | 4060 | 235 | 578.82 | 2 | 4.93 | 0.85 | 10.4 | 11.25 | -0.85 | 0.85 | -0.05 | 0.05 |
| gene3793 | https://www.ncbi.nlm.nih.gov/gene/?term=LOC105276429 | 2901 | 112 | 386.07 | 1 | 3.45 | 0.89 | 25.16 | 26.39 | -1.23 | 1.23 | -0.05 | 0.05 |
| gene3727 | https://www.ncbi.nlm.nih.gov/gene/?term=LOC105276367 | 2484 | 63 | 253.62 | 1 | 4.03 | 1.59 | 45.38 | 47.24 | -1.87 | 1.87 | -0.05 | 0.05 |
| gene3599 | https://www.ncbi.nlm.nih.gov/gene/?term=LOC105276196 | 2955 | 59 | 199.66 | 1 | 3.38 | 1.69 | 37.76 | 40.32 | -2.56 | 2.56 | -0.05 | 0.05 |
| gene6237 | https://www.ncbi.nlm.nih.gov/gene/?term=LOC105279244 | 1484 | 25 | 168.46 | 1 | 6.74 | 4 | 41.13 | 44.31 | -3.18 | 3.18 | -0.05 | 0.05 |
| gene1008 | https://www.ncbi.nlm.nih.gov/gene/?term=LOC105283492 | 1479 | 10 | 67.61 | 1 | 6.76 | 10 | 34.7 | 42 | -7.3 | 7.3 | -0.05 | 0.05 |
| gene393 | https://www.ncbi.nlm.nih.gov/gene/?term=LOC105276566 | 1123 | 2 | 17.81 | 1 | 8.9 | 50 | 47.22 | 76.78 | -29.56 | 29.56 | -0.05 | 0.05 |
| gene1518 | https://www.ncbi.nlm.nih.gov/gene/?term=LOC105287648 | 2195 | 25 | 113.9 | 1 | 4.56 | 4 | 44.43 | 41.18 | 3.25 | 3.25 | 0.04 | 0.04 |
| gene2844 | https://www.ncbi.nlm.nih.gov/gene/?term=LOC105275363 | 4330 | 190 | 438.8 | 1 | 2.31 | 0.53 | 14.79 | 13.86 | 0.93 | 0.93 | 0.04 | 0.04 |
| gene10678 | https://www.ncbi.nlm.nih.gov/gene/?term=LOC105284355 | 4031 | 166 | 411.81 | 1 | 2.48 | 0.6 | 19.71 | 18.79 | 0.92 | 0.92 | 0.04 | 0.04 |
| gene4034 | https://www.ncbi.nlm.nih.gov/gene/?term=LOC105276704 | 2065 | 87 | 421.31 | 1 | 4.84 | 1.15 | 14.97 | 14.08 | 0.89 | 0.89 | 0.04 | 0.04 |
| gene5831 | https://www.ncbi.nlm.nih.gov/gene/?term=LOC105278761 | 2506 | 110 | 438.95 | 1 | 3.99 | 0.91 | 23.47 | 22.6 | 0.88 | 0.88 | 0.04 | 0.04 |
| gene11401 | https://www.ncbi.nlm.nih.gov/gene/?term=LOC105285202 | 1841 | 79 | 429.11 | 1 | 5.43 | 1.27 | 24.33 | 23.48 | 0.85 | 0.85 | 0.04 | 0.04 |
| gene9740 | https://www.ncbi.nlm.nih.gov/gene/?term=LOC105283277 | 2887 | 137 | 474.54 | 1 | 3.46 | 0.73 | 19.95 | 20.71 | -0.76 | 0.76 | -0.04 | 0.04 |
| gene10213 | https://www.ncbi.nlm.nih.gov/gene/?term=LOC105283826 | 1503 | 54 | 359.28 | 1 | 6.65 | 1.85 | 14.52 | 15.73 | -1.21 | 1.21 | -0.04 | 0.04 |
| gene11340 | https://www.ncbi.nlm.nih.gov/gene/?term=LOC105285131 | 2918 | 88 | 301.58 | 1 | 3.43 | 1.14 | 10.13 | 11.45 | -1.32 | 1.32 | -0.04 | 0.04 |
| gene1147 | https://www.ncbi.nlm.nih.gov/gene/?term=LOC105284954 | 4022 | 90 | 223.77 | 1 | 2.49 | 1.11 | 16.68 | 18.39 | -1.71 | 1.71 | -0.04 | 0.04 |
| gene2413 | https://www.ncbi.nlm.nih.gov/gene/?term=LOC105274863 | 2006 | 49 | 244.27 | 1 | 4.99 | 2.04 | 53.75 | 55.47 | -1.72 | 1.72 | -0.04 | 0.04 |
| gene10107 | https://www.ncbi.nlm.nih.gov/gene/?term=LOC105283712 | 1883 | 35 | 185.87 | 1 | 5.31 | 2.86 | 26.3 | 28.27 | -1.97 | 1.97 | -0.04 | 0.04 |
| gene1505 | https://www.ncbi.nlm.nih.gov/gene/?term=LOC105287633 | 1693 | 27 | 159.48 | 1 | 5.91 | 3.7 | 42.55 | 45.35 | -2.8 | 2.8 | -0.04 | 0.04 |
| gene5647 | https://www.ncbi.nlm.nih.gov/gene/?term=LOC105278567 | 3118 | 36 | 115.46 | 1 | 3.21 | 2.78 | 16.65 | 14.27 | 2.38 | 2.38 | 0.03 | 0.03 |
| gene5273 | https://www.ncbi.nlm.nih.gov/gene/?term=LOC105278108 | 4067 | 46 | 113.11 | 1 | 2.46 | 2.17 | 27.18 | 24.92 | 2.26 | 2.26 | 0.03 | 0.03 |
| gene7734 | https://www.ncbi.nlm.nih.gov/gene/?term=LOC105280972 | 1528 | 30 | 196.34 | 1 | 6.54 | 3.33 | 20.49 | 18.79 | 1.71 | 1.71 | 0.03 | 0.03 |
| gene8661 | https://www.ncbi.nlm.nih.gov/gene/?term=LOC105282063 | 1926 | 34 | 176.53 | 1 | 5.19 | 2.94 | 30.6 | 28.98 | 1.62 | 1.62 | 0.03 | 0.03 |
| gene4213 | https://www.ncbi.nlm.nih.gov/gene/?term=LOC105276901 | 3257 | 81 | 248.7 | 1 | 3.07 | 1.23 | 28.04 | 26.96 | 1.09 | 1.09 | 0.03 | 0.03 |
| gene8274 | https://www.ncbi.nlm.nih.gov/gene/?term=LOC105281624 | 5425 | 211 | 388.94 | 1 | 1.84 | 0.47 | 9.95 | 9.18 | 0.77 | 0.77 | 0.03 | 0.03 |
| gene12265 | https://www.ncbi.nlm.nih.gov/gene/?term=LOC105286166 | 3050 | 110 | 360.66 | 1 | 3.28 | 0.91 | 22.97 | 23.8 | -0.82 | 0.82 | -0.03 | 0.03 |
| gene10606 | https://www.ncbi.nlm.nih.gov/gene/?term=LOC105284273 | 6119 | 248 | 405.29 | 1 | 1.63 | 0.4 | 17.15 | 17.98 | -0.83 | 0.83 | -0.03 | 0.03 |
| gene1220 | https://www.ncbi.nlm.nih.gov/gene/?term=LOC105285742 | 3453 | 111 | 321.46 | 1 | 2.9 | 0.9 | 21.24 | 22.19 | -0.96 | 0.96 | -0.03 | 0.03 |
| gene6031 | https://www.ncbi.nlm.nih.gov/gene/?term=LOC105279000 | 5691 | 179 | 314.53 | 1 | 1.76 | 0.56 | 13.44 | 14.41 | -0.97 | 0.97 | -0.03 | 0.03 |
| gene7211 | https://www.ncbi.nlm.nih.gov/gene/?term=LOC105280377 | 2024 | 67 | 331.03 | 1 | 4.94 | 1.49 | 44.45 | 45.48 | -1.03 | 1.03 | -0.03 | 0.03 |
| gene12981 | https://www.ncbi.nlm.nih.gov/gene/?term=LOC105286999 | 2063 | 46 | 222.98 | 1 | 4.85 | 2.17 | 31.8 | 32.93 | -1.13 | 1.13 | -0.03 | 0.03 |
| gene12406 | https://www.ncbi.nlm.nih.gov/gene/?term=LOC105286316 | 1386 | 33 | 238.1 | 1 | 7.22 | 3.03 | 40.83 | 41.98 | -1.15 | 1.15 | -0.03 | 0.03 |
| gene388 | https://www.ncbi.nlm.nih.gov/gene/?term=LOC105276538 | 1558 | 18 | 115.53 | 2 | 12.84 | 11.11 | 66.09 | 68.41 | -2.32 | 2.32 | -0.03 | 0.03 |
| gene3168 | https://www.ncbi.nlm.nih.gov/gene/?term=LOC105275711 | 6150 | 69 | 112.2 | 1 | 1.63 | 1.45 | 12.54 | 15.09 | -2.54 | 2.54 | -0.03 | 0.03 |
| gene7199 | https://www.ncbi.nlm.nih.gov/gene/?term=LOC105280366 | 1414 | 22 | 155.59 | 1 | 7.07 | 4.55 | 43.24 | 41.76 | 1.47 | 1.47 | 0.02 | 0.02 |
| gene2748 | https://www.ncbi.nlm.nih.gov/gene/?term=LOC105275261 | 3845 | 54 | 140.44 | 1 | 2.6 | 1.85 | 40.26 | 39.17 | 1.09 | 1.09 | 0.02 | 0.02 |
| gene10600 | https://www.ncbi.nlm.nih.gov/gene/?term=LOC105284270 | 4837 | 119 | 246.02 | 1 | 2.07 | 0.84 | 10.98 | 10.08 | 0.9 | 0.9 | 0.02 | 0.02 |
| gene9336 | https://www.ncbi.nlm.nih.gov/gene/?term=LOC105282834 | 5993 | 124 | 206.91 | 1 | 1.67 | 0.81 | 16.5 | 15.65 | 0.85 | 0.85 | 0.02 | 0.02 |
| gene303 | https://www.ncbi.nlm.nih.gov/gene/?term=LOC105275584 | 3081 | 86 | 279.13 | 1 | 3.25 | 1.16 | 20.08 | 19.37 | 0.71 | 0.71 | 0.02 | 0.02 |
| gene12516 | https://www.ncbi.nlm.nih.gov/gene/?term=LOC105286452 | 1529 | 38 | 248.53 | 1 | 6.54 | 2.63 | 50.05 | 49.37 | 0.68 | 0.68 | 0.02 | 0.02 |
| gene11568 | https://www.ncbi.nlm.nih.gov/gene/?term=LOC105285395 | 8086 | 190 | 234.97 | 1 | 1.24 | 0.53 | 16.67 | 16 | 0.67 | 0.67 | 0.02 | 0.02 |
| gene4892 | https://www.ncbi.nlm.nih.gov/gene/?term=LOC105277667 | 3898 | 126 | 323.24 | 1 | 2.57 | 0.79 | 27.81 | 27.16 | 0.65 | 0.65 | 0.02 | 0.02 |
| gene10520 | https://www.ncbi.nlm.nih.gov/gene/?term=LOC105284184 | 2451 | 84 | 342.72 | 1 | 4.08 | 1.19 | 33.8 | 33.16 | 0.64 | 0.64 | 0.02 | 0.02 |
| gene473 | https://www.ncbi.nlm.nih.gov/gene/?term=LOC105277423 | 2342 | 106 | 452.6 | 1 | 4.27 | 0.94 | 28.94 | 28.4 | 0.54 | 0.54 | 0.02 | 0.02 |

| | | | | | | | | | | | | | | |
|---|---|---|---|---|---|---|---|---|---|---|---|---|---|---|
| gene10540 | https://www.ncbi.nlm.nih.gov/gene/?term=LOC105284200 | 5267 | 220 | 417.7 | 1 | 1.9 | 0.45 | 12.54 | 12.08 | 0.45 | 0.45 | 0.02 | 0.02 | |
| gene6583 | https://www.ncbi.nlm.nih.gov/gene/?term=LOC105279649 | 7444 | 320 | 429.88 | | 1 | 1.34 | 0.31 | 11.91 | 11.5 | 0.41 | 0.41 | 0.02 | 0.02 |
| gene298 | https://www.ncbi.nlm.nih.gov/gene/?term=LOC105275460 | 2473 | 117 | 473.11 | | 1 | 4.04 | 0.85 | 13.71 | 13.31 | 0.39 | 0.39 | 0.02 | 0.02 |
| gene13114 | https://www.ncbi.nlm.nih.gov/gene/?term=LOC105287147 | 7609 | 398 | 523.06 | | 1 | 1.31 | 0.25 | 9.72 | 9.4 | 0.33 | 0.33 | 0.02 | 0.02 |
| gene920 | https://www.ncbi.nlm.nih.gov/gene/?term=LOC105282518 | 1197 | 52 | 434.42 | | 1 | 8.35 | 1.92 | 30.27 | 30.77 | -0.49 | 0.49 | -0.02 | 0.02 |
| gene10549 | https://www.ncbi.nlm.nih.gov/gene/?term=LOC105284210 | 7145 | 290 | 405.88 | | 2 | 2.8 | 0.69 | 11.49 | 12 | -0.51 | 0.51 | -0.02 | 0.02 |
| gene7249 | https://www.ncbi.nlm.nih.gov/gene/?term=LOC105280422 | 1775 | 59 | 332.39 | | 1 | 5.63 | 1.69 | 16.45 | 17.08 | -0.63 | 0.63 | -0.02 | 0.02 |
| gene35 | https://www.ncbi.nlm.nih.gov/gene/?term=LOC105275197 | 2643 | 50 | 189.18 | | 1 | 3.78 | 2 | 37.01 | 37.84 | -0.83 | 0.83 | -0.02 | 0.02 |
| gene4376 | https://www.ncbi.nlm.nih.gov/gene/?term=LOC105277083 | 4307 | 102 | 236.82 | | 1 | 2.32 | 0.98 | 24.25 | 25.14 | -0.89 | 0.89 | -0.02 | 0.02 |
| gene3385 | https://www.ncbi.nlm.nih.gov/gene/?term=LOC105275966 | 3305 | 58 | 175.49 | | 1 | 3.03 | 1.72 | 49.09 | 50.19 | -1.1 | 1.1 | -0.02 | 0.02 |
| gene322 | https://www.ncbi.nlm.nih.gov/gene/?term=LOC105275777 | 1606 | 28 | 174.35 | | 1 | 6.23 | 3.57 | 42.75 | 44.07 | -1.32 | 1.32 | -0.02 | 0.02 |
| gene7248 | https://www.ncbi.nlm.nih.gov/gene/?term=LOC105280408 | 5175 | 52 | 100.48 | | 1 | 1.93 | 1.92 | 16.62 | 18.28 | -1.65 | 1.65 | -0.02 | 0.02 |
| gene8343 | https://www.ncbi.nlm.nih.gov/gene/?term=LOC105281700 | 2172 | 32 | 147.33 | | 1 | 4.6 | 3.12 | 37.31 | 38.97 | -1.66 | 1.66 | -0.02 | 0.02 |
| gene13235 | https://www.ncbi.nlm.nih.gov/gene/?term=LOC105287278 | 2679 | 34 | 126.91 | | 1 | 3.73 | 2.94 | 22.53 | 24.28 | -1.75 | 1.75 | -0.02 | 0.02 |
| gene564 | https://www.ncbi.nlm.nih.gov/gene/?term=LOC105278478 | 2802 | 17 | 60.67 | 1 | 3.57 | 5.88 | 14.04 | 17.34 | -3.31 | 3.31 | -0.02 | 0.02 | |
| gene8178 | https://www.ncbi.nlm.nih.gov/gene/?term=LOC105281515 | 8647 | 44 | 50.88 | 1 | 1.16 | 2.27 | 25.92 | 24.04 | 1.88 | 1.88 | 0.01 | 0.01 | |
| gene6455 | https://www.ncbi.nlm.nih.gov/gene/?term=LOC105279500 | 2612 | 35 | 134 | 1 | 3.83 | 2.86 | 49.33 | 48.29 | 1.04 | 1.04 | 0.01 | 0.01 | |
| gene7398 | https://www.ncbi.nlm.nih.gov/gene/?term=LOC105280589 | 3452 | 33 | 95.6 | 1 | 2.9 | 3.03 | 21.34 | 20.45 | 0.9 | 0.9 | 0.01 | 0.01 | |
| gene3470 | https://www.ncbi.nlm.nih.gov/gene/?term=LOC105276059 | 8716 | 157 | 180.13 | | 1 | 1.15 | 0.64 | 8.72 | 7.97 | 0.76 | 0.76 | 0.01 | 0.01 |
| gene8252 | https://www.ncbi.nlm.nih.gov/gene/?term=LOC105281599 | 3813 | 49 | 128.51 | | 1 | 2.62 | 2.04 | 4.72 | 4.12 | 0.6 | 0.6 | 0.01 | 0.01 |
| gene7884 | https://www.ncbi.nlm.nih.gov/gene/?term=LOC105281266 | 3390 | 76 | 224.19 | | 1 | 2.95 | 1.32 | 16.63 | 16.11 | 0.52 | 0.52 | 0.01 | 0.01 |
| gene11579 | https://www.ncbi.nlm.nih.gov/gene/?term=LOC105285406 | 13571 | 226 | 166.53 | | 1 | 0.74 | 0.44 | 7.69 | 7.18 | 0.51 | 0.51 | 0.01 | 0.01 |
| gene5877 | https://www.ncbi.nlm.nih.gov/gene/?term=LOC105278807 | 1719 | 49 | 285.05 | | 1 | 5.82 | 2.04 | 27.11 | 26.65 | 0.46 | 0.46 | 0.01 | 0.01 |
| gene11082 | https://www.ncbi.nlm.nih.gov/gene/?term=LOC105284824 | 7436 | 200 | 268.96 | | 1 | 1.34 | 0.5 | 7.89 | 7.46 | 0.43 | 0.43 | 0.01 | 0.01 |
| gene7552 | https://www.ncbi.nlm.nih.gov/gene/?term=LOC105280769 | 8566 | 287 | 335.05 | | 1 | 1.17 | 0.35 | 2.06 | 1.63 | 0.43 | 0.43 | 0.01 | 0.01 |
| gene10531 | https://www.ncbi.nlm.nih.gov/gene/?term=LOC105284190 | 6881 | 186 | 270.31 | | 1 | 1.45 | 0.54 | 12.46 | 12.07 | 0.39 | 0.39 | 0.01 | 0.01 |
| gene7556 | https://www.ncbi.nlm.nih.gov/gene/?term=LOC105280774 | 2792 | 97 | 347.42 | | 1 | 3.58 | 1.03 | 33.33 | 32.94 | 0.39 | 0.39 | 0.01 | 0.01 |
| gene12806 | https://www.ncbi.nlm.nih.gov/gene/?term=LOC105286803 | 4196 | 141 | 336.03 | | 1 | 2.38 | 0.71 | 9.78 | 9.4 | 0.38 | 0.38 | 0.01 | 0.01 |
| gene10844 | https://www.ncbi.nlm.nih.gov/gene/?term=LOC105284535 | 9086 | 231 | 254.24 | | 1 | 1.1 | 0.43 | 15.03 | 14.66 | 0.37 | 0.37 | 0.01 | 0.01 |
| gene292 | https://www.ncbi.nlm.nih.gov/gene/?term=LOC105275398 | 2825 | 81 | 286.73 | | 1 | 3.54 | 1.23 | 24.98 | 24.69 | 0.3 | 0.3 | 0.01 | 0.01 |
| gene5524 | https://www.ncbi.nlm.nih.gov/gene/?term=LOC105278397 | 5177 | 198 | 382.46 | | 1 | 1.93 | 0.51 | 24.98 | 24.69 | 0.29 | 0.29 | 0.01 | 0.01 |
| gene2271 | https://www.ncbi.nlm.nih.gov/gene/?term=LOC105274706 | 1448 | 51 | 352.21 | | 1 | 6.91 | 1.96 | 40.12 | 39.86 | 0.26 | 0.26 | 0.01 | 0.01 |
| gene4216 | https://www.ncbi.nlm.nih.gov/gene/?term=LOC105276906 | 2127 | 90 | 423.13 | | 1 | 4.7 | 1.11 | 21.35 | 21.12 | 0.23 | 0.23 | 0.01 | 0.01 |
| gene10018 | https://www.ncbi.nlm.nih.gov/gene/?term=LOC105283607 | 4414 | 207 | 468.96 | | 1 | 2.27 | 0.48 | 15.29 | 15.09 | 0.2 | 0.2 | 0.01 | 0.01 |
| gene10596 | https://www.ncbi.nlm.nih.gov/gene/?term=LOC105284262 | 6520 | 208 | 319.02 | | 1 | 1.53 | 0.48 | 4.3 | 4.14 | 0.16 | 0.16 | 0.01 | 0.01 |
| gene3206 | https://www.ncbi.nlm.nih.gov/gene/?term=LOC105275752 | 3474 | 118 | 339.67 | | 1 | 2.88 | 0.85 | 32.15 | 31.99 | 0.16 | 0.16 | 0.01 | 0.01 |
| gene570 | https://www.ncbi.nlm.nih.gov/gene/?term=LOC105278549 | 1673 | 63 | 376.57 | | 1 | 5.98 | 1.59 | 43.38 | 43.23 | 0.15 | 0.15 | 0.01 | 0.01 |
| gene5525 | https://www.ncbi.nlm.nih.gov/gene/?term=LOC105278396 | 3090 | 182 | 589 | 1 | 3.24 | 0.55 | 24.88 | 24.76 | 0.12 | 0.12 | 0.01 | 0.01 | |
| gene5746 | https://www.ncbi.nlm.nih.gov/gene/?term=LOC105278666 | 10343 | 441 | 426.38 | | 1 | 0.97 | 0.23 | 6.88 | 6.76 | 0.12 | 0.12 | 0.01 | 0.01 |
| gene1470 | https://www.ncbi.nlm.nih.gov/gene/?term=LOC105287597 | 2974 | 186 | 625.42 | | 1 | 3.36 | 0.54 | 18.48 | 18.39 | 0.09 | 0.09 | 0.01 | 0.01 |
| gene4190 | https://www.ncbi.nlm.nih.gov/gene/?term=LOC105276875 | 3150 | 117 | 371.43 | | 1 | 3.17 | 0.85 | 21.8 | 21.95 | -0.15 | 0.15 | -0.01 | 0.01 |
| gene12802 | https://www.ncbi.nlm.nih.gov/gene/?term=LOC105286799 | 3183 | 164 | 515.24 | | 1 | 3.14 | 0.61 | 17.26 | 17.43 | -0.17 | 0.17 | -0.01 | 0.01 |
| gene1430 | https://www.ncbi.nlm.nih.gov/gene/?term=LOC105287551 | 6433 | 223 | 346.65 | | 1 | 1.55 | 0.45 | 8.23 | 8.41 | -0.18 | 0.18 | -0.01 | 0.01 |
| gene6286 | https://www.ncbi.nlm.nih.gov/gene/?term=LOC105279298 | 2861 | 129 | 450.89 | | 2 | 6.99 | 1.55 | 23.63 | 23.86 | -0.23 | 0.23 | -0.01 | 0.01 |
| gene4962 | https://www.ncbi.nlm.nih.gov/gene/?term=LOC105277761 | 4645 | 144 | 310.01 | | 1 | 2.15 | 0.69 | 13.44 | 13.75 | -0.31 | 0.31 | -0.01 | 0.01 |
| gene9327 | https://www.ncbi.nlm.nih.gov/gene/?term=LOC105282821 | 7793 | 145 | 186.06 | | 1 | 1.28 | 0.69 | 12.57 | 12.89 | -0.32 | 0.32 | -0.01 | 0.01 |
| gene2957 | https://www.ncbi.nlm.nih.gov/gene/?term=LOC105275500 | 10926 | 240 | 219.66 | | 1 | 0.92 | 0.42 | 6.99 | 7.33 | -0.34 | 0.34 | -0.01 | 0.01 |
| gene12307 | https://www.ncbi.nlm.nih.gov/gene/?term=LOC105286216 | 10313 | 187 | 181.32 | | 1 | 0.97 | 0.53 | 14.69 | 15.04 | -0.35 | 0.35 | -0.01 | 0.01 |
| gene10155 | https://www.ncbi.nlm.nih.gov/gene/?term=LOC105283756 | 7030 | 112 | 159.32 | | 1 | 1.42 | 0.89 | 5.92 | 6.29 | -0.37 | 0.37 | -0.01 | 0.01 |
| gene4023 | https://www.ncbi.nlm.nih.gov/gene/?term=LOC105276692 | 1759 | 31 | 176.24 | | 1 | 5.69 | 3.23 | 23.25 | 23.63 | -0.38 | 0.38 | -0.01 | 0.01 |
| gene1310 | https://www.ncbi.nlm.nih.gov/gene/?term=LOC105286661 | 2582 | 71 | 274.98 | | 1 | 3.87 | 1.41 | 21.29 | 21.68 | -0.39 | 0.39 | -0.01 | 0.01 |
| gene12573 | https://www.ncbi.nlm.nih.gov/gene/?term=LOC105286515 | 3647 | 100 | 274.2 | 1 | 2.74 | 1 | 13.03 | 13.44 | -0.42 | 0.42 | -0.01 | 0.01 | |
| gene13036 | https://www.ncbi.nlm.nih.gov/gene/?term=LOC105287059 | 4959 | 160 | 322.65 | | 1 | 2.02 | 0.62 | 19.69 | 20.15 | -0.46 | 0.46 | -0.01 | 0.01 |
| gene1199 | https://www.ncbi.nlm.nih.gov/gene/?term=LOC105285498 | 2220 | 28 | 126.13 | | 1 | 4.5 | 3.57 | 27.44 | 27.99 | -0.55 | 0.55 | -0.01 | 0.01 |
| gene657 | https://www.ncbi.nlm.nih.gov/gene/?term=LOC105279667 | 6421 | 81 | 126.15 | | 1 | 1.56 | 1.23 | 2.49 | 3.12 | -0.63 | 0.63 | -0.01 | 0.01 |
| gene7183 | https://www.ncbi.nlm.nih.gov/gene/?term=LOC105280348 | 3925 | 70 | 178.34 | | 1 | 2.55 | 1.43 | 20.85 | 21.48 | -0.63 | 0.63 | -0.01 | 0.01 |
| gene12021 | https://www.ncbi.nlm.nih.gov/gene/?term=LOC105285905 | 1399 | 15 | 107.22 | | 1 | 7.15 | 6.67 | 50 | 50.68 | -0.68 | 0.68 | -0.01 | 0.01 |
| gene7247 | https://www.ncbi.nlm.nih.gov/gene/?term=LOC105280407 | 10486 | 206 | 196.45 | | 1 | 0.95 | 0.49 | 16.14 | 16.82 | -0.68 | 0.68 | -0.01 | 0.01 |
| gene11561 | https://www.ncbi.nlm.nih.gov/gene/?term=LOC105285387 | 11528 | 178 | 154.41 | | 1 | 0.87 | 0.56 | 9.89 | 10.61 | -0.72 | 0.72 | -0.01 | 0.01 |
| gene12203 | https://www.ncbi.nlm.nih.gov/gene/?term=LOC105286099 | 5967 | 75 | 125.69 | | 1 | 1.68 | 1.33 | 9.56 | 10.52 | -0.96 | 0.96 | -0.01 | 0.01 |
| gene4033 | https://www.ncbi.nlm.nih.gov/gene/?term=LOC105276705 | 10546 | 142 | 134.65 | | 1 | 0.95 | 0.7 | 7.83 | 8.89 | -1.06 | 1.06 | -0.01 | 0.01 |
| gene10345 | https://www.ncbi.nlm.nih.gov/gene/?term=LOC105283989 | 4444 | 27 | 60.76 | 1 | 2.25 | 3.7 | 36.46 | 38.69 | -2.24 | 2.24 | -0.01 | 0.01 | |
| gene2173 | https://www.ncbi.nlm.nih.gov/gene/?term=LOC105274590 | 4387 | 70 | 159.56 | | 2 | 4.56 | 2.86 | 26.31 | 26.05 | 0.27 | 0.27 | 0 | 0 |
| gene6639 | https://www.ncbi.nlm.nih.gov/gene/?term=LOC105279705 | 7656 | 67 | 87.51 | 1 | 1.31 | 1.49 | 22.85 | 22.67 | 0.17 | 0.17 | 0 | 0 | |

| | | | | | | | | | | | | | |
|---|---|---|---|---|---|---|---|---|---|---|---|---|---|
| gene1632 | https://www.ncbi.nlm.nih.gov/gene/?term=LOC105287764 | 18061 | 391 | 216.49 | 2 | 1.11 | 0.51 | 3.3 | 3.14 | 0.16 | 0.16 | 0 | 0 |
| gene862 | https://www.ncbi.nlm.nih.gov/gene/?term=LOC105281802 | 73922 | 1907 | 257.97 | 1 | 0.14 | 0.05 | 2.33 | 2.19 | 0.14 | 0.14 | 0 | 0 |
| gene7888 | https://www.ncbi.nlm.nih.gov/gene/?term=LOC105281141 | 4357 | 153 | 351.16 | 1 | 2.3 | 0.65 | 32.3 | 32.17 | 0.13 | 0.13 | 0 | 0 |
| gene10392 | https://www.ncbi.nlm.nih.gov/gene/?term=LOC105284036 | 3216 | 99 | 307.84 | 2 | 6.22 | 2.02 | 22.54 | 22.43 | 0.11 | 0.11 | 0 | 0 |
| gene7368 | https://www.ncbi.nlm.nih.gov/gene/?term=LOC105280556 | 4105 | 41 | 99.88 1 | 2.44 | 2.44 | 16.06 | 15.95 | 0.11 | 0.11 | 0 | 0 | |
| gene2806 | https://www.ncbi.nlm.nih.gov/gene/?term=LOC105275327 | 14257 | 473 | 331.77 | 1 | 0.7 | 0.21 | 7.14 | 7.04 | 0.1 | 0.1 | 0 | 0 |
| gene3885 | https://www.ncbi.nlm.nih.gov/gene/?term=LOC105276527 | 7928 | 318 | 401.11 | 1 | 1.26 | 0.31 | 9.68 | 9.6 | 0.09 | 0.09 | 0 | 0 |
| gene9101 | https://www.ncbi.nlm.nih.gov/gene/?term=LOC105282553 | 3068 | 104 | 338.98 | 1 | 3.26 | 0.96 | 23.44 | 23.39 | 0.06 | 0.06 | 0 | 0 |
| gene1408 | https://www.ncbi.nlm.nih.gov/gene/?term=LOC109611520 | 8645 | 306 | 353.96 | 1 | 1.16 | 0.33 | 6.02 | 5.98 | 0.05 | 0.05 | 0 | 0 |
| gene2633 | https://www.ncbi.nlm.nih.gov/gene/?term=LOC105275127 | 7457 | 252 | 337.94 | 1 | 1.34 | 0.4 | 8.36 | 8.33 | 0.03 | 0.03 | 0 | 0 |
| gene10326 | https://www.ncbi.nlm.nih.gov/gene/?term=LOC105283962 | 3042 | 124 | 407.63 | 1 | 3.29 | 0.81 | 8.49 | 8.5 | -0.01 | 0.01 | 0 | 0 |
| gene11275 | https://www.ncbi.nlm.nih.gov/gene/?term=LOC105285055 | 4119 | 252 | 611.8 1 | 2.43 | 0.4 | 11.84 | 11.86 | -0.03 | 0.03 | 0 | 0 | |
| gene6620 | https://www.ncbi.nlm.nih.gov/gene/?term=LOC105279687 | 17998 | 355 | 197.24 | 1 | 0.56 | 0.28 | 5.15 | 5.19 | -0.04 | 0.04 | 0 | 0 |
| gene7915 | https://www.ncbi.nlm.nih.gov/gene/?term=LOC105281173 | 1597 | 39 | 244.21 | 1 | 6.26 | 2.56 | 59.75 | 59.8 | -0.05 | 0.05 | 0 | 0 |
| gene12572 | https://www.ncbi.nlm.nih.gov/gene/?term=LOC105286514 | 2374 | 49 | 206.4 1 | 4.21 | 2.04 | 35.44 | 35.51 | -0.07 | 0.07 | 0 | 0 | |
| gene2683 | https://www.ncbi.nlm.nih.gov/gene/?term=LOC105275187 | 5492 | 70 | 127.46 | 1 | 1.82 | 1.43 | 12.3 | 12.38 | -0.07 | 0.07 | 0 | 0 |
| gene5387 | https://www.ncbi.nlm.nih.gov/gene/?term=LOC105278245 | 6451 | 112 | 173.62 | 1 | 1.55 | 0.89 | 22.85 | 22.92 | -0.07 | 0.07 | 0 | 0 |
| gene8814 | https://www.ncbi.nlm.nih.gov/gene/?term=LOC105282240 | 6910 | 176 | 254.7 1 | 1.45 | 0.57 | 9.18 | 9.27 | -0.08 | 0.08 | 0 | 0 | |
| gene8600 | https://www.ncbi.nlm.nih.gov/gene/?term=LOC105282001 | 8513 | 151 | 177.38 | 1 | 1.17 | 0.66 | 10.34 | 10.43 | -0.09 | 0.09 | 0 | 0 |
| gene8729 | https://www.ncbi.nlm.nih.gov/gene/?term=LOC105282137 | 5005 | 211 | 421.58 | 1 | 2 | 0.47 | 15.81 | 15.91 | -0.09 | 0.09 | 0 | 0 |
| gene2404 | https://www.ncbi.nlm.nih.gov/gene/?term=LOC105274853 | 1767 | 35 | 198.08 | 1 | 5.66 | 2.86 | 33.43 | 33.53 | -0.1 | 0.1 | 0 | 0 |
| gene5101 | https://www.ncbi.nlm.nih.gov/gene/?term=LOC105277903 | 1813 | 35 | 193.05 | 1 | 5.52 | 2.86 | 51.16 | 51.27 | -0.1 | 0.1 | 0 | 0 |
| gene53 | https://www.ncbi.nlm.nih.gov/gene/?term=LOC105277148 | 19283 | 295 | 152.98 | 1 | 0.52 | 0.34 | 3.8 | 3.9 | -0.1 | 0.1 | 0 | 0 |
| gene5646 | https://www.ncbi.nlm.nih.gov/gene/?term=LOC105278534 | 7475 | 187 | 250.17 | 1 | 1.34 | 0.53 | 18.3 | 18.4 | -0.1 | 0.1 | 0 | 0 |
| gene2846 | https://www.ncbi.nlm.nih.gov/gene/?term=LOC105275374 | 11246 | 477 | 424.15 | 1 | 0.89 | 0.21 | 10.56 | 10.68 | -0.11 | 0.11 | 0 | 0 |
| gene34 | https://www.ncbi.nlm.nih.gov/gene/?term=LOC105278416 | 7120 | 140 | 196.63 | 1 | 1.4 | 0.71 | 13.56 | 13.67 | -0.11 | 0.11 | 0 | 0 |
| gene7166 | https://www.ncbi.nlm.nih.gov/gene/?term=LOC105280326 | 6117 | 243 | 397.25 | 1 | 1.63 | 0.41 | 14.54 | 14.65 | -0.11 | 0.11 | 0 | 0 |
| gene9326 | https://www.ncbi.nlm.nih.gov/gene/?term=LOC105282818 | 16612 | 178 | 107.15 | 1 | 0.6 | 0.56 | 13.41 | 13.58 | -0.18 | 0.18 | 0 | 0 |

Directory: OGLall File: rnk-dmg-Cb_LI_BCphaseB2.vs.RphaseA1.txt

| gene_ID | gene_link | gwidth | #Sites | #per10Kb | #dmSites | #dmsp10kb | %dmSites | %pSite1 | %pSite2 | DMpSite | ADMpSite | DMpNucl | ADMpNucl |
|---|---|---|---|---|---|---|---|---|---|---|---|---|---|
| gene3165 | https://www.ncbi.nlm.nih.gov/gene/?term=LOC105275706 | 2374 | 60 | 252.74 | 3 | 12.64 | 5 | 22.21 | 32.53 | -10.32 | 10.32 | -0.26 | 0.26 |
| gene2593 | https://www.ncbi.nlm.nih.gov/gene/?term=LOC105275070 | 1774 | 5 | 28.18 | 5 | 28.18 | 100 | 10.96 | 79.22 | -68.26 | 68.26 | -0.19 | 0.19 |
| gene5714 | https://www.ncbi.nlm.nih.gov/gene/?term=LOC105278729 | 710 | 21 | 295.77 | 1 | 14.08 | 4.76 | 56.35 | 60.77 | -4.42 | 4.42 | -0.13 | 0.13 |
| gene10714 | https://www.ncbi.nlm.nih.gov/gene/?term=LOC105284396 | 1966 | 62 | 315.36 | 1 | 5.09 | 1.61 | 36.35 | 33.21 | 3.14 | 3.14 | 0.1 | 0.1 |
| gene12021 | https://www.ncbi.nlm.nih.gov/gene/?term=LOC105285905 | 1399 | 21 | 150.11 | 1 | 7.15 | 4.76 | 36.22 | 31.01 | 5.22 | 5.22 | 0.08 | 0.08 |
| gene6652 | https://www.ncbi.nlm.nih.gov/gene/?term=LOC105279719 | 2831 | 86 | 303.78 | 1 | 3.53 | 1.16 | 23.08 | 20.39 | 2.69 | 2.69 | 0.08 | 0.08 |
| gene9101 | https://www.ncbi.nlm.nih.gov/gene/?term=LOC105282553 | 3068 | 106 | 345.5 | 1 | 3.26 | 0.94 | 23.35 | 21.05 | 2.29 | 2.29 | 0.08 | 0.08 |
| gene4498 | https://www.ncbi.nlm.nih.gov/gene/?term=LOC105277239 | 2831 | 130 | 459.2 | 1 | 3.53 | 0.77 | 12.47 | 10.69 | 1.78 | 1.78 | 0.08 | 0.08 |
| gene2271 | https://www.ncbi.nlm.nih.gov/gene/?term=LOC105274706 | 1448 | 51 | 352.21 | 1 | 6.91 | 1.96 | 39.73 | 37.8 | 1.94 | 1.94 | 0.07 | 0.07 |
| gene2856 | https://www.ncbi.nlm.nih.gov/gene/?term=LOC105275384 | 1987 | 87 | 437.85 | 1 | 5.03 | 1.15 | 34.24 | 35.77 | -1.53 | 1.53 | -0.07 | 0.07 |
| gene7568 | https://www.ncbi.nlm.nih.gov/gene/?term=LOC105280786 | 2525 | 109 | 431.68 | 1 | 3.96 | 0.92 | 28.7 | 30.24 | -1.54 | 1.54 | -0.07 | 0.07 |
| gene3817 | https://www.ncbi.nlm.nih.gov/gene/?term=LOC105276449 | 1230 | 45 | 365.85 | 1 | 8.13 | 2.22 | 27.32 | 29.29 | -1.97 | 1.97 | -0.07 | 0.07 |
| gene12406 | https://www.ncbi.nlm.nih.gov/gene/?term=LOC105286316 | 1386 | 31 | 223.67 | 1 | 7.22 | 3.23 | 46.61 | 43.84 | 2.77 | 2.77 | 0.06 | 0.06 |
| gene7550 | https://www.ncbi.nlm.nih.gov/gene/?term=LOC105280763 | 2173 | 98 | 450.99 | 1 | 4.6 | 1.02 | 27.13 | 25.7 | 1.43 | 1.43 | 0.06 | 0.06 |
| gene2071 | https://www.ncbi.nlm.nih.gov/gene/?term=LOC105274471 | 2446 | 121 | 494.69 | 1 | 4.09 | 0.83 | 19.71 | 20.9 | -1.19 | 1.19 | -0.06 | 0.06 |
| gene5877 | https://www.ncbi.nlm.nih.gov/gene/?term=LOC105278807 | 1719 | 53 | 308.32 | 1 | 5.82 | 1.89 | 22.63 | 24.74 | -2.11 | 2.11 | -0.06 | 0.06 |
| gene5101 | https://www.ncbi.nlm.nih.gov/gene/?term=LOC105277903 | 1813 | 36 | 198.57 | 1 | 5.52 | 2.78 | 53.23 | 56.32 | -3.09 | 3.09 | -0.06 | 0.06 |
| gene5647 | https://www.ncbi.nlm.nih.gov/gene/?term=LOC105278567 | 3118 | 29 | 93.01 | 1 | 3.21 | 3.45 | 13.21 | 19.43 | -6.21 | 6.21 | -0.06 | 0.06 |
| gene7199 | https://www.ncbi.nlm.nih.gov/gene/?term=LOC105280366 | 1414 | 33 | 233.38 | 1 | 7.07 | 3.03 | 43.27 | 41 | 2.28 | 2.28 | 0.05 | 0.05 |
| gene2404 | https://www.ncbi.nlm.nih.gov/gene/?term=LOC105274853 | 1767 | 43 | 243.35 | 1 | 5.66 | 2.33 | 31.85 | 29.62 | 2.23 | 2.23 | 0.05 | 0.05 |
| gene1907 | https://www.ncbi.nlm.nih.gov/gene/?term=LOC105288099 | 1238 | 35 | 282.71 | 1 | 8.08 | 2.86 | 34.89 | 33.16 | 1.73 | 1.73 | 0.05 | 0.05 |
| gene920 | https://www.ncbi.nlm.nih.gov/gene/?term=LOC105282518 | 1197 | 41 | 342.52 | 1 | 8.35 | 2.44 | 39.23 | 37.74 | 1.49 | 1.49 | 0.05 | 0.05 |
| gene4892 | https://www.ncbi.nlm.nih.gov/gene/?term=LOC105277667 | 3898 | 120 | 307.85 | 1 | 2.57 | 0.83 | 27.15 | 28.74 | -1.59 | 1.59 | -0.05 | 0.05 |
| gene393 | https://www.ncbi.nlm.nih.gov/gene/?term=LOC105276566 | 1123 | 2 | 17.81 | 1 | 8.9 | 50 | 70.57 | 50 | 20.57 | 20.57 | 0.04 | 0.04 |
| gene1199 | https://www.ncbi.nlm.nih.gov/gene/?term=LOC105285498 | 2220 | 25 | 112.61 | 1 | 4.5 | 4 | 37.37 | 33.76 | 3.61 | 3.61 | 0.04 | 0.04 |
| gene388 | https://www.ncbi.nlm.nih.gov/gene/?term=LOC105276538 | 1558 | 20 | 128.37 | 2 | 12.84 | 10 | 66.14 | 62.93 | 3.2 | 3.2 | 0.04 | 0.04 |
| gene322 | https://www.ncbi.nlm.nih.gov/gene/?term=LOC105275777 | 1606 | 25 | 155.67 | 1 | 6.23 | 4 | 44.37 | 41.49 | 2.89 | 2.89 | 0.04 | 0.04 |
| gene6422 | https://www.ncbi.nlm.nih.gov/gene/?term=LOC105279458 | 2419 | 37 | 152.96 | 1 | 4.13 | 2.7 | 52.5 | 49.97 | 2.53 | 2.53 | 0.04 | 0.04 |
| gene6286 | https://www.ncbi.nlm.nih.gov/gene/?term=LOC105279298 | 2861 | 131 | 457.88 | 2 | 6.99 | 1.53 | 24.99 | 24.15 | 0.84 | 0.84 | 0.04 | 0.04 |
| gene1430 | https://www.ncbi.nlm.nih.gov/gene/?term=LOC105287551 | 6433 | 235 | 365.3 | 1 | 1.55 | 0.43 | 8.56 | 9.58 | -1.02 | 1.02 | -0.04 | 0.04 |
| gene11401 | https://www.ncbi.nlm.nih.gov/gene/?term=LOC105285202 | 1841 | 75 | 407.39 | 1 | 5.43 | 1.33 | 24.08 | 25.12 | -1.03 | 1.03 | -0.04 | 0.04 |
| gene4720 | https://www.ncbi.nlm.nih.gov/gene/?term=LOC105277479 | 1697 | 50 | 294.64 | 1 | 5.89 | 2 | 35.87 | 37.27 | -1.4 | 1.4 | -0.04 | 0.04 |
| gene1884 | https://www.ncbi.nlm.nih.gov/gene/?term=LOC105288072 | 4169 | 78 | 187.1 | 1 | 2.4 | 1.28 | 22.55 | 24.51 | -1.96 | 1.96 | -0.04 | 0.04 |
| gene8661 | https://www.ncbi.nlm.nih.gov/gene/?term=LOC105282063 | 1926 | 30 | 155.76 | 1 | 5.19 | 3.33 | 28.06 | 30.33 | -2.28 | 2.28 | -0.04 | 0.04 |
| gene9411 | https://www.ncbi.nlm.nih.gov/gene/?term=LOC105282914 | 5534 | 41 | 74.09 | 1 | 1.81 | 2.44 | 52.21 | 48.58 | 3.63 | 3.63 | 0.03 | 0.03 |
| gene3168 | https://www.ncbi.nlm.nih.gov/gene/?term=LOC105275711 | 6150 | 73 | 118.7 | 1 | 1.63 | 1.37 | 14.69 | 12.05 | 2.64 | 2.64 | 0.03 | 0.03 |
| gene6455 | https://www.ncbi.nlm.nih.gov/gene/?term=LOC105279500 | 2612 | 30 | 114.85 | 1 | 3.83 | 3.33 | 48.51 | 46 | 2.5 | 2.5 | 0.03 | 0.03 |
| gene6237 | https://www.ncbi.nlm.nih.gov/gene/?term=LOC105279244 | 1484 | 23 | 154.99 | 1 | 6.74 | 4.35 | 50.08 | 47.92 | 2.16 | 2.16 | 0.03 | 0.03 |
| gene2748 | https://www.ncbi.nlm.nih.gov/gene/?term=LOC105275261 | 3845 | 54 | 140.44 | 1 | 2.6 | 1.85 | 39.59 | 37.65 | 1.94 | 1.94 | 0.03 | 0.03 |
| gene10107 | https://www.ncbi.nlm.nih.gov/gene/?term=LOC105283712 | 1883 | 35 | 185.87 | 1 | 5.31 | 2.86 | 27.16 | 25.31 | 1.85 | 1.85 | 0.03 | 0.03 |
| gene3671 | https://www.ncbi.nlm.nih.gov/gene/?term=LOC105276272 | 1966 | 46 | 233.98 | 1 | 5.09 | 2.17 | 27.89 | 26.72 | 1.16 | 1.16 | 0.03 | 0.03 |
| gene5646 | https://www.ncbi.nlm.nih.gov/gene/?term=LOC105278534 | 7475 | 185 | 247.49 | 1 | 1.34 | 0.54 | 18.24 | 17.21 | 1.03 | 1.03 | 0.03 | 0.03 |
| gene10392 | https://www.ncbi.nlm.nih.gov/gene/?term=LOC105284036 | 3216 | 106 | 329.6 | 2 | 6.22 | 1.89 | 21.2 | 20.33 | 0.87 | 0.87 | 0.03 | 0.03 |
| gene11340 | https://www.ncbi.nlm.nih.gov/gene/?term=LOC105285131 | 2918 | 105 | 359.84 | 1 | 3.43 | 0.95 | 14.38 | 13.54 | 0.84 | 0.84 | 0.03 | 0.03 |
| gene8729 | https://www.ncbi.nlm.nih.gov/gene/?term=LOC105282137 | 5005 | 200 | 399.6 | 1 | 2 | 0.5 | 16.14 | 15.41 | 0.72 | 0.72 | 0.03 | 0.03 |
| gene10018 | https://www.ncbi.nlm.nih.gov/gene/?term=LOC105283607 | 4414 | 201 | 455.37 | 1 | 2.27 | 0.5 | 15.61 | 14.98 | 0.63 | 0.63 | 0.03 | 0.03 |
| gene5298 | https://www.ncbi.nlm.nih.gov/gene/?term=LOC105278145 | 4060 | 237 | 583.74 | 3 | 7.39 | 1.27 | 11.65 | 11.08 | 0.57 | 0.57 | 0.03 | 0.03 |
| gene298 | https://www.ncbi.nlm.nih.gov/gene/?term=LOC105275460 | 2473 | 120 | 485.24 | 1 | 4.04 | 0.83 | 13.22 | 13.75 | -0.53 | 0.53 | -0.03 | 0.03 |
| gene6583 | https://www.ncbi.nlm.nih.gov/gene/?term=LOC105279649 | 7444 | 340 | 456.74 | 1 | 1.34 | 0.29 | 10.48 | 11.08 | -0.59 | 0.59 | -0.03 | 0.03 |
| gene7514 | https://www.ncbi.nlm.nih.gov/gene/?term=LOC105280723 | 4471 | 208 | 465.22 | 1 | 2.24 | 0.48 | 13.49 | 14.1 | -0.6 | 0.6 | -0.03 | 0.03 |
| gene2056 | https://www.ncbi.nlm.nih.gov/gene/?term=LOC105274452 | 6584 | 243 | 369.08 | 1 | 1.52 | 0.41 | 13.17 | 13.88 | -0.71 | 0.71 | -0.03 | 0.03 |
| gene7884 | https://www.ncbi.nlm.nih.gov/gene/?term=LOC105281266 | 3390 | 81 | 238.94 | 1 | 2.95 | 1.23 | 16.21 | 17.33 | -1.12 | 1.12 | -0.03 | 0.03 |
| gene1518 | https://www.ncbi.nlm.nih.gov/gene/?term=LOC105287648 | 2195 | 35 | 159.45 | 1 | 4.56 | 2.86 | 35.74 | 37.41 | -1.67 | 1.67 | -0.03 | 0.03 |
| gene9478 | https://www.ncbi.nlm.nih.gov/gene/?term=LOC105282991 | 3726 | 58 | 155.66 | 1 | 2.68 | 1.72 | 33.94 | 35.62 | -1.68 | 1.68 | -0.03 | 0.03 |
| gene927 | https://www.ncbi.nlm.nih.gov/gene/?term=LOC105282600 | 3727 | 52 | 139.52 | 1 | 2.68 | 1.92 | 31.26 | 33.17 | -1.91 | 1.91 | -0.03 | 0.03 |
| gene9648 | https://www.ncbi.nlm.nih.gov/gene/?term=LOC105283183 | 2602 | 33 | 126.83 | 1 | 3.84 | 3.03 | 40.53 | 42.61 | -2.08 | 2.08 | -0.03 | 0.03 |
| gene564 | https://www.ncbi.nlm.nih.gov/gene/?term=LOC105278478 | 2802 | 18 | 64.24 | 1 | 3.57 | 5.56 | 17.32 | 14.97 | 2.35 | 2.35 | 0.02 | 0.02 |
| gene6806 | https://www.ncbi.nlm.nih.gov/gene/?term=LOC105279894 | 3649 | 35 | 95.92 | 1 | 2.74 | 2.86 | 40.58 | 38.26 | 2.32 | 2.32 | 0.02 | 0.02 |
| gene6639 | https://www.ncbi.nlm.nih.gov/gene/?term=LOC105279705 | 7656 | 68 | 88.82 | 1 | 1.31 | 1.47 | 23.58 | 21.76 | 1.82 | 1.82 | 0.02 | 0.02 |
| gene3046 | https://www.ncbi.nlm.nih.gov/gene/?term=LOC105275587 | 2394 | 58 | 242.27 | 1 | 4.18 | 1.72 | 36.72 | 35.74 | 0.98 | 0.98 | 0.02 | 0.02 |

| | | | | | | | | | | | | | |
|---|---|---|---|---|---|---|---|---|---|---|---|---|---|
| gene7915 | https://www.ncbi.nlm.nih.gov/gene/?term=LOC105281173 | 1597 | 38 | 237.95 | 1 | 6.26 | 2.63 | 60.79 | 59.81 | 0.98 | 0.98 | 0.02 | 0.02 |
| gene8015 | https://www.ncbi.nlm.nih.gov/gene/?term=LOC105281312 | 2998 | 65 | 216.81 | 1 | 3.34 | 1.54 | 33.24 | 32.3 | 0.93 | 0.93 | 0.02 | 0.02 |
| gene1310 | https://www.ncbi.nlm.nih.gov/gene/?term=LOC105286661 | 2582 | 62 | 240.12 | 1 | 3.87 | 1.61 | 22.65 | 21.78 | 0.87 | 0.87 | 0.02 | 0.02 |
| gene9462 | https://www.ncbi.nlm.nih.gov/gene/?term=LOC105282970 | 1469 | 41 | 279.1 1 | 6.81 | 2.44 | 50 | 49.15 | 0.85 | 0.85 | 0.02 | 0.02 | |
| gene1220 | https://www.ncbi.nlm.nih.gov/gene/?term=LOC105285742 | 3453 | 106 | 306.98 | 1 | 2.9 | 0.94 | 18.53 | 17.74 | 0.79 | 0.79 | 0.02 | 0.02 |
| gene4190 | https://www.ncbi.nlm.nih.gov/gene/?term=LOC105276875 | 3150 | 111 | 352.38 | 1 | 3.17 | 0.9 | 22.75 | 22.14 | 0.61 | 0.61 | 0.02 | 0.02 |
| gene4962 | https://www.ncbi.nlm.nih.gov/gene/?term=LOC105277761 | 4645 | 147 | 316.47 | 1 | 2.15 | 0.68 | 14.43 | 13.91 | 0.52 | 0.52 | 0.02 | 0.02 |
| gene10520 | https://www.ncbi.nlm.nih.gov/gene/?term=LOC105284184 | 2451 | 93 | 379.44 | 1 | 4.08 | 1.08 | 31.28 | 30.8 | 0.49 | 0.49 | 0.02 | 0.02 |
| gene10326 | https://www.ncbi.nlm.nih.gov/gene/?term=LOC105283962 | 3042 | 126 | 414.2 1 | 3.29 | 0.79 | 9.42 | 8.95 | 0.47 | 0.47 | 0.02 | 0.02 | |
| gene6031 | https://www.ncbi.nlm.nih.gov/gene/?term=LOC105279000 | 5691 | 195 | 342.65 | 1 | 1.76 | 0.51 | 13.76 | 13.29 | 0.47 | 0.47 | 0.02 | 0.02 |
| gene3885 | https://www.ncbi.nlm.nih.gov/gene/?term=LOC105276527 | 7928 | 318 | 401.11 | 1 | 1.26 | 0.31 | 10.14 | 9.68 | 0.46 | 0.46 | 0.02 | 0.02 |
| gene8274 | https://www.ncbi.nlm.nih.gov/gene/?term=LOC105281624 | 5425 | 219 | 403.69 | 1 | 1.84 | 0.46 | 10.56 | 10.14 | 0.42 | 0.42 | 0.02 | 0.02 |
| gene9740 | https://www.ncbi.nlm.nih.gov/gene/?term=LOC105283277 | 2887 | 136 | 471.08 | 1 | 3.46 | 0.74 | 19.63 | 19.22 | 0.41 | 0.41 | 0.02 | 0.02 |
| gene10213 | https://www.ncbi.nlm.nih.gov/gene/?term=LOC105283826 | 1503 | 64 | 425.82 | 1 | 6.65 | 1.56 | 16.48 | 16.1 | 0.38 | 0.38 | 0.02 | 0.02 |
| gene1470 | https://www.ncbi.nlm.nih.gov/gene/?term=LOC105287597 | 2974 | 195 | 655.68 | 1 | 3.36 | 0.51 | 18.15 | 17.88 | 0.27 | 0.27 | 0.02 | 0.02 |
| gene738 | https://www.ncbi.nlm.nih.gov/gene/?term=LOC105280421 | 4927 | 216 | 438.4 1 | 2.03 | 0.46 | 14.22 | 14.61 | -0.39 | 0.39 | -0.02 | 0.02 | |
| gene3206 | https://www.ncbi.nlm.nih.gov/gene/?term=LOC105275752 | 3474 | 129 | 371.33 | 1 | 2.88 | 0.78 | 29.67 | 30.17 | -0.5 | 0.5 | -0.02 | 0.02 |
| gene4376 | https://www.ncbi.nlm.nih.gov/gene/?term=LOC105277083 | 4307 | 114 | 264.69 | 1 | 2.32 | 0.88 | 20.15 | 20.76 | -0.61 | 0.61 | -0.02 | 0.02 |
| gene13033 | https://www.ncbi.nlm.nih.gov/gene/?term=LOC105287063 | 1877 | 51 | 271.71 | 1 | 5.33 | 1.96 | 30.35 | 31.05 | -0.7 | 0.7 | -0.02 | 0.02 |
| gene12439 | https://www.ncbi.nlm.nih.gov/gene/?term=LOC105286357 | 4761 | 81 | 170.13 | 1 | 2.1 | 1.23 | 29.49 | 30.52 | -1.03 | 1.03 | -0.02 | 0.02 |
| gene4023 | https://www.ncbi.nlm.nih.gov/gene/?term=LOC105276692 | 1759 | 36 | 204.66 | 1 | 5.69 | 2.78 | 20.19 | 21.39 | -1.2 | 1.2 | -0.02 | 0.02 |
| gene7368 | https://www.ncbi.nlm.nih.gov/gene/?term=LOC105280556 | 4105 | 50 | 121.8 1 | 2.44 | 2 | 14.06 | 15.36 | -1.3 | 1.3 | -0.02 | 0.02 | |
| gene10654 | https://www.ncbi.nlm.nih.gov/gene/?term=LOC105284329 | 4027 | 51 | 126.65 | 1 | 2.48 | 1.96 | 17.3 | 18.75 | -1.45 | 1.45 | -0.02 | 0.02 |
| gene9647 | https://www.ncbi.nlm.nih.gov/gene/?term=LOC109611233 | 3094 | 35 | 113.12 | 1 | 3.23 | 2.86 | 38.21 | 40.18 | -1.97 | 1.97 | -0.02 | 0.02 |
| gene8178 | https://www.ncbi.nlm.nih.gov/gene/?term=LOC105281515 | 8647 | 63 | 72.86 1 | 1.16 | 1.59 | 24.53 | 26.83 | -2.3 | 2.3 | -0.02 | 0.02 | |
| gene10602 | https://www.ncbi.nlm.nih.gov/gene/?term=LOC105284268 | 2122 | 21 | 98.96 1 | 4.71 | 4.76 | 25.67 | 24.45 | 1.22 | 1.22 | 0.01 | 0.01 | |
| gene7398 | https://www.ncbi.nlm.nih.gov/gene/?term=LOC105280589 | 3452 | 36 | 104.29 | 1 | 2.9 | 2.78 | 20.14 | 19.24 | 0.9 | 0.9 | 0.01 | 0.01 |
| gene12203 | https://www.ncbi.nlm.nih.gov/gene/?term=LOC105286099 | 5967 | 72 | 120.66 | 1 | 1.68 | 1.39 | 10.57 | 10 | 0.57 | 0.57 | 0.01 | 0.01 |
| gene10844 | https://www.ncbi.nlm.nih.gov/gene/?term=LOC105284535 | 9086 | 238 | 261.94 | 1 | 1.1 | 0.42 | 14.5 | 14.01 | 0.49 | 0.49 | 0.01 | 0.01 |
| gene2173 | https://www.ncbi.nlm.nih.gov/gene/?term=LOC105274590 | 4387 | 80 | 182.36 | 1 | 2.28 | 1.25 | 27.39 | 26.91 | 0.48 | 0.48 | 0.01 | 0.01 |
| gene10600 | https://www.ncbi.nlm.nih.gov/gene/?term=LOC105284270 | 4837 | 115 | 237.75 | 1 | 2.07 | 0.87 | 11.71 | 11.26 | 0.46 | 0.46 | 0.01 | 0.01 |
| gene2257 | https://www.ncbi.nlm.nih.gov/gene/?term=LOC105274689 | 3658 | 73 | 199.56 | 1 | 2.73 | 1.37 | 16.2 | 15.75 | 0.46 | 0.46 | 0.01 | 0.01 |
| gene12572 | https://www.ncbi.nlm.nih.gov/gene/?term=LOC105286514 | 2374 | 44 | 185.34 | 1 | 4.21 | 2.27 | 36.23 | 35.78 | 0.45 | 0.45 | 0.01 | 0.01 |
| gene7600 | https://www.ncbi.nlm.nih.gov/gene/?term=LOC105280819 | 1615 | 45 | 278.64 | 1 | 6.19 | 2.22 | 37.23 | 36.8 | 0.43 | 0.43 | 0.01 | 0.01 |
| gene3599 | https://www.ncbi.nlm.nih.gov/gene/?term=LOC105276196 | 2955 | 57 | 192.89 | 1 | 3.38 | 1.75 | 38.84 | 38.45 | 0.38 | 0.38 | 0.01 | 0.01 |
| gene3107 | https://www.ncbi.nlm.nih.gov/gene/?term=LOC105275650 | 5103 | 174 | 340.98 | 1 | 1.96 | 0.57 | 16.52 | 16.19 | 0.33 | 0.33 | 0.01 | 0.01 |
| gene6966 | https://www.ncbi.nlm.nih.gov/gene/?term=LOC105280104 | 3747 | 121 | 322.93 | 1 | 2.67 | 0.83 | 16.24 | 15.96 | 0.28 | 0.28 | 0.01 | 0.01 |
| gene9445 | https://www.ncbi.nlm.nih.gov/gene/?term=LOC105282951 | 2285 | 114 | 498.91 | 1 | 4.38 | 0.88 | 20.31 | 20.03 | 0.28 | 0.28 | 0.01 | 0.01 |
| gene11454 | https://www.ncbi.nlm.nih.gov/gene/?term=LOC105285256 | 1906 | 51 | 267.58 | 1 | 5.25 | 1.96 | 42.03 | 41.77 | 0.26 | 0.26 | 0.01 | 0.01 |
| gene5524 | https://www.ncbi.nlm.nih.gov/gene/?term=LOC105278397 | 5177 | 202 | 390.19 | 1 | 1.93 | 0.5 | 25.27 | 25.04 | 0.22 | 0.22 | 0.01 | 0.01 |
| gene570 | https://www.ncbi.nlm.nih.gov/gene/?term=LOC105278549 | 1673 | 69 | 412.43 | 1 | 5.98 | 1.45 | 43.78 | 43.57 | 0.21 | 0.21 | 0.01 | 0.01 |
| gene7547 | https://www.ncbi.nlm.nih.gov/gene/?term=LOC105280767 | 24576 | 785 | 319.42 | 1 | 0.41 | 0.13 | 4.48 | 4.27 | 0.21 | 0.21 | 0.01 | 0.01 |
| gene9177 | https://www.ncbi.nlm.nih.gov/gene/?term=LOC105282646 | 3320 | 101 | 304.22 | 1 | 3.01 | 0.99 | 21.24 | 21.05 | 0.2 | 0.2 | 0.01 | 0.01 |
| gene10886 | https://www.ncbi.nlm.nih.gov/gene/?term=LOC105284582 | 1475 | 49 | 332.2 1 | 6.78 | 2.04 | 22 | 21.8 | 0.19 | 0.19 | 0.01 | 0.01 | |
| gene5525 | https://www.ncbi.nlm.nih.gov/gene/?term=LOC105278396 | 3090 | 186 | 601.94 | 1 | 3.24 | 0.54 | 25.06 | 24.87 | 0.18 | 0.18 | 0.01 | 0.01 |
| gene7249 | https://www.ncbi.nlm.nih.gov/gene/?term=LOC105280422 | 1775 | 63 | 354.93 | 1 | 5.63 | 1.59 | 17.74 | 17.55 | 0.18 | 0.18 | 0.01 | 0.01 |
| gene13036 | https://www.ncbi.nlm.nih.gov/gene/?term=LOC105287059 | 4959 | 165 | 332.73 | 1 | 2.02 | 0.61 | 17.15 | 16.98 | 0.17 | 0.17 | 0.01 | 0.01 |
| gene10606 | https://www.ncbi.nlm.nih.gov/gene/?term=LOC105284273 | 6119 | 255 | 416.73 | 1 | 1.63 | 0.39 | 19.24 | 19.09 | 0.16 | 0.16 | 0.01 | 0.01 |
| gene5777 | https://www.ncbi.nlm.nih.gov/gene/?term=LOC105278691 | 3883 | 152 | 391.45 | 1 | 2.58 | 0.66 | 14.89 | 14.73 | 0.16 | 0.16 | 0.01 | 0.01 |
| gene7166 | https://www.ncbi.nlm.nih.gov/gene/?term=LOC105280326 | 6117 | 238 | 389.08 | 1 | 1.63 | 0.42 | 15.04 | 14.89 | 0.15 | 0.15 | 0.01 | 0.01 |
| gene13114 | https://www.ncbi.nlm.nih.gov/gene/?term=LOC105287147 | 7609 | 419 | 550.66 | 1 | 1.31 | 0.24 | 8.81 | 8.67 | 0.14 | 0.14 | 0.01 | 0.01 |
| gene2846 | https://www.ncbi.nlm.nih.gov/gene/?term=LOC105275374 | 11246 | 465 | 413.48 | 1 | 0.89 | 0.22 | 10.6 | 10.73 | -0.13 | 0.13 | -0.01 | 0.01 |
| gene12802 | https://www.ncbi.nlm.nih.gov/gene/?term=LOC105286799 | 3183 | 190 | 596.92 | 1 | 3.14 | 0.53 | 15.49 | 15.63 | -0.14 | 0.14 | -0.01 | 0.01 |
| gene10678 | https://www.ncbi.nlm.nih.gov/gene/?term=LOC105284355 | 4031 | 182 | 451.5 1 | 2.48 | 0.55 | 17.39 | 17.62 | -0.23 | 0.23 | -0.01 | 0.01 | |
| gene7556 | https://www.ncbi.nlm.nih.gov/gene/?term=LOC105280774 | 2792 | 118 | 422.64 | 1 | 3.58 | 0.85 | 27.01 | 27.27 | -0.26 | 0.26 | -0.01 | 0.01 |
| gene8600 | https://www.ncbi.nlm.nih.gov/gene/?term=LOC105282001 | 8513 | 186 | 218.49 | 1 | 1.17 | 0.54 | 10 | 10.27 | -0.27 | 0.27 | -0.01 | 0.01 |
| gene2806 | https://www.ncbi.nlm.nih.gov/gene/?term=LOC105275327 | 14257 | 511 | 358.42 | 1 | 0.7 | 0.2 | 6.7 | 7.03 | -0.33 | 0.33 | -0.01 | 0.01 |
| gene35 | https://www.ncbi.nlm.nih.gov/gene/?term=LOC105275197 | 2643 | 48 | 181.61 | 1 | 3.78 | 2.08 | 38.22 | 38.55 | -0.33 | 0.33 | -0.01 | 0.01 |
| gene2668 | https://www.ncbi.nlm.nih.gov/gene/?term=LOC105275170 | 12496 | 279 | 223.27 | 1 | 0.8 | 0.36 | 1.97 | 2.32 | -0.35 | 0.35 | -0.01 | 0.01 |
| gene1855 | https://www.ncbi.nlm.nih.gov/gene/?term=LOC105288036 | 2387 | 84 | 351.91 | 1 | 4.19 | 1.19 | 29.18 | 29.59 | -0.41 | 0.41 | -0.01 | 0.01 |
| gene6620 | https://www.ncbi.nlm.nih.gov/gene/?term=LOC105279687 | 17998 | 368 | 204.47 | 1 | 0.56 | 0.27 | 4.64 | 5.1 | -0.46 | 0.46 | -0.01 | 0.01 |
| gene7247 | https://www.ncbi.nlm.nih.gov/gene/?term=LOC105280407 | 10486 | 218 | 207.9 1 | 0.95 | 0.46 | 16.54 | 17.03 | -0.49 | 0.49 | -0.01 | 0.01 | |
| gene9336 | https://www.ncbi.nlm.nih.gov/gene/?term=LOC105282834 | 5993 | 129 | 215.25 | 1 | 1.67 | 0.78 | 14.53 | 15.02 | -0.49 | 0.49 | -0.01 | 0.01 |
| gene940 | https://www.ncbi.nlm.nih.gov/gene/?term=LOC105282794 | 3578 | 85 | 237.56 | 1 | 2.79 | 1.18 | 13.96 | 14.47 | -0.51 | 0.51 | -0.01 | 0.01 |

| | | | | | | | | | | | | | |
|---|---|---|---|---|---|---|---|---|---|---|---|---|---|
| gene3470 | https://www.ncbi.nlm.nih.gov/gene/?term=LOC105276059 | 8716 | 152 | 174.39 | 1 | 1.15 | 0.66 | 7.9 | 8.44 | -0.54 | 0.54 | -0.01 | 0.01 |
| gene12516 | https://www.ncbi.nlm.nih.gov/gene/?term=LOC105286452 | 1529 | 35 | 228.91 | 1 | 6.54 | 2.86 | 51.89 | 52.54 | -0.65 | 0.65 | -0.01 | 0.01 |
| gene5733 | https://www.ncbi.nlm.nih.gov/gene/?term=LOC105278649 | 5221 | 91 | 174.3 1 | 1.92 | 1.1 | 19.36 | 20.1 | -0.74 | 0.74 | -0.01 | 0.01 | |
| gene8742 | https://www.ncbi.nlm.nih.gov/gene/?term=LOC105282151 | 2563 | 42 | 163.87 | 1 | 3.9 | 2.38 | 34.83 | 35.57 | -0.75 | 0.75 | -0.01 | 0.01 |
| gene2683 | https://www.ncbi.nlm.nih.gov/gene/?term=LOC105275187 | 5492 | 70 | 127.46 | 1 | 1.82 | 1.43 | 13.71 | 14.76 | -1.05 | 1.05 | -0.01 | 0.01 |
| gene6528 | https://www.ncbi.nlm.nih.gov/gene/?term=LOC105279591 | 7222 | 59 | 81.69 1 | 1.38 | 1.69 | 26.22 | 27.32 | -1.1 | 1.1 | -0.01 | 0.01 | |
| gene10155 | https://www.ncbi.nlm.nih.gov/gene/?term=LOC105283756 | 7030 | 104 | 147.94 | 1 | 1.42 | 0.96 | 5.54 | 5.23 | 0.31 | 0.31 | 0 | 0 |
| gene12307 | https://www.ncbi.nlm.nih.gov/gene/?term=LOC105286216 | 10313 | 160 | 155.14 | 1 | 0.97 | 0.62 | 16.94 | 16.73 | 0.22 | 0.22 | 0 | 0 |
| gene7183 | https://www.ncbi.nlm.nih.gov/gene/?term=LOC105280348 | 3925 | 70 | 178.34 | 1 | 2.55 | 1.43 | 18.6 | 18.42 | 0.18 | 0.18 | 0 | 0 |
| gene10531 | https://www.ncbi.nlm.nih.gov/gene/?term=LOC105284190 | 6881 | 175 | 254.32 | 1 | 1.45 | 0.57 | 12.25 | 12.09 | 0.16 | 0.16 | 0 | 0 |
| gene11082 | https://www.ncbi.nlm.nih.gov/gene/?term=LOC105284824 | 7436 | 212 | 285.1 1 | 1.34 | 0.47 | 6.95 | 6.8 | 0.15 | 0.15 | 0 | 0 | |
| gene8343 | https://www.ncbi.nlm.nih.gov/gene/?term=LOC105281700 | 2172 | 37 | 170.35 | 1 | 4.6 | 2.7 | 39.63 | 39.48 | 0.15 | 0.15 | 0 | 0 |
| gene53 | https://www.ncbi.nlm.nih.gov/gene/?term=LOC105277148 | 19283 | 291 | 150.91 | 1 | 0.52 | 0.34 | 4.8 | 4.67 | 0.13 | 0.13 | 0 | 0 |
| gene1632 | https://www.ncbi.nlm.nih.gov/gene/?term=LOC105287764 | 18061 | 379 | 209.84 | 1 | 0.55 | 0.26 | 3.62 | 3.51 | 0.11 | 0.11 | 0 | 0 |
| gene10549 | https://www.ncbi.nlm.nih.gov/gene/?term=LOC105284210 | 7145 | 335 | 468.86 | 2 | 2.8 | 0.6 | 10.35 | 10.25 | 0.09 | 0.09 | 0 | 0 |
| gene2413 | https://www.ncbi.nlm.nih.gov/gene/?term=LOC105274863 | 2006 | 53 | 264.21 | 1 | 4.99 | 1.89 | 52.76 | 52.69 | 0.07 | 0.07 | 0 | 0 |
| gene6805 | https://www.ncbi.nlm.nih.gov/gene/?term=LOC105279895 | 87888 | 973 | 110.71 | 1 | 0.11 | 0.1 | 1.87 | 1.81 | 0.06 | 0.06 | 0 | 0 |
| gene11275 | https://www.ncbi.nlm.nih.gov/gene/?term=LOC105285055 | 4119 | 252 | 611.8 1 | 2.43 | 0.4 | 10.71 | 10.67 | 0.05 | 0.05 | 0 | 0 | |
| gene862 | https://www.ncbi.nlm.nih.gov/gene/?term=LOC105281802 | 73922 | 1969 | 266.36 | 1 | 0.14 | 0.05 | 2.17 | 2.12 | 0.05 | 0.05 | 0 | 0 |
| gene1332 | https://www.ncbi.nlm.nih.gov/gene/?term=LOC105286872 | 2460 | 158 | 642.28 | 1 | 4.07 | 0.63 | 20.39 | 20.35 | 0.04 | 0.04 | 0 | 0 |
| gene3856 | https://www.ncbi.nlm.nih.gov/gene/?term=LOC105276499 | 5236 | 196 | 374.33 | 1 | 1.91 | 0.51 | 10.18 | 10.15 | 0.03 | 0.03 | 0 | 0 |
| gene1147 | https://www.ncbi.nlm.nih.gov/gene/?term=LOC105284954 | 4022 | 111 | 275.98 | 1 | 2.49 | 0.9 | 11.93 | 11.9 | 0.02 | 0.02 | 0 | 0 |
| gene5197 | https://www.ncbi.nlm.nih.gov/gene/?term=LOC105278018 | 89233 | 3377 | 378.45 | 1 | 0.11 | 0.03 | 0.69 | 0.68 | 0 | 0 | 0 | 0 |
| gene11583 | https://www.ncbi.nlm.nih.gov/gene/?term=LOC105285414 | 8192 | 178 | 217.29 | 1 | 1.22 | 0.56 | 9.44 | 9.48 | -0.04 | 0.04 | 0 | 0 |
| gene5375 | https://www.ncbi.nlm.nih.gov/gene/?term=LOC105278228 | 8640 | 154 | 178.24 | 1 | 1.16 | 0.65 | 24.99 | 25.04 | -0.05 | 0.05 | 0 | 0 |
| gene5746 | https://www.ncbi.nlm.nih.gov/gene/?term=LOC105278666 | 10343 | 477 | 461.18 | 1 | 0.97 | 0.21 | 6.71 | 6.77 | -0.05 | 0.05 | 0 | 0 |
| gene2844 | https://www.ncbi.nlm.nih.gov/gene/?term=LOC105275363 | 4330 | 197 | 454.97 | 1 | 2.31 | 0.51 | 14.26 | 14.32 | -0.06 | 0.06 | 0 | 0 |
| gene7319 | https://www.ncbi.nlm.nih.gov/gene/?term=LOC105280513 | 68997 | 1336 | 193.63 | 1 | 0.14 | 0.07 | 0.68 | 0.75 | -0.06 | 0.06 | 0 | 0 |
| gene10345 | https://www.ncbi.nlm.nih.gov/gene/?term=LOC105283989 | 4444 | 26 | 58.51 1 | 2.25 | 3.85 | 49.48 | 49.58 | -0.09 | 0.09 | 0 | 0 | |
| gene11579 | https://www.ncbi.nlm.nih.gov/gene/?term=LOC105285406 | 13571 | 280 | 206.32 | 1 | 0.74 | 0.36 | 6.48 | 6.65 | -0.17 | 0.17 | 0 | 0 |
| gene8814 | https://www.ncbi.nlm.nih.gov/gene/?term=LOC105282240 | 6910 | 185 | 267.73 | 1 | 1.45 | 0.54 | 9.48 | 9.66 | -0.18 | 0.18 | 0 | 0 |
| gene11561 | https://www.ncbi.nlm.nih.gov/gene/?term=LOC105285387 | 11528 | 207 | 179.56 | 1 | 0.87 | 0.48 | 8.9 | 9.12 | -0.22 | 0.22 | 0 | 0 |
| gene34 | https://www.ncbi.nlm.nih.gov/gene/?term=LOC105278416 | 7120 | 132 | 185.39 | 1 | 1.4 | 0.76 | 14.08 | 14.34 | -0.26 | 0.26 | 0 | 0 |
| gene5052 | https://www.ncbi.nlm.nih.gov/gene/?term=LOC105277856 | 3167 | 33 | 104.2 1 | 3.16 | 3.03 | 39.08 | 39.37 | -0.29 | 0.29 | 0 | 0 | |
| gene7248 | https://www.ncbi.nlm.nih.gov/gene/?term=LOC105280408 | 5175 | 61 | 117.87 | 1 | 1.93 | 1.64 | 20.52 | 20.93 | -0.41 | 0.41 | 0 | 0 |

Directory: OGLall File: rnk-dmg-Cb_LI_BCphaseB2.vs.RphaseA2.txt

| gene_ID | gene_link | gwidth | #Sites | #per10Kb | #dmSites | #dmsp10kb | %dmSites | %pSite1 | %pSite2 | DMpSite | ADMpSite | DMpNucl | ADMpNucl |
|---|---|---|---|---|---|---|---|---|---|---|---|---|---|
| gene2824 | https://www.ncbi.nlm.nih.gov/gene/?term=LOC105275337 | 1772 | 64 | 361.17 | 2 | 11.29 | 3.12 | 29.09 | 37.7 | -8.61 | 8.61 | -0.31 | 0.31 |
| gene3165 | https://www.ncbi.nlm.nih.gov/gene/?term=LOC105275706 | 2374 | 69 | 290.65 | 5 | 21.06 | 7.25 | 23.81 | 34.06 | -10.25 | 10.25 | -0.3 | 0.3 |
| gene10963 | https://www.ncbi.nlm.nih.gov/gene/?term=LOC105284677 | 1576 | 30 | 190.36 | 2 | 12.69 | 6.67 | 38.74 | 23.8 | 14.95 | 14.95 | 0.28 | 0.28 |
| gene2593 | https://www.ncbi.nlm.nih.gov/gene/?term=LOC105275070 | 1774 | 5 | 28.18 | 5 | 28.18 | 100 | 10.96 | 81.03 | -70.07 | 70.07 | -0.2 | 0.2 |
| gene5714 | https://www.ncbi.nlm.nih.gov/gene/?term=LOC105278729 | 710 | 24 | 338.03 | 1 | 14.08 | 4.17 | 56.79 | 61.08 | -4.29 | 4.29 | -0.14 | 0.14 |
| gene7568 | https://www.ncbi.nlm.nih.gov/gene/?term=LOC105280786 | 2525 | 119 | 471.29 | 1 | 3.96 | 0.84 | 29.84 | 32.5 | -2.66 | 2.66 | -0.13 | 0.13 |
| gene7165 | https://www.ncbi.nlm.nih.gov/gene/?term=LOC105280327 | 2688 | 109 | 405.51 | 1 | 3.72 | 0.92 | 26.1 | 29.06 | -2.96 | 2.96 | -0.12 | 0.12 |
| gene12961 | https://www.ncbi.nlm.nih.gov/gene/?term=LOC105286983 | 3094 | 69 | 223.01 | 2 | 6.46 | 2.9 | 0.97 | 5.84 | -4.87 | 4.87 | -0.11 | 0.11 |
| gene3817 | https://www.ncbi.nlm.nih.gov/gene/?term=LOC105276449 | 1230 | 57 | 463.41 | 1 | 8.13 | 1.75 | 31.59 | 33.84 | -2.25 | 2.25 | -0.1 | 0.1 |
| gene920 | https://www.ncbi.nlm.nih.gov/gene/?term=LOC105282518 | 1197 | 42 | 350.88 | 1 | 8.35 | 2.38 | 38.3 | 35.67 | 2.62 | 2.62 | 0.09 | 0.09 |
| gene5525 | https://www.ncbi.nlm.nih.gov/gene/?term=LOC105278396 | 3090 | 190 | 614.89 | 1 | 3.24 | 0.53 | 25.02 | 26.4 | -1.38 | 1.38 | -0.08 | 0.08 |
| gene10864 | https://www.ncbi.nlm.nih.gov/gene/?term=LOC105284557 | 1666 | 84 | 504.2 | 1 | 6 | 1.19 | 28.16 | 29.73 | -1.56 | 1.56 | -0.08 | 0.08 |
| gene3200 | https://www.ncbi.nlm.nih.gov/gene/?term=LOC105275743 | 1786 | 76 | 425.53 | 1 | 5.6 | 1.32 | 23.84 | 22.24 | 1.6 | 1.6 | 0.07 | 0.07 |
| gene4498 | https://www.ncbi.nlm.nih.gov/gene/?term=LOC105277239 | 2831 | 141 | 498.06 | 1 | 3.53 | 0.71 | 15.21 | 13.89 | 1.32 | 1.32 | 0.07 | 0.07 |
| gene12572 | https://www.ncbi.nlm.nih.gov/gene/?term=LOC105286514 | 2374 | 47 | 197.98 | 1 | 4.21 | 2.13 | 34.72 | 38.06 | -3.34 | 3.34 | -0.07 | 0.07 |
| gene11619 | https://www.ncbi.nlm.nih.gov/gene/?term=LOC105285454 | 1649 | 29 | 175.86 | 1 | 6.06 | 3.45 | 26.22 | 30.26 | -4.03 | 4.03 | -0.07 | 0.07 |
| gene10816 | https://www.ncbi.nlm.nih.gov/gene/?term=LOC105284502 | 1974 | 21 | 106.38 | 1 | 5.07 | 4.76 | 45.81 | 40.14 | 5.67 | 5.67 | 0.06 | 0.06 |
| gene9101 | https://www.ncbi.nlm.nih.gov/gene/?term=LOC105282553 | 3068 | 115 | 374.84 | 1 | 3.26 | 0.87 | 23.82 | 22.25 | 1.57 | 1.57 | 0.06 | 0.06 |
| gene12802 | https://www.ncbi.nlm.nih.gov/gene/?term=LOC105286799 | 3183 | 193 | 606.35 | 2 | 6.28 | 1.04 | 15.46 | 16.4 | -0.94 | 0.94 | -0.06 | 0.06 |
| gene4034 | https://www.ncbi.nlm.nih.gov/gene/?term=LOC105276704 | 2065 | 106 | 513.32 | 2 | 9.69 | 1.89 | 12.65 | 13.91 | -1.26 | 1.26 | -0.06 | 0.06 |
| gene2907 | https://www.ncbi.nlm.nih.gov/gene/?term=LOC105275447 | 2539 | 43 | 169.36 | 1 | 3.94 | 2.33 | 3.37 | 7.11 | -3.74 | 3.74 | -0.06 | 0.06 |
| gene1199 | https://www.ncbi.nlm.nih.gov/gene/?term=LOC105285498 | 2220 | 26 | 117.12 | 1 | 4.5 | 3.85 | 37.03 | 32.91 | 4.12 | 4.12 | 0.05 | 0.05 |
| gene6652 | https://www.ncbi.nlm.nih.gov/gene/?term=LOC105279719 | 2831 | 82 | 289.65 | 1 | 3.53 | 1.22 | 24.05 | 22.34 | 1.72 | 1.72 | 0.05 | 0.05 |
| gene10714 | https://www.ncbi.nlm.nih.gov/gene/?term=LOC105284396 | 1966 | 66 | 335.71 | 1 | 5.09 | 1.52 | 34.65 | 33.2 | 1.45 | 1.45 | 0.05 | 0.05 |

...

... only first 25 lines shown ...

Directory: OGLall File: rnk-dmg-Cb_LI_BCphaseB2.vs.RphaseB1.txt

| gene_ID | gene_link | gwidth | #Sites | #per10Kb | #dmSites | #dmsp10kb | %dmSites | %pSite1 | %pSite2 | DMpSite | ADMpSite | DMpNucl | ADMpNucl |
|---|---|---|---|---|---|---|---|---|---|---|---|---|---|
| gene1470 | https://www.ncbi.nlm.nih.gov/gene/?term=LOC105287597 | 2974 | 190 | 638.87 | 1 | 3.36 | 0.53 | 18.6 | 19.2 | -0.6 | 0.6 | -0.04 | 0.04 |
| gene5877 | https://www.ncbi.nlm.nih.gov/gene/?term=LOC105278807 | 1719 | 50 | 290.87 | 1 | 5.82 | 2 | 23.49 | 23.85 | -0.36 | 0.36 | -0.01 | 0.01 |

Directory: OGLall File: rnk-dmg-Cb_LI_BCphaseB2.vs.RphaseB2.txt

```
gene_ID     gene_link   gwidth      #Sites      #per10Kb    #dmSites    #dmsp10kb   %dmSites    %pSite1     %pSite2     DMpSite     ADMpSite    DMpNucl     ADMpNucl
gene558     https://www.ncbi.nlm.nih.gov/gene/?term=LOC105278458 216   3     138.89      1     46.3  33.33 10.44 29.54 -19.1 19.1  -0.27 0.27
gene10219   https://www.ncbi.nlm.nih.gov/gene/?term=LOC105283830 235   4     170.21      1     42.55 25    29.84 43.16 -13.31      13.31 -0.23 0.23
gene11399   https://www.ncbi.nlm.nih.gov/gene/?term=LOC105285203 1029  28    272.11      2     19.44 7.14  30.21 24.31 5.9   5.9   0.16  0.16
gene3334    https://www.ncbi.nlm.nih.gov/gene/?term=LOC105275897 1689  72    426.29      2     11.84 2.78  36.11 39.55 -3.44 3.44  -0.15 0.15
gene10278   https://www.ncbi.nlm.nih.gov/gene/?term=LOC105283908 902   14    155.21      4     44.35 28.57 48.14 38.93 9.21  9.21  0.14  0.14
gene5987    https://www.ncbi.nlm.nih.gov/gene/?term=LOC105278938 1882  110   584.48      1     5.31  0.91  21.13 23.32 -2.19 2.19  -0.13 0.13
gene1562    https://www.ncbi.nlm.nih.gov/gene/?term=LOC105287692 1491  89    596.91      4     26.83 4.49  31.38 29.32 2.06  2.06  0.12  0.12
gene3910    https://www.ncbi.nlm.nih.gov/gene/?term=LOC105276563 1957  112   572.3 6     30.66 5.36  19.61 17.59 2.02  2.02  0.12  0.12
gene762     https://www.ncbi.nlm.nih.gov/gene/?term=LOC105280710 1173  60    511.51      2     17.05 3.33  23.52 25.93 -2.41 2.41  -0.12 0.12
gene5191    https://www.ncbi.nlm.nih.gov/gene/?term=LOC105278012 411   6     145.99      1     24.33 16.67 44.15 52.1  -7.95 7.95  -0.12 0.12
gene9965    https://www.ncbi.nlm.nih.gov/gene/?term=LOC105283552 1563  37    236.72      1     6.4   2.7   55.71 50.95 4.76  4.76  0.11  0.11
gene5809    https://www.ncbi.nlm.nih.gov/gene/?term=LOC105278737 1159  91    785.16      1     8.63  1.1   17.55 16.13 1.42  1.42  0.11  0.11
gene12660   https://www.ncbi.nlm.nih.gov/gene/?term=LOC105286633 1293  47    363.5 2     15.47 4.26  13.29 16.18 -2.89 2.89  -0.11 0.11
gene9019    https://www.ncbi.nlm.nih.gov/gene/?term=LOC105282481 1743  57    327.02      1     5.74  1.75  34.58 37.81 -3.22 3.22  -0.11 0.11
gene10743   https://www.ncbi.nlm.nih.gov/gene/?term=LOC105284456 1028  30    291.83      2     19.46 6.67  47.46 51.1  -3.64 3.64  -0.11 0.11
gene10644   https://www.ncbi.nlm.nih.gov/gene/?term=LOC105284320 358   9     251.4 1     27.93 11.11 24.22 28.77 -4.55 4.55  -0.11 0.11
gene12033   https://www.ncbi.nlm.nih.gov/gene/?term=LOC105285914 844   28    331.75      2     23.7  7.14  16.23 13.21 3.02  3.02  0.1   0.1
gene8463    https://www.ncbi.nlm.nih.gov/gene/?term=LOC105281837 2474  106   428.46      1     4.04  0.94  22.3  19.98 2.31  2.31  0.1   0.1
gene6075    https://www.ncbi.nlm.nih.gov/gene/?term=LOC105279049 1285  90    700.39      2     15.56 2.22  17.35 18.81 -1.46 1.46  -0.1  0.1
gene5412    https://www.ncbi.nlm.nih.gov/gene/?term=LOC105278266 1868  115   615.63      1     5.35  0.87  29.91 31.54 -1.63 1.63  -0.1  0.1
gene3268    https://www.ncbi.nlm.nih.gov/gene/?term=LOC105275833 1040  42    403.85      2     19.23 4.76  32.43 34.9  -2.47 2.47  -0.1  0.1
gene10584   https://www.ncbi.nlm.nih.gov/gene/?term=LOC105284253 1255  41    326.69      1     7.97  2.44  28.64 31.83 -3.18 3.18  -0.1  0.1
gene2481    https://www.ncbi.nlm.nih.gov/gene/?term=LOC105274938 1937  54    278.78      5     25.81 9.26  26.94 30.66 -3.73 3.73  -0.1  0.1
gene2709    https://www.ncbi.nlm.nih.gov/gene/?term=LOC105275219 999   18    180.18      1     10.01 5.56  37.14 42.54 -5.4  5.4   -0.1  0.1
...
... only first 25 lines shown ...
```

Directory: OGLall File: rnk-dmg-Cb_LI_RphaseA1.vs.RphaseA2.txt

| gene_ID | gene_link | gwidth | #Sites | #per10Kb | #dmSites | #dmsp10kb | %dmSites | %pSite1 | %pSite2 | DMpSite | ADMpSite | DMpNucl | ADMpNucl |
|---|---|---|---|---|---|---|---|---|---|---|---|---|---|
| gene10232 | https://www.ncbi.nlm.nih.gov/gene/?term=LOC105283847 | 1591 | 72 | 452.55 | 2 | 12.57 | 2.78 | 32.03 | 35.7 | -3.67 | 3.67 | -0.17 | 0.17 |
| gene12411 | https://www.ncbi.nlm.nih.gov/gene/?term=LOC105286320 | 1660 | 64 | 385.54 | 4 | 24.1 | 6.25 | 38.65 | 42.95 | -4.3 | 4.3 | -0.17 | 0.17 |
| gene3268 | https://www.ncbi.nlm.nih.gov/gene/?term=LOC105275833 | 1040 | 41 | 394.23 | 1 | 9.62 | 2.44 | 35.12 | 39.5 | -4.38 | 4.38 | -0.17 | 0.17 |
| gene3719 | https://www.ncbi.nlm.nih.gov/gene/?term=LOC105276360 | 1471 | 107 | 727.4 | 1 | 6.8 | 0.93 | 30.24 | 32.33 | -2.08 | 2.08 | -0.15 | 0.15 |
| gene12925 | https://www.ncbi.nlm.nih.gov/gene/?term=LOC109611487 | 391 | 27 | 690.54 | 1 | 25.58 | 3.7 | 15.18 | 17.34 | -2.15 | 2.15 | -0.15 | 0.15 |
| gene3803 | https://www.ncbi.nlm.nih.gov/gene/?term=LOC105276438 | 2613 | 162 | 619.98 | 2 | 7.65 | 1.23 | 19.62 | 21.97 | -2.35 | 2.35 | -0.15 | 0.15 |
| gene3910 | https://www.ncbi.nlm.nih.gov/gene/?term=LOC105276563 | 1957 | 118 | 602.96 | 7 | 35.77 | 5.93 | 16.25 | 18.68 | -2.43 | 2.43 | -0.15 | 0.15 |
| gene11566 | https://www.ncbi.nlm.nih.gov/gene/?term=LOC105285392 | 1603 | 91 | 567.69 | 4 | 24.95 | 4.4 | 31.49 | 34.19 | -2.7 | 2.7 | -0.15 | 0.15 |
| gene2837 | https://www.ncbi.nlm.nih.gov/gene/?term=LOC105275355 | 2872 | 161 | 560.58 | 8 | 27.86 | 4.97 | 27.89 | 25.3 | 2.58 | 2.58 | 0.14 | 0.14 |
| gene6781 | https://www.ncbi.nlm.nih.gov/gene/?term=LOC105279857 | 241 | 12 | 497.93 | 1 | 41.49 | 8.33 | 7.9 | 10.68 | -2.78 | 2.78 | -0.14 | 0.14 |
| gene5027 | https://www.ncbi.nlm.nih.gov/gene/?term=LOC105277833 | 905 | 61 | 674.03 | 1 | 11.05 | 1.64 | 2.47 | 4.36 | -1.89 | 1.89 | -0.13 | 0.13 |
| gene10176 | https://www.ncbi.nlm.nih.gov/gene/?term=LOC105283785 | 1637 | 42 | 256.57 | 3 | 18.33 | 7.14 | 12.82 | 18.05 | -5.23 | 5.23 | -0.13 | 0.13 |
| gene5828 | https://www.ncbi.nlm.nih.gov/gene/?term=LOC105278756 | 4586 | 413 | 900.57 | 3 | 6.54 | 0.73 | 9.45 | 10.77 | -1.33 | 1.33 | -0.12 | 0.12 |
| gene9183 | https://www.ncbi.nlm.nih.gov/gene/?term=LOC105282640 | 2752 | 206 | 748.55 | 3 | 10.9 | 1.46 | 21.54 | 23.12 | -1.58 | 1.58 | -0.12 | 0.12 |
| gene12652 | https://www.ncbi.nlm.nih.gov/gene/?term=LOC109610640 | 1122 | 45 | 401.07 | 4 | 35.65 | 8.89 | 47.28 | 50.3 | -3.02 | 3.02 | -0.12 | 0.12 |
| gene2274 | https://www.ncbi.nlm.nih.gov/gene/?term=LOC109610680 | 1467 | 31 | 211.32 | 2 | 13.63 | 6.45 | 65.47 | 71.37 | -5.89 | 5.89 | -0.12 | 0.12 |
| gene8798 | https://www.ncbi.nlm.nih.gov/gene/?term=LOC105282220 | 2300 | 232 | 1008.7 | 2 | 8.7 | 0.86 | 7.56 | 8.63 | -1.07 | 1.07 | -0.11 | 0.11 |
| gene12416 | https://www.ncbi.nlm.nih.gov/gene/?term=LOC105286327 | 3689 | 219 | 593.66 | 2 | 5.42 | 0.91 | 9 | 10.78 | -1.78 | 1.78 | -0.11 | 0.11 |
| gene5447 | https://www.ncbi.nlm.nih.gov/gene/?term=LOC105278314 | 1482 | 95 | 641.03 | 2 | 13.5 | 2.11 | 21.76 | 23.54 | -1.78 | 1.78 | -0.11 | 0.11 |
| gene3314 | https://www.ncbi.nlm.nih.gov/gene/?term=LOC105275878 | 2314 | 133 | 574.76 | 2 | 8.64 | 1.5 | 29.12 | 30.97 | -1.84 | 1.84 | -0.11 | 0.11 |
| gene10084 | https://www.ncbi.nlm.nih.gov/gene/?term=LOC105283683 | 1448 | 36 | 248.62 | 4 | 27.62 | 11.11 | 29.43 | 33.99 | -4.56 | 4.56 | -0.11 | 0.11 |
| gene8333 | https://www.ncbi.nlm.nih.gov/gene/?term=LOC105281691 | 1264 | 28 | 221.52 | 3 | 23.73 | 10.71 | 52.62 | 57.6 | -4.98 | 4.98 | -0.11 | 0.11 |
| gene1024 | https://www.ncbi.nlm.nih.gov/gene/?term=LOC105283693 | 1307 | 16 | 122.42 | 4 | 30.6 | 25 | 60.96 | 69.71 | -8.75 | 8.75 | -0.11 | 0.11 |
| gene13341 | https://www.ncbi.nlm.nih.gov/gene/?term=LOC105287399 | 534 | 6 | 112.36 | 1 | 18.73 | 16.67 | 23.22 | 33.07 | -9.85 | 9.85 | -0.11 | 0.11 |

...

... only first 25 lines shown ...

Directory: OGLall File: rnk-dmg-Cb_LI_RphaseA1.vs.RphaseB1.txt

| gene_ID | gene_link | gwidth | #Sites | #per10Kb | #dmSites | #dmsp10kb | %dmSites | %pSite1 | %pSite2 | DMpSite | ADMpSite | DMpNucl | ADMpNucl |
|---|---|---|---|---|---|---|---|---|---|---|---|---|---|
| gene2593 | https://www.ncbi.nlm.nih.gov/gene/?term=LOC105275070 | 1774 | 11 | 62.01 | 3 | 16.91 | 27.27 | 69.15 | 25.28 | 43.86 | 43.86 | 0.27 | 0.27 |
| gene2837 | https://www.ncbi.nlm.nih.gov/gene/?term=LOC105275355 | 2872 | 164 | 571.03 | 1 | 3.48 | 0.61 | 27.39 | 23.06 | 4.33 | 4.33 | 0.25 | 0.25 |
| gene3165 | https://www.ncbi.nlm.nih.gov/gene/?term=LOC105275706 | 2374 | 63 | 265.37 | 3 | 12.64 | 4.76 | 31.1 | 21.89 | 9.2 | 9.2 | 0.24 | 0.24 |
| gene2856 | https://www.ncbi.nlm.nih.gov/gene/?term=LOC105275384 | 1987 | 88 | 442.88 | 1 | 5.03 | 1.14 | 36.35 | 32.79 | 3.56 | 3.56 | 0.16 | 0.16 |
| gene5714 | https://www.ncbi.nlm.nih.gov/gene/?term=LOC105278729 | 710 | 16 | 225.35 | 1 | 14.08 | 6.25 | 55.81 | 51.9 | 3.91 | 3.91 | 0.09 | 0.09 |
| gene7550 | https://www.ncbi.nlm.nih.gov/gene/?term=LOC105280763 | 2173 | 101 | 464.8 | 1 | 4.6 | 0.99 | 26.35 | 28.2 | -1.85 | 1.85 | -0.09 | 0.09 |
| gene1855 | https://www.ncbi.nlm.nih.gov/gene/?term=LOC105288036 | 2387 | 90 | 377.04 | 1 | 4.19 | 1.11 | 28.76 | 26.75 | 2.02 | 2.02 | 0.08 | 0.08 |
| gene13033 | https://www.ncbi.nlm.nih.gov/gene/?term=LOC105287063 | 1877 | 49 | 261.05 | 1 | 5.33 | 2.04 | 32.74 | 30.05 | 2.68 | 2.68 | 0.07 | 0.07 |
| gene5877 | https://www.ncbi.nlm.nih.gov/gene/?term=LOC105278807 | 1719 | 51 | 296.68 | 1 | 5.82 | 1.96 | 27.19 | 24.91 | 2.27 | 2.27 | 0.07 | 0.07 |
| gene11235 | https://www.ncbi.nlm.nih.gov/gene/?term=LOC105285004 | 1339 | 38 | 283.79 | 1 | 7.47 | 2.63 | 16.34 | 18.94 | -2.6 | 2.6 | -0.07 | 0.07 |
| gene9478 | https://www.ncbi.nlm.nih.gov/gene/?term=LOC105282991 | 3726 | 58 | 155.66 | 1 | 2.68 | 1.72 | 31.92 | 28.27 | 3.65 | 3.65 | 0.06 | 0.06 |
| gene1198 | https://www.ncbi.nlm.nih.gov/gene/?term=LOC105285507 | 2738 | 98 | 357.93 | 1 | 3.65 | 1.02 | 19.77 | 21.55 | -1.78 | 1.78 | -0.06 | 0.06 |
| gene12406 | https://www.ncbi.nlm.nih.gov/gene/?term=LOC105286316 | 1386 | 35 | 252.53 | 1 | 7.22 | 2.86 | 40.6 | 43.17 | -2.57 | 2.57 | -0.06 | 0.06 |
| gene12021 | https://www.ncbi.nlm.nih.gov/gene/?term=LOC105285905 | 1399 | 30 | 214.44 | 1 | 7.15 | 3.33 | 30.6 | 33.22 | -2.62 | 2.62 | -0.06 | 0.06 |
| gene388 | https://www.ncbi.nlm.nih.gov/gene/?term=LOC105276538 | 1558 | 22 | 141.21 | 2 | 12.84 | 9.09 | 54.75 | 58.78 | -4.03 | 4.03 | -0.06 | 0.06 |
| gene3671 | https://www.ncbi.nlm.nih.gov/gene/?term=LOC105276272 | 1966 | 44 | 223.8 | 1 | 5.09 | 2.27 | 27.22 | 24.96 | 2.26 | 2.26 | 0.05 | 0.05 |
| gene11819 | https://www.ncbi.nlm.nih.gov/gene/?term=LOC105285671 | 2366 | 100 | 422.65 | 1 | 4.23 | 1 | 31.41 | 30.12 | 1.29 | 1.29 | 0.05 | 0.05 |
| gene4641 | https://www.ncbi.nlm.nih.gov/gene/?term=LOC105277398 | 1937 | 73 | 376.87 | 1 | 5.16 | 1.37 | 36.14 | 34.87 | 1.27 | 1.27 | 0.05 | 0.05 |
| gene298 | https://www.ncbi.nlm.nih.gov/gene/?term=LOC105275460 | 2473 | 126 | 509.5 | 1 | 4.04 | 0.79 | 13.41 | 12.43 | 0.98 | 0.98 | 0.05 | 0.05 |
| gene1470 | https://www.ncbi.nlm.nih.gov/gene/?term=LOC105287597 | 2974 | 182 | 611.97 | 1 | 3.36 | 0.55 | 19.45 | 20.26 | -0.81 | 0.81 | -0.05 | 0.05 |
| gene9101 | https://www.ncbi.nlm.nih.gov/gene/?term=LOC105282553 | 3068 | 106 | 345.5 | 1 | 3.26 | 0.94 | 21.02 | 22.34 | -1.33 | 1.33 | -0.05 | 0.05 |
| gene11340 | https://www.ncbi.nlm.nih.gov/gene/?term=LOC105285131 | 2918 | 98 | 335.85 | 1 | 3.43 | 1.02 | 13.77 | 15.33 | -1.56 | 1.56 | -0.05 | 0.05 |
| gene2413 | https://www.ncbi.nlm.nih.gov/gene/?term=LOC105274863 | 2006 | 60 | 299.1 | 1 | 4.99 | 1.67 | 50.89 | 52.54 | -1.65 | 1.65 | -0.05 | 0.05 |
| gene7199 | https://www.ncbi.nlm.nih.gov/gene/?term=LOC105280366 | 1414 | 37 | 261.67 | 1 | 7.07 | 2.7 | 44.35 | 46.38 | -2.03 | 2.03 | -0.05 | 0.05 |
| gene12516 | https://www.ncbi.nlm.nih.gov/gene/?term=LOC105286452 | 1529 | 38 | 248.53 | 1 | 6.54 | 2.63 | 49.76 | 48.05 | 1.71 | 1.71 | 0.04 | 0.04 |
| gene3817 | https://www.ncbi.nlm.nih.gov/gene/?term=LOC105276449 | 1230 | 48 | 390.24 | 1 | 8.13 | 2.08 | 25.64 | 24.6 | 1.04 | 1.04 | 0.04 | 0.04 |
| gene9740 | https://www.ncbi.nlm.nih.gov/gene/?term=LOC105283277 | 2887 | 138 | 478 | 1 | 3.46 | 0.72 | 17.69 | 18.47 | -0.78 | 0.78 | -0.04 | 0.04 |
| gene8729 | https://www.ncbi.nlm.nih.gov/gene/?term=LOC105282137 | 5005 | 225 | 449.55 | 1 | 2 | 0.44 | 13.97 | 14.84 | -0.86 | 0.86 | -0.04 | 0.04 |
| gene570 | https://www.ncbi.nlm.nih.gov/gene/?term=LOC105278549 | 1673 | 70 | 418.41 | 1 | 5.98 | 1.43 | 42.95 | 43.99 | -1.04 | 1.04 | -0.04 | 0.04 |
| gene920 | https://www.ncbi.nlm.nih.gov/gene/?term=LOC105282518 | 1197 | 44 | 367.59 | 1 | 8.35 | 2.27 | 32.92 | 33.98 | -1.06 | 1.06 | -0.04 | 0.04 |
| gene4962 | https://www.ncbi.nlm.nih.gov/gene/?term=LOC105277761 | 4645 | 160 | 344.46 | 1 | 2.15 | 0.62 | 12.8 | 13.96 | -1.16 | 1.16 | -0.04 | 0.04 |
| gene3335 | https://www.ncbi.nlm.nih.gov/gene/?term=LOC105275898 | 4665 | 157 | 336.55 | 1 | 2.14 | 0.64 | 12.01 | 13.22 | -1.21 | 1.21 | -0.04 | 0.04 |
| gene10392 | https://www.ncbi.nlm.nih.gov/gene/?term=LOC105284036 | 3216 | 98 | 304.73 | 1 | 3.11 | 1.02 | 22.15 | 23.39 | -1.25 | 1.25 | -0.04 | 0.04 |
| gene1310 | https://www.ncbi.nlm.nih.gov/gene/?term=LOC105286661 | 2582 | 67 | 259.49 | 1 | 3.87 | 1.49 | 17.48 | 18.89 | -1.41 | 1.41 | -0.04 | 0.04 |
| gene2404 | https://www.ncbi.nlm.nih.gov/gene/?term=LOC105274853 | 1767 | 42 | 237.69 | 1 | 5.66 | 2.38 | 30.33 | 32.12 | -1.8 | 1.8 | -0.04 | 0.04 |
| gene12572 | https://www.ncbi.nlm.nih.gov/gene/?term=LOC105286514 | 2374 | 51 | 214.83 | 1 | 4.21 | 1.96 | 34.55 | 36.64 | -2.09 | 2.09 | -0.04 | 0.04 |
| gene11584 | https://www.ncbi.nlm.nih.gov/gene/?term=LOC105285416 | 3709 | 13 | 35.05 | 2 | 5.39 | 15.38 | 23.21 | 33.23 | -10.02 | 10.02 | -0.04 | 0.04 |
| gene393 | https://www.ncbi.nlm.nih.gov/gene/?term=LOC105276566 | 1123 | 2 | 17.81 | 1 | 8.9 | 50 | 50 | 74.3 | -24.3 | 24.3 | -0.04 | 0.04 |
| gene9189 | https://www.ncbi.nlm.nih.gov/gene/?term=LOC105282652 | 4699 | 84 | 178.76 | 1 | 2.13 | 1.19 | 16.38 | 14.61 | 1.78 | 1.78 | 0.03 | 0.03 |
| gene4023 | https://www.ncbi.nlm.nih.gov/gene/?term=LOC105276692 | 1759 | 36 | 204.66 | 1 | 5.69 | 2.78 | 23.9 | 22.4 | 1.5 | 1.5 | 0.03 | 0.03 |
| gene1884 | https://www.ncbi.nlm.nih.gov/gene/?term=LOC105288072 | 4169 | 82 | 196.69 | 1 | 2.4 | 1.22 | 24.39 | 23.08 | 1.31 | 1.31 | 0.03 | 0.03 |
| gene10091 | https://www.ncbi.nlm.nih.gov/gene/?term=LOC105283697 | 4988 | 154 | 308.74 | 1 | 2 | 0.65 | 16.48 | 15.36 | 1.12 | 1.12 | 0.03 | 0.03 |
| gene5831 | https://www.ncbi.nlm.nih.gov/gene/?term=LOC105278761 | 2506 | 115 | 458.9 | 2 | 7.98 | 1.74 | 22.22 | 21.51 | 0.71 | 0.71 | 0.03 | 0.03 |
| gene10540 | https://www.ncbi.nlm.nih.gov/gene/?term=LOC105284200 | 5267 | 209 | 396.81 | 1 | 1.9 | 0.48 | 13.12 | 12.47 | 0.65 | 0.65 | 0.03 | 0.03 |
| gene5298 | https://www.ncbi.nlm.nih.gov/gene/?term=LOC105278145 | 4060 | 253 | 623.15 | 3 | 7.39 | 1.19 | 10.61 | 11.09 | -0.49 | 0.49 | -0.03 | 0.03 |
| gene10326 | https://www.ncbi.nlm.nih.gov/gene/?term=LOC105283962 | 3042 | 132 | 433.93 | 1 | 3.29 | 0.76 | 8.06 | 8.65 | -0.59 | 0.59 | -0.03 | 0.03 |
| gene7166 | https://www.ncbi.nlm.nih.gov/gene/?term=LOC105280326 | 6117 | 249 | 407.06 | 1 | 1.63 | 0.4 | 14.56 | 15.24 | -0.68 | 0.68 | -0.03 | 0.03 |
| gene4190 | https://www.ncbi.nlm.nih.gov/gene/?term=LOC105276875 | 3150 | 125 | 396.83 | 1 | 3.17 | 0.8 | 19.56 | 20.35 | -0.79 | 0.79 | -0.03 | 0.03 |
| gene6652 | https://www.ncbi.nlm.nih.gov/gene/?term=LOC105279719 | 2831 | 87 | 307.31 | 1 | 3.53 | 1.15 | 20.49 | 21.33 | -0.83 | 0.83 | -0.03 | 0.03 |
| gene7915 | https://www.ncbi.nlm.nih.gov/gene/?term=LOC105281173 | 1597 | 38 | 237.95 | 1 | 6.26 | 2.63 | 59.81 | 61.01 | -1.2 | 1.2 | -0.03 | 0.03 |
| gene3599 | https://www.ncbi.nlm.nih.gov/gene/?term=LOC105276196 | 2955 | 57 | 192.89 | 1 | 3.38 | 1.75 | 39.57 | 40.89 | -1.32 | 1.32 | -0.03 | 0.03 |
| gene12203 | https://www.ncbi.nlm.nih.gov/gene/?term=LOC105286099 | 5967 | 80 | 134.07 | 1 | 1.68 | 1.25 | 10.4 | 12.39 | -2 | 2 | -0.03 | 0.03 |
| gene3028 | https://www.ncbi.nlm.nih.gov/gene/?term=LOC105275571 | 1924 | 30 | 155.93 | 1 | 5.2 | 3.33 | 42.66 | 44.75 | -2.09 | 2.09 | -0.03 | 0.03 |
| gene3168 | https://www.ncbi.nlm.nih.gov/gene/?term=LOC105275711 | 6150 | 87 | 141.46 | 1 | 1.63 | 1.15 | 10.39 | 12.71 | -2.33 | 2.33 | -0.03 | 0.03 |
| gene9411 | https://www.ncbi.nlm.nih.gov/gene/?term=LOC105282914 | 5534 | 42 | 75.89 | 1 | 1.81 | 2.38 | 47.45 | 50.92 | -3.47 | 3.47 | -0.03 | 0.03 |
| gene927 | https://www.ncbi.nlm.nih.gov/gene/?term=LOC105282600 | 3727 | 60 | 160.99 | 1 | 2.68 | 1.67 | 27.81 | 26.32 | 1.49 | 1.49 | 0.02 | 0.02 |
| gene9336 | https://www.ncbi.nlm.nih.gov/gene/?term=LOC105282834 | 5993 | 134 | 223.59 | 1 | 1.67 | 0.75 | 14.48 | 13.37 | 1.1 | 1.1 | 0.02 | 0.02 |
| gene940 | https://www.ncbi.nlm.nih.gov/gene/?term=LOC105282794 | 3578 | 93 | 259.92 | 1 | 2.79 | 1.08 | 14.3 | 13.63 | 0.67 | 0.67 | 0.02 | 0.02 |
| gene4892 | https://www.ncbi.nlm.nih.gov/gene/?term=LOC105277667 | 3898 | 121 | 310.42 | 1 | 2.57 | 0.83 | 29.02 | 28.4 | 0.62 | 0.62 | 0.02 | 0.02 |
| gene13036 | https://www.ncbi.nlm.nih.gov/gene/?term=LOC105287059 | 4959 | 180 | 362.98 | 1 | 2.02 | 0.56 | 17.65 | 17.23 | 0.42 | 0.42 | 0.02 | 0.02 |

| | | | | | | | | | | | | | |
|---|---|---|---|---|---|---|---|---|---|---|---|---|---|
| gene12441 | https://www.ncbi.nlm.nih.gov/gene/?term=LOC105286355 | 3252 | 149 | 458.18 | 1 | 3.08 | 0.67 | 21.45 | 21.08 | 0.38 | 0.38 | 0.02 | 0.02 |
| gene2844 | https://www.ncbi.nlm.nih.gov/gene/?term=LOC105275363 | 4330 | 218 | 503.46 | 1 | 2.31 | 0.46 | 13.23 | 12.92 | 0.3 | 0.3 | 0.02 | 0.02 |
| gene12802 | https://www.ncbi.nlm.nih.gov/gene/?term=LOC105286799 | 3183 | 178 | 559.22 | 1 | 3.14 | 0.56 | 16.8 | 17.1 | -0.29 | 0.29 | -0.02 | 0.02 |
| gene5525 | https://www.ncbi.nlm.nih.gov/gene/?term=LOC105278396 | 3090 | 186 | 601.94 | 1 | 3.24 | 0.54 | 24.12 | 24.45 | -0.33 | 0.33 | -0.02 | 0.02 |
| gene5524 | https://www.ncbi.nlm.nih.gov/gene/?term=LOC105278397 | 5177 | 205 | 395.98 | 1 | 1.93 | 0.49 | 24.01 | 24.4 | -0.39 | 0.39 | -0.02 | 0.02 |
| gene6583 | https://www.ncbi.nlm.nih.gov/gene/?term=LOC105279649 | 7444 | 364 | 488.98 | 1 | 1.34 | 0.27 | 10.46 | 10.89 | -0.43 | 0.43 | -0.02 | 0.02 |
| gene10678 | https://www.ncbi.nlm.nih.gov/gene/?term=LOC105284355 | 4031 | 173 | 429.17 | 1 | 2.48 | 0.58 | 18.28 | 18.72 | -0.44 | 0.44 | -0.02 | 0.02 |
| gene3107 | https://www.ncbi.nlm.nih.gov/gene/?term=LOC105275650 | 5103 | 180 | 352.73 | 1 | 1.96 | 0.56 | 15.71 | 16.3 | -0.59 | 0.59 | -0.02 | 0.02 |
| gene5646 | https://www.ncbi.nlm.nih.gov/gene/?term=LOC105278534 | 7475 | 198 | 264.88 | 1 | 1.34 | 0.51 | 17.1 | 17.7 | -0.6 | 0.6 | -0.02 | 0.02 |
| gene10844 | https://www.ncbi.nlm.nih.gov/gene/?term=LOC105284535 | 9086 | 258 | 283.95 | 1 | 1.1 | 0.39 | 13.99 | 14.72 | -0.73 | 0.73 | -0.02 | 0.02 |
| gene11454 | https://www.ncbi.nlm.nih.gov/gene/?term=LOC105285256 | 1906 | 50 | 262.33 | 1 | 5.25 | 2 | 43.22 | 44.08 | -0.87 | 0.87 | -0.02 | 0.02 |
| gene6422 | https://www.ncbi.nlm.nih.gov/gene/?term=LOC105279458 | 2419 | 43 | 177.76 | 1 | 4.13 | 2.33 | 44.62 | 45.68 | -1.05 | 1.05 | -0.02 | 0.02 |
| gene2173 | https://www.ncbi.nlm.nih.gov/gene/?term=LOC105274590 | 4387 | 84 | 191.47 | 1 | 2.28 | 1.19 | 23.89 | 24.96 | -1.07 | 1.07 | -0.02 | 0.02 |
| gene7248 | https://www.ncbi.nlm.nih.gov/gene/?term=LOC105280408 | 5175 | 60 | 115.94 | 1 | 1.93 | 1.67 | 15.92 | 17.41 | -1.49 | 1.49 | -0.02 | 0.02 |
| gene1575 | https://www.ncbi.nlm.nih.gov/gene/?term=LOC105287706 | 3783 | 44 | 116.31 | 1 | 2.64 | 2.27 | 12.72 | 14.57 | -1.85 | 1.85 | -0.02 | 0.02 |
| gene7398 | https://www.ncbi.nlm.nih.gov/gene/?term=LOC105280589 | 3452 | 41 | 118.77 | 1 | 2.9 | 2.44 | 24.33 | 26.31 | -1.98 | 1.98 | -0.02 | 0.02 |
| gene564 | https://www.ncbi.nlm.nih.gov/gene/?term=LOC105278478 | 2802 | 17 | 60.67 1 | 3.57 | 5.88 | 15.85 | 19.36 | -3.51 | 3.51 | -0.02 | 0.02 | |
| gene8178 | https://www.ncbi.nlm.nih.gov/gene/?term=LOC105281515 | 8647 | 60 | 69.39 1 | 1.16 | 1.67 | 29.73 | 28.04 | 1.69 | 1.69 | 0.01 | 0.01 | |
| gene2683 | https://www.ncbi.nlm.nih.gov/gene/?term=LOC105275187 | 5492 | 73 | 132.92 | 1 | 1.82 | 1.37 | 14.7 | 13.62 | 1.08 | 1.08 | 0.01 | 0.01 |
| gene652 | https://www.ncbi.nlm.nih.gov/gene/?term=LOC105279528 | 5806 | 39 | 67.17 1 | 1.72 | 2.56 | 22.38 | 21.58 | 0.8 | 0.8 | 0.01 | 0.01 | |
| gene10107 | https://www.ncbi.nlm.nih.gov/gene/?term=LOC105283712 | 1883 | 36 | 191.18 | 1 | 5.31 | 2.78 | 27.63 | 26.89 | 0.74 | 0.74 | 0.01 | 0.01 |
| gene7600 | https://www.ncbi.nlm.nih.gov/gene/?term=LOC105280819 | 1615 | 41 | 253.87 | 1 | 6.19 | 2.44 | 39.36 | 38.88 | 0.48 | 0.48 | 0.01 | 0.01 |
| gene1147 | https://www.ncbi.nlm.nih.gov/gene/?term=LOC105284954 | 4022 | 118 | 293.39 | 1 | 2.49 | 0.85 | 11.22 | 10.78 | 0.44 | 0.44 | 0.01 | 0.01 |
| gene10222 | https://www.ncbi.nlm.nih.gov/gene/?term=LOC105283838 | 1748 | 30 | 171.62 | 1 | 5.72 | 3.33 | 33.06 | 32.69 | 0.37 | 0.37 | 0.01 | 0.01 |
| gene5101 | https://www.ncbi.nlm.nih.gov/gene/?term=LOC105277903 | 1813 | 36 | 198.57 | 1 | 5.52 | 2.78 | 56.09 | 55.74 | 0.35 | 0.35 | 0.01 | 0.01 |
| gene7247 | https://www.ncbi.nlm.nih.gov/gene/?term=LOC105280407 | 10486 | 226 | 215.53 | 1 | 0.95 | 0.44 | 15.61 | 15.31 | 0.3 | 0.3 | 0.01 | 0.01 |
| gene10520 | https://www.ncbi.nlm.nih.gov/gene/?term=LOC105284184 | 2451 | 92 | 375.36 | 1 | 4.08 | 1.09 | 31.61 | 31.4 | 0.21 | 0.21 | 0.01 | 0.01 |
| gene2846 | https://www.ncbi.nlm.nih.gov/gene/?term=LOC105275374 | 11246 | 493 | 438.38 | 1 | 0.89 | 0.2 | 9.9 | 9.75 | 0.16 | 0.16 | 0.01 | 0.01 |
| gene12978 | https://www.ncbi.nlm.nih.gov/gene/?term=LOC105286997 | 2858 | 142 | 496.85 | 1 | 3.5 | 0.7 | 11.15 | 11.02 | 0.14 | 0.14 | 0.01 | 0.01 |
| gene2271 | https://www.ncbi.nlm.nih.gov/gene/?term=LOC105274706 | 1448 | 51 | 352.21 | 1 | 6.91 | 1.96 | 37.35 | 37.52 | -0.17 | 0.17 | -0.01 | 0.01 |
| gene3885 | https://www.ncbi.nlm.nih.gov/gene/?term=LOC105276527 | 7928 | 336 | 423.81 | 1 | 1.26 | 0.3 | 8.31 | 8.48 | -0.17 | 0.17 | -0.01 | 0.01 |
| gene8274 | https://www.ncbi.nlm.nih.gov/gene/?term=LOC105281624 | 5425 | 216 | 398.16 | 1 | 1.84 | 0.46 | 9.81 | 9.99 | -0.18 | 0.18 | -0.01 | 0.01 |
| gene7547 | https://www.ncbi.nlm.nih.gov/gene/?term=LOC105280767 | 24576 | 808 | 328.78 | 1 | 0.41 | 0.12 | 4.37 | 4.58 | -0.21 | 0.21 | -0.01 | 0.01 |
| gene8814 | https://www.ncbi.nlm.nih.gov/gene/?term=LOC105282240 | 6910 | 188 | 272.07 | 1 | 1.45 | 0.53 | 8.71 | 8.93 | -0.22 | 0.22 | -0.01 | 0.01 |
| gene10018 | https://www.ncbi.nlm.nih.gov/gene/?term=LOC105283607 | 4414 | 219 | 496.15 | 1 | 2.27 | 0.46 | 13.97 | 14.21 | -0.23 | 0.23 | -0.01 | 0.01 |
| gene13114 | https://www.ncbi.nlm.nih.gov/gene/?term=LOC105287147 | 7609 | 415 | 545.41 | 1 | 1.31 | 0.24 | 8.59 | 8.85 | -0.26 | 0.26 | -0.01 | 0.01 |
| gene6286 | https://www.ncbi.nlm.nih.gov/gene/?term=LOC105279298 | 2861 | 125 | 436.91 | 2 | 6.99 | 1.6 | 23.63 | 23.91 | -0.29 | 0.29 | -0.01 | 0.01 |
| gene10549 | https://www.ncbi.nlm.nih.gov/gene/?term=LOC105284210 | 7145 | 348 | 487.05 | 2 | 2.8 | 0.57 | 9.98 | 10.29 | -0.31 | 0.31 | -0.01 | 0.01 |
| gene9330 | https://www.ncbi.nlm.nih.gov/gene/?term=LOC105282825 | 3936 | 96 | 243.9 1 | 2.54 | 1.04 | 14.55 | 14.91 | -0.36 | 0.36 | -0.01 | 0.01 | |
| gene4376 | https://www.ncbi.nlm.nih.gov/gene/?term=LOC105277083 | 4307 | 120 | 278.62 | 1 | 2.32 | 0.83 | 19.12 | 19.51 | -0.39 | 0.39 | -0.01 | 0.01 |
| gene6031 | https://www.ncbi.nlm.nih.gov/gene/?term=LOC105279000 | 5691 | 210 | 369 1 | 1.76 | 0.48 | 11.92 | 12.31 | -0.4 | 0.4 | -0.01 | 0.01 | |
| gene12307 | https://www.ncbi.nlm.nih.gov/gene/?term=LOC105286216 | 10313 | 186 | 180.35 | 1 | 0.97 | 0.54 | 14.88 | 15.35 | -0.47 | 0.47 | -0.01 | 0.01 |
| gene9647 | https://www.ncbi.nlm.nih.gov/gene/?term=LOC109611233 | 3094 | 33 | 106.66 | 1 | 3.23 | 3.03 | 41.64 | 42.16 | -0.52 | 0.52 | -0.01 | 0.01 |
| gene9648 | https://www.ncbi.nlm.nih.gov/gene/?term=LOC105283183 | 2602 | 31 | 119.14 | 1 | 3.84 | 3.23 | 44.33 | 44.88 | -0.55 | 0.55 | -0.01 | 0.01 |
| gene6237 | https://www.ncbi.nlm.nih.gov/gene/?term=LOC105279244 | 1484 | 25 | 168.46 | 1 | 6.74 | 4 | 45.17 | 45.86 | -0.69 | 0.69 | -0.01 | 0.01 |
| gene10155 | https://www.ncbi.nlm.nih.gov/gene/?term=LOC105283756 | 7030 | 112 | 159.32 | 1 | 1.42 | 0.89 | 7.35 | 8.24 | -0.89 | 0.89 | -0.01 | 0.01 |
| gene6639 | https://www.ncbi.nlm.nih.gov/gene/?term=LOC105279705 | 7656 | 81 | 105.8 1 | 1.31 | 1.23 | 19.09 | 20.33 | -1.25 | 1.25 | -0.01 | 0.01 | |
| gene10243 | https://www.ncbi.nlm.nih.gov/gene/?term=LOC105283862 | 3239 | 43 | 132.76 | 1 | 3.09 | 2.33 | 29.28 | 28.91 | 0.37 | 0.37 | 0 | 0 |
| gene10654 | https://www.ncbi.nlm.nih.gov/gene/?term=LOC105284329 | 4027 | 71 | 176.31 | 1 | 2.48 | 1.41 | 19.25 | 18.99 | 0.26 | 0.26 | 0 | 0 |
| gene6455 | https://www.ncbi.nlm.nih.gov/gene/?term=LOC105279500 | 2612 | 31 | 118.68 | 1 | 3.83 | 3.23 | 47.76 | 47.54 | 0.22 | 0.22 | 0 | 0 |
| gene10345 | https://www.ncbi.nlm.nih.gov/gene/?term=LOC105283989 | 4444 | 29 | 65.26 1 | 2.25 | 3.45 | 49.14 | 48.92 | 0.21 | 0.21 | 0 | 0 | |
| gene7183 | https://www.ncbi.nlm.nih.gov/gene/?term=LOC105280348 | 3925 | 79 | 201.27 | 1 | 2.55 | 1.27 | 17.02 | 16.83 | 0.19 | 0.19 | 0 | 0 |
| gene2668 | https://www.ncbi.nlm.nih.gov/gene/?term=LOC105275170 | 12496 | 348 | 278.49 | 1 | 0.8 | 0.29 | 1.95 | 1.78 | 0.18 | 0.18 | 0 | 0 |
| gene9327 | https://www.ncbi.nlm.nih.gov/gene/?term=LOC105282821 | 7793 | 162 | 207.88 | 1 | 1.28 | 0.62 | 13.28 | 13.13 | 0.16 | 0.16 | 0 | 0 |
| gene7556 | https://www.ncbi.nlm.nih.gov/gene/?term=LOC105280774 | 2792 | 132 | 472.78 | 1 | 3.58 | 0.76 | 25.2 | 25.1 | 0.1 | 0.1 | 0 | 0 |
| gene10606 | https://www.ncbi.nlm.nih.gov/gene/?term=LOC105284273 | 6119 | 261 | 426.54 | 1 | 1.63 | 0.38 | 19.09 | 19 | 0.08 | 0.08 | 0 | 0 |
| gene8600 | https://www.ncbi.nlm.nih.gov/gene/?term=LOC105282001 | 8513 | 189 | 222.01 | 1 | 1.17 | 0.53 | 10.4 | 10.32 | 0.08 | 0.08 | 0 | 0 |
| gene2806 | https://www.ncbi.nlm.nih.gov/gene/?term=LOC105275327 | 14257 | 529 | 371.05 | 1 | 0.7 | 0.19 | 6.66 | 6.63 | 0.03 | 0.03 | 0 | 0 |
| gene2748 | https://www.ncbi.nlm.nih.gov/gene/?term=LOC105275261 | 3845 | 55 | 143.04 | 1 | 2.6 | 1.82 | 38.35 | 38.32 | 0.02 | 0.02 | 0 | 0 |
| gene7368 | https://www.ncbi.nlm.nih.gov/gene/?term=LOC105280556 | 4105 | 65 | 158.34 | 1 | 2.44 | 1.54 | 14.12 | 14.11 | 0.01 | 0.01 | 0 | 0 |
| gene11583 | https://www.ncbi.nlm.nih.gov/gene/?term=LOC105285414 | 8192 | 189 | 230.71 | 1 | 1.22 | 0.53 | 8.96 | 8.96 | 0 | 0 | 0 | 0 |
| gene10879 | https://www.ncbi.nlm.nih.gov/gene/?term=LOC105284574 | 2423 | 50 | 206.36 | 1 | 4.13 | 2 | 52.18 | 52.2 | -0.01 | 0.01 | 0 | 0 |
| gene3470 | https://www.ncbi.nlm.nih.gov/gene/?term=LOC105276059 | 8716 | 161 | 184.72 | 1 | 1.15 | 0.62 | 9.23 | 9.23 | -0.01 | 0.01 | 0 | 0 |
| gene5746 | https://www.ncbi.nlm.nih.gov/gene/?term=LOC105278666 | 10343 | 473 | 457.31 | 1 | 0.97 | 0.21 | 6.89 | 6.9 | -0.01 | 0.01 | 0 | 0 |

| | | | | | | | | | | | | | |
|---|---|---|---|---|---|---|---|---|---|---|---|---|---|
| gene5375 | https://www.ncbi.nlm.nih.gov/gene/?term=LOC105278228 | 8640 | 170 | 196.76 | 1 | 1.16 | 0.59 | 23.53 | 23.55 | -0.02 | 0.02 | 0 | 0 |
| gene9326 | https://www.ncbi.nlm.nih.gov/gene/?term=LOC105282818 | 16612 | 210 | 126.41 | 1 | 0.6 | 0.48 | 13.96 | 13.99 | -0.03 | 0.03 | 0 | 0 |
| gene12573 | https://www.ncbi.nlm.nih.gov/gene/?term=LOC105286515 | 3647 | 112 | 307.1 | 1 | 2.74 | 0.89 | 11.12 | 11.16 | -0.04 | 0.04 | 0 | 0 |
| gene2443 | https://www.ncbi.nlm.nih.gov/gene/?term=LOC105274895 | 10433 | 409 | 392.03 | 1 | 0.96 | 0.24 | 4.52 | 4.57 | -0.04 | 0.04 | 0 | 0 |
| gene7249 | https://www.ncbi.nlm.nih.gov/gene/?term=LOC105280422 | 1775 | 66 | 371.83 | 1 | 5.63 | 1.52 | 15.99 | 16.04 | -0.05 | 0.05 | 0 | 0 |
| gene11275 | https://www.ncbi.nlm.nih.gov/gene/?term=LOC105285055 | 4119 | 251 | 609.37 | 1 | 2.43 | 0.4 | 10.55 | 10.62 | -0.07 | 0.07 | 0 | 0 |
| gene5539 | https://www.ncbi.nlm.nih.gov/gene/?term=LOC105278415 | 25024 | 724 | 289.32 | 1 | 0.4 | 0.14 | 3.42 | 3.48 | -0.07 | 0.07 | 0 | 0 |
| gene5197 | https://www.ncbi.nlm.nih.gov/gene/?term=LOC105278018 | 89233 | 3396 | 380.58 | 1 | 0.11 | 0.03 | 0.66 | 0.73 | -0.08 | 0.08 | 0 | 0 |
| gene1220 | https://www.ncbi.nlm.nih.gov/gene/?term=LOC105285742 | 3453 | 109 | 315.67 | 1 | 2.9 | 0.92 | 16.97 | 17.05 | -0.09 | 0.09 | 0 | 0 |
| gene6620 | https://www.ncbi.nlm.nih.gov/gene/?term=LOC105279687 | 17998 | 433 | 240.58 | 1 | 0.56 | 0.23 | 4.48 | 4.58 | -0.1 | 0.1 | 0 | 0 |
| gene53 | https://www.ncbi.nlm.nih.gov/gene/?term=LOC105277148 | 19283 | 305 | 158.17 | 1 | 0.52 | 0.33 | 4.34 | 4.47 | -0.13 | 0.13 | 0 | 0 |
| gene5052 | https://www.ncbi.nlm.nih.gov/gene/?term=LOC105277856 | 3167 | 33 | 104.2 | 1 | 3.16 | 3.03 | 39.38 | 39.53 | -0.15 | 0.15 | 0 | 0 |
| gene862 | https://www.ncbi.nlm.nih.gov/gene/?term=LOC105281802 | 73922 | 2087 | 282.32 | 1 | 0.14 | 0.05 | 2.14 | 2.3 | -0.15 | 0.15 | 0 | 0 |
| gene11561 | https://www.ncbi.nlm.nih.gov/gene/?term=LOC105285387 | 11528 | 219 | 189.97 | 1 | 0.87 | 0.46 | 9.39 | 9.54 | -0.16 | 0.16 | 0 | 0 |
| gene292 | https://www.ncbi.nlm.nih.gov/gene/?term=LOC105275398 | 2825 | 83 | 293.81 | 1 | 3.54 | 1.2 | 20.15 | 20.32 | -0.16 | 0.16 | 0 | 0 |
| gene1632 | https://www.ncbi.nlm.nih.gov/gene/?term=LOC105287764 | 18061 | 398 | 220.36 | 1 | 0.55 | 0.25 | 3.49 | 3.69 | -0.19 | 0.19 | 0 | 0 |
| gene10600 | https://www.ncbi.nlm.nih.gov/gene/?term=LOC105284270 | 4837 | 103 | 212.94 | 1 | 2.07 | 0.97 | 12.43 | 12.66 | -0.23 | 0.23 | 0 | 0 |

Directory: OGLall File: rnk-dmg-Cb_LI_RphaseA1.vs.RphaseB2.txt

| gene_ID | gene_link | gwidth | #Sites | #per10Kb | #dmSites | #dmsp10kb | %dmSites | %pSite1 | %pSite2 | DMpSite | ADMpSite | DMpNucl | ADMpNucl |
|---|---|---|---|---|---|---|---|---|---|---|---|---|---|
| gene2824 | https://www.ncbi.nlm.nih.gov/gene/?term=LOC105275337 | 1772 | 58 | 327.31 | 1 | 5.64 | 1.72 | 38.96 | 30.77 | 8.18 | 8.18 | 0.27 | 0.27 |
| gene2593 | https://www.ncbi.nlm.nih.gov/gene/?term=LOC105275070 | 1774 | 7 | 39.46 | 5 | 28.18 | 71.43 | 68.01 | 4.09 | 63.92 | 63.92 | 0.25 | 0.25 |
| gene3165 | https://www.ncbi.nlm.nih.gov/gene/?term=LOC105275706 | 2374 | 65 | 273.8 | 5 | 21.06 | 7.69 | 30.28 | 21 | 9.28 | 9.28 | 0.25 | 0.25 |
| gene11819 | https://www.ncbi.nlm.nih.gov/gene/?term=LOC105285671 | 2366 | 98 | 414.2 | 1 | 4.23 | 1.02 | 29.1 | 26.97 | 2.13 | 2.13 | 0.09 | 0.09 |
| gene6334 | https://www.ncbi.nlm.nih.gov/gene/?term=LOC105279356 | 1471 | 45 | 305.91 | 1 | 6.8 | 2.22 | 31.17 | 28.75 | 2.42 | 2.42 | 0.07 | 0.07 |
| gene11401 | https://www.ncbi.nlm.nih.gov/gene/?term=LOC105285202 | 1841 | 78 | 423.68 | 1 | 5.43 | 1.28 | 26.41 | 24.78 | 1.62 | 1.62 | 0.07 | 0.07 |
| gene7406 | https://www.ncbi.nlm.nih.gov/gene/?term=LOC105280596 | 3432 | 106 | 308.86 | 1 | 2.91 | 0.94 | 17.02 | 19.17 | -2.15 | 2.15 | -0.07 | 0.07 |
| gene7568 | https://www.ncbi.nlm.nih.gov/gene/?term=LOC105280786 | 2525 | 114 | 451.49 | 1 | 3.96 | 0.88 | 28.64 | 27.23 | 1.41 | 1.41 | 0.06 | 0.06 |
| gene9101 | https://www.ncbi.nlm.nih.gov/gene/?term=LOC105282553 | 3068 | 102 | 332.46 | 1 | 3.26 | 0.98 | 21.53 | 23.25 | -1.72 | 1.72 | -0.06 | 0.06 |
| gene12406 | https://www.ncbi.nlm.nih.gov/gene/?term=LOC105286316 | 1386 | 31 | 223.67 | 1 | 7.22 | 3.23 | 42.03 | 44.54 | -2.51 | 2.51 | -0.06 | 0.06 |
| gene322 | https://www.ncbi.nlm.nih.gov/gene/?term=LOC105275777 | 1606 | 27 | 168.12 | 1 | 6.23 | 3.7 | 38.42 | 42 | -3.59 | 3.59 | -0.06 | 0.06 |
| gene5877 | https://www.ncbi.nlm.nih.gov/gene/?term=LOC105278807 | 1719 | 48 | 279.23 | 1 | 5.82 | 2.08 | 26.96 | 25.13 | 1.83 | 1.83 | 0.05 | 0.05 |
| gene1855 | https://www.ncbi.nlm.nih.gov/gene/?term=LOC105288036 | 2387 | 77 | 322.58 | 1 | 4.19 | 1.3 | 32.99 | 31.48 | 1.51 | 1.51 | 0.05 | 0.05 |
| gene4641 | https://www.ncbi.nlm.nih.gov/gene/?term=LOC105277398 | 1937 | 71 | 366.55 | 1 | 5.16 | 1.41 | 38.26 | 39.69 | -1.43 | 1.43 | -0.05 | 0.05 |
| gene12981 | https://www.ncbi.nlm.nih.gov/gene/?term=LOC105286999 | 2063 | 42 | 203.59 | 1 | 4.85 | 2.38 | 33.61 | 35.96 | -2.34 | 2.34 | -0.05 | 0.05 |
| gene10107 | https://www.ncbi.nlm.nih.gov/gene/?term=LOC105283712 | 1883 | 35 | 185.87 | 1 | 5.31 | 2.86 | 23.29 | 25.95 | -2.66 | 2.66 | -0.05 | 0.05 |
| gene3599 | https://www.ncbi.nlm.nih.gov/gene/?term=LOC105276196 | 2955 | 58 | 196.28 | 1 | 3.38 | 1.72 | 37.85 | 40.55 | -2.7 | 2.7 | -0.05 | 0.05 |
| gene6422 | https://www.ncbi.nlm.nih.gov/gene/?term=LOC105279458 | 2419 | 38 | 157.09 | 1 | 4.13 | 2.63 | 47.43 | 50.37 | -2.94 | 2.94 | -0.05 | 0.05 |
| gene388 | https://www.ncbi.nlm.nih.gov/gene/?term=LOC105276538 | 1558 | 19 | 121.95 | 2 | 12.84 | 10.53 | 61.29 | 65.13 | -3.85 | 3.85 | -0.05 | 0.05 |
| gene10602 | https://www.ncbi.nlm.nih.gov/gene/?term=LOC105284268 | 2122 | 19 | 89.54 | 1 | 4.71 | 5.26 | 27.9 | 32.99 | -5.08 | 5.08 | -0.05 | 0.05 |
| gene393 | https://www.ncbi.nlm.nih.gov/gene/?term=LOC105276566 | 1123 | 2 | 17.81 | 1 | 8.9 | 50 | 50 | 76.78 | -26.78 | 26.78 | -0.05 | 0.05 |
| gene1884 | https://www.ncbi.nlm.nih.gov/gene/?term=LOC105288072 | 4169 | 70 | 167.91 | 1 | 2.4 | 1.43 | 26.72 | 24.36 | 2.36 | 2.36 | 0.04 | 0.04 |
| gene7884 | https://www.ncbi.nlm.nih.gov/gene/?term=LOC105281266 | 3390 | 76 | 224.19 | 1 | 2.95 | 1.32 | 18.91 | 17.21 | 1.7 | 1.7 | 0.04 | 0.04 |
| gene940 | https://www.ncbi.nlm.nih.gov/gene/?term=LOC105282794 | 3578 | 85 | 237.56 | 1 | 2.79 | 1.18 | 14.76 | 13.28 | 1.48 | 1.48 | 0.04 | 0.04 |
| gene2844 | https://www.ncbi.nlm.nih.gov/gene/?term=LOC105275363 | 4330 | 186 | 429.56 | 1 | 2.31 | 0.54 | 15.2 | 14.36 | 0.85 | 0.85 | 0.04 | 0.04 |
| gene8729 | https://www.ncbi.nlm.nih.gov/gene/?term=LOC105282137 | 5005 | 213 | 425.57 | 1 | 2 | 0.47 | 14.62 | 15.67 | -1.05 | 1.05 | -0.04 | 0.04 |
| gene10213 | https://www.ncbi.nlm.nih.gov/gene/?term=LOC105283826 | 1503 | 53 | 352.63 | 1 | 6.65 | 1.89 | 14.78 | 15.95 | -1.17 | 1.17 | -0.04 | 0.04 |
| gene6031 | https://www.ncbi.nlm.nih.gov/gene/?term=LOC105279000 | 5691 | 180 | 316.29 | 1 | 1.76 | 0.56 | 14.61 | 15.92 | -1.31 | 1.31 | -0.04 | 0.04 |
| gene2413 | https://www.ncbi.nlm.nih.gov/gene/?term=LOC105274863 | 2006 | 49 | 244.27 | 1 | 4.99 | 2.04 | 53.65 | 55.47 | -1.81 | 1.81 | -0.04 | 0.04 |
| gene5101 | https://www.ncbi.nlm.nih.gov/gene/?term=LOC105277903 | 1813 | 35 | 193.05 | 1 | 5.52 | 2.86 | 52.77 | 51.27 | 1.51 | 1.51 | 0.03 | 0.03 |
| gene7915 | https://www.ncbi.nlm.nih.gov/gene/?term=LOC105281173 | 1597 | 38 | 237.95 | 1 | 6.26 | 2.63 | 59.81 | 58.74 | 1.06 | 1.06 | 0.03 | 0.03 |
| gene10540 | https://www.ncbi.nlm.nih.gov/gene/?term=LOC105284200 | 5267 | 215 | 408.2 | 1 | 1.9 | 0.47 | 12.8 | 12.05 | 0.75 | 0.75 | 0.03 | 0.03 |
| gene298 | https://www.ncbi.nlm.nih.gov/gene/?term=LOC105275460 | 2473 | 118 | 477.15 | 1 | 4.04 | 0.85 | 13.95 | 13.28 | 0.67 | 0.67 | 0.03 | 0.03 |
| gene1332 | https://www.ncbi.nlm.nih.gov/gene/?term=LOC105286872 | 2460 | 156 | 634.15 | 1 | 4.07 | 0.64 | 20.52 | 20.12 | 0.41 | 0.41 | 0.03 | 0.03 |
| gene12802 | https://www.ncbi.nlm.nih.gov/gene/?term=LOC105286799 | 3183 | 159 | 499.53 | 1 | 3.14 | 0.63 | 17.45 | 17.99 | -0.54 | 0.54 | -0.03 | 0.03 |
| gene5298 | https://www.ncbi.nlm.nih.gov/gene/?term=LOC105278145 | 4060 | 235 | 578.82 | 2 | 4.93 | 0.85 | 10.98 | 11.56 | -0.58 | 0.58 | -0.03 | 0.03 |
| gene920 | https://www.ncbi.nlm.nih.gov/gene/?term=LOC105282518 | 1197 | 48 | 401 | 1 | 8.35 | 2.08 | 32.24 | 33.02 | -0.78 | 0.78 | -0.03 | 0.03 |
| gene11340 | https://www.ncbi.nlm.nih.gov/gene/?term=LOC105285131 | 2918 | 87 | 298.15 | 1 | 3.43 | 1.15 | 10.66 | 11.58 | -0.92 | 0.92 | -0.03 | 0.03 |
| gene1310 | https://www.ncbi.nlm.nih.gov/gene/?term=LOC105286661 | 2582 | 68 | 263.36 | 1 | 3.87 | 1.47 | 19.86 | 20.85 | -1 | 1 | -0.03 | 0.03 |
| gene7600 | https://www.ncbi.nlm.nih.gov/gene/?term=LOC105280819 | 1615 | 37 | 229.1 | 1 | 6.19 | 2.7 | 36.99 | 38.16 | -1.16 | 1.16 | -0.03 | 0.03 |
| gene7183 | https://www.ncbi.nlm.nih.gov/gene/?term=LOC105280348 | 3925 | 70 | 178.34 | 1 | 2.55 | 1.43 | 19.96 | 21.48 | -1.52 | 1.52 | -0.03 | 0.03 |
| gene7199 | https://www.ncbi.nlm.nih.gov/gene/?term=LOC105280366 | 1414 | 24 | 169.73 | 1 | 7.07 | 4.17 | 43.55 | 45.17 | -1.62 | 1.62 | -0.03 | 0.03 |
| gene12572 | https://www.ncbi.nlm.nih.gov/gene/?term=LOC105286514 | 2374 | 47 | 197.98 | 1 | 4.21 | 2.13 | 35.17 | 36.86 | -1.69 | 1.69 | -0.03 | 0.03 |
| gene6237 | https://www.ncbi.nlm.nih.gov/gene/?term=LOC105279244 | 1484 | 22 | 148.25 | 1 | 6.74 | 4.55 | 40.19 | 41.99 | -1.8 | 1.8 | -0.03 | 0.03 |
| gene3168 | https://www.ncbi.nlm.nih.gov/gene/?term=LOC105275711 | 6150 | 69 | 112.2 | 1 | 1.63 | 1.45 | 12.17 | 15.18 | -3.01 | 3.01 | -0.03 | 0.03 |
| gene1008 | https://www.ncbi.nlm.nih.gov/gene/?term=LOC105283492 | 1479 | 9 | 60.85 | 1 | 6.76 | 11.11 | 30.57 | 36.08 | -5.51 | 5.51 | -0.03 | 0.03 |
| gene10654 | https://www.ncbi.nlm.nih.gov/gene/?term=LOC105284329 | 4027 | 51 | 126.65 | 1 | 2.48 | 1.96 | 20.19 | 18.78 | 1.41 | 1.41 | 0.02 | 0.02 |
| gene12516 | https://www.ncbi.nlm.nih.gov/gene/?term=LOC105286452 | 1529 | 38 | 248.53 | 1 | 6.54 | 2.63 | 50.31 | 49.37 | 0.94 | 0.94 | 0.02 | 0.02 |
| gene11454 | https://www.ncbi.nlm.nih.gov/gene/?term=LOC105285256 | 1906 | 45 | 236.1 | 1 | 5.25 | 2.22 | 47.83 | 47.02 | 0.81 | 0.81 | 0.02 | 0.02 |
| gene4892 | https://www.ncbi.nlm.nih.gov/gene/?term=LOC105277667 | 3898 | 119 | 305.28 | 1 | 2.57 | 0.84 | 28.98 | 28.19 | 0.8 | 0.8 | 0.02 | 0.02 |
| gene4213 | https://www.ncbi.nlm.nih.gov/gene/?term=LOC105276901 | 3257 | 83 | 254.84 | 1 | 3.07 | 1.2 | 27.23 | 26.49 | 0.74 | 0.74 | 0.02 | 0.02 |
| gene10520 | https://www.ncbi.nlm.nih.gov/gene/?term=LOC105284184 | 2451 | 79 | 322.32 | 1 | 4.08 | 1.27 | 35.68 | 34.99 | 0.69 | 0.69 | 0.02 | 0.02 |
| gene10600 | https://www.ncbi.nlm.nih.gov/gene/?term=LOC105284270 | 4837 | 115 | 237.75 | 1 | 2.07 | 0.87 | 11.06 | 10.43 | 0.63 | 0.63 | 0.02 | 0.02 |
| gene6966 | https://www.ncbi.nlm.nih.gov/gene/?term=LOC105280104 | 3747 | 124 | 330.93 | 1 | 2.67 | 0.81 | 17.23 | 16.6 | 0.63 | 0.63 | 0.02 | 0.02 |
| gene1430 | https://www.ncbi.nlm.nih.gov/gene/?term=LOC105287551 | 6433 | 216 | 335.77 | 1 | 1.55 | 0.46 | 9.29 | 8.67 | 0.62 | 0.62 | 0.02 | 0.02 |
| gene570 | https://www.ncbi.nlm.nih.gov/gene/?term=LOC105278549 | 1673 | 62 | 370.59 | 1 | 5.98 | 1.61 | 43.95 | 43.37 | 0.58 | 0.58 | 0.02 | 0.02 |
| gene6583 | https://www.ncbi.nlm.nih.gov/gene/?term=LOC105279649 | 7444 | 315 | 423.16 | 1 | 1.34 | 0.32 | 11.55 | 11.04 | 0.51 | 0.51 | 0.02 | 0.02 |
| gene12978 | https://www.ncbi.nlm.nih.gov/gene/?term=LOC105286997 | 2858 | 106 | 370.89 | 1 | 3.5 | 0.94 | 12 | 11.5 | 0.5 | 0.5 | 0.02 | 0.02 |
| gene5831 | https://www.ncbi.nlm.nih.gov/gene/?term=LOC105278761 | 2506 | 105 | 418.99 | 1 | 3.99 | 0.95 | 24.55 | 24.07 | 0.48 | 0.48 | 0.02 | 0.02 |
| gene8274 | https://www.ncbi.nlm.nih.gov/gene/?term=LOC105281624 | 5425 | 198 | 364.98 | 1 | 1.84 | 0.51 | 10.15 | 9.67 | 0.48 | 0.48 | 0.02 | 0.02 |

| | | | | | | | | | | | | | |
|---|---|---|---|---|---|---|---|---|---|---|---|---|---|
| gene2846 | https://www.ncbi.nlm.nih.gov/gene/?term=LOC105275374 | 11246 | 432 | 384.14 | 1 | 0.89 | 0.23 | 11.15 | 10.76 | 0.39 | 0.39 | 0.02 | 0.02 |
| gene7211 | https://www.ncbi.nlm.nih.gov/gene/?term=LOC105280377 | 2024 | 63 | 311.26 | 1 | 4.94 | 1.59 | 48.25 | 48.74 | -0.49 | 0.49 | -0.02 | 0.02 |
| gene7550 | https://www.ncbi.nlm.nih.gov/gene/?term=LOC105280763 | 2173 | 76 | 349.75 | 1 | 4.6 | 1.32 | 32.2 | 32.7 | -0.49 | 0.49 | -0.02 | 0.02 |
| gene292 | https://www.ncbi.nlm.nih.gov/gene/?term=LOC105275398 | 2825 | 78 | 276.11 | 1 | 3.54 | 1.28 | 22.03 | 22.57 | -0.55 | 0.55 | -0.02 | 0.02 |
| gene4216 | https://www.ncbi.nlm.nih.gov/gene/?term=LOC105276906 | 2127 | 86 | 404.33 | 1 | 4.7 | 1.16 | 19.06 | 19.63 | -0.57 | 0.57 | -0.02 | 0.02 |
| gene13036 | https://www.ncbi.nlm.nih.gov/gene/?term=LOC105287059 | 4959 | 148 | 298.45 | 1 | 2.02 | 0.68 | 20.35 | 20.94 | -0.59 | 0.59 | -0.02 | 0.02 |
| gene3107 | https://www.ncbi.nlm.nih.gov/gene/?term=LOC105275650 | 5103 | 152 | 297.86 | 1 | 1.96 | 0.66 | 16.19 | 16.78 | -0.59 | 0.59 | -0.02 | 0.02 |
| gene4190 | https://www.ncbi.nlm.nih.gov/gene/?term=LOC105276875 | 3150 | 105 | 333.33 | 1 | 3.17 | 0.95 | 22.87 | 23.46 | -0.6 | 0.6 | -0.02 | 0.02 |
| gene7249 | https://www.ncbi.nlm.nih.gov/gene/?term=LOC105280422 | 1775 | 60 | 338.03 | 1 | 5.63 | 1.67 | 16.46 | 17.06 | -0.6 | 0.6 | -0.02 | 0.02 |
| gene4376 | https://www.ncbi.nlm.nih.gov/gene/?term=LOC105277083 | 4307 | 97 | 225.21 | 1 | 2.32 | 1.03 | 24.28 | 25.05 | -0.77 | 0.77 | -0.02 | 0.02 |
| gene10392 | https://www.ncbi.nlm.nih.gov/gene/?term=LOC105284036 | 3216 | 94 | 292.29 | 2 | 6.22 | 2.13 | 22.41 | 23.26 | -0.84 | 0.84 | -0.02 | 0.02 |
| gene35 | https://www.ncbi.nlm.nih.gov/gene/?term=LOC105275197 | 2643 | 45 | 170.26 | 1 | 3.78 | 2.22 | 38.91 | 40.15 | -1.23 | 1.23 | -0.02 | 0.02 |
| gene8343 | https://www.ncbi.nlm.nih.gov/gene/?term=LOC105281700 | 2172 | 32 | 147.33 | 1 | 4.6 | 3.12 | 37.43 | 38.97 | -1.55 | 1.55 | -0.02 | 0.02 |
| gene10345 | https://www.ncbi.nlm.nih.gov/gene/?term=LOC105283989 | 4444 | 22 | 49.5 1 | 2.25 | 4.55 | 37.68 | 41.66 | -3.99 | 3.99 | -0.02 | 0.02 | |
| gene8178 | https://www.ncbi.nlm.nih.gov/gene/?term=LOC105281515 | 8647 | 43 | 49.73 1 | 1.16 | 2.33 | 27.02 | 24.75 | 2.27 | 2.27 | 0.01 | 0.01 | |
| gene8661 | https://www.ncbi.nlm.nih.gov/gene/?term=LOC105282063 | 1926 | 26 | 134.99 | 1 | 5.19 | 3.85 | 38.79 | 37.9 | 0.89 | 0.89 | 0.01 | 0.01 |
| gene4023 | https://www.ncbi.nlm.nih.gov/gene/?term=LOC105276692 | 1759 | 31 | 176.24 | 1 | 5.69 | 3.23 | 24.44 | 23.63 | 0.81 | 0.81 | 0.01 | 0.01 |
| gene3470 | https://www.ncbi.nlm.nih.gov/gene/?term=LOC105276059 | 8716 | 146 | 167.51 | 1 | 1.15 | 0.68 | 9.33 | 8.54 | 0.79 | 0.79 | 0.01 | 0.01 |
| gene9336 | https://www.ncbi.nlm.nih.gov/gene/?term=LOC105282834 | 5993 | 123 | 205.24 | 1 | 1.67 | 0.81 | 15.27 | 14.61 | 0.66 | 0.66 | 0.01 | 0.01 |
| gene5273 | https://www.ncbi.nlm.nih.gov/gene/?term=LOC105278108 | 4067 | 43 | 105.73 | 1 | 2.46 | 2.33 | 26.66 | 26.12 | 0.55 | 0.55 | 0.01 | 0.01 |
| gene11579 | https://www.ncbi.nlm.nih.gov/gene/?term=LOC105285406 | 13571 | 229 | 168.74 | 1 | 0.74 | 0.44 | 7.6 | 7.22 | 0.38 | 0.38 | 0.01 | 0.01 |
| gene12439 | https://www.ncbi.nlm.nih.gov/gene/?term=LOC105286357 | 4761 | 87 | 182.73 | 1 | 2.1 | 1.15 | 25.3 | 24.92 | 0.38 | 0.38 | 0.01 | 0.01 |
| gene7247 | https://www.ncbi.nlm.nih.gov/gene/?term=LOC105280407 | 10486 | 208 | 198.36 | 1 | 0.95 | 0.48 | 16.58 | 16.25 | 0.33 | 0.33 | 0.01 | 0.01 |
| gene2266 | https://www.ncbi.nlm.nih.gov/gene/?term=LOC105274699 | 2391 | 49 | 204.94 | 1 | 4.18 | 2.04 | 23.32 | 23 | 0.32 | 0.32 | 0.01 | 0.01 |
| gene13114 | https://www.ncbi.nlm.nih.gov/gene/?term=LOC105287147 | 7609 | 393 | 516.49 | 1 | 1.31 | 0.25 | 9.24 | 9.02 | 0.23 | 0.23 | 0.01 | 0.01 |
| gene5525 | https://www.ncbi.nlm.nih.gov/gene/?term=LOC105278396 | 3090 | 180 | 582.52 | 1 | 3.24 | 0.56 | 24.99 | 24.77 | 0.22 | 0.22 | 0.01 | 0.01 |
| gene11082 | https://www.ncbi.nlm.nih.gov/gene/?term=LOC105284824 | 7436 | 189 | 254.17 | 1 | 1.34 | 0.53 | 8.04 | 7.84 | 0.2 | 0.2 | 0.01 | 0.01 |
| gene11055 | https://www.ncbi.nlm.nih.gov/gene/?term=LOC105284796 | 2283 | 121 | 530 1 | 4.38 | 0.83 | 34.82 | 34.65 | 0.17 | 0.17 | 0.01 | 0.01 | |
| gene1470 | https://www.ncbi.nlm.nih.gov/gene/?term=LOC105287597 | 2974 | 181 | 608.61 | 1 | 3.36 | 0.55 | 19.06 | 18.9 | 0.16 | 0.16 | 0.01 | 0.01 |
| gene7556 | https://www.ncbi.nlm.nih.gov/gene/?term=LOC105280774 | 2792 | 98 | 351 1 | 3.58 | 1.02 | 32.45 | 32.61 | -0.17 | 0.17 | -0.01 | 0.01 | |
| gene10018 | https://www.ncbi.nlm.nih.gov/gene/?term=LOC105283607 | 4414 | 202 | 457.63 | 1 | 2.27 | 0.5 | 14.99 | 15.23 | -0.24 | 0.24 | -0.01 | 0.01 |
| gene6286 | https://www.ncbi.nlm.nih.gov/gene/?term=LOC105279298 | 2861 | 130 | 454.39 | 2 | 6.99 | 1.54 | 23.66 | 23.9 | -0.24 | 0.24 | -0.01 | 0.01 |
| gene10606 | https://www.ncbi.nlm.nih.gov/gene/?term=LOC105284273 | 6119 | 244 | 398.76 | 1 | 1.63 | 0.41 | 18.11 | 18.39 | -0.29 | 0.29 | -0.01 | 0.01 |
| gene34 | https://www.ncbi.nlm.nih.gov/gene/?term=LOC105278416 | 7120 | 124 | 174.16 | 1 | 1.4 | 0.81 | 14.41 | 14.72 | -0.3 | 0.3 | -0.01 | 0.01 |
| gene12573 | https://www.ncbi.nlm.nih.gov/gene/?term=LOC105286515 | 3647 | 105 | 287.91 | 1 | 2.74 | 0.95 | 12.53 | 12.84 | -0.31 | 0.31 | -0.01 | 0.01 |
| gene3885 | https://www.ncbi.nlm.nih.gov/gene/?term=LOC105276527 | 7928 | 299 | 377.14 | 1 | 1.26 | 0.33 | 9.83 | 10.14 | -0.31 | 0.31 | -0.01 | 0.01 |
| gene7166 | https://www.ncbi.nlm.nih.gov/gene/?term=LOC105280326 | 6117 | 226 | 369.46 | 1 | 1.63 | 0.44 | 14.91 | 15.21 | -0.31 | 0.31 | -0.01 | 0.01 |
| gene10549 | https://www.ncbi.nlm.nih.gov/gene/?term=LOC105284210 | 7145 | 284 | 397.48 | 2 | 2.8 | 0.7 | 11.85 | 12.19 | -0.33 | 0.33 | -0.01 | 0.01 |
| gene5646 | https://www.ncbi.nlm.nih.gov/gene/?term=LOC105278534 | 7475 | 176 | 235.45 | 1 | 1.34 | 0.57 | 18.67 | 19 | -0.33 | 0.33 | -0.01 | 0.01 |
| gene2404 | https://www.ncbi.nlm.nih.gov/gene/?term=LOC105274853 | 1767 | 41 | 232.03 | 1 | 5.66 | 2.44 | 30.51 | 30.87 | -0.36 | 0.36 | -0.01 | 0.01 |
| gene9327 | https://www.ncbi.nlm.nih.gov/gene/?term=LOC105282821 | 7793 | 140 | 179.65 | 1 | 1.28 | 0.71 | 12.3 | 12.71 | -0.41 | 0.41 | -0.01 | 0.01 |
| gene5375 | https://www.ncbi.nlm.nih.gov/gene/?term=LOC105278228 | 8640 | 152 | 175.93 | 1 | 1.16 | 0.66 | 22.96 | 23.39 | -0.42 | 0.42 | -0.01 | 0.01 |
| gene10879 | https://www.ncbi.nlm.nih.gov/gene/?term=LOC105284574 | 2423 | 50 | 206.36 | 1 | 4.13 | 2 | 52.82 | 53.31 | -0.49 | 0.49 | -0.01 | 0.01 |
| gene12307 | https://www.ncbi.nlm.nih.gov/gene/?term=LOC105286216 | 10313 | 164 | 159.02 | 1 | 0.97 | 0.61 | 16.27 | 16.77 | -0.5 | 0.5 | -0.01 | 0.01 |
| gene1147 | https://www.ncbi.nlm.nih.gov/gene/?term=LOC105284954 | 4022 | 91 | 226.26 | 1 | 2.49 | 1.1 | 14.14 | 14.77 | -0.63 | 0.63 | -0.01 | 0.01 |
| gene9647 | https://www.ncbi.nlm.nih.gov/gene/?term=LOC109611233 | 3094 | 32 | 103.43 | 1 | 3.23 | 3.12 | 37.53 | 38.18 | -0.65 | 0.65 | -0.01 | 0.01 |
| gene9648 | https://www.ncbi.nlm.nih.gov/gene/?term=LOC105283183 | 2602 | 30 | 115.3 1 | 3.84 | 3.33 | 40.03 | 40.73 | -0.7 | 0.7 | -0.01 | 0.01 | |
| gene12203 | https://www.ncbi.nlm.nih.gov/gene/?term=LOC105286099 | 5967 | 70 | 117.31 | 1 | 1.68 | 1.43 | 9.91 | 10.79 | -0.88 | 0.88 | -0.01 | 0.01 |
| gene7248 | https://www.ncbi.nlm.nih.gov/gene/?term=LOC105280408 | 5175 | 52 | 100.48 | 1 | 1.93 | 1.92 | 17.59 | 18.54 | -0.95 | 0.95 | -0.01 | 0.01 |
| gene10155 | https://www.ncbi.nlm.nih.gov/gene/?term=LOC105283756 | 7030 | 108 | 153.63 | 1 | 1.42 | 0.93 | 5.58 | 6.54 | -0.96 | 0.96 | -0.01 | 0.01 |
| gene12021 | https://www.ncbi.nlm.nih.gov/gene/?term=LOC105285905 | 1399 | 16 | 114.37 | 1 | 7.15 | 6.25 | 46.76 | 47.75 | -0.98 | 0.98 | -0.01 | 0.01 |
| gene6639 | https://www.ncbi.nlm.nih.gov/gene/?term=LOC105279705 | 7656 | 69 | 90.13 1 | 1.31 | 1.45 | 20.74 | 21.96 | -1.21 | 1.21 | -0.01 | 0.01 | |
| gene564 | https://www.ncbi.nlm.nih.gov/gene/?term=LOC105278478 | 2802 | 18 | 64.24 1 | 3.57 | 5.56 | 14.97 | 16.38 | -1.41 | 1.41 | -0.01 | 0.01 | |
| gene2683 | https://www.ncbi.nlm.nih.gov/gene/?term=LOC105275187 | 5492 | 65 | 118.35 | 1 | 1.82 | 1.54 | 11.46 | 11.25 | 0.21 | 0.21 | 0 | 0 |
| gene5052 | https://www.ncbi.nlm.nih.gov/gene/?term=LOC105277856 | 3167 | 28 | 88.41 1 | 3.16 | 3.57 | 33.66 | 33.45 | 0.21 | 0.21 | 0 | 0 | |
| gene2806 | https://www.ncbi.nlm.nih.gov/gene/?term=LOC105275327 | 14257 | 456 | 319.84 | 1 | 0.7 | 0.22 | 7.38 | 7.24 | 0.14 | 0.14 | 0 | 0 |
| gene1220 | https://www.ncbi.nlm.nih.gov/gene/?term=LOC105285742 | 3453 | 100 | 289.6 1 | 2.9 | 1 | 18.13 | 18.02 | 0.11 | 0.11 | 0 | 0 | |
| gene5733 | https://www.ncbi.nlm.nih.gov/gene/?term=LOC105278649 | 5221 | 92 | 176.21 | 1 | 1.92 | 1.09 | 19.86 | 19.75 | 0.11 | 0.11 | 0 | 0 |
| gene5746 | https://www.ncbi.nlm.nih.gov/gene/?term=LOC105278666 | 10343 | 436 | 421.54 | 1 | 0.97 | 0.23 | 6.97 | 6.87 | 0.11 | 0.11 | 0 | 0 |
| gene8600 | https://www.ncbi.nlm.nih.gov/gene/?term=LOC105282001 | 8513 | 151 | 177.38 | 1 | 1.17 | 0.66 | 10.5 | 10.38 | 0.11 | 0.11 | 0 | 0 |
| gene10844 | https://www.ncbi.nlm.nih.gov/gene/?term=LOC105284535 | 9086 | 222 | 244.33 | 1 | 1.1 | 0.45 | 15.15 | 15.05 | 0.1 | 0.1 | 0 | 0 |
| gene1632 | https://www.ncbi.nlm.nih.gov/gene/?term=LOC105287764 | 18061 | 367 | 203.2 1 | 0.55 | 0.27 | 3.44 | 3.36 | 0.08 | 0.08 | 0 | 0 | |
| gene5524 | https://www.ncbi.nlm.nih.gov/gene/?term=LOC105278397 | 5177 | 196 | 378.6 1 | 1.93 | 0.51 | 24.77 | 24.69 | 0.07 | 0.07 | 0 | 0 | |
| gene1408 | https://www.ncbi.nlm.nih.gov/gene/?term=LOC109611520 | 8645 | 289 | 334.3 1 | 1.16 | 0.35 | 6.21 | 6.16 | 0.05 | 0.05 | 0 | 0 | |

| | | | | | | | | | | | | | |
|---|---|---|---|---|---|---|---|---|---|---|---|---|---|
| gene7368 | https://www.ncbi.nlm.nih.gov/gene/?term=LOC105280556 | 4105 | 38 | 92.57 | 1 | 2.44 | 2.63 | 16.79 | 16.73 | 0.05 | 0.05 | 0 | 0 |
| gene9740 | https://www.ncbi.nlm.nih.gov/gene/?term=LOC105283277 | 2887 | 131 | 453.76 | 1 | 3.46 | 0.76 | 19.31 | 19.27 | 0.04 | 0.04 | 0 | 0 |
| gene8814 | https://www.ncbi.nlm.nih.gov/gene/?term=LOC105282240 | 6910 | 163 | 235.89 | 1 | 1.45 | 0.61 | 9.99 | 9.96 | 0.03 | 0.03 | 0 | 0 |
| gene2271 | https://www.ncbi.nlm.nih.gov/gene/?term=LOC105274706 | 1448 | 50 | 345.3 | 1 | 6.91 | 2 | 38.1 | 38.09 | 0.01 | 0.01 | 0 | 0 |
| gene2173 | https://www.ncbi.nlm.nih.gov/gene/?term=LOC105274590 | 4387 | 68 | 155 | 1 | 2.28 | 1.47 | 26.72 | 26.74 | -0.02 | 0.02 | 0 | 0 |
| gene7547 | https://www.ncbi.nlm.nih.gov/gene/?term=LOC105280767 | 24576 | 671 | 273.03 | 1 | 0.41 | 0.15 | 4.75 | 4.77 | -0.02 | 0.02 | 0 | 0 |
| gene10678 | https://www.ncbi.nlm.nih.gov/gene/?term=LOC105284355 | 4031 | 168 | 416.77 | 1 | 2.48 | 0.6 | 18.14 | 18.16 | -0.03 | 0.03 | 0 | 0 |
| gene10984 | https://www.ncbi.nlm.nih.gov/gene/?term=LOC105284704 | 1987 | 58 | 291.9 | 1 | 5.03 | 1.72 | 23.02 | 23.06 | -0.04 | 0.04 | 0 | 0 |
| gene862 | https://www.ncbi.nlm.nih.gov/gene/?term=LOC105281802 | 73922 | 1763 | 238.49 | 1 | 0.14 | 0.06 | 2.22 | 2.27 | -0.05 | 0.05 | 0 | 0 |
| gene10326 | https://www.ncbi.nlm.nih.gov/gene/?term=LOC105283962 | 3042 | 119 | 391.19 | 1 | 3.29 | 0.84 | 8.91 | 8.98 | -0.07 | 0.07 | 0 | 0 |
| gene11275 | https://www.ncbi.nlm.nih.gov/gene/?term=LOC105285055 | 4119 | 257 | 623.94 | 1 | 2.43 | 0.39 | 10.45 | 10.51 | -0.07 | 0.07 | 0 | 0 |
| gene4962 | https://www.ncbi.nlm.nih.gov/gene/?term=LOC105277761 | 4645 | 142 | 305.71 | 1 | 2.15 | 0.7 | 13.6 | 13.69 | -0.09 | 0.09 | 0 | 0 |
| gene7398 | https://www.ncbi.nlm.nih.gov/gene/?term=LOC105280589 | 3452 | 33 | 95.6 | 1 | 2.9 | 3.03 | 20.49 | 20.58 | -0.09 | 0.09 | 0 | 0 |
| gene6620 | https://www.ncbi.nlm.nih.gov/gene/?term=LOC105279687 | 17998 | 344 | 191.13 | 1 | 0.56 | 0.29 | 5.2 | 5.3 | -0.1 | 0.1 | 0 | 0 |
| gene53 | https://www.ncbi.nlm.nih.gov/gene/?term=LOC105277148 | 19283 | 268 | 138.98 | 1 | 0.52 | 0.37 | 3.71 | 3.95 | -0.23 | 0.23 | 0 | 0 |
| gene9326 | https://www.ncbi.nlm.nih.gov/gene/?term=LOC105282818 | 16612 | 169 | 101.73 | 1 | 0.6 | 0.59 | 13.49 | 13.76 | -0.26 | 0.26 | 0 | 0 |
| gene2748 | https://www.ncbi.nlm.nih.gov/gene/?term=LOC105275261 | 3845 | 53 | 137.84 | 1 | 2.6 | 1.89 | 38.36 | 38.68 | -0.33 | 0.33 | 0 | 0 |

Directory: OGLall File: rnk-dmg-Cb_LI_RphaseA2.vs.RphaseB1.txt

| gene_ID | gene_link | gwidth | #Sites | #per10Kb | #dmSites | #dmsp10kb | %dmSites | %pSite1 | %pSite2 | DMpSite | ADMpSite | DMpNucl | ADMpNucl |
|---|---|---|---|---|---|---|---|---|---|---|---|---|---|
| gene2824 | https://www.ncbi.nlm.nih.gov/gene/?term=LOC105275337 | 1772 | 63 | 355.53 | 2 | 11.29 | 3.17 | 38.53 | 29.97 | 8.56 | 8.56 | 0.3 | 0.3 |
| gene2593 | https://www.ncbi.nlm.nih.gov/gene/?term=LOC105275070 | 1774 | 11 | 62.01 6 | 33.82 | 54.55 | 67.77 | 25.28 | 42.49 | 42.49 | 0.26 | 0.26 | |
| gene3165 | https://www.ncbi.nlm.nih.gov/gene/?term=LOC105275706 | 2374 | 70 | 294.86 | 5 | 21.06 | 7.14 | 33.14 | 24.38 | 8.76 | 8.76 | 0.26 | 0.26 |
| gene12961 | https://www.ncbi.nlm.nih.gov/gene/?term=LOC105286983 | 3094 | 71 | 229.48 | 1 | 3.23 | 1.41 | 6.3 | 1.17 | 5.12 | 5.12 | 0.12 | 0.12 |
| gene5714 | https://www.ncbi.nlm.nih.gov/gene/?term=LOC105278729 | 710 | 16 | 225.35 | 1 | 14.08 | 6.25 | 56.86 | 51.9 | 4.96 | 4.96 | 0.11 | 0.11 |
| gene7568 | https://www.ncbi.nlm.nih.gov/gene/?term=LOC105280786 | 2525 | 143 | 566.34 | 1 | 3.96 | 0.7 | 27.63 | 25.63 | 2 | 2 | 0.11 | 0.11 |
| gene13033 | https://www.ncbi.nlm.nih.gov/gene/?term=LOC105287063 | 1877 | 55 | 293.02 | 1 | 5.33 | 1.82 | 30.5 | 27.04 | 3.46 | 3.46 | 0.1 | 0.1 |
| gene2413 | https://www.ncbi.nlm.nih.gov/gene/?term=LOC105274863 | 2006 | 65 | 324.03 | 1 | 4.99 | 1.54 | 49.74 | 52.89 | -3.14 | 3.14 | -0.1 | 0.1 |
| gene1007 | https://www.ncbi.nlm.nih.gov/gene/?term=LOC105283457 | 1878 | 49 | 260.92 | 1 | 5.32 | 2.04 | 35.2 | 31.9 | 3.3 | 3.3 | 0.09 | 0.09 |
| gene2271 | https://www.ncbi.nlm.nih.gov/gene/?term=LOC105274706 | 1448 | 57 | 393.65 | 1 | 6.91 | 1.75 | 39.53 | 37.37 | 2.16 | 2.16 | 0.09 | 0.09 |
| gene4034 | https://www.ncbi.nlm.nih.gov/gene/?term=LOC105276704 | 2065 | 92 | 445.52 | 5 | 24.21 | 5.43 | 14.65 | 12.65 | 2 | 2 | 0.09 | 0.09 |
| gene10984 | https://www.ncbi.nlm.nih.gov/gene/?term=LOC105284704 | 1987 | 66 | 332.16 | 1 | 5.03 | 1.52 | 25.73 | 23.31 | 2.42 | 2.42 | 0.08 | 0.08 |
| gene5525 | https://www.ncbi.nlm.nih.gov/gene/?term=LOC105278396 | 3090 | 189 | 611.65 | 1 | 3.24 | 0.53 | 25.87 | 24.52 | 1.35 | 1.35 | 0.08 | 0.08 |
| gene920 | https://www.ncbi.nlm.nih.gov/gene/?term=LOC105282518 | 1197 | 46 | 384.29 | 1 | 8.35 | 2.17 | 30.45 | 32.64 | -2.19 | 2.19 | -0.08 | 0.08 |
| gene1768 | https://www.ncbi.nlm.nih.gov/gene/?term=LOC105287938 | 2792 | 89 | 318.77 | 1 | 3.58 | 1.12 | 25.55 | 28.06 | -2.51 | 2.51 | -0.08 | 0.08 |
| gene3817 | https://www.ncbi.nlm.nih.gov/gene/?term=LOC105276449 | 1230 | 55 | 447.15 | 1 | 8.13 | 1.82 | 24.78 | 23.21 | 1.57 | 1.57 | 0.07 | 0.07 |
| gene10552 | https://www.ncbi.nlm.nih.gov/gene/?term=LOC105284213 | 878 | 23 | 261.96 | 1 | 11.39 | 4.35 | 9.72 | 12.3 | -2.58 | 2.58 | -0.07 | 0.07 |
| gene3774 | https://www.ncbi.nlm.nih.gov/gene/?term=LOC105276553 | 2716 | 84 | 309.28 | 1 | 3.68 | 1.19 | 43.69 | 41.82 | 1.87 | 1.87 | 0.06 | 0.06 |
| gene7493 | https://www.ncbi.nlm.nih.gov/gene/?term=LOC105280704 | 4225 | 155 | 366.86 | 1 | 2.37 | 0.65 | 17.59 | 16.02 | 1.58 | 1.58 | 0.06 | 0.06 |
| gene935 | https://www.ncbi.nlm.nih.gov/gene/?term=LOC105282682 | 4368 | 240 | 549.45 | 1 | 2.29 | 0.42 | 13.64 | 12.5 | 1.14 | 1.14 | 0.06 | 0.06 |
| gene11340 | https://www.ncbi.nlm.nih.gov/gene/?term=LOC105285131 | 2918 | 100 | 342.7 1 | 3.43 | 1 | 13.44 | 15.08 | -1.64 | 1.64 | -0.06 | 0.06 | |
| gene274 | https://www.ncbi.nlm.nih.gov/gene/?term=LOC105275190 | 3488 | 74 | 212.16 | 1 | 2.87 | 1.35 | 22.52 | 20.11 | 2.4 | 2.4 | 0.05 | 0.05 |
| gene3671 | https://www.ncbi.nlm.nih.gov/gene/?term=LOC105276272 | 1966 | 47 | 239.06 | 1 | 5.09 | 2.13 | 25.43 | 23.44 | 1.98 | 1.98 | 0.05 | 0.05 |
| gene5831 | https://www.ncbi.nlm.nih.gov/gene/?term=LOC105278761 | 2506 | 119 | 474.86 | 1 | 3.99 | 0.84 | 22.74 | 21.63 | 1.11 | 1.11 | 0.05 | 0.05 |
| gene298 | https://www.ncbi.nlm.nih.gov/gene/?term=LOC105275460 | 2473 | 123 | 497.37 | 1 | 4.04 | 0.81 | 13.29 | 12.33 | 0.96 | 0.96 | 0.05 | 0.05 |
| gene7556 | https://www.ncbi.nlm.nih.gov/gene/?term=LOC105280774 | 2792 | 135 | 483.52 | 1 | 3.58 | 0.74 | 25.37 | 24.41 | 0.96 | 0.96 | 0.05 | 0.05 |
| gene1505 | https://www.ncbi.nlm.nih.gov/gene/?term=LOC105287633 | 1693 | 24 | 141.76 | 1 | 5.91 | 4.17 | 48.26 | 51.48 | -3.22 | 3.22 | -0.05 | 0.05 |
| gene3166 | https://www.ncbi.nlm.nih.gov/gene/?term=LOC105275707 | 2393 | 12 | 50.15 1 | 4.18 | 8.33 | 62.45 | 73.29 | -10.83 | | 10.83 | -0.05 | 0.05 |
| gene393 | https://www.ncbi.nlm.nih.gov/gene/?term=LOC105276566 | 1123 | 2 | 17.81 1 | 8.9 | 50 | 47.62 | 74.3 | -26.68 | | 26.68 | -0.05 | 0.05 |
| gene10107 | https://www.ncbi.nlm.nih.gov/gene/?term=LOC105283712 | 1883 | 34 | 180.56 | 1 | 5.31 | 2.94 | 31.9 | 29.48 | 2.42 | 2.42 | 0.04 | 0.04 |
| gene9336 | https://www.ncbi.nlm.nih.gov/gene/?term=LOC105282834 | 5993 | 137 | 228.6 1 | 1.67 | 0.73 | 15.64 | 13.69 | 1.95 | 1.95 | 0.04 | 0.04 | |
| gene10879 | https://www.ncbi.nlm.nih.gov/gene/?term=LOC105284574 | 2423 | 52 | 214.61 | 1 | 4.13 | 1.92 | 56.31 | 54.67 | 1.63 | 1.63 | 0.04 | 0.04 |
| gene940 | https://www.ncbi.nlm.nih.gov/gene/?term=LOC105282794 | 3578 | 95 | 265.51 | 1 | 2.79 | 1.05 | 15.95 | 14.34 | 1.61 | 1.61 | 0.04 | 0.04 |
| gene2241 | https://www.ncbi.nlm.nih.gov/gene/?term=LOC105274666 | 2598 | 64 | 246.34 | 1 | 3.85 | 1.56 | 28.81 | 27.27 | 1.54 | 1.54 | 0.04 | 0.04 |
| gene1147 | https://www.ncbi.nlm.nih.gov/gene/?term=LOC105284954 | 4022 | 114 | 283.44 | 1 | 2.49 | 0.88 | 14.88 | 13.52 | 1.36 | 1.36 | 0.04 | 0.04 |
| gene5524 | https://www.ncbi.nlm.nih.gov/gene/?term=LOC105278397 | 5177 | 206 | 397.91 | 1 | 1.93 | 0.49 | 25.82 | 24.7 | 1.12 | 1.12 | 0.04 | 0.04 |
| gene12406 | https://www.ncbi.nlm.nih.gov/gene/?term=LOC105286316 | 1386 | 52 | 375.18 | 1 | 7.22 | 1.92 | 33.87 | 32.77 | 1.1 | 1.1 | 0.04 | 0.04 |
| gene3773 | https://www.ncbi.nlm.nih.gov/gene/?term=LOC105276408 | 5522 | 200 | 362.19 | 1 | 1.81 | 0.5 | 21.6 | 20.62 | 0.98 | 0.98 | 0.04 | 0.04 |
| gene11819 | https://www.ncbi.nlm.nih.gov/gene/?term=LOC105285671 | 2366 | 103 | 435.33 | 1 | 4.23 | 0.97 | 30.19 | 29.25 | 0.95 | 0.95 | 0.04 | 0.04 |
| gene10540 | https://www.ncbi.nlm.nih.gov/gene/?term=LOC105284200 | 5267 | 219 | 415.8 1 | 1.9 | 0.46 | 13.98 | 13.1 | 0.88 | 0.88 | 0.04 | 0.04 | |
| gene6286 | https://www.ncbi.nlm.nih.gov/gene/?term=LOC105279298 | 2861 | 133 | 464.87 | 2 | 6.99 | 1.5 | 23.83 | 22.97 | 0.86 | 0.86 | 0.04 | 0.04 |
| gene10678 | https://www.ncbi.nlm.nih.gov/gene/?term=LOC105284355 | 4031 | 172 | 426.69 | 1 | 2.48 | 0.58 | 20.12 | 19.3 | 0.83 | 0.83 | 0.04 | 0.04 |
| gene10606 | https://www.ncbi.nlm.nih.gov/gene/?term=LOC105284273 | 6119 | 286 | 467.4 1 | 1.63 | 0.35 | 18.29 | 17.53 | 0.75 | 0.75 | 0.04 | 0.04 | |
| gene10864 | https://www.ncbi.nlm.nih.gov/gene/?term=LOC105284557 | 1666 | 91 | 546.22 | 1 | 6 | 1.1 | 27.88 | 27.16 | 0.72 | 0.72 | 0.04 | 0.04 |
| gene12802 | https://www.ncbi.nlm.nih.gov/gene/?term=LOC105286799 | 3183 | 184 | 578.07 | 1 | 3.14 | 0.54 | 17.39 | 16.7 | 0.69 | 0.69 | 0.04 | 0.04 |
| gene5298 | https://www.ncbi.nlm.nih.gov/gene/?term=LOC105278145 | 4060 | 275 | 677.34 | 2 | 4.93 | 0.73 | 9.89 | 10.42 | -0.53 | 0.53 | -0.04 | 0.04 |
| gene7550 | https://www.ncbi.nlm.nih.gov/gene/?term=LOC105280763 | 2173 | 109 | 501.61 | 1 | 4.6 | 0.92 | 27.44 | 28.14 | -0.7 | 0.7 | -0.04 | 0.04 |
| gene11401 | https://www.ncbi.nlm.nih.gov/gene/?term=LOC105285202 | 1841 | 89 | 483.43 | 1 | 5.43 | 1.12 | 23.71 | 24.59 | -0.89 | 0.89 | -0.04 | 0.04 |
| gene9101 | https://www.ncbi.nlm.nih.gov/gene/?term=LOC105282553 | 3068 | 110 | 358.54 | 1 | 3.26 | 0.91 | 21.48 | 22.61 | -1.13 | 1.13 | -0.04 | 0.04 |
| gene1199 | https://www.ncbi.nlm.nih.gov/gene/?term=LOC105285498 | 2220 | 29 | 130.63 | 1 | 4.5 | 3.45 | 32.78 | 35.99 | -3.21 | 3.21 | -0.04 | 0.04 |
| gene12395 | https://www.ncbi.nlm.nih.gov/gene/?term=LOC105286302 | 4093 | 24 | 58.64 1 | 2.44 | 4.17 | 20.4 | 15.15 | 5.24 | 5.24 | 0.03 | 0.03 | |
| gene2970 | https://www.ncbi.nlm.nih.gov/gene/?term=LOC105275518 | 2340 | 28 | 119.66 | 1 | 4.27 | 3.57 | 41.75 | 39.36 | 2.39 | 2.39 | 0.03 | 0.03 |
| gene388 | https://www.ncbi.nlm.nih.gov/gene/?term=LOC105276538 | 1558 | 24 | 154.04 | 2 | 12.84 | 8.33 | 66.6 | 64.8 | 1.8 | 1.8 | 0.03 | 0.03 |
| gene8742 | https://www.ncbi.nlm.nih.gov/gene/?term=LOC105282151 | 2563 | 44 | 171.67 | 1 | 3.9 | 2.27 | 43.82 | 42.21 | 1.61 | 1.61 | 0.03 | 0.03 |
| gene2825 | https://www.ncbi.nlm.nih.gov/gene/?term=LOC105275343 | 5545 | 98 | 176.74 | 1 | 1.8 | 1.02 | 9.51 | 8.1 | 1.42 | 1.42 | 0.03 | 0.03 |
| gene12572 | https://www.ncbi.nlm.nih.gov/gene/?term=LOC105286514 | 2374 | 56 | 235.89 | 1 | 4.21 | 1.79 | 36.49 | 35.22 | 1.27 | 1.27 | 0.03 | 0.03 |
| gene5877 | https://www.ncbi.nlm.nih.gov/gene/?term=LOC105278807 | 1719 | 53 | 308.32 | 1 | 5.82 | 1.89 | 27.62 | 26.53 | 1.08 | 1.08 | 0.03 | 0.03 |
| gene4892 | https://www.ncbi.nlm.nih.gov/gene/?term=LOC105277667 | 3898 | 127 | 325.81 | 1 | 2.57 | 0.79 | 28.13 | 27.36 | 0.77 | 0.77 | 0.03 | 0.03 |
| gene289 | https://www.ncbi.nlm.nih.gov/gene/?term=LOC105275347 | 2533 | 90 | 355.31 | 1 | 3.95 | 1.11 | 24.42 | 23.69 | 0.73 | 0.73 | 0.03 | 0.03 |
| gene13036 | https://www.ncbi.nlm.nih.gov/gene/?term=LOC105287059 | 4959 | 200 | 403.31 | 1 | 2.02 | 0.5 | 17.56 | 16.85 | 0.71 | 0.71 | 0.03 | 0.03 |

| | | | | | | | | | | | | | |
|---|---|---|---|---|---|---|---|---|---|---|---|---|---|
| gene1573 | https://www.ncbi.nlm.nih.gov/gene/?term=LOC105287704 | 3030 | 143 | 471.95 | 1 | 3.3 | 0.7 | 19.92 | 19.38 | 0.54 | 0.54 | 0.03 | 0.03 |
| gene1470 | https://www.ncbi.nlm.nih.gov/gene/?term=LOC105287597 | 2974 | 200 | 672.49 | 1 | 3.36 | 0.5 | 18.11 | 18.56 | -0.46 | 0.46 | -0.03 | 0.03 |
| gene12377 | https://www.ncbi.nlm.nih.gov/gene/?term=LOC105286279 | 5846 | 278 | 475.54 | 1 | 1.71 | 0.36 | 2.15 | 2.82 | -0.67 | 0.67 | -0.03 | 0.03 |
| gene10213 | https://www.ncbi.nlm.nih.gov/gene/?term=LOC105283826 | 1503 | 62 | 412.51 | 1 | 6.65 | 1.61 | 14.74 | 15.54 | -0.8 | 0.8 | -0.03 | 0.03 |
| gene10392 | https://www.ncbi.nlm.nih.gov/gene/?term=LOC105284036 | 3216 | 103 | 320.27 | 2 | 6.22 | 1.94 | 24.04 | 24.86 | -0.81 | 0.81 | -0.03 | 0.03 |
| gene2404 | https://www.ncbi.nlm.nih.gov/gene/?term=LOC105274853 | 1767 | 45 | 254.67 | 1 | 5.66 | 2.22 | 30.61 | 31.94 | -1.33 | 1.33 | -0.03 | 0.03 |
| gene10654 | https://www.ncbi.nlm.nih.gov/gene/?term=LOC105284329 | 4027 | 72 | 178.79 | 1 | 2.48 | 1.39 | 18.86 | 20.42 | -1.56 | 1.56 | -0.03 | 0.03 |
| gene6237 | https://www.ncbi.nlm.nih.gov/gene/?term=LOC105279244 | 1484 | 30 | 202.16 | 1 | 6.74 | 3.33 | 49.36 | 51.05 | -1.69 | 1.69 | -0.03 | 0.03 |
| gene7398 | https://www.ncbi.nlm.nih.gov/gene/?term=LOC105280589 | 3452 | 42 | 121.67 | 1 | 2.9 | 2.38 | 23.2 | 25.91 | -2.71 | 2.71 | -0.03 | 0.03 |
| gene8691 | https://www.ncbi.nlm.nih.gov/gene/?term=LOC105282094 | 2028 | 6 | 29.59 1 | 4.93 | 16.67 | 86.17 | 97.58 | -11.41 | | 11.41 | -0.03 | 0.03 |
| gene8178 | https://www.ncbi.nlm.nih.gov/gene/?term=LOC105281515 | 8647 | 59 | 68.23 1 | 1.16 | 1.69 | 30.95 | 28.43 | 2.52 | 2.52 | 0.02 | 0.02 | |
| gene4996 | https://www.ncbi.nlm.nih.gov/gene/?term=LOC105277798 | 5617 | 74 | 131.74 | 1 | 1.78 | 1.35 | 11.47 | 9.83 | 1.64 | 1.64 | 0.02 | 0.02 |
| gene2748 | https://www.ncbi.nlm.nih.gov/gene/?term=LOC105275261 | 3845 | 55 | 143.04 | 1 | 2.6 | 1.82 | 41.25 | 39.71 | 1.54 | 1.54 | 0.02 | 0.02 |
| gene5273 | https://www.ncbi.nlm.nih.gov/gene/?term=LOC105278108 | 4067 | 57 | 140.15 | 1 | 2.46 | 1.75 | 25.93 | 24.39 | 1.54 | 1.54 | 0.02 | 0.02 |
| gene322 | https://www.ncbi.nlm.nih.gov/gene/?term=LOC105275777 | 1606 | 29 | 180.57 | 1 | 6.23 | 3.45 | 36.14 | 34.8 | 1.35 | 1.35 | 0.02 | 0.02 |
| gene927 | https://www.ncbi.nlm.nih.gov/gene/?term=LOC105282600 | 3727 | 67 | 179.77 | 1 | 2.68 | 1.49 | 27.6 | 26.32 | 1.28 | 1.28 | 0.02 | 0.02 |
| gene6455 | https://www.ncbi.nlm.nih.gov/gene/?term=LOC105279500 | 2612 | 36 | 137.83 | 1 | 3.83 | 2.78 | 48.83 | 47.56 | 1.26 | 1.26 | 0.02 | 0.02 |
| gene4048 | https://www.ncbi.nlm.nih.gov/gene/?term=LOC105276724 | 2955 | 47 | 159.05 | 1 | 3.38 | 2.13 | 12.29 | 11.14 | 1.14 | 1.14 | 0.02 | 0.02 |
| gene10222 | https://www.ncbi.nlm.nih.gov/gene/?term=LOC105283838 | 1748 | 33 | 188.79 | 1 | 5.72 | 3.03 | 33.58 | 32.49 | 1.1 | 1.1 | 0.02 | 0.02 |
| gene12021 | https://www.ncbi.nlm.nih.gov/gene/?term=LOC105285905 | 1399 | 27 | 192.99 | 1 | 7.15 | 3.7 | 37.97 | 36.9 | 1.06 | 1.06 | 0.02 | 0.02 |
| gene7183 | https://www.ncbi.nlm.nih.gov/gene/?term=LOC105280348 | 3925 | 79 | 201.27 | 1 | 2.55 | 1.27 | 18.1 | 17.08 | 1.01 | 1.01 | 0.02 | 0.02 |
| gene1884 | https://www.ncbi.nlm.nih.gov/gene/?term=LOC105288072 | 4169 | 88 | 211.08 | 1 | 2.4 | 1.14 | 22.65 | 21.71 | 0.94 | 0.94 | 0.02 | 0.02 |
| gene12516 | https://www.ncbi.nlm.nih.gov/gene/?term=LOC105286452 | 1529 | 42 | 274.69 | 1 | 6.54 | 2.38 | 47.45 | 46.6 | 0.86 | 0.86 | 0.02 | 0.02 |
| gene10600 | https://www.ncbi.nlm.nih.gov/gene/?term=LOC105284270 | 4837 | 114 | 235.68 | 1 | 2.07 | 0.88 | 12.4 | 11.65 | 0.75 | 0.75 | 0.02 | 0.02 |
| gene8015 | https://www.ncbi.nlm.nih.gov/gene/?term=LOC105281312 | 2998 | 69 | 230.15 | 1 | 3.34 | 1.45 | 31.17 | 30.43 | 0.74 | 0.74 | 0.02 | 0.02 |
| gene9330 | https://www.ncbi.nlm.nih.gov/gene/?term=LOC105282825 | 3936 | 102 | 259.15 | 1 | 2.54 | 0.98 | 14.91 | 14.23 | 0.68 | 0.68 | 0.02 | 0.02 |
| gene6031 | https://www.ncbi.nlm.nih.gov/gene/?term=LOC105279000 | 5691 | 226 | 397.12 | 1 | 1.76 | 0.44 | 12.2 | 11.72 | 0.48 | 0.48 | 0.02 | 0.02 |
| gene3206 | https://www.ncbi.nlm.nih.gov/gene/?term=LOC105275752 | 3474 | 138 | 397.24 | 1 | 2.88 | 0.72 | 31.47 | 31.01 | 0.46 | 0.46 | 0.02 | 0.02 |
| gene7888 | https://www.ncbi.nlm.nih.gov/gene/?term=LOC105281141 | 4357 | 180 | 413.13 | 1 | 2.3 | 0.56 | 27.03 | 26.57 | 0.45 | 0.45 | 0.02 | 0.02 |
| gene4641 | https://www.ncbi.nlm.nih.gov/gene/?term=LOC105277398 | 1937 | 74 | 382.03 | 1 | 5.16 | 1.35 | 35.22 | 34.78 | 0.44 | 0.44 | 0.02 | 0.02 |
| gene7249 | https://www.ncbi.nlm.nih.gov/gene/?term=LOC105280422 | 1775 | 66 | 371.83 | 1 | 5.63 | 1.52 | 16.48 | 16.04 | 0.44 | 0.44 | 0.02 | 0.02 |
| gene10520 | https://www.ncbi.nlm.nih.gov/gene/?term=LOC105284184 | 2451 | 105 | 428.4 1 | 4.08 | 0.95 | 28.89 | 28.49 | 0.4 | 0.4 | 0.02 | 0.02 | |
| gene2846 | https://www.ncbi.nlm.nih.gov/gene/?term=LOC105275374 | 11246 | 566 | 503.29 | 1 | 0.89 | 0.18 | 9.56 | 9.16 | 0.4 | 0.4 | 0.02 | 0.02 |
| gene570 | https://www.ncbi.nlm.nih.gov/gene/?term=LOC105278549 | 1673 | 74 | 442.32 | 1 | 5.98 | 1.35 | 44.93 | 44.55 | 0.38 | 0.38 | 0.02 | 0.02 |
| gene3885 | https://www.ncbi.nlm.nih.gov/gene/?term=LOC105276527 | 7928 | 380 | 479.31 | 1 | 1.26 | 0.26 | 8.1 | 7.75 | 0.34 | 0.34 | 0.02 | 0.02 |
| gene2844 | https://www.ncbi.nlm.nih.gov/gene/?term=LOC105275363 | 4330 | 258 | 595.84 | 1 | 2.31 | 0.39 | 11.65 | 11.32 | 0.33 | 0.33 | 0.02 | 0.02 |
| gene6583 | https://www.ncbi.nlm.nih.gov/gene/?term=LOC105279649 | 7444 | 393 | 527.94 | 1 | 1.34 | 0.25 | 10.67 | 11.14 | -0.47 | 0.47 | -0.02 | 0.02 |
| gene3848 | https://www.ncbi.nlm.nih.gov/gene/?term=LOC105276486 | 5899 | 176 | 298.36 | 1 | 1.7 | 0.57 | 23.6 | 24.17 | -0.57 | 0.57 | -0.02 | 0.02 |
| gene4962 | https://www.ncbi.nlm.nih.gov/gene/?term=LOC105277761 | 4645 | 175 | 376.75 | 1 | 2.15 | 0.57 | 13.58 | 14.2 | -0.62 | 0.62 | -0.02 | 0.02 |
| gene7207 | https://www.ncbi.nlm.nih.gov/gene/?term=LOC105280376 | 4659 | 124 | 266.15 | 1 | 2.15 | 0.81 | 26.54 | 27.17 | -0.63 | 0.63 | -0.02 | 0.02 |
| gene3385 | https://www.ncbi.nlm.nih.gov/gene/?term=LOC105275966 | 3305 | 75 | 226.93 | 1 | 3.03 | 1.33 | 43.86 | 44.6 | -0.75 | 0.75 | -0.02 | 0.02 |
| gene11454 | https://www.ncbi.nlm.nih.gov/gene/?term=LOC105285256 | 1906 | 50 | 262.33 | 1 | 5.25 | 2 | 43.28 | 44.08 | -0.8 | 0.8 | -0.02 | 0.02 |
| gene7915 | https://www.ncbi.nlm.nih.gov/gene/?term=LOC105281173 | 1597 | 39 | 244.21 | 1 | 6.26 | 2.56 | 61.13 | 62.01 | -0.88 | 0.88 | -0.02 | 0.02 |
| gene11447 | https://www.ncbi.nlm.nih.gov/gene/?term=LOC105285252 | 7437 | 150 | 201.69 | 1 | 1.34 | 0.67 | 11.78 | 12.78 | -0.99 | 0.99 | -0.02 | 0.02 |
| gene3168 | https://www.ncbi.nlm.nih.gov/gene/?term=LOC105275711 | 6150 | 95 | 154.47 | 1 | 1.63 | 1.05 | 10.97 | 11.98 | -1 | 1 | -0.02 | 0.02 |
| gene7657 | https://www.ncbi.nlm.nih.gov/gene/?term=LOC105280890 | 1109 | 23 | 207.39 | 1 | 9.02 | 4.35 | 23.17 | 24.26 | -1.09 | 1.09 | -0.02 | 0.02 |
| gene10155 | https://www.ncbi.nlm.nih.gov/gene/?term=LOC105283756 | 7030 | 119 | 169.27 | 1 | 1.42 | 0.84 | 6.89 | 8.19 | -1.3 | 1.3 | -0.02 | 0.02 |
| gene9411 | https://www.ncbi.nlm.nih.gov/gene/?term=LOC105282914 | 5534 | 42 | 75.89 1 | 1.81 | 2.38 | 48.54 | 50.92 | -2.38 | 2.38 | -0.02 | 0.02 | |
| gene9109 | https://www.ncbi.nlm.nih.gov/gene/?term=LOC105282561 | 2400 | 17 | 70.83 1 | 4.17 | 5.88 | 28.52 | 31.86 | -3.34 | 3.34 | -0.02 | 0.02 | |
| gene11584 | https://www.ncbi.nlm.nih.gov/gene/?term=LOC105285416 | 3709 | 15 | 40.44 2 | 5.39 | 13.33 | 24.1 | 29.33 | -5.23 | 5.23 | -0.02 | 0.02 | |
| gene12106 | https://www.ncbi.nlm.nih.gov/gene/?term=LOC105286002 | 8556 | 83 | 97.01 1 | 1.17 | 1.2 | 3.39 | 2.1 | 1.29 | 1.29 | 0.01 | 0.01 | |
| gene5052 | https://www.ncbi.nlm.nih.gov/gene/?term=LOC105277856 | 3167 | 36 | 113.67 | 1 | 3.16 | 2.78 | 37.89 | 36.75 | 1.14 | 1.14 | 0.01 | 0.01 |
| gene10243 | https://www.ncbi.nlm.nih.gov/gene/?term=LOC105283862 | 3239 | 45 | 138.93 | 1 | 3.09 | 2.22 | 29.81 | 28.76 | 1.05 | 1.05 | 0.01 | 0.01 |
| gene2833 | https://www.ncbi.nlm.nih.gov/gene/?term=LOC105275352 | 5528 | 43 | 77.79 1 | 1.81 | 2.33 | 22.51 | 21.53 | 0.98 | 0.98 | 0.01 | 0.01 | |
| gene3912 | https://www.ncbi.nlm.nih.gov/gene/?term=LOC105276560 | 17900 | 373 | 208.38 | 3 | 1.68 | 0.8 | 2.56 | 1.89 | 0.67 | 0.67 | 0.01 | 0.01 |
| gene814 | https://www.ncbi.nlm.nih.gov/gene/?term=LOC105281296 | 2722 | 57 | 209.4 1 | 3.67 | 1.75 | 23.86 | 23.2 | 0.65 | 0.65 | 0.01 | 0.01 | |
| gene8343 | https://www.ncbi.nlm.nih.gov/gene/?term=LOC105281700 | 2172 | 43 | 197.97 | 1 | 4.6 | 2.33 | 36.96 | 36.39 | 0.57 | 0.57 | 0.01 | 0.01 |
| gene2257 | https://www.ncbi.nlm.nih.gov/gene/?term=LOC105274689 | 3658 | 90 | 246.04 | 1 | 2.73 | 1.11 | 15.54 | 15.05 | 0.49 | 0.49 | 0.01 | 0.01 |
| gene9648 | https://www.ncbi.nlm.nih.gov/gene/?term=LOC105283183 | 2602 | 33 | 126.83 | 1 | 3.84 | 3.03 | 45.18 | 44.75 | 0.44 | 0.44 | 0.01 | 0.01 |
| gene2330 | https://www.ncbi.nlm.nih.gov/gene/?term=LOC105274773 | 13104 | 208 | 158.73 | 1 | 0.76 | 0.48 | 15.28 | 14.85 | 0.43 | 0.43 | 0.01 | 0.01 |
| gene8814 | https://www.ncbi.nlm.nih.gov/gene/?term=LOC105282240 | 6910 | 209 | 302.46 | 1 | 1.45 | 0.48 | 8.7 | 8.29 | 0.4 | 0.4 | 0.01 | 0.01 |
| gene9029 | https://www.ncbi.nlm.nih.gov/gene/?term=LOC105282491 | 4762 | 190 | 398.99 | 1 | 2.1 | 0.53 | 8.59 | 8.25 | 0.34 | 0.34 | 0.01 | 0.01 |
| gene2056 | https://www.ncbi.nlm.nih.gov/gene/?term=LOC105274452 | 6584 | 268 | 407.05 | 1 | 1.52 | 0.37 | 13.37 | 13.03 | 0.33 | 0.33 | 0.01 | 0.01 |
| gene11568 | https://www.ncbi.nlm.nih.gov/gene/?term=LOC105285395 | 8086 | 257 | 317.83 | 1 | 1.24 | 0.39 | 13.19 | 12.93 | 0.27 | 0.27 | 0.01 | 0.01 |

| | | | | | | | | | | | | | |
|---|---|---|---|---|---|---|---|---|---|---|---|---|---|
| gene10018 | https://www.ncbi.nlm.nih.gov/gene/?term=LOC105283607 | 4414 | 222 | 502.95 | 1 | 2.27 | 0.45 | 14.44 | 14.18 | 0.26 | 0.26 | 0.01 | 0.01 |
| gene6620 | https://www.ncbi.nlm.nih.gov/gene/?term=LOC105279687 | 17998 | 456 | 253.36 | 1 | 0.56 | 0.22 | 4.79 | 4.53 | 0.26 | 0.26 | 0.01 | 0.01 |
| gene8600 | https://www.ncbi.nlm.nih.gov/gene/?term=LOC105282001 | 8513 | 211 | 247.86 | 1 | 1.17 | 0.47 | 9.85 | 9.6 | 0.25 | 0.25 | 0.01 | 0.01 |
| gene1310 | https://www.ncbi.nlm.nih.gov/gene/?term=LOC105286661 | 2582 | 72 | 278.85 | 1 | 3.87 | 1.39 | 20.69 | 20.45 | 0.24 | 0.24 | 0.01 | 0.01 |
| gene292 | https://www.ncbi.nlm.nih.gov/gene/?term=LOC105275398 | 2825 | 92 | 325.66 | 1 | 3.54 | 1.09 | 21.85 | 21.62 | 0.23 | 0.23 | 0.01 | 0.01 |
| gene11275 | https://www.ncbi.nlm.nih.gov/gene/?term=LOC105285055 | 4119 | 262 | 636.08 | 1 | 2.43 | 0.38 | 11.65 | 11.43 | 0.22 | 0.22 | 0.01 | 0.01 |
| gene8729 | https://www.ncbi.nlm.nih.gov/gene/?term=LOC105282137 | 5005 | 236 | 471.53 | 1 | 2 | 0.42 | 14.47 | 14.26 | 0.21 | 0.21 | 0.01 | 0.01 |
| gene2806 | https://www.ncbi.nlm.nih.gov/gene/?term=LOC105275327 | 14257 | 550 | 385.78 | 1 | 0.7 | 0.18 | 6.58 | 6.38 | 0.2 | 0.2 | 0.01 | 0.01 |
| gene3747 | https://www.ncbi.nlm.nih.gov/gene/?term=LOC105276387 | 3736 | 198 | 529.98 | 1 | 2.68 | 0.51 | 14.52 | 14.33 | 0.19 | 0.19 | 0.01 | 0.01 |
| gene5746 | https://www.ncbi.nlm.nih.gov/gene/?term=LOC105278666 | 10343 | 491 | 474.72 | 1 | 0.97 | 0.2 | 6.89 | 6.71 | 0.19 | 0.19 | 0.01 | 0.01 |
| gene10596 | https://www.ncbi.nlm.nih.gov/gene/?term=LOC105284262 | 6520 | 247 | 378.83 | 1 | 1.53 | 0.4 | 4.51 | 4.34 | 0.18 | 0.18 | 0.01 | 0.01 |
| gene13114 | https://www.ncbi.nlm.nih.gov/gene/?term=LOC105287147 | 7609 | 447 | 587.46 | 1 | 1.31 | 0.22 | 9.07 | 9.17 | -0.1 | 0.1 | -0.01 | 0.01 |
| gene8274 | https://www.ncbi.nlm.nih.gov/gene/?term=LOC105281624 | 5425 | 244 | 449.77 | 1 | 1.84 | 0.41 | 9.34 | 9.46 | -0.12 | 0.12 | -0.01 | 0.01 |
| gene12674 | https://www.ncbi.nlm.nih.gov/gene/?term=LOC105286649 | 18283 | 836 | 457.26 | 1 | 0.55 | 0.12 | 1.34 | 1.5 | -0.16 | 0.16 | -0.01 | 0.01 |
| gene5539 | https://www.ncbi.nlm.nih.gov/gene/?term=LOC105278415 | 25024 | 835 | 333.68 | 1 | 0.4 | 0.12 | 3.01 | 3.18 | -0.17 | 0.17 | -0.01 | 0.01 |
| gene11082 | https://www.ncbi.nlm.nih.gov/gene/?term=LOC105284824 | 7436 | 245 | 329.48 | 1 | 1.34 | 0.41 | 7.32 | 7.57 | -0.25 | 0.25 | -0.01 | 0.01 |
| gene1198 | https://www.ncbi.nlm.nih.gov/gene/?term=LOC105285507 | 2738 | 98 | 357.93 | 1 | 3.65 | 1.02 | 21.45 | 21.7 | -0.25 | 0.25 | -0.01 | 0.01 |
| gene12407 | https://www.ncbi.nlm.nih.gov/gene/?term=LOC105286315 | 8593 | 227 | 264.17 | 1 | 1.16 | 0.44 | 13.44 | 13.72 | -0.28 | 0.28 | -0.01 | 0.01 |
| gene11583 | https://www.ncbi.nlm.nih.gov/gene/?term=LOC105285414 | 8192 | 204 | 249.02 | 1 | 1.22 | 0.49 | 7.78 | 8.09 | -0.31 | 0.31 | -0.01 | 0.01 |
| gene12573 | https://www.ncbi.nlm.nih.gov/gene/?term=LOC105286515 | 3647 | 128 | 350.97 | 1 | 2.74 | 0.78 | 10.64 | 10.97 | -0.33 | 0.33 | -0.01 | 0.01 |
| gene9598 | https://www.ncbi.nlm.nih.gov/gene/?term=LOC105283132 | 1769 | 56 | 316.56 | 1 | 5.65 | 1.79 | 39.81 | 40.18 | -0.38 | 0.38 | -0.01 | 0.01 |
| gene1220 | https://www.ncbi.nlm.nih.gov/gene/?term=LOC105285742 | 3453 | 123 | 356.21 | 1 | 2.9 | 0.81 | 16.77 | 17.17 | -0.4 | 0.4 | -0.01 | 0.01 |
| gene657 | https://www.ncbi.nlm.nih.gov/gene/?term=LOC105279667 | 6421 | 106 | 165.08 | 1 | 1.56 | 0.94 | 2.17 | 2.57 | -0.4 | 0.4 | -0.01 | 0.01 |
| gene5646 | https://www.ncbi.nlm.nih.gov/gene/?term=LOC105278534 | 7475 | 214 | 286.29 | 1 | 1.34 | 0.47 | 16.15 | 16.57 | -0.41 | 0.41 | -0.01 | 0.01 |
| gene7148 | https://www.ncbi.nlm.nih.gov/gene/?term=LOC105280304 | 8920 | 143 | 160.31 | 1 | 1.12 | 0.7 | 6.95 | 7.37 | -0.42 | 0.42 | -0.01 | 0.01 |
| gene7600 | https://www.ncbi.nlm.nih.gov/gene/?term=LOC105280819 | 1615 | 43 | 266.25 | 1 | 6.19 | 2.33 | 37.46 | 37.88 | -0.42 | 0.42 | -0.01 | 0.01 |
| gene10819 | https://www.ncbi.nlm.nih.gov/gene/?term=LOC105284507 | 5767 | 122 | 211.55 | 1 | 1.73 | 0.82 | 18.54 | 19.25 | -0.71 | 0.71 | -0.01 | 0.01 |
| gene12203 | https://www.ncbi.nlm.nih.gov/gene/?term=LOC105286099 | 5967 | 87 | 145.8 1 | 1.68 | 1.15 | 11.05 | 12.06 | -1.01 | 1.01 | -0.01 | 0.01 | |
| gene652 | https://www.ncbi.nlm.nih.gov/gene/?term=LOC105279528 | 5806 | 39 | 67.17 1 | 1.72 | 2.56 | 19.31 | 20.42 | -1.11 | 1.11 | -0.01 | 0.01 | |
| gene6639 | https://www.ncbi.nlm.nih.gov/gene/?term=LOC105279705 | 7656 | 88 | 114.94 | 1 | 1.31 | 1.14 | 19.96 | 21.09 | -1.13 | 1.13 | -0.01 | 0.01 |
| gene564 | https://www.ncbi.nlm.nih.gov/gene/?term=LOC105278478 | 2802 | 22 | 78.52 1 | 3.57 | 4.55 | 23.15 | 24.78 | -1.63 | 1.63 | -0.01 | 0.01 | |
| gene9647 | https://www.ncbi.nlm.nih.gov/gene/?term=LOC109611233 | 3094 | 35 | 113.12 | 1 | 3.23 | 2.86 | 42.6 | 42.19 | 0.41 | 0.41 | 0 | 0 |
| gene2683 | https://www.ncbi.nlm.nih.gov/gene/?term=LOC105275187 | 5492 | 87 | 158.41 | 1 | 1.82 | 1.15 | 15.13 | 14.88 | 0.25 | 0.25 | 0 | 0 |
| gene7245 | https://www.ncbi.nlm.nih.gov/gene/?term=LOC105280410 | 12478 | 319 | 255.65 | 1 | 0.8 | 0.31 | 17.86 | 17.71 | 0.15 | 0.15 | 0 | 0 |
| gene2173 | https://www.ncbi.nlm.nih.gov/gene/?term=LOC105274590 | 4387 | 91 | 207.43 | 2 | 4.56 | 2.2 | 24.1 | 23.96 | 0.14 | 0.14 | 0 | 0 |
| gene4225 | https://www.ncbi.nlm.nih.gov/gene/?term=LOC105276917 | 2443 | 40 | 163.73 | 1 | 4.09 | 2.5 | 29.99 | 29.86 | 0.14 | 0.14 | 0 | 0 |
| gene6966 | https://www.ncbi.nlm.nih.gov/gene/?term=LOC105280104 | 3747 | 133 | 354.95 | 1 | 2.67 | 0.75 | 17.09 | 16.96 | 0.13 | 0.13 | 0 | 0 |
| gene6652 | https://www.ncbi.nlm.nih.gov/gene/?term=LOC105279719 | 2831 | 83 | 293.18 | 1 | 3.53 | 1.2 | 23.57 | 23.44 | 0.12 | 0.12 | 0 | 0 |
| gene9722 | https://www.ncbi.nlm.nih.gov/gene/?term=LOC105283264 | 3951 | 94 | 237.91 | 1 | 2.53 | 1.06 | 22.96 | 22.87 | 0.1 | 0.1 | 0 | 0 |
| gene4033 | https://www.ncbi.nlm.nih.gov/gene/?term=LOC105276705 | 10546 | 157 | 148.87 | 1 | 0.95 | 0.64 | 6.45 | 6.38 | 0.06 | 0.06 | 0 | 0 |
| gene3727 | https://www.ncbi.nlm.nih.gov/gene/?term=LOC105276367 | 2484 | 76 | 305.96 | 1 | 4.03 | 1.32 | 41.02 | 40.98 | 0.04 | 0.04 | 0 | 0 |
| gene7319 | https://www.ncbi.nlm.nih.gov/gene/?term=LOC105280513 | 68997 | 1592 | 230.73 | 1 | 0.14 | 0.06 | 0.75 | 0.72 | 0.03 | 0.03 | 0 | 0 |
| gene12307 | https://www.ncbi.nlm.nih.gov/gene/?term=LOC105286216 | 10313 | 209 | 202.66 | 1 | 0.97 | 0.48 | 14.07 | 14.04 | 0.02 | 0.02 | 0 | 0 |
| gene3470 | https://www.ncbi.nlm.nih.gov/gene/?term=LOC105276059 | 8716 | 186 | 213.4 1 | 1.15 | 0.54 | 9.64 | 9.62 | 0.02 | 0.02 | 0 | 0 | |
| gene3107 | https://www.ncbi.nlm.nih.gov/gene/?term=LOC105275650 | 5103 | 187 | 366.45 | 1 | 1.96 | 0.53 | 16.49 | 16.48 | 0.01 | 0.01 | 0 | 0 |
| gene3335 | https://www.ncbi.nlm.nih.gov/gene/?term=LOC105275898 | 4665 | 172 | 368.7 1 | 2.14 | 0.58 | 12.53 | 12.51 | 0.01 | 0.01 | 0 | 0 | |
| gene9326 | https://www.ncbi.nlm.nih.gov/gene/?term=LOC105282818 | 16612 | 251 | 151.1 1 | 0.6 | 0.4 | 12.99 | 12.99 | 0 | 0 | 0 | 0 | |
| gene11561 | https://www.ncbi.nlm.nih.gov/gene/?term=LOC105285387 | 11528 | 226 | 196.04 | 1 | 0.87 | 0.44 | 9.46 | 9.47 | -0.01 | 0.01 | 0 | 0 |
| gene13039 | https://www.ncbi.nlm.nih.gov/gene/?term=LOC105287065 | 4902 | 263 | 536.52 | 1 | 2.04 | 0.38 | 11.98 | 12.01 | -0.02 | 0.02 | 0 | 0 |
| gene4209 | https://www.ncbi.nlm.nih.gov/gene/?term=LOC105277125 | 50550 | 2067 | 408.9 1 | 0.2 | 0.05 | 1.64 | 1.66 | -0.02 | 0.02 | 0 | 0 | |
| gene7166 | https://www.ncbi.nlm.nih.gov/gene/?term=LOC105280326 | 6117 | 258 | 421.78 | 1 | 1.63 | 0.39 | 14.79 | 14.83 | -0.04 | 0.04 | 0 | 0 |
| gene6649 | https://www.ncbi.nlm.nih.gov/gene/?term=LOC105279717 | 6017 | 181 | 300.81 | 1 | 1.66 | 0.55 | 11.29 | 11.35 | -0.06 | 0.06 | 0 | 0 |
| gene3856 | https://www.ncbi.nlm.nih.gov/gene/?term=LOC105276499 | 5236 | 206 | 393.43 | 1 | 1.91 | 0.49 | 10.22 | 10.29 | -0.07 | 0.07 | 0 | 0 |
| gene10326 | https://www.ncbi.nlm.nih.gov/gene/?term=LOC105283962 | 3042 | 145 | 476.66 | 1 | 3.29 | 0.69 | 9.18 | 9.26 | -0.08 | 0.08 | 0 | 0 |
| gene10549 | https://www.ncbi.nlm.nih.gov/gene/?term=LOC105284210 | 7145 | 374 | 523.44 | 2 | 2.8 | 0.53 | 9.66 | 9.75 | -0.09 | 0.09 | 0 | 0 |
| gene2835 | https://www.ncbi.nlm.nih.gov/gene/?term=LOC105275357 | 2492 | 122 | 489.57 | 1 | 4.01 | 0.82 | 14.14 | 14.23 | -0.09 | 0.09 | 0 | 0 |
| gene4190 | https://www.ncbi.nlm.nih.gov/gene/?term=LOC105276875 | 3150 | 135 | 428.57 | 1 | 3.17 | 0.74 | 20.47 | 20.57 | -0.09 | 0.09 | 0 | 0 |
| gene5613 | https://www.ncbi.nlm.nih.gov/gene/?term=LOC105278504 | 8524 | 274 | 321.45 | 1 | 1.17 | 0.36 | 5.89 | 5.98 | -0.09 | 0.09 | 0 | 0 |
| gene9755 | https://www.ncbi.nlm.nih.gov/gene/?term=LOC105283296 | 2177 | 97 | 445.57 | 1 | 4.59 | 1.03 | 14.1 | 14.18 | -0.09 | 0.09 | 0 | 0 |
| gene4376 | https://www.ncbi.nlm.nih.gov/gene/?term=LOC105277083 | 4307 | 140 | 325.05 | 1 | 2.32 | 0.71 | 17.67 | 17.76 | -0.1 | 0.1 | 0 | 0 |
| gene53 | https://www.ncbi.nlm.nih.gov/gene/?term=LOC105277148 | 19283 | 339 | 175.8 1 | 0.52 | 0.29 | 4.53 | 4.65 | -0.11 | 0.11 | 0 | 0 | |
| gene1632 | https://www.ncbi.nlm.nih.gov/gene/?term=LOC105287764 | 18061 | 462 | 255.8 1 | 0.55 | 0.22 | 3.51 | 3.63 | -0.12 | 0.12 | 0 | 0 | |
| gene7547 | https://www.ncbi.nlm.nih.gov/gene/?term=LOC105280767 | 24576 | 950 | 386.56 | 1 | 0.41 | 0.11 | 4.24 | 4.36 | -0.12 | 0.12 | 0 | 0 |
| gene3599 | https://www.ncbi.nlm.nih.gov/gene/?term=LOC105276196 | 2955 | 61 | 206.43 | 1 | 3.38 | 1.64 | 38.99 | 39.11 | -0.13 | 0.13 | 0 | 0 |

| | | | | | | | | | | | | | |
|---|---|---|---|---|---|---|---|---|---|---|---|---|---|
| gene9327 | https://www.ncbi.nlm.nih.gov/gene/?term=LOC105282821 | 7793 | 184 | 236.11 | 1 | 1.28 | 0.54 | 12.73 | 12.87 | -0.14 | 0.14 | 0 | 0 |
| gene11579 | https://www.ncbi.nlm.nih.gov/gene/?term=LOC105285406 | 13571 | 341 | 251.27 | 1 | 0.74 | 0.29 | 6.04 | 6.2 | -0.15 | 0.15 | 0 | 0 |
| gene7368 | https://www.ncbi.nlm.nih.gov/gene/?term=LOC105280556 | 4105 | 76 | 185.14 | 1 | 2.44 | 1.32 | 15.19 | 15.44 | -0.25 | 0.25 | 0 | 0 |
| gene9464 | https://www.ncbi.nlm.nih.gov/gene/?term=LOC105282975 | 1450 | 25 | 172.41 | 1 | 6.9 | 4 | 40.46 | 40.7 | -0.25 | 0.25 | 0 | 0 |
| gene10345 | https://www.ncbi.nlm.nih.gov/gene/?term=LOC105283989 | 4444 | 40 | 90.01 | 1 | 2.25 | 2.5 | 37.64 | 38.15 | -0.5 | 0.5 | 0 | 0 |

Directory: OGLall File: rnk-dmg-Cb_LI_RphaseA2.vs.RphaseB2.txt

| gene_ID | gene_link | gwidth | #Sites | #per10Kb | #dmSites | #dmsp10kb | %dmSites | %pSite1 | %pSite2 | DMpSite | ADMpSite | DMpNucl | ADMpNucl |
|---|---|---|---|---|---|---|---|---|---|---|---|---|---|
| gene3165 | https://www.ncbi.nlm.nih.gov/gene/?term=LOC105275706 | 2374 | 74 | 311.71 | 6 | 25.27 | 8.11 | 32.18 | 22.24 | 9.94 | 9.94 | 0.31 | 0.31 |
| gene2824 | https://www.ncbi.nlm.nih.gov/gene/?term=LOC105275337 | 1772 | 60 | 338.6 | 2 | 11.29 | 3.33 | 38.84 | 29.82 | 9.02 | 9.02 | 0.31 | 0.31 |
| gene10963 | https://www.ncbi.nlm.nih.gov/gene/?term=LOC105284677 | 1576 | 30 | 190.36 | 1 | 6.35 | 3.33 | 25.25 | 41.23 | -15.97 | 15.97 | -0.3 | 0.3 |
| gene2593 | https://www.ncbi.nlm.nih.gov/gene/?term=LOC105275070 | 1774 | 7 | 39.46 | 5 | 28.18 | 71.43 | 65.68 | 4.09 | 61.59 | 61.59 | 0.24 | 0.24 |
| gene7568 | https://www.ncbi.nlm.nih.gov/gene/?term=LOC105280786 | 2525 | 126 | 499.01 | 1 | 3.96 | 0.79 | 30.35 | 27.57 | 2.78 | 2.78 | 0.14 | 0.14 |
| gene5714 | https://www.ncbi.nlm.nih.gov/gene/?term=LOC105278729 | 710 | 17 | 239.44 | 1 | 14.08 | 5.88 | 59.17 | 53.54 | 5.62 | 5.62 | 0.13 | 0.13 |
| gene8943 | https://www.ncbi.nlm.nih.gov/gene/?term=LOC105282388 | 1220 | 33 | 270.49 | 1 | 8.2 | 3.03 | 49.82 | 45.11 | 4.71 | 4.71 | 0.13 | 0.13 |
| gene5525 | https://www.ncbi.nlm.nih.gov/gene/?term=LOC105278396 | 3090 | 184 | 595.47 | 1 | 3.24 | 0.54 | 26.64 | 24.83 | 1.81 | 1.81 | 0.11 | 0.11 |
| gene2271 | https://www.ncbi.nlm.nih.gov/gene/?term=LOC105274706 | 1448 | 56 | 386.74 | 1 | 6.91 | 1.79 | 38.93 | 36.4 | 2.53 | 2.53 | 0.1 | 0.1 |
| gene4641 | https://www.ncbi.nlm.nih.gov/gene/?term=LOC105277398 | 1937 | 72 | 371.71 | 1 | 5.16 | 1.39 | 36.58 | 39.26 | -2.68 | 2.68 | -0.1 | 0.1 |
| gene4034 | https://www.ncbi.nlm.nih.gov/gene/?term=LOC105276704 | 2065 | 88 | 426.15 | 1 | 4.84 | 1.14 | 16.14 | 13.92 | 2.22 | 2.22 | 0.09 | 0.09 |
| gene11819 | https://www.ncbi.nlm.nih.gov/gene/?term=LOC105285671 | 2366 | 103 | 435.33 | 1 | 4.23 | 0.97 | 27.63 | 25.66 | 1.97 | 1.97 | 0.09 | 0.09 |
| gene2907 | https://www.ncbi.nlm.nih.gov/gene/?term=LOC105275447 | 2539 | 37 | 145.73 | 1 | 3.94 | 2.7 | 8.22 | 2.41 | 5.8 | 5.8 | 0.08 | 0.08 |
| gene920 | https://www.ncbi.nlm.nih.gov/gene/?term=LOC105282518 | 1197 | 52 | 434.42 | 1 | 8.35 | 1.92 | 29.01 | 30.77 | -1.76 | 1.76 | -0.08 | 0.08 |
| gene9598 | https://www.ncbi.nlm.nih.gov/gene/?term=LOC105283132 | 1769 | 57 | 322.22 | 1 | 5.65 | 1.75 | 39.92 | 37.87 | 2.05 | 2.05 | 0.07 | 0.07 |
| gene940 | https://www.ncbi.nlm.nih.gov/gene/?term=LOC105282794 | 3578 | 88 | 245.95 | 1 | 2.79 | 1.14 | 16.61 | 14.17 | 2.44 | 2.44 | 0.06 | 0.06 |
| gene13033 | https://www.ncbi.nlm.nih.gov/gene/?term=LOC105287063 | 1877 | 48 | 255.73 | 1 | 5.33 | 2.08 | 32.8 | 30.37 | 2.43 | 2.43 | 0.06 | 0.06 |
| gene7211 | https://www.ncbi.nlm.nih.gov/gene/?term=LOC105280377 | 2024 | 69 | 340.91 | 1 | 4.94 | 1.45 | 48.01 | 46.12 | 1.89 | 1.89 | 0.06 | 0.06 |
| gene5524 | https://www.ncbi.nlm.nih.gov/gene/?term=LOC105278397 | 5177 | 201 | 388.26 | 1 | 1.93 | 0.5 | 26.23 | 24.63 | 1.61 | 1.61 | 0.06 | 0.06 |
| gene9646 | https://www.ncbi.nlm.nih.gov/gene/?term=LOC105283182 | 3588 | 133 | 370.68 | 1 | 2.79 | 0.75 | 13.97 | 12.43 | 1.54 | 1.54 | 0.06 | 0.06 |
| gene3817 | https://www.ncbi.nlm.nih.gov/gene/?term=LOC105276449 | 1230 | 52 | 422.76 | 1 | 8.13 | 1.92 | 31.62 | 30.14 | 1.48 | 1.48 | 0.06 | 0.06 |
| gene10678 | https://www.ncbi.nlm.nih.gov/gene/?term=LOC105284355 | 4031 | 169 | 419.25 | 1 | 2.48 | 0.59 | 19.82 | 18.64 | 1.18 | 1.18 | 0.05 | 0.05 |
| gene1993 | https://www.ncbi.nlm.nih.gov/gene/?term=LOC105288193 | 1449 | 77 | 531.4 | 1 | 6.9 | 1.3 | 34.64 | 33.62 | 1.02 | 1.02 | 0.05 | 0.05 |
| gene935 | https://www.ncbi.nlm.nih.gov/gene/?term=LOC105282682 | 4368 | 222 | 508.24 | 1 | 2.29 | 0.45 | 13.71 | 12.73 | 0.98 | 0.98 | 0.05 | 0.05 |

...
... only first 25 lines shown ...

Directory: OGLall File: rnk-dmg-Cb_LI_RphaseB1.vs.RphaseB2.txt

| gene_ID | gene_link | gwidth | #Sites | #per10Kb | #dmSites | #dmsp10kb | %dmSites | %pSite1 | %pSite2 | DMpSite | ADMpSite | DMpNucl | ADMpNucl |
|---|---|---|---|---|---|---|---|---|---|---|---|---|---|
| gene10442 | https://www.ncbi.nlm.nih.gov/gene/?term=LOC105284093 | 809 | 61 | 754.02 | 2 | 24.72 | 3.28 | 10.04 | 7.71 | 2.34 | 2.34 | 0.18 | 0.18 |
| gene10584 | https://www.ncbi.nlm.nih.gov/gene/?term=LOC105284253 | 1255 | 41 | 326.69 | 5 | 39.84 | 12.2 | 26.29 | 31.83 | -5.54 | 5.54 | -0.18 | 0.18 |
| gene5917 | https://www.ncbi.nlm.nih.gov/gene/?term=LOC105278849 | 823 | 80 | 972.05 | 2 | 24.3 | 2.5 | 21.53 | 23.29 | -1.76 | 1.76 | -0.17 | 0.17 |
| gene10644 | https://www.ncbi.nlm.nih.gov/gene/?term=LOC105284320 | 358 | 10 | 279.33 | 2 | 55.87 | 20 | 24.37 | 30.4 | -6.02 | 6.02 | -0.17 | 0.17 |
| gene4515 | https://www.ncbi.nlm.nih.gov/gene/?term=LOC109610839 | 335 | 7 | 208.96 | 1 | 29.85 | 14.29 | 66.57 | 58.9 | 7.67 | 7.67 | 0.16 | 0.16 |
| gene1993 | https://www.ncbi.nlm.nih.gov/gene/?term=LOC105288193 | 1449 | 77 | 531.4 | 5 | 34.51 | 6.49 | 30.7 | 33.62 | -2.92 | 2.92 | -0.16 | 0.16 |
| gene9019 | https://www.ncbi.nlm.nih.gov/gene/?term=LOC105282481 | 1743 | 58 | 332.76 | 4 | 22.95 | 6.9 | 32.56 | 37.33 | -4.77 | 4.77 | -0.16 | 0.16 |
| gene2636 | https://www.ncbi.nlm.nih.gov/gene/?term=LOC105275141 | 947 | 11 | 116.16 | 4 | 42.24 | 36.36 | 32.51 | 19.93 | 12.58 | 12.58 | 0.15 | 0.15 |
| gene9291 | https://www.ncbi.nlm.nih.gov/gene/?term=LOC105282787 | 1045 | 42 | 401.91 | 3 | 28.71 | 7.14 | 37.93 | 34.07 | 3.85 | 3.85 | 0.15 | 0.15 |
| gene5309 | https://www.ncbi.nlm.nih.gov/gene/?term=LOC105278160 | 1704 | 91 | 534.04 | 1 | 5.87 | 1.1 | 34.43 | 37.28 | -2.85 | 2.85 | -0.15 | 0.15 |
| gene10529 | https://www.ncbi.nlm.nih.gov/gene/?term=LOC105284189 | 2602 | 133 | 511.15 | 3 | 11.53 | 2.26 | 17.44 | 14.96 | 2.49 | 2.49 | 0.13 | 0.13 |
| gene1589 | https://www.ncbi.nlm.nih.gov/gene/?term=LOC105287721 | 1407 | 41 | 291.4 | 2 | 14.21 | 4.88 | 25.63 | 21.54 | 4.09 | 4.09 | 0.12 | 0.12 |
| gene4597 | https://www.ncbi.nlm.nih.gov/gene/?term=LOC105277345 | 2173 | 78 | 358.95 | 3 | 13.81 | 3.85 | 18.24 | 14.85 | 3.38 | 3.38 | 0.12 | 0.12 |
| gene6324 | https://www.ncbi.nlm.nih.gov/gene/?term=LOC105279346 | 4650 | 253 | 544.09 | 7 | 15.05 | 2.77 | 14.36 | 16.48 | -2.12 | 2.12 | -0.12 | 0.12 |
| gene558 | https://www.ncbi.nlm.nih.gov/gene/?term=LOC105278458 | 216 | 3 | 138.89 | 1 | 46.3 | 33.33 | 20.77 | 29.54 | -8.77 | 8.77 | -0.12 | 0.12 |
| gene11399 | https://www.ncbi.nlm.nih.gov/gene/?term=LOC105285203 | 1029 | 30 | 291.55 | 2 | 19.44 | 6.67 | 30.87 | 26.95 | 3.93 | 3.93 | 0.11 | 0.11 |
| gene2559 | https://www.ncbi.nlm.nih.gov/gene/?term=LOC105275030 | 1897 | 119 | 627.31 | 6 | 31.63 | 5.04 | 19.12 | 20.91 | -1.79 | 1.79 | -0.11 | 0.11 |
| gene1223 | https://www.ncbi.nlm.nih.gov/gene/?term=LOC105285713 | 1048 | 38 | 362.6 | 2 | 19.08 | 5.26 | 46.12 | 43.33 | 2.79 | 2.79 | 0.1 | 0.1 |
| gene2130 | https://www.ncbi.nlm.nih.gov/gene/?term=LOC105274542 | 1254 | 52 | 414.67 | 3 | 23.92 | 5.77 | 30.14 | 27.83 | 2.32 | 2.32 | 0.1 | 0.1 |
| gene6441 | https://www.ncbi.nlm.nih.gov/gene/?term=LOC105279487 | 2117 | 100 | 472.37 | 2 | 9.45 | 2 | 15.08 | 13.01 | 2.07 | 2.07 | 0.1 | 0.1 |
| gene3910 | https://www.ncbi.nlm.nih.gov/gene/?term=LOC105276563 | 1957 | 112 | 572.3 | 4 | 20.44 | 3.57 | 18.76 | 17 | 1.77 | 1.77 | 0.1 | 0.1 |
| gene12842 | https://www.ncbi.nlm.nih.gov/gene/?term=LOC105286845 | 1157 | 78 | 674.16 | 2 | 17.29 | 2.56 | 11.01 | 12.53 | -1.53 | 1.53 | -0.1 | 0.1 |
| gene5987 | https://www.ncbi.nlm.nih.gov/gene/?term=LOC105278938 | 1882 | 109 | 579.17 | 4 | 21.25 | 3.67 | 19.75 | 21.46 | -1.71 | 1.71 | -0.1 | 0.1 |
| gene6781 | https://www.ncbi.nlm.nih.gov/gene/?term=LOC105279857 | 241 | 10 | 414.94 | 1 | 41.49 | 10 | 8.85 | 11.17 | -2.32 | 2.32 | -0.1 | 0.1 |

...

... only first 25 lines shown ...

```
Directory: OGLall File: wrt-Cb_LI.txt

... comparing  BCphaseA1.vs.BCphaseA2 ...

Number of genes with #dmsites >=  1 :  139

sample1     sum of percent methylation:  3385.72     average:    24.36
sample2     sum of percent methylation:  3419.48     average:    24.60

sum difference:   -33.73     average difference:  -0.2427

      Wilcoxon signed rank test with continuity correction

data:  pw_summary$`%pSite1` and pw_summary$`%pSite2`
V = 4962, p-value = 0.7
alternative hypothesis: true location shift is not equal to 0

... comparing  BCphaseA1.vs.BCphaseB1 ...

Number of genes with #dmsites >=  1 :  141

sample1     sum of percent methylation:  3284.84     average:    23.30
sample2     sum of percent methylation:  3301.47     average:    23.41

sum difference:   -16.58     average difference:  -0.1176
...
... only first 25 lines shown ...
```

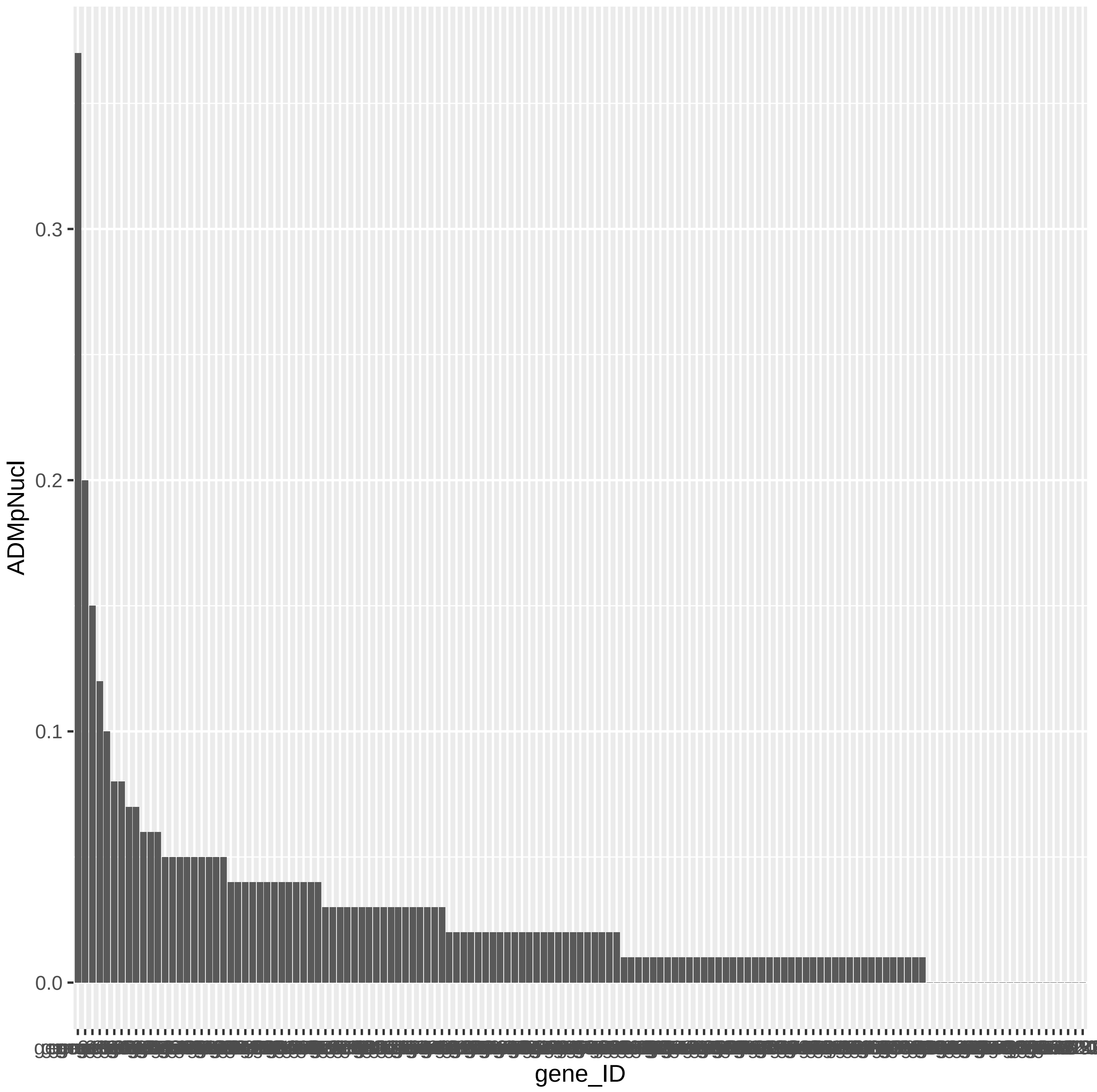

ADMpNucl
0.3
0.2
0.1
0.0
gene_ID

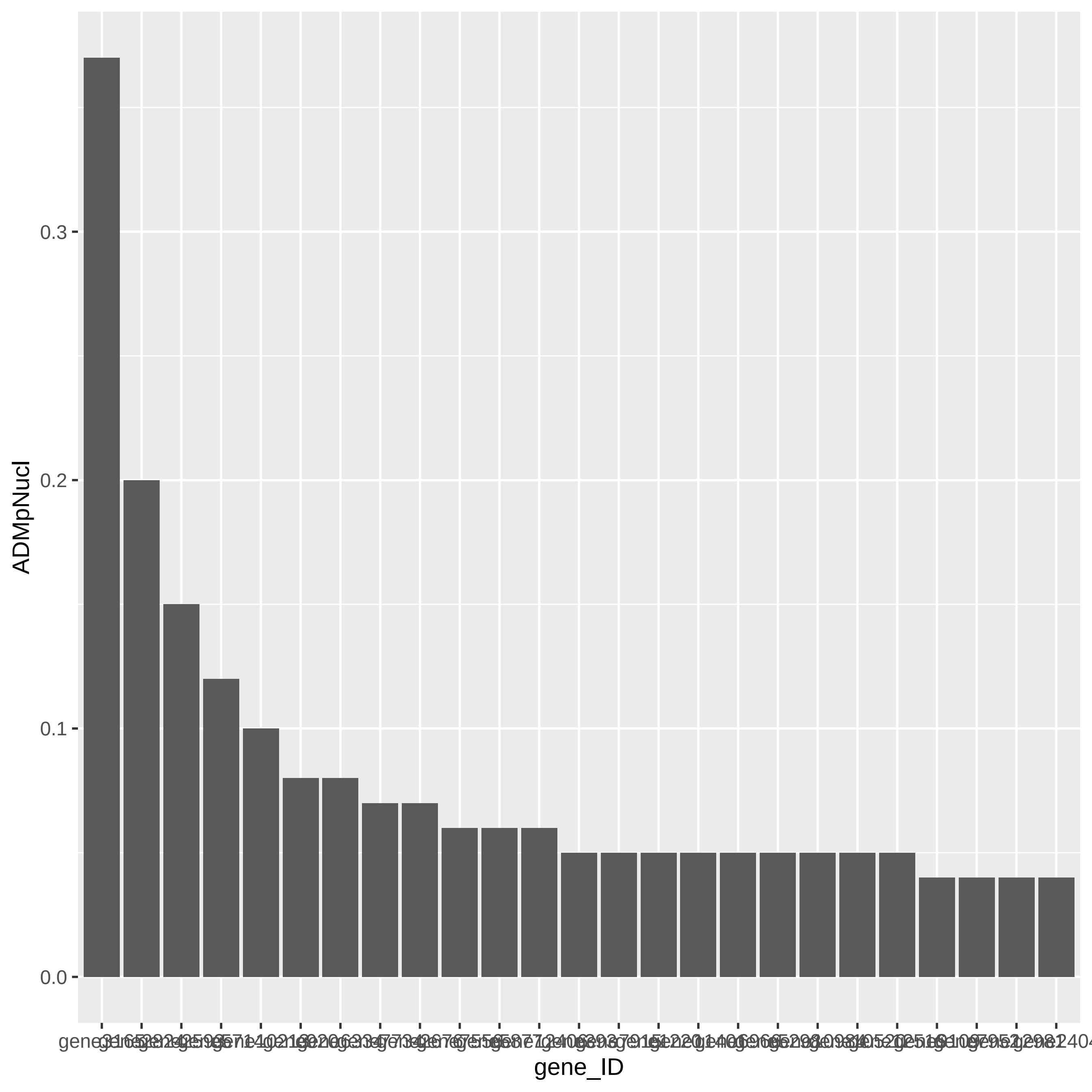

0.3
0.2
0.1
0.0
ADMpNucl
gene_ID

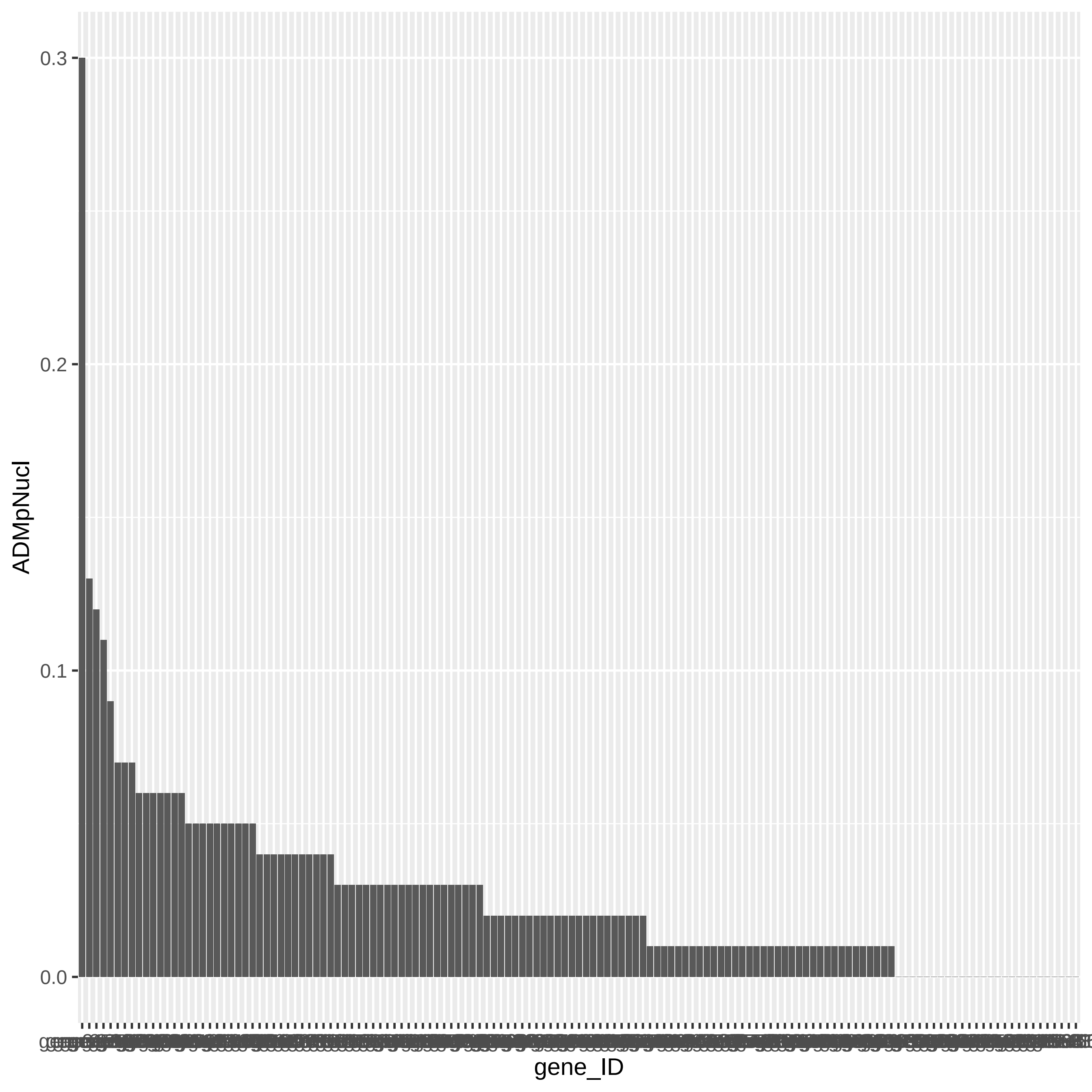

0.3
0.2
0.1
0.0
ADMpNucl
gene_ID

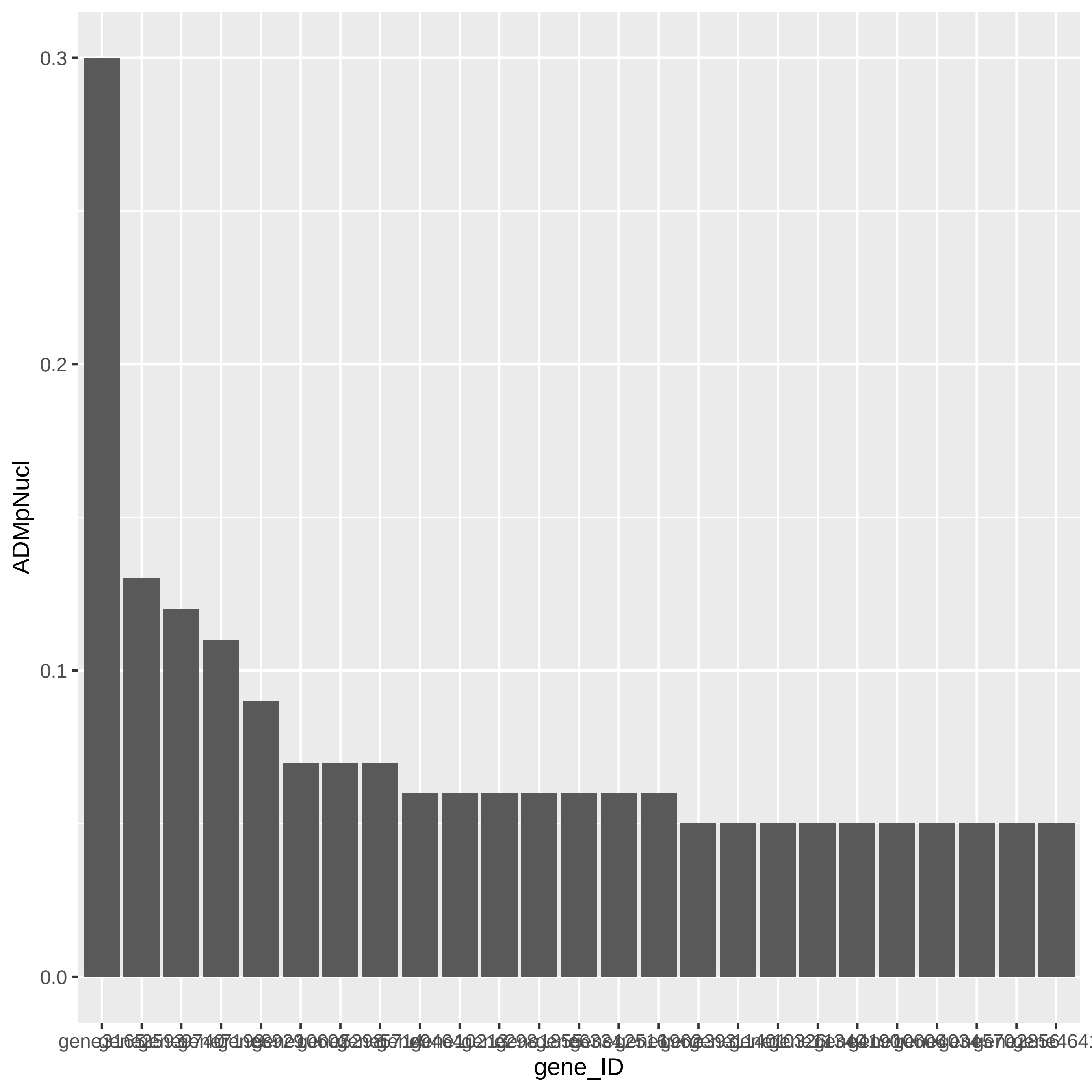

0.3
0.2
0.1
0.0
ADMpNucl
gene_ID

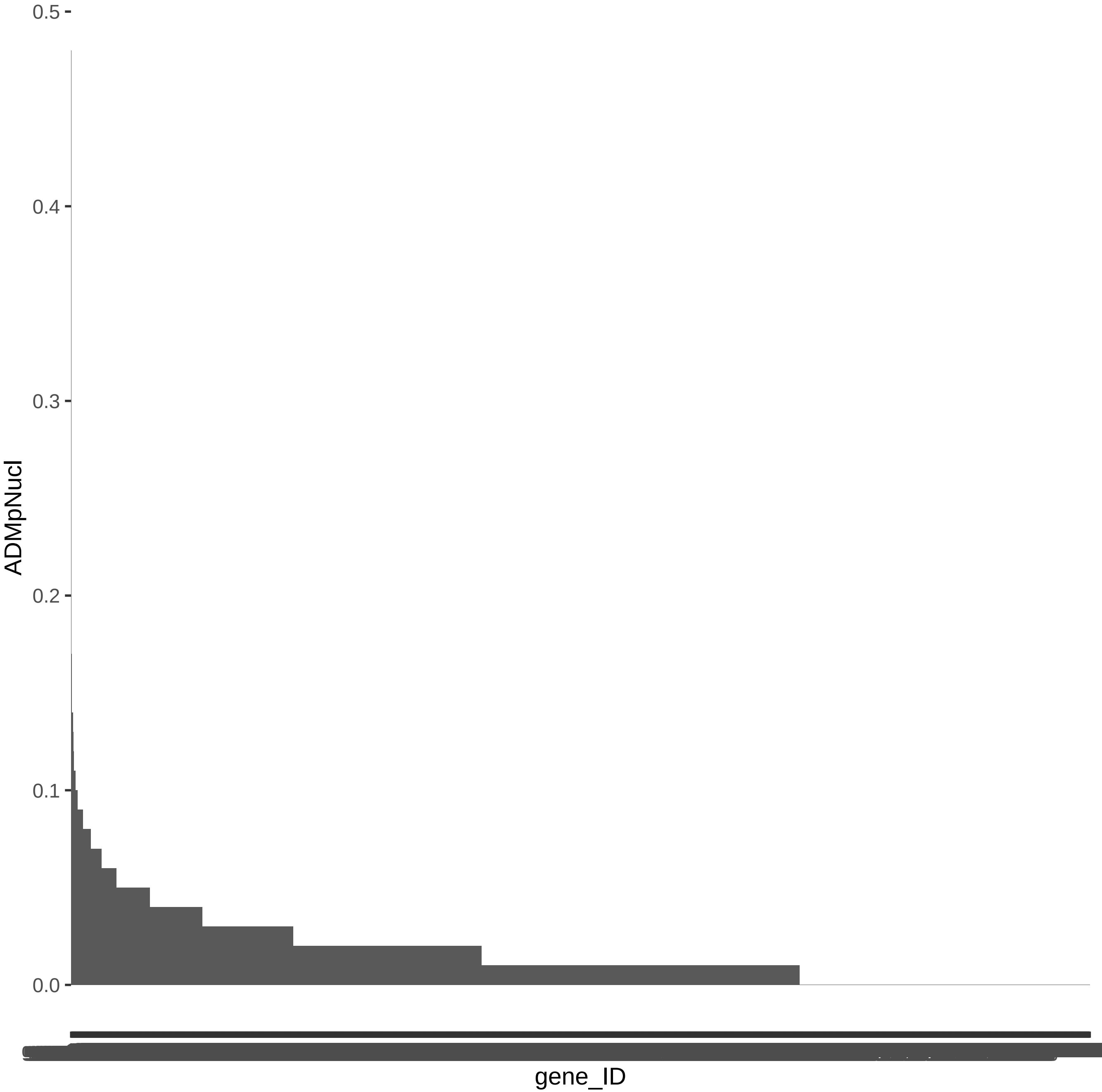

0.5
0.4
0.3
0.2
0.1
0.0
ADMpNucl
gene_ID

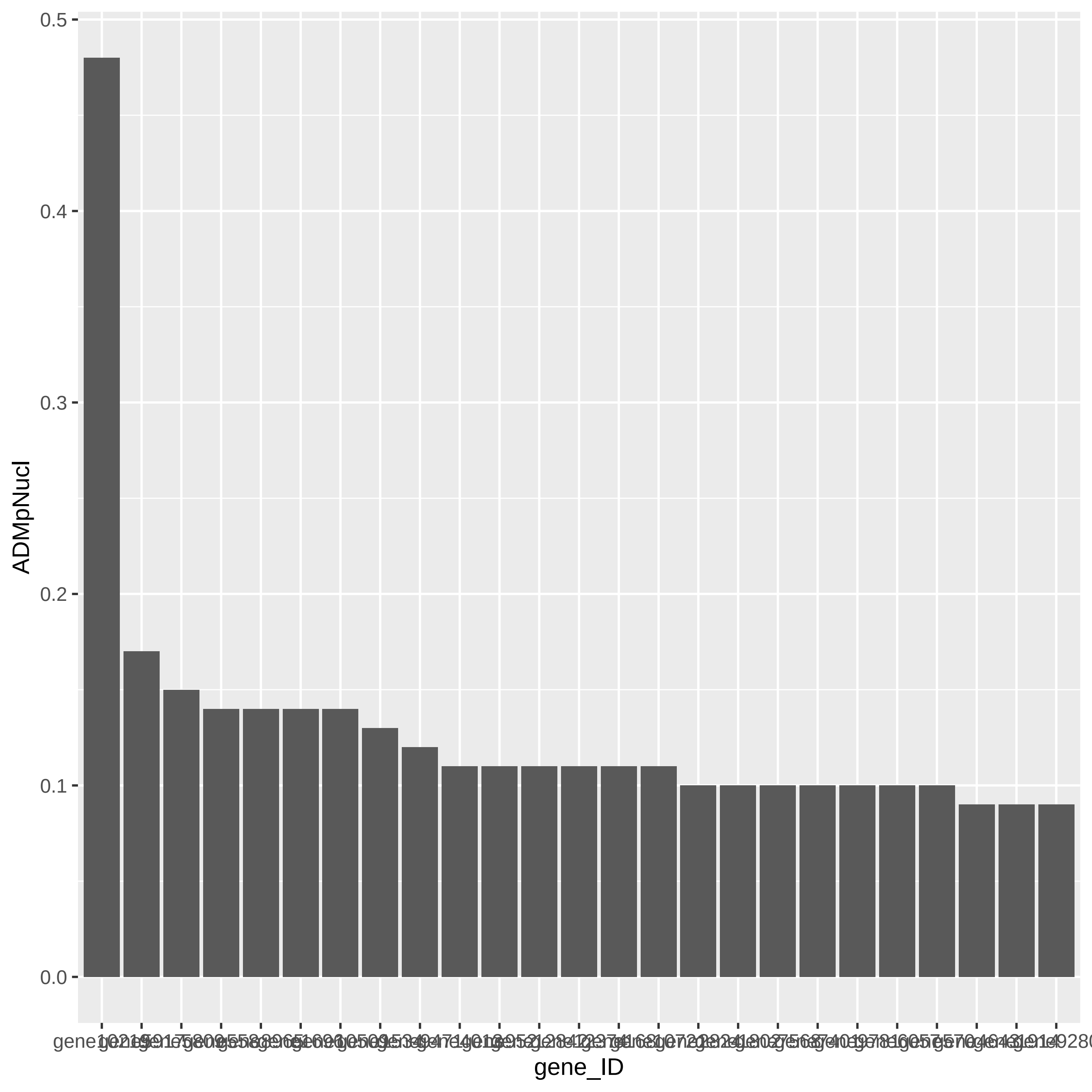

0.5
0.4
0.3
0.2
0.1
0.0
ADMpNucl
gene_ID

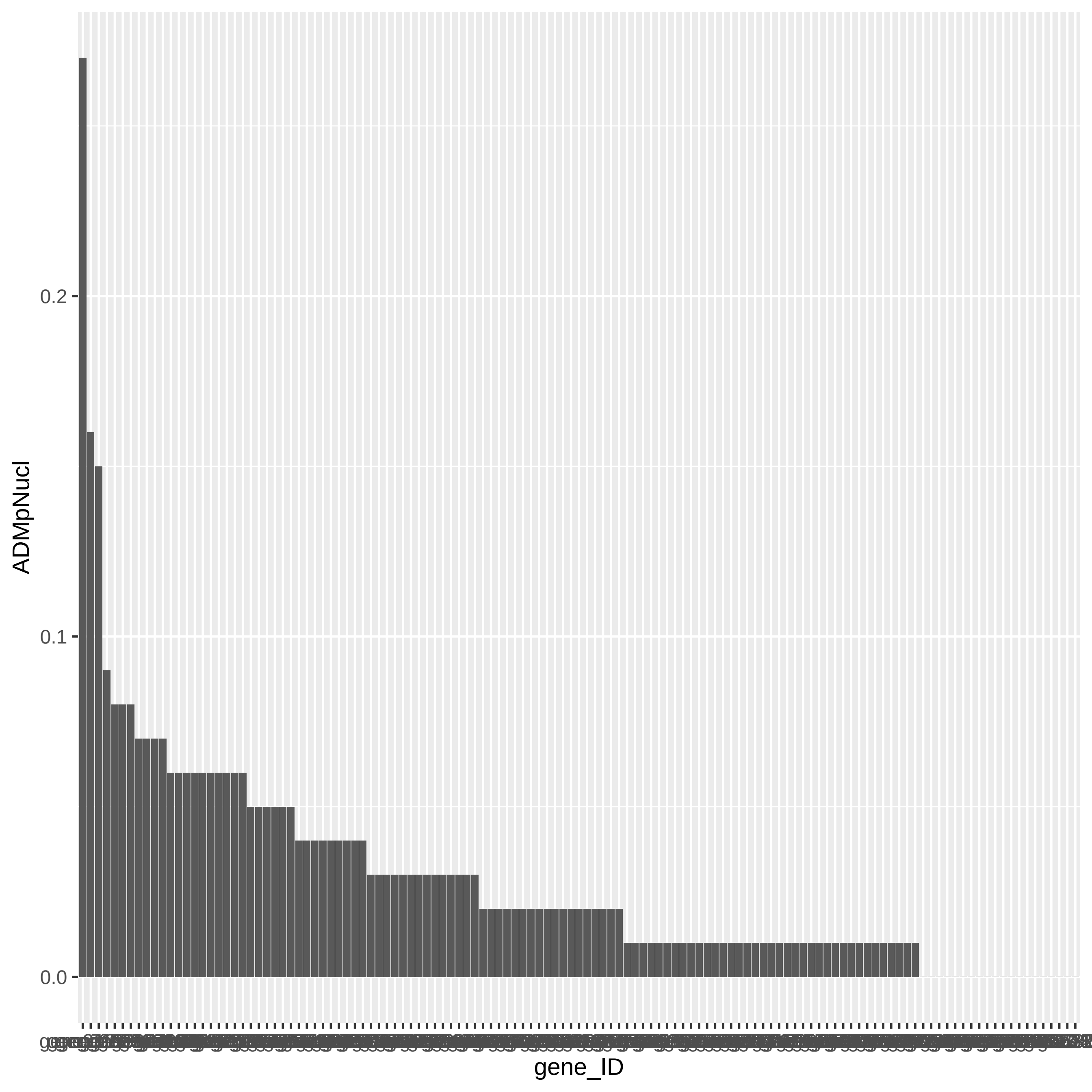
ADMpNucl
0.2
0.1
0.0
gene_ID

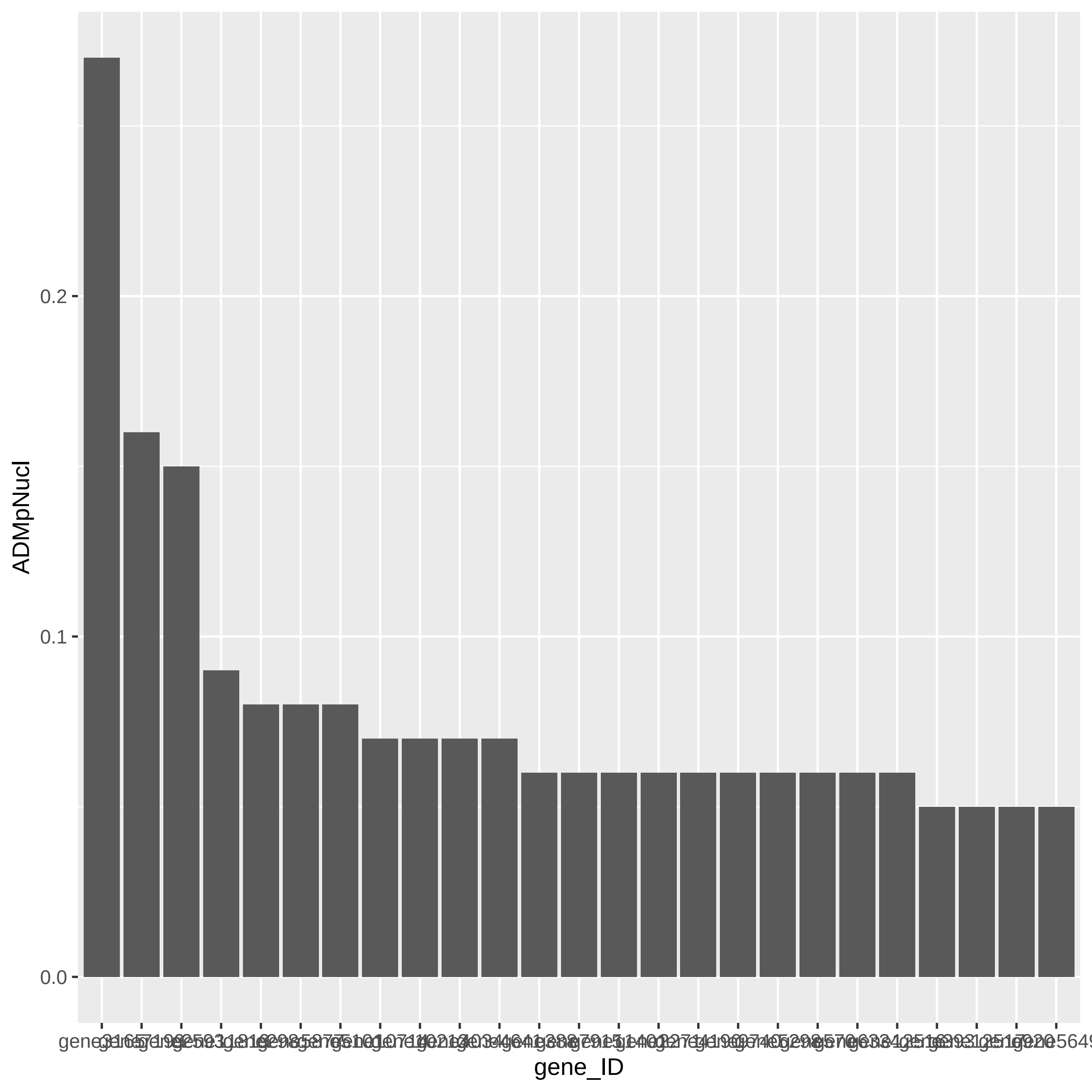

ADMpNucl
0.2
0.1
0.0
gene_ID

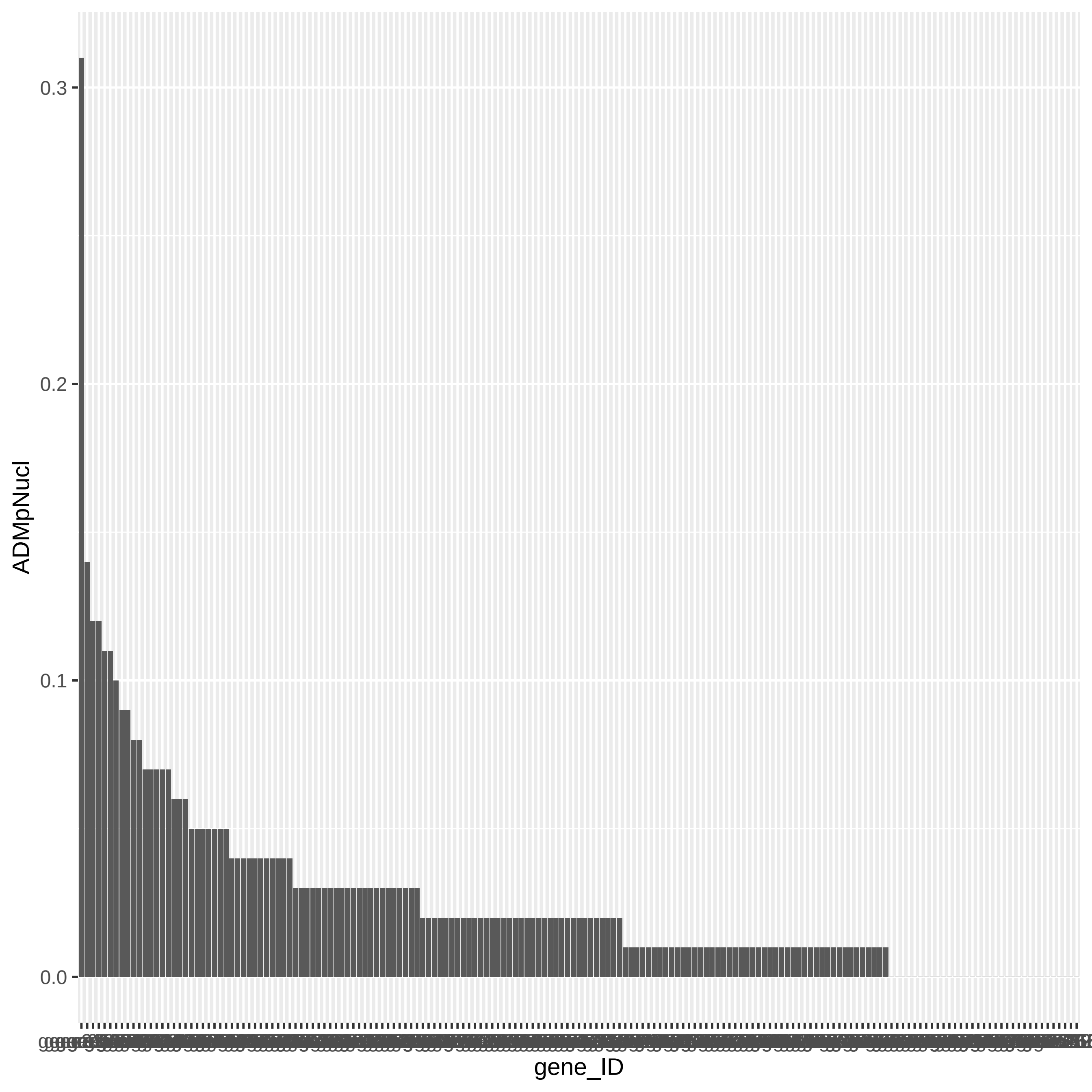

0.3
0.2
0.1
0.0
ADMpNucl
gene_ID

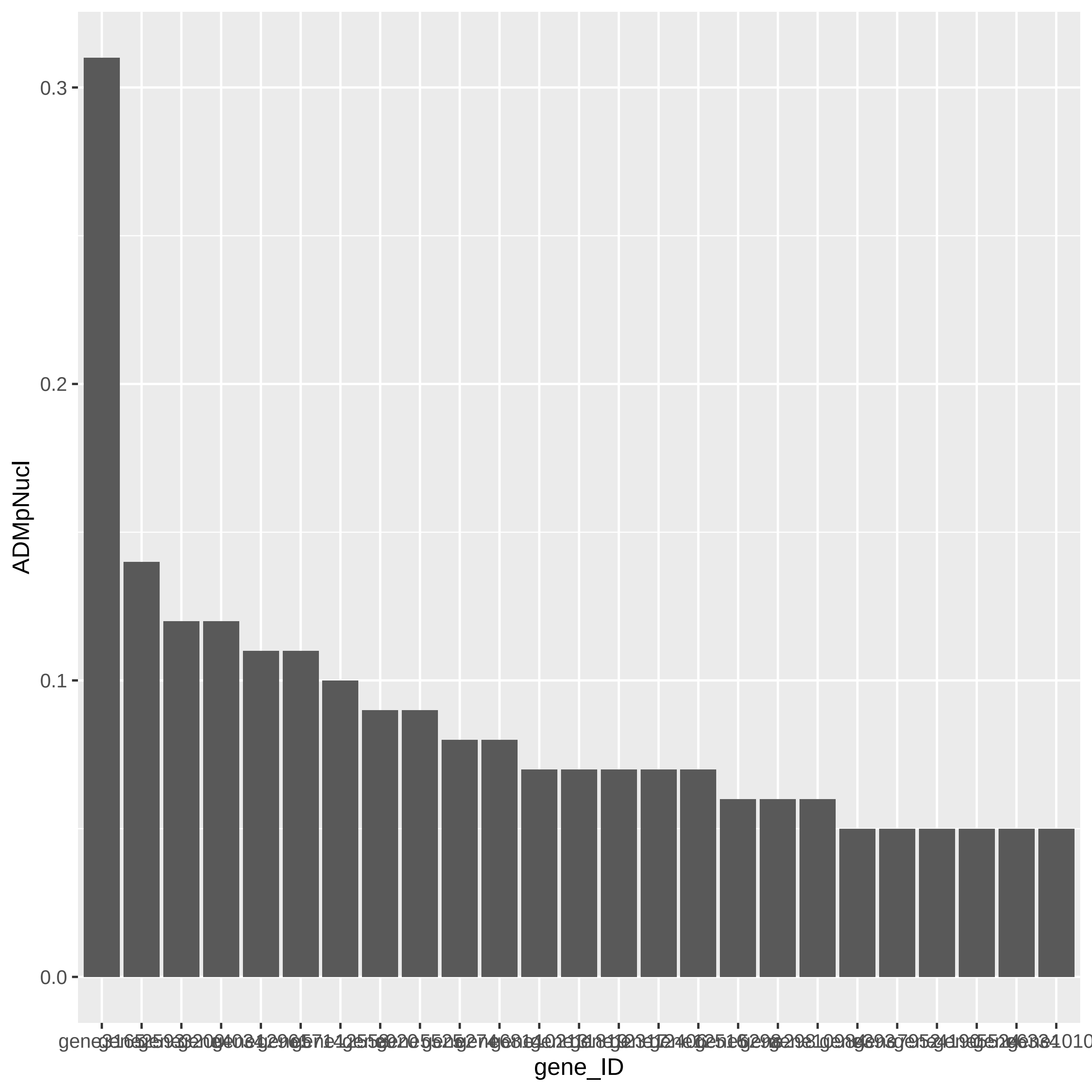
0.3
0.2
0.1
0.0
ADMpNucl
gene_ID

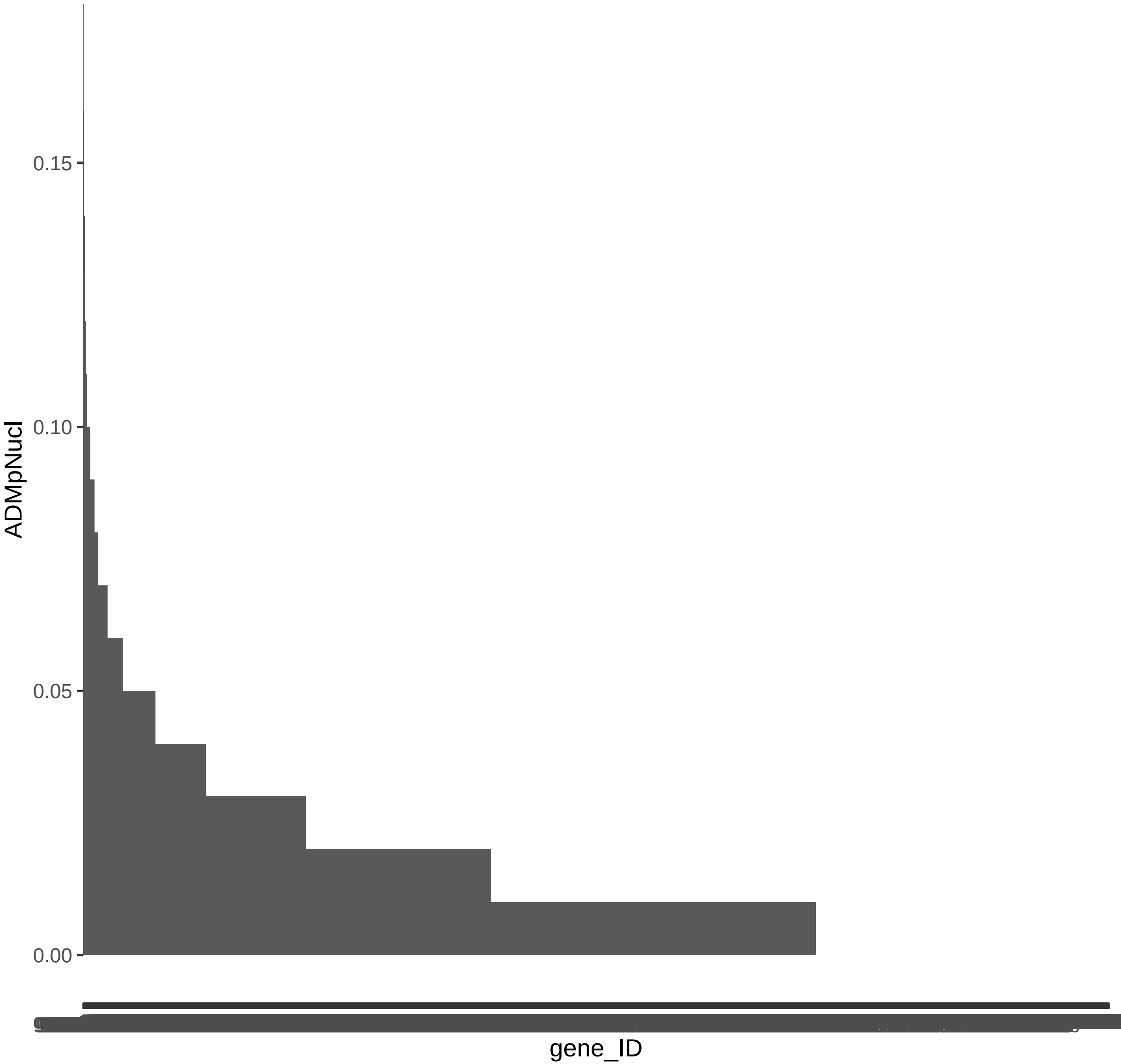

ADMpNucl
0.15
0.10
0.05
0.00
gene_ID

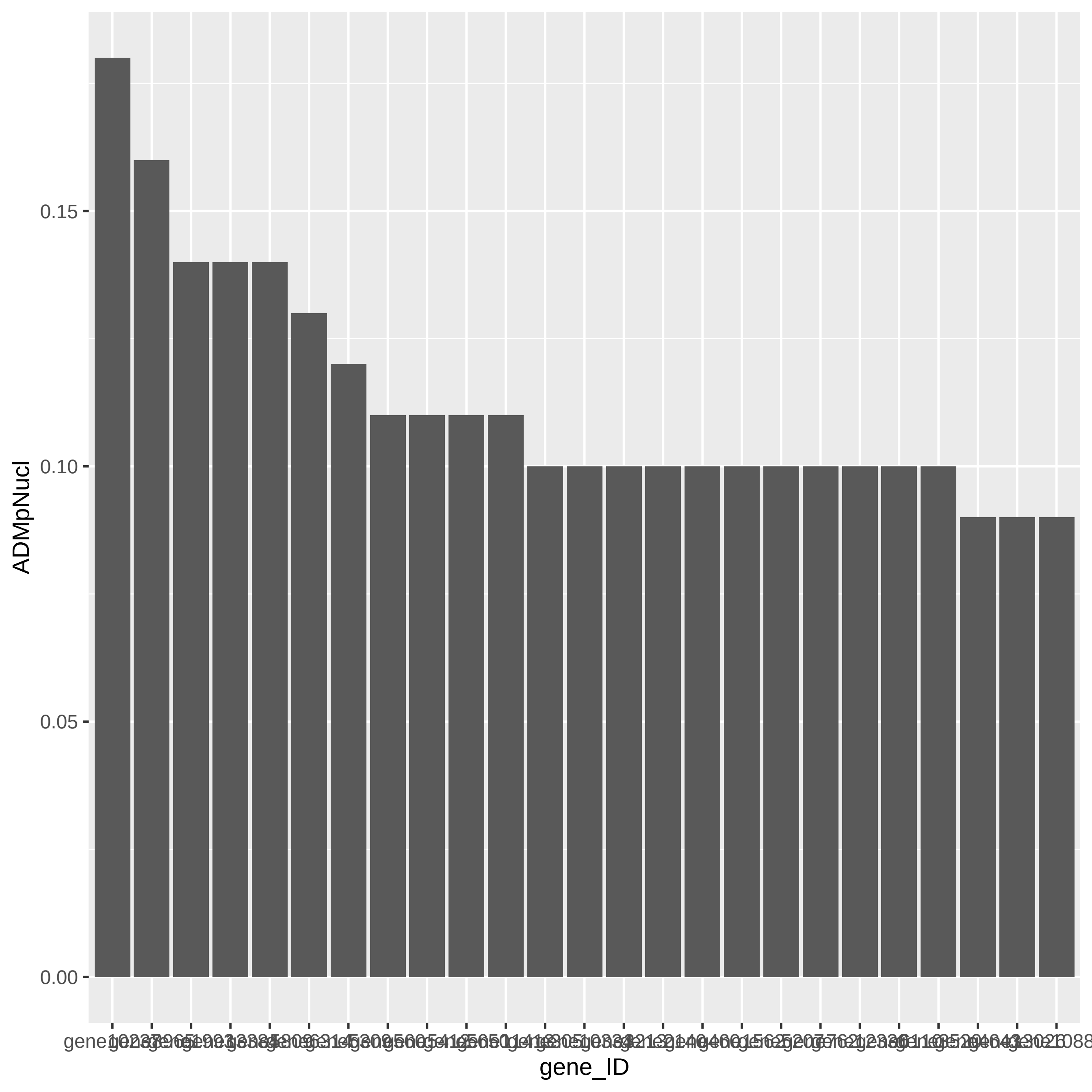
ADMpNucl
0.15
0.10
0.05
0.00
gene_ID

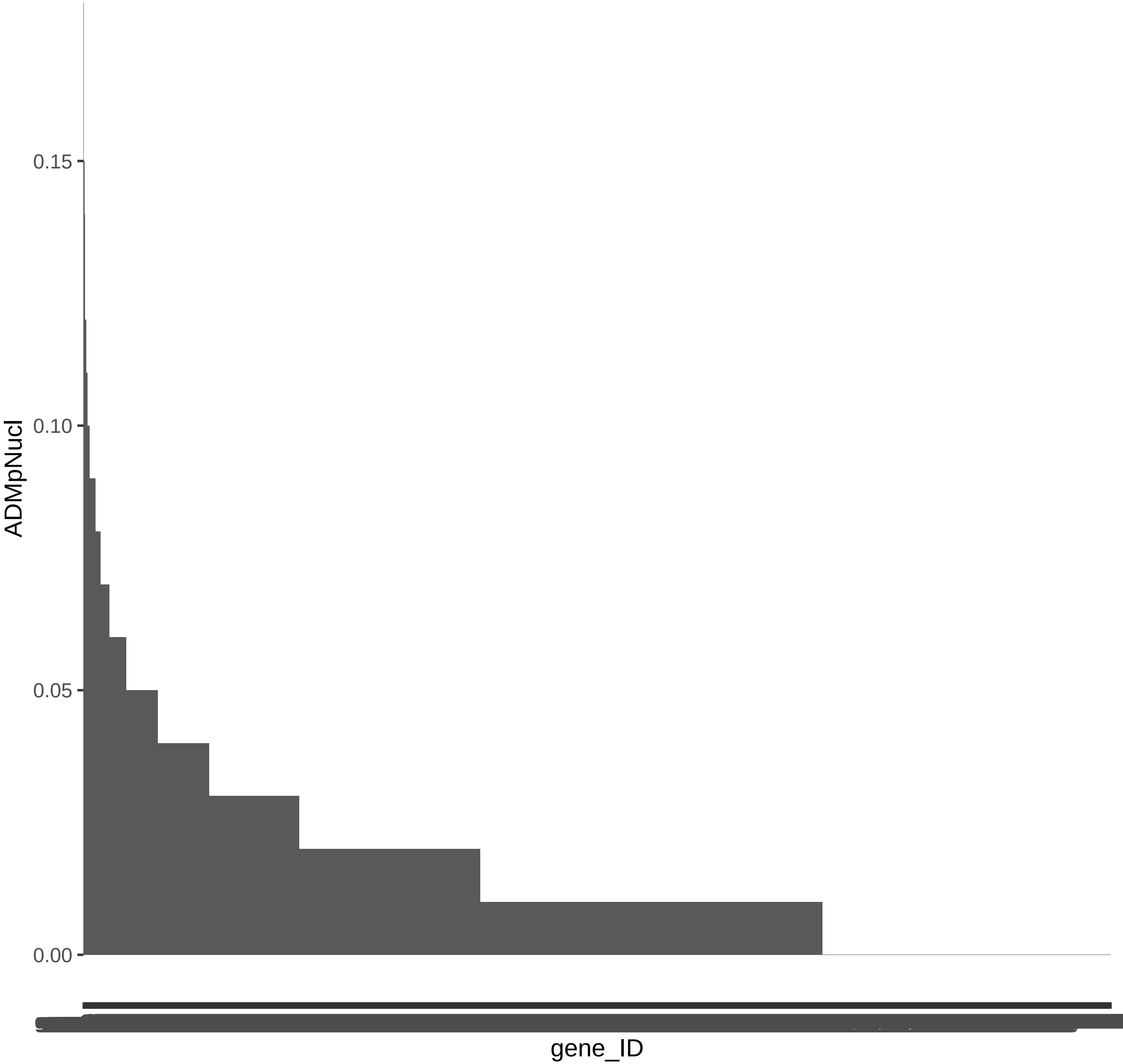

ADMpNucl
0.15
0.10
0.05
0.00
gene_ID

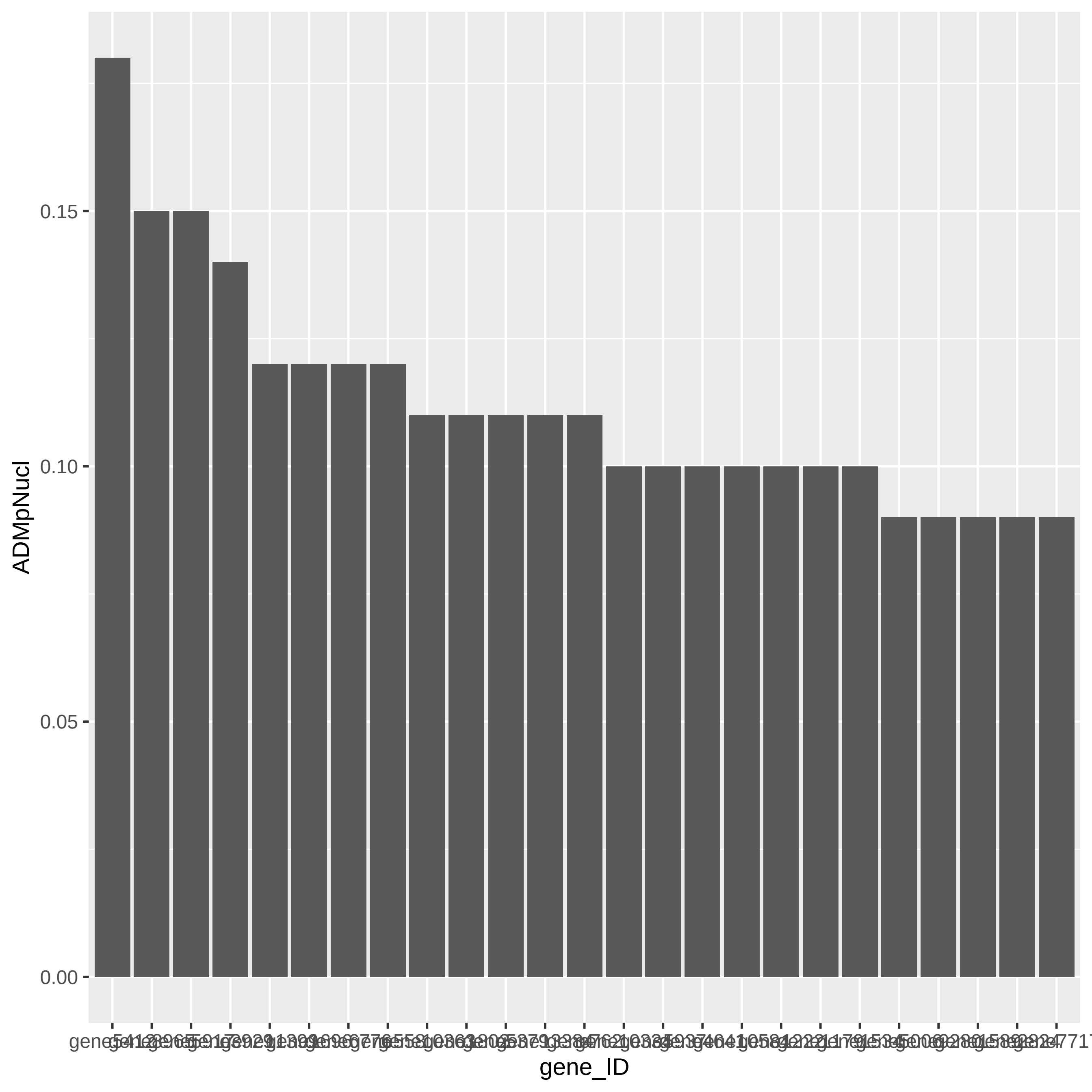
0.15
0.10
0.05
0.00
ADMpNucl
gene_ID

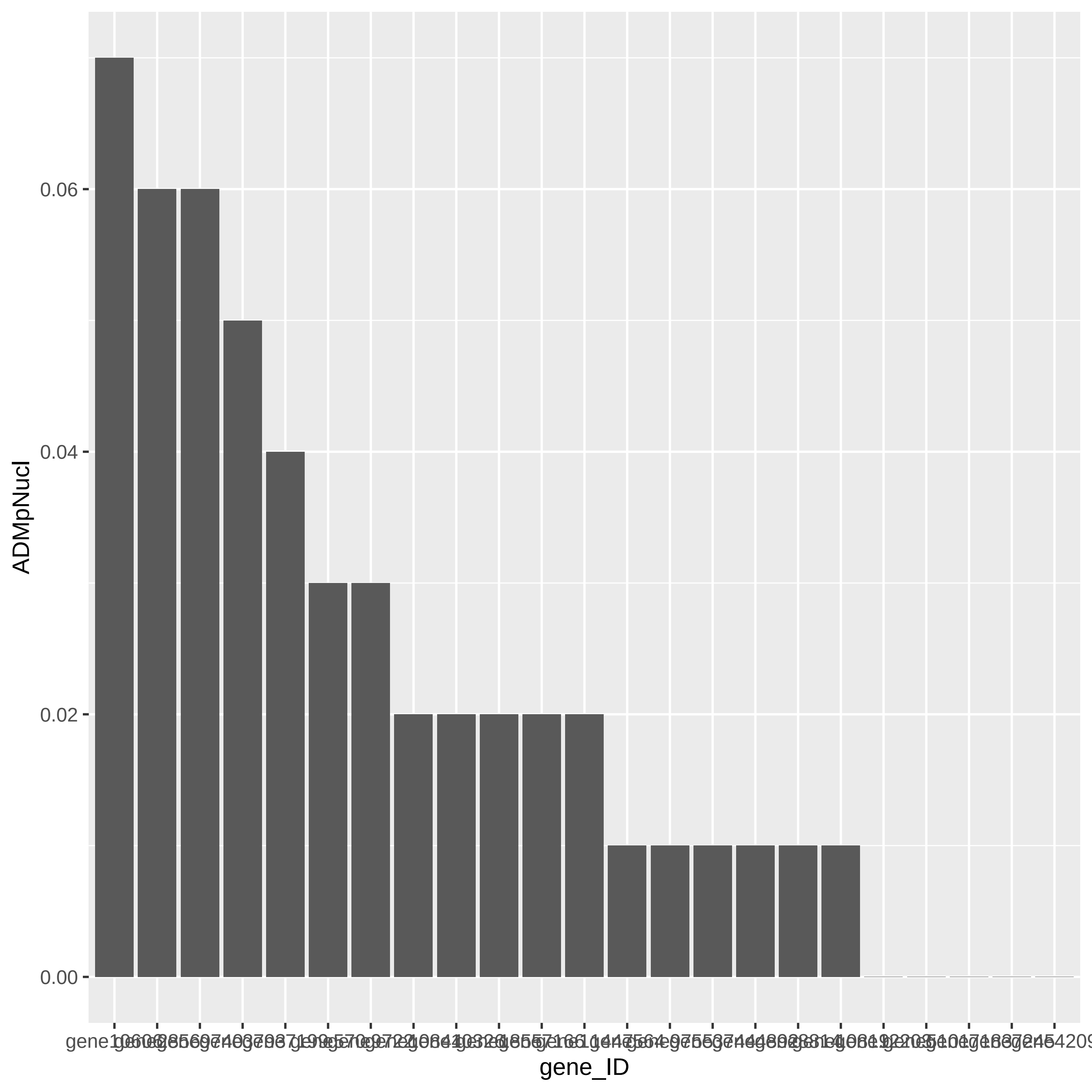
ADMpNucl
0.06
0.04
0.02
0.00
gene_ID

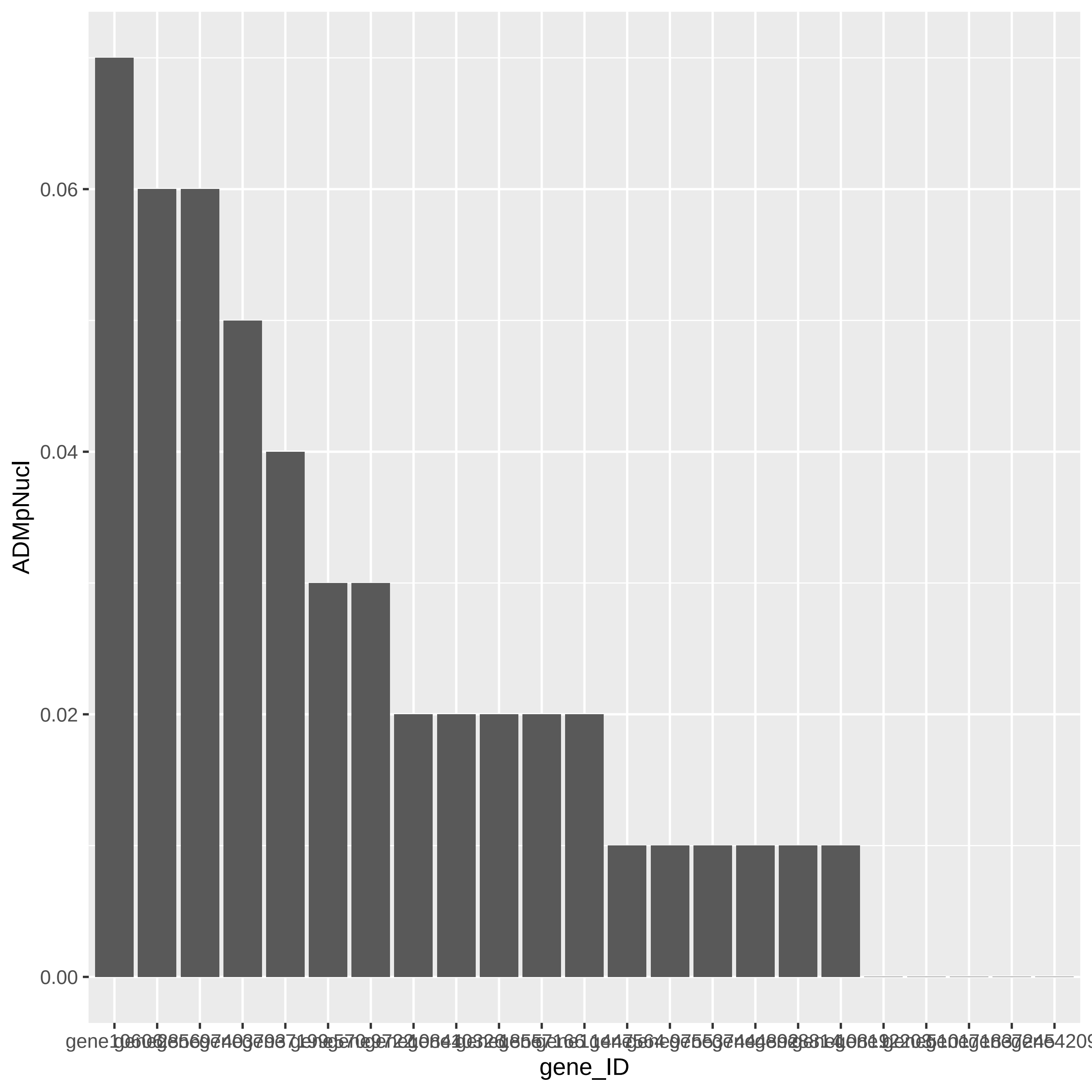

0.06
0.04
0.02
0.00
ADMpNucl
gene_ID

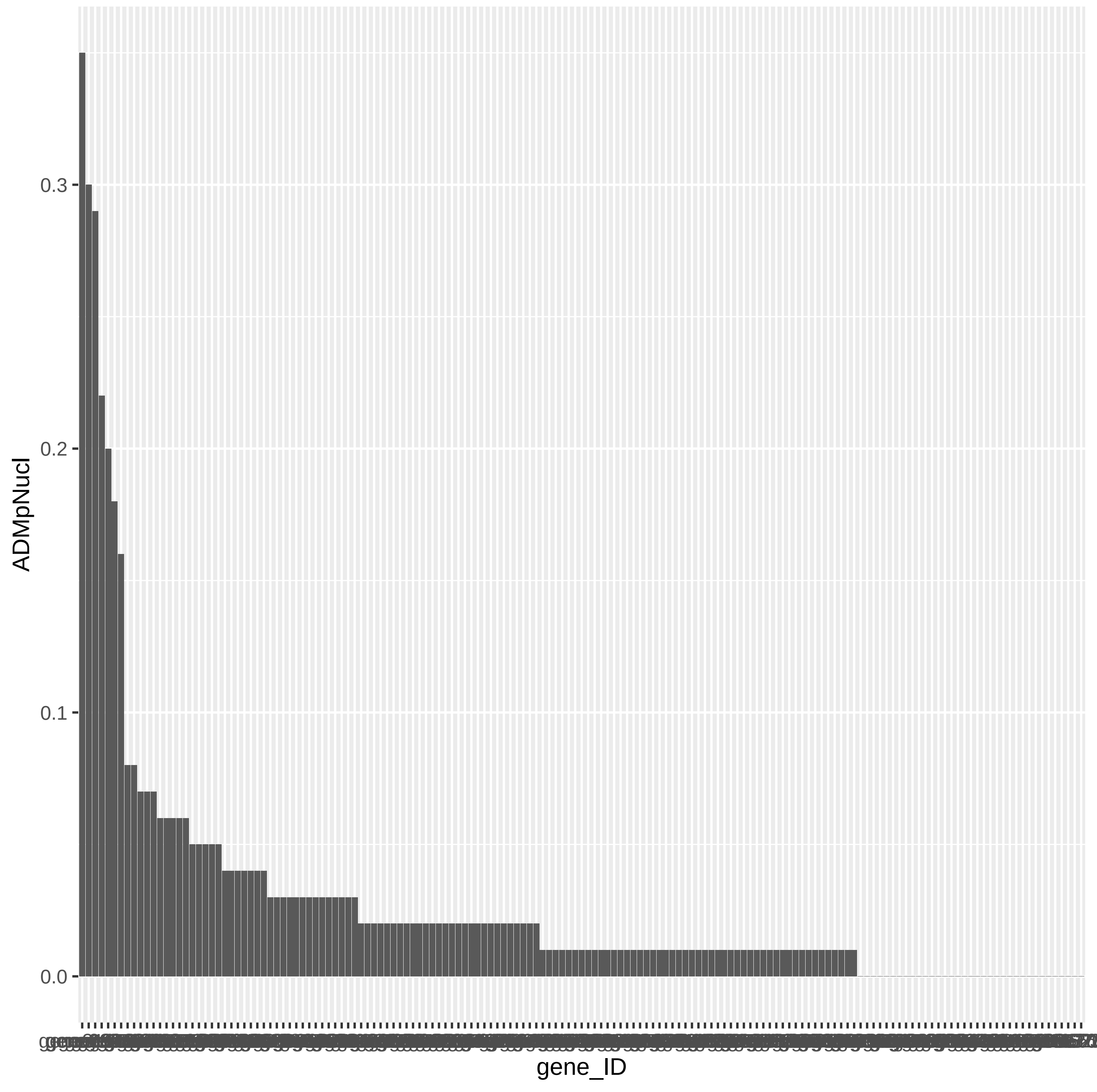
0.3
0.2
0.1
0.0
ADMpNucl
gene_ID

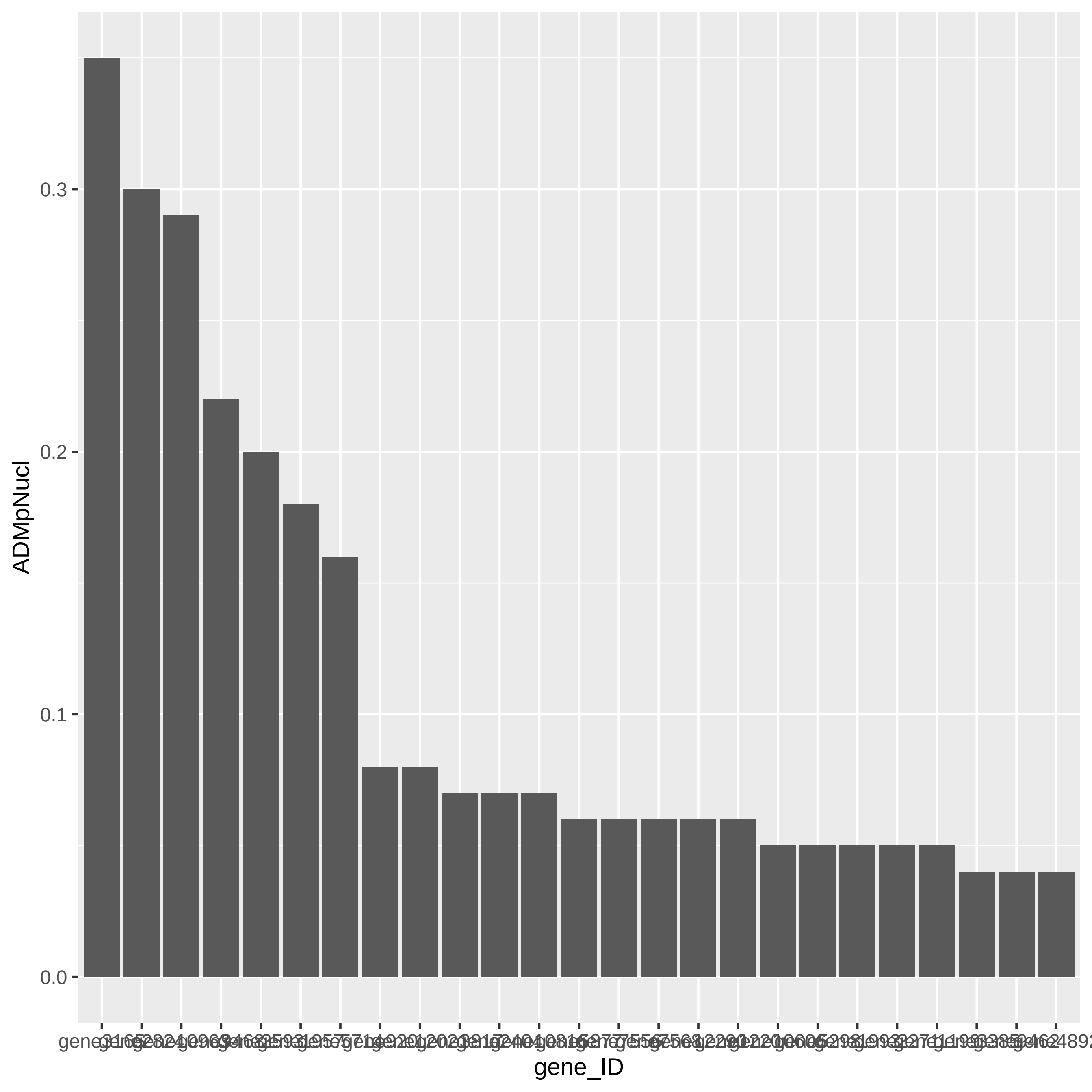
ADMpNucl
0.0
0.1
0.2
0.3
gene_ID

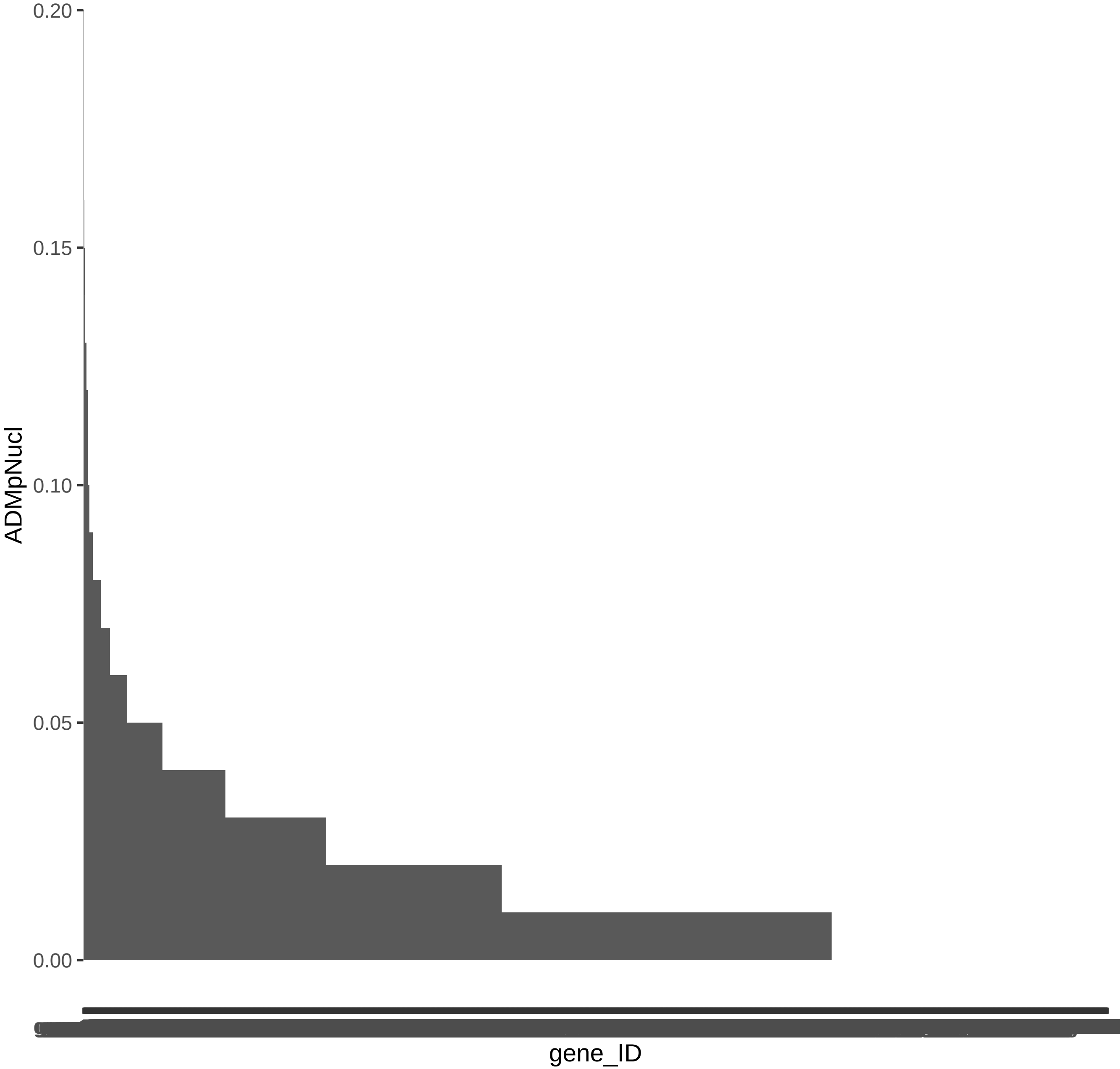
ADMpNucl
0.20
0.15
0.10
0.05
0.00
gene_ID

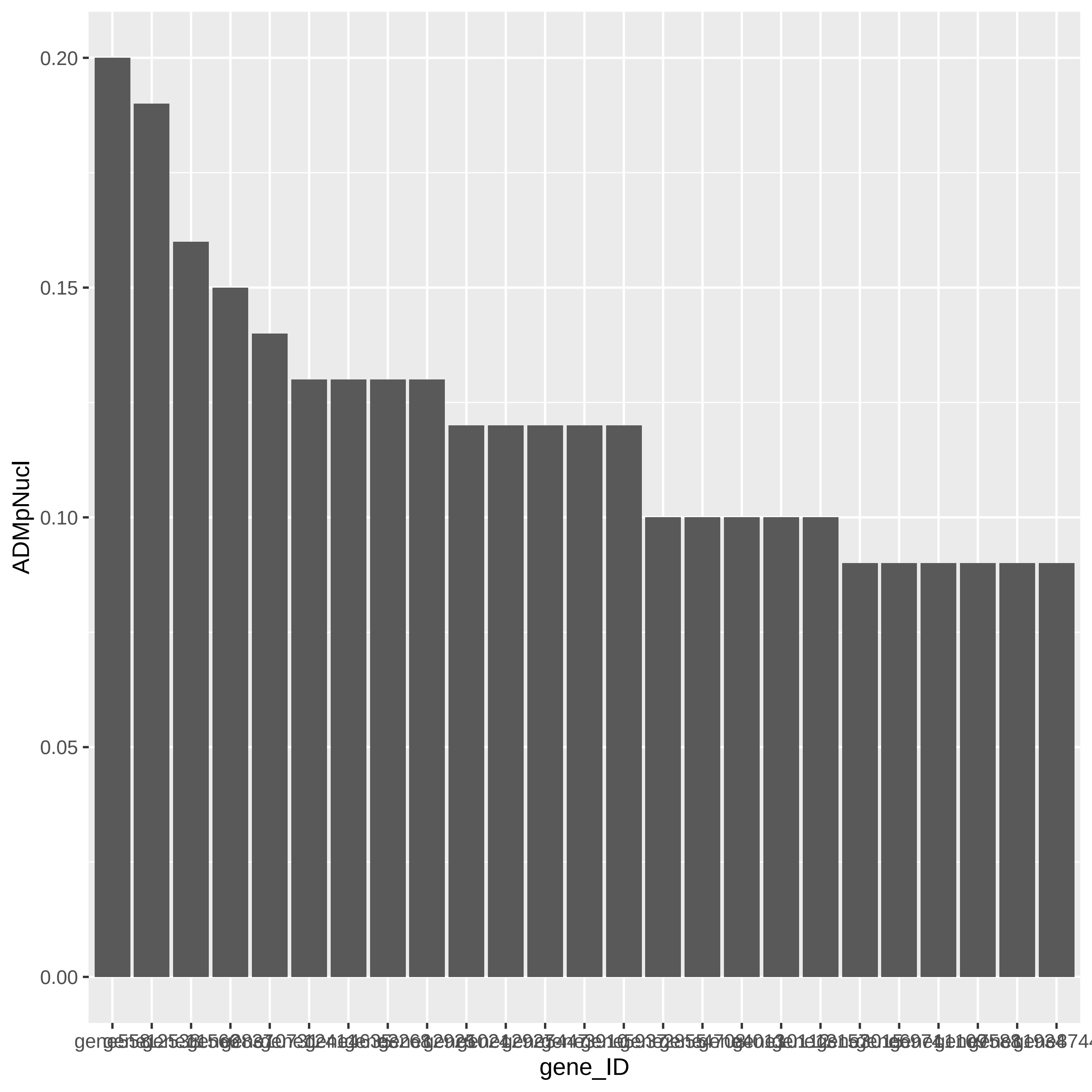

0.20
0.15
0.10
0.05
0.00
ADMpNucl
gene_ID

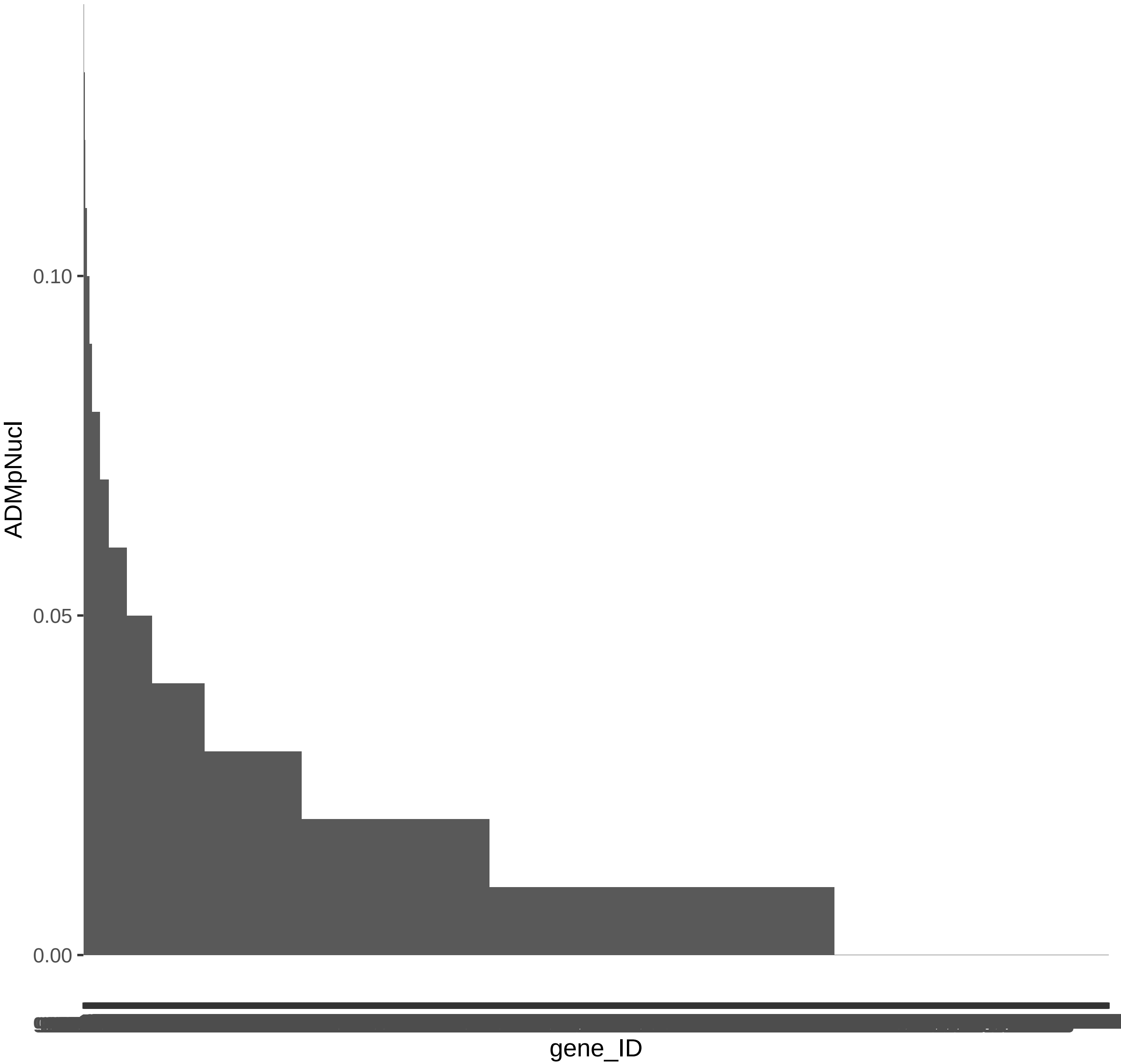

ADMpNucl
0.10
0.05
0.00
gene_ID

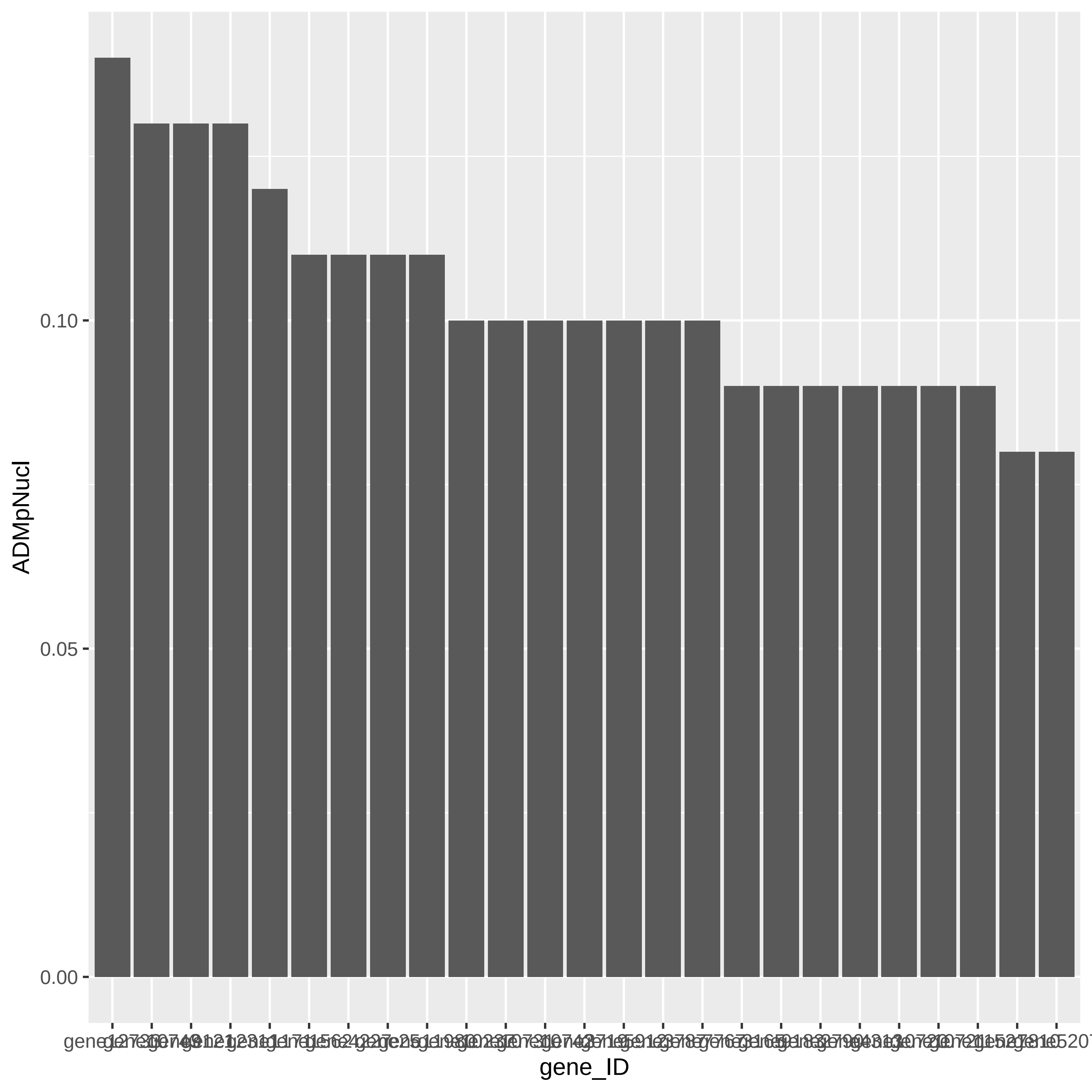

0.10
0.05
0.00
ADMpNucl
gene_ID

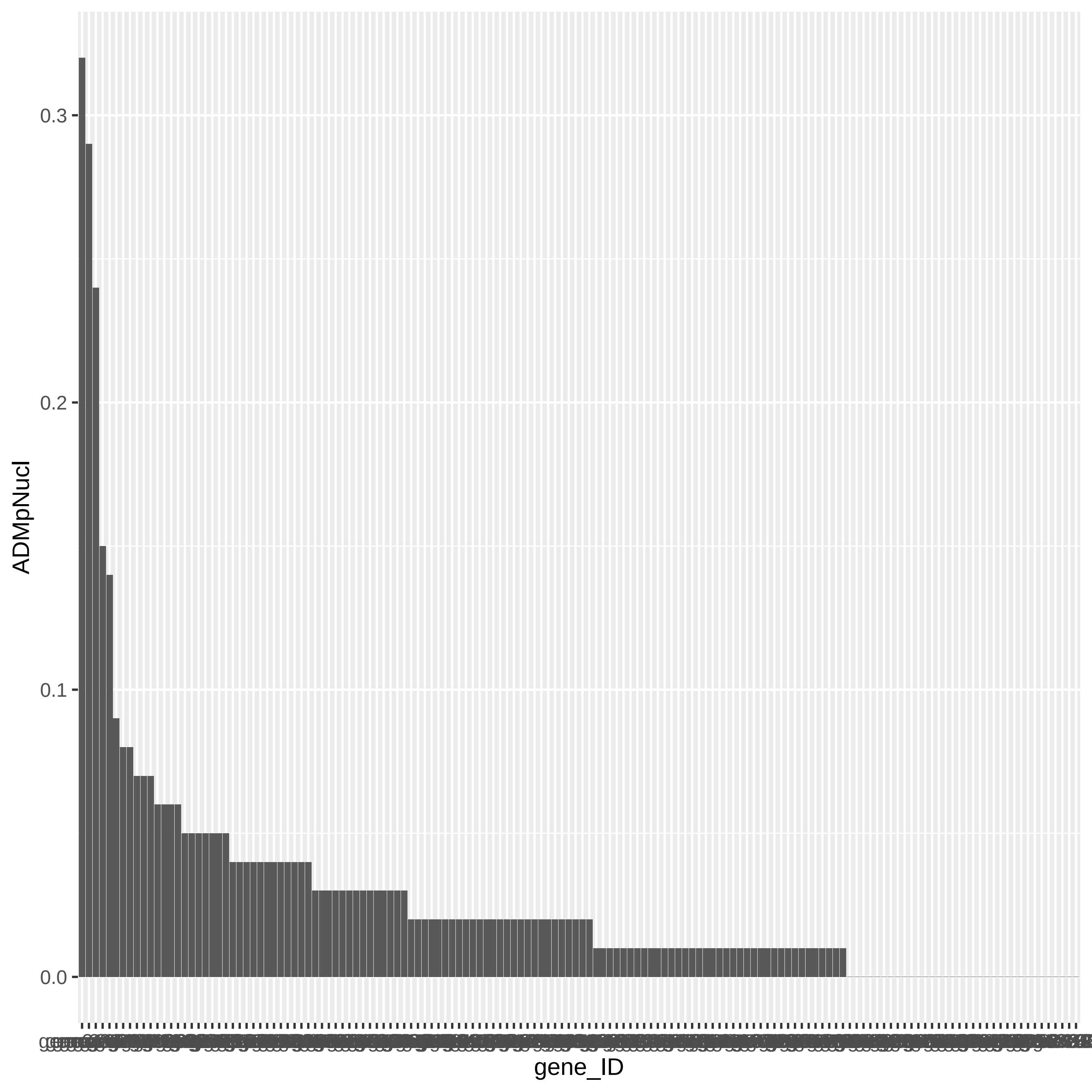
ADMpNucl
0.3
0.2
0.1
0.0
gene_ID

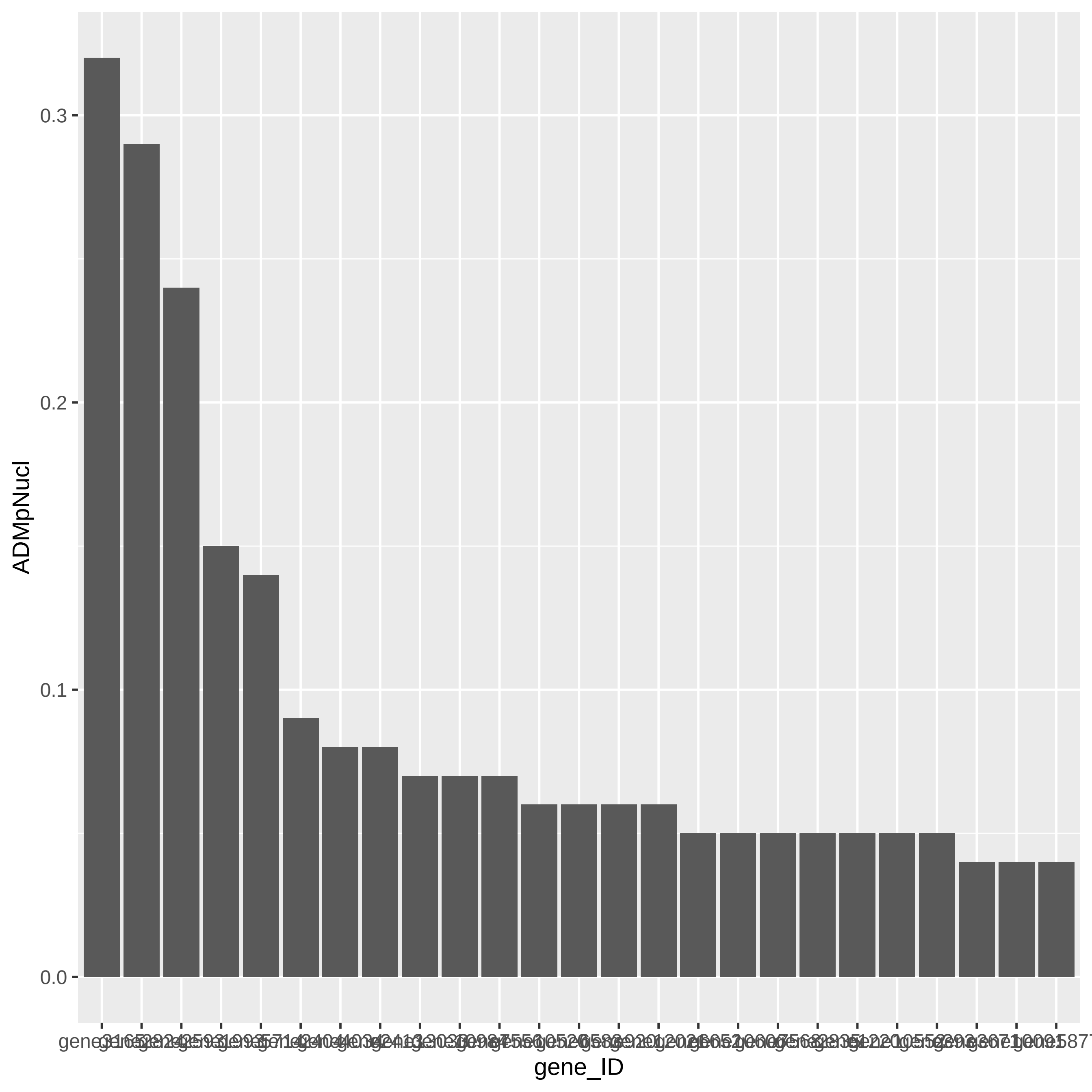

0.3
0.2
0.1
0.0
ADMpNucl
gene_ID

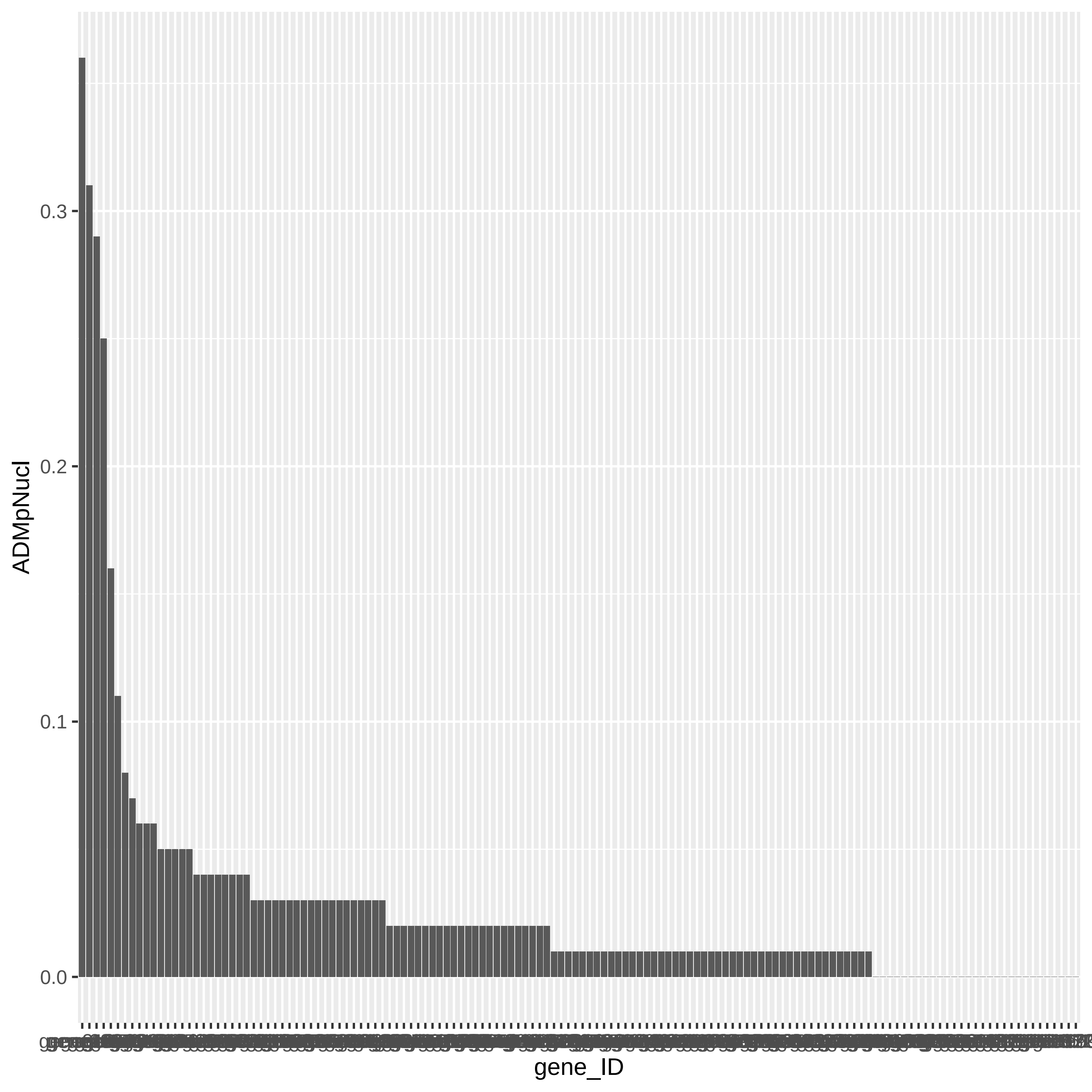

ADMpNucl
0.3
0.2
0.1
0.0
gene_ID

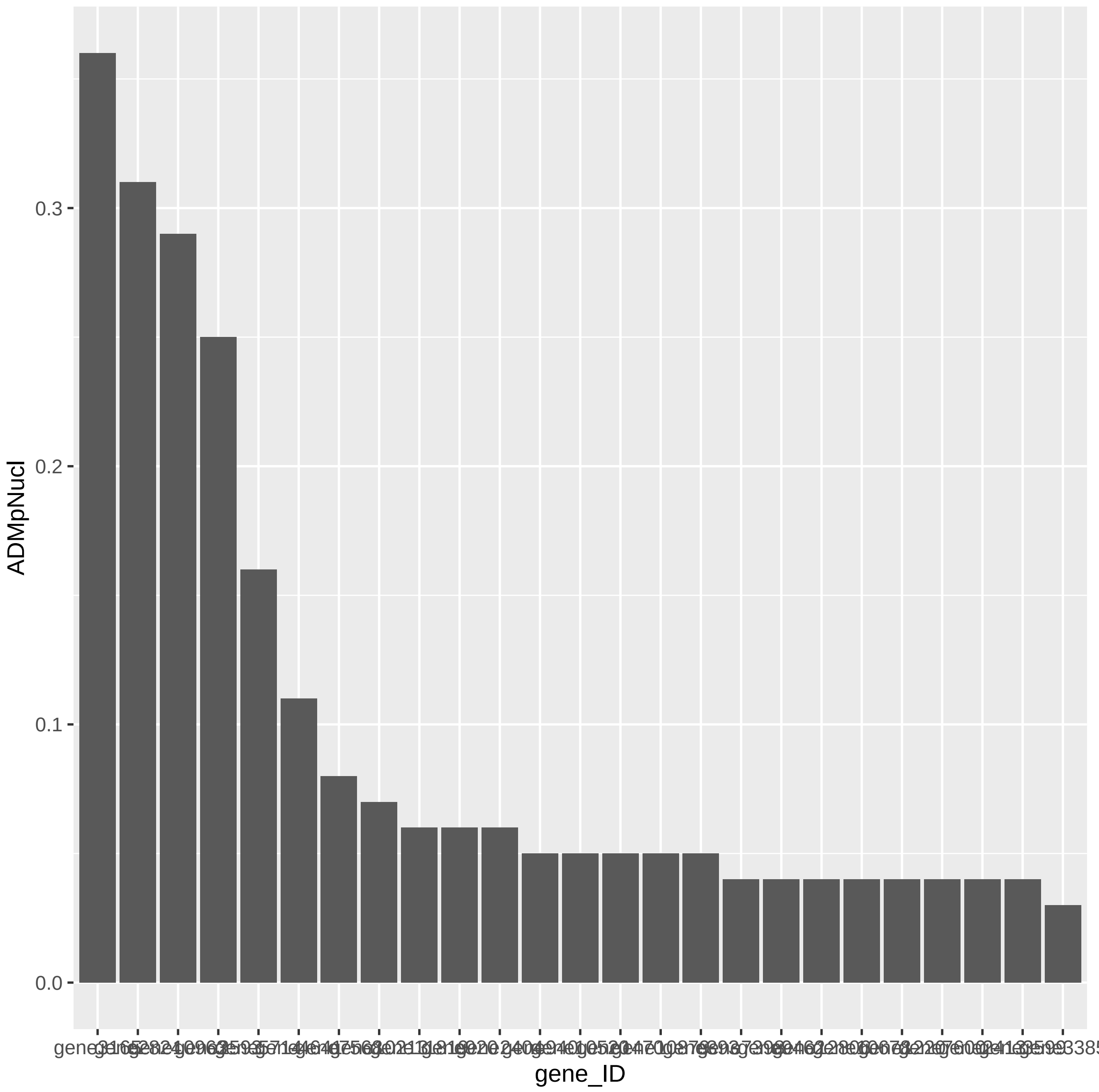
ADMpNucl
0.3
0.2
0.1
0.0
gene_ID

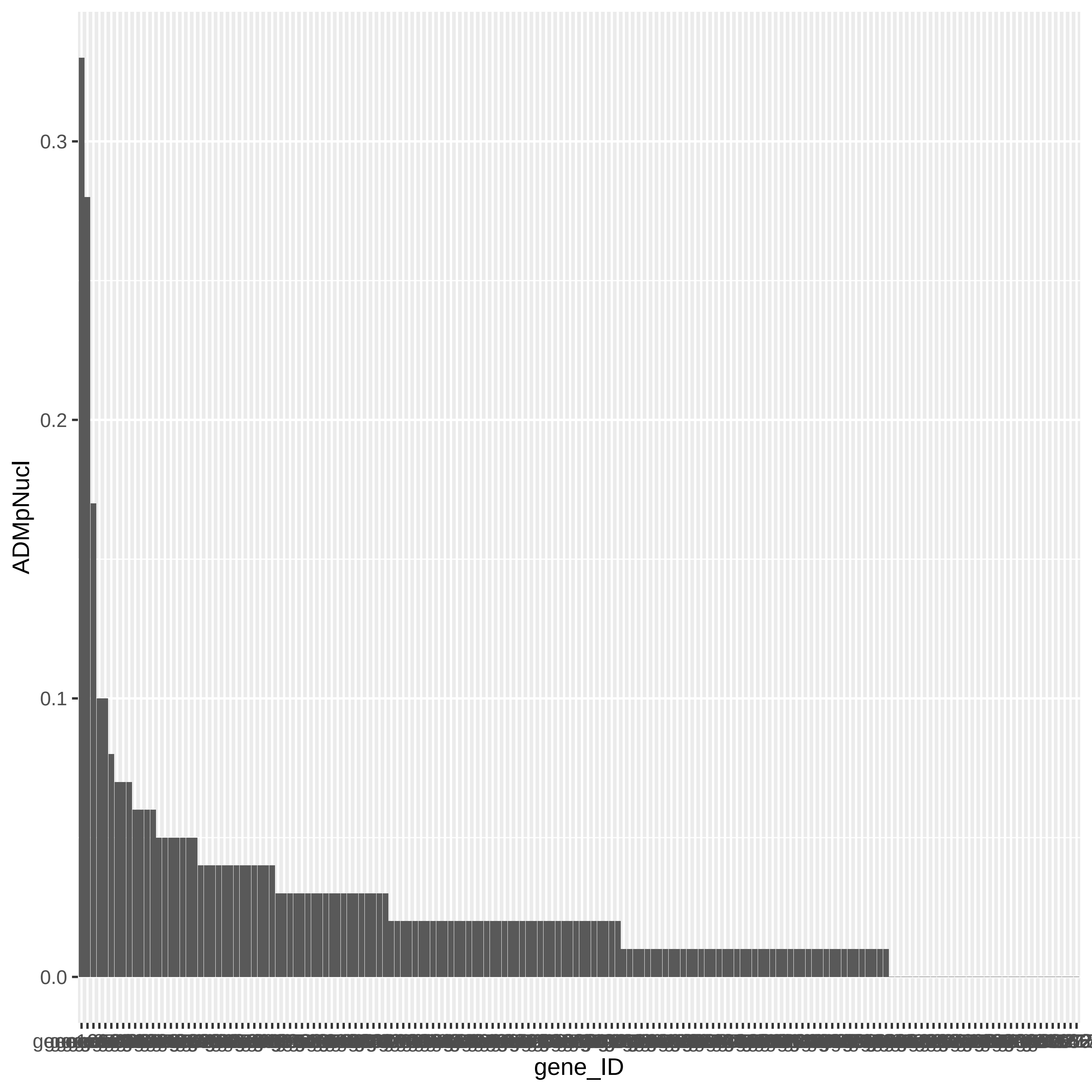

0.3
0.2
0.1
0.0
ADMpNucl
gene_ID

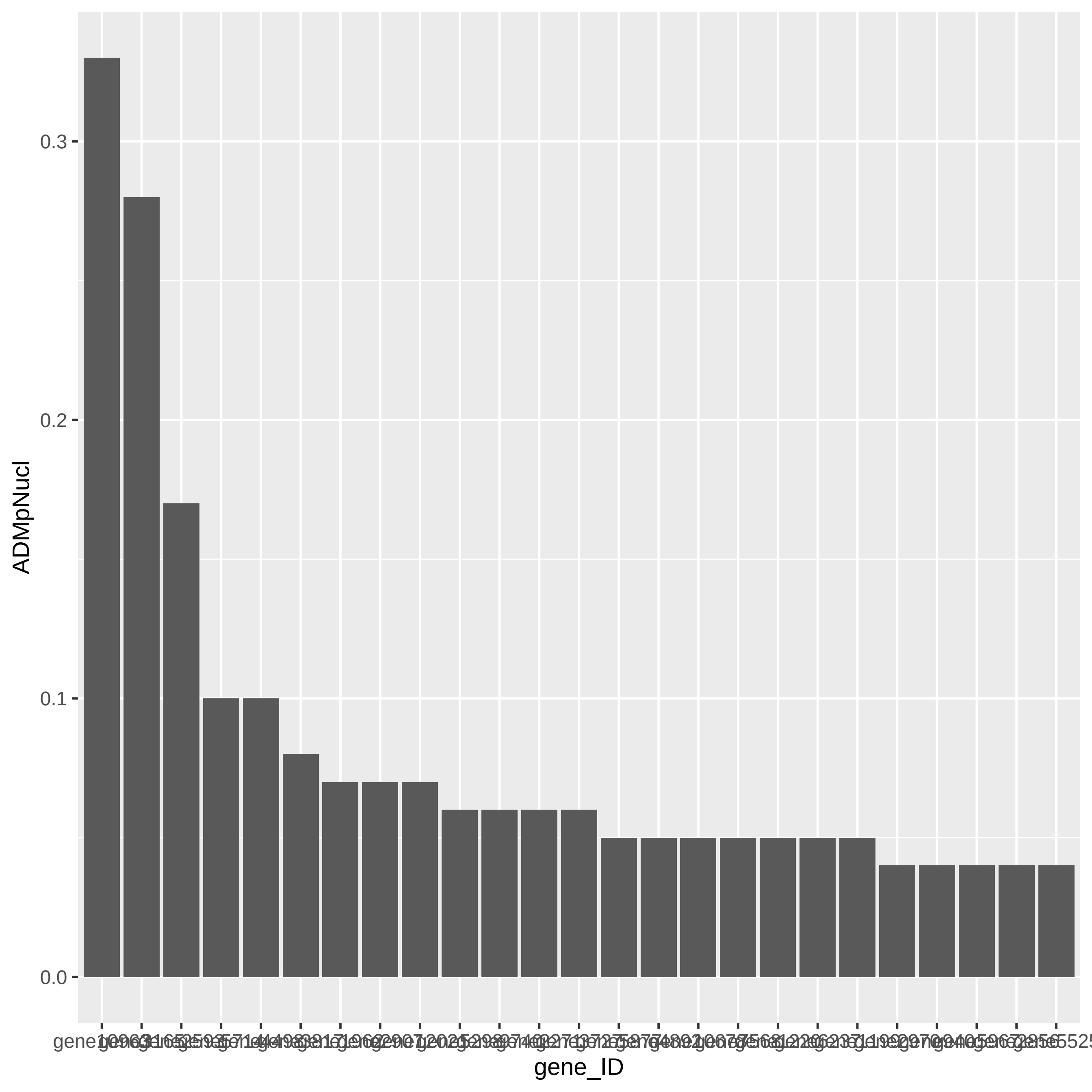

ADMpNucl
0.3
0.2
0.1
0.0
gene_ID

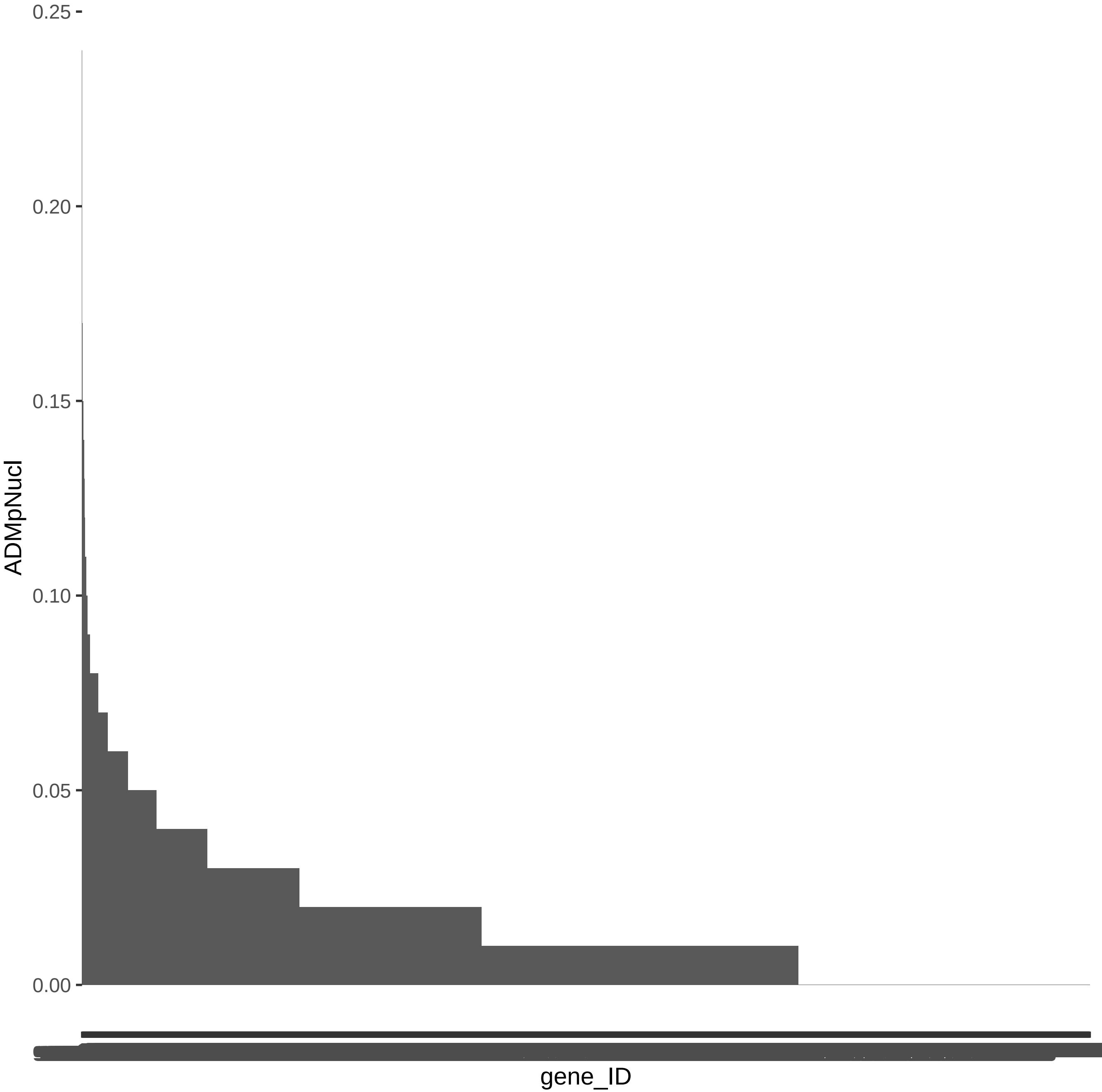

0.25
0.20
0.15
0.10
0.05
0.00
ADMpNucl
gene_ID

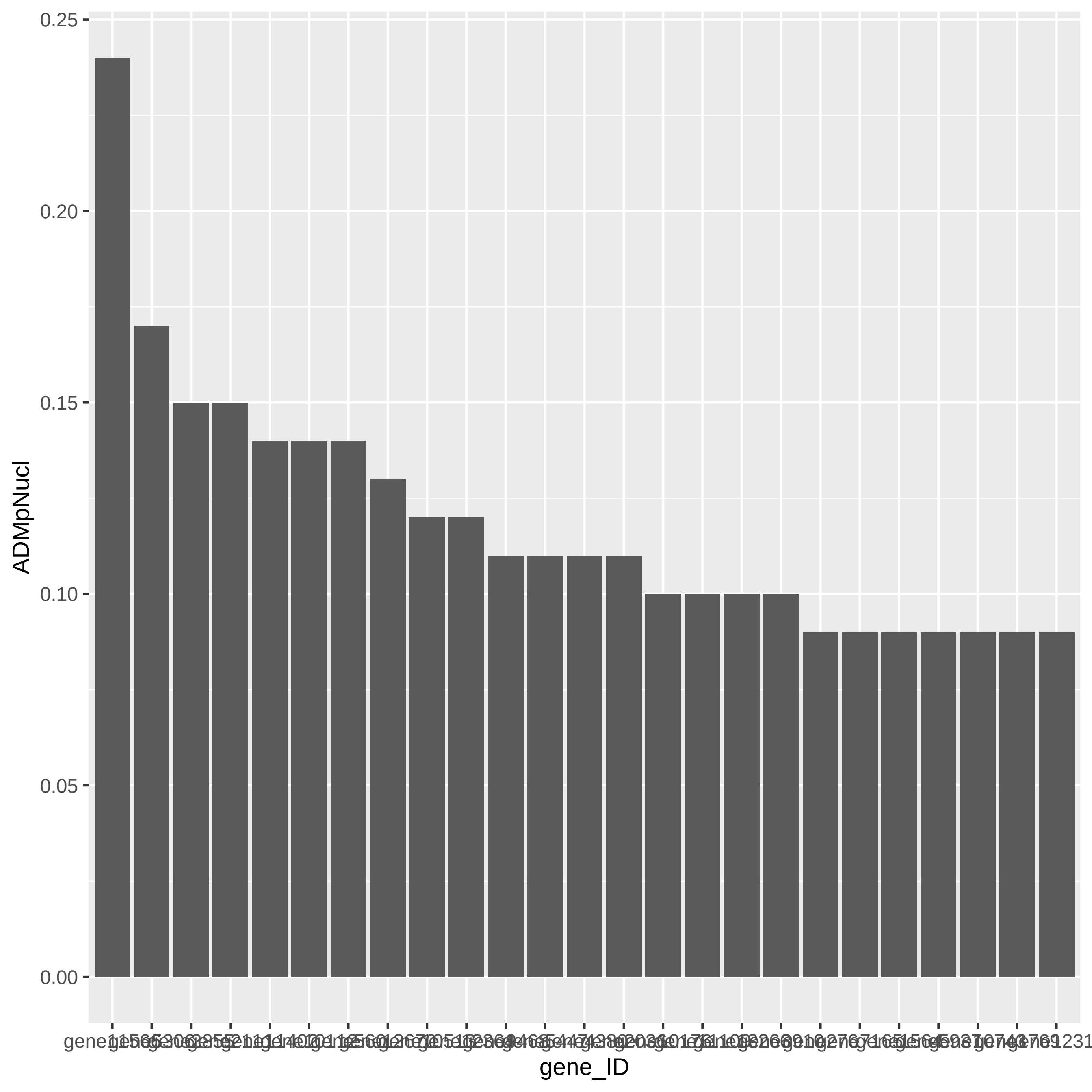
0.25
0.20
0.15
0.10
0.05
0.00
ADMpNucl
gene_ID

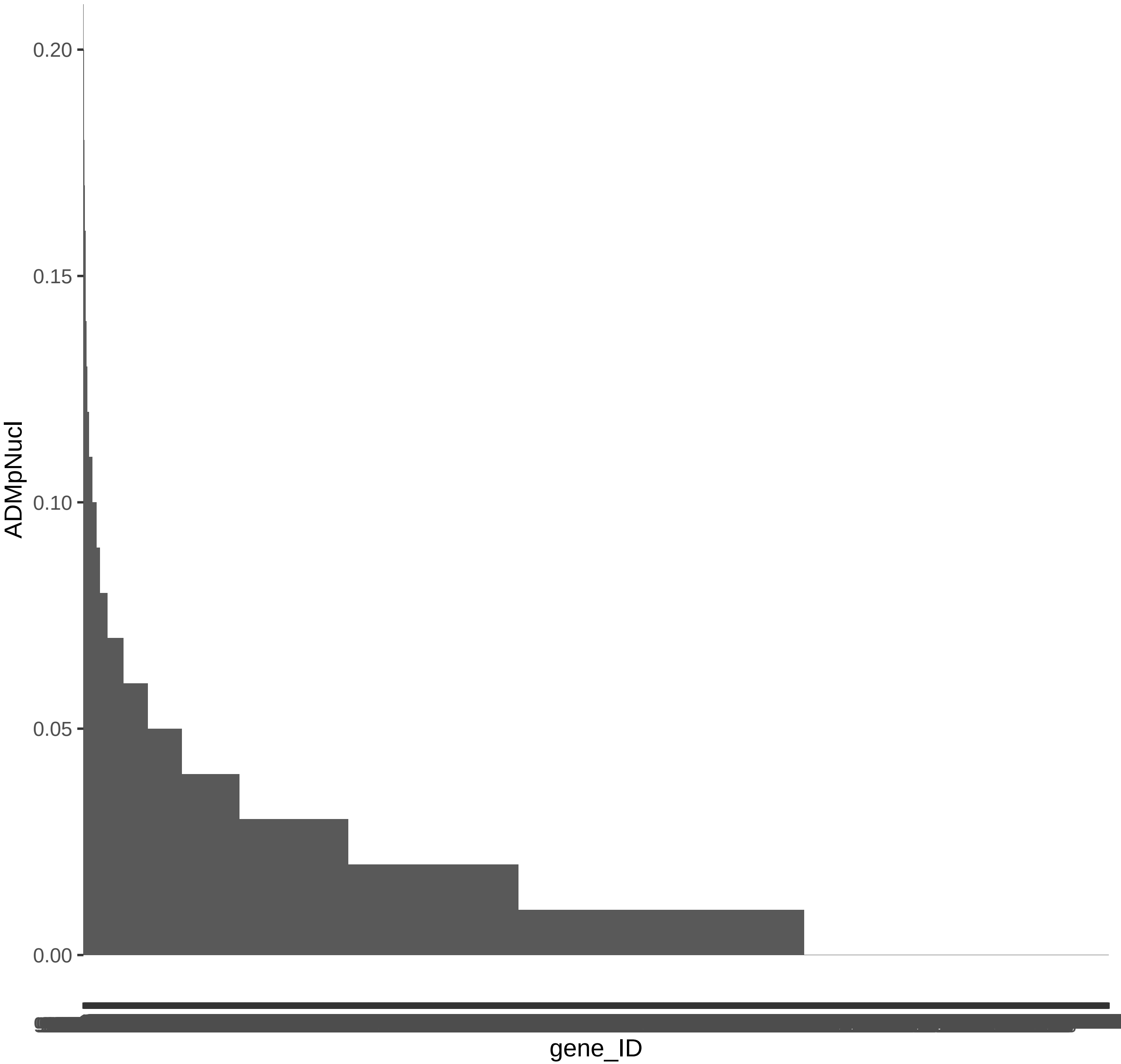
0.20
0.15
0.10
0.05
0.00
ADMpNucl
gene_ID

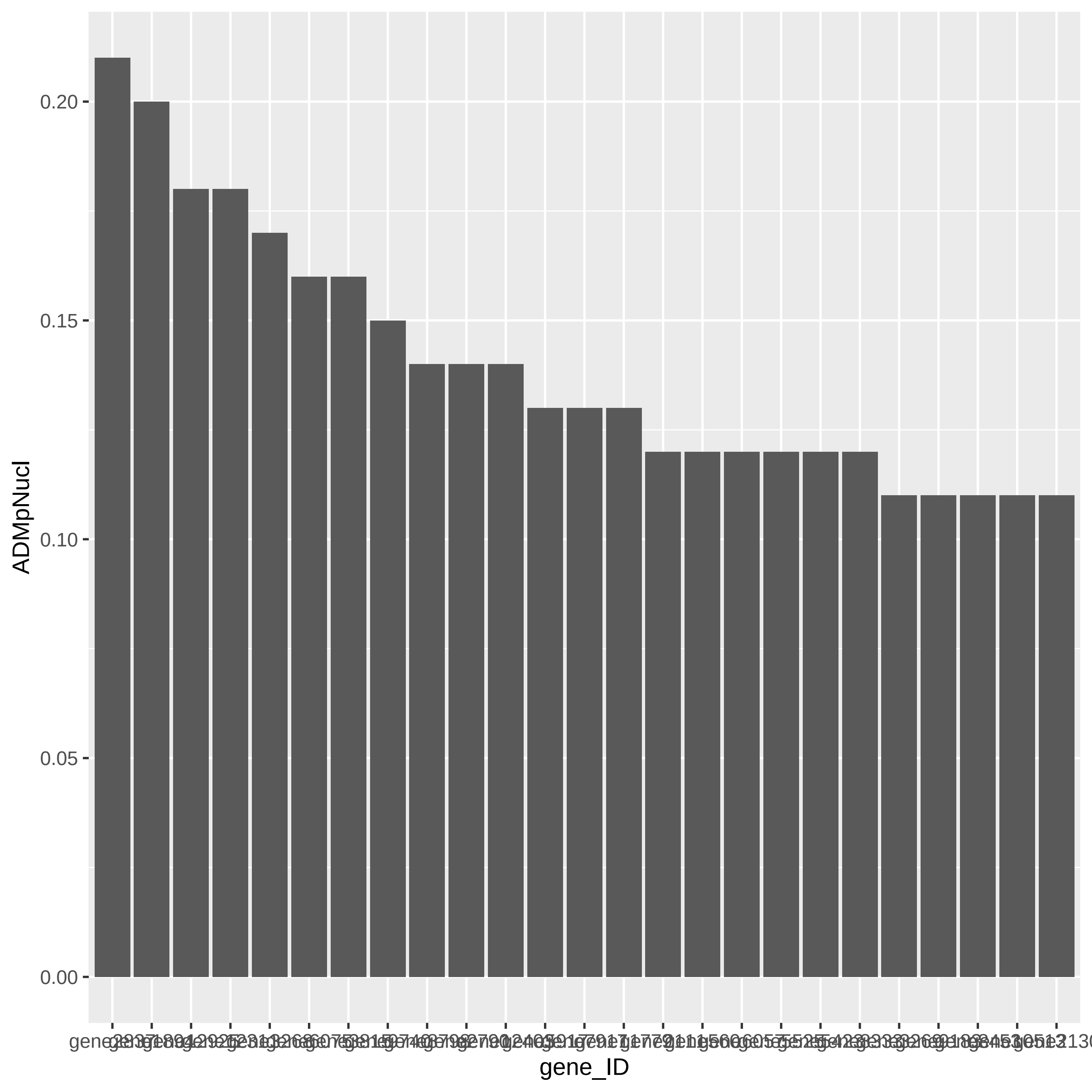

ADMpNucl
0.20
0.15
0.10
0.05
0.00
gene_ID

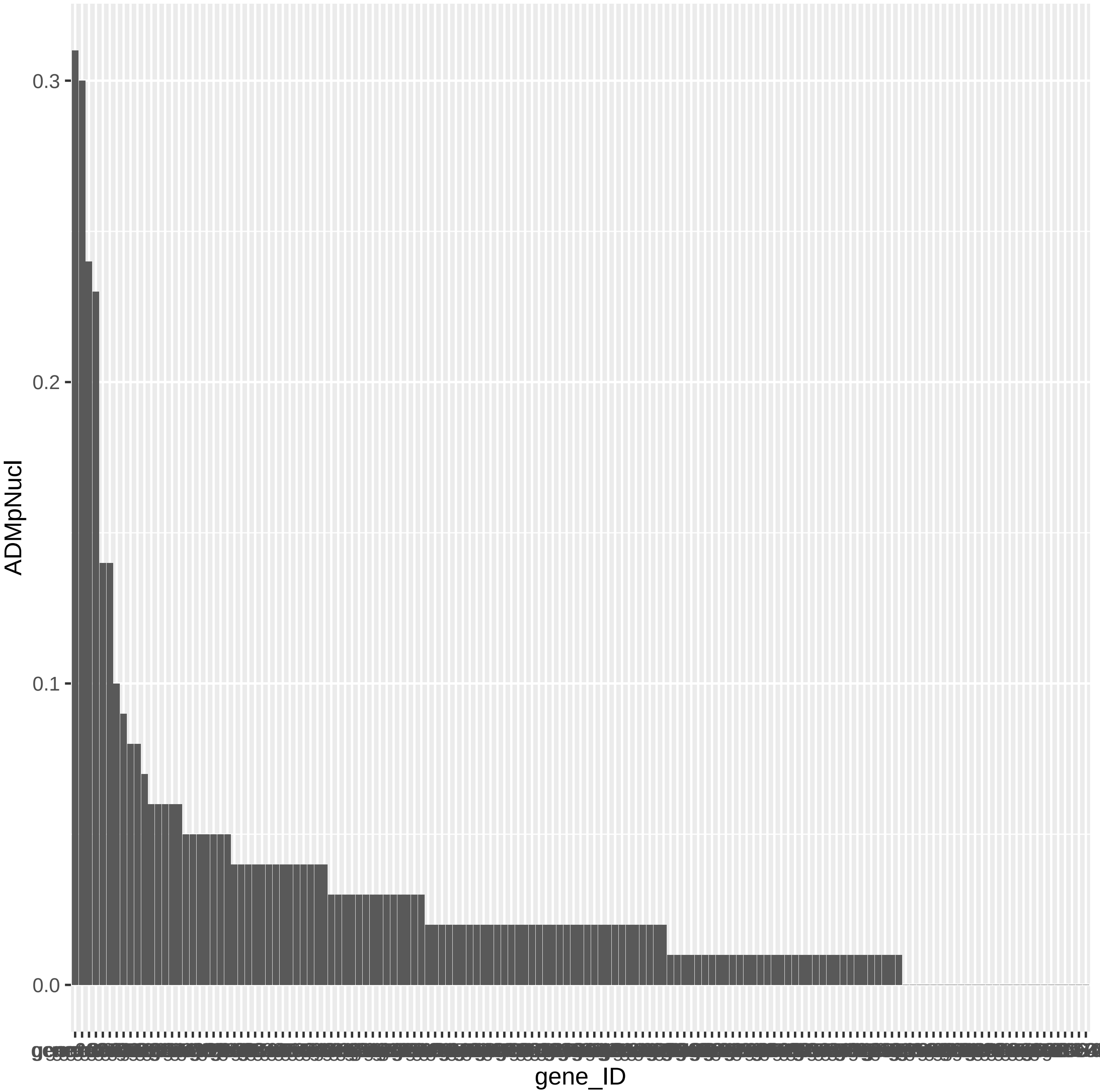
0.3
0.2
0.1
0.0
ADMpNucl
gene_ID

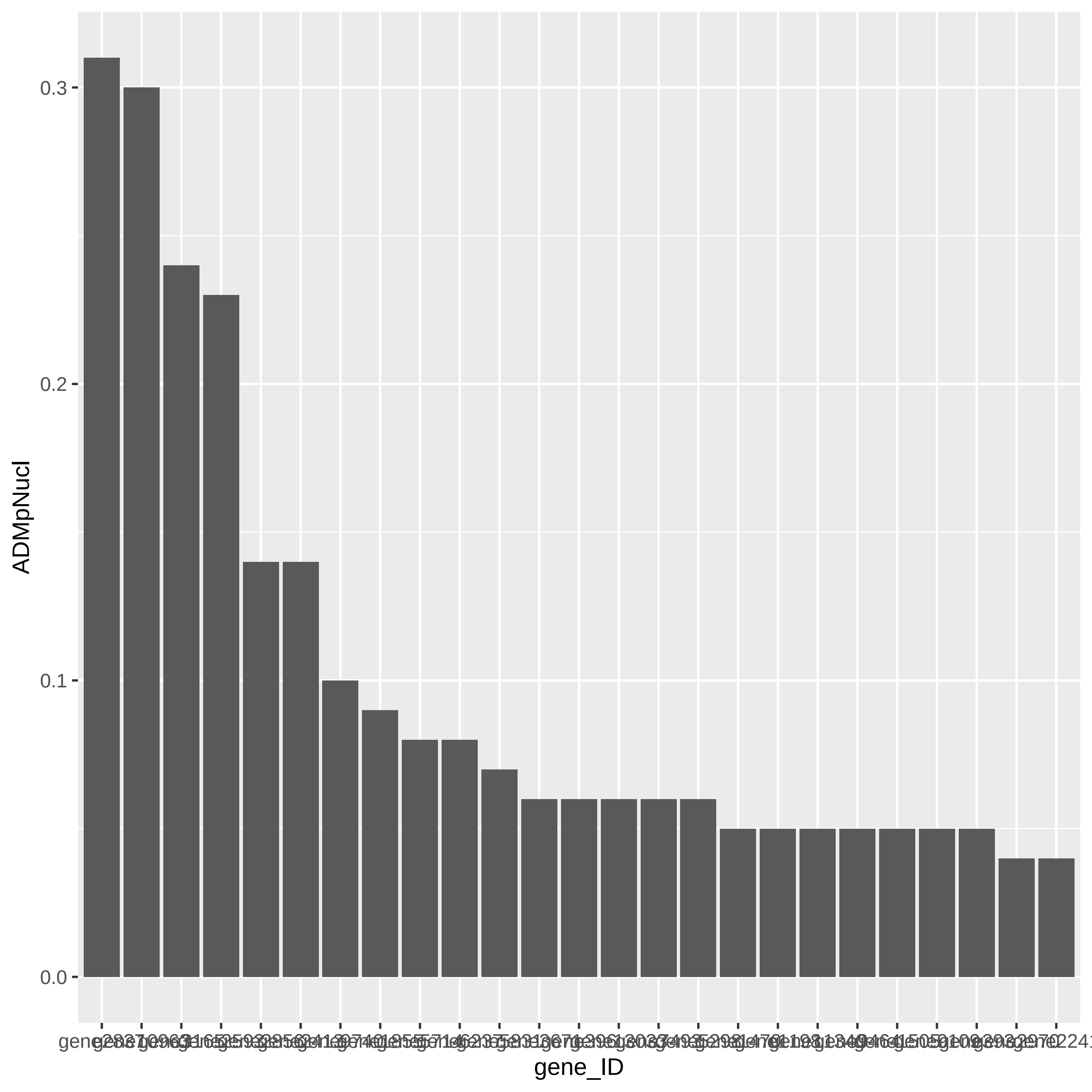
ADMpNucl
0.3
0.2
0.1
0.0
gene_ID

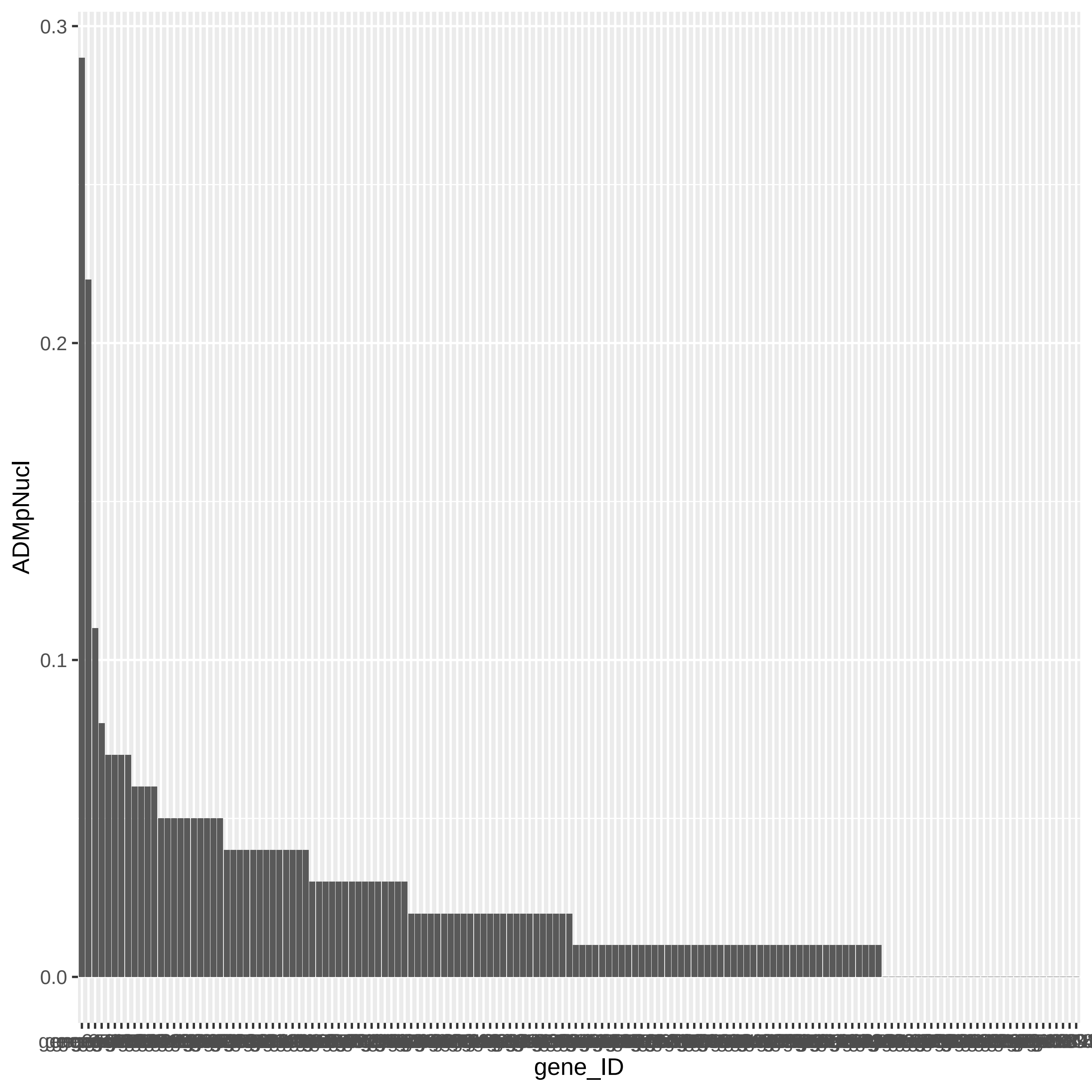
0.3
0.2
0.1
0.0
ADMpNucl
gene_ID

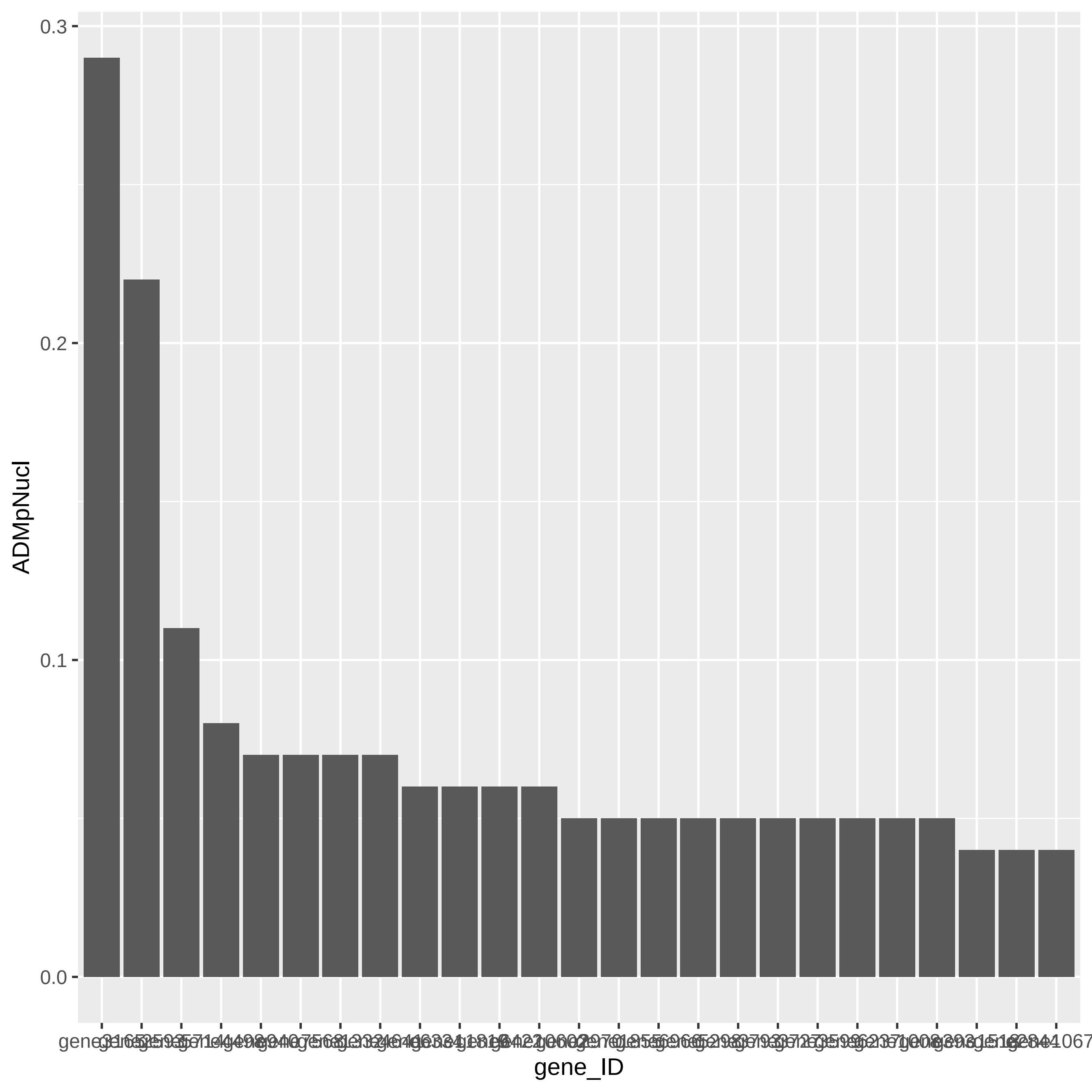
0.3
0.2
0.1
0.0
ADMpNucl
gene_ID

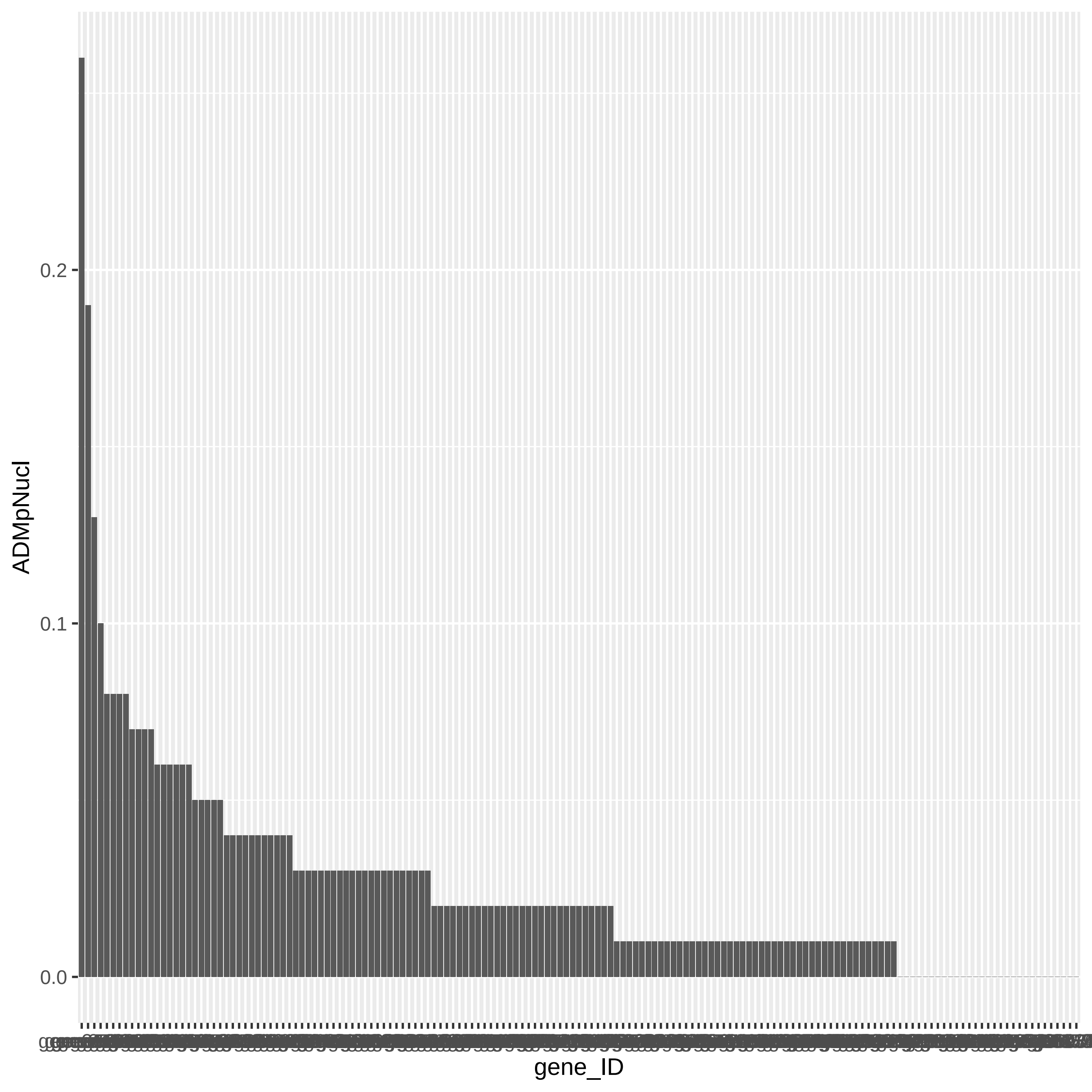

ADMpNucl
0.2
0.1
0.0
gene_ID

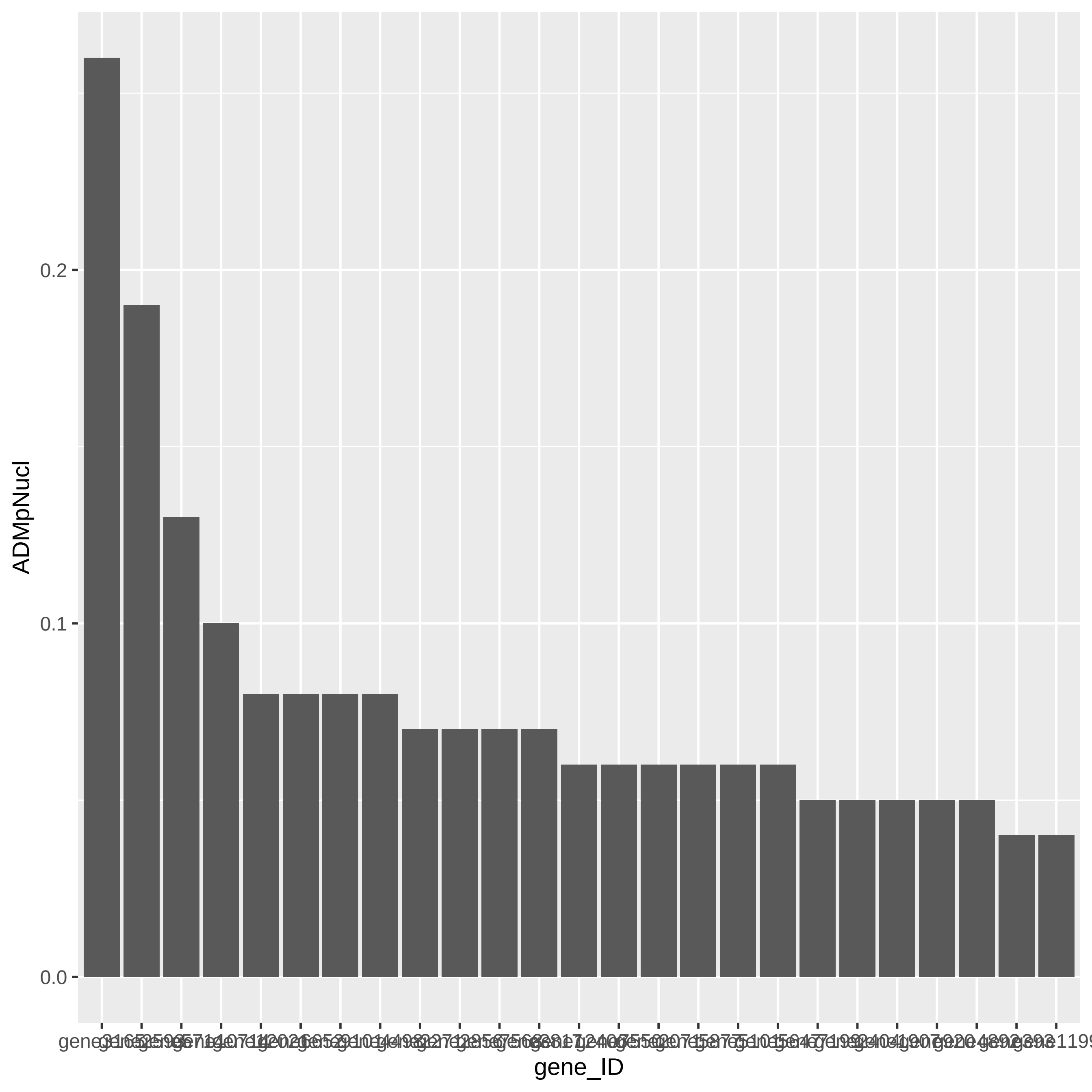
ADMpNucl
0.2
0.1
0.0
gene_ID

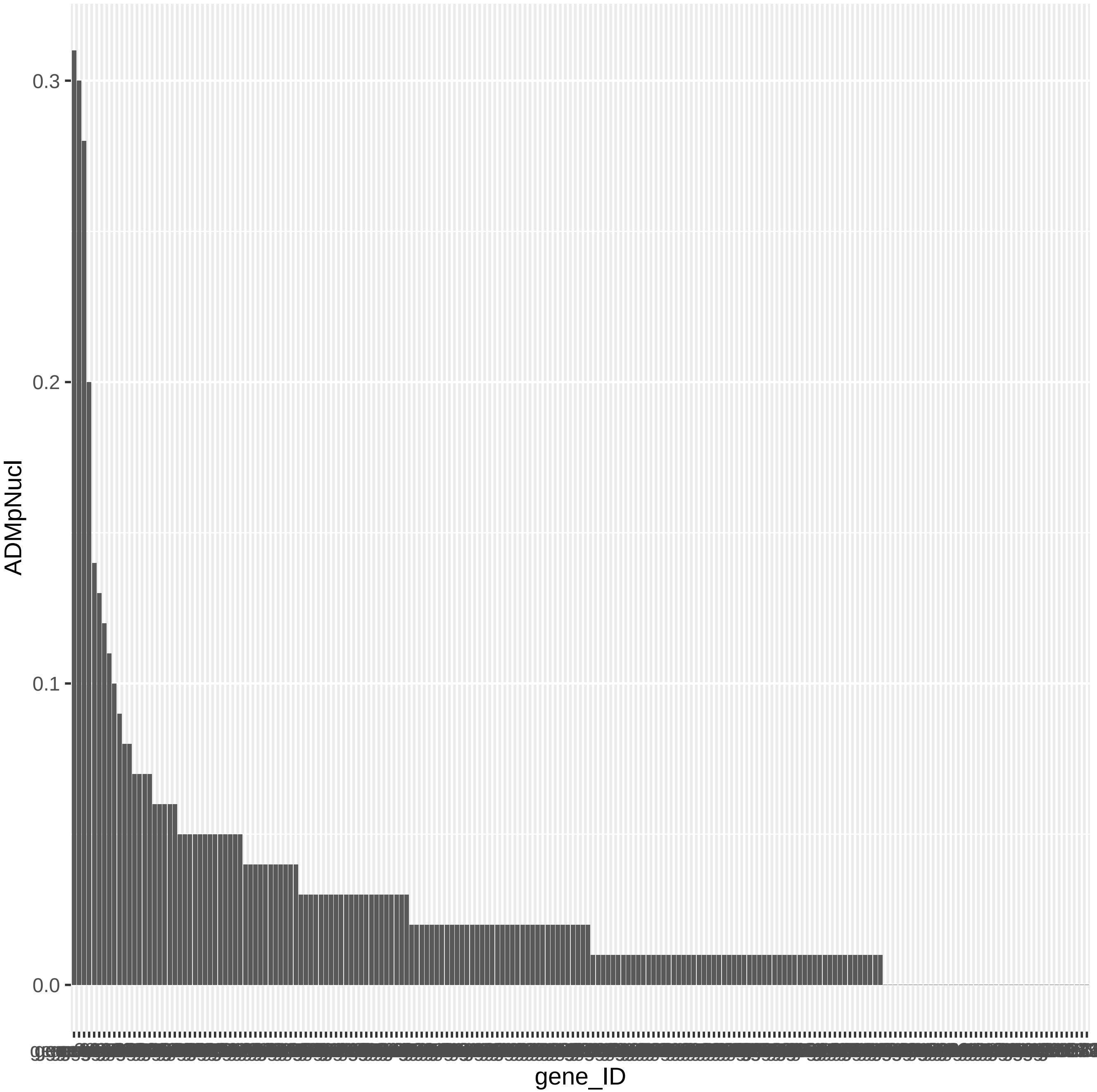
ADMpNucl
0.3
0.2
0.1
0.0
gene_ID

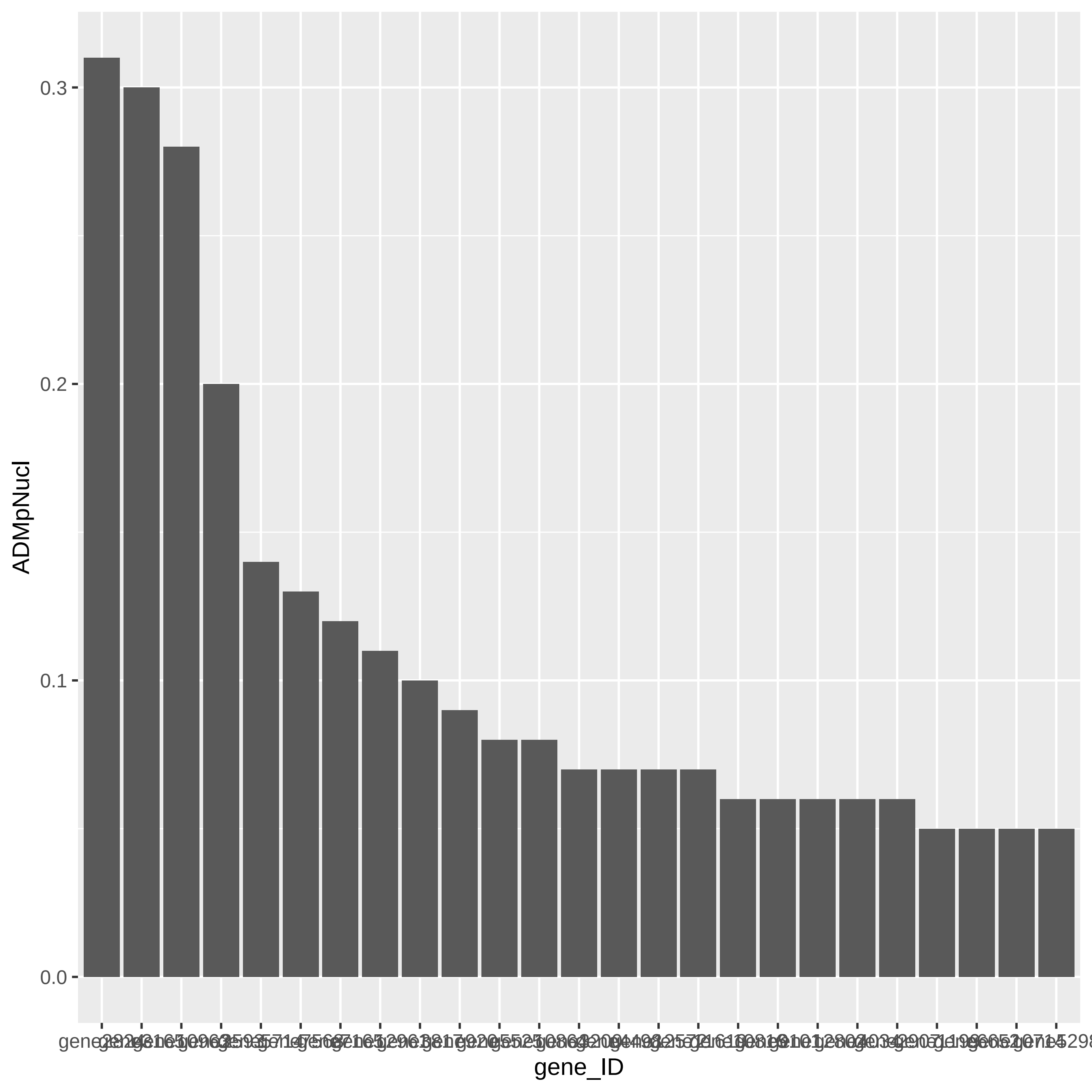

ADMpNucl
0.0
0.1
0.2
0.3
gene_ID

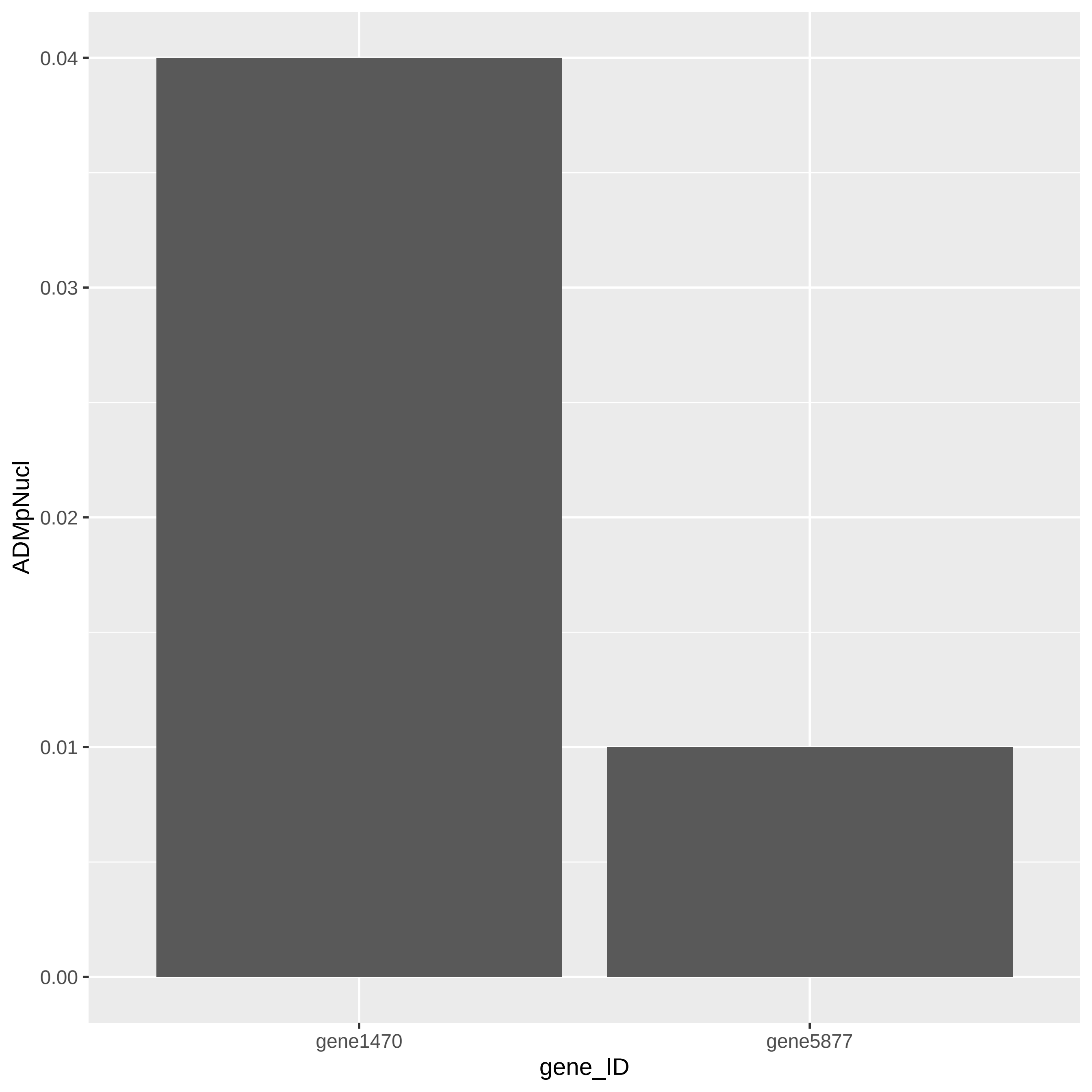
0.04
0.03
0.02
0.01
0.00
ADMpNucl
gene1470
gene5877
gene_ID

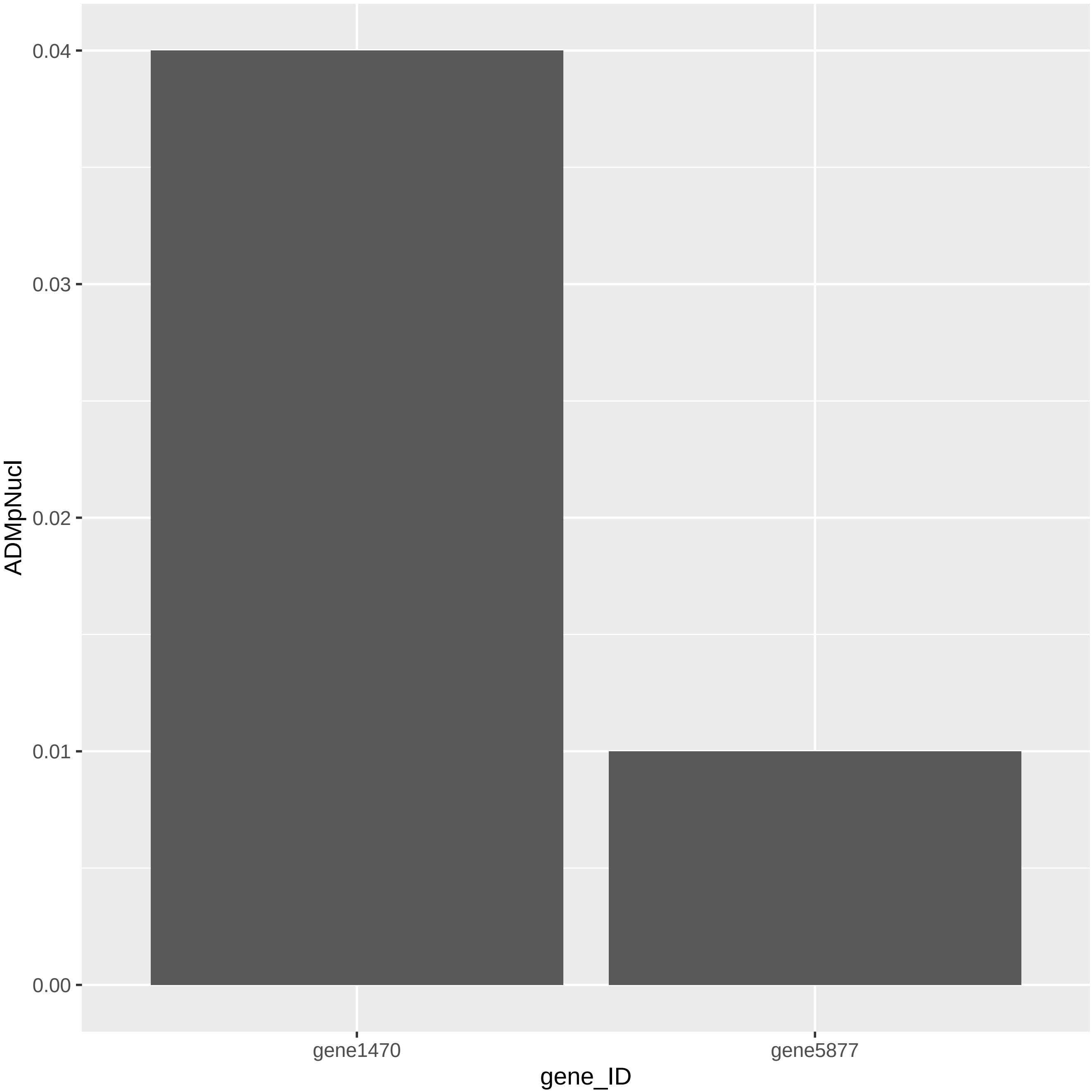
0.04
0.03
0.02
0.01
0.00
ADMpNucl
gene1470
gene5877
gene_ID

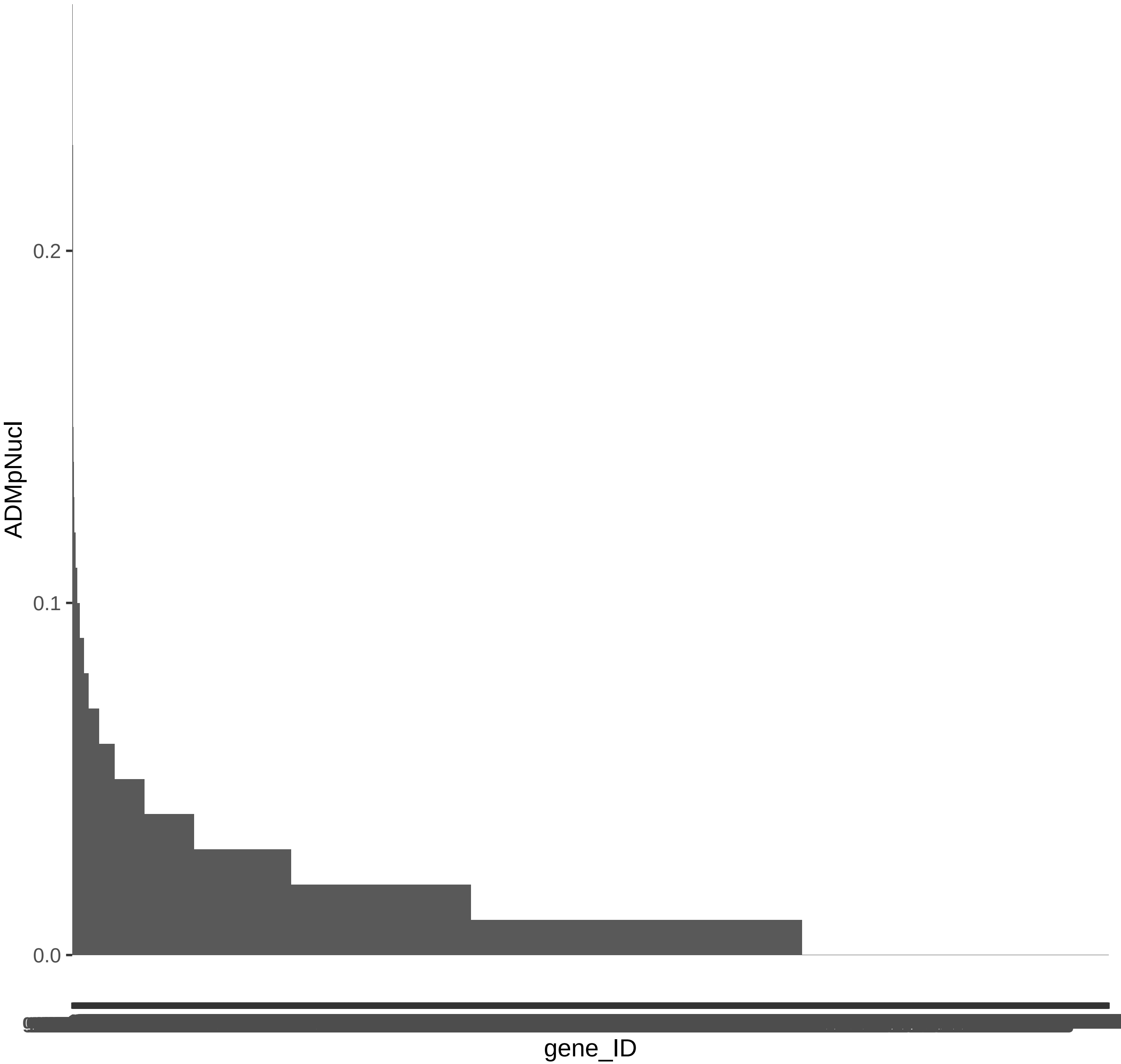
ADMpNucl
0.2
0.1
0.0
gene_ID

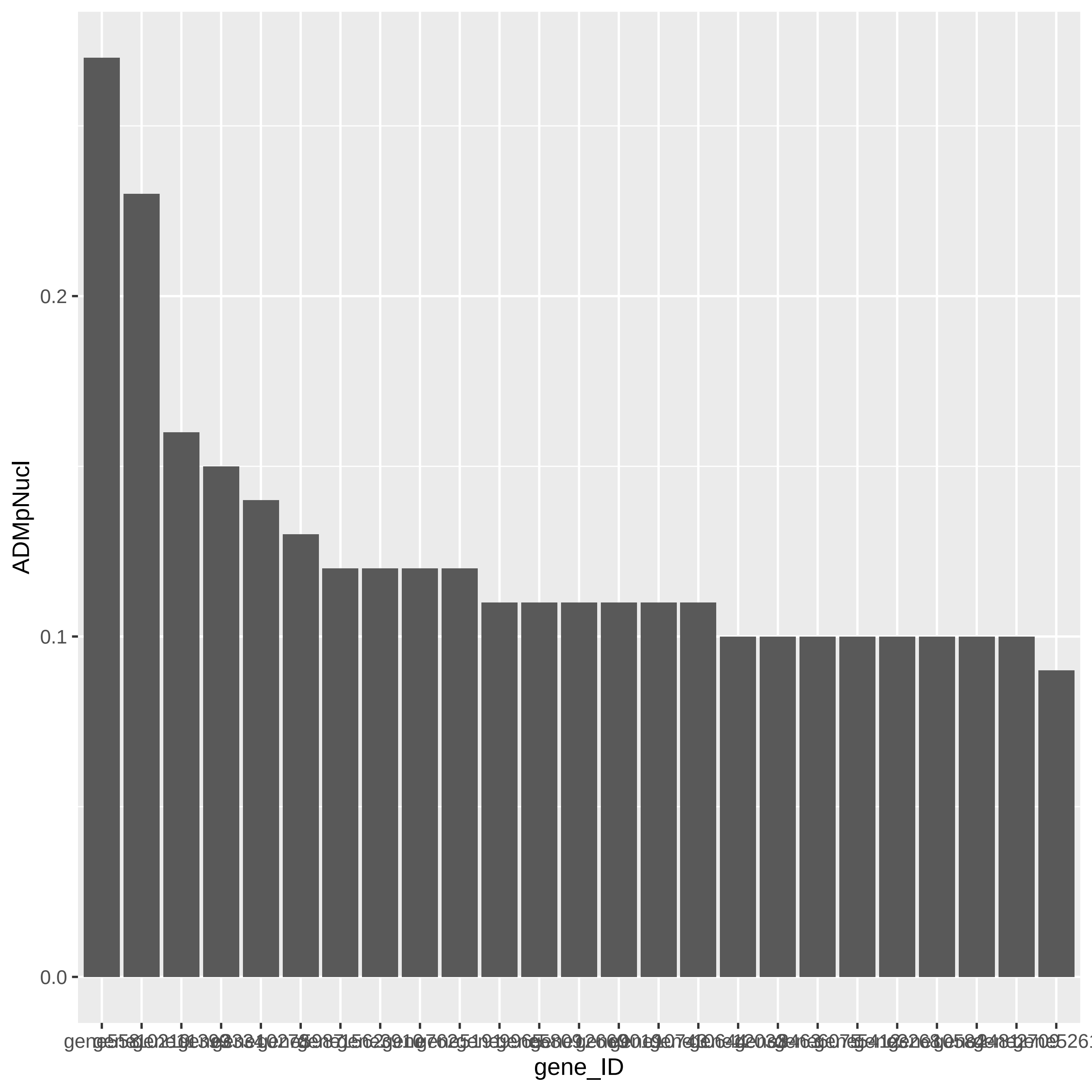
ADMpNucl
0.2
0.1
0.0
gene_ID

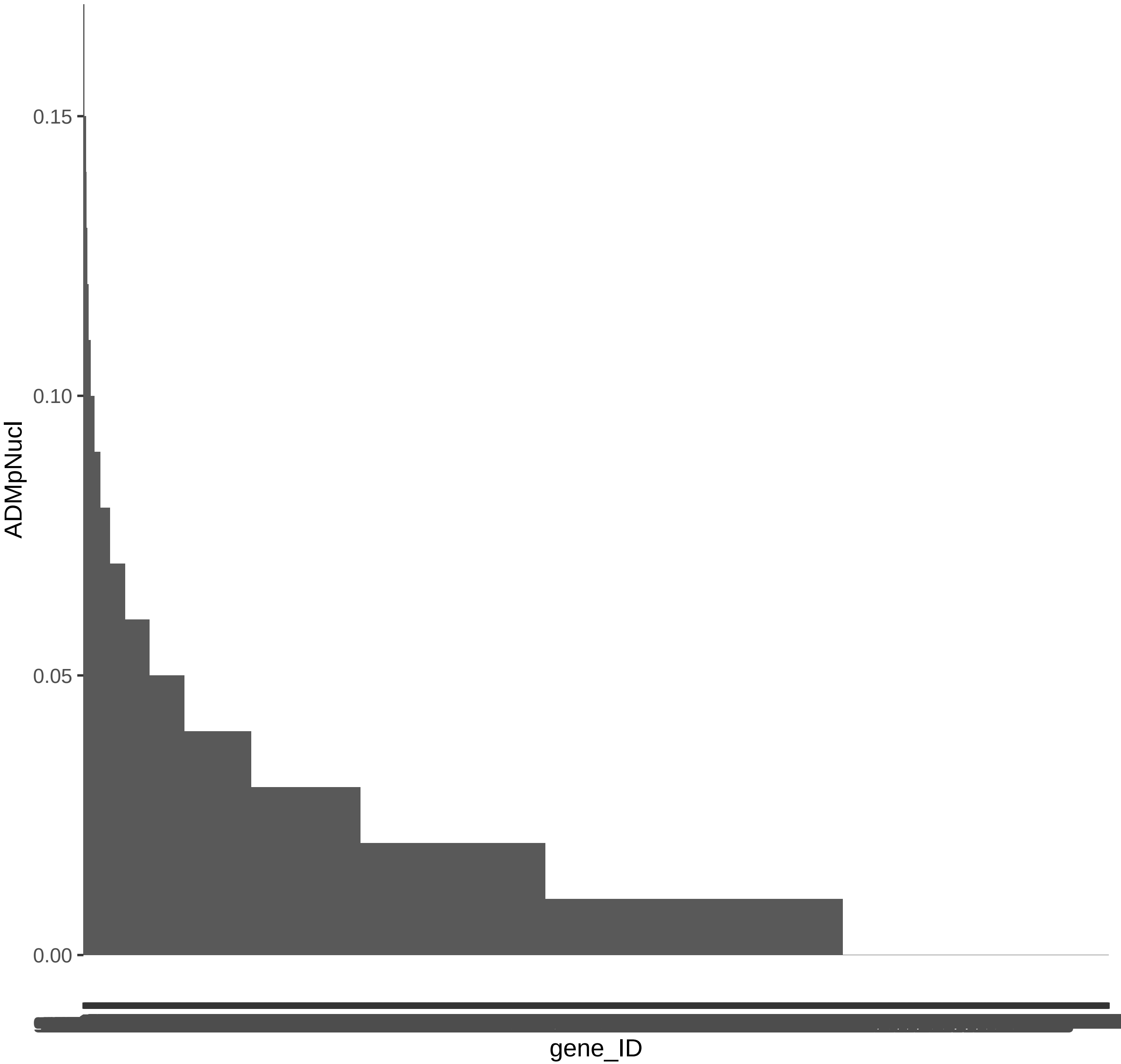
0.15
0.10
0.05
0.00
ADMpNucl
gene_ID

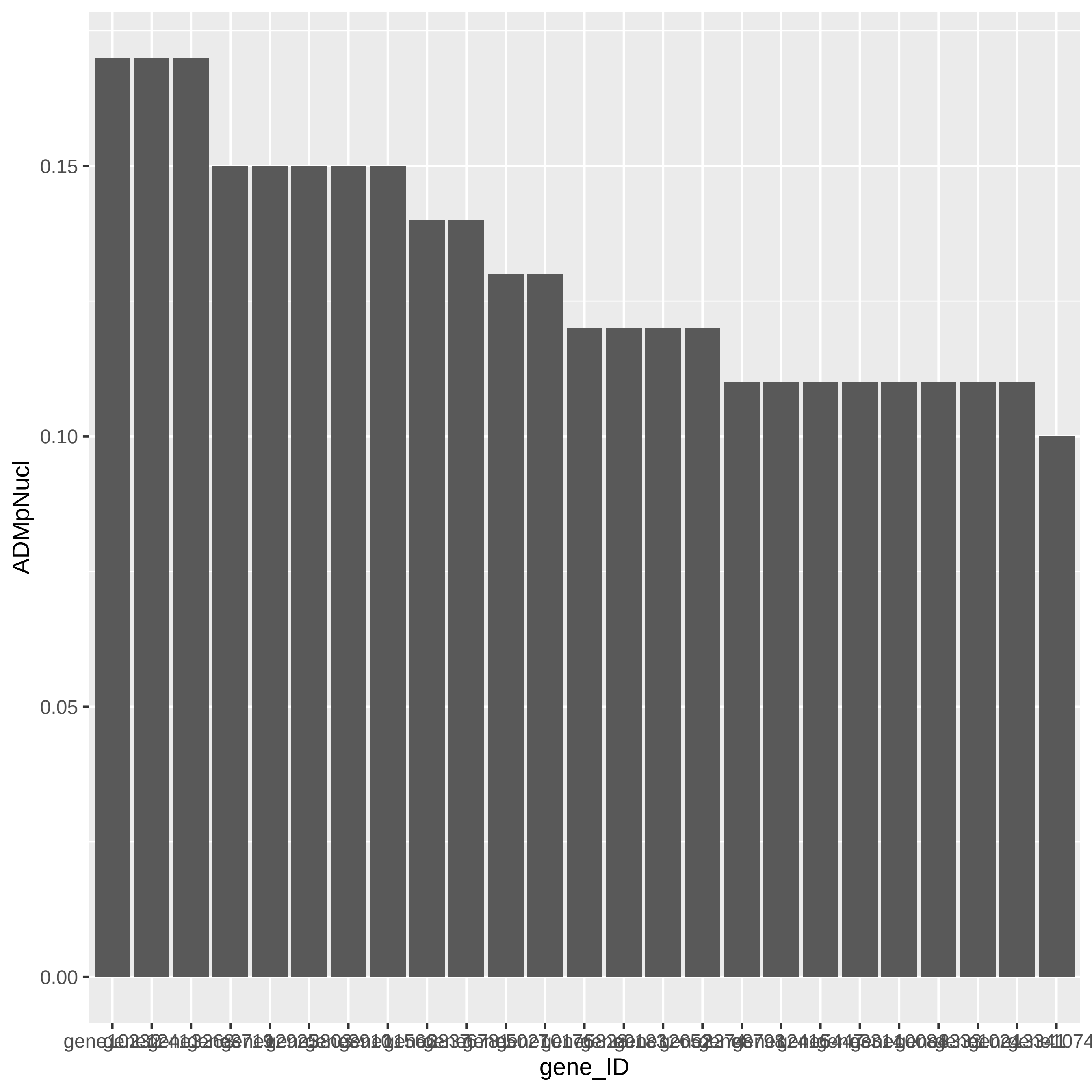

ADMpNucl
0.15
0.10
0.05
0.00
gene_ID

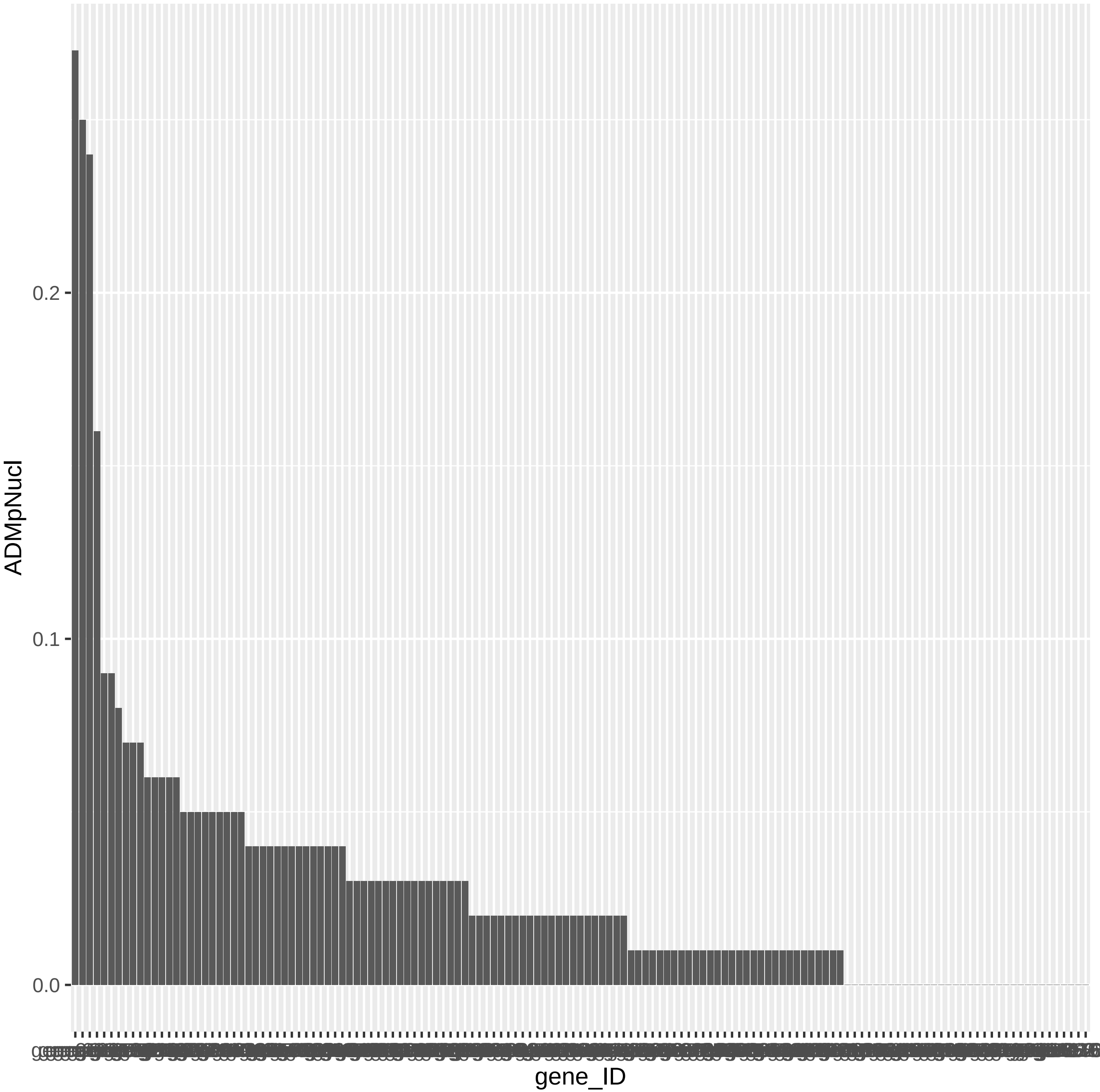
ADMpNucl
0.2
0.1
0.0
gene_ID

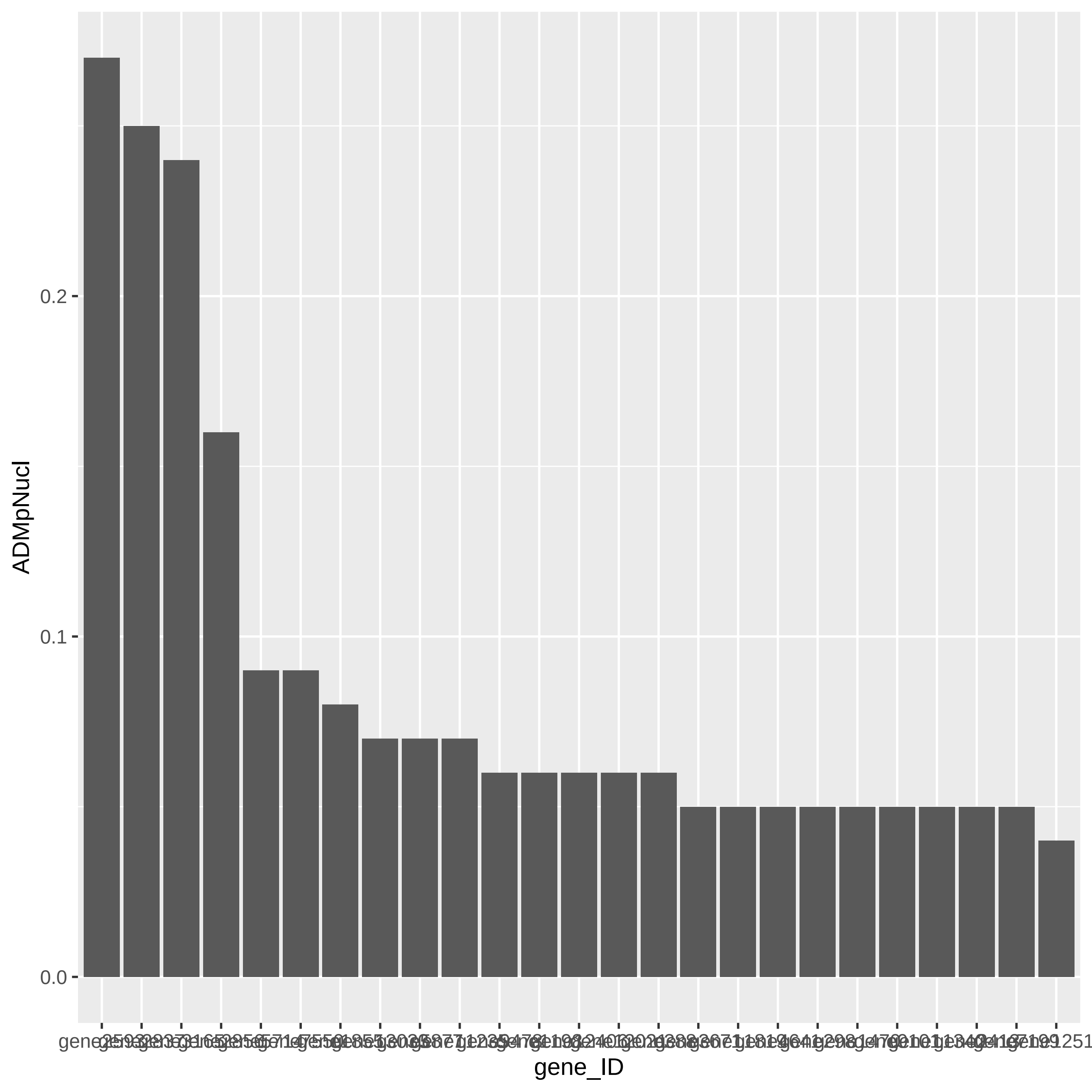
ADMpNucl
0.0
0.1
0.2
gene_ID

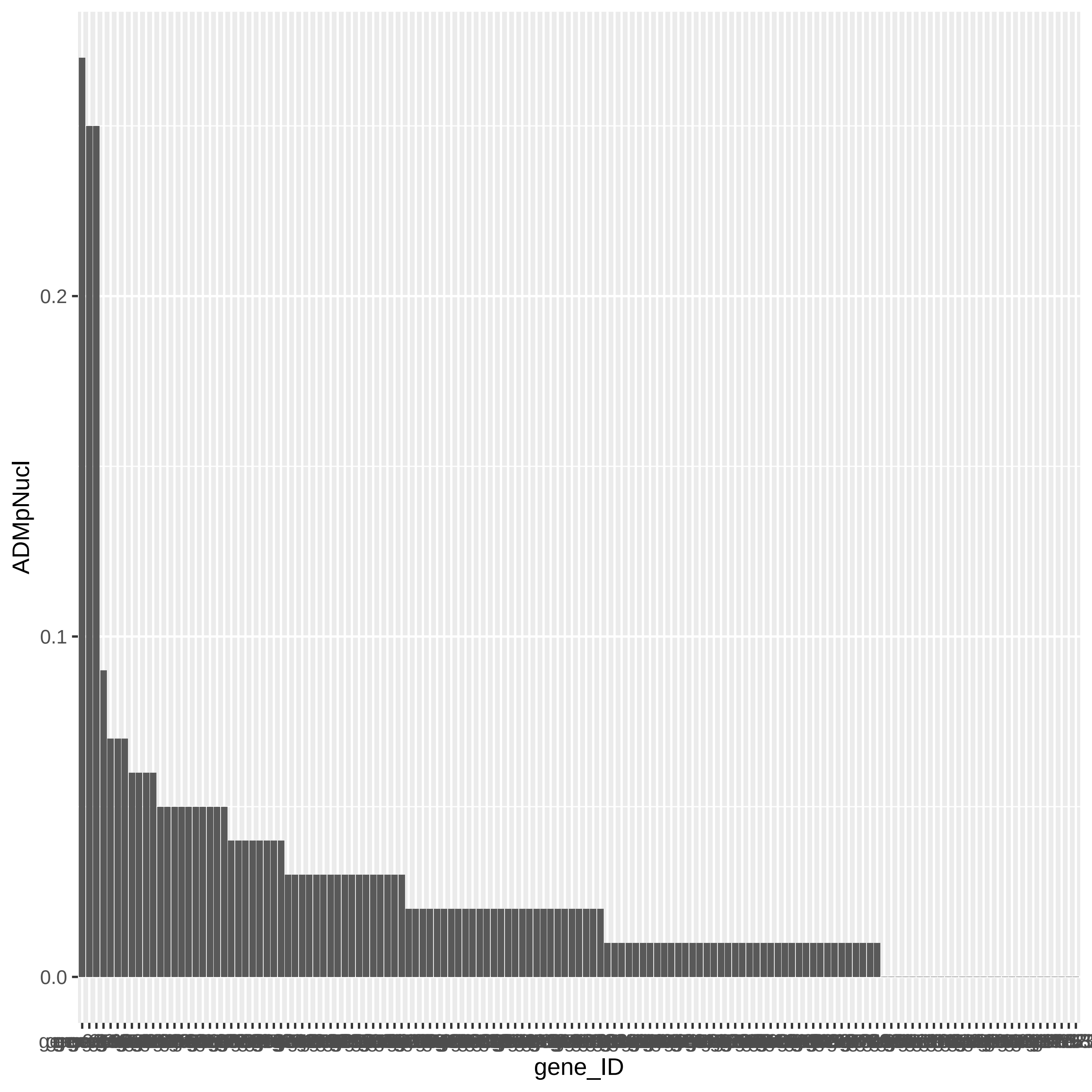
ADMpNucl
0.2
0.1
0.0
gene_ID

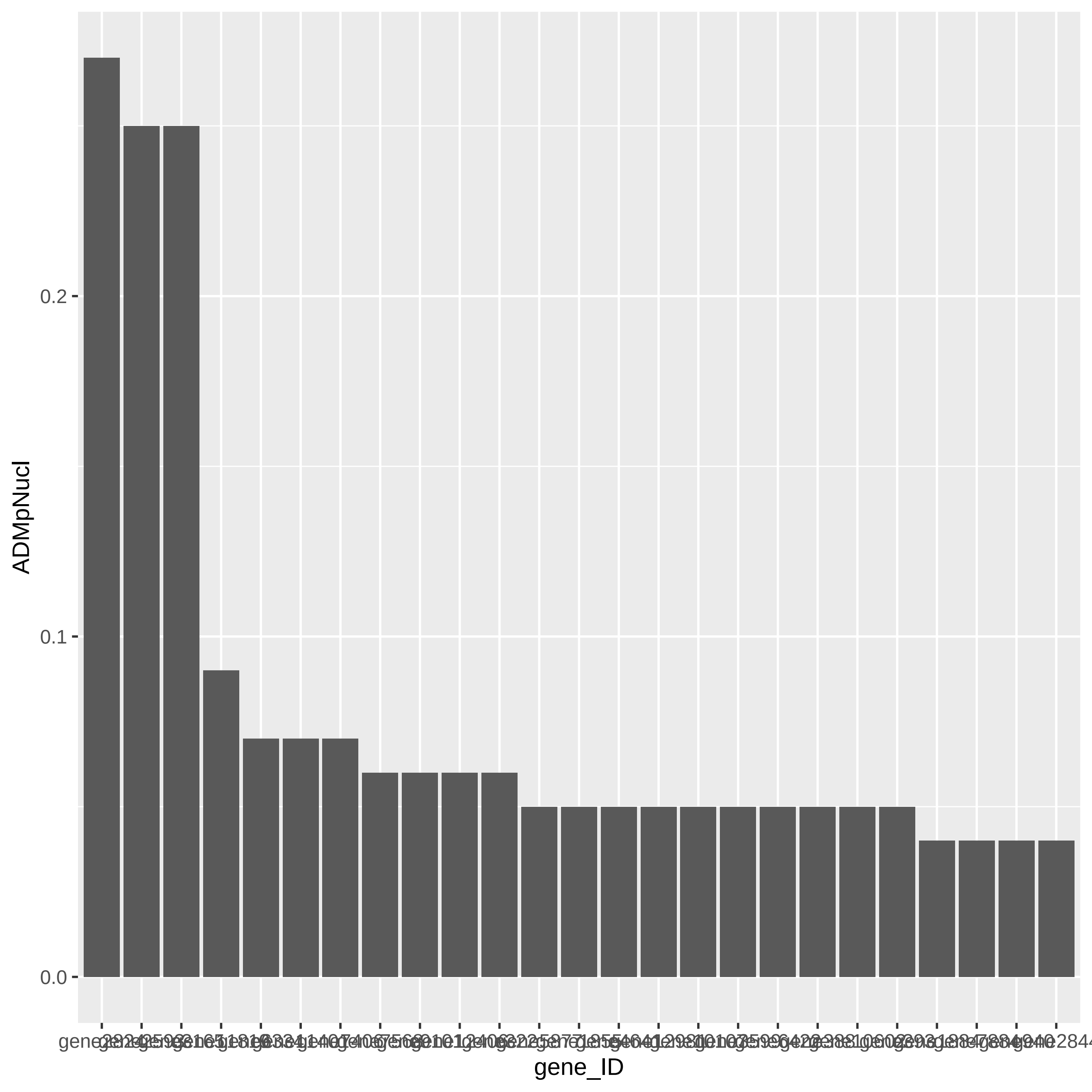
ADMpNucl
0.2
0.1
0.0
gene_ID

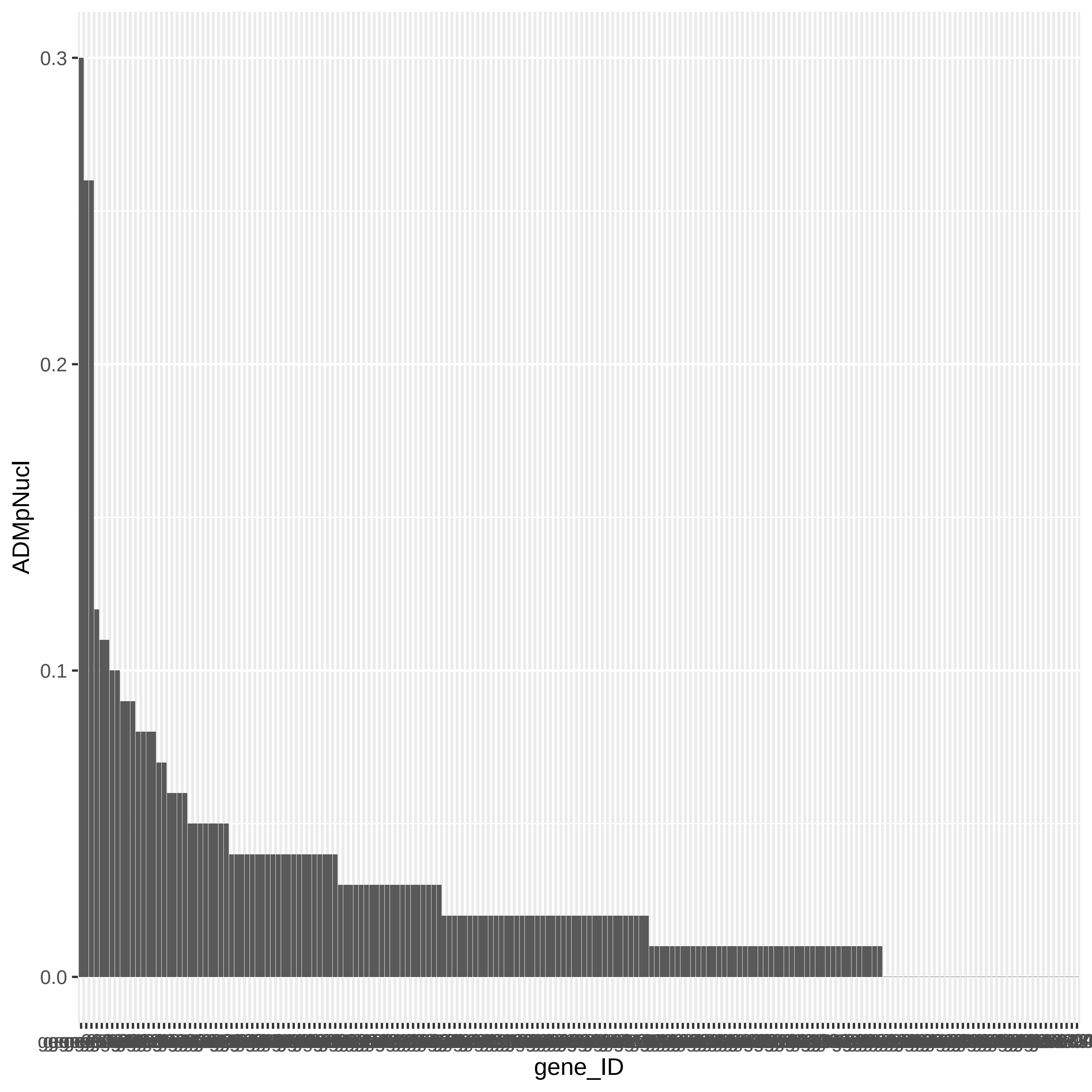

0.3
0.2
0.1
0.0
ADMpNucl
gene_ID

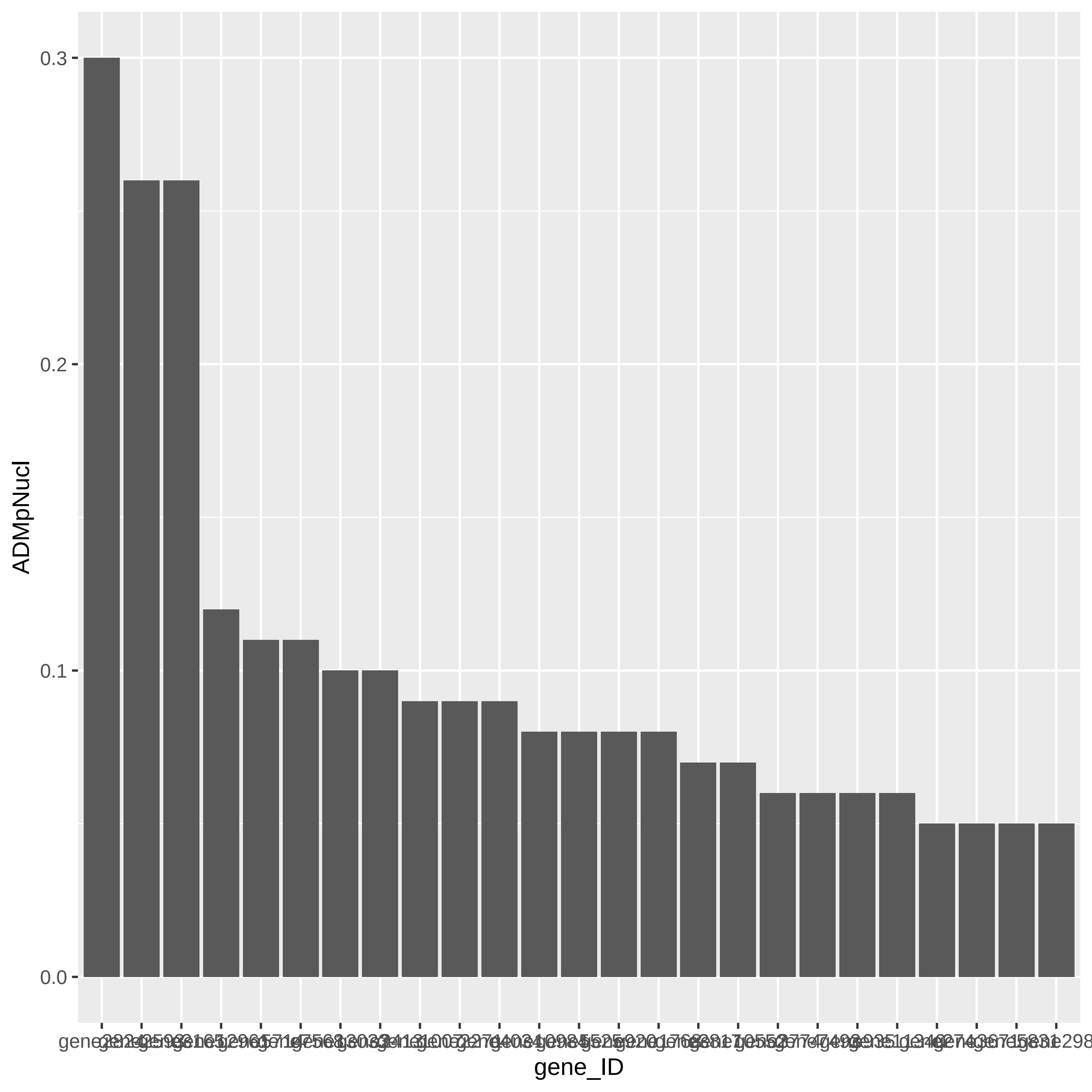

0.3
0.2
0.1
0.0
ADMpNucl
gene_ID

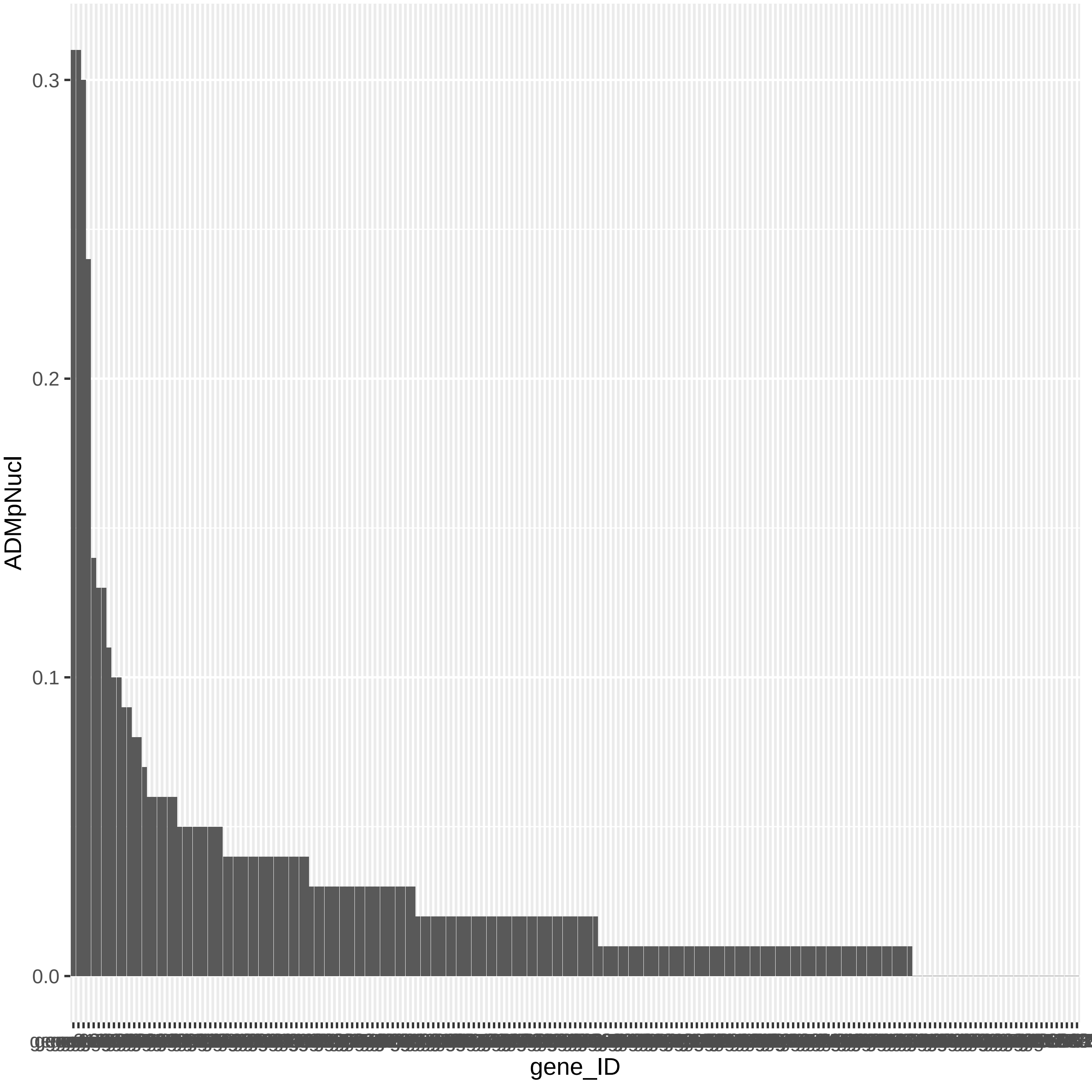

ADMpNucl
0.3
0.2
0.1
0.0
gene_ID

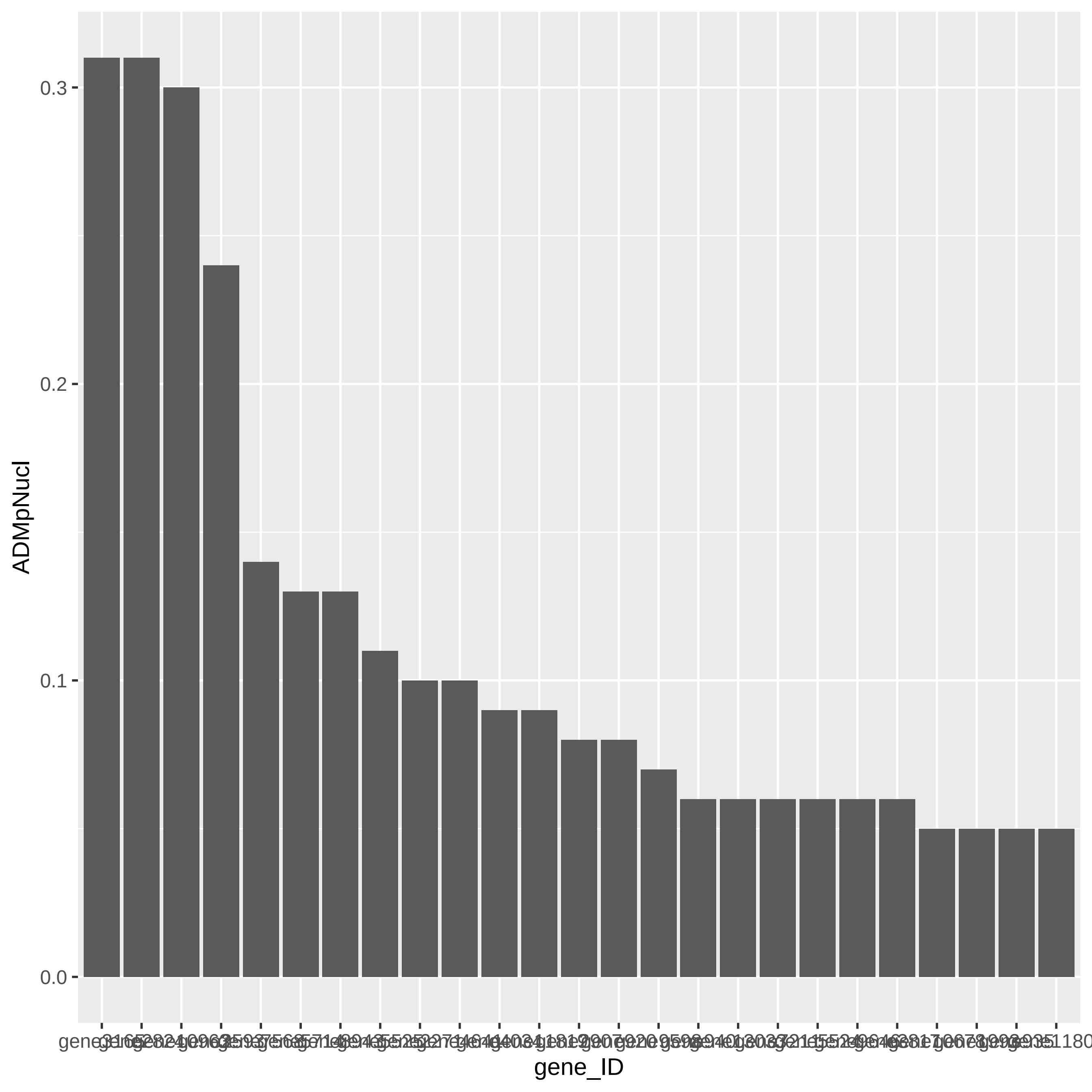
ADMpNucl
0.0
0.1
0.2
0.3
gene_ID

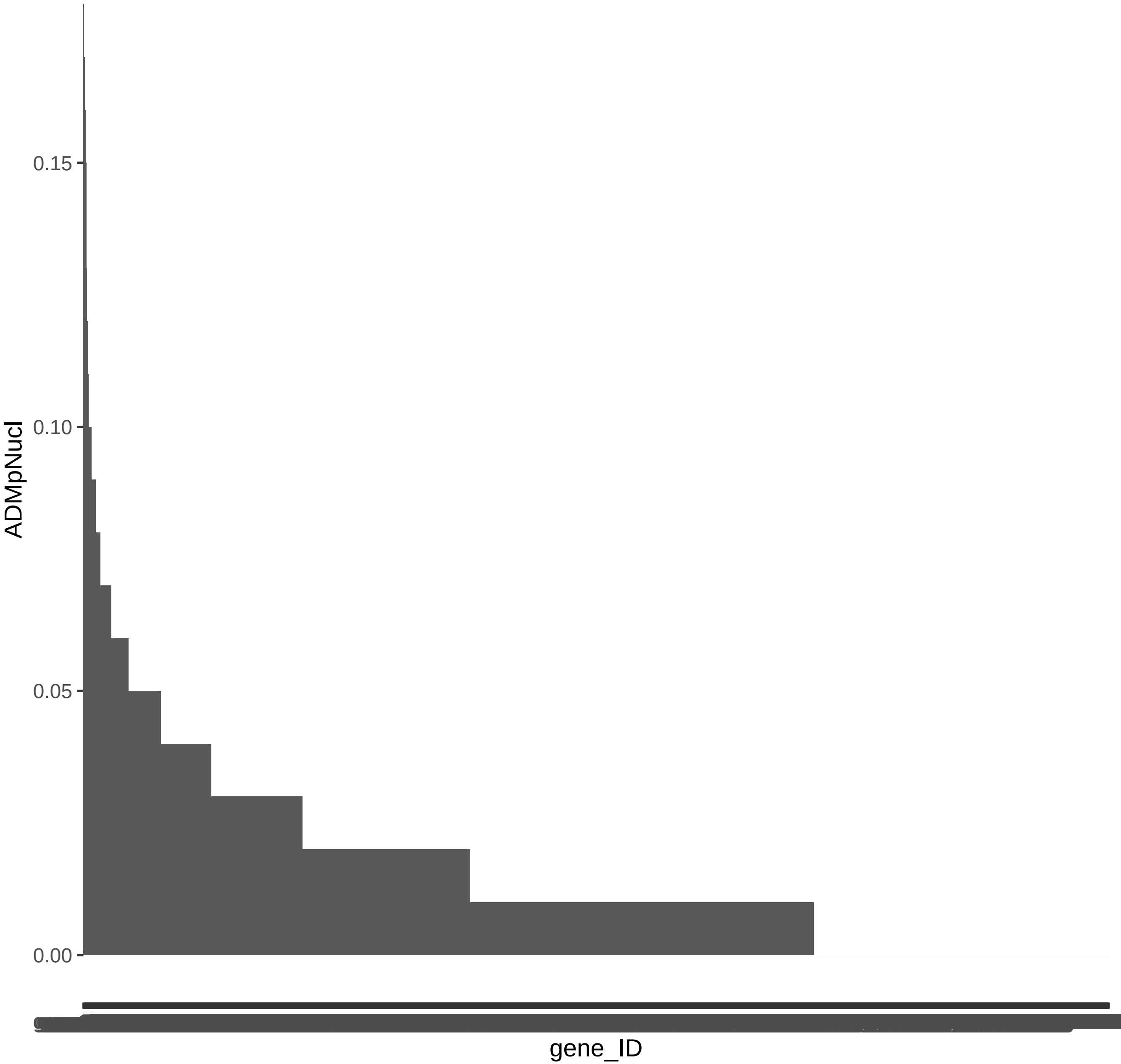
0.15
0.10
0.05
0.00
ADMpNucl
gene_ID

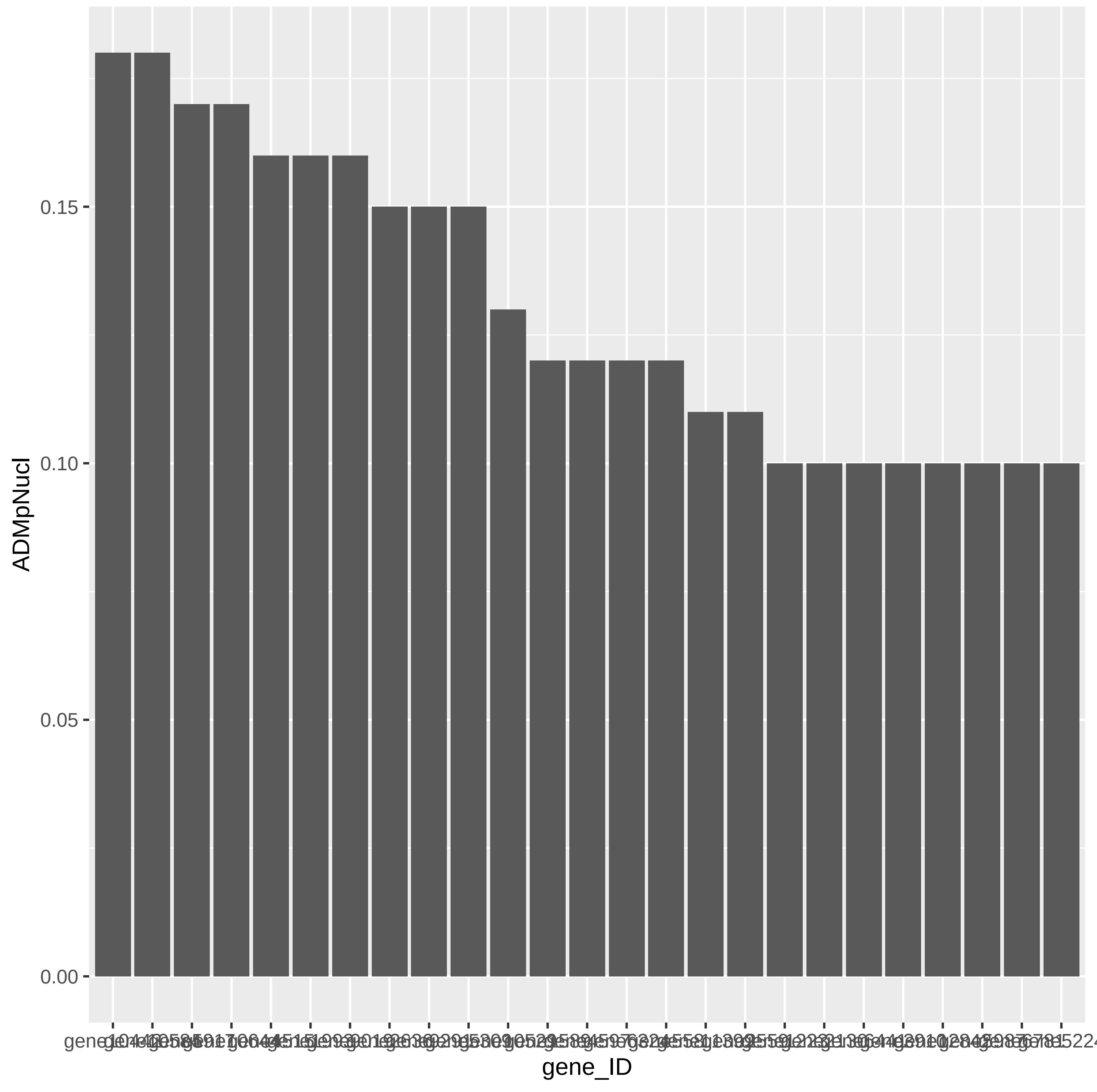
0.15
0.10
0.05
0.00
ADMpNucl
gene_ID
