## Supplementary material for "Tools and applications for integrative analysis of DNA methylation in social insects": Dataset S20

Ob_LI.conf

```
#Customize variables here:#####################################################
##############################################################################

#Load the files:
#
infiles       <- setup_BWASPR(datafile="./Ob.dat",
                              parfile="./Ob.par")

#Set the study and samples in the study:
#
species       <- "Ob"
study         <- "LI"
samplelist    <- list("BCphaseA1", "BCphaseA2", "BCphaseB1", "BCphaseB2", "RphaseA1", "RphaseA2", "RphaseB1", "RphaseB2")
## The following two variables are used for output file labeling:
studyLabel    <- "LI"
sampleLabels  <- list("Ob_LI_BCphaseA1", "Ob_LI_BCphaseA2", "Ob_LI_BCphaseB1", "Ob_LI_BCphaseB2", "Ob_LI_RphaseA1", "Ob_LI_RphaseA2", "Ob_LI_RphaseB1", "Ob_LI_RphaseB2")

hasreplicates <- FALSE
type          <- "CpG"
destrand      <- TRUE
covlist       <- c(6,20,50)
locount       <- 100
hicount       <- 1000
repcovlist    <- c(6,10,15)
replocount    <- 10
rephicount    <- 100
hheight       <- 0.035
nbrpnts       <- 5000

## The following four variables are sent to rank_rbm() and determine what data
#  points get plotted:
minnbrmsprm   <- 5      # minimum number of methylation sites per promoter
mingenewidth  <- 500    # minimum gene width
maxgenewidth  <- 5000   # maximum gene width
minnbrmsgene  <- 5      # minimum number of methylation sites per gene

## Other parameters (see Rscript.BWASPR for usage notes):
#
highcoverage  <- 20     # high read coverage threshold for studyhc methylRawList object
threshold     <- 20.0   # "difference" threshold for getMethylDiff(), called by det_dmsg()
qvalue        <- 0.05   # "qvalue" setting for getMethylDiff(), called by det_dmsg()
wsize         <- 500    # "win.size" parameter for tileMethylCounts() in det_dmt()
stepsize      <- 500    # "step.size" parameter for tileMethylCounts() in det_dmt()

minNsites     <-  1     # minimum number of hc sites in a gene to be heatmapped in show_dmsg()
maxNsites     <- 60     # maximum number of hc sites in a gene to be heatmapped in show_dmsg()
minPdmsites   <- 10     # minimum ratio of dm/hc (in %) in a gene to be heatmapped in show_dmsg()

maxgwidth     <- 20000  # maximal gene width for a gene to be considered by explore_dmsg()
minnbrdmsites <- 1      # minimum number of differentially methylated sites for a gene to be
                        #  considered in sample comparisons by explore_dmsg()
glink         <- "NCBIgene"   # URL to show in explore_dmsg(); options: "" or "NCBIgene"

#Set the number of processors to use:
#
numprc <- 6

#Determine what analyses to run:
#
RUNload    <- FALSE
```

```
RUNcms     <- TRUE
RUNpwc     <- TRUE
RUNcrl     <- TRUE

RUNrepcms  <- FALSE
RUNrepcrl  <- FALSE

RUNmmp     <- TRUE
RUNacs     <- TRUE
RUNrnk     <- TRUE
RUNmrpr    <- TRUE

RUNdmt     <- TRUE
RUNdmsg    <- TRUE
RUNdmgdtls <- TRUE
RUNogl     <- TRUE

RUNsave    <- TRUE

mymessage <- sprintf("\nAnalyzing %s study %s for type %s\n\n",species,studyLabel,type)
message(mymessage)

################################################################################
#End of typical customization.##################################################
```

Ob.par

```
SPECIESNAME Ooceraea biroi
TOTALNBRPMSITES   28317694
ASSEMBLYVERSION   Obir_v5.4
GENOMESIZE  223876465
SPECIESGFF3DIR    ./MCALLS/Obir/genome/GFF3DIR
GENELISTGFF3      Obir.gene.gff3
EXONLISTGFF3      Obir.exon.gff3
PCGEXNLISTGFF3    Obir.pcg-exon.gff3
PROMOTRLISTGFF3   Obir.promoter.gff3
CDSLISTGFF3 Obir.pcg-CDS.gff3
UTRFLAGSET  1
5UTRLISTGFF3      Obir.pcg-5pUTR.gff3
3UTRLISTGFF3      Obir.pcg-3pUTR.gff3
```

Ob.dat

```
# Samples from Libbrecht et al. (2016) Current Biology:
#

### 1. Comparisons between all 8 data sets (no replicates) ###
#

Ob    LI    BCphaseA1   0     CpGhsm      ./MCALLS/Obir/Libbrecht2016/BCphaseA/replicate1/BCphaseA1.CpGhsm.mcalls
Ob    LI    BCphaseA1   0     CpGscd      ./MCALLS/Obir/Libbrecht2016/BCphaseA/replicate1/BCphaseA1.CpGscd.mcalls
Ob    LI    BCphaseA2   0     CpGhsm      ./MCALLS/Obir/Libbrecht2016/BCphaseA/replicate2/BCphaseA2.CpGhsm.mcalls
Ob    LI    BCphaseA2   0     CpGscd      ./MCALLS/Obir/Libbrecht2016/BCphaseA/replicate2/BCphaseA2.CpGscd.mcalls

Ob    LI    BCphaseB1   0     CpGhsm      ./MCALLS/Obir/Libbrecht2016/BCphaseB/replicate1/BCphaseB1.CpGhsm.mcalls
Ob    LI    BCphaseB1   0     CpGscd      ./MCALLS/Obir/Libbrecht2016/BCphaseB/replicate1/BCphaseB1.CpGscd.mcalls
Ob    LI    BCphaseB2   0     CpGhsm      ./MCALLS/Obir/Libbrecht2016/BCphaseB/replicate2/BCphaseB2.CpGhsm.mcalls
Ob    LI    BCphaseB2   0     CpGscd      ./MCALLS/Obir/Libbrecht2016/BCphaseB/replicate2/BCphaseB2.CpGscd.mcalls

Ob    LI    RphaseA1    0     CpGhsm      ./MCALLS/Obir/Libbrecht2016/RphaseA/replicate1/RphaseA1.CpGhsm.mcalls
Ob    LI    RphaseA1    0     CpGscd      ./MCALLS/Obir/Libbrecht2016/RphaseA/replicate1/RphaseA1.CpGscd.mcalls
Ob    LI    RphaseA2    0     CpGhsm      ./MCALLS/Obir/Libbrecht2016/RphaseA/replicate2/RphaseA2.CpGhsm.mcalls
Ob    LI    RphaseA2    0     CpGscd      ./MCALLS/Obir/Libbrecht2016/RphaseA/replicate2/RphaseA2.CpGscd.mcalls

Ob    LI    RphaseB1    0     CpGhsm      ./MCALLS/Obir/Libbrecht2016/RphaseB/replicate1/RphaseB1.CpGhsm.mcalls
Ob    LI    RphaseB1    0     CpGscd      ./MCALLS/Obir/Libbrecht2016/RphaseB/replicate1/RphaseB1.CpGscd.mcalls
Ob    LI    RphaseB2    0     CpGhsm      ./MCALLS/Obir/Libbrecht2016/RphaseB/replicate2/RphaseB2.CpGhsm.mcalls
Ob    LI    RphaseB2    0     CpGscd      ./MCALLS/Obir/Libbrecht2016/RphaseB/replicate2/RphaseB2.CpGscd.mcalls

### 2. Pairwise omparisons between data sets split accorinding to phase, or line, or batch (2 samples, each with 4 replicates) ###
### 2.1 Comparing BCphase versus Rphase ###
#
# BCphase
Ob    LI    BCphase     0     CpGhsm      ./MCALLS/Obir/Libbrecht2016/PhaseBCvsR/PhaseBC/PhaseBC.CpGhsm.mcalls
Ob    LI    BCphase     0     CpGscd      ./MCALLS/Obir/Libbrecht2016/PhaseBCvsR/PhaseBC/PhaseBC.CpGscd.mcalls

# ...with replicates
Ob    LI    BCphase     1     CpGhsm      ./MCALLS/Obir/Libbrecht2016/BCphaseA/replicate1/BCphaseA1.CpGhsm.mcalls
Ob    LI    BCphase     1     CpGscd      ./MCALLS/Obir/Libbrecht2016/BCphaseA/replicate1/BCphaseA1.CpGscd.mcalls
Ob    LI    BCphase     2     CpGhsm      ./MCALLS/Obir/Libbrecht2016/BCphaseA/replicate2/BCphaseA2.CpGhsm.mcalls
Ob    LI    BCphase     2     CpGscd      ./MCALLS/Obir/Libbrecht2016/BCphaseA/replicate2/BCphaseA2.CpGscd.mcalls
Ob    LI    BCphase     3     CpGhsm      ./MCALLS/Obir/Libbrecht2016/BCphaseB/replicate1/BCphaseB1.CpGhsm.mcalls
Ob    LI    BCphase     3     CpGscd      ./MCALLS/Obir/Libbrecht2016/BCphaseB/replicate1/BCphaseB1.CpGscd.mcalls
Ob    LI    BCphase     4     CpGhsm      ./MCALLS/Obir/Libbrecht2016/BCphaseB/replicate2/BCphaseB2.CpGhsm.mcalls
Ob    LI    BCphase     4     CpGscd      ./MCALLS/Obir/Libbrecht2016/BCphaseB/replicate2/BCphaseB2.CpGscd.mcalls

# Rphase
Ob    LI    Rphase      0     CpGhsm      ./MCALLS/Obir/Libbrecht2016/PhaseBCvsR/PhaseR/PhaseR.CpGhsm.mcalls
Ob    LI    Rphase      0     CpGscd      ./MCALLS/Obir/Libbrecht2016/PhaseBCvsR/PhaseR/PhaseR.CpGscd.mcalls

# ...with replicates
Ob    LI    Rphase      1     CpGhsm      ./MCALLS/Obir/Libbrecht2016/RphaseA/replicate1/RphaseA1.CpGhsm.mcalls
Ob    LI    Rphase      1     CpGscd      ./MCALLS/Obir/Libbrecht2016/RphaseA/replicate1/RphaseA1.CpGscd.mcalls
Ob    LI    Rphase      2     CpGhsm      ./MCALLS/Obir/Libbrecht2016/RphaseA/replicate2/RphaseA2.CpGhsm.mcalls
Ob    LI    Rphase      2     CpGscd      ./MCALLS/Obir/Libbrecht2016/RphaseA/replicate2/RphaseA2.CpGscd.mcalls
Ob    LI    Rphase      3     CpGhsm      ./MCALLS/Obir/Libbrecht2016/RphaseB/replicate1/RphaseB1.CpGhsm.mcalls
Ob    LI    Rphase      3     CpGscd      ./MCALLS/Obir/Libbrecht2016/RphaseB/replicate1/RphaseB1.CpGscd.mcalls
Ob    LI    Rphase      4     CpGhsm      ./MCALLS/Obir/Libbrecht2016/RphaseB/replicate2/RphaseB2.CpGhsm.mcalls
Ob    LI    Rphase      4     CpGscd      ./MCALLS/Obir/Libbrecht2016/RphaseB/replicate2/RphaseB2.CpGscd.mcalls

### 2.2 Comparing LineA versus LineB ###
```

```
#
# LineA
Ob    LI    LineA 0     CpGhsm      ./MCALLS/Obir/Libbrecht2016/LineAvsB/LineA/LineA.CpGhsm.mcalls
Ob    LI    LineA 0     CpGscd      ./MCALLS/Obir/Libbrecht2016/LineAvsB/LineA/LineA.CpGscd.mcalls

# ...with replicates
Ob    LI    LineA 1     CpGhsm      ./MCALLS/Obir/Libbrecht2016/BCphaseA/replicate1/BCphaseA1.CpGhsm.mcalls
Ob    LI    LineA 1     CpGscd      ./MCALLS/Obir/Libbrecht2016/BCphaseA/replicate1/BCphaseA1.CpGscd.mcalls
Ob    LI    LineA 2     CpGhsm      ./MCALLS/Obir/Libbrecht2016/BCphaseA/replicate2/BCphaseA2.CpGhsm.mcalls
Ob    LI    LineA 2     CpGscd      ./MCALLS/Obir/Libbrecht2016/BCphaseA/replicate2/BCphaseA2.CpGscd.mcalls
Ob    LI    LineA 3     CpGhsm      ./MCALLS/Obir/Libbrecht2016/RphaseA/replicate1/RphaseA1.CpGhsm.mcalls
Ob    LI    LineA 3     CpGscd      ./MCALLS/Obir/Libbrecht2016/RphaseA/replicate1/RphaseA1.CpGscd.mcalls
Ob    LI    LineA 4     CpGhsm      ./MCALLS/Obir/Libbrecht2016/RphaseA/replicate2/RphaseA2.CpGhsm.mcalls
Ob    LI    LineA 4     CpGscd      ./MCALLS/Obir/Libbrecht2016/RphaseA/replicate2/RphaseA2.CpGscd.mcalls

# LineB
Ob    LI    LineB 0     CpGhsm      ./MCALLS/Obir/Libbrecht2016/LineAvsB/LineB/LineB.CpGhsm.mcalls
Ob    LI    LineB 0     CpGscd      ./MCALLS/Obir/Libbrecht2016/LineAvsB/LineB/LineB.CpGscd.mcalls

# ...with replicates
Ob    LI    LineB 1     CpGhsm      ./MCALLS/Obir/Libbrecht2016/BCphaseB/replicate1/BCphaseB1.CpGhsm.mcalls
Ob    LI    LineB 1     CpGscd      ./MCALLS/Obir/Libbrecht2016/BCphaseB/replicate1/BCphaseB1.CpGscd.mcalls
Ob    LI    LineB 2     CpGhsm      ./MCALLS/Obir/Libbrecht2016/BCphaseB/replicate2/BCphaseB2.CpGhsm.mcalls
Ob    LI    LineB 2     CpGscd      ./MCALLS/Obir/Libbrecht2016/BCphaseB/replicate2/BCphaseB2.CpGscd.mcalls
Ob    LI    LineB 3     CpGhsm      ./MCALLS/Obir/Libbrecht2016/RphaseB/replicate1/RphaseB1.CpGhsm.mcalls
Ob    LI    LineB 3     CpGscd      ./MCALLS/Obir/Libbrecht2016/RphaseB/replicate1/RphaseB1.CpGscd.mcalls
Ob    LI    LineB 4     CpGhsm      ./MCALLS/Obir/Libbrecht2016/RphaseB/replicate2/RphaseB2.CpGhsm.mcalls
Ob    LI    LineB 4     CpGscd      ./MCALLS/Obir/Libbrecht2016/RphaseB/replicate2/RphaseB2.CpGscd.mcalls

### 2.2 2.2 Comparing Batch1 versus Batch2 ###
#
# Batch1
Ob    LI    Batch1      0     CpGhsm      ./MCALLS/Obir/Libbrecht2016/Batch1vs2/Batch1/Batch1.CpGhsm.mcalls
Ob    LI    Batch1      0     CpGscd      ./MCALLS/Obir/Libbrecht2016/Batch1vs2/Batch1/Batch1.CpGscd.mcalls

# ...with replicates
Ob    LI    Batch1      1     CpGhsm      ./MCALLS/Obir/Libbrecht2016/BCphaseA/replicate1/BCphaseA1.CpGhsm.mcalls
Ob    LI    Batch1      1     CpGscd      ./MCALLS/Obir/Libbrecht2016/BCphaseA/replicate1/BCphaseA1.CpGscd.mcalls
Ob    LI    Batch1      2     CpGhsm      ./MCALLS/Obir/Libbrecht2016/BCphaseB/replicate1/BCphaseB1.CpGhsm.mcalls
Ob    LI    Batch1      2     CpGscd      ./MCALLS/Obir/Libbrecht2016/BCphaseB/replicate1/BCphaseB1.CpGscd.mcalls
Ob    LI    Batch1      3     CpGhsm      ./MCALLS/Obir/Libbrecht2016/RphaseA/replicate1/RphaseA1.CpGhsm.mcalls
Ob    LI    Batch1      3     CpGscd      ./MCALLS/Obir/Libbrecht2016/RphaseA/replicate1/RphaseA1.CpGscd.mcalls
Ob    LI    Batch1      4     CpGhsm      ./MCALLS/Obir/Libbrecht2016/RphaseB/replicate1/RphaseB1.CpGhsm.mcalls
Ob    LI    Batch1      4     CpGscd      ./MCALLS/Obir/Libbrecht2016/RphaseB/replicate1/RphaseB1.CpGscd.mcalls

# Batch2
Ob    LI    Batch2      0     CpGhsm      ./MCALLS/Obir/Libbrecht2016/Batch1vs2/Batch2/Batch2.CpGhsm.mcalls
Ob    LI    Batch2      0     CpGscd      ./MCALLS/Obir/Libbrecht2016/Batch1vs2/Batch2/Batch2.CpGscd.mcalls

# ...with replicates
Ob    LI    Batch2      1     CpGhsm      ./MCALLS/Obir/Libbrecht2016/BCphaseA/replicate2/BCphaseA2.CpGhsm.mcalls
Ob    LI    Batch2      1     CpGscd      ./MCALLS/Obir/Libbrecht2016/BCphaseA/replicate2/BCphaseA2.CpGscd.mcalls
Ob    LI    Batch2      2     CpGhsm      ./MCALLS/Obir/Libbrecht2016/BCphaseB/replicate2/BCphaseB2.CpGhsm.mcalls
Ob    LI    Batch2      2     CpGscd      ./MCALLS/Obir/Libbrecht2016/BCphaseB/replicate2/BCphaseB2.CpGscd.mcalls
Ob    LI    Batch2      3     CpGhsm      ./MCALLS/Obir/Libbrecht2016/RphaseA/replicate2/RphaseA2.CpGhsm.mcalls
Ob    LI    Batch2      3     CpGscd      ./MCALLS/Obir/Libbrecht2016/RphaseA/replicate2/RphaseA2.CpGscd.mcalls
Ob    LI    Batch2      4     CpGhsm      ./MCALLS/Obir/Libbrecht2016/RphaseB/replicate2/RphaseB2.CpGhsm.mcalls
Ob    LI    Batch2      4     CpGscd      ./MCALLS/Obir/Libbrecht2016/RphaseB/replicate2/RphaseB2.CpGscd.mcalls

### 3. Pairwise comparisons between BCphaseA, BCphaseB, RphaseA, and RphaseB (2 replicates each, corresponding to batches 1 and 2) ###
#
```

```
# BCphaseA
Ob    LI    BCphaseA    0     CpGhsm      ./MCALLS/Obir/Libbrecht2016/BCphaseA/BCphaseA.CpGhsm.mcalls
Ob    LI    BCphaseA    0     CpGscd      ./MCALLS/Obir/Libbrecht2016/BCphaseA/BCphaseA.CpGscd.mcalls

# ... with replicates:
Ob    LI    BCphaseA    1     CpGhsm      ./MCALLS/Obir/Libbrecht2016/BCphaseA/replicate1/BCphaseA1.CpGhsm.mcalls
Ob    LI    BCphaseA    1     CpGscd      ./MCALLS/Obir/Libbrecht2016/BCphaseA/replicate1/BCphaseA1.CpGscd.mcalls
Ob    LI    BCphaseA    2     CpGhsm      ./MCALLS/Obir/Libbrecht2016/BCphaseA/replicate2/BCphaseA2.CpGhsm.mcalls
Ob    LI    BCphaseA    2     CpGscd      ./MCALLS/Obir/Libbrecht2016/BCphaseA/replicate2/BCphaseA2.CpGscd.mcalls

# BCphaseB
Ob    LI    BCphaseB    0     CpGhsm      ./MCALLS/Obir/Libbrecht2016/BCphaseB/BCphaseB.CpGhsm.mcalls
Ob    LI    BCphaseB    0     CpGscd      ./MCALLS/Obir/Libbrecht2016/BCphaseB/BCphaseB.CpGscd.mcalls

# ... with replicates:
Ob    LI    BCphaseB    1     CpGhsm      ./MCALLS/Obir/Libbrecht2016/BCphaseB/replicate1/BCphaseB1.CpGhsm.mcalls
Ob    LI    BCphaseB    1     CpGscd      ./MCALLS/Obir/Libbrecht2016/BCphaseB/replicate1/BCphaseB1.CpGscd.mcalls
Ob    LI    BCphaseB    2     CpGhsm      ./MCALLS/Obir/Libbrecht2016/BCphaseB/replicate2/BCphaseB2.CpGhsm.mcalls
Ob    LI    BCphaseB    2     CpGscd      ./MCALLS/Obir/Libbrecht2016/BCphaseB/replicate2/BCphaseB2.CpGscd.mcalls

# RphaseA
Ob    LI    RphaseA     0     CpGhsm      ./MCALLS/Obir/Libbrecht2016/RphaseA/RphaseA.CpGhsm.mcalls
Ob    LI    RphaseA     0     CpGscd      ./MCALLS/Obir/Libbrecht2016/RphaseA/RphaseA.CpGscd.mcalls

# ... with replicates:
Ob    LI    RphaseA     1     CpGhsm      ./MCALLS/Obir/Libbrecht2016/RphaseA/replicate1/RphaseA1.CpGhsm.mcalls
Ob    LI    RphaseA     1     CpGscd      ./MCALLS/Obir/Libbrecht2016/RphaseA/replicate1/RphaseA1.CpGscd.mcalls
Ob    LI    RphaseA     2     CpGhsm      ./MCALLS/Obir/Libbrecht2016/RphaseA/replicate2/RphaseA2.CpGhsm.mcalls
Ob    LI    RphaseA     2     CpGscd      ./MCALLS/Obir/Libbrecht2016/RphaseA/replicate2/RphaseA2.CpGscd.mcalls

# RphaseB
Ob    LI    RphaseB     0     CpGhsm      ./MCALLS/Obir/Libbrecht2016/RphaseB/RphaseB.CpGhsm.mcalls
Ob    LI    RphaseB     0     CpGscd      ./MCALLS/Obir/Libbrecht2016/RphaseB/RphaseB.CpGscd.mcalls

# ... with replicates:
Ob    LI    RphaseB     1     CpGhsm      ./MCALLS/Obir/Libbrecht2016/RphaseB/replicate1/RphaseB1.CpGhsm.mcalls
Ob    LI    RphaseB     1     CpGscd      ./MCALLS/Obir/Libbrecht2016/RphaseB/replicate1/RphaseB1.CpGscd.mcalls
Ob    LI    RphaseB     2     CpGhsm      ./MCALLS/Obir/Libbrecht2016/RphaseB/replicate2/RphaseB2.CpGhsm.mcalls
Ob    LI    RphaseB     2     CpGscd      ./MCALLS/Obir/Libbrecht2016/RphaseB/replicate2/RphaseB2.CpGscd.mcalls
```

```
0README

Rscript.BWASPR output for .conf file Ob_LI.conf.

Data input are the specified *.mcalls files and parameters set in Ob_LI.conf.
The *.mcalls input data are saved in the following data structures:

studymk - a methylKit methylRaw(List) object storing the CpGhsm site data for all samples
studymc - a methylKit methylRaw(List) object storing the CpGscd site data for all samples
studyhc - a methylKit methylRaw(List) object storing the CpGscd site data for all samples,
           restricted to sites with high coverage (here set to 20)
mkrd    - a methylKit methylRawList object storing the CpGhsm site data for all
           replicates of a given sample

Output of the analysis is stored in the following subdirectories (if the corresponding
 function calls were specified in Ob_LI.conf; please consult the 0README
 files in the subdirectories for details on the output files):

CMS - Coverage and methylation statistics for aggregate samples
PWC - Pairwise comparisons between all samples
CRL - Correlations between aggregate samples
REPCMS - Coverage and methylation statistics for replicate samples
REPCRL - Correlations between replicates
MMP - Mapping of methylation sites on genome annotation
ACS - Annotation of conserved methylation sites
RNK - Genomic feature regions ranked by CpGhsm statistics
MRPR - Methylation-rich and -poor regions
DMSG - Differentially methylateds sites and genes
OGL - Ordered gene lists
```

```
Directory: CMS File: 0READMEcms

CMS - Coverage and methylation statistics for aggregate samples.

Input: studymk, covlist, locount, hicount

Output: files cms-*.txt and cms-*.pdf

Notes: The statistics are given for CpGhsm sites restricted to minimum or
        higher level coverage.  At the minimum level, the number of sites will be
        the number of lines of the corresponding *CpGhsm.mcalls file minus one
        (consistency check).  The distribution of methylation levels is really only
        interesting for high coverage because at low coverage there will be a strong
        bias towards high methylation levels (by definition of hsm sites).
        The same bias at high coverage would suggest that the sample includes a
        preponderance of consistently methylated genomic sites.

        locount and hicount set bounds on the coverage to exclude sites with too few
        or too many covering reads to provide statistics on a typical range.
```

```
Directory: CMS File: cms-Ob_LI_BCphaseA1.txt

Number of "BCphaseA1" CpGhsm-sites with minimal and higher level coverage:

  number of "BCphaseA1" CpGhsm-sites with coverage >=  5: 197029
  number of "BCphaseA1" CpGhsm-sites with coverage >=  6: 193888
  number of "BCphaseA1" CpGhsm-sites with coverage >= 20: 112061
  number of "BCphaseA1" CpGhsm-sites with coverage >= 50:  10742

Coverage and methylation statistics for "BCphaseA1" CpGhsm-sites at different levels of minimum coverage:

methylKit::getCoverageStats output for "BCphaseA1" CpGhsm-sites (#: 197029) at minimum coverage 5    - read coverage statistics per base
summary:
   Min. 1st Qu.  Median    Mean 3rd Qu.    Max.
      5      14      22      25      33     601
percentiles:
   0%   10%   20%   30%   40%   50%   60%   70%   80%   90%   95%   99% 99.5% 99.9%  100%
    5     9    12    15    18    22    26    31    36    44    50    63    68    80   601

methylKit::getMethylationStats output for "BCphaseA1" CpGhsm-sites (#: 197029) at minimum coverage 5 - methylation statistics per base
summary:
   Min. 1st Qu.  Median    Mean 3rd Qu.    Max.
    5.3    46.7    71.4    68.0    90.6   100.0
percentiles:
    0%    10%    20%    30%    40%    50%    60%    70%    80%    90%    95%    99%  99.5%  99.9%   100%
  5.32  31.82  41.67  52.17  61.70  71.43  80.00  87.50  93.55 100.00 100.00 100.00 100.00 100.00 100.00

methylKit::getCoverageStats output for "BCphaseA1" CpGhsm-sites (#: 193888) at minimum coverage 6    - read coverage statistics per base
summary:
   Min. 1st Qu.  Median    Mean 3rd Qu.    Max.
      6      14      23      25      33     601
percentiles:
   0%   10%   20%   30%   40%   50%   60%   70%   80%   90%   95%   99% 99.5% 99.9%  100%
    6     9    12    16    19    23    27    31    36    44    50    63    68    80   601

methylKit::getMethylationStats output for "BCphaseA1" CpGhsm-sites (#: 193888) at minimum coverage 6 - methylation statistics per base
summary:
   Min. 1st Qu.  Median    Mean 3rd Qu.    Max.
    5.3    46.2    70.6    67.5    90.0   100.0
percentiles:
    0%    10%    20%    30%    40%    50%    60%    70%    80%    90%    95%    99%  99.5%  99.9%   100%
  5.32  31.63  41.18  51.52  61.11  70.59  79.41  86.67  92.86 100.00 100.00 100.00 100.00 100.00 100.00

methylKit::getCoverageStats output for "BCphaseA1" CpGhsm-sites (#: 112061) at minimum coverage 20   - read coverage statistics per base
summary:
   Min. 1st Qu.  Median    Mean 3rd Qu.    Max.
     20      25      31      34      40     601
percentiles:
   0%   10%   20%   30%   40%   50%   60%   70%   80%   90%   95%   99% 99.5% 99.9%  100%
   20    22    24    26    29    31    34    38    42    49    55    67    72    85   601

methylKit::getMethylationStats output for "BCphaseA1" CpGhsm-sites (#: 112061) at minimum coverage 20      - methylation statistics per base
summary:
   Min. 1st Qu.  Median    Mean 3rd Qu.    Max.
    5.3    37.5    60.5    61.1    85.7   100.0
percentiles:
    0%    10%    20%    30%    40%    50%    60%    70%    80%    90%    95%    99%  99.5%  99.9%   100%
```

```
  5.32  26.92  33.33  41.18  50.00  60.47  71.15  81.48  90.00  95.83  98.41 100.00 100.00 100.00 100.00
```

```
methylKit::getCoverageStats output for "BCphaseA1" CpGhsm-sites (#: 10742) at minimum coverage 50    - read coverage statistics per base
summary:
   Min. 1st Qu.  Median    Mean 3rd Qu.    Max.
     50      52      55      58      61     601
percentiles:
   0%   10%   20%   30%   40%   50%   60%   70%   80%   90%   95%   99% 99.5% 99.9%  100%
   50    50    51    53    54    55    57    59    63    68    72    85    91   168   601

methylKit::getMethylationStats output for "BCphaseA1" CpGhsm-sites (#: 10742) at minimum coverage 50 - methylation statistics per base
summary:
   Min. 1st Qu.  Median    Mean 3rd Qu.    Max.
    5.3    33.9    61.8    60.0    86.8   100.0
percentiles:
    0%    10%    20%    30%    40%    50%    60%    70%    80%    90%    95%    99%  99.5%  99.9%   100%
  5.32  21.05  29.31  39.22  50.00  61.82  72.87  82.69  90.57  96.00  98.08 100.00 100.00 100.00 100.00
```

```
methylKit::getCoverageStats output for "BCphaseA1" CpGhsm-sites (#: 28) in coverage range [100-1000] - read coverage statistics per base
summary:
   Min. 1st Qu.  Median    Mean 3rd Qu.    Max.
    100     110     154     165     178     601
percentiles:
   0%   10%   20%   30%   40%   50%   60%   70%   80%   90%   95%   99% 99.5% 99.9%  100%
  100   101   105   114   132   154   167   176   180   204   238   507   554   592   601

methylKit::getMethylationStats output for "BCphaseA1" CpGhsm-sites (#: 28) in coverage range [100-1000]    - methylation statistics per base
summary:
   Min. 1st Qu.  Median    Mean 3rd Qu.    Max.
    5.3    19.1    44.7    41.2    55.8    99.0
percentiles:
   0%   10%   20%   30%   40%   50%   60%   70%   80%   90%   95%   99% 99.5% 99.9%  100%
 5.32  8.49 15.82 24.72 40.38 44.66 49.98 55.10 57.61 62.81 76.50 94.42 96.72 98.56 99.02
```

```
Directory: CMS File: cms-Ob_LI_BCphaseA2.txt

Number of "BCphaseA2" CpGhsm-sites with minimal and higher level coverage:

  number of "BCphaseA2" CpGhsm-sites with coverage >=  5: 213136
  number of "BCphaseA2" CpGhsm-sites with coverage >=  6: 210280
  number of "BCphaseA2" CpGhsm-sites with coverage >= 20: 137168
  number of "BCphaseA2" CpGhsm-sites with coverage >= 50:  39055

Coverage and methylation statistics for "BCphaseA2" CpGhsm-sites at different levels of minimum coverage:

methylKit::getCoverageStats output for "BCphaseA2" CpGhsm-sites (#: 213136) at minimum coverage 5    - read coverage statistics per base
summary:
   Min. 1st Qu.  Median    Mean 3rd Qu.    Max.
    5.0    15.0    27.0    31.4    43.0   203.0
percentiles:
   0%   10%   20%   30%   40%   50%   60%   70%   80%   90%   95%   99% 99.5% 99.9%  100%
    5     9    13    17    22    27    32    39    48    61    72    95   105   126   203

methylKit::getMethylationStats output for "BCphaseA2" CpGhsm-sites (#: 213136) at minimum coverage 5 - methylation statistics per base
summary:
   Min. 1st Qu.  Median    Mean 3rd Qu.    Max.
    7.7    42.9    69.0    65.7    90.0   100.0
percentiles:
    0%    10%    20%    30%    40%    50%    60%    70%    80%    90%    95%    99%  99.5%  99.9%   100%
  7.75  27.66  37.50  48.00  58.33  69.05  78.57  86.67  92.98 100.00 100.00 100.00 100.00 100.00 100.00

methylKit::getCoverageStats output for "BCphaseA2" CpGhsm-sites (#: 210280) at minimum coverage 6    - read coverage statistics per base
summary:
   Min. 1st Qu.  Median    Mean 3rd Qu.    Max.
    6.0    16.0    27.0    31.8    43.0   203.0
percentiles:
   0%   10%   20%   30%   40%   50%   60%   70%   80%   90%   95%   99% 99.5% 99.9%  100%
    6    10    14    18    22    27    33    39    48    61    72    95   105   126   203

methylKit::getMethylationStats output for "BCphaseA2" CpGhsm-sites (#: 210280) at minimum coverage 6 - methylation statistics per base
summary:
   Min. 1st Qu.  Median    Mean 3rd Qu.    Max.
    7.7    42.6    68.2    65.2    89.3   100.0
percentiles:
    0%    10%    20%    30%    40%    50%    60%    70%    80%    90%    95%    99%  99.5%  99.9%   100%
  7.75  27.59  37.50  47.62  57.92  68.18  77.78  85.71  92.31 100.00 100.00 100.00 100.00 100.00 100.00

methylKit::getCoverageStats output for "BCphaseA2" CpGhsm-sites (#: 137168) at minimum coverage 20   - read coverage statistics per base
summary:
   Min. 1st Qu.  Median    Mean 3rd Qu.    Max.
     20      27      37      42      52     203
percentiles:
   0%   10%   20%   30%   40%   50%   60%   70%   80%   90%   95%   99% 99.5% 99.9%  100%
   20    23    26    29    33    37    42    48    56    68    79   101   111   132   203

methylKit::getMethylationStats output for "BCphaseA2" CpGhsm-sites (#: 137168) at minimum coverage 20      - methylation statistics per base
summary:
   Min. 1st Qu.  Median    Mean 3rd Qu.    Max.
    7.7    34.5    57.9    59.1    85.2   100.0
percentiles:
    0%    10%    20%    30%    40%    50%    60%    70%    80%    90%    95%    99%  99.5%  99.9%   100%
```

```
  7.75  23.53  30.77  38.27  47.62  57.89  69.57  80.36  89.66  95.65  97.96 100.00 100.00 100.00 100.00
```

```
methylKit::getCoverageStats output for "BCphaseA2" CpGhsm-sites (#: 39055) at minimum coverage 50   - read coverage statistics per base
summary:
   Min. 1st Qu.  Median    Mean 3rd Qu.    Max.
   50.0    55.0    62.0    66.4    73.0   203.0
percentiles:
   0%   10%   20%   30%   40%   50%   60%   70%   80%   90%   95%   99% 99.5% 99.9%  100%
   50    52    54    56    59    62    66    71    77    87    96   118   128   147   203

methylKit::getMethylationStats output for "BCphaseA2" CpGhsm-sites (#: 39055) at minimum coverage 50 - methylation statistics per base
summary:
   Min. 1st Qu.  Median    Mean 3rd Qu.    Max.
    7.7    27.5    52.6    54.7    82.7   100.0
percentiles:
    0%    10%    20%    30%    40%    50%    60%    70%    80%    90%    95%    99%  99.5%  99.9%   100%
  7.75  17.65  23.81  31.58  41.27  52.63  64.71  77.19  87.50  94.64  97.06 100.00 100.00 100.00 100.00
```

```
methylKit::getCoverageStats output for "BCphaseA2" CpGhsm-sites (#: 1546) in coverage range [100-1000]    - read coverage statistics per base
summary:
   Min. 1st Qu.  Median    Mean 3rd Qu.    Max.
    100     103     109     113     119     203
percentiles:
   0%   10%   20%   30%   40%   50%   60%   70%   80%   90%   95%   99% 99.5% 99.9%  100%
  100   101   102   104   106   109   112   116   121   131   139   163   175   188   203

methylKit::getMethylationStats output for "BCphaseA2" CpGhsm-sites (#: 1546) in coverage range [100-1000] - methylation statistics per base
summary:
   Min. 1st Qu.  Median    Mean 3rd Qu.    Max.
    7.7    31.8    59.5    57.6    85.5   100.0
percentiles:
    0%    10%    20%    30%    40%    50%    60%    70%    80%    90%    95%    99%  99.5%  99.9%   100%
  7.75  17.21  26.61  36.48  47.86  59.46  69.52  80.75  89.42  94.92  97.14  99.08  99.25 100.00 100.00
```

```
Directory: CMS File: cms-Ob_LI_BCphaseB1.txt

Number of "BCphaseB1" CpGhsm-sites with minimal and higher level coverage:

  number of "BCphaseB1" CpGhsm-sites with coverage >=  5: 226855
  number of "BCphaseB1" CpGhsm-sites with coverage >=  6: 224253
  number of "BCphaseB1" CpGhsm-sites with coverage >= 20: 149882
  number of "BCphaseB1" CpGhsm-sites with coverage >= 50:  29431

Coverage and methylation statistics for "BCphaseB1" CpGhsm-sites at different levels of minimum coverage:

methylKit::getCoverageStats output for "BCphaseB1" CpGhsm-sites (#: 226855) at minimum coverage 5    - read coverage statistics per base
summary:
   Min. 1st Qu.  Median    Mean 3rd Qu.    Max.
    5.0    16.0    27.0    29.5    40.0   247.0
percentiles:
   0%   10%   20%   30%   40%   50%   60%   70%   80%   90%   95%   99% 99.5% 99.9%  100%
    5    10    14    18    22    27    32    37    44    53    60    76    82    96   247

methylKit::getMethylationStats output for "BCphaseB1" CpGhsm-sites (#: 226855) at minimum coverage 5 - methylation statistics per base
summary:
   Min. 1st Qu.  Median    Mean 3rd Qu.    Max.
    6.9    42.1    67.4    65.1    89.5   100.0
percentiles:
    0%    10%    20%    30%    40%    50%    60%    70%    80%    90%    95%    99%  99.5%  99.9%   100%
  6.88  28.00  37.50  47.06  57.14  67.39  77.50  85.71  92.59 100.00 100.00 100.00 100.00 100.00 100.00

methylKit::getCoverageStats output for "BCphaseB1" CpGhsm-sites (#: 224253) at minimum coverage 6    - read coverage statistics per base
summary:
   Min. 1st Qu.  Median    Mean 3rd Qu.    Max.
    6.0    16.0    27.0    29.7    40.0   247.0
percentiles:
   0%   10%   20%   30%   40%   50%   60%   70%   80%   90%   95%   99% 99.5% 99.9%  100%
    6    10    14    18    23    27    32    37    44    53    60    76    82    96   247

methylKit::getMethylationStats output for "BCphaseB1" CpGhsm-sites (#: 224253) at minimum coverage 6 - methylation statistics per base
summary:
   Min. 1st Qu.  Median    Mean 3rd Qu.    Max.
    6.9    41.7    66.7    64.7    88.9   100.0
percentiles:
    0%    10%    20%    30%    40%    50%    60%    70%    80%    90%    95%    99%  99.5%  99.9%   100%
  6.88  27.78  37.04  46.67  56.67  66.67  76.81  85.71  92.00 100.00 100.00 100.00 100.00 100.00 100.00

methylKit::getCoverageStats output for "BCphaseB1" CpGhsm-sites (#: 149882) at minimum coverage 20   - read coverage statistics per base
summary:
   Min. 1st Qu.  Median    Mean 3rd Qu.    Max.
   20.0    27.0    35.0    38.1    46.0   247.0
percentiles:
   0%   10%   20%   30%   40%   50%   60%   70%   80%   90%   95%   99% 99.5% 99.9%  100%
   20    22    26    29    32    35    39    44    49    57    65    79    85    99   247

methylKit::getMethylationStats output for "BCphaseB1" CpGhsm-sites (#: 149882) at minimum coverage 20      - methylation statistics per base
summary:
   Min. 1st Qu.  Median    Mean 3rd Qu.    Max.
    6.9    34.6    57.5    59.1    85.0   100.0
percentiles:
    0%    10%    20%    30%    40%    50%    60%    70%    80%    90%    95%    99%  99.5%  99.9%   100%
```

```
  6.88  24.32  31.15  38.46  47.27  57.50  68.97  80.00  89.47  95.65  98.00 100.00 100.00 100.00 100.00

methylKit::getCoverageStats output for "BCphaseB1" CpGhsm-sites (#: 29431) at minimum coverage 50    - read coverage statistics per base
summary:
   Min. 1st Qu.  Median    Mean 3rd Qu.    Max.
   50.0    53.0    58.0    60.3    65.0   247.0
percentiles:
   0%   10%   20%   30%   40%   50%   60%   70%   80%   90%   95%   99% 99.5% 99.9%  100%
   50    51    52    54    56    58    60    63    67    74    80    94    99   110   247

methylKit::getMethylationStats output for "BCphaseB1" CpGhsm-sites (#: 29431) at minimum coverage 50 - methylation statistics per base
summary:
   Min. 1st Qu.  Median    Mean 3rd Qu.    Max.
    6.9    29.5    55.4    56.4    84.3   100.0
percentiles:
    0%    10%    20%    30%    40%    50%    60%    70%    80%    90%    95%    99%  99.5%  99.9%   100%
  6.88  18.87  25.45  33.90  43.86  55.36  67.27  79.25  88.68  94.92  97.40 100.00 100.00 100.00 100.00

methylKit::getCoverageStats output for "BCphaseB1" CpGhsm-sites (#: 138) in coverage range [100-1000]      - read coverage statistics per base
summary:
   Min. 1st Qu.  Median    Mean 3rd Qu.    Max.
    100     102     105     109     109     247
percentiles:
   0%   10%   20%   30%   40%   50%   60%   70%   80%   90%   95%   99% 99.5% 99.9%  100%
  100   101   101   102   103   105   106   108   111   121   129   158   196   237   247

methylKit::getMethylationStats output for "BCphaseB1" CpGhsm-sites (#: 138) in coverage range [100-1000]   - methylation statistics per base
summary:
   Min. 1st Qu.  Median    Mean 3rd Qu.    Max.
    6.9    38.0    66.0    61.8    88.9    99.0
percentiles:
   0%   10%   20%   30%   40%   50%   60%   70%   80%   90%   95%   99% 99.5% 99.9%  100%
 6.88 20.09 30.53 45.31 56.87 66.01 74.87 85.51 90.46 94.02 96.39 98.10 98.42 98.89 99.01
```

```
Directory: CMS File: cms-Ob_LI_BCphaseB2.txt

Number of "BCphaseB2" CpGhsm-sites with minimal and higher level coverage:

  number of "BCphaseB2" CpGhsm-sites with coverage >=  5: 209549
  number of "BCphaseB2" CpGhsm-sites with coverage >=  6: 206680
  number of "BCphaseB2" CpGhsm-sites with coverage >= 20: 133461
  number of "BCphaseB2" CpGhsm-sites with coverage >= 50:  30958

Coverage and methylation statistics for "BCphaseB2" CpGhsm-sites at different levels of minimum coverage:

methylKit::getCoverageStats output for "BCphaseB2" CpGhsm-sites (#: 209549) at minimum coverage 5    - read coverage statistics per base
summary:
   Min. 1st Qu.  Median    Mean 3rd Qu.    Max.
      5      15      26      30      40     582
percentiles:
   0%   10%   20%   30%   40%   50%   60%   70%   80%   90%   95%   99% 99.5% 99.9%  100%
    5     9    13    17    21    26    31    37    44    56    65    85    93   111   582

methylKit::getMethylationStats output for "BCphaseB2" CpGhsm-sites (#: 209549) at minimum coverage 5 - methylation statistics per base
summary:
   Min. 1st Qu.  Median    Mean 3rd Qu.    Max.
    4.0    42.9    68.8    65.8    89.7   100.0
percentiles:
    0%    10%    20%    30%    40%    50%    60%    70%    80%    90%    95%    99%  99.5%  99.9%   100%
  3.95  28.57  38.10  48.28  58.46  68.75  78.05  86.21  92.86 100.00 100.00 100.00 100.00 100.00 100.00

methylKit::getCoverageStats output for "BCphaseB2" CpGhsm-sites (#: 206680) at minimum coverage 6    - read coverage statistics per base
summary:
   Min. 1st Qu.  Median    Mean 3rd Qu.    Max.
      6      16      26      30      41     582
percentiles:
   0%   10%   20%   30%   40%   50%   60%   70%   80%   90%   95%   99% 99.5% 99.9%  100%
    6    10    14    17    21    26    31    37    45    56    65    85    93   111   582

methylKit::getMethylationStats output for "BCphaseB2" CpGhsm-sites (#: 206680) at minimum coverage 6 - methylation statistics per base
summary:
   Min. 1st Qu.  Median    Mean 3rd Qu.    Max.
    4.0    42.9    67.9    65.3    88.9   100.0
percentiles:
    0%    10%    20%    30%    40%    50%    60%    70%    80%    90%    95%    99%  99.5%  99.9%   100%
  3.95  28.21  37.84  47.83  58.06  67.86  77.55  85.71  92.31 100.00 100.00 100.00 100.00 100.00 100.00

methylKit::getCoverageStats output for "BCphaseB2" CpGhsm-sites (#: 133461) at minimum coverage 20   - read coverage statistics per base
summary:
   Min. 1st Qu.  Median    Mean 3rd Qu.    Max.
     20      27      36      40      48     582
percentiles:
   0%   10%   20%   30%   40%   50%   60%   70%   80%   90%   95%   99% 99.5% 99.9%  100%
   20    22    25    28    32    36    40    45    52    62    71    90    98   117   582

methylKit::getMethylationStats output for "BCphaseB2" CpGhsm-sites (#: 133461) at minimum coverage 20      - methylation statistics per base
summary:
   Min. 1st Qu.  Median    Mean 3rd Qu.    Max.
    4.0    35.0    58.2    59.3    85.0   100.0
percentiles:
    0%    10%    20%    30%    40%    50%    60%    70%    80%    90%    95%    99%  99.5%  99.9%   100%
```

```
  3.95  24.39  31.48  38.89  47.83  58.23  69.23  80.00  89.29  95.56  97.96 100.00 100.00 100.00 100.00

methylKit::getCoverageStats output for "BCphaseB2" CpGhsm-sites (#: 30958) at minimum coverage 50   - read coverage statistics per base
summary:
   Min. 1st Qu.  Median    Mean 3rd Qu.    Max.
     50      54      60      63      69     582
percentiles:
   0%   10%   20%   30%   40%   50%   60%   70%   80%   90%   95%   99% 99.5% 99.9%  100%
   50    51    53    55    57    60    63    67    72    80    88   107   115   138   582

methylKit::getMethylationStats output for "BCphaseB2" CpGhsm-sites (#: 30958) at minimum coverage 50 - methylation statistics per base
summary:
   Min. 1st Qu.  Median    Mean 3rd Qu.    Max.
    4.0    28.9    54.4    55.9    83.6   100.0
percentiles:
    0%    10%    20%    30%    40%    50%    60%    70%    80%    90%    95%    99%  99.5%  99.9%   100%
  3.95  18.47  25.00  33.33  43.14  54.39  66.28  78.33  88.00  94.83  97.26 100.00 100.00 100.00 100.00

methylKit::getCoverageStats output for "BCphaseB2" CpGhsm-sites (#: 577) in coverage range [100-1000]     - read coverage statistics per base
summary:
   Min. 1st Qu.  Median    Mean 3rd Qu.    Max.
    100     102     107     114     116     582
percentiles:
   0%   10%   20%   30%   40%   50%   60%   70%   80%   90%   95%   99% 99.5% 99.9%  100%
  100   101   102   103   105   107   110   114   118   126   140   223   231   401   582

methylKit::getMethylationStats output for "BCphaseB2" CpGhsm-sites (#: 577) in coverage range [100-1000]  - methylation statistics per base
summary:
   Min. 1st Qu.  Median    Mean 3rd Qu.    Max.
    4.0    34.3    59.4    58.1    86.0   100.0
percentiles:
    0%    10%    20%    30%    40%    50%    60%    70%    80%    90%    95%    99%  99.5%  99.9%   100%
  3.95  18.09  27.56  38.99  47.08  59.43  71.27  81.18  88.36  95.00  97.03  99.04  99.15 100.00 100.00
```

```
Directory: CMS File: cms-Ob_LI_RphaseA1.txt

Number of "RphaseA1" CpGhsm-sites with minimal and higher level coverage:

  number of "RphaseA1" CpGhsm-sites with coverage >=  5: 213360
  number of "RphaseA1" CpGhsm-sites with coverage >=  6: 210617
  number of "RphaseA1" CpGhsm-sites with coverage >= 20: 133209
  number of "RphaseA1" CpGhsm-sites with coverage >= 50:  18034

Coverage and methylation statistics for "RphaseA1" CpGhsm-sites at different levels of minimum coverage:

methylKit::getCoverageStats output for "RphaseA1" CpGhsm-sites (#: 213360) at minimum coverage 5     - read coverage statistics per base
summary:
   Min. 1st Qu.  Median    Mean 3rd Qu.    Max.
      5      15      25      27      37     133
percentiles:
   0%   10%   20%   30%   40%   50%   60%   70%   80%   90%   95%   99% 99.5% 99.9%  100%
    5     9    13    17    21    25    29    34    40    48    54    68    73    86   133

methylKit::getMethylationStats output for "RphaseA1" CpGhsm-sites (#: 213360) at minimum coverage 5  - methylation statistics per base
summary:
   Min. 1st Qu.  Median    Mean 3rd Qu.    Max.
    8.3    44.4    70.0    66.7    90.0   100.0
percentiles:
    0%    10%    20%    30%    40%    50%    60%    70%    80%    90%    95%    99%  99.5%  99.9%   100%
  8.27  30.00  39.53  50.00  60.00  70.00  78.95  87.04  93.33 100.00 100.00 100.00 100.00 100.00 100.00

methylKit::getCoverageStats output for "RphaseA1" CpGhsm-sites (#: 210617) at minimum coverage 6     - read coverage statistics per base
summary:
   Min. 1st Qu.  Median    Mean 3rd Qu.    Max.
    6.0    15.0    25.0    27.3    37.0   133.0
percentiles:
   0%   10%   20%   30%   40%   50%   60%   70%   80%   90%   95%   99% 99.5% 99.9%  100%
    6    10    13    17    21    25    29    34    40    48    54    68    73    86   133

methylKit::getMethylationStats output for "RphaseA1" CpGhsm-sites (#: 210617) at minimum coverage 6  - methylation statistics per base
summary:
   Min. 1st Qu.  Median    Mean 3rd Qu.    Max.
    8.3    44.0    69.2    66.2    89.7   100.0
percentiles:
    0%    10%    20%    30%    40%    50%    60%    70%    80%    90%    95%    99%  99.5%  99.9%   100%
  8.27  29.69  39.29  50.00  59.09  69.23  78.26  86.27  92.59 100.00 100.00 100.00 100.00 100.00 100.00

methylKit::getCoverageStats output for "RphaseA1" CpGhsm-sites (#: 133209) at minimum coverage 20    - read coverage statistics per base
summary:
   Min. 1st Qu.  Median    Mean 3rd Qu.    Max.
   20.0    26.0    33.0    35.7    43.0   133.0
percentiles:
   0%   10%   20%   30%   40%   50%   60%   70%   80%   90%   95%   99% 99.5% 99.9%  100%
   20    22    25    28    30    33    37    41    45    52    58    72    77    90   133

methylKit::getMethylationStats output for "RphaseA1" CpGhsm-sites (#: 133209) at minimum coverage 20 - methylation statistics per base
summary:
   Min. 1st Qu.  Median    Mean 3rd Qu.    Max.
    8.3    36.0    59.3    60.3    85.7   100.0
percentiles:
    0%    10%    20%    30%    40%    50%    60%    70%    80%    90%    95%    99%  99.5%  99.9%   100%
```

```
  8.27  25.71  32.50  40.00  48.94  59.26  70.37  80.95  90.00  95.83  98.18 100.00 100.00 100.00 100.00

methylKit::getCoverageStats output for "RphaseA1" CpGhsm-sites (#: 18034) at minimum coverage 50    - read coverage statistics per base
summary:
   Min. 1st Qu.  Median    Mean 3rd Qu.    Max. 
   50.0    52.0    56.0    58.3    62.0   133.0 
percentiles:
   0%   10%   20%   30%   40%   50%   60%   70%   80%   90%   95%   99% 99.5% 99.9%  100% 
 50.0  50.0  52.0  53.0  54.0  56.0  58.0  60.0  64.0  69.0  74.0  87.7  93.0 106.0 133.0 

methylKit::getMethylationStats output for "RphaseA1" CpGhsm-sites (#: 18034) at minimum coverage 50  - methylation statistics per base
summary:
   Min. 1st Qu.  Median    Mean 3rd Qu.    Max. 
    8.3    32.1    58.5    58.5    86.0   100.0 
percentiles:
    0%    10%    20%    30%    40%    50%    60%    70%    80%    90%    95%    99%  99.5%  99.9%   100% 
  8.27  20.00  27.87  36.84  47.46  58.49  70.00  81.13  89.86  95.52  98.04 100.00 100.00 100.00 100.00 

methylKit::getCoverageStats output for "RphaseA1" CpGhsm-sites (#: 33) in coverage range [100-1000]  - read coverage statistics per base
summary:
   Min. 1st Qu.  Median    Mean 3rd Qu.    Max. 
    100     102     106     108     112     133 
percentiles:
   0%   10%   20%   30%   40%   50%   60%   70%   80%   90%   95%   99% 99.5% 99.9%  100% 
  100   100   101   103   106   106   110   111   113   116   124   130   132   133   133 

methylKit::getMethylationStats output for "RphaseA1" CpGhsm-sites (#: 33) in coverage range [100-1000]    - methylation statistics per base
summary:
   Min. 1st Qu.  Median    Mean 3rd Qu.    Max. 
    8.3    36.2    59.4    60.1    83.1    98.1 
percentiles:
   0%   10%   20%   30%   40%   50%   60%   70%   80%   90%   95%   99% 99.5% 99.9%  100% 
 8.27 21.60 35.75 47.18 56.06 59.43 68.55 79.14 87.14 94.97 96.20 97.86 97.99 98.09 98.11 
```

```
Directory: CMS File: cms-Ob_LI_RphaseA2.txt

Number of "RphaseA2" CpGhsm-sites with minimal and higher level coverage:

  number of "RphaseA2" CpGhsm-sites with coverage >=  5: 265465
  number of "RphaseA2" CpGhsm-sites with coverage >=  6: 263149
  number of "RphaseA2" CpGhsm-sites with coverage >= 20: 200126
  number of "RphaseA2" CpGhsm-sites with coverage >= 50:  88014

Coverage and methylation statistics for "RphaseA2" CpGhsm-sites at different levels of minimum coverage:

methylKit::getCoverageStats output for "RphaseA2" CpGhsm-sites (#: 265465) at minimum coverage 5     - read coverage statistics per base
summary:
   Min. 1st Qu.  Median    Mean 3rd Qu.    Max.
    5.0    20.0    36.0    42.2    58.0   281.0
percentiles:
   0%   10%   20%   30%   40%   50%   60%   70%   80%   90%   95%   99% 99.5% 99.9%  100%
    5    11    17    23    29    36    43    53    65    83    98   129   141   167   281

methylKit::getMethylationStats output for "RphaseA2" CpGhsm-sites (#: 265465) at minimum coverage 5  - methylation statistics per base
summary:
   Min. 1st Qu.  Median    Mean 3rd Qu.    Max.
    5.0    35.6    62.5    61.1    87.5   100.0
percentiles:
    0%    10%    20%    30%    40%    50%    60%    70%    80%    90%    95%    99%  99.5%  99.9%   100%
  4.98  22.50  31.37  40.54  51.16  62.50  73.33  83.33  91.30  97.44 100.00 100.00 100.00 100.00 100.00

methylKit::getCoverageStats output for "RphaseA2" CpGhsm-sites (#: 263149) at minimum coverage 6     - read coverage statistics per base
summary:
   Min. 1st Qu.  Median    Mean 3rd Qu.    Max.
    6.0    20.0    36.0    42.5    58.0   281.0
percentiles:
   0%   10%   20%   30%   40%   50%   60%   70%   80%   90%   95%   99% 99.5% 99.9%  100%
    6    12    17    23    29    36    44    53    65    83    98   129   141   167   281

methylKit::getMethylationStats output for "RphaseA2" CpGhsm-sites (#: 263149) at minimum coverage 6  - methylation statistics per base
summary:
   Min. 1st Qu.  Median    Mean 3rd Qu.    Max.
    5.0    35.3    61.8    60.7    87.5   100.0
percentiles:
    0%    10%    20%    30%    40%    50%    60%    70%    80%    90%    95%    99%  99.5%  99.9%   100%
  4.98  22.45  31.17  40.00  50.00  61.76  72.73  83.33  90.91  96.88 100.00 100.00 100.00 100.00 100.00

methylKit::getCoverageStats output for "RphaseA2" CpGhsm-sites (#: 200126) at minimum coverage 20    - read coverage statistics per base
summary:
   Min. 1st Qu.  Median    Mean 3rd Qu.    Max.
   20.0    31.0    45.0    51.9    66.0   281.0
percentiles:
   0%   10%   20%   30%   40%   50%   60%   70%   80%   90%   95%   99% 99.5% 99.9%  100%
   20    24    29    34    39    45    52    61    72    89   104   134   146   173   281

methylKit::getMethylationStats output for "RphaseA2" CpGhsm-sites (#: 200126) at minimum coverage 20 - methylation statistics per base
summary:
   Min. 1st Qu.  Median    Mean 3rd Qu.    Max.
    5.0    30.3    53.3    56.0    83.3   100.0
percentiles:
    0%    10%    20%    30%    40%    50%    60%    70%    80%    90%    95%    99%  99.5%  99.9%   100%
```

```
  4.98  20.45  26.83  33.85  42.59  53.26  65.31  77.78  88.18  95.24  97.53 100.00 100.00 100.00 100.00

methylKit::getCoverageStats output for "RphaseA2" CpGhsm-sites (#: 88014) at minimum coverage 50    - read coverage statistics per base
summary:
   Min. 1st Qu.  Median    Mean 3rd Qu.    Max.
   50.0    58.0    70.0    75.6    87.0   281.0
percentiles:
   0%   10%   20%   30%   40%   50%   60%   70%   80%   90%   95%   99% 99.5% 99.9%  100%
   50    53    56    60    65    70    76    83    92   106   119   148   160   190   281

methylKit::getMethylationStats output for "RphaseA2" CpGhsm-sites (#: 88014) at minimum coverage 50 - methylation statistics per base
summary:
   Min. 1st Qu.  Median    Mean 3rd Qu.    Max.
    5.0    24.4    48.4    52.3    80.8   100.0
percentiles:
    0%    10%    20%    30%    40%    50%    60%    70%    80%    90%    95%    99%  99.5%  99.9%   100%
  4.98  16.07  21.31  28.07  37.25  48.39  61.36  74.60  86.41  94.44  96.97 100.00 100.00 100.00 100.00

methylKit::getCoverageStats output for "RphaseA2" CpGhsm-sites (#: 12480) in coverage range [100-1000]    - read coverage statistics per base
summary:
   Min. 1st Qu.  Median    Mean 3rd Qu.    Max.
    100     105     113     118     126     281
percentiles:
   0%   10%   20%   30%   40%   50%   60%   70%   80%   90%   95%   99% 99.5% 99.9%  100%
  100   102   104   106   109   113   117   122   130   142   154   182   196   227   281

methylKit::getMethylationStats output for "RphaseA2" CpGhsm-sites (#: 12480) in coverage range [100-1000] - methylation statistics per base
summary:
   Min. 1st Qu.  Median    Mean 3rd Qu.    Max.
    5.0    25.6    52.7    53.9    83.2   100.0
percentiles:
    0%    10%    20%    30%    40%    50%    60%    70%    80%    90%    95%    99%  99.5%  99.9%   100%
  4.98  14.12  21.17  30.08  40.97  52.65  65.35  77.97  87.81  94.64  97.03  99.03  99.20 100.00 100.00
```

```
Directory: CMS File: cms-Ob_LI_RphaseB1.txt

Number of "RphaseB1" CpGhsm-sites with minimal and higher level coverage:

  number of "RphaseB1" CpGhsm-sites with coverage >=  5: 217571
  number of "RphaseB1" CpGhsm-sites with coverage >=  6: 214719
  number of "RphaseB1" CpGhsm-sites with coverage >= 20: 138237
  number of "RphaseB1" CpGhsm-sites with coverage >= 50:  22908

Coverage and methylation statistics for "RphaseB1" CpGhsm-sites at different levels of minimum coverage:

methylKit::getCoverageStats output for "RphaseB1" CpGhsm-sites (#: 217571) at minimum coverage 5    - read coverage statistics per base
summary:
   Min. 1st Qu.  Median    Mean 3rd Qu.    Max.
      5      15      25      28      38     692
percentiles:
   0%   10%   20%   30%   40%   50%   60%   70%   80%   90%   95%   99% 99.5% 99.9%  100%
    5     9    13    17    21    25    30    35    41    50    57    73    79    94   692

methylKit::getMethylationStats output for "RphaseB1" CpGhsm-sites (#: 217571) at minimum coverage 5  - methylation statistics per base
summary:
   Min. 1st Qu.  Median    Mean 3rd Qu.    Max.
    4.0    43.5    68.8    66.0    90.0   100.0
percentiles:
    0%    10%    20%    30%    40%    50%    60%    70%    80%    90%    95%    99%  99.5%  99.9%   100%
  4.05  29.17  38.71  48.65  58.33  68.75  77.78  86.36  92.86 100.00 100.00 100.00 100.00 100.00 100.00

methylKit::getCoverageStats output for "RphaseB1" CpGhsm-sites (#: 214719) at minimum coverage 6    - read coverage statistics per base
summary:
   Min. 1st Qu.  Median    Mean 3rd Qu.    Max.
      6      16      26      28      38     692
percentiles:
   0%   10%   20%   30%   40%   50%   60%   70%   80%   90%   95%   99% 99.5% 99.9%  100%
    6    10    14    17    21    26    30    35    42    50    58    73    79    94   692

methylKit::getMethylationStats output for "RphaseB1" CpGhsm-sites (#: 214719) at minimum coverage 6  - methylation statistics per base
summary:
   Min. 1st Qu.  Median    Mean 3rd Qu.    Max.
    4.0    43.1    67.9    65.5    88.9   100.0
percentiles:
    0%    10%    20%    30%    40%    50%    60%    70%    80%    90%    95%    99%  99.5%  99.9%   100%
  4.05  29.03  38.46  48.08  58.06  67.86  77.50  85.71  92.31 100.00 100.00 100.00 100.00 100.00 100.00

methylKit::getCoverageStats output for "RphaseB1" CpGhsm-sites (#: 138237) at minimum coverage 20   - read coverage statistics per base
summary:
   Min. 1st Qu.  Median    Mean 3rd Qu.    Max.
     20      27      34      37      44     692
percentiles:
   0%   10%   20%   30%   40%   50%   60%   70%   80%   90%   95%   99% 99.5% 99.9%  100%
   20    22    25    28    31    34    38    42    47    55    62    77    83    98   692

methylKit::getMethylationStats output for "RphaseB1" CpGhsm-sites (#: 138237) at minimum coverage 20 - methylation statistics per base
summary:
   Min. 1st Qu.  Median    Mean 3rd Qu.    Max.
    4.0    35.5    58.3    59.7    85.2   100.0
percentiles:
    0%    10%    20%    30%    40%    50%    60%    70%    80%    90%    95%    99%  99.5%  99.9%   100%
```

```
  4.05  25.00  32.08  39.39  48.28  58.33  69.57  80.00  89.58  95.74  98.08 100.00 100.00 100.00 100.00

methylKit::getCoverageStats output for "RphaseB1" CpGhsm-sites (#: 22908) at minimum coverage 50    - read coverage statistics per base
summary:
   Min. 1st Qu.  Median    Mean 3rd Qu.    Max. 
     50      53      57      60      64     692 
percentiles:
   0%   10%   20%   30%   40%   50%   60%   70%   80%   90%   95%   99% 99.5% 99.9%  100% 
   50    51    52    53    55    57    59    62    66    72    78    93    99   131   692 

methylKit::getMethylationStats output for "RphaseB1" CpGhsm-sites (#: 22908) at minimum coverage 50  - methylation statistics per base
summary:
   Min. 1st Qu.  Median    Mean 3rd Qu.    Max. 
    4.0    31.0    57.4    57.6    84.8   100.0 
percentiles:
    0%    10%    20%    30%    40%    50%    60%    70%    80%    90%    95%    99%  99.5%  99.9%   100% 
  4.05  19.64  26.79  35.71  46.15  57.41  69.23  80.00  88.89  95.16  98.00 100.00 100.00 100.00 100.00

methylKit::getCoverageStats output for "RphaseB1" CpGhsm-sites (#: 107) in coverage range [100-1000] - read coverage statistics per base
summary:
   Min. 1st Qu.  Median    Mean 3rd Qu.    Max. 
    100     103     110     132     124     692 
percentiles:
   0%   10%   20%   30%   40%   50%   60%   70%   80%   90%   95%   99% 99.5% 99.9%  100% 
  100   101   103   104   106   110   113   120   137   197   217   390   535   661   692 

methylKit::getMethylationStats output for "RphaseB1" CpGhsm-sites (#: 107) in coverage range [100-1000]   - methylation statistics per base
summary:
   Min. 1st Qu.  Median    Mean 3rd Qu.    Max. 
    4.0    33.6    54.1    55.3    79.1    98.4 
percentiles:
   0%   10%   20%   30%   40%   50%   60%   70%   80%   90%   95%   99% 99.5% 99.9%  100% 
 4.05 16.58 26.78 37.60 49.13 54.07 63.87 73.94 83.66 95.32 96.29 98.04 98.24 98.37 98.40 
```

```
Directory: CMS File: cms-Ob_LI_RphaseB2.txt

Number of "RphaseB2" CpGhsm-sites with minimal and higher level coverage:

  number of "RphaseB2" CpGhsm-sites with coverage >=  5: 201205
  number of "RphaseB2" CpGhsm-sites with coverage >=  6: 198266
  number of "RphaseB2" CpGhsm-sites with coverage >= 20: 122498
  number of "RphaseB2" CpGhsm-sites with coverage >= 50:  25596

Coverage and methylation statistics for "RphaseB2" CpGhsm-sites at different levels of minimum coverage:

methylKit::getCoverageStats output for "RphaseB2" CpGhsm-sites (#: 201205) at minimum coverage 5    - read coverage statistics per base
summary:
   Min. 1st Qu.  Median    Mean 3rd Qu.    Max.
      5      14      24      28      38     541
percentiles:
   0%   10%   20%   30%   40%   50%   60%   70%   80%   90%   95%   99% 99.5% 99.9%  100%
    5     9    12    16    20    24    29    35    42    53    63    82    90   107   541

methylKit::getMethylationStats output for "RphaseB2" CpGhsm-sites (#: 201205) at minimum coverage 5  - methylation statistics per base
summary:
   Min. 1st Qu.  Median    Mean 3rd Qu.    Max.
    3.9    44.8    70.0    66.8    90.0   100.0
percentiles:
    0%    10%    20%    30%    40%    50%    60%    70%    80%    90%    95%    99%  99.5%  99.9%   100%
  3.88  29.73  40.00  50.00  60.00  70.00  79.31  86.96  93.33 100.00 100.00 100.00 100.00 100.00 100.00

methylKit::getCoverageStats output for "RphaseB2" CpGhsm-sites (#: 198266) at minimum coverage 6    - read coverage statistics per base
summary:
   Min. 1st Qu.  Median    Mean 3rd Qu.    Max.
      6      15      25      28      38     541
percentiles:
   0%   10%   20%   30%   40%   50%   60%   70%   80%   90%   95%   99% 99.5% 99.9%  100%
    6     9    13    16    20    25    29    35    42    53    63    82    90   107   541

methylKit::getMethylationStats output for "RphaseB2" CpGhsm-sites (#: 198266) at minimum coverage 6  - methylation statistics per base
summary:
   Min. 1st Qu.  Median    Mean 3rd Qu.    Max.
    3.9    44.4    69.4    66.3    89.5   100.0
percentiles:
    0%    10%    20%    30%    40%    50%    60%    70%    80%    90%    95%    99%  99.5%  99.9%   100%
  3.88  29.63  39.47  50.00  60.00  69.44  78.38  86.11  92.31 100.00 100.00 100.00 100.00 100.00 100.00

methylKit::getCoverageStats output for "RphaseB2" CpGhsm-sites (#: 122498) at minimum coverage 20   - read coverage statistics per base
summary:
   Min. 1st Qu.  Median    Mean 3rd Qu.    Max.
     20      26      34      38      47     541
percentiles:
   0%   10%   20%   30%   40%   50%   60%   70%   80%   90%   95%   99% 99.5% 99.9%  100%
   20    22    25    28    31    34    39    44    50    60    69    87    95   114   541

methylKit::getMethylationStats output for "RphaseB2" CpGhsm-sites (#: 122498) at minimum coverage 20 - methylation statistics per base
summary:
   Min. 1st Qu.  Median    Mean 3rd Qu.    Max.
    3.9    35.9    59.3    60.0    85.7   100.0
percentiles:
    0%    10%    20%    30%    40%    50%    60%    70%    80%    90%    95%    99%  99.5%  99.9%   100%
```

```
  3.88  25.00  32.26  40.00  48.84  59.26  70.18  80.77  89.66  95.65  98.00 100.00 100.00 100.00 100.00
```

```
methylKit::getCoverageStats output for "RphaseB2" CpGhsm-sites (#: 25596) at minimum coverage 50    - read coverage statistics per base
summary:
   Min. 1st Qu.  Median    Mean 3rd Qu.    Max. 
     50      54      59      63      68     541 
percentiles:
   0%   10%   20%   30%   40%   50%   60%   70%   80%   90%   95%   99% 99.5% 99.9%  100% 
   50    51    53    55    57    59    62    66    71    79    87   104   113   139   541 

methylKit::getMethylationStats output for "RphaseB2" CpGhsm-sites (#: 25596) at minimum coverage 50 - methylation statistics per base
summary:
   Min. 1st Qu.  Median    Mean 3rd Qu.    Max. 
    3.9    30.1    55.7    56.6    84.0   100.0 
percentiles:
    0%    10%    20%    30%    40%    50%    60%    70%    80%    90%    95%    99%  99.5%  99.9%   100% 
  3.88  18.97  25.93  34.55  44.64  55.74  67.27  78.85  88.41  94.83  97.33 100.00 100.00 100.00 100.00
```

```
methylKit::getCoverageStats output for "RphaseB2" CpGhsm-sites (#: 417) in coverage range [100-1000]- read coverage statistics per base
summary:
   Min. 1st Qu.  Median    Mean 3rd Qu.    Max. 
    100     103     106     115     116     541 
percentiles:
   0%   10%   20%   30%   40%   50%   60%   70%   80%   90%   95%   99% 99.5% 99.9%  100% 
  100   101   102   103   105   106   110   113   119   130   155   221   228   425   541 

methylKit::getMethylationStats output for "RphaseB2" CpGhsm-sites (#: 417) in coverage range [100-1000]   - methylation statistics per base
summary:
   Min. 1st Qu.  Median    Mean 3rd Qu.    Max. 
    3.9    38.1    66.7    61.6    88.3   100.0 
percentiles:
    0%    10%    20%    30%    40%    50%    60%    70%    80%    90%    95%    99%  99.5%  99.9%   100% 
  3.88  22.58  32.86  42.13  53.98  66.67  74.68  83.29  91.18  95.67  97.20  99.10  99.27 100.00 100.00
```

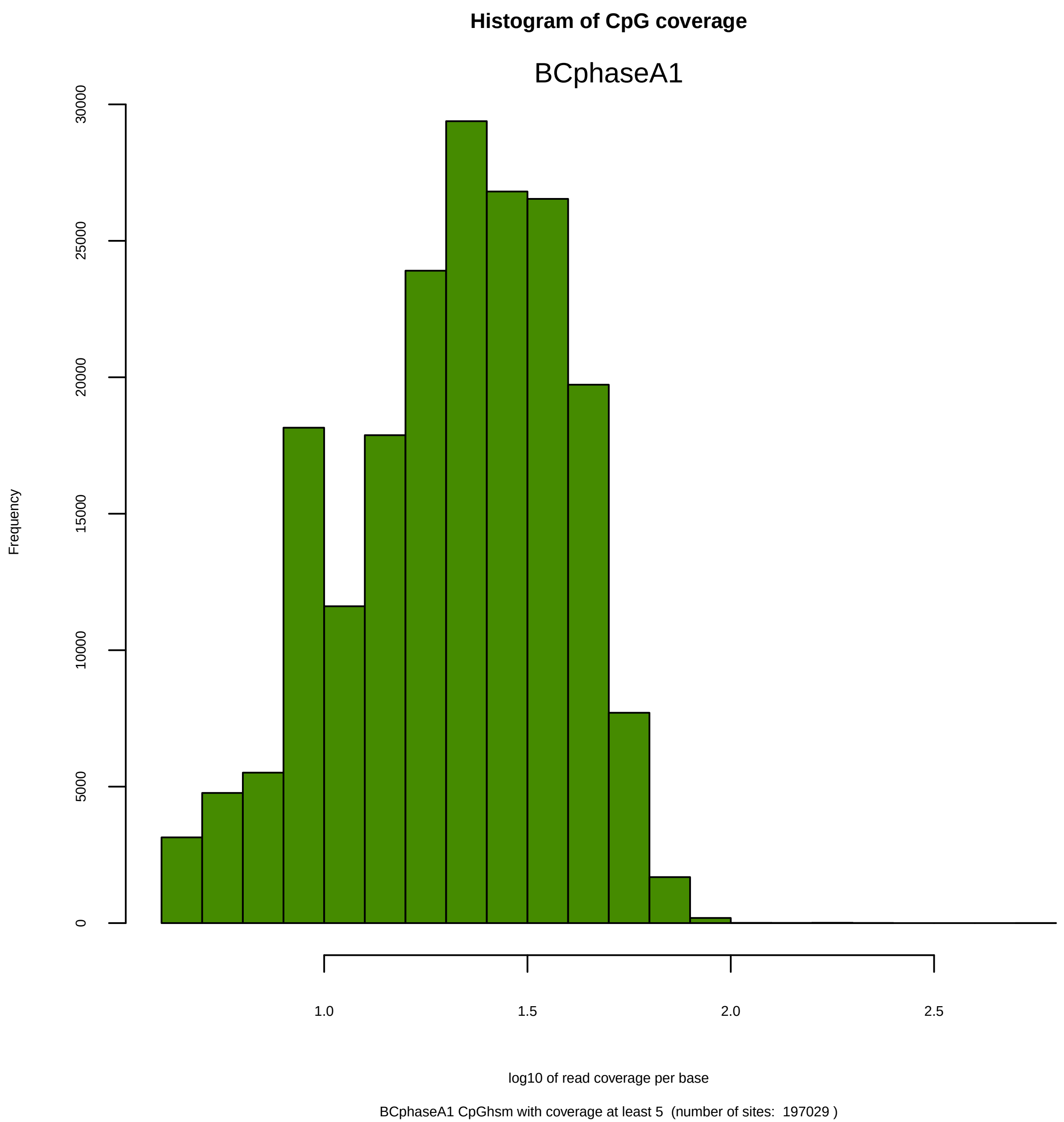
Histogram of CpG coverage
BCphaseA1
Frequency
0
5000
10000
15000
20000
25000
30000
1.0
1.5
2.0
2.5
log10 of read coverage per base
BCphaseA1 CpGhsm with coverage at least 5 (number of sites: 197029 )

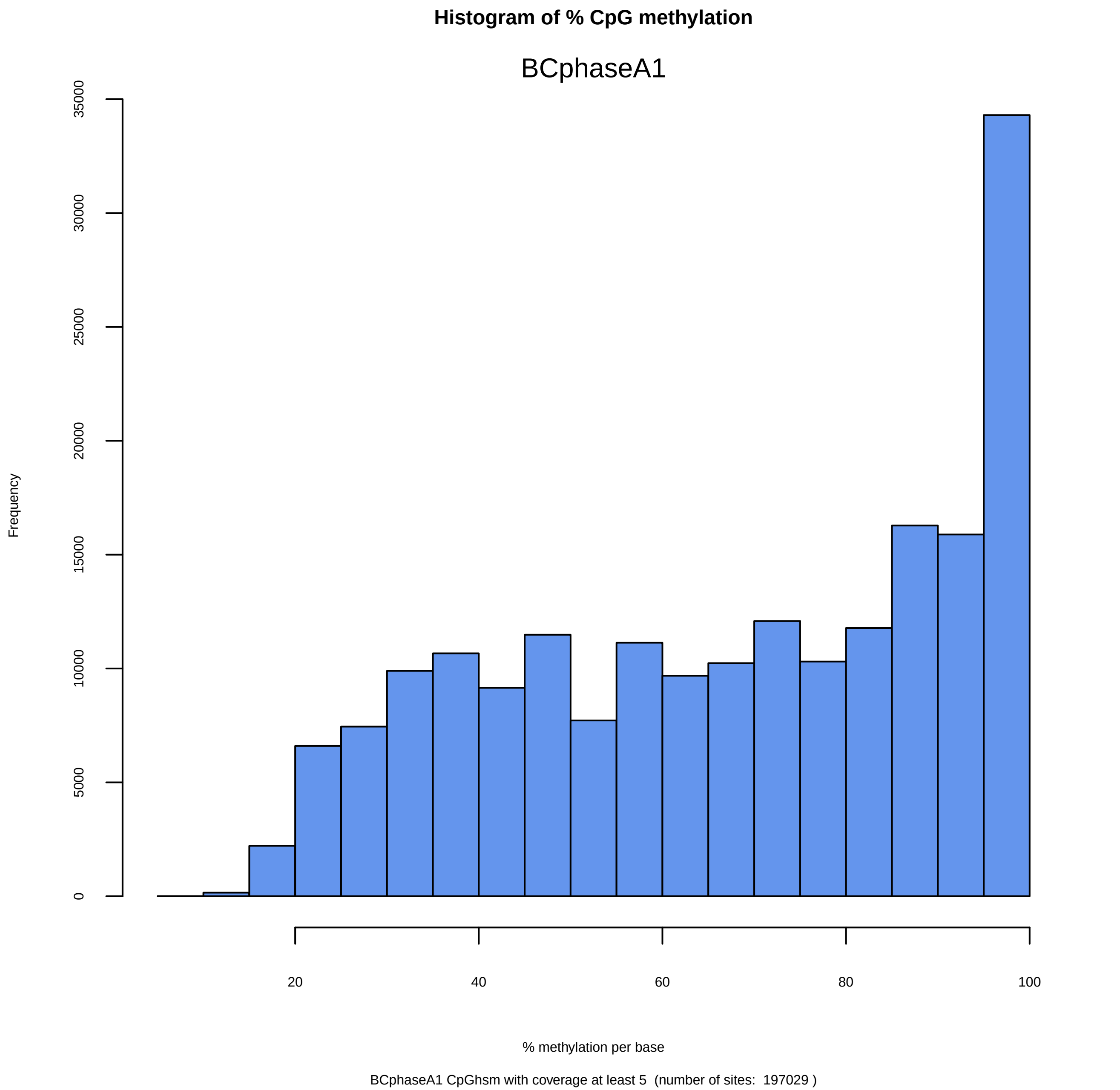
Histogram of % CpG methylation
BCphaseA1
Frequency
0
5000
10000
15000
20000
25000
30000
35000
20
40
60
80
100
% methylation per base
BCphaseA1 CpGhsm with coverage at least 5  (number of sites:  197029 )

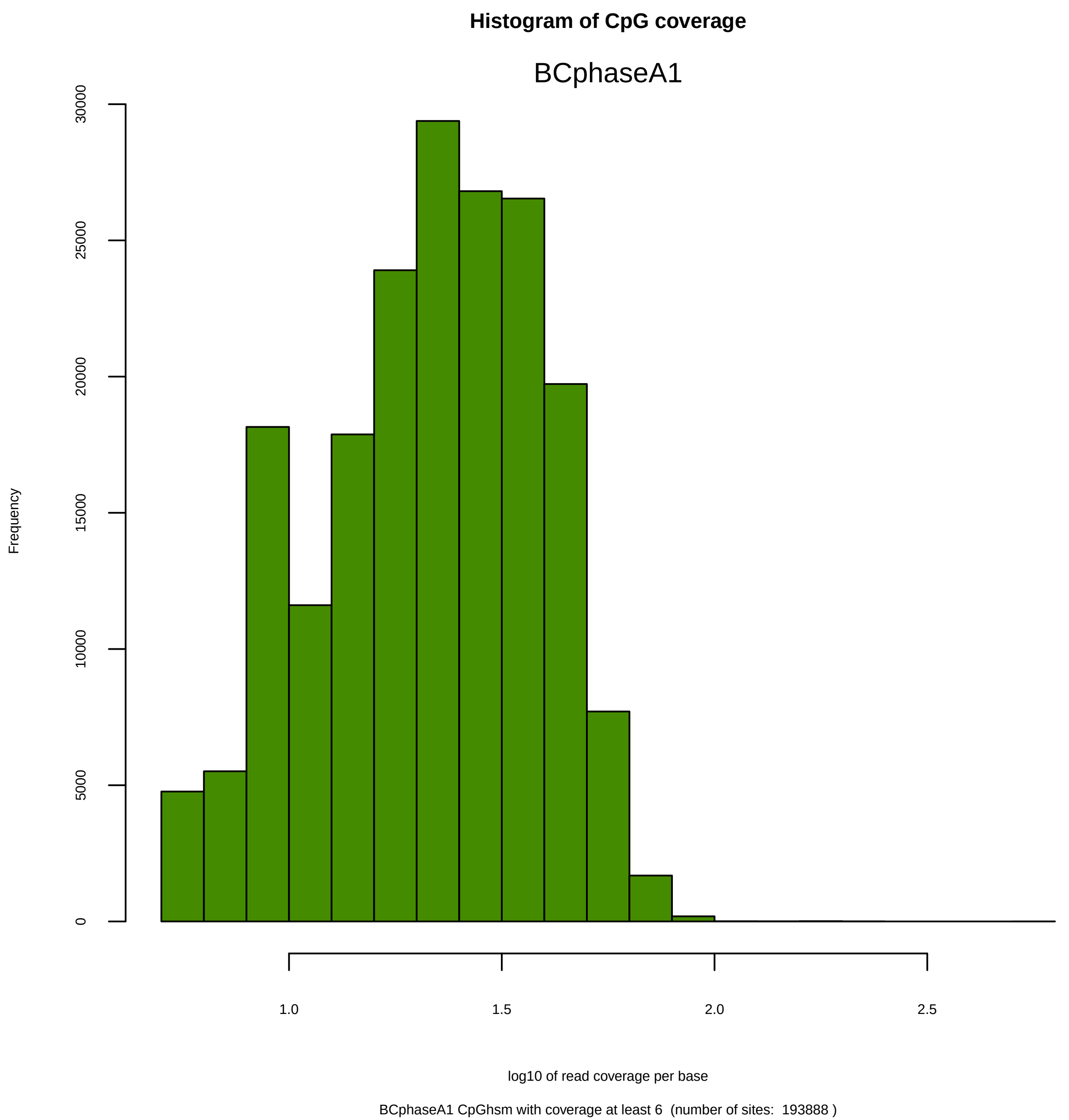
Histogram of CpG coverage
BCphaseA1
Frequency
0
5000
10000
15000
20000
25000
30000
1.0
1.5
2.0
2.5
log10 of read coverage per base
BCphaseA1 CpGhsm with coverage at least 6 (number of sites: 193888 )

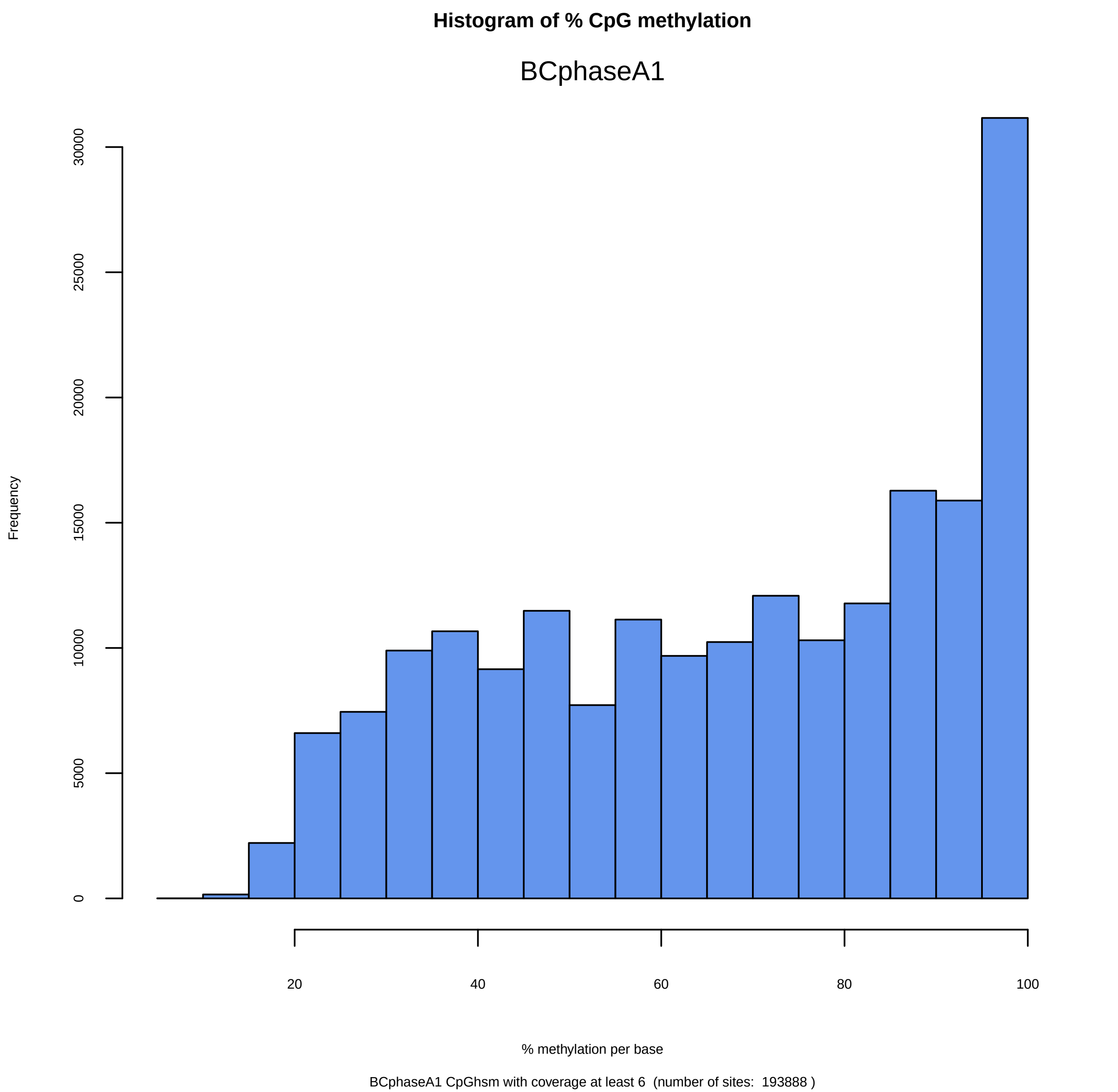
Histogram of % CpG methylation
BCphaseA1
Frequency
0
5000
10000
15000
20000
25000
30000
20
40
60
80
100
% methylation per base
BCphaseA1 CpGhsm with coverage at least 6 (number of sites: 193888 )

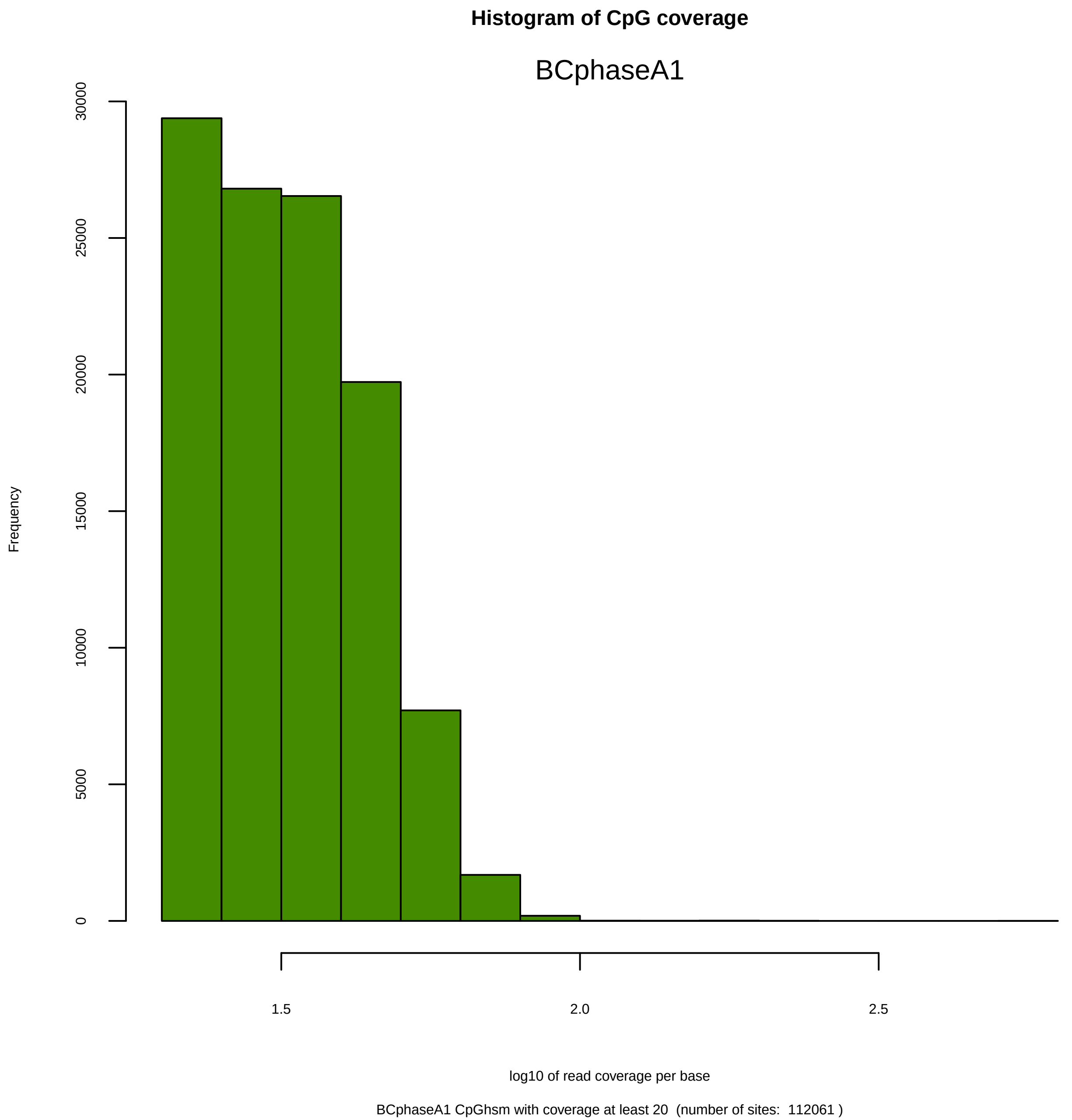

Histogram of CpG coverage
BCphaseA1
Frequency
0
5000
10000
15000
20000
25000
30000
1.5
2.0
2.5
log10 of read coverage per base
BCphaseA1 CpGhsm with coverage at least 20  (number of sites:  112061 )

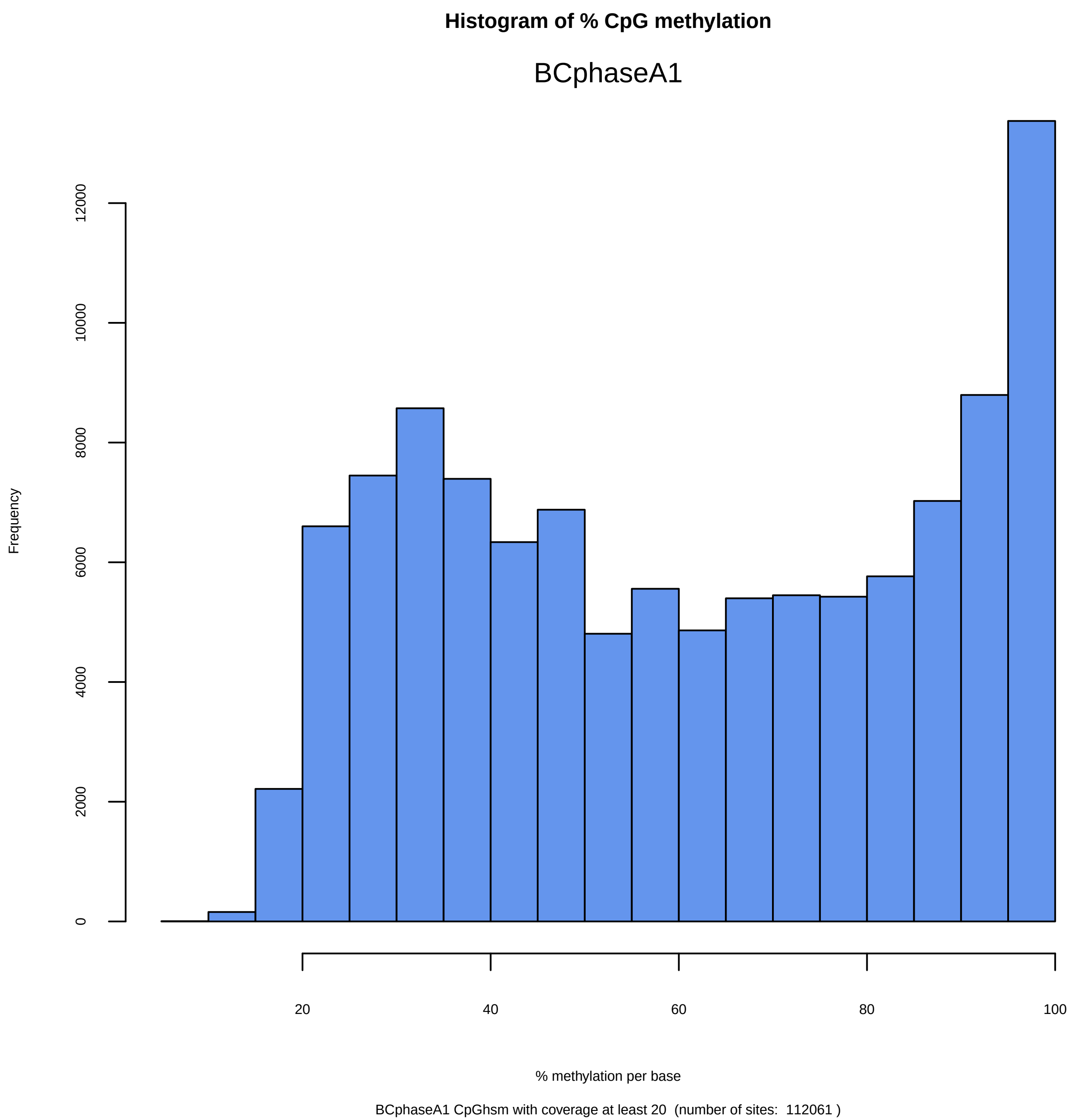

Histogram of % CpG methylation
BCphaseA1
Frequency
0
2000
4000
6000
8000
10000
12000
20
40
60
80
100
% methylation per base
BCphaseA1 CpGhsm with coverage at least 20 (number of sites: 112061 )

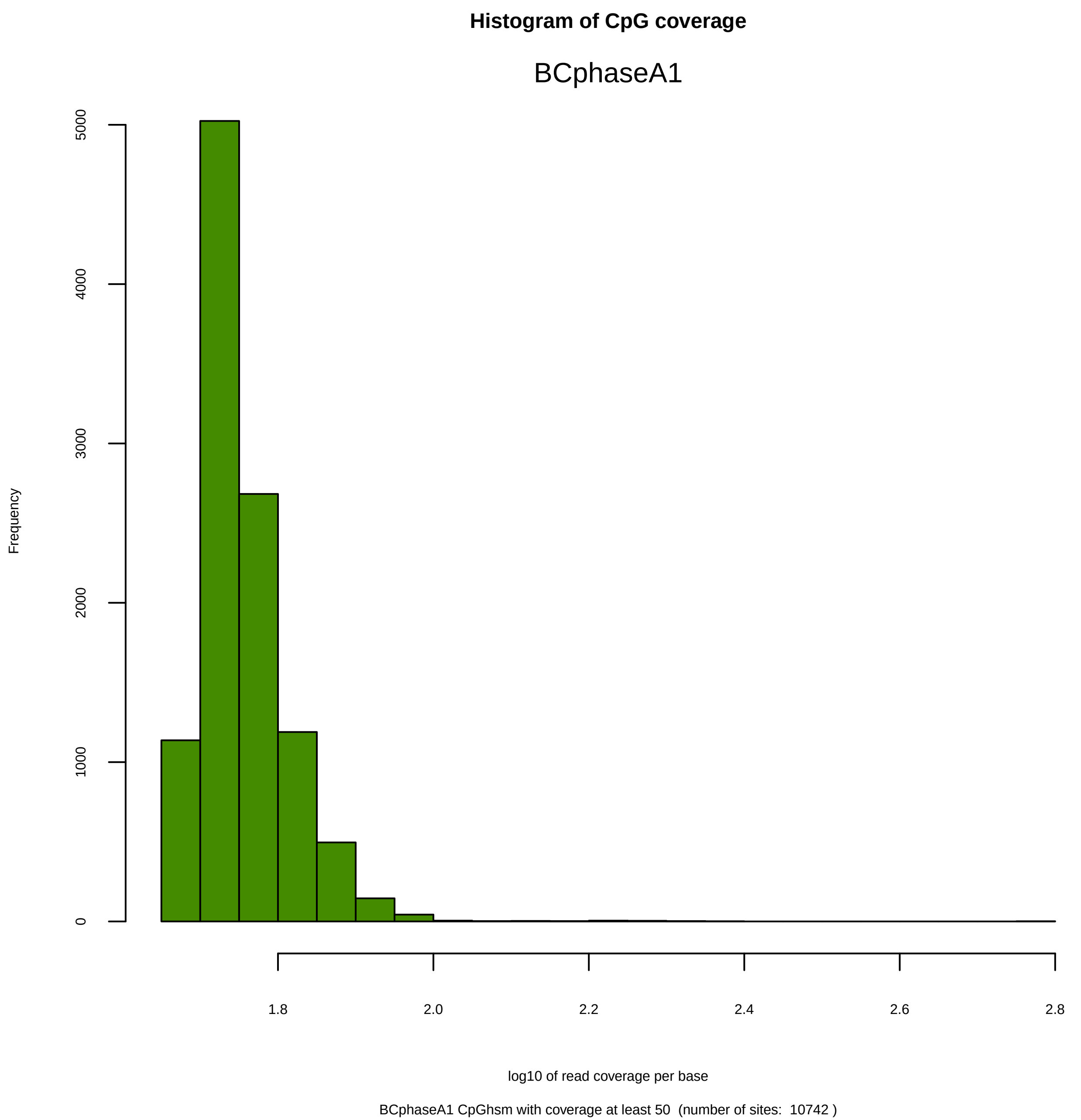

Histogram of CpG coverage
BCphaseA1
Frequency
0
1000
2000
3000
4000
5000
1.8
2.0
2.2
2.4
2.6
2.8
log10 of read coverage per base
BCphaseA1 CpGhsm with coverage at least 50 (number of sites: 10742 )

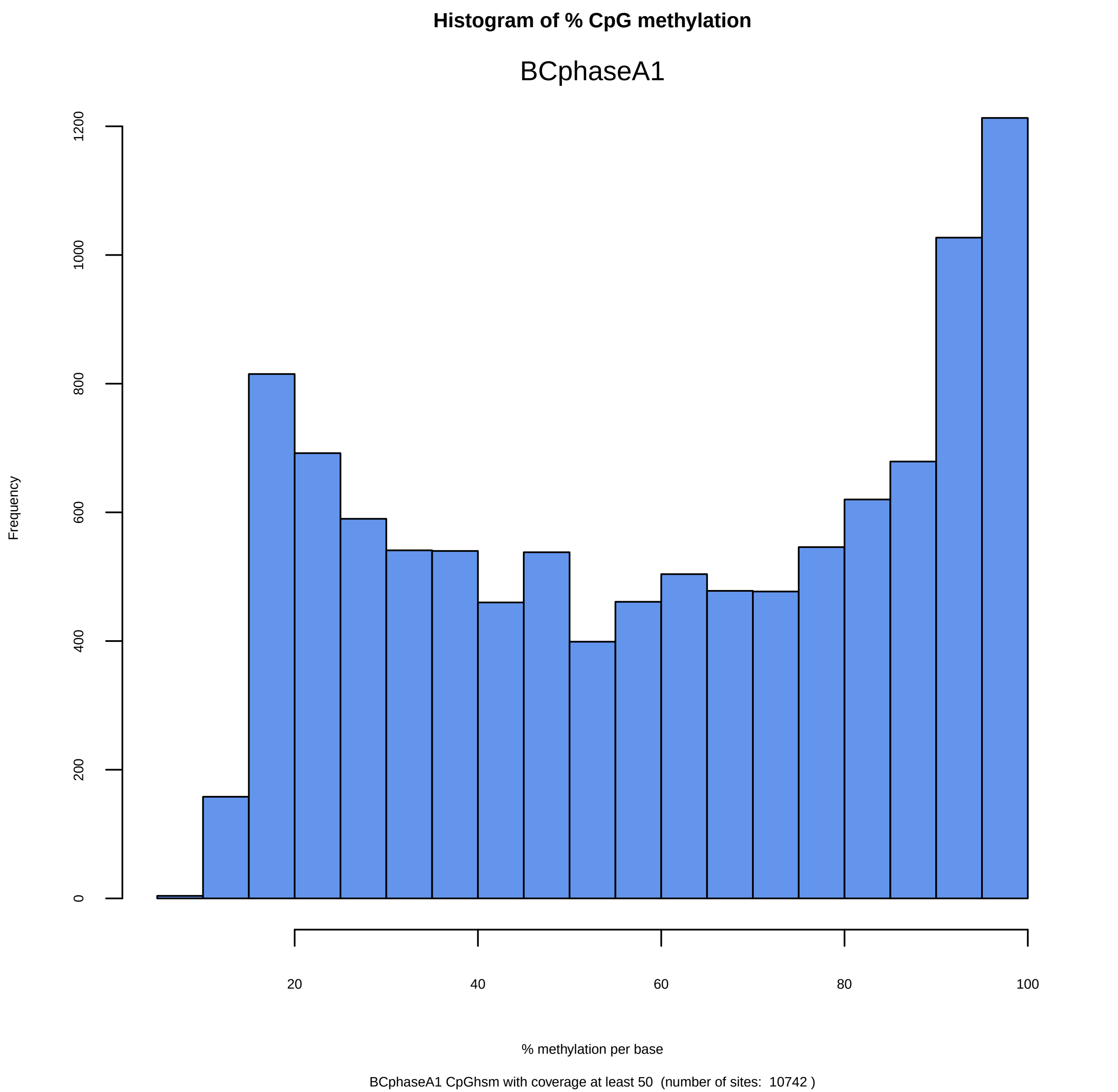

Histogram of % CpG methylation
BCphaseA1
Frequency
0
200
400
600
800
1000
1200
20
40
60
80
100
% methylation per base
BCphaseA1 CpGhsm with coverage at least 50  (number of sites:  10742 )

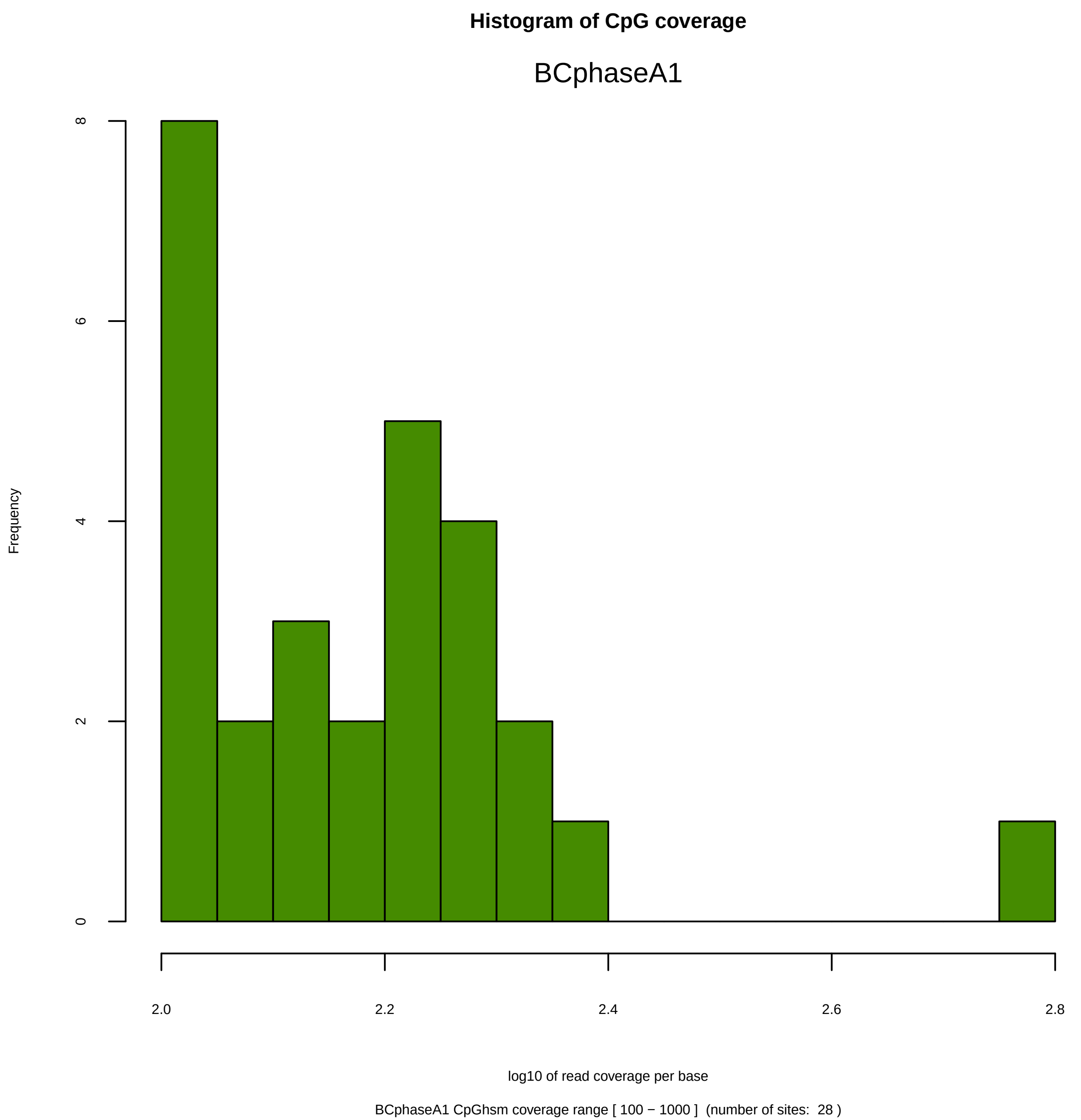
Histogram of CpG coverage
BCphaseA1
Frequency
0
2
4
6
8
2.0
2.2
2.4
2.6
2.8
log10 of read coverage per base
BCphaseA1 CpGhsm coverage range [ 100 − 1000 ]  (number of sites:  28 )

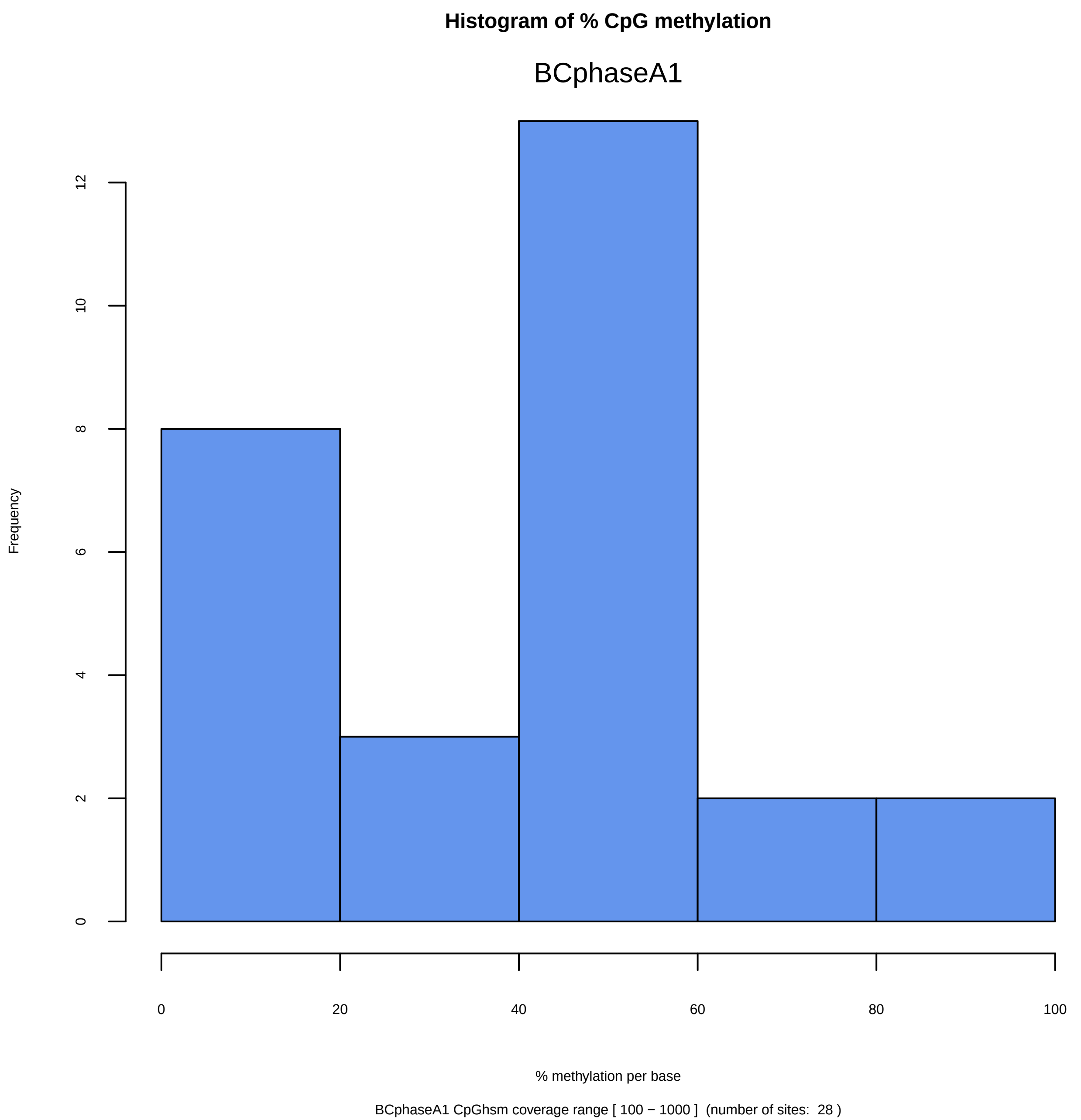
Histogram of % CpG methylation
BCphaseA1
Frequency
0
2
4
6
8
10
12
0
20
40
60
80
100
% methylation per base
BCphaseA1 CpGhsm coverage range [ 100 − 1000 ] (number of sites: 28 )

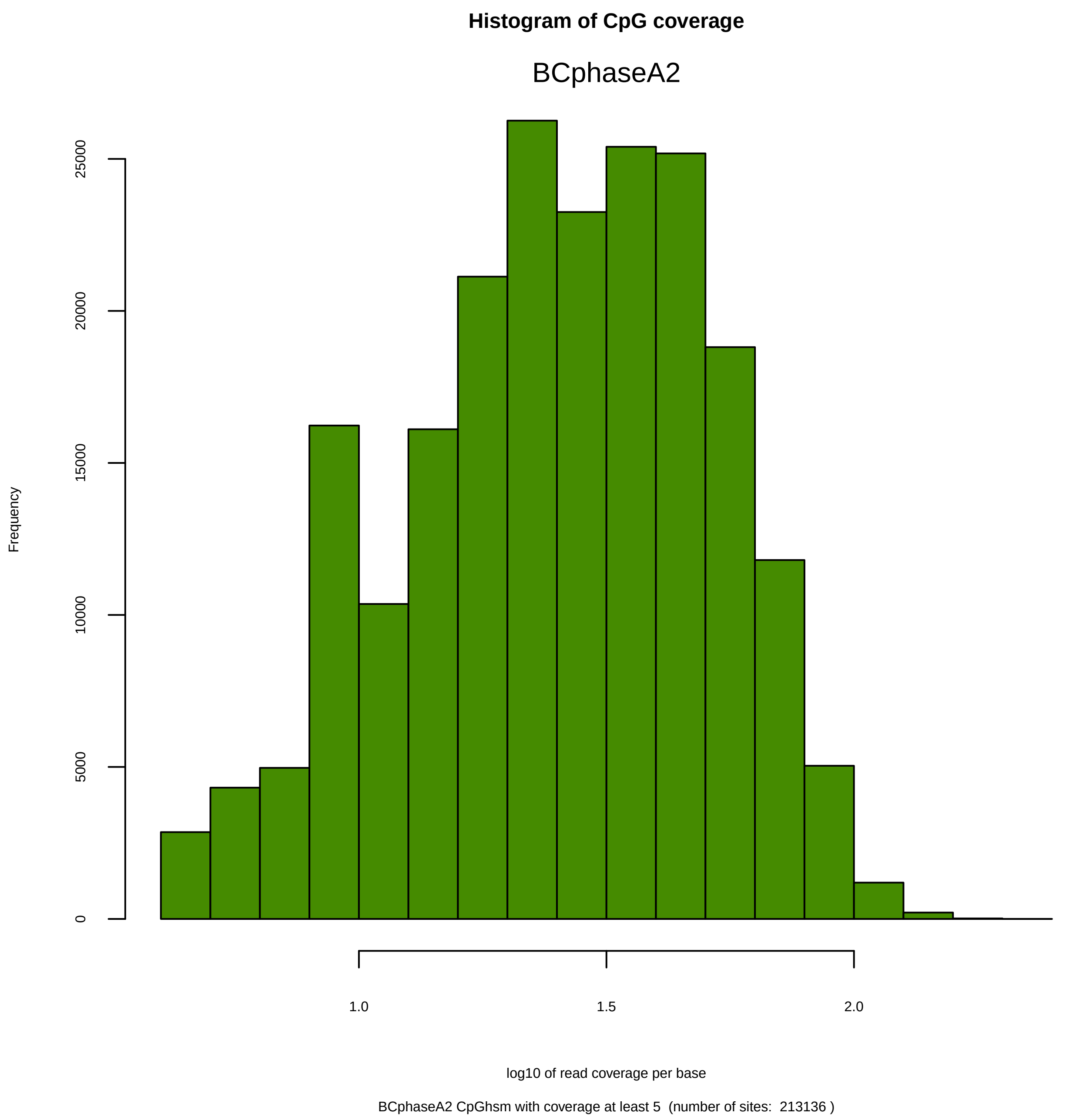

Histogram of CpG coverage
BCphaseA2
Frequency
0
5000
10000
15000
20000
25000
1.0
1.5
2.0
log10 of read coverage per base
BCphaseA2 CpGhsm with coverage at least 5  (number of sites:  213136 )

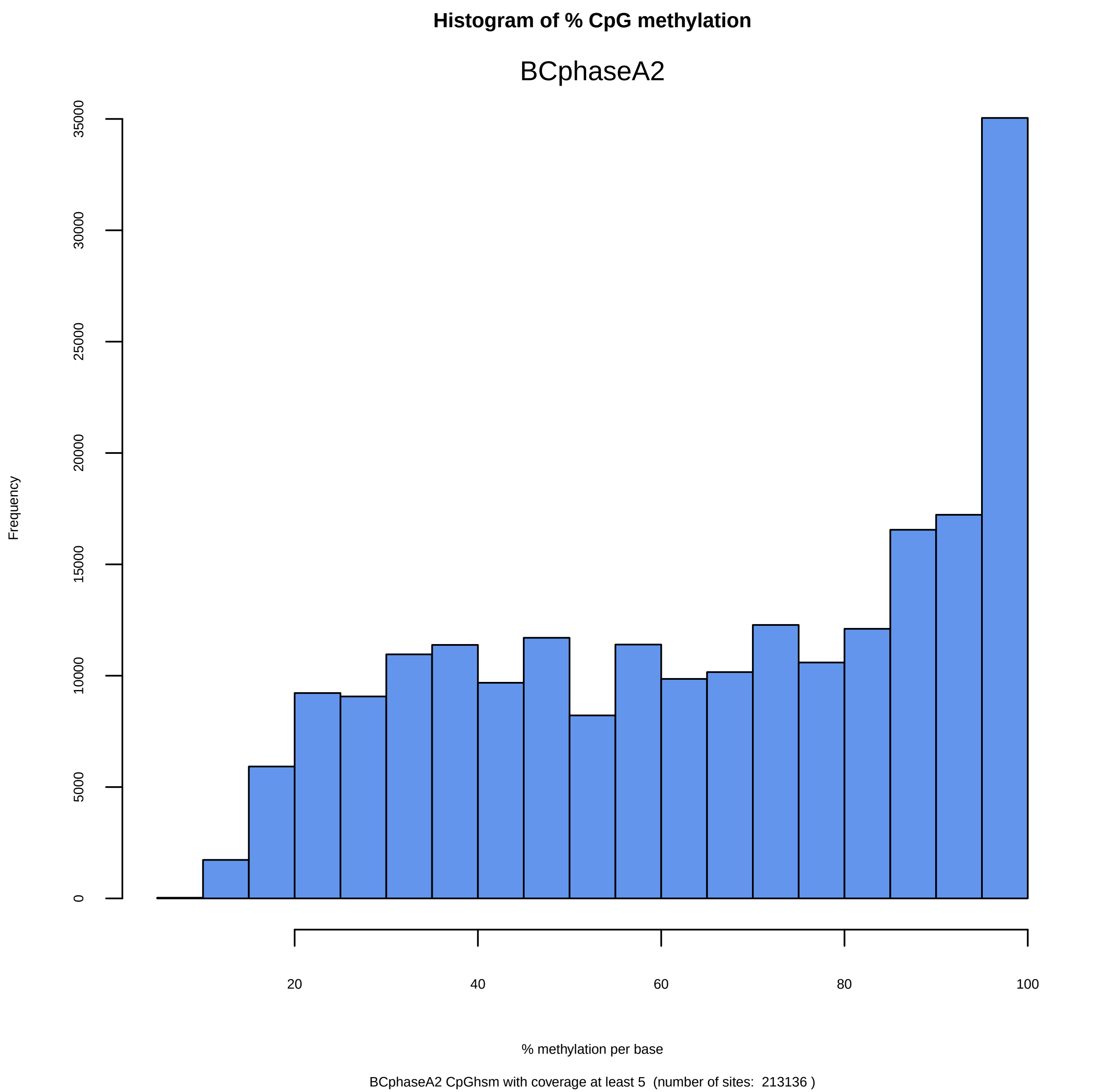

Histogram of % CpG methylation
BCphaseA2
Frequency
0
5000
10000
15000
20000
25000
30000
35000
20
40
60
80
100
% methylation per base
BCphaseA2 CpGhsm with coverage at least 5 (number of sites: 213136 )

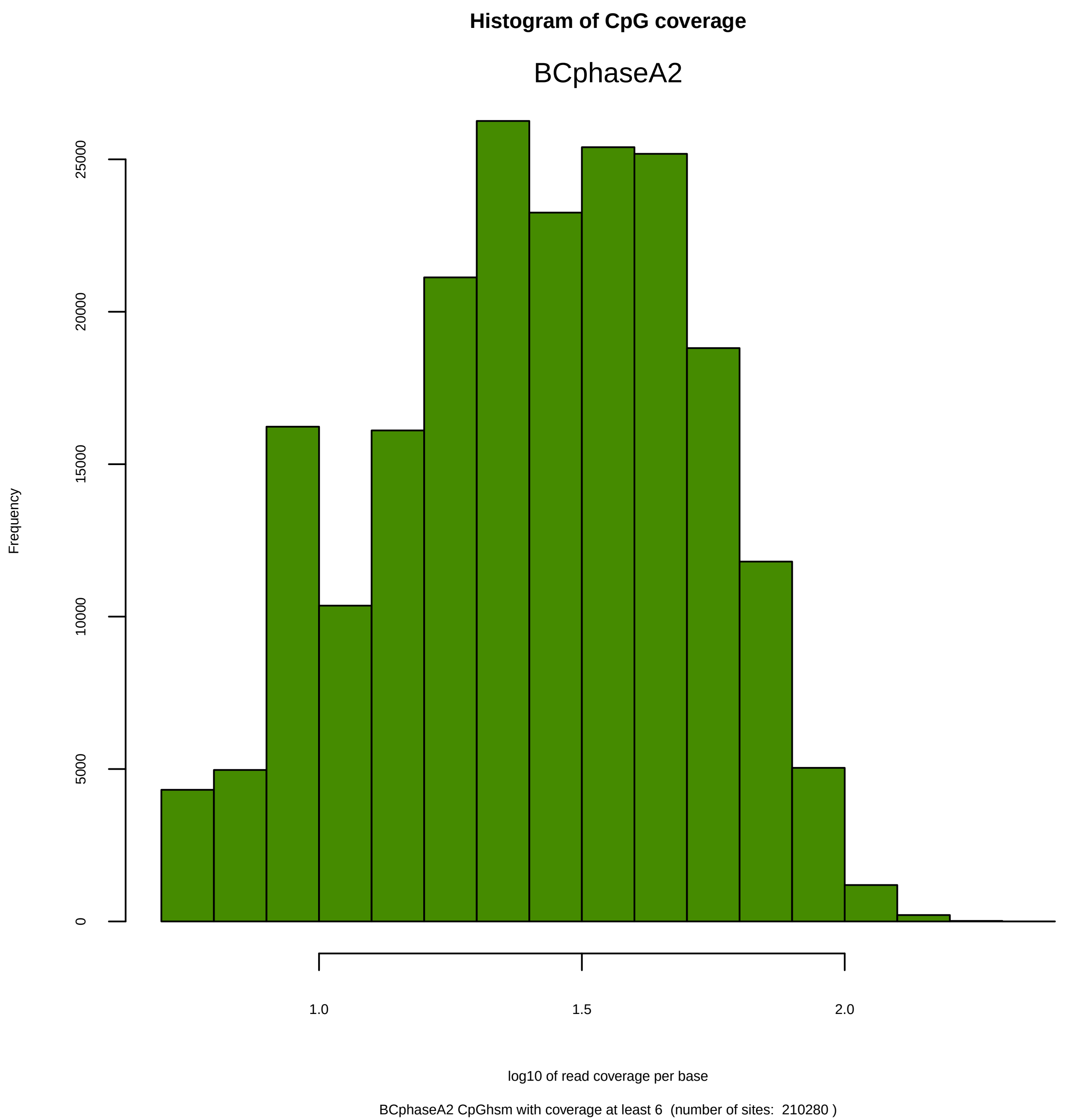
Histogram of CpG coverage
BCphaseA2
Frequency
0
5000
10000
15000
20000
25000
1.0
1.5
2.0
log10 of read coverage per base
BCphaseA2 CpGhsm with coverage at least 6 (number of sites: 210280 )

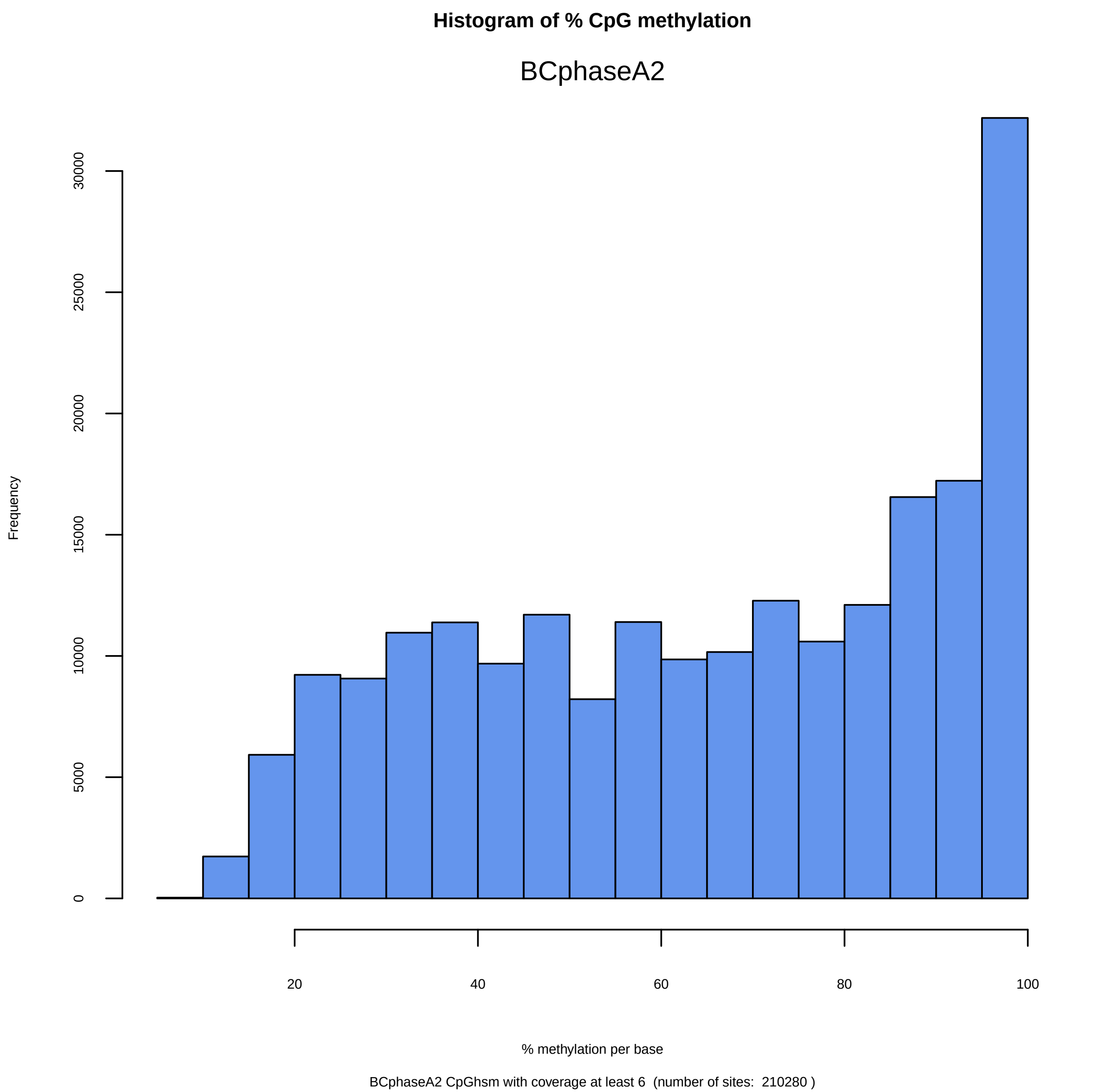
Histogram of % CpG methylation
BCphaseA2
Frequency
0
5000
10000
15000
20000
25000
30000
20
40
60
80
100
% methylation per base
BCphaseA2 CpGhsm with coverage at least 6 (number of sites: 210280 )

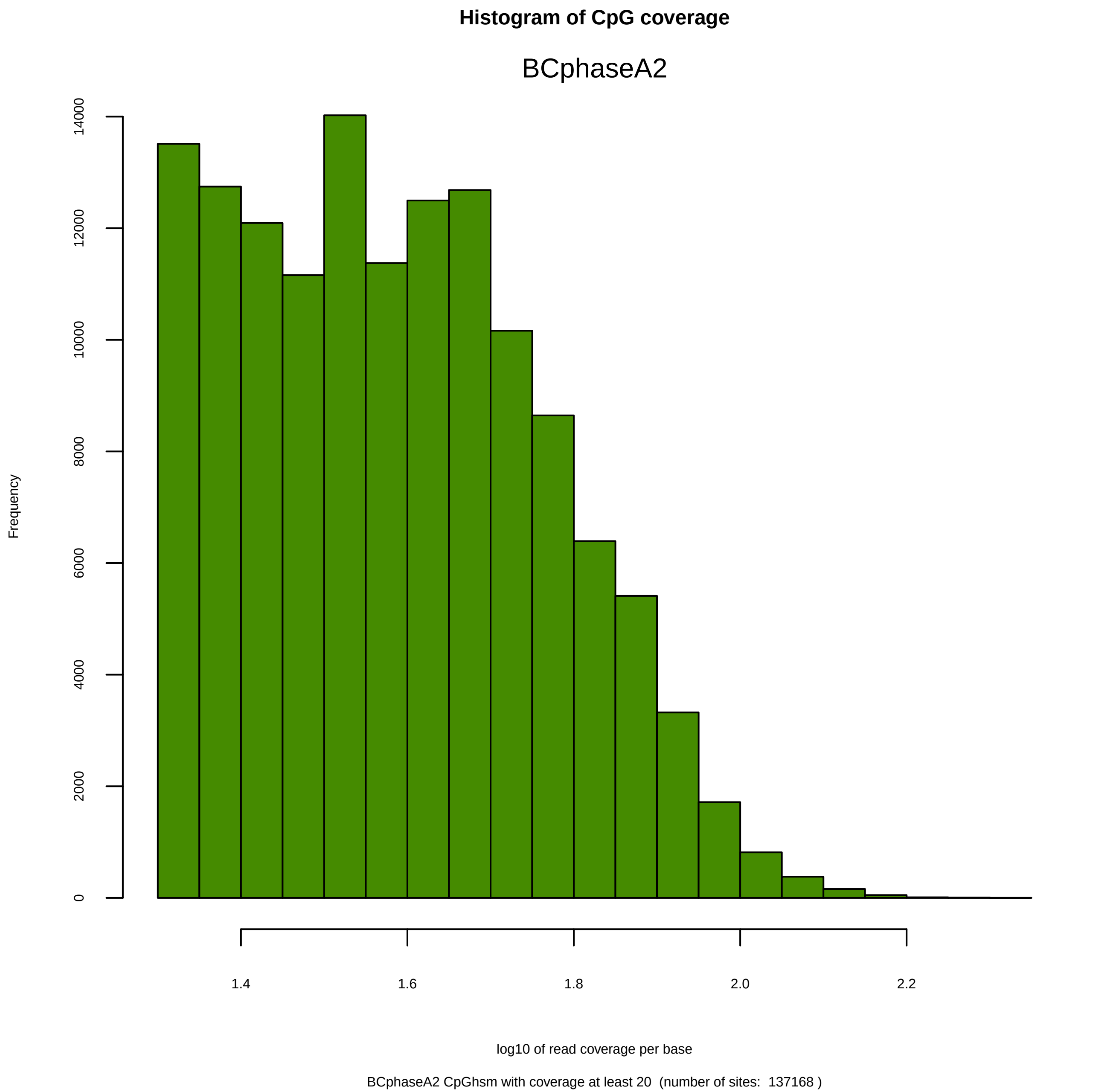

Histogram of CpG coverage
BCphaseA2
Frequency
0
2000
4000
6000
8000
10000
12000
14000
1.4
1.6
1.8
2.0
2.2
log10 of read coverage per base
BCphaseA2 CpGhsm with coverage at least 20  (number of sites:  137168 )

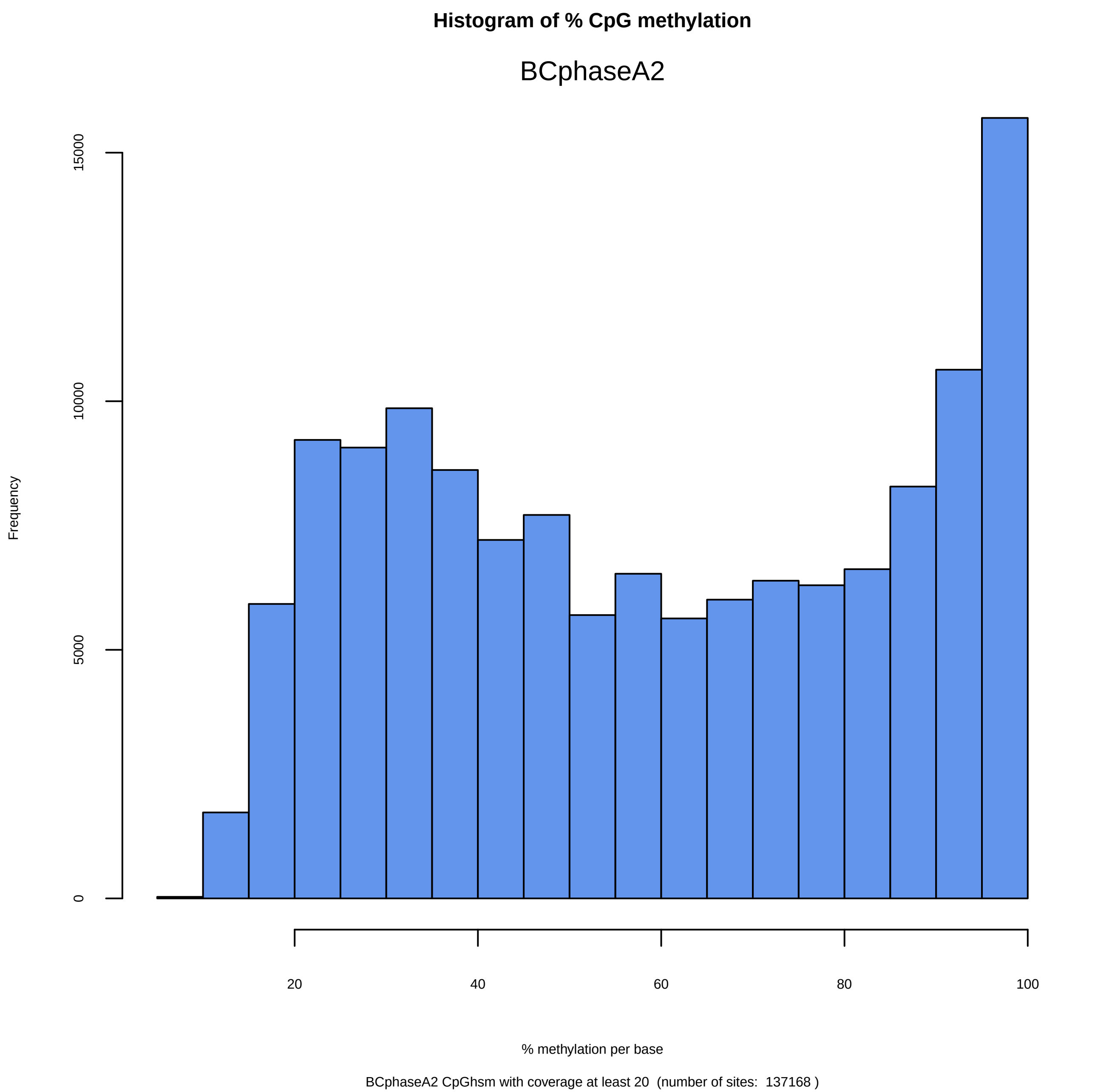

Histogram of % CpG methylation
BCphaseA2
Frequency
0
5000
10000
15000
20
40
60
80
100
% methylation per base
BCphaseA2 CpGhsm with coverage at least 20 (number of sites: 137168 )

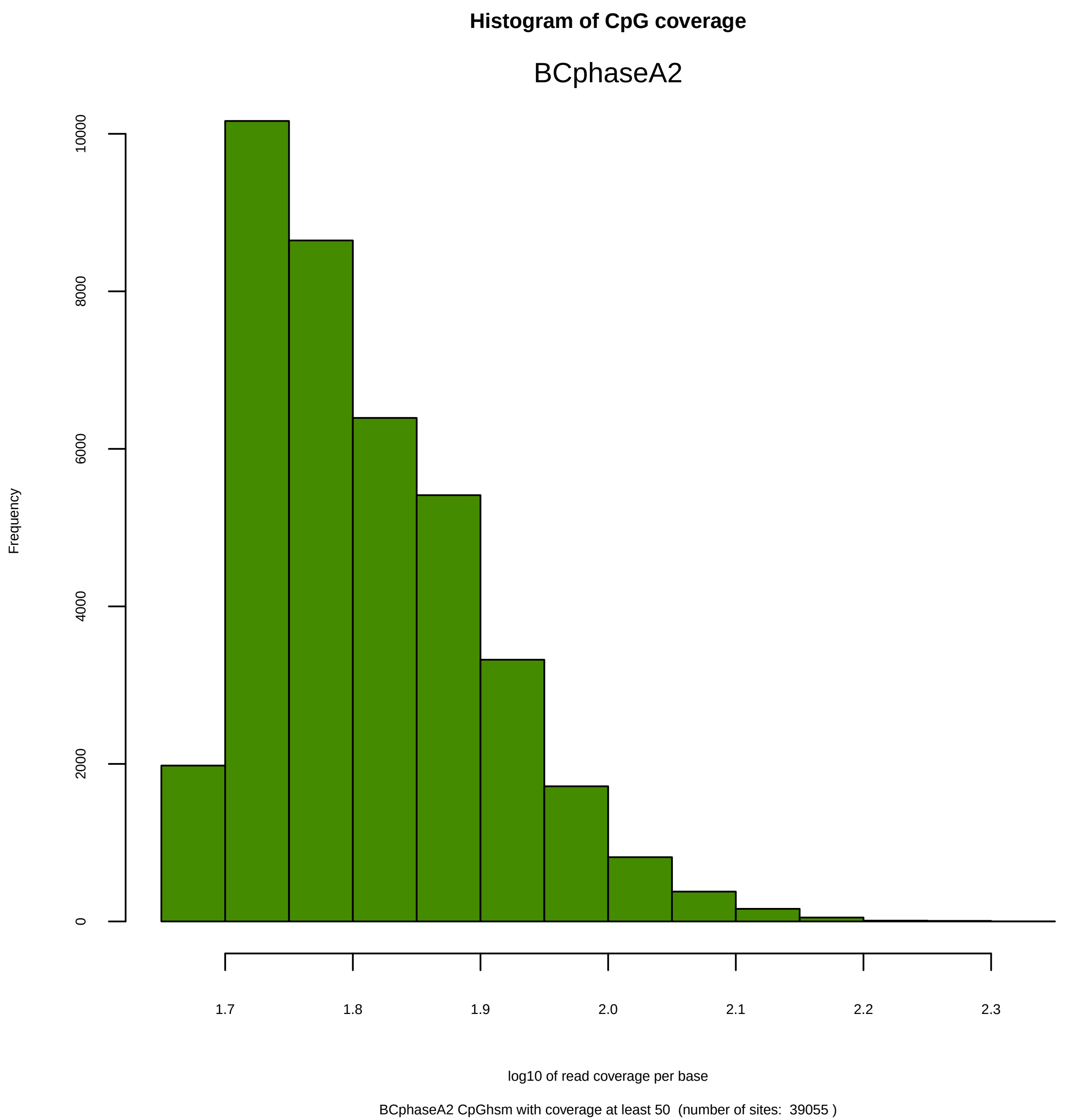
Histogram of CpG coverage
BCphaseA2
Frequency
0
2000
4000
6000
8000
10000
1.7
1.8
1.9
2.0
2.1
2.2
2.3
log10 of read coverage per base
BCphaseA2 CpGhsm with coverage at least 50 (number of sites: 39055 )

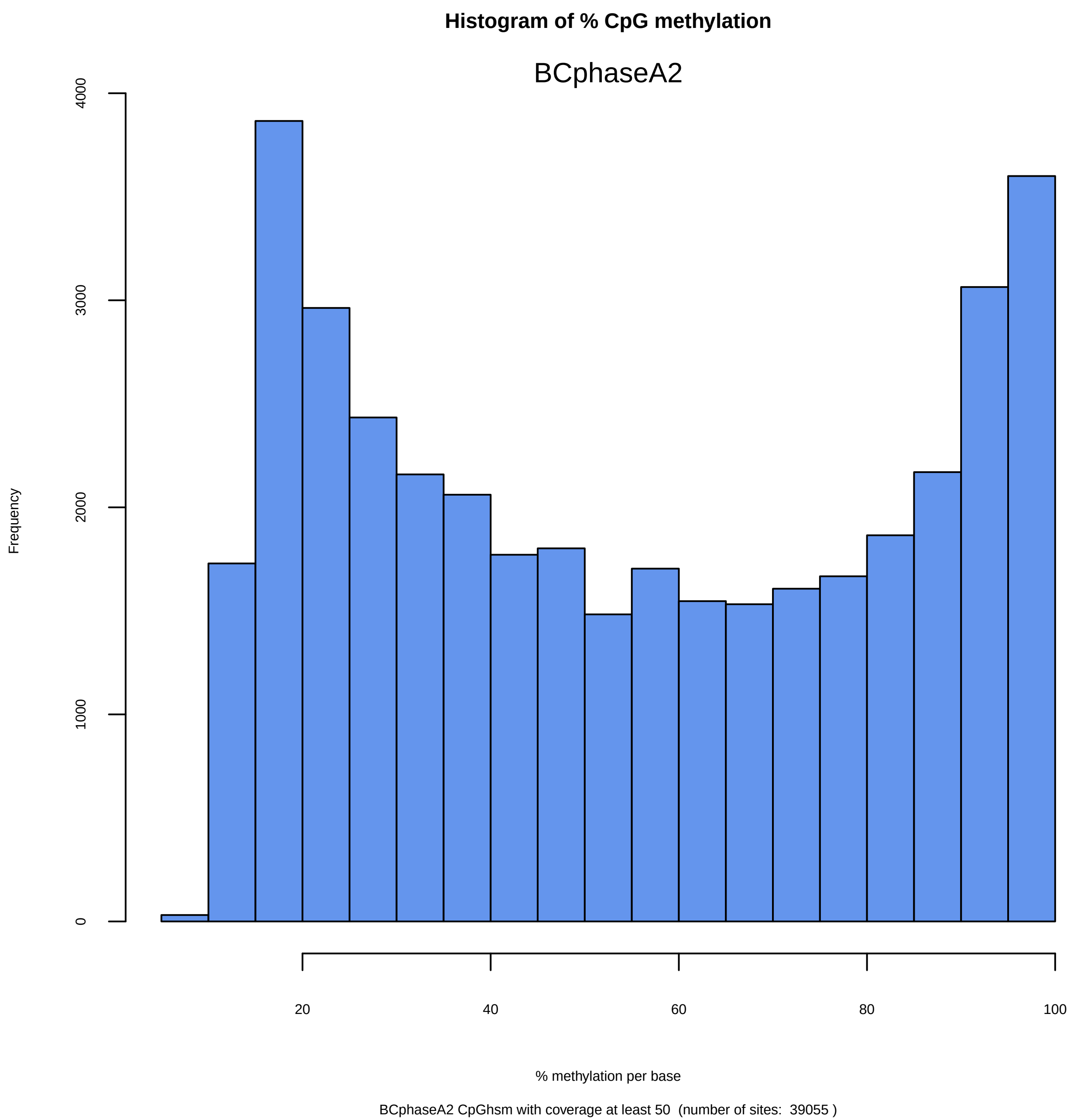
Histogram of % CpG methylation
BCphaseA2
Frequency
0
1000
2000
3000
4000
20
40
60
80
100
% methylation per base
BCphaseA2 CpGhsm with coverage at least 50  (number of sites:  39055 )

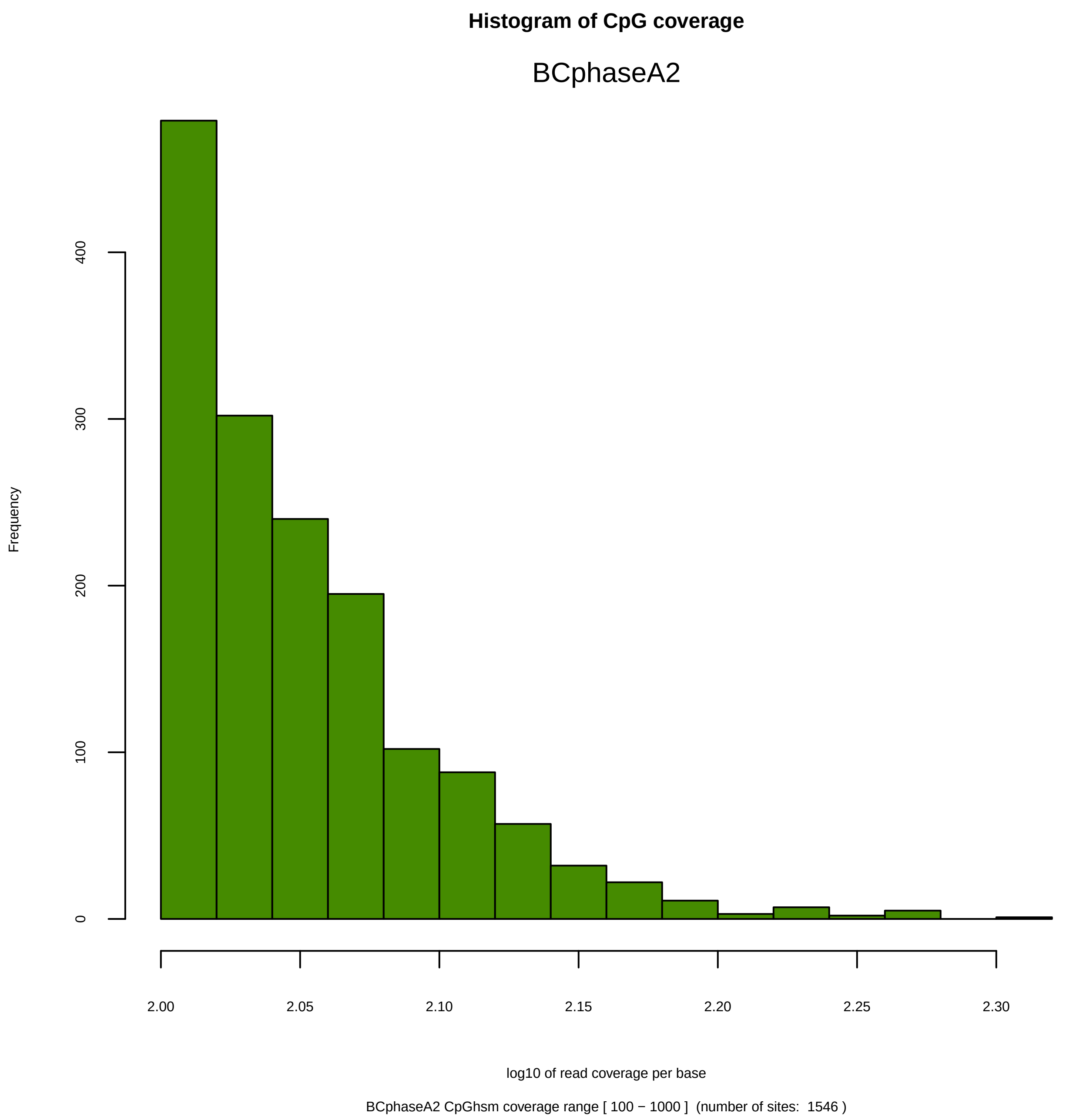
Histogram of CpG coverage
BCphaseA2
Frequency
0
100
200
300
400
2.00
2.05
2.10
2.15
2.20
2.25
2.30
log10 of read coverage per base
BCphaseA2 CpGhsm coverage range [ 100 − 1000 ]  (number of sites:  1546 )

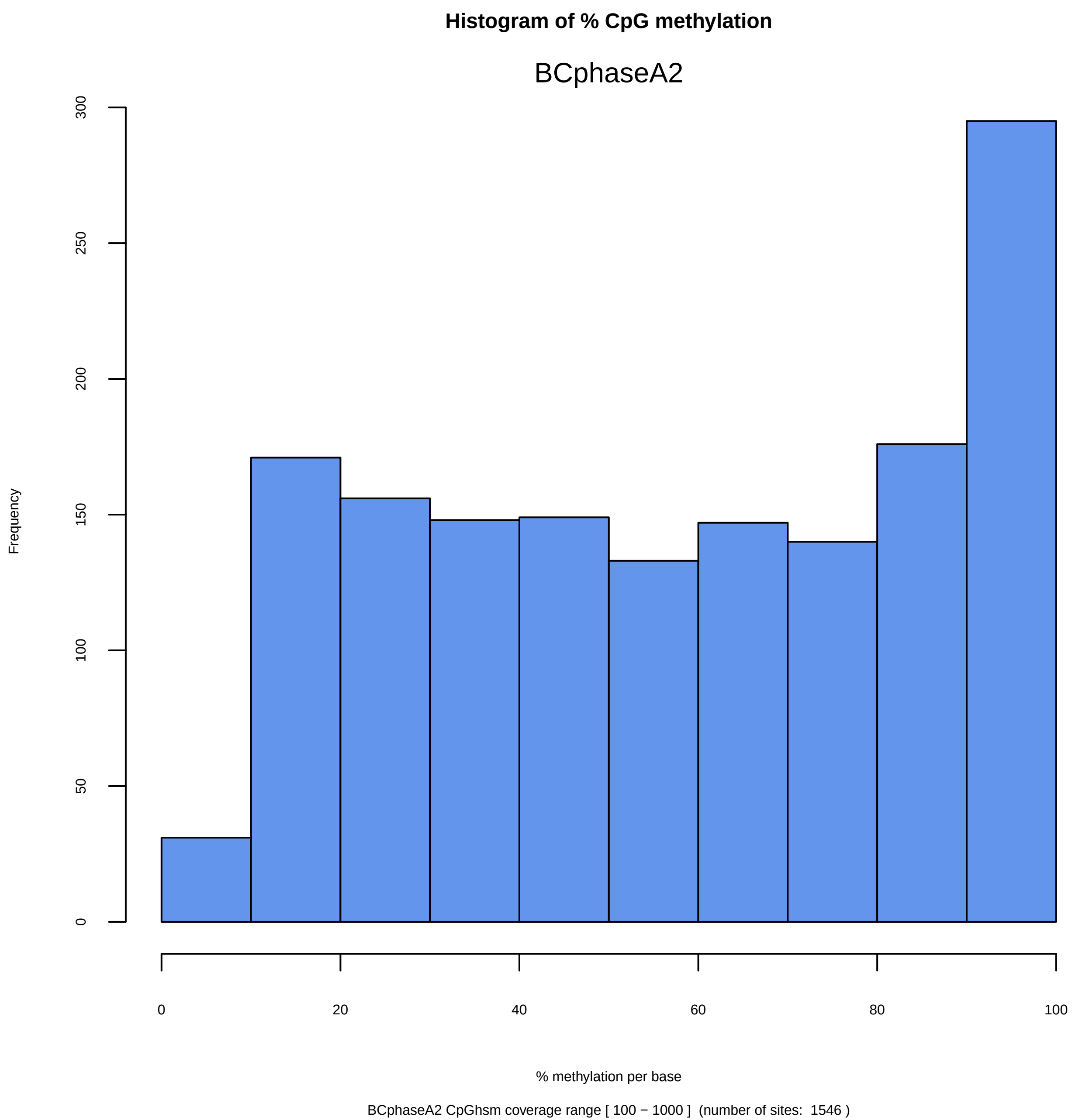
Histogram of % CpG methylation
BCphaseA2
Frequency
0
50
100
150
200
250
300
0
20
40
60
80
100
% methylation per base
BCphaseA2 CpGhsm coverage range [ 100 − 1000 ]  (number of sites:  1546 )

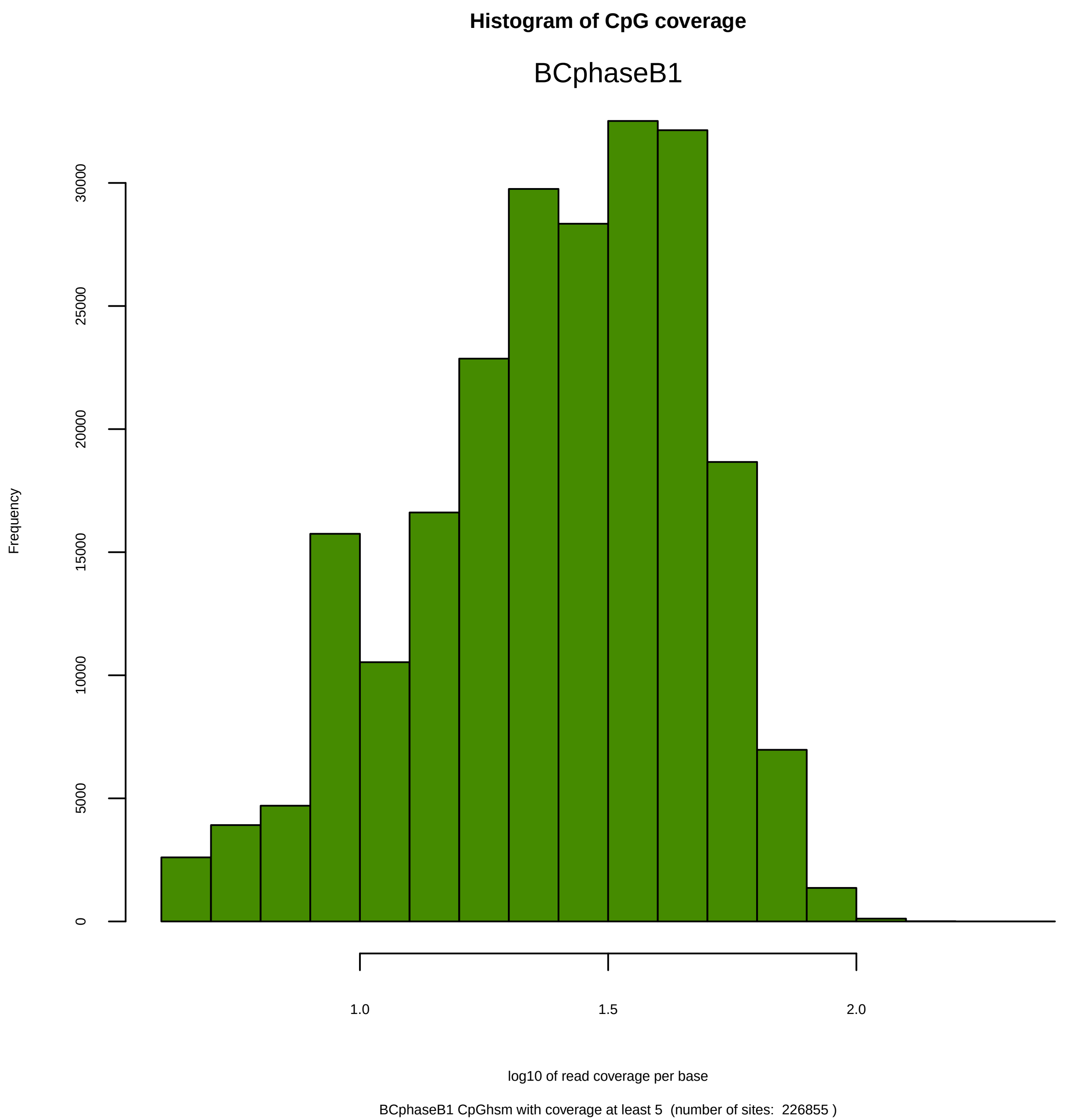

Histogram of CpG coverage
BCphaseB1
Frequency
0
5000
10000
15000
20000
25000
30000
1.0
1.5
2.0
log10 of read coverage per base
BCphaseB1 CpGhsm with coverage at least 5 (number of sites: 226855 )

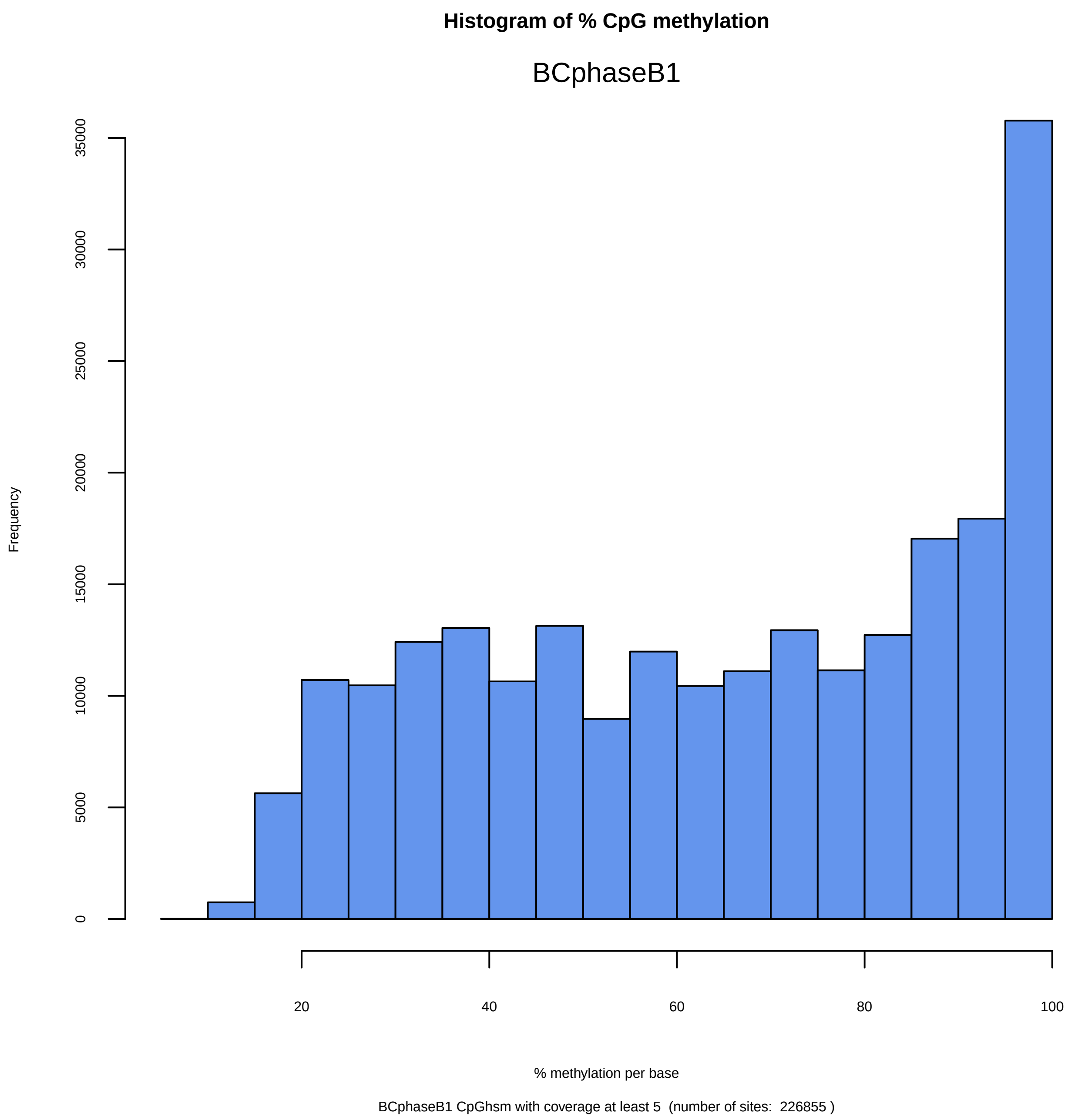

Histogram of % CpG methylation
BCphaseB1
Frequency
0
5000
10000
15000
20000
25000
30000
35000
20
40
60
80
100
% methylation per base
BCphaseB1 CpGhsm with coverage at least 5 (number of sites: 226855 )

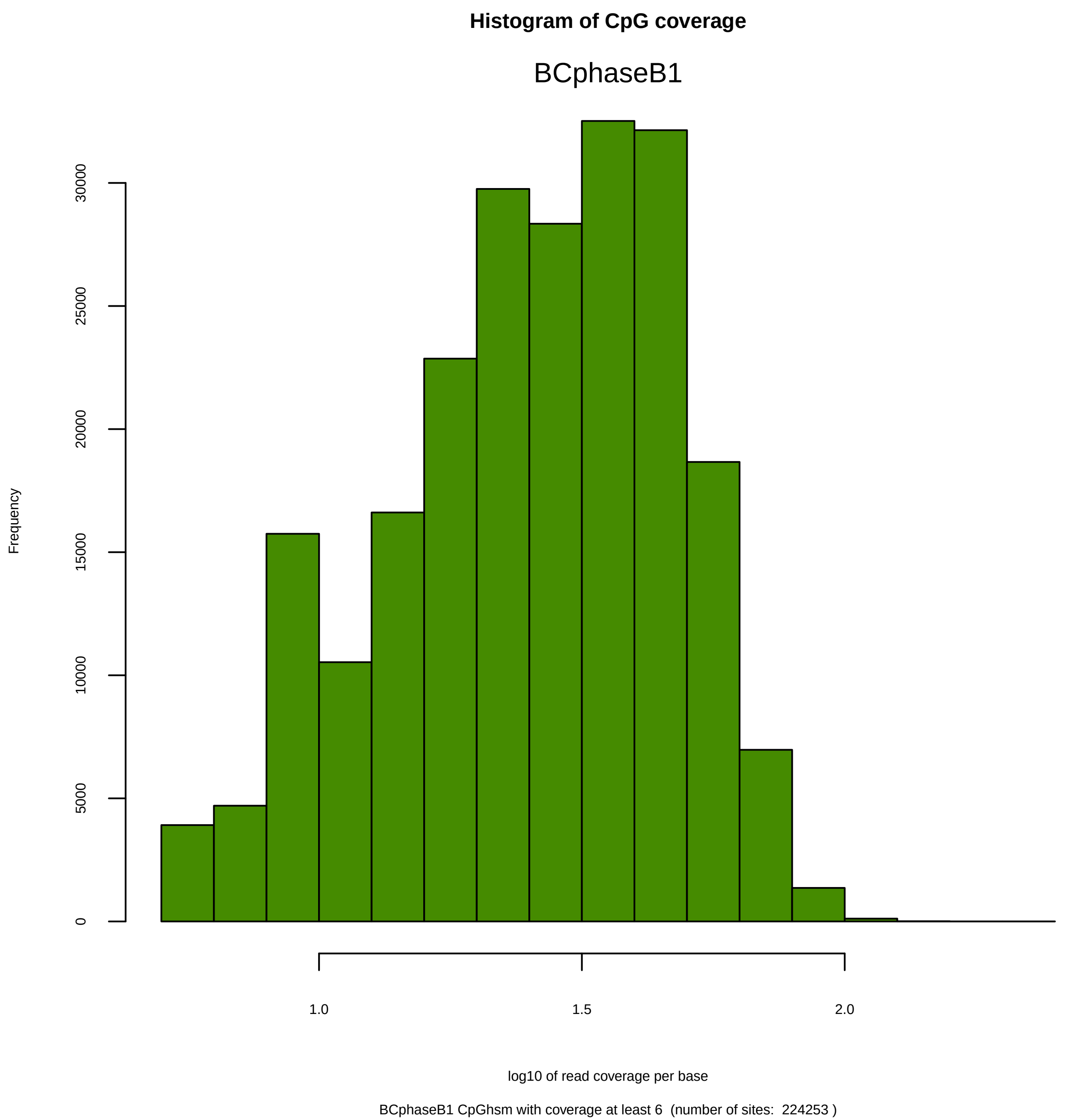

Histogram of CpG coverage
BCphaseB1
Frequency
0
5000
10000
15000
20000
25000
30000
1.0
1.5
2.0
log10 of read coverage per base
BCphaseB1 CpGhsm with coverage at least 6  (number of sites:  224253 )

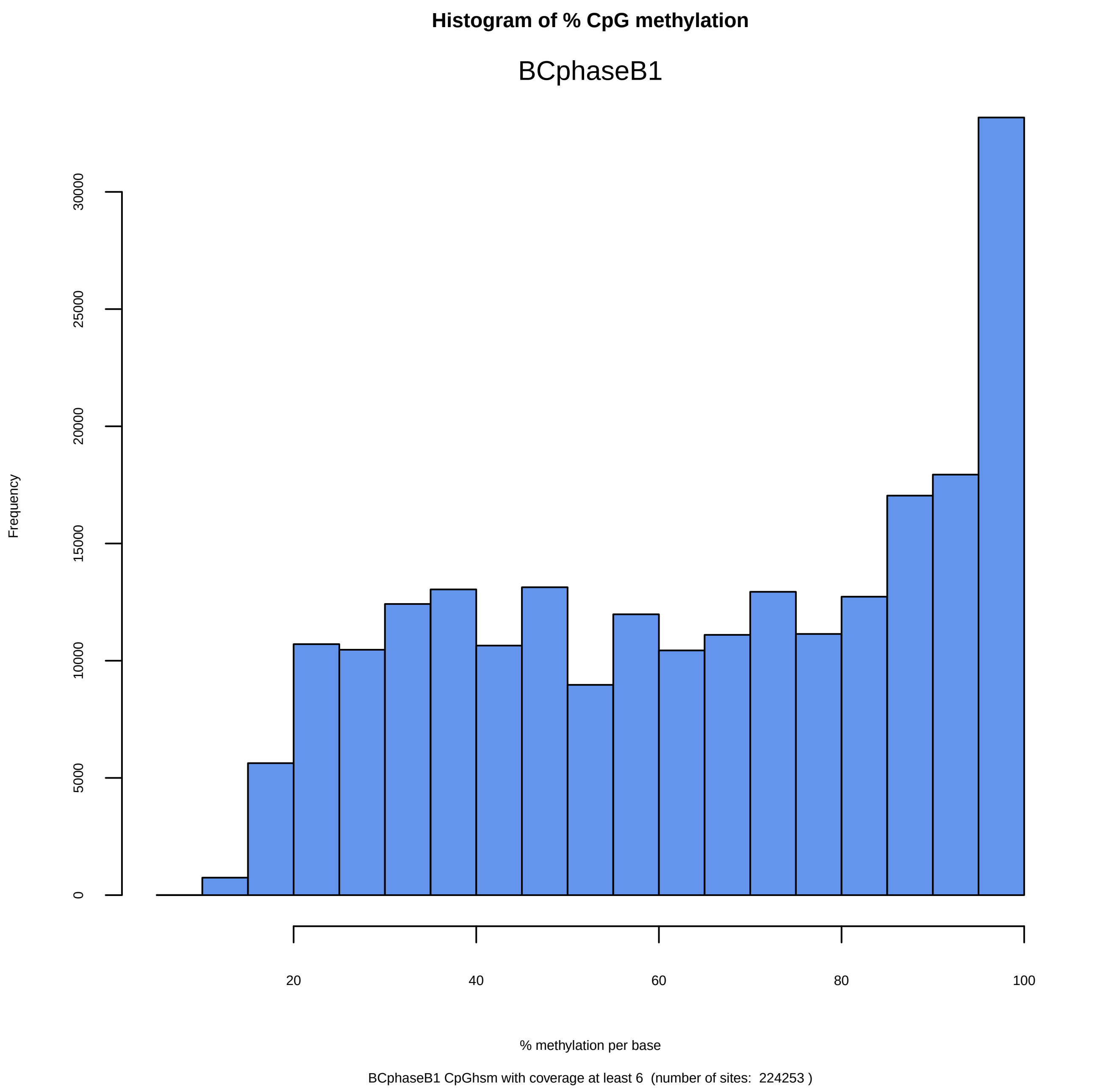
Histogram of % CpG methylation
BCphaseB1
Frequency
0
5000
10000
15000
20000
25000
30000
20
40
60
80
100
% methylation per base
BCphaseB1 CpGhsm with coverage at least 6 (number of sites: 224253 )

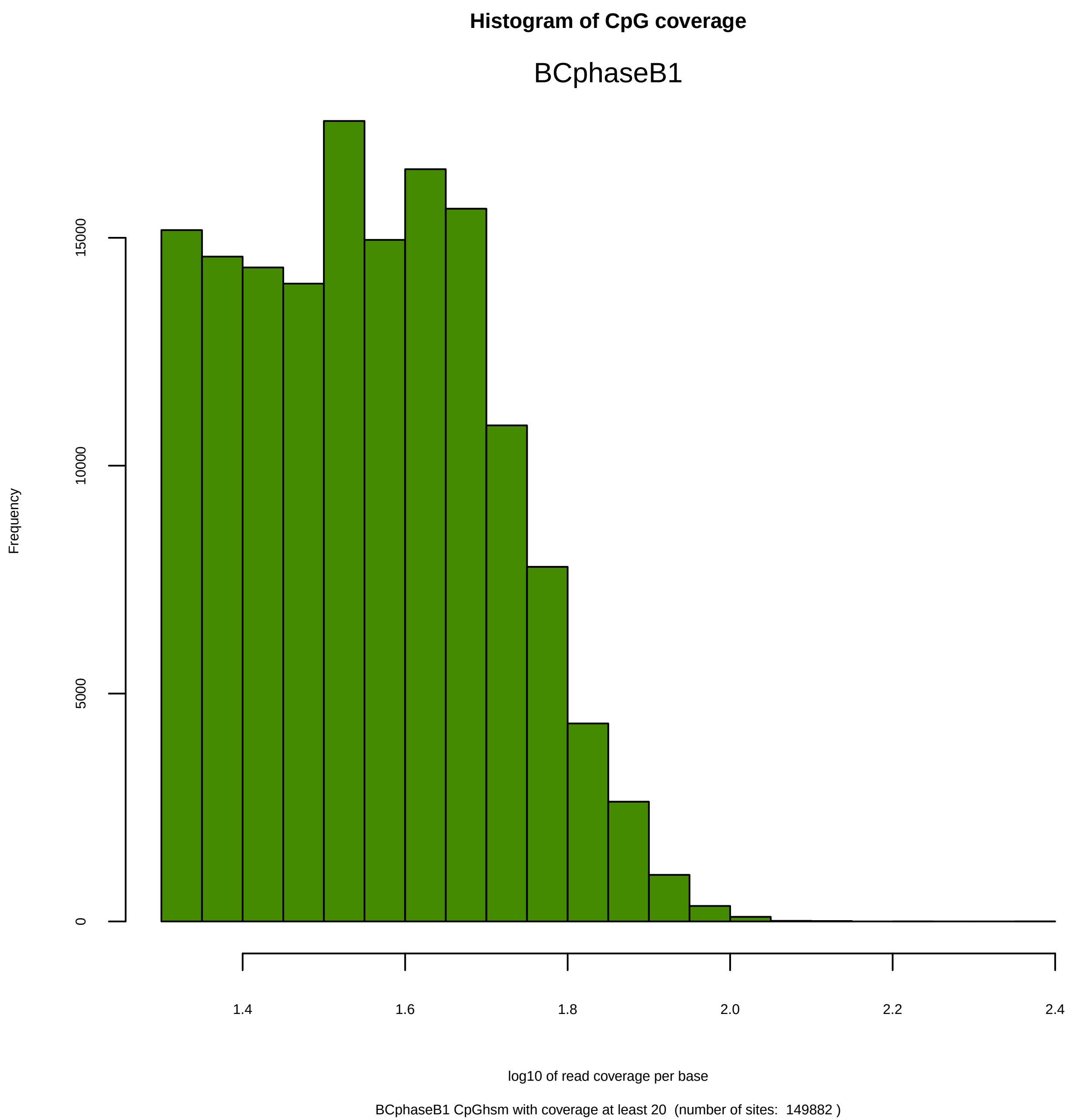
Histogram of CpG coverage
BCphaseB1
Frequency
0
5000
10000
15000
1.4
1.6
1.8
2.0
2.2
2.4
log10 of read coverage per base
BCphaseB1 CpGhsm with coverage at least 20 (number of sites: 149882 )

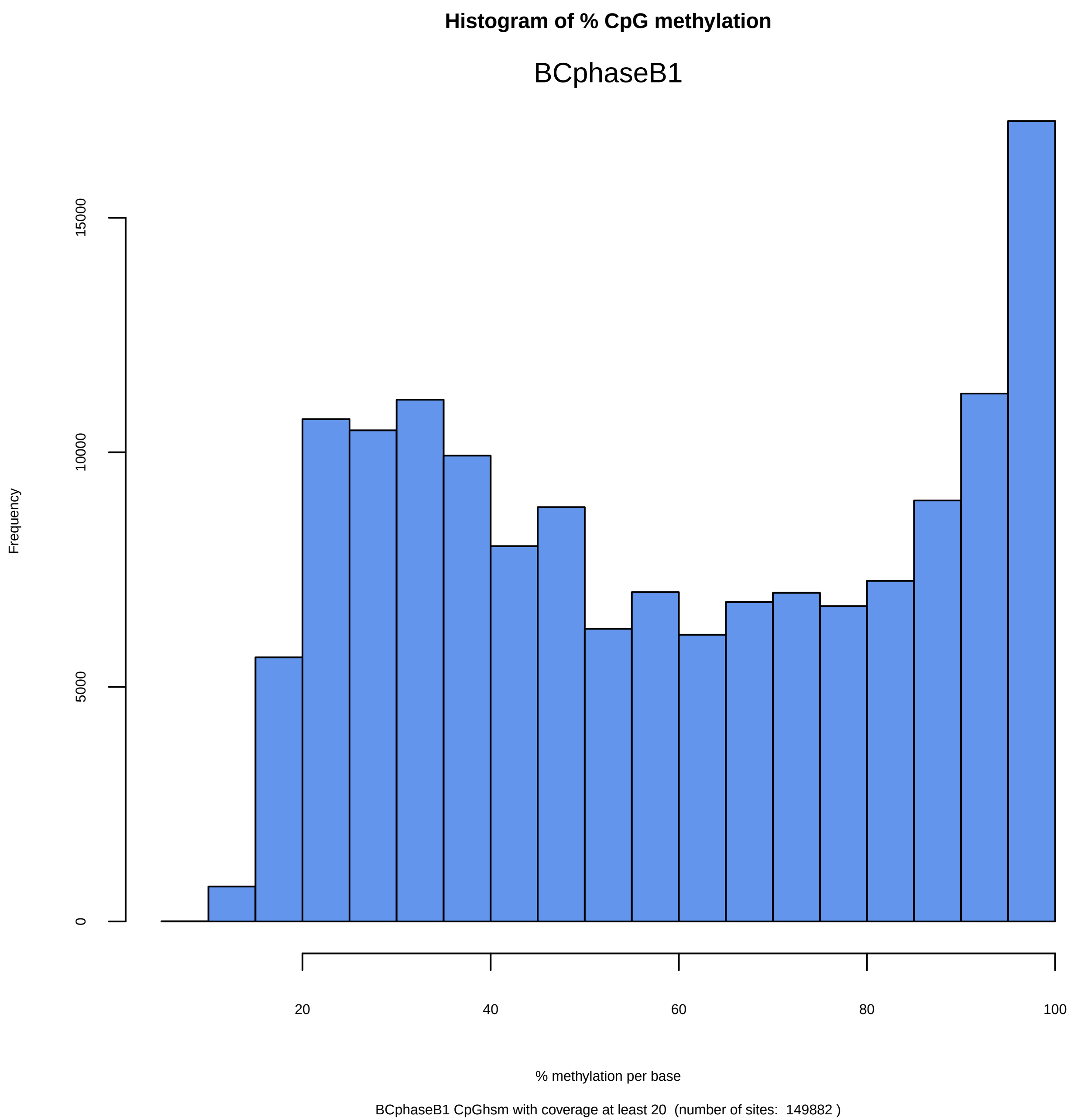
Histogram of % CpG methylation
BCphaseB1
Frequency
0
5000
10000
15000
20
40
60
80
100
% methylation per base
BCphaseB1 CpGhsm with coverage at least 20  (number of sites:  149882 )

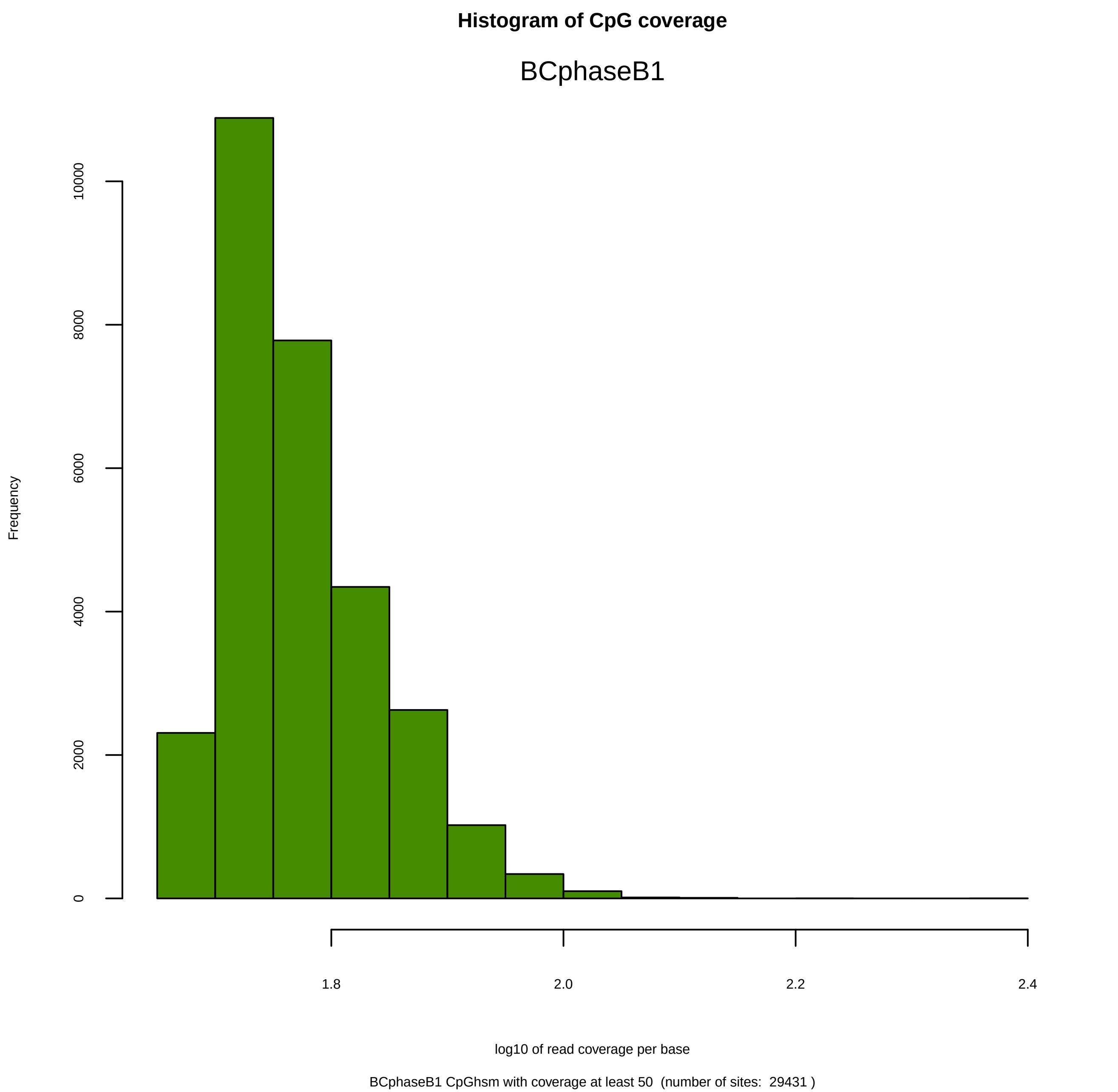

Histogram of CpG coverage
BCphaseB1
Frequency
0
2000
4000
6000
8000
10000
1.8
2.0
2.2
2.4
log10 of read coverage per base
BCphaseB1 CpGhsm with coverage at least 50  (number of sites:  29431 )

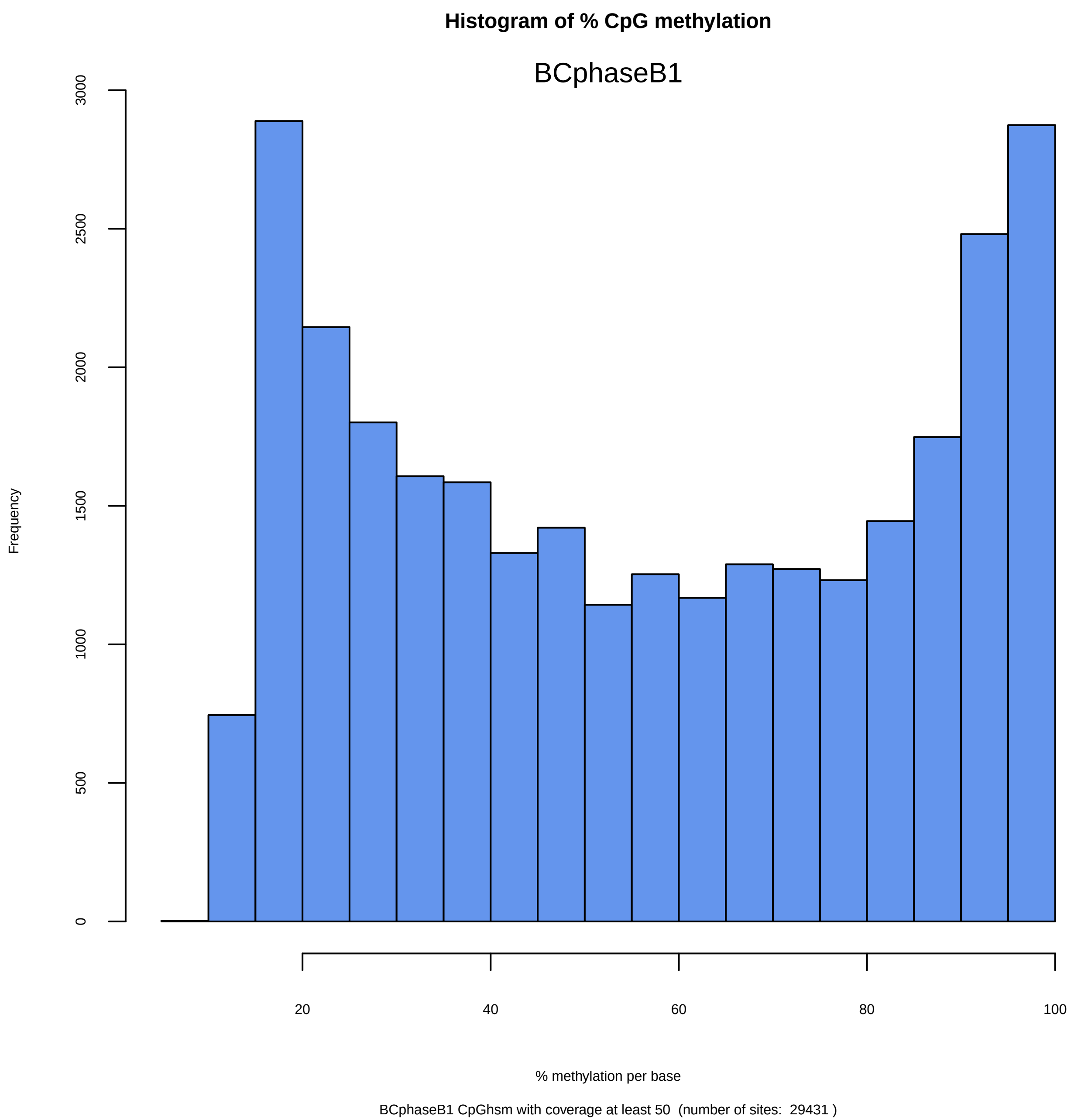
Histogram of % CpG methylation
BCphaseB1
Frequency
0
500
1000
1500
2000
2500
3000
20
40
60
80
100
% methylation per base
BCphaseB1 CpGhsm with coverage at least 50 (number of sites: 29431 )

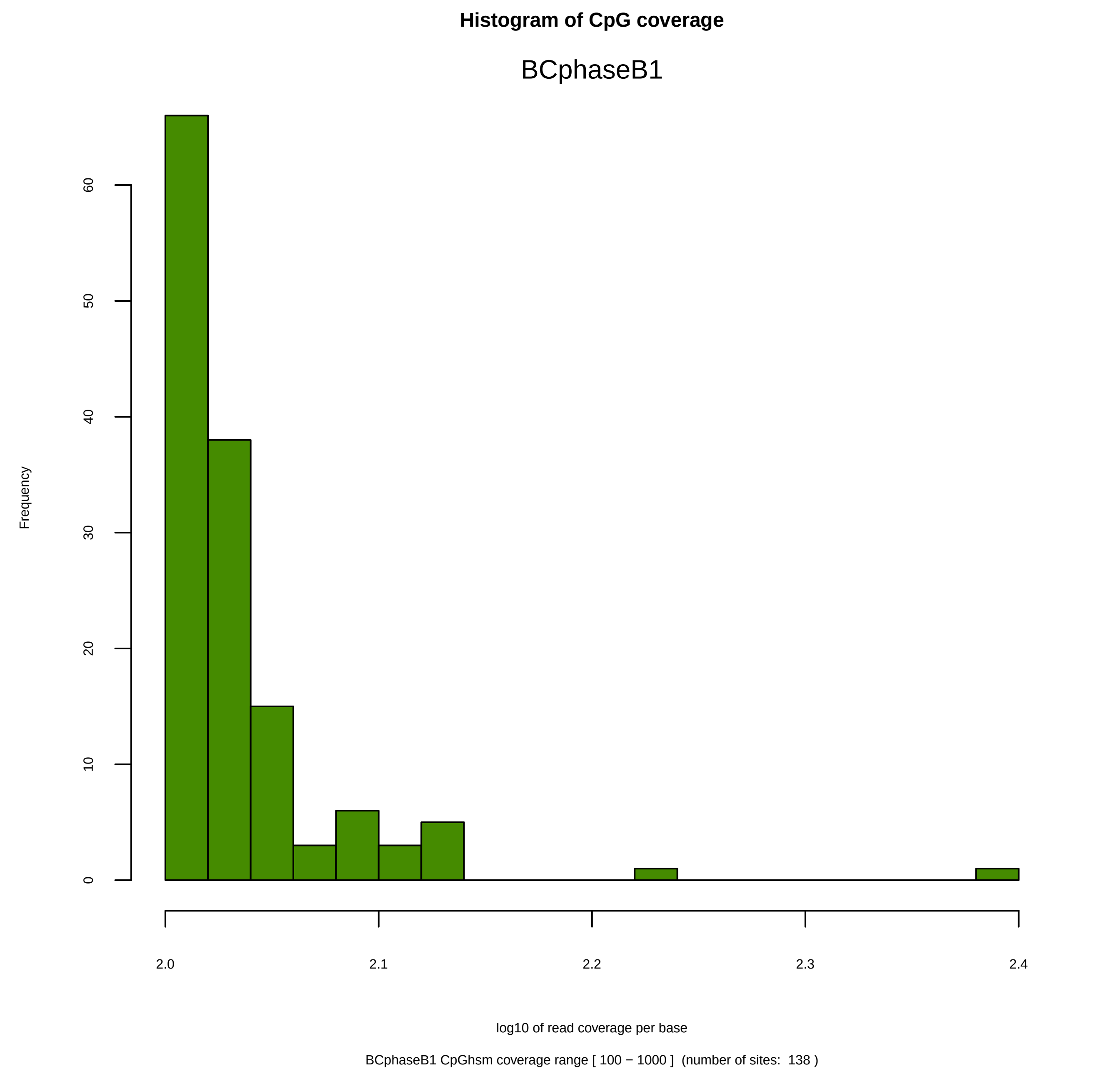
Histogram of CpG coverage
BCphaseB1
Frequency
0
10
20
30
40
50
60
2.0
2.1
2.2
2.3
2.4
log10 of read coverage per base
BCphaseB1 CpGhsm coverage range [ 100 − 1000 ]  (number of sites:  138 )

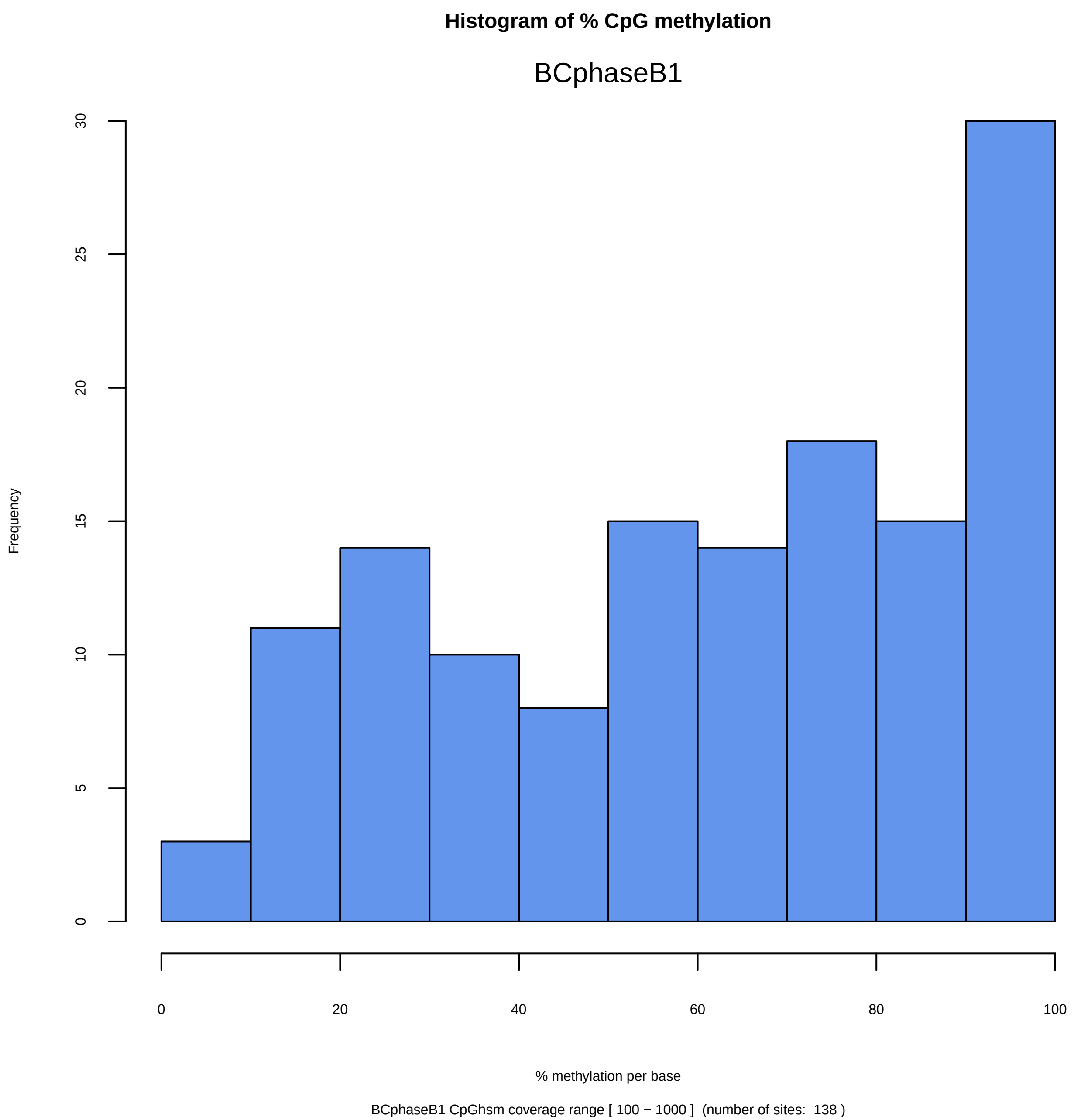
Histogram of % CpG methylation
BCphaseB1
Frequency
0
5
10
15
20
25
30
0
20
40
60
80
100
% methylation per base
BCphaseB1 CpGhsm coverage range [ 100 − 1000 ] (number of sites: 138 )

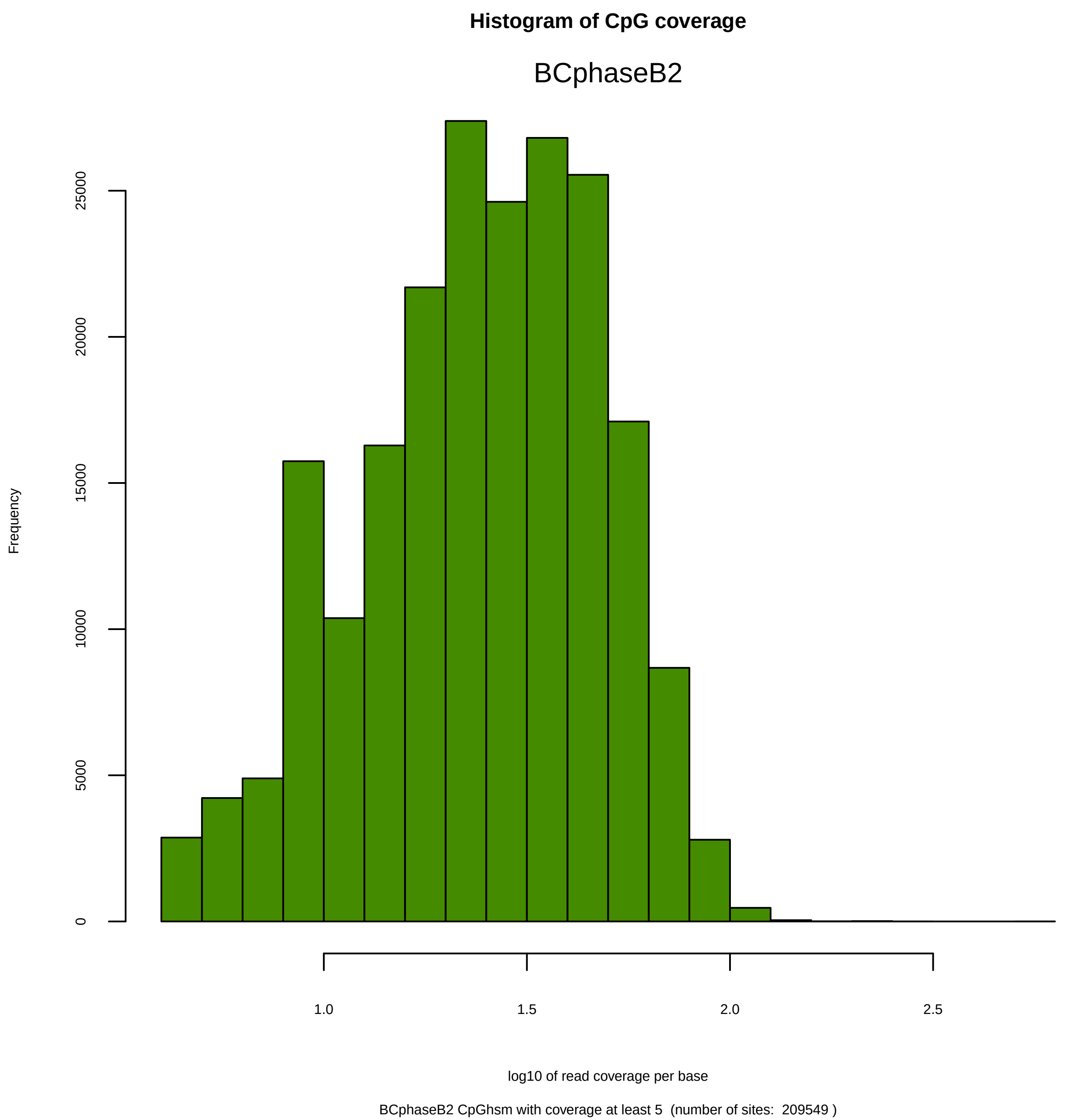
Histogram of CpG coverage
BCphaseB2
Frequency
0
5000
10000
15000
20000
25000
1.0
1.5
2.0
2.5
log10 of read coverage per base
BCphaseB2 CpGhsm with coverage at least 5  (number of sites:  209549 )

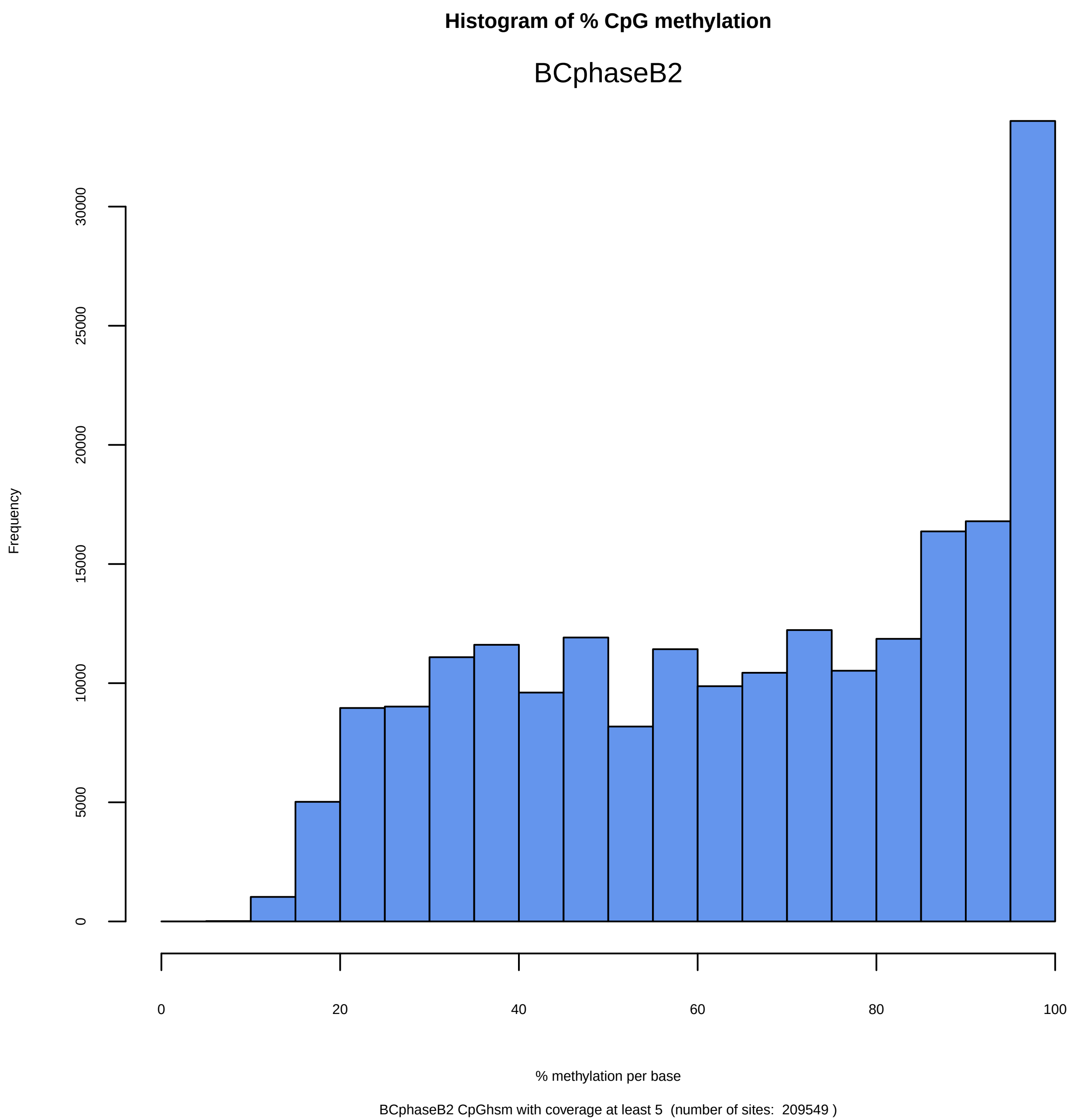

Histogram of % CpG methylation
BCphaseB2
Frequency
0
5000
10000
15000
20000
25000
30000
0
20
40
60
80
100
% methylation per base
BCphaseB2 CpGhsm with coverage at least 5 (number of sites: 209549 )

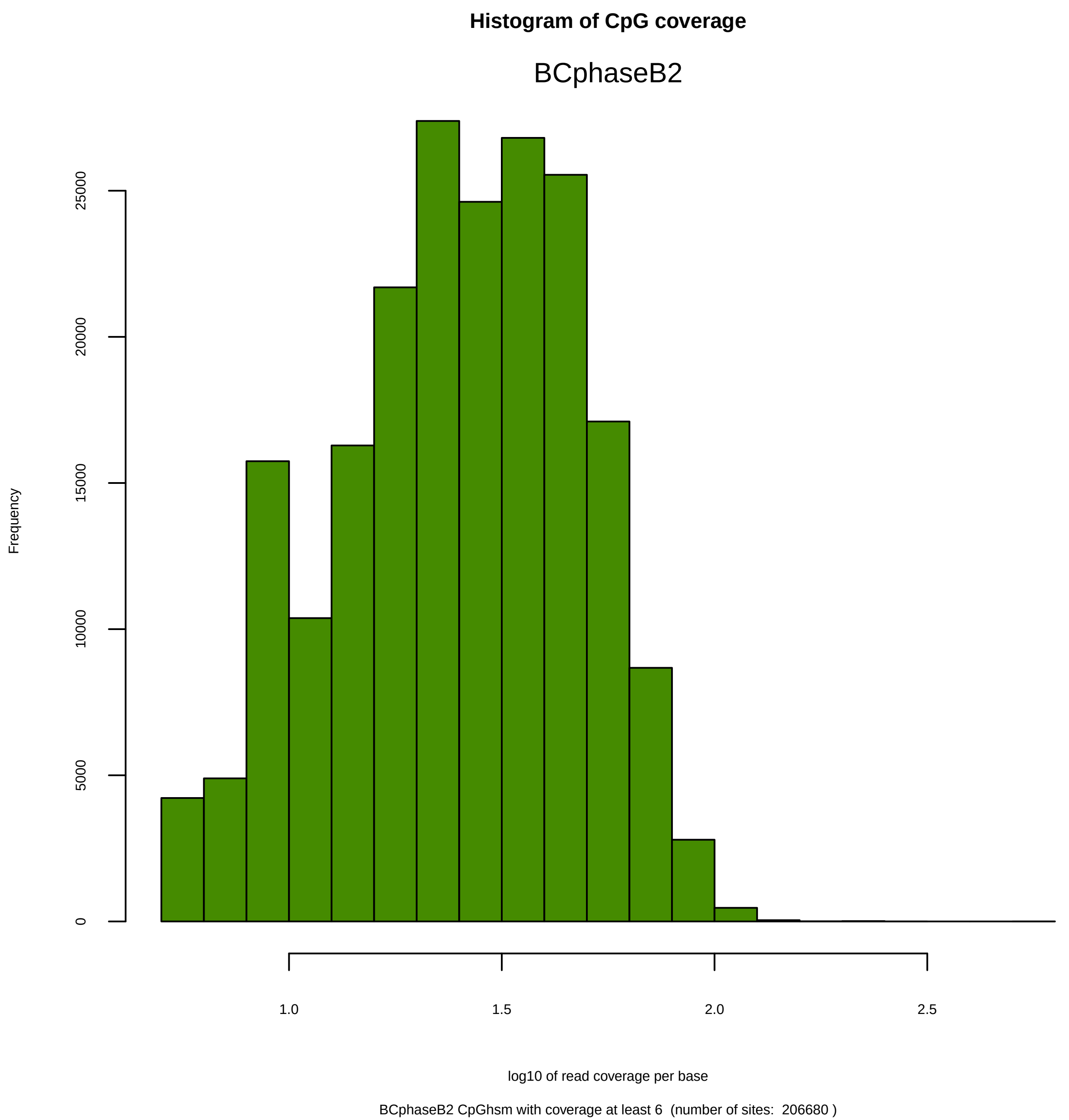
Histogram of CpG coverage
BCphaseB2
Frequency
0
5000
10000
15000
20000
25000
1.0
1.5
2.0
2.5
log10 of read coverage per base
BCphaseB2 CpGhsm with coverage at least 6  (number of sites:  206680 )

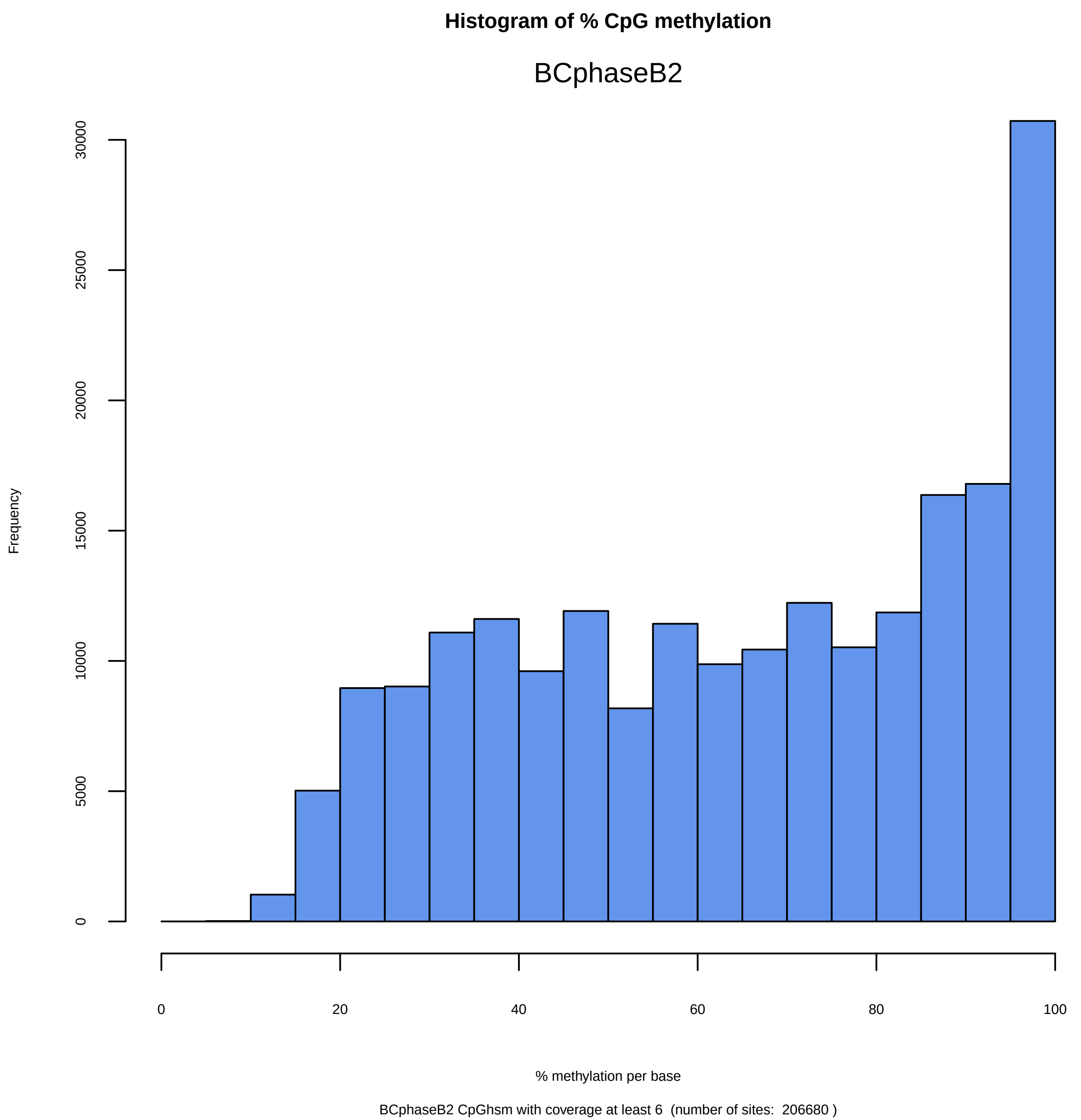
Histogram of % CpG methylation
BCphaseB2
Frequency
0
5000
10000
15000
20000
25000
30000
0
20
40
60
80
100
% methylation per base
BCphaseB2 CpGhsm with coverage at least 6  (number of sites:  206680 )

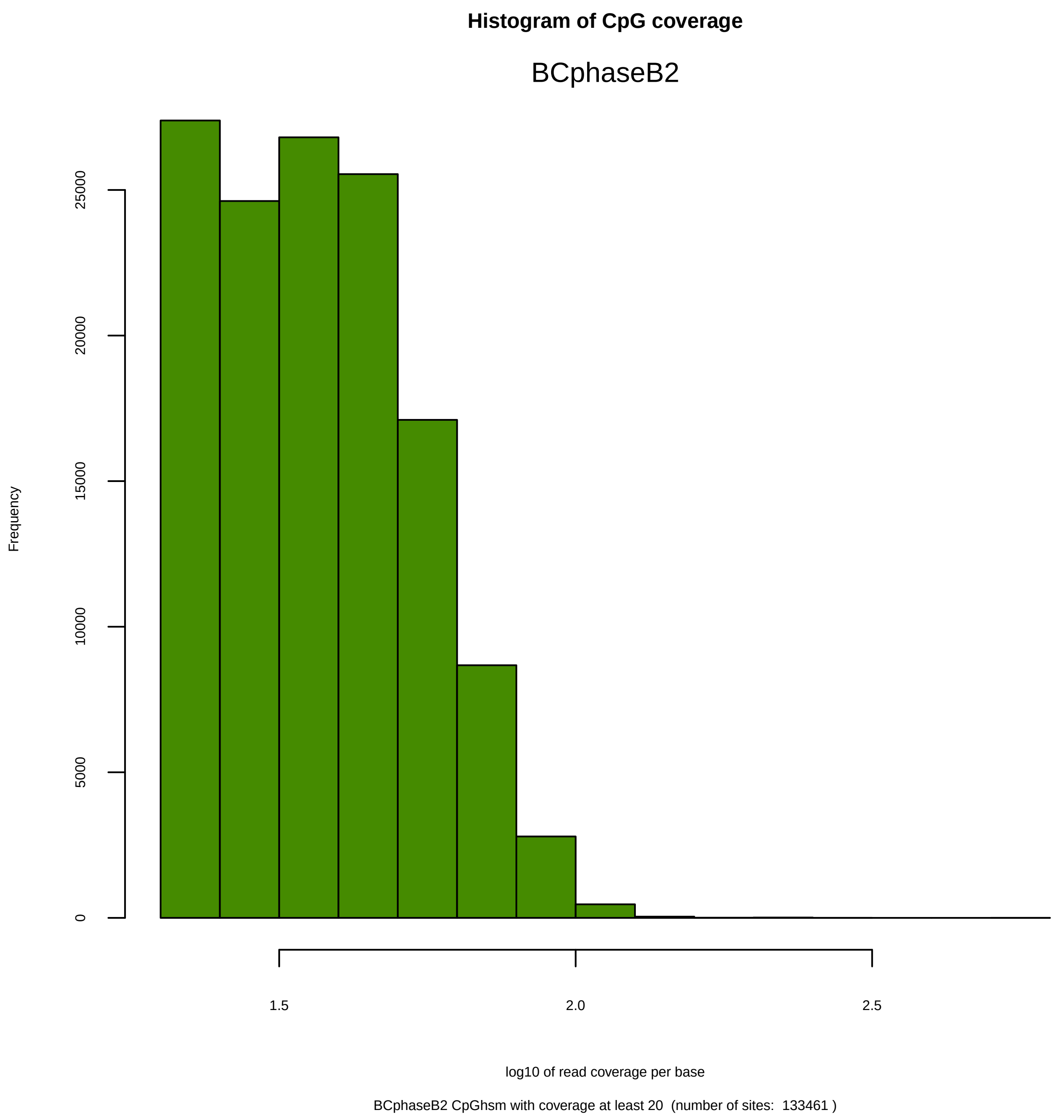

Histogram of CpG coverage
BCphaseB2
Frequency
0
5000
10000
15000
20000
25000
1.5
2.0
2.5
log10 of read coverage per base
BCphaseB2 CpGhsm with coverage at least 20 (number of sites: 133461 )

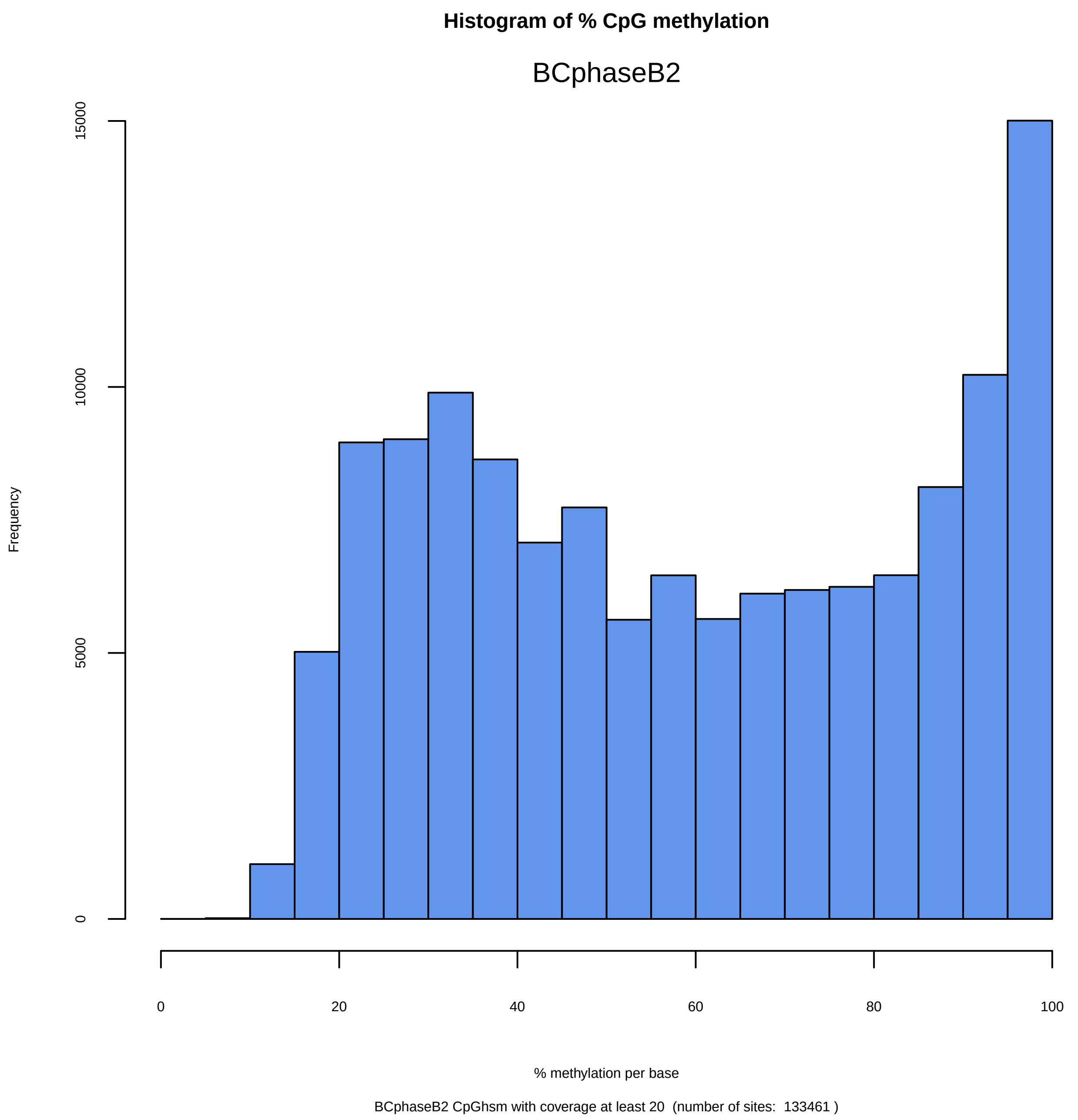
Histogram of % CpG methylation
BCphaseB2
Frequency
0
5000
10000
15000
0
20
40
60
80
100
% methylation per base
BCphaseB2 CpGhsm with coverage at least 20  (number of sites:  133461 )

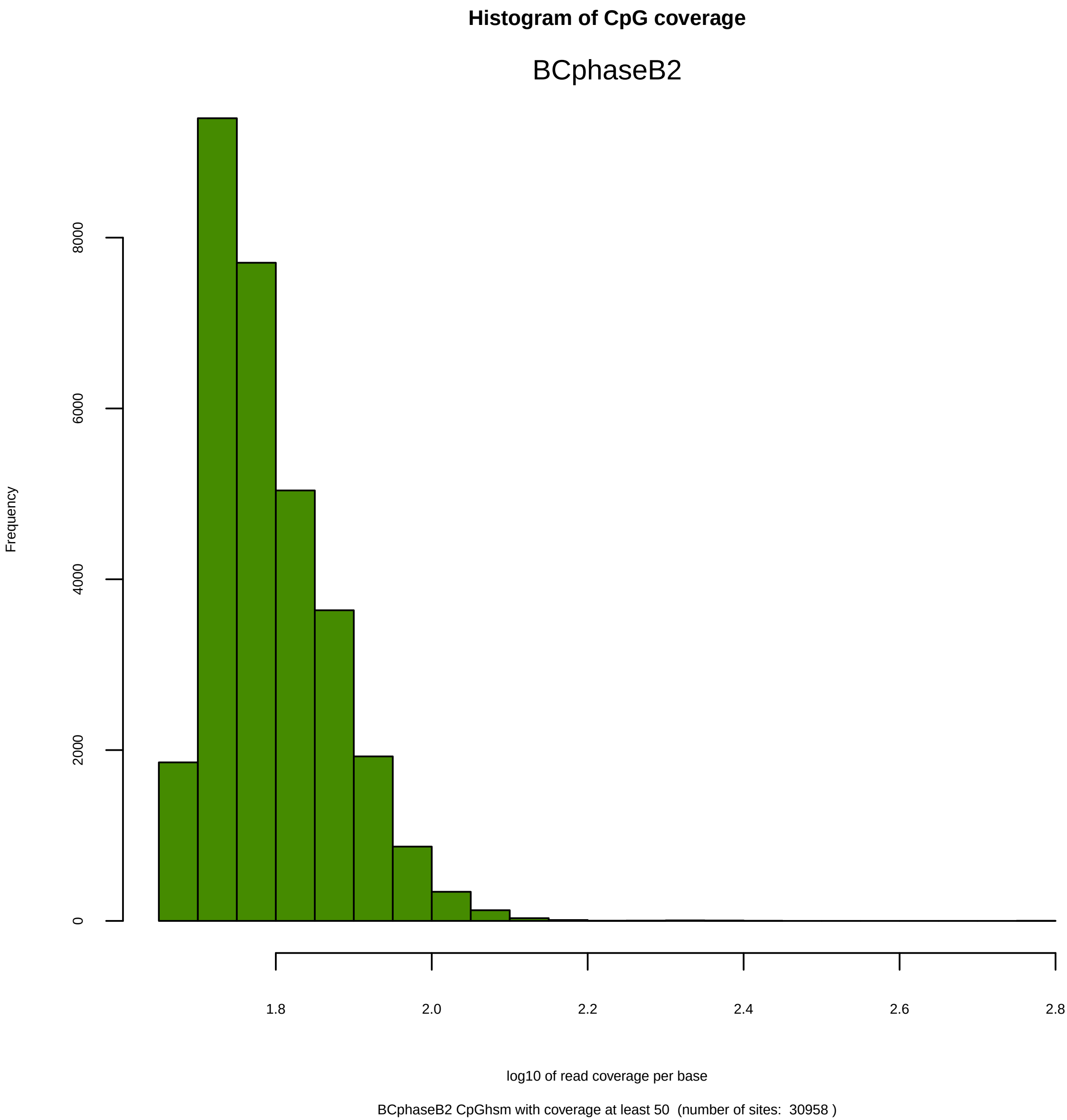
Histogram of CpG coverage
BCphaseB2
Frequency
0
2000
4000
6000
8000
1.8
2.0
2.2
2.4
2.6
2.8
log10 of read coverage per base
BCphaseB2 CpGhsm with coverage at least 50  (number of sites:  30958 )

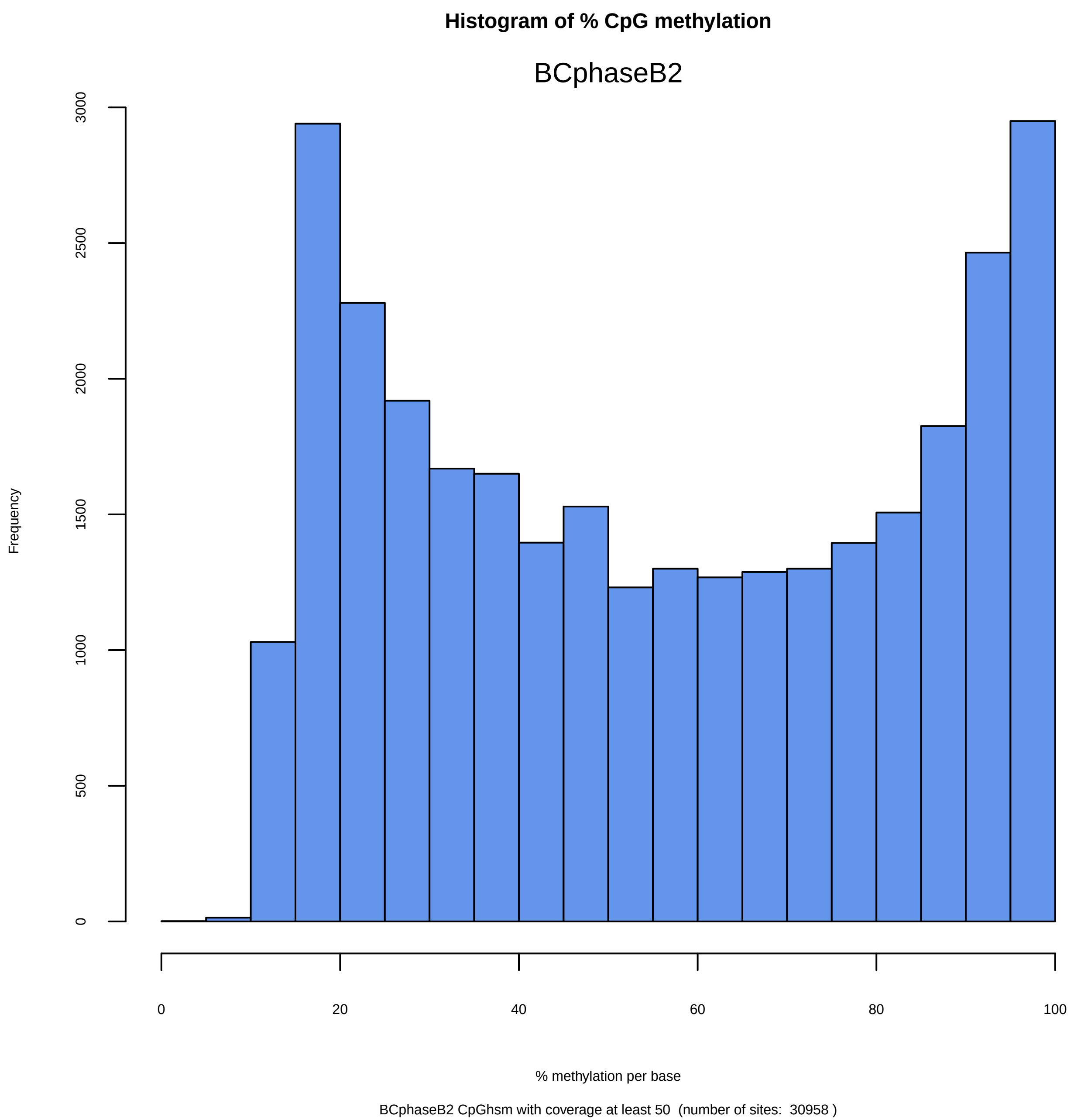
Histogram of % CpG methylation
BCphaseB2
Frequency
0
500
1000
1500
2000
2500
3000
0
20
40
60
80
100
% methylation per base
BCphaseB2 CpGhsm with coverage at least 50 (number of sites: 30958 )

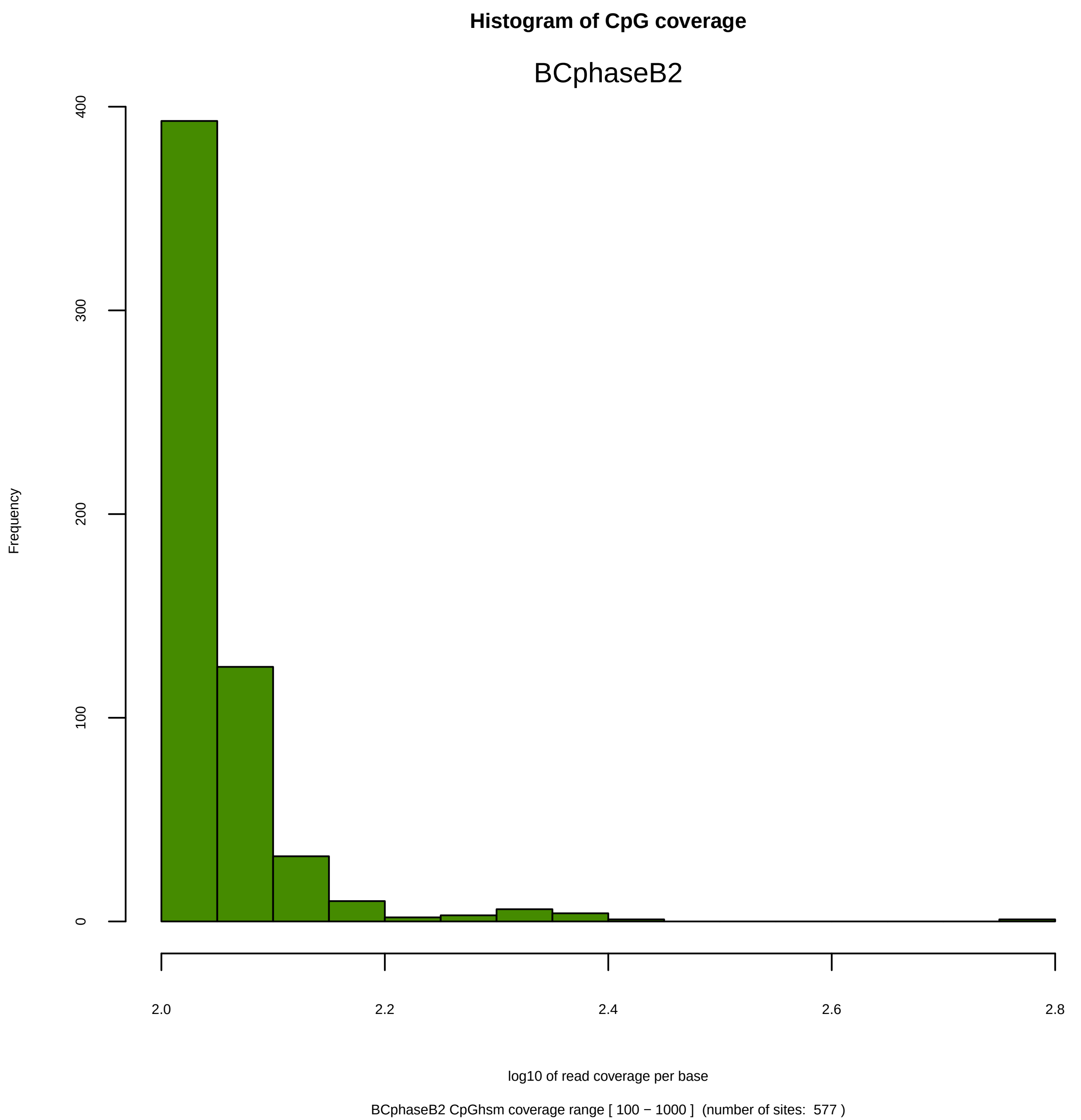

Histogram of CpG coverage
BCphaseB2
Frequency
0
100
200
300
400
2.0
2.2
2.4
2.6
2.8
log10 of read coverage per base
BCphaseB2 CpGhsm coverage range [ 100 − 1000 ]  (number of sites:  577 )

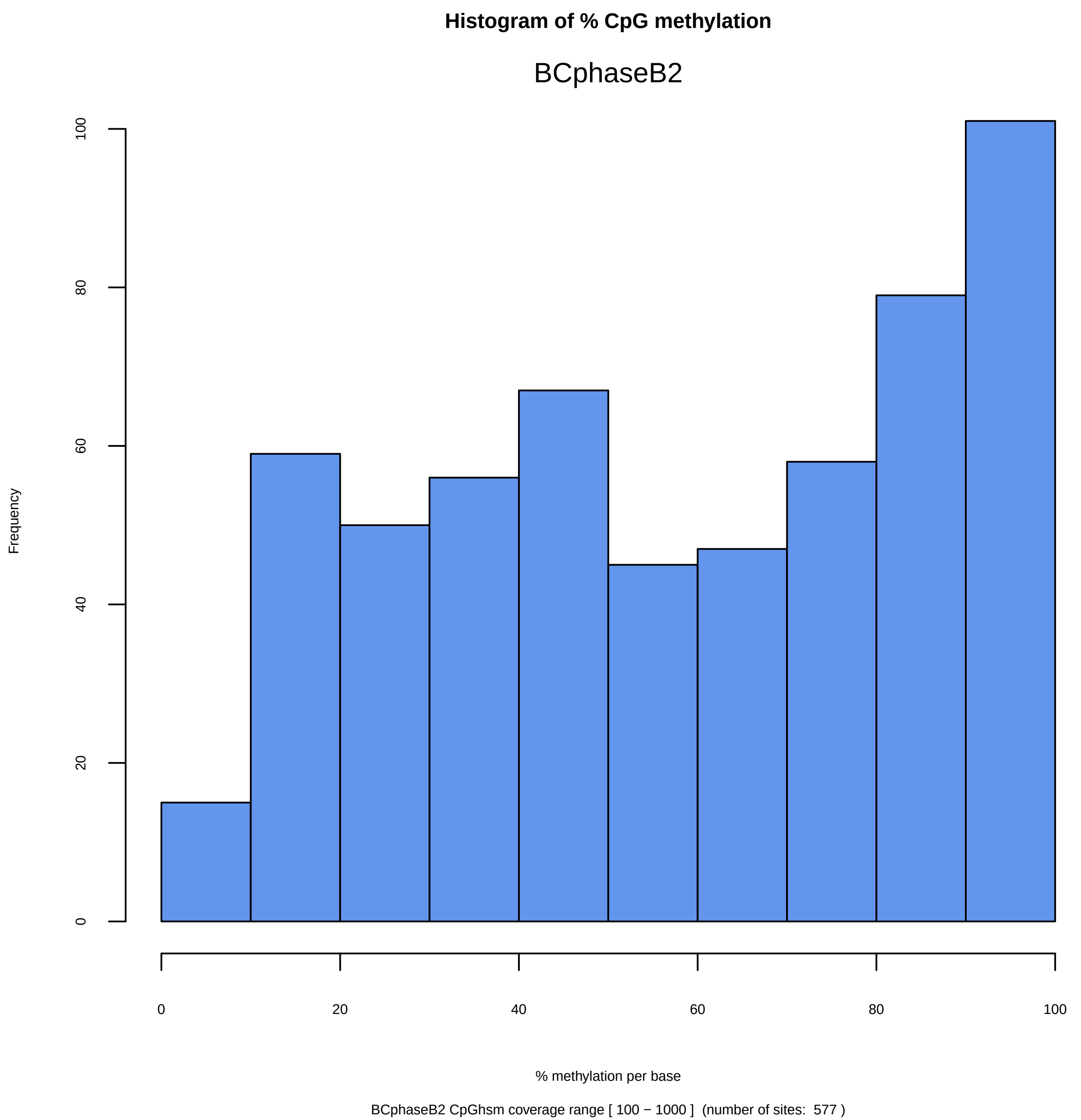

Histogram of % CpG methylation
BCphaseB2
Frequency
0
20
40
60
80
100
0
20
40
60
80
100
% methylation per base
BCphaseB2 CpGhsm coverage range [ 100 − 1000 ]  (number of sites:  577 )

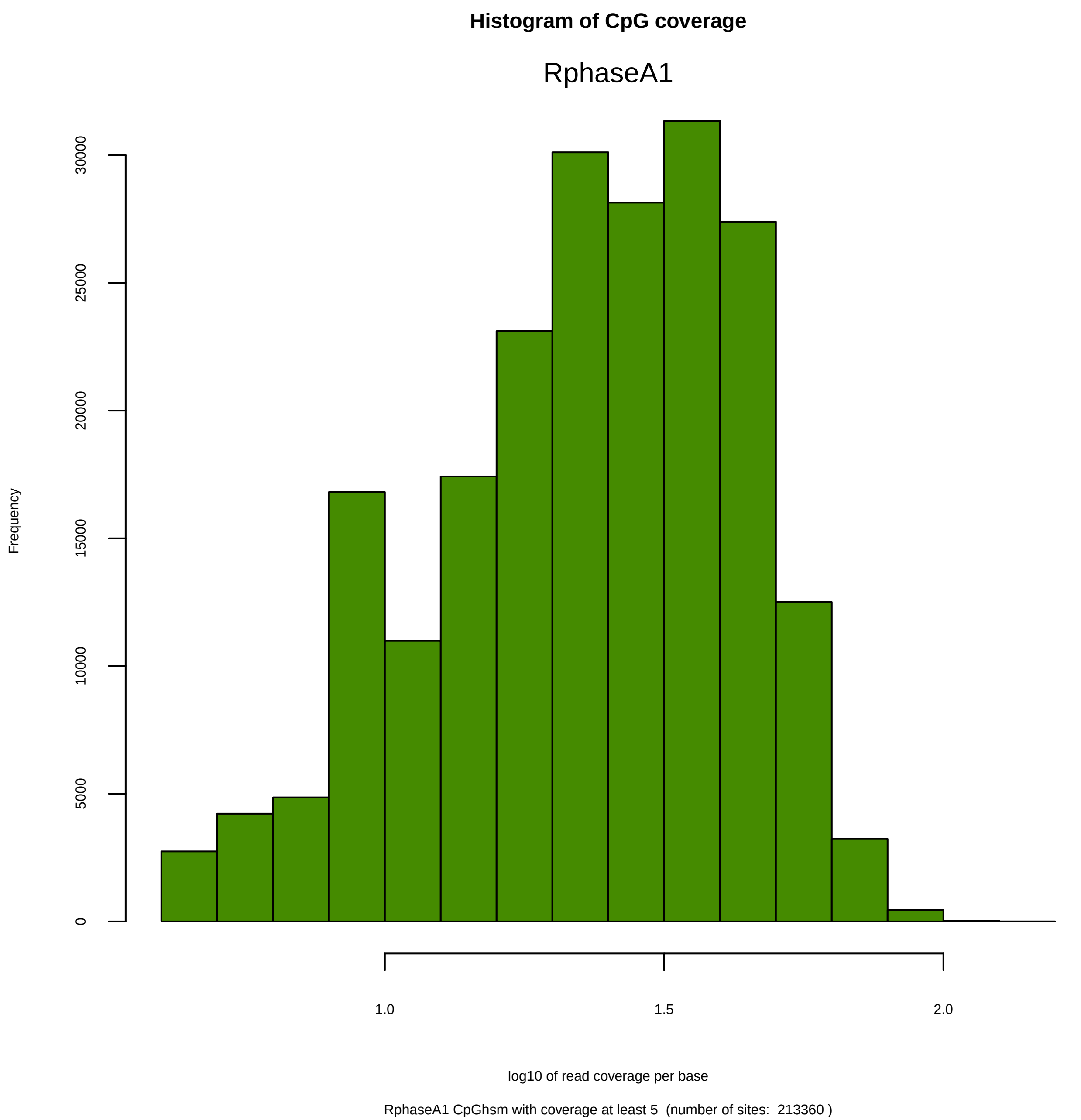
Histogram of CpG coverage
RphaseA1
Frequency
0
5000
10000
15000
20000
25000
30000
1.0
1.5
2.0
log10 of read coverage per base
RphaseA1 CpGhsm with coverage at least 5 (number of sites: 213360 )

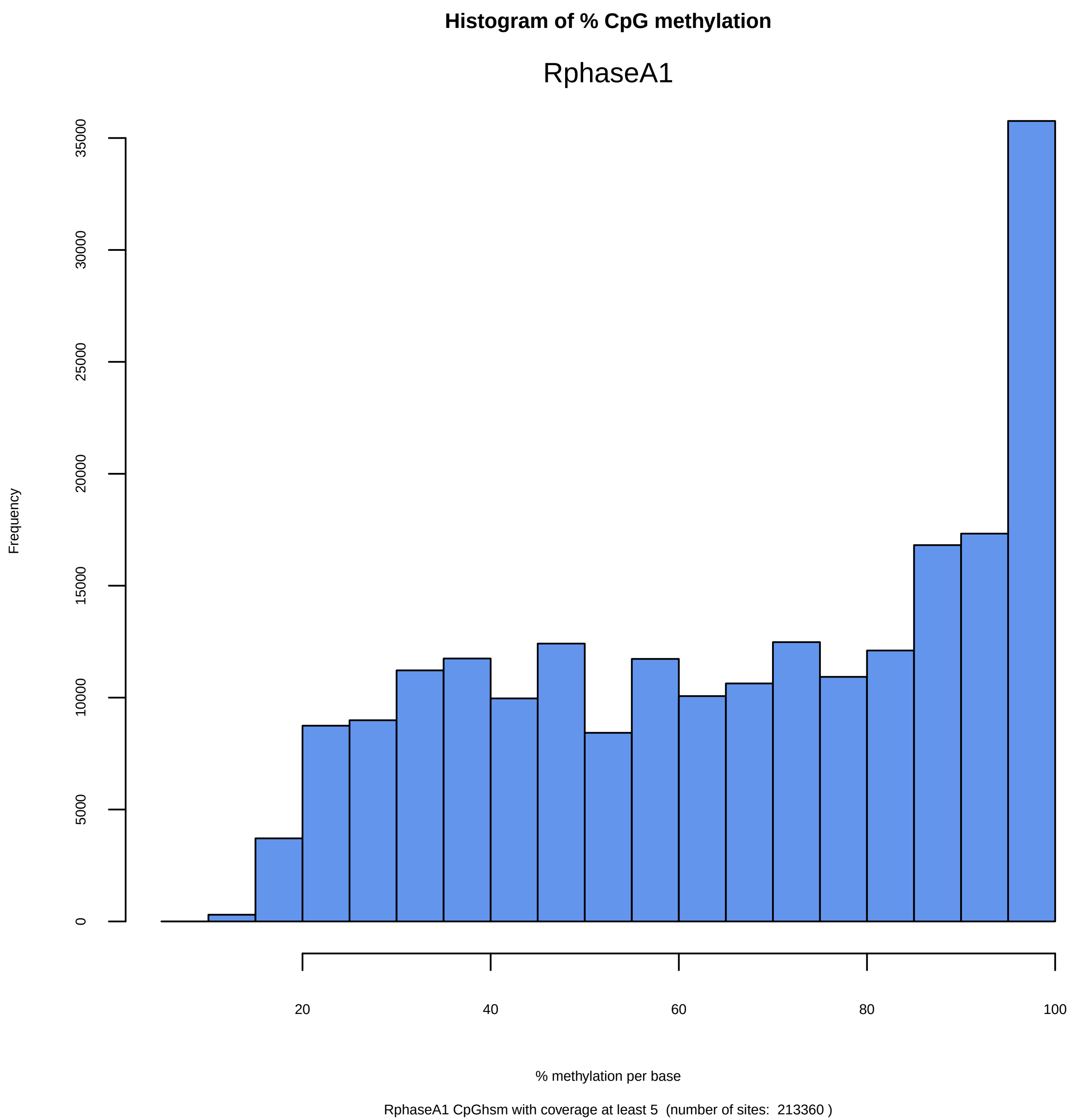
Histogram of % CpG methylation
RphaseA1
Frequency
0
5000
10000
15000
20000
25000
30000
35000
20
40
60
80
100
% methylation per base
RphaseA1 CpGhsm with coverage at least 5  (number of sites:  213360 )

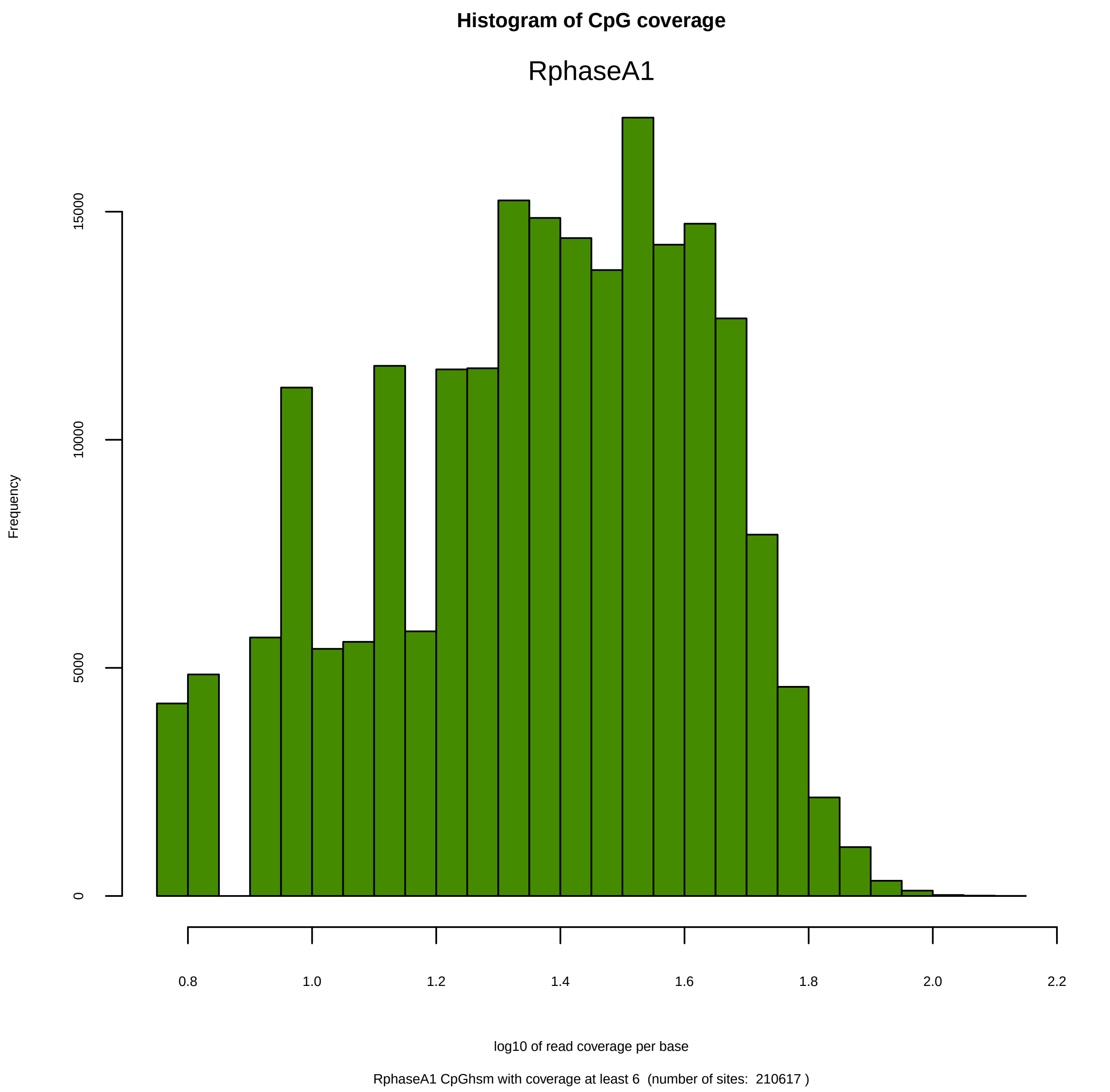
Histogram of CpG coverage
RphaseA1
Frequency
0
5000
10000
15000
0.8
1.0
1.2
1.4
1.6
1.8
2.0
2.2
log10 of read coverage per base
RphaseA1 CpGhsm with coverage at least 6 (number of sites: 210617 )

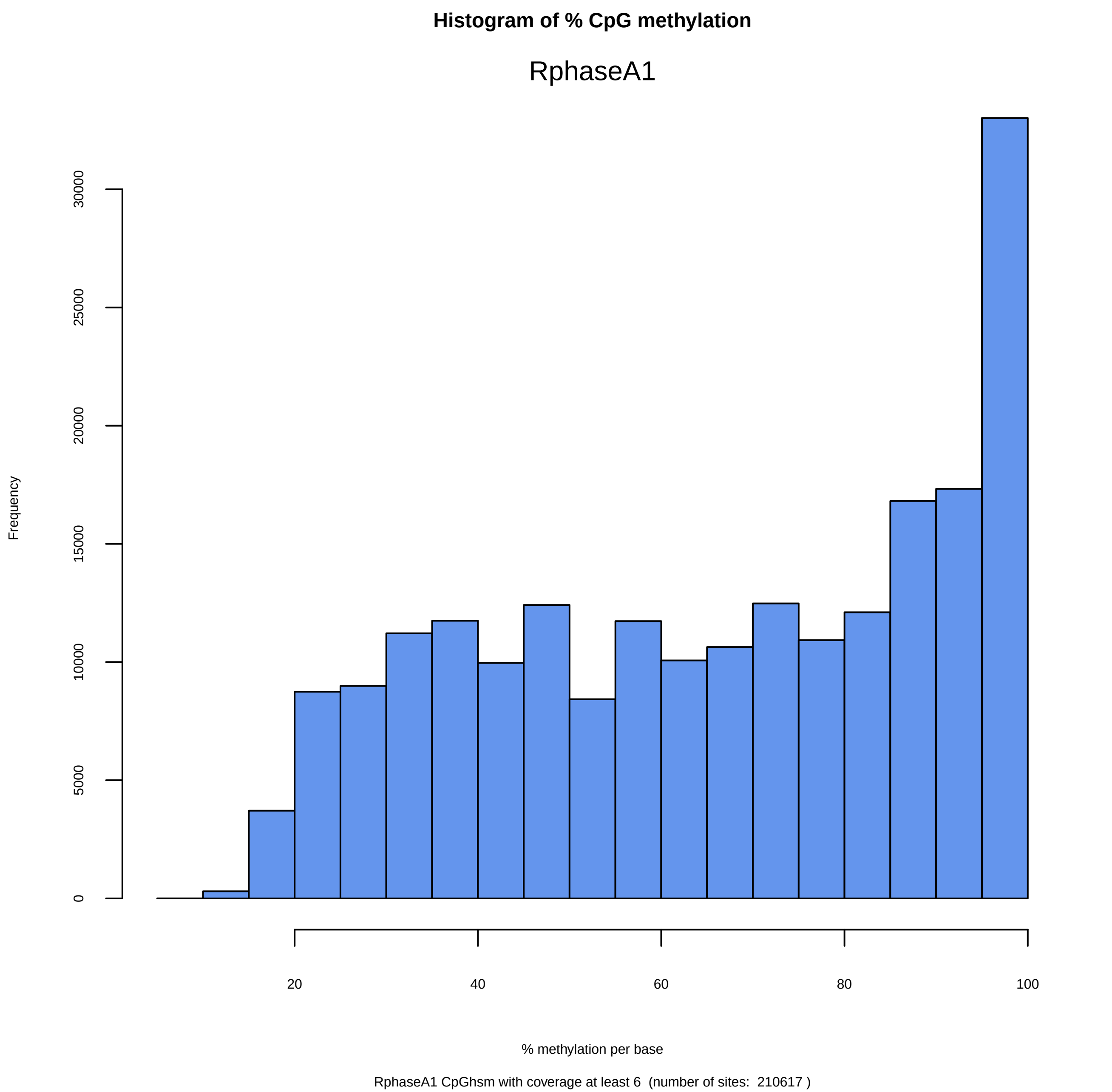

Histogram of % CpG methylation
RphaseA1
Frequency
0
5000
10000
15000
20000
25000
30000
20
40
60
80
100
% methylation per base
RphaseA1 CpGhsm with coverage at least 6  (number of sites:  210617 )

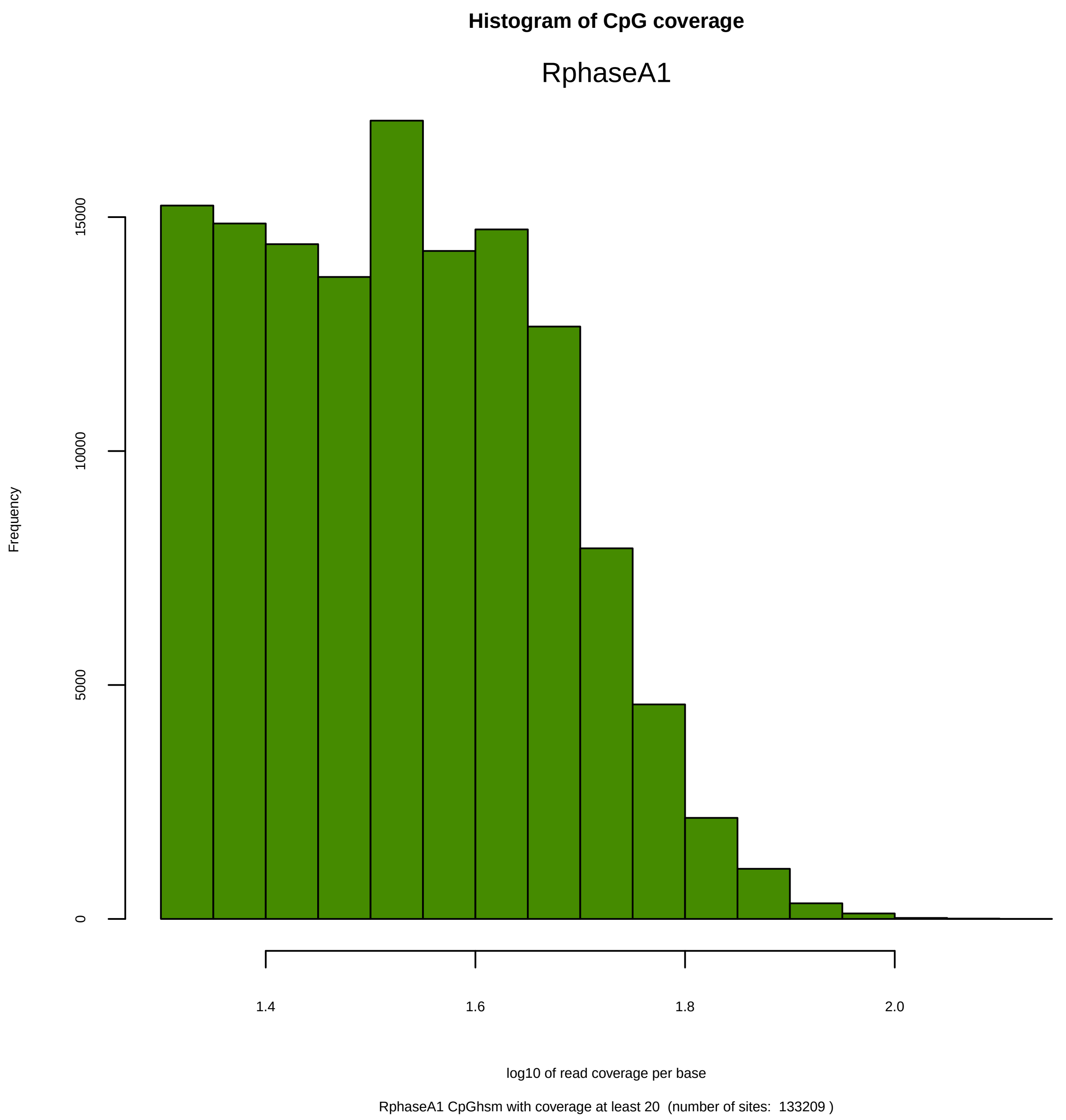
Histogram of CpG coverage
RphaseA1
Frequency
0
5000
10000
15000
1.4
1.6
1.8
2.0
log10 of read coverage per base
RphaseA1 CpGhsm with coverage at least 20 (number of sites: 133209 )

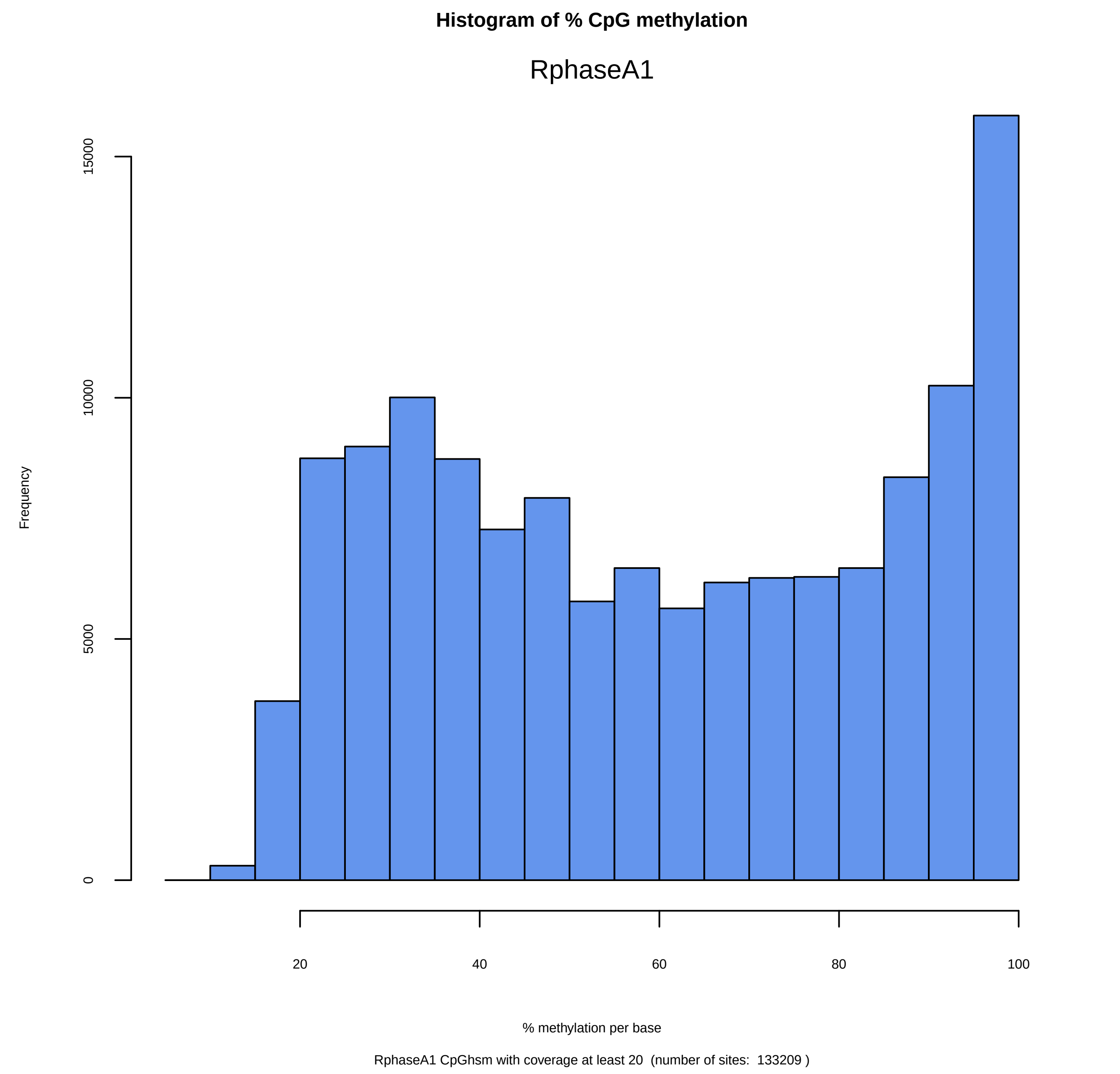
Histogram of % CpG methylation
RphaseA1
Frequency
0
5000
10000
15000
20
40
60
80
100
% methylation per base
RphaseA1 CpGhsm with coverage at least 20 (number of sites: 133209 )

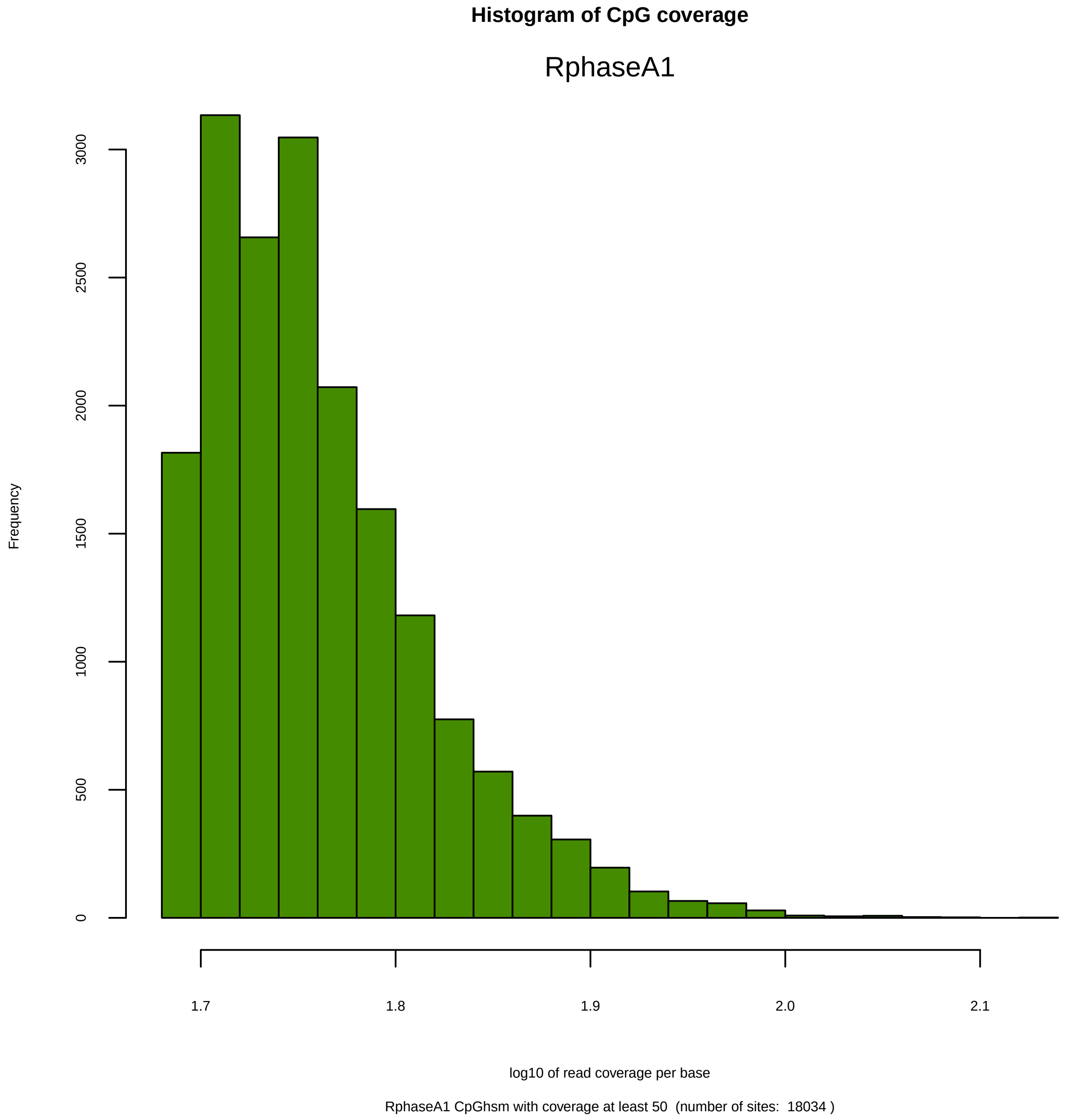
Histogram of CpG coverage
RphaseA1
Frequency
0
500
1000
1500
2000
2500
3000
1.7
1.8
1.9
2.0
2.1
log10 of read coverage per base
RphaseA1 CpGhsm with coverage at least 50  (number of sites:  18034 )

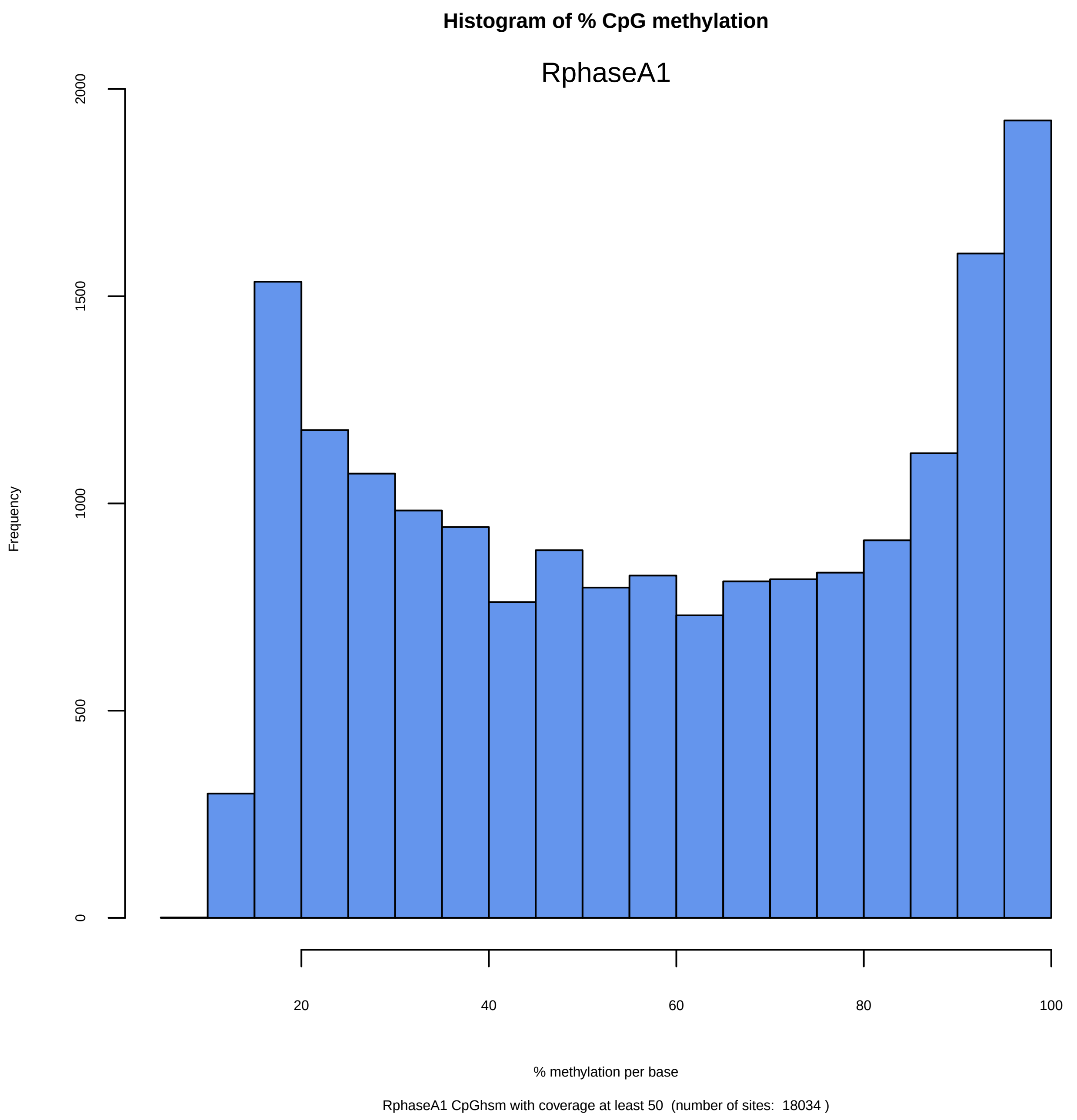

Histogram of % CpG methylation
RphaseA1
Frequency
0
500
1000
1500
2000
20
40
60
80
100
% methylation per base
RphaseA1 CpGhsm with coverage at least 50 (number of sites: 18034 )

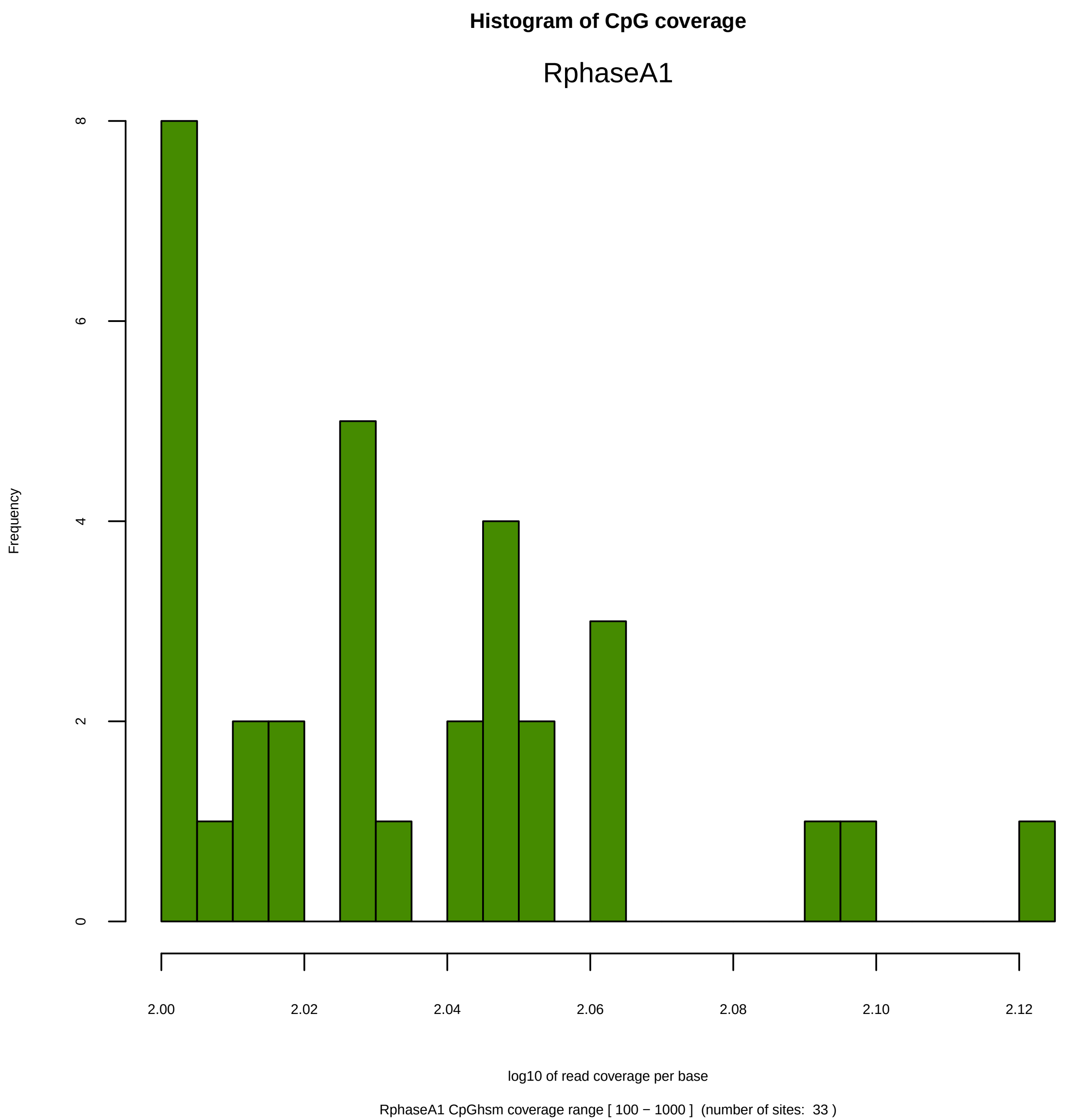
Histogram of CpG coverage
RphaseA1
Frequency
0
2
4
6
8
2.00
2.02
2.04
2.06
2.08
2.10
2.12
log10 of read coverage per base
RphaseA1 CpGhsm coverage range [ 100 − 1000 ] (number of sites: 33 )

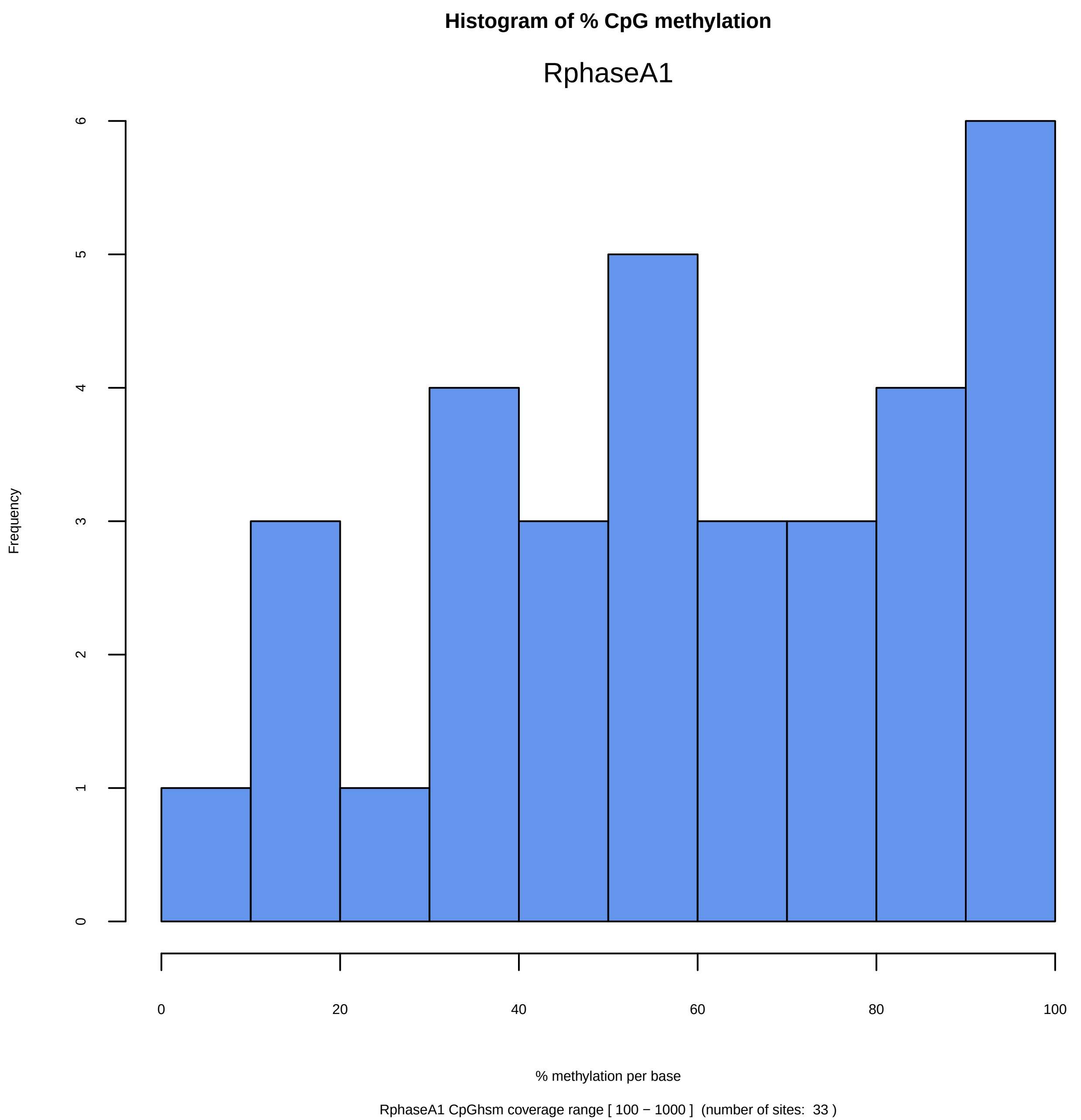

Histogram of % CpG methylation
RphaseA1
Frequency
0
1
2
3
4
5
6
0
20
40
60
80
100
% methylation per base
RphaseA1 CpGhsm coverage range [ 100 − 1000 ]  (number of sites:  33 )

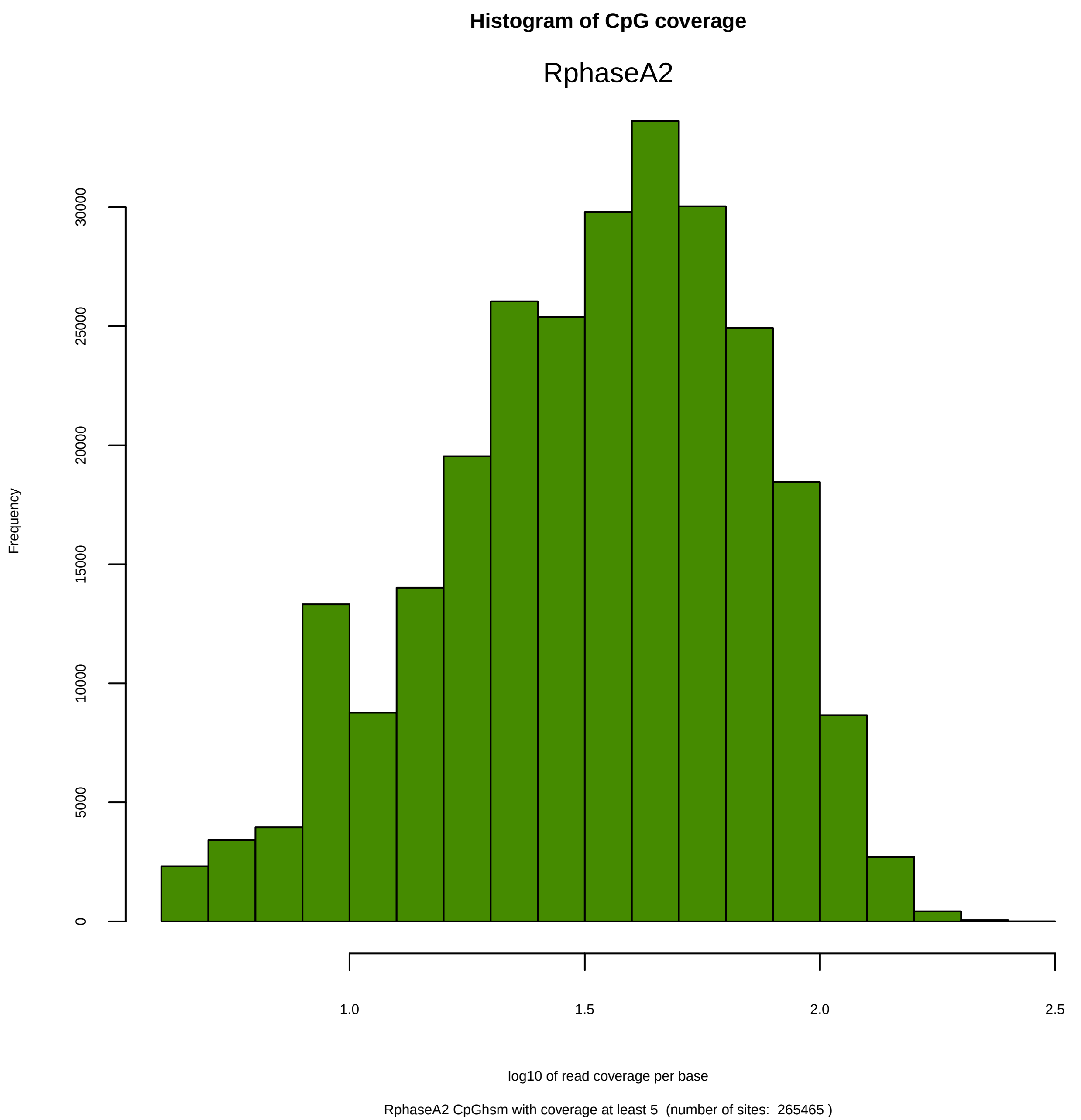
Histogram of CpG coverage
RphaseA2
Frequency
0
5000
10000
15000
20000
25000
30000
1.0
1.5
2.0
2.5
log10 of read coverage per base
RphaseA2 CpGhsm with coverage at least 5  (number of sites:  265465 )

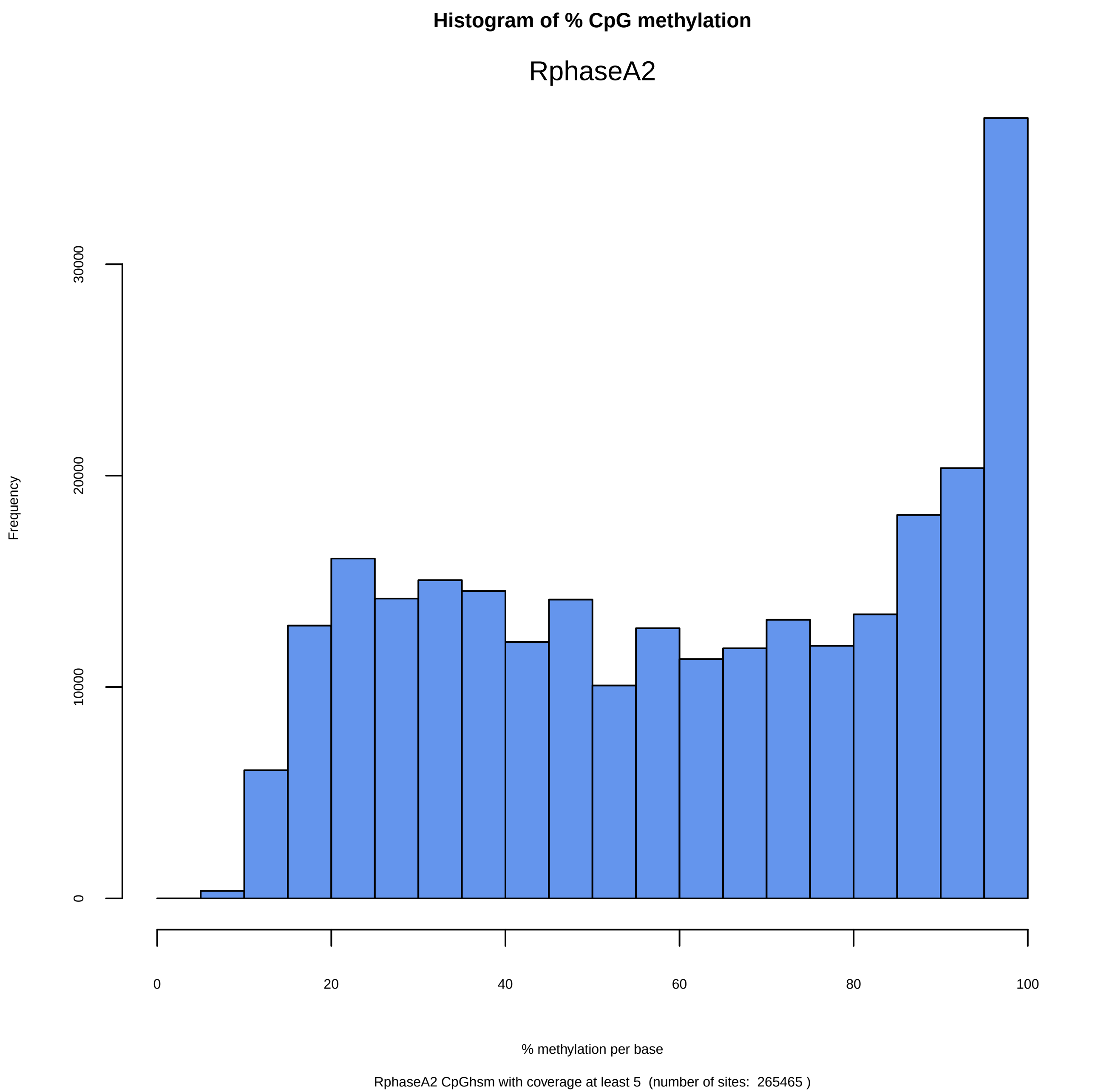
Histogram of % CpG methylation
RphaseA2
Frequency
0
10000
20000
30000
0
20
40
60
80
100
% methylation per base
RphaseA2 CpGhsm with coverage at least 5 (number of sites: 265465 )

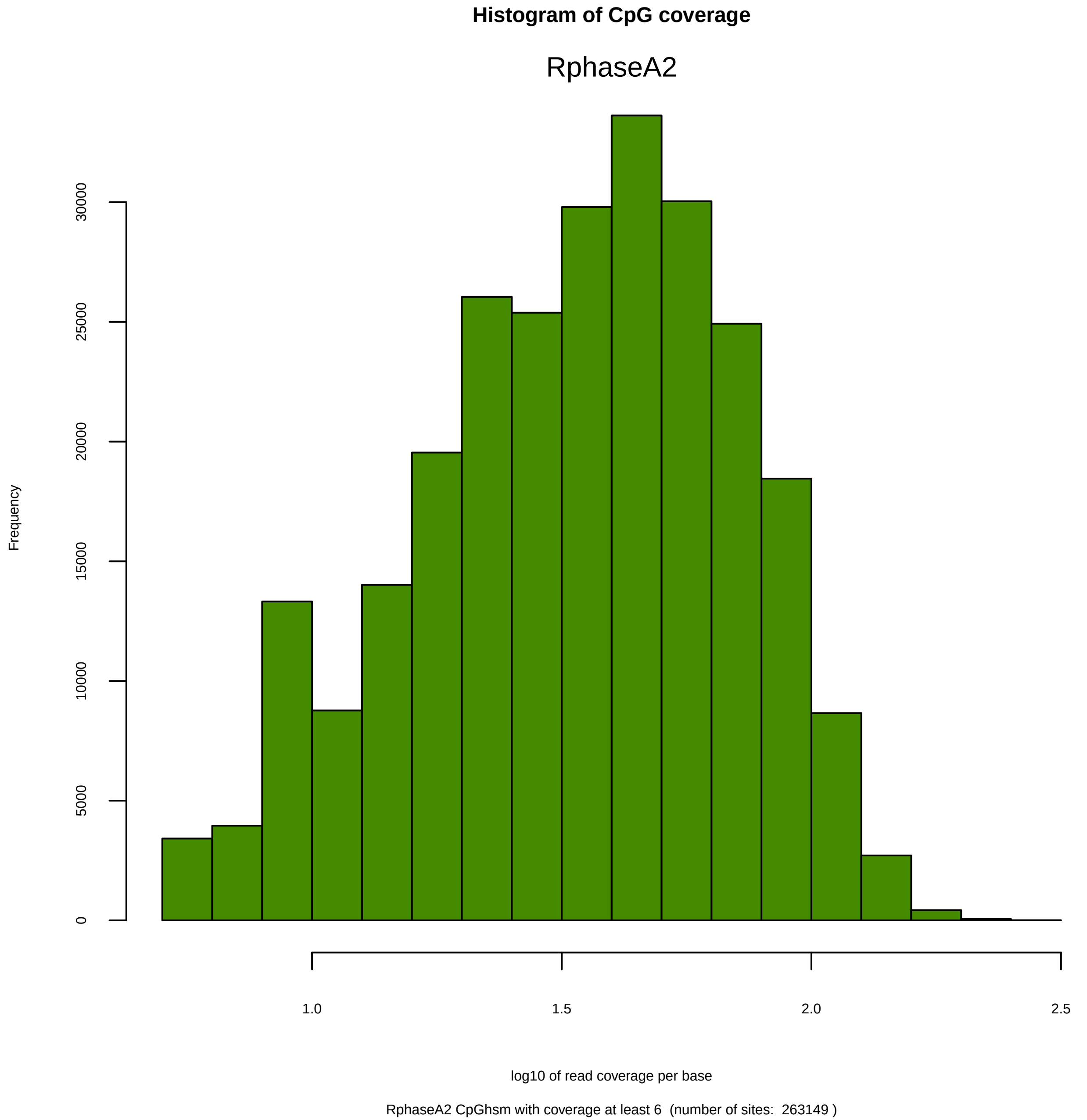
Histogram of CpG coverage
RphaseA2
Frequency
0
5000
10000
15000
20000
25000
30000
1.0
1.5
2.0
2.5
log10 of read coverage per base
RphaseA2 CpGhsm with coverage at least 6  (number of sites:  263149 )

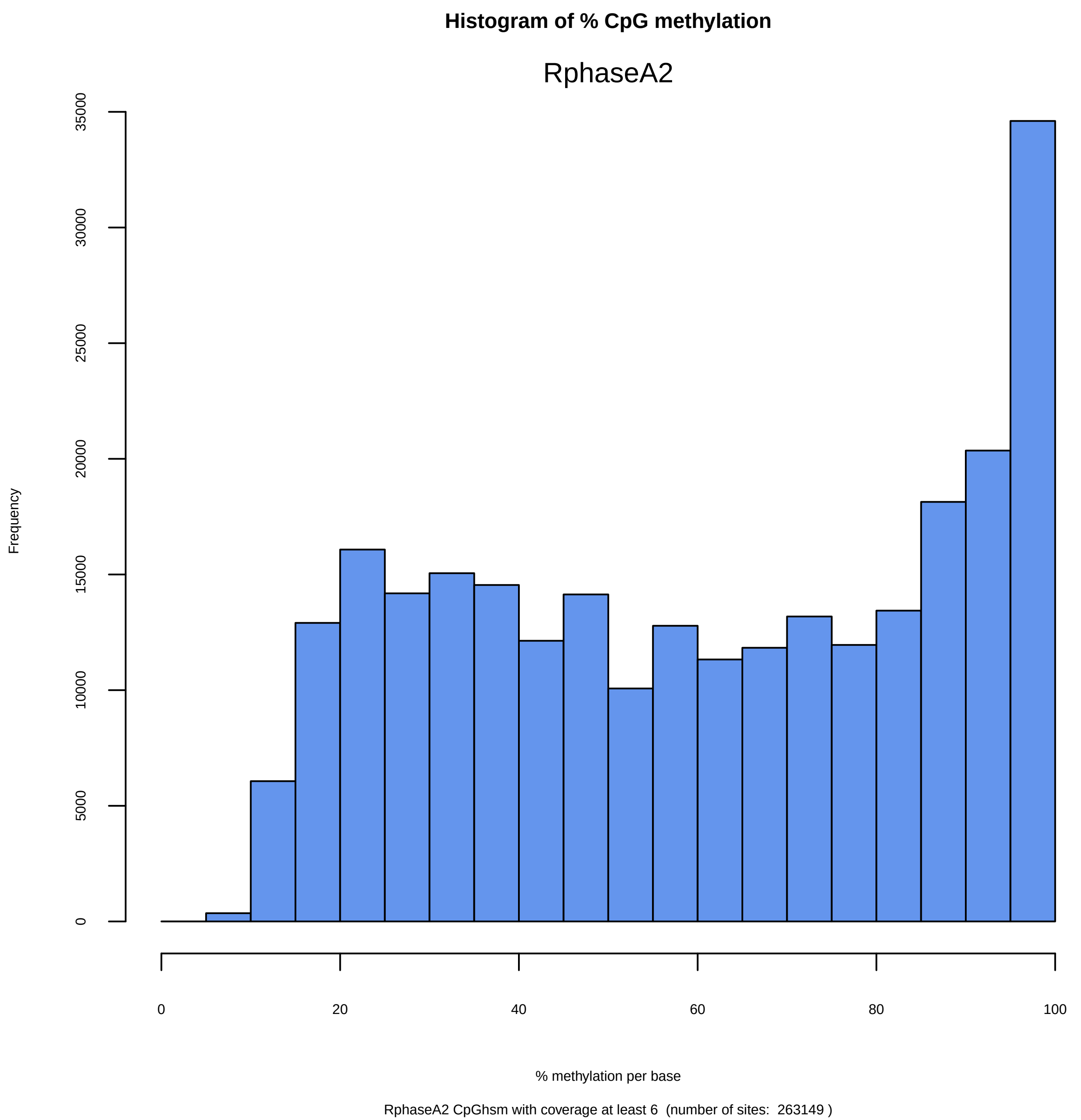
Histogram of % CpG methylation
RphaseA2
Frequency
0
5000
10000
15000
20000
25000
30000
35000
0
20
40
60
80
100
% methylation per base
RphaseA2 CpGhsm with coverage at least 6  (number of sites:  263149 )

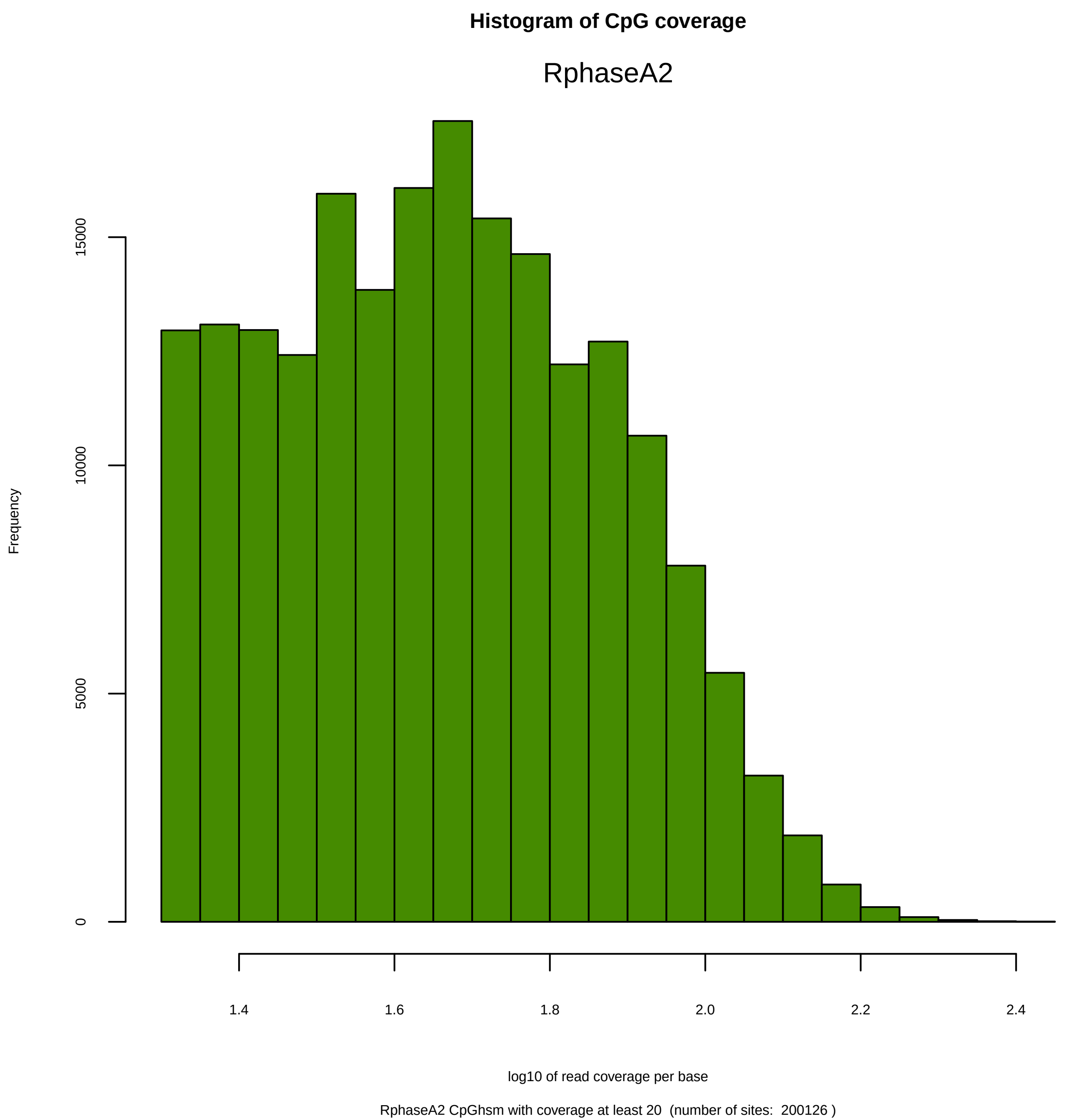
Histogram of CpG coverage
RphaseA2
Frequency
0
5000
10000
15000
1.4
1.6
1.8
2.0
2.2
2.4
log10 of read coverage per base
RphaseA2 CpGhsm with coverage at least 20 (number of sites: 200126 )

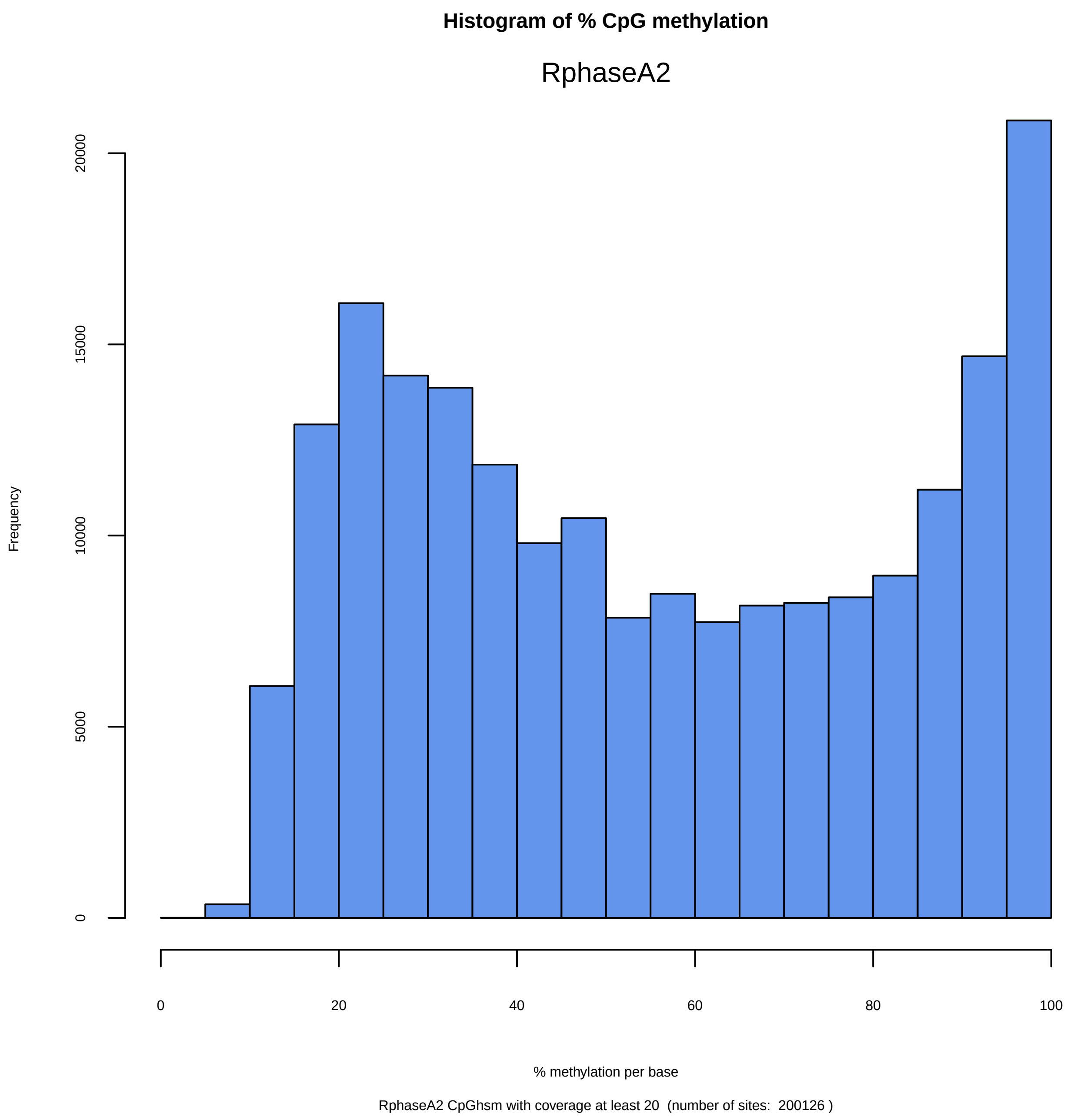

Histogram of % CpG methylation
RphaseA2
Frequency
0
5000
10000
15000
20000
0
20
40
60
80
100
% methylation per base
RphaseA2 CpGhsm with coverage at least 20 (number of sites: 200126 )

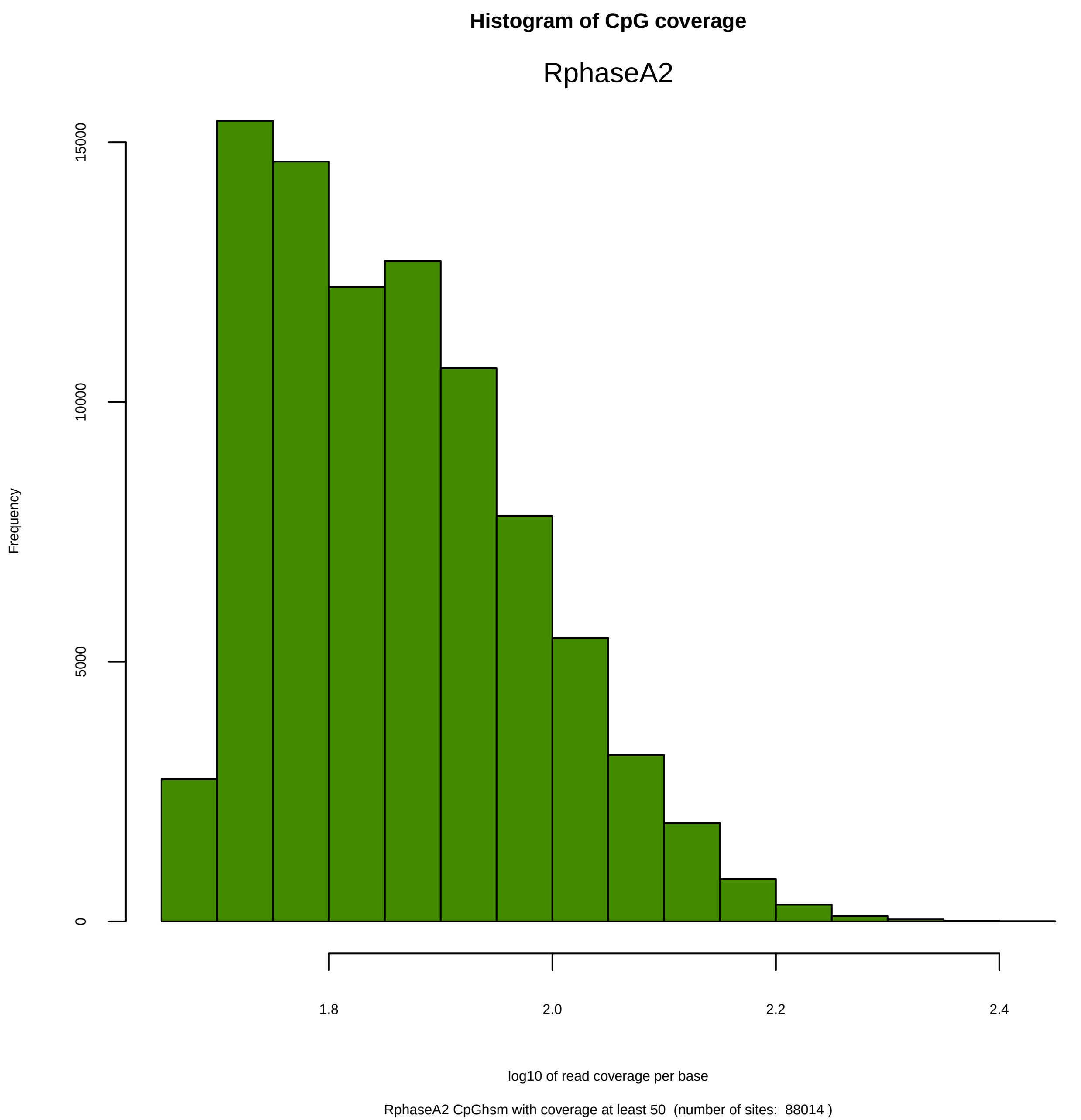

Histogram of CpG coverage
RphaseA2
Frequency
0
5000
10000
15000
1.8
2.0
2.2
2.4
log10 of read coverage per base
RphaseA2 CpGhsm with coverage at least 50  (number of sites:  88014 )

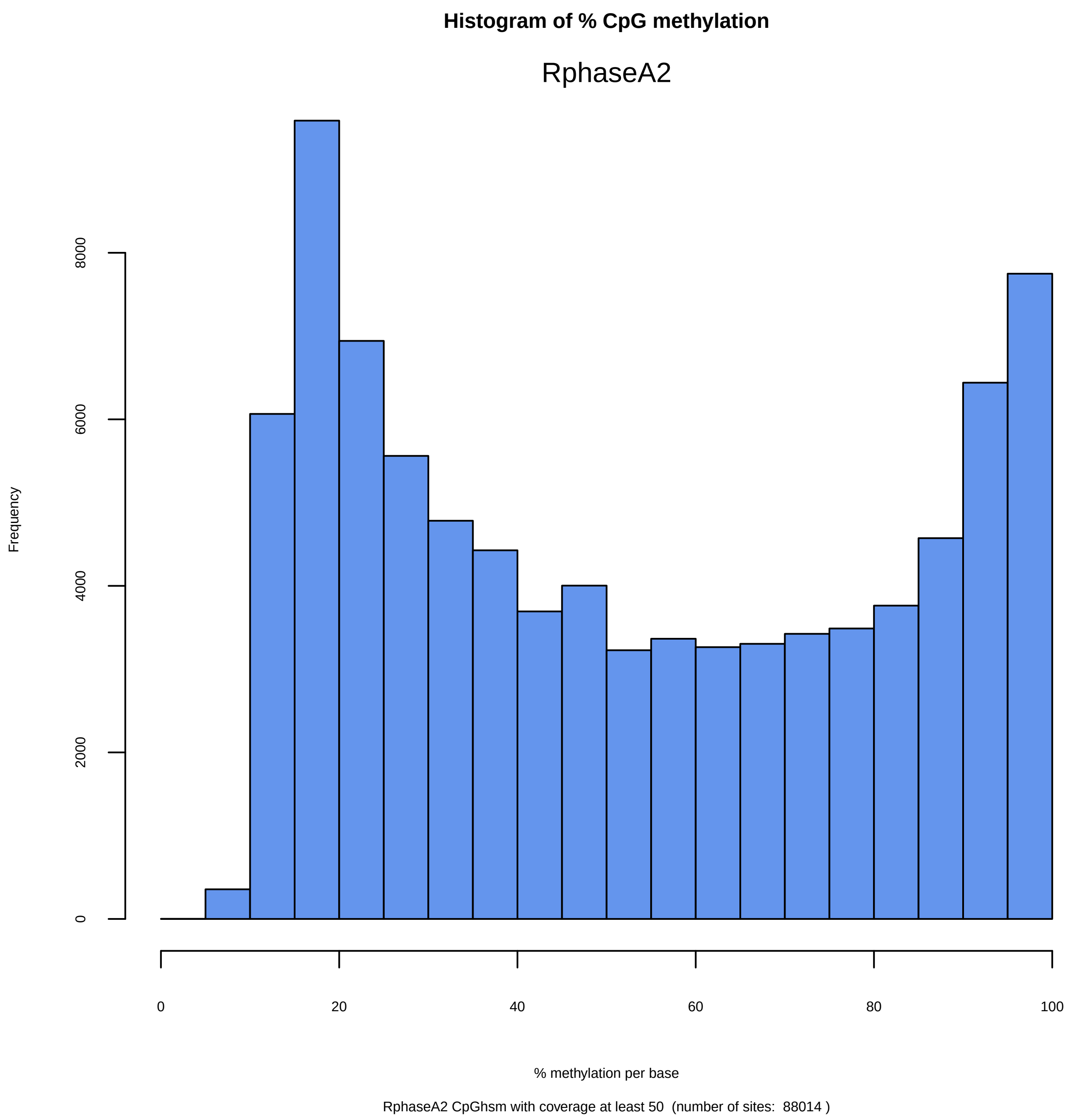

Histogram of % CpG methylation
RphaseA2
Frequency
0
2000
4000
6000
8000
0
20
40
60
80
100
% methylation per base
RphaseA2 CpGhsm with coverage at least 50 (number of sites: 88014 )

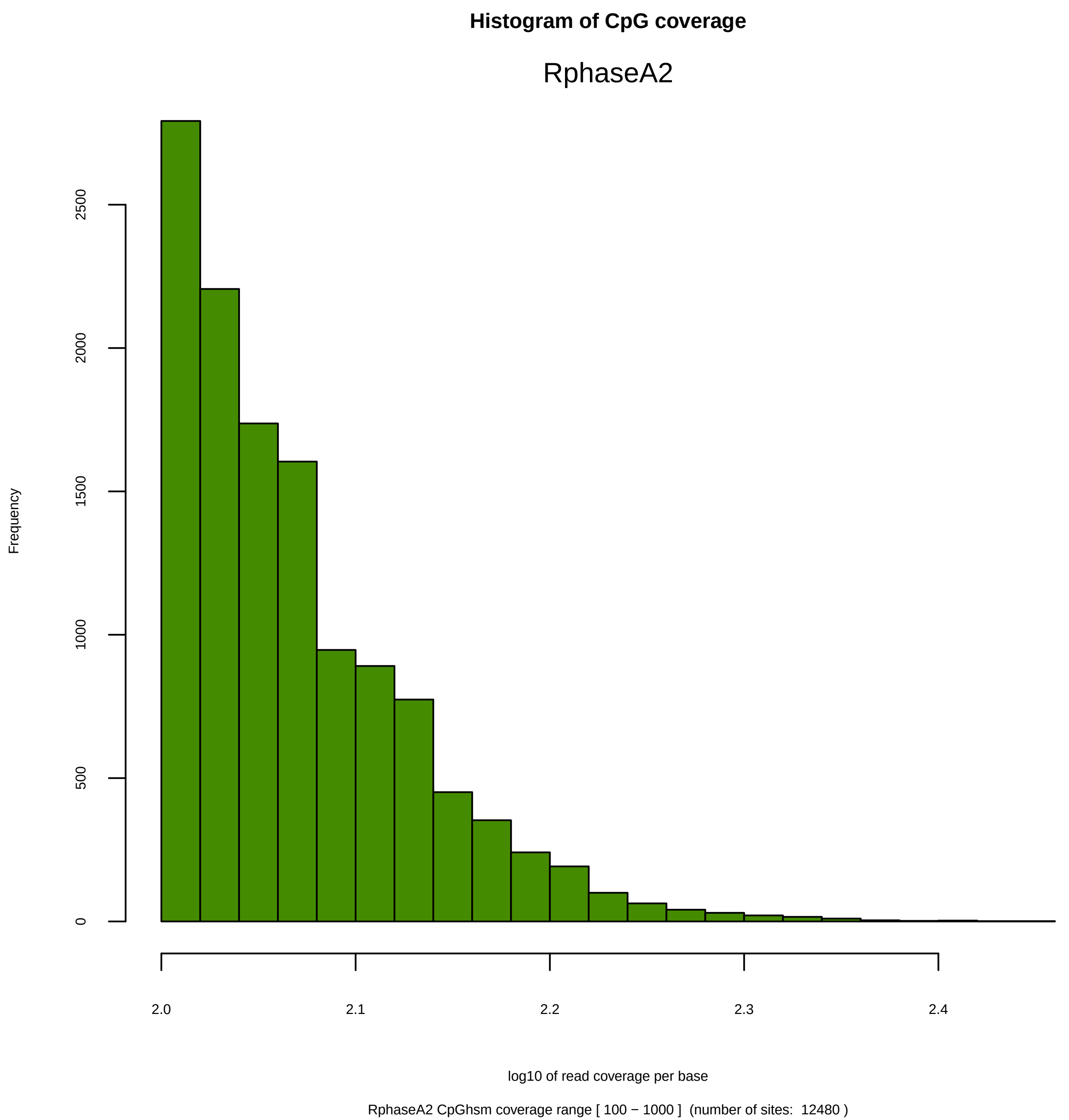
Histogram of CpG coverage
RphaseA2
Frequency
0
500
1000
1500
2000
2500
2.0
2.1
2.2
2.3
2.4
log10 of read coverage per base
RphaseA2 CpGhsm coverage range [ 100 − 1000 ]  (number of sites:  12480 )

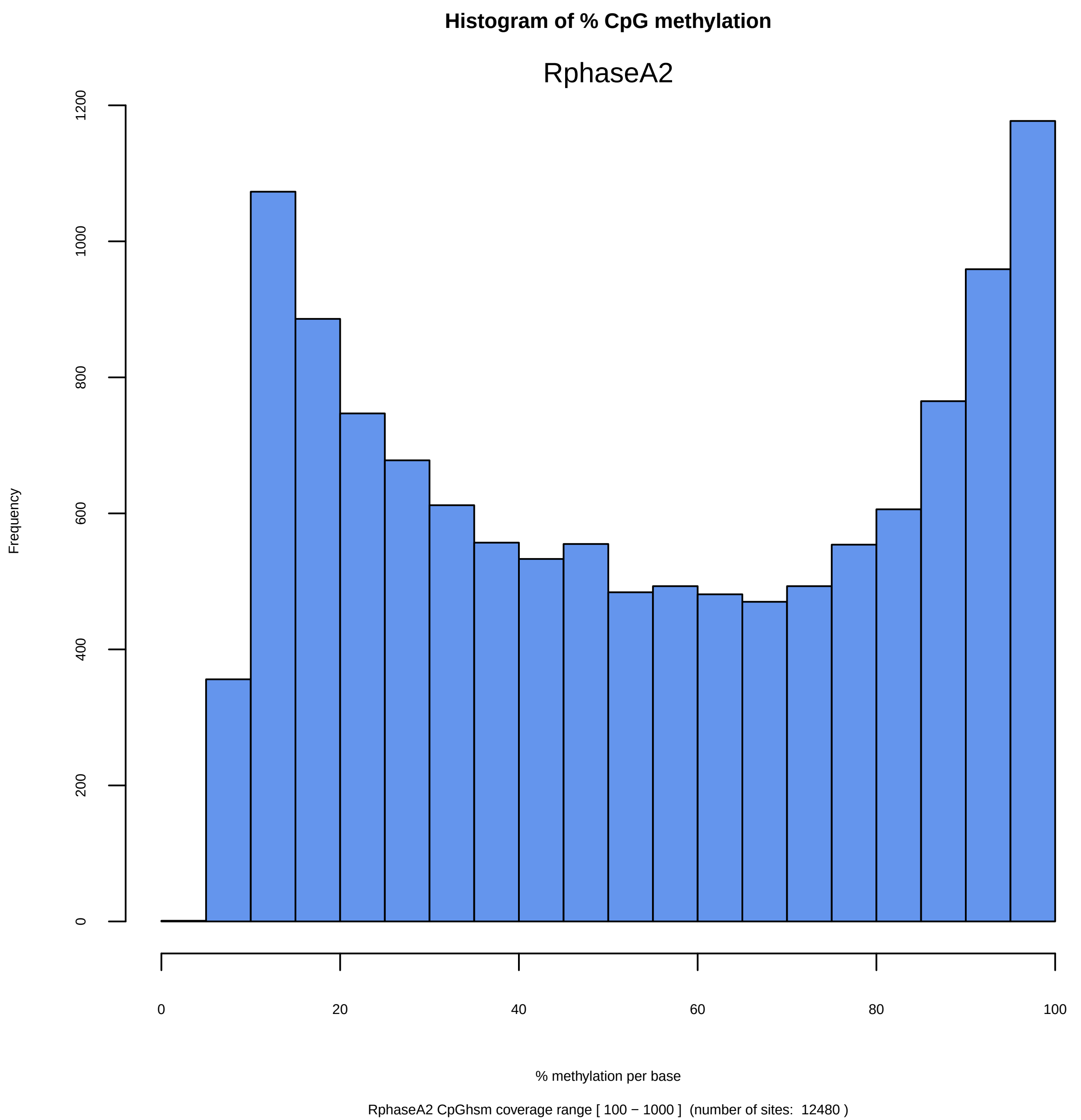

Histogram of % CpG methylation
RphaseA2
Frequency
0
200
400
600
800
1000
1200
0
20
40
60
80
100
% methylation per base
RphaseA2 CpGhsm coverage range [ 100 − 1000 ]  (number of sites:  12480 )

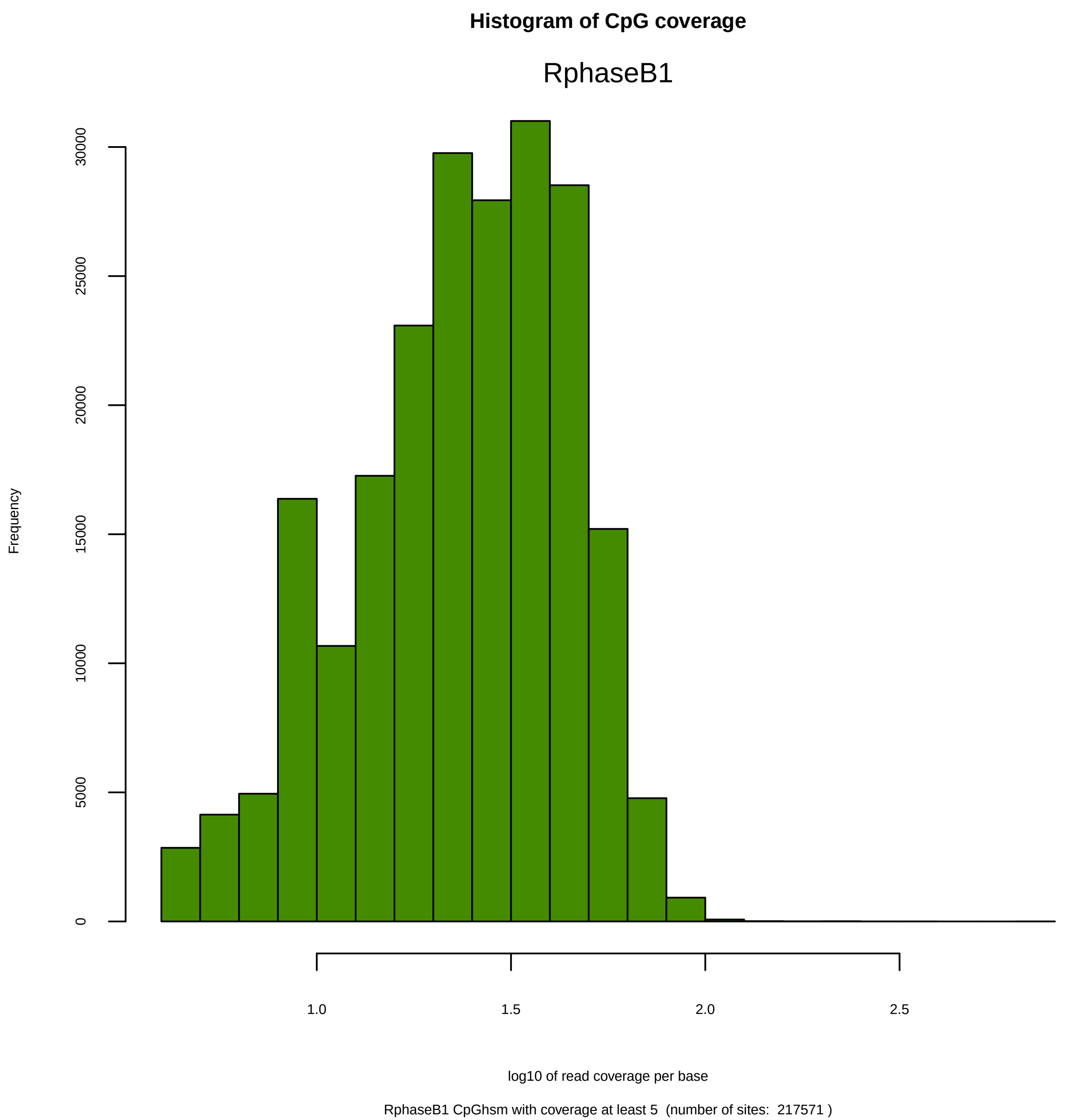
Histogram of CpG coverage
RphaseB1
Frequency
0
5000
10000
15000
20000
25000
30000
1.0
1.5
2.0
2.5
log10 of read coverage per base
RphaseB1 CpGhsm with coverage at least 5 (number of sites: 217571 )

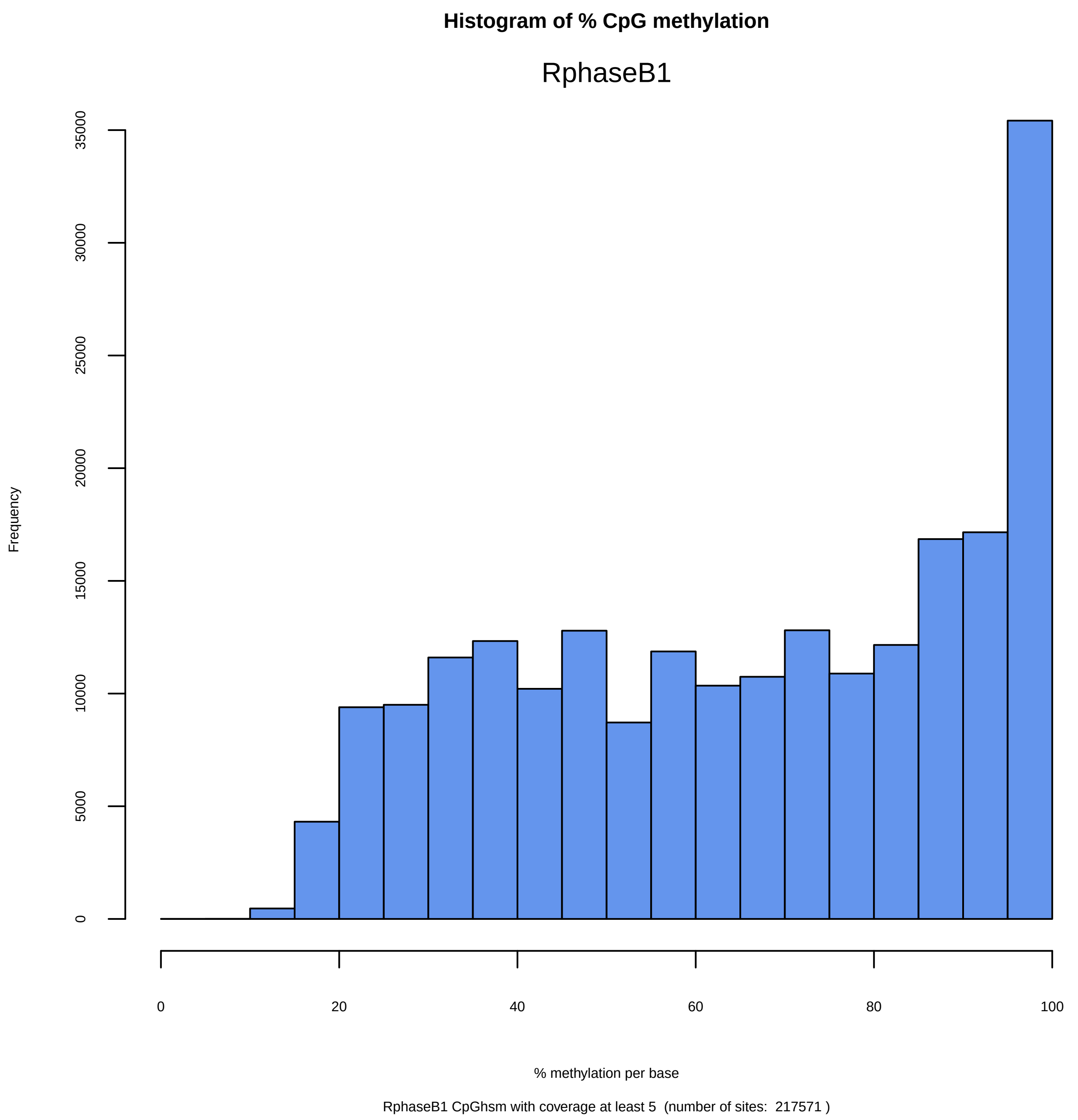

Histogram of % CpG methylation
RphaseB1
Frequency
0
5000
10000
15000
20000
25000
30000
35000
0
20
40
60
80
100
% methylation per base
RphaseB1 CpGhsm with coverage at least 5 (number of sites: 217571 )

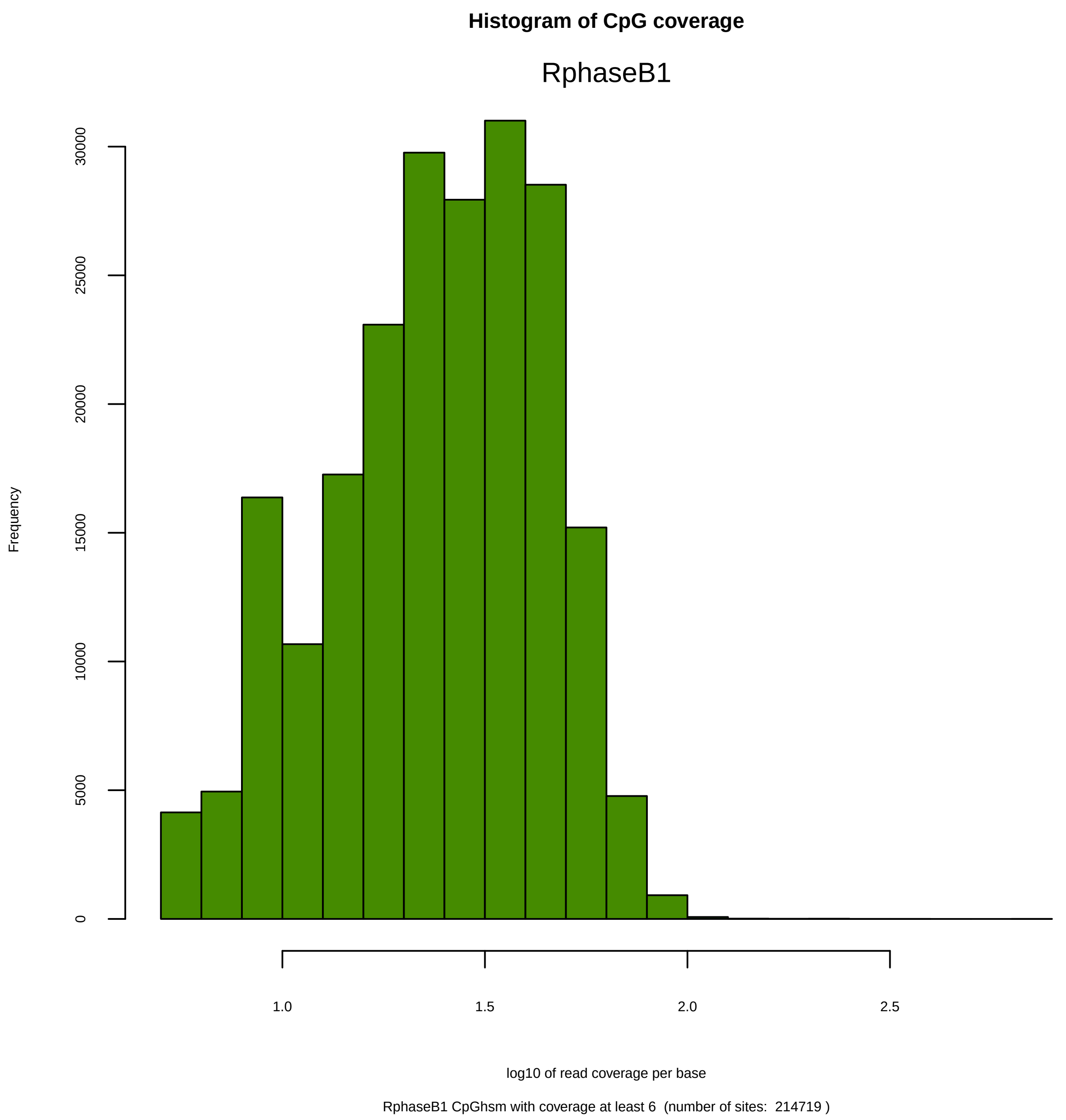

Histogram of CpG coverage
RphaseB1
Frequency
0
5000
10000
15000
20000
25000
30000
1.0
1.5
2.0
2.5
log10 of read coverage per base
RphaseB1 CpGhsm with coverage at least 6  (number of sites:  214719 )

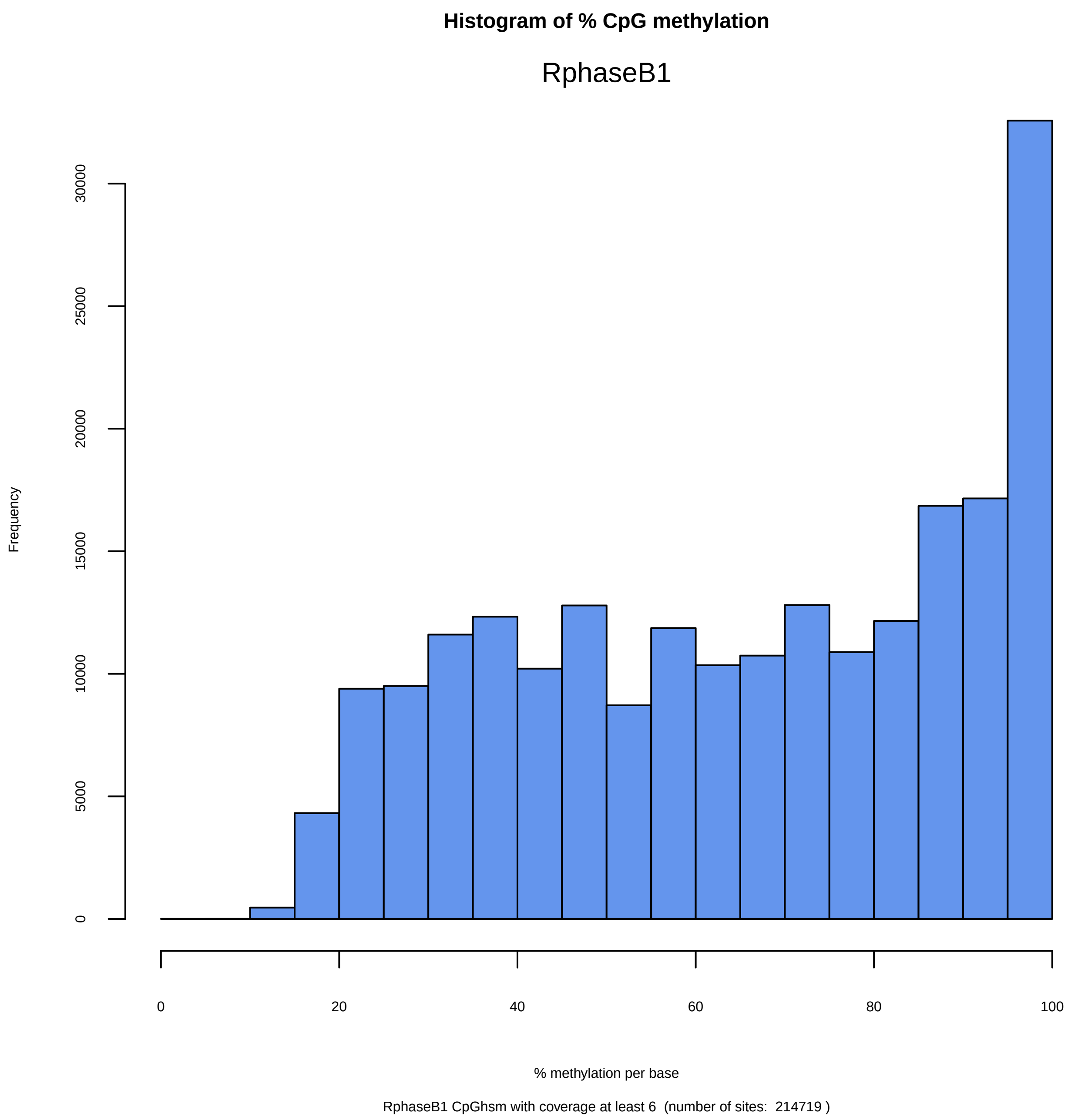
Histogram of % CpG methylation
RphaseB1
Frequency
0
5000
10000
15000
20000
25000
30000
0
20
40
60
80
100
% methylation per base
RphaseB1 CpGhsm with coverage at least 6 (number of sites: 214719 )

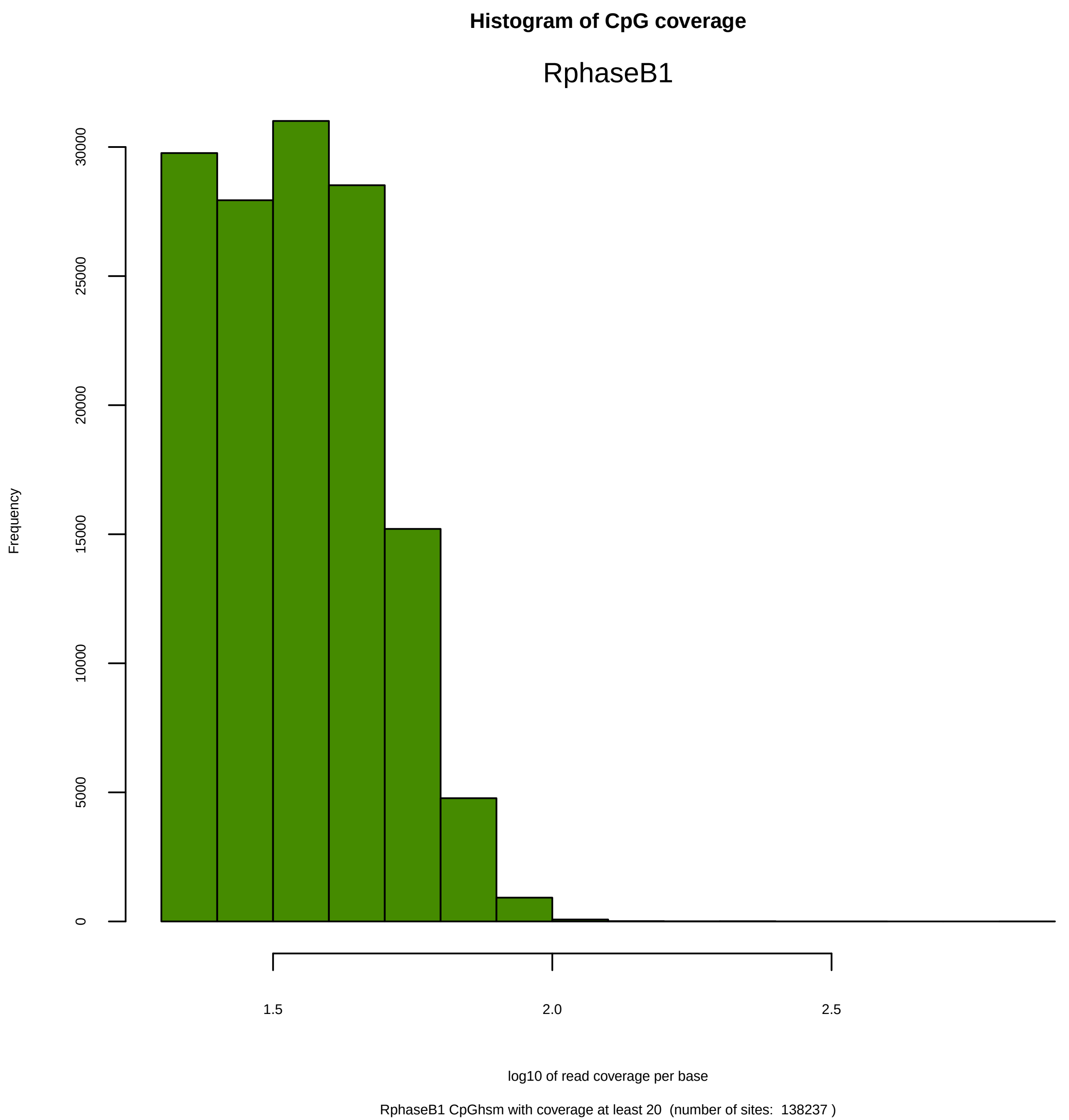
Histogram of CpG coverage
RphaseB1
Frequency
0
5000
10000
15000
20000
25000
30000
1.5
2.0
2.5
log10 of read coverage per base
RphaseB1 CpGhsm with coverage at least 20  (number of sites:  138237 )

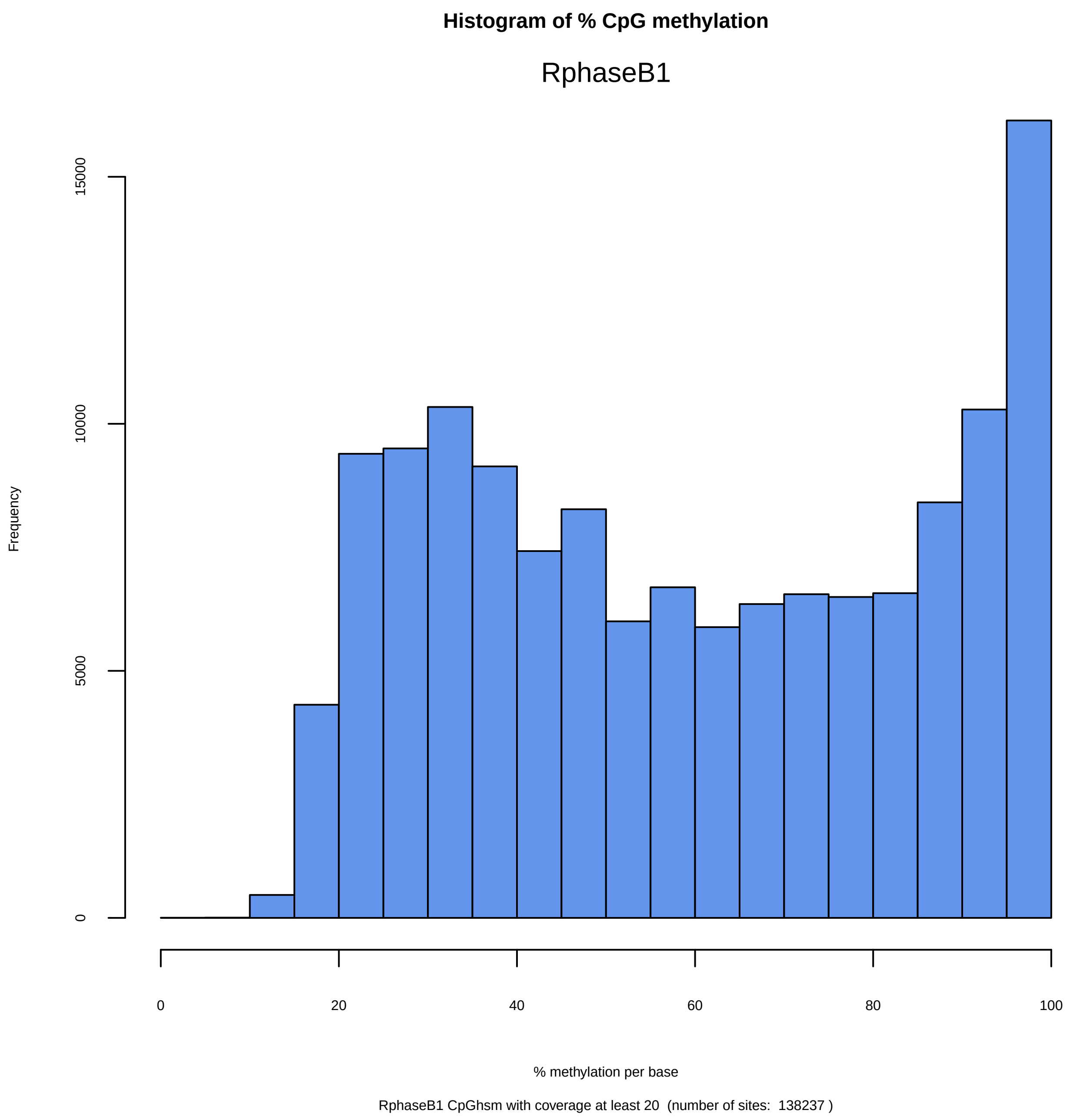
Histogram of % CpG methylation
RphaseB1
Frequency
0
5000
10000
15000
0
20
40
60
80
100
% methylation per base
RphaseB1 CpGhsm with coverage at least 20  (number of sites:  138237 )

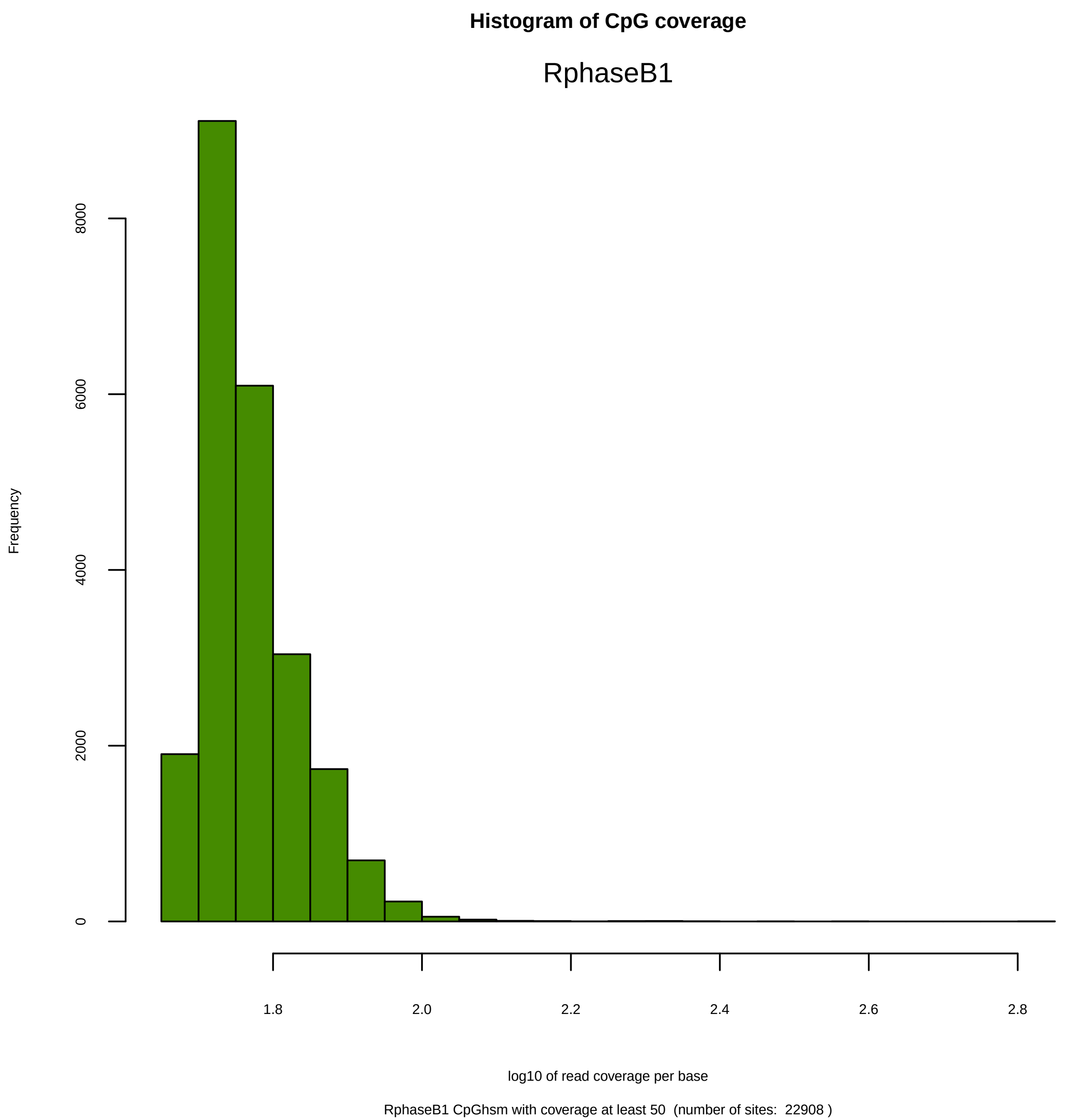
Histogram of CpG coverage
RphaseB1
Frequency
0
2000
4000
6000
8000
1.8
2.0
2.2
2.4
2.6
2.8
log10 of read coverage per base
RphaseB1 CpGhsm with coverage at least 50  (number of sites:  22908 )

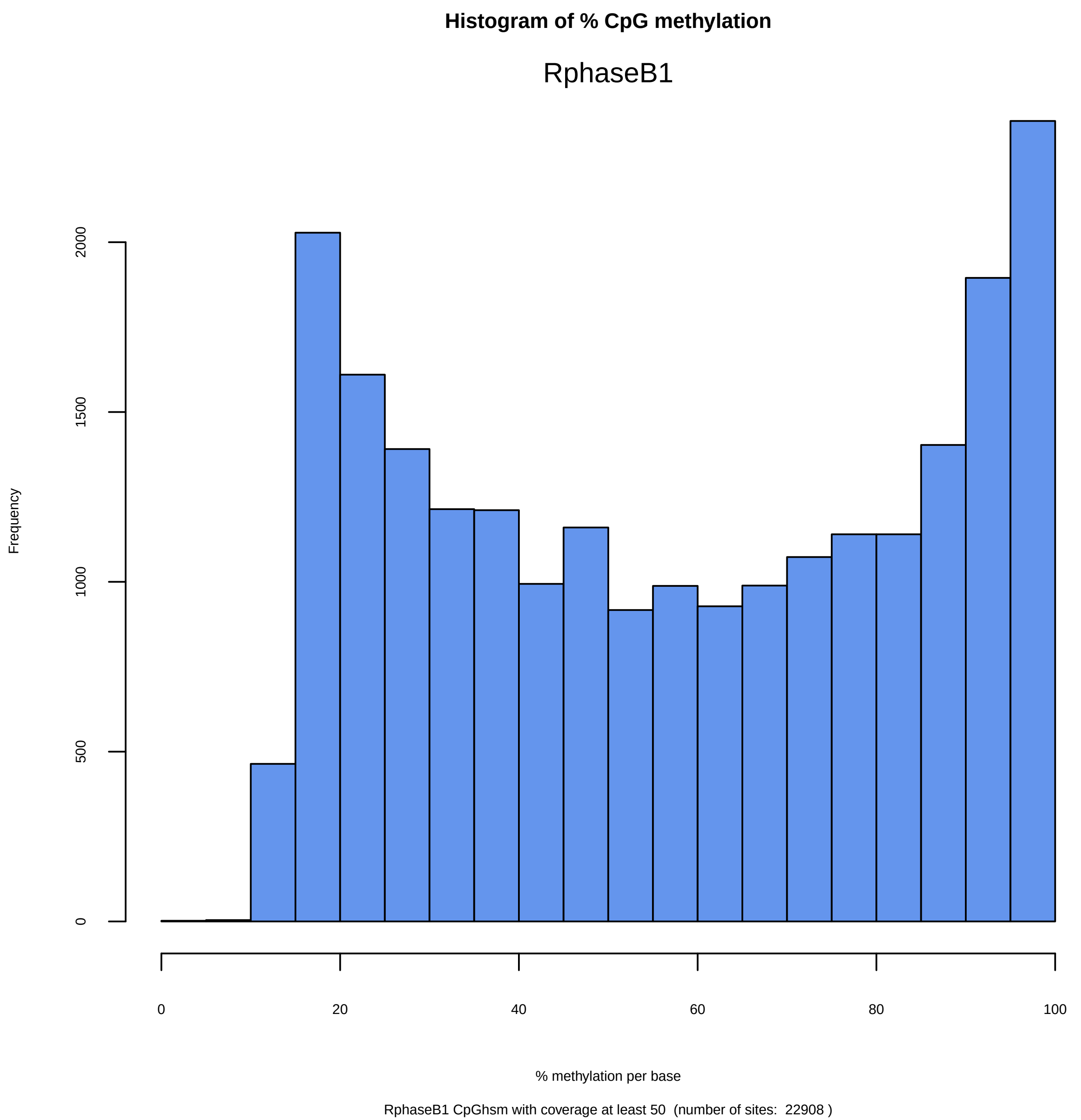
Histogram of % CpG methylation
RphaseB1
Frequency
0
500
1000
1500
2000
0
20
40
60
80
100
% methylation per base
RphaseB1 CpGhsm with coverage at least 50  (number of sites:  22908 )

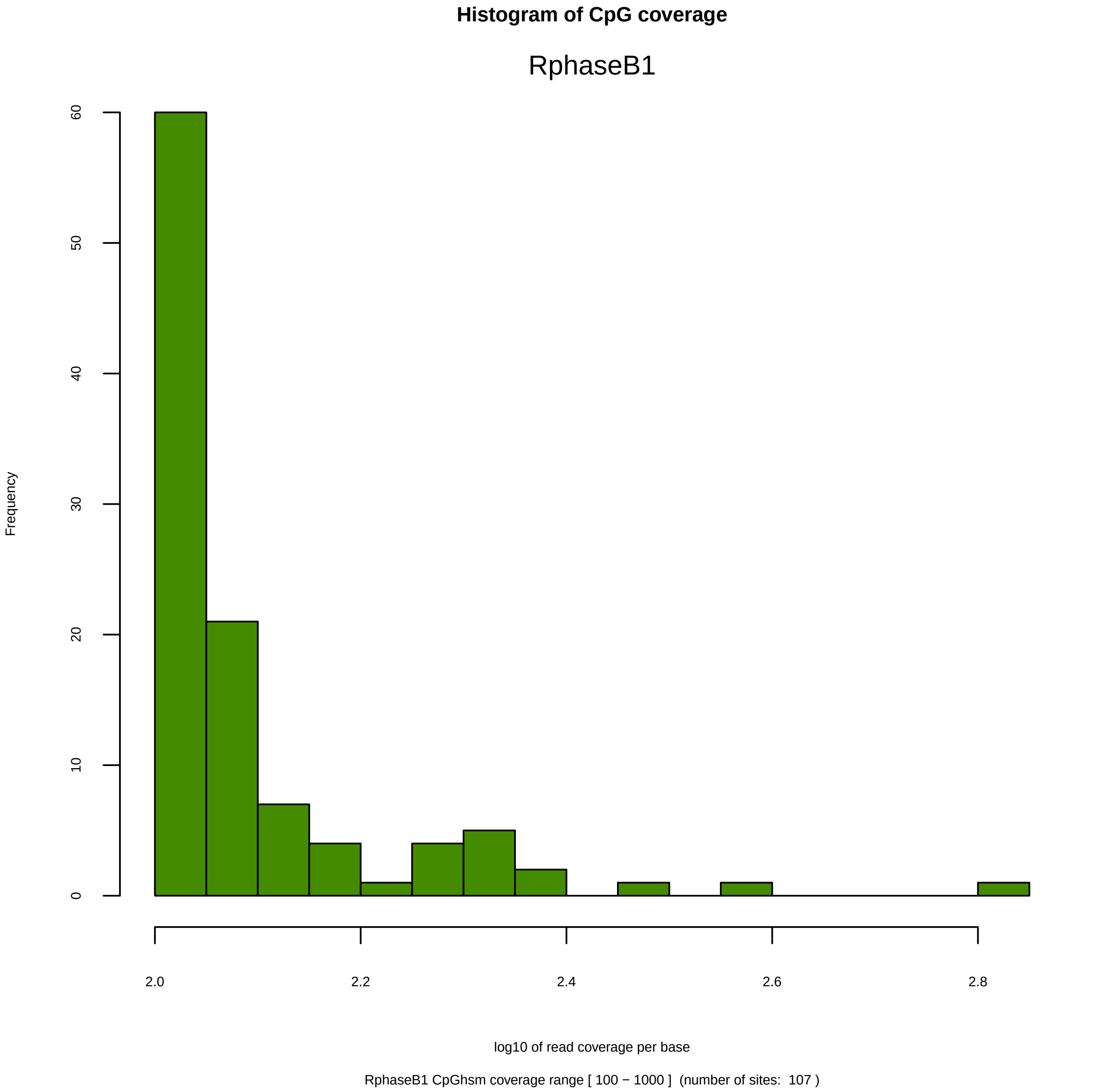
Histogram of CpG coverage
RphaseB1
Frequency
0
10
20
30
40
50
60
2.0
2.2
2.4
2.6
2.8
log10 of read coverage per base
RphaseB1 CpGhsm coverage range [ 100 − 1000 ]  (number of sites:  107 )

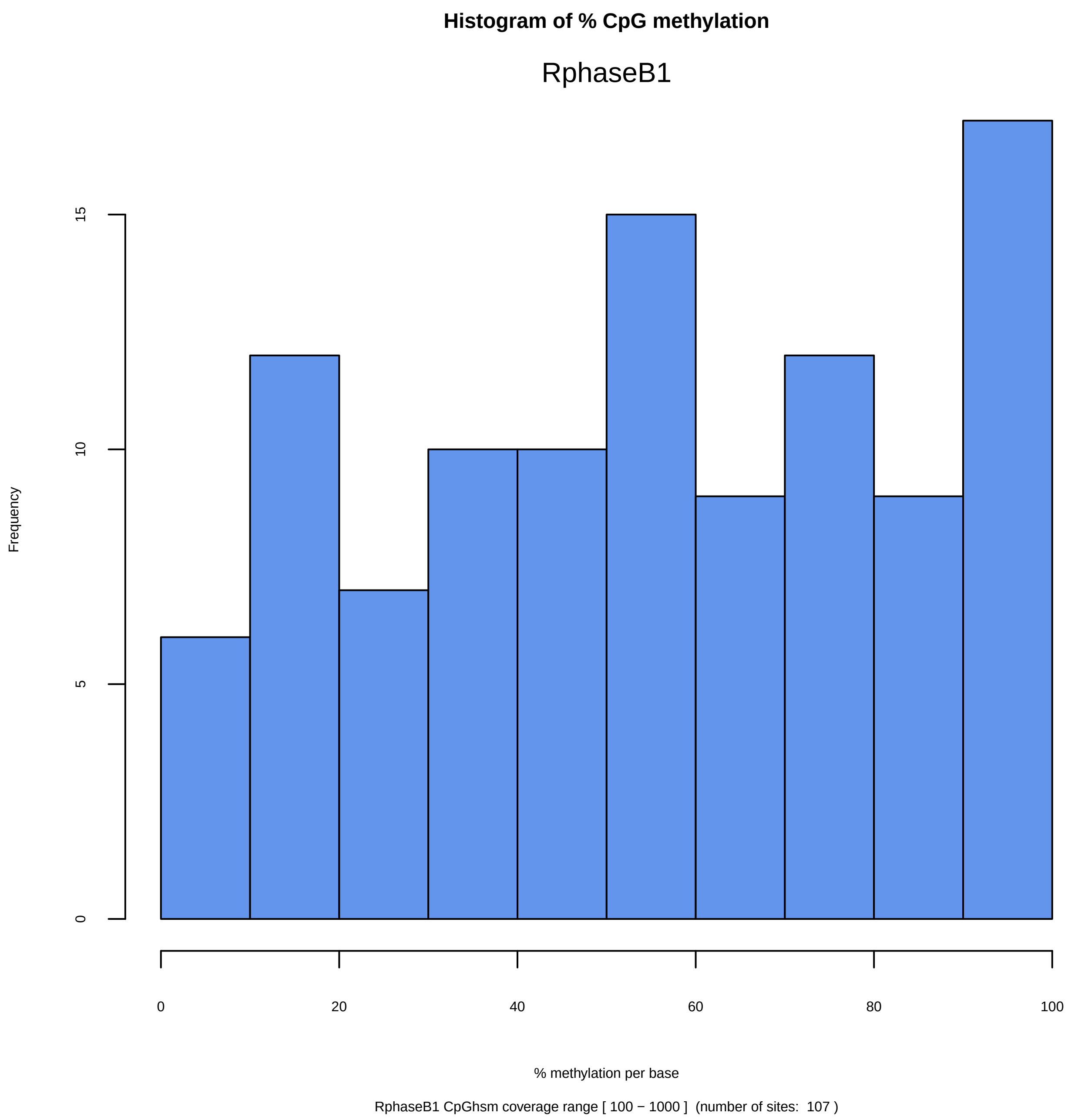
Histogram of % CpG methylation
RphaseB1
Frequency
0
5
10
15
0
20
40
60
80
100
% methylation per base
RphaseB1 CpGhsm coverage range [ 100 − 1000 ]  (number of sites:  107 )

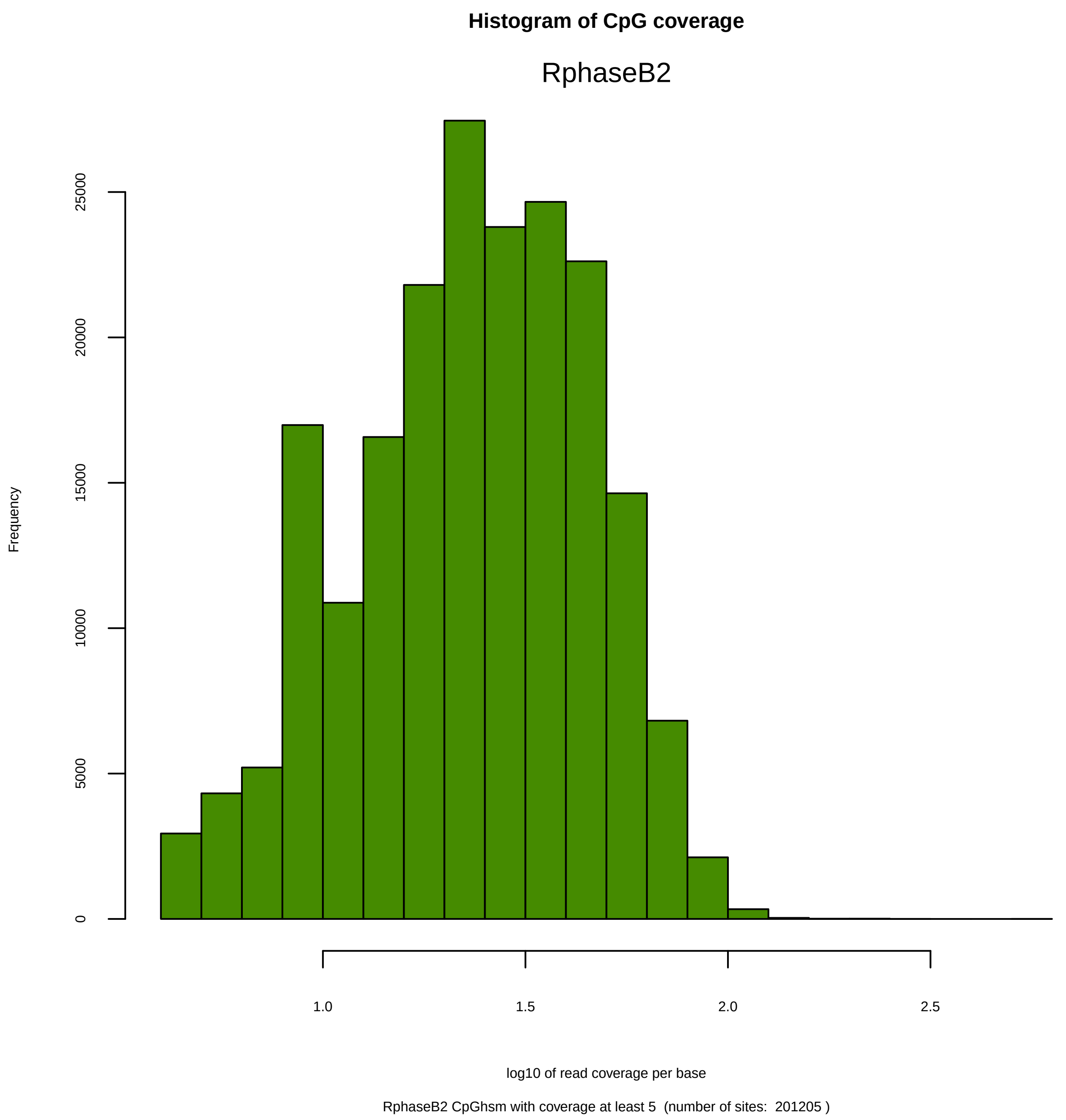

Histogram of CpG coverage
RphaseB2
Frequency
0
5000
10000
15000
20000
25000
1.0
1.5
2.0
2.5
log10 of read coverage per base
RphaseB2 CpGhsm with coverage at least 5  (number of sites:  201205 )

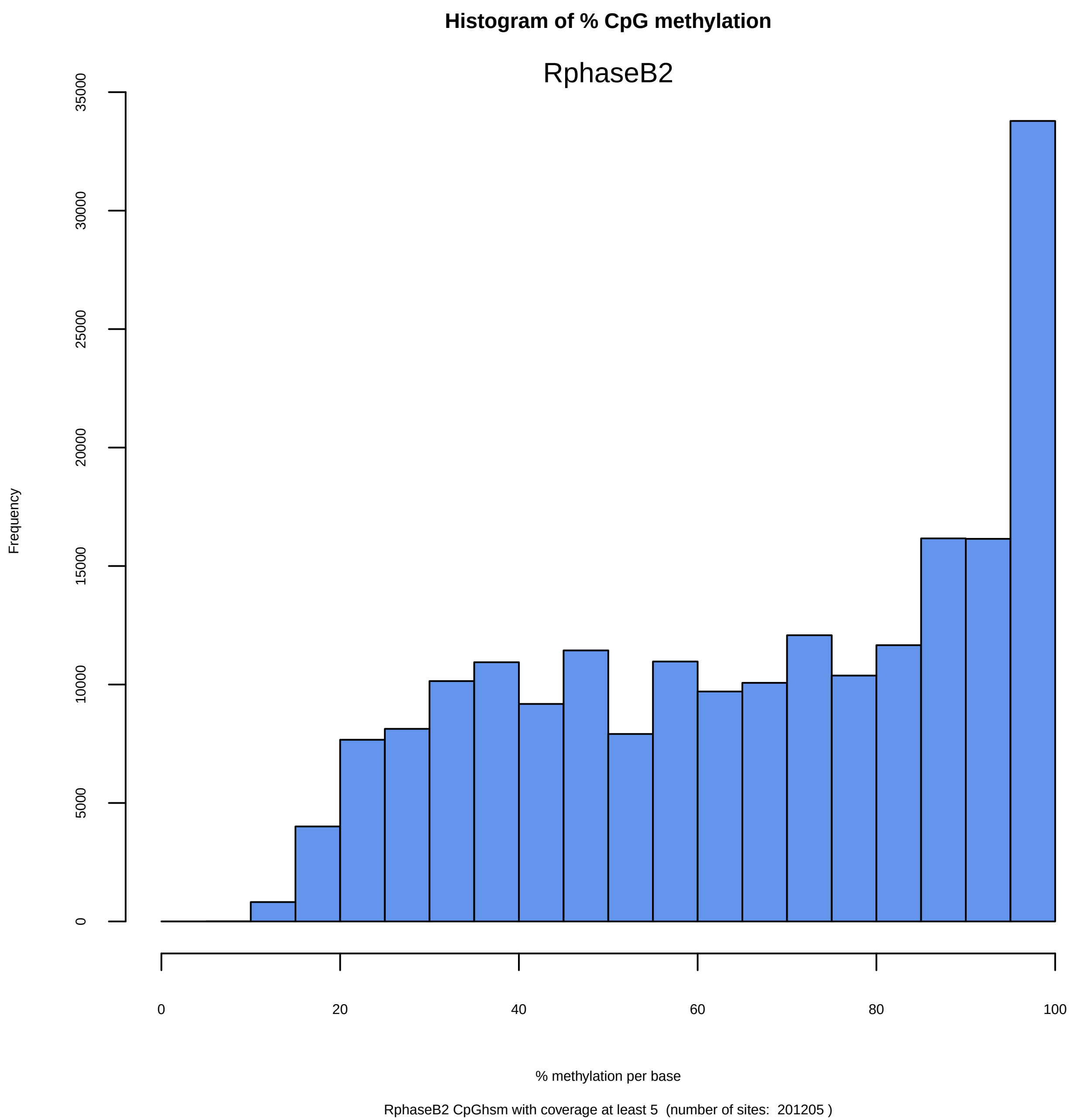
Histogram of % CpG methylation
RphaseB2
Frequency
0
5000
10000
15000
20000
25000
30000
35000
0
20
40
60
80
100
% methylation per base
RphaseB2 CpGhsm with coverage at least 5  (number of sites:  201205 )

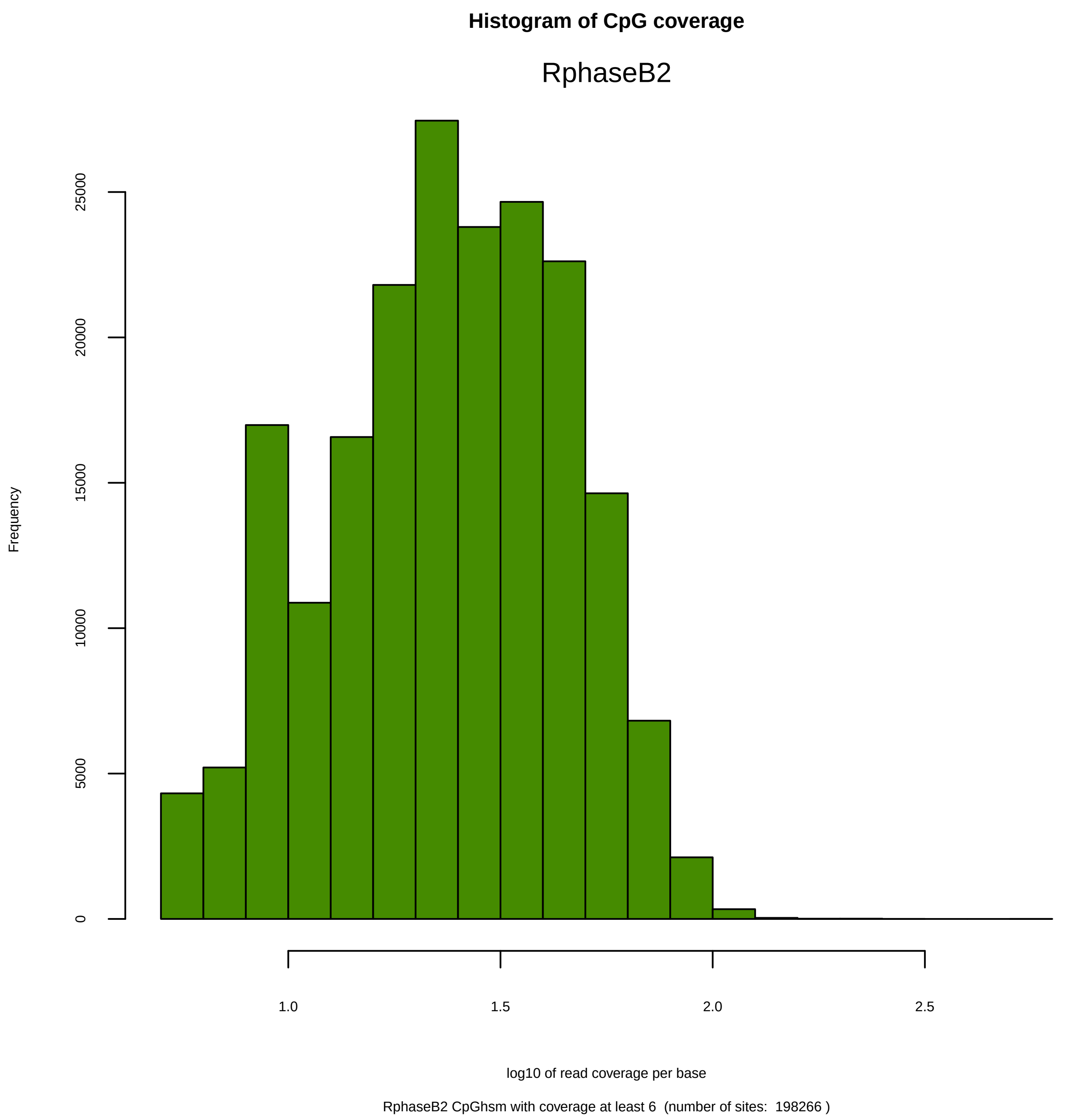

Histogram of CpG coverage
RphaseB2
Frequency
0
5000
10000
15000
20000
25000
1.0
1.5
2.0
2.5
log10 of read coverage per base
RphaseB2 CpGhsm with coverage at least 6  (number of sites:  198266 )

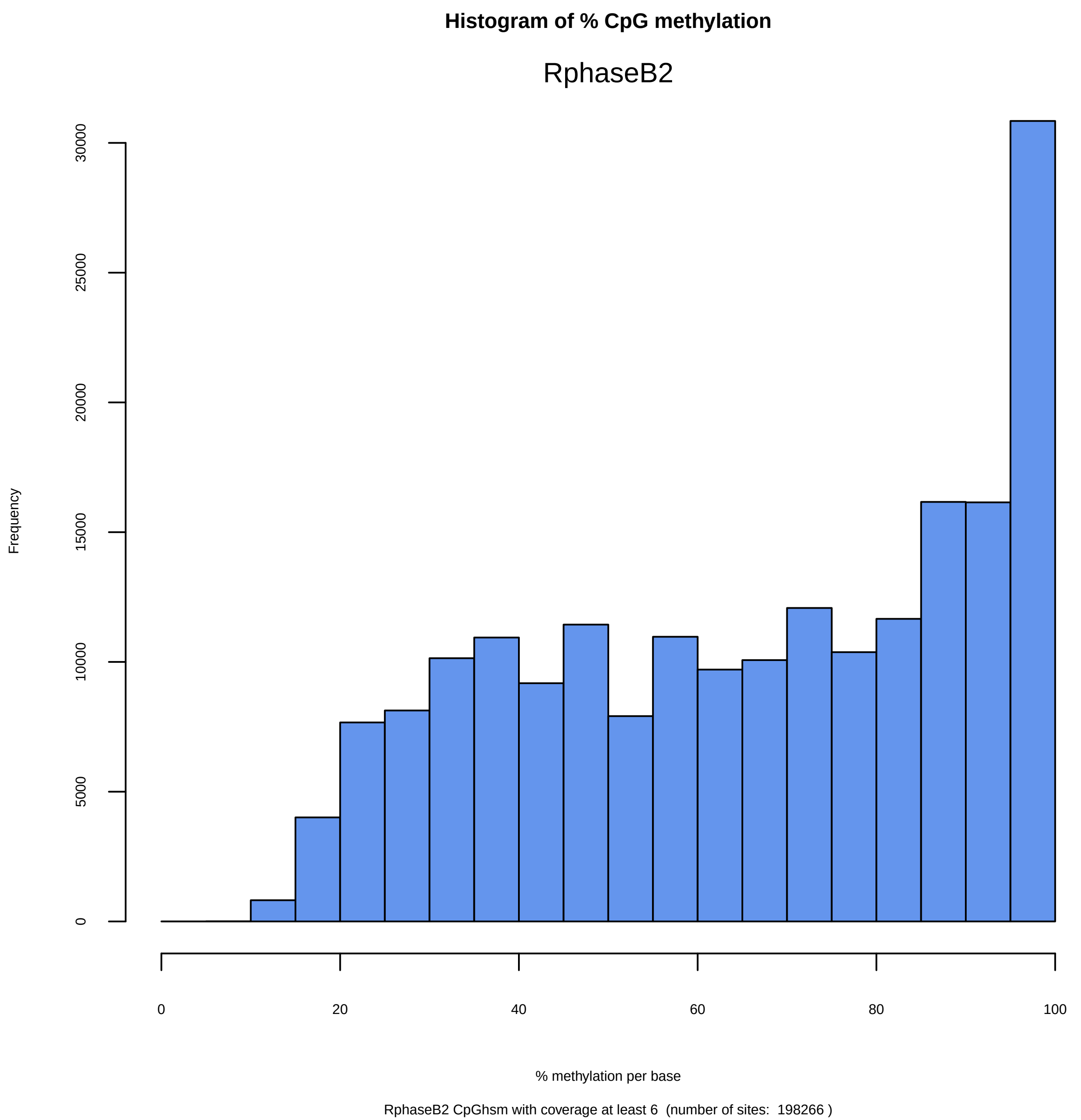

Histogram of % CpG methylation
RphaseB2
Frequency
0
5000
10000
15000
20000
25000
30000
0
20
40
60
80
100
% methylation per base
RphaseB2 CpGhsm with coverage at least 6 (number of sites: 198266 )

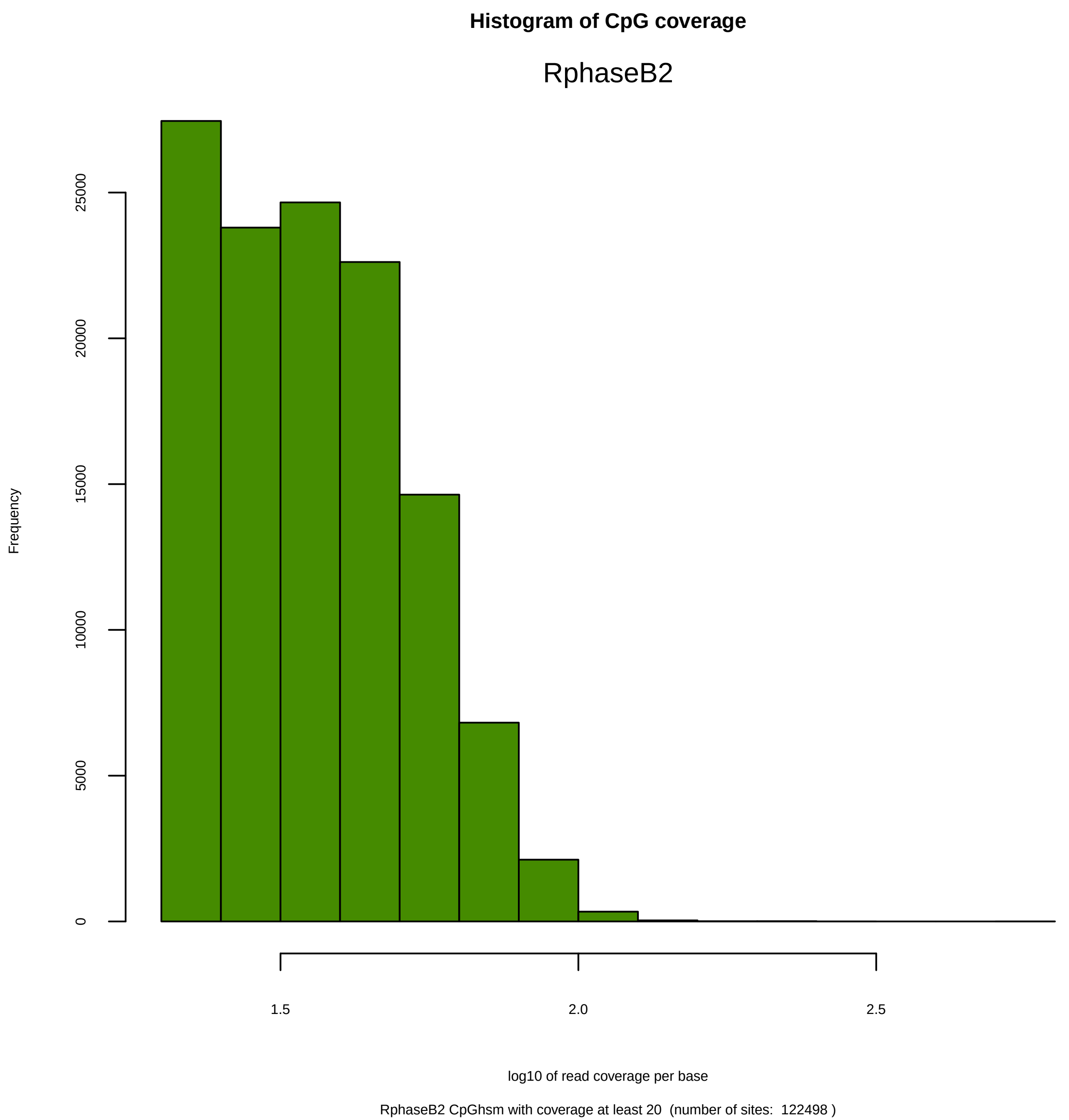

Histogram of CpG coverage
RphaseB2
Frequency
0
5000
10000
15000
20000
25000
1.5
2.0
2.5
log10 of read coverage per base
RphaseB2 CpGhsm with coverage at least 20 (number of sites: 122498 )

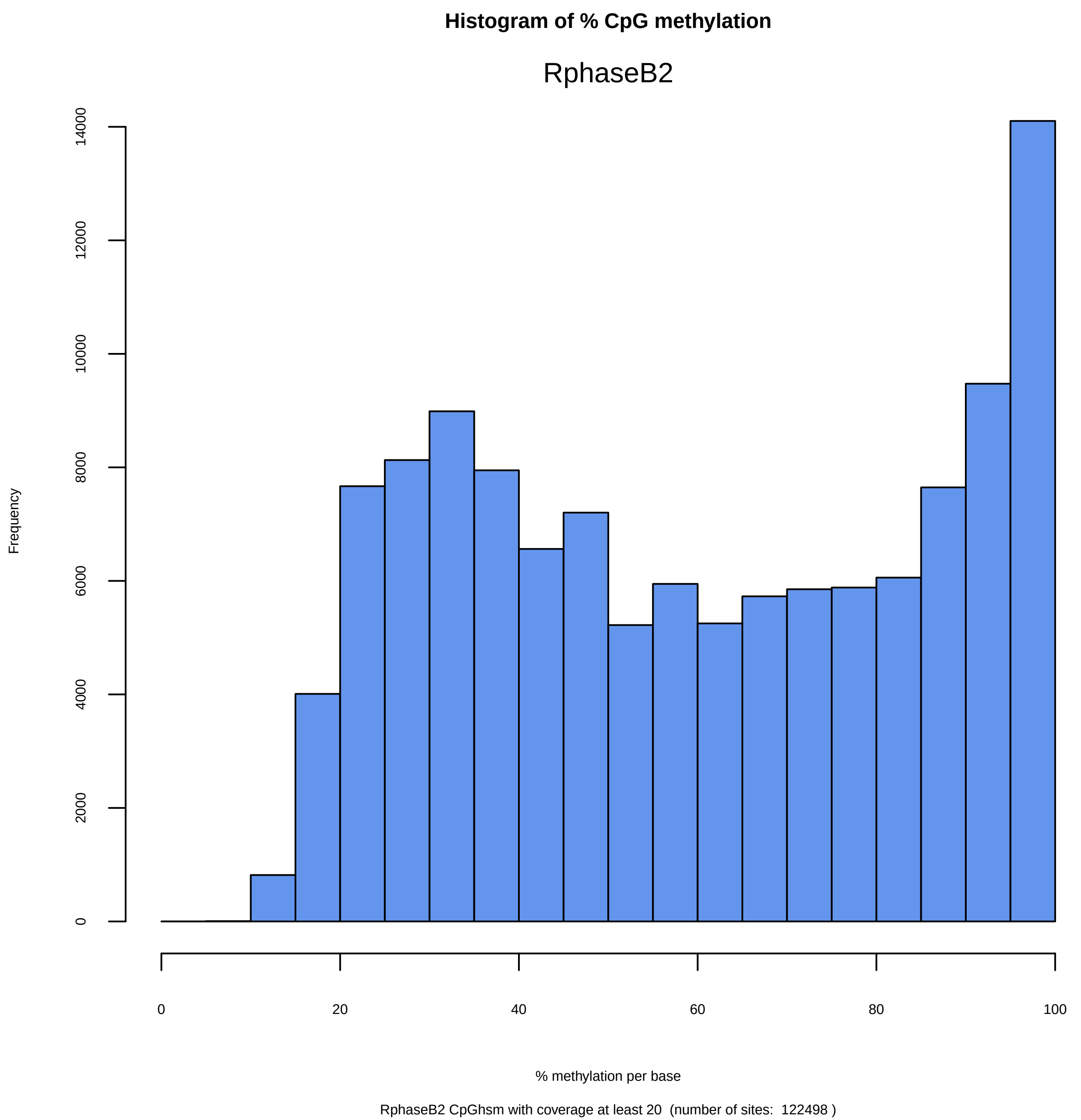
Histogram of % CpG methylation
RphaseB2
Frequency
0
2000
4000
6000
8000
10000
12000
14000
0
20
40
60
80
100
% methylation per base
RphaseB2 CpGhsm with coverage at least 20 (number of sites: 122498 )

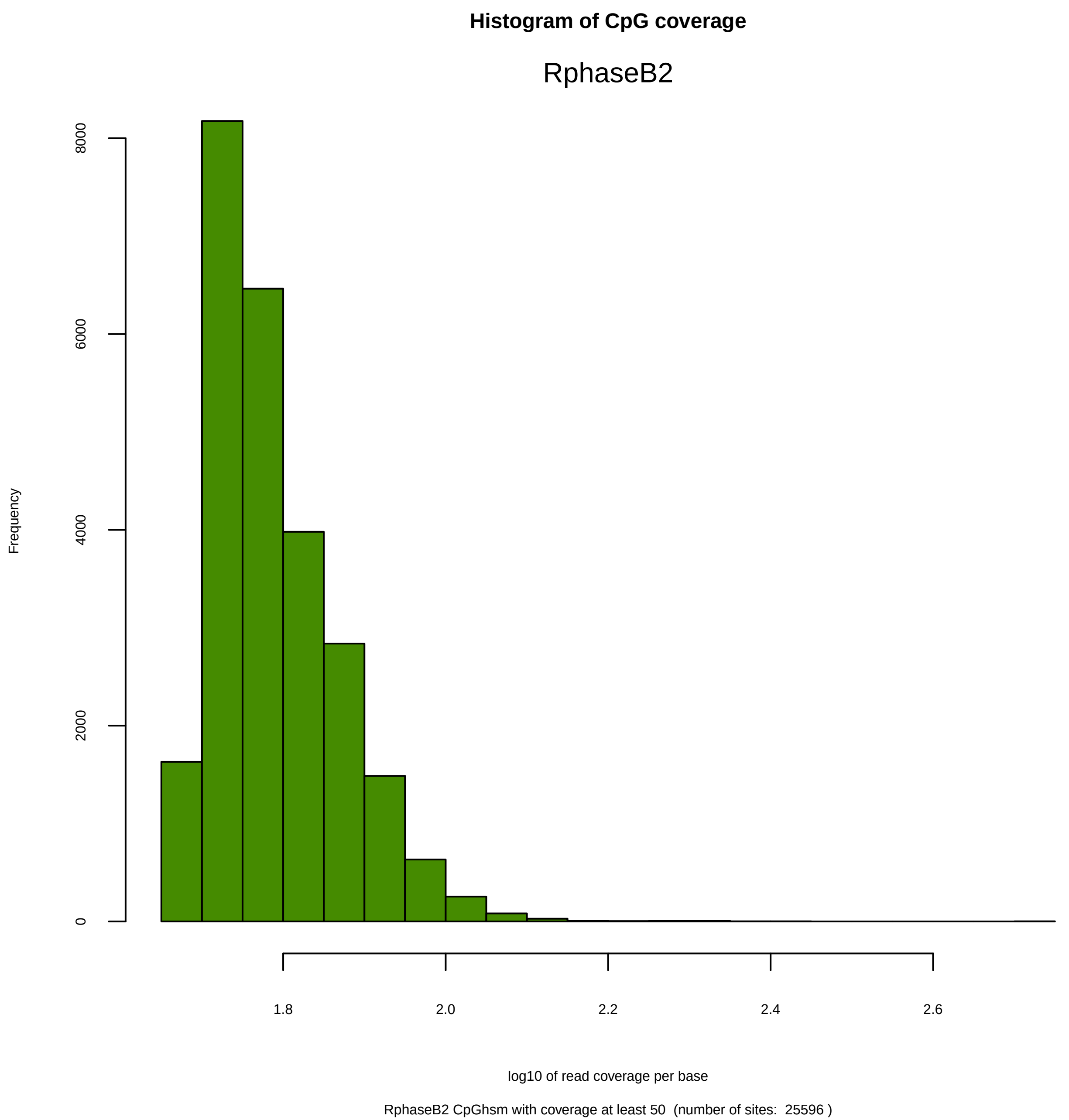
Histogram of CpG coverage
RphaseB2
Frequency
0
2000
4000
6000
8000
1.8
2.0
2.2
2.4
2.6
log10 of read coverage per base
RphaseB2 CpGhsm with coverage at least 50  (number of sites:  25596 )

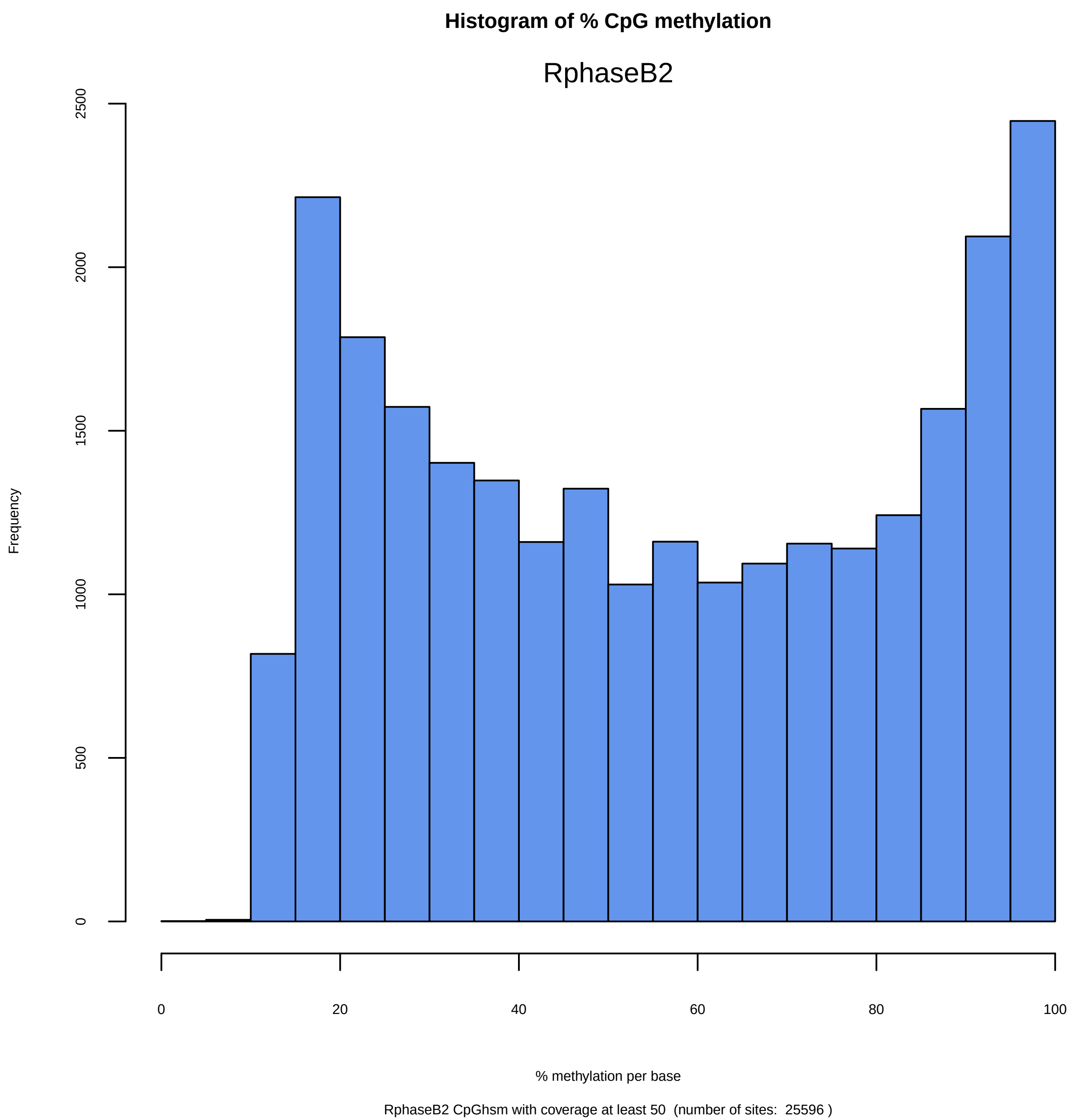
Histogram of % CpG methylation
RphaseB2
Frequency
0
500
1000
1500
2000
2500
0
20
40
60
80
100
% methylation per base
RphaseB2 CpGhsm with coverage at least 50  (number of sites:  25596 )

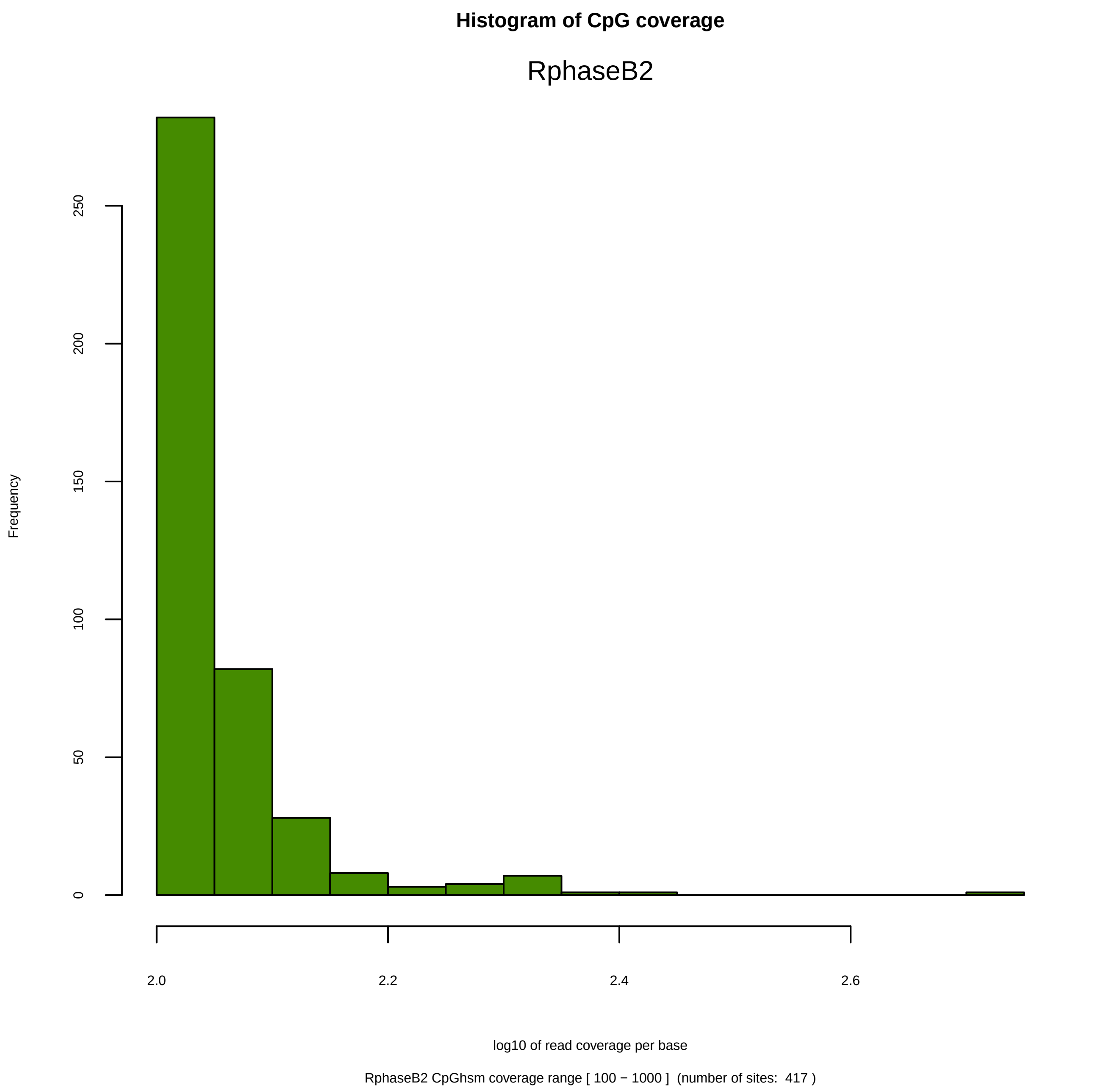

Histogram of CpG coverage
RphaseB2
Frequency
0
50
100
150
200
250
2.0
2.2
2.4
2.6
log10 of read coverage per base
RphaseB2 CpGhsm coverage range [ 100 − 1000 ]  (number of sites:  417 )

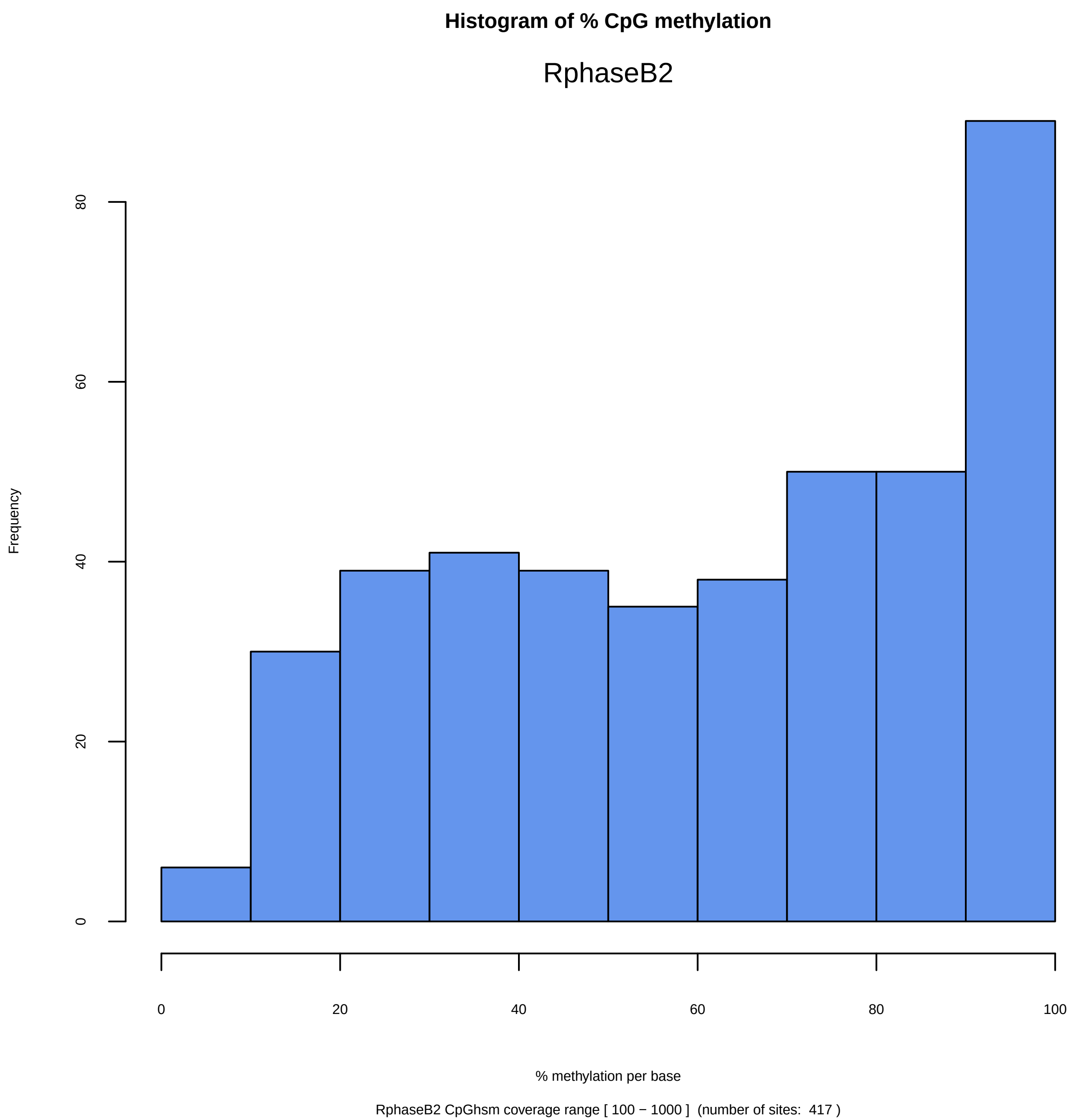
Histogram of % CpG methylation
RphaseB2
Frequency
0
20
40
60
80
0
20
40
60
80
100
% methylation per base
RphaseB2 CpGhsm coverage range [ 100 − 1000 ] (number of sites: 417 )

```
Directory: PWC File: 0READMEpwc

PWC - Pairwise comparisons between all samples.

Input: studymk, studymc, nbrpms, hheight, nbrpnts

Output: files pwc-*.txt pwc-.*pdf

Notes: The output is generated by BWASPR::cmpSites(), which determines for each pairwise
        comparison the numbers of common and distinct sites.  The function also calculates
        the overlap index between the samples and estimates the size of the common pool of
        potential methylation sites.

       The CpGscd data input (studymc) is necessary to determine sites that are detectable
        in both samples, and nbrpms (listed in the *.par file specified in the Ob_LI.conf
        configuration file as argument to TOTALNBRPMSITES) provides the total number of
        potential methylation sites (typically, all CpG sites), a necessary parameter for
        the estimations.
       hheight (default: 0.10) specifies the y-axis limit in the histogram plots.
       nbrpnts (default: 5000) specifies the number of common sites to be plotted in the
        methylation levels scatter plot.
```

```
Directory: PWC File: pwc-Ob_LI_BCphaseA1.vs.Ob_LI_BCphaseA2.txt

Numbers of common and distinct sites comparing Ob_LI_BCphaseA1 versus Ob_LI_BCphaseA2
================================================================================

  total number of potential sites:  28317694

  number of "Ob_LI_BCphaseA1_hsm"         sites:     197029
  number of "Ob_LI_BCphaseA2_hsm"         sites:     213136
    number of "Ob_LI_BCphaseA1_hsm"-unique sites:     28042
    number of common sites:        168987
    number of "Ob_LI_BCphaseA2_hsm"-unique sites:     44149
  total number of "Ob_LI_BCphaseA1_hsm+Ob_LI_BCphaseA2_hsm"-sites observed:  241178

  number of "Ob_LI_BCphaseA1_scd"        sites: 18563480 ( 65.55% of total)
  number of "Ob_LI_BCphaseA2_scd"        sites: 18737737 ( 66.17% of total)
    number of "Ob_LI_BCphaseA1_scd"-unique sites:      1644221
    number of sites in common:      16919259   (Expected:  12283402; O/E:       1.4)
    number of "Ob_LI_BCphaseA2_scd"-unique sites:      1818478
  total number of "Ob_LI_BCphaseA1_scd+Ob_LI_BCphaseA2_scd"-sites observed:   20381958

  number of sites in "Ob_LI_BCphaseA1_hsm" that are not  detectable in "Ob_LI_BCphaseA2_hsm":    6815
  number of sites in "Ob_LI_BCphaseA1_hsm" that are also detectable in "Ob_LI_BCphaseA2_hsm":  190214
    number of sites unique to "Ob_LI_BCphaseA1_hsm" although detectable in "Ob_LI_BCphaseA2_hsm":          21227
    number of "Ob_LI_BCphaseA1_hsm" / "Ob_LI_BCphaseA2_hsm" common sites                        :         168987      (Expected:   2314; O/E:      73.0)

  number of sites in "Ob_LI_BCphaseA2_hsm" that are not  detectable in "Ob_LI_BCphaseA1_hsm":    7323
  number of sites in "Ob_LI_BCphaseA2_hsm" that are also detectable in "Ob_LI_BCphaseA1_hsm":  205813
    number of sites unique to "Ob_LI_BCphaseA2_hsm" although detectable in "Ob_LI_BCphaseA1_hsm":          36826
    number of "Ob_LI_BCphaseA2_hsm" / "Ob_LI_BCphaseA1_hsm" common sites                        :         168987      (Expected:   2314; O/E:      73.0)

  Overlap index of "Ob_LI_BCphaseA1_hsm" with "Ob_LI_BCphaseA2_hsm":         0.805
  Estimated number of "Ob_LI_BCphaseA1_hsm" = "Ob_LI_BCphaseA2_hsm" sites (assuming sampling from one population):             231666
  Adjusted population size of "Ob_LI_BCphaseA1_hsm" = "Ob_LI_BCphaseA2_hsm" sites (assuming all sites detectable):             387738 (  1.61x of observed)
```

```
Directory: PWC File: pwc-Ob_LI_BCphaseA1.vs.Ob_LI_BCphaseB1.txt

Numbers of common and distinct sites comparing Ob_LI_BCphaseA1 versus Ob_LI_BCphaseB1
================================================================================

  total number of potential sites:  28317694

  number of "Ob_LI_BCphaseA1_hsm"         sites:     197029
  number of "Ob_LI_BCphaseB1_hsm"         sites:     226855
    number of "Ob_LI_BCphaseA1_hsm"-unique sites:     21999
    number of common sites:        175030
    number of "Ob_LI_BCphaseB1_hsm"-unique sites:     51825
  total number of "Ob_LI_BCphaseA1_hsm+Ob_LI_BCphaseB1_hsm"-sites observed:  248854

  number of "Ob_LI_BCphaseA1_scd"        sites: 18563480 ( 65.55% of total)
  number of "Ob_LI_BCphaseB1_scd"        sites: 20546412 ( 72.56% of total)
    number of "Ob_LI_BCphaseA1_scd"-unique sites:       723732
    number of sites in common:      17839748   (Expected:  13469067; O/E:       1.3)
    number of "Ob_LI_BCphaseB1_scd"-unique sites:      2706664
  total number of "Ob_LI_BCphaseA1_scd+Ob_LI_BCphaseB1_scd"-sites observed:   21270144

  number of sites in "Ob_LI_BCphaseA1_hsm" that are not  detectable in "Ob_LI_BCphaseB1_hsm":    3189
  number of sites in "Ob_LI_BCphaseA1_hsm" that are also detectable in "Ob_LI_BCphaseB1_hsm":  193840
    number of sites unique to "Ob_LI_BCphaseA1_hsm" although detectable in "Ob_LI_BCphaseB1_hsm":          18810
    number of "Ob_LI_BCphaseA1_hsm" / "Ob_LI_BCphaseB1_hsm" common sites                        :         175030      (Expected:   2350; O/E:      74.5)

  number of sites in "Ob_LI_BCphaseB1_hsm" that are not  detectable in "Ob_LI_BCphaseA1_hsm":   10599
  number of sites in "Ob_LI_BCphaseB1_hsm" that are also detectable in "Ob_LI_BCphaseA1_hsm":  216256
    number of sites unique to "Ob_LI_BCphaseB1_hsm" although detectable in "Ob_LI_BCphaseA1_hsm":          41226
    number of "Ob_LI_BCphaseB1_hsm" / "Ob_LI_BCphaseA1_hsm" common sites                        :         175030      (Expected:   2350; O/E:      74.5)

  Overlap index of "Ob_LI_BCphaseA1_hsm" with "Ob_LI_BCphaseB1_hsm":         0.831
  Estimated number of "Ob_LI_BCphaseA1_hsm" = "Ob_LI_BCphaseB1_hsm" sites (assuming sampling from one population):             239496
  Adjusted population size of "Ob_LI_BCphaseA1_hsm" = "Ob_LI_BCphaseB1_hsm" sites (assuming all sites detectable):             380162 (  1.53x of observed)
```

```
Directory: PWC File: pwc-Ob_LI_BCphaseA1.vs.Ob_LI_BCphaseB2.txt

Numbers of common and distinct sites comparing Ob_LI_BCphaseA1 versus Ob_LI_BCphaseB2
================================================================================

  total number of potential sites:  28317694

  number of "Ob_LI_BCphaseA1_hsm"         sites:     197029
  number of "Ob_LI_BCphaseB2_hsm"         sites:     209549
    number of "Ob_LI_BCphaseA1_hsm"-unique sites:     28873
    number of common sites:        168156
    number of "Ob_LI_BCphaseB2_hsm"-unique sites:     41393
  total number of "Ob_LI_BCphaseA1_hsm+Ob_LI_BCphaseB2_hsm"-sites observed:  238422

  number of "Ob_LI_BCphaseA1_scd"        sites: 18563480 ( 65.55% of total)
  number of "Ob_LI_BCphaseB2_scd"        sites: 19220462 ( 67.87% of total)
    number of "Ob_LI_BCphaseA1_scd"-unique sites:      1351985
    number of sites in common:      17211495   (Expected:  12599849; O/E:       1.4)
    number of "Ob_LI_BCphaseB2_scd"-unique sites:      2008967
  total number of "Ob_LI_BCphaseA1_scd+Ob_LI_BCphaseB2_scd"-sites observed:   20572447

  number of sites in "Ob_LI_BCphaseA1_hsm" that are not  detectable in "Ob_LI_BCphaseB2_hsm":    5872
  number of sites in "Ob_LI_BCphaseA1_hsm" that are also detectable in "Ob_LI_BCphaseB2_hsm":  191157
    number of sites unique to "Ob_LI_BCphaseA1_hsm" although detectable in "Ob_LI_BCphaseB2_hsm":          23001
    number of "Ob_LI_BCphaseA1_hsm" / "Ob_LI_BCphaseB2_hsm" common sites                        :         168156      (Expected:   2240; O/E:      75.1)

  number of sites in "Ob_LI_BCphaseB2_hsm" that are not  detectable in "Ob_LI_BCphaseA1_hsm":    7821
  number of sites in "Ob_LI_BCphaseB2_hsm" that are also detectable in "Ob_LI_BCphaseA1_hsm":  201728
    number of sites unique to "Ob_LI_BCphaseB2_hsm" although detectable in "Ob_LI_BCphaseA1_hsm":          33572
    number of "Ob_LI_BCphaseB2_hsm" / "Ob_LI_BCphaseA1_hsm" common sites                        :         168156      (Expected:   2240; O/E:      75.1)

  Overlap index of "Ob_LI_BCphaseA1_hsm" with "Ob_LI_BCphaseB2_hsm":         0.790
  Estimated number of "Ob_LI_BCphaseA1_hsm" = "Ob_LI_BCphaseB2_hsm" sites (assuming sampling from one population):             229321
  Adjusted population size of "Ob_LI_BCphaseA1_hsm" = "Ob_LI_BCphaseB2_hsm" sites (assuming all sites detectable):             377297 (  1.58x of observed)
```

```
Directory: PWC File: pwc-Ob_LI_BCphaseA1.vs.Ob_LI_RphaseA1.txt

Numbers of common and distinct sites comparing Ob_LI_BCphaseA1 versus Ob_LI_RphaseA1
================================================================================

  total number of potential sites:  28317694

  number of "Ob_LI_BCphaseA1_hsm"         sites:     197029
  number of "Ob_LI_RphaseA1_hsm"         sites:213360
    number of "Ob_LI_BCphaseA1_hsm"-unique sites:     26850
    number of common sites:        170179
    number of "Ob_LI_RphaseA1_hsm"-unique sites:      43181
  total number of "Ob_LI_BCphaseA1_hsm+Ob_LI_RphaseA1_hsm"-sites observed:   240210

  number of "Ob_LI_BCphaseA1_scd"        sites: 18563480 ( 65.55% of total)
  number of "Ob_LI_RphaseA1_scd"        sites:  19848503 ( 70.09% of total)
    number of "Ob_LI_BCphaseA1_scd"-unique sites:      1023490
    number of sites in common:      17539990   (Expected:  13011557; O/E:       1.3)
    number of "Ob_LI_RphaseA1_scd"-unique sites:       2308513
  total number of "Ob_LI_BCphaseA1_scd+Ob_LI_RphaseA1_scd"-sites observed:    20871993

  number of sites in "Ob_LI_BCphaseA1_hsm" that are not  detectable in "Ob_LI_RphaseA1_hsm":     4267
  number of sites in "Ob_LI_BCphaseA1_hsm" that are also detectable in "Ob_LI_RphaseA1_hsm":   192762
    number of sites unique to "Ob_LI_BCphaseA1_hsm" although detectable in "Ob_LI_RphaseA1_hsm":           22583
    number of "Ob_LI_BCphaseA1_hsm" / "Ob_LI_RphaseA1_hsm" common sites                        :          170179      (Expected:   2245; O/E:      75.8)

  number of sites in "Ob_LI_RphaseA1_hsm" that are not  detectable in "Ob_LI_BCphaseA1_hsm":     9039
  number of sites in "Ob_LI_RphaseA1_hsm" that are also detectable in "Ob_LI_BCphaseA1_hsm":   204321
    number of sites unique to "Ob_LI_RphaseA1_hsm" although detectable in "Ob_LI_BCphaseA1_hsm":           34142
    number of "Ob_LI_RphaseA1_hsm" / "Ob_LI_BCphaseA1_hsm" common sites                        :          170179      (Expected:   2245; O/E:      75.8)

  Overlap index of "Ob_LI_BCphaseA1_hsm" with "Ob_LI_RphaseA1_hsm":          0.795
  Estimated number of "Ob_LI_BCphaseA1_hsm" = "Ob_LI_RphaseA1_hsm" sites (assuming sampling from one population):              231435
  Adjusted population size of "Ob_LI_BCphaseA1_hsm" = "Ob_LI_RphaseA1_hsm" sites (assuming all sites detectable):              373643 (  1.56x of observed)
```

```
Directory: PWC File: pwc-Ob_LI_BCphaseA1.vs.Ob_LI_RphaseA2.txt

Numbers of common and distinct sites comparing Ob_LI_BCphaseA1 versus Ob_LI_RphaseA2
================================================================================

  total number of potential sites:   28317694

  number of "Ob_LI_BCphaseA1_hsm"         sites:     197029
  number of "Ob_LI_RphaseA2_hsm"         sites: 265465
    number of "Ob_LI_BCphaseA1_hsm"-unique sites:     13337
    number of common sites:        183692
    number of "Ob_LI_RphaseA2_hsm"-unique sites:      81773
  total number of "Ob_LI_BCphaseA1_hsm+Ob_LI_RphaseA2_hsm"-sites observed:   278802

  number of "Ob_LI_BCphaseA1_scd"        sites: 18563480 ( 65.55% of total)
  number of "Ob_LI_RphaseA2_scd"        sites:  20631688 ( 72.86% of total)
    number of "Ob_LI_BCphaseA1_scd"-unique sites:       733178
    number of sites in common:      17830302   (Expected:  13524969; O/E:       1.3)
    number of "Ob_LI_RphaseA2_scd"-unique sites:       2801386
  total number of "Ob_LI_BCphaseA1_scd+Ob_LI_RphaseA2_scd"-sites observed:    21364866

  number of sites in "Ob_LI_BCphaseA1_hsm" that are not  detectable in "Ob_LI_RphaseA2_hsm":     2192
  number of sites in "Ob_LI_BCphaseA1_hsm" that are also detectable in "Ob_LI_RphaseA2_hsm":   194837
    number of sites unique to "Ob_LI_BCphaseA1_hsm" although detectable in "Ob_LI_RphaseA2_hsm":           11145
    number of "Ob_LI_BCphaseA1_hsm" / "Ob_LI_RphaseA2_hsm" common sites                        :          183692      (Expected:   2743; O/E:      67.0)

  number of sites in "Ob_LI_RphaseA2_hsm" that are not  detectable in "Ob_LI_BCphaseA1_hsm":    14485
  number of sites in "Ob_LI_RphaseA2_hsm" that are also detectable in "Ob_LI_BCphaseA1_hsm":   250980
    number of sites unique to "Ob_LI_RphaseA2_hsm" although detectable in "Ob_LI_BCphaseA1_hsm":           67288
    number of "Ob_LI_RphaseA2_hsm" / "Ob_LI_BCphaseA1_hsm" common sites                        :          183692      (Expected:   2743; O/E:      67.0)

  Overlap index of "Ob_LI_BCphaseA1_hsm" with "Ob_LI_RphaseA2_hsm":          0.903
  Estimated number of "Ob_LI_BCphaseA1_hsm" = "Ob_LI_RphaseA2_hsm" sites (assuming sampling from one population):              266208
  Adjusted population size of "Ob_LI_BCphaseA1_hsm" = "Ob_LI_RphaseA2_hsm" sites (assuming all sites detectable):              422785 (  1.52x of observed)
```

```
Directory: PWC File: pwc-Ob_LI_BCphaseA1.vs.Ob_LI_RphaseB1.txt

Numbers of common and distinct sites comparing Ob_LI_BCphaseA1 versus Ob_LI_RphaseB1
================================================================================

  total number of potential sites:  28317694

  number of "Ob_LI_BCphaseA1_hsm"         sites:     197029
  number of "Ob_LI_RphaseB1_hsm"         sites:217571
    number of "Ob_LI_BCphaseA1_hsm"-unique sites:     24914
    number of common sites:        172115
    number of "Ob_LI_RphaseB1_hsm"-unique sites:      45456
  total number of "Ob_LI_BCphaseA1_hsm+Ob_LI_RphaseB1_hsm"-sites observed:   242485

  number of "Ob_LI_BCphaseA1_scd"        sites: 18563480 ( 65.55% of total)
  number of "Ob_LI_RphaseB1_scd"        sites:  19708064 ( 69.60% of total)
    number of "Ob_LI_BCphaseA1_scd"-unique sites:      1039227
    number of sites in common:      17524253   (Expected:  12919493; O/E:       1.4)
    number of "Ob_LI_RphaseB1_scd"-unique sites:       2183811
  total number of "Ob_LI_BCphaseA1_scd+Ob_LI_RphaseB1_scd"-sites observed:    20747291

  number of sites in "Ob_LI_BCphaseA1_hsm" that are not  detectable in "Ob_LI_RphaseB1_hsm":     3902
  number of sites in "Ob_LI_BCphaseA1_hsm" that are also detectable in "Ob_LI_RphaseB1_hsm":   193127
    number of sites unique to "Ob_LI_BCphaseA1_hsm" although detectable in "Ob_LI_RphaseB1_hsm":           21012
    number of "Ob_LI_BCphaseA1_hsm" / "Ob_LI_RphaseB1_hsm" common sites                        :          172115      (Expected:   2297; O/E:      74.9)

  number of sites in "Ob_LI_RphaseB1_hsm" that are not  detectable in "Ob_LI_BCphaseA1_hsm":     9124
  number of sites in "Ob_LI_RphaseB1_hsm" that are also detectable in "Ob_LI_BCphaseA1_hsm":   208447
    number of sites unique to "Ob_LI_RphaseB1_hsm" although detectable in "Ob_LI_BCphaseA1_hsm":           36332
    number of "Ob_LI_RphaseB1_hsm" / "Ob_LI_BCphaseA1_hsm" common sites                        :          172115      (Expected:   2297; O/E:      74.9)

  Overlap index of "Ob_LI_BCphaseA1_hsm" with "Ob_LI_RphaseB1_hsm":          0.810
  Estimated number of "Ob_LI_BCphaseA1_hsm" = "Ob_LI_RphaseB1_hsm" sites (assuming sampling from one population):              233894
  Adjusted population size of "Ob_LI_BCphaseA1_hsm" = "Ob_LI_RphaseB1_hsm" sites (assuming all sites detectable):              377953 (  1.56x of observed)
```

```
Directory: PWC File: pwc-Ob_LI_BCphaseA1.vs.Ob_LI_RphaseB2.txt

Numbers of common and distinct sites comparing Ob_LI_BCphaseA1 versus Ob_LI_RphaseB2
================================================================================

  total number of potential sites:  28317694

  number of "Ob_LI_BCphaseA1_hsm"         sites:     197029
  number of "Ob_LI_RphaseB2_hsm"         sites:201205
    number of "Ob_LI_BCphaseA1_hsm"-unique sites:     32249
    number of common sites:        164780
    number of "Ob_LI_RphaseB2_hsm"-unique sites:      36425
  total number of "Ob_LI_BCphaseA1_hsm+Ob_LI_RphaseB2_hsm"-sites observed:   233454

  number of "Ob_LI_BCphaseA1_scd"        sites: 18563480 ( 65.55% of total)
  number of "Ob_LI_RphaseB2_scd"        sites:  18531805 ( 65.44% of total)
    number of "Ob_LI_BCphaseA1_scd"-unique sites:      1737175
    number of sites in common:      16826305   (Expected:  12148404; O/E:       1.4)
    number of "Ob_LI_RphaseB2_scd"-unique sites:       1705500
  total number of "Ob_LI_BCphaseA1_scd+Ob_LI_RphaseB2_scd"-sites observed:    20268980

  number of sites in "Ob_LI_BCphaseA1_hsm" that are not  detectable in "Ob_LI_RphaseB2_hsm":     6912
  number of sites in "Ob_LI_BCphaseA1_hsm" that are also detectable in "Ob_LI_RphaseB2_hsm":   190117
    number of sites unique to "Ob_LI_BCphaseA1_hsm" although detectable in "Ob_LI_RphaseB2_hsm":           25337
    number of "Ob_LI_BCphaseA1_hsm" / "Ob_LI_RphaseB2_hsm" common sites                        :          164780      (Expected:   2195; O/E:      75.1)

  number of sites in "Ob_LI_RphaseB2_hsm" that are not  detectable in "Ob_LI_BCphaseA1_hsm":     6938
  number of sites in "Ob_LI_RphaseB2_hsm" that are also detectable in "Ob_LI_BCphaseA1_hsm":   194267
    number of sites unique to "Ob_LI_RphaseB2_hsm" although detectable in "Ob_LI_BCphaseA1_hsm":           29487
    number of "Ob_LI_RphaseB2_hsm" / "Ob_LI_BCphaseA1_hsm" common sites                        :          164780      (Expected:   2195; O/E:      75.1)

  Overlap index of "Ob_LI_BCphaseA1_hsm" with "Ob_LI_RphaseB2_hsm":          0.767
  Estimated number of "Ob_LI_BCphaseA1_hsm" = "Ob_LI_RphaseB2_hsm" sites (assuming sampling from one population):              224138
  Adjusted population size of "Ob_LI_BCphaseA1_hsm" = "Ob_LI_RphaseB2_hsm" sites (assuming all sites detectable):              377211 (  1.62x of observed)
```

```
Directory: PWC File: pwc-Ob_LI_BCphaseA2.vs.Ob_LI_BCphaseB1.txt

Numbers of common and distinct sites comparing Ob_LI_BCphaseA2 versus Ob_LI_BCphaseB1
================================================================================

  total number of potential sites:   28317694

  number of "Ob_LI_BCphaseA2_hsm"         sites:     213136
  number of "Ob_LI_BCphaseB1_hsm"         sites:     226855
    number of "Ob_LI_BCphaseA2_hsm"-unique sites:     29337
    number of common sites:        183799
    number of "Ob_LI_BCphaseB1_hsm"-unique sites:     43056
  total number of "Ob_LI_BCphaseA2_hsm+Ob_LI_BCphaseB1_hsm"-sites observed:  256192

  number of "Ob_LI_BCphaseA2_scd"        sites: 18737737 ( 66.17% of total)
  number of "Ob_LI_BCphaseB1_scd"        sites: 20546412 ( 72.56% of total)
    number of "Ob_LI_BCphaseA2_scd"-unique sites:       798024
    number of sites in common:       17939713   (Expected:  13595502; O/E:       1.3)
    number of "Ob_LI_BCphaseB1_scd"-unique sites:      2606699
  total number of "Ob_LI_BCphaseA2_scd+Ob_LI_BCphaseB1_scd"-sites observed:   21344436

  number of sites in "Ob_LI_BCphaseA2_hsm" that are not  detectable in "Ob_LI_BCphaseB1_hsm":    3481
  number of sites in "Ob_LI_BCphaseA2_hsm" that are also detectable in "Ob_LI_BCphaseB1_hsm":  209655
    number of sites unique to "Ob_LI_BCphaseA2_hsm" although detectable in "Ob_LI_BCphaseB1_hsm":          25856
    number of "Ob_LI_BCphaseA2_hsm" / "Ob_LI_BCphaseB1_hsm" common sites                        :         183799      (Expected:   2527; O/E:      72.7)

  number of sites in "Ob_LI_BCphaseB1_hsm" that are not  detectable in "Ob_LI_BCphaseA2_hsm":   10623
  number of sites in "Ob_LI_BCphaseB1_hsm" that are also detectable in "Ob_LI_BCphaseA2_hsm":  216232
    number of sites unique to "Ob_LI_BCphaseB1_hsm" although detectable in "Ob_LI_BCphaseA2_hsm":          32433
    number of "Ob_LI_BCphaseB1_hsm" / "Ob_LI_BCphaseA2_hsm" common sites                        :         183799      (Expected:   2527; O/E:      72.7)

  Overlap index of "Ob_LI_BCphaseA2_hsm" with "Ob_LI_BCphaseB1_hsm":         0.783
  Estimated number of "Ob_LI_BCphaseA2_hsm" = "Ob_LI_BCphaseB1_hsm" sites (assuming sampling from one population):             246651
  Adjusted population size of "Ob_LI_BCphaseA2_hsm" = "Ob_LI_BCphaseB1_hsm" sites (assuming all sites detectable):             389336 (  1.52x of observed)
```

```
Directory: PWC File: pwc-Ob_LI_BCphaseA2.vs.Ob_LI_BCphaseB2.txt

Numbers of common and distinct sites comparing Ob_LI_BCphaseA2 versus Ob_LI_BCphaseB2
================================================================================

  total number of potential sites:  28317694

  number of "Ob_LI_BCphaseA2_hsm"         sites:     213136
  number of "Ob_LI_BCphaseB2_hsm"         sites:     209549
    number of "Ob_LI_BCphaseA2_hsm"-unique sites:     35753
    number of common sites:        177383
    number of "Ob_LI_BCphaseB2_hsm"-unique sites:     32166
  total number of "Ob_LI_BCphaseA2_hsm+Ob_LI_BCphaseB2_hsm"-sites observed:  245302

  number of "Ob_LI_BCphaseA2_scd"        sites: 18737737 ( 66.17% of total)
  number of "Ob_LI_BCphaseB2_scd"        sites: 19220462 ( 67.87% of total)
    number of "Ob_LI_BCphaseA2_scd"-unique sites:      1336307
    number of sites in common:      17401430   (Expected:  12718125; O/E:       1.4)
    number of "Ob_LI_BCphaseB2_scd"-unique sites:      1819032
  total number of "Ob_LI_BCphaseA2_scd+Ob_LI_BCphaseB2_scd"-sites observed:   20556769

  number of sites in "Ob_LI_BCphaseA2_hsm" that are not  detectable in "Ob_LI_BCphaseB2_hsm":    5470
  number of sites in "Ob_LI_BCphaseA2_hsm" that are also detectable in "Ob_LI_BCphaseB2_hsm":  207666
    number of sites unique to "Ob_LI_BCphaseA2_hsm" although detectable in "Ob_LI_BCphaseB2_hsm":          30283
    number of "Ob_LI_BCphaseA2_hsm" / "Ob_LI_BCphaseB2_hsm" common sites                        :         177383      (Expected:   2418; O/E:      73.4)

  number of sites in "Ob_LI_BCphaseB2_hsm" that are not  detectable in "Ob_LI_BCphaseA2_hsm":    6908
  number of sites in "Ob_LI_BCphaseB2_hsm" that are also detectable in "Ob_LI_BCphaseA2_hsm":  202641
    number of sites unique to "Ob_LI_BCphaseB2_hsm" although detectable in "Ob_LI_BCphaseA2_hsm":          25258
    number of "Ob_LI_BCphaseB2_hsm" / "Ob_LI_BCphaseA2_hsm" common sites                        :         177383      (Expected:   2418; O/E:      73.4)

  Overlap index of "Ob_LI_BCphaseA2_hsm" with "Ob_LI_BCphaseB2_hsm":         0.780
  Estimated number of "Ob_LI_BCphaseA2_hsm" = "Ob_LI_BCphaseB2_hsm" sites (assuming sampling from one population):             237236
  Adjusted population size of "Ob_LI_BCphaseA2_hsm" = "Ob_LI_BCphaseB2_hsm" sites (assuming all sites detectable):             386059 (  1.57x of observed)
```

```
Directory: PWC File: pwc-Ob_LI_BCphaseA2.vs.Ob_LI_RphaseA1.txt

Numbers of common and distinct sites comparing Ob_LI_BCphaseA2 versus Ob_LI_RphaseA1
================================================================================

  total number of potential sites:  28317694

  number of "Ob_LI_BCphaseA2_hsm"         sites:     213136
  number of "Ob_LI_RphaseA1_hsm"         sites:213360
    number of "Ob_LI_BCphaseA2_hsm"-unique sites:     35229
    number of common sites:        177907
    number of "Ob_LI_RphaseA1_hsm"-unique sites:      35453
  total number of "Ob_LI_BCphaseA2_hsm+Ob_LI_RphaseA1_hsm"-sites observed:   248589

  number of "Ob_LI_BCphaseA2_scd"        sites: 18737737 ( 66.17% of total)
  number of "Ob_LI_RphaseA1_scd"        sites:  19848503 ( 70.09% of total)
    number of "Ob_LI_BCphaseA2_scd"-unique sites:      1115586
    number of sites in common:      17622151   (Expected:  13133698; O/E:       1.3)
    number of "Ob_LI_RphaseA1_scd"-unique sites:       2226352
  total number of "Ob_LI_BCphaseA2_scd+Ob_LI_RphaseA1_scd"-sites observed:    20964089

  number of sites in "Ob_LI_BCphaseA2_hsm" that are not  detectable in "Ob_LI_RphaseA1_hsm":     4732
  number of sites in "Ob_LI_BCphaseA2_hsm" that are also detectable in "Ob_LI_RphaseA1_hsm":   208404
    number of sites unique to "Ob_LI_BCphaseA2_hsm" although detectable in "Ob_LI_RphaseA1_hsm":           30497
    number of "Ob_LI_BCphaseA2_hsm" / "Ob_LI_RphaseA1_hsm" common sites                        :          177907      (Expected:   2415; O/E:      73.7)

  number of sites in "Ob_LI_RphaseA1_hsm" that are not  detectable in "Ob_LI_BCphaseA2_hsm":     9118
  number of sites in "Ob_LI_RphaseA1_hsm" that are also detectable in "Ob_LI_BCphaseA2_hsm":   204242
    number of sites unique to "Ob_LI_RphaseA1_hsm" although detectable in "Ob_LI_BCphaseA2_hsm":           26335
    number of "Ob_LI_RphaseA1_hsm" / "Ob_LI_BCphaseA2_hsm" common sites                        :          177907      (Expected:   2415; O/E:      73.7)

  Overlap index of "Ob_LI_BCphaseA2_hsm" with "Ob_LI_RphaseA1_hsm":          0.773
  Estimated number of "Ob_LI_BCphaseA2_hsm" = "Ob_LI_RphaseA1_hsm" sites (assuming sampling from one population):              239253
  Adjusted population size of "Ob_LI_BCphaseA2_hsm" = "Ob_LI_RphaseA1_hsm" sites (assuming all sites detectable):              384465 (  1.55x of observed)
```

```
Directory: PWC File: pwc-Ob_LI_BCphaseA2.vs.Ob_LI_RphaseA2.txt

Numbers of common and distinct sites comparing Ob_LI_BCphaseA2 versus Ob_LI_RphaseA2
================================================================================

  total number of potential sites:  28317694

  number of "Ob_LI_BCphaseA2_hsm"         sites:     213136
  number of "Ob_LI_RphaseA2_hsm"         sites:265465
    number of "Ob_LI_BCphaseA2_hsm"-unique sites:     15194
    number of common sites:        197942
    number of "Ob_LI_RphaseA2_hsm"-unique sites:      67523
  total number of "Ob_LI_BCphaseA2_hsm+Ob_LI_RphaseA2_hsm"-sites observed:   280659

  number of "Ob_LI_BCphaseA2_scd"        sites: 18737737 ( 66.17% of total)
  number of "Ob_LI_RphaseA2_scd"        sites:  20631688 ( 72.86% of total)
    number of "Ob_LI_BCphaseA2_scd"-unique sites:       662118
    number of sites in common:      18075619   (Expected:  13651929; O/E:       1.3)
    number of "Ob_LI_RphaseA2_scd"-unique sites:       2556069
  total number of "Ob_LI_BCphaseA2_scd+Ob_LI_RphaseA2_scd"-sites observed:    21293806

  number of sites in "Ob_LI_BCphaseA2_hsm" that are not  detectable in "Ob_LI_RphaseA2_hsm":     1802
  number of sites in "Ob_LI_BCphaseA2_hsm" that are also detectable in "Ob_LI_RphaseA2_hsm":   211334
    number of sites unique to "Ob_LI_BCphaseA2_hsm" although detectable in "Ob_LI_RphaseA2_hsm":           13392
    number of "Ob_LI_BCphaseA2_hsm" / "Ob_LI_RphaseA2_hsm" common sites                        :          197942      (Expected:   2949; O/E:      67.1)

  number of sites in "Ob_LI_RphaseA2_hsm" that are not  detectable in "Ob_LI_BCphaseA2_hsm":    13236
  number of sites in "Ob_LI_RphaseA2_hsm" that are also detectable in "Ob_LI_BCphaseA2_hsm":   252229
    number of sites unique to "Ob_LI_RphaseA2_hsm" although detectable in "Ob_LI_BCphaseA2_hsm":           54287
    number of "Ob_LI_RphaseA2_hsm" / "Ob_LI_BCphaseA2_hsm" common sites                        :          197942      (Expected:   2949; O/E:      67.1)

  Overlap index of "Ob_LI_BCphaseA2_hsm" with "Ob_LI_RphaseA2_hsm":          0.889
  Estimated number of "Ob_LI_BCphaseA2_hsm" = "Ob_LI_RphaseA2_hsm" sites (assuming sampling from one population):              269294
  Adjusted population size of "Ob_LI_BCphaseA2_hsm" = "Ob_LI_RphaseA2_hsm" sites (assuming all sites detectable):              421882 (  1.50x of observed)
```

```
Directory: PWC File: pwc-Ob_LI_BCphaseA2.vs.Ob_LI_RphaseB1.txt

Numbers of common and distinct sites comparing Ob_LI_BCphaseA2 versus Ob_LI_RphaseB1
================================================================================

  total number of potential sites:  28317694

  number of "Ob_LI_BCphaseA2_hsm"         sites:     213136
  number of "Ob_LI_RphaseB1_hsm"         sites:217571
    number of "Ob_LI_BCphaseA2_hsm"-unique sites:     34157
    number of common sites:        178979
    number of "Ob_LI_RphaseB1_hsm"-unique sites:      38592
  total number of "Ob_LI_BCphaseA2_hsm+Ob_LI_RphaseB1_hsm"-sites observed:   251728

  number of "Ob_LI_BCphaseA2_scd"        sites: 18737737 ( 66.17% of total)
  number of "Ob_LI_RphaseB1_scd"        sites:  19708064 ( 69.60% of total)
    number of "Ob_LI_BCphaseA2_scd"-unique sites:      1188793
    number of sites in common:      17548944   (Expected:  13040769; O/E:       1.3)
    number of "Ob_LI_RphaseB1_scd"-unique sites:       2159120
  total number of "Ob_LI_BCphaseA2_scd+Ob_LI_RphaseB1_scd"-sites observed:    20896857

  number of sites in "Ob_LI_BCphaseA2_hsm" that are not  detectable in "Ob_LI_RphaseB1_hsm":     4606
  number of sites in "Ob_LI_BCphaseA2_hsm" that are also detectable in "Ob_LI_RphaseB1_hsm":   208530
    number of sites unique to "Ob_LI_BCphaseA2_hsm" although detectable in "Ob_LI_RphaseB1_hsm":           29551
    number of "Ob_LI_BCphaseA2_hsm" / "Ob_LI_RphaseB1_hsm" common sites                        :          178979      (Expected:   2474; O/E:      72.3)

  number of sites in "Ob_LI_RphaseB1_hsm" that are not  detectable in "Ob_LI_BCphaseA2_hsm":     9373
  number of sites in "Ob_LI_RphaseB1_hsm" that are also detectable in "Ob_LI_BCphaseA2_hsm":   208198
    number of sites unique to "Ob_LI_RphaseB1_hsm" although detectable in "Ob_LI_BCphaseA2_hsm":           29219
    number of "Ob_LI_RphaseB1_hsm" / "Ob_LI_BCphaseA2_hsm" common sites                        :          178979      (Expected:   2474; O/E:      72.3)

  Overlap index of "Ob_LI_BCphaseA2_hsm" with "Ob_LI_RphaseB1_hsm":          0.754
  Estimated number of "Ob_LI_BCphaseA2_hsm" = "Ob_LI_RphaseB1_hsm" sites (assuming sampling from one population):              242573
  Adjusted population size of "Ob_LI_BCphaseA2_hsm" = "Ob_LI_RphaseB1_hsm" sites (assuming all sites detectable):              391426 (  1.55x of observed)
```

```
Directory: PWC File: pwc-Ob_LI_BCphaseA2.vs.Ob_LI_RphaseB2.txt

Numbers of common and distinct sites comparing Ob_LI_BCphaseA2 versus Ob_LI_RphaseB2
================================================================================

  total number of potential sites:   28317694

  number of "Ob_LI_BCphaseA2_hsm"         sites:     213136
  number of "Ob_LI_RphaseB2_hsm"         sites:201205
    number of "Ob_LI_BCphaseA2_hsm"-unique sites:     39617
    number of common sites:        173519
    number of "Ob_LI_RphaseB2_hsm"-unique sites:      27686
  total number of "Ob_LI_BCphaseA2_hsm+Ob_LI_RphaseB2_hsm"-sites observed:   240822

  number of "Ob_LI_BCphaseA2_scd"        sites: 18737737 ( 66.17% of total)
  number of "Ob_LI_RphaseB2_scd"        sites:  18531805 ( 65.44% of total)
    number of "Ob_LI_BCphaseA2_scd"-unique sites:      1726145
    number of sites in common:      17011592   (Expected:  12262442; O/E:       1.4)
    number of "Ob_LI_RphaseB2_scd"-unique sites:       1520213
  total number of "Ob_LI_BCphaseA2_scd+Ob_LI_RphaseB2_scd"-sites observed:    20257950

  number of sites in "Ob_LI_BCphaseA2_hsm" that are not  detectable in "Ob_LI_RphaseB2_hsm":     6620
  number of sites in "Ob_LI_BCphaseA2_hsm" that are also detectable in "Ob_LI_RphaseB2_hsm":   206516
    number of sites unique to "Ob_LI_BCphaseA2_hsm" although detectable in "Ob_LI_RphaseB2_hsm":           32997
    number of "Ob_LI_BCphaseA2_hsm" / "Ob_LI_RphaseB2_hsm" common sites                        :          173519      (Expected:   2370; O/E:      73.2)

  number of sites in "Ob_LI_RphaseB2_hsm" that are not  detectable in "Ob_LI_BCphaseA2_hsm":     5991
  number of sites in "Ob_LI_RphaseB2_hsm" that are also detectable in "Ob_LI_BCphaseA2_hsm":   195214
    number of sites unique to "Ob_LI_RphaseB2_hsm" although detectable in "Ob_LI_BCphaseA2_hsm":           21695
    number of "Ob_LI_RphaseB2_hsm" / "Ob_LI_BCphaseA2_hsm" common sites                        :          173519      (Expected:   2370; O/E:      73.2)

  Overlap index of "Ob_LI_BCphaseA2_hsm" with "Ob_LI_RphaseB2_hsm":          0.804
  Estimated number of "Ob_LI_BCphaseA2_hsm" = "Ob_LI_RphaseB2_hsm" sites (assuming sampling from one population):              232337
  Adjusted population size of "Ob_LI_BCphaseA2_hsm" = "Ob_LI_RphaseB2_hsm" sites (assuming all sites detectable):              386750 (  1.61x of observed)
```

```
Directory: PWC File: pwc-Ob_LI_BCphaseB1.vs.Ob_LI_BCphaseB2.txt

Numbers of common and distinct sites comparing Ob_LI_BCphaseB1 versus Ob_LI_BCphaseB2
================================================================================

  total number of potential sites:  28317694

  number of "Ob_LI_BCphaseB1_hsm"         sites:     226855
  number of "Ob_LI_BCphaseB2_hsm"         sites:     209549
    number of "Ob_LI_BCphaseB1_hsm"-unique sites:     45450
    number of common sites:        181405
    number of "Ob_LI_BCphaseB2_hsm"-unique sites:     28144
  total number of "Ob_LI_BCphaseB1_hsm+Ob_LI_BCphaseB2_hsm"-sites observed:  254999

  number of "Ob_LI_BCphaseB1_scd"        sites: 20546412 ( 72.56% of total)
  number of "Ob_LI_BCphaseB2_scd"        sites: 19220462 ( 67.87% of total)
    number of "Ob_LI_BCphaseB1_scd"-unique sites:      2269606
    number of sites in common:      18276806   (Expected:  13945752; O/E:       1.3)
    number of "Ob_LI_BCphaseB2_scd"-unique sites:       943656
  total number of "Ob_LI_BCphaseB1_scd+Ob_LI_BCphaseB2_scd"-sites observed:   21490068

  number of sites in "Ob_LI_BCphaseB1_hsm" that are not  detectable in "Ob_LI_BCphaseB2_hsm":    9632
  number of sites in "Ob_LI_BCphaseB1_hsm" that are also detectable in "Ob_LI_BCphaseB2_hsm":  217223
    number of sites unique to "Ob_LI_BCphaseB1_hsm" although detectable in "Ob_LI_BCphaseB2_hsm":          35818
    number of "Ob_LI_BCphaseB1_hsm" / "Ob_LI_BCphaseB2_hsm" common sites                        :         181405      (Expected:   2442; O/E:      74.3)

  number of sites in "Ob_LI_BCphaseB2_hsm" that are not  detectable in "Ob_LI_BCphaseB1_hsm":    4047
  number of sites in "Ob_LI_BCphaseB2_hsm" that are also detectable in "Ob_LI_BCphaseB1_hsm":  205502
    number of sites unique to "Ob_LI_BCphaseB2_hsm" although detectable in "Ob_LI_BCphaseB1_hsm":          24097
    number of "Ob_LI_BCphaseB2_hsm" / "Ob_LI_BCphaseB1_hsm" common sites                        :         181405      (Expected:   2442; O/E:      74.3)

  Overlap index of "Ob_LI_BCphaseB1_hsm" with "Ob_LI_BCphaseB2_hsm":         0.795
  Estimated number of "Ob_LI_BCphaseB1_hsm" = "Ob_LI_BCphaseB2_hsm" sites (assuming sampling from one population):             246078
  Adjusted population size of "Ob_LI_BCphaseB1_hsm" = "Ob_LI_BCphaseB2_hsm" sites (assuming all sites detectable):             381268 (  1.50x of observed)
```

Directory: PWC File: pwc-Ob_LI_BCphaseB1.vs.Ob_LI_RphaseA1.txt

Numbers of common and distinct sites comparing Ob_LI_BCphaseB1 versus Ob_LI_RphaseA1
================================================================================

```
  total number of potential sites:   28317694

  number of "Ob_LI_BCphaseB1_hsm"         sites:     226855
  number of "Ob_LI_RphaseA1_hsm"         sites: 213360
    number of "Ob_LI_BCphaseB1_hsm"-unique sites:     41654
    number of common sites:        185201
    number of "Ob_LI_RphaseA1_hsm"-unique sites:      28159
  total number of "Ob_LI_BCphaseB1_hsm+Ob_LI_RphaseA1_hsm"-sites observed:   255014

  number of "Ob_LI_BCphaseB1_scd"        sites: 20546412 ( 72.56% of total)
  number of "Ob_LI_RphaseA1_scd"        sites:  19848503 ( 70.09% of total)
    number of "Ob_LI_BCphaseB1_scd"-unique sites:      1762851
    number of sites in common:      18783561   (Expected:  14401438; O/E:       1.3)
    number of "Ob_LI_RphaseA1_scd"-unique sites:       1064942
  total number of "Ob_LI_BCphaseB1_scd+Ob_LI_RphaseA1_scd"-sites observed:    21611354

  number of sites in "Ob_LI_BCphaseB1_hsm" that are not  detectable in "Ob_LI_RphaseA1_hsm":     7141
  number of sites in "Ob_LI_BCphaseB1_hsm" that are also detectable in "Ob_LI_RphaseA1_hsm":   219714
    number of sites unique to "Ob_LI_BCphaseB1_hsm" although detectable in "Ob_LI_RphaseA1_hsm":           34513
    number of "Ob_LI_BCphaseB1_hsm" / "Ob_LI_RphaseA1_hsm" common sites                        :          185201      (Expected:   2442; O/E:      75.8)

  number of sites in "Ob_LI_RphaseA1_hsm" that are not  detectable in "Ob_LI_BCphaseB1_hsm":     4617
  number of sites in "Ob_LI_RphaseA1_hsm" that are also detectable in "Ob_LI_BCphaseB1_hsm":   208743
    number of sites unique to "Ob_LI_RphaseA1_hsm" although detectable in "Ob_LI_BCphaseB1_hsm":           23542
    number of "Ob_LI_RphaseA1_hsm" / "Ob_LI_BCphaseB1_hsm" common sites                        :          185201      (Expected:   2442; O/E:      75.8)

  Overlap index of "Ob_LI_BCphaseB1_hsm" with "Ob_LI_RphaseA1_hsm":          0.801
  Estimated number of "Ob_LI_BCphaseB1_hsm" = "Ob_LI_RphaseA1_hsm" sites (assuming sampling from one population):              247643
  Adjusted population size of "Ob_LI_BCphaseB1_hsm" = "Ob_LI_RphaseA1_hsm" sites (assuming all sites detectable):              373342 (  1.46x of observed)
```

```
Directory: PWC File: pwc-Ob_LI_BCphaseB1.vs.Ob_LI_RphaseA2.txt

Numbers of common and distinct sites comparing Ob_LI_BCphaseB1 versus Ob_LI_RphaseA2
================================================================================

  total number of potential sites:  28317694

  number of "Ob_LI_BCphaseB1_hsm"         sites:     226855
  number of "Ob_LI_RphaseA2_hsm"         sites:265465
    number of "Ob_LI_BCphaseB1_hsm"-unique sites:     22494
    number of common sites:        204361
    number of "Ob_LI_RphaseA2_hsm"-unique sites:      61104
  total number of "Ob_LI_BCphaseB1_hsm+Ob_LI_RphaseA2_hsm"-sites observed:   287959

  number of "Ob_LI_BCphaseB1_scd"        sites: 20546412 ( 72.56% of total)
  number of "Ob_LI_RphaseA2_scd"        sites:  20631688 ( 72.86% of total)
    number of "Ob_LI_BCphaseB1_scd"-unique sites:      1326699
    number of sites in common:      19219713   (Expected:  14969692; O/E:       1.3)
    number of "Ob_LI_RphaseA2_scd"-unique sites:       1411975
  total number of "Ob_LI_BCphaseB1_scd+Ob_LI_RphaseA2_scd"-sites observed:    21958387

  number of sites in "Ob_LI_BCphaseB1_hsm" that are not  detectable in "Ob_LI_RphaseA2_hsm":     3953
  number of sites in "Ob_LI_BCphaseB1_hsm" that are also detectable in "Ob_LI_RphaseA2_hsm":   222902
    number of sites unique to "Ob_LI_BCphaseB1_hsm" although detectable in "Ob_LI_RphaseA2_hsm":           18541
    number of "Ob_LI_BCphaseB1_hsm" / "Ob_LI_RphaseA2_hsm" common sites                        :          204361      (Expected:   2985; O/E:      68.5)

  number of sites in "Ob_LI_RphaseA2_hsm" that are not  detectable in "Ob_LI_BCphaseB1_hsm":     8061
  number of sites in "Ob_LI_RphaseA2_hsm" that are also detectable in "Ob_LI_BCphaseB1_hsm":   257404
    number of sites unique to "Ob_LI_RphaseA2_hsm" although detectable in "Ob_LI_BCphaseB1_hsm":           53043
    number of "Ob_LI_RphaseA2_hsm" / "Ob_LI_BCphaseB1_hsm" common sites                        :          204361      (Expected:   2985; O/E:      68.5)

  Overlap index of "Ob_LI_BCphaseB1_hsm" with "Ob_LI_RphaseA2_hsm":          0.855
  Estimated number of "Ob_LI_BCphaseB1_hsm" = "Ob_LI_RphaseA2_hsm" sites (assuming sampling from one population):              280757
  Adjusted population size of "Ob_LI_BCphaseB1_hsm" = "Ob_LI_RphaseA2_hsm" sites (assuming all sites detectable):              413659 (  1.44x of observed)
```

```
Directory: PWC File: pwc-Ob_LI_BCphaseB1.vs.Ob_LI_RphaseB1.txt

Numbers of common and distinct sites comparing Ob_LI_BCphaseB1 versus Ob_LI_RphaseB1
================================================================================

  total number of potential sites:  28317694

  number of "Ob_LI_BCphaseB1_hsm"         sites:     226855
  number of "Ob_LI_RphaseB1_hsm"         sites:217571
    number of "Ob_LI_BCphaseB1_hsm"-unique sites:     40329
    number of common sites:        186526
    number of "Ob_LI_RphaseB1_hsm"-unique sites:      31045
  total number of "Ob_LI_BCphaseB1_hsm+Ob_LI_RphaseB1_hsm"-sites observed:   257900

  number of "Ob_LI_BCphaseB1_scd"        sites: 20546412 ( 72.56% of total)
  number of "Ob_LI_RphaseB1_scd"        sites:  19708064 ( 69.60% of total)
    number of "Ob_LI_BCphaseB1_scd"-unique sites:      1868551
    number of sites in common:      18677861   (Expected:  14299540; O/E:       1.3)
    number of "Ob_LI_RphaseB1_scd"-unique sites:       1030203
  total number of "Ob_LI_BCphaseB1_scd+Ob_LI_RphaseB1_scd"-sites observed:    21576615

  number of sites in "Ob_LI_BCphaseB1_hsm" that are not  detectable in "Ob_LI_RphaseB1_hsm":     6999
  number of sites in "Ob_LI_BCphaseB1_hsm" that are also detectable in "Ob_LI_RphaseB1_hsm":   219856
    number of sites unique to "Ob_LI_BCphaseB1_hsm" although detectable in "Ob_LI_RphaseB1_hsm":           33330
    number of "Ob_LI_BCphaseB1_hsm" / "Ob_LI_RphaseB1_hsm" common sites                        :          186526      (Expected:   2505; O/E:      74.5)

  number of sites in "Ob_LI_RphaseB1_hsm" that are not  detectable in "Ob_LI_BCphaseB1_hsm":     4720
  number of sites in "Ob_LI_RphaseB1_hsm" that are also detectable in "Ob_LI_BCphaseB1_hsm":   212851
    number of sites unique to "Ob_LI_RphaseB1_hsm" although detectable in "Ob_LI_BCphaseB1_hsm":           26325
    number of "Ob_LI_RphaseB1_hsm" / "Ob_LI_BCphaseB1_hsm" common sites                        :          186526      (Expected:   2505; O/E:      74.5)

  Overlap index of "Ob_LI_BCphaseB1_hsm" with "Ob_LI_RphaseB1_hsm":          0.783
  Estimated number of "Ob_LI_BCphaseB1_hsm" = "Ob_LI_RphaseB1_hsm" sites (assuming sampling from one population):              250885
  Adjusted population size of "Ob_LI_BCphaseB1_hsm" = "Ob_LI_RphaseB1_hsm" sites (assuming all sites detectable):              380369 (  1.47x of observed)
```

```
Directory: PWC File: pwc-Ob_LI_BCphaseB1.vs.Ob_LI_RphaseB2.txt

Numbers of common and distinct sites comparing Ob_LI_BCphaseB1 versus Ob_LI_RphaseB2
================================================================================

  total number of potential sites:  28317694

  number of "Ob_LI_BCphaseB1_hsm"         sites:     226855
  number of "Ob_LI_RphaseB2_hsm"         sites:201205
    number of "Ob_LI_BCphaseB1_hsm"-unique sites:     49873
    number of common sites:        176982
    number of "Ob_LI_RphaseB2_hsm"-unique sites:      24223
  total number of "Ob_LI_BCphaseB1_hsm+Ob_LI_RphaseB2_hsm"-sites observed:   251078

  number of "Ob_LI_BCphaseB1_scd"        sites: 20546412 ( 72.56% of total)
  number of "Ob_LI_RphaseB2_scd"        sites:  18531805 ( 65.44% of total)
    number of "Ob_LI_BCphaseB1_scd"-unique sites:      2790278
    number of sites in common:      17756134   (Expected:  13446084; O/E:       1.3)
    number of "Ob_LI_RphaseB2_scd"-unique sites:        775671
  total number of "Ob_LI_BCphaseB1_scd+Ob_LI_RphaseB2_scd"-sites observed:    21322083

  number of sites in "Ob_LI_BCphaseB1_hsm" that are not  detectable in "Ob_LI_RphaseB2_hsm":    11141
  number of sites in "Ob_LI_BCphaseB1_hsm" that are also detectable in "Ob_LI_RphaseB2_hsm":   215714
    number of sites unique to "Ob_LI_BCphaseB1_hsm" although detectable in "Ob_LI_RphaseB2_hsm":           38732
    number of "Ob_LI_BCphaseB1_hsm" / "Ob_LI_RphaseB2_hsm" common sites                        :          176982      (Expected:   2402; O/E:      73.7)

  number of sites in "Ob_LI_RphaseB2_hsm" that are not  detectable in "Ob_LI_BCphaseB1_hsm":     3495
  number of sites in "Ob_LI_RphaseB2_hsm" that are also detectable in "Ob_LI_BCphaseB1_hsm":   197710
    number of sites unique to "Ob_LI_RphaseB2_hsm" although detectable in "Ob_LI_BCphaseB1_hsm":           20728
    number of "Ob_LI_RphaseB2_hsm" / "Ob_LI_BCphaseB1_hsm" common sites                        :          176982      (Expected:   2402; O/E:      73.7)

  Overlap index of "Ob_LI_BCphaseB1_hsm" with "Ob_LI_RphaseB2_hsm":          0.817
  Estimated number of "Ob_LI_BCphaseB1_hsm" = "Ob_LI_RphaseB2_hsm" sites (assuming sampling from one population):              240978
  Adjusted population size of "Ob_LI_BCphaseB1_hsm" = "Ob_LI_RphaseB2_hsm" sites (assuming all sites detectable):              384315 (  1.53x of observed)
```

```
Directory: PWC File: pwc-Ob_LI_BCphaseB2.vs.Ob_LI_RphaseA1.txt

Numbers of common and distinct sites comparing Ob_LI_BCphaseB2 versus Ob_LI_RphaseA1
================================================================================

  total number of potential sites:  28317694

  number of "Ob_LI_BCphaseB2_hsm"         sites:     209549
  number of "Ob_LI_RphaseA1_hsm"         sites: 213360
    number of "Ob_LI_BCphaseB2_hsm"-unique sites:     33788
    number of common sites:        175761
    number of "Ob_LI_RphaseA1_hsm"-unique sites:      37599
  total number of "Ob_LI_BCphaseB2_hsm+Ob_LI_RphaseA1_hsm"-sites observed:   247148

  number of "Ob_LI_BCphaseB2_scd"        sites: 19220462 ( 67.87% of total)
  number of "Ob_LI_RphaseA1_scd"        sites:  19848503 ( 70.09% of total)
    number of "Ob_LI_BCphaseB2_scd"-unique sites:      1297828
    number of sites in common:      17922634   (Expected:  13472050; O/E:       1.3)
    number of "Ob_LI_RphaseA1_scd"-unique sites:       1925869
  total number of "Ob_LI_BCphaseB2_scd+Ob_LI_RphaseA1_scd"-sites observed:    21146331

  number of sites in "Ob_LI_BCphaseB2_hsm" that are not  detectable in "Ob_LI_RphaseA1_hsm":     5362
  number of sites in "Ob_LI_BCphaseB2_hsm" that are also detectable in "Ob_LI_RphaseA1_hsm":   204187
    number of sites unique to "Ob_LI_BCphaseB2_hsm" although detectable in "Ob_LI_RphaseA1_hsm":           28426
    number of "Ob_LI_BCphaseB2_hsm" / "Ob_LI_RphaseA1_hsm" common sites                        :          175761      (Expected:   2337; O/E:      75.2)

  number of sites in "Ob_LI_RphaseA1_hsm" that are not  detectable in "Ob_LI_BCphaseB2_hsm":     8212
  number of sites in "Ob_LI_RphaseA1_hsm" that are also detectable in "Ob_LI_BCphaseB2_hsm":   205148
    number of sites unique to "Ob_LI_RphaseA1_hsm" although detectable in "Ob_LI_BCphaseB2_hsm":           29387
    number of "Ob_LI_RphaseA1_hsm" / "Ob_LI_BCphaseB2_hsm" common sites                        :          175761      (Expected:   2337; O/E:      75.2)

  Overlap index of "Ob_LI_BCphaseB2_hsm" with "Ob_LI_RphaseA1_hsm":          0.756
  Estimated number of "Ob_LI_BCphaseB2_hsm" = "Ob_LI_RphaseA1_hsm" sites (assuming sampling from one population):              238327
  Adjusted population size of "Ob_LI_BCphaseB2_hsm" = "Ob_LI_RphaseA1_hsm" sites (assuming all sites detectable):              376555 (  1.52x of observed)
```

```
Directory: PWC File: pwc-Ob_LI_BCphaseB2.vs.Ob_LI_RphaseA2.txt

Numbers of common and distinct sites comparing Ob_LI_BCphaseB2 versus Ob_LI_RphaseA2
================================================================================

  total number of potential sites:   28317694

  number of "Ob_LI_BCphaseB2_hsm"         sites:     209549
  number of "Ob_LI_RphaseA2_hsm"         sites:265465
    number of "Ob_LI_BCphaseB2_hsm"-unique sites:     14928
    number of common sites:        194621
    number of "Ob_LI_RphaseA2_hsm"-unique sites:      70844
  total number of "Ob_LI_BCphaseB2_hsm+Ob_LI_RphaseA2_hsm"-sites observed:   280393

  number of "Ob_LI_BCphaseB2_scd"        sites: 19220462 ( 67.87% of total)
  number of "Ob_LI_RphaseA2_scd"        sites:  20631688 ( 72.86% of total)
    number of "Ob_LI_BCphaseB2_scd"-unique sites:       803352
    number of sites in common:      18417110   (Expected:  14003632; O/E:       1.3)
    number of "Ob_LI_RphaseA2_scd"-unique sites:       2214578
  total number of "Ob_LI_BCphaseB2_scd+Ob_LI_RphaseA2_scd"-sites observed:    21435040

  number of sites in "Ob_LI_BCphaseB2_hsm" that are not  detectable in "Ob_LI_RphaseA2_hsm":     1991
  number of sites in "Ob_LI_BCphaseB2_hsm" that are also detectable in "Ob_LI_RphaseA2_hsm":   207558
    number of sites unique to "Ob_LI_BCphaseB2_hsm" although detectable in "Ob_LI_RphaseA2_hsm":           12937
    number of "Ob_LI_BCphaseB2_hsm" / "Ob_LI_RphaseA2_hsm" common sites                        :          194621      (Expected:   2861; O/E:      68.0)

  number of sites in "Ob_LI_RphaseA2_hsm" that are not  detectable in "Ob_LI_BCphaseB2_hsm":    11621
  number of sites in "Ob_LI_RphaseA2_hsm" that are also detectable in "Ob_LI_BCphaseB2_hsm":   253844
    number of sites unique to "Ob_LI_RphaseA2_hsm" although detectable in "Ob_LI_BCphaseB2_hsm":           59223
    number of "Ob_LI_RphaseA2_hsm" / "Ob_LI_BCphaseB2_hsm" common sites                        :          194621      (Expected:   2861; O/E:      68.0)

  Overlap index of "Ob_LI_BCphaseB2_hsm" with "Ob_LI_RphaseA2_hsm":          0.892
  Estimated number of "Ob_LI_BCphaseB2_hsm" = "Ob_LI_RphaseA2_hsm" sites (assuming sampling from one population):              270718
  Adjusted population size of "Ob_LI_BCphaseB2_hsm" = "Ob_LI_RphaseA2_hsm" sites (assuming all sites detectable):              416249 (  1.48x of observed)
```

```
Directory: PWC File: pwc-Ob_LI_BCphaseB2.vs.Ob_LI_RphaseB1.txt

Numbers of common and distinct sites comparing Ob_LI_BCphaseB2 versus Ob_LI_RphaseB1
================================================================================

  total number of potential sites:   28317694

  number of "Ob_LI_BCphaseB2_hsm"         sites:     209549
  number of "Ob_LI_RphaseB1_hsm"         sites: 217571
    number of "Ob_LI_BCphaseB2_hsm"-unique sites:     31446
    number of common sites:        178103
    number of "Ob_LI_RphaseB1_hsm"-unique sites:      39468
  total number of "Ob_LI_BCphaseB2_hsm+Ob_LI_RphaseB1_hsm"-sites observed:   249017

  number of "Ob_LI_BCphaseB2_scd"        sites: 19220462 ( 67.87% of total)
  number of "Ob_LI_RphaseB1_scd"        sites:  19708064 ( 69.60% of total)
    number of "Ob_LI_BCphaseB2_scd"-unique sites:      1323425
    number of sites in common:      17897037   (Expected:  13376728; O/E:       1.3)
    number of "Ob_LI_RphaseB1_scd"-unique sites:       1811027
  total number of "Ob_LI_BCphaseB2_scd+Ob_LI_RphaseB1_scd"-sites observed:    21031489

  number of sites in "Ob_LI_BCphaseB2_hsm" that are not  detectable in "Ob_LI_RphaseB1_hsm":     4876
  number of sites in "Ob_LI_BCphaseB2_hsm" that are also detectable in "Ob_LI_RphaseB1_hsm":   204673
    number of sites unique to "Ob_LI_BCphaseB2_hsm" although detectable in "Ob_LI_RphaseB1_hsm":           26570
    number of "Ob_LI_BCphaseB2_hsm" / "Ob_LI_RphaseB1_hsm" common sites                        :          178103      (Expected:   2396; O/E:      74.3)

  number of sites in "Ob_LI_RphaseB1_hsm" that are not  detectable in "Ob_LI_BCphaseB2_hsm":     8024
  number of sites in "Ob_LI_RphaseB1_hsm" that are also detectable in "Ob_LI_BCphaseB2_hsm":   209547
    number of sites unique to "Ob_LI_RphaseB1_hsm" although detectable in "Ob_LI_BCphaseB2_hsm":           31444
    number of "Ob_LI_RphaseB1_hsm" / "Ob_LI_BCphaseB2_hsm" common sites                        :          178103      (Expected:   2396; O/E:      74.3)

  Overlap index of "Ob_LI_BCphaseB2_hsm" with "Ob_LI_RphaseB1_hsm":          0.772
  Estimated number of "Ob_LI_BCphaseB2_hsm" = "Ob_LI_RphaseB1_hsm" sites (assuming sampling from one population):              240808
  Adjusted population size of "Ob_LI_BCphaseB2_hsm" = "Ob_LI_RphaseB1_hsm" sites (assuming all sites detectable):              381020 (  1.53x of observed)
```

Directory: PWC File: pwc-Ob_LI_BCphaseB2.vs.Ob_LI_RphaseB2.txt

Numbers of common and distinct sites comparing Ob_LI_BCphaseB2 versus Ob_LI_RphaseB2
================================================================================

```
  total number of potential sites:  28317694

  number of "Ob_LI_BCphaseB2_hsm"        sites:     209549
  number of "Ob_LI_RphaseB2_hsm"        sites:201205
    number of "Ob_LI_BCphaseB2_hsm"-unique sites:     36882
    number of common sites:        172667
    number of "Ob_LI_RphaseB2_hsm"-unique sites:      28538
  total number of "Ob_LI_BCphaseB2_hsm+Ob_LI_RphaseB2_hsm"-sites observed:   238087

  number of "Ob_LI_BCphaseB2_scd"       sites: 19220462 ( 67.87% of total)
  number of "Ob_LI_RphaseB2_scd"       sites:  18531805 ( 65.44% of total)
    number of "Ob_LI_BCphaseB2_scd"-unique sites:      1928347
    number of sites in common:      17292115   (Expected:  12578350; O/E:       1.4)
    number of "Ob_LI_RphaseB2_scd"-unique sites:       1239690
  total number of "Ob_LI_BCphaseB2_scd+Ob_LI_RphaseB2_scd"-sites observed:    20460152

  number of sites in "Ob_LI_BCphaseB2_hsm" that are not  detectable in "Ob_LI_RphaseB2_hsm":     6927
  number of sites in "Ob_LI_BCphaseB2_hsm" that are also detectable in "Ob_LI_RphaseB2_hsm":   202622
    number of sites unique to "Ob_LI_BCphaseB2_hsm" although detectable in "Ob_LI_RphaseB2_hsm":           29955
    number of "Ob_LI_BCphaseB2_hsm" / "Ob_LI_RphaseB2_hsm" common sites                        :          172667      (Expected:   2299; O/E:      75.1)

  number of sites in "Ob_LI_RphaseB2_hsm" that are not  detectable in "Ob_LI_BCphaseB2_hsm":     4992
  number of sites in "Ob_LI_RphaseB2_hsm" that are also detectable in "Ob_LI_BCphaseB2_hsm":   196213
    number of sites unique to "Ob_LI_RphaseB2_hsm" although detectable in "Ob_LI_BCphaseB2_hsm":           23546
    number of "Ob_LI_RphaseB2_hsm" / "Ob_LI_BCphaseB2_hsm" common sites                        :          172667      (Expected:   2299; O/E:      75.1)

  Overlap index of "Ob_LI_BCphaseB2_hsm" with "Ob_LI_RphaseB2_hsm":          0.788
  Estimated number of "Ob_LI_BCphaseB2_hsm" = "Ob_LI_RphaseB2_hsm" sites (assuming sampling from one population):              230253
  Adjusted population size of "Ob_LI_BCphaseB2_hsm" = "Ob_LI_RphaseB2_hsm" sites (assuming all sites detectable):              377064 (  1.58x of observed)
```

```
Directory: PWC File: pwc-Ob_LI_RphaseA1.vs.Ob_LI_RphaseA2.txt

Numbers of common and distinct sites comparing Ob_LI_RphaseA1 versus Ob_LI_RphaseA2
================================================================================

  total number of potential sites:   28317694

  number of "Ob_LI_RphaseA1_hsm"         sites:213360
  number of "Ob_LI_RphaseA2_hsm"         sites:265465
    number of "Ob_LI_RphaseA1_hsm"-unique sites:      17708
    number of common sites:        195652
    number of "Ob_LI_RphaseA2_hsm"-unique sites:      69813
  total number of "Ob_LI_RphaseA1_hsm+Ob_LI_RphaseA2_hsm"-sites observed:    283173

  number of "Ob_LI_RphaseA1_scd"        sites:  19848503 ( 70.09% of total)
  number of "Ob_LI_RphaseA2_scd"        sites:  20631688 ( 72.86% of total)
    number of "Ob_LI_RphaseA1_scd"-unique sites:       1085838
    number of sites in common:      18762665   (Expected:  14461210; O/E:       1.3)
    number of "Ob_LI_RphaseA2_scd"-unique sites:       1869023
  total number of "Ob_LI_RphaseA1_scd+Ob_LI_RphaseA2_scd"-sites observed:     21717526

  number of sites in "Ob_LI_RphaseA1_hsm" that are not  detectable in "Ob_LI_RphaseA2_hsm":      3328
  number of sites in "Ob_LI_RphaseA1_hsm" that are also detectable in "Ob_LI_RphaseA2_hsm":    210032
    number of sites unique to "Ob_LI_RphaseA1_hsm" although detectable in "Ob_LI_RphaseA2_hsm":            14380
    number of "Ob_LI_RphaseA1_hsm" / "Ob_LI_RphaseA2_hsm" common sites                        :           195652      (Expected:   2856; O/E:      68.5)

  number of sites in "Ob_LI_RphaseA2_hsm" that are not  detectable in "Ob_LI_RphaseA1_hsm":     10367
  number of sites in "Ob_LI_RphaseA2_hsm" that are also detectable in "Ob_LI_RphaseA1_hsm":    255098
    number of sites unique to "Ob_LI_RphaseA2_hsm" although detectable in "Ob_LI_RphaseA1_hsm":            59446
    number of "Ob_LI_RphaseA2_hsm" / "Ob_LI_RphaseA1_hsm" common sites                        :           195652      (Expected:   2856; O/E:      68.5)

  Overlap index of "Ob_LI_RphaseA1_hsm" with "Ob_LI_RphaseA2_hsm":           0.882
  Estimated number of "Ob_LI_RphaseA1_hsm" = "Ob_LI_RphaseA2_hsm" sites (assuming sampling from one population):         273847
  Adjusted population size of "Ob_LI_RphaseA1_hsm" = "Ob_LI_RphaseA2_hsm" sites (assuming all sites detectable):         413306 (  1.46x of observed)
```

```
Directory: PWC File: pwc-Ob_LI_RphaseA1.vs.Ob_LI_RphaseB1.txt

Numbers of common and distinct sites comparing Ob_LI_RphaseA1 versus Ob_LI_RphaseB1
================================================================================

  total number of potential sites:  28317694

  number of "Ob_LI_RphaseA1_hsm"         sites:213360
  number of "Ob_LI_RphaseB1_hsm"         sites:217571
    number of "Ob_LI_RphaseA1_hsm"-unique sites:      32772
    number of common sites:        180588
    number of "Ob_LI_RphaseB1_hsm"-unique sites:      36983
  total number of "Ob_LI_RphaseA1_hsm+Ob_LI_RphaseB1_hsm"-sites observed:    250343

  number of "Ob_LI_RphaseA1_scd"        sites:  19848503 ( 70.09% of total)
  number of "Ob_LI_RphaseB1_scd"        sites:  19708064 ( 69.60% of total)
    number of "Ob_LI_RphaseA1_scd"-unique sites:       1555796
    number of sites in common:      18292707   (Expected:  13813821; O/E:       1.3)
    number of "Ob_LI_RphaseB1_scd"-unique sites:       1415357
  total number of "Ob_LI_RphaseA1_scd+Ob_LI_RphaseB1_scd"-sites observed:     21263860

  number of sites in "Ob_LI_RphaseA1_hsm" that are not  detectable in "Ob_LI_RphaseB1_hsm":      5826
  number of sites in "Ob_LI_RphaseA1_hsm" that are also detectable in "Ob_LI_RphaseB1_hsm":    207534
    number of sites unique to "Ob_LI_RphaseA1_hsm" although detectable in "Ob_LI_RphaseB1_hsm":            26946
    number of "Ob_LI_RphaseA1_hsm" / "Ob_LI_RphaseB1_hsm" common sites                        :           180588      (Expected:   2398; O/E:      75.3)

  number of sites in "Ob_LI_RphaseB1_hsm" that are not  detectable in "Ob_LI_RphaseA1_hsm":      6186
  number of sites in "Ob_LI_RphaseB1_hsm" that are also detectable in "Ob_LI_RphaseA1_hsm":    211385
    number of sites unique to "Ob_LI_RphaseB1_hsm" although detectable in "Ob_LI_RphaseA1_hsm":            30797
    number of "Ob_LI_RphaseB1_hsm" / "Ob_LI_RphaseA1_hsm" common sites                        :           180588      (Expected:   2398; O/E:      75.3)

  Overlap index of "Ob_LI_RphaseA1_hsm" with "Ob_LI_RphaseB1_hsm":           0.772
  Estimated number of "Ob_LI_RphaseA1_hsm" = "Ob_LI_RphaseB1_hsm" sites (assuming sampling from one population):         242926
  Adjusted population size of "Ob_LI_RphaseA1_hsm" = "Ob_LI_RphaseB1_hsm" sites (assuming all sites detectable):         376058 (  1.50x of observed)
```

```
Directory: PWC File: pwc-Ob_LI_RphaseA1.vs.Ob_LI_RphaseB2.txt

Numbers of common and distinct sites comparing Ob_LI_RphaseA1 versus Ob_LI_RphaseB2
================================================================================

  total number of potential sites:  28317694

  number of "Ob_LI_RphaseA1_hsm"         sites:213360
  number of "Ob_LI_RphaseB2_hsm"         sites:201205
    number of "Ob_LI_RphaseA1_hsm"-unique sites:      41675
    number of common sites:        171685
    number of "Ob_LI_RphaseB2_hsm"-unique sites:      29520
  total number of "Ob_LI_RphaseA1_hsm+Ob_LI_RphaseB2_hsm"-sites observed:    242880

  number of "Ob_LI_RphaseA1_scd"        sites:  19848503 ( 70.09% of total)
  number of "Ob_LI_RphaseB2_scd"        sites:  18531805 ( 65.44% of total)
    number of "Ob_LI_RphaseA1_scd"-unique sites:       2403512
    number of sites in common:      17444991   (Expected:  12989355; O/E:       1.3)
    number of "Ob_LI_RphaseB2_scd"-unique sites:       1086814
  total number of "Ob_LI_RphaseA1_scd+Ob_LI_RphaseB2_scd"-sites observed:     20935317

  number of sites in "Ob_LI_RphaseA1_hsm" that are not  detectable in "Ob_LI_RphaseB2_hsm":      9616
  number of sites in "Ob_LI_RphaseA1_hsm" that are also detectable in "Ob_LI_RphaseB2_hsm":    203744
    number of sites unique to "Ob_LI_RphaseA1_hsm" although detectable in "Ob_LI_RphaseB2_hsm":            32059
    number of "Ob_LI_RphaseA1_hsm" / "Ob_LI_RphaseB2_hsm" common sites                        :           171685      (Expected:   2296; O/E:      74.8)

  number of sites in "Ob_LI_RphaseB2_hsm" that are not  detectable in "Ob_LI_RphaseA1_hsm":      4647
  number of sites in "Ob_LI_RphaseB2_hsm" that are also detectable in "Ob_LI_RphaseA1_hsm":    196558
    number of sites unique to "Ob_LI_RphaseB2_hsm" although detectable in "Ob_LI_RphaseA1_hsm":            24873
    number of "Ob_LI_RphaseB2_hsm" / "Ob_LI_RphaseA1_hsm" common sites                        :           171685      (Expected:   2296; O/E:      74.8)

  Overlap index of "Ob_LI_RphaseA1_hsm" with "Ob_LI_RphaseB2_hsm":           0.778
  Estimated number of "Ob_LI_RphaseA1_hsm" = "Ob_LI_RphaseB2_hsm" sites (assuming sampling from one population):         233262
  Adjusted population size of "Ob_LI_RphaseA1_hsm" = "Ob_LI_RphaseB2_hsm" sites (assuming all sites detectable):         378643 (  1.56x of observed)
```

```
Directory: PWC File: pwc-Ob_LI_RphaseA2.vs.Ob_LI_RphaseB1.txt

Numbers of common and distinct sites comparing Ob_LI_RphaseA2 versus Ob_LI_RphaseB1
================================================================================

  total number of potential sites:  28317694

  number of "Ob_LI_RphaseA2_hsm"         sites:265465
  number of "Ob_LI_RphaseB1_hsm"         sites:217571
    number of "Ob_LI_RphaseA2_hsm"-unique sites:      67532
    number of common sites:        197933
    number of "Ob_LI_RphaseB1_hsm"-unique sites:      19638
  total number of "Ob_LI_RphaseA2_hsm+Ob_LI_RphaseB1_hsm"-sites observed:    285103

  number of "Ob_LI_RphaseA2_scd"        sites:  20631688 ( 72.86% of total)
  number of "Ob_LI_RphaseB1_scd"        sites:  19708064 ( 69.60% of total)
    number of "Ob_LI_RphaseA2_scd"-unique sites:       1958959
    number of sites in common:      18672729   (Expected:  14358889; O/E:       1.3)
    number of "Ob_LI_RphaseB1_scd"-unique sites:       1035335
  total number of "Ob_LI_RphaseA2_scd+Ob_LI_RphaseB1_scd"-sites observed:     21667023

  number of sites in "Ob_LI_RphaseA2_hsm" that are not  detectable in "Ob_LI_RphaseB1_hsm":      9829
  number of sites in "Ob_LI_RphaseA2_hsm" that are also detectable in "Ob_LI_RphaseB1_hsm":    255636
    number of sites unique to "Ob_LI_RphaseA2_hsm" although detectable in "Ob_LI_RphaseB1_hsm":            57703
    number of "Ob_LI_RphaseA2_hsm" / "Ob_LI_RphaseB1_hsm" common sites                        :           197933      (Expected:   2934; O/E:      67.5)

  number of sites in "Ob_LI_RphaseB1_hsm" that are not  detectable in "Ob_LI_RphaseA2_hsm":      3285
  number of sites in "Ob_LI_RphaseB1_hsm" that are also detectable in "Ob_LI_RphaseA2_hsm":    214286
    number of sites unique to "Ob_LI_RphaseB1_hsm" although detectable in "Ob_LI_RphaseA2_hsm":            16353
    number of "Ob_LI_RphaseB1_hsm" / "Ob_LI_RphaseA2_hsm" common sites                        :           197933      (Expected:   2934; O/E:      67.5)

  Overlap index of "Ob_LI_RphaseA2_hsm" with "Ob_LI_RphaseB1_hsm":           0.868
  Estimated number of "Ob_LI_RphaseA2_hsm" = "Ob_LI_RphaseB1_hsm" sites (assuming sampling from one population):         276756
  Adjusted population size of "Ob_LI_RphaseA2_hsm" = "Ob_LI_RphaseB1_hsm" sites (assuming all sites detectable):         419708 (  1.47x of observed)
```

Directory: PWC File: pwc-Ob_LI_RphaseA2.vs.Ob_LI_RphaseB2.txt

Numbers of common and distinct sites comparing Ob_LI_RphaseA2 versus Ob_LI_RphaseB2
================================================================================

```
  total number of potential sites:  28317694

  number of "Ob_LI_RphaseA2_hsm"        sites:265465
  number of "Ob_LI_RphaseB2_hsm"        sites:201205
    number of "Ob_LI_RphaseA2_hsm"-unique sites:      76723
    number of common sites:        188742
    number of "Ob_LI_RphaseB2_hsm"-unique sites:      12463
  total number of "Ob_LI_RphaseA2_hsm+Ob_LI_RphaseB2_hsm"-sites observed:    277928

  number of "Ob_LI_RphaseA2_scd"       sites:  20631688 ( 72.86% of total)
  number of "Ob_LI_RphaseB2_scd"       sites:  18531805 ( 65.44% of total)
    number of "Ob_LI_RphaseA2_scd"-unique sites:       2742140
    number of sites in common:      17889548   (Expected:  13501891; O/E:       1.3)
    number of "Ob_LI_RphaseB2_scd"-unique sites:        642257
  total number of "Ob_LI_RphaseA2_scd+Ob_LI_RphaseB2_scd"-sites observed:     21273945

  number of sites in "Ob_LI_RphaseA2_hsm" that are not  detectable in "Ob_LI_RphaseB2_hsm":     13641
  number of sites in "Ob_LI_RphaseA2_hsm" that are also detectable in "Ob_LI_RphaseB2_hsm":    251824
    number of sites unique to "Ob_LI_RphaseA2_hsm" although detectable in "Ob_LI_RphaseB2_hsm":            63082
    number of "Ob_LI_RphaseA2_hsm" / "Ob_LI_RphaseB2_hsm" common sites                        :           188742      (Expected:   2808; O/E:      67.2)

  number of sites in "Ob_LI_RphaseB2_hsm" that are not  detectable in "Ob_LI_RphaseA2_hsm":      1732
  number of sites in "Ob_LI_RphaseB2_hsm" that are also detectable in "Ob_LI_RphaseA2_hsm":    199473
    number of sites unique to "Ob_LI_RphaseB2_hsm" although detectable in "Ob_LI_RphaseA2_hsm":            10731
    number of "Ob_LI_RphaseB2_hsm" / "Ob_LI_RphaseA2_hsm" common sites                        :           188742      (Expected:   2808; O/E:      67.2)

  Overlap index of "Ob_LI_RphaseA2_hsm" with "Ob_LI_RphaseB2_hsm":           0.908
  Estimated number of "Ob_LI_RphaseA2_hsm" = "Ob_LI_RphaseB2_hsm" sites (assuming sampling from one population):         266142
  Adjusted population size of "Ob_LI_RphaseA2_hsm" = "Ob_LI_RphaseB2_hsm" sites (assuming all sites detectable):         421280 (  1.52x of observed)
```

```
Directory: PWC File: pwc-Ob_LI_RphaseB1.vs.Ob_LI_RphaseB2.txt

Numbers of common and distinct sites comparing Ob_LI_RphaseB1 versus Ob_LI_RphaseB2
================================================================================

  total number of potential sites:  28317694

  number of "Ob_LI_RphaseB1_hsm"         sites: 217571
  number of "Ob_LI_RphaseB2_hsm"         sites: 201205
    number of "Ob_LI_RphaseB1_hsm"-unique sites:      43647
    number of common sites:        173924
    number of "Ob_LI_RphaseB2_hsm"-unique sites:      27281
  total number of "Ob_LI_RphaseB1_hsm+Ob_LI_RphaseB2_hsm"-sites observed:    244852

  number of "Ob_LI_RphaseB1_scd"        sites:  19708064 ( 69.60% of total)
  number of "Ob_LI_RphaseB2_scd"        sites:  18531805 ( 65.44% of total)
    number of "Ob_LI_RphaseB1_scd"-unique sites:       2276363
    number of sites in common:      17431701   (Expected:  12897448; O/E:       1.4)
    number of "Ob_LI_RphaseB2_scd"-unique sites:       1100104
  total number of "Ob_LI_RphaseB1_scd+Ob_LI_RphaseB2_scd"-sites observed:     20808168

  number of sites in "Ob_LI_RphaseB1_hsm" that are not  detectable in "Ob_LI_RphaseB2_hsm":      9411
  number of sites in "Ob_LI_RphaseB1_hsm" that are also detectable in "Ob_LI_RphaseB2_hsm":    208160
    number of sites unique to "Ob_LI_RphaseB1_hsm" although detectable in "Ob_LI_RphaseB2_hsm":            34236
    number of "Ob_LI_RphaseB1_hsm" / "Ob_LI_RphaseB2_hsm" common sites                        :           173924      (Expected:   2352; O/E:      73.9)

  number of sites in "Ob_LI_RphaseB2_hsm" that are not  detectable in "Ob_LI_RphaseB1_hsm":      4213
  number of sites in "Ob_LI_RphaseB2_hsm" that are also detectable in "Ob_LI_RphaseB1_hsm":    196992
    number of sites unique to "Ob_LI_RphaseB2_hsm" although detectable in "Ob_LI_RphaseB1_hsm":            23068
    number of "Ob_LI_RphaseB2_hsm" / "Ob_LI_RphaseB1_hsm" common sites                        :           173924      (Expected:   2352; O/E:      73.9)

  Overlap index of "Ob_LI_RphaseB1_hsm" with "Ob_LI_RphaseB2_hsm":           0.795
  Estimated number of "Ob_LI_RphaseB1_hsm" = "Ob_LI_RphaseB2_hsm" sites (assuming sampling from one population):         235769
  Adjusted population size of "Ob_LI_RphaseB1_hsm" = "Ob_LI_RphaseB2_hsm" sites (assuming all sites detectable):         383005 (  1.56x of observed)
```

```
Directory: CRL File: 0READMEcrl

CRL - Correlations between aggregate samples.

Input: studymk

Output: files crl-*.txt crl-*.pdf

Notes: The output is generated by BWASPR::cmpSamples(), which uses methylKit::unite()
        to determine the sites common to all samples and applies methylKit::getCorrelation()
        and methylKit::PCASamples().  The *.txt files show the correlations in a table, and
        the *.pdf files show graphics.

       Beware that the number of common sites (shown at the bottom of the *.txt files) may be
        small, which may make the correlations less informative.
```

Directory: CRL File: crl-Ob_LI.txt

| | BCphaseA1 | BCphaseA2 | BCphaseB1 | BCphaseB2 | RphaseA1 | RphaseA2 | RphaseB1 | RphaseB2 |
|---|---|---|---|---|---|---|---|---|
| BCphaseA1 | 1.000 | 0.903 | 0.906 | 0.912 | 0.903 | 0.913 | 0.913 | 0.909 |
| BCphaseA2 | 0.903 | 1.000 | 0.921 | 0.911 | 0.918 | 0.933 | 0.910 | 0.909 |
| BCphaseB1 | 0.906 | 0.921 | 1.000 | 0.913 | 0.923 | 0.933 | 0.913 | 0.910 |
| BCphaseB2 | 0.912 | 0.911 | 0.913 | 1.000 | 0.908 | 0.921 | 0.918 | 0.919 |
| RphaseA1 | 0.903 | 0.918 | 0.923 | 0.908 | 1.000 | 0.929 | 0.910 | 0.906 |
| RphaseA2 | 0.913 | 0.933 | 0.933 | 0.921 | 0.929 | 1.000 | 0.920 | 0.918 |
| RphaseB1 | 0.913 | 0.910 | 0.913 | 0.918 | 0.910 | 0.920 | 1.000 | 0.915 |
| RphaseB2 | 0.909 | 0.909 | 0.910 | 0.919 | 0.906 | 0.918 | 0.915 | 1.000 |

The number of conserved sites is 104327.

#### CpG base pearson cor.

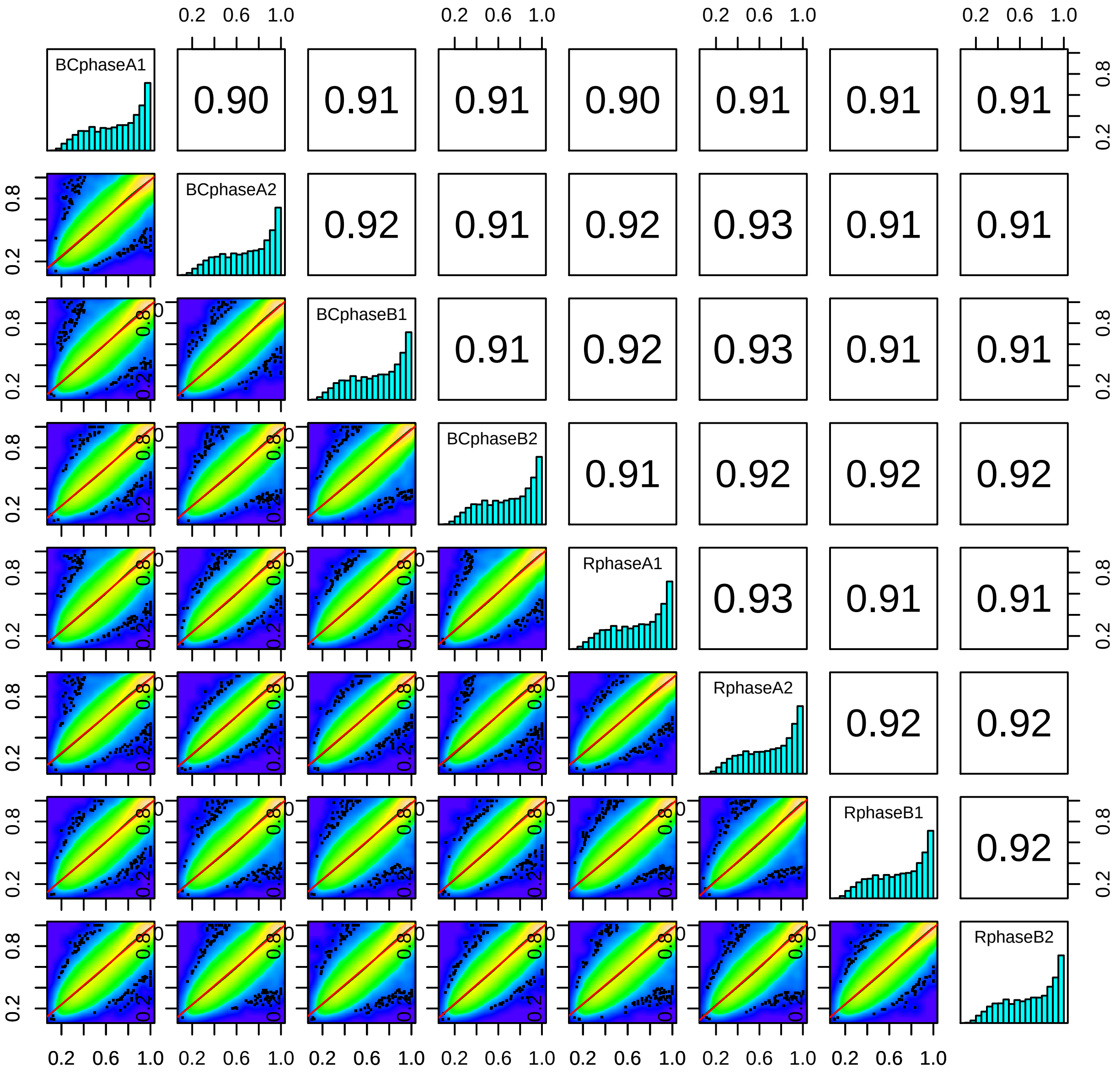

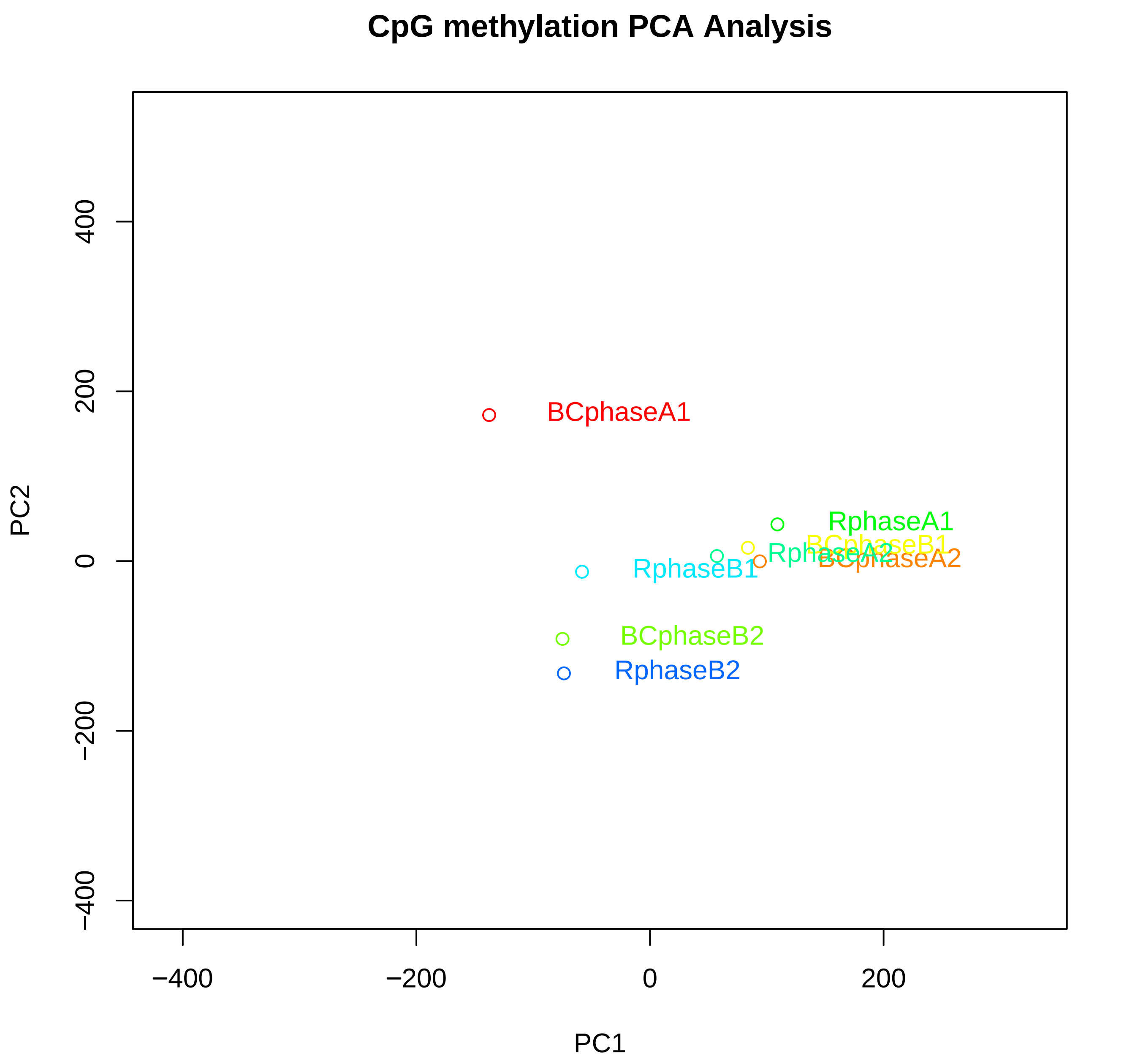
CpG methylation PCA Analysis
BCphaseA1
RphaseA1
BCphaseB1
RphaseA2
BCphaseA2
RphaseB1
BCphaseB2
RphaseB2
PC1
PC2
−400
−200
0
200
400

Directory: MMP File: 0READMEmmp

MMP - Mapping of methylation sites on genome annotation.

Input: studymk, studymc, genome_ann (a list of GRanges objects providing annotated region labels
and boundaries; output of BWASPR::get_genome_annotation())

Output: files mmp-*.txt

Notes: The output is generated by BWASPR::map_methylome() and gives an accounting for every sample
as to where the CpGhsm and CpGscd (control) sites reside relative to the genome annotation.
Abbreviations used: O/E, observed over expected
(expected percentages are based on the respective feature proportions
in the genome, as annotated).

Directory: MMP File: mmp-Ob_LI_BCphaseA1.txt

================================================================================
Genomic composition in terms of feature regions
================================================================================

| | | |
|---|---|---|
| Ob genome size: | 223876465 bp | |
| Ob genic region size: | 166752117 bp | ( 74.5%) |
| Ob exon region size: | 44907373 bp | ( 20.1%) |
| Ob CDS region size: | 20651162 bp | ( 46.0%) |
| Ob five-prime UTR region size: | 5434886 bp | ( 12.1%) |
| Ob three-prime UTR region size: | 10723012 bp | ( 23.9%) |
| Ob other exon region size: | 8098313 bp | ( 18.0%) |
| Ob intron region size: | 121844744 bp | ( 54.4%) |
| Ob intergenic region size: | 57124348 bp | ( 25.5%) |
| Ob promoter region size: | 6874701 bp | ( 3.1%) |
| Ob other intergenic region size: | 50249647 bp | ( 22.4%) |

================================================================================
Methylation sites in genomic feature regions
================================================================================

Number and density (per 10kb) of sites in Ob_LI_BCphaseA1_scd

| | | | | |
|---|---|---|---|---|
| Number of sites identified in Ob Ob_LI_BCphaseA1_scd: | 18563480 | ( 829.18) | | |
| Number of sites identified in Ob Ob_LI_BCphaseA1_scd genic regions: | 13518904 | ( 810.72) | 72.83% | ( 0.98 O/E) |
| Number of sites identified in Ob Ob_LI_BCphaseA1_scd exon regions: | 3833476 | ( 853.64) | 20.65% | ( 1.03 O/E) |
| Number of sites identified in Ob Ob_LI_BCphaseA1_scd CDS regions: | 2206242 | ( 1068.34) | 57.55% | ( 1.25 O/E) |
| Number of sites identified in Ob Ob_LI_BCphaseA1_scd five-prime UTR regions: | 605259 | ( 1113.66) | 15.79% | ( 1.30 O/E) |
| Number of sites identified in Ob Ob_LI_BCphaseA1_scd three-prime UTR regions: | 657786 | ( 613.43) | 17.16% | ( 0.72 O/E) |
| Number of sites identified in Ob Ob_LI_BCphaseA1_scd other exon regions: | 364189 | ( 449.71) | 9.50% | ( 0.53 O/E) |
| Number of sites identified in Ob Ob_LI_BCphaseA1_scd intron regions: | 9685428 | ( 794.90) | 52.17% | ( 0.96 O/E) |
| Number of sites identified in Ob Ob_LI_BCphaseA1_scd intergenic regions: | 5044576 | ( 883.09) | 27.17% | ( 1.07 O/E) |
| Number of sites identified in Ob Ob_LI_BCphaseA1_scd promoter regions: | 306195 | ( 445.39) | 1.65% | ( 0.54 O/E) |
| Number of sites identified in Ob Ob_LI_BCphaseA1_scd other intergenic regions: | 4738381 | ( 942.97) | 25.53% | ( 1.14 O/E) |

Number and density (per 10kb) of sites in Ob_LI_BCphaseA1_hsm

| | | | | | |
|---|---|---|---|---|---|
| Number of sites identified in Ob Ob_LI_BCphaseA1_hsm: | 197029 | ( 8.80) | ( 1.06% of control) | | |
| Number of sites identified in Ob Ob_LI_BCphaseA1_hsm genic regions: | 194706 | ( 11.68) | ( 1.44% of control) | 98.82% | ( 1.33 O/E) |
| Number of sites identified in Ob Ob_LI_BCphaseA1_hsm exon regions: | 159604 | ( 35.54) | ( 4.16% of control) | 81.01% | ( 4.04 O/E) |
| Number of sites identified in Ob Ob_LI_BCphaseA1_hsm CDS regions: | 152119 | ( 73.66) | ( 6.89% of control) | 95.31% | ( 2.07 O/E) |
| Number of sites identified in Ob Ob_LI_BCphaseA1_hsm five-prime UTR regions: | 2388 | ( 4.39) | ( 0.39% of control) | 1.50% | ( 0.12 O/E) |
| Number of sites identified in Ob Ob_LI_BCphaseA1_hsm three-prime UTR regions: | 3912 | ( 3.65) | ( 0.59% of control) | 2.45% | ( 0.10 O/E) |
| Number of sites identified in Ob Ob_LI_BCphaseA1_hsm other exon regions: | 1185 | ( 1.46) | ( 0.33% of control) | 0.74% | ( 0.04 O/E) |
| Number of sites identified in Ob Ob_LI_BCphaseA1_hsm intron regions: | 35102 | ( 2.88) | ( 0.36% of control) | 17.82% | ( 0.33 O/E) |
| Number of sites identified in Ob Ob_LI_BCphaseA1_hsm intergenic regions: | 2323 | ( 0.41) | ( 0.05% of control) | 1.18% | ( 0.05 O/E) |
| Number of sites identified in Ob Ob_LI_BCphaseA1_hsm promoter regions: | 382 | ( 0.56) | ( 0.12% of control) | 0.19% | ( 0.06 O/E) |
| Number of sites identified in Ob Ob_LI_BCphaseA1_hsm other intergenic regions: | 1941 | ( 0.39) | ( 0.04% of control) | 0.99% | ( 0.04 O/E) |

```
Directory: MMP File: mmp-Ob_LI_BCphaseA2.txt

================================================================================
Genomic composition in terms of feature regions
================================================================================

Ob genome size:                        223876465 bp
  Ob genic region size:                  166752117 bp ( 74.5%)
    Ob exon region size:                    44907373 bp ( 20.1%)
      Ob CDS region size:                     20651162 bp ( 46.0%)
      Ob five-prime UTR region size:           5434886 bp ( 12.1%)
      Ob three-prime UTR region size:         10723012 bp ( 23.9%)
      Ob other exon region size:               8098313 bp ( 18.0%)
    Ob intron region size:                 121844744 bp ( 54.4%)
  Ob intergenic region size:              57124348 bp ( 25.5%)
    Ob promoter region size:                 6874701 bp (  3.1%)
    Ob other intergenic region size:        50249647 bp ( 22.4%)

================================================================================
Methylation sites in genomic feature regions
================================================================================

Number and density (per 10kb) of sites in Ob_LI_BCphaseA2_scd

Number of sites identified in Ob Ob_LI_BCphaseA2_scd:                            18737737     (  836.97)                            
Number of sites identified in Ob Ob_LI_BCphaseA2_scd genic regions:              13638053     (  817.86)                        72.78%     ( 0.98 O/E)
Number of sites identified in Ob Ob_LI_BCphaseA2_scd   exon regions:               3846901   (  856.63)                          20.53%   ( 1.02 O/E)
Number of sites identified in Ob Ob_LI_BCphaseA2_scd     CDS regions:                2212439 ( 1071.34)                            57.51% ( 1.25 O/E)
Number of sites identified in Ob Ob_LI_BCphaseA2_scd     five-prime UTR regions:      611552 ( 1125.23)                            15.90% ( 1.31 O/E)
Number of sites identified in Ob Ob_LI_BCphaseA2_scd     three-prime UTR regions:     655149 (  610.97)                            17.03% ( 0.71 O/E)
Number of sites identified in Ob Ob_LI_BCphaseA2_scd     other exon regions:          367761 (  454.12)                             9.56% ( 0.53 O/E)
Number of sites identified in Ob Ob_LI_BCphaseA2_scd   intron regions:             9791152   (  803.58)                          52.25%   ( 0.96 O/E)
Number of sites identified in Ob Ob_LI_BCphaseA2_scd intergenic regions:          5099684     (  892.73)                        27.22%     ( 1.07 O/E)
Number of sites identified in Ob Ob_LI_BCphaseA2_scd   promoter regions:            311225   (  452.71)                           1.66%   ( 0.54 O/E)
Number of sites identified in Ob Ob_LI_BCphaseA2_scd   other intergenic regions:   4788459   (  952.93)                          25.56%   ( 1.14 O/E)

Number and density (per 10kb) of sites in Ob_LI_BCphaseA2_hsm

Number of sites identified in Ob Ob_LI_BCphaseA2_hsm:                              213136     (    9.52) (  1.14% of control)
Number of sites identified in Ob Ob_LI_BCphaseA2_hsm genic regions:                210748     (   12.64) (  1.55% of control)  98.88%     ( 1.33 O/E)
Number of sites identified in Ob Ob_LI_BCphaseA2_hsm   exon regions:                 172993   (   38.52) (  4.50% of control)   81.17%   ( 4.05 O/E)
Number of sites identified in Ob Ob_LI_BCphaseA2_hsm     CDS regions:                  164849 (   79.83) (  7.45% of control)     95.29% ( 2.07 O/E)
Number of sites identified in Ob Ob_LI_BCphaseA2_hsm     five-prime UTR regions:         2660 (    4.89) (  0.43% of control)      1.54% ( 0.13 O/E)
Number of sites identified in Ob Ob_LI_BCphaseA2_hsm     three-prime UTR regions:        4227 (    3.94) (  0.65% of control)      2.44% ( 0.10 O/E)
Number of sites identified in Ob Ob_LI_BCphaseA2_hsm     other exon regions:             1257 (    1.55) (  0.34% of control)      0.73% ( 0.04 O/E)
Number of sites identified in Ob Ob_LI_BCphaseA2_hsm   intron regions:                37755   (    3.10) (  0.39% of control)   17.71%   ( 0.33 O/E)
Number of sites identified in Ob Ob_LI_BCphaseA2_hsm intergenic regions:             2388     (    0.42) (  0.05% of control)  1.12%      ( 0.04 O/E)
Number of sites identified in Ob Ob_LI_BCphaseA2_hsm   promoter regions:                366   (    0.53) (  0.12% of control)    0.17%   ( 0.06 O/E)
Number of sites identified in Ob Ob_LI_BCphaseA2_hsm   other intergenic regions:       2022   (    0.40) (  0.04% of control)    0.95%   ( 0.04 O/E)
```

Directory: MMP File: mmp-Ob_LI_BCphaseB1.txt

================================================================================
Genomic composition in terms of feature regions
================================================================================

| | | |
|---|---|---|
| Ob genome size: | 223876465 bp | |
| Ob genic region size: | 166752117 bp | ( 74.5%) |
| Ob exon region size: | 44907373 bp | ( 20.1%) |
| Ob CDS region size: | 20651162 bp | ( 46.0%) |
| Ob five-prime UTR region size: | 5434886 bp | ( 12.1%) |
| Ob three-prime UTR region size: | 10723012 bp | ( 23.9%) |
| Ob other exon region size: | 8098313 bp | ( 18.0%) |
| Ob intron region size: | 121844744 bp | ( 54.4%) |
| Ob intergenic region size: | 57124348 bp | ( 25.5%) |
| Ob promoter region size: | 6874701 bp | ( 3.1%) |
| Ob other intergenic region size: | 50249647 bp | ( 22.4%) |

================================================================================
Methylation sites in genomic feature regions
================================================================================

Number and density (per 10kb) of sites in Ob_LI_BCphaseB1_scd

| | | | | |
|---|---|---|---|---|
| Number of sites identified in Ob Ob_LI_BCphaseB1_scd: | 20546412 | ( 917.76) | | |
| Number of sites identified in Ob Ob_LI_BCphaseB1_scd genic regions: | 14927576 | ( 895.20) | 72.65% | ( 0.98 O/E) |
| Number of sites identified in Ob Ob_LI_BCphaseB1_scd exon regions: | 4166778 | ( 927.86) | 20.28% | ( 1.01 O/E) |
| Number of sites identified in Ob Ob_LI_BCphaseB1_scd CDS regions: | 2363689 | ( 1144.58) | 56.73% | ( 1.23 O/E) |
| Number of sites identified in Ob Ob_LI_BCphaseB1_scd five-prime UTR regions: | 655764 | ( 1206.58) | 15.74% | ( 1.30 O/E) |
| Number of sites identified in Ob Ob_LI_BCphaseB1_scd three-prime UTR regions: | 741205 | ( 691.23) | 17.79% | ( 0.74 O/E) |
| Number of sites identified in Ob Ob_LI_BCphaseB1_scd other exon regions: | 406120 | ( 501.49) | 9.75% | ( 0.54 O/E) |
| Number of sites identified in Ob Ob_LI_BCphaseB1_scd intron regions: | 10760798 | ( 883.16) | 52.37% | ( 0.96 O/E) |
| Number of sites identified in Ob Ob_LI_BCphaseB1_scd intergenic regions: | 5618836 | ( 983.61) | 27.35% | ( 1.07 O/E) |
| Number of sites identified in Ob Ob_LI_BCphaseB1_scd promoter regions: | 337873 | ( 491.47) | 1.64% | ( 0.54 O/E) |
| Number of sites identified in Ob Ob_LI_BCphaseB1_scd other intergenic regions: | 5280963 | ( 1050.95) | 25.70% | ( 1.15 O/E) |

Number and density (per 10kb) of sites in Ob_LI_BCphaseB1_hsm

| | | | | | |
|---|---|---|---|---|---|
| Number of sites identified in Ob Ob_LI_BCphaseB1_hsm: | 226855 | ( 10.13) | ( 1.10% of control) | | |
| Number of sites identified in Ob Ob_LI_BCphaseB1_hsm genic regions: | 223811 | ( 13.42) | ( 1.50% of control) | 98.66% | ( 1.32 O/E) |
| Number of sites identified in Ob Ob_LI_BCphaseB1_hsm exon regions: | 182647 | ( 40.67) | ( 4.38% of control) | 80.51% | ( 4.01 O/E) |
| Number of sites identified in Ob Ob_LI_BCphaseB1_hsm CDS regions: | 173532 | ( 84.03) | ( 7.34% of control) | 95.01% | ( 2.07 O/E) |
| Number of sites identified in Ob Ob_LI_BCphaseB1_hsm five-prime UTR regions: | 2813 | ( 5.18) | ( 0.43% of control) | 1.54% | ( 0.13 O/E) |
| Number of sites identified in Ob Ob_LI_BCphaseB1_hsm three-prime UTR regions: | 4904 | ( 4.57) | ( 0.66% of control) | 2.68% | ( 0.11 O/E) |
| Number of sites identified in Ob Ob_LI_BCphaseB1_hsm other exon regions: | 1398 | ( 1.73) | ( 0.34% of control) | 0.77% | ( 0.04 O/E) |
| Number of sites identified in Ob Ob_LI_BCphaseB1_hsm intron regions: | 41164 | ( 3.38) | ( 0.38% of control) | 18.15% | ( 0.33 O/E) |
| Number of sites identified in Ob Ob_LI_BCphaseB1_hsm intergenic regions: | 3044 | ( 0.53) | ( 0.05% of control) | 1.34% | ( 0.05 O/E) |
| Number of sites identified in Ob Ob_LI_BCphaseB1_hsm promoter regions: | 454 | ( 0.66) | ( 0.13% of control) | 0.20% | ( 0.07 O/E) |
| Number of sites identified in Ob Ob_LI_BCphaseB1_hsm other intergenic regions: | 2590 | ( 0.52) | ( 0.05% of control) | 1.14% | ( 0.05 O/E) |

Directory: MMP File: mmp-Ob_LI_BCphaseB2.txt

================================================================================
#### Genomic composition in terms of feature regions
================================================================================

| | | |
|---|---|---|
| Ob genome size: | 223876465 bp | |
| Ob genic region size: | 166752117 bp | ( 74.5%) |
| Ob exon region size: | 44907373 bp | ( 20.1%) |
| Ob CDS region size: | 20651162 bp | ( 46.0%) |
| Ob five-prime UTR region size: | 5434886 bp | ( 12.1%) |
| Ob three-prime UTR region size: | 10723012 bp | ( 23.9%) |
| Ob other exon region size: | 8098313 bp | ( 18.0%) |
| Ob intron region size: | 121844744 bp | ( 54.4%) |
| Ob intergenic region size: | 57124348 bp | ( 25.5%) |
| Ob promoter region size: | 6874701 bp | ( 3.1%) |
| Ob other intergenic region size: | 50249647 bp | ( 22.4%) |

================================================================================
#### Methylation sites in genomic feature regions
================================================================================

Number and density (per 10kb) of sites in Ob_LI_BCphaseB2_scd

| | | | | |
|---|---|---|---|---|
| Number of sites identified in Ob Ob_LI_BCphaseB2_scd: | 19220462 | ( 858.53) | | |
| Number of sites identified in Ob Ob_LI_BCphaseB2_scd genic regions: | 13990235 | ( 838.98) | 72.79% | ( 0.98 O/E) |
| Number of sites identified in Ob Ob_LI_BCphaseB2_scd exon regions: | 3920615 | ( 873.04) | 20.40% | ( 1.02 O/E) |
| Number of sites identified in Ob Ob_LI_BCphaseB2_scd CDS regions: | 2242331 | ( 1085.81) | 57.19% | ( 1.24 O/E) |
| Number of sites identified in Ob Ob_LI_BCphaseB2_scd five-prime UTR regions: | 622073 | ( 1144.59) | 15.87% | ( 1.31 O/E) |
| Number of sites identified in Ob Ob_LI_BCphaseB2_scd three-prime UTR regions: | 679013 | ( 633.23) | 17.32% | ( 0.73 O/E) |
| Number of sites identified in Ob Ob_LI_BCphaseB2_scd other exon regions: | 377198 | ( 465.77) | 9.62% | ( 0.53 O/E) |
| Number of sites identified in Ob Ob_LI_BCphaseB2_scd intron regions: | 10069620 | ( 826.43) | 52.39% | ( 0.96 O/E) |
| Number of sites identified in Ob Ob_LI_BCphaseB2_scd intergenic regions: | 5230227 | ( 915.59) | 27.21% | ( 1.07 O/E) |
| Number of sites identified in Ob Ob_LI_BCphaseB2_scd promoter regions: | 316405 | ( 460.25) | 1.65% | ( 0.54 O/E) |
| Number of sites identified in Ob Ob_LI_BCphaseB2_scd other intergenic regions: | 4913822 | ( 977.88) | 25.57% | ( 1.14 O/E) |

Number and density (per 10kb) of sites in Ob_LI_BCphaseB2_hsm

| | | | | | |
|---|---|---|---|---|---|
| Number of sites identified in Ob Ob_LI_BCphaseB2_hsm: | 209549 | ( 9.36) | ( 1.09% of control) | | |
| Number of sites identified in Ob Ob_LI_BCphaseB2_hsm genic regions: | 206993 | ( 12.41) | ( 1.48% of control) | 98.78% | ( 1.33 O/E) |
| Number of sites identified in Ob Ob_LI_BCphaseB2_hsm exon regions: | 169594 | ( 37.77) | ( 4.33% of control) | 80.93% | ( 4.03 O/E) |
| Number of sites identified in Ob Ob_LI_BCphaseB2_hsm CDS regions: | 161468 | ( 78.19) | ( 7.20% of control) | 95.21% | ( 2.07 O/E) |
| Number of sites identified in Ob Ob_LI_BCphaseB2_hsm five-prime UTR regions: | 2516 | ( 4.63) | ( 0.40% of control) | 1.48% | ( 0.12 O/E) |
| Number of sites identified in Ob Ob_LI_BCphaseB2_hsm three-prime UTR regions: | 4340 | ( 4.05) | ( 0.64% of control) | 2.56% | ( 0.11 O/E) |
| Number of sites identified in Ob Ob_LI_BCphaseB2_hsm other exon regions: | 1270 | ( 1.57) | ( 0.34% of control) | 0.75% | ( 0.04 O/E) |
| Number of sites identified in Ob Ob_LI_BCphaseB2_hsm intron regions: | 37399 | ( 3.07) | ( 0.37% of control) | 17.85% | ( 0.33 O/E) |
| Number of sites identified in Ob Ob_LI_BCphaseB2_hsm intergenic regions: | 2556 | ( 0.45) | ( 0.05% of control) | 1.22% | ( 0.05 O/E) |
| Number of sites identified in Ob Ob_LI_BCphaseB2_hsm promoter regions: | 370 | ( 0.54) | ( 0.12% of control) | 0.18% | ( 0.06 O/E) |
| Number of sites identified in Ob Ob_LI_BCphaseB2_hsm other intergenic regions: | 2186 | ( 0.44) | ( 0.04% of control) | 1.04% | ( 0.05 O/E) |

```
Directory: MMP File: mmp-Ob_LI_RphaseA1.txt

================================================================================
Genomic composition in terms of feature regions
================================================================================

Ob genome size:                        223876465 bp
  Ob genic region size:                  166752117 bp ( 74.5%)
    Ob exon region size:                    44907373 bp ( 20.1%)
      Ob CDS region size:                     20651162 bp ( 46.0%)
      Ob five-prime UTR region size:           5434886 bp ( 12.1%)
      Ob three-prime UTR region size:         10723012 bp ( 23.9%)
      Ob other exon region size:               8098313 bp ( 18.0%)
    Ob intron region size:                 121844744 bp ( 54.4%)
  Ob intergenic region size:              57124348 bp ( 25.5%)
    Ob promoter region size:                 6874701 bp (  3.1%)
    Ob other intergenic region size:        50249647 bp ( 22.4%)

================================================================================
Methylation sites in genomic feature regions
================================================================================

Number and density (per 10kb) of sites in Ob_LI_RphaseA1_scd

Number of sites identified in Ob Ob_LI_RphaseA1_scd:                            19848503     (  886.58)
Number of sites identified in Ob Ob_LI_RphaseA1_scd genic regions:              14434391     (  865.62)                         72.72%     ( 0.98 O/E)
Number of sites identified in Ob Ob_LI_RphaseA1_scd   exon regions:                4058745   (  903.80)                           20.45%   ( 1.02 O/E)
Number of sites identified in Ob Ob_LI_RphaseA1_scd     CDS regions:                 2316763 ( 1121.86)                             57.08% ( 1.24 O/E)
Number of sites identified in Ob Ob_LI_RphaseA1_scd     five-prime UTR regions:       638850 ( 1175.46)                             15.74% ( 1.30 O/E)
Number of sites identified in Ob Ob_LI_RphaseA1_scd     three-prime UTR regions:      711142 (  663.19)                             17.52% ( 0.73 O/E)
Number of sites identified in Ob Ob_LI_RphaseA1_scd     other exon regions:           391990 (  484.04)                              9.66% ( 0.54 O/E)
Number of sites identified in Ob Ob_LI_RphaseA1_scd   intron regions:             10375646   (  851.55)                           52.27%   ( 0.96 O/E)
Number of sites identified in Ob Ob_LI_RphaseA1_scd intergenic regions:          5414112     (  947.78)                         27.28%     ( 1.07 O/E)
Number of sites identified in Ob Ob_LI_RphaseA1_scd   promoter regions:             327416   (  476.26)                            1.65%   ( 0.54 O/E)
Number of sites identified in Ob Ob_LI_RphaseA1_scd   other intergenic regions:    5086696   ( 1012.28)                           25.63%   ( 1.14 O/E)

Number and density (per 10kb) of sites in Ob_LI_RphaseA1_hsm

Number of sites identified in Ob Ob_LI_RphaseA1_hsm:                              213360     (    9.53) (  1.07% of control)
Number of sites identified in Ob Ob_LI_RphaseA1_hsm genic regions:                210878     (   12.65) (  1.46% of control)  98.84%     ( 1.33 O/E)
Number of sites identified in Ob Ob_LI_RphaseA1_hsm   exon regions:                 172334   (   38.38) (  4.25% of control)    80.77%   ( 4.03 O/E)
Number of sites identified in Ob Ob_LI_RphaseA1_hsm     CDS regions:                  163932 (   79.38) (  7.08% of control)      95.12% ( 2.07 O/E)
Number of sites identified in Ob Ob_LI_RphaseA1_hsm     five-prime UTR regions:         2719 (    5.00) (  0.43% of control)       1.58% ( 0.13 O/E)
Number of sites identified in Ob Ob_LI_RphaseA1_hsm     three-prime UTR regions:        4387 (    4.09) (  0.62% of control)       2.55% ( 0.11 O/E)
Number of sites identified in Ob Ob_LI_RphaseA1_hsm     other exon regions:             1296 (    1.60) (  0.33% of control)       0.75% ( 0.04 O/E)
Number of sites identified in Ob Ob_LI_RphaseA1_hsm   intron regions:                38544   (    3.16) (  0.37% of control)    18.07%   ( 0.33 O/E)
Number of sites identified in Ob Ob_LI_RphaseA1_hsm intergenic regions:             2482     (    0.43) (  0.05% of control)   1.16%     ( 0.05 O/E)
Number of sites identified in Ob Ob_LI_RphaseA1_hsm   promoter regions:                389   (    0.57) (  0.12% of control)     0.18%   ( 0.06 O/E)
Number of sites identified in Ob Ob_LI_RphaseA1_hsm   other intergenic regions:       2093   (    0.42) (  0.04% of control)     0.98%   ( 0.04 O/E)
```

```
Directory: MMP File: mmp-Ob_LI_RphaseA2.txt

================================================================================
Genomic composition in terms of feature regions
================================================================================

Ob genome size:                        223876465 bp
  Ob genic region size:                  166752117 bp ( 74.5%)
    Ob exon region size:                    44907373 bp ( 20.1%)
      Ob CDS region size:                     20651162 bp ( 46.0%)
      Ob five-prime UTR region size:           5434886 bp ( 12.1%)
      Ob three-prime UTR region size:         10723012 bp ( 23.9%)
      Ob other exon region size:               8098313 bp ( 18.0%)
    Ob intron region size:                 121844744 bp ( 54.4%)
  Ob intergenic region size:              57124348 bp ( 25.5%)
    Ob promoter region size:                 6874701 bp (  3.1%)
    Ob other intergenic region size:        50249647 bp ( 22.4%)

================================================================================
Methylation sites in genomic feature regions
================================================================================

Number and density (per 10kb) of sites in Ob_LI_RphaseA2_scd

Number of sites identified in Ob Ob_LI_RphaseA2_scd:                          20631688     (  921.57)
Number of sites identified in Ob Ob_LI_RphaseA2_scd genic regions:            15023408     (  900.94)                       72.82%     ( 0.98 O/E)
Number of sites identified in Ob Ob_LI_RphaseA2_scd   exon regions:             4195856   (  934.34)                         20.34%   ( 1.01 O/E)
Number of sites identified in Ob Ob_LI_RphaseA2_scd     CDS regions:              2372508 ( 1148.85)                           56.54% ( 1.23 O/E)
Number of sites identified in Ob Ob_LI_RphaseA2_scd     five-prime UTR regions:    661757 ( 1217.61)                           15.77% ( 1.30 O/E)
Number of sites identified in Ob Ob_LI_RphaseA2_scd     three-prime UTR regions:   752458 (  701.72)                           17.93% ( 0.75 O/E)
Number of sites identified in Ob Ob_LI_RphaseA2_scd     other exon regions:        409133 (  505.21)                            9.75% ( 0.54 O/E)
Number of sites identified in Ob Ob_LI_RphaseA2_scd   intron regions:          10827552   (  888.64)                         52.48%   ( 0.96 O/E)
Number of sites identified in Ob Ob_LI_RphaseA2_scd intergenic regions:        5608280     (  981.77)                       27.18%     ( 1.07 O/E)
Number of sites identified in Ob Ob_LI_RphaseA2_scd   promoter regions:          341904   (  497.34)                          1.66%   ( 0.54 O/E)
Number of sites identified in Ob Ob_LI_RphaseA2_scd   other intergenic regions:     5266376   ( 1048.04)                         25.53%   ( 1.14 O/E)

Number and density (per 10kb) of sites in Ob_LI_RphaseA2_hsm

Number of sites identified in Ob Ob_LI_RphaseA2_hsm:                            265465     (   11.86) (  1.29% of control)
Number of sites identified in Ob Ob_LI_RphaseA2_hsm genic regions:              261090     (   15.66) (  1.74% of control)  98.35%     ( 1.32 O/E)
Number of sites identified in Ob Ob_LI_RphaseA2_hsm   exon regions:               211275   (   47.05) (  5.04% of control)   79.59%   ( 3.97 O/E)
Number of sites identified in Ob Ob_LI_RphaseA2_hsm     CDS regions:                199735 (   96.72) (  8.42% of control)     94.54% ( 2.06 O/E)
Number of sites identified in Ob Ob_LI_RphaseA2_hsm     five-prime UTR regions:       3358 (    6.18) (  0.51% of control)      1.59% ( 0.13 O/E)
Number of sites identified in Ob Ob_LI_RphaseA2_hsm     three-prime UTR regions:      6450 (    6.02) (  0.86% of control)      3.05% ( 0.13 O/E)
Number of sites identified in Ob Ob_LI_RphaseA2_hsm     other exon regions:           1732 (    2.14) (  0.42% of control)      0.82% ( 0.05 O/E)
Number of sites identified in Ob Ob_LI_RphaseA2_hsm   intron regions:              49815   (    4.09) (  0.46% of control)   18.77%   ( 0.34 O/E)
Number of sites identified in Ob Ob_LI_RphaseA2_hsm intergenic regions:            4375     (    0.77) (  0.08% of control)  1.65%     ( 0.06 O/E)
Number of sites identified in Ob Ob_LI_RphaseA2_hsm   promoter regions:              602   (    0.88) (  0.18% of control)    0.23%   ( 0.07 O/E)
Number of sites identified in Ob Ob_LI_RphaseA2_hsm   other intergenic regions:        3773   (    0.75) (  0.07% of control)    1.42%   ( 0.06 O/E)
```

Directory: MMP File: mmp-Ob_LI_RphaseB1.txt

================================================================================
#### Genomic composition in terms of feature regions
================================================================================

| | | |
|---|---|---|
| Ob genome size: | 223876465 bp | |
| Ob genic region size: | 166752117 bp | ( 74.5%) |
| Ob exon region size: | 44907373 bp | ( 20.1%) |
| Ob CDS region size: | 20651162 bp | ( 46.0%) |
| Ob five-prime UTR region size: | 5434886 bp | ( 12.1%) |
| Ob three-prime UTR region size: | 10723012 bp | ( 23.9%) |
| Ob other exon region size: | 8098313 bp | ( 18.0%) |
| Ob intron region size: | 121844744 bp | ( 54.4%) |
| Ob intergenic region size: | 57124348 bp | ( 25.5%) |
| Ob promoter region size: | 6874701 bp | ( 3.1%) |
| Ob other intergenic region size: | 50249647 bp | ( 22.4%) |

================================================================================
#### Methylation sites in genomic feature regions
================================================================================

Number and density (per 10kb) of sites in Ob_LI_RphaseB1_scd

| | | | | |
|---|---|---|---|---|
| Number of sites identified in Ob Ob_LI_RphaseB1_scd: | 19708064 | ( 880.31) | | |
| Number of sites identified in Ob Ob_LI_RphaseB1_scd genic regions: | 14345119 | ( 860.27) | 72.79% | ( 0.98 O/E) |
| Number of sites identified in Ob Ob_LI_RphaseB1_scd exon regions: | 4036886 | ( 898.94) | 20.48% | ( 1.02 O/E) |
| Number of sites identified in Ob Ob_LI_RphaseB1_scd CDS regions: | 2305080 | ( 1116.20) | 57.10% | ( 1.24 O/E) |
| Number of sites identified in Ob Ob_LI_RphaseB1_scd five-prime UTR regions: | 636146 | ( 1170.49) | 15.76% | ( 1.30 O/E) |
| Number of sites identified in Ob Ob_LI_RphaseB1_scd three-prime UTR regions: | 706621 | ( 658.98) | 17.50% | ( 0.73 O/E) |
| Number of sites identified in Ob Ob_LI_RphaseB1_scd other exon regions: | 389039 | ( 480.40) | 9.64% | ( 0.53 O/E) |
| Number of sites identified in Ob Ob_LI_RphaseB1_scd intron regions: | 10308233 | ( 846.01) | 52.30% | ( 0.96 O/E) |
| Number of sites identified in Ob Ob_LI_RphaseB1_scd intergenic regions: | 5362945 | ( 938.82) | 27.21% | ( 1.07 O/E) |
| Number of sites identified in Ob Ob_LI_RphaseB1_scd promoter regions: | 325070 | ( 472.85) | 1.65% | ( 0.54 O/E) |
| Number of sites identified in Ob Ob_LI_RphaseB1_scd other intergenic regions: | 5037875 | ( 1002.57) | 25.56% | ( 1.14 O/E) |

Number and density (per 10kb) of sites in Ob_LI_RphaseB1_hsm

| | | | | | |
|---|---|---|---|---|---|
| Number of sites identified in Ob Ob_LI_RphaseB1_hsm: | 217571 | ( 9.72) | ( 1.10% of control) | | |
| Number of sites identified in Ob Ob_LI_RphaseB1_hsm genic regions: | 214832 | ( 12.88) | ( 1.50% of control) | 98.74% | ( 1.33 O/E) |
| Number of sites identified in Ob Ob_LI_RphaseB1_hsm exon regions: | 175410 | ( 39.06) | ( 4.35% of control) | 80.62% | ( 4.02 O/E) |
| Number of sites identified in Ob Ob_LI_RphaseB1_hsm CDS regions: | 166834 | ( 80.79) | ( 7.24% of control) | 95.11% | ( 2.07 O/E) |
| Number of sites identified in Ob Ob_LI_RphaseB1_hsm five-prime UTR regions: | 2645 | ( 4.87) | ( 0.42% of control) | 1.51% | ( 0.12 O/E) |
| Number of sites identified in Ob Ob_LI_RphaseB1_hsm three-prime UTR regions: | 4572 | ( 4.26) | ( 0.65% of control) | 2.61% | ( 0.11 O/E) |
| Number of sites identified in Ob Ob_LI_RphaseB1_hsm other exon regions: | 1359 | ( 1.68) | ( 0.35% of control) | 0.77% | ( 0.04 O/E) |
| Number of sites identified in Ob Ob_LI_RphaseB1_hsm intron regions: | 39422 | ( 3.24) | ( 0.38% of control) | 18.12% | ( 0.33 O/E) |
| Number of sites identified in Ob Ob_LI_RphaseB1_hsm intergenic regions: | 2739 | ( 0.48) | ( 0.05% of control) | 1.26% | ( 0.05 O/E) |
| Number of sites identified in Ob Ob_LI_RphaseB1_hsm promoter regions: | 411 | ( 0.60) | ( 0.13% of control) | 0.19% | ( 0.06 O/E) |
| Number of sites identified in Ob Ob_LI_RphaseB1_hsm other intergenic regions: | 2328 | ( 0.46) | ( 0.05% of control) | 1.07% | ( 0.05 O/E) |

```
Directory: MMP File: mmp-Ob_LI_RphaseB2.txt

================================================================================
Genomic composition in terms of feature regions
================================================================================

Ob genome size:                        223876465 bp
  Ob genic region size:                  166752117 bp ( 74.5%)
    Ob exon region size:                    44907373 bp ( 20.1%)
      Ob CDS region size:                     20651162 bp ( 46.0%)
      Ob five-prime UTR region size:           5434886 bp ( 12.1%)
      Ob three-prime UTR region size:         10723012 bp ( 23.9%)
      Ob other exon region size:               8098313 bp ( 18.0%)
    Ob intron region size:                 121844744 bp ( 54.4%)
  Ob intergenic region size:              57124348 bp ( 25.5%)
    Ob promoter region size:                 6874701 bp (  3.1%)
    Ob other intergenic region size:        50249647 bp ( 22.4%)

================================================================================
Methylation sites in genomic feature regions
================================================================================

Number and density (per 10kb) of sites in Ob_LI_RphaseB2_scd

Number of sites identified in Ob Ob_LI_RphaseB2_scd:                             18531805     (  827.77)
Number of sites identified in Ob Ob_LI_RphaseB2_scd genic regions:               13501538     (  809.68)                          72.86%     ( 0.98 O/E)
Number of sites identified in Ob Ob_LI_RphaseB2_scd   exon regions:                 3807651   (  847.89)                            20.55%   ( 1.02 O/E)
Number of sites identified in Ob Ob_LI_RphaseB2_scd     CDS regions:                  2186801 ( 1058.92)                              57.43% ( 1.25 O/E)
Number of sites identified in Ob Ob_LI_RphaseB2_scd     five-prime UTR regions:        604660 ( 1112.55)                              15.88% ( 1.31 O/E)
Number of sites identified in Ob Ob_LI_RphaseB2_scd     three-prime UTR regions:       652553 (  608.55)                              17.14% ( 0.72 O/E)
Number of sites identified in Ob Ob_LI_RphaseB2_scd     other exon regions:            363637 (  449.03)                               9.55% ( 0.53 O/E)
Number of sites identified in Ob Ob_LI_RphaseB2_scd   intron regions:               9693887   (  795.59)                            52.31%   ( 0.96 O/E)
Number of sites identified in Ob Ob_LI_RphaseB2_scd intergenic regions:           5030267     (  880.58)                          27.14%     ( 1.06 O/E)
Number of sites identified in Ob Ob_LI_RphaseB2_scd   promoter regions:              306706   (  446.14)                             1.66%   ( 0.54 O/E)
Number of sites identified in Ob Ob_LI_RphaseB2_scd   other intergenic regions:     4723561   (  940.02)                            25.49%   ( 1.14 O/E)

Number and density (per 10kb) of sites in Ob_LI_RphaseB2_hsm

Number of sites identified in Ob Ob_LI_RphaseB2_hsm:                               201205     (    8.99) (  1.09% of control)
Number of sites identified in Ob Ob_LI_RphaseB2_hsm genic regions:                 198904     (   11.93) (  1.47% of control)  98.86%     ( 1.33 O/E)
Number of sites identified in Ob Ob_LI_RphaseB2_hsm   exon regions:                  162884   (   36.27) (  4.28% of control)    80.95%   ( 4.04 O/E)
Number of sites identified in Ob Ob_LI_RphaseB2_hsm     CDS regions:                   155250 (   75.18) (  7.10% of control)      95.31% ( 2.07 O/E)
Number of sites identified in Ob Ob_LI_RphaseB2_hsm     five-prime UTR regions:          2376 (    4.37) (  0.39% of control)       1.46% ( 0.12 O/E)
Number of sites identified in Ob Ob_LI_RphaseB2_hsm     three-prime UTR regions:         4062 (    3.79) (  0.62% of control)       2.49% ( 0.10 O/E)
Number of sites identified in Ob Ob_LI_RphaseB2_hsm     other exon regions:              1196 (    1.48) (  0.33% of control)       0.73% ( 0.04 O/E)
Number of sites identified in Ob Ob_LI_RphaseB2_hsm   intron regions:                 36020   (    2.96) (  0.37% of control)    17.90%   ( 0.33 O/E)
Number of sites identified in Ob Ob_LI_RphaseB2_hsm intergenic regions:             2301      (    0.40) (  0.05% of control)   1.14%     ( 0.04 O/E)
Number of sites identified in Ob Ob_LI_RphaseB2_hsm   promoter regions:                 361   (    0.53) (  0.12% of control)     0.18%   ( 0.06 O/E)
Number of sites identified in Ob Ob_LI_RphaseB2_hsm   other intergenic regions:        1940   (    0.39) (  0.04% of control)     0.96%   ( 0.04 O/E)
```

```
Directory: ACS File: 0READMEacs

ACS - Annotation of conserved methylation sites.

Input: studymk, genome_ann (a list of GRanges objects providing annotated region labels and bounds;
                            output of BWASPR::get_genome_annotation())

Output: file acs-*.txt

Notes: The output is generated by BWASPR::annotate_methylome(), which uses methylKit::unite()
        to determine the sites common to all experiments and applies genomation::annotateWithFeature().
        The acs-*.txt files show the common sites with methylation levels in all samples
        and columns with booleans indicating whether the site fits a specific annotation feature.

       Abbreviations used: pc, protein coding; nc, non-coding; fp, five prime; tp, three prime.
                           UTR, untranslated region; unique, not overlapping with other categories.
```

Directory: ACS File: acs-Ob_LI.txt

| BCphaseA1 | BCphaseA2 | BCphaseB1 | BCphaseB2 | RphaseA1 | RphaseA2 | RphaseB1 | RphaseB2 | chr | start | end | strand | gene | exon | pcexon | promoter | CDS | fpUTR | tpUTR | fpUTRnotCDS | tpUTRnotCDS | tpUTRunique | ncexon |
|---|---|---|---|---|---|---|---|---|---|---|---|---|---|---|---|---|---|---|---|---|---|---|
| 38.46 | 34.29 | 30 | 44 | 59.26 | 44.68 | 27.27 | 36 | NC_039506.1 | 1097588 | 1097588 | + | 1 | 1 | 1 | 0 | 1 | 0 | 0 | 0 | 0 | 0 | 0 |
| 69.44 | 43.75 | 54.35 | 51.85 | 51.61 | 63.93 | 50 | 66.67 | NC_039506.1 | 1097598 | 1097598 | + | 1 | 1 | 1 | 0 | 1 | 0 | 0 | 0 | 0 | 0 | 0 |
| 92.31 | 97.78 | 96.77 | 96.88 | 97.22 | 93.75 | 91.67 | 97.37 | NC_039506.1 | 1097773 | 1097773 | + | 1 | 1 | 1 | 0 | 1 | 0 | 0 | 0 | 0 | 0 | 0 |
| 100 | 96.88 | 100 | 95.45 | 96.3 | 94.59 | 96.67 | 100 | NC_039506.1 | 1098055 | 1098055 | + | 1 | 1 | 1 | 0 | 1 | 0 | 0 | 0 | 0 | 0 | 0 |
| 100 | 96.77 | 95.45 | 85.71 | 100 | 100 | 100 | 93.33 | NC_039506.1 | 1098069 | 1098069 | + | 1 | 1 | 1 | 0 | 1 | 0 | 0 | 0 | 0 | 0 | 0 |
| 58.82 | 82.86 | 52.17 | 52.38 | 70.83 | 61.9 | 73.33 | 61.11 | NC_039506.1 | 1098123 | 1098123 | + | 1 | 1 | 1 | 0 | 1 | 0 | 0 | 0 | 0 | 0 | 0 |
| 91.67 | 100 | 94.74 | 100 | 91.67 | 84 | 95.65 | 100 | NC_039506.1 | 1098144 | 1098144 | + | 1 | 1 | 1 | 0 | 1 | 0 | 0 | 0 | 0 | 0 | 0 |
| 84.62 | 100 | 85.71 | 92.31 | 76.19 | 82.14 | 86.21 | 100 | NC_039506.1 | 1098163 | 1098163 | + | 1 | 1 | 1 | 0 | 1 | 0 | 0 | 0 | 0 | 0 | 0 |
| 65.38 | 76.47 | 67.65 | 58.62 | 78.38 | 62.79 | 66.67 | 60.71 | NC_039506.1 | 1098206 | 1098206 | + | 1 | 1 | 1 | 0 | 1 | 0 | 0 | 0 | 0 | 0 | 0 |
| 51.22 | 58.14 | 54 | 57.41 | 56.14 | 61.43 | 56.25 | 60 | NC_039506.1 | 1098266 | 1098266 | + | 1 | 1 | 1 | 0 | 1 | 0 | 0 | 0 | 0 | 0 | 0 |
| 52.38 | 38.3 | 39.58 | 45.76 | 30.36 | 45.83 | 45.1 | 36.96 | NC_039506.1 | 1098279 | 1098279 | + | 1 | 1 | 1 | 0 | 1 | 0 | 0 | 0 | 0 | 0 | 0 |
| 40 | 42.31 | 33.33 | 25.71 | 38.71 | 41.86 | 24.32 | 37.93 | NC_039506.1 | 1098293 | 1098293 | + | 1 | 1 | 1 | 0 | 1 | 0 | 0 | 0 | 0 | 0 | 0 |
| 73.33 | 60.87 | 76.92 | 57.14 | 53.12 | 52.78 | 61.7 | 69.57 | NC_039506.1 | 1098301 | 1098301 | + | 1 | 1 | 1 | 0 | 1 | 0 | 0 | 0 | 0 | 0 | 0 |
| 34.21 | 44.44 | 17.02 | 19.57 | 19.61 | 35.62 | 40 | 32.14 | NC_039506.1 | 1098330 | 1098330 | + | 1 | 1 | 1 | 0 | 1 | 0 | 0 | 0 | 0 | 0 | 0 |
| 64.86 | 60.34 | 54 | 60 | 43.4 | 61.33 | 56.86 | 68.97 | NC_039506.1 | 1098334 | 1098334 | + | 1 | 1 | 1 | 0 | 1 | 0 | 0 | 0 | 0 | 0 | 0 |
| 41.67 | 49.12 | 22.95 | 47.27 | 34.38 | 40.48 | 28.57 | 32.79 | NC_039506.1 | 1098377 | 1098377 | + | 1 | 1 | 1 | 0 | 1 | 0 | 0 | 0 | 0 | 0 | 0 |
| 53.49 | 56.67 | 50 | 54.55 | 53.7 | 51.14 | 52.73 | 43.86 | NC_039506.1 | 1098387 | 1098387 | + | 1 | 1 | 1 | 0 | 1 | 0 | 0 | 0 | 0 | 0 | 0 |
| 28.57 | 53.06 | 44.74 | 46.67 | 44.44 | 59.21 | 41.86 | 58.7 | NC_039506.1 | 1098400 | 1098400 | + | 1 | 1 | 1 | 0 | 1 | 0 | 0 | 0 | 0 | 0 | 0 |
| 41.67 | 40 | 41.18 | 31.03 | 48.57 | 37.74 | 48.84 | 51.85 | NC_039506.1 | 1098420 | 1098420 | + | 1 | 1 | 1 | 0 | 1 | 0 | 0 | 0 | 0 | 0 | 0 |
| 52.78 | 44.44 | 50 | 54.84 | 50 | 46.43 | 63.04 | 62.07 | NC_039506.1 | 1098427 | 1098427 | + | 1 | 1 | 1 | 0 | 1 | 0 | 0 | 0 | 0 | 0 | 0 |
| 58.82 | 54.55 | 65.71 | 42.86 | 33.33 | 46.67 | 63.16 | 60 | NC_039506.1 | 1113185 | 1113185 | + | 1 | 0 | 0 | 0 | 0 | 0 | 0 | 0 | 0 | 0 | 0 |
| 44.19 | 35.42 | 59.46 | 56.9 | 46 | 36.07 | 36.36 | 45.16 | NC_039506.1 | 1113298 | 1113298 | + | 1 | 1 | 1 | 0 | 1 | 0 | 0 | 0 | 0 | 0 | 0 |
| 33.33 | 41.67 | 39.13 | 43.14 | 37.5 | 25.64 | 37.5 | 37.04 | NC_039506.1 | 1122757 | 1122757 | + | 1 | 1 | 1 | 0 | 1 | 0 | 0 | 0 | 0 | 0 | 0 |
| 63.64 | 72.22 | 71.88 | 65.38 | 70.59 | 75 | 76 | 83.33 | NC_039506.1 | 1214614 | 1214614 | + | 1 | 1 | 1 | 0 | 1 | 0 | 0 | 0 | 0 | 0 | 0 |

...
... only first 25 lines shown ...

```
Directory: RNK File: 0READMErnk

RNK - Genomic feature regions ranked by CpG methylation and hsm statistics.

Input: studymc, studymk, region.gr (GRanges object providing annotated region labels and bounds)

Output: files ranked-genes-*.txt sites-in-genes-*.txt
              ranked-promoters-*.txt sites-in-promoters-*.txt
              plot*.pdf

Notes: The output is generated by BWASPR::rank_rbm().
       The files ranked-genes-*.txt provide ranked lists of genes based on
        overall methylation percentage or the occurrence of CpGhms sites
        within the annotated gene bounds.  The output columns include
          region_ID    (= region/gene name)
          rwidth       (= region/gene width)
          nbrsites     (= number of CpGhsm sites in the region/gene)
          nbrper10kb   (= number of CpGhsm sites in the region/gene normalized to 10kb width)
          pmrpersite   (= average % methylation per CpGhsm site in the region/gene)
          pmrpernucl   (= average % CpGhsm methylation per nucleotide in the region/gene)
          prcntM       (= overall % CpGhsm methylation in the region/gene)

       The tables are sorted by prcntM (*byPrcntM* files) or by nbper10kb
         (*bySiteDensity* files) in descending order.

       The ranked-promoters-*.txt files provide analogous tables for promoter regions.
        If the promoter annotation was derived with BWASP, then these regions are simply
        defined here as 500 nucleotides upstream of the 5'-end of the gene (shorter if the
        scaffold ends before).

       The files sites-*.txt provide details for the CpGhsm sites in the respective
        regions.
       The files plot-*.pdf explore concordance of the two methylation measures.
       The optional parameters minrwidth, maxrwidth, and minnbrsites specify
       the length range and minimal number of methylation sites for regions
       to be included in the plot.
```

Directory: RNK File: ranked-genes-byPrcntM-Ob_LI_BCphaseA1.txt

| region_ID | seqnames | start | end | width | strand | coverage | numCs | numTs | prcntM |
|---|---|---|---|---|---|---|---|---|---|
| gene-LOC113562209 | NC_039511.1 | 14632766 | 14633743 | 978 | - | 13 | 13 | 0 | 100 |
| gene-LOC105278868 | NC_039517.1 | 529826 | 532361 | 2536 | + | 5 | 5 | 0 | 100 |
| gene-LOC113561532 | NC_039507.1 | 7278071 | 7278142 | 72 | + | 112 | 108 | 4 | 96.43 |
| gene-LOC113563560 | NC_039506.1 | 6691659 | 6696603 | 4945 | + | 200 | 176 | 24 | 88 |
| gene-LOC113562300 | NC_039512.1 | 5854372 | 5855269 | 898 | - | 101 | 87 | 14 | 86.14 |
| gene-LOC105277804 | NC_039506.1 | 6089683 | 6090856 | 1174 | + | 431 | 344 | 87 | 79.81 |
| gene-LOC113563324 | NC_039518.1 | 9465393 | 9469066 | 3674 | + | 325 | 254 | 71 | 78.15 |
| gene-LOC105276706 | NC_039511.1 | 13248978 | 13250417 | 1440 | - | 957 | 726 | 231 | 75.86 |
| gene-LOC105281831 | NC_039511.1 | 2540409 | 2541476 | 1068 | - | 511 | 385 | 126 | 75.34 |
| gene-LOC105283535 | NC_039509.1 | 110345 | 111564 | 1220 | + | 551 | 412 | 139 | 74.77 |
| gene-LOC105275557 | NC_039513.1 | 10333074 | 10333884 | 811 | + | 607 | 453 | 154 | 74.63 |
| gene-LOC105286755 | NC_039506.1 | 6803777 | 6805333 | 1557 | - | 431 | 320 | 111 | 74.25 |
| gene-LOC105281669 | NC_039507.1 | 5181562 | 5182434 | 873 | - | 479 | 354 | 125 | 73.9 |
| gene-LOC105283045 | NC_039519.1 | 7797834 | 7799515 | 1682 | + | 841 | 606 | 235 | 72.06 |
| gene-LOC105283854 | NC_039509.1 | 2130793 | 2133136 | 2344 | - | 253 | 180 | 73 | 71.15 |
| gene-LOC105275946 | NC_039506.1 | 12729899 | 12732494 | 2596 | + | 339 | 241 | 98 | 71.09 |
| gene-LOC113562627 | NC_039514.1 | 15577696 | 15578092 | 397 | + | 657 | 466 | 191 | 70.93 |
| gene-LOC105277649 | NC_039512.1 | 2275901 | 2276899 | 999 | - | 523 | 369 | 154 | 70.55 |
| gene-LOC105281226 | NC_039510.1 | 10003292 | 10004599 | 1308 | + | 535 | 377 | 158 | 70.47 |
| gene-LOC105278227 | NC_039515.1 | 4632903 | 4635548 | 2646 | + | 256 | 180 | 76 | 70.31 |
| gene-LOC105281543 | NC_039517.1 | 3028824 | 3030314 | 1491 | - | 421 | 290 | 131 | 68.88 |
| gene-LOC105284008 | NC_039507.1 | 14582710 | 14583816 | 1107 | + | 494 | 336 | 158 | 68.02 |
| gene-LOC105275270 | NC_039516.1 | 12307603 | 12308894 | 1292 | + | 1239 | 836 | 403 | 67.47 |
| gene-LOC105277325 | NC_039514.1 | 13671986 | 13674003 | 2018 | + | 681 | 459 | 222 | 67.4 |

...

... only first 25 lines shown ...

Directory: RNK File: ranked-genes-byPrcntM-Ob_LI_BCphaseA2.txt

| region_ID | seqnames | start | end | width | strand | coverage | numCs | numTs | prcntM |
|---|---|---|---|---|---|---|---|---|---|
| gene-LOC113561532 | NC_039507.1 | 7278071 | 7278142 | 72 | + | 137 | 131 | 6 | 95.62 |
| gene-LOC113563560 | NC_039506.1 | 6691659 | 6696603 | 4945 | + | 131 | 121 | 10 | 92.37 |
| gene-LOC105274682 | NC_039516.1 | 737277 | 738341 | 1065 | + | 30 | 26 | 4 | 86.67 |
| gene-LOC105283998 | NC_039507.1 | 14515051 | 14517463 | 2413 | + | 135 | 110 | 25 | 81.48 |
| gene-LOC113563324 | NC_039518.1 | 9465393 | 9469066 | 3674 | + | 417 | 334 | 83 | 80.1 |
| gene-LOC105277804 | NC_039506.1 | 6089683 | 6090856 | 1174 | + | 401 | 314 | 87 | 78.3 |
| gene-LOC105275133 | NC_039509.1 | 10713340 | 10714785 | 1446 | + | 160 | 123 | 37 | 76.88 |
| gene-LOC105281688 | NC_039507.1 | 5097593 | 5098771 | 1179 | - | 233 | 179 | 54 | 76.82 |
| gene-LOC105287835 | NC_039513.1 | 7295631 | 7296689 | 1059 | + | 141 | 108 | 33 | 76.6 |
| gene-LOC105281543 | NC_039517.1 | 3028824 | 3030314 | 1491 | - | 546 | 416 | 130 | 76.19 |
| gene-LOC105286755 | NC_039506.1 | 6803777 | 6805333 | 1557 | - | 616 | 466 | 150 | 75.65 |
| gene-LOC105275557 | NC_039513.1 | 10333074 | 10333884 | 811 | + | 602 | 448 | 154 | 74.42 |
| gene-LOC105276706 | NC_039511.1 | 13248978 | 13250417 | 1440 | - | 1167 | 859 | 308 | 73.61 |
| gene-LOC105283535 | NC_039509.1 | 110345 | 111564 | 1220 | + | 515 | 371 | 144 | 72.04 |
| gene-LOC105287395 | NC_039517.1 | 5748085 | 5749071 | 987 | + | 196 | 141 | 55 | 71.94 |
| gene-LOC105275498 | NC_039513.1 | 10737439 | 10738859 | 1421 | - | 520 | 374 | 146 | 71.92 |
| gene-LOC105275946 | NC_039506.1 | 12729899 | 12732494 | 2596 | + | 263 | 189 | 74 | 71.86 |
| gene-LOC105278227 | NC_039515.1 | 4632903 | 4635548 | 2646 | + | 290 | 208 | 82 | 71.72 |
| gene-LOC105281669 | NC_039507.1 | 5181562 | 5182434 | 873 | - | 675 | 481 | 194 | 71.26 |
| gene-LOC105282802 | NC_039512.1 | 16231639 | 16232693 | 1055 | + | 229 | 162 | 67 | 70.74 |
| gene-LOC105277649 | NC_039512.1 | 2275901 | 2276899 | 999 | - | 467 | 328 | 139 | 70.24 |
| gene-LOC105283045 | NC_039519.1 | 7797834 | 7799515 | 1682 | + | 758 | 532 | 226 | 70.18 |
| gene-LOC105283854 | NC_039509.1 | 2130793 | 2133136 | 2344 | - | 357 | 249 | 108 | 69.75 |
| gene-LOC105281226 | NC_039510.1 | 10003292 | 10004599 | 1308 | + | 598 | 417 | 181 | 69.73 |

...
... only first 25 lines shown ...

Directory: RNK File: ranked-genes-byPrcntM-Ob_LI_BCphaseB1.txt

| region_ID | seqnames | start | end | width | strand | coverage | numCs | numTs | prcntM |
|---|---|---|---|---|---|---|---|---|---|
| gene-LOC113562209 | NC_039511.1 | 14632766 | 14633743 | 978 | - | 26 | 26 | 0 | 100 |
| gene-LOC113561532 | NC_039507.1 | 7278071 | 7278142 | 72 | + | 144 | 137 | 7 | 95.14 |
| gene-LOC113563560 | NC_039506.1 | 6691659 | 6696603 | 4945 | + | 300 | 264 | 36 | 88 |
| gene-LOC105283998 | NC_039507.1 | 14515051 | 14517463 | 2413 | + | 184 | 157 | 27 | 85.33 |
| gene-LOC105286755 | NC_039506.1 | 6803777 | 6805333 | 1557 | - | 729 | 563 | 166 | 77.23 |
| gene-LOC105277804 | NC_039506.1 | 6089683 | 6090856 | 1174 | + | 486 | 375 | 111 | 77.16 |
| gene-LOC113563324 | NC_039518.1 | 9465393 | 9469066 | 3674 | + | 411 | 316 | 95 | 76.89 |
| gene-LOC105275557 | NC_039513.1 | 10333074 | 10333884 | 811 | + | 859 | 628 | 231 | 73.11 |
| gene-LOC105283535 | NC_039509.1 | 110345 | 111564 | 1220 | + | 719 | 522 | 197 | 72.6 |
| gene-LOC105276706 | NC_039511.1 | 13248978 | 13250417 | 1440 | - | 1457 | 1054 | 403 | 72.34 |
| gene-LOC105281669 | NC_039507.1 | 5181562 | 5182434 | 873 | - | 661 | 475 | 186 | 71.86 |
| gene-LOC105281543 | NC_039517.1 | 3028824 | 3030314 | 1491 | - | 506 | 361 | 145 | 71.34 |
| gene-LOC105282913 | NC_039507.1 | 725648 | 727283 | 1636 | - | 699 | 489 | 210 | 69.96 |
| gene-LOC113562627 | NC_039514.1 | 15577696 | 15578092 | 397 | + | 955 | 666 | 289 | 69.74 |
| gene-LOC105283997 | NC_039507.1 | 14517342 | 14519998 | 2657 | - | 36 | 25 | 11 | 69.44 |
| gene-LOC105277649 | NC_039512.1 | 2275901 | 2276899 | 999 | - | 717 | 497 | 220 | 69.32 |
| gene-LOC105283854 | NC_039509.1 | 2130793 | 2133136 | 2344 | - | 411 | 281 | 130 | 68.37 |
| gene-LOC113562396 | NC_039513.1 | 5906271 | 5907921 | 1651 | + | 104 | 71 | 33 | 68.27 |
| gene-LOC113561389 | NC_039516.1 | 7634243 | 7635142 | 900 | + | 22 | 15 | 7 | 68.18 |
| gene-LOC113562959 | NC_039516.1 | 7634915 | 7636446 | 1532 | - | 22 | 15 | 7 | 68.18 |
| gene-LOC105283045 | NC_039519.1 | 7797834 | 7799515 | 1682 | + | 1037 | 699 | 338 | 67.41 |
| gene-LOC105286178 | NC_039509.1 | 7797927 | 7799516 | 1590 | + | 204 | 136 | 68 | 66.67 |
| gene-LOC105286857 | NC_039506.1 | 3999764 | 4001339 | 1576 | - | 214 | 142 | 72 | 66.36 |
| gene-LOC109610680 | NC_039515.1 | 1495098 | 1496589 | 1492 | - | 2216 | 1467 | 749 | 66.2 |

...

... only first 25 lines shown ...

Directory: RNK File: ranked-genes-byPrcntM-Ob_LI_BCphaseB2.txt

| region_ID | seqnames | start | end | width | strand | coverage | numCs | numTs | prcntM |
|---|---|---|---|---|---|---|---|---|---|
| gene-LOC113561389 | NC_039516.1 | 7634243 | 7635142 | 900 | + | 10 | 10 | 0 | 100 |
| gene-LOC113562959 | NC_039516.1 | 7634915 | 7636446 | 1532 | - | 10 | 10 | 0 | 100 |
| gene-LOC113561532 | NC_039507.1 | 7278071 | 7278142 | 72 | + | 113 | 109 | 4 | 96.46 |
| gene-LOC105283998 | NC_039507.1 | 14515051 | 14517463 | 2413 | + | 150 | 130 | 20 | 86.67 |
| gene-LOC113563560 | NC_039506.1 | 6691659 | 6696603 | 4945 | + | 206 | 176 | 30 | 85.44 |
| gene-LOC113562533 | NC_039514.1 | 1299355 | 1299805 | 451 | + | 25 | 21 | 4 | 84 |
| gene-LOC105283997 | NC_039507.1 | 14517342 | 14519998 | 2657 | - | 5 | 4 | 1 | 80 |
| gene-LOC105283854 | NC_039509.1 | 2130793 | 2133136 | 2344 | - | 286 | 227 | 59 | 79.37 |
| gene-LOC105286755 | NC_039506.1 | 6803777 | 6805333 | 1557 | - | 457 | 360 | 97 | 78.77 |
| gene-LOC113563324 | NC_039518.1 | 9465393 | 9469066 | 3674 | + | 249 | 196 | 53 | 78.71 |
| gene-LOC105281543 | NC_039517.1 | 3028824 | 3030314 | 1491 | - | 561 | 429 | 132 | 76.47 |
| gene-LOC113562300 | NC_039512.1 | 5854372 | 5855269 | 898 | - | 113 | 85 | 28 | 75.22 |
| gene-LOC105275557 | NC_039513.1 | 10333074 | 10333884 | 811 | + | 673 | 506 | 167 | 75.19 |
| gene-LOC105283045 | NC_039519.1 | 7797834 | 7799515 | 1682 | + | 903 | 678 | 225 | 75.08 |
| gene-LOC105277804 | NC_039506.1 | 6089683 | 6090856 | 1174 | + | 450 | 332 | 118 | 73.78 |
| gene-LOC105275498 | NC_039513.1 | 10737439 | 10738859 | 1421 | - | 631 | 458 | 173 | 72.58 |
| gene-LOC105281831 | NC_039511.1 | 2540409 | 2541476 | 1068 | - | 595 | 426 | 169 | 71.6 |
| gene-LOC105276706 | NC_039511.1 | 13248978 | 13250417 | 1440 | - | 1187 | 827 | 360 | 69.67 |
| gene-LOC105286178 | NC_039509.1 | 7797927 | 7799516 | 1590 | + | 153 | 106 | 47 | 69.28 |
| gene-LOC105287955 | NC_039507.1 | 1428396 | 1430881 | 2486 | - | 986 | 682 | 304 | 69.17 |
| gene-LOC105283535 | NC_039509.1 | 110345 | 111564 | 1220 | + | 514 | 355 | 159 | 69.07 |
| gene-LOC105277649 | NC_039512.1 | 2275901 | 2276899 | 999 | - | 506 | 349 | 157 | 68.97 |
| gene-LOC105282390 | NC_039512.1 | 8801935 | 8803557 | 1623 | + | 1919 | 1322 | 597 | 68.89 |
| gene-LOC109610680 | NC_039515.1 | 1495098 | 1496589 | 1492 | - | 2034 | 1392 | 642 | 68.44 |

...
... only first 25 lines shown ...

Directory: RNK File: ranked-genes-byPrcntM-Ob_LI_RphaseA1.txt

| region_ID | seqnames | start | end | width | strand | coverage | numCs | numTs | prcntM |
|---|---|---|---|---|---|---|---|---|---|
| gene-LOC113561532 | NC_039507.1 | 7278071 | 7278142 | 72 | + | 135 | 132 | 3 | 97.78 |
| gene-LOC113562209 | NC_039511.1 | 14632766 | 14633743 | 978 | - | 16 | 15 | 1 | 93.75 |
| gene-LOC113563560 | NC_039506.1 | 6691659 | 6696603 | 4945 | + | 260 | 227 | 33 | 87.31 |
| gene-LOC105286755 | NC_039506.1 | 6803777 | 6805333 | 1557 | - | 503 | 407 | 96 | 80.91 |
| gene-LOC113563324 | NC_039518.1 | 9465393 | 9469066 | 3674 | + | 546 | 435 | 111 | 79.67 |
| gene-LOC105283998 | NC_039507.1 | 14515051 | 14517463 | 2413 | + | 191 | 147 | 44 | 76.96 |
| gene-LOC105277649 | NC_039512.1 | 2275901 | 2276899 | 999 | - | 594 | 442 | 152 | 74.41 |
| gene-LOC105277804 | NC_039506.1 | 6089683 | 6090856 | 1174 | + | 365 | 267 | 98 | 73.15 |
| gene-LOC105275557 | NC_039513.1 | 10333074 | 10333884 | 811 | + | 657 | 478 | 179 | 72.75 |
| gene-LOC105281669 | NC_039507.1 | 5181562 | 5182434 | 873 | - | 691 | 498 | 193 | 72.07 |
| gene-LOC105278227 | NC_039515.1 | 4632903 | 4635548 | 2646 | + | 334 | 230 | 104 | 68.86 |
| gene-LOC105276706 | NC_039511.1 | 13248978 | 13250417 | 1440 | - | 1298 | 893 | 405 | 68.8 |
| gene-LOC113562959 | NC_039516.1 | 7634915 | 7636446 | 1532 | - | 119 | 81 | 38 | 68.07 |
| gene-LOC105275946 | NC_039506.1 | 12729899 | 12732494 | 2596 | + | 323 | 219 | 104 | 67.8 |
| gene-LOC105281543 | NC_039517.1 | 3028824 | 3030314 | 1491 | - | 552 | 373 | 179 | 67.57 |
| gene-LOC105281226 | NC_039510.1 | 10003292 | 10004599 | 1308 | + | 555 | 374 | 181 | 67.39 |
| gene-LOC113562396 | NC_039513.1 | 5906271 | 5907921 | 1651 | + | 101 | 68 | 33 | 67.33 |
| gene-LOC105283854 | NC_039509.1 | 2130793 | 2133136 | 2344 | - | 355 | 239 | 116 | 67.32 |
| gene-LOC105283045 | NC_039519.1 | 7797834 | 7799515 | 1682 | + | 715 | 475 | 240 | 66.43 |
| gene-LOC105283535 | NC_039509.1 | 110345 | 111564 | 1220 | + | 750 | 497 | 253 | 66.27 |
| gene-LOC113562627 | NC_039514.1 | 15577696 | 15578092 | 397 | + | 790 | 519 | 271 | 65.7 |
| gene-LOC105287395 | NC_039517.1 | 5748085 | 5749071 | 987 | + | 192 | 126 | 66 | 65.62 |
| gene-LOC105277650 | NC_039512.1 | 2275742 | 2277091 | 1350 | + | 779 | 509 | 270 | 65.34 |
| gene-LOC113563090 | NC_039506.1 | 11568116 | 11569844 | 1729 | - | 406 | 265 | 141 | 65.27 |

...
... only first 25 lines shown ...

Directory: RNK File: ranked-genes-byPrcntM-Ob_LI_RphaseA2.txt

| region_ID | seqnames | start | end | width | strand | coverage | numCs | numTs | prcntM |
|---|---|---|---|---|---|---|---|---|---|
| gene-LOC113561532 | NC_039507.1 | 7278071 | 7278142 | 72 | + | 203 | 194 | 9 | 95.57 |
| gene-LOC113563560 | NC_039506.1 | 6691659 | 6696603 | 4945 | + | 239 | 217 | 22 | 90.79 |
| gene-LOC113563370 | NC_039519.1 | 415372 | 417073 | 1702 | - | 10 | 9 | 1 | 90 |
| gene-LOC105286755 | NC_039506.1 | 6803777 | 6805333 | 1557 | - | 701 | 552 | 149 | 78.74 |
| gene-LOC113563324 | NC_039518.1 | 9465393 | 9469066 | 3674 | + | 490 | 382 | 108 | 77.96 |
| gene-LOC105283854 | NC_039509.1 | 2130793 | 2133136 | 2344 | - | 519 | 398 | 121 | 76.69 |
| gene-LOC105275557 | NC_039513.1 | 10333074 | 10333884 | 811 | + | 866 | 659 | 207 | 76.1 |
| gene-LOC105277804 | NC_039506.1 | 6089683 | 6090856 | 1174 | + | 689 | 523 | 166 | 75.91 |
| gene-LOC113562300 | NC_039512.1 | 5854372 | 5855269 | 898 | - | 220 | 166 | 54 | 75.45 |
| gene-LOC105276706 | NC_039511.1 | 13248978 | 13250417 | 1440 | - | 2010 | 1513 | 497 | 75.27 |
| gene-LOC105283998 | NC_039507.1 | 14515051 | 14517463 | 2413 | + | 261 | 194 | 67 | 74.33 |
| gene-LOC105281669 | NC_039507.1 | 5181562 | 5182434 | 873 | - | 906 | 659 | 247 | 72.74 |
| gene-LOC105277649 | NC_039512.1 | 2275901 | 2276899 | 999 | - | 795 | 576 | 219 | 72.45 |
| gene-LOC105281543 | NC_039517.1 | 3028824 | 3030314 | 1491 | - | 823 | 594 | 229 | 72.17 |
| gene-LOC105275498 | NC_039513.1 | 10737439 | 10738859 | 1421 | - | 945 | 680 | 265 | 71.96 |
| gene-LOC105283045 | NC_039519.1 | 7797834 | 7799515 | 1682 | + | 1287 | 925 | 362 | 71.87 |
| gene-LOC113563090 | NC_039506.1 | 11568116 | 11569844 | 1729 | - | 514 | 363 | 151 | 70.62 |
| gene-LOC105281831 | NC_039511.1 | 2540409 | 2541476 | 1068 | - | 955 | 669 | 286 | 70.05 |
| gene-LOC105287955 | NC_039507.1 | 1428396 | 1430881 | 2486 | - | 1348 | 944 | 404 | 70.03 |
| gene-LOC109610680 | NC_039515.1 | 1495098 | 1496589 | 1492 | - | 3492 | 2445 | 1047 | 70.02 |
| gene-LOC105283997 | NC_039507.1 | 14517342 | 14519998 | 2657 | - | 33 | 23 | 10 | 69.7 |
| gene-LOC113562627 | NC_039514.1 | 15577696 | 15578092 | 397 | + | 1297 | 901 | 396 | 69.47 |
| gene-LOC105281226 | NC_039510.1 | 10003292 | 10004599 | 1308 | + | 877 | 607 | 270 | 69.21 |
| gene-LOC105276123 | NC_039514.1 | 349087 | 351144 | 2058 | + | 1390 | 945 | 445 | 67.99 |

...

... only first 25 lines shown ...

Directory: RNK File: ranked-genes-byPrcntM-Ob_LI_RphaseB1.txt

| region_ID | seqnames | start | end | width | strand | coverage | numCs | numTs | prcntM |
|---|---|---|---|---|---|---|---|---|---|
| gene-LOC113562209 | NC_039511.1 | 14632766 | 14633743 | 978 | - | 15 | 15 | 0 | 100 |
| gene-LOC113561532 | NC_039507.1 | 7278071 | 7278142 | 72 | + | 155 | 148 | 7 | 95.48 |
| gene-LOC113563560 | NC_039506.1 | 6691659 | 6696603 | 4945 | + | 217 | 190 | 27 | 87.56 |
| gene-LOC105286755 | NC_039506.1 | 6803777 | 6805333 | 1557 | - | 656 | 527 | 129 | 80.34 |
| gene-LOC113563324 | NC_039518.1 | 9465393 | 9469066 | 3674 | + | 568 | 448 | 120 | 78.87 |
| gene-LOC105283535 | NC_039509.1 | 110345 | 111564 | 1220 | + | 621 | 468 | 153 | 75.36 |
| gene-LOC105277804 | NC_039506.1 | 6089683 | 6090856 | 1174 | + | 407 | 306 | 101 | 75.18 |
| gene-LOC105281543 | NC_039517.1 | 3028824 | 3030314 | 1491 | - | 532 | 397 | 135 | 74.62 |
| gene-LOC105281688 | NC_039507.1 | 5097593 | 5098771 | 1179 | - | 264 | 194 | 70 | 73.48 |
| gene-LOC105283854 | NC_039509.1 | 2130793 | 2133136 | 2344 | - | 320 | 235 | 85 | 73.44 |
| gene-LOC105275557 | NC_039513.1 | 10333074 | 10333884 | 811 | + | 727 | 532 | 195 | 73.18 |
| gene-LOC105278227 | NC_039515.1 | 4632903 | 4635548 | 2646 | + | 302 | 215 | 87 | 71.19 |
| gene-LOC105283045 | NC_039519.1 | 7797834 | 7799515 | 1682 | + | 816 | 575 | 241 | 70.47 |
| gene-LOC105276706 | NC_039511.1 | 13248978 | 13250417 | 1440 | - | 1244 | 873 | 371 | 70.18 |
| gene-LOC105282802 | NC_039512.1 | 16231639 | 16232693 | 1055 | + | 325 | 228 | 97 | 70.15 |
| gene-LOC105275133 | NC_039509.1 | 10713340 | 10714785 | 1446 | + | 156 | 109 | 47 | 69.87 |
| gene-LOC105277649 | NC_039512.1 | 2275901 | 2276899 | 999 | - | 535 | 371 | 164 | 69.35 |
| gene-LOC105281669 | NC_039507.1 | 5181562 | 5182434 | 873 | - | 633 | 434 | 199 | 68.56 |
| gene-LOC105284656 | NC_039514.1 | 9403937 | 9405248 | 1312 | - | 940 | 636 | 304 | 67.66 |
| gene-LOC105283997 | NC_039507.1 | 14517342 | 14519998 | 2657 | - | 37 | 25 | 12 | 67.57 |
| gene-LOC105287955 | NC_039507.1 | 1428396 | 1430881 | 2486 | - | 1037 | 698 | 339 | 67.31 |
| gene-LOC105275498 | NC_039513.1 | 10737439 | 10738859 | 1421 | - | 706 | 473 | 233 | 67 |
| gene-LOC105282913 | NC_039507.1 | 725648 | 727283 | 1636 | - | 630 | 422 | 208 | 66.98 |
| gene-LOC113562627 | NC_039514.1 | 15577696 | 15578092 | 397 | + | 873 | 581 | 292 | 66.55 |

...

... only first 25 lines shown ...

Directory: RNK File: ranked-genes-byPrcntM-Ob_LI_RphaseB2.txt

| region_ID | seqnames | start | end | width | strand | coverage | numCs | numTs | prcntM |
|---|---|---|---|---|---|---|---|---|---|
| gene-LOC113561532 | NC_039507.1 | 7278071 | 7278142 | 72 | + | 150 | 143 | 7 | 95.33 |
| gene-LOC113563560 | NC_039506.1 | 6691659 | 6696603 | 4945 | + | 169 | 151 | 18 | 89.35 |
| gene-LOC113563324 | NC_039518.1 | 9465393 | 9469066 | 3674 | + | 444 | 364 | 80 | 81.98 |
| gene-LOC105277804 | NC_039506.1 | 6089683 | 6090856 | 1174 | + | 390 | 310 | 80 | 79.49 |
| gene-LOC105283854 | NC_039509.1 | 2130793 | 2133136 | 2344 | - | 282 | 222 | 60 | 78.72 |
| gene-LOC105281543 | NC_039517.1 | 3028824 | 3030314 | 1491 | - | 538 | 413 | 125 | 76.77 |
| gene-LOC105283045 | NC_039519.1 | 7797834 | 7799515 | 1682 | + | 787 | 603 | 184 | 76.62 |
| gene-LOC105282802 | NC_039512.1 | 16231639 | 16232693 | 1055 | + | 225 | 171 | 54 | 76 |
| gene-LOC105275498 | NC_039513.1 | 10737439 | 10738859 | 1421 | - | 535 | 400 | 135 | 74.77 |
| gene-LOC105275557 | NC_039513.1 | 10333074 | 10333884 | 811 | + | 663 | 495 | 168 | 74.66 |
| gene-LOC113561389 | NC_039516.1 | 7634243 | 7635142 | 900 | + | 15 | 11 | 4 | 73.33 |
| gene-LOC113562959 | NC_039516.1 | 7634915 | 7636446 | 1532 | - | 15 | 11 | 4 | 73.33 |
| gene-LOC105277649 | NC_039512.1 | 2275901 | 2276899 | 999 | - | 547 | 401 | 146 | 73.31 |
| gene-LOC113563090 | NC_039506.1 | 11568116 | 11569844 | 1729 | - | 301 | 217 | 84 | 72.09 |
| gene-LOC105283535 | NC_039509.1 | 110345 | 111564 | 1220 | + | 488 | 351 | 137 | 71.93 |
| gene-LOC105286755 | NC_039506.1 | 6803777 | 6805333 | 1557 | - | 472 | 339 | 133 | 71.82 |
| gene-LOC105281193 | NC_039510.1 | 10533970 | 10535137 | 1168 | + | 808 | 571 | 237 | 70.67 |
| gene-LOC113562300 | NC_039512.1 | 5854372 | 5855269 | 898 | - | 134 | 94 | 40 | 70.15 |
| gene-LOC109610680 | NC_039515.1 | 1495098 | 1496589 | 1492 | - | 2076 | 1454 | 622 | 70.04 |
| gene-LOC105276706 | NC_039511.1 | 13248978 | 13250417 | 1440 | - | 1044 | 731 | 313 | 70.02 |
| gene-LOC105278227 | NC_039515.1 | 4632903 | 4635548 | 2646 | + | 293 | 205 | 88 | 69.97 |
| gene-LOC105282770 | NC_039506.1 | 9064989 | 9066028 | 1040 | - | 135 | 94 | 41 | 69.63 |
| gene-LOC105281226 | NC_039510.1 | 10003292 | 10004599 | 1308 | + | 516 | 357 | 159 | 69.19 |
| gene-LOC105282390 | NC_039512.1 | 8801935 | 8803557 | 1623 | + | 1876 | 1285 | 591 | 68.5 |

...
... only first 25 lines shown ...

Directory: RNK File: ranked-genes-bySiteDensity-Ob_LI_BCphaseA1.txt

| region_ID | rwidth | nbrsites | nbrper10kb | pmpersite | pmpernucl | pglink | seqnames | start | end | strand | coverage | numCs | numTs | prcntM |
|---|---|---|---|---|---|---|---|---|---|---|---|---|---|---|
| gene-LOC113561532 | 72 | 7 | 972.22 | 96.55 | 9.39 | https://www.ncbi.nlm.nih.gov/gene/?term=LOC113561532 | NC_039507.1 | 7278071 | 7278142 | + | 112 | 108 | 4 | 96.43 |
| gene-LOC105283830 | 252 | 18 | 714.29 | 61.3 | 4.38 | https://www.ncbi.nlm.nih.gov/gene/?term=LOC105283830 | NC_039509.1 | 2003101 | 2003352 | - | 787 | 305 | 482 | 38.75 |
| gene-LOC105285394 | 1307 | 92 | 703.9 | 74.88 | 5.27 | https://www.ncbi.nlm.nih.gov/gene/?term=LOC105285394 | NC_039518.1 | 2681489 | 2682795 | - | 2807 | 1684 | 1123 | 59.99 |
| gene-LOC113562627 | 397 | 26 | 654.91 | 81.27 | 5.32 | https://www.ncbi.nlm.nih.gov/gene/?term=LOC113562627 | NC_039514.1 | 15577696 | 15578092 | + | 657 | 466 | 191 | 70.93 |
| gene-LOC105280739 | 1991 | 129 | 647.92 | 62.7 | 4.06 | https://www.ncbi.nlm.nih.gov/gene/?term=LOC105280739 | NC_039513.1 | 11674366 | 11676356 | - | 5288 | 2807 | 2481 | 53.08 |
| gene-LOC105278849 | 776 | 49 | 631.44 | 42.31 | 2.67 | https://www.ncbi.nlm.nih.gov/gene/?term=LOC105278849 | NC_039517.1 | 644964 | 645739 | - | 3950 | 932 | 3018 | 23.59 |
| gene-LOC105282149 | 1787 | 112 | 626.75 | 68.18 | 4.27 | https://www.ncbi.nlm.nih.gov/gene/?term=LOC105282149 | NC_039515.1 | 12586764 | 12588550 | + | 5453 | 2252 | 3201 | 41.3 |
| gene-LOC105274711 | 1655 | 101 | 610.27 | 62.08 | 3.79 | https://www.ncbi.nlm.nih.gov/gene/?term=LOC105274711 | NC_039515.1 | 1527980 | 1529634 | + | 4443 | 2088 | 2355 | 47 |
| gene-LOC105274534 | 1329 | 79 | 594.43 | 63.38 | 3.77 | https://www.ncbi.nlm.nih.gov/gene/?term=LOC105274534 | NC_039512.1 | 15726645 | 15727973 | + | 3825 | 1589 | 2236 | 41.54 |
| gene-LOC105286269 | 1401 | 83 | 592.43 | 67.38 | 3.99 | https://www.ncbi.nlm.nih.gov/gene/?term=LOC105286269 | NC_039518.1 | 8735257 | 8736657 | - | 3478 | 1680 | 1798 | 48.3 |
| gene-LOC105278160 | 1872 | 107 | 571.58 | 53.56 | 3.06 | https://www.ncbi.nlm.nih.gov/gene/?term=LOC105278160 | NC_039507.1 | 17010609 | 17012480 | - | 5727 | 2152 | 3575 | 37.58 |
| gene-LOC105287692 | 1505 | 85 | 564.78 | 58.01 | 3.28 | https://www.ncbi.nlm.nih.gov/gene/?term=LOC105287692 | NC_039510.1 | 3132542 | 3134046 | + | 5107 | 1541 | 3566 | 30.17 |
| gene-LOC105287811 | 2155 | 121 | 561.48 | 65.71 | 3.69 | https://www.ncbi.nlm.nih.gov/gene/?term=LOC105287811 | NC_039513.1 | 3140959 | 3143113 | - | 5847 | 2722 | 3125 | 46.55 |
| gene-LOC105285635 | 1175 | 65 | 553.19 | 60.85 | 3.37 | https://www.ncbi.nlm.nih.gov/gene/?term=LOC105285635 | NC_039513.1 | 15171298 | 15172472 | + | 4387 | 1318 | 3069 | 30.04 |
| gene-LOC105276360 | 1466 | 81 | 552.52 | 69.88 | 3.86 | https://www.ncbi.nlm.nih.gov/gene/?term=LOC105276360 | NC_039513.1 | 12169421 | 12170886 | - | 5250 | 1698 | 3552 | 32.34 |
| gene-LOC105276371 | 871 | 48 | 551.09 | 59.85 | 3.3 | https://www.ncbi.nlm.nih.gov/gene/?term=LOC105276371 | NC_039513.1 | 12529590 | 12530460 | - | 2337 | 989 | 1348 | 42.32 |
| gene-LOC105275314 | 1489 | 81 | 543.99 | 65.71 | 3.57 | https://www.ncbi.nlm.nih.gov/gene/?term=LOC105275314 | NC_039516.1 | 12883875 | 12885363 | + | 3816 | 1543 | 2273 | 40.44 |
| gene-LOC105285392 | 1601 | 87 | 543.41 | 58.06 | 3.16 | https://www.ncbi.nlm.nih.gov/gene/?term=LOC105285392 | NC_039518.1 | 2657034 | 2658634 | + | 4173 | 1486 | 2687 | 35.61 |
| gene-LOC105286531 | 372 | 20 | 537.63 | 66.91 | 3.6 | https://www.ncbi.nlm.nih.gov/gene/?term=LOC105286531 | NC_039510.1 | 15744784 | 15745155 | - | 1214 | 422 | 792 | 34.76 |
| gene-LOC105286801 | 1102 | 59 | 535.39 | 73.93 | 3.96 | https://www.ncbi.nlm.nih.gov/gene/?term=LOC105286801 | NC_039508.1 | 14384322 | 14385423 | - | 3181 | 1283 | 1898 | 40.33 |
| gene-LOC105277648 | 722 | 38 | 526.32 | 78.47 | 4.13 | https://www.ncbi.nlm.nih.gov/gene/?term=LOC105277648 | NC_039512.1 | 2277115 | 2277836 | + | 1230 | 746 | 484 | 60.65 |
| gene-LOC105285713 | 1048 | 55 | 524.81 | 65.25 | 3.42 | https://www.ncbi.nlm.nih.gov/gene/?term=LOC105285713 | NC_039509.1 | 17676015 | 17677062 | + | 2098 | 983 | 1115 | 46.85 |
| gene-LOC105278917 | 2176 | 114 | 523.9 | 59.8 | 3.13 | https://www.ncbi.nlm.nih.gov/gene/?term=LOC105278917 | NC_039514.1 | 8611043 | 8613218 | + | 6600 | 2218 | 4382 | 33.61 |
| gene-LOC105286235 | 1510 | 78 | 516.56 | 65.49 | 3.38 | https://www.ncbi.nlm.nih.gov/gene/?term=LOC105286235 | NC_039518.1 | 8581721 | 8583230 | - | 3900 | 1650 | 2250 | 42.31 |

...

... only first 25 lines shown ...

Directory: RNK File: ranked-genes-bySiteDensity-Ob_LI_BCphaseA2.txt

| region_ID | rwidth | nbrsites | nbrper10kb | pmpersite | pmpernucl | pglink | seqnames | start | end | strand | coverage | numCs | numTs | prcntM |
|---|---|---|---|---|---|---|---|---|---|---|---|---|---|---|
| gene-LOC113561532 | 72 | 7 | 972.22 | 96.6 | 9.39 | https://www.ncbi.nlm.nih.gov/gene/?term=LOC113561532 | NC_039507.1 | 7278071 | 7278142 | + | 137 | 131 | 6 | 95.62 |
| gene-LOC105278849 | 776 | 70 | 902.06 | 40.39 | 3.64 | https://www.ncbi.nlm.nih.gov/gene/?term=LOC105278849 | NC_039517.1 | 644964 | 645739 | - | 6569 | 1556 | 5013 | 23.69 |
| gene-LOC105285635 | 1175 | 87 | 740.43 | 53.48 | 3.96 | https://www.ncbi.nlm.nih.gov/gene/?term=LOC105285635 | NC_039513.1 | 15171298 | 15172472 | + | 6330 | 2115 | 4215 | 33.41 |
| gene-LOC113562627 | 397 | 29 | 730.48 | 78.38 | 5.73 | https://www.ncbi.nlm.nih.gov/gene/?term=LOC113562627 | NC_039514.1 | 15577696 | 15578092 | + | 923 | 635 | 288 | 68.8 |
| gene-LOC105282149 | 1787 | 128 | 716.28 | 65.5 | 4.69 | https://www.ncbi.nlm.nih.gov/gene/?term=LOC105282149 | NC_039515.1 | 12586764 | 12588550 | + | 8310 | 3802 | 4508 | 45.75 |
| gene-LOC105285394 | 1307 | 92 | 703.9 | 77.64 | 5.47 | https://www.ncbi.nlm.nih.gov/gene/?term=LOC105285394 | NC_039518.1 | 2681489 | 2682795 | - | 3715 | 2333 | 1382 | 62.8 |
| gene-LOC105278160 | 1872 | 127 | 678.42 | 53.05 | 3.6 | https://www.ncbi.nlm.nih.gov/gene/?term=LOC105278160 | NC_039507.1 | 17010609 | 17012480 | - | 8439 | 3340 | 5099 | 39.58 |
| gene-LOC105280739 | 1991 | 135 | 678.05 | 62.99 | 4.27 | https://www.ncbi.nlm.nih.gov/gene/?term=LOC105280739 | NC_039513.1 | 11674366 | 11676356 | - | 7488 | 4116 | 3372 | 54.97 |
| gene-LOC105286531 | 372 | 25 | 672.04 | 61.42 | 4.13 | https://www.ncbi.nlm.nih.gov/gene/?term=LOC105286531 | NC_039510.1 | 15744784 | 15745155 | - | 1839 | 685 | 1154 | 37.25 |
| gene-LOC105285392 | 1601 | 105 | 655.84 | 55.03 | 3.61 | https://www.ncbi.nlm.nih.gov/gene/?term=LOC105285392 | NC_039518.1 | 2657034 | 2658634 | + | 5989 | 2272 | 3717 | 37.94 |
| gene-LOC105286269 | 1401 | 91 | 649.54 | 66.37 | 4.31 | https://www.ncbi.nlm.nih.gov/gene/?term=LOC105286269 | NC_039518.1 | 8735257 | 8736657 | - | 4276 | 2147 | 2129 | 50.21 |
| gene-LOC105274711 | 1655 | 107 | 646.53 | 61.53 | 3.98 | https://www.ncbi.nlm.nih.gov/gene/?term=LOC105274711 | NC_039515.1 | 1527980 | 1529634 | + | 6158 | 2860 | 3298 | 46.44 |
| gene-LOC105274534 | 1329 | 83 | 624.53 | 62.45 | 3.9 | https://www.ncbi.nlm.nih.gov/gene/?term=LOC105274534 | NC_039512.1 | 15726645 | 15727973 | + | 4523 | 2012 | 2511 | 44.48 |
| gene-LOC105287677 | 1588 | 98 | 617.13 | 57.98 | 3.58 | https://www.ncbi.nlm.nih.gov/gene/?term=LOC105287677 | NC_039510.1 | 3210462 | 3212049 | - | 5439 | 2283 | 3156 | 41.97 |
| gene-LOC105287811 | 2155 | 130 | 603.25 | 66.29 | 4 | https://www.ncbi.nlm.nih.gov/gene/?term=LOC105287811 | NC_039513.1 | 3140959 | 3143113 | - | 7678 | 3693 | 3985 | 48.1 |
| gene-LOC105275314 | 1489 | 89 | 597.72 | 64.84 | 3.88 | https://www.ncbi.nlm.nih.gov/gene/?term=LOC105275314 | NC_039516.1 | 12883875 | 12885363 | + | 5031 | 2279 | 2752 | 45.3 |
| gene-LOC105286235 | 1510 | 90 | 596.03 | 62.31 | 3.71 | https://www.ncbi.nlm.nih.gov/gene/?term=LOC105286235 | NC_039518.1 | 8581721 | 8583230 | - | 5153 | 2321 | 2832 | 45.04 |
| gene-LOC105278938 | 1818 | 105 | 577.56 | 49.33 | 2.85 | https://www.ncbi.nlm.nih.gov/gene/?term=LOC105278938 | NC_039507.1 | 12939006 | 12940823 | - | 7658 | 1898 | 5760 | 24.78 |
| gene-LOC105276371 | 871 | 50 | 574.05 | 59.38 | 3.41 | https://www.ncbi.nlm.nih.gov/gene/?term=LOC105276371 | NC_039513.1 | 12529590 | 12530460 | - | 2407 | 1050 | 1357 | 43.62 |
| gene-LOC105276360 | 1466 | 84 | 572.99 | 70.56 | 4.04 | https://www.ncbi.nlm.nih.gov/gene/?term=LOC105276360 | NC_039513.1 | 12169421 | 12170886 | - | 6753 | 2253 | 4500 | 33.36 |
| gene-LOC105283847 | 1590 | 90 | 566.04 | 48.5 | 2.75 | https://www.ncbi.nlm.nih.gov/gene/?term=LOC105283847 | NC_039509.1 | 1750527 | 1752116 | - | 6350 | 2176 | 4174 | 34.27 |
| gene-LOC105278917 | 2176 | 123 | 565.26 | 59.94 | 3.39 | https://www.ncbi.nlm.nih.gov/gene/?term=LOC105278917 | NC_039514.1 | 8611043 | 8613218 | + | 9759 | 3550 | 6209 | 36.38 |
| gene-LOC105287692 | 1505 | 85 | 564.78 | 56.74 | 3.2 | https://www.ncbi.nlm.nih.gov/gene/?term=LOC105287692 | NC_039510.1 | 3132542 | 3134046 | + | 6338 | 2059 | 4279 | 32.49 |
| gene-LOC105286801 | 1102 | 62 | 562.61 | 75.61 | 4.25 | https://www.ncbi.nlm.nih.gov/gene/?term=LOC105286801 | NC_039508.1 | 14384322 | 14385423 | - | 4801 | 2115 | 2686 | 44.05 |

...

... only first 25 lines shown ...

Directory: RNK File: ranked-genes-bySiteDensity-Ob_LI_BCphaseB1.txt

| region_ID | rwidth | nbrsites | nbrper10kb | pmpersite | pmpernucl | pglink | seqnames | start | end | strand | coverage | numCs | numTs | prcntM |
|---|---|---|---|---|---|---|---|---|---|---|---|---|---|---|
| gene-LOC105278849 | 776 | 69 | 889.18 | 43.6 | 3.88 | https://www.ncbi.nlm.nih.gov/gene/?term=LOC105278849 | NC_039517.1 | 644964 | 645739 | - | 5093 | 1243 | 3850 | 24.41 |
| gene-LOC113562627 | 397 | 34 | 856.42 | 74.92 | 6.42 | https://www.ncbi.nlm.nih.gov/gene/?term=LOC113562627 | NC_039514.1 | 15577696 | 15578092 | + | 955 | 666 | 289 | 69.74 |
| gene-LOC113561532 | 72 | 6 | 833.33 | 95.37 | 7.95 | https://www.ncbi.nlm.nih.gov/gene/?term=LOC113561532 | NC_039507.1 | 7278071 | 7278142 | + | 144 | 137 | 7 | 95.14 |
| gene-LOC105285394 | 1307 | 91 | 696.25 | 76.92 | 5.36 | https://www.ncbi.nlm.nih.gov/gene/?term=LOC105285394 | NC_039518.1 | 2681489 | 2682795 | - | 3231 | 1960 | 1271 | 60.66 |
| gene-LOC105278160 | 1872 | 128 | 683.76 | 53.98 | 3.69 | https://www.ncbi.nlm.nih.gov/gene/?term=LOC105278160 | NC_039507.1 | 17010609 | 17012480 | - | 7441 | 3006 | 4435 | 40.4 |
| gene-LOC105280739 | 1991 | 136 | 683.07 | 63.36 | 4.33 | https://www.ncbi.nlm.nih.gov/gene/?term=LOC105280739 | NC_039513.1 | 11674366 | 11676356 | - | 7153 | 3781 | 3372 | 52.86 |
| gene-LOC105274711 | 1655 | 113 | 682.78 | 63.34 | 4.32 | https://www.ncbi.nlm.nih.gov/gene/?term=LOC105274711 | NC_039515.1 | 1527980 | 1529634 | + | 5363 | 2575 | 2788 | 48.01 |
| gene-LOC105286531 | 372 | 25 | 672.04 | 64.86 | 4.36 | https://www.ncbi.nlm.nih.gov/gene/?term=LOC105286531 | NC_039510.1 | 15744784 | 15745155 | - | 1776 | 646 | 1130 | 36.37 |
| gene-LOC105285635 | 1175 | 77 | 655.32 | 57.17 | 3.75 | https://www.ncbi.nlm.nih.gov/gene/?term=LOC105285635 | NC_039513.1 | 15171298 | 15172472 | + | 6096 | 2044 | 4052 | 33.53 |
| gene-LOC105282149 | 1787 | 117 | 654.73 | 68.17 | 4.46 | https://www.ncbi.nlm.nih.gov/gene/?term=LOC105282149 | NC_039515.1 | 12586764 | 12588550 | + | 7328 | 2939 | 4389 | 40.11 |
| gene-LOC105274534 | 1329 | 85 | 639.58 | 59.73 | 3.82 | https://www.ncbi.nlm.nih.gov/gene/?term=LOC105274534 | NC_039512.1 | 15726645 | 15727973 | + | 4780 | 1892 | 2888 | 39.58 |
| gene-LOC105286269 | 1401 | 88 | 628.12 | 67.05 | 4.21 | https://www.ncbi.nlm.nih.gov/gene/?term=LOC105286269 | NC_039518.1 | 8735257 | 8736657 | - | 4076 | 1971 | 2105 | 48.36 |
| gene-LOC105283847 | 1590 | 98 | 616.35 | 49.13 | 3.03 | https://www.ncbi.nlm.nih.gov/gene/?term=LOC105283847 | NC_039509.1 | 1750527 | 1752116 | - | 6027 | 2033 | 3994 | 33.73 |
| gene-LOC105286235 | 1510 | 93 | 615.89 | 62.47 | 3.85 | https://www.ncbi.nlm.nih.gov/gene/?term=LOC105286235 | NC_039518.1 | 8581721 | 8583230 | - | 5536 | 2331 | 3205 | 42.11 |
| gene-LOC105276360 | 1466 | 90 | 613.92 | 66.81 | 4.1 | https://www.ncbi.nlm.nih.gov/gene/?term=LOC105276360 | NC_039513.1 | 12169421 | 12170886 | - | 7028 | 2153 | 4875 | 30.63 |
| gene-LOC105285392 | 1601 | 95 | 593.38 | 58.07 | 3.45 | https://www.ncbi.nlm.nih.gov/gene/?term=LOC105285392 | NC_039518.1 | 2657034 | 2658634 | + | 6015 | 2095 | 3920 | 34.83 |
| gene-LOC105286562 | 1484 | 88 | 592.99 | 71.24 | 4.22 | https://www.ncbi.nlm.nih.gov/gene/?term=LOC105286562 | NC_039512.1 | 13281118 | 13282601 | + | 3010 | 1559 | 1451 | 51.79 |
| gene-LOC105287692 | 1505 | 88 | 584.72 | 55.37 | 3.24 | https://www.ncbi.nlm.nih.gov/gene/?term=LOC105287692 | NC_039510.1 | 3132542 | 3134046 | + | 5936 | 1885 | 4051 | 31.76 |
| gene-LOC105277648 | 722 | 42 | 581.72 | 79.38 | 4.62 | https://www.ncbi.nlm.nih.gov/gene/?term=LOC105277648 | NC_039512.1 | 2277115 | 2277836 | + | 1772 | 1061 | 711 | 59.88 |
| gene-LOC105287811 | 2155 | 125 | 580.05 | 64.64 | 3.75 | https://www.ncbi.nlm.nih.gov/gene/?term=LOC105287811 | NC_039513.1 | 3140959 | 3143113 | - | 7232 | 3384 | 3848 | 46.79 |
| gene-LOC105278549 | 1771 | 101 | 570.3 | 60.8 | 3.47 | https://www.ncbi.nlm.nih.gov/gene/?term=LOC105278549 | NC_039509.1 | 13031833 | 13033603 | - | 5612 | 2283 | 3329 | 40.68 |
| gene-LOC105275314 | 1489 | 84 | 564.14 | 64.47 | 3.64 | https://www.ncbi.nlm.nih.gov/gene/?term=LOC105275314 | NC_039516.1 | 12883875 | 12885363 | + | 4917 | 1967 | 2950 | 40 |
| gene-LOC105286801 | 1102 | 62 | 562.61 | 76.41 | 4.3 | https://www.ncbi.nlm.nih.gov/gene/?term=LOC105286801 | NC_039508.1 | 14384322 | 14385423 | - | 4293 | 1913 | 2380 | 44.56 |
| gene-LOC105275231 | 1270 | 71 | 559.06 | 72.68 | 4.06 | https://www.ncbi.nlm.nih.gov/gene/?term=LOC105275231 | NC_039516.1 | 12121205 | 12122474 | - | 3668 | 1459 | 2209 | 39.78 |

...
... only first 25 lines shown ...

Directory: RNK File: ranked-genes-bySiteDensity-Ob_LI_BCphaseB2.txt

| region_ID | rwidth | nbrsites | nbrper10kb | pmpersite | pmpernucl | pglink | seqnames | start | end | strand | coverage | numCs | numTs | prcntM |
|---|---|---|---|---|---|---|---|---|---|---|---|---|---|---|
| gene-LOC113561532 | 72 | 6 | 833.33 | 96.1 | 8.01 | https://www.ncbi.nlm.nih.gov/gene/?term=LOC113561532 | NC_039507.1 | 7278071 | 7278142 | + | 113 | 109 | 4 | 96.46 |
| gene-LOC105278849 | 776 | 62 | 798.97 | 38.26 | 3.06 | https://www.ncbi.nlm.nih.gov/gene/?term=LOC105278849 | NC_039517.1 | 644964 | 645739 | - | 5668 | 1223 | 4445 | 21.58 |
| gene-LOC113562627 | 397 | 30 | 755.67 | 74.51 | 5.63 | https://www.ncbi.nlm.nih.gov/gene/?term=LOC113562627 | NC_039514.1 | 15577696 | 15578092 | + | 962 | 645 | 317 | 67.05 |
| gene-LOC105282149 | 1787 | 129 | 721.88 | 66.65 | 4.81 | https://www.ncbi.nlm.nih.gov/gene/?term=LOC105282149 | NC_039515.1 | 12586764 | 12588550 | + | 8162 | 3501 | 4661 | 42.89 |
| gene-LOC105285635 | 1175 | 82 | 697.87 | 54.62 | 3.81 | https://www.ncbi.nlm.nih.gov/gene/?term=LOC105285635 | NC_039513.1 | 15171298 | 15172472 | + | 5695 | 1800 | 3895 | 31.61 |
| gene-LOC105285394 | 1307 | 89 | 680.95 | 74.6 | 5.08 | https://www.ncbi.nlm.nih.gov/gene/?term=LOC105285394 | NC_039518.1 | 2681489 | 2682795 | - | 3369 | 2041 | 1328 | 60.58 |
| gene-LOC105280739 | 1991 | 132 | 662.98 | 60.45 | 4.01 | https://www.ncbi.nlm.nih.gov/gene/?term=LOC105280739 | NC_039513.1 | 11674366 | 11676356 | - | 6859 | 3616 | 3243 | 52.72 |
| gene-LOC105274711 | 1655 | 109 | 658.61 | 60.24 | 3.97 | https://www.ncbi.nlm.nih.gov/gene/?term=LOC105274711 | NC_039515.1 | 1527980 | 1529634 | + | 5665 | 2561 | 3104 | 45.21 |
| gene-LOC105278160 | 1872 | 117 | 625 | 51.54 | 3.22 | https://www.ncbi.nlm.nih.gov/gene/?term=LOC105278160 | NC_039507.1 | 17010609 | 17012480 | - | 7227 | 2748 | 4479 | 38.02 |
| gene-LOC105286531 | 372 | 23 | 618.28 | 63.3 | 3.91 | https://www.ncbi.nlm.nih.gov/gene/?term=LOC105286531 | NC_039510.1 | 15744784 | 15745155 | - | 1785 | 629 | 1156 | 35.24 |
| gene-LOC105287692 | 1505 | 93 | 617.94 | 56.07 | 3.47 | https://www.ncbi.nlm.nih.gov/gene/?term=LOC105287692 | NC_039510.1 | 3132542 | 3134046 | + | 5852 | 1994 | 3858 | 34.07 |
| gene-LOC105274534 | 1329 | 81 | 609.48 | 64.25 | 3.92 | https://www.ncbi.nlm.nih.gov/gene/?term=LOC105274534 | NC_039512.1 | 15726645 | 15727973 | + | 4778 | 2030 | 2748 | 42.49 |
| gene-LOC105276360 | 1466 | 89 | 607.09 | 68.79 | 4.18 | https://www.ncbi.nlm.nih.gov/gene/?term=LOC105276360 | NC_039513.1 | 12169421 | 12170886 | - | 7036 | 2196 | 4840 | 31.21 |
| gene-LOC105286269 | 1401 | 85 | 606.71 | 67.14 | 4.07 | https://www.ncbi.nlm.nih.gov/gene/?term=LOC105286269 | NC_039518.1 | 8735257 | 8736657 | - | 4015 | 1917 | 2098 | 47.75 |
| gene-LOC105275314 | 1489 | 89 | 597.72 | 64.76 | 3.87 | https://www.ncbi.nlm.nih.gov/gene/?term=LOC105275314 | NC_039516.1 | 12883875 | 12885363 | + | 4837 | 1972 | 2865 | 40.77 |
| gene-LOC105287811 | 2155 | 128 | 593.97 | 65.25 | 3.88 | https://www.ncbi.nlm.nih.gov/gene/?term=LOC105287811 | NC_039513.1 | 3140959 | 3143113 | - | 7280 | 3302 | 3978 | 45.36 |
| gene-LOC105285392 | 1601 | 90 | 562.15 | 57.81 | 3.25 | https://www.ncbi.nlm.nih.gov/gene/?term=LOC105285392 | NC_039518.1 | 2657034 | 2658634 | + | 5792 | 2023 | 3769 | 34.93 |
| gene-LOC105283830 | 252 | 14 | 555.56 | 64.03 | 3.56 | https://www.ncbi.nlm.nih.gov/gene/?term=LOC105283830 | NC_039509.1 | 2003101 | 2003352 | - | 861 | 280 | 581 | 32.52 |
| gene-LOC105287677 | 1588 | 88 | 554.16 | 59.79 | 3.31 | https://www.ncbi.nlm.nih.gov/gene/?term=LOC105287677 | NC_039510.1 | 3210462 | 3212049 | - | 4866 | 2010 | 2856 | 41.31 |
| gene-LOC105286562 | 1484 | 82 | 552.56 | 73.27 | 4.05 | https://www.ncbi.nlm.nih.gov/gene/?term=LOC105286562 | NC_039512.1 | 13281118 | 13282601 | + | 2836 | 1479 | 1357 | 52.15 |
| gene-LOC105276371 | 871 | 48 | 551.09 | 58.26 | 3.21 | https://www.ncbi.nlm.nih.gov/gene/?term=LOC105276371 | NC_039513.1 | 12529590 | 12530460 | - | 2428 | 943 | 1485 | 38.84 |
| gene-LOC105286235 | 1510 | 83 | 549.67 | 63.71 | 3.5 | https://www.ncbi.nlm.nih.gov/gene/?term=LOC105286235 | NC_039518.1 | 8581721 | 8583230 | - | 4653 | 1969 | 2684 | 42.32 |
| gene-LOC105277648 | 722 | 39 | 540.17 | 75.08 | 4.06 | https://www.ncbi.nlm.nih.gov/gene/?term=LOC105277648 | NC_039512.1 | 2277115 | 2277836 | + | 1391 | 787 | 604 | 56.58 |
| gene-LOC105278844 | 1101 | 59 | 535.88 | 60.74 | 3.25 | https://www.ncbi.nlm.nih.gov/gene/?term=LOC105278844 | NC_039517.1 | 685051 | 686151 | + | 5659 | 1938 | 3721 | 34.25 |

...

... only first 25 lines shown ...

Directory: RNK File: ranked-genes-bySiteDensity-Ob_LI_RphaseA1.txt

| region_ID | rwidth | nbrsites | nbrper10kb | pmpersite | pmpernucl | pglink | seqnames | start | end | strand | coverage | numCs | numTs | prcntM |
|---|---|---|---|---|---|---|---|---|---|---|---|---|---|---|
| gene-LOC113561532 | 72 | 7 | 972.22 | 97.45 | 9.47 | https://www.ncbi.nlm.nih.gov/gene/?term=LOC113561532 | NC_039507.1 | 7278071 | 7278142 | + | 135 | 132 | 3 | 97.78 |
| gene-LOC113562627 | 397 | 29 | 730.48 | 77.11 | 5.63 | https://www.ncbi.nlm.nih.gov/gene/?term=LOC113562627 | NC_039514.1 | 15577696 | 15578092 | + | 790 | 519 | 271 | 65.7 |
| gene-LOC105282149 | 1787 | 124 | 693.9 | 67.58 | 4.69 | https://www.ncbi.nlm.nih.gov/gene/?term=LOC105282149 | NC_039515.1 | 12586764 | 12588550 | + | 6573 | 2758 | 3815 | 41.96 |
| gene-LOC105280739 | 1991 | 131 | 657.96 | 62.84 | 4.13 | https://www.ncbi.nlm.nih.gov/gene/?term=LOC105280739 | NC_039513.1 | 11674366 | 11676356 | - | 6740 | 3531 | 3209 | 52.39 |
| gene-LOC105285394 | 1307 | 84 | 642.69 | 77.72 | 5 | https://www.ncbi.nlm.nih.gov/gene/?term=LOC105285394 | NC_039518.1 | 2681489 | 2682795 | - | 3308 | 2003 | 1305 | 60.55 |
| gene-LOC105285635 | 1175 | 75 | 638.3 | 56.66 | 3.62 | https://www.ncbi.nlm.nih.gov/gene/?term=LOC105285635 | NC_039513.1 | 15171298 | 15172472 | + | 5315 | 1675 | 3640 | 31.51 |
| gene-LOC105278849 | 776 | 49 | 631.44 | 44.94 | 2.84 | https://www.ncbi.nlm.nih.gov/gene/?term=LOC105278849 | NC_039517.1 | 644964 | 645739 | - | 4082 | 912 | 3170 | 22.34 |
| gene-LOC105286269 | 1401 | 88 | 628.12 | 66.88 | 4.2 | https://www.ncbi.nlm.nih.gov/gene/?term=LOC105286269 | NC_039518.1 | 8735257 | 8736657 | - | 4029 | 1958 | 2071 | 48.6 |
| gene-LOC105286531 | 372 | 23 | 618.28 | 64.99 | 4.02 | https://www.ncbi.nlm.nih.gov/gene/?term=LOC105286531 | NC_039510.1 | 15744784 | 15745155 | - | 1418 | 504 | 914 | 35.54 |
| gene-LOC105274534 | 1329 | 82 | 617.01 | 64.63 | 3.99 | https://www.ncbi.nlm.nih.gov/gene/?term=LOC105274534 | NC_039512.1 | 15726645 | 15727973 | + | 4400 | 1879 | 2521 | 42.7 |
| gene-LOC105278160 | 1872 | 115 | 614.32 | 55.25 | 3.39 | https://www.ncbi.nlm.nih.gov/gene/?term=LOC105278160 | NC_039507.1 | 17010609 | 17012480 | - | 6863 | 2576 | 4287 | 37.53 |
| gene-LOC105274711 | 1655 | 100 | 604.23 | 62.97 | 3.81 | https://www.ncbi.nlm.nih.gov/gene/?term=LOC105274711 | NC_039515.1 | 1527980 | 1529634 | + | 4784 | 2212 | 2572 | 46.24 |
| gene-LOC105283830 | 252 | 15 | 595.24 | 65.68 | 3.91 | https://www.ncbi.nlm.nih.gov/gene/?term=LOC105283830 | NC_039509.1 | 2003101 | 2003352 | - | 779 | 273 | 506 | 35.04 |
| gene-LOC105285392 | 1601 | 95 | 593.38 | 54.33 | 3.22 | https://www.ncbi.nlm.nih.gov/gene/?term=LOC105285392 | NC_039518.1 | 2657034 | 2658634 | + | 5744 | 1933 | 3811 | 33.65 |
| gene-LOC105287811 | 2155 | 126 | 584.69 | 65.05 | 3.8 | https://www.ncbi.nlm.nih.gov/gene/?term=LOC105287811 | NC_039513.1 | 3140959 | 3143113 | - | 6943 | 3208 | 3735 | 46.2 |
| gene-LOC105286235 | 1510 | 86 | 569.54 | 64.27 | 3.66 | https://www.ncbi.nlm.nih.gov/gene/?term=LOC105286235 | NC_039518.1 | 8581721 | 8583230 | - | 4624 | 1945 | 2679 | 42.06 |
| gene-LOC105287677 | 1588 | 89 | 560.45 | 62.66 | 3.51 | https://www.ncbi.nlm.nih.gov/gene/?term=LOC105287677 | NC_039510.1 | 3210462 | 3212049 | - | 4565 | 1920 | 2645 | 42.06 |
| gene-LOC105276360 | 1466 | 82 | 559.35 | 68.3 | 3.82 | https://www.ncbi.nlm.nih.gov/gene/?term=LOC105276360 | NC_039513.1 | 12169421 | 12170886 | - | 6306 | 1906 | 4400 | 30.23 |
| gene-LOC105275314 | 1489 | 83 | 557.42 | 65.53 | 3.65 | https://www.ncbi.nlm.nih.gov/gene/?term=LOC105275314 | NC_039516.1 | 12883875 | 12885363 | + | 4368 | 1859 | 2509 | 42.56 |
| gene-LOC105278917 | 2176 | 119 | 546.88 | 59.81 | 3.27 | https://www.ncbi.nlm.nih.gov/gene/?term=LOC105278917 | NC_039514.1 | 8611043 | 8613218 | + | 7845 | 2677 | 5168 | 34.12 |
| gene-LOC105286562 | 1484 | 81 | 545.82 | 71.25 | 3.89 | https://www.ncbi.nlm.nih.gov/gene/?term=LOC105286562 | NC_039512.1 | 13281118 | 13282601 | + | 2543 | 1251 | 1292 | 49.19 |
| gene-LOC105287692 | 1505 | 82 | 544.85 | 59.4 | 3.24 | https://www.ncbi.nlm.nih.gov/gene/?term=LOC105287692 | NC_039510.1 | 3132542 | 3134046 | + | 5202 | 1689 | 3513 | 32.47 |
| gene-LOC105288193 | 1495 | 81 | 541.81 | 52.75 | 2.86 | https://www.ncbi.nlm.nih.gov/gene/?term=LOC105288193 | NC_039516.1 | 3100198 | 3101692 | + | 5048 | 1393 | 3655 | 27.6 |
| gene-LOC105283847 | 1590 | 86 | 540.88 | 47.58 | 2.57 | https://www.ncbi.nlm.nih.gov/gene/?term=LOC105283847 | NC_039509.1 | 1750527 | 1752116 | - | 5122 | 1637 | 3485 | 31.96 |

...

... only first 25 lines shown ...

Directory: RNK File: ranked-genes-bySiteDensity-Ob_LI_RphaseA2.txt

| region_ID | rwidth | nbrsites | nbrper10kb | pmpersite | pmpernucl | pglink | seqnames | start | end | strand | coverage | numCs | numTs | prcntM |
|---|---|---|---|---|---|---|---|---|---|---|---|---|---|---|
| gene-LOC105278849 | 776 | 90 | 1159.79 | 33.97 | 3.94 | https://www.ncbi.nlm.nih.gov/gene/?term=LOC105278849 | NC_039517.1 | 644964 | 645739 | - | 9798 | 2273 | 7525 | 23.2 |
| gene-LOC113561532 | 72 | 7 | 972.22 | 96.26 | 9.36 | https://www.ncbi.nlm.nih.gov/gene/?term=LOC113561532 | NC_039507.1 | 7278071 | 7278142 | + | 203 | 194 | 9 | 95.57 |
| gene-LOC113562627 | 397 | 34 | 856.42 | 78.47 | 6.72 | https://www.ncbi.nlm.nih.gov/gene/?term=LOC113562627 | NC_039514.1 | 15577696 | 15578092 | + | 1297 | 901 | 396 | 69.47 |
| gene-LOC105283830 | 252 | 20 | 793.65 | 57 | 4.52 | https://www.ncbi.nlm.nih.gov/gene/?term=LOC105283830 | NC_039509.1 | 2003101 | 2003352 | - | 1162 | 436 | 726 | 37.52 |
| gene-LOC105285394 | 1307 | 102 | 780.41 | 75.91 | 5.92 | https://www.ncbi.nlm.nih.gov/gene/?term=LOC105285394 | NC_039518.1 | 2681489 | 2682795 | - | 5137 | 3333 | 1804 | 64.88 |
| gene-LOC105282149 | 1787 | 138 | 772.24 | 64.71 | 5 | https://www.ncbi.nlm.nih.gov/gene/?term=LOC105282149 | NC_039515.1 | 12586764 | 12588550 | + | 12440 | 5469 | 6971 | 43.96 |
| gene-LOC105285635 | 1175 | 90 | 765.96 | 52.39 | 4.01 | https://www.ncbi.nlm.nih.gov/gene/?term=LOC105285635 | NC_039513.1 | 15171298 | 15172472 | + | 9027 | 3026 | 6001 | 33.52 |
| gene-LOC105278160 | 1872 | 135 | 721.15 | 50.21 | 3.62 | https://www.ncbi.nlm.nih.gov/gene/?term=LOC105278160 | NC_039507.1 | 17010609 | 17012480 | - | 11687 | 4548 | 7139 | 38.92 |
| gene-LOC105278873 | 1897 | 135 | 711.65 | 39.88 | 2.84 | https://www.ncbi.nlm.nih.gov/gene/?term=LOC105278873 | NC_039517.1 | 502319 | 504215 | + | 16432 | 3551 | 12881 | 21.61 |
| gene-LOC105280739 | 1991 | 141 | 708.19 | 61.19 | 4.33 | https://www.ncbi.nlm.nih.gov/gene/?term=LOC105280739 | NC_039513.1 | 11674366 | 11676356 | - | 11590 | 6325 | 5265 | 54.57 |
| gene-LOC105276360 | 1466 | 103 | 702.59 | 64.53 | 4.53 | https://www.ncbi.nlm.nih.gov/gene/?term=LOC105276360 | NC_039513.1 | 12169421 | 12170886 | - | 10577 | 3521 | 7056 | 33.29 |
| gene-LOC105274534 | 1329 | 93 | 699.77 | 59.98 | 4.2 | https://www.ncbi.nlm.nih.gov/gene/?term=LOC105274534 | NC_039512.1 | 15726645 | 15727973 | + | 7322 | 3225 | 4097 | 44.05 |
| gene-LOC105286269 | 1401 | 98 | 699.5 | 63.68 | 4.45 | https://www.ncbi.nlm.nih.gov/gene/?term=LOC105286269 | NC_039518.1 | 8735257 | 8736657 | - | 6098 | 3089 | 3009 | 50.66 |
| gene-LOC105287692 | 1505 | 103 | 684.39 | 53.06 | 3.63 | https://www.ncbi.nlm.nih.gov/gene/?term=LOC105287692 | NC_039510.1 | 3132542 | 3134046 | + | 9944 | 3469 | 6475 | 34.89 |
| gene-LOC105287811 | 2155 | 147 | 682.13 | 60.4 | 4.12 | https://www.ncbi.nlm.nih.gov/gene/?term=LOC105287811 | NC_039513.1 | 3140959 | 3143113 | - | 11451 | 5316 | 6135 | 46.42 |
| gene-LOC105288193 | 1495 | 101 | 675.59 | 46.16 | 3.12 | https://www.ncbi.nlm.nih.gov/gene/?term=LOC105288193 | NC_039516.1 | 3100198 | 3101692 | + | 8645 | 2573 | 6072 | 29.76 |
| gene-LOC105285392 | 1601 | 108 | 674.58 | 52.56 | 3.55 | https://www.ncbi.nlm.nih.gov/gene/?term=LOC105285392 | NC_039518.1 | 2657034 | 2658634 | + | 8693 | 3278 | 5415 | 37.71 |
| gene-LOC105283847 | 1590 | 107 | 672.96 | 46.23 | 3.11 | https://www.ncbi.nlm.nih.gov/gene/?term=LOC105283847 | NC_039509.1 | 1750527 | 1752116 | - | 8503 | 3102 | 5401 | 36.48 |
| gene-LOC105286531 | 372 | 25 | 672.04 | 61.6 | 4.14 | https://www.ncbi.nlm.nih.gov/gene/?term=LOC105286531 | NC_039510.1 | 15744784 | 15745155 | - | 2425 | 869 | 1556 | 35.84 |
| gene-LOC105274711 | 1655 | 111 | 670.69 | 60.57 | 4.06 | https://www.ncbi.nlm.nih.gov/gene/?term=LOC105274711 | NC_039515.1 | 1527980 | 1529634 | + | 8638 | 4017 | 4621 | 46.5 |
| gene-LOC105287677 | 1588 | 106 | 667.51 | 53.31 | 3.56 | https://www.ncbi.nlm.nih.gov/gene/?term=LOC105287677 | NC_039510.1 | 3210462 | 3212049 | - | 7499 | 3046 | 4453 | 40.62 |
| gene-LOC105275314 | 1489 | 96 | 644.73 | 62.31 | 4.02 | https://www.ncbi.nlm.nih.gov/gene/?term=LOC105275314 | NC_039516.1 | 12883875 | 12885363 | + | 7455 | 3164 | 4291 | 42.44 |
| gene-LOC105286562 | 1484 | 95 | 640.16 | 72.09 | 4.61 | https://www.ncbi.nlm.nih.gov/gene/?term=LOC105286562 | NC_039512.1 | 13281118 | 13282601 | + | 4191 | 2324 | 1867 | 55.45 |
| gene-LOC105286801 | 1102 | 69 | 626.13 | 74.75 | 4.68 | https://www.ncbi.nlm.nih.gov/gene/?term=LOC105286801 | NC_039508.1 | 14384322 | 14385423 | - | 6903 | 2977 | 3926 | 43.13 |

...
... only first 25 lines shown ...

Directory: RNK File: ranked-genes-bySiteDensity-Ob_LI_RphaseB1.txt

| region_ID | rwidth | nbrsites | nbrper10kb | pmpersite | pmpernucl | pglink | seqnames | start | end | strand | coverage | numCs | numTs | prcntM |
|---|---|---|---|---|---|---|---|---|---|---|---|---|---|---|
| gene-LOC113561532 | 72 | 7 | 972.22 | 94.78 | 9.21 | https://www.ncbi.nlm.nih.gov/gene/?term=LOC113561532 | NC_039507.1 | 7278071 | 7278142 | + | 155 | 148 | 7 | 95.48 |
| gene-LOC113562627 | 397 | 33 | 831.23 | 74.18 | 6.17 | https://www.ncbi.nlm.nih.gov/gene/?term=LOC113562627 | NC_039514.1 | 15577696 | 15578092 | + | 873 | 581 | 292 | 66.55 |
| gene-LOC105278849 | 776 | 57 | 734.54 | 39.01 | 2.87 | https://www.ncbi.nlm.nih.gov/gene/?term=LOC105278849 | NC_039517.1 | 644964 | 645739 | - | 4532 | 943 | 3589 | 20.81 |
| gene-LOC105280739 | 1991 | 136 | 683.07 | 62.16 | 4.25 | https://www.ncbi.nlm.nih.gov/gene/?term=LOC105280739 | NC_039513.1 | 11674366 | 11676356 | - | 6505 | 3471 | 3034 | 53.36 |
| gene-LOC105285394 | 1307 | 89 | 680.95 | 75.99 | 5.17 | https://www.ncbi.nlm.nih.gov/gene/?term=LOC105285394 | NC_039518.1 | 2681489 | 2682795 | - | 3127 | 1889 | 1238 | 60.41 |
| gene-LOC105286269 | 1401 | 95 | 678.09 | 63.53 | 4.31 | https://www.ncbi.nlm.nih.gov/gene/?term=LOC105286269 | NC_039518.1 | 8735257 | 8736657 | - | 4205 | 2105 | 2100 | 50.06 |
| gene-LOC105285635 | 1175 | 79 | 672.34 | 55.29 | 3.72 | https://www.ncbi.nlm.nih.gov/gene/?term=LOC105285635 | NC_039513.1 | 15171298 | 15172472 | + | 5269 | 1657 | 3612 | 31.45 |
| gene-LOC105282149 | 1787 | 116 | 649.13 | 66.18 | 4.3 | https://www.ncbi.nlm.nih.gov/gene/?term=LOC105282149 | NC_039515.1 | 12586764 | 12588550 | + | 6424 | 2598 | 3826 | 40.44 |
| gene-LOC105283830 | 252 | 16 | 634.92 | 68.55 | 4.35 | https://www.ncbi.nlm.nih.gov/gene/?term=LOC105283830 | NC_039509.1 | 2003101 | 2003352 | - | 753 | 295 | 458 | 39.18 |
| gene-LOC105274534 | 1329 | 84 | 632.05 | 62.4 | 3.94 | https://www.ncbi.nlm.nih.gov/gene/?term=LOC105274534 | NC_039512.1 | 15726645 | 15727973 | + | 4591 | 1955 | 2636 | 42.58 |
| gene-LOC105286531 | 372 | 23 | 618.28 | 62.34 | 3.85 | https://www.ncbi.nlm.nih.gov/gene/?term=LOC105286531 | NC_039510.1 | 15744784 | 15745155 | - | 1632 | 551 | 1081 | 33.76 |
| gene-LOC105278160 | 1872 | 115 | 614.32 | 50.7 | 3.11 | https://www.ncbi.nlm.nih.gov/gene/?term=LOC105278160 | NC_039507.1 | 17010609 | 17012480 | - | 7123 | 2543 | 4580 | 35.7 |
| gene-LOC105276360 | 1466 | 88 | 600.27 | 68.28 | 4.1 | https://www.ncbi.nlm.nih.gov/gene/?term=LOC105276360 | NC_039513.1 | 12169421 | 12170886 | - | 6588 | 2091 | 4497 | 31.74 |
| gene-LOC105286801 | 1102 | 65 | 589.84 | 72.61 | 4.28 | https://www.ncbi.nlm.nih.gov/gene/?term=LOC105286801 | NC_039508.1 | 14384322 | 14385423 | - | 4010 | 1634 | 2376 | 40.75 |
| gene-LOC105286562 | 1484 | 86 | 579.51 | 72.6 | 4.21 | https://www.ncbi.nlm.nih.gov/gene/?term=LOC105286562 | NC_039512.1 | 13281118 | 13282601 | + | 2614 | 1309 | 1305 | 50.08 |
| gene-LOC105287811 | 2155 | 124 | 575.41 | 65.33 | 3.76 | https://www.ncbi.nlm.nih.gov/gene/?term=LOC105287811 | NC_039513.1 | 3140959 | 3143113 | - | 6779 | 3266 | 3513 | 48.18 |
| gene-LOC105287677 | 1588 | 91 | 573.05 | 59.98 | 3.44 | https://www.ncbi.nlm.nih.gov/gene/?term=LOC105287677 | NC_039510.1 | 3210462 | 3212049 | - | 4720 | 2038 | 2682 | 43.18 |
| gene-LOC105287692 | 1505 | 86 | 571.43 | 56.52 | 3.23 | https://www.ncbi.nlm.nih.gov/gene/?term=LOC105287692 | NC_039510.1 | 3132542 | 3134046 | + | 6064 | 1905 | 4159 | 31.41 |
| gene-LOC105275314 | 1489 | 85 | 570.85 | 64.88 | 3.7 | https://www.ncbi.nlm.nih.gov/gene/?term=LOC105275314 | NC_039516.1 | 12883875 | 12885363 | + | 4719 | 1967 | 2752 | 41.68 |
| gene-LOC105286235 | 1510 | 86 | 569.54 | 63.93 | 3.64 | https://www.ncbi.nlm.nih.gov/gene/?term=LOC105286235 | NC_039518.1 | 8581721 | 8583230 | - | 4577 | 2008 | 2569 | 43.87 |
| gene-LOC105274711 | 1655 | 94 | 567.98 | 63.01 | 3.58 | https://www.ncbi.nlm.nih.gov/gene/?term=LOC105274711 | NC_039515.1 | 1527980 | 1529634 | + | 5176 | 2333 | 2843 | 45.07 |
| gene-LOC105278549 | 1771 | 99 | 559.01 | 60.4 | 3.38 | https://www.ncbi.nlm.nih.gov/gene/?term=LOC105278549 | NC_039509.1 | 13031833 | 13033603 | - | 5383 | 2190 | 3193 | 40.68 |
| gene-LOC105285392 | 1601 | 87 | 543.41 | 58.13 | 3.16 | https://www.ncbi.nlm.nih.gov/gene/?term=LOC105285392 | NC_039518.1 | 2657034 | 2658634 | + | 5456 | 2005 | 3451 | 36.75 |
| gene-LOC105278917 | 2176 | 118 | 542.28 | 61.85 | 3.35 | https://www.ncbi.nlm.nih.gov/gene/?term=LOC105278917 | NC_039514.1 | 8611043 | 8613218 | + | 7414 | 2704 | 4710 | 36.47 |

...

... only first 25 lines shown ...

Directory: RNK File: ranked-genes-bySiteDensity-Ob_LI_RphaseB2.txt

| region_ID | rwidth | nbrsites | nbrper10kb | pmpersite | pmpernucl | pglink | seqnames | start | end | strand | coverage | numCs | numTs | prcntM |
|---|---|---|---|---|---|---|---|---|---|---|---|---|---|---|
| gene-LOC113561532 | 72 | 7 | 972.22 | 96.12 | 9.34 | https://www.ncbi.nlm.nih.gov/gene/?term=LOC113561532 | NC_039507.1 | 7278071 | 7278142 | + | 150 | 143 | 7 | 95.33 |
| gene-LOC113562627 | 397 | 30 | 755.67 | 75.94 | 5.74 | https://www.ncbi.nlm.nih.gov/gene/?term=LOC113562627 | NC_039514.1 | 15577696 | 15578092 | + | 789 | 537 | 252 | 68.06 |
| gene-LOC105278849 | 776 | 55 | 708.76 | 42.12 | 2.99 | https://www.ncbi.nlm.nih.gov/gene/?term=LOC105278849 | NC_039517.1 | 644964 | 645739 | - | 4809 | 1094 | 3715 | 22.75 |
| gene-LOC105282149 | 1787 | 125 | 699.5 | 66.43 | 4.65 | https://www.ncbi.nlm.nih.gov/gene/?term=LOC105282149 | NC_039515.1 | 12586764 | 12588550 | + | 7523 | 3324 | 4199 | 44.18 |
| gene-LOC105280739 | 1991 | 133 | 668.01 | 60.94 | 4.07 | https://www.ncbi.nlm.nih.gov/gene/?term=LOC105280739 | NC_039513.1 | 11674366 | 11676356 | - | 6288 | 3300 | 2988 | 52.48 |
| gene-LOC105285394 | 1307 | 86 | 658 | 75.08 | 4.94 | https://www.ncbi.nlm.nih.gov/gene/?term=LOC105285394 | NC_039518.1 | 2681489 | 2682795 | - | 3169 | 2009 | 1160 | 63.4 |
| gene-LOC105274711 | 1655 | 107 | 646.53 | 62.09 | 4.01 | https://www.ncbi.nlm.nih.gov/gene/?term=LOC105274711 | NC_039515.1 | 1527980 | 1529634 | + | 5601 | 2586 | 3015 | 46.17 |
| gene-LOC105278160 | 1872 | 119 | 635.68 | 53.28 | 3.39 | https://www.ncbi.nlm.nih.gov/gene/?term=LOC105278160 | NC_039507.1 | 17010609 | 17012480 | - | 7499 | 2935 | 4564 | 39.14 |
| gene-LOC105283830 | 252 | 16 | 634.92 | 67.3 | 4.27 | https://www.ncbi.nlm.nih.gov/gene/?term=LOC105283830 | NC_039509.1 | 2003101 | 2003352 | - | 539 | 214 | 325 | 39.7 |
| gene-LOC105285635 | 1175 | 74 | 629.79 | 56.19 | 3.54 | https://www.ncbi.nlm.nih.gov/gene/?term=LOC105285635 | NC_039513.1 | 15171298 | 15172472 | + | 5428 | 1730 | 3698 | 31.87 |
| gene-LOC105274534 | 1329 | 83 | 624.53 | 62.19 | 3.88 | https://www.ncbi.nlm.nih.gov/gene/?term=LOC105274534 | NC_039512.1 | 15726645 | 15727973 | + | 4570 | 1996 | 2574 | 43.68 |
| gene-LOC105286531 | 372 | 23 | 618.28 | 63.72 | 3.94 | https://www.ncbi.nlm.nih.gov/gene/?term=LOC105286531 | NC_039510.1 | 15744784 | 15745155 | - | 1572 | 547 | 1025 | 34.8 |
| gene-LOC105275314 | 1489 | 90 | 604.43 | 63.51 | 3.84 | https://www.ncbi.nlm.nih.gov/gene/?term=LOC105275314 | NC_039516.1 | 12883875 | 12885363 | + | 4754 | 2115 | 2639 | 44.49 |
| gene-LOC105287811 | 2155 | 130 | 603.25 | 64.39 | 3.88 | https://www.ncbi.nlm.nih.gov/gene/?term=LOC105287811 | NC_039513.1 | 3140959 | 3143113 | - | 6928 | 3244 | 3684 | 46.82 |
| gene-LOC105286801 | 1102 | 65 | 589.84 | 74.27 | 4.38 | https://www.ncbi.nlm.nih.gov/gene/?term=LOC105286801 | NC_039508.1 | 14384322 | 14385423 | - | 4192 | 1849 | 2343 | 44.11 |
| gene-LOC105276360 | 1466 | 85 | 579.81 | 69.15 | 4.01 | https://www.ncbi.nlm.nih.gov/gene/?term=LOC105276360 | NC_039513.1 | 12169421 | 12170886 | - | 6071 | 2007 | 4064 | 33.06 |
| gene-LOC105285392 | 1601 | 91 | 568.39 | 55.68 | 3.16 | https://www.ncbi.nlm.nih.gov/gene/?term=LOC105285392 | NC_039518.1 | 2657034 | 2658634 | + | 5529 | 2026 | 3503 | 36.64 |
| gene-LOC105287677 | 1588 | 90 | 566.75 | 58.52 | 3.32 | https://www.ncbi.nlm.nih.gov/gene/?term=LOC105287677 | NC_039510.1 | 3210462 | 3212049 | - | 4923 | 2059 | 2864 | 41.82 |
| gene-LOC105287692 | 1505 | 85 | 564.78 | 56.23 | 3.18 | https://www.ncbi.nlm.nih.gov/gene/?term=LOC105287692 | NC_039510.1 | 3132542 | 3134046 | + | 5976 | 2037 | 3939 | 34.09 |
| gene-LOC105283847 | 1590 | 89 | 559.75 | 46.75 | 2.62 | https://www.ncbi.nlm.nih.gov/gene/?term=LOC105283847 | NC_039509.1 | 1750527 | 1752116 | - | 4847 | 1641 | 3206 | 33.86 |
| gene-LOC105276371 | 871 | 48 | 551.09 | 58.59 | 3.23 | https://www.ncbi.nlm.nih.gov/gene/?term=LOC105276371 | NC_039513.1 | 12529590 | 12530460 | - | 2456 | 1030 | 1426 | 41.94 |
| gene-LOC105286269 | 1401 | 77 | 549.61 | 71.26 | 3.92 | https://www.ncbi.nlm.nih.gov/gene/?term=LOC105286269 | NC_039518.1 | 8735257 | 8736657 | - | 3420 | 1697 | 1723 | 49.62 |
| gene-LOC105286235 | 1510 | 81 | 536.42 | 64.08 | 3.44 | https://www.ncbi.nlm.nih.gov/gene/?term=LOC105286235 | NC_039518.1 | 8581721 | 8583230 | - | 4332 | 1926 | 2406 | 44.46 |
| gene-LOC105275231 | 1270 | 67 | 527.56 | 69.1 | 3.65 | https://www.ncbi.nlm.nih.gov/gene/?term=LOC105275231 | NC_039516.1 | 12121205 | 12122474 | - | 3083 | 1422 | 1661 | 46.12 |

...
... only first 25 lines shown ...

Directory: RNK File: ranked-promoters-byPrcntM-Ob_LI_BCphaseA1.txt

| region_ID | seqnames | start | end | width | strand | coverage | numCs | numTs | prcntM |
|---|---|---|---|---|---|---|---|---|---|
| gene-LOC105285355 | NC_039509.1 | 8035614 | 8036113 | 500 | - | 24 | 23 | 1 | 95.83 |
| gene-LOC105276002 | NC_039511.1 | 12550061 | 12550560 | 500 | + | 52 | 47 | 5 | 90.38 |
| gene-LOC105276685 | NC_039511.1 | 14299458 | 14299957 | 500 | - | 752 | 674 | 78 | 89.63 |
| gene-LOC105280917 | NC_039506.1 | 11578183 | 11578682 | 500 | - | 83 | 74 | 9 | 89.16 |
| gene-LOC105276139 | NC_039514.1 | 452530 | 453029 | 500 | - | 64 | 57 | 7 | 89.06 |
| gene-LOC105277073 | NC_039508.1 | 12946948 | 12947447 | 500 | + | 17 | 15 | 2 | 88.24 |
| gene-LOC105279930 | NC_039512.1 | 11167002 | 11167501 | 500 | + | 92 | 81 | 11 | 88.04 |
| gene-LOC105285429 | NC_039517.1 | 10183111 | 10183610 | 500 | + | 307 | 268 | 39 | 87.3 |
| gene-LOC113561391 | NC_039507.1 | 999503 | 1000002 | 500 | + | 116 | 101 | 15 | 87.07 |
| gene-LOC113561532 | NC_039507.1 | 7277571 | 7278070 | 500 | + | 815 | 708 | 107 | 86.87 |
| gene-LOC105282078 | NC_039512.1 | 6125187 | 6125686 | 500 | + | 65 | 56 | 9 | 86.15 |
| gene-LOC105282801 | NC_039512.1 | 16232132 | 16232631 | 500 | + | 176 | 151 | 25 | 85.8 |
| gene-LOC113562329 | NC_039506.1 | 5740607 | 5741106 | 500 | + | 28 | 24 | 4 | 85.71 |
| gene-LOC105281689 | NC_039507.1 | 5097604 | 5098103 | 500 | - | 117 | 99 | 18 | 84.62 |
| gene-LOC105286418 | NC_039514.1 | 3321606 | 3322105 | 500 | - | 13 | 11 | 2 | 84.62 |
| gene-LOC105276074 | NC_039514.1 | 15118229 | 15118728 | 500 | - | 12 | 10 | 2 | 83.33 |
| gene-LOC105281627 | NC_039514.1 | 11099400 | 11099899 | 500 | + | 304 | 240 | 64 | 78.95 |
| gene-LOC105284331 | NC_039511.1 | 5357461 | 5357960 | 500 | + | 237 | 187 | 50 | 78.9 |
| gene-LOC113561934 | NC_039506.1 | 15618494 | 15618993 | 500 | + | 803 | 627 | 176 | 78.08 |
| gene-LOC105281345 | NC_039508.1 | 2649490 | 2649989 | 500 | - | 70 | 53 | 17 | 75.71 |
| gene-LOC105288107 | NC_039515.1 | 1102618 | 1103117 | 500 | + | 841 | 633 | 208 | 75.27 |
| gene-LOC105281671 | NC_039507.1 | 5181504 | 5182003 | 500 | - | 479 | 354 | 125 | 73.9 |
| gene-LOC105287833 | NC_039513.1 | 7297618 | 7298117 | 500 | + | 254 | 187 | 67 | 73.62 |
| gene-LOC105276585 | NC_039517.1 | 4571850 | 4572349 | 500 | - | 496 | 357 | 139 | 71.98 |

...
... only first 25 lines shown ...

Directory: RNK File: ranked-promoters-byPrcntM-Ob_LI_BCphaseA2.txt

| region_ID | seqnames | start | end | width | strand | coverage | numCs | numTs | prcntM |
|---|---|---|---|---|---|---|---|---|---|
| gene-LOC113562329 | NC_039506.1 | 5740607 | 5741106 | 500 | + | 9 | 9 | 0 | 100 |
| gene-LOC105276139 | NC_039514.1 | 452530 | 453029 | 500 | - | 49 | 48 | 1 | 97.96 |
| gene-LOC105285355 | NC_039509.1 | 8035614 | 8036113 | 500 | - | 36 | 34 | 2 | 94.44 |
| gene-LOC105276002 | NC_039511.1 | 12550061 | 12550560 | 500 | + | 88 | 82 | 6 | 93.18 |
| gene-LOC105276685 | NC_039511.1 | 14299458 | 14299957 | 500 | - | 925 | 857 | 68 | 92.65 |
| gene-LOC105285429 | NC_039517.1 | 10183111 | 10183610 | 500 | + | 271 | 243 | 28 | 89.67 |
| gene-LOC105281689 | NC_039507.1 | 5097604 | 5098103 | 500 | - | 204 | 179 | 25 | 87.75 |
| gene-LOC105277073 | NC_039508.1 | 12946948 | 12947447 | 500 | + | 8 | 7 | 1 | 87.5 |
| gene-LOC113561532 | NC_039507.1 | 7277571 | 7278070 | 500 | + | 1134 | 985 | 149 | 86.86 |
| gene-LOC113561391 | NC_039507.1 | 999503 | 1000002 | 500 | + | 107 | 92 | 15 | 85.98 |
| gene-LOC105279917 | NC_039512.1 | 13122631 | 13123130 | 500 | - | 228 | 196 | 32 | 85.96 |
| gene-LOC105282801 | NC_039512.1 | 16232132 | 16232631 | 500 | + | 189 | 158 | 31 | 83.6 |
| gene-LOC105282379 | NC_039512.1 | 8812640 | 8813139 | 500 | + | 36 | 30 | 6 | 83.33 |
| gene-LOC105281345 | NC_039508.1 | 2649490 | 2649989 | 500 | - | 45 | 37 | 8 | 82.22 |
| gene-LOC105274533 | NC_039516.1 | 1349957 | 1350456 | 500 | - | 73 | 59 | 14 | 80.82 |
| gene-LOC105284331 | NC_039511.1 | 5357461 | 5357960 | 500 | + | 254 | 203 | 51 | 79.92 |
| gene-LOC113561934 | NC_039506.1 | 15618494 | 15618993 | 500 | + | 1114 | 887 | 227 | 79.62 |
| gene-LOC105282078 | NC_039512.1 | 6125187 | 6125686 | 500 | + | 68 | 54 | 14 | 79.41 |
| gene-LOC105279930 | NC_039512.1 | 11167002 | 11167501 | 500 | + | 45 | 35 | 10 | 77.78 |
| gene-LOC105282822 | NC_039512.1 | 16435014 | 16435513 | 500 | + | 418 | 323 | 95 | 77.27 |
| gene-LOC105287837 | NC_039513.1 | 7295612 | 7296111 | 500 | - | 78 | 60 | 18 | 76.92 |
| gene-LOC105287833 | NC_039513.1 | 7297618 | 7298117 | 500 | + | 371 | 284 | 87 | 76.55 |
| gene-LOC113562533 | NC_039514.1 | 1298855 | 1299354 | 500 | + | 409 | 308 | 101 | 75.31 |
| gene-LOC105286944 | NC_039506.1 | 13094778 | 13095277 | 500 | + | 795 | 592 | 203 | 74.47 |

...
... only first 25 lines shown ...

Directory: RNK File: ranked-promoters-byPrcntM-Ob_LI_BCphaseB1.txt

| region_ID | seqnames | start | end | width | strand | coverage | numCs | numTs | prcntM |
|---|---|---|---|---|---|---|---|---|---|
| gene-LOC105285355 | NC_039509.1 | 8035614 | 8036113 | 500 | - | 54 | 51 | 3 | 94.44 |
| gene-LOC105276139 | NC_039514.1 | 452530 | 453029 | 500 | - | 65 | 60 | 5 | 92.31 |
| gene-LOC105276685 | NC_039511.1 | 14299458 | 14299957 | 500 | - | 911 | 822 | 89 | 90.23 |
| gene-LOC105282078 | NC_039512.1 | 6125187 | 6125686 | 500 | + | 107 | 96 | 11 | 89.72 |
| gene-LOC105285429 | NC_039517.1 | 10183111 | 10183610 | 500 | + | 386 | 344 | 42 | 89.12 |
| gene-LOC113561391 | NC_039507.1 | 999503 | 1000002 | 500 | + | 174 | 155 | 19 | 89.08 |
| gene-LOC113562329 | NC_039506.1 | 5740607 | 5741106 | 500 | + | 41 | 36 | 5 | 87.8 |
| gene-LOC105277073 | NC_039508.1 | 12946948 | 12947447 | 500 | + | 39 | 34 | 5 | 87.18 |
| gene-LOC105279930 | NC_039512.1 | 11167002 | 11167501 | 500 | + | 128 | 108 | 20 | 84.38 |
| gene-LOC113561532 | NC_039507.1 | 7277571 | 7278070 | 500 | + | 1041 | 869 | 172 | 83.48 |
| gene-LOC105276002 | NC_039511.1 | 12550061 | 12550560 | 500 | + | 90 | 75 | 15 | 83.33 |
| gene-LOC113561934 | NC_039506.1 | 15618494 | 15618993 | 500 | + | 1050 | 842 | 208 | 80.19 |
| gene-LOC105280917 | NC_039506.1 | 11578183 | 11578682 | 500 | - | 118 | 94 | 24 | 79.66 |
| gene-LOC105281040 | NC_039510.1 | 12238683 | 12239182 | 500 | + | 168 | 132 | 36 | 78.57 |
| gene-LOC105281689 | NC_039507.1 | 5097604 | 5098103 | 500 | - | 223 | 175 | 48 | 78.48 |
| gene-LOC105281627 | NC_039514.1 | 11099400 | 11099899 | 500 | + | 342 | 267 | 75 | 78.07 |
| gene-LOC105282801 | NC_039512.1 | 16232132 | 16232631 | 500 | + | 239 | 184 | 55 | 76.99 |
| gene-LOC105287833 | NC_039513.1 | 7297618 | 7298117 | 500 | + | 325 | 250 | 75 | 76.92 |
| gene-LOC105276408 | NC_039511.1 | 2016594 | 2017093 | 500 | + | 33 | 25 | 8 | 75.76 |
| gene-LOC105282822 | NC_039512.1 | 16435014 | 16435513 | 500 | + | 540 | 407 | 133 | 75.37 |
| gene-LOC105281671 | NC_039507.1 | 5181504 | 5182003 | 500 | - | 644 | 475 | 169 | 73.76 |
| gene-LOC113562533 | NC_039514.1 | 1298855 | 1299354 | 500 | + | 572 | 410 | 162 | 71.68 |
| gene-LOC109610674 | NC_039516.1 | 731871 | 732370 | 500 | + | 742 | 529 | 213 | 71.29 |
| gene-LOC105282379 | NC_039512.1 | 8812640 | 8813139 | 500 | + | 72 | 51 | 21 | 70.83 |

...
... only first 25 lines shown ...

Directory: RNK File: ranked-promoters-byPrcntM-Ob_LI_BCphaseB2.txt

| region_ID | seqnames | start | end | width | strand | coverage | numCs | numTs | prcntM |
|---|---|---|---|---|---|---|---|---|---|
| gene-LOC113562329 | NC_039506.1 | 5740607 | 5741106 | 500 | + | 16 | 16 | 0 | 100 |
| gene-LOC105276685 | NC_039511.1 | 14299458 | 14299957 | 500 | - | 833 | 760 | 73 | 91.24 |
| gene-LOC105276139 | NC_039514.1 | 452530 | 453029 | 500 | - | 66 | 60 | 6 | 90.91 |
| gene-LOC105285429 | NC_039517.1 | 10183111 | 10183610 | 500 | + | 229 | 208 | 21 | 90.83 |
| gene-LOC105285355 | NC_039509.1 | 8035614 | 8036113 | 500 | - | 45 | 40 | 5 | 88.89 |
| gene-LOC105276002 | NC_039511.1 | 12550061 | 12550560 | 500 | + | 68 | 60 | 8 | 88.24 |
| gene-LOC113561391 | NC_039507.1 | 999503 | 1000002 | 500 | + | 99 | 87 | 12 | 87.88 |
| gene-LOC105286418 | NC_039514.1 | 3321606 | 3322105 | 500 | - | 33 | 29 | 4 | 87.88 |
| gene-LOC105280917 | NC_039506.1 | 11578183 | 11578682 | 500 | - | 81 | 70 | 11 | 86.42 |
| gene-LOC113561532 | NC_039507.1 | 7277571 | 7278070 | 500 | + | 1155 | 990 | 165 | 85.71 |
| gene-LOC113563070 | NC_039517.1 | 3179512 | 3180011 | 500 | - | 7 | 6 | 1 | 85.71 |
| gene-LOC105281689 | NC_039507.1 | 5097604 | 5098103 | 500 | - | 179 | 152 | 27 | 84.92 |
| gene-LOC105282078 | NC_039512.1 | 6125187 | 6125686 | 500 | + | 66 | 56 | 10 | 84.85 |
| gene-LOC105282801 | NC_039512.1 | 16232132 | 16232631 | 500 | + | 188 | 153 | 35 | 81.38 |
| gene-LOC113561934 | NC_039506.1 | 15618494 | 15618993 | 500 | + | 1048 | 835 | 213 | 79.68 |
| gene-LOC105276074 | NC_039514.1 | 15118229 | 15118728 | 500 | - | 33 | 26 | 7 | 78.79 |
| gene-LOC105288107 | NC_039515.1 | 1102618 | 1103117 | 500 | + | 800 | 625 | 175 | 78.12 |
| gene-LOC105277073 | NC_039508.1 | 12946948 | 12947447 | 500 | + | 18 | 14 | 4 | 77.78 |
| gene-LOC113562533 | NC_039514.1 | 1298855 | 1299354 | 500 | + | 390 | 297 | 93 | 76.15 |
| gene-LOC105284331 | NC_039511.1 | 5357461 | 5357960 | 500 | + | 336 | 254 | 82 | 75.6 |
| gene-LOC105281627 | NC_039514.1 | 11099400 | 11099899 | 500 | + | 274 | 207 | 67 | 75.55 |
| gene-LOC105276585 | NC_039517.1 | 4571850 | 4572349 | 500 | - | 682 | 509 | 173 | 74.63 |
| gene-LOC105279930 | NC_039512.1 | 11167002 | 11167501 | 500 | + | 76 | 56 | 20 | 73.68 |
| gene-LOC105282822 | NC_039512.1 | 16435014 | 16435513 | 500 | + | 432 | 318 | 114 | 73.61 |

...
... only first 25 lines shown ...

Directory: RNK File: ranked-promoters-byPrcntM-Ob_LI_RphaseA1.txt

| region_ID | seqnames | start | end | width | strand | coverage | numCs | numTs | prcntM |
|---|---|---|---|---|---|---|---|---|---|
| gene-LOC105278219 | NC_039509.1 | 14123230 | 14123729 | 500 | + | 5 | 5 | 0 | 100 |
| gene-LOC105276139 | NC_039514.1 | 452530 | 453029 | 500 | - | 75 | 72 | 3 | 96 |
| gene-LOC105285355 | NC_039509.1 | 8035614 | 8036113 | 500 | - | 43 | 41 | 2 | 95.35 |
| gene-LOC105276685 | NC_039511.1 | 14299458 | 14299957 | 500 | - | 877 | 796 | 81 | 90.76 |
| gene-LOC105285429 | NC_039517.1 | 10183111 | 10183610 | 500 | + | 295 | 265 | 30 | 89.83 |
| gene-LOC105282078 | NC_039512.1 | 6125187 | 6125686 | 500 | + | 76 | 68 | 8 | 89.47 |
| gene-LOC113561391 | NC_039507.1 | 999503 | 1000002 | 500 | + | 127 | 113 | 14 | 88.98 |
| gene-LOC105280917 | NC_039506.1 | 11578183 | 11578682 | 500 | - | 106 | 91 | 15 | 85.85 |
| gene-LOC113562329 | NC_039506.1 | 5740607 | 5741106 | 500 | + | 20 | 17 | 3 | 85 |
| gene-LOC113561934 | NC_039506.1 | 15618494 | 15618993 | 500 | + | 683 | 570 | 113 | 83.46 |
| gene-LOC105276074 | NC_039514.1 | 15118229 | 15118728 | 500 | - | 36 | 30 | 6 | 83.33 |
| gene-LOC105277073 | NC_039508.1 | 12946948 | 12947447 | 500 | + | 29 | 24 | 5 | 82.76 |
| gene-LOC113561532 | NC_039507.1 | 7277571 | 7278070 | 500 | + | 907 | 746 | 161 | 82.25 |
| gene-LOC105276002 | NC_039511.1 | 12550061 | 12550560 | 500 | + | 93 | 75 | 18 | 80.65 |
| gene-LOC105279917 | NC_039512.1 | 13122631 | 13123130 | 500 | - | 188 | 151 | 37 | 80.32 |
| gene-LOC105281689 | NC_039507.1 | 5097604 | 5098103 | 500 | - | 191 | 151 | 40 | 79.06 |
| gene-LOC105282822 | NC_039512.1 | 16435014 | 16435513 | 500 | + | 475 | 369 | 106 | 77.68 |
| gene-LOC105276585 | NC_039517.1 | 4571850 | 4572349 | 500 | - | 640 | 497 | 143 | 77.66 |
| gene-LOC105282801 | NC_039512.1 | 16232132 | 16232631 | 500 | + | 204 | 158 | 46 | 77.45 |
| gene-LOC105279930 | NC_039512.1 | 11167002 | 11167501 | 500 | + | 113 | 87 | 26 | 76.99 |
| gene-LOC113562533 | NC_039514.1 | 1298855 | 1299354 | 500 | + | 538 | 410 | 128 | 76.21 |
| gene-LOC105286944 | NC_039506.1 | 13094778 | 13095277 | 500 | + | 590 | 447 | 143 | 75.76 |
| gene-LOC105281040 | NC_039510.1 | 12238683 | 12239182 | 500 | + | 122 | 92 | 30 | 75.41 |
| gene-LOC105281345 | NC_039508.1 | 2649490 | 2649989 | 500 | - | 110 | 81 | 29 | 73.64 |

...
... only first 25 lines shown ...

Directory: RNK File: ranked-promoters-byPrcntM-Ob_LI_RphaseA2.txt

| region_ID | seqnames | start | end | width | strand | coverage | numCs | numTs | prcntM |
|---|---|---|---|---|---|---|---|---|---|
| gene-LOC105285355 | NC_039509.1 | 8035614 | 8036113 | 500 | - | 66 | 65 | 1 | 98.48 |
| gene-LOC105276139 | NC_039514.1 | 452530 | 453029 | 500 | - | 61 | 57 | 4 | 93.44 |
| gene-LOC105276685 | NC_039511.1 | 14299458 | 14299957 | 500 | - | 1485 | 1385 | 100 | 93.27 |
| gene-LOC113561391 | NC_039507.1 | 999503 | 1000002 | 500 | + | 118 | 109 | 9 | 92.37 |
| gene-LOC113563070 | NC_039517.1 | 3179512 | 3180011 | 500 | - | 12 | 11 | 1 | 91.67 |
| gene-LOC113562329 | NC_039506.1 | 5740607 | 5741106 | 500 | + | 40 | 36 | 4 | 90 |
| gene-LOC105280917 | NC_039506.1 | 11578183 | 11578682 | 500 | - | 79 | 71 | 8 | 89.87 |
| gene-LOC105285429 | NC_039517.1 | 10183111 | 10183610 | 500 | + | 416 | 372 | 44 | 89.42 |
| gene-LOC105282379 | NC_039512.1 | 8812640 | 8813139 | 500 | + | 58 | 51 | 7 | 87.93 |
| gene-LOC105279917 | NC_039512.1 | 13122631 | 13123130 | 500 | - | 252 | 221 | 31 | 87.7 |
| gene-LOC105276074 | NC_039514.1 | 15118229 | 15118728 | 500 | - | 21 | 18 | 3 | 85.71 |
| gene-LOC113561532 | NC_039507.1 | 7277571 | 7278070 | 500 | + | 1604 | 1361 | 243 | 84.85 |
| gene-LOC105276002 | NC_039511.1 | 12550061 | 12550560 | 500 | + | 126 | 106 | 20 | 84.13 |
| gene-LOC105282801 | NC_039512.1 | 16232132 | 16232631 | 500 | + | 297 | 248 | 49 | 83.5 |
| gene-LOC105277073 | NC_039508.1 | 12946948 | 12947447 | 500 | + | 42 | 35 | 7 | 83.33 |
| gene-LOC105279451 | NC_039514.1 | 9158109 | 9158608 | 500 | + | 6 | 5 | 1 | 83.33 |
| gene-LOC105276408 | NC_039511.1 | 2016594 | 2017093 | 500 | + | 29 | 24 | 5 | 82.76 |
| gene-LOC113561934 | NC_039506.1 | 15618494 | 15618993 | 500 | + | 1497 | 1238 | 259 | 82.7 |
| gene-LOC105282078 | NC_039512.1 | 6125187 | 6125686 | 500 | + | 120 | 97 | 23 | 80.83 |
| gene-LOC105275802 | NC_039516.1 | 7330402 | 7330901 | 500 | - | 54 | 43 | 11 | 79.63 |
| gene-LOC105281689 | NC_039507.1 | 5097604 | 5098103 | 500 | - | 303 | 240 | 63 | 79.21 |
| gene-LOC105282822 | NC_039512.1 | 16435014 | 16435513 | 500 | + | 728 | 576 | 152 | 79.12 |
| gene-LOC105287833 | NC_039513.1 | 7297618 | 7298117 | 500 | + | 489 | 385 | 104 | 78.73 |
| gene-LOC113562533 | NC_039514.1 | 1298855 | 1299354 | 500 | + | 849 | 668 | 181 | 78.68 |

...

... only first 25 lines shown ...

Directory: RNK File: ranked-promoters-byPrcntM-Ob_LI_RphaseB1.txt

| region_ID | seqnames | start | end | width | strand | coverage | numCs | numTs | prcntM |
|---|---|---|---|---|---|---|---|---|---|
| gene-LOC113563070 | NC_039517.1 | 3179512 | 3180011 | 500 | - | 5 | 5 | 0 | 100 |
| gene-LOC105285355 | NC_039509.1 | 8035614 | 8036113 | 500 | - | 45 | 43 | 2 | 95.56 |
| gene-LOC105280917 | NC_039506.1 | 11578183 | 11578682 | 500 | - | 98 | 91 | 7 | 92.86 |
| gene-LOC105276685 | NC_039511.1 | 14299458 | 14299957 | 500 | - | 898 | 827 | 71 | 92.09 |
| gene-LOC105277264 | NC_039518.1 | 2605800 | 2606299 | 500 | - | 10 | 9 | 1 | 90 |
| gene-LOC105276139 | NC_039514.1 | 452530 | 453029 | 500 | - | 51 | 45 | 6 | 88.24 |
| gene-LOC105285429 | NC_039517.1 | 10183111 | 10183610 | 500 | + | 379 | 334 | 45 | 88.13 |
| gene-LOC105282078 | NC_039512.1 | 6125187 | 6125686 | 500 | + | 92 | 81 | 11 | 88.04 |
| gene-LOC113561391 | NC_039507.1 | 999503 | 1000002 | 500 | + | 139 | 120 | 19 | 86.33 |
| gene-LOC105281689 | NC_039507.1 | 5097604 | 5098103 | 500 | - | 229 | 194 | 35 | 84.72 |
| gene-LOC113561532 | NC_039507.1 | 7277571 | 7278070 | 500 | + | 999 | 840 | 159 | 84.08 |
| gene-LOC105276002 | NC_039511.1 | 12550061 | 12550560 | 500 | + | 91 | 74 | 17 | 81.32 |
| gene-LOC105275802 | NC_039516.1 | 7330402 | 7330901 | 500 | - | 50 | 40 | 10 | 80 |
| gene-LOC105276074 | NC_039514.1 | 15118229 | 15118728 | 500 | - | 39 | 31 | 8 | 79.49 |
| gene-LOC105279930 | NC_039512.1 | 11167002 | 11167501 | 500 | + | 100 | 79 | 21 | 79 |
| gene-LOC105283432 | NC_039506.1 | 23395368 | 23395867 | 500 | - | 28 | 22 | 6 | 78.57 |
| gene-LOC113562533 | NC_039514.1 | 1298855 | 1299354 | 500 | + | 529 | 410 | 119 | 77.5 |
| gene-LOC105282801 | NC_039512.1 | 16232132 | 16232631 | 500 | + | 282 | 217 | 65 | 76.95 |
| gene-LOC105276585 | NC_039517.1 | 4571850 | 4572349 | 500 | - | 655 | 502 | 153 | 76.64 |
| gene-LOC105288107 | NC_039515.1 | 1102618 | 1103117 | 500 | + | 802 | 614 | 188 | 76.56 |
| gene-LOC105276408 | NC_039511.1 | 2016594 | 2017093 | 500 | + | 21 | 16 | 5 | 76.19 |
| gene-LOC113561934 | NC_039506.1 | 15618494 | 15618993 | 500 | + | 874 | 665 | 209 | 76.09 |
| gene-LOC105284215 | NC_039517.1 | 7397076 | 7397575 | 500 | + | 242 | 182 | 60 | 75.21 |
| gene-LOC105281627 | NC_039514.1 | 11099400 | 11099899 | 500 | + | 351 | 260 | 91 | 74.07 |

...

... only first 25 lines shown ...

Directory: RNK File: ranked-promoters-byPrcntM-Ob_LI_RphaseB2.txt

| region_ID | seqnames | start | end | width | strand | coverage | numCs | numTs | prcntM |
|---|---|---|---|---|---|---|---|---|---|
| gene-LOC105285429 | NC_039517.1 | 10183111 | 10183610 | 500 | + | 260 | 243 | 17 | 93.46 |
| gene-LOC105276139 | NC_039514.1 | 452530 | 453029 | 500 | - | 44 | 41 | 3 | 93.18 |
| gene-LOC105285355 | NC_039509.1 | 8035614 | 8036113 | 500 | - | 42 | 39 | 3 | 92.86 |
| gene-LOC105276002 | NC_039511.1 | 12550061 | 12550560 | 500 | + | 70 | 65 | 5 | 92.86 |
| gene-LOC105276685 | NC_039511.1 | 14299458 | 14299957 | 500 | - | 804 | 733 | 71 | 91.17 |
| gene-LOC113561391 | NC_039507.1 | 999503 | 1000002 | 500 | + | 109 | 99 | 10 | 90.83 |
| gene-LOC113562329 | NC_039506.1 | 5740607 | 5741106 | 500 | + | 30 | 27 | 3 | 90 |
| gene-LOC113561532 | NC_039507.1 | 7277571 | 7278070 | 500 | + | 1137 | 966 | 171 | 84.96 |
| gene-LOC105282801 | NC_039512.1 | 16232132 | 16232631 | 500 | + | 199 | 169 | 30 | 84.92 |
| gene-LOC105282078 | NC_039512.1 | 6125187 | 6125686 | 500 | + | 49 | 40 | 9 | 81.63 |
| gene-LOC105281627 | NC_039514.1 | 11099400 | 11099899 | 500 | + | 274 | 219 | 55 | 79.93 |
| gene-LOC105281689 | NC_039507.1 | 5097604 | 5098103 | 500 | - | 178 | 141 | 37 | 79.21 |
| gene-LOC105284331 | NC_039511.1 | 5357461 | 5357960 | 500 | + | 368 | 291 | 77 | 79.08 |
| gene-LOC105281345 | NC_039508.1 | 2649490 | 2649989 | 500 | - | 85 | 67 | 18 | 78.82 |
| gene-LOC105279917 | NC_039512.1 | 13122631 | 13123130 | 500 | - | 212 | 166 | 46 | 78.3 |
| gene-LOC113561934 | NC_039506.1 | 15618494 | 15618993 | 500 | + | 828 | 636 | 192 | 76.81 |
| gene-LOC105279502 | NC_039514.1 | 15553326 | 15553825 | 500 | - | 267 | 204 | 63 | 76.4 |
| gene-LOC105280917 | NC_039506.1 | 11578183 | 11578682 | 500 | - | 75 | 57 | 18 | 76 |
| gene-LOC105274987 | NC_039506.1 | 17638152 | 17638651 | 500 | - | 143 | 108 | 35 | 75.52 |
| gene-LOC105282822 | NC_039512.1 | 16435014 | 16435513 | 500 | + | 474 | 354 | 120 | 74.68 |
| gene-LOC105287833 | NC_039513.1 | 7297618 | 7298117 | 500 | + | 298 | 222 | 76 | 74.5 |
| gene-LOC105288107 | NC_039515.1 | 1102618 | 1103117 | 500 | + | 869 | 639 | 230 | 73.53 |
| gene-LOC105286944 | NC_039506.1 | 13094778 | 13095277 | 500 | + | 866 | 636 | 230 | 73.44 |
| gene-Trnav-aac-5 | NC_039517.1 | 7367422 | 7367921 | 500 | + | 222 | 163 | 59 | 73.42 |

...

... only first 25 lines shown ...

Directory: RNK File: ranked-promoters-bySiteDensity-Ob_LI_BCphaseA1.txt

| region_ID | rwidth | nbrsites | nbrper10kb | pmpersite | pmpernucl | pglink | seqnames | start | end | strand | coverage | numCs | numTs | prcntM |
|---|---|---|---|---|---|---|---|---|---|---|---|---|---|---|
| gene-LOC105288107 | 500 | 40 | 800 | 80.61 | 6.45 | https://www.ncbi.nlm.nih.gov/gene/?term=LOC105288107 | NC_039515.1 | 1102618 | 1103117 | + | 841 | 633 | 208 | 75.27 |
| gene-LOC113561934 | 500 | 37 | 740 | 84.19 | 6.23 | https://www.ncbi.nlm.nih.gov/gene/?term=LOC113561934 | NC_039506.1 | 15618494 | 15618993 | + | 803 | 627 | 176 | 78.08 |
| gene-LOC105282404 | 500 | 36 | 720 | 60.78 | 4.38 | https://www.ncbi.nlm.nih.gov/gene/?term=LOC105282404 | NC_039512.1 | 8476210 | 8476709 | + | 1281 | 571 | 710 | 44.57 |
| gene-LOC105277808 | 500 | 34 | 680 | 59.31 | 4.03 | https://www.ncbi.nlm.nih.gov/gene/?term=LOC105277808 | NC_039516.1 | 13518519 | 13519018 | - | 1893 | 565 | 1328 | 29.85 |
| gene-LOC105278673 | 500 | 33 | 660 | 75.64 | 4.99 | https://www.ncbi.nlm.nih.gov/gene/?term=LOC105278673 | NC_039516.1 | 11703885 | 11704384 | - | 1073 | 718 | 355 | 66.92 |
| gene-LOC105280727 | 500 | 33 | 660 | 67.1 | 4.43 | https://www.ncbi.nlm.nih.gov/gene/?term=LOC105280727 | NC_039513.1 | 11520171 | 11520670 | + | 1306 | 578 | 728 | 44.26 |
| gene-LOC105280861 | 500 | 32 | 640 | 73.61 | 4.71 | https://www.ncbi.nlm.nih.gov/gene/?term=LOC105280861 | NC_039506.1 | 15371187 | 15371686 | + | 1163 | 648 | 515 | 55.72 |
| gene-LOC105277304 | 500 | 31 | 620 | 70.26 | 4.36 | https://www.ncbi.nlm.nih.gov/gene/?term=LOC105277304 | NC_039516.1 | 9356205 | 9356704 | - | 1145 | 611 | 534 | 53.36 |
| gene-LOC105283833 | 500 | 31 | 620 | 66.65 | 4.13 | https://www.ncbi.nlm.nih.gov/gene/?term=LOC105283833 | NC_039509.1 | 2003067 | 2003566 | - | 1268 | 523 | 745 | 41.25 |
| gene-LOC105286944 | 500 | 31 | 620 | 80.81 | 5.01 | https://www.ncbi.nlm.nih.gov/gene/?term=LOC105286944 | NC_039506.1 | 13094778 | 13095277 | + | 681 | 460 | 221 | 67.55 |
| gene-LOC105279250 | 500 | 30 | 600 | 75.36 | 4.52 | https://www.ncbi.nlm.nih.gov/gene/?term=LOC105279250 | NC_039512.1 | 7901112 | 7901611 | - | 865 | 537 | 328 | 62.08 |
| gene-LOC105283749 | 500 | 29 | 580 | 84.37 | 4.89 | https://www.ncbi.nlm.nih.gov/gene/?term=LOC105283749 | NC_039515.1 | 7648626 | 7649125 | - | 1062 | 600 | 462 | 56.5 |
| gene-LOC105286234 | 500 | 28 | 560 | 57.23 | 3.2 | https://www.ncbi.nlm.nih.gov/gene/?term=LOC105286234 | NC_039518.1 | 8582509 | 8583008 | + | 1568 | 534 | 1034 | 34.06 |
| gene-LOC113561532 | 500 | 28 | 560 | 91.49 | 5.12 | https://www.ncbi.nlm.nih.gov/gene/?term=LOC113561532 | NC_039507.1 | 7277571 | 7278070 | + | 815 | 708 | 107 | 86.87 |
| gene-LOC105278632 | 500 | 27 | 540 | 88.01 | 4.75 | https://www.ncbi.nlm.nih.gov/gene/?term=LOC105278632 | NC_039516.1 | 11911628 | 11912127 | + | 602 | 313 | 289 | 51.99 |
| gene-LOC105286681 | 500 | 27 | 540 | 56.79 | 3.07 | https://www.ncbi.nlm.nih.gov/gene/?term=LOC105286681 | NC_039512.1 | 14397556 | 14398055 | + | 1014 | 450 | 564 | 44.38 |
| gene-LOC113562689 | 500 | 27 | 540 | 70.32 | 3.8 | https://www.ncbi.nlm.nih.gov/gene/?term=LOC113562689 | NC_039515.1 | 5223138 | 5223637 | - | 1066 | 450 | 616 | 42.21 |
| gene-LOC105276685 | 500 | 26 | 520 | 89.6 | 4.66 | https://www.ncbi.nlm.nih.gov/gene/?term=LOC105276685 | NC_039511.1 | 14299458 | 14299957 | - | 752 | 674 | 78 | 89.63 |
| gene-LOC105283965 | 500 | 26 | 520 | 58.45 | 3.04 | https://www.ncbi.nlm.nih.gov/gene/?term=LOC105283965 | NC_039506.1 | 15529793 | 15530292 | - | 801 | 363 | 438 | 45.32 |
| gene-LOC109611487 | 500 | 26 | 520 | 53.71 | 2.79 | https://www.ncbi.nlm.nih.gov/gene/?term=LOC109611487 | NC_039506.1 | 13096374 | 13096873 | + | 1569 | 409 | 1160 | 26.07 |
| gene-LOC105278635 | 500 | 25 | 500 | 87.05 | 4.35 | https://www.ncbi.nlm.nih.gov/gene/?term=LOC105278635 | NC_039516.1 | 11911609 | 11912108 | - | 584 | 295 | 289 | 50.51 |
| gene-LOC105286266 | 500 | 24 | 480 | 66.34 | 3.18 | https://www.ncbi.nlm.nih.gov/gene/?term=LOC105286266 | NC_039518.1 | 8735110 | 8735609 | - | 774 | 298 | 476 | 38.5 |
| gene-LOC113562533 | 500 | 24 | 480 | 79.52 | 3.82 | https://www.ncbi.nlm.nih.gov/gene/?term=LOC113562533 | NC_039514.1 | 1298855 | 1299354 | + | 416 | 298 | 118 | 71.63 |
| gene-LOC105276585 | 500 | 23 | 460 | 80.01 | 3.68 | https://www.ncbi.nlm.nih.gov/gene/?term=LOC105276585 | NC_039517.1 | 4571850 | 4572349 | - | 496 | 357 | 139 | 71.98 |

...

... only first 25 lines shown ...

Directory: RNK File: ranked-promoters-bySiteDensity-Ob_LI_BCphaseA2.txt

| region_ID | rwidth | nbrsites | nbrper10kb | pmpersite | pmpernucl | pglink | seqnames | start | end | strand | coverage | numCs | numTs | prcntM |
|---|---|---|---|---|---|---|---|---|---|---|---|---|---|---|
| gene-LOC105277808 | 500 | 39 | 780 | 57.54 | 4.49 | https://www.ncbi.nlm.nih.gov/gene/?term=LOC105277808 | NC_039516.1 | 13518519 | 13519018 | - | 3102 | 984 | 2118 | 31.72 |
| gene-LOC105286234 | 500 | 39 | 780 | 53.74 | 4.19 | https://www.ncbi.nlm.nih.gov/gene/?term=LOC105286234 | NC_039518.1 | 8582509 | 8583008 | + | 2460 | 1007 | 1453 | 40.93 |
| gene-LOC105280727 | 500 | 38 | 760 | 63.63 | 4.84 | https://www.ncbi.nlm.nih.gov/gene/?term=LOC105280727 | NC_039513.1 | 11520171 | 11520670 | + | 1939 | 908 | 1031 | 46.83 |
| gene-LOC105288107 | 500 | 36 | 720 | 78.61 | 5.66 | https://www.ncbi.nlm.nih.gov/gene/?term=LOC105288107 | NC_039515.1 | 1102618 | 1103117 | + | 874 | 604 | 270 | 69.11 |
| gene-LOC113561934 | 500 | 35 | 700 | 80.98 | 5.67 | https://www.ncbi.nlm.nih.gov/gene/?term=LOC113561934 | NC_039506.1 | 15618494 | 15618993 | + | 1114 | 887 | 227 | 79.62 |
| gene-LOC105282404 | 500 | 34 | 680 | 59.28 | 4.03 | https://www.ncbi.nlm.nih.gov/gene/?term=LOC105282404 | NC_039512.1 | 8476210 | 8476709 | + | 1614 | 705 | 909 | 43.68 |
| gene-LOC105279250 | 500 | 33 | 660 | 71.55 | 4.72 | https://www.ncbi.nlm.nih.gov/gene/?term=LOC105279250 | NC_039512.1 | 7901112 | 7901611 | - | 1132 | 778 | 354 | 68.73 |
| gene-LOC105280861 | 500 | 32 | 640 | 74.81 | 4.79 | https://www.ncbi.nlm.nih.gov/gene/?term=LOC105280861 | NC_039506.1 | 15371187 | 15371686 | + | 1208 | 684 | 524 | 56.62 |
| gene-LOC105277304 | 500 | 31 | 620 | 72.61 | 4.5 | https://www.ncbi.nlm.nih.gov/gene/?term=LOC105277304 | NC_039516.1 | 9356205 | 9356704 | - | 1417 | 670 | 747 | 47.28 |
| gene-LOC113561532 | 500 | 31 | 620 | 88.84 | 5.51 | https://www.ncbi.nlm.nih.gov/gene/?term=LOC113561532 | NC_039507.1 | 7277571 | 7278070 | + | 1134 | 985 | 149 | 86.86 |
| gene-LOC105278673 | 500 | 30 | 600 | 78.37 | 4.7 | https://www.ncbi.nlm.nih.gov/gene/?term=LOC105278673 | NC_039516.1 | 11703885 | 11704384 | - | 1113 | 800 | 313 | 71.88 |
| gene-LOC105283749 | 500 | 29 | 580 | 85.6 | 4.96 | https://www.ncbi.nlm.nih.gov/gene/?term=LOC105283749 | NC_039515.1 | 7648626 | 7649125 | - | 1392 | 742 | 650 | 53.3 |
| gene-LOC109611487 | 500 | 29 | 580 | 54.43 | 3.16 | https://www.ncbi.nlm.nih.gov/gene/?term=LOC109611487 | NC_039506.1 | 13096374 | 13096873 | + | 2349 | 637 | 1712 | 27.12 |
| gene-LOC105283833 | 500 | 28 | 560 | 68.07 | 3.81 | https://www.ncbi.nlm.nih.gov/gene/?term=LOC105283833 | NC_039509.1 | 2003067 | 2003566 | - | 1208 | 488 | 720 | 40.4 |
| gene-LOC105284094 | 500 | 28 | 560 | 39.48 | 2.21 | https://www.ncbi.nlm.nih.gov/gene/?term=LOC105284094 | NC_039511.1 | 3031639 | 3032138 | + | 2602 | 532 | 2070 | 20.45 |
| gene-LOC113562689 | 500 | 28 | 560 | 69.97 | 3.92 | https://www.ncbi.nlm.nih.gov/gene/?term=LOC113562689 | NC_039515.1 | 5223138 | 5223637 | - | 1214 | 494 | 720 | 40.69 |
| gene-LOC105286681 | 500 | 27 | 540 | 57.61 | 3.11 | https://www.ncbi.nlm.nih.gov/gene/?term=LOC105286681 | NC_039512.1 | 14397556 | 14398055 | + | 1120 | 482 | 638 | 43.04 |
| gene-LOC105276585 | 500 | 26 | 520 | 86.03 | 4.47 | https://www.ncbi.nlm.nih.gov/gene/?term=LOC105276585 | NC_039517.1 | 4571850 | 4572349 | - | 705 | 506 | 199 | 71.77 |
| gene-LOC105276685 | 500 | 26 | 520 | 91.92 | 4.78 | https://www.ncbi.nlm.nih.gov/gene/?term=LOC105276685 | NC_039511.1 | 14299458 | 14299957 | - | 925 | 857 | 68 | 92.65 |
| gene-LOC105286944 | 500 | 26 | 520 | 87.54 | 4.55 | https://www.ncbi.nlm.nih.gov/gene/?term=LOC105286944 | NC_039506.1 | 13094778 | 13095277 | + | 795 | 592 | 203 | 74.47 |
| gene-LOC105277395 | 500 | 25 | 500 | 79.79 | 3.99 | https://www.ncbi.nlm.nih.gov/gene/?term=LOC105277395 | NC_039508.1 | 1340463 | 1340962 | - | 485 | 314 | 171 | 64.74 |
| gene-LOC105278632 | 500 | 25 | 500 | 76.43 | 3.82 | https://www.ncbi.nlm.nih.gov/gene/?term=LOC105278632 | NC_039516.1 | 11911628 | 11912127 | + | 807 | 439 | 368 | 54.4 |
| gene-LOC105283965 | 500 | 25 | 500 | 61.2 | 3.06 | https://www.ncbi.nlm.nih.gov/gene/?term=LOC105283965 | NC_039506.1 | 15529793 | 15530292 | - | 792 | 385 | 407 | 48.61 |
| gene-LOC105286479 | 500 | 25 | 500 | 61.42 | 3.07 | https://www.ncbi.nlm.nih.gov/gene/?term=LOC105286479 | NC_039510.1 | 15744783 | 15745282 | - | 2365 | 695 | 1670 | 29.39 |

...
... only first 25 lines shown ...

Directory: RNK File: ranked-promoters-bySiteDensity-Ob_LI_BCphaseB1.txt

| region_ID | rwidth | nbrsites | nbrper10kb | pmpersite | pmpernucl | pglink | seqnames | start | end | strand | coverage | numCs | numTs | prcntM |
|---|---|---|---|---|---|---|---|---|---|---|---|---|---|---|
| gene-LOC109611487 | 500 | 41 | 820 | 44.19 | 3.62 | https://www.ncbi.nlm.nih.gov/gene/?term=LOC109611487 | NC_039506.1 | 13096374 | 13096873 | + | 2819 | 719 | 2100 | 25.51 |
| gene-LOC105277808 | 500 | 39 | 780 | 58.66 | 4.58 | https://www.ncbi.nlm.nih.gov/gene/?term=LOC105277808 | NC_039516.1 | 13518519 | 13519018 | - | 2573 | 845 | 1728 | 32.84 |
| gene-LOC105286234 | 500 | 39 | 780 | 54.61 | 4.26 | https://www.ncbi.nlm.nih.gov/gene/?term=LOC105286234 | NC_039518.1 | 8582509 | 8583008 | + | 2503 | 926 | 1577 | 37 |
| gene-LOC105280727 | 500 | 38 | 760 | 62.3 | 4.74 | https://www.ncbi.nlm.nih.gov/gene/?term=LOC105280727 | NC_039513.1 | 11520171 | 11520670 | + | 1812 | 790 | 1022 | 43.6 |
| gene-LOC113561934 | 500 | 38 | 760 | 85.1 | 6.47 | https://www.ncbi.nlm.nih.gov/gene/?term=LOC113561934 | NC_039506.1 | 15618494 | 15618993 | + | 1050 | 842 | 208 | 80.19 |
| gene-LOC105282404 | 500 | 37 | 740 | 50.28 | 3.72 | https://www.ncbi.nlm.nih.gov/gene/?term=LOC105282404 | NC_039512.1 | 8476210 | 8476709 | + | 1744 | 656 | 1088 | 37.61 |
| gene-LOC105288107 | 500 | 37 | 740 | 78.54 | 5.81 | https://www.ncbi.nlm.nih.gov/gene/?term=LOC105288107 | NC_039515.1 | 1102618 | 1103117 | + | 990 | 696 | 294 | 70.3 |
| gene-LOC105277304 | 500 | 36 | 720 | 72.21 | 5.2 | https://www.ncbi.nlm.nih.gov/gene/?term=LOC105277304 | NC_039516.1 | 9356205 | 9356704 | - | 1387 | 671 | 716 | 48.38 |
| gene-LOC105280861 | 500 | 36 | 720 | 72.53 | 5.22 | https://www.ncbi.nlm.nih.gov/gene/?term=LOC105280861 | NC_039506.1 | 15371187 | 15371686 | + | 1357 | 725 | 632 | 53.43 |
| gene-LOC105278673 | 500 | 34 | 680 | 77.03 | 5.24 | https://www.ncbi.nlm.nih.gov/gene/?term=LOC105278673 | NC_039516.1 | 11703885 | 11704384 | - | 1132 | 754 | 378 | 66.61 |
| gene-LOC105279250 | 500 | 33 | 660 | 72.85 | 4.81 | https://www.ncbi.nlm.nih.gov/gene/?term=LOC105279250 | NC_039512.1 | 7901112 | 7901611 | - | 1189 | 802 | 387 | 67.45 |
| gene-LOC105280668 | 500 | 33 | 660 | 49.19 | 3.25 | https://www.ncbi.nlm.nih.gov/gene/?term=LOC105280668 | NC_039508.1 | 18012855 | 18013354 | - | 1508 | 525 | 983 | 34.81 |
| gene-LOC113561532 | 500 | 33 | 660 | 91.41 | 6.03 | https://www.ncbi.nlm.nih.gov/gene/?term=LOC113561532 | NC_039507.1 | 7277571 | 7278070 | + | 1041 | 869 | 172 | 83.48 |
| gene-LOC105274855 | 500 | 31 | 620 | 40.39 | 2.5 | https://www.ncbi.nlm.nih.gov/gene/?term=LOC105274855 | NC_039513.1 | 3348900 | 3349399 | + | 1883 | 440 | 1443 | 23.37 |
| gene-LOC113562689 | 500 | 31 | 620 | 67.8 | 4.2 | https://www.ncbi.nlm.nih.gov/gene/?term=LOC113562689 | NC_039515.1 | 5223138 | 5223637 | - | 1363 | 590 | 773 | 43.29 |
| gene-LOC105283749 | 500 | 29 | 580 | 84.41 | 4.9 | https://www.ncbi.nlm.nih.gov/gene/?term=LOC105283749 | NC_039515.1 | 7648626 | 7649125 | - | 1444 | 797 | 647 | 55.19 |
| gene-LOC105283760 | 500 | 29 | 580 | 69.71 | 4.04 | https://www.ncbi.nlm.nih.gov/gene/?term=LOC105283760 | NC_039514.1 | 3324673 | 3325172 | + | 698 | 450 | 248 | 64.47 |
| gene-LOC105286944 | 500 | 28 | 560 | 84.56 | 4.74 | https://www.ncbi.nlm.nih.gov/gene/?term=LOC105286944 | NC_039506.1 | 13094778 | 13095277 | + | 600 | 421 | 179 | 70.17 |
| gene-LOC105276585 | 500 | 27 | 540 | 77.05 | 4.16 | https://www.ncbi.nlm.nih.gov/gene/?term=LOC105276585 | NC_039517.1 | 4571850 | 4572349 | - | 757 | 507 | 250 | 66.97 |
| gene-LOC105277395 | 500 | 27 | 540 | 75.49 | 4.08 | https://www.ncbi.nlm.nih.gov/gene/?term=LOC105277395 | NC_039508.1 | 1340463 | 1340962 | - | 552 | 361 | 191 | 65.4 |
| gene-LOC105277648 | 500 | 27 | 540 | 78.42 | 4.23 | https://www.ncbi.nlm.nih.gov/gene/?term=LOC105277648 | NC_039512.1 | 2276615 | 2277114 | + | 648 | 441 | 207 | 68.06 |
| gene-LOC105283833 | 500 | 27 | 540 | 70.81 | 3.82 | https://www.ncbi.nlm.nih.gov/gene/?term=LOC105283833 | NC_039509.1 | 2003067 | 2003566 | - | 1397 | 554 | 843 | 39.66 |
| gene-LOC105276685 | 500 | 26 | 520 | 90.71 | 4.72 | https://www.ncbi.nlm.nih.gov/gene/?term=LOC105276685 | NC_039511.1 | 14299458 | 14299957 | - | 911 | 822 | 89 | 90.23 |
| gene-LOC105278632 | 500 | 26 | 520 | 80.95 | 4.21 | https://www.ncbi.nlm.nih.gov/gene/?term=LOC105278632 | NC_039516.1 | 11911628 | 11912127 | + | 857 | 457 | 400 | 53.33 |

...
... only first 25 lines shown ...

Directory: RNK File: ranked-promoters-bySiteDensity-Ob_LI_BCphaseB2.txt

| region_ID | rwidth | nbrsites | nbrper10kb | pmpersite | pmpernucl | pglink | seqnames | start | end | strand | coverage | numCs | numTs | prcntM |
|---|---|---|---|---|---|---|---|---|---|---|---|---|---|---|
| gene-LOC105280861 | 500 | 39 | 780 | 68.7 | 5.36 | https://www.ncbi.nlm.nih.gov/gene/?term=LOC105280861 | NC_039506.1 | 15371187 | 15371686 | + | 1424 | 780 | 644 | 54.78 |
| gene-LOC105288107 | 500 | 39 | 780 | 83.59 | 6.52 | https://www.ncbi.nlm.nih.gov/gene/?term=LOC105288107 | NC_039515.1 | 1102618 | 1103117 | + | 800 | 625 | 175 | 78.12 |
| gene-LOC113561934 | 500 | 37 | 740 | 84.31 | 6.24 | https://www.ncbi.nlm.nih.gov/gene/?term=LOC113561934 | NC_039506.1 | 15618494 | 15618993 | + | 1048 | 835 | 213 | 79.68 |
| gene-LOC105277808 | 500 | 36 | 720 | 55.62 | 4 | https://www.ncbi.nlm.nih.gov/gene/?term=LOC105277808 | NC_039516.1 | 13518519 | 13519018 | - | 3149 | 944 | 2205 | 29.98 |
| gene-LOC105278673 | 500 | 36 | 720 | 77.48 | 5.58 | https://www.ncbi.nlm.nih.gov/gene/?term=LOC105278673 | NC_039516.1 | 11703885 | 11704384 | - | 1050 | 731 | 319 | 69.62 |
| gene-LOC105286234 | 500 | 35 | 700 | 54.94 | 3.85 | https://www.ncbi.nlm.nih.gov/gene/?term=LOC105286234 | NC_039518.1 | 8582509 | 8583008 | + | 2019 | 734 | 1285 | 36.35 |
| gene-LOC105280727 | 500 | 34 | 680 | 63.01 | 4.28 | https://www.ncbi.nlm.nih.gov/gene/?term=LOC105280727 | NC_039513.1 | 11520171 | 11520670 | + | 1808 | 764 | 1044 | 42.26 |
| gene-LOC109611487 | 500 | 34 | 680 | 47.47 | 3.23 | https://www.ncbi.nlm.nih.gov/gene/?term=LOC109611487 | NC_039506.1 | 13096374 | 13096873 | + | 2330 | 601 | 1729 | 25.79 |
| gene-LOC105279250 | 500 | 33 | 660 | 75.29 | 4.97 | https://www.ncbi.nlm.nih.gov/gene/?term=LOC105279250 | NC_039512.1 | 7901112 | 7901611 | - | 1076 | 654 | 422 | 60.78 |
| gene-LOC105282404 | 500 | 33 | 660 | 55.41 | 3.66 | https://www.ncbi.nlm.nih.gov/gene/?term=LOC105282404 | NC_039512.1 | 8476210 | 8476709 | + | 1742 | 710 | 1032 | 40.76 |
| gene-LOC105286681 | 500 | 33 | 660 | 61.11 | 4.03 | https://www.ncbi.nlm.nih.gov/gene/?term=LOC105286681 | NC_039512.1 | 14397556 | 14398055 | + | 1100 | 498 | 602 | 45.27 |
| gene-LOC113561532 | 500 | 32 | 640 | 88.45 | 5.66 | https://www.ncbi.nlm.nih.gov/gene/?term=LOC113561532 | NC_039507.1 | 7277571 | 7278070 | + | 1155 | 990 | 165 | 85.71 |
| gene-LOC105276585 | 500 | 30 | 600 | 80.71 | 4.84 | https://www.ncbi.nlm.nih.gov/gene/?term=LOC105276585 | NC_039517.1 | 4571850 | 4572349 | - | 682 | 509 | 173 | 74.63 |
| gene-LOC105283749 | 500 | 30 | 600 | 81.82 | 4.91 | https://www.ncbi.nlm.nih.gov/gene/?term=LOC105283749 | NC_039515.1 | 7648626 | 7649125 | - | 1315 | 714 | 601 | 54.3 |
| gene-LOC105277304 | 500 | 28 | 560 | 75.3 | 4.22 | https://www.ncbi.nlm.nih.gov/gene/?term=LOC105277304 | NC_039516.1 | 9356205 | 9356704 | - | 1271 | 603 | 668 | 47.44 |
| gene-LOC113562689 | 500 | 28 | 560 | 67.43 | 3.78 | https://www.ncbi.nlm.nih.gov/gene/?term=LOC113562689 | NC_039515.1 | 5223138 | 5223637 | - | 1265 | 505 | 760 | 39.92 |
| gene-LOC105274855 | 500 | 27 | 540 | 41.26 | 2.23 | https://www.ncbi.nlm.nih.gov/gene/?term=LOC105274855 | NC_039513.1 | 3348900 | 3349399 | + | 1798 | 434 | 1364 | 24.14 |
| gene-LOC105276685 | 500 | 27 | 540 | 90.73 | 4.9 | https://www.ncbi.nlm.nih.gov/gene/?term=LOC105276685 | NC_039511.1 | 14299458 | 14299957 | - | 833 | 760 | 73 | 91.24 |
| gene-LOC105286944 | 500 | 27 | 540 | 83.6 | 4.51 | https://www.ncbi.nlm.nih.gov/gene/?term=LOC105286944 | NC_039506.1 | 13094778 | 13095277 | + | 771 | 543 | 228 | 70.43 |
| gene-LOC105278632 | 500 | 26 | 520 | 80.6 | 4.19 | https://www.ncbi.nlm.nih.gov/gene/?term=LOC105278632 | NC_039516.1 | 11911628 | 11912127 | + | 814 | 434 | 380 | 53.32 |
| gene-LOC105286266 | 500 | 26 | 520 | 61.51 | 3.2 | https://www.ncbi.nlm.nih.gov/gene/?term=LOC105286266 | NC_039518.1 | 8735110 | 8735609 | - | 979 | 395 | 584 | 40.35 |
| gene-LOC105277395 | 500 | 25 | 500 | 71.32 | 3.57 | https://www.ncbi.nlm.nih.gov/gene/?term=LOC105277395 | NC_039508.1 | 1340463 | 1340962 | - | 488 | 302 | 186 | 61.89 |
| gene-LOC105288013 | 500 | 25 | 500 | 59.55 | 2.98 | https://www.ncbi.nlm.nih.gov/gene/?term=LOC105288013 | NC_039512.1 | 130325 | 130824 | + | 659 | 277 | 382 | 42.03 |
| gene-LOC105276797 | 500 | 24 | 480 | 72.06 | 3.46 | https://www.ncbi.nlm.nih.gov/gene/?term=LOC105276797 | NC_039507.1 | 8906378 | 8906877 | - | 714 | 404 | 310 | 56.58 |

...
... only first 25 lines shown ...

Directory: RNK File: ranked-promoters-bySiteDensity-Ob_LI_RphaseA1.txt

| region_ID | rwidth | nbrsites | nbrper10kb | pmpersite | pmpernucl | pglink | seqnames | start | end | strand | coverage | numCs | numTs | prcntM |
|---|---|---|---|---|---|---|---|---|---|---|---|---|---|---|
| gene-LOC105280727 | 500 | 39 | 780 | 66.14 | 5.16 | https://www.ncbi.nlm.nih.gov/gene/?term=LOC105280727 | NC_039513.1 | 11520171 | 11520670 | + | 1668 | 731 | 937 | 43.82 |
| gene-LOC105277808 | 500 | 37 | 740 | 60.57 | 4.48 | https://www.ncbi.nlm.nih.gov/gene/?term=LOC105277808 | NC_039516.1 | 13518519 | 13519018 | - | 2685 | 872 | 1813 | 32.48 |
| gene-LOC105286234 | 500 | 37 | 740 | 53.48 | 3.96 | https://www.ncbi.nlm.nih.gov/gene/?term=LOC105286234 | NC_039518.1 | 8582509 | 8583008 | + | 2018 | 711 | 1307 | 35.23 |
| gene-LOC105288107 | 500 | 36 | 720 | 78.73 | 5.67 | https://www.ncbi.nlm.nih.gov/gene/?term=LOC105288107 | NC_039515.1 | 1102618 | 1103117 | + | 935 | 687 | 248 | 73.48 |
| gene-LOC109611487 | 500 | 36 | 720 | 51.6 | 3.71 | https://www.ncbi.nlm.nih.gov/gene/?term=LOC109611487 | NC_039506.1 | 13096374 | 13096873 | + | 2153 | 648 | 1505 | 30.1 |
| gene-LOC105278673 | 500 | 35 | 700 | 80.46 | 5.63 | https://www.ncbi.nlm.nih.gov/gene/?term=LOC105278673 | NC_039516.1 | 11703885 | 11704384 | - | 1081 | 749 | 332 | 69.29 |
| gene-LOC105280861 | 500 | 35 | 700 | 73.61 | 5.15 | https://www.ncbi.nlm.nih.gov/gene/?term=LOC105280861 | NC_039506.1 | 15371187 | 15371686 | + | 1103 | 569 | 534 | 51.59 |
| gene-LOC105282404 | 500 | 33 | 660 | 51.35 | 3.39 | https://www.ncbi.nlm.nih.gov/gene/?term=LOC105282404 | NC_039512.1 | 8476210 | 8476709 | + | 1232 | 438 | 794 | 35.55 |
| gene-LOC113561532 | 500 | 32 | 640 | 88.16 | 5.64 | https://www.ncbi.nlm.nih.gov/gene/?term=LOC113561532 | NC_039507.1 | 7277571 | 7278070 | + | 907 | 746 | 161 | 82.25 |
| gene-LOC113561934 | 500 | 32 | 640 | 85.97 | 5.5 | https://www.ncbi.nlm.nih.gov/gene/?term=LOC113561934 | NC_039506.1 | 15618494 | 15618993 | + | 683 | 570 | 113 | 83.46 |
| gene-LOC105276585 | 500 | 31 | 620 | 81.46 | 5.05 | https://www.ncbi.nlm.nih.gov/gene/?term=LOC105276585 | NC_039517.1 | 4571850 | 4572349 | - | 640 | 497 | 143 | 77.66 |
| gene-LOC105279250 | 500 | 31 | 620 | 71.71 | 4.45 | https://www.ncbi.nlm.nih.gov/gene/?term=LOC105279250 | NC_039512.1 | 7901112 | 7901611 | - | 1107 | 680 | 427 | 61.43 |
| gene-LOC105283833 | 500 | 31 | 620 | 66.41 | 4.12 | https://www.ncbi.nlm.nih.gov/gene/?term=LOC105283833 | NC_039509.1 | 2003067 | 2003566 | - | 1301 | 545 | 756 | 41.89 |
| gene-LOC113562689 | 500 | 31 | 620 | 63.06 | 3.91 | https://www.ncbi.nlm.nih.gov/gene/?term=LOC113562689 | NC_039515.1 | 5223138 | 5223637 | - | 1626 | 617 | 1009 | 37.95 |
| gene-LOC105283749 | 500 | 29 | 580 | 82.32 | 4.77 | https://www.ncbi.nlm.nih.gov/gene/?term=LOC105283749 | NC_039515.1 | 7648626 | 7649125 | - | 1316 | 761 | 555 | 57.83 |
| gene-LOC105286681 | 500 | 28 | 560 | 57.95 | 3.25 | https://www.ncbi.nlm.nih.gov/gene/?term=LOC105286681 | NC_039512.1 | 14397556 | 14398055 | + | 1011 | 426 | 585 | 42.14 |
| gene-LOC105286944 | 500 | 28 | 560 | 84.86 | 4.75 | https://www.ncbi.nlm.nih.gov/gene/?term=LOC105286944 | NC_039506.1 | 13094778 | 13095277 | + | 590 | 447 | 143 | 75.76 |
| gene-LOC105278632 | 500 | 27 | 540 | 82.57 | 4.46 | https://www.ncbi.nlm.nih.gov/gene/?term=LOC105278632 | NC_039516.1 | 11911628 | 11912127 | + | 871 | 450 | 421 | 51.66 |
| gene-LOC105276685 | 500 | 26 | 520 | 91.2 | 4.74 | https://www.ncbi.nlm.nih.gov/gene/?term=LOC105276685 | NC_039511.1 | 14299458 | 14299957 | - | 877 | 796 | 81 | 90.76 |
| gene-LOC105283760 | 500 | 26 | 520 | 65.79 | 3.42 | https://www.ncbi.nlm.nih.gov/gene/?term=LOC105283760 | NC_039514.1 | 3324673 | 3325172 | + | 664 | 423 | 241 | 63.7 |
| gene-LOC105277304 | 500 | 25 | 500 | 72.36 | 3.62 | https://www.ncbi.nlm.nih.gov/gene/?term=LOC105277304 | NC_039516.1 | 9356205 | 9356704 | - | 1242 | 591 | 651 | 47.58 |
| gene-LOC105278635 | 500 | 25 | 500 | 81.18 | 4.06 | https://www.ncbi.nlm.nih.gov/gene/?term=LOC105278635 | NC_039516.1 | 11911609 | 11912108 | - | 851 | 430 | 421 | 50.53 |
| gene-LOC113562533 | 500 | 25 | 500 | 79.96 | 4 | https://www.ncbi.nlm.nih.gov/gene/?term=LOC113562533 | NC_039514.1 | 1298855 | 1299354 | + | 538 | 410 | 128 | 76.21 |
| gene-LOC105274855 | 500 | 24 | 480 | 43.29 | 2.08 | https://www.ncbi.nlm.nih.gov/gene/?term=LOC105274855 | NC_039513.1 | 3348900 | 3349399 | + | 1855 | 449 | 1406 | 24.2 |

...

... only first 25 lines shown ...

Directory: RNK File: ranked-promoters-bySiteDensity-Ob_LI_RphaseA2.txt

| region_ID | rwidth | nbrsites | nbrper10kb | pmpersite | pmpernucl | pglink | seqnames | start | end | strand | coverage | numCs | numTs | prcntM |
|---|---|---|---|---|---|---|---|---|---|---|---|---|---|---|
| gene-LOC105277808 | 500 | 46 | 920 | 56.11 | 5.16 | https://www.ncbi.nlm.nih.gov/gene/?term=LOC105277808 | NC_039516.1 | 13518519 | 13519018 | - | 4284 | 1498 | 2786 | 34.97 |
| gene-LOC109611487 | 500 | 44 | 880 | 47.2 | 4.15 | https://www.ncbi.nlm.nih.gov/gene/?term=LOC109611487 | NC_039506.1 | 13096374 | 13096873 | + | 3868 | 1086 | 2782 | 28.08 |
| gene-LOC105282404 | 500 | 43 | 860 | 50.09 | 4.31 | https://www.ncbi.nlm.nih.gov/gene/?term=LOC105282404 | NC_039512.1 | 8476210 | 8476709 | + | 2463 | 1029 | 1434 | 41.78 |
| gene-LOC105280861 | 500 | 42 | 840 | 69.06 | 5.8 | https://www.ncbi.nlm.nih.gov/gene/?term=LOC105280861 | NC_039506.1 | 15371187 | 15371686 | + | 1966 | 1075 | 891 | 54.68 |
| gene-LOC105274855 | 500 | 41 | 820 | 38.89 | 3.19 | https://www.ncbi.nlm.nih.gov/gene/?term=LOC105274855 | NC_039513.1 | 3348900 | 3349399 | + | 3084 | 828 | 2256 | 26.85 |
| gene-LOC105279250 | 500 | 41 | 820 | 72.55 | 5.95 | https://www.ncbi.nlm.nih.gov/gene/?term=LOC105279250 | NC_039512.1 | 7901112 | 7901611 | - | 1981 | 1322 | 659 | 66.73 |
| gene-LOC105280727 | 500 | 41 | 820 | 62.82 | 5.15 | https://www.ncbi.nlm.nih.gov/gene/?term=LOC105280727 | NC_039513.1 | 11520171 | 11520670 | + | 2993 | 1510 | 1483 | 50.45 |
| gene-LOC105278673 | 500 | 40 | 800 | 75.69 | 6.06 | https://www.ncbi.nlm.nih.gov/gene/?term=LOC105278673 | NC_039516.1 | 11703885 | 11704384 | - | 1592 | 1058 | 534 | 66.46 |
| gene-LOC105286234 | 500 | 40 | 800 | 51.71 | 4.14 | https://www.ncbi.nlm.nih.gov/gene/?term=LOC105286234 | NC_039518.1 | 8582509 | 8583008 | + | 3325 | 1331 | 1994 | 40.03 |
| gene-LOC113561934 | 500 | 39 | 780 | 86.12 | 6.72 | https://www.ncbi.nlm.nih.gov/gene/?term=LOC113561934 | NC_039506.1 | 15618494 | 15618993 | + | 1497 | 1238 | 259 | 82.7 |
| gene-LOC105277304 | 500 | 37 | 740 | 68.24 | 5.05 | https://www.ncbi.nlm.nih.gov/gene/?term=LOC105277304 | NC_039516.1 | 9356205 | 9356704 | - | 2020 | 1028 | 992 | 50.89 |
| gene-LOC105288107 | 500 | 37 | 740 | 79.74 | 5.9 | https://www.ncbi.nlm.nih.gov/gene/?term=LOC105288107 | NC_039515.1 | 1102618 | 1103117 | + | 1113 | 804 | 309 | 72.24 |
| gene-LOC105286681 | 500 | 36 | 720 | 58.87 | 4.24 | https://www.ncbi.nlm.nih.gov/gene/?term=LOC105286681 | NC_039512.1 | 14397556 | 14398055 | + | 1514 | 677 | 837 | 44.72 |
| gene-LOC105286944 | 500 | 36 | 720 | 81.52 | 5.87 | https://www.ncbi.nlm.nih.gov/gene/?term=LOC105286944 | NC_039506.1 | 13094778 | 13095277 | + | 1091 | 814 | 277 | 74.61 |
| gene-LOC105276585 | 500 | 34 | 680 | 80.62 | 5.48 | https://www.ncbi.nlm.nih.gov/gene/?term=LOC105276585 | NC_039517.1 | 4571850 | 4572349 | - | 1118 | 848 | 270 | 75.85 |
| gene-LOC105278632 | 500 | 33 | 660 | 73.9 | 4.88 | https://www.ncbi.nlm.nih.gov/gene/?term=LOC105278632 | NC_039516.1 | 11911628 | 11912127 | + | 1128 | 627 | 501 | 55.59 |
| gene-LOC105283833 | 500 | 33 | 660 | 63.8 | 4.21 | https://www.ncbi.nlm.nih.gov/gene/?term=LOC105283833 | NC_039509.1 | 2003067 | 2003566 | - | 1872 | 784 | 1088 | 41.88 |
| gene-LOC113561532 | 500 | 33 | 660 | 87.86 | 5.8 | https://www.ncbi.nlm.nih.gov/gene/?term=LOC113561532 | NC_039507.1 | 7277571 | 7278070 | + | 1604 | 1361 | 243 | 84.85 |
| gene-LOC113562689 | 500 | 33 | 660 | 63.22 | 4.17 | https://www.ncbi.nlm.nih.gov/gene/?term=LOC113562689 | NC_039515.1 | 5223138 | 5223637 | - | 2025 | 814 | 1211 | 40.2 |
| gene-LOC105283965 | 500 | 32 | 640 | 58.88 | 3.77 | https://www.ncbi.nlm.nih.gov/gene/?term=LOC105283965 | NC_039506.1 | 15529793 | 15530292 | - | 1254 | 586 | 668 | 46.73 |
| gene-LOC105278635 | 500 | 31 | 620 | 72.34 | 4.49 | https://www.ncbi.nlm.nih.gov/gene/?term=LOC105278635 | NC_039516.1 | 11911609 | 11912108 | - | 1066 | 567 | 499 | 53.19 |
| gene-LOC105284094 | 500 | 31 | 620 | 34.6 | 2.15 | https://www.ncbi.nlm.nih.gov/gene/?term=LOC105284094 | NC_039511.1 | 3031639 | 3032138 | + | 3565 | 706 | 2859 | 19.8 |
| gene-LOC105277648 | 500 | 29 | 580 | 77.28 | 4.48 | https://www.ncbi.nlm.nih.gov/gene/?term=LOC105277648 | NC_039512.1 | 2276615 | 2277114 | + | 735 | 498 | 237 | 67.76 |
| gene-LOC105283749 | 500 | 29 | 580 | 84.39 | 4.89 | https://www.ncbi.nlm.nih.gov/gene/?term=LOC105283749 | NC_039515.1 | 7648626 | 7649125 | - | 1713 | 955 | 758 | 55.75 |

...

... only first 25 lines shown ...

Directory: RNK File: ranked-promoters-bySiteDensity-Ob_LI_RphaseB1.txt

| region_ID | rwidth | nbrsites | nbrper10kb | pmpersite | pmpernucl | pglink | seqnames | start | end | strand | coverage | numCs | numTs | prcntM |
|---|---|---|---|---|---|---|---|---|---|---|---|---|---|---|
| gene-LOC105288107 | 500 | 40 | 800 | 82.1 | 6.57 | https://www.ncbi.nlm.nih.gov/gene/?term=LOC105288107 | NC_039515.1 | 1102618 | 1103117 | + | 802 | 614 | 188 | 76.56 |
| gene-LOC105277808 | 500 | 39 | 780 | 56.66 | 4.42 | https://www.ncbi.nlm.nih.gov/gene/?term=LOC105277808 | NC_039516.1 | 13518519 | 13519018 | - | 2814 | 869 | 1945 | 30.88 |
| gene-LOC105280727 | 500 | 39 | 780 | 61.77 | 4.82 | https://www.ncbi.nlm.nih.gov/gene/?term=LOC105280727 | NC_039513.1 | 11520171 | 11520670 | + | 1729 | 694 | 1035 | 40.14 |
| gene-LOC105277304 | 500 | 38 | 760 | 68.4 | 5.2 | https://www.ncbi.nlm.nih.gov/gene/?term=LOC105277304 | NC_039516.1 | 9356205 | 9356704 | - | 1526 | 751 | 775 | 49.21 |
| gene-LOC105282404 | 500 | 36 | 720 | 56.74 | 4.09 | https://www.ncbi.nlm.nih.gov/gene/?term=LOC105282404 | NC_039512.1 | 8476210 | 8476709 | + | 1640 | 641 | 999 | 39.09 |
| gene-LOC105286234 | 500 | 34 | 680 | 53.14 | 3.61 | https://www.ncbi.nlm.nih.gov/gene/?term=LOC105286234 | NC_039518.1 | 8582509 | 8583008 | + | 1979 | 708 | 1271 | 35.78 |
| gene-LOC109611487 | 500 | 33 | 660 | 47.55 | 3.14 | https://www.ncbi.nlm.nih.gov/gene/?term=LOC109611487 | NC_039506.1 | 13096374 | 13096873 | + | 2454 | 635 | 1819 | 25.88 |
| gene-LOC113561532 | 500 | 33 | 660 | 88.54 | 5.84 | https://www.ncbi.nlm.nih.gov/gene/?term=LOC113561532 | NC_039507.1 | 7277571 | 7278070 | + | 999 | 840 | 159 | 84.08 |
| gene-LOC113561934 | 500 | 33 | 660 | 81.57 | 5.38 | https://www.ncbi.nlm.nih.gov/gene/?term=LOC113561934 | NC_039506.1 | 15618494 | 15618993 | + | 874 | 665 | 209 | 76.09 |
| gene-LOC113562689 | 500 | 32 | 640 | 65.65 | 4.2 | https://www.ncbi.nlm.nih.gov/gene/?term=LOC113562689 | NC_039515.1 | 5223138 | 5223637 | - | 1292 | 583 | 709 | 45.12 |
| gene-LOC105278673 | 500 | 30 | 600 | 72.2 | 4.33 | https://www.ncbi.nlm.nih.gov/gene/?term=LOC105278673 | NC_039516.1 | 11703885 | 11704384 | - | 1002 | 663 | 339 | 66.17 |
| gene-LOC105279250 | 500 | 30 | 600 | 74.53 | 4.47 | https://www.ncbi.nlm.nih.gov/gene/?term=LOC105279250 | NC_039512.1 | 7901112 | 7901611 | - | 1048 | 629 | 419 | 60.02 |
| gene-LOC105286681 | 500 | 30 | 600 | 58.02 | 3.48 | https://www.ncbi.nlm.nih.gov/gene/?term=LOC105286681 | NC_039512.1 | 14397556 | 14398055 | + | 1223 | 509 | 714 | 41.62 |
| gene-LOC105276585 | 500 | 29 | 580 | 82.1 | 4.76 | https://www.ncbi.nlm.nih.gov/gene/?term=LOC105276585 | NC_039517.1 | 4571850 | 4572349 | - | 655 | 502 | 153 | 76.64 |
| gene-LOC105283833 | 500 | 29 | 580 | 69.45 | 4.03 | https://www.ncbi.nlm.nih.gov/gene/?term=LOC105283833 | NC_039509.1 | 2003067 | 2003566 | - | 1219 | 522 | 697 | 42.82 |
| gene-LOC105286944 | 500 | 29 | 580 | 82.28 | 4.77 | https://www.ncbi.nlm.nih.gov/gene/?term=LOC105286944 | NC_039506.1 | 13094778 | 13095277 | + | 603 | 417 | 186 | 69.15 |
| gene-LOC105280861 | 500 | 28 | 560 | 74.12 | 4.15 | https://www.ncbi.nlm.nih.gov/gene/?term=LOC105280861 | NC_039506.1 | 15371187 | 15371686 | + | 1116 | 553 | 563 | 49.55 |
| gene-LOC105283749 | 500 | 28 | 560 | 85.27 | 4.78 | https://www.ncbi.nlm.nih.gov/gene/?term=LOC105283749 | NC_039515.1 | 7648626 | 7649125 | - | 1388 | 746 | 642 | 53.75 |
| gene-LOC105276685 | 500 | 27 | 540 | 91.97 | 4.97 | https://www.ncbi.nlm.nih.gov/gene/?term=LOC105276685 | NC_039511.1 | 14299458 | 14299957 | - | 898 | 827 | 71 | 92.09 |
| gene-LOC113562533 | 500 | 27 | 540 | 80.64 | 4.35 | https://www.ncbi.nlm.nih.gov/gene/?term=LOC113562533 | NC_039514.1 | 1298855 | 1299354 | + | 529 | 410 | 119 | 77.5 |
| gene-LOC105276797 | 500 | 25 | 500 | 74.29 | 3.71 | https://www.ncbi.nlm.nih.gov/gene/?term=LOC105276797 | NC_039507.1 | 8906378 | 8906877 | - | 933 | 523 | 410 | 56.06 |
| gene-LOC105278632 | 500 | 25 | 500 | 84.81 | 4.24 | https://www.ncbi.nlm.nih.gov/gene/?term=LOC105278632 | NC_039516.1 | 11911628 | 11912127 | + | 815 | 399 | 416 | 48.96 |
| gene-LOC105283965 | 500 | 25 | 500 | 58.53 | 2.93 | https://www.ncbi.nlm.nih.gov/gene/?term=LOC105283965 | NC_039506.1 | 15529793 | 15530292 | - | 698 | 316 | 382 | 45.27 |
| gene-LOC105277395 | 500 | 24 | 480 | 72.69 | 3.49 | https://www.ncbi.nlm.nih.gov/gene/?term=LOC105277395 | NC_039508.1 | 1340463 | 1340962 | - | 536 | 321 | 215 | 59.89 |

...

... only first 25 lines shown ...

Directory: RNK File: ranked-promoters-bySiteDensity-Ob_LI_RphaseB2.txt

| region_ID | rwidth | nbrsites | nbrper10kb | pmpersite | pmpernucl | pglink | seqnames | start | end | strand | coverage | numCs | numTs | prcntM |
|---|---|---|---|---|---|---|---|---|---|---|---|---|---|---|
| gene-LOC113561934 | 500 | 37 | 740 | 82.55 | 6.11 | https://www.ncbi.nlm.nih.gov/gene/?term=LOC113561934 | NC_039506.1 | 15618494 | 15618993 | + | 828 | 636 | 192 | 76.81 |
| gene-LOC105277808 | 500 | 36 | 720 | 58.17 | 4.19 | https://www.ncbi.nlm.nih.gov/gene/?term=LOC105277808 | NC_039516.1 | 13518519 | 13519018 | - | 2553 | 783 | 1770 | 30.67 |
| gene-LOC105288107 | 500 | 36 | 720 | 79.61 | 5.73 | https://www.ncbi.nlm.nih.gov/gene/?term=LOC105288107 | NC_039515.1 | 1102618 | 1103117 | + | 869 | 639 | 230 | 73.53 |
| gene-LOC105280727 | 500 | 34 | 680 | 61.18 | 4.16 | https://www.ncbi.nlm.nih.gov/gene/?term=LOC105280727 | NC_039513.1 | 11520171 | 11520670 | + | 1820 | 775 | 1045 | 42.58 |
| gene-LOC105286234 | 500 | 33 | 660 | 51.79 | 3.42 | https://www.ncbi.nlm.nih.gov/gene/?term=LOC105286234 | NC_039518.1 | 8582509 | 8583008 | + | 1878 | 713 | 1165 | 37.97 |
| gene-LOC105280861 | 500 | 32 | 640 | 74.16 | 4.75 | https://www.ncbi.nlm.nih.gov/gene/?term=LOC105280861 | NC_039506.1 | 15371187 | 15371686 | + | 1195 | 641 | 554 | 53.64 |
| gene-LOC109611487 | 500 | 31 | 620 | 50.13 | 3.11 | https://www.ncbi.nlm.nih.gov/gene/?term=LOC109611487 | NC_039506.1 | 13096374 | 13096873 | + | 2322 | 595 | 1727 | 25.62 |
| gene-LOC113561532 | 500 | 31 | 620 | 89.77 | 5.57 | https://www.ncbi.nlm.nih.gov/gene/?term=LOC113561532 | NC_039507.1 | 7277571 | 7278070 | + | 1137 | 966 | 171 | 84.96 |
| gene-LOC105277304 | 500 | 30 | 600 | 78.09 | 4.69 | https://www.ncbi.nlm.nih.gov/gene/?term=LOC105277304 | NC_039516.1 | 9356205 | 9356704 | - | 1225 | 617 | 608 | 50.37 |
| gene-LOC105282404 | 500 | 30 | 600 | 55.77 | 3.35 | https://www.ncbi.nlm.nih.gov/gene/?term=LOC105282404 | NC_039512.1 | 8476210 | 8476709 | + | 1387 | 566 | 821 | 40.81 |
| gene-LOC105286944 | 500 | 28 | 560 | 86.7 | 4.86 | https://www.ncbi.nlm.nih.gov/gene/?term=LOC105286944 | NC_039506.1 | 13094778 | 13095277 | + | 866 | 636 | 230 | 73.44 |
| gene-LOC105276685 | 500 | 27 | 540 | 88.55 | 4.78 | https://www.ncbi.nlm.nih.gov/gene/?term=LOC105276685 | NC_039511.1 | 14299458 | 14299957 | - | 804 | 733 | 71 | 91.17 |
| gene-LOC105278673 | 500 | 27 | 540 | 78.87 | 4.26 | https://www.ncbi.nlm.nih.gov/gene/?term=LOC105278673 | NC_039516.1 | 11703885 | 11704384 | - | 983 | 648 | 335 | 65.92 |
| gene-LOC105279250 | 500 | 27 | 540 | 71.13 | 3.84 | https://www.ncbi.nlm.nih.gov/gene/?term=LOC105279250 | NC_039512.1 | 7901112 | 7901611 | - | 1092 | 682 | 410 | 62.45 |
| gene-LOC105283749 | 500 | 27 | 540 | 83.39 | 4.5 | https://www.ncbi.nlm.nih.gov/gene/?term=LOC105283749 | NC_039515.1 | 7648626 | 7649125 | - | 1147 | 639 | 508 | 55.71 |
| gene-LOC105283833 | 500 | 27 | 540 | 70.39 | 3.8 | https://www.ncbi.nlm.nih.gov/gene/?term=LOC105283833 | NC_039509.1 | 2003067 | 2003566 | - | 964 | 407 | 557 | 42.22 |
| gene-LOC105283965 | 500 | 27 | 540 | 62.24 | 3.36 | https://www.ncbi.nlm.nih.gov/gene/?term=LOC105283965 | NC_039506.1 | 15529793 | 15530292 | - | 780 | 388 | 392 | 49.74 |
| gene-LOC105276585 | 500 | 26 | 520 | 79.1 | 4.11 | https://www.ncbi.nlm.nih.gov/gene/?term=LOC105276585 | NC_039517.1 | 4571850 | 4572349 | - | 681 | 498 | 183 | 73.13 |
| gene-LOC105286681 | 500 | 26 | 520 | 57.86 | 3.01 | https://www.ncbi.nlm.nih.gov/gene/?term=LOC105286681 | NC_039512.1 | 14397556 | 14398055 | + | 1005 | 419 | 586 | 41.69 |
| gene-LOC105276797 | 500 | 24 | 480 | 76.95 | 3.69 | https://www.ncbi.nlm.nih.gov/gene/?term=LOC105276797 | NC_039507.1 | 8906378 | 8906877 | - | 654 | 401 | 253 | 61.31 |
| gene-LOC109610674 | 500 | 24 | 480 | 82.93 | 3.98 | https://www.ncbi.nlm.nih.gov/gene/?term=LOC109610674 | NC_039516.1 | 731871 | 732370 | + | 729 | 494 | 235 | 67.76 |
| gene-LOC105274855 | 500 | 23 | 460 | 40.29 | 1.85 | https://www.ncbi.nlm.nih.gov/gene/?term=LOC105274855 | NC_039513.1 | 3348900 | 3349399 | + | 1873 | 432 | 1441 | 23.06 |
| gene-LOC105277648 | 500 | 23 | 460 | 76.94 | 3.54 | https://www.ncbi.nlm.nih.gov/gene/?term=LOC105277648 | NC_039512.1 | 2276615 | 2277114 | + | 459 | 318 | 141 | 69.28 |
| gene-LOC105278632 | 500 | 23 | 460 | 80.74 | 3.71 | https://www.ncbi.nlm.nih.gov/gene/?term=LOC105278632 | NC_039516.1 | 11911628 | 11912127 | + | 657 | 348 | 309 | 52.97 |

...

... only first 25 lines shown ...

Directory: RNK File: sites-in-genes-Ob_LI_BCphaseA1.txt

```
seqnames	start	end	width	strand	coverage	numCs	numTs	perc_meth	region_seqnames	region_start	region_end	region_width	region_strand	region_source
	region_type	region_score	region_phase	region_ID	region_Dbxref	region_Name	region_gbkey	region_gene	region_gene_biotype	region_end_range	region_partial
	region_start_range
NC_039506.1	956202	956202	1	+	32	7	25	21.875	NC_039506.1	926117	968035	41919	-	Gnomon	gene	NA	NA	gene-LOC113561445
	GeneID:113561445	LOC113561445	Gene	LOC113561445	protein_coding	character(0)	NA	character(0)
NC_039506.1	1097588	1097588	1	+	31	10	21	32.258064516129	NC_039506.1	1095614	1100480	4867	+	Gnomon	gene	NA	NA	gene-LOC105286750
	GeneID:105286750	LOC105286750	Gene	LOC105286750	protein_coding	character(0)	NA	character(0)
NC_039506.1	1097589	1097589	1	-	8	5	3	62.5	NC_039506.1	1095614	1100480	4867	+	Gnomon	gene	NA	NA	gene-LOC105286750
	GeneID:105286750	LOC105286750	Gene	LOC105286750	protein_coding	character(0)	NA	character(0)
NC_039506.1	1097598	1097598	1	+	36	25	11	69.4444444444444	NC_039506.1	1095614	1100480	4867	+	Gnomon	gene	NA	NA	gene-LOC105286750
	GeneID:105286750	LOC105286750	Gene	LOC105286750	protein_coding	character(0)	NA	character(0)
NC_039506.1	1097702	1097702	1	+	52	8	44	15.3846153846154	NC_039506.1	1095614	1100480	4867	+	Gnomon	gene	NA	NA	gene-LOC105286750
	GeneID:105286750	LOC105286750	Gene	LOC105286750	protein_coding	character(0)	NA	character(0)
NC_039506.1	1097773	1097773	1	+	31	29	2	93.5483870967742	NC_039506.1	1095614	1100480	4867	+	Gnomon	gene	NA	NA	gene-LOC105286750
	GeneID:105286750	LOC105286750	Gene	LOC105286750	protein_coding	character(0)	NA	character(0)
NC_039506.1	1097774	1097774	1	-	8	7	1	87.5	NC_039506.1	1095614	1100480	4867	+	Gnomon	gene	NA	NA	gene-LOC105286750
	GeneID:105286750	LOC105286750	Gene	LOC105286750	protein_coding	character(0)	NA	character(0)
NC_039506.1	1097925	1097925	1	-	5	5	0	100	NC_039506.1	1095614	1100480	4867	+	Gnomon	gene	NA	NA	gene-LOC105286750
	GeneID:105286750	LOC105286750	Gene	LOC105286750	protein_coding	character(0)	NA	character(0)
NC_039506.1	1097943	1097943	1	+	6	6	0	100	NC_039506.1	1095614	1100480	4867	+	Gnomon	gene	NA	NA	gene-LOC105286750
	GeneID:105286750	LOC105286750	Gene	LOC105286750	protein_coding	character(0)	NA	character(0)
NC_039506.1	1097944	1097944	1	-	6	6	0	100	NC_039506.1	1095614	1100480	4867	+	Gnomon	gene	NA	NA	gene-LOC105286750
	GeneID:105286750	LOC105286750	Gene	LOC105286750	protein_coding	character(0)	NA	character(0)
NC_039506.1	1097981	1097981	1	+	9	8	1	88.8888888888889	NC_039506.1	1095614	1100480	4867	+	Gnomon	gene	NA	NA	gene-LOC105286750
	GeneID:105286750	LOC105286750	Gene	LOC105286750	protein_coding	character(0)	NA	character(0)
NC_039506.1	1098055	1098055	1	+	13	13	0	100	NC_039506.1	1095614	1100480	4867	+	Gnomon	gene	NA	NA	gene-LOC105286750
	GeneID:105286750	LOC105286750	Gene	LOC105286750	protein_coding	character(0)	NA	character(0)
NC_039506.1	1098056	1098056	1	-	11	11	0	100	NC_039506.1	1095614	1100480	4867	+	Gnomon	gene	NA	NA	gene-LOC105286750
	GeneID:105286750	LOC105286750	Gene	LOC105286750	protein_coding	character(0)	NA	character(0)
NC_039506.1	1098069	1098069	1	+	14	14	0	100	NC_039506.1	1095614	1100480	4867	+	Gnomon	gene	NA	NA	gene-LOC105286750
	GeneID:105286750	LOC105286750	Gene	LOC105286750	protein_coding	character(0)	NA	character(0)
NC_039506.1	1098070	1098070	1	-	10	10	0	100	NC_039506.1	1095614	1100480	4867	+	Gnomon	gene	NA	NA	gene-LOC105286750
	GeneID:105286750	LOC105286750	Gene	LOC105286750	protein_coding	character(0)	NA	character(0)
NC_039506.1	1098123	1098123	1	+	17	10	7	58.8235294117647	NC_039506.1	1095614	1100480	4867	+	Gnomon	gene	NA	NA	gene-LOC105286750
	GeneID:105286750	LOC105286750	Gene	LOC105286750	protein_coding	character(0)	NA	character(0)
NC_039506.1	1098144	1098144	1	+	12	11	1	91.6666666666667	NC_039506.1	1095614	1100480	4867	+	Gnomon	gene	NA	NA	gene-LOC105286750
	GeneID:105286750	LOC105286750	Gene	LOC105286750	protein_coding	character(0)	NA	character(0)
NC_039506.1	1098163	1098163	1	+	13	11	2	84.6153846153846	NC_039506.1	1095614	1100480	4867	+	Gnomon	gene	NA	NA	gene-LOC105286750
	GeneID:105286750	LOC105286750	Gene	LOC105286750	protein_coding	character(0)	NA	character(0)
NC_039506.1	1098206	1098206	1	+	26	17	9	65.3846153846154	NC_039506.1	1095614	1100480	4867	+	Gnomon	gene	NA	NA	gene-LOC105286750
	GeneID:105286750	LOC105286750	Gene	LOC105286750	protein_coding	character(0)	NA	character(0)
NC_039506.1	1098266	1098266	1	+	41	21	20	51.219512195122	NC_039506.1	1095614	1100480	4867	+	Gnomon	gene	NA	NA	gene-LOC105286750
	GeneID:105286750	LOC105286750	Gene	LOC105286750	protein_coding	character(0)	NA	character(0)
NC_039506.1	1098279	1098279	1	+	42	22	20	52.3809523809524	NC_039506.1	1095614	1100480	4867	+	Gnomon	gene	NA	NA	gene-LOC105286750
	GeneID:105286750	LOC105286750	Gene	LOC105286750	protein_coding	character(0)	NA	character(0)
NC_039506.1	1098293	1098293	1	+	25	10	15	40	NC_039506.1	1095614	1100480	4867	+	Gnomon	gene	NA	NA	gene-LOC105286750
	GeneID:105286750	LOC105286750	Gene	LOC105286750	protein_coding	character(0)	NA	character(0)
NC_039506.1	1098301	1098301	1	+	30	22	8	73.3333333333333	NC_039506.1	1095614	1100480	4867	+	Gnomon	gene	NA	NA	gene-LOC105286750
	GeneID:105286750	LOC105286750	Gene	LOC105286750	protein_coding	character(0)	NA	character(0)
NC_039506.1	1098330	1098330	1	+	38	13	25	34.2105263157895	NC_039506.1	1095614	1100480	4867	+	Gnomon	gene	NA	NA	gene-LOC105286750
	GeneID:105286750	LOC105286750	Gene	LOC105286750	protein_coding	character(0)	NA	character(0)
...
... only first 25 lines shown ...
```

Directory: RNK File: sites-in-genes-Ob_LI_BCphaseA2.txt

| seqnames | start | end | width | strand | coverage | numCs | numTs | perc_meth | region_seqnames | region_start | region_end | region_width | region_strand | region_source | region_type | region_score | region_phase | region_ID | region_Dbxref | region_Name | region_gbkey | region_gene | region_gene_biotype | region_end_range | region_partial | region_start_range |
|---|---|---|---|---|---|---|---|---|---|---|---|---|---|---|---|---|---|---|---|---|---|---|---|---|---|---|
| NC_039506.1 | 1097588 | 1097588 | 1 | + | 35 | 12 | 23 | 34.2857142857143 | NC_039506.1 | 1095614 | 1100480 | 4867 | + | Gnomon | gene | NA | NA | gene-LOC105286750 | GeneID:105286750 | LOC105286750 | Gene | LOC105286750 | protein_coding | character(0) | NA | character(0) |
| NC_039506.1 | 1097598 | 1097598 | 1 | + | 38 | 15 | 23 | 39.4736842105263 | NC_039506.1 | 1095614 | 1100480 | 4867 | + | Gnomon | gene | NA | NA | gene-LOC105286750 | GeneID:105286750 | LOC105286750 | Gene | LOC105286750 | protein_coding | character(0) | NA | character(0) |
| NC_039506.1 | 1097599 | 1097599 | 1 | - | 10 | 6 | 4 | 60 | NC_039506.1 | 1095614 | 1100480 | 4867 | + | Gnomon | gene | NA | NA | gene-LOC105286750 | GeneID:105286750 | LOC105286750 | Gene | LOC105286750 | protein_coding | character(0) | NA | character(0) |
| NC_039506.1 | 1097702 | 1097702 | 1 | + | 50 | 8 | 42 | 16 | NC_039506.1 | 1095614 | 1100480 | 4867 | + | Gnomon | gene | NA | NA | gene-LOC105286750 | GeneID:105286750 | LOC105286750 | Gene | LOC105286750 | protein_coding | character(0) | NA | character(0) |
| NC_039506.1 | 1097773 | 1097773 | 1 | + | 45 | 44 | 1 | 97.7777777777778 | NC_039506.1 | 1095614 | 1100480 | 4867 | + | Gnomon | gene | NA | NA | gene-LOC105286750 | GeneID:105286750 | LOC105286750 | Gene | LOC105286750 | protein_coding | character(0) | NA | character(0) |
| NC_039506.1 | 1097943 | 1097943 | 1 | + | 5 | 5 | 0 | 100 | NC_039506.1 | 1095614 | 1100480 | 4867 | + | Gnomon | gene | NA | NA | gene-LOC105286750 | GeneID:105286750 | LOC105286750 | Gene | LOC105286750 | protein_coding | character(0) | NA | character(0) |
| NC_039506.1 | 1097981 | 1097981 | 1 | + | 6 | 6 | 0 | 100 | NC_039506.1 | 1095614 | 1100480 | 4867 | + | Gnomon | gene | NA | NA | gene-LOC105286750 | GeneID:105286750 | LOC105286750 | Gene | LOC105286750 | protein_coding | character(0) | NA | character(0) |
| NC_039506.1 | 1098055 | 1098055 | 1 | + | 16 | 15 | 1 | 93.75 | NC_039506.1 | 1095614 | 1100480 | 4867 | + | Gnomon | gene | NA | NA | gene-LOC105286750 | GeneID:105286750 | LOC105286750 | Gene | LOC105286750 | protein_coding | character(0) | NA | character(0) |
| NC_039506.1 | 1098056 | 1098056 | 1 | - | 16 | 16 | 0 | 100 | NC_039506.1 | 1095614 | 1100480 | 4867 | + | Gnomon | gene | NA | NA | gene-LOC105286750 | GeneID:105286750 | LOC105286750 | Gene | LOC105286750 | protein_coding | character(0) | NA | character(0) |
| NC_039506.1 | 1098069 | 1098069 | 1 | + | 17 | 16 | 1 | 94.1176470588235 | NC_039506.1 | 1095614 | 1100480 | 4867 | + | Gnomon | gene | NA | NA | gene-LOC105286750 | GeneID:105286750 | LOC105286750 | Gene | LOC105286750 | protein_coding | character(0) | NA | character(0) |
| NC_039506.1 | 1098070 | 1098070 | 1 | - | 14 | 14 | 0 | 100 | NC_039506.1 | 1095614 | 1100480 | 4867 | + | Gnomon | gene | NA | NA | gene-LOC105286750 | GeneID:105286750 | LOC105286750 | Gene | LOC105286750 | protein_coding | character(0) | NA | character(0) |
| NC_039506.1 | 1098123 | 1098123 | 1 | + | 29 | 23 | 6 | 79.3103448275862 | NC_039506.1 | 1095614 | 1100480 | 4867 | + | Gnomon | gene | NA | NA | gene-LOC105286750 | GeneID:105286750 | LOC105286750 | Gene | LOC105286750 | protein_coding | character(0) | NA | character(0) |
| NC_039506.1 | 1098124 | 1098124 | 1 | - | 6 | 6 | 0 | 100 | NC_039506.1 | 1095614 | 1100480 | 4867 | + | Gnomon | gene | NA | NA | gene-LOC105286750 | GeneID:105286750 | LOC105286750 | Gene | LOC105286750 | protein_coding | character(0) | NA | character(0) |
| NC_039506.1 | 1098144 | 1098144 | 1 | + | 25 | 25 | 0 | 100 | NC_039506.1 | 1095614 | 1100480 | 4867 | + | Gnomon | gene | NA | NA | gene-LOC105286750 | GeneID:105286750 | LOC105286750 | Gene | LOC105286750 | protein_coding | character(0) | NA | character(0) |
| NC_039506.1 | 1098145 | 1098145 | 1 | - | 5 | 5 | 0 | 100 | NC_039506.1 | 1095614 | 1100480 | 4867 | + | Gnomon | gene | NA | NA | gene-LOC105286750 | GeneID:105286750 | LOC105286750 | Gene | LOC105286750 | protein_coding | character(0) | NA | character(0) |
| NC_039506.1 | 1098163 | 1098163 | 1 | + | 15 | 15 | 0 | 100 | NC_039506.1 | 1095614 | 1100480 | 4867 | + | Gnomon | gene | NA | NA | gene-LOC105286750 | GeneID:105286750 | LOC105286750 | Gene | LOC105286750 | protein_coding | character(0) | NA | character(0) |
| NC_039506.1 | 1098206 | 1098206 | 1 | + | 34 | 26 | 8 | 76.4705882352941 | NC_039506.1 | 1095614 | 1100480 | 4867 | + | Gnomon | gene | NA | NA | gene-LOC105286750 | GeneID:105286750 | LOC105286750 | Gene | LOC105286750 | protein_coding | character(0) | NA | character(0) |
| NC_039506.1 | 1098266 | 1098266 | 1 | + | 43 | 25 | 18 | 58.1395348837209 | NC_039506.1 | 1095614 | 1100480 | 4867 | + | Gnomon | gene | NA | NA | gene-LOC105286750 | GeneID:105286750 | LOC105286750 | Gene | LOC105286750 | protein_coding | character(0) | NA | character(0) |
| NC_039506.1 | 1098279 | 1098279 | 1 | + | 47 | 18 | 29 | 38.2978723404255 | NC_039506.1 | 1095614 | 1100480 | 4867 | + | Gnomon | gene | NA | NA | gene-LOC105286750 | GeneID:105286750 | LOC105286750 | Gene | LOC105286750 | protein_coding | character(0) | NA | character(0) |
| NC_039506.1 | 1098293 | 1098293 | 1 | + | 26 | 11 | 15 | 42.3076923076923 | NC_039506.1 | 1095614 | 1100480 | 4867 | + | Gnomon | gene | NA | NA | gene-LOC105286750 | GeneID:105286750 | LOC105286750 | Gene | LOC105286750 | protein_coding | character(0) | NA | character(0) |
| NC_039506.1 | 1098301 | 1098301 | 1 | + | 23 | 14 | 9 | 60.8695652173913 | NC_039506.1 | 1095614 | 1100480 | 4867 | + | Gnomon | gene | NA | NA | gene-LOC105286750 | GeneID:105286750 | LOC105286750 | Gene | LOC105286750 | protein_coding | character(0) | NA | character(0) |
| NC_039506.1 | 1098330 | 1098330 | 1 | + | 54 | 24 | 30 | 44.4444444444444 | NC_039506.1 | 1095614 | 1100480 | 4867 | + | Gnomon | gene | NA | NA | gene-LOC105286750 | GeneID:105286750 | LOC105286750 | Gene | LOC105286750 | protein_coding | character(0) | NA | character(0) |
| NC_039506.1 | 1098334 | 1098334 | 1 | + | 58 | 35 | 23 | 60.3448275862069 | NC_039506.1 | 1095614 | 1100480 | 4867 | + | Gnomon | gene | NA | NA | gene-LOC105286750 | GeneID:105286750 | LOC105286750 | Gene | LOC105286750 | protein_coding | character(0) | NA | character(0) |
| NC_039506.1 | 1098362 | 1098362 | 1 | + | 58 | 9 | 49 | 15.5172413793103 | NC_039506.1 | 1095614 | 1100480 | 4867 | + | Gnomon | gene | NA | NA | gene-LOC105286750 | GeneID:105286750 | LOC105286750 | Gene | LOC105286750 | protein_coding | character(0) | NA | character(0) |

...
... only first 25 lines shown ...

Directory: RNK File: sites-in-genes-Ob_LI_BCphaseB1.txt

| seqnames | start | end | width | strand | coverage | numCs | numTs | perc_meth | region_seqnames | region_start | region_end | region_width | region_strand | region_source | region_type | region_score | region_phase | region_ID | region_Dbxref | region_Name | region_gbkey | region_gene | region_gene_biotype | region_end_range | region_partial | region_start_range |
|---|---|---|---|---|---|---|---|---|---|---|---|---|---|---|---|---|---|---|---|---|---|---|---|---|---|---|
| NC_039506.1 | 137598 | 137598 | 1 | - | 31 | 7 | 24 | 22.5806451612903 | NC_039506.1 | 134260 | 142204 | 7945 | - | Gnomon | gene | NA | NA | gene-LOC105288138 | GeneID:105288138 | LOC105288138 | Gene | LOC105288138 | protein_coding | character(0) | NA | character(0) |
| NC_039506.1 | 1058205 | 1058205 | 1 | + | 32 | 9 | 23 | 28.125 | NC_039506.1 | 1056151 | 1064257 | 8107 | + | Gnomon | gene | NA | NA | gene-LOC105286747 | GeneID:105286747 | LOC105286747 | Gene | LOC105286747 | lncRNA | character(0) | NA | character(0) |
| NC_039506.1 | 1097588 | 1097588 | 1 | + | 30 | 9 | 21 | 30 | NC_039506.1 | 1095614 | 1100480 | 4867 | + | Gnomon | gene | NA | NA | gene-LOC105286750 | GeneID:105286750 | LOC105286750 | Gene | LOC105286750 | protein_coding | character(0) | NA | character(0) |
| NC_039506.1 | 1097598 | 1097598 | 1 | + | 33 | 17 | 16 | 51.5151515151515 | NC_039506.1 | 1095614 | 1100480 | 4867 | + | Gnomon | gene | NA | NA | gene-LOC105286750 | GeneID:105286750 | LOC105286750 | Gene | LOC105286750 | protein_coding | character(0) | NA | character(0) |
| NC_039506.1 | 1097599 | 1097599 | 1 | - | 13 | 8 | 5 | 61.5384615384615 | NC_039506.1 | 1095614 | 1100480 | 4867 | + | Gnomon | gene | NA | NA | gene-LOC105286750 | GeneID:105286750 | LOC105286750 | Gene | LOC105286750 | protein_coding | character(0) | NA | character(0) |
| NC_039506.1 | 1097647 | 1097647 | 1 | + | 46 | 8 | 38 | 17.3913043478261 | NC_039506.1 | 1095614 | 1100480 | 4867 | + | Gnomon | gene | NA | NA | gene-LOC105286750 | GeneID:105286750 | LOC105286750 | Gene | LOC105286750 | protein_coding | character(0) | NA | character(0) |
| NC_039506.1 | 1097773 | 1097773 | 1 | + | 24 | 23 | 1 | 95.8333333333333 | NC_039506.1 | 1095614 | 1100480 | 4867 | + | Gnomon | gene | NA | NA | gene-LOC105286750 | GeneID:105286750 | LOC105286750 | Gene | LOC105286750 | protein_coding | character(0) | NA | character(0) |
| NC_039506.1 | 1097774 | 1097774 | 1 | - | 7 | 7 | 0 | 100 | NC_039506.1 | 1095614 | 1100480 | 4867 | + | Gnomon | gene | NA | NA | gene-LOC105286750 | GeneID:105286750 | LOC105286750 | Gene | LOC105286750 | protein_coding | character(0) | NA | character(0) |
| NC_039506.1 | 1097925 | 1097925 | 1 | - | 7 | 6 | 1 | 85.7142857142857 | NC_039506.1 | 1095614 | 1100480 | 4867 | + | Gnomon | gene | NA | NA | gene-LOC105286750 | GeneID:105286750 | LOC105286750 | Gene | LOC105286750 | protein_coding | character(0) | NA | character(0) |
| NC_039506.1 | 1097943 | 1097943 | 1 | + | 6 | 6 | 0 | 100 | NC_039506.1 | 1095614 | 1100480 | 4867 | + | Gnomon | gene | NA | NA | gene-LOC105286750 | GeneID:105286750 | LOC105286750 | Gene | LOC105286750 | protein_coding | character(0) | NA | character(0) |
| NC_039506.1 | 1097944 | 1097944 | 1 | - | 7 | 7 | 0 | 100 | NC_039506.1 | 1095614 | 1100480 | 4867 | + | Gnomon | gene | NA | NA | gene-LOC105286750 | GeneID:105286750 | LOC105286750 | Gene | LOC105286750 | protein_coding | character(0) | NA | character(0) |
| NC_039506.1 | 1097981 | 1097981 | 1 | + | 10 | 10 | 0 | 100 | NC_039506.1 | 1095614 | 1100480 | 4867 | + | Gnomon | gene | NA | NA | gene-LOC105286750 | GeneID:105286750 | LOC105286750 | Gene | LOC105286750 | protein_coding | character(0) | NA | character(0) |
| NC_039506.1 | 1098055 | 1098055 | 1 | + | 14 | 14 | 0 | 100 | NC_039506.1 | 1095614 | 1100480 | 4867 | + | Gnomon | gene | NA | NA | gene-LOC105286750 | GeneID:105286750 | LOC105286750 | Gene | LOC105286750 | protein_coding | character(0) | NA | character(0) |
| NC_039506.1 | 1098056 | 1098056 | 1 | - | 8 | 8 | 0 | 100 | NC_039506.1 | 1095614 | 1100480 | 4867 | + | Gnomon | gene | NA | NA | gene-LOC105286750 | GeneID:105286750 | LOC105286750 | Gene | LOC105286750 | protein_coding | character(0) | NA | character(0) |
| NC_039506.1 | 1098069 | 1098069 | 1 | + | 16 | 16 | 0 | 100 | NC_039506.1 | 1095614 | 1100480 | 4867 | + | Gnomon | gene | NA | NA | gene-LOC105286750 | GeneID:105286750 | LOC105286750 | Gene | LOC105286750 | protein_coding | character(0) | NA | character(0) |
| NC_039506.1 | 1098070 | 1098070 | 1 | - | 6 | 5 | 1 | 83.3333333333333 | NC_039506.1 | 1095614 | 1100480 | 4867 | + | Gnomon | gene | NA | NA | gene-LOC105286750 | GeneID:105286750 | LOC105286750 | Gene | LOC105286750 | protein_coding | character(0) | NA | character(0) |
| NC_039506.1 | 1098123 | 1098123 | 1 | + | 23 | 12 | 11 | 52.1739130434783 | NC_039506.1 | 1095614 | 1100480 | 4867 | + | Gnomon | gene | NA | NA | gene-LOC105286750 | GeneID:105286750 | LOC105286750 | Gene | LOC105286750 | protein_coding | character(0) | NA | character(0) |
| NC_039506.1 | 1098144 | 1098144 | 1 | + | 19 | 18 | 1 | 94.7368421052632 | NC_039506.1 | 1095614 | 1100480 | 4867 | + | Gnomon | gene | NA | NA | gene-LOC105286750 | GeneID:105286750 | LOC105286750 | Gene | LOC105286750 | protein_coding | character(0) | NA | character(0) |
| NC_039506.1 | 1098163 | 1098163 | 1 | + | 21 | 18 | 3 | 85.7142857142857 | NC_039506.1 | 1095614 | 1100480 | 4867 | + | Gnomon | gene | NA | NA | gene-LOC105286750 | GeneID:105286750 | LOC105286750 | Gene | LOC105286750 | protein_coding | character(0) | NA | character(0) |
| NC_039506.1 | 1098206 | 1098206 | 1 | + | 34 | 23 | 11 | 67.6470588235294 | NC_039506.1 | 1095614 | 1100480 | 4867 | + | Gnomon | gene | NA | NA | gene-LOC105286750 | GeneID:105286750 | LOC105286750 | Gene | LOC105286750 | protein_coding | character(0) | NA | character(0) |
| NC_039506.1 | 1098266 | 1098266 | 1 | + | 50 | 27 | 23 | 54 | NC_039506.1 | 1095614 | 1100480 | 4867 | + | Gnomon | gene | NA | NA | gene-LOC105286750 | GeneID:105286750 | LOC105286750 | Gene | LOC105286750 | protein_coding | character(0) | NA | character(0) |
| NC_039506.1 | 1098279 | 1098279 | 1 | + | 48 | 19 | 29 | 39.5833333333333 | NC_039506.1 | 1095614 | 1100480 | 4867 | + | Gnomon | gene | NA | NA | gene-LOC105286750 | GeneID:105286750 | LOC105286750 | Gene | LOC105286750 | protein_coding | character(0) | NA | character(0) |
| NC_039506.1 | 1098293 | 1098293 | 1 | + | 21 | 7 | 14 | 33.3333333333333 | NC_039506.1 | 1095614 | 1100480 | 4867 | + | Gnomon | gene | NA | NA | gene-LOC105286750 | GeneID:105286750 | LOC105286750 | Gene | LOC105286750 | protein_coding | character(0) | NA | character(0) |
| NC_039506.1 | 1098301 | 1098301 | 1 | + | 26 | 20 | 6 | 76.9230769230769 | NC_039506.1 | 1095614 | 1100480 | 4867 | + | Gnomon | gene | NA | NA | gene-LOC105286750 | GeneID:105286750 | LOC105286750 | Gene | LOC105286750 | protein_coding | character(0) | NA | character(0) |

...
... only first 25 lines shown ...

Directory: RNK File: sites-in-genes-Ob_LI_BCphaseB2.txt

| seqnames | start | end | width | strand | coverage | numCs | numTs | perc_meth | region_seqnames | region_start | region_end | region_width | region_strand | region_source | region_type | region_score | region_phase | region_ID | region_Dbxref | region_Name | region_gbkey | region_gene | region_gene_biotype | region_end_range | region_partial | region_start_range |
|---|---|---|---|---|---|---|---|---|---|---|---|---|---|---|---|---|---|---|---|---|---|---|---|---|---|---|
| NC_039506.1 | 1097588 | 1097588 | 1 | + | 25 | 11 | 14 | 44 | NC_039506.1 | 1095614 | 1100480 | 4867 | + | Gnomon | gene | NA | NA | gene-LOC105286750 | GeneID:105286750 | LOC105286750 | Gene | LOC105286750 | protein_coding | character(0) | NA | character(0) |
| NC_039506.1 | 1097598 | 1097598 | 1 | + | 27 | 14 | 13 | 51.8518518518518 | NC_039506.1 | 1095614 | 1100480 | 4867 | + | Gnomon | gene | NA | NA | gene-LOC105286750 | GeneID:105286750 | LOC105286750 | Gene | LOC105286750 | protein_coding | character(0) | NA | character(0) |
| NC_039506.1 | 1097702 | 1097702 | 1 | + | 33 | 8 | 25 | 24.2424242424242 | NC_039506.1 | 1095614 | 1100480 | 4867 | + | Gnomon | gene | NA | NA | gene-LOC105286750 | GeneID:105286750 | LOC105286750 | Gene | LOC105286750 | protein_coding | character(0) | NA | character(0) |
| NC_039506.1 | 1097773 | 1097773 | 1 | + | 32 | 31 | 1 | 96.875 | NC_039506.1 | 1095614 | 1100480 | 4867 | + | Gnomon | gene | NA | NA | gene-LOC105286750 | GeneID:105286750 | LOC105286750 | Gene | LOC105286750 | protein_coding | character(0) | NA | character(0) |
| NC_039506.1 | 1097981 | 1097981 | 1 | + | 7 | 7 | 0 | 100 | NC_039506.1 | 1095614 | 1100480 | 4867 | + | Gnomon | gene | NA | NA | gene-LOC105286750 | GeneID:105286750 | LOC105286750 | Gene | LOC105286750 | protein_coding | character(0) | NA | character(0) |
| NC_039506.1 | 1098055 | 1098055 | 1 | + | 11 | 10 | 1 | 90.9090909090909 | NC_039506.1 | 1095614 | 1100480 | 4867 | + | Gnomon | gene | NA | NA | gene-LOC105286750 | GeneID:105286750 | LOC105286750 | Gene | LOC105286750 | protein_coding | character(0) | NA | character(0) |
| NC_039506.1 | 1098056 | 1098056 | 1 | - | 11 | 11 | 0 | 100 | NC_039506.1 | 1095614 | 1100480 | 4867 | + | Gnomon | gene | NA | NA | gene-LOC105286750 | GeneID:105286750 | LOC105286750 | Gene | LOC105286750 | protein_coding | character(0) | NA | character(0) |
| NC_039506.1 | 1098069 | 1098069 | 1 | + | 11 | 10 | 1 | 90.9090909090909 | NC_039506.1 | 1095614 | 1100480 | 4867 | + | Gnomon | gene | NA | NA | gene-LOC105286750 | GeneID:105286750 | LOC105286750 | Gene | LOC105286750 | protein_coding | character(0) | NA | character(0) |
| NC_039506.1 | 1098070 | 1098070 | 1 | - | 10 | 8 | 2 | 80 | NC_039506.1 | 1095614 | 1100480 | 4867 | + | Gnomon | gene | NA | NA | gene-LOC105286750 | GeneID:105286750 | LOC105286750 | Gene | LOC105286750 | protein_coding | character(0) | NA | character(0) |
| NC_039506.1 | 1098123 | 1098123 | 1 | + | 21 | 11 | 10 | 52.3809523809524 | NC_039506.1 | 1095614 | 1100480 | 4867 | + | Gnomon | gene | NA | NA | gene-LOC105286750 | GeneID:105286750 | LOC105286750 | Gene | LOC105286750 | protein_coding | character(0) | NA | character(0) |
| NC_039506.1 | 1098144 | 1098144 | 1 | + | 18 | 18 | 0 | 100 | NC_039506.1 | 1095614 | 1100480 | 4867 | + | Gnomon | gene | NA | NA | gene-LOC105286750 | GeneID:105286750 | LOC105286750 | Gene | LOC105286750 | protein_coding | character(0) | NA | character(0) |
| NC_039506.1 | 1098163 | 1098163 | 1 | + | 13 | 12 | 1 | 92.3076923076923 | NC_039506.1 | 1095614 | 1100480 | 4867 | + | Gnomon | gene | NA | NA | gene-LOC105286750 | GeneID:105286750 | LOC105286750 | Gene | LOC105286750 | protein_coding | character(0) | NA | character(0) |
| NC_039506.1 | 1098206 | 1098206 | 1 | + | 29 | 17 | 12 | 58.6206896551724 | NC_039506.1 | 1095614 | 1100480 | 4867 | + | Gnomon | gene | NA | NA | gene-LOC105286750 | GeneID:105286750 | LOC105286750 | Gene | LOC105286750 | protein_coding | character(0) | NA | character(0) |
| NC_039506.1 | 1098266 | 1098266 | 1 | + | 54 | 31 | 23 | 57.4074074074074 | NC_039506.1 | 1095614 | 1100480 | 4867 | + | Gnomon | gene | NA | NA | gene-LOC105286750 | GeneID:105286750 | LOC105286750 | Gene | LOC105286750 | protein_coding | character(0) | NA | character(0) |
| NC_039506.1 | 1098279 | 1098279 | 1 | + | 59 | 27 | 32 | 45.7627118644068 | NC_039506.1 | 1095614 | 1100480 | 4867 | + | Gnomon | gene | NA | NA | gene-LOC105286750 | GeneID:105286750 | LOC105286750 | Gene | LOC105286750 | protein_coding | character(0) | NA | character(0) |
| NC_039506.1 | 1098293 | 1098293 | 1 | + | 35 | 9 | 26 | 25.7142857142857 | NC_039506.1 | 1095614 | 1100480 | 4867 | + | Gnomon | gene | NA | NA | gene-LOC105286750 | GeneID:105286750 | LOC105286750 | Gene | LOC105286750 | protein_coding | character(0) | NA | character(0) |
| NC_039506.1 | 1098301 | 1098301 | 1 | + | 28 | 16 | 12 | 57.1428571428571 | NC_039506.1 | 1095614 | 1100480 | 4867 | + | Gnomon | gene | NA | NA | gene-LOC105286750 | GeneID:105286750 | LOC105286750 | Gene | LOC105286750 | protein_coding | character(0) | NA | character(0) |
| NC_039506.1 | 1098330 | 1098330 | 1 | + | 46 | 9 | 37 | 19.5652173913043 | NC_039506.1 | 1095614 | 1100480 | 4867 | + | Gnomon | gene | NA | NA | gene-LOC105286750 | GeneID:105286750 | LOC105286750 | Gene | LOC105286750 | protein_coding | character(0) | NA | character(0) |
| NC_039506.1 | 1098334 | 1098334 | 1 | + | 45 | 27 | 18 | 60 | NC_039506.1 | 1095614 | 1100480 | 4867 | + | Gnomon | gene | NA | NA | gene-LOC105286750 | GeneID:105286750 | LOC105286750 | Gene | LOC105286750 | protein_coding | character(0) | NA | character(0) |
| NC_039506.1 | 1098377 | 1098377 | 1 | + | 55 | 26 | 29 | 47.2727272727273 | NC_039506.1 | 1095614 | 1100480 | 4867 | + | Gnomon | gene | NA | NA | gene-LOC105286750 | GeneID:105286750 | LOC105286750 | Gene | LOC105286750 | protein_coding | character(0) | NA | character(0) |
| NC_039506.1 | 1098387 | 1098387 | 1 | + | 55 | 30 | 25 | 54.5454545454545 | NC_039506.1 | 1095614 | 1100480 | 4867 | + | Gnomon | gene | NA | NA | gene-LOC105286750 | GeneID:105286750 | LOC105286750 | Gene | LOC105286750 | protein_coding | character(0) | NA | character(0) |
| NC_039506.1 | 1098400 | 1098400 | 1 | + | 45 | 21 | 24 | 46.6666666666667 | NC_039506.1 | 1095614 | 1100480 | 4867 | + | Gnomon | gene | NA | NA | gene-LOC105286750 | GeneID:105286750 | LOC105286750 | Gene | LOC105286750 | protein_coding | character(0) | NA | character(0) |
| NC_039506.1 | 1098420 | 1098420 | 1 | + | 29 | 9 | 20 | 31.0344827586207 | NC_039506.1 | 1095614 | 1100480 | 4867 | + | Gnomon | gene | NA | NA | gene-LOC105286750 | GeneID:105286750 | LOC105286750 | Gene | LOC105286750 | protein_coding | character(0) | NA | character(0) |
| NC_039506.1 | 1098427 | 1098427 | 1 | + | 31 | 17 | 14 | 54.8387096774194 | NC_039506.1 | 1095614 | 1100480 | 4867 | + | Gnomon | gene | NA | NA | gene-LOC105286750 | GeneID:105286750 | LOC105286750 | Gene | LOC105286750 | protein_coding | character(0) | NA | character(0) |

...
... only first 25 lines shown ...

Directory: RNK File: sites-in-genes-Ob_LI_RphaseA1.txt

```
seqnames	start	end	width	strand	coverage	numCs	numTs	perc_meth	region_seqnames	region_start	region_end	region_width	region_strand	region_source
	region_type	region_score	region_phase	region_ID	region_Dbxref	region_Name	region_gbkey	region_gene	region_gene_biotype	region_end_range	region_partial
	region_start_range
NC_039506.1	1097588	1097588	1	+	27	16	11	59.2592592592593	NC_039506.1	1095614	1100480	4867	+	Gnomon	gene	NA	NA	gene-LOC105286750
	GeneID:105286750	LOC105286750	Gene	LOC105286750	protein_coding	character(0)	NA	character(0)
NC_039506.1	1097598	1097598	1	+	31	16	15	51.6129032258064	NC_039506.1	1095614	1100480	4867	+	Gnomon	gene	NA	NA	gene-LOC105286750
	GeneID:105286750	LOC105286750	Gene	LOC105286750	protein_coding	character(0)	NA	character(0)
NC_039506.1	1097702	1097702	1	+	50	15	35	30	NC_039506.1	1095614	1100480	4867	+	Gnomon	gene	NA	NA	gene-LOC105286750
	GeneID:105286750	LOC105286750	Gene	LOC105286750	protein_coding	character(0)	NA	character(0)
NC_039506.1	1097703	1097703	1	-	30	11	19	36.6666666666667	NC_039506.1	1095614	1100480	4867	+	Gnomon	gene	NA	NA	gene-LOC105286750
	GeneID:105286750	LOC105286750	Gene	LOC105286750	protein_coding	character(0)	NA	character(0)
NC_039506.1	1097773	1097773	1	+	29	28	1	96.551724137931	NC_039506.1	1095614	1100480	4867	+	Gnomon	gene	NA	NA	gene-LOC105286750
	GeneID:105286750	LOC105286750	Gene	LOC105286750	protein_coding	character(0)	NA	character(0)
NC_039506.1	1097774	1097774	1	-	7	7	0	100	NC_039506.1	1095614	1100480	4867	+	Gnomon	gene	NA	NA	gene-LOC105286750
	GeneID:105286750	LOC105286750	Gene	LOC105286750	protein_coding	character(0)	NA	character(0)
NC_039506.1	1097925	1097925	1	-	6	6	0	100	NC_039506.1	1095614	1100480	4867	+	Gnomon	gene	NA	NA	gene-LOC105286750
	GeneID:105286750	LOC105286750	Gene	LOC105286750	protein_coding	character(0)	NA	character(0)
NC_039506.1	1097944	1097944	1	-	6	6	0	100	NC_039506.1	1095614	1100480	4867	+	Gnomon	gene	NA	NA	gene-LOC105286750
	GeneID:105286750	LOC105286750	Gene	LOC105286750	protein_coding	character(0)	NA	character(0)
NC_039506.1	1098055	1098055	1	+	11	11	0	100	NC_039506.1	1095614	1100480	4867	+	Gnomon	gene	NA	NA	gene-LOC105286750
	GeneID:105286750	LOC105286750	Gene	LOC105286750	protein_coding	character(0)	NA	character(0)
NC_039506.1	1098056	1098056	1	-	16	15	1	93.75	NC_039506.1	1095614	1100480	4867	+	Gnomon	gene	NA	NA	gene-LOC105286750
	GeneID:105286750	LOC105286750	Gene	LOC105286750	protein_coding	character(0)	NA	character(0)
NC_039506.1	1098069	1098069	1	+	14	14	0	100	NC_039506.1	1095614	1100480	4867	+	Gnomon	gene	NA	NA	gene-LOC105286750
	GeneID:105286750	LOC105286750	Gene	LOC105286750	protein_coding	character(0)	NA	character(0)
NC_039506.1	1098070	1098070	1	-	13	13	0	100	NC_039506.1	1095614	1100480	4867	+	Gnomon	gene	NA	NA	gene-LOC105286750
	GeneID:105286750	LOC105286750	Gene	LOC105286750	protein_coding	character(0)	NA	character(0)
NC_039506.1	1098123	1098123	1	+	24	17	7	70.8333333333333	NC_039506.1	1095614	1100480	4867	+	Gnomon	gene	NA	NA	gene-LOC105286750
	GeneID:105286750	LOC105286750	Gene	LOC105286750	protein_coding	character(0)	NA	character(0)
NC_039506.1	1098144	1098144	1	+	24	22	2	91.6666666666667	NC_039506.1	1095614	1100480	4867	+	Gnomon	gene	NA	NA	gene-LOC105286750
	GeneID:105286750	LOC105286750	Gene	LOC105286750	protein_coding	character(0)	NA	character(0)
NC_039506.1	1098163	1098163	1	+	21	16	5	76.1904761904762	NC_039506.1	1095614	1100480	4867	+	Gnomon	gene	NA	NA	gene-LOC105286750
	GeneID:105286750	LOC105286750	Gene	LOC105286750	protein_coding	character(0)	NA	character(0)
NC_039506.1	1098206	1098206	1	+	37	29	8	78.3783783783784	NC_039506.1	1095614	1100480	4867	+	Gnomon	gene	NA	NA	gene-LOC105286750
	GeneID:105286750	LOC105286750	Gene	LOC105286750	protein_coding	character(0)	NA	character(0)
NC_039506.1	1098266	1098266	1	+	57	32	25	56.140350877193	NC_039506.1	1095614	1100480	4867	+	Gnomon	gene	NA	NA	gene-LOC105286750
	GeneID:105286750	LOC105286750	Gene	LOC105286750	protein_coding	character(0)	NA	character(0)
NC_039506.1	1098279	1098279	1	+	56	17	39	30.3571428571429	NC_039506.1	1095614	1100480	4867	+	Gnomon	gene	NA	NA	gene-LOC105286750
	GeneID:105286750	LOC105286750	Gene	LOC105286750	protein_coding	character(0)	NA	character(0)
NC_039506.1	1098293	1098293	1	+	31	12	19	38.7096774193548	NC_039506.1	1095614	1100480	4867	+	Gnomon	gene	NA	NA	gene-LOC105286750
	GeneID:105286750	LOC105286750	Gene	LOC105286750	protein_coding	character(0)	NA	character(0)
NC_039506.1	1098301	1098301	1	+	32	17	15	53.125	NC_039506.1	1095614	1100480	4867	+	Gnomon	gene	NA	NA	gene-LOC105286750
	GeneID:105286750	LOC105286750	Gene	LOC105286750	protein_coding	character(0)	NA	character(0)
NC_039506.1	1098330	1098330	1	+	51	10	41	19.6078431372549	NC_039506.1	1095614	1100480	4867	+	Gnomon	gene	NA	NA	gene-LOC105286750
	GeneID:105286750	LOC105286750	Gene	LOC105286750	protein_coding	character(0)	NA	character(0)
NC_039506.1	1098334	1098334	1	+	53	23	30	43.3962264150943	NC_039506.1	1095614	1100480	4867	+	Gnomon	gene	NA	NA	gene-LOC105286750
	GeneID:105286750	LOC105286750	Gene	LOC105286750	protein_coding	character(0)	NA	character(0)
NC_039506.1	1098377	1098377	1	+	64	22	42	34.375	NC_039506.1	1095614	1100480	4867	+	Gnomon	gene	NA	NA	gene-LOC105286750
	GeneID:105286750	LOC105286750	Gene	LOC105286750	protein_coding	character(0)	NA	character(0)
NC_039506.1	1098387	1098387	1	+	54	29	25	53.7037037037037	NC_039506.1	1095614	1100480	4867	+	Gnomon	gene	NA	NA	gene-LOC105286750
	GeneID:105286750	LOC105286750	Gene	LOC105286750	protein_coding	character(0)	NA	character(0)
...
... only first 25 lines shown ...
```

Directory: RNK File: sites-in-genes-Ob_LI_RphaseA2.txt

```
seqnames    start end   width strand      coverage    numCs numTs perc_meth   region_seqnames   region_start      region_end  region_width      region_strand     region_source
      region_type region_score      region_phase      region_ID   region_Dbxref     region_Name region_gbkey      region_gene region_gene_biotype     region_end_range  region_partial
      region_start_range
NC_039506.1 138205      138205      1     +     30    8     22    26.6666666666667  NC_039506.1 134260      142204      7945  -     Gnomon      gene  NA    NA    gene-LOC105288138
      GeneID:105288138  LOC105288138      Gene  LOC105288138      protein_coding    character(0)      NA    character(0)
NC_039506.1 1097588     1097588     1     +     47    21    26    44.6808510638298  NC_039506.1 1095614     1100480     4867  +     Gnomon      gene  NA    NA    gene-LOC105286750
      GeneID:105286750  LOC105286750      Gene  LOC105286750      protein_coding    character(0)      NA    character(0)
NC_039506.1 1097598     1097598     1     +     50    29    21    58    NC_039506.1 1095614     1100480     4867  +     Gnomon      gene  NA    NA    gene-LOC105286750
      GeneID:105286750  LOC105286750      Gene  LOC105286750      protein_coding    character(0)      NA    character(0)
NC_039506.1 1097599     1097599     1     -     11    10    1     90.9090909090909  NC_039506.1 1095614     1100480     4867  +     Gnomon      gene  NA    NA    gene-LOC105286750
      GeneID:105286750  LOC105286750      Gene  LOC105286750      protein_coding    character(0)      NA    character(0)
NC_039506.1 1097647     1097647     1     +     72    13    59    18.0555555555556  NC_039506.1 1095614     1100480     4867  +     Gnomon      gene  NA    NA    gene-LOC105286750
      GeneID:105286750  LOC105286750      Gene  LOC105286750      protein_coding    character(0)      NA    character(0)
NC_039506.1 1097702     1097702     1     +     87    10    77    11.4942528735632  NC_039506.1 1095614     1100480     4867  +     Gnomon      gene  NA    NA    gene-LOC105286750
      GeneID:105286750  LOC105286750      Gene  LOC105286750      protein_coding    character(0)      NA    character(0)
NC_039506.1 1097703     1097703     1     -     21    7     14    33.3333333333333  NC_039506.1 1095614     1100480     4867  +     Gnomon      gene  NA    NA    gene-LOC105286750
      GeneID:105286750  LOC105286750      Gene  LOC105286750      protein_coding    character(0)      NA    character(0)
NC_039506.1 1097773     1097773     1     +     74    69    5     93.2432432432432  NC_039506.1 1095614     1100480     4867  +     Gnomon      gene  NA    NA    gene-LOC105286750
      GeneID:105286750  LOC105286750      Gene  LOC105286750      protein_coding    character(0)      NA    character(0)
NC_039506.1 1097774     1097774     1     -     6     6     0     100   NC_039506.1 1095614     1100480     4867  +     Gnomon      gene  NA    NA    gene-LOC105286750
      GeneID:105286750  LOC105286750      Gene  LOC105286750      protein_coding    character(0)      NA    character(0)
NC_039506.1 1097924     1097924     1     +     6     6     0     100   NC_039506.1 1095614     1100480     4867  +     Gnomon      gene  NA    NA    gene-LOC105286750
      GeneID:105286750  LOC105286750      Gene  LOC105286750      protein_coding    character(0)      NA    character(0)
NC_039506.1 1097943     1097943     1     +     6     6     0     100   NC_039506.1 1095614     1100480     4867  +     Gnomon      gene  NA    NA    gene-LOC105286750
      GeneID:105286750  LOC105286750      Gene  LOC105286750      protein_coding    character(0)      NA    character(0)
NC_039506.1 1097981     1097981     1     +     12    12    0     100   NC_039506.1 1095614     1100480     4867  +     Gnomon      gene  NA    NA    gene-LOC105286750
      GeneID:105286750  LOC105286750      Gene  LOC105286750      protein_coding    character(0)      NA    character(0)
NC_039506.1 1098055     1098055     1     +     26    25    1     96.1538461538462  NC_039506.1 1095614     1100480     4867  +     Gnomon      gene  NA    NA    gene-LOC105286750
      GeneID:105286750  LOC105286750      Gene  LOC105286750      protein_coding    character(0)      NA    character(0)
NC_039506.1 1098056     1098056     1     -     11    10    1     90.9090909090909  NC_039506.1 1095614     1100480     4867  +     Gnomon      gene  NA    NA    gene-LOC105286750
      GeneID:105286750  LOC105286750      Gene  LOC105286750      protein_coding    character(0)      NA    character(0)
NC_039506.1 1098069     1098069     1     +     29    29    0     100   NC_039506.1 1095614     1100480     4867  +     Gnomon      gene  NA    NA    gene-LOC105286750
      GeneID:105286750  LOC105286750      Gene  LOC105286750      protein_coding    character(0)      NA    character(0)
NC_039506.1 1098070     1098070     1     -     9     9     0     100   NC_039506.1 1095614     1100480     4867  +     Gnomon      gene  NA    NA    gene-LOC105286750
      GeneID:105286750  LOC105286750      Gene  LOC105286750      protein_coding    character(0)      NA    character(0)
NC_039506.1 1098123     1098123     1     +     42    26    16    61.9047619047619  NC_039506.1 1095614     1100480     4867  +     Gnomon      gene  NA    NA    gene-LOC105286750
      GeneID:105286750  LOC105286750      Gene  LOC105286750      protein_coding    character(0)      NA    character(0)
NC_039506.1 1098144     1098144     1     +     25    21    4     84    NC_039506.1 1095614     1100480     4867  +     Gnomon      gene  NA    NA    gene-LOC105286750
      GeneID:105286750  LOC105286750      Gene  LOC105286750      protein_coding    character(0)      NA    character(0)
NC_039506.1 1098163     1098163     1     +     28    23    5     82.1428571428571  NC_039506.1 1095614     1100480     4867  +     Gnomon      gene  NA    NA    gene-LOC105286750
      GeneID:105286750  LOC105286750      Gene  LOC105286750      protein_coding    character(0)      NA    character(0)
NC_039506.1 1098206     1098206     1     +     43    27    16    62.7906976744186  NC_039506.1 1095614     1100480     4867  +     Gnomon      gene  NA    NA    gene-LOC105286750
      GeneID:105286750  LOC105286750      Gene  LOC105286750      protein_coding    character(0)      NA    character(0)
NC_039506.1 1098266     1098266     1     +     70    43    27    61.4285714285714  NC_039506.1 1095614     1100480     4867  +     Gnomon      gene  NA    NA    gene-LOC105286750
      GeneID:105286750  LOC105286750      Gene  LOC105286750      protein_coding    character(0)      NA    character(0)
NC_039506.1 1098279     1098279     1     +     72    33    39    45.8333333333333  NC_039506.1 1095614     1100480     4867  +     Gnomon      gene  NA    NA    gene-LOC105286750
      GeneID:105286750  LOC105286750      Gene  LOC105286750      protein_coding    character(0)      NA    character(0)
NC_039506.1 1098293     1098293     1     +     43    18    25    41.8604651162791  NC_039506.1 1095614     1100480     4867  +     Gnomon      gene  NA    NA    gene-LOC105286750
      GeneID:105286750  LOC105286750      Gene  LOC105286750      protein_coding    character(0)      NA    character(0)
NC_039506.1 1098301     1098301     1     +     36    19    17    52.7777777777778  NC_039506.1 1095614     1100480     4867  +     Gnomon      gene  NA    NA    gene-LOC105286750
      GeneID:105286750  LOC105286750      Gene  LOC105286750      protein_coding    character(0)      NA    character(0)
...
... only first 25 lines shown ...
```

Directory: RNK File: sites-in-genes-Ob_LI_RphaseB1.txt

```
seqnames	start	end	width	strand	coverage	numCs	numTs	perc_meth	region_seqnames	region_start	region_end	region_width	region_strand	region_source
	region_type	region_score	region_phase	region_ID	region_Dbxref	region_Name	region_gbkey	region_gene	region_gene_biotype	region_end_range	region_partial
	region_start_range
NC_039506.1	463932	463932	1	-	12	7	5	58.3333333333333	NC_039506.1	458174	503061	44888	-	Gnomon	gene	NA	NA	gene-LOC105285445
	GeneID:105285445	LOC105285445	Gene	LOC105285445	protein_coding	character(0)	NA	character(0)
NC_039506.1	1058205	1058205	1	+	21	9	12	42.8571428571429	NC_039506.1	1056151	1064257	8107	+	Gnomon	gene	NA	NA	gene-LOC105286747
	GeneID:105286747	LOC105286747	Gene	LOC105286747	lncRNA	character(0)	NA	character(0)
NC_039506.1	1097588	1097588	1	+	33	9	24	27.2727272727273	NC_039506.1	1095614	1100480	4867	+	Gnomon	gene	NA	NA	gene-LOC105286750
	GeneID:105286750	LOC105286750	Gene	LOC105286750	protein_coding	character(0)	NA	character(0)
NC_039506.1	1097598	1097598	1	+	38	19	19	50	NC_039506.1	1095614	1100480	4867	+	Gnomon	gene	NA	NA	gene-LOC105286750
	GeneID:105286750	LOC105286750	Gene	LOC105286750	protein_coding	character(0)	NA	character(0)
NC_039506.1	1097702	1097702	1	+	43	11	32	25.5813953488372	NC_039506.1	1095614	1100480	4867	+	Gnomon	gene	NA	NA	gene-LOC105286750
	GeneID:105286750	LOC105286750	Gene	LOC105286750	protein_coding	character(0)	NA	character(0)
NC_039506.1	1097773	1097773	1	+	30	28	2	93.3333333333333	NC_039506.1	1095614	1100480	4867	+	Gnomon	gene	NA	NA	gene-LOC105286750
	GeneID:105286750	LOC105286750	Gene	LOC105286750	protein_coding	character(0)	NA	character(0)
NC_039506.1	1097774	1097774	1	-	6	5	1	83.3333333333333	NC_039506.1	1095614	1100480	4867	+	Gnomon	gene	NA	NA	gene-LOC105286750
	GeneID:105286750	LOC105286750	Gene	LOC105286750	protein_coding	character(0)	NA	character(0)
NC_039506.1	1097924	1097924	1	+	6	6	0	100	NC_039506.1	1095614	1100480	4867	+	Gnomon	gene	NA	NA	gene-LOC105286750
	GeneID:105286750	LOC105286750	Gene	LOC105286750	protein_coding	character(0)	NA	character(0)
NC_039506.1	1097943	1097943	1	+	6	5	1	83.3333333333333	NC_039506.1	1095614	1100480	4867	+	Gnomon	gene	NA	NA	gene-LOC105286750
	GeneID:105286750	LOC105286750	Gene	LOC105286750	protein_coding	character(0)	NA	character(0)
NC_039506.1	1097944	1097944	1	-	5	5	0	100	NC_039506.1	1095614	1100480	4867	+	Gnomon	gene	NA	NA	gene-LOC105286750
	GeneID:105286750	LOC105286750	Gene	LOC105286750	protein_coding	character(0)	NA	character(0)
NC_039506.1	1097981	1097981	1	+	8	8	0	100	NC_039506.1	1095614	1100480	4867	+	Gnomon	gene	NA	NA	gene-LOC105286750
	GeneID:105286750	LOC105286750	Gene	LOC105286750	protein_coding	character(0)	NA	character(0)
NC_039506.1	1098055	1098055	1	+	11	11	0	100	NC_039506.1	1095614	1100480	4867	+	Gnomon	gene	NA	NA	gene-LOC105286750
	GeneID:105286750	LOC105286750	Gene	LOC105286750	protein_coding	character(0)	NA	character(0)
NC_039506.1	1098056	1098056	1	-	19	18	1	94.7368421052632	NC_039506.1	1095614	1100480	4867	+	Gnomon	gene	NA	NA	gene-LOC105286750
	GeneID:105286750	LOC105286750	Gene	LOC105286750	protein_coding	character(0)	NA	character(0)
NC_039506.1	1098069	1098069	1	+	12	12	0	100	NC_039506.1	1095614	1100480	4867	+	Gnomon	gene	NA	NA	gene-LOC105286750
	GeneID:105286750	LOC105286750	Gene	LOC105286750	protein_coding	character(0)	NA	character(0)
NC_039506.1	1098070	1098070	1	-	17	17	0	100	NC_039506.1	1095614	1100480	4867	+	Gnomon	gene	NA	NA	gene-LOC105286750
	GeneID:105286750	LOC105286750	Gene	LOC105286750	protein_coding	character(0)	NA	character(0)
NC_039506.1	1098123	1098123	1	+	21	14	7	66.6666666666667	NC_039506.1	1095614	1100480	4867	+	Gnomon	gene	NA	NA	gene-LOC105286750
	GeneID:105286750	LOC105286750	Gene	LOC105286750	protein_coding	character(0)	NA	character(0)
NC_039506.1	1098124	1098124	1	-	9	8	1	88.8888888888889	NC_039506.1	1095614	1100480	4867	+	Gnomon	gene	NA	NA	gene-LOC105286750
	GeneID:105286750	LOC105286750	Gene	LOC105286750	protein_coding	character(0)	NA	character(0)
NC_039506.1	1098144	1098144	1	+	17	16	1	94.1176470588235	NC_039506.1	1095614	1100480	4867	+	Gnomon	gene	NA	NA	gene-LOC105286750
	GeneID:105286750	LOC105286750	Gene	LOC105286750	protein_coding	character(0)	NA	character(0)
NC_039506.1	1098145	1098145	1	-	6	6	0	100	NC_039506.1	1095614	1100480	4867	+	Gnomon	gene	NA	NA	gene-LOC105286750
	GeneID:105286750	LOC105286750	Gene	LOC105286750	protein_coding	character(0)	NA	character(0)
NC_039506.1	1098163	1098163	1	+	22	18	4	81.8181818181818	NC_039506.1	1095614	1100480	4867	+	Gnomon	gene	NA	NA	gene-LOC105286750
	GeneID:105286750	LOC105286750	Gene	LOC105286750	protein_coding	character(0)	NA	character(0)
NC_039506.1	1098164	1098164	1	-	7	7	0	100	NC_039506.1	1095614	1100480	4867	+	Gnomon	gene	NA	NA	gene-LOC105286750
	GeneID:105286750	LOC105286750	Gene	LOC105286750	protein_coding	character(0)	NA	character(0)
NC_039506.1	1098206	1098206	1	+	24	16	8	66.6666666666667	NC_039506.1	1095614	1100480	4867	+	Gnomon	gene	NA	NA	gene-LOC105286750
	GeneID:105286750	LOC105286750	Gene	LOC105286750	protein_coding	character(0)	NA	character(0)
NC_039506.1	1098266	1098266	1	+	48	27	21	56.25	NC_039506.1	1095614	1100480	4867	+	Gnomon	gene	NA	NA	gene-LOC105286750
	GeneID:105286750	LOC105286750	Gene	LOC105286750	protein_coding	character(0)	NA	character(0)
NC_039506.1	1098279	1098279	1	+	51	23	28	45.0980392156863	NC_039506.1	1095614	1100480	4867	+	Gnomon	gene	NA	NA	gene-LOC105286750
	GeneID:105286750	LOC105286750	Gene	LOC105286750	protein_coding	character(0)	NA	character(0)
...
... only first 25 lines shown ...
```

Directory: RNK File: sites-in-genes-Ob_LI_RphaseB2.txt

| seqnames | start | end | width | strand | coverage | numCs | numTs | perc_meth | region_seqnames | region_start | region_end | region_width | region_strand | region_source | region_type | region_score | region_phase | region_ID | region_Dbxref | region_Name | region_gbkey | region_gene | region_gene_biotype | region_end_range | region_partial | region_start_range |
|---|---|---|---|---|---|---|---|---|---|---|---|---|---|---|---|---|---|---|---|---|---|---|---|---|---|---|
| NC_039506.1 | 1097588 | 1097588 | 1 | + | 25 | 9 | 16 | 36 | NC_039506.1 | 1095614 | 1100480 | 4867 | + | Gnomon | gene | NA | NA | gene-LOC105286750 | GeneID:105286750 | LOC105286750 | Gene | LOC105286750 | protein_coding | character(0) | NA | character(0) |
| NC_039506.1 | 1097598 | 1097598 | 1 | + | 27 | 18 | 9 | 66.6666666666667 | NC_039506.1 | 1095614 | 1100480 | 4867 | + | Gnomon | gene | NA | NA | gene-LOC105286750 | GeneID:105286750 | LOC105286750 | Gene | LOC105286750 | protein_coding | character(0) | NA | character(0) |
| NC_039506.1 | 1097703 | 1097703 | 1 | - | 15 | 6 | 9 | 40 | NC_039506.1 | 1095614 | 1100480 | 4867 | + | Gnomon | gene | NA | NA | gene-LOC105286750 | GeneID:105286750 | LOC105286750 | Gene | LOC105286750 | protein_coding | character(0) | NA | character(0) |
| NC_039506.1 | 1097773 | 1097773 | 1 | + | 38 | 37 | 1 | 97.3684210526316 | NC_039506.1 | 1095614 | 1100480 | 4867 | + | Gnomon | gene | NA | NA | gene-LOC105286750 | GeneID:105286750 | LOC105286750 | Gene | LOC105286750 | protein_coding | character(0) | NA | character(0) |
| NC_039506.1 | 1098055 | 1098055 | 1 | + | 9 | 9 | 0 | 100 | NC_039506.1 | 1095614 | 1100480 | 4867 | + | Gnomon | gene | NA | NA | gene-LOC105286750 | GeneID:105286750 | LOC105286750 | Gene | LOC105286750 | protein_coding | character(0) | NA | character(0) |
| NC_039506.1 | 1098056 | 1098056 | 1 | - | 7 | 7 | 0 | 100 | NC_039506.1 | 1095614 | 1100480 | 4867 | + | Gnomon | gene | NA | NA | gene-LOC105286750 | GeneID:105286750 | LOC105286750 | Gene | LOC105286750 | protein_coding | character(0) | NA | character(0) |
| NC_039506.1 | 1098069 | 1098069 | 1 | + | 10 | 9 | 1 | 90 | NC_039506.1 | 1095614 | 1100480 | 4867 | + | Gnomon | gene | NA | NA | gene-LOC105286750 | GeneID:105286750 | LOC105286750 | Gene | LOC105286750 | protein_coding | character(0) | NA | character(0) |
| NC_039506.1 | 1098070 | 1098070 | 1 | - | 5 | 5 | 0 | 100 | NC_039506.1 | 1095614 | 1100480 | 4867 | + | Gnomon | gene | NA | NA | gene-LOC105286750 | GeneID:105286750 | LOC105286750 | Gene | LOC105286750 | protein_coding | character(0) | NA | character(0) |
| NC_039506.1 | 1098123 | 1098123 | 1 | + | 18 | 11 | 7 | 61.1111111111111 | NC_039506.1 | 1095614 | 1100480 | 4867 | + | Gnomon | gene | NA | NA | gene-LOC105286750 | GeneID:105286750 | LOC105286750 | Gene | LOC105286750 | protein_coding | character(0) | NA | character(0) |
| NC_039506.1 | 1098144 | 1098144 | 1 | + | 16 | 16 | 0 | 100 | NC_039506.1 | 1095614 | 1100480 | 4867 | + | Gnomon | gene | NA | NA | gene-LOC105286750 | GeneID:105286750 | LOC105286750 | Gene | LOC105286750 | protein_coding | character(0) | NA | character(0) |
| NC_039506.1 | 1098163 | 1098163 | 1 | + | 15 | 15 | 0 | 100 | NC_039506.1 | 1095614 | 1100480 | 4867 | + | Gnomon | gene | NA | NA | gene-LOC105286750 | GeneID:105286750 | LOC105286750 | Gene | LOC105286750 | protein_coding | character(0) | NA | character(0) |
| NC_039506.1 | 1098206 | 1098206 | 1 | + | 28 | 17 | 11 | 60.7142857142857 | NC_039506.1 | 1095614 | 1100480 | 4867 | + | Gnomon | gene | NA | NA | gene-LOC105286750 | GeneID:105286750 | LOC105286750 | Gene | LOC105286750 | protein_coding | character(0) | NA | character(0) |
| NC_039506.1 | 1098266 | 1098266 | 1 | + | 45 | 27 | 18 | 60 | NC_039506.1 | 1095614 | 1100480 | 4867 | + | Gnomon | gene | NA | NA | gene-LOC105286750 | GeneID:105286750 | LOC105286750 | Gene | LOC105286750 | protein_coding | character(0) | NA | character(0) |
| NC_039506.1 | 1098279 | 1098279 | 1 | + | 46 | 17 | 29 | 36.9565217391304 | NC_039506.1 | 1095614 | 1100480 | 4867 | + | Gnomon | gene | NA | NA | gene-LOC105286750 | GeneID:105286750 | LOC105286750 | Gene | LOC105286750 | protein_coding | character(0) | NA | character(0) |
| NC_039506.1 | 1098293 | 1098293 | 1 | + | 29 | 11 | 18 | 37.9310344827586 | NC_039506.1 | 1095614 | 1100480 | 4867 | + | Gnomon | gene | NA | NA | gene-LOC105286750 | GeneID:105286750 | LOC105286750 | Gene | LOC105286750 | protein_coding | character(0) | NA | character(0) |
| NC_039506.1 | 1098301 | 1098301 | 1 | + | 23 | 16 | 7 | 69.5652173913043 | NC_039506.1 | 1095614 | 1100480 | 4867 | + | Gnomon | gene | NA | NA | gene-LOC105286750 | GeneID:105286750 | LOC105286750 | Gene | LOC105286750 | protein_coding | character(0) | NA | character(0) |
| NC_039506.1 | 1098330 | 1098330 | 1 | + | 56 | 18 | 38 | 32.1428571428571 | NC_039506.1 | 1095614 | 1100480 | 4867 | + | Gnomon | gene | NA | NA | gene-LOC105286750 | GeneID:105286750 | LOC105286750 | Gene | LOC105286750 | protein_coding | character(0) | NA | character(0) |
| NC_039506.1 | 1098334 | 1098334 | 1 | + | 58 | 40 | 18 | 68.9655172413793 | NC_039506.1 | 1095614 | 1100480 | 4867 | + | Gnomon | gene | NA | NA | gene-LOC105286750 | GeneID:105286750 | LOC105286750 | Gene | LOC105286750 | protein_coding | character(0) | NA | character(0) |
| NC_039506.1 | 1098377 | 1098377 | 1 | + | 61 | 20 | 41 | 32.7868852459016 | NC_039506.1 | 1095614 | 1100480 | 4867 | + | Gnomon | gene | NA | NA | gene-LOC105286750 | GeneID:105286750 | LOC105286750 | Gene | LOC105286750 | protein_coding | character(0) | NA | character(0) |
| NC_039506.1 | 1098387 | 1098387 | 1 | + | 57 | 25 | 32 | 43.859649122807 | NC_039506.1 | 1095614 | 1100480 | 4867 | + | Gnomon | gene | NA | NA | gene-LOC105286750 | GeneID:105286750 | LOC105286750 | Gene | LOC105286750 | protein_coding | character(0) | NA | character(0) |
| NC_039506.1 | 1098400 | 1098400 | 1 | + | 46 | 27 | 19 | 58.695652173913 | NC_039506.1 | 1095614 | 1100480 | 4867 | + | Gnomon | gene | NA | NA | gene-LOC105286750 | GeneID:105286750 | LOC105286750 | Gene | LOC105286750 | protein_coding | character(0) | NA | character(0) |
| NC_039506.1 | 1098420 | 1098420 | 1 | + | 27 | 14 | 13 | 51.8518518518518 | NC_039506.1 | 1095614 | 1100480 | 4867 | + | Gnomon | gene | NA | NA | gene-LOC105286750 | GeneID:105286750 | LOC105286750 | Gene | LOC105286750 | protein_coding | character(0) | NA | character(0) |
| NC_039506.1 | 1098427 | 1098427 | 1 | + | 29 | 18 | 11 | 62.0689655172414 | NC_039506.1 | 1095614 | 1100480 | 4867 | + | Gnomon | gene | NA | NA | gene-LOC105286750 | GeneID:105286750 | LOC105286750 | Gene | LOC105286750 | protein_coding | character(0) | NA | character(0) |
| NC_039506.1 | 1098639 | 1098639 | 1 | + | 12 | 7 | 5 | 58.3333333333333 | NC_039506.1 | 1095614 | 1100480 | 4867 | + | Gnomon | gene | NA | NA | gene-LOC105286750 | GeneID:105286750 | LOC105286750 | Gene | LOC105286750 | protein_coding | character(0) | NA | character(0) |

...
... only first 25 lines shown ...

Directory: RNK File: sites-in-promoters-Ob_LI_BCphaseA1.txt

| seqnames | start | end | width | strand | coverage | numCs | numTs | perc_meth | region_seqnames | region_start | region_end | region_width | region_strand | region_source | region_type | region_score | region_phase | region_ID | region_Dbxref | region_Name | region_gbkey | region_gene | region_gene_biotype | region_end_range | region_partial | region_start_range |
|---|---|---|---|---|---|---|---|---|---|---|---|---|---|---|---|---|---|---|---|---|---|---|---|---|---|---|
| NC_039506.1 | 1664198 | 1664198 | 1 | + | 31 | 8 | 23 | 25.8064516129032 | NC_039506.1 | 1664107 | 1664606 | 500 | + | Gnomon | promoter | NA | NA | gene-LOC105280261 | GeneID:105280261 | LOC105280261 | Gene | LOC105280261 | protein_coding | character(0) | NA | character(0) |
| NC_039506.1 | 1664400 | 1664400 | 1 | + | 16 | 7 | 9 | 43.75 | NC_039506.1 | 1664107 | 1664606 | 500 | + | Gnomon | promoter | NA | NA | gene-LOC105280261 | GeneID:105280261 | LOC105280261 | Gene | LOC105280261 | protein_coding | character(0) | NA | character(0) |
| NC_039506.1 | 1673111 | 1673111 | 1 | - | 15 | 13 | 2 | 86.6666666666667 | NC_039506.1 | 1673027 | 1673526 | 500 | + | Gnomon | promoter | NA | NA | gene-LOC109611032 | GeneID:109611032 | LOC109611032 | Gene | LOC109611032 | lncRNA | character(0) | NA | character(0) |
| NC_039506.1 | 1673135 | 1673135 | 1 | - | 14 | 12 | 2 | 85.7142857142857 | NC_039506.1 | 1673027 | 1673526 | 500 | + | Gnomon | promoter | NA | NA | gene-LOC109611032 | GeneID:109611032 | LOC109611032 | Gene | LOC109611032 | lncRNA | character(0) | NA | character(0) |
| NC_039506.1 | 1673160 | 1673160 | 1 | - | 11 | 8 | 3 | 72.7272727272727 | NC_039506.1 | 1673027 | 1673526 | 500 | + | Gnomon | promoter | NA | NA | gene-LOC109611032 | GeneID:109611032 | LOC109611032 | Gene | LOC109611032 | lncRNA | character(0) | NA | character(0) |
| NC_039506.1 | 1673189 | 1673189 | 1 | - | 19 | 15 | 4 | 78.9473684210526 | NC_039506.1 | 1673027 | 1673526 | 500 | + | Gnomon | promoter | NA | NA | gene-LOC109611032 | GeneID:109611032 | LOC109611032 | Gene | LOC109611032 | lncRNA | character(0) | NA | character(0) |
| NC_039506.1 | 1673197 | 1673197 | 1 | - | 22 | 8 | 14 | 36.3636363636364 | NC_039506.1 | 1673027 | 1673526 | 500 | + | Gnomon | promoter | NA | NA | gene-LOC109611032 | GeneID:109611032 | LOC109611032 | Gene | LOC109611032 | lncRNA | character(0) | NA | character(0) |
| NC_039506.1 | 1673205 | 1673205 | 1 | - | 31 | 29 | 2 | 93.5483870967742 | NC_039506.1 | 1673027 | 1673526 | 500 | + | Gnomon | promoter | NA | NA | gene-LOC109611032 | GeneID:109611032 | LOC109611032 | Gene | LOC109611032 | lncRNA | character(0) | NA | character(0) |
| NC_039506.1 | 1673219 | 1673219 | 1 | - | 46 | 12 | 34 | 26.0869565217391 | NC_039506.1 | 1673027 | 1673526 | 500 | + | Gnomon | promoter | NA | NA | gene-LOC109611032 | GeneID:109611032 | LOC109611032 | Gene | LOC109611032 | lncRNA | character(0) | NA | character(0) |
| NC_039506.1 | 1673225 | 1673225 | 1 | - | 44 | 32 | 12 | 72.7272727272727 | NC_039506.1 | 1673027 | 1673526 | 500 | + | Gnomon | promoter | NA | NA | gene-LOC109611032 | GeneID:109611032 | LOC109611032 | Gene | LOC109611032 | lncRNA | character(0) | NA | character(0) |
| NC_039506.1 | 1673234 | 1673234 | 1 | - | 47 | 28 | 19 | 59.5744680851064 | NC_039506.1 | 1673027 | 1673526 | 500 | + | Gnomon | promoter | NA | NA | gene-LOC109611032 | GeneID:109611032 | LOC109611032 | Gene | LOC109611032 | lncRNA | character(0) | NA | character(0) |
| NC_039506.1 | 1673241 | 1673241 | 1 | - | 47 | 30 | 17 | 63.8297872340426 | NC_039506.1 | 1673027 | 1673526 | 500 | + | Gnomon | promoter | NA | NA | gene-LOC109611032 | GeneID:109611032 | LOC109611032 | Gene | LOC109611032 | lncRNA | character(0) | NA | character(0) |
| NC_039506.1 | 1673284 | 1673284 | 1 | - | 38 | 31 | 7 | 81.5789473684211 | NC_039506.1 | 1673027 | 1673526 | 500 | + | Gnomon | promoter | NA | NA | gene-LOC109611032 | GeneID:109611032 | LOC109611032 | Gene | LOC109611032 | lncRNA | character(0) | NA | character(0) |
| NC_039506.1 | 1673353 | 1673353 | 1 | - | 21 | 21 | 0 | 100 | NC_039506.1 | 1673027 | 1673526 | 500 | + | Gnomon | promoter | NA | NA | gene-LOC109611032 | GeneID:109611032 | LOC109611032 | Gene | LOC109611032 | lncRNA | character(0) | NA | character(0) |
| NC_039506.1 | 1673389 | 1673389 | 1 | - | 69 | 68 | 1 | 98.5507246376812 | NC_039506.1 | 1673027 | 1673526 | 500 | + | Gnomon | promoter | NA | NA | gene-LOC109611032 | GeneID:109611032 | LOC109611032 | Gene | LOC109611032 | lncRNA | character(0) | NA | character(0) |
| NC_039506.1 | 1673463 | 1673463 | 1 | + | 6 | 6 | 0 | 100 | NC_039506.1 | 1673027 | 1673526 | 500 | + | Gnomon | promoter | NA | NA | gene-LOC109611032 | GeneID:109611032 | LOC109611032 | Gene | LOC109611032 | lncRNA | character(0) | NA | character(0) |
| NC_039506.1 | 1673464 | 1673464 | 1 | - | 47 | 39 | 8 | 82.9787234042553 | NC_039506.1 | 1673027 | 1673526 | 500 | + | Gnomon | promoter | NA | NA | gene-LOC109611032 | GeneID:109611032 | LOC109611032 | Gene | LOC109611032 | lncRNA | character(0) | NA | character(0) |
| NC_039506.1 | 1673488 | 1673488 | 1 | - | 37 | 30 | 7 | 81.0810810810811 | NC_039506.1 | 1673027 | 1673526 | 500 | + | Gnomon | promoter | NA | NA | gene-LOC109611032 | GeneID:109611032 | LOC109611032 | Gene | LOC109611032 | lncRNA | character(0) | NA | character(0) |
| NC_039506.1 | 1673511 | 1673511 | 1 | - | 30 | 26 | 4 | 86.6666666666667 | NC_039506.1 | 1673027 | 1673526 | 500 | + | Gnomon | promoter | NA | NA | gene-LOC109611032 | GeneID:109611032 | LOC109611032 | Gene | LOC109611032 | lncRNA | character(0) | NA | character(0) |
| NC_039506.1 | 2172293 | 2172293 | 1 | - | 12 | 7 | 5 | 58.3333333333333 | NC_039506.1 | 2172291 | 2172790 | 500 | + | Gnomon | promoter | NA | NA | gene-LOC105280304 | GeneID:105280304 | LOC105280304 | Gene | LOC105280304 | protein_coding | character(0) | NA | character(0) |
| NC_039506.1 | 2187052 | 2187052 | 1 | + | 7 | 5 | 2 | 71.4285714285714 | NC_039506.1 | 2187023 | 2187522 | 500 | + | Gnomon | promoter | NA | NA | gene-LOC105280309 | GeneID:105280309 | LOC105280309 | Gene | LOC105280309 | protein_coding | character(0) | NA | character(0) |
| NC_039506.1 | 2187053 | 2187053 | 1 | - | 6 | 5 | 1 | 83.3333333333333 | NC_039506.1 | 2187023 | 2187522 | 500 | + | Gnomon | promoter | NA | NA | gene-LOC105280309 | GeneID:105280309 | LOC105280309 | Gene | LOC105280309 | protein_coding | character(0) | NA | character(0) |
| NC_039506.1 | 2200872 | 2200872 | 1 | - | 8 | 5 | 3 | 62.5 | NC_039506.1 | 2200872 | 2201371 | 500 | + | Gnomon | promoter | NA | NA | gene-LOC105280312 | GeneID:105280312 | LOC105280312 | Gene | LOC105280312 | protein_coding | character(0) | NA | character(0) |
| NC_039506.1 | 3094044 | 3094044 | 1 | + | 19 | 9 | 10 | 47.3684210526316 | NC_039506.1 | 3094025 | 3094524 | 500 | + | Gnomon | promoter | NA | NA | gene-LOC105280352 | GeneID:105280352 | LOC105280352 | Gene | LOC105280352 | protein_coding | character(0) | NA | character(0) |

...

... only first 25 lines shown ...

Directory: RNK File: sites-in-promoters-Ob_LI_BCphaseA2.txt

seqnames start end width strand coverage numCs numTs perc_meth region_seqnames region_start region_end region_width region_strand region_source
region_type region_score region_phase region_ID region_Dbxref region_Name region_gbkey region_gene region_gene_biotype region_end_range region_partial
region_start_range
NC_039506.1 1664198 1664198 1 + 30 11 19 36.6666666666667 NC_039506.1 1664107 1664606 500 + Gnomon promoter NA NA gene-
LOC105280261 GeneID:105280261 LOC105280261 Gene LOC105280261 protein_coding character(0) NA character(0)
NC_039506.1 1664241 1664241 1 + 35 7 28 20 NC_039506.1 1664107 1664606 500 + Gnomon promoter NA NA gene-LOC105280261
GeneID:105280261 LOC105280261 Gene LOC105280261 protein_coding character(0) NA character(0)
NC_039506.1 1664392 1664392 1 + 16 6 10 37.5 NC_039506.1 1664107 1664606 500 + Gnomon promoter NA NA gene-LOC105280261
GeneID:105280261 LOC105280261 Gene LOC105280261 protein_coding character(0) NA character(0)
NC_039506.1 1673110 1673110 1 + 5 5 0 100 NC_039506.1 1673027 1673526 500 + Gnomon promoter NA NA gene-LOC109611032
GeneID:109611032 LOC109611032 Gene LOC109611032 lncRNA character(0) NA character(0)
NC_039506.1 1673111 1673111 1 - 19 18 1 94.7368421052632 NC_039506.1 1673027 1673526 500 + Gnomon promoter NA NA gene-
LOC109611032 GeneID:109611032 LOC109611032 Gene LOC109611032 lncRNA character(0) NA character(0)
NC_039506.1 1673135 1673135 1 - 17 9 8 52.9411764705882 NC_039506.1 1673027 1673526 500 + Gnomon promoter NA NA gene-
LOC109611032 GeneID:109611032 LOC109611032 Gene LOC109611032 lncRNA character(0) NA character(0)
NC_039506.1 1673160 1673160 1 - 11 11 0 100 NC_039506.1 1673027 1673526 500 + Gnomon promoter NA NA gene-LOC109611032
GeneID:109611032 LOC109611032 Gene LOC109611032 lncRNA character(0) NA character(0)
NC_039506.1 1673189 1673189 1 - 29 23 6 79.3103448275862 NC_039506.1 1673027 1673526 500 + Gnomon promoter NA NA gene-
LOC109611032 GeneID:109611032 LOC109611032 Gene LOC109611032 lncRNA character(0) NA character(0)
NC_039506.1 1673197 1673197 1 - 32 23 9 71.875 NC_039506.1 1673027 1673526 500 + Gnomon promoter NA NA gene-LOC109611032
GeneID:109611032 LOC109611032 Gene LOC109611032 lncRNA character(0) NA character(0)
NC_039506.1 1673205 1673205 1 - 46 43 3 93.4782608695652 NC_039506.1 1673027 1673526 500 + Gnomon promoter NA NA gene-
LOC109611032 GeneID:109611032 LOC109611032 Gene LOC109611032 lncRNA character(0) NA character(0)
NC_039506.1 1673219 1673219 1 - 66 25 41 37.8787878787879 NC_039506.1 1673027 1673526 500 + Gnomon promoter NA NA gene-
LOC109611032 GeneID:109611032 LOC109611032 Gene LOC109611032 lncRNA character(0) NA character(0)
NC_039506.1 1673225 1673225 1 - 64 54 10 84.375 NC_039506.1 1673027 1673526 500 + Gnomon promoter NA NA gene-LOC109611032
GeneID:109611032 LOC109611032 Gene LOC109611032 lncRNA character(0) NA character(0)
NC_039506.1 1673234 1673234 1 - 64 39 25 60.9375 NC_039506.1 1673027 1673526 500 + Gnomon promoter NA NA gene-LOC109611032
GeneID:109611032 LOC109611032 Gene LOC109611032 lncRNA character(0) NA character(0)
NC_039506.1 1673241 1673241 1 - 63 41 22 65.0793650793651 NC_039506.1 1673027 1673526 500 + Gnomon promoter NA NA gene-
LOC109611032 GeneID:109611032 LOC109611032 Gene LOC109611032 lncRNA character(0) NA character(0)
NC_039506.1 1673284 1673284 1 - 49 45 4 91.8367346938776 NC_039506.1 1673027 1673526 500 + Gnomon promoter NA NA gene-
LOC109611032 GeneID:109611032 LOC109611032 Gene LOC109611032 lncRNA character(0) NA character(0)
NC_039506.1 1673353 1673353 1 - 22 21 1 95.4545454545455 NC_039506.1 1673027 1673526 500 + Gnomon promoter NA NA gene-
LOC109611032 GeneID:109611032 LOC109611032 Gene LOC109611032 lncRNA character(0) NA character(0)
NC_039506.1 1673389 1673389 1 - 74 72 2 97.2972972972973 NC_039506.1 1673027 1673526 500 + Gnomon promoter NA NA gene-
LOC109611032 GeneID:109611032 LOC109611032 Gene LOC109611032 lncRNA character(0) NA character(0)
NC_039506.1 1673463 1673463 1 + 5 5 0 100 NC_039506.1 1673027 1673526 500 + Gnomon promoter NA NA gene-LOC109611032
GeneID:109611032 LOC109611032 Gene LOC109611032 lncRNA character(0) NA character(0)
NC_039506.1 1673464 1673464 1 - 28 27 1 96.4285714285714 NC_039506.1 1673027 1673526 500 + Gnomon promoter NA NA gene-
LOC109611032 GeneID:109611032 LOC109611032 Gene LOC109611032 lncRNA character(0) NA character(0)
NC_039506.1 1673488 1673488 1 - 32 27 5 84.375 NC_039506.1 1673027 1673526 500 + Gnomon promoter NA NA gene-LOC109611032
GeneID:109611032 LOC109611032 Gene LOC109611032 lncRNA character(0) NA character(0)
NC_039506.1 1673511 1673511 1 - 28 23 5 82.1428571428571 NC_039506.1 1673027 1673526 500 + Gnomon promoter NA NA gene-
LOC109611032 GeneID:109611032 LOC109611032 Gene LOC109611032 lncRNA character(0) NA character(0)
NC_039506.1 3094044 3094044 1 + 18 9 9 50 NC_039506.1 3094025 3094524 500 + Gnomon promoter NA NA gene-LOC105280352
GeneID:105280352 LOC105280352 Gene LOC105280352 protein_coding character(0) NA character(0)
NC_039506.1 3094045 3094045 1 - 16 11 5 68.75 NC_039506.1 3094025 3094524 500 + Gnomon promoter NA NA gene-LOC105280352
GeneID:105280352 LOC105280352 Gene LOC105280352 protein_coding character(0) NA character(0)
NC_039506.1 3094067 3094067 1 + 11 8 3 72.7272727272727 NC_039506.1 3094025 3094524 500 + Gnomon promoter NA NA gene-
LOC105280352 GeneID:105280352 LOC105280352 Gene LOC105280352 protein_coding character(0) NA character(0)
...
... only first 25 lines shown ...

Directory: RNK File: sites-in-promoters-Ob_LI_BCphaseB1.txt

| seqnames | start | end | width | strand | coverage | numCs | numTs | perc_meth | region_seqnames | region_start | region_end | region_width | region_strand | region_source | region_type | region_score | region_phase | region_ID | region_Dbxref | region_Name | region_gbkey | region_gene | region_gene_biotype | region_end_range | region_partial | region_start_range |
|---|---|---|---|---|---|---|---|---|---|---|---|---|---|---|---|---|---|---|---|---|---|---|---|---|---|---|
| NC_039506.1 | 1664198 | 1664198 | 1 | + | 39 | 13 | 26 | 33.3333333333333 | NC_039506.1 | 1664107 | 1664606 | 500 | + | Gnomon | promoter | NA | NA | gene-LOC105280261 | GeneID:105280261 | LOC105280261 | Gene | LOC105280261 | protein_coding | character(0) | NA | character(0) |
| NC_039506.1 | 1673111 | 1673111 | 1 | - | 12 | 12 | 0 | 100 | NC_039506.1 | 1673027 | 1673526 | 500 | + | Gnomon | promoter | NA | NA | gene-LOC109611032 | GeneID:109611032 | LOC109611032 | Gene | LOC109611032 | lncRNA | character(0) | NA | character(0) |
| NC_039506.1 | 1673135 | 1673135 | 1 | - | 15 | 11 | 4 | 73.3333333333333 | NC_039506.1 | 1673027 | 1673526 | 500 | + | Gnomon | promoter | NA | NA | gene-LOC109611032 | GeneID:109611032 | LOC109611032 | Gene | LOC109611032 | lncRNA | character(0) | NA | character(0) |
| NC_039506.1 | 1673160 | 1673160 | 1 | - | 13 | 9 | 4 | 69.2307692307692 | NC_039506.1 | 1673027 | 1673526 | 500 | + | Gnomon | promoter | NA | NA | gene-LOC109611032 | GeneID:109611032 | LOC109611032 | Gene | LOC109611032 | lncRNA | character(0) | NA | character(0) |
| NC_039506.1 | 1673189 | 1673189 | 1 | - | 27 | 21 | 6 | 77.7777777777778 | NC_039506.1 | 1673027 | 1673526 | 500 | + | Gnomon | promoter | NA | NA | gene-LOC109611032 | GeneID:109611032 | LOC109611032 | Gene | LOC109611032 | lncRNA | character(0) | NA | character(0) |
| NC_039506.1 | 1673197 | 1673197 | 1 | - | 34 | 17 | 17 | 50 | NC_039506.1 | 1673027 | 1673526 | 500 | + | Gnomon | promoter | NA | NA | gene-LOC109611032 | GeneID:109611032 | LOC109611032 | Gene | LOC109611032 | lncRNA | character(0) | NA | character(0) |
| NC_039506.1 | 1673205 | 1673205 | 1 | - | 44 | 34 | 10 | 77.2727272727273 | NC_039506.1 | 1673027 | 1673526 | 500 | + | Gnomon | promoter | NA | NA | gene-LOC109611032 | GeneID:109611032 | LOC109611032 | Gene | LOC109611032 | lncRNA | character(0) | NA | character(0) |
| NC_039506.1 | 1673219 | 1673219 | 1 | - | 61 | 21 | 40 | 34.4262295081967 | NC_039506.1 | 1673027 | 1673526 | 500 | + | Gnomon | promoter | NA | NA | gene-LOC109611032 | GeneID:109611032 | LOC109611032 | Gene | LOC109611032 | lncRNA | character(0) | NA | character(0) |
| NC_039506.1 | 1673225 | 1673225 | 1 | - | 54 | 40 | 14 | 74.0740740740741 | NC_039506.1 | 1673027 | 1673526 | 500 | + | Gnomon | promoter | NA | NA | gene-LOC109611032 | GeneID:109611032 | LOC109611032 | Gene | LOC109611032 | lncRNA | character(0) | NA | character(0) |
| NC_039506.1 | 1673234 | 1673234 | 1 | - | 57 | 29 | 28 | 50.8771929824561 | NC_039506.1 | 1673027 | 1673526 | 500 | + | Gnomon | promoter | NA | NA | gene-LOC109611032 | GeneID:109611032 | LOC109611032 | Gene | LOC109611032 | lncRNA | character(0) | NA | character(0) |
| NC_039506.1 | 1673241 | 1673241 | 1 | - | 56 | 33 | 23 | 58.9285714285714 | NC_039506.1 | 1673027 | 1673526 | 500 | + | Gnomon | promoter | NA | NA | gene-LOC109611032 | GeneID:109611032 | LOC109611032 | Gene | LOC109611032 | lncRNA | character(0) | NA | character(0) |
| NC_039506.1 | 1673284 | 1673284 | 1 | - | 48 | 37 | 11 | 77.0833333333333 | NC_039506.1 | 1673027 | 1673526 | 500 | + | Gnomon | promoter | NA | NA | gene-LOC109611032 | GeneID:109611032 | LOC109611032 | Gene | LOC109611032 | lncRNA | character(0) | NA | character(0) |
| NC_039506.1 | 1673353 | 1673353 | 1 | - | 21 | 19 | 2 | 90.4761904761905 | NC_039506.1 | 1673027 | 1673526 | 500 | + | Gnomon | promoter | NA | NA | gene-LOC109611032 | GeneID:109611032 | LOC109611032 | Gene | LOC109611032 | lncRNA | character(0) | NA | character(0) |
| NC_039506.1 | 1673389 | 1673389 | 1 | - | 71 | 70 | 1 | 98.5915492957747 | NC_039506.1 | 1673027 | 1673526 | 500 | + | Gnomon | promoter | NA | NA | gene-LOC109611032 | GeneID:109611032 | LOC109611032 | Gene | LOC109611032 | lncRNA | character(0) | NA | character(0) |
| NC_039506.1 | 1673464 | 1673464 | 1 | - | 38 | 33 | 5 | 86.8421052631579 | NC_039506.1 | 1673027 | 1673526 | 500 | + | Gnomon | promoter | NA | NA | gene-LOC109611032 | GeneID:109611032 | LOC109611032 | Gene | LOC109611032 | lncRNA | character(0) | NA | character(0) |
| NC_039506.1 | 1673488 | 1673488 | 1 | - | 34 | 28 | 6 | 82.3529411764706 | NC_039506.1 | 1673027 | 1673526 | 500 | + | Gnomon | promoter | NA | NA | gene-LOC109611032 | GeneID:109611032 | LOC109611032 | Gene | LOC109611032 | lncRNA | character(0) | NA | character(0) |
| NC_039506.1 | 1673510 | 1673510 | 1 | + | 8 | 6 | 2 | 75 | NC_039506.1 | 1673027 | 1673526 | 500 | + | Gnomon | promoter | NA | NA | gene-LOC109611032 | GeneID:109611032 | LOC109611032 | Gene | LOC109611032 | lncRNA | character(0) | NA | character(0) |
| NC_039506.1 | 1673511 | 1673511 | 1 | - | 33 | 20 | 13 | 60.6060606060606 | NC_039506.1 | 1673027 | 1673526 | 500 | + | Gnomon | promoter | NA | NA | gene-LOC109611032 | GeneID:109611032 | LOC109611032 | Gene | LOC109611032 | lncRNA | character(0) | NA | character(0) |
| NC_039506.1 | 1811800 | 1811800 | 1 | + | 23 | 8 | 15 | 34.7826086956522 | NC_039506.1 | 1811643 | 1812142 | 500 | + | Gnomon | promoter | NA | NA | gene-LOC105280272 | GeneID:105280272 | LOC105280272 | Gene | LOC105280272 | protein_coding | character(0) | NA | character(0) |
| NC_039506.1 | 2172293 | 2172293 | 1 | - | 22 | 8 | 14 | 36.3636363636364 | NC_039506.1 | 2172291 | 2172790 | 500 | + | Gnomon | promoter | NA | NA | gene-LOC105280304 | GeneID:105280304 | LOC105280304 | Gene | LOC105280304 | protein_coding | character(0) | NA | character(0) |
| NC_039506.1 | 2187053 | 2187053 | 1 | - | 11 | 7 | 4 | 63.6363636363636 | NC_039506.1 | 2187023 | 2187522 | 500 | + | Gnomon | promoter | NA | NA | gene-LOC105280309 | GeneID:105280309 | LOC105280309 | Gene | LOC105280309 | protein_coding | character(0) | NA | character(0) |
| NC_039506.1 | 2187086 | 2187086 | 1 | + | 9 | 7 | 2 | 77.7777777777778 | NC_039506.1 | 2187023 | 2187522 | 500 | + | Gnomon | promoter | NA | NA | gene-LOC105280309 | GeneID:105280309 | LOC105280309 | Gene | LOC105280309 | protein_coding | character(0) | NA | character(0) |
| NC_039506.1 | 2187087 | 2187087 | 1 | - | 8 | 6 | 2 | 75 | NC_039506.1 | 2187023 | 2187522 | 500 | + | Gnomon | promoter | NA | NA | gene-LOC105280309 | GeneID:105280309 | LOC105280309 | Gene | LOC105280309 | protein_coding | character(0) | NA | character(0) |
| NC_039506.1 | 2200872 | 2200872 | 1 | - | 10 | 6 | 4 | 60 | NC_039506.1 | 2200872 | 2201371 | 500 | + | Gnomon | promoter | NA | NA | gene-LOC105280312 | GeneID:105280312 | LOC105280312 | Gene | LOC105280312 | protein_coding | character(0) | NA | character(0) |

...
... only first 25 lines shown ...

Directory: RNK File: sites-in-promoters-Ob_LI_BCphaseB2.txt

| seqnames | start | end | width | strand | coverage | numCs | numTs | perc_meth | region_seqnames | region_start | region_end | region_width | region_strand | region_source | region_type | region_score | region_phase | region_ID | region_Dbxref | region_Name | region_gbkey | region_gene | region_gene_biotype | region_end_range | region_partial | region_start_range |
|---|---|---|---|---|---|---|---|---|---|---|---|---|---|---|---|---|---|---|---|---|---|---|---|---|---|---|
| NC_039506.1 | 1664198 | 1664198 | 1 | + | 37 | 14 | 23 | 37.8378378378378 | NC_039506.1 | 1664107 | 1664606 | 500 | + | Gnomon | promoter | NA | NA | gene-LOC105280261 | GeneID:105280261 | LOC105280261 | Gene | LOC105280261 | protein_coding | character(0) | NA | character(0) |
| NC_039506.1 | 1673111 | 1673111 | 1 | - | 13 | 11 | 2 | 84.6153846153846 | NC_039506.1 | 1673027 | 1673526 | 500 | + | Gnomon | promoter | NA | NA | gene-LOC109611032 | GeneID:109611032 | LOC109611032 | Gene | LOC109611032 | lncRNA | character(0) | NA | character(0) |
| NC_039506.1 | 1673135 | 1673135 | 1 | - | 12 | 9 | 3 | 75 | NC_039506.1 | 1673027 | 1673526 | 500 | + | Gnomon | promoter | NA | NA | gene-LOC109611032 | GeneID:109611032 | LOC109611032 | Gene | LOC109611032 | lncRNA | character(0) | NA | character(0) |
| NC_039506.1 | 1673160 | 1673160 | 1 | - | 12 | 12 | 0 | 100 | NC_039506.1 | 1673027 | 1673526 | 500 | + | Gnomon | promoter | NA | NA | gene-LOC109611032 | GeneID:109611032 | LOC109611032 | Gene | LOC109611032 | lncRNA | character(0) | NA | character(0) |
| NC_039506.1 | 1673189 | 1673189 | 1 | - | 36 | 29 | 7 | 80.5555555555556 | NC_039506.1 | 1673027 | 1673526 | 500 | + | Gnomon | promoter | NA | NA | gene-LOC109611032 | GeneID:109611032 | LOC109611032 | Gene | LOC109611032 | lncRNA | character(0) | NA | character(0) |
| NC_039506.1 | 1673197 | 1673197 | 1 | - | 37 | 20 | 17 | 54.0540540540541 | NC_039506.1 | 1673027 | 1673526 | 500 | + | Gnomon | promoter | NA | NA | gene-LOC109611032 | GeneID:109611032 | LOC109611032 | Gene | LOC109611032 | lncRNA | character(0) | NA | character(0) |
| NC_039506.1 | 1673205 | 1673205 | 1 | - | 53 | 40 | 13 | 75.4716981132076 | NC_039506.1 | 1673027 | 1673526 | 500 | + | Gnomon | promoter | NA | NA | gene-LOC109611032 | GeneID:109611032 | LOC109611032 | Gene | LOC109611032 | lncRNA | character(0) | NA | character(0) |
| NC_039506.1 | 1673219 | 1673219 | 1 | - | 64 | 9 | 55 | 14.0625 | NC_039506.1 | 1673027 | 1673526 | 500 | + | Gnomon | promoter | NA | NA | gene-LOC109611032 | GeneID:109611032 | LOC109611032 | Gene | LOC109611032 | lncRNA | character(0) | NA | character(0) |
| NC_039506.1 | 1673225 | 1673225 | 1 | - | 61 | 48 | 13 | 78.6885245901639 | NC_039506.1 | 1673027 | 1673526 | 500 | + | Gnomon | promoter | NA | NA | gene-LOC109611032 | GeneID:109611032 | LOC109611032 | Gene | LOC109611032 | lncRNA | character(0) | NA | character(0) |
| NC_039506.1 | 1673234 | 1673234 | 1 | - | 60 | 35 | 25 | 58.3333333333333 | NC_039506.1 | 1673027 | 1673526 | 500 | + | Gnomon | promoter | NA | NA | gene-LOC109611032 | GeneID:109611032 | LOC109611032 | Gene | LOC109611032 | lncRNA | character(0) | NA | character(0) |
| NC_039506.1 | 1673241 | 1673241 | 1 | - | 58 | 38 | 20 | 65.5172413793103 | NC_039506.1 | 1673027 | 1673526 | 500 | + | Gnomon | promoter | NA | NA | gene-LOC109611032 | GeneID:109611032 | LOC109611032 | Gene | LOC109611032 | lncRNA | character(0) | NA | character(0) |
| NC_039506.1 | 1673284 | 1673284 | 1 | - | 44 | 35 | 9 | 79.5454545454545 | NC_039506.1 | 1673027 | 1673526 | 500 | + | Gnomon | promoter | NA | NA | gene-LOC109611032 | GeneID:109611032 | LOC109611032 | Gene | LOC109611032 | lncRNA | character(0) | NA | character(0) |
| NC_039506.1 | 1673353 | 1673353 | 1 | - | 24 | 24 | 0 | 100 | NC_039506.1 | 1673027 | 1673526 | 500 | + | Gnomon | promoter | NA | NA | gene-LOC109611032 | GeneID:109611032 | LOC109611032 | Gene | LOC109611032 | lncRNA | character(0) | NA | character(0) |
| NC_039506.1 | 1673389 | 1673389 | 1 | - | 73 | 72 | 1 | 98.6301369863014 | NC_039506.1 | 1673027 | 1673526 | 500 | + | Gnomon | promoter | NA | NA | gene-LOC109611032 | GeneID:109611032 | LOC109611032 | Gene | LOC109611032 | lncRNA | character(0) | NA | character(0) |
| NC_039506.1 | 1673464 | 1673464 | 1 | - | 31 | 30 | 1 | 96.7741935483871 | NC_039506.1 | 1673027 | 1673526 | 500 | + | Gnomon | promoter | NA | NA | gene-LOC109611032 | GeneID:109611032 | LOC109611032 | Gene | LOC109611032 | lncRNA | character(0) | NA | character(0) |
| NC_039506.1 | 1673488 | 1673488 | 1 | - | 37 | 30 | 7 | 81.0810810810811 | NC_039506.1 | 1673027 | 1673526 | 500 | + | Gnomon | promoter | NA | NA | gene-LOC109611032 | GeneID:109611032 | LOC109611032 | Gene | LOC109611032 | lncRNA | character(0) | NA | character(0) |
| NC_039506.1 | 1673511 | 1673511 | 1 | - | 37 | 29 | 8 | 78.3783783783784 | NC_039506.1 | 1673027 | 1673526 | 500 | + | Gnomon | promoter | NA | NA | gene-LOC109611032 | GeneID:109611032 | LOC109611032 | Gene | LOC109611032 | lncRNA | character(0) | NA | character(0) |
| NC_039506.1 | 2114196 | 2114196 | 1 | + | 14 | 7 | 7 | 50 | NC_039506.1 | 2114158 | 2114657 | 500 | + | Gnomon | promoter | NA | NA | gene-LOC105280298 | GeneID:105280298 | LOC105280298 | Gene | LOC105280298 | protein_coding | character(0) | NA | character(0) |
| NC_039506.1 | 2187052 | 2187052 | 1 | + | 5 | 5 | 0 | 100 | NC_039506.1 | 2187023 | 2187522 | 500 | + | Gnomon | promoter | NA | NA | gene-LOC105280309 | GeneID:105280309 | LOC105280309 | Gene | LOC105280309 | protein_coding | character(0) | NA | character(0) |
| NC_039506.1 | 2187053 | 2187053 | 1 | - | 7 | 5 | 2 | 71.4285714285714 | NC_039506.1 | 2187023 | 2187522 | 500 | + | Gnomon | promoter | NA | NA | gene-LOC105280309 | GeneID:105280309 | LOC105280309 | Gene | LOC105280309 | protein_coding | character(0) | NA | character(0) |
| NC_039506.1 | 2187087 | 2187087 | 1 | - | 7 | 5 | 2 | 71.4285714285714 | NC_039506.1 | 2187023 | 2187522 | 500 | + | Gnomon | promoter | NA | NA | gene-LOC105280309 | GeneID:105280309 | LOC105280309 | Gene | LOC105280309 | protein_coding | character(0) | NA | character(0) |
| NC_039506.1 | 2200872 | 2200872 | 1 | - | 10 | 7 | 3 | 70 | NC_039506.1 | 2200872 | 2201371 | 500 | + | Gnomon | promoter | NA | NA | gene-LOC105280312 | GeneID:105280312 | LOC105280312 | Gene | LOC105280312 | protein_coding | character(0) | NA | character(0) |
| NC_039506.1 | 2274299 | 2274299 | 1 | - | 10 | 6 | 4 | 60 | NC_039506.1 | 2273836 | 2274335 | 500 | - | Gnomon | promoter | NA | NA | gene-LOC105280317 | GeneID:105280317 | LOC105280317 | Gene | LOC105280317 | protein_coding | character(0) | NA | character(0) |
| NC_039506.1 | 3094044 | 3094044 | 1 | + | 20 | 7 | 13 | 35 | NC_039506.1 | 3094025 | 3094524 | 500 | + | Gnomon | promoter | NA | NA | gene-LOC105280352 | GeneID:105280352 | LOC105280352 | Gene | LOC105280352 | protein_coding | character(0) | NA | character(0) |

...
... only first 25 lines shown ...

Directory: RNK File: sites-in-promoters-Ob_LI_RphaseA1.txt

| seqnames | start | end | width | strand | coverage | numCs | numTs | perc_meth | region_seqnames | region_start | region_end | region_width | region_strand | region_source | region_type | region_score | region_phase | region_ID | region_Dbxref | region_Name | region_gbkey | region_gene | region_gene_biotype | region_end_range | region_partial | region_start_range |
|---|---|---|---|---|---|---|---|---|---|---|---|---|---|---|---|---|---|---|---|---|---|---|---|---|---|---|
| NC_039506.1 | 1664198 | 1664198 | 1 | + | 41 | 15 | 26 | 36.5853658536585 | NC_039506.1 | 1664107 | 1664606 | 500 | + | Gnomon | promoter | NA | NA | gene-LOC105280261 | GeneID:105280261 | LOC105280261 | Gene | LOC105280261 | protein_coding | character(0) | NA | character(0) |
| NC_039506.1 | 1664400 | 1664400 | 1 | + | 13 | 6 | 7 | 46.1538461538462 | NC_039506.1 | 1664107 | 1664606 | 500 | + | Gnomon | promoter | NA | NA | gene-LOC105280261 | GeneID:105280261 | LOC105280261 | Gene | LOC105280261 | protein_coding | character(0) | NA | character(0) |
| NC_039506.1 | 1673111 | 1673111 | 1 | - | 14 | 13 | 1 | 92.8571428571429 | NC_039506.1 | 1673027 | 1673526 | 500 | + | Gnomon | promoter | NA | NA | gene-LOC109611032 | GeneID:109611032 | LOC109611032 | Gene | LOC109611032 | lncRNA | character(0) | NA | character(0) |
| NC_039506.1 | 1673135 | 1673135 | 1 | - | 12 | 9 | 3 | 75 | NC_039506.1 | 1673027 | 1673526 | 500 | + | Gnomon | promoter | NA | NA | gene-LOC109611032 | GeneID:109611032 | LOC109611032 | Gene | LOC109611032 | lncRNA | character(0) | NA | character(0) |
| NC_039506.1 | 1673160 | 1673160 | 1 | - | 12 | 9 | 3 | 75 | NC_039506.1 | 1673027 | 1673526 | 500 | + | Gnomon | promoter | NA | NA | gene-LOC109611032 | GeneID:109611032 | LOC109611032 | Gene | LOC109611032 | lncRNA | character(0) | NA | character(0) |
| NC_039506.1 | 1673189 | 1673189 | 1 | - | 24 | 19 | 5 | 79.1666666666667 | NC_039506.1 | 1673027 | 1673526 | 500 | + | Gnomon | promoter | NA | NA | gene-LOC109611032 | GeneID:109611032 | LOC109611032 | Gene | LOC109611032 | lncRNA | character(0) | NA | character(0) |
| NC_039506.1 | 1673197 | 1673197 | 1 | - | 28 | 23 | 5 | 82.1428571428571 | NC_039506.1 | 1673027 | 1673526 | 500 | + | Gnomon | promoter | NA | NA | gene-LOC109611032 | GeneID:109611032 | LOC109611032 | Gene | LOC109611032 | lncRNA | character(0) | NA | character(0) |
| NC_039506.1 | 1673205 | 1673205 | 1 | - | 33 | 28 | 5 | 84.8484848484848 | NC_039506.1 | 1673027 | 1673526 | 500 | + | Gnomon | promoter | NA | NA | gene-LOC109611032 | GeneID:109611032 | LOC109611032 | Gene | LOC109611032 | lncRNA | character(0) | NA | character(0) |
| NC_039506.1 | 1673219 | 1673219 | 1 | - | 57 | 16 | 41 | 28.0701754385965 | NC_039506.1 | 1673027 | 1673526 | 500 | + | Gnomon | promoter | NA | NA | gene-LOC109611032 | GeneID:109611032 | LOC109611032 | Gene | LOC109611032 | lncRNA | character(0) | NA | character(0) |
| NC_039506.1 | 1673225 | 1673225 | 1 | - | 57 | 43 | 14 | 75.4385964912281 | NC_039506.1 | 1673027 | 1673526 | 500 | + | Gnomon | promoter | NA | NA | gene-LOC109611032 | GeneID:109611032 | LOC109611032 | Gene | LOC109611032 | lncRNA | character(0) | NA | character(0) |
| NC_039506.1 | 1673234 | 1673234 | 1 | - | 59 | 36 | 23 | 61.0169491525424 | NC_039506.1 | 1673027 | 1673526 | 500 | + | Gnomon | promoter | NA | NA | gene-LOC109611032 | GeneID:109611032 | LOC109611032 | Gene | LOC109611032 | lncRNA | character(0) | NA | character(0) |
| NC_039506.1 | 1673241 | 1673241 | 1 | - | 60 | 37 | 23 | 61.6666666666667 | NC_039506.1 | 1673027 | 1673526 | 500 | + | Gnomon | promoter | NA | NA | gene-LOC109611032 | GeneID:109611032 | LOC109611032 | Gene | LOC109611032 | lncRNA | character(0) | NA | character(0) |
| NC_039506.1 | 1673284 | 1673284 | 1 | - | 57 | 50 | 7 | 87.719298245614 | NC_039506.1 | 1673027 | 1673526 | 500 | + | Gnomon | promoter | NA | NA | gene-LOC109611032 | GeneID:109611032 | LOC109611032 | Gene | LOC109611032 | lncRNA | character(0) | NA | character(0) |
| NC_039506.1 | 1673353 | 1673353 | 1 | - | 25 | 25 | 0 | 100 | NC_039506.1 | 1673027 | 1673526 | 500 | + | Gnomon | promoter | NA | NA | gene-LOC109611032 | GeneID:109611032 | LOC109611032 | Gene | LOC109611032 | lncRNA | character(0) | NA | character(0) |
| NC_039506.1 | 1673388 | 1673388 | 1 | + | 6 | 6 | 0 | 100 | NC_039506.1 | 1673027 | 1673526 | 500 | + | Gnomon | promoter | NA | NA | gene-LOC109611032 | GeneID:109611032 | LOC109611032 | Gene | LOC109611032 | lncRNA | character(0) | NA | character(0) |
| NC_039506.1 | 1673389 | 1673389 | 1 | - | 56 | 54 | 2 | 96.4285714285714 | NC_039506.1 | 1673027 | 1673526 | 500 | + | Gnomon | promoter | NA | NA | gene-LOC109611032 | GeneID:109611032 | LOC109611032 | Gene | LOC109611032 | lncRNA | character(0) | NA | character(0) |
| NC_039506.1 | 1673464 | 1673464 | 1 | - | 45 | 40 | 5 | 88.8888888888889 | NC_039506.1 | 1673027 | 1673526 | 500 | + | Gnomon | promoter | NA | NA | gene-LOC109611032 | GeneID:109611032 | LOC109611032 | Gene | LOC109611032 | lncRNA | character(0) | NA | character(0) |
| NC_039506.1 | 1673488 | 1673488 | 1 | - | 34 | 29 | 5 | 85.2941176470588 | NC_039506.1 | 1673027 | 1673526 | 500 | + | Gnomon | promoter | NA | NA | gene-LOC109611032 | GeneID:109611032 | LOC109611032 | Gene | LOC109611032 | lncRNA | character(0) | NA | character(0) |
| NC_039506.1 | 1673511 | 1673511 | 1 | - | 30 | 27 | 3 | 90 | NC_039506.1 | 1673027 | 1673526 | 500 | + | Gnomon | promoter | NA | NA | gene-LOC109611032 | GeneID:109611032 | LOC109611032 | Gene | LOC109611032 | lncRNA | character(0) | NA | character(0) |
| NC_039506.1 | 2187052 | 2187052 | 1 | + | 8 | 6 | 2 | 75 | NC_039506.1 | 2187023 | 2187522 | 500 | + | Gnomon | promoter | NA | NA | gene-LOC105280309 | GeneID:105280309 | LOC105280309 | Gene | LOC105280309 | protein_coding | character(0) | NA | character(0) |
| NC_039506.1 | 2187053 | 2187053 | 1 | - | 6 | 6 | 0 | 100 | NC_039506.1 | 2187023 | 2187522 | 500 | + | Gnomon | promoter | NA | NA | gene-LOC105280309 | GeneID:105280309 | LOC105280309 | Gene | LOC105280309 | protein_coding | character(0) | NA | character(0) |
| NC_039506.1 | 2187086 | 2187086 | 1 | + | 8 | 5 | 3 | 62.5 | NC_039506.1 | 2187023 | 2187522 | 500 | + | Gnomon | promoter | NA | NA | gene-LOC105280309 | GeneID:105280309 | LOC105280309 | Gene | LOC105280309 | protein_coding | character(0) | NA | character(0) |
| NC_039506.1 | 3094044 | 3094044 | 1 | + | 16 | 6 | 10 | 37.5 | NC_039506.1 | 3094025 | 3094524 | 500 | + | Gnomon | promoter | NA | NA | gene-LOC105280352 | GeneID:105280352 | LOC105280352 | Gene | LOC105280352 | protein_coding | character(0) | NA | character(0) |
| NC_039506.1 | 3094045 | 3094045 | 1 | - | 18 | 9 | 9 | 50 | NC_039506.1 | 3094025 | 3094524 | 500 | + | Gnomon | promoter | NA | NA | gene-LOC105280352 | GeneID:105280352 | LOC105280352 | Gene | LOC105280352 | protein_coding | character(0) | NA | character(0) |

...
... only first 25 lines shown ...

Directory: RNK File: sites-in-promoters-Ob_LI_RphaseA2.txt

| seqnames | start | end | width | strand | coverage | numCs | numTs | perc_meth | region_seqnames | region_start | region_end | region_width | region_strand | region_source | region_type | region_score | region_phase | region_ID | region_Dbxref | region_Name | region_gbkey | region_gene | region_gene_biotype | region_end_range | region_partial | region_start_range |
|---|---|---|---|---|---|---|---|---|---|---|---|---|---|---|---|---|---|---|---|---|---|---|---|---|---|---|
| NC_039506.1 | 1487355 | 1487355 | 1 | + | 53 | 8 | 45 | 15.0943396226415 | NC_039506.1 | 1487279 | 1487778 | 500 | + | Gnomon | promoter | NA | NA | gene-LOC105280236 | GeneID:105280236 | LOC105280236 | Gene | LOC105280236 | protein_coding | character(0) | NA | character(0) |
| NC_039506.1 | 1664198 | 1664198 | 1 | + | 45 | 10 | 35 | 22.2222222222222 | NC_039506.1 | 1664107 | 1664606 | 500 | + | Gnomon | promoter | NA | NA | gene-LOC105280261 | GeneID:105280261 | LOC105280261 | Gene | LOC105280261 | protein_coding | character(0) | NA | character(0) |
| NC_039506.1 | 1673083 | 1673083 | 1 | + | 45 | 9 | 36 | 20 | NC_039506.1 | 1673027 | 1673526 | 500 | + | Gnomon | promoter | NA | NA | gene-LOC109611032 | GeneID:109611032 | LOC109611032 | Gene | LOC109611032 | lncRNA | character(0) | NA | character(0) |
| NC_039506.1 | 1673110 | 1673110 | 1 | + | 11 | 10 | 1 | 90.9090909090909 | NC_039506.1 | 1673027 | 1673526 | 500 | + | Gnomon | promoter | NA | NA | gene-LOC109611032 | GeneID:109611032 | LOC109611032 | Gene | LOC109611032 | lncRNA | character(0) | NA | character(0) |
| NC_039506.1 | 1673111 | 1673111 | 1 | - | 27 | 26 | 1 | 96.2962962962963 | NC_039506.1 | 1673027 | 1673526 | 500 | + | Gnomon | promoter | NA | NA | gene-LOC109611032 | GeneID:109611032 | LOC109611032 | Gene | LOC109611032 | lncRNA | character(0) | NA | character(0) |
| NC_039506.1 | 1673135 | 1673135 | 1 | - | 26 | 22 | 4 | 84.6153846153846 | NC_039506.1 | 1673027 | 1673526 | 500 | + | Gnomon | promoter | NA | NA | gene-LOC109611032 | GeneID:109611032 | LOC109611032 | Gene | LOC109611032 | lncRNA | character(0) | NA | character(0) |
| NC_039506.1 | 1673160 | 1673160 | 1 | - | 18 | 16 | 2 | 88.8888888888889 | NC_039506.1 | 1673027 | 1673526 | 500 | + | Gnomon | promoter | NA | NA | gene-LOC109611032 | GeneID:109611032 | LOC109611032 | Gene | LOC109611032 | lncRNA | character(0) | NA | character(0) |
| NC_039506.1 | 1673189 | 1673189 | 1 | - | 34 | 28 | 6 | 82.3529411764706 | NC_039506.1 | 1673027 | 1673526 | 500 | + | Gnomon | promoter | NA | NA | gene-LOC109611032 | GeneID:109611032 | LOC109611032 | Gene | LOC109611032 | lncRNA | character(0) | NA | character(0) |
| NC_039506.1 | 1673197 | 1673197 | 1 | - | 35 | 21 | 14 | 60 | NC_039506.1 | 1673027 | 1673526 | 500 | + | Gnomon | promoter | NA | NA | gene-LOC109611032 | GeneID:109611032 | LOC109611032 | Gene | LOC109611032 | lncRNA | character(0) | NA | character(0) |
| NC_039506.1 | 1673205 | 1673205 | 1 | - | 53 | 45 | 8 | 84.9056603773585 | NC_039506.1 | 1673027 | 1673526 | 500 | + | Gnomon | promoter | NA | NA | gene-LOC109611032 | GeneID:109611032 | LOC109611032 | Gene | LOC109611032 | lncRNA | character(0) | NA | character(0) |
| NC_039506.1 | 1673219 | 1673219 | 1 | - | 85 | 18 | 67 | 21.1764705882353 | NC_039506.1 | 1673027 | 1673526 | 500 | + | Gnomon | promoter | NA | NA | gene-LOC109611032 | GeneID:109611032 | LOC109611032 | Gene | LOC109611032 | lncRNA | character(0) | NA | character(0) |
| NC_039506.1 | 1673225 | 1673225 | 1 | - | 84 | 62 | 22 | 73.8095238095238 | NC_039506.1 | 1673027 | 1673526 | 500 | + | Gnomon | promoter | NA | NA | gene-LOC109611032 | GeneID:109611032 | LOC109611032 | Gene | LOC109611032 | lncRNA | character(0) | NA | character(0) |
| NC_039506.1 | 1673234 | 1673234 | 1 | - | 82 | 43 | 39 | 52.4390243902439 | NC_039506.1 | 1673027 | 1673526 | 500 | + | Gnomon | promoter | NA | NA | gene-LOC109611032 | GeneID:109611032 | LOC109611032 | Gene | LOC109611032 | lncRNA | character(0) | NA | character(0) |
| NC_039506.1 | 1673241 | 1673241 | 1 | - | 79 | 42 | 37 | 53.1645569620253 | NC_039506.1 | 1673027 | 1673526 | 500 | + | Gnomon | promoter | NA | NA | gene-LOC109611032 | GeneID:109611032 | LOC109611032 | Gene | LOC109611032 | lncRNA | character(0) | NA | character(0) |
| NC_039506.1 | 1673284 | 1673284 | 1 | - | 63 | 52 | 11 | 82.5396825396825 | NC_039506.1 | 1673027 | 1673526 | 500 | + | Gnomon | promoter | NA | NA | gene-LOC109611032 | GeneID:109611032 | LOC109611032 | Gene | LOC109611032 | lncRNA | character(0) | NA | character(0) |
| NC_039506.1 | 1673353 | 1673353 | 1 | - | 37 | 34 | 3 | 91.8918918918919 | NC_039506.1 | 1673027 | 1673526 | 500 | + | Gnomon | promoter | NA | NA | gene-LOC109611032 | GeneID:109611032 | LOC109611032 | Gene | LOC109611032 | lncRNA | character(0) | NA | character(0) |
| NC_039506.1 | 1673388 | 1673388 | 1 | + | 9 | 9 | 0 | 100 | NC_039506.1 | 1673027 | 1673526 | 500 | + | Gnomon | promoter | NA | NA | gene-LOC109611032 | GeneID:109611032 | LOC109611032 | Gene | LOC109611032 | lncRNA | character(0) | NA | character(0) |
| NC_039506.1 | 1673389 | 1673389 | 1 | - | 123 | 121 | 2 | 98.3739837398374 | NC_039506.1 | 1673027 | 1673526 | 500 | + | Gnomon | promoter | NA | NA | gene-LOC109611032 | GeneID:109611032 | LOC109611032 | Gene | LOC109611032 | lncRNA | character(0) | NA | character(0) |
| NC_039506.1 | 1673463 | 1673463 | 1 | + | 11 | 9 | 2 | 81.8181818181818 | NC_039506.1 | 1673027 | 1673526 | 500 | + | Gnomon | promoter | NA | NA | gene-LOC109611032 | GeneID:109611032 | LOC109611032 | Gene | LOC109611032 | lncRNA | character(0) | NA | character(0) |
| NC_039506.1 | 1673464 | 1673464 | 1 | - | 58 | 57 | 1 | 98.2758620689655 | NC_039506.1 | 1673027 | 1673526 | 500 | + | Gnomon | promoter | NA | NA | gene-LOC109611032 | GeneID:109611032 | LOC109611032 | Gene | LOC109611032 | lncRNA | character(0) | NA | character(0) |
| NC_039506.1 | 1673488 | 1673488 | 1 | - | 55 | 43 | 12 | 78.1818181818182 | NC_039506.1 | 1673027 | 1673526 | 500 | + | Gnomon | promoter | NA | NA | gene-LOC109611032 | GeneID:109611032 | LOC109611032 | Gene | LOC109611032 | lncRNA | character(0) | NA | character(0) |
| NC_039506.1 | 1673511 | 1673511 | 1 | - | 48 | 41 | 7 | 85.4166666666667 | NC_039506.1 | 1673027 | 1673526 | 500 | + | Gnomon | promoter | NA | NA | gene-LOC109611032 | GeneID:109611032 | LOC109611032 | Gene | LOC109611032 | lncRNA | character(0) | NA | character(0) |
| NC_039506.1 | 1828389 | 1828389 | 1 | + | 70 | 10 | 60 | 14.2857142857143 | NC_039506.1 | 1827907 | 1828406 | 500 | - | Gnomon | promoter | NA | NA | gene-LOC105280274 | GeneID:105280274 | LOC105280274 | Gene | LOC105280274 | protein_coding | character(0) | NA | character(0) |
| NC_039506.1 | 2172293 | 2172293 | 1 | - | 17 | 12 | 5 | 70.5882352941177 | NC_039506.1 | 2172291 | 2172790 | 500 | + | Gnomon | promoter | NA | NA | gene-LOC105280304 | GeneID:105280304 | LOC105280304 | Gene | LOC105280304 | protein_coding | character(0) | NA | character(0) |

...
... only first 25 lines shown ...

Directory: RNK File: sites-in-promoters-Ob_LI_RphaseB1.txt

| seqnames | start | end | width | strand | coverage | numCs | numTs | perc_meth | region_seqnames | region_start | region_end | region_width | region_strand | region_source | region_type | region_score | region_phase | region_ID | region_Dbxref | region_Name | region_gbkey | region_gene | region_gene_biotype | region_end_range | region_partial | region_start_range |
|---|---|---|---|---|---|---|---|---|---|---|---|---|---|---|---|---|---|---|---|---|---|---|---|---|---|---|
| NC_039506.1 | 1664198 | 1664198 | 1 | + | 37 | 9 | 28 | 24.3243243243243 | NC_039506.1 | 1664107 | 1664606 | 500 | + | Gnomon | promoter | NA | NA | gene-LOC105280261 | GeneID:105280261 | LOC105280261 | Gene | LOC105280261 | protein_coding | character(0) | NA | character(0) |
| NC_039506.1 | 1673111 | 1673111 | 1 | - | 15 | 14 | 1 | 93.3333333333333 | NC_039506.1 | 1673027 | 1673526 | 500 | + | Gnomon | promoter | NA | NA | gene-LOC109611032 | GeneID:109611032 | LOC109611032 | Gene | LOC109611032 | lncRNA | character(0) | NA | character(0) |
| NC_039506.1 | 1673135 | 1673135 | 1 | - | 13 | 11 | 2 | 84.6153846153846 | NC_039506.1 | 1673027 | 1673526 | 500 | + | Gnomon | promoter | NA | NA | gene-LOC109611032 | GeneID:109611032 | LOC109611032 | Gene | LOC109611032 | lncRNA | character(0) | NA | character(0) |
| NC_039506.1 | 1673160 | 1673160 | 1 | - | 12 | 8 | 4 | 66.6666666666667 | NC_039506.1 | 1673027 | 1673526 | 500 | + | Gnomon | promoter | NA | NA | gene-LOC109611032 | GeneID:109611032 | LOC109611032 | Gene | LOC109611032 | lncRNA | character(0) | NA | character(0) |
| NC_039506.1 | 1673189 | 1673189 | 1 | - | 31 | 24 | 7 | 77.4193548387097 | NC_039506.1 | 1673027 | 1673526 | 500 | + | Gnomon | promoter | NA | NA | gene-LOC109611032 | GeneID:109611032 | LOC109611032 | Gene | LOC109611032 | lncRNA | character(0) | NA | character(0) |
| NC_039506.1 | 1673197 | 1673197 | 1 | - | 39 | 12 | 27 | 30.7692307692308 | NC_039506.1 | 1673027 | 1673526 | 500 | + | Gnomon | promoter | NA | NA | gene-LOC109611032 | GeneID:109611032 | LOC109611032 | Gene | LOC109611032 | lncRNA | character(0) | NA | character(0) |
| NC_039506.1 | 1673205 | 1673205 | 1 | - | 42 | 27 | 15 | 64.2857142857143 | NC_039506.1 | 1673027 | 1673526 | 500 | + | Gnomon | promoter | NA | NA | gene-LOC109611032 | GeneID:109611032 | LOC109611032 | Gene | LOC109611032 | lncRNA | character(0) | NA | character(0) |
| NC_039506.1 | 1673219 | 1673219 | 1 | - | 54 | 15 | 39 | 27.7777777777778 | NC_039506.1 | 1673027 | 1673526 | 500 | + | Gnomon | promoter | NA | NA | gene-LOC109611032 | GeneID:109611032 | LOC109611032 | Gene | LOC109611032 | lncRNA | character(0) | NA | character(0) |
| NC_039506.1 | 1673225 | 1673225 | 1 | - | 55 | 42 | 13 | 76.3636363636364 | NC_039506.1 | 1673027 | 1673526 | 500 | + | Gnomon | promoter | NA | NA | gene-LOC109611032 | GeneID:109611032 | LOC109611032 | Gene | LOC109611032 | lncRNA | character(0) | NA | character(0) |
| NC_039506.1 | 1673234 | 1673234 | 1 | - | 55 | 31 | 24 | 56.3636363636364 | NC_039506.1 | 1673027 | 1673526 | 500 | + | Gnomon | promoter | NA | NA | gene-LOC109611032 | GeneID:109611032 | LOC109611032 | Gene | LOC109611032 | lncRNA | character(0) | NA | character(0) |
| NC_039506.1 | 1673241 | 1673241 | 1 | - | 53 | 25 | 28 | 47.1698113207547 | NC_039506.1 | 1673027 | 1673526 | 500 | + | Gnomon | promoter | NA | NA | gene-LOC109611032 | GeneID:109611032 | LOC109611032 | Gene | LOC109611032 | lncRNA | character(0) | NA | character(0) |
| NC_039506.1 | 1673284 | 1673284 | 1 | - | 45 | 38 | 7 | 84.4444444444444 | NC_039506.1 | 1673027 | 1673526 | 500 | + | Gnomon | promoter | NA | NA | gene-LOC109611032 | GeneID:109611032 | LOC109611032 | Gene | LOC109611032 | lncRNA | character(0) | NA | character(0) |
| NC_039506.1 | 1673353 | 1673353 | 1 | - | 24 | 24 | 0 | 100 | NC_039506.1 | 1673027 | 1673526 | 500 | + | Gnomon | promoter | NA | NA | gene-LOC109611032 | GeneID:109611032 | LOC109611032 | Gene | LOC109611032 | lncRNA | character(0) | NA | character(0) |
| NC_039506.1 | 1673389 | 1673389 | 1 | - | 64 | 62 | 2 | 96.875 | NC_039506.1 | 1673027 | 1673526 | 500 | + | Gnomon | promoter | NA | NA | gene-LOC109611032 | GeneID:109611032 | LOC109611032 | Gene | LOC109611032 | lncRNA | character(0) | NA | character(0) |
| NC_039506.1 | 1673463 | 1673463 | 1 | + | 6 | 6 | 0 | 100 | NC_039506.1 | 1673027 | 1673526 | 500 | + | Gnomon | promoter | NA | NA | gene-LOC109611032 | GeneID:109611032 | LOC109611032 | Gene | LOC109611032 | lncRNA | character(0) | NA | character(0) |
| NC_039506.1 | 1673464 | 1673464 | 1 | - | 52 | 47 | 5 | 90.3846153846154 | NC_039506.1 | 1673027 | 1673526 | 500 | + | Gnomon | promoter | NA | NA | gene-LOC109611032 | GeneID:109611032 | LOC109611032 | Gene | LOC109611032 | lncRNA | character(0) | NA | character(0) |
| NC_039506.1 | 1673488 | 1673488 | 1 | - | 31 | 24 | 7 | 77.4193548387097 | NC_039506.1 | 1673027 | 1673526 | 500 | + | Gnomon | promoter | NA | NA | gene-LOC109611032 | GeneID:109611032 | LOC109611032 | Gene | LOC109611032 | lncRNA | character(0) | NA | character(0) |
| NC_039506.1 | 1673511 | 1673511 | 1 | - | 22 | 21 | 1 | 95.4545454545455 | NC_039506.1 | 1673027 | 1673526 | 500 | + | Gnomon | promoter | NA | NA | gene-LOC109611032 | GeneID:109611032 | LOC109611032 | Gene | LOC109611032 | lncRNA | character(0) | NA | character(0) |
| NC_039506.1 | 2172293 | 2172293 | 1 | - | 18 | 9 | 9 | 50 | NC_039506.1 | 2172291 | 2172790 | 500 | + | Gnomon | promoter | NA | NA | gene-LOC105280304 | GeneID:105280304 | LOC105280304 | Gene | LOC105280304 | protein_coding | character(0) | NA | character(0) |
| NC_039506.1 | 2187053 | 2187053 | 1 | - | 8 | 5 | 3 | 62.5 | NC_039506.1 | 2187023 | 2187522 | 500 | + | Gnomon | promoter | NA | NA | gene-LOC105280309 | GeneID:105280309 | LOC105280309 | Gene | LOC105280309 | protein_coding | character(0) | NA | character(0) |
| NC_039506.1 | 2187086 | 2187086 | 1 | + | 10 | 7 | 3 | 70 | NC_039506.1 | 2187023 | 2187522 | 500 | + | Gnomon | promoter | NA | NA | gene-LOC105280309 | GeneID:105280309 | LOC105280309 | Gene | LOC105280309 | protein_coding | character(0) | NA | character(0) |
| NC_039506.1 | 2187087 | 2187087 | 1 | - | 7 | 6 | 1 | 85.7142857142857 | NC_039506.1 | 2187023 | 2187522 | 500 | + | Gnomon | promoter | NA | NA | gene-LOC105280309 | GeneID:105280309 | LOC105280309 | Gene | LOC105280309 | protein_coding | character(0) | NA | character(0) |
| NC_039506.1 | 2187097 | 2187097 | 1 | + | 14 | 8 | 6 | 57.1428571428571 | NC_039506.1 | 2187023 | 2187522 | 500 | + | Gnomon | promoter | NA | NA | gene-LOC105280309 | GeneID:105280309 | LOC105280309 | Gene | LOC105280309 | protein_coding | character(0) | NA | character(0) |
| NC_039506.1 | 3094044 | 3094044 | 1 | + | 18 | 10 | 8 | 55.5555555555556 | NC_039506.1 | 3094025 | 3094524 | 500 | + | Gnomon | promoter | NA | NA | gene-LOC105280352 | GeneID:105280352 | LOC105280352 | Gene | LOC105280352 | protein_coding | character(0) | NA | character(0) |

...
... only first 25 lines shown ...

Directory: RNK File: sites-in-promoters-Ob_LI_RphaseB2.txt

| seqnames | start | end | width | strand | coverage | numCs | numTs | perc_meth | region_seqnames | region_start | region_end | region_width | region_strand | region_source | region_type | region_score | region_phase | region_ID | region_Dbxref | region_Name | region_gbkey | region_gene | region_gene_biotype | region_end_range | region_partial | region_start_range |
|---|---|---|---|---|---|---|---|---|---|---|---|---|---|---|---|---|---|---|---|---|---|---|---|---|---|---|
| NC_039506.1 | 1673111 | 1673111 | 1 | - | 19 | 14 | 5 | 73.6842105263158 | NC_039506.1 | 1673027 | 1673526 | 500 | + | Gnomon | promoter | NA | NA | gene-LOC109611032 | GeneID:109611032 | LOC109611032 | Gene | LOC109611032 | lncRNA | character(0) | NA | character(0) |
| NC_039506.1 | 1673135 | 1673135 | 1 | - | 21 | 14 | 7 | 66.6666666666667 | NC_039506.1 | 1673027 | 1673526 | 500 | + | Gnomon | promoter | NA | NA | gene-LOC109611032 | GeneID:109611032 | LOC109611032 | Gene | LOC109611032 | lncRNA | character(0) | NA | character(0) |
| NC_039506.1 | 1673160 | 1673160 | 1 | - | 14 | 12 | 2 | 85.7142857142857 | NC_039506.1 | 1673027 | 1673526 | 500 | + | Gnomon | promoter | NA | NA | gene-LOC109611032 | GeneID:109611032 | LOC109611032 | Gene | LOC109611032 | lncRNA | character(0) | NA | character(0) |
| NC_039506.1 | 1673189 | 1673189 | 1 | - | 21 | 16 | 5 | 76.1904761904762 | NC_039506.1 | 1673027 | 1673526 | 500 | + | Gnomon | promoter | NA | NA | gene-LOC109611032 | GeneID:109611032 | LOC109611032 | Gene | LOC109611032 | lncRNA | character(0) | NA | character(0) |
| NC_039506.1 | 1673197 | 1673197 | 1 | - | 19 | 7 | 12 | 36.8421052631579 | NC_039506.1 | 1673027 | 1673526 | 500 | + | Gnomon | promoter | NA | NA | gene-LOC109611032 | GeneID:109611032 | LOC109611032 | Gene | LOC109611032 | lncRNA | character(0) | NA | character(0) |
| NC_039506.1 | 1673205 | 1673205 | 1 | - | 34 | 23 | 11 | 67.6470588235294 | NC_039506.1 | 1673027 | 1673526 | 500 | + | Gnomon | promoter | NA | NA | gene-LOC109611032 | GeneID:109611032 | LOC109611032 | Gene | LOC109611032 | lncRNA | character(0) | NA | character(0) |
| NC_039506.1 | 1673219 | 1673219 | 1 | - | 60 | 15 | 45 | 25 | NC_039506.1 | 1673027 | 1673526 | 500 | + | Gnomon | promoter | NA | NA | gene-LOC109611032 | GeneID:109611032 | LOC109611032 | Gene | LOC109611032 | lncRNA | character(0) | NA | character(0) |
| NC_039506.1 | 1673225 | 1673225 | 1 | - | 58 | 42 | 16 | 72.4137931034483 | NC_039506.1 | 1673027 | 1673526 | 500 | + | Gnomon | promoter | NA | NA | gene-LOC109611032 | GeneID:109611032 | LOC109611032 | Gene | LOC109611032 | lncRNA | character(0) | NA | character(0) |
| NC_039506.1 | 1673234 | 1673234 | 1 | - | 61 | 32 | 29 | 52.4590163934426 | NC_039506.1 | 1673027 | 1673526 | 500 | + | Gnomon | promoter | NA | NA | gene-LOC109611032 | GeneID:109611032 | LOC109611032 | Gene | LOC109611032 | lncRNA | character(0) | NA | character(0) |
| NC_039506.1 | 1673241 | 1673241 | 1 | - | 57 | 37 | 20 | 64.9122807017544 | NC_039506.1 | 1673027 | 1673526 | 500 | + | Gnomon | promoter | NA | NA | gene-LOC109611032 | GeneID:109611032 | LOC109611032 | Gene | LOC109611032 | lncRNA | character(0) | NA | character(0) |
| NC_039506.1 | 1673284 | 1673284 | 1 | - | 42 | 33 | 9 | 78.5714285714286 | NC_039506.1 | 1673027 | 1673526 | 500 | + | Gnomon | promoter | NA | NA | gene-LOC109611032 | GeneID:109611032 | LOC109611032 | Gene | LOC109611032 | lncRNA | character(0) | NA | character(0) |
| NC_039506.1 | 1673353 | 1673353 | 1 | - | 29 | 28 | 1 | 96.551724137931 | NC_039506.1 | 1673027 | 1673526 | 500 | + | Gnomon | promoter | NA | NA | gene-LOC109611032 | GeneID:109611032 | LOC109611032 | Gene | LOC109611032 | lncRNA | character(0) | NA | character(0) |
| NC_039506.1 | 1673389 | 1673389 | 1 | - | 77 | 74 | 3 | 96.1038961038961 | NC_039506.1 | 1673027 | 1673526 | 500 | + | Gnomon | promoter | NA | NA | gene-LOC109611032 | GeneID:109611032 | LOC109611032 | Gene | LOC109611032 | lncRNA | character(0) | NA | character(0) |
| NC_039506.1 | 1673463 | 1673463 | 1 | + | 5 | 5 | 0 | 100 | NC_039506.1 | 1673027 | 1673526 | 500 | + | Gnomon | promoter | NA | NA | gene-LOC109611032 | GeneID:109611032 | LOC109611032 | Gene | LOC109611032 | lncRNA | character(0) | NA | character(0) |
| NC_039506.1 | 1673464 | 1673464 | 1 | - | 32 | 25 | 7 | 78.125 | NC_039506.1 | 1673027 | 1673526 | 500 | + | Gnomon | promoter | NA | NA | gene-LOC109611032 | GeneID:109611032 | LOC109611032 | Gene | LOC109611032 | lncRNA | character(0) | NA | character(0) |
| NC_039506.1 | 1673487 | 1673487 | 1 | + | 5 | 5 | 0 | 100 | NC_039506.1 | 1673027 | 1673526 | 500 | + | Gnomon | promoter | NA | NA | gene-LOC109611032 | GeneID:109611032 | LOC109611032 | Gene | LOC109611032 | lncRNA | character(0) | NA | character(0) |
| NC_039506.1 | 1673488 | 1673488 | 1 | - | 31 | 24 | 7 | 77.4193548387097 | NC_039506.1 | 1673027 | 1673526 | 500 | + | Gnomon | promoter | NA | NA | gene-LOC109611032 | GeneID:109611032 | LOC109611032 | Gene | LOC109611032 | lncRNA | character(0) | NA | character(0) |
| NC_039506.1 | 1673510 | 1673510 | 1 | + | 7 | 7 | 0 | 100 | NC_039506.1 | 1673027 | 1673526 | 500 | + | Gnomon | promoter | NA | NA | gene-LOC109611032 | GeneID:109611032 | LOC109611032 | Gene | LOC109611032 | lncRNA | character(0) | NA | character(0) |
| NC_039506.1 | 1673511 | 1673511 | 1 | - | 28 | 20 | 8 | 71.4285714285714 | NC_039506.1 | 1673027 | 1673526 | 500 | + | Gnomon | promoter | NA | NA | gene-LOC109611032 | GeneID:109611032 | LOC109611032 | Gene | LOC109611032 | lncRNA | character(0) | NA | character(0) |
| NC_039506.1 | 2187052 | 2187052 | 1 | + | 5 | 5 | 0 | 100 | NC_039506.1 | 2187023 | 2187522 | 500 | + | Gnomon | promoter | NA | NA | gene-LOC105280309 | GeneID:105280309 | LOC105280309 | Gene | LOC105280309 | protein_coding | character(0) | NA | character(0) |
| NC_039506.1 | 2200872 | 2200872 | 1 | - | 9 | 6 | 3 | 66.6666666666667 | NC_039506.1 | 2200872 | 2201371 | 500 | + | Gnomon | promoter | NA | NA | gene-LOC105280312 | GeneID:105280312 | LOC105280312 | Gene | LOC105280312 | protein_coding | character(0) | NA | character(0) |
| NC_039506.1 | 3094044 | 3094044 | 1 | + | 20 | 9 | 11 | 45 | NC_039506.1 | 3094025 | 3094524 | 500 | + | Gnomon | promoter | NA | NA | gene-LOC105280352 | GeneID:105280352 | LOC105280352 | Gene | LOC105280352 | protein_coding | character(0) | NA | character(0) |
| NC_039506.1 | 3094045 | 3094045 | 1 | - | 15 | 8 | 7 | 53.3333333333333 | NC_039506.1 | 3094025 | 3094524 | 500 | + | Gnomon | promoter | NA | NA | gene-LOC105280352 | GeneID:105280352 | LOC105280352 | Gene | LOC105280352 | protein_coding | character(0) | NA | character(0) |
| NC_039506.1 | 3094067 | 3094067 | 1 | + | 14 | 11 | 3 | 78.5714285714286 | NC_039506.1 | 3094025 | 3094524 | 500 | + | Gnomon | promoter | NA | NA | gene-LOC105280352 | GeneID:105280352 | LOC105280352 | Gene | LOC105280352 | protein_coding | character(0) | NA | character(0) |

...

... only first 25 lines shown ...

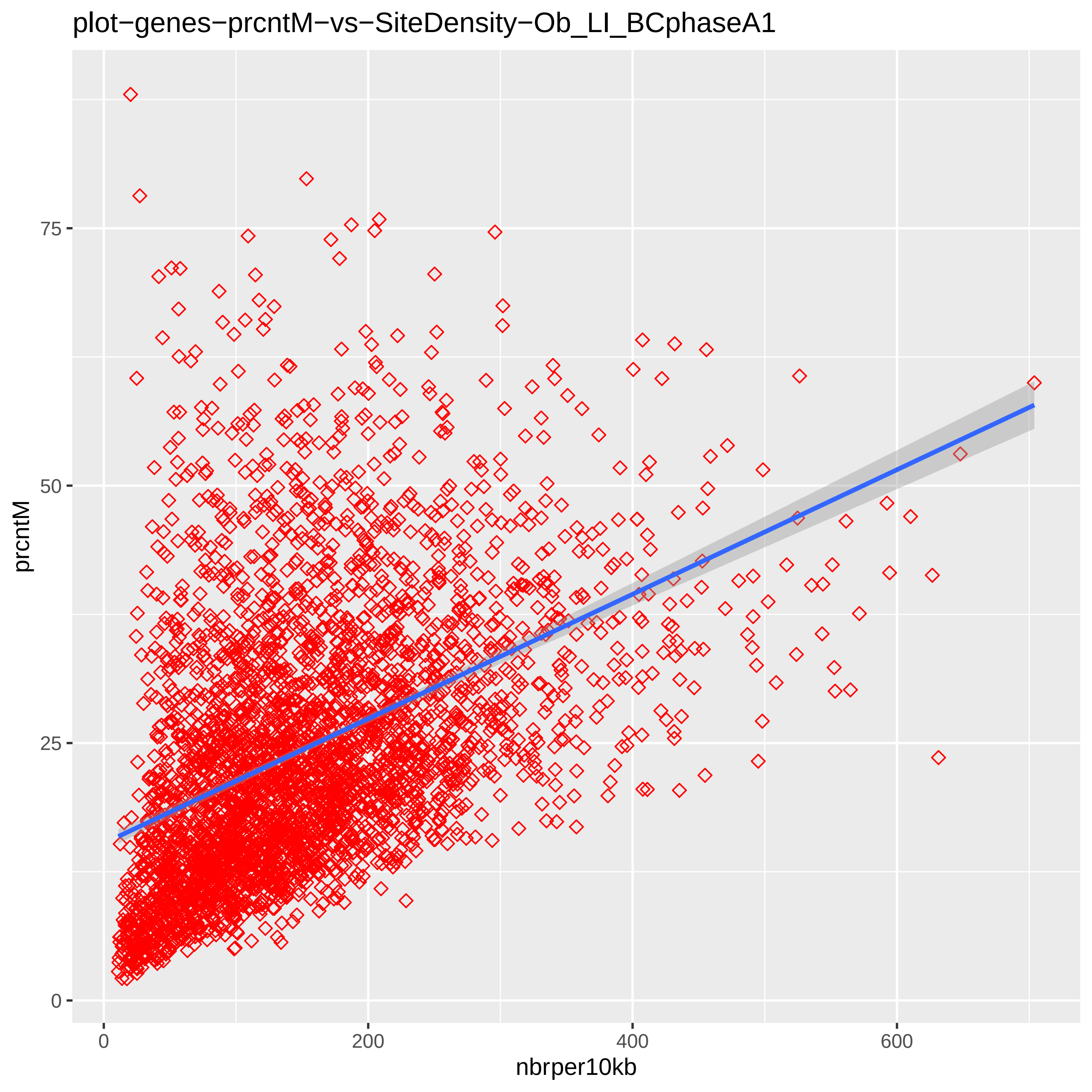
plot−genes−prcntM−vs−SiteDensity−Ob_LI_BCphaseA1
prcntM
75
50
25
0
nbrper10kb
0
200
400
600

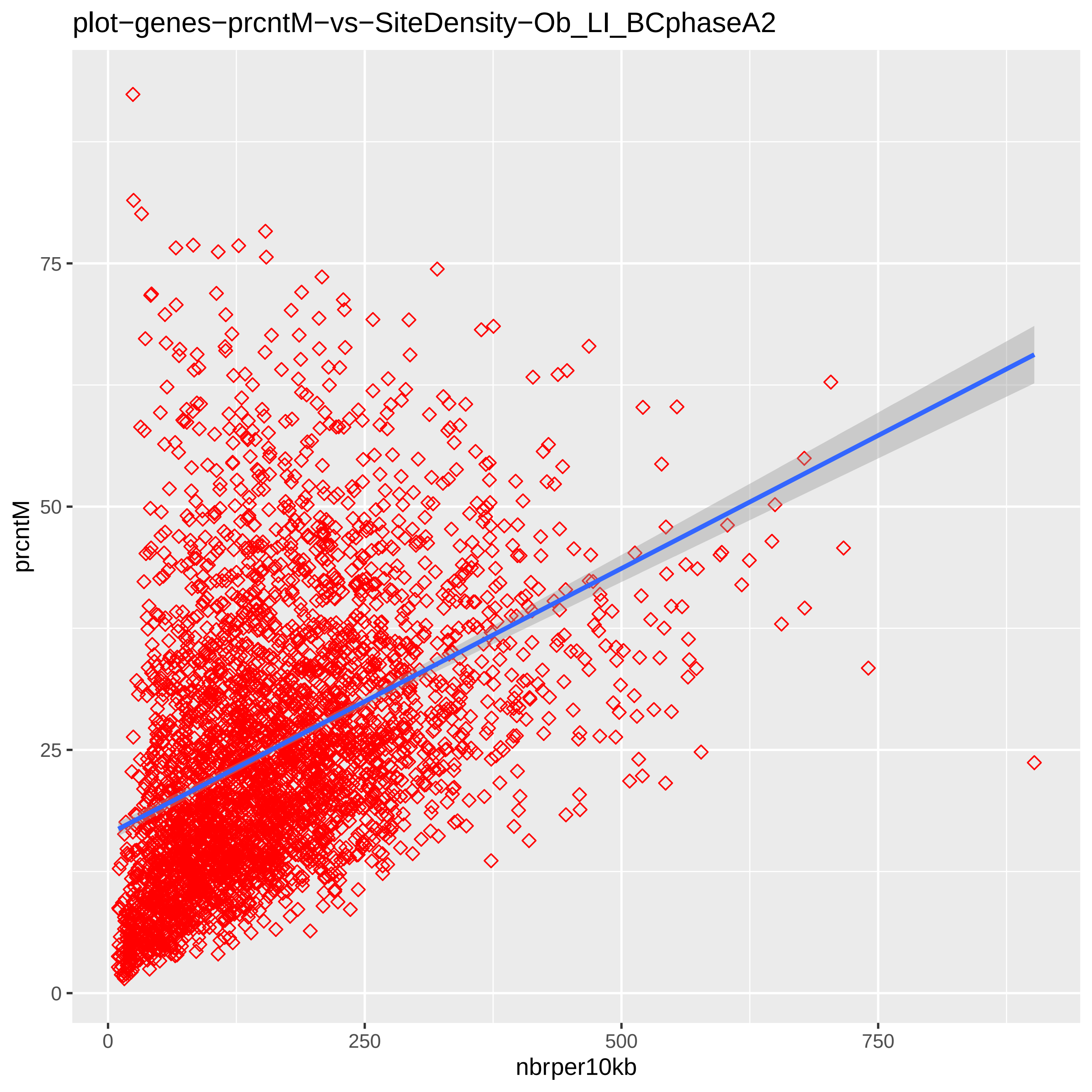
plot−genes−prcntM−vs−SiteDensity−Ob_LI_BCphaseA2
prcntM
nbrper10kb
0
25
50
75
0
250
500
750

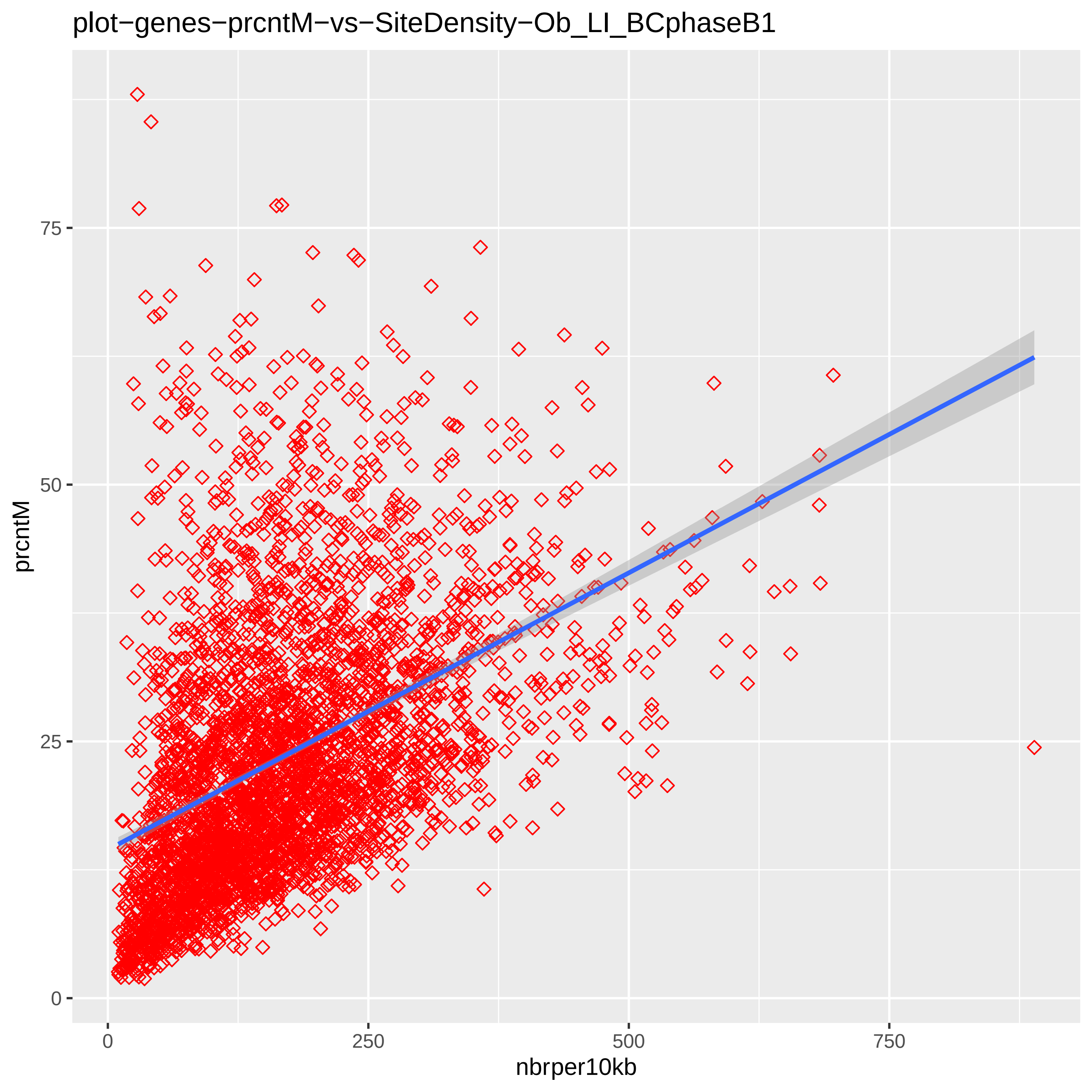

plot−genes−prcntM−vs−SiteDensity−Ob_LI_BCphaseB1
prcntM
75
50
25
0
0
250
500
750
nbrper10kb

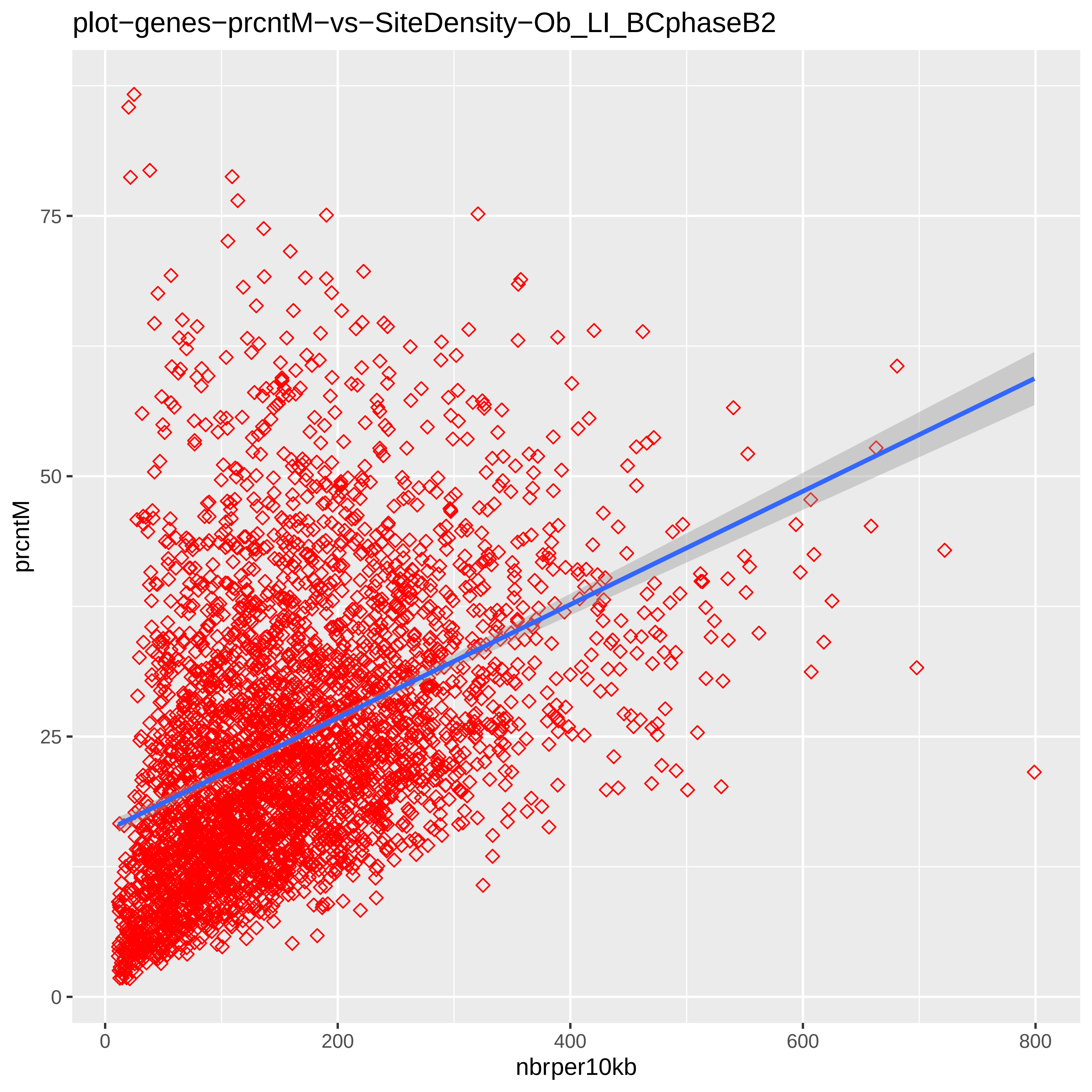
plot−genes−prcntM−vs−SiteDensity−Ob_LI_BCphaseB2
prcntM
75
50
25
0
0
200
400
600
800
nbrper10kb

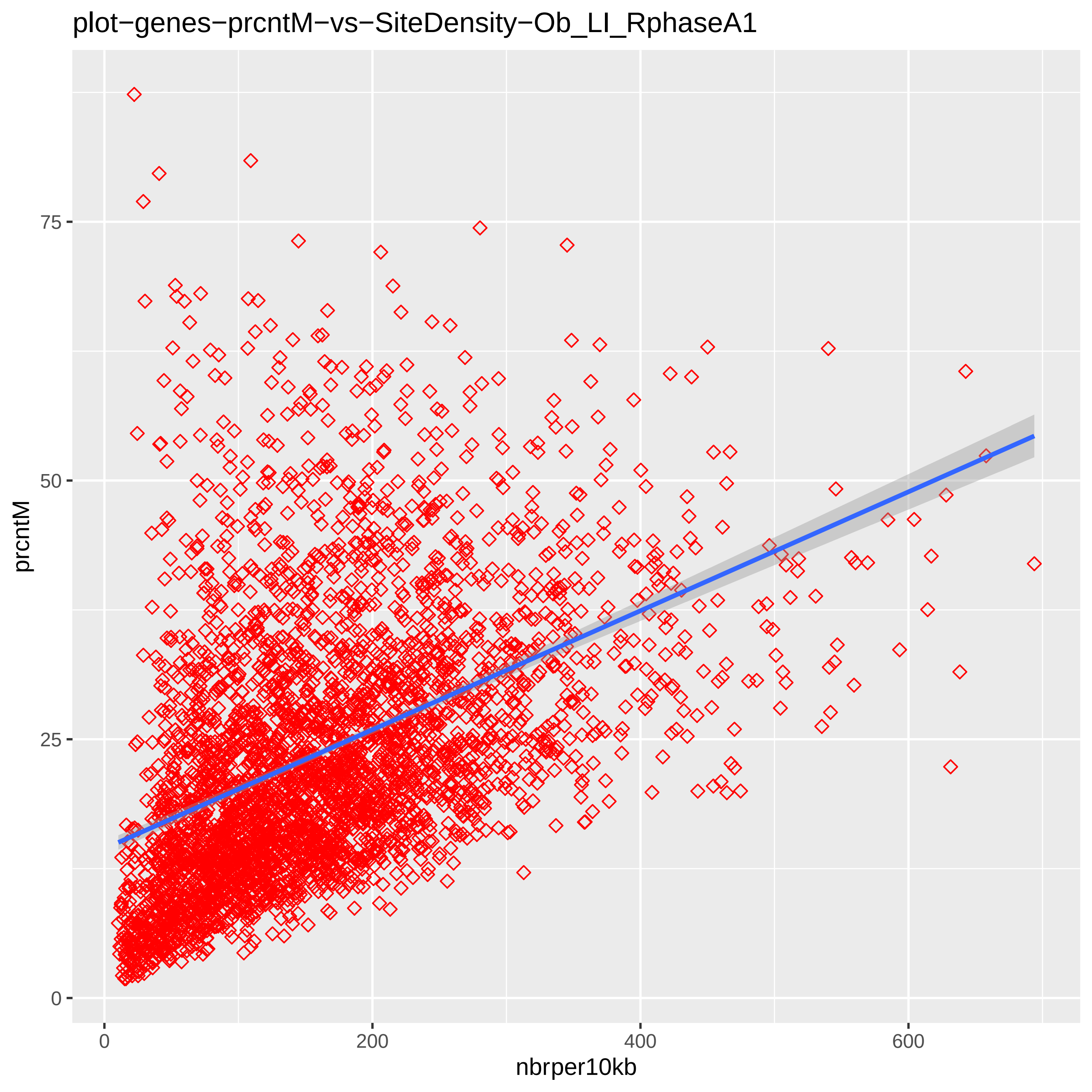
plot−genes−prcntM−vs−SiteDensity−Ob_LI_RphaseA1
prcntM
0
25
50
75
nbrper10kb
0
200
400
600

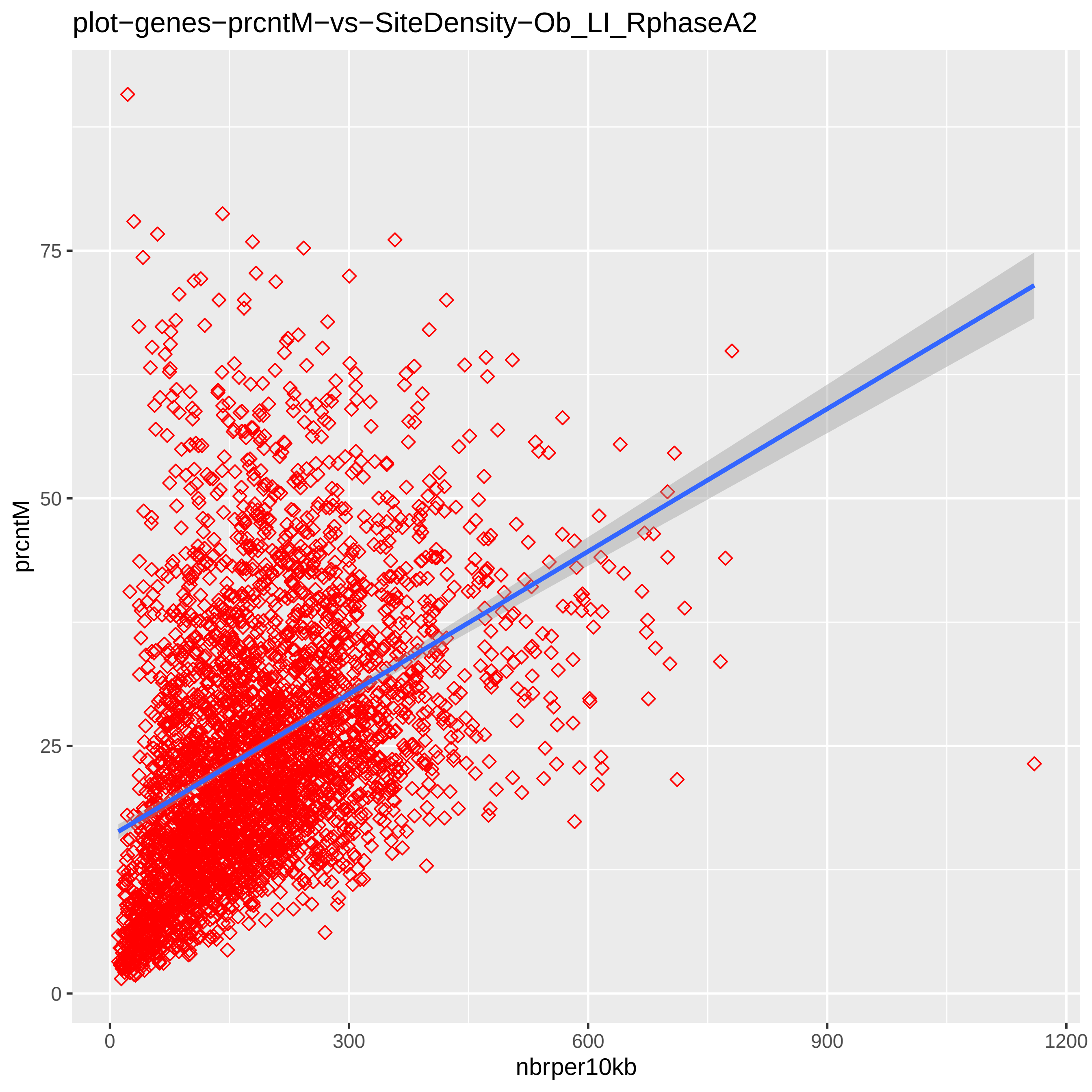
plot−genes−prcntM−vs−SiteDensity−Ob_LI_RphaseA2
prcntM
nbrper10kb
0
25
50
75
0
300
600
900
1200

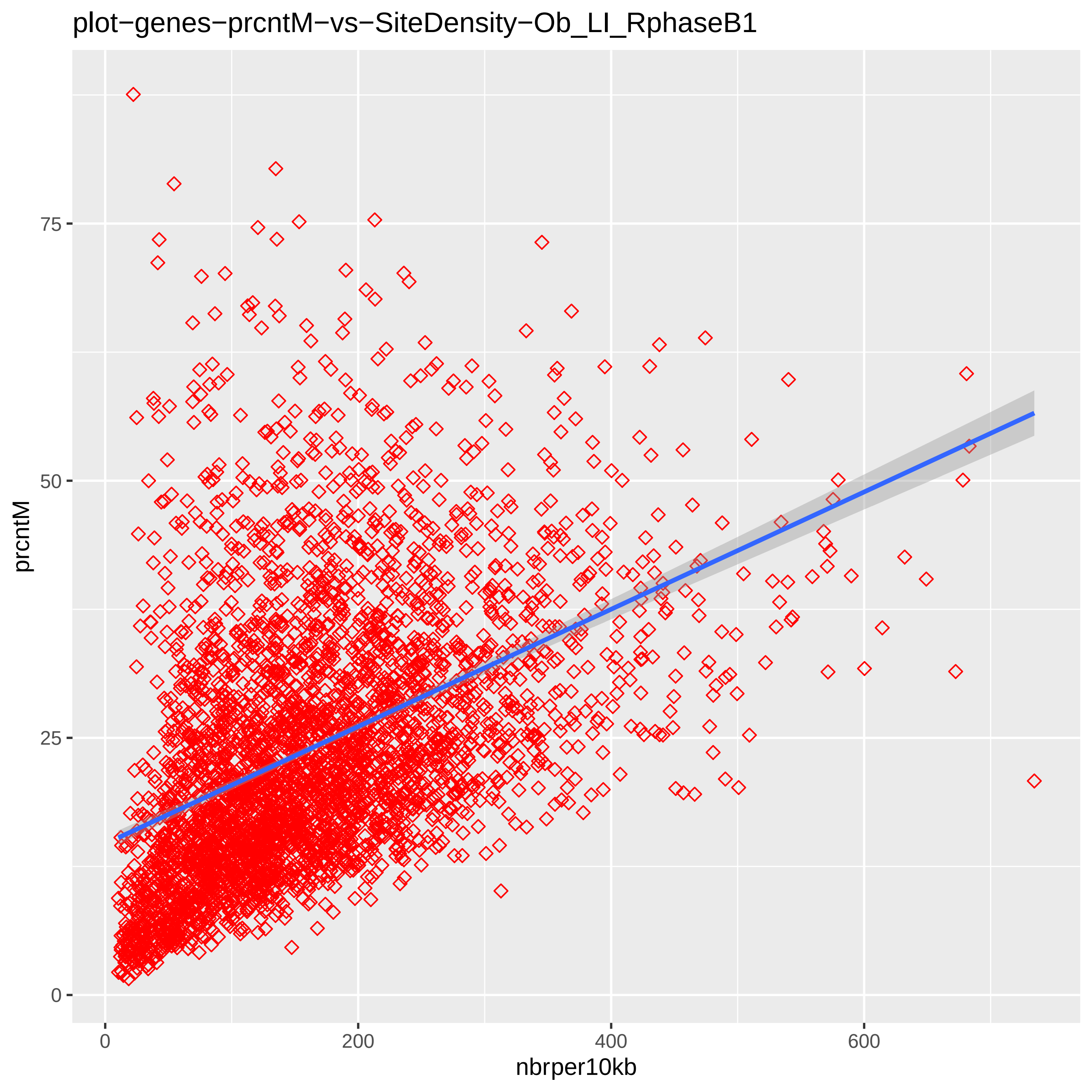
plot−genes−prcntM−vs−SiteDensity−Ob_LI_RphaseB1
prcntM
nbrper10kb
0
25
50
75
0
200
400
600

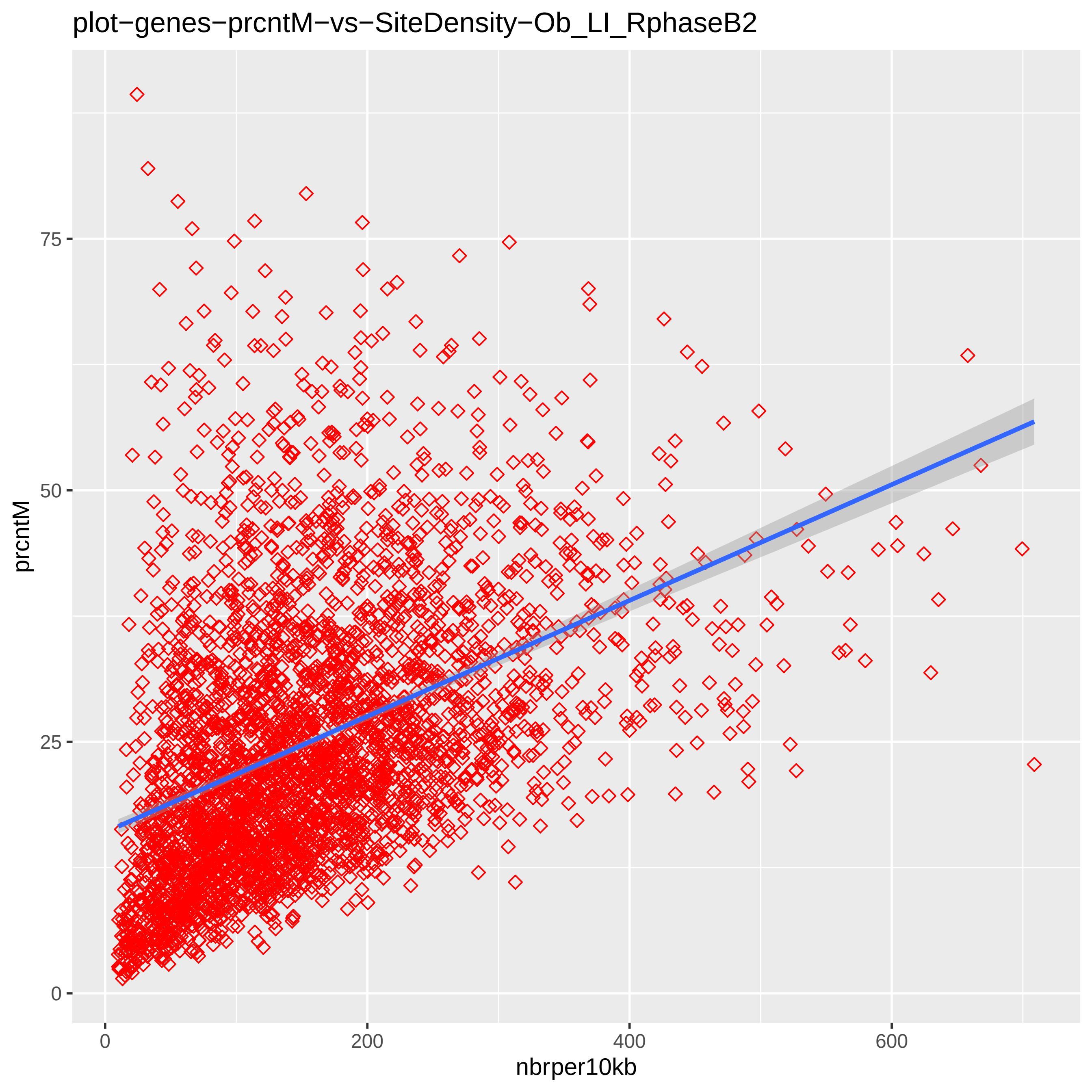
plot−genes−prcntM−vs−SiteDensity−Ob_LI_RphaseB2
prcntM
nbrper10kb
0
25
50
75
0
200
400
600

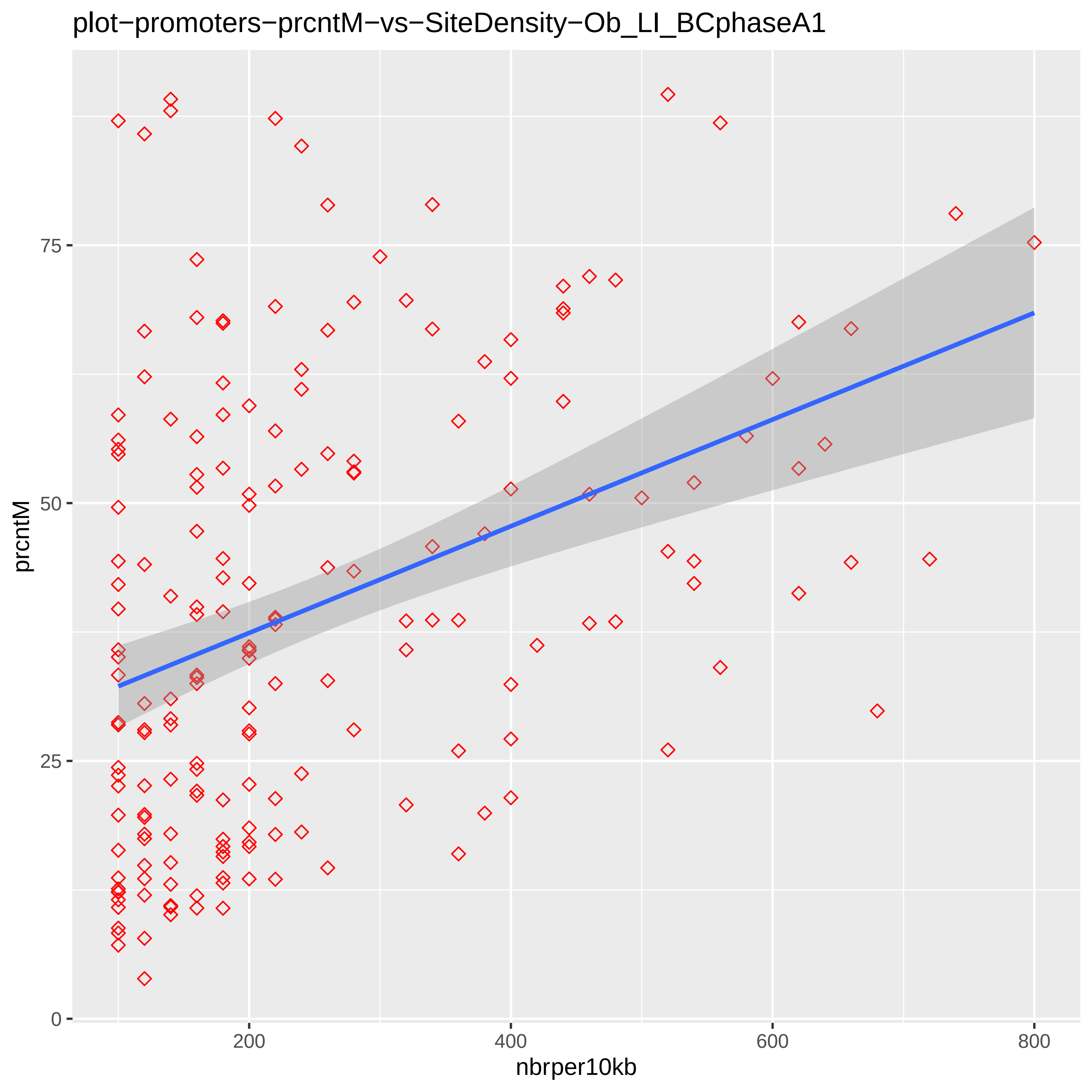

plot−promoters−prcntM−vs−SiteDensity−Ob_LI_BCphaseA1
prcntM
0
25
50
75
200
400
600
800
nbrper10kb

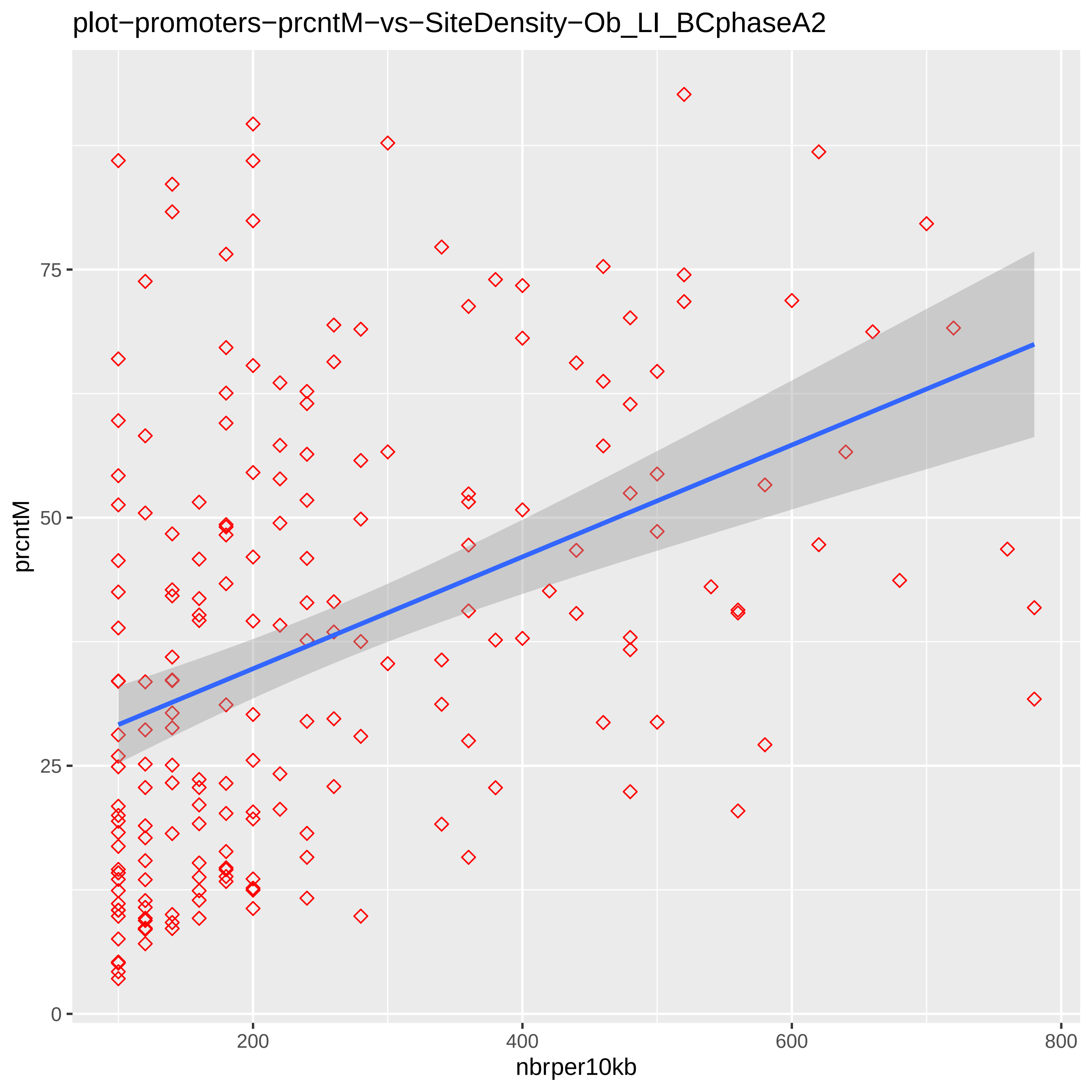

plot−promoters−prcntM−vs−SiteDensity−Ob_LI_BCphaseA2
prcntM
0
25
50
75
200
400
600
800
nbrper10kb

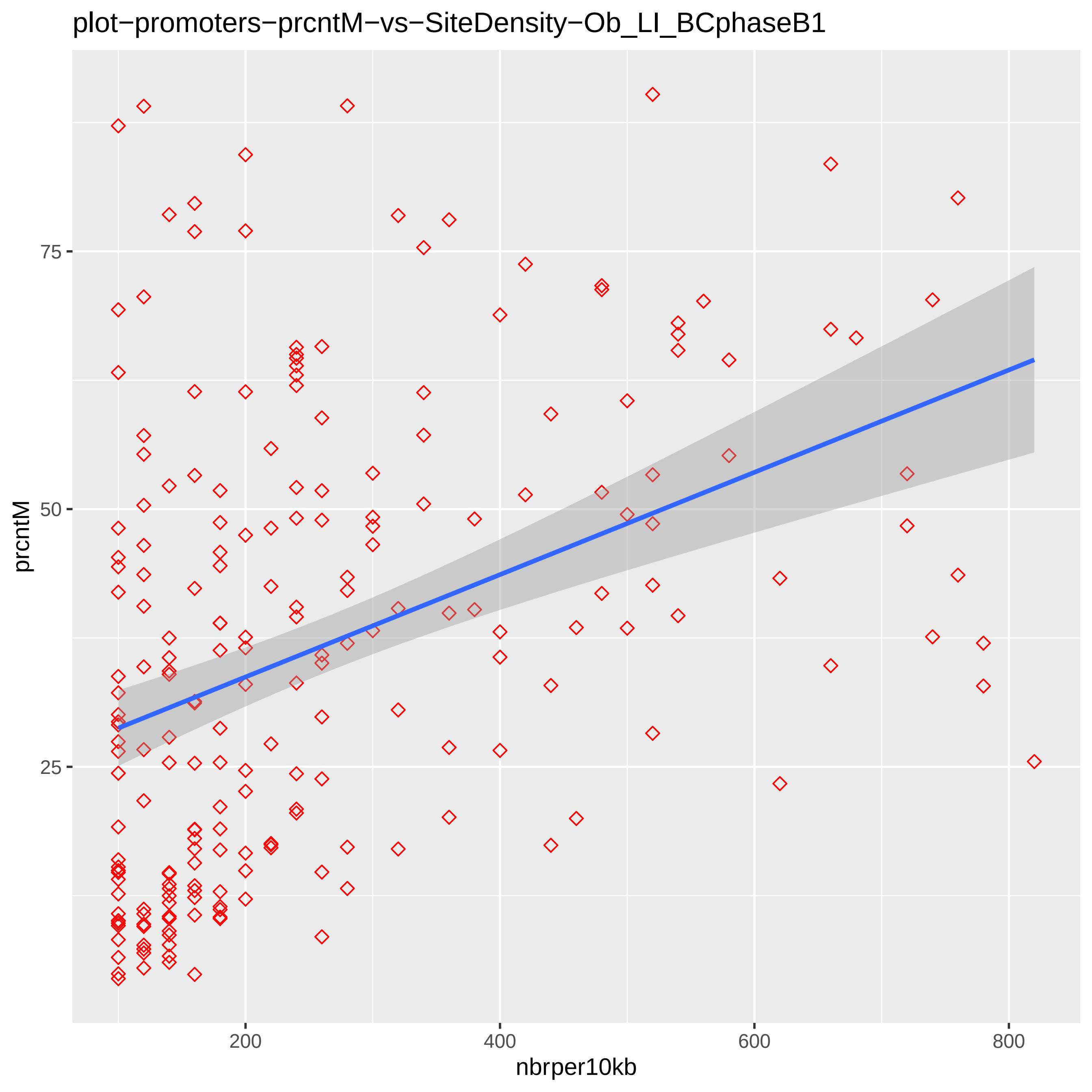
plot−promoters−prcntM−vs−SiteDensity−Ob_LI_BCphaseB1
prcntM
75
50
25
200
400
600
800
nbrper10kb

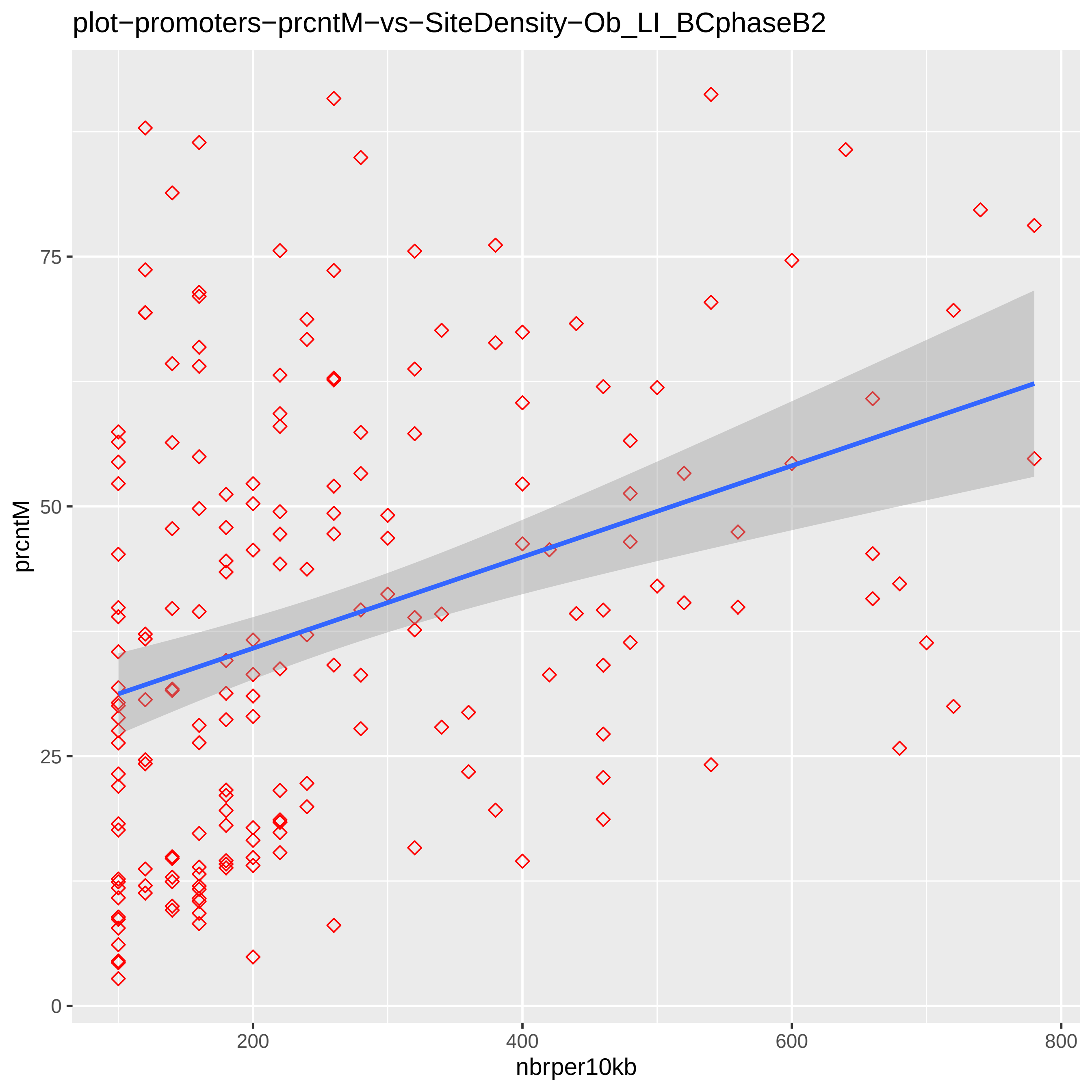
plot−promoters−prcntM−vs−SiteDensity−Ob_LI_BCphaseB2
prcntM
nbrper10kb
0
25
50
75
200
400
600
800

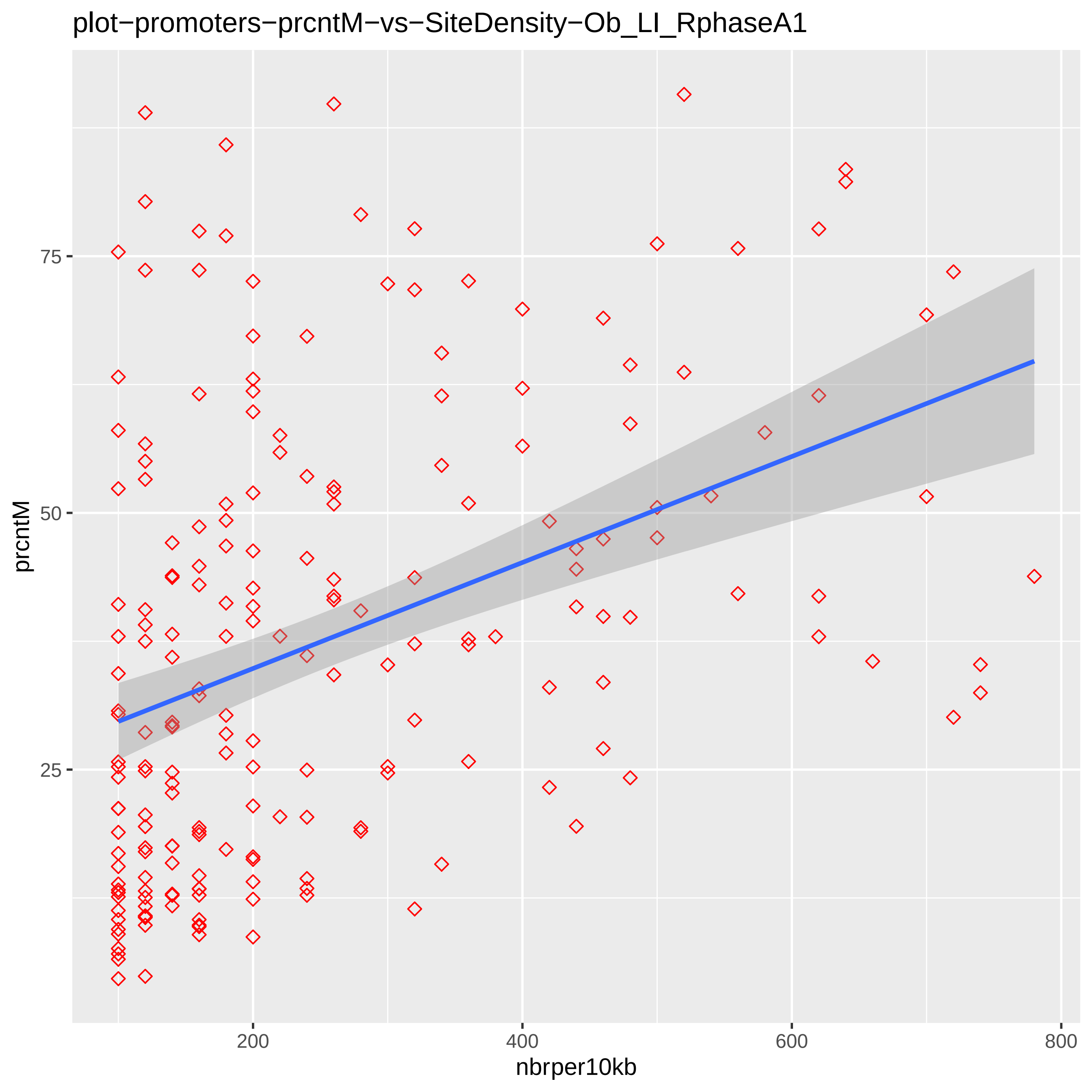
plot−promoters−prcntM−vs−SiteDensity−Ob_LI_RphaseA1
prcntM
75
50
25
200
400
600
800
nbrper10kb

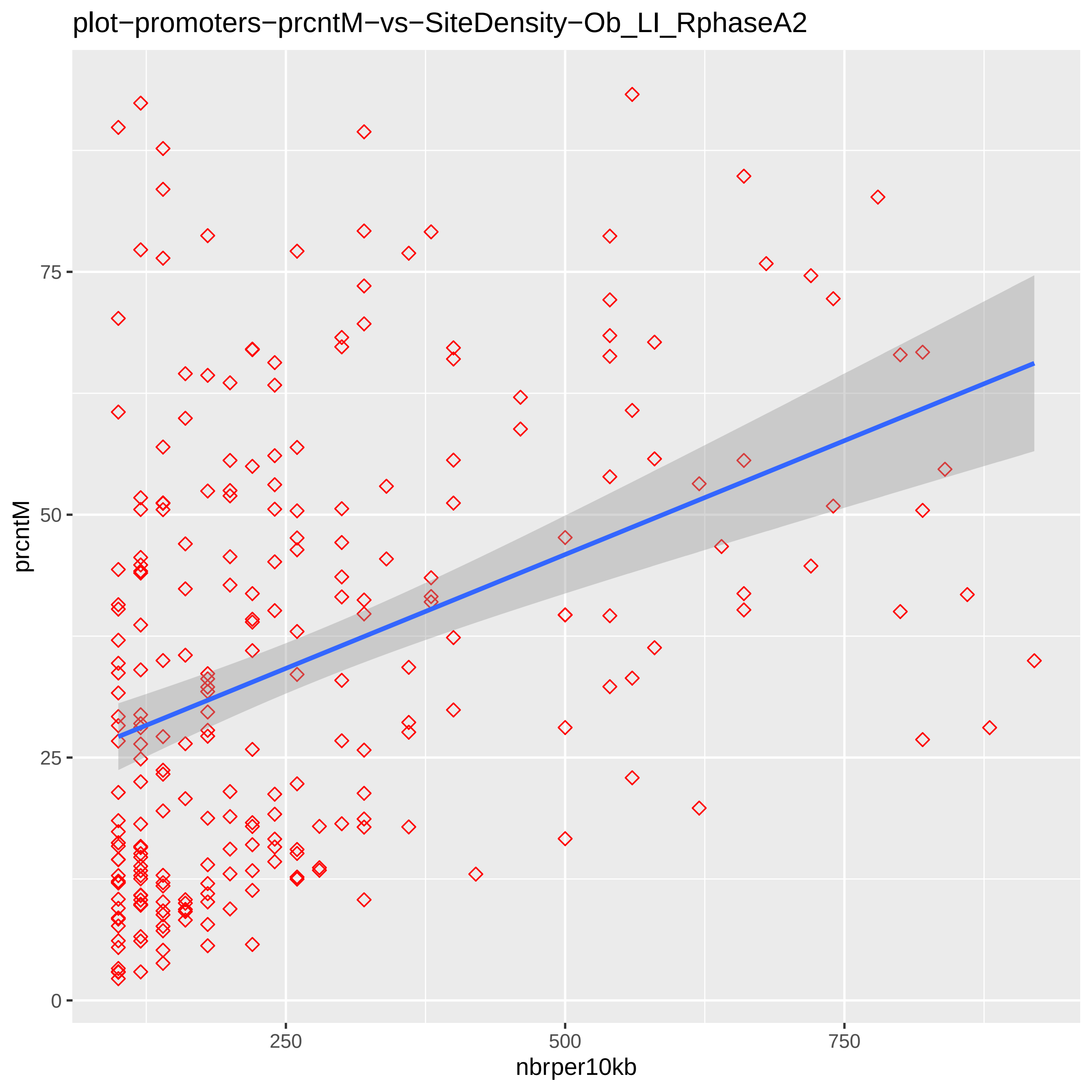
plot−promoters−prcntM−vs−SiteDensity−Ob_LI_RphaseA2
prcntM
75
50
25
0
250
500
750
nbrper10kb

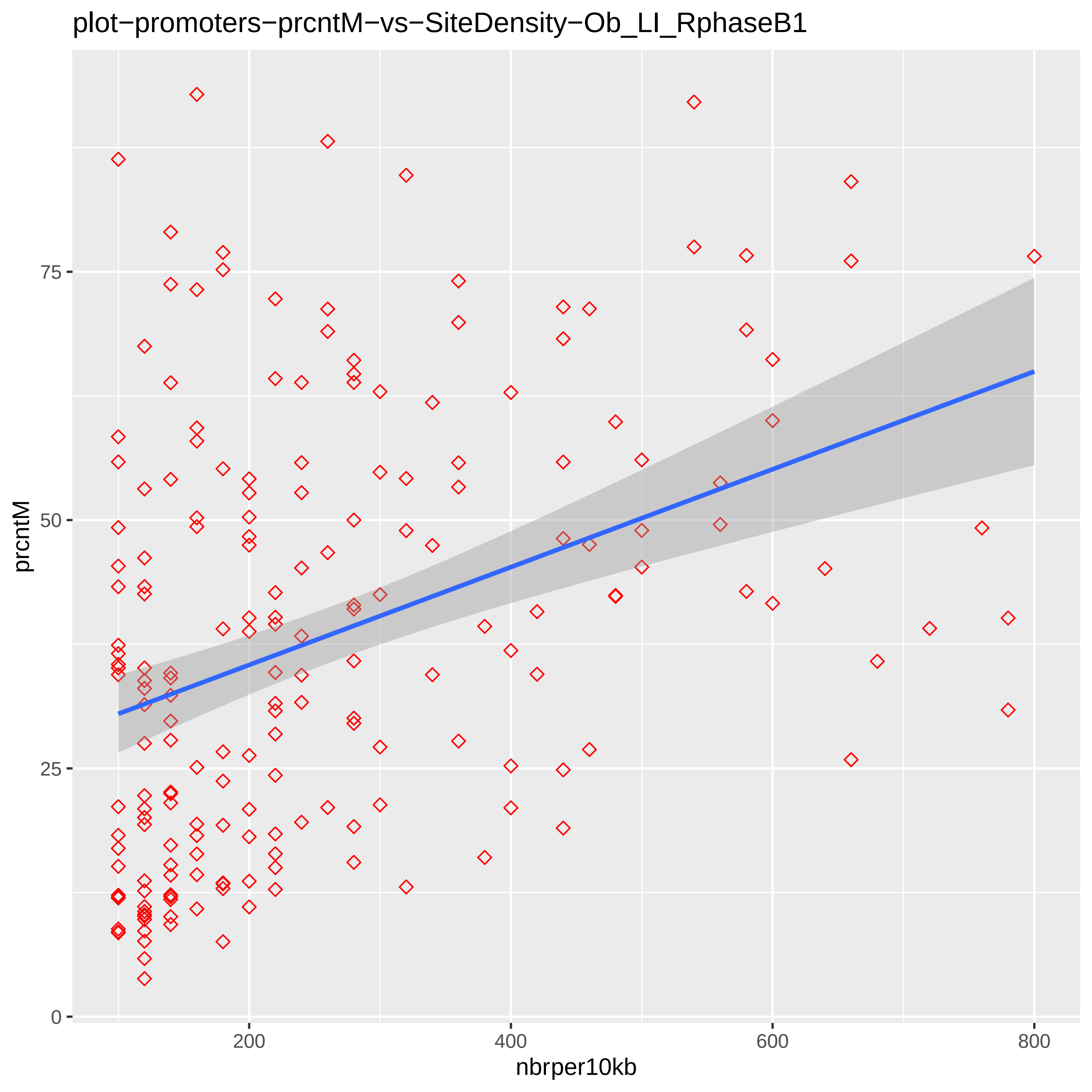

plot−promoters−prcntM−vs−SiteDensity−Ob_LI_RphaseB1
prcntM
nbrper10kb
0
25
50
75
200
400
600
800

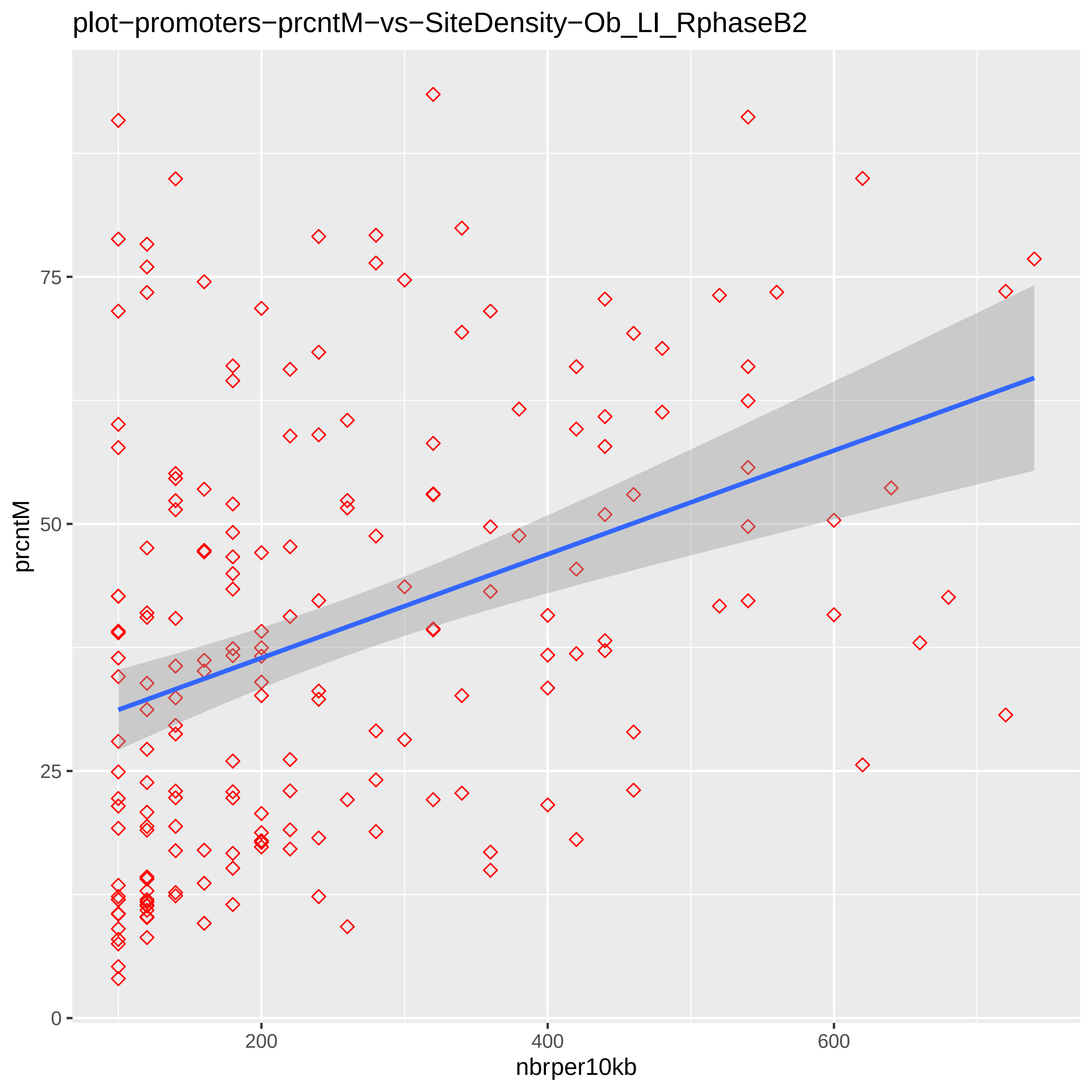
plot−promoters−prcntM−vs−SiteDensity−Ob_LI_RphaseB2
prcntM
75
50
25
0
200
400
600
nbrper10kb

```
Directory: MRPR File: 0READMEmrpr

MRPR - Methylation-rich and -poor regions.

Input: studymk, genome_ann (a list of GRanges objects providing annotated region labels and bounds;
                           output of BWASPR::get_genome_annotation())
       nbrxtrms (specifies the number of richest and poorest regions to be displayed; default: 100)

Output: files dst-*.txt 1ds-*.pdf 5ds-*.pdf    mdr-*.bed   rmp-* gwp-* gwr-*

Notes: The output is generated by BWASPR functions det_mrpr() and map_mrpr().
       Methylation-rich and -poor regions are determined based on the spacing between
        neighboring CpGhsm sites.  If sites occur at positions a, b, c, d, e, and f
        (and not in between), then b-a, c-b, ... are 1-distances, and f-a is a 5-distance.
        The file dst-*.txt shows the empirical distribution of d-distances for the sample.
        Methylation-poor regions are determined as long 1-distances (in the top <nbrxtrms>; merged
        if adjacent 1-distances are both in the top <nbrxtrms>).  Methylation-rich regions are determined
        as short 5-distances (in the low <nbrxtrms>; merged if adjacent 5-distances are both in the low
        <nbrxtrms>).  Parts of the distribution are plotted in the 1ds-*.pdf and 5ds-*.pdf files.

       The methylation-rich and -poor regions are listed in files dst-*.txt and mdr-*.bed,
        the latter in BED format for potential display in genome browsers.

       The overlap of methylation regions with genome features is summarized in files rmp-*.txt.
        Files gwr-*.txt show genes overlapping with methylation-rich regions, ordered by site
        density in the methylation-rich region.
        Files gwp-*.txt show genes overlapping with methylation-poor regions, ordered by site
        density in the methylation-poor region.
```

Directory: MRPR File: dst-Ob_LI_BCphaseA1.txt

Analysis of 1-distances for sample "Ob_LI_BCphaseA1" (197029 sites):

Total number of 1-distances: 197010
Median: 11.0 Mean: 1113.3 Std: 14938.4
Quantiles:

| 0% | 5% | 10% | 15% | 20% | 25% | 30% | 35% | 40% | 45% | 50% | 55% | 60% | 65% | 70% | 75% | 80% | 85% | 90% | 95% | 100% |
|---|---|---|---|---|---|---|---|---|---|---|---|---|---|---|---|---|---|---|---|---|
| 1 | 1 | 1 | 1 | 1 | 1 | 1 | 3 | 6 | 8 | 11 | 14 | 18 | 22 | 28 | 36 | 47 | 69 | 116 | 328 | 1333649 |

Low density region distance cutoff (lowest 100): 301082
High density region distance cutoff (highest 100): 1

Ordered lists of methylation-poor regions:

Top 100 (out of 92) methylation-poor regions in "Ob_LI_BCphaseA1":
(Sdnsty = sites per 1kb)

| | Rtype | Sample | SeqID | From | To | Rlgth | NbrSites | Sdnsty |
|---|---|---|---|---|---|---|---|---|
| 1 | Poor | BCphaseA1 | NC_039518.1 | 5545094 | 6878743 | 1333650 | 2 | 0.00150 |
| 2 | Poor | BCphaseA1 | NC_039510.1 | 6991940 | 8277430 | 1285491 | 2 | 0.00156 |
| 3 | Poor | BCphaseA1 | NC_039513.1 | 49855 | 1653046 | 1603192 | 3 | 0.00187 |
| 4 | Poor | BCphaseA1 | NC_039516.1 | 7633246 | 8594200 | 960955 | 2 | 0.00208 |
| 5 | Poor | BCphaseA1 | NC_039511.1 | 6611048 | 8304235 | 1693188 | 4 | 0.00236 |
| 6 | Poor | BCphaseA1 | NC_039506.1 | 6804709 | 7641314 | 836606 | 2 | 0.00239 |

...
... only first 25 lines shown ...

Directory: MRPR File: dst-Ob_LI_BCphaseA2.txt

Analysis of 1-distances for sample "Ob_LI_BCphaseA2" (213136 sites):

Total number of 1-distances: 213121
Median: 10.0 Mean: 1017.5 Std: 16381.0
Quantiles:

| 0% | 5% | 10% | 15% | 20% | 25% | 30% | 35% | 40% | 45% | 50% | 55% | 60% | 65% | 70% | 75% | 80% | 85% | 90% | 95% | 100% |
|---|---|---|---|---|---|---|---|---|---|---|---|---|---|---|---|---|---|---|---|---|
| 1 | 1 | 1 | 1 | 1 | 1 | 1 | 3 | 5 | 8 | 10 | 13 | 17 | 21 | 26 | 33 | 43 | 61 | 102 | 262 | 1403739 |

Low density region distance cutoff (lowest 100): 356501
High density region distance cutoff (highest 100): 1

Ordered lists of methylation-poor regions:

Top 100 (out of 96) methylation-poor regions in "Ob_LI_BCphaseA2":
(Sdnsty = sites per 1kb)

| | Rtype | Sample | SeqID | From | To | Rlgth | NbrSites | Sdnsty |
|---|---|---|---|---|---|---|---|---|
| 1 | Poor | BCphaseA2 | NC_039510.1 | 6992643 | 8396382 | 1403740 | 2 | 0.00142 |
| 2 | Poor | BCphaseA2 | NC_039508.1 | 7797312 | 9041543 | 1244232 | 2 | 0.00161 |
| 3 | Poor | BCphaseA2 | NC_039517.1 | 10919759 | 12683795 | 1764037 | 3 | 0.00170 |
| 4 | Poor | BCphaseA2 | NC_039511.1 | 6586955 | 8305089 | 1718135 | 3 | 0.00175 |
| 5 | Poor | BCphaseA2 | NC_039514.1 | 4377103 | 5490389 | 1113287 | 2 | 0.00180 |
| 6 | Poor | BCphaseA2 | NC_039519.1 | 923043 | 1956245 | 1033203 | 2 | 0.00194 |

...
... only first 25 lines shown ...

Directory: MRPR File: dst-Ob_LI_BCphaseB1.txt

Analysis of 1-distances for sample "Ob_LI_BCphaseB1" (226855 sites):

Total number of 1-distances: 226838
Median: 10.0 Mean: 969.3 Std: 12209.5
Quantiles:

| 0% | 5% | 10% | 15% | 20% | 25% | 30% | 35% | 40% | 45% | 50% | 55% | 60% | 65% | 70% | 75% | 80% | 85% | 90% | 95% | 100% |
|---|---|---|---|---|---|---|---|---|---|---|---|---|---|---|---|---|---|---|---|---|
| 1 | 1 | 1 | 1 | 1 | 1 | 1 | 2 | 5 | 7 | 10 | 13 | 17 | 21 | 26 | 34 | 45 | 65 | 111 | 307 | 1098092 |

Low density region distance cutoff (lowest 100): 258964
High density region distance cutoff (highest 100): 1

Ordered lists of methylation-poor regions:

Top 100 (out of 96) methylation-poor regions in "Ob_LI_BCphaseB1":
(Sdnsty = sites per 1kb)

| | Rtype | Sample | SeqID | From | To | Rlgth | NbrSites | Sdnsty |
|---|---|---|---|---|---|---|---|---|
| 1 | Poor | BCphaseB1 | NC_039518.1 | 5621056 | 6719148 | 1098093 | 2 | 0.00182 |
| 2 | Poor | BCphaseB1 | NC_039506.1 | 137598 | 1058205 | 920608 | 2 | 0.00217 |
| 3 | Poor | BCphaseB1 | NC_039506.1 | 6804755 | 7641314 | 836560 | 2 | 0.00239 |
| 4 | Poor | BCphaseB1 | NC_039519.1 | 4759443 | 5455854 | 696412 | 2 | 0.00287 |
| 5 | Poor | BCphaseB1 | NC_039518.1 | 3077899 | 3765933 | 688035 | 2 | 0.00291 |
| 6 | Poor | BCphaseB1 | NC_039515.1 | 12841060 | 13515447 | 674388 | 2 | 0.00297 |

...
... only first 25 lines shown ...

Directory: MRPR File: dst-Ob_LI_BCphaseB2.txt

Analysis of 1-distances for sample "Ob_LI_BCphaseB2" (209549 sites):

Total number of 1-distances: 209533
Median: 11.0 Mean: 1046.0 Std: 13359.6
Quantiles:

| 0% | 5% | 10% | 15% | 20% | 25% | 30% | 35% | 40% | 45% | 50% | 55% | 60% | 65% | 70% | 75% | 80% | 85% | 90% | 95% | 100% |
|---|---|---|---|---|---|---|---|---|---|---|---|---|---|---|---|---|---|---|---|---|
| 1 | 1 | 1 | 1 | 1 | 1 | 1 | 3 | 5 | 8 | 11 | 14 | 17 | 21 | 27 | 34 | 45 | 65 | 111 | 316 | 1123560 |

Low density region distance cutoff (lowest 100): 270841
High density region distance cutoff (highest 100): 1

Ordered lists of methylation-poor regions:

Top 100 (out of 90) methylation-poor regions in "Ob_LI_BCphaseB2":
(Sdnsty = sites per 1kb)

| | Rtype | Sample | SeqID | From | To | Rlgth | NbrSites | Sdnsty |
|---|---|---|---|---|---|---|---|---|
| 1 | Poor | BCphaseB2 | NC_039518.1 | 5974026 | 7049520 | 1075495 | 2 | 0.00186 |
| 2 | Poor | BCphaseB2 | NC_039511.1 | 6715713 | 8126204 | 1410492 | 3 | 0.00213 |
| 3 | Poor | BCphaseB2 | NC_039516.1 | 7715620 | 8594453 | 878834 | 2 | 0.00228 |
| 4 | Poor | BCphaseB2 | NC_039510.1 | 7068682 | 7937215 | 868534 | 2 | 0.00230 |
| 5 | Poor | BCphaseB2 | NC_039513.1 | 12988 | 871185 | 858198 | 2 | 0.00233 |
| 6 | Poor | BCphaseB2 | NC_039506.1 | 6804709 | 7641130 | 836422 | 2 | 0.00239 |

...
... only first 25 lines shown ...

Directory: MRPR File: dst-Ob_LI_RphaseA1.txt

Analysis of 1-distances for sample "Ob_LI_RphaseA1" (213360 sites):

Total number of 1-distances: 213341
Median: 10.0 Mean: 1022.3 Std: 15817.2
Quantiles:

| 0% | 5% | 10% | 15% | 20% | 25% | 30% | 35% | 40% | 45% | 50% | 55% | 60% | 65% | 70% | 75% | 80% | 85% | 90% | 95% | 100% |
|---|---|---|---|---|---|---|---|---|---|---|---|---|---|---|---|---|---|---|---|---|
| 1 | 1 | 1 | 1 | 1 | 1 | 1 | 2 | 5 | 7 | 10 | 13 | 17 | 21 | 27 | 34 | 45 | 64 | 109 | 276 | 1668843 |

Low density region distance cutoff (lowest 100): 355548
High density region distance cutoff (highest 100): 1

Ordered lists of methylation-poor regions:

Top 100 (out of 94) methylation-poor regions in "Ob_LI_RphaseA1":
(Sdnsty = sites per 1kb)

| | Rtype | Sample | SeqID | From | To | Rlgth | NbrSites | Sdnsty |
|---|---|---|---|---|---|---|---|---|
| 1 | Poor | RphaseA1 | NC_039517.1 | 11083779 | 12752622 | 1668844 | 2 | 0.00120 |
| 2 | Poor | RphaseA1 | NC_039511.1 | 6586955 | 8305089 | 1718135 | 3 | 0.00175 |
| 3 | Poor | RphaseA1 | NC_039514.1 | 4485672 | 5550310 | 1064639 | 2 | 0.00188 |
| 4 | Poor | RphaseA1 | NC_039518.1 | 5545094 | 7051390 | 1506297 | 3 | 0.00199 |
| 5 | Poor | RphaseA1 | NC_039510.1 | 7063270 | 8396382 | 1333113 | 3 | 0.00225 |
| 6 | Poor | RphaseA1 | NC_039506.1 | 8173736 | 9049014 | 875279 | 2 | 0.00228 |

...
... only first 25 lines shown ...

Directory: MRPR File: dst-Ob_LI_RphaseA2.txt

Analysis of 1-distances for sample "Ob_LI_RphaseA2" (265465 sites):

Total number of 1-distances: 265444
Median: 10.0 Mean: 831.0 Std: 8066.3
Quantiles:

| 0% | 5% | 10% | 15% | 20% | 25% | 30% | 35% | 40% | 45% | 50% | 55% | 60% | 65% | 70% | 75% | 80% | 85% | 90% | 95% | 100% |
|---|---|---|---|---|---|---|---|---|---|---|---|---|---|---|---|---|---|---|---|---|
| 1 | 1 | 1 | 1 | 1 | 1 | 1 | 2 | 5 | 7 | 10 | 13 | 16 | 20 | 26 | 33 | 44 | 65 | 116 | 432 | 710164 |

Low density region distance cutoff (lowest 100): 162764
High density region distance cutoff (highest 100): 1

Ordered lists of methylation-poor regions:

Top 100 (out of 87) methylation-poor regions in "Ob_LI_RphaseA2":
(Sdnsty = sites per 1kb)

| | Rtype | Sample | SeqID | From | To | Rlgth | NbrSites | Sdnsty |
|---|---|---|---|---|---|---|---|---|
| 1 | Poor | RphaseA2 | NC_039506.1 | 11672856 | 12383020 | 710165 | 2 | 0.00282 |
| 2 | Poor | RphaseA2 | NC_039506.1 | 138205 | 1097588 | 959384 | 3 | 0.00313 |
| 3 | Poor | RphaseA2 | NC_039517.1 | 12407713 | 13005470 | 597758 | 2 | 0.00335 |
| 4 | Poor | RphaseA2 | NC_039506.1 | 6804709 | 7641269 | 836561 | 3 | 0.00359 |
| 5 | Poor | RphaseA2 | NC_039508.1 | 8376285 | 8880732 | 504448 | 2 | 0.00396 |
| 6 | Poor | RphaseA2 | NC_039511.1 | 6981230 | 7735053 | 753824 | 3 | 0.00398 |

...
... only first 25 lines shown ...

Directory: MRPR File: dst-Ob_LI_RphaseB1.txt

Analysis of 1-distances for sample "Ob_LI_RphaseB1" (217571 sites):

Total number of 1-distances: 217552
Median: 10.0 Mean: 1011.2 Std: 12131.7
Quantiles:

| 0% | 5% | 10% | 15% | 20% | 25% | 30% | 35% | 40% | 45% | 50% | 55% | 60% | 65% | 70% | 75% | 80% | 85% | 90% | 95% | 100% |
|---|---|---|---|---|---|---|---|---|---|---|---|---|---|---|---|---|---|---|---|---|
| 1 | 1 | 1 | 1 | 1 | 1 | 1 | 3 | 5 | 8 | 10 | 14 | 17 | 21 | 27 | 35 | 46 | 66 | 113 | 320 | 836605 |

Low density region distance cutoff (lowest 100): 259714
High density region distance cutoff (highest 100): 1

Ordered lists of methylation-poor regions:

Top 100 (out of 87) methylation-poor regions in "Ob_LI_RphaseB1":
(Sdnsty = sites per 1kb)

| | Rtype | Sample | SeqID | From | To | Rlgth | NbrSites | Sdnsty |
|---|---|---|---|---|---|---|---|---|
| 1 | Poor | RphaseB1 | NC_039506.1 | 6804709 | 7641314 | 836606 | 2 | 0.00239 |
| 2 | Poor | RphaseB1 | NC_039513.1 | 144218 | 1283069 | 1138852 | 3 | 0.00263 |
| 3 | Poor | RphaseB1 | NC_039517.1 | 1492996 | 2206478 | 713483 | 2 | 0.00280 |
| 4 | Poor | RphaseB1 | NC_039514.1 | 3518826 | 4530760 | 1011935 | 3 | 0.00296 |
| 5 | Poor | RphaseB1 | NC_039508.1 | 7825906 | 9039560 | 1213655 | 4 | 0.00330 |
| 6 | Poor | RphaseB1 | NC_039511.1 | 7750741 | 8304235 | 553495 | 2 | 0.00361 |

...
... only first 25 lines shown ...

Directory: MRPR File: dst-Ob_LI_RphaseB2.txt

Analysis of 1-distances for sample "Ob_LI_RphaseB2" (201205 sites):

Total number of 1-distances: 201187
Median: 11.0 Mean: 1083.0 Std: 14663.1
Quantiles:

| 0% | 5% | 10% | 15% | 20% | 25% | 30% | 35% | 40% | 45% | 50% | 55% | 60% | 65% | 70% | 75% | 80% | 85% | 90% | 95% | 100% |
|---|---|---|---|---|---|---|---|---|---|---|---|---|---|---|---|---|---|---|---|---|
| 1 | 1 | 1 | 1 | 1 | 1 | 1 | 3 | 6 | 8 | 11 | 14 | 17 | 22 | 27 | 34 | 45 | 65 | 110 | 299 | 1230897 |

Low density region distance cutoff (lowest 100): 300259
High density region distance cutoff (highest 100): 1

Ordered lists of methylation-poor regions:

Top 100 (out of 96) methylation-poor regions in "Ob_LI_RphaseB2":
(Sdnsty = sites per 1kb)

| | Rtype | Sample | SeqID | From | To | Rlgth | NbrSites | Sdnsty |
|---|---|---|---|---|---|---|---|---|
| 1 | Poor | RphaseB2 | NC_039518.1 | 5647846 | 6878743 | 1230898 | 2 | 0.00162 |
| 2 | Poor | RphaseB2 | NC_039519.1 | 832234 | 2001608 | 1169375 | 2 | 0.00171 |
| 3 | Poor | RphaseB2 | NC_039517.1 | 11350528 | 13017550 | 1667023 | 3 | 0.00180 |
| 4 | Poor | RphaseB2 | NC_039515.1 | 11635493 | 12529093 | 893601 | 2 | 0.00224 |
| 5 | Poor | RphaseB2 | NC_039515.1 | 12652577 | 13515447 | 862871 | 2 | 0.00232 |
| 6 | Poor | RphaseB2 | NC_039506.1 | 6804669 | 7641269 | 836601 | 2 | 0.00239 |

...
... only first 25 lines shown ...

Directory: MRPR File: gwp-Ob_LI_BCphaseA1.txt

```
seqnames    start end   width strand      source      type  score phase ID    Dbxref      Name  gbkey gene  gene_biotype      end_range   partial     start_range seqnames    start
      end   width strand      NbrSites    Sdnsty
NC_039518.1 5540575     5545590     5016  -     Gnomon      gene  NA    NA    gene-LOC105281899 GeneID:105281899  LOC105281899      Gene  LOC105281899      protein_coding
      character(0)      NA    character(0)      NC_039518.1 5545094     6878743     1333650     *     2     0.00149964383458928
NC_039518.1 5654291     5654596     306   -     Gnomon      gene  NA    NA    gene-LOC113563312 GeneID:113563312  LOC113563312      Gene  LOC113563312      protein_coding
      c("5654596", ".") true  c(".", "5654291") NC_039518.1 5545094     6878743     1333650     *     2     0.00149964383458928
NC_039518.1 5666846     5667151     306   -     Gnomon      gene  NA    NA    gene-LOC113563174 GeneID:113563174  LOC113563174      Gene  LOC113563174      protein_coding
      c("5667151", ".") true  c(".", "5666846") NC_039518.1 5545094     6878743     1333650     *     2     0.00149964383458928
NC_039518.1 5831582     5835441     3860  -     Gnomon      gene  NA    NA    gene-LOC105284858 GeneID:105284858  LOC105284858      Gene  LOC105284858      protein_coding
      character(0)      NA    character(0)      NC_039518.1 5545094     6878743     1333650     *     2     0.00149964383458928
NC_039518.1 5903338     5903643     306   -     Gnomon      gene  NA    NA    gene-LOC113563176 GeneID:113563176  LOC113563176      Gene  LOC113563176      protein_coding
      c("5903643", ".") true  c(".", "5903338") NC_039518.1 5545094     6878743     1333650     *     2     0.00149964383458928
NC_039518.1 5955002     5956241     1240  +     Gnomon      gene  NA    NA    gene-LOC105275939 GeneID:105275939  LOC105275939      Gene  LOC105275939      protein_coding
      character(0)      NA    character(0)      NC_039518.1 5545094     6878743     1333650     *     2     0.00149964383458928
NC_039518.1 6147897     6148202     306   +     Gnomon      gene  NA    NA    gene-LOC113563311 GeneID:113563311  LOC113563311      Gene  LOC113563311      protein_coding
      c("6148202", ".") true  c(".", "6147897") NC_039518.1 5545094     6878743     1333650     *     2     0.00149964383458928
NC_039518.1 6168060     6171567     3508  +     Gnomon      gene  NA    NA    gene-LOC105287193 GeneID:105287193  LOC105287193      Gene  LOC105287193      protein_coding
      character(0)      NA    character(0)      NC_039518.1 5545094     6878743     1333650     *     2     0.00149964383458928
NC_039518.1 6191135     6191440     306   -     Gnomon      gene  NA    NA    gene-LOC113563173 GeneID:113563173  LOC113563173      Gene  LOC113563173      protein_coding
      c("6191440", ".") true  c(".", "6191135") NC_039518.1 5545094     6878743     1333650     *     2     0.00149964383458928
NC_039518.1 6192330     6192635     306   -     Gnomon      gene  NA    NA    gene-LOC113563175 GeneID:113563175  LOC113563175      Gene  LOC113563175      protein_coding
      c("6192635", ".") true  c(".", "6192330") NC_039518.1 5545094     6878743     1333650     *     2     0.00149964383458928
NC_039518.1 6246480     6248316     1837  +     Gnomon      gene  NA    NA    gene-LOC105286058 GeneID:105286058  LOC105286058      Gene  LOC105286058      protein_coding
      character(0)      NA    character(0)      NC_039518.1 5545094     6878743     1333650     *     2     0.00149964383458928
NC_039518.1 6372000     6375086     3087  -     Gnomon      gene  NA    NA    gene-LOC113563235 GeneID:113563235  LOC113563235      Gene  LOC113563235      protein_coding
      character(0)      NA    character(0)      NC_039518.1 5545094     6878743     1333650     *     2     0.00149964383458928
NC_039518.1 6768698     6873781     105084      -     Gnomon      gene  NA    NA    gene-LOC105285730 GeneID:105285730  LOC105285730      Gene  LOC105285730      protein_coding
      character(0)      NA    character(0)      NC_039518.1 5545094     6878743     1333650     *     2     0.00149964383458928
NC_039510.1 6990084     6994039     3956  +     Gnomon      gene  NA    NA    gene-LOC105287545 GeneID:105287545  LOC105287545      Gene  LOC105287545      protein_coding
      character(0)      NA    character(0)      NC_039510.1 6991940     8277430     1285491     *     2     0.00155582575062758
NC_039510.1 6994301     7009423     15123 -     Gnomon      gene  NA    NA    gene-LOC105287546 GeneID:105287546  LOC105287546      Gene  LOC105287546      protein_coding
      character(0)      NA    character(0)      NC_039510.1 6991940     8277430     1285491     *     2     0.00155582575062758
NC_039510.1 7057641     7118044     60404 -     Gnomon      gene  NA    NA    gene-LOC105287547 GeneID:105287547  LOC105287547      Gene  LOC105287547      protein_coding
      character(0)      NA    character(0)      NC_039510.1 6991940     8277430     1285491     *     2     0.00155582575062758
NC_039510.1 7151619     7153950     2332  -     Gnomon      gene  NA    NA    gene-LOC105286892 GeneID:105286892  LOC105286892      Gene  LOC105286892      protein_coding
      character(0)      NA    character(0)      NC_039510.1 6991940     8277430     1285491     *     2     0.00155582575062758
NC_039510.1 7164083     7165989     1907  +     Gnomon      gene  NA    NA    gene-LOC105286891 GeneID:105286891  LOC105286891      Gene  LOC105286891      protein_coding
      character(0)      NA    character(0)      NC_039510.1 6991940     8277430     1285491     *     2     0.00155582575062758
NC_039510.1 7166460     7168400     1941  -     Gnomon      gene  NA    NA    gene-LOC105286895 GeneID:105286895  LOC105286895      Gene  LOC105286895      protein_coding
      character(0)      NA    character(0)      NC_039510.1 6991940     8277430     1285491     *     2     0.00155582575062758
NC_039510.1 7171986     7173486     1501  -     Gnomon      gene  NA    NA    gene-LOC109611483 GeneID:109611483  LOC109611483      Gene  LOC109611483      protein_coding
      character(0)      NA    character(0)      NC_039510.1 6991940     8277430     1285491     *     2     0.00155582575062758
NC_039510.1 7183155     7192107     8953  -     Gnomon      gene  NA    NA    gene-LOC105286890 GeneID:105286890  LOC105286890      Gene  LOC105286890      protein_coding
      character(0)      NA    character(0)      NC_039510.1 6991940     8277430     1285491     *     2     0.00155582575062758
NC_039510.1 7200311     7207829     7519  -     Gnomon      gene  NA    NA    gene-LOC105286888 GeneID:105286888  LOC105286888      Gene  LOC105286888      protein_coding
      character(0)      NA    character(0)      NC_039510.1 6991940     8277430     1285491     *     2     0.00155582575062758
NC_039510.1 7214122     7283344     69223 -     Gnomon      gene  NA    NA    gene-LOC105286415 GeneID:105286415  LOC105286415      Gene  LOC105286415      protein_coding
      character(0)      NA    character(0)      NC_039510.1 6991940     8277430     1285491     *     2     0.00155582575062758
NC_039510.1 7239495     7290328     50834 +     Gnomon      gene  NA    NA    gene-LOC105286414 GeneID:105286414  LOC105286414      Gene  LOC105286414      lncRNA      character(0)
      NA    character(0)      NC_039510.1 6991940     8277430     1285491     *     2     0.00155582575062758
...
... only first 25 lines shown ...
```

Directory: MRPR File: gwp-Ob_LI_BCphaseA2.txt

| seqnames | start | end | width | strand | source | type | score | phase | ID | Dbxref | Name | gbkey | gene | gene_biotype | end_range | partial | start_range | seqnames | start | end | width | strand | NbrSites | Sdnsty |
|---|---|---|---|---|---|---|---|---|---|---|---|---|---|---|---|---|---|---|---|---|---|---|---|---|
| NC_039510.1 | 6990084 | 6994039 | 3956 | + | Gnomon | gene | NA | NA | gene-LOC105287545 | GeneID:105287545 | LOC105287545 | Gene | LOC105287545 | protein_coding | character(0) | NA | character(0) | NC_039510.1 | 6992643 | 8396382 | 1403740 | * | 2 | 0.00142476526992178 |
| NC_039510.1 | 6994301 | 7009423 | 15123 | - | Gnomon | gene | NA | NA | gene-LOC105287546 | GeneID:105287546 | LOC105287546 | Gene | LOC105287546 | protein_coding | character(0) | NA | character(0) | NC_039510.1 | 6992643 | 8396382 | 1403740 | * | 2 | 0.00142476526992178 |
| NC_039510.1 | 7057641 | 7118044 | 60404 | - | Gnomon | gene | NA | NA | gene-LOC105287547 | GeneID:105287547 | LOC105287547 | Gene | LOC105287547 | protein_coding | character(0) | NA | character(0) | NC_039510.1 | 6992643 | 8396382 | 1403740 | * | 2 | 0.00142476526992178 |
| NC_039510.1 | 7151619 | 7153950 | 2332 | - | Gnomon | gene | NA | NA | gene-LOC105286892 | GeneID:105286892 | LOC105286892 | Gene | LOC105286892 | protein_coding | character(0) | NA | character(0) | NC_039510.1 | 6992643 | 8396382 | 1403740 | * | 2 | 0.00142476526992178 |
| NC_039510.1 | 7164083 | 7165989 | 1907 | + | Gnomon | gene | NA | NA | gene-LOC105286891 | GeneID:105286891 | LOC105286891 | Gene | LOC105286891 | protein_coding | character(0) | NA | character(0) | NC_039510.1 | 6992643 | 8396382 | 1403740 | * | 2 | 0.00142476526992178 |
| NC_039510.1 | 7166460 | 7168400 | 1941 | - | Gnomon | gene | NA | NA | gene-LOC105286895 | GeneID:105286895 | LOC105286895 | Gene | LOC105286895 | protein_coding | character(0) | NA | character(0) | NC_039510.1 | 6992643 | 8396382 | 1403740 | * | 2 | 0.00142476526992178 |
| NC_039510.1 | 7171986 | 7173486 | 1501 | - | Gnomon | gene | NA | NA | gene-LOC109611483 | GeneID:109611483 | LOC109611483 | Gene | LOC109611483 | protein_coding | character(0) | NA | character(0) | NC_039510.1 | 6992643 | 8396382 | 1403740 | * | 2 | 0.00142476526992178 |
| NC_039510.1 | 7183155 | 7192107 | 8953 | - | Gnomon | gene | NA | NA | gene-LOC105286890 | GeneID:105286890 | LOC105286890 | Gene | LOC105286890 | protein_coding | character(0) | NA | character(0) | NC_039510.1 | 6992643 | 8396382 | 1403740 | * | 2 | 0.00142476526992178 |
| NC_039510.1 | 7200311 | 7207829 | 7519 | - | Gnomon | gene | NA | NA | gene-LOC105286888 | GeneID:105286888 | LOC105286888 | Gene | LOC105286888 | protein_coding | character(0) | NA | character(0) | NC_039510.1 | 6992643 | 8396382 | 1403740 | * | 2 | 0.00142476526992178 |
| NC_039510.1 | 7214122 | 7283344 | 69223 | - | Gnomon | gene | NA | NA | gene-LOC105286415 | GeneID:105286415 | LOC105286415 | Gene | LOC105286415 | protein_coding | character(0) | NA | character(0) | NC_039510.1 | 6992643 | 8396382 | 1403740 | * | 2 | 0.00142476526992178 |
| NC_039510.1 | 7239495 | 7290328 | 50834 | + | Gnomon | gene | NA | NA | gene-LOC105286414 | GeneID:105286414 | LOC105286414 | Gene | LOC105286414 | lncRNA | character(0) | NA | character(0) | NC_039510.1 | 6992643 | 8396382 | 1403740 | * | 2 | 0.00142476526992178 |
| NC_039510.1 | 7289171 | 7341309 | 52139 | - | Gnomon | gene | NA | NA | gene-LOC105286413 | GeneID:105286413 | LOC105286413 | Gene | LOC105286413 | protein_coding | character(0) | NA | character(0) | NC_039510.1 | 6992643 | 8396382 | 1403740 | * | 2 | 0.00142476526992178 |
| NC_039510.1 | 7325553 | 7329075 | 3523 | + | Gnomon | gene | NA | NA | gene-LOC105286411 | GeneID:105286411 | LOC105286411 | Gene | LOC105286411 | lncRNA | character(0) | NA | character(0) | NC_039510.1 | 6992643 | 8396382 | 1403740 | * | 2 | 0.00142476526992178 |
| NC_039510.1 | 7361053 | 7362045 | 993 | - | Gnomon | gene | NA | NA | gene-LOC113561386 | GeneID:113561386 | LOC113561386 | Gene | LOC113561386 | protein_coding | character(0) | NA | character(0) | NC_039510.1 | 6992643 | 8396382 | 1403740 | * | 2 | 0.00142476526992178 |
| NC_039510.1 | 7363117 | 7365842 | 2726 | + | Gnomon | gene | NA | NA | gene-LOC105275937 | GeneID:105275937 | LOC105275937 | Gene | LOC105275937 | protein_coding | character(0) | NA | character(0) | NC_039510.1 | 6992643 | 8396382 | 1403740 | * | 2 | 0.00142476526992178 |
| NC_039510.1 | 7367002 | 7368518 | 1517 | - | Gnomon | gene | NA | NA | gene-LOC113561942 | GeneID:113561942 | LOC113561942 | Gene | LOC113561942 | protein_coding | character(0) | NA | character(0) | NC_039510.1 | 6992643 | 8396382 | 1403740 | * | 2 | 0.00142476526992178 |
| NC_039510.1 | 7368467 | 7371064 | 2598 | + | Gnomon | gene | NA | NA | gene-LOC113561924 | GeneID:113561924 | LOC113561924 | Gene | LOC113561924 | protein_coding | character(0) | NA | character(0) | NC_039510.1 | 6992643 | 8396382 | 1403740 | * | 2 | 0.00142476526992178 |
| NC_039510.1 | 7370125 | 7373894 | 3770 | + | Gnomon | gene | NA | NA | gene-LOC113561943 | GeneID:113561943 | LOC113561943 | Gene | LOC113561943 | protein_coding | character(0) | NA | character(0) | NC_039510.1 | 6992643 | 8396382 | 1403740 | * | 2 | 0.00142476526992178 |
| NC_039510.1 | 7375933 | 7376857 | 925 | + | Gnomon | gene | NA | NA | gene-LOC105275941 | GeneID:105275941 | LOC105275941 | Gene | LOC105275941 | protein_coding | character(0) | NA | character(0) | NC_039510.1 | 6992643 | 8396382 | 1403740 | * | 2 | 0.00142476526992178 |
| NC_039510.1 | 7381993 | 7382640 | 648 | + | Gnomon | gene | NA | NA | gene-LOC105282200 | GeneID:105282200 | LOC105282200 | Gene | LOC105282200 | protein_coding | character(0) | NA | character(0) | NC_039510.1 | 6992643 | 8396382 | 1403740 | * | 2 | 0.00142476526992178 |
| NC_039510.1 | 7409207 | 7409851 | 645 | - | Gnomon | gene | NA | NA | gene-LOC113561944 | GeneID:113561944 | LOC113561944 | Gene | LOC113561944 | protein_coding | character(0) | NA | character(0) | NC_039510.1 | 6992643 | 8396382 | 1403740 | * | 2 | 0.00142476526992178 |
| NC_039510.1 | 7460558 | 7461490 | 933 | + | Gnomon | gene | NA | NA | gene-LOC113561899 | GeneID:113561899 | LOC113561899 | Gene | LOC113561899 | protein_coding | character(0) | NA | character(0) | NC_039510.1 | 6992643 | 8396382 | 1403740 | * | 2 | 0.00142476526992178 |
| NC_039510.1 | 7469479 | 7471917 | 2439 | + | Gnomon | gene | NA | NA | gene-LOC105275943 | GeneID:105275943 | LOC105275943 | Gene | LOC105275943 | protein_coding | character(0) | NA | character(0) | NC_039510.1 | 6992643 | 8396382 | 1403740 | * | 2 | 0.00142476526992178 |
| NC_039510.1 | 7480666 | 7484337 | 3672 | + | Gnomon | gene | NA | NA | gene-LOC105287483 | GeneID:105287483 | LOC105287483 | Gene | LOC105287483 | protein_coding | character(0) | NA | character(0) | NC_039510.1 | 6992643 | 8396382 | 1403740 | * | 2 | 0.00142476526992178 |

...
... only first 25 lines shown ...

Directory: MRPR File: gwp-Ob_LI_BCphaseB1.txt

```
seqnames    start end   width strand      source      type  score phase ID    Dbxref      Name  gbkey gene  gene_biotype      end_range   partial     start_range seqnames    start
      end   width strand      NbrSites    Sdnsty
NC_039518.1 5654291     5654596     306   -     Gnomon      gene  NA    NA    gene-LOC113563312 GeneID:113563312  LOC113563312      Gene  LOC113563312      protein_coding
      c("5654596", ".") true  c(".", "5654291") NC_039518.1 5621056     6719148     1098093     *     2     0.00182133935832393
NC_039518.1 5666846     5667151     306   -     Gnomon      gene  NA    NA    gene-LOC113563174 GeneID:113563174  LOC113563174      Gene  LOC113563174      protein_coding
      c("5667151", ".") true  c(".", "5666846") NC_039518.1 5621056     6719148     1098093     *     2     0.00182133935832393
NC_039518.1 5831582     5835441     3860  -     Gnomon      gene  NA    NA    gene-LOC105284858 GeneID:105284858  LOC105284858      Gene  LOC105284858      protein_coding
      character(0)      NA    character(0)      NC_039518.1 5621056     6719148     1098093     *     2     0.00182133935832393
NC_039518.1 5903338     5903643     306   -     Gnomon      gene  NA    NA    gene-LOC113563176 GeneID:113563176  LOC113563176      Gene  LOC113563176      protein_coding
      c("5903643", ".") true  c(".", "5903338") NC_039518.1 5621056     6719148     1098093     *     2     0.00182133935832393
NC_039518.1 5955002     5956241     1240  +     Gnomon      gene  NA    NA    gene-LOC105275939 GeneID:105275939  LOC105275939      Gene  LOC105275939      protein_coding
      character(0)      NA    character(0)      NC_039518.1 5621056     6719148     1098093     *     2     0.00182133935832393
NC_039518.1 6147897     6148202     306   +     Gnomon      gene  NA    NA    gene-LOC113563311 GeneID:113563311  LOC113563311      Gene  LOC113563311      protein_coding
      c("6148202", ".") true  c(".", "6147897") NC_039518.1 5621056     6719148     1098093     *     2     0.00182133935832393
NC_039518.1 6168060     6171567     3508  +     Gnomon      gene  NA    NA    gene-LOC105287193 GeneID:105287193  LOC105287193      Gene  LOC105287193      protein_coding
      character(0)      NA    character(0)      NC_039518.1 5621056     6719148     1098093     *     2     0.00182133935832393
NC_039518.1 6191135     6191440     306   -     Gnomon      gene  NA    NA    gene-LOC113563173 GeneID:113563173  LOC113563173      Gene  LOC113563173      protein_coding
      c("6191440", ".") true  c(".", "6191135") NC_039518.1 5621056     6719148     1098093     *     2     0.00182133935832393
NC_039518.1 6192330     6192635     306   -     Gnomon      gene  NA    NA    gene-LOC113563175 GeneID:113563175  LOC113563175      Gene  LOC113563175      protein_coding
      c("6192635", ".") true  c(".", "6192330") NC_039518.1 5621056     6719148     1098093     *     2     0.00182133935832393
NC_039518.1 6246480     6248316     1837  +     Gnomon      gene  NA    NA    gene-LOC105286058 GeneID:105286058  LOC105286058      Gene  LOC105286058      protein_coding
      character(0)      NA    character(0)      NC_039518.1 5621056     6719148     1098093     *     2     0.00182133935832393
NC_039518.1 6372000     6375086     3087  -     Gnomon      gene  NA    NA    gene-LOC113563235 GeneID:113563235  LOC113563235      Gene  LOC113563235      protein_coding
      character(0)      NA    character(0)      NC_039518.1 5621056     6719148     1098093     *     2     0.00182133935832393
NC_039506.1 134260      142204      7945  -     Gnomon      gene  NA    NA    gene-LOC105288138 GeneID:105288138  LOC105288138      Gene  LOC105288138      protein_coding
      character(0)      NA    character(0)      NC_039506.1 137598      1058205     920608      *     2     0.00217247731933679
NC_039506.1 143775      267542      123768      -     Gnomon      gene  NA    NA    gene-LOC105288140 GeneID:105288140  LOC105288140      Gene  LOC105288140      protein_coding
      character(0)      NA    character(0)      NC_039506.1 137598      1058205     920608      *     2     0.00217247731933679
NC_039506.1 260227      263340      3114  +     Gnomon      gene  NA    NA    gene-LOC105288139 GeneID:105288139  LOC105288139      Gene  LOC105288139      lncRNA      character(0)
      NA    character(0)      NC_039506.1 137598      1058205     920608      *     2     0.00217247731933679
NC_039506.1 271139      273181      2043  -     Gnomon      gene  NA    NA    gene-LOC113562194 GeneID:113562194  LOC113562194      Gene  LOC113562194      protein_coding
      character(0)      NA    character(0)      NC_039506.1 137598      1058205     920608      *     2     0.00217247731933679
NC_039506.1 273665      275760      2096  +     Gnomon      gene  NA    NA    gene-LOC105288142 GeneID:105288142  LOC105288142      Gene  LOC105288142      protein_coding
      character(0)      NA    character(0)      NC_039506.1 137598      1058205     920608      *     2     0.00217247731933679
NC_039506.1 279299      297434      18136 -     Gnomon      gene  NA    NA    gene-LOC105288150 GeneID:105288150  LOC105288150      Gene  LOC105288150      protein_coding
      character(0)      NA    character(0)      NC_039506.1 137598      1058205     920608      *     2     0.00217247731933679
NC_039506.1 299178      309425      10248 -     Gnomon      gene  NA    NA    gene-LOC105288144 GeneID:105288144  LOC105288144      Gene  LOC105288144      protein_coding
      character(0)      NA    character(0)      NC_039506.1 137598      1058205     920608      *     2     0.00217247731933679
NC_039506.1 310674      322599      11926 -     Gnomon      gene  NA    NA    gene-LOC105287009 GeneID:105287009  LOC105287009      Gene  LOC105287009      protein_coding
      character(0)      NA    character(0)      NC_039506.1 137598      1058205     920608      *     2     0.00217247731933679
NC_039506.1 324544      334991      10448 -     Gnomon      gene  NA    NA    gene-LOC105287258 GeneID:105287258  LOC105287258      Gene  LOC105287258      protein_coding
      character(0)      NA    character(0)      NC_039506.1 137598      1058205     920608      *     2     0.00217247731933679
NC_039506.1 337710      351091      13382 -     Gnomon      gene  NA    NA    gene-LOC105287298 GeneID:105287298  LOC105287298      Gene  LOC105287298      protein_coding
      character(0)      NA    character(0)      NC_039506.1 137598      1058205     920608      *     2     0.00217247731933679
NC_039506.1 352522      364599      12078 -     Gnomon      gene  NA    NA    gene-LOC105285568 GeneID:105285568  LOC105285568      Gene  LOC105285568      protein_coding
      character(0)      NA    character(0)      NC_039506.1 137598      1058205     920608      *     2     0.00217247731933679
NC_039506.1 364796      369330      4535  -     Gnomon      gene  NA    NA    gene-LOC105285437 GeneID:105285437  LOC105285437      Gene  LOC105285437      protein_coding
      character(0)      NA    character(0)      NC_039506.1 137598      1058205     920608      *     2     0.00217247731933679
NC_039506.1 370197      377142      6946  +     Gnomon      gene  NA    NA    gene-LOC105285439 GeneID:105285439  LOC105285439      Gene  LOC105285439      protein_coding
      character(0)      NA    character(0)      NC_039506.1 137598      1058205     920608      *     2     0.00217247731933679
...
... only first 25 lines shown ...
```

Directory: MRPR File: gwp-Ob_LI_BCphaseB2.txt

```
seqnames    start end   width strand      source      type  score phase ID    Dbxref      Name  gbkey gene  gene_biotype      end_range   partial     start_range seqnames    start
      end   width strand      NbrSites    Sdnsty
NC_039518.1 6147897     6148202     306   +     Gnomon      gene  NA    NA    gene-LOC113563311 GeneID:113563311  LOC113563311      Gene  LOC113563311      protein_coding
      c("6148202", ".") true  c(".", "6147897") NC_039518.1 5974026     7049520     1075495     *     2     0.00185960883128234
NC_039518.1 6168060     6171567     3508  +     Gnomon      gene  NA    NA    gene-LOC105287193 GeneID:105287193  LOC105287193      Gene  LOC105287193      protein_coding
      character(0)      NA    character(0)      NC_039518.1 5974026     7049520     1075495     *     2     0.00185960883128234
NC_039518.1 6191135     6191440     306   -     Gnomon      gene  NA    NA    gene-LOC113563173 GeneID:113563173  LOC113563173      Gene  LOC113563173      protein_coding
      c("6191440", ".") true  c(".", "6191135") NC_039518.1 5974026     7049520     1075495     *     2     0.00185960883128234
NC_039518.1 6192330     6192635     306   -     Gnomon      gene  NA    NA    gene-LOC113563175 GeneID:113563175  LOC113563175      Gene  LOC113563175      protein_coding
      c("6192635", ".") true  c(".", "6192330") NC_039518.1 5974026     7049520     1075495     *     2     0.00185960883128234
NC_039518.1 6246480     6248316     1837  +     Gnomon      gene  NA    NA    gene-LOC105286058 GeneID:105286058  LOC105286058      Gene  LOC105286058      protein_coding
      character(0)      NA    character(0)      NC_039518.1 5974026     7049520     1075495     *     2     0.00185960883128234
NC_039518.1 6372000     6375086     3087  -     Gnomon      gene  NA    NA    gene-LOC113563235 GeneID:113563235  LOC113563235      Gene  LOC113563235      protein_coding
      character(0)      NA    character(0)      NC_039518.1 5974026     7049520     1075495     *     2     0.00185960883128234
NC_039518.1 6768698     6873781     105084      -     Gnomon      gene  NA    NA    gene-LOC105285730 GeneID:105285730  LOC105285730      Gene  LOC105285730      protein_coding
      character(0)      NA    character(0)      NC_039518.1 5974026     7049520     1075495     *     2     0.00185960883128234
NC_039518.1 6915302     6948947     33646 +     Gnomon      gene  NA    NA    gene-LOC113563191 GeneID:113563191  LOC113563191      Gene  LOC113563191      protein_coding
      character(0)      NA    character(0)      NC_039518.1 5974026     7049520     1075495     *     2     0.00185960883128234
NC_039518.1 6959377     6962879     3503  -     Gnomon      gene  NA    NA    gene-LOC105280842 GeneID:105280842  LOC105280842      Gene  LOC105280842      protein_coding
      character(0)      NA    character(0)      NC_039518.1 5974026     7049520     1075495     *     2     0.00185960883128234
NC_039518.1 6964879     6966366     1488  -     Gnomon      gene  NA    NA    gene-LOC109611063 GeneID:109611063  LOC109611063      Gene  LOC109611063      protein_coding
      character(0)      NA    character(0)      NC_039518.1 5974026     7049520     1075495     *     2     0.00185960883128234
NC_039518.1 6988547     7033055     44509 +     Gnomon      gene  NA    NA    gene-LOC105280616 GeneID:105280616  LOC105280616      Gene  LOC105280616      protein_coding
      character(0)      NA    character(0)      NC_039518.1 5974026     7049520     1075495     *     2     0.00185960883128234
NC_039518.1 7042945     7056012     13068 -     Gnomon      gene  NA    NA    gene-LOC105280615 GeneID:105280615  LOC105280615      Gene  LOC105280615      protein_coding
      character(0)      NA    character(0)      NC_039518.1 5974026     7049520     1075495     *     2     0.00185960883128234
NC_039511.1 6640532     6993146     352615      +     Gnomon      gene  NA    NA    gene-LOC105276776 GeneID:105276776  LOC105276776      Gene  LOC105276776      protein_coding
      character(0)      NA    character(0)      NC_039511.1 6715713     8126204     1410492     *     3     0.00212691741605057
NC_039511.1 6736497     6737777     1281  -     Gnomon      gene  NA    NA    gene-LOC105276775 GeneID:105276775  LOC105276775      Gene  LOC105276775      lncRNA      character(0)
      NA    character(0)      NC_039511.1 6715713     8126204     1410492     *     3     0.00212691741605057
NC_039511.1 6867576     6869339     1764  -     Gnomon      gene  NA    NA    gene-LOC113562075 GeneID:113562075  LOC113562075      Gene  LOC113562075      protein_coding
      character(0)      NA    character(0)      NC_039511.1 6715713     8126204     1410492     *     3     0.00212691741605057
NC_039511.1 6932472     6934628     2157  -     Gnomon      gene  NA    NA    gene-LOC113562074 GeneID:113562074  LOC113562074      Gene  LOC113562074      protein_coding
      character(0)      NA    character(0)      NC_039511.1 6715713     8126204     1410492     *     3     0.00212691741605057
NC_039511.1 6965860     6970599     4740  +     Gnomon      gene  NA    NA    gene-LOC109610815 GeneID:109610815  LOC109610815      Gene  LOC109610815      protein_coding
      character(0)      NA    character(0)      NC_039511.1 6715713     8126204     1410492     *     3     0.00212691741605057
NC_039511.1 7013281     7013736     456   +     Gnomon      gene  NA    NA    gene-LOC105280903 GeneID:105280903  LOC105280903      Gene  LOC105280903      protein_coding
      character(0)      NA    character(0)      NC_039511.1 6715713     8126204     1410492     *     3     0.00212691741605057
NC_039511.1 7015323     7016008     686   +     Gnomon      gene  NA    NA    gene-LOC113562136 GeneID:113562136  LOC113562136      Gene  LOC113562136      protein_coding
      character(0)      NA    character(0)      NC_039511.1 6715713     8126204     1410492     *     3     0.00212691741605057
NC_039511.1 7030342     7031429     1088  -     Gnomon      gene  NA    NA    gene-LOC113562137 GeneID:113562137  LOC113562137      Gene  LOC113562137      protein_coding
      character(0)      NA    character(0)      NC_039511.1 6715713     8126204     1410492     *     3     0.00212691741605057
NC_039511.1 7033382     7034155     774   -     Gnomon      gene  NA    NA    gene-LOC105280906 GeneID:105280906  LOC105280906      Gene  LOC105280906      protein_coding
      character(0)      NA    character(0)      NC_039511.1 6715713     8126204     1410492     *     3     0.00212691741605057
NC_039511.1 7042111     7042790     680   -     Gnomon      gene  NA    NA    gene-LOC113562138 GeneID:113562138  LOC113562138      Gene  LOC113562138      protein_coding
      character(0)      NA    character(0)      NC_039511.1 6715713     8126204     1410492     *     3     0.00212691741605057
NC_039511.1 7042793     7050794     8002  +     Gnomon      gene  NA    NA    gene-LOC105280907 GeneID:105280907  LOC105280907      Gene  LOC105280907      protein_coding
      character(0)      NA    character(0)      NC_039511.1 6715713     8126204     1410492     *     3     0.00212691741605057
NC_039511.1 7056909     7057787     879   -     Gnomon      gene  NA    NA    gene-LOC113562139 GeneID:113562139  LOC113562139      Gene  LOC113562139      protein_coding
      character(0)      NA    character(0)      NC_039511.1 6715713     8126204     1410492     *     3     0.00212691741605057
...
... only first 25 lines shown ...
```

Directory: MRPR File: gwp-Ob_LI_RphaseA1.txt

| seqnames | start | end | width | strand | source | type | score | phase | ID | Dbxref | Name | gbkey | gene | gene_biotype | end_range | partial | start_range | seqnames | start | end | width | strand | NbrSites | Sdnsty |
|---|---|---|---|---|---|---|---|---|---|---|---|---|---|---|---|---|---|---|---|---|---|---|---|---|
| NC_039517.1 | 11057436 | 11454413 | 396978 | + | Gnomon | gene | NA | NA | gene-LOC105283053 | GeneID:105283053 | LOC105283053 | Gene | LOC105283053 | protein_coding | character(0) | NA | character(0) | NC_039517.1 | 11083779 | 12752622 | 1668844 | * | 2 | 0.00119843436534511 |
| NC_039517.1 | 11082922 | 11084374 | 1453 | + | Gnomon | gene | NA | NA | gene-LOC113563160 | GeneID:113563160 | LOC113563160 | Gene | LOC113563160 | protein_coding | character(0) | NA | character(0) | NC_039517.1 | 11083779 | 12752622 | 1668844 | * | 2 | 0.00119843436534511 |
| NC_039517.1 | 11083896 | 11087902 | 4007 | - | Gnomon | gene | NA | NA | gene-LOC113563156 | GeneID:113563156 | LOC113563156 | Gene | LOC113563156 | protein_coding | character(0) | NA | character(0) | NC_039517.1 | 11083779 | 12752622 | 1668844 | * | 2 | 0.00119843436534511 |
| NC_039517.1 | 11117694 | 11119099 | 1406 | - | Gnomon | gene | NA | NA | gene-LOC113563163 | GeneID:113563163 | LOC113563163 | Gene | LOC113563163 | lncRNA | character(0) | NA | character(0) | NC_039517.1 | 11083779 | 12752622 | 1668844 | * | 2 | 0.00119843436534511 |
| NC_039517.1 | 11121241 | 11122313 | 1073 | + | Gnomon | gene | NA | NA | gene-LOC113563165 | GeneID:113563165 | LOC113563165 | Gene | LOC113563165 | lncRNA | character(0) | NA | character(0) | NC_039517.1 | 11083779 | 12752622 | 1668844 | * | 2 | 0.00119843436534511 |
| NC_039517.1 | 11232974 | 11237884 | 4911 | - | Gnomon | gene | NA | NA | gene-LOC105286061 | GeneID:105286061 | LOC105286061 | Gene | LOC105286061 | lncRNA | character(0) | NA | character(0) | NC_039517.1 | 11083779 | 12752622 | 1668844 | * | 2 | 0.00119843436534511 |
| NC_039517.1 | 11373912 | 11375270 | 1359 | - | Gnomon | gene | NA | NA | gene-LOC113563161 | GeneID:113563161 | LOC113563161 | Gene | LOC113563161 | protein_coding | character(0) | NA | character(0) | NC_039517.1 | 11083779 | 12752622 | 1668844 | * | 2 | 0.00119843436534511 |
| NC_039517.1 | 11461130 | 11462225 | 1096 | - | Gnomon | gene | NA | NA | gene-LOC113563057 | GeneID:113563057 | LOC113563057 | Gene | LOC113563057 | protein_coding | character(0) | NA | character(0) | NC_039517.1 | 11083779 | 12752622 | 1668844 | * | 2 | 0.00119843436534511 |
| NC_039517.1 | 11474062 | 11474517 | 456 | + | Gnomon | gene | NA | NA | gene-LOC113563139 | GeneID:113563139 | LOC113563139 | Gene | LOC113563139 | protein_coding | character(0) | NA | character(0) | NC_039517.1 | 11083779 | 12752622 | 1668844 | * | 2 | 0.00119843436534511 |
| NC_039517.1 | 11555820 | 11556868 | 1049 | + | Gnomon | gene | NA | NA | gene-LOC113563140 | GeneID:113563140 | LOC113563140 | Gene | LOC113563140 | protein_coding | character(0) | NA | character(0) | NC_039517.1 | 11083779 | 12752622 | 1668844 | * | 2 | 0.00119843436534511 |
| NC_039517.1 | 11567773 | 11568537 | 765 | - | Gnomon | gene | NA | NA | gene-LOC105279241 | GeneID:105279241 | LOC105279241 | Gene | LOC105279241 | protein_coding | character(0) | NA | character(0) | NC_039517.1 | 11083779 | 12752622 | 1668844 | * | 2 | 0.00119843436534511 |
| NC_039517.1 | 11569360 | 11570148 | 789 | - | Gnomon | gene | NA | NA | gene-LOC113563141 | GeneID:113563141 | LOC113563141 | Gene | LOC113563141 | protein_coding | character(0) | NA | character(0) | NC_039517.1 | 11083779 | 12752622 | 1668844 | * | 2 | 0.00119843436534511 |
| NC_039517.1 | 11613047 | 11615117 | 2071 | - | Gnomon | gene | NA | NA | gene-LOC105282194 | GeneID:105282194 | LOC105282194 | Gene | LOC105282194 | protein_coding | character(0) | NA | character(0) | NC_039517.1 | 11083779 | 12752622 | 1668844 | * | 2 | 0.00119843436534511 |
| NC_039517.1 | 11619801 | 11620895 | 1095 | - | Gnomon | gene | NA | NA | gene-LOC105282197 | GeneID:105282197 | LOC105282197 | Gene | LOC105282197 | protein_coding | character(0) | NA | character(0) | NC_039517.1 | 11083779 | 12752622 | 1668844 | * | 2 | 0.00119843436534511 |
| NC_039517.1 | 11642648 | 11643659 | 1012 | - | Gnomon | gene | NA | NA | gene-LOC105282195 | GeneID:105282195 | LOC105282195 | Gene | LOC105282195 | lncRNA | character(0) | NA | character(0) | NC_039517.1 | 11083779 | 12752622 | 1668844 | * | 2 | 0.00119843436534511 |
| NC_039517.1 | 11648482 | 11650734 | 2253 | - | Gnomon | gene | NA | NA | gene-LOC113563142 | GeneID:113563142 | LOC113563142 | Gene | LOC113563142 | protein_coding | character(0) | NA | character(0) | NC_039517.1 | 11083779 | 12752622 | 1668844 | * | 2 | 0.00119843436534511 |
| NC_039517.1 | 11652796 | 11653626 | 831 | - | Gnomon | gene | NA | NA | gene-LOC105282198 | GeneID:105282198 | LOC105282198 | Gene | LOC105282198 | protein_coding | character(0) | NA | character(0) | NC_039517.1 | 11083779 | 12752622 | 1668844 | * | 2 | 0.00119843436534511 |
| NC_039517.1 | 11655626 | 11659685 | 4060 | - | Gnomon | gene | NA | NA | gene-LOC113563143 | GeneID:113563143 | LOC113563143 | Gene | LOC113563143 | protein_coding | character(0) | NA | character(0) | NC_039517.1 | 11083779 | 12752622 | 1668844 | * | 2 | 0.00119843436534511 |
| NC_039517.1 | 11663971 | 11666022 | 2052 | + | Gnomon | gene | NA | NA | gene-LOC113563144 | GeneID:113563144 | LOC113563144 | Gene | LOC113563144 | protein_coding | character(0) | NA | character(0) | NC_039517.1 | 11083779 | 12752622 | 1668844 | * | 2 | 0.00119843436534511 |
| NC_039517.1 | 11685616 | 11686938 | 1323 | + | Gnomon | gene | NA | NA | gene-LOC113563072 | GeneID:113563072 | LOC113563072 | Gene | LOC113563072 | protein_coding | character(0) | NA | character(0) | NC_039517.1 | 11083779 | 12752622 | 1668844 | * | 2 | 0.00119843436534511 |
| NC_039517.1 | 11748481 | 11749879 | 1399 | + | Gnomon | gene | NA | NA | gene-LOC109611302 | GeneID:109611302 | LOC109611302 | Gene | LOC109611302 | protein_coding | character(0) | NA | character(0) | NC_039517.1 | 11083779 | 12752622 | 1668844 | * | 2 | 0.00119843436534511 |
| NC_039517.1 | 11782362 | 11782837 | 476 | - | Gnomon | gene | NA | NA | gene-LOC105284345 | GeneID:105284345 | LOC105284345 | Gene | LOC105284345 | protein_coding | character(0) | NA | character(0) | NC_039517.1 | 11083779 | 12752622 | 1668844 | * | 2 | 0.00119843436534511 |
| NC_039517.1 | 11797846 | 11798385 | 540 | + | Gnomon | gene | NA | NA | gene-LOC113563145 | GeneID:113563145 | LOC113563145 | Gene | LOC113563145 | protein_coding | character(0) | NA | character(0) | NC_039517.1 | 11083779 | 12752622 | 1668844 | * | 2 | 0.00119843436534511 |
| NC_039517.1 | 11798437 | 11799051 | 615 | + | Gnomon | gene | NA | NA | gene-LOC113563146 | GeneID:113563146 | LOC113563146 | Gene | LOC113563146 | protein_coding | character(0) | NA | character(0) | NC_039517.1 | 11083779 | 12752622 | 1668844 | * | 2 | 0.00119843436534511 |

...
... only first 25 lines shown ...

Directory: MRPR File: gwp-Ob_LI_RphaseA2.txt

```
seqnames	start	end	width	strand	source	type	score	phase	ID	Dbxref	Name	gbkey	gene	gene_biotype	end_range	partial	start_range	seqnames	start
	end	width	strand	NbrSites	Sdnsty
NC_039506.1	11665249	11676616	11368	+	Gnomon	gene	NA	NA	gene-LOC105284629	GeneID:105284629	LOC105284629	Gene	LOC105284629	protein_coding
	character(0)	NA	character(0)	NC_039506.1	11672856	12383020	710165	*	2	0.00281624692853069
NC_039506.1	11684976	11687683	2708	+	Gnomon	gene	NA	NA	gene-LOC105284630	GeneID:105284630	LOC105284630	Gene	LOC105284630	protein_coding
	character(0)	NA	character(0)	NC_039506.1	11672856	12383020	710165	*	2	0.00281624692853069
NC_039506.1	11688099	11691089	2991	-	Gnomon	gene	NA	NA	gene-LOC105284632	GeneID:105284632	LOC105284632	Gene	LOC105284632	protein_coding
	character(0)	NA	character(0)	NC_039506.1	11672856	12383020	710165	*	2	0.00281624692853069
NC_039506.1	11719215	11720420	1206	-	Gnomon	gene	NA	NA	gene-LOC105284634	GeneID:105284634	LOC105284634	Gene	LOC105284634	protein_coding
	character(0)	NA	character(0)	NC_039506.1	11672856	12383020	710165	*	2	0.00281624692853069
NC_039506.1	11723141	11726318	3178	+	Gnomon	gene	NA	NA	gene-LOC113562735	GeneID:113562735	LOC113562735	Gene	LOC113562735	protein_coding
	character(0)	NA	character(0)	NC_039506.1	11672856	12383020	710165	*	2	0.00281624692853069
NC_039506.1	11766307	11766708	402	+	Gnomon	gene	NA	NA	gene-LOC113562904	GeneID:113562904	LOC113562904	Gene	LOC113562904	protein_coding
	character(0)	NA	character(0)	NC_039506.1	11672856	12383020	710165	*	2	0.00281624692853069
NC_039506.1	11810234	11811670	1437	+	Gnomon	gene	NA	NA	gene-LOC113563583	GeneID:113563583	LOC113563583	Gene	LOC113563583	protein_coding
	character(0)	NA	character(0)	NC_039506.1	11672856	12383020	710165	*	2	0.00281624692853069
NC_039506.1	11836294	11837625	1332	-	Gnomon	gene	NA	NA	gene-LOC113562554	GeneID:113562554	LOC113562554	Gene	LOC113562554	protein_coding
	character(0)	NA	character(0)	NC_039506.1	11672856	12383020	710165	*	2	0.00281624692853069
NC_039506.1	11850743	11852397	1655	-	Gnomon	gene	NA	NA	gene-LOC113562061	GeneID:113562061	LOC113562061	Gene	LOC113562061	lncRNA	character(0)
	NA	character(0)	NC_039506.1	11672856	12383020	710165	*	2	0.00281624692853069
NC_039506.1	11874101	11876912	2812	+	Gnomon	gene	NA	NA	gene-LOC113561801	GeneID:113561801	LOC113561801	Gene	LOC113561801	protein_coding
	character(0)	NA	character(0)	NC_039506.1	11672856	12383020	710165	*	2	0.00281624692853069
NC_039506.1	11938701	11939905	1205	+	Gnomon	gene	NA	NA	gene-LOC113563653	GeneID:113563653	LOC113563653	Gene	LOC113563653	protein_coding
	character(0)	NA	character(0)	NC_039506.1	11672856	12383020	710165	*	2	0.00281624692853069
NC_039506.1	11947131	11949811	2681	-	Gnomon	gene	NA	NA	gene-LOC113561452	GeneID:113561452	LOC113561452	Gene	LOC113561452	protein_coding
	character(0)	NA	character(0)	NC_039506.1	11672856	12383020	710165	*	2	0.00281624692853069
NC_039506.1	12029120	12030495	1376	+	Gnomon	gene	NA	NA	gene-LOC113562163	GeneID:113562163	LOC113562163	Gene	LOC113562163	protein_coding
	character(0)	NA	character(0)	NC_039506.1	11672856	12383020	710165	*	2	0.00281624692853069
NC_039506.1	12055523	12056340	818	+	Gnomon	gene	NA	NA	gene-LOC113561894	GeneID:113561894	LOC113561894	Gene	LOC113561894	protein_coding
	character(0)	NA	character(0)	NC_039506.1	11672856	12383020	710165	*	2	0.00281624692853069
NC_039506.1	12061930	12067335	5406	+	Gnomon	gene	NA	NA	gene-LOC105279196	GeneID:105279196	LOC105279196	Gene	LOC105279196	protein_coding
	character(0)	true	c(".", "12061930")	NC_039506.1	11672856	12383020	710165	*	2	0.00281624692853069
NC_039506.1	12075579	12077637	2059	+	Gnomon	gene	NA	NA	gene-LOC105279194	GeneID:105279194	LOC105279194	Gene	LOC105279194	protein_coding
	character(0)	NA	character(0)	NC_039506.1	11672856	12383020	710165	*	2	0.00281624692853069
NC_039506.1	12077867	12080487	2621	-	Gnomon	gene	NA	NA	gene-LOC105281922	GeneID:105281922	LOC105281922	Gene	LOC105281922	protein_coding
	character(0)	NA	character(0)	NC_039506.1	11672856	12383020	710165	*	2	0.00281624692853069
NC_039506.1	12099807	12100963	1157	+	Gnomon	gene	NA	NA	gene-LOC113562999	GeneID:113562999	LOC113562999	Gene	LOC113562999	protein_coding
	character(0)	NA	character(0)	NC_039506.1	11672856	12383020	710165	*	2	0.00281624692853069
NC_039506.1	12101958	12103371	1414	-	Gnomon	gene	NA	NA	gene-LOC113562265	GeneID:113562265	LOC113562265	Gene	LOC113562265	protein_coding
	character(0)	NA	character(0)	NC_039506.1	11672856	12383020	710165	*	2	0.00281624692853069
NC_039506.1	12103416	12104604	1189	+	Gnomon	gene	NA	NA	gene-LOC113562315	GeneID:113562315	LOC113562315	Gene	LOC113562315	protein_coding
	character(0)	NA	character(0)	NC_039506.1	11672856	12383020	710165	*	2	0.00281624692853069
NC_039506.1	12122903	12125538	2636	+	Gnomon	gene	NA	NA	gene-LOC113561897	GeneID:113561897	LOC113561897	Gene	LOC113561897	protein_coding
	character(0)	NA	character(0)	NC_039506.1	11672856	12383020	710165	*	2	0.00281624692853069
NC_039506.1	12197722	12222696	24975	+	Gnomon	gene	NA	NA	gene-LOC105283216	GeneID:105283216	LOC105283216	Gene	LOC105283216	protein_coding
	character(0)	NA	character(0)	NC_039506.1	11672856	12383020	710165	*	2	0.00281624692853069
NC_039506.1	12206578	12207252	675	-	Gnomon	gene	NA	NA	gene-LOC113561698	GeneID:113561698	LOC113561698	Gene	LOC113561698	lncRNA	character(0)
	NA	character(0)	NC_039506.1	11672856	12383020	710165	*	2	0.00281624692853069
NC_039506.1	12243999	12251051	7053	-	Gnomon	gene	NA	NA	gene-LOC105283217	GeneID:105283217	LOC105283217	Gene	LOC105283217	protein_coding
	character(0)	NA	character(0)	NC_039506.1	11672856	12383020	710165	*	2	0.00281624692853069
...
... only first 25 lines shown ...
```

Directory: MRPR File: gwp-Ob_LI_RphaseB1.txt

| seqnames | start | end | width | strand | source | type | score | phase | ID | Dbxref | Name | gbkey | gene | gene_biotype | end_range | partial | start_range | seqnames | start | end | width | strand | NbrSites | Sdnsty |
|---|---|---|---|---|---|---|---|---|---|---|---|---|---|---|---|---|---|---|---|---|---|---|---|---|
| NC_039506.1 | 6803777 | 6805333 | 1557 | - | Gnomon | gene | NA | NA | gene-LOC105286755 | GeneID:105286755 | LOC105286755 | Gene | LOC105286755 | protein_coding | character(0) | NA | character(0) | NC_039506.1 | 6804709 | 7641314 | 836606 | * | 2 | 0.00239061159016311 |
| NC_039506.1 | 6806386 | 6809461 | 3076 | + | Gnomon | gene | NA | NA | gene-LOC113562021 | GeneID:113562021 | LOC113562021 | Gene | LOC113562021 | protein_coding | character(0) | NA | character(0) | NC_039506.1 | 6804709 | 7641314 | 836606 | * | 2 | 0.00239061159016311 |
| NC_039506.1 | 6811328 | 6813556 | 2229 | + | Gnomon | gene | NA | NA | gene-LOC113562182 | GeneID:113562182 | LOC113562182 | Gene | LOC113562182 | protein_coding | character(0) | NA | character(0) | NC_039506.1 | 6804709 | 7641314 | 836606 | * | 2 | 0.00239061159016311 |
| NC_039506.1 | 6818791 | 6827588 | 8798 | - | Gnomon | gene | NA | NA | gene-LOC105286743 | GeneID:105286743 | LOC105286743 | Gene | LOC105286743 | protein_coding | character(0) | NA | character(0) | NC_039506.1 | 6804709 | 7641314 | 836606 | * | 2 | 0.00239061159016311 |
| NC_039506.1 | 6827443 | 6828619 | 1177 | - | Gnomon | gene | NA | NA | gene-LOC113563549 | GeneID:113563549 | LOC113563549 | Gene | LOC113563549 | protein_coding | character(0) | NA | character(0) | NC_039506.1 | 6804709 | 7641314 | 836606 | * | 2 | 0.00239061159016311 |
| NC_039506.1 | 6830926 | 6842053 | 11128 | - | Gnomon | gene | NA | NA | gene-LOC113563542 | GeneID:113563542 | LOC113563542 | Gene | LOC113563542 | protein_coding | character(0) | NA | character(0) | NC_039506.1 | 6804709 | 7641314 | 836606 | * | 2 | 0.00239061159016311 |
| NC_039506.1 | 6853633 | 6856026 | 2394 | - | Gnomon | gene | NA | NA | gene-LOC113563551 | GeneID:113563551 | LOC113563551 | Gene | LOC113563551 | lncRNA | character(0) | NA | character(0) | NC_039506.1 | 6804709 | 7641314 | 836606 | * | 2 | 0.00239061159016311 |
| NC_039506.1 | 6864750 | 6880262 | 15513 | - | Gnomon | gene | NA | NA | gene-LOC113563550 | GeneID:113563550 | LOC113563550 | Gene | LOC113563550 | protein_coding | character(0) | NA | character(0) | NC_039506.1 | 6804709 | 7641314 | 836606 | * | 2 | 0.00239061159016311 |
| NC_039506.1 | 6886485 | 6893071 | 6587 | + | Gnomon | gene | NA | NA | gene-LOC113562173 | GeneID:113562173 | LOC113562173 | Gene | LOC113562173 | protein_coding | character(0) | NA | character(0) | NC_039506.1 | 6804709 | 7641314 | 836606 | * | 2 | 0.00239061159016311 |
| NC_039506.1 | 6893206 | 6895055 | 1850 | - | Gnomon | gene | NA | NA | gene-LOC113562172 | GeneID:113562172 | LOC113562172 | Gene | LOC113562172 | protein_coding | character(0) | NA | character(0) | NC_039506.1 | 6804709 | 7641314 | 836606 | * | 2 | 0.00239061159016311 |
| NC_039506.1 | 6894837 | 6904965 | 10129 | - | Gnomon | gene | NA | NA | gene-LOC105285469 | GeneID:105285469 | LOC105285469 | Gene | LOC105285469 | protein_coding | character(0) | NA | character(0) | NC_039506.1 | 6804709 | 7641314 | 836606 | * | 2 | 0.00239061159016311 |
| NC_039506.1 | 6911997 | 6929826 | 17830 | + | Gnomon | gene | NA | NA | gene-LOC105285467 | GeneID:105285467 | LOC105285467 | Gene | LOC105285467 | protein_coding | character(0) | NA | character(0) | NC_039506.1 | 6804709 | 7641314 | 836606 | * | 2 | 0.00239061159016311 |
| NC_039506.1 | 6930430 | 6935483 | 5054 | + | Gnomon | gene | NA | NA | gene-LOC105285466 | GeneID:105285466 | LOC105285466 | Gene | LOC105285466 | protein_coding | character(0) | NA | character(0) | NC_039506.1 | 6804709 | 7641314 | 836606 | * | 2 | 0.00239061159016311 |
| NC_039506.1 | 6936103 | 6939754 | 3652 | + | Gnomon | gene | NA | NA | gene-LOC113563155 | GeneID:113563155 | LOC113563155 | Gene | LOC113563155 | protein_coding | character(0) | NA | character(0) | NC_039506.1 | 6804709 | 7641314 | 836606 | * | 2 | 0.00239061159016311 |
| NC_039506.1 | 6940841 | 6984364 | 43524 | + | Gnomon | gene | NA | NA | gene-LOC105285464 | GeneID:105285464 | LOC105285464 | Gene | LOC105285464 | protein_coding | character(0) | NA | character(0) | NC_039506.1 | 6804709 | 7641314 | 836606 | * | 2 | 0.00239061159016311 |
| NC_039506.1 | 6991269 | 7032087 | 40819 | - | Gnomon | gene | NA | NA | gene-LOC105285462 | GeneID:105285462 | LOC105285462 | Gene | LOC105285462 | protein_coding | character(0) | NA | character(0) | NC_039506.1 | 6804709 | 7641314 | 836606 | * | 2 | 0.00239061159016311 |
| NC_039506.1 | 7068352 | 7072795 | 4444 | + | Gnomon | gene | NA | NA | gene-LOC105285461 | GeneID:105285461 | LOC105285461 | Gene | LOC105285461 | protein_coding | character(0) | NA | character(0) | NC_039506.1 | 6804709 | 7641314 | 836606 | * | 2 | 0.00239061159016311 |
| NC_039506.1 | 7087928 | 7195955 | 108028 | + | Gnomon | gene | NA | NA | gene-LOC105285459 | GeneID:105285459 | LOC105285459 | Gene | LOC105285459 | protein_coding | character(0) | NA | character(0) | NC_039506.1 | 6804709 | 7641314 | 836606 | * | 2 | 0.00239061159016311 |
| NC_039506.1 | 7200050 | 7204119 | 4070 | - | Gnomon | gene | NA | NA | gene-LOC105285458 | GeneID:105285458 | LOC105285458 | Gene | LOC105285458 | protein_coding | character(0) | NA | character(0) | NC_039506.1 | 6804709 | 7641314 | 836606 | * | 2 | 0.00239061159016311 |
| NC_039506.1 | 7218049 | 7225481 | 7433 | + | Gnomon | gene | NA | NA | gene-LOC105285457 | GeneID:105285457 | LOC105285457 | Gene | LOC105285457 | protein_coding | character(0) | NA | character(0) | NC_039506.1 | 6804709 | 7641314 | 836606 | * | 2 | 0.00239061159016311 |
| NC_039506.1 | 7233376 | 7276873 | 43498 | + | Gnomon | gene | NA | NA | gene-LOC113562292 | GeneID:113562292 | LOC113562292 | Gene | LOC113562292 | protein_coding | character(0) | NA | character(0) | NC_039506.1 | 6804709 | 7641314 | 836606 | * | 2 | 0.00239061159016311 |
| NC_039506.1 | 7277811 | 7279366 | 1556 | + | Gnomon | gene | NA | NA | gene-LOC113562297 | GeneID:113562297 | LOC113562297 | Gene | LOC113562297 | protein_coding | character(0) | NA | character(0) | NC_039506.1 | 6804709 | 7641314 | 836606 | * | 2 | 0.00239061159016311 |
| NC_039506.1 | 7279744 | 7280766 | 1023 | + | Gnomon | gene | NA | NA | gene-LOC105285452 | GeneID:105285452 | LOC105285452 | Gene | LOC105285452 | protein_coding | character(0) | NA | character(0) | NC_039506.1 | 6804709 | 7641314 | 836606 | * | 2 | 0.00239061159016311 |
| NC_039506.1 | 7288143 | 7290085 | 1943 | - | Gnomon | gene | NA | NA | gene-LOC113562235 | GeneID:113562235 | LOC113562235 | Gene | LOC113562235 | protein_coding | character(0) | NA | character(0) | NC_039506.1 | 6804709 | 7641314 | 836606 | * | 2 | 0.00239061159016311 |

...
... only first 25 lines shown ...

Directory: MRPR File: gwp-Ob_LI_RphaseB2.txt

| seqnames | start | end | width | strand | source | type | score | phase | ID | Dbxref | Name | gbkey | gene | gene_biotype | end_range | partial | start_range | seqnames | start | end | width | strand | NbrSites | Sdnsty |
|---|---|---|---|---|---|---|---|---|---|---|---|---|---|---|---|---|---|---|---|---|---|---|---|---|
| NC_039518.1 | 5654291 | 5654596 | 306 | - | Gnomon | gene | NA | NA | gene-LOC113563312 | GeneID:113563312 | LOC113563312 | Gene | LOC113563312 | protein_coding | c("5654596", ".") | true | c(".", "5654291") | NC_039518.1 | 5647846 | 6878743 | 1230898 | * | 2 | 0.00162483000216102 |
| NC_039518.1 | 5666846 | 5667151 | 306 | - | Gnomon | gene | NA | NA | gene-LOC113563174 | GeneID:113563174 | LOC113563174 | Gene | LOC113563174 | protein_coding | c("5667151", ".") | true | c(".", "5666846") | NC_039518.1 | 5647846 | 6878743 | 1230898 | * | 2 | 0.00162483000216102 |
| NC_039518.1 | 5831582 | 5835441 | 3860 | - | Gnomon | gene | NA | NA | gene-LOC105284858 | GeneID:105284858 | LOC105284858 | Gene | LOC105284858 | protein_coding | character(0) | NA | character(0) | NC_039518.1 | 5647846 | 6878743 | 1230898 | * | 2 | 0.00162483000216102 |
| NC_039518.1 | 5903338 | 5903643 | 306 | - | Gnomon | gene | NA | NA | gene-LOC113563176 | GeneID:113563176 | LOC113563176 | Gene | LOC113563176 | protein_coding | c("5903643", ".") | true | c(".", "5903338") | NC_039518.1 | 5647846 | 6878743 | 1230898 | * | 2 | 0.00162483000216102 |
| NC_039518.1 | 5955002 | 5956241 | 1240 | + | Gnomon | gene | NA | NA | gene-LOC105275939 | GeneID:105275939 | LOC105275939 | Gene | LOC105275939 | protein_coding | character(0) | NA | character(0) | NC_039518.1 | 5647846 | 6878743 | 1230898 | * | 2 | 0.00162483000216102 |
| NC_039518.1 | 6147897 | 6148202 | 306 | + | Gnomon | gene | NA | NA | gene-LOC113563311 | GeneID:113563311 | LOC113563311 | Gene | LOC113563311 | protein_coding | c("6148202", ".") | true | c(".", "6147897") | NC_039518.1 | 5647846 | 6878743 | 1230898 | * | 2 | 0.00162483000216102 |
| NC_039518.1 | 6168060 | 6171567 | 3508 | + | Gnomon | gene | NA | NA | gene-LOC105287193 | GeneID:105287193 | LOC105287193 | Gene | LOC105287193 | protein_coding | character(0) | NA | character(0) | NC_039518.1 | 5647846 | 6878743 | 1230898 | * | 2 | 0.00162483000216102 |
| NC_039518.1 | 6191135 | 6191440 | 306 | - | Gnomon | gene | NA | NA | gene-LOC113563173 | GeneID:113563173 | LOC113563173 | Gene | LOC113563173 | protein_coding | c("6191440", ".") | true | c(".", "6191135") | NC_039518.1 | 5647846 | 6878743 | 1230898 | * | 2 | 0.00162483000216102 |
| NC_039518.1 | 6192330 | 6192635 | 306 | - | Gnomon | gene | NA | NA | gene-LOC113563175 | GeneID:113563175 | LOC113563175 | Gene | LOC113563175 | protein_coding | c("6192635", ".") | true | c(".", "6192330") | NC_039518.1 | 5647846 | 6878743 | 1230898 | * | 2 | 0.00162483000216102 |
| NC_039518.1 | 6246480 | 6248316 | 1837 | + | Gnomon | gene | NA | NA | gene-LOC105286058 | GeneID:105286058 | LOC105286058 | Gene | LOC105286058 | protein_coding | character(0) | NA | character(0) | NC_039518.1 | 5647846 | 6878743 | 1230898 | * | 2 | 0.00162483000216102 |
| NC_039518.1 | 6372000 | 6375086 | 3087 | - | Gnomon | gene | NA | NA | gene-LOC113563235 | GeneID:113563235 | LOC113563235 | Gene | LOC113563235 | protein_coding | character(0) | NA | character(0) | NC_039518.1 | 5647846 | 6878743 | 1230898 | * | 2 | 0.00162483000216102 |
| NC_039518.1 | 6768698 | 6873781 | 105084 | - | Gnomon | gene | NA | NA | gene-LOC105285730 | GeneID:105285730 | LOC105285730 | Gene | LOC105285730 | protein_coding | character(0) | NA | character(0) | NC_039518.1 | 5647846 | 6878743 | 1230898 | * | 2 | 0.00162483000216102 |
| NC_039519.1 | 827002 | 834196 | 7195 | + | Gnomon | gene | NA | NA | gene-LOC105285516 | GeneID:105285516 | LOC105285516 | Gene | LOC105285516 | protein_coding | character(0) | NA | character(0) | NC_039519.1 | 832234 | 2001608 | 1169375 | * | 2 | 0.0017103153393907 |
| NC_039519.1 | 834519 | 835972 | 1454 | - | Gnomon | gene | NA | NA | gene-LOC105285515 | GeneID:105285515 | LOC105285515 | Gene | LOC105285515 | protein_coding | character(0) | NA | character(0) | NC_039519.1 | 832234 | 2001608 | 1169375 | * | 2 | 0.0017103153393907 |
| NC_039519.1 | 836527 | 851534 | 15008 | + | Gnomon | gene | NA | NA | gene-LOC109611484 | GeneID:109611484 | LOC109611484 | Gene | LOC109611484 | lncRNA | character(0) | NA | character(0) | NC_039519.1 | 832234 | 2001608 | 1169375 | * | 2 | 0.0017103153393907 |
| NC_039519.1 | 872044 | 1040040 | 167997 | + | Gnomon | gene | NA | NA | gene-LOC105286206 | GeneID:105286206 | LOC105286206 | Gene | LOC105286206 | protein_coding | character(0) | NA | character(0) | NC_039519.1 | 832234 | 2001608 | 1169375 | * | 2 | 0.0017103153393907 |
| NC_039519.1 | 1045374 | 1049821 | 4448 | - | Gnomon | gene | NA | NA | gene-LOC109611448 | GeneID:109611448 | LOC109611448 | Gene | LOC109611448 | lncRNA | character(0) | NA | character(0) | NC_039519.1 | 832234 | 2001608 | 1169375 | * | 2 | 0.0017103153393907 |
| NC_039519.1 | 1092913 | 1097563 | 4651 | - | Gnomon | gene | NA | NA | gene-LOC105286916 | GeneID:105286916 | LOC105286916 | Gene | LOC105286916 | protein_coding | character(0) | true | c(".", "1092913") | NC_039519.1 | 832234 | 2001608 | 1169375 | * | 2 | 0.0017103153393907 |
| NC_039519.1 | 1111369 | 1112262 | 894 | - | Gnomon | gene | NA | NA | gene-LOC113563418 | GeneID:113563418 | LOC113563418 | Gene | LOC113563418 | protein_coding | character(0) | NA | character(0) | NC_039519.1 | 832234 | 2001608 | 1169375 | * | 2 | 0.0017103153393907 |
| NC_039519.1 | 1163945 | 1164493 | 549 | - | Gnomon | gene | NA | NA | gene-LOC113563419 | GeneID:113563419 | LOC113563419 | Gene | LOC113563419 | protein_coding | character(0) | NA | character(0) | NC_039519.1 | 832234 | 2001608 | 1169375 | * | 2 | 0.0017103153393907 |
| NC_039519.1 | 1170163 | 1171363 | 1201 | + | Gnomon | gene | NA | NA | gene-LOC113563420 | GeneID:113563420 | LOC113563420 | Gene | LOC113563420 | protein_coding | character(0) | NA | character(0) | NC_039519.1 | 832234 | 2001608 | 1169375 | * | 2 | 0.0017103153393907 |
| NC_039519.1 | 1192872 | 1193651 | 780 | + | Gnomon | gene | NA | NA | gene-LOC113563421 | GeneID:113563421 | LOC113563421 | Gene | LOC113563421 | protein_coding | character(0) | NA | character(0) | NC_039519.1 | 832234 | 2001608 | 1169375 | * | 2 | 0.0017103153393907 |
| NC_039519.1 | 1248210 | 1283537 | 35328 | + | Gnomon | gene | NA | NA | gene-LOC105282746 | GeneID:105282746 | LOC105282746 | Gene | LOC105282746 | protein_coding | character(0) | NA | character(0) | NC_039519.1 | 832234 | 2001608 | 1169375 | * | 2 | 0.0017103153393907 |
| NC_039519.1 | 1301835 | 1314231 | 12397 | + | Gnomon | gene | NA | NA | gene-LOC113563379 | GeneID:113563379 | LOC113563379 | Gene | LOC113563379 | lncRNA | character(0) | NA | character(0) | NC_039519.1 | 832234 | 2001608 | 1169375 | * | 2 | 0.0017103153393907 |

...
... only first 25 lines shown ...

Directory: MRPR File: gwr-Ob_LI_BCphaseA1.txt

| seqnames | start | end | width | strand | source | type | score | phase | ID | Dbxref | Name | gbkey | gene | gene_biotype | end_range | partial | start_range | seqnames | start | end | width | strand | NbrSites | Sdnsty |
|---|---|---|---|---|---|---|---|---|---|---|---|---|---|---|---|---|---|---|---|---|---|---|---|---|
| NC_039506.1 | 6089683 | 6090856 | 1174 | + | Gnomon | gene | NA | NA | gene-LOC105277804 | GeneID:105277804 | LOC105277804 | Gene | LOC105277804 | protein_coding | character(0) | NA | character(0) | NC_039506.1 | 6090624 | 6090629 | 6 | * | 6 | 1000 |
| NC_039507.1 | 7276866 | 7278961 | 2096 | + | Gnomon | gene | NA | NA | gene-LOC105283373 | GeneID:105283373 | LOC105283373 | Gene | LOC105283373 | protein_coding | character(0) | NA | character(0) | NC_039507.1 | 7277947 | 7277952 | 6 | * | 6 | 1000 |
| NC_039507.1 | 18333035 | 18336377 | 3343 | - | Gnomon | gene | NA | NA | gene-LOC105281577 | GeneID:105281577 | LOC105281577 | Gene | LOC105281577 | protein_coding | character(0) | NA | character(0) | NC_039507.1 | 18335809 | 18335814 | 6 | * | 6 | 1000 |
| NC_039508.1 | 2590841 | 2594093 | 3253 | + | Gnomon | gene | NA | NA | gene-LOC105281341 | GeneID:105281341 | LOC105281341 | Gene | LOC105281341 | protein_coding | character(0) | NA | character(0) | NC_039508.1 | 2591612 | 2591617 | 6 | * | 6 | 1000 |
| NC_039508.1 | 10274002 | 10277440 | 3439 | + | Gnomon | gene | NA | NA | gene-LOC105276881 | GeneID:105276881 | LOC105276881 | Gene | LOC105276881 | protein_coding | character(0) | NA | character(0) | NC_039508.1 | 10275212 | 10275217 | 6 | * | 6 | 1000 |
| NC_039508.1 | 17548373 | 17550348 | 1976 | - | Gnomon | gene | NA | NA | gene-LOC105284396 | GeneID:105284396 | LOC105284396 | Gene | LOC105284396 | protein_coding | character(0) | NA | character(0) | NC_039508.1 | 17549290 | 17549295 | 6 | * | 6 | 1000 |
| NC_039509.1 | 1002697 | 1006107 | 3411 | - | Gnomon | gene | NA | NA | gene-LOC105283620 | GeneID:105283620 | LOC105283620 | Gene | LOC105283620 | protein_coding | character(0) | NA | character(0) | NC_039509.1 | 1004827 | 1004832 | 6 | * | 6 | 1000 |
| NC_039509.1 | 1135464 | 1137481 | 2018 | - | Gnomon | gene | NA | NA | gene-LOC105283634 | GeneID:105283634 | LOC105283634 | Gene | LOC105283634 | protein_coding | character(0) | NA | character(0) | NC_039509.1 | 1136266 | 1136271 | 6 | * | 6 | 1000 |
| NC_039509.1 | 11068956 | 11086837 | 17882 | - | Gnomon | gene | NA | NA | gene-LOC105275770 | GeneID:105275770 | LOC105275770 | Gene | LOC105275770 | protein_coding | character(0) | NA | character(0) | NC_039509.1 | 11085457 | 11085462 | 6 | * | 6 | 1000 |
| NC_039509.1 | 13033738 | 13035966 | 2229 | + | Gnomon | gene | NA | NA | gene-LOC105278527 | GeneID:105278527 | LOC105278527 | Gene | LOC105278527 | protein_coding | character(0) | NA | character(0) | NC_039509.1 | 13034669 | 13034674 | 6 | * | 6 | 1000 |
| NC_039509.1 | 13067024 | 13068370 | 1347 | + | Gnomon | gene | NA | NA | gene-LOC105278478 | GeneID:105278478 | LOC105278478 | Gene | LOC105278478 | protein_coding | character(0) | NA | character(0) | NC_039509.1 | 13067776 | 13067781 | 6 | * | 6 | 1000 |
| NC_039511.1 | 480353 | 488275 | 7923 | + | Gnomon | gene | NA | NA | gene-LOC105276527 | GeneID:105276527 | LOC105276527 | Gene | LOC105276527 | protein_coding | character(0) | NA | character(0) | NC_039511.1 | 481652 | 481657 | 6 | * | 6 | 1000 |
| NC_039511.1 | 2234830 | 2236789 | 1960 | - | Gnomon | gene | NA | NA | gene-LOC105276382 | GeneID:105276382 | LOC105276382 | Gene | LOC105276382 | protein_coding | character(0) | NA | character(0) | NC_039511.1 | 2236016 | 2236021 | 6 | * | 6 | 1000 |
| NC_039511.1 | 3614241 | 3616939 | 2699 | + | Gnomon | gene | NA | NA | gene-LOC105284057 | GeneID:105284057 | LOC105284057 | Gene | LOC105284057 | protein_coding | character(0) | NA | character(0) | NC_039511.1 | 3615462 | 3615467 | 6 | * | 6 | 1000 |
| NC_039512.1 | 10275466 | 10280187 | 4722 | + | Gnomon | gene | NA | NA | gene-LOC105282847 | GeneID:105282847 | LOC105282847 | Gene | LOC105282847 | protein_coding | character(0) | NA | character(0) | NC_039512.1 | 10276481 | 10276486 | 6 | * | 6 | 1000 |
| NC_039512.1 | 15979275 | 15981529 | 2255 | - | Gnomon | gene | NA | NA | gene-LOC105274775 | GeneID:105274775 | LOC105274775 | Gene | LOC105274775 | protein_coding | character(0) | NA | character(0) | NC_039512.1 | 15980634 | 15980639 | 6 | * | 6 | 1000 |
| NC_039513.1 | 10401827 | 10404590 | 2764 | - | Gnomon | gene | NA | NA | gene-LOC105275550 | GeneID:105275550 | LOC105275550 | Gene | LOC105275550 | protein_coding | character(0) | NA | character(0) | NC_039513.1 | 10403469 | 10403474 | 6 | * | 6 | 1000 |
| NC_039513.1 | 12028675 | 12030941 | 2267 | - | Gnomon | gene | NA | NA | gene-LOC105280791 | GeneID:105280791 | LOC105280791 | Gene | LOC105280791 | protein_coding | character(0) | NA | character(0) | NC_039513.1 | 12029791 | 12029796 | 6 | * | 6 | 1000 |
| NC_039514.1 | 6622839 | 6626782 | 3944 | - | Gnomon | gene | NA | NA | gene-LOC105285534 | GeneID:105285534 | LOC105285534 | Gene | LOC105285534 | protein_coding | character(0) | NA | character(0) | NC_039514.1 | 6625853 | 6625858 | 6 | * | 6 | 1000 |
| NC_039514.1 | 9336154 | 9343762 | 7609 | + | Gnomon | gene | NA | NA | gene-LOC105284644 | GeneID:105284644 | LOC105284644 | Gene | LOC105284644 | protein_coding | character(0) | NA | character(0) | NC_039514.1 | 9337043 | 9337048 | 6 | * | 6 | 1000 |
| NC_039514.1 | 11109536 | 11117116 | 7581 | - | Gnomon | gene | NA | NA | gene-LOC105281601 | GeneID:105281601 | LOC105281601 | Gene | LOC105281601 | protein_coding | character(0) | NA | character(0) | NC_039514.1 | 11112955 | 11112960 | 6 | * | 6 | 1000 |
| NC_039514.1 | 11111779 | 11115572 | 3794 | + | Gnomon | gene | NA | NA | gene-LOC105281603 | GeneID:105281603 | LOC105281603 | Gene | LOC105281603 | protein_coding | character(0) | NA | character(0) | NC_039514.1 | 11112955 | 11112960 | 6 | * | 6 | 1000 |
| NC_039515.1 | 7098785 | 7103508 | 4724 | - | Gnomon | gene | NA | NA | gene-LOC105286003 | GeneID:105286003 | LOC105286003 | Gene | LOC105286003 | protein_coding | character(0) | NA | character(0) | NC_039515.1 | 7102685 | 7102690 | 6 | * | 6 | 1000 |
| NC_039515.1 | 12582469 | 12585642 | 3174 | - | Gnomon | gene | NA | NA | gene-LOC105282150 | GeneID:105282150 | LOC105282150 | Gene | LOC105282150 | protein_coding | character(0) | NA | character(0) | NC_039515.1 | 12584748 | 12584753 | 6 | * | 6 | 1000 |
| NC_039516.1 | 12889538 | 12895772 | 6235 | + | Gnomon | gene | NA | NA | gene-LOC105275316 | GeneID:105275316 | LOC105275316 | Gene | LOC105275316 | protein_coding | character(0) | NA | character(0) | NC_039516.1 | 12890326 | 12890331 | 6 | * | 6 | 1000 |
| NC_039516.1 | 13065193 | 13068389 | 3197 | + | Gnomon | gene | NA | NA | gene-LOC105275355 | GeneID:105275355 | LOC105275355 | Gene | LOC105275355 | protein_coding | character(0) | NA | character(0) | NC_039516.1 | 13066190 | 13066195 | 6 | * | 6 | 1000 |
| NC_039517.1 | 1360424 | 1367346 | 6923 | + | Gnomon | gene | NA | NA | gene-LOC105278812 | GeneID:105278812 | LOC105278812 | Gene | LOC105278812 | protein_coding | character(0) | NA | character(0) | NC_039517.1 | 1361411 | 1361416 | 6 | * | 6 | 1000 |
| NC_039517.1 | 8175046 | 8181714 | 6669 | - | Gnomon | gene | NA | NA | gene-LOC105284262 | GeneID:105284262 | LOC105284262 | Gene | LOC105284262 | protein_coding | character(0) | NA | character(0) | NC_039517.1 | 8175507 | 8175512 | 6 | * | 6 | 1000 |
| NC_039518.1 | 5496870 | 5510429 | 13560 | - | Gnomon | gene | NA | NA | gene-LOC105281906 | GeneID:105281906 | LOC105281906 | Gene | LOC105281906 | protein_coding | character(0) | NA | character(0) | NC_039518.1 | 5508860 | 5508865 | 6 | * | 6 | 1000 |
| NC_039506.1 | 23567682 | 23574351 | 6670 | + | Gnomon | gene | NA | NA | gene-LOC105279333 | GeneID:105279333 | LOC105279333 | Gene | LOC105279333 | protein_coding | | | | | | | | | | |

character(0) NA character(0) NC_039506.1 23568887 23568897 11 * 10 909.090909090909
NC_039513.1 13434303 13438575 4273 - Gnomon gene NA NA gene-LOC105275036 GeneID:105275036 LOC105275036 Gene LOC105275036 protein_coding
character(0) NA character(0) NC_039513.1 13436991 13437000 10 * 9 900
NC_039509.1 17207204 17209527 2324 + Gnomon gene NA NA gene-LOC105283016 GeneID:105283016 LOC105283016 Gene LOC105283016 protein_coding
character(0) NA character(0) NC_039509.1 17208071 17208079 9 * 8 888.888888888889
NC_039510.1 8568121 8569477 1357 - Gnomon gene NA NA gene-LOC105275137 GeneID:105275137 LOC105275137 Gene LOC105275137 protein_coding
character(0) NA character(0) NC_039510.1 8568914 8568922 9 * 8 888.888888888889
NC_039513.1 3303685 3306319 2635 + Gnomon gene NA NA gene-LOC105275761 GeneID:105275761 LOC105275761 Gene LOC105275761 protein_coding
character(0) NA character(0) NC_039513.1 3304807 3304814 8 * 7 875
NC_039518.1 8474988 8476009 1022 + Gnomon gene NA NA gene-LOC105286212 GeneID:105286212 LOC105286212 Gene LOC105286212 protein_coding
character(0) NA character(0) NC_039518.1 8475762 8475769 8 * 7 875
NC_039506.1 1487779 1489683 1905 + Gnomon gene NA NA gene-LOC105280236 GeneID:105280236 LOC105280236 Gene LOC105280236 protein_coding
character(0) NA character(0) NC_039506.1 1488926 1488932 7 * 6 857.142857142857
NC_039506.1 2274108 2276090 1983 - Gnomon gene NA NA gene-LOC105280319 GeneID:105280319 LOC105280319 Gene LOC105280319 protein_coding
character(0) NA character(0) NC_039506.1 2275368 2275374 7 * 6 857.142857142857
NC_039506.1 4874444 4880252 5809 - Gnomon gene NA NA gene-LOC105277909 GeneID:105277909 LOC105277909 Gene LOC105277909 protein_coding
character(0) NA character(0) NC_039506.1 4878628 4878634 7 * 6 857.142857142857
NC_039506.1 15509387 15513034 3648 + Gnomon gene NA NA gene-LOC105283961 GeneID:105283961 LOC105283961 Gene LOC105283961 protein_coding
character(0) NA character(0) NC_039506.1 15510126 15510132 7 * 6 857.142857142857
NC_039506.1 15517083 15520247 3165 - Gnomon gene NA NA gene-LOC105283964 GeneID:105283964 LOC105283964 Gene LOC105283964 protein_coding
character(0) NA character(0) NC_039506.1 15519235 15519241 7 * 6 857.142857142857
NC_039506.1 20553263 20559098 5836 - Gnomon gene NA NA gene-LOC105278544 GeneID:105278544 LOC105278544 Gene LOC105278544 protein_coding
character(0) NA character(0) NC_039506.1 20557629 20557635 7 * 6 857.142857142857
NC_039506.1 22941360 22946330 4971 - Gnomon gene NA NA gene-LOC105283484 GeneID:105283484 LOC105283484 Gene LOC105283484 protein_coding
character(0) NA character(0) NC_039506.1 22945036 22945042 7 * 6 857.142857142857
NC_039506.1 23689672 23694320 4649 - Gnomon gene NA NA gene-LOC105279346 GeneID:105279346 LOC105279346 Gene LOC105279346 protein_coding
character(0) NA character(0) NC_039506.1 23693180 23693186 7 * 6 857.142857142857
NC_039507.1 790243 798925 8683 + Gnomon gene NA NA gene-LOC105282921 GeneID:105282921 LOC105282921 Gene LOC105282921 protein_coding
character(0) NA character(0) NC_039507.1 791692 791698 7 * 6 857.142857142857
NC_039507.1 5492373 5502473 10101 + Gnomon gene NA NA gene-LOC105281645 GeneID:105281645 LOC105281645 Gene LOC105281645 protein_coding
character(0) NA character(0) NC_039507.1 5493751 5493757 7 * 6 857.142857142857
NC_039507.1 6347235 6350936 3702 + Gnomon gene NA NA gene-LOC105275401 GeneID:105275401 LOC105275401 Gene LOC105275401 protein_coding
character(0) NA character(0) NC_039507.1 6348012 6348018 7 * 6 857.142857142857
NC_039507.1 6560548 6563391 2844 - Gnomon gene NA NA gene-LOC105285345 GeneID:105285345 LOC105285345 Gene LOC105285345 protein_coding
character(0) NA character(0) NC_039507.1 6562518 6562524 7 * 6 857.142857142857
NC_039507.1 15481857 15484141 2285 + Gnomon gene NA NA gene-LOC105280092 GeneID:105280092 LOC105280092 Gene LOC105280092 protein_coding
character(0) NA character(0) NC_039507.1 15482876 15482882 7 * 6 857.142857142857
NC_039507.1 15506083 15508702 2620 + Gnomon gene NA NA gene-LOC105280102 GeneID:105280102 LOC105280102 Gene LOC105280102 protein_coding
character(0) NA character(0) NC_039507.1 15507322 15507328 7 * 6 857.142857142857
NC_039507.1 18333035 18336377 3343 - Gnomon gene NA NA gene-LOC105281577 GeneID:105281577 LOC105281577 Gene LOC105281577 protein_coding
character(0) NA character(0) NC_039507.1 18335506 18335512 7 * 6 857.142857142857
NC_039508.1 1386723 1389150 2428 - Gnomon gene NA NA gene-LOC105277389 GeneID:105277389 LOC105277389 Gene LOC105277389 protein_coding
character(0) NA character(0) NC_039508.1 1387947 1387953 7 * 6 857.142857142857
NC_039508.1 2598434 2600165 1732 - Gnomon gene NA NA gene-LOC105281339 GeneID:105281339 LOC105281339 Gene LOC105281339 protein_coding
character(0) NA character(0) NC_039508.1 2599031 2599037 7 * 6 857.142857142857
NC_039508.1 5353027 5357392 4366 - Gnomon gene NA NA gene-LOC105277468 GeneID:105277468 LOC105277468 Gene LOC105277468 protein_coding
character(0) NA character(0) NC_039508.1 5356681 5356687 7 * 6 857.142857142857
NC_039508.1 9041025 9045259 4235 + Gnomon gene NA NA gene-LOC105278106 GeneID:105278106 LOC105278106 Gene LOC105278106 protein_coding
character(0) NA character(0) NC_039508.1 9042110 9042116 7 * 6 857.142857142857
NC_039508.1 10763110 10766843 3734 + Gnomon gene NA NA gene-LOC105276948 GeneID:105276948 LOC105276948 Gene LOC105276948 protein_coding
character(0) NA character(0) NC_039508.1 10764758 10764764 7 * 6 857.142857142857
NC_039508.1 10797665 10799797 2133 + Gnomon gene NA NA gene-LOC105276949 GeneID:105276949 LOC105276949 Gene LOC105276949 protein_coding
character(0) NA character(0) NC_039508.1 10798605 10798611 7 * 6 857.142857142857
NC_039508.1 13116146 13120560 4415 - Gnomon gene NA NA gene-LOC105277128 GeneID:105277128 LOC105277128 Gene LOC105277128 protein_coding
character(0) NA character(0) NC_039508.1 13116245 13116251 7 * 6 857.142857142857
NC_039508.1 14019414 14022018 2605 + Gnomon gene NA NA gene-LOC105278684 GeneID:105278684 LOC105278684 Gene LOC105278684 protein_coding
character(0) NA character(0) NC_039508.1 14021521 14021527 7 * 6 857.142857142857
NC_039508.1 15763225 15782851 19627 + Gnomon gene NA NA gene-LOC105277234 GeneID:105277234 LOC105277234 Gene LOC105277234 protein_coding
character(0) NA character(0) NC_039508.1 15764452 15764458 7 * 6 857.142857142857
NC_039508.1 17548373 17550348 1976 - Gnomon gene NA NA gene-LOC105284396 GeneID:105284396 LOC105284396 Gene LOC105284396 protein_coding
character(0) NA character(0) NC_039508.1 17549403 17549409 7 * 6 857.142857142857
NC_039509.1 89280 90451 1172 + Gnomon gene NA NA gene-LOC113561814 GeneID:113561814 LOC113561814 Gene LOC113561814 protein_coding character(0) NA
character(0) NC_039509.1 90329 90335 7 * 6 857.142857142857
NC_039509.1 2072593 2082307 9715 - Gnomon gene NA NA gene-LOC105283818 GeneID:105283818 LOC105283818 Gene LOC105283818 protein_coding

character(0) NA character(0) NC_039509.1 2081387 2081393 7 * 6 857.142857142857
NC_039509.1 2365507 2377666 12160 + Gnomon gene NA NA gene-LOC105285865 GeneID:105285865 LOC105285865 Gene LOC105285865 protein_coding
character(0) NA character(0) NC_039509.1 2366218 2366224 7 * 6 857.142857142857
NC_039509.1 5250307 5257914 7608 - Gnomon gene NA NA gene-LOC105287147 GeneID:105287147 LOC105287147 Gene LOC105287147 protein_coding
character(0) NA character(0) NC_039509.1 5256794 5256800 7 * 6 857.142857142857
NC_039509.1 7092641 7097742 5102 + Gnomon gene NA NA gene-LOC105282559 GeneID:105282559 LOC105282559 Gene LOC105282559 protein_coding
character(0) NA character(0) NC_039509.1 7093578 7093584 7 * 6 857.142857142857
NC_039509.1 10232323 10244464 12142 + Gnomon gene NA NA gene-LOC105282655 GeneID:105282655 LOC105282655 Gene LOC105282655 protein_coding
character(0) NA character(0) NC_039509.1 10233310 10233316 7 * 6 857.142857142857
NC_039509.1 10710016 10711124 1109 + Gnomon gene NA NA gene-LOC105275153 GeneID:105275153 LOC105275153 Gene LOC105275153 protein_coding
character(0) NA character(0) NC_039509.1 10710730 10710736 7 * 6 857.142857142857
NC_039509.1 13909149 13915606 6458 + Gnomon gene NA NA gene-LOC105278288 GeneID:105278288 LOC105278288 Gene LOC105278288 protein_coding
character(0) NA character(0) NC_039509.1 13909987 13909993 7 * 6 857.142857142857
NC_039509.1 17375981 17382001 6021 + Gnomon gene NA NA gene-LOC105282459 GeneID:105282459 LOC105282459 Gene LOC105282459 protein_coding
character(0) NA character(0) NC_039509.1 17378741 17378747 7 * 6 857.142857142857
NC_039510.1 624625 628259 3635 + Gnomon gene NA NA gene-LOC105279313 GeneID:105279313 LOC105279313 Gene LOC105279313 protein_coding
character(0) NA character(0) NC_039510.1 625494 625500 7 * 6 857.142857142857
NC_039510.1 3040660 3042070 1411 + Gnomon gene NA NA gene-LOC105287710 GeneID:105287710 LOC105287710 Gene LOC105287710 protein_coding
character(0) NA character(0) NC_039510.1 3041562 3041568 7 * 6 857.142857142857
NC_039510.1 3110492 3117379 6888 - Gnomon gene NA NA gene-LOC105287697 GeneID:105287697 LOC105287697 Gene LOC105287697 protein_coding
character(0) NA character(0) NC_039510.1 3116602 3116608 7 * 6 857.142857142857
NC_039510.1 5067676 5072600 4925 - Gnomon gene NA NA gene-LOC105287597 GeneID:105287597 LOC105287597 Gene LOC105287597 protein_coding
character(0) NA character(0) NC_039510.1 5071200 5071206 7 * 6 857.142857142857
NC_039510.1 5537223 5539036 1814 - Gnomon gene NA NA gene-LOC105287569 GeneID:105287569 LOC105287569 Gene LOC105287569 protein_coding
character(0) NA character(0) NC_039510.1 5538139 5538145 7 * 6 857.142857142857
NC_039510.1 6122701 6125649 2949 - Gnomon gene NA NA gene-LOC105287279 GeneID:105287279 LOC105287279 Gene LOC105287279 protein_coding
character(0) NA character(0) NC_039510.1 6124458 6124464 7 * 6 857.142857142857
NC_039510.1 6163214 6171437 8224 - Gnomon gene NA NA gene-LOC105287371 GeneID:105287371 LOC105287371 Gene LOC105287371 protein_coding
character(0) NA character(0) NC_039510.1 6170155 6170161 7 * 6 857.142857142857
NC_039510.1 6269006 6275115 6110 + Gnomon gene NA NA gene-LOC105287511 GeneID:105287511 LOC105287511 Gene LOC105287511 protein_coding
character(0) NA character(0) NC_039510.1 6270460 6270466 7 * 6 857.142857142857
NC_039510.1 8487238 8489293 2056 - Gnomon gene NA NA gene-LOC105275119 GeneID:105275119 LOC105275119 Gene LOC105275119 misc_RNA character(0)
NA character(0) NC_039510.1 8488481 8488487 7 * 6 857.142857142857
NC_039510.1 8568121 8569477 1357 - Gnomon gene NA NA gene-LOC105275137 GeneID:105275137 LOC105275137 Gene LOC105275137 protein_coding
character(0) NA character(0) NC_039510.1 8568885 8568891 7 * 6 857.142857142857
NC_039510.1 9996113 9998708 2596 - Gnomon gene NA NA gene-LOC105281227 GeneID:105281227 LOC105281227 Gene LOC105281227 protein_coding
character(0) NA character(0) NC_039510.1 9997644 9997650 7 * 6 857.142857142857
NC_039510.1 13531247 13535548 4302 - Gnomon gene NA NA gene-LOC105280964 GeneID:105280964 LOC105280964 Gene LOC105280964 protein_coding
character(0) NA character(0) NC_039510.1 13533348 13533354 7 * 6 857.142857142857
NC_039510.1 15582075 15587881 5807 - Gnomon gene NA NA gene-LOC105286461 GeneID:105286461 LOC105286461 Gene LOC105286461 protein_coding
character(0) NA character(0) NC_039510.1 15586562 15586568 7 * 6 857.142857142857
NC_039511.1 1662842 1668395 5554 + Gnomon gene NA NA gene-LOC105276437 GeneID:105276437 LOC105276437 Gene LOC105276437 protein_coding
character(0) NA character(0) NC_039511.1 1664306 1664312 7 * 6 857.142857142857
NC_039511.1 2004889 2015973 11085 - Gnomon gene NA NA gene-LOC105276410 GeneID:105276410 LOC105276410 Gene LOC105276410 protein_coding
character(0) NA character(0) NC_039511.1 2014683 2014689 7 * 6 857.142857142857
NC_039511.1 2004889 2015973 11085 - Gnomon gene NA NA gene-LOC105276410 GeneID:105276410 LOC105276410 Gene LOC105276410 protein_coding
character(0) NA character(0) NC_039511.1 2014740 2014746 7 * 6 857.142857142857
NC_039511.1 3635459 3642537 7079 - Gnomon gene NA NA gene-LOC105284052 GeneID:105284052 LOC105284052 Gene LOC105284052 protein_coding
character(0) NA character(0) NC_039511.1 3639538 3639544 7 * 6 857.142857142857
NC_039511.1 4554146 4561340 7195 - Gnomon gene NA NA gene-LOC105281750 GeneID:105281750 LOC105281750 Gene LOC105281750 protein_coding
character(0) NA character(0) NC_039511.1 4560502 4560508 7 * 6 857.142857142857
NC_039511.1 12403841 12413518 9678 - Gnomon gene NA NA gene-LOC105285135 GeneID:105285135 LOC105285135 Gene LOC105285135 protein_coding
character(0) NA character(0) NC_039511.1 12411122 12411128 7 * 6 857.142857142857
NC_039511.1 14704311 14706938 2628 - Gnomon gene NA NA gene-LOC105285889 GeneID:105285889 LOC105285889 Gene LOC105285889 protein_coding
character(0) NA character(0) NC_039511.1 14705900 14705906 7 * 6 857.142857142857
NC_039511.1 15983428 15985754 2327 + Gnomon gene NA NA gene-LOC105284576 GeneID:105284576 LOC105284576 Gene LOC105284576 protein_coding
character(0) NA character(0) NC_039511.1 15984100 15984106 7 * 6 857.142857142857
NC_039512.1 11140867 11151152 10286 - Gnomon gene NA NA gene-LOC105279854 GeneID:105279854 LOC105279854 Gene LOC105279854 protein_coding
character(0) NA character(0) NC_039512.1 11150305 11150311 7 * 6 857.142857142857
NC_039512.1 12469542 12475093 5552 - Gnomon gene NA NA gene-LOC105279748 GeneID:105279748 LOC105279748 Gene LOC105279748 protein_coding
character(0) NA character(0) NC_039512.1 12474009 12474015 7 * 6 857.142857142857
NC_039512.1 12621899 12625222 3324 + Gnomon gene NA NA gene-LOC105279740 GeneID:105279740 LOC105279740 Gene LOC105279740 protein_coding
character(0) NA character(0) NC_039512.1 12623117 12623123 7 * 6 857.142857142857
NC_039512.1 12739826 12742653 2828 + Gnomon gene NA NA gene-LOC105279719 GeneID:105279719 LOC105279719 Gene LOC105279719 protein_coding

| | | | | | | | | | | | | | | |
|---|---|---|---|---|---|---|---|---|---|---|---|---|---|---|
| character(0) | NA | character(0) | NC_039512.1 | 12740699 | 12740705 | 7 | * | 6 | 857.142857142857 | | | | | |
| NC_039512.1 | 12814910 | 12819293 | 4384 | - | Gnomon | gene | NA | NA | gene-LOC105279709 | GeneID:105279709 | LOC105279709 | Gene | LOC105279709 | protein_coding |
| character(0) | NA | character(0) | NC_039512.1 | 12818052 | 12818058 | 7 | * | 6 | 857.142857142857 | | | | | |
| NC_039512.1 | 13089540 | 13093562 | 4023 | - | Gnomon | gene | NA | NA | gene-LOC105279664 | GeneID:105279664 | LOC105279664 | Gene | LOC105279664 | protein_coding |
| character(0) | NA | character(0) | NC_039512.1 | 13092263 | 13092269 | 7 | * | 6 | 857.142857142857 | | | | | |
| NC_039512.1 | 14720220 | 14725286 | 5067 | + | Gnomon | gene | NA | NA | gene-LOC105287687 | GeneID:105287687 | LOC105287687 | Gene | LOC105287687 | protein_coding |
| character(0) | NA | character(0) | NC_039512.1 | 14721247 | 14721253 | 7 | * | 6 | 857.142857142857 | | | | | |
| NC_039513.1 | 3171172 | 3175402 | 4231 | + | Gnomon | gene | NA | NA | gene-LOC105287820 | GeneID:105287820 | LOC105287820 | Gene | LOC105287820 | protein_coding |
| character(0) | NA | character(0) | NC_039513.1 | 3172807 | 3172813 | 7 | * | 6 | 857.142857142857 | | | | | |
| NC_039513.1 | 3253321 | 3257474 | 4154 | - | Gnomon | gene | NA | NA | gene-LOC105275753 | GeneID:105275753 | LOC105275753 | Gene | LOC105275753 | protein_coding |
| character(0) | NA | character(0) | NC_039513.1 | 3256149 | 3256155 | 7 | * | 6 | 857.142857142857 | | | | | |
| NC_039513.1 | 3253321 | 3257474 | 4154 | - | Gnomon | gene | NA | NA | gene-LOC105275753 | GeneID:105275753 | LOC105275753 | Gene | LOC105275753 | protein_coding |
| character(0) | NA | character(0) | NC_039513.1 | 3256358 | 3256364 | 7 | * | 6 | 857.142857142857 | | | | | |
| NC_039513.1 | 3257374 | 3260835 | 3462 | - | Gnomon | gene | NA | NA | gene-LOC105275752 | GeneID:105275752 | LOC105275752 | Gene | LOC105275752 | protein_coding |
| character(0) | NA | character(0) | NC_039513.1 | 3259768 | 3259774 | 7 | * | 6 | 857.142857142857 | | | | | |
| NC_039513.1 | 3373854 | 3375485 | 1632 | + | Gnomon | gene | NA | NA | gene-LOC105274850 | GeneID:105274850 | LOC105274850 | Gene | LOC105274850 | protein_coding |
| character(0) | NA | character(0) | NC_039513.1 | 3374674 | 3374680 | 7 | * | 6 | 857.142857142857 | | | | | |
| NC_039513.1 | 8483464 | 8491634 | 8171 | + | Gnomon | gene | NA | NA | gene-LOC105275689 | GeneID:105275689 | LOC105275689 | Gene | LOC105275689 | protein_coding |
| character(0) | NA | character(0) | NC_039513.1 | 8484406 | 8484412 | 7 | * | 6 | 857.142857142857 | | | | | |
| NC_039513.1 | 9603616 | 9607153 | 3538 | + | Gnomon | gene | NA | NA | gene-LOC105275615 | GeneID:105275615 | LOC105275615 | Gene | LOC105275615 | protein_coding |
| character(0) | NA | character(0) | NC_039513.1 | 9605113 | 9605119 | 7 | * | 6 | 857.142857142857 | | | | | |
| NC_039513.1 | 9603616 | 9607153 | 3538 | + | Gnomon | gene | NA | NA | gene-LOC105275615 | GeneID:105275615 | LOC105275615 | Gene | LOC105275615 | protein_coding |
| character(0) | NA | character(0) | NC_039513.1 | 9605414 | 9605420 | 7 | * | 6 | 857.142857142857 | | | | | |
| NC_039513.1 | 11674366 | 11676356 | 1991 | - | Gnomon | gene | NA | NA | gene-LOC105280739 | GeneID:105280739 | LOC105280739 | Gene | LOC105280739 | protein_coding |
| character(0) | NA | character(0) | NC_039513.1 | 11675069 | 11675075 | 7 | * | 6 | 857.142857142857 | | | | | |
| NC_039513.1 | 11674366 | 11676356 | 1991 | - | Gnomon | gene | NA | NA | gene-LOC105280739 | GeneID:105280739 | LOC105280739 | Gene | LOC105280739 | protein_coding |
| character(0) | NA | character(0) | NC_039513.1 | 11675195 | 11675201 | 7 | * | 6 | 857.142857142857 | | | | | |
| NC_039513.1 | 13452183 | 13456265 | 4083 | + | Gnomon | gene | NA | NA | gene-LOC105275031 | GeneID:105275031 | LOC105275031 | Gene | LOC105275031 | protein_coding |
| character(0) | NA | character(0) | NC_039513.1 | 13453683 | 13453689 | 7 | * | 6 | 857.142857142857 | | | | | |
| NC_039513.1 | 15035458 | 15043240 | 7783 | + | Gnomon | gene | NA | NA | gene-LOC105285648 | GeneID:105285648 | LOC105285648 | Gene | LOC105285648 | protein_coding |
| character(0) | NA | character(0) | NC_039513.1 | 15037033 | 15037039 | 7 | * | 6 | 857.142857142857 | | | | | |
| NC_039514.1 | 1286674 | 1297576 | 10903 | - | Gnomon | gene | NA | NA | gene-LOC105276233 | GeneID:105276233 | LOC105276233 | Gene | LOC105276233 | protein_coding |
| character(0) | NA | character(0) | NC_039514.1 | 1295968 | 1295974 | 7 | * | 6 | 857.142857142857 | | | | | |
| NC_039514.1 | 8629561 | 8648221 | 18661 | + | Gnomon | gene | NA | NA | gene-LOC105278994 | GeneID:105278994 | LOC105278994 | Gene | LOC105278994 | protein_coding |
| character(0) | NA | character(0) | NC_039514.1 | 8630645 | 8630651 | 7 | * | 6 | 857.142857142857 | | | | | |
| NC_039514.1 | 9413016 | 9415003 | 1988 | - | Gnomon | gene | NA | NA | gene-LOC105284659 | GeneID:105284659 | LOC105284659 | Gene | LOC105284659 | protein_coding |
| character(0) | NA | character(0) | NC_039514.1 | 9414116 | 9414122 | 7 | * | 6 | 857.142857142857 | | | | | |
| NC_039514.1 | 11109536 | 11117116 | 7581 | - | Gnomon | gene | NA | NA | gene-LOC105281601 | GeneID:105281601 | LOC105281601 | Gene | LOC105281601 | protein_coding |
| character(0) | NA | character(0) | NC_039514.1 | 11113139 | 11113145 | 7 | * | 6 | 857.142857142857 | | | | | |
| NC_039514.1 | 11111779 | 11115572 | 3794 | + | Gnomon | gene | NA | NA | gene-LOC105281603 | GeneID:105281603 | LOC105281603 | Gene | LOC105281603 | protein_coding |
| character(0) | NA | character(0) | NC_039514.1 | 11113139 | 11113145 | 7 | * | 6 | 857.142857142857 | | | | | |
| NC_039515.1 | 1527980 | 1529634 | 1655 | + | Gnomon | gene | NA | NA | gene-LOC105274711 | GeneID:105274711 | LOC105274711 | Gene | LOC105274711 | protein_coding |
| character(0) | NA | character(0) | NC_039515.1 | 1528293 | 1528299 | 7 | * | 6 | 857.142857142857 | | | | | |
| NC_039515.1 | 2215137 | 2216986 | 1850 | - | Gnomon | gene | NA | NA | gene-LOC105278431 | GeneID:105278431 | LOC105278431 | Gene | LOC105278431 | protein_coding |
| character(0) | NA | character(0) | NC_039515.1 | 2215927 | 2215933 | 7 | * | 6 | 857.142857142857 | | | | | |
| NC_039515.1 | 5537006 | 5539696 | 2691 | + | Gnomon | gene | NA | NA | gene-LOC105277256 | GeneID:105277256 | LOC105277256 | Gene | LOC105277256 | protein_coding |
| character(0) | NA | character(0) | NC_039515.1 | 5537966 | 5537972 | 7 | * | 6 | 857.142857142857 | | | | | |
| NC_039515.1 | 6480766 | 6491993 | 11228 | + | Gnomon | gene | NA | NA | gene-LOC105282298 | GeneID:105282298 | LOC105282298 | Gene | LOC105282298 | protein_coding |
| character(0) | NA | character(0) | NC_039515.1 | 6481705 | 6481711 | 7 | * | 6 | 857.142857142857 | | | | | |
| NC_039515.1 | 6604872 | 6605723 | 852 | + | Gnomon | gene | NA | NA | gene-LOC105282280 | GeneID:105282280 | LOC105282280 | Gene | LOC105282280 | protein_coding |
| character(0) | NA | character(0) | NC_039515.1 | 6605539 | 6605545 | 7 | * | 6 | 857.142857142857 | | | | | |
| NC_039515.1 | 6615586 | 6618213 | 2628 | - | Gnomon | gene | NA | NA | gene-LOC105282278 | GeneID:105282278 | LOC105282278 | Gene | LOC105282278 | protein_coding |
| character(0) | NA | character(0) | NC_039515.1 | 6617273 | 6617279 | 7 | * | 6 | 857.142857142857 | | | | | |
| NC_039515.1 | 7926262 | 7929466 | 3205 | + | Gnomon | gene | NA | NA | gene-LOC105283741 | GeneID:105283741 | LOC105283741 | Gene | LOC105283741 | protein_coding |
| character(0) | NA | character(0) | NC_039515.1 | 7927531 | 7927537 | 7 | * | 6 | 857.142857142857 | | | | | |
| NC_039516.1 | 756663 | 760199 | 3537 | + | Gnomon | gene | NA | NA | gene-LOC105274638 | GeneID:105274638 | LOC105274638 | Gene | LOC105274638 | protein_coding |
| character(0) | NA | character(0) | NC_039516.1 | 757725 | 757731 | 7 | * | 6 | 857.142857142857 | | | | | |
| NC_039516.1 | 842233 | 846795 | 4563 | + | Gnomon | gene | NA | NA | gene-LOC105274618 | GeneID:105274618 | LOC105274618 | Gene | LOC105274618 | protein_coding |
| character(0) | NA | character(0) | NC_039516.1 | 844061 | 844067 | 7 | * | 6 | 857.142857142857 | | | | | |
| NC_039516.1 | 985951 | 990466 | 4516 | + | Gnomon | gene | NA | NA | gene-LOC105274590 | GeneID:105274590 | LOC105274590 | Gene | LOC105274590 | protein_coding |
| character(0) | NA | character(0) | NC_039516.1 | 987063 | 987069 | 7 | * | 6 | 857.142857142857 | | | | | |
| NC_039516.1 | 3100198 | 3101692 | 1495 | + | Gnomon | gene | NA | NA | gene-LOC105288193 | GeneID:105288193 | LOC105288193 | Gene | LOC105288193 | protein_coding |
| character(0) | NA | character(0) | NC_039516.1 | 3101202 | 3101208 | 7 | * | 6 | 857.142857142857 | | | | | |
| NC_039516.1 | 4713084 | 4714767 | 1684 | - | Gnomon | gene | NA | NA | gene-LOC105275897 | GeneID:105275897 | LOC105275897 | Gene | LOC105275897 | protein_coding |

| | | | | | | | | | | | | | | |
|---|---|---|---|---|---|---|---|---|---|---|---|---|---|---|
| character(0) | NA | character(0) | NC_039516.1 | 4713795 | 4713801 | 7 | * | 6 | 857.142857142857 | | | | | |
| NC_039516.1 | 11790047 | 11792531 | 2485 | + | Gnomon | gene | NA | NA | gene-LOC105278656 | GeneID:105278656 | LOC105278656 | Gene | LOC105278656 | protein_coding |
| character(0) | NA | character(0) | NC_039516.1 | 11791250 | 11791256 | 7 | * | 6 | 857.142857142857 | | | | | |
| NC_039516.1 | 12834895 | 12836314 | 1420 | - | Gnomon | gene | NA | NA | gene-LOC105275307 | GeneID:105275307 | LOC105275307 | Gene | LOC105275307 | protein_coding |
| character(0) | NA | character(0) | NC_039516.1 | 12835552 | 12835558 | 7 | * | 6 | 857.142857142857 | | | | | |
| NC_039516.1 | 13386687 | 13390618 | 3932 | + | Gnomon | gene | NA | NA | gene-LOC105278022 | GeneID:105278022 | LOC105278022 | Gene | LOC105278022 | protein_coding |
| character(0) | NA | character(0) | NC_039516.1 | 13387740 | 13387746 | 7 | * | 6 | 857.142857142857 | | | | | |
| NC_039516.1 | 13386687 | 13390618 | 3932 | + | Gnomon | gene | NA | NA | gene-LOC105278022 | GeneID:105278022 | LOC105278022 | Gene | LOC105278022 | protein_coding |
| character(0) | NA | character(0) | NC_039516.1 | 13387869 | 13387875 | 7 | * | 6 | 857.142857142857 | | | | | |
| NC_039516.1 | 13518014 | 13520692 | 2679 | + | Gnomon | gene | NA | NA | gene-LOC105277817 | GeneID:105277817 | LOC105277817 | Gene | LOC105277817 | protein_coding |
| character(0) | NA | character(0) | NC_039516.1 | 13518940 | 13518946 | 7 | * | 6 | 857.142857142857 | | | | | |
| NC_039517.1 | 867927 | 869156 | 1230 | - | Gnomon | gene | NA | NA | gene-LOC105278824 | GeneID:105278824 | LOC105278824 | Gene | LOC105278824 | protein_coding |
| character(0) | NA | character(0) | NC_039517.1 | 868363 | 868369 | 7 | * | 6 | 857.142857142857 | | | | | |
| NC_039517.1 | 2307792 | 2312395 | 4604 | - | Gnomon | gene | NA | NA | gene-LOC105281943 | GeneID:105281943 | LOC105281943 | Gene | LOC105281943 | protein_coding |
| character(0) | NA | character(0) | NC_039517.1 | 2309012 | 2309018 | 7 | * | 6 | 857.142857142857 | | | | | |
| NC_039517.1 | 2393025 | 2396928 | 3904 | - | Gnomon | gene | NA | NA | gene-LOC105281949 | GeneID:105281949 | LOC105281949 | Gene | LOC105281949 | protein_coding |
| character(0) | NA | character(0) | NC_039517.1 | 2395845 | 2395851 | 7 | * | 6 | 857.142857142857 | | | | | |
| NC_039517.1 | 7306074 | 7309288 | 3215 | + | Gnomon | gene | NA | NA | gene-LOC105284202 | GeneID:105284202 | LOC105284202 | Gene | LOC105284202 | protein_coding |
| character(0) | NA | character(0) | NC_039517.1 | 7307471 | 7307477 | 7 | * | 6 | 857.142857142857 | | | | | |
| NC_039517.1 | 9368753 | 9371050 | 2298 | - | Gnomon | gene | NA | NA | gene-LOC105282220 | GeneID:105282220 | LOC105282220 | Gene | LOC105282220 | protein_coding |
| character(0) | NA | character(0) | NC_039517.1 | 9370300 | 9370306 | 7 | * | 6 | 857.142857142857 | | | | | |
| NC_039518.1 | 2681489 | 2682795 | 1307 | - | Gnomon | gene | NA | NA | gene-LOC105285394 | GeneID:105285394 | LOC105285394 | Gene | LOC105285394 | protein_coding |
| character(0) | NA | character(0) | NC_039518.1 | 2681945 | 2681951 | 7 | * | 6 | 857.142857142857 | | | | | |
| NC_039518.1 | 5288531 | 5294826 | 6296 | - | Gnomon | gene | NA | NA | gene-LOC105285410 | GeneID:105285410 | LOC105285410 | Gene | LOC105285410 | protein_coding |
| character(0) | NA | character(0) | NC_039518.1 | 5293748 | 5293754 | 7 | * | 6 | 857.142857142857 | | | | | |
| NC_039518.1 | 9430521 | 9433765 | 3245 | - | Gnomon | gene | NA | NA | gene-LOC105286355 | GeneID:105286355 | LOC105286355 | Gene | LOC105286355 | protein_coding |
| character(0) | NA | character(0) | NC_039518.1 | 9432671 | 9432677 | 7 | * | 6 | 857.142857142857 | | | | | |
| NC_039519.1 | 197513 | 200423 | 2911 | + | Gnomon | gene | NA | NA | gene-LOC105282793 | GeneID:105282793 | LOC105282793 | Gene | LOC105282793 | protein_coding |
| character(0) | NA | character(0) | NC_039519.1 | 198095 | 198101 | 7 | * | 6 | 857.142857142857 | | | | | |
| NC_039519.1 | 8239374 | 8248322 | 8949 | - | Gnomon | gene | NA | NA | gene-LOC105287985 | GeneID:105287985 | LOC105287985 | Gene | LOC105287985 | protein_coding |
| character(0) | NA | character(0) | NC_039519.1 | 8247911 | 8247917 | 7 | * | 6 | 857.142857142857 | | | | | |
| NC_039508.1 | 13115312 | 13116213 | 902 | + | Gnomon | gene | NA | NA | gene-LOC105277091 | GeneID:105277091 | LOC105277091 | Gene | LOC105277091 | protein_coding |
| character(0) | NA | character(0) | NC_039508.1 | 13116053 | 13116062 | 10 | * | 8 | 800 | | | | | |
| NC_039513.1 | 5176546 | 5178099 | 1554 | + | Gnomon | gene | NA | NA | gene-LOC105279718 | GeneID:105279718 | LOC105279718 | Gene | LOC105279718 | protein_coding |
| character(0) | NA | character(0) | NC_039513.1 | 5177245 | 5177254 | 10 | * | 8 | 800 | | | | | |
| NC_039519.1 | 8084633 | 8088546 | 3914 | + | Gnomon | gene | NA | NA | gene-LOC105274767 | GeneID:105274767 | LOC105274767 | Gene | LOC105274767 | protein_coding |
| character(0) | NA | character(0) | NC_039519.1 | 8085717 | 8085726 | 10 | * | 8 | 800 | | | | | |

Directory: MRPR File: gwr-Ob_LI_BCphaseA2.txt

| seqnames | start | end | width | strand | source | type | score | phase | ID | Dbxref | Name | gbkey | gene | gene_biotype | end_range | partial | start_range | seqnames | start | end | width | strand | NbrSites | Sdnsty |
|---|---|---|---|---|---|---|---|---|---|---|---|---|---|---|---|---|---|---|---|---|---|---|---|---|
| NC_039506.1 | 3342881 | 3348391 | 5511 | + | Gnomon | gene | NA | NA | gene-LOC105280395 | GeneID:105280395 | LOC105280395 | Gene | LOC105280395 | protein_coding | character(0) | NA | character(0) | NC_039506.1 | 3344178 | 3344183 | 6 | * | 6 | 1000 |
| NC_039506.1 | 6089683 | 6090856 | 1174 | + | Gnomon | gene | NA | NA | gene-LOC105277804 | GeneID:105277804 | LOC105277804 | Gene | LOC105277804 | protein_coding | character(0) | NA | character(0) | NC_039506.1 | 6090624 | 6090629 | 6 | * | 6 | 1000 |
| NC_039507.1 | 5093844 | 5095945 | 2102 | + | Gnomon | gene | NA | NA | gene-LOC105281690 | GeneID:105281690 | LOC105281690 | Gene | LOC105281690 | protein_coding | character(0) | NA | character(0) | NC_039507.1 | 5094466 | 5094471 | 6 | * | 6 | 1000 |
| NC_039507.1 | 7181199 | 7188522 | 7324 | + | Gnomon | gene | NA | NA | gene-LOC105283383 | GeneID:105283383 | LOC105283383 | Gene | LOC105283383 | protein_coding | character(0) | NA | character(0) | NC_039507.1 | 7182290 | 7182295 | 6 | * | 6 | 1000 |
| NC_039507.1 | 7276866 | 7278961 | 2096 | + | Gnomon | gene | NA | NA | gene-LOC105283373 | GeneID:105283373 | LOC105283373 | Gene | LOC105283373 | protein_coding | character(0) | NA | character(0) | NC_039507.1 | 7277947 | 7277952 | 6 | * | 6 | 1000 |
| NC_039507.1 | 18333035 | 18336377 | 3343 | - | Gnomon | gene | NA | NA | gene-LOC105281577 | GeneID:105281577 | LOC105281577 | Gene | LOC105281577 | protein_coding | character(0) | NA | character(0) | NC_039507.1 | 18335809 | 18335814 | 6 | * | 6 | 1000 |
| NC_039508.1 | 10274002 | 10277440 | 3439 | + | Gnomon | gene | NA | NA | gene-LOC105276881 | GeneID:105276881 | LOC105276881 | Gene | LOC105276881 | protein_coding | character(0) | NA | character(0) | NC_039508.1 | 10275212 | 10275217 | 6 | * | 6 | 1000 |
| NC_039508.1 | 10716085 | 10732040 | 15956 | - | Gnomon | gene | NA | NA | gene-LOC105276941 | GeneID:105276941 | LOC105276941 | Gene | LOC105276941 | protein_coding | character(0) | NA | character(0) | NC_039508.1 | 10730559 | 10730564 | 6 | * | 6 | 1000 |
| NC_039509.1 | 1002697 | 1006107 | 3411 | - | Gnomon | gene | NA | NA | gene-LOC105283620 | GeneID:105283620 | LOC105283620 | Gene | LOC105283620 | protein_coding | character(0) | NA | character(0) | NC_039509.1 | 1004827 | 1004832 | 6 | * | 6 | 1000 |
| NC_039509.1 | 1135464 | 1137481 | 2018 | - | Gnomon | gene | NA | NA | gene-LOC105283634 | GeneID:105283634 | LOC105283634 | Gene | LOC105283634 | protein_coding | character(0) | NA | character(0) | NC_039509.1 | 1136266 | 1136271 | 6 | * | 6 | 1000 |
| NC_039509.1 | 2825666 | 2828757 | 3092 | + | Gnomon | gene | NA | NA | gene-LOC105282589 | GeneID:105282589 | LOC105282589 | Gene | LOC105282589 | protein_coding | character(0) | NA | character(0) | NC_039509.1 | 2826537 | 2826542 | 6 | * | 6 | 1000 |
| NC_039509.1 | 11068956 | 11086837 | 17882 | - | Gnomon | gene | NA | NA | gene-LOC105275770 | GeneID:105275770 | LOC105275770 | Gene | LOC105275770 | protein_coding | character(0) | NA | character(0) | NC_039509.1 | 11085457 | 11085462 | 6 | * | 6 | 1000 |
| NC_039509.1 | 13033738 | 13035966 | 2229 | + | Gnomon | gene | NA | NA | gene-LOC105278527 | GeneID:105278527 | LOC105278527 | Gene | LOC105278527 | protein_coding | character(0) | NA | character(0) | NC_039509.1 | 13034669 | 13034674 | 6 | * | 6 | 1000 |
| NC_039510.1 | 2607810 | 2610599 | 2790 | - | Gnomon | gene | NA | NA | gene-LOC105287754 | GeneID:105287754 | LOC105287754 | Gene | LOC105287754 | protein_coding | character(0) | NA | character(0) | NC_039510.1 | 2609179 | 2609184 | 6 | * | 6 | 1000 |
| NC_039510.1 | 3319355 | 3322927 | 3573 | + | Gnomon | gene | NA | NA | gene-LOC105287663 | GeneID:105287663 | LOC105287663 | Gene | LOC105287663 | protein_coding | character(0) | NA | character(0) | NC_039510.1 | 3320255 | 3320260 | 6 | * | 6 | 1000 |
| NC_039510.1 | 16287375 | 16291357 | 3983 | - | Gnomon | gene | NA | NA | gene-LOC105286509 | GeneID:105286509 | LOC105286509 | Gene | LOC105286509 | protein_coding | character(0) | NA | character(0) | NC_039510.1 | 16290598 | 16290603 | 6 | * | 6 | 1000 |
| NC_039511.1 | 2234830 | 2236789 | 1960 | - | Gnomon | gene | NA | NA | gene-LOC105276382 | GeneID:105276382 | LOC105276382 | Gene | LOC105276382 | protein_coding | character(0) | NA | character(0) | NC_039511.1 | 2236016 | 2236021 | 6 | * | 6 | 1000 |
| NC_039511.1 | 3614241 | 3616939 | 2699 | + | Gnomon | gene | NA | NA | gene-LOC105284057 | GeneID:105284057 | LOC105284057 | Gene | LOC105284057 | protein_coding | character(0) | NA | character(0) | NC_039511.1 | 3615462 | 3615467 | 6 | * | 6 | 1000 |
| NC_039511.1 | 3625986 | 3631991 | 6006 | - | Gnomon | gene | NA | NA | gene-LOC105284053 | GeneID:105284053 | LOC105284053 | Gene | LOC105284053 | protein_coding | character(0) | NA | character(0) | NC_039511.1 | 3630771 | 3630776 | 6 | * | 6 | 1000 |
| NC_039511.1 | 8407553 | 8425032 | 17480 | + | Gnomon | gene | NA | NA | gene-LOC105284845 | GeneID:105284845 | LOC105284845 | Gene | LOC105284845 | protein_coding | character(0) | NA | character(0) | NC_039511.1 | 8409706 | 8409711 | 6 | * | 6 | 1000 |
| NC_039512.1 | 384950 | 392996 | 8047 | - | Gnomon | gene | NA | NA | gene-LOC105288044 | GeneID:105288044 | LOC105288044 | Gene | LOC105288044 | protein_coding | character(0) | NA | character(0) | NC_039512.1 | 391510 | 391515 | 6 | * | 6 | 1000 |
| NC_039512.1 | 10275466 | 10280187 | 4722 | + | Gnomon | gene | NA | NA | gene-LOC105282847 | GeneID:105282847 | LOC105282847 | Gene | LOC105282847 | protein_coding | character(0) | NA | character(0) | NC_039512.1 | 10276481 | 10276486 | 6 | * | 6 | 1000 |
| NC_039512.1 | 12092666 | 12097773 | 5108 | + | Gnomon | gene | NA | NA | gene-LOC105279779 | GeneID:105279779 | LOC105279779 | Gene | LOC105279779 | protein_coding | character(0) | NA | character(0) | NC_039512.1 | 12094200 | 12094205 | 6 | * | 6 | 1000 |
| NC_039512.1 | 15707994 | 15714445 | 6452 | + | Gnomon | gene | NA | NA | gene-LOC105274475 | GeneID:105274475 | LOC105274475 | Gene | LOC105274475 | protein_coding | character(0) | NA | character(0) | NC_039512.1 | 15711477 | 15711482 | 6 | * | 6 | 1000 |
| NC_039512.1 | 15979275 | 15981529 | 2255 | - | Gnomon | gene | NA | NA | gene-LOC105274775 | GeneID:105274775 | LOC105274775 | Gene | LOC105274775 | protein_coding | character(0) | NA | character(0) | NC_039512.1 | 15980634 | 15980639 | 6 | * | 6 | 1000 |
| NC_039513.1 | 10401827 | 10404590 | 2764 | - | Gnomon | gene | NA | NA | gene-LOC105275550 | GeneID:105275550 | LOC105275550 | Gene | LOC105275550 | protein_coding | character(0) | NA | character(0) | NC_039513.1 | 10403469 | 10403474 | 6 | * | 6 | 1000 |
| NC_039513.1 | 11029007 | 11033701 | 4695 | - | Gnomon | gene | NA | NA | gene-LOC105275763 | GeneID:105275763 | LOC105275763 | Gene | LOC105275763 | protein_coding | character(0) | NA | character(0) | NC_039513.1 | 11030126 | 11030131 | 6 | * | 6 | 1000 |
| NC_039513.1 | 12028675 | 12030941 | 2267 | - | Gnomon | gene | NA | NA | gene-LOC105280791 | GeneID:105280791 | LOC105280791 | Gene | LOC105280791 | protein_coding | character(0) | NA | character(0) | NC_039513.1 | 12029791 | 12029796 | 6 | * | 6 | 1000 |
| NC_039513.1 | 13434303 | 13438575 | 4273 | - | Gnomon | gene | NA | NA | gene-LOC105275036 | GeneID:105275036 | LOC105275036 | Gene | LOC105275036 | protein_coding | character(0) | NA | character(0) | NC_039513.1 | 13436991 | 13437000 | 10 | * | 10 | 1000 |
| NC_039513.1 | 14931340 | 14932816 | 1477 | + | Gnomon | gene | NA | NA | gene-LOC105285670 | GeneID:105285670 | LOC105285670 | Gene | LOC105285670 | protein_coding | | | | | | | | | | |

character(0) NA character(0) NC_039513.1 14932042 14932047 6 * 6 1000
NC_039514.1 6622839 6626782 3944 - Gnomon gene NA NA gene-LOC105285534 GeneID:105285534 LOC105285534 Gene LOC105285534 protein_coding
character(0) NA character(0) NC_039514.1 6625853 6625858 6 * 6 1000
NC_039514.1 9336154 9343762 7609 + Gnomon gene NA NA gene-LOC105284644 GeneID:105284644 LOC105284644 Gene LOC105284644 protein_coding
character(0) NA character(0) NC_039514.1 9337043 9337048 6 * 6 1000
NC_039514.1 11109536 11117116 7581 - Gnomon gene NA NA gene-LOC105281601 GeneID:105281601 LOC105281601 Gene LOC105281601 protein_coding
character(0) NA character(0) NC_039514.1 11112955 11112960 6 * 6 1000
NC_039514.1 11111779 11115572 3794 + Gnomon gene NA NA gene-LOC105281603 GeneID:105281603 LOC105281603 Gene LOC105281603 protein_coding
character(0) NA character(0) NC_039514.1 11112955 11112960 6 * 6 1000
NC_039515.1 7098785 7103508 4724 - Gnomon gene NA NA gene-LOC105286003 GeneID:105286003 LOC105286003 Gene LOC105286003 protein_coding
character(0) NA character(0) NC_039515.1 7102685 7102690 6 * 6 1000
NC_039515.1 12582469 12585642 3174 - Gnomon gene NA NA gene-LOC105282150 GeneID:105282150 LOC105282150 Gene LOC105282150 protein_coding
character(0) NA character(0) NC_039515.1 12584748 12584753 6 * 6 1000
NC_039515.1 12596735 12605489 8755 - Gnomon gene NA NA gene-LOC105282142 GeneID:105282142 LOC105282142 Gene LOC105282142 protein_coding
character(0) NA character(0) NC_039515.1 12604757 12604762 6 * 6 1000
NC_039515.1 12601969 12605663 3695 + Gnomon gene NA NA gene-LOC105282144 GeneID:105282144 LOC105282144 Gene LOC105282144 protein_coding
character(0) NA character(0) NC_039515.1 12604757 12604762 6 * 6 1000
NC_039516.1 2372548 2378885 6338 + Gnomon gene NA NA gene-LOC105274446 GeneID:105274446 LOC105274446 Gene LOC105274446 protein_coding
character(0) NA character(0) NC_039516.1 2377394 2377399 6 * 6 1000
NC_039516.1 4713084 4714767 1684 - Gnomon gene NA NA gene-LOC105275897 GeneID:105275897 LOC105275897 Gene LOC105275897 protein_coding
character(0) NA character(0) NC_039516.1 4713694 4713699 6 * 6 1000
NC_039516.1 10251143 10256122 4980 - Gnomon gene NA NA gene-LOC105287447 GeneID:105287447 LOC105287447 Gene LOC105287447 protein_coding
character(0) NA character(0) NC_039516.1 10254626 10254631 6 * 6 1000
NC_039516.1 12889538 12895772 6235 + Gnomon gene NA NA gene-LOC105275316 GeneID:105275316 LOC105275316 Gene LOC105275316 protein_coding
character(0) NA character(0) NC_039516.1 12890326 12890331 6 * 6 1000
NC_039516.1 13065193 13068389 3197 + Gnomon gene NA NA gene-LOC105275355 GeneID:105275355 LOC105275355 Gene LOC105275355 protein_coding
character(0) NA character(0) NC_039516.1 13066190 13066195 6 * 6 1000
NC_039517.1 1360424 1367346 6923 + Gnomon gene NA NA gene-LOC105278812 GeneID:105278812 LOC105278812 Gene LOC105278812 protein_coding
character(0) NA character(0) NC_039517.1 1361411 1361416 6 * 6 1000
NC_039517.1 7309114 7314427 5314 - Gnomon gene NA NA gene-LOC105284201 GeneID:105284201 LOC105284201 Gene LOC105284201 protein_coding
character(0) NA character(0) NC_039517.1 7312519 7312524 6 * 6 1000
NC_039517.1 8175046 8181714 6669 - Gnomon gene NA NA gene-LOC105284262 GeneID:105284262 LOC105284262 Gene LOC105284262 protein_coding
character(0) NA character(0) NC_039517.1 8175507 8175512 6 * 6 1000
NC_039518.1 5496870 5510429 13560 - Gnomon gene NA NA gene-LOC105281906 GeneID:105281906 LOC105281906 Gene LOC105281906 protein_coding
character(0) NA character(0) NC_039518.1 5508860 5508865 6 * 6 1000
NC_039518.1 8474988 8476009 1022 + Gnomon gene NA NA gene-LOC105286212 GeneID:105286212 LOC105286212 Gene LOC105286212 protein_coding
character(0) NA character(0) NC_039518.1 8475762 8475769 8 * 8 1000
NC_039518.1 8721748 8727127 5380 + Gnomon gene NA NA gene-LOC105286364 GeneID:105286364 LOC105286364 Gene LOC105286364 protein_coding
character(0) NA character(0) NC_039518.1 8722922 8722927 6 * 6 1000
NC_039518.1 9164450 9166108 1659 - Gnomon gene NA NA gene-LOC105286320 GeneID:105286320 LOC105286320 Gene LOC105286320 protein_coding
character(0) NA character(0) NC_039518.1 9164960 9164965 6 * 6 1000
NC_039506.1 23567682 23574351 6670 + Gnomon gene NA NA gene-LOC105279333 GeneID:105279333 LOC105279333 Gene LOC105279333 protein_coding
character(0) NA character(0) NC_039506.1 23568887 23568897 11 * 10 909.090909090909
NC_039507.1 9161485 9166239 4755 + Gnomon gene NA NA gene-LOC105282461 GeneID:105282461 LOC105282461 Gene LOC105282461 protein_coding
character(0) NA character(0) NC_039507.1 9162374 9162382 9 * 8 888.888888888889
NC_039509.1 17207204 17209527 2324 + Gnomon gene NA NA gene-LOC105283016 GeneID:105283016 LOC105283016 Gene LOC105283016 protein_coding
character(0) NA character(0) NC_039509.1 17208071 17208079 9 * 8 888.888888888889
NC_039510.1 8568121 8569477 1357 - Gnomon gene NA NA gene-LOC105275137 GeneID:105275137 LOC105275137 Gene LOC105275137 protein_coding
character(0) NA character(0) NC_039510.1 8568914 8568922 9 * 8 888.888888888889
NC_039513.1 3303685 3306319 2635 + Gnomon gene NA NA gene-LOC105275761 GeneID:105275761 LOC105275761 Gene LOC105275761 protein_coding
character(0) NA character(0) NC_039513.1 3304807 3304814 8 * 7 875
NC_039516.1 5472871 5475154 2284 - Gnomon gene NA NA gene-LOC105275878 GeneID:105275878 LOC105275878 Gene LOC105275878 protein_coding
character(0) NA character(0) NC_039516.1 5474293 5474300 8 * 7 875
NC_039506.1 1487779 1489683 1905 + Gnomon gene NA NA gene-LOC105280236 GeneID:105280236 LOC105280236 Gene LOC105280236 protein_coding
character(0) NA character(0) NC_039506.1 1488926 1488932 7 * 6 857.142857142857
NC_039506.1 2274108 2276090 1983 - Gnomon gene NA NA gene-LOC105280319 GeneID:105280319 LOC105280319 Gene LOC105280319 protein_coding
character(0) NA character(0) NC_039506.1 2275368 2275374 7 * 6 857.142857142857
NC_039506.1 3247135 3248424 1290 + Gnomon gene NA NA gene-LOC105280378 GeneID:105280378 LOC105280378 Gene LOC105280378 protein_coding
character(0) NA character(0) NC_039506.1 3248091 3248097 7 * 6 857.142857142857
NC_039506.1 4874444 4880252 5809 - Gnomon gene NA NA gene-LOC105277909 GeneID:105277909 LOC105277909 Gene LOC105277909 protein_coding
character(0) NA character(0) NC_039506.1 4878628 4878634 7 * 6 857.142857142857
NC_039506.1 5594174 5596919 2746 - Gnomon gene NA NA gene-LOC105277849 GeneID:105277849 LOC105277849 Gene LOC105277849 protein_coding
character(0) NA character(0) NC_039506.1 5596093 5596099 7 * 6 857.142857142857
NC_039506.1 11102693 11121131 18439 - Gnomon gene NA NA gene-LOC105284754 GeneID:105284754 LOC105284754 Gene LOC105284754 protein_coding

character(0) NA character(0) NC_039506.1 11119816 11119822 7 * 6 857.142857142857
NC_039506.1 13552800 13986605 433806 - Gnomon gene NA NA gene-LOC105280980 GeneID:105280980 LOC105280980 Gene LOC105280980 protein_coding
character(0) NA character(0) NC_039506.1 13985875 13985881 7 * 6 857.142857142857
NC_039506.1 15506447 15508913 2467 - Gnomon gene NA NA gene-LOC105283960 GeneID:105283960 LOC105283960 Gene LOC105283960 protein_coding
character(0) NA character(0) NC_039506.1 15508062 15508068 7 * 6 857.142857142857
NC_039506.1 15509387 15513034 3648 + Gnomon gene NA NA gene-LOC105283961 GeneID:105283961 LOC105283961 Gene LOC105283961 protein_coding
character(0) NA character(0) NC_039506.1 15510126 15510132 7 * 6 857.142857142857
NC_039506.1 15517083 15520247 3165 - Gnomon gene NA NA gene-LOC105283964 GeneID:105283964 LOC105283964 Gene LOC105283964 protein_coding
character(0) NA character(0) NC_039506.1 15519235 15519241 7 * 6 857.142857142857
NC_039506.1 16116359 16120578 4220 - Gnomon gene NA NA gene-LOC105274912 GeneID:105274912 LOC105274912 Gene LOC105274912 protein_coding
character(0) NA character(0) NC_039506.1 16119072 16119078 7 * 6 857.142857142857
NC_039506.1 19093260 19098014 4755 - Gnomon gene NA NA gene-LOC105278005 GeneID:105278005 LOC105278005 Gene LOC105278005 protein_coding
character(0) NA character(0) NC_039506.1 19096860 19096866 7 * 6 857.142857142857
NC_039506.1 20553263 20559098 5836 - Gnomon gene NA NA gene-LOC105278544 GeneID:105278544 LOC105278544 Gene LOC105278544 protein_coding
character(0) NA character(0) NC_039506.1 20557629 20557635 7 * 6 857.142857142857
NC_039506.1 22941360 22946330 4971 - Gnomon gene NA NA gene-LOC105283484 GeneID:105283484 LOC105283484 Gene LOC105283484 protein_coding
character(0) NA character(0) NC_039506.1 22945036 22945042 7 * 6 857.142857142857
NC_039506.1 23689672 23694320 4649 - Gnomon gene NA NA gene-LOC105279346 GeneID:105279346 LOC105279346 Gene LOC105279346 protein_coding
character(0) NA character(0) NC_039506.1 23693180 23693186 7 * 6 857.142857142857
NC_039507.1 124714 126752 2039 + Gnomon gene NA NA gene-LOC105279088 GeneID:105279088 LOC105279088 Gene LOC105279088 protein_coding
character(0) NA character(0) NC_039507.1 125332 125338 7 * 6 857.142857142857
NC_039507.1 124714 126752 2039 + Gnomon gene NA NA gene-LOC105279088 GeneID:105279088 LOC105279088 Gene LOC105279088 protein_coding
character(0) NA character(0) NC_039507.1 125663 125669 7 * 6 857.142857142857
NC_039507.1 790243 798925 8683 + Gnomon gene NA NA gene-LOC105282921 GeneID:105282921 LOC105282921 Gene LOC105282921 protein_coding
character(0) NA character(0) NC_039507.1 791692 791698 7 * 6 857.142857142857
NC_039507.1 6347235 6350936 3702 + Gnomon gene NA NA gene-LOC105275401 GeneID:105275401 LOC105275401 Gene LOC105275401 protein_coding
character(0) NA character(0) NC_039507.1 6348012 6348018 7 * 6 857.142857142857
NC_039507.1 15481857 15484141 2285 + Gnomon gene NA NA gene-LOC105280092 GeneID:105280092 LOC105280092 Gene LOC105280092 protein_coding
character(0) NA character(0) NC_039507.1 15482876 15482882 7 * 6 857.142857142857
NC_039507.1 16969693 16973301 3609 - Gnomon gene NA NA gene-LOC105278147 GeneID:105278147 LOC105278147 Gene LOC105278147 protein_coding
character(0) NA character(0) NC_039507.1 16972138 16972144 7 * 6 857.142857142857
NC_039507.1 18333035 18336377 3343 - Gnomon gene NA NA gene-LOC105281577 GeneID:105281577 LOC105281577 Gene LOC105281577 protein_coding
character(0) NA character(0) NC_039507.1 18335506 18335512 7 * 6 857.142857142857
NC_039508.1 192252 200875 8624 + Gnomon gene NA NA gene-LOC105277452 GeneID:105277452 LOC105277452 Gene LOC105277452 protein_coding
character(0) NA character(0) NC_039508.1 193513 193519 7 * 6 857.142857142857
NC_039508.1 1384214 1386820 2607 + Gnomon gene NA NA gene-LOC105277388 GeneID:105277388 LOC105277388 Gene LOC105277388 protein_coding
character(0) NA character(0) NC_039508.1 1384932 1384938 7 * 6 857.142857142857
NC_039508.1 1386723 1389150 2428 - Gnomon gene NA NA gene-LOC105277389 GeneID:105277389 LOC105277389 Gene LOC105277389 protein_coding
character(0) NA character(0) NC_039508.1 1387947 1387953 7 * 6 857.142857142857
NC_039508.1 10763110 10766843 3734 + Gnomon gene NA NA gene-LOC105276948 GeneID:105276948 LOC105276948 Gene LOC105276948 protein_coding
character(0) NA character(0) NC_039508.1 10764758 10764764 7 * 6 857.142857142857
NC_039508.1 10797665 10799797 2133 + Gnomon gene NA NA gene-LOC105276949 GeneID:105276949 LOC105276949 Gene LOC105276949 protein_coding
character(0) NA character(0) NC_039508.1 10798605 10798611 7 * 6 857.142857142857
NC_039508.1 13116146 13120560 4415 - Gnomon gene NA NA gene-LOC105277128 GeneID:105277128 LOC105277128 Gene LOC105277128 protein_coding
character(0) NA character(0) NC_039508.1 13116245 13116251 7 * 6 857.142857142857
NC_039508.1 15763225 15782851 19627 + Gnomon gene NA NA gene-LOC105277234 GeneID:105277234 LOC105277234 Gene LOC105277234 protein_coding
character(0) NA character(0) NC_039508.1 15764452 15764458 7 * 6 857.142857142857
NC_039508.1 17368821 17424169 55349 + Gnomon gene NA NA gene-LOC105284404 GeneID:105284404 LOC105284404 Gene LOC105284404 protein_coding
character(0) NA character(0) NC_039508.1 17370215 17370221 7 * 6 857.142857142857
NC_039508.1 17548373 17550348 1976 - Gnomon gene NA NA gene-LOC105284396 GeneID:105284396 LOC105284396 Gene LOC105284396 protein_coding
character(0) NA character(0) NC_039508.1 17549403 17549409 7 * 6 857.142857142857
NC_039509.1 89280 90451 1172 + Gnomon gene NA NA gene-LOC113561814 GeneID:113561814 LOC113561814 Gene LOC113561814 protein_coding character(0) NA
character(0) NC_039509.1 90329 90335 7 * 6 857.142857142857
NC_039509.1 2365507 2377666 12160 + Gnomon gene NA NA gene-LOC105285865 GeneID:105285865 LOC105285865 Gene LOC105285865 protein_coding
character(0) NA character(0) NC_039509.1 2366218 2366224 7 * 6 857.142857142857
NC_039509.1 10232323 10244464 12142 + Gnomon gene NA NA gene-LOC105282655 GeneID:105282655 LOC105282655 Gene LOC105282655 protein_coding
character(0) NA character(0) NC_039509.1 10233310 10233316 7 * 6 857.142857142857
NC_039509.1 10710016 10711124 1109 + Gnomon gene NA NA gene-LOC105275153 GeneID:105275153 LOC105275153 Gene LOC105275153 protein_coding
character(0) NA character(0) NC_039509.1 10710730 10710736 7 * 6 857.142857142857
NC_039509.1 13909149 13915606 6458 + Gnomon gene NA NA gene-LOC105278288 GeneID:105278288 LOC105278288 Gene LOC105278288 protein_coding
character(0) NA character(0) NC_039509.1 13909987 13909993 7 * 6 857.142857142857
NC_039509.1 15295604 15304160 8557 + Gnomon gene NA NA gene-LOC105284317 GeneID:105284317 LOC105284317 Gene LOC105284317 protein_coding
character(0) NA character(0) NC_039509.1 15296358 15296364 7 * 6 857.142857142857
NC_039510.1 624625 628259 3635 + Gnomon gene NA NA gene-LOC105279313 GeneID:105279313 LOC105279313 Gene LOC105279313 protein_coding

character(0) NA character(0) NC_039510.1 625494 625500 7 * 6 857.142857142857
NC_039510.1 3040660 3042070 1411 + Gnomon gene NA NA gene-LOC105287710 GeneID:105287710 LOC105287710 Gene LOC105287710 protein_coding
character(0) NA character(0) NC_039510.1 3041562 3041568 7 * 6 857.142857142857
NC_039510.1 3110492 3117379 6888 - Gnomon gene NA NA gene-LOC105287697 GeneID:105287697 LOC105287697 Gene LOC105287697 protein_coding
character(0) NA character(0) NC_039510.1 3116602 3116608 7 * 6 857.142857142857
NC_039510.1 3167251 3170185 2935 - Gnomon gene NA NA gene-LOC105287684 GeneID:105287684 LOC105287684 Gene LOC105287684 protein_coding
character(0) NA character(0) NC_039510.1 3169234 3169240 7 * 6 857.142857142857
NC_039510.1 4220534 4227851 7318 + Gnomon gene NA NA gene-LOC105287620 GeneID:105287620 LOC105287620 Gene LOC105287620 protein_coding
character(0) NA character(0) NC_039510.1 4221389 4221395 7 * 6 857.142857142857
NC_039510.1 5067676 5072600 4925 - Gnomon gene NA NA gene-LOC105287597 GeneID:105287597 LOC105287597 Gene LOC105287597 protein_coding
character(0) NA character(0) NC_039510.1 5071200 5071206 7 * 6 857.142857142857
NC_039510.1 5537223 5539036 1814 - Gnomon gene NA NA gene-LOC105287569 GeneID:105287569 LOC105287569 Gene LOC105287569 protein_coding
character(0) NA character(0) NC_039510.1 5538139 5538145 7 * 6 857.142857142857
NC_039510.1 6038459 6040963 2505 + Gnomon gene NA NA gene-LOC105287185 GeneID:105287185 LOC105287185 Gene LOC105287185 protein_coding
character(0) NA character(0) NC_039510.1 6040036 6040042 7 * 6 857.142857142857
NC_039510.1 6122701 6125649 2949 - Gnomon gene NA NA gene-LOC105287279 GeneID:105287279 LOC105287279 Gene LOC105287279 protein_coding
character(0) NA character(0) NC_039510.1 6124458 6124464 7 * 6 857.142857142857
NC_039510.1 6163214 6171437 8224 - Gnomon gene NA NA gene-LOC105287371 GeneID:105287371 LOC105287371 Gene LOC105287371 protein_coding
character(0) NA character(0) NC_039510.1 6170155 6170161 7 * 6 857.142857142857
NC_039510.1 6212141 6215222 3082 + Gnomon gene NA NA gene-LOC105287473 GeneID:105287473 LOC105287473 Gene LOC105287473 protein_coding
character(0) NA character(0) NC_039510.1 6213095 6213101 7 * 6 857.142857142857
NC_039510.1 6269006 6275115 6110 + Gnomon gene NA NA gene-LOC105287511 GeneID:105287511 LOC105287511 Gene LOC105287511 protein_coding
character(0) NA character(0) NC_039510.1 6270460 6270466 7 * 6 857.142857142857
NC_039510.1 8487238 8489293 2056 - Gnomon gene NA NA gene-LOC105275119 GeneID:105275119 LOC105275119 Gene LOC105275119 misc_RNA character(0)
NA character(0) NC_039510.1 8488481 8488487 7 * 6 857.142857142857
NC_039510.1 8568121 8569477 1357 - Gnomon gene NA NA gene-LOC105275137 GeneID:105275137 LOC105275137 Gene LOC105275137 protein_coding
character(0) NA character(0) NC_039510.1 8568885 8568891 7 * 6 857.142857142857
NC_039510.1 9996113 9998708 2596 - Gnomon gene NA NA gene-LOC105281227 GeneID:105281227 LOC105281227 Gene LOC105281227 protein_coding
character(0) NA character(0) NC_039510.1 9997644 9997650 7 * 6 857.142857142857
NC_039510.1 13531247 13535548 4302 - Gnomon gene NA NA gene-LOC105280964 GeneID:105280964 LOC105280964 Gene LOC105280964 protein_coding
character(0) NA character(0) NC_039510.1 13533348 13533354 7 * 6 857.142857142857
NC_039510.1 15582075 15587881 5807 - Gnomon gene NA NA gene-LOC105286461 GeneID:105286461 LOC105286461 Gene LOC105286461 protein_coding
character(0) NA character(0) NC_039510.1 15586562 15586568 7 * 6 857.142857142857
NC_039510.1 15745071 15751836 6766 + Gnomon gene NA NA gene-LOC105286478 GeneID:105286478 LOC105286478 Gene LOC105286478 protein_coding
character(0) NA character(0) NC_039510.1 15746500 15746506 7 * 6 857.142857142857
NC_039510.1 16500898 16505659 4762 - Gnomon gene NA NA gene-LOC105286526 GeneID:105286526 LOC105286526 Gene LOC105286526 protein_coding
character(0) NA character(0) NC_039510.1 16504460 16504466 7 * 6 857.142857142857
NC_039511.1 517195 530396 13202 + Gnomon gene NA NA gene-LOC105276519 GeneID:105276519 LOC105276519 Gene LOC105276519 protein_coding
character(0) NA character(0) NC_039511.1 518659 518665 7 * 6 857.142857142857
NC_039511.1 1376414 1379330 2917 - Gnomon gene NA NA gene-LOC105276452 GeneID:105276452 LOC105276452 Gene LOC105276452 protein_coding
character(0) NA character(0) NC_039511.1 1377342 1377348 7 * 6 857.142857142857
NC_039511.1 1856026 1859536 3511 + Gnomon gene NA NA gene-LOC105276422 GeneID:105276422 LOC105276422 Gene LOC105276422 protein_coding
character(0) NA character(0) NC_039511.1 1857678 1857684 7 * 6 857.142857142857
NC_039511.1 1861744 1869078 7335 + Gnomon gene NA NA gene-LOC105276417 GeneID:105276417 LOC105276417 Gene LOC105276417 protein_coding
character(0) NA character(0) NC_039511.1 1867899 1867905 7 * 6 857.142857142857
NC_039511.1 2004889 2015973 11085 - Gnomon gene NA NA gene-LOC105276410 GeneID:105276410 LOC105276410 Gene LOC105276410 protein_coding
character(0) NA character(0) NC_039511.1 2014683 2014689 7 * 6 857.142857142857
NC_039511.1 2004889 2015973 11085 - Gnomon gene NA NA gene-LOC105276410 GeneID:105276410 LOC105276410 Gene LOC105276410 protein_coding
character(0) NA character(0) NC_039511.1 2014740 2014746 7 * 6 857.142857142857
NC_039511.1 3635459 3642537 7079 - Gnomon gene NA NA gene-LOC105284052 GeneID:105284052 LOC105284052 Gene LOC105284052 protein_coding
character(0) NA character(0) NC_039511.1 3639538 3639544 7 * 6 857.142857142857
NC_039511.1 4554146 4561340 7195 - Gnomon gene NA NA gene-LOC105281750 GeneID:105281750 LOC105281750 Gene LOC105281750 protein_coding
character(0) NA character(0) NC_039511.1 4560502 4560508 7 * 6 857.142857142857
NC_039511.1 12403841 12413518 9678 - Gnomon gene NA NA gene-LOC105285135 GeneID:105285135 LOC105285135 Gene LOC105285135 protein_coding
character(0) NA character(0) NC_039511.1 12411122 12411128 7 * 6 857.142857142857
NC_039511.1 14704311 14706938 2628 - Gnomon gene NA NA gene-LOC105285889 GeneID:105285889 LOC105285889 Gene LOC105285889 protein_coding
character(0) NA character(0) NC_039511.1 14705900 14705906 7 * 6 857.142857142857
NC_039511.1 15983428 15985754 2327 + Gnomon gene NA NA gene-LOC105284576 GeneID:105284576 LOC105284576 Gene LOC105284576 protein_coding
character(0) NA character(0) NC_039511.1 15984100 15984106 7 * 6 857.142857142857
NC_039512.1 2097714 2101611 3898 - Gnomon gene NA NA gene-LOC105277667 GeneID:105277667 LOC105277667 Gene LOC105277667 protein_coding
character(0) NA character(0) NC_039512.1 2100585 2100591 7 * 6 857.142857142857
NC_039512.1 4385041 4390283 5243 - Gnomon gene NA NA gene-LOC105277552 GeneID:105277552 LOC105277552 Gene LOC105277552 protein_coding
character(0) NA character(0) NC_039512.1 4389120 4389126 7 * 6 857.142857142857
NC_039512.1 11140867 11151152 10286 - Gnomon gene NA NA gene-LOC105279854 GeneID:105279854 LOC105279854 Gene LOC105279854 protein_coding

| | | | | | | | | | | | | | | |
|---|---|---|---|---|---|---|---|---|---|---|---|---|---|---|
| character(0) | NA | character(0) | NC_039512.1 | 11150305 | 11150311 | 7 | * | 6 | 857.142857142857 | | | | | |
| NC_039512.1 | 12469542 | 12475093 | 5552 | - | Gnomon | gene | NA | NA | gene-LOC105279748 | GeneID:105279748 | LOC105279748 | Gene | LOC105279748 | protein_coding |
| character(0) | NA | character(0) | NC_039512.1 | 12474009 | 12474015 | 7 | * | 6 | 857.142857142857 | | | | | |
| NC_039512.1 | 12621899 | 12625222 | 3324 | + | Gnomon | gene | NA | NA | gene-LOC105279740 | GeneID:105279740 | LOC105279740 | Gene | LOC105279740 | protein_coding |
| character(0) | NA | character(0) | NC_039512.1 | 12623117 | 12623123 | 7 | * | 6 | 857.142857142857 | | | | | |
| NC_039512.1 | 12739826 | 12742653 | 2828 | + | Gnomon | gene | NA | NA | gene-LOC105279719 | GeneID:105279719 | LOC105279719 | Gene | LOC105279719 | protein_coding |
| character(0) | NA | character(0) | NC_039512.1 | 12740699 | 12740705 | 7 | * | 6 | 857.142857142857 | | | | | |
| NC_039512.1 | 12814910 | 12819293 | 4384 | - | Gnomon | gene | NA | NA | gene-LOC105279709 | GeneID:105279709 | LOC105279709 | Gene | LOC105279709 | protein_coding |
| character(0) | NA | character(0) | NC_039512.1 | 12818052 | 12818058 | 7 | * | 6 | 857.142857142857 | | | | | |
| NC_039512.1 | 13089540 | 13093562 | 4023 | - | Gnomon | gene | NA | NA | gene-LOC105279664 | GeneID:105279664 | LOC105279664 | Gene | LOC105279664 | protein_coding |
| character(0) | NA | character(0) | NC_039512.1 | 13092263 | 13092269 | 7 | * | 6 | 857.142857142857 | | | | | |
| NC_039513.1 | 3253321 | 3257474 | 4154 | - | Gnomon | gene | NA | NA | gene-LOC105275753 | GeneID:105275753 | LOC105275753 | Gene | LOC105275753 | protein_coding |
| character(0) | NA | character(0) | NC_039513.1 | 3256149 | 3256155 | 7 | * | 6 | 857.142857142857 | | | | | |
| NC_039513.1 | 3253321 | 3257474 | 4154 | - | Gnomon | gene | NA | NA | gene-LOC105275753 | GeneID:105275753 | LOC105275753 | Gene | LOC105275753 | protein_coding |
| character(0) | NA | character(0) | NC_039513.1 | 3256358 | 3256364 | 7 | * | 6 | 857.142857142857 | | | | | |
| NC_039513.1 | 3257374 | 3260835 | 3462 | - | Gnomon | gene | NA | NA | gene-LOC105275752 | GeneID:105275752 | LOC105275752 | Gene | LOC105275752 | protein_coding |
| character(0) | NA | character(0) | NC_039513.1 | 3259768 | 3259774 | 7 | * | 6 | 857.142857142857 | | | | | |
| NC_039513.1 | 6784235 | 6788101 | 3867 | - | Gnomon | gene | NA | NA | gene-LOC105287855 | GeneID:105287855 | LOC105287855 | Gene | LOC105287855 | protein_coding |
| character(0) | NA | character(0) | NC_039513.1 | 6787239 | 6787245 | 7 | * | 6 | 857.142857142857 | | | | | |
| NC_039513.1 | 9603616 | 9607153 | 3538 | + | Gnomon | gene | NA | NA | gene-LOC105275615 | GeneID:105275615 | LOC105275615 | Gene | LOC105275615 | protein_coding |
| character(0) | NA | character(0) | NC_039513.1 | 9605113 | 9605119 | 7 | * | 6 | 857.142857142857 | | | | | |
| NC_039513.1 | 9603616 | 9607153 | 3538 | + | Gnomon | gene | NA | NA | gene-LOC105275615 | GeneID:105275615 | LOC105275615 | Gene | LOC105275615 | protein_coding |
| character(0) | NA | character(0) | NC_039513.1 | 9605414 | 9605420 | 7 | * | 6 | 857.142857142857 | | | | | |
| NC_039513.1 | 11674366 | 11676356 | 1991 | - | Gnomon | gene | NA | NA | gene-LOC105280739 | GeneID:105280739 | LOC105280739 | Gene | LOC105280739 | protein_coding |
| character(0) | NA | character(0) | NC_039513.1 | 11675069 | 11675075 | 7 | * | 6 | 857.142857142857 | | | | | |
| NC_039513.1 | 11674366 | 11676356 | 1991 | - | Gnomon | gene | NA | NA | gene-LOC105280739 | GeneID:105280739 | LOC105280739 | Gene | LOC105280739 | protein_coding |
| character(0) | NA | character(0) | NC_039513.1 | 11675195 | 11675201 | 7 | * | 6 | 857.142857142857 | | | | | |
| NC_039513.1 | 13452183 | 13456265 | 4083 | + | Gnomon | gene | NA | NA | gene-LOC105275031 | GeneID:105275031 | LOC105275031 | Gene | LOC105275031 | protein_coding |
| character(0) | NA | character(0) | NC_039513.1 | 13453683 | 13453689 | 7 | * | 6 | 857.142857142857 | | | | | |
| NC_039513.1 | 13482189 | 13486570 | 4382 | - | Gnomon | gene | NA | NA | gene-LOC105275064 | GeneID:105275064 | LOC105275064 | Gene | LOC105275064 | protein_coding |
| character(0) | NA | character(0) | NC_039513.1 | 13485760 | 13485766 | 7 | * | 6 | 857.142857142857 | | | | | |
| NC_039513.1 | 15725203 | 15744087 | 18885 | + | Gnomon | gene | NA | NA | gene-LOC105280058 | GeneID:105280058 | LOC105280058 | Gene | LOC105280058 | protein_coding |
| character(0) | NA | character(0) | NC_039513.1 | 15726271 | 15726277 | 7 | * | 6 | 857.142857142857 | | | | | |
| NC_039514.1 | 873449 | 876660 | 3212 | + | Gnomon | gene | NA | NA | gene-LOC105276205 | GeneID:105276205 | LOC105276205 | Gene | LOC105276205 | protein_coding |
| character(0) | NA | character(0) | NC_039514.1 | 874466 | 874472 | 7 | * | 6 | 857.142857142857 | | | | | |
| NC_039514.1 | 1286674 | 1297576 | 10903 | - | Gnomon | gene | NA | NA | gene-LOC105276233 | GeneID:105276233 | LOC105276233 | Gene | LOC105276233 | protein_coding |
| character(0) | NA | character(0) | NC_039514.1 | 1295968 | 1295974 | 7 | * | 6 | 857.142857142857 | | | | | |
| NC_039514.1 | 5919621 | 5920916 | 1296 | - | Gnomon | gene | NA | NA | gene-LOC105279626 | GeneID:105279626 | LOC105279626 | Gene | LOC105279626 | protein_coding |
| character(0) | NA | character(0) | NC_039514.1 | 5920192 | 5920198 | 7 | * | 6 | 857.142857142857 | | | | | |
| NC_039514.1 | 6622839 | 6626782 | 3944 | - | Gnomon | gene | NA | NA | gene-LOC105285534 | GeneID:105285534 | LOC105285534 | Gene | LOC105285534 | protein_coding |
| character(0) | NA | character(0) | NC_039514.1 | 6625801 | 6625807 | 7 | * | 6 | 857.142857142857 | | | | | |
| NC_039514.1 | 8629561 | 8648221 | 18661 | + | Gnomon | gene | NA | NA | gene-LOC105278994 | GeneID:105278994 | LOC105278994 | Gene | LOC105278994 | protein_coding |
| character(0) | NA | character(0) | NC_039514.1 | 8630645 | 8630651 | 7 | * | 6 | 857.142857142857 | | | | | |
| NC_039514.1 | 8629561 | 8648221 | 18661 | + | Gnomon | gene | NA | NA | gene-LOC105278994 | GeneID:105278994 | LOC105278994 | Gene | LOC105278994 | protein_coding |
| character(0) | NA | character(0) | NC_039514.1 | 8630804 | 8630810 | 7 | * | 6 | 857.142857142857 | | | | | |
| NC_039514.1 | 9413016 | 9415003 | 1988 | - | Gnomon | gene | NA | NA | gene-LOC105284659 | GeneID:105284659 | LOC105284659 | Gene | LOC105284659 | protein_coding |
| character(0) | NA | character(0) | NC_039514.1 | 9414116 | 9414122 | 7 | * | 6 | 857.142857142857 | | | | | |
| NC_039514.1 | 14398821 | 14400325 | 1505 | - | Gnomon | gene | NA | NA | gene-LOC105275991 | GeneID:105275991 | LOC105275991 | Gene | LOC105275991 | protein_coding |
| character(0) | NA | character(0) | NC_039514.1 | 14399289 | 14399295 | 7 | * | 6 | 857.142857142857 | | | | | |
| NC_039514.1 | 15117749 | 15130344 | 12596 | + | Gnomon | gene | NA | NA | gene-LOC105276073 | GeneID:105276073 | LOC105276073 | Gene | LOC105276073 | protein_coding |
| character(0) | NA | character(0) | NC_039514.1 | 15123389 | 15123395 | 7 | * | 6 | 857.142857142857 | | | | | |
| NC_039514.1 | 15122114 | 15127165 | 5052 | + | Gnomon | gene | NA | NA | gene-LOC105276070 | GeneID:105276070 | LOC105276070 | Gene | LOC105276070 | protein_coding |
| character(0) | NA | character(0) | NC_039514.1 | 15123389 | 15123395 | 7 | * | 6 | 857.142857142857 | | | | | |
| NC_039515.1 | 1472003 | 1475695 | 3693 | - | Gnomon | gene | NA | NA | gene-LOC105274705 | GeneID:105274705 | LOC105274705 | Gene | LOC105274705 | protein_coding |
| character(0) | NA | character(0) | NC_039515.1 | 1475067 | 1475073 | 7 | * | 6 | 857.142857142857 | | | | | |
| NC_039515.1 | 1527980 | 1529634 | 1655 | + | Gnomon | gene | NA | NA | gene-LOC105274711 | GeneID:105274711 | LOC105274711 | Gene | LOC105274711 | protein_coding |
| character(0) | NA | character(0) | NC_039515.1 | 1528293 | 1528299 | 7 | * | 6 | 857.142857142857 | | | | | |
| NC_039515.1 | 2215137 | 2216986 | 1850 | - | Gnomon | gene | NA | NA | gene-LOC105278431 | GeneID:105278431 | LOC105278431 | Gene | LOC105278431 | protein_coding |
| character(0) | NA | character(0) | NC_039515.1 | 2215927 | 2215933 | 7 | * | 6 | 857.142857142857 | | | | | |
| NC_039515.1 | 3498019 | 3503642 | 5624 | + | Gnomon | gene | NA | NA | gene-LOC105278309 | GeneID:105278309 | LOC105278309 | Gene | LOC105278309 | protein_coding |
| character(0) | NA | character(0) | NC_039515.1 | 3499095 | 3499101 | 7 | * | 6 | 857.142857142857 | | | | | |
| NC_039515.1 | 4407130 | 4431402 | 24273 | - | Gnomon | gene | NA | NA | gene-LOC105278244 | GeneID:105278244 | LOC105278244 | Gene | LOC105278244 | protein_coding |
| character(0) | NA | character(0) | NC_039515.1 | 4428992 | 4428998 | 7 | * | 6 | 857.142857142857 | | | | | |
| NC_039515.1 | 5537006 | 5539696 | 2691 | + | Gnomon | gene | NA | NA | gene-LOC105277256 | GeneID:105277256 | LOC105277256 | Gene | LOC105277256 | protein_coding |

| | | | | | | | | | | | | | | |
|---|---|---|---|---|---|---|---|---|---|---|---|---|---|---|
| character(0) | NA | character(0) | NC_039515.1 | 5537966 | 5537972 | 7 | * | 6 | 857.142857142857 | | | | | |
| NC_039515.1 | 6529779 | 6536130 | 6352 | + | Gnomon | gene | NA | NA | gene-LOC105282286 | GeneID:105282286 | LOC105282286 | Gene | LOC105282286 | protein_coding |
| character(0) | NA | character(0) | NC_039515.1 | 6530893 | 6530899 | 7 | * | 6 | 857.142857142857 | | | | | |
| NC_039515.1 | 6604872 | 6605723 | 852 | + | Gnomon | gene | NA | NA | gene-LOC105282280 | GeneID:105282280 | LOC105282280 | Gene | LOC105282280 | protein_coding |
| character(0) | NA | character(0) | NC_039515.1 | 6605539 | 6605545 | 7 | * | 6 | 857.142857142857 | | | | | |
| NC_039515.1 | 6615586 | 6618213 | 2628 | - | Gnomon | gene | NA | NA | gene-LOC105282278 | GeneID:105282278 | LOC105282278 | Gene | LOC105282278 | protein_coding |
| character(0) | NA | character(0) | NC_039515.1 | 6617273 | 6617279 | 7 | * | 6 | 857.142857142857 | | | | | |
| NC_039515.1 | 7926262 | 7929466 | 3205 | + | Gnomon | gene | NA | NA | gene-LOC105283741 | GeneID:105283741 | LOC105283741 | Gene | LOC105283741 | protein_coding |
| character(0) | NA | character(0) | NC_039515.1 | 7927531 | 7927537 | 7 | * | 6 | 857.142857142857 | | | | | |
| NC_039515.1 | 8973009 | 8978309 | 5301 | + | Gnomon | gene | NA | NA | gene-LOC105283697 | GeneID:105283697 | LOC105283697 | Gene | LOC105283697 | protein_coding |
| character(0) | NA | character(0) | NC_039515.1 | 8974548 | 8974554 | 7 | * | 6 | 857.142857142857 | | | | | |
| NC_039516.1 | 756663 | 760199 | 3537 | + | Gnomon | gene | NA | NA | gene-LOC105274638 | GeneID:105274638 | LOC105274638 | Gene | LOC105274638 | protein_coding |
| character(0) | NA | character(0) | NC_039516.1 | 757725 | 757731 | 7 | * | 6 | 857.142857142857 | | | | | |
| NC_039516.1 | 985951 | 990466 | 4516 | + | Gnomon | gene | NA | NA | gene-LOC105274590 | GeneID:105274590 | LOC105274590 | Gene | LOC105274590 | protein_coding |
| character(0) | NA | character(0) | NC_039516.1 | 987063 | 987069 | 7 | * | 6 | 857.142857142857 | | | | | |
| NC_039516.1 | 985951 | 990466 | 4516 | + | Gnomon | gene | NA | NA | gene-LOC105274590 | GeneID:105274590 | LOC105274590 | Gene | LOC105274590 | protein_coding |
| character(0) | NA | character(0) | NC_039516.1 | 987547 | 987553 | 7 | * | 6 | 857.142857142857 | | | | | |
| NC_039516.1 | 1549696 | 1551173 | 1478 | + | Gnomon | gene | NA | NA | gene-LOC105274497 | GeneID:105274497 | LOC105274497 | Gene | LOC105274497 | protein_coding |
| character(0) | NA | character(0) | NC_039516.1 | 1550744 | 1550750 | 7 | * | 6 | 857.142857142857 | | | | | |
| NC_039516.1 | 2241904 | 2245378 | 3475 | + | Gnomon | gene | NA | NA | gene-LOC105274464 | GeneID:105274464 | LOC105274464 | Gene | LOC105274464 | protein_coding |
| character(0) | NA | character(0) | NC_039516.1 | 2242648 | 2242654 | 7 | * | 6 | 857.142857142857 | | | | | |
| NC_039516.1 | 3088715 | 3091076 | 2362 | + | Gnomon | gene | NA | NA | gene-LOC105288196 | GeneID:105288196 | LOC105288196 | Gene | LOC105288196 | protein_coding |
| character(0) | NA | character(0) | NC_039516.1 | 3089812 | 3089818 | 7 | * | 6 | 857.142857142857 | | | | | |
| NC_039516.1 | 4713084 | 4714767 | 1684 | - | Gnomon | gene | NA | NA | gene-LOC105275897 | GeneID:105275897 | LOC105275897 | Gene | LOC105275897 | protein_coding |
| character(0) | NA | character(0) | NC_039516.1 | 4713795 | 4713801 | 7 | * | 6 | 857.142857142857 | | | | | |
| NC_039516.1 | 7248249 | 7252272 | 4024 | + | Gnomon | gene | NA | NA | gene-LOC105275812 | GeneID:105275812 | LOC105275812 | Gene | LOC105275812 | protein_coding |
| character(0) | NA | character(0) | NC_039516.1 | 7251402 | 7251408 | 7 | * | 6 | 857.142857142857 | | | | | |
| NC_039516.1 | 10251143 | 10256122 | 4980 | - | Gnomon | gene | NA | NA | gene-LOC105287447 | GeneID:105287447 | LOC105287447 | Gene | LOC105287447 | protein_coding |
| character(0) | NA | character(0) | NC_039516.1 | 10255095 | 10255101 | 7 | * | 6 | 857.142857142857 | | | | | |
| NC_039516.1 | 11747670 | 11758010 | 10341 | - | Gnomon | gene | NA | NA | gene-LOC105278666 | GeneID:105278666 | LOC105278666 | Gene | LOC105278666 | protein_coding |
| character(0) | NA | character(0) | NC_039516.1 | 11756589 | 11756595 | 7 | * | 6 | 857.142857142857 | | | | | |
| NC_039516.1 | 11790047 | 11792531 | 2485 | + | Gnomon | gene | NA | NA | gene-LOC105278656 | GeneID:105278656 | LOC105278656 | Gene | LOC105278656 | protein_coding |
| character(0) | NA | character(0) | NC_039516.1 | 11791250 | 11791256 | 7 | * | 6 | 857.142857142857 | | | | | |
| NC_039516.1 | 12834895 | 12836314 | 1420 | - | Gnomon | gene | NA | NA | gene-LOC105275307 | GeneID:105275307 | LOC105275307 | Gene | LOC105275307 | protein_coding |
| character(0) | NA | character(0) | NC_039516.1 | 12835552 | 12835558 | 7 | * | 6 | 857.142857142857 | | | | | |
| NC_039516.1 | 13386687 | 13390618 | 3932 | + | Gnomon | gene | NA | NA | gene-LOC105278022 | GeneID:105278022 | LOC105278022 | Gene | LOC105278022 | protein_coding |
| character(0) | NA | character(0) | NC_039516.1 | 13387740 | 13387746 | 7 | * | 6 | 857.142857142857 | | | | | |
| NC_039516.1 | 13386687 | 13390618 | 3932 | + | Gnomon | gene | NA | NA | gene-LOC105278022 | GeneID:105278022 | LOC105278022 | Gene | LOC105278022 | protein_coding |
| character(0) | NA | character(0) | NC_039516.1 | 13387869 | 13387875 | 7 | * | 6 | 857.142857142857 | | | | | |
| NC_039516.1 | 13497458 | 13503461 | 6004 | - | Gnomon | gene | NA | NA | gene-LOC105277915 | GeneID:105277915 | LOC105277915 | Gene | LOC105277915 | protein_coding |
| character(0) | NA | character(0) | NC_039516.1 | 13502711 | 13502717 | 7 | * | 6 | 857.142857142857 | | | | | |
| NC_039516.1 | 13518014 | 13520692 | 2679 | + | Gnomon | gene | NA | NA | gene-LOC105277817 | GeneID:105277817 | LOC105277817 | Gene | LOC105277817 | protein_coding |
| character(0) | NA | character(0) | NC_039516.1 | 13518940 | 13518946 | 7 | * | 6 | 857.142857142857 | | | | | |
| NC_039517.1 | 867927 | 869156 | 1230 | - | Gnomon | gene | NA | NA | gene-LOC105278824 | GeneID:105278824 | LOC105278824 | Gene | LOC105278824 | protein_coding |
| character(0) | NA | character(0) | NC_039517.1 | 868363 | 868369 | 7 | * | 6 | 857.142857142857 | | | | | |
| NC_039517.1 | 2307792 | 2312395 | 4604 | - | Gnomon | gene | NA | NA | gene-LOC105281943 | GeneID:105281943 | LOC105281943 | Gene | LOC105281943 | protein_coding |
| character(0) | NA | character(0) | NC_039517.1 | 2309012 | 2309018 | 7 | * | 6 | 857.142857142857 | | | | | |
| NC_039517.1 | 3225755 | 3318595 | 92841 | - | Gnomon | gene | NA | NA | gene-LOC105280521 | GeneID:105280521 | LOC105280521 | Gene | LOC105280521 | protein_coding |
| character(0) | NA | character(0) | NC_039517.1 | 3227153 | 3227159 | 7 | * | 6 | 857.142857142857 | | | | | |
| NC_039517.1 | 7306074 | 7309288 | 3215 | + | Gnomon | gene | NA | NA | gene-LOC105284202 | GeneID:105284202 | LOC105284202 | Gene | LOC105284202 | protein_coding |
| character(0) | NA | character(0) | NC_039517.1 | 7307471 | 7307477 | 7 | * | 6 | 857.142857142857 | | | | | |
| NC_039517.1 | 7446985 | 7449823 | 2839 | + | Gnomon | gene | NA | NA | gene-LOC105284223 | GeneID:105284223 | LOC105284223 | Gene | LOC105284223 | protein_coding |
| character(0) | NA | character(0) | NC_039517.1 | 7448190 | 7448196 | 7 | * | 6 | 857.142857142857 | | | | | |
| NC_039517.1 | 9368753 | 9371050 | 2298 | - | Gnomon | gene | NA | NA | gene-LOC105282220 | GeneID:105282220 | LOC105282220 | Gene | LOC105282220 | protein_coding |
| character(0) | NA | character(0) | NC_039517.1 | 9370300 | 9370306 | 7 | * | 6 | 857.142857142857 | | | | | |
| NC_039517.1 | 10161586 | 10164793 | 3208 | - | Gnomon | gene | NA | NA | gene-LOC105285418 | GeneID:105285418 | LOC105285418 | Gene | LOC105285418 | protein_coding |
| character(0) | NA | character(0) | NC_039517.1 | 10163396 | 10163402 | 7 | * | 6 | 857.142857142857 | | | | | |
| NC_039518.1 | 673305 | 686962 | 13658 | + | Gnomon | gene | NA | NA | gene-LOC105287313 | GeneID:105287313 | LOC105287313 | Gene | LOC105287313 | protein_coding |
| character(0) | NA | character(0) | NC_039518.1 | 674158 | 674164 | 7 | * | 6 | 857.142857142857 | | | | | |
| NC_039518.1 | 2232165 | 2249269 | 17105 | + | Gnomon | gene | NA | NA | gene-LOC105283878 | GeneID:105283878 | LOC105283878 | Gene | LOC105283878 | protein_coding |
| character(0) | NA | character(0) | NC_039518.1 | 2233357 | 2233363 | 7 | * | 6 | 857.142857142857 | | | | | |
| NC_039518.1 | 2657034 | 2658634 | 1601 | + | Gnomon | gene | NA | NA | gene-LOC105285392 | GeneID:105285392 | LOC105285392 | Gene | LOC105285392 | protein_coding |
| character(0) | NA | character(0) | NC_039518.1 | 2658047 | 2658053 | 7 | * | 6 | 857.142857142857 | | | | | |
| NC_039518.1 | 2681489 | 2682795 | 1307 | - | Gnomon | gene | NA | NA | gene-LOC105285394 | GeneID:105285394 | LOC105285394 | Gene | LOC105285394 | protein_coding |

| | | | | | | | | | | | | | | |
|---|---|---|---|---|---|---|---|---|---|---|---|---|---|---|
| | character(0) | NA | character(0) | NC_039518.1 | 2681945 | 2681951 | 7 | * | 6 | 857.142857142857 | | | | |
| NC_039518.1 | 5288531 | 5294826 | 6296 | - | Gnomon | gene | NA | NA | gene-LOC105285410 | GeneID:105285410 | LOC105285410 | Gene | LOC105285410 | protein_coding |
| | character(0) | NA | character(0) | NC_039518.1 | 5293748 | 5293754 | 7 | * | 6 | 857.142857142857 | | | | |
| NC_039519.1 | 197513 | 200423 | 2911 | + | Gnomon | gene | NA | NA | gene-LOC105282793 | GeneID:105282793 | LOC105282793 | Gene | LOC105282793 | protein_coding |
| | character(0) | NA | character(0) | NC_039519.1 | 198095 | 198101 | 7 | * | 6 | 857.142857142857 | | | | |
| NC_039519.1 | 8239374 | 8248322 | 8949 | - | Gnomon | gene | NA | NA | gene-LOC105287985 | GeneID:105287985 | LOC105287985 | Gene | LOC105287985 | protein_coding |
| | character(0) | NA | character(0) | NC_039519.1 | 8247911 | 8247917 | 7 | * | 6 | 857.142857142857 | | | | |
| NC_039508.1 | 13115312 | 13116213 | 902 | + | Gnomon | gene | NA | NA | gene-LOC105277091 | GeneID:105277091 | LOC105277091 | Gene | LOC105277091 | protein_coding |
| | character(0) | NA | character(0) | NC_039508.1 | 13116053 | 13116062 | 10 | * | 8 | 800 | | | | |
| NC_039509.1 | 17375981 | 17382001 | 6021 | + | Gnomon | gene | NA | NA | gene-LOC105282459 | GeneID:105282459 | LOC105282459 | Gene | LOC105282459 | protein_coding |
| | character(0) | NA | character(0) | NC_039509.1 | 17378741 | 17378750 | 10 | * | 8 | 800 | | | | |
| NC_039513.1 | 5176546 | 5178099 | 1554 | + | Gnomon | gene | NA | NA | gene-LOC105279718 | GeneID:105279718 | LOC105279718 | Gene | LOC105279718 | protein_coding |
| | character(0) | NA | character(0) | NC_039513.1 | 5177245 | 5177254 | 10 | * | 8 | 800 | | | | |
| NC_039516.1 | 12104816 | 12108596 | 3781 | + | Gnomon | gene | NA | NA | gene-LOC105275223 | GeneID:105275223 | LOC105275223 | Gene | LOC105275223 | protein_coding |
| | character(0) | NA | character(0) | NC_039516.1 | 12106134 | 12106143 | 10 | * | 8 | 800 | | | | |
| NC_039519.1 | 8084633 | 8088546 | 3914 | + | Gnomon | gene | NA | NA | gene-LOC105274767 | GeneID:105274767 | LOC105274767 | Gene | LOC105274767 | protein_coding |
| | character(0) | NA | character(0) | NC_039519.1 | 8085717 | 8085726 | 10 | * | 8 | 800 | | | | |

Directory: MRPR File: gwr-Ob_LI_BCphaseB1.txt

```
seqnames    start end   width strand      source      type  score phase ID    Dbxref      Name  gbkey gene  gene_biotype      end_range   partial     start_range seqnames    start
      end   width strand      NbrSites    Sdnsty
NC_039506.1 3342881     3348391     5511  +     Gnomon      gene  NA    NA    gene-LOC105280395 GeneID:105280395  LOC105280395      Gene  LOC105280395      protein_coding
      character(0)      NA    character(0)      NC_039506.1 3344178     3344183     6     *     6     1000
NC_039506.1 6089683     6090856     1174  +     Gnomon      gene  NA    NA    gene-LOC105277804 GeneID:105277804  LOC105277804      Gene  LOC105277804      protein_coding
      character(0)      NA    character(0)      NC_039506.1 6090624     6090629     6     *     6     1000
NC_039506.1 15370515    15371538    1024  +     Gnomon      gene  NA    NA    gene-LOC105280860 GeneID:105280860  LOC105280860      Gene  LOC105280860      protein_coding
      character(0)      NA    character(0)      NC_039506.1 15371422    15371427    6     *     6     1000
NC_039507.1 7276866     7278961     2096  +     Gnomon      gene  NA    NA    gene-LOC105283373 GeneID:105283373  LOC105283373      Gene  LOC105283373      protein_coding
      character(0)      NA    character(0)      NC_039507.1 7277947     7277952     6     *     6     1000
NC_039507.1 18333035    18336377    3343  -     Gnomon      gene  NA    NA    gene-LOC105281577 GeneID:105281577  LOC105281577      Gene  LOC105281577      protein_coding
      character(0)      NA    character(0)      NC_039507.1 18335809    18335814    6     *     6     1000
NC_039508.1 2590841     2594093     3253  +     Gnomon      gene  NA    NA    gene-LOC105281341 GeneID:105281341  LOC105281341      Gene  LOC105281341      protein_coding
      character(0)      NA    character(0)      NC_039508.1 2591612     2591617     6     *     6     1000
NC_039508.1 10274002    10277440    3439  +     Gnomon      gene  NA    NA    gene-LOC105276881 GeneID:105276881  LOC105276881      Gene  LOC105276881      protein_coding
      character(0)      NA    character(0)      NC_039508.1 10275212    10275217    6     *     6     1000
NC_039509.1 1002697     1006107     3411  -     Gnomon      gene  NA    NA    gene-LOC105283620 GeneID:105283620  LOC105283620      Gene  LOC105283620      protein_coding
      character(0)      NA    character(0)      NC_039509.1 1004827     1004832     6     *     6     1000
NC_039509.1 1135464     1137481     2018  -     Gnomon      gene  NA    NA    gene-LOC105283634 GeneID:105283634  LOC105283634      Gene  LOC105283634      protein_coding
      character(0)      NA    character(0)      NC_039509.1 1136266     1136271     6     *     6     1000
NC_039509.1 11068956    11086837    17882 -     Gnomon      gene  NA    NA    gene-LOC105275770 GeneID:105275770  LOC105275770      Gene  LOC105275770      protein_coding
      character(0)      NA    character(0)      NC_039509.1 11085457    11085462    6     *     6     1000
NC_039509.1 13033738    13035966    2229  +     Gnomon      gene  NA    NA    gene-LOC105278527 GeneID:105278527  LOC105278527      Gene  LOC105278527      protein_coding
      character(0)      NA    character(0)      NC_039509.1 13034669    13034674    6     *     6     1000
NC_039510.1 2607810     2610599     2790  -     Gnomon      gene  NA    NA    gene-LOC105287754 GeneID:105287754  LOC105287754      Gene  LOC105287754      protein_coding
      character(0)      NA    character(0)      NC_039510.1 2609179     2609184     6     *     6     1000
NC_039510.1 16287375    16291357    3983  -     Gnomon      gene  NA    NA    gene-LOC105286509 GeneID:105286509  LOC105286509      Gene  LOC105286509      protein_coding
      character(0)      NA    character(0)      NC_039510.1 16290598    16290603    6     *     6     1000
NC_039511.1 480353      488275      7923  +     Gnomon      gene  NA    NA    gene-LOC105276527 GeneID:105276527  LOC105276527      Gene  LOC105276527      protein_coding
      character(0)      NA    character(0)      NC_039511.1 481652      481657      6     *     6     1000
NC_039511.1 2234830     2236789     1960  -     Gnomon      gene  NA    NA    gene-LOC105276382 GeneID:105276382  LOC105276382      Gene  LOC105276382      protein_coding
      character(0)      NA    character(0)      NC_039511.1 2236016     2236021     6     *     6     1000
NC_039511.1 3614241     3616939     2699  +     Gnomon      gene  NA    NA    gene-LOC105284057 GeneID:105284057  LOC105284057      Gene  LOC105284057      protein_coding
      character(0)      NA    character(0)      NC_039511.1 3615462     3615467     6     *     6     1000
NC_039511.1 3625986     3631991     6006  -     Gnomon      gene  NA    NA    gene-LOC105284053 GeneID:105284053  LOC105284053      Gene  LOC105284053      protein_coding
      character(0)      NA    character(0)      NC_039511.1 3630771     3630776     6     *     6     1000
NC_039511.1 8407553     8425032     17480 +     Gnomon      gene  NA    NA    gene-LOC105284845 GeneID:105284845  LOC105284845      Gene  LOC105284845      protein_coding
      character(0)      NA    character(0)      NC_039511.1 8409706     8409711     6     *     6     1000
NC_039511.1 8695181     8698086     2906  +     Gnomon      gene  NA    NA    gene-LOC105280856 GeneID:105280856  LOC105280856      Gene  LOC105280856      protein_coding
      character(0)      NA    character(0)      NC_039511.1 8695844     8695849     6     *     6     1000
NC_039512.1 10275466    10280187    4722  +     Gnomon      gene  NA    NA    gene-LOC105282847 GeneID:105282847  LOC105282847      Gene  LOC105282847      protein_coding
      character(0)      NA    character(0)      NC_039512.1 10276481    10276486    6     *     6     1000
NC_039512.1 15979275    15981529    2255  -     Gnomon      gene  NA    NA    gene-LOC105274775 GeneID:105274775  LOC105274775      Gene  LOC105274775      protein_coding
      character(0)      NA    character(0)      NC_039512.1 15980634    15980639    6     *     6     1000
NC_039513.1 10401827    10404590    2764  -     Gnomon      gene  NA    NA    gene-LOC105275550 GeneID:105275550  LOC105275550      Gene  LOC105275550      protein_coding
      character(0)      NA    character(0)      NC_039513.1 10403469    10403474    6     *     6     1000
NC_039513.1 11029007    11033701    4695  -     Gnomon      gene  NA    NA    gene-LOC105275763 GeneID:105275763  LOC105275763      Gene  LOC105275763      protein_coding
      character(0)      NA    character(0)      NC_039513.1 11030126    11030131    6     *     6     1000
NC_039513.1 12028675    12030941    2267  -     Gnomon      gene  NA    NA    gene-LOC105280791 GeneID:105280791  LOC105280791      Gene  LOC105280791      protein_coding
      character(0)      NA    character(0)      NC_039513.1 12029791    12029796    6     *     6     1000
...
... only first 25 lines shown ...
```

Directory: MRPR File: gwr-Ob_LI_BCphaseB2.txt

| seqnames | start | end | width | strand | source | type | score | phase | ID | Dbxref | Name | gbkey | gene | gene_biotype | end_range | partial | start_range | seqnames | start | end | width | strand | NbrSites | Sdnsty |
|---|---|---|---|---|---|---|---|---|---|---|---|---|---|---|---|---|---|---|---|---|---|---|---|---|
| NC_039506.1 | 3342881 | 3348391 | 5511 | + | Gnomon | gene | NA | NA | gene-LOC105280395 | GeneID:105280395 | LOC105280395 | Gene | LOC105280395 | protein_coding | character(0) | NA | character(0) | NC_039506.1 | 3344178 | 3344183 | 6 | * | 6 | 1000 |
| NC_039506.1 | 6089683 | 6090856 | 1174 | + | Gnomon | gene | NA | NA | gene-LOC105277804 | GeneID:105277804 | LOC105277804 | Gene | LOC105277804 | protein_coding | character(0) | NA | character(0) | NC_039506.1 | 6090624 | 6090629 | 6 | * | 6 | 1000 |
| NC_039506.1 | 15370515 | 15371538 | 1024 | + | Gnomon | gene | NA | NA | gene-LOC105280860 | GeneID:105280860 | LOC105280860 | Gene | LOC105280860 | protein_coding | character(0) | NA | character(0) | NC_039506.1 | 15371422 | 15371427 | 6 | * | 6 | 1000 |
| NC_039506.1 | 24043149 | 24047806 | 4658 | + | Gnomon | gene | NA | NA | gene-LOC105279367 | GeneID:105279367 | LOC105279367 | Gene | LOC105279367 | protein_coding | character(0) | NA | character(0) | NC_039506.1 | 24044697 | 24044702 | 6 | * | 6 | 1000 |
| NC_039507.1 | 7181199 | 7188522 | 7324 | + | Gnomon | gene | NA | NA | gene-LOC105283383 | GeneID:105283383 | LOC105283383 | Gene | LOC105283383 | protein_coding | character(0) | NA | character(0) | NC_039507.1 | 7182290 | 7182295 | 6 | * | 6 | 1000 |
| NC_039507.1 | 7276866 | 7278961 | 2096 | + | Gnomon | gene | NA | NA | gene-LOC105283373 | GeneID:105283373 | LOC105283373 | Gene | LOC105283373 | protein_coding | character(0) | NA | character(0) | NC_039507.1 | 7277947 | 7277952 | 6 | * | 6 | 1000 |
| NC_039507.1 | 18333035 | 18336377 | 3343 | - | Gnomon | gene | NA | NA | gene-LOC105281577 | GeneID:105281577 | LOC105281577 | Gene | LOC105281577 | protein_coding | character(0) | NA | character(0) | NC_039507.1 | 18335809 | 18335814 | 6 | * | 6 | 1000 |
| NC_039508.1 | 2590841 | 2594093 | 3253 | + | Gnomon | gene | NA | NA | gene-LOC105281341 | GeneID:105281341 | LOC105281341 | Gene | LOC105281341 | protein_coding | character(0) | NA | character(0) | NC_039508.1 | 2591612 | 2591617 | 6 | * | 6 | 1000 |
| NC_039508.1 | 10716085 | 10732040 | 15956 | - | Gnomon | gene | NA | NA | gene-LOC105276941 | GeneID:105276941 | LOC105276941 | Gene | LOC105276941 | protein_coding | character(0) | NA | character(0) | NC_039508.1 | 10730559 | 10730564 | 6 | * | 6 | 1000 |
| NC_039508.1 | 17548373 | 17550348 | 1976 | - | Gnomon | gene | NA | NA | gene-LOC105284396 | GeneID:105284396 | LOC105284396 | Gene | LOC105284396 | protein_coding | character(0) | NA | character(0) | NC_039508.1 | 17549290 | 17549295 | 6 | * | 6 | 1000 |
| NC_039509.1 | 1002697 | 1006107 | 3411 | - | Gnomon | gene | NA | NA | gene-LOC105283620 | GeneID:105283620 | LOC105283620 | Gene | LOC105283620 | protein_coding | character(0) | NA | character(0) | NC_039509.1 | 1004827 | 1004832 | 6 | * | 6 | 1000 |
| NC_039509.1 | 1135464 | 1137481 | 2018 | - | Gnomon | gene | NA | NA | gene-LOC105283634 | GeneID:105283634 | LOC105283634 | Gene | LOC105283634 | protein_coding | character(0) | NA | character(0) | NC_039509.1 | 1136266 | 1136271 | 6 | * | 6 | 1000 |
| NC_039509.1 | 2825666 | 2828757 | 3092 | + | Gnomon | gene | NA | NA | gene-LOC105282589 | GeneID:105282589 | LOC105282589 | Gene | LOC105282589 | protein_coding | character(0) | NA | character(0) | NC_039509.1 | 2826537 | 2826542 | 6 | * | 6 | 1000 |
| NC_039509.1 | 11068956 | 11086837 | 17882 | - | Gnomon | gene | NA | NA | gene-LOC105275770 | GeneID:105275770 | LOC105275770 | Gene | LOC105275770 | protein_coding | character(0) | NA | character(0) | NC_039509.1 | 11085457 | 11085462 | 6 | * | 6 | 1000 |
| NC_039509.1 | 13033738 | 13035966 | 2229 | + | Gnomon | gene | NA | NA | gene-LOC105278527 | GeneID:105278527 | LOC105278527 | Gene | LOC105278527 | protein_coding | character(0) | NA | character(0) | NC_039509.1 | 13034669 | 13034674 | 6 | * | 6 | 1000 |
| NC_039510.1 | 2607810 | 2610599 | 2790 | - | Gnomon | gene | NA | NA | gene-LOC105287754 | GeneID:105287754 | LOC105287754 | Gene | LOC105287754 | protein_coding | character(0) | NA | character(0) | NC_039510.1 | 2609179 | 2609184 | 6 | * | 6 | 1000 |
| NC_039510.1 | 16287375 | 16291357 | 3983 | - | Gnomon | gene | NA | NA | gene-LOC105286509 | GeneID:105286509 | LOC105286509 | Gene | LOC105286509 | protein_coding | character(0) | NA | character(0) | NC_039510.1 | 16290598 | 16290603 | 6 | * | 6 | 1000 |
| NC_039511.1 | 480353 | 488275 | 7923 | + | Gnomon | gene | NA | NA | gene-LOC105276527 | GeneID:105276527 | LOC105276527 | Gene | LOC105276527 | protein_coding | character(0) | NA | character(0) | NC_039511.1 | 481652 | 481657 | 6 | * | 6 | 1000 |
| NC_039511.1 | 2234830 | 2236789 | 1960 | - | Gnomon | gene | NA | NA | gene-LOC105276382 | GeneID:105276382 | LOC105276382 | Gene | LOC105276382 | protein_coding | character(0) | NA | character(0) | NC_039511.1 | 2236016 | 2236021 | 6 | * | 6 | 1000 |
| NC_039511.1 | 3614241 | 3616939 | 2699 | + | Gnomon | gene | NA | NA | gene-LOC105284057 | GeneID:105284057 | LOC105284057 | Gene | LOC105284057 | protein_coding | character(0) | NA | character(0) | NC_039511.1 | 3615462 | 3615467 | 6 | * | 6 | 1000 |
| NC_039511.1 | 8407553 | 8425032 | 17480 | + | Gnomon | gene | NA | NA | gene-LOC105284845 | GeneID:105284845 | LOC105284845 | Gene | LOC105284845 | protein_coding | character(0) | NA | character(0) | NC_039511.1 | 8409706 | 8409711 | 6 | * | 6 | 1000 |
| NC_039511.1 | 8695181 | 8698086 | 2906 | + | Gnomon | gene | NA | NA | gene-LOC105280856 | GeneID:105280856 | LOC105280856 | Gene | LOC105280856 | protein_coding | character(0) | NA | character(0) | NC_039511.1 | 8695844 | 8695849 | 6 | * | 6 | 1000 |
| NC_039512.1 | 12092666 | 12097773 | 5108 | + | Gnomon | gene | NA | NA | gene-LOC105279779 | GeneID:105279779 | LOC105279779 | Gene | LOC105279779 | protein_coding | character(0) | NA | character(0) | NC_039512.1 | 12094200 | 12094205 | 6 | * | 6 | 1000 |
| NC_039512.1 | 15979275 | 15981529 | 2255 | - | Gnomon | gene | NA | NA | gene-LOC105274775 | GeneID:105274775 | LOC105274775 | Gene | LOC105274775 | protein_coding | character(0) | NA | character(0) | NC_039512.1 | 15980634 | 15980639 | 6 | * | 6 | 1000 |
| NC_039513.1 | 10401827 | 10404590 | 2764 | - | Gnomon | gene | NA | NA | gene-LOC105275550 | GeneID:105275550 | LOC105275550 | Gene | LOC105275550 | protein_coding | character(0) | NA | character(0) | NC_039513.1 | 10403469 | 10403474 | 6 | * | 6 | 1000 |
| NC_039513.1 | 12028675 | 12030941 | 2267 | - | Gnomon | gene | NA | NA | gene-LOC105280791 | GeneID:105280791 | LOC105280791 | Gene | LOC105280791 | protein_coding | character(0) | NA | character(0) | NC_039513.1 | 12029791 | 12029796 | 6 | * | 6 | 1000 |
| NC_039513.1 | 13434303 | 13438575 | 4273 | - | Gnomon | gene | NA | NA | gene-LOC105275036 | GeneID:105275036 | LOC105275036 | Gene | LOC105275036 | protein_coding | character(0) | NA | character(0) | NC_039513.1 | 13436991 | 13437000 | 10 | * | 10 | 1000 |
| NC_039514.1 | 6622839 | 6626782 | 3944 | - | Gnomon | gene | NA | NA | gene-LOC105285534 | GeneID:105285534 | LOC105285534 | Gene | LOC105285534 | protein_coding | character(0) | NA | character(0) | NC_039514.1 | 6625853 | 6625858 | 6 | * | 6 | 1000 |
| NC_039514.1 | 9336154 | 9343762 | 7609 | + | Gnomon | gene | NA | NA | gene-LOC105284644 | GeneID:105284644 | LOC105284644 | Gene | LOC105284644 | protein_coding | character(0) | NA | character(0) | NC_039514.1 | 9337043 | 9337048 | 6 | * | 6 | 1000 |
| NC_039514.1 | 11109536 | 11117116 | 7581 | - | Gnomon | gene | NA | NA | gene-LOC105281601 | GeneID:105281601 | LOC105281601 | Gene | LOC105281601 | protein_coding | | | | | | | | | | |

| | | | | | | | | | | | | | | |
|---|---|---|---|---|---|---|---|---|---|---|---|---|---|---|
| character(0) | NA | character(0) | NC_039514.1 | 11112955 | 11112960 | 6 | * | 6 | 1000 | | | | | |
| NC_039514.1 | 11111779 | 11115572 | 3794 | + | Gnomon | gene | NA | NA | gene-LOC105281603 | GeneID:105281603 | LOC105281603 | Gene | LOC105281603 | protein_coding |
| character(0) | NA | character(0) | NC_039514.1 | 11112955 | 11112960 | 6 | * | 6 | 1000 | | | | | |
| NC_039515.1 | 7098785 | 7103508 | 4724 | - | Gnomon | gene | NA | NA | gene-LOC105286003 | GeneID:105286003 | LOC105286003 | Gene | LOC105286003 | protein_coding |
| character(0) | NA | character(0) | NC_039515.1 | 7102685 | 7102690 | 6 | * | 6 | 1000 | | | | | |
| NC_039515.1 | 12582469 | 12585642 | 3174 | - | Gnomon | gene | NA | NA | gene-LOC105282150 | GeneID:105282150 | LOC105282150 | Gene | LOC105282150 | protein_coding |
| character(0) | NA | character(0) | NC_039515.1 | 12584748 | 12584753 | 6 | * | 6 | 1000 | | | | | |
| NC_039516.1 | 12889538 | 12895772 | 6235 | + | Gnomon | gene | NA | NA | gene-LOC105275316 | GeneID:105275316 | LOC105275316 | Gene | LOC105275316 | protein_coding |
| character(0) | NA | character(0) | NC_039516.1 | 12890326 | 12890331 | 6 | * | 6 | 1000 | | | | | |
| NC_039516.1 | 13065193 | 13068389 | 3197 | + | Gnomon | gene | NA | NA | gene-LOC105275355 | GeneID:105275355 | LOC105275355 | Gene | LOC105275355 | protein_coding |
| character(0) | NA | character(0) | NC_039516.1 | 13066190 | 13066195 | 6 | * | 6 | 1000 | | | | | |
| NC_039517.1 | 1360424 | 1367346 | 6923 | + | Gnomon | gene | NA | NA | gene-LOC105278812 | GeneID:105278812 | LOC105278812 | Gene | LOC105278812 | protein_coding |
| character(0) | NA | character(0) | NC_039517.1 | 1361411 | 1361416 | 6 | * | 6 | 1000 | | | | | |
| NC_039517.1 | 8175046 | 8181714 | 6669 | - | Gnomon | gene | NA | NA | gene-LOC105284262 | GeneID:105284262 | LOC105284262 | Gene | LOC105284262 | protein_coding |
| character(0) | NA | character(0) | NC_039517.1 | 8175507 | 8175512 | 6 | * | 6 | 1000 | | | | | |
| NC_039518.1 | 5496870 | 5510429 | 13560 | - | Gnomon | gene | NA | NA | gene-LOC105281906 | GeneID:105281906 | LOC105281906 | Gene | LOC105281906 | protein_coding |
| character(0) | NA | character(0) | NC_039518.1 | 5508860 | 5508865 | 6 | * | 6 | 1000 | | | | | |
| NC_039518.1 | 8721748 | 8727127 | 5380 | + | Gnomon | gene | NA | NA | gene-LOC105286364 | GeneID:105286364 | LOC105286364 | Gene | LOC105286364 | protein_coding |
| character(0) | NA | character(0) | NC_039518.1 | 8722922 | 8722927 | 6 | * | 6 | 1000 | | | | | |
| NC_039506.1 | 23567682 | 23574351 | 6670 | + | Gnomon | gene | NA | NA | gene-LOC105279333 | GeneID:105279333 | LOC105279333 | Gene | LOC105279333 | protein_coding |
| character(0) | NA | character(0) | NC_039506.1 | 23568887 | 23568897 | 11 | * | 10 | 909.090909090909 | | | | | |
| NC_039507.1 | 9161485 | 9166239 | 4755 | + | Gnomon | gene | NA | NA | gene-LOC105282461 | GeneID:105282461 | LOC105282461 | Gene | LOC105282461 | protein_coding |
| character(0) | NA | character(0) | NC_039507.1 | 9162374 | 9162382 | 9 | * | 8 | 888.888888888889 | | | | | |
| NC_039509.1 | 17207204 | 17209527 | 2324 | + | Gnomon | gene | NA | NA | gene-LOC105283016 | GeneID:105283016 | LOC105283016 | Gene | LOC105283016 | protein_coding |
| character(0) | NA | character(0) | NC_039509.1 | 17208071 | 17208079 | 9 | * | 8 | 888.888888888889 | | | | | |
| NC_039510.1 | 8568121 | 8569477 | 1357 | - | Gnomon | gene | NA | NA | gene-LOC105275137 | GeneID:105275137 | LOC105275137 | Gene | LOC105275137 | protein_coding |
| character(0) | NA | character(0) | NC_039510.1 | 8568914 | 8568922 | 9 | * | 8 | 888.888888888889 | | | | | |
| NC_039516.1 | 5472871 | 5475154 | 2284 | - | Gnomon | gene | NA | NA | gene-LOC105275878 | GeneID:105275878 | LOC105275878 | Gene | LOC105275878 | protein_coding |
| character(0) | NA | character(0) | NC_039516.1 | 5474292 | 5474300 | 9 | * | 8 | 888.888888888889 | | | | | |
| NC_039513.1 | 3303685 | 3306319 | 2635 | + | Gnomon | gene | NA | NA | gene-LOC105275761 | GeneID:105275761 | LOC105275761 | Gene | LOC105275761 | protein_coding |
| character(0) | NA | character(0) | NC_039513.1 | 3304807 | 3304814 | 8 | * | 7 | 875 | | | | | |
| NC_039506.1 | 1487779 | 1489683 | 1905 | + | Gnomon | gene | NA | NA | gene-LOC105280236 | GeneID:105280236 | LOC105280236 | Gene | LOC105280236 | protein_coding |
| character(0) | NA | character(0) | NC_039506.1 | 1488926 | 1488932 | 7 | * | 6 | 857.142857142857 | | | | | |
| NC_039506.1 | 1728863 | 1739284 | 10422 | - | Gnomon | gene | NA | NA | gene-LOC105280269 | GeneID:105280269 | LOC105280269 | Gene | LOC105280269 | protein_coding |
| character(0) | NA | character(0) | NC_039506.1 | 1737091 | 1737097 | 7 | * | 6 | 857.142857142857 | | | | | |
| NC_039506.1 | 2274108 | 2276090 | 1983 | - | Gnomon | gene | NA | NA | gene-LOC105280319 | GeneID:105280319 | LOC105280319 | Gene | LOC105280319 | protein_coding |
| character(0) | NA | character(0) | NC_039506.1 | 2275368 | 2275374 | 7 | * | 6 | 857.142857142857 | | | | | |
| NC_039506.1 | 3929216 | 3931981 | 2766 | + | Gnomon | gene | NA | NA | gene-LOC105280469 | GeneID:105280469 | LOC105280469 | Gene | LOC105280469 | protein_coding |
| character(0) | NA | character(0) | NC_039506.1 | 3930638 | 3930644 | 7 | * | 6 | 857.142857142857 | | | | | |
| NC_039506.1 | 4874444 | 4880252 | 5809 | - | Gnomon | gene | NA | NA | gene-LOC105277909 | GeneID:105277909 | LOC105277909 | Gene | LOC105277909 | protein_coding |
| character(0) | NA | character(0) | NC_039506.1 | 4878628 | 4878634 | 7 | * | 6 | 857.142857142857 | | | | | |
| NC_039506.1 | 13552800 | 13986605 | 433806 | - | Gnomon | gene | NA | NA | gene-LOC105280980 | GeneID:105280980 | LOC105280980 | Gene | LOC105280980 | protein_coding |
| character(0) | NA | character(0) | NC_039506.1 | 13985875 | 13985881 | 7 | * | 6 | 857.142857142857 | | | | | |
| NC_039506.1 | 15509387 | 15513034 | 3648 | + | Gnomon | gene | NA | NA | gene-LOC105283961 | GeneID:105283961 | LOC105283961 | Gene | LOC105283961 | protein_coding |
| character(0) | NA | character(0) | NC_039506.1 | 15510126 | 15510132 | 7 | * | 6 | 857.142857142857 | | | | | |
| NC_039506.1 | 16116359 | 16120578 | 4220 | - | Gnomon | gene | NA | NA | gene-LOC105274912 | GeneID:105274912 | LOC105274912 | Gene | LOC105274912 | protein_coding |
| character(0) | NA | character(0) | NC_039506.1 | 16119072 | 16119078 | 7 | * | 6 | 857.142857142857 | | | | | |
| NC_039506.1 | 19093260 | 19098014 | 4755 | - | Gnomon | gene | NA | NA | gene-LOC105278005 | GeneID:105278005 | LOC105278005 | Gene | LOC105278005 | protein_coding |
| character(0) | NA | character(0) | NC_039506.1 | 19096860 | 19096866 | 7 | * | 6 | 857.142857142857 | | | | | |
| NC_039506.1 | 20553263 | 20559098 | 5836 | - | Gnomon | gene | NA | NA | gene-LOC105278544 | GeneID:105278544 | LOC105278544 | Gene | LOC105278544 | protein_coding |
| character(0) | NA | character(0) | NC_039506.1 | 20557629 | 20557635 | 7 | * | 6 | 857.142857142857 | | | | | |
| NC_039506.1 | 23689672 | 23694320 | 4649 | - | Gnomon | gene | NA | NA | gene-LOC105279346 | GeneID:105279346 | LOC105279346 | Gene | LOC105279346 | protein_coding |
| character(0) | NA | character(0) | NC_039506.1 | 23693180 | 23693186 | 7 | * | 6 | 857.142857142857 | | | | | |
| NC_039507.1 | 790243 | 798925 | 8683 | + | Gnomon | gene | NA | NA | gene-LOC105282921 | GeneID:105282921 | LOC105282921 | Gene | LOC105282921 | protein_coding |
| character(0) | NA | character(0) | NC_039507.1 | 791692 | 791698 | 7 | * | 6 | 857.142857142857 | | | | | |
| NC_039507.1 | 6347235 | 6350936 | 3702 | + | Gnomon | gene | NA | NA | gene-LOC105275401 | GeneID:105275401 | LOC105275401 | Gene | LOC105275401 | protein_coding |
| character(0) | NA | character(0) | NC_039507.1 | 6348012 | 6348018 | 7 | * | 6 | 857.142857142857 | | | | | |
| NC_039507.1 | 6560548 | 6563391 | 2844 | - | Gnomon | gene | NA | NA | gene-LOC105285345 | GeneID:105285345 | LOC105285345 | Gene | LOC105285345 | protein_coding |
| character(0) | NA | character(0) | NC_039507.1 | 6562518 | 6562524 | 7 | * | 6 | 857.142857142857 | | | | | |
| NC_039507.1 | 15481857 | 15484141 | 2285 | + | Gnomon | gene | NA | NA | gene-LOC105280092 | GeneID:105280092 | LOC105280092 | Gene | LOC105280092 | protein_coding |
| character(0) | NA | character(0) | NC_039507.1 | 15482876 | 15482882 | 7 | * | 6 | 857.142857142857 | | | | | |
| NC_039507.1 | 15506083 | 15508702 | 2620 | + | Gnomon | gene | NA | NA | gene-LOC105280102 | GeneID:105280102 | LOC105280102 | Gene | LOC105280102 | protein_coding |
| character(0) | NA | character(0) | NC_039507.1 | 15507322 | 15507328 | 7 | * | 6 | 857.142857142857 | | | | | |
| NC_039507.1 | 18333035 | 18336377 | 3343 | - | Gnomon | gene | NA | NA | gene-LOC105281577 | GeneID:105281577 | LOC105281577 | Gene | LOC105281577 | protein_coding |

character(0) NA character(0) NC_039507.1 18335506 18335512 7 * 6 857.142857142857
NC_039508.1 1384214 1386820 2607 + Gnomon gene NA NA gene-LOC105277388 GeneID:105277388 LOC105277388 Gene LOC105277388 protein_coding
character(0) NA character(0) NC_039508.1 1384932 1384938 7 * 6 857.142857142857
NC_039508.1 1386723 1389150 2428 - Gnomon gene NA NA gene-LOC105277389 GeneID:105277389 LOC105277389 Gene LOC105277389 protein_coding
character(0) NA character(0) NC_039508.1 1387947 1387953 7 * 6 857.142857142857
NC_039508.1 9041025 9045259 4235 + Gnomon gene NA NA gene-LOC105278106 GeneID:105278106 LOC105278106 Gene LOC105278106 protein_coding
character(0) NA character(0) NC_039508.1 9042110 9042116 7 * 6 857.142857142857
NC_039508.1 9080743 9083827 3085 - Gnomon gene NA NA gene-LOC105278098 GeneID:105278098 LOC105278098 Gene LOC105278098 protein_coding
character(0) NA character(0) NC_039508.1 9082000 9082006 7 * 6 857.142857142857
NC_039508.1 10763110 10766843 3734 + Gnomon gene NA NA gene-LOC105276948 GeneID:105276948 LOC105276948 Gene LOC105276948 protein_coding
character(0) NA character(0) NC_039508.1 10764758 10764764 7 * 6 857.142857142857
NC_039508.1 10797665 10799797 2133 + Gnomon gene NA NA gene-LOC105276949 GeneID:105276949 LOC105276949 Gene LOC105276949 protein_coding
character(0) NA character(0) NC_039508.1 10798605 10798611 7 * 6 857.142857142857
NC_039508.1 13116146 13120560 4415 - Gnomon gene NA NA gene-LOC105277128 GeneID:105277128 LOC105277128 Gene LOC105277128 protein_coding
character(0) NA character(0) NC_039508.1 13116245 13116251 7 * 6 857.142857142857
NC_039508.1 15763225 15782851 19627 + Gnomon gene NA NA gene-LOC105277234 GeneID:105277234 LOC105277234 Gene LOC105277234 protein_coding
character(0) NA character(0) NC_039508.1 15764452 15764458 7 * 6 857.142857142857
NC_039508.1 17548373 17550348 1976 - Gnomon gene NA NA gene-LOC105284396 GeneID:105284396 LOC105284396 Gene LOC105284396 protein_coding
character(0) NA character(0) NC_039508.1 17549403 17549409 7 * 6 857.142857142857
NC_039509.1 89280 90451 1172 + Gnomon gene NA NA gene-LOC113561814 GeneID:113561814 LOC113561814 Gene LOC113561814 protein_coding character(0) NA
character(0) NC_039509.1 90329 90335 7 * 6 857.142857142857
NC_039509.1 2072593 2082307 9715 - Gnomon gene NA NA gene-LOC105283818 GeneID:105283818 LOC105283818 Gene LOC105283818 protein_coding
character(0) NA character(0) NC_039509.1 2081387 2081393 7 * 6 857.142857142857
NC_039509.1 2365507 2377666 12160 + Gnomon gene NA NA gene-LOC105285865 GeneID:105285865 LOC105285865 Gene LOC105285865 protein_coding
character(0) NA character(0) NC_039509.1 2366218 2366224 7 * 6 857.142857142857
NC_039509.1 2385054 2391699 6646 - Gnomon gene NA NA gene-LOC105285867 GeneID:105285867 LOC105285867 Gene LOC105285867 protein_coding
character(0) NA character(0) NC_039509.1 2389203 2389209 7 * 6 857.142857142857
NC_039509.1 5250307 5257914 7608 - Gnomon gene NA NA gene-LOC105287147 GeneID:105287147 LOC105287147 Gene LOC105287147 protein_coding
character(0) NA character(0) NC_039509.1 5256794 5256800 7 * 6 857.142857142857
NC_039509.1 10232323 10244464 12142 + Gnomon gene NA NA gene-LOC105282655 GeneID:105282655 LOC105282655 Gene LOC105282655 protein_coding
character(0) NA character(0) NC_039509.1 10233310 10233316 7 * 6 857.142857142857
NC_039509.1 10710016 10711124 1109 + Gnomon gene NA NA gene-LOC105275153 GeneID:105275153 LOC105275153 Gene LOC105275153 protein_coding
character(0) NA character(0) NC_039509.1 10710730 10710736 7 * 6 857.142857142857
NC_039509.1 13909149 13915606 6458 + Gnomon gene NA NA gene-LOC105278288 GeneID:105278288 LOC105278288 Gene LOC105278288 protein_coding
character(0) NA character(0) NC_039509.1 13909987 13909993 7 * 6 857.142857142857
NC_039510.1 3040660 3042070 1411 + Gnomon gene NA NA gene-LOC105287710 GeneID:105287710 LOC105287710 Gene LOC105287710 protein_coding
character(0) NA character(0) NC_039510.1 3041562 3041568 7 * 6 857.142857142857
NC_039510.1 3110492 3117379 6888 - Gnomon gene NA NA gene-LOC105287697 GeneID:105287697 LOC105287697 Gene LOC105287697 protein_coding
character(0) NA character(0) NC_039510.1 3116602 3116608 7 * 6 857.142857142857
NC_039510.1 3167251 3170185 2935 - Gnomon gene NA NA gene-LOC105287684 GeneID:105287684 LOC105287684 Gene LOC105287684 protein_coding
character(0) NA character(0) NC_039510.1 3169234 3169240 7 * 6 857.142857142857
NC_039510.1 4220534 4227851 7318 + Gnomon gene NA NA gene-LOC105287620 GeneID:105287620 LOC105287620 Gene LOC105287620 protein_coding
character(0) NA character(0) NC_039510.1 4221389 4221395 7 * 6 857.142857142857
NC_039510.1 5067676 5072600 4925 - Gnomon gene NA NA gene-LOC105287597 GeneID:105287597 LOC105287597 Gene LOC105287597 protein_coding
character(0) NA character(0) NC_039510.1 5071200 5071206 7 * 6 857.142857142857
NC_039510.1 5537223 5539036 1814 - Gnomon gene NA NA gene-LOC105287569 GeneID:105287569 LOC105287569 Gene LOC105287569 protein_coding
character(0) NA character(0) NC_039510.1 5538139 5538145 7 * 6 857.142857142857
NC_039510.1 6043706 6047166 3461 + Gnomon gene NA NA gene-LOC105287208 GeneID:105287208 LOC105287208 Gene LOC105287208 protein_coding
character(0) NA character(0) NC_039510.1 6045163 6045169 7 * 6 857.142857142857
NC_039510.1 6163214 6171437 8224 - Gnomon gene NA NA gene-LOC105287371 GeneID:105287371 LOC105287371 Gene LOC105287371 protein_coding
character(0) NA character(0) NC_039510.1 6170155 6170161 7 * 6 857.142857142857
NC_039510.1 6212141 6215222 3082 + Gnomon gene NA NA gene-LOC105287473 GeneID:105287473 LOC105287473 Gene LOC105287473 protein_coding
character(0) NA character(0) NC_039510.1 6213095 6213101 7 * 6 857.142857142857
NC_039510.1 8487238 8489293 2056 - Gnomon gene NA NA gene-LOC105275119 GeneID:105275119 LOC105275119 Gene LOC105275119 misc_RNA character(0)
NA character(0) NC_039510.1 8488481 8488487 7 * 6 857.142857142857
NC_039510.1 8568121 8569477 1357 - Gnomon gene NA NA gene-LOC105275137 GeneID:105275137 LOC105275137 Gene LOC105275137 protein_coding
character(0) NA character(0) NC_039510.1 8568885 8568891 7 * 6 857.142857142857
NC_039510.1 9996113 9998708 2596 - Gnomon gene NA NA gene-LOC105281227 GeneID:105281227 LOC105281227 Gene LOC105281227 protein_coding
character(0) NA character(0) NC_039510.1 9997644 9997650 7 * 6 857.142857142857
NC_039510.1 13531247 13535548 4302 - Gnomon gene NA NA gene-LOC105280964 GeneID:105280964 LOC105280964 Gene LOC105280964 protein_coding
character(0) NA character(0) NC_039510.1 13533348 13533354 7 * 6 857.142857142857
NC_039510.1 15582075 15587881 5807 - Gnomon gene NA NA gene-LOC105286461 GeneID:105286461 LOC105286461 Gene LOC105286461 protein_coding
character(0) NA character(0) NC_039510.1 15586562 15586568 7 * 6 857.142857142857
NC_039510.1 15745071 15751836 6766 + Gnomon gene NA NA gene-LOC105286478 GeneID:105286478 LOC105286478 Gene LOC105286478 protein_coding

character(0) NA character(0) NC_039510.1 15746500 15746506 7 * 6 857.142857142857
NC_039511.1 517195 530396 13202 + Gnomon gene NA NA gene-LOC105276519 GeneID:105276519 LOC105276519 Gene LOC105276519 protein_coding
character(0) NA character(0) NC_039511.1 518659 518665 7 * 6 857.142857142857
NC_039511.1 1856026 1859536 3511 + Gnomon gene NA NA gene-LOC105276422 GeneID:105276422 LOC105276422 Gene LOC105276422 protein_coding
character(0) NA character(0) NC_039511.1 1857678 1857684 7 * 6 857.142857142857
NC_039511.1 2004889 2015973 11085 - Gnomon gene NA NA gene-LOC105276410 GeneID:105276410 LOC105276410 Gene LOC105276410 protein_coding
character(0) NA character(0) NC_039511.1 2014683 2014689 7 * 6 857.142857142857
NC_039511.1 2004889 2015973 11085 - Gnomon gene NA NA gene-LOC105276410 GeneID:105276410 LOC105276410 Gene LOC105276410 protein_coding
character(0) NA character(0) NC_039511.1 2014740 2014746 7 * 6 857.142857142857
NC_039511.1 3635459 3642537 7079 - Gnomon gene NA NA gene-LOC105284052 GeneID:105284052 LOC105284052 Gene LOC105284052 protein_coding
character(0) NA character(0) NC_039511.1 3639538 3639544 7 * 6 857.142857142857
NC_039511.1 4554146 4561340 7195 - Gnomon gene NA NA gene-LOC105281750 GeneID:105281750 LOC105281750 Gene LOC105281750 protein_coding
character(0) NA character(0) NC_039511.1 4560502 4560508 7 * 6 857.142857142857
NC_039511.1 12403841 12413518 9678 - Gnomon gene NA NA gene-LOC105285135 GeneID:105285135 LOC105285135 Gene LOC105285135 protein_coding
character(0) NA character(0) NC_039511.1 12411122 12411128 7 * 6 857.142857142857
NC_039511.1 14704311 14706938 2628 - Gnomon gene NA NA gene-LOC105285889 GeneID:105285889 LOC105285889 Gene LOC105285889 protein_coding
character(0) NA character(0) NC_039511.1 14705900 14705906 7 * 6 857.142857142857
NC_039511.1 15978904 15980446 1543 + Gnomon gene NA NA gene-LOC105284579 GeneID:105284579 LOC105284579 Gene LOC105284579 protein_coding
character(0) NA character(0) NC_039511.1 15979535 15979541 7 * 6 857.142857142857
NC_039511.1 15983428 15985754 2327 + Gnomon gene NA NA gene-LOC105284576 GeneID:105284576 LOC105284576 Gene LOC105284576 protein_coding
character(0) NA character(0) NC_039511.1 15984100 15984106 7 * 6 857.142857142857
NC_039512.1 4385041 4390283 5243 - Gnomon gene NA NA gene-LOC105277552 GeneID:105277552 LOC105277552 Gene LOC105277552 protein_coding
character(0) NA character(0) NC_039512.1 4389120 4389126 7 * 6 857.142857142857
NC_039512.1 11140867 11151152 10286 - Gnomon gene NA NA gene-LOC105279854 GeneID:105279854 LOC105279854 Gene LOC105279854 protein_coding
character(0) NA character(0) NC_039512.1 11150305 11150311 7 * 6 857.142857142857
NC_039512.1 12469542 12475093 5552 - Gnomon gene NA NA gene-LOC105279748 GeneID:105279748 LOC105279748 Gene LOC105279748 protein_coding
character(0) NA character(0) NC_039512.1 12474009 12474015 7 * 6 857.142857142857
NC_039512.1 12621899 12625222 3324 + Gnomon gene NA NA gene-LOC105279740 GeneID:105279740 LOC105279740 Gene LOC105279740 protein_coding
character(0) NA character(0) NC_039512.1 12623117 12623123 7 * 6 857.142857142857
NC_039512.1 12739826 12742653 2828 + Gnomon gene NA NA gene-LOC105279719 GeneID:105279719 LOC105279719 Gene LOC105279719 protein_coding
character(0) NA character(0) NC_039512.1 12740699 12740705 7 * 6 857.142857142857
NC_039512.1 12814910 12819293 4384 - Gnomon gene NA NA gene-LOC105279709 GeneID:105279709 LOC105279709 Gene LOC105279709 protein_coding
character(0) NA character(0) NC_039512.1 12818052 12818058 7 * 6 857.142857142857
NC_039512.1 13089540 13093562 4023 - Gnomon gene NA NA gene-LOC105279664 GeneID:105279664 LOC105279664 Gene LOC105279664 protein_coding
character(0) NA character(0) NC_039512.1 13092263 13092269 7 * 6 857.142857142857
NC_039512.1 16555603 16557649 2047 - Gnomon gene NA NA gene-LOC105282841 GeneID:105282841 LOC105282841 Gene LOC105282841 protein_coding
character(0) NA character(0) NC_039512.1 16556845 16556851 7 * 6 857.142857142857
NC_039513.1 3253321 3257474 4154 - Gnomon gene NA NA gene-LOC105275753 GeneID:105275753 LOC105275753 Gene LOC105275753 protein_coding
character(0) NA character(0) NC_039513.1 3256149 3256155 7 * 6 857.142857142857
NC_039513.1 3253321 3257474 4154 - Gnomon gene NA NA gene-LOC105275753 GeneID:105275753 LOC105275753 Gene LOC105275753 protein_coding
character(0) NA character(0) NC_039513.1 3256358 3256364 7 * 6 857.142857142857
NC_039513.1 3257374 3260835 3462 - Gnomon gene NA NA gene-LOC105275752 GeneID:105275752 LOC105275752 Gene LOC105275752 protein_coding
character(0) NA character(0) NC_039513.1 3259768 3259774 7 * 6 857.142857142857
NC_039513.1 3373854 3375485 1632 + Gnomon gene NA NA gene-LOC105274850 GeneID:105274850 LOC105274850 Gene LOC105274850 protein_coding
character(0) NA character(0) NC_039513.1 3374674 3374680 7 * 6 857.142857142857
NC_039513.1 3472678 3520828 48151 - Gnomon gene NA NA gene-LOC105274836 GeneID:105274836 LOC105274836 Gene LOC105274836 protein_coding
character(0) NA character(0) NC_039513.1 3480047 3480053 7 * 6 857.142857142857
NC_039513.1 5176546 5178099 1554 + Gnomon gene NA NA gene-LOC105279718 GeneID:105279718 LOC105279718 Gene LOC105279718 protein_coding
character(0) NA character(0) NC_039513.1 5177248 5177254 7 * 6 857.142857142857
NC_039513.1 6784235 6788101 3867 - Gnomon gene NA NA gene-LOC105287855 GeneID:105287855 LOC105287855 Gene LOC105287855 protein_coding
character(0) NA character(0) NC_039513.1 6787239 6787245 7 * 6 857.142857142857
NC_039513.1 8483464 8491634 8171 + Gnomon gene NA NA gene-LOC105275689 GeneID:105275689 LOC105275689 Gene LOC105275689 protein_coding
character(0) NA character(0) NC_039513.1 8484406 8484412 7 * 6 857.142857142857
NC_039513.1 9227749 9230551 2803 - Gnomon gene NA NA gene-LOC105275656 GeneID:105275656 LOC105275656 Gene LOC105275656 protein_coding
character(0) NA character(0) NC_039513.1 9229018 9229024 7 * 6 857.142857142857
NC_039513.1 9603616 9607153 3538 + Gnomon gene NA NA gene-LOC105275615 GeneID:105275615 LOC105275615 Gene LOC105275615 protein_coding
character(0) NA character(0) NC_039513.1 9605113 9605119 7 * 6 857.142857142857
NC_039513.1 11674366 11676356 1991 - Gnomon gene NA NA gene-LOC105280739 GeneID:105280739 LOC105280739 Gene LOC105280739 protein_coding
character(0) NA character(0) NC_039513.1 11675195 11675201 7 * 6 857.142857142857
NC_039513.1 13452183 13456265 4083 + Gnomon gene NA NA gene-LOC105275031 GeneID:105275031 LOC105275031 Gene LOC105275031 protein_coding
character(0) NA character(0) NC_039513.1 13453683 13453689 7 * 6 857.142857142857
NC_039513.1 15035458 15043240 7783 + Gnomon gene NA NA gene-LOC105285648 GeneID:105285648 LOC105285648 Gene LOC105285648 protein_coding
character(0) NA character(0) NC_039513.1 15037033 15037039 7 * 6 857.142857142857
NC_039513.1 15725203 15744087 18885 + Gnomon gene NA NA gene-LOC105280058 GeneID:105280058 LOC105280058 Gene LOC105280058 protein_coding

| | | | | | | | | | | | | | | |
|---|---|---|---|---|---|---|---|---|---|---|---|---|---|---|
| | character(0) | NA | character(0) | NC_039513.1 | 15726271 | 15726277 | 7 | * | 6 | 857.142857142857 | | | | |
| NC_039514.1 | 873449 | 876660 | 3212 | + | Gnomon | gene | NA | NA | gene-LOC105276205 | GeneID:105276205 | LOC105276205 | Gene | LOC105276205 | protein_coding |
| | character(0) | NA | character(0) | NC_039514.1 | 874466 | 874472 | 7 | * | 6 | 857.142857142857 | | | | |
| NC_039514.1 | 1286674 | 1297576 | 10903 | - | Gnomon | gene | NA | NA | gene-LOC105276233 | GeneID:105276233 | LOC105276233 | Gene | LOC105276233 | protein_coding |
| | character(0) | NA | character(0) | NC_039514.1 | 1295968 | 1295974 | 7 | * | 6 | 857.142857142857 | | | | |
| NC_039514.1 | 6551602 | 6556437 | 4836 | - | Gnomon | gene | NA | NA | gene-LOC105285546 | GeneID:105285546 | LOC105285546 | Gene | LOC105285546 | protein_coding |
| | character(0) | NA | character(0) | NC_039514.1 | 6555389 | 6555395 | 7 | * | 6 | 857.142857142857 | | | | |
| NC_039514.1 | 8629561 | 8648221 | 18661 | + | Gnomon | gene | NA | NA | gene-LOC105278994 | GeneID:105278994 | LOC105278994 | Gene | LOC105278994 | protein_coding |
| | character(0) | NA | character(0) | NC_039514.1 | 8630645 | 8630651 | 7 | * | 6 | 857.142857142857 | | | | |
| NC_039514.1 | 8629561 | 8648221 | 18661 | + | Gnomon | gene | NA | NA | gene-LOC105278994 | GeneID:105278994 | LOC105278994 | Gene | LOC105278994 | protein_coding |
| | character(0) | NA | character(0) | NC_039514.1 | 8630804 | 8630810 | 7 | * | 6 | 857.142857142857 | | | | |
| NC_039514.1 | 9413016 | 9415003 | 1988 | - | Gnomon | gene | NA | NA | gene-LOC105284659 | GeneID:105284659 | LOC105284659 | Gene | LOC105284659 | protein_coding |
| | character(0) | NA | character(0) | NC_039514.1 | 9414116 | 9414122 | 7 | * | 6 | 857.142857142857 | | | | |
| NC_039514.1 | 11109536 | 11117116 | 7581 | - | Gnomon | gene | NA | NA | gene-LOC105281601 | GeneID:105281601 | LOC105281601 | Gene | LOC105281601 | protein_coding |
| | character(0) | NA | character(0) | NC_039514.1 | 11113139 | 11113145 | 7 | * | 6 | 857.142857142857 | | | | |
| NC_039514.1 | 11111779 | 11115572 | 3794 | + | Gnomon | gene | NA | NA | gene-LOC105281603 | GeneID:105281603 | LOC105281603 | Gene | LOC105281603 | protein_coding |
| | character(0) | NA | character(0) | NC_039514.1 | 11113139 | 11113145 | 7 | * | 6 | 857.142857142857 | | | | |
| NC_039514.1 | 12976430 | 12980725 | 4296 | - | Gnomon | gene | NA | NA | gene-LOC105277376 | GeneID:105277376 | LOC105277376 | Gene | LOC105277376 | protein_coding |
| | character(0) | NA | character(0) | NC_039514.1 | 12979531 | 12979537 | 7 | * | 6 | 857.142857142857 | | | | |
| NC_039514.1 | 14398821 | 14400325 | 1505 | - | Gnomon | gene | NA | NA | gene-LOC105275991 | GeneID:105275991 | LOC105275991 | Gene | LOC105275991 | protein_coding |
| | character(0) | NA | character(0) | NC_039514.1 | 14399289 | 14399295 | 7 | * | 6 | 857.142857142857 | | | | |
| NC_039514.1 | 15117749 | 15130344 | 12596 | + | Gnomon | gene | NA | NA | gene-LOC105276073 | GeneID:105276073 | LOC105276073 | Gene | LOC105276073 | protein_coding |
| | character(0) | NA | character(0) | NC_039514.1 | 15123389 | 15123395 | 7 | * | 6 | 857.142857142857 | | | | |
| NC_039514.1 | 15122114 | 15127165 | 5052 | + | Gnomon | gene | NA | NA | gene-LOC105276070 | GeneID:105276070 | LOC105276070 | Gene | LOC105276070 | protein_coding |
| | character(0) | NA | character(0) | NC_039514.1 | 15123389 | 15123395 | 7 | * | 6 | 857.142857142857 | | | | |
| NC_039515.1 | 1527980 | 1529634 | 1655 | + | Gnomon | gene | NA | NA | gene-LOC105274711 | GeneID:105274711 | LOC105274711 | Gene | LOC105274711 | protein_coding |
| | character(0) | NA | character(0) | NC_039515.1 | 1528293 | 1528299 | 7 | * | 6 | 857.142857142857 | | | | |
| NC_039515.1 | 2215137 | 2216986 | 1850 | - | Gnomon | gene | NA | NA | gene-LOC105278431 | GeneID:105278431 | LOC105278431 | Gene | LOC105278431 | protein_coding |
| | character(0) | NA | character(0) | NC_039515.1 | 2215927 | 2215933 | 7 | * | 6 | 857.142857142857 | | | | |
| NC_039515.1 | 3498019 | 3503642 | 5624 | + | Gnomon | gene | NA | NA | gene-LOC105278309 | GeneID:105278309 | LOC105278309 | Gene | LOC105278309 | protein_coding |
| | character(0) | NA | character(0) | NC_039515.1 | 3499095 | 3499101 | 7 | * | 6 | 857.142857142857 | | | | |
| NC_039515.1 | 3819603 | 3823750 | 4148 | + | Gnomon | gene | NA | NA | gene-LOC105278277 | GeneID:105278277 | LOC105278277 | Gene | LOC105278277 | protein_coding |
| | character(0) | NA | character(0) | NC_039515.1 | 3820723 | 3820729 | 7 | * | 6 | 857.142857142857 | | | | |
| NC_039515.1 | 3819837 | 3821619 | 1783 | + | Gnomon | gene | NA | NA | gene-LOC105278275 | GeneID:105278275 | LOC105278275 | Gene | LOC105278275 | protein_coding |
| | character(0) | NA | character(0) | NC_039515.1 | 3820723 | 3820729 | 7 | * | 6 | 857.142857142857 | | | | |
| NC_039515.1 | 5537006 | 5539696 | 2691 | + | Gnomon | gene | NA | NA | gene-LOC105277256 | GeneID:105277256 | LOC105277256 | Gene | LOC105277256 | protein_coding |
| | character(0) | NA | character(0) | NC_039515.1 | 5537966 | 5537972 | 7 | * | 6 | 857.142857142857 | | | | |
| NC_039515.1 | 6014495 | 6167113 | 152619 | + | Gnomon | gene | NA | NA | gene-LOC105286194 | GeneID:105286194 | LOC105286194 | Gene | LOC105286194 | protein_coding |
| | character(0) | NA | character(0) | NC_039515.1 | 6066909 | 6066915 | 7 | * | 6 | 857.142857142857 | | | | |
| NC_039515.1 | 6065547 | 6067835 | 2289 | + | Gnomon | gene | NA | NA | gene-LOC105285808 | GeneID:105285808 | LOC105285808 | Gene | LOC105285808 | protein_coding |
| | character(0) | NA | character(0) | NC_039515.1 | 6066909 | 6066915 | 7 | * | 6 | 857.142857142857 | | | | |
| NC_039515.1 | 6480766 | 6491993 | 11228 | + | Gnomon | gene | NA | NA | gene-LOC105282298 | GeneID:105282298 | LOC105282298 | Gene | LOC105282298 | protein_coding |
| | character(0) | NA | character(0) | NC_039515.1 | 6481705 | 6481711 | 7 | * | 6 | 857.142857142857 | | | | |
| NC_039515.1 | 6604872 | 6605723 | 852 | + | Gnomon | gene | NA | NA | gene-LOC105282280 | GeneID:105282280 | LOC105282280 | Gene | LOC105282280 | protein_coding |
| | character(0) | NA | character(0) | NC_039515.1 | 6605539 | 6605545 | 7 | * | 6 | 857.142857142857 | | | | |
| NC_039515.1 | 6615586 | 6618213 | 2628 | - | Gnomon | gene | NA | NA | gene-LOC105282278 | GeneID:105282278 | LOC105282278 | Gene | LOC105282278 | protein_coding |
| | character(0) | NA | character(0) | NC_039515.1 | 6617273 | 6617279 | 7 | * | 6 | 857.142857142857 | | | | |
| NC_039515.1 | 7926262 | 7929466 | 3205 | + | Gnomon | gene | NA | NA | gene-LOC105283741 | GeneID:105283741 | LOC105283741 | Gene | LOC105283741 | protein_coding |
| | character(0) | NA | character(0) | NC_039515.1 | 7927531 | 7927537 | 7 | * | 6 | 857.142857142857 | | | | |
| NC_039515.1 | 8973009 | 8978309 | 5301 | + | Gnomon | gene | NA | NA | gene-LOC105283697 | GeneID:105283697 | LOC105283697 | Gene | LOC105283697 | protein_coding |
| | character(0) | NA | character(0) | NC_039515.1 | 8974548 | 8974554 | 7 | * | 6 | 857.142857142857 | | | | |
| NC_039515.1 | 12611158 | 12612372 | 1215 | - | Gnomon | gene | NA | NA | gene-LOC105282147 | GeneID:105282147 | LOC105282147 | Gene | LOC105282147 | protein_coding |
| | character(0) | NA | character(0) | NC_039515.1 | 12611603 | 12611609 | 7 | * | 6 | 857.142857142857 | | | | |
| NC_039516.1 | 756663 | 760199 | 3537 | + | Gnomon | gene | NA | NA | gene-LOC105274638 | GeneID:105274638 | LOC105274638 | Gene | LOC105274638 | protein_coding |
| | character(0) | NA | character(0) | NC_039516.1 | 757725 | 757731 | 7 | * | 6 | 857.142857142857 | | | | |
| NC_039516.1 | 985951 | 990466 | 4516 | + | Gnomon | gene | NA | NA | gene-LOC105274590 | GeneID:105274590 | LOC105274590 | Gene | LOC105274590 | protein_coding |
| | character(0) | NA | character(0) | NC_039516.1 | 987063 | 987069 | 7 | * | 6 | 857.142857142857 | | | | |
| NC_039516.1 | 1549696 | 1551173 | 1478 | + | Gnomon | gene | NA | NA | gene-LOC105274497 | GeneID:105274497 | LOC105274497 | Gene | LOC105274497 | protein_coding |
| | character(0) | NA | character(0) | NC_039516.1 | 1550744 | 1550750 | 7 | * | 6 | 857.142857142857 | | | | |
| NC_039516.1 | 2241904 | 2245378 | 3475 | + | Gnomon | gene | NA | NA | gene-LOC105274464 | GeneID:105274464 | LOC105274464 | Gene | LOC105274464 | protein_coding |
| | character(0) | NA | character(0) | NC_039516.1 | 2242648 | 2242654 | 7 | * | 6 | 857.142857142857 | | | | |
| NC_039516.1 | 4713084 | 4714767 | 1684 | - | Gnomon | gene | NA | NA | gene-LOC105275897 | GeneID:105275897 | LOC105275897 | Gene | LOC105275897 | protein_coding |
| | character(0) | NA | character(0) | NC_039516.1 | 4713795 | 4713801 | 7 | * | 6 | 857.142857142857 | | | | |
| NC_039516.1 | 7248249 | 7252272 | 4024 | + | Gnomon | gene | NA | NA | gene-LOC105275812 | GeneID:105275812 | LOC105275812 | Gene | LOC105275812 | protein_coding |

| | | | | | | | | | | | | | | |
|---|---|---|---|---|---|---|---|---|---|---|---|---|---|---|
| | character(0) | NA | character(0) | NC_039516.1 | 7251402 | 7251408 | 7 | * | 6 | 857.142857142857 | | | | |
| NC_039516.1 | 12834895 | 12836314 | 1420 | - | Gnomon | gene | NA | NA | gene-LOC105275307 | GeneID:105275307 | LOC105275307 | Gene | LOC105275307 | protein_coding |
| | character(0) | NA | character(0) | NC_039516.1 | 12835552 | 12835558 | 7 | * | 6 | 857.142857142857 | | | | |
| NC_039516.1 | 13497458 | 13503461 | 6004 | - | Gnomon | gene | NA | NA | gene-LOC105277915 | GeneID:105277915 | LOC105277915 | Gene | LOC105277915 | protein_coding |
| | character(0) | NA | character(0) | NC_039516.1 | 13502711 | 13502717 | 7 | * | 6 | 857.142857142857 | | | | |
| NC_039516.1 | 13518014 | 13520692 | 2679 | + | Gnomon | gene | NA | NA | gene-LOC105277817 | GeneID:105277817 | LOC105277817 | Gene | LOC105277817 | protein_coding |
| | character(0) | NA | character(0) | NC_039516.1 | 13518940 | 13518946 | 7 | * | 6 | 857.142857142857 | | | | |
| NC_039517.1 | 867927 | 869156 | 1230 | - | Gnomon | gene | NA | NA | gene-LOC105278824 | GeneID:105278824 | LOC105278824 | Gene | LOC105278824 | protein_coding |
| | character(0) | NA | character(0) | NC_039517.1 | 868363 | 868369 | 7 | * | 6 | 857.142857142857 | | | | |
| NC_039517.1 | 2307792 | 2312395 | 4604 | - | Gnomon | gene | NA | NA | gene-LOC105281943 | GeneID:105281943 | LOC105281943 | Gene | LOC105281943 | protein_coding |
| | character(0) | NA | character(0) | NC_039517.1 | 2309012 | 2309018 | 7 | * | 6 | 857.142857142857 | | | | |
| NC_039517.1 | 2393025 | 2396928 | 3904 | - | Gnomon | gene | NA | NA | gene-LOC105281949 | GeneID:105281949 | LOC105281949 | Gene | LOC105281949 | protein_coding |
| | character(0) | NA | character(0) | NC_039517.1 | 2395845 | 2395851 | 7 | * | 6 | 857.142857142857 | | | | |
| NC_039517.1 | 2399031 | 2400965 | 1935 | + | Gnomon | gene | NA | NA | gene-LOC105281950 | GeneID:105281950 | LOC105281950 | Gene | LOC105281950 | protein_coding |
| | character(0) | NA | character(0) | NC_039517.1 | 2400125 | 2400131 | 7 | * | 6 | 857.142857142857 | | | | |
| NC_039517.1 | 7306074 | 7309288 | 3215 | + | Gnomon | gene | NA | NA | gene-LOC105284202 | GeneID:105284202 | LOC105284202 | Gene | LOC105284202 | protein_coding |
| | character(0) | NA | character(0) | NC_039517.1 | 7307471 | 7307477 | 7 | * | 6 | 857.142857142857 | | | | |
| NC_039517.1 | 9368753 | 9371050 | 2298 | - | Gnomon | gene | NA | NA | gene-LOC105282220 | GeneID:105282220 | LOC105282220 | Gene | LOC105282220 | protein_coding |
| | character(0) | NA | character(0) | NC_039517.1 | 9370300 | 9370306 | 7 | * | 6 | 857.142857142857 | | | | |
| NC_039517.1 | 9921720 | 9925663 | 3944 | + | Gnomon | gene | NA | NA | gene-LOC105287104 | GeneID:105287104 | LOC105287104 | Gene | LOC105287104 | protein_coding |
| | character(0) | NA | character(0) | NC_039517.1 | 9922442 | 9922448 | 7 | * | 6 | 857.142857142857 | | | | |
| NC_039517.1 | 10161586 | 10164793 | 3208 | - | Gnomon | gene | NA | NA | gene-LOC105285418 | GeneID:105285418 | LOC105285418 | Gene | LOC105285418 | protein_coding |
| | character(0) | NA | character(0) | NC_039517.1 | 10163396 | 10163402 | 7 | * | 6 | 857.142857142857 | | | | |
| NC_039518.1 | 2232165 | 2249269 | 17105 | + | Gnomon | gene | NA | NA | gene-LOC105283878 | GeneID:105283878 | LOC105283878 | Gene | LOC105283878 | protein_coding |
| | character(0) | NA | character(0) | NC_039518.1 | 2233357 | 2233363 | 7 | * | 6 | 857.142857142857 | | | | |
| NC_039518.1 | 2657034 | 2658634 | 1601 | + | Gnomon | gene | NA | NA | gene-LOC105285392 | GeneID:105285392 | LOC105285392 | Gene | LOC105285392 | protein_coding |
| | character(0) | NA | character(0) | NC_039518.1 | 2658047 | 2658053 | 7 | * | 6 | 857.142857142857 | | | | |
| NC_039518.1 | 2681489 | 2682795 | 1307 | - | Gnomon | gene | NA | NA | gene-LOC105285394 | GeneID:105285394 | LOC105285394 | Gene | LOC105285394 | protein_coding |
| | character(0) | NA | character(0) | NC_039518.1 | 2681945 | 2681951 | 7 | * | 6 | 857.142857142857 | | | | |
| NC_039518.1 | 5288531 | 5294826 | 6296 | - | Gnomon | gene | NA | NA | gene-LOC105285410 | GeneID:105285410 | LOC105285410 | Gene | LOC105285410 | protein_coding |
| | character(0) | NA | character(0) | NC_039518.1 | 5293748 | 5293754 | 7 | * | 6 | 857.142857142857 | | | | |
| NC_039518.1 | 8632464 | 8637212 | 4749 | + | Gnomon | gene | NA | NA | gene-LOC105286242 | GeneID:105286242 | LOC105286242 | Gene | LOC105286242 | protein_coding |
| | character(0) | NA | character(0) | NC_039518.1 | 8633518 | 8633524 | 7 | * | 6 | 857.142857142857 | | | | |
| NC_039519.1 | 197513 | 200423 | 2911 | + | Gnomon | gene | NA | NA | gene-LOC105282793 | GeneID:105282793 | LOC105282793 | Gene | LOC105282793 | protein_coding |
| | character(0) | NA | character(0) | NC_039519.1 | 198095 | 198101 | 7 | * | 6 | 857.142857142857 | | | | |
| NC_039519.1 | 3406510 | 3418101 | 11592 | - | Gnomon | gene | NA | NA | gene-LOC105286844 | GeneID:105286844 | LOC105286844 | Gene | LOC105286844 | protein_coding |
| | character(0) | NA | character(0) | NC_039519.1 | 3416621 | 3416627 | 7 | * | 6 | 857.142857142857 | | | | |
| NC_039519.1 | 8239374 | 8248322 | 8949 | - | Gnomon | gene | NA | NA | gene-LOC105287985 | GeneID:105287985 | LOC105287985 | Gene | LOC105287985 | protein_coding |
| | character(0) | NA | character(0) | NC_039519.1 | 8247911 | 8247917 | 7 | * | 6 | 857.142857142857 | | | | |
| NC_039508.1 | 13115312 | 13116213 | 902 | + | Gnomon | gene | NA | NA | gene-LOC105277091 | GeneID:105277091 | LOC105277091 | Gene | LOC105277091 | protein_coding |
| | character(0) | NA | character(0) | NC_039508.1 | 13116053 | 13116062 | 10 | * | 8 | 800 | | | | |
| NC_039509.1 | 17375981 | 17382001 | 6021 | + | Gnomon | gene | NA | NA | gene-LOC105282459 | GeneID:105282459 | LOC105282459 | Gene | LOC105282459 | protein_coding |
| | character(0) | NA | character(0) | NC_039509.1 | 17378741 | 17378750 | 10 | * | 8 | 800 | | | | |
| NC_039519.1 | 8084633 | 8088546 | 3914 | + | Gnomon | gene | NA | NA | gene-LOC105274767 | GeneID:105274767 | LOC105274767 | Gene | LOC105274767 | protein_coding |
| | character(0) | NA | character(0) | NC_039519.1 | 8085717 | 8085726 | 10 | * | 8 | 800 | | | | |

Directory: MRPR File: gwr-Ob_LI_RphaseA1.txt

| seqnames | start | end | width | strand | source | type | score | phase | ID | Dbxref | Name | gbkey | gene | gene_biotype | end_range | partial | start_range | seqnames | start | end | width | strand | NbrSites | Sdnsty |
|---|---|---|---|---|---|---|---|---|---|---|---|---|---|---|---|---|---|---|---|---|---|---|---|---|
| NC_039506.1 | 3342881 | 3348391 | 5511 | + | Gnomon | gene | NA | NA | gene-LOC105280395 | GeneID:105280395 | LOC105280395 | Gene | LOC105280395 | protein_coding | character(0) | NA | character(0) | NC_039506.1 | 3344178 | 3344183 | 6 | * | 6 | 1000 |
| NC_039506.1 | 6089683 | 6090856 | 1174 | + | Gnomon | gene | NA | NA | gene-LOC105277804 | GeneID:105277804 | LOC105277804 | Gene | LOC105277804 | protein_coding | character(0) | NA | character(0) | NC_039506.1 | 6090624 | 6090629 | 6 | * | 6 | 1000 |
| NC_039506.1 | 13177256 | 13183827 | 6572 | + | Gnomon | gene | NA | NA | gene-LOC105284041 | GeneID:105284041 | LOC105284041 | Gene | LOC105284041 | protein_coding | character(0) | NA | character(0) | NC_039506.1 | 13181314 | 13181319 | 6 | * | 6 | 1000 |
| NC_039506.1 | 15370515 | 15371538 | 1024 | + | Gnomon | gene | NA | NA | gene-LOC105280860 | GeneID:105280860 | LOC105280860 | Gene | LOC105280860 | protein_coding | character(0) | NA | character(0) | NC_039506.1 | 15371422 | 15371427 | 6 | * | 6 | 1000 |
| NC_039507.1 | 7181199 | 7188522 | 7324 | + | Gnomon | gene | NA | NA | gene-LOC105283383 | GeneID:105283383 | LOC105283383 | Gene | LOC105283383 | protein_coding | character(0) | NA | character(0) | NC_039507.1 | 7182290 | 7182295 | 6 | * | 6 | 1000 |
| NC_039507.1 | 7276866 | 7278961 | 2096 | + | Gnomon | gene | NA | NA | gene-LOC105283373 | GeneID:105283373 | LOC105283373 | Gene | LOC105283373 | protein_coding | character(0) | NA | character(0) | NC_039507.1 | 7277947 | 7277952 | 6 | * | 6 | 1000 |
| NC_039507.1 | 18333035 | 18336377 | 3343 | - | Gnomon | gene | NA | NA | gene-LOC105281577 | GeneID:105281577 | LOC105281577 | Gene | LOC105281577 | protein_coding | character(0) | NA | character(0) | NC_039507.1 | 18335809 | 18335814 | 6 | * | 6 | 1000 |
| NC_039508.1 | 10274002 | 10277440 | 3439 | + | Gnomon | gene | NA | NA | gene-LOC105276881 | GeneID:105276881 | LOC105276881 | Gene | LOC105276881 | protein_coding | character(0) | NA | character(0) | NC_039508.1 | 10275212 | 10275217 | 6 | * | 6 | 1000 |
| NC_039509.1 | 1135464 | 1137481 | 2018 | - | Gnomon | gene | NA | NA | gene-LOC105283634 | GeneID:105283634 | LOC105283634 | Gene | LOC105283634 | protein_coding | character(0) | NA | character(0) | NC_039509.1 | 1136266 | 1136271 | 6 | * | 6 | 1000 |
| NC_039509.1 | 10756577 | 10757875 | 1299 | - | Gnomon | gene | NA | NA | gene-LOC105275214 | GeneID:105275214 | LOC105275214 | Gene | LOC105275214 | protein_coding | character(0) | NA | character(0) | NC_039509.1 | 10756803 | 10756808 | 6 | * | 6 | 1000 |
| NC_039509.1 | 11068956 | 11086837 | 17882 | - | Gnomon | gene | NA | NA | gene-LOC105275770 | GeneID:105275770 | LOC105275770 | Gene | LOC105275770 | protein_coding | character(0) | NA | character(0) | NC_039509.1 | 11085457 | 11085462 | 6 | * | 6 | 1000 |
| NC_039509.1 | 13033738 | 13035966 | 2229 | + | Gnomon | gene | NA | NA | gene-LOC105278527 | GeneID:105278527 | LOC105278527 | Gene | LOC105278527 | protein_coding | character(0) | NA | character(0) | NC_039509.1 | 13034669 | 13034674 | 6 | * | 6 | 1000 |
| NC_039509.1 | 13067024 | 13068370 | 1347 | + | Gnomon | gene | NA | NA | gene-LOC105278478 | GeneID:105278478 | LOC105278478 | Gene | LOC105278478 | protein_coding | character(0) | NA | character(0) | NC_039509.1 | 13067776 | 13067781 | 6 | * | 6 | 1000 |
| NC_039510.1 | 2607810 | 2610599 | 2790 | - | Gnomon | gene | NA | NA | gene-LOC105287754 | GeneID:105287754 | LOC105287754 | Gene | LOC105287754 | protein_coding | character(0) | NA | character(0) | NC_039510.1 | 2609179 | 2609184 | 6 | * | 6 | 1000 |
| NC_039510.1 | 16287375 | 16291357 | 3983 | - | Gnomon | gene | NA | NA | gene-LOC105286509 | GeneID:105286509 | LOC105286509 | Gene | LOC105286509 | protein_coding | character(0) | NA | character(0) | NC_039510.1 | 16290598 | 16290603 | 6 | * | 6 | 1000 |
| NC_039511.1 | 2234830 | 2236789 | 1960 | - | Gnomon | gene | NA | NA | gene-LOC105276382 | GeneID:105276382 | LOC105276382 | Gene | LOC105276382 | protein_coding | character(0) | NA | character(0) | NC_039511.1 | 2236016 | 2236021 | 6 | * | 6 | 1000 |
| NC_039511.1 | 3614241 | 3616939 | 2699 | + | Gnomon | gene | NA | NA | gene-LOC105284057 | GeneID:105284057 | LOC105284057 | Gene | LOC105284057 | protein_coding | character(0) | NA | character(0) | NC_039511.1 | 3615462 | 3615467 | 6 | * | 6 | 1000 |
| NC_039511.1 | 3625986 | 3631991 | 6006 | - | Gnomon | gene | NA | NA | gene-LOC105284053 | GeneID:105284053 | LOC105284053 | Gene | LOC105284053 | protein_coding | character(0) | NA | character(0) | NC_039511.1 | 3630771 | 3630776 | 6 | * | 6 | 1000 |
| NC_039511.1 | 8695181 | 8698086 | 2906 | + | Gnomon | gene | NA | NA | gene-LOC105280856 | GeneID:105280856 | LOC105280856 | Gene | LOC105280856 | protein_coding | character(0) | NA | character(0) | NC_039511.1 | 8695844 | 8695849 | 6 | * | 6 | 1000 |
| NC_039512.1 | 10275466 | 10280187 | 4722 | + | Gnomon | gene | NA | NA | gene-LOC105282847 | GeneID:105282847 | LOC105282847 | Gene | LOC105282847 | protein_coding | character(0) | NA | character(0) | NC_039512.1 | 10276481 | 10276486 | 6 | * | 6 | 1000 |
| NC_039512.1 | 15979275 | 15981529 | 2255 | - | Gnomon | gene | NA | NA | gene-LOC105274775 | GeneID:105274775 | LOC105274775 | Gene | LOC105274775 | protein_coding | character(0) | NA | character(0) | NC_039512.1 | 15980634 | 15980639 | 6 | * | 6 | 1000 |
| NC_039513.1 | 10401827 | 10404590 | 2764 | - | Gnomon | gene | NA | NA | gene-LOC105275550 | GeneID:105275550 | LOC105275550 | Gene | LOC105275550 | protein_coding | character(0) | NA | character(0) | NC_039513.1 | 10403469 | 10403474 | 6 | * | 6 | 1000 |
| NC_039513.1 | 11029007 | 11033701 | 4695 | - | Gnomon | gene | NA | NA | gene-LOC105275763 | GeneID:105275763 | LOC105275763 | Gene | LOC105275763 | protein_coding | character(0) | NA | character(0) | NC_039513.1 | 11030126 | 11030131 | 6 | * | 6 | 1000 |
| NC_039513.1 | 12028675 | 12030941 | 2267 | - | Gnomon | gene | NA | NA | gene-LOC105280791 | GeneID:105280791 | LOC105280791 | Gene | LOC105280791 | protein_coding | character(0) | NA | character(0) | NC_039513.1 | 12029791 | 12029796 | 6 | * | 6 | 1000 |

...
... only first 25 lines shown ...

Directory: MRPR File: gwr-Ob_LI_RphaseA2.txt

| seqnames | start | end | width | strand | source | type | score | phase | ID | Dbxref | Name | gbkey | gene | gene_biotype | end_range | partial | start_range | seqnames | start | end | width | strand | NbrSites | Sdnsty |
|---|---|---|---|---|---|---|---|---|---|---|---|---|---|---|---|---|---|---|---|---|---|---|---|---|
| NC_039506.1 | 3342881 | 3348391 | 5511 | + | Gnomon | gene | NA | NA | gene-LOC105280395 | GeneID:105280395 | LOC105280395 | Gene | LOC105280395 | protein_coding | character(0) | NA | character(0) | NC_039506.1 | 3344178 | 3344183 | 6 | * | 6 | 1000 |
| NC_039506.1 | 6089683 | 6090856 | 1174 | + | Gnomon | gene | NA | NA | gene-LOC105277804 | GeneID:105277804 | LOC105277804 | Gene | LOC105277804 | protein_coding | character(0) | NA | character(0) | NC_039506.1 | 6090624 | 6090629 | 6 | * | 6 | 1000 |
| NC_039506.1 | 15370515 | 15371538 | 1024 | + | Gnomon | gene | NA | NA | gene-LOC105280860 | GeneID:105280860 | LOC105280860 | Gene | LOC105280860 | protein_coding | character(0) | NA | character(0) | NC_039506.1 | 15371422 | 15371427 | 6 | * | 6 | 1000 |
| NC_039506.1 | 24043149 | 24047806 | 4658 | + | Gnomon | gene | NA | NA | gene-LOC105279367 | GeneID:105279367 | LOC105279367 | Gene | LOC105279367 | protein_coding | character(0) | NA | character(0) | NC_039506.1 | 24044697 | 24044702 | 6 | * | 6 | 1000 |
| NC_039507.1 | 5093844 | 5095945 | 2102 | + | Gnomon | gene | NA | NA | gene-LOC105281690 | GeneID:105281690 | LOC105281690 | Gene | LOC105281690 | protein_coding | character(0) | NA | character(0) | NC_039507.1 | 5094466 | 5094471 | 6 | * | 6 | 1000 |
| NC_039507.1 | 7276866 | 7278961 | 2096 | + | Gnomon | gene | NA | NA | gene-LOC105283373 | GeneID:105283373 | LOC105283373 | Gene | LOC105283373 | protein_coding | character(0) | NA | character(0) | NC_039507.1 | 7277947 | 7277952 | 6 | * | 6 | 1000 |
| NC_039507.1 | 18333035 | 18336377 | 3343 | - | Gnomon | gene | NA | NA | gene-LOC105281577 | GeneID:105281577 | LOC105281577 | Gene | LOC105281577 | protein_coding | character(0) | NA | character(0) | NC_039507.1 | 18335809 | 18335814 | 6 | * | 6 | 1000 |
| NC_039508.1 | 2590841 | 2594093 | 3253 | + | Gnomon | gene | NA | NA | gene-LOC105281341 | GeneID:105281341 | LOC105281341 | Gene | LOC105281341 | protein_coding | character(0) | NA | character(0) | NC_039508.1 | 2591612 | 2591617 | 6 | * | 6 | 1000 |
| NC_039508.1 | 10274002 | 10277440 | 3439 | + | Gnomon | gene | NA | NA | gene-LOC105276881 | GeneID:105276881 | LOC105276881 | Gene | LOC105276881 | protein_coding | character(0) | NA | character(0) | NC_039508.1 | 10275212 | 10275217 | 6 | * | 6 | 1000 |
| NC_039508.1 | 10799899 | 10803333 | 3435 | + | Gnomon | gene | NA | NA | gene-LOC105276950 | GeneID:105276950 | LOC105276950 | Gene | LOC105276950 | protein_coding | character(0) | NA | character(0) | NC_039508.1 | 10801049 | 10801054 | 6 | * | 6 | 1000 |
| NC_039508.1 | 17548373 | 17550348 | 1976 | - | Gnomon | gene | NA | NA | gene-LOC105284396 | GeneID:105284396 | LOC105284396 | Gene | LOC105284396 | protein_coding | character(0) | NA | character(0) | NC_039508.1 | 17549290 | 17549295 | 6 | * | 6 | 1000 |
| NC_039509.1 | 1002697 | 1006107 | 3411 | - | Gnomon | gene | NA | NA | gene-LOC105283620 | GeneID:105283620 | LOC105283620 | Gene | LOC105283620 | protein_coding | character(0) | NA | character(0) | NC_039509.1 | 1004827 | 1004832 | 6 | * | 6 | 1000 |
| NC_039509.1 | 1135464 | 1137481 | 2018 | - | Gnomon | gene | NA | NA | gene-LOC105283634 | GeneID:105283634 | LOC105283634 | Gene | LOC105283634 | protein_coding | character(0) | NA | character(0) | NC_039509.1 | 1136266 | 1136271 | 6 | * | 6 | 1000 |
| NC_039509.1 | 2825666 | 2828757 | 3092 | + | Gnomon | gene | NA | NA | gene-LOC105282589 | GeneID:105282589 | LOC105282589 | Gene | LOC105282589 | protein_coding | character(0) | NA | character(0) | NC_039509.1 | 2826537 | 2826542 | 6 | * | 6 | 1000 |
| NC_039509.1 | 10756577 | 10757875 | 1299 | - | Gnomon | gene | NA | NA | gene-LOC105275214 | GeneID:105275214 | LOC105275214 | Gene | LOC105275214 | protein_coding | character(0) | NA | character(0) | NC_039509.1 | 10756803 | 10756808 | 6 | * | 6 | 1000 |
| NC_039509.1 | 11068956 | 11086837 | 17882 | - | Gnomon | gene | NA | NA | gene-LOC105275770 | GeneID:105275770 | LOC105275770 | Gene | LOC105275770 | protein_coding | character(0) | NA | character(0) | NC_039509.1 | 11085457 | 11085462 | 6 | * | 6 | 1000 |
| NC_039509.1 | 13033738 | 13035966 | 2229 | + | Gnomon | gene | NA | NA | gene-LOC105278527 | GeneID:105278527 | LOC105278527 | Gene | LOC105278527 | protein_coding | character(0) | NA | character(0) | NC_039509.1 | 13034669 | 13034674 | 6 | * | 6 | 1000 |
| NC_039510.1 | 2607810 | 2610599 | 2790 | - | Gnomon | gene | NA | NA | gene-LOC105287754 | GeneID:105287754 | LOC105287754 | Gene | LOC105287754 | protein_coding | character(0) | NA | character(0) | NC_039510.1 | 2609179 | 2609184 | 6 | * | 6 | 1000 |
| NC_039510.1 | 3126910 | 3147548 | 20639 | - | Gnomon | gene | NA | NA | gene-LOC105287694 | GeneID:105287694 | LOC105287694 | Gene | LOC105287694 | protein_coding | character(0) | NA | character(0) | NC_039510.1 | 3133558 | 3133563 | 6 | * | 6 | 1000 |
| NC_039510.1 | 3132542 | 3134046 | 1505 | + | Gnomon | gene | NA | NA | gene-LOC105287692 | GeneID:105287692 | LOC105287692 | Gene | LOC105287692 | protein_coding | character(0) | NA | character(0) | NC_039510.1 | 3133558 | 3133563 | 6 | * | 6 | 1000 |
| NC_039510.1 | 6135149 | 6137854 | 2706 | + | Gnomon | gene | NA | NA | gene-LOC105287333 | GeneID:105287333 | LOC105287333 | Gene | LOC105287333 | protein_coding | character(0) | NA | character(0) | NC_039510.1 | 6136456 | 6136461 | 6 | * | 6 | 1000 |
| NC_039510.1 | 12000027 | 12025662 | 25636 | - | Gnomon | gene | NA | NA | gene-LOC105281066 | GeneID:105281066 | LOC105281066 | Gene | LOC105281066 | protein_coding | character(0) | NA | character(0) | NC_039510.1 | 12024411 | 12024416 | 6 | * | 6 | 1000 |
| NC_039510.1 | 12023589 | 12029249 | 5661 | + | Gnomon | gene | NA | NA | gene-LOC105281067 | GeneID:105281067 | LOC105281067 | Gene | LOC105281067 | protein_coding | character(0) | NA | character(0) | NC_039510.1 | 12024411 | 12024416 | 6 | * | 6 | 1000 |
| NC_039510.1 | 16287375 | 16291357 | 3983 | - | Gnomon | gene | NA | NA | gene-LOC105286509 | GeneID:105286509 | LOC105286509 | Gene | LOC105286509 | protein_coding | character(0) | NA | character(0) | NC_039510.1 | 16290598 | 16290603 | 6 | * | 6 | 1000 |

...
... only first 25 lines shown ...

Directory: MRPR File: gwr-Ob_LI_RphaseB1.txt

| seqnames | start | end | width | strand | source | type | score | phase | ID | Dbxref | Name | gbkey | gene | gene_biotype | end_range | partial | start_range | seqnames | start | end | width | strand | NbrSites | Sdnsty |
|---|---|---|---|---|---|---|---|---|---|---|---|---|---|---|---|---|---|---|---|---|---|---|---|---|
| NC_039506.1 | 3342881 | 3348391 | 5511 | + | Gnomon | gene | NA | NA | gene-LOC105280395 | GeneID:105280395 | LOC105280395 | Gene | LOC105280395 | protein_coding | character(0) | NA | character(0) | NC_039506.1 | 3344178 | 3344183 | 6 | * | 6 | 1000 |
| NC_039506.1 | 6089683 | 6090856 | 1174 | + | Gnomon | gene | NA | NA | gene-LOC105277804 | GeneID:105277804 | LOC105277804 | Gene | LOC105277804 | protein_coding | character(0) | NA | character(0) | NC_039506.1 | 6090624 | 6090629 | 6 | * | 6 | 1000 |
| NC_039506.1 | 24043149 | 24047806 | 4658 | + | Gnomon | gene | NA | NA | gene-LOC105279367 | GeneID:105279367 | LOC105279367 | Gene | LOC105279367 | protein_coding | character(0) | NA | character(0) | NC_039506.1 | 24044697 | 24044702 | 6 | * | 6 | 1000 |
| NC_039507.1 | 7181199 | 7188522 | 7324 | + | Gnomon | gene | NA | NA | gene-LOC105283383 | GeneID:105283383 | LOC105283383 | Gene | LOC105283383 | protein_coding | character(0) | NA | character(0) | NC_039507.1 | 7182290 | 7182295 | 6 | * | 6 | 1000 |
| NC_039507.1 | 7276866 | 7278961 | 2096 | + | Gnomon | gene | NA | NA | gene-LOC105283373 | GeneID:105283373 | LOC105283373 | Gene | LOC105283373 | protein_coding | character(0) | NA | character(0) | NC_039507.1 | 7277947 | 7277952 | 6 | * | 6 | 1000 |
| NC_039507.1 | 18333035 | 18336377 | 3343 | - | Gnomon | gene | NA | NA | gene-LOC105281577 | GeneID:105281577 | LOC105281577 | Gene | LOC105281577 | protein_coding | character(0) | NA | character(0) | NC_039507.1 | 18335809 | 18335814 | 6 | * | 6 | 1000 |
| NC_039508.1 | 2590841 | 2594093 | 3253 | + | Gnomon | gene | NA | NA | gene-LOC105281341 | GeneID:105281341 | LOC105281341 | Gene | LOC105281341 | protein_coding | character(0) | NA | character(0) | NC_039508.1 | 2591612 | 2591617 | 6 | * | 6 | 1000 |
| NC_039509.1 | 1135464 | 1137481 | 2018 | - | Gnomon | gene | NA | NA | gene-LOC105283634 | GeneID:105283634 | LOC105283634 | Gene | LOC105283634 | protein_coding | character(0) | NA | character(0) | NC_039509.1 | 1136266 | 1136271 | 6 | * | 6 | 1000 |
| NC_039509.1 | 2825666 | 2828757 | 3092 | + | Gnomon | gene | NA | NA | gene-LOC105282589 | GeneID:105282589 | LOC105282589 | Gene | LOC105282589 | protein_coding | character(0) | NA | character(0) | NC_039509.1 | 2826537 | 2826542 | 6 | * | 6 | 1000 |
| NC_039509.1 | 10756577 | 10757875 | 1299 | - | Gnomon | gene | NA | NA | gene-LOC105275214 | GeneID:105275214 | LOC105275214 | Gene | LOC105275214 | protein_coding | character(0) | NA | character(0) | NC_039509.1 | 10756803 | 10756808 | 6 | * | 6 | 1000 |
| NC_039509.1 | 11068956 | 11086837 | 17882 | - | Gnomon | gene | NA | NA | gene-LOC105275770 | GeneID:105275770 | LOC105275770 | Gene | LOC105275770 | protein_coding | character(0) | NA | character(0) | NC_039509.1 | 11085457 | 11085462 | 6 | * | 6 | 1000 |
| NC_039509.1 | 13033738 | 13035966 | 2229 | + | Gnomon | gene | NA | NA | gene-LOC105278527 | GeneID:105278527 | LOC105278527 | Gene | LOC105278527 | protein_coding | character(0) | NA | character(0) | NC_039509.1 | 13034669 | 13034674 | 6 | * | 6 | 1000 |
| NC_039510.1 | 2607810 | 2610599 | 2790 | - | Gnomon | gene | NA | NA | gene-LOC105287754 | GeneID:105287754 | LOC105287754 | Gene | LOC105287754 | protein_coding | character(0) | NA | character(0) | NC_039510.1 | 2609179 | 2609184 | 6 | * | 6 | 1000 |
| NC_039510.1 | 3319355 | 3322927 | 3573 | + | Gnomon | gene | NA | NA | gene-LOC105287663 | GeneID:105287663 | LOC105287663 | Gene | LOC105287663 | protein_coding | character(0) | NA | character(0) | NC_039510.1 | 3320255 | 3320260 | 6 | * | 6 | 1000 |
| NC_039510.1 | 16287375 | 16291357 | 3983 | - | Gnomon | gene | NA | NA | gene-LOC105286509 | GeneID:105286509 | LOC105286509 | Gene | LOC105286509 | protein_coding | character(0) | NA | character(0) | NC_039510.1 | 16290598 | 16290603 | 6 | * | 6 | 1000 |
| NC_039511.1 | 480353 | 488275 | 7923 | + | Gnomon | gene | NA | NA | gene-LOC105276527 | GeneID:105276527 | LOC105276527 | Gene | LOC105276527 | protein_coding | character(0) | NA | character(0) | NC_039511.1 | 481652 | 481657 | 6 | * | 6 | 1000 |
| NC_039511.1 | 2234830 | 2236789 | 1960 | - | Gnomon | gene | NA | NA | gene-LOC105276382 | GeneID:105276382 | LOC105276382 | Gene | LOC105276382 | protein_coding | character(0) | NA | character(0) | NC_039511.1 | 2236016 | 2236021 | 6 | * | 6 | 1000 |
| NC_039511.1 | 3614241 | 3616939 | 2699 | + | Gnomon | gene | NA | NA | gene-LOC105284057 | GeneID:105284057 | LOC105284057 | Gene | LOC105284057 | protein_coding | character(0) | NA | character(0) | NC_039511.1 | 3615462 | 3615467 | 6 | * | 6 | 1000 |
| NC_039511.1 | 8407553 | 8425032 | 17480 | + | Gnomon | gene | NA | NA | gene-LOC105284845 | GeneID:105284845 | LOC105284845 | Gene | LOC105284845 | protein_coding | character(0) | NA | character(0) | NC_039511.1 | 8409706 | 8409711 | 6 | * | 6 | 1000 |
| NC_039511.1 | 8695181 | 8698086 | 2906 | + | Gnomon | gene | NA | NA | gene-LOC105280856 | GeneID:105280856 | LOC105280856 | Gene | LOC105280856 | protein_coding | character(0) | NA | character(0) | NC_039511.1 | 8695844 | 8695849 | 6 | * | 6 | 1000 |
| NC_039512.1 | 10275466 | 10280187 | 4722 | + | Gnomon | gene | NA | NA | gene-LOC105282847 | GeneID:105282847 | LOC105282847 | Gene | LOC105282847 | protein_coding | character(0) | NA | character(0) | NC_039512.1 | 10276481 | 10276486 | 6 | * | 6 | 1000 |
| NC_039512.1 | 12092666 | 12097773 | 5108 | + | Gnomon | gene | NA | NA | gene-LOC105279779 | GeneID:105279779 | LOC105279779 | Gene | LOC105279779 | protein_coding | character(0) | NA | character(0) | NC_039512.1 | 12094200 | 12094205 | 6 | * | 6 | 1000 |
| NC_039512.1 | 15707994 | 15714445 | 6452 | + | Gnomon | gene | NA | NA | gene-LOC105274475 | GeneID:105274475 | LOC105274475 | Gene | LOC105274475 | protein_coding | character(0) | NA | character(0) | NC_039512.1 | 15711477 | 15711482 | 6 | * | 6 | 1000 |
| NC_039512.1 | 15979275 | 15981529 | 2255 | - | Gnomon | gene | NA | NA | gene-LOC105274775 | GeneID:105274775 | LOC105274775 | Gene | LOC105274775 | protein_coding | character(0) | NA | character(0) | NC_039512.1 | 15980634 | 15980639 | 6 | * | 6 | 1000 |
| NC_039513.1 | 10401827 | 10404590 | 2764 | - | Gnomon | gene | NA | NA | gene-LOC105275550 | GeneID:105275550 | LOC105275550 | Gene | LOC105275550 | protein_coding | character(0) | NA | character(0) | NC_039513.1 | 10403469 | 10403474 | 6 | * | 6 | 1000 |
| NC_039513.1 | 11029007 | 11033701 | 4695 | - | Gnomon | gene | NA | NA | gene-LOC105275763 | GeneID:105275763 | LOC105275763 | Gene | LOC105275763 | protein_coding | character(0) | NA | character(0) | NC_039513.1 | 11030126 | 11030131 | 6 | * | 6 | 1000 |
| NC_039513.1 | 12028675 | 12030941 | 2267 | - | Gnomon | gene | NA | NA | gene-LOC105280791 | GeneID:105280791 | LOC105280791 | Gene | LOC105280791 | protein_coding | character(0) | NA | character(0) | NC_039513.1 | 12029791 | 12029796 | 6 | * | 6 | 1000 |
| NC_039514.1 | 9336154 | 9343762 | 7609 | + | Gnomon | gene | NA | NA | gene-LOC105284644 | GeneID:105284644 | LOC105284644 | Gene | LOC105284644 | protein_coding | character(0) | NA | character(0) | NC_039514.1 | 9337043 | 9337048 | 6 | * | 6 | 1000 |
| NC_039515.1 | 7098785 | 7103508 | 4724 | - | Gnomon | gene | NA | NA | gene-LOC105286003 | GeneID:105286003 | LOC105286003 | Gene | LOC105286003 | protein_coding | character(0) | NA | character(0) | NC_039515.1 | 7102685 | 7102690 | 6 | * | 6 | 1000 |
| NC_039515.1 | 12582469 | 12585642 | 3174 | - | Gnomon | gene | NA | NA | gene-LOC105282150 | GeneID:105282150 | LOC105282150 | Gene | LOC105282150 | protein_coding | | | | | | | | | | |

| | | | | | | | | | | | | | | |
|---|---|---|---|---|---|---|---|---|---|---|---|---|---|---|
| character(0) | | NA | character(0) | | NC_039515.1 | 12584748 | | 12584753 | 6 | * | 6 | 1000 | | |
| NC_039515.1 | 12596735 | 12605489 | 8755 | - | Gnomon | gene | NA | NA | gene-LOC105282142 | GeneID:105282142 | LOC105282142 | Gene | LOC105282142 | protein_coding |
| character(0) | | NA | character(0) | | NC_039515.1 | 12604757 | | 12604762 | 6 | * | 6 | 1000 | | |
| NC_039515.1 | 12601969 | 12605663 | 3695 | + | Gnomon | gene | NA | NA | gene-LOC105282144 | GeneID:105282144 | LOC105282144 | Gene | LOC105282144 | protein_coding |
| character(0) | | NA | character(0) | | NC_039515.1 | 12604757 | | 12604762 | 6 | * | 6 | 1000 | | |
| NC_039516.1 | 7444132 | 7447943 | 3812 | - | Gnomon | gene | NA | NA | gene-LOC105275781 | GeneID:105275781 | LOC105275781 | Gene | LOC105275781 | protein_coding |
| character(0) | | NA | character(0) | | NC_039516.1 | 7446427 | | 7446432 | 6 | * | 6 | 1000 | | |
| NC_039516.1 | 10251143 | 10256122 | 4980 | - | Gnomon | gene | NA | NA | gene-LOC105287447 | GeneID:105287447 | LOC105287447 | Gene | LOC105287447 | protein_coding |
| character(0) | | NA | character(0) | | NC_039516.1 | 10254626 | | 10254631 | 6 | * | 6 | 1000 | | |
| NC_039516.1 | 12307603 | 12308894 | 1292 | + | Gnomon | gene | NA | NA | gene-LOC105275270 | GeneID:105275270 | LOC105275270 | Gene | LOC105275270 | protein_coding |
| character(0) | | NA | character(0) | | NC_039516.1 | 12308308 | | 12308313 | 6 | * | 6 | 1000 | | |
| NC_039516.1 | 12889538 | 12895772 | 6235 | + | Gnomon | gene | NA | NA | gene-LOC105275316 | GeneID:105275316 | LOC105275316 | Gene | LOC105275316 | protein_coding |
| character(0) | | NA | character(0) | | NC_039516.1 | 12890326 | | 12890331 | 6 | * | 6 | 1000 | | |
| NC_039516.1 | 12996090 | 13006535 | 10446 | + | Gnomon | gene | NA | NA | gene-LOC105275342 | GeneID:105275342 | LOC105275342 | Gene | LOC105275342 | protein_coding |
| character(0) | | NA | character(0) | | NC_039516.1 | 12997443 | | 12997448 | 6 | * | 6 | 1000 | | |
| NC_039516.1 | 13065193 | 13068389 | 3197 | + | Gnomon | gene | NA | NA | gene-LOC105275355 | GeneID:105275355 | LOC105275355 | Gene | LOC105275355 | protein_coding |
| character(0) | | NA | character(0) | | NC_039516.1 | 13066190 | | 13066195 | 6 | * | 6 | 1000 | | |
| NC_039517.1 | 1360424 | 1367346 | 6923 | + | Gnomon | gene | NA | NA | gene-LOC105278812 | GeneID:105278812 | LOC105278812 | Gene | LOC105278812 | protein_coding |
| character(0) | | NA | character(0) | | NC_039517.1 | 1361411 | | 1361416 | 6 | * | 6 | 1000 | | |
| NC_039517.1 | 2989996 | 2992325 | 2330 | + | Gnomon | gene | NA | NA | gene-LOC105281549 | GeneID:105281549 | LOC105281549 | Gene | LOC105281549 | protein_coding |
| character(0) | | NA | character(0) | | NC_039517.1 | 2990626 | | 2990631 | 6 | * | 6 | 1000 | | |
| NC_039517.1 | 8175046 | 8181714 | 6669 | - | Gnomon | gene | NA | NA | gene-LOC105284262 | GeneID:105284262 | LOC105284262 | Gene | LOC105284262 | protein_coding |
| character(0) | | NA | character(0) | | NC_039517.1 | 8175507 | | 8175512 | 6 | * | 6 | 1000 | | |
| NC_039518.1 | 5496870 | 5510429 | 13560 | - | Gnomon | gene | NA | NA | gene-LOC105281906 | GeneID:105281906 | LOC105281906 | Gene | LOC105281906 | protein_coding |
| character(0) | | NA | character(0) | | NC_039518.1 | 5508860 | | 5508865 | 6 | * | 6 | 1000 | | |
| NC_039518.1 | 8721748 | 8727127 | 5380 | + | Gnomon | gene | NA | NA | gene-LOC105286364 | GeneID:105286364 | LOC105286364 | Gene | LOC105286364 | protein_coding |
| character(0) | | NA | character(0) | | NC_039518.1 | 8722922 | | 8722927 | 6 | * | 6 | 1000 | | |
| NC_039506.1 | 23567682 | 23574351 | 6670 | + | Gnomon | gene | NA | NA | gene-LOC105279333 | GeneID:105279333 | LOC105279333 | Gene | LOC105279333 | protein_coding |
| character(0) | | NA | character(0) | | NC_039506.1 | 23568887 | | 23568897 | 11 | * | 10 | 909.090909090909 | | |
| NC_039513.1 | 13434303 | 13438575 | 4273 | - | Gnomon | gene | NA | NA | gene-LOC105275036 | GeneID:105275036 | LOC105275036 | Gene | LOC105275036 | protein_coding |
| character(0) | | NA | character(0) | | NC_039513.1 | 13436991 | | 13437000 | 10 | * | 9 | 900 | | |
| NC_039507.1 | 5492373 | 5502473 | 10101 | + | Gnomon | gene | NA | NA | gene-LOC105281645 | GeneID:105281645 | LOC105281645 | Gene | LOC105281645 | protein_coding |
| character(0) | | NA | character(0) | | NC_039507.1 | 5493751 | | 5493759 | 9 | * | 8 | 888.888888888889 | | |
| NC_039507.1 | 9161485 | 9166239 | 4755 | + | Gnomon | gene | NA | NA | gene-LOC105282461 | GeneID:105282461 | LOC105282461 | Gene | LOC105282461 | protein_coding |
| character(0) | | NA | character(0) | | NC_039507.1 | 9162374 | | 9162382 | 9 | * | 8 | 888.888888888889 | | |
| NC_039509.1 | 17207204 | 17209527 | 2324 | + | Gnomon | gene | NA | NA | gene-LOC105283016 | GeneID:105283016 | LOC105283016 | Gene | LOC105283016 | protein_coding |
| character(0) | | NA | character(0) | | NC_039509.1 | 17208071 | | 17208079 | 9 | * | 8 | 888.888888888889 | | |
| NC_039510.1 | 8568121 | 8569477 | 1357 | - | Gnomon | gene | NA | NA | gene-LOC105275137 | GeneID:105275137 | LOC105275137 | Gene | LOC105275137 | protein_coding |
| character(0) | | NA | character(0) | | NC_039510.1 | 8568914 | | 8568922 | 9 | * | 8 | 888.888888888889 | | |
| NC_039513.1 | 3303685 | 3306319 | 2635 | + | Gnomon | gene | NA | NA | gene-LOC105275761 | GeneID:105275761 | LOC105275761 | Gene | LOC105275761 | protein_coding |
| character(0) | | NA | character(0) | | NC_039513.1 | 3304806 | | 3304814 | 9 | * | 8 | 888.888888888889 | | |
| NC_039512.1 | 11140867 | 11151152 | 10286 | - | Gnomon | gene | NA | NA | gene-LOC105279854 | GeneID:105279854 | LOC105279854 | Gene | LOC105279854 | protein_coding |
| character(0) | | NA | character(0) | | NC_039512.1 | 11150038 | | 11150045 | 8 | * | 7 | 875 | | |
| NC_039517.1 | 9921720 | 9925663 | 3944 | + | Gnomon | gene | NA | NA | gene-LOC105287104 | GeneID:105287104 | LOC105287104 | Gene | LOC105287104 | protein_coding |
| character(0) | | NA | character(0) | | NC_039517.1 | 9922442 | | 9922449 | 8 | * | 7 | 875 | | |
| NC_039518.1 | 8474988 | 8476009 | 1022 | + | Gnomon | gene | NA | NA | gene-LOC105286212 | GeneID:105286212 | LOC105286212 | Gene | LOC105286212 | protein_coding |
| character(0) | | NA | character(0) | | NC_039518.1 | 8475762 | | 8475769 | 8 | * | 7 | 875 | | |
| NC_039506.1 | 1487779 | 1489683 | 1905 | + | Gnomon | gene | NA | NA | gene-LOC105280236 | GeneID:105280236 | LOC105280236 | Gene | LOC105280236 | protein_coding |
| character(0) | | NA | character(0) | | NC_039506.1 | 1488926 | | 1488932 | 7 | * | 6 | 857.142857142857 | | |
| NC_039506.1 | 2183297 | 2187485 | 4189 | - | Gnomon | gene | NA | NA | gene-LOC105280306 | GeneID:105280306 | LOC105280306 | Gene | LOC105280306 | protein_coding |
| character(0) | | NA | character(0) | | NC_039506.1 | 2186928 | | 2186934 | 7 | * | 6 | 857.142857142857 | | |
| NC_039506.1 | 2274108 | 2276090 | 1983 | - | Gnomon | gene | NA | NA | gene-LOC105280319 | GeneID:105280319 | LOC105280319 | Gene | LOC105280319 | protein_coding |
| character(0) | | NA | character(0) | | NC_039506.1 | 2275368 | | 2275374 | 7 | * | 6 | 857.142857142857 | | |
| NC_039506.1 | 11102693 | 11121131 | 18439 | - | Gnomon | gene | NA | NA | gene-LOC105284754 | GeneID:105284754 | LOC105284754 | Gene | LOC105284754 | protein_coding |
| character(0) | | NA | character(0) | | NC_039506.1 | 11119816 | | 11119822 | 7 | * | 6 | 857.142857142857 | | |
| NC_039506.1 | 15506447 | 15508913 | 2467 | - | Gnomon | gene | NA | NA | gene-LOC105283960 | GeneID:105283960 | LOC105283960 | Gene | LOC105283960 | protein_coding |
| character(0) | | NA | character(0) | | NC_039506.1 | 15508062 | | 15508068 | 7 | * | 6 | 857.142857142857 | | |
| NC_039506.1 | 15509387 | 15513034 | 3648 | + | Gnomon | gene | NA | NA | gene-LOC105283961 | GeneID:105283961 | LOC105283961 | Gene | LOC105283961 | protein_coding |
| character(0) | | NA | character(0) | | NC_039506.1 | 15510126 | | 15510132 | 7 | * | 6 | 857.142857142857 | | |
| NC_039506.1 | 19093260 | 19098014 | 4755 | - | Gnomon | gene | NA | NA | gene-LOC105278005 | GeneID:105278005 | LOC105278005 | Gene | LOC105278005 | protein_coding |
| character(0) | | NA | character(0) | | NC_039506.1 | 19096860 | | 19096866 | 7 | * | 6 | 857.142857142857 | | |
| NC_039506.1 | 20553263 | 20559098 | 5836 | - | Gnomon | gene | NA | NA | gene-LOC105278544 | GeneID:105278544 | LOC105278544 | Gene | LOC105278544 | protein_coding |
| character(0) | | NA | character(0) | | NC_039506.1 | 20557629 | | 20557635 | 7 | * | 6 | 857.142857142857 | | |
| NC_039506.1 | 22941360 | 22946330 | 4971 | - | Gnomon | gene | NA | NA | gene-LOC105283484 | GeneID:105283484 | LOC105283484 | Gene | LOC105283484 | protein_coding |

character(0) NA character(0) NC_039506.1 22945036 22945042 7 * 6 857.142857142857
NC_039506.1 23689672 23694320 4649 - Gnomon gene NA NA gene-LOC105279346 GeneID:105279346 LOC105279346 Gene LOC105279346 protein_coding
character(0) NA character(0) NC_039506.1 23693180 23693186 7 * 6 857.142857142857
NC_039507.1 124714 126752 2039 + Gnomon gene NA NA gene-LOC105279088 GeneID:105279088 LOC105279088 Gene LOC105279088 protein_coding
character(0) NA character(0) NC_039507.1 125663 125669 7 * 6 857.142857142857
NC_039507.1 790243 798925 8683 + Gnomon gene NA NA gene-LOC105282921 GeneID:105282921 LOC105282921 Gene LOC105282921 protein_coding
character(0) NA character(0) NC_039507.1 791692 791698 7 * 6 857.142857142857
NC_039507.1 6347235 6350936 3702 + Gnomon gene NA NA gene-LOC105275401 GeneID:105275401 LOC105275401 Gene LOC105275401 protein_coding
character(0) NA character(0) NC_039507.1 6348012 6348018 7 * 6 857.142857142857
NC_039507.1 15481857 15484141 2285 + Gnomon gene NA NA gene-LOC105280092 GeneID:105280092 LOC105280092 Gene LOC105280092 protein_coding
character(0) NA character(0) NC_039507.1 15482876 15482882 7 * 6 857.142857142857
NC_039507.1 15506083 15508702 2620 + Gnomon gene NA NA gene-LOC105280102 GeneID:105280102 LOC105280102 Gene LOC105280102 protein_coding
character(0) NA character(0) NC_039507.1 15507322 15507328 7 * 6 857.142857142857
NC_039507.1 17006579 17010876 4298 + Gnomon gene NA NA gene-LOC105278157 GeneID:105278157 LOC105278157 Gene LOC105278157 protein_coding
character(0) NA character(0) NC_039507.1 17008850 17008856 7 * 6 857.142857142857
NC_039507.1 18333035 18336377 3343 - Gnomon gene NA NA gene-LOC105281577 GeneID:105281577 LOC105281577 Gene LOC105281577 protein_coding
character(0) NA character(0) NC_039507.1 18335506 18335512 7 * 6 857.142857142857
NC_039508.1 192252 200875 8624 + Gnomon gene NA NA gene-LOC105277452 GeneID:105277452 LOC105277452 Gene LOC105277452 protein_coding
character(0) NA character(0) NC_039508.1 193513 193519 7 * 6 857.142857142857
NC_039508.1 1384214 1386820 2607 + Gnomon gene NA NA gene-LOC105277388 GeneID:105277388 LOC105277388 Gene LOC105277388 protein_coding
character(0) NA character(0) NC_039508.1 1384932 1384938 7 * 6 857.142857142857
NC_039508.1 1386723 1389150 2428 - Gnomon gene NA NA gene-LOC105277389 GeneID:105277389 LOC105277389 Gene LOC105277389 protein_coding
character(0) NA character(0) NC_039508.1 1387947 1387953 7 * 6 857.142857142857
NC_039508.1 3996551 4002196 5646 - Gnomon gene NA NA gene-LOC105285719 GeneID:105285719 LOC105285719 Gene LOC105285719 protein_coding
character(0) NA character(0) NC_039508.1 4001313 4001319 7 * 6 857.142857142857
NC_039508.1 5353027 5357392 4366 - Gnomon gene NA NA gene-LOC105277468 GeneID:105277468 LOC105277468 Gene LOC105277468 protein_coding
character(0) NA character(0) NC_039508.1 5356681 5356687 7 * 6 857.142857142857
NC_039508.1 9041025 9045259 4235 + Gnomon gene NA NA gene-LOC105278106 GeneID:105278106 LOC105278106 Gene LOC105278106 protein_coding
character(0) NA character(0) NC_039508.1 9042110 9042116 7 * 6 857.142857142857
NC_039508.1 10763110 10766843 3734 + Gnomon gene NA NA gene-LOC105276948 GeneID:105276948 LOC105276948 Gene LOC105276948 protein_coding
character(0) NA character(0) NC_039508.1 10764758 10764764 7 * 6 857.142857142857
NC_039508.1 10797665 10799797 2133 + Gnomon gene NA NA gene-LOC105276949 GeneID:105276949 LOC105276949 Gene LOC105276949 protein_coding
character(0) NA character(0) NC_039508.1 10798605 10798611 7 * 6 857.142857142857
NC_039508.1 15763225 15782851 19627 + Gnomon gene NA NA gene-LOC105277234 GeneID:105277234 LOC105277234 Gene LOC105277234 protein_coding
character(0) NA character(0) NC_039508.1 15764452 15764458 7 * 6 857.142857142857
NC_039508.1 17368821 17424169 55349 + Gnomon gene NA NA gene-LOC105284404 GeneID:105284404 LOC105284404 Gene LOC105284404 protein_coding
character(0) NA character(0) NC_039508.1 17370215 17370221 7 * 6 857.142857142857
NC_039508.1 17548373 17550348 1976 - Gnomon gene NA NA gene-LOC105284396 GeneID:105284396 LOC105284396 Gene LOC105284396 protein_coding
character(0) NA character(0) NC_039508.1 17549403 17549409 7 * 6 857.142857142857
NC_039509.1 89280 90451 1172 + Gnomon gene NA NA gene-LOC113561814 GeneID:113561814 LOC113561814 Gene LOC113561814 protein_coding character(0) NA
character(0) NC_039509.1 90329 90335 7 * 6 857.142857142857
NC_039509.1 2365507 2377666 12160 + Gnomon gene NA NA gene-LOC105285865 GeneID:105285865 LOC105285865 Gene LOC105285865 protein_coding
character(0) NA character(0) NC_039509.1 2366218 2366224 7 * 6 857.142857142857
NC_039509.1 5250307 5257914 7608 - Gnomon gene NA NA gene-LOC105287147 GeneID:105287147 LOC105287147 Gene LOC105287147 protein_coding
character(0) NA character(0) NC_039509.1 5256794 5256800 7 * 6 857.142857142857
NC_039509.1 7092641 7097742 5102 + Gnomon gene NA NA gene-LOC105282559 GeneID:105282559 LOC105282559 Gene LOC105282559 protein_coding
character(0) NA character(0) NC_039509.1 7093578 7093584 7 * 6 857.142857142857
NC_039509.1 7141680 7152824 11145 - Gnomon gene NA NA gene-LOC105282568 GeneID:105282568 LOC105282568 Gene LOC105282568 protein_coding
character(0) NA character(0) NC_039509.1 7145645 7145651 7 * 6 857.142857142857
NC_039509.1 10232323 10244464 12142 + Gnomon gene NA NA gene-LOC105282655 GeneID:105282655 LOC105282655 Gene LOC105282655 protein_coding
character(0) NA character(0) NC_039509.1 10233310 10233316 7 * 6 857.142857142857
NC_039509.1 10710016 10711124 1109 + Gnomon gene NA NA gene-LOC105275153 GeneID:105275153 LOC105275153 Gene LOC105275153 protein_coding
character(0) NA character(0) NC_039509.1 10710730 10710736 7 * 6 857.142857142857
NC_039509.1 13909149 13915606 6458 + Gnomon gene NA NA gene-LOC105278288 GeneID:105278288 LOC105278288 Gene LOC105278288 protein_coding
character(0) NA character(0) NC_039509.1 13909987 13909993 7 * 6 857.142857142857
NC_039510.1 624625 628259 3635 + Gnomon gene NA NA gene-LOC105279313 GeneID:105279313 LOC105279313 Gene LOC105279313 protein_coding
character(0) NA character(0) NC_039510.1 625494 625500 7 * 6 857.142857142857
NC_039510.1 3040660 3042070 1411 + Gnomon gene NA NA gene-LOC105287710 GeneID:105287710 LOC105287710 Gene LOC105287710 protein_coding
character(0) NA character(0) NC_039510.1 3041562 3041568 7 * 6 857.142857142857
NC_039510.1 3110492 3117379 6888 - Gnomon gene NA NA gene-LOC105287697 GeneID:105287697 LOC105287697 Gene LOC105287697 protein_coding
character(0) NA character(0) NC_039510.1 3116602 3116608 7 * 6 857.142857142857
NC_039510.1 3167251 3170185 2935 - Gnomon gene NA NA gene-LOC105287684 GeneID:105287684 LOC105287684 Gene LOC105287684 protein_coding
character(0) NA character(0) NC_039510.1 3169234 3169240 7 * 6 857.142857142857
NC_039510.1 4220534 4227851 7318 + Gnomon gene NA NA gene-LOC105287620 GeneID:105287620 LOC105287620 Gene LOC105287620 protein_coding

character(0) NA character(0) NC_039510.1 4221389 4221395 7 * 6 857.142857142857
NC_039510.1 5067676 5072600 4925 - Gnomon gene NA NA gene-LOC105287597 GeneID:105287597 LOC105287597 Gene LOC105287597 protein_coding
character(0) NA character(0) NC_039510.1 5071200 5071206 7 * 6 857.142857142857
NC_039510.1 5537223 5539036 1814 - Gnomon gene NA NA gene-LOC105287569 GeneID:105287569 LOC105287569 Gene LOC105287569 protein_coding
character(0) NA character(0) NC_039510.1 5538139 5538145 7 * 6 857.142857142857
NC_039510.1 6038459 6040963 2505 + Gnomon gene NA NA gene-LOC105287185 GeneID:105287185 LOC105287185 Gene LOC105287185 protein_coding
character(0) NA character(0) NC_039510.1 6040036 6040042 7 * 6 857.142857142857
NC_039510.1 6122701 6125649 2949 - Gnomon gene NA NA gene-LOC105287279 GeneID:105287279 LOC105287279 Gene LOC105287279 protein_coding
character(0) NA character(0) NC_039510.1 6124458 6124464 7 * 6 857.142857142857
NC_039510.1 6163214 6171437 8224 - Gnomon gene NA NA gene-LOC105287371 GeneID:105287371 LOC105287371 Gene LOC105287371 protein_coding
character(0) NA character(0) NC_039510.1 6170155 6170161 7 * 6 857.142857142857
NC_039510.1 6212141 6215222 3082 + Gnomon gene NA NA gene-LOC105287473 GeneID:105287473 LOC105287473 Gene LOC105287473 protein_coding
character(0) NA character(0) NC_039510.1 6213095 6213101 7 * 6 857.142857142857
NC_039510.1 6269006 6275115 6110 + Gnomon gene NA NA gene-LOC105287511 GeneID:105287511 LOC105287511 Gene LOC105287511 protein_coding
character(0) NA character(0) NC_039510.1 6270460 6270466 7 * 6 857.142857142857
NC_039510.1 8487238 8489293 2056 - Gnomon gene NA NA gene-LOC105275119 GeneID:105275119 LOC105275119 Gene LOC105275119 misc_RNA character(0)
NA character(0) NC_039510.1 8488481 8488487 7 * 6 857.142857142857
NC_039510.1 9996113 9998708 2596 - Gnomon gene NA NA gene-LOC105281227 GeneID:105281227 LOC105281227 Gene LOC105281227 protein_coding
character(0) NA character(0) NC_039510.1 9997644 9997650 7 * 6 857.142857142857
NC_039510.1 13531247 13535548 4302 - Gnomon gene NA NA gene-LOC105280964 GeneID:105280964 LOC105280964 Gene LOC105280964 protein_coding
character(0) NA character(0) NC_039510.1 13533348 13533354 7 * 6 857.142857142857
NC_039510.1 15582075 15587881 5807 - Gnomon gene NA NA gene-LOC105286461 GeneID:105286461 LOC105286461 Gene LOC105286461 protein_coding
character(0) NA character(0) NC_039510.1 15586562 15586568 7 * 6 857.142857142857
NC_039510.1 15745071 15751836 6766 + Gnomon gene NA NA gene-LOC105286478 GeneID:105286478 LOC105286478 Gene LOC105286478 protein_coding
character(0) NA character(0) NC_039510.1 15746500 15746506 7 * 6 857.142857142857
NC_039510.1 16500898 16505659 4762 - Gnomon gene NA NA gene-LOC105286526 GeneID:105286526 LOC105286526 Gene LOC105286526 protein_coding
character(0) NA character(0) NC_039510.1 16504460 16504466 7 * 6 857.142857142857
NC_039511.1 517195 530396 13202 + Gnomon gene NA NA gene-LOC105276519 GeneID:105276519 LOC105276519 Gene LOC105276519 protein_coding
character(0) NA character(0) NC_039511.1 518659 518665 7 * 6 857.142857142857
NC_039511.1 1662842 1668395 5554 + Gnomon gene NA NA gene-LOC105276437 GeneID:105276437 LOC105276437 Gene LOC105276437 protein_coding
character(0) NA character(0) NC_039511.1 1664306 1664312 7 * 6 857.142857142857
NC_039511.1 1861744 1869078 7335 + Gnomon gene NA NA gene-LOC105276417 GeneID:105276417 LOC105276417 Gene LOC105276417 protein_coding
character(0) NA character(0) NC_039511.1 1867899 1867905 7 * 6 857.142857142857
NC_039511.1 2004889 2015973 11085 - Gnomon gene NA NA gene-LOC105276410 GeneID:105276410 LOC105276410 Gene LOC105276410 protein_coding
character(0) NA character(0) NC_039511.1 2014683 2014689 7 * 6 857.142857142857
NC_039511.1 2004889 2015973 11085 - Gnomon gene NA NA gene-LOC105276410 GeneID:105276410 LOC105276410 Gene LOC105276410 protein_coding
character(0) NA character(0) NC_039511.1 2014740 2014746 7 * 6 857.142857142857
NC_039511.1 3034592 3043570 8979 + Gnomon gene NA NA gene-LOC105284096 GeneID:105284096 LOC105284096 Gene LOC105284096 protein_coding
character(0) NA character(0) NC_039511.1 3035910 3035916 7 * 6 857.142857142857
NC_039511.1 3635459 3642537 7079 - Gnomon gene NA NA gene-LOC105284052 GeneID:105284052 LOC105284052 Gene LOC105284052 protein_coding
character(0) NA character(0) NC_039511.1 3639538 3639544 7 * 6 857.142857142857
NC_039511.1 4554146 4561340 7195 - Gnomon gene NA NA gene-LOC105281750 GeneID:105281750 LOC105281750 Gene LOC105281750 protein_coding
character(0) NA character(0) NC_039511.1 4560502 4560508 7 * 6 857.142857142857
NC_039511.1 12403841 12413518 9678 - Gnomon gene NA NA gene-LOC105285135 GeneID:105285135 LOC105285135 Gene LOC105285135 protein_coding
character(0) NA character(0) NC_039511.1 12411122 12411128 7 * 6 857.142857142857
NC_039511.1 14704311 14706938 2628 - Gnomon gene NA NA gene-LOC105285889 GeneID:105285889 LOC105285889 Gene LOC105285889 protein_coding
character(0) NA character(0) NC_039511.1 14705900 14705906 7 * 6 857.142857142857
NC_039511.1 15983428 15985754 2327 + Gnomon gene NA NA gene-LOC105284576 GeneID:105284576 LOC105284576 Gene LOC105284576 protein_coding
character(0) NA character(0) NC_039511.1 15984100 15984106 7 * 6 857.142857142857
NC_039511.1 16541849 16544005 2157 + Gnomon gene NA NA gene-LOC105284502 GeneID:105284502 LOC105284502 Gene LOC105284502 protein_coding
character(0) NA character(0) NC_039511.1 16542944 16542950 7 * 6 857.142857142857
NC_039512.1 4385041 4390283 5243 - Gnomon gene NA NA gene-LOC105277552 GeneID:105277552 LOC105277552 Gene LOC105277552 protein_coding
character(0) NA character(0) NC_039512.1 4389120 4389126 7 * 6 857.142857142857
NC_039512.1 11140867 11151152 10286 - Gnomon gene NA NA gene-LOC105279854 GeneID:105279854 LOC105279854 Gene LOC105279854 protein_coding
character(0) NA character(0) NC_039512.1 11150305 11150311 7 * 6 857.142857142857
NC_039512.1 12621899 12625222 3324 + Gnomon gene NA NA gene-LOC105279740 GeneID:105279740 LOC105279740 Gene LOC105279740 protein_coding
character(0) NA character(0) NC_039512.1 12623117 12623123 7 * 6 857.142857142857
NC_039512.1 12739826 12742653 2828 + Gnomon gene NA NA gene-LOC105279719 GeneID:105279719 LOC105279719 Gene LOC105279719 protein_coding
character(0) NA character(0) NC_039512.1 12740699 12740705 7 * 6 857.142857142857
NC_039512.1 12814910 12819293 4384 - Gnomon gene NA NA gene-LOC105279709 GeneID:105279709 LOC105279709 Gene LOC105279709 protein_coding
character(0) NA character(0) NC_039512.1 12818052 12818058 7 * 6 857.142857142857
NC_039512.1 13089540 13093562 4023 - Gnomon gene NA NA gene-LOC105279664 GeneID:105279664 LOC105279664 Gene LOC105279664 protein_coding
character(0) NA character(0) NC_039512.1 13092263 13092269 7 * 6 857.142857142857
NC_039512.1 14720220 14725286 5067 + Gnomon gene NA NA gene-LOC105287687 GeneID:105287687 LOC105287687 Gene LOC105287687 protein_coding

character(0) NA character(0) NC_039512.1 14721247 14721253 7 * 6 857.142857142857
NC_039512.1 16555603 16557649 2047 - Gnomon gene NA NA gene-LOC105282841 GeneID:105282841 LOC105282841 Gene LOC105282841 protein_coding
character(0) NA character(0) NC_039512.1 16556845 16556851 7 * 6 857.142857142857
NC_039513.1 3253321 3257474 4154 - Gnomon gene NA NA gene-LOC105275753 GeneID:105275753 LOC105275753 Gene LOC105275753 protein_coding
character(0) NA character(0) NC_039513.1 3256149 3256155 7 * 6 857.142857142857
NC_039513.1 3253321 3257474 4154 - Gnomon gene NA NA gene-LOC105275753 GeneID:105275753 LOC105275753 Gene LOC105275753 protein_coding
character(0) NA character(0) NC_039513.1 3256358 3256364 7 * 6 857.142857142857
NC_039513.1 3257374 3260835 3462 - Gnomon gene NA NA gene-LOC105275752 GeneID:105275752 LOC105275752 Gene LOC105275752 protein_coding
character(0) NA character(0) NC_039513.1 3259768 3259774 7 * 6 857.142857142857
NC_039513.1 6784235 6788101 3867 - Gnomon gene NA NA gene-LOC105287855 GeneID:105287855 LOC105287855 Gene LOC105287855 protein_coding
character(0) NA character(0) NC_039513.1 6787239 6787245 7 * 6 857.142857142857
NC_039513.1 8483464 8491634 8171 + Gnomon gene NA NA gene-LOC105275689 GeneID:105275689 LOC105275689 Gene LOC105275689 protein_coding
character(0) NA character(0) NC_039513.1 8484406 8484412 7 * 6 857.142857142857
NC_039513.1 9603616 9607153 3538 + Gnomon gene NA NA gene-LOC105275615 GeneID:105275615 LOC105275615 Gene LOC105275615 protein_coding
character(0) NA character(0) NC_039513.1 9605113 9605119 7 * 6 857.142857142857
NC_039513.1 9603616 9607153 3538 + Gnomon gene NA NA gene-LOC105275615 GeneID:105275615 LOC105275615 Gene LOC105275615 protein_coding
character(0) NA character(0) NC_039513.1 9605414 9605420 7 * 6 857.142857142857
NC_039513.1 11674366 11676356 1991 - Gnomon gene NA NA gene-LOC105280739 GeneID:105280739 LOC105280739 Gene LOC105280739 protein_coding
character(0) NA character(0) NC_039513.1 11675069 11675075 7 * 6 857.142857142857
NC_039513.1 11674366 11676356 1991 - Gnomon gene NA NA gene-LOC105280739 GeneID:105280739 LOC105280739 Gene LOC105280739 protein_coding
character(0) NA character(0) NC_039513.1 11675195 11675201 7 * 6 857.142857142857
NC_039513.1 12032461 12036421 3961 + Gnomon gene NA NA gene-LOC105280793 GeneID:105280793 LOC105280793 Gene LOC105280793 protein_coding
character(0) NA character(0) NC_039513.1 12033291 12033297 7 * 6 857.142857142857
NC_039513.1 13452183 13456265 4083 + Gnomon gene NA NA gene-LOC105275031 GeneID:105275031 LOC105275031 Gene LOC105275031 protein_coding
character(0) NA character(0) NC_039513.1 13453683 13453689 7 * 6 857.142857142857
NC_039513.1 15035458 15043240 7783 + Gnomon gene NA NA gene-LOC105285648 GeneID:105285648 LOC105285648 Gene LOC105285648 protein_coding
character(0) NA character(0) NC_039513.1 15037033 15037039 7 * 6 857.142857142857
NC_039514.1 1286674 1297576 10903 - Gnomon gene NA NA gene-LOC105276233 GeneID:105276233 LOC105276233 Gene LOC105276233 protein_coding
character(0) NA character(0) NC_039514.1 1295968 1295974 7 * 6 857.142857142857
NC_039514.1 6551602 6556437 4836 - Gnomon gene NA NA gene-LOC105285546 GeneID:105285546 LOC105285546 Gene LOC105285546 protein_coding
character(0) NA character(0) NC_039514.1 6555389 6555395 7 * 6 857.142857142857
NC_039514.1 8629561 8648221 18661 + Gnomon gene NA NA gene-LOC105278994 GeneID:105278994 LOC105278994 Gene LOC105278994 protein_coding
character(0) NA character(0) NC_039514.1 8630645 8630651 7 * 6 857.142857142857
NC_039514.1 8629561 8648221 18661 + Gnomon gene NA NA gene-LOC105278994 GeneID:105278994 LOC105278994 Gene LOC105278994 protein_coding
character(0) NA character(0) NC_039514.1 8630804 8630810 7 * 6 857.142857142857
NC_039514.1 8729983 8735648 5666 + Gnomon gene NA NA gene-LOC105279000 GeneID:105279000 LOC105279000 Gene LOC105279000 protein_coding
character(0) NA character(0) NC_039514.1 8731123 8731129 7 * 6 857.142857142857
NC_039514.1 9413016 9415003 1988 - Gnomon gene NA NA gene-LOC105284659 GeneID:105284659 LOC105284659 Gene LOC105284659 protein_coding
character(0) NA character(0) NC_039514.1 9414116 9414122 7 * 6 857.142857142857
NC_039514.1 11109536 11117116 7581 - Gnomon gene NA NA gene-LOC105281601 GeneID:105281601 LOC105281601 Gene LOC105281601 protein_coding
character(0) NA character(0) NC_039514.1 11113139 11113145 7 * 6 857.142857142857
NC_039514.1 11111779 11115572 3794 + Gnomon gene NA NA gene-LOC105281603 GeneID:105281603 LOC105281603 Gene LOC105281603 protein_coding
character(0) NA character(0) NC_039514.1 11113139 11113145 7 * 6 857.142857142857
NC_039514.1 14398821 14400325 1505 - Gnomon gene NA NA gene-LOC105275991 GeneID:105275991 LOC105275991 Gene LOC105275991 protein_coding
character(0) NA character(0) NC_039514.1 14399289 14399295 7 * 6 857.142857142857
NC_039515.1 1472003 1475695 3693 - Gnomon gene NA NA gene-LOC105274705 GeneID:105274705 LOC105274705 Gene LOC105274705 protein_coding
character(0) NA character(0) NC_039515.1 1475067 1475073 7 * 6 857.142857142857
NC_039515.1 1527980 1529634 1655 + Gnomon gene NA NA gene-LOC105274711 GeneID:105274711 LOC105274711 Gene LOC105274711 protein_coding
character(0) NA character(0) NC_039515.1 1528293 1528299 7 * 6 857.142857142857
NC_039515.1 2215137 2216986 1850 - Gnomon gene NA NA gene-LOC105278431 GeneID:105278431 LOC105278431 Gene LOC105278431 protein_coding
character(0) NA character(0) NC_039515.1 2215927 2215933 7 * 6 857.142857142857
NC_039515.1 3498019 3503642 5624 + Gnomon gene NA NA gene-LOC105278309 GeneID:105278309 LOC105278309 Gene LOC105278309 protein_coding
character(0) NA character(0) NC_039515.1 3499095 3499101 7 * 6 857.142857142857
NC_039515.1 6014495 6167113 152619 + Gnomon gene NA NA gene-LOC105286194 GeneID:105286194 LOC105286194 Gene LOC105286194 protein_coding
character(0) NA character(0) NC_039515.1 6066909 6066915 7 * 6 857.142857142857
NC_039515.1 6065547 6067835 2289 + Gnomon gene NA NA gene-LOC105285808 GeneID:105285808 LOC105285808 Gene LOC105285808 protein_coding
character(0) NA character(0) NC_039515.1 6066909 6066915 7 * 6 857.142857142857
NC_039515.1 6604872 6605723 852 + Gnomon gene NA NA gene-LOC105282280 GeneID:105282280 LOC105282280 Gene LOC105282280 protein_coding
character(0) NA character(0) NC_039515.1 6605539 6605545 7 * 6 857.142857142857
NC_039515.1 6615586 6618213 2628 - Gnomon gene NA NA gene-LOC105282278 GeneID:105282278 LOC105282278 Gene LOC105282278 protein_coding
character(0) NA character(0) NC_039515.1 6617273 6617279 7 * 6 857.142857142857
NC_039515.1 7926262 7929466 3205 + Gnomon gene NA NA gene-LOC105283741 GeneID:105283741 LOC105283741 Gene LOC105283741 protein_coding
character(0) NA character(0) NC_039515.1 7927531 7927537 7 * 6 857.142857142857
NC_039515.1 8973009 8978309 5301 + Gnomon gene NA NA gene-LOC105283697 GeneID:105283697 LOC105283697 Gene LOC105283697 protein_coding

| | | | | | | | | | | | | | | |
|---|---|---|---|---|---|---|---|---|---|---|---|---|---|---|
| character(0) | NA | character(0) | NC_039515.1 | 8974548 | 8974554 | 7 | * | 6 | 857.142857142857 | | | | | |
| NC_039515.1 | 12611158 | 12612372 | 1215 | - | Gnomon | gene | NA | NA | gene-LOC105282147 | GeneID:105282147 | LOC105282147 | Gene | LOC105282147 | protein_coding |
| character(0) | NA | character(0) | NC_039515.1 | 12611603 | 12611609 | 7 | * | 6 | 857.142857142857 | | | | | |
| NC_039516.1 | 756663 | 760199 | 3537 | + | Gnomon | gene | NA | NA | gene-LOC105274638 | GeneID:105274638 | LOC105274638 | Gene | LOC105274638 | protein_coding |
| character(0) | NA | character(0) | NC_039516.1 | 757725 | 757731 | 7 | * | 6 | 857.142857142857 | | | | | |
| NC_039516.1 | 985951 | 990466 | 4516 | + | Gnomon | gene | NA | NA | gene-LOC105274590 | GeneID:105274590 | LOC105274590 | Gene | LOC105274590 | protein_coding |
| character(0) | NA | character(0) | NC_039516.1 | 987063 | 987069 | 7 | * | 6 | 857.142857142857 | | | | | |
| NC_039516.1 | 985951 | 990466 | 4516 | + | Gnomon | gene | NA | NA | gene-LOC105274590 | GeneID:105274590 | LOC105274590 | Gene | LOC105274590 | protein_coding |
| character(0) | NA | character(0) | NC_039516.1 | 987547 | 987553 | 7 | * | 6 | 857.142857142857 | | | | | |
| NC_039516.1 | 1549696 | 1551173 | 1478 | + | Gnomon | gene | NA | NA | gene-LOC105274497 | GeneID:105274497 | LOC105274497 | Gene | LOC105274497 | protein_coding |
| character(0) | NA | character(0) | NC_039516.1 | 1550744 | 1550750 | 7 | * | 6 | 857.142857142857 | | | | | |
| NC_039516.1 | 4713084 | 4714767 | 1684 | - | Gnomon | gene | NA | NA | gene-LOC105275897 | GeneID:105275897 | LOC105275897 | Gene | LOC105275897 | protein_coding |
| character(0) | NA | character(0) | NC_039516.1 | 4713795 | 4713801 | 7 | * | 6 | 857.142857142857 | | | | | |
| NC_039516.1 | 7248249 | 7252272 | 4024 | + | Gnomon | gene | NA | NA | gene-LOC105275812 | GeneID:105275812 | LOC105275812 | Gene | LOC105275812 | protein_coding |
| character(0) | NA | character(0) | NC_039516.1 | 7251402 | 7251408 | 7 | * | 6 | 857.142857142857 | | | | | |
| NC_039516.1 | 12834895 | 12836314 | 1420 | - | Gnomon | gene | NA | NA | gene-LOC105275307 | GeneID:105275307 | LOC105275307 | Gene | LOC105275307 | protein_coding |
| character(0) | NA | character(0) | NC_039516.1 | 12835552 | 12835558 | 7 | * | 6 | 857.142857142857 | | | | | |
| NC_039516.1 | 13386687 | 13390618 | 3932 | + | Gnomon | gene | NA | NA | gene-LOC105278022 | GeneID:105278022 | LOC105278022 | Gene | LOC105278022 | protein_coding |
| character(0) | NA | character(0) | NC_039516.1 | 13387740 | 13387746 | 7 | * | 6 | 857.142857142857 | | | | | |
| NC_039516.1 | 13386687 | 13390618 | 3932 | + | Gnomon | gene | NA | NA | gene-LOC105278022 | GeneID:105278022 | LOC105278022 | Gene | LOC105278022 | protein_coding |
| character(0) | NA | character(0) | NC_039516.1 | 13387911 | 13387917 | 7 | * | 6 | 857.142857142857 | | | | | |
| NC_039516.1 | 13497458 | 13503461 | 6004 | - | Gnomon | gene | NA | NA | gene-LOC105277915 | GeneID:105277915 | LOC105277915 | Gene | LOC105277915 | protein_coding |
| character(0) | NA | character(0) | NC_039516.1 | 13502711 | 13502717 | 7 | * | 6 | 857.142857142857 | | | | | |
| NC_039516.1 | 13518014 | 13520692 | 2679 | + | Gnomon | gene | NA | NA | gene-LOC105277817 | GeneID:105277817 | LOC105277817 | Gene | LOC105277817 | protein_coding |
| character(0) | NA | character(0) | NC_039516.1 | 13518940 | 13518946 | 7 | * | 6 | 857.142857142857 | | | | | |
| NC_039517.1 | 867927 | 869156 | 1230 | - | Gnomon | gene | NA | NA | gene-LOC105278824 | GeneID:105278824 | LOC105278824 | Gene | LOC105278824 | protein_coding |
| character(0) | NA | character(0) | NC_039517.1 | 868363 | 868369 | 7 | * | 6 | 857.142857142857 | | | | | |
| NC_039517.1 | 2307792 | 2312395 | 4604 | - | Gnomon | gene | NA | NA | gene-LOC105281943 | GeneID:105281943 | LOC105281943 | Gene | LOC105281943 | protein_coding |
| character(0) | NA | character(0) | NC_039517.1 | 2309012 | 2309018 | 7 | * | 6 | 857.142857142857 | | | | | |
| NC_039517.1 | 2393025 | 2396928 | 3904 | - | Gnomon | gene | NA | NA | gene-LOC105281949 | GeneID:105281949 | LOC105281949 | Gene | LOC105281949 | protein_coding |
| character(0) | NA | character(0) | NC_039517.1 | 2395845 | 2395851 | 7 | * | 6 | 857.142857142857 | | | | | |
| NC_039517.1 | 3018758 | 3022375 | 3618 | + | Gnomon | gene | NA | NA | gene-LOC105281545 | GeneID:105281545 | LOC105281545 | Gene | LOC105281545 | protein_coding |
| character(0) | NA | character(0) | NC_039517.1 | 3019985 | 3019991 | 7 | * | 6 | 857.142857142857 | | | | | |
| NC_039517.1 | 7306074 | 7309288 | 3215 | + | Gnomon | gene | NA | NA | gene-LOC105284202 | GeneID:105284202 | LOC105284202 | Gene | LOC105284202 | protein_coding |
| character(0) | NA | character(0) | NC_039517.1 | 7307471 | 7307477 | 7 | * | 6 | 857.142857142857 | | | | | |
| NC_039518.1 | 2232165 | 2249269 | 17105 | + | Gnomon | gene | NA | NA | gene-LOC105283878 | GeneID:105283878 | LOC105283878 | Gene | LOC105283878 | protein_coding |
| character(0) | NA | character(0) | NC_039518.1 | 2233357 | 2233363 | 7 | * | 6 | 857.142857142857 | | | | | |
| NC_039518.1 | 2657034 | 2658634 | 1601 | + | Gnomon | gene | NA | NA | gene-LOC105285392 | GeneID:105285392 | LOC105285392 | Gene | LOC105285392 | protein_coding |
| character(0) | NA | character(0) | NC_039518.1 | 2658047 | 2658053 | 7 | * | 6 | 857.142857142857 | | | | | |
| NC_039518.1 | 2681489 | 2682795 | 1307 | - | Gnomon | gene | NA | NA | gene-LOC105285394 | GeneID:105285394 | LOC105285394 | Gene | LOC105285394 | protein_coding |
| character(0) | NA | character(0) | NC_039518.1 | 2681945 | 2681951 | 7 | * | 6 | 857.142857142857 | | | | | |
| NC_039518.1 | 5288531 | 5294826 | 6296 | - | Gnomon | gene | NA | NA | gene-LOC105285410 | GeneID:105285410 | LOC105285410 | Gene | LOC105285410 | protein_coding |
| character(0) | NA | character(0) | NC_039518.1 | 5293748 | 5293754 | 7 | * | 6 | 857.142857142857 | | | | | |
| NC_039518.1 | 8632464 | 8637212 | 4749 | + | Gnomon | gene | NA | NA | gene-LOC105286242 | GeneID:105286242 | LOC105286242 | Gene | LOC105286242 | protein_coding |
| character(0) | NA | character(0) | NC_039518.1 | 8633518 | 8633524 | 7 | * | 6 | 857.142857142857 | | | | | |
| NC_039519.1 | 197513 | 200423 | 2911 | + | Gnomon | gene | NA | NA | gene-LOC105282793 | GeneID:105282793 | LOC105282793 | Gene | LOC105282793 | protein_coding |
| character(0) | NA | character(0) | NC_039519.1 | 198095 | 198101 | 7 | * | 6 | 857.142857142857 | | | | | |
| NC_039519.1 | 3406510 | 3418101 | 11592 | - | Gnomon | gene | NA | NA | gene-LOC105286844 | GeneID:105286844 | LOC105286844 | Gene | LOC105286844 | protein_coding |
| character(0) | NA | character(0) | NC_039519.1 | 3416621 | 3416627 | 7 | * | 6 | 857.142857142857 | | | | | |
| NC_039519.1 | 6006305 | 6010966 | 4662 | + | Gnomon | gene | NA | NA | gene-LOC105274778 | GeneID:105274778 | LOC105274778 | Gene | LOC105274778 | protein_coding |
| character(0) | NA | character(0) | NC_039519.1 | 6007366 | 6007372 | 7 | * | 6 | 857.142857142857 | | | | | |
| NC_039519.1 | 8084633 | 8088546 | 3914 | + | Gnomon | gene | NA | NA | gene-LOC105274767 | GeneID:105274767 | LOC105274767 | Gene | LOC105274767 | protein_coding |
| character(0) | NA | character(0) | NC_039519.1 | 8085720 | 8085726 | 7 | * | 6 | 857.142857142857 | | | | | |
| NC_039519.1 | 8239374 | 8248322 | 8949 | - | Gnomon | gene | NA | NA | gene-LOC105287985 | GeneID:105287985 | LOC105287985 | Gene | LOC105287985 | protein_coding |
| character(0) | NA | character(0) | NC_039519.1 | 8247911 | 8247917 | 7 | * | 6 | 857.142857142857 | | | | | |
| NC_039508.1 | 13115312 | 13116213 | 902 | + | Gnomon | gene | NA | NA | gene-LOC105277091 | GeneID:105277091 | LOC105277091 | Gene | LOC105277091 | protein_coding |
| character(0) | NA | character(0) | NC_039508.1 | 13116053 | 13116062 | 10 | * | 8 | 800 | | | | | |
| NC_039513.1 | 5176546 | 5178099 | 1554 | + | Gnomon | gene | NA | NA | gene-LOC105279718 | GeneID:105279718 | LOC105279718 | Gene | LOC105279718 | protein_coding |
| character(0) | NA | character(0) | NC_039513.1 | 5177245 | 5177254 | 10 | * | 8 | 800 | | | | | |
| NC_039519.1 | 333719 | 337798 | 4080 | - | Gnomon | gene | NA | NA | gene-LOC105285496 | GeneID:105285496 | LOC105285496 | Gene | LOC105285496 | protein_coding |
| character(0) | NA | character(0) | NC_039519.1 | 337077 | 337086 | 10 | * | 8 | 800 | | | | | |

Directory: MRPR File: gwr-Ob_LI_RphaseB2.txt

| seqnames | start | end | width | strand | source | type | score | phase | ID | Dbxref | Name | gbkey | gene | gene_biotype | end_range | partial | start_range | seqnames | start | end | width | strand | NbrSites | Sdnsty |
|---|---|---|---|---|---|---|---|---|---|---|---|---|---|---|---|---|---|---|---|---|---|---|---|---|
| NC_039506.1 | 6089683 | 6090856 | 1174 | + | Gnomon | gene | NA | NA | gene-LOC105277804 | GeneID:105277804 | LOC105277804 | Gene | LOC105277804 | protein_coding | character(0) | NA | character(0) | NC_039506.1 | 6090624 | 6090629 | 6 | * | 6 | 1000 |
| NC_039506.1 | 15370515 | 15371538 | 1024 | + | Gnomon | gene | NA | NA | gene-LOC105280860 | GeneID:105280860 | LOC105280860 | Gene | LOC105280860 | protein_coding | character(0) | NA | character(0) | NC_039506.1 | 15371422 | 15371427 | 6 | * | 6 | 1000 |
| NC_039507.1 | 5093844 | 5095945 | 2102 | + | Gnomon | gene | NA | NA | gene-LOC105281690 | GeneID:105281690 | LOC105281690 | Gene | LOC105281690 | protein_coding | character(0) | NA | character(0) | NC_039507.1 | 5094466 | 5094471 | 6 | * | 6 | 1000 |
| NC_039507.1 | 7181199 | 7188522 | 7324 | + | Gnomon | gene | NA | NA | gene-LOC105283383 | GeneID:105283383 | LOC105283383 | Gene | LOC105283383 | protein_coding | character(0) | NA | character(0) | NC_039507.1 | 7182290 | 7182295 | 6 | * | 6 | 1000 |
| NC_039507.1 | 7276866 | 7278961 | 2096 | + | Gnomon | gene | NA | NA | gene-LOC105283373 | GeneID:105283373 | LOC105283373 | Gene | LOC105283373 | protein_coding | character(0) | NA | character(0) | NC_039507.1 | 7277947 | 7277952 | 6 | * | 6 | 1000 |
| NC_039507.1 | 18333035 | 18336377 | 3343 | - | Gnomon | gene | NA | NA | gene-LOC105281577 | GeneID:105281577 | LOC105281577 | Gene | LOC105281577 | protein_coding | character(0) | NA | character(0) | NC_039507.1 | 18335809 | 18335814 | 6 | * | 6 | 1000 |
| NC_039508.1 | 2590841 | 2594093 | 3253 | + | Gnomon | gene | NA | NA | gene-LOC105281341 | GeneID:105281341 | LOC105281341 | Gene | LOC105281341 | protein_coding | character(0) | NA | character(0) | NC_039508.1 | 2591612 | 2591617 | 6 | * | 6 | 1000 |
| NC_039508.1 | 17548373 | 17550348 | 1976 | - | Gnomon | gene | NA | NA | gene-LOC105284396 | GeneID:105284396 | LOC105284396 | Gene | LOC105284396 | protein_coding | character(0) | NA | character(0) | NC_039508.1 | 17549290 | 17549295 | 6 | * | 6 | 1000 |
| NC_039509.1 | 1135464 | 1137481 | 2018 | - | Gnomon | gene | NA | NA | gene-LOC105283634 | GeneID:105283634 | LOC105283634 | Gene | LOC105283634 | protein_coding | character(0) | NA | character(0) | NC_039509.1 | 1136266 | 1136271 | 6 | * | 6 | 1000 |
| NC_039509.1 | 10756577 | 10757875 | 1299 | - | Gnomon | gene | NA | NA | gene-LOC105275214 | GeneID:105275214 | LOC105275214 | Gene | LOC105275214 | protein_coding | character(0) | NA | character(0) | NC_039509.1 | 10756803 | 10756808 | 6 | * | 6 | 1000 |
| NC_039509.1 | 11068956 | 11086837 | 17882 | - | Gnomon | gene | NA | NA | gene-LOC105275770 | GeneID:105275770 | LOC105275770 | Gene | LOC105275770 | protein_coding | character(0) | NA | character(0) | NC_039509.1 | 11085457 | 11085462 | 6 | * | 6 | 1000 |
| NC_039509.1 | 13033738 | 13035966 | 2229 | + | Gnomon | gene | NA | NA | gene-LOC105278527 | GeneID:105278527 | LOC105278527 | Gene | LOC105278527 | protein_coding | character(0) | NA | character(0) | NC_039509.1 | 13034669 | 13034674 | 6 | * | 6 | 1000 |
| NC_039509.1 | 13067024 | 13068370 | 1347 | + | Gnomon | gene | NA | NA | gene-LOC105278478 | GeneID:105278478 | LOC105278478 | Gene | LOC105278478 | protein_coding | character(0) | NA | character(0) | NC_039509.1 | 13067776 | 13067781 | 6 | * | 6 | 1000 |
| NC_039510.1 | 2607810 | 2610599 | 2790 | - | Gnomon | gene | NA | NA | gene-LOC105287754 | GeneID:105287754 | LOC105287754 | Gene | LOC105287754 | protein_coding | character(0) | NA | character(0) | NC_039510.1 | 2609179 | 2609184 | 6 | * | 6 | 1000 |
| NC_039510.1 | 16287375 | 16291357 | 3983 | - | Gnomon | gene | NA | NA | gene-LOC105286509 | GeneID:105286509 | LOC105286509 | Gene | LOC105286509 | protein_coding | character(0) | NA | character(0) | NC_039510.1 | 16290598 | 16290603 | 6 | * | 6 | 1000 |
| NC_039511.1 | 2234830 | 2236789 | 1960 | - | Gnomon | gene | NA | NA | gene-LOC105276382 | GeneID:105276382 | LOC105276382 | Gene | LOC105276382 | protein_coding | character(0) | NA | character(0) | NC_039511.1 | 2236016 | 2236021 | 6 | * | 6 | 1000 |
| NC_039511.1 | 8407553 | 8425032 | 17480 | + | Gnomon | gene | NA | NA | gene-LOC105284845 | GeneID:105284845 | LOC105284845 | Gene | LOC105284845 | protein_coding | character(0) | NA | character(0) | NC_039511.1 | 8409706 | 8409711 | 6 | * | 6 | 1000 |
| NC_039511.1 | 8695181 | 8698086 | 2906 | + | Gnomon | gene | NA | NA | gene-LOC105280856 | GeneID:105280856 | LOC105280856 | Gene | LOC105280856 | protein_coding | character(0) | NA | character(0) | NC_039511.1 | 8695844 | 8695849 | 6 | * | 6 | 1000 |
| NC_039512.1 | 15979275 | 15981529 | 2255 | - | Gnomon | gene | NA | NA | gene-LOC105274775 | GeneID:105274775 | LOC105274775 | Gene | LOC105274775 | protein_coding | character(0) | NA | character(0) | NC_039512.1 | 15980634 | 15980639 | 6 | * | 6 | 1000 |
| NC_039513.1 | 10401827 | 10404590 | 2764 | - | Gnomon | gene | NA | NA | gene-LOC105275550 | GeneID:105275550 | LOC105275550 | Gene | LOC105275550 | protein_coding | character(0) | NA | character(0) | NC_039513.1 | 10403469 | 10403474 | 6 | * | 6 | 1000 |
| NC_039513.1 | 12028675 | 12030941 | 2267 | - | Gnomon | gene | NA | NA | gene-LOC105280791 | GeneID:105280791 | LOC105280791 | Gene | LOC105280791 | protein_coding | character(0) | NA | character(0) | NC_039513.1 | 12029791 | 12029796 | 6 | * | 6 | 1000 |
| NC_039513.1 | 13434303 | 13438575 | 4273 | - | Gnomon | gene | NA | NA | gene-LOC105275036 | GeneID:105275036 | LOC105275036 | Gene | LOC105275036 | protein_coding | character(0) | NA | character(0) | NC_039513.1 | 13436991 | 13437000 | 10 | * | 10 | 1000 |
| NC_039514.1 | 6622839 | 6626782 | 3944 | - | Gnomon | gene | NA | NA | gene-LOC105285534 | GeneID:105285534 | LOC105285534 | Gene | LOC105285534 | protein_coding | character(0) | NA | character(0) | NC_039514.1 | 6625853 | 6625858 | 6 | * | 6 | 1000 |
| NC_039514.1 | 9336154 | 9343762 | 7609 | + | Gnomon | gene | NA | NA | gene-LOC105284644 | GeneID:105284644 | LOC105284644 | Gene | LOC105284644 | protein_coding | character(0) | NA | character(0) | NC_039514.1 | 9337043 | 9337048 | 6 | * | 6 | 1000 |
| NC_039515.1 | 7098785 | 7103508 | 4724 | - | Gnomon | gene | NA | NA | gene-LOC105286003 | GeneID:105286003 | LOC105286003 | Gene | LOC105286003 | protein_coding | character(0) | NA | character(0) | NC_039515.1 | 7102685 | 7102690 | 6 | * | 6 | 1000 |
| NC_039515.1 | 12582469 | 12585642 | 3174 | - | Gnomon | gene | NA | NA | gene-LOC105282150 | GeneID:105282150 | LOC105282150 | Gene | LOC105282150 | protein_coding | character(0) | NA | character(0) | NC_039515.1 | 12584748 | 12584753 | 6 | * | 6 | 1000 |
| NC_039516.1 | 1062627 | 1071280 | 8654 | + | Gnomon | gene | NA | NA | gene-LOC105274578 | GeneID:105274578 | LOC105274578 | Gene | LOC105274578 | protein_coding | character(0) | NA | character(0) | NC_039516.1 | 1064627 | 1064632 | 6 | * | 6 | 1000 |
| NC_039516.1 | 2464828 | 2468017 | 3190 | + | Gnomon | gene | NA | NA | gene-LOC105288248 | GeneID:105288248 | LOC105288248 | Gene | LOC105288248 | protein_coding | character(0) | NA | character(0) | NC_039516.1 | 2465766 | 2465771 | 6 | * | 6 | 1000 |
| NC_039516.1 | 12996090 | 13006535 | 10446 | + | Gnomon | gene | NA | NA | gene-LOC105275342 | GeneID:105275342 | LOC105275342 | Gene | LOC105275342 | protein_coding | character(0) | NA | character(0) | NC_039516.1 | 12997443 | 12997448 | 6 | * | 6 | 1000 |
| NC_039516.1 | 13065193 | 13068389 | 3197 | + | Gnomon | gene | NA | NA | gene-LOC105275355 | GeneID:105275355 | LOC105275355 | Gene | LOC105275355 | protein_coding | | | | | | | | | | |

| | | | | | | | | | | | | | | |
|---|---|---|---|---|---|---|---|---|---|---|---|---|---|---|
| character(0) | NA | character(0) | NC_039516.1 | 13066190 | 13066195 | 6 | * | 6 | 1000 | | | | | |
| NC_039517.1 | 1360424 | 1367346 | 6923 | + | Gnomon | gene | NA | NA | gene-LOC105278812 | GeneID:105278812 | LOC105278812 | Gene | LOC105278812 | protein_coding |
| character(0) | NA | character(0) | NC_039517.1 | 1361411 | 1361416 | 6 | * | 6 | 1000 | | | | | |
| NC_039517.1 | 7309114 | 7314427 | 5314 | - | Gnomon | gene | NA | NA | gene-LOC105284201 | GeneID:105284201 | LOC105284201 | Gene | LOC105284201 | protein_coding |
| character(0) | NA | character(0) | NC_039517.1 | 7312519 | 7312524 | 6 | * | 6 | 1000 | | | | | |
| NC_039517.1 | 8175046 | 8181714 | 6669 | - | Gnomon | gene | NA | NA | gene-LOC105284262 | GeneID:105284262 | LOC105284262 | Gene | LOC105284262 | protein_coding |
| character(0) | NA | character(0) | NC_039517.1 | 8175507 | 8175512 | 6 | * | 6 | 1000 | | | | | |
| NC_039518.1 | 5496870 | 5510429 | 13560 | - | Gnomon | gene | NA | NA | gene-LOC105281906 | GeneID:105281906 | LOC105281906 | Gene | LOC105281906 | protein_coding |
| character(0) | NA | character(0) | NC_039518.1 | 5508860 | 5508865 | 6 | * | 6 | 1000 | | | | | |
| NC_039518.1 | 8474988 | 8476009 | 1022 | + | Gnomon | gene | NA | NA | gene-LOC105286212 | GeneID:105286212 | LOC105286212 | Gene | LOC105286212 | protein_coding |
| character(0) | NA | character(0) | NC_039518.1 | 8475762 | 8475769 | 8 | * | 8 | 1000 | | | | | |
| NC_039518.1 | 8721748 | 8727127 | 5380 | + | Gnomon | gene | NA | NA | gene-LOC105286364 | GeneID:105286364 | LOC105286364 | Gene | LOC105286364 | protein_coding |
| character(0) | NA | character(0) | NC_039518.1 | 8722922 | 8722927 | 6 | * | 6 | 1000 | | | | | |
| NC_039518.1 | 9164450 | 9166108 | 1659 | - | Gnomon | gene | NA | NA | gene-LOC105286320 | GeneID:105286320 | LOC105286320 | Gene | LOC105286320 | protein_coding |
| character(0) | NA | character(0) | NC_039518.1 | 9164960 | 9164965 | 6 | * | 6 | 1000 | | | | | |
| NC_039506.1 | 23567682 | 23574351 | 6670 | + | Gnomon | gene | NA | NA | gene-LOC105279333 | GeneID:105279333 | LOC105279333 | Gene | LOC105279333 | protein_coding |
| character(0) | NA | character(0) | NC_039506.1 | 23568887 | 23568897 | 11 | * | 10 | 909.090909090909 | | | | | |
| NC_039507.1 | 9161485 | 9166239 | 4755 | + | Gnomon | gene | NA | NA | gene-LOC105282461 | GeneID:105282461 | LOC105282461 | Gene | LOC105282461 | protein_coding |
| character(0) | NA | character(0) | NC_039507.1 | 9162374 | 9162382 | 9 | * | 8 | 888.888888888889 | | | | | |
| NC_039509.1 | 17207204 | 17209527 | 2324 | + | Gnomon | gene | NA | NA | gene-LOC105283016 | GeneID:105283016 | LOC105283016 | Gene | LOC105283016 | protein_coding |
| character(0) | NA | character(0) | NC_039509.1 | 17208071 | 17208079 | 9 | * | 8 | 888.888888888889 | | | | | |
| NC_039510.1 | 8568121 | 8569477 | 1357 | - | Gnomon | gene | NA | NA | gene-LOC105275137 | GeneID:105275137 | LOC105275137 | Gene | LOC105275137 | protein_coding |
| character(0) | NA | character(0) | NC_039510.1 | 8568914 | 8568922 | 9 | * | 8 | 888.888888888889 | | | | | |
| NC_039516.1 | 5472871 | 5475154 | 2284 | - | Gnomon | gene | NA | NA | gene-LOC105275878 | GeneID:105275878 | LOC105275878 | Gene | LOC105275878 | protein_coding |
| character(0) | NA | character(0) | NC_039516.1 | 5474292 | 5474300 | 9 | * | 8 | 888.888888888889 | | | | | |
| NC_039512.1 | 15373917 | 15376509 | 2593 | + | Gnomon | gene | NA | NA | gene-LOC105288114 | GeneID:105288114 | LOC105288114 | Gene | LOC105288114 | protein_coding |
| character(0) | NA | character(0) | NC_039512.1 | 15375336 | 15375343 | 8 | * | 7 | 875 | | | | | |
| NC_039513.1 | 3303685 | 3306319 | 2635 | + | Gnomon | gene | NA | NA | gene-LOC105275761 | GeneID:105275761 | LOC105275761 | Gene | LOC105275761 | protein_coding |
| character(0) | NA | character(0) | NC_039513.1 | 3304807 | 3304814 | 8 | * | 7 | 875 | | | | | |
| NC_039506.1 | 1487779 | 1489683 | 1905 | + | Gnomon | gene | NA | NA | gene-LOC105280236 | GeneID:105280236 | LOC105280236 | Gene | LOC105280236 | protein_coding |
| character(0) | NA | character(0) | NC_039506.1 | 1488926 | 1488932 | 7 | * | 6 | 857.142857142857 | | | | | |
| NC_039506.1 | 2274108 | 2276090 | 1983 | - | Gnomon | gene | NA | NA | gene-LOC105280319 | GeneID:105280319 | LOC105280319 | Gene | LOC105280319 | protein_coding |
| character(0) | NA | character(0) | NC_039506.1 | 2275368 | 2275374 | 7 | * | 6 | 857.142857142857 | | | | | |
| NC_039506.1 | 13552800 | 13986605 | 433806 | - | Gnomon | gene | NA | NA | gene-LOC105280980 | GeneID:105280980 | LOC105280980 | Gene | LOC105280980 | protein_coding |
| character(0) | NA | character(0) | NC_039506.1 | 13985875 | 13985881 | 7 | * | 6 | 857.142857142857 | | | | | |
| NC_039506.1 | 15506447 | 15508913 | 2467 | - | Gnomon | gene | NA | NA | gene-LOC105283960 | GeneID:105283960 | LOC105283960 | Gene | LOC105283960 | protein_coding |
| character(0) | NA | character(0) | NC_039506.1 | 15508062 | 15508068 | 7 | * | 6 | 857.142857142857 | | | | | |
| NC_039506.1 | 15509387 | 15513034 | 3648 | + | Gnomon | gene | NA | NA | gene-LOC105283961 | GeneID:105283961 | LOC105283961 | Gene | LOC105283961 | protein_coding |
| character(0) | NA | character(0) | NC_039506.1 | 15510126 | 15510132 | 7 | * | 6 | 857.142857142857 | | | | | |
| NC_039506.1 | 16116359 | 16120578 | 4220 | - | Gnomon | gene | NA | NA | gene-LOC105274912 | GeneID:105274912 | LOC105274912 | Gene | LOC105274912 | protein_coding |
| character(0) | NA | character(0) | NC_039506.1 | 16119072 | 16119078 | 7 | * | 6 | 857.142857142857 | | | | | |
| NC_039506.1 | 19093260 | 19098014 | 4755 | - | Gnomon | gene | NA | NA | gene-LOC105278005 | GeneID:105278005 | LOC105278005 | Gene | LOC105278005 | protein_coding |
| character(0) | NA | character(0) | NC_039506.1 | 19096860 | 19096866 | 7 | * | 6 | 857.142857142857 | | | | | |
| NC_039506.1 | 20553263 | 20559098 | 5836 | - | Gnomon | gene | NA | NA | gene-LOC105278544 | GeneID:105278544 | LOC105278544 | Gene | LOC105278544 | protein_coding |
| character(0) | NA | character(0) | NC_039506.1 | 20557629 | 20557635 | 7 | * | 6 | 857.142857142857 | | | | | |
| NC_039506.1 | 22941360 | 22946330 | 4971 | - | Gnomon | gene | NA | NA | gene-LOC105283484 | GeneID:105283484 | LOC105283484 | Gene | LOC105283484 | protein_coding |
| character(0) | NA | character(0) | NC_039506.1 | 22945036 | 22945042 | 7 | * | 6 | 857.142857142857 | | | | | |
| NC_039506.1 | 23689672 | 23694320 | 4649 | - | Gnomon | gene | NA | NA | gene-LOC105279346 | GeneID:105279346 | LOC105279346 | Gene | LOC105279346 | protein_coding |
| character(0) | NA | character(0) | NC_039506.1 | 23693180 | 23693186 | 7 | * | 6 | 857.142857142857 | | | | | |
| NC_039507.1 | 124714 | 126752 | 2039 | + | Gnomon | gene | NA | NA | gene-LOC105279088 | GeneID:105279088 | LOC105279088 | Gene | LOC105279088 | protein_coding |
| character(0) | NA | character(0) | NC_039507.1 | 125663 | 125669 | 7 | * | 6 | 857.142857142857 | | | | | |
| NC_039507.1 | 790243 | 798925 | 8683 | + | Gnomon | gene | NA | NA | gene-LOC105282921 | GeneID:105282921 | LOC105282921 | Gene | LOC105282921 | protein_coding |
| character(0) | NA | character(0) | NC_039507.1 | 791692 | 791698 | 7 | * | 6 | 857.142857142857 | | | | | |
| NC_039507.1 | 6347235 | 6350936 | 3702 | + | Gnomon | gene | NA | NA | gene-LOC105275401 | GeneID:105275401 | LOC105275401 | Gene | LOC105275401 | protein_coding |
| character(0) | NA | character(0) | NC_039507.1 | 6348012 | 6348018 | 7 | * | 6 | 857.142857142857 | | | | | |
| NC_039507.1 | 15481857 | 15484141 | 2285 | + | Gnomon | gene | NA | NA | gene-LOC105280092 | GeneID:105280092 | LOC105280092 | Gene | LOC105280092 | protein_coding |
| character(0) | NA | character(0) | NC_039507.1 | 15482876 | 15482882 | 7 | * | 6 | 857.142857142857 | | | | | |
| NC_039507.1 | 15506083 | 15508702 | 2620 | + | Gnomon | gene | NA | NA | gene-LOC105280102 | GeneID:105280102 | LOC105280102 | Gene | LOC105280102 | protein_coding |
| character(0) | NA | character(0) | NC_039507.1 | 15507322 | 15507328 | 7 | * | 6 | 857.142857142857 | | | | | |
| NC_039507.1 | 18333035 | 18336377 | 3343 | - | Gnomon | gene | NA | NA | gene-LOC105281577 | GeneID:105281577 | LOC105281577 | Gene | LOC105281577 | protein_coding |
| character(0) | NA | character(0) | NC_039507.1 | 18335506 | 18335512 | 7 | * | 6 | 857.142857142857 | | | | | |
| NC_039508.1 | 192252 | 200875 | 8624 | + | Gnomon | gene | NA | NA | gene-LOC105277452 | GeneID:105277452 | LOC105277452 | Gene | LOC105277452 | protein_coding |
| character(0) | NA | character(0) | NC_039508.1 | 193513 | 193519 | 7 | * | 6 | 857.142857142857 | | | | | |
| NC_039508.1 | 1384214 | 1386820 | 2607 | + | Gnomon | gene | NA | NA | gene-LOC105277388 | GeneID:105277388 | LOC105277388 | Gene | LOC105277388 | protein_coding |

character(0) NA character(0) NC_039508.1 1384932 1384938 7 * 6 857.142857142857
NC_039508.1 1386723 1389150 2428 - Gnomon gene NA NA gene-LOC105277389 GeneID:105277389 LOC105277389 Gene LOC105277389 protein_coding
character(0) NA character(0) NC_039508.1 1387947 1387953 7 * 6 857.142857142857
NC_039508.1 3996551 4002196 5646 - Gnomon gene NA NA gene-LOC105285719 GeneID:105285719 LOC105285719 Gene LOC105285719 protein_coding
character(0) NA character(0) NC_039508.1 4001396 4001402 7 * 6 857.142857142857
NC_039508.1 9041025 9045259 4235 + Gnomon gene NA NA gene-LOC105278106 GeneID:105278106 LOC105278106 Gene LOC105278106 protein_coding
character(0) NA character(0) NC_039508.1 9042110 9042116 7 * 6 857.142857142857
NC_039508.1 10763110 10766843 3734 + Gnomon gene NA NA gene-LOC105276948 GeneID:105276948 LOC105276948 Gene LOC105276948 protein_coding
character(0) NA character(0) NC_039508.1 10764758 10764764 7 * 6 857.142857142857
NC_039508.1 10797665 10799797 2133 + Gnomon gene NA NA gene-LOC105276949 GeneID:105276949 LOC105276949 Gene LOC105276949 protein_coding
character(0) NA character(0) NC_039508.1 10798605 10798611 7 * 6 857.142857142857
NC_039508.1 13116146 13120560 4415 - Gnomon gene NA NA gene-LOC105277128 GeneID:105277128 LOC105277128 Gene LOC105277128 protein_coding
character(0) NA character(0) NC_039508.1 13116245 13116251 7 * 6 857.142857142857
NC_039508.1 15763225 15782851 19627 + Gnomon gene NA NA gene-LOC105277234 GeneID:105277234 LOC105277234 Gene LOC105277234 protein_coding
character(0) NA character(0) NC_039508.1 15764452 15764458 7 * 6 857.142857142857
NC_039508.1 15807211 15809918 2708 - Gnomon gene NA NA gene-LOC105277238 GeneID:105277238 LOC105277238 Gene LOC105277238 protein_coding
character(0) NA character(0) NC_039508.1 15808870 15808876 7 * 6 857.142857142857
NC_039508.1 17548373 17550348 1976 - Gnomon gene NA NA gene-LOC105284396 GeneID:105284396 LOC105284396 Gene LOC105284396 protein_coding
character(0) NA character(0) NC_039508.1 17549403 17549409 7 * 6 857.142857142857
NC_039509.1 2385054 2391699 6646 - Gnomon gene NA NA gene-LOC105285867 GeneID:105285867 LOC105285867 Gene LOC105285867 protein_coding
character(0) NA character(0) NC_039509.1 2389203 2389209 7 * 6 857.142857142857
NC_039509.1 5250307 5257914 7608 - Gnomon gene NA NA gene-LOC105287147 GeneID:105287147 LOC105287147 Gene LOC105287147 protein_coding
character(0) NA character(0) NC_039509.1 5256794 5256800 7 * 6 857.142857142857
NC_039509.1 7092641 7097742 5102 + Gnomon gene NA NA gene-LOC105282559 GeneID:105282559 LOC105282559 Gene LOC105282559 protein_coding
character(0) NA character(0) NC_039509.1 7093578 7093584 7 * 6 857.142857142857
NC_039509.1 7141680 7152824 11145 - Gnomon gene NA NA gene-LOC105282568 GeneID:105282568 LOC105282568 Gene LOC105282568 protein_coding
character(0) NA character(0) NC_039509.1 7145645 7145651 7 * 6 857.142857142857
NC_039509.1 10232323 10244464 12142 + Gnomon gene NA NA gene-LOC105282655 GeneID:105282655 LOC105282655 Gene LOC105282655 protein_coding
character(0) NA character(0) NC_039509.1 10233310 10233316 7 * 6 857.142857142857
NC_039509.1 10710016 10711124 1109 + Gnomon gene NA NA gene-LOC105275153 GeneID:105275153 LOC105275153 Gene LOC105275153 protein_coding
character(0) NA character(0) NC_039509.1 10710730 10710736 7 * 6 857.142857142857
NC_039509.1 13909149 13915606 6458 + Gnomon gene NA NA gene-LOC105278288 GeneID:105278288 LOC105278288 Gene LOC105278288 protein_coding
character(0) NA character(0) NC_039509.1 13909987 13909993 7 * 6 857.142857142857
NC_039509.1 15295604 15304160 8557 + Gnomon gene NA NA gene-LOC105284317 GeneID:105284317 LOC105284317 Gene LOC105284317 protein_coding
character(0) NA character(0) NC_039509.1 15296358 15296364 7 * 6 857.142857142857
NC_039510.1 3040660 3042070 1411 + Gnomon gene NA NA gene-LOC105287710 GeneID:105287710 LOC105287710 Gene LOC105287710 protein_coding
character(0) NA character(0) NC_039510.1 3041562 3041568 7 * 6 857.142857142857
NC_039510.1 3110492 3117379 6888 - Gnomon gene NA NA gene-LOC105287697 GeneID:105287697 LOC105287697 Gene LOC105287697 protein_coding
character(0) NA character(0) NC_039510.1 3116602 3116608 7 * 6 857.142857142857
NC_039510.1 3167251 3170185 2935 - Gnomon gene NA NA gene-LOC105287684 GeneID:105287684 LOC105287684 Gene LOC105287684 protein_coding
character(0) NA character(0) NC_039510.1 3169234 3169240 7 * 6 857.142857142857
NC_039510.1 4220534 4227851 7318 + Gnomon gene NA NA gene-LOC105287620 GeneID:105287620 LOC105287620 Gene LOC105287620 protein_coding
character(0) NA character(0) NC_039510.1 4221389 4221395 7 * 6 857.142857142857
NC_039510.1 5067676 5072600 4925 - Gnomon gene NA NA gene-LOC105287597 GeneID:105287597 LOC105287597 Gene LOC105287597 protein_coding
character(0) NA character(0) NC_039510.1 5071200 5071206 7 * 6 857.142857142857
NC_039510.1 5537223 5539036 1814 - Gnomon gene NA NA gene-LOC105287569 GeneID:105287569 LOC105287569 Gene LOC105287569 protein_coding
character(0) NA character(0) NC_039510.1 5538139 5538145 7 * 6 857.142857142857
NC_039510.1 6043706 6047166 3461 + Gnomon gene NA NA gene-LOC105287208 GeneID:105287208 LOC105287208 Gene LOC105287208 protein_coding
character(0) NA character(0) NC_039510.1 6045163 6045169 7 * 6 857.142857142857
NC_039510.1 6122701 6125649 2949 - Gnomon gene NA NA gene-LOC105287279 GeneID:105287279 LOC105287279 Gene LOC105287279 protein_coding
character(0) NA character(0) NC_039510.1 6124458 6124464 7 * 6 857.142857142857
NC_039510.1 6163214 6171437 8224 - Gnomon gene NA NA gene-LOC105287371 GeneID:105287371 LOC105287371 Gene LOC105287371 protein_coding
character(0) NA character(0) NC_039510.1 6170155 6170161 7 * 6 857.142857142857
NC_039510.1 6212141 6215222 3082 + Gnomon gene NA NA gene-LOC105287473 GeneID:105287473 LOC105287473 Gene LOC105287473 protein_coding
character(0) NA character(0) NC_039510.1 6213095 6213101 7 * 6 857.142857142857
NC_039510.1 8487238 8489293 2056 - Gnomon gene NA NA gene-LOC105275119 GeneID:105275119 LOC105275119 Gene LOC105275119 misc_RNA character(0)
NA character(0) NC_039510.1 8488481 8488487 7 * 6 857.142857142857
NC_039510.1 8568121 8569477 1357 - Gnomon gene NA NA gene-LOC105275137 GeneID:105275137 LOC105275137 Gene LOC105275137 protein_coding
character(0) NA character(0) NC_039510.1 8568885 8568891 7 * 6 857.142857142857
NC_039510.1 9996113 9998708 2596 - Gnomon gene NA NA gene-LOC105281227 GeneID:105281227 LOC105281227 Gene LOC105281227 protein_coding
character(0) NA character(0) NC_039510.1 9997644 9997650 7 * 6 857.142857142857
NC_039510.1 13531247 13535548 4302 - Gnomon gene NA NA gene-LOC105280964 GeneID:105280964 LOC105280964 Gene LOC105280964 protein_coding
character(0) NA character(0) NC_039510.1 13533348 13533354 7 * 6 857.142857142857
NC_039510.1 15582075 15587881 5807 - Gnomon gene NA NA gene-LOC105286461 GeneID:105286461 LOC105286461 Gene LOC105286461 protein_coding

| | | | | | | | | | | | | | | |
|---|---|---|---|---|---|---|---|---|---|---|---|---|---|---|
| character(0) | NA | character(0) | NC_039510.1 | 15586562 | 15586568 | 7 | * | 6 | 857.142857142857 | | | | | |
| NC_039510.1 | 15745071 | 15751836 | 6766 | + | Gnomon | gene | NA | NA | gene-LOC105286478 | GeneID:105286478 | LOC105286478 | Gene | LOC105286478 | protein_coding |
| character(0) | NA | character(0) | NC_039510.1 | 15746500 | 15746506 | 7 | * | 6 | 857.142857142857 | | | | | |
| NC_039510.1 | 16500898 | 16505659 | 4762 | - | Gnomon | gene | NA | NA | gene-LOC105286526 | GeneID:105286526 | LOC105286526 | Gene | LOC105286526 | protein_coding |
| character(0) | NA | character(0) | NC_039510.1 | 16504460 | 16504466 | 7 | * | 6 | 857.142857142857 | | | | | |
| NC_039511.1 | 517195 | 530396 | 13202 | + | Gnomon | gene | NA | NA | gene-LOC105276519 | GeneID:105276519 | LOC105276519 | Gene | LOC105276519 | protein_coding |
| character(0) | NA | character(0) | NC_039511.1 | 518659 | 518665 | 7 | * | 6 | 857.142857142857 | | | | | |
| NC_039511.1 | 1662842 | 1668395 | 5554 | + | Gnomon | gene | NA | NA | gene-LOC105276437 | GeneID:105276437 | LOC105276437 | Gene | LOC105276437 | protein_coding |
| character(0) | NA | character(0) | NC_039511.1 | 1664306 | 1664312 | 7 | * | 6 | 857.142857142857 | | | | | |
| NC_039511.1 | 1856026 | 1859536 | 3511 | + | Gnomon | gene | NA | NA | gene-LOC105276422 | GeneID:105276422 | LOC105276422 | Gene | LOC105276422 | protein_coding |
| character(0) | NA | character(0) | NC_039511.1 | 1857678 | 1857684 | 7 | * | 6 | 857.142857142857 | | | | | |
| NC_039511.1 | 2004889 | 2015973 | 11085 | - | Gnomon | gene | NA | NA | gene-LOC105276410 | GeneID:105276410 | LOC105276410 | Gene | LOC105276410 | protein_coding |
| character(0) | NA | character(0) | NC_039511.1 | 2014683 | 2014689 | 7 | * | 6 | 857.142857142857 | | | | | |
| NC_039511.1 | 2004889 | 2015973 | 11085 | - | Gnomon | gene | NA | NA | gene-LOC105276410 | GeneID:105276410 | LOC105276410 | Gene | LOC105276410 | protein_coding |
| character(0) | NA | character(0) | NC_039511.1 | 2014740 | 2014746 | 7 | * | 6 | 857.142857142857 | | | | | |
| NC_039511.1 | 3635459 | 3642537 | 7079 | - | Gnomon | gene | NA | NA | gene-LOC105284052 | GeneID:105284052 | LOC105284052 | Gene | LOC105284052 | protein_coding |
| character(0) | NA | character(0) | NC_039511.1 | 3639538 | 3639544 | 7 | * | 6 | 857.142857142857 | | | | | |
| NC_039511.1 | 4554146 | 4561340 | 7195 | - | Gnomon | gene | NA | NA | gene-LOC105281750 | GeneID:105281750 | LOC105281750 | Gene | LOC105281750 | protein_coding |
| character(0) | NA | character(0) | NC_039511.1 | 4560502 | 4560508 | 7 | * | 6 | 857.142857142857 | | | | | |
| NC_039511.1 | 12403841 | 12413518 | 9678 | - | Gnomon | gene | NA | NA | gene-LOC105285135 | GeneID:105285135 | LOC105285135 | Gene | LOC105285135 | protein_coding |
| character(0) | NA | character(0) | NC_039511.1 | 12411122 | 12411128 | 7 | * | 6 | 857.142857142857 | | | | | |
| NC_039511.1 | 14704311 | 14706938 | 2628 | - | Gnomon | gene | NA | NA | gene-LOC105285889 | GeneID:105285889 | LOC105285889 | Gene | LOC105285889 | protein_coding |
| character(0) | NA | character(0) | NC_039511.1 | 14705900 | 14705906 | 7 | * | 6 | 857.142857142857 | | | | | |
| NC_039511.1 | 15983428 | 15985754 | 2327 | + | Gnomon | gene | NA | NA | gene-LOC105284576 | GeneID:105284576 | LOC105284576 | Gene | LOC105284576 | protein_coding |
| character(0) | NA | character(0) | NC_039511.1 | 15984100 | 15984106 | 7 | * | 6 | 857.142857142857 | | | | | |
| NC_039511.1 | 16541849 | 16544005 | 2157 | + | Gnomon | gene | NA | NA | gene-LOC105284502 | GeneID:105284502 | LOC105284502 | Gene | LOC105284502 | protein_coding |
| character(0) | NA | character(0) | NC_039511.1 | 16542944 | 16542950 | 7 | * | 6 | 857.142857142857 | | | | | |
| NC_039512.1 | 4385041 | 4390283 | 5243 | - | Gnomon | gene | NA | NA | gene-LOC105277552 | GeneID:105277552 | LOC105277552 | Gene | LOC105277552 | protein_coding |
| character(0) | NA | character(0) | NC_039512.1 | 4389120 | 4389126 | 7 | * | 6 | 857.142857142857 | | | | | |
| NC_039512.1 | 11140867 | 11151152 | 10286 | - | Gnomon | gene | NA | NA | gene-LOC105279854 | GeneID:105279854 | LOC105279854 | Gene | LOC105279854 | protein_coding |
| character(0) | NA | character(0) | NC_039512.1 | 11150039 | 11150045 | 7 | * | 6 | 857.142857142857 | | | | | |
| NC_039512.1 | 11140867 | 11151152 | 10286 | - | Gnomon | gene | NA | NA | gene-LOC105279854 | GeneID:105279854 | LOC105279854 | Gene | LOC105279854 | protein_coding |
| character(0) | NA | character(0) | NC_039512.1 | 11150305 | 11150311 | 7 | * | 6 | 857.142857142857 | | | | | |
| NC_039512.1 | 12469542 | 12475093 | 5552 | - | Gnomon | gene | NA | NA | gene-LOC105279748 | GeneID:105279748 | LOC105279748 | Gene | LOC105279748 | protein_coding |
| character(0) | NA | character(0) | NC_039512.1 | 12474009 | 12474015 | 7 | * | 6 | 857.142857142857 | | | | | |
| NC_039512.1 | 12621899 | 12625222 | 3324 | + | Gnomon | gene | NA | NA | gene-LOC105279740 | GeneID:105279740 | LOC105279740 | Gene | LOC105279740 | protein_coding |
| character(0) | NA | character(0) | NC_039512.1 | 12623117 | 12623123 | 7 | * | 6 | 857.142857142857 | | | | | |
| NC_039512.1 | 12739826 | 12742653 | 2828 | + | Gnomon | gene | NA | NA | gene-LOC105279719 | GeneID:105279719 | LOC105279719 | Gene | LOC105279719 | protein_coding |
| character(0) | NA | character(0) | NC_039512.1 | 12740699 | 12740705 | 7 | * | 6 | 857.142857142857 | | | | | |
| NC_039512.1 | 12814910 | 12819293 | 4384 | - | Gnomon | gene | NA | NA | gene-LOC105279709 | GeneID:105279709 | LOC105279709 | Gene | LOC105279709 | protein_coding |
| character(0) | NA | character(0) | NC_039512.1 | 12818052 | 12818058 | 7 | * | 6 | 857.142857142857 | | | | | |
| NC_039512.1 | 13089540 | 13093562 | 4023 | - | Gnomon | gene | NA | NA | gene-LOC105279664 | GeneID:105279664 | LOC105279664 | Gene | LOC105279664 | protein_coding |
| character(0) | NA | character(0) | NC_039512.1 | 13092263 | 13092269 | 7 | * | 6 | 857.142857142857 | | | | | |
| NC_039512.1 | 14920902 | 14923325 | 2424 | - | Gnomon | gene | NA | NA | gene-LOC105287895 | GeneID:105287895 | LOC105287895 | Gene | LOC105287895 | protein_coding |
| character(0) | NA | character(0) | NC_039512.1 | 14922255 | 14922261 | 7 | * | 6 | 857.142857142857 | | | | | |
| NC_039512.1 | 16555603 | 16557649 | 2047 | - | Gnomon | gene | NA | NA | gene-LOC105282841 | GeneID:105282841 | LOC105282841 | Gene | LOC105282841 | protein_coding |
| character(0) | NA | character(0) | NC_039512.1 | 16556845 | 16556851 | 7 | * | 6 | 857.142857142857 | | | | | |
| NC_039513.1 | 3140959 | 3143113 | 2155 | - | Gnomon | gene | NA | NA | gene-LOC105287811 | GeneID:105287811 | LOC105287811 | Gene | LOC105287811 | protein_coding |
| character(0) | NA | character(0) | NC_039513.1 | 3142573 | 3142579 | 7 | * | 6 | 857.142857142857 | | | | | |
| NC_039513.1 | 3253321 | 3257474 | 4154 | - | Gnomon | gene | NA | NA | gene-LOC105275753 | GeneID:105275753 | LOC105275753 | Gene | LOC105275753 | protein_coding |
| character(0) | NA | character(0) | NC_039513.1 | 3256149 | 3256155 | 7 | * | 6 | 857.142857142857 | | | | | |
| NC_039513.1 | 3253321 | 3257474 | 4154 | - | Gnomon | gene | NA | NA | gene-LOC105275753 | GeneID:105275753 | LOC105275753 | Gene | LOC105275753 | protein_coding |
| character(0) | NA | character(0) | NC_039513.1 | 3256358 | 3256364 | 7 | * | 6 | 857.142857142857 | | | | | |
| NC_039513.1 | 3257374 | 3260835 | 3462 | - | Gnomon | gene | NA | NA | gene-LOC105275752 | GeneID:105275752 | LOC105275752 | Gene | LOC105275752 | protein_coding |
| character(0) | NA | character(0) | NC_039513.1 | 3259768 | 3259774 | 7 | * | 6 | 857.142857142857 | | | | | |
| NC_039513.1 | 3373854 | 3375485 | 1632 | + | Gnomon | gene | NA | NA | gene-LOC105274850 | GeneID:105274850 | LOC105274850 | Gene | LOC105274850 | protein_coding |
| character(0) | NA | character(0) | NC_039513.1 | 3374674 | 3374680 | 7 | * | 6 | 857.142857142857 | | | | | |
| NC_039513.1 | 8483464 | 8491634 | 8171 | + | Gnomon | gene | NA | NA | gene-LOC105275689 | GeneID:105275689 | LOC105275689 | Gene | LOC105275689 | protein_coding |
| character(0) | NA | character(0) | NC_039513.1 | 8484406 | 8484412 | 7 | * | 6 | 857.142857142857 | | | | | |
| NC_039513.1 | 9603616 | 9607153 | 3538 | + | Gnomon | gene | NA | NA | gene-LOC105275615 | GeneID:105275615 | LOC105275615 | Gene | LOC105275615 | protein_coding |
| character(0) | NA | character(0) | NC_039513.1 | 9605113 | 9605119 | 7 | * | 6 | 857.142857142857 | | | | | |
| NC_039513.1 | 9603616 | 9607153 | 3538 | + | Gnomon | gene | NA | NA | gene-LOC105275615 | GeneID:105275615 | LOC105275615 | Gene | LOC105275615 | protein_coding |
| character(0) | NA | character(0) | NC_039513.1 | 9605414 | 9605420 | 7 | * | 6 | 857.142857142857 | | | | | |
| NC_039513.1 | 11674366 | 11676356 | 1991 | - | Gnomon | gene | NA | NA | gene-LOC105280739 | GeneID:105280739 | LOC105280739 | Gene | LOC105280739 | protein_coding |

| | | | | | | | | | | | | | | |
|---|---|---|---|---|---|---|---|---|---|---|---|---|---|---|
| character(0) | NA | character(0) | NC_039513.1 | 11675069 | 11675075 | 7 | * | 6 | 857.142857142857 | | | | | |
| NC_039513.1 | 11674366 | 11676356 | 1991 | - | Gnomon | gene | NA | NA | gene-LOC105280739 | GeneID:105280739 | LOC105280739 | Gene | LOC105280739 | protein_coding |
| character(0) | NA | character(0) | NC_039513.1 | 11675195 | 11675201 | 7 | * | 6 | 857.142857142857 | | | | | |
| NC_039513.1 | 13452183 | 13456265 | 4083 | + | Gnomon | gene | NA | NA | gene-LOC105275031 | GeneID:105275031 | LOC105275031 | Gene | LOC105275031 | protein_coding |
| character(0) | NA | character(0) | NC_039513.1 | 13453683 | 13453689 | 7 | * | 6 | 857.142857142857 | | | | | |
| NC_039513.1 | 13482189 | 13486570 | 4382 | - | Gnomon | gene | NA | NA | gene-LOC105275064 | GeneID:105275064 | LOC105275064 | Gene | LOC105275064 | protein_coding |
| character(0) | NA | character(0) | NC_039513.1 | 13485760 | 13485766 | 7 | * | 6 | 857.142857142857 | | | | | |
| NC_039513.1 | 15035458 | 15043240 | 7783 | + | Gnomon | gene | NA | NA | gene-LOC105285648 | GeneID:105285648 | LOC105285648 | Gene | LOC105285648 | protein_coding |
| character(0) | NA | character(0) | NC_039513.1 | 15037033 | 15037039 | 7 | * | 6 | 857.142857142857 | | | | | |
| NC_039513.1 | 15725203 | 15744087 | 18885 | + | Gnomon | gene | NA | NA | gene-LOC105280058 | GeneID:105280058 | LOC105280058 | Gene | LOC105280058 | protein_coding |
| character(0) | NA | character(0) | NC_039513.1 | 15726271 | 15726277 | 7 | * | 6 | 857.142857142857 | | | | | |
| NC_039514.1 | 5919621 | 5920916 | 1296 | - | Gnomon | gene | NA | NA | gene-LOC105279626 | GeneID:105279626 | LOC105279626 | Gene | LOC105279626 | protein_coding |
| character(0) | NA | character(0) | NC_039514.1 | 5920192 | 5920198 | 7 | * | 6 | 857.142857142857 | | | | | |
| NC_039514.1 | 6551602 | 6556437 | 4836 | - | Gnomon | gene | NA | NA | gene-LOC105285546 | GeneID:105285546 | LOC105285546 | Gene | LOC105285546 | protein_coding |
| character(0) | NA | character(0) | NC_039514.1 | 6555389 | 6555395 | 7 | * | 6 | 857.142857142857 | | | | | |
| NC_039514.1 | 8629561 | 8648221 | 18661 | + | Gnomon | gene | NA | NA | gene-LOC105278994 | GeneID:105278994 | LOC105278994 | Gene | LOC105278994 | protein_coding |
| character(0) | NA | character(0) | NC_039514.1 | 8630645 | 8630651 | 7 | * | 6 | 857.142857142857 | | | | | |
| NC_039514.1 | 8729983 | 8735648 | 5666 | + | Gnomon | gene | NA | NA | gene-LOC105279000 | GeneID:105279000 | LOC105279000 | Gene | LOC105279000 | protein_coding |
| character(0) | NA | character(0) | NC_039514.1 | 8731123 | 8731129 | 7 | * | 6 | 857.142857142857 | | | | | |
| NC_039514.1 | 9413016 | 9415003 | 1988 | - | Gnomon | gene | NA | NA | gene-LOC105284659 | GeneID:105284659 | LOC105284659 | Gene | LOC105284659 | protein_coding |
| character(0) | NA | character(0) | NC_039514.1 | 9414116 | 9414122 | 7 | * | 6 | 857.142857142857 | | | | | |
| NC_039514.1 | 11109536 | 11117116 | 7581 | - | Gnomon | gene | NA | NA | gene-LOC105281601 | GeneID:105281601 | LOC105281601 | Gene | LOC105281601 | protein_coding |
| character(0) | NA | character(0) | NC_039514.1 | 11113139 | 11113145 | 7 | * | 6 | 857.142857142857 | | | | | |
| NC_039514.1 | 11111779 | 11115572 | 3794 | + | Gnomon | gene | NA | NA | gene-LOC105281603 | GeneID:105281603 | LOC105281603 | Gene | LOC105281603 | protein_coding |
| character(0) | NA | character(0) | NC_039514.1 | 11113139 | 11113145 | 7 | * | 6 | 857.142857142857 | | | | | |
| NC_039515.1 | 1527980 | 1529634 | 1655 | + | Gnomon | gene | NA | NA | gene-LOC105274711 | GeneID:105274711 | LOC105274711 | Gene | LOC105274711 | protein_coding |
| character(0) | NA | character(0) | NC_039515.1 | 1528293 | 1528299 | 7 | * | 6 | 857.142857142857 | | | | | |
| NC_039515.1 | 2215137 | 2216986 | 1850 | - | Gnomon | gene | NA | NA | gene-LOC105278431 | GeneID:105278431 | LOC105278431 | Gene | LOC105278431 | protein_coding |
| character(0) | NA | character(0) | NC_039515.1 | 2215927 | 2215933 | 7 | * | 6 | 857.142857142857 | | | | | |
| NC_039515.1 | 3819603 | 3823750 | 4148 | + | Gnomon | gene | NA | NA | gene-LOC105278277 | GeneID:105278277 | LOC105278277 | Gene | LOC105278277 | protein_coding |
| character(0) | NA | character(0) | NC_039515.1 | 3820723 | 3820729 | 7 | * | 6 | 857.142857142857 | | | | | |
| NC_039515.1 | 3819837 | 3821619 | 1783 | + | Gnomon | gene | NA | NA | gene-LOC105278275 | GeneID:105278275 | LOC105278275 | Gene | LOC105278275 | protein_coding |
| character(0) | NA | character(0) | NC_039515.1 | 3820723 | 3820729 | 7 | * | 6 | 857.142857142857 | | | | | |
| NC_039515.1 | 5537006 | 5539696 | 2691 | + | Gnomon | gene | NA | NA | gene-LOC105277256 | GeneID:105277256 | LOC105277256 | Gene | LOC105277256 | protein_coding |
| character(0) | NA | character(0) | NC_039515.1 | 5537966 | 5537972 | 7 | * | 6 | 857.142857142857 | | | | | |
| NC_039515.1 | 6480766 | 6491993 | 11228 | + | Gnomon | gene | NA | NA | gene-LOC105282298 | GeneID:105282298 | LOC105282298 | Gene | LOC105282298 | protein_coding |
| character(0) | NA | character(0) | NC_039515.1 | 6481705 | 6481711 | 7 | * | 6 | 857.142857142857 | | | | | |
| NC_039515.1 | 6604872 | 6605723 | 852 | + | Gnomon | gene | NA | NA | gene-LOC105282280 | GeneID:105282280 | LOC105282280 | Gene | LOC105282280 | protein_coding |
| character(0) | NA | character(0) | NC_039515.1 | 6605539 | 6605545 | 7 | * | 6 | 857.142857142857 | | | | | |
| NC_039515.1 | 8113916 | 8116205 | 2290 | - | Gnomon | gene | NA | NA | gene-LOC105283727 | GeneID:105283727 | LOC105283727 | Gene | LOC105283727 | protein_coding |
| character(0) | NA | character(0) | NC_039515.1 | 8115487 | 8115493 | 7 | * | 6 | 857.142857142857 | | | | | |
| NC_039515.1 | 8973009 | 8978309 | 5301 | + | Gnomon | gene | NA | NA | gene-LOC105283697 | GeneID:105283697 | LOC105283697 | Gene | LOC105283697 | protein_coding |
| character(0) | NA | character(0) | NC_039515.1 | 8974548 | 8974554 | 7 | * | 6 | 857.142857142857 | | | | | |
| NC_039515.1 | 12611158 | 12612372 | 1215 | - | Gnomon | gene | NA | NA | gene-LOC105282147 | GeneID:105282147 | LOC105282147 | Gene | LOC105282147 | protein_coding |
| character(0) | NA | character(0) | NC_039515.1 | 12611603 | 12611609 | 7 | * | 6 | 857.142857142857 | | | | | |
| NC_039516.1 | 756663 | 760199 | 3537 | + | Gnomon | gene | NA | NA | gene-LOC105274638 | GeneID:105274638 | LOC105274638 | Gene | LOC105274638 | protein_coding |
| character(0) | NA | character(0) | NC_039516.1 | 757725 | 757731 | 7 | * | 6 | 857.142857142857 | | | | | |
| NC_039516.1 | 985951 | 990466 | 4516 | + | Gnomon | gene | NA | NA | gene-LOC105274590 | GeneID:105274590 | LOC105274590 | Gene | LOC105274590 | protein_coding |
| character(0) | NA | character(0) | NC_039516.1 | 987063 | 987069 | 7 | * | 6 | 857.142857142857 | | | | | |
| NC_039516.1 | 985951 | 990466 | 4516 | + | Gnomon | gene | NA | NA | gene-LOC105274590 | GeneID:105274590 | LOC105274590 | Gene | LOC105274590 | protein_coding |
| character(0) | NA | character(0) | NC_039516.1 | 987547 | 987553 | 7 | * | 6 | 857.142857142857 | | | | | |
| NC_039516.1 | 1549696 | 1551173 | 1478 | + | Gnomon | gene | NA | NA | gene-LOC105274497 | GeneID:105274497 | LOC105274497 | Gene | LOC105274497 | protein_coding |
| character(0) | NA | character(0) | NC_039516.1 | 1550744 | 1550750 | 7 | * | 6 | 857.142857142857 | | | | | |
| NC_039516.1 | 2241904 | 2245378 | 3475 | + | Gnomon | gene | NA | NA | gene-LOC105274464 | GeneID:105274464 | LOC105274464 | Gene | LOC105274464 | protein_coding |
| character(0) | NA | character(0) | NC_039516.1 | 2242648 | 2242654 | 7 | * | 6 | 857.142857142857 | | | | | |
| NC_039516.1 | 3088715 | 3091076 | 2362 | + | Gnomon | gene | NA | NA | gene-LOC105288196 | GeneID:105288196 | LOC105288196 | Gene | LOC105288196 | protein_coding |
| character(0) | NA | character(0) | NC_039516.1 | 3089812 | 3089818 | 7 | * | 6 | 857.142857142857 | | | | | |
| NC_039516.1 | 10251143 | 10256122 | 4980 | - | Gnomon | gene | NA | NA | gene-LOC105287447 | GeneID:105287447 | LOC105287447 | Gene | LOC105287447 | protein_coding |
| character(0) | NA | character(0) | NC_039516.1 | 10255095 | 10255101 | 7 | * | 6 | 857.142857142857 | | | | | |
| NC_039516.1 | 11790047 | 11792531 | 2485 | + | Gnomon | gene | NA | NA | gene-LOC105278656 | GeneID:105278656 | LOC105278656 | Gene | LOC105278656 | protein_coding |
| character(0) | NA | character(0) | NC_039516.1 | 11791250 | 11791256 | 7 | * | 6 | 857.142857142857 | | | | | |
| NC_039516.1 | 12834895 | 12836314 | 1420 | - | Gnomon | gene | NA | NA | gene-LOC105275307 | GeneID:105275307 | LOC105275307 | Gene | LOC105275307 | protein_coding |
| character(0) | NA | character(0) | NC_039516.1 | 12835552 | 12835558 | 7 | * | 6 | 857.142857142857 | | | | | |
| NC_039516.1 | 13054997 | 13057481 | 2485 | + | Gnomon | gene | NA | NA | gene-LOC105275352 | GeneID:105275352 | LOC105275352 | Gene | LOC105275352 | protein_coding |

| | | | | | | | | | | | | | | |
|---|---|---|---|---|---|---|---|---|---|---|---|---|---|---|
| character(0) | NA | character(0) | NC_039516.1 | 13055949 | 13055955 | 7 | * | 6 | 857.142857142857 | | | | | |
| NC_039516.1 | 13386687 | 13390618 | 3932 | + | Gnomon | gene | NA | NA | gene-LOC105278022 | GeneID:105278022 | LOC105278022 | Gene | LOC105278022 | protein_coding |
| character(0) | NA | character(0) | NC_039516.1 | 13387869 | 13387875 | 7 | * | 6 | 857.142857142857 | | | | | |
| NC_039516.1 | 13497458 | 13503461 | 6004 | - | Gnomon | gene | NA | NA | gene-LOC105277915 | GeneID:105277915 | LOC105277915 | Gene | LOC105277915 | protein_coding |
| character(0) | NA | character(0) | NC_039516.1 | 13502711 | 13502717 | 7 | * | 6 | 857.142857142857 | | | | | |
| NC_039516.1 | 13518014 | 13520692 | 2679 | + | Gnomon | gene | NA | NA | gene-LOC105277817 | GeneID:105277817 | LOC105277817 | Gene | LOC105277817 | protein_coding |
| character(0) | NA | character(0) | NC_039516.1 | 13518940 | 13518946 | 7 | * | 6 | 857.142857142857 | | | | | |
| NC_039517.1 | 502319 | 504215 | 1897 | + | Gnomon | gene | NA | NA | gene-LOC105278873 | GeneID:105278873 | LOC105278873 | Gene | LOC105278873 | protein_coding |
| character(0) | NA | character(0) | NC_039517.1 | 503110 | 503116 | 7 | * | 6 | 857.142857142857 | | | | | |
| NC_039517.1 | 867927 | 869156 | 1230 | - | Gnomon | gene | NA | NA | gene-LOC105278824 | GeneID:105278824 | LOC105278824 | Gene | LOC105278824 | protein_coding |
| character(0) | NA | character(0) | NC_039517.1 | 868363 | 868369 | 7 | * | 6 | 857.142857142857 | | | | | |
| NC_039517.1 | 2307792 | 2312395 | 4604 | - | Gnomon | gene | NA | NA | gene-LOC105281943 | GeneID:105281943 | LOC105281943 | Gene | LOC105281943 | protein_coding |
| character(0) | NA | character(0) | NC_039517.1 | 2309012 | 2309018 | 7 | * | 6 | 857.142857142857 | | | | | |
| NC_039517.1 | 7306074 | 7309288 | 3215 | + | Gnomon | gene | NA | NA | gene-LOC105284202 | GeneID:105284202 | LOC105284202 | Gene | LOC105284202 | protein_coding |
| character(0) | NA | character(0) | NC_039517.1 | 7307471 | 7307477 | 7 | * | 6 | 857.142857142857 | | | | | |
| NC_039517.1 | 7446985 | 7449823 | 2839 | + | Gnomon | gene | NA | NA | gene-LOC105284223 | GeneID:105284223 | LOC105284223 | Gene | LOC105284223 | protein_coding |
| character(0) | NA | character(0) | NC_039517.1 | 7448190 | 7448196 | 7 | * | 6 | 857.142857142857 | | | | | |
| NC_039517.1 | 9368753 | 9371050 | 2298 | - | Gnomon | gene | NA | NA | gene-LOC105282220 | GeneID:105282220 | LOC105282220 | Gene | LOC105282220 | protein_coding |
| character(0) | NA | character(0) | NC_039517.1 | 9370300 | 9370306 | 7 | * | 6 | 857.142857142857 | | | | | |
| NC_039518.1 | 2232165 | 2249269 | 17105 | + | Gnomon | gene | NA | NA | gene-LOC105283878 | GeneID:105283878 | LOC105283878 | Gene | LOC105283878 | protein_coding |
| character(0) | NA | character(0) | NC_039518.1 | 2233357 | 2233363 | 7 | * | 6 | 857.142857142857 | | | | | |
| NC_039518.1 | 2657034 | 2658634 | 1601 | + | Gnomon | gene | NA | NA | gene-LOC105285392 | GeneID:105285392 | LOC105285392 | Gene | LOC105285392 | protein_coding |
| character(0) | NA | character(0) | NC_039518.1 | 2658047 | 2658053 | 7 | * | 6 | 857.142857142857 | | | | | |
| NC_039518.1 | 2681489 | 2682795 | 1307 | - | Gnomon | gene | NA | NA | gene-LOC105285394 | GeneID:105285394 | LOC105285394 | Gene | LOC105285394 | protein_coding |
| character(0) | NA | character(0) | NC_039518.1 | 2681945 | 2681951 | 7 | * | 6 | 857.142857142857 | | | | | |
| NC_039518.1 | 5288531 | 5294826 | 6296 | - | Gnomon | gene | NA | NA | gene-LOC105285410 | GeneID:105285410 | LOC105285410 | Gene | LOC105285410 | protein_coding |
| character(0) | NA | character(0) | NC_039518.1 | 5293748 | 5293754 | 7 | * | 6 | 857.142857142857 | | | | | |
| NC_039519.1 | 197513 | 200423 | 2911 | + | Gnomon | gene | NA | NA | gene-LOC105282793 | GeneID:105282793 | LOC105282793 | Gene | LOC105282793 | protein_coding |
| character(0) | NA | character(0) | NC_039519.1 | 198095 | 198101 | 7 | * | 6 | 857.142857142857 | | | | | |
| NC_039519.1 | 8239374 | 8248322 | 8949 | - | Gnomon | gene | NA | NA | gene-LOC105287985 | GeneID:105287985 | LOC105287985 | Gene | LOC105287985 | protein_coding |
| character(0) | NA | character(0) | NC_039519.1 | 8247911 | 8247917 | 7 | * | 6 | 857.142857142857 | | | | | |
| NC_039513.1 | 5176546 | 5178099 | 1554 | + | Gnomon | gene | NA | NA | gene-LOC105279718 | GeneID:105279718 | LOC105279718 | Gene | LOC105279718 | protein_coding |
| character(0) | NA | character(0) | NC_039513.1 | 5177245 | 5177254 | 10 | * | 8 | 800 | | | | | |
| NC_039519.1 | 333719 | 337798 | 4080 | - | Gnomon | gene | NA | NA | gene-LOC105285496 | GeneID:105285496 | LOC105285496 | Gene | LOC105285496 | protein_coding |
| character(0) | NA | character(0) | NC_039519.1 | 337077 | 337086 | 10 | * | 8 | 800 | | | | | |
| NC_039519.1 | 8084633 | 8088546 | 3914 | + | Gnomon | gene | NA | NA | gene-LOC105274767 | GeneID:105274767 | LOC105274767 | Gene | LOC105274767 | protein_coding |
| character(0) | NA | character(0) | NC_039519.1 | 8085717 | 8085726 | 10 | * | 8 | 800 | | | | | |

```
Directory: MRPR File: rmp-Ob_LI_BCphaseA1.txt

Overlap of methylation-rich regions with genome features for Ob sample Ob_LI_BCphaseA1

Total size of the methylation-rich regions of Ob_LI_BCphaseA1:                                       1009 bp (  0.0% of genome)
Size of overlap of methylation-rich regions of Ob Ob_LI_BCphaseA1 with genic regions:                    988 bp ( 97.9% of total size)   ( 1.31 O/E)
Size of overlap of methylation-rich regions of Ob Ob_LI_BCphaseA1 with   exon regions:                     916 bp ( 90.8% of total size) ( 0.36 O/E)
Size of overlap of methylation-rich regions of Ob Ob_LI_BCphaseA1 with   intron regions:                    72 bp (  7.1% of total size) ( 0.13 O/E)
Size of overlap of methylation-rich regions of Ob Ob_LI_BCphaseA1 with intergenic regions:               21 bp (  2.1% of total size)   ( 3.84 O/E)
Size of overlap of methylation-rich regions of Ob Ob_LI_BCphaseA1 with   promoter regions:                   0 bp (  0.0% of total size) ( 0.00 O/E)
Size of overlap of methylation-rich regions of Ob Ob_LI_BCphaseA1 with   other intergenic regions:          21 bp (  2.1% of total size) ( 0.09 O/E)

Overlap of methylation-poor regions with genome features for Ob sample Ob_LI_BCphaseA1

Total size of the methylation-poor regions of Ob_LI_BCphaseA1:                                   46352207 bp ( 20.7% of genome)
Size of overlap of methylation-poor regions of Ob Ob_LI_BCphaseA1 with genic regions:               24456610 bp ( 52.8% of total size)   ( 0.71 O/E)
Size of overlap of methylation-poor regions of Ob Ob_LI_BCphaseA1 with   exon regions:                3907596 bp (  8.4% of total size) ( 2.21 O/E)
Size of overlap of methylation-poor regions of Ob Ob_LI_BCphaseA1 with   intron regions:             20549014 bp ( 44.3% of total size) ( 0.81 O/E)
Size of overlap of methylation-poor regions of Ob Ob_LI_BCphaseA1 with intergenic regions:          21895597 bp ( 47.2% of total size)   ( 2.07 O/E)
Size of overlap of methylation-poor regions of Ob Ob_LI_BCphaseA1 with   promoter regions:             473787 bp (  1.0% of total size) ( 0.33 O/E)
Size of overlap of methylation-poor regions of Ob Ob_LI_BCphaseA1 with   other intergenic regions:    21421810 bp ( 46.2% of total size) ( 2.06 O/E)
```

```
Directory: MRPR File: rmp-Ob_LI_BCphaseA2.txt

Overlap of methylation-rich regions with genome features for Ob sample Ob_LI_BCphaseA2

Total size of the methylation-rich regions of Ob_LI_BCphaseA2:                                       1364 bp (  0.0% of genome)
Size of overlap of methylation-rich regions of Ob Ob_LI_BCphaseA2 with genic regions:                   1350 bp ( 99.0% of total size)   ( 1.33 O/E)
Size of overlap of methylation-rich regions of Ob Ob_LI_BCphaseA2 with   exon regions:                    1262 bp ( 92.5% of total size) ( 0.32 O/E)
Size of overlap of methylation-rich regions of Ob Ob_LI_BCphaseA2 with   intron regions:                    88 bp (  6.5% of total size) ( 0.12 O/E)
Size of overlap of methylation-rich regions of Ob Ob_LI_BCphaseA2 with intergenic regions:                14 bp (  1.0% of total size)   ( 3.88 O/E)
Size of overlap of methylation-rich regions of Ob Ob_LI_BCphaseA2 with   promoter regions:                   0 bp (  0.0% of total size) ( 0.00 O/E)
Size of overlap of methylation-rich regions of Ob Ob_LI_BCphaseA2 with   other intergenic regions:          14 bp (  1.0% of total size) ( 0.05 O/E)

Overlap of methylation-poor regions with genome features for Ob sample Ob_LI_BCphaseA2

Total size of the methylation-poor regions of Ob_LI_BCphaseA2:                                   56747838 bp ( 25.4% of genome)
Size of overlap of methylation-poor regions of Ob Ob_LI_BCphaseA2 with genic regions:               33175031 bp ( 58.5% of total size)   ( 0.78 O/E)
Size of overlap of methylation-poor regions of Ob Ob_LI_BCphaseA2 with   exon regions:                4981976 bp (  8.8% of total size) ( 2.48 O/E)
Size of overlap of methylation-poor regions of Ob Ob_LI_BCphaseA2 with   intron regions:             28193055 bp ( 49.7% of total size) ( 0.91 O/E)
Size of overlap of methylation-poor regions of Ob Ob_LI_BCphaseA2 with intergenic regions:          23572807 bp ( 41.5% of total size)   ( 2.29 O/E)
Size of overlap of methylation-poor regions of Ob Ob_LI_BCphaseA2 with   promoter regions:             555757 bp (  1.0% of total size) ( 0.32 O/E)
Size of overlap of methylation-poor regions of Ob Ob_LI_BCphaseA2 with   other intergenic regions:    23017050 bp ( 40.6% of total size) ( 1.81 O/E)
```

```
Directory: MRPR File: rmp-Ob_LI_BCphaseB1.txt

Overlap of methylation-rich regions with genome features for Ob sample Ob_LI_BCphaseB1

Total size of the methylation-rich regions of Ob_LI_BCphaseB1:                                       1436 bp (  0.0% of genome)
Size of overlap of methylation-rich regions of Ob Ob_LI_BCphaseB1 with genic regions:                   1422 bp ( 99.0% of total size)   ( 1.33 O/E)
Size of overlap of methylation-rich regions of Ob Ob_LI_BCphaseB1 with   exon regions:                    1307 bp ( 91.0% of total size) ( 0.40 O/E)
Size of overlap of methylation-rich regions of Ob Ob_LI_BCphaseB1 with   intron regions:                   115 bp (  8.0% of total size) ( 0.15 O/E)
Size of overlap of methylation-rich regions of Ob Ob_LI_BCphaseB1 with intergenic regions:                14 bp (  1.0% of total size)   ( 3.88 O/E)
Size of overlap of methylation-rich regions of Ob Ob_LI_BCphaseB1 with   promoter regions:                   0 bp (  0.0% of total size) ( 0.00 O/E)
Size of overlap of methylation-rich regions of Ob Ob_LI_BCphaseB1 with   other intergenic regions:          14 bp (  1.0% of total size) ( 0.04 O/E)

Overlap of methylation-poor regions with genome features for Ob sample Ob_LI_BCphaseB1

Total size of the methylation-poor regions of Ob_LI_BCphaseB1:                                   38634147 bp ( 17.3% of genome)
Size of overlap of methylation-poor regions of Ob Ob_LI_BCphaseB1 with genic regions:               20933073 bp ( 54.2% of total size)   ( 0.73 O/E)
Size of overlap of methylation-poor regions of Ob Ob_LI_BCphaseB1 with   exon regions:                 3580330 bp (  9.3% of total size) ( 2.24 O/E)
Size of overlap of methylation-poor regions of Ob Ob_LI_BCphaseB1 with   intron regions:              17352743 bp ( 44.9% of total size) ( 0.83 O/E)
Size of overlap of methylation-poor regions of Ob Ob_LI_BCphaseB1 with intergenic regions:          17701074 bp ( 45.8% of total size)   ( 2.12 O/E)
Size of overlap of methylation-poor regions of Ob Ob_LI_BCphaseB1 with   promoter regions:              428247 bp (  1.1% of total size) ( 0.36 O/E)
Size of overlap of methylation-poor regions of Ob Ob_LI_BCphaseB1 with   other intergenic regions:    17272827 bp ( 44.7% of total size) ( 1.99 O/E)
```

Directory: MRPR File: rmp-Ob_LI_BCphaseB2.txt

Overlap of methylation-rich regions with genome features for Ob sample Ob_LI_BCphaseB2

```
Total size of the methylation-rich regions of Ob_LI_BCphaseB2:                                       1234 bp (  0.0% of genome)
Size of overlap of methylation-rich regions of Ob Ob_LI_BCphaseB2 with genic regions:                   1213 bp ( 98.3% of total size)   ( 1.32 O/E)
Size of overlap of methylation-rich regions of Ob Ob_LI_BCphaseB2 with   exon regions:                    1127 bp ( 91.3% of total size) ( 0.35 O/E)
Size of overlap of methylation-rich regions of Ob Ob_LI_BCphaseB2 with   intron regions:                    86 bp (  7.0% of total size) ( 0.13 O/E)
Size of overlap of methylation-rich regions of Ob Ob_LI_BCphaseB2 with intergenic regions:                21 bp (  1.7% of total size)   ( 3.85 O/E)
Size of overlap of methylation-rich regions of Ob Ob_LI_BCphaseB2 with   promoter regions:                   0 bp (  0.0% of total size) ( 0.00 O/E)
Size of overlap of methylation-rich regions of Ob Ob_LI_BCphaseB2 with   other intergenic regions:          21 bp (  1.7% of total size) ( 0.08 O/E)
```

Overlap of methylation-poor regions with genome features for Ob sample Ob_LI_BCphaseB2

```
Total size of the methylation-poor regions of Ob_LI_BCphaseB2:                                   41432691 bp ( 18.5% of genome)
Size of overlap of methylation-poor regions of Ob Ob_LI_BCphaseB2 with genic regions:               20999154 bp ( 50.7% of total size)   ( 0.68 O/E)
Size of overlap of methylation-poor regions of Ob Ob_LI_BCphaseB2 with   exon regions:                 3615052 bp (  8.7% of total size) ( 2.09 O/E)
Size of overlap of methylation-poor regions of Ob Ob_LI_BCphaseB2 with   intron regions:              17384102 bp ( 42.0% of total size) ( 0.77 O/E)
Size of overlap of methylation-poor regions of Ob Ob_LI_BCphaseB2 with intergenic regions:          20433537 bp ( 49.3% of total size)   ( 1.99 O/E)
Size of overlap of methylation-poor regions of Ob Ob_LI_BCphaseB2 with   promoter regions:             483141 bp (  1.2% of total size) ( 0.38 O/E)
Size of overlap of methylation-poor regions of Ob Ob_LI_BCphaseB2 with   other intergenic regions:    19950396 bp ( 48.1% of total size) ( 2.15 O/E)
```

Directory: MRPR File: rmp-Ob_LI_RphaseA1.txt

```
Overlap of methylation-rich regions with genome features for Ob sample Ob_LI_RphaseA1

Total size of the methylation-rich regions of Ob_LI_RphaseA1:                                   1372 bp (  0.0% of genome)
Size of overlap of methylation-rich regions of Ob Ob_LI_RphaseA1 with genic regions:                1358 bp ( 99.0% of total size)   ( 1.33 O/E)
Size of overlap of methylation-rich regions of Ob Ob_LI_RphaseA1 with   exon regions:                 1277 bp ( 93.1% of total size) ( 0.29 O/E)
Size of overlap of methylation-rich regions of Ob Ob_LI_RphaseA1 with   intron regions:                 81 bp (  5.9% of total size) ( 0.11 O/E)
Size of overlap of methylation-rich regions of Ob Ob_LI_RphaseA1 with intergenic regions:             14 bp (  1.0% of total size)   ( 3.88 O/E)
Size of overlap of methylation-rich regions of Ob Ob_LI_RphaseA1 with   promoter regions:                0 bp (  0.0% of total size) ( 0.00 O/E)
Size of overlap of methylation-rich regions of Ob Ob_LI_RphaseA1 with   other intergenic regions:       14 bp (  1.0% of total size) ( 0.05 O/E)

Overlap of methylation-poor regions with genome features for Ob sample Ob_LI_RphaseA1

Total size of the methylation-poor regions of Ob_LI_RphaseA1:                               53142440 bp ( 23.7% of genome)
Size of overlap of methylation-poor regions of Ob Ob_LI_RphaseA1 with genic regions:            31324203 bp ( 58.9% of total size)   ( 0.79 O/E)
Size of overlap of methylation-poor regions of Ob Ob_LI_RphaseA1 with   exon regions:              4619707 bp (  8.7% of total size) ( 2.51 O/E)
Size of overlap of methylation-poor regions of Ob Ob_LI_RphaseA1 with   intron regions:           26704496 bp ( 50.2% of total size) ( 0.92 O/E)
Size of overlap of methylation-poor regions of Ob Ob_LI_RphaseA1 with intergenic regions:       21818237 bp ( 41.1% of total size)   ( 2.31 O/E)
Size of overlap of methylation-poor regions of Ob Ob_LI_RphaseA1 with   promoter regions:            537988 bp (  1.0% of total size) ( 0.33 O/E)
Size of overlap of methylation-poor regions of Ob Ob_LI_RphaseA1 with   other intergenic regions:  21280249 bp ( 40.0% of total size) ( 1.78 O/E)
```

Directory: MRPR File: rmp-Ob_LI_RphaseA2.txt

Overlap of methylation-rich regions with genome features for Ob sample Ob_LI_RphaseA2

```
Total size of the methylation-rich regions of Ob_LI_RphaseA2:                                    1991 bp (  0.0% of genome)
Size of overlap of methylation-rich regions of Ob Ob_LI_RphaseA2 with genic regions:                  1970 bp ( 99.0% of total size)   ( 1.33 O/E)
Size of overlap of methylation-rich regions of Ob Ob_LI_RphaseA2 with   exon regions:                 1829 bp ( 91.9% of total size) ( 0.35 O/E)
Size of overlap of methylation-rich regions of Ob Ob_LI_RphaseA2 with   intron regions:                141 bp (  7.1% of total size) ( 0.13 O/E)
Size of overlap of methylation-rich regions of Ob Ob_LI_RphaseA2 with intergenic regions:              21 bp (  1.1% of total size)   ( 3.88 O/E)
Size of overlap of methylation-rich regions of Ob Ob_LI_RphaseA2 with   promoter regions:                0 bp (  0.0% of total size) ( 0.00 O/E)
Size of overlap of methylation-rich regions of Ob Ob_LI_RphaseA2 with   other intergenic regions:       21 bp (  1.1% of total size) ( 0.05 O/E)
```

Overlap of methylation-poor regions with genome features for Ob sample Ob_LI_RphaseA2

```
Total size of the methylation-poor regions of Ob_LI_RphaseA2:                                24695357 bp ( 11.0% of genome)
Size of overlap of methylation-poor regions of Ob Ob_LI_RphaseA2 with genic regions:             11052036 bp ( 44.8% of total size)   ( 0.60 O/E)
Size of overlap of methylation-poor regions of Ob Ob_LI_RphaseA2 with   exon regions:             2466755 bp ( 10.0% of total size) ( 1.73 O/E)
Size of overlap of methylation-poor regions of Ob Ob_LI_RphaseA2 with   intron regions:           8585281 bp ( 34.8% of total size) ( 0.64 O/E)
Size of overlap of methylation-poor regions of Ob Ob_LI_RphaseA2 with intergenic regions:        13643321 bp ( 55.2% of total size)   ( 1.75 O/E)
Size of overlap of methylation-poor regions of Ob Ob_LI_RphaseA2 with   promoter regions:          350356 bp (  1.4% of total size) ( 0.46 O/E)
Size of overlap of methylation-poor regions of Ob Ob_LI_RphaseA2 with   other intergenic regions:  13292965 bp ( 53.8% of total size) ( 2.40 O/E)
```

```
Directory: MRPR File: rmp-Ob_LI_RphaseB1.txt

Overlap of methylation-rich regions with genome features for Ob sample Ob_LI_RphaseB1

Total size of the methylation-rich regions of Ob_LI_RphaseB1:                                       1296 bp (  0.0% of genome)
Size of overlap of methylation-rich regions of Ob Ob_LI_RphaseB1 with genic regions:                   1282 bp ( 98.9% of total size)   ( 1.33 O/E)
Size of overlap of methylation-rich regions of Ob Ob_LI_RphaseB1 with   exon regions:                    1195 bp ( 92.2% of total size) ( 0.33 O/E)
Size of overlap of methylation-rich regions of Ob Ob_LI_RphaseB1 with   intron regions:                    87 bp (  6.7% of total size) ( 0.12 O/E)
Size of overlap of methylation-rich regions of Ob Ob_LI_RphaseB1 with intergenic regions:                14 bp (  1.1% of total size)   ( 3.88 O/E)
Size of overlap of methylation-rich regions of Ob Ob_LI_RphaseB1 with   promoter regions:                   0 bp (  0.0% of total size) ( 0.00 O/E)
Size of overlap of methylation-rich regions of Ob Ob_LI_RphaseB1 with   other intergenic regions:          14 bp (  1.1% of total size) ( 0.05 O/E)

Overlap of methylation-poor regions with genome features for Ob sample Ob_LI_RphaseB1

Total size of the methylation-poor regions of Ob_LI_RphaseB1:                                   37055002 bp ( 16.6% of genome)
Size of overlap of methylation-poor regions of Ob Ob_LI_RphaseB1 with genic regions:               18531884 bp ( 50.0% of total size)   ( 0.67 O/E)
Size of overlap of methylation-poor regions of Ob Ob_LI_RphaseB1 with   exon regions:                 3177874 bp (  8.6% of total size) ( 2.07 O/E)
Size of overlap of methylation-poor regions of Ob Ob_LI_RphaseB1 with   intron regions:              15354010 bp ( 41.4% of total size) ( 0.76 O/E)
Size of overlap of methylation-poor regions of Ob Ob_LI_RphaseB1 with intergenic regions:          18523118 bp ( 50.0% of total size)   ( 1.96 O/E)
Size of overlap of methylation-poor regions of Ob Ob_LI_RphaseB1 with   promoter regions:              417582 bp (  1.1% of total size) ( 0.37 O/E)
Size of overlap of methylation-poor regions of Ob Ob_LI_RphaseB1 with   other intergenic regions:    18105536 bp ( 48.9% of total size) ( 2.18 O/E)
```

Directory: MRPR File: rmp-Ob_LI_RphaseB2.txt

Overlap of methylation-rich regions with genome features for Ob sample Ob_LI_RphaseB2

```
Total size of the methylation-rich regions of Ob_LI_RphaseB2:                                       1224 bp (  0.0% of genome)
Size of overlap of methylation-rich regions of Ob Ob_LI_RphaseB2 with genic regions:                   1210 bp ( 98.9% of total size)   ( 1.33 O/E)
Size of overlap of methylation-rich regions of Ob Ob_LI_RphaseB2 with   exon regions:                    1144 bp ( 93.5% of total size) ( 0.27 O/E)
Size of overlap of methylation-rich regions of Ob Ob_LI_RphaseB2 with   intron regions:                    66 bp (  5.4% of total size) ( 0.10 O/E)
Size of overlap of methylation-rich regions of Ob Ob_LI_RphaseB2 with intergenic regions:                14 bp (  1.1% of total size)   ( 3.87 O/E)
Size of overlap of methylation-rich regions of Ob Ob_LI_RphaseB2 with   promoter regions:                   0 bp (  0.0% of total size) ( 0.00 O/E)
Size of overlap of methylation-rich regions of Ob Ob_LI_RphaseB2 with   other intergenic regions:          14 bp (  1.1% of total size) ( 0.05 O/E)
```

Overlap of methylation-poor regions with genome features for Ob sample Ob_LI_RphaseB2

```
Total size of the methylation-poor regions of Ob_LI_RphaseB2:                                   46161627 bp ( 20.6% of genome)
Size of overlap of methylation-poor regions of Ob Ob_LI_RphaseB2 with genic regions:               25118523 bp ( 54.4% of total size)   ( 0.73 O/E)
Size of overlap of methylation-poor regions of Ob Ob_LI_RphaseB2 with   exon regions:                4030708 bp (  8.7% of total size) ( 2.28 O/E)
Size of overlap of methylation-poor regions of Ob Ob_LI_RphaseB2 with   intron regions:             21087815 bp ( 45.7% of total size) ( 0.84 O/E)
Size of overlap of methylation-poor regions of Ob Ob_LI_RphaseB2 with intergenic regions:          21043104 bp ( 45.6% of total size)   ( 2.13 O/E)
Size of overlap of methylation-poor regions of Ob Ob_LI_RphaseB2 with   promoter regions:              483863 bp (  1.1% of total size) ( 0.34 O/E)
Size of overlap of methylation-poor regions of Ob Ob_LI_RphaseB2 with   other intergenic regions:    20559241 bp ( 44.5% of total size) ( 1.98 O/E)
```

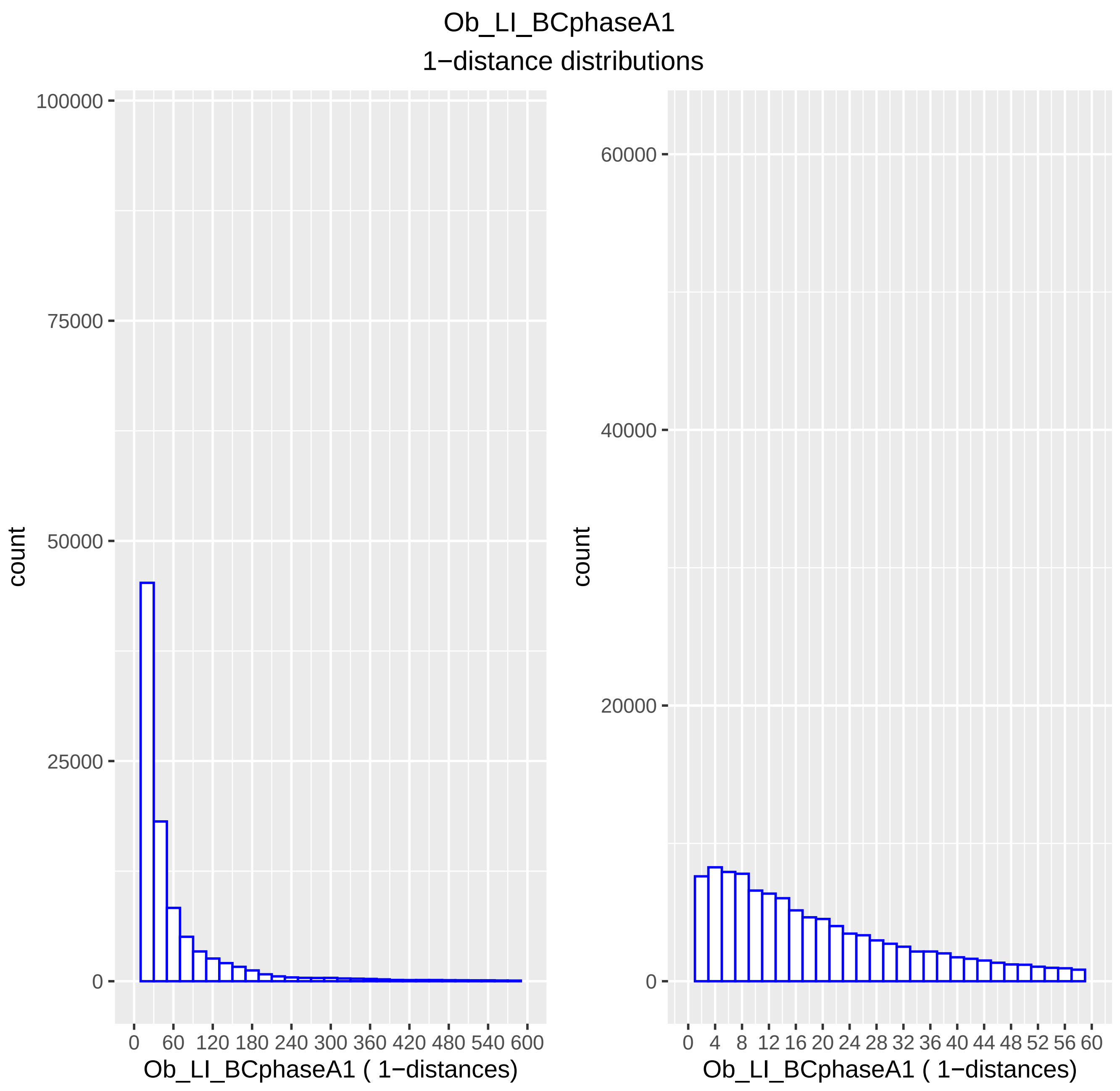
Ob_LI_BCphaseA1
1−distance distributions
count
100000
75000
50000
25000
0
0 60 120 180 240 300 360 420 480 540 600
Ob_LI_BCphaseA1 ( 1−distances)
count
60000
40000
20000
0
0 4 8 12 16 20 24 28 32 36 40 44 48 52 56 60
Ob_LI_BCphaseA1 ( 1−distances)

Ob_LI_BCphaseA2
1−distance distributions

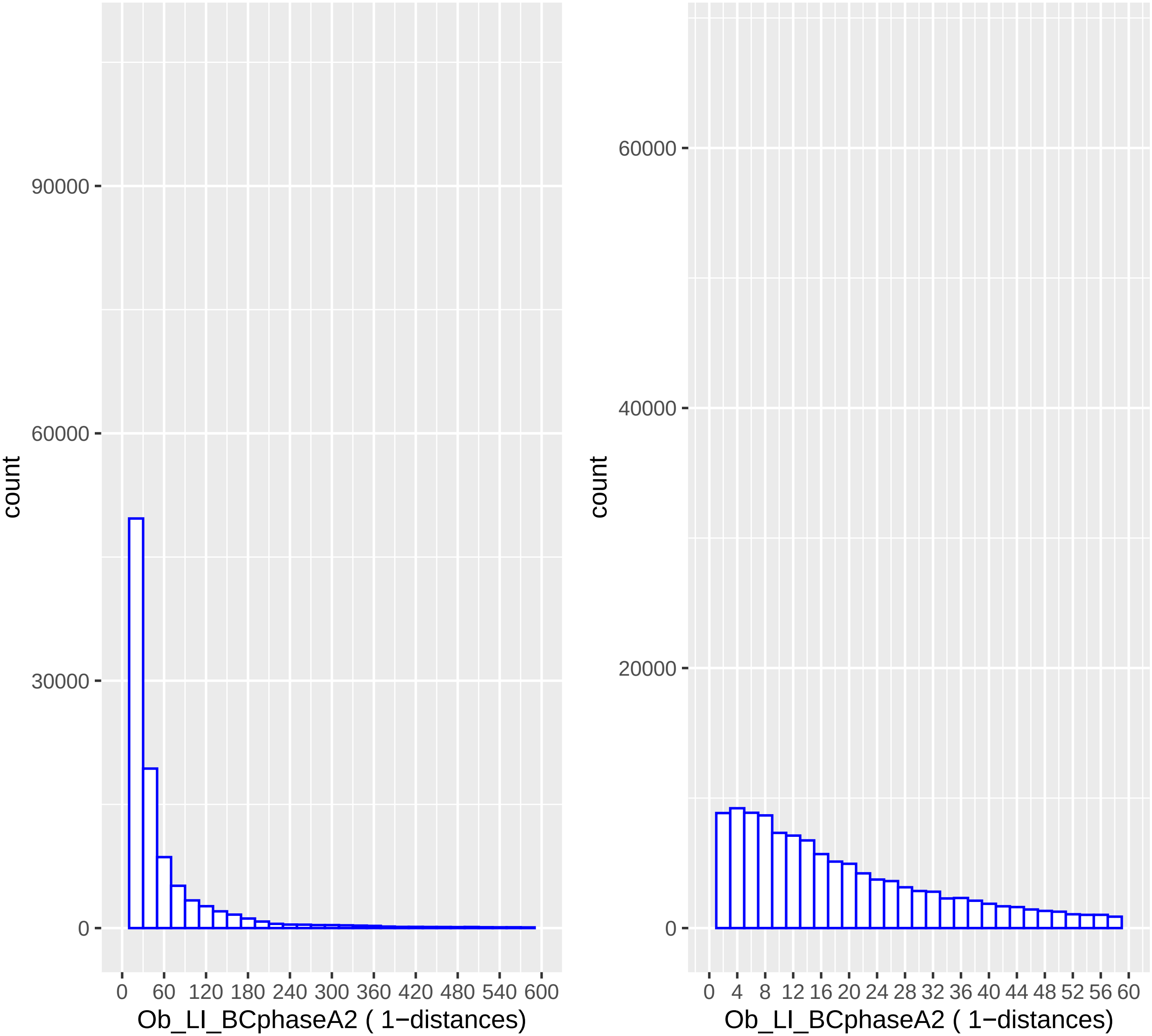

Ob_LI_BCphaseB1
1−distance distributions

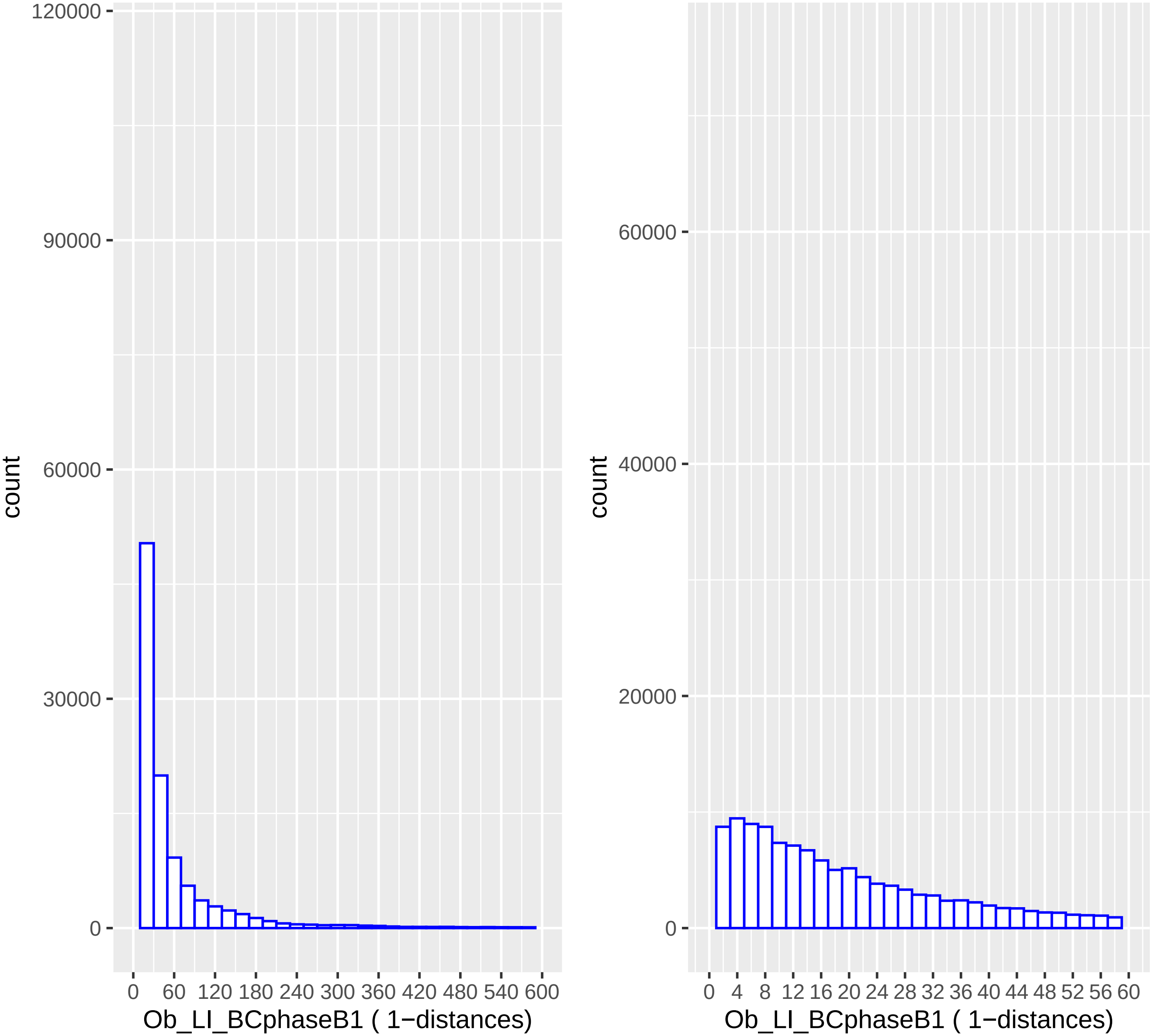

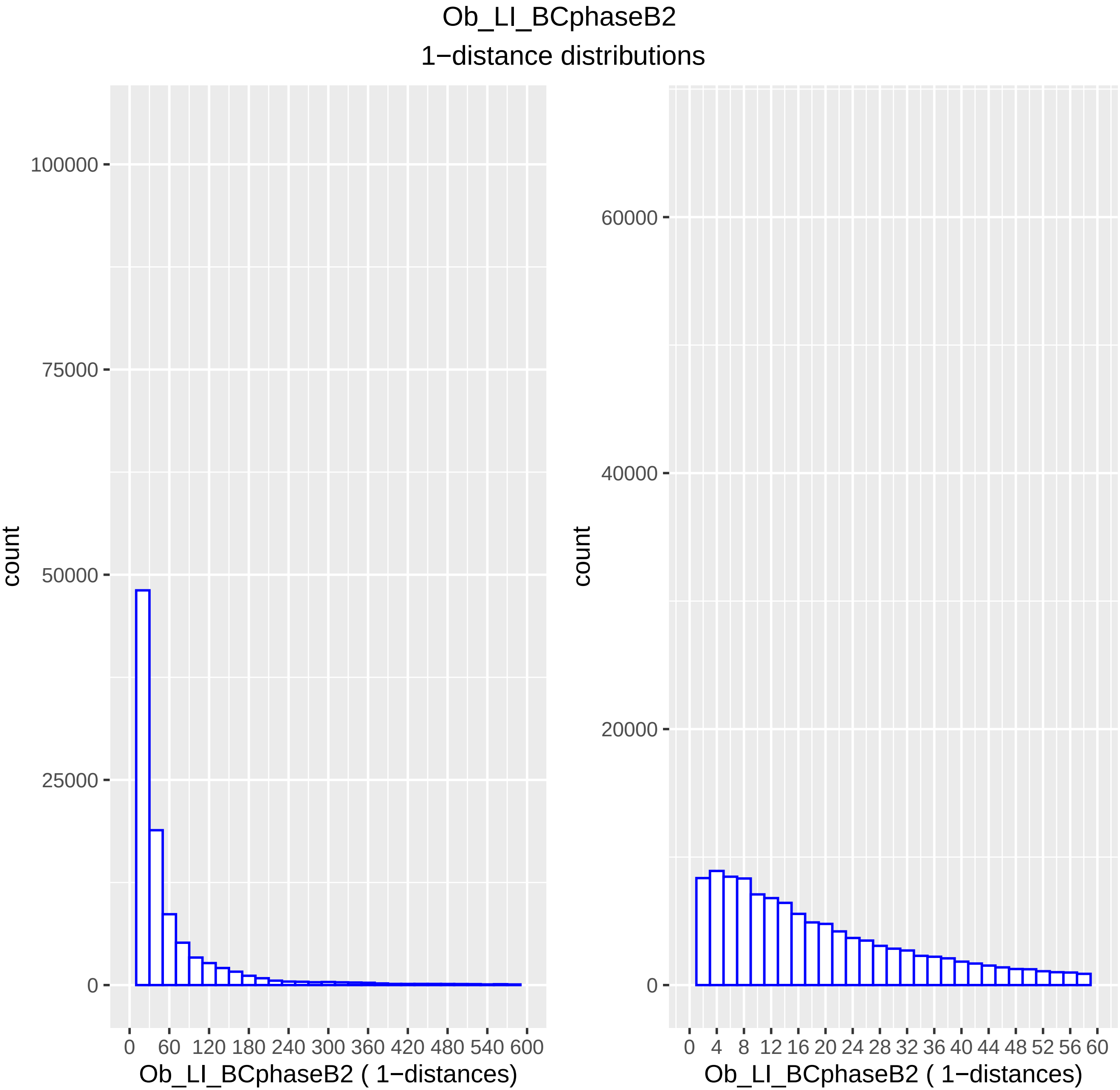
Ob_LI_BCphaseB2
1−distance distributions
count
100000
75000
50000
25000
0
0 60 120 180 240 300 360 420 480 540 600
Ob_LI_BCphaseB2 ( 1−distances)
count
60000
40000
20000
0
0 4 8 12 16 20 24 28 32 36 40 44 48 52 56 60
Ob_LI_BCphaseB2 ( 1−distances)

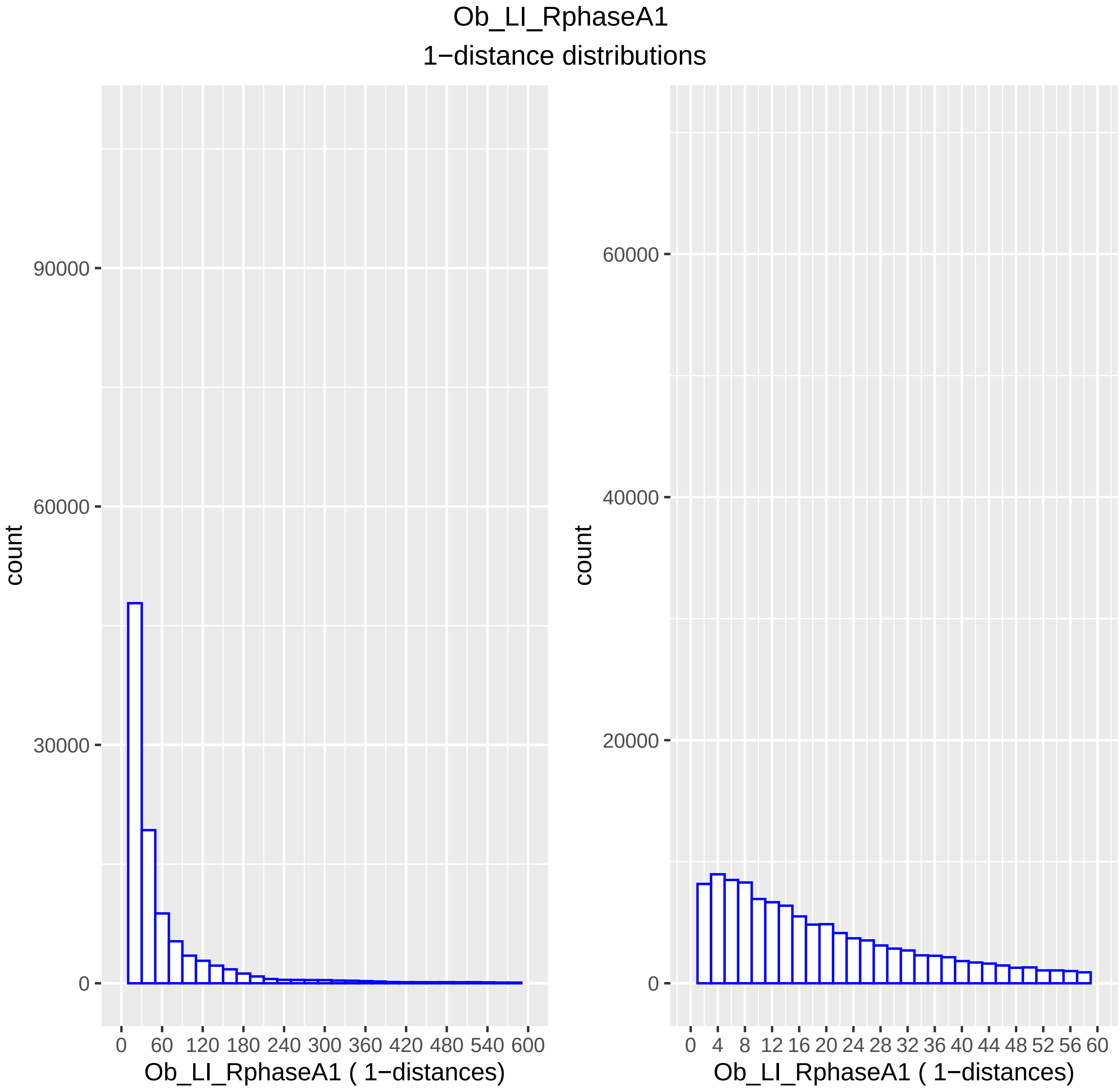

Ob_LI_RphaseA1
1−distance distributions
count
90000
60000
30000
0
0 60 120 180 240 300 360 420 480 540 600
Ob_LI_RphaseA1 ( 1−distances)
count
60000
40000
20000
0
0 4 8 12 16 20 24 28 32 36 40 44 48 52 56 60
Ob_LI_RphaseA1 ( 1−distances)

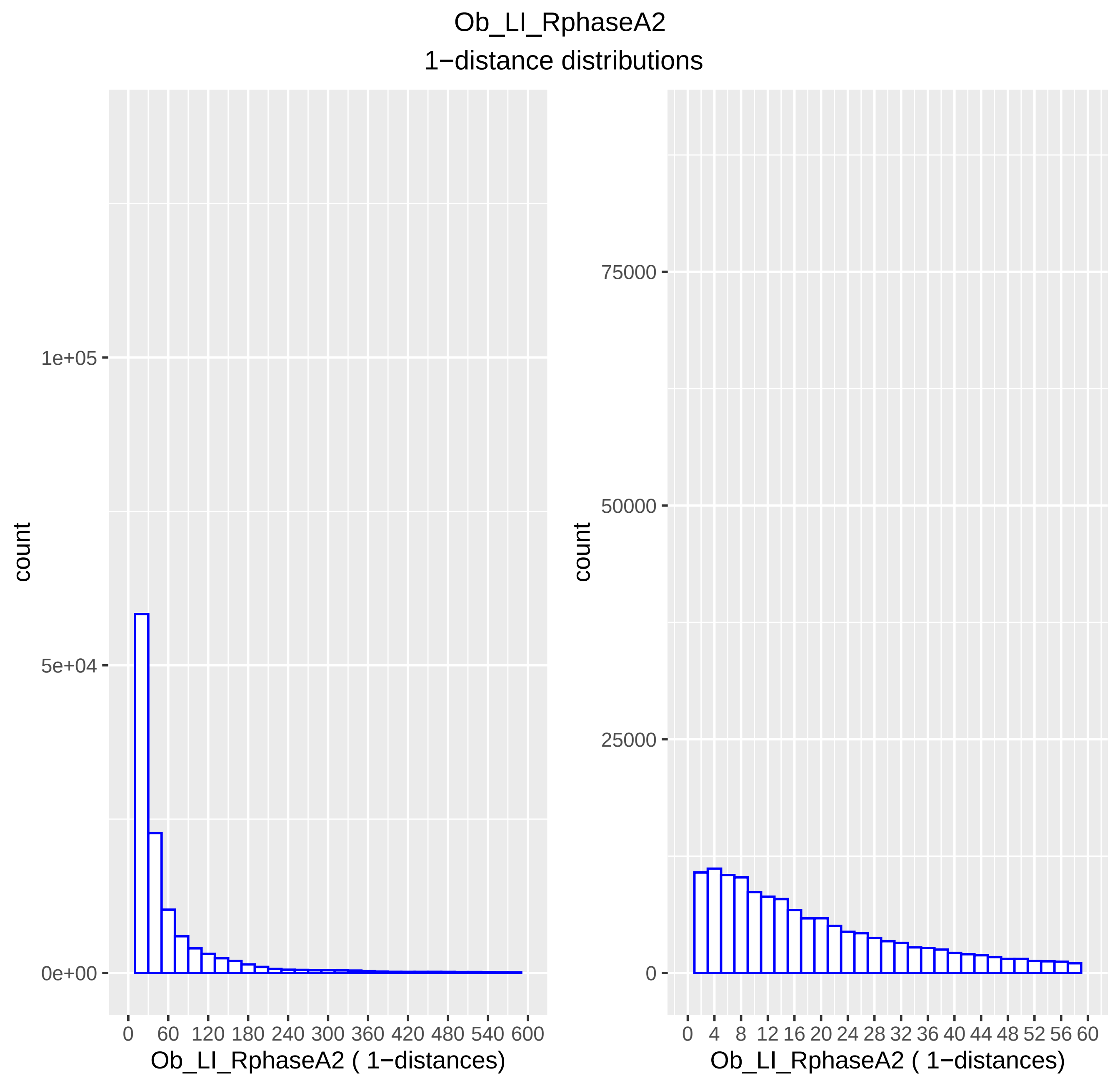
Ob_LI_RphaseA2
1−distance distributions
count
1e+05
5e+04
0e+00
0
60
120
180
240
300
360
420
480
540
600
Ob_LI_RphaseA2 ( 1−distances)
count
75000
50000
25000
0
0
4
8
12
16
20
24
28
32
36
40
44
48
52
56
60
Ob_LI_RphaseA2 ( 1−distances)

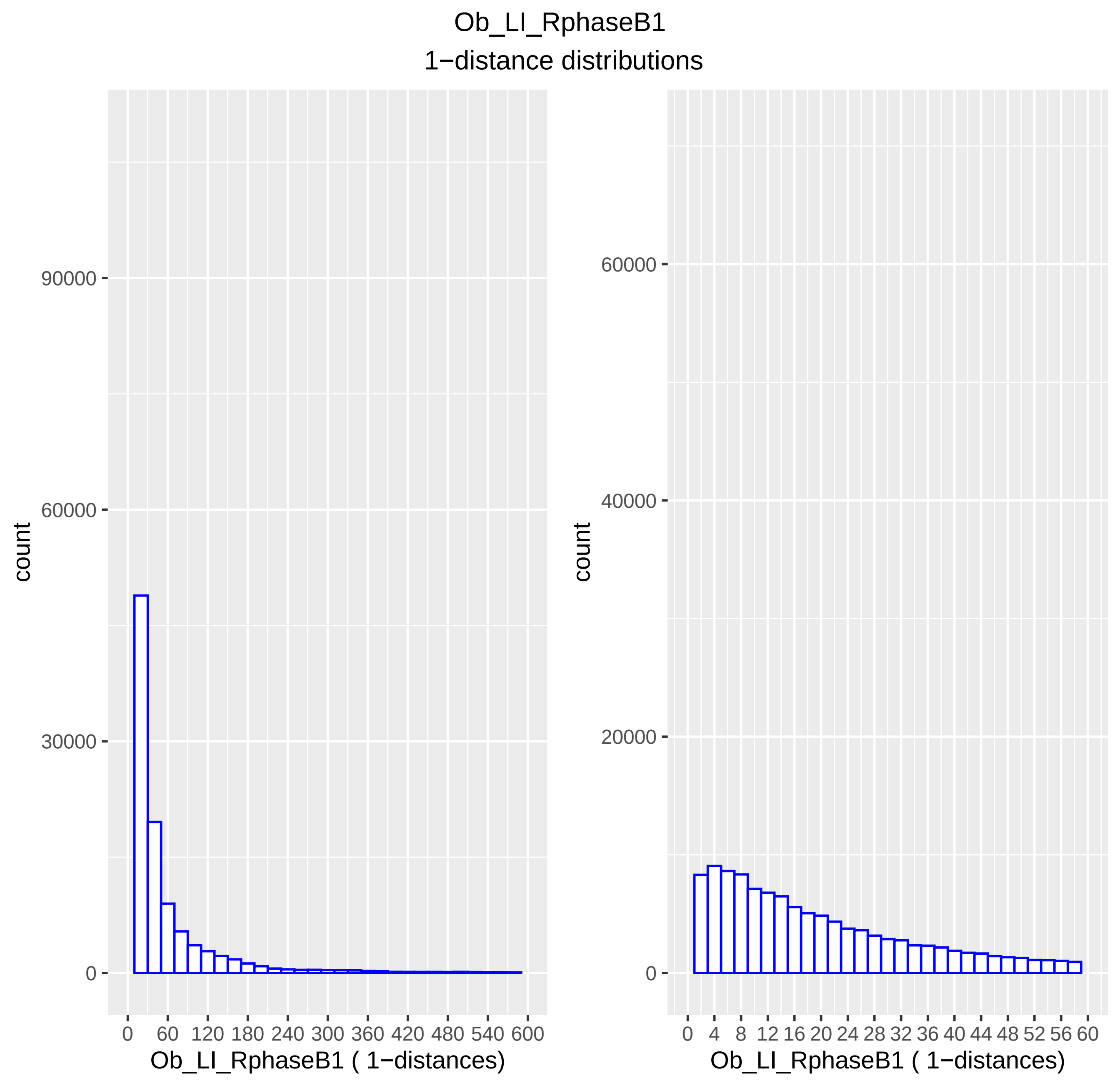

Ob_LI_RphaseB1
1−distance distributions
count
90000
60000
30000
0
0 60 120 180 240 300 360 420 480 540 600
Ob_LI_RphaseB1 ( 1−distances)
count
60000
40000
20000
0
0 4 8 12 16 20 24 28 32 36 40 44 48 52 56 60
Ob_LI_RphaseB1 ( 1−distances)

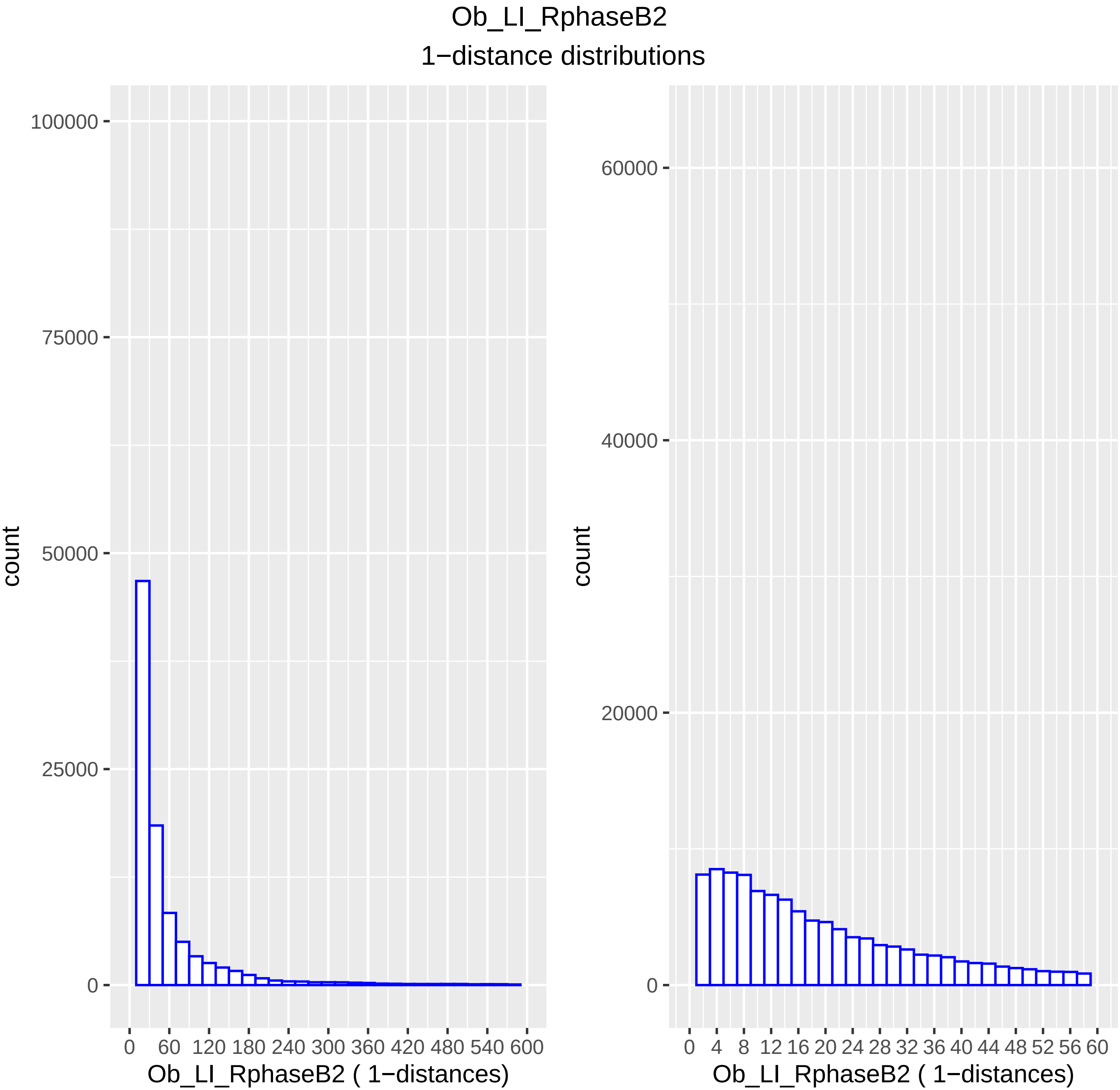

Ob_LI_RphaseB2
1−distance distributions
count
100000
75000
50000
25000
0
0 60 120 180 240 300 360 420 480 540 600
Ob_LI_RphaseB2 ( 1−distances)
count
60000
40000
20000
0
0 4 8 12 16 20 24 28 32 36 40 44 48 52 56 60
Ob_LI_RphaseB2 ( 1−distances)

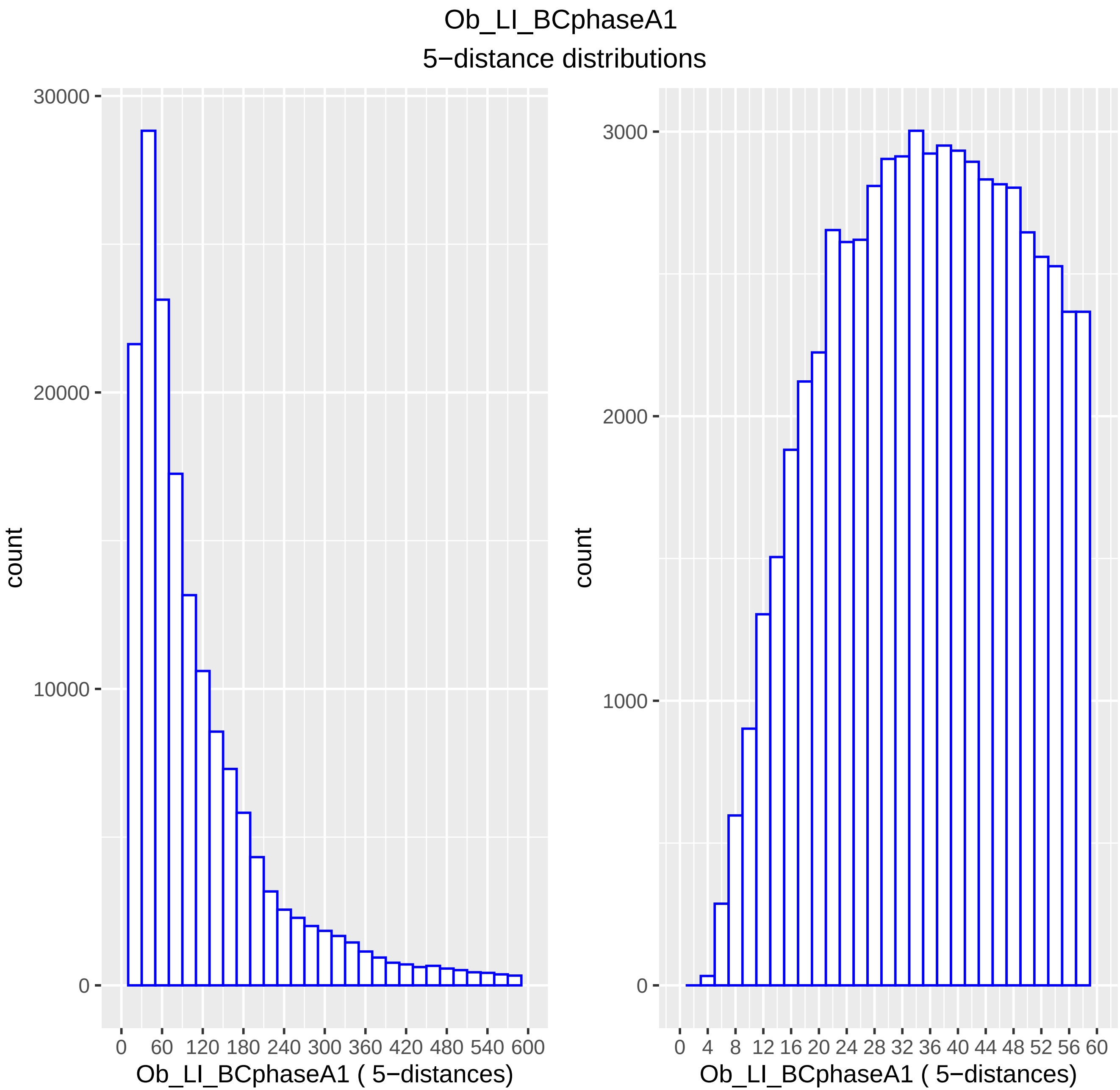
Ob_LI_BCphaseA1
5−distance distributions
count
0
10000
20000
30000
0 60 120 180 240 300 360 420 480 540 600
Ob_LI_BCphaseA1 ( 5−distances)
count
0
1000
2000
3000
0 4 8 12 16 20 24 28 32 36 40 44 48 52 56 60
Ob_LI_BCphaseA1 ( 5−distances)

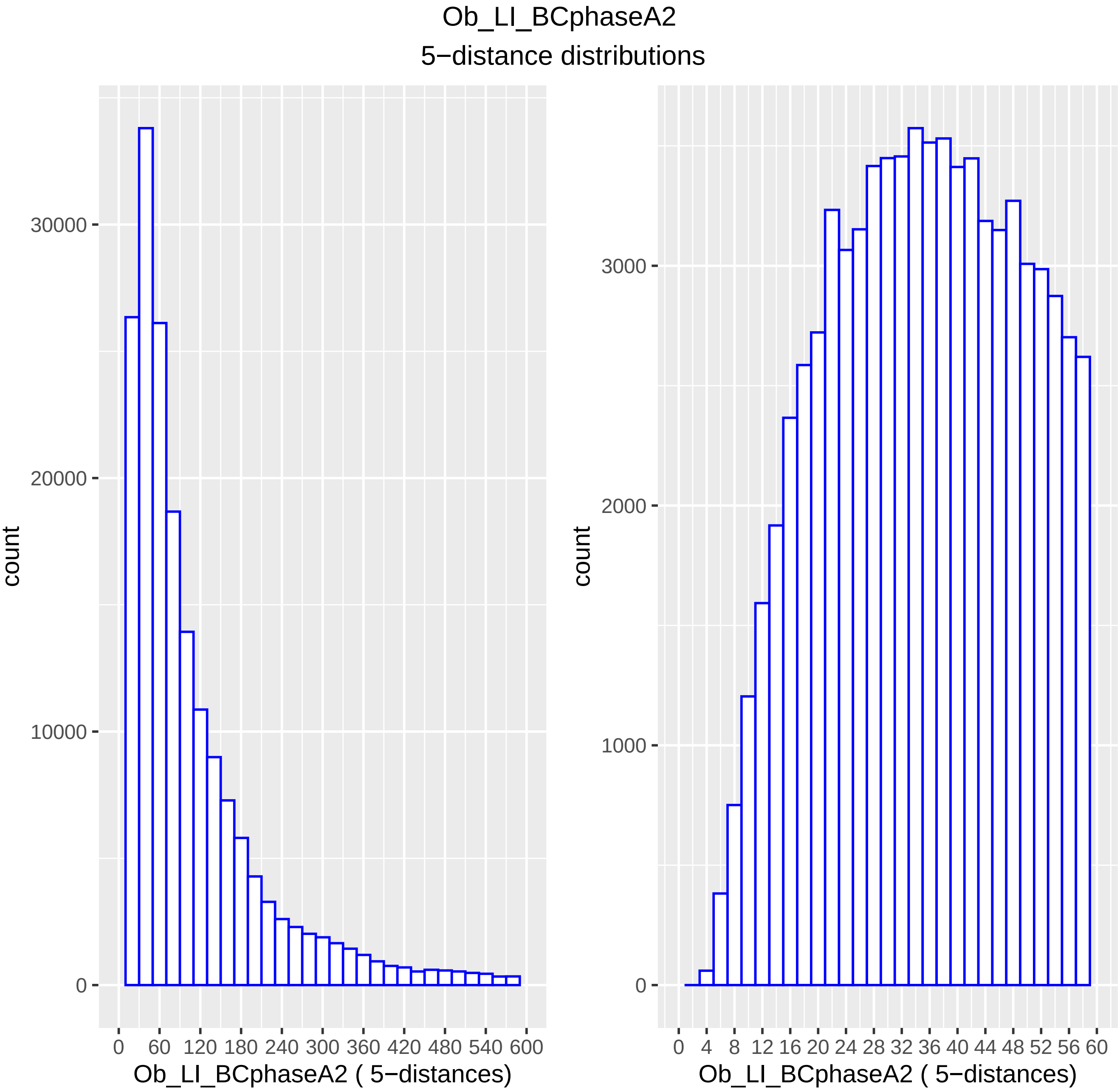
Ob_LI_BCphaseA2
5−distance distributions
count
30000
20000
10000
0
0 60 120 180 240 300 360 420 480 540 600
Ob_LI_BCphaseA2 ( 5−distances)
count
3000
2000
1000
0
0 4 8 12 16 20 24 28 32 36 40 44 48 52 56 60
Ob_LI_BCphaseA2 ( 5−distances)

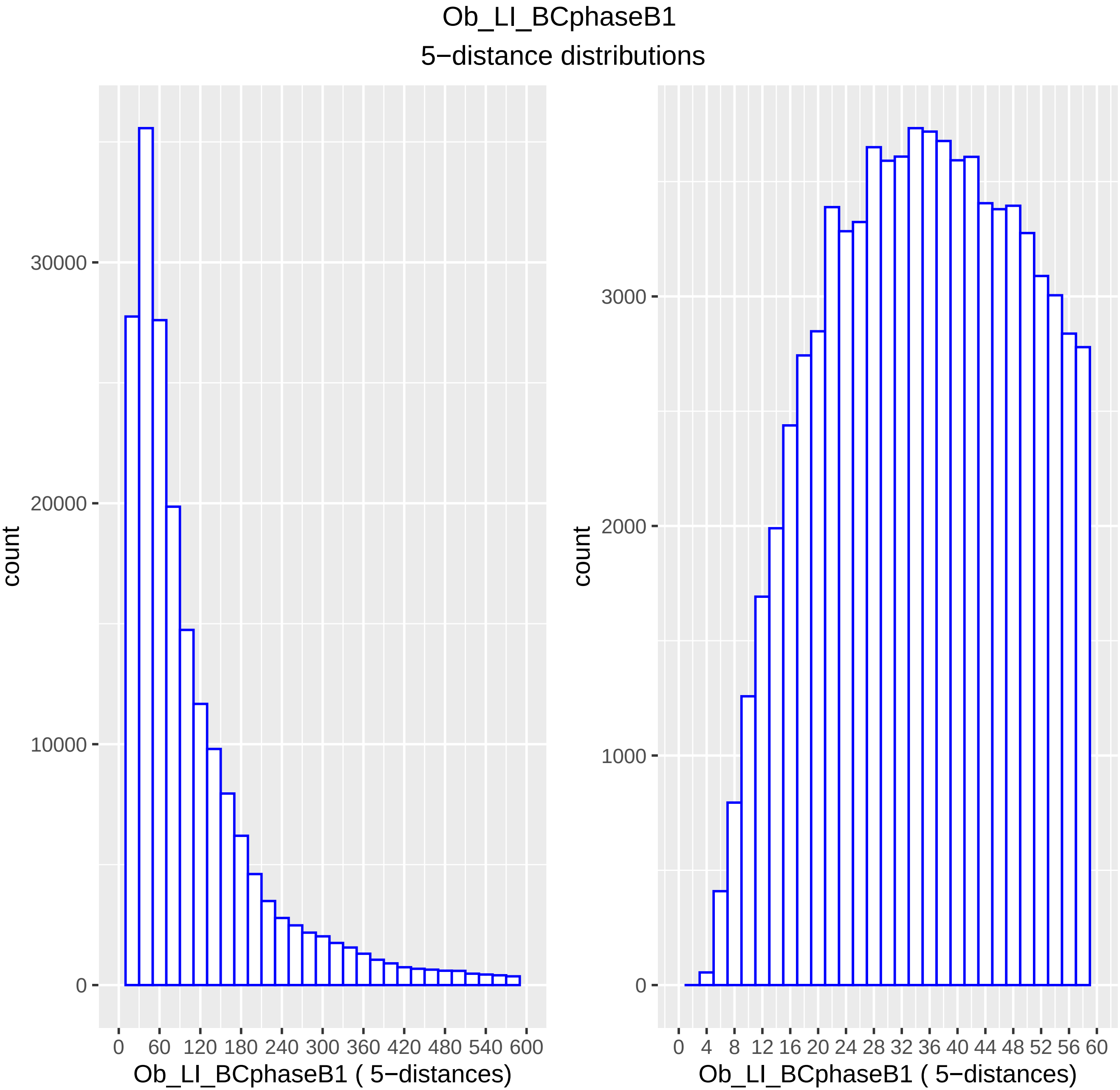

Ob_LI_BCphaseB1
5−distance distributions
30000
20000
10000
0
count
0 60 120 180 240 300 360 420 480 540 600
Ob_LI_BCphaseB1 ( 5−distances)
3000
2000
1000
0
count
0 4 8 12 16 20 24 28 32 36 40 44 48 52 56 60
Ob_LI_BCphaseB1 ( 5−distances)

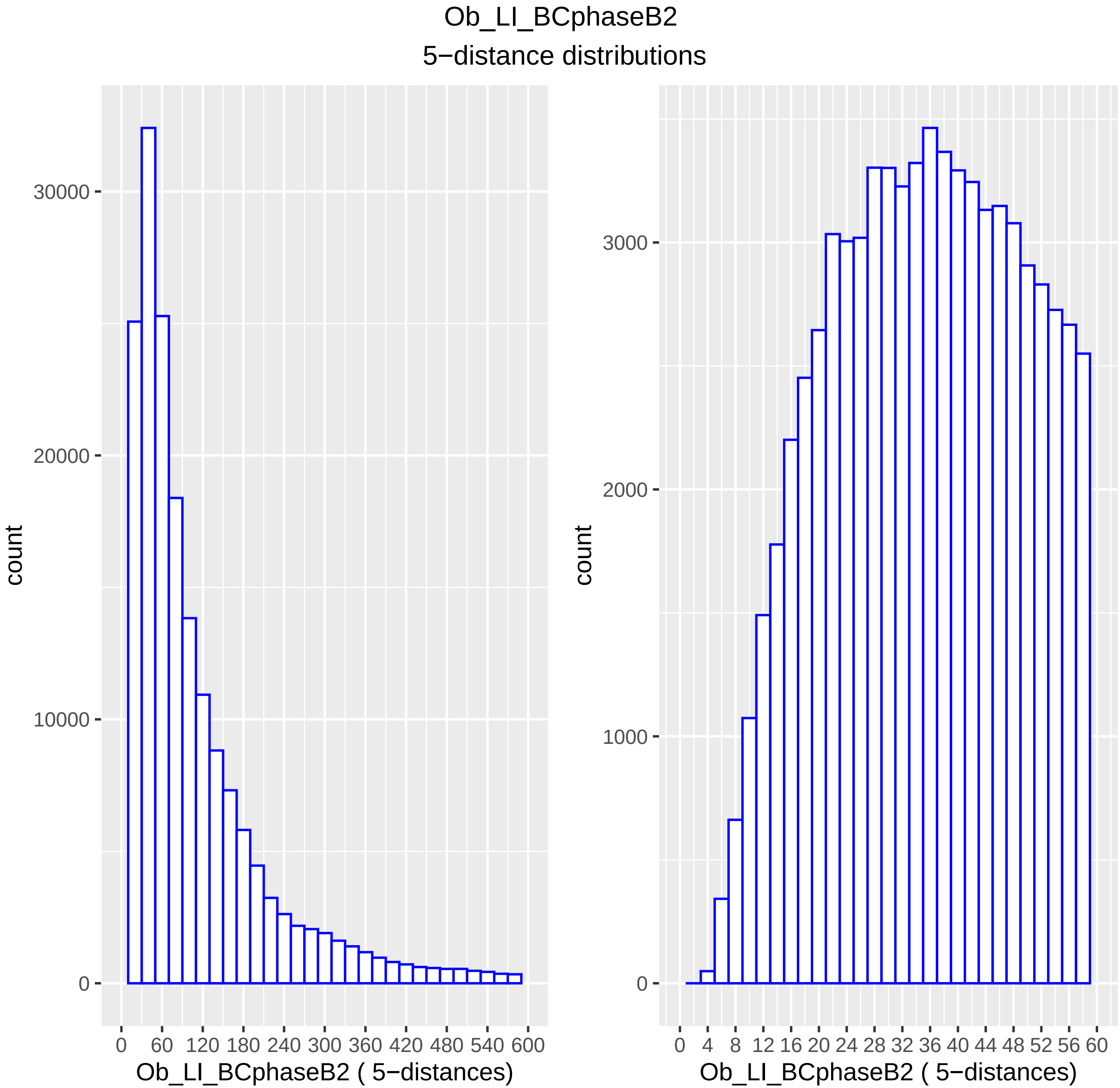

Ob_LI_BCphaseB2
5−distance distributions
count
30000
20000
10000
0
0 60 120 180 240 300 360 420 480 540 600
Ob_LI_BCphaseB2 ( 5−distances)
count
3000
2000
1000
0
0 4 8 12 16 20 24 28 32 36 40 44 48 52 56 60
Ob_LI_BCphaseB2 ( 5−distances)

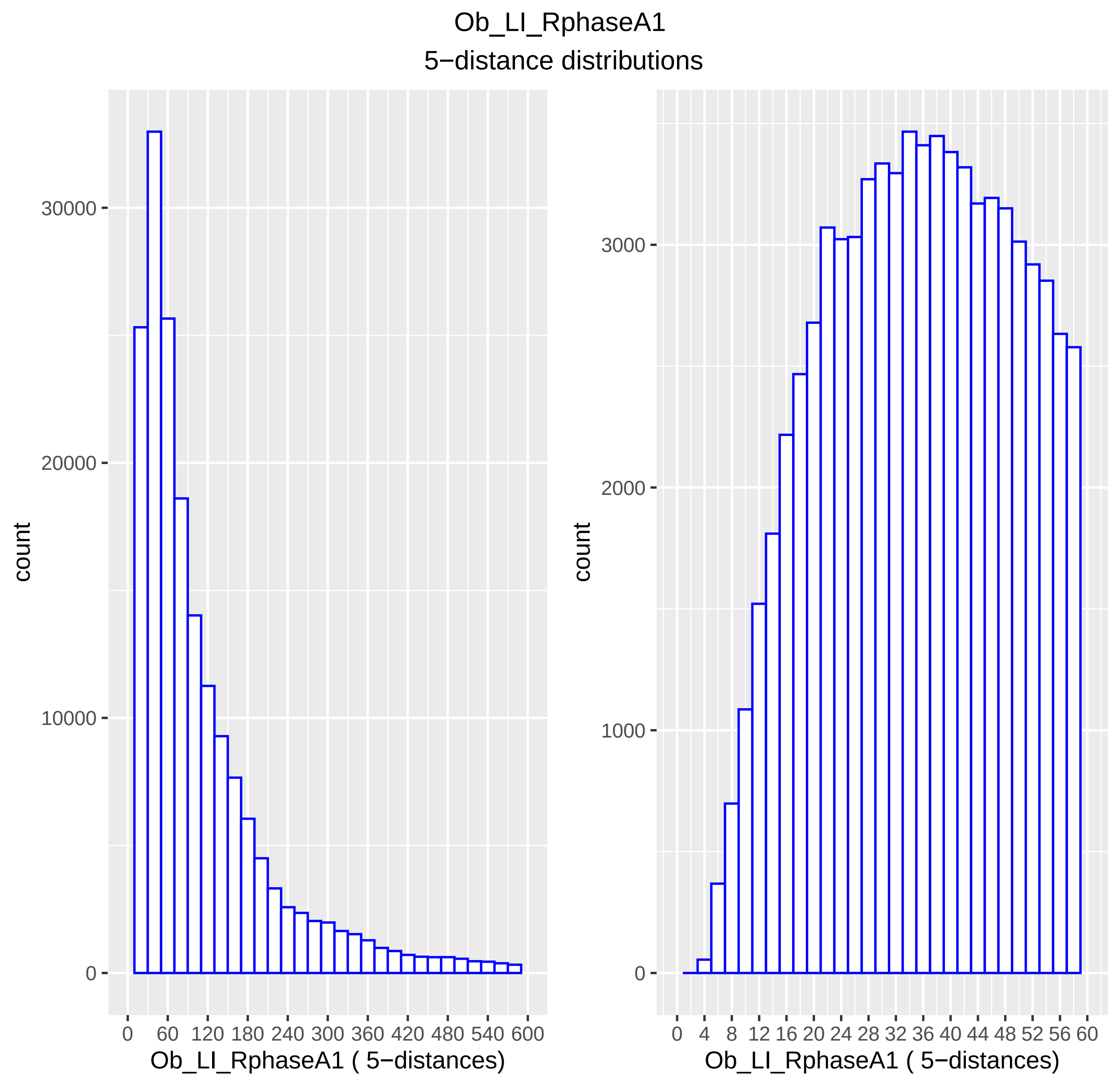
Ob_LI_RphaseA1
5−distance distributions
count
30000
20000
10000
0
0 60 120 180 240 300 360 420 480 540 600
Ob_LI_RphaseA1 ( 5−distances)
count
3000
2000
1000
0
0 4 8 12 16 20 24 28 32 36 40 44 48 52 56 60
Ob_LI_RphaseA1 ( 5−distances)

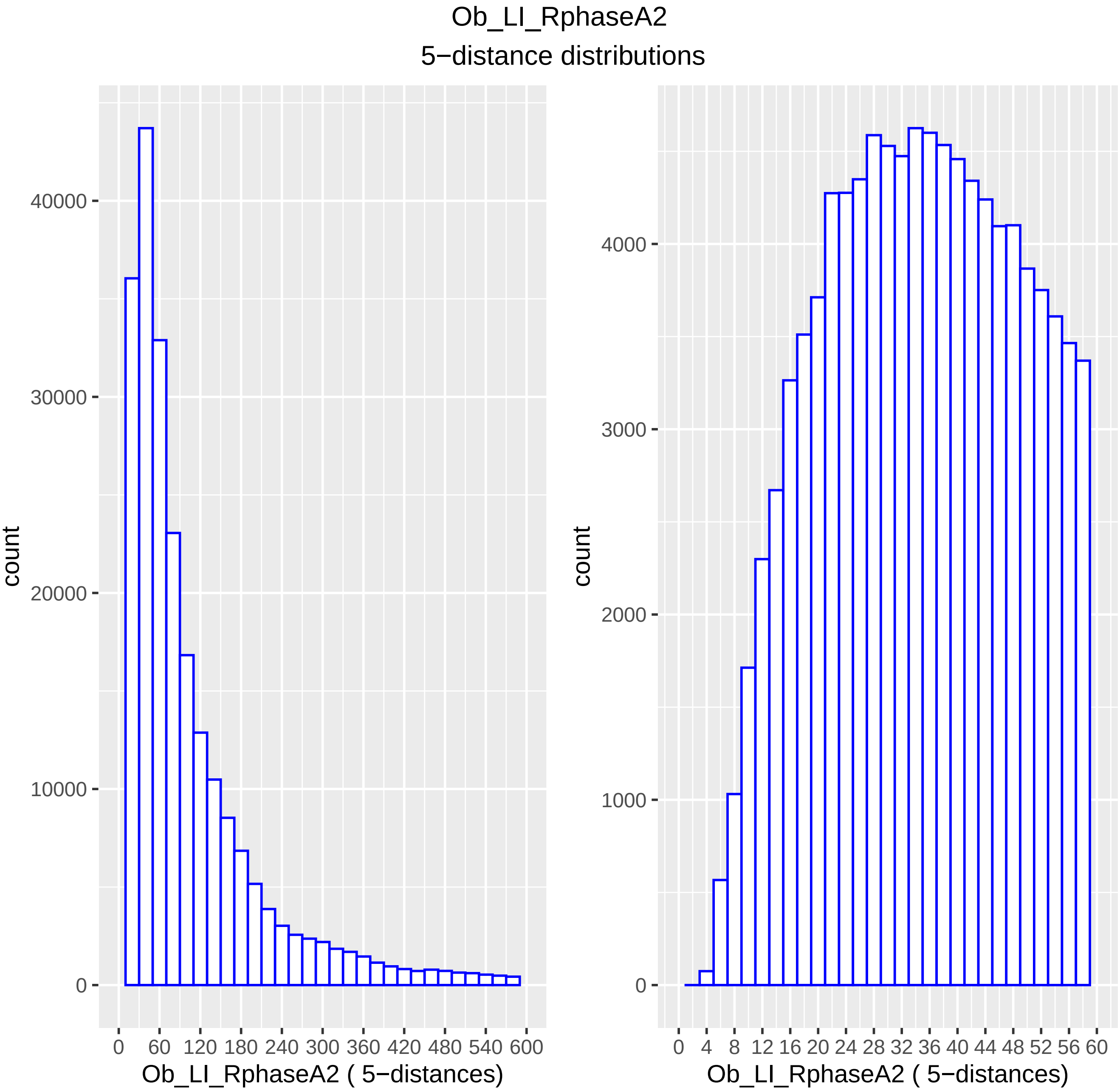

Ob_LI_RphaseA2
5−distance distributions
count
40000
30000
20000
10000
0
0
60
120
180
240
300
360
420
480
540
600
Ob_LI_RphaseA2 ( 5−distances)
count
4000
3000
2000
1000
0
0
4
8
12
16
20
24
28
32
36
40
44
48
52
56
60
Ob_LI_RphaseA2 ( 5−distances)

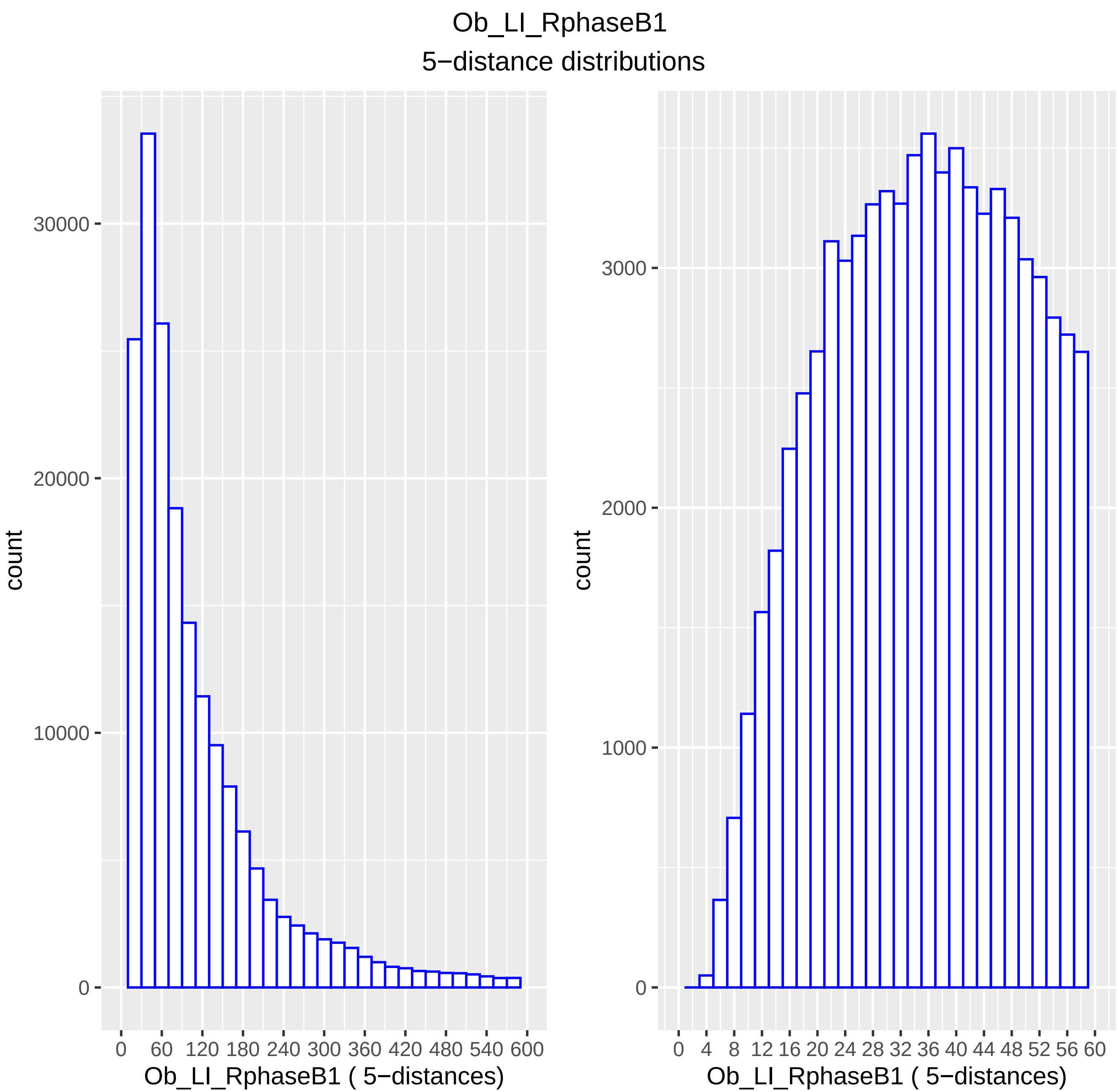

Ob_LI_RphaseB1
5−distance distributions
count
30000
20000
10000
0
0 60 120 180 240 300 360 420 480 540 600
Ob_LI_RphaseB1 ( 5−distances)
count
3000
2000
1000
0
0 4 8 12 16 20 24 28 32 36 40 44 48 52 56 60
Ob_LI_RphaseB1 ( 5−distances)

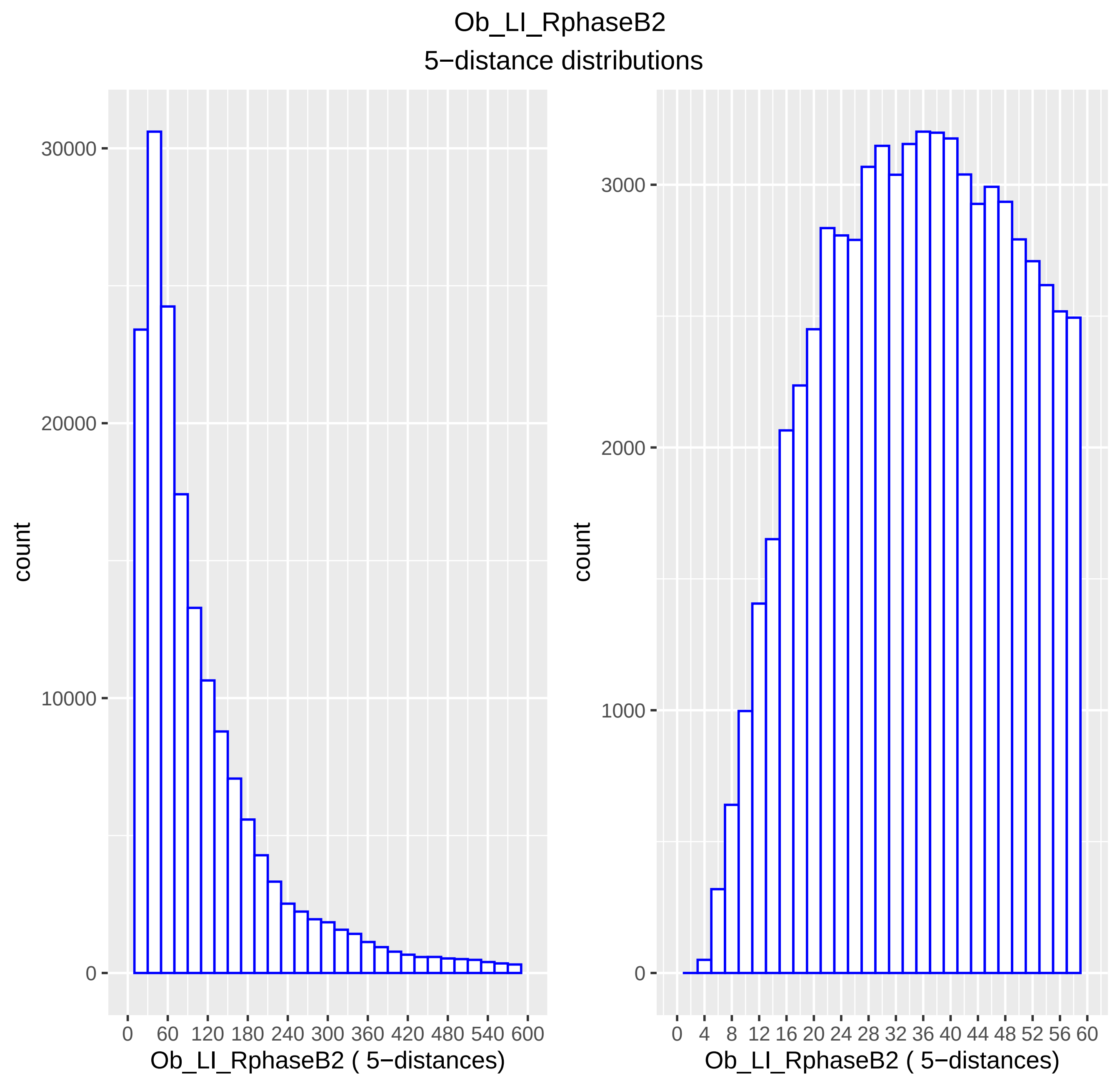
Ob_LI_RphaseB2
5−distance distributions
count
30000
20000
10000
0
0 60 120 180 240 300 360 420 480 540 600
Ob_LI_RphaseB2 ( 5−distances)
count
3000
2000
1000
0
0 4 8 12 16 20 24 28 32 36 40 44 48 52 56 60
Ob_LI_RphaseB2 ( 5−distances)

```
Directory: DMT File: 0READMEdmt

DMT - Differentially methylated tiles.

Input: studymc wsize stepsize threshold qvalue
Output: files dmt-*.txt dmg-*.txt

Notes: Output files dmt-*.txt and dmg-*.txt are generated by BWASPR::det_dmt() and show the
        differentially methylated tiles and genes as determined by methylKit::getMethylDiff with
        parameters difference=threshold and qvalue, as provided in the Ob_LI.conf
        configuration file.  Tiles refer to sliding windows along the genome within which
        methylation calls are cumulated by methylKit::tileMethylCounts.
        The dmg-*.txt files show all genes with at least one differentially methylated tile.

        A positive meth.diff value in comparison A.vs.B means that the B methylation percentage
        in that tile is higher than the A methylation percentage.
```

Directory: DMT File: dmg-Ob_LI.txt

| seqnames | start | end | width | strand | source | type | score | phase | ID | Dbxref | Name | gbkey | gene | gene_biotype | end_range | partial | start_range | comparison |
|---|---|---|---|---|---|---|---|---|---|---|---|---|---|---|---|---|---|---|
| NC_039506.1 | 10584230 | 10588663 | 4434 | - | Gnomon | gene | NA | NA | gene-LOC105286711 | GeneID:105286711 | LOC105286711 | Gene | LOC105286711 | protein_coding | character(0) | NA | | BCphaseA1.vs.BCphaseA2 |
| NC_039506.1 | 12495232 | 12500898 | 5667 | + | Gnomon | gene | NA | NA | gene-LOC105275070 | GeneID:105275070 | LOC105275070 | Gene | LOC105275070 | protein_coding | character(0) | NA | character(0) | BCphaseA1.vs.BCphaseA2 |
| NC_039507.1 | 9148975 | 9159660 | 10686 | + | Gnomon | gene | NA | NA | gene-LOC105278822 | GeneID:105278822 | LOC105278822 | Gene | LOC105278822 | protein_coding | character(0) | NA | character(0) | BCphaseA1.vs.BCphaseA2 |
| NC_039507.1 | 12942711 | 12945953 | 3243 | - | Gnomon | gene | NA | NA | gene-LOC105278934 | GeneID:105278934 | LOC105278934 | Gene | LOC105278934 | protein_coding | character(0) | NA | character(0) | BCphaseA1.vs.BCphaseA2 |
| NC_039507.1 | 14505339 | 14508190 | 2852 | + | Gnomon | gene | NA | NA | gene-LOC105283991 | GeneID:105283991 | LOC105283991 | Gene | LOC105283991 | protein_coding | character(0) | NA | character(0) | BCphaseA1.vs.BCphaseA2 |
| NC_039508.1 | 1201156 | 1209530 | 8375 | + | Gnomon | gene | NA | NA | gene-LOC105277409 | GeneID:105277409 | LOC105277409 | Gene | LOC105277409 | protein_coding | character(0) | NA | character(0) | BCphaseA1.vs.BCphaseA2 |
| NC_039508.1 | 3915820 | 3917712 | 1893 | + | Gnomon | gene | NA | NA | gene-LOC105281418 | GeneID:105281418 | LOC105281418 | Gene | LOC105281418 | protein_coding | character(0) | NA | character(0) | BCphaseA1.vs.BCphaseA2 |
| NC_039508.1 | 18076454 | 18078162 | 1709 | - | Gnomon | gene | NA | NA | gene-LOC105280656 | GeneID:105280656 | LOC105280656 | Gene | LOC105280656 | protein_coding | character(0) | NA | character(0) | BCphaseA1.vs.BCphaseA2 |
| NC_039509.1 | 8320869 | 8324118 | 3250 | - | Gnomon | gene | NA | NA | gene-LOC105281282 | GeneID:105281282 | LOC105281282 | Gene | LOC105281282 | protein_coding | character(0) | NA | character(0) | BCphaseA1.vs.BCphaseA2 |
| NC_039509.1 | 8327307 | 8443148 | 115842 | + | Gnomon | gene | NA | NA | gene-LOC105280645 | GeneID:105280645 | LOC105280645 | Gene | LOC105280645 | protein_coding | character(0) | NA | character(0) | BCphaseA1.vs.BCphaseA2 |
| NC_039509.1 | 8981313 | 8983857 | 2545 | - | Gnomon | gene | NA | NA | gene-LOC105283091 | GeneID:105283091 | LOC105283091 | Gene | LOC105283091 | protein_coding | character(0) | NA | character(0) | BCphaseA1.vs.BCphaseA2 |
| NC_039509.1 | 11068956 | 11086837 | 17882 | - | Gnomon | gene | NA | NA | gene-LOC105275770 | GeneID:105275770 | LOC105275770 | Gene | LOC105275770 | protein_coding | character(0) | NA | character(0) | BCphaseA1.vs.BCphaseA2 |
| NC_039509.1 | 16643753 | 16645287 | 1535 | - | Gnomon | gene | NA | NA | gene-LOC105283492 | GeneID:105283492 | LOC105283492 | Gene | LOC105283492 | protein_coding | character(0) | NA | character(0) | BCphaseA1.vs.BCphaseA2 |
| NC_039510.1 | 234125 | 236987 | 2863 | + | Gnomon | gene | NA | NA | gene-LOC105283071 | GeneID:105283071 | LOC105283071 | Gene | LOC105283071 | protein_coding | character(0) | NA | character(0) | BCphaseA1.vs.BCphaseA2 |
| NC_039510.1 | 16331362 | 16334383 | 3022 | - | Gnomon | gene | NA | NA | gene-LOC105286521 | GeneID:105286521 | LOC105286521 | Gene | LOC105286521 | protein_coding | character(0) | NA | character(0) | BCphaseA1.vs.BCphaseA2 |
| NC_039511.1 | 13140135 | 13149418 | 9284 | + | Gnomon | gene | NA | NA | gene-LOC105276599 | GeneID:105276599 | LOC105276599 | Gene | LOC105276599 | protein_coding | character(0) | NA | character(0) | BCphaseA1.vs.BCphaseA2 |
| NC_039511.1 | 16402468 | 16408726 | 6259 | - | Gnomon | gene | NA | NA | gene-LOC105284528 | GeneID:105284528 | LOC105284528 | Gene | LOC105284528 | protein_coding | character(0) | NA | character(0) | BCphaseA1.vs.BCphaseA2 |
| NC_039512.1 | 4289608 | 4290786 | 1179 | + | Gnomon | gene | NA | NA | gene-LOC105287276 | GeneID:105287276 | LOC105287276 | Gene | LOC105287276 | protein_coding | character(0) | NA | character(0) | BCphaseA1.vs.BCphaseA2 |
| NC_039512.1 | 4290708 | 4298417 | 7710 | - | Gnomon | gene | NA | NA | gene-LOC105277554 | GeneID:105277554 | LOC105277554 | Gene | LOC105277554 | protein_coding | character(0) | NA | character(0) | BCphaseA1.vs.BCphaseA2 |
| NC_039512.1 | 5367891 | 5406727 | 38837 | - | Gnomon | gene | NA | NA | gene-LOC105282015 | GeneID:105282015 | LOC105282015 | Gene | LOC105282015 | protein_coding | character(0) | NA | character(0) | BCphaseA1.vs.BCphaseA2 |
| NC_039512.1 | 5854372 | 5855269 | 898 | - | Gnomon | gene | NA | NA | gene-LOC113562300 | GeneID:113562300 | LOC113562300 | Gene | LOC113562300 | protein_coding | character(0) | NA | character(0) | BCphaseA1.vs.BCphaseA2 |
| NC_039512.1 | 8846758 | 8848479 | 1722 | + | Gnomon | gene | NA | NA | gene-LOC105282371 | GeneID:105282371 | LOC105282371 | Gene | LOC105282371 | protein_coding | character(0) | NA | character(0) | BCphaseA1.vs.BCphaseA2 |
| NC_039512.1 | 9796572 | 9800489 | 3918 | + | Gnomon | gene | NA | NA | gene-LOC105279899 | GeneID:105279899 | LOC105279899 | Gene | LOC105279899 | protein_coding | character(0) | NA | character(0) | BCphaseA1.vs.BCphaseA2 |
| NC_039512.1 | 14393197 | 14394878 | 1682 | + | Gnomon | gene | NA | NA | gene-LOC105286831 | GeneID:105286831 | LOC105286831 | Gene | LOC105286831 | protein_coding | character(0) | NA | character(0) | BCphaseA1.vs.BCphaseA2 |

...
... only first 25 lines shown ...

Directory: DMT File: dmt-Ob_LI.txt

| seqnames | start | end | width | strand | pvalue | qvalue | meth.diff | comparison |
|---|---|---|---|---|---|---|---|---|
| NC_039506.1 | 10588001 | 10588500 | 500 | * | 7.56008533707562e-06 | 0.014 | 28.99 | BCphaseA1.vs.BCphaseA2 |
| NC_039506.1 | 12496001 | 12496500 | 500 | * | 9.94525184496602e-43 | 0 | 55.23 | BCphaseA1.vs.BCphaseA2 |
| NC_039506.1 | 12496501 | 12497000 | 500 | * | 2.06532892392705e-20 | 0 | 28.36 | BCphaseA1.vs.BCphaseA2 |
| NC_039507.1 | 9151501 | 9152000 | 500 | * | 4.05083413316457e-06 | 0.009 | -62.53 | BCphaseA1.vs.BCphaseA2 |
| NC_039507.1 | 12945501 | 12946000 | 500 | * | 6.02827802233786e-07 | 0.002 | 22.53 | BCphaseA1.vs.BCphaseA2 |
| NC_039507.1 | 14505501 | 14506000 | 500 | * | 2.55367842421072e-05 | 0.034 | 29.16 | BCphaseA1.vs.BCphaseA2 |
| NC_039508.1 | 1205001 | 1205500 | 500 | * | 2.26172959742878e-06 | 0.006 | 30.25 | BCphaseA1.vs.BCphaseA2 |
| NC_039508.1 | 3916001 | 3916500 | 500 | * | 9.96175785983529e-09 | 0 | 29.99 | BCphaseA1.vs.BCphaseA2 |
| NC_039508.1 | 7771501 | 7772000 | 500 | * | 3.10624124108235e-05 | 0.038 | 35.2 | BCphaseA1.vs.BCphaseA2 |
| NC_039508.1 | 18077001 | 18077500 | 500 | * | 1.77390330035201e-12 | 0 | 31.16 | BCphaseA1.vs.BCphaseA2 |
| NC_039509.1 | 8321501 | 8322000 | 500 | * | 5.16376754306592e-06 | 0.011 | 22.36 | BCphaseA1.vs.BCphaseA2 |
| NC_039509.1 | 8328001 | 8328500 | 500 | * | 2.36016737772857e-05 | 0.032 | 66.46 | BCphaseA1.vs.BCphaseA2 |
| NC_039509.1 | 8328501 | 8329000 | 500 | * | 2.50474554639287e-07 | 0.001 | 23.74 | BCphaseA1.vs.BCphaseA2 |
| NC_039509.1 | 8983001 | 8983500 | 500 | * | 1.91352306459505e-13 | 0 | 27.33 | BCphaseA1.vs.BCphaseA2 |
| NC_039509.1 | 9291001 | 9291500 | 500 | * | 1.68897957952752e-43 | 0 | 29.4 | BCphaseA1.vs.BCphaseA2 |
| NC_039509.1 | 11086001 | 11086500 | 500 | * | 2.89305140139684e-06 | 0.008 | 25.02 | BCphaseA1.vs.BCphaseA2 |
| NC_039509.1 | 16644501 | 16645000 | 500 | * | 1.78225560447143e-06 | 0.005 | 22.36 | BCphaseA1.vs.BCphaseA2 |
| NC_039510.1 | 234001 | 234500 | 500 | * | 1.54644117492415e-06 | 0.005 | -100 | BCphaseA1.vs.BCphaseA2 |
| NC_039510.1 | 16333001 | 16333500 | 500 | * | 7.70696977295299e-08 | 0 | -51.58 | BCphaseA1.vs.BCphaseA2 |
| NC_039511.1 | 13140501 | 13141000 | 500 | * | 3.20596477263989e-08 | 0 | 34.61 | BCphaseA1.vs.BCphaseA2 |
| NC_039511.1 | 16405501 | 16406000 | 500 | * | 1.08982375062791e-06 | 0.004 | 23.74 | BCphaseA1.vs.BCphaseA2 |
| NC_039512.1 | 4290501 | 4291000 | 500 | * | 4.10764936582692e-08 | 0 | 80.65 | BCphaseA1.vs.BCphaseA2 |
| NC_039512.1 | 5405001 | 5405500 | 500 | * | 8.4868855011857e-10 | 0 | -43.37 | BCphaseA1.vs.BCphaseA2 |
| NC_039512.1 | 5854501 | 5855000 | 500 | * | 9.9655719763823e-07 | 0.003 | -29.94 | BCphaseA1.vs.BCphaseA2 |

...
... only first 25 lines shown ...

```
Directory: DMSG File: 0READMEdmsg

DMSG - Differentially methylated sites and genes.

Input: studyhc threshold qvalue
       The studyhc methylKit raw object contains all CpGscd sites with coverage at least
        20 reads.
Output: files dms-*.txt dmg-*.txt  dmg-*details.txt dmg-*heatmaps.pdf

Notes: Output files dms-*.txt and dmg-*.txt are generated by BWASPR::det_dmsg() and show the
        differentially methylated sites and genes as determined by methylKit::getMethylDiff with
        parameters difference=threshold and qvalue, as provided in the Ob_LI.conf
        configuration file.  The table of differentially methylated genes contains all genes
        with at least one differentially methylated site.

       The files dmg-*details.txt, generated by BWASPR::show_dmsg(), show all CpGscd sites in
        the differentially methylated genes, with coverage numbers and methylation percentages.

       The files dmg-*heatmaps.pdf, generated by BWASPR::show_dmsg(), give heatmap displays for
        genes meeting the following criteria: 1) there are between minNsites and maxNsites common
        CpGscd sites; 2) at least minPdmsites % of these sites are differentially methylated sites.
        The parameters are set in the Ob_LI.conf configuration file.
```

Directory: DMSG File: dmg-Ob_LI.txt

| seqnames | start | end | width | strand | source | type | score | phase | ID | Dbxref | Name | gbkey | gene | gene_biotype | end_range | partial | start_range | comparison |
|---|---|---|---|---|---|---|---|---|---|---|---|---|---|---|---|---|---|---|
| NC_039506.1 | 2172791 | 2181709 | 8919 | + | Gnomon | gene | NA | NA | gene-LOC105280304 | GeneID:105280304 | LOC105280304 | Gene | LOC105280304 | protein_coding | character(0) | NA | character(0) | BCphaseA1.vs.BCphaseA2 |
| NC_039506.1 | 2980439 | 2984235 | 3797 | + | Gnomon | gene | NA | NA | gene-LOC105280348 | GeneID:105280348 | LOC105280348 | Gene | LOC105280348 | protein_coding | character(0) | NA | character(0) | BCphaseA1.vs.BCphaseA2 |
| NC_039506.1 | 3575547 | 3588126 | 12580 | - | Gnomon | gene | NA | NA | gene-LOC105280410 | GeneID:105280410 | LOC105280410 | Gene | LOC105280410 | protein_coding | character(0) | NA | character(0) | BCphaseA1.vs.BCphaseA2 |
| NC_039506.1 | 5466806 | 5499850 | 33045 | - | Gnomon | gene | NA | NA | gene-LOC105277866 | GeneID:105277866 | LOC105277866 | Gene | LOC105277866 | protein_coding | character(0) | NA | character(0) | BCphaseA1.vs.BCphaseA2 |
| NC_039506.1 | 10972299 | 10984387 | 12089 | - | Gnomon | gene | NA | NA | gene-LOC105284774 | GeneID:105284774 | LOC105284774 | Gene | LOC105284774 | protein_coding | character(0) | NA | character(0) | BCphaseA1.vs.BCphaseA2 |
| NC_039506.1 | 12495232 | 12500898 | 5667 | + | Gnomon | gene | NA | NA | gene-LOC105275070 | GeneID:105275070 | LOC105275070 | Gene | LOC105275070 | protein_coding | character(0) | NA | character(0) | BCphaseA1.vs.BCphaseA2 |
| NC_039506.1 | 13173343 | 13174793 | 1451 | + | Gnomon | gene | NA | NA | gene-LOC109611283 | GeneID:109611283 | LOC109611283 | Gene | LOC109611283 | protein_coding | character(0) | NA | character(0) | BCphaseA1.vs.BCphaseA2 |
| NC_039506.1 | 15935415 | 15940975 | 5561 | - | Gnomon | gene | NA | NA | gene-LOC105274900 | GeneID:105274900 | LOC105274900 | Gene | LOC105274900 | protein_coding | character(0) | NA | character(0) | BCphaseA1.vs.BCphaseA2 |
| NC_039506.1 | 19217974 | 19220222 | 2249 | - | Gnomon | gene | NA | NA | gene-LOC113561621 | GeneID:113561621 | LOC113561621 | Gene | LOC113561621 | protein_coding | character(0) | NA | character(0) | BCphaseA1.vs.BCphaseA2 |
| NC_039506.1 | 20580521 | 20582670 | 2150 | - | Gnomon | gene | NA | NA | gene-LOC105278538 | GeneID:105278538 | LOC105278538 | Gene | LOC105278538 | protein_coding | character(0) | NA | character(0) | BCphaseA1.vs.BCphaseA2 |
| NC_039506.1 | 20610062 | 20617425 | 7364 | + | Gnomon | gene | NA | NA | gene-LOC105278534 | GeneID:105278534 | LOC105278534 | Gene | LOC105278534 | protein_coding | character(0) | NA | character(0) | BCphaseA1.vs.BCphaseA2 |
| NC_039506.1 | 23928483 | 23929796 | 1314 | - | Gnomon | gene | NA | NA | gene-LOC105279356 | GeneID:105279356 | LOC105279356 | Gene | LOC105279356 | protein_coding | character(0) | NA | character(0) | BCphaseA1.vs.BCphaseA2 |
| NC_039507.1 | 5014963 | 5017170 | 2208 | - | Gnomon | gene | NA | NA | gene-LOC105281700 | GeneID:105281700 | LOC105281700 | Gene | LOC105281700 | protein_coding | character(0) | NA | character(0) | BCphaseA1.vs.BCphaseA2 |
| NC_039507.1 | 14490394 | 14495234 | 4841 | - | Gnomon | gene | NA | NA | gene-LOC105283987 | GeneID:105283987 | LOC105283987 | Gene | LOC105283987 | protein_coding | character(0) | NA | character(0) | BCphaseA1.vs.BCphaseA2 |
| NC_039507.1 | 14495724 | 14500671 | 4948 | + | Gnomon | gene | NA | NA | gene-LOC105283989 | GeneID:105283989 | LOC105283989 | Gene | LOC105283989 | protein_coding | character(0) | NA | character(0) | BCphaseA1.vs.BCphaseA2 |
| NC_039507.1 | 16959121 | 16963590 | 4470 | + | Gnomon | gene | NA | NA | gene-LOC105278145 | GeneID:105278145 | LOC105278145 | Gene | LOC105278145 | protein_coding | character(0) | NA | character(0) | BCphaseA1.vs.BCphaseA2 |
| NC_039508.1 | 4118007 | 4119377 | 1371 | - | Gnomon | gene | NA | NA | gene-LOC105285695 | GeneID:105285695 | LOC105285695 | Gene | LOC105285695 | protein_coding | character(0) | NA | character(0) | BCphaseA1.vs.BCphaseA2 |
| NC_039508.1 | 7055657 | 7059685 | 4029 | + | Gnomon | gene | NA | NA | gene-LOC105278109 | GeneID:105278109 | LOC105278109 | Gene | LOC105278109 | protein_coding | character(0) | NA | character(0) | BCphaseA1.vs.BCphaseA2 |
| NC_039508.1 | 10474651 | 10525729 | 51079 | - | Gnomon | gene | NA | NA | gene-LOC105277125 | GeneID:105277125 | LOC105277125 | Gene | LOC105277125 | protein_coding | character(0) | NA | character(0) | BCphaseA1.vs.BCphaseA2 |
| NC_039508.1 | 10716085 | 10732040 | 15956 | - | Gnomon | gene | NA | NA | gene-LOC105276941 | GeneID:105276941 | LOC105276941 | Gene | LOC105276941 | protein_coding | character(0) | NA | character(0) | BCphaseA1.vs.BCphaseA2 |
| NC_039508.1 | 15147286 | 15154912 | 7627 | - | Gnomon | gene | NA | NA | gene-LOC105277216 | GeneID:105277216 | LOC105277216 | Gene | LOC105277216 | lncRNA | character(0) | NA | character(0) | BCphaseA1.vs.BCphaseA2 |
| NC_039508.1 | 15151829 | 15154447 | 2619 | + | Gnomon | gene | NA | NA | gene-LOC105277215 | GeneID:105277215 | LOC105277215 | Gene | LOC105277215 | protein_coding | character(0) | NA | character(0) | BCphaseA1.vs.BCphaseA2 |
| NC_039509.1 | 184477 | 186038 | 1562 | + | Gnomon | gene | NA | NA | gene-LOC105283552 | GeneID:105283552 | LOC105283552 | Gene | LOC105283552 | protein_coding | character(0) | NA | character(0) | BCphaseA1.vs.BCphaseA2 |
| NC_039509.1 | 607985 | 611463 | 3479 | + | Gnomon | gene | NA | NA | gene-LOC105283592 | GeneID:105283592 | LOC105283592 | Gene | LOC105283592 | protein_coding | character(0) | NA | character(0) | BCphaseA1.vs.BCphaseA2 |

...
... only first 25 lines shown ...

Directory: DMSG File: dmg-Ob_LI_BCphaseA1.vs.BCphaseA2_details.txt

| seqnames | start | end | width | strand | coverage1 | numCs1 | numTs1 | coverage2 | numCs2 | numTs2 | BCphaseA1 | BCphaseA2 | gene_seqnames | gene_start | gene_end | gene_width | gene_strand | gene_source | gene_type | gene_ID | gene_Dbxref | gene_Name | gene_gbkey | gene_gene | gene_gene_biotype | gene_end_range | gene_start_range | gene_comparison | is.dm |
|---|---|---|---|---|---|---|---|---|---|---|---|---|---|---|---|---|---|---|---|---|---|---|---|---|---|---|---|---|---|
| NC_039506.1 | 2173614 | 2173614 | 1 | + | 34 | 0 | 34 | 22 | 1 | 21 | 0 | 4.55 | NC_039506.1 | 2172791 | 2181709 | 8919 | + | Gnomon | gene | gene-LOC105280304 | GeneID:105280304 | LOC105280304 | Gene | LOC105280304 | protein_coding | character(0) | character(0) | BCphaseA1.vs.BCphaseA2 | FALSE |
| NC_039506.1 | 2173709 | 2173709 | 1 | + | 34 | 0 | 34 | 41 | 0 | 41 | 0 | 0 | NC_039506.1 | 2172791 | 2181709 | 8919 | + | Gnomon | gene | gene-LOC105280304 | GeneID:105280304 | LOC105280304 | Gene | LOC105280304 | protein_coding | character(0) | character(0) | BCphaseA1.vs.BCphaseA2 | FALSE |
| NC_039506.1 | 2173733 | 2173733 | 1 | + | 46 | 1 | 45 | 33 | 1 | 32 | 2.17 | 3.03 | NC_039506.1 | 2172791 | 2181709 | 8919 | + | Gnomon | gene | gene-LOC105280304 | GeneID:105280304 | LOC105280304 | Gene | LOC105280304 | protein_coding | character(0) | character(0) | BCphaseA1.vs.BCphaseA2 | FALSE |
| NC_039506.1 | 2173742 | 2173742 | 1 | + | 46 | 0 | 46 | 41 | 0 | 41 | 0 | 0 | NC_039506.1 | 2172791 | 2181709 | 8919 | + | Gnomon | gene | gene-LOC105280304 | GeneID:105280304 | LOC105280304 | Gene | LOC105280304 | protein_coding | character(0) | character(0) | BCphaseA1.vs.BCphaseA2 | FALSE |
| NC_039506.1 | 2173746 | 2173746 | 1 | + | 46 | 1 | 45 | 42 | 0 | 42 | 2.17 | 0 | NC_039506.1 | 2172791 | 2181709 | 8919 | + | Gnomon | gene | gene-LOC105280304 | GeneID:105280304 | LOC105280304 | Gene | LOC105280304 | protein_coding | character(0) | character(0) | BCphaseA1.vs.BCphaseA2 | FALSE |
| NC_039506.1 | 2173753 | 2173753 | 1 | + | 46 | 1 | 45 | 44 | 0 | 44 | 2.17 | 0 | NC_039506.1 | 2172791 | 2181709 | 8919 | + | Gnomon | gene | gene-LOC105280304 | GeneID:105280304 | LOC105280304 | Gene | LOC105280304 | protein_coding | character(0) | character(0) | BCphaseA1.vs.BCphaseA2 | FALSE |
| NC_039506.1 | 2173759 | 2173759 | 1 | + | 47 | 0 | 47 | 46 | 0 | 46 | 0 | 0 | NC_039506.1 | 2172791 | 2181709 | 8919 | + | Gnomon | gene | gene-LOC105280304 | GeneID:105280304 | LOC105280304 | Gene | LOC105280304 | protein_coding | character(0) | character(0) | BCphaseA1.vs.BCphaseA2 | FALSE |
| NC_039506.1 | 2173764 | 2173764 | 1 | + | 49 | 0 | 49 | 50 | 0 | 50 | 0 | 0 | NC_039506.1 | 2172791 | 2181709 | 8919 | + | Gnomon | gene | gene-LOC105280304 | GeneID:105280304 | LOC105280304 | Gene | LOC105280304 | protein_coding | character(0) | character(0) | BCphaseA1.vs.BCphaseA2 | FALSE |
| NC_039506.1 | 2173768 | 2173768 | 1 | + | 50 | 1 | 49 | 49 | 1 | 48 | 2 | 2.04 | NC_039506.1 | 2172791 | 2181709 | 8919 | + | Gnomon | gene | gene-LOC105280304 | GeneID:105280304 | LOC105280304 | Gene | LOC105280304 | protein_coding | character(0) | character(0) | BCphaseA1.vs.BCphaseA2 | FALSE |
| NC_039506.1 | 2173779 | 2173779 | 1 | + | 56 | 1 | 55 | 53 | 0 | 53 | 1.79 | 0 | NC_039506.1 | 2172791 | 2181709 | 8919 | + | Gnomon | gene | gene-LOC105280304 | GeneID:105280304 | LOC105280304 | Gene | LOC105280304 | protein_coding | character(0) | character(0) | BCphaseA1.vs.BCphaseA2 | FALSE |
| NC_039506.1 | 2173789 | 2173789 | 1 | + | 54 | 0 | 54 | 57 | 0 | 57 | 0 | 0 | NC_039506.1 | 2172791 | 2181709 | 8919 | + | Gnomon | gene | gene-LOC105280304 | GeneID:105280304 | LOC105280304 | Gene | LOC105280304 | protein_coding | character(0) | character(0) | BCphaseA1.vs.BCphaseA2 | FALSE |
| NC_039506.1 | 2173798 | 2173798 | 1 | + | 48 | 0 | 48 | 55 | 0 | 55 | 0 | 0 | NC_039506.1 | 2172791 | 2181709 | 8919 | + | Gnomon | gene | gene-LOC105280304 | GeneID:105280304 | LOC105280304 | Gene | LOC105280304 | protein_coding | character(0) | character(0) | BCphaseA1.vs.BCphaseA2 | FALSE |
| NC_039506.1 | 2173801 | 2173801 | 1 | + | 46 | 0 | 46 | 45 | 0 | 45 | 0 | 0 | NC_039506.1 | 2172791 | 2181709 | 8919 | + | Gnomon | gene | gene-LOC105280304 | GeneID:105280304 | LOC105280304 | Gene | LOC105280304 | protein_coding | character(0) | character(0) | BCphaseA1.vs.BCphaseA2 | FALSE |
| NC_039506.1 | 2173810 | 2173810 | 1 | + | 36 | 2 | 34 | 39 | 1 | 38 | 5.56 | 2.56 | NC_039506.1 | 2172791 | 2181709 | 8919 | + | Gnomon | gene | gene-LOC105280304 | GeneID:105280304 | LOC105280304 | Gene | LOC105280304 | protein_coding | character(0) | character(0) | BCphaseA1.vs.BCphaseA2 | FALSE |
| NC_039506.1 | 2173812 | 2173812 | 1 | + | 38 | 2 | 36 | 39 | 1 | 38 | 5.26 | 2.56 | NC_039506.1 | 2172791 | 2181709 | 8919 | + | Gnomon | gene | gene-LOC105280304 | GeneID:105280304 | LOC105280304 | Gene | LOC105280304 | protein_coding | character(0) | character(0) | BCphaseA1.vs.BCphaseA2 | FALSE |
| NC_039506.1 | 2173823 | 2173823 | 1 | + | 31 | 0 | 31 | 32 | 0 | 32 | 0 | 0 | NC_039506.1 | 2172791 | 2181709 | 8919 | + | Gnomon | gene | gene-LOC105280304 | GeneID:105280304 | LOC105280304 | Gene | LOC105280304 | protein_coding | character(0) | character(0) | BCphaseA1.vs.BCphaseA2 | FALSE |
| NC_039506.1 | 2173831 | 2173831 | 1 | + | 30 | 0 | 30 | 32 | 0 | 32 | 0 | 0 | NC_039506.1 | 2172791 | 2181709 | 8919 | + | Gnomon | gene | gene-LOC105280304 | GeneID:105280304 | LOC105280304 | Gene | LOC105280304 | protein_coding | character(0) | character(0) | BCphaseA1.vs.BCphaseA2 | FALSE |
| NC_039506.1 | 2174095 | 2174095 | 1 | + | 23 | 0 | 23 | 22 | 0 | 22 | 0 | 0 | NC_039506.1 | 2172791 | 2181709 | 8919 | + | Gnomon | gene | gene-LOC105280304 | GeneID:105280304 | LOC105280304 | Gene | LOC105280304 | protein_coding | character(0) | character(0) | BCphaseA1.vs.BCphaseA2 | FALSE |
| NC_039506.1 | 2174105 | 2174105 | 1 | + | 25 | 0 | 25 | 23 | 0 | 23 | 0 | 0 | NC_039506.1 | 2172791 | 2181709 | 8919 | + | Gnomon | gene | gene-LOC105280304 | GeneID:105280304 | LOC105280304 | Gene | LOC105280304 | protein_coding | character(0) | character(0) | BCphaseA1.vs.BCphaseA2 | FALSE |
| NC_039506.1 | 2174110 | 2174110 | 1 | + | 23 | 0 | 23 | 24 | 0 | 24 | 0 | 0 | NC_039506.1 | 2172791 | 2181709 | 8919 | + | Gnomon | gene | gene-LOC105280304 | GeneID:105280304 | LOC105280304 | Gene | LOC105280304 | protein_coding | character(0) | character(0) | BCphaseA1.vs.BCphaseA2 | FALSE |
| NC_039506.1 | 2175207 | 2175207 | 1 | + | 25 | 13 | 12 | 44 | 17 | 27 | 52 | 38.64 | NC_039506.1 | 2172791 | 2181709 | 8919 | + | Gnomon | gene | gene-LOC105280304 | GeneID:105280304 | LOC105280304 | Gene | LOC105280304 | protein_coding | character(0) | character(0) | BCphaseA1.vs.BCphaseA2 | FALSE |
| NC_039506.1 | 2175399 | 2175399 | 1 | + | 34 | 34 | 0 | 25 | 11 | 14 | 100 | 44 | NC_039506.1 | 2172791 | 2181709 | 8919 | + | Gnomon | gene | gene-LOC105280304 | GeneID:105280304 | LOC105280304 | Gene | LOC105280304 | protein_coding | character(0) | character(0) | BCphaseA1.vs.BCphaseA2 | TRUE |
| NC_039506.1 | 2175432 | 2175432 | 1 | + | 41 | 28 | 13 | 38 | 31 | 7 | 68.29 | 81.58 | NC_039506.1 | 2172791 | 2181709 | 8919 | + | Gnomon | gene | gene-LOC105280304 | GeneID:105280304 | LOC105280304 | Gene | LOC105280304 | protein_coding | character(0) | character(0) | BCphaseA1.vs.BCphaseA2 | FALSE |
| NC_039506.1 | 2175438 | 2175438 | 1 | + | 40 | 26 | 14 | 38 | 28 | 10 | 65 | 73.68 | NC_039506.1 | 2172791 | 2181709 | 8919 | + | Gnomon | gene | gene-LOC105280304 | GeneID:105280304 | LOC105280304 | Gene | LOC105280304 | protein_coding | character(0) | character(0) | BCphaseA1.vs.BCphaseA2 | FALSE |

...
... only first 25 lines shown ...

Directory: DMSG File: dmg-Ob_LI_BCphaseA1.vs.BCphaseB1_details.txt

| seqnames | start | end | width | strand | coverage1 | numCs1 | numTs1 | coverage2 | numCs2 | numTs2 | BCphaseA1 | BCphaseB1 | gene_seqnames | gene_start | gene_end | gene_width | gene_strand | gene_source | gene_type | gene_ID | gene_Dbxref | gene_Name | gene_gbkey | gene_gene | gene_gene_biotype | gene_end_range | gene_start_range | gene_comparison | is.dm |
|---|---|---|---|---|---|---|---|---|---|---|---|---|---|---|---|---|---|---|---|---|---|---|---|---|---|---|---|---|---|
| NC_039506.1 | 2980750 | 2980750 | 1 | + | 21 | 0 | 21 | 29 | 0 | 29 | 0 | 0 | NC_039506.1 | 2980439 | 2984235 | 3797 | + | Gnomon | gene | gene-LOC105280348 | GeneID:105280348 | LOC105280348 | Gene | LOC105280348 | protein_coding | character(0) | character(0) | BCphaseA1.vs.BCphaseB1 | FALSE |
| NC_039506.1 | 2980761 | 2980761 | 1 | + | 22 | 1 | 21 | 32 | 0 | 32 | 4.55 | 0 | NC_039506.1 | 2980439 | 2984235 | 3797 | + | Gnomon | gene | gene-LOC105280348 | GeneID:105280348 | LOC105280348 | Gene | LOC105280348 | protein_coding | character(0) | character(0) | BCphaseA1.vs.BCphaseB1 | FALSE |
| NC_039506.1 | 2980767 | 2980767 | 1 | + | 23 | 0 | 23 | 34 | 0 | 34 | 0 | 0 | NC_039506.1 | 2980439 | 2984235 | 3797 | + | Gnomon | gene | gene-LOC105280348 | GeneID:105280348 | LOC105280348 | Gene | LOC105280348 | protein_coding | character(0) | character(0) | BCphaseA1.vs.BCphaseB1 | FALSE |
| NC_039506.1 | 2980989 | 2980989 | 1 | + | 33 | 3 | 30 | 29 | 3 | 26 | 9.09 | 10.34 | NC_039506.1 | 2980439 | 2984235 | 3797 | + | Gnomon | gene | gene-LOC105280348 | GeneID:105280348 | LOC105280348 | Gene | LOC105280348 | protein_coding | character(0) | character(0) | BCphaseA1.vs.BCphaseB1 | FALSE |
| NC_039506.1 | 2980993 | 2980993 | 1 | + | 31 | 10 | 21 | 29 | 12 | 17 | 32.26 | 41.38 | NC_039506.1 | 2980439 | 2984235 | 3797 | + | Gnomon | gene | gene-LOC105280348 | GeneID:105280348 | LOC105280348 | Gene | LOC105280348 | protein_coding | character(0) | character(0) | BCphaseA1.vs.BCphaseB1 | FALSE |
| NC_039506.1 | 2980998 | 2980998 | 1 | + | 32 | 6 | 26 | 28 | 8 | 20 | 18.75 | 28.57 | NC_039506.1 | 2980439 | 2984235 | 3797 | + | Gnomon | gene | gene-LOC105280348 | GeneID:105280348 | LOC105280348 | Gene | LOC105280348 | protein_coding | character(0) | character(0) | BCphaseA1.vs.BCphaseB1 | FALSE |
| NC_039506.1 | 2981030 | 2981030 | 1 | + | 55 | 46 | 9 | 56 | 50 | 6 | 83.64 | 89.29 | NC_039506.1 | 2980439 | 2984235 | 3797 | + | Gnomon | gene | gene-LOC105280348 | GeneID:105280348 | LOC105280348 | Gene | LOC105280348 | protein_coding | character(0) | character(0) | BCphaseA1.vs.BCphaseB1 | FALSE |
| NC_039506.1 | 2981077 | 2981077 | 1 | + | 34 | 18 | 16 | 49 | 25 | 24 | 52.94 | 51.02 | NC_039506.1 | 2980439 | 2984235 | 3797 | + | Gnomon | gene | gene-LOC105280348 | GeneID:105280348 | LOC105280348 | Gene | LOC105280348 | protein_coding | character(0) | character(0) | BCphaseA1.vs.BCphaseB1 | FALSE |
| NC_039506.1 | 2981125 | 2981125 | 1 | + | 29 | 18 | 11 | 32 | 19 | 13 | 62.07 | 59.38 | NC_039506.1 | 2980439 | 2984235 | 3797 | + | Gnomon | gene | gene-LOC105280348 | GeneID:105280348 | LOC105280348 | Gene | LOC105280348 | protein_coding | character(0) | character(0) | BCphaseA1.vs.BCphaseB1 | FALSE |
| NC_039506.1 | 2981146 | 2981146 | 1 | + | 22 | 21 | 1 | 42 | 38 | 4 | 95.45 | 90.48 | NC_039506.1 | 2980439 | 2984235 | 3797 | + | Gnomon | gene | gene-LOC105280348 | GeneID:105280348 | LOC105280348 | Gene | LOC105280348 | protein_coding | character(0) | character(0) | BCphaseA1.vs.BCphaseB1 | FALSE |
| NC_039506.1 | 2981183 | 2981183 | 1 | + | 56 | 25 | 31 | 83 | 78 | 5 | 44.64 | 93.98 | NC_039506.1 | 2980439 | 2984235 | 3797 | + | Gnomon | gene | gene-LOC105280348 | GeneID:105280348 | LOC105280348 | Gene | LOC105280348 | protein_coding | character(0) | character(0) | BCphaseA1.vs.BCphaseB1 | TRUE |
| NC_039506.1 | 2981225 | 2981225 | 1 | + | 25 | 17 | 8 | 37 | 23 | 14 | 68 | 62.16 | NC_039506.1 | 2980439 | 2984235 | 3797 | + | Gnomon | gene | gene-LOC105280348 | GeneID:105280348 | LOC105280348 | Gene | LOC105280348 | protein_coding | character(0) | character(0) | BCphaseA1.vs.BCphaseB1 | FALSE |
| NC_039506.1 | 2981245 | 2981245 | 1 | + | 31 | 15 | 16 | 40 | 18 | 22 | 48.39 | 45 | NC_039506.1 | 2980439 | 2984235 | 3797 | + | Gnomon | gene | gene-LOC105280348 | GeneID:105280348 | LOC105280348 | Gene | LOC105280348 | protein_coding | character(0) | character(0) | BCphaseA1.vs.BCphaseB1 | FALSE |
| NC_039506.1 | 2981292 | 2981292 | 1 | + | 31 | 26 | 5 | 68 | 64 | 4 | 83.87 | 94.12 | NC_039506.1 | 2980439 | 2984235 | 3797 | + | Gnomon | gene | gene-LOC105280348 | GeneID:105280348 | LOC105280348 | Gene | LOC105280348 | protein_coding | character(0) | character(0) | BCphaseA1.vs.BCphaseB1 | FALSE |
| NC_039506.1 | 2981308 | 2981308 | 1 | + | 38 | 17 | 21 | 78 | 52 | 26 | 44.74 | 66.67 | NC_039506.1 | 2980439 | 2984235 | 3797 | + | Gnomon | gene | gene-LOC105280348 | GeneID:105280348 | LOC105280348 | Gene | LOC105280348 | protein_coding | character(0) | character(0) | BCphaseA1.vs.BCphaseB1 | FALSE |
| NC_039506.1 | 2981315 | 2981315 | 1 | + | 39 | 28 | 11 | 77 | 69 | 8 | 71.79 | 89.61 | NC_039506.1 | 2980439 | 2984235 | 3797 | + | Gnomon | gene | gene-LOC105280348 | GeneID:105280348 | LOC105280348 | Gene | LOC105280348 | protein_coding | character(0) | character(0) | BCphaseA1.vs.BCphaseB1 | FALSE |
| NC_039506.1 | 2981335 | 2981335 | 1 | + | 35 | 34 | 1 | 56 | 54 | 2 | 97.14 | 96.43 | NC_039506.1 | 2980439 | 2984235 | 3797 | + | Gnomon | gene | gene-LOC105280348 | GeneID:105280348 | LOC105280348 | Gene | LOC105280348 | protein_coding | character(0) | character(0) | BCphaseA1.vs.BCphaseB1 | FALSE |
| NC_039506.1 | 2981350 | 2981350 | 1 | + | 51 | 50 | 1 | 58 | 57 | 1 | 98.04 | 98.28 | NC_039506.1 | 2980439 | 2984235 | 3797 | + | Gnomon | gene | gene-LOC105280348 | GeneID:105280348 | LOC105280348 | Gene | LOC105280348 | protein_coding | character(0) | character(0) | BCphaseA1.vs.BCphaseB1 | FALSE |
| NC_039506.1 | 2981391 | 2981391 | 1 | + | 63 | 58 | 5 | 60 | 49 | 11 | 92.06 | 81.67 | NC_039506.1 | 2980439 | 2984235 | 3797 | + | Gnomon | gene | gene-LOC105280348 | GeneID:105280348 | LOC105280348 | Gene | LOC105280348 | protein_coding | character(0) | character(0) | BCphaseA1.vs.BCphaseB1 | FALSE |
| NC_039506.1 | 2981402 | 2981402 | 1 | + | 69 | 63 | 6 | 66 | 60 | 6 | 91.3 | 90.91 | NC_039506.1 | 2980439 | 2984235 | 3797 | + | Gnomon | gene | gene-LOC105280348 | GeneID:105280348 | LOC105280348 | Gene | LOC105280348 | protein_coding | character(0) | character(0) | BCphaseA1.vs.BCphaseB1 | FALSE |
| NC_039506.1 | 2981411 | 2981411 | 1 | + | 73 | 69 | 4 | 68 | 65 | 3 | 94.52 | 95.59 | NC_039506.1 | 2980439 | 2984235 | 3797 | + | Gnomon | gene | gene-LOC105280348 | GeneID:105280348 | LOC105280348 | Gene | LOC105280348 | protein_coding | character(0) | character(0) | BCphaseA1.vs.BCphaseB1 | FALSE |
| NC_039506.1 | 2981975 | 2981975 | 1 | + | 23 | 9 | 14 | 25 | 7 | 18 | 39.13 | 28 | NC_039506.1 | 2980439 | 2984235 | 3797 | + | Gnomon | gene | gene-LOC105280348 | GeneID:105280348 | LOC105280348 | Gene | LOC105280348 | protein_coding | character(0) | character(0) | BCphaseA1.vs.BCphaseB1 | FALSE |
| NC_039506.1 | 2982097 | 2982097 | 1 | + | 31 | 0 | 31 | 34 | 1 | 33 | 0 | 2.94 | NC_039506.1 | 2980439 | 2984235 | 3797 | + | Gnomon | gene | gene-LOC105280348 | GeneID:105280348 | LOC105280348 | Gene | LOC105280348 | protein_coding | character(0) | character(0) | BCphaseA1.vs.BCphaseB1 | FALSE |
| NC_039506.1 | 2982133 | 2982133 | 1 | + | 38 | 0 | 38 | 47 | 1 | 46 | 0 | 2.13 | NC_039506.1 | 2980439 | 2984235 | 3797 | + | Gnomon | gene | gene-LOC105280348 | GeneID:105280348 | LOC105280348 | Gene | LOC105280348 | protein_coding | character(0) | character(0) | BCphaseA1.vs.BCphaseB1 | FALSE |

...

... only first 25 lines shown ...

Directory: DMSG File: dmg-Ob_LI_BCphaseA1.vs.BCphaseB2_details.txt

| seqnames | start | end | width | strand | coverage1 | numCs1 | numTs1 | coverage2 | numCs2 | numTs2 | BCphaseA1 | BCphaseB2 | gene_seqnames | gene_start | gene_end | gene_width | gene_strand | gene_source | gene_type | gene_ID | gene_Dbxref | gene_Name | gene_gbkey | gene_gene | gene_gene_biotype | gene_end_range | gene_partial | gene_start_range | gene_comparison | is.dm |
|---|---|---|---|---|---|---|---|---|---|---|---|---|---|---|---|---|---|---|---|---|---|---|---|---|---|---|---|---|---|---|
| NC_039506.1 | 1120656 | 1120656 | 1 | + | 20 | 0 | 20 | 26 | 0 | 26 | 0 | 0 | NC_039506.1 | 1116271 | 1134722 | 18452 | - | Gnomon | gene | gene-LOC113561443 | GeneID:113561443 | LOC113561443 | Gene | LOC113561443 | protein_coding | character(0) | NA | character(0) | BCphaseA1.vs.BCphaseB2 | FALSE |
| NC_039506.1 | 1120664 | 1120664 | 1 | + | 20 | 0 | 20 | 26 | 0 | 26 | 0 | 0 | NC_039506.1 | 1116271 | 1134722 | 18452 | - | Gnomon | gene | gene-LOC113561443 | GeneID:113561443 | LOC113561443 | Gene | LOC113561443 | protein_coding | character(0) | NA | character(0) | BCphaseA1.vs.BCphaseB2 | FALSE |
| NC_039506.1 | 1122355 | 1122355 | 1 | + | 42 | 1 | 41 | 42 | 10 | 32 | 2.38 | 23.81 | NC_039506.1 | 1116271 | 1134722 | 18452 | - | Gnomon | gene | gene-LOC113561443 | GeneID:113561443 | LOC113561443 | Gene | LOC113561443 | protein_coding | character(0) | NA | character(0) | BCphaseA1.vs.BCphaseB2 | TRUE |
| NC_039506.1 | 1122394 | 1122394 | 1 | + | 40 | 2 | 38 | 42 | 0 | 42 | 5 | 0 | NC_039506.1 | 1116271 | 1134722 | 18452 | - | Gnomon | gene | gene-LOC113561443 | GeneID:113561443 | LOC113561443 | Gene | LOC113561443 | protein_coding | character(0) | NA | character(0) | BCphaseA1.vs.BCphaseB2 | FALSE |
| NC_039506.1 | 1122397 | 1122397 | 1 | + | 38 | 1 | 37 | 40 | 2 | 38 | 2.63 | 5 | NC_039506.1 | 1116271 | 1134722 | 18452 | - | Gnomon | gene | gene-LOC113561443 | GeneID:113561443 | LOC113561443 | Gene | LOC113561443 | protein_coding | character(0) | NA | character(0) | BCphaseA1.vs.BCphaseB2 | FALSE |
| NC_039506.1 | 1122400 | 1122400 | 1 | + | 38 | 3 | 35 | 39 | 0 | 39 | 7.89 | 0 | NC_039506.1 | 1116271 | 1134722 | 18452 | - | Gnomon | gene | gene-LOC113561443 | GeneID:113561443 | LOC113561443 | Gene | LOC113561443 | protein_coding | character(0) | NA | character(0) | BCphaseA1.vs.BCphaseB2 | FALSE |
| NC_039506.1 | 1122426 | 1122426 | 1 | + | 33 | 0 | 33 | 31 | 1 | 30 | 0 | 3.23 | NC_039506.1 | 1116271 | 1134722 | 18452 | - | Gnomon | gene | gene-LOC113561443 | GeneID:113561443 | LOC113561443 | Gene | LOC113561443 | protein_coding | character(0) | NA | character(0) | BCphaseA1.vs.BCphaseB2 | FALSE |
| NC_039506.1 | 1122677 | 1122677 | 1 | + | 48 | 1 | 47 | 20 | 1 | 19 | 2.08 | 5 | NC_039506.1 | 1116271 | 1134722 | 18452 | - | Gnomon | gene | gene-LOC113561443 | GeneID:113561443 | LOC113561443 | Gene | LOC113561443 | protein_coding | character(0) | NA | character(0) | BCphaseA1.vs.BCphaseB2 | FALSE |
| NC_039506.1 | 1122707 | 1122707 | 1 | + | 28 | 0 | 28 | 24 | 0 | 24 | 0 | 0 | NC_039506.1 | 1116271 | 1134722 | 18452 | - | Gnomon | gene | gene-LOC113561443 | GeneID:113561443 | LOC113561443 | Gene | LOC113561443 | protein_coding | character(0) | NA | character(0) | BCphaseA1.vs.BCphaseB2 | FALSE |
| NC_039506.1 | 1122719 | 1122719 | 1 | + | 58 | 3 | 55 | 60 | 0 | 60 | 5.17 | 0 | NC_039506.1 | 1116271 | 1134722 | 18452 | - | Gnomon | gene | gene-LOC113561443 | GeneID:113561443 | LOC113561443 | Gene | LOC113561443 | protein_coding | character(0) | NA | character(0) | BCphaseA1.vs.BCphaseB2 | FALSE |
| NC_039506.1 | 1122757 | 1122757 | 1 | + | 45 | 13 | 32 | 51 | 22 | 29 | 28.89 | 43.14 | NC_039506.1 | 1116271 | 1134722 | 18452 | - | Gnomon | gene | gene-LOC113561443 | GeneID:113561443 | LOC113561443 | Gene | LOC113561443 | protein_coding | character(0) | NA | character(0) | BCphaseA1.vs.BCphaseB2 | FALSE |
| NC_039506.1 | 1122768 | 1122768 | 1 | + | 23 | 2 | 21 | 22 | 1 | 21 | 8.7 | 4.55 | NC_039506.1 | 1116271 | 1134722 | 18452 | - | Gnomon | gene | gene-LOC113561443 | GeneID:113561443 | LOC113561443 | Gene | LOC113561443 | protein_coding | character(0) | NA | character(0) | BCphaseA1.vs.BCphaseB2 | FALSE |
| NC_039506.1 | 1122778 | 1122778 | 1 | + | 21 | 1 | 20 | 22 | 0 | 22 | 4.76 | 0 | NC_039506.1 | 1116271 | 1134722 | 18452 | - | Gnomon | gene | gene-LOC113561443 | GeneID:113561443 | LOC113561443 | Gene | LOC113561443 | protein_coding | character(0) | NA | character(0) | BCphaseA1.vs.BCphaseB2 | FALSE |
| NC_039506.1 | 1122786 | 1122786 | 1 | + | 21 | 0 | 21 | 20 | 0 | 20 | 0 | 0 | NC_039506.1 | 1116271 | 1134722 | 18452 | - | Gnomon | gene | gene-LOC113561443 | GeneID:113561443 | LOC113561443 | Gene | LOC113561443 | protein_coding | character(0) | NA | character(0) | BCphaseA1.vs.BCphaseB2 | FALSE |
| NC_039506.1 | 1125825 | 1125825 | 1 | + | 23 | 3 | 20 | 20 | 4 | 16 | 13.04 | 20 | NC_039506.1 | 1116271 | 1134722 | 18452 | - | Gnomon | gene | gene-LOC113561443 | GeneID:113561443 | LOC113561443 | Gene | LOC113561443 | protein_coding | character(0) | NA | character(0) | BCphaseA1.vs.BCphaseB2 | FALSE |
| NC_039506.1 | 1127447 | 1127447 | 1 | + | 31 | 1 | 30 | 29 | 2 | 27 | 3.23 | 6.9 | NC_039506.1 | 1116271 | 1134722 | 18452 | - | Gnomon | gene | gene-LOC113561443 | GeneID:113561443 | LOC113561443 | Gene | LOC113561443 | protein_coding | character(0) | NA | character(0) | BCphaseA1.vs.BCphaseB2 | FALSE |
| NC_039506.1 | 1540915 | 1540915 | 1 | + | 35 | 0 | 35 | 25 | 0 | 25 | 0 | 0 | NC_039506.1 | 1540640 | 1547998 | 7359 | - | Gnomon | gene | gene-LOC105280244 | GeneID:105280244 | LOC105280244 | Gene | LOC105280244 | protein_coding | character(0) | NA | character(0) | BCphaseA1.vs.BCphaseB2 | FALSE |
| NC_039506.1 | 1540917 | 1540917 | 1 | + | 35 | 0 | 35 | 26 | 0 | 26 | 0 | 0 | NC_039506.1 | 1540640 | 1547998 | 7359 | - | Gnomon | gene | gene-LOC105280244 | GeneID:105280244 | LOC105280244 | Gene | LOC105280244 | protein_coding | character(0) | NA | character(0) | BCphaseA1.vs.BCphaseB2 | FALSE |
| NC_039506.1 | 1540928 | 1540928 | 1 | + | 33 | 0 | 33 | 27 | 0 | 27 | 0 | 0 | NC_039506.1 | 1540640 | 1547998 | 7359 | - | Gnomon | gene | gene-LOC105280244 | GeneID:105280244 | LOC105280244 | Gene | LOC105280244 | protein_coding | character(0) | NA | character(0) | BCphaseA1.vs.BCphaseB2 | FALSE |
| NC_039506.1 | 1542484 | 1542484 | 1 | + | 20 | 0 | 20 | 20 | 0 | 20 | 0 | 0 | NC_039506.1 | 1540640 | 1547998 | 7359 | - | Gnomon | gene | gene-LOC105280244 | GeneID:105280244 | LOC105280244 | Gene | LOC105280244 | protein_coding | character(0) | NA | character(0) | BCphaseA1.vs.BCphaseB2 | FALSE |
| NC_039506.1 | 1542627 | 1542627 | 1 | + | 36 | 0 | 36 | 33 | 0 | 33 | 0 | 0 | NC_039506.1 | 1540640 | 1547998 | 7359 | - | Gnomon | gene | gene-LOC105280244 | GeneID:105280244 | LOC105280244 | Gene | LOC105280244 | protein_coding | character(0) | NA | character(0) | BCphaseA1.vs.BCphaseB2 | FALSE |
| NC_039506.1 | 1542653 | 1542653 | 1 | + | 28 | 1 | 27 | 30 | 0 | 30 | 3.57 | 0 | NC_039506.1 | 1540640 | 1547998 | 7359 | - | Gnomon | gene | gene-LOC105280244 | GeneID:105280244 | LOC105280244 | Gene | LOC105280244 | protein_coding | character(0) | NA | character(0) | BCphaseA1.vs.BCphaseB2 | FALSE |
| NC_039506.1 | 1542672 | 1542672 | 1 | + | 27 | 0 | 27 | 23 | 1 | 22 | 0 | 4.35 | NC_039506.1 | 1540640 | 1547998 | 7359 | - | Gnomon | gene | gene-LOC105280244 | GeneID:105280244 | LOC105280244 | Gene | LOC105280244 | protein_coding | character(0) | NA | character(0) | BCphaseA1.vs.BCphaseB2 | FALSE |
| NC_039506.1 | 1542674 | 1542674 | 1 | + | 26 | 0 | 26 | 23 | 0 | 23 | 0 | 0 | NC_039506.1 | 1540640 | 1547998 | 7359 | - | Gnomon | gene | gene-LOC105280244 | GeneID:105280244 | LOC105280244 | Gene | LOC105280244 | protein_coding | character(0) | NA | character(0) | BCphaseA1.vs.BCphaseB2 | FALSE |

...
... only first 25 lines shown ...

Directory: DMSG File: dmg-Ob_LI_BCphaseA1.vs.RphaseA1_details.txt

| seqnames | start | end | width | strand | coverage1 | numCs1 | numTs1 | coverage2 | numCs2 | numTs2 | BCphaseA1 | RphaseA1 | gene_seqnames | gene_start | gene_end | gene_width | gene_strand | gene_source | gene_type | gene_ID | gene_Dbxref | gene_Name | gene_gbkey | gene_gene | gene_gene_biotype | gene_end_range | gene_start_range | gene_comparison | is.dm |
|---|---|---|---|---|---|---|---|---|---|---|---|---|---|---|---|---|---|---|---|---|---|---|---|---|---|---|---|---|---|
| NC_039506.1 | 2980750 | 2980750 | 1 | + | 21 | 0 | 21 | 28 | 0 | 28 | 0 | 0 | NC_039506.1 | 2980439 | 2984235 | 3797 | + | Gnomon | gene | gene-LOC105280348 | GeneID:105280348 | LOC105280348 | Gene | LOC105280348 | protein_coding | character(0) | character(0) | BCphaseA1.vs.RphaseA1 | FALSE |
| NC_039506.1 | 2980761 | 2980761 | 1 | + | 22 | 1 | 21 | 31 | 0 | 31 | 4.55 | 0 | NC_039506.1 | 2980439 | 2984235 | 3797 | + | Gnomon | gene | gene-LOC105280348 | GeneID:105280348 | LOC105280348 | Gene | LOC105280348 | protein_coding | character(0) | character(0) | BCphaseA1.vs.RphaseA1 | FALSE |
| NC_039506.1 | 2980767 | 2980767 | 1 | + | 23 | 0 | 23 | 32 | 1 | 31 | 0 | 3.12 | NC_039506.1 | 2980439 | 2984235 | 3797 | + | Gnomon | gene | gene-LOC105280348 | GeneID:105280348 | LOC105280348 | Gene | LOC105280348 | protein_coding | character(0) | character(0) | BCphaseA1.vs.RphaseA1 | FALSE |
| NC_039506.1 | 2980989 | 2980989 | 1 | + | 33 | 3 | 30 | 28 | 3 | 25 | 9.09 | 10.71 | NC_039506.1 | 2980439 | 2984235 | 3797 | + | Gnomon | gene | gene-LOC105280348 | GeneID:105280348 | LOC105280348 | Gene | LOC105280348 | protein_coding | character(0) | character(0) | BCphaseA1.vs.RphaseA1 | FALSE |
| NC_039506.1 | 2980993 | 2980993 | 1 | + | 31 | 10 | 21 | 29 | 15 | 14 | 32.26 | 51.72 | NC_039506.1 | 2980439 | 2984235 | 3797 | + | Gnomon | gene | gene-LOC105280348 | GeneID:105280348 | LOC105280348 | Gene | LOC105280348 | protein_coding | character(0) | character(0) | BCphaseA1.vs.RphaseA1 | FALSE |
| NC_039506.1 | 2980998 | 2980998 | 1 | + | 32 | 6 | 26 | 28 | 7 | 21 | 18.75 | 25 | NC_039506.1 | 2980439 | 2984235 | 3797 | + | Gnomon | gene | gene-LOC105280348 | GeneID:105280348 | LOC105280348 | Gene | LOC105280348 | protein_coding | character(0) | character(0) | BCphaseA1.vs.RphaseA1 | FALSE |
| NC_039506.1 | 2981030 | 2981030 | 1 | + | 55 | 46 | 9 | 28 | 25 | 3 | 83.64 | 89.29 | NC_039506.1 | 2980439 | 2984235 | 3797 | + | Gnomon | gene | gene-LOC105280348 | GeneID:105280348 | LOC105280348 | Gene | LOC105280348 | protein_coding | character(0) | character(0) | BCphaseA1.vs.RphaseA1 | FALSE |
| NC_039506.1 | 2981077 | 2981077 | 1 | + | 34 | 18 | 16 | 30 | 11 | 19 | 52.94 | 36.67 | NC_039506.1 | 2980439 | 2984235 | 3797 | + | Gnomon | gene | gene-LOC105280348 | GeneID:105280348 | LOC105280348 | Gene | LOC105280348 | protein_coding | character(0) | character(0) | BCphaseA1.vs.RphaseA1 | FALSE |
| NC_039506.1 | 2981125 | 2981125 | 1 | + | 29 | 18 | 11 | 20 | 10 | 10 | 62.07 | 50 | NC_039506.1 | 2980439 | 2984235 | 3797 | + | Gnomon | gene | gene-LOC105280348 | GeneID:105280348 | LOC105280348 | Gene | LOC105280348 | protein_coding | character(0) | character(0) | BCphaseA1.vs.RphaseA1 | FALSE |
| NC_039506.1 | 2981146 | 2981146 | 1 | + | 22 | 21 | 1 | 31 | 28 | 3 | 95.45 | 90.32 | NC_039506.1 | 2980439 | 2984235 | 3797 | + | Gnomon | gene | gene-LOC105280348 | GeneID:105280348 | LOC105280348 | Gene | LOC105280348 | protein_coding | character(0) | character(0) | BCphaseA1.vs.RphaseA1 | FALSE |
| NC_039506.1 | 2981183 | 2981183 | 1 | + | 56 | 25 | 31 | 60 | 59 | 1 | 44.64 | 98.33 | NC_039506.1 | 2980439 | 2984235 | 3797 | + | Gnomon | gene | gene-LOC105280348 | GeneID:105280348 | LOC105280348 | Gene | LOC105280348 | protein_coding | character(0) | character(0) | BCphaseA1.vs.RphaseA1 | TRUE |
| NC_039506.1 | 2981225 | 2981225 | 1 | + | 25 | 17 | 8 | 46 | 29 | 17 | 68 | 63.04 | NC_039506.1 | 2980439 | 2984235 | 3797 | + | Gnomon | gene | gene-LOC105280348 | GeneID:105280348 | LOC105280348 | Gene | LOC105280348 | protein_coding | character(0) | character(0) | BCphaseA1.vs.RphaseA1 | FALSE |
| NC_039506.1 | 2981245 | 2981245 | 1 | + | 31 | 15 | 16 | 38 | 15 | 23 | 48.39 | 39.47 | NC_039506.1 | 2980439 | 2984235 | 3797 | + | Gnomon | gene | gene-LOC105280348 | GeneID:105280348 | LOC105280348 | Gene | LOC105280348 | protein_coding | character(0) | character(0) | BCphaseA1.vs.RphaseA1 | FALSE |
| NC_039506.1 | 2981292 | 2981292 | 1 | + | 31 | 26 | 5 | 42 | 38 | 4 | 83.87 | 90.48 | NC_039506.1 | 2980439 | 2984235 | 3797 | + | Gnomon | gene | gene-LOC105280348 | GeneID:105280348 | LOC105280348 | Gene | LOC105280348 | protein_coding | character(0) | character(0) | BCphaseA1.vs.RphaseA1 | FALSE |
| NC_039506.1 | 2981308 | 2981308 | 1 | + | 38 | 17 | 21 | 73 | 49 | 24 | 44.74 | 67.12 | NC_039506.1 | 2980439 | 2984235 | 3797 | + | Gnomon | gene | gene-LOC105280348 | GeneID:105280348 | LOC105280348 | Gene | LOC105280348 | protein_coding | character(0) | character(0) | BCphaseA1.vs.RphaseA1 | FALSE |
| NC_039506.1 | 2981315 | 2981315 | 1 | + | 39 | 28 | 11 | 52 | 48 | 4 | 71.79 | 92.31 | NC_039506.1 | 2980439 | 2984235 | 3797 | + | Gnomon | gene | gene-LOC105280348 | GeneID:105280348 | LOC105280348 | Gene | LOC105280348 | protein_coding | character(0) | character(0) | BCphaseA1.vs.RphaseA1 | FALSE |
| NC_039506.1 | 2981335 | 2981335 | 1 | + | 35 | 34 | 1 | 40 | 38 | 2 | 97.14 | 95 | NC_039506.1 | 2980439 | 2984235 | 3797 | + | Gnomon | gene | gene-LOC105280348 | GeneID:105280348 | LOC105280348 | Gene | LOC105280348 | protein_coding | character(0) | character(0) | BCphaseA1.vs.RphaseA1 | FALSE |
| NC_039506.1 | 2981350 | 2981350 | 1 | + | 51 | 50 | 1 | 52 | 51 | 1 | 98.04 | 98.08 | NC_039506.1 | 2980439 | 2984235 | 3797 | + | Gnomon | gene | gene-LOC105280348 | GeneID:105280348 | LOC105280348 | Gene | LOC105280348 | protein_coding | character(0) | character(0) | BCphaseA1.vs.RphaseA1 | FALSE |
| NC_039506.1 | 2981391 | 2981391 | 1 | + | 63 | 58 | 5 | 60 | 49 | 11 | 92.06 | 81.67 | NC_039506.1 | 2980439 | 2984235 | 3797 | + | Gnomon | gene | gene-LOC105280348 | GeneID:105280348 | LOC105280348 | Gene | LOC105280348 | protein_coding | character(0) | character(0) | BCphaseA1.vs.RphaseA1 | FALSE |
| NC_039506.1 | 2981402 | 2981402 | 1 | + | 69 | 63 | 6 | 73 | 68 | 5 | 91.3 | 93.15 | NC_039506.1 | 2980439 | 2984235 | 3797 | + | Gnomon | gene | gene-LOC105280348 | GeneID:105280348 | LOC105280348 | Gene | LOC105280348 | protein_coding | character(0) | character(0) | BCphaseA1.vs.RphaseA1 | FALSE |
| NC_039506.1 | 2981411 | 2981411 | 1 | + | 73 | 69 | 4 | 73 | 70 | 3 | 94.52 | 95.89 | NC_039506.1 | 2980439 | 2984235 | 3797 | + | Gnomon | gene | gene-LOC105280348 | GeneID:105280348 | LOC105280348 | Gene | LOC105280348 | protein_coding | character(0) | character(0) | BCphaseA1.vs.RphaseA1 | FALSE |
| NC_039506.1 | 2981975 | 2981975 | 1 | + | 23 | 9 | 14 | 29 | 14 | 15 | 39.13 | 48.28 | NC_039506.1 | 2980439 | 2984235 | 3797 | + | Gnomon | gene | gene-LOC105280348 | GeneID:105280348 | LOC105280348 | Gene | LOC105280348 | protein_coding | character(0) | character(0) | BCphaseA1.vs.RphaseA1 | FALSE |
| NC_039506.1 | 2982097 | 2982097 | 1 | + | 31 | 0 | 31 | 35 | 0 | 35 | 0 | 0 | NC_039506.1 | 2980439 | 2984235 | 3797 | + | Gnomon | gene | gene-LOC105280348 | GeneID:105280348 | LOC105280348 | Gene | LOC105280348 | protein_coding | character(0) | character(0) | BCphaseA1.vs.RphaseA1 | FALSE |
| NC_039506.1 | 2982133 | 2982133 | 1 | + | 38 | 0 | 38 | 40 | 0 | 40 | 0 | 0 | NC_039506.1 | 2980439 | 2984235 | 3797 | + | Gnomon | gene | gene-LOC105280348 | GeneID:105280348 | LOC105280348 | Gene | LOC105280348 | protein_coding | character(0) | character(0) | BCphaseA1.vs.RphaseA1 | FALSE |

...
... only first 25 lines shown ...

Directory: DMSG File: dmg-Ob_LI_BCphaseA1.vs.RphaseA2_details.txt

| seqnames | start | end | width | strand | coverage1 | numCs1 | numTs1 | coverage2 | numCs2 | numTs2 | BCphaseA1 | RphaseA2 | gene_seqnames | gene_start | gene_end | gene_width | gene_strand | gene_source | gene_type | gene_ID | gene_Dbxref | gene_Name | gene_gbkey | gene_gene | gene_gene_biotype | gene_end_range | gene_start_range | gene_comparison | is.dm |
|---|---|---|---|---|---|---|---|---|---|---|---|---|---|---|---|---|---|---|---|---|---|---|---|---|---|---|---|---|---|
| NC_039506.1 | 2173614 | 2173614 | 1 | + | 34 | 0 | 34 | 63 | 0 | 63 | 0 | 0 | NC_039506.1 | 2172791 | 2181709 | 8919 | + | Gnomon | gene | gene-LOC105280304 | GeneID:105280304 | LOC105280304 | Gene | LOC105280304 | protein_coding | character(0) | character(0) | BCphaseA1.vs.RphaseA2 | FALSE |
| NC_039506.1 | 2173709 | 2173709 | 1 | + | 34 | 0 | 34 | 64 | 0 | 64 | 0 | 0 | NC_039506.1 | 2172791 | 2181709 | 8919 | + | Gnomon | gene | gene-LOC105280304 | GeneID:105280304 | LOC105280304 | Gene | LOC105280304 | protein_coding | character(0) | character(0) | BCphaseA1.vs.RphaseA2 | FALSE |
| NC_039506.1 | 2173733 | 2173733 | 1 | + | 46 | 1 | 45 | 61 | 1 | 60 | 2.17 | 1.64 | NC_039506.1 | 2172791 | 2181709 | 8919 | + | Gnomon | gene | gene-LOC105280304 | GeneID:105280304 | LOC105280304 | Gene | LOC105280304 | protein_coding | character(0) | character(0) | BCphaseA1.vs.RphaseA2 | FALSE |
| NC_039506.1 | 2173742 | 2173742 | 1 | + | 46 | 0 | 46 | 66 | 0 | 66 | 0 | 0 | NC_039506.1 | 2172791 | 2181709 | 8919 | + | Gnomon | gene | gene-LOC105280304 | GeneID:105280304 | LOC105280304 | Gene | LOC105280304 | protein_coding | character(0) | character(0) | BCphaseA1.vs.RphaseA2 | FALSE |
| NC_039506.1 | 2173746 | 2173746 | 1 | + | 46 | 1 | 45 | 67 | 0 | 67 | 2.17 | 0 | NC_039506.1 | 2172791 | 2181709 | 8919 | + | Gnomon | gene | gene-LOC105280304 | GeneID:105280304 | LOC105280304 | Gene | LOC105280304 | protein_coding | character(0) | character(0) | BCphaseA1.vs.RphaseA2 | FALSE |
| NC_039506.1 | 2173753 | 2173753 | 1 | + | 46 | 1 | 45 | 69 | 0 | 69 | 2.17 | 0 | NC_039506.1 | 2172791 | 2181709 | 8919 | + | Gnomon | gene | gene-LOC105280304 | GeneID:105280304 | LOC105280304 | Gene | LOC105280304 | protein_coding | character(0) | character(0) | BCphaseA1.vs.RphaseA2 | FALSE |
| NC_039506.1 | 2173759 | 2173759 | 1 | + | 47 | 0 | 47 | 70 | 1 | 69 | 0 | 1.43 | NC_039506.1 | 2172791 | 2181709 | 8919 | + | Gnomon | gene | gene-LOC105280304 | GeneID:105280304 | LOC105280304 | Gene | LOC105280304 | protein_coding | character(0) | character(0) | BCphaseA1.vs.RphaseA2 | FALSE |
| NC_039506.1 | 2173764 | 2173764 | 1 | + | 49 | 0 | 49 | 75 | 0 | 75 | 0 | 0 | NC_039506.1 | 2172791 | 2181709 | 8919 | + | Gnomon | gene | gene-LOC105280304 | GeneID:105280304 | LOC105280304 | Gene | LOC105280304 | protein_coding | character(0) | character(0) | BCphaseA1.vs.RphaseA2 | FALSE |
| NC_039506.1 | 2173768 | 2173768 | 1 | + | 50 | 1 | 49 | 75 | 0 | 75 | 2 | 0 | NC_039506.1 | 2172791 | 2181709 | 8919 | + | Gnomon | gene | gene-LOC105280304 | GeneID:105280304 | LOC105280304 | Gene | LOC105280304 | protein_coding | character(0) | character(0) | BCphaseA1.vs.RphaseA2 | FALSE |
| NC_039506.1 | 2173779 | 2173779 | 1 | + | 56 | 1 | 55 | 82 | 0 | 82 | 1.79 | 0 | NC_039506.1 | 2172791 | 2181709 | 8919 | + | Gnomon | gene | gene-LOC105280304 | GeneID:105280304 | LOC105280304 | Gene | LOC105280304 | protein_coding | character(0) | character(0) | BCphaseA1.vs.RphaseA2 | FALSE |
| NC_039506.1 | 2173789 | 2173789 | 1 | + | 54 | 0 | 54 | 84 | 0 | 84 | 0 | 0 | NC_039506.1 | 2172791 | 2181709 | 8919 | + | Gnomon | gene | gene-LOC105280304 | GeneID:105280304 | LOC105280304 | Gene | LOC105280304 | protein_coding | character(0) | character(0) | BCphaseA1.vs.RphaseA2 | FALSE |
| NC_039506.1 | 2173798 | 2173798 | 1 | + | 48 | 0 | 48 | 74 | 0 | 74 | 0 | 0 | NC_039506.1 | 2172791 | 2181709 | 8919 | + | Gnomon | gene | gene-LOC105280304 | GeneID:105280304 | LOC105280304 | Gene | LOC105280304 | protein_coding | character(0) | character(0) | BCphaseA1.vs.RphaseA2 | FALSE |
| NC_039506.1 | 2173801 | 2173801 | 1 | + | 46 | 0 | 46 | 68 | 1 | 67 | 0 | 1.47 | NC_039506.1 | 2172791 | 2181709 | 8919 | + | Gnomon | gene | gene-LOC105280304 | GeneID:105280304 | LOC105280304 | Gene | LOC105280304 | protein_coding | character(0) | character(0) | BCphaseA1.vs.RphaseA2 | FALSE |
| NC_039506.1 | 2173810 | 2173810 | 1 | + | 36 | 2 | 34 | 54 | 1 | 53 | 5.56 | 1.85 | NC_039506.1 | 2172791 | 2181709 | 8919 | + | Gnomon | gene | gene-LOC105280304 | GeneID:105280304 | LOC105280304 | Gene | LOC105280304 | protein_coding | character(0) | character(0) | BCphaseA1.vs.RphaseA2 | FALSE |
| NC_039506.1 | 2173812 | 2173812 | 1 | + | 38 | 2 | 36 | 54 | 3 | 51 | 5.26 | 5.56 | NC_039506.1 | 2172791 | 2181709 | 8919 | + | Gnomon | gene | gene-LOC105280304 | GeneID:105280304 | LOC105280304 | Gene | LOC105280304 | protein_coding | character(0) | character(0) | BCphaseA1.vs.RphaseA2 | FALSE |
| NC_039506.1 | 2173823 | 2173823 | 1 | + | 31 | 0 | 31 | 48 | 1 | 47 | 0 | 2.08 | NC_039506.1 | 2172791 | 2181709 | 8919 | + | Gnomon | gene | gene-LOC105280304 | GeneID:105280304 | LOC105280304 | Gene | LOC105280304 | protein_coding | character(0) | character(0) | BCphaseA1.vs.RphaseA2 | FALSE |
| NC_039506.1 | 2173831 | 2173831 | 1 | + | 30 | 0 | 30 | 48 | 0 | 48 | 0 | 0 | NC_039506.1 | 2172791 | 2181709 | 8919 | + | Gnomon | gene | gene-LOC105280304 | GeneID:105280304 | LOC105280304 | Gene | LOC105280304 | protein_coding | character(0) | character(0) | BCphaseA1.vs.RphaseA2 | FALSE |
| NC_039506.1 | 2174095 | 2174095 | 1 | + | 23 | 0 | 23 | 36 | 0 | 36 | 0 | 0 | NC_039506.1 | 2172791 | 2181709 | 8919 | + | Gnomon | gene | gene-LOC105280304 | GeneID:105280304 | LOC105280304 | Gene | LOC105280304 | protein_coding | character(0) | character(0) | BCphaseA1.vs.RphaseA2 | FALSE |
| NC_039506.1 | 2174105 | 2174105 | 1 | + | 25 | 0 | 25 | 39 | 0 | 39 | 0 | 0 | NC_039506.1 | 2172791 | 2181709 | 8919 | + | Gnomon | gene | gene-LOC105280304 | GeneID:105280304 | LOC105280304 | Gene | LOC105280304 | protein_coding | character(0) | character(0) | BCphaseA1.vs.RphaseA2 | FALSE |
| NC_039506.1 | 2174110 | 2174110 | 1 | + | 23 | 0 | 23 | 39 | 0 | 39 | 0 | 0 | NC_039506.1 | 2172791 | 2181709 | 8919 | + | Gnomon | gene | gene-LOC105280304 | GeneID:105280304 | LOC105280304 | Gene | LOC105280304 | protein_coding | character(0) | character(0) | BCphaseA1.vs.RphaseA2 | FALSE |
| NC_039506.1 | 2174115 | 2174115 | 1 | + | 24 | 0 | 24 | 37 | 0 | 37 | 0 | 0 | NC_039506.1 | 2172791 | 2181709 | 8919 | + | Gnomon | gene | gene-LOC105280304 | GeneID:105280304 | LOC105280304 | Gene | LOC105280304 | protein_coding | character(0) | character(0) | BCphaseA1.vs.RphaseA2 | FALSE |
| NC_039506.1 | 2174125 | 2174125 | 1 | + | 23 | 0 | 23 | 36 | 0 | 36 | 0 | 0 | NC_039506.1 | 2172791 | 2181709 | 8919 | + | Gnomon | gene | gene-LOC105280304 | GeneID:105280304 | LOC105280304 | Gene | LOC105280304 | protein_coding | character(0) | character(0) | BCphaseA1.vs.RphaseA2 | FALSE |
| NC_039506.1 | 2174139 | 2174139 | 1 | + | 21 | 0 | 21 | 25 | 0 | 25 | 0 | 0 | NC_039506.1 | 2172791 | 2181709 | 8919 | + | Gnomon | gene | gene-LOC105280304 | GeneID:105280304 | LOC105280304 | Gene | LOC105280304 | protein_coding | character(0) | character(0) | BCphaseA1.vs.RphaseA2 | FALSE |
| NC_039506.1 | 2174148 | 2174148 | 1 | + | 22 | 0 | 22 | 21 | 0 | 21 | 0 | 0 | NC_039506.1 | 2172791 | 2181709 | 8919 | + | Gnomon | gene | gene-LOC105280304 | GeneID:105280304 | LOC105280304 | Gene | LOC105280304 | protein_coding | character(0) | character(0) | BCphaseA1.vs.RphaseA2 | FALSE |

...

... only first 25 lines shown ...

Directory: DMSG File: dmg-Ob_LI_BCphaseA1.vs.RphaseB1_details.txt

| seqnames | start | end | width | strand | coverage1 | numCs1 | numTs1 | coverage2 | numCs2 | numTs2 | BCphaseA1 | RphaseB1 | gene_seqnames | gene_start | gene_end | gene_width | gene_strand | gene_source | gene_type | gene_ID | gene_Dbxref | gene_Name | gene_gbkey | gene_gene | gene_gene_biotype | gene_end_range | gene_partial | gene_start_range | gene_comparison | is.dm |
|---|---|---|---|---|---|---|---|---|---|---|---|---|---|---|---|---|---|---|---|---|---|---|---|---|---|---|---|---|---|---|
| NC_039506.1 | 1540915 | 1540915 | 1 | + | 35 | 0 | 35 | 24 | 0 | 24 | 0 | 0 | NC_039506.1 | 1540640 | 1547998 | 7359 | - | Gnomon | gene | gene-LOC105280244 | GeneID:105280244 | LOC105280244 | Gene | LOC105280244 | protein_coding | character(0) | NA | character(0) | BCphaseA1.vs.RphaseB1 | FALSE |
| NC_039506.1 | 1540917 | 1540917 | 1 | + | 35 | 0 | 35 | 24 | 0 | 24 | 0 | 0 | NC_039506.1 | 1540640 | 1547998 | 7359 | - | Gnomon | gene | gene-LOC105280244 | GeneID:105280244 | LOC105280244 | Gene | LOC105280244 | protein_coding | character(0) | NA | character(0) | BCphaseA1.vs.RphaseB1 | FALSE |
| NC_039506.1 | 1540928 | 1540928 | 1 | + | 33 | 0 | 33 | 24 | 0 | 24 | 0 | 0 | NC_039506.1 | 1540640 | 1547998 | 7359 | - | Gnomon | gene | gene-LOC105280244 | GeneID:105280244 | LOC105280244 | Gene | LOC105280244 | protein_coding | character(0) | NA | character(0) | BCphaseA1.vs.RphaseB1 | FALSE |
| NC_039506.1 | 1542484 | 1542484 | 1 | + | 20 | 0 | 20 | 21 | 0 | 21 | 0 | 0 | NC_039506.1 | 1540640 | 1547998 | 7359 | - | Gnomon | gene | gene-LOC105280244 | GeneID:105280244 | LOC105280244 | Gene | LOC105280244 | protein_coding | character(0) | NA | character(0) | BCphaseA1.vs.RphaseB1 | FALSE |
| NC_039506.1 | 1542627 | 1542627 | 1 | + | 36 | 0 | 36 | 31 | 0 | 31 | 0 | 0 | NC_039506.1 | 1540640 | 1547998 | 7359 | - | Gnomon | gene | gene-LOC105280244 | GeneID:105280244 | LOC105280244 | Gene | LOC105280244 | protein_coding | character(0) | NA | character(0) | BCphaseA1.vs.RphaseB1 | FALSE |
| NC_039506.1 | 1542653 | 1542653 | 1 | + | 28 | 1 | 27 | 24 | 0 | 24 | 3.57 | 0 | NC_039506.1 | 1540640 | 1547998 | 7359 | - | Gnomon | gene | gene-LOC105280244 | GeneID:105280244 | LOC105280244 | Gene | LOC105280244 | protein_coding | character(0) | NA | character(0) | BCphaseA1.vs.RphaseB1 | FALSE |
| NC_039506.1 | 1542672 | 1542672 | 1 | + | 27 | 0 | 27 | 22 | 0 | 22 | 0 | 0 | NC_039506.1 | 1540640 | 1547998 | 7359 | - | Gnomon | gene | gene-LOC105280244 | GeneID:105280244 | LOC105280244 | Gene | LOC105280244 | protein_coding | character(0) | NA | character(0) | BCphaseA1.vs.RphaseB1 | FALSE |
| NC_039506.1 | 1542674 | 1542674 | 1 | + | 26 | 0 | 26 | 22 | 1 | 21 | 0 | 4.55 | NC_039506.1 | 1540640 | 1547998 | 7359 | - | Gnomon | gene | gene-LOC105280244 | GeneID:105280244 | LOC105280244 | Gene | LOC105280244 | protein_coding | character(0) | NA | character(0) | BCphaseA1.vs.RphaseB1 | FALSE |
| NC_039506.1 | 1543020 | 1543020 | 1 | + | 20 | 1 | 19 | 23 | 0 | 23 | 5 | 0 | NC_039506.1 | 1540640 | 1547998 | 7359 | - | Gnomon | gene | gene-LOC105280244 | GeneID:105280244 | LOC105280244 | Gene | LOC105280244 | protein_coding | character(0) | NA | character(0) | BCphaseA1.vs.RphaseB1 | FALSE |
| NC_039506.1 | 1543060 | 1543060 | 1 | + | 41 | 1 | 40 | 57 | 0 | 57 | 2.44 | 0 | NC_039506.1 | 1540640 | 1547998 | 7359 | - | Gnomon | gene | gene-LOC105280244 | GeneID:105280244 | LOC105280244 | Gene | LOC105280244 | protein_coding | character(0) | NA | character(0) | BCphaseA1.vs.RphaseB1 | FALSE |
| NC_039506.1 | 1543103 | 1543103 | 1 | + | 26 | 1 | 25 | 39 | 1 | 38 | 3.85 | 2.56 | NC_039506.1 | 1540640 | 1547998 | 7359 | - | Gnomon | gene | gene-LOC105280244 | GeneID:105280244 | LOC105280244 | Gene | LOC105280244 | protein_coding | character(0) | NA | character(0) | BCphaseA1.vs.RphaseB1 | FALSE |
| NC_039506.1 | 1543108 | 1543108 | 1 | + | 25 | 1 | 24 | 39 | 1 | 38 | 4 | 2.56 | NC_039506.1 | 1540640 | 1547998 | 7359 | - | Gnomon | gene | gene-LOC105280244 | GeneID:105280244 | LOC105280244 | Gene | LOC105280244 | protein_coding | character(0) | NA | character(0) | BCphaseA1.vs.RphaseB1 | FALSE |
| NC_039506.1 | 1543137 | 1543137 | 1 | + | 21 | 0 | 21 | 24 | 1 | 23 | 0 | 4.17 | NC_039506.1 | 1540640 | 1547998 | 7359 | - | Gnomon | gene | gene-LOC105280244 | GeneID:105280244 | LOC105280244 | Gene | LOC105280244 | protein_coding | character(0) | NA | character(0) | BCphaseA1.vs.RphaseB1 | FALSE |
| NC_039506.1 | 1543240 | 1543240 | 1 | + | 31 | 0 | 31 | 22 | 0 | 22 | 0 | 0 | NC_039506.1 | 1540640 | 1547998 | 7359 | - | Gnomon | gene | gene-LOC105280244 | GeneID:105280244 | LOC105280244 | Gene | LOC105280244 | protein_coding | character(0) | NA | character(0) | BCphaseA1.vs.RphaseB1 | FALSE |
| NC_039506.1 | 1543341 | 1543341 | 1 | + | 26 | 0 | 26 | 25 | 0 | 25 | 0 | 0 | NC_039506.1 | 1540640 | 1547998 | 7359 | - | Gnomon | gene | gene-LOC105280244 | GeneID:105280244 | LOC105280244 | Gene | LOC105280244 | protein_coding | character(0) | NA | character(0) | BCphaseA1.vs.RphaseB1 | FALSE |
| NC_039506.1 | 1543415 | 1543415 | 1 | + | 56 | 1 | 55 | 49 | 0 | 49 | 1.79 | 0 | NC_039506.1 | 1540640 | 1547998 | 7359 | - | Gnomon | gene | gene-LOC105280244 | GeneID:105280244 | LOC105280244 | Gene | LOC105280244 | protein_coding | character(0) | NA | character(0) | BCphaseA1.vs.RphaseB1 | FALSE |
| NC_039506.1 | 1543443 | 1543443 | 1 | + | 47 | 0 | 47 | 48 | 0 | 48 | 0 | 0 | NC_039506.1 | 1540640 | 1547998 | 7359 | - | Gnomon | gene | gene-LOC105280244 | GeneID:105280244 | LOC105280244 | Gene | LOC105280244 | protein_coding | character(0) | NA | character(0) | BCphaseA1.vs.RphaseB1 | FALSE |
| NC_039506.1 | 1543445 | 1543445 | 1 | + | 46 | 0 | 46 | 48 | 1 | 47 | 0 | 2.08 | NC_039506.1 | 1540640 | 1547998 | 7359 | - | Gnomon | gene | gene-LOC105280244 | GeneID:105280244 | LOC105280244 | Gene | LOC105280244 | protein_coding | character(0) | NA | character(0) | BCphaseA1.vs.RphaseB1 | FALSE |
| NC_039506.1 | 1543451 | 1543451 | 1 | + | 45 | 1 | 44 | 40 | 1 | 39 | 2.22 | 2.5 | NC_039506.1 | 1540640 | 1547998 | 7359 | - | Gnomon | gene | gene-LOC105280244 | GeneID:105280244 | LOC105280244 | Gene | LOC105280244 | protein_coding | character(0) | NA | character(0) | BCphaseA1.vs.RphaseB1 | FALSE |
| NC_039506.1 | 1543458 | 1543458 | 1 | + | 45 | 0 | 45 | 41 | 1 | 40 | 0 | 2.44 | NC_039506.1 | 1540640 | 1547998 | 7359 | - | Gnomon | gene | gene-LOC105280244 | GeneID:105280244 | LOC105280244 | Gene | LOC105280244 | protein_coding | character(0) | NA | character(0) | BCphaseA1.vs.RphaseB1 | FALSE |
| NC_039506.1 | 1543470 | 1543470 | 1 | + | 34 | 0 | 34 | 36 | 0 | 36 | 0 | 0 | NC_039506.1 | 1540640 | 1547998 | 7359 | - | Gnomon | gene | gene-LOC105280244 | GeneID:105280244 | LOC105280244 | Gene | LOC105280244 | protein_coding | character(0) | NA | character(0) | BCphaseA1.vs.RphaseB1 | FALSE |
| NC_039506.1 | 1543476 | 1543476 | 1 | + | 30 | 1 | 29 | 32 | 0 | 32 | 3.33 | 0 | NC_039506.1 | 1540640 | 1547998 | 7359 | - | Gnomon | gene | gene-LOC105280244 | GeneID:105280244 | LOC105280244 | Gene | LOC105280244 | protein_coding | character(0) | NA | character(0) | BCphaseA1.vs.RphaseB1 | FALSE |
| NC_039506.1 | 1543479 | 1543479 | 1 | + | 32 | 1 | 31 | 32 | 0 | 32 | 3.12 | 0 | NC_039506.1 | 1540640 | 1547998 | 7359 | - | Gnomon | gene | gene-LOC105280244 | GeneID:105280244 | LOC105280244 | Gene | LOC105280244 | protein_coding | character(0) | NA | character(0) | BCphaseA1.vs.RphaseB1 | FALSE |
| NC_039506.1 | 1543482 | 1543482 | 1 | + | 29 | 0 | 29 | 32 | 0 | 32 | 0 | 0 | NC_039506.1 | 1540640 | 1547998 | 7359 | - | Gnomon | gene | gene-LOC105280244 | GeneID:105280244 | LOC105280244 | Gene | LOC105280244 | protein_coding | character(0) | NA | character(0) | BCphaseA1.vs.RphaseB1 | FALSE |

...

... only first 25 lines shown ...

Directory: DMSG File: dmg-Ob_LI_BCphaseA1.vs.RphaseB2_details.txt

| seqnames | start | end | width | strand | coverage1 | numCs1 | numTs1 | coverage2 | numCs2 | numTs2 | BCphaseA1 | RphaseB2 | gene_seqnames | gene_start | gene_end | gene_width | gene_strand | gene_source | gene_type | gene_ID | gene_Dbxref | gene_Name | gene_gbkey | gene_gene | gene_gene_biotype | gene_end_range | gene_partial | gene_start_range | gene_comparison | is.dm |
|---|---|---|---|---|---|---|---|---|---|---|---|---|---|---|---|---|---|---|---|---|---|---|---|---|---|---|---|---|---|---|
| NC_039506.1 | 416303 | 416303 | 1 | + | 23 | 0 | 23 | 24 | 0 | 24 | 0 | 0 | NC_039506.1 | 412793 | 450860 | 38068 | - | Gnomon | gene | gene-LOC105285468 | GeneID:105285468 | LOC105285468 | Gene | LOC105285468 | protein_coding | character(0) | NA | character(0) | BCphaseA1.vs.RphaseB2 | FALSE |
| NC_039506.1 | 424882 | 424882 | 1 | + | 36 | 0 | 36 | 36 | 0 | 36 | 0 | 0 | NC_039506.1 | 412793 | 450860 | 38068 | - | Gnomon | gene | gene-LOC105285468 | GeneID:105285468 | LOC105285468 | Gene | LOC105285468 | protein_coding | character(0) | NA | character(0) | BCphaseA1.vs.RphaseB2 | FALSE |
| NC_039506.1 | 424887 | 424887 | 1 | + | 37 | 0 | 37 | 36 | 1 | 35 | 0 | 2.78 | NC_039506.1 | 412793 | 450860 | 38068 | - | Gnomon | gene | gene-LOC105285468 | GeneID:105285468 | LOC105285468 | Gene | LOC105285468 | protein_coding | character(0) | NA | character(0) | BCphaseA1.vs.RphaseB2 | FALSE |
| NC_039506.1 | 424889 | 424889 | 1 | + | 37 | 0 | 37 | 36 | 0 | 36 | 0 | 0 | NC_039506.1 | 412793 | 450860 | 38068 | - | Gnomon | gene | gene-LOC105285468 | GeneID:105285468 | LOC105285468 | Gene | LOC105285468 | protein_coding | character(0) | NA | character(0) | BCphaseA1.vs.RphaseB2 | FALSE |
| NC_039506.1 | 424891 | 424891 | 1 | + | 37 | 0 | 37 | 36 | 0 | 36 | 0 | 0 | NC_039506.1 | 412793 | 450860 | 38068 | - | Gnomon | gene | gene-LOC105285468 | GeneID:105285468 | LOC105285468 | Gene | LOC105285468 | protein_coding | character(0) | NA | character(0) | BCphaseA1.vs.RphaseB2 | FALSE |
| NC_039506.1 | 424919 | 424919 | 1 | + | 30 | 0 | 30 | 24 | 0 | 24 | 0 | 0 | NC_039506.1 | 412793 | 450860 | 38068 | - | Gnomon | gene | gene-LOC105285468 | GeneID:105285468 | LOC105285468 | Gene | LOC105285468 | protein_coding | character(0) | NA | character(0) | BCphaseA1.vs.RphaseB2 | FALSE |
| NC_039506.1 | 424927 | 424927 | 1 | + | 27 | 0 | 27 | 22 | 0 | 22 | 0 | 0 | NC_039506.1 | 412793 | 450860 | 38068 | - | Gnomon | gene | gene-LOC105285468 | GeneID:105285468 | LOC105285468 | Gene | LOC105285468 | protein_coding | character(0) | NA | character(0) | BCphaseA1.vs.RphaseB2 | FALSE |
| NC_039506.1 | 426085 | 426085 | 1 | + | 27 | 0 | 27 | 31 | 0 | 31 | 0 | 0 | NC_039506.1 | 412793 | 450860 | 38068 | - | Gnomon | gene | gene-LOC105285468 | GeneID:105285468 | LOC105285468 | Gene | LOC105285468 | protein_coding | character(0) | NA | character(0) | BCphaseA1.vs.RphaseB2 | FALSE |
| NC_039506.1 | 429092 | 429092 | 1 | + | 21 | 0 | 21 | 21 | 2 | 19 | 0 | 9.52 | NC_039506.1 | 412793 | 450860 | 38068 | - | Gnomon | gene | gene-LOC105285468 | GeneID:105285468 | LOC105285468 | Gene | LOC105285468 | protein_coding | character(0) | NA | character(0) | BCphaseA1.vs.RphaseB2 | FALSE |
| NC_039506.1 | 429120 | 429120 | 1 | + | 30 | 0 | 30 | 39 | 0 | 39 | 0 | 0 | NC_039506.1 | 412793 | 450860 | 38068 | - | Gnomon | gene | gene-LOC105285468 | GeneID:105285468 | LOC105285468 | Gene | LOC105285468 | protein_coding | character(0) | NA | character(0) | BCphaseA1.vs.RphaseB2 | FALSE |
| NC_039506.1 | 429141 | 429141 | 1 | + | 32 | 0 | 32 | 41 | 0 | 41 | 0 | 0 | NC_039506.1 | 412793 | 450860 | 38068 | - | Gnomon | gene | gene-LOC105285468 | GeneID:105285468 | LOC105285468 | Gene | LOC105285468 | protein_coding | character(0) | NA | character(0) | BCphaseA1.vs.RphaseB2 | FALSE |
| NC_039506.1 | 429144 | 429144 | 1 | + | 30 | 1 | 29 | 37 | 1 | 36 | 3.33 | 2.7 | NC_039506.1 | 412793 | 450860 | 38068 | - | Gnomon | gene | gene-LOC105285468 | GeneID:105285468 | LOC105285468 | Gene | LOC105285468 | protein_coding | character(0) | NA | character(0) | BCphaseA1.vs.RphaseB2 | FALSE |
| NC_039506.1 | 432540 | 432540 | 1 | + | 22 | 0 | 22 | 28 | 0 | 28 | 0 | 0 | NC_039506.1 | 412793 | 450860 | 38068 | - | Gnomon | gene | gene-LOC105285468 | GeneID:105285468 | LOC105285468 | Gene | LOC105285468 | protein_coding | character(0) | NA | character(0) | BCphaseA1.vs.RphaseB2 | FALSE |
| NC_039506.1 | 432561 | 432561 | 1 | + | 22 | 0 | 22 | 30 | 0 | 30 | 0 | 0 | NC_039506.1 | 412793 | 450860 | 38068 | - | Gnomon | gene | gene-LOC105285468 | GeneID:105285468 | LOC105285468 | Gene | LOC105285468 | protein_coding | character(0) | NA | character(0) | BCphaseA1.vs.RphaseB2 | FALSE |
| NC_039506.1 | 432566 | 432566 | 1 | + | 22 | 0 | 22 | 30 | 0 | 30 | 0 | 0 | NC_039506.1 | 412793 | 450860 | 38068 | - | Gnomon | gene | gene-LOC105285468 | GeneID:105285468 | LOC105285468 | Gene | LOC105285468 | protein_coding | character(0) | NA | character(0) | BCphaseA1.vs.RphaseB2 | FALSE |
| NC_039506.1 | 432596 | 432596 | 1 | + | 21 | 0 | 21 | 26 | 0 | 26 | 0 | 0 | NC_039506.1 | 412793 | 450860 | 38068 | - | Gnomon | gene | gene-LOC105285468 | GeneID:105285468 | LOC105285468 | Gene | LOC105285468 | protein_coding | character(0) | NA | character(0) | BCphaseA1.vs.RphaseB2 | FALSE |
| NC_039506.1 | 432599 | 432599 | 1 | + | 21 | 0 | 21 | 26 | 0 | 26 | 0 | 0 | NC_039506.1 | 412793 | 450860 | 38068 | - | Gnomon | gene | gene-LOC105285468 | GeneID:105285468 | LOC105285468 | Gene | LOC105285468 | protein_coding | character(0) | NA | character(0) | BCphaseA1.vs.RphaseB2 | FALSE |
| NC_039506.1 | 432604 | 432604 | 1 | + | 23 | 0 | 23 | 26 | 0 | 26 | 0 | 0 | NC_039506.1 | 412793 | 450860 | 38068 | - | Gnomon | gene | gene-LOC105285468 | GeneID:105285468 | LOC105285468 | Gene | LOC105285468 | protein_coding | character(0) | NA | character(0) | BCphaseA1.vs.RphaseB2 | FALSE |
| NC_039506.1 | 432607 | 432607 | 1 | + | 20 | 0 | 20 | 26 | 0 | 26 | 0 | 0 | NC_039506.1 | 412793 | 450860 | 38068 | - | Gnomon | gene | gene-LOC105285468 | GeneID:105285468 | LOC105285468 | Gene | LOC105285468 | protein_coding | character(0) | NA | character(0) | BCphaseA1.vs.RphaseB2 | FALSE |
| NC_039506.1 | 432613 | 432613 | 1 | + | 20 | 0 | 20 | 25 | 0 | 25 | 0 | 0 | NC_039506.1 | 412793 | 450860 | 38068 | - | Gnomon | gene | gene-LOC105285468 | GeneID:105285468 | LOC105285468 | Gene | LOC105285468 | protein_coding | character(0) | NA | character(0) | BCphaseA1.vs.RphaseB2 | FALSE |
| NC_039506.1 | 432631 | 432631 | 1 | + | 22 | 0 | 22 | 22 | 0 | 22 | 0 | 0 | NC_039506.1 | 412793 | 450860 | 38068 | - | Gnomon | gene | gene-LOC105285468 | GeneID:105285468 | LOC105285468 | Gene | LOC105285468 | protein_coding | character(0) | NA | character(0) | BCphaseA1.vs.RphaseB2 | FALSE |
| NC_039506.1 | 432635 | 432635 | 1 | + | 23 | 0 | 23 | 24 | 0 | 24 | 0 | 0 | NC_039506.1 | 412793 | 450860 | 38068 | - | Gnomon | gene | gene-LOC105285468 | GeneID:105285468 | LOC105285468 | Gene | LOC105285468 | protein_coding | character(0) | NA | character(0) | BCphaseA1.vs.RphaseB2 | FALSE |
| NC_039506.1 | 432640 | 432640 | 1 | + | 23 | 0 | 23 | 24 | 0 | 24 | 0 | 0 | NC_039506.1 | 412793 | 450860 | 38068 | - | Gnomon | gene | gene-LOC105285468 | GeneID:105285468 | LOC105285468 | Gene | LOC105285468 | protein_coding | character(0) | NA | character(0) | BCphaseA1.vs.RphaseB2 | FALSE |
| NC_039506.1 | 432647 | 432647 | 1 | + | 22 | 0 | 22 | 23 | 0 | 23 | 0 | 0 | NC_039506.1 | 412793 | 450860 | 38068 | - | Gnomon | gene | gene-LOC105285468 | GeneID:105285468 | LOC105285468 | Gene | LOC105285468 | protein_coding | character(0) | NA | character(0) | BCphaseA1.vs.RphaseB2 | FALSE |

...
... only first 25 lines shown ...

Directory: DMSG File: dmg-Ob_LI_BCphaseA2.vs.BCphaseB1_details.txt

| seqnames | start | end | width | strand | coverage1 | numCs1 | numTs1 | coverage2 | numCs2 | numTs2 | BCphaseA2 | BCphaseB1 | gene_seqnames | gene_start | gene_end | gene_width | gene_strand | gene_source | gene_type | gene_ID | gene_Dbxref | gene_Name | gene_gbkey | gene_gene | gene_gene_biotype | gene_end_range | gene_start_range | gene_comparison | is.dm |
|---|---|---|---|---|---|---|---|---|---|---|---|---|---|---|---|---|---|---|---|---|---|---|---|---|---|---|---|---|---|
| NC_039506.1 | 3575883 | 3575883 | 1 | + | 20 | 0 | 20 | 23 | 1 | 22 | 0 | 4.35 | NC_039506.1 | 3575547 | 3588126 | 12580 | - | Gnomon | gene | gene-LOC105280410 | GeneID:105280410 | LOC105280410 | Gene | LOC105280410 | protein_coding | character(0) | character(0) | BCphaseA2.vs.BCphaseB1 | FALSE |
| NC_039506.1 | 3575907 | 3575907 | 1 | + | 20 | 0 | 20 | 22 | 1 | 21 | 0 | 4.55 | NC_039506.1 | 3575547 | 3588126 | 12580 | - | Gnomon | gene | gene-LOC105280410 | GeneID:105280410 | LOC105280410 | Gene | LOC105280410 | protein_coding | character(0) | character(0) | BCphaseA2.vs.BCphaseB1 | FALSE |
| NC_039506.1 | 3575910 | 3575910 | 1 | + | 21 | 0 | 21 | 24 | 0 | 24 | 0 | 0 | NC_039506.1 | 3575547 | 3588126 | 12580 | - | Gnomon | gene | gene-LOC105280410 | GeneID:105280410 | LOC105280410 | Gene | LOC105280410 | protein_coding | character(0) | character(0) | BCphaseA2.vs.BCphaseB1 | FALSE |
| NC_039506.1 | 3575921 | 3575921 | 1 | + | 23 | 0 | 23 | 28 | 0 | 28 | 0 | 0 | NC_039506.1 | 3575547 | 3588126 | 12580 | - | Gnomon | gene | gene-LOC105280410 | GeneID:105280410 | LOC105280410 | Gene | LOC105280410 | protein_coding | character(0) | character(0) | BCphaseA2.vs.BCphaseB1 | FALSE |
| NC_039506.1 | 3576234 | 3576234 | 1 | + | 23 | 0 | 23 | 52 | 0 | 52 | 0 | 0 | NC_039506.1 | 3575547 | 3588126 | 12580 | - | Gnomon | gene | gene-LOC105280410 | GeneID:105280410 | LOC105280410 | Gene | LOC105280410 | protein_coding | character(0) | character(0) | BCphaseA2.vs.BCphaseB1 | FALSE |
| NC_039506.1 | 3576241 | 3576241 | 1 | + | 23 | 0 | 23 | 55 | 0 | 55 | 0 | 0 | NC_039506.1 | 3575547 | 3588126 | 12580 | - | Gnomon | gene | gene-LOC105280410 | GeneID:105280410 | LOC105280410 | Gene | LOC105280410 | protein_coding | character(0) | character(0) | BCphaseA2.vs.BCphaseB1 | FALSE |
| NC_039506.1 | 3576243 | 3576243 | 1 | + | 23 | 0 | 23 | 54 | 1 | 53 | 0 | 1.85 | NC_039506.1 | 3575547 | 3588126 | 12580 | - | Gnomon | gene | gene-LOC105280410 | GeneID:105280410 | LOC105280410 | Gene | LOC105280410 | protein_coding | character(0) | character(0) | BCphaseA2.vs.BCphaseB1 | FALSE |
| NC_039506.1 | 3576265 | 3576265 | 1 | + | 20 | 1 | 19 | 26 | 2 | 24 | 5 | 7.69 | NC_039506.1 | 3575547 | 3588126 | 12580 | - | Gnomon | gene | gene-LOC105280410 | GeneID:105280410 | LOC105280410 | Gene | LOC105280410 | protein_coding | character(0) | character(0) | BCphaseA2.vs.BCphaseB1 | FALSE |
| NC_039506.1 | 3576431 | 3576431 | 1 | + | 33 | 1 | 32 | 41 | 0 | 41 | 3.03 | 0 | NC_039506.1 | 3575547 | 3588126 | 12580 | - | Gnomon | gene | gene-LOC105280410 | GeneID:105280410 | LOC105280410 | Gene | LOC105280410 | protein_coding | character(0) | character(0) | BCphaseA2.vs.BCphaseB1 | FALSE |
| NC_039506.1 | 3576447 | 3576447 | 1 | + | 32 | 1 | 31 | 40 | 3 | 37 | 3.12 | 7.5 | NC_039506.1 | 3575547 | 3588126 | 12580 | - | Gnomon | gene | gene-LOC105280410 | GeneID:105280410 | LOC105280410 | Gene | LOC105280410 | protein_coding | character(0) | character(0) | BCphaseA2.vs.BCphaseB1 | FALSE |
| NC_039506.1 | 3576449 | 3576449 | 1 | + | 32 | 1 | 31 | 40 | 0 | 40 | 3.12 | 0 | NC_039506.1 | 3575547 | 3588126 | 12580 | - | Gnomon | gene | gene-LOC105280410 | GeneID:105280410 | LOC105280410 | Gene | LOC105280410 | protein_coding | character(0) | character(0) | BCphaseA2.vs.BCphaseB1 | FALSE |
| NC_039506.1 | 3576461 | 3576461 | 1 | + | 32 | 0 | 32 | 41 | 0 | 41 | 0 | 0 | NC_039506.1 | 3575547 | 3588126 | 12580 | - | Gnomon | gene | gene-LOC105280410 | GeneID:105280410 | LOC105280410 | Gene | LOC105280410 | protein_coding | character(0) | character(0) | BCphaseA2.vs.BCphaseB1 | FALSE |
| NC_039506.1 | 3576470 | 3576470 | 1 | + | 32 | 1 | 31 | 41 | 1 | 40 | 3.12 | 2.44 | NC_039506.1 | 3575547 | 3588126 | 12580 | - | Gnomon | gene | gene-LOC105280410 | GeneID:105280410 | LOC105280410 | Gene | LOC105280410 | protein_coding | character(0) | character(0) | BCphaseA2.vs.BCphaseB1 | FALSE |
| NC_039506.1 | 3576482 | 3576482 | 1 | + | 25 | 0 | 25 | 41 | 0 | 41 | 0 | 0 | NC_039506.1 | 3575547 | 3588126 | 12580 | - | Gnomon | gene | gene-LOC105280410 | GeneID:105280410 | LOC105280410 | Gene | LOC105280410 | protein_coding | character(0) | character(0) | BCphaseA2.vs.BCphaseB1 | FALSE |
| NC_039506.1 | 3576494 | 3576494 | 1 | + | 21 | 0 | 21 | 32 | 1 | 31 | 0 | 3.12 | NC_039506.1 | 3575547 | 3588126 | 12580 | - | Gnomon | gene | gene-LOC105280410 | GeneID:105280410 | LOC105280410 | Gene | LOC105280410 | protein_coding | character(0) | character(0) | BCphaseA2.vs.BCphaseB1 | FALSE |
| NC_039506.1 | 3576515 | 3576515 | 1 | + | 20 | 1 | 19 | 24 | 3 | 21 | 5 | 12.5 | NC_039506.1 | 3575547 | 3588126 | 12580 | - | Gnomon | gene | gene-LOC105280410 | GeneID:105280410 | LOC105280410 | Gene | LOC105280410 | protein_coding | character(0) | character(0) | BCphaseA2.vs.BCphaseB1 | FALSE |
| NC_039506.1 | 3576540 | 3576540 | 1 | + | 22 | 0 | 22 | 29 | 0 | 29 | 0 | 0 | NC_039506.1 | 3575547 | 3588126 | 12580 | - | Gnomon | gene | gene-LOC105280410 | GeneID:105280410 | LOC105280410 | Gene | LOC105280410 | protein_coding | character(0) | character(0) | BCphaseA2.vs.BCphaseB1 | FALSE |
| NC_039506.1 | 3576549 | 3576549 | 1 | + | 24 | 2 | 22 | 29 | 0 | 29 | 8.33 | 0 | NC_039506.1 | 3575547 | 3588126 | 12580 | - | Gnomon | gene | gene-LOC105280410 | GeneID:105280410 | LOC105280410 | Gene | LOC105280410 | protein_coding | character(0) | character(0) | BCphaseA2.vs.BCphaseB1 | FALSE |
| NC_039506.1 | 3576578 | 3576578 | 1 | + | 20 | 1 | 19 | 26 | 3 | 23 | 5 | 11.54 | NC_039506.1 | 3575547 | 3588126 | 12580 | - | Gnomon | gene | gene-LOC105280410 | GeneID:105280410 | LOC105280410 | Gene | LOC105280410 | protein_coding | character(0) | character(0) | BCphaseA2.vs.BCphaseB1 | FALSE |
| NC_039506.1 | 3576645 | 3576645 | 1 | + | 30 | 1 | 29 | 27 | 1 | 26 | 3.33 | 3.7 | NC_039506.1 | 3575547 | 3588126 | 12580 | - | Gnomon | gene | gene-LOC105280410 | GeneID:105280410 | LOC105280410 | Gene | LOC105280410 | protein_coding | character(0) | character(0) | BCphaseA2.vs.BCphaseB1 | FALSE |
| NC_039506.1 | 3576652 | 3576652 | 1 | + | 28 | 0 | 28 | 28 | 0 | 28 | 0 | 0 | NC_039506.1 | 3575547 | 3588126 | 12580 | - | Gnomon | gene | gene-LOC105280410 | GeneID:105280410 | LOC105280410 | Gene | LOC105280410 | protein_coding | character(0) | character(0) | BCphaseA2.vs.BCphaseB1 | FALSE |
| NC_039506.1 | 3576659 | 3576659 | 1 | + | 30 | 0 | 30 | 28 | 0 | 28 | 0 | 0 | NC_039506.1 | 3575547 | 3588126 | 12580 | - | Gnomon | gene | gene-LOC105280410 | GeneID:105280410 | LOC105280410 | Gene | LOC105280410 | protein_coding | character(0) | character(0) | BCphaseA2.vs.BCphaseB1 | FALSE |
| NC_039506.1 | 3576679 | 3576679 | 1 | + | 26 | 0 | 26 | 27 | 0 | 27 | 0 | 0 | NC_039506.1 | 3575547 | 3588126 | 12580 | - | Gnomon | gene | gene-LOC105280410 | GeneID:105280410 | LOC105280410 | Gene | LOC105280410 | protein_coding | character(0) | character(0) | BCphaseA2.vs.BCphaseB1 | FALSE |
| NC_039506.1 | 3576691 | 3576691 | 1 | + | 25 | 0 | 25 | 22 | 0 | 22 | 0 | 0 | NC_039506.1 | 3575547 | 3588126 | 12580 | - | Gnomon | gene | gene-LOC105280410 | GeneID:105280410 | LOC105280410 | Gene | LOC105280410 | protein_coding | character(0) | character(0) | BCphaseA2.vs.BCphaseB1 | FALSE |

...

... only first 25 lines shown ...

Directory: DMSG File: dmg-Ob_LI_BCphaseA2.vs.BCphaseB2_details.txt

| seqnames | start | end | width | strand | coverage1 | numCs1 | numTs1 | coverage2 | numCs2 | numTs2 | BCphaseA2 | BCphaseB2 | gene_seqnames | gene_start | gene_end | gene_width | gene_strand | gene_source | gene_type | gene_ID | gene_Dbxref | gene_Name | gene_gbkey | gene_gene | gene_gene_biotype | gene_end_range | gene_start_range | gene_comparison | is.dm |
|---|---|---|---|---|---|---|---|---|---|---|---|---|---|---|---|---|---|---|---|---|---|---|---|---|---|---|---|---|---|
| NC_039506.1 | 2980750 | 2980750 | 1 | + | 23 | 0 | 23 | 32 | 0 | 32 | 0 | 0 | NC_039506.1 | 2980439 | 2984235 | 3797 | + | Gnomon | gene | gene-LOC105280348 | GeneID:105280348 | LOC105280348 | Gene | LOC105280348 | protein_coding | character(0) | character(0) | BCphaseA2.vs.BCphaseB2 | FALSE |
| NC_039506.1 | 2980761 | 2980761 | 1 | + | 22 | 0 | 22 | 28 | 0 | 28 | 0 | 0 | NC_039506.1 | 2980439 | 2984235 | 3797 | + | Gnomon | gene | gene-LOC105280348 | GeneID:105280348 | LOC105280348 | Gene | LOC105280348 | protein_coding | character(0) | character(0) | BCphaseA2.vs.BCphaseB2 | FALSE |
| NC_039506.1 | 2980767 | 2980767 | 1 | + | 22 | 0 | 22 | 28 | 0 | 28 | 0 | 0 | NC_039506.1 | 2980439 | 2984235 | 3797 | + | Gnomon | gene | gene-LOC105280348 | GeneID:105280348 | LOC105280348 | Gene | LOC105280348 | protein_coding | character(0) | character(0) | BCphaseA2.vs.BCphaseB2 | FALSE |
| NC_039506.1 | 2980928 | 2980928 | 1 | + | 22 | 6 | 16 | 23 | 6 | 17 | 27.27 | 26.09 | NC_039506.1 | 2980439 | 2984235 | 3797 | + | Gnomon | gene | gene-LOC105280348 | GeneID:105280348 | LOC105280348 | Gene | LOC105280348 | protein_coding | character(0) | character(0) | BCphaseA2.vs.BCphaseB2 | FALSE |
| NC_039506.1 | 2980949 | 2980949 | 1 | + | 20 | 6 | 14 | 22 | 8 | 14 | 30 | 36.36 | NC_039506.1 | 2980439 | 2984235 | 3797 | + | Gnomon | gene | gene-LOC105280348 | GeneID:105280348 | LOC105280348 | Gene | LOC105280348 | protein_coding | character(0) | character(0) | BCphaseA2.vs.BCphaseB2 | FALSE |
| NC_039506.1 | 2980989 | 2980989 | 1 | + | 27 | 3 | 24 | 29 | 7 | 22 | 11.11 | 24.14 | NC_039506.1 | 2980439 | 2984235 | 3797 | + | Gnomon | gene | gene-LOC105280348 | GeneID:105280348 | LOC105280348 | Gene | LOC105280348 | protein_coding | character(0) | character(0) | BCphaseA2.vs.BCphaseB2 | FALSE |
| NC_039506.1 | 2980993 | 2980993 | 1 | + | 28 | 17 | 11 | 29 | 10 | 19 | 60.71 | 34.48 | NC_039506.1 | 2980439 | 2984235 | 3797 | + | Gnomon | gene | gene-LOC105280348 | GeneID:105280348 | LOC105280348 | Gene | LOC105280348 | protein_coding | character(0) | character(0) | BCphaseA2.vs.BCphaseB2 | FALSE |
| NC_039506.1 | 2980998 | 2980998 | 1 | + | 27 | 5 | 22 | 30 | 11 | 19 | 18.52 | 36.67 | NC_039506.1 | 2980439 | 2984235 | 3797 | + | Gnomon | gene | gene-LOC105280348 | GeneID:105280348 | LOC105280348 | Gene | LOC105280348 | protein_coding | character(0) | character(0) | BCphaseA2.vs.BCphaseB2 | FALSE |
| NC_039506.1 | 2981030 | 2981030 | 1 | + | 22 | 20 | 2 | 43 | 35 | 8 | 90.91 | 81.4 | NC_039506.1 | 2980439 | 2984235 | 3797 | + | Gnomon | gene | gene-LOC105280348 | GeneID:105280348 | LOC105280348 | Gene | LOC105280348 | protein_coding | character(0) | character(0) | BCphaseA2.vs.BCphaseB2 | FALSE |
| NC_039506.1 | 2981077 | 2981077 | 1 | + | 33 | 16 | 17 | 36 | 13 | 23 | 48.48 | 36.11 | NC_039506.1 | 2980439 | 2984235 | 3797 | + | Gnomon | gene | gene-LOC105280348 | GeneID:105280348 | LOC105280348 | Gene | LOC105280348 | protein_coding | character(0) | character(0) | BCphaseA2.vs.BCphaseB2 | FALSE |
| NC_039506.1 | 2981183 | 2981183 | 1 | + | 85 | 78 | 7 | 49 | 25 | 24 | 91.76 | 51.02 | NC_039506.1 | 2980439 | 2984235 | 3797 | + | Gnomon | gene | gene-LOC105280348 | GeneID:105280348 | LOC105280348 | Gene | LOC105280348 | protein_coding | character(0) | character(0) | BCphaseA2.vs.BCphaseB2 | TRUE |
| NC_039506.1 | 2981225 | 2981225 | 1 | + | 45 | 31 | 14 | 32 | 24 | 8 | 68.89 | 75 | NC_039506.1 | 2980439 | 2984235 | 3797 | + | Gnomon | gene | gene-LOC105280348 | GeneID:105280348 | LOC105280348 | Gene | LOC105280348 | protein_coding | character(0) | character(0) | BCphaseA2.vs.BCphaseB2 | FALSE |
| NC_039506.1 | 2981245 | 2981245 | 1 | + | 35 | 12 | 23 | 38 | 14 | 24 | 34.29 | 36.84 | NC_039506.1 | 2980439 | 2984235 | 3797 | + | Gnomon | gene | gene-LOC105280348 | GeneID:105280348 | LOC105280348 | Gene | LOC105280348 | protein_coding | character(0) | character(0) | BCphaseA2.vs.BCphaseB2 | FALSE |
| NC_039506.1 | 2981292 | 2981292 | 1 | + | 32 | 32 | 0 | 42 | 42 | 0 | 100 | 100 | NC_039506.1 | 2980439 | 2984235 | 3797 | + | Gnomon | gene | gene-LOC105280348 | GeneID:105280348 | LOC105280348 | Gene | LOC105280348 | protein_coding | character(0) | character(0) | BCphaseA2.vs.BCphaseB2 | FALSE |
| NC_039506.1 | 2981308 | 2981308 | 1 | + | 39 | 22 | 17 | 49 | 34 | 15 | 56.41 | 69.39 | NC_039506.1 | 2980439 | 2984235 | 3797 | + | Gnomon | gene | gene-LOC105280348 | GeneID:105280348 | LOC105280348 | Gene | LOC105280348 | protein_coding | character(0) | character(0) | BCphaseA2.vs.BCphaseB2 | FALSE |
| NC_039506.1 | 2981315 | 2981315 | 1 | + | 36 | 30 | 6 | 49 | 40 | 9 | 83.33 | 81.63 | NC_039506.1 | 2980439 | 2984235 | 3797 | + | Gnomon | gene | gene-LOC105280348 | GeneID:105280348 | LOC105280348 | Gene | LOC105280348 | protein_coding | character(0) | character(0) | BCphaseA2.vs.BCphaseB2 | FALSE |
| NC_039506.1 | 2981335 | 2981335 | 1 | + | 29 | 28 | 1 | 41 | 39 | 2 | 96.55 | 95.12 | NC_039506.1 | 2980439 | 2984235 | 3797 | + | Gnomon | gene | gene-LOC105280348 | GeneID:105280348 | LOC105280348 | Gene | LOC105280348 | protein_coding | character(0) | character(0) | BCphaseA2.vs.BCphaseB2 | FALSE |
| NC_039506.1 | 2981350 | 2981350 | 1 | + | 59 | 57 | 2 | 72 | 68 | 4 | 96.61 | 94.44 | NC_039506.1 | 2980439 | 2984235 | 3797 | + | Gnomon | gene | gene-LOC105280348 | GeneID:105280348 | LOC105280348 | Gene | LOC105280348 | protein_coding | character(0) | character(0) | BCphaseA2.vs.BCphaseB2 | FALSE |
| NC_039506.1 | 2981391 | 2981391 | 1 | + | 72 | 64 | 8 | 87 | 69 | 18 | 88.89 | 79.31 | NC_039506.1 | 2980439 | 2984235 | 3797 | + | Gnomon | gene | gene-LOC105280348 | GeneID:105280348 | LOC105280348 | Gene | LOC105280348 | protein_coding | character(0) | character(0) | BCphaseA2.vs.BCphaseB2 | FALSE |
| NC_039506.1 | 2981402 | 2981402 | 1 | + | 71 | 69 | 2 | 88 | 85 | 3 | 97.18 | 96.59 | NC_039506.1 | 2980439 | 2984235 | 3797 | + | Gnomon | gene | gene-LOC105280348 | GeneID:105280348 | LOC105280348 | Gene | LOC105280348 | protein_coding | character(0) | character(0) | BCphaseA2.vs.BCphaseB2 | FALSE |
| NC_039506.1 | 2981411 | 2981411 | 1 | + | 76 | 72 | 4 | 101 | 94 | 7 | 94.74 | 93.07 | NC_039506.1 | 2980439 | 2984235 | 3797 | + | Gnomon | gene | gene-LOC105280348 | GeneID:105280348 | LOC105280348 | Gene | LOC105280348 | protein_coding | character(0) | character(0) | BCphaseA2.vs.BCphaseB2 | FALSE |
| NC_039506.1 | 2981688 | 2981688 | 1 | + | 20 | 4 | 16 | 43 | 16 | 27 | 20 | 37.21 | NC_039506.1 | 2980439 | 2984235 | 3797 | + | Gnomon | gene | gene-LOC105280348 | GeneID:105280348 | LOC105280348 | Gene | LOC105280348 | protein_coding | character(0) | character(0) | BCphaseA2.vs.BCphaseB2 | FALSE |
| NC_039506.1 | 2981802 | 2981802 | 1 | + | 27 | 8 | 19 | 24 | 9 | 15 | 29.63 | 37.5 | NC_039506.1 | 2980439 | 2984235 | 3797 | + | Gnomon | gene | gene-LOC105280348 | GeneID:105280348 | LOC105280348 | Gene | LOC105280348 | protein_coding | character(0) | character(0) | BCphaseA2.vs.BCphaseB2 | FALSE |
| NC_039506.1 | 2982097 | 2982097 | 1 | + | 34 | 1 | 33 | 22 | 0 | 22 | 2.94 | 0 | NC_039506.1 | 2980439 | 2984235 | 3797 | + | Gnomon | gene | gene-LOC105280348 | GeneID:105280348 | LOC105280348 | Gene | LOC105280348 | protein_coding | character(0) | character(0) | BCphaseA2.vs.BCphaseB2 | FALSE |

...
... only first 25 lines shown ...

Directory: DMSG File: dmg-Ob_LI_BCphaseA2.vs.RphaseA1_details.txt

| seqnames | start | end | width | strand | coverage1 | numCs1 | numTs1 | coverage2 | numCs2 | numTs2 | BCphaseA2 | RphaseA1 | gene_seqnames | gene_start | gene_end | gene_width | gene_strand | gene_source | gene_type | gene_ID | gene_Dbxref | gene_Name | gene_gbkey | gene_gene | gene_gene_biotype | gene_end_range | gene_partial | gene_start_range | gene_comparison | is.dm |
|---|---|---|---|---|---|---|---|---|---|---|---|---|---|---|---|---|---|---|---|---|---|---|---|---|---|---|---|---|---|---|
| NC_039506.1 | 137017 | 137017 | 1 | + | 26 | 1 | 25 | 33 | 1 | 32 | 3.85 | 3.03 | NC_039506.1 | 134260 | 142204 | 7945 | - | Gnomon | gene | gene-LOC105288138 | GeneID:105288138 | LOC105288138 | Gene | LOC105288138 | protein_coding | character(0) | NA | character(0) | BCphaseA2.vs.RphaseA1 | FALSE |
| NC_039506.1 | 137087 | 137087 | 1 | + | 21 | 0 | 21 | 37 | 5 | 32 | 0 | 13.51 | NC_039506.1 | 134260 | 142204 | 7945 | - | Gnomon | gene | gene-LOC105288138 | GeneID:105288138 | LOC105288138 | Gene | LOC105288138 | protein_coding | character(0) | NA | character(0) | BCphaseA2.vs.RphaseA1 | FALSE |
| NC_039506.1 | 137597 | 137597 | 1 | + | 22 | 2 | 20 | 24 | 2 | 22 | 9.09 | 8.33 | NC_039506.1 | 134260 | 142204 | 7945 | - | Gnomon | gene | gene-LOC105288138 | GeneID:105288138 | LOC105288138 | Gene | LOC105288138 | protein_coding | character(0) | NA | character(0) | BCphaseA2.vs.RphaseA1 | FALSE |
| NC_039506.1 | 137683 | 137683 | 1 | + | 39 | 0 | 39 | 26 | 6 | 20 | 0 | 23.08 | NC_039506.1 | 134260 | 142204 | 7945 | - | Gnomon | gene | gene-LOC105288138 | GeneID:105288138 | LOC105288138 | Gene | LOC105288138 | protein_coding | character(0) | NA | character(0) | BCphaseA2.vs.RphaseA1 | TRUE |
| NC_039506.1 | 138104 | 138104 | 1 | + | 22 | 0 | 22 | 20 | 0 | 20 | 0 | 0 | NC_039506.1 | 134260 | 142204 | 7945 | - | Gnomon | gene | gene-LOC105288138 | GeneID:105288138 | LOC105288138 | Gene | LOC105288138 | protein_coding | character(0) | NA | character(0) | BCphaseA2.vs.RphaseA1 | FALSE |
| NC_039506.1 | 138118 | 138118 | 1 | + | 29 | 1 | 28 | 26 | 0 | 26 | 3.45 | 0 | NC_039506.1 | 134260 | 142204 | 7945 | - | Gnomon | gene | gene-LOC105288138 | GeneID:105288138 | LOC105288138 | Gene | LOC105288138 | protein_coding | character(0) | NA | character(0) | BCphaseA2.vs.RphaseA1 | FALSE |
| NC_039506.1 | 138130 | 138130 | 1 | + | 33 | 0 | 33 | 29 | 0 | 29 | 0 | 0 | NC_039506.1 | 134260 | 142204 | 7945 | - | Gnomon | gene | gene-LOC105288138 | GeneID:105288138 | LOC105288138 | Gene | LOC105288138 | protein_coding | character(0) | NA | character(0) | BCphaseA2.vs.RphaseA1 | FALSE |
| NC_039506.1 | 138133 | 138133 | 1 | + | 34 | 0 | 34 | 29 | 2 | 27 | 0 | 6.9 | NC_039506.1 | 134260 | 142204 | 7945 | - | Gnomon | gene | gene-LOC105288138 | GeneID:105288138 | LOC105288138 | Gene | LOC105288138 | protein_coding | character(0) | NA | character(0) | BCphaseA2.vs.RphaseA1 | FALSE |
| NC_039506.1 | 138146 | 138146 | 1 | + | 36 | 0 | 36 | 37 | 0 | 37 | 0 | 0 | NC_039506.1 | 134260 | 142204 | 7945 | - | Gnomon | gene | gene-LOC105288138 | GeneID:105288138 | LOC105288138 | Gene | LOC105288138 | protein_coding | character(0) | NA | character(0) | BCphaseA2.vs.RphaseA1 | FALSE |
| NC_039506.1 | 138154 | 138154 | 1 | + | 43 | 0 | 43 | 38 | 1 | 37 | 0 | 2.63 | NC_039506.1 | 134260 | 142204 | 7945 | - | Gnomon | gene | gene-LOC105288138 | GeneID:105288138 | LOC105288138 | Gene | LOC105288138 | protein_coding | character(0) | NA | character(0) | BCphaseA2.vs.RphaseA1 | FALSE |
| NC_039506.1 | 138160 | 138160 | 1 | + | 43 | 1 | 42 | 39 | 0 | 39 | 2.33 | 0 | NC_039506.1 | 134260 | 142204 | 7945 | - | Gnomon | gene | gene-LOC105288138 | GeneID:105288138 | LOC105288138 | Gene | LOC105288138 | protein_coding | character(0) | NA | character(0) | BCphaseA2.vs.RphaseA1 | FALSE |
| NC_039506.1 | 138173 | 138173 | 1 | + | 50 | 1 | 49 | 40 | 0 | 40 | 2 | 0 | NC_039506.1 | 134260 | 142204 | 7945 | - | Gnomon | gene | gene-LOC105288138 | GeneID:105288138 | LOC105288138 | Gene | LOC105288138 | protein_coding | character(0) | NA | character(0) | BCphaseA2.vs.RphaseA1 | FALSE |
| NC_039506.1 | 138185 | 138185 | 1 | + | 44 | 4 | 40 | 34 | 2 | 32 | 9.09 | 5.88 | NC_039506.1 | 134260 | 142204 | 7945 | - | Gnomon | gene | gene-LOC105288138 | GeneID:105288138 | LOC105288138 | Gene | LOC105288138 | protein_coding | character(0) | NA | character(0) | BCphaseA2.vs.RphaseA1 | FALSE |
| NC_039506.1 | 138555 | 138555 | 1 | + | 23 | 0 | 23 | 22 | 2 | 20 | 0 | 9.09 | NC_039506.1 | 134260 | 142204 | 7945 | - | Gnomon | gene | gene-LOC105288138 | GeneID:105288138 | LOC105288138 | Gene | LOC105288138 | protein_coding | character(0) | NA | character(0) | BCphaseA2.vs.RphaseA1 | FALSE |
| NC_039506.1 | 138568 | 138568 | 1 | + | 20 | 2 | 18 | 27 | 1 | 26 | 10 | 3.7 | NC_039506.1 | 134260 | 142204 | 7945 | - | Gnomon | gene | gene-LOC105288138 | GeneID:105288138 | LOC105288138 | Gene | LOC105288138 | protein_coding | character(0) | NA | character(0) | BCphaseA2.vs.RphaseA1 | FALSE |
| NC_039506.1 | 138578 | 138578 | 1 | + | 22 | 1 | 21 | 30 | 6 | 24 | 4.55 | 20 | NC_039506.1 | 134260 | 142204 | 7945 | - | Gnomon | gene | gene-LOC105288138 | GeneID:105288138 | LOC105288138 | Gene | LOC105288138 | protein_coding | character(0) | NA | character(0) | BCphaseA2.vs.RphaseA1 | FALSE |
| NC_039506.1 | 138592 | 138592 | 1 | + | 24 | 2 | 22 | 30 | 6 | 24 | 8.33 | 20 | NC_039506.1 | 134260 | 142204 | 7945 | - | Gnomon | gene | gene-LOC105288138 | GeneID:105288138 | LOC105288138 | Gene | LOC105288138 | protein_coding | character(0) | NA | character(0) | BCphaseA2.vs.RphaseA1 | FALSE |
| NC_039506.1 | 138597 | 138597 | 1 | + | 23 | 1 | 22 | 30 | 2 | 28 | 4.35 | 6.67 | NC_039506.1 | 134260 | 142204 | 7945 | - | Gnomon | gene | gene-LOC105288138 | GeneID:105288138 | LOC105288138 | Gene | LOC105288138 | protein_coding | character(0) | NA | character(0) | BCphaseA2.vs.RphaseA1 | FALSE |
| NC_039506.1 | 1097536 | 1097536 | 1 | + | 26 | 0 | 26 | 32 | 0 | 32 | 0 | 0 | NC_039506.1 | 1095614 | 1100480 | 4867 | + | Gnomon | gene | gene-LOC105286750 | GeneID:105286750 | LOC105286750 | Gene | LOC105286750 | protein_coding | character(0) | NA | character(0) | BCphaseA2.vs.RphaseA1 | FALSE |
| NC_039506.1 | 1097543 | 1097543 | 1 | + | 26 | 0 | 26 | 31 | 1 | 30 | 0 | 3.23 | NC_039506.1 | 1095614 | 1100480 | 4867 | + | Gnomon | gene | gene-LOC105286750 | GeneID:105286750 | LOC105286750 | Gene | LOC105286750 | protein_coding | character(0) | NA | character(0) | BCphaseA2.vs.RphaseA1 | FALSE |
| NC_039506.1 | 1097546 | 1097546 | 1 | + | 27 | 0 | 27 | 32 | 0 | 32 | 0 | 0 | NC_039506.1 | 1095614 | 1100480 | 4867 | + | Gnomon | gene | gene-LOC105286750 | GeneID:105286750 | LOC105286750 | Gene | LOC105286750 | protein_coding | character(0) | NA | character(0) | BCphaseA2.vs.RphaseA1 | FALSE |
| NC_039506.1 | 1097588 | 1097588 | 1 | + | 35 | 12 | 23 | 27 | 16 | 11 | 34.29 | 59.26 | NC_039506.1 | 1095614 | 1100480 | 4867 | + | Gnomon | gene | gene-LOC105286750 | GeneID:105286750 | LOC105286750 | Gene | LOC105286750 | protein_coding | character(0) | NA | character(0) | BCphaseA2.vs.RphaseA1 | TRUE |
| NC_039506.1 | 1097598 | 1097598 | 1 | + | 38 | 15 | 23 | 31 | 16 | 15 | 39.47 | 51.61 | NC_039506.1 | 1095614 | 1100480 | 4867 | + | Gnomon | gene | gene-LOC105286750 | GeneID:105286750 | LOC105286750 | Gene | LOC105286750 | protein_coding | character(0) | NA | character(0) | BCphaseA2.vs.RphaseA1 | FALSE |
| NC_039506.1 | 1097647 | 1097647 | 1 | + | 46 | 4 | 42 | 45 | 4 | 41 | 8.7 | 8.89 | NC_039506.1 | 1095614 | 1100480 | 4867 | + | Gnomon | gene | gene-LOC105286750 | GeneID:105286750 | LOC105286750 | Gene | LOC105286750 | protein_coding | character(0) | NA | character(0) | BCphaseA2.vs.RphaseA1 | FALSE |

...
... only first 25 lines shown ...

Directory: DMSG File: dmg-Ob_LI_BCphaseA2.vs.RphaseA2_details.txt

| seqnames | start | end | width | strand | coverage1 | numCs1 | numTs1 | coverage2 | numCs2 | numTs2 | BCphaseA2 | RphaseA2 | gene_seqnames | gene_start | gene_end | gene_width | gene_strand | gene_source | gene_type | gene_ID | gene_Dbxref | gene_Name | gene_gbkey | gene_gene | gene_gene_biotype | gene_end_range | gene_partial | gene_start_range | gene_comparison | is.dm |
|---|---|---|---|---|---|---|---|---|---|---|---|---|---|---|---|---|---|---|---|---|---|---|---|---|---|---|---|---|---|---|
| NC_039506.1 | 1097536 | 1097536 | 1 | + | 26 | 0 | 26 | 37 | 4 | 33 | 0 | 10.81 | NC_039506.1 | 1095614 | 1100480 | 4867 | + | Gnomon | gene | gene-LOC105286750 | GeneID:105286750 | LOC105286750 | Gene | LOC105286750 | protein_coding | character(0) | NA | character(0) | BCphaseA2.vs.RphaseA2 | FALSE |
| NC_039506.1 | 1097543 | 1097543 | 1 | + | 26 | 0 | 26 | 38 | 4 | 34 | 0 | 10.53 | NC_039506.1 | 1095614 | 1100480 | 4867 | + | Gnomon | gene | gene-LOC105286750 | GeneID:105286750 | LOC105286750 | Gene | LOC105286750 | protein_coding | character(0) | NA | character(0) | BCphaseA2.vs.RphaseA2 | FALSE |
| NC_039506.1 | 1097546 | 1097546 | 1 | + | 27 | 0 | 27 | 41 | 4 | 37 | 0 | 9.76 | NC_039506.1 | 1095614 | 1100480 | 4867 | + | Gnomon | gene | gene-LOC105286750 | GeneID:105286750 | LOC105286750 | Gene | LOC105286750 | protein_coding | character(0) | NA | character(0) | BCphaseA2.vs.RphaseA2 | FALSE |
| NC_039506.1 | 1097588 | 1097588 | 1 | + | 35 | 12 | 23 | 47 | 21 | 26 | 34.29 | 44.68 | NC_039506.1 | 1095614 | 1100480 | 4867 | + | Gnomon | gene | gene-LOC105286750 | GeneID:105286750 | LOC105286750 | Gene | LOC105286750 | protein_coding | character(0) | NA | character(0) | BCphaseA2.vs.RphaseA2 | FALSE |
| NC_039506.1 | 1097598 | 1097598 | 1 | + | 38 | 15 | 23 | 50 | 29 | 21 | 39.47 | 58 | NC_039506.1 | 1095614 | 1100480 | 4867 | + | Gnomon | gene | gene-LOC105286750 | GeneID:105286750 | LOC105286750 | Gene | LOC105286750 | protein_coding | character(0) | NA | character(0) | BCphaseA2.vs.RphaseA2 | FALSE |
| NC_039506.1 | 1097647 | 1097647 | 1 | + | 46 | 4 | 42 | 72 | 13 | 59 | 8.7 | 18.06 | NC_039506.1 | 1095614 | 1100480 | 4867 | + | Gnomon | gene | gene-LOC105286750 | GeneID:105286750 | LOC105286750 | Gene | LOC105286750 | protein_coding | character(0) | NA | character(0) | BCphaseA2.vs.RphaseA2 | FALSE |
| NC_039506.1 | 1097653 | 1097653 | 1 | + | 47 | 2 | 45 | 94 | 4 | 90 | 4.26 | 4.26 | NC_039506.1 | 1095614 | 1100480 | 4867 | + | Gnomon | gene | gene-LOC105286750 | GeneID:105286750 | LOC105286750 | Gene | LOC105286750 | protein_coding | character(0) | NA | character(0) | BCphaseA2.vs.RphaseA2 | FALSE |
| NC_039506.1 | 1097702 | 1097702 | 1 | + | 50 | 8 | 42 | 108 | 17 | 91 | 16 | 15.74 | NC_039506.1 | 1095614 | 1100480 | 4867 | + | Gnomon | gene | gene-LOC105286750 | GeneID:105286750 | LOC105286750 | Gene | LOC105286750 | protein_coding | character(0) | NA | character(0) | BCphaseA2.vs.RphaseA2 | FALSE |
| NC_039506.1 | 1097773 | 1097773 | 1 | + | 45 | 44 | 1 | 74 | 69 | 5 | 97.78 | 93.24 | NC_039506.1 | 1095614 | 1100480 | 4867 | + | Gnomon | gene | gene-LOC105286750 | GeneID:105286750 | LOC105286750 | Gene | LOC105286750 | protein_coding | character(0) | NA | character(0) | BCphaseA2.vs.RphaseA2 | FALSE |
| NC_039506.1 | 1098123 | 1098123 | 1 | + | 29 | 23 | 6 | 42 | 26 | 16 | 79.31 | 61.9 | NC_039506.1 | 1095614 | 1100480 | 4867 | + | Gnomon | gene | gene-LOC105286750 | GeneID:105286750 | LOC105286750 | Gene | LOC105286750 | protein_coding | character(0) | NA | character(0) | BCphaseA2.vs.RphaseA2 | FALSE |
| NC_039506.1 | 1098144 | 1098144 | 1 | + | 25 | 25 | 0 | 25 | 21 | 4 | 100 | 84 | NC_039506.1 | 1095614 | 1100480 | 4867 | + | Gnomon | gene | gene-LOC105286750 | GeneID:105286750 | LOC105286750 | Gene | LOC105286750 | protein_coding | character(0) | NA | character(0) | BCphaseA2.vs.RphaseA2 | FALSE |
| NC_039506.1 | 1098206 | 1098206 | 1 | + | 34 | 26 | 8 | 43 | 27 | 16 | 76.47 | 62.79 | NC_039506.1 | 1095614 | 1100480 | 4867 | + | Gnomon | gene | gene-LOC105286750 | GeneID:105286750 | LOC105286750 | Gene | LOC105286750 | protein_coding | character(0) | NA | character(0) | BCphaseA2.vs.RphaseA2 | FALSE |
| NC_039506.1 | 1098266 | 1098266 | 1 | + | 43 | 25 | 18 | 70 | 43 | 27 | 58.14 | 61.43 | NC_039506.1 | 1095614 | 1100480 | 4867 | + | Gnomon | gene | gene-LOC105286750 | GeneID:105286750 | LOC105286750 | Gene | LOC105286750 | protein_coding | character(0) | NA | character(0) | BCphaseA2.vs.RphaseA2 | FALSE |
| NC_039506.1 | 1098279 | 1098279 | 1 | + | 47 | 18 | 29 | 72 | 33 | 39 | 38.3 | 45.83 | NC_039506.1 | 1095614 | 1100480 | 4867 | + | Gnomon | gene | gene-LOC105286750 | GeneID:105286750 | LOC105286750 | Gene | LOC105286750 | protein_coding | character(0) | NA | character(0) | BCphaseA2.vs.RphaseA2 | FALSE |
| NC_039506.1 | 1098293 | 1098293 | 1 | + | 26 | 11 | 15 | 43 | 18 | 25 | 42.31 | 41.86 | NC_039506.1 | 1095614 | 1100480 | 4867 | + | Gnomon | gene | gene-LOC105286750 | GeneID:105286750 | LOC105286750 | Gene | LOC105286750 | protein_coding | character(0) | NA | character(0) | BCphaseA2.vs.RphaseA2 | FALSE |
| NC_039506.1 | 1098301 | 1098301 | 1 | + | 23 | 14 | 9 | 36 | 19 | 17 | 60.87 | 52.78 | NC_039506.1 | 1095614 | 1100480 | 4867 | + | Gnomon | gene | gene-LOC105286750 | GeneID:105286750 | LOC105286750 | Gene | LOC105286750 | protein_coding | character(0) | NA | character(0) | BCphaseA2.vs.RphaseA2 | FALSE |
| NC_039506.1 | 1098330 | 1098330 | 1 | + | 54 | 24 | 30 | 73 | 26 | 47 | 44.44 | 35.62 | NC_039506.1 | 1095614 | 1100480 | 4867 | + | Gnomon | gene | gene-LOC105286750 | GeneID:105286750 | LOC105286750 | Gene | LOC105286750 | protein_coding | character(0) | NA | character(0) | BCphaseA2.vs.RphaseA2 | FALSE |
| NC_039506.1 | 1098334 | 1098334 | 1 | + | 58 | 35 | 23 | 75 | 46 | 29 | 60.34 | 61.33 | NC_039506.1 | 1095614 | 1100480 | 4867 | + | Gnomon | gene | gene-LOC105286750 | GeneID:105286750 | LOC105286750 | Gene | LOC105286750 | protein_coding | character(0) | NA | character(0) | BCphaseA2.vs.RphaseA2 | FALSE |
| NC_039506.1 | 1098362 | 1098362 | 1 | + | 58 | 9 | 49 | 83 | 6 | 77 | 15.52 | 7.23 | NC_039506.1 | 1095614 | 1100480 | 4867 | + | Gnomon | gene | gene-LOC105286750 | GeneID:105286750 | LOC105286750 | Gene | LOC105286750 | protein_coding | character(0) | NA | character(0) | BCphaseA2.vs.RphaseA2 | FALSE |
| NC_039506.1 | 1098377 | 1098377 | 1 | + | 57 | 28 | 29 | 84 | 34 | 50 | 49.12 | 40.48 | NC_039506.1 | 1095614 | 1100480 | 4867 | + | Gnomon | gene | gene-LOC105286750 | GeneID:105286750 | LOC105286750 | Gene | LOC105286750 | protein_coding | character(0) | NA | character(0) | BCphaseA2.vs.RphaseA2 | FALSE |
| NC_039506.1 | 1098387 | 1098387 | 1 | + | 60 | 34 | 26 | 83 | 40 | 43 | 56.67 | 48.19 | NC_039506.1 | 1095614 | 1100480 | 4867 | + | Gnomon | gene | gene-LOC105286750 | GeneID:105286750 | LOC105286750 | Gene | LOC105286750 | protein_coding | character(0) | NA | character(0) | BCphaseA2.vs.RphaseA2 | FALSE |
| NC_039506.1 | 1098400 | 1098400 | 1 | + | 49 | 26 | 23 | 76 | 45 | 31 | 53.06 | 59.21 | NC_039506.1 | 1095614 | 1100480 | 4867 | + | Gnomon | gene | gene-LOC105286750 | GeneID:105286750 | LOC105286750 | Gene | LOC105286750 | protein_coding | character(0) | NA | character(0) | BCphaseA2.vs.RphaseA2 | FALSE |
| NC_039506.1 | 1098420 | 1098420 | 1 | + | 25 | 10 | 15 | 53 | 20 | 33 | 40 | 37.74 | NC_039506.1 | 1095614 | 1100480 | 4867 | + | Gnomon | gene | gene-LOC105286750 | GeneID:105286750 | LOC105286750 | Gene | LOC105286750 | protein_coding | character(0) | NA | character(0) | BCphaseA2.vs.RphaseA2 | FALSE |
| NC_039506.1 | 1098427 | 1098427 | 1 | + | 27 | 12 | 15 | 56 | 26 | 30 | 44.44 | 46.43 | NC_039506.1 | 1095614 | 1100480 | 4867 | + | Gnomon | gene | gene-LOC105286750 | GeneID:105286750 | LOC105286750 | Gene | LOC105286750 | protein_coding | character(0) | NA | character(0) | BCphaseA2.vs.RphaseA2 | FALSE |

...
... only first 25 lines shown ...

Directory: DMSG File: dmg-Ob_LI_BCphaseA2.vs.RphaseB1_details.txt

| seqnames | start | end | width | strand | coverage1 | numCs1 | numTs1 | coverage2 | numCs2 | numTs2 | BCphaseA2 | RphaseB1 | gene_seqnames | gene_start | gene_end | gene_width | gene_strand | gene_source | gene_type | gene_ID | gene_Dbxref | gene_Name | gene_gbkey | gene_gene | gene_gene_biotype | gene_end_range | gene_start_range | gene_comparison | is.dm |
|---|---|---|---|---|---|---|---|---|---|---|---|---|---|---|---|---|---|---|---|---|---|---|---|---|---|---|---|---|---|
| NC_039506.1 | 2980750 | 2980750 | 1 | + | 23 | 0 | 23 | 33 | 0 | 33 | 0 | 0 | NC_039506.1 | 2980439 | 2984235 | 3797 | + | Gnomon | gene | gene-LOC105280348 | GeneID:105280348 | LOC105280348 | Gene | LOC105280348 | protein_coding | character(0) | character(0) | BCphaseA2.vs.RphaseB1 | FALSE |
| NC_039506.1 | 2980761 | 2980761 | 1 | + | 22 | 0 | 22 | 34 | 0 | 34 | 0 | 0 | NC_039506.1 | 2980439 | 2984235 | 3797 | + | Gnomon | gene | gene-LOC105280348 | GeneID:105280348 | LOC105280348 | Gene | LOC105280348 | protein_coding | character(0) | character(0) | BCphaseA2.vs.RphaseB1 | FALSE |
| NC_039506.1 | 2980767 | 2980767 | 1 | + | 22 | 0 | 22 | 38 | 0 | 38 | 0 | 0 | NC_039506.1 | 2980439 | 2984235 | 3797 | + | Gnomon | gene | gene-LOC105280348 | GeneID:105280348 | LOC105280348 | Gene | LOC105280348 | protein_coding | character(0) | character(0) | BCphaseA2.vs.RphaseB1 | FALSE |
| NC_039506.1 | 2980928 | 2980928 | 1 | + | 22 | 6 | 16 | 21 | 4 | 17 | 27.27 | 19.05 | NC_039506.1 | 2980439 | 2984235 | 3797 | + | Gnomon | gene | gene-LOC105280348 | GeneID:105280348 | LOC105280348 | Gene | LOC105280348 | protein_coding | character(0) | character(0) | BCphaseA2.vs.RphaseB1 | FALSE |
| NC_039506.1 | 2980949 | 2980949 | 1 | + | 20 | 6 | 14 | 20 | 5 | 15 | 30 | 25 | NC_039506.1 | 2980439 | 2984235 | 3797 | + | Gnomon | gene | gene-LOC105280348 | GeneID:105280348 | LOC105280348 | Gene | LOC105280348 | protein_coding | character(0) | character(0) | BCphaseA2.vs.RphaseB1 | FALSE |
| NC_039506.1 | 2980989 | 2980989 | 1 | + | 27 | 3 | 24 | 32 | 2 | 30 | 11.11 | 6.25 | NC_039506.1 | 2980439 | 2984235 | 3797 | + | Gnomon | gene | gene-LOC105280348 | GeneID:105280348 | LOC105280348 | Gene | LOC105280348 | protein_coding | character(0) | character(0) | BCphaseA2.vs.RphaseB1 | FALSE |
| NC_039506.1 | 2980993 | 2980993 | 1 | + | 28 | 17 | 11 | 32 | 16 | 16 | 60.71 | 50 | NC_039506.1 | 2980439 | 2984235 | 3797 | + | Gnomon | gene | gene-LOC105280348 | GeneID:105280348 | LOC105280348 | Gene | LOC105280348 | protein_coding | character(0) | character(0) | BCphaseA2.vs.RphaseB1 | FALSE |
| NC_039506.1 | 2980998 | 2980998 | 1 | + | 27 | 5 | 22 | 31 | 13 | 18 | 18.52 | 41.94 | NC_039506.1 | 2980439 | 2984235 | 3797 | + | Gnomon | gene | gene-LOC105280348 | GeneID:105280348 | LOC105280348 | Gene | LOC105280348 | protein_coding | character(0) | character(0) | BCphaseA2.vs.RphaseB1 | FALSE |
| NC_039506.1 | 2981030 | 2981030 | 1 | + | 22 | 20 | 2 | 61 | 54 | 7 | 90.91 | 88.52 | NC_039506.1 | 2980439 | 2984235 | 3797 | + | Gnomon | gene | gene-LOC105280348 | GeneID:105280348 | LOC105280348 | Gene | LOC105280348 | protein_coding | character(0) | character(0) | BCphaseA2.vs.RphaseB1 | FALSE |
| NC_039506.1 | 2981077 | 2981077 | 1 | + | 33 | 16 | 17 | 43 | 27 | 16 | 48.48 | 62.79 | NC_039506.1 | 2980439 | 2984235 | 3797 | + | Gnomon | gene | gene-LOC105280348 | GeneID:105280348 | LOC105280348 | Gene | LOC105280348 | protein_coding | character(0) | character(0) | BCphaseA2.vs.RphaseB1 | FALSE |
| NC_039506.1 | 2981125 | 2981125 | 1 | + | 24 | 14 | 10 | 24 | 11 | 13 | 58.33 | 45.83 | NC_039506.1 | 2980439 | 2984235 | 3797 | + | Gnomon | gene | gene-LOC105280348 | GeneID:105280348 | LOC105280348 | Gene | LOC105280348 | protein_coding | character(0) | character(0) | BCphaseA2.vs.RphaseB1 | FALSE |
| NC_039506.1 | 2981183 | 2981183 | 1 | + | 85 | 78 | 7 | 51 | 24 | 27 | 91.76 | 47.06 | NC_039506.1 | 2980439 | 2984235 | 3797 | + | Gnomon | gene | gene-LOC105280348 | GeneID:105280348 | LOC105280348 | Gene | LOC105280348 | protein_coding | character(0) | character(0) | BCphaseA2.vs.RphaseB1 | TRUE |
| NC_039506.1 | 2981225 | 2981225 | 1 | + | 45 | 31 | 14 | 27 | 16 | 11 | 68.89 | 59.26 | NC_039506.1 | 2980439 | 2984235 | 3797 | + | Gnomon | gene | gene-LOC105280348 | GeneID:105280348 | LOC105280348 | Gene | LOC105280348 | protein_coding | character(0) | character(0) | BCphaseA2.vs.RphaseB1 | FALSE |
| NC_039506.1 | 2981245 | 2981245 | 1 | + | 35 | 12 | 23 | 27 | 13 | 14 | 34.29 | 48.15 | NC_039506.1 | 2980439 | 2984235 | 3797 | + | Gnomon | gene | gene-LOC105280348 | GeneID:105280348 | LOC105280348 | Gene | LOC105280348 | protein_coding | character(0) | character(0) | BCphaseA2.vs.RphaseB1 | FALSE |
| NC_039506.1 | 2981292 | 2981292 | 1 | + | 32 | 32 | 0 | 57 | 51 | 6 | 100 | 89.47 | NC_039506.1 | 2980439 | 2984235 | 3797 | + | Gnomon | gene | gene-LOC105280348 | GeneID:105280348 | LOC105280348 | Gene | LOC105280348 | protein_coding | character(0) | character(0) | BCphaseA2.vs.RphaseB1 | FALSE |
| NC_039506.1 | 2981308 | 2981308 | 1 | + | 39 | 22 | 17 | 64 | 36 | 28 | 56.41 | 56.25 | NC_039506.1 | 2980439 | 2984235 | 3797 | + | Gnomon | gene | gene-LOC105280348 | GeneID:105280348 | LOC105280348 | Gene | LOC105280348 | protein_coding | character(0) | character(0) | BCphaseA2.vs.RphaseB1 | FALSE |
| NC_039506.1 | 2981315 | 2981315 | 1 | + | 36 | 30 | 6 | 67 | 50 | 17 | 83.33 | 74.63 | NC_039506.1 | 2980439 | 2984235 | 3797 | + | Gnomon | gene | gene-LOC105280348 | GeneID:105280348 | LOC105280348 | Gene | LOC105280348 | protein_coding | character(0) | character(0) | BCphaseA2.vs.RphaseB1 | FALSE |
| NC_039506.1 | 2981335 | 2981335 | 1 | + | 29 | 28 | 1 | 65 | 63 | 2 | 96.55 | 96.92 | NC_039506.1 | 2980439 | 2984235 | 3797 | + | Gnomon | gene | gene-LOC105280348 | GeneID:105280348 | LOC105280348 | Gene | LOC105280348 | protein_coding | character(0) | character(0) | BCphaseA2.vs.RphaseB1 | FALSE |
| NC_039506.1 | 2981350 | 2981350 | 1 | + | 59 | 57 | 2 | 55 | 55 | 0 | 96.61 | 100 | NC_039506.1 | 2980439 | 2984235 | 3797 | + | Gnomon | gene | gene-LOC105280348 | GeneID:105280348 | LOC105280348 | Gene | LOC105280348 | protein_coding | character(0) | character(0) | BCphaseA2.vs.RphaseB1 | FALSE |
| NC_039506.1 | 2981391 | 2981391 | 1 | + | 72 | 64 | 8 | 65 | 53 | 12 | 88.89 | 81.54 | NC_039506.1 | 2980439 | 2984235 | 3797 | + | Gnomon | gene | gene-LOC105280348 | GeneID:105280348 | LOC105280348 | Gene | LOC105280348 | protein_coding | character(0) | character(0) | BCphaseA2.vs.RphaseB1 | FALSE |
| NC_039506.1 | 2981402 | 2981402 | 1 | + | 71 | 69 | 2 | 74 | 67 | 7 | 97.18 | 90.54 | NC_039506.1 | 2980439 | 2984235 | 3797 | + | Gnomon | gene | gene-LOC105280348 | GeneID:105280348 | LOC105280348 | Gene | LOC105280348 | protein_coding | character(0) | character(0) | BCphaseA2.vs.RphaseB1 | FALSE |
| NC_039506.1 | 2981411 | 2981411 | 1 | + | 76 | 72 | 4 | 81 | 71 | 10 | 94.74 | 87.65 | NC_039506.1 | 2980439 | 2984235 | 3797 | + | Gnomon | gene | gene-LOC105280348 | GeneID:105280348 | LOC105280348 | Gene | LOC105280348 | protein_coding | character(0) | character(0) | BCphaseA2.vs.RphaseB1 | FALSE |
| NC_039506.1 | 2982097 | 2982097 | 1 | + | 34 | 1 | 33 | 28 | 0 | 28 | 2.94 | 0 | NC_039506.1 | 2980439 | 2984235 | 3797 | + | Gnomon | gene | gene-LOC105280348 | GeneID:105280348 | LOC105280348 | Gene | LOC105280348 | protein_coding | character(0) | character(0) | BCphaseA2.vs.RphaseB1 | FALSE |
| NC_039506.1 | 2982133 | 2982133 | 1 | + | 35 | 1 | 34 | 33 | 0 | 33 | 2.86 | 0 | NC_039506.1 | 2980439 | 2984235 | 3797 | + | Gnomon | gene | gene-LOC105280348 | GeneID:105280348 | LOC105280348 | Gene | LOC105280348 | protein_coding | character(0) | character(0) | BCphaseA2.vs.RphaseB1 | FALSE |

...
... only first 25 lines shown ...

Directory: DMSG File: dmg-Ob_LI_BCphaseA2.vs.RphaseB2_details.txt

| seqnames | start | end | width | strand | coverage1 | numCs1 | numTs1 | coverage2 | numCs2 | numTs2 | BCphaseA2 | RphaseB2 | gene_seqnames | gene_start | gene_end | gene_width | gene_strand | gene_source | gene_type | gene_ID | gene_Dbxref | gene_Name | gene_gbkey | gene_gene | gene_gene_biotype | gene_end_range | gene_start_range | gene_comparison | is.dm |
|---|---|---|---|---|---|---|---|---|---|---|---|---|---|---|---|---|---|---|---|---|---|---|---|---|---|---|---|---|---|
| NC_039506.1 | 3248474 | 3248474 | 1 | + | 30 | 7 | 23 | 51 | 14 | 37 | 23.33 | 27.45 | NC_039506.1 | 3248415 | 3250438 | 2024 | - | Gnomon | gene | gene-LOC105280377 | GeneID:105280377 | LOC105280377 | Gene | LOC105280377 | protein_coding | character(0) | character(0) | BCphaseA2.vs.RphaseB2 | FALSE |
| NC_039506.1 | 3248483 | 3248483 | 1 | + | 31 | 15 | 16 | 57 | 27 | 30 | 48.39 | 47.37 | NC_039506.1 | 3248415 | 3250438 | 2024 | - | Gnomon | gene | gene-LOC105280377 | GeneID:105280377 | LOC105280377 | Gene | LOC105280377 | protein_coding | character(0) | character(0) | BCphaseA2.vs.RphaseB2 | FALSE |
| NC_039506.1 | 3248495 | 3248495 | 1 | + | 53 | 10 | 43 | 65 | 13 | 52 | 18.87 | 20 | NC_039506.1 | 3248415 | 3250438 | 2024 | - | Gnomon | gene | gene-LOC105280377 | GeneID:105280377 | LOC105280377 | Gene | LOC105280377 | protein_coding | character(0) | character(0) | BCphaseA2.vs.RphaseB2 | FALSE |
| NC_039506.1 | 3248534 | 3248534 | 1 | + | 52 | 14 | 38 | 69 | 22 | 47 | 26.92 | 31.88 | NC_039506.1 | 3248415 | 3250438 | 2024 | - | Gnomon | gene | gene-LOC105280377 | GeneID:105280377 | LOC105280377 | Gene | LOC105280377 | protein_coding | character(0) | character(0) | BCphaseA2.vs.RphaseB2 | FALSE |
| NC_039506.1 | 3248539 | 3248539 | 1 | + | 50 | 3 | 47 | 63 | 5 | 58 | 6 | 7.94 | NC_039506.1 | 3248415 | 3250438 | 2024 | - | Gnomon | gene | gene-LOC105280377 | GeneID:105280377 | LOC105280377 | Gene | LOC105280377 | protein_coding | character(0) | character(0) | BCphaseA2.vs.RphaseB2 | FALSE |
| NC_039506.1 | 3248551 | 3248551 | 1 | + | 26 | 1 | 25 | 48 | 3 | 45 | 3.85 | 6.25 | NC_039506.1 | 3248415 | 3250438 | 2024 | - | Gnomon | gene | gene-LOC105280377 | GeneID:105280377 | LOC105280377 | Gene | LOC105280377 | protein_coding | character(0) | character(0) | BCphaseA2.vs.RphaseB2 | FALSE |
| NC_039506.1 | 3248553 | 3248553 | 1 | + | 25 | 1 | 24 | 44 | 0 | 44 | 4 | 0 | NC_039506.1 | 3248415 | 3250438 | 2024 | - | Gnomon | gene | gene-LOC105280377 | GeneID:105280377 | LOC105280377 | Gene | LOC105280377 | protein_coding | character(0) | character(0) | BCphaseA2.vs.RphaseB2 | FALSE |
| NC_039506.1 | 3248556 | 3248556 | 1 | + | 22 | 2 | 20 | 46 | 8 | 38 | 9.09 | 17.39 | NC_039506.1 | 3248415 | 3250438 | 2024 | - | Gnomon | gene | gene-LOC105280377 | GeneID:105280377 | LOC105280377 | Gene | LOC105280377 | protein_coding | character(0) | character(0) | BCphaseA2.vs.RphaseB2 | FALSE |
| NC_039506.1 | 3248598 | 3248598 | 1 | + | 26 | 3 | 23 | 24 | 2 | 22 | 11.54 | 8.33 | NC_039506.1 | 3248415 | 3250438 | 2024 | - | Gnomon | gene | gene-LOC105280377 | GeneID:105280377 | LOC105280377 | Gene | LOC105280377 | protein_coding | character(0) | character(0) | BCphaseA2.vs.RphaseB2 | FALSE |
| NC_039506.1 | 3248631 | 3248631 | 1 | + | 28 | 2 | 26 | 38 | 2 | 36 | 7.14 | 5.26 | NC_039506.1 | 3248415 | 3250438 | 2024 | - | Gnomon | gene | gene-LOC105280377 | GeneID:105280377 | LOC105280377 | Gene | LOC105280377 | protein_coding | character(0) | character(0) | BCphaseA2.vs.RphaseB2 | FALSE |
| NC_039506.1 | 3248698 | 3248698 | 1 | + | 42 | 5 | 37 | 52 | 3 | 49 | 11.9 | 5.77 | NC_039506.1 | 3248415 | 3250438 | 2024 | - | Gnomon | gene | gene-LOC105280377 | GeneID:105280377 | LOC105280377 | Gene | LOC105280377 | protein_coding | character(0) | character(0) | BCphaseA2.vs.RphaseB2 | FALSE |
| NC_039506.1 | 3248700 | 3248700 | 1 | + | 42 | 0 | 42 | 52 | 1 | 51 | 0 | 1.92 | NC_039506.1 | 3248415 | 3250438 | 2024 | - | Gnomon | gene | gene-LOC105280377 | GeneID:105280377 | LOC105280377 | Gene | LOC105280377 | protein_coding | character(0) | character(0) | BCphaseA2.vs.RphaseB2 | FALSE |
| NC_039506.1 | 3248715 | 3248715 | 1 | + | 23 | 0 | 23 | 26 | 1 | 25 | 0 | 3.85 | NC_039506.1 | 3248415 | 3250438 | 2024 | - | Gnomon | gene | gene-LOC105280377 | GeneID:105280377 | LOC105280377 | Gene | LOC105280377 | protein_coding | character(0) | character(0) | BCphaseA2.vs.RphaseB2 | FALSE |
| NC_039506.1 | 3248760 | 3248760 | 1 | + | 21 | 0 | 21 | 23 | 1 | 22 | 0 | 4.35 | NC_039506.1 | 3248415 | 3250438 | 2024 | - | Gnomon | gene | gene-LOC105280377 | GeneID:105280377 | LOC105280377 | Gene | LOC105280377 | protein_coding | character(0) | character(0) | BCphaseA2.vs.RphaseB2 | FALSE |
| NC_039506.1 | 3248793 | 3248793 | 1 | + | 46 | 9 | 37 | 35 | 6 | 29 | 19.57 | 17.14 | NC_039506.1 | 3248415 | 3250438 | 2024 | - | Gnomon | gene | gene-LOC105280377 | GeneID:105280377 | LOC105280377 | Gene | LOC105280377 | protein_coding | character(0) | character(0) | BCphaseA2.vs.RphaseB2 | FALSE |
| NC_039506.1 | 3248801 | 3248801 | 1 | + | 46 | 5 | 41 | 55 | 11 | 44 | 10.87 | 20 | NC_039506.1 | 3248415 | 3250438 | 2024 | - | Gnomon | gene | gene-LOC105280377 | GeneID:105280377 | LOC105280377 | Gene | LOC105280377 | protein_coding | character(0) | character(0) | BCphaseA2.vs.RphaseB2 | FALSE |
| NC_039506.1 | 3248827 | 3248827 | 1 | + | 85 | 8 | 77 | 69 | 12 | 57 | 9.41 | 17.39 | NC_039506.1 | 3248415 | 3250438 | 2024 | - | Gnomon | gene | gene-LOC105280377 | GeneID:105280377 | LOC105280377 | Gene | LOC105280377 | protein_coding | character(0) | character(0) | BCphaseA2.vs.RphaseB2 | FALSE |
| NC_039506.1 | 3248834 | 3248834 | 1 | + | 89 | 2 | 87 | 69 | 2 | 67 | 2.25 | 2.9 | NC_039506.1 | 3248415 | 3250438 | 2024 | - | Gnomon | gene | gene-LOC105280377 | GeneID:105280377 | LOC105280377 | Gene | LOC105280377 | protein_coding | character(0) | character(0) | BCphaseA2.vs.RphaseB2 | FALSE |
| NC_039506.1 | 3248849 | 3248849 | 1 | + | 52 | 7 | 45 | 24 | 2 | 22 | 13.46 | 8.33 | NC_039506.1 | 3248415 | 3250438 | 2024 | - | Gnomon | gene | gene-LOC105280377 | GeneID:105280377 | LOC105280377 | Gene | LOC105280377 | protein_coding | character(0) | character(0) | BCphaseA2.vs.RphaseB2 | FALSE |
| NC_039506.1 | 3248853 | 3248853 | 1 | + | 31 | 1 | 30 | 23 | 0 | 23 | 3.23 | 0 | NC_039506.1 | 3248415 | 3250438 | 2024 | - | Gnomon | gene | gene-LOC105280377 | GeneID:105280377 | LOC105280377 | Gene | LOC105280377 | protein_coding | character(0) | character(0) | BCphaseA2.vs.RphaseB2 | FALSE |
| NC_039506.1 | 3248860 | 3248860 | 1 | + | 39 | 10 | 29 | 34 | 8 | 26 | 25.64 | 23.53 | NC_039506.1 | 3248415 | 3250438 | 2024 | - | Gnomon | gene | gene-LOC105280377 | GeneID:105280377 | LOC105280377 | Gene | LOC105280377 | protein_coding | character(0) | character(0) | BCphaseA2.vs.RphaseB2 | FALSE |
| NC_039506.1 | 3248865 | 3248865 | 1 | + | 37 | 1 | 36 | 34 | 2 | 32 | 2.7 | 5.88 | NC_039506.1 | 3248415 | 3250438 | 2024 | - | Gnomon | gene | gene-LOC105280377 | GeneID:105280377 | LOC105280377 | Gene | LOC105280377 | protein_coding | character(0) | character(0) | BCphaseA2.vs.RphaseB2 | FALSE |
| NC_039506.1 | 3248875 | 3248875 | 1 | + | 67 | 1 | 66 | 51 | 1 | 50 | 1.49 | 1.96 | NC_039506.1 | 3248415 | 3250438 | 2024 | - | Gnomon | gene | gene-LOC105280377 | GeneID:105280377 | LOC105280377 | Gene | LOC105280377 | protein_coding | character(0) | character(0) | BCphaseA2.vs.RphaseB2 | FALSE |
| NC_039506.1 | 3248893 | 3248893 | 1 | + | 49 | 8 | 41 | 37 | 3 | 34 | 16.33 | 8.11 | NC_039506.1 | 3248415 | 3250438 | 2024 | - | Gnomon | gene | gene-LOC105280377 | GeneID:105280377 | LOC105280377 | Gene | LOC105280377 | protein_coding | character(0) | character(0) | BCphaseA2.vs.RphaseB2 | FALSE |

...
... only first 25 lines shown ...

Directory: DMSG File: dmg-Ob_LI_BCphaseB1.vs.BCphaseB2_details.txt

| seqnames | start | end | width | strand | coverage1 | numCs1 | numTs1 | coverage2 | numCs2 | numTs2 | BCphaseB1 | BCphaseB2 | gene_seqnames | gene_start | gene_end | gene_width | gene_strand | gene_source | gene_type | gene_ID | gene_Dbxref | gene_Name | gene_gbkey | gene_gene | gene_gene_biotype | gene_end_range | gene_start_range | gene_comparison | is.dm |
|---|---|---|---|---|---|---|---|---|---|---|---|---|---|---|---|---|---|---|---|---|---|---|---|---|---|---|---|---|---|
| NC_039506.1 | 2980668 | 2980668 | 1 | + | 28 | 0 | 28 | 29 | 1 | 28 | 0 | 3.45 | NC_039506.1 | 2980439 | 2984235 | 3797 | + | Gnomon | gene | gene-LOC105280348 | GeneID:105280348 | LOC105280348 | Gene | LOC105280348 | protein_coding | character(0) | character(0) | BCphaseB1.vs.BCphaseB2 | FALSE |
| NC_039506.1 | 2980692 | 2980692 | 1 | + | 24 | 1 | 23 | 22 | 2 | 20 | 4.17 | 9.09 | NC_039506.1 | 2980439 | 2984235 | 3797 | + | Gnomon | gene | gene-LOC105280348 | GeneID:105280348 | LOC105280348 | Gene | LOC105280348 | protein_coding | character(0) | character(0) | BCphaseB1.vs.BCphaseB2 | FALSE |
| NC_039506.1 | 2980750 | 2980750 | 1 | + | 29 | 0 | 29 | 32 | 0 | 32 | 0 | 0 | NC_039506.1 | 2980439 | 2984235 | 3797 | + | Gnomon | gene | gene-LOC105280348 | GeneID:105280348 | LOC105280348 | Gene | LOC105280348 | protein_coding | character(0) | character(0) | BCphaseB1.vs.BCphaseB2 | FALSE |
| NC_039506.1 | 2980761 | 2980761 | 1 | + | 32 | 0 | 32 | 28 | 0 | 28 | 0 | 0 | NC_039506.1 | 2980439 | 2984235 | 3797 | + | Gnomon | gene | gene-LOC105280348 | GeneID:105280348 | LOC105280348 | Gene | LOC105280348 | protein_coding | character(0) | character(0) | BCphaseB1.vs.BCphaseB2 | FALSE |
| NC_039506.1 | 2980767 | 2980767 | 1 | + | 34 | 0 | 34 | 28 | 0 | 28 | 0 | 0 | NC_039506.1 | 2980439 | 2984235 | 3797 | + | Gnomon | gene | gene-LOC105280348 | GeneID:105280348 | LOC105280348 | Gene | LOC105280348 | protein_coding | character(0) | character(0) | BCphaseB1.vs.BCphaseB2 | FALSE |
| NC_039506.1 | 2980928 | 2980928 | 1 | + | 23 | 8 | 15 | 23 | 6 | 17 | 34.78 | 26.09 | NC_039506.1 | 2980439 | 2984235 | 3797 | + | Gnomon | gene | gene-LOC105280348 | GeneID:105280348 | LOC105280348 | Gene | LOC105280348 | protein_coding | character(0) | character(0) | BCphaseB1.vs.BCphaseB2 | FALSE |
| NC_039506.1 | 2980989 | 2980989 | 1 | + | 29 | 3 | 26 | 29 | 7 | 22 | 10.34 | 24.14 | NC_039506.1 | 2980439 | 2984235 | 3797 | + | Gnomon | gene | gene-LOC105280348 | GeneID:105280348 | LOC105280348 | Gene | LOC105280348 | protein_coding | character(0) | character(0) | BCphaseB1.vs.BCphaseB2 | FALSE |
| NC_039506.1 | 2980993 | 2980993 | 1 | + | 29 | 12 | 17 | 29 | 10 | 19 | 41.38 | 34.48 | NC_039506.1 | 2980439 | 2984235 | 3797 | + | Gnomon | gene | gene-LOC105280348 | GeneID:105280348 | LOC105280348 | Gene | LOC105280348 | protein_coding | character(0) | character(0) | BCphaseB1.vs.BCphaseB2 | FALSE |
| NC_039506.1 | 2980998 | 2980998 | 1 | + | 28 | 8 | 20 | 30 | 11 | 19 | 28.57 | 36.67 | NC_039506.1 | 2980439 | 2984235 | 3797 | + | Gnomon | gene | gene-LOC105280348 | GeneID:105280348 | LOC105280348 | Gene | LOC105280348 | protein_coding | character(0) | character(0) | BCphaseB1.vs.BCphaseB2 | FALSE |
| NC_039506.1 | 2981030 | 2981030 | 1 | + | 56 | 50 | 6 | 43 | 35 | 8 | 89.29 | 81.4 | NC_039506.1 | 2980439 | 2984235 | 3797 | + | Gnomon | gene | gene-LOC105280348 | GeneID:105280348 | LOC105280348 | Gene | LOC105280348 | protein_coding | character(0) | character(0) | BCphaseB1.vs.BCphaseB2 | FALSE |
| NC_039506.1 | 2981077 | 2981077 | 1 | + | 49 | 25 | 24 | 36 | 13 | 23 | 51.02 | 36.11 | NC_039506.1 | 2980439 | 2984235 | 3797 | + | Gnomon | gene | gene-LOC105280348 | GeneID:105280348 | LOC105280348 | Gene | LOC105280348 | protein_coding | character(0) | character(0) | BCphaseB1.vs.BCphaseB2 | FALSE |
| NC_039506.1 | 2981183 | 2981183 | 1 | + | 83 | 78 | 5 | 49 | 25 | 24 | 93.98 | 51.02 | NC_039506.1 | 2980439 | 2984235 | 3797 | + | Gnomon | gene | gene-LOC105280348 | GeneID:105280348 | LOC105280348 | Gene | LOC105280348 | protein_coding | character(0) | character(0) | BCphaseB1.vs.BCphaseB2 | TRUE |
| NC_039506.1 | 2981225 | 2981225 | 1 | + | 37 | 23 | 14 | 32 | 24 | 8 | 62.16 | 75 | NC_039506.1 | 2980439 | 2984235 | 3797 | + | Gnomon | gene | gene-LOC105280348 | GeneID:105280348 | LOC105280348 | Gene | LOC105280348 | protein_coding | character(0) | character(0) | BCphaseB1.vs.BCphaseB2 | FALSE |
| NC_039506.1 | 2981245 | 2981245 | 1 | + | 40 | 18 | 22 | 38 | 14 | 24 | 45 | 36.84 | NC_039506.1 | 2980439 | 2984235 | 3797 | + | Gnomon | gene | gene-LOC105280348 | GeneID:105280348 | LOC105280348 | Gene | LOC105280348 | protein_coding | character(0) | character(0) | BCphaseB1.vs.BCphaseB2 | FALSE |
| NC_039506.1 | 2981292 | 2981292 | 1 | + | 68 | 64 | 4 | 42 | 42 | 0 | 94.12 | 100 | NC_039506.1 | 2980439 | 2984235 | 3797 | + | Gnomon | gene | gene-LOC105280348 | GeneID:105280348 | LOC105280348 | Gene | LOC105280348 | protein_coding | character(0) | character(0) | BCphaseB1.vs.BCphaseB2 | FALSE |
| NC_039506.1 | 2981308 | 2981308 | 1 | + | 78 | 52 | 26 | 49 | 34 | 15 | 66.67 | 69.39 | NC_039506.1 | 2980439 | 2984235 | 3797 | + | Gnomon | gene | gene-LOC105280348 | GeneID:105280348 | LOC105280348 | Gene | LOC105280348 | protein_coding | character(0) | character(0) | BCphaseB1.vs.BCphaseB2 | FALSE |
| NC_039506.1 | 2981315 | 2981315 | 1 | + | 77 | 69 | 8 | 49 | 40 | 9 | 89.61 | 81.63 | NC_039506.1 | 2980439 | 2984235 | 3797 | + | Gnomon | gene | gene-LOC105280348 | GeneID:105280348 | LOC105280348 | Gene | LOC105280348 | protein_coding | character(0) | character(0) | BCphaseB1.vs.BCphaseB2 | FALSE |
| NC_039506.1 | 2981335 | 2981335 | 1 | + | 56 | 54 | 2 | 41 | 39 | 2 | 96.43 | 95.12 | NC_039506.1 | 2980439 | 2984235 | 3797 | + | Gnomon | gene | gene-LOC105280348 | GeneID:105280348 | LOC105280348 | Gene | LOC105280348 | protein_coding | character(0) | character(0) | BCphaseB1.vs.BCphaseB2 | FALSE |
| NC_039506.1 | 2981350 | 2981350 | 1 | + | 58 | 57 | 1 | 72 | 68 | 4 | 98.28 | 94.44 | NC_039506.1 | 2980439 | 2984235 | 3797 | + | Gnomon | gene | gene-LOC105280348 | GeneID:105280348 | LOC105280348 | Gene | LOC105280348 | protein_coding | character(0) | character(0) | BCphaseB1.vs.BCphaseB2 | FALSE |
| NC_039506.1 | 2981391 | 2981391 | 1 | + | 60 | 49 | 11 | 87 | 69 | 18 | 81.67 | 79.31 | NC_039506.1 | 2980439 | 2984235 | 3797 | + | Gnomon | gene | gene-LOC105280348 | GeneID:105280348 | LOC105280348 | Gene | LOC105280348 | protein_coding | character(0) | character(0) | BCphaseB1.vs.BCphaseB2 | FALSE |
| NC_039506.1 | 2981402 | 2981402 | 1 | + | 66 | 60 | 6 | 88 | 85 | 3 | 90.91 | 96.59 | NC_039506.1 | 2980439 | 2984235 | 3797 | + | Gnomon | gene | gene-LOC105280348 | GeneID:105280348 | LOC105280348 | Gene | LOC105280348 | protein_coding | character(0) | character(0) | BCphaseB1.vs.BCphaseB2 | FALSE |
| NC_039506.1 | 2981411 | 2981411 | 1 | + | 68 | 65 | 3 | 101 | 94 | 7 | 95.59 | 93.07 | NC_039506.1 | 2980439 | 2984235 | 3797 | + | Gnomon | gene | gene-LOC105280348 | GeneID:105280348 | LOC105280348 | Gene | LOC105280348 | protein_coding | character(0) | character(0) | BCphaseB1.vs.BCphaseB2 | FALSE |
| NC_039506.1 | 2981645 | 2981645 | 1 | + | 23 | 12 | 11 | 21 | 14 | 7 | 52.17 | 66.67 | NC_039506.1 | 2980439 | 2984235 | 3797 | + | Gnomon | gene | gene-LOC105280348 | GeneID:105280348 | LOC105280348 | Gene | LOC105280348 | protein_coding | character(0) | character(0) | BCphaseB1.vs.BCphaseB2 | FALSE |
| NC_039506.1 | 2981661 | 2981661 | 1 | + | 21 | 14 | 7 | 24 | 15 | 9 | 66.67 | 62.5 | NC_039506.1 | 2980439 | 2984235 | 3797 | + | Gnomon | gene | gene-LOC105280348 | GeneID:105280348 | LOC105280348 | Gene | LOC105280348 | protein_coding | character(0) | character(0) | BCphaseB1.vs.BCphaseB2 | FALSE |

...
... only first 25 lines shown ...

Directory: DMSG File: dmg-Ob_LI_BCphaseB1.vs.RphaseA1_details.txt

| seqnames | start | end | width | strand | coverage1 | numCs1 | numTs1 | coverage2 | numCs2 | numTs2 | BCphaseB1 | RphaseA1 | gene_seqnames | gene_start | gene_end | gene_width | gene_strand | gene_source | gene_type | gene_ID | gene_Dbxref | gene_Name | gene_gbkey | gene_gene | gene_gene_biotype | gene_end_range | gene_partial | gene_start_range | gene_comparison | is.dm |
|---|---|---|---|---|---|---|---|---|---|---|---|---|---|---|---|---|---|---|---|---|---|---|---|---|---|---|---|---|---|---|
| NC_039506.1 | 1097103 | 1097103 | 1 | + | 26 | 0 | 26 | 21 | 0 | 21 | 0 | 0 | NC_039506.1 | 1095614 | 1100480 | 4867 | + | Gnomon | gene | gene-LOC105286750 | GeneID:105286750 | LOC105286750 | Gene | LOC105286750 | protein_coding | character(0) | NA | character(0) | BCphaseB1.vs.RphaseA1 | FALSE |
| NC_039506.1 | 1097536 | 1097536 | 1 | + | 27 | 1 | 26 | 32 | 0 | 32 | 3.7 | 0 | NC_039506.1 | 1095614 | 1100480 | 4867 | + | Gnomon | gene | gene-LOC105286750 | GeneID:105286750 | LOC105286750 | Gene | LOC105286750 | protein_coding | character(0) | NA | character(0) | BCphaseB1.vs.RphaseA1 | FALSE |
| NC_039506.1 | 1097543 | 1097543 | 1 | + | 28 | 3 | 25 | 31 | 1 | 30 | 10.71 | 3.23 | NC_039506.1 | 1095614 | 1100480 | 4867 | + | Gnomon | gene | gene-LOC105286750 | GeneID:105286750 | LOC105286750 | Gene | LOC105286750 | protein_coding | character(0) | NA | character(0) | BCphaseB1.vs.RphaseA1 | FALSE |
| NC_039506.1 | 1097546 | 1097546 | 1 | + | 28 | 3 | 25 | 32 | 0 | 32 | 10.71 | 0 | NC_039506.1 | 1095614 | 1100480 | 4867 | + | Gnomon | gene | gene-LOC105286750 | GeneID:105286750 | LOC105286750 | Gene | LOC105286750 | protein_coding | character(0) | NA | character(0) | BCphaseB1.vs.RphaseA1 | FALSE |
| NC_039506.1 | 1097588 | 1097588 | 1 | + | 30 | 9 | 21 | 27 | 16 | 11 | 30 | 59.26 | NC_039506.1 | 1095614 | 1100480 | 4867 | + | Gnomon | gene | gene-LOC105286750 | GeneID:105286750 | LOC105286750 | Gene | LOC105286750 | protein_coding | character(0) | NA | character(0) | BCphaseB1.vs.RphaseA1 | TRUE |
| NC_039506.1 | 1097598 | 1097598 | 1 | + | 33 | 17 | 16 | 31 | 16 | 15 | 51.52 | 51.61 | NC_039506.1 | 1095614 | 1100480 | 4867 | + | Gnomon | gene | gene-LOC105286750 | GeneID:105286750 | LOC105286750 | Gene | LOC105286750 | protein_coding | character(0) | NA | character(0) | BCphaseB1.vs.RphaseA1 | FALSE |
| NC_039506.1 | 1097647 | 1097647 | 1 | + | 46 | 8 | 38 | 45 | 4 | 41 | 17.39 | 8.89 | NC_039506.1 | 1095614 | 1100480 | 4867 | + | Gnomon | gene | gene-LOC105286750 | GeneID:105286750 | LOC105286750 | Gene | LOC105286750 | protein_coding | character(0) | NA | character(0) | BCphaseB1.vs.RphaseA1 | FALSE |
| NC_039506.1 | 1097653 | 1097653 | 1 | + | 46 | 2 | 44 | 45 | 1 | 44 | 4.35 | 2.22 | NC_039506.1 | 1095614 | 1100480 | 4867 | + | Gnomon | gene | gene-LOC105286750 | GeneID:105286750 | LOC105286750 | Gene | LOC105286750 | protein_coding | character(0) | NA | character(0) | BCphaseB1.vs.RphaseA1 | FALSE |
| NC_039506.1 | 1097702 | 1097702 | 1 | + | 68 | 11 | 57 | 80 | 26 | 54 | 16.18 | 32.5 | NC_039506.1 | 1095614 | 1100480 | 4867 | + | Gnomon | gene | gene-LOC105286750 | GeneID:105286750 | LOC105286750 | Gene | LOC105286750 | protein_coding | character(0) | NA | character(0) | BCphaseB1.vs.RphaseA1 | FALSE |
| NC_039506.1 | 1097773 | 1097773 | 1 | + | 24 | 23 | 1 | 29 | 28 | 1 | 95.83 | 96.55 | NC_039506.1 | 1095614 | 1100480 | 4867 | + | Gnomon | gene | gene-LOC105286750 | GeneID:105286750 | LOC105286750 | Gene | LOC105286750 | protein_coding | character(0) | NA | character(0) | BCphaseB1.vs.RphaseA1 | FALSE |
| NC_039506.1 | 1098123 | 1098123 | 1 | + | 23 | 12 | 11 | 24 | 17 | 7 | 52.17 | 70.83 | NC_039506.1 | 1095614 | 1100480 | 4867 | + | Gnomon | gene | gene-LOC105286750 | GeneID:105286750 | LOC105286750 | Gene | LOC105286750 | protein_coding | character(0) | NA | character(0) | BCphaseB1.vs.RphaseA1 | FALSE |
| NC_039506.1 | 1098163 | 1098163 | 1 | + | 21 | 18 | 3 | 21 | 16 | 5 | 85.71 | 76.19 | NC_039506.1 | 1095614 | 1100480 | 4867 | + | Gnomon | gene | gene-LOC105286750 | GeneID:105286750 | LOC105286750 | Gene | LOC105286750 | protein_coding | character(0) | NA | character(0) | BCphaseB1.vs.RphaseA1 | FALSE |
| NC_039506.1 | 1098206 | 1098206 | 1 | + | 34 | 23 | 11 | 37 | 29 | 8 | 67.65 | 78.38 | NC_039506.1 | 1095614 | 1100480 | 4867 | + | Gnomon | gene | gene-LOC105286750 | GeneID:105286750 | LOC105286750 | Gene | LOC105286750 | protein_coding | character(0) | NA | character(0) | BCphaseB1.vs.RphaseA1 | FALSE |
| NC_039506.1 | 1098266 | 1098266 | 1 | + | 50 | 27 | 23 | 57 | 32 | 25 | 54 | 56.14 | NC_039506.1 | 1095614 | 1100480 | 4867 | + | Gnomon | gene | gene-LOC105286750 | GeneID:105286750 | LOC105286750 | Gene | LOC105286750 | protein_coding | character(0) | NA | character(0) | BCphaseB1.vs.RphaseA1 | FALSE |
| NC_039506.1 | 1098279 | 1098279 | 1 | + | 48 | 19 | 29 | 56 | 17 | 39 | 39.58 | 30.36 | NC_039506.1 | 1095614 | 1100480 | 4867 | + | Gnomon | gene | gene-LOC105286750 | GeneID:105286750 | LOC105286750 | Gene | LOC105286750 | protein_coding | character(0) | NA | character(0) | BCphaseB1.vs.RphaseA1 | FALSE |
| NC_039506.1 | 1098293 | 1098293 | 1 | + | 21 | 7 | 14 | 31 | 12 | 19 | 33.33 | 38.71 | NC_039506.1 | 1095614 | 1100480 | 4867 | + | Gnomon | gene | gene-LOC105286750 | GeneID:105286750 | LOC105286750 | Gene | LOC105286750 | protein_coding | character(0) | NA | character(0) | BCphaseB1.vs.RphaseA1 | FALSE |
| NC_039506.1 | 1098301 | 1098301 | 1 | + | 26 | 20 | 6 | 32 | 17 | 15 | 76.92 | 53.12 | NC_039506.1 | 1095614 | 1100480 | 4867 | + | Gnomon | gene | gene-LOC105286750 | GeneID:105286750 | LOC105286750 | Gene | LOC105286750 | protein_coding | character(0) | NA | character(0) | BCphaseB1.vs.RphaseA1 | TRUE |
| NC_039506.1 | 1098330 | 1098330 | 1 | + | 47 | 8 | 39 | 51 | 10 | 41 | 17.02 | 19.61 | NC_039506.1 | 1095614 | 1100480 | 4867 | + | Gnomon | gene | gene-LOC105286750 | GeneID:105286750 | LOC105286750 | Gene | LOC105286750 | protein_coding | character(0) | NA | character(0) | BCphaseB1.vs.RphaseA1 | FALSE |
| NC_039506.1 | 1098334 | 1098334 | 1 | + | 50 | 27 | 23 | 53 | 23 | 30 | 54 | 43.4 | NC_039506.1 | 1095614 | 1100480 | 4867 | + | Gnomon | gene | gene-LOC105286750 | GeneID:105286750 | LOC105286750 | Gene | LOC105286750 | protein_coding | character(0) | NA | character(0) | BCphaseB1.vs.RphaseA1 | FALSE |
| NC_039506.1 | 1098362 | 1098362 | 1 | + | 61 | 4 | 57 | 64 | 3 | 61 | 6.56 | 4.69 | NC_039506.1 | 1095614 | 1100480 | 4867 | + | Gnomon | gene | gene-LOC105286750 | GeneID:105286750 | LOC105286750 | Gene | LOC105286750 | protein_coding | character(0) | NA | character(0) | BCphaseB1.vs.RphaseA1 | FALSE |
| NC_039506.1 | 1098377 | 1098377 | 1 | + | 61 | 14 | 47 | 64 | 22 | 42 | 22.95 | 34.38 | NC_039506.1 | 1095614 | 1100480 | 4867 | + | Gnomon | gene | gene-LOC105286750 | GeneID:105286750 | LOC105286750 | Gene | LOC105286750 | protein_coding | character(0) | NA | character(0) | BCphaseB1.vs.RphaseA1 | FALSE |
| NC_039506.1 | 1098387 | 1098387 | 1 | + | 56 | 28 | 28 | 54 | 29 | 25 | 50 | 53.7 | NC_039506.1 | 1095614 | 1100480 | 4867 | + | Gnomon | gene | gene-LOC105286750 | GeneID:105286750 | LOC105286750 | Gene | LOC105286750 | protein_coding | character(0) | NA | character(0) | BCphaseB1.vs.RphaseA1 | FALSE |
| NC_039506.1 | 1098400 | 1098400 | 1 | + | 38 | 17 | 21 | 36 | 16 | 20 | 44.74 | 44.44 | NC_039506.1 | 1095614 | 1100480 | 4867 | + | Gnomon | gene | gene-LOC105286750 | GeneID:105286750 | LOC105286750 | Gene | LOC105286750 | protein_coding | character(0) | NA | character(0) | BCphaseB1.vs.RphaseA1 | FALSE |
| NC_039506.1 | 1098420 | 1098420 | 1 | + | 34 | 14 | 20 | 35 | 17 | 18 | 41.18 | 48.57 | NC_039506.1 | 1095614 | 1100480 | 4867 | + | Gnomon | gene | gene-LOC105286750 | GeneID:105286750 | LOC105286750 | Gene | LOC105286750 | protein_coding | character(0) | NA | character(0) | BCphaseB1.vs.RphaseA1 | FALSE |

...
... only first 25 lines shown ...

Directory: DMSG File: dmg-Ob_LI_BCphaseB1.vs.RphaseA2_details.txt

| seqnames | start | end | width | strand | coverage1 | numCs1 | numTs1 | coverage2 | numCs2 | numTs2 | BCphaseB1 | RphaseA2 | gene_seqnames | gene_start | gene_end | gene_width | gene_strand | gene_source | gene_type | gene_ID | gene_Dbxref | gene_Name | gene_gbkey | gene_gene | gene_gene_biotype | gene_end_range | gene_partial | gene_start_range | gene_comparison | is.dm |
|---|---|---|---|---|---|---|---|---|---|---|---|---|---|---|---|---|---|---|---|---|---|---|---|---|---|---|---|---|---|---|
| NC_039506.1 | 1057034 | 1057034 | 1 | + | 24 | 0 | 24 | 40 | 0 | 40 | 0 | 0 | NC_039506.1 | 1056151 | 1064257 | 8107 | + | Gnomon | gene | gene-LOC105286747 | GeneID:105286747 | LOC105286747 | Gene | LOC105286747 | lncRNA | character(0) | NA | character(0) | BCphaseB1.vs.RphaseA2 | FALSE |
| NC_039506.1 | 1057043 | 1057043 | 1 | + | 28 | 0 | 28 | 44 | 0 | 44 | 0 | 0 | NC_039506.1 | 1056151 | 1064257 | 8107 | + | Gnomon | gene | gene-LOC105286747 | GeneID:105286747 | LOC105286747 | Gene | LOC105286747 | lncRNA | character(0) | NA | character(0) | BCphaseB1.vs.RphaseA2 | FALSE |
| NC_039506.1 | 1057047 | 1057047 | 1 | + | 27 | 0 | 27 | 44 | 0 | 44 | 0 | 0 | NC_039506.1 | 1056151 | 1064257 | 8107 | + | Gnomon | gene | gene-LOC105286747 | GeneID:105286747 | LOC105286747 | Gene | LOC105286747 | lncRNA | character(0) | NA | character(0) | BCphaseB1.vs.RphaseA2 | FALSE |
| NC_039506.1 | 1057061 | 1057061 | 1 | + | 28 | 0 | 28 | 44 | 1 | 43 | 0 | 2.27 | NC_039506.1 | 1056151 | 1064257 | 8107 | + | Gnomon | gene | gene-LOC105286747 | GeneID:105286747 | LOC105286747 | Gene | LOC105286747 | lncRNA | character(0) | NA | character(0) | BCphaseB1.vs.RphaseA2 | FALSE |
| NC_039506.1 | 1057064 | 1057064 | 1 | + | 28 | 0 | 28 | 44 | 0 | 44 | 0 | 0 | NC_039506.1 | 1056151 | 1064257 | 8107 | + | Gnomon | gene | gene-LOC105286747 | GeneID:105286747 | LOC105286747 | Gene | LOC105286747 | lncRNA | character(0) | NA | character(0) | BCphaseB1.vs.RphaseA2 | FALSE |
| NC_039506.1 | 1057110 | 1057110 | 1 | + | 31 | 1 | 30 | 55 | 0 | 55 | 3.23 | 0 | NC_039506.1 | 1056151 | 1064257 | 8107 | + | Gnomon | gene | gene-LOC105286747 | GeneID:105286747 | LOC105286747 | Gene | LOC105286747 | lncRNA | character(0) | NA | character(0) | BCphaseB1.vs.RphaseA2 | FALSE |
| NC_039506.1 | 1057112 | 1057112 | 1 | + | 37 | 0 | 37 | 53 | 0 | 53 | 0 | 0 | NC_039506.1 | 1056151 | 1064257 | 8107 | + | Gnomon | gene | gene-LOC105286747 | GeneID:105286747 | LOC105286747 | Gene | LOC105286747 | lncRNA | character(0) | NA | character(0) | BCphaseB1.vs.RphaseA2 | FALSE |
| NC_039506.1 | 1057128 | 1057128 | 1 | + | 31 | 0 | 31 | 49 | 0 | 49 | 0 | 0 | NC_039506.1 | 1056151 | 1064257 | 8107 | + | Gnomon | gene | gene-LOC105286747 | GeneID:105286747 | LOC105286747 | Gene | LOC105286747 | lncRNA | character(0) | NA | character(0) | BCphaseB1.vs.RphaseA2 | FALSE |
| NC_039506.1 | 1057134 | 1057134 | 1 | + | 31 | 0 | 31 | 48 | 2 | 46 | 0 | 4.17 | NC_039506.1 | 1056151 | 1064257 | 8107 | + | Gnomon | gene | gene-LOC105286747 | GeneID:105286747 | LOC105286747 | Gene | LOC105286747 | lncRNA | character(0) | NA | character(0) | BCphaseB1.vs.RphaseA2 | FALSE |
| NC_039506.1 | 1057145 | 1057145 | 1 | + | 33 | 0 | 33 | 47 | 0 | 47 | 0 | 0 | NC_039506.1 | 1056151 | 1064257 | 8107 | + | Gnomon | gene | gene-LOC105286747 | GeneID:105286747 | LOC105286747 | Gene | LOC105286747 | lncRNA | character(0) | NA | character(0) | BCphaseB1.vs.RphaseA2 | FALSE |
| NC_039506.1 | 1057157 | 1057157 | 1 | + | 31 | 0 | 31 | 43 | 0 | 43 | 0 | 0 | NC_039506.1 | 1056151 | 1064257 | 8107 | + | Gnomon | gene | gene-LOC105286747 | GeneID:105286747 | LOC105286747 | Gene | LOC105286747 | lncRNA | character(0) | NA | character(0) | BCphaseB1.vs.RphaseA2 | FALSE |
| NC_039506.1 | 1057169 | 1057169 | 1 | + | 31 | 0 | 31 | 38 | 2 | 36 | 0 | 5.26 | NC_039506.1 | 1056151 | 1064257 | 8107 | + | Gnomon | gene | gene-LOC105286747 | GeneID:105286747 | LOC105286747 | Gene | LOC105286747 | lncRNA | character(0) | NA | character(0) | BCphaseB1.vs.RphaseA2 | FALSE |
| NC_039506.1 | 1057181 | 1057181 | 1 | + | 30 | 0 | 30 | 31 | 0 | 31 | 0 | 0 | NC_039506.1 | 1056151 | 1064257 | 8107 | + | Gnomon | gene | gene-LOC105286747 | GeneID:105286747 | LOC105286747 | Gene | LOC105286747 | lncRNA | character(0) | NA | character(0) | BCphaseB1.vs.RphaseA2 | FALSE |
| NC_039506.1 | 1057290 | 1057290 | 1 | + | 20 | 0 | 20 | 26 | 0 | 26 | 0 | 0 | NC_039506.1 | 1056151 | 1064257 | 8107 | + | Gnomon | gene | gene-LOC105286747 | GeneID:105286747 | LOC105286747 | Gene | LOC105286747 | lncRNA | character(0) | NA | character(0) | BCphaseB1.vs.RphaseA2 | FALSE |
| NC_039506.1 | 1057294 | 1057294 | 1 | + | 20 | 1 | 19 | 29 | 0 | 29 | 5 | 0 | NC_039506.1 | 1056151 | 1064257 | 8107 | + | Gnomon | gene | gene-LOC105286747 | GeneID:105286747 | LOC105286747 | Gene | LOC105286747 | lncRNA | character(0) | NA | character(0) | BCphaseB1.vs.RphaseA2 | FALSE |
| NC_039506.1 | 1057300 | 1057300 | 1 | + | 20 | 1 | 19 | 29 | 1 | 28 | 5 | 3.45 | NC_039506.1 | 1056151 | 1064257 | 8107 | + | Gnomon | gene | gene-LOC105286747 | GeneID:105286747 | LOC105286747 | Gene | LOC105286747 | lncRNA | character(0) | NA | character(0) | BCphaseB1.vs.RphaseA2 | FALSE |
| NC_039506.1 | 1057302 | 1057302 | 1 | + | 20 | 1 | 19 | 30 | 0 | 30 | 5 | 0 | NC_039506.1 | 1056151 | 1064257 | 8107 | + | Gnomon | gene | gene-LOC105286747 | GeneID:105286747 | LOC105286747 | Gene | LOC105286747 | lncRNA | character(0) | NA | character(0) | BCphaseB1.vs.RphaseA2 | FALSE |
| NC_039506.1 | 1057308 | 1057308 | 1 | + | 23 | 1 | 22 | 31 | 0 | 31 | 4.35 | 0 | NC_039506.1 | 1056151 | 1064257 | 8107 | + | Gnomon | gene | gene-LOC105286747 | GeneID:105286747 | LOC105286747 | Gene | LOC105286747 | lncRNA | character(0) | NA | character(0) | BCphaseB1.vs.RphaseA2 | FALSE |
| NC_039506.1 | 1057314 | 1057314 | 1 | + | 47 | 0 | 47 | 64 | 1 | 63 | 0 | 1.56 | NC_039506.1 | 1056151 | 1064257 | 8107 | + | Gnomon | gene | gene-LOC105286747 | GeneID:105286747 | LOC105286747 | Gene | LOC105286747 | lncRNA | character(0) | NA | character(0) | BCphaseB1.vs.RphaseA2 | FALSE |
| NC_039506.1 | 1057316 | 1057316 | 1 | + | 47 | 0 | 47 | 66 | 0 | 66 | 0 | 0 | NC_039506.1 | 1056151 | 1064257 | 8107 | + | Gnomon | gene | gene-LOC105286747 | GeneID:105286747 | LOC105286747 | Gene | LOC105286747 | lncRNA | character(0) | NA | character(0) | BCphaseB1.vs.RphaseA2 | FALSE |
| NC_039506.1 | 1057320 | 1057320 | 1 | + | 48 | 1 | 47 | 70 | 0 | 70 | 2.08 | 0 | NC_039506.1 | 1056151 | 1064257 | 8107 | + | Gnomon | gene | gene-LOC105286747 | GeneID:105286747 | LOC105286747 | Gene | LOC105286747 | lncRNA | character(0) | NA | character(0) | BCphaseB1.vs.RphaseA2 | FALSE |
| NC_039506.1 | 1057331 | 1057331 | 1 | + | 78 | 0 | 78 | 109 | 2 | 107 | 0 | 1.83 | NC_039506.1 | 1056151 | 1064257 | 8107 | + | Gnomon | gene | gene-LOC105286747 | GeneID:105286747 | LOC105286747 | Gene | LOC105286747 | lncRNA | character(0) | NA | character(0) | BCphaseB1.vs.RphaseA2 | FALSE |
| NC_039506.1 | 1057337 | 1057337 | 1 | + | 76 | 0 | 76 | 107 | 0 | 107 | 0 | 0 | NC_039506.1 | 1056151 | 1064257 | 8107 | + | Gnomon | gene | gene-LOC105286747 | GeneID:105286747 | LOC105286747 | Gene | LOC105286747 | lncRNA | character(0) | NA | character(0) | BCphaseB1.vs.RphaseA2 | FALSE |
| NC_039506.1 | 1057344 | 1057344 | 1 | + | 70 | 0 | 70 | 100 | 0 | 100 | 0 | 0 | NC_039506.1 | 1056151 | 1064257 | 8107 | + | Gnomon | gene | gene-LOC105286747 | GeneID:105286747 | LOC105286747 | Gene | LOC105286747 | lncRNA | character(0) | NA | character(0) | BCphaseB1.vs.RphaseA2 | FALSE |

...

... only first 25 lines shown ...

Directory: DMSG File: dmg-Ob_LI_BCphaseB1.vs.RphaseB1_details.txt

| seqnames | start | end | width | strand | coverage1 | numCs1 | numTs1 | coverage2 | numCs2 | numTs2 | BCphaseB1 | RphaseB1 | gene_seqnames | gene_start | gene_end | gene_width | gene_strand | gene_source | gene_type | gene_ID | gene_Dbxref | gene_Name | gene_gbkey | gene_gene | gene_gene_biotype | gene_end_range | gene_start_range | gene_comparison | is.dm |
|---|---|---|---|---|---|---|---|---|---|---|---|---|---|---|---|---|---|---|---|---|---|---|---|---|---|---|---|---|---|
| NC_039506.1 | 2980627 | 2980627 | 1 | + | 21 | 1 | 20 | 22 | 0 | 22 | 4.76 | 0 | NC_039506.1 | 2980439 | 2984235 | 3797 | + | Gnomon | gene | gene-LOC105280348 | GeneID:105280348 | LOC105280348 | Gene | LOC105280348 | protein_coding | character(0) | character(0) | BCphaseB1.vs.RphaseB1 | FALSE |
| NC_039506.1 | 2980668 | 2980668 | 1 | + | 28 | 0 | 28 | 29 | 0 | 29 | 0 | 0 | NC_039506.1 | 2980439 | 2984235 | 3797 | + | Gnomon | gene | gene-LOC105280348 | GeneID:105280348 | LOC105280348 | Gene | LOC105280348 | protein_coding | character(0) | character(0) | BCphaseB1.vs.RphaseB1 | FALSE |
| NC_039506.1 | 2980710 | 2980710 | 1 | + | 21 | 0 | 21 | 20 | 1 | 19 | 0 | 5 | NC_039506.1 | 2980439 | 2984235 | 3797 | + | Gnomon | gene | gene-LOC105280348 | GeneID:105280348 | LOC105280348 | Gene | LOC105280348 | protein_coding | character(0) | character(0) | BCphaseB1.vs.RphaseB1 | FALSE |
| NC_039506.1 | 2980750 | 2980750 | 1 | + | 29 | 0 | 29 | 33 | 0 | 33 | 0 | 0 | NC_039506.1 | 2980439 | 2984235 | 3797 | + | Gnomon | gene | gene-LOC105280348 | GeneID:105280348 | LOC105280348 | Gene | LOC105280348 | protein_coding | character(0) | character(0) | BCphaseB1.vs.RphaseB1 | FALSE |
| NC_039506.1 | 2980761 | 2980761 | 1 | + | 32 | 0 | 32 | 34 | 0 | 34 | 0 | 0 | NC_039506.1 | 2980439 | 2984235 | 3797 | + | Gnomon | gene | gene-LOC105280348 | GeneID:105280348 | LOC105280348 | Gene | LOC105280348 | protein_coding | character(0) | character(0) | BCphaseB1.vs.RphaseB1 | FALSE |
| NC_039506.1 | 2980767 | 2980767 | 1 | + | 34 | 0 | 34 | 38 | 0 | 38 | 0 | 0 | NC_039506.1 | 2980439 | 2984235 | 3797 | + | Gnomon | gene | gene-LOC105280348 | GeneID:105280348 | LOC105280348 | Gene | LOC105280348 | protein_coding | character(0) | character(0) | BCphaseB1.vs.RphaseB1 | FALSE |
| NC_039506.1 | 2980928 | 2980928 | 1 | + | 23 | 8 | 15 | 21 | 4 | 17 | 34.78 | 19.05 | NC_039506.1 | 2980439 | 2984235 | 3797 | + | Gnomon | gene | gene-LOC105280348 | GeneID:105280348 | LOC105280348 | Gene | LOC105280348 | protein_coding | character(0) | character(0) | BCphaseB1.vs.RphaseB1 | FALSE |
| NC_039506.1 | 2980989 | 2980989 | 1 | + | 29 | 3 | 26 | 32 | 2 | 30 | 10.34 | 6.25 | NC_039506.1 | 2980439 | 2984235 | 3797 | + | Gnomon | gene | gene-LOC105280348 | GeneID:105280348 | LOC105280348 | Gene | LOC105280348 | protein_coding | character(0) | character(0) | BCphaseB1.vs.RphaseB1 | FALSE |
| NC_039506.1 | 2980993 | 2980993 | 1 | + | 29 | 12 | 17 | 32 | 16 | 16 | 41.38 | 50 | NC_039506.1 | 2980439 | 2984235 | 3797 | + | Gnomon | gene | gene-LOC105280348 | GeneID:105280348 | LOC105280348 | Gene | LOC105280348 | protein_coding | character(0) | character(0) | BCphaseB1.vs.RphaseB1 | FALSE |
| NC_039506.1 | 2980998 | 2980998 | 1 | + | 28 | 8 | 20 | 31 | 13 | 18 | 28.57 | 41.94 | NC_039506.1 | 2980439 | 2984235 | 3797 | + | Gnomon | gene | gene-LOC105280348 | GeneID:105280348 | LOC105280348 | Gene | LOC105280348 | protein_coding | character(0) | character(0) | BCphaseB1.vs.RphaseB1 | FALSE |
| NC_039506.1 | 2981030 | 2981030 | 1 | + | 56 | 50 | 6 | 61 | 54 | 7 | 89.29 | 88.52 | NC_039506.1 | 2980439 | 2984235 | 3797 | + | Gnomon | gene | gene-LOC105280348 | GeneID:105280348 | LOC105280348 | Gene | LOC105280348 | protein_coding | character(0) | character(0) | BCphaseB1.vs.RphaseB1 | FALSE |
| NC_039506.1 | 2981077 | 2981077 | 1 | + | 49 | 25 | 24 | 43 | 27 | 16 | 51.02 | 62.79 | NC_039506.1 | 2980439 | 2984235 | 3797 | + | Gnomon | gene | gene-LOC105280348 | GeneID:105280348 | LOC105280348 | Gene | LOC105280348 | protein_coding | character(0) | character(0) | BCphaseB1.vs.RphaseB1 | FALSE |
| NC_039506.1 | 2981125 | 2981125 | 1 | + | 32 | 19 | 13 | 24 | 11 | 13 | 59.38 | 45.83 | NC_039506.1 | 2980439 | 2984235 | 3797 | + | Gnomon | gene | gene-LOC105280348 | GeneID:105280348 | LOC105280348 | Gene | LOC105280348 | protein_coding | character(0) | character(0) | BCphaseB1.vs.RphaseB1 | FALSE |
| NC_039506.1 | 2981183 | 2981183 | 1 | + | 83 | 78 | 5 | 51 | 24 | 27 | 93.98 | 47.06 | NC_039506.1 | 2980439 | 2984235 | 3797 | + | Gnomon | gene | gene-LOC105280348 | GeneID:105280348 | LOC105280348 | Gene | LOC105280348 | protein_coding | character(0) | character(0) | BCphaseB1.vs.RphaseB1 | TRUE |
| NC_039506.1 | 2981225 | 2981225 | 1 | + | 37 | 23 | 14 | 27 | 16 | 11 | 62.16 | 59.26 | NC_039506.1 | 2980439 | 2984235 | 3797 | + | Gnomon | gene | gene-LOC105280348 | GeneID:105280348 | LOC105280348 | Gene | LOC105280348 | protein_coding | character(0) | character(0) | BCphaseB1.vs.RphaseB1 | FALSE |
| NC_039506.1 | 2981245 | 2981245 | 1 | + | 40 | 18 | 22 | 27 | 13 | 14 | 45 | 48.15 | NC_039506.1 | 2980439 | 2984235 | 3797 | + | Gnomon | gene | gene-LOC105280348 | GeneID:105280348 | LOC105280348 | Gene | LOC105280348 | protein_coding | character(0) | character(0) | BCphaseB1.vs.RphaseB1 | FALSE |
| NC_039506.1 | 2981292 | 2981292 | 1 | + | 68 | 64 | 4 | 57 | 51 | 6 | 94.12 | 89.47 | NC_039506.1 | 2980439 | 2984235 | 3797 | + | Gnomon | gene | gene-LOC105280348 | GeneID:105280348 | LOC105280348 | Gene | LOC105280348 | protein_coding | character(0) | character(0) | BCphaseB1.vs.RphaseB1 | FALSE |
| NC_039506.1 | 2981308 | 2981308 | 1 | + | 78 | 52 | 26 | 64 | 36 | 28 | 66.67 | 56.25 | NC_039506.1 | 2980439 | 2984235 | 3797 | + | Gnomon | gene | gene-LOC105280348 | GeneID:105280348 | LOC105280348 | Gene | LOC105280348 | protein_coding | character(0) | character(0) | BCphaseB1.vs.RphaseB1 | FALSE |
| NC_039506.1 | 2981315 | 2981315 | 1 | + | 77 | 69 | 8 | 67 | 50 | 17 | 89.61 | 74.63 | NC_039506.1 | 2980439 | 2984235 | 3797 | + | Gnomon | gene | gene-LOC105280348 | GeneID:105280348 | LOC105280348 | Gene | LOC105280348 | protein_coding | character(0) | character(0) | BCphaseB1.vs.RphaseB1 | FALSE |
| NC_039506.1 | 2981335 | 2981335 | 1 | + | 56 | 54 | 2 | 65 | 63 | 2 | 96.43 | 96.92 | NC_039506.1 | 2980439 | 2984235 | 3797 | + | Gnomon | gene | gene-LOC105280348 | GeneID:105280348 | LOC105280348 | Gene | LOC105280348 | protein_coding | character(0) | character(0) | BCphaseB1.vs.RphaseB1 | FALSE |
| NC_039506.1 | 2981350 | 2981350 | 1 | + | 58 | 57 | 1 | 55 | 55 | 0 | 98.28 | 100 | NC_039506.1 | 2980439 | 2984235 | 3797 | + | Gnomon | gene | gene-LOC105280348 | GeneID:105280348 | LOC105280348 | Gene | LOC105280348 | protein_coding | character(0) | character(0) | BCphaseB1.vs.RphaseB1 | FALSE |
| NC_039506.1 | 2981391 | 2981391 | 1 | + | 60 | 49 | 11 | 65 | 53 | 12 | 81.67 | 81.54 | NC_039506.1 | 2980439 | 2984235 | 3797 | + | Gnomon | gene | gene-LOC105280348 | GeneID:105280348 | LOC105280348 | Gene | LOC105280348 | protein_coding | character(0) | character(0) | BCphaseB1.vs.RphaseB1 | FALSE |
| NC_039506.1 | 2981402 | 2981402 | 1 | + | 66 | 60 | 6 | 74 | 67 | 7 | 90.91 | 90.54 | NC_039506.1 | 2980439 | 2984235 | 3797 | + | Gnomon | gene | gene-LOC105280348 | GeneID:105280348 | LOC105280348 | Gene | LOC105280348 | protein_coding | character(0) | character(0) | BCphaseB1.vs.RphaseB1 | FALSE |
| NC_039506.1 | 2981411 | 2981411 | 1 | + | 68 | 65 | 3 | 81 | 71 | 10 | 95.59 | 87.65 | NC_039506.1 | 2980439 | 2984235 | 3797 | + | Gnomon | gene | gene-LOC105280348 | GeneID:105280348 | LOC105280348 | Gene | LOC105280348 | protein_coding | character(0) | character(0) | BCphaseB1.vs.RphaseB1 | FALSE |

...

... only first 25 lines shown ...

Directory: DMSG File: dmg-Ob_LI_BCphaseB1.vs.RphaseB2_details.txt

| seqnames | start | end | width | strand | coverage1 | numCs1 | numTs1 | coverage2 | numCs2 | numTs2 | BCphaseB1 | RphaseB2 | gene_seqnames | gene_start | gene_end | gene_width | gene_strand | gene_source | gene_type | gene_ID | gene_Dbxref | gene_Name | gene_gbkey | gene_gene | gene_gene_biotype | gene_end_range | gene_start_range | gene_comparison | is.dm |
|---|---|---|---|---|---|---|---|---|---|---|---|---|---|---|---|---|---|---|---|---|---|---|---|---|---|---|---|---|---|
| NC_039506.1 | 3248474 | 3248474 | 1 | + | 49 | 13 | 36 | 51 | 14 | 37 | 26.53 | 27.45 | NC_039506.1 | 3248415 | 3250438 | 2024 | - | Gnomon | gene | gene-LOC105280377 | GeneID:105280377 | LOC105280377 | Gene | LOC105280377 | protein_coding | character(0) | character(0) | BCphaseB1.vs.RphaseB2 | FALSE |
| NC_039506.1 | 3248483 | 3248483 | 1 | + | 71 | 32 | 39 | 57 | 27 | 30 | 45.07 | 47.37 | NC_039506.1 | 3248415 | 3250438 | 2024 | - | Gnomon | gene | gene-LOC105280377 | GeneID:105280377 | LOC105280377 | Gene | LOC105280377 | protein_coding | character(0) | character(0) | BCphaseB1.vs.RphaseB2 | FALSE |
| NC_039506.1 | 3248495 | 3248495 | 1 | + | 75 | 13 | 62 | 65 | 13 | 52 | 17.33 | 20 | NC_039506.1 | 3248415 | 3250438 | 2024 | - | Gnomon | gene | gene-LOC105280377 | GeneID:105280377 | LOC105280377 | Gene | LOC105280377 | protein_coding | character(0) | character(0) | BCphaseB1.vs.RphaseB2 | FALSE |
| NC_039506.1 | 3248534 | 3248534 | 1 | + | 76 | 20 | 56 | 69 | 22 | 47 | 26.32 | 31.88 | NC_039506.1 | 3248415 | 3250438 | 2024 | - | Gnomon | gene | gene-LOC105280377 | GeneID:105280377 | LOC105280377 | Gene | LOC105280377 | protein_coding | character(0) | character(0) | BCphaseB1.vs.RphaseB2 | FALSE |
| NC_039506.1 | 3248539 | 3248539 | 1 | + | 64 | 4 | 60 | 63 | 5 | 58 | 6.25 | 7.94 | NC_039506.1 | 3248415 | 3250438 | 2024 | - | Gnomon | gene | gene-LOC105280377 | GeneID:105280377 | LOC105280377 | Gene | LOC105280377 | protein_coding | character(0) | character(0) | BCphaseB1.vs.RphaseB2 | FALSE |
| NC_039506.1 | 3248551 | 3248551 | 1 | + | 58 | 5 | 53 | 48 | 3 | 45 | 8.62 | 6.25 | NC_039506.1 | 3248415 | 3250438 | 2024 | - | Gnomon | gene | gene-LOC105280377 | GeneID:105280377 | LOC105280377 | Gene | LOC105280377 | protein_coding | character(0) | character(0) | BCphaseB1.vs.RphaseB2 | FALSE |
| NC_039506.1 | 3248553 | 3248553 | 1 | + | 56 | 0 | 56 | 44 | 0 | 44 | 0 | 0 | NC_039506.1 | 3248415 | 3250438 | 2024 | - | Gnomon | gene | gene-LOC105280377 | GeneID:105280377 | LOC105280377 | Gene | LOC105280377 | protein_coding | character(0) | character(0) | BCphaseB1.vs.RphaseB2 | FALSE |
| NC_039506.1 | 3248556 | 3248556 | 1 | + | 52 | 3 | 49 | 46 | 8 | 38 | 5.77 | 17.39 | NC_039506.1 | 3248415 | 3250438 | 2024 | - | Gnomon | gene | gene-LOC105280377 | GeneID:105280377 | LOC105280377 | Gene | LOC105280377 | protein_coding | character(0) | character(0) | BCphaseB1.vs.RphaseB2 | FALSE |
| NC_039506.1 | 3248560 | 3248560 | 1 | + | 49 | 2 | 47 | 21 | 1 | 20 | 4.08 | 4.76 | NC_039506.1 | 3248415 | 3250438 | 2024 | - | Gnomon | gene | gene-LOC105280377 | GeneID:105280377 | LOC105280377 | Gene | LOC105280377 | protein_coding | character(0) | character(0) | BCphaseB1.vs.RphaseB2 | FALSE |
| NC_039506.1 | 3248578 | 3248578 | 1 | + | 46 | 9 | 37 | 20 | 1 | 19 | 19.57 | 5 | NC_039506.1 | 3248415 | 3250438 | 2024 | - | Gnomon | gene | gene-LOC105280377 | GeneID:105280377 | LOC105280377 | Gene | LOC105280377 | protein_coding | character(0) | character(0) | BCphaseB1.vs.RphaseB2 | FALSE |
| NC_039506.1 | 3248598 | 3248598 | 1 | + | 28 | 4 | 24 | 24 | 2 | 22 | 14.29 | 8.33 | NC_039506.1 | 3248415 | 3250438 | 2024 | - | Gnomon | gene | gene-LOC105280377 | GeneID:105280377 | LOC105280377 | Gene | LOC105280377 | protein_coding | character(0) | character(0) | BCphaseB1.vs.RphaseB2 | FALSE |
| NC_039506.1 | 3248631 | 3248631 | 1 | + | 39 | 1 | 38 | 38 | 2 | 36 | 2.56 | 5.26 | NC_039506.1 | 3248415 | 3250438 | 2024 | - | Gnomon | gene | gene-LOC105280377 | GeneID:105280377 | LOC105280377 | Gene | LOC105280377 | protein_coding | character(0) | character(0) | BCphaseB1.vs.RphaseB2 | FALSE |
| NC_039506.1 | 3248698 | 3248698 | 1 | + | 53 | 4 | 49 | 52 | 3 | 49 | 7.55 | 5.77 | NC_039506.1 | 3248415 | 3250438 | 2024 | - | Gnomon | gene | gene-LOC105280377 | GeneID:105280377 | LOC105280377 | Gene | LOC105280377 | protein_coding | character(0) | character(0) | BCphaseB1.vs.RphaseB2 | FALSE |
| NC_039506.1 | 3248700 | 3248700 | 1 | + | 53 | 0 | 53 | 52 | 1 | 51 | 0 | 1.92 | NC_039506.1 | 3248415 | 3250438 | 2024 | - | Gnomon | gene | gene-LOC105280377 | GeneID:105280377 | LOC105280377 | Gene | LOC105280377 | protein_coding | character(0) | character(0) | BCphaseB1.vs.RphaseB2 | FALSE |
| NC_039506.1 | 3248715 | 3248715 | 1 | + | 53 | 6 | 47 | 26 | 1 | 25 | 11.32 | 3.85 | NC_039506.1 | 3248415 | 3250438 | 2024 | - | Gnomon | gene | gene-LOC105280377 | GeneID:105280377 | LOC105280377 | Gene | LOC105280377 | protein_coding | character(0) | character(0) | BCphaseB1.vs.RphaseB2 | FALSE |
| NC_039506.1 | 3248724 | 3248724 | 1 | + | 57 | 1 | 56 | 21 | 1 | 20 | 1.75 | 4.76 | NC_039506.1 | 3248415 | 3250438 | 2024 | - | Gnomon | gene | gene-LOC105280377 | GeneID:105280377 | LOC105280377 | Gene | LOC105280377 | protein_coding | character(0) | character(0) | BCphaseB1.vs.RphaseB2 | FALSE |
| NC_039506.1 | 3248733 | 3248733 | 1 | + | 65 | 1 | 64 | 26 | 0 | 26 | 1.54 | 0 | NC_039506.1 | 3248415 | 3250438 | 2024 | - | Gnomon | gene | gene-LOC105280377 | GeneID:105280377 | LOC105280377 | Gene | LOC105280377 | protein_coding | character(0) | character(0) | BCphaseB1.vs.RphaseB2 | FALSE |
| NC_039506.1 | 3248748 | 3248748 | 1 | + | 62 | 2 | 60 | 24 | 0 | 24 | 3.23 | 0 | NC_039506.1 | 3248415 | 3250438 | 2024 | - | Gnomon | gene | gene-LOC105280377 | GeneID:105280377 | LOC105280377 | Gene | LOC105280377 | protein_coding | character(0) | character(0) | BCphaseB1.vs.RphaseB2 | FALSE |
| NC_039506.1 | 3248750 | 3248750 | 1 | + | 62 | 2 | 60 | 24 | 0 | 24 | 3.23 | 0 | NC_039506.1 | 3248415 | 3250438 | 2024 | - | Gnomon | gene | gene-LOC105280377 | GeneID:105280377 | LOC105280377 | Gene | LOC105280377 | protein_coding | character(0) | character(0) | BCphaseB1.vs.RphaseB2 | FALSE |
| NC_039506.1 | 3248755 | 3248755 | 1 | + | 57 | 0 | 57 | 23 | 0 | 23 | 0 | 0 | NC_039506.1 | 3248415 | 3250438 | 2024 | - | Gnomon | gene | gene-LOC105280377 | GeneID:105280377 | LOC105280377 | Gene | LOC105280377 | protein_coding | character(0) | character(0) | BCphaseB1.vs.RphaseB2 | FALSE |
| NC_039506.1 | 3248760 | 3248760 | 1 | + | 65 | 3 | 62 | 23 | 1 | 22 | 4.62 | 4.35 | NC_039506.1 | 3248415 | 3250438 | 2024 | - | Gnomon | gene | gene-LOC105280377 | GeneID:105280377 | LOC105280377 | Gene | LOC105280377 | protein_coding | character(0) | character(0) | BCphaseB1.vs.RphaseB2 | FALSE |
| NC_039506.1 | 3248793 | 3248793 | 1 | + | 76 | 18 | 58 | 35 | 6 | 29 | 23.68 | 17.14 | NC_039506.1 | 3248415 | 3250438 | 2024 | - | Gnomon | gene | gene-LOC105280377 | GeneID:105280377 | LOC105280377 | Gene | LOC105280377 | protein_coding | character(0) | character(0) | BCphaseB1.vs.RphaseB2 | FALSE |
| NC_039506.1 | 3248801 | 3248801 | 1 | + | 79 | 8 | 71 | 55 | 11 | 44 | 10.13 | 20 | NC_039506.1 | 3248415 | 3250438 | 2024 | - | Gnomon | gene | gene-LOC105280377 | GeneID:105280377 | LOC105280377 | Gene | LOC105280377 | protein_coding | character(0) | character(0) | BCphaseB1.vs.RphaseB2 | FALSE |
| NC_039506.1 | 3248827 | 3248827 | 1 | + | 85 | 2 | 83 | 69 | 12 | 57 | 2.35 | 17.39 | NC_039506.1 | 3248415 | 3250438 | 2024 | - | Gnomon | gene | gene-LOC105280377 | GeneID:105280377 | LOC105280377 | Gene | LOC105280377 | protein_coding | character(0) | character(0) | BCphaseB1.vs.RphaseB2 | FALSE |

...
... only first 25 lines shown ...

Directory: DMSG File: dmg-Ob_LI_BCphaseB2.vs.RphaseA1_details.txt

| seqnames | start | end | width | strand | coverage1 | numCs1 | numTs1 | coverage2 | numCs2 | numTs2 | BCphaseB2 | RphaseA1 | gene_seqnames | gene_start | gene_end | gene_width | gene_strand | gene_source | gene_type | gene_ID | gene_Dbxref | gene_Name | gene_gbkey | gene_gene | gene_gene_biotype | gene_end_range | gene_start_range | gene_comparison | is.dm |
|---|---|---|---|---|---|---|---|---|---|---|---|---|---|---|---|---|---|---|---|---|---|---|---|---|---|---|---|---|---|
| NC_039506.1 | 2980668 | 2980668 | 1 | + | 29 | 1 | 28 | 21 | 2 | 19 | 3.45 | 9.52 | NC_039506.1 | 2980439 | 2984235 | 3797 | + | Gnomon | gene | gene-LOC105280348 | GeneID:105280348 | LOC105280348 | Gene | LOC105280348 | protein_coding | character(0) | character(0) | BCphaseB2.vs.RphaseA1 | FALSE |
| NC_039506.1 | 2980750 | 2980750 | 1 | + | 32 | 0 | 32 | 28 | 0 | 28 | 0 | 0 | NC_039506.1 | 2980439 | 2984235 | 3797 | + | Gnomon | gene | gene-LOC105280348 | GeneID:105280348 | LOC105280348 | Gene | LOC105280348 | protein_coding | character(0) | character(0) | BCphaseB2.vs.RphaseA1 | FALSE |
| NC_039506.1 | 2980761 | 2980761 | 1 | + | 28 | 0 | 28 | 31 | 0 | 31 | 0 | 0 | NC_039506.1 | 2980439 | 2984235 | 3797 | + | Gnomon | gene | gene-LOC105280348 | GeneID:105280348 | LOC105280348 | Gene | LOC105280348 | protein_coding | character(0) | character(0) | BCphaseB2.vs.RphaseA1 | FALSE |
| NC_039506.1 | 2980767 | 2980767 | 1 | + | 28 | 0 | 28 | 32 | 1 | 31 | 0 | 3.12 | NC_039506.1 | 2980439 | 2984235 | 3797 | + | Gnomon | gene | gene-LOC105280348 | GeneID:105280348 | LOC105280348 | Gene | LOC105280348 | protein_coding | character(0) | character(0) | BCphaseB2.vs.RphaseA1 | FALSE |
| NC_039506.1 | 2980928 | 2980928 | 1 | + | 23 | 6 | 17 | 20 | 8 | 12 | 26.09 | 40 | NC_039506.1 | 2980439 | 2984235 | 3797 | + | Gnomon | gene | gene-LOC105280348 | GeneID:105280348 | LOC105280348 | Gene | LOC105280348 | protein_coding | character(0) | character(0) | BCphaseB2.vs.RphaseA1 | FALSE |
| NC_039506.1 | 2980989 | 2980989 | 1 | + | 29 | 7 | 22 | 28 | 3 | 25 | 24.14 | 10.71 | NC_039506.1 | 2980439 | 2984235 | 3797 | + | Gnomon | gene | gene-LOC105280348 | GeneID:105280348 | LOC105280348 | Gene | LOC105280348 | protein_coding | character(0) | character(0) | BCphaseB2.vs.RphaseA1 | FALSE |
| NC_039506.1 | 2980993 | 2980993 | 1 | + | 29 | 10 | 19 | 29 | 15 | 14 | 34.48 | 51.72 | NC_039506.1 | 2980439 | 2984235 | 3797 | + | Gnomon | gene | gene-LOC105280348 | GeneID:105280348 | LOC105280348 | Gene | LOC105280348 | protein_coding | character(0) | character(0) | BCphaseB2.vs.RphaseA1 | FALSE |
| NC_039506.1 | 2980998 | 2980998 | 1 | + | 30 | 11 | 19 | 28 | 7 | 21 | 36.67 | 25 | NC_039506.1 | 2980439 | 2984235 | 3797 | + | Gnomon | gene | gene-LOC105280348 | GeneID:105280348 | LOC105280348 | Gene | LOC105280348 | protein_coding | character(0) | character(0) | BCphaseB2.vs.RphaseA1 | FALSE |
| NC_039506.1 | 2981030 | 2981030 | 1 | + | 43 | 35 | 8 | 28 | 25 | 3 | 81.4 | 89.29 | NC_039506.1 | 2980439 | 2984235 | 3797 | + | Gnomon | gene | gene-LOC105280348 | GeneID:105280348 | LOC105280348 | Gene | LOC105280348 | protein_coding | character(0) | character(0) | BCphaseB2.vs.RphaseA1 | FALSE |
| NC_039506.1 | 2981077 | 2981077 | 1 | + | 36 | 13 | 23 | 30 | 11 | 19 | 36.11 | 36.67 | NC_039506.1 | 2980439 | 2984235 | 3797 | + | Gnomon | gene | gene-LOC105280348 | GeneID:105280348 | LOC105280348 | Gene | LOC105280348 | protein_coding | character(0) | character(0) | BCphaseB2.vs.RphaseA1 | FALSE |
| NC_039506.1 | 2981183 | 2981183 | 1 | + | 49 | 25 | 24 | 60 | 59 | 1 | 51.02 | 98.33 | NC_039506.1 | 2980439 | 2984235 | 3797 | + | Gnomon | gene | gene-LOC105280348 | GeneID:105280348 | LOC105280348 | Gene | LOC105280348 | protein_coding | character(0) | character(0) | BCphaseB2.vs.RphaseA1 | TRUE |
| NC_039506.1 | 2981225 | 2981225 | 1 | + | 32 | 24 | 8 | 46 | 29 | 17 | 75 | 63.04 | NC_039506.1 | 2980439 | 2984235 | 3797 | + | Gnomon | gene | gene-LOC105280348 | GeneID:105280348 | LOC105280348 | Gene | LOC105280348 | protein_coding | character(0) | character(0) | BCphaseB2.vs.RphaseA1 | FALSE |
| NC_039506.1 | 2981245 | 2981245 | 1 | + | 38 | 14 | 24 | 38 | 15 | 23 | 36.84 | 39.47 | NC_039506.1 | 2980439 | 2984235 | 3797 | + | Gnomon | gene | gene-LOC105280348 | GeneID:105280348 | LOC105280348 | Gene | LOC105280348 | protein_coding | character(0) | character(0) | BCphaseB2.vs.RphaseA1 | FALSE |
| NC_039506.1 | 2981292 | 2981292 | 1 | + | 42 | 42 | 0 | 42 | 38 | 4 | 100 | 90.48 | NC_039506.1 | 2980439 | 2984235 | 3797 | + | Gnomon | gene | gene-LOC105280348 | GeneID:105280348 | LOC105280348 | Gene | LOC105280348 | protein_coding | character(0) | character(0) | BCphaseB2.vs.RphaseA1 | FALSE |
| NC_039506.1 | 2981308 | 2981308 | 1 | + | 49 | 34 | 15 | 73 | 49 | 24 | 69.39 | 67.12 | NC_039506.1 | 2980439 | 2984235 | 3797 | + | Gnomon | gene | gene-LOC105280348 | GeneID:105280348 | LOC105280348 | Gene | LOC105280348 | protein_coding | character(0) | character(0) | BCphaseB2.vs.RphaseA1 | FALSE |
| NC_039506.1 | 2981315 | 2981315 | 1 | + | 49 | 40 | 9 | 52 | 48 | 4 | 81.63 | 92.31 | NC_039506.1 | 2980439 | 2984235 | 3797 | + | Gnomon | gene | gene-LOC105280348 | GeneID:105280348 | LOC105280348 | Gene | LOC105280348 | protein_coding | character(0) | character(0) | BCphaseB2.vs.RphaseA1 | FALSE |
| NC_039506.1 | 2981335 | 2981335 | 1 | + | 41 | 39 | 2 | 40 | 38 | 2 | 95.12 | 95 | NC_039506.1 | 2980439 | 2984235 | 3797 | + | Gnomon | gene | gene-LOC105280348 | GeneID:105280348 | LOC105280348 | Gene | LOC105280348 | protein_coding | character(0) | character(0) | BCphaseB2.vs.RphaseA1 | FALSE |
| NC_039506.1 | 2981350 | 2981350 | 1 | + | 72 | 68 | 4 | 52 | 51 | 1 | 94.44 | 98.08 | NC_039506.1 | 2980439 | 2984235 | 3797 | + | Gnomon | gene | gene-LOC105280348 | GeneID:105280348 | LOC105280348 | Gene | LOC105280348 | protein_coding | character(0) | character(0) | BCphaseB2.vs.RphaseA1 | FALSE |
| NC_039506.1 | 2981391 | 2981391 | 1 | + | 87 | 69 | 18 | 60 | 49 | 11 | 79.31 | 81.67 | NC_039506.1 | 2980439 | 2984235 | 3797 | + | Gnomon | gene | gene-LOC105280348 | GeneID:105280348 | LOC105280348 | Gene | LOC105280348 | protein_coding | character(0) | character(0) | BCphaseB2.vs.RphaseA1 | FALSE |
| NC_039506.1 | 2981402 | 2981402 | 1 | + | 88 | 85 | 3 | 73 | 68 | 5 | 96.59 | 93.15 | NC_039506.1 | 2980439 | 2984235 | 3797 | + | Gnomon | gene | gene-LOC105280348 | GeneID:105280348 | LOC105280348 | Gene | LOC105280348 | protein_coding | character(0) | character(0) | BCphaseB2.vs.RphaseA1 | FALSE |
| NC_039506.1 | 2981411 | 2981411 | 1 | + | 101 | 94 | 7 | 73 | 70 | 3 | 93.07 | 95.89 | NC_039506.1 | 2980439 | 2984235 | 3797 | + | Gnomon | gene | gene-LOC105280348 | GeneID:105280348 | LOC105280348 | Gene | LOC105280348 | protein_coding | character(0) | character(0) | BCphaseB2.vs.RphaseA1 | FALSE |
| NC_039506.1 | 2981802 | 2981802 | 1 | + | 24 | 9 | 15 | 26 | 9 | 17 | 37.5 | 34.62 | NC_039506.1 | 2980439 | 2984235 | 3797 | + | Gnomon | gene | gene-LOC105280348 | GeneID:105280348 | LOC105280348 | Gene | LOC105280348 | protein_coding | character(0) | character(0) | BCphaseB2.vs.RphaseA1 | FALSE |
| NC_039506.1 | 2981975 | 2981975 | 1 | + | 22 | 9 | 13 | 29 | 14 | 15 | 40.91 | 48.28 | NC_039506.1 | 2980439 | 2984235 | 3797 | + | Gnomon | gene | gene-LOC105280348 | GeneID:105280348 | LOC105280348 | Gene | LOC105280348 | protein_coding | character(0) | character(0) | BCphaseB2.vs.RphaseA1 | FALSE |
| NC_039506.1 | 2982097 | 2982097 | 1 | + | 22 | 0 | 22 | 35 | 0 | 35 | 0 | 0 | NC_039506.1 | 2980439 | 2984235 | 3797 | + | Gnomon | gene | gene-LOC105280348 | GeneID:105280348 | LOC105280348 | Gene | LOC105280348 | protein_coding | character(0) | character(0) | BCphaseB2.vs.RphaseA1 | FALSE |

...

... only first 25 lines shown ...

Directory: DMSG File: dmg-Ob_LI_BCphaseB2.vs.RphaseA2_details.txt

| seqnames | start | end | width | strand | coverage1 | numCs1 | numTs1 | coverage2 | numCs2 | numTs2 | BCphaseB2 | RphaseA2 | gene_seqnames | gene_start | gene_end | gene_width | gene_strand | gene_source | gene_type | gene_ID | gene_Dbxref | gene_Name | gene_gbkey | gene_gene | gene_gene_biotype | gene_end_range | gene_start_range | gene_comparison | is.dm |
|---|---|---|---|---|---|---|---|---|---|---|---|---|---|---|---|---|---|---|---|---|---|---|---|---|---|---|---|---|---|
| NC_039506.1 | 2173709 | 2173709 | 1 | + | 44 | 1 | 43 | 64 | 0 | 64 | 2.27 | 0 | NC_039506.1 | 2172791 | 2181709 | 8919 | + | Gnomon | gene | gene-LOC105280304 | GeneID:105280304 | LOC105280304 | Gene | LOC105280304 | protein_coding | character(0) | character(0) | BCphaseB2.vs.RphaseA2 | FALSE |
| NC_039506.1 | 2173733 | 2173733 | 1 | + | 35 | 0 | 35 | 61 | 1 | 60 | 0 | 1.64 | NC_039506.1 | 2172791 | 2181709 | 8919 | + | Gnomon | gene | gene-LOC105280304 | GeneID:105280304 | LOC105280304 | Gene | LOC105280304 | protein_coding | character(0) | character(0) | BCphaseB2.vs.RphaseA2 | FALSE |
| NC_039506.1 | 2173742 | 2173742 | 1 | + | 39 | 0 | 39 | 66 | 0 | 66 | 0 | 0 | NC_039506.1 | 2172791 | 2181709 | 8919 | + | Gnomon | gene | gene-LOC105280304 | GeneID:105280304 | LOC105280304 | Gene | LOC105280304 | protein_coding | character(0) | character(0) | BCphaseB2.vs.RphaseA2 | FALSE |
| NC_039506.1 | 2173746 | 2173746 | 1 | + | 38 | 0 | 38 | 67 | 0 | 67 | 0 | 0 | NC_039506.1 | 2172791 | 2181709 | 8919 | + | Gnomon | gene | gene-LOC105280304 | GeneID:105280304 | LOC105280304 | Gene | LOC105280304 | protein_coding | character(0) | character(0) | BCphaseB2.vs.RphaseA2 | FALSE |
| NC_039506.1 | 2173753 | 2173753 | 1 | + | 38 | 0 | 38 | 69 | 0 | 69 | 0 | 0 | NC_039506.1 | 2172791 | 2181709 | 8919 | + | Gnomon | gene | gene-LOC105280304 | GeneID:105280304 | LOC105280304 | Gene | LOC105280304 | protein_coding | character(0) | character(0) | BCphaseB2.vs.RphaseA2 | FALSE |
| NC_039506.1 | 2173759 | 2173759 | 1 | + | 39 | 0 | 39 | 70 | 1 | 69 | 0 | 1.43 | NC_039506.1 | 2172791 | 2181709 | 8919 | + | Gnomon | gene | gene-LOC105280304 | GeneID:105280304 | LOC105280304 | Gene | LOC105280304 | protein_coding | character(0) | character(0) | BCphaseB2.vs.RphaseA2 | FALSE |
| NC_039506.1 | 2173764 | 2173764 | 1 | + | 42 | 0 | 42 | 75 | 0 | 75 | 0 | 0 | NC_039506.1 | 2172791 | 2181709 | 8919 | + | Gnomon | gene | gene-LOC105280304 | GeneID:105280304 | LOC105280304 | Gene | LOC105280304 | protein_coding | character(0) | character(0) | BCphaseB2.vs.RphaseA2 | FALSE |
| NC_039506.1 | 2173768 | 2173768 | 1 | + | 42 | 0 | 42 | 75 | 0 | 75 | 0 | 0 | NC_039506.1 | 2172791 | 2181709 | 8919 | + | Gnomon | gene | gene-LOC105280304 | GeneID:105280304 | LOC105280304 | Gene | LOC105280304 | protein_coding | character(0) | character(0) | BCphaseB2.vs.RphaseA2 | FALSE |
| NC_039506.1 | 2173779 | 2173779 | 1 | + | 50 | 0 | 50 | 82 | 0 | 82 | 0 | 0 | NC_039506.1 | 2172791 | 2181709 | 8919 | + | Gnomon | gene | gene-LOC105280304 | GeneID:105280304 | LOC105280304 | Gene | LOC105280304 | protein_coding | character(0) | character(0) | BCphaseB2.vs.RphaseA2 | FALSE |
| NC_039506.1 | 2173789 | 2173789 | 1 | + | 52 | 1 | 51 | 84 | 0 | 84 | 1.92 | 0 | NC_039506.1 | 2172791 | 2181709 | 8919 | + | Gnomon | gene | gene-LOC105280304 | GeneID:105280304 | LOC105280304 | Gene | LOC105280304 | protein_coding | character(0) | character(0) | BCphaseB2.vs.RphaseA2 | FALSE |
| NC_039506.1 | 2173798 | 2173798 | 1 | + | 47 | 0 | 47 | 74 | 0 | 74 | 0 | 0 | NC_039506.1 | 2172791 | 2181709 | 8919 | + | Gnomon | gene | gene-LOC105280304 | GeneID:105280304 | LOC105280304 | Gene | LOC105280304 | protein_coding | character(0) | character(0) | BCphaseB2.vs.RphaseA2 | FALSE |
| NC_039506.1 | 2173801 | 2173801 | 1 | + | 40 | 0 | 40 | 68 | 1 | 67 | 0 | 1.47 | NC_039506.1 | 2172791 | 2181709 | 8919 | + | Gnomon | gene | gene-LOC105280304 | GeneID:105280304 | LOC105280304 | Gene | LOC105280304 | protein_coding | character(0) | character(0) | BCphaseB2.vs.RphaseA2 | FALSE |
| NC_039506.1 | 2173810 | 2173810 | 1 | + | 34 | 0 | 34 | 54 | 1 | 53 | 0 | 1.85 | NC_039506.1 | 2172791 | 2181709 | 8919 | + | Gnomon | gene | gene-LOC105280304 | GeneID:105280304 | LOC105280304 | Gene | LOC105280304 | protein_coding | character(0) | character(0) | BCphaseB2.vs.RphaseA2 | FALSE |
| NC_039506.1 | 2173812 | 2173812 | 1 | + | 33 | 1 | 32 | 54 | 3 | 51 | 3.03 | 5.56 | NC_039506.1 | 2172791 | 2181709 | 8919 | + | Gnomon | gene | gene-LOC105280304 | GeneID:105280304 | LOC105280304 | Gene | LOC105280304 | protein_coding | character(0) | character(0) | BCphaseB2.vs.RphaseA2 | FALSE |
| NC_039506.1 | 2173823 | 2173823 | 1 | + | 48 | 0 | 48 | 48 | 1 | 47 | 0 | 2.08 | NC_039506.1 | 2172791 | 2181709 | 8919 | + | Gnomon | gene | gene-LOC105280304 | GeneID:105280304 | LOC105280304 | Gene | LOC105280304 | protein_coding | character(0) | character(0) | BCphaseB2.vs.RphaseA2 | FALSE |
| NC_039506.1 | 2173831 | 2173831 | 1 | + | 49 | 0 | 49 | 48 | 0 | 48 | 0 | 0 | NC_039506.1 | 2172791 | 2181709 | 8919 | + | Gnomon | gene | gene-LOC105280304 | GeneID:105280304 | LOC105280304 | Gene | LOC105280304 | protein_coding | character(0) | character(0) | BCphaseB2.vs.RphaseA2 | FALSE |
| NC_039506.1 | 2173849 | 2173849 | 1 | + | 20 | 0 | 20 | 28 | 0 | 28 | 0 | 0 | NC_039506.1 | 2172791 | 2181709 | 8919 | + | Gnomon | gene | gene-LOC105280304 | GeneID:105280304 | LOC105280304 | Gene | LOC105280304 | protein_coding | character(0) | character(0) | BCphaseB2.vs.RphaseA2 | FALSE |
| NC_039506.1 | 2174051 | 2174051 | 1 | + | 23 | 0 | 23 | 36 | 0 | 36 | 0 | 0 | NC_039506.1 | 2172791 | 2181709 | 8919 | + | Gnomon | gene | gene-LOC105280304 | GeneID:105280304 | LOC105280304 | Gene | LOC105280304 | protein_coding | character(0) | character(0) | BCphaseB2.vs.RphaseA2 | FALSE |
| NC_039506.1 | 2174072 | 2174072 | 1 | + | 25 | 0 | 25 | 40 | 0 | 40 | 0 | 0 | NC_039506.1 | 2172791 | 2181709 | 8919 | + | Gnomon | gene | gene-LOC105280304 | GeneID:105280304 | LOC105280304 | Gene | LOC105280304 | protein_coding | character(0) | character(0) | BCphaseB2.vs.RphaseA2 | FALSE |
| NC_039506.1 | 2174088 | 2174088 | 1 | + | 25 | 0 | 25 | 36 | 0 | 36 | 0 | 0 | NC_039506.1 | 2172791 | 2181709 | 8919 | + | Gnomon | gene | gene-LOC105280304 | GeneID:105280304 | LOC105280304 | Gene | LOC105280304 | protein_coding | character(0) | character(0) | BCphaseB2.vs.RphaseA2 | FALSE |
| NC_039506.1 | 2174095 | 2174095 | 1 | + | 23 | 0 | 23 | 36 | 0 | 36 | 0 | 0 | NC_039506.1 | 2172791 | 2181709 | 8919 | + | Gnomon | gene | gene-LOC105280304 | GeneID:105280304 | LOC105280304 | Gene | LOC105280304 | protein_coding | character(0) | character(0) | BCphaseB2.vs.RphaseA2 | FALSE |
| NC_039506.1 | 2174105 | 2174105 | 1 | + | 25 | 0 | 25 | 39 | 0 | 39 | 0 | 0 | NC_039506.1 | 2172791 | 2181709 | 8919 | + | Gnomon | gene | gene-LOC105280304 | GeneID:105280304 | LOC105280304 | Gene | LOC105280304 | protein_coding | character(0) | character(0) | BCphaseB2.vs.RphaseA2 | FALSE |
| NC_039506.1 | 2174110 | 2174110 | 1 | + | 25 | 0 | 25 | 39 | 0 | 39 | 0 | 0 | NC_039506.1 | 2172791 | 2181709 | 8919 | + | Gnomon | gene | gene-LOC105280304 | GeneID:105280304 | LOC105280304 | Gene | LOC105280304 | protein_coding | character(0) | character(0) | BCphaseB2.vs.RphaseA2 | FALSE |
| NC_039506.1 | 2174115 | 2174115 | 1 | + | 26 | 1 | 25 | 37 | 0 | 37 | 3.85 | 0 | NC_039506.1 | 2172791 | 2181709 | 8919 | + | Gnomon | gene | gene-LOC105280304 | GeneID:105280304 | LOC105280304 | Gene | LOC105280304 | protein_coding | character(0) | character(0) | BCphaseB2.vs.RphaseA2 | FALSE |

...
... only first 25 lines shown ...

Directory: DMSG File: dmg-Ob_LI_BCphaseB2.vs.RphaseB1_details.txt

| seqnames | start | end | width | strand | coverage1 | numCs1 | numTs1 | coverage2 | numCs2 | numTs2 | BCphaseB2 | RphaseB1 | gene_seqnames | gene_start | gene_end | gene_width | gene_strand | gene_source | gene_type | gene_ID | gene_Dbxref | gene_Name | gene_gbkey | gene_gene | gene_gene_biotype | gene_end_range | gene_partial | gene_start_range | gene_comparison | is.dm |
|---|---|---|---|---|---|---|---|---|---|---|---|---|---|---|---|---|---|---|---|---|---|---|---|---|---|---|---|---|---|---|
| NC_039506.1 | 1057034 | 1057034 | 1 | + | 39 | 0 | 39 | 25 | 0 | 25 | 0 | 0 | NC_039506.1 | 1056151 | 1064257 | 8107 | + | Gnomon | gene | gene-LOC105286747 | GeneID:105286747 | LOC105286747 | Gene | LOC105286747 | lncRNA | character(0) | NA | character(0) | BCphaseB2.vs.RphaseB1 | FALSE |
| NC_039506.1 | 1057043 | 1057043 | 1 | + | 37 | 0 | 37 | 29 | 0 | 29 | 0 | 0 | NC_039506.1 | 1056151 | 1064257 | 8107 | + | Gnomon | gene | gene-LOC105286747 | GeneID:105286747 | LOC105286747 | Gene | LOC105286747 | lncRNA | character(0) | NA | character(0) | BCphaseB2.vs.RphaseB1 | FALSE |
| NC_039506.1 | 1057047 | 1057047 | 1 | + | 38 | 0 | 38 | 30 | 0 | 30 | 0 | 0 | NC_039506.1 | 1056151 | 1064257 | 8107 | + | Gnomon | gene | gene-LOC105286747 | GeneID:105286747 | LOC105286747 | Gene | LOC105286747 | lncRNA | character(0) | NA | character(0) | BCphaseB2.vs.RphaseB1 | FALSE |
| NC_039506.1 | 1057061 | 1057061 | 1 | + | 39 | 0 | 39 | 30 | 0 | 30 | 0 | 0 | NC_039506.1 | 1056151 | 1064257 | 8107 | + | Gnomon | gene | gene-LOC105286747 | GeneID:105286747 | LOC105286747 | Gene | LOC105286747 | lncRNA | character(0) | NA | character(0) | BCphaseB2.vs.RphaseB1 | FALSE |
| NC_039506.1 | 1057064 | 1057064 | 1 | + | 39 | 0 | 39 | 30 | 0 | 30 | 0 | 0 | NC_039506.1 | 1056151 | 1064257 | 8107 | + | Gnomon | gene | gene-LOC105286747 | GeneID:105286747 | LOC105286747 | Gene | LOC105286747 | lncRNA | character(0) | NA | character(0) | BCphaseB2.vs.RphaseB1 | FALSE |
| NC_039506.1 | 1057110 | 1057110 | 1 | + | 38 | 0 | 38 | 37 | 0 | 37 | 0 | 0 | NC_039506.1 | 1056151 | 1064257 | 8107 | + | Gnomon | gene | gene-LOC105286747 | GeneID:105286747 | LOC105286747 | Gene | LOC105286747 | lncRNA | character(0) | NA | character(0) | BCphaseB2.vs.RphaseB1 | FALSE |
| NC_039506.1 | 1057112 | 1057112 | 1 | + | 38 | 0 | 38 | 40 | 0 | 40 | 0 | 0 | NC_039506.1 | 1056151 | 1064257 | 8107 | + | Gnomon | gene | gene-LOC105286747 | GeneID:105286747 | LOC105286747 | Gene | LOC105286747 | lncRNA | character(0) | NA | character(0) | BCphaseB2.vs.RphaseB1 | FALSE |
| NC_039506.1 | 1057128 | 1057128 | 1 | + | 37 | 0 | 37 | 30 | 0 | 30 | 0 | 0 | NC_039506.1 | 1056151 | 1064257 | 8107 | + | Gnomon | gene | gene-LOC105286747 | GeneID:105286747 | LOC105286747 | Gene | LOC105286747 | lncRNA | character(0) | NA | character(0) | BCphaseB2.vs.RphaseB1 | FALSE |
| NC_039506.1 | 1057134 | 1057134 | 1 | + | 35 | 1 | 34 | 30 | 0 | 30 | 2.86 | 0 | NC_039506.1 | 1056151 | 1064257 | 8107 | + | Gnomon | gene | gene-LOC105286747 | GeneID:105286747 | LOC105286747 | Gene | LOC105286747 | lncRNA | character(0) | NA | character(0) | BCphaseB2.vs.RphaseB1 | FALSE |
| NC_039506.1 | 1057145 | 1057145 | 1 | + | 37 | 0 | 37 | 32 | 0 | 32 | 0 | 0 | NC_039506.1 | 1056151 | 1064257 | 8107 | + | Gnomon | gene | gene-LOC105286747 | GeneID:105286747 | LOC105286747 | Gene | LOC105286747 | lncRNA | character(0) | NA | character(0) | BCphaseB2.vs.RphaseB1 | FALSE |
| NC_039506.1 | 1057157 | 1057157 | 1 | + | 37 | 0 | 37 | 30 | 1 | 29 | 0 | 3.33 | NC_039506.1 | 1056151 | 1064257 | 8107 | + | Gnomon | gene | gene-LOC105286747 | GeneID:105286747 | LOC105286747 | Gene | LOC105286747 | lncRNA | character(0) | NA | character(0) | BCphaseB2.vs.RphaseB1 | FALSE |
| NC_039506.1 | 1057169 | 1057169 | 1 | + | 35 | 0 | 35 | 28 | 0 | 28 | 0 | 0 | NC_039506.1 | 1056151 | 1064257 | 8107 | + | Gnomon | gene | gene-LOC105286747 | GeneID:105286747 | LOC105286747 | Gene | LOC105286747 | lncRNA | character(0) | NA | character(0) | BCphaseB2.vs.RphaseB1 | FALSE |
| NC_039506.1 | 1057181 | 1057181 | 1 | + | 30 | 0 | 30 | 26 | 0 | 26 | 0 | 0 | NC_039506.1 | 1056151 | 1064257 | 8107 | + | Gnomon | gene | gene-LOC105286747 | GeneID:105286747 | LOC105286747 | Gene | LOC105286747 | lncRNA | character(0) | NA | character(0) | BCphaseB2.vs.RphaseB1 | FALSE |
| NC_039506.1 | 1057308 | 1057308 | 1 | + | 20 | 1 | 19 | 58 | 0 | 58 | 5 | 0 | NC_039506.1 | 1056151 | 1064257 | 8107 | + | Gnomon | gene | gene-LOC105286747 | GeneID:105286747 | LOC105286747 | Gene | LOC105286747 | lncRNA | character(0) | NA | character(0) | BCphaseB2.vs.RphaseB1 | FALSE |
| NC_039506.1 | 1057314 | 1057314 | 1 | + | 36 | 0 | 36 | 61 | 4 | 57 | 0 | 6.56 | NC_039506.1 | 1056151 | 1064257 | 8107 | + | Gnomon | gene | gene-LOC105286747 | GeneID:105286747 | LOC105286747 | Gene | LOC105286747 | lncRNA | character(0) | NA | character(0) | BCphaseB2.vs.RphaseB1 | FALSE |
| NC_039506.1 | 1057316 | 1057316 | 1 | + | 36 | 0 | 36 | 61 | 0 | 61 | 0 | 0 | NC_039506.1 | 1056151 | 1064257 | 8107 | + | Gnomon | gene | gene-LOC105286747 | GeneID:105286747 | LOC105286747 | Gene | LOC105286747 | lncRNA | character(0) | NA | character(0) | BCphaseB2.vs.RphaseB1 | FALSE |
| NC_039506.1 | 1057320 | 1057320 | 1 | + | 56 | 0 | 56 | 58 | 0 | 58 | 0 | 0 | NC_039506.1 | 1056151 | 1064257 | 8107 | + | Gnomon | gene | gene-LOC105286747 | GeneID:105286747 | LOC105286747 | Gene | LOC105286747 | lncRNA | character(0) | NA | character(0) | BCphaseB2.vs.RphaseB1 | FALSE |
| NC_039506.1 | 1057331 | 1057331 | 1 | + | 80 | 1 | 79 | 77 | 0 | 77 | 1.25 | 0 | NC_039506.1 | 1056151 | 1064257 | 8107 | + | Gnomon | gene | gene-LOC105286747 | GeneID:105286747 | LOC105286747 | Gene | LOC105286747 | lncRNA | character(0) | NA | character(0) | BCphaseB2.vs.RphaseB1 | FALSE |
| NC_039506.1 | 1057337 | 1057337 | 1 | + | 75 | 0 | 75 | 74 | 0 | 74 | 0 | 0 | NC_039506.1 | 1056151 | 1064257 | 8107 | + | Gnomon | gene | gene-LOC105286747 | GeneID:105286747 | LOC105286747 | Gene | LOC105286747 | lncRNA | character(0) | NA | character(0) | BCphaseB2.vs.RphaseB1 | FALSE |
| NC_039506.1 | 1057344 | 1057344 | 1 | + | 74 | 0 | 74 | 72 | 0 | 72 | 0 | 0 | NC_039506.1 | 1056151 | 1064257 | 8107 | + | Gnomon | gene | gene-LOC105286747 | GeneID:105286747 | LOC105286747 | Gene | LOC105286747 | lncRNA | character(0) | NA | character(0) | BCphaseB2.vs.RphaseB1 | FALSE |
| NC_039506.1 | 1057373 | 1057373 | 1 | + | 43 | 0 | 43 | 32 | 0 | 32 | 0 | 0 | NC_039506.1 | 1056151 | 1064257 | 8107 | + | Gnomon | gene | gene-LOC105286747 | GeneID:105286747 | LOC105286747 | Gene | LOC105286747 | lncRNA | character(0) | NA | character(0) | BCphaseB2.vs.RphaseB1 | FALSE |
| NC_039506.1 | 1057386 | 1057386 | 1 | + | 36 | 0 | 36 | 29 | 1 | 28 | 0 | 3.45 | NC_039506.1 | 1056151 | 1064257 | 8107 | + | Gnomon | gene | gene-LOC105286747 | GeneID:105286747 | LOC105286747 | Gene | LOC105286747 | lncRNA | character(0) | NA | character(0) | BCphaseB2.vs.RphaseB1 | FALSE |
| NC_039506.1 | 1058047 | 1058047 | 1 | + | 20 | 0 | 20 | 22 | 0 | 22 | 0 | 0 | NC_039506.1 | 1056151 | 1064257 | 8107 | + | Gnomon | gene | gene-LOC105286747 | GeneID:105286747 | LOC105286747 | Gene | LOC105286747 | lncRNA | character(0) | NA | character(0) | BCphaseB2.vs.RphaseB1 | FALSE |
| NC_039506.1 | 1058053 | 1058053 | 1 | + | 20 | 0 | 20 | 22 | 0 | 22 | 0 | 0 | NC_039506.1 | 1056151 | 1064257 | 8107 | + | Gnomon | gene | gene-LOC105286747 | GeneID:105286747 | LOC105286747 | Gene | LOC105286747 | lncRNA | character(0) | NA | character(0) | BCphaseB2.vs.RphaseB1 | FALSE |

...
... only first 25 lines shown ...

Directory: DMSG File: dmg-Ob_LI_BCphaseB2.vs.RphaseB2_details.txt

| seqnames | start | end | width | strand | coverage1 | numCs1 | numTs1 | coverage2 | numCs2 | numTs2 | BCphaseB2 | RphaseB2 | gene_seqnames | gene_start | gene_end | gene_width | gene_strand | gene_source | gene_type | gene_ID | gene_Dbxref | gene_Name | gene_gbkey | gene_gene | gene_gene_biotype | gene_end_range | gene_partial | gene_start_range | gene_comparison | is.dm |
|---|---|---|---|---|---|---|---|---|---|---|---|---|---|---|---|---|---|---|---|---|---|---|---|---|---|---|---|---|---|---|
| NC_039506.1 | 1097588 | 1097588 | 1 | + | 25 | 11 | 14 | 25 | 9 | 16 | 44 | 36 | NC_039506.1 | 1095614 | 1100480 | 4867 | + | Gnomon | gene | gene-LOC105286750 | GeneID:105286750 | LOC105286750 | Gene | LOC105286750 | protein_coding | character(0) | NA | character(0) | BCphaseB2.vs.RphaseB2 | FALSE |
| NC_039506.1 | 1097598 | 1097598 | 1 | + | 27 | 14 | 13 | 27 | 18 | 9 | 51.85 | 66.67 | NC_039506.1 | 1095614 | 1100480 | 4867 | + | Gnomon | gene | gene-LOC105286750 | GeneID:105286750 | LOC105286750 | Gene | LOC105286750 | protein_coding | character(0) | NA | character(0) | BCphaseB2.vs.RphaseB2 | FALSE |
| NC_039506.1 | 1097647 | 1097647 | 1 | + | 42 | 6 | 36 | 38 | 4 | 34 | 14.29 | 10.53 | NC_039506.1 | 1095614 | 1100480 | 4867 | + | Gnomon | gene | gene-LOC105286750 | GeneID:105286750 | LOC105286750 | Gene | LOC105286750 | protein_coding | character(0) | NA | character(0) | BCphaseB2.vs.RphaseB2 | FALSE |
| NC_039506.1 | 1097653 | 1097653 | 1 | + | 42 | 2 | 40 | 38 | 1 | 37 | 4.76 | 2.63 | NC_039506.1 | 1095614 | 1100480 | 4867 | + | Gnomon | gene | gene-LOC105286750 | GeneID:105286750 | LOC105286750 | Gene | LOC105286750 | protein_coding | character(0) | NA | character(0) | BCphaseB2.vs.RphaseB2 | FALSE |
| NC_039506.1 | 1097702 | 1097702 | 1 | + | 33 | 8 | 25 | 39 | 7 | 32 | 24.24 | 17.95 | NC_039506.1 | 1095614 | 1100480 | 4867 | + | Gnomon | gene | gene-LOC105286750 | GeneID:105286750 | LOC105286750 | Gene | LOC105286750 | protein_coding | character(0) | NA | character(0) | BCphaseB2.vs.RphaseB2 | FALSE |
| NC_039506.1 | 1097773 | 1097773 | 1 | + | 32 | 31 | 1 | 38 | 37 | 1 | 96.88 | 97.37 | NC_039506.1 | 1095614 | 1100480 | 4867 | + | Gnomon | gene | gene-LOC105286750 | GeneID:105286750 | LOC105286750 | Gene | LOC105286750 | protein_coding | character(0) | NA | character(0) | BCphaseB2.vs.RphaseB2 | FALSE |
| NC_039506.1 | 1098206 | 1098206 | 1 | + | 29 | 17 | 12 | 28 | 17 | 11 | 58.62 | 60.71 | NC_039506.1 | 1095614 | 1100480 | 4867 | + | Gnomon | gene | gene-LOC105286750 | GeneID:105286750 | LOC105286750 | Gene | LOC105286750 | protein_coding | character(0) | NA | character(0) | BCphaseB2.vs.RphaseB2 | FALSE |
| NC_039506.1 | 1098266 | 1098266 | 1 | + | 54 | 31 | 23 | 45 | 27 | 18 | 57.41 | 60 | NC_039506.1 | 1095614 | 1100480 | 4867 | + | Gnomon | gene | gene-LOC105286750 | GeneID:105286750 | LOC105286750 | Gene | LOC105286750 | protein_coding | character(0) | NA | character(0) | BCphaseB2.vs.RphaseB2 | FALSE |
| NC_039506.1 | 1098279 | 1098279 | 1 | + | 59 | 27 | 32 | 46 | 17 | 29 | 45.76 | 36.96 | NC_039506.1 | 1095614 | 1100480 | 4867 | + | Gnomon | gene | gene-LOC105286750 | GeneID:105286750 | LOC105286750 | Gene | LOC105286750 | protein_coding | character(0) | NA | character(0) | BCphaseB2.vs.RphaseB2 | FALSE |
| NC_039506.1 | 1098293 | 1098293 | 1 | + | 35 | 9 | 26 | 29 | 11 | 18 | 25.71 | 37.93 | NC_039506.1 | 1095614 | 1100480 | 4867 | + | Gnomon | gene | gene-LOC105286750 | GeneID:105286750 | LOC105286750 | Gene | LOC105286750 | protein_coding | character(0) | NA | character(0) | BCphaseB2.vs.RphaseB2 | FALSE |
| NC_039506.1 | 1098301 | 1098301 | 1 | + | 28 | 16 | 12 | 23 | 16 | 7 | 57.14 | 69.57 | NC_039506.1 | 1095614 | 1100480 | 4867 | + | Gnomon | gene | gene-LOC105286750 | GeneID:105286750 | LOC105286750 | Gene | LOC105286750 | protein_coding | character(0) | NA | character(0) | BCphaseB2.vs.RphaseB2 | FALSE |
| NC_039506.1 | 1098330 | 1098330 | 1 | + | 46 | 9 | 37 | 56 | 18 | 38 | 19.57 | 32.14 | NC_039506.1 | 1095614 | 1100480 | 4867 | + | Gnomon | gene | gene-LOC105286750 | GeneID:105286750 | LOC105286750 | Gene | LOC105286750 | protein_coding | character(0) | NA | character(0) | BCphaseB2.vs.RphaseB2 | FALSE |
| NC_039506.1 | 1098334 | 1098334 | 1 | + | 45 | 27 | 18 | 58 | 40 | 18 | 60 | 68.97 | NC_039506.1 | 1095614 | 1100480 | 4867 | + | Gnomon | gene | gene-LOC105286750 | GeneID:105286750 | LOC105286750 | Gene | LOC105286750 | protein_coding | character(0) | NA | character(0) | BCphaseB2.vs.RphaseB2 | FALSE |
| NC_039506.1 | 1098362 | 1098362 | 1 | + | 52 | 5 | 47 | 60 | 6 | 54 | 9.62 | 10 | NC_039506.1 | 1095614 | 1100480 | 4867 | + | Gnomon | gene | gene-LOC105286750 | GeneID:105286750 | LOC105286750 | Gene | LOC105286750 | protein_coding | character(0) | NA | character(0) | BCphaseB2.vs.RphaseB2 | FALSE |
| NC_039506.1 | 1098377 | 1098377 | 1 | + | 55 | 26 | 29 | 61 | 20 | 41 | 47.27 | 32.79 | NC_039506.1 | 1095614 | 1100480 | 4867 | + | Gnomon | gene | gene-LOC105286750 | GeneID:105286750 | LOC105286750 | Gene | LOC105286750 | protein_coding | character(0) | NA | character(0) | BCphaseB2.vs.RphaseB2 | FALSE |
| NC_039506.1 | 1098387 | 1098387 | 1 | + | 55 | 30 | 25 | 57 | 25 | 32 | 54.55 | 43.86 | NC_039506.1 | 1095614 | 1100480 | 4867 | + | Gnomon | gene | gene-LOC105286750 | GeneID:105286750 | LOC105286750 | Gene | LOC105286750 | protein_coding | character(0) | NA | character(0) | BCphaseB2.vs.RphaseB2 | FALSE |
| NC_039506.1 | 1098400 | 1098400 | 1 | + | 45 | 21 | 24 | 46 | 27 | 19 | 46.67 | 58.7 | NC_039506.1 | 1095614 | 1100480 | 4867 | + | Gnomon | gene | gene-LOC105286750 | GeneID:105286750 | LOC105286750 | Gene | LOC105286750 | protein_coding | character(0) | NA | character(0) | BCphaseB2.vs.RphaseB2 | FALSE |
| NC_039506.1 | 1098420 | 1098420 | 1 | + | 29 | 9 | 20 | 27 | 14 | 13 | 31.03 | 51.85 | NC_039506.1 | 1095614 | 1100480 | 4867 | + | Gnomon | gene | gene-LOC105286750 | GeneID:105286750 | LOC105286750 | Gene | LOC105286750 | protein_coding | character(0) | NA | character(0) | BCphaseB2.vs.RphaseB2 | TRUE |
| NC_039506.1 | 1098427 | 1098427 | 1 | + | 31 | 17 | 14 | 29 | 18 | 11 | 54.84 | 62.07 | NC_039506.1 | 1095614 | 1100480 | 4867 | + | Gnomon | gene | gene-LOC105286750 | GeneID:105286750 | LOC105286750 | Gene | LOC105286750 | protein_coding | character(0) | NA | character(0) | BCphaseB2.vs.RphaseB2 | FALSE |
| NC_039506.1 | 1098994 | 1098994 | 1 | + | 22 | 1 | 21 | 31 | 1 | 30 | 4.55 | 3.23 | NC_039506.1 | 1095614 | 1100480 | 4867 | + | Gnomon | gene | gene-LOC105286750 | GeneID:105286750 | LOC105286750 | Gene | LOC105286750 | protein_coding | character(0) | NA | character(0) | BCphaseB2.vs.RphaseB2 | FALSE |
| NC_039506.1 | 1099002 | 1099002 | 1 | + | 22 | 1 | 21 | 32 | 1 | 31 | 4.55 | 3.12 | NC_039506.1 | 1095614 | 1100480 | 4867 | + | Gnomon | gene | gene-LOC105286750 | GeneID:105286750 | LOC105286750 | Gene | LOC105286750 | protein_coding | character(0) | NA | character(0) | BCphaseB2.vs.RphaseB2 | FALSE |
| NC_039506.1 | 1099012 | 1099012 | 1 | + | 22 | 1 | 21 | 31 | 3 | 28 | 4.55 | 9.68 | NC_039506.1 | 1095614 | 1100480 | 4867 | + | Gnomon | gene | gene-LOC105286750 | GeneID:105286750 | LOC105286750 | Gene | LOC105286750 | protein_coding | character(0) | NA | character(0) | BCphaseB2.vs.RphaseB2 | FALSE |
| NC_039506.1 | 1099051 | 1099051 | 1 | + | 33 | 1 | 32 | 40 | 0 | 40 | 3.03 | 0 | NC_039506.1 | 1095614 | 1100480 | 4867 | + | Gnomon | gene | gene-LOC105286750 | GeneID:105286750 | LOC105286750 | Gene | LOC105286750 | protein_coding | character(0) | NA | character(0) | BCphaseB2.vs.RphaseB2 | FALSE |
| NC_039506.1 | 1099061 | 1099061 | 1 | + | 34 | 6 | 28 | 44 | 1 | 43 | 17.65 | 2.27 | NC_039506.1 | 1095614 | 1100480 | 4867 | + | Gnomon | gene | gene-LOC105286750 | GeneID:105286750 | LOC105286750 | Gene | LOC105286750 | protein_coding | character(0) | NA | character(0) | BCphaseB2.vs.RphaseB2 | FALSE |

...
... only first 25 lines shown ...

Directory: DMSG File: dmg-Ob_LI_RphaseA1.vs.RphaseA2_details.txt

| seqnames | start | end | width | strand | coverage1 | numCs1 | numTs1 | coverage2 | numCs2 | numTs2 | RphaseA1 | RphaseA2 | gene_seqnames | gene_start | gene_end | gene_width | gene_strand | gene_source | gene_type | gene_ID | gene_Dbxref | gene_Name | gene_gbkey | gene_gene | gene_gene_biotype | gene_end_range | gene_start_range | gene_comparison | is.dm |
|---|---|---|---|---|---|---|---|---|---|---|---|---|---|---|---|---|---|---|---|---|---|---|---|---|---|---|---|---|---|
| NC_039506.1 | 2173614 | 2173614 | 1 | + | 30 | 0 | 30 | 63 | 0 | 63 | 0 | 0 | NC_039506.1 | 2172791 | 2181709 | 8919 | + | Gnomon | gene | gene-LOC105280304 | GeneID:105280304 | LOC105280304 | Gene | LOC105280304 | protein_coding | character(0) | character(0) | RphaseA1.vs.RphaseA2 | FALSE |
| NC_039506.1 | 2173709 | 2173709 | 1 | + | 46 | 0 | 46 | 64 | 0 | 64 | 0 | 0 | NC_039506.1 | 2172791 | 2181709 | 8919 | + | Gnomon | gene | gene-LOC105280304 | GeneID:105280304 | LOC105280304 | Gene | LOC105280304 | protein_coding | character(0) | character(0) | RphaseA1.vs.RphaseA2 | FALSE |
| NC_039506.1 | 2173733 | 2173733 | 1 | + | 52 | 2 | 50 | 61 | 1 | 60 | 3.85 | 1.64 | NC_039506.1 | 2172791 | 2181709 | 8919 | + | Gnomon | gene | gene-LOC105280304 | GeneID:105280304 | LOC105280304 | Gene | LOC105280304 | protein_coding | character(0) | character(0) | RphaseA1.vs.RphaseA2 | FALSE |
| NC_039506.1 | 2173742 | 2173742 | 1 | + | 57 | 0 | 57 | 66 | 0 | 66 | 0 | 0 | NC_039506.1 | 2172791 | 2181709 | 8919 | + | Gnomon | gene | gene-LOC105280304 | GeneID:105280304 | LOC105280304 | Gene | LOC105280304 | protein_coding | character(0) | character(0) | RphaseA1.vs.RphaseA2 | FALSE |
| NC_039506.1 | 2173746 | 2173746 | 1 | + | 57 | 0 | 57 | 67 | 0 | 67 | 0 | 0 | NC_039506.1 | 2172791 | 2181709 | 8919 | + | Gnomon | gene | gene-LOC105280304 | GeneID:105280304 | LOC105280304 | Gene | LOC105280304 | protein_coding | character(0) | character(0) | RphaseA1.vs.RphaseA2 | FALSE |
| NC_039506.1 | 2173753 | 2173753 | 1 | + | 57 | 0 | 57 | 69 | 0 | 69 | 0 | 0 | NC_039506.1 | 2172791 | 2181709 | 8919 | + | Gnomon | gene | gene-LOC105280304 | GeneID:105280304 | LOC105280304 | Gene | LOC105280304 | protein_coding | character(0) | character(0) | RphaseA1.vs.RphaseA2 | FALSE |
| NC_039506.1 | 2173759 | 2173759 | 1 | + | 58 | 0 | 58 | 70 | 1 | 69 | 0 | 1.43 | NC_039506.1 | 2172791 | 2181709 | 8919 | + | Gnomon | gene | gene-LOC105280304 | GeneID:105280304 | LOC105280304 | Gene | LOC105280304 | protein_coding | character(0) | character(0) | RphaseA1.vs.RphaseA2 | FALSE |
| NC_039506.1 | 2173764 | 2173764 | 1 | + | 60 | 1 | 59 | 75 | 0 | 75 | 1.67 | 0 | NC_039506.1 | 2172791 | 2181709 | 8919 | + | Gnomon | gene | gene-LOC105280304 | GeneID:105280304 | LOC105280304 | Gene | LOC105280304 | protein_coding | character(0) | character(0) | RphaseA1.vs.RphaseA2 | FALSE |
| NC_039506.1 | 2173768 | 2173768 | 1 | + | 59 | 0 | 59 | 75 | 0 | 75 | 0 | 0 | NC_039506.1 | 2172791 | 2181709 | 8919 | + | Gnomon | gene | gene-LOC105280304 | GeneID:105280304 | LOC105280304 | Gene | LOC105280304 | protein_coding | character(0) | character(0) | RphaseA1.vs.RphaseA2 | FALSE |
| NC_039506.1 | 2173779 | 2173779 | 1 | + | 64 | 0 | 64 | 82 | 0 | 82 | 0 | 0 | NC_039506.1 | 2172791 | 2181709 | 8919 | + | Gnomon | gene | gene-LOC105280304 | GeneID:105280304 | LOC105280304 | Gene | LOC105280304 | protein_coding | character(0) | character(0) | RphaseA1.vs.RphaseA2 | FALSE |
| NC_039506.1 | 2173789 | 2173789 | 1 | + | 56 | 0 | 56 | 84 | 0 | 84 | 0 | 0 | NC_039506.1 | 2172791 | 2181709 | 8919 | + | Gnomon | gene | gene-LOC105280304 | GeneID:105280304 | LOC105280304 | Gene | LOC105280304 | protein_coding | character(0) | character(0) | RphaseA1.vs.RphaseA2 | FALSE |
| NC_039506.1 | 2173798 | 2173798 | 1 | + | 51 | 1 | 50 | 74 | 0 | 74 | 1.96 | 0 | NC_039506.1 | 2172791 | 2181709 | 8919 | + | Gnomon | gene | gene-LOC105280304 | GeneID:105280304 | LOC105280304 | Gene | LOC105280304 | protein_coding | character(0) | character(0) | RphaseA1.vs.RphaseA2 | FALSE |
| NC_039506.1 | 2173801 | 2173801 | 1 | + | 49 | 1 | 48 | 68 | 1 | 67 | 2.04 | 1.47 | NC_039506.1 | 2172791 | 2181709 | 8919 | + | Gnomon | gene | gene-LOC105280304 | GeneID:105280304 | LOC105280304 | Gene | LOC105280304 | protein_coding | character(0) | character(0) | RphaseA1.vs.RphaseA2 | FALSE |
| NC_039506.1 | 2173810 | 2173810 | 1 | + | 48 | 3 | 45 | 54 | 1 | 53 | 6.25 | 1.85 | NC_039506.1 | 2172791 | 2181709 | 8919 | + | Gnomon | gene | gene-LOC105280304 | GeneID:105280304 | LOC105280304 | Gene | LOC105280304 | protein_coding | character(0) | character(0) | RphaseA1.vs.RphaseA2 | FALSE |
| NC_039506.1 | 2173812 | 2173812 | 1 | + | 46 | 0 | 46 | 54 | 3 | 51 | 0 | 5.56 | NC_039506.1 | 2172791 | 2181709 | 8919 | + | Gnomon | gene | gene-LOC105280304 | GeneID:105280304 | LOC105280304 | Gene | LOC105280304 | protein_coding | character(0) | character(0) | RphaseA1.vs.RphaseA2 | FALSE |
| NC_039506.1 | 2173823 | 2173823 | 1 | + | 60 | 0 | 60 | 48 | 1 | 47 | 0 | 2.08 | NC_039506.1 | 2172791 | 2181709 | 8919 | + | Gnomon | gene | gene-LOC105280304 | GeneID:105280304 | LOC105280304 | Gene | LOC105280304 | protein_coding | character(0) | character(0) | RphaseA1.vs.RphaseA2 | FALSE |
| NC_039506.1 | 2173831 | 2173831 | 1 | + | 61 | 0 | 61 | 48 | 0 | 48 | 0 | 0 | NC_039506.1 | 2172791 | 2181709 | 8919 | + | Gnomon | gene | gene-LOC105280304 | GeneID:105280304 | LOC105280304 | Gene | LOC105280304 | protein_coding | character(0) | character(0) | RphaseA1.vs.RphaseA2 | FALSE |
| NC_039506.1 | 2173849 | 2173849 | 1 | + | 23 | 0 | 23 | 28 | 0 | 28 | 0 | 0 | NC_039506.1 | 2172791 | 2181709 | 8919 | + | Gnomon | gene | gene-LOC105280304 | GeneID:105280304 | LOC105280304 | Gene | LOC105280304 | protein_coding | character(0) | character(0) | RphaseA1.vs.RphaseA2 | FALSE |
| NC_039506.1 | 2174072 | 2174072 | 1 | + | 26 | 0 | 26 | 40 | 0 | 40 | 0 | 0 | NC_039506.1 | 2172791 | 2181709 | 8919 | + | Gnomon | gene | gene-LOC105280304 | GeneID:105280304 | LOC105280304 | Gene | LOC105280304 | protein_coding | character(0) | character(0) | RphaseA1.vs.RphaseA2 | FALSE |
| NC_039506.1 | 2174088 | 2174088 | 1 | + | 26 | 0 | 26 | 36 | 0 | 36 | 0 | 0 | NC_039506.1 | 2172791 | 2181709 | 8919 | + | Gnomon | gene | gene-LOC105280304 | GeneID:105280304 | LOC105280304 | Gene | LOC105280304 | protein_coding | character(0) | character(0) | RphaseA1.vs.RphaseA2 | FALSE |
| NC_039506.1 | 2174095 | 2174095 | 1 | + | 50 | 1 | 49 | 36 | 0 | 36 | 2 | 0 | NC_039506.1 | 2172791 | 2181709 | 8919 | + | Gnomon | gene | gene-LOC105280304 | GeneID:105280304 | LOC105280304 | Gene | LOC105280304 | protein_coding | character(0) | character(0) | RphaseA1.vs.RphaseA2 | FALSE |
| NC_039506.1 | 2174105 | 2174105 | 1 | + | 51 | 0 | 51 | 39 | 0 | 39 | 0 | 0 | NC_039506.1 | 2172791 | 2181709 | 8919 | + | Gnomon | gene | gene-LOC105280304 | GeneID:105280304 | LOC105280304 | Gene | LOC105280304 | protein_coding | character(0) | character(0) | RphaseA1.vs.RphaseA2 | FALSE |
| NC_039506.1 | 2174110 | 2174110 | 1 | + | 48 | 0 | 48 | 39 | 0 | 39 | 0 | 0 | NC_039506.1 | 2172791 | 2181709 | 8919 | + | Gnomon | gene | gene-LOC105280304 | GeneID:105280304 | LOC105280304 | Gene | LOC105280304 | protein_coding | character(0) | character(0) | RphaseA1.vs.RphaseA2 | FALSE |
| NC_039506.1 | 2174115 | 2174115 | 1 | + | 49 | 0 | 49 | 37 | 0 | 37 | 0 | 0 | NC_039506.1 | 2172791 | 2181709 | 8919 | + | Gnomon | gene | gene-LOC105280304 | GeneID:105280304 | LOC105280304 | Gene | LOC105280304 | protein_coding | character(0) | character(0) | RphaseA1.vs.RphaseA2 | FALSE |

...
... only first 25 lines shown ...

Directory: DMSG File: dmg-Ob_LI_RphaseA1.vs.RphaseB1_details.txt

| seqnames | start | end | width | strand | coverage1 | numCs1 | numTs1 | coverage2 | numCs2 | numTs2 | RphaseA1 | RphaseB1 | gene_seqnames | gene_start | gene_end | gene_width | gene_strand | gene_source | gene_type | gene_ID | gene_Dbxref | gene_Name | gene_gbkey | gene_gene | gene_gene_biotype | gene_end_range | gene_start_range | gene_comparison | is.dm |
|---|---|---|---|---|---|---|---|---|---|---|---|---|---|---|---|---|---|---|---|---|---|---|---|---|---|---|---|---|---|
| NC_039506.1 | 2980627 | 2980627 | 1 | + | 21 | 0 | 21 | 22 | 0 | 22 | 0 | 0 | NC_039506.1 | 2980439 | 2984235 | 3797 | + | Gnomon | gene | gene-LOC105280348 | GeneID:105280348 | LOC105280348 | Gene | LOC105280348 | protein_coding | character(0) | character(0) | RphaseA1.vs.RphaseB1 | FALSE |
| NC_039506.1 | 2980668 | 2980668 | 1 | + | 21 | 2 | 19 | 29 | 0 | 29 | 9.52 | 0 | NC_039506.1 | 2980439 | 2984235 | 3797 | + | Gnomon | gene | gene-LOC105280348 | GeneID:105280348 | LOC105280348 | Gene | LOC105280348 | protein_coding | character(0) | character(0) | RphaseA1.vs.RphaseB1 | FALSE |
| NC_039506.1 | 2980710 | 2980710 | 1 | + | 21 | 0 | 21 | 20 | 1 | 19 | 0 | 5 | NC_039506.1 | 2980439 | 2984235 | 3797 | + | Gnomon | gene | gene-LOC105280348 | GeneID:105280348 | LOC105280348 | Gene | LOC105280348 | protein_coding | character(0) | character(0) | RphaseA1.vs.RphaseB1 | FALSE |
| NC_039506.1 | 2980750 | 2980750 | 1 | + | 28 | 0 | 28 | 33 | 0 | 33 | 0 | 0 | NC_039506.1 | 2980439 | 2984235 | 3797 | + | Gnomon | gene | gene-LOC105280348 | GeneID:105280348 | LOC105280348 | Gene | LOC105280348 | protein_coding | character(0) | character(0) | RphaseA1.vs.RphaseB1 | FALSE |
| NC_039506.1 | 2980761 | 2980761 | 1 | + | 31 | 0 | 31 | 34 | 0 | 34 | 0 | 0 | NC_039506.1 | 2980439 | 2984235 | 3797 | + | Gnomon | gene | gene-LOC105280348 | GeneID:105280348 | LOC105280348 | Gene | LOC105280348 | protein_coding | character(0) | character(0) | RphaseA1.vs.RphaseB1 | FALSE |
| NC_039506.1 | 2980767 | 2980767 | 1 | + | 32 | 1 | 31 | 38 | 0 | 38 | 3.12 | 0 | NC_039506.1 | 2980439 | 2984235 | 3797 | + | Gnomon | gene | gene-LOC105280348 | GeneID:105280348 | LOC105280348 | Gene | LOC105280348 | protein_coding | character(0) | character(0) | RphaseA1.vs.RphaseB1 | FALSE |
| NC_039506.1 | 2980928 | 2980928 | 1 | + | 20 | 8 | 12 | 21 | 4 | 17 | 40 | 19.05 | NC_039506.1 | 2980439 | 2984235 | 3797 | + | Gnomon | gene | gene-LOC105280348 | GeneID:105280348 | LOC105280348 | Gene | LOC105280348 | protein_coding | character(0) | character(0) | RphaseA1.vs.RphaseB1 | FALSE |
| NC_039506.1 | 2980989 | 2980989 | 1 | + | 28 | 3 | 25 | 32 | 2 | 30 | 10.71 | 6.25 | NC_039506.1 | 2980439 | 2984235 | 3797 | + | Gnomon | gene | gene-LOC105280348 | GeneID:105280348 | LOC105280348 | Gene | LOC105280348 | protein_coding | character(0) | character(0) | RphaseA1.vs.RphaseB1 | FALSE |
| NC_039506.1 | 2980993 | 2980993 | 1 | + | 29 | 15 | 14 | 32 | 16 | 16 | 51.72 | 50 | NC_039506.1 | 2980439 | 2984235 | 3797 | + | Gnomon | gene | gene-LOC105280348 | GeneID:105280348 | LOC105280348 | Gene | LOC105280348 | protein_coding | character(0) | character(0) | RphaseA1.vs.RphaseB1 | FALSE |
| NC_039506.1 | 2980998 | 2980998 | 1 | + | 28 | 7 | 21 | 31 | 13 | 18 | 25 | 41.94 | NC_039506.1 | 2980439 | 2984235 | 3797 | + | Gnomon | gene | gene-LOC105280348 | GeneID:105280348 | LOC105280348 | Gene | LOC105280348 | protein_coding | character(0) | character(0) | RphaseA1.vs.RphaseB1 | FALSE |
| NC_039506.1 | 2981030 | 2981030 | 1 | + | 28 | 25 | 3 | 61 | 54 | 7 | 89.29 | 88.52 | NC_039506.1 | 2980439 | 2984235 | 3797 | + | Gnomon | gene | gene-LOC105280348 | GeneID:105280348 | LOC105280348 | Gene | LOC105280348 | protein_coding | character(0) | character(0) | RphaseA1.vs.RphaseB1 | FALSE |
| NC_039506.1 | 2981077 | 2981077 | 1 | + | 30 | 11 | 19 | 43 | 27 | 16 | 36.67 | 62.79 | NC_039506.1 | 2980439 | 2984235 | 3797 | + | Gnomon | gene | gene-LOC105280348 | GeneID:105280348 | LOC105280348 | Gene | LOC105280348 | protein_coding | character(0) | character(0) | RphaseA1.vs.RphaseB1 | FALSE |
| NC_039506.1 | 2981125 | 2981125 | 1 | + | 20 | 10 | 10 | 24 | 11 | 13 | 50 | 45.83 | NC_039506.1 | 2980439 | 2984235 | 3797 | + | Gnomon | gene | gene-LOC105280348 | GeneID:105280348 | LOC105280348 | Gene | LOC105280348 | protein_coding | character(0) | character(0) | RphaseA1.vs.RphaseB1 | FALSE |
| NC_039506.1 | 2981183 | 2981183 | 1 | + | 60 | 59 | 1 | 51 | 24 | 27 | 98.33 | 47.06 | NC_039506.1 | 2980439 | 2984235 | 3797 | + | Gnomon | gene | gene-LOC105280348 | GeneID:105280348 | LOC105280348 | Gene | LOC105280348 | protein_coding | character(0) | character(0) | RphaseA1.vs.RphaseB1 | TRUE |
| NC_039506.1 | 2981225 | 2981225 | 1 | + | 46 | 29 | 17 | 27 | 16 | 11 | 63.04 | 59.26 | NC_039506.1 | 2980439 | 2984235 | 3797 | + | Gnomon | gene | gene-LOC105280348 | GeneID:105280348 | LOC105280348 | Gene | LOC105280348 | protein_coding | character(0) | character(0) | RphaseA1.vs.RphaseB1 | FALSE |
| NC_039506.1 | 2981245 | 2981245 | 1 | + | 38 | 15 | 23 | 27 | 13 | 14 | 39.47 | 48.15 | NC_039506.1 | 2980439 | 2984235 | 3797 | + | Gnomon | gene | gene-LOC105280348 | GeneID:105280348 | LOC105280348 | Gene | LOC105280348 | protein_coding | character(0) | character(0) | RphaseA1.vs.RphaseB1 | FALSE |
| NC_039506.1 | 2981292 | 2981292 | 1 | + | 42 | 38 | 4 | 57 | 51 | 6 | 90.48 | 89.47 | NC_039506.1 | 2980439 | 2984235 | 3797 | + | Gnomon | gene | gene-LOC105280348 | GeneID:105280348 | LOC105280348 | Gene | LOC105280348 | protein_coding | character(0) | character(0) | RphaseA1.vs.RphaseB1 | FALSE |
| NC_039506.1 | 2981308 | 2981308 | 1 | + | 73 | 49 | 24 | 64 | 36 | 28 | 67.12 | 56.25 | NC_039506.1 | 2980439 | 2984235 | 3797 | + | Gnomon | gene | gene-LOC105280348 | GeneID:105280348 | LOC105280348 | Gene | LOC105280348 | protein_coding | character(0) | character(0) | RphaseA1.vs.RphaseB1 | FALSE |
| NC_039506.1 | 2981315 | 2981315 | 1 | + | 52 | 48 | 4 | 67 | 50 | 17 | 92.31 | 74.63 | NC_039506.1 | 2980439 | 2984235 | 3797 | + | Gnomon | gene | gene-LOC105280348 | GeneID:105280348 | LOC105280348 | Gene | LOC105280348 | protein_coding | character(0) | character(0) | RphaseA1.vs.RphaseB1 | FALSE |
| NC_039506.1 | 2981335 | 2981335 | 1 | + | 40 | 38 | 2 | 65 | 63 | 2 | 95 | 96.92 | NC_039506.1 | 2980439 | 2984235 | 3797 | + | Gnomon | gene | gene-LOC105280348 | GeneID:105280348 | LOC105280348 | Gene | LOC105280348 | protein_coding | character(0) | character(0) | RphaseA1.vs.RphaseB1 | FALSE |
| NC_039506.1 | 2981350 | 2981350 | 1 | + | 52 | 51 | 1 | 55 | 55 | 0 | 98.08 | 100 | NC_039506.1 | 2980439 | 2984235 | 3797 | + | Gnomon | gene | gene-LOC105280348 | GeneID:105280348 | LOC105280348 | Gene | LOC105280348 | protein_coding | character(0) | character(0) | RphaseA1.vs.RphaseB1 | FALSE |
| NC_039506.1 | 2981391 | 2981391 | 1 | + | 60 | 49 | 11 | 65 | 53 | 12 | 81.67 | 81.54 | NC_039506.1 | 2980439 | 2984235 | 3797 | + | Gnomon | gene | gene-LOC105280348 | GeneID:105280348 | LOC105280348 | Gene | LOC105280348 | protein_coding | character(0) | character(0) | RphaseA1.vs.RphaseB1 | FALSE |
| NC_039506.1 | 2981402 | 2981402 | 1 | + | 73 | 68 | 5 | 74 | 67 | 7 | 93.15 | 90.54 | NC_039506.1 | 2980439 | 2984235 | 3797 | + | Gnomon | gene | gene-LOC105280348 | GeneID:105280348 | LOC105280348 | Gene | LOC105280348 | protein_coding | character(0) | character(0) | RphaseA1.vs.RphaseB1 | FALSE |
| NC_039506.1 | 2981411 | 2981411 | 1 | + | 73 | 70 | 3 | 81 | 71 | 10 | 95.89 | 87.65 | NC_039506.1 | 2980439 | 2984235 | 3797 | + | Gnomon | gene | gene-LOC105280348 | GeneID:105280348 | LOC105280348 | Gene | LOC105280348 | protein_coding | character(0) | character(0) | RphaseA1.vs.RphaseB1 | FALSE |

...
... only first 25 lines shown ...

Directory: DMSG File: dmg-Ob_LI_RphaseA1.vs.RphaseB2_details.txt

| seqnames | start | end | width | strand | coverage1 | numCs1 | numTs1 | coverage2 | numCs2 | numTs2 | RphaseA1 | RphaseB2 | gene_seqnames | gene_start | gene_end | gene_width | gene_strand | gene_source | gene_type | gene_ID | gene_Dbxref | gene_Name | gene_gbkey | gene_gene | gene_gene_biotype | gene_end_range | gene_start_range | gene_comparison | is.dm |
|---|---|---|---|---|---|---|---|---|---|---|---|---|---|---|---|---|---|---|---|---|---|---|---|---|---|---|---|---|---|
| NC_039506.1 | 2980668 | 2980668 | 1 | + | 21 | 2 | 19 | 28 | 0 | 28 | 9.52 | 0 | NC_039506.1 | 2980439 | 2984235 | 3797 | + | Gnomon | gene | gene-LOC105280348 | GeneID:105280348 | LOC105280348 | Gene | LOC105280348 | protein_coding | character(0) | character(0) | RphaseA1.vs.RphaseB2 | FALSE |
| NC_039506.1 | 2980750 | 2980750 | 1 | + | 28 | 0 | 28 | 25 | 1 | 24 | 0 | 4 | NC_039506.1 | 2980439 | 2984235 | 3797 | + | Gnomon | gene | gene-LOC105280348 | GeneID:105280348 | LOC105280348 | Gene | LOC105280348 | protein_coding | character(0) | character(0) | RphaseA1.vs.RphaseB2 | FALSE |
| NC_039506.1 | 2980761 | 2980761 | 1 | + | 31 | 0 | 31 | 26 | 1 | 25 | 0 | 3.85 | NC_039506.1 | 2980439 | 2984235 | 3797 | + | Gnomon | gene | gene-LOC105280348 | GeneID:105280348 | LOC105280348 | Gene | LOC105280348 | protein_coding | character(0) | character(0) | RphaseA1.vs.RphaseB2 | FALSE |
| NC_039506.1 | 2980767 | 2980767 | 1 | + | 32 | 1 | 31 | 26 | 0 | 26 | 3.12 | 0 | NC_039506.1 | 2980439 | 2984235 | 3797 | + | Gnomon | gene | gene-LOC105280348 | GeneID:105280348 | LOC105280348 | Gene | LOC105280348 | protein_coding | character(0) | character(0) | RphaseA1.vs.RphaseB2 | FALSE |
| NC_039506.1 | 2980989 | 2980989 | 1 | + | 28 | 3 | 25 | 27 | 3 | 24 | 10.71 | 11.11 | NC_039506.1 | 2980439 | 2984235 | 3797 | + | Gnomon | gene | gene-LOC105280348 | GeneID:105280348 | LOC105280348 | Gene | LOC105280348 | protein_coding | character(0) | character(0) | RphaseA1.vs.RphaseB2 | FALSE |
| NC_039506.1 | 2980993 | 2980993 | 1 | + | 29 | 15 | 14 | 27 | 9 | 18 | 51.72 | 33.33 | NC_039506.1 | 2980439 | 2984235 | 3797 | + | Gnomon | gene | gene-LOC105280348 | GeneID:105280348 | LOC105280348 | Gene | LOC105280348 | protein_coding | character(0) | character(0) | RphaseA1.vs.RphaseB2 | FALSE |
| NC_039506.1 | 2980998 | 2980998 | 1 | + | 28 | 7 | 21 | 27 | 7 | 20 | 25 | 25.93 | NC_039506.1 | 2980439 | 2984235 | 3797 | + | Gnomon | gene | gene-LOC105280348 | GeneID:105280348 | LOC105280348 | Gene | LOC105280348 | protein_coding | character(0) | character(0) | RphaseA1.vs.RphaseB2 | FALSE |
| NC_039506.1 | 2981030 | 2981030 | 1 | + | 28 | 25 | 3 | 22 | 19 | 3 | 89.29 | 86.36 | NC_039506.1 | 2980439 | 2984235 | 3797 | + | Gnomon | gene | gene-LOC105280348 | GeneID:105280348 | LOC105280348 | Gene | LOC105280348 | protein_coding | character(0) | character(0) | RphaseA1.vs.RphaseB2 | FALSE |
| NC_039506.1 | 2981077 | 2981077 | 1 | + | 30 | 11 | 19 | 37 | 22 | 15 | 36.67 | 59.46 | NC_039506.1 | 2980439 | 2984235 | 3797 | + | Gnomon | gene | gene-LOC105280348 | GeneID:105280348 | LOC105280348 | Gene | LOC105280348 | protein_coding | character(0) | character(0) | RphaseA1.vs.RphaseB2 | FALSE |
| NC_039506.1 | 2981125 | 2981125 | 1 | + | 20 | 10 | 10 | 29 | 19 | 10 | 50 | 65.52 | NC_039506.1 | 2980439 | 2984235 | 3797 | + | Gnomon | gene | gene-LOC105280348 | GeneID:105280348 | LOC105280348 | Gene | LOC105280348 | protein_coding | character(0) | character(0) | RphaseA1.vs.RphaseB2 | FALSE |
| NC_039506.1 | 2981146 | 2981146 | 1 | + | 31 | 28 | 3 | 24 | 23 | 1 | 90.32 | 95.83 | NC_039506.1 | 2980439 | 2984235 | 3797 | + | Gnomon | gene | gene-LOC105280348 | GeneID:105280348 | LOC105280348 | Gene | LOC105280348 | protein_coding | character(0) | character(0) | RphaseA1.vs.RphaseB2 | FALSE |
| NC_039506.1 | 2981183 | 2981183 | 1 | + | 60 | 59 | 1 | 59 | 39 | 20 | 98.33 | 66.1 | NC_039506.1 | 2980439 | 2984235 | 3797 | + | Gnomon | gene | gene-LOC105280348 | GeneID:105280348 | LOC105280348 | Gene | LOC105280348 | protein_coding | character(0) | character(0) | RphaseA1.vs.RphaseB2 | TRUE |
| NC_039506.1 | 2981225 | 2981225 | 1 | + | 46 | 29 | 17 | 39 | 29 | 10 | 63.04 | 74.36 | NC_039506.1 | 2980439 | 2984235 | 3797 | + | Gnomon | gene | gene-LOC105280348 | GeneID:105280348 | LOC105280348 | Gene | LOC105280348 | protein_coding | character(0) | character(0) | RphaseA1.vs.RphaseB2 | FALSE |
| NC_039506.1 | 2981245 | 2981245 | 1 | + | 38 | 15 | 23 | 36 | 21 | 15 | 39.47 | 58.33 | NC_039506.1 | 2980439 | 2984235 | 3797 | + | Gnomon | gene | gene-LOC105280348 | GeneID:105280348 | LOC105280348 | Gene | LOC105280348 | protein_coding | character(0) | character(0) | RphaseA1.vs.RphaseB2 | FALSE |
| NC_039506.1 | 2981292 | 2981292 | 1 | + | 42 | 38 | 4 | 42 | 40 | 2 | 90.48 | 95.24 | NC_039506.1 | 2980439 | 2984235 | 3797 | + | Gnomon | gene | gene-LOC105280348 | GeneID:105280348 | LOC105280348 | Gene | LOC105280348 | protein_coding | character(0) | character(0) | RphaseA1.vs.RphaseB2 | FALSE |
| NC_039506.1 | 2981308 | 2981308 | 1 | + | 73 | 49 | 24 | 46 | 20 | 26 | 67.12 | 43.48 | NC_039506.1 | 2980439 | 2984235 | 3797 | + | Gnomon | gene | gene-LOC105280348 | GeneID:105280348 | LOC105280348 | Gene | LOC105280348 | protein_coding | character(0) | character(0) | RphaseA1.vs.RphaseB2 | FALSE |
| NC_039506.1 | 2981315 | 2981315 | 1 | + | 52 | 48 | 4 | 49 | 39 | 10 | 92.31 | 79.59 | NC_039506.1 | 2980439 | 2984235 | 3797 | + | Gnomon | gene | gene-LOC105280348 | GeneID:105280348 | LOC105280348 | Gene | LOC105280348 | protein_coding | character(0) | character(0) | RphaseA1.vs.RphaseB2 | FALSE |
| NC_039506.1 | 2981335 | 2981335 | 1 | + | 40 | 38 | 2 | 40 | 38 | 2 | 95 | 95 | NC_039506.1 | 2980439 | 2984235 | 3797 | + | Gnomon | gene | gene-LOC105280348 | GeneID:105280348 | LOC105280348 | Gene | LOC105280348 | protein_coding | character(0) | character(0) | RphaseA1.vs.RphaseB2 | FALSE |
| NC_039506.1 | 2981350 | 2981350 | 1 | + | 52 | 51 | 1 | 74 | 73 | 1 | 98.08 | 98.65 | NC_039506.1 | 2980439 | 2984235 | 3797 | + | Gnomon | gene | gene-LOC105280348 | GeneID:105280348 | LOC105280348 | Gene | LOC105280348 | protein_coding | character(0) | character(0) | RphaseA1.vs.RphaseB2 | FALSE |
| NC_039506.1 | 2981391 | 2981391 | 1 | + | 60 | 49 | 11 | 80 | 65 | 15 | 81.67 | 81.25 | NC_039506.1 | 2980439 | 2984235 | 3797 | + | Gnomon | gene | gene-LOC105280348 | GeneID:105280348 | LOC105280348 | Gene | LOC105280348 | protein_coding | character(0) | character(0) | RphaseA1.vs.RphaseB2 | FALSE |
| NC_039506.1 | 2981402 | 2981402 | 1 | + | 73 | 68 | 5 | 81 | 79 | 2 | 93.15 | 97.53 | NC_039506.1 | 2980439 | 2984235 | 3797 | + | Gnomon | gene | gene-LOC105280348 | GeneID:105280348 | LOC105280348 | Gene | LOC105280348 | protein_coding | character(0) | character(0) | RphaseA1.vs.RphaseB2 | FALSE |
| NC_039506.1 | 2981411 | 2981411 | 1 | + | 73 | 70 | 3 | 88 | 84 | 4 | 95.89 | 95.45 | NC_039506.1 | 2980439 | 2984235 | 3797 | + | Gnomon | gene | gene-LOC105280348 | GeneID:105280348 | LOC105280348 | Gene | LOC105280348 | protein_coding | character(0) | character(0) | RphaseA1.vs.RphaseB2 | FALSE |
| NC_039506.1 | 2981802 | 2981802 | 1 | + | 26 | 9 | 17 | 23 | 10 | 13 | 34.62 | 43.48 | NC_039506.1 | 2980439 | 2984235 | 3797 | + | Gnomon | gene | gene-LOC105280348 | GeneID:105280348 | LOC105280348 | Gene | LOC105280348 | protein_coding | character(0) | character(0) | RphaseA1.vs.RphaseB2 | FALSE |
| NC_039506.1 | 2981975 | 2981975 | 1 | + | 29 | 14 | 15 | 20 | 13 | 7 | 48.28 | 65 | NC_039506.1 | 2980439 | 2984235 | 3797 | + | Gnomon | gene | gene-LOC105280348 | GeneID:105280348 | LOC105280348 | Gene | LOC105280348 | protein_coding | character(0) | character(0) | RphaseA1.vs.RphaseB2 | FALSE |

...

... only first 25 lines shown ...

Directory: DMSG File: dmg-Ob_LI_RphaseA2.vs.RphaseB1_details.txt

| seqnames | start | end | width | strand | coverage1 | numCs1 | numTs1 | coverage2 | numCs2 | numTs2 | RphaseA2 | RphaseB1 | gene_seqnames | gene_start | gene_end | gene_width | gene_strand | gene_source | gene_type | gene_ID | gene_Dbxref | gene_Name | gene_gbkey | gene_gene | gene_gene_biotype | gene_end_range | gene_start_range | gene_comparison | is.dm |
|---|---|---|---|---|---|---|---|---|---|---|---|---|---|---|---|---|---|---|---|---|---|---|---|---|---|---|---|---|---|
| NC_039506.1 | 2173251 | 2173251 | 1 | + | 22 | 0 | 22 | 20 | 0 | 20 | 0 | 0 | NC_039506.1 | 2172791 | 2181709 | 8919 | + | Gnomon | gene | gene-LOC105280304 | GeneID:105280304 | LOC105280304 | Gene | LOC105280304 | protein_coding | character(0) | character(0) | RphaseA2.vs.RphaseB1 | FALSE |
| NC_039506.1 | 2173602 | 2173602 | 1 | + | 50 | 2 | 48 | 22 | 2 | 20 | 4 | 9.09 | NC_039506.1 | 2172791 | 2181709 | 8919 | + | Gnomon | gene | gene-LOC105280304 | GeneID:105280304 | LOC105280304 | Gene | LOC105280304 | protein_coding | character(0) | character(0) | RphaseA2.vs.RphaseB1 | FALSE |
| NC_039506.1 | 2173608 | 2173608 | 1 | + | 49 | 1 | 48 | 21 | 0 | 21 | 2.04 | 0 | NC_039506.1 | 2172791 | 2181709 | 8919 | + | Gnomon | gene | gene-LOC105280304 | GeneID:105280304 | LOC105280304 | Gene | LOC105280304 | protein_coding | character(0) | character(0) | RphaseA2.vs.RphaseB1 | FALSE |
| NC_039506.1 | 2173614 | 2173614 | 1 | + | 63 | 0 | 63 | 28 | 0 | 28 | 0 | 0 | NC_039506.1 | 2172791 | 2181709 | 8919 | + | Gnomon | gene | gene-LOC105280304 | GeneID:105280304 | LOC105280304 | Gene | LOC105280304 | protein_coding | character(0) | character(0) | RphaseA2.vs.RphaseB1 | FALSE |
| NC_039506.1 | 2173709 | 2173709 | 1 | + | 64 | 0 | 64 | 46 | 0 | 46 | 0 | 0 | NC_039506.1 | 2172791 | 2181709 | 8919 | + | Gnomon | gene | gene-LOC105280304 | GeneID:105280304 | LOC105280304 | Gene | LOC105280304 | protein_coding | character(0) | character(0) | RphaseA2.vs.RphaseB1 | FALSE |
| NC_039506.1 | 2173733 | 2173733 | 1 | + | 61 | 1 | 60 | 52 | 0 | 52 | 1.64 | 0 | NC_039506.1 | 2172791 | 2181709 | 8919 | + | Gnomon | gene | gene-LOC105280304 | GeneID:105280304 | LOC105280304 | Gene | LOC105280304 | protein_coding | character(0) | character(0) | RphaseA2.vs.RphaseB1 | FALSE |
| NC_039506.1 | 2173742 | 2173742 | 1 | + | 66 | 0 | 66 | 51 | 1 | 50 | 0 | 1.96 | NC_039506.1 | 2172791 | 2181709 | 8919 | + | Gnomon | gene | gene-LOC105280304 | GeneID:105280304 | LOC105280304 | Gene | LOC105280304 | protein_coding | character(0) | character(0) | RphaseA2.vs.RphaseB1 | FALSE |
| NC_039506.1 | 2173746 | 2173746 | 1 | + | 67 | 0 | 67 | 53 | 1 | 52 | 0 | 1.89 | NC_039506.1 | 2172791 | 2181709 | 8919 | + | Gnomon | gene | gene-LOC105280304 | GeneID:105280304 | LOC105280304 | Gene | LOC105280304 | protein_coding | character(0) | character(0) | RphaseA2.vs.RphaseB1 | FALSE |
| NC_039506.1 | 2173753 | 2173753 | 1 | + | 69 | 0 | 69 | 54 | 0 | 54 | 0 | 0 | NC_039506.1 | 2172791 | 2181709 | 8919 | + | Gnomon | gene | gene-LOC105280304 | GeneID:105280304 | LOC105280304 | Gene | LOC105280304 | protein_coding | character(0) | character(0) | RphaseA2.vs.RphaseB1 | FALSE |
| NC_039506.1 | 2173759 | 2173759 | 1 | + | 70 | 1 | 69 | 54 | 0 | 54 | 1.43 | 0 | NC_039506.1 | 2172791 | 2181709 | 8919 | + | Gnomon | gene | gene-LOC105280304 | GeneID:105280304 | LOC105280304 | Gene | LOC105280304 | protein_coding | character(0) | character(0) | RphaseA2.vs.RphaseB1 | FALSE |
| NC_039506.1 | 2173764 | 2173764 | 1 | + | 75 | 0 | 75 | 61 | 0 | 61 | 0 | 0 | NC_039506.1 | 2172791 | 2181709 | 8919 | + | Gnomon | gene | gene-LOC105280304 | GeneID:105280304 | LOC105280304 | Gene | LOC105280304 | protein_coding | character(0) | character(0) | RphaseA2.vs.RphaseB1 | FALSE |
| NC_039506.1 | 2173768 | 2173768 | 1 | + | 75 | 0 | 75 | 59 | 1 | 58 | 0 | 1.69 | NC_039506.1 | 2172791 | 2181709 | 8919 | + | Gnomon | gene | gene-LOC105280304 | GeneID:105280304 | LOC105280304 | Gene | LOC105280304 | protein_coding | character(0) | character(0) | RphaseA2.vs.RphaseB1 | FALSE |
| NC_039506.1 | 2173779 | 2173779 | 1 | + | 82 | 0 | 82 | 66 | 0 | 66 | 0 | 0 | NC_039506.1 | 2172791 | 2181709 | 8919 | + | Gnomon | gene | gene-LOC105280304 | GeneID:105280304 | LOC105280304 | Gene | LOC105280304 | protein_coding | character(0) | character(0) | RphaseA2.vs.RphaseB1 | FALSE |
| NC_039506.1 | 2173789 | 2173789 | 1 | + | 84 | 0 | 84 | 62 | 1 | 61 | 0 | 1.61 | NC_039506.1 | 2172791 | 2181709 | 8919 | + | Gnomon | gene | gene-LOC105280304 | GeneID:105280304 | LOC105280304 | Gene | LOC105280304 | protein_coding | character(0) | character(0) | RphaseA2.vs.RphaseB1 | FALSE |
| NC_039506.1 | 2173798 | 2173798 | 1 | + | 74 | 0 | 74 | 56 | 0 | 56 | 0 | 0 | NC_039506.1 | 2172791 | 2181709 | 8919 | + | Gnomon | gene | gene-LOC105280304 | GeneID:105280304 | LOC105280304 | Gene | LOC105280304 | protein_coding | character(0) | character(0) | RphaseA2.vs.RphaseB1 | FALSE |
| NC_039506.1 | 2173801 | 2173801 | 1 | + | 68 | 1 | 67 | 50 | 0 | 50 | 1.47 | 0 | NC_039506.1 | 2172791 | 2181709 | 8919 | + | Gnomon | gene | gene-LOC105280304 | GeneID:105280304 | LOC105280304 | Gene | LOC105280304 | protein_coding | character(0) | character(0) | RphaseA2.vs.RphaseB1 | FALSE |
| NC_039506.1 | 2173810 | 2173810 | 1 | + | 54 | 1 | 53 | 40 | 1 | 39 | 1.85 | 2.5 | NC_039506.1 | 2172791 | 2181709 | 8919 | + | Gnomon | gene | gene-LOC105280304 | GeneID:105280304 | LOC105280304 | Gene | LOC105280304 | protein_coding | character(0) | character(0) | RphaseA2.vs.RphaseB1 | FALSE |
| NC_039506.1 | 2173812 | 2173812 | 1 | + | 54 | 3 | 51 | 40 | 2 | 38 | 5.56 | 5 | NC_039506.1 | 2172791 | 2181709 | 8919 | + | Gnomon | gene | gene-LOC105280304 | GeneID:105280304 | LOC105280304 | Gene | LOC105280304 | protein_coding | character(0) | character(0) | RphaseA2.vs.RphaseB1 | FALSE |
| NC_039506.1 | 2173823 | 2173823 | 1 | + | 48 | 1 | 47 | 58 | 0 | 58 | 2.08 | 0 | NC_039506.1 | 2172791 | 2181709 | 8919 | + | Gnomon | gene | gene-LOC105280304 | GeneID:105280304 | LOC105280304 | Gene | LOC105280304 | protein_coding | character(0) | character(0) | RphaseA2.vs.RphaseB1 | FALSE |
| NC_039506.1 | 2173831 | 2173831 | 1 | + | 48 | 0 | 48 | 56 | 1 | 55 | 0 | 1.79 | NC_039506.1 | 2172791 | 2181709 | 8919 | + | Gnomon | gene | gene-LOC105280304 | GeneID:105280304 | LOC105280304 | Gene | LOC105280304 | protein_coding | character(0) | character(0) | RphaseA2.vs.RphaseB1 | FALSE |
| NC_039506.1 | 2173849 | 2173849 | 1 | + | 28 | 0 | 28 | 43 | 0 | 43 | 0 | 0 | NC_039506.1 | 2172791 | 2181709 | 8919 | + | Gnomon | gene | gene-LOC105280304 | GeneID:105280304 | LOC105280304 | Gene | LOC105280304 | protein_coding | character(0) | character(0) | RphaseA2.vs.RphaseB1 | FALSE |
| NC_039506.1 | 2174095 | 2174095 | 1 | + | 36 | 0 | 36 | 21 | 0 | 21 | 0 | 0 | NC_039506.1 | 2172791 | 2181709 | 8919 | + | Gnomon | gene | gene-LOC105280304 | GeneID:105280304 | LOC105280304 | Gene | LOC105280304 | protein_coding | character(0) | character(0) | RphaseA2.vs.RphaseB1 | FALSE |
| NC_039506.1 | 2174105 | 2174105 | 1 | + | 39 | 0 | 39 | 25 | 0 | 25 | 0 | 0 | NC_039506.1 | 2172791 | 2181709 | 8919 | + | Gnomon | gene | gene-LOC105280304 | GeneID:105280304 | LOC105280304 | Gene | LOC105280304 | protein_coding | character(0) | character(0) | RphaseA2.vs.RphaseB1 | FALSE |
| NC_039506.1 | 2174115 | 2174115 | 1 | + | 37 | 0 | 37 | 21 | 0 | 21 | 0 | 0 | NC_039506.1 | 2172791 | 2181709 | 8919 | + | Gnomon | gene | gene-LOC105280304 | GeneID:105280304 | LOC105280304 | Gene | LOC105280304 | protein_coding | character(0) | character(0) | RphaseA2.vs.RphaseB1 | FALSE |

...
... only first 25 lines shown ...

Directory: DMSG File: dmg-Ob_LI_RphaseA2.vs.RphaseB2_details.txt

| seqnames | start | end | width | strand | coverage1 | numCs1 | numTs1 | coverage2 | numCs2 | numTs2 | RphaseA2 | RphaseB2 | gene_seqnames | gene_start | gene_end | gene_width | gene_strand | gene_source | gene_type | gene_ID | gene_Dbxref | gene_Name | gene_gbkey | gene_gene | gene_gene_biotype | gene_end_range | gene_start_range | gene_comparison | is.dm |
|---|---|---|---|---|---|---|---|---|---|---|---|---|---|---|---|---|---|---|---|---|---|---|---|---|---|---|---|---|---|
| NC_039506.1 | 2980668 | 2980668 | 1 | + | 35 | 0 | 35 | 28 | 0 | 28 | 0 | 0 | NC_039506.1 | 2980439 | 2984235 | 3797 | + | Gnomon | gene | gene-LOC105280348 | GeneID:105280348 | LOC105280348 | Gene | LOC105280348 | protein_coding | character(0) | character(0) | RphaseA2.vs.RphaseB2 | FALSE |
| NC_039506.1 | 2980750 | 2980750 | 1 | + | 31 | 1 | 30 | 25 | 1 | 24 | 3.23 | 4 | NC_039506.1 | 2980439 | 2984235 | 3797 | + | Gnomon | gene | gene-LOC105280348 | GeneID:105280348 | LOC105280348 | Gene | LOC105280348 | protein_coding | character(0) | character(0) | RphaseA2.vs.RphaseB2 | FALSE |
| NC_039506.1 | 2980761 | 2980761 | 1 | + | 33 | 0 | 33 | 26 | 1 | 25 | 0 | 3.85 | NC_039506.1 | 2980439 | 2984235 | 3797 | + | Gnomon | gene | gene-LOC105280348 | GeneID:105280348 | LOC105280348 | Gene | LOC105280348 | protein_coding | character(0) | character(0) | RphaseA2.vs.RphaseB2 | FALSE |
| NC_039506.1 | 2980767 | 2980767 | 1 | + | 35 | 0 | 35 | 26 | 0 | 26 | 0 | 0 | NC_039506.1 | 2980439 | 2984235 | 3797 | + | Gnomon | gene | gene-LOC105280348 | GeneID:105280348 | LOC105280348 | Gene | LOC105280348 | protein_coding | character(0) | character(0) | RphaseA2.vs.RphaseB2 | FALSE |
| NC_039506.1 | 2980989 | 2980989 | 1 | + | 52 | 11 | 41 | 27 | 3 | 24 | 21.15 | 11.11 | NC_039506.1 | 2980439 | 2984235 | 3797 | + | Gnomon | gene | gene-LOC105280348 | GeneID:105280348 | LOC105280348 | Gene | LOC105280348 | protein_coding | character(0) | character(0) | RphaseA2.vs.RphaseB2 | FALSE |
| NC_039506.1 | 2980993 | 2980993 | 1 | + | 52 | 23 | 29 | 27 | 9 | 18 | 44.23 | 33.33 | NC_039506.1 | 2980439 | 2984235 | 3797 | + | Gnomon | gene | gene-LOC105280348 | GeneID:105280348 | LOC105280348 | Gene | LOC105280348 | protein_coding | character(0) | character(0) | RphaseA2.vs.RphaseB2 | FALSE |
| NC_039506.1 | 2980998 | 2980998 | 1 | + | 51 | 24 | 27 | 27 | 7 | 20 | 47.06 | 25.93 | NC_039506.1 | 2980439 | 2984235 | 3797 | + | Gnomon | gene | gene-LOC105280348 | GeneID:105280348 | LOC105280348 | Gene | LOC105280348 | protein_coding | character(0) | character(0) | RphaseA2.vs.RphaseB2 | FALSE |
| NC_039506.1 | 2981030 | 2981030 | 1 | + | 81 | 74 | 7 | 22 | 19 | 3 | 91.36 | 86.36 | NC_039506.1 | 2980439 | 2984235 | 3797 | + | Gnomon | gene | gene-LOC105280348 | GeneID:105280348 | LOC105280348 | Gene | LOC105280348 | protein_coding | character(0) | character(0) | RphaseA2.vs.RphaseB2 | FALSE |
| NC_039506.1 | 2981077 | 2981077 | 1 | + | 75 | 30 | 45 | 37 | 22 | 15 | 40 | 59.46 | NC_039506.1 | 2980439 | 2984235 | 3797 | + | Gnomon | gene | gene-LOC105280348 | GeneID:105280348 | LOC105280348 | Gene | LOC105280348 | protein_coding | character(0) | character(0) | RphaseA2.vs.RphaseB2 | FALSE |
| NC_039506.1 | 2981125 | 2981125 | 1 | + | 30 | 16 | 14 | 29 | 19 | 10 | 53.33 | 65.52 | NC_039506.1 | 2980439 | 2984235 | 3797 | + | Gnomon | gene | gene-LOC105280348 | GeneID:105280348 | LOC105280348 | Gene | LOC105280348 | protein_coding | character(0) | character(0) | RphaseA2.vs.RphaseB2 | FALSE |
| NC_039506.1 | 2981146 | 2981146 | 1 | + | 51 | 48 | 3 | 24 | 23 | 1 | 94.12 | 95.83 | NC_039506.1 | 2980439 | 2984235 | 3797 | + | Gnomon | gene | gene-LOC105280348 | GeneID:105280348 | LOC105280348 | Gene | LOC105280348 | protein_coding | character(0) | character(0) | RphaseA2.vs.RphaseB2 | FALSE |
| NC_039506.1 | 2981183 | 2981183 | 1 | + | 108 | 105 | 3 | 59 | 39 | 20 | 97.22 | 66.1 | NC_039506.1 | 2980439 | 2984235 | 3797 | + | Gnomon | gene | gene-LOC105280348 | GeneID:105280348 | LOC105280348 | Gene | LOC105280348 | protein_coding | character(0) | character(0) | RphaseA2.vs.RphaseB2 | TRUE |
| NC_039506.1 | 2981225 | 2981225 | 1 | + | 83 | 51 | 32 | 39 | 29 | 10 | 61.45 | 74.36 | NC_039506.1 | 2980439 | 2984235 | 3797 | + | Gnomon | gene | gene-LOC105280348 | GeneID:105280348 | LOC105280348 | Gene | LOC105280348 | protein_coding | character(0) | character(0) | RphaseA2.vs.RphaseB2 | FALSE |
| NC_039506.1 | 2981245 | 2981245 | 1 | + | 69 | 28 | 41 | 36 | 21 | 15 | 40.58 | 58.33 | NC_039506.1 | 2980439 | 2984235 | 3797 | + | Gnomon | gene | gene-LOC105280348 | GeneID:105280348 | LOC105280348 | Gene | LOC105280348 | protein_coding | character(0) | character(0) | RphaseA2.vs.RphaseB2 | FALSE |
| NC_039506.1 | 2981292 | 2981292 | 1 | + | 92 | 86 | 6 | 42 | 40 | 2 | 93.48 | 95.24 | NC_039506.1 | 2980439 | 2984235 | 3797 | + | Gnomon | gene | gene-LOC105280348 | GeneID:105280348 | LOC105280348 | Gene | LOC105280348 | protein_coding | character(0) | character(0) | RphaseA2.vs.RphaseB2 | FALSE |
| NC_039506.1 | 2981308 | 2981308 | 1 | + | 104 | 70 | 34 | 46 | 20 | 26 | 67.31 | 43.48 | NC_039506.1 | 2980439 | 2984235 | 3797 | + | Gnomon | gene | gene-LOC105280348 | GeneID:105280348 | LOC105280348 | Gene | LOC105280348 | protein_coding | character(0) | character(0) | RphaseA2.vs.RphaseB2 | FALSE |
| NC_039506.1 | 2981315 | 2981315 | 1 | + | 108 | 90 | 18 | 49 | 39 | 10 | 83.33 | 79.59 | NC_039506.1 | 2980439 | 2984235 | 3797 | + | Gnomon | gene | gene-LOC105280348 | GeneID:105280348 | LOC105280348 | Gene | LOC105280348 | protein_coding | character(0) | character(0) | RphaseA2.vs.RphaseB2 | FALSE |
| NC_039506.1 | 2981335 | 2981335 | 1 | + | 68 | 67 | 1 | 40 | 38 | 2 | 98.53 | 95 | NC_039506.1 | 2980439 | 2984235 | 3797 | + | Gnomon | gene | gene-LOC105280348 | GeneID:105280348 | LOC105280348 | Gene | LOC105280348 | protein_coding | character(0) | character(0) | RphaseA2.vs.RphaseB2 | FALSE |
| NC_039506.1 | 2981350 | 2981350 | 1 | + | 97 | 95 | 2 | 74 | 73 | 1 | 97.94 | 98.65 | NC_039506.1 | 2980439 | 2984235 | 3797 | + | Gnomon | gene | gene-LOC105280348 | GeneID:105280348 | LOC105280348 | Gene | LOC105280348 | protein_coding | character(0) | character(0) | RphaseA2.vs.RphaseB2 | FALSE |
| NC_039506.1 | 2981391 | 2981391 | 1 | + | 102 | 84 | 18 | 80 | 65 | 15 | 82.35 | 81.25 | NC_039506.1 | 2980439 | 2984235 | 3797 | + | Gnomon | gene | gene-LOC105280348 | GeneID:105280348 | LOC105280348 | Gene | LOC105280348 | protein_coding | character(0) | character(0) | RphaseA2.vs.RphaseB2 | FALSE |
| NC_039506.1 | 2981402 | 2981402 | 1 | + | 115 | 102 | 13 | 81 | 79 | 2 | 88.7 | 97.53 | NC_039506.1 | 2980439 | 2984235 | 3797 | + | Gnomon | gene | gene-LOC105280348 | GeneID:105280348 | LOC105280348 | Gene | LOC105280348 | protein_coding | character(0) | character(0) | RphaseA2.vs.RphaseB2 | FALSE |
| NC_039506.1 | 2981411 | 2981411 | 1 | + | 132 | 127 | 5 | 88 | 84 | 4 | 96.21 | 95.45 | NC_039506.1 | 2980439 | 2984235 | 3797 | + | Gnomon | gene | gene-LOC105280348 | GeneID:105280348 | LOC105280348 | Gene | LOC105280348 | protein_coding | character(0) | character(0) | RphaseA2.vs.RphaseB2 | FALSE |
| NC_039506.1 | 2981975 | 2981975 | 1 | + | 26 | 10 | 16 | 20 | 13 | 7 | 38.46 | 65 | NC_039506.1 | 2980439 | 2984235 | 3797 | + | Gnomon | gene | gene-LOC105280348 | GeneID:105280348 | LOC105280348 | Gene | LOC105280348 | protein_coding | character(0) | character(0) | RphaseA2.vs.RphaseB2 | FALSE |
| NC_039506.1 | 2982097 | 2982097 | 1 | + | 51 | 2 | 49 | 31 | 3 | 28 | 3.92 | 9.68 | NC_039506.1 | 2980439 | 2984235 | 3797 | + | Gnomon | gene | gene-LOC105280348 | GeneID:105280348 | LOC105280348 | Gene | LOC105280348 | protein_coding | character(0) | character(0) | RphaseA2.vs.RphaseB2 | FALSE |

...

... only first 25 lines shown ...

Directory: DMSG File: dmg-Ob_LI_RphaseB1.vs.RphaseB2_details.txt

| seqnames | start | end | width | strand | coverage1 | numCs1 | numTs1 | coverage2 | numCs2 | numTs2 | RphaseB1 | RphaseB2 | gene_seqnames | gene_start | gene_end | gene_width | gene_strand | gene_source | gene_type | gene_ID | gene_Dbxref | gene_Name | gene_gbkey | gene_gene | gene_gene_biotype | gene_end_range | gene_partial | gene_start_range | gene_comparison | is.dm |
|---|---|---|---|---|---|---|---|---|---|---|---|---|---|---|---|---|---|---|---|---|---|---|---|---|---|---|---|---|---|---|
| NC_039506.1 | 1057034 | 1057034 | 1 | + | 25 | 0 | 25 | 31 | 0 | 31 | 0 | 0 | NC_039506.1 | 1056151 | 1064257 | 8107 | + | Gnomon | gene | gene-LOC105286747 | GeneID:105286747 | LOC105286747 | Gene | LOC105286747 | lncRNA | character(0) | NA | character(0) | RphaseB1.vs.RphaseB2 | FALSE |
| NC_039506.1 | 1057043 | 1057043 | 1 | + | 29 | 0 | 29 | 31 | 0 | 31 | 0 | 0 | NC_039506.1 | 1056151 | 1064257 | 8107 | + | Gnomon | gene | gene-LOC105286747 | GeneID:105286747 | LOC105286747 | Gene | LOC105286747 | lncRNA | character(0) | NA | character(0) | RphaseB1.vs.RphaseB2 | FALSE |
| NC_039506.1 | 1057047 | 1057047 | 1 | + | 30 | 0 | 30 | 31 | 0 | 31 | 0 | 0 | NC_039506.1 | 1056151 | 1064257 | 8107 | + | Gnomon | gene | gene-LOC105286747 | GeneID:105286747 | LOC105286747 | Gene | LOC105286747 | lncRNA | character(0) | NA | character(0) | RphaseB1.vs.RphaseB2 | FALSE |
| NC_039506.1 | 1057061 | 1057061 | 1 | + | 30 | 0 | 30 | 33 | 0 | 33 | 0 | 0 | NC_039506.1 | 1056151 | 1064257 | 8107 | + | Gnomon | gene | gene-LOC105286747 | GeneID:105286747 | LOC105286747 | Gene | LOC105286747 | lncRNA | character(0) | NA | character(0) | RphaseB1.vs.RphaseB2 | FALSE |
| NC_039506.1 | 1057064 | 1057064 | 1 | + | 30 | 0 | 30 | 34 | 0 | 34 | 0 | 0 | NC_039506.1 | 1056151 | 1064257 | 8107 | + | Gnomon | gene | gene-LOC105286747 | GeneID:105286747 | LOC105286747 | Gene | LOC105286747 | lncRNA | character(0) | NA | character(0) | RphaseB1.vs.RphaseB2 | FALSE |
| NC_039506.1 | 1057110 | 1057110 | 1 | + | 37 | 0 | 37 | 35 | 1 | 34 | 0 | 2.86 | NC_039506.1 | 1056151 | 1064257 | 8107 | + | Gnomon | gene | gene-LOC105286747 | GeneID:105286747 | LOC105286747 | Gene | LOC105286747 | lncRNA | character(0) | NA | character(0) | RphaseB1.vs.RphaseB2 | FALSE |
| NC_039506.1 | 1057112 | 1057112 | 1 | + | 40 | 0 | 40 | 35 | 0 | 35 | 0 | 0 | NC_039506.1 | 1056151 | 1064257 | 8107 | + | Gnomon | gene | gene-LOC105286747 | GeneID:105286747 | LOC105286747 | Gene | LOC105286747 | lncRNA | character(0) | NA | character(0) | RphaseB1.vs.RphaseB2 | FALSE |
| NC_039506.1 | 1057128 | 1057128 | 1 | + | 30 | 0 | 30 | 30 | 0 | 30 | 0 | 0 | NC_039506.1 | 1056151 | 1064257 | 8107 | + | Gnomon | gene | gene-LOC105286747 | GeneID:105286747 | LOC105286747 | Gene | LOC105286747 | lncRNA | character(0) | NA | character(0) | RphaseB1.vs.RphaseB2 | FALSE |
| NC_039506.1 | 1057134 | 1057134 | 1 | + | 30 | 0 | 30 | 32 | 0 | 32 | 0 | 0 | NC_039506.1 | 1056151 | 1064257 | 8107 | + | Gnomon | gene | gene-LOC105286747 | GeneID:105286747 | LOC105286747 | Gene | LOC105286747 | lncRNA | character(0) | NA | character(0) | RphaseB1.vs.RphaseB2 | FALSE |
| NC_039506.1 | 1057145 | 1057145 | 1 | + | 32 | 0 | 32 | 30 | 0 | 30 | 0 | 0 | NC_039506.1 | 1056151 | 1064257 | 8107 | + | Gnomon | gene | gene-LOC105286747 | GeneID:105286747 | LOC105286747 | Gene | LOC105286747 | lncRNA | character(0) | NA | character(0) | RphaseB1.vs.RphaseB2 | FALSE |
| NC_039506.1 | 1057157 | 1057157 | 1 | + | 30 | 1 | 29 | 29 | 0 | 29 | 3.33 | 0 | NC_039506.1 | 1056151 | 1064257 | 8107 | + | Gnomon | gene | gene-LOC105286747 | GeneID:105286747 | LOC105286747 | Gene | LOC105286747 | lncRNA | character(0) | NA | character(0) | RphaseB1.vs.RphaseB2 | FALSE |
| NC_039506.1 | 1057169 | 1057169 | 1 | + | 28 | 0 | 28 | 27 | 0 | 27 | 0 | 0 | NC_039506.1 | 1056151 | 1064257 | 8107 | + | Gnomon | gene | gene-LOC105286747 | GeneID:105286747 | LOC105286747 | Gene | LOC105286747 | lncRNA | character(0) | NA | character(0) | RphaseB1.vs.RphaseB2 | FALSE |
| NC_039506.1 | 1057181 | 1057181 | 1 | + | 26 | 0 | 26 | 26 | 0 | 26 | 0 | 0 | NC_039506.1 | 1056151 | 1064257 | 8107 | + | Gnomon | gene | gene-LOC105286747 | GeneID:105286747 | LOC105286747 | Gene | LOC105286747 | lncRNA | character(0) | NA | character(0) | RphaseB1.vs.RphaseB2 | FALSE |
| NC_039506.1 | 1057294 | 1057294 | 1 | + | 50 | 0 | 50 | 21 | 0 | 21 | 0 | 0 | NC_039506.1 | 1056151 | 1064257 | 8107 | + | Gnomon | gene | gene-LOC105286747 | GeneID:105286747 | LOC105286747 | Gene | LOC105286747 | lncRNA | character(0) | NA | character(0) | RphaseB1.vs.RphaseB2 | FALSE |
| NC_039506.1 | 1057300 | 1057300 | 1 | + | 53 | 2 | 51 | 24 | 0 | 24 | 3.77 | 0 | NC_039506.1 | 1056151 | 1064257 | 8107 | + | Gnomon | gene | gene-LOC105286747 | GeneID:105286747 | LOC105286747 | Gene | LOC105286747 | lncRNA | character(0) | NA | character(0) | RphaseB1.vs.RphaseB2 | FALSE |
| NC_039506.1 | 1057302 | 1057302 | 1 | + | 52 | 0 | 52 | 26 | 0 | 26 | 0 | 0 | NC_039506.1 | 1056151 | 1064257 | 8107 | + | Gnomon | gene | gene-LOC105286747 | GeneID:105286747 | LOC105286747 | Gene | LOC105286747 | lncRNA | character(0) | NA | character(0) | RphaseB1.vs.RphaseB2 | FALSE |
| NC_039506.1 | 1057308 | 1057308 | 1 | + | 58 | 0 | 58 | 26 | 0 | 26 | 0 | 0 | NC_039506.1 | 1056151 | 1064257 | 8107 | + | Gnomon | gene | gene-LOC105286747 | GeneID:105286747 | LOC105286747 | Gene | LOC105286747 | lncRNA | character(0) | NA | character(0) | RphaseB1.vs.RphaseB2 | FALSE |
| NC_039506.1 | 1057314 | 1057314 | 1 | + | 61 | 4 | 57 | 53 | 0 | 53 | 6.56 | 0 | NC_039506.1 | 1056151 | 1064257 | 8107 | + | Gnomon | gene | gene-LOC105286747 | GeneID:105286747 | LOC105286747 | Gene | LOC105286747 | lncRNA | character(0) | NA | character(0) | RphaseB1.vs.RphaseB2 | FALSE |
| NC_039506.1 | 1057316 | 1057316 | 1 | + | 61 | 0 | 61 | 55 | 0 | 55 | 0 | 0 | NC_039506.1 | 1056151 | 1064257 | 8107 | + | Gnomon | gene | gene-LOC105286747 | GeneID:105286747 | LOC105286747 | Gene | LOC105286747 | lncRNA | character(0) | NA | character(0) | RphaseB1.vs.RphaseB2 | FALSE |
| NC_039506.1 | 1057320 | 1057320 | 1 | + | 58 | 0 | 58 | 52 | 1 | 51 | 0 | 1.92 | NC_039506.1 | 1056151 | 1064257 | 8107 | + | Gnomon | gene | gene-LOC105286747 | GeneID:105286747 | LOC105286747 | Gene | LOC105286747 | lncRNA | character(0) | NA | character(0) | RphaseB1.vs.RphaseB2 | FALSE |
| NC_039506.1 | 1057331 | 1057331 | 1 | + | 77 | 0 | 77 | 83 | 0 | 83 | 0 | 0 | NC_039506.1 | 1056151 | 1064257 | 8107 | + | Gnomon | gene | gene-LOC105286747 | GeneID:105286747 | LOC105286747 | Gene | LOC105286747 | lncRNA | character(0) | NA | character(0) | RphaseB1.vs.RphaseB2 | FALSE |
| NC_039506.1 | 1057337 | 1057337 | 1 | + | 74 | 0 | 74 | 80 | 0 | 80 | 0 | 0 | NC_039506.1 | 1056151 | 1064257 | 8107 | + | Gnomon | gene | gene-LOC105286747 | GeneID:105286747 | LOC105286747 | Gene | LOC105286747 | lncRNA | character(0) | NA | character(0) | RphaseB1.vs.RphaseB2 | FALSE |
| NC_039506.1 | 1057344 | 1057344 | 1 | + | 72 | 0 | 72 | 79 | 0 | 79 | 0 | 0 | NC_039506.1 | 1056151 | 1064257 | 8107 | + | Gnomon | gene | gene-LOC105286747 | GeneID:105286747 | LOC105286747 | Gene | LOC105286747 | lncRNA | character(0) | NA | character(0) | RphaseB1.vs.RphaseB2 | FALSE |
| NC_039506.1 | 1057373 | 1057373 | 1 | + | 32 | 0 | 32 | 46 | 1 | 45 | 0 | 2.17 | NC_039506.1 | 1056151 | 1064257 | 8107 | + | Gnomon | gene | gene-LOC105286747 | GeneID:105286747 | LOC105286747 | Gene | LOC105286747 | lncRNA | character(0) | NA | character(0) | RphaseB1.vs.RphaseB2 | FALSE |

...
... only first 25 lines shown ...

Directory: DMSG File: dms-Ob_LI.txt

| seqnames | start | end | width | strand | pvalue | qvalue | meth.diff | comparison |
|---|---|---|---|---|---|---|---|---|
| NC_039506.1 | 2175399 | 2175399 | 1 | + | 3.3518009746166e-07 | 0.006 | -56 | BCphaseA1.vs.BCphaseA2 |
| NC_039506.1 | 2981183 | 2981183 | 1 | + | 8.62545898096015e-10 | 0 | 47.12 | BCphaseA1.vs.BCphaseA2 |
| NC_039506.1 | 3585329 | 3585329 | 1 | + | 1.35015770592042e-09 | 0 | -49.1 | BCphaseA1.vs.BCphaseA2 |
| NC_039506.1 | 5498915 | 5498915 | 1 | + | 2.15282553701382e-07 | 0.004 | -44.93 | BCphaseA1.vs.BCphaseA2 |
| NC_039506.1 | 10974955 | 10974955 | 1 | + | 2.96668037346733e-06 | 0.042 | 54.17 | BCphaseA1.vs.BCphaseA2 |
| NC_039506.1 | 12496370 | 12496370 | 1 | + | 7.38217163087816e-08 | 0.002 | 55.92 | BCphaseA1.vs.BCphaseA2 |
| NC_039506.1 | 12496715 | 12496715 | 1 | + | 2.29671619364023e-09 | 0 | 60.32 | BCphaseA1.vs.BCphaseA2 |
| NC_039506.1 | 13174267 | 13174267 | 1 | + | 2.10914456271352e-07 | 0.004 | -45.68 | BCphaseA1.vs.BCphaseA2 |
| NC_039506.1 | 15940056 | 15940056 | 1 | + | 1.57083538711083e-06 | 0.024 | 51.6 | BCphaseA1.vs.BCphaseA2 |
| NC_039506.1 | 15940061 | 15940061 | 1 | + | 1.33072107109468e-06 | 0.021 | 48.8 | BCphaseA1.vs.BCphaseA2 |
| NC_039506.1 | 19219354 | 19219354 | 1 | + | 1.19617597079373e-06 | 0.02 | 55.8 | BCphaseA1.vs.BCphaseA2 |
| NC_039506.1 | 20581616 | 20581616 | 1 | + | 1.47438643715683e-07 | 0.003 | -49.74 | BCphaseA1.vs.BCphaseA2 |
| NC_039506.1 | 20611384 | 20611384 | 1 | + | 1.1996962005141e-12 | 0 | 66.06 | BCphaseA1.vs.BCphaseA2 |
| NC_039506.1 | 23929017 | 23929017 | 1 | + | 2.27979698997725e-08 | 0.001 | -47.73 | BCphaseA1.vs.BCphaseA2 |
| NC_039507.1 | 5016313 | 5016313 | 1 | + | 9.02588105279279e-10 | 0 | 45.99 | BCphaseA1.vs.BCphaseA2 |
| NC_039507.1 | 14494249 | 14494249 | 1 | + | 6.35683529022794e-07 | 0.011 | 45.86 | BCphaseA1.vs.BCphaseA2 |
| NC_039507.1 | 14496529 | 14496529 | 1 | + | 1.15901290479277e-06 | 0.019 | -42.26 | BCphaseA1.vs.BCphaseA2 |
| NC_039507.1 | 16959867 | 16959867 | 1 | + | 9.48818831692176e-23 | 0 | 91.3 | BCphaseA1.vs.BCphaseA2 |
| NC_039508.1 | 4118752 | 4118752 | 1 | + | 1.37959967115003e-06 | 0.022 | 58.65 | BCphaseA1.vs.BCphaseA2 |
| NC_039508.1 | 7056837 | 7056837 | 1 | + | 2.78417501597611e-09 | 0 | 53.04 | BCphaseA1.vs.BCphaseA2 |
| NC_039508.1 | 10480425 | 10480425 | 1 | + | 2.22150230629451e-10 | 0 | -46.97 | BCphaseA1.vs.BCphaseA2 |
| NC_039508.1 | 10729874 | 10729874 | 1 | + | 1.4725955689355e-07 | 0.003 | 62.6 | BCphaseA1.vs.BCphaseA2 |
| NC_039508.1 | 15152165 | 15152165 | 1 | + | 1.66615172720232e-10 | 0 | 75 | BCphaseA1.vs.BCphaseA2 |
| NC_039509.1 | 185576 | 185576 | 1 | + | 1.50241302590709e-06 | 0.023 | 37.21 | BCphaseA1.vs.BCphaseA2 |

...
... only first 25 lines shown ...

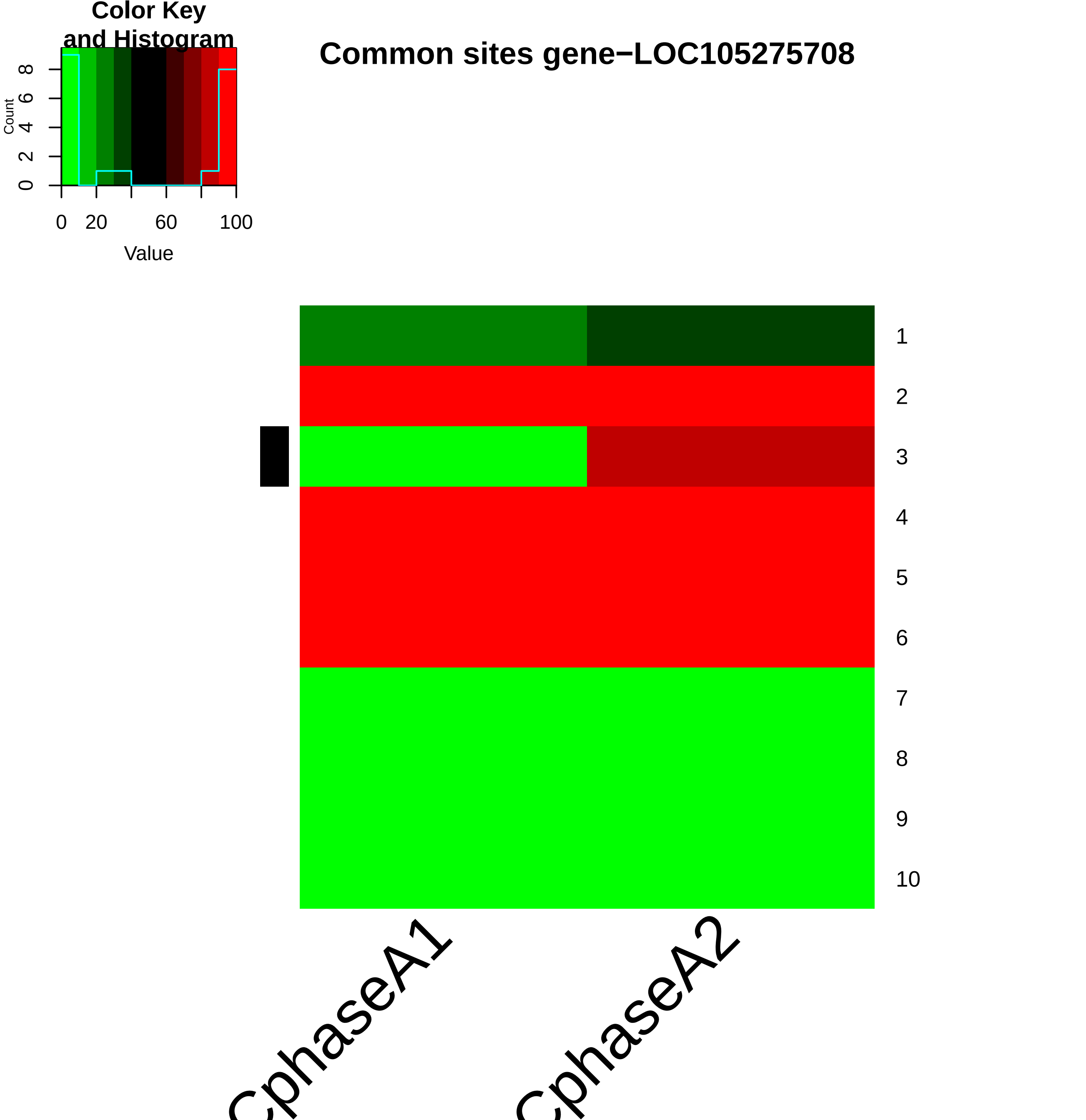
Color Key
and Histogram
Count
0
2
4
6
8
0
20
60
100
Value
Common sites gene−LOC105275708
1
2
3
4
5
6
7
8
9
10
CphaseA1
CphaseA2

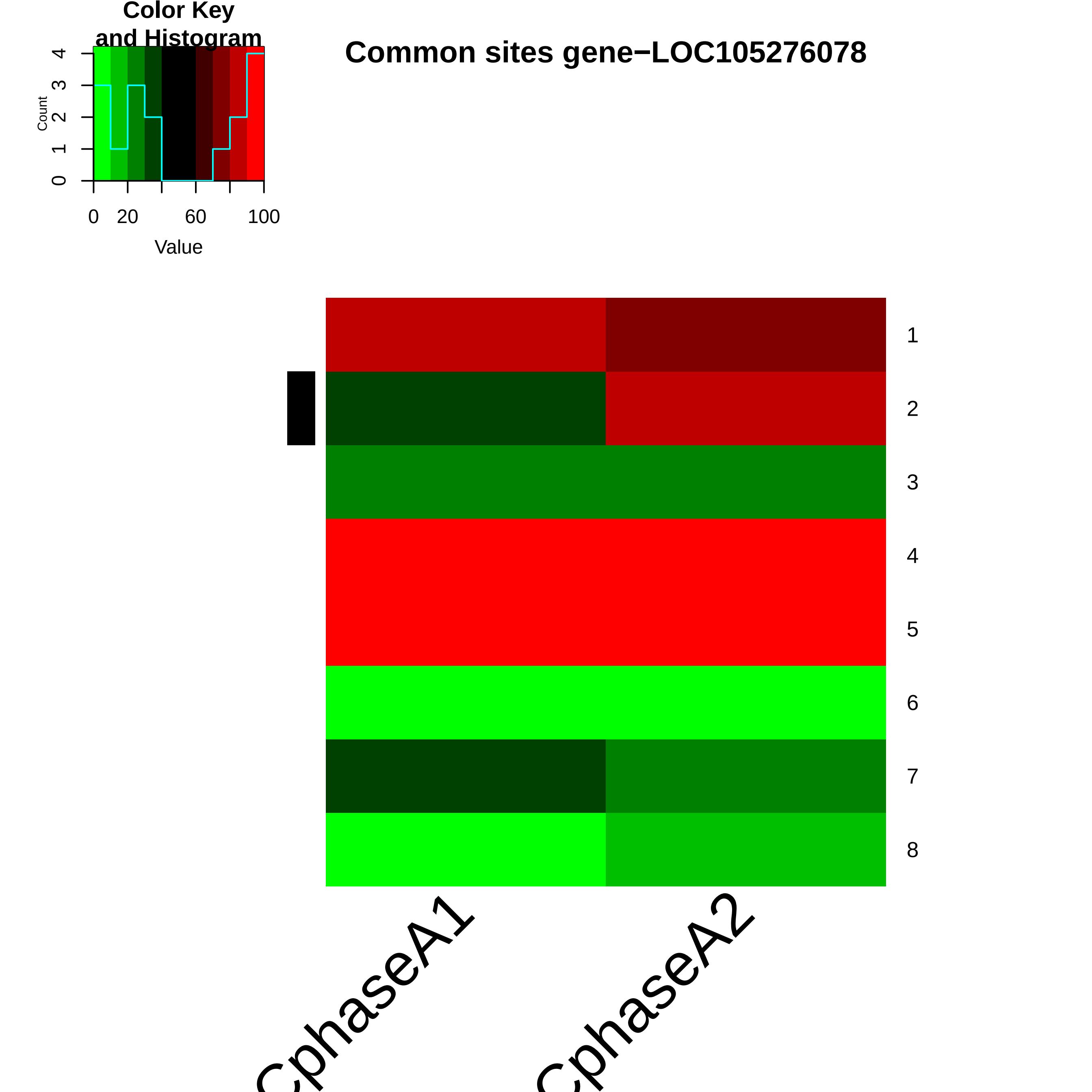
Color Key
and Histogram
Count
0
1
2
3
4
0
20
60
100
Value
Common sites gene−LOC105276078
1
2
3
4
5
6
7
8
CphaseA1
CphaseA2

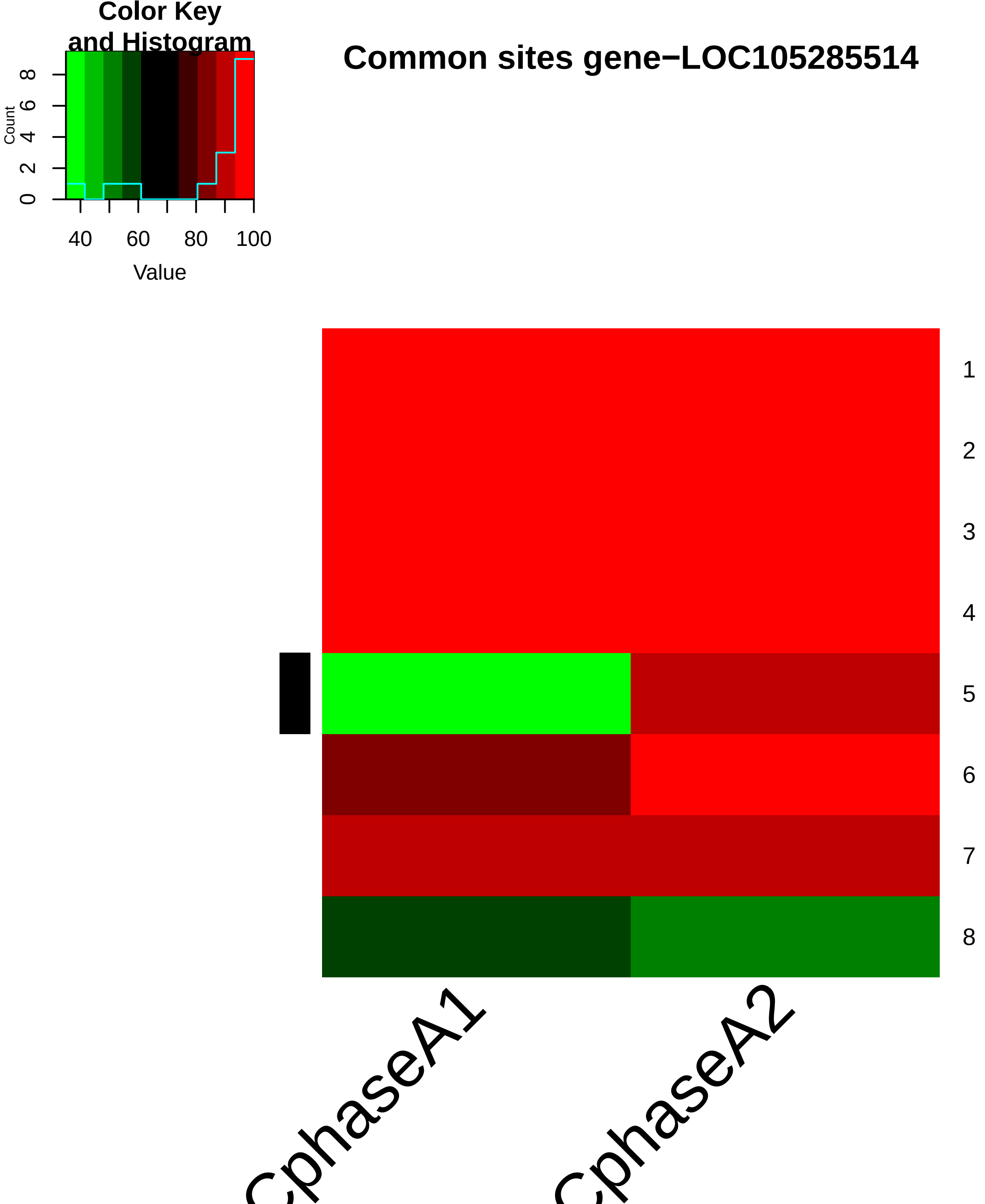

Color Key
and Histogram
Count
0 2 4 6 8
40 60 80 100
Value
Common sites gene−LOC105285514
1
2
3
4
5
6
7
8
CphaseA1
CphaseA2

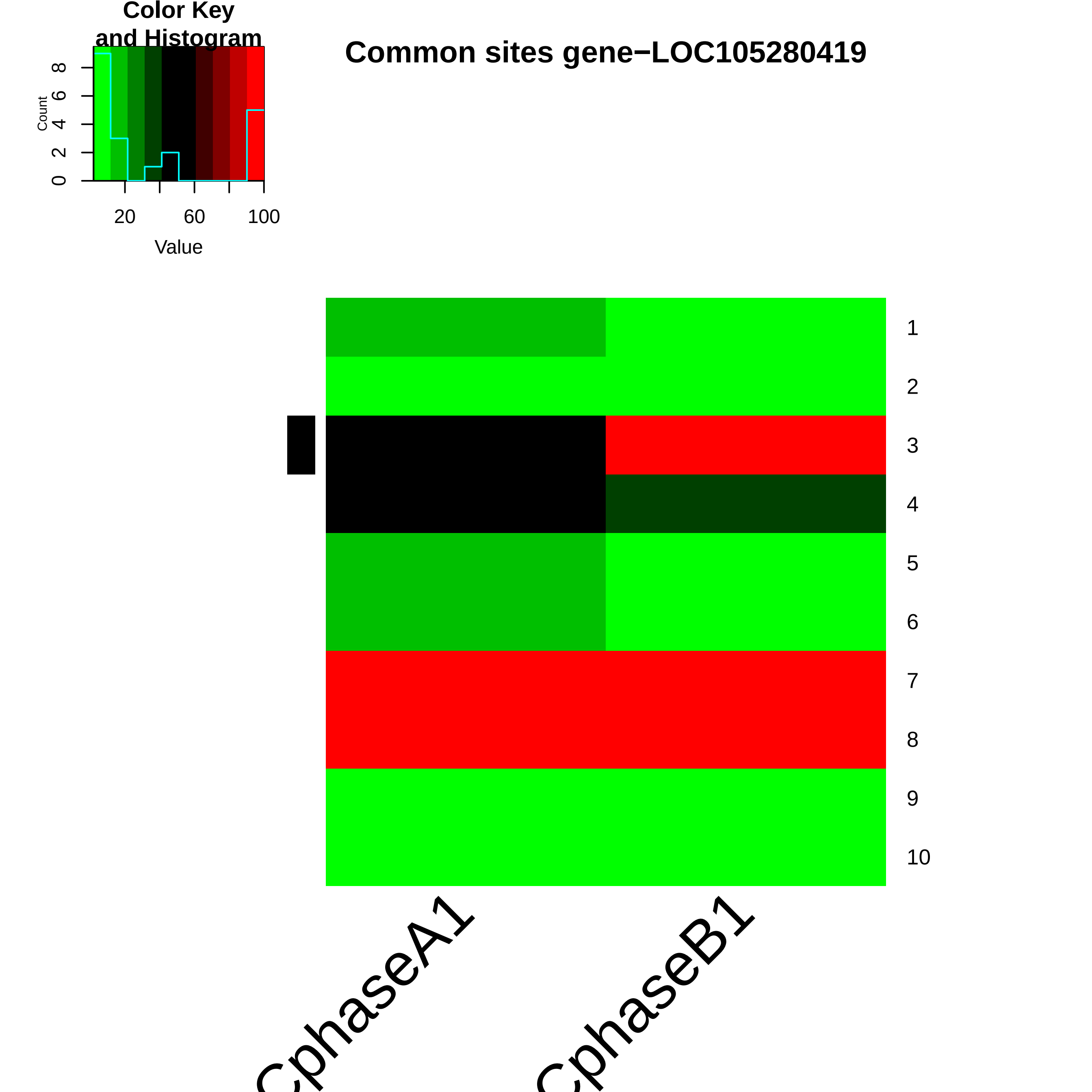
Color Key
and Histogram
Count
Value
Common sites gene−LOC105280419
1
2
3
4
5
6
7
8
9
10
CphaseA1
CphaseB1

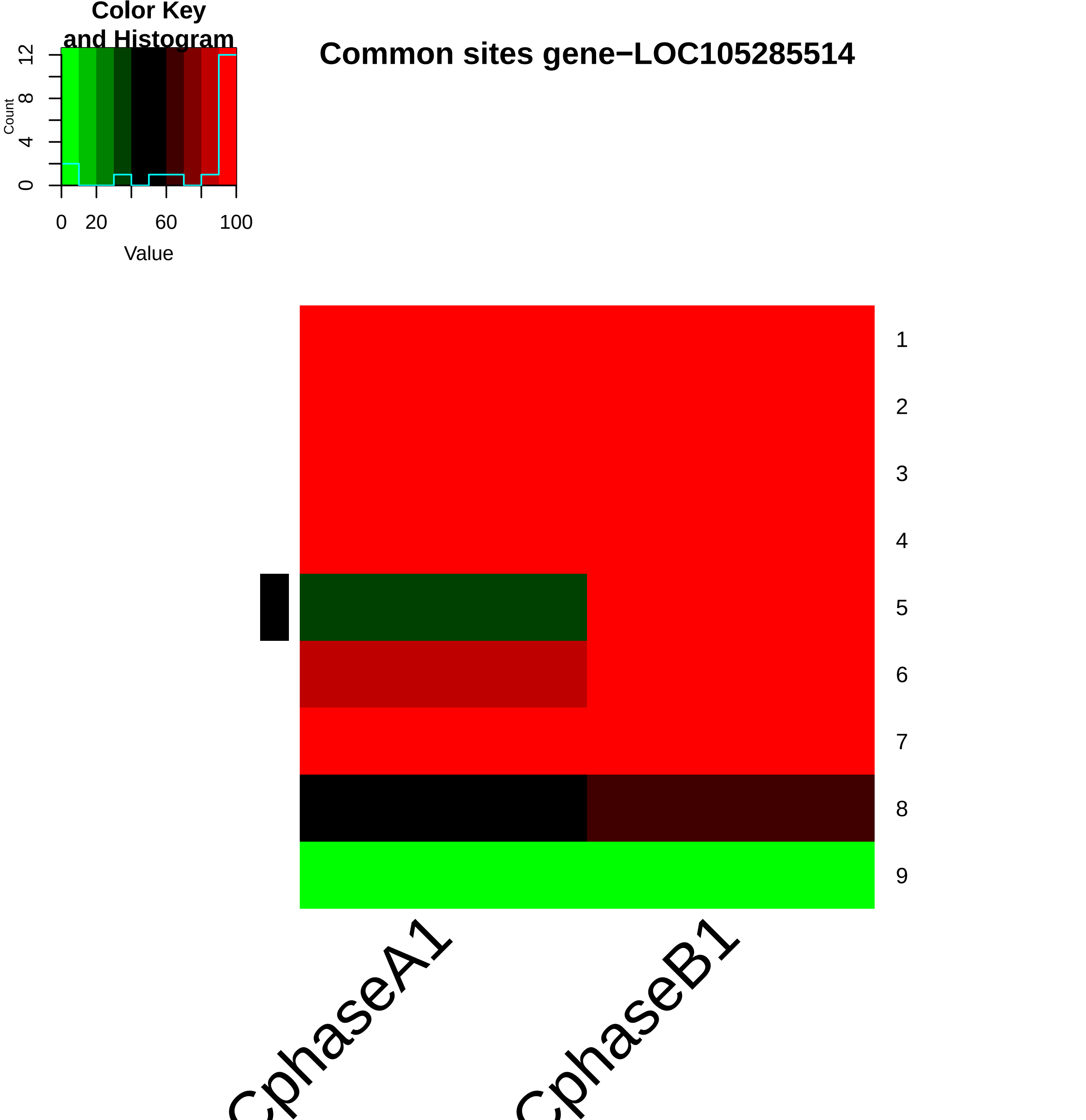
Color Key
and Histogram
Count
0
4
8
12
0
20
60
100
Value
Common sites gene−LOC105285514
1
2
3
4
5
6
7
8
9
CphaseA1
CphaseB1

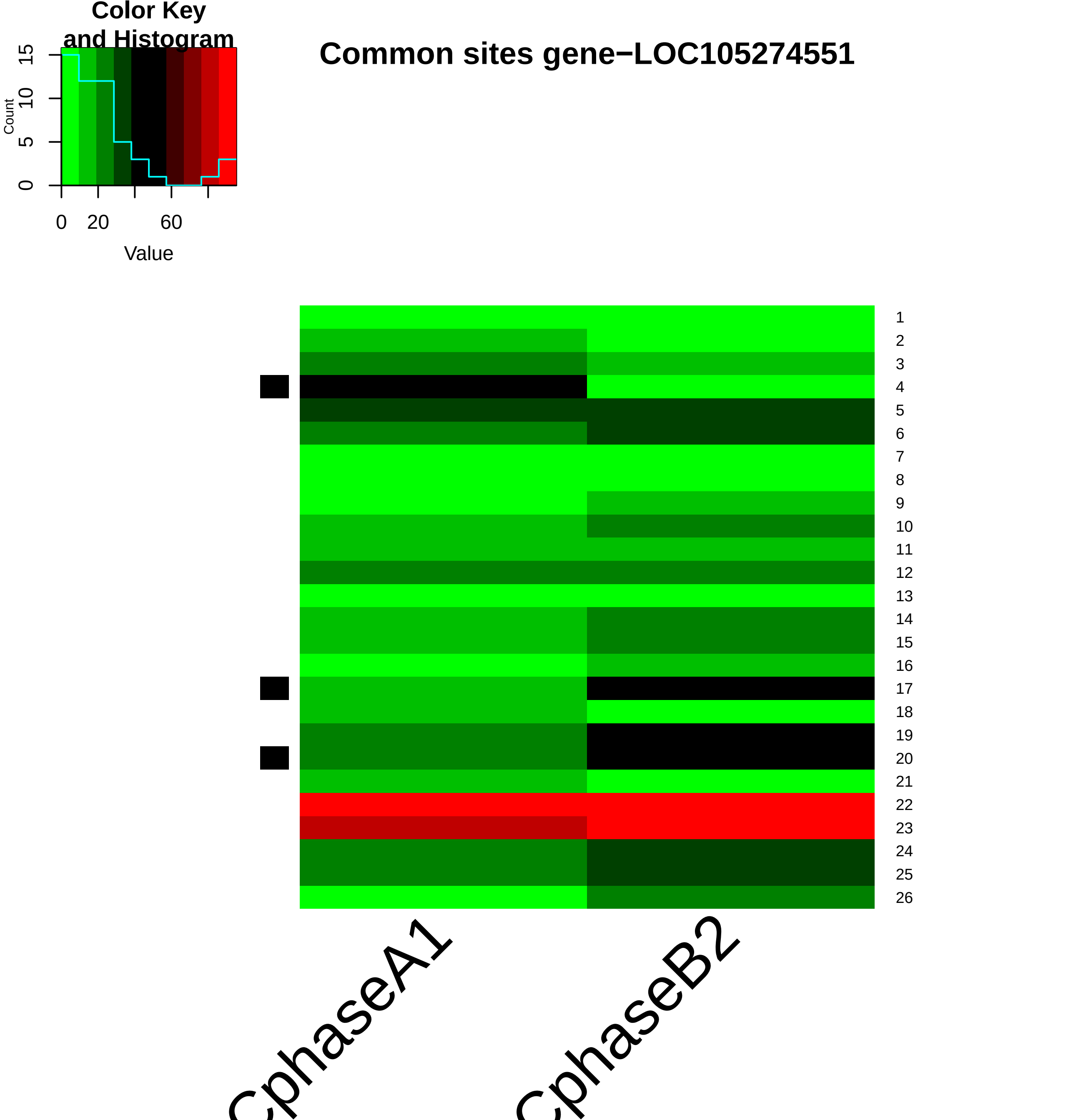
Color Key
and Histogram
Count
Value
0
5
10
15
0
20
60
Common sites gene−LOC105274551
CphaseA1
CphaseB2

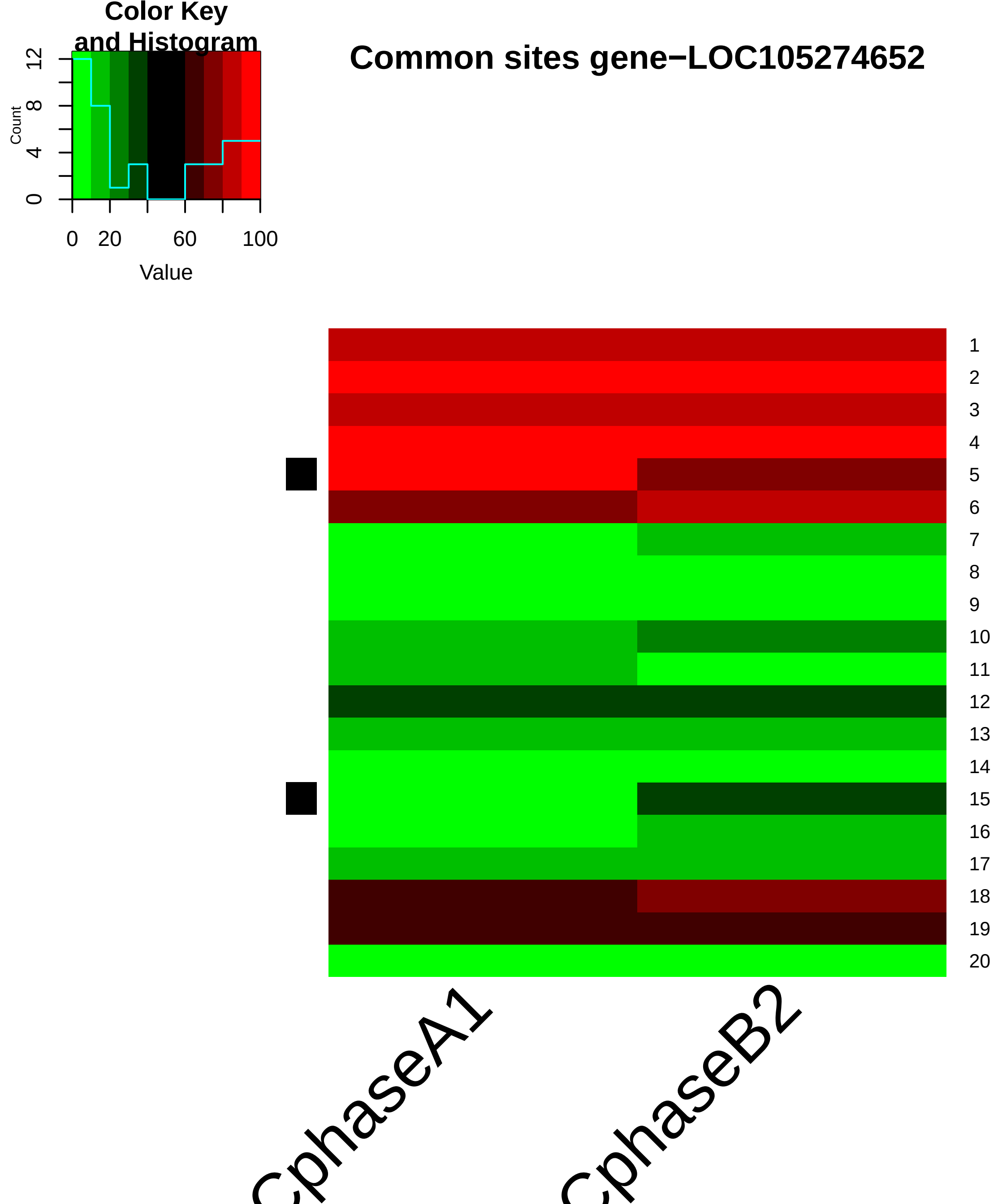
Color Key
and Histogram
Count
0
4
8
12
0
20
60
100
Value
Common sites gene−LOC105274652
CphaseA1
CphaseB2

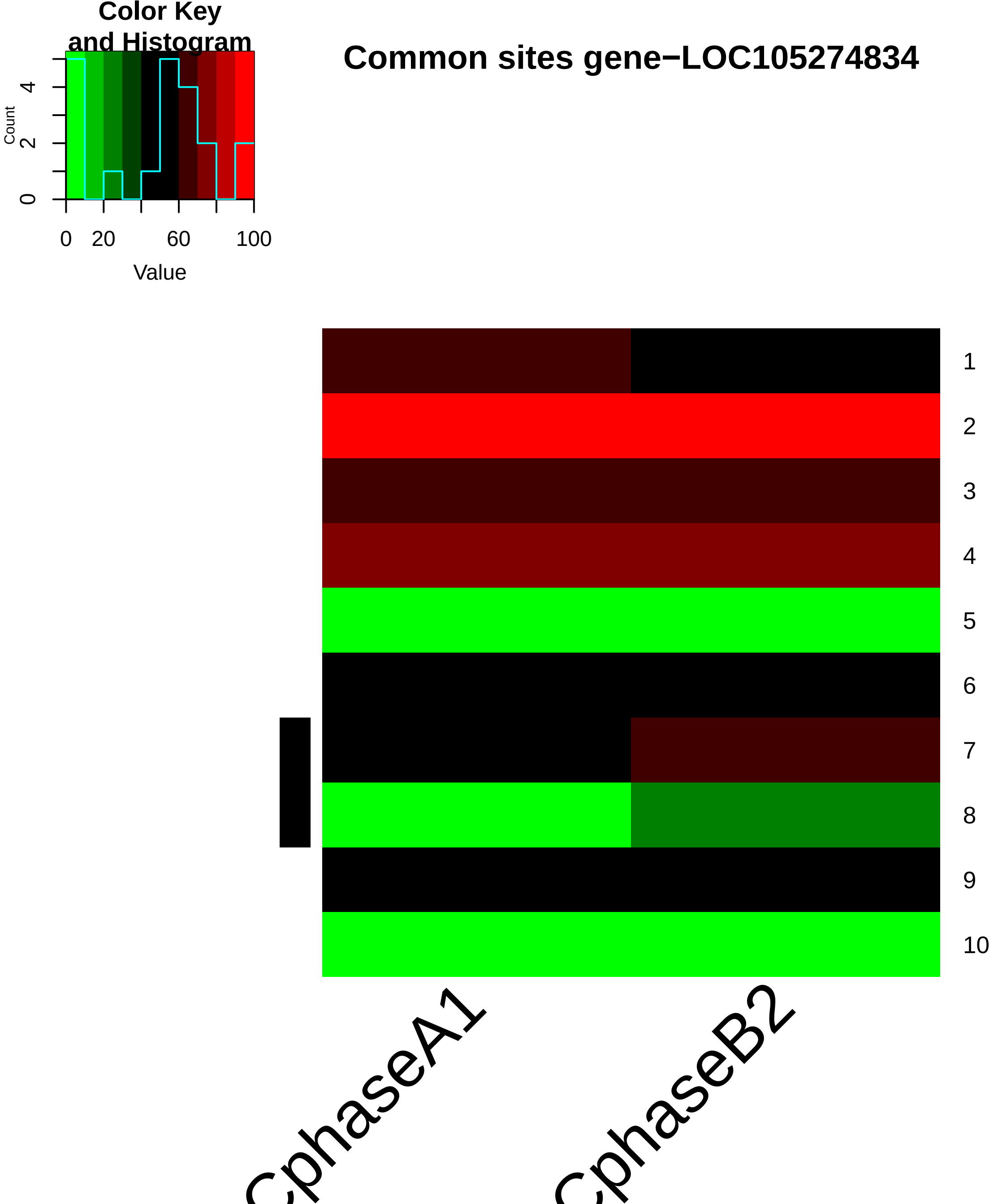
Color Key
and Histogram
Count
0
2
4
0
20
60
100
Value
Common sites gene−LOC105274834
1
2
3
4
5
6
7
8
9
10

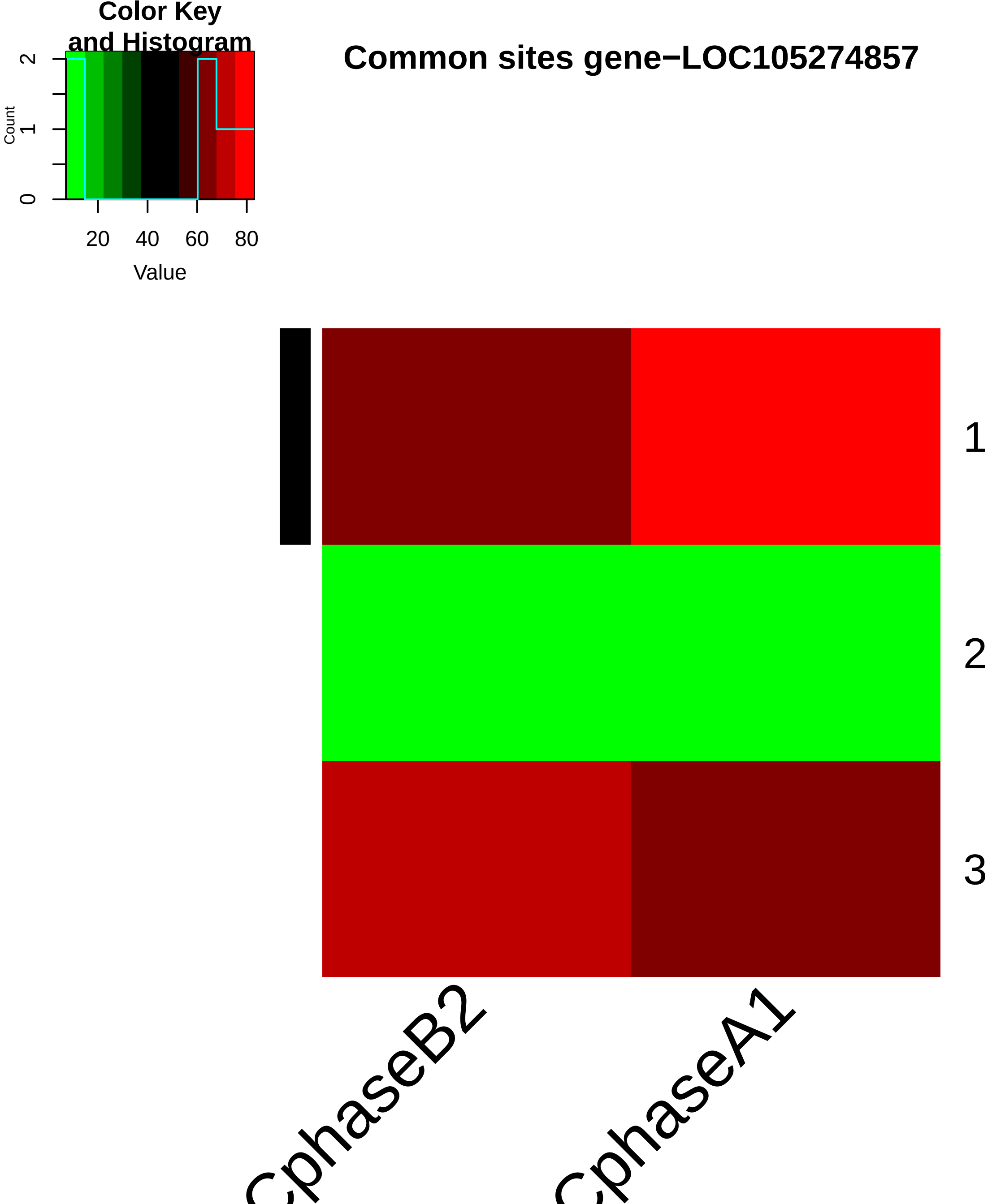

Color Key
and Histogram
Count
0
1
2
20
40
60
80
Value
Common sites gene−LOC105274857
1
2
3
CphaseB2
CphaseA1

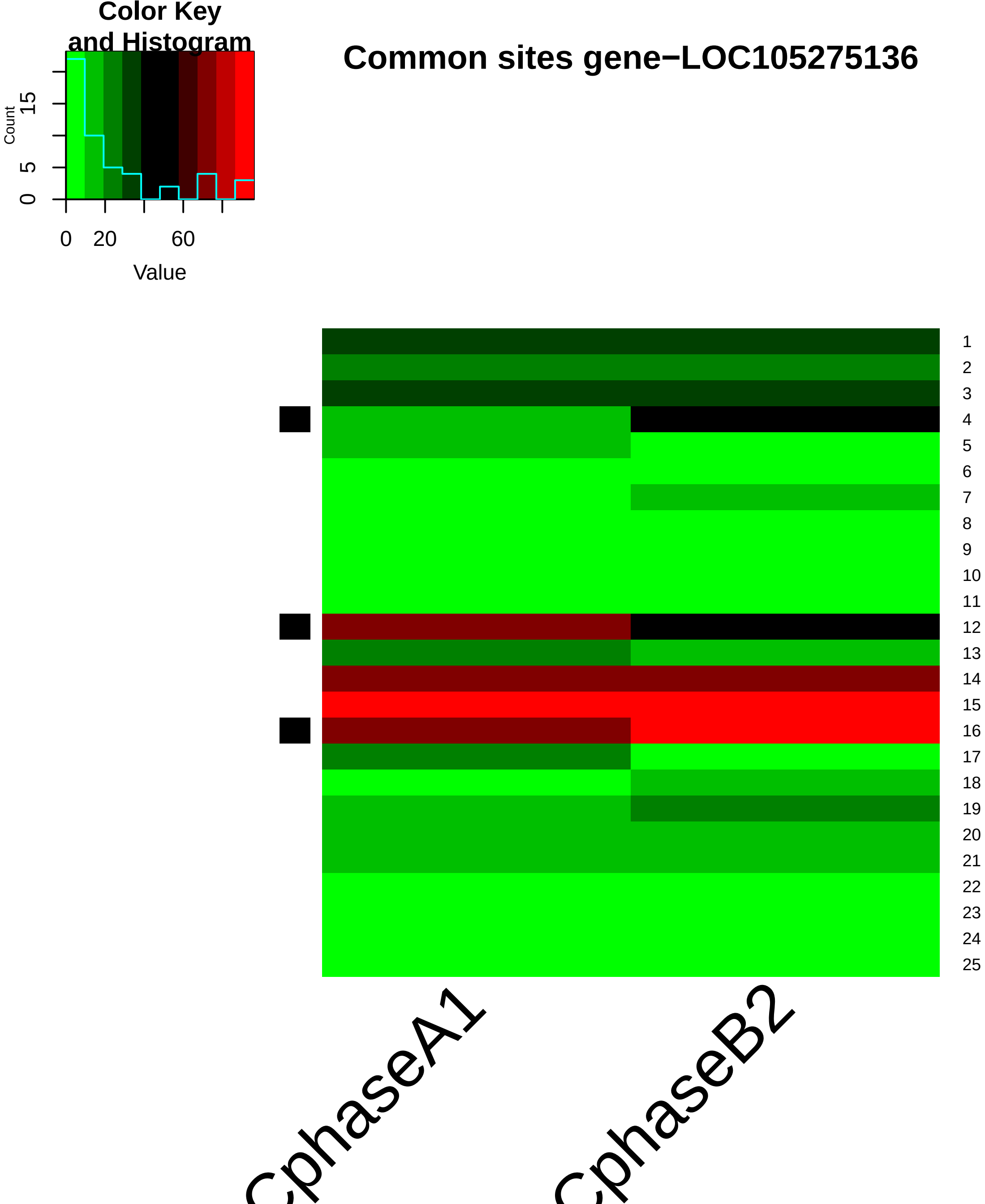
Color Key
and Histogram
Count
0
5
15
0
20
60
Value
Common sites gene−LOC105275136
CphaseA1
CphaseB2

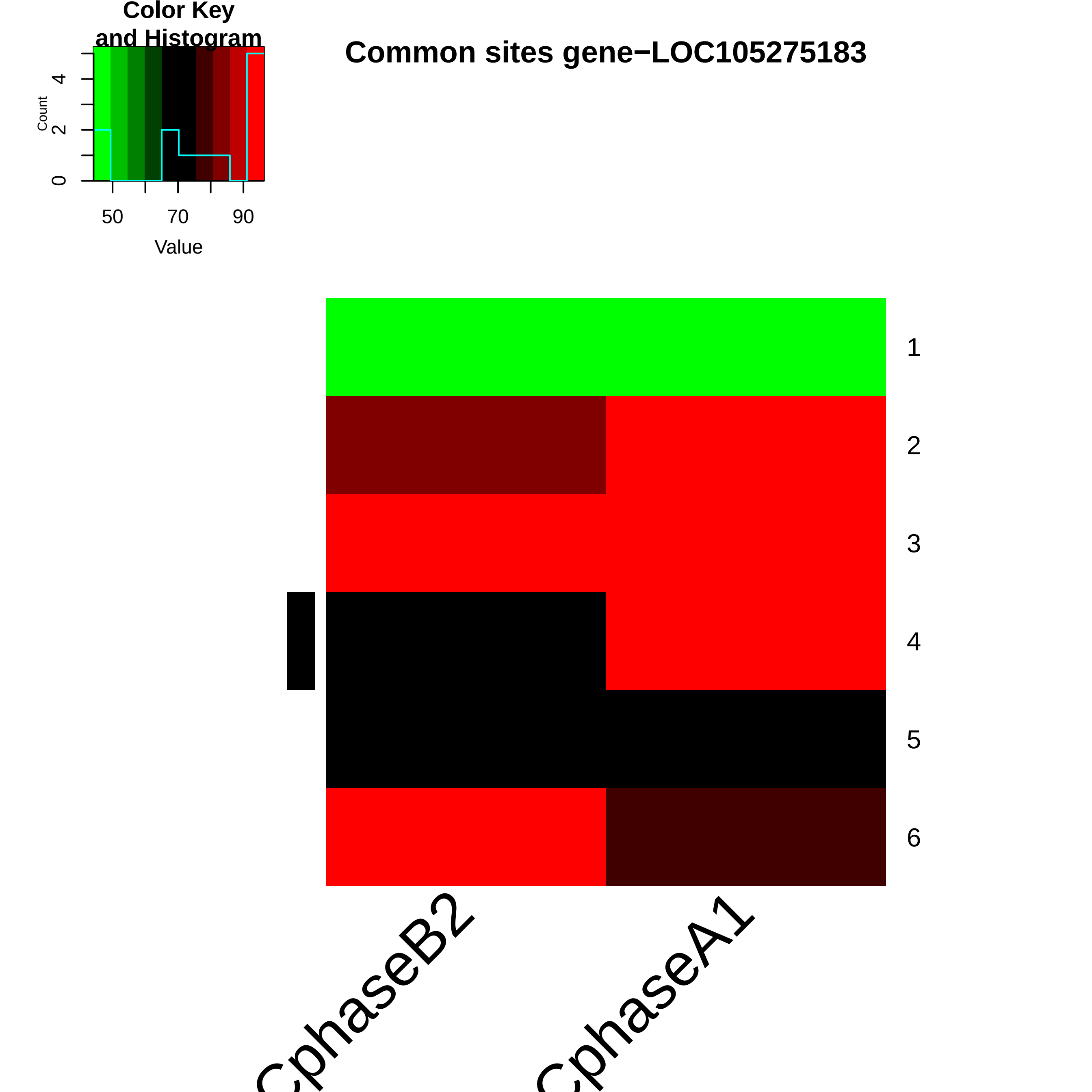
Color Key
and Histogram
Count
0
2
4
50
70
90
Value
Common sites gene−LOC105275183
1
2
3
4
5
6
CphaseB2
CphaseA1

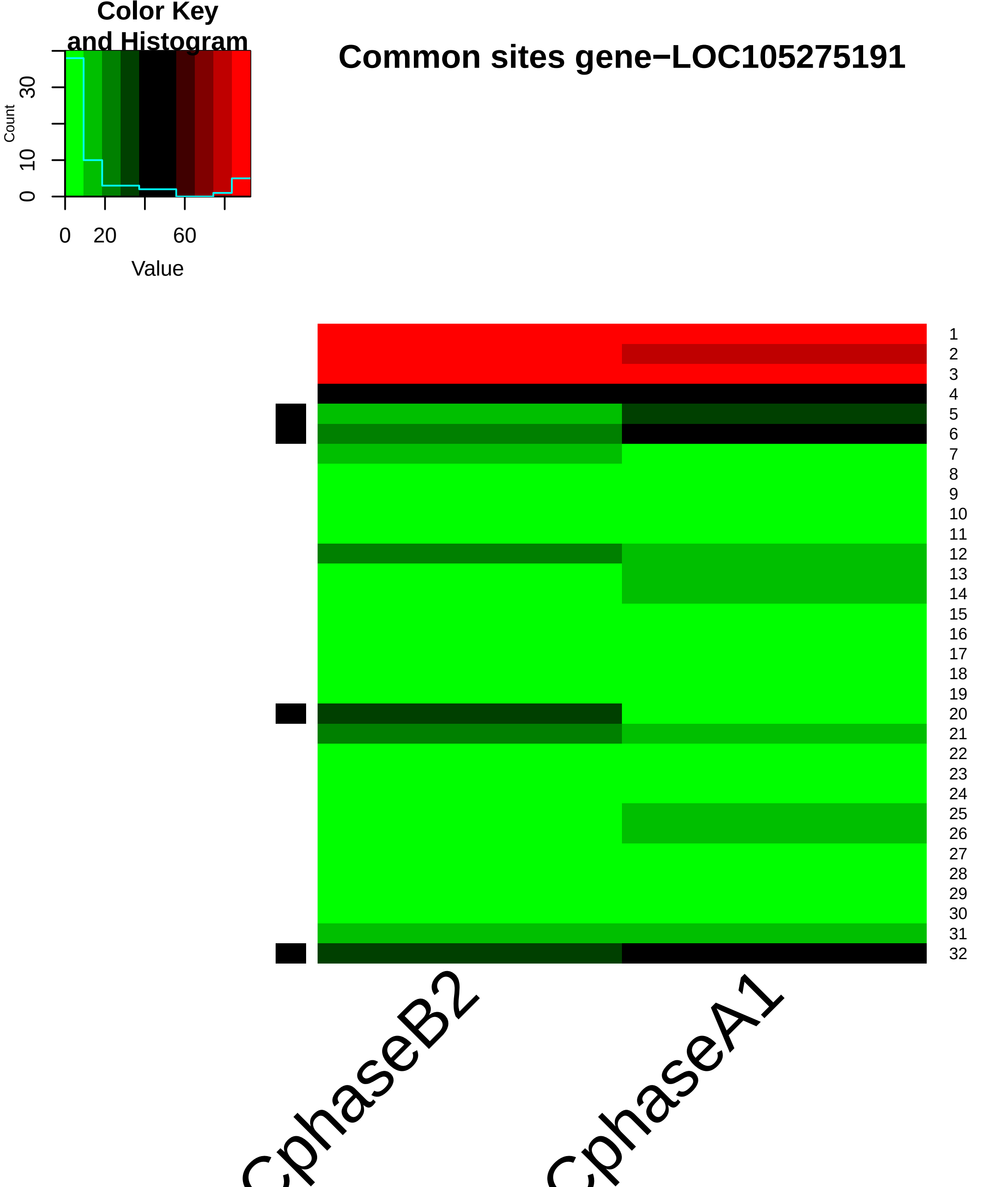
Color Key
and Histogram
Count
0
10
30
0
20
60
Value
Common sites gene−LOC105275191
CphaseB2
CphaseA1

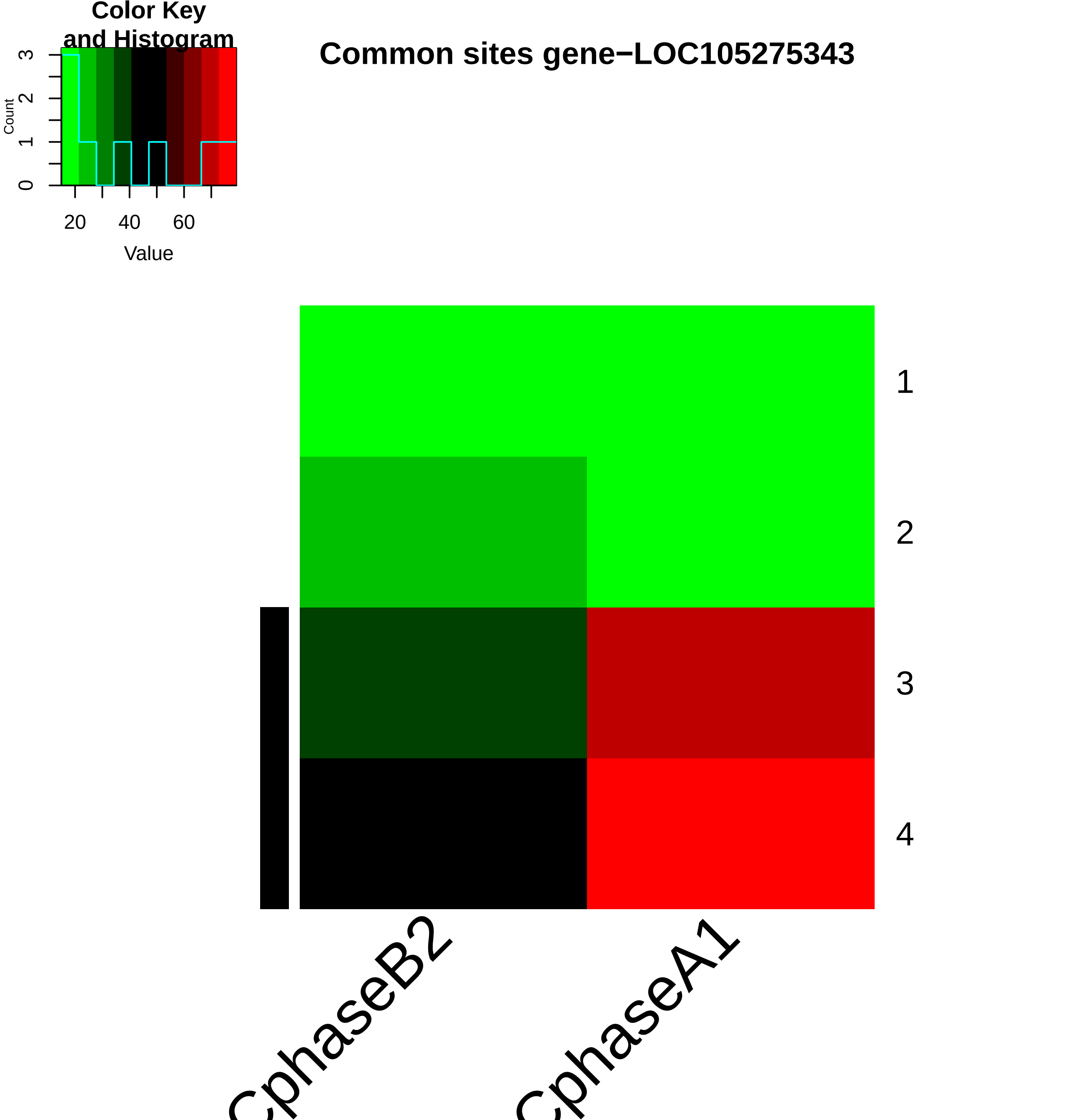

Color Key
and Histogram
Count
0
1
2
3
20
40
60
Value
Common sites gene−LOC105275343
1
2
3
4
CphaseB2
CphaseA1

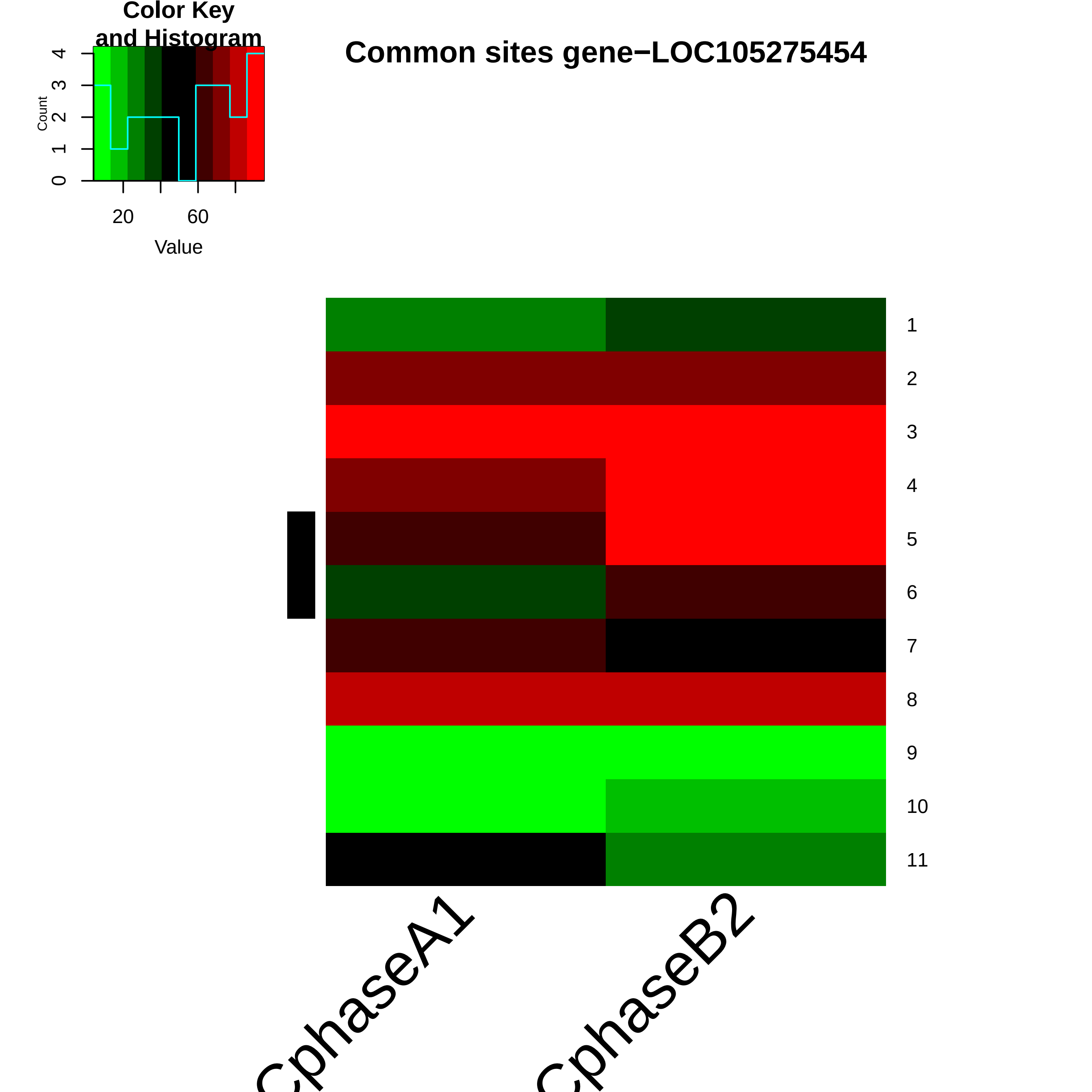

Color Key
and Histogram
Count
0
1
2
3
4
20
60
Value
Common sites gene−LOC105275454
1
2
3
4
5
6
7
8
9
10
11
CphaseA1
CphaseB2

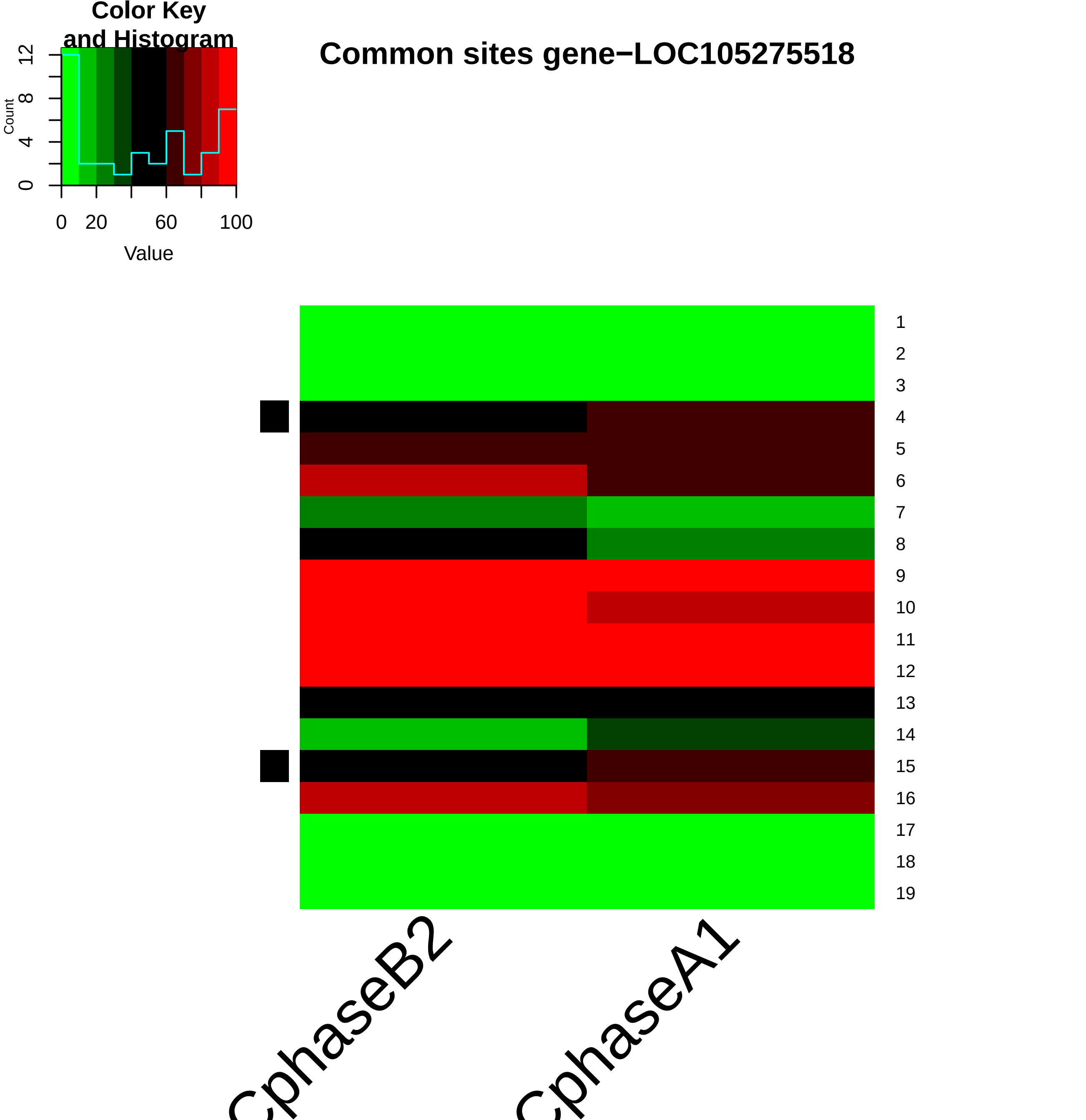
Color Key
and Histogram
Count
0
4
8
12
0
20
60
100
Value
Common sites gene−LOC105275518
CphaseB2
CphaseA1

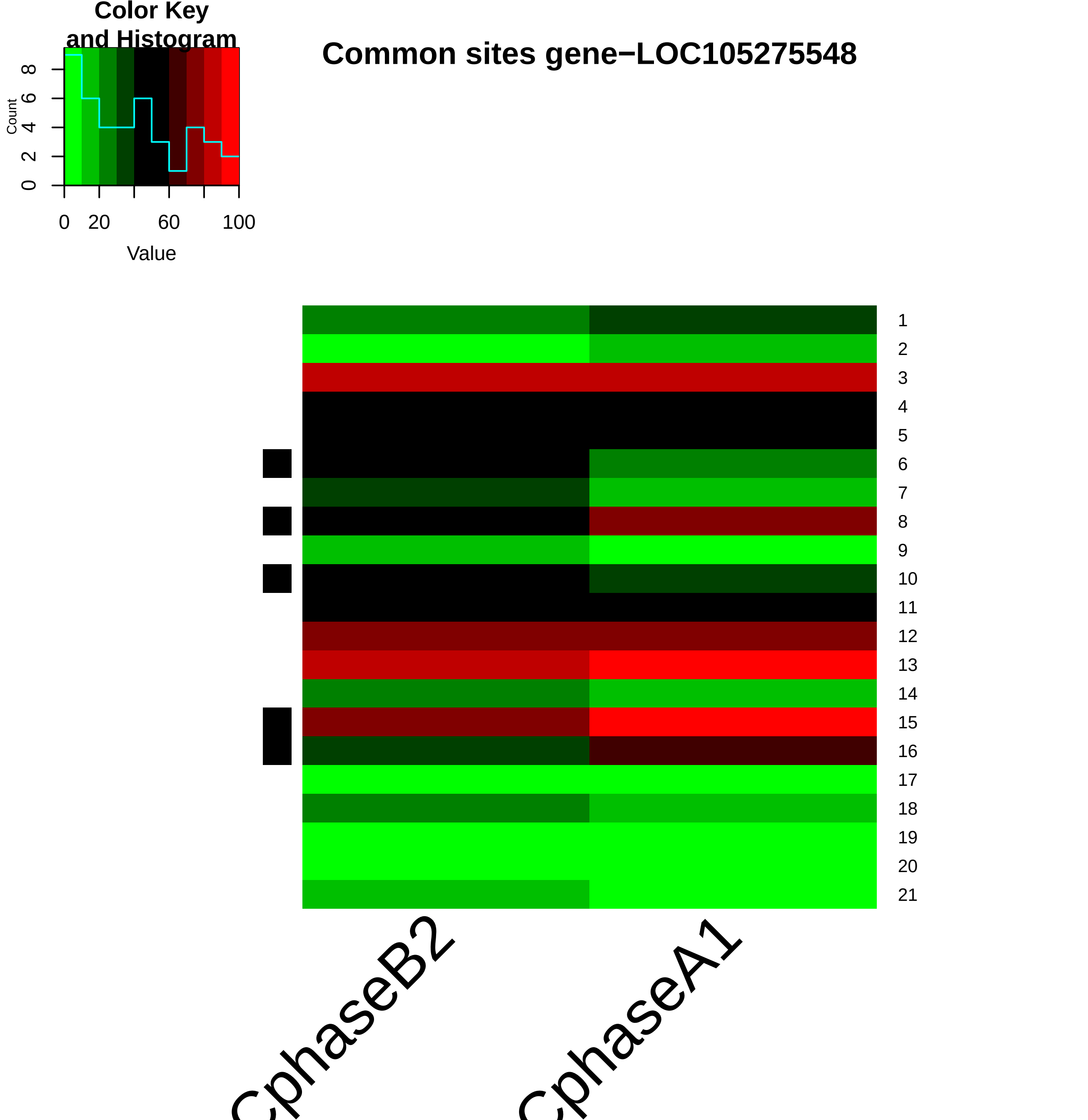

Color Key
and Histogram
Count
0
2
4
6
8
0
20
60
100
Value
Common sites gene−LOC105275548
CphaseB2
CphaseA1

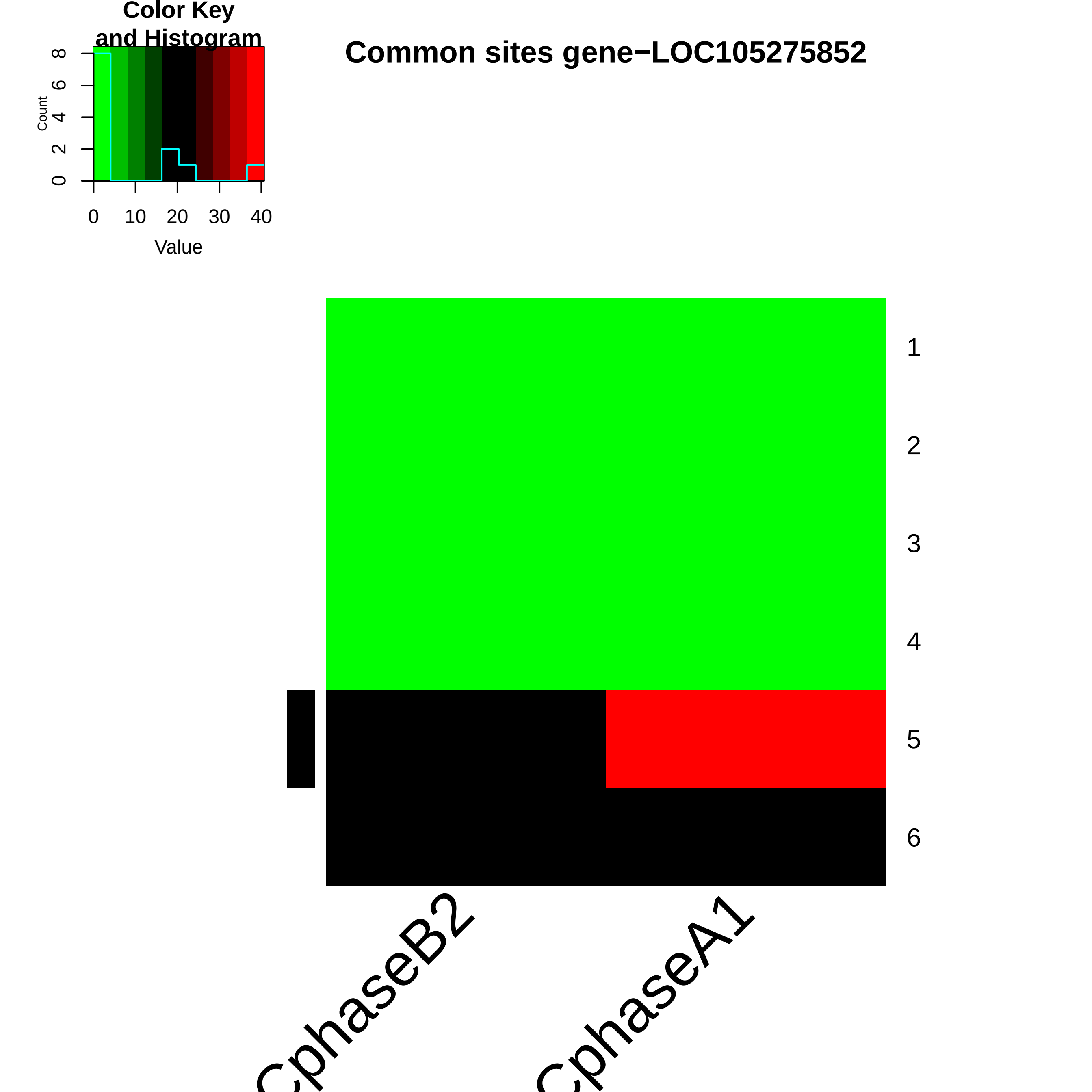
Color Key
and Histogram
Count
Value
0
10
20
30
40
0
2
4
6
8
Common sites gene−LOC105275852
1
2
3
4
5
6
CphaseB2
CphaseA1

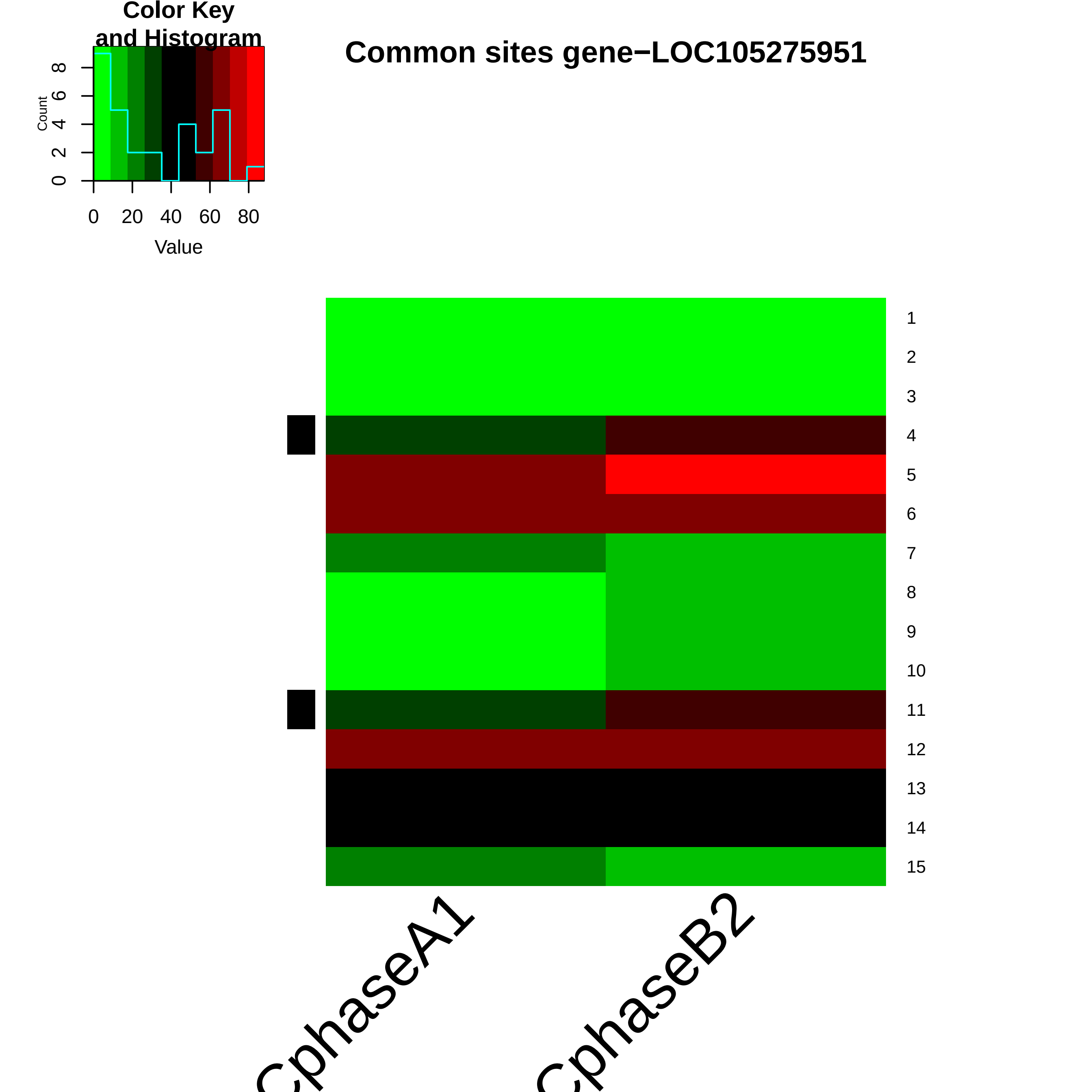
Color Key
and Histogram
Count
0
2
4
6
8
0
20
40
60
80
Value
Common sites gene−LOC105275951
CphaseA1
CphaseB2

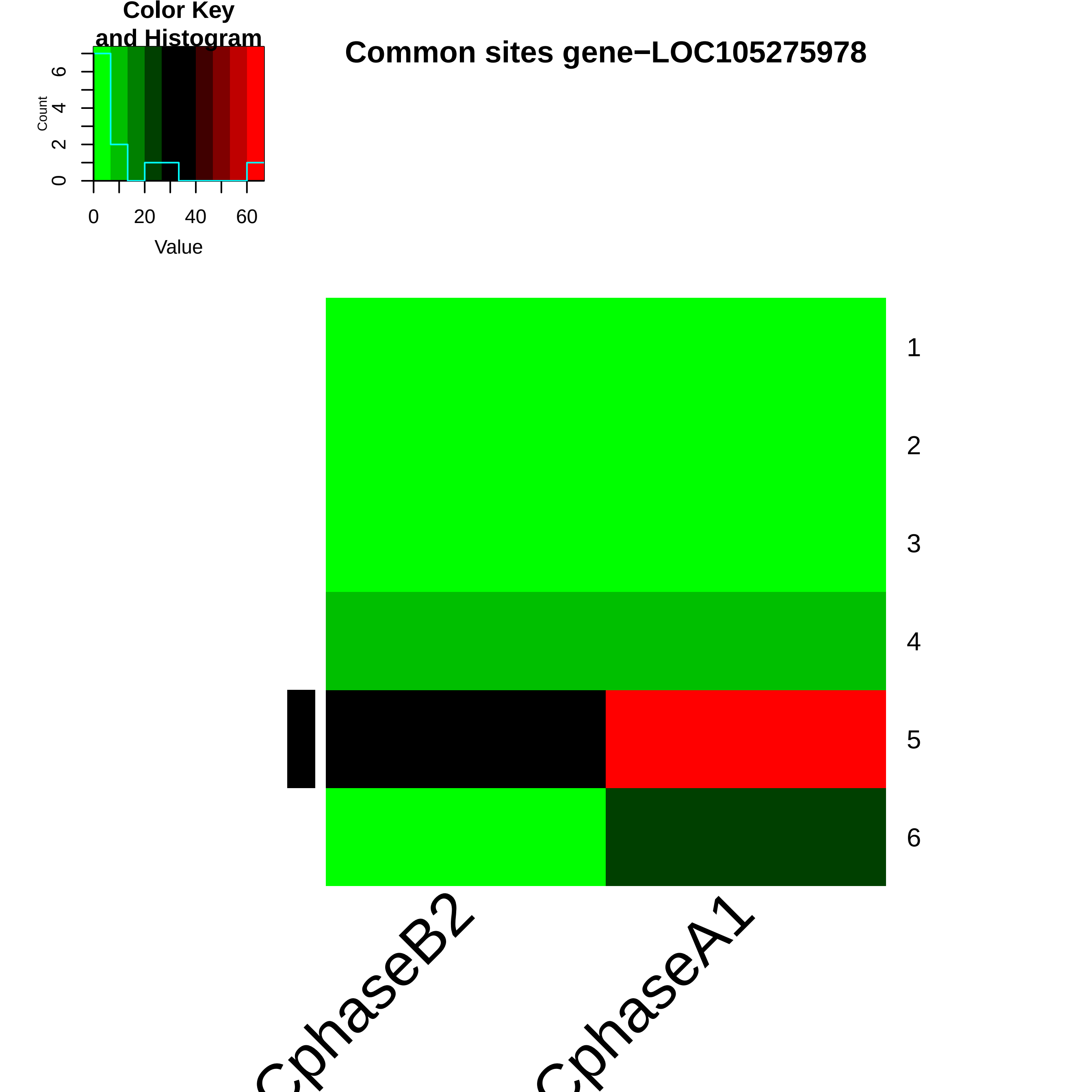
Color Key
and Histogram
Count
0
2
4
6
0
20
40
60
Value
Common sites gene−LOC105275978
1
2
3
4
5
6
CphaseB2
CphaseA1

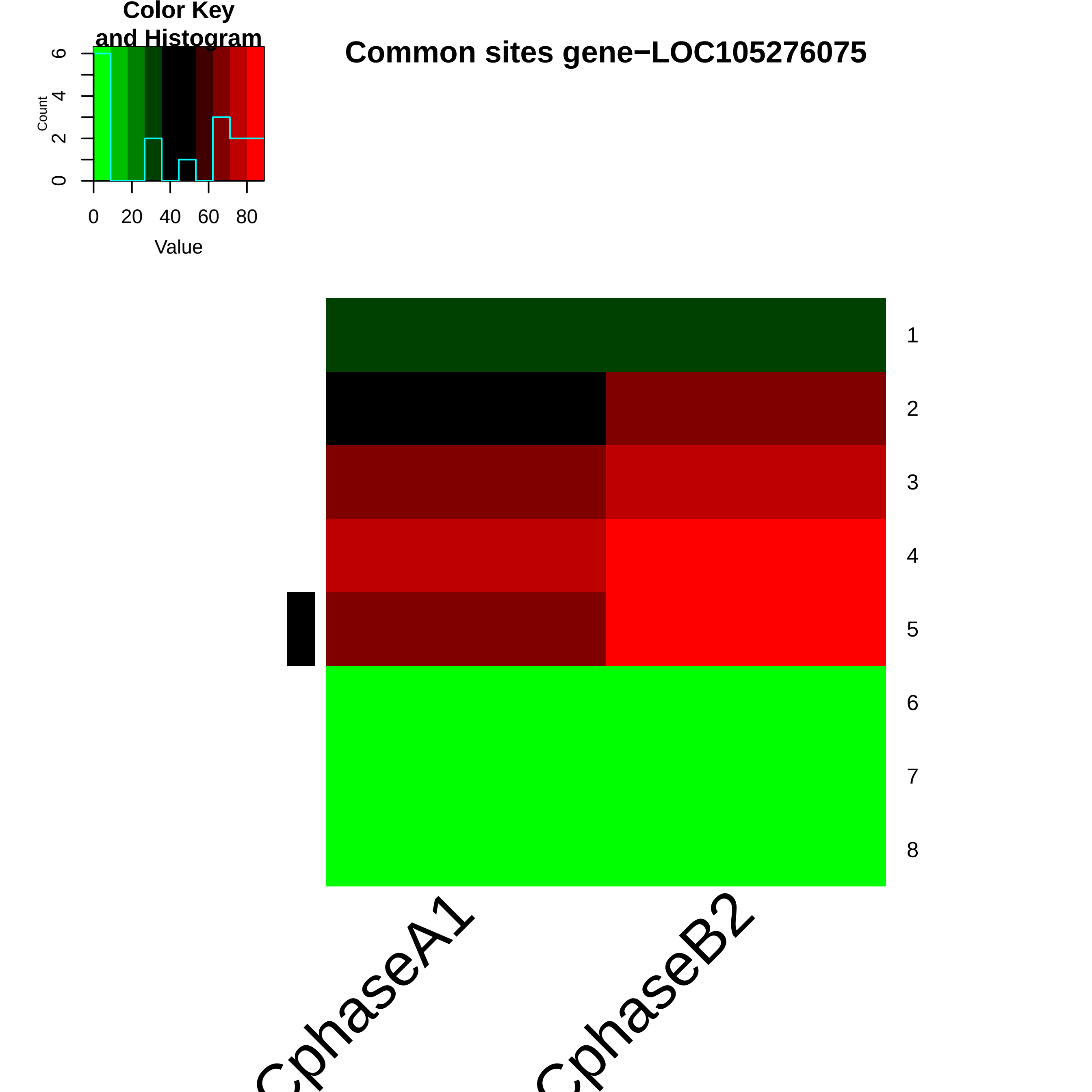
Color Key
and Histogram
Count
0
2
4
6
0
20
40
60
80
Value
Common sites gene−LOC105276075
1
2
3
4
5
6
7
8
CphaseA1
CphaseB2

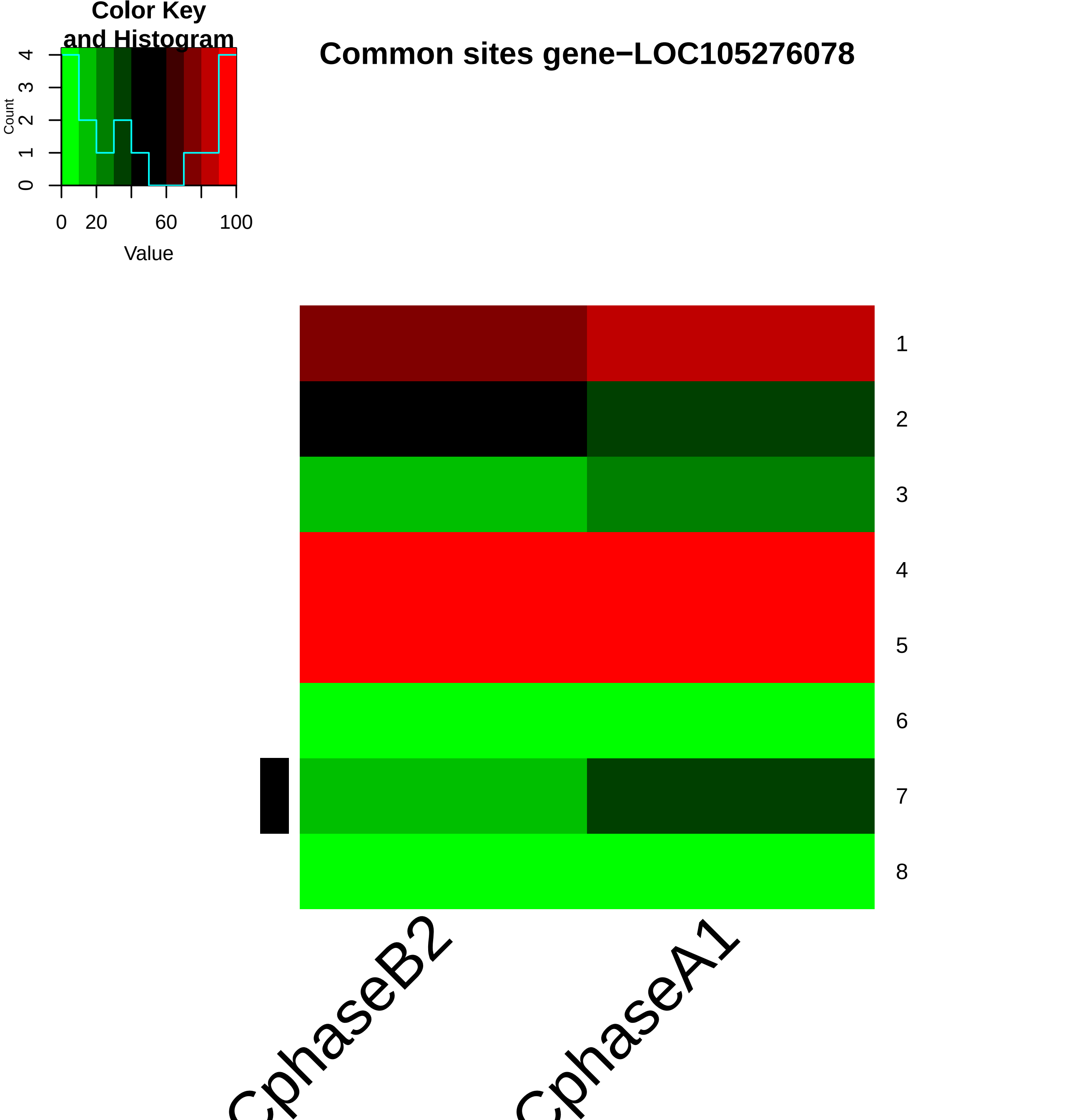

Color Key
and Histogram
Count
0
1
2
3
4
0
20
60
100
Value
Common sites gene−LOC105276078
1
2
3
4
5
6
7
8
CphaseB2
CphaseA1

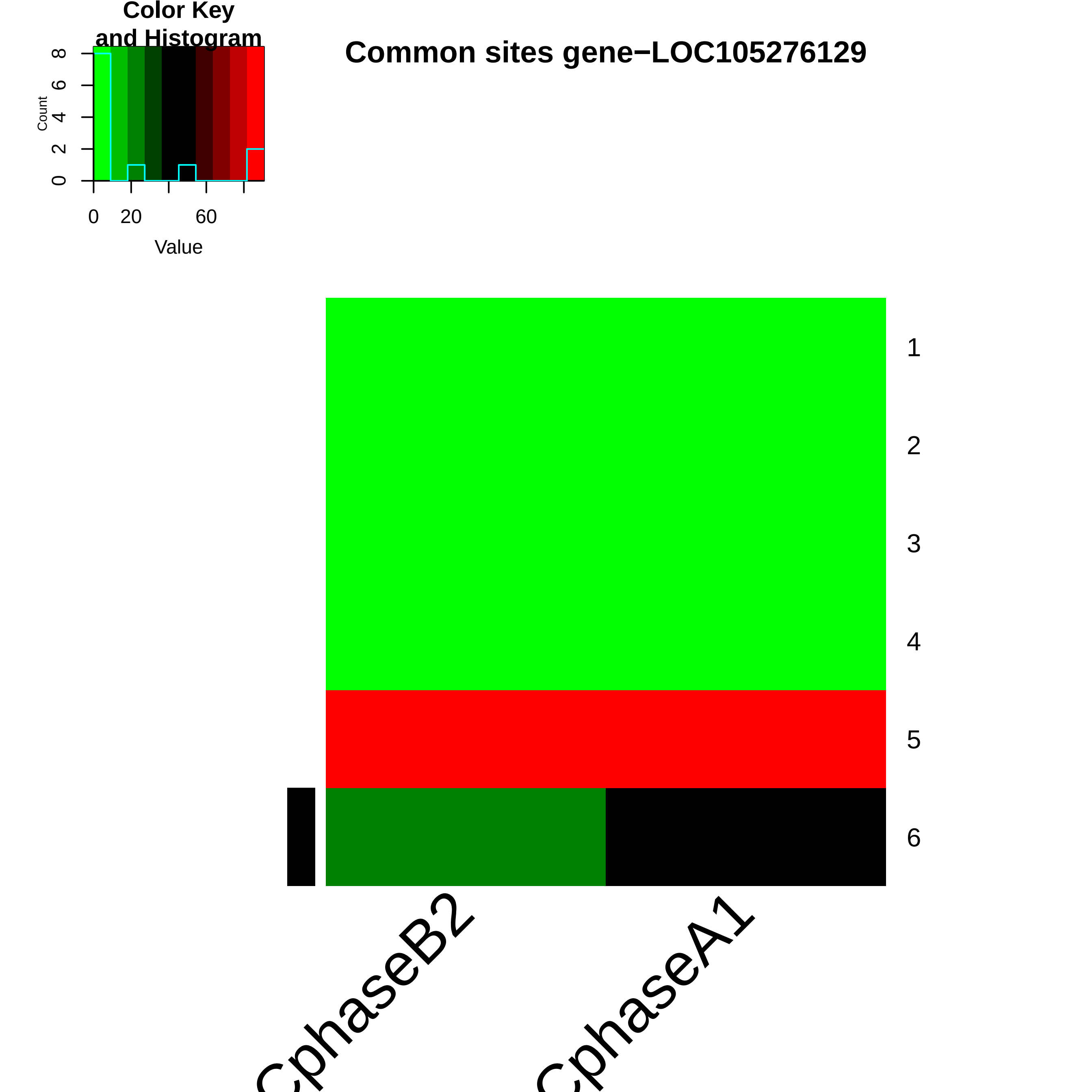

Color Key
and Histogram
Count
0
2
4
6
8
0
20
60
Value
Common sites gene−LOC105276129
1
2
3
4
5
6
CphaseB2
CphaseA1

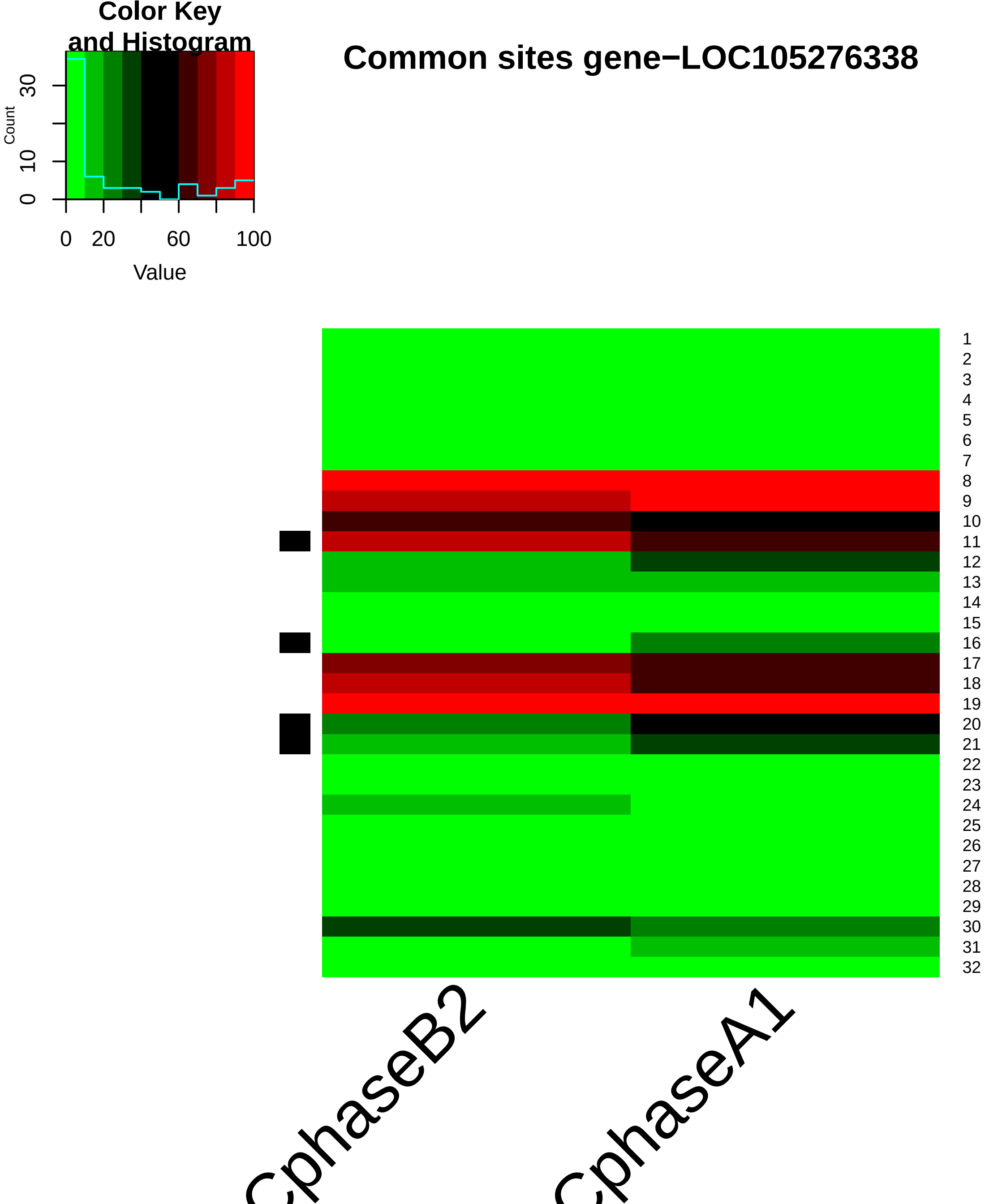
Color Key
and Histogram
Count
0
10
30
0
20
60
100
Value
Common sites gene−LOC105276338
CphaseB2
CphaseA1

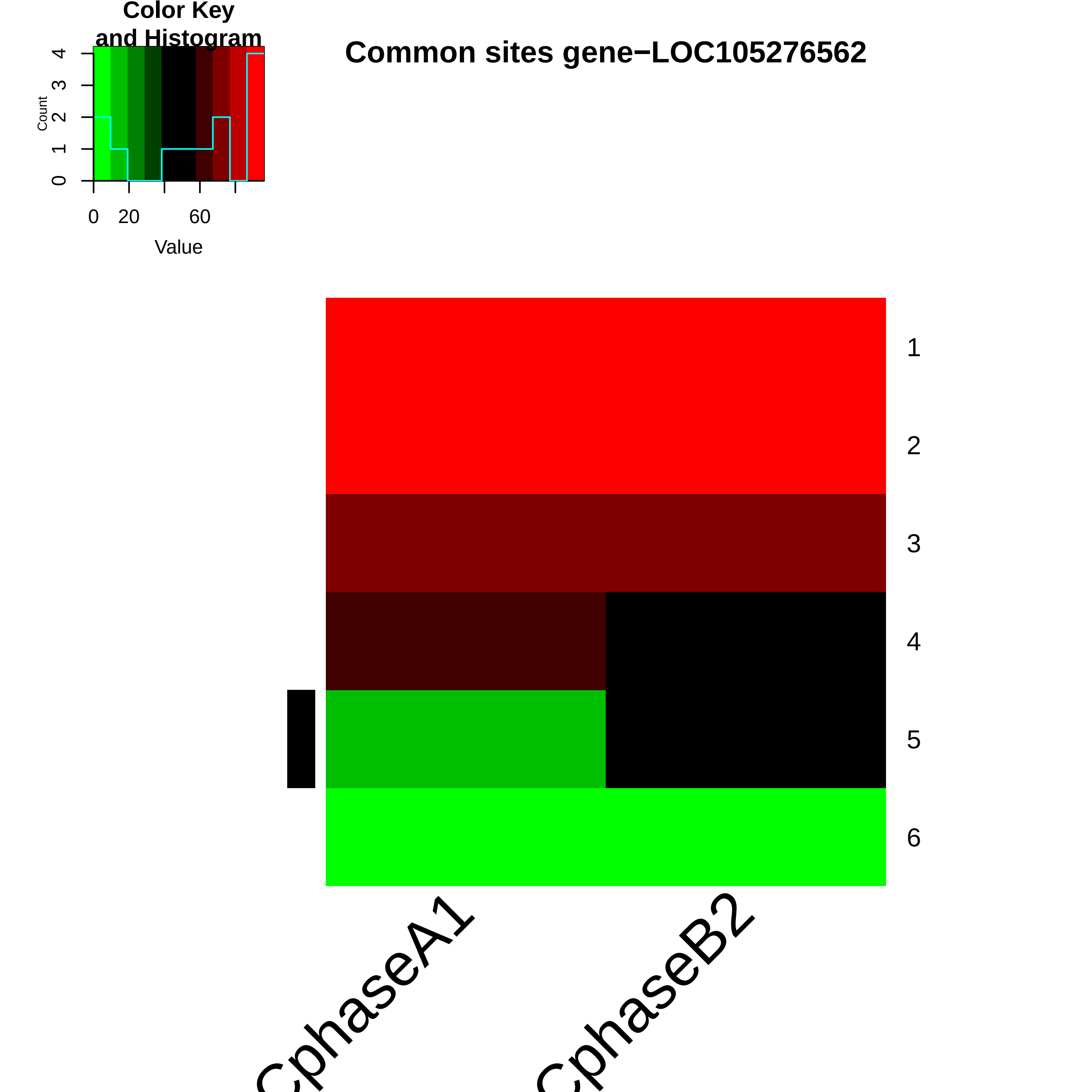
Color Key
and Histogram
Count
0
1
2
3
4
0
20
60
Value
Common sites gene−LOC105276562
1
2
3
4
5
6
CphaseA1
CphaseB2

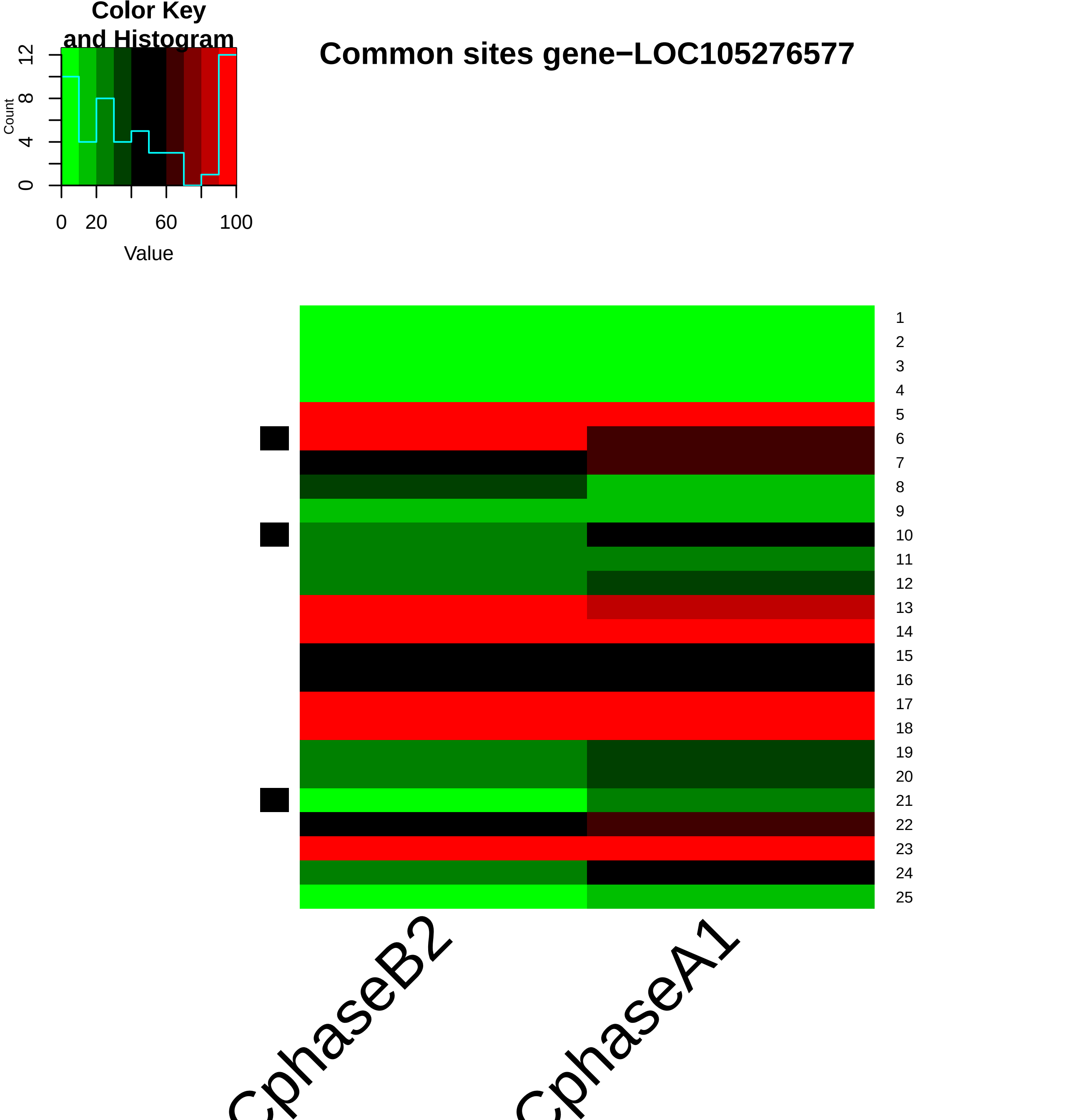
Color Key
and Histogram
Count
0
4
8
12
0
20
60
100
Value
Common sites gene−LOC105276577
CphaseB2
CphaseA1

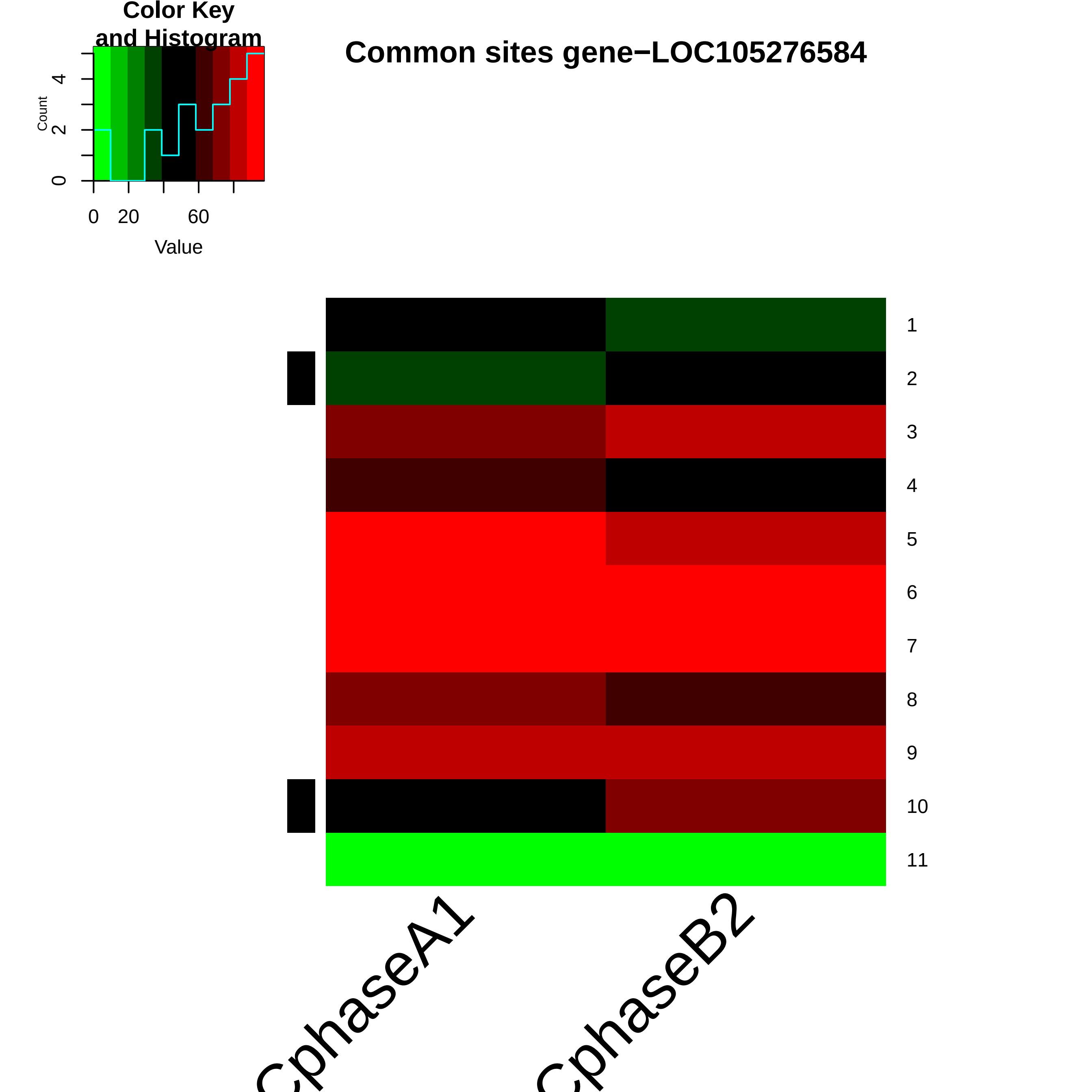

Color Key
and Histogram
Count
0
2
4
0
20
60
Value
Common sites gene−LOC105276584
1
2
3
4
5
6
7
8
9
10
11
CphaseA1
CphaseB2

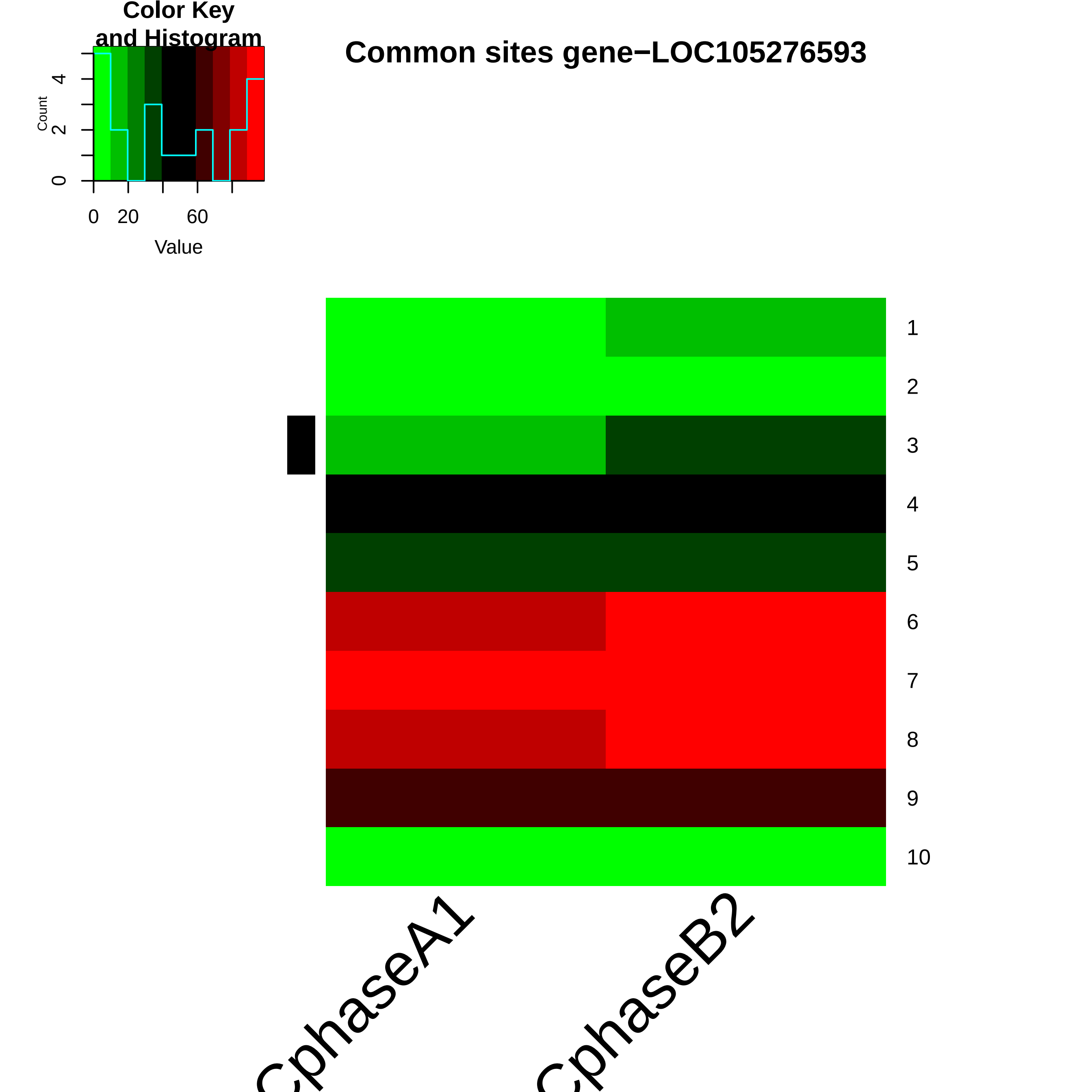
Color Key
and Histogram
Count
0
2
4
0
20
60
Value
Common sites gene−LOC105276593
1
2
3
4
5
6
7
8
9
10
CphaseA1
CphaseB2

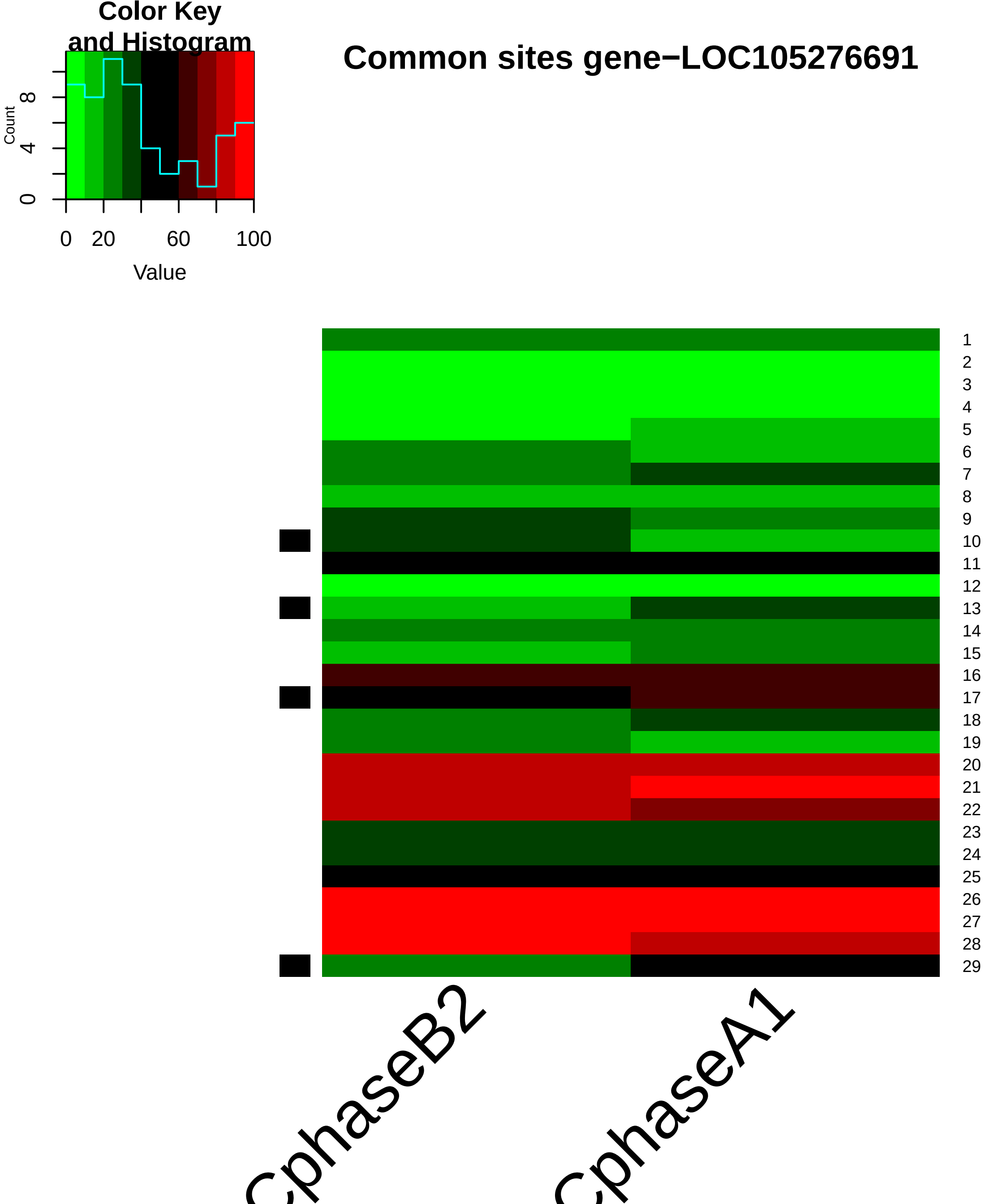

Color Key
and Histogram
Count
Value
0
4
8
0
20
60
100
Common sites gene−LOC105276691
CphaseB2
CphaseA1

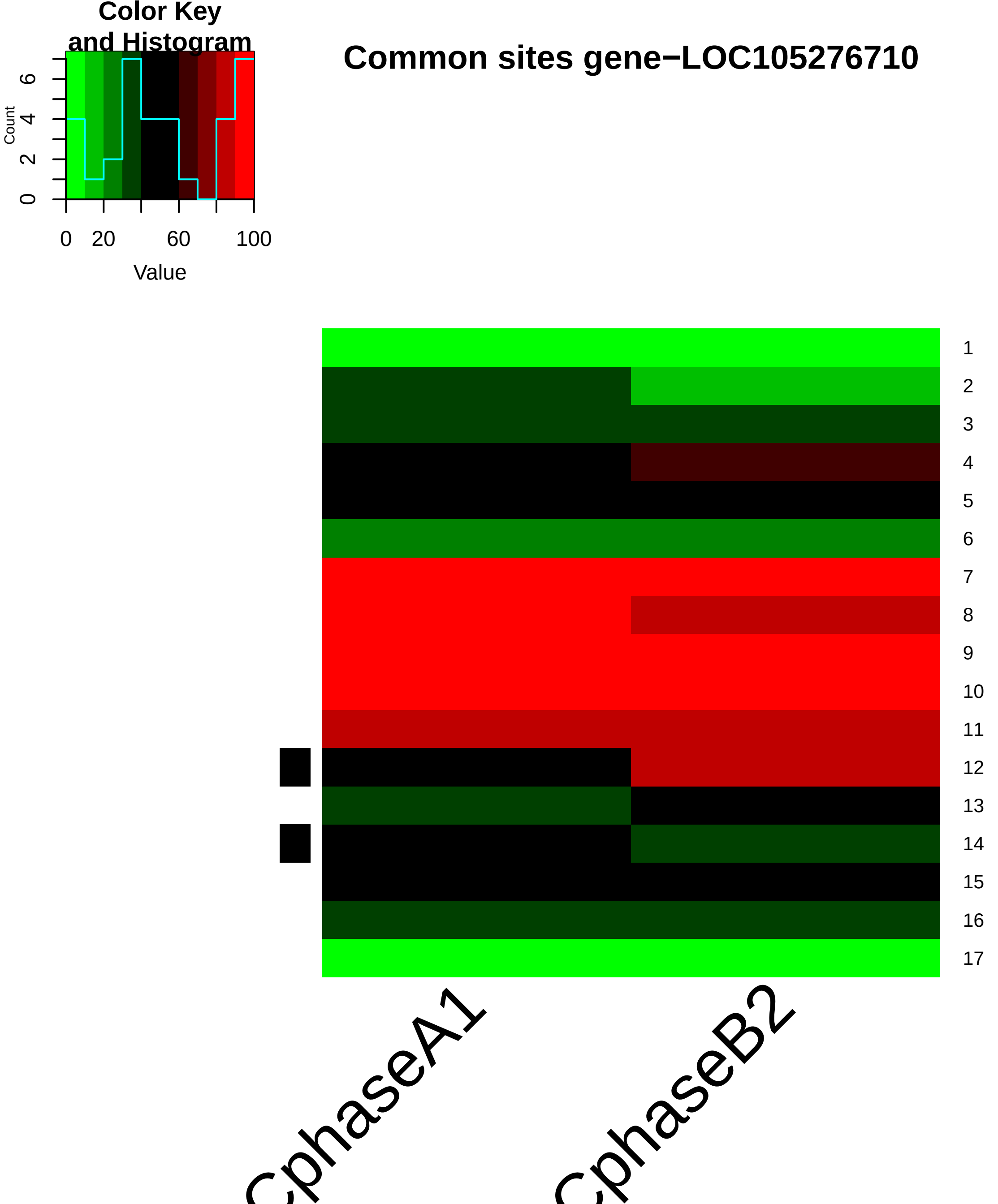

Color Key and Histogram
Count
Value
0
20
60
100
0
2
4
6
Common sites gene−LOC105276710
CphaseA1
CphaseB2

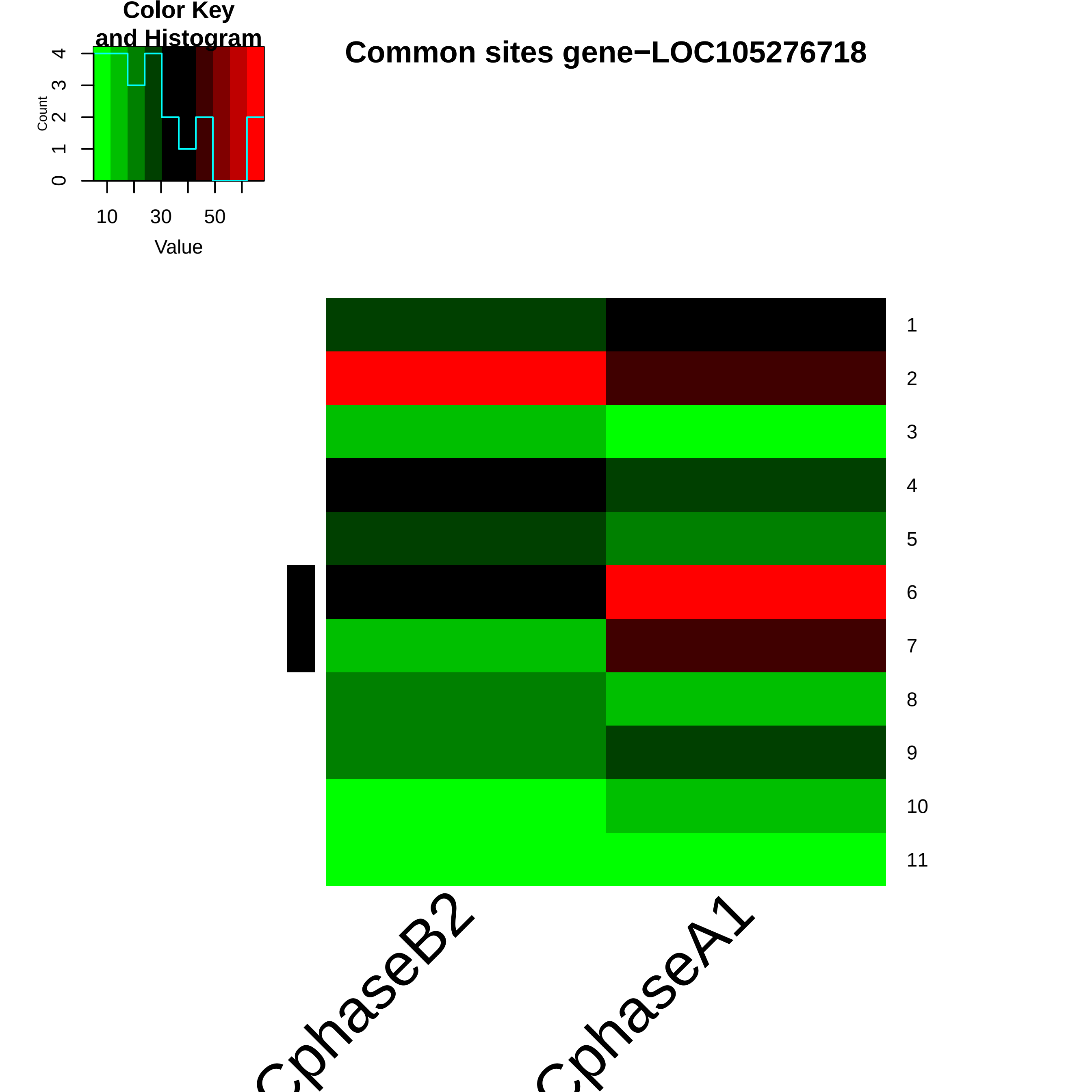

Color Key
and Histogram
Count
0
1
2
3
4
10
30
50
Value
Common sites gene−LOC105276718
1
2
3
4
5
6
7
8
9
10
11
CphaseB2
CphaseA1

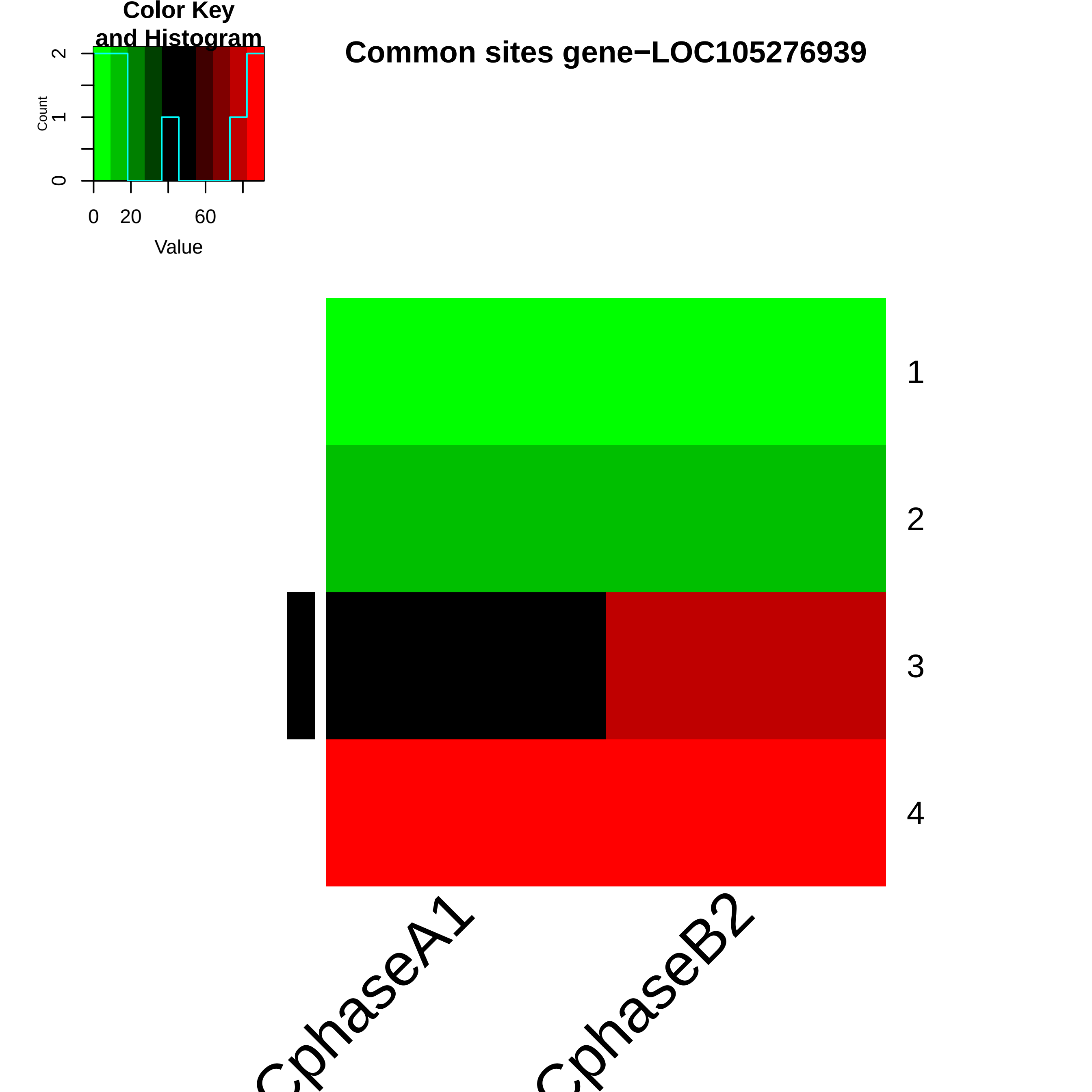

Color Key
and Histogram
Count
0
1
2
0
20
60
Value
Common sites gene−LOC105276939
1
2
3
4
CphaseA1
CphaseB2

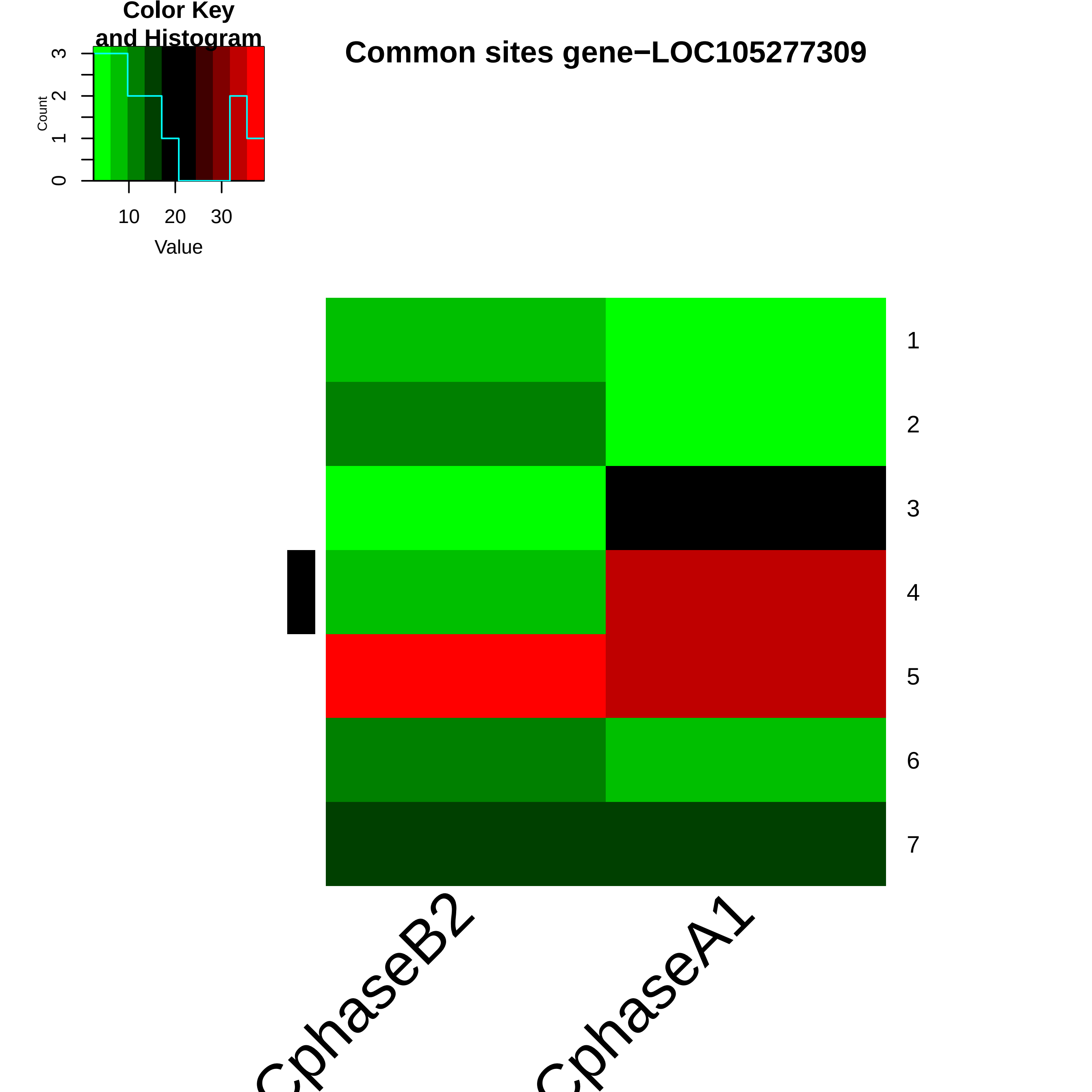

Color Key
and Histogram
Count
0
1
2
3
10
20
30
Value
Common sites gene−LOC105277309
1
2
3
4
5
6
7
CphaseB2
CphaseA1

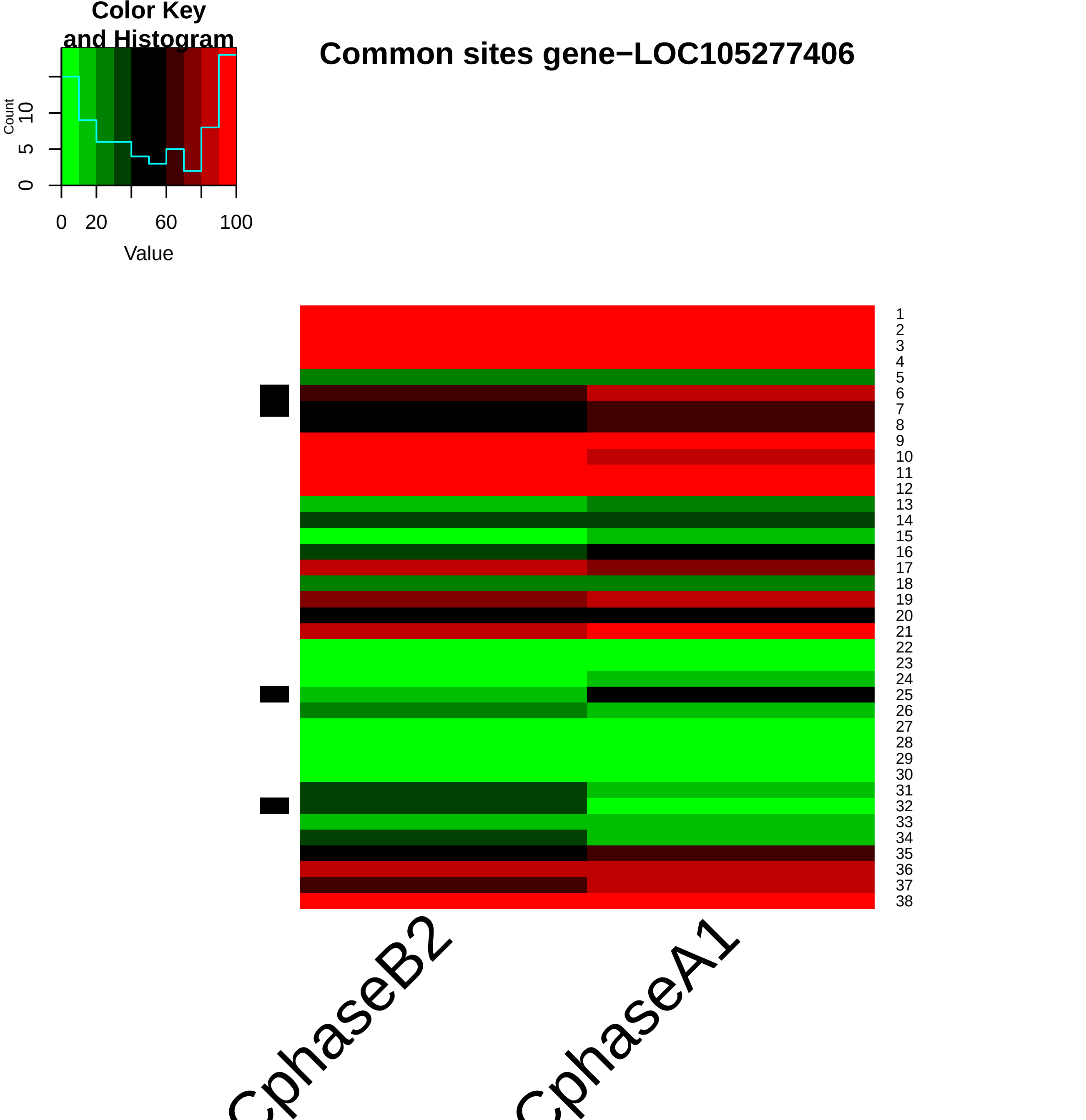
Color Key
and Histogram
Count
0
5
10
0
20
60
100
Value
Common sites gene−LOC105277406
CphaseB2
CphaseA1

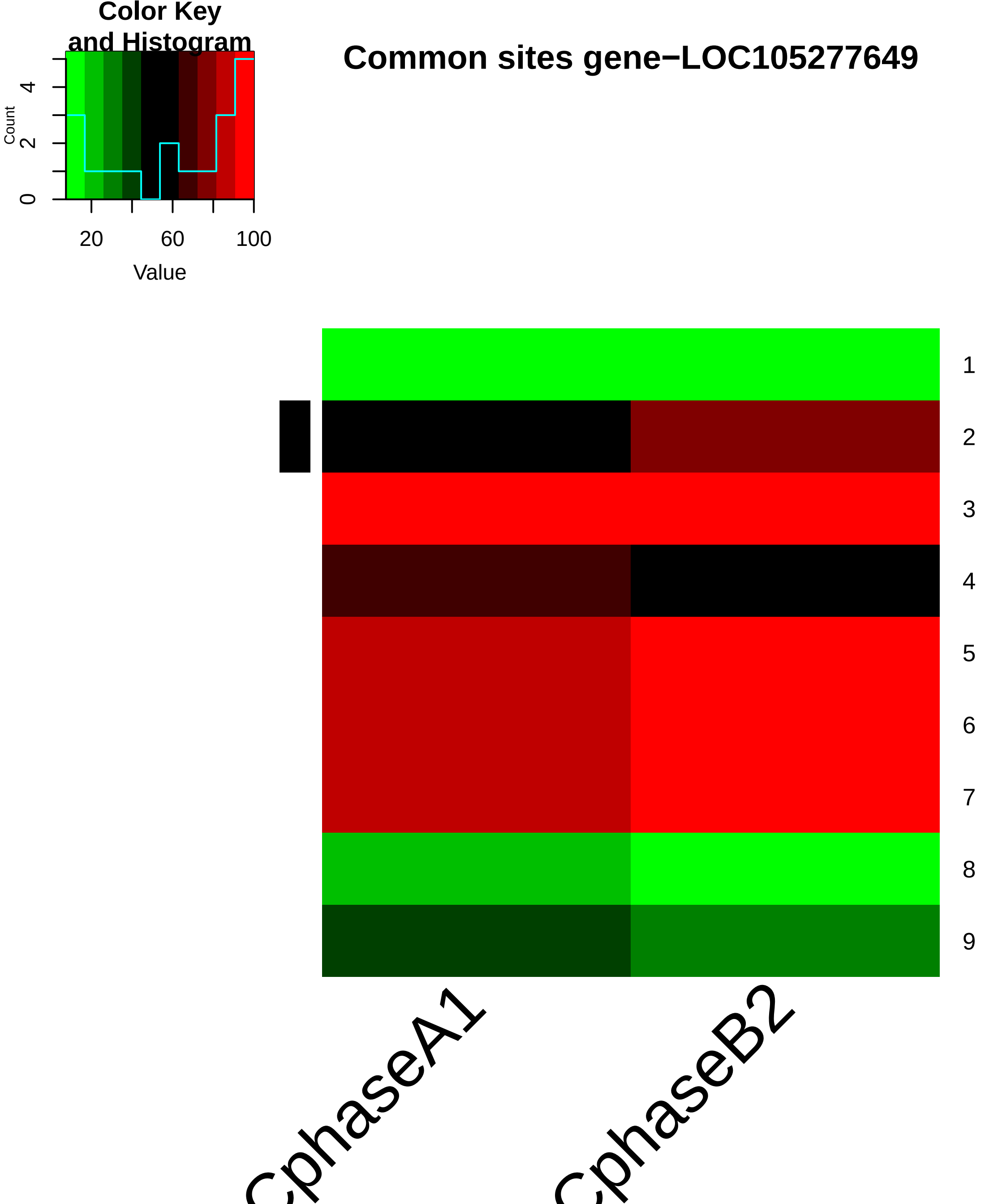

Color Key
and Histogram
Count
Value
0
2
4
20
60
100
Common sites gene−LOC105277649
1
2
3
4
5
6
7
8
9
CphaseA1
CphaseB2

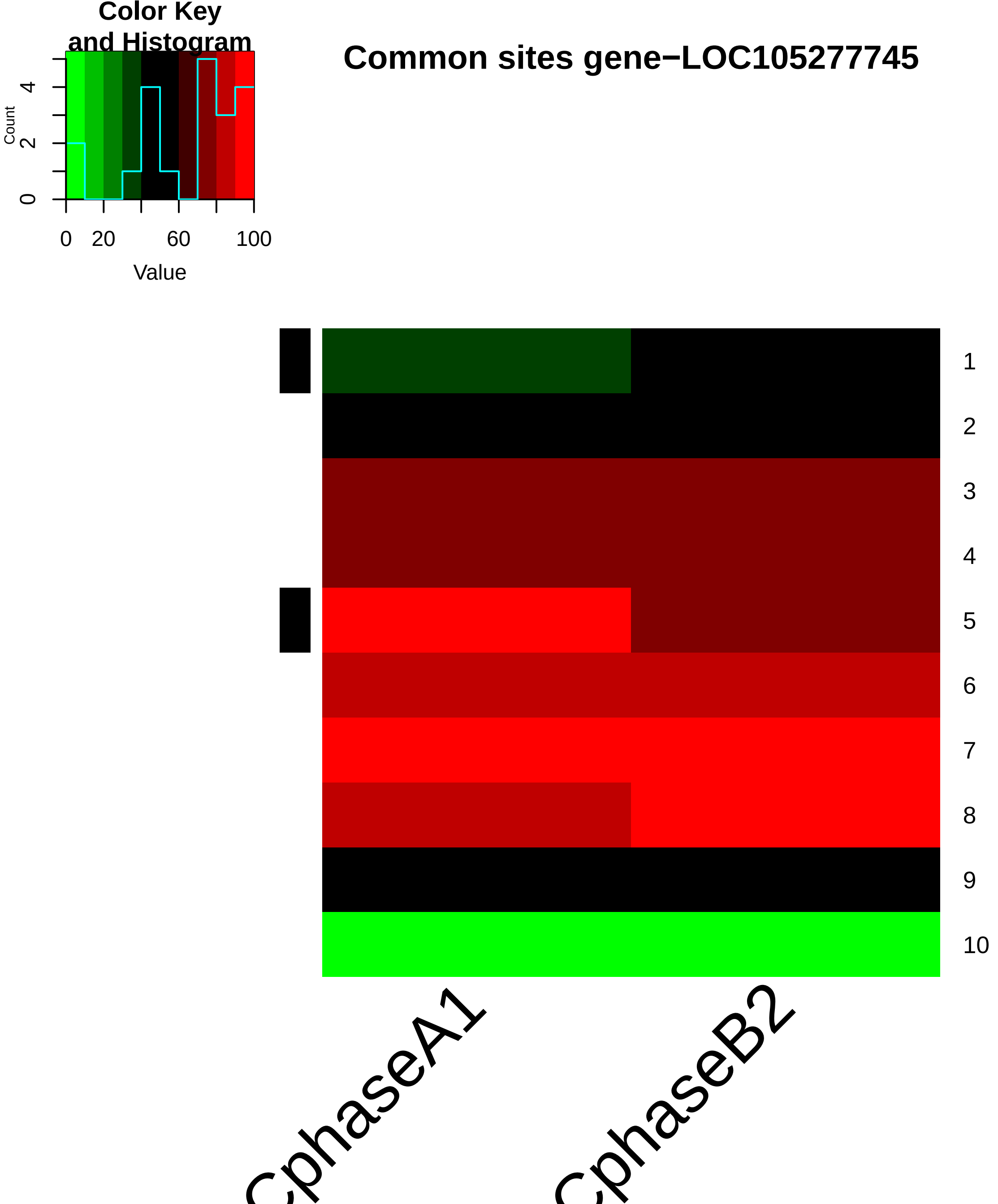
Color Key
and Histogram
Count
0
2
4
0
20
60
100
Value
Common sites gene−LOC105277745
1
2
3
4
5
6
7
8
9
10
CphaseA1
CphaseB2

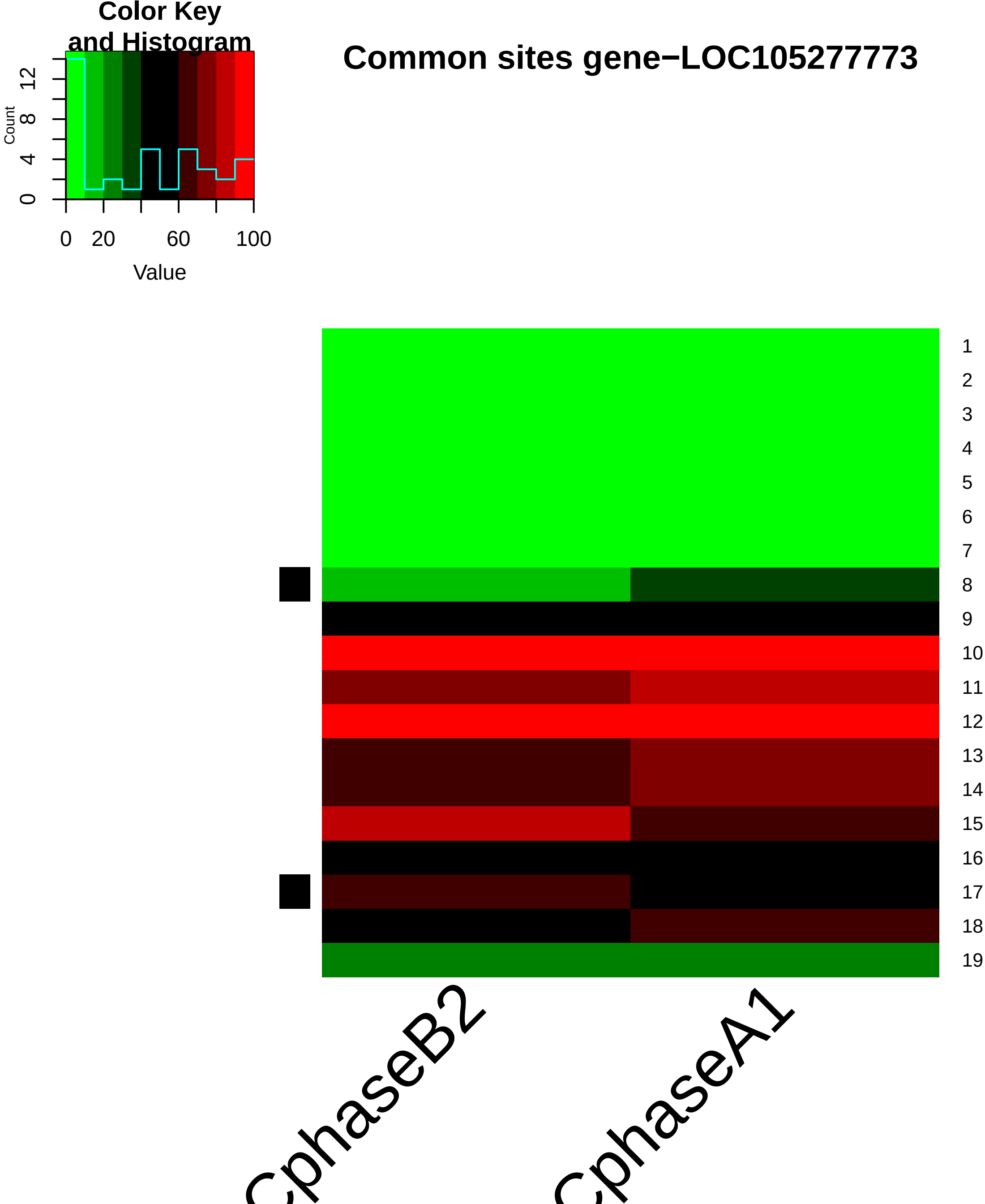
Color Key
and Histogram
Count
0
4
8
12
0
20
60
100
Value
Common sites gene−LOC105277773
CphaseB2
CphaseA1

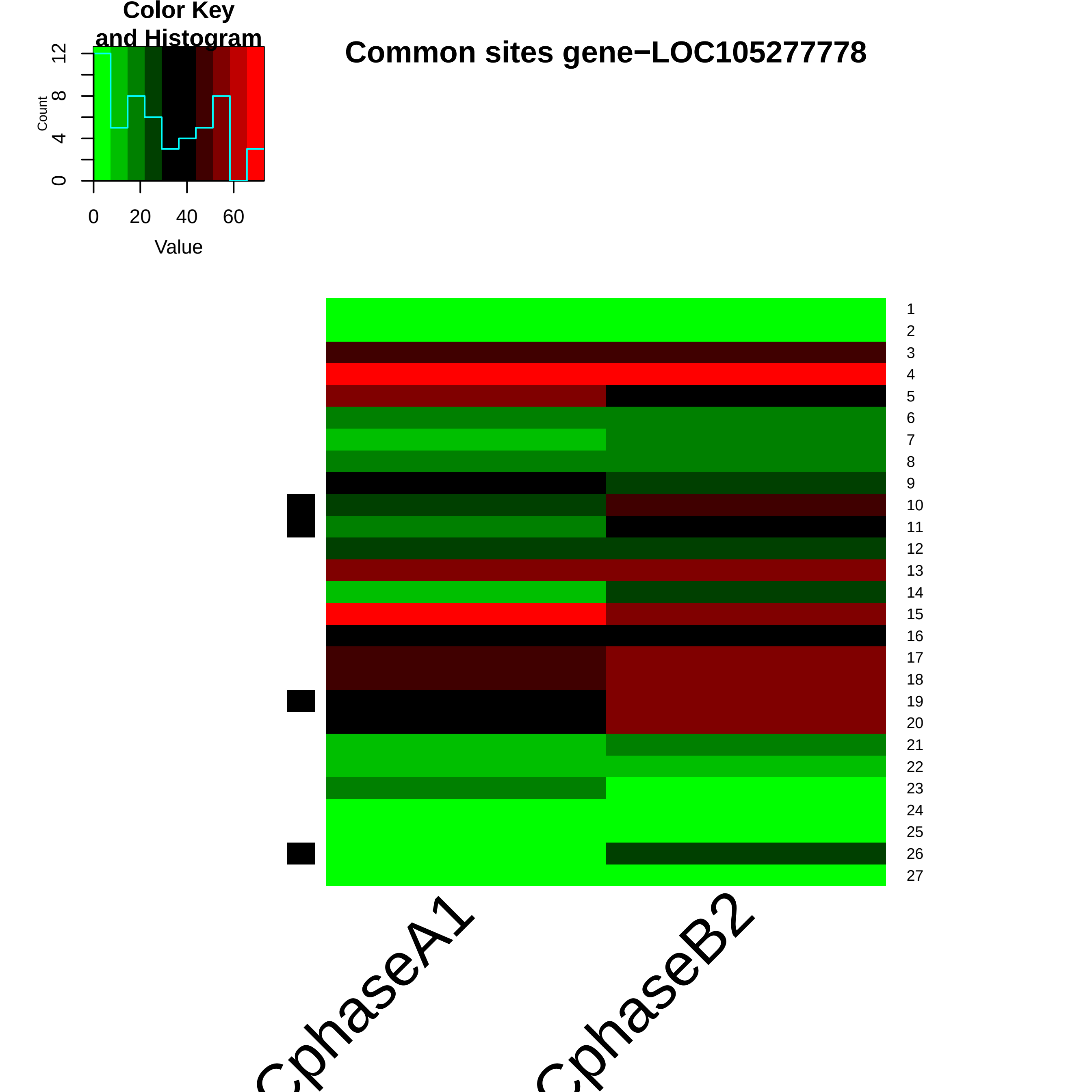
Color Key
and Histogram
Count
Value
0
4
8
12
0
20
40
60
Common sites gene−LOC105277778
CphaseA1
CphaseB2

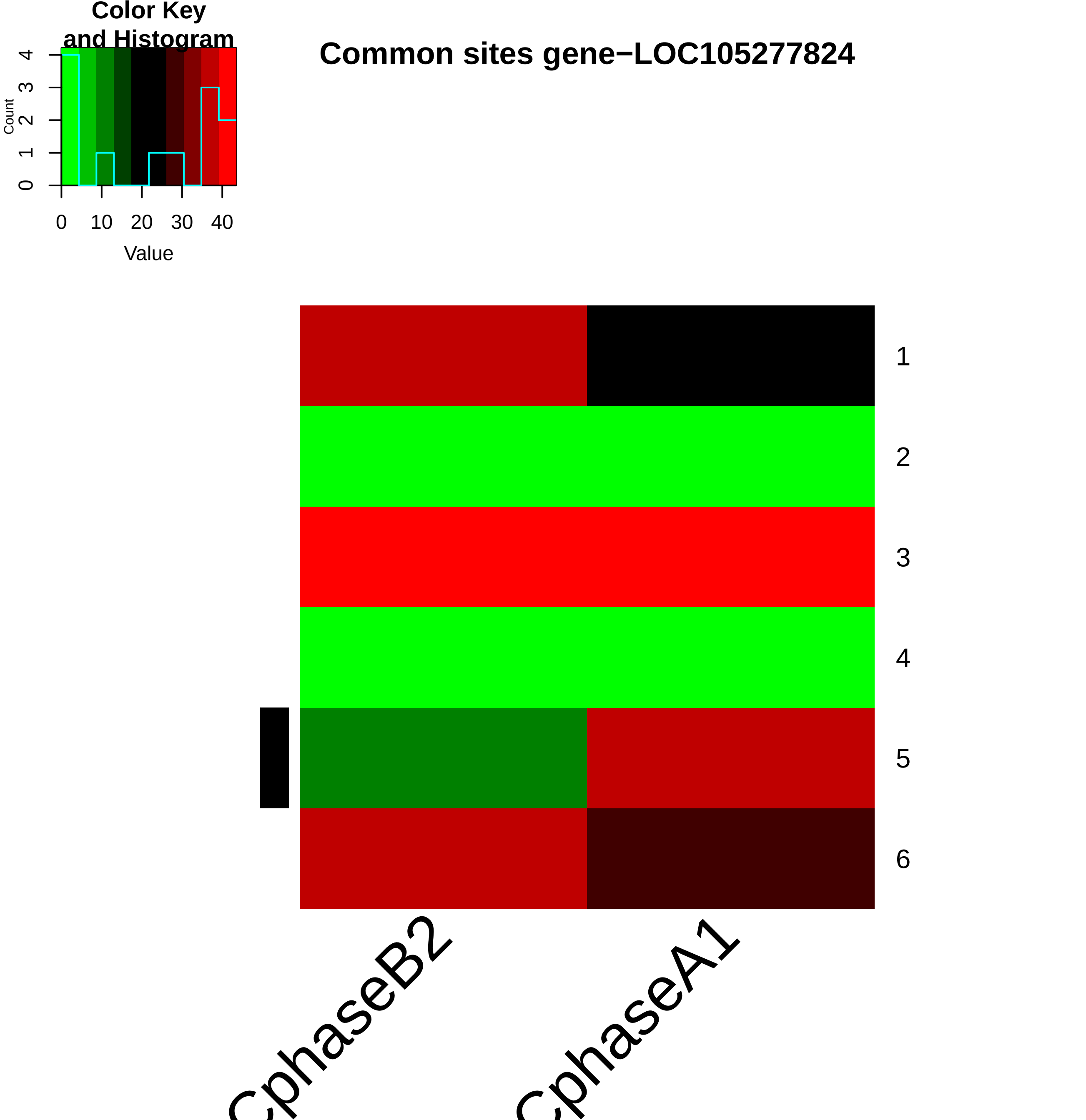
Color Key
and Histogram
Count
0
1
2
3
4
0
10
20
30
40
Value
Common sites gene−LOC105277824
1
2
3
4
5
6
CphaseB2
CphaseA1

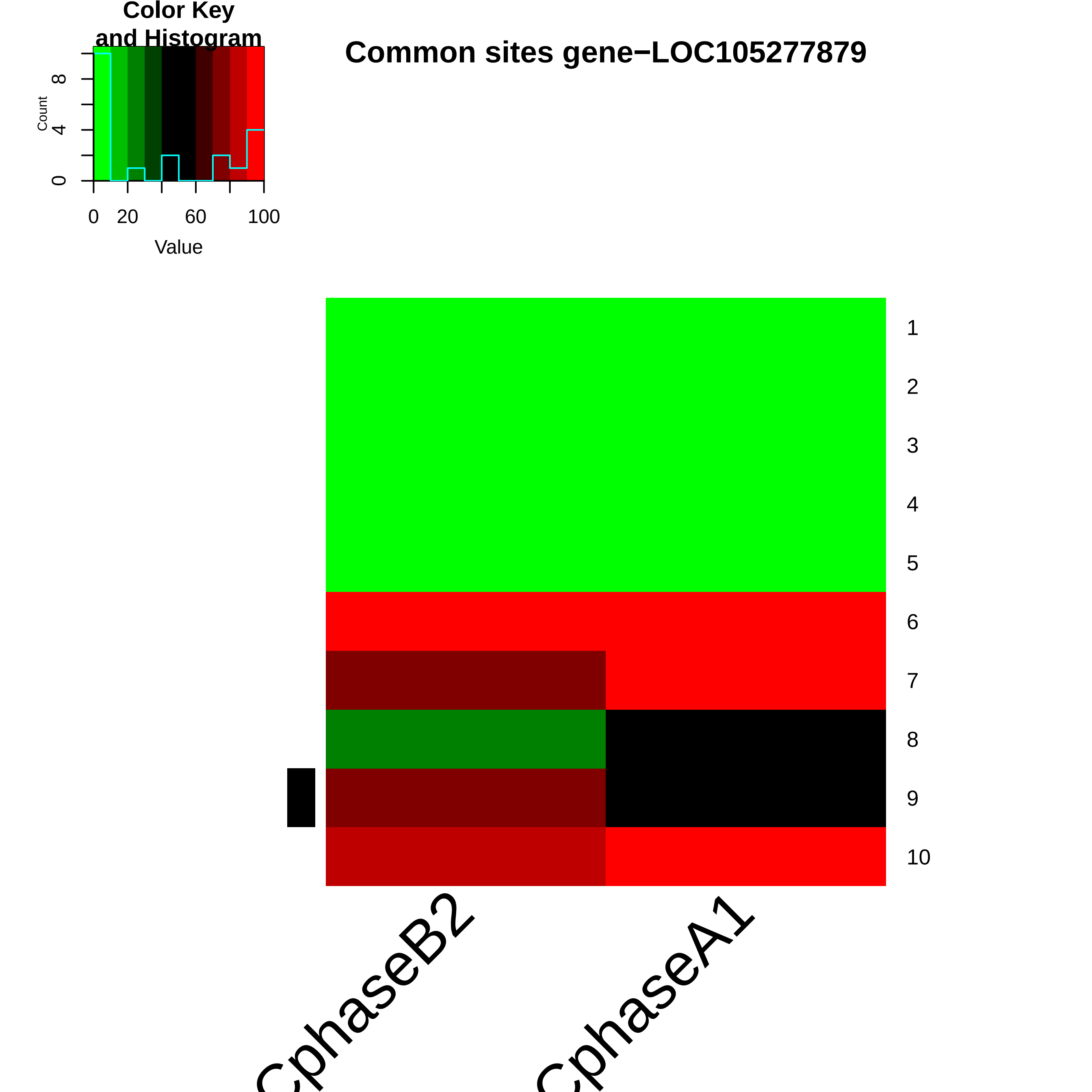
Color Key
and Histogram
Count
0
4
8
0
20
60
100
Value
Common sites gene−LOC105277879
1
2
3
4
5
6
7
8
9
10
CphaseB2
CphaseA1

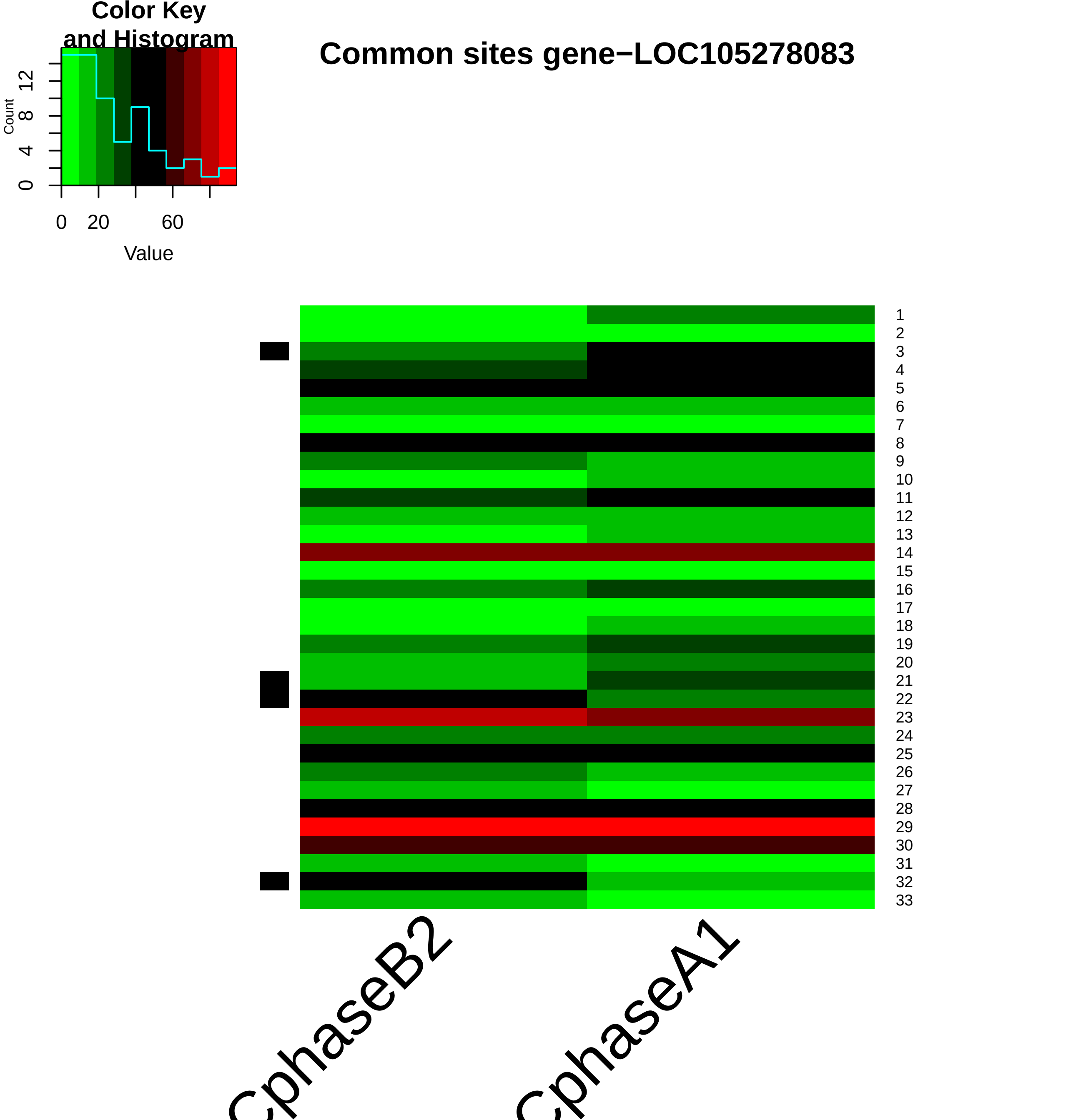
Color Key
and Histogram
Count
0
4
8
12
0
20
60
Value
Common sites gene−LOC105278083
CphaseB2
CphaseA1

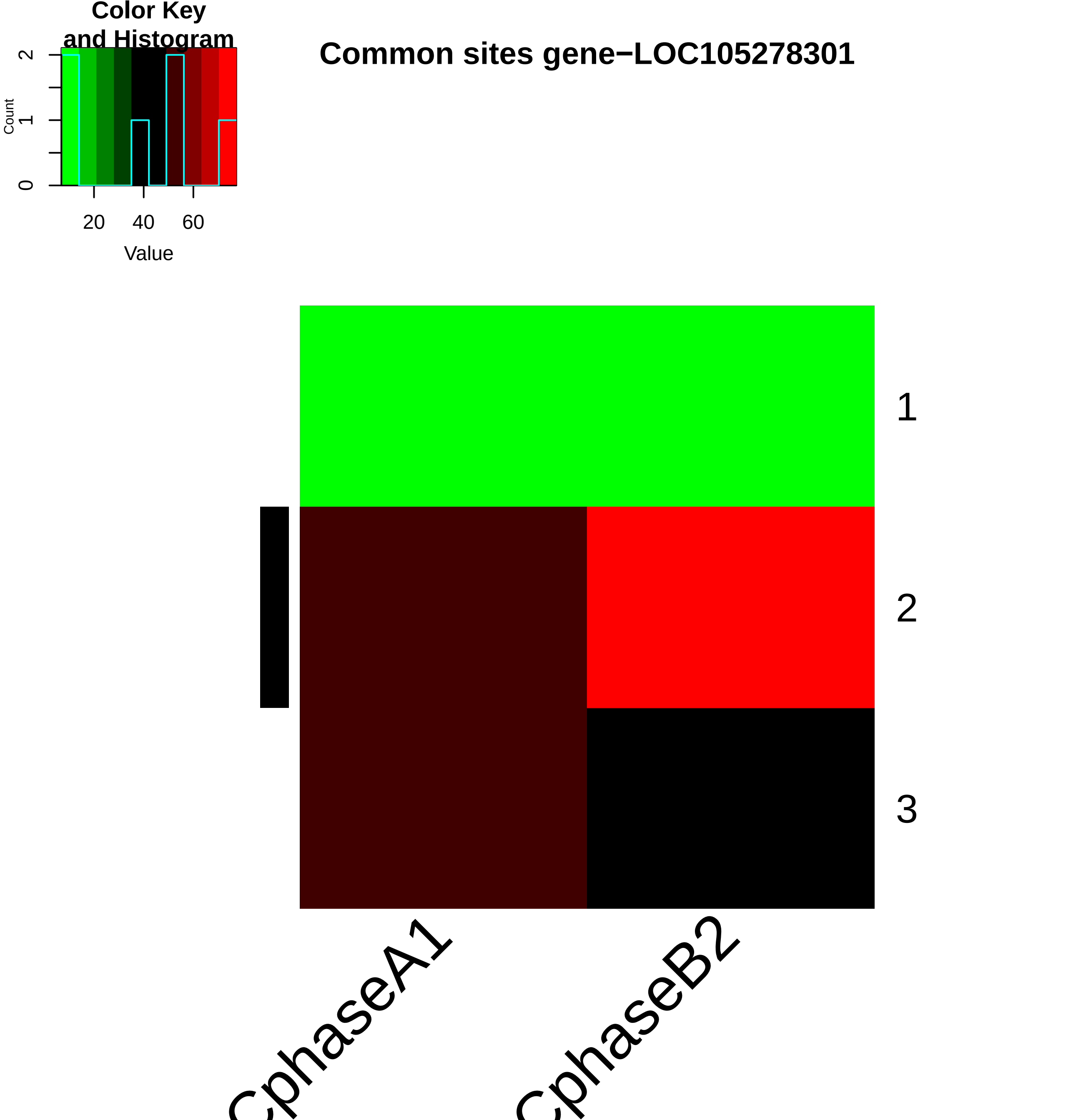

Color Key
and Histogram
Count
0
1
2
20
40
60
Value
Common sites gene−LOC105278301
1
2
3
CphaseA1
CphaseB2

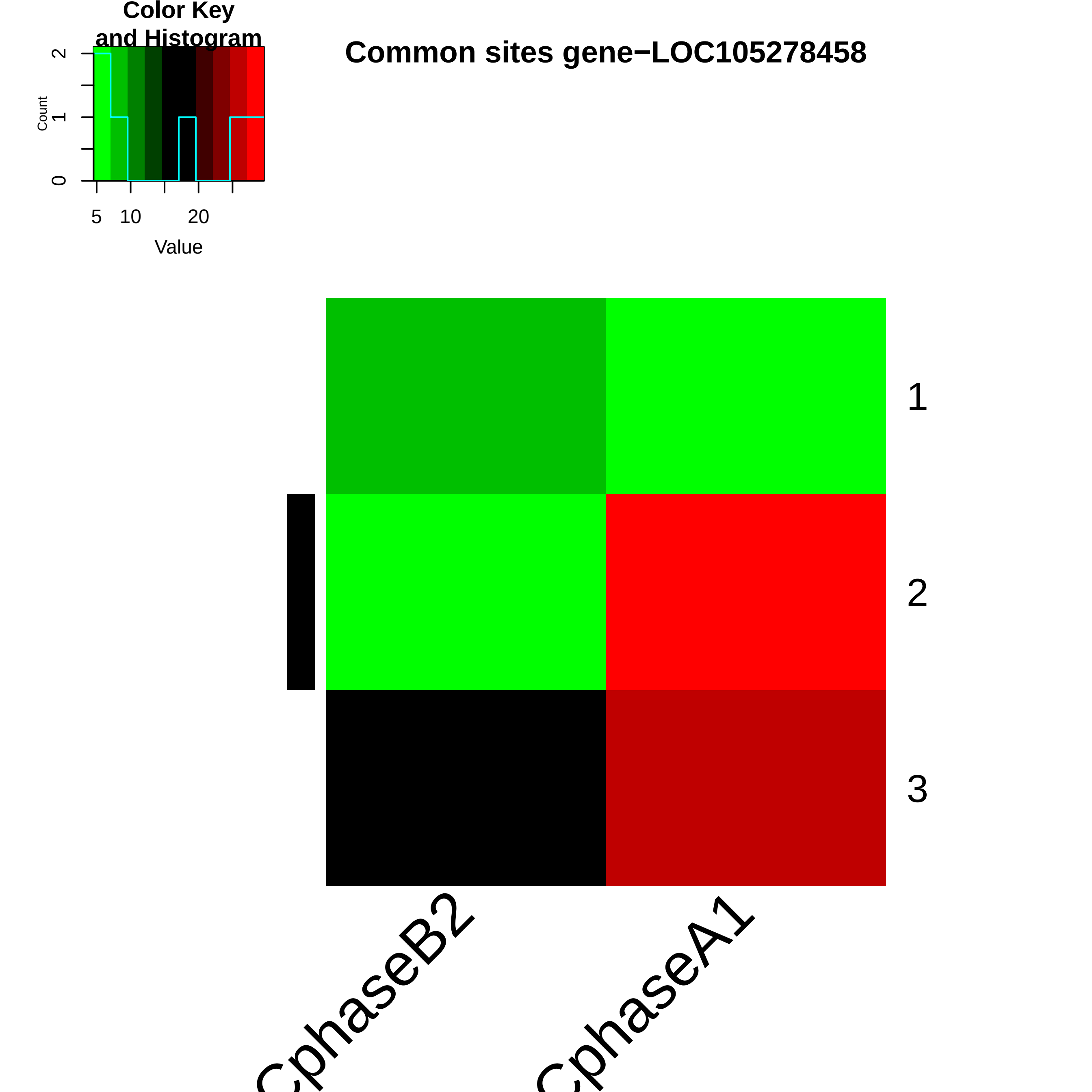
Color Key
and Histogram
Count
0
1
2
5
10
20
Value
Common sites gene−LOC105278458
1
2
3
CphaseB2
CphaseA1

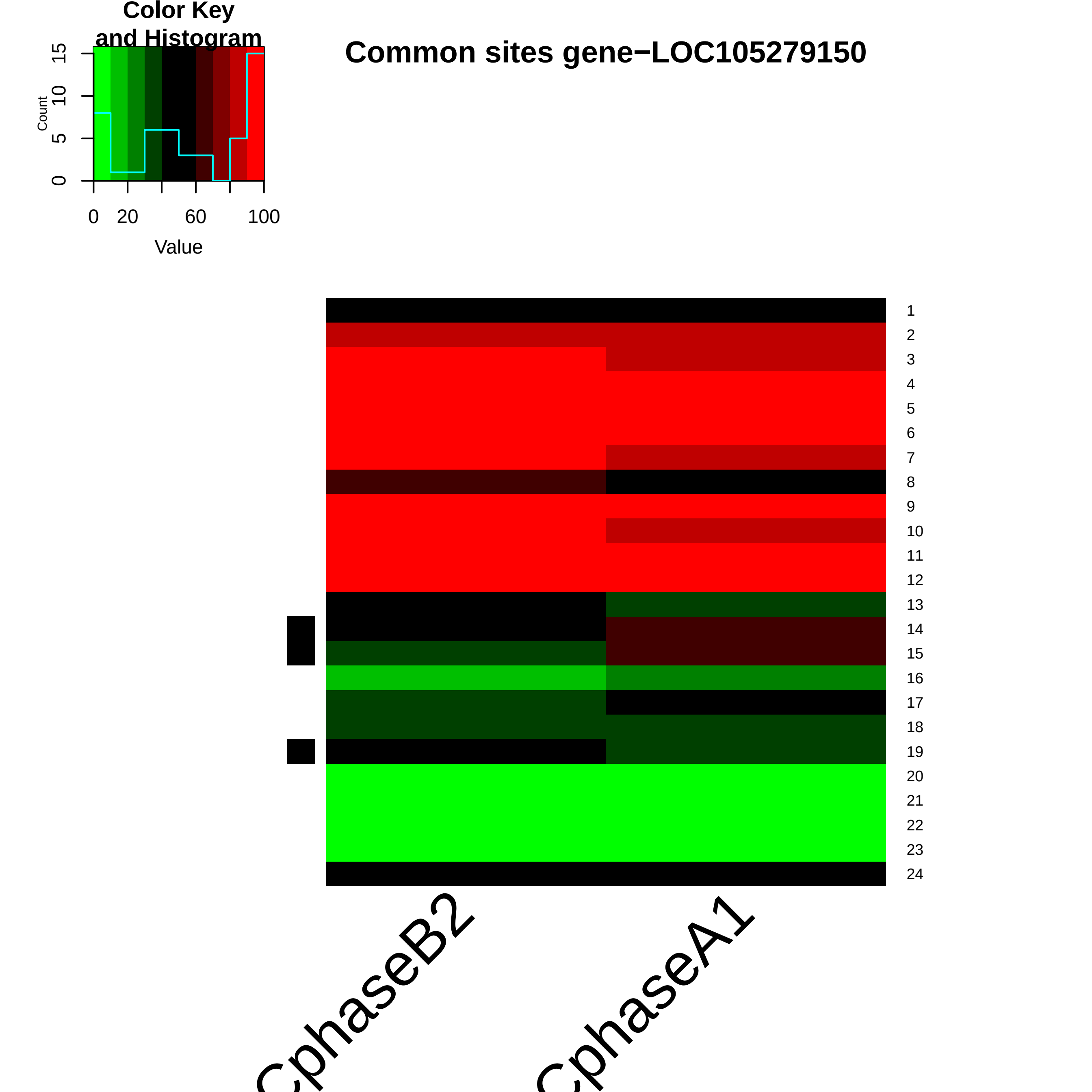

Color Key
and Histogram
Count
0
5
10
15
0
20
60
100
Value
Common sites gene−LOC105279150
CphaseB2
CphaseA1

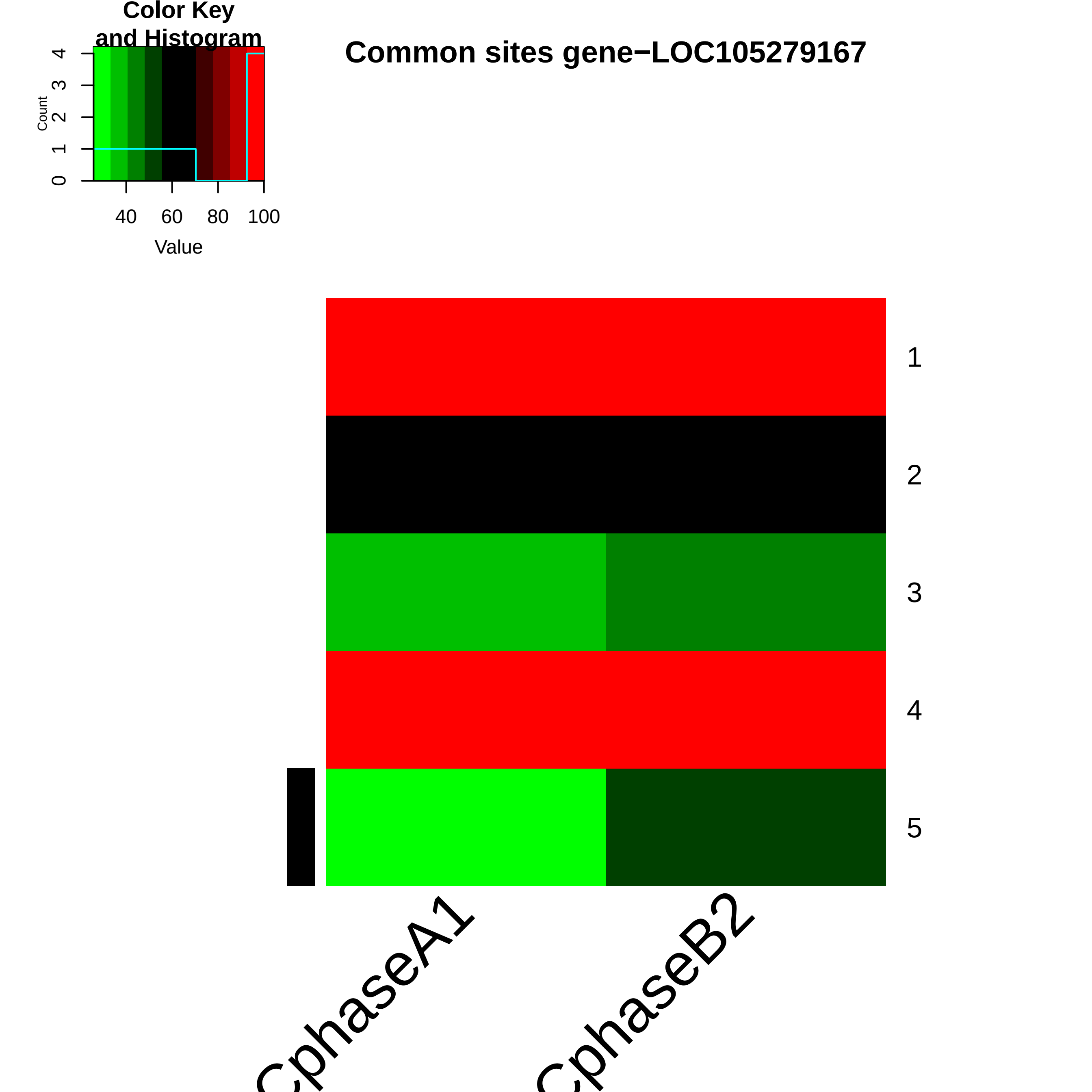

Color Key
and Histogram
Count
0
1
2
3
4
40
60
80
100
Value
Common sites gene−LOC105279167
1
2
3
4
5
CphaseA1
CphaseB2

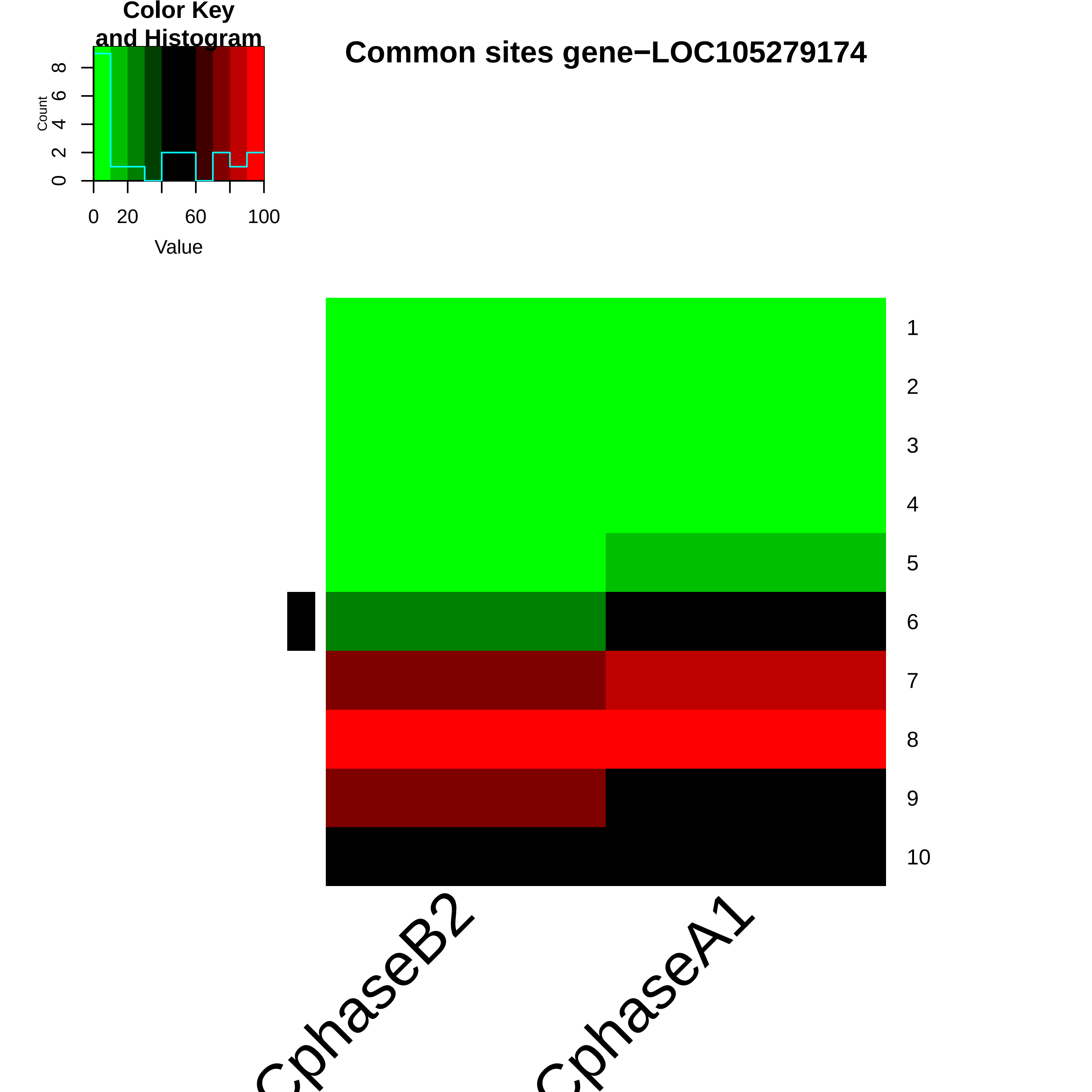
Color Key
and Histogram
Count
Value
0
20
60
100
0
2
4
6
8
Common sites gene−LOC105279174
1
2
3
4
5
6
7
8
9
10
CphaseB2
CphaseA1

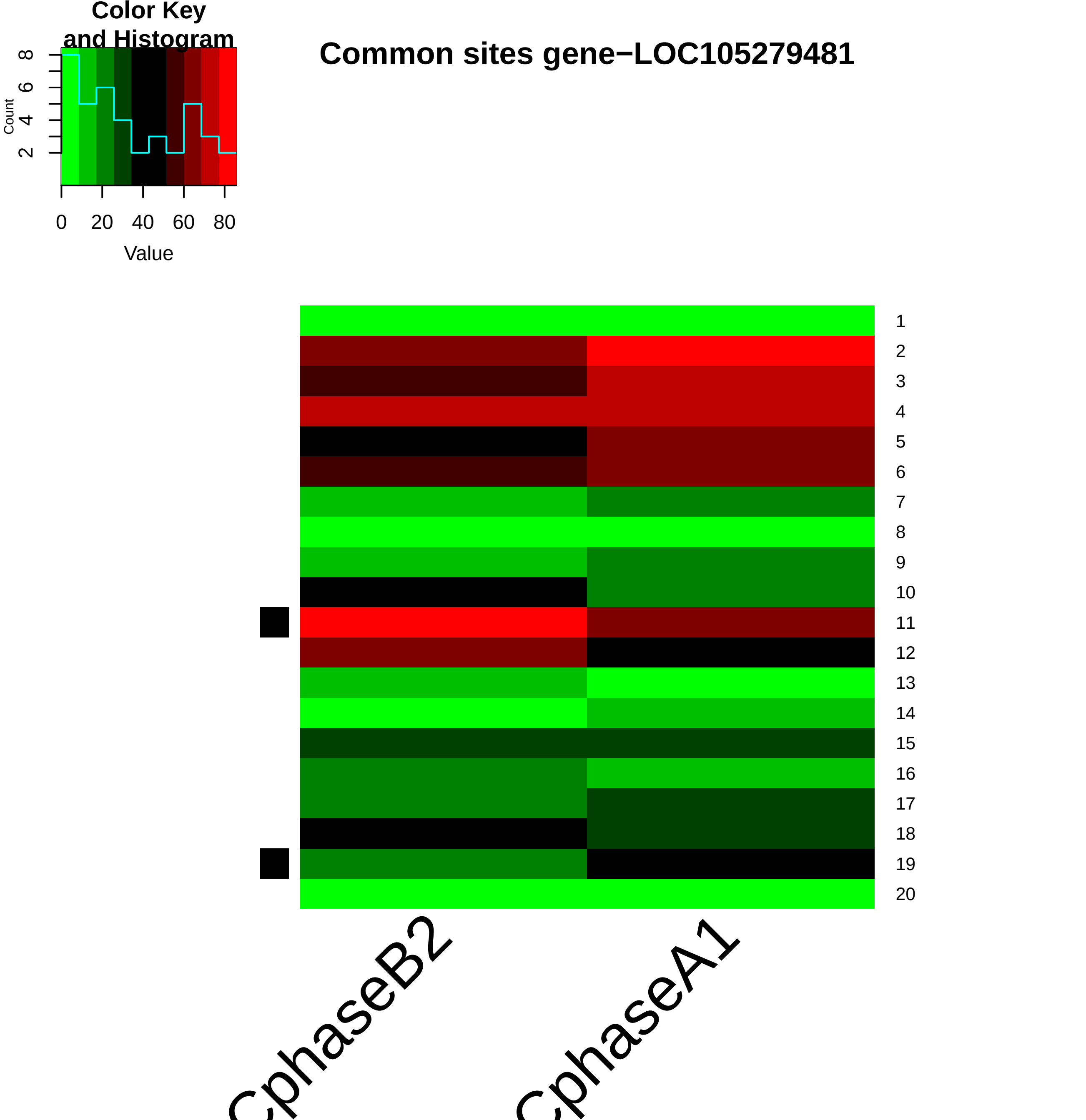
Color Key
and Histogram
Count
Value
Common sites gene−LOC105279481
CphaseB2
CphaseA1

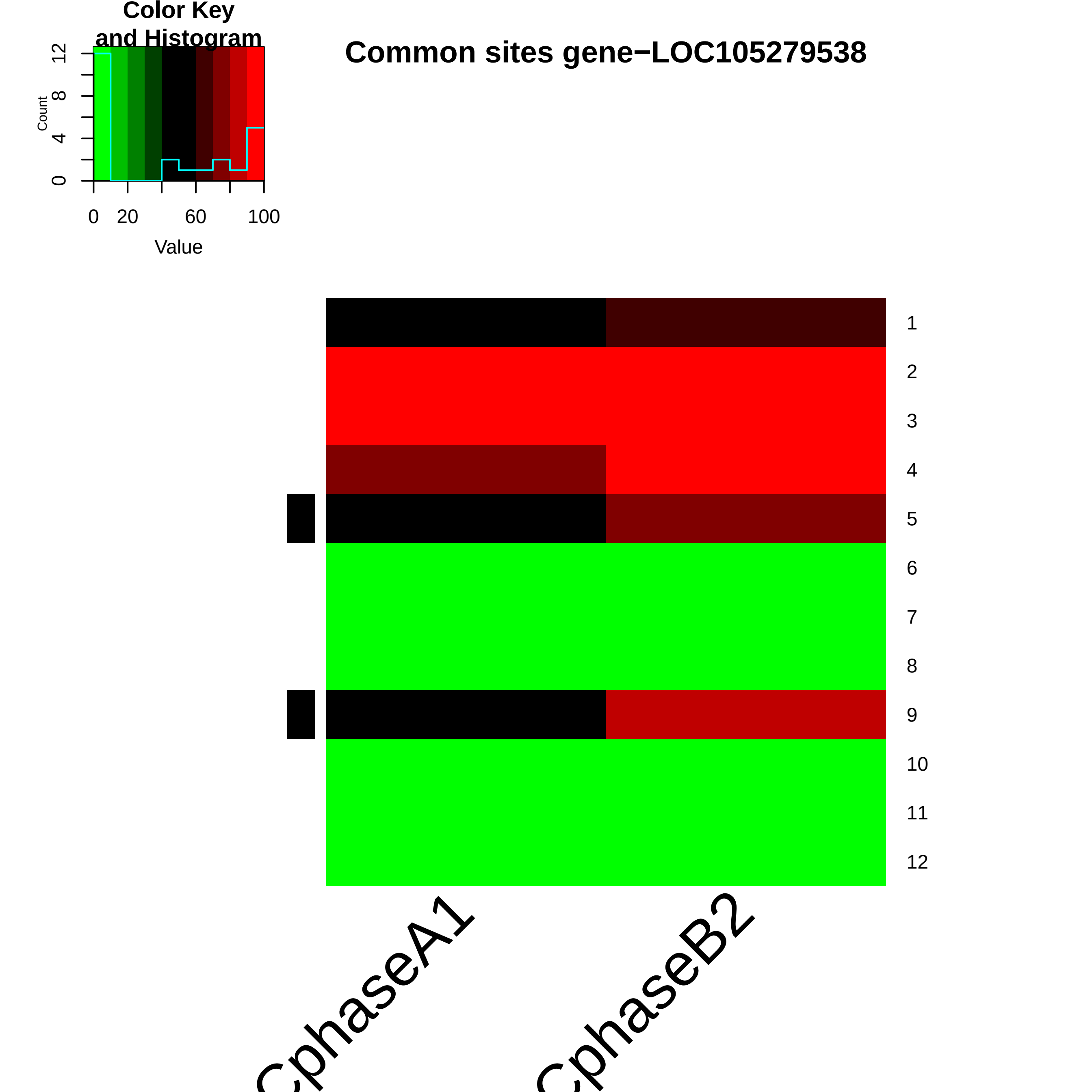
Color Key
and Histogram
Count
0
4
8
12
0
20
60
100
Value
Common sites gene−LOC105279538
1
2
3
4
5
6
7
8
9
10
11
12
CphaseA1
CphaseB2

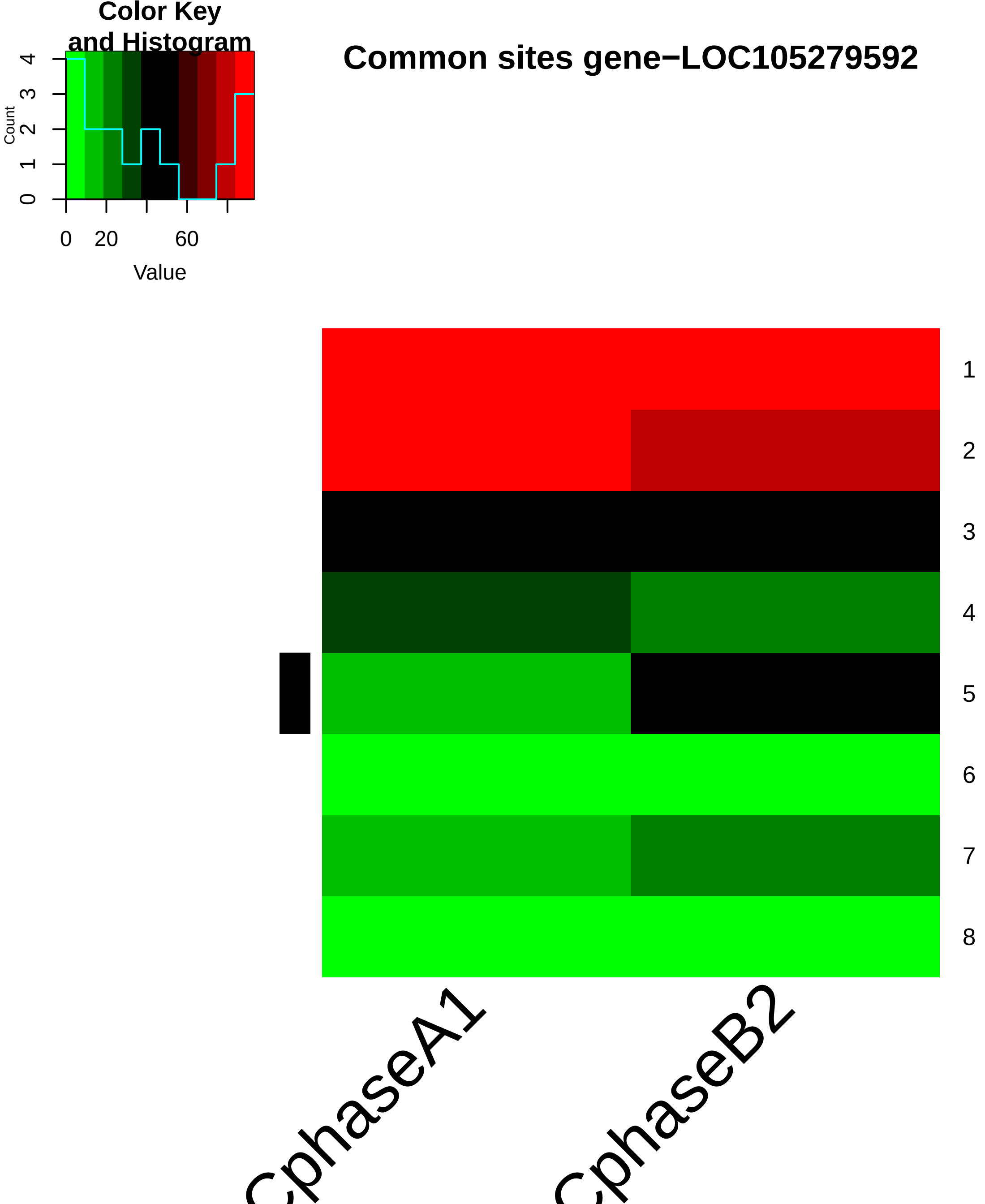

Color Key
and Histogram
Count
0 1 2 3 4
0 20 60
Value
Common sites gene−LOC105279592
1
2
3
4
5
6
7
8
CphaseA1
CphaseB2

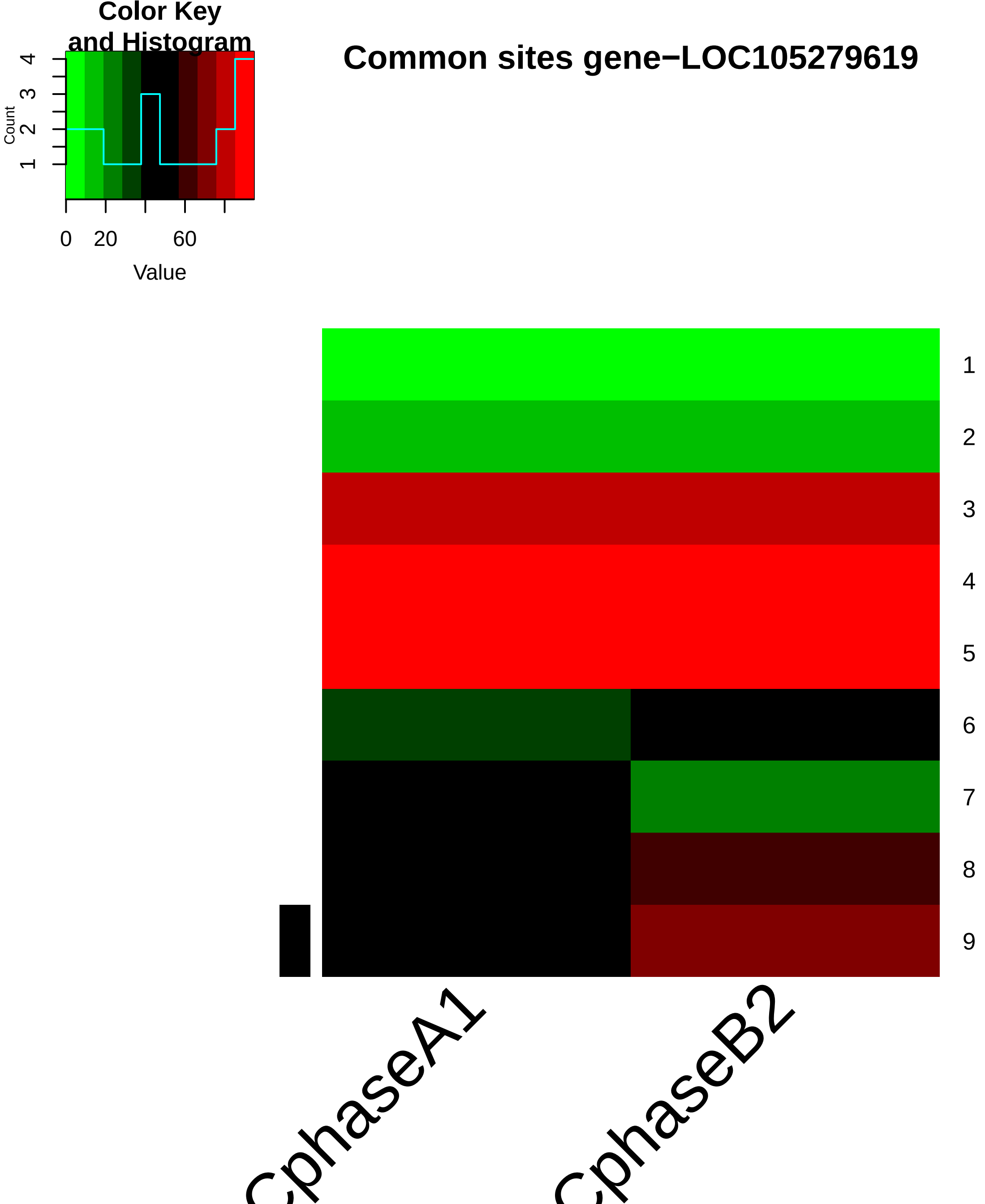

Color Key
and Histogram
Count
4
3
2
1
0
20
60
Value
Common sites gene−LOC105279619
1
2
3
4
5
6
7
8
9
CphaseA1
CphaseB2

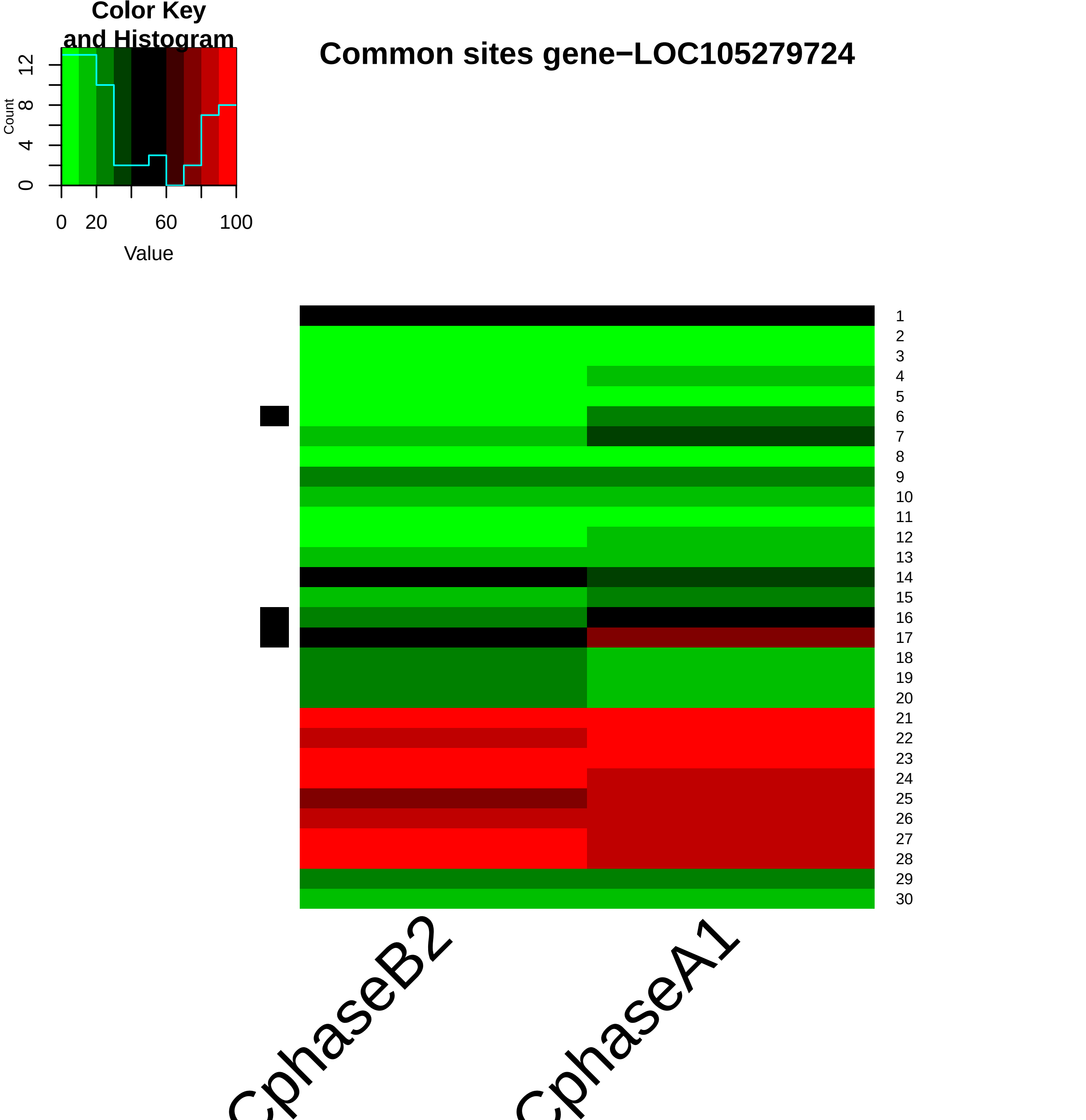
Color Key
and Histogram
Count
0
4
8
12
0
20
60
100
Value
Common sites gene−LOC105279724
CphaseB2
CphaseA1

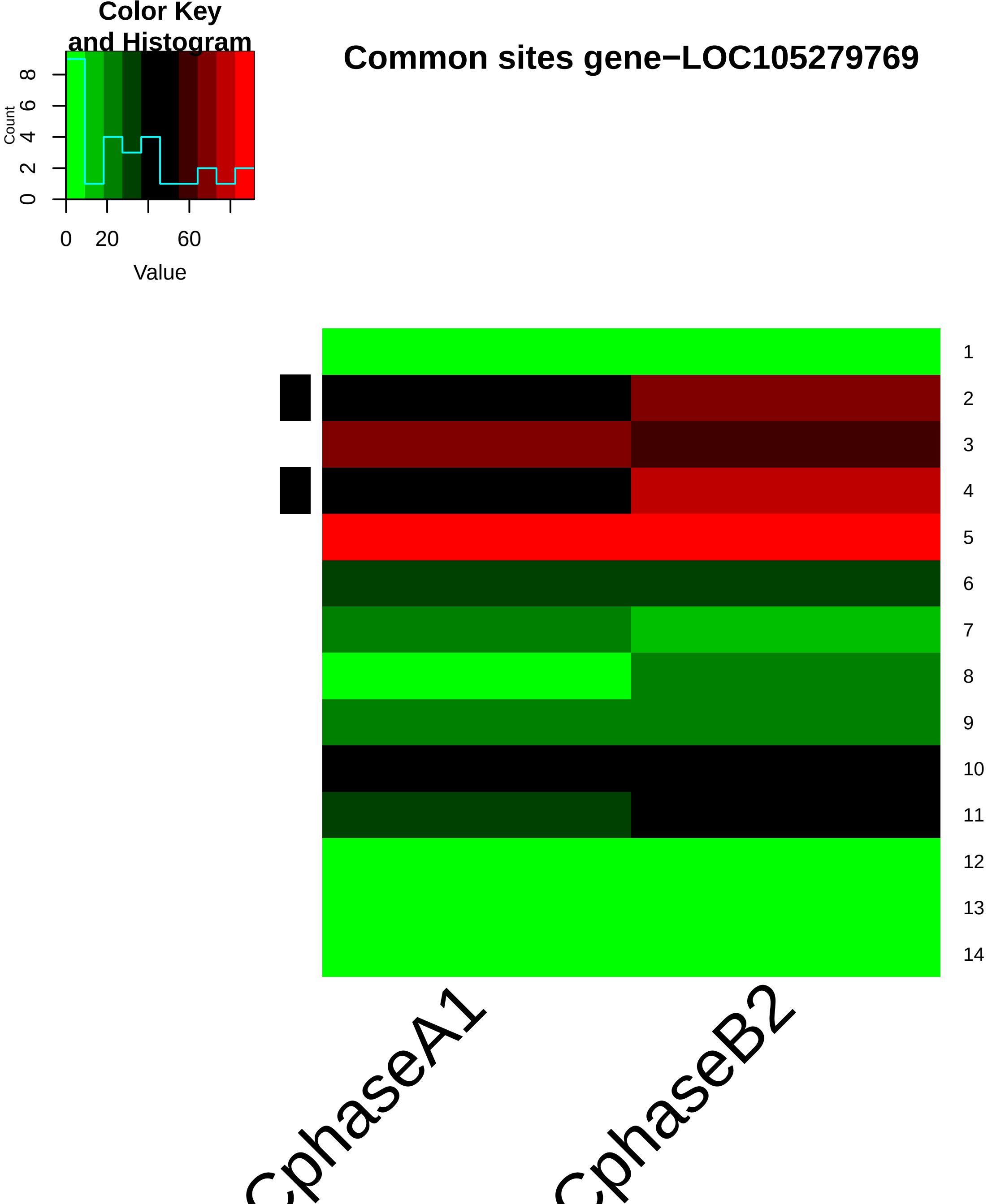
Color Key
and Histogram
Count
0
2
4
6
8
0
20
60
Value
Common sites gene−LOC105279769
1
2
3
4
5
6
7
8
9
10
11
12
13
14
CphaseA1
CphaseB2

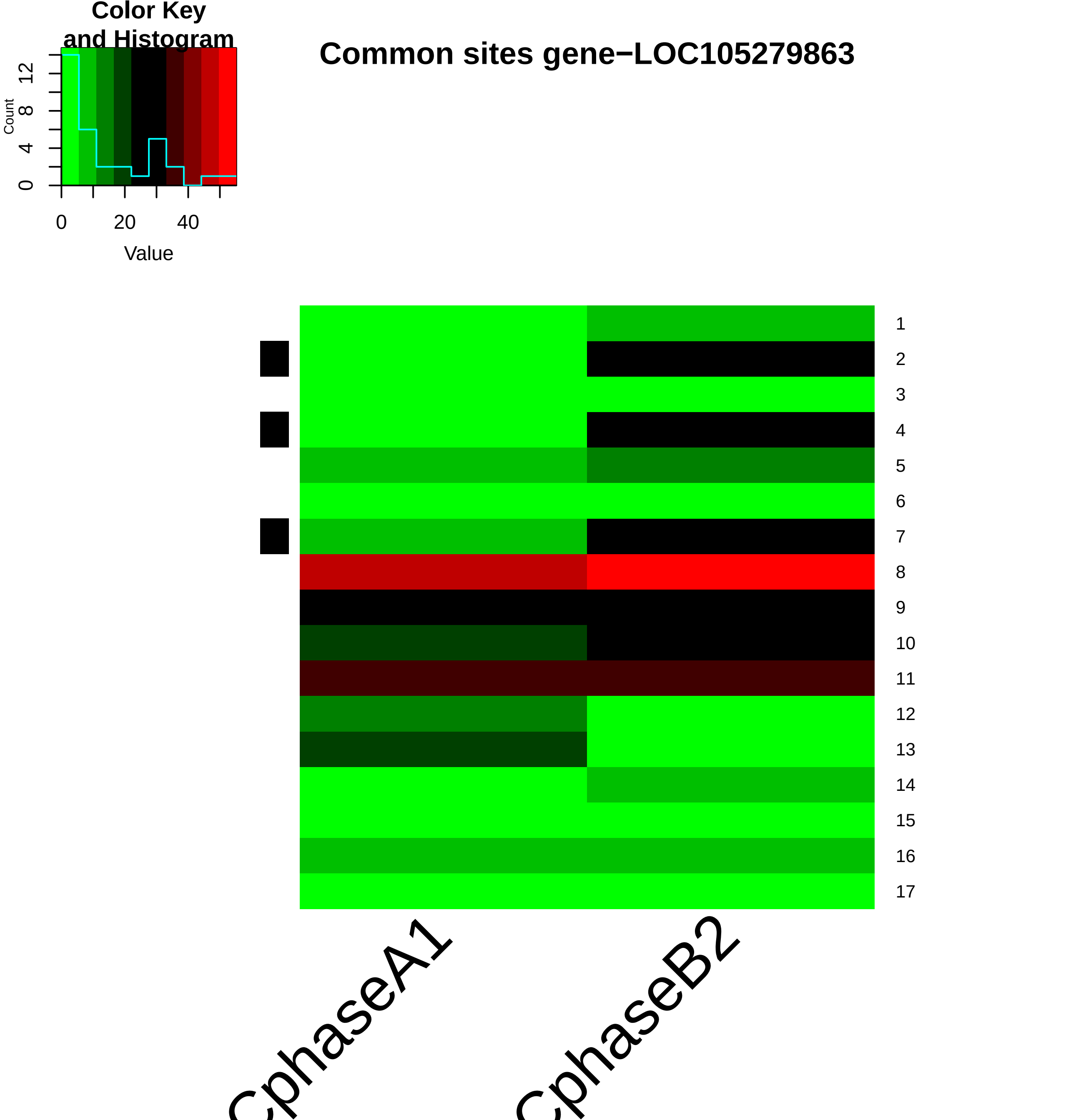
Color Key
and Histogram
Count
0
4
8
12
0
20
40
Value
Common sites gene−LOC105279863
CphaseA1
CphaseB2

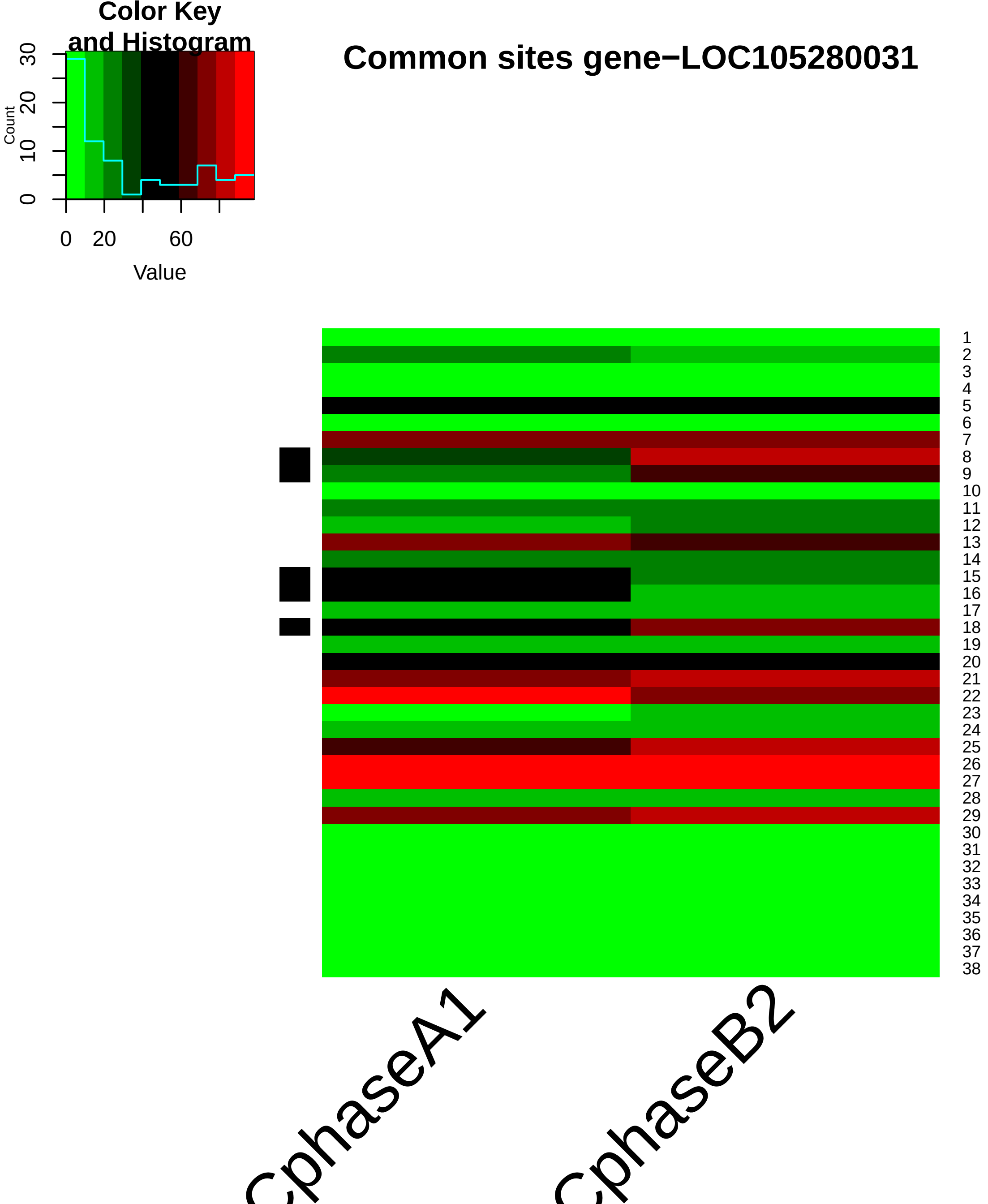
Color Key
and Histogram
Count
0
10
20
30
0
20
60
Value
Common sites gene−LOC105280031
CphaseA1
CphaseB2

### Common sites gene−LOC105280220

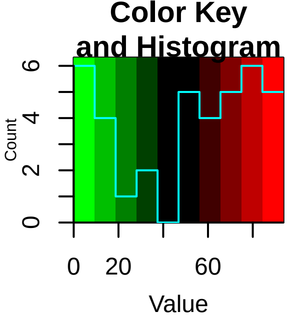

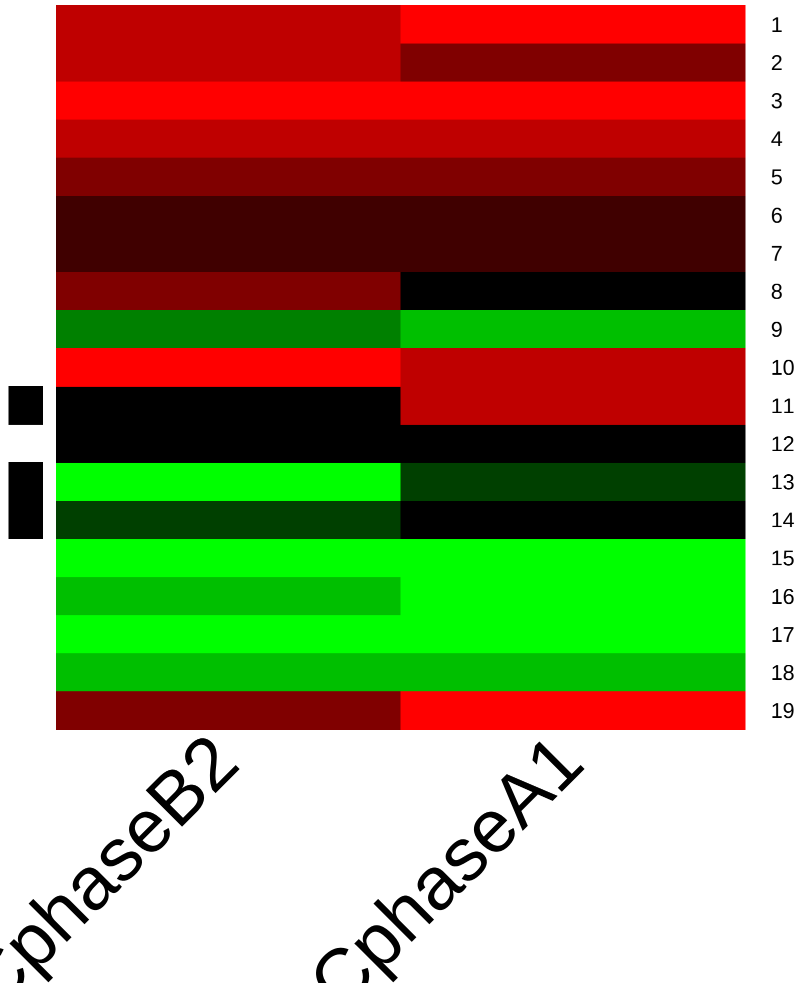

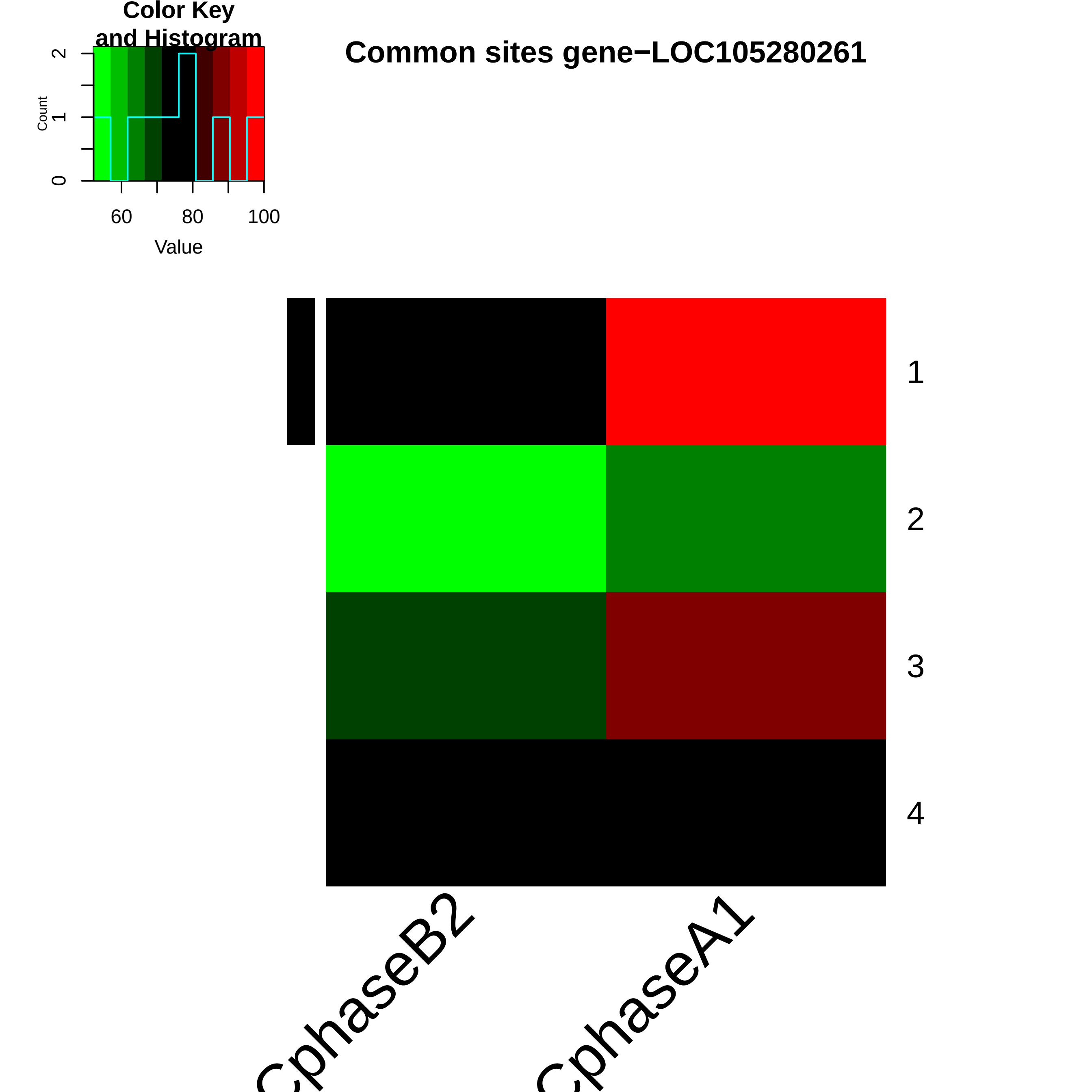

Color Key
and Histogram
Count
0
1
2
60
80
100
Value
Common sites gene−LOC105280261
1
2
3
4
CphaseB2
CphaseA1

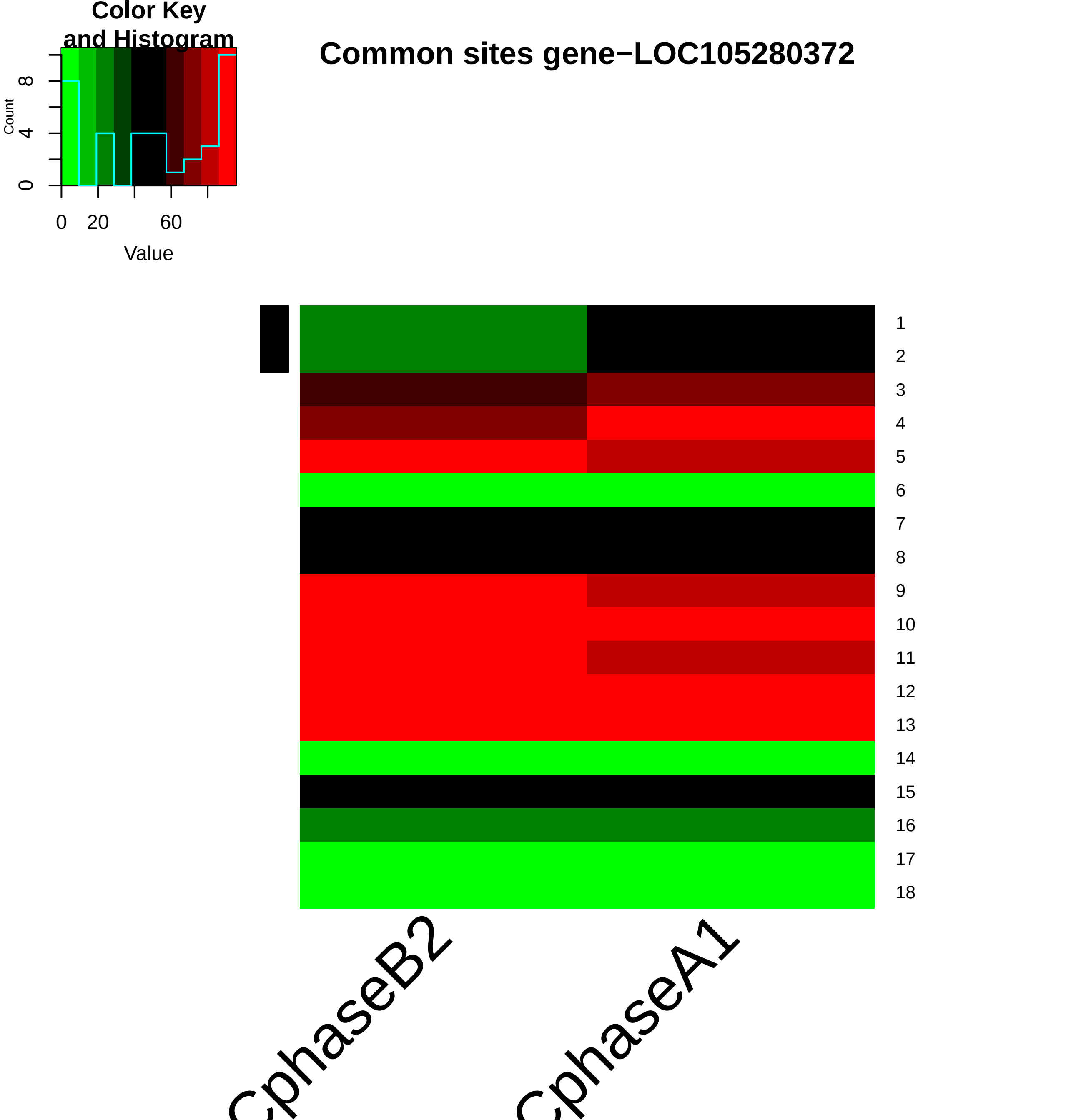

Color Key
and Histogram
Count
0
4
8
0
20
60
Value
Common sites gene−LOC105280372
CphaseB2
CphaseA1

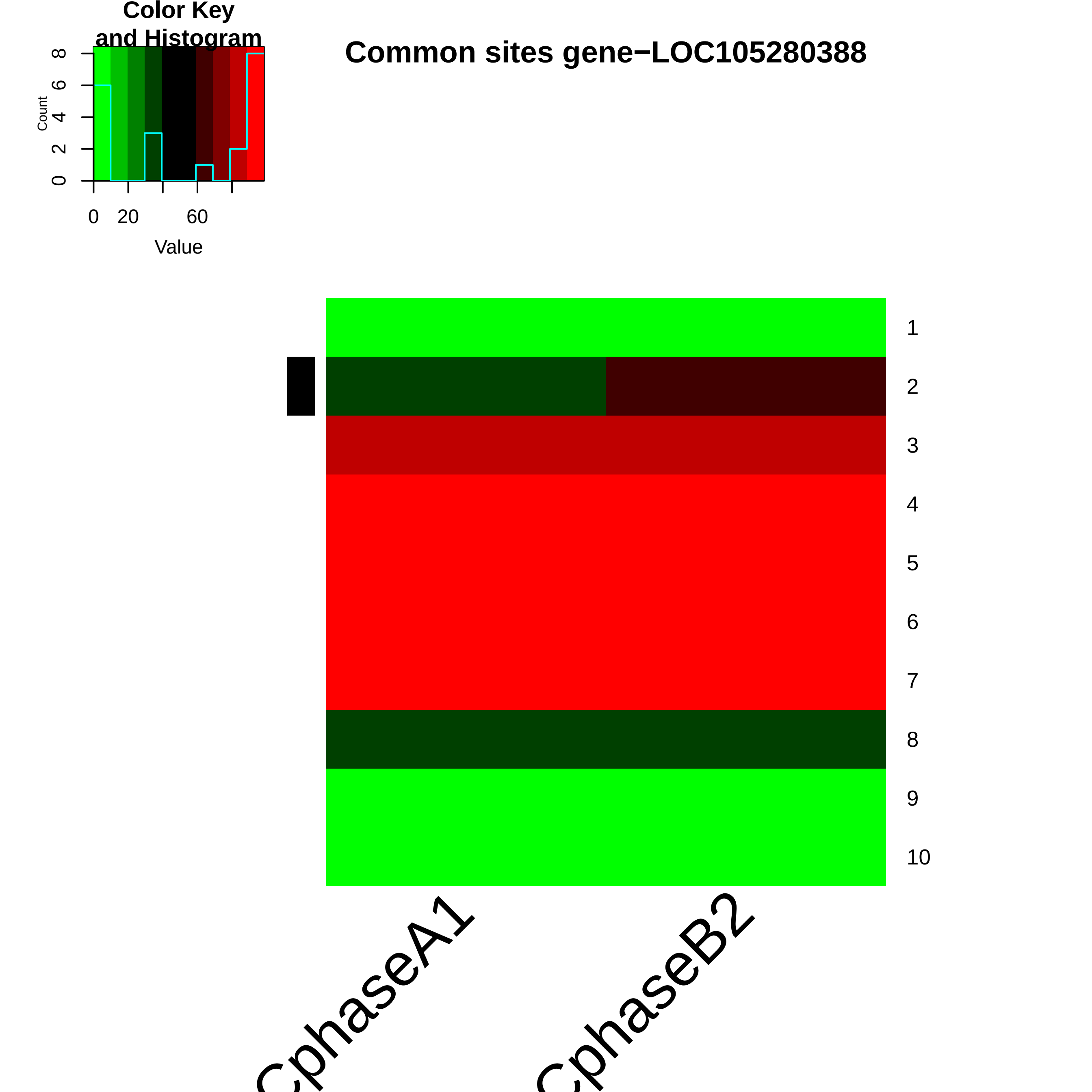

Color Key
and Histogram
Count
0
2
4
6
8
0
20
60
Value
Common sites gene−LOC105280388
1
2
3
4
5
6
7
8
9
10
CphaseA1
CphaseB2

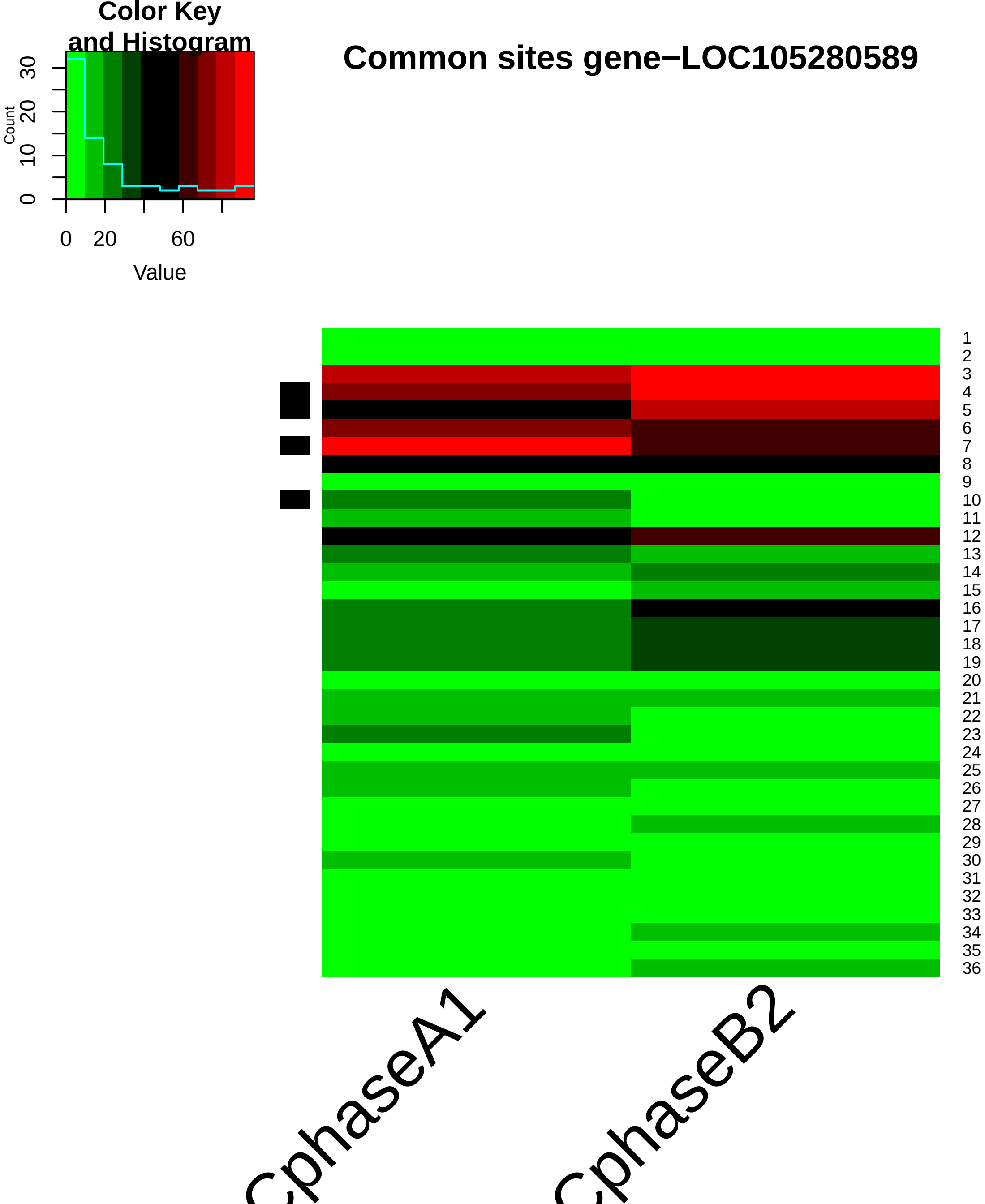
Color Key
and Histogram
Count
0
10
20
30
0
20
60
Value
Common sites gene−LOC105280589
CphaseA1
CphaseB2

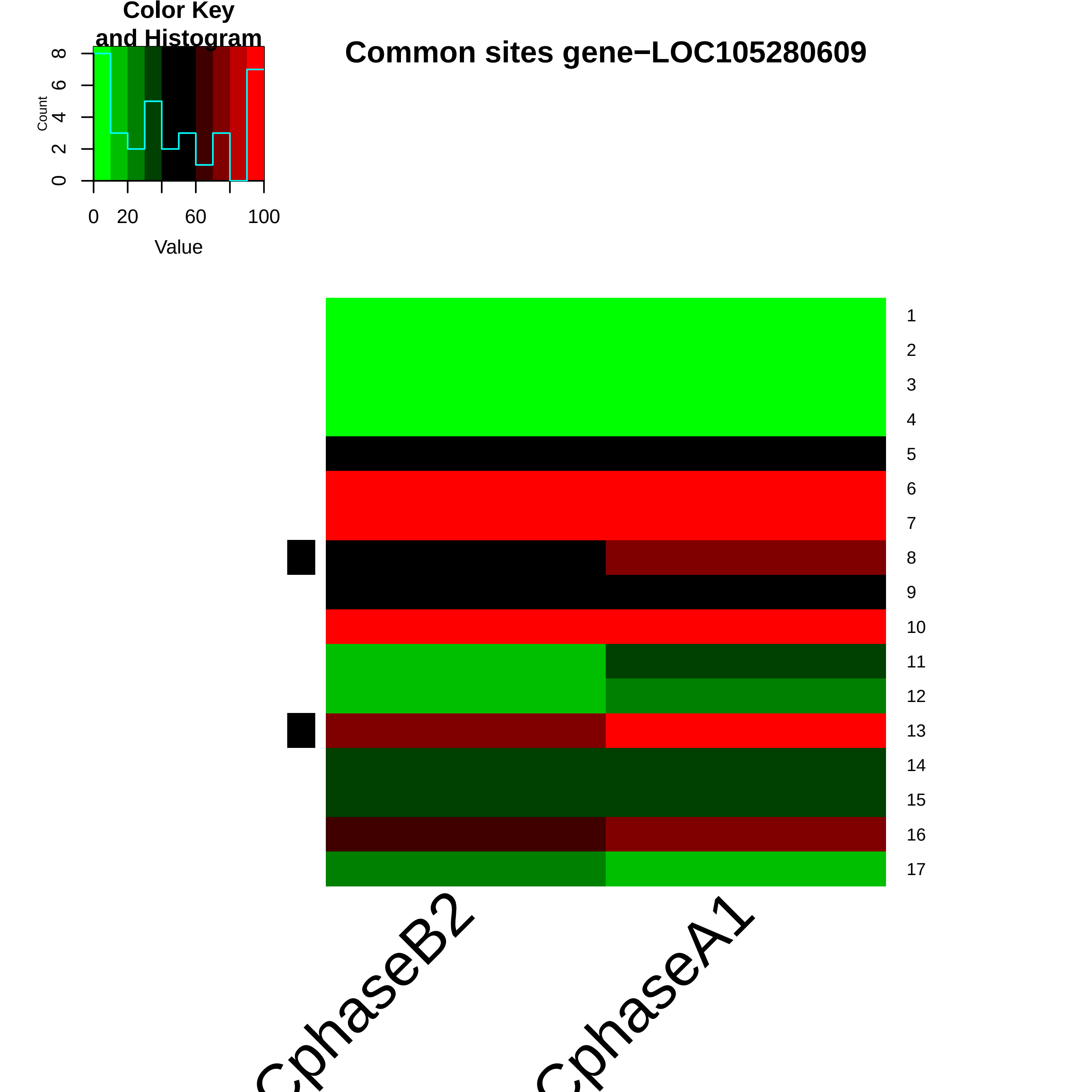
Color Key
and Histogram
Count
0
2
4
6
8
0
20
60
100
Value
Common sites gene−LOC105280609
CphaseB2
CphaseA1

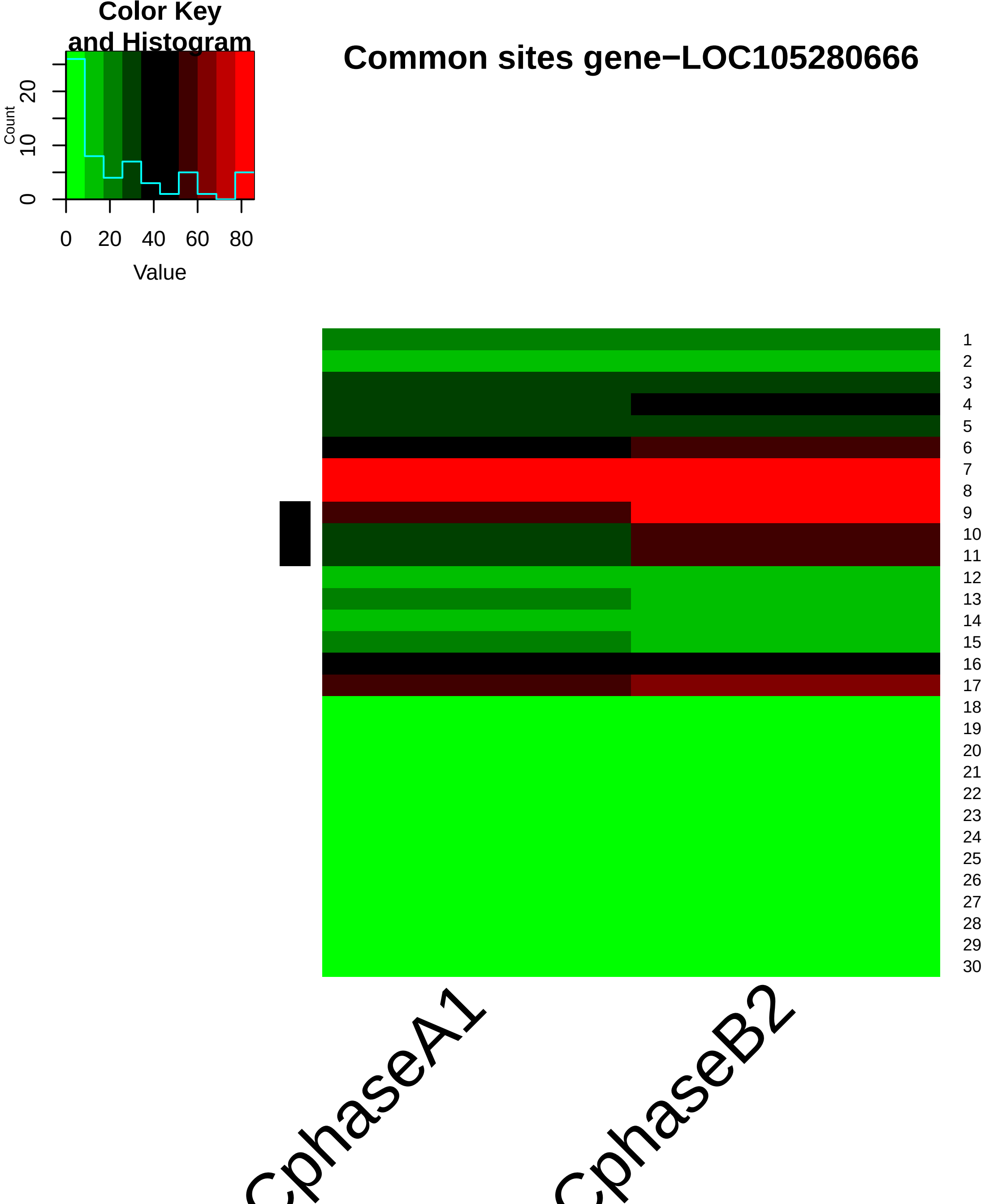

Color Key
and Histogram
Count
0
10
20
0
20
40
60
80
Value
Common sites gene−LOC105280666
CphaseA1
CphaseB2

### Common sites gene−LOC105280891

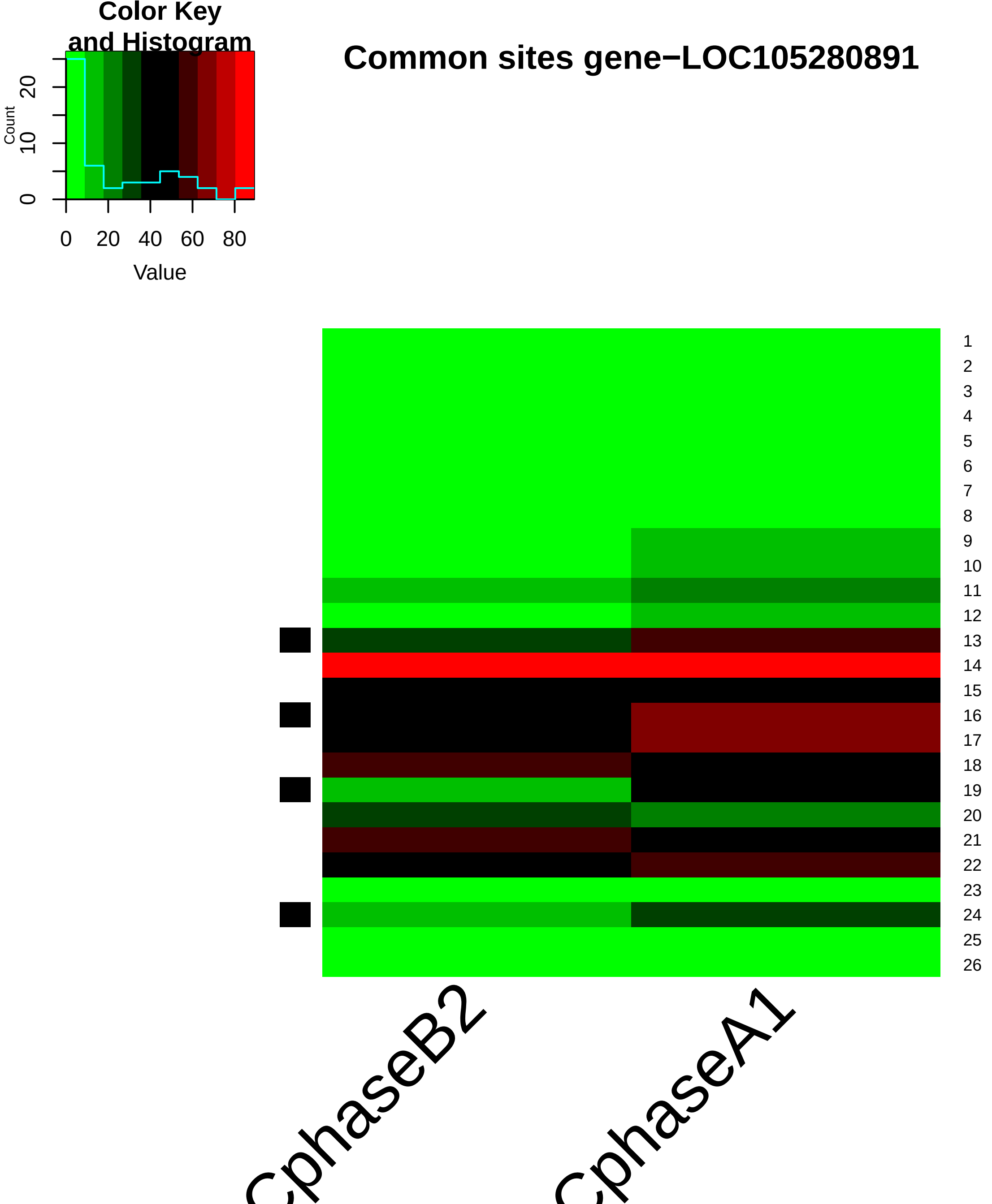

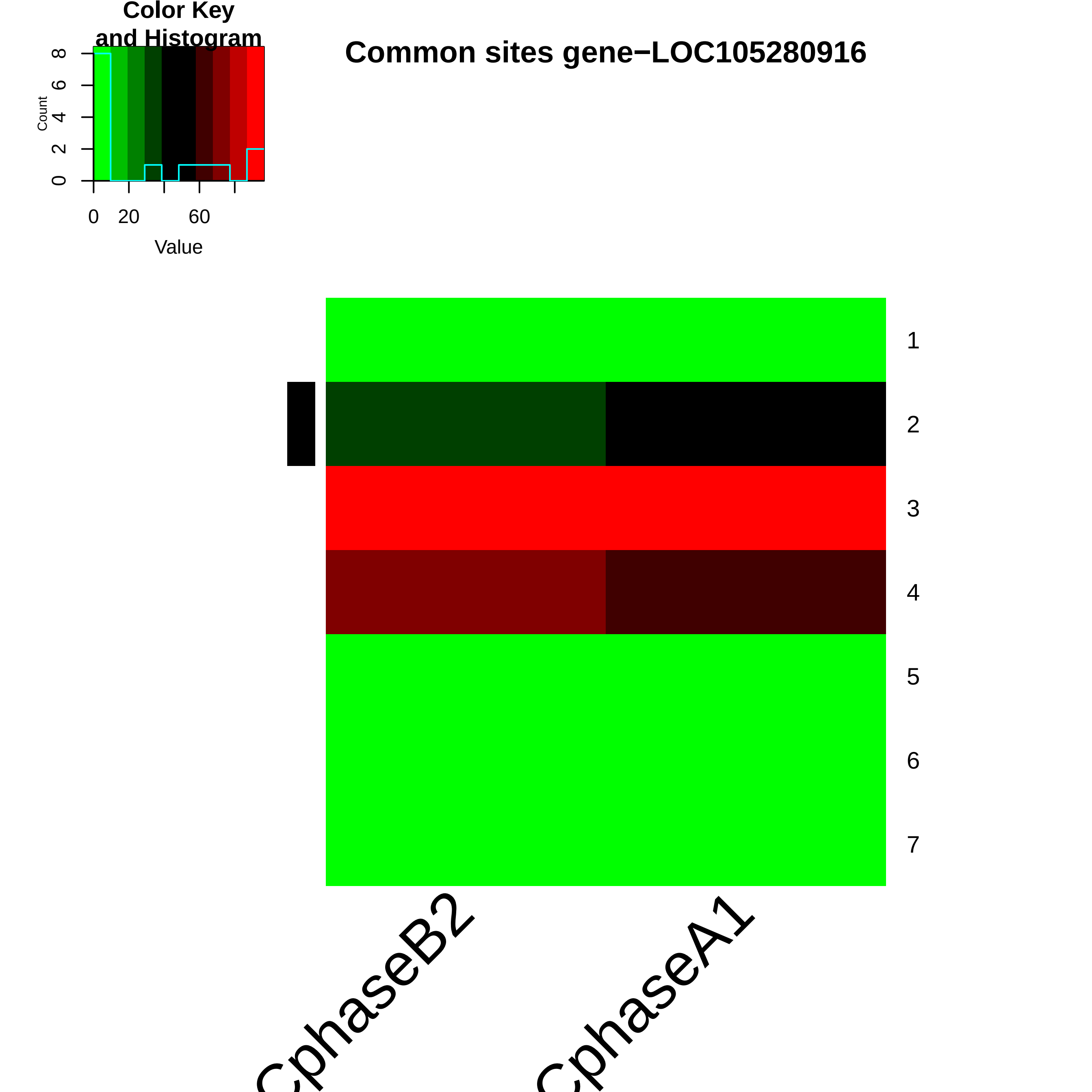
Color Key
and Histogram
Count
Value
Common sites gene−LOC105280916
1
2
3
4
5
6
7
CphaseB2
CphaseA1

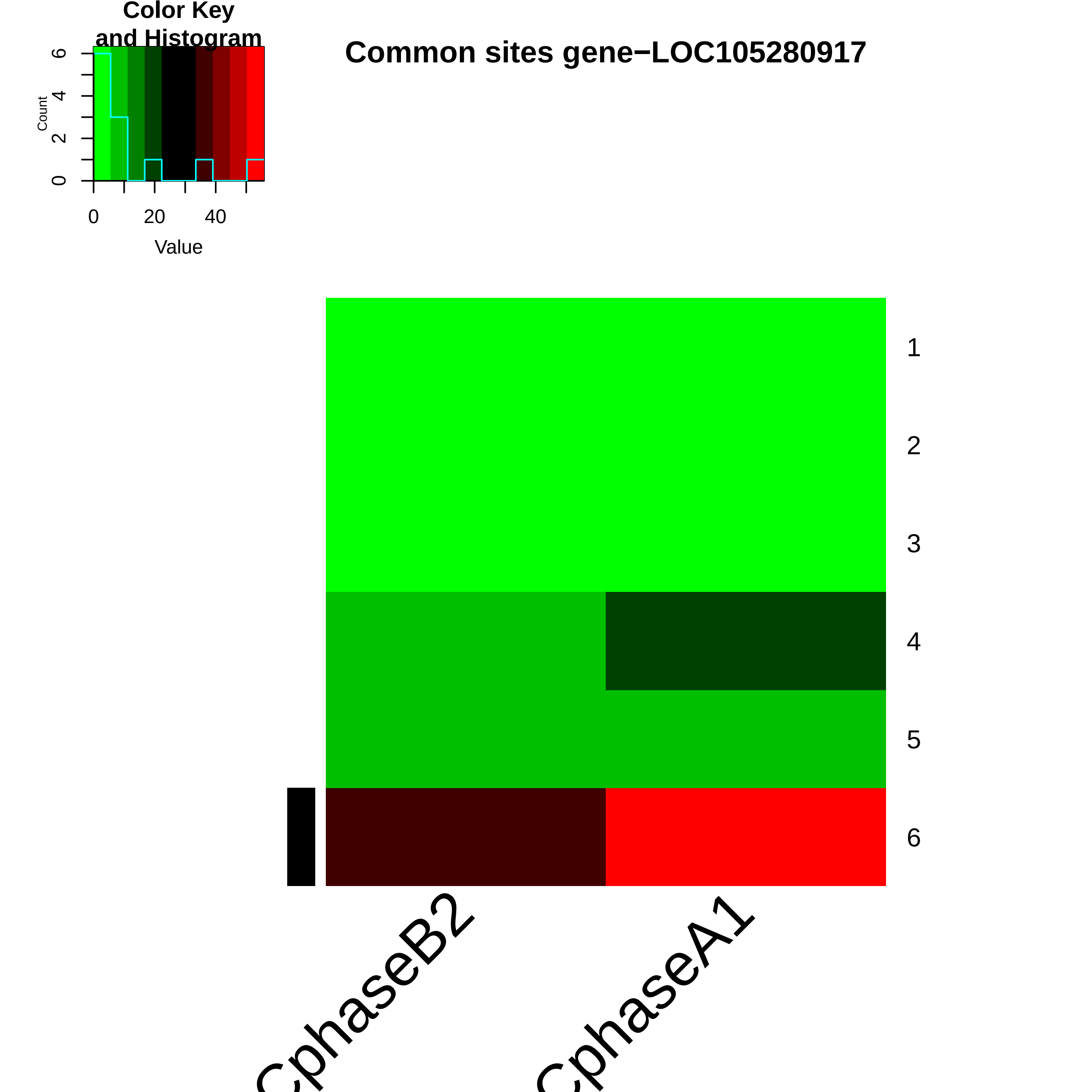

Color Key
and Histogram
Count
0
2
4
6
0
20
40
Value
Common sites gene−LOC105280917
1
2
3
4
5
6
CphaseB2
CphaseA1

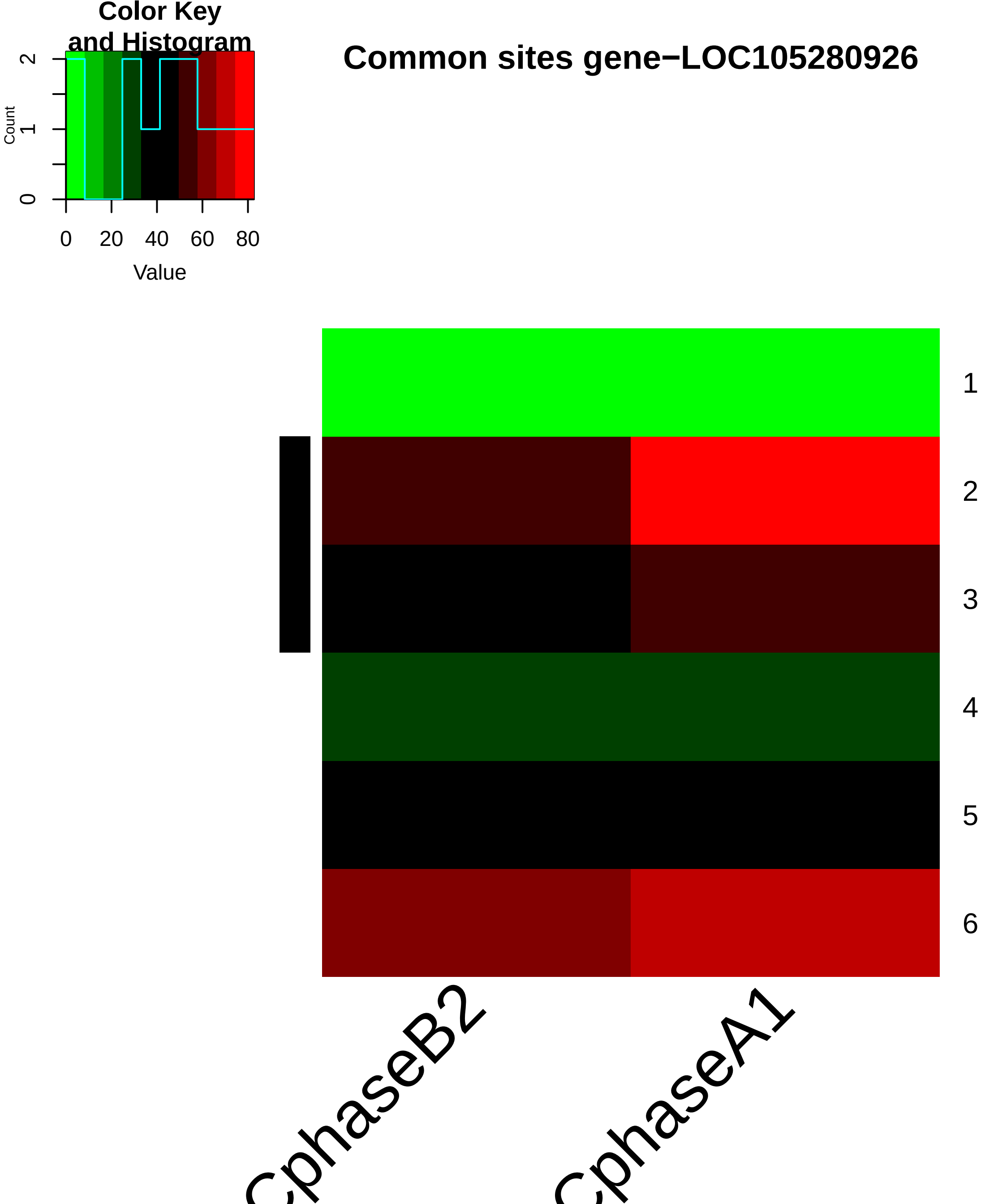
Color Key
and Histogram
Count
Value
0
1
2
0
20
40
60
80
Common sites gene−LOC105280926
1
2
3
4
5
6
CphaseB2
CphaseA1

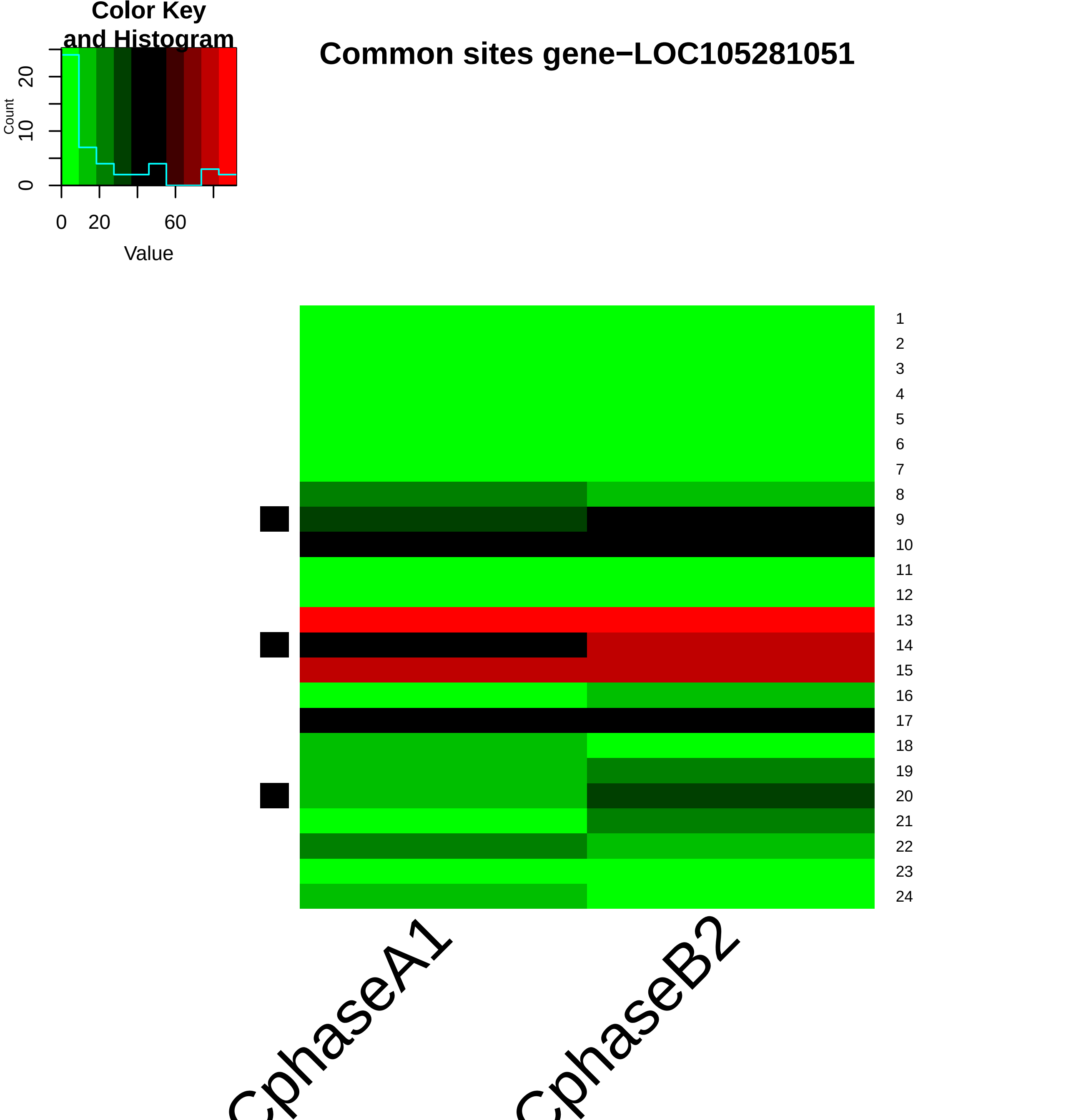
Color Key
and Histogram
Count
0
10
20
0
20
60
Value
Common sites gene−LOC105281051
CphaseA1
CphaseB2

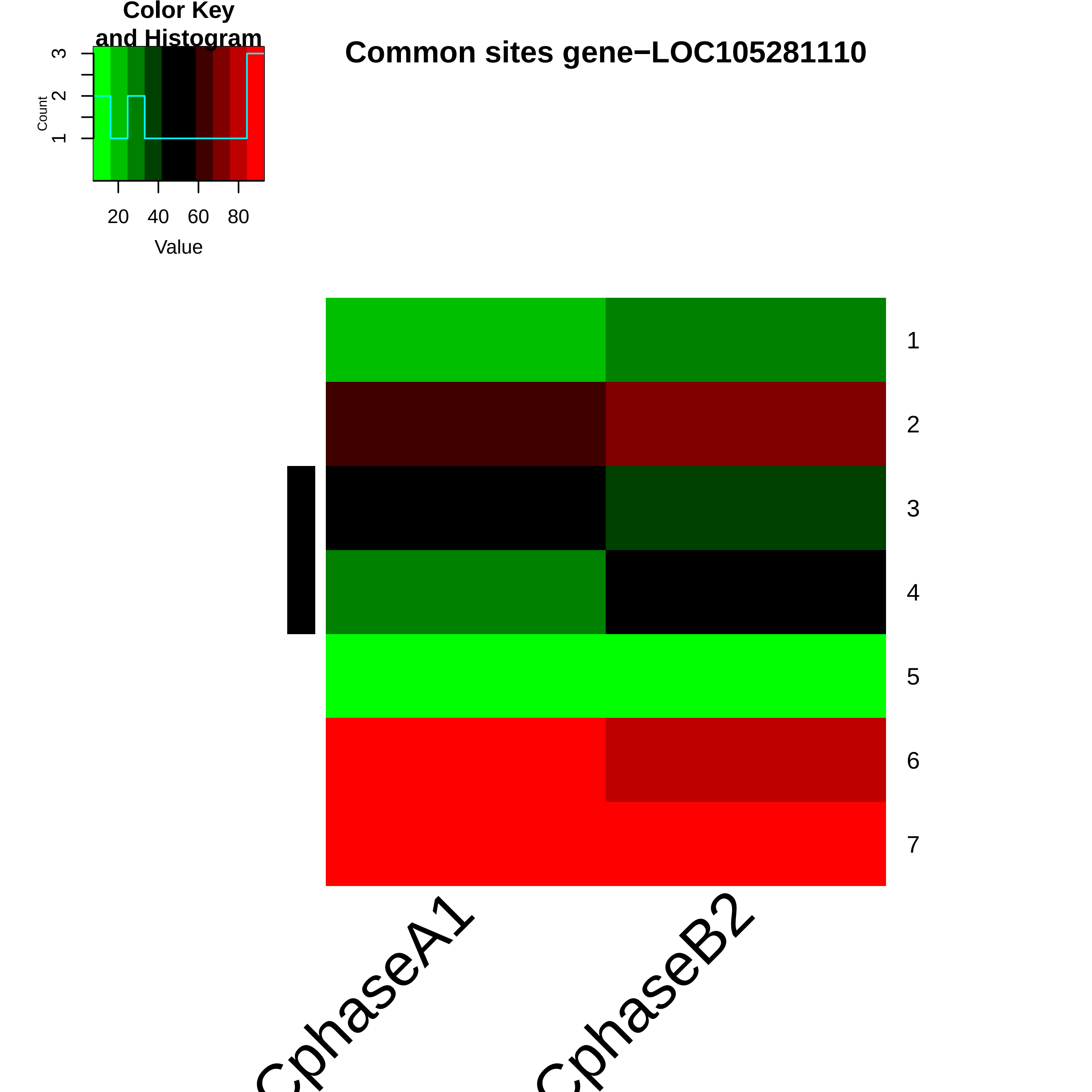

Color Key
and Histogram
Count
Value
Common sites gene−LOC105281110
1
2
3
4
5
6
7
CphaseA1
CphaseB2

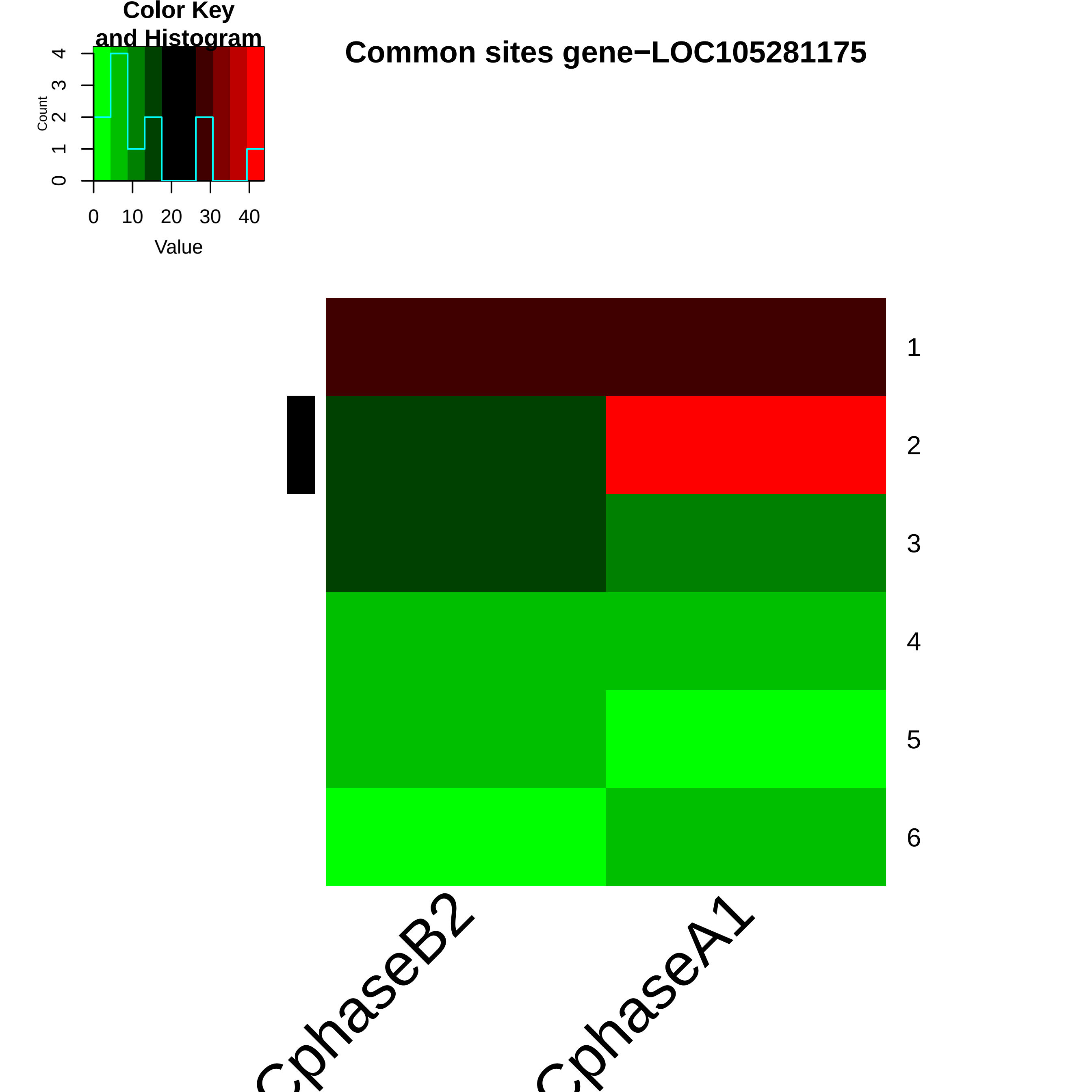
Color Key
and Histogram
Count
0
1
2
3
4
0
10
20
30
40
Value
Common sites gene−LOC105281175
1
2
3
4
5
6
CphaseB2
CphaseA1

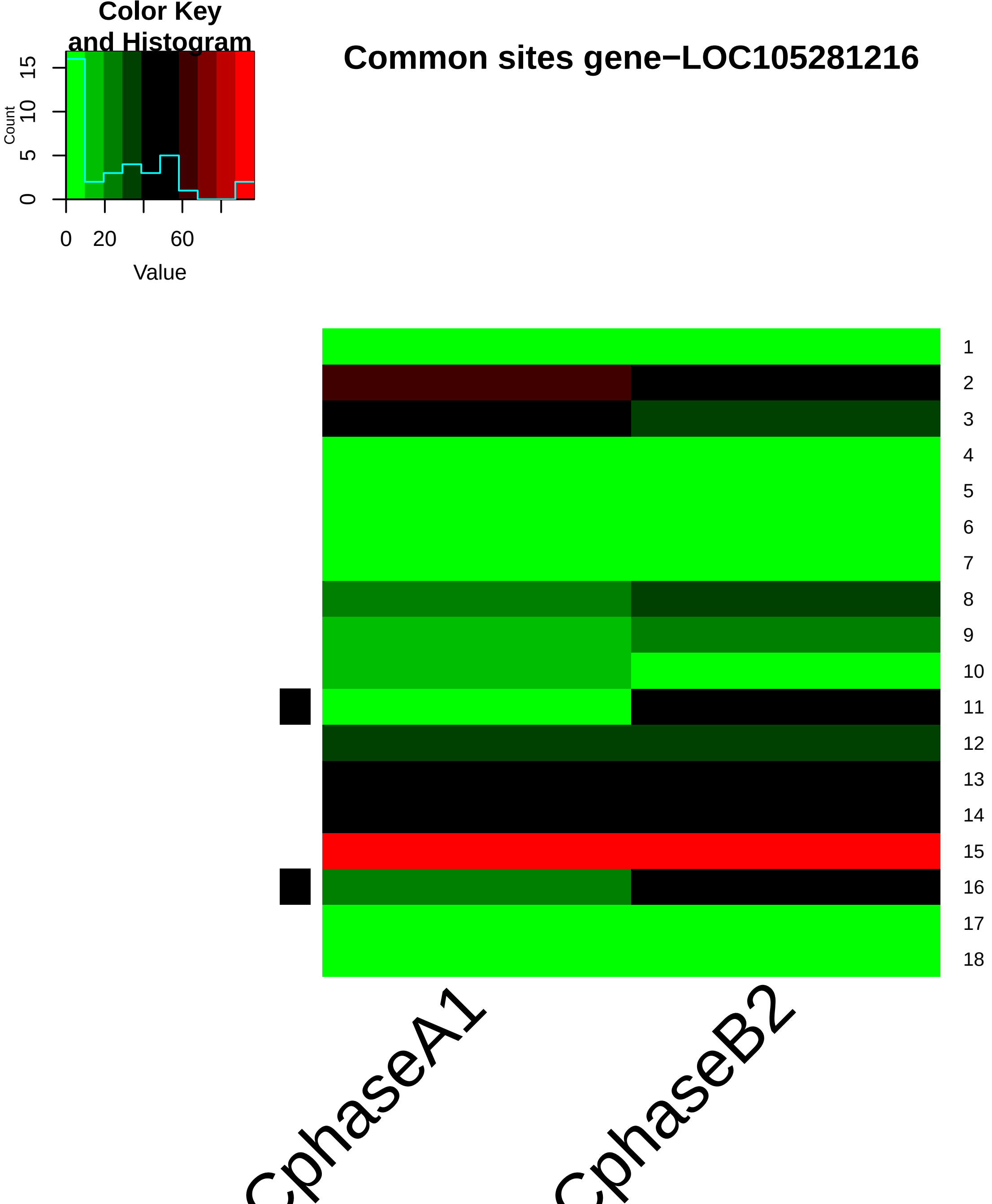
Color Key
and Histogram
Count
0
5
10
15
0
20
60
Value
Common sites gene−LOC105281216
CphaseA1
CphaseB2

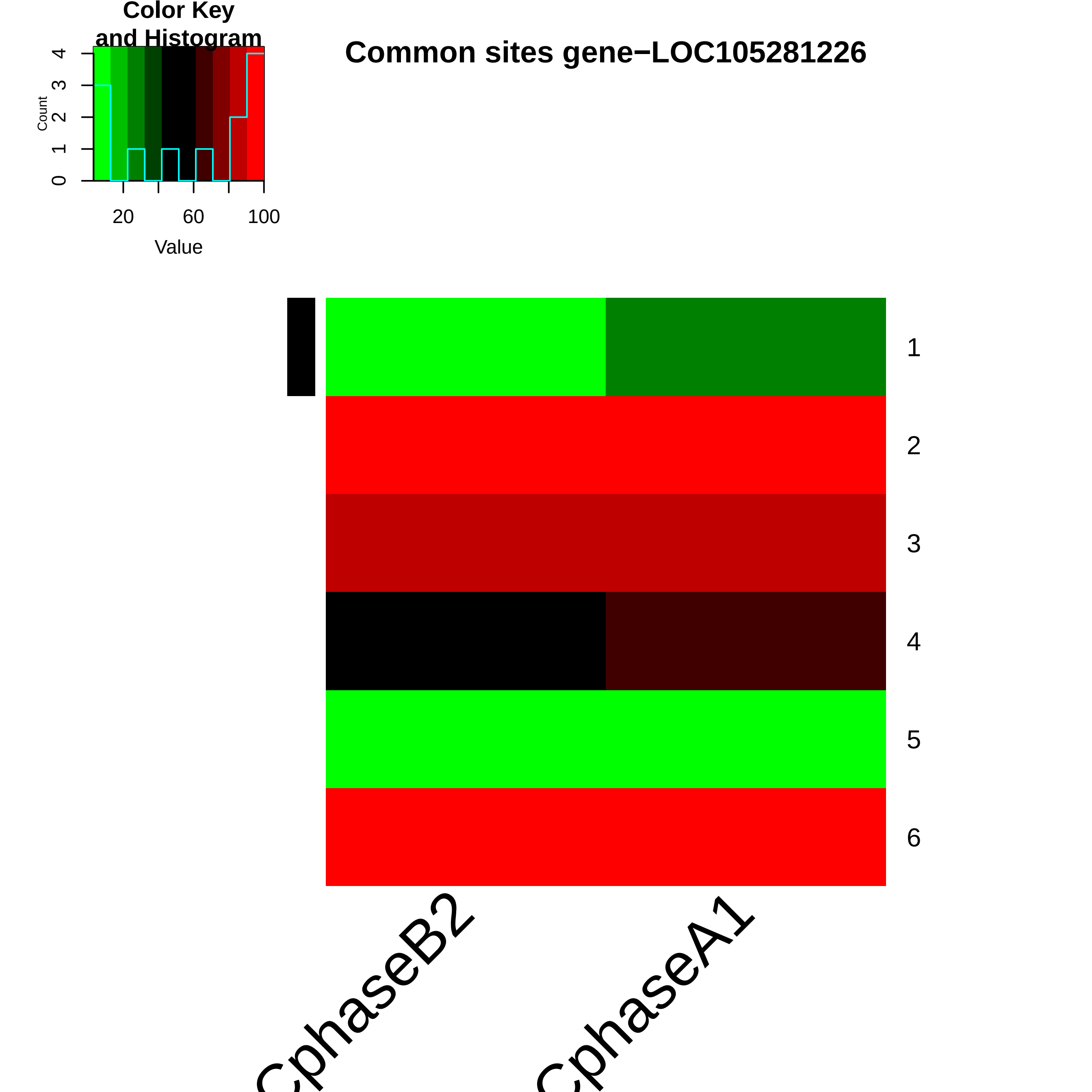
Color Key
and Histogram
Count
Value
20
60
100
0
1
2
3
4
Common sites gene−LOC105281226
1
2
3
4
5
6
CphaseB2
CphaseA1

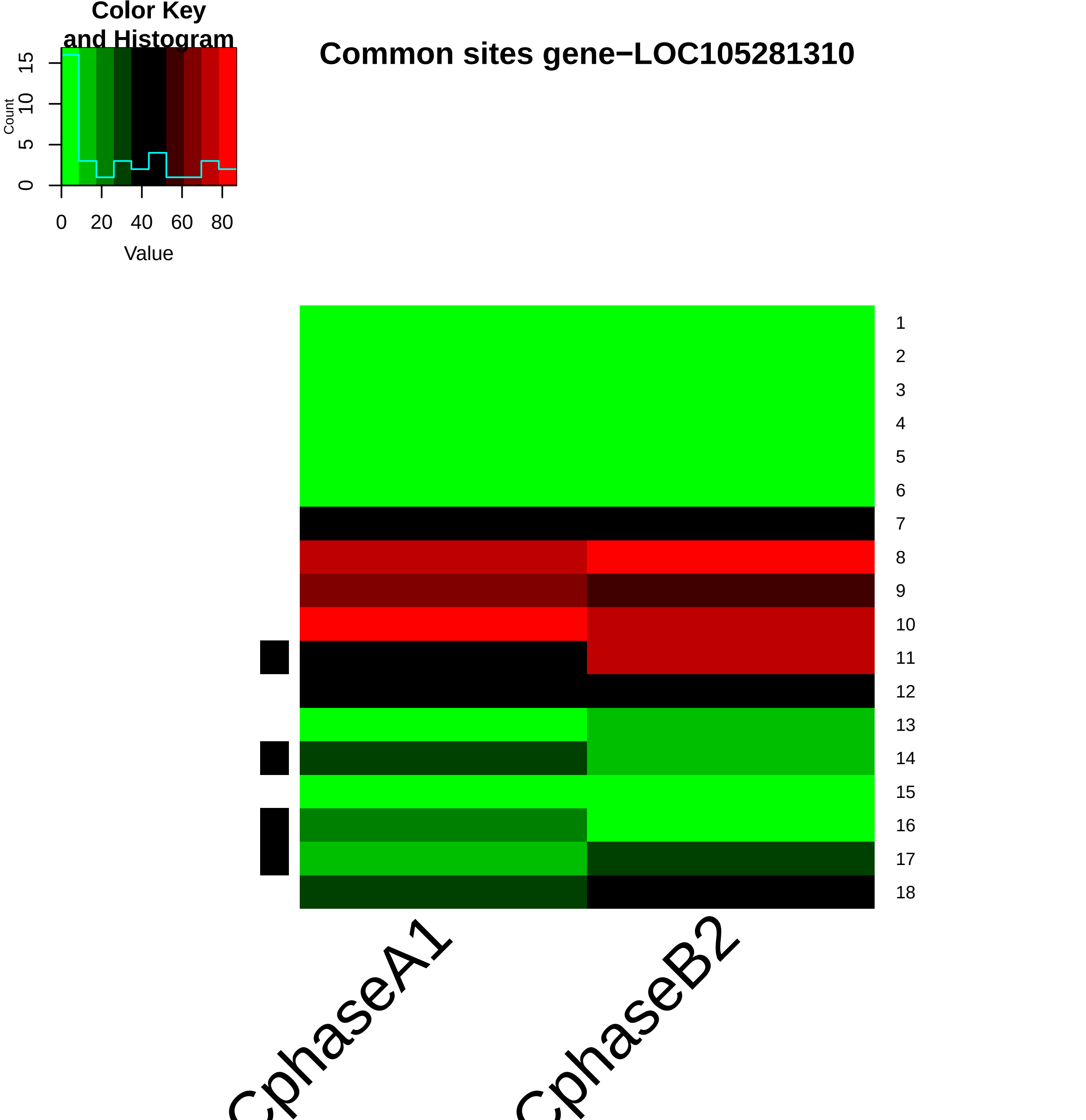
Color Key
and Histogram
Count
0
5
10
15
0
20
40
60
80
Value
Common sites gene−LOC105281310
CphaseA1
CphaseB2

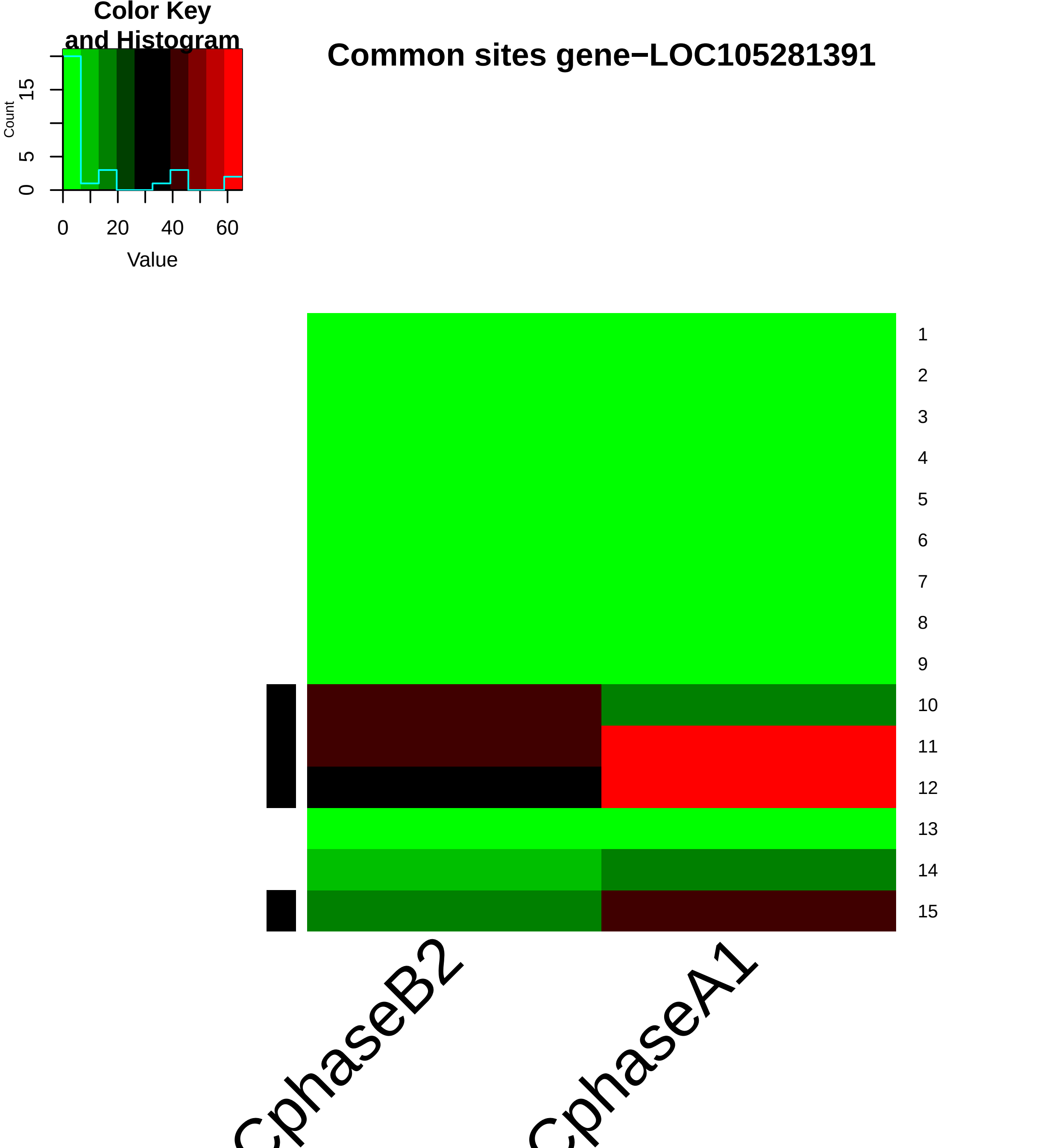

Color Key
and Histogram
Count
0
5
15
0
20
40
60
Value
Common sites gene−LOC105281391
CphaseB2
CphaseA1

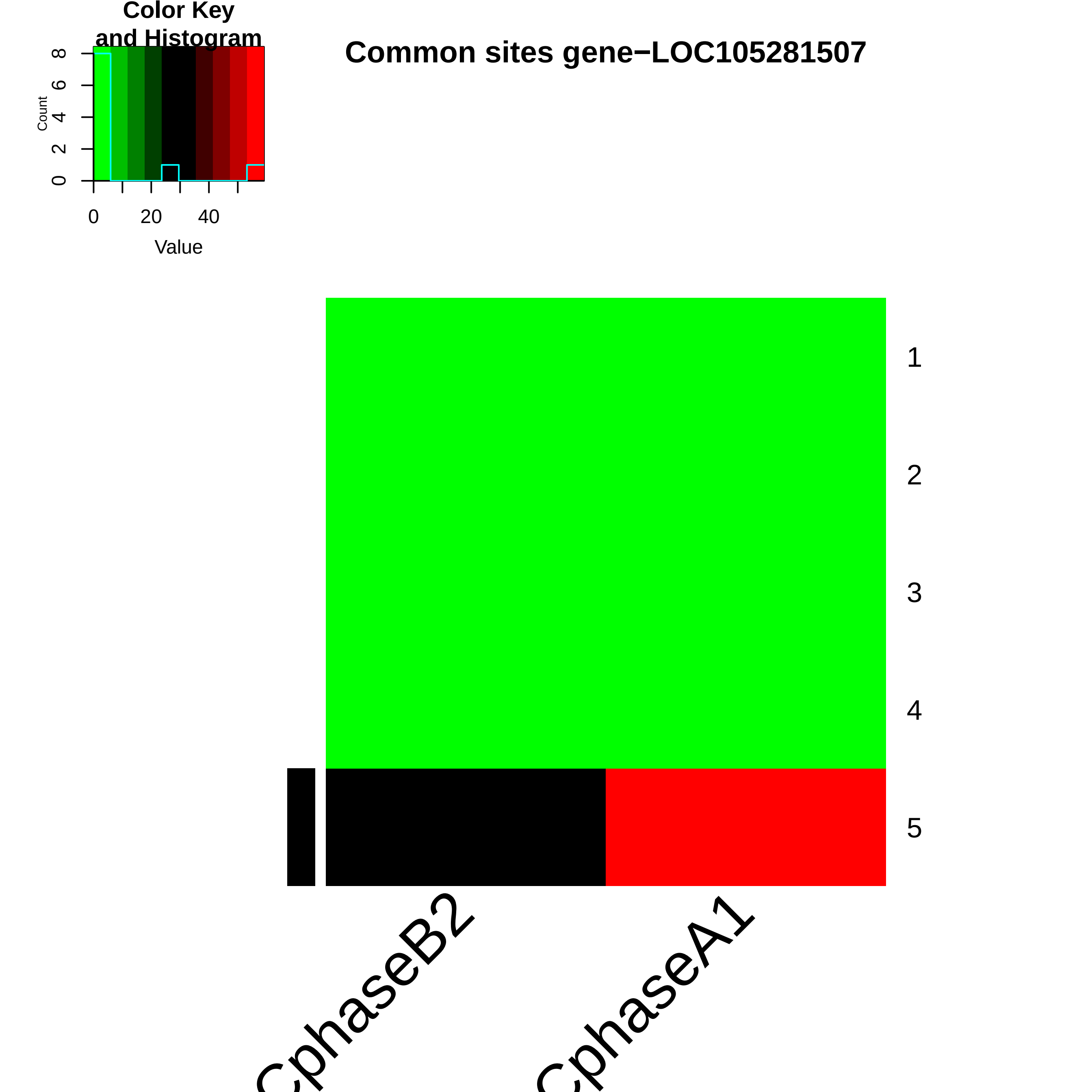
Color Key
and Histogram
Count
0
2
4
6
8
0
20
40
Value
Common sites gene−LOC105281507
1
2
3
4
5
CphaseB2
CphaseA1

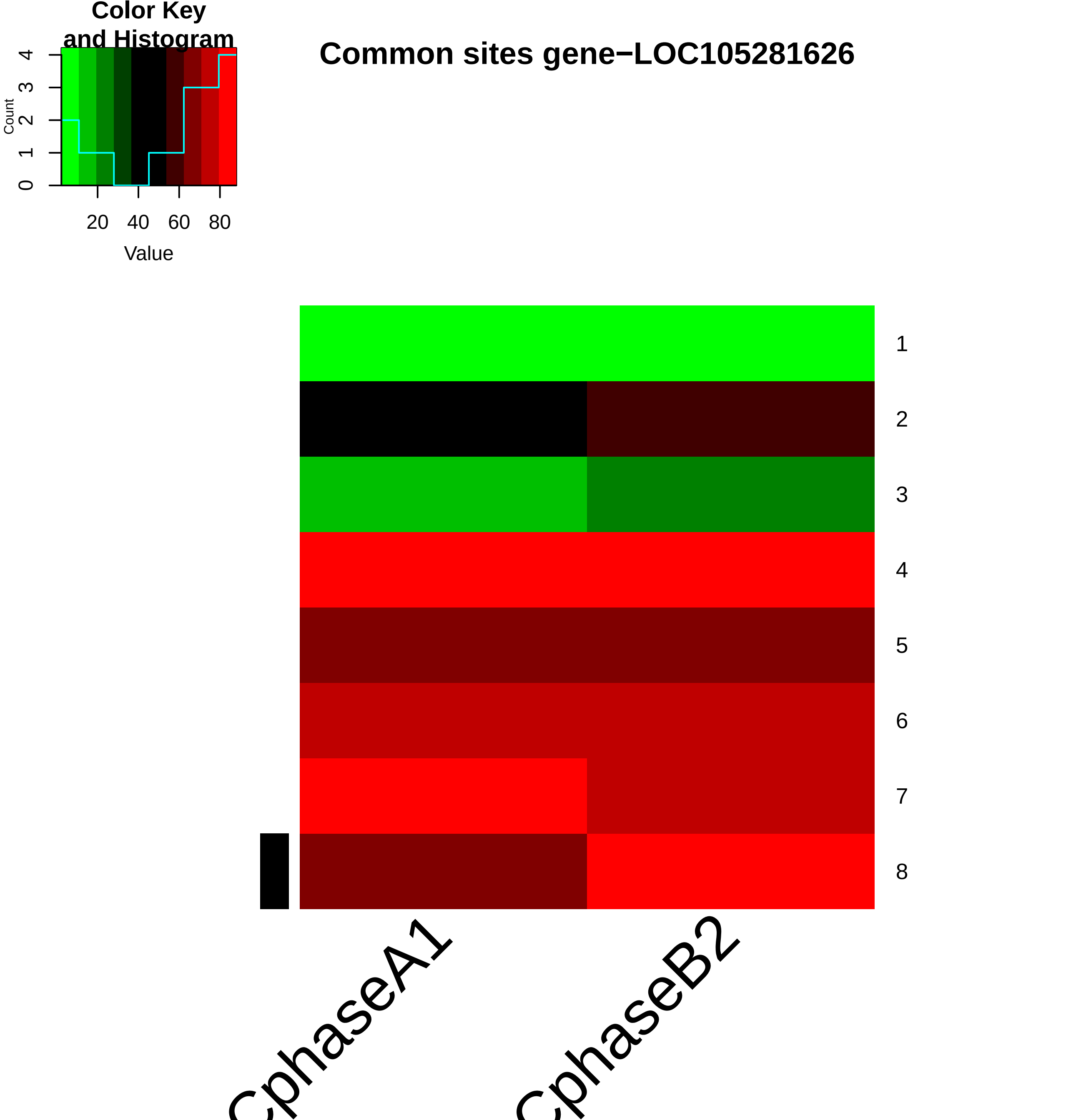
Color Key
and Histogram
Count
0
1
2
3
4
20
40
60
80
Value
Common sites gene−LOC105281626
1
2
3
4
5
6
7
8
CphaseA1
CphaseB2

### Common sites gene−LOC105281628

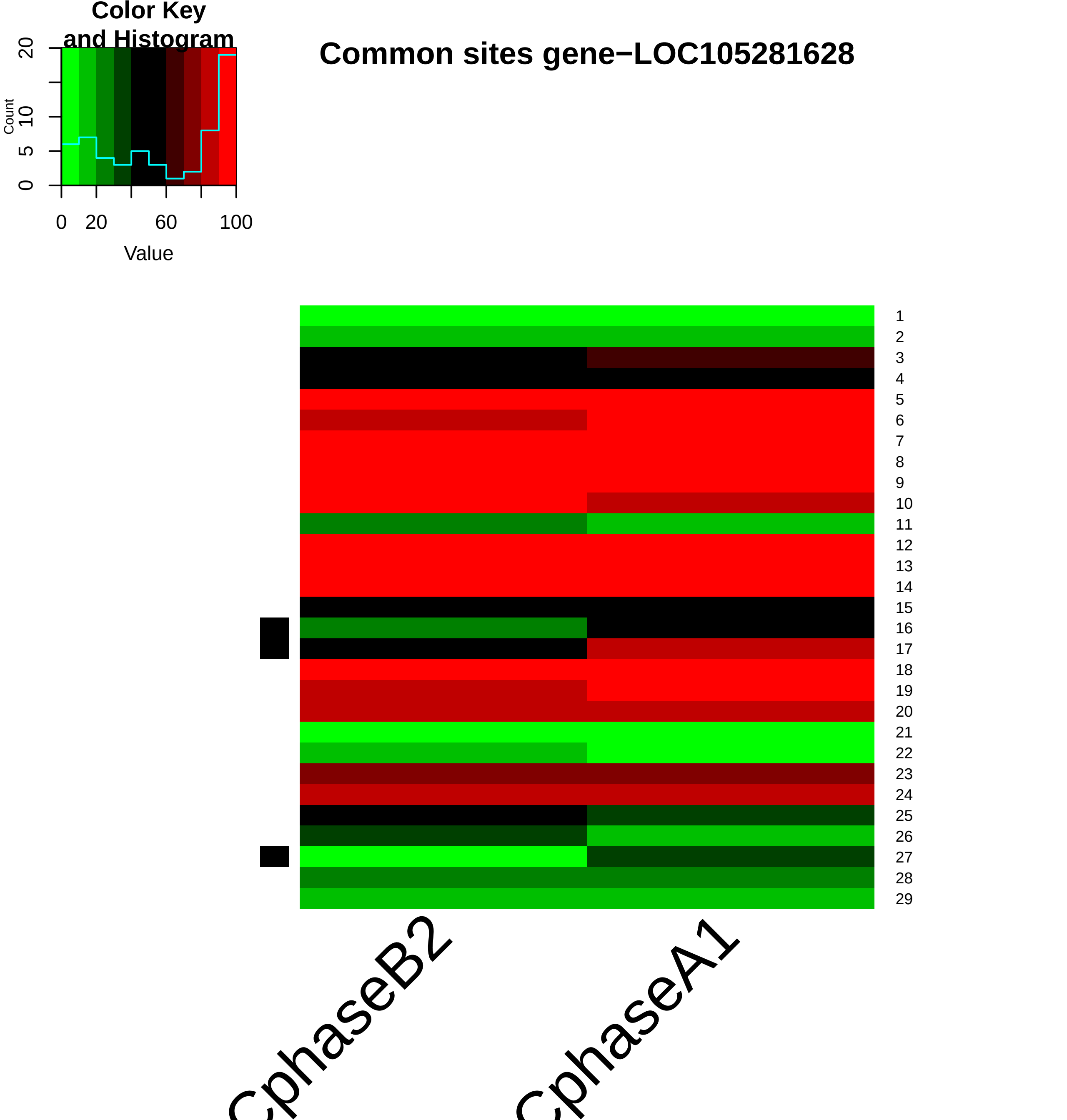

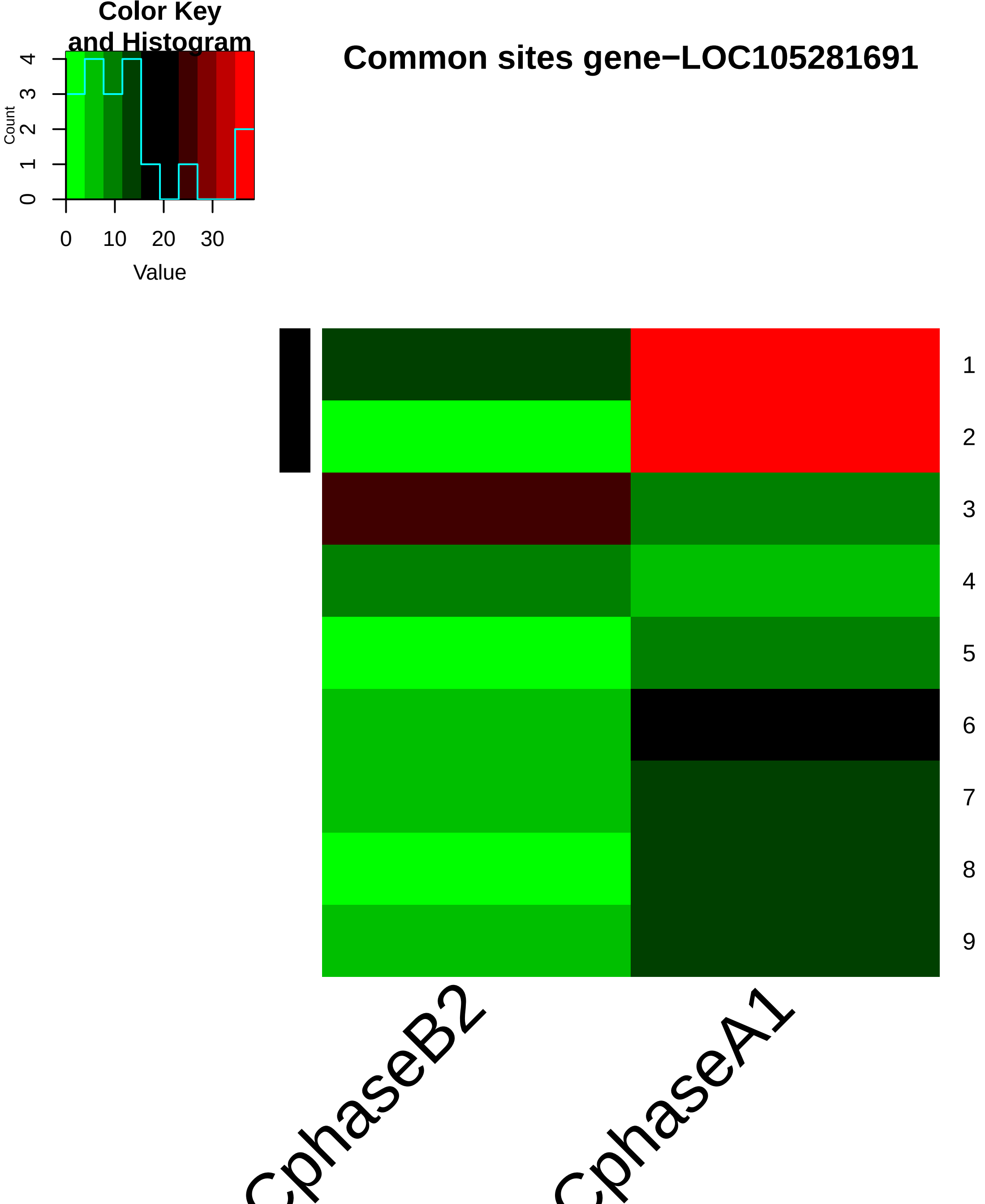

Color Key
and Histogram
Count
0
1
2
3
4
0
10
20
30
Value
Common sites gene−LOC105281691
1
2
3
4
5
6
7
8
9
CphaseB2
CphaseA1

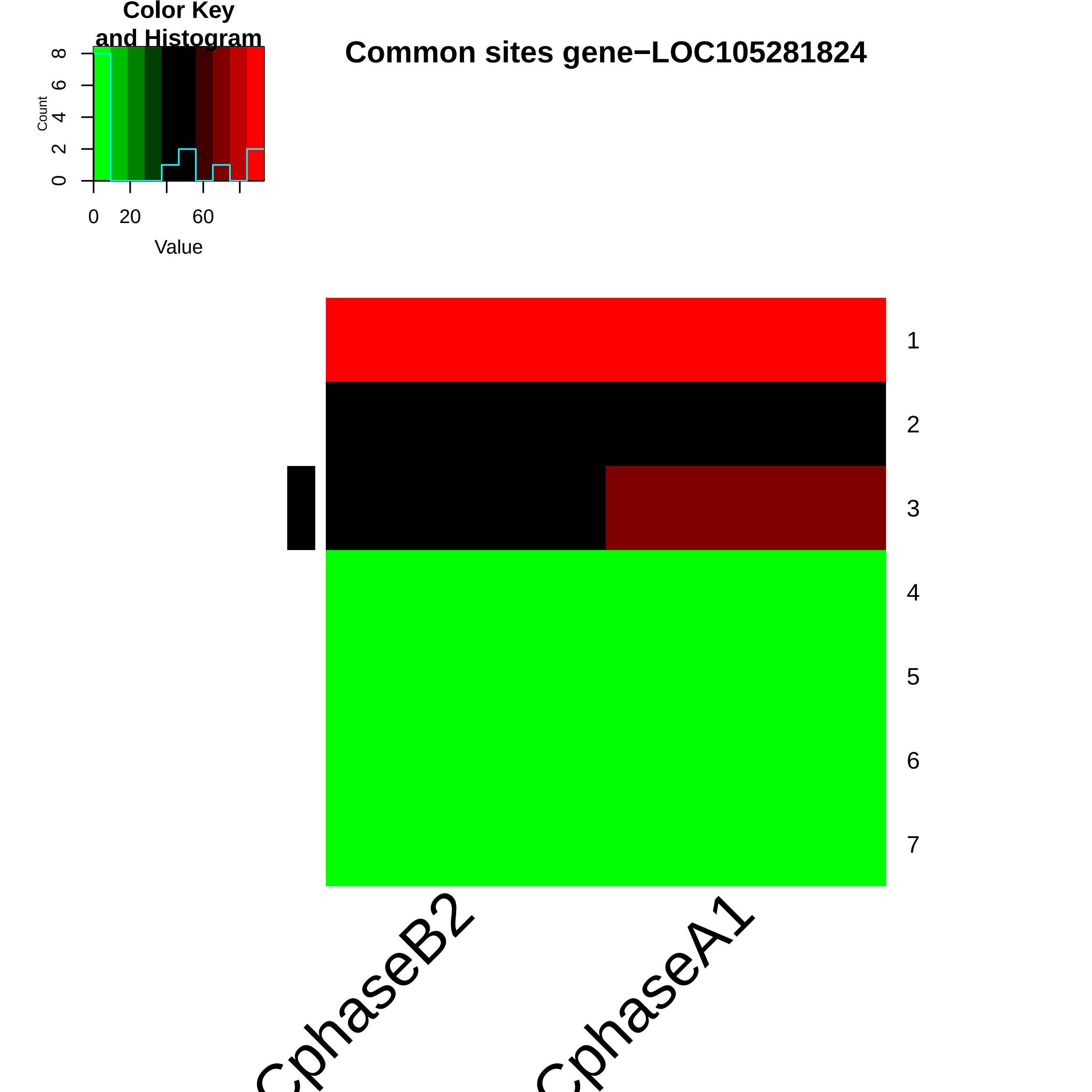

Color Key
and Histogram
Count
0
2
4
6
8
0
20
60
Value
Common sites gene−LOC105281824
1
2
3
4
5
6
7
CphaseB2
CphaseA1

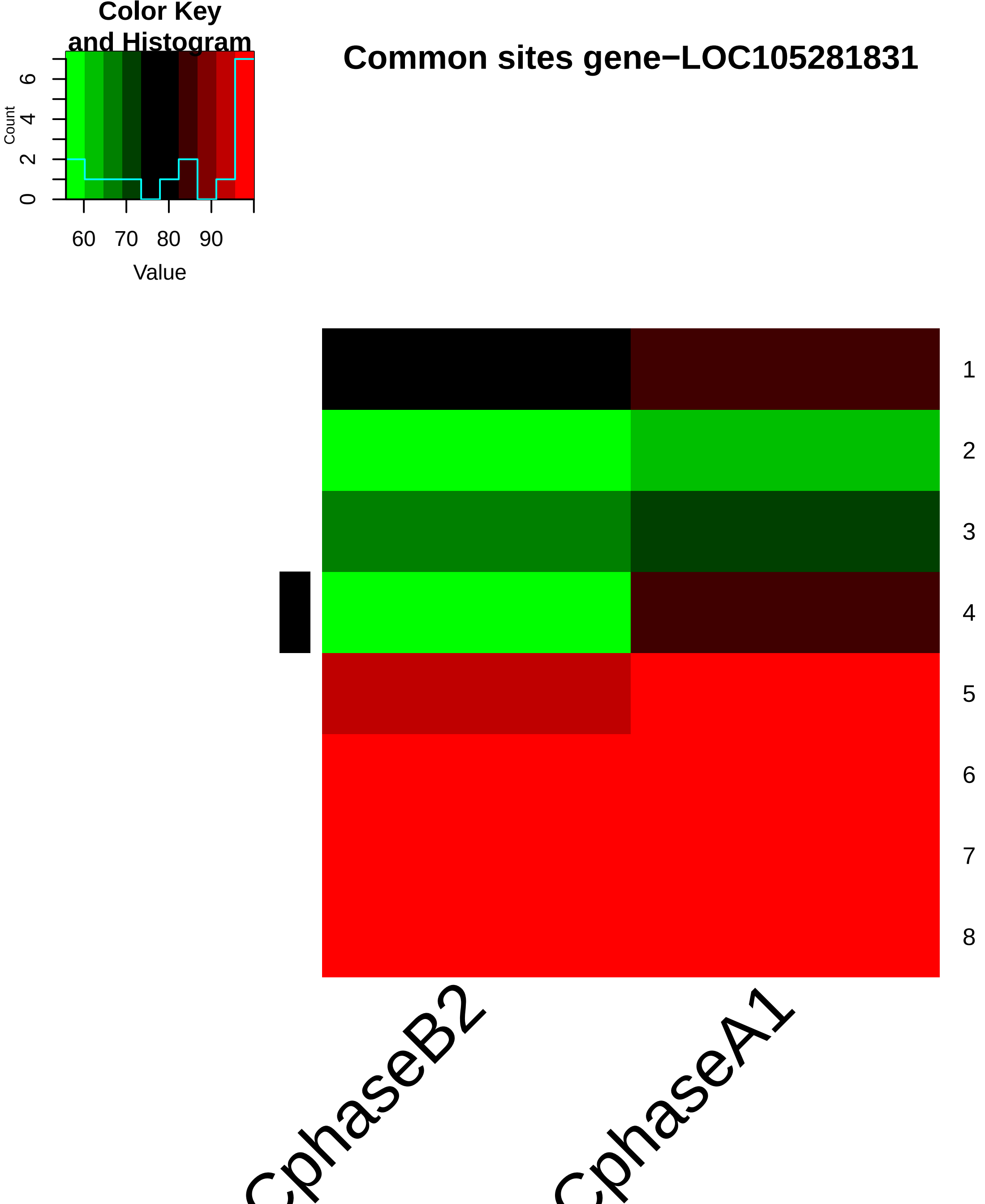
Color Key
and Histogram
Count
0
2
4
6
60
70
80
90
Value
Common sites gene−LOC105281831
1
2
3
4
5
6
7
8
CphaseB2
CphaseA1

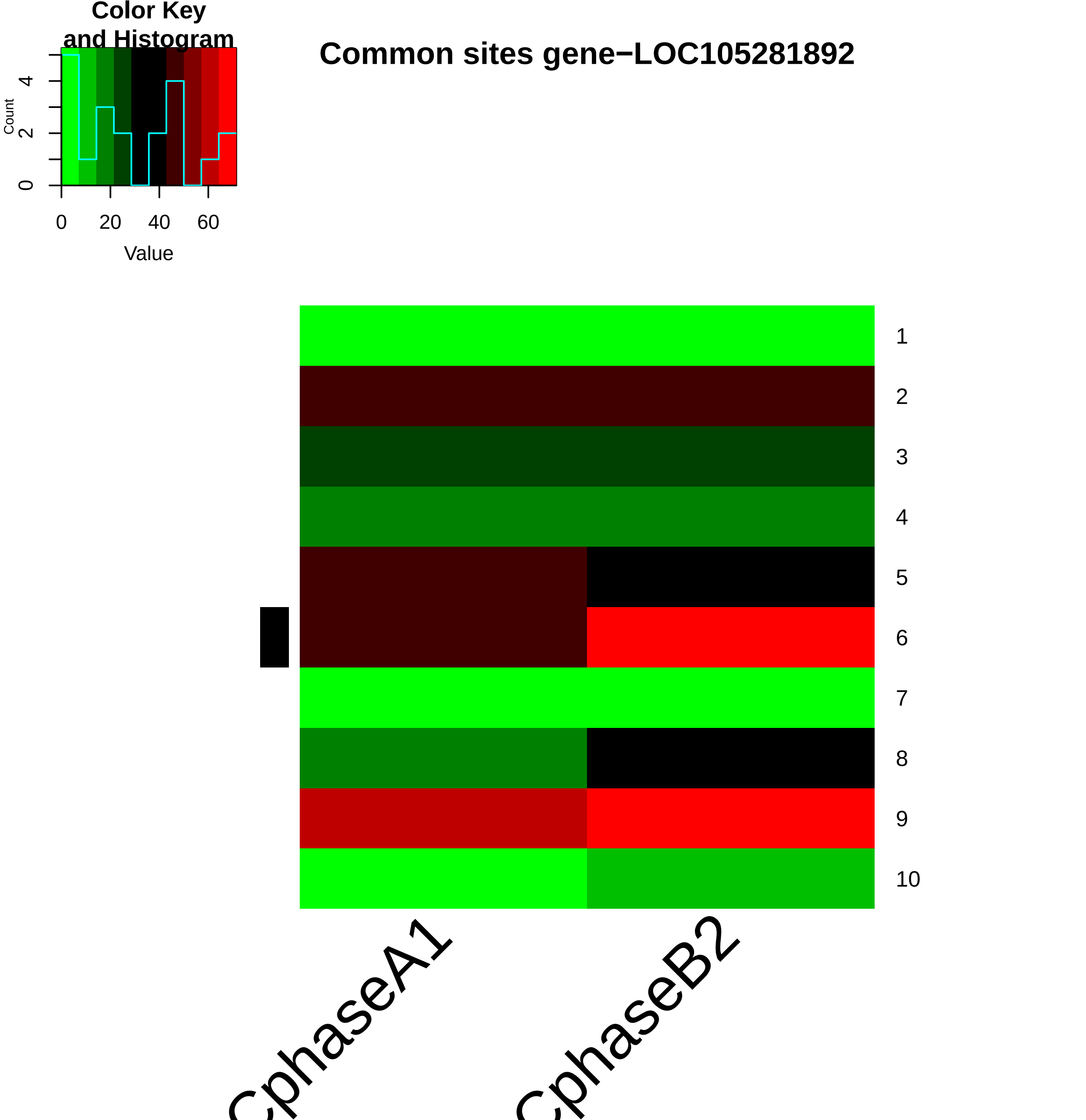

Color Key
and Histogram
Count
0
2
4
0
20
40
60
Value
Common sites gene−LOC105281892
1
2
3
4
5
6
7
8
9
10
CphaseA1
CphaseB2

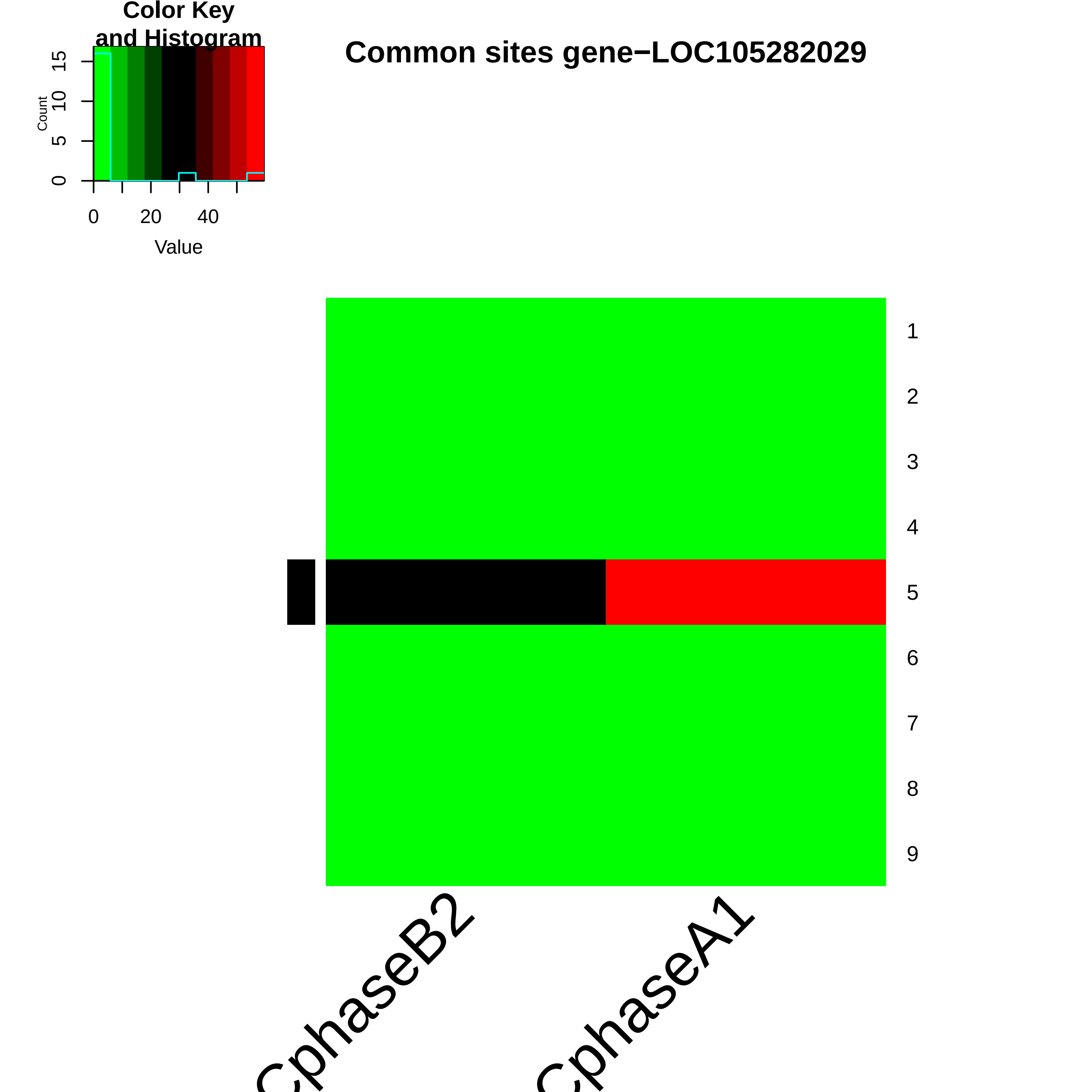

Color Key
and Histogram
Count
0
5
10
15
0
20
40
Value
Common sites gene−LOC105282029
1
2
3
4
5
6
7
8
9
CphaseB2
CphaseA1

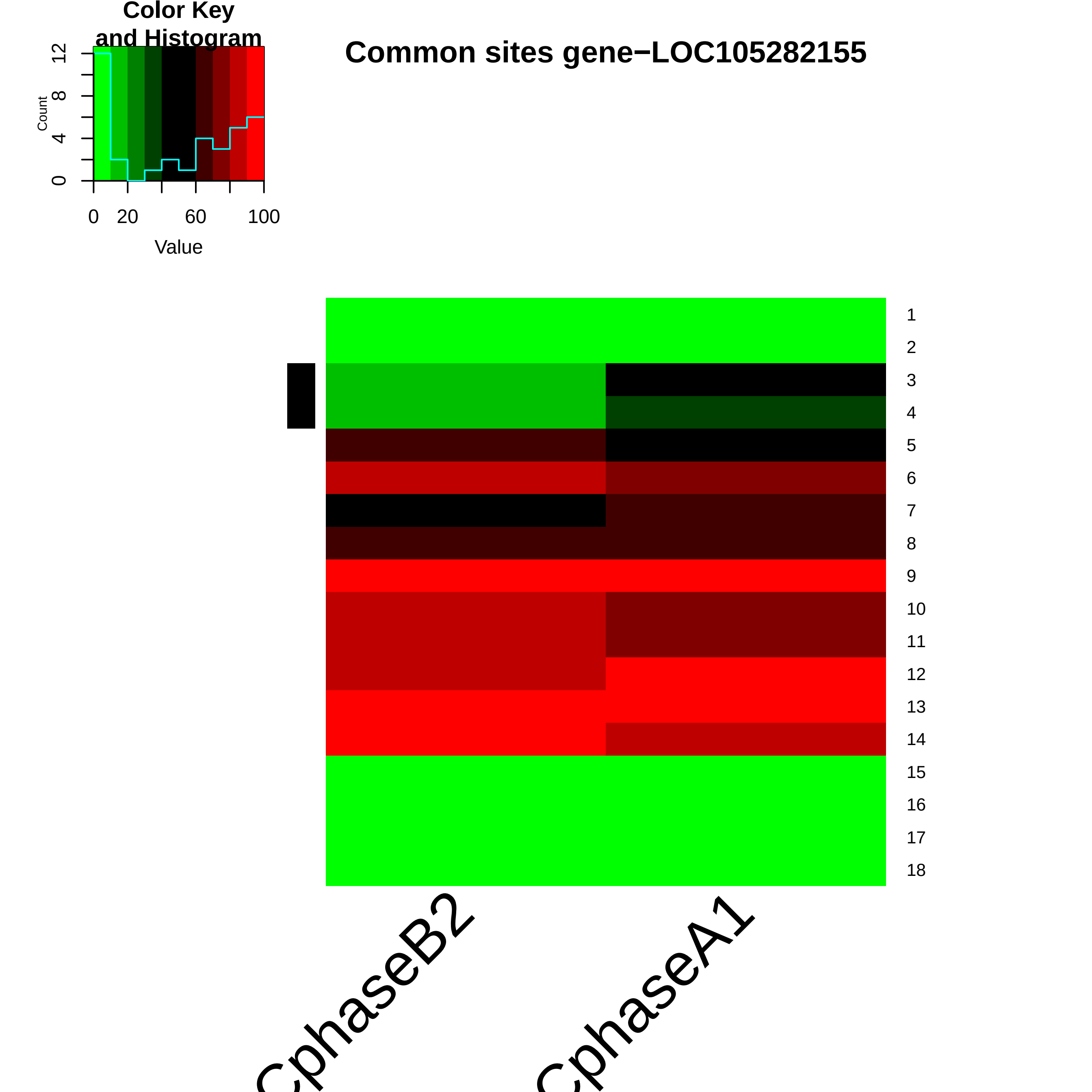

Color Key
and Histogram
Count
0
4
8
12
0
20
60
100
Value
Common sites gene−LOC105282155
CphaseB2
CphaseA1

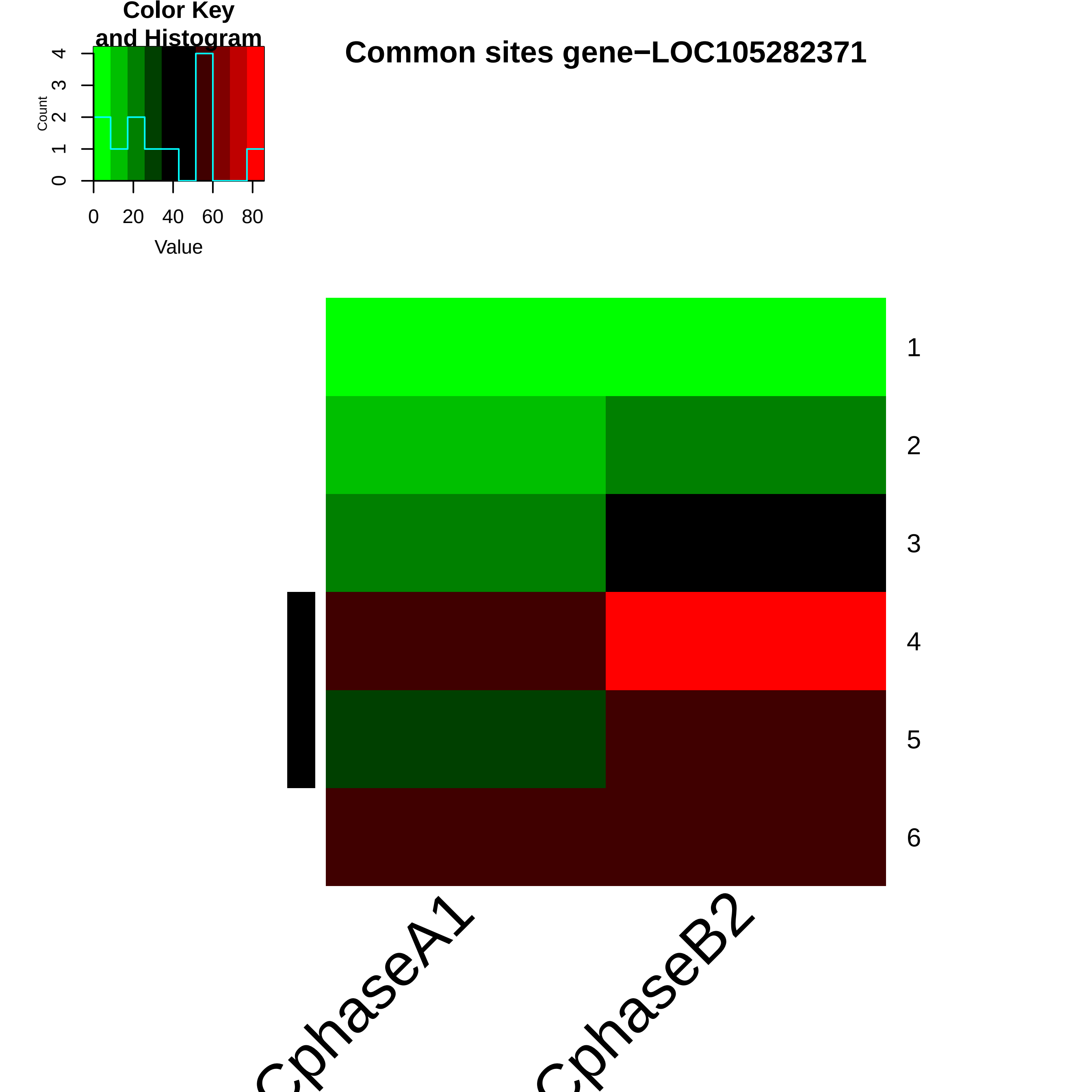

Color Key
and Histogram
Count
0
1
2
3
4
0
20
40
60
80
Value
Common sites gene−LOC105282371
1
2
3
4
5
6
CphaseA1
CphaseB2

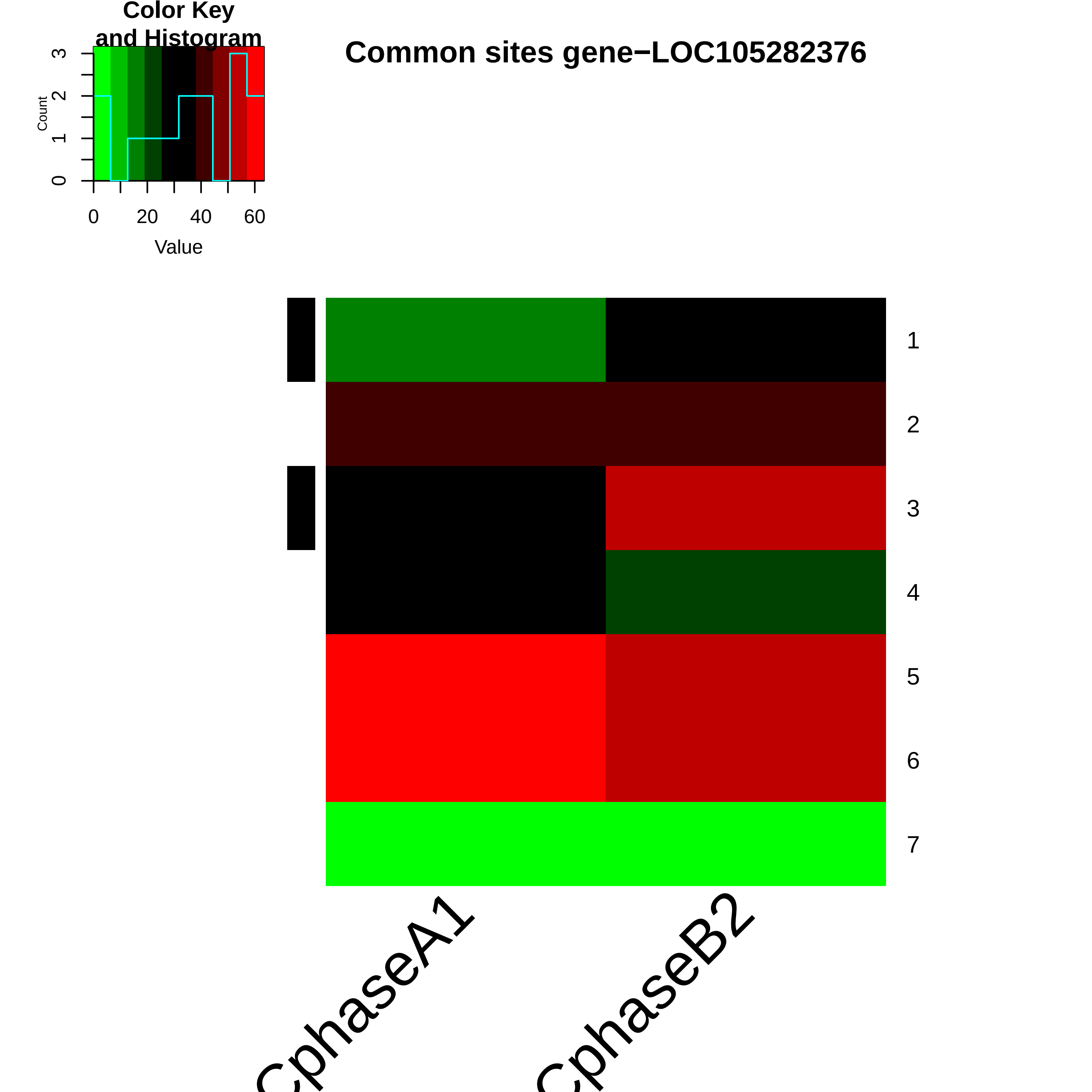
Color Key
and Histogram
Count
Value
0
20
40
60
0
1
2
3
Common sites gene−LOC105282376
1
2
3
4
5
6
7
CphaseA1
CphaseB2

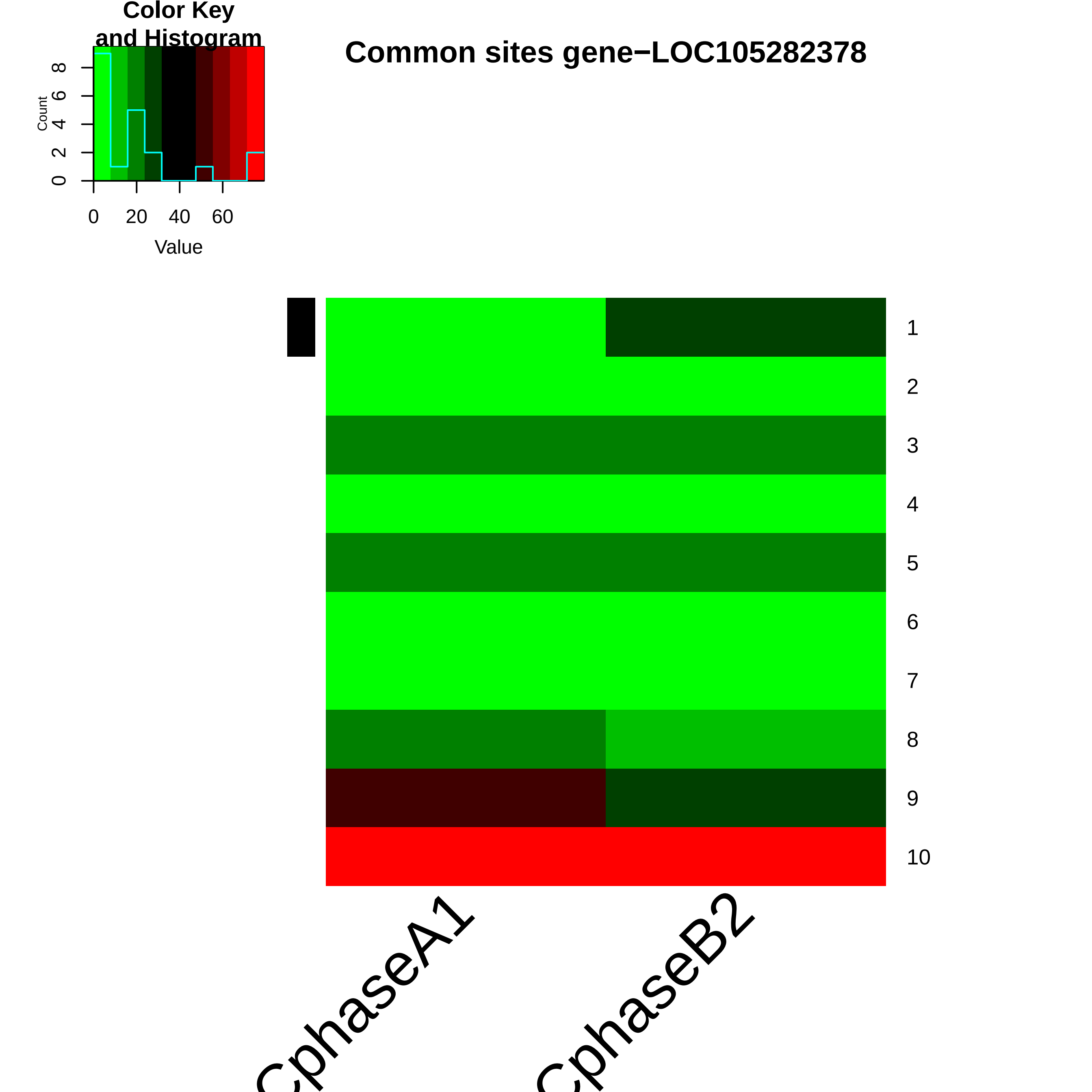
Color Key
and Histogram
Count
0
2
4
6
8
0
20
40
60
Value
Common sites gene−LOC105282378
1
2
3
4
5
6
7
8
9
10
CphaseA1
CphaseB2

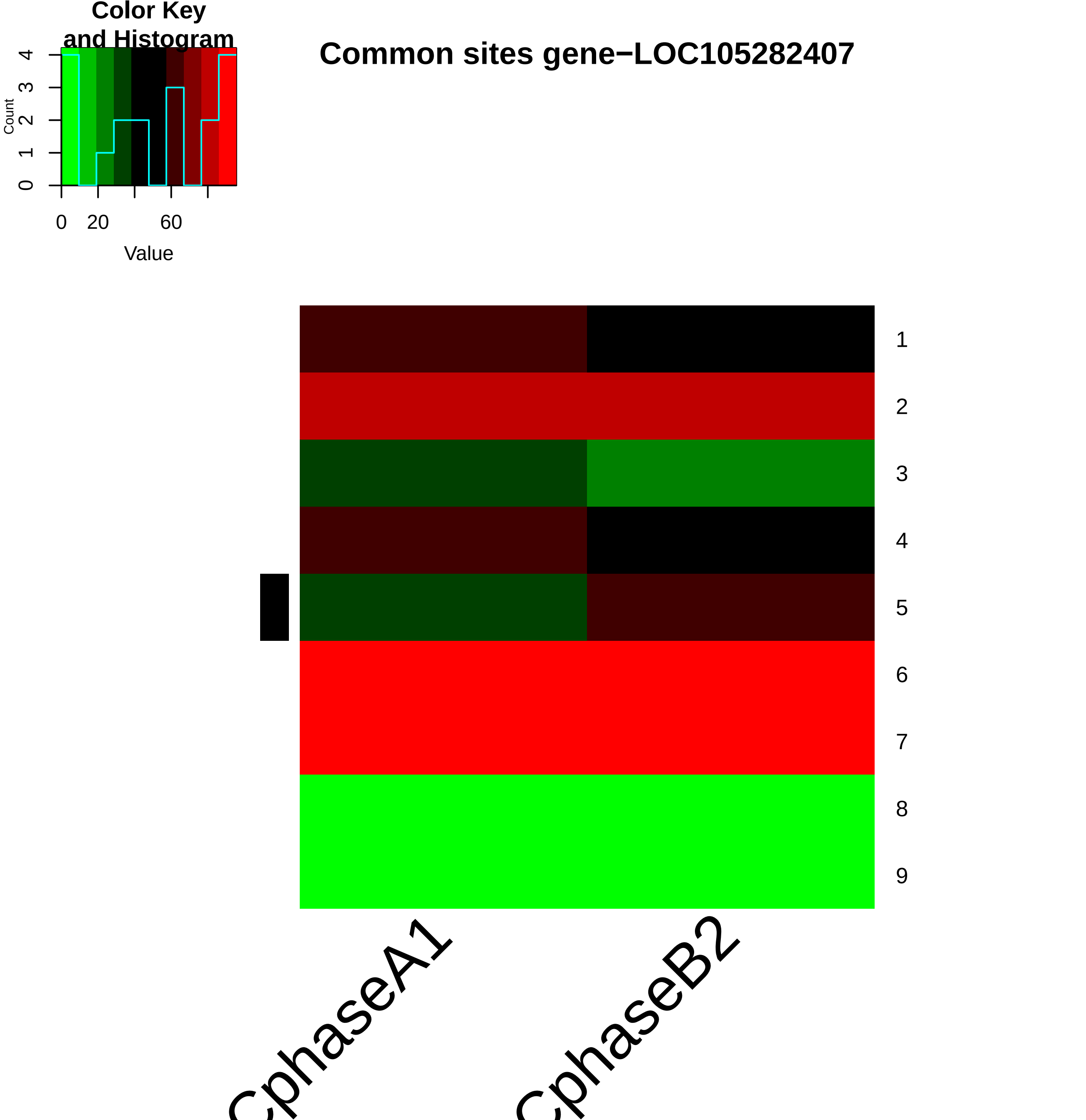

Color Key
and Histogram
Count
Value
0
20
60
0
1
2
3
4
Common sites gene−LOC105282407
1
2
3
4
5
6
7
8
9
CphaseA1
CphaseB2

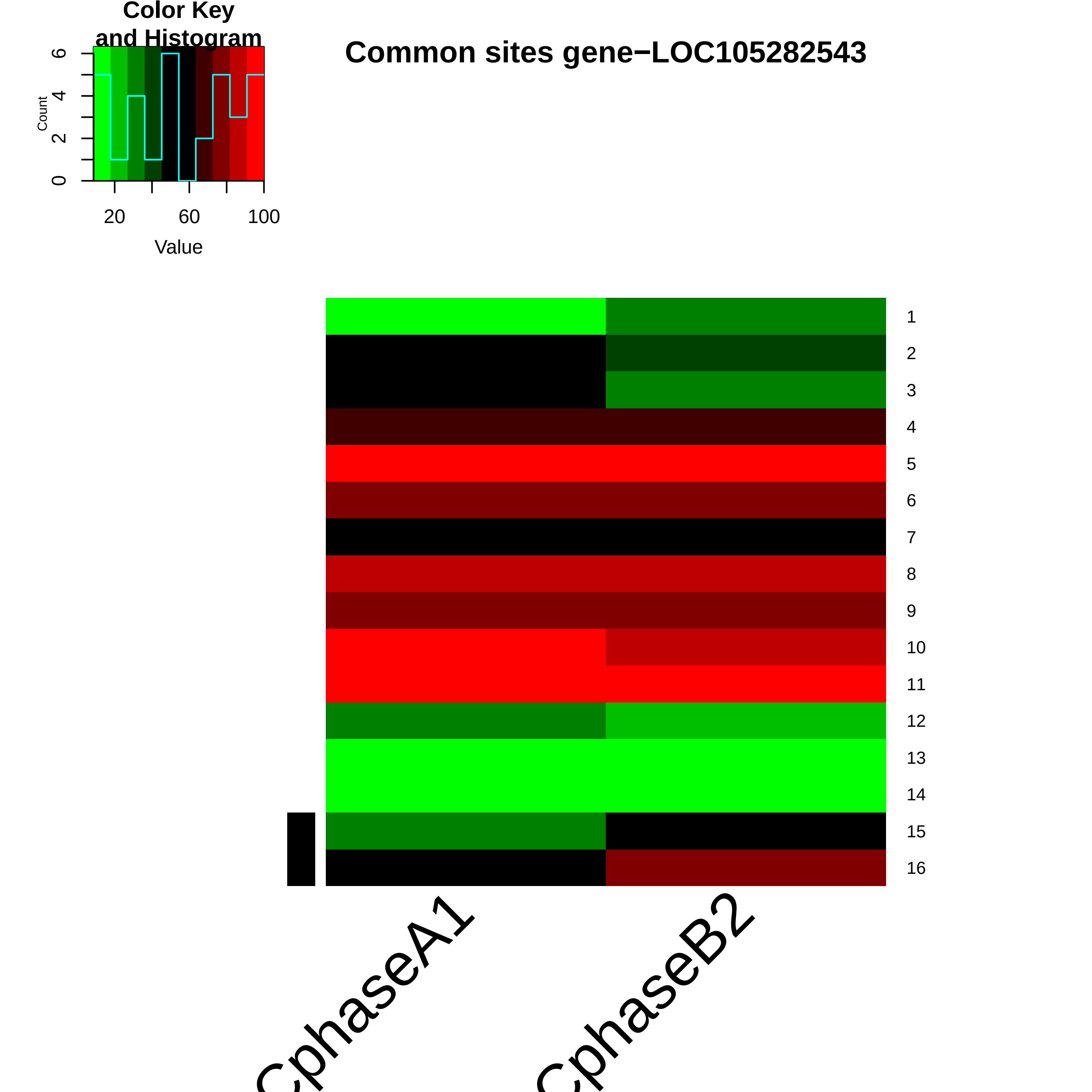

Color Key
and Histogram
Count
Value
20
60
100
0
2
4
6
Common sites gene−LOC105282543
CphaseA1
CphaseB2

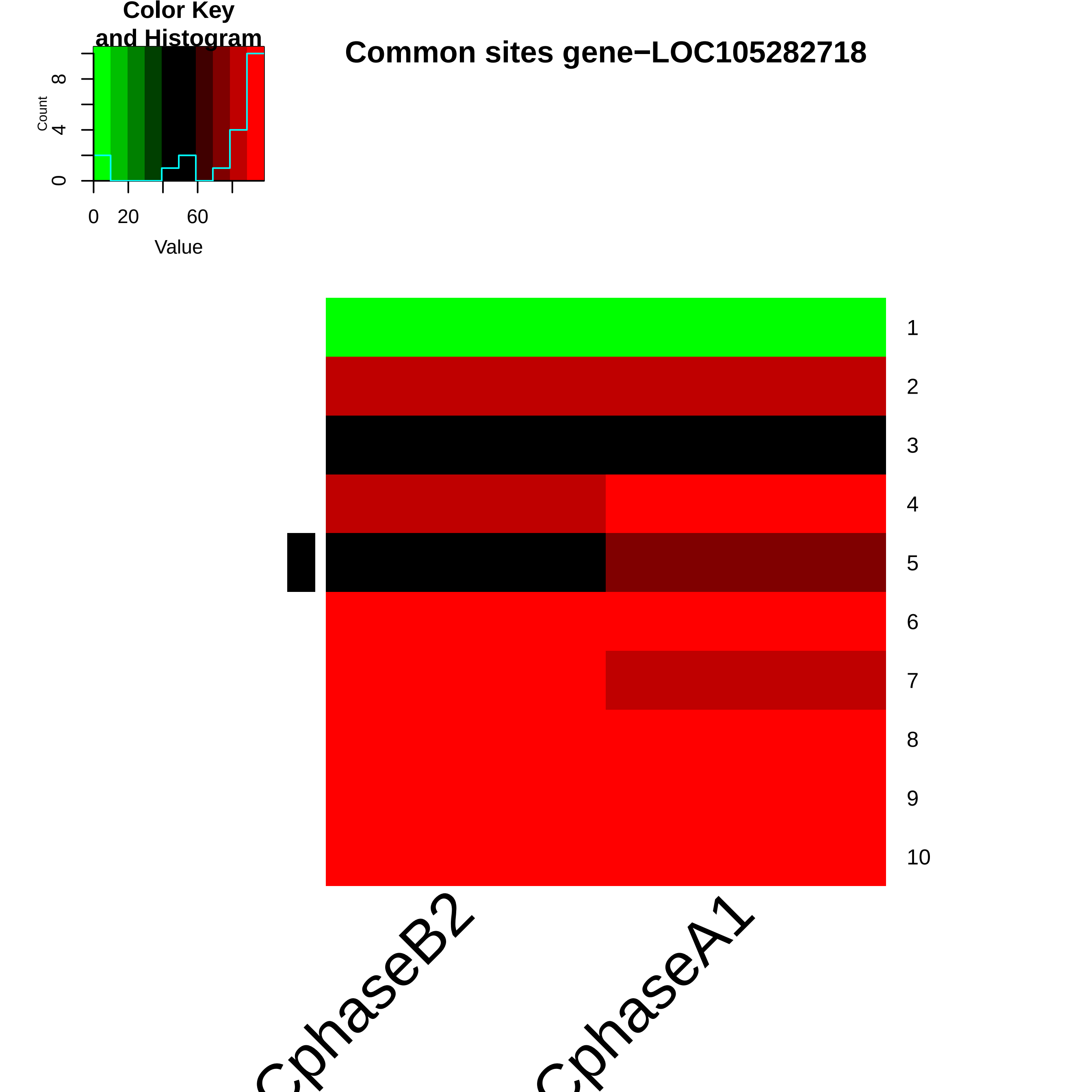
Color Key
and Histogram
Count
0
4
8
0
20
60
Value
Common sites gene−LOC105282718
1
2
3
4
5
6
7
8
9
10
CphaseB2
CphaseA1

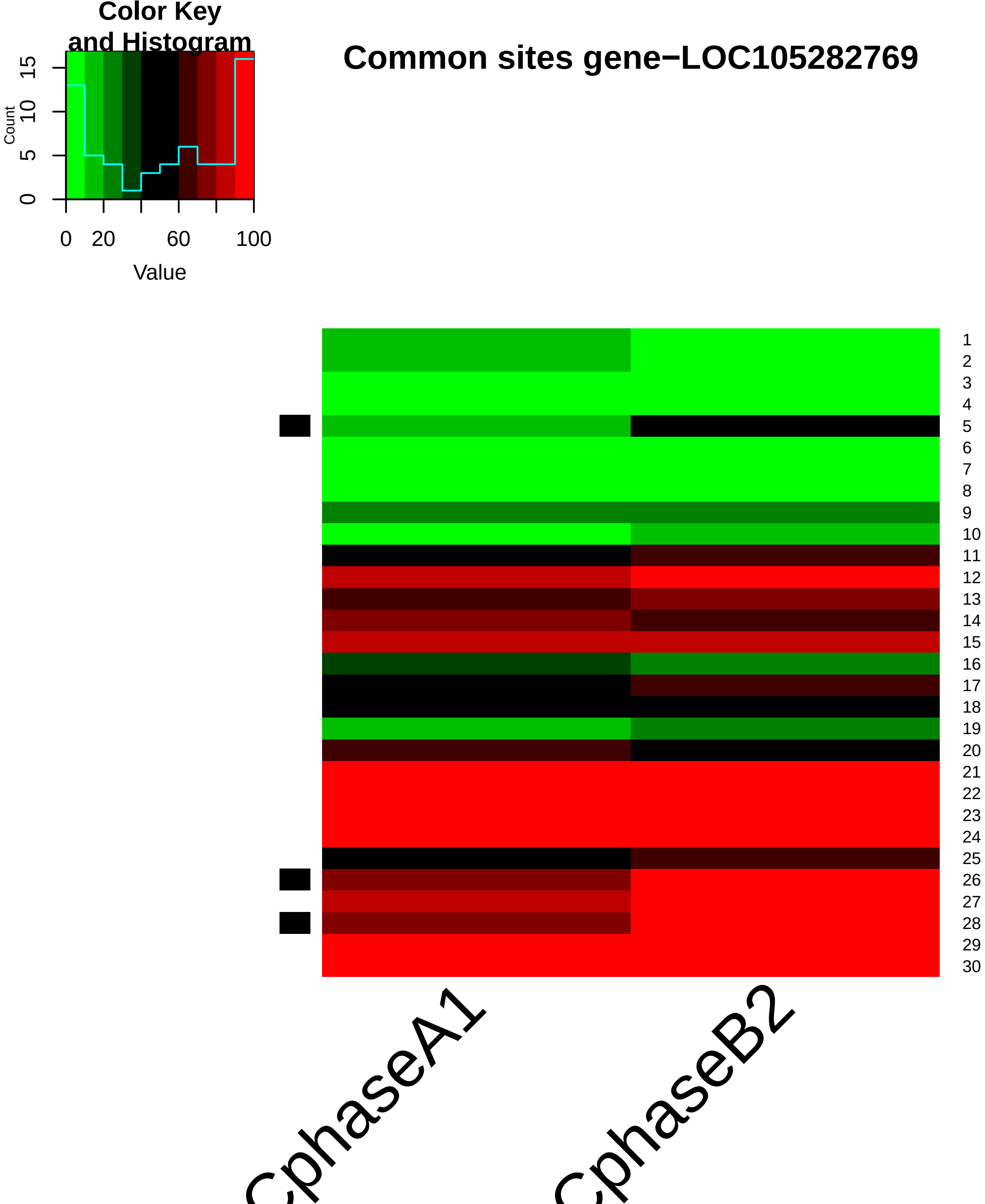
Color Key
and Histogram
Count
0
5
10
15
0
20
60
100
Value
Common sites gene−LOC105282769
CphaseA1
CphaseB2

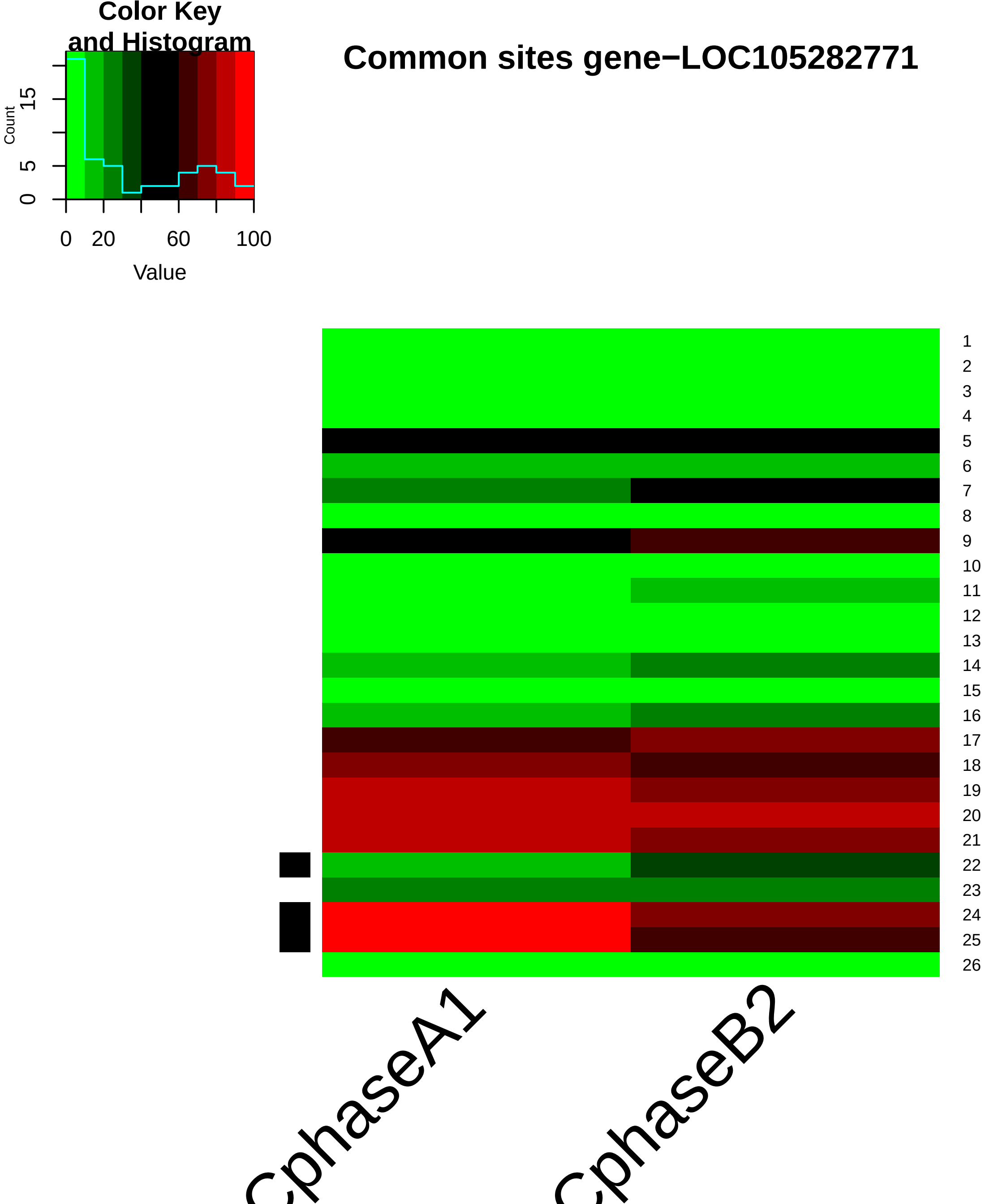
Color Key
and Histogram
Count
0
5
15
0
20
60
100
Value
Common sites gene−LOC105282771
CphaseA1
CphaseB2

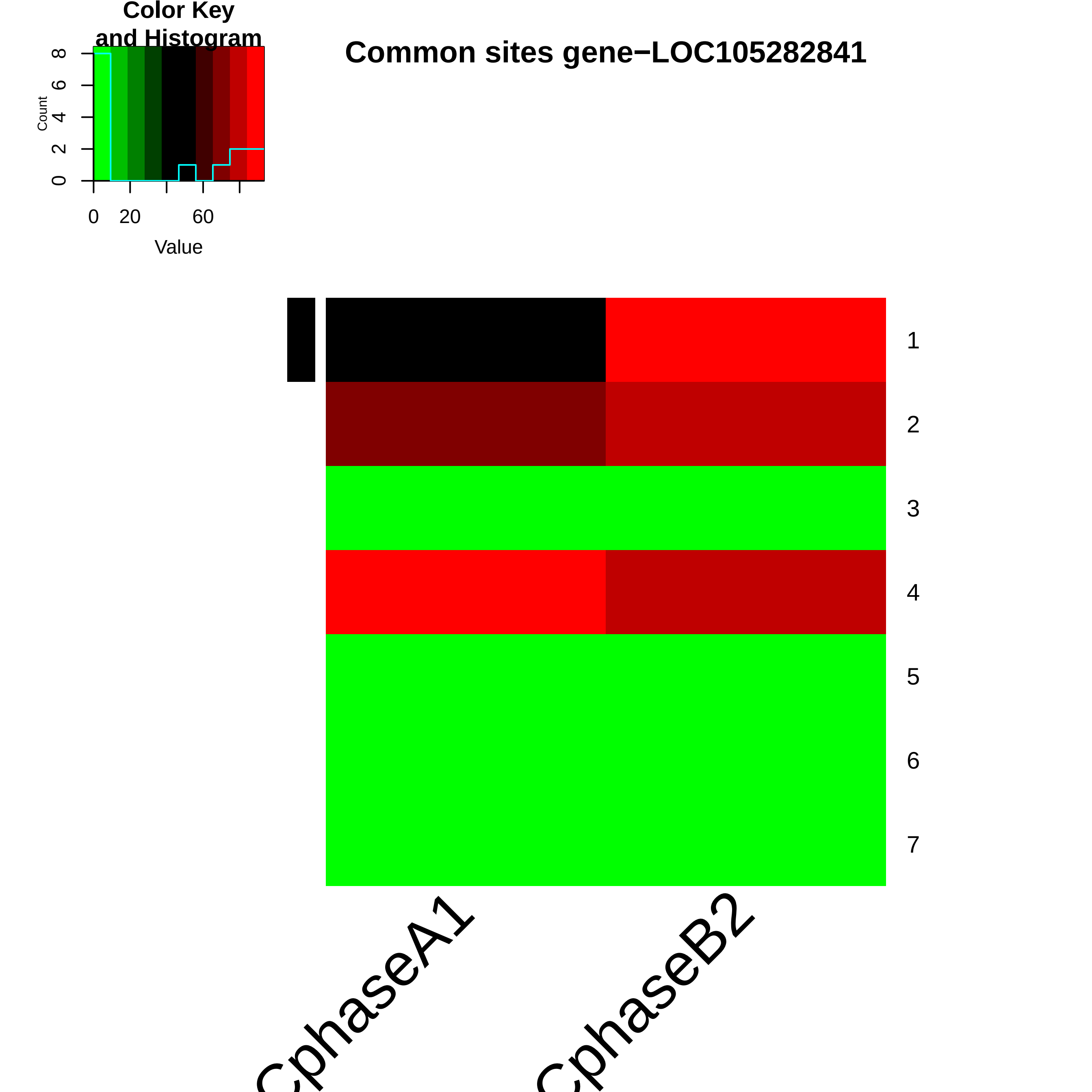

Color Key
and Histogram
Count
0
2
4
6
8
0
20
60
Value
Common sites gene−LOC105282841
1
2
3
4
5
6
7
CphaseA1
CphaseB2

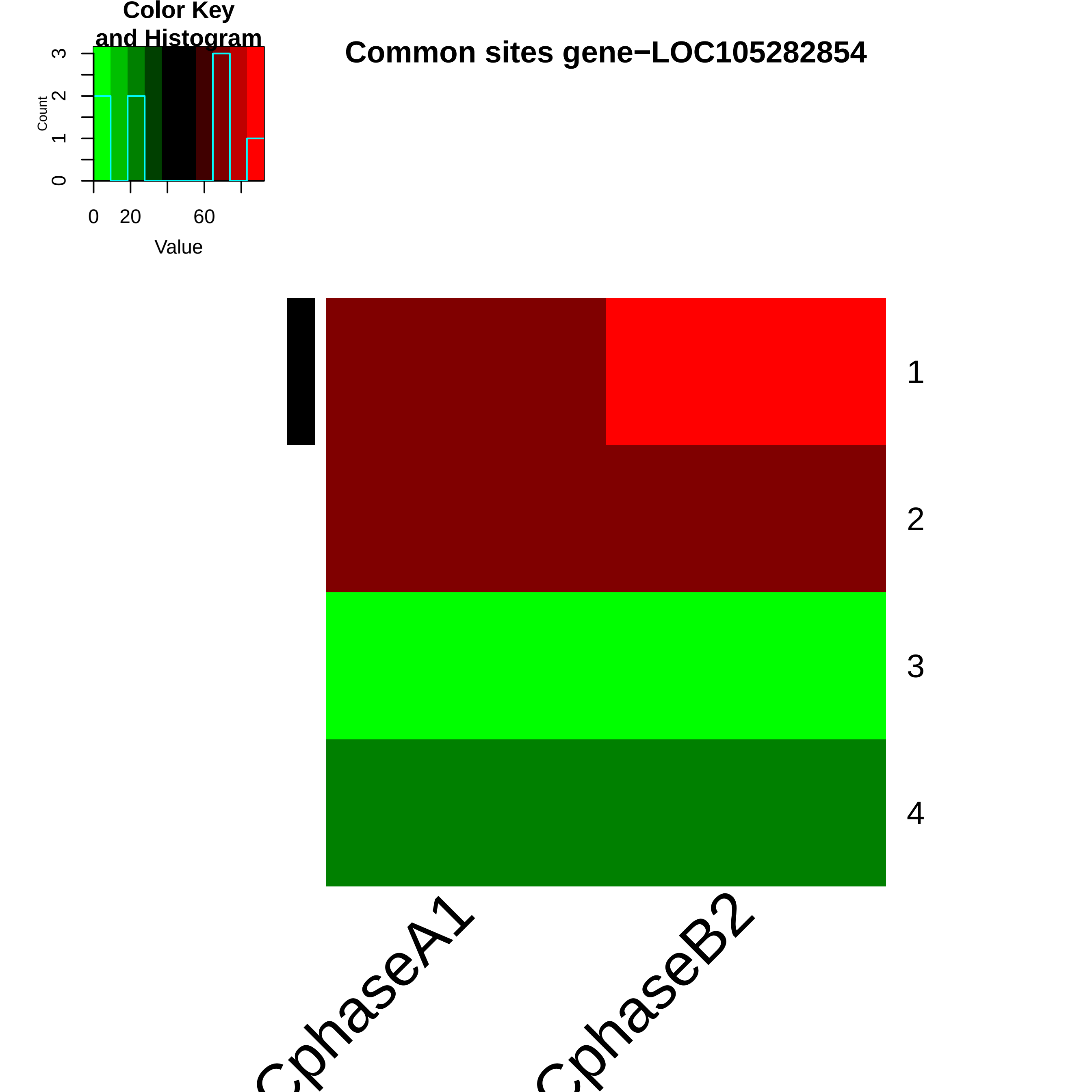

Color Key
and Histogram
Count
0
1
2
3
0
20
60
Value
Common sites gene−LOC105282854
1
2
3
4
CphaseA1
CphaseB2

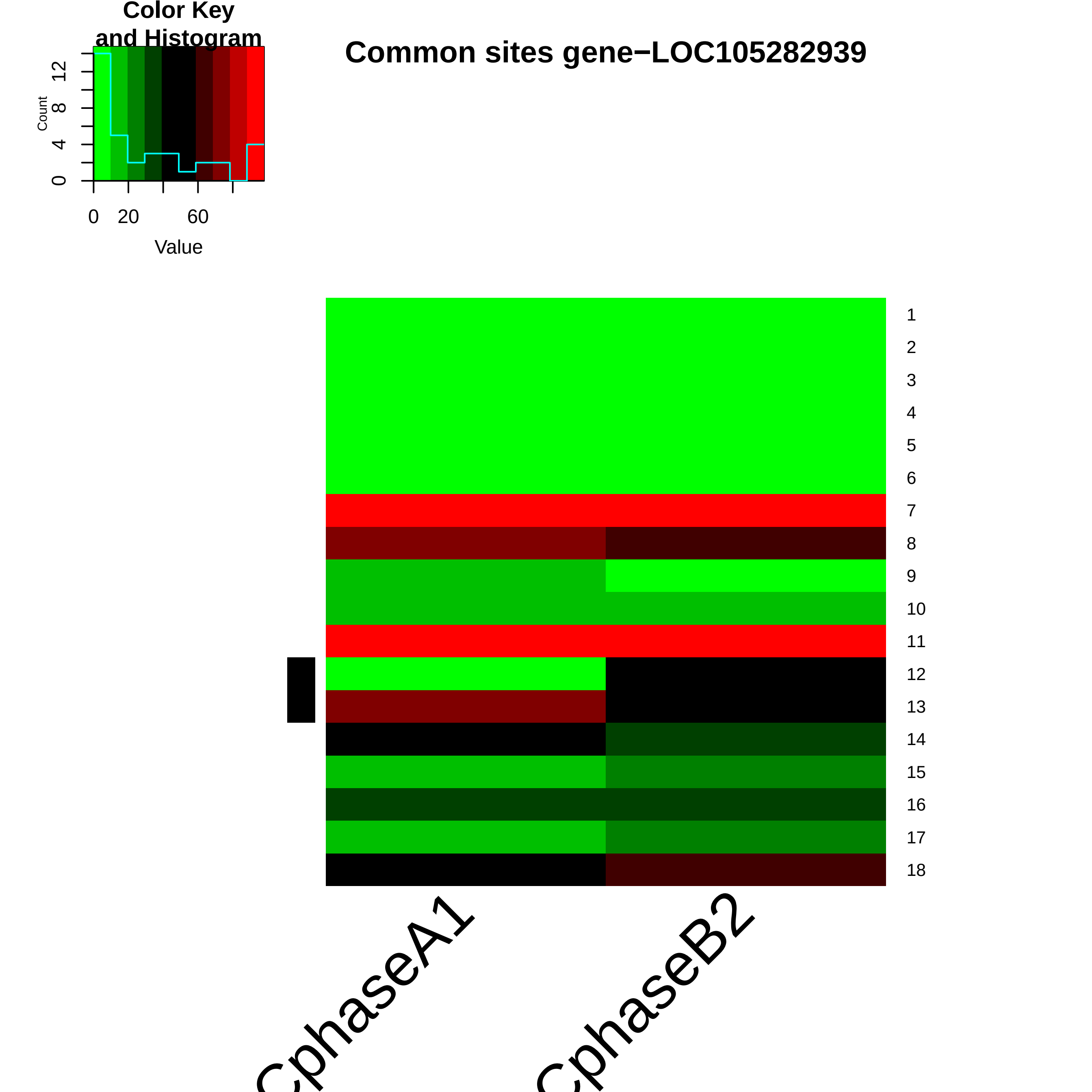

Color Key
and Histogram
Count
0
4
8
12
0
20
60
Value
Common sites gene−LOC105282939
CphaseA1
CphaseB2

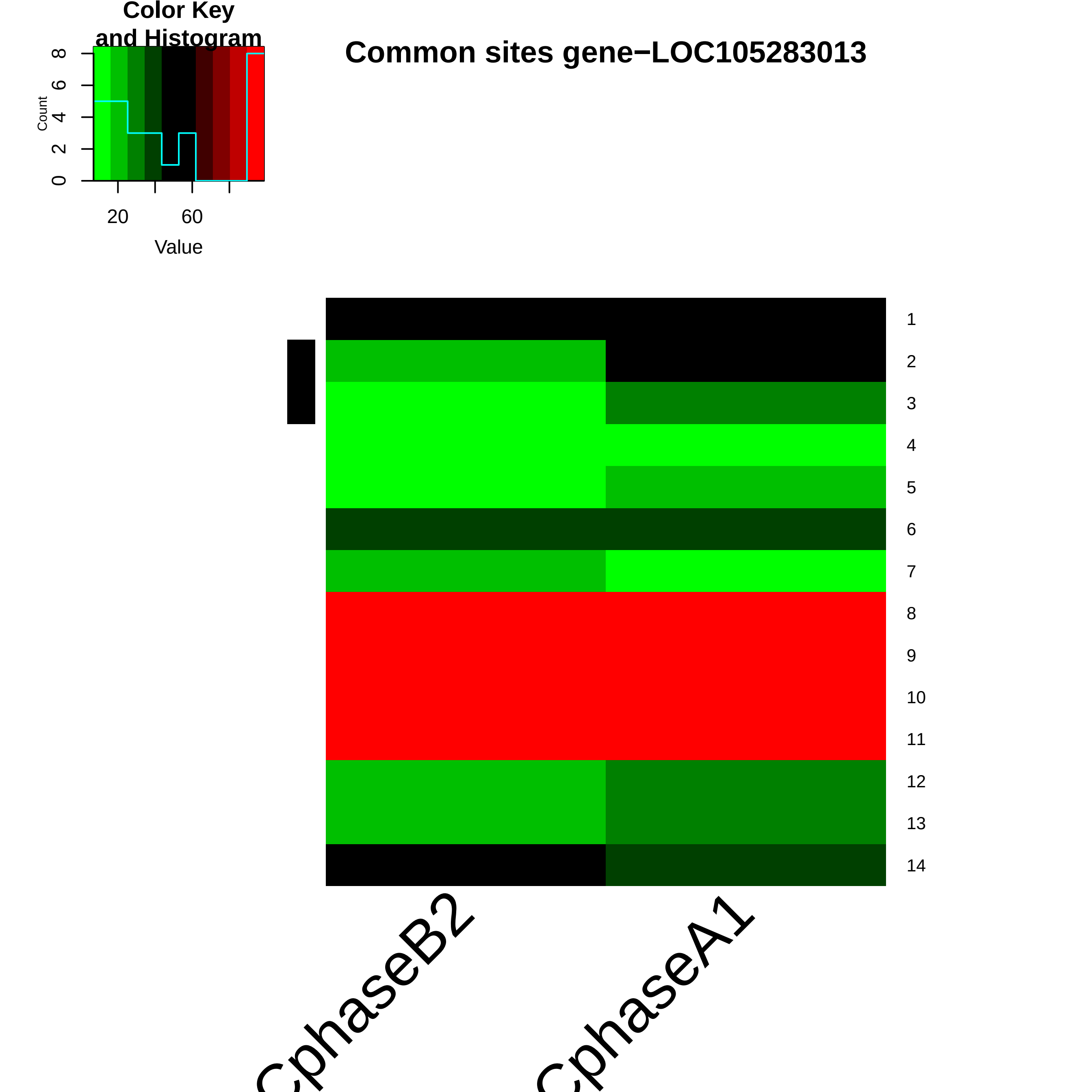

Color Key
and Histogram
Count
0
2
4
6
8
20
60
Value
Common sites gene−LOC105283013
1
2
3
4
5
6
7
8
9
10
11
12
13
14
CphaseB2
CphaseA1

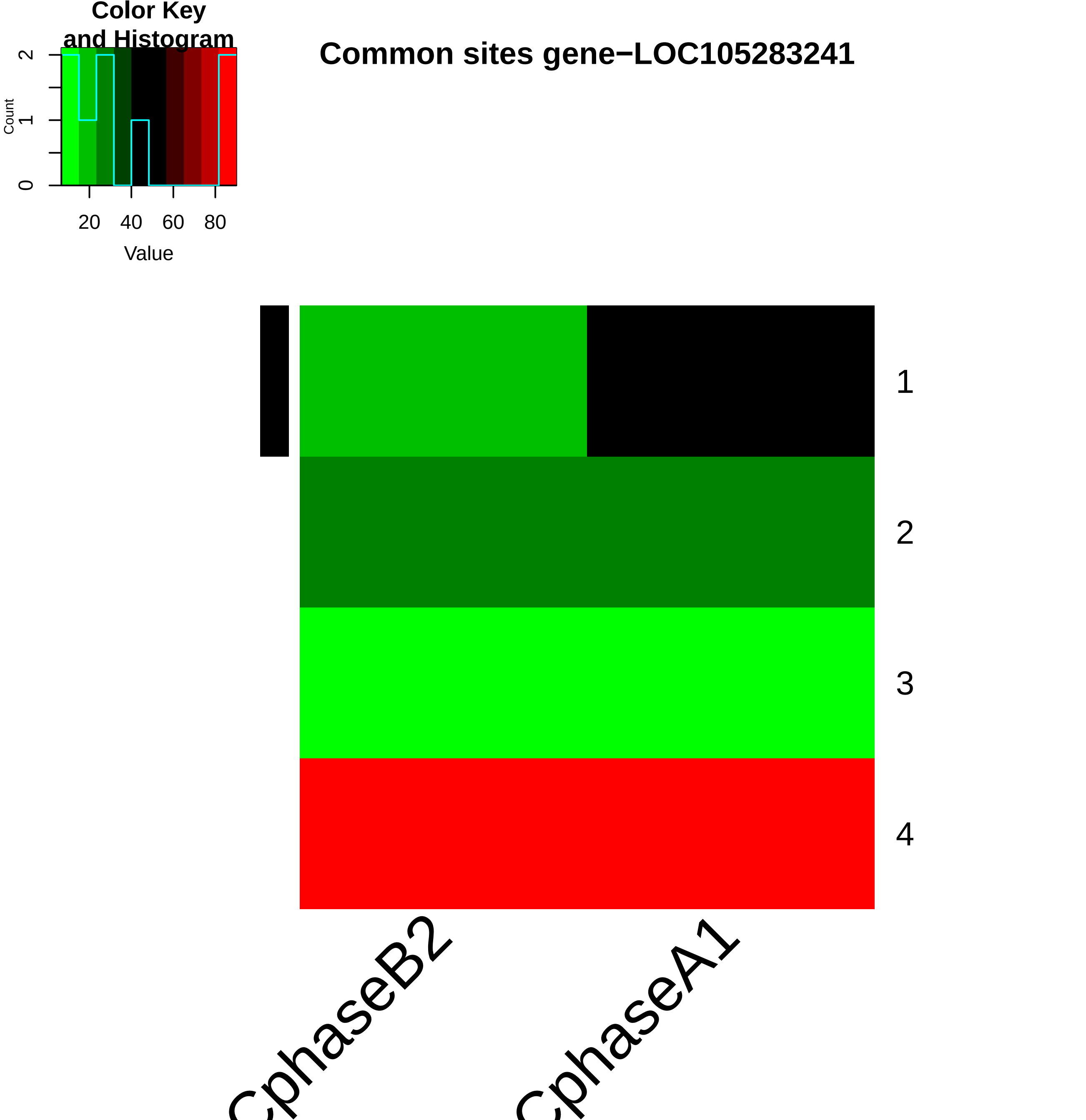
Color Key
and Histogram
Count
0
1
2
20
40
60
80
Value
Common sites gene−LOC105283241
1
2
3
4
CphaseB2
CphaseA1

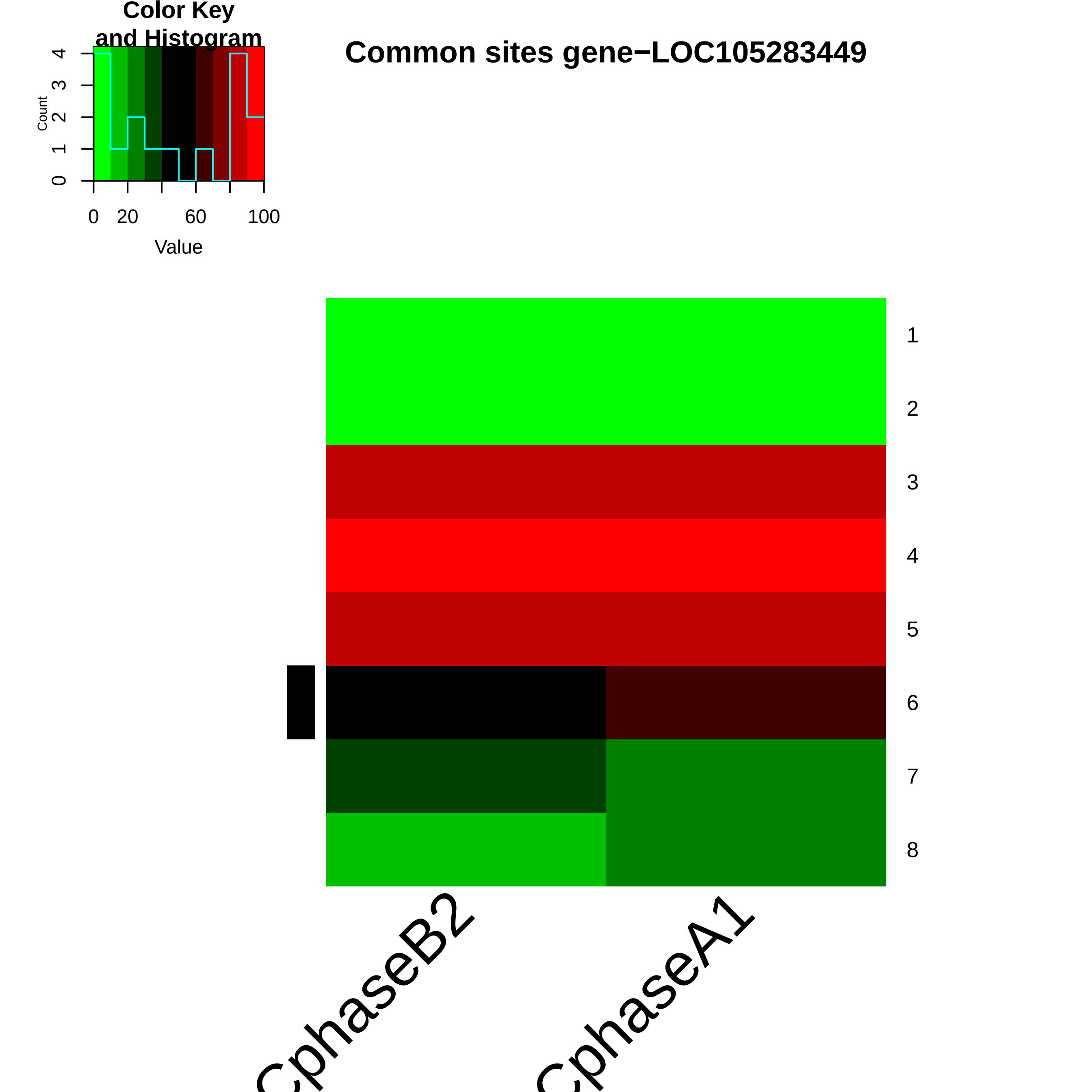
Color Key
and Histogram
Count
0
1
2
3
4
0
20
60
100
Value
Common sites gene−LOC105283449
1
2
3
4
5
6
7
8
CphaseB2
CphaseA1

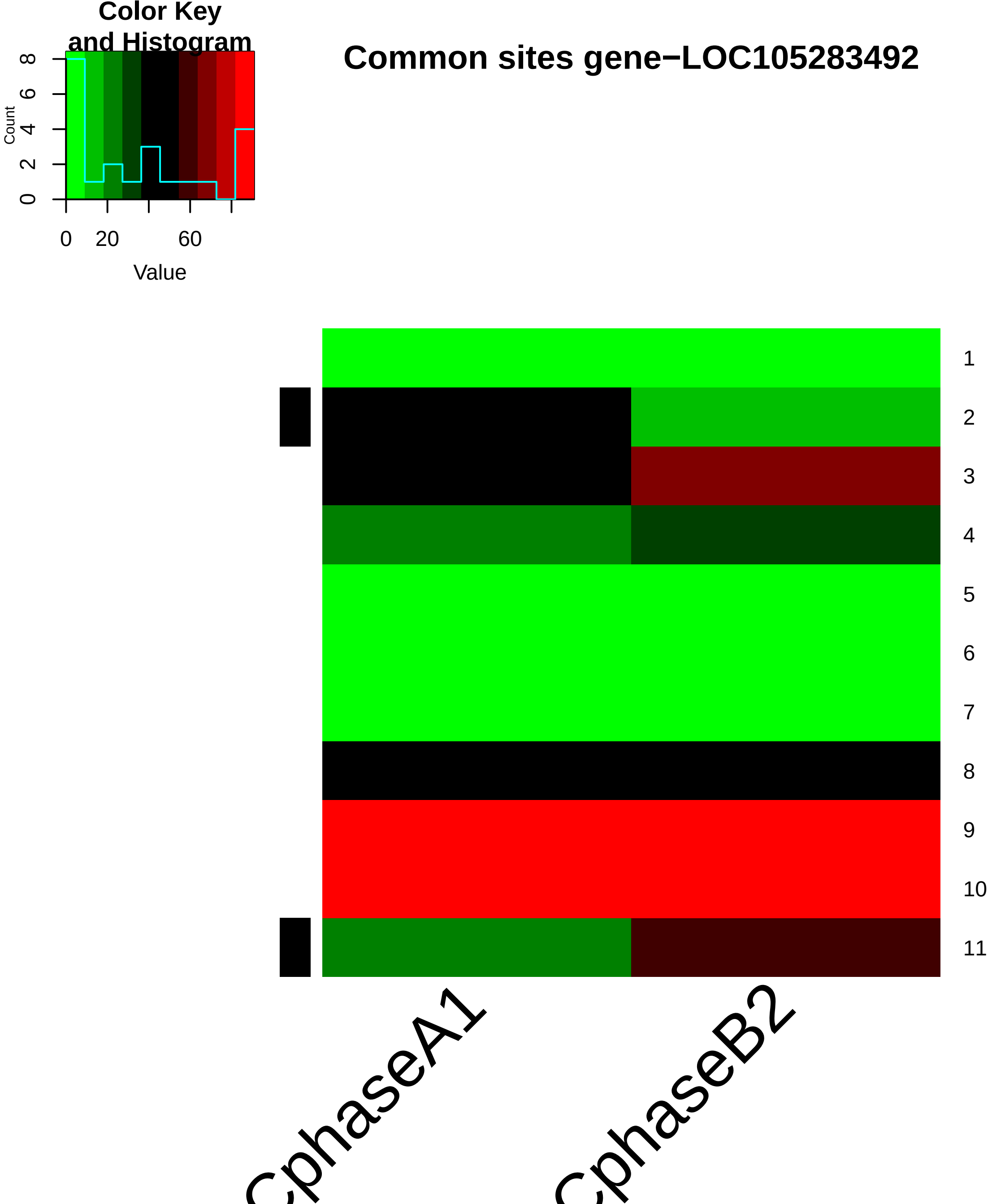
Color Key
and Histogram
Count
0
2
4
6
8
0
20
60
Value
Common sites gene−LOC105283492
1
2
3
4
5
6
7
8
9
10
11
CphaseA1
CphaseB2

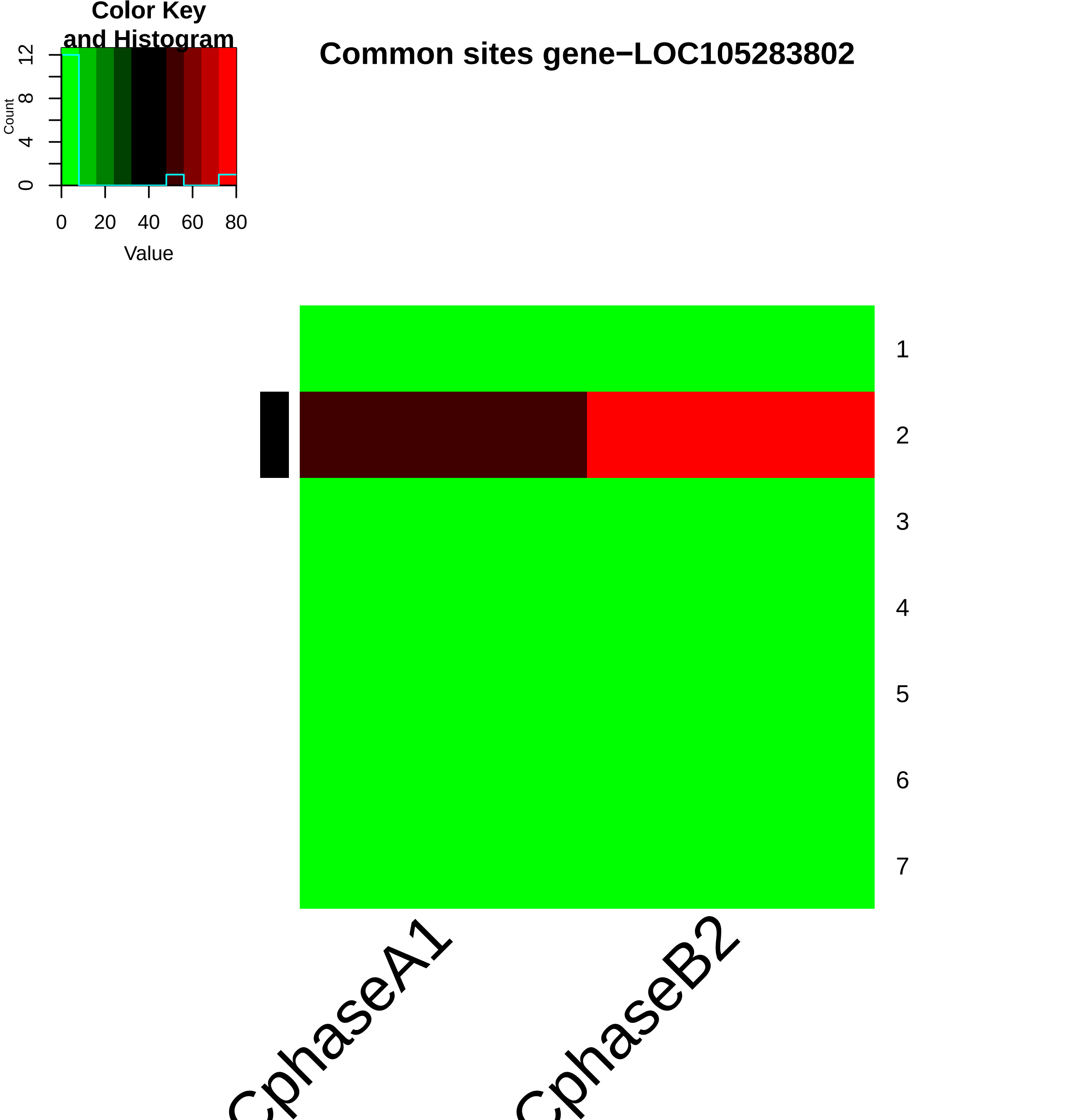
Color Key
and Histogram
Count
0
4
8
12
0
20
40
60
80
Value
Common sites gene−LOC105283802
1
2
3
4
5
6
7

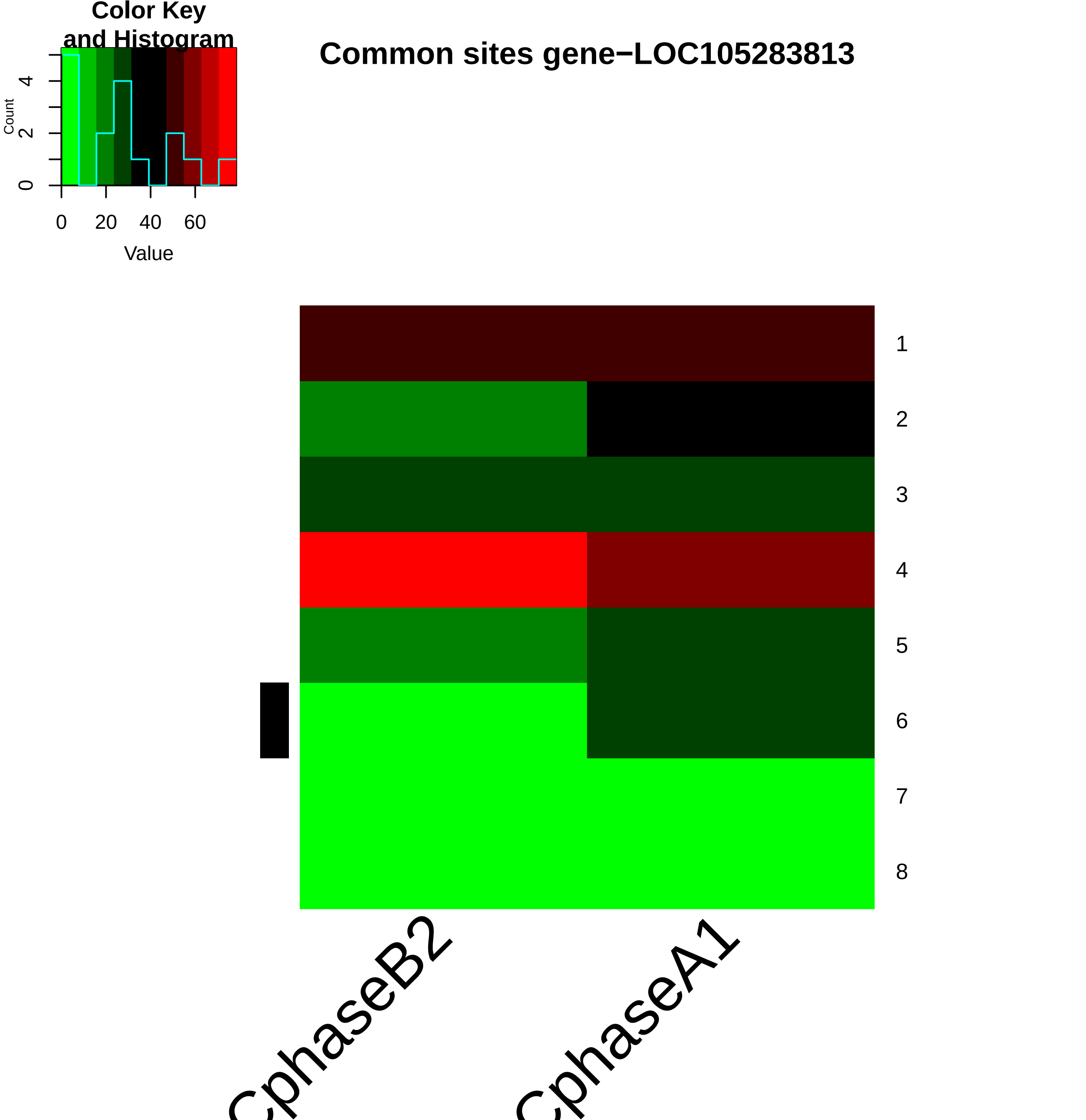

Color Key
and Histogram
Count
0
2
4
0
20
40
60
Value
Common sites gene−LOC105283813
1
2
3
4
5
6
7
8
CphaseB2
CphaseA1

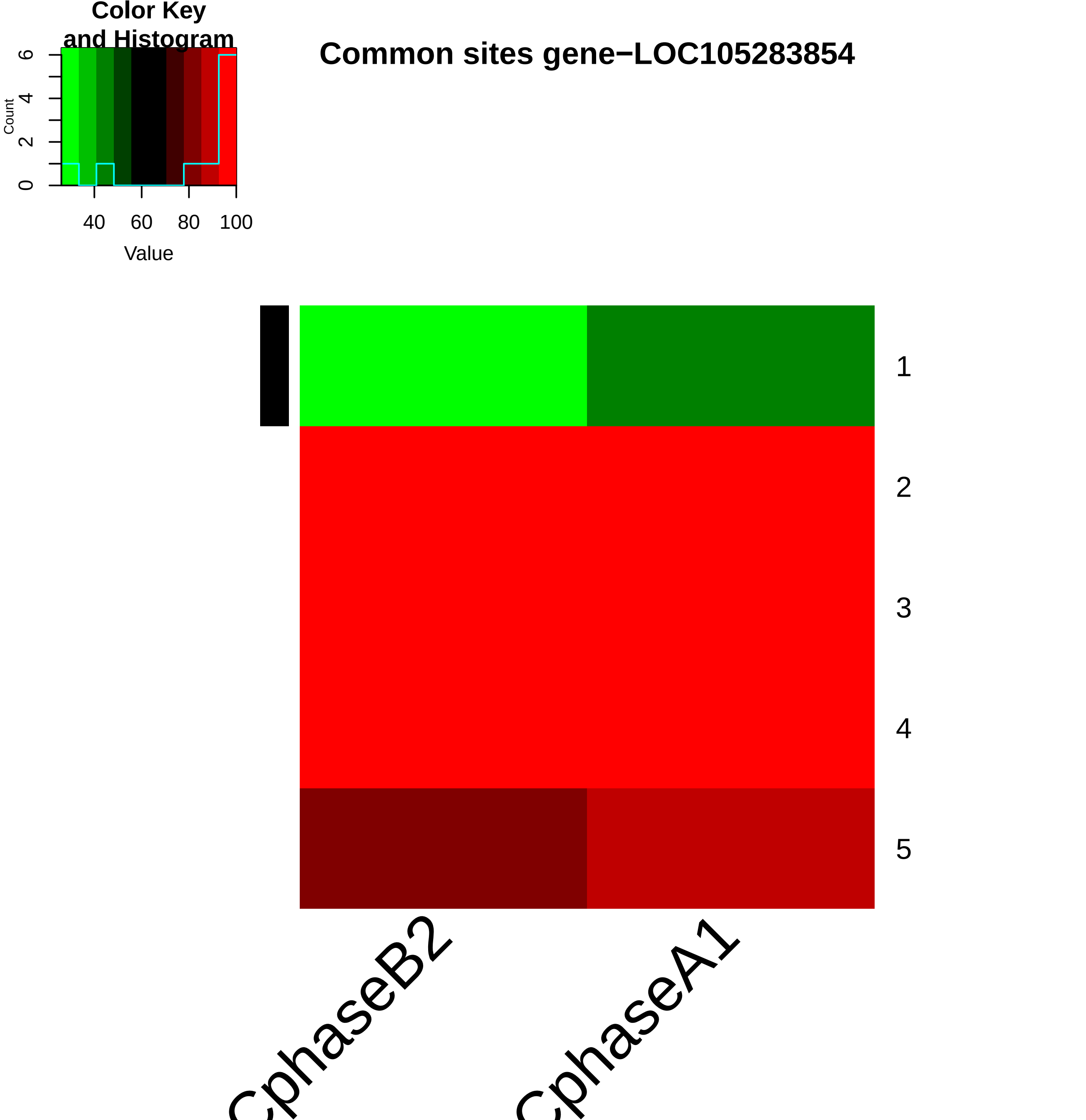

Color Key
and Histogram
Count
0
2
4
6
40
60
80
100
Value
Common sites gene−LOC105283854
1
2
3
4
5
CphaseB2
CphaseA1

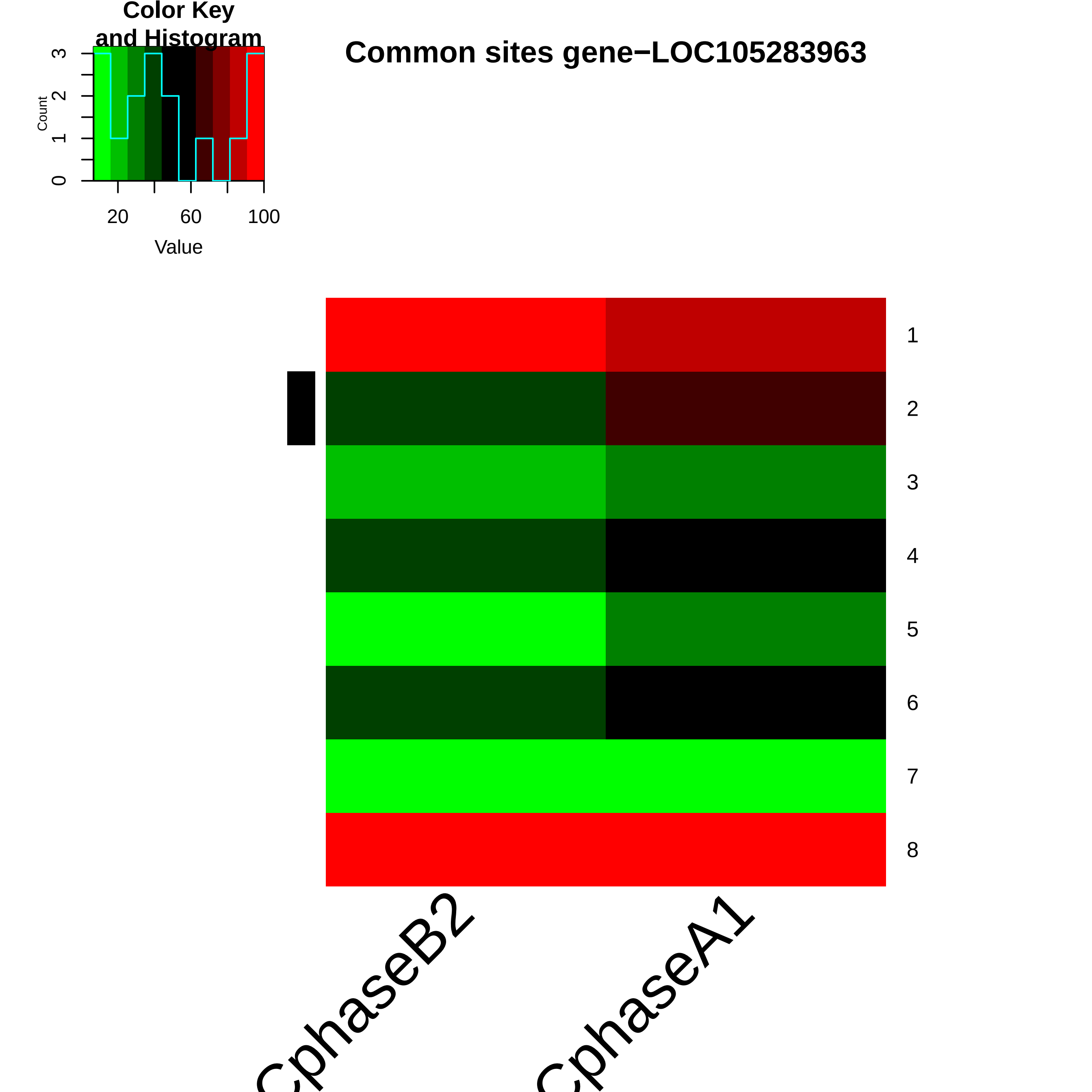
Color Key
and Histogram
Count
0
1
2
3
20
60
100
Value
Common sites gene−LOC105283963
1
2
3
4
5
6
7
8
CphaseB2
CphaseA1

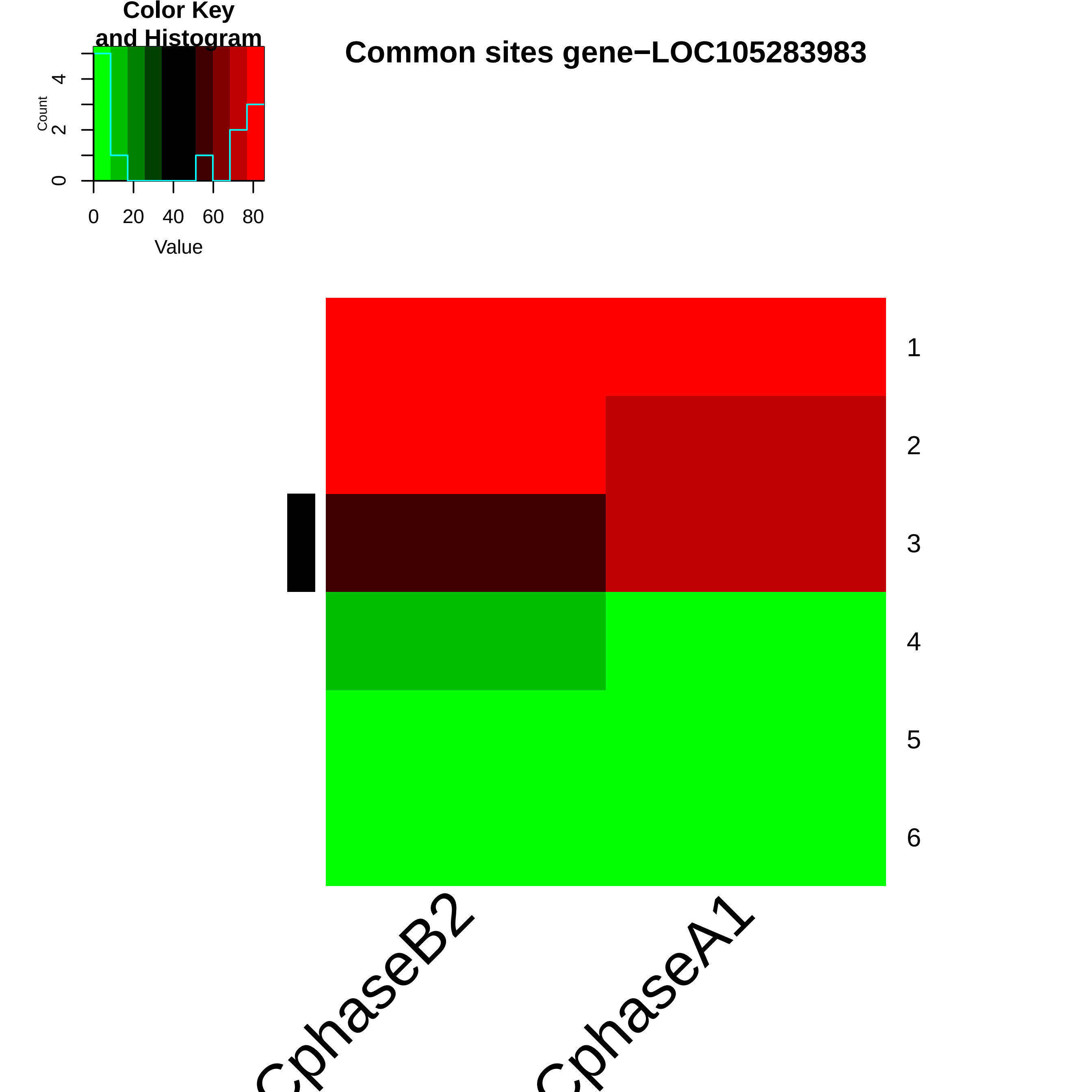

Color Key
and Histogram
Count
0
2
4
0
20
40
60
80
Value
Common sites gene−LOC105283983
1
2
3
4
5
6
CphaseB2
CphaseA1

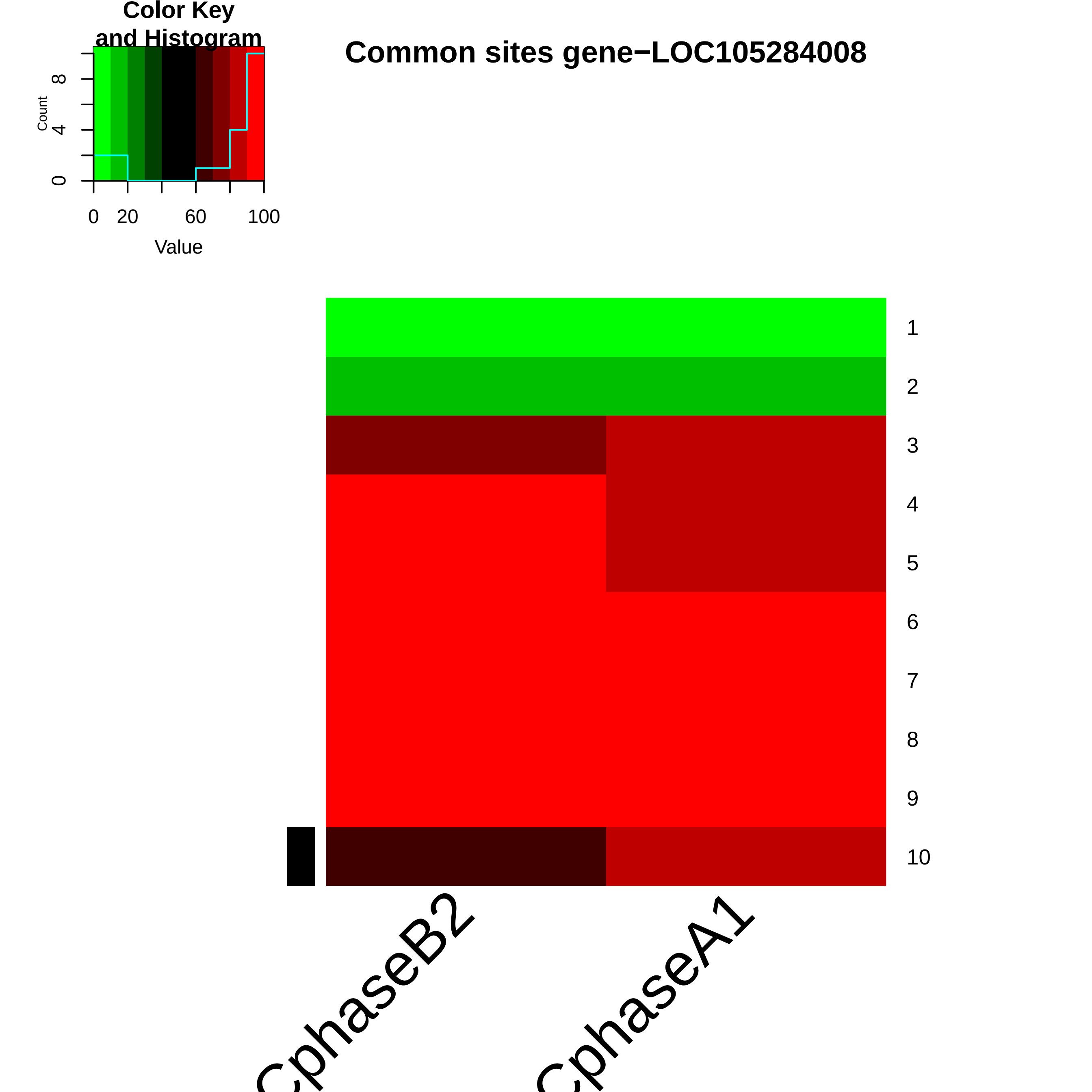

Color Key and Histogram
Count
0
4
8
0
20
60
100
Value
Common sites gene−LOC105284008
1
2
3
4
5
6
7
8
9
10
CphaseB2
CphaseA1

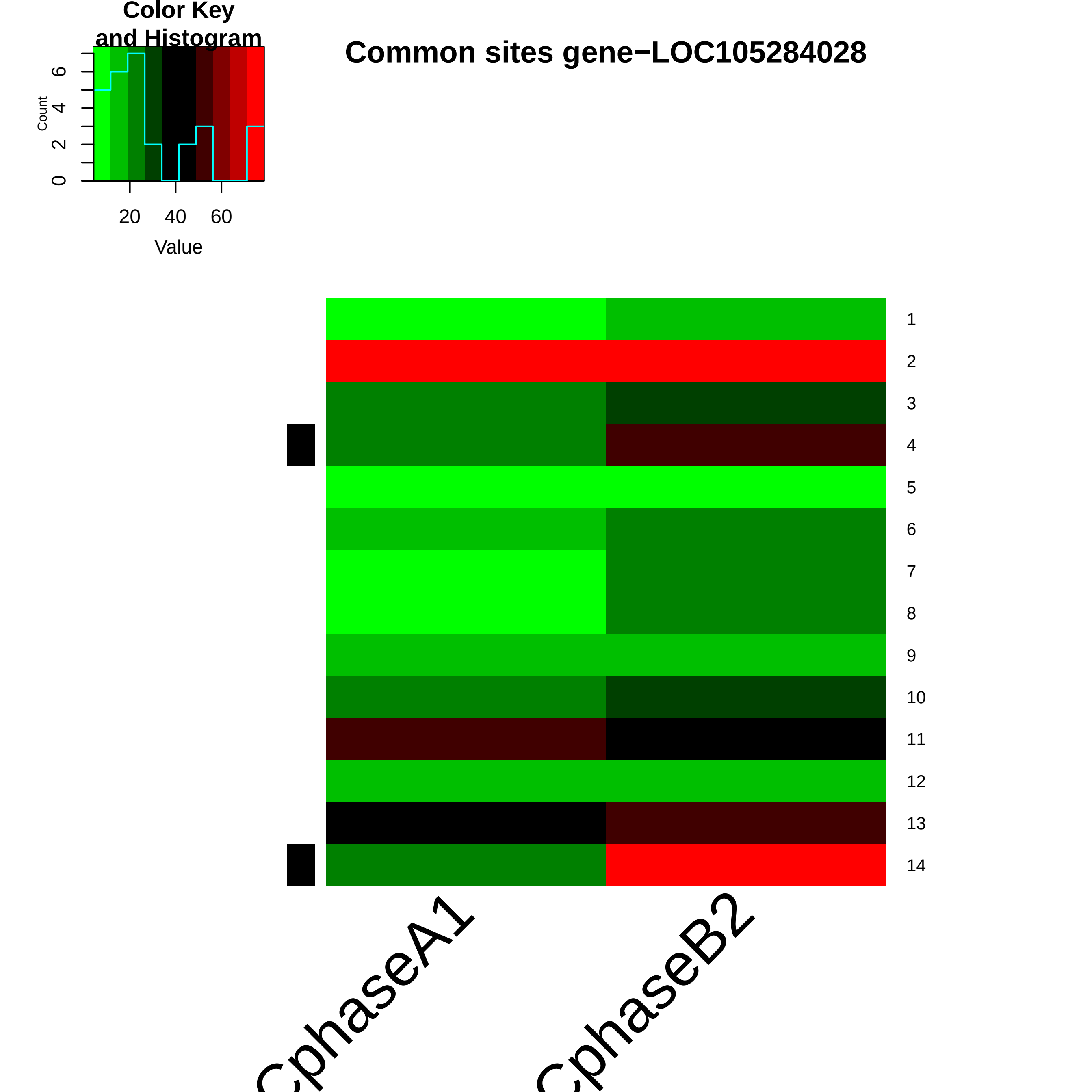
Color Key
and Histogram
Count
0
2
4
6
20
40
60
Value
Common sites gene−LOC105284028
1
2
3
4
5
6
7
8
9
10
11
12
13
14
CphaseA1
CphaseB2

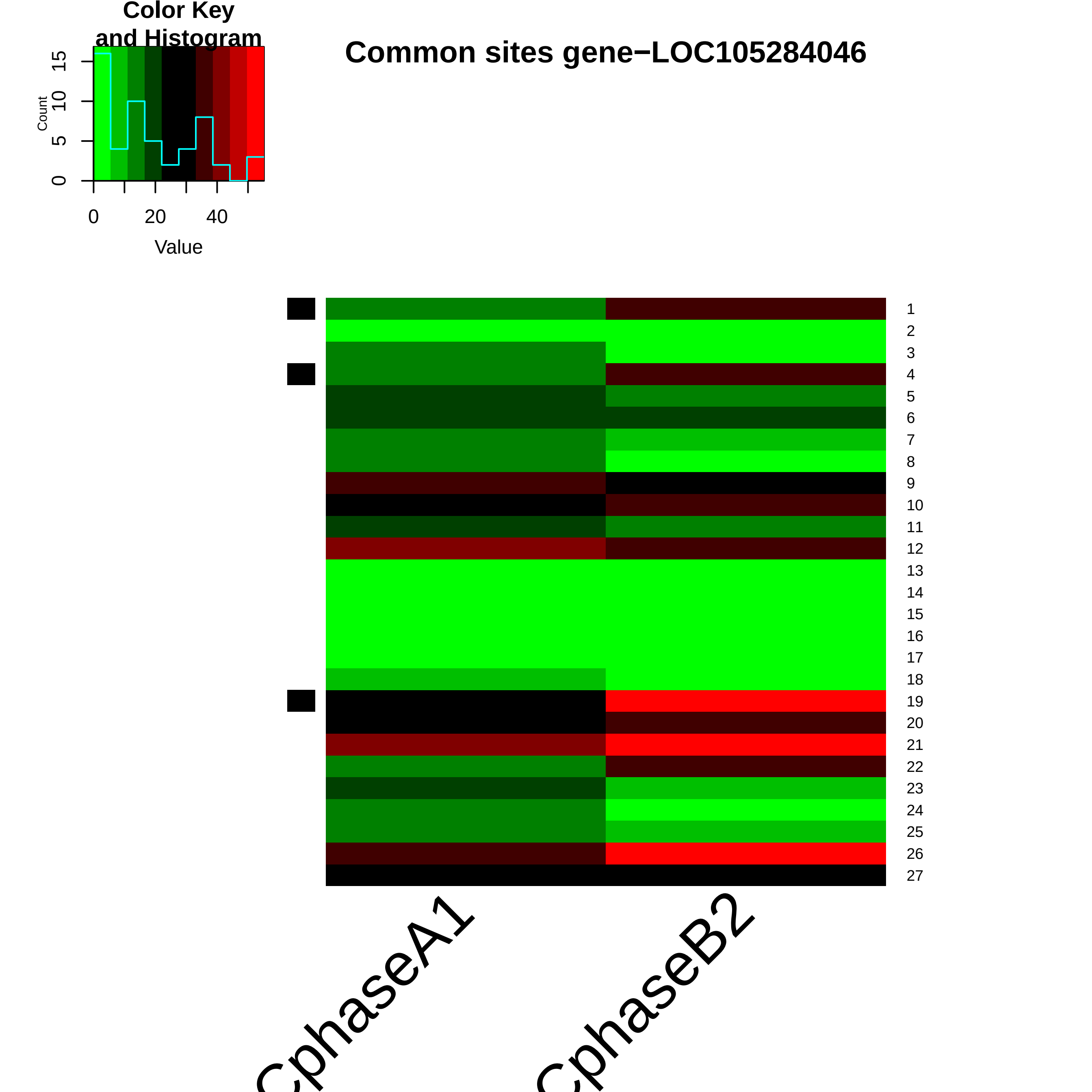
Color Key
and Histogram
Count
0
5
10
15
0
20
40
Value
Common sites gene−LOC105284046
CphaseA1
CphaseB2

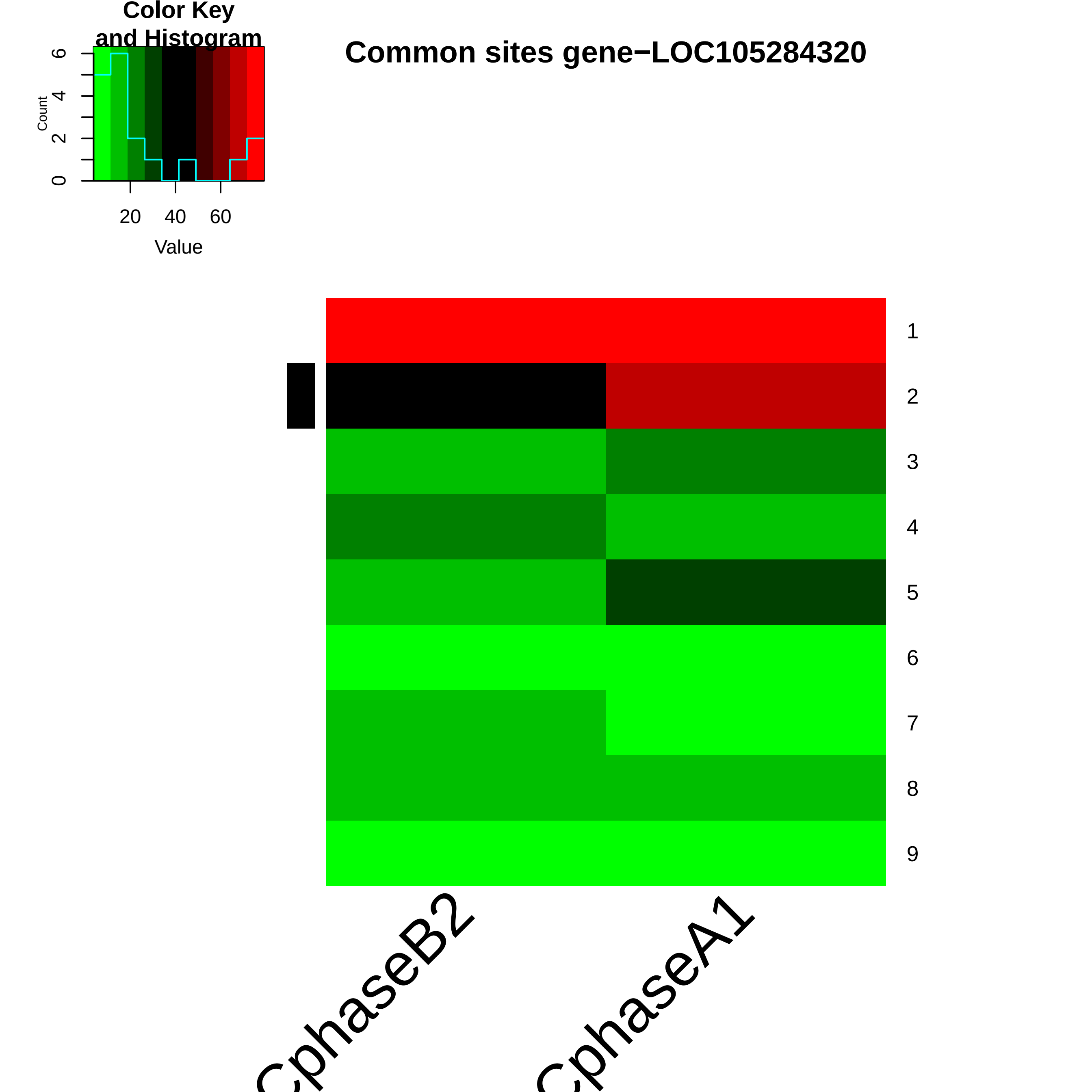
Color Key
and Histogram
Count
0
2
4
6
20
40
60
Value
Common sites gene−LOC105284320
1
2
3
4
5
6
7
8
9
CphaseB2
CphaseA1

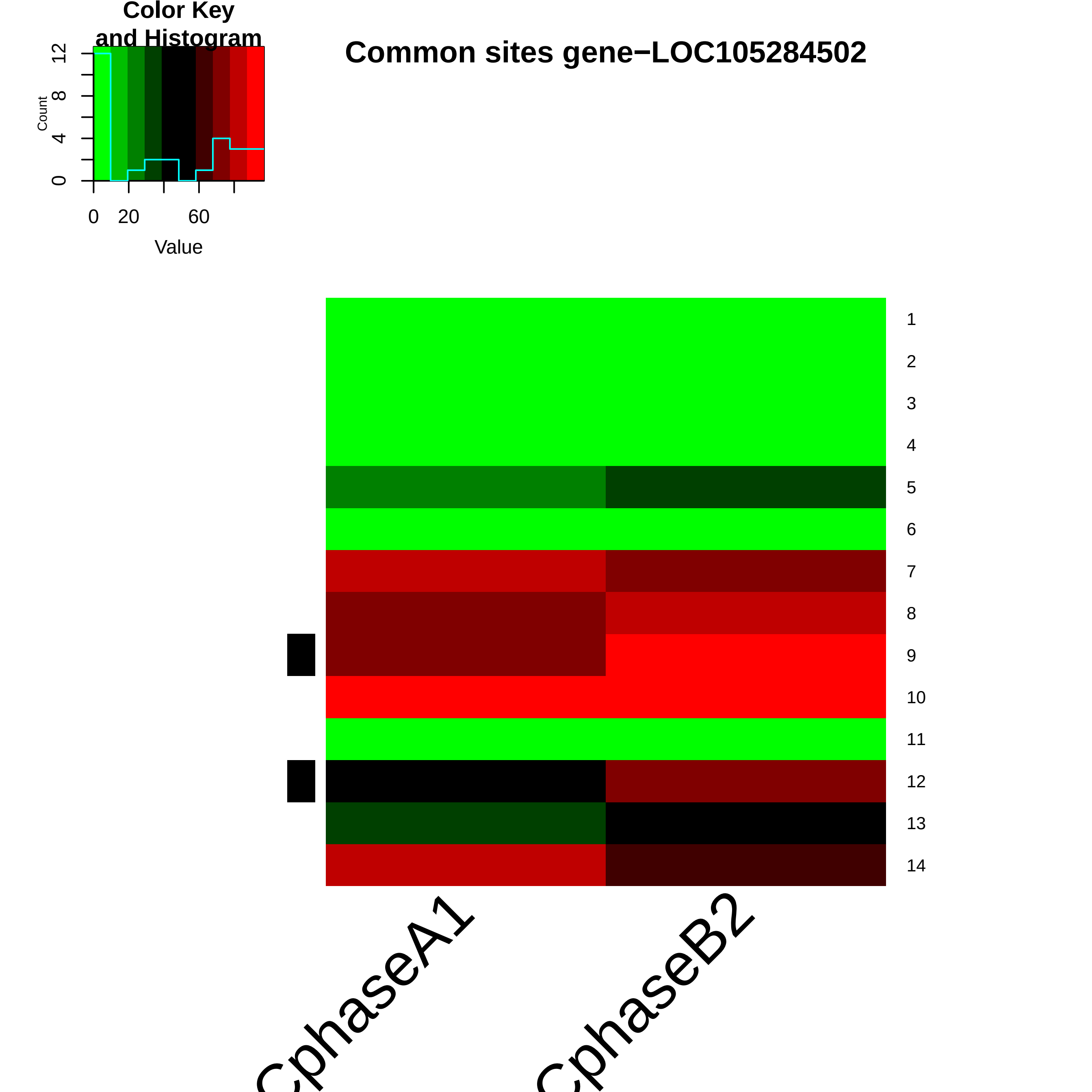

Color Key
and Histogram
Count
0
4
8
12
0
20
60
Value
Common sites gene−LOC105284502
1
2
3
4
5
6
7
8
9
10
11
12
13
14
CphaseA1
CphaseB2

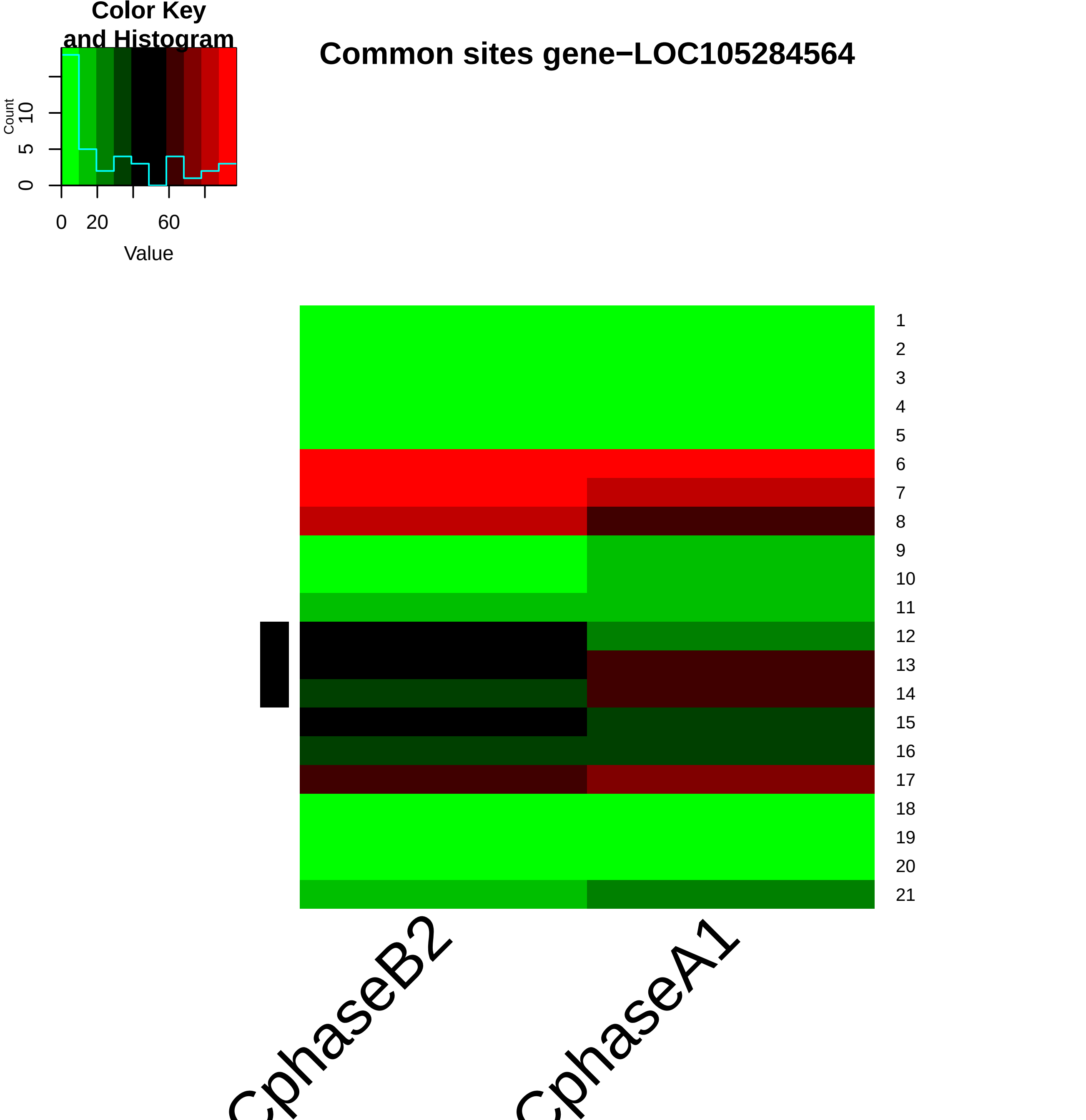
Color Key
and Histogram
Count
0
5
10
0
20
60
Value
Common sites gene−LOC105284564
CphaseB2
CphaseA1

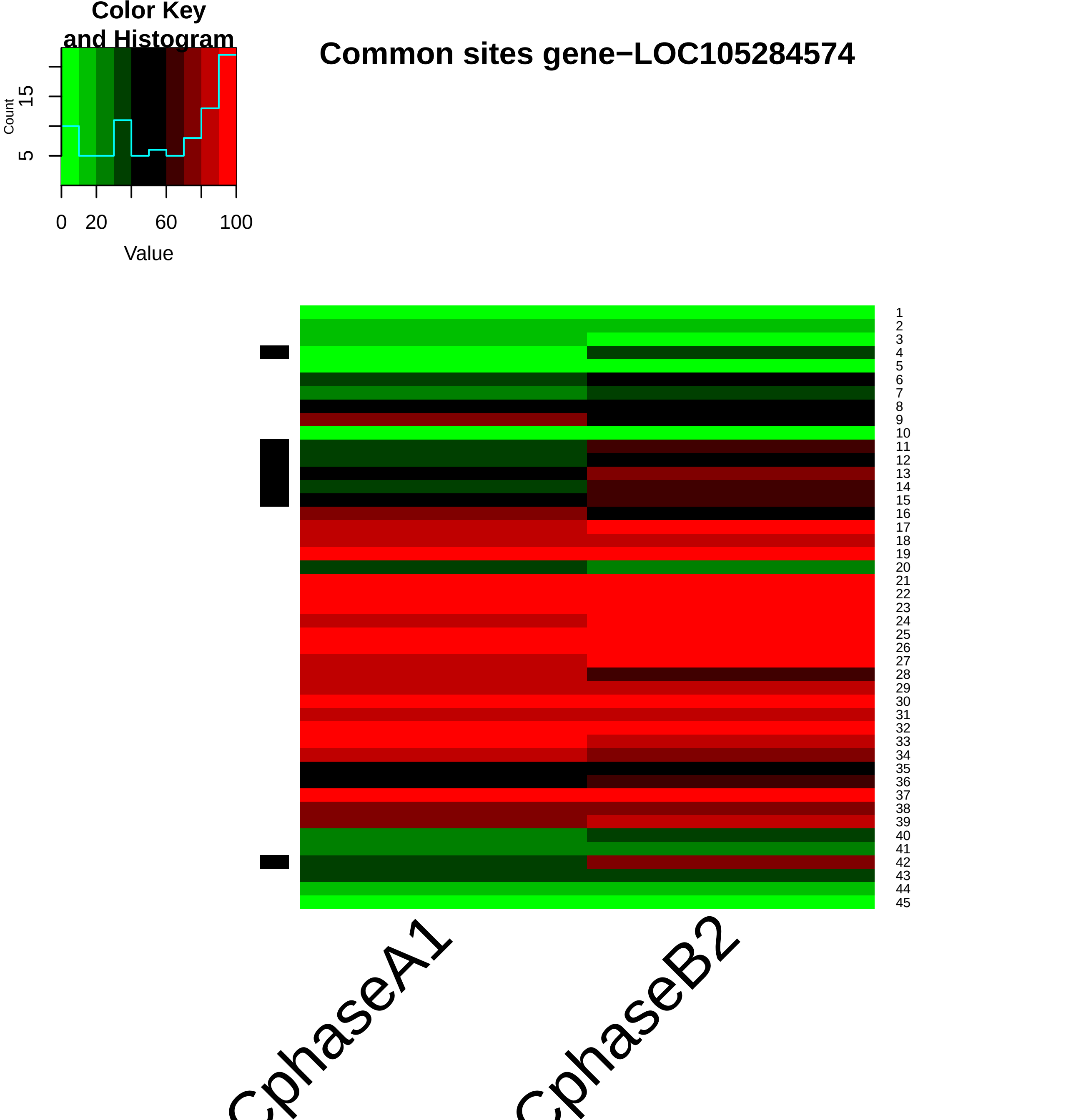
Color Key
and Histogram
Count
5
15
0
20
60
100
Value
Common sites gene−LOC105284574
CphaseA1
CphaseB2

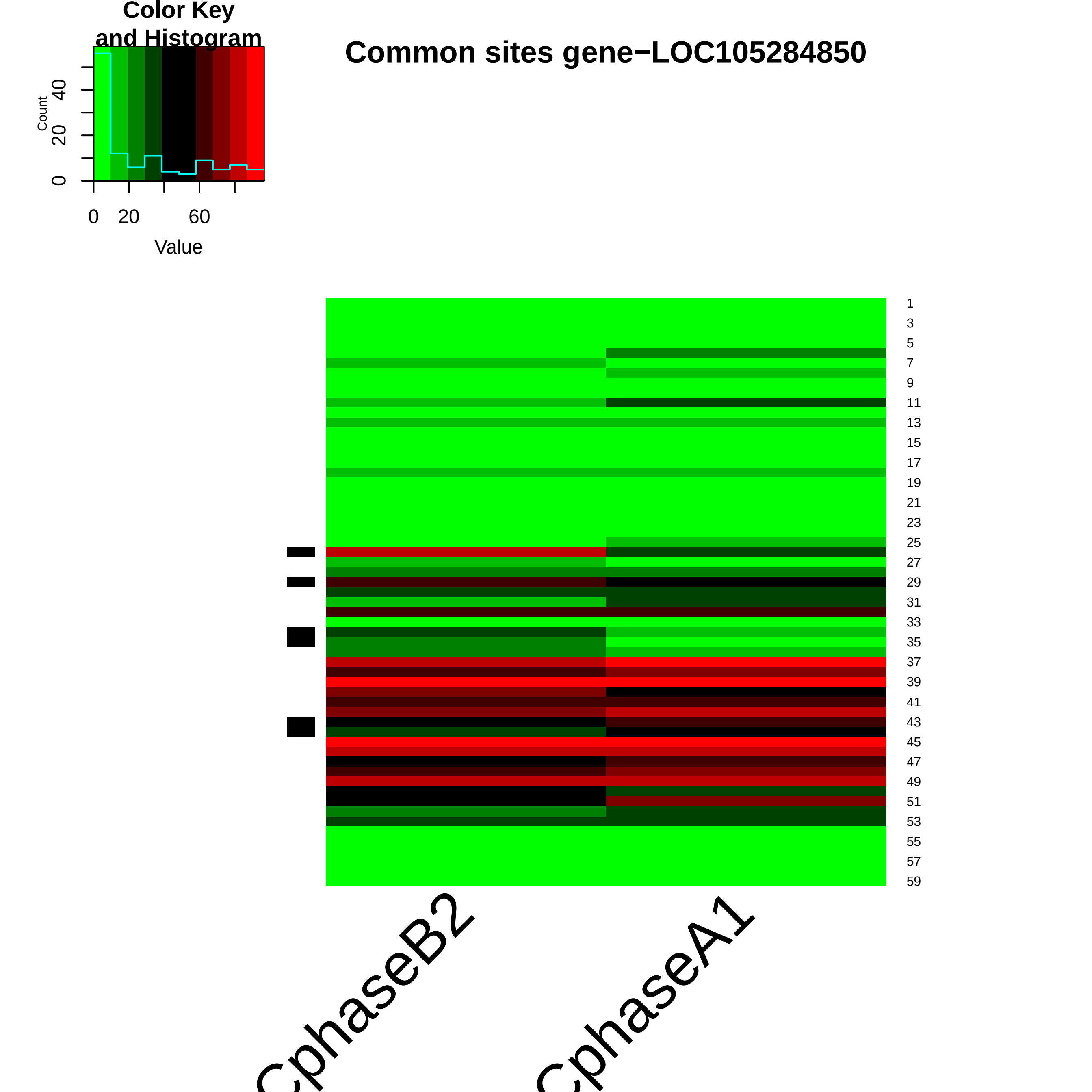
Color Key
and Histogram
Count
0
20
40
0
20
60
Value
Common sites gene−LOC105284850
1
3
5
7
9
11
13
15
17
19
21
23
25
27
29
31
33
35
37
39
41
43
45
47
49
51
53
55
57
59
CphaseB2
CphaseA1

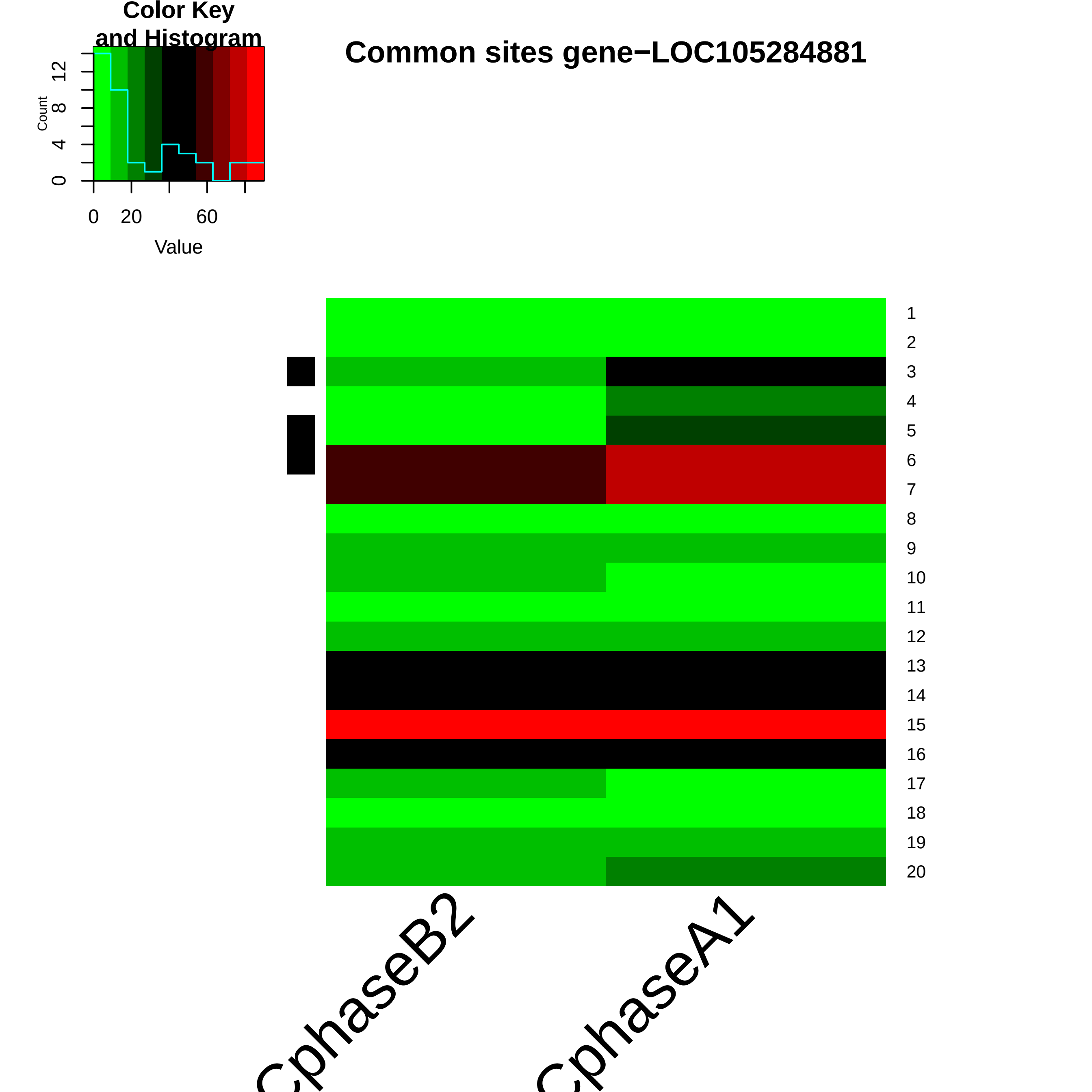

Color Key
and Histogram
Count
0
4
8
12
0
20
60
Value
Common sites gene−LOC105284881
CphaseB2
CphaseA1

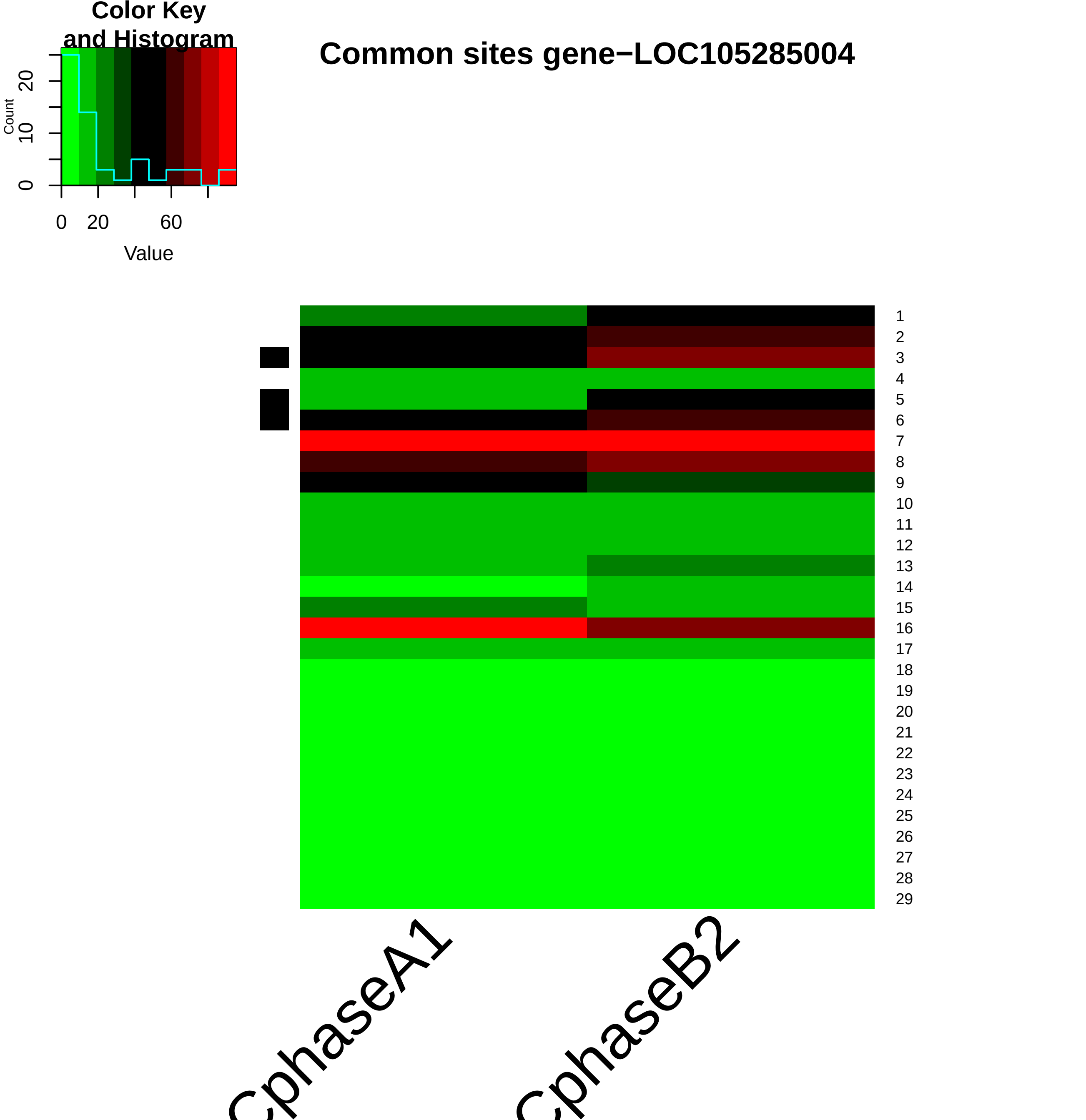
Color Key
and Histogram
Count
0
10
20
0
20
60
Value
Common sites gene−LOC105285004
CphaseA1
CphaseB2

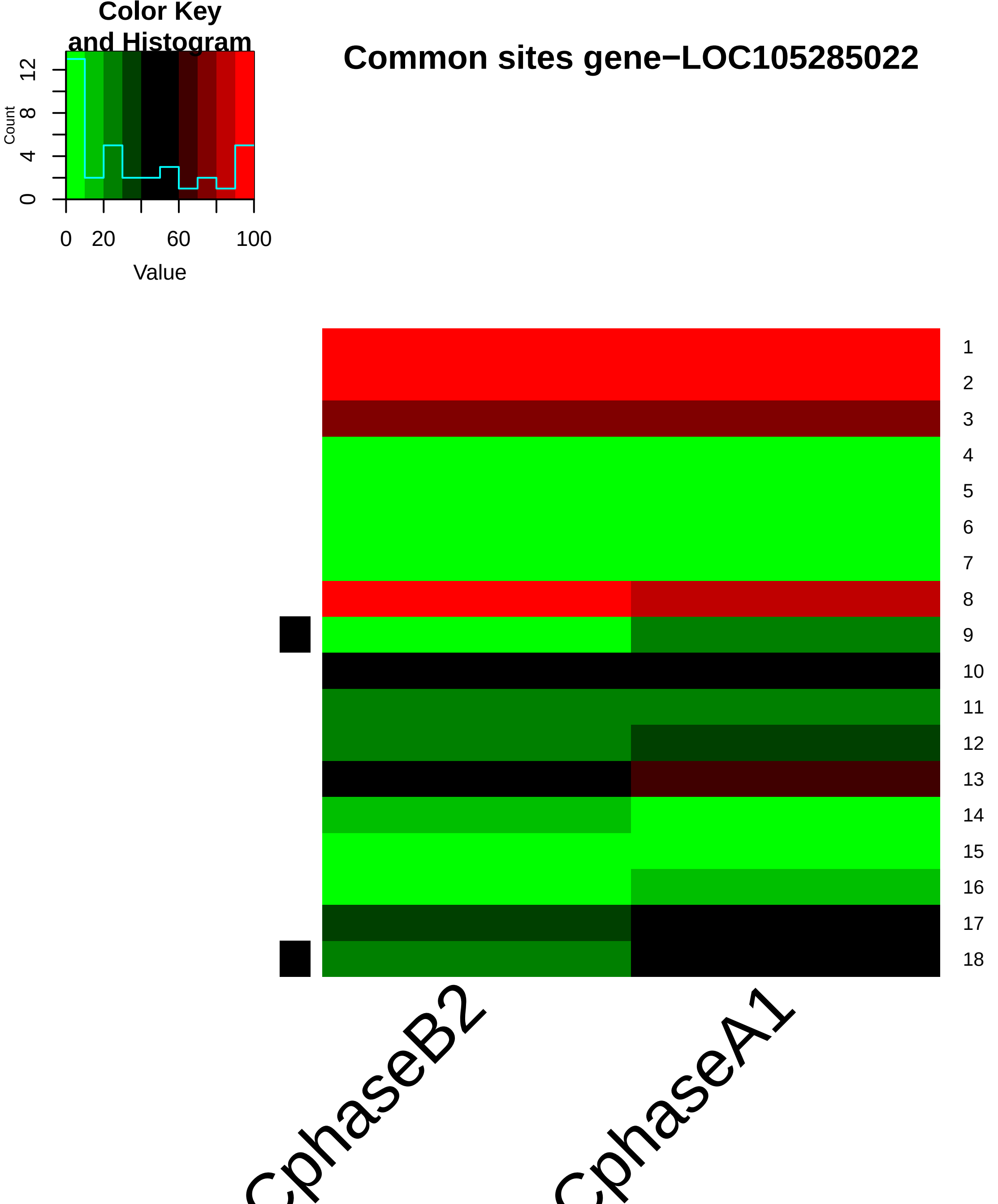

Color Key
and Histogram
Count
0
4
8
12
0
20
60
100
Value
Common sites gene−LOC105285022
CphaseB2
CphaseA1

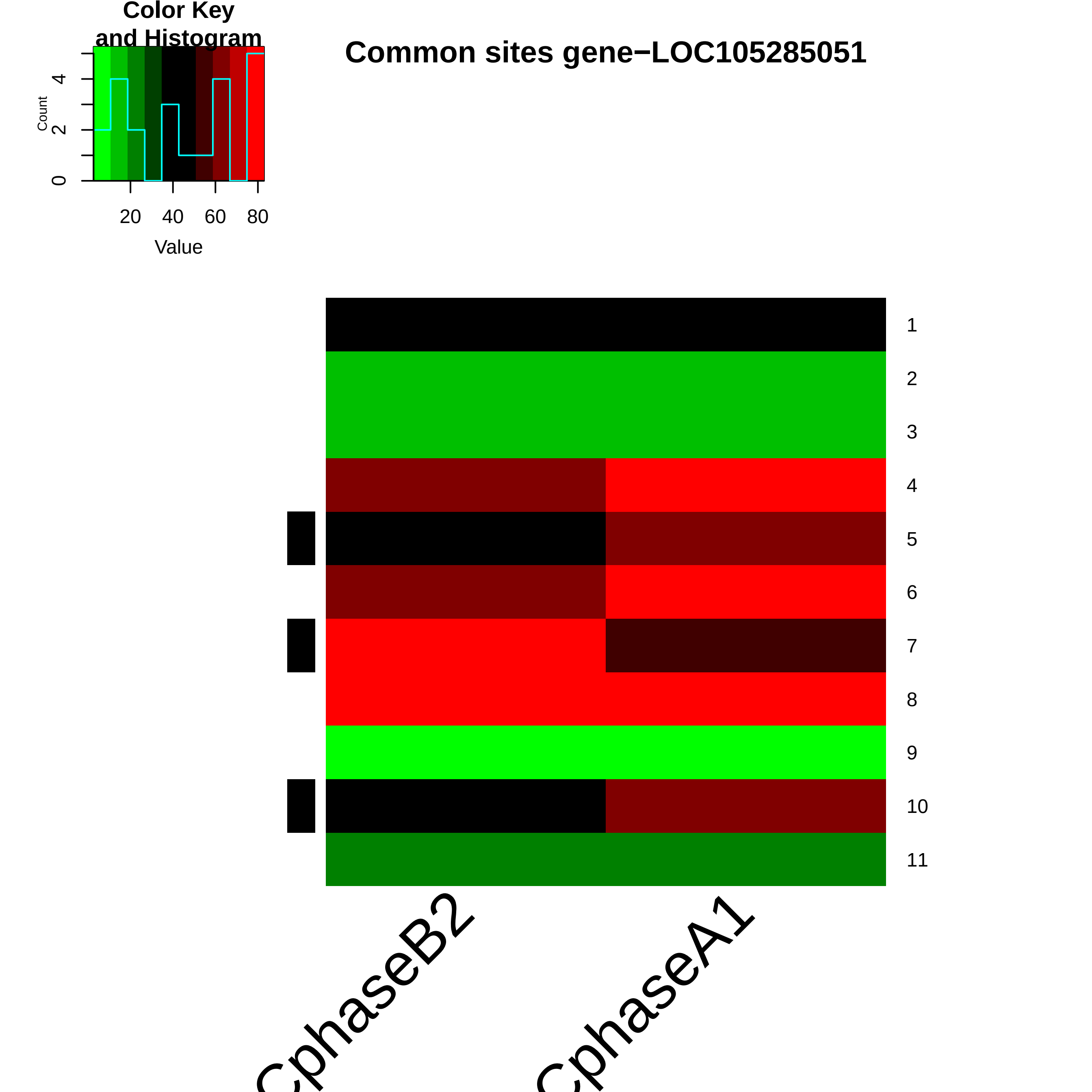

Color Key
and Histogram
Count
0
2
4
20
40
60
80
Value
Common sites gene−LOC105285051
1
2
3
4
5
6
7
8
9
10
11
CphaseB2
CphaseA1

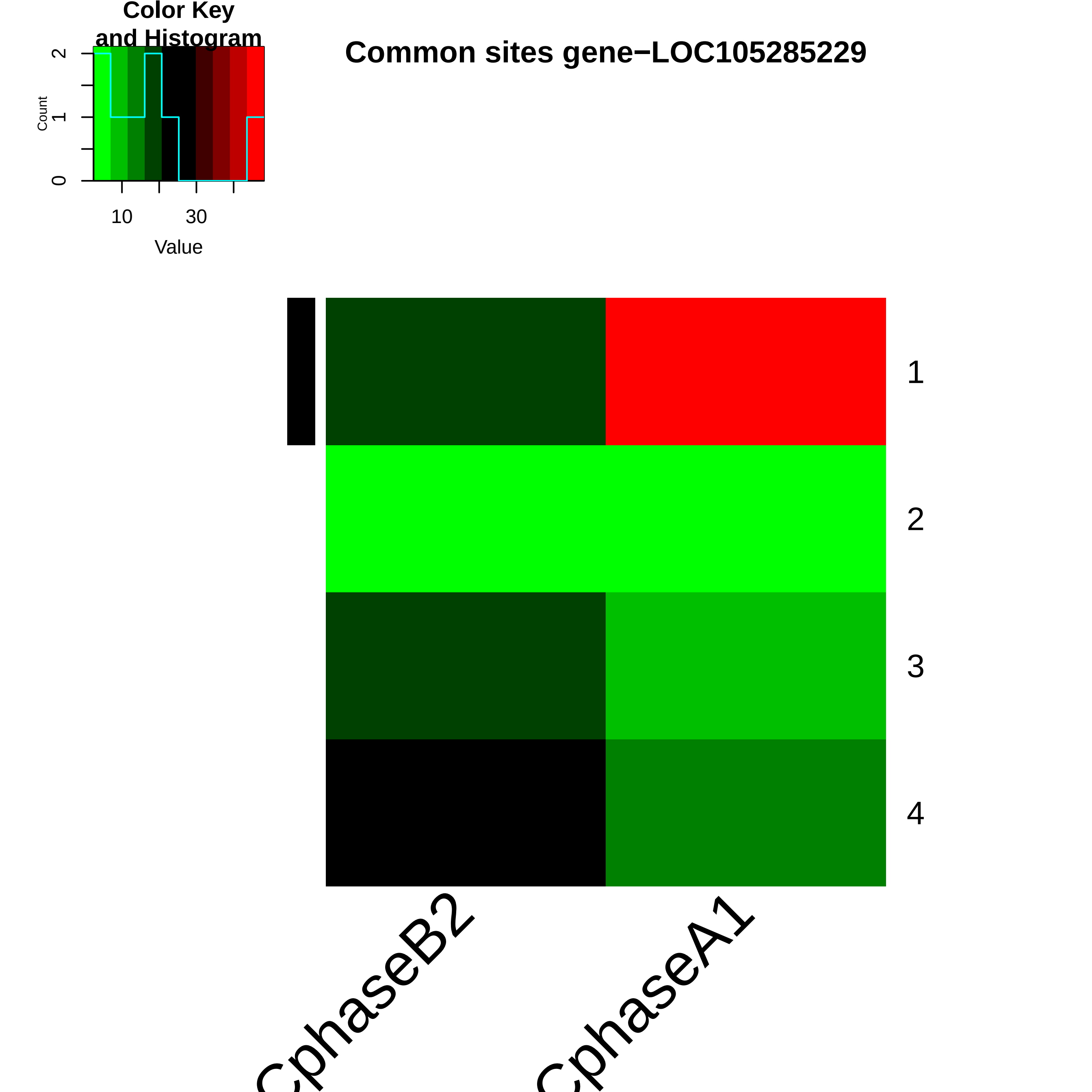

Color Key
and Histogram
Count
0
1
2
10
30
Value
Common sites gene−LOC105285229
1
2
3
4
CphaseB2
CphaseA1

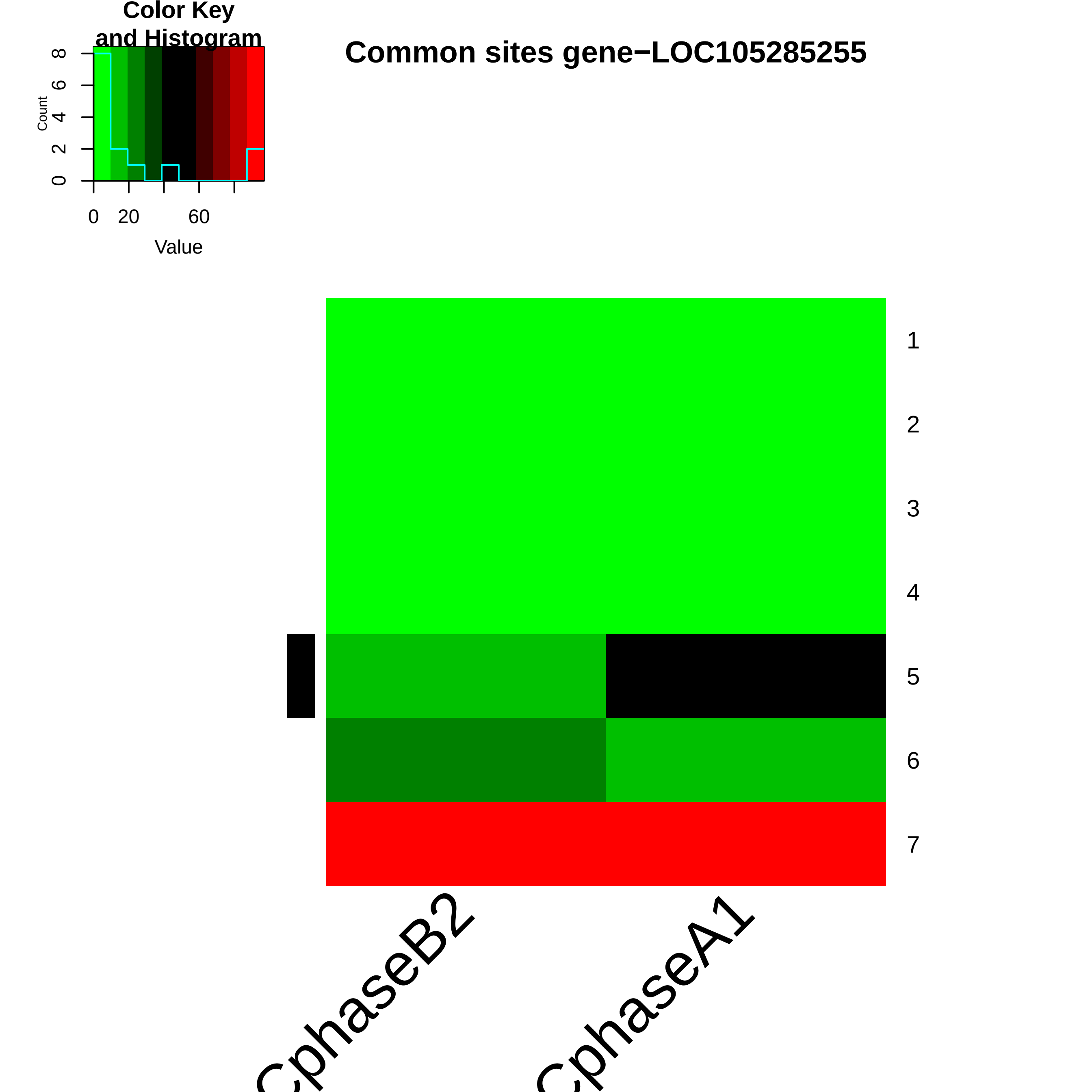
Color Key
and Histogram
Count
0
2
4
6
8
0
20
60
Value
Common sites gene−LOC105285255
1
2
3
4
5
6
7
CphaseB2
CphaseA1

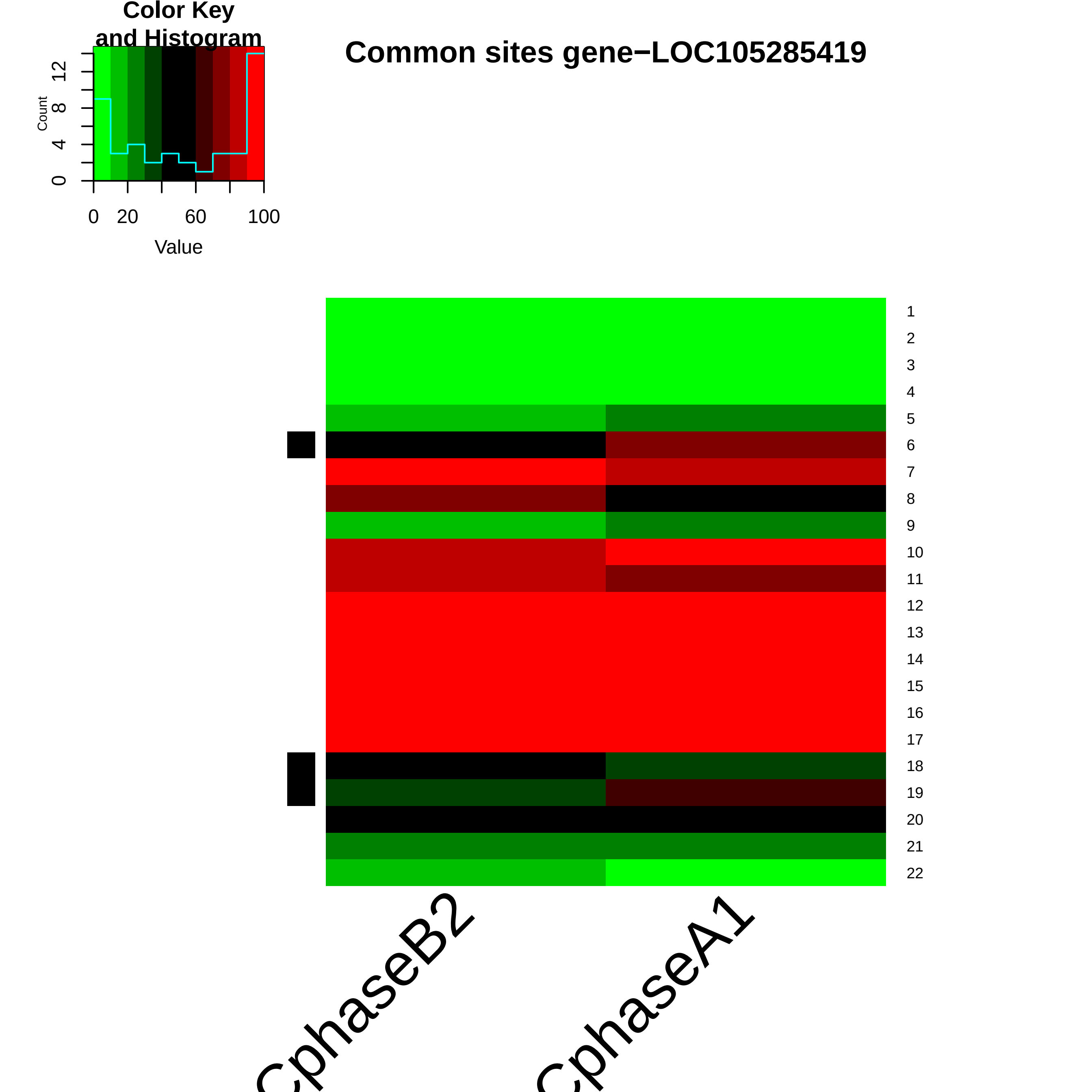

Color Key
and Histogram
Count
0
4
8
12
0
20
60
100
Value
Common sites gene−LOC105285419
CphaseB2
CphaseA1

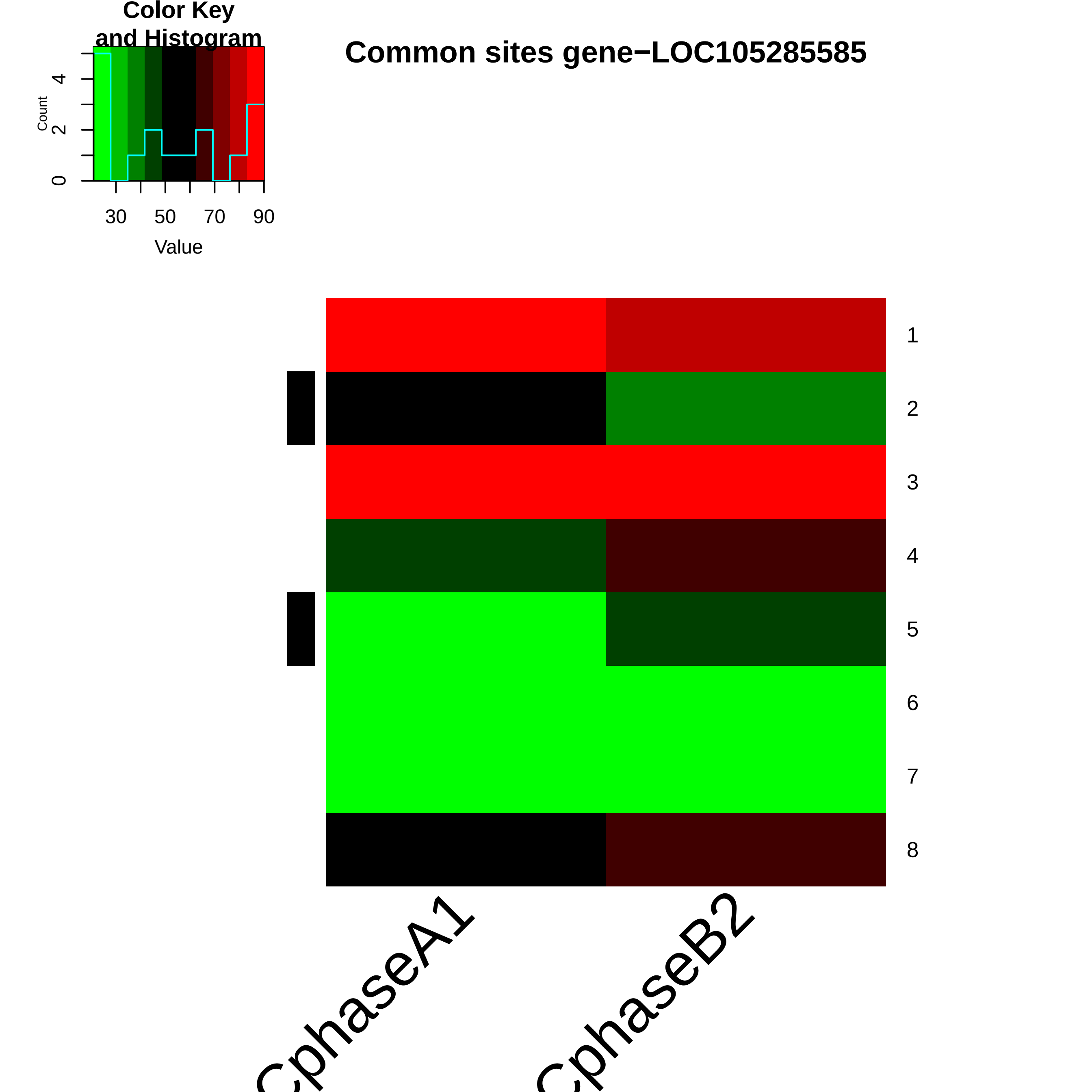
Color Key
and Histogram
Count
0
2
4
30
50
70
90
Value
Common sites gene−LOC105285585
1
2
3
4
5
6
7
8

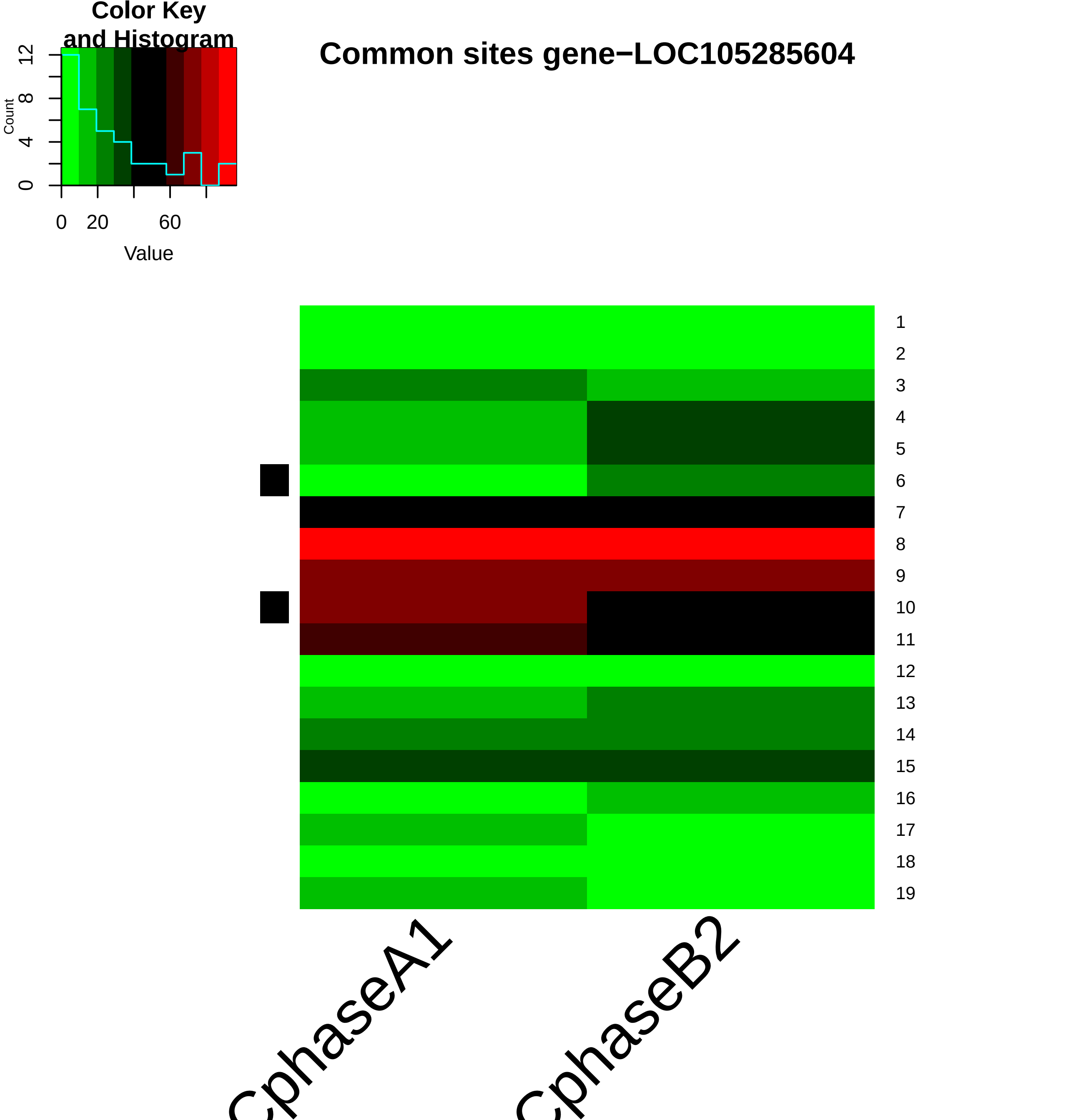
Color Key
and Histogram
Count
0
4
8
12
0
20
60
Value
Common sites gene−LOC105285604
CphaseA1
CphaseB2

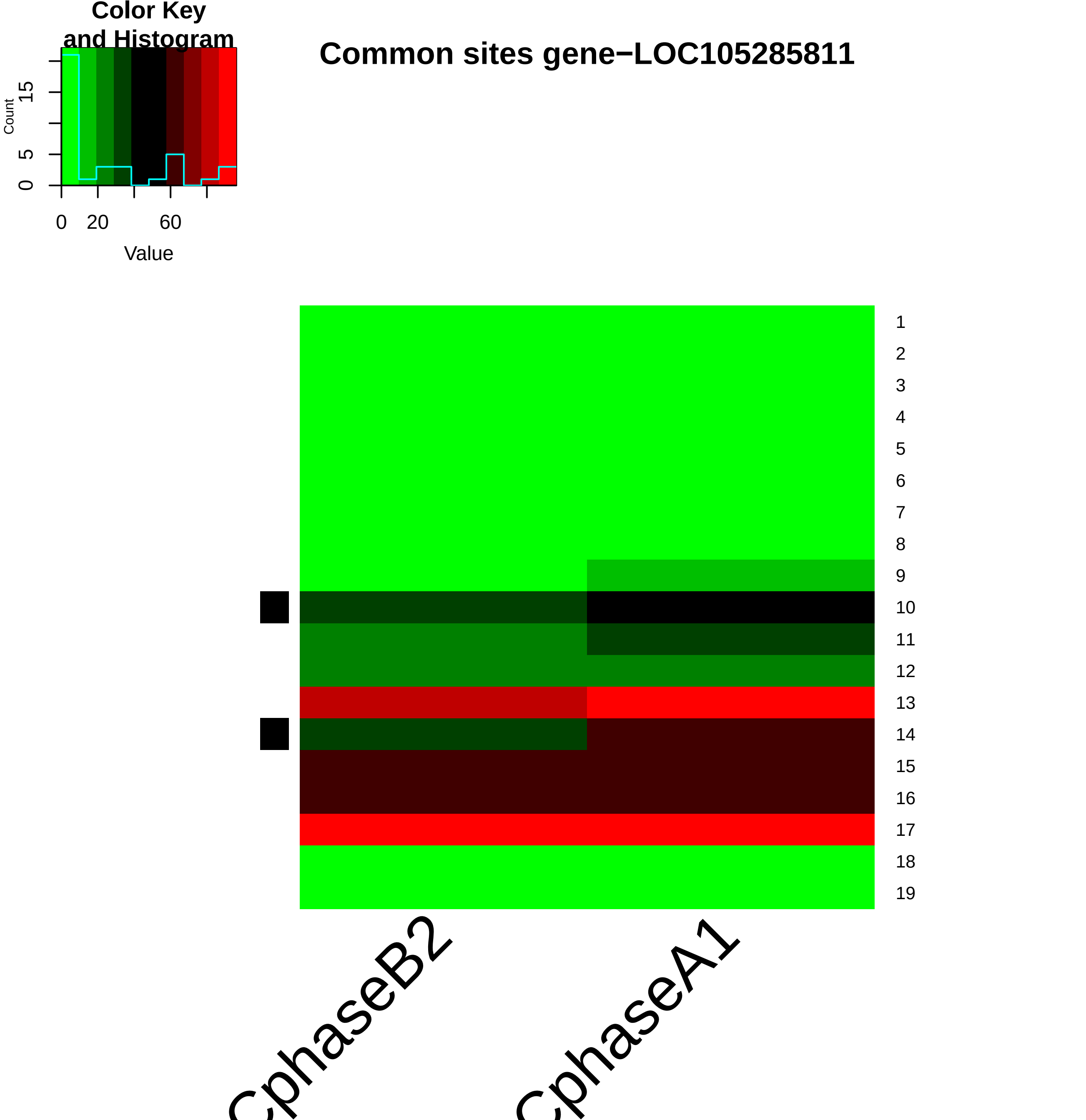

Color Key
and Histogram
Count
0
5
15
0
20
60
Value
Common sites gene−LOC105285811
CphaseB2
CphaseA1

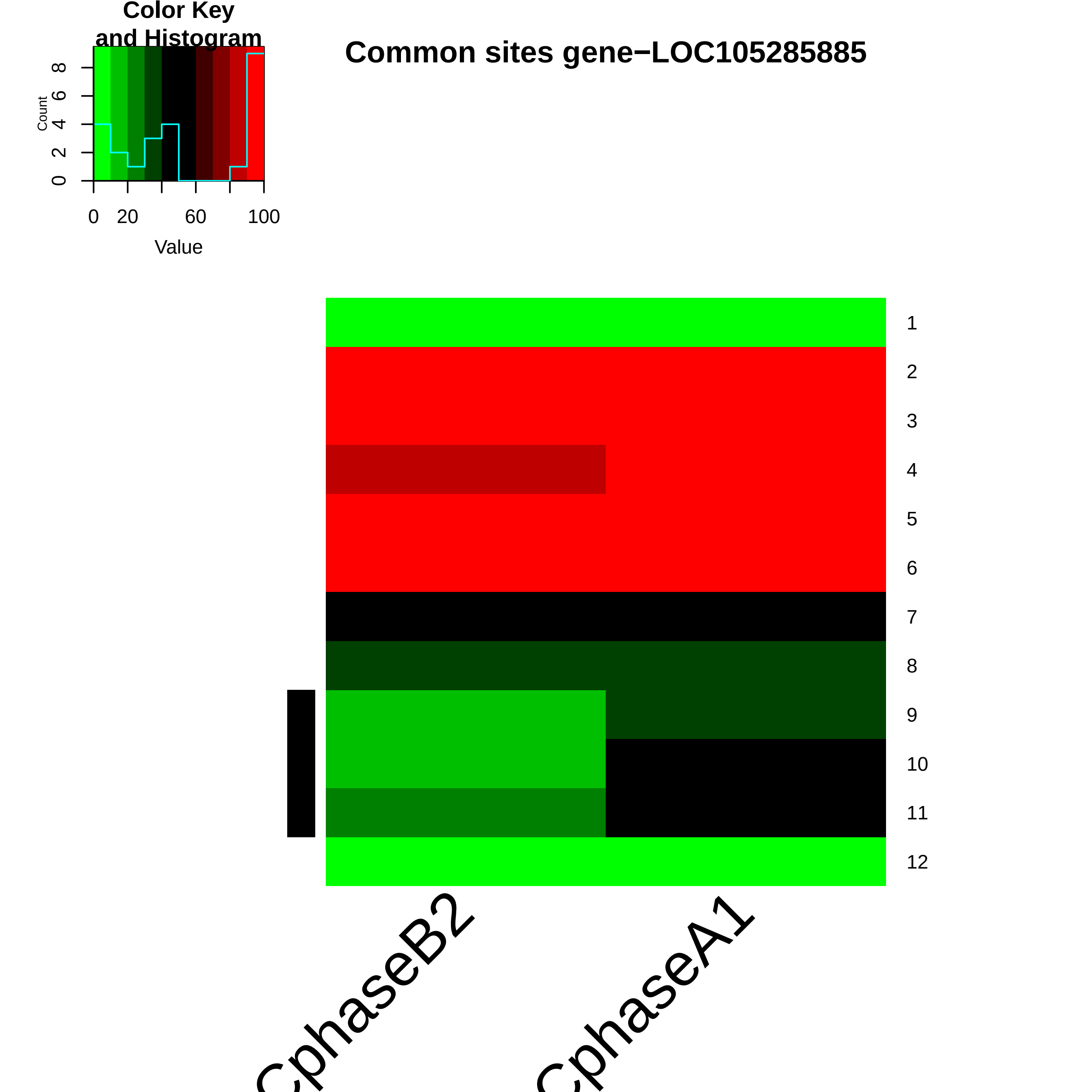

Color Key
and Histogram
Count
0
2
4
6
8
0
20
60
100
Value
Common sites gene−LOC105285885
1
2
3
4
5
6
7
8
9
10
11
12
CphaseB2
CphaseA1

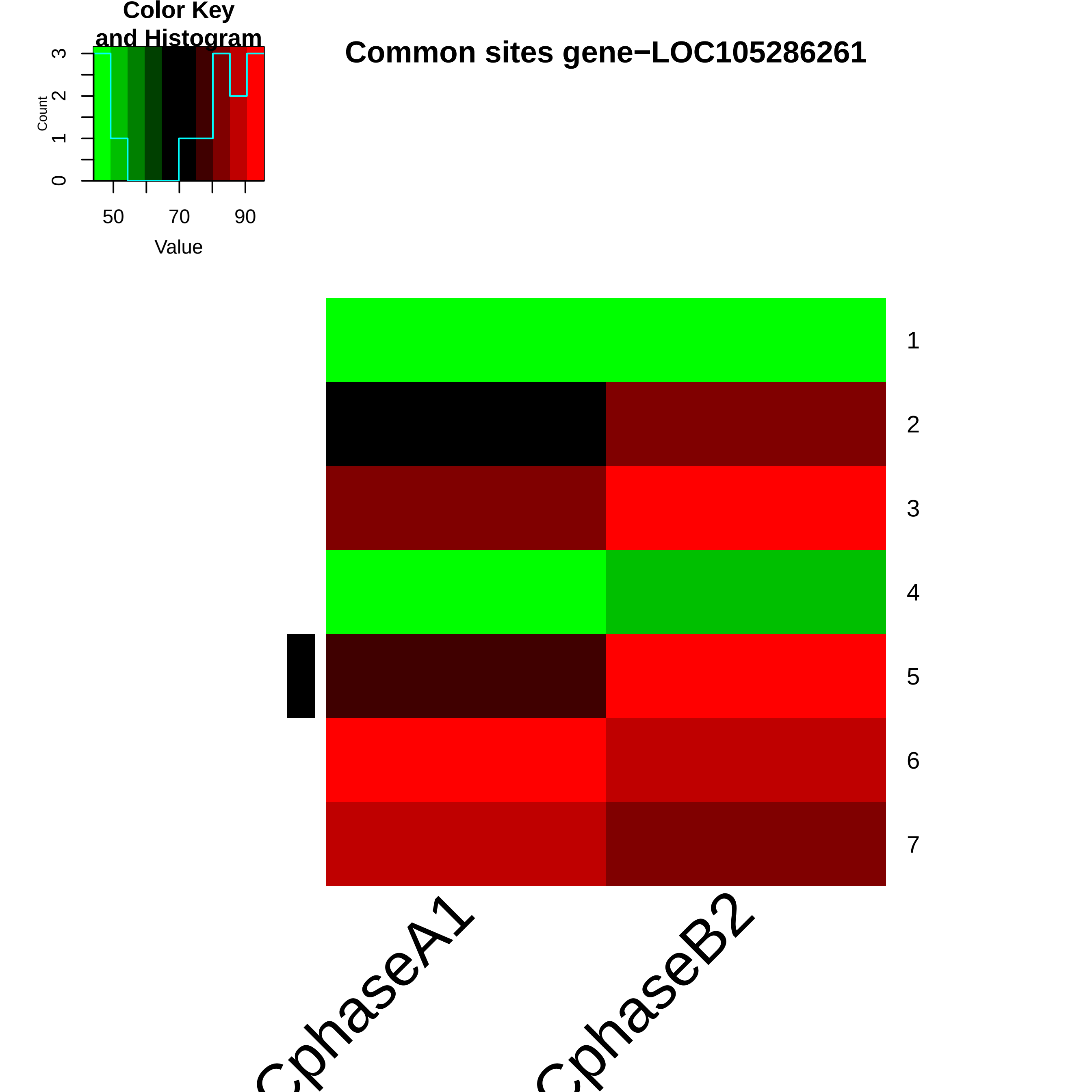

Color Key
and Histogram
Count
0
1
2
3
50
70
90
Value
Common sites gene−LOC105286261
1
2
3
4
5
6
7
CphaseA1
CphaseB2

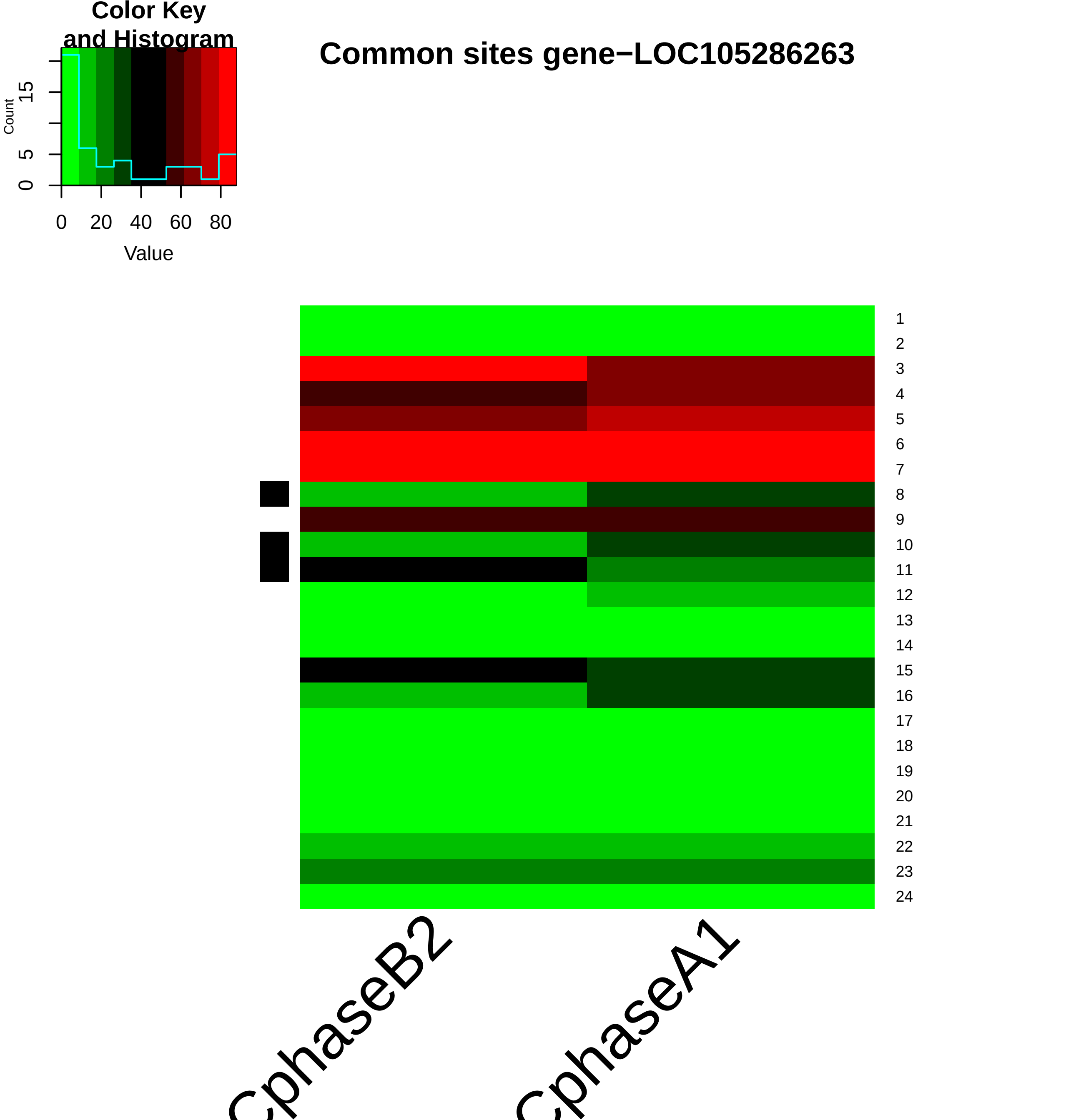
Color Key
and Histogram
Count
0
5
15
0
20
40
60
80
Value
Common sites gene−LOC105286263
CphaseB2
CphaseA1

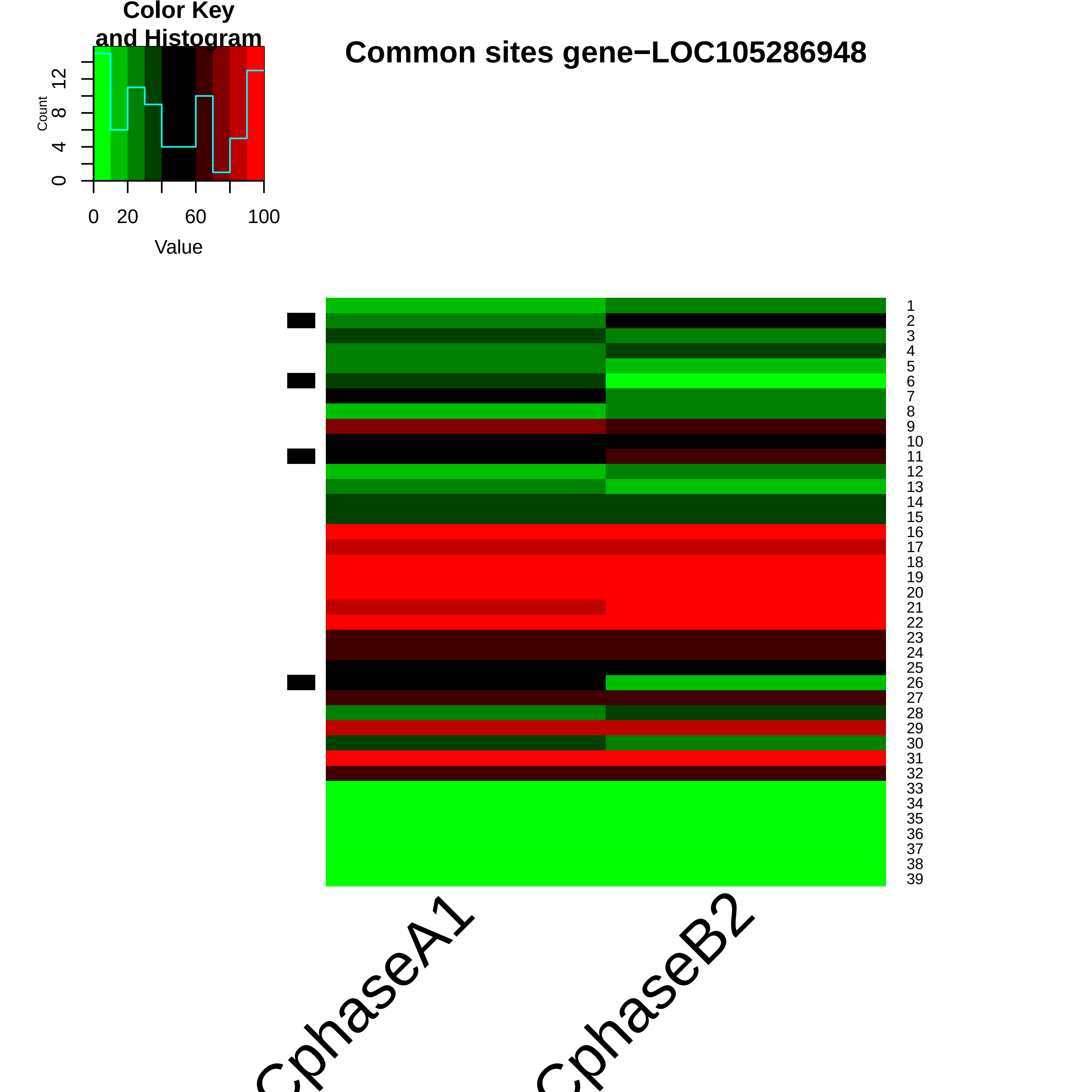

Color Key
and Histogram
Count
0
4
8
12
0
20
60
100
Value
Common sites gene−LOC105286948
CphaseA1
CphaseB2

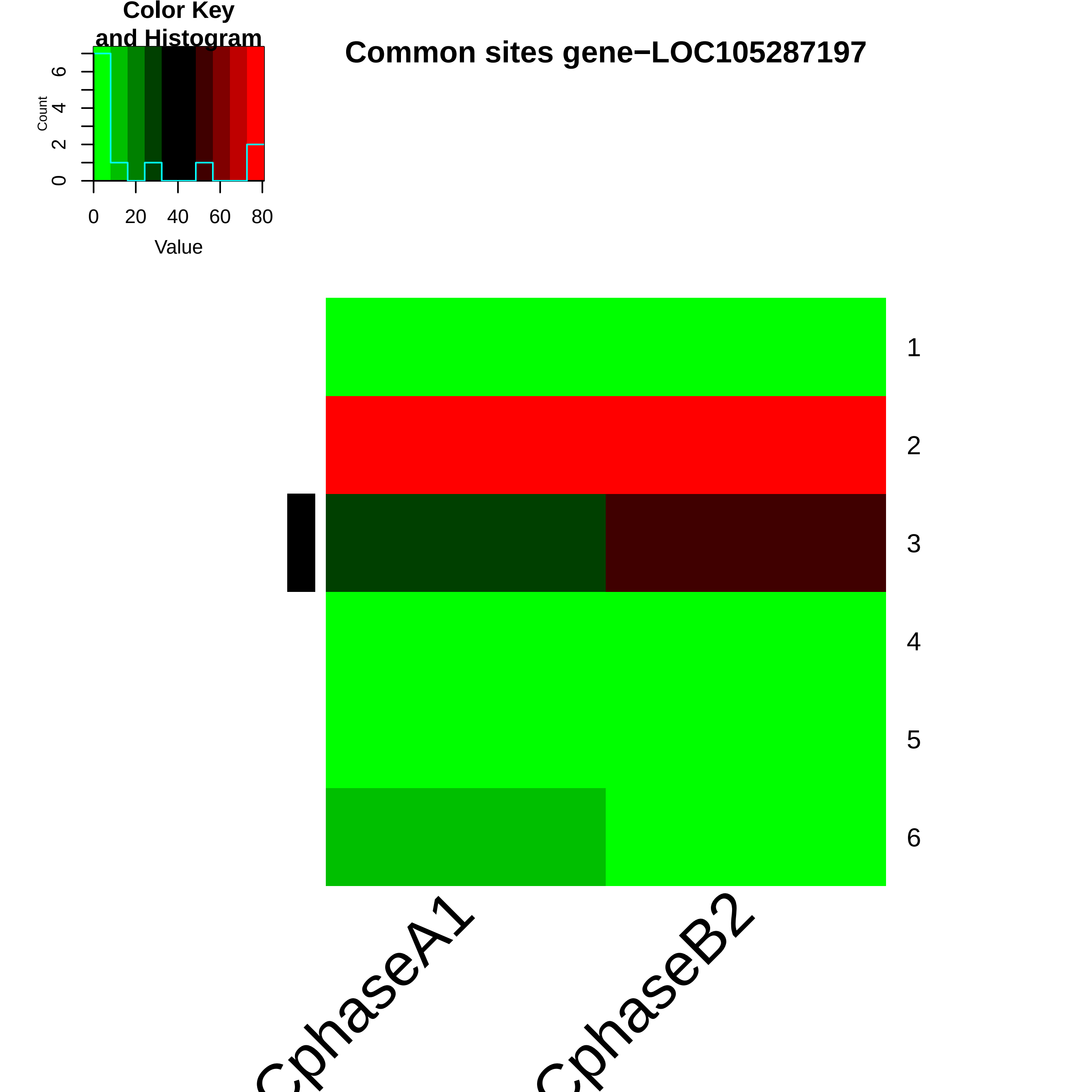
Color Key
and Histogram
Count
0
2
4
6
0
20
40
60
80
Value
Common sites gene−LOC105287197
1
2
3
4
5
6
CphaseA1
CphaseB2

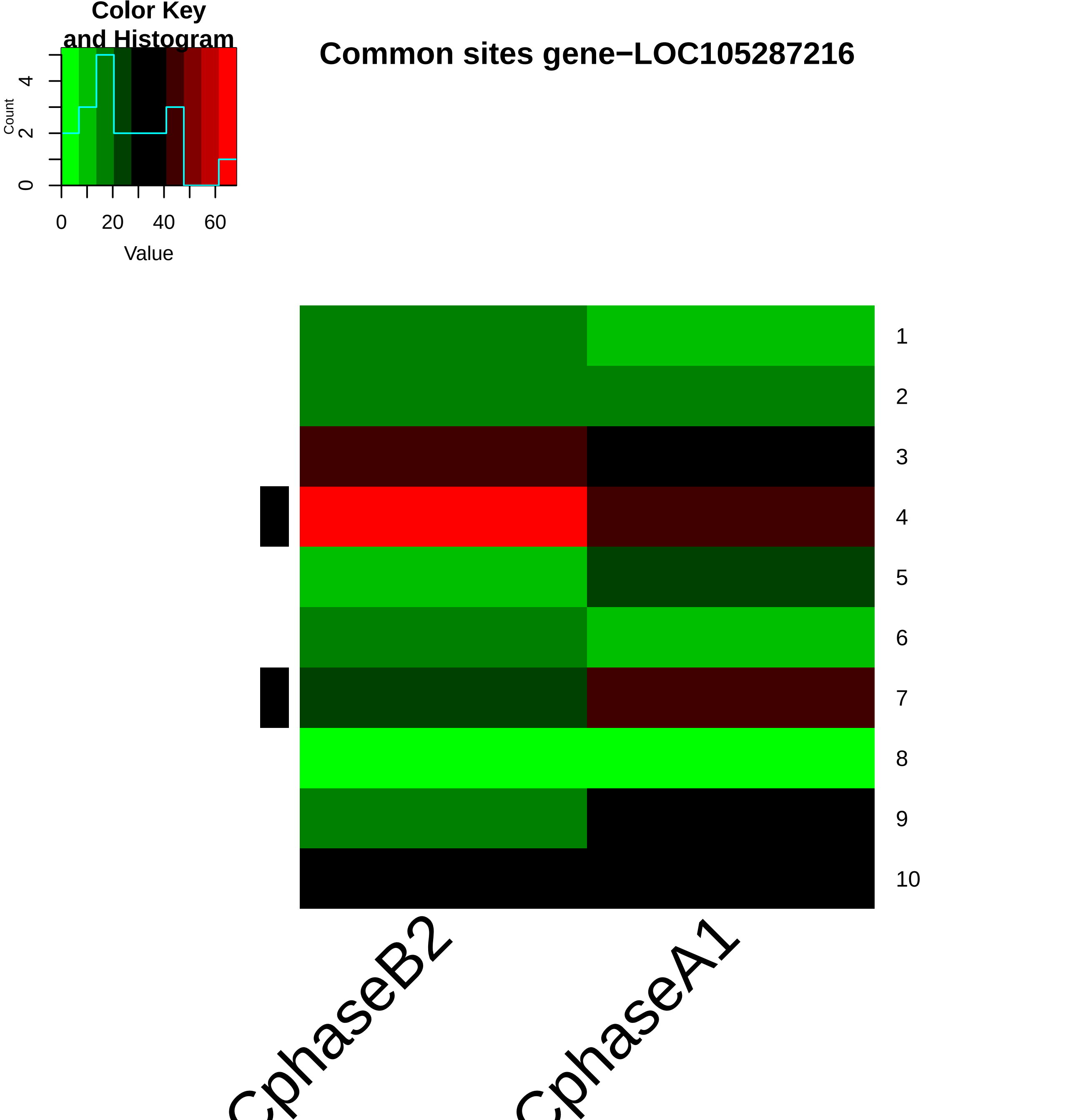

Color Key
and Histogram
Count
0
2
4
0
20
40
60
Value
Common sites gene−LOC105287216
1
2
3
4
5
6
7
8
9
10
CphaseB2
CphaseA1

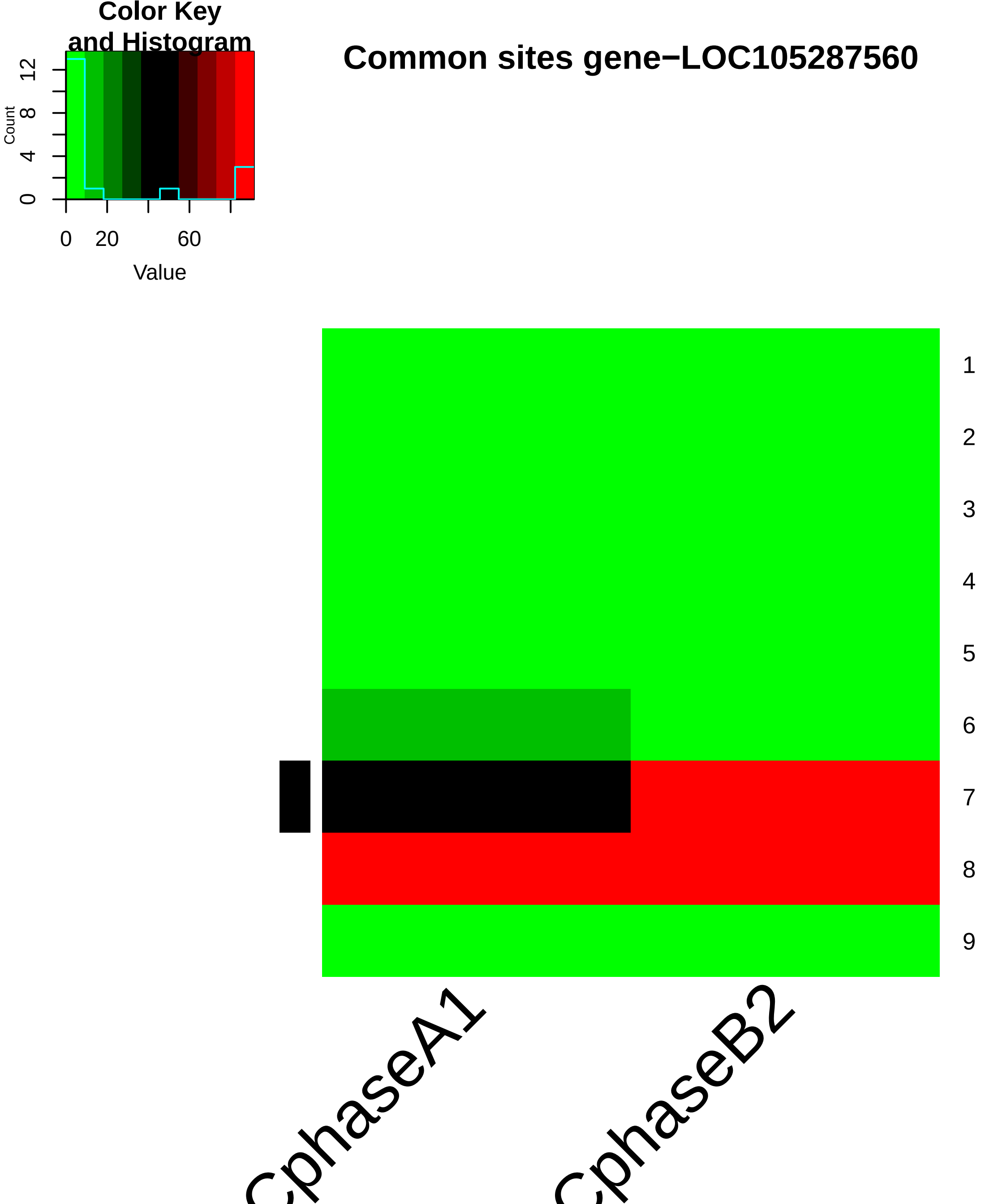
Color Key
and Histogram
Count
0
4
8
12
0
20
60
Value
Common sites gene−LOC105287560
1
2
3
4
5
6
7
8
9

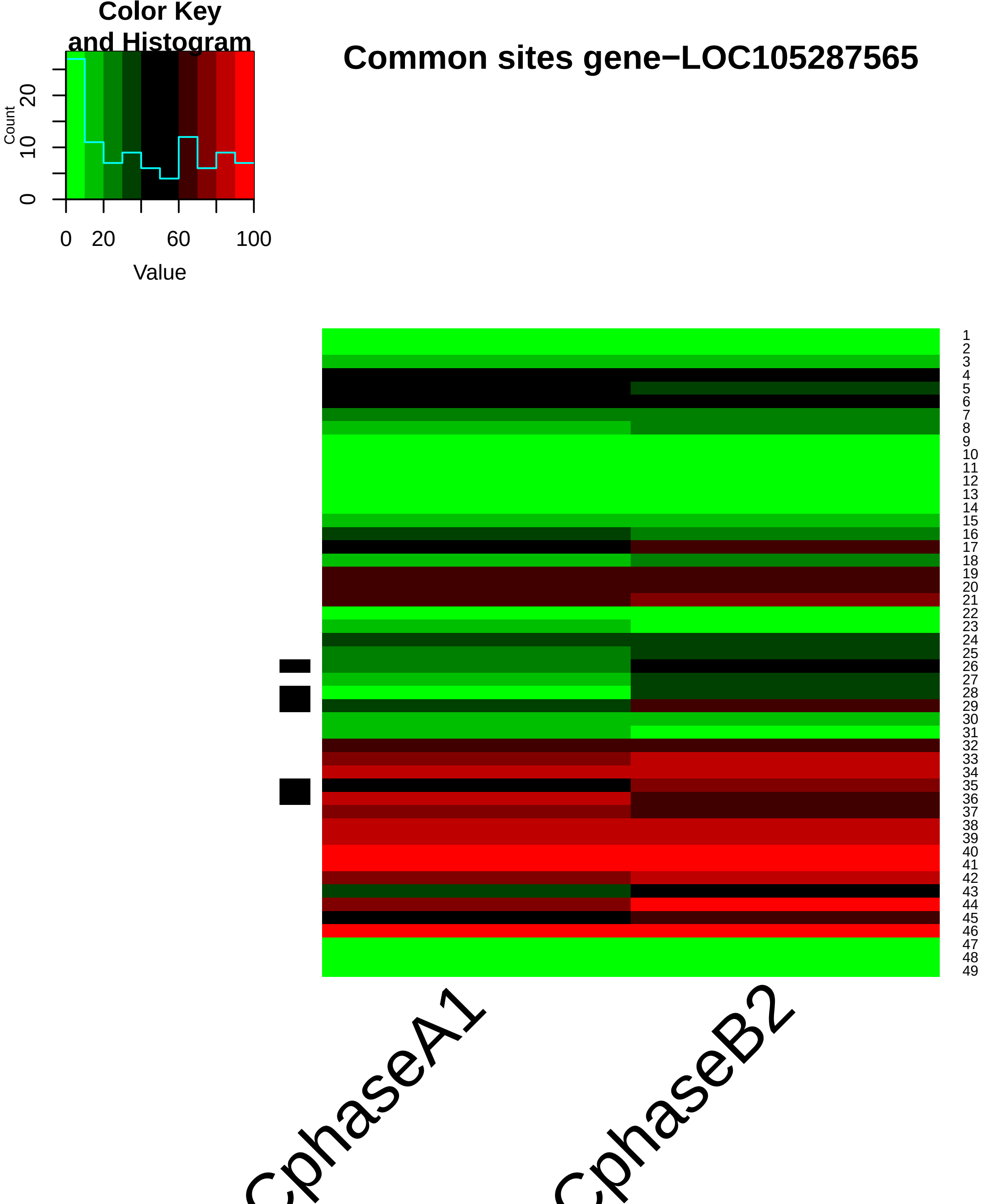
Color Key
and Histogram
Count
0
10
20
0
20
60
100
Value
Common sites gene−LOC105287565
1 2 3 4 5 6 7 8 9 10 11 12 13 14 15 16 17 18 19 20 21 22 23 24 25 26 27 28 29 30 31 32 33 34 35 36 37 38 39 40 41 42 43 44 45 46 47 48 49
CphaseA1
CphaseB2

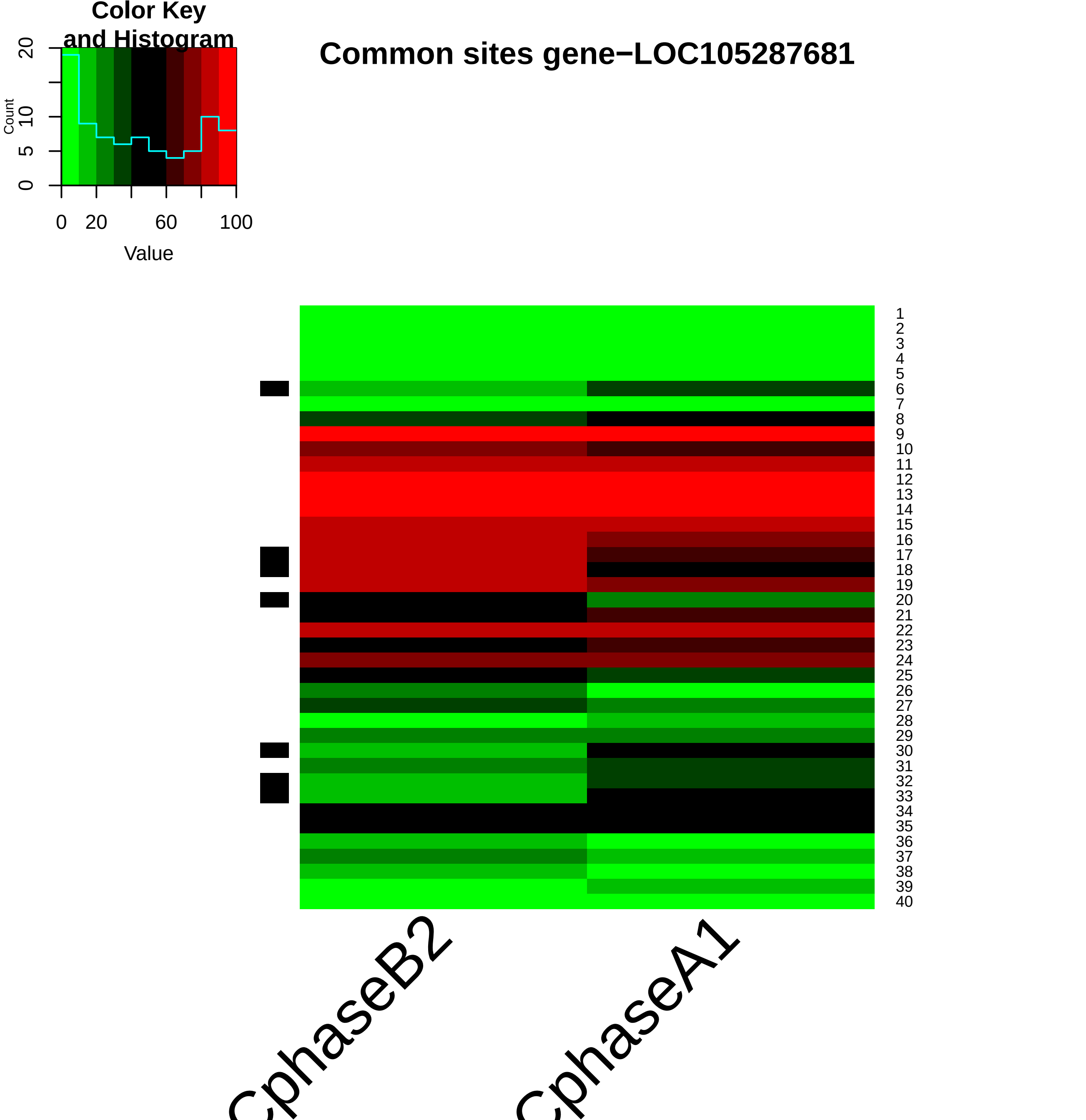

Color Key
and Histogram
Common sites gene−LOC105287681
Count
Value
0
20
60
100
0
5
10
20

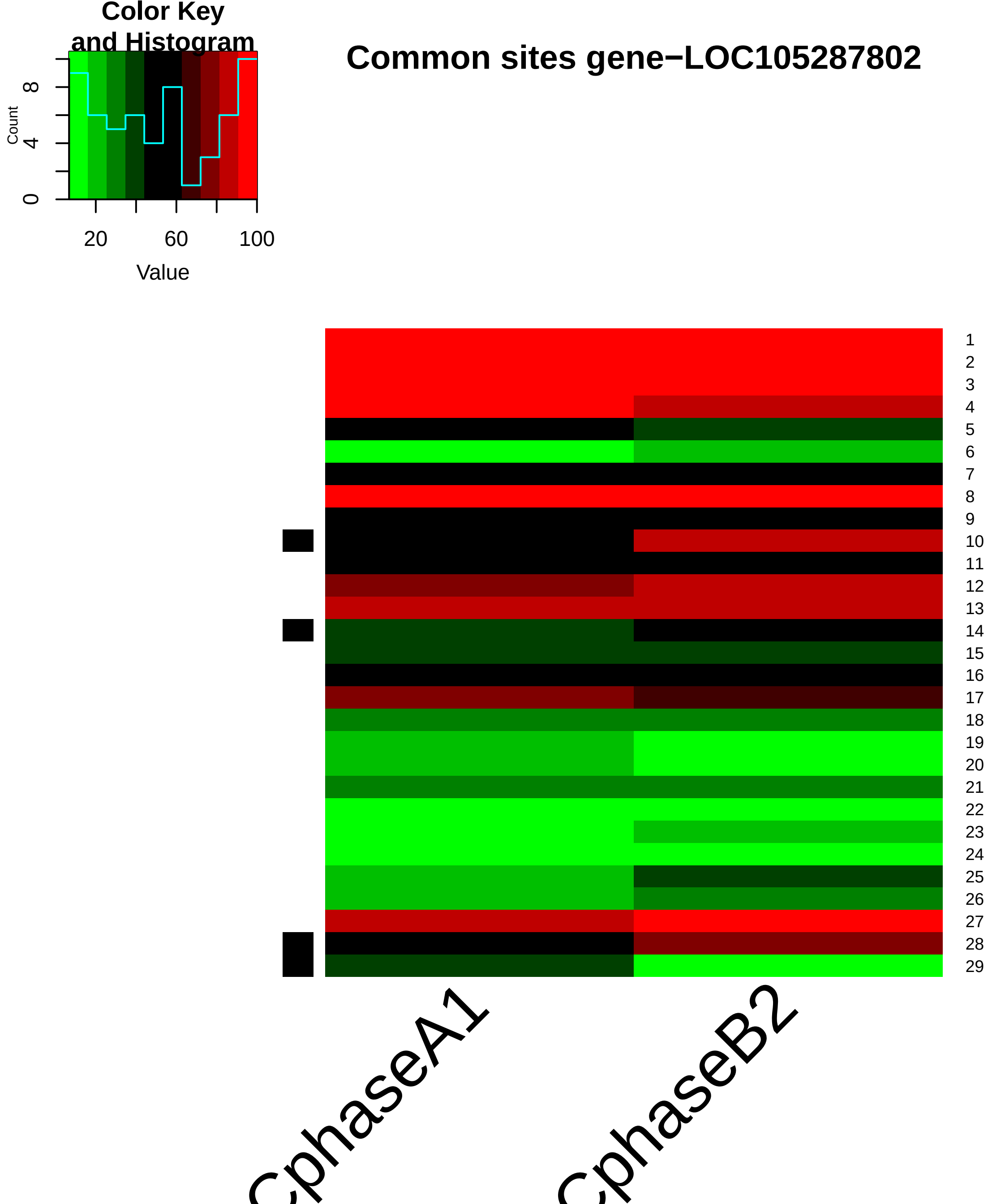
Color Key
and Histogram
Count
0
4
8
20
60
100
Value
Common sites gene−LOC105287802
CphaseA1
CphaseB2

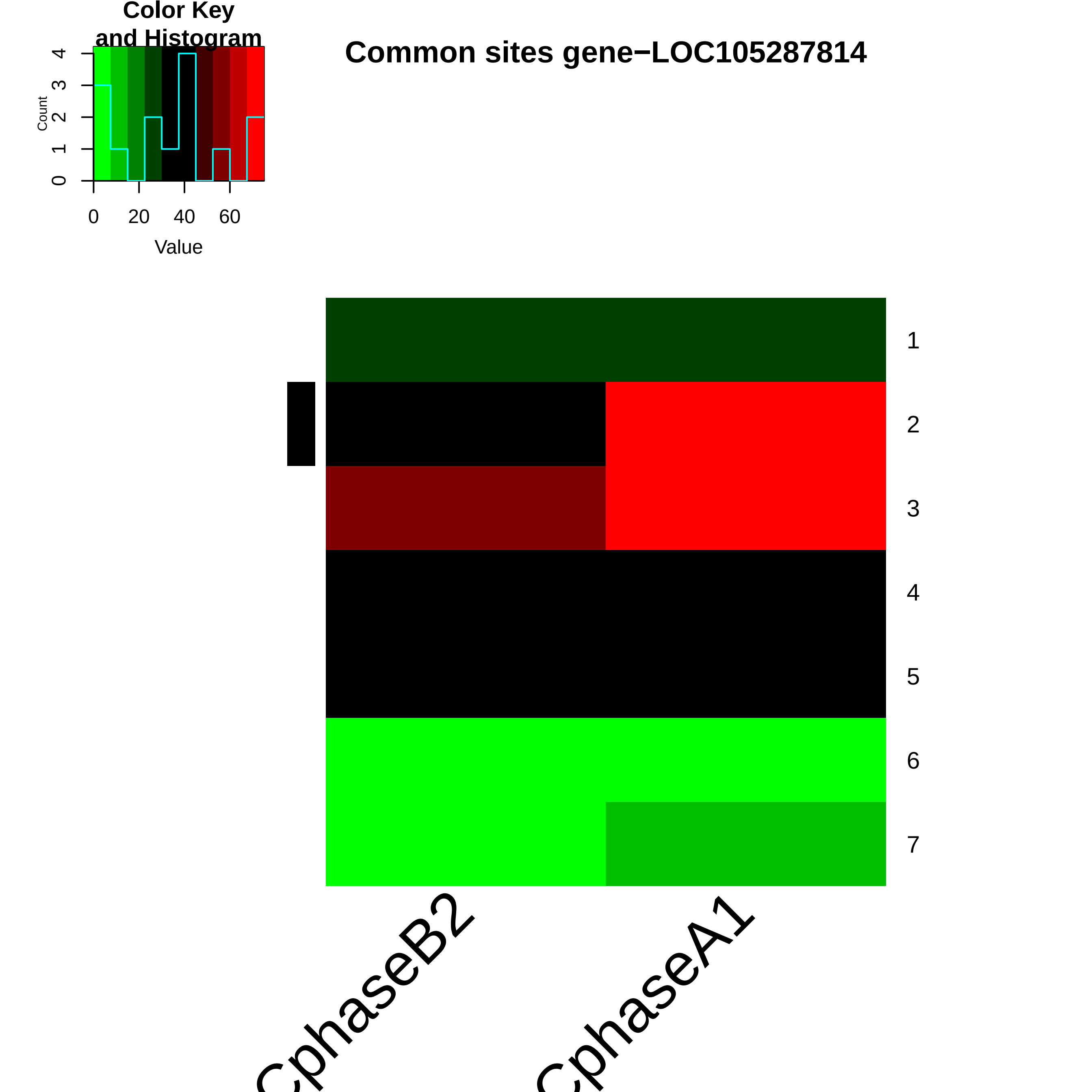
Color Key
and Histogram
Count
Value
0
20
40
60
0
1
2
3
4
Common sites gene−LOC105287814
1
2
3
4
5
6
7
CphaseB2
CphaseA1

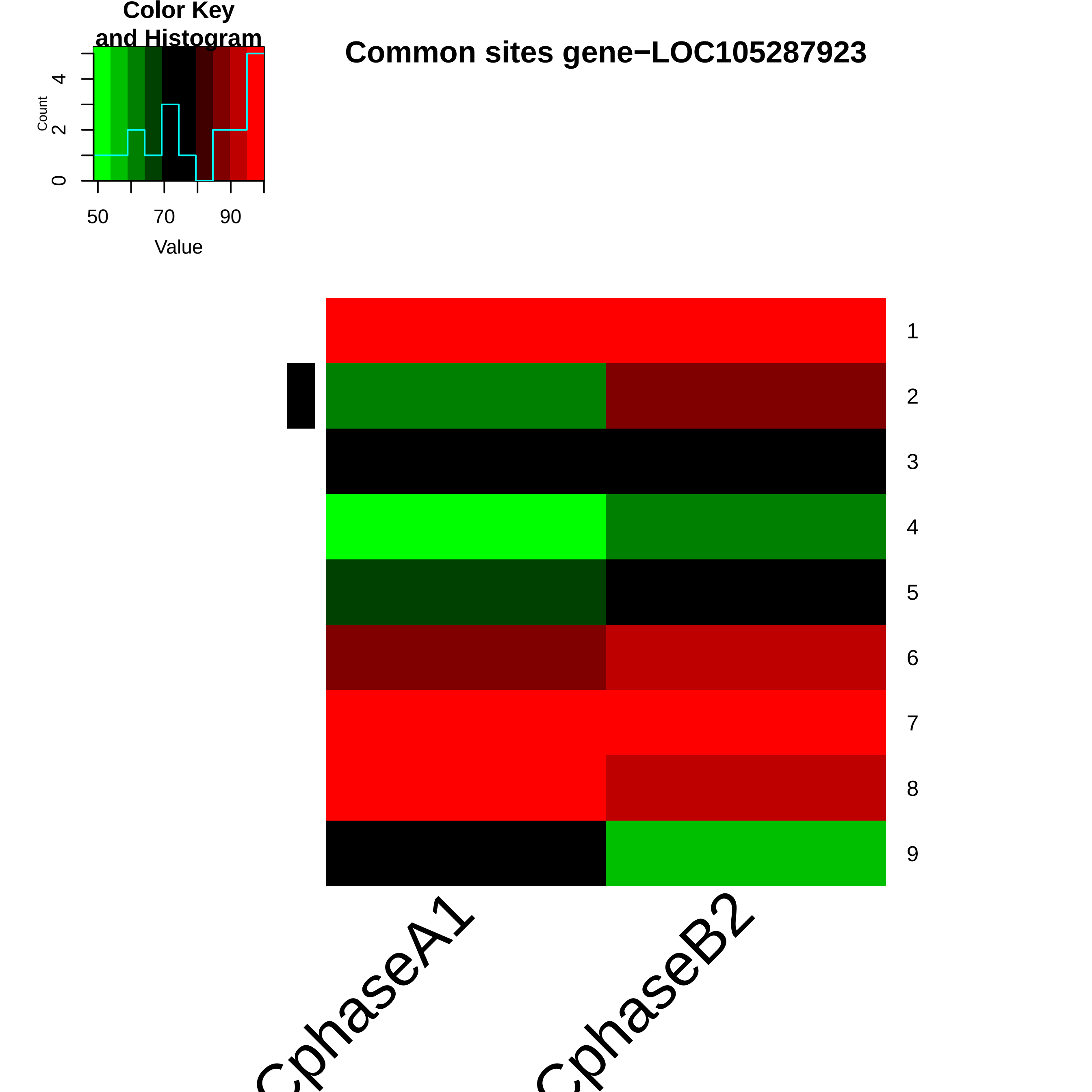
Color Key
and Histogram
Count
0
2
4
50
70
90
Value
Common sites gene−LOC105287923
1
2
3
4
5
6
7
8
9
CphaseA1
CphaseB2

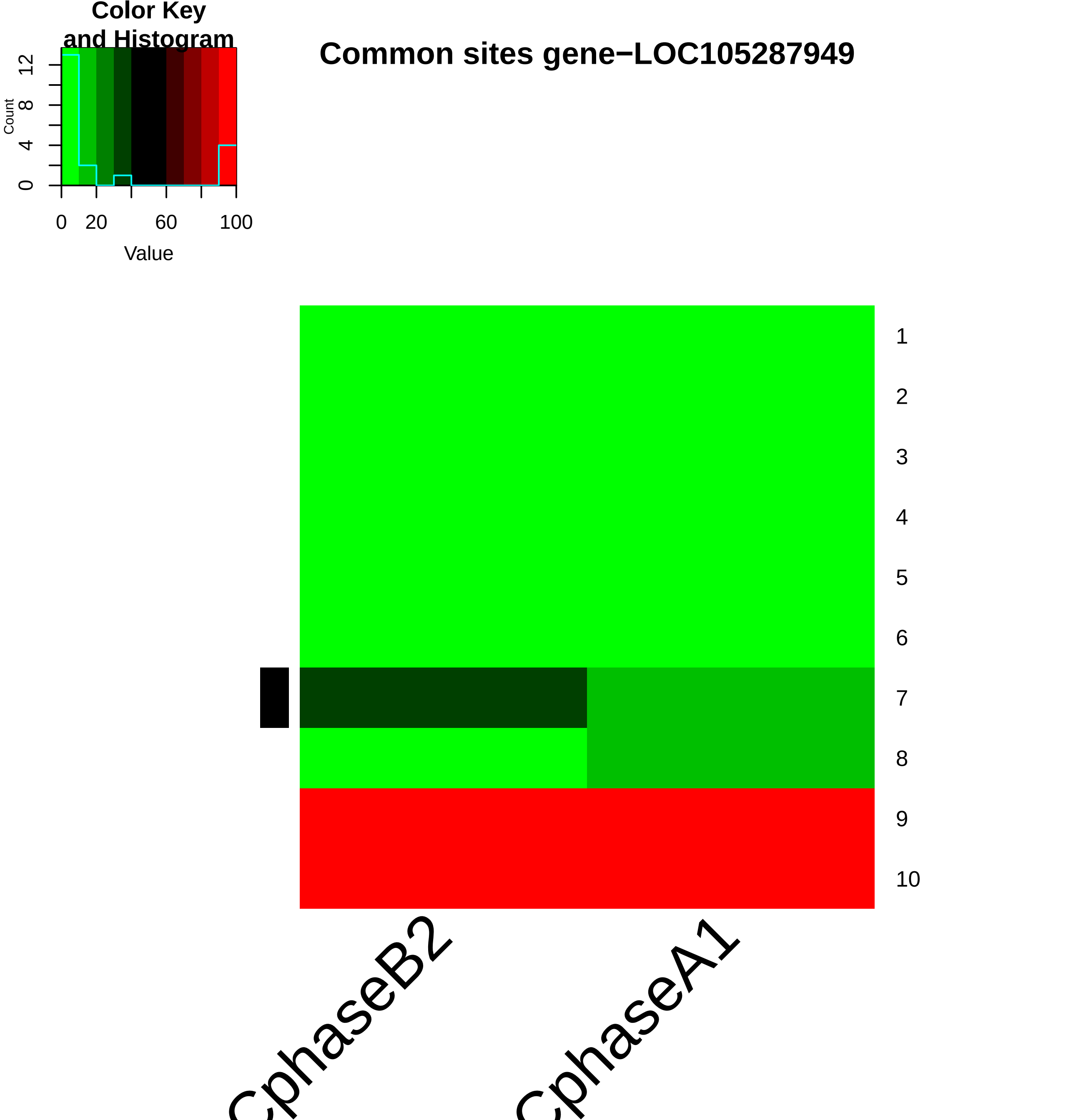

Color Key
and Histogram
Count
0
4
8
12
0
20
60
100
Value
Common sites gene−LOC105287949
1
2
3
4
5
6
7
8
9
10
CphaseB2
CphaseA1

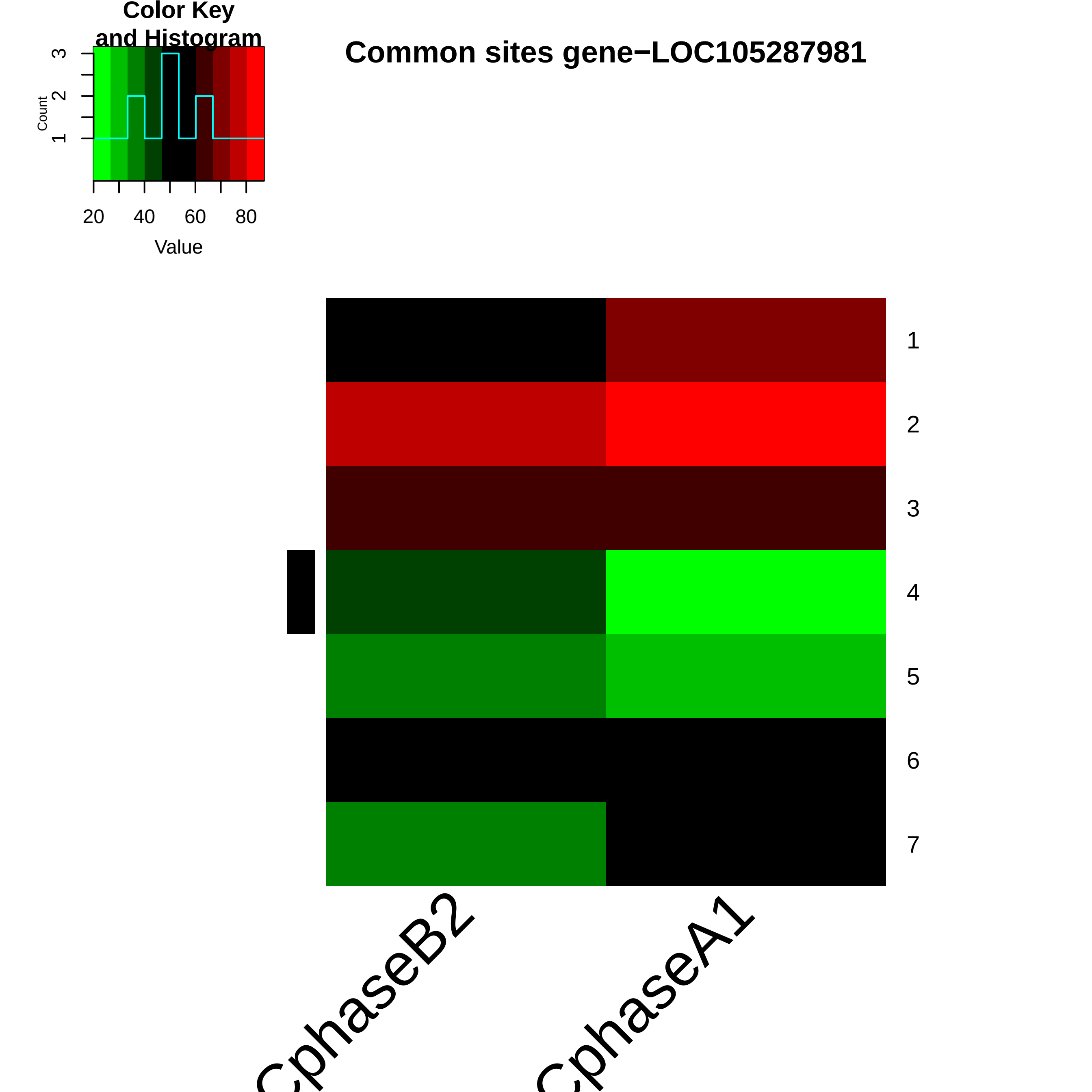

Color Key
and Histogram
Count
3
2
1
20
40
60
80
Value
Common sites gene−LOC105287981
1
2
3
4
5
6
7
CphaseB2
CphaseA1

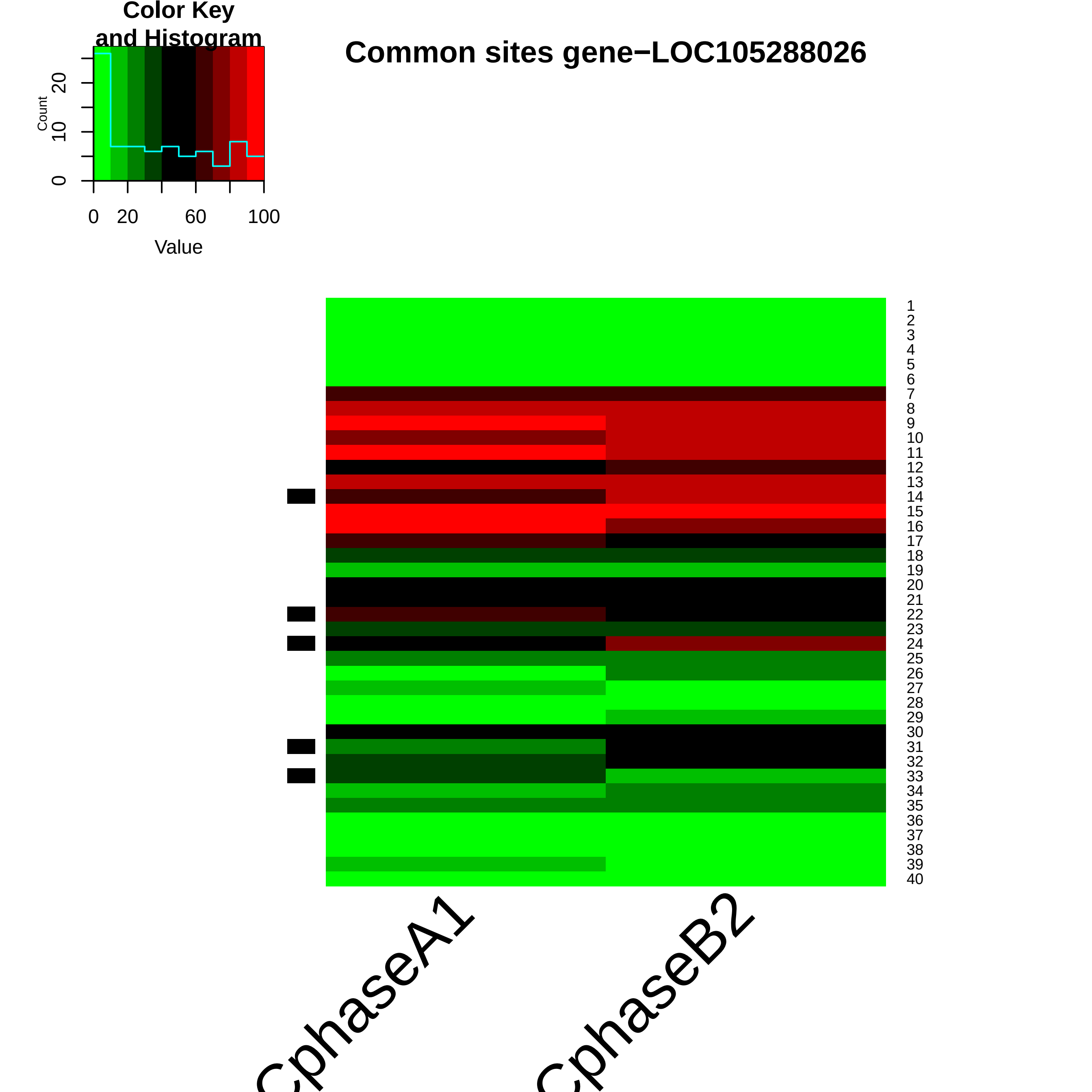

Color Key
and Histogram
Count
0
10
20
0
20
60
100
Value
Common sites gene−LOC105288026
CphaseA1
CphaseB2

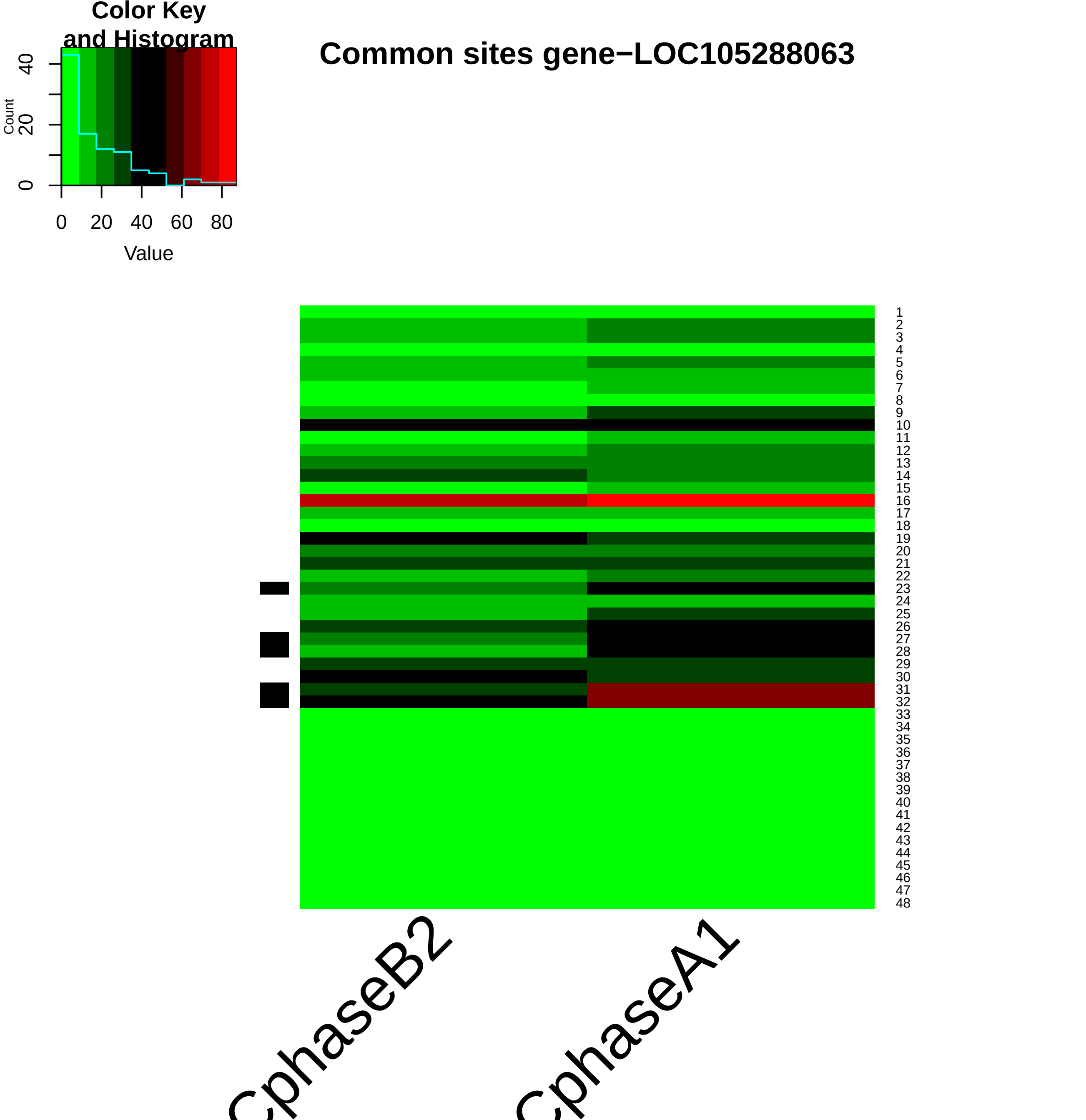
Color Key
and Histogram
Count
0
20
40
0
20
40
60
80
Value
Common sites gene−LOC105288063
1 2 3 4 5 6 7 8 9 10 11 12 13 14 15 16 17 18 19 20 21 22 23 24 25 26 27 28 29 30 31 32 33 34 35 36 37 38 39 40 41 42 43 44 45 46 47 48
CphaseB2
CphaseA1

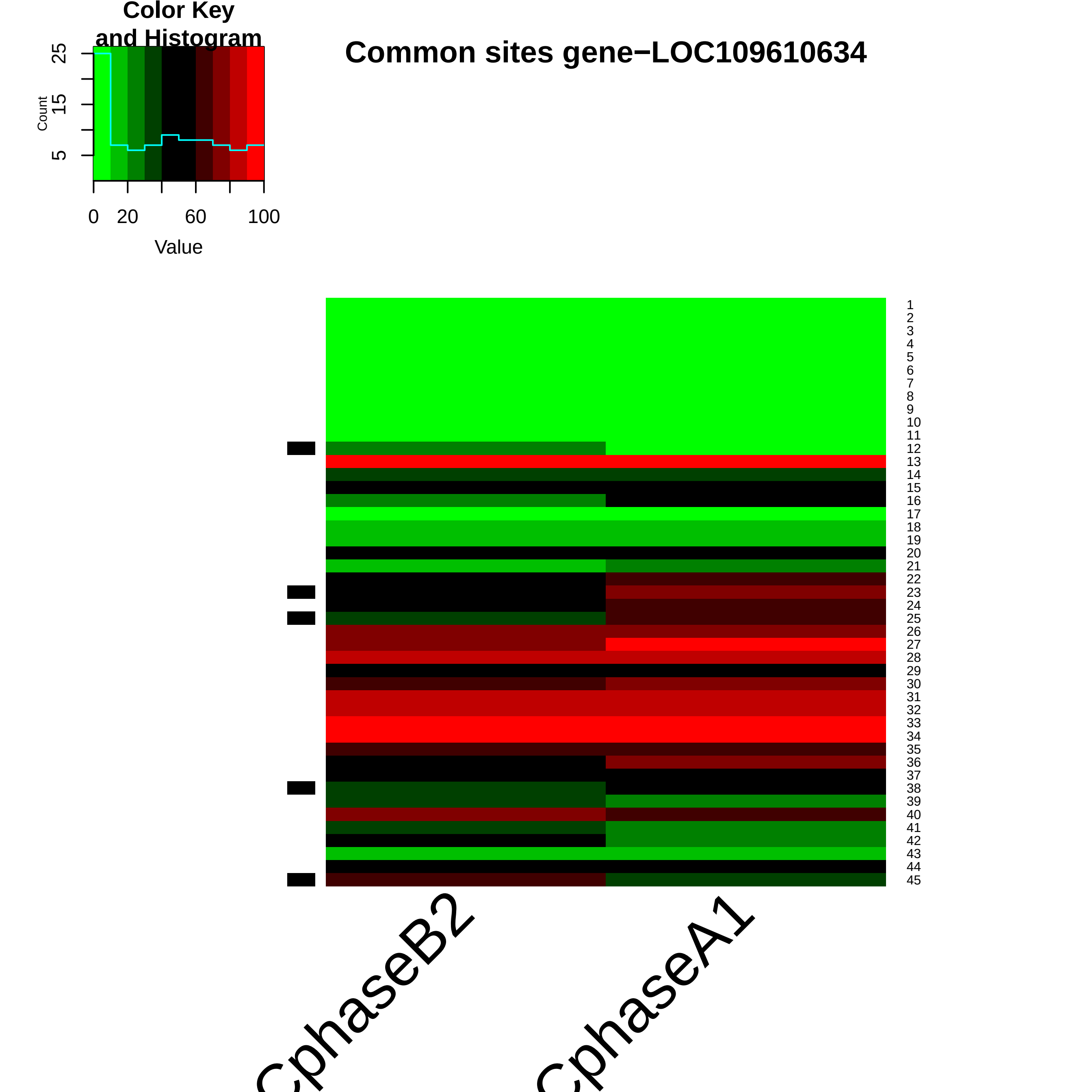

Common sites gene−LOC109610634
Color Key and Histogram
Count
25
15
5
0
20
60
100
Value
CphaseB2
CphaseA1

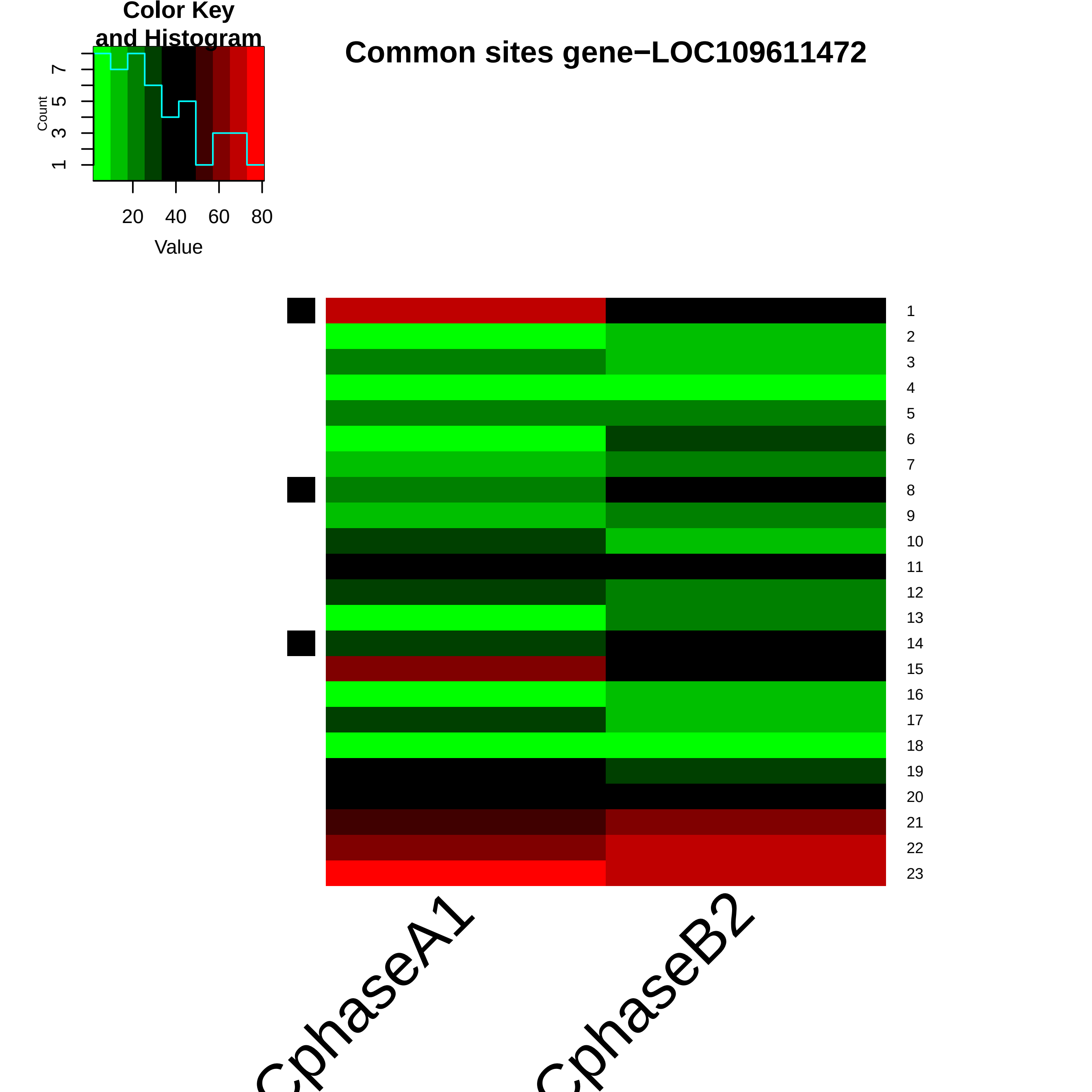
Color Key
and Histogram
Count
Value
Common sites gene−LOC109611472
CphaseA1
CphaseB2

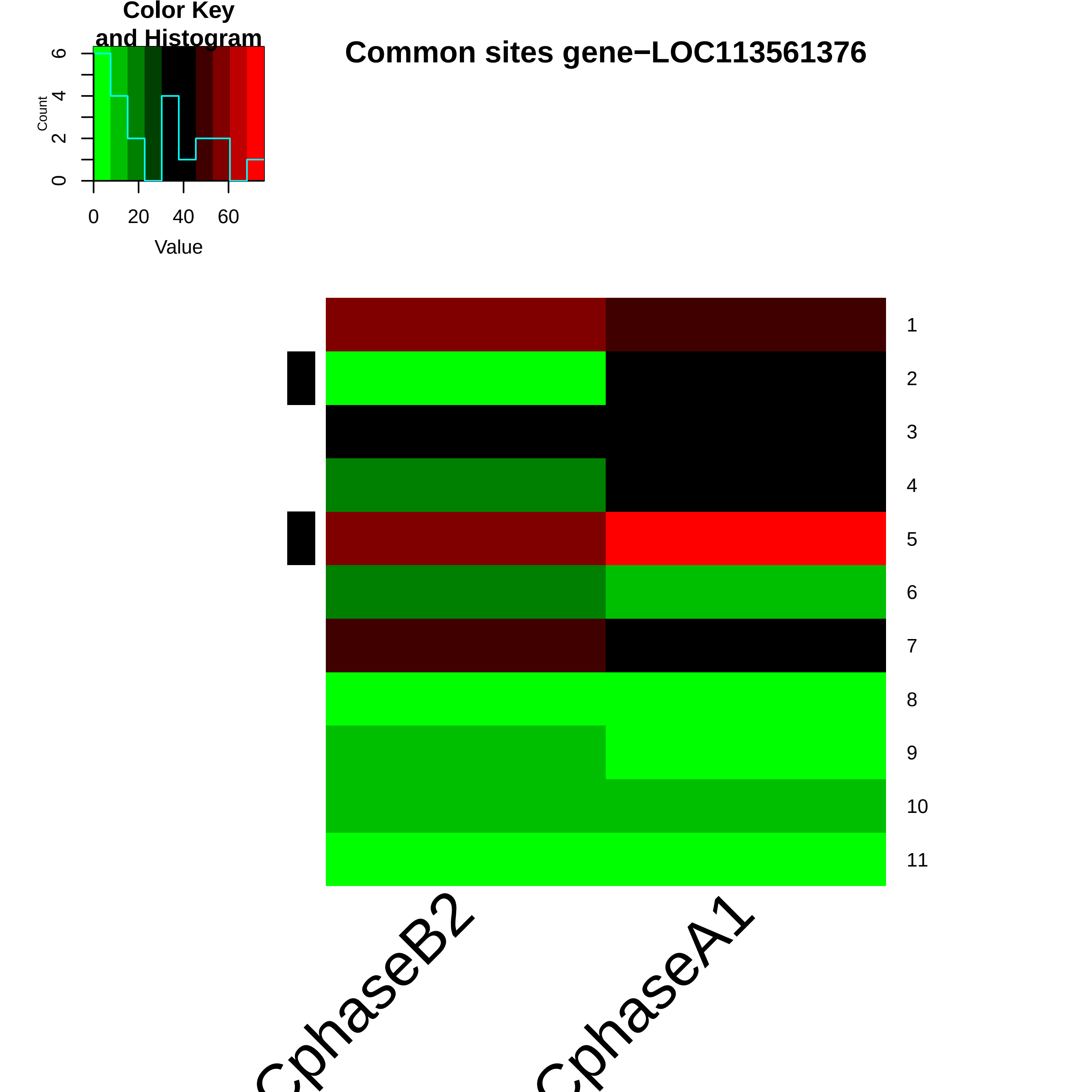

Color Key
and Histogram
Count
0
2
4
6
0
20
40
60
Value
Common sites gene−LOC113561376
1
2
3
4
5
6
7
8
9
10
11
CphaseB2
CphaseA1

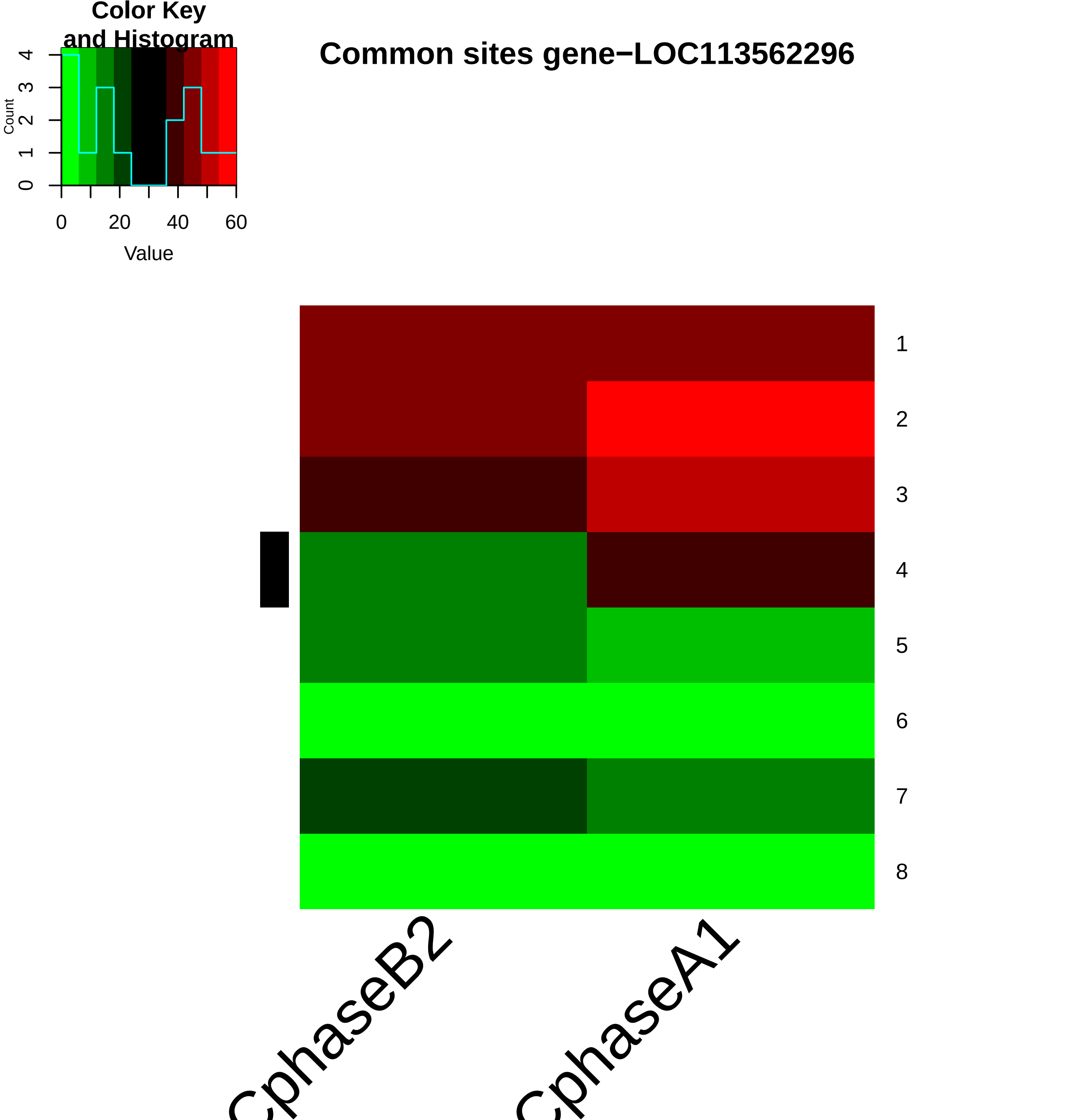
Color Key
and Histogram
Count
Value
0
20
40
60
0
1
2
3
4
Common sites gene−LOC113562296
1
2
3
4
5
6
7
8
CphaseB2
CphaseA1

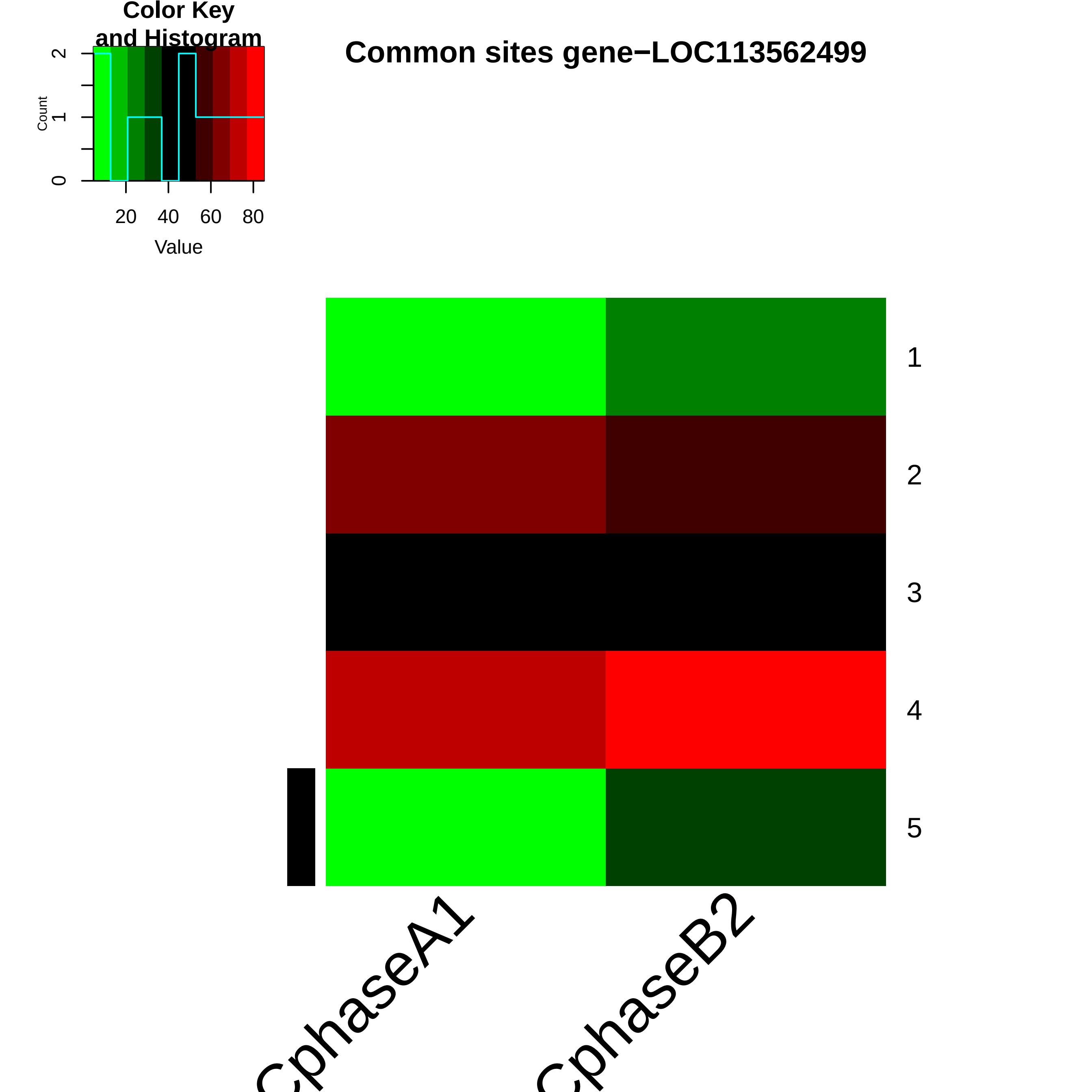

Color Key
and Histogram
Count
0
1
2
20
40
60
80
Value
Common sites gene−LOC113562499
1
2
3
4
5
CphaseA1
CphaseB2

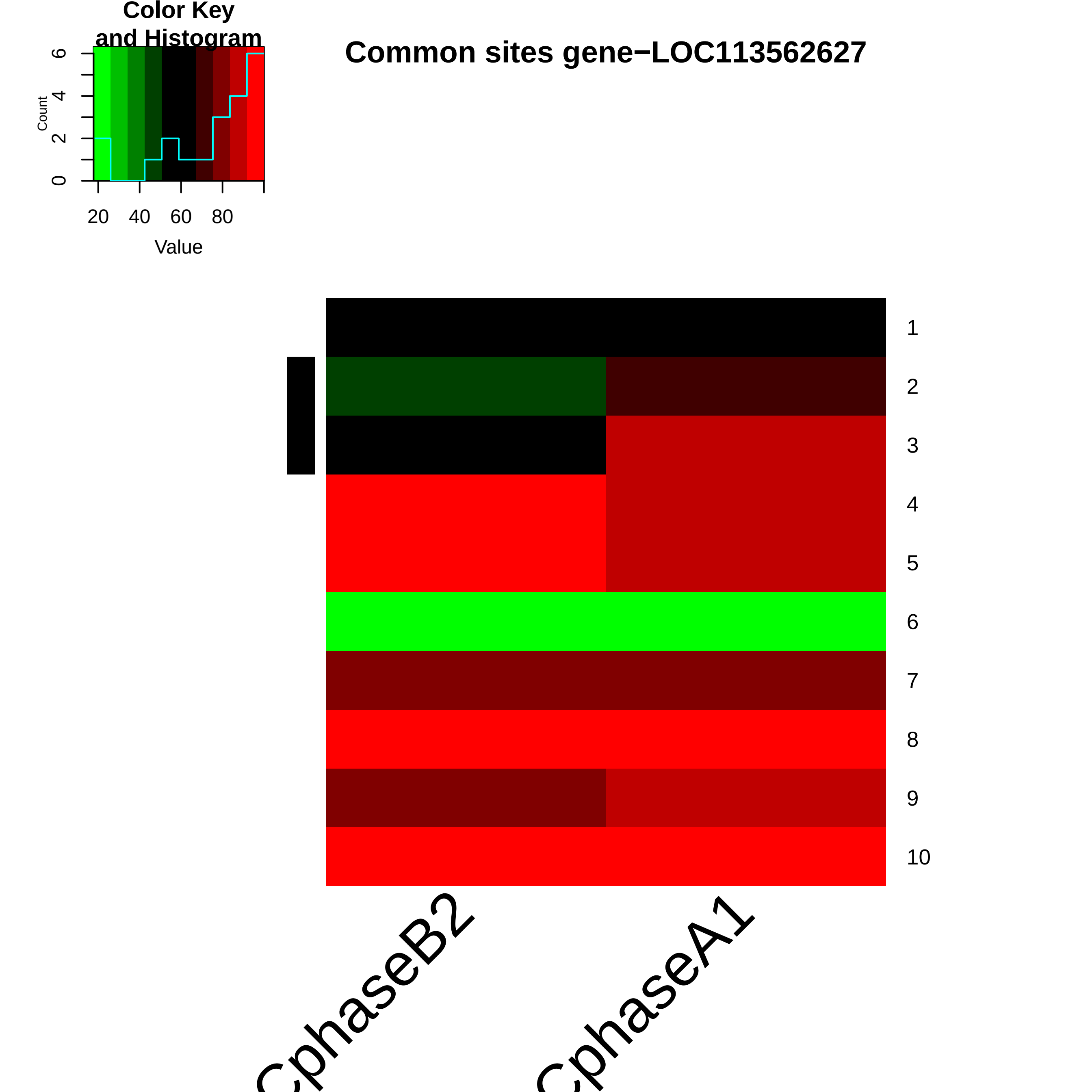
Color Key
and Histogram
Count
0
2
4
6
20
40
60
80
Value
Common sites gene−LOC113562627
1
2
3
4
5
6
7
8
9
10
CphaseB2
CphaseA1

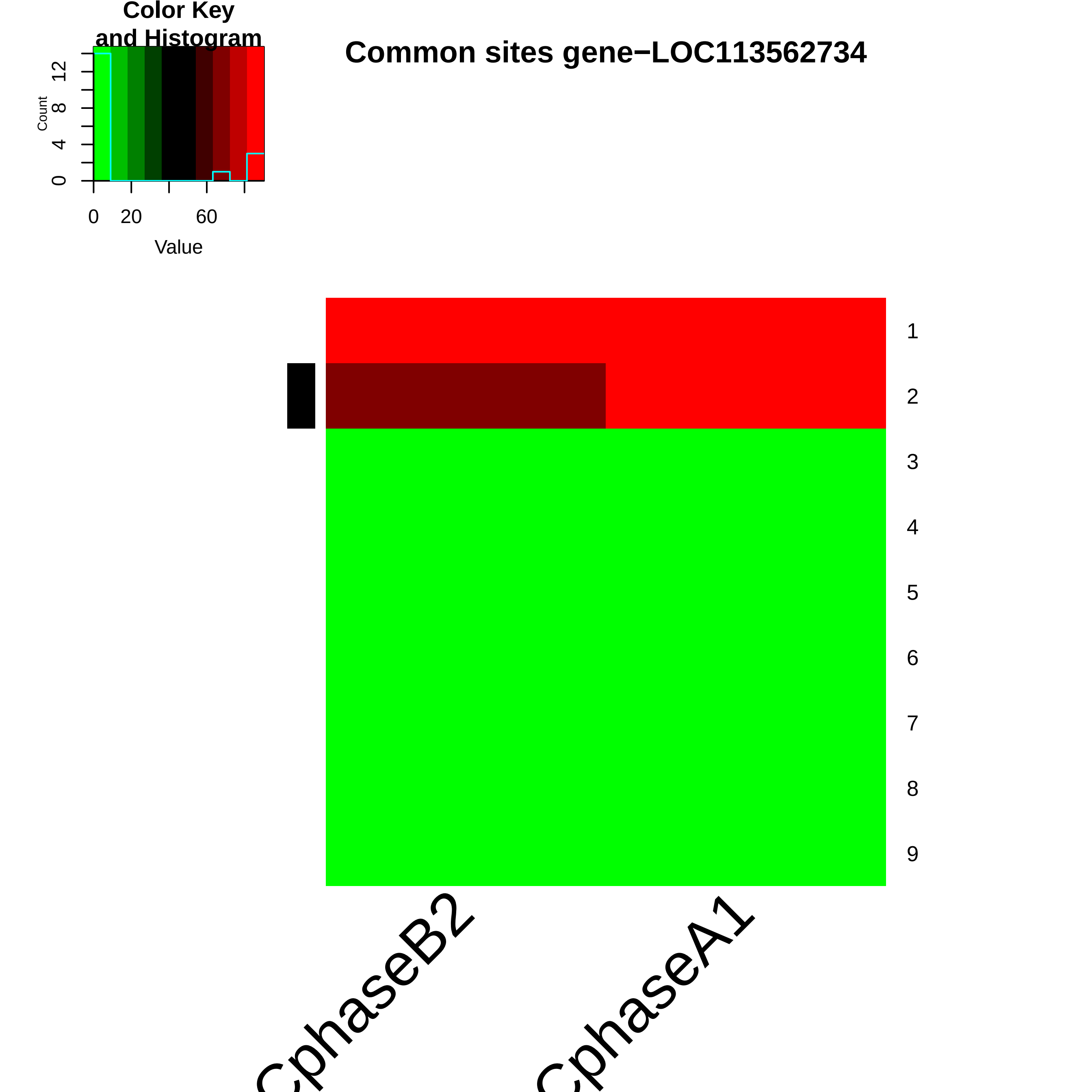

Color Key
and Histogram
Count
0
4
8
12
0
20
60
Value
Common sites gene−LOC113562734
1
2
3
4
5
6
7
8
9
CphaseB2
CphaseA1

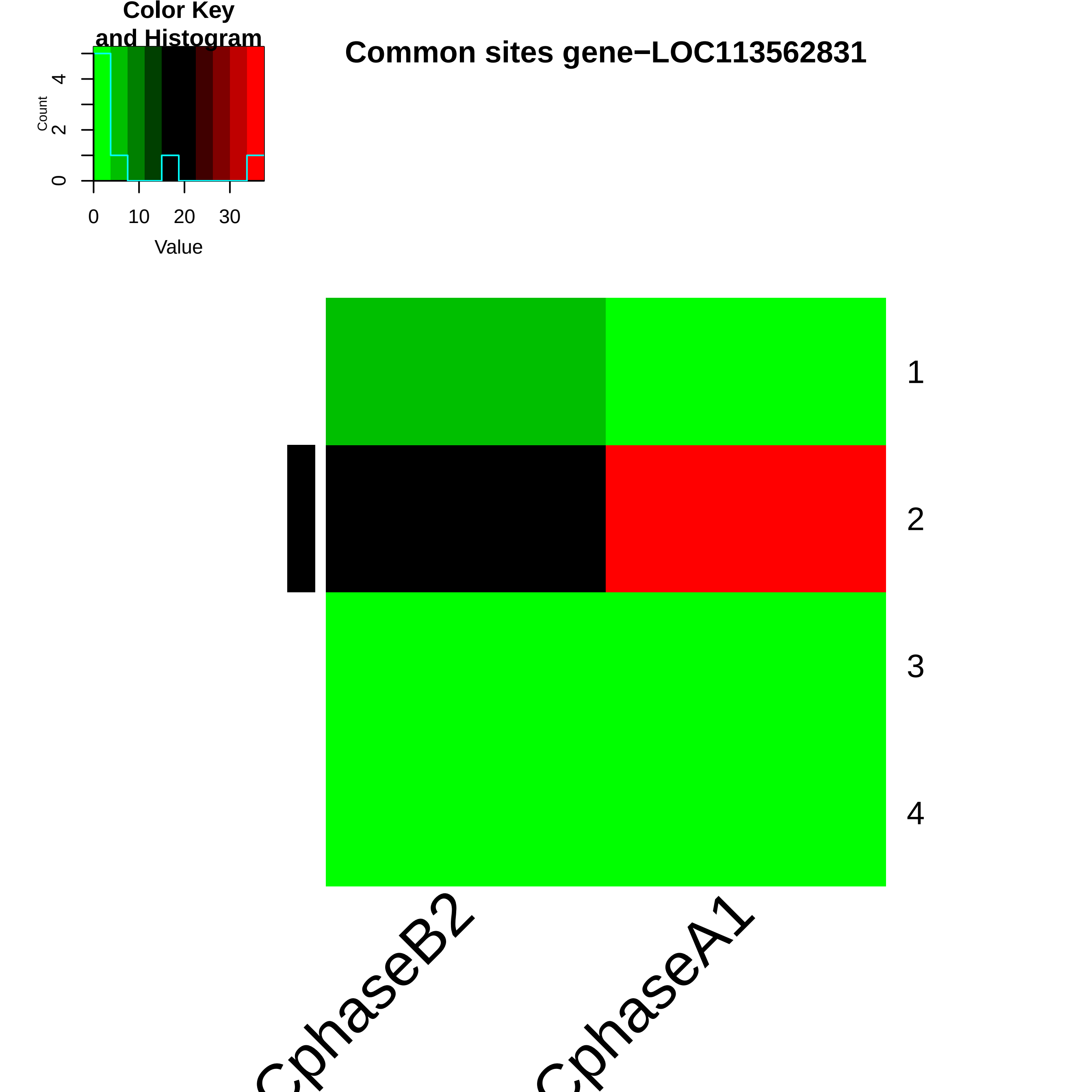

Color Key
and Histogram
Count
0
2
4
0
10
20
30
Value
Common sites gene−LOC113562831
1
2
3
4
CphaseB2
CphaseA1

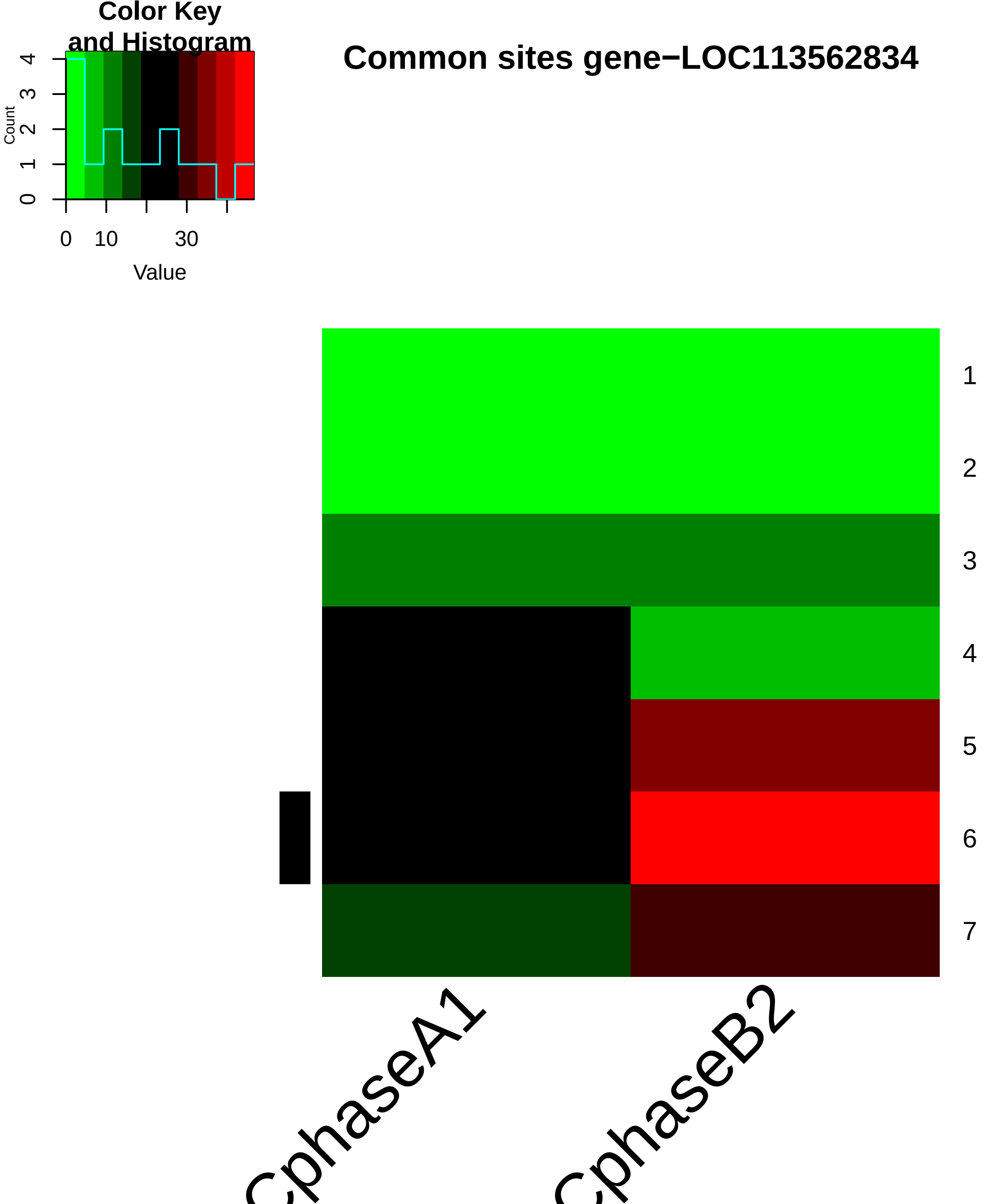
Color Key
and Histogram
Count
0
1
2
3
4
0
10
30
Value
Common sites gene−LOC113562834
1
2
3
4
5
6
7
CphaseA1
CphaseB2

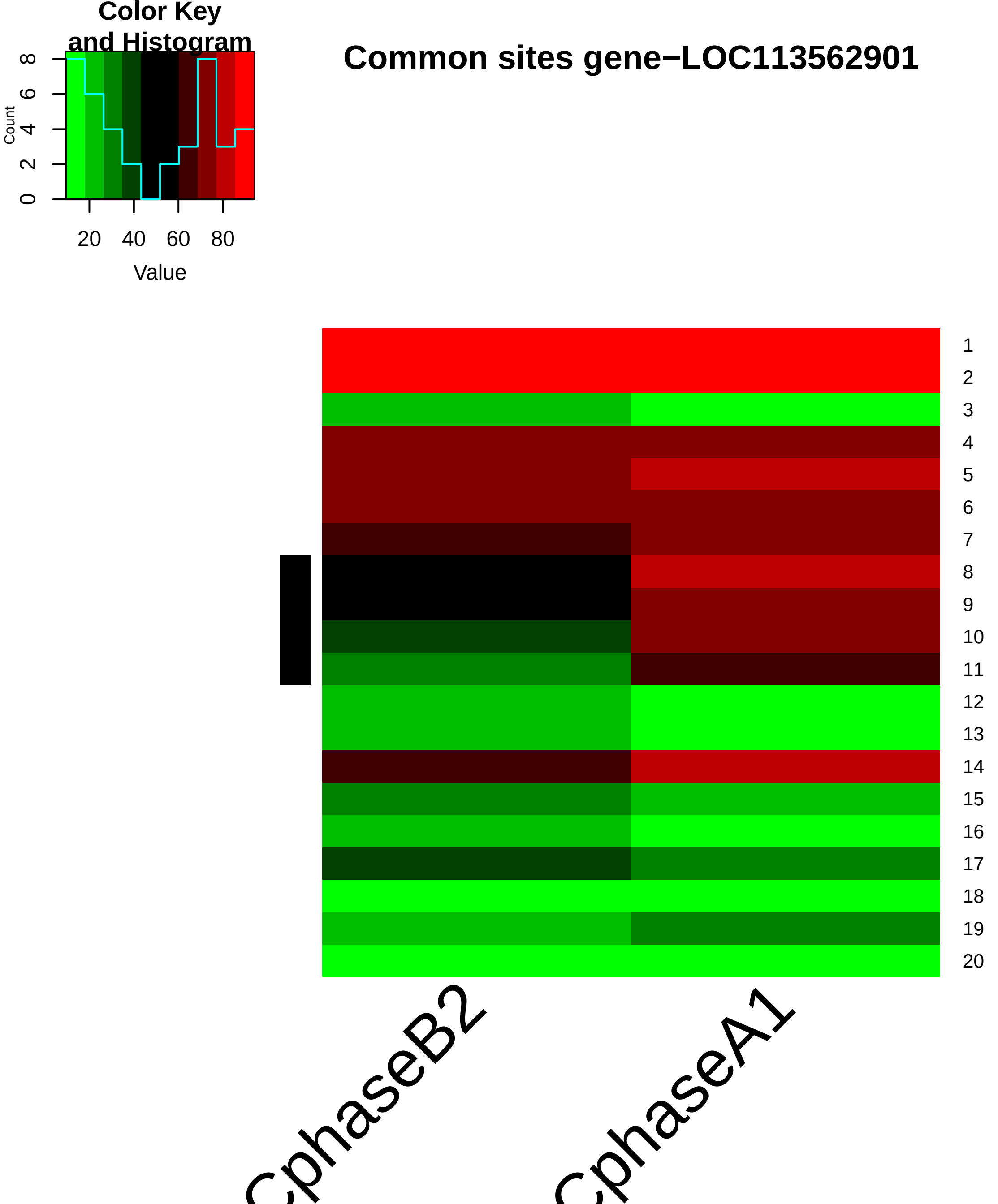

Color Key
and Histogram
Count
Value
0
2
4
6
8
20
40
60
80
Common sites gene−LOC113562901
CphaseB2
CphaseA1

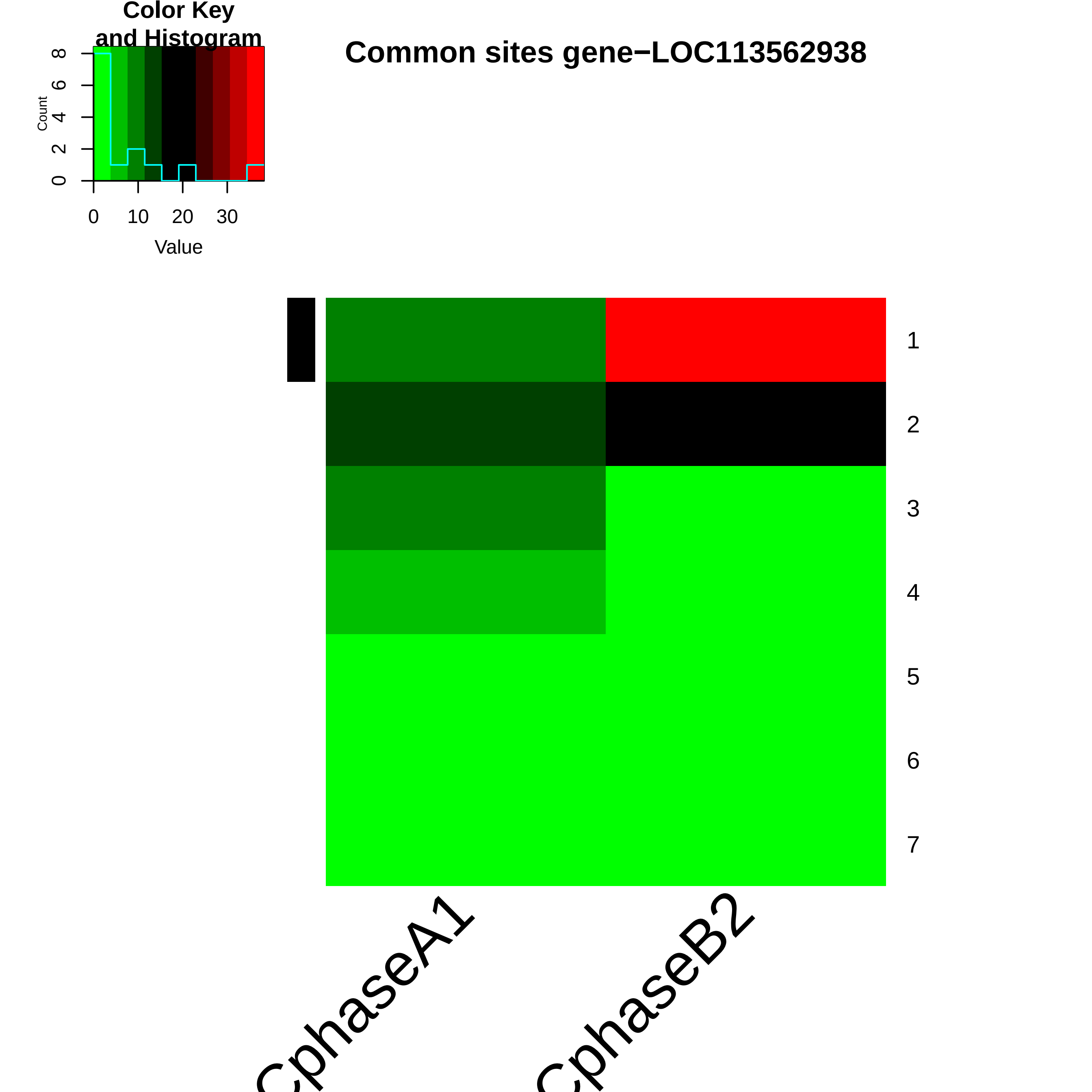

Color Key
and Histogram
Count
0
2
4
6
8
0
10
20
30
Value
Common sites gene−LOC113562938
1
2
3
4
5
6
7
CphaseA1
CphaseB2

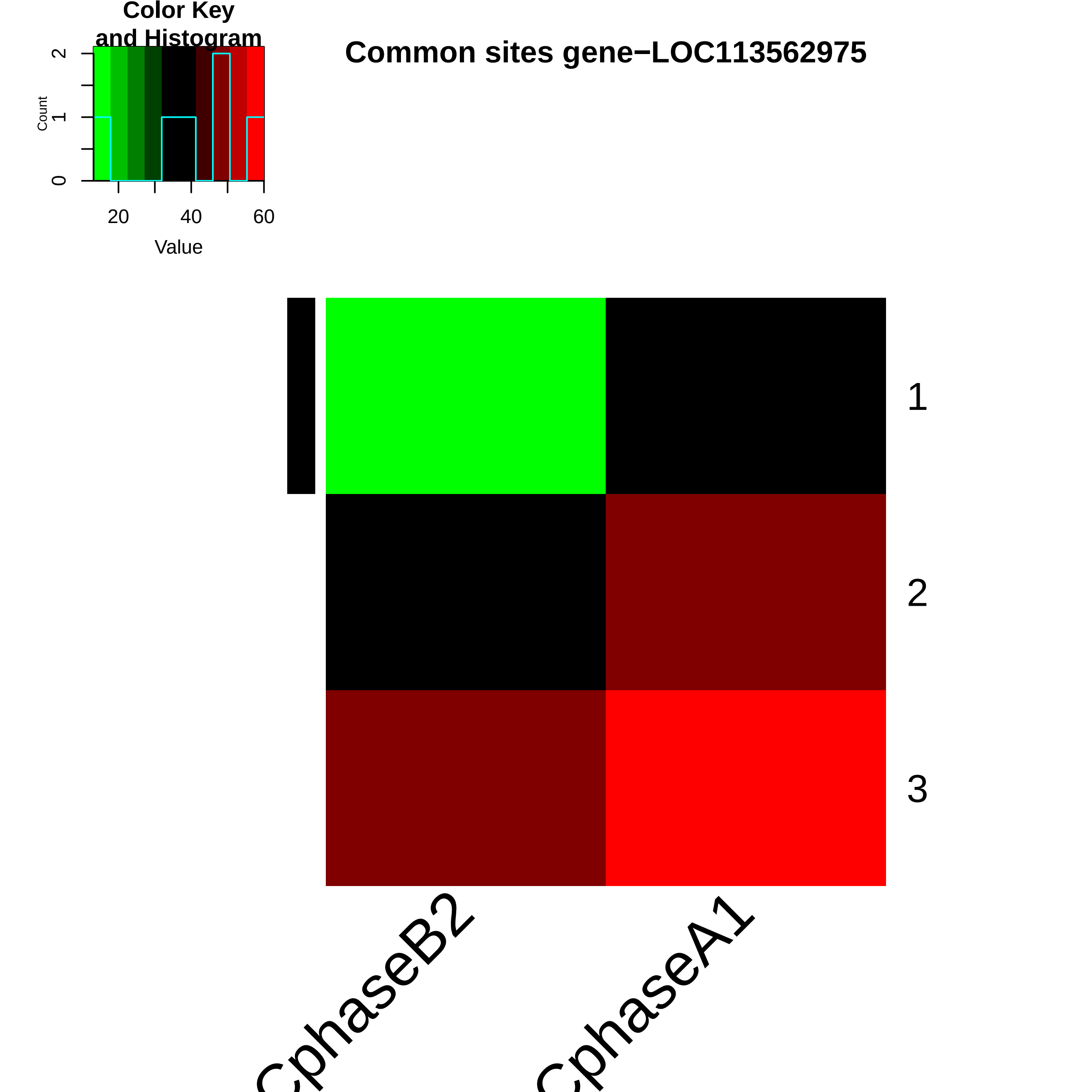
Color Key
and Histogram
Count
0
1
2
20
40
60
Value
Common sites gene−LOC113562975
1
2
3
CphaseB2
CphaseA1

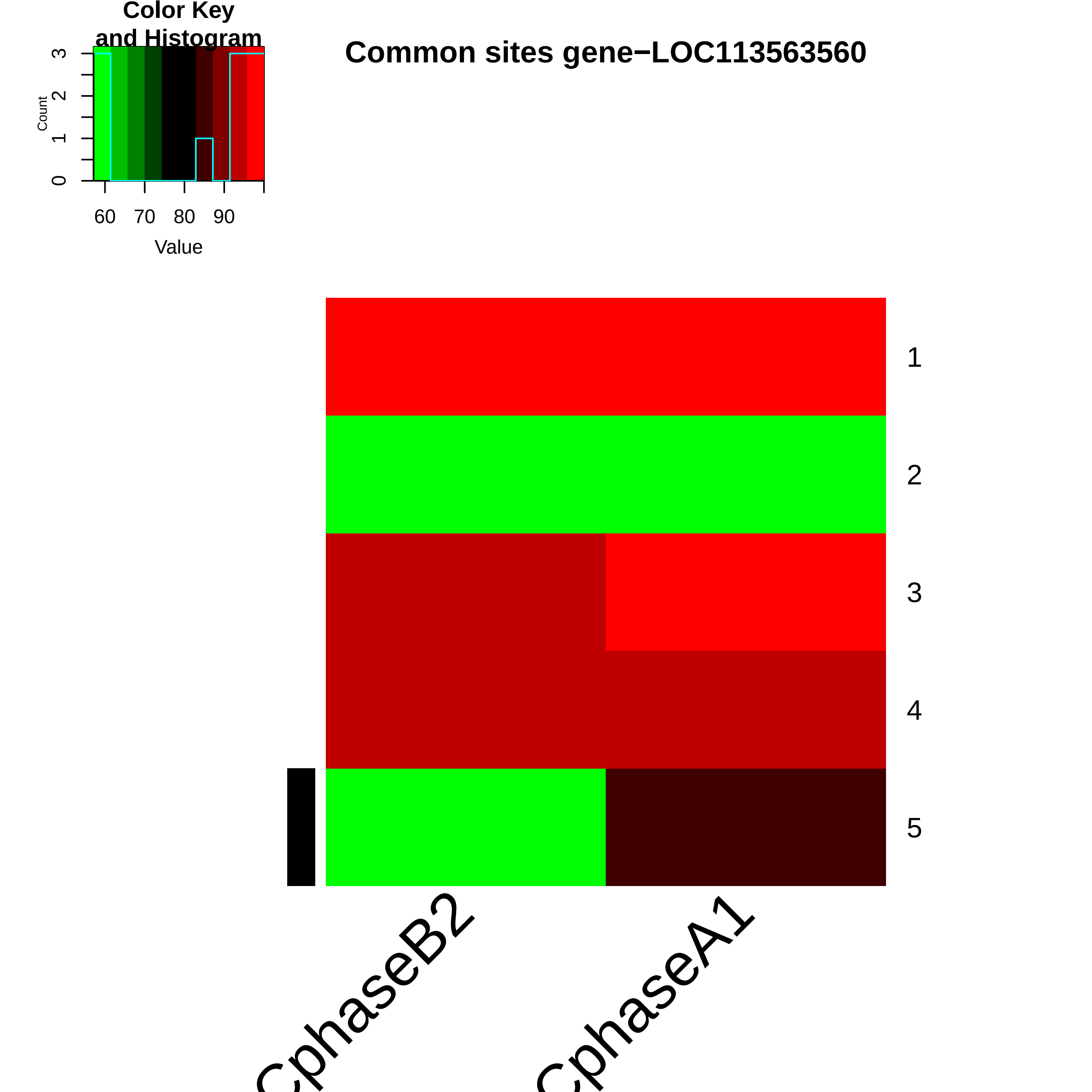
Color Key
and Histogram
Count
0
1
2
3
60
70
80
90
Value
Common sites gene−LOC113563560
1
2
3
4
5
CphaseB2
CphaseA1

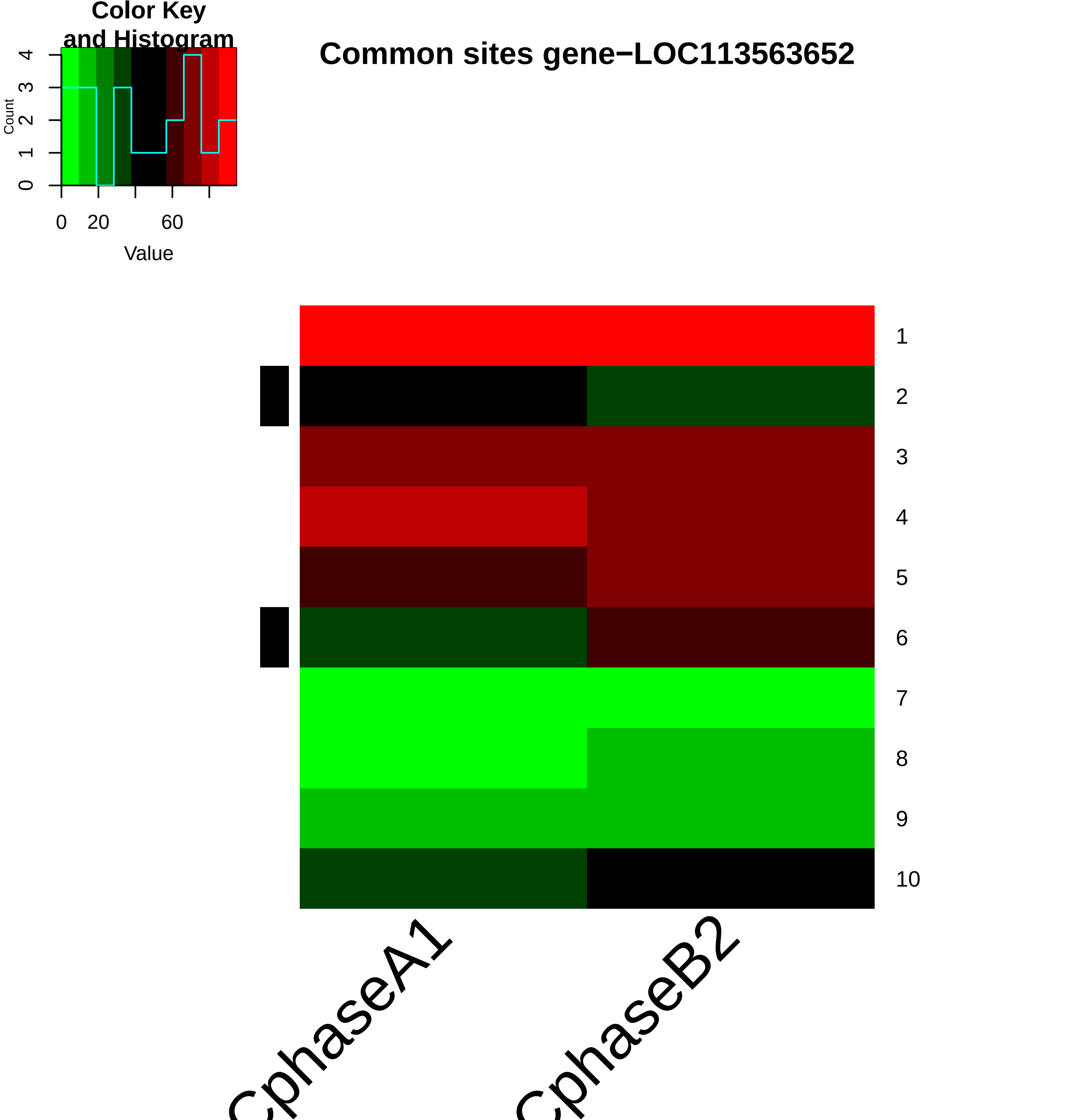
Color Key
and Histogram
Count
Value
0
20
60
0
1
2
3
4
Common sites gene−LOC113563652
1
2
3
4
5
6
7
8
9
10
CphaseA1
CphaseB2

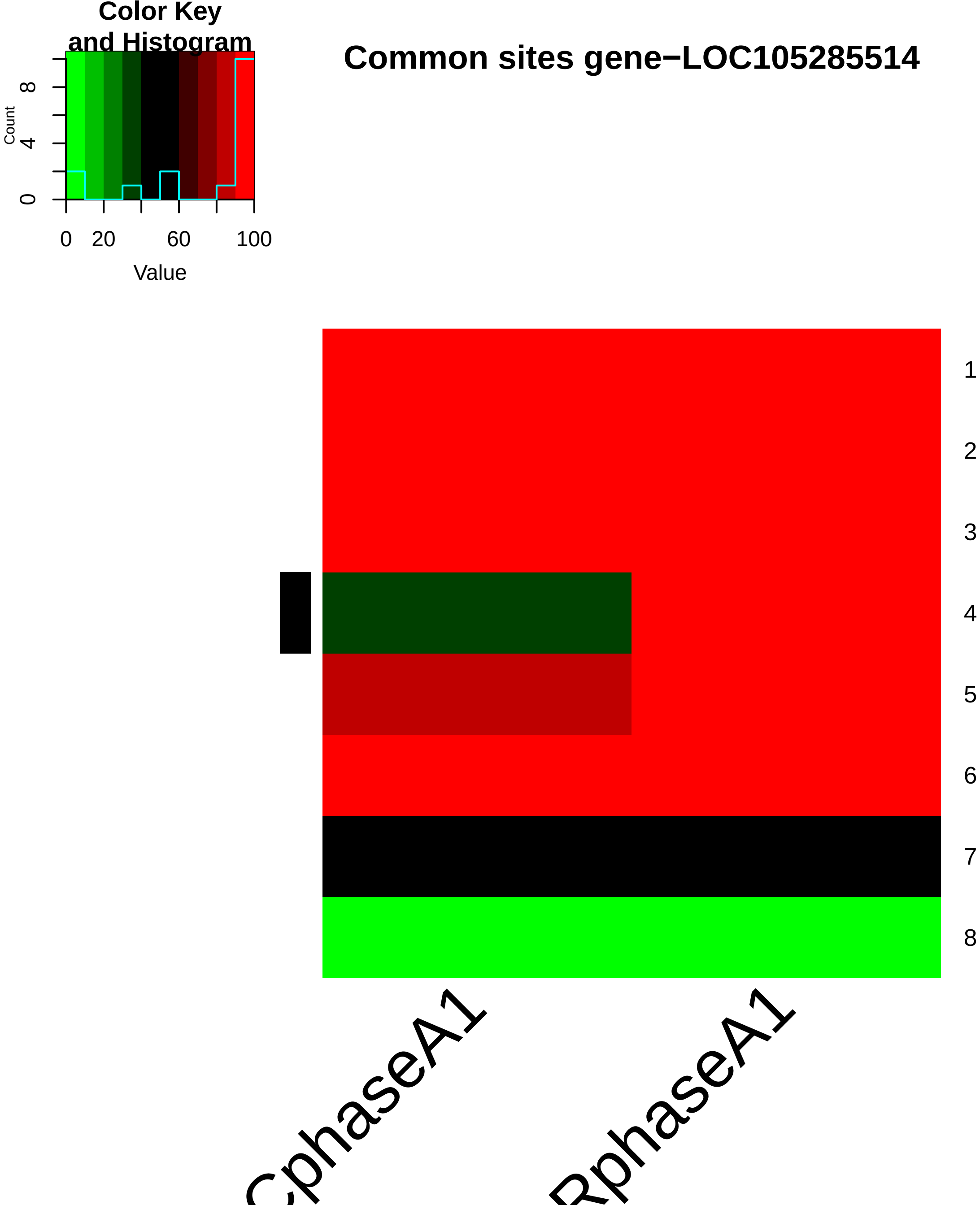
Color Key
and Histogram
Count
0
4
8
0
20
60
100
Value
Common sites gene−LOC105285514
1
2
3
4
5
6
7
8
CphaseA1
RphaseA1

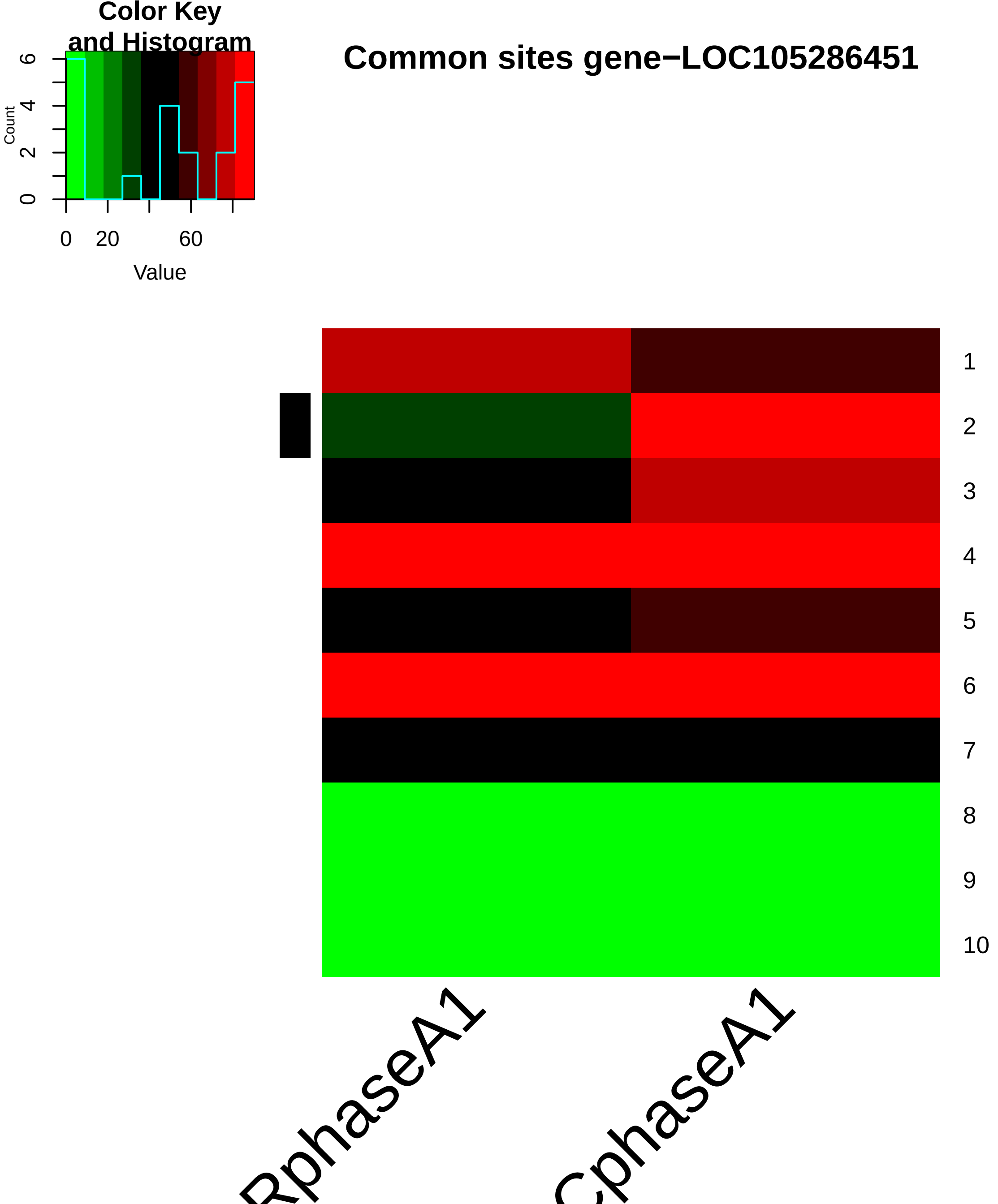
Color Key
and Histogram
Count
0
2
4
6
0
20
60
Value
Common sites gene−LOC105286451
1
2
3
4
5
6
7
8
9
10
RphaseA1
CphaseA1

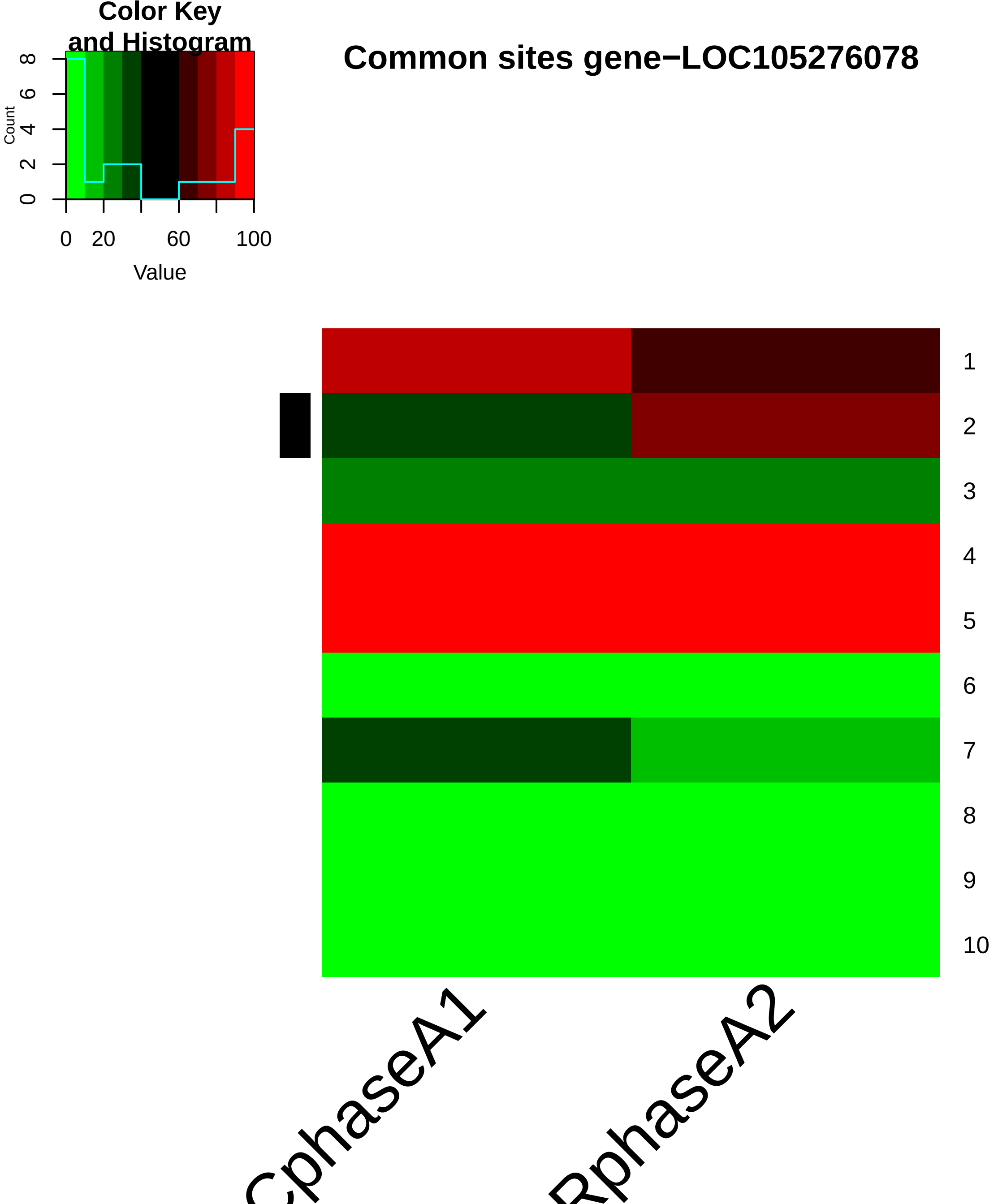
Color Key
and Histogram
Count
0
2
4
6
8
0
20
60
100
Value
Common sites gene−LOC105276078
1
2
3
4
5
6
7
8
9
10
CphaseA1
RphaseA2

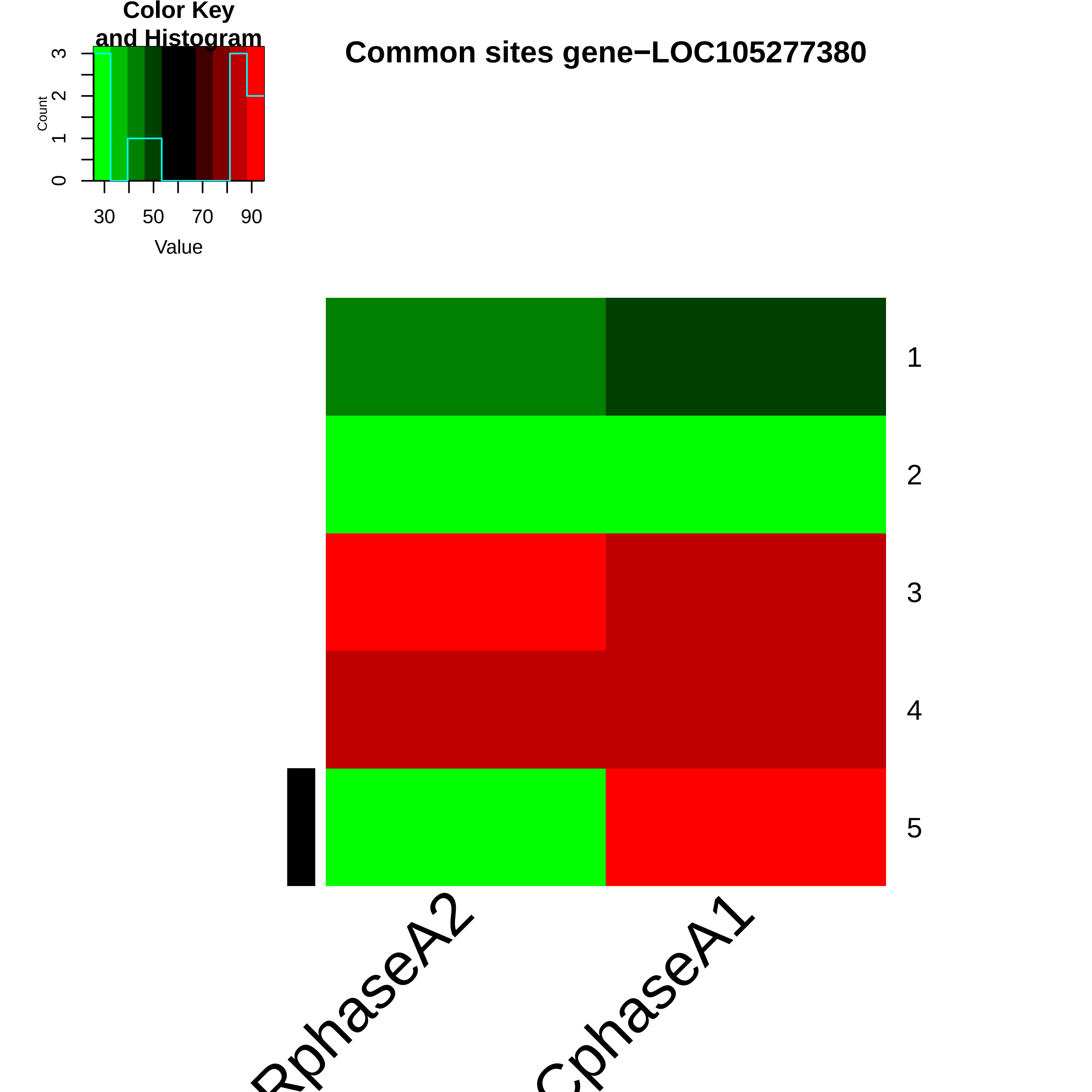

Color Key
and Histogram
Count
0
1
2
3
30
50
70
90
Value
Common sites gene−LOC105277380
1
2
3
4
5
RphaseA2
CphaseA1

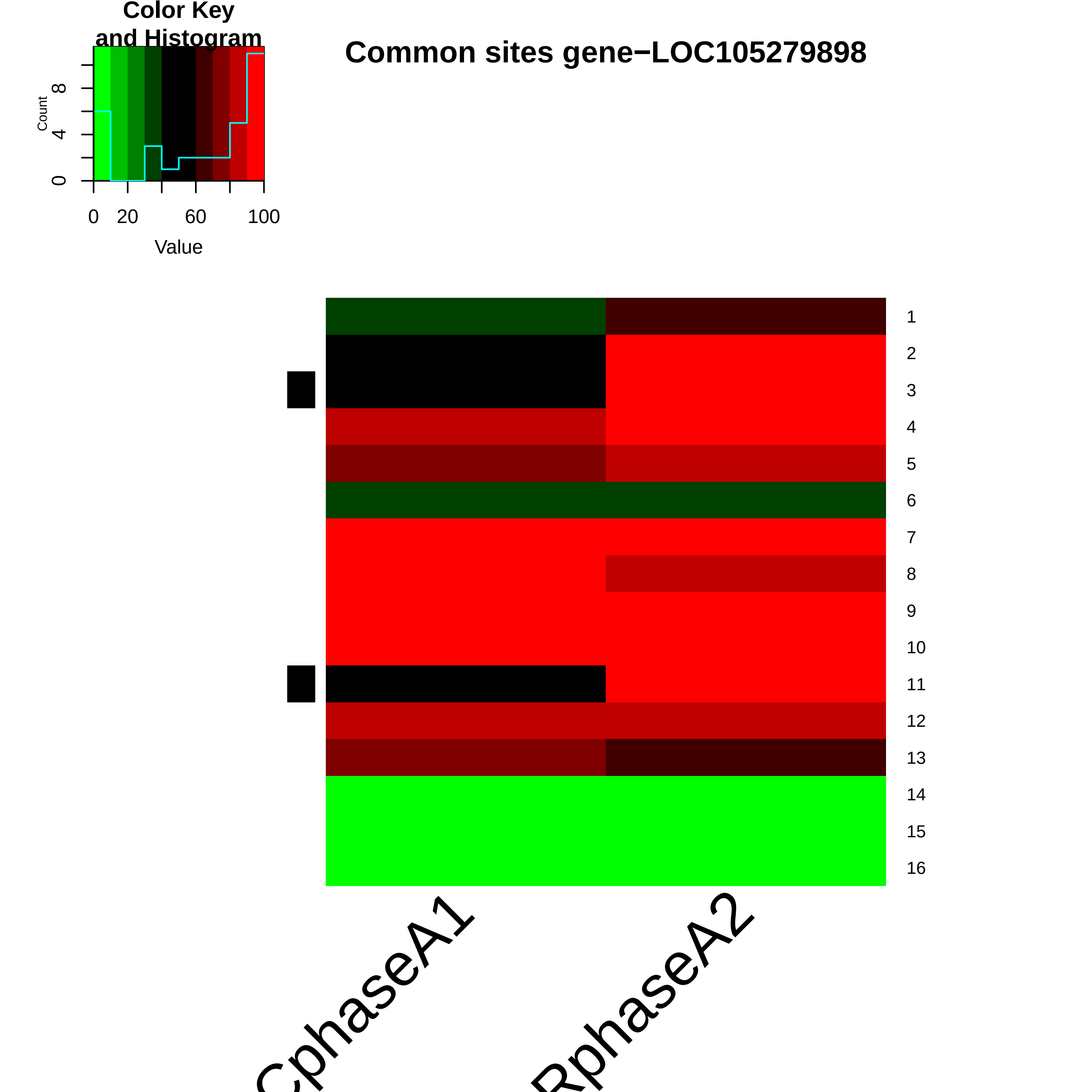
Color Key
and Histogram
Count
0
4
8
0
20
60
100
Value
Common sites gene−LOC105279898
CphaseA1
RphaseA2

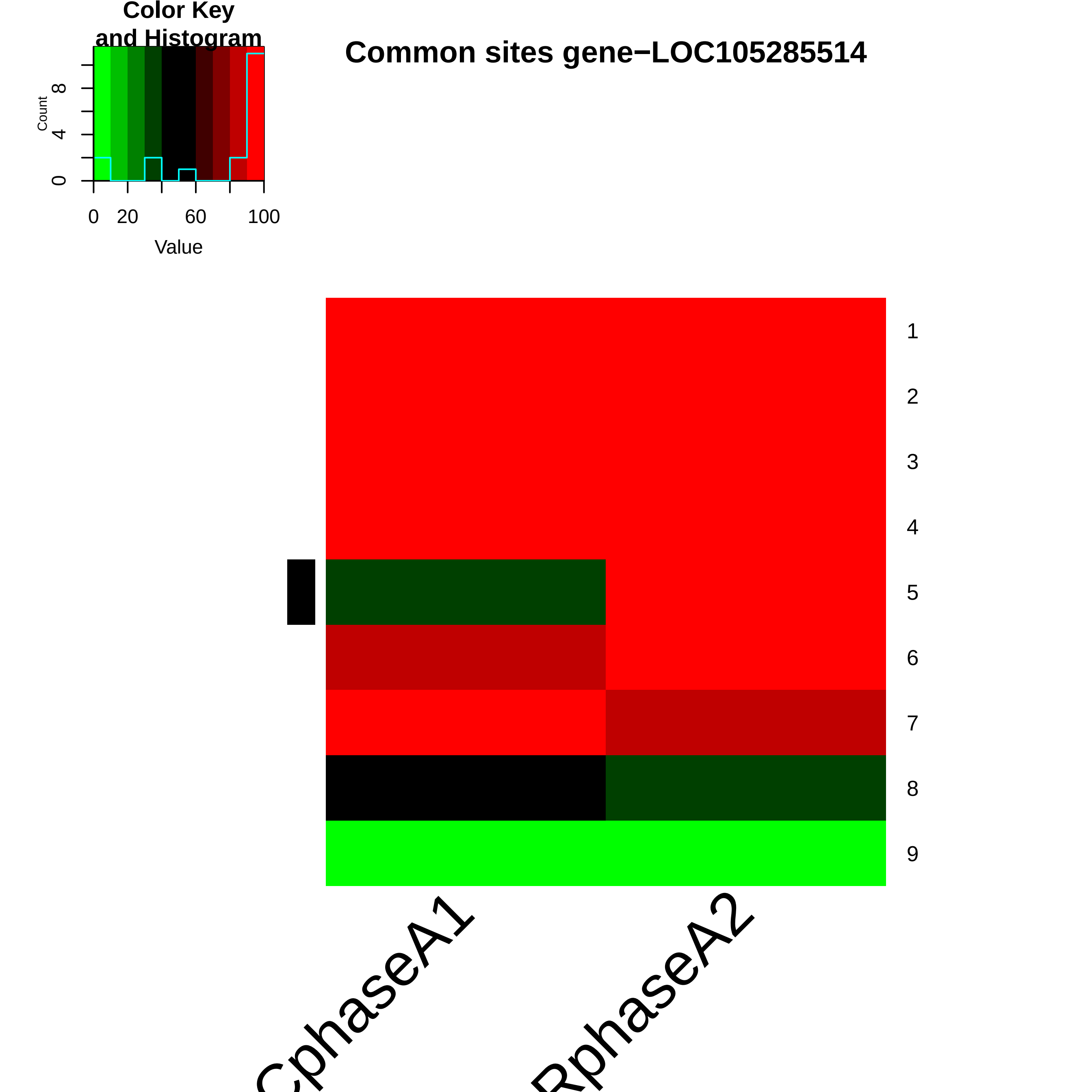
Color Key
and Histogram
Count
0
4
8
0
20
60
100
Value
Common sites gene−LOC105285514
1
2
3
4
5
6
7
8
9
CphaseA1
RphaseA2

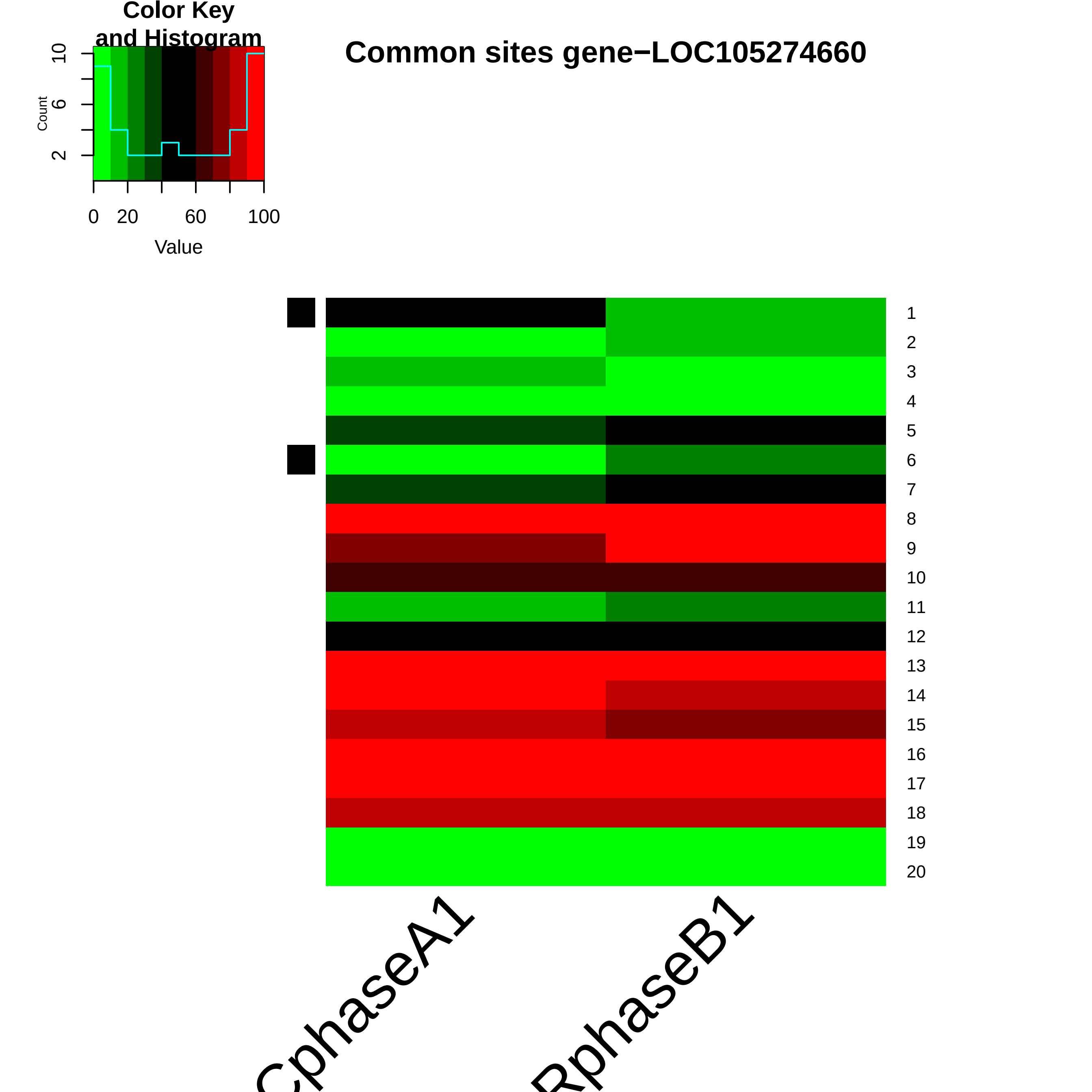
Color Key
and Histogram
Count
2
6
10
0
20
60
100
Value
Common sites gene−LOC105274660
CphaseA1
RphaseB1

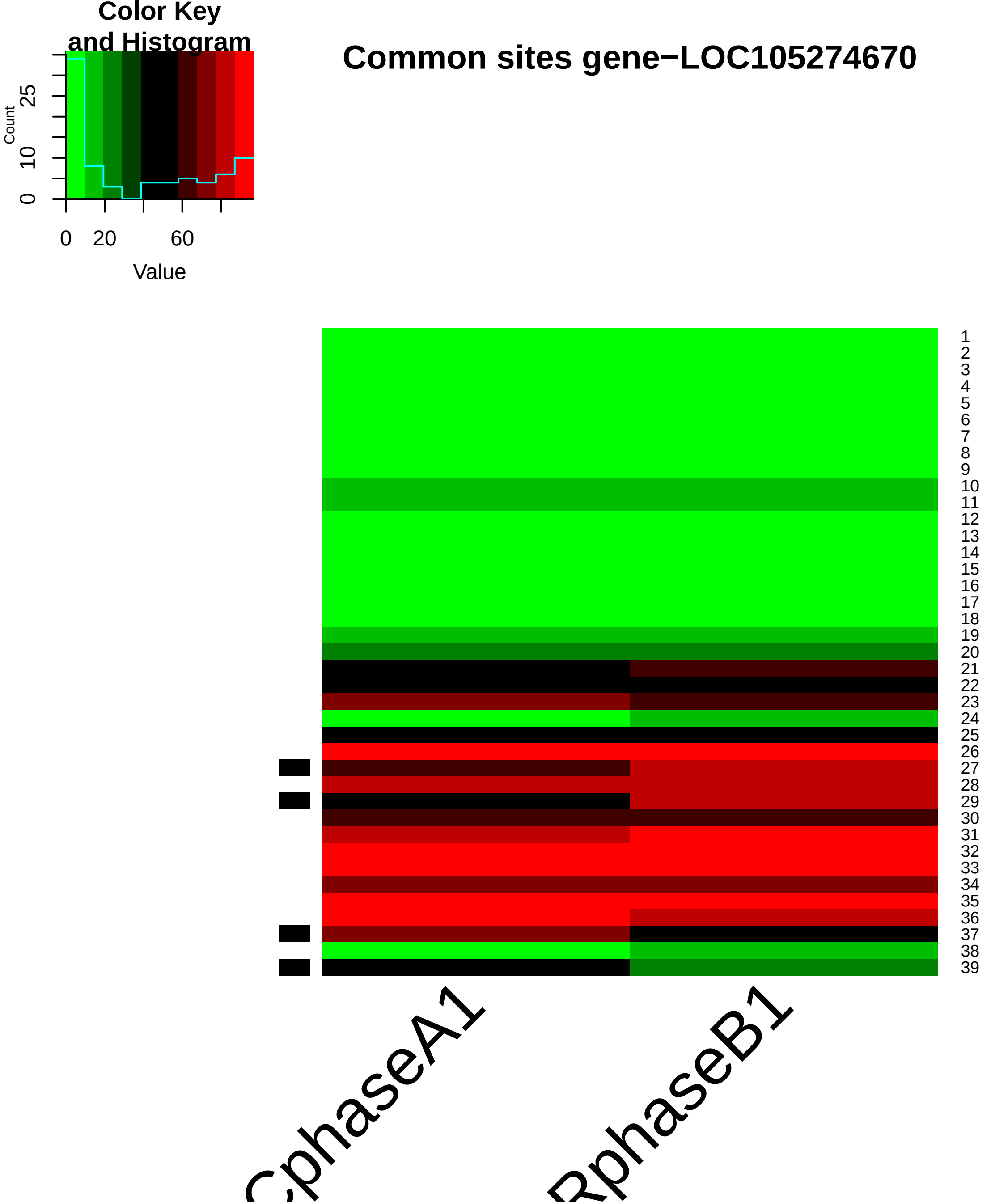
Color Key
and Histogram
Count
0
10
25
0
20
60
Value
Common sites gene−LOC105274670
CphaseA1
RphaseB1

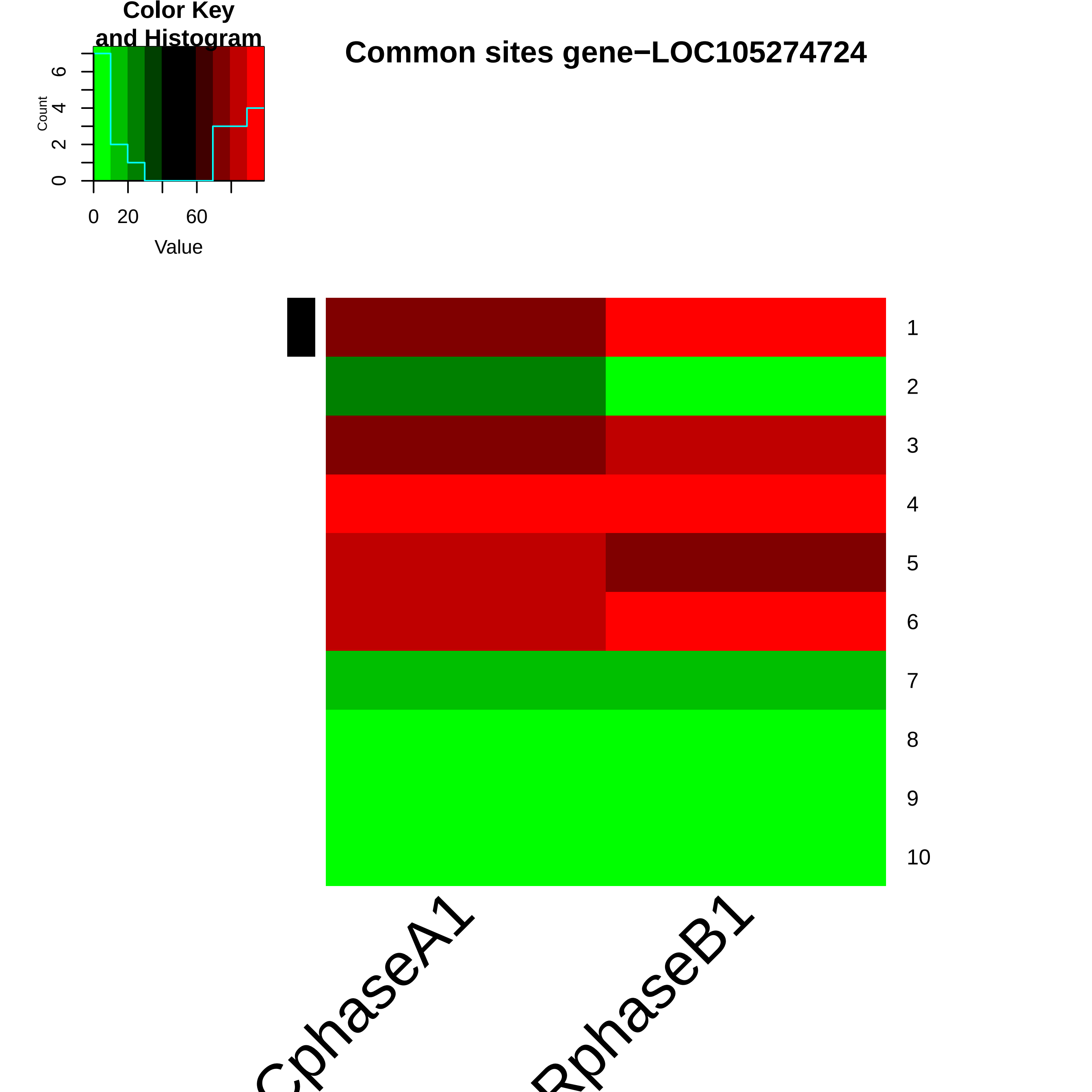
Color Key
and Histogram
Count
0
2
4
6
0
20
60
Value
Common sites gene−LOC105274724
1
2
3
4
5
6
7
8
9
10
CphaseA1
RphaseB1

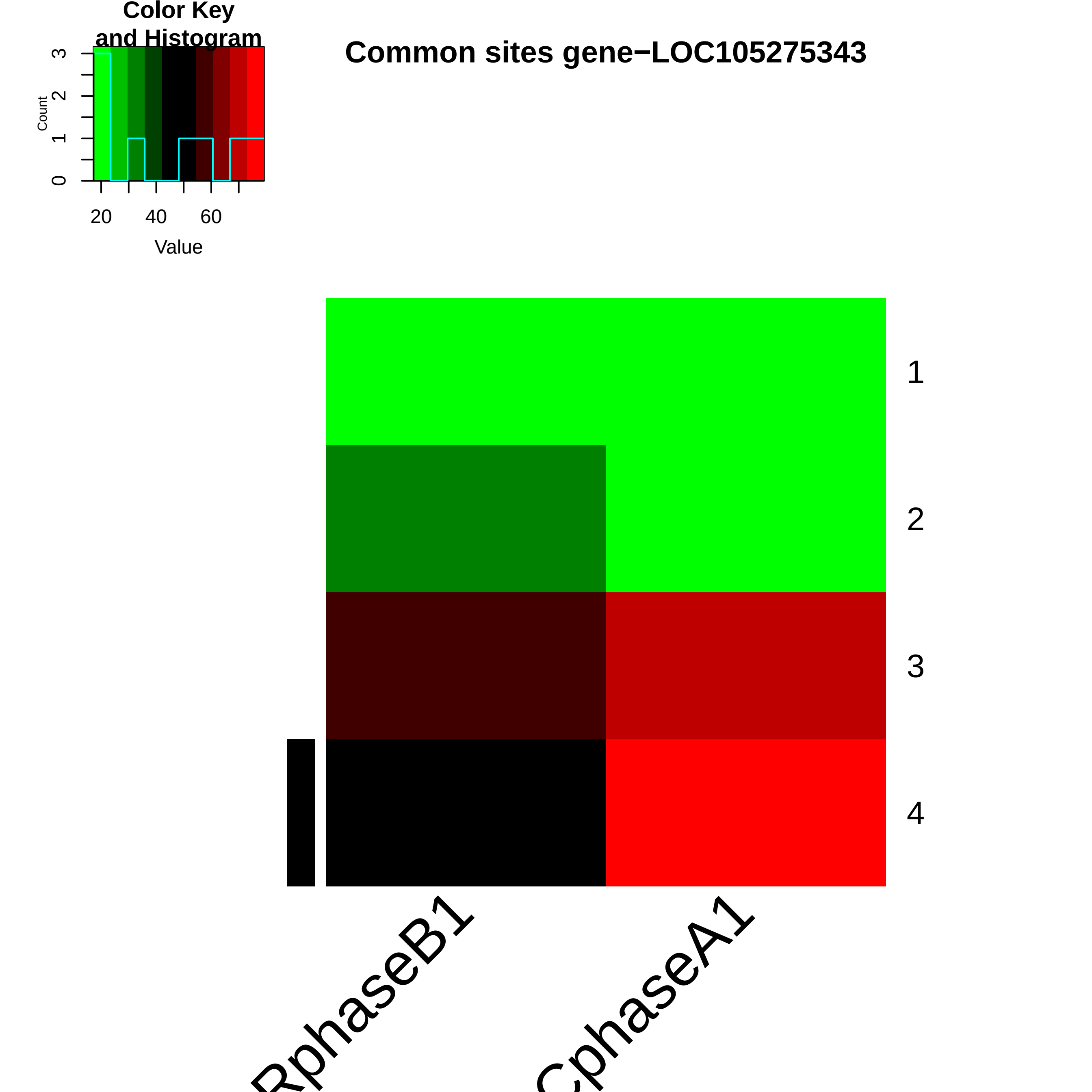
Color Key
and Histogram
Count
Value
20
40
60
0
1
2
3
Common sites gene−LOC105275343
1
2
3
4
RphaseB1
CphaseA1

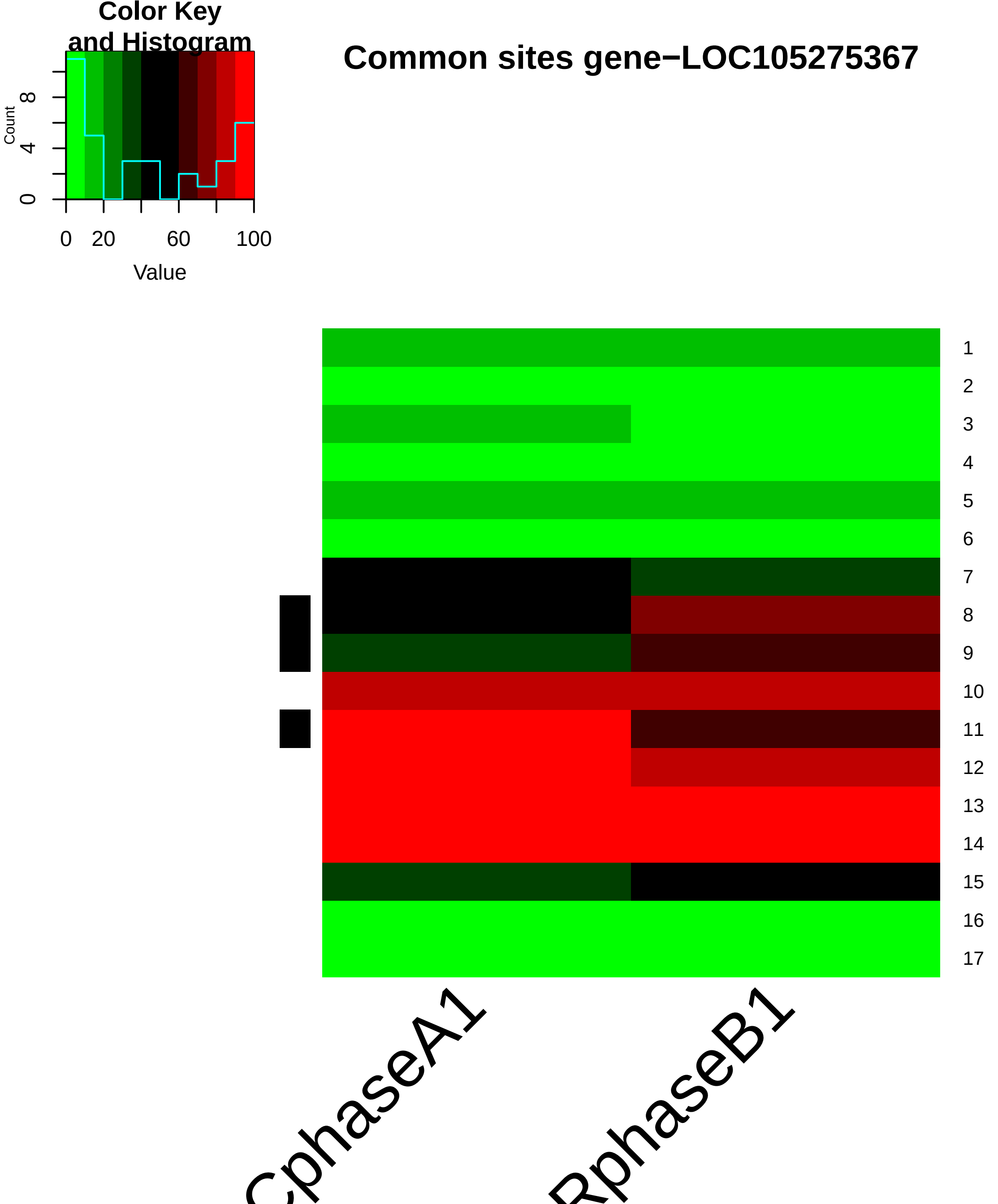
Color Key
and Histogram
Count
0
4
8
0
20
60
100
Value
Common sites gene−LOC105275367
CphaseA1
RphaseB1

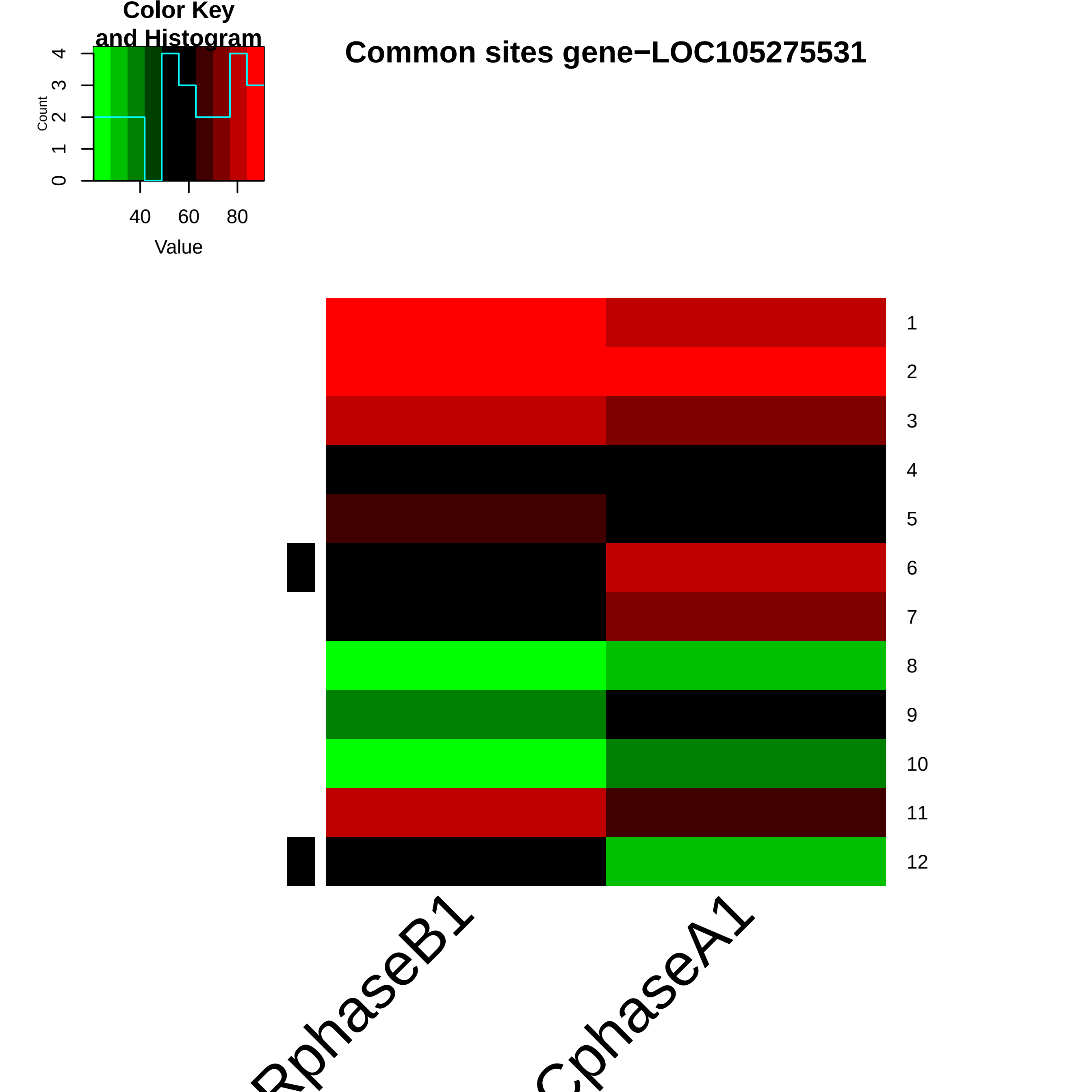

Color Key
and Histogram
Count
0
1
2
3
4
40
60
80
Value
Common sites gene−LOC105275531
1
2
3
4
5
6
7
8
9
10
11
12
RphaseB1
CphaseA1

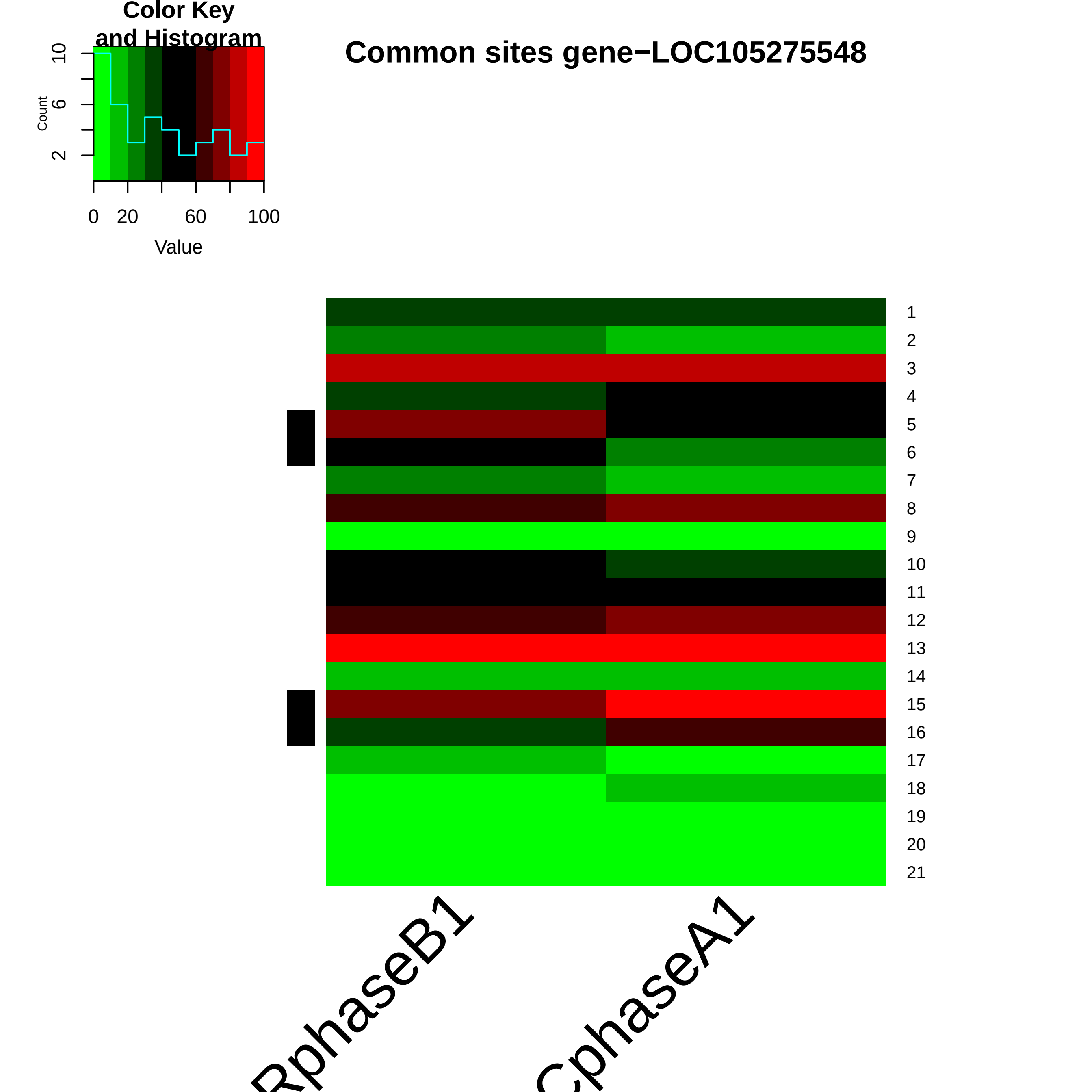

Color Key
and Histogram
Count
10
6
2
0
20
60
100
Value
Common sites gene−LOC105275548
RphaseB1
CphaseA1

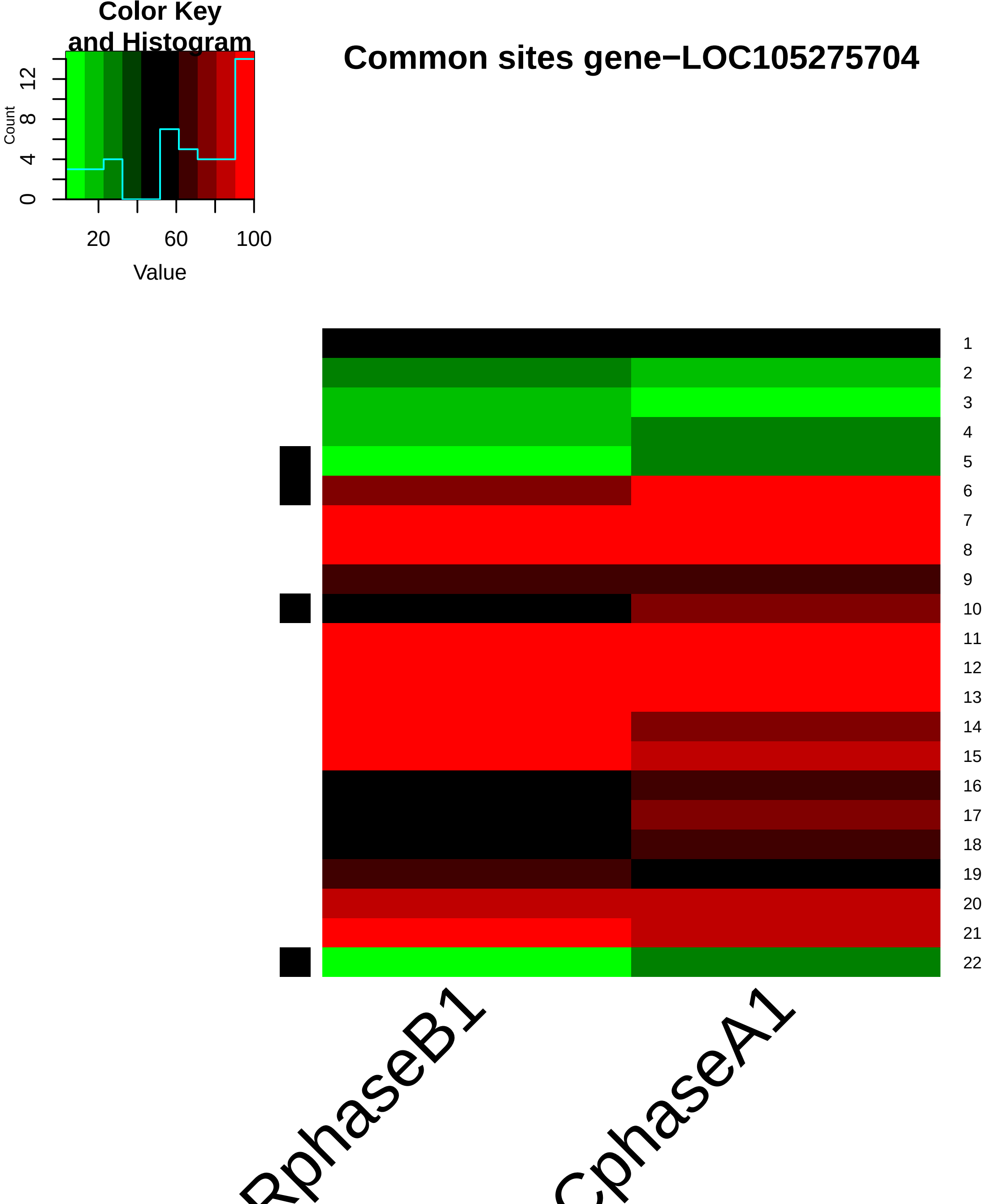
Color Key
and Histogram
Count
0
4
8
12
20
60
100
Value
Common sites gene−LOC105275704
RphaseB1
CphaseA1

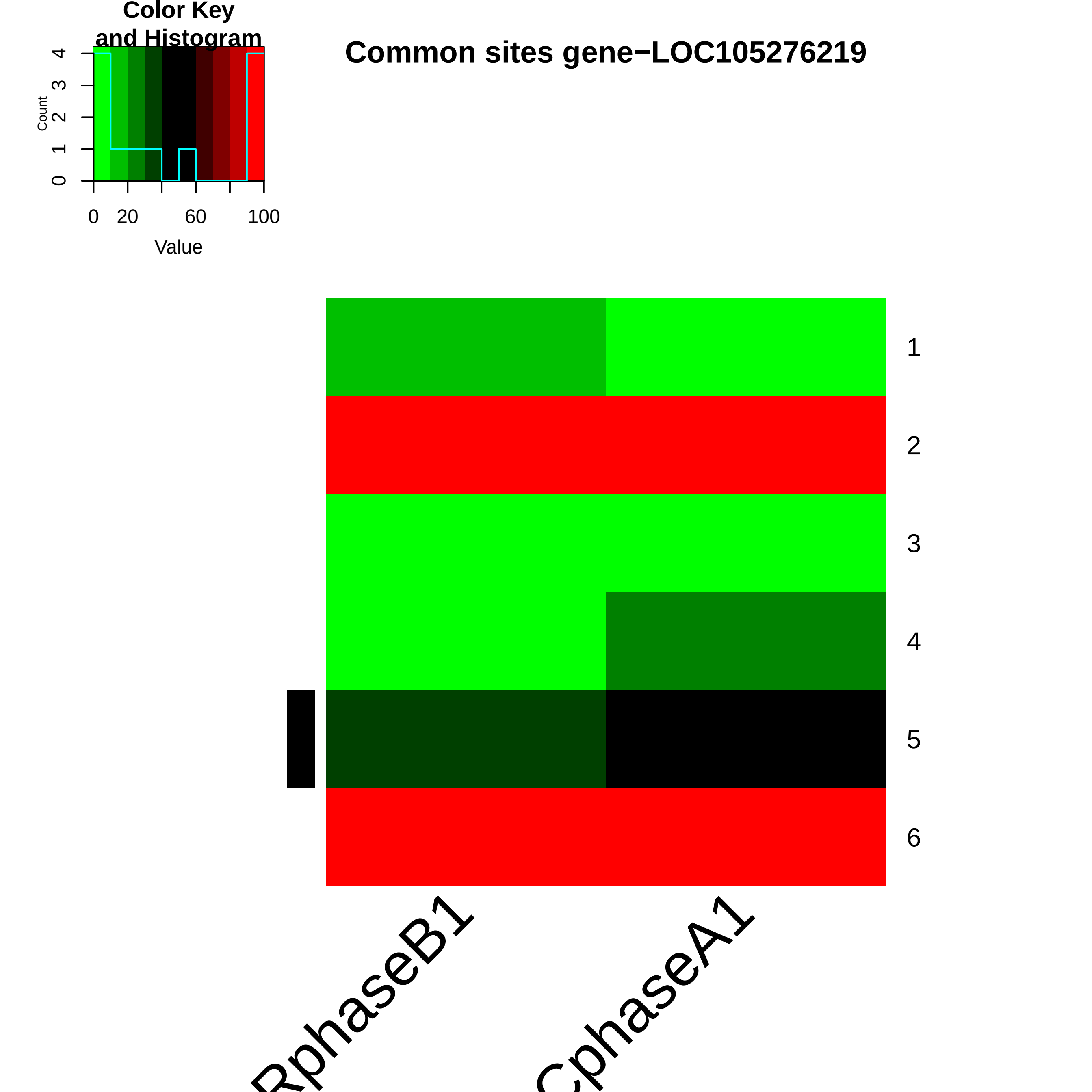
Color Key
and Histogram
Count
Value
0
20
60
100
0
1
2
3
4
Common sites gene−LOC105276219
1
2
3
4
5
6
RphaseB1
CphaseA1

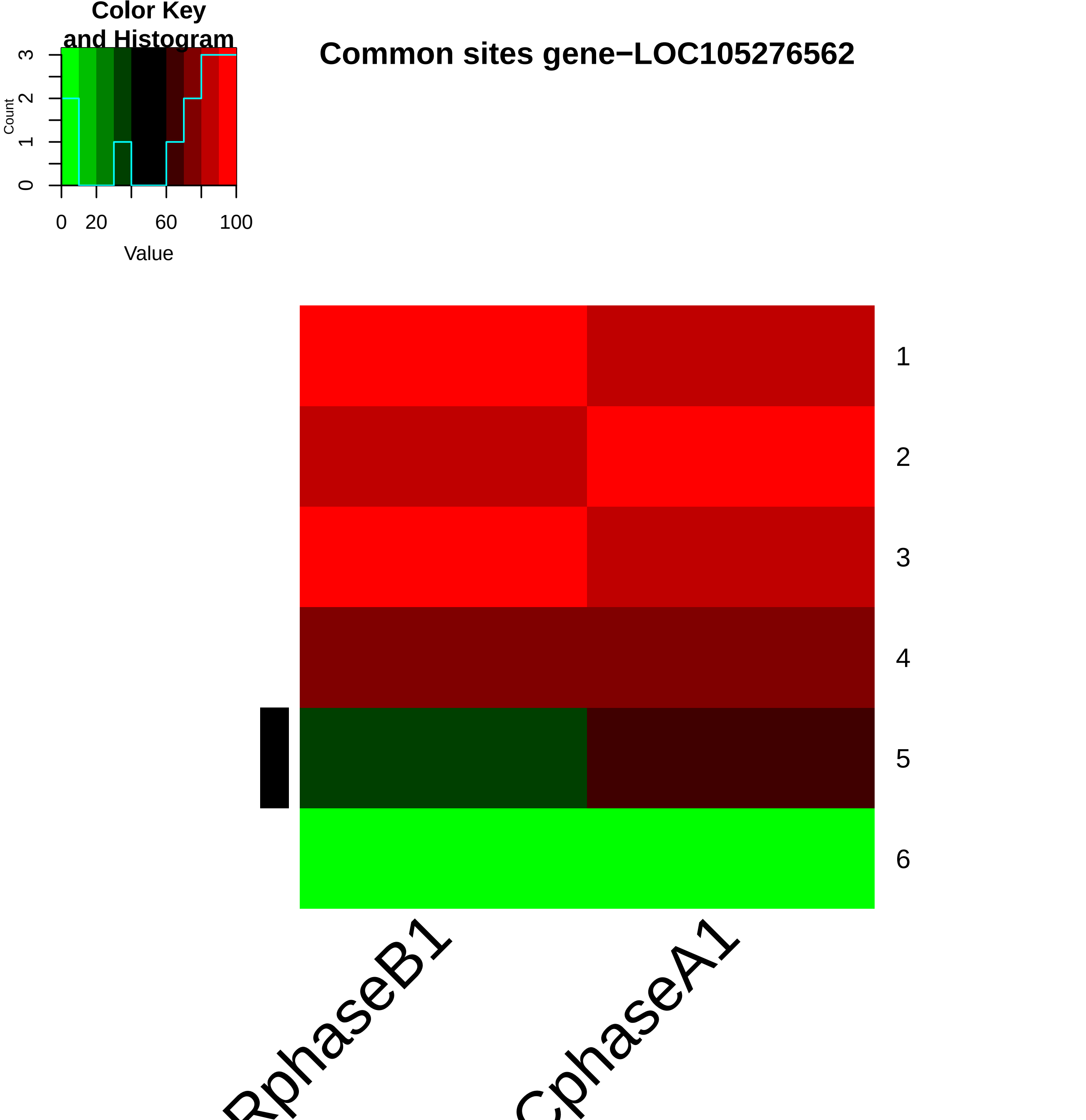
Color Key
and Histogram
Count
Value
0
20
60
100
0
1
2
3
Common sites gene−LOC105276562
1
2
3
4
5
6
RphaseB1
CphaseA1

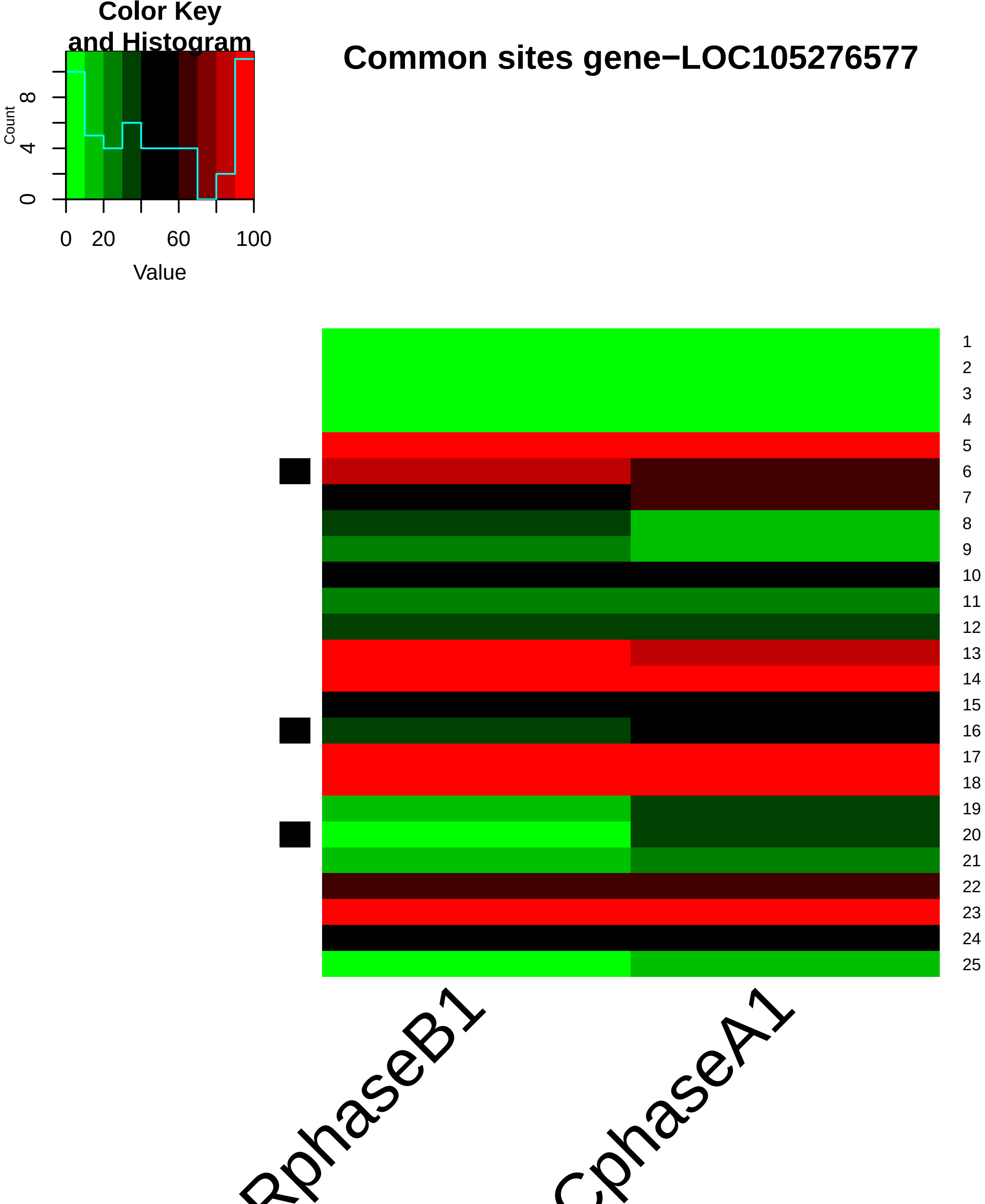

Color Key and Histogram
Count
0
4
8
0
20
60
100
Value
Common sites gene−LOC105276577
RphaseB1
CphaseA1

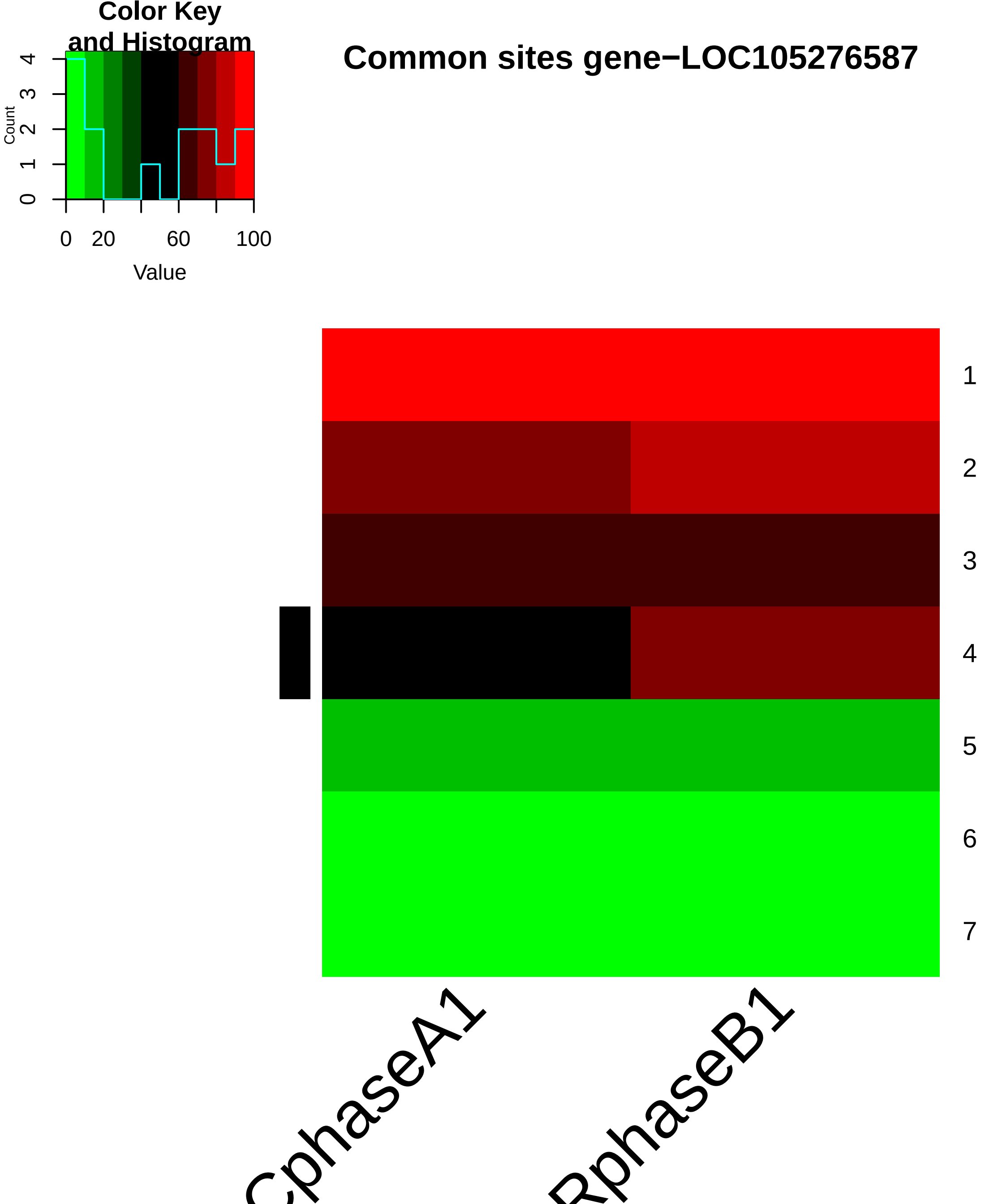

Color Key
and Histogram
Count
0
1
2
3
4
0
20
60
100
Value
Common sites gene−LOC105276587
1
2
3
4
5
6
7
CphaseA1
RphaseB1

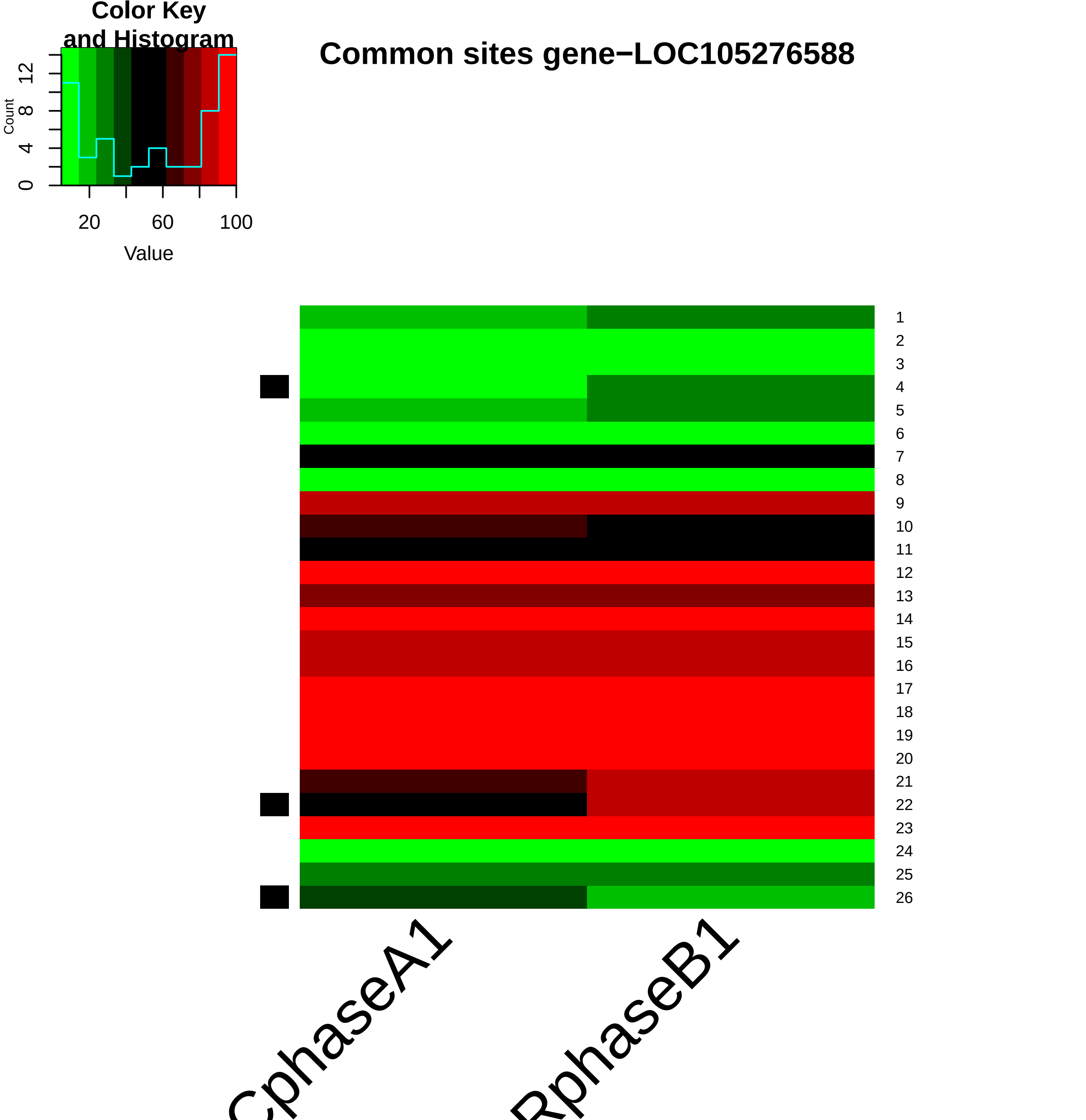

Color Key
and Histogram
Count
0
4
8
12
20
60
100
Value
Common sites gene−LOC105276588

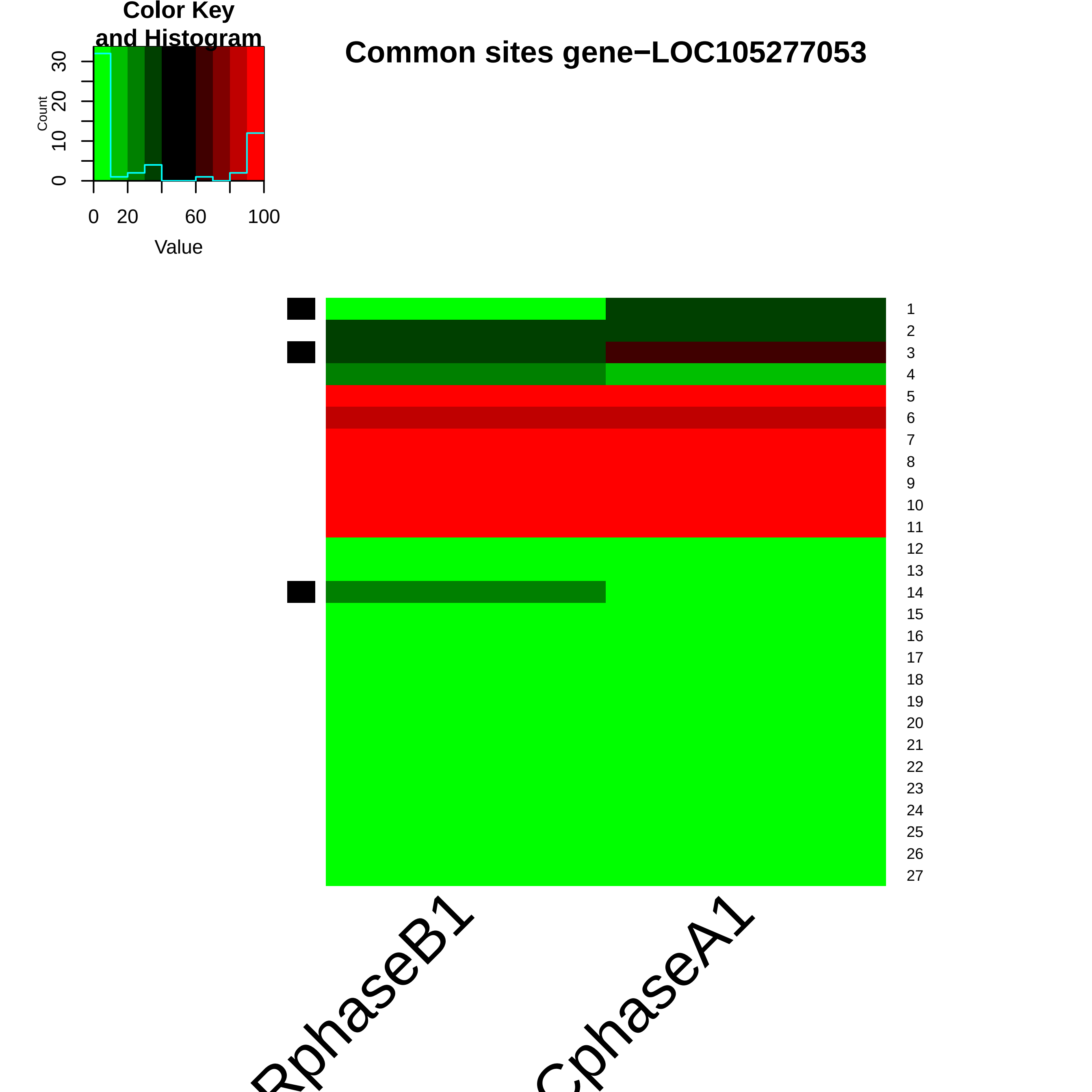

Color Key
and Histogram
Count
0
10
20
30
0
20
60
100
Value
Common sites gene−LOC105277053
RphaseB1
CphaseA1

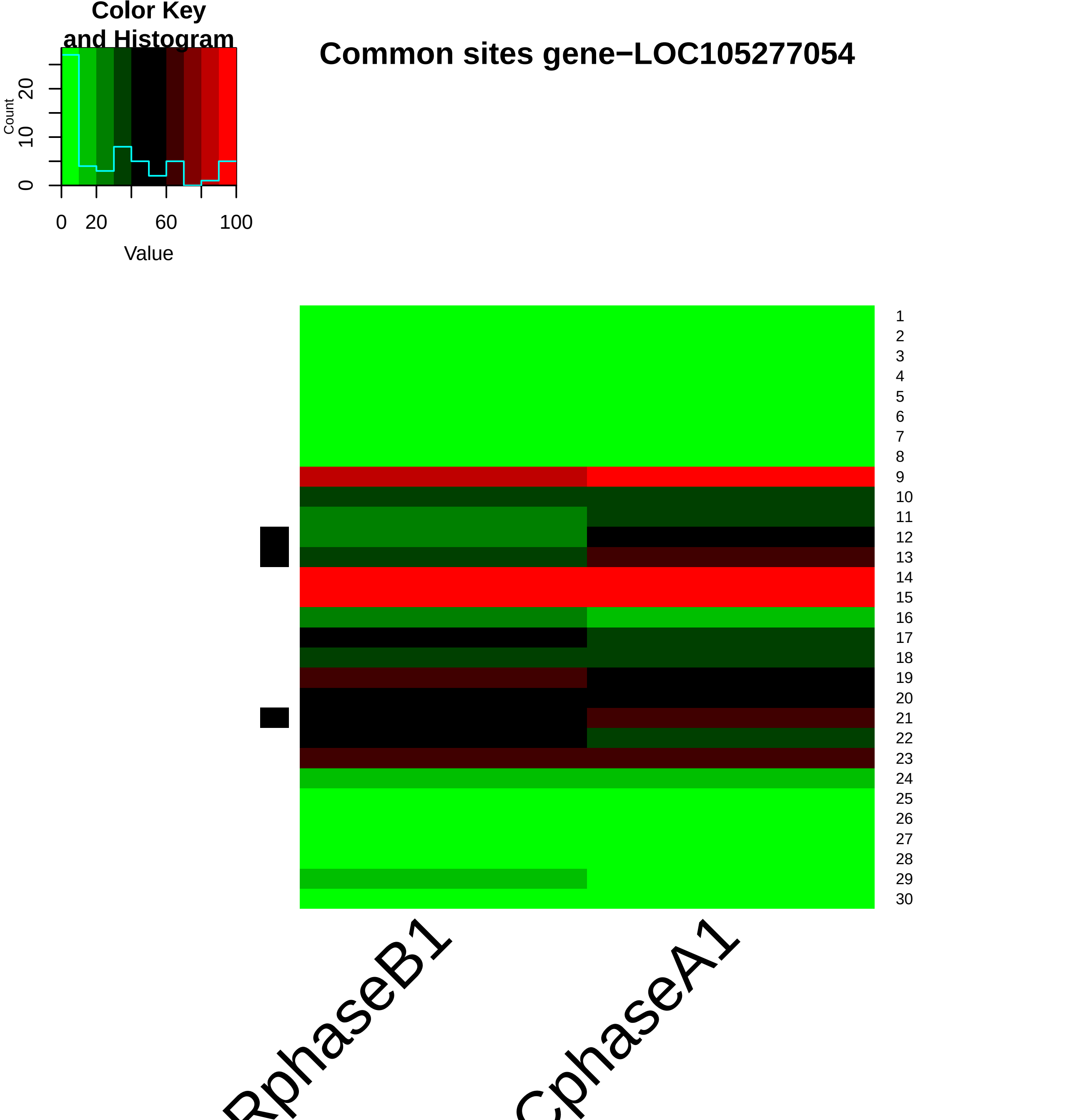

Color Key
and Histogram
Count
0
10
20
0
20
60
100
Value
Common sites gene−LOC105277054
RphaseB1
CphaseA1

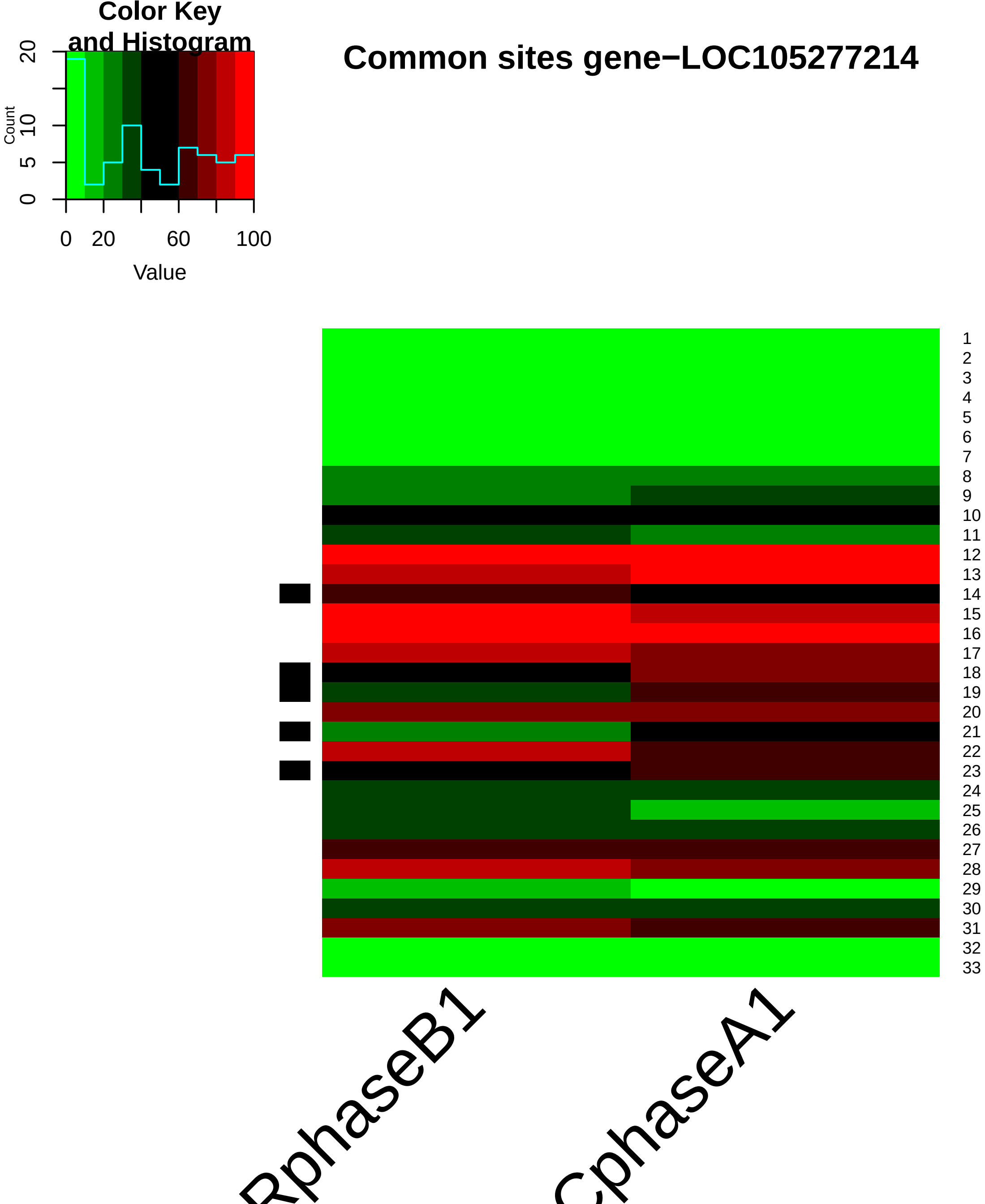

Color Key
and Histogram
Count
0
5
10
20
0
20
60
100
Value
Common sites gene−LOC105277214
RphaseB1
CphaseA1

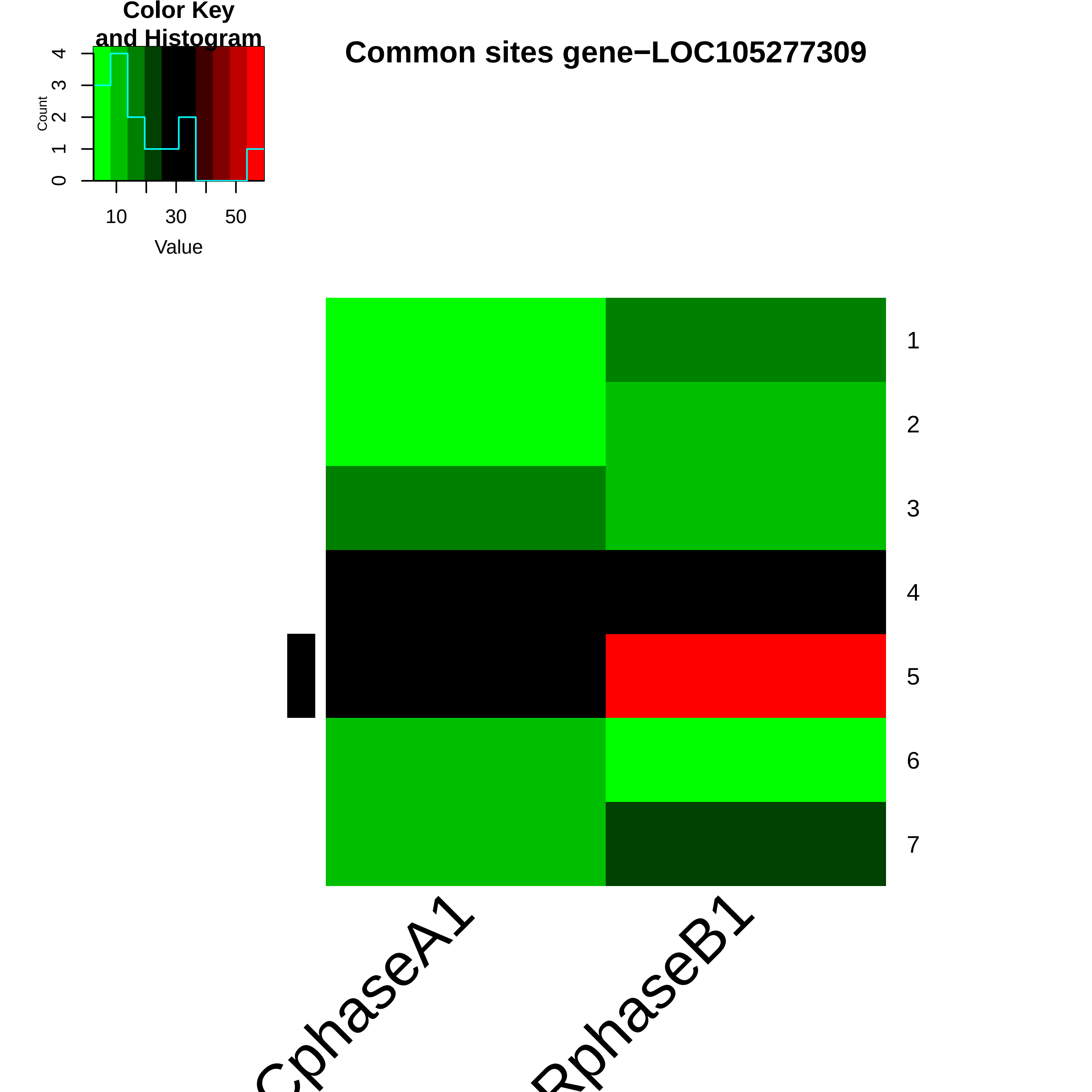

Color Key
and Histogram
Count
0
1
2
3
4
10
30
50
Value
Common sites gene−LOC105277309
1
2
3
4
5
6
7
CphaseA1
RphaseB1

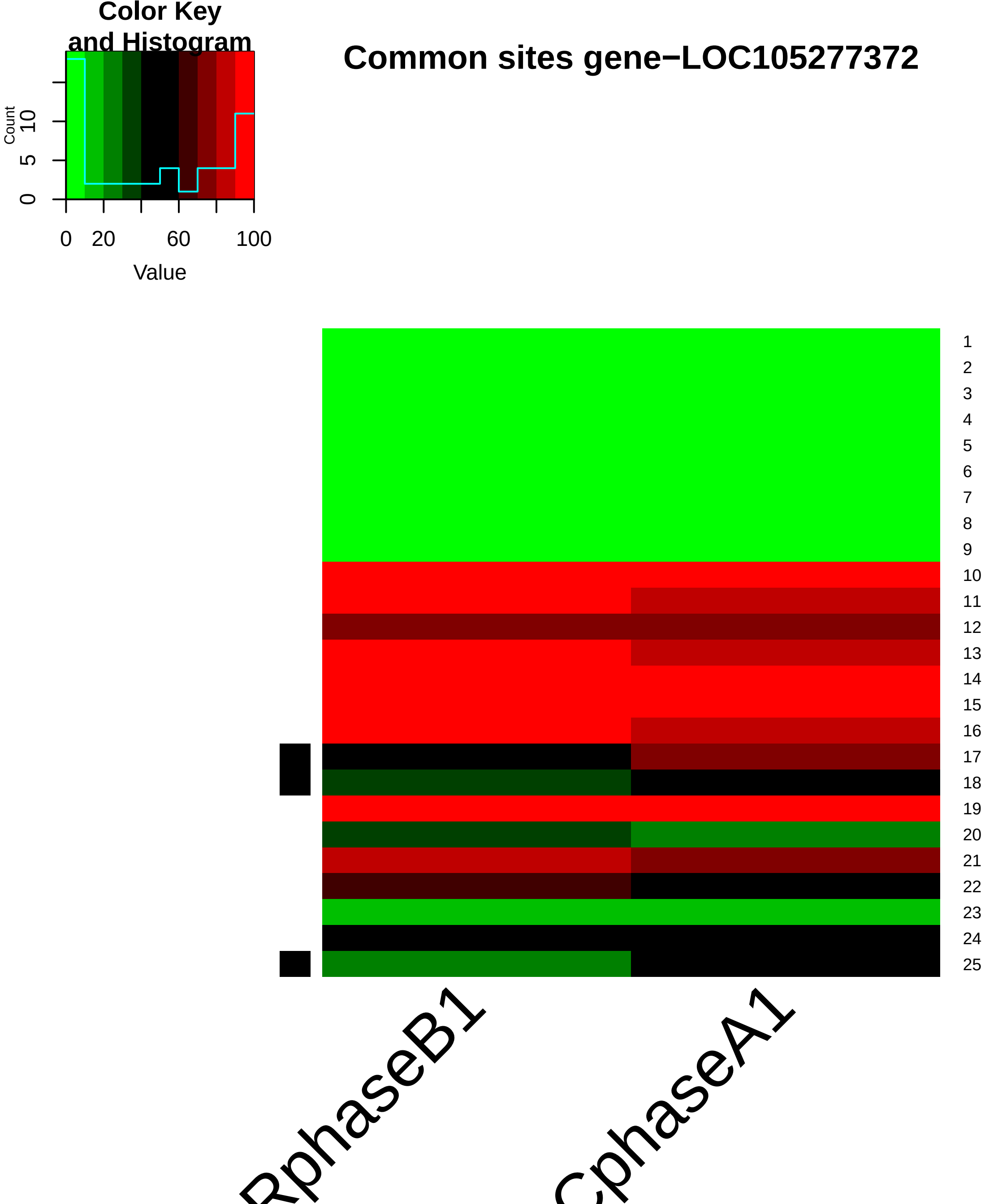

Color Key and Histogram
Count
0
5
10
0
20
60
100
Value
Common sites gene−LOC105277372
RphaseB1
CphaseA1

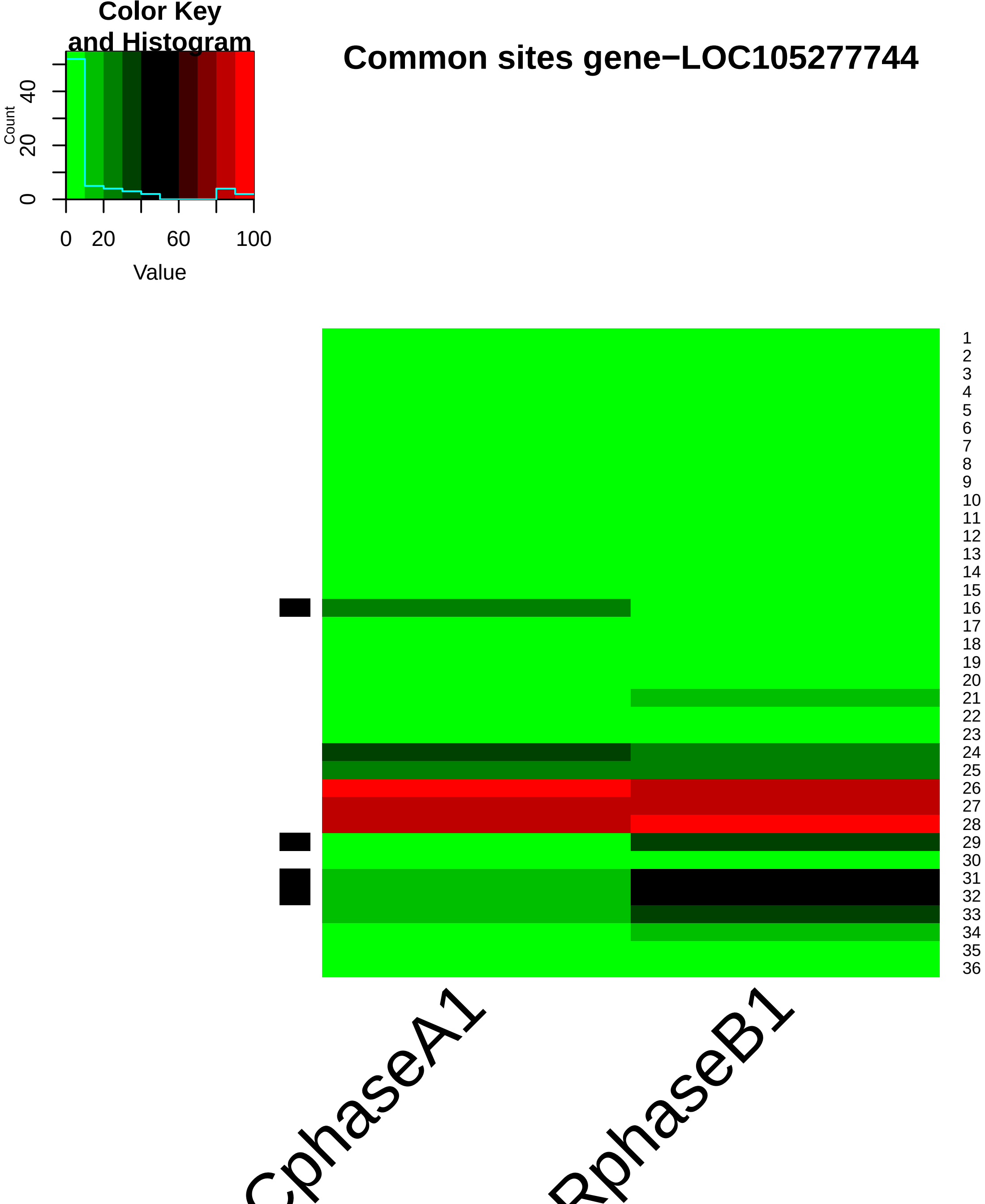

Color Key
and Histogram
Count
0
20
40
0
20
60
100
Value
Common sites gene−LOC105277744
CphaseA1
RphaseB1

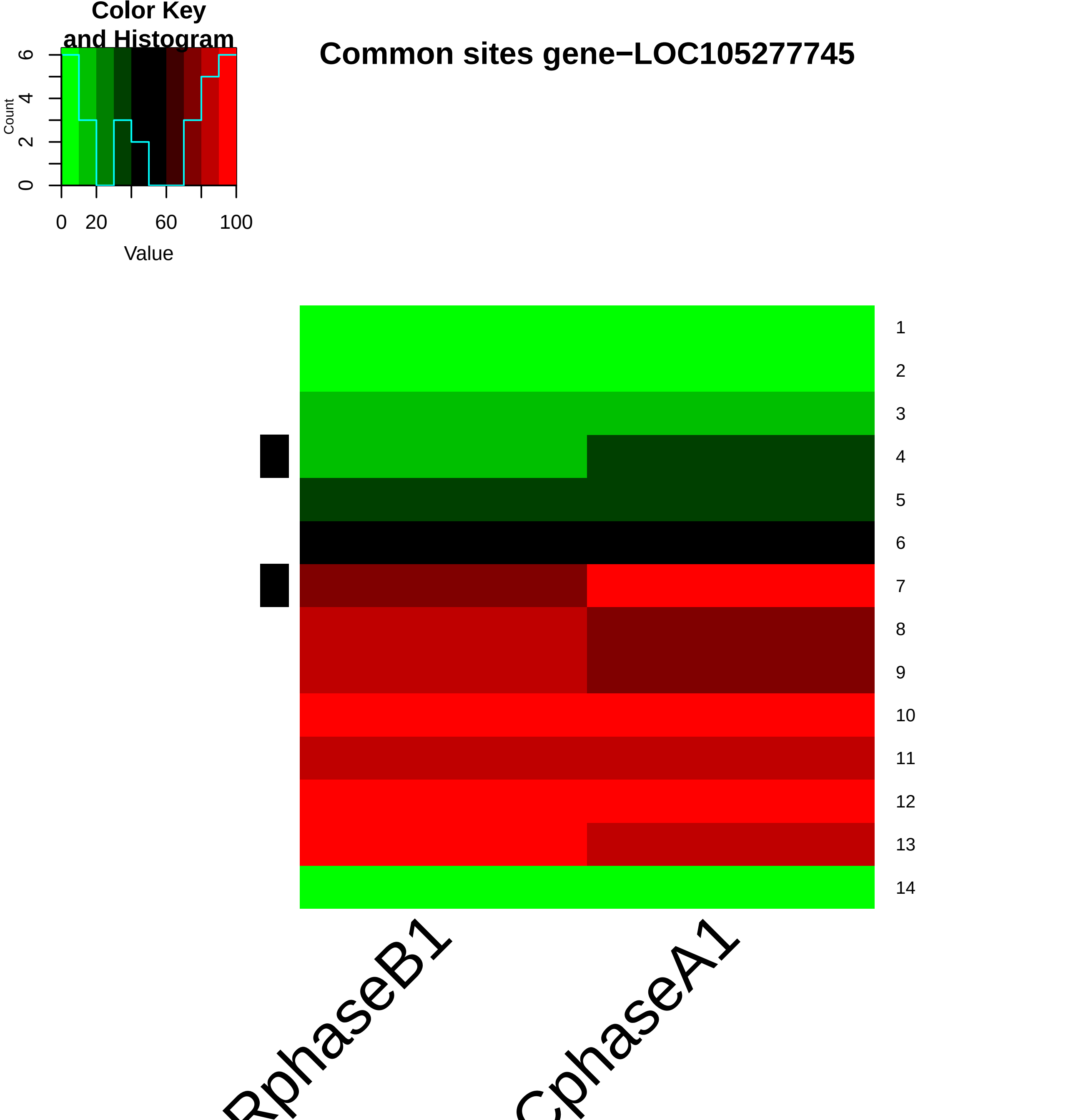

Color Key
and Histogram
Count
0
2
4
6
0
20
60
100
Value
Common sites gene−LOC105277745
1
2
3
4
5
6
7
8
9
10
11
12
13
14
RphaseB1
CphaseA1

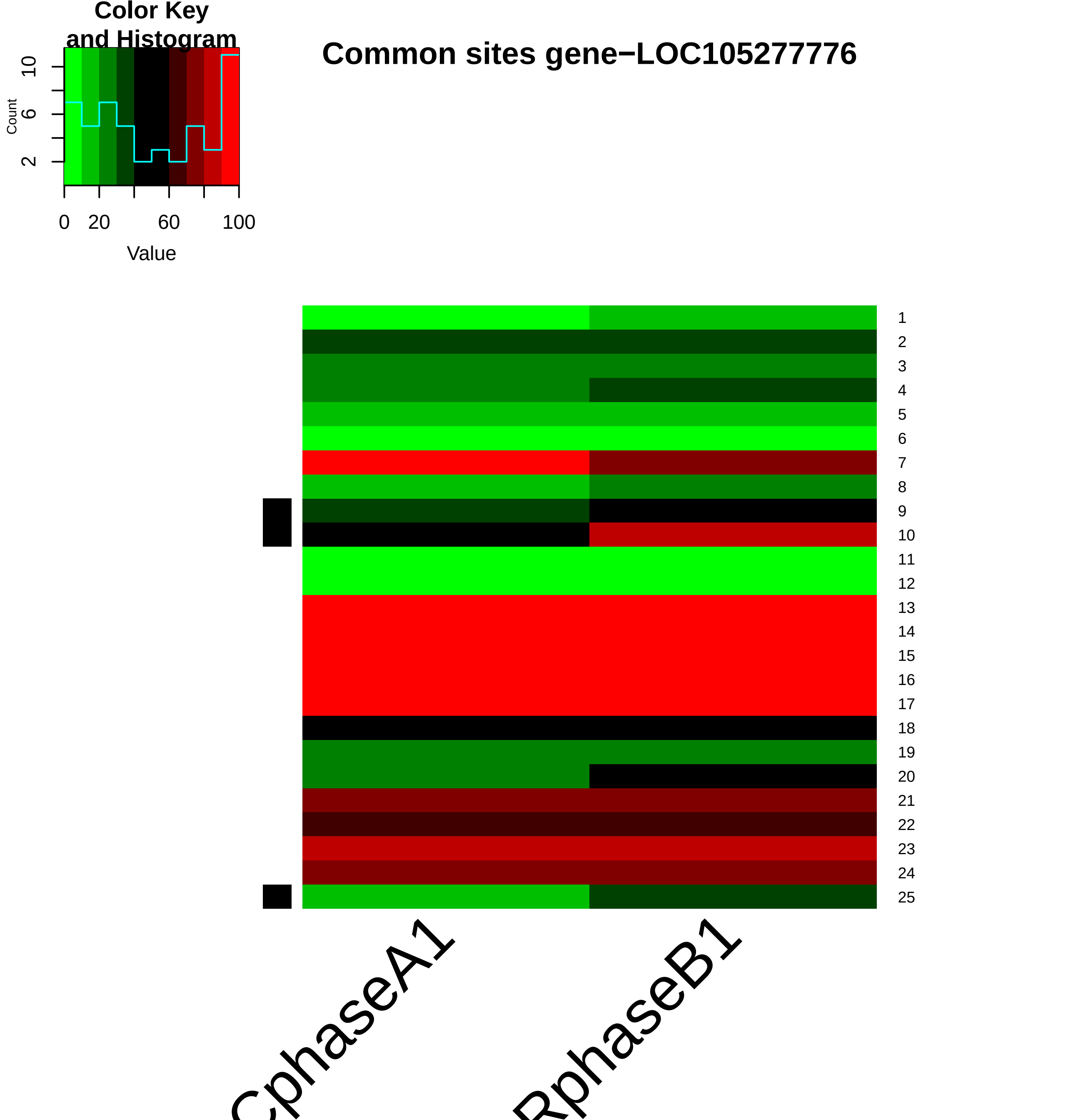

Color Key
and Histogram
Count
2
6
10
0
20
60
100
Value
Common sites gene−LOC105277776
CphaseA1
RphaseB1

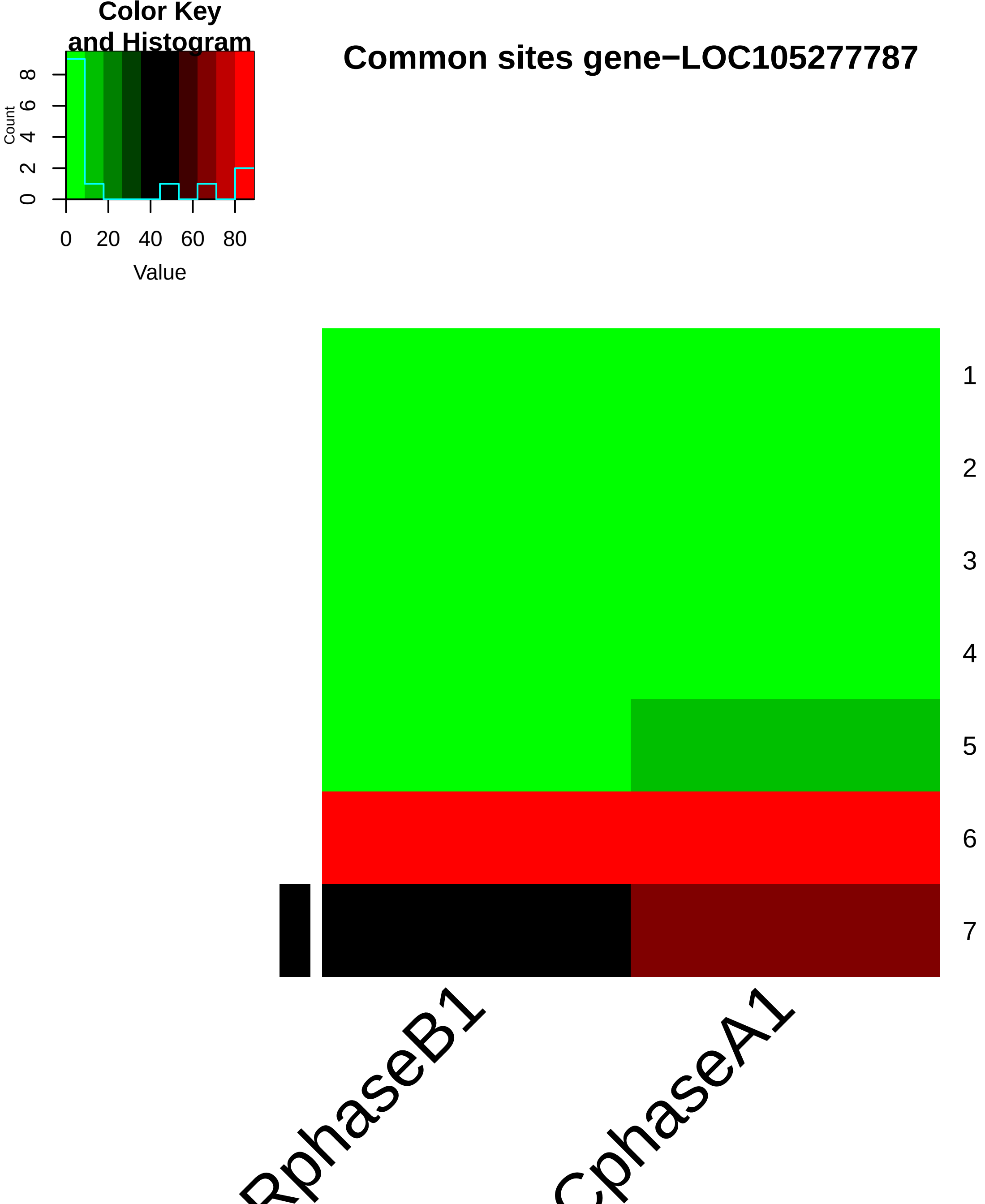

Color Key
and Histogram
Count
Value
0 20 40 60 80
0 2 4 6 8
Common sites gene−LOC105277787
1
2
3
4
5
6
7
RphaseB1
CphaseA1

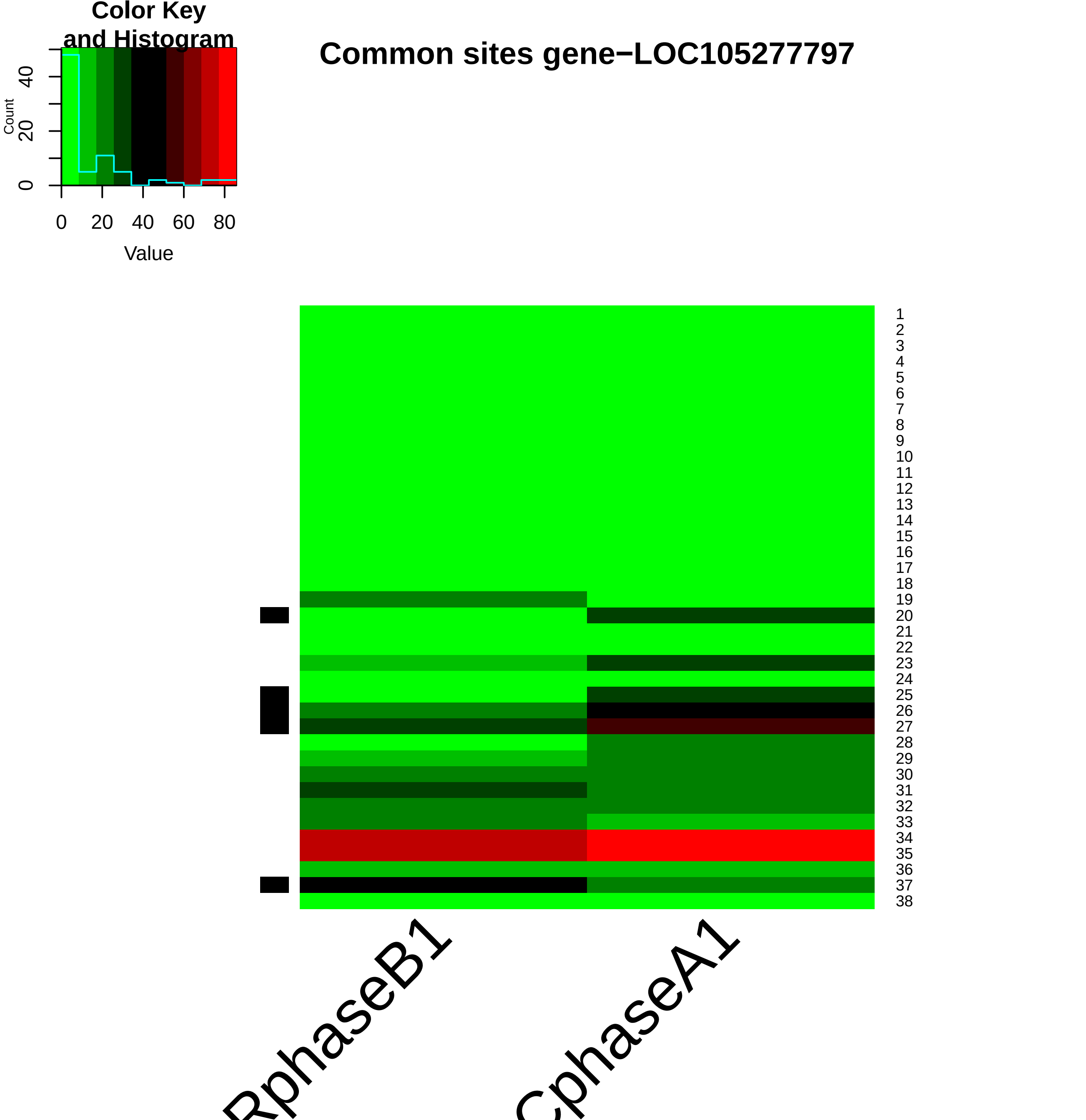
Color Key
and Histogram
Count
0
20
40
0
20
40
60
80
Value
Common sites gene−LOC105277797
RphaseB1
CphaseA1

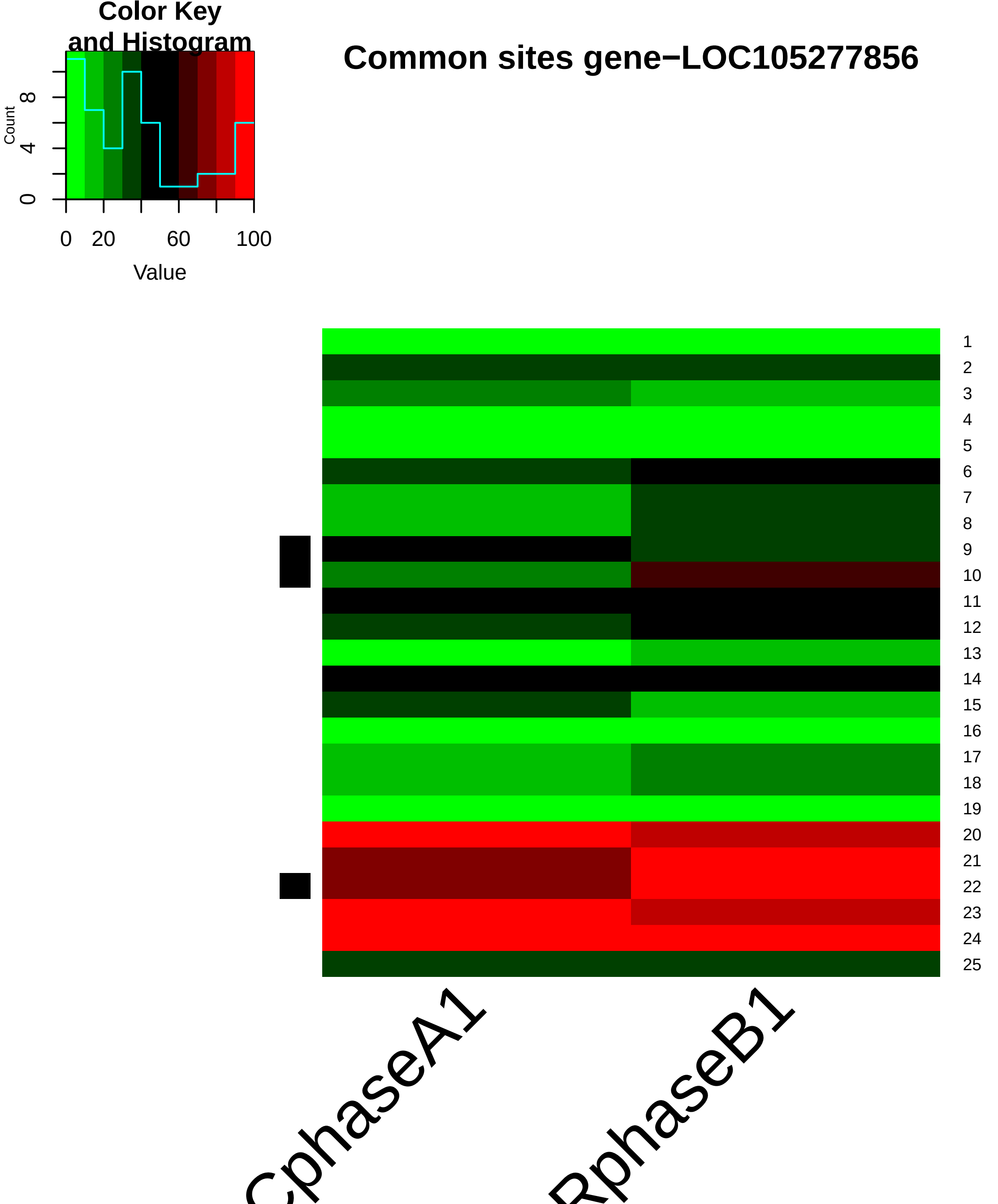
Color Key
and Histogram
Count
0
4
8
0
20
60
100
Value
Common sites gene−LOC105277856
CphaseA1
RphaseB1

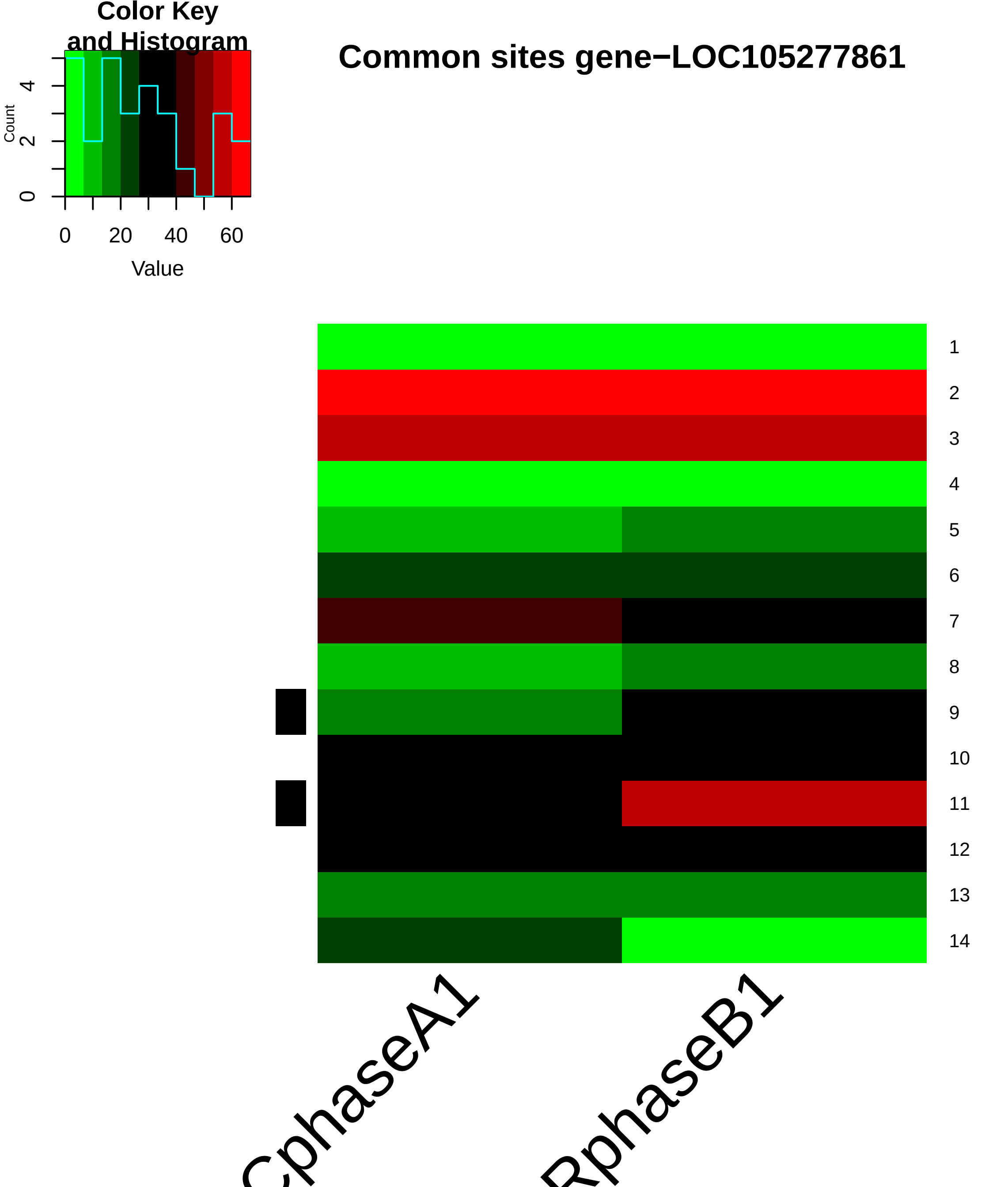
Color Key
and Histogram
Count
0
2
4
0
20
40
60
Value
Common sites gene−LOC105277861
1
2
3
4
5
6
7
8
9
10
11
12
13
14
CphaseA1
RphaseB1

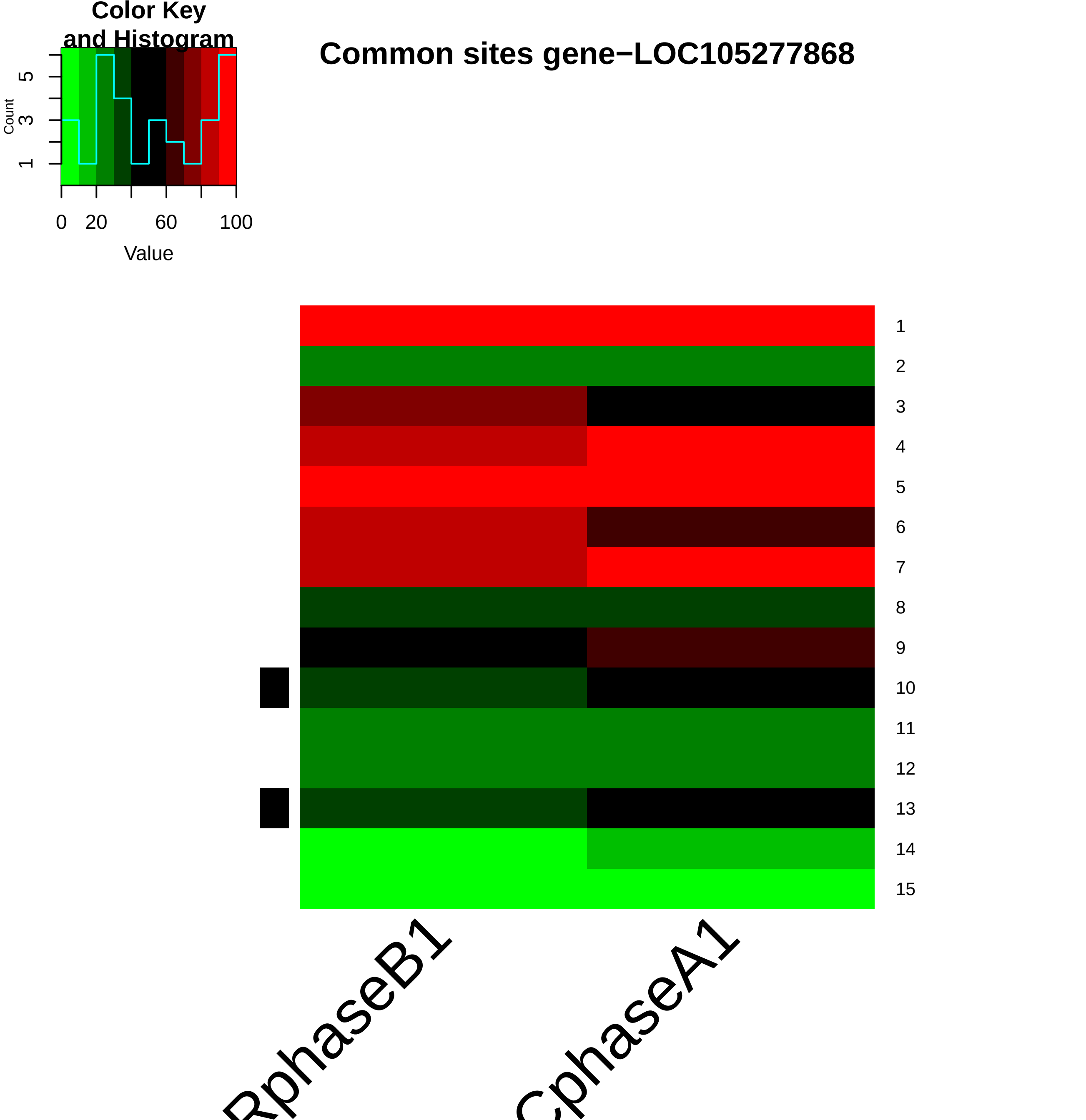
Color Key
and Histogram
Count
1
3
5
0
20
60
100
Value
Common sites gene−LOC105277868
RphaseB1
CphaseA1

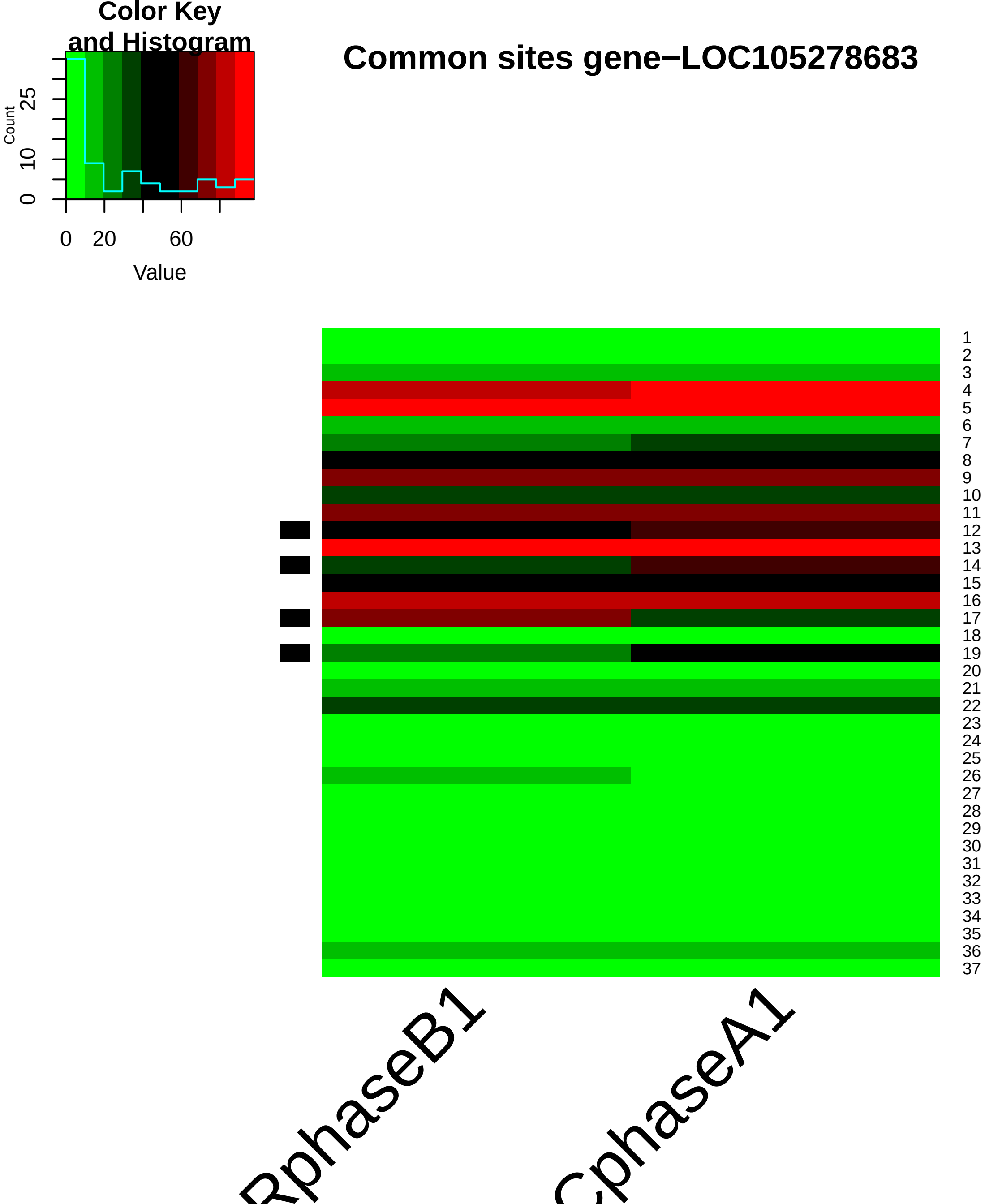
Color Key
and Histogram
Count
0
10
25
0
20
60
Value
Common sites gene−LOC105278683
RphaseB1
CphaseA1

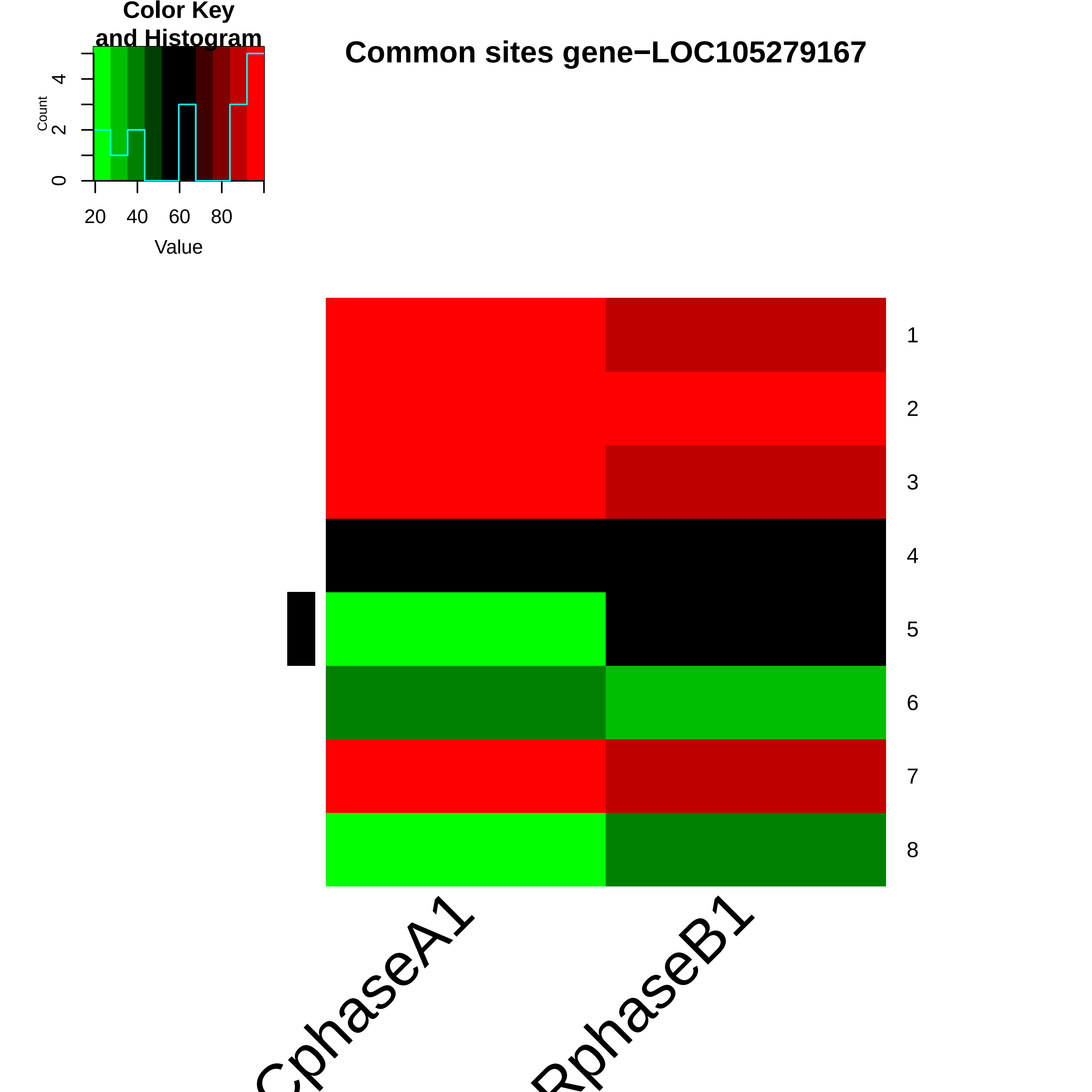

Color Key
and Histogram
Count
0
2
4
20
40
60
80
Value
Common sites gene−LOC105279167
1
2
3
4
5
6
7
8
CphaseA1
RphaseB1

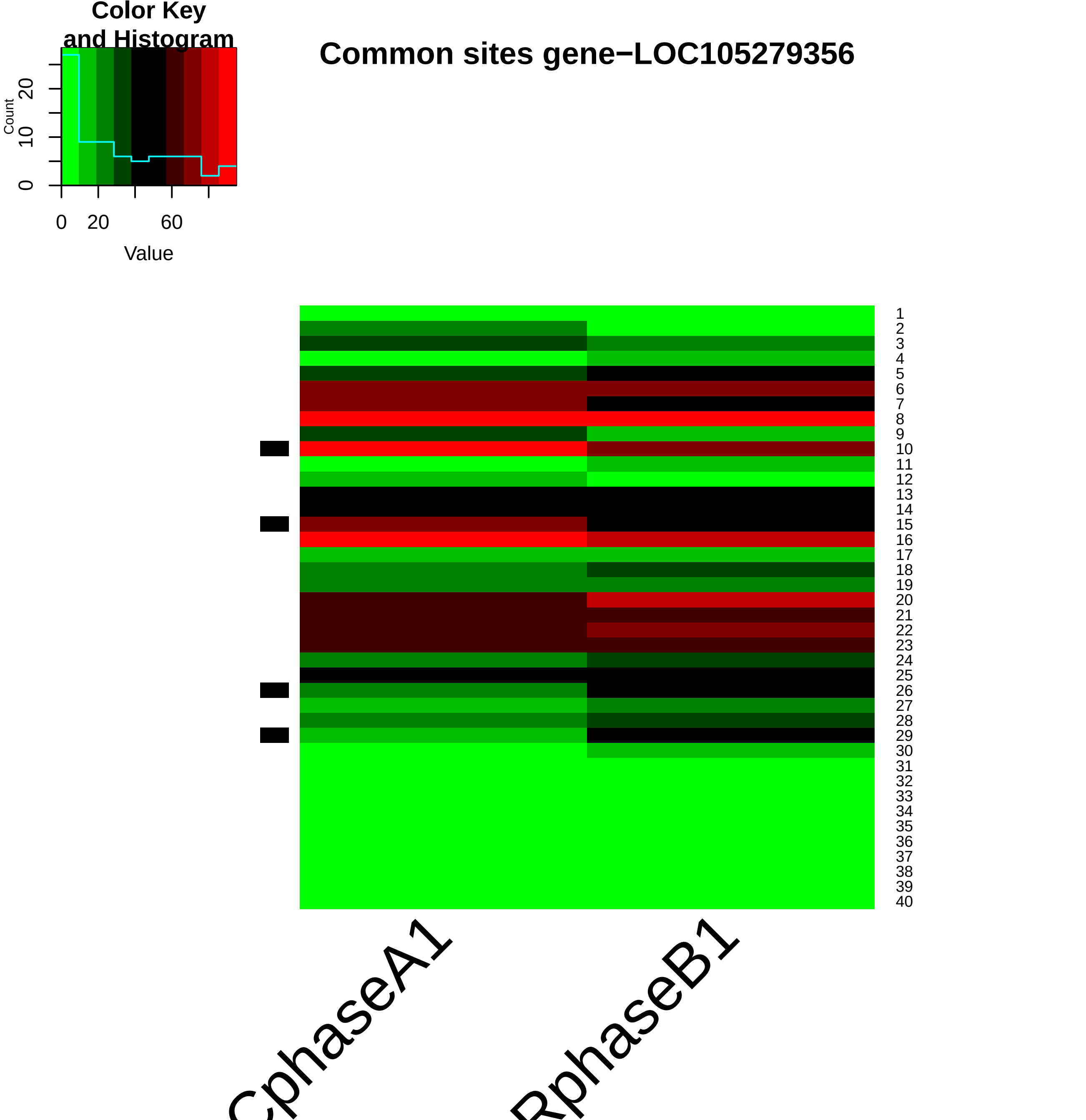
Color Key
and Histogram
Count
0
10
20
0
20
60
Value
Common sites gene−LOC105279356
CphaseA1
RphaseB1

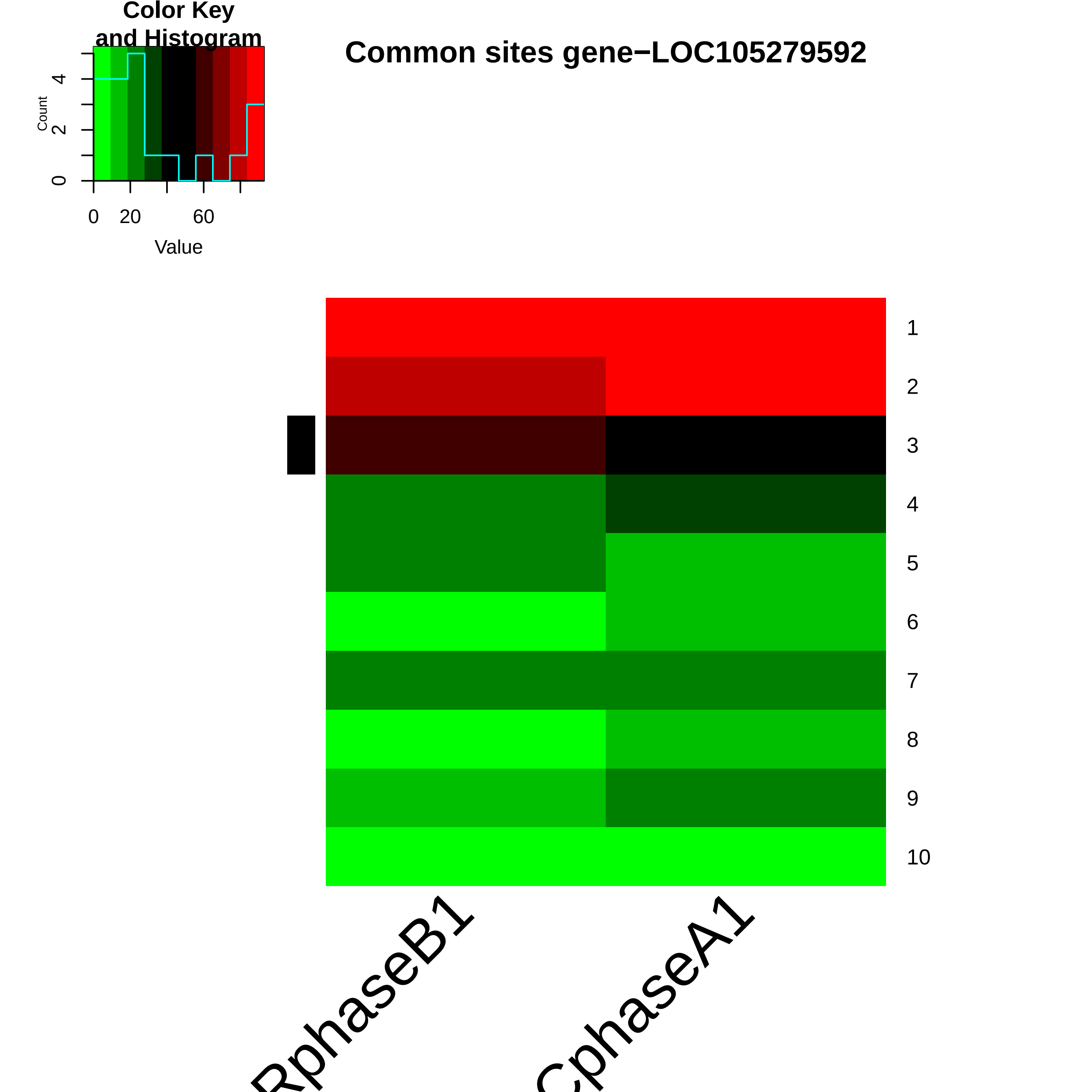

Color Key
and Histogram
Count
Value
0
20
60
0
2
4
Common sites gene−LOC105279592
1
2
3
4
5
6
7
8
9
10
RphaseB1
CphaseA1

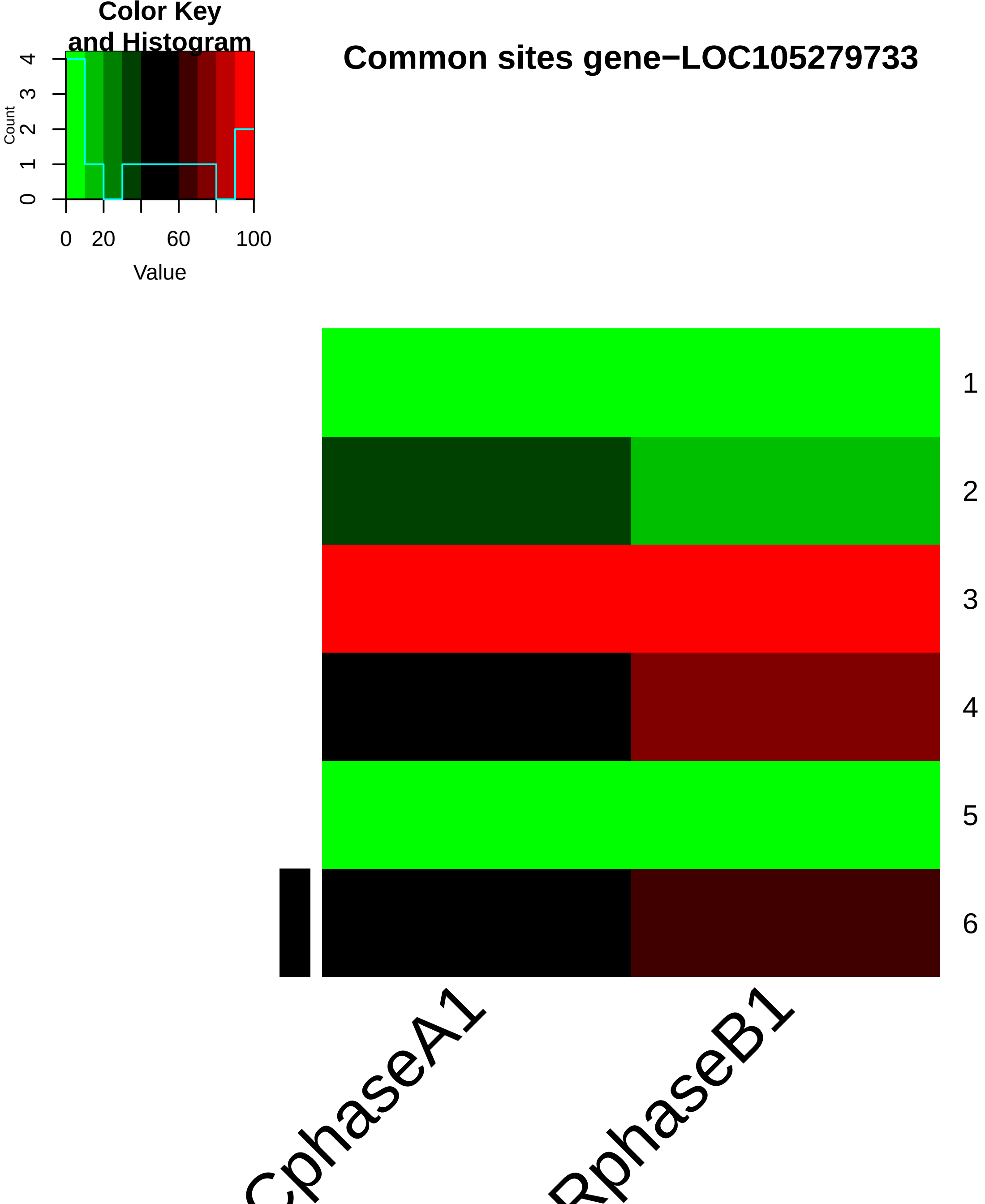
Color Key and Histogram
Count
0
1
2
3
4
0
20
60
100
Value
Common sites gene−LOC105279733
1
2
3
4
5
6
CphaseA1
RphaseB1

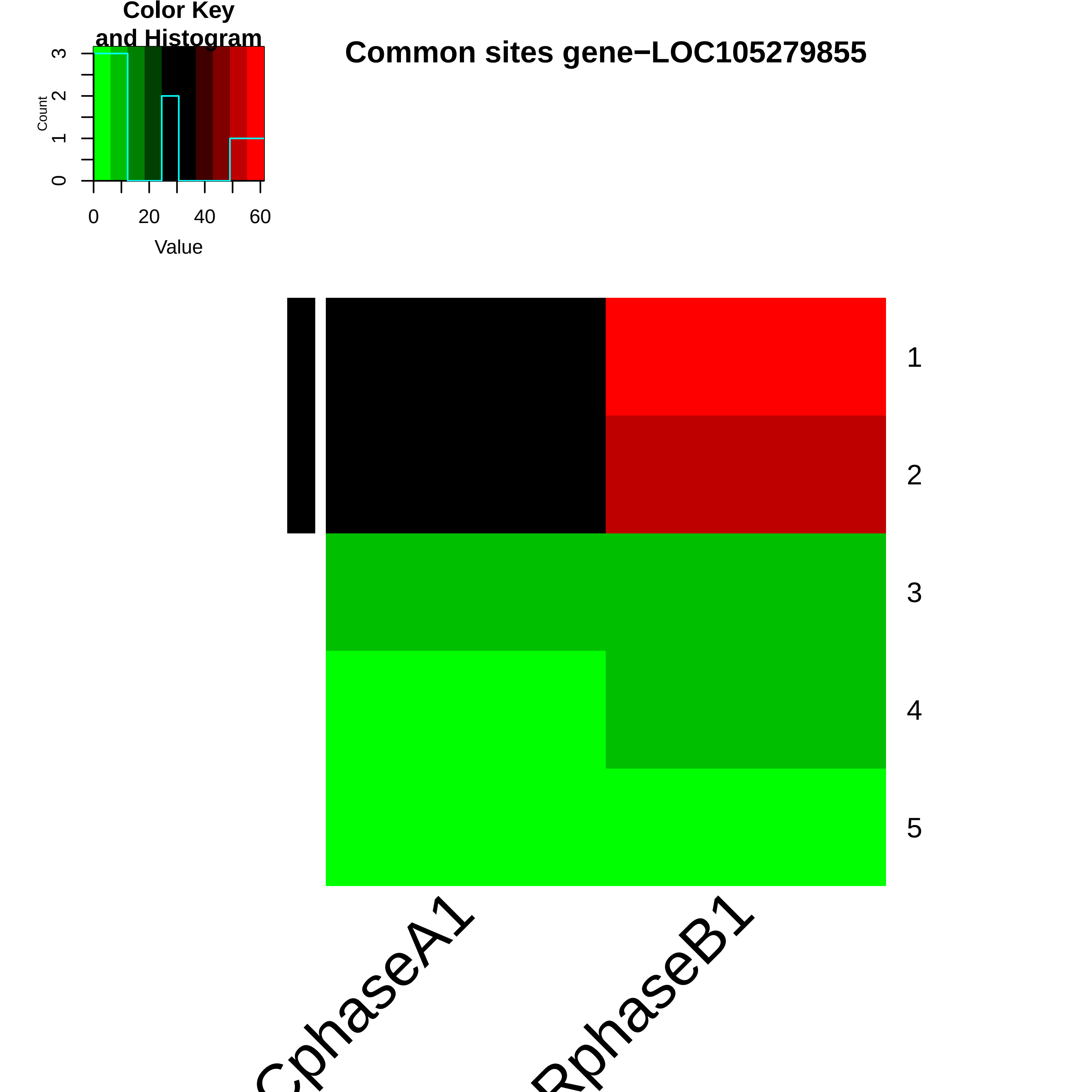

Color Key
and Histogram
Count
0
1
2
3
0
20
40
60
Value
Common sites gene−LOC105279855
1
2
3
4
5
CphaseA1
RphaseB1

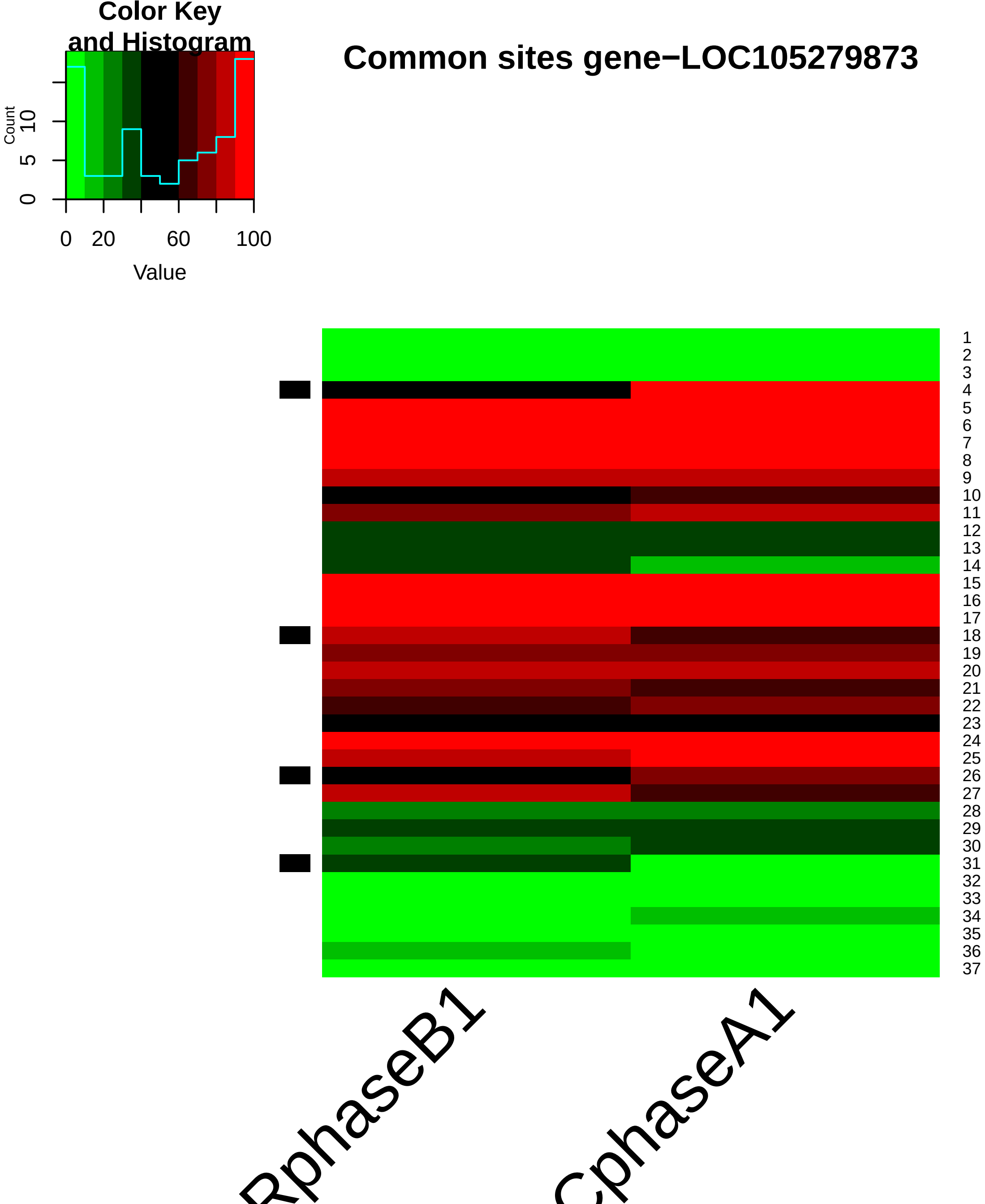

Color Key
and Histogram
Count
0
5
10
0
20
60
100
Value
Common sites gene−LOC105279873
RphaseB1
CphaseA1

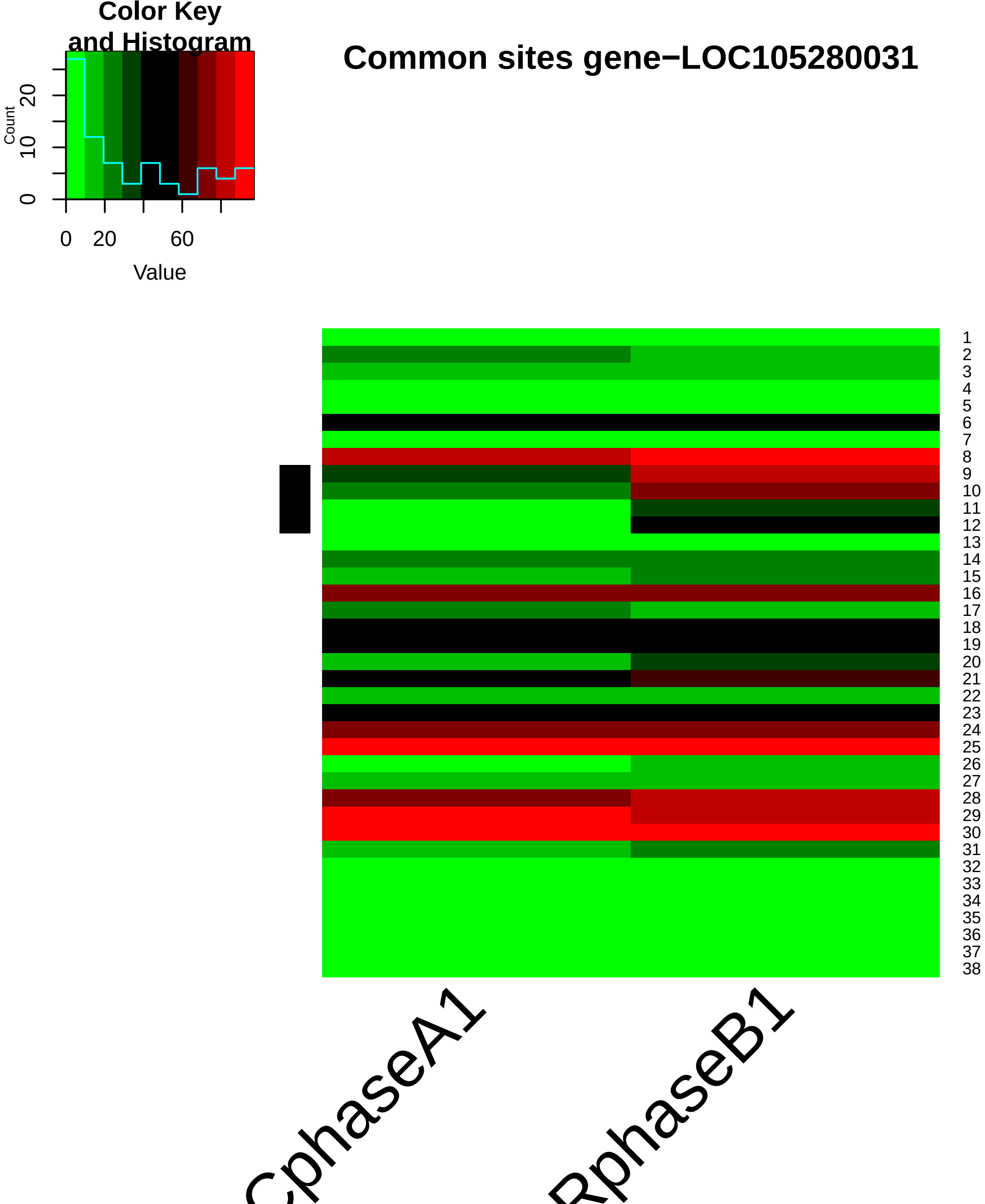

Color Key
and Histogram
Count
0
10
20
0
20
60
Value
Common sites gene−LOC105280031
CphaseA1
RphaseB1

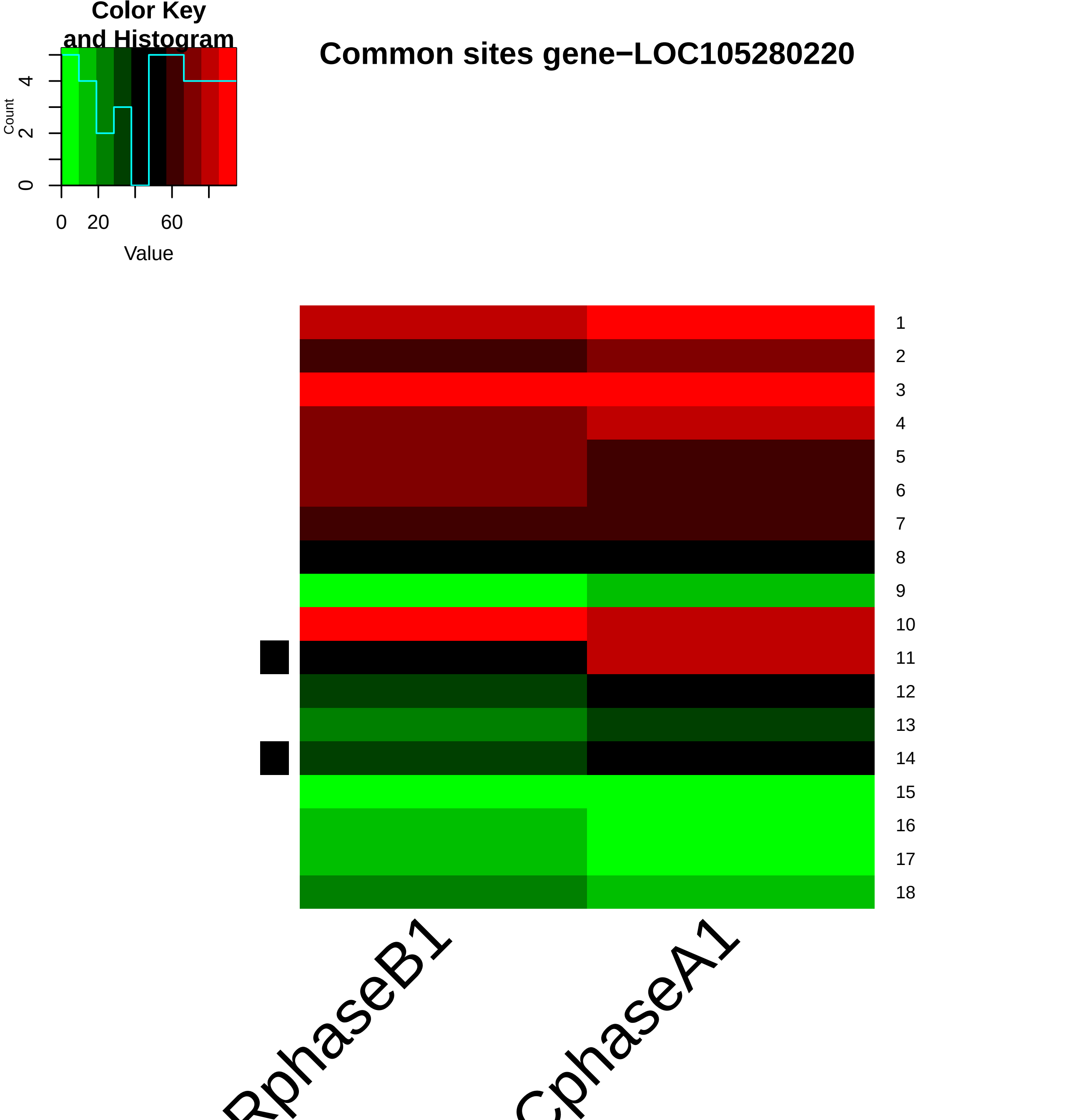
Color Key
and Histogram
Count
0
2
4
0
20
60
Value
Common sites gene−LOC105280220
RphaseB1
CphaseA1

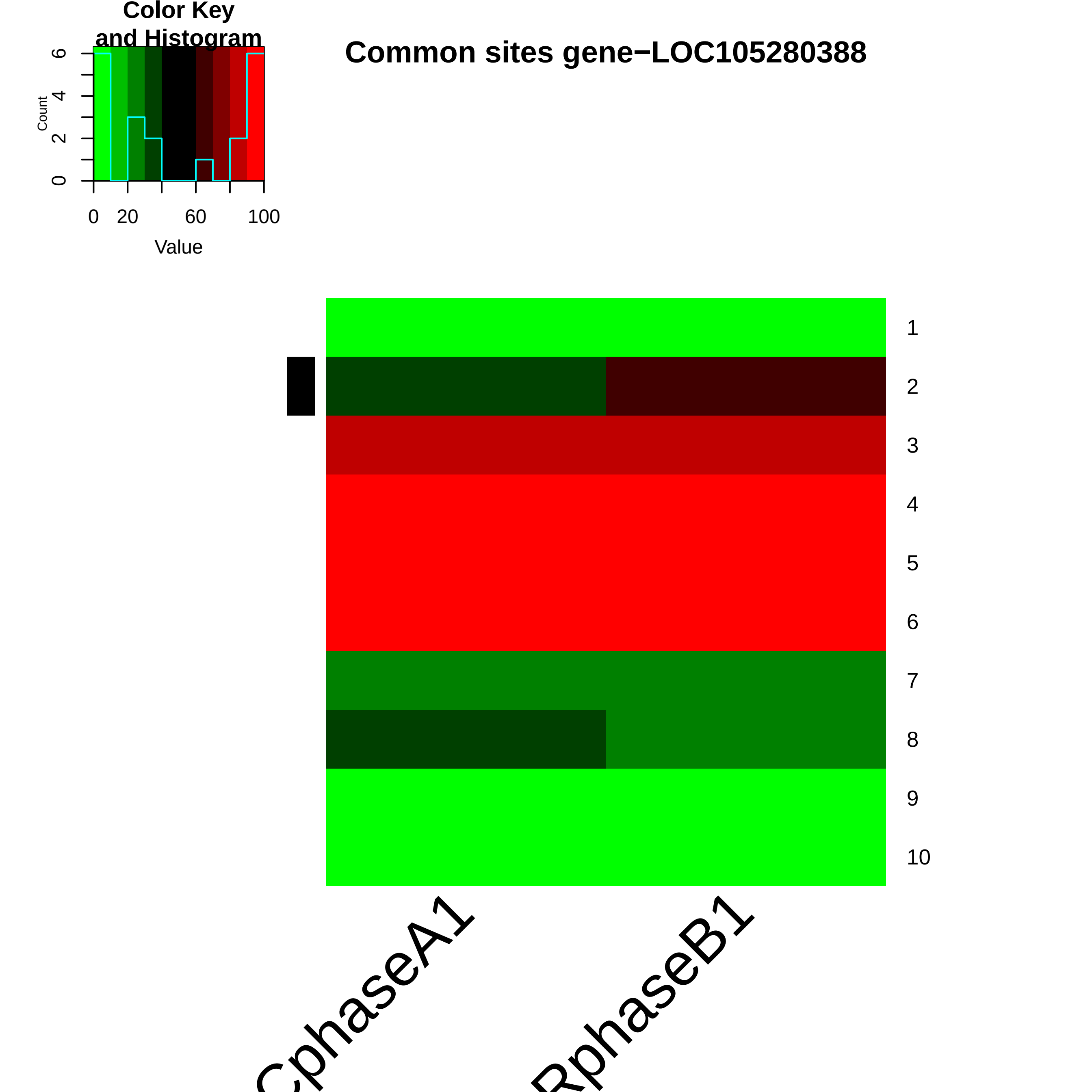

Color Key
and Histogram
Count
0
2
4
6
0
20
60
100
Value
Common sites gene−LOC105280388
1
2
3
4
5
6
7
8
9
10
CphaseA1
RphaseB1

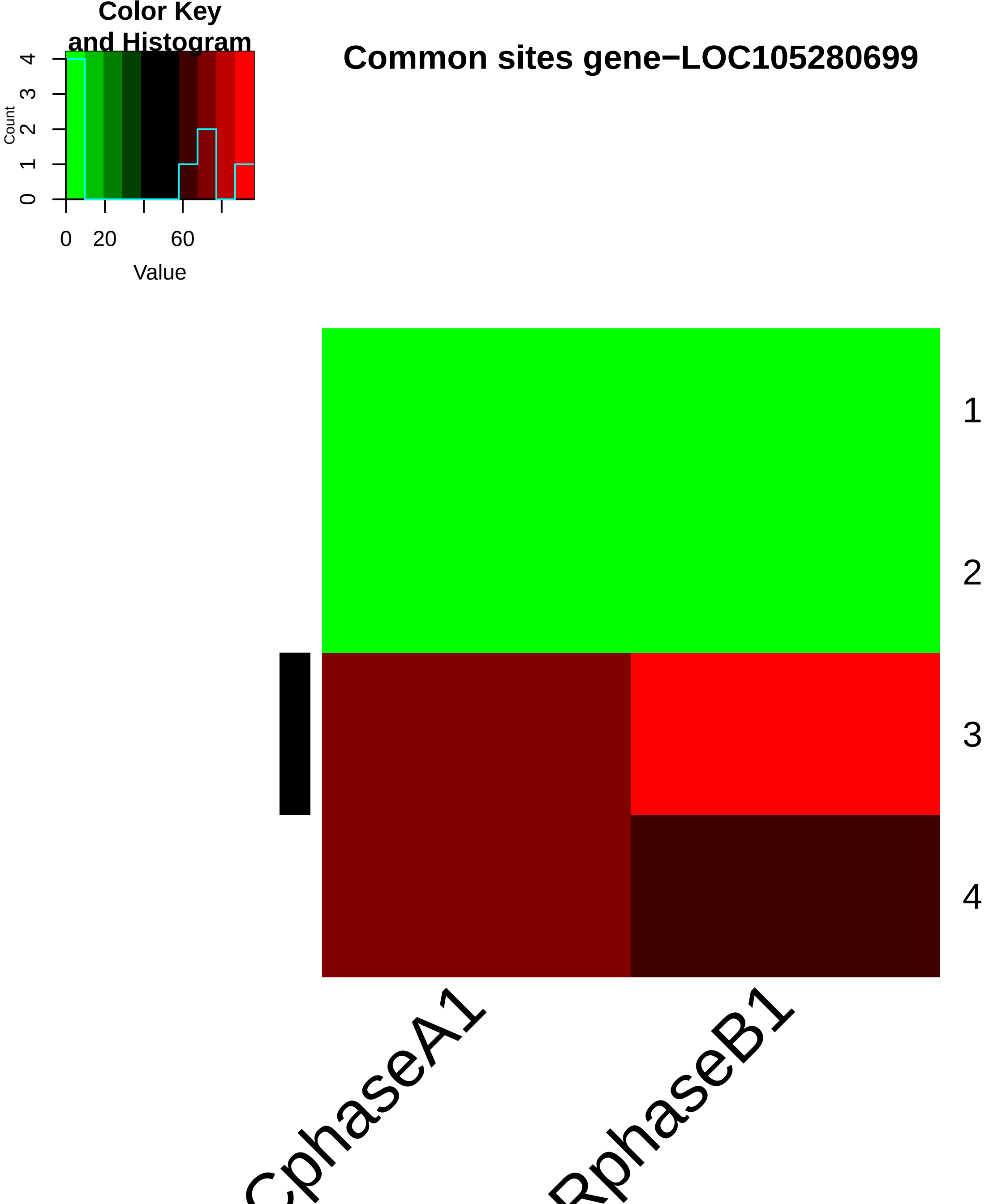

Color Key
and Histogram
Count
0
1
2
3
4
0
20
60
Value
Common sites gene−LOC105280699
1
2
3
4
CphaseA1
RphaseB1

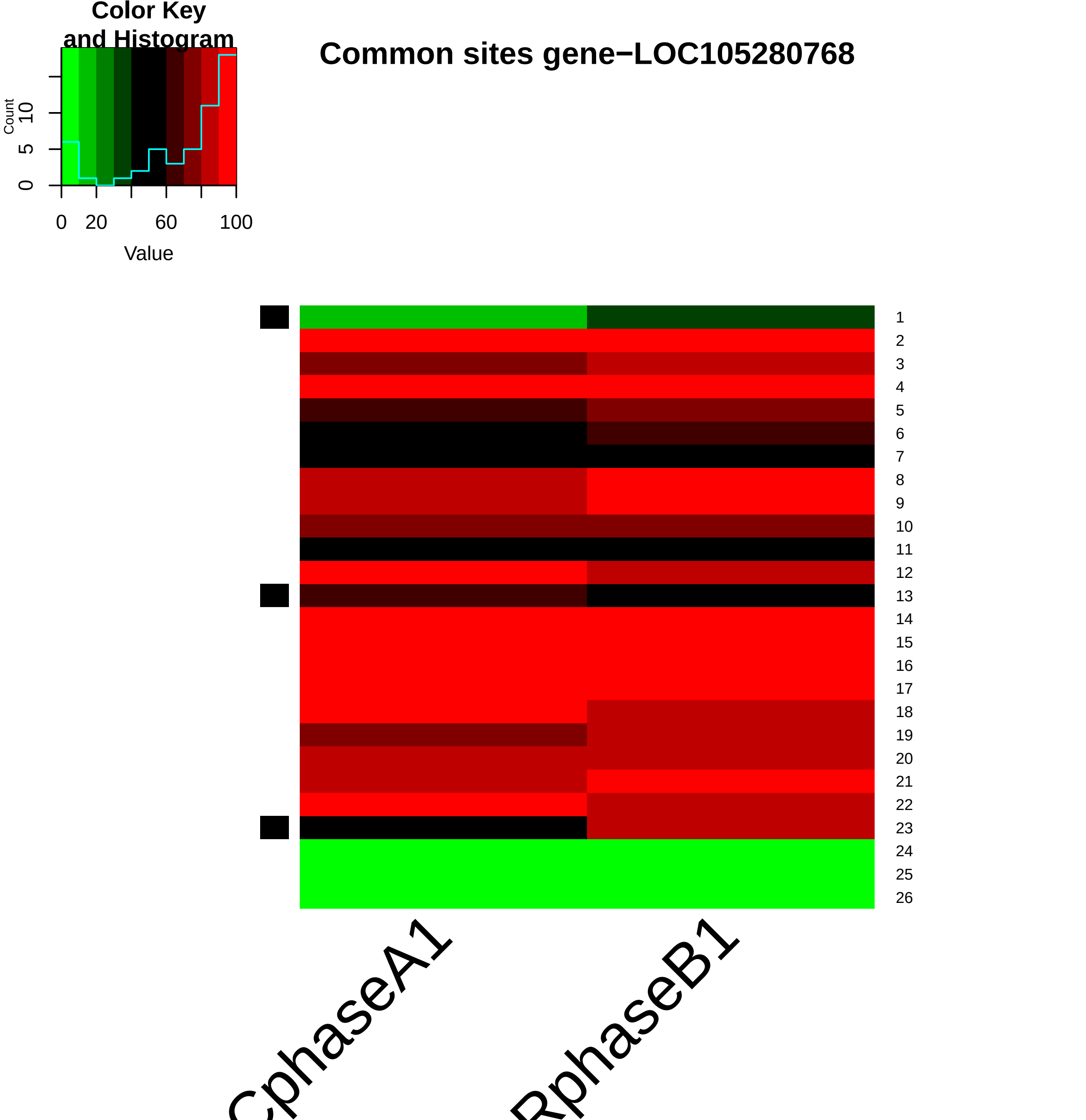
Color Key
and Histogram
Count
0
5
10
0
20
60
100
Value
Common sites gene−LOC105280768

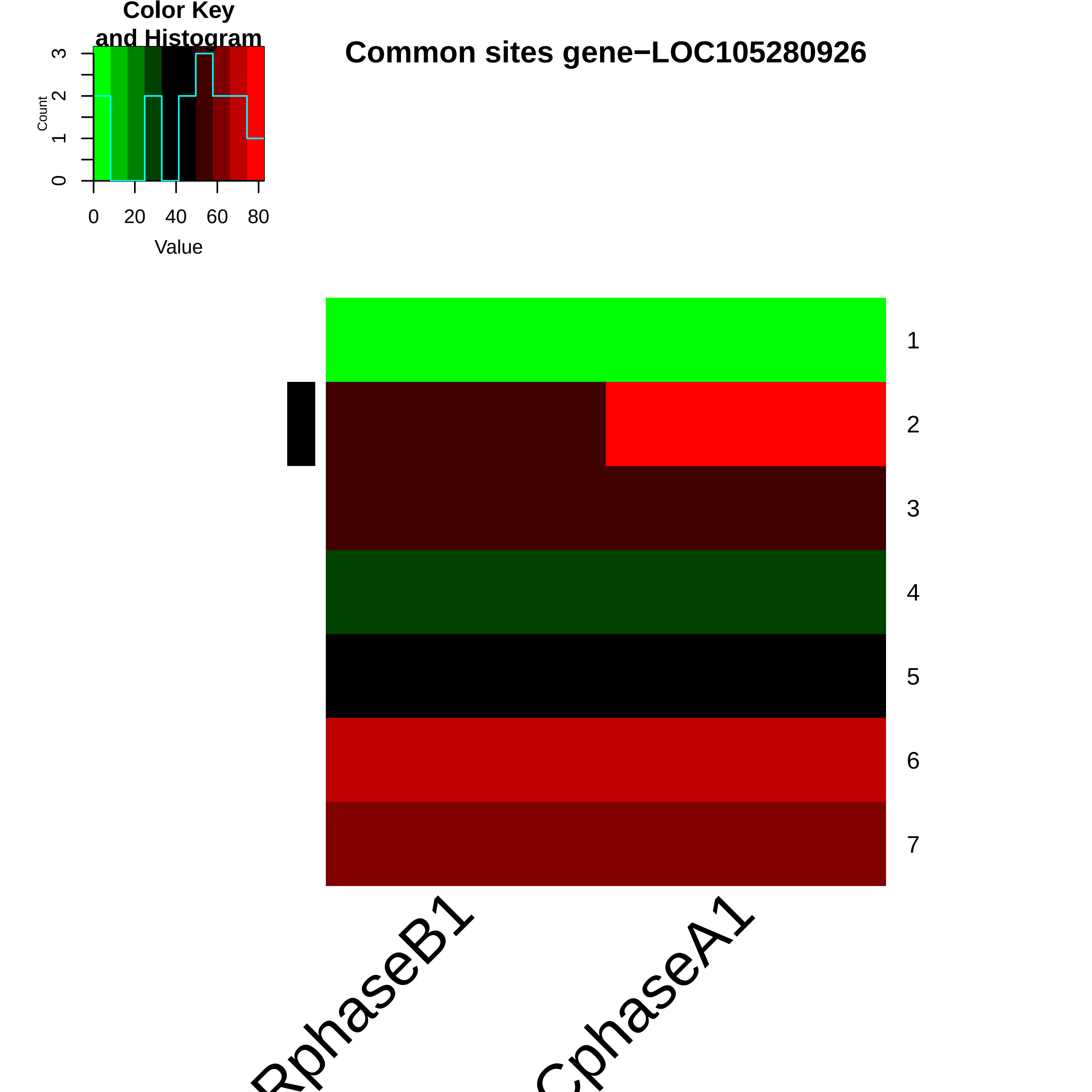
Color Key
and Histogram
Count
0
1
2
3
0
20
40
60
80
Value
Common sites gene−LOC105280926
1
2
3
4
5
6
7
RphaseB1
CphaseA1

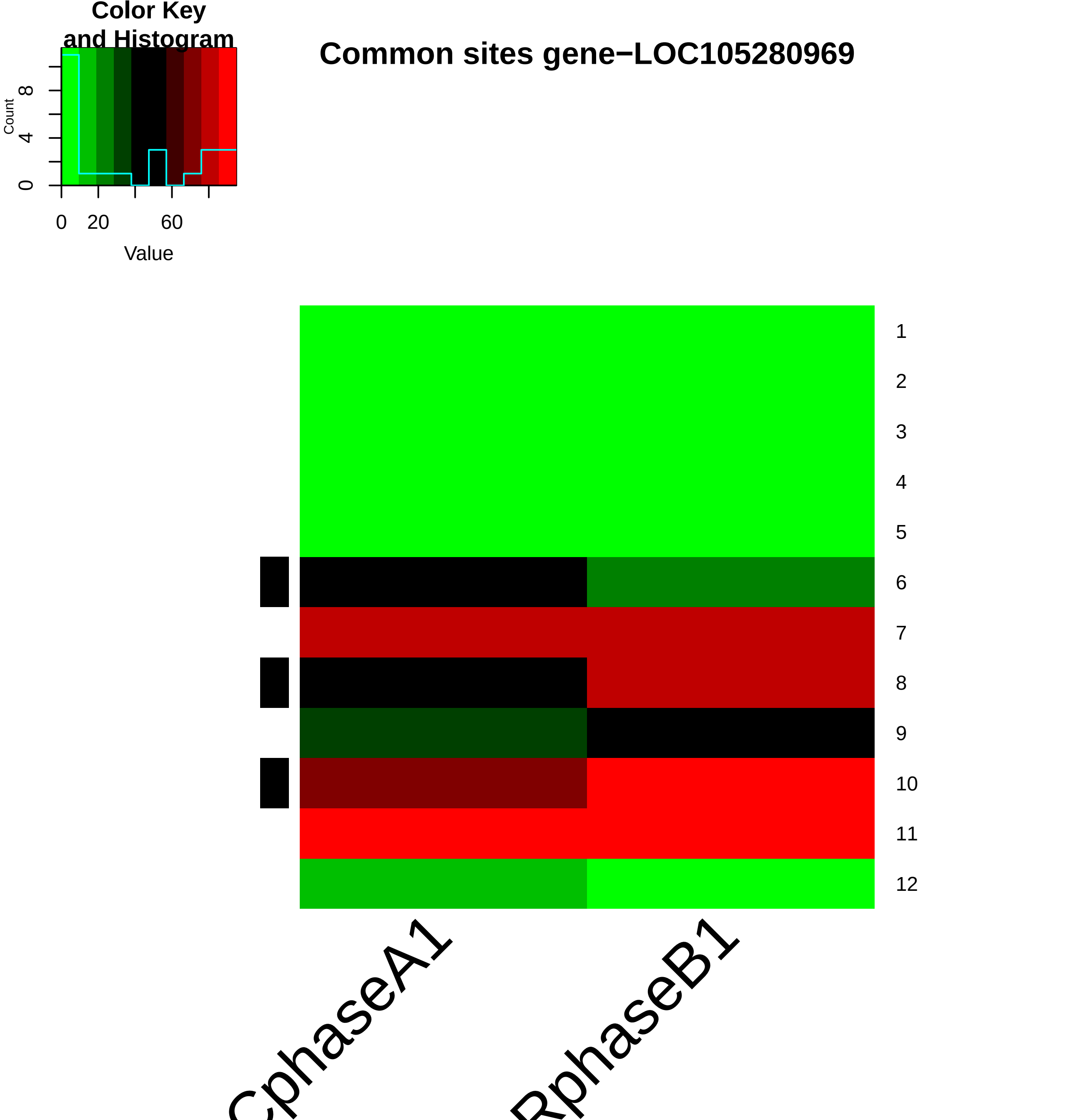
Color Key
and Histogram
Count
0
4
8
0
20
60
Value
Common sites gene−LOC105280969
1
2
3
4
5
6
7
8
9
10
11
12
CphaseA1
RphaseB1

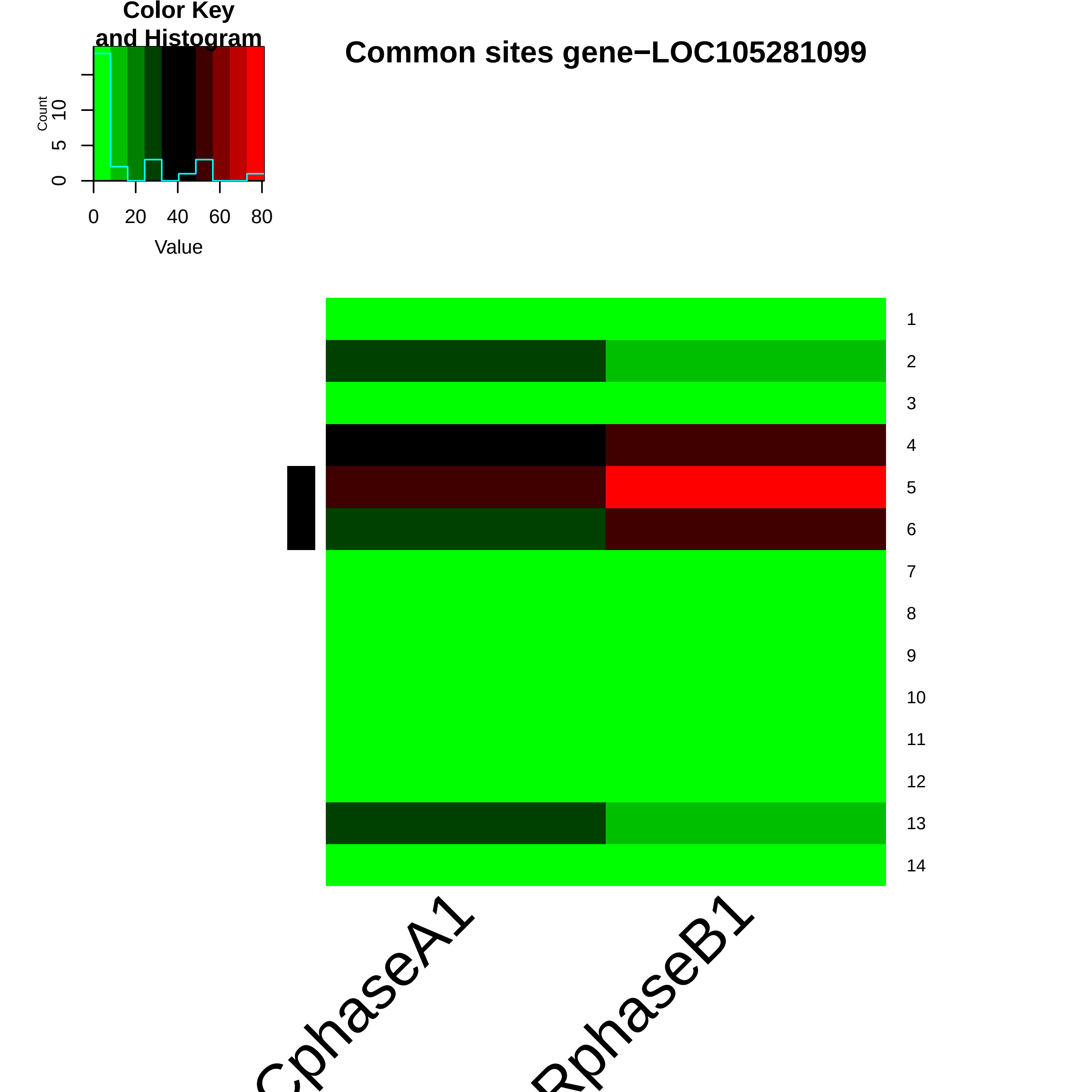
Color Key
and Histogram
Count
0
5
10
0
20
40
60
80
Value
Common sites gene−LOC105281099
1
2
3
4
5
6
7
8
9
10
11
12
13
14

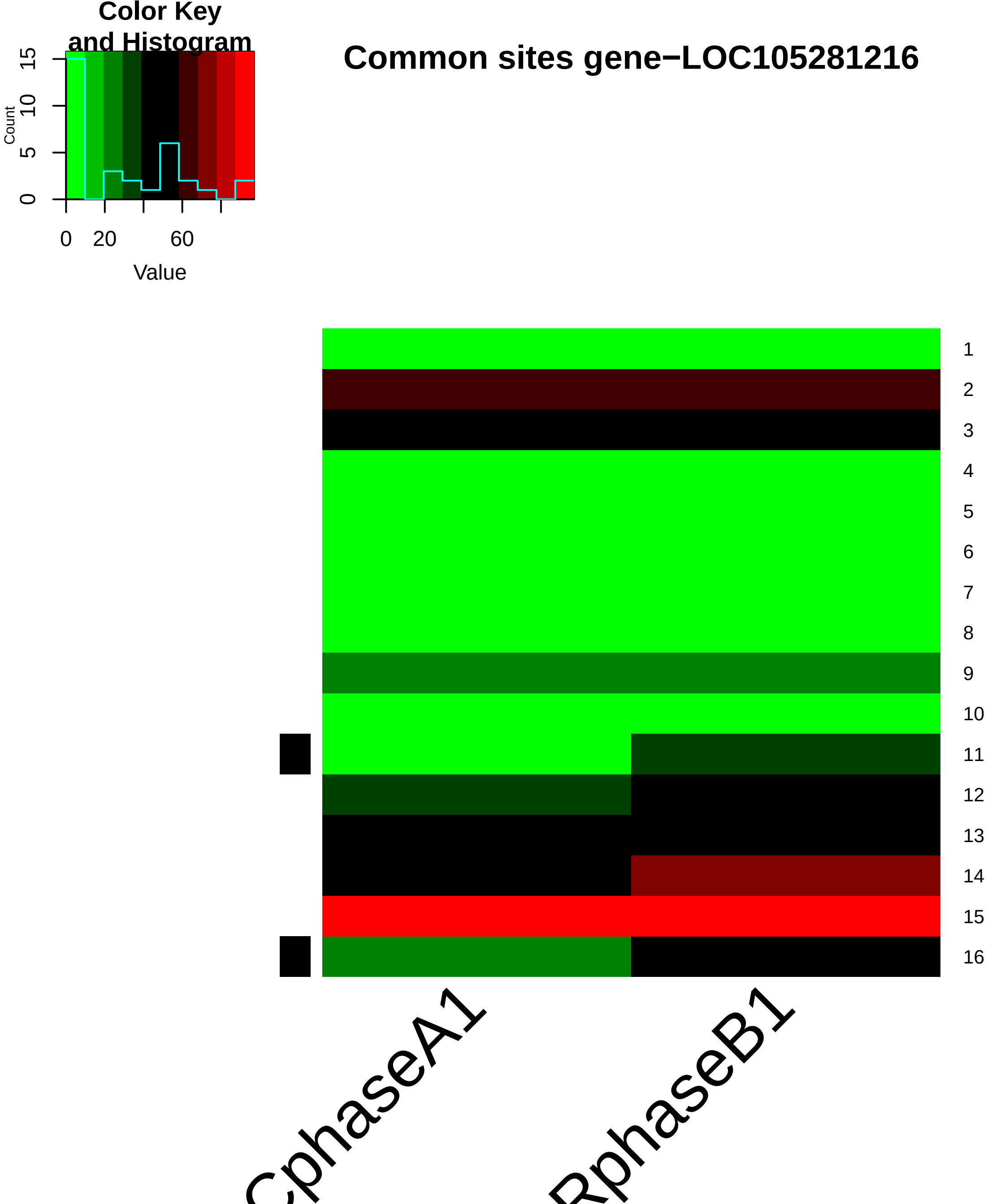
Color Key
and Histogram
Count
0
5
10
15
0
20
60
Value
Common sites gene−LOC105281216
CphaseA1
RphaseB1

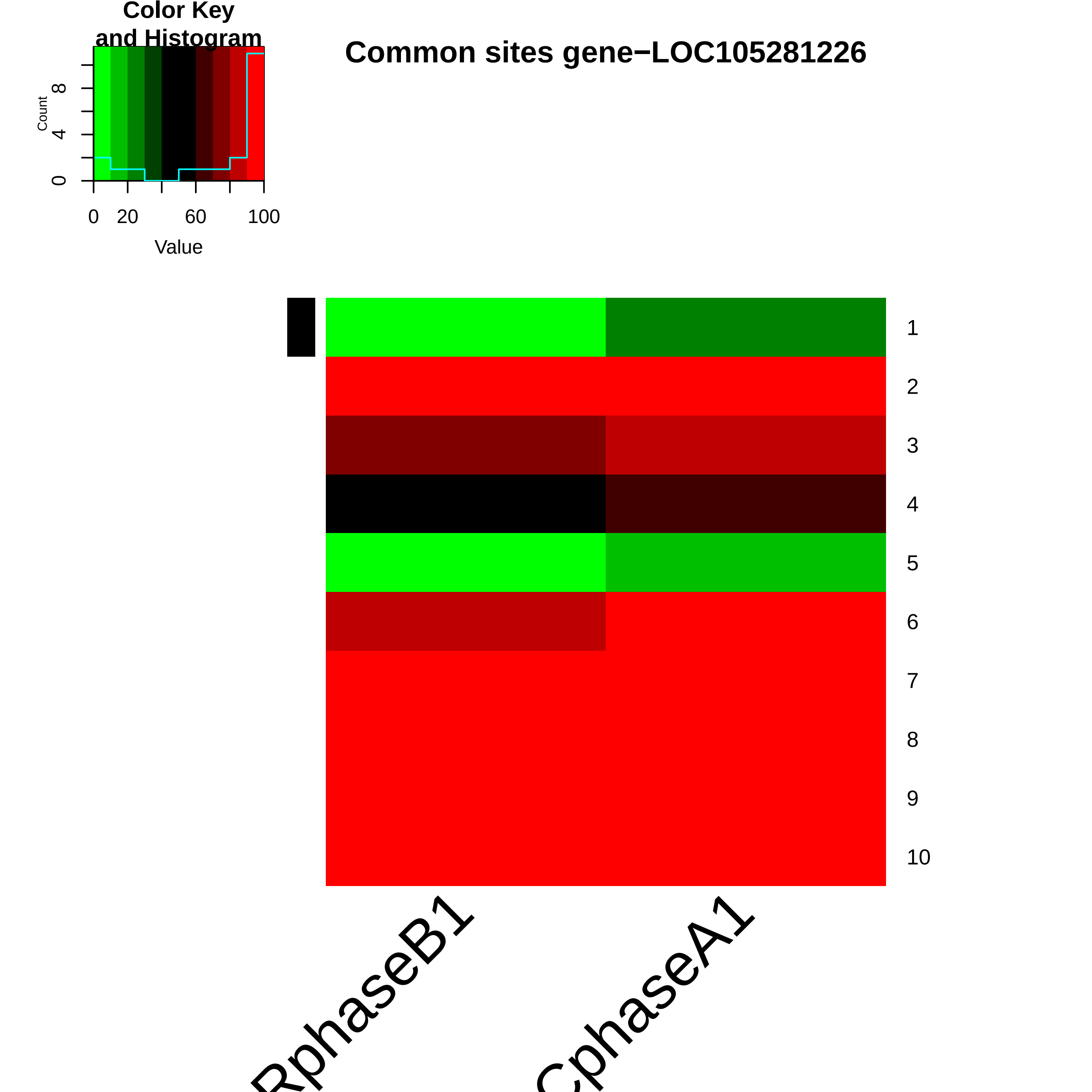
Color Key
and Histogram
Count
0
4
8
0
20
60
100
Value
Common sites gene−LOC105281226
1
2
3
4
5
6
7
8
9
10
RphaseB1
CphaseA1

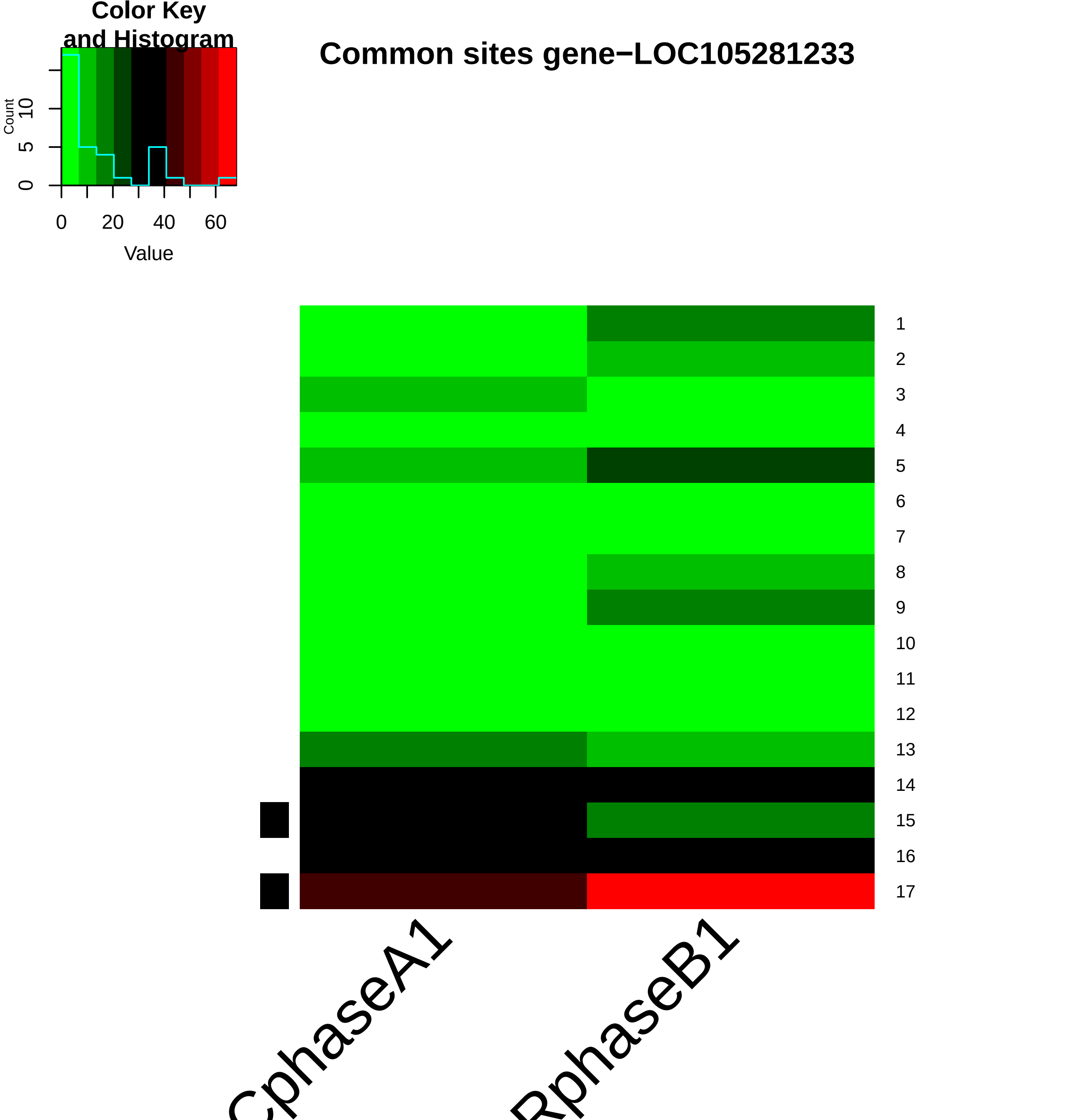

Color Key
and Histogram
Count
0
5
10
0
20
40
60
Value
Common sites gene−LOC105281233
CphaseA1
RphaseB1

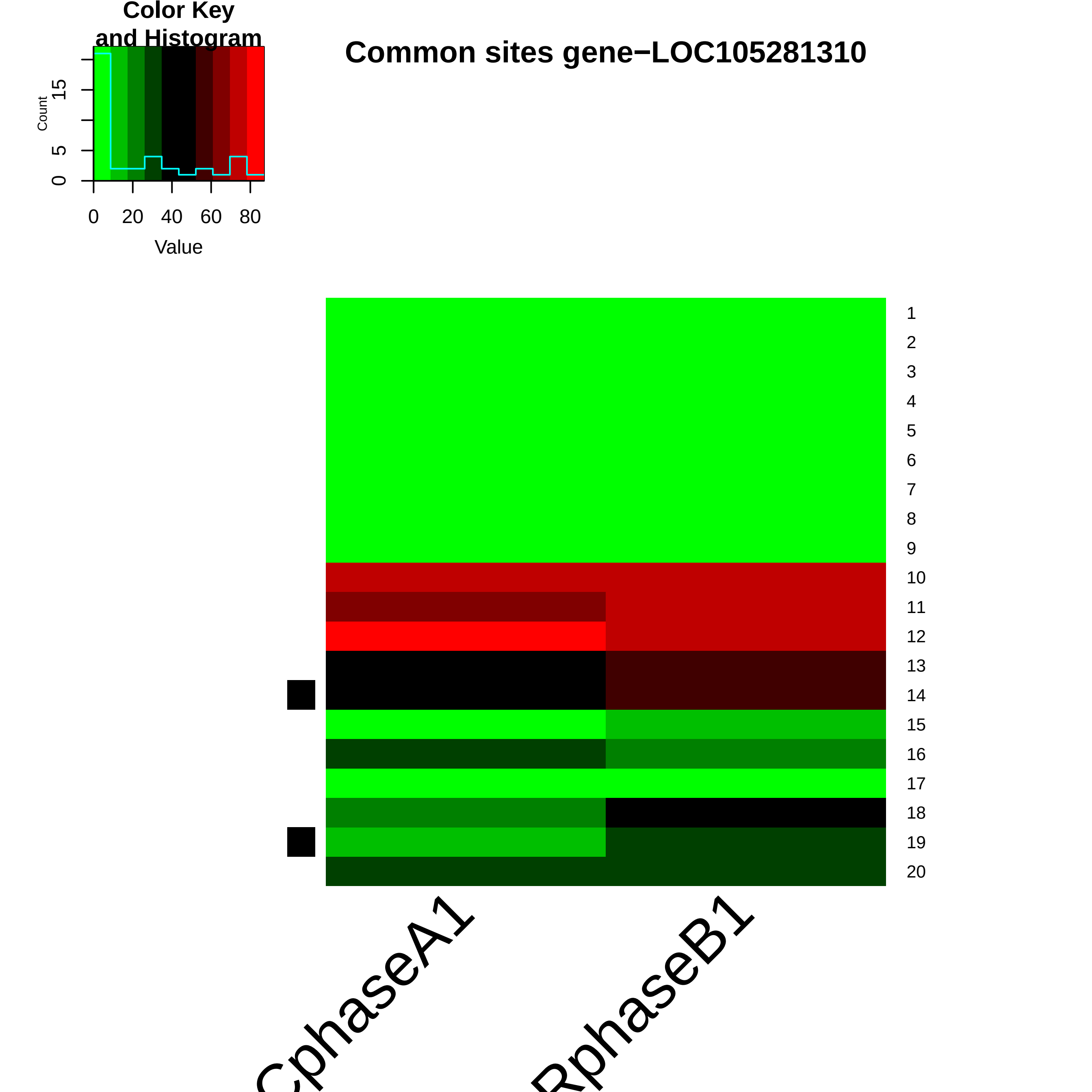

Color Key
and Histogram
Count
0
5
15
0
20
40
60
80
Value
Common sites gene−LOC105281310
CphaseA1
RphaseB1

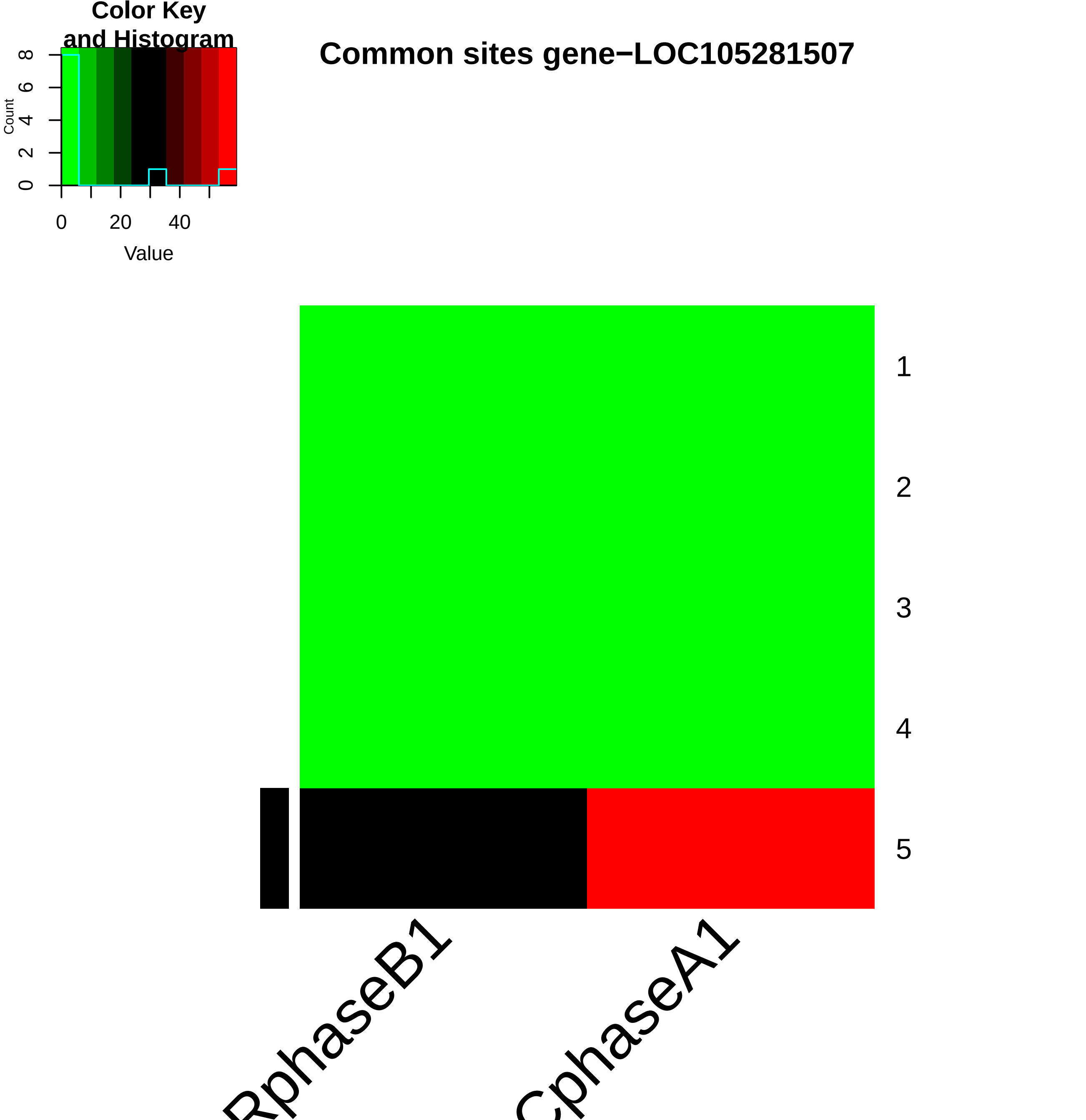

Color Key
and Histogram
Count
Value
Common sites gene−LOC105281507
1
2
3
4
5
RphaseB1
CphaseA1

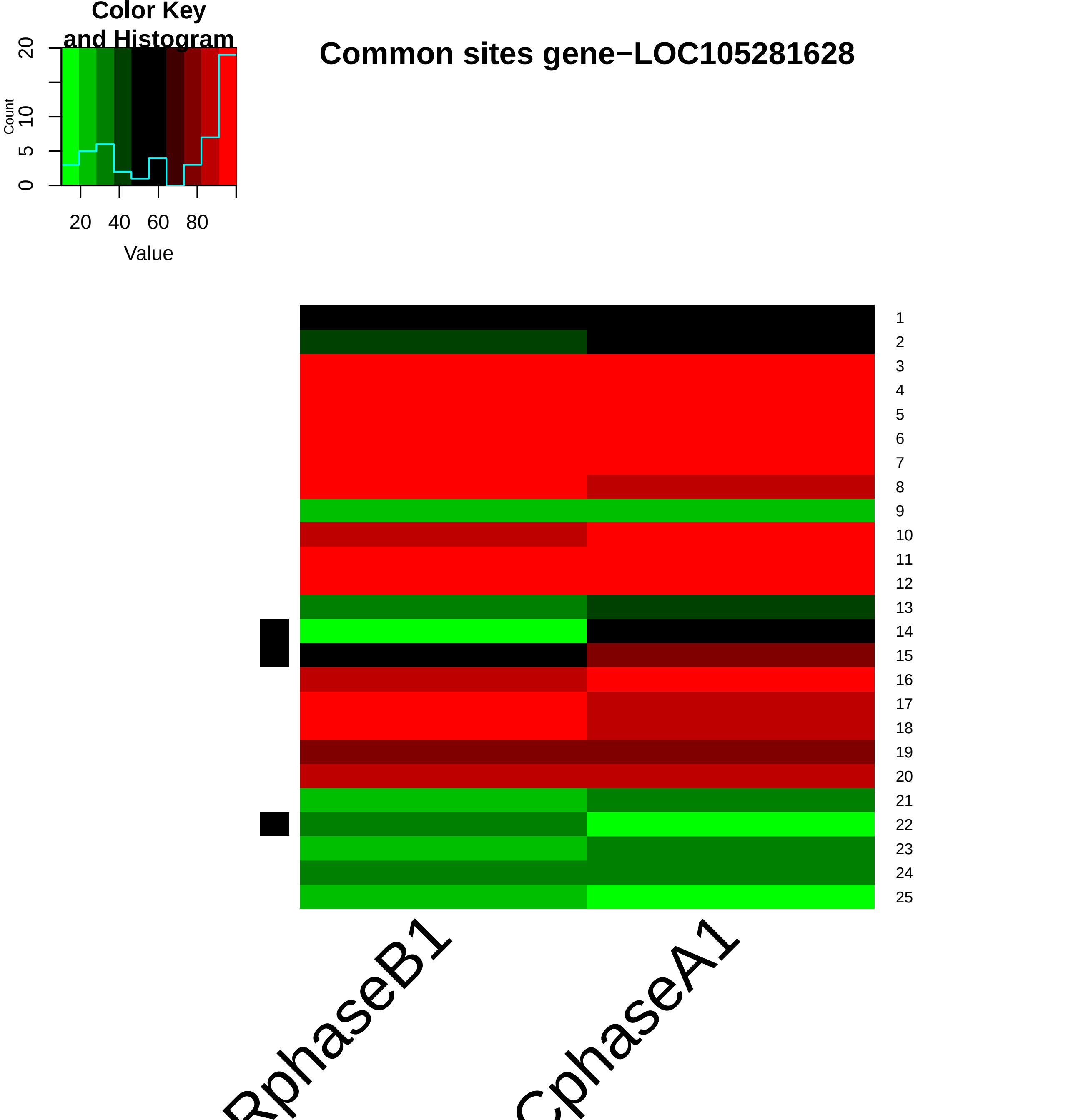
Color Key
and Histogram
Count
0
5
10
20
20
40
60
80
Value
Common sites gene−LOC105281628
RphaseB1
CphaseA1

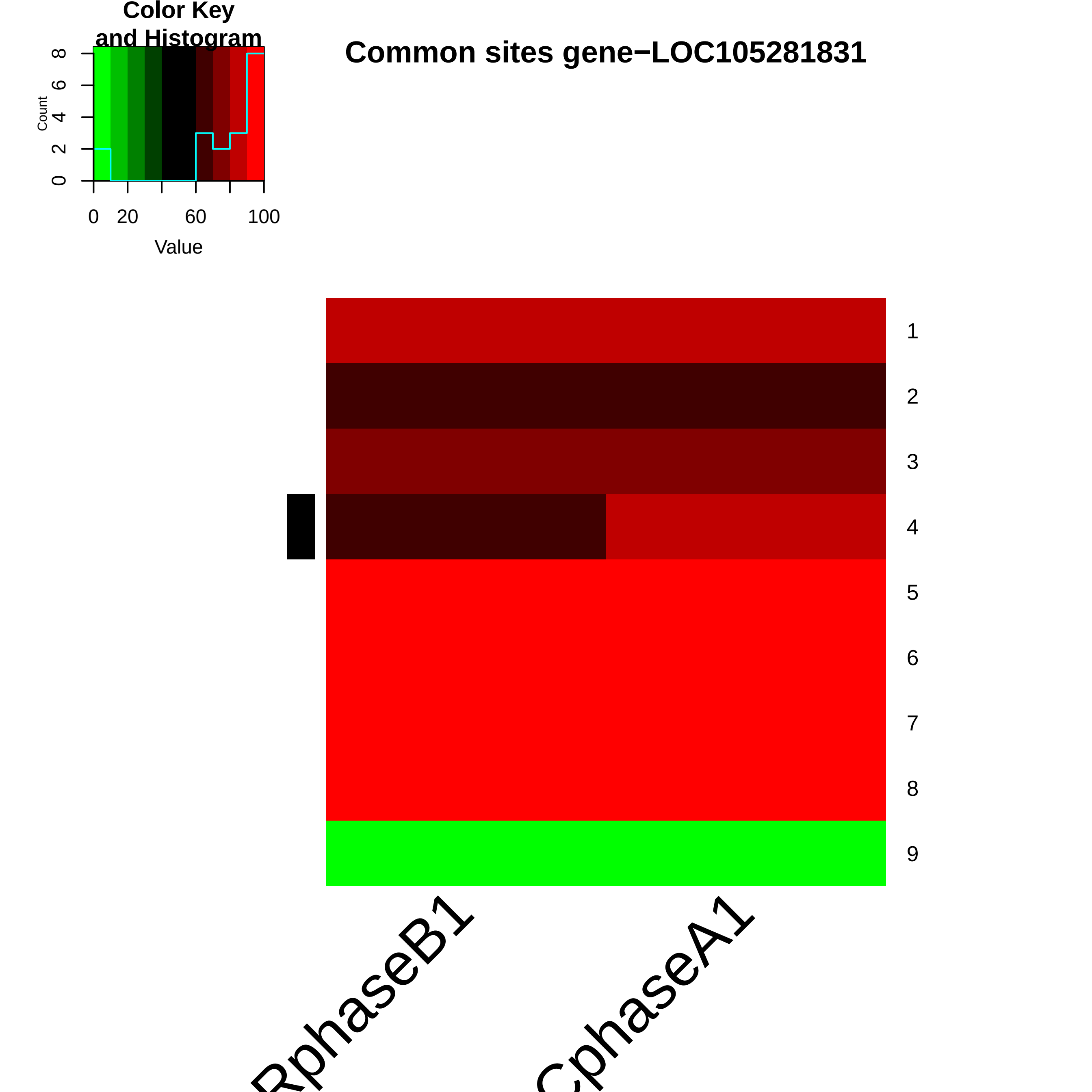
Color Key
and Histogram
Count
Value
0
20
60
100
0
2
4
6
8
Common sites gene−LOC105281831
1
2
3
4
5
6
7
8
9
RphaseB1
CphaseA1

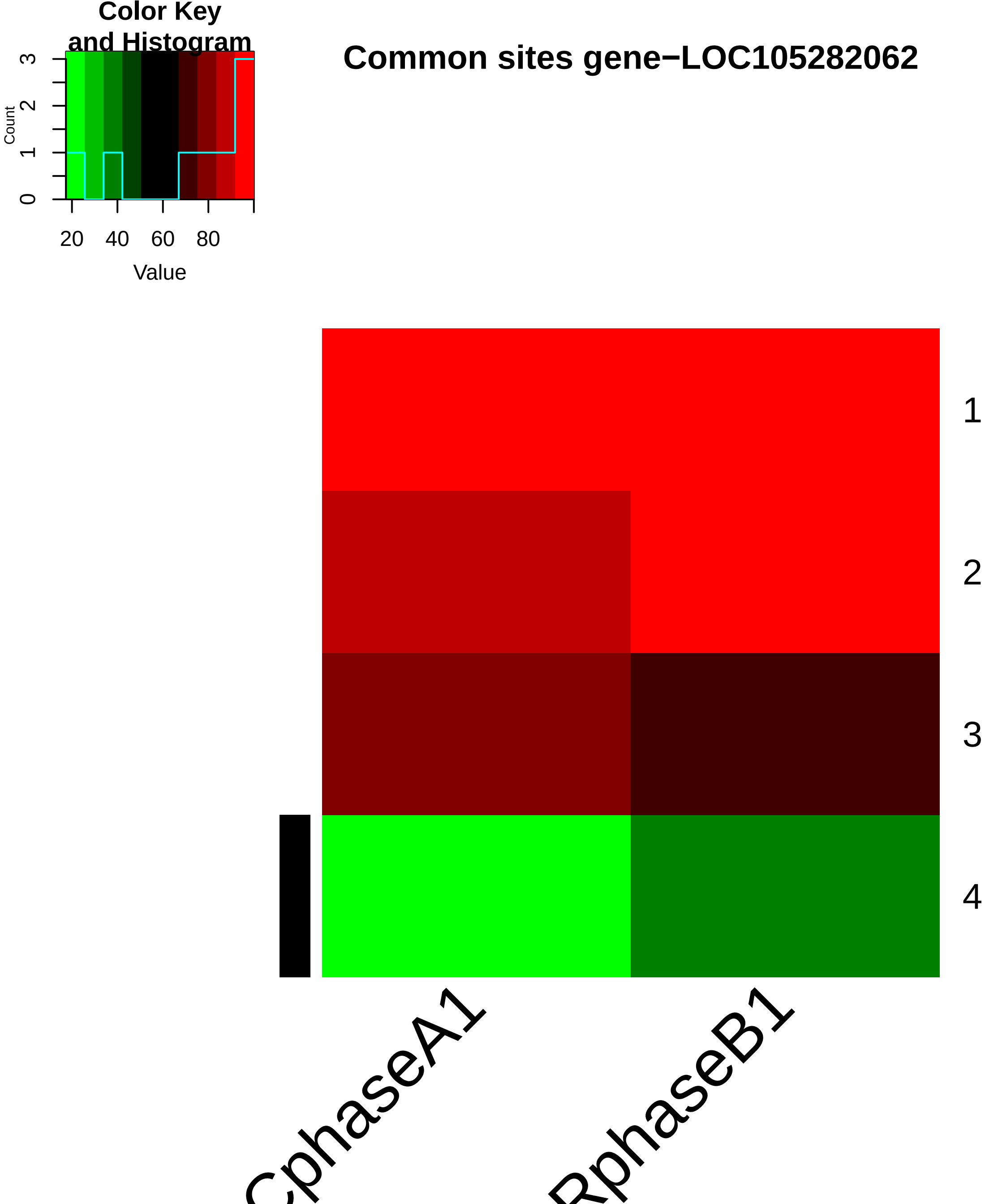

Color Key
and Histogram
Count
0
1
2
3
20
40
60
80
Value
Common sites gene−LOC105282062
1
2
3
4
CphaseA1
RphaseB1

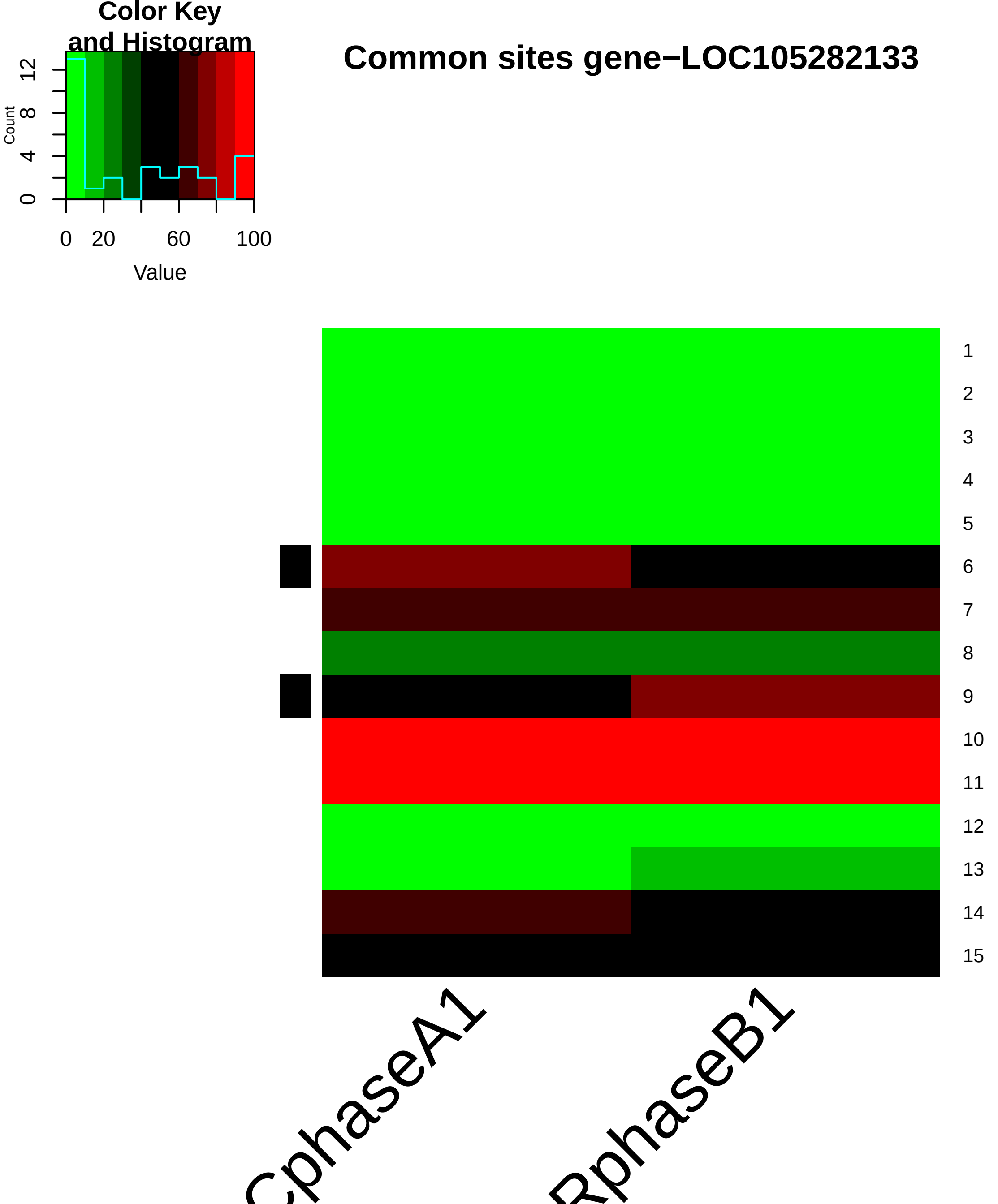
Color Key
and Histogram
Count
0
4
8
12
0
20
60
100
Value
Common sites gene−LOC105282133
CphaseA1
RphaseB1

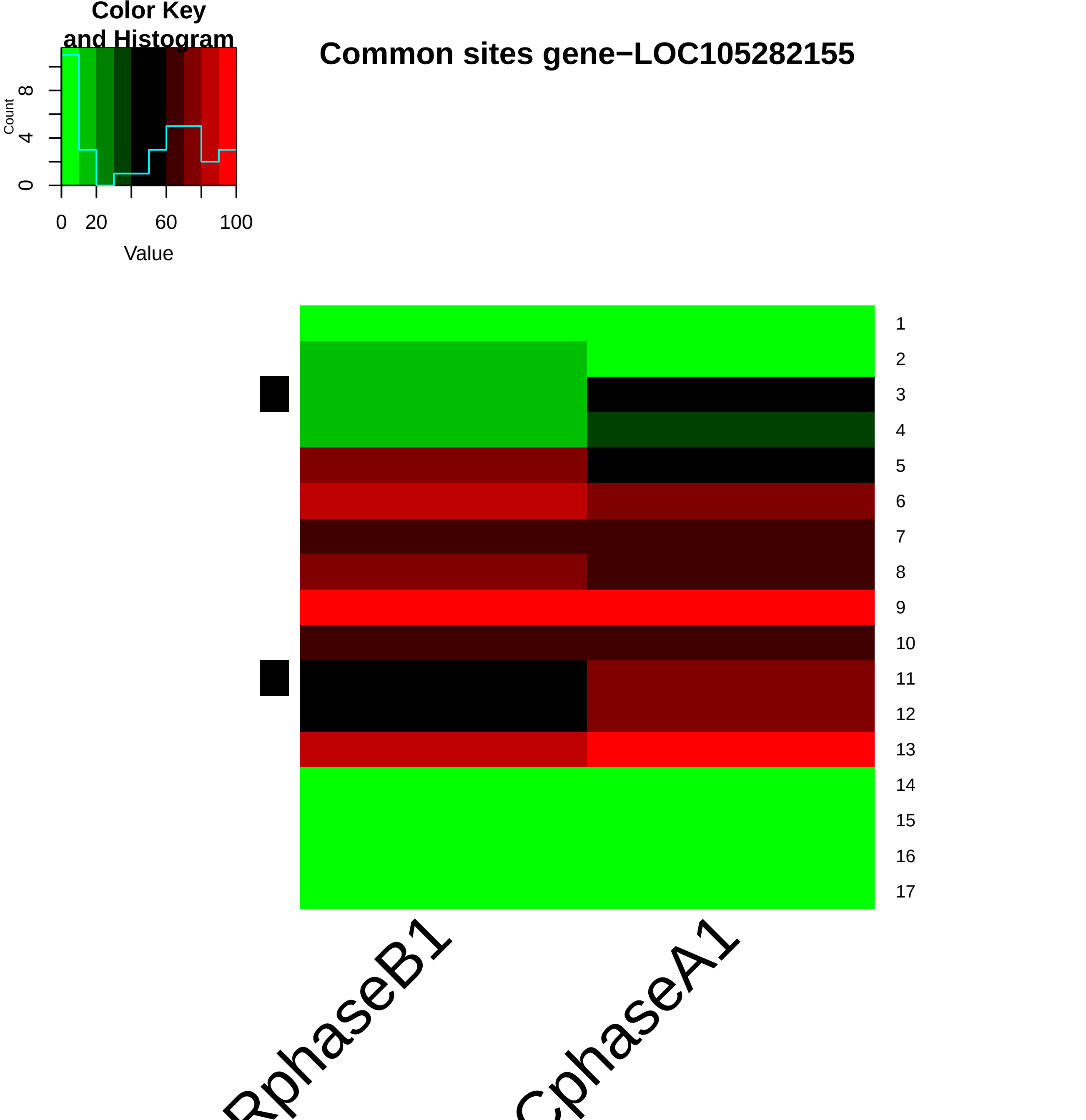

Color Key
and Histogram
Count
0
4
8
0
20
60
100
Value
Common sites gene−LOC105282155
RphaseB1
CphaseA1

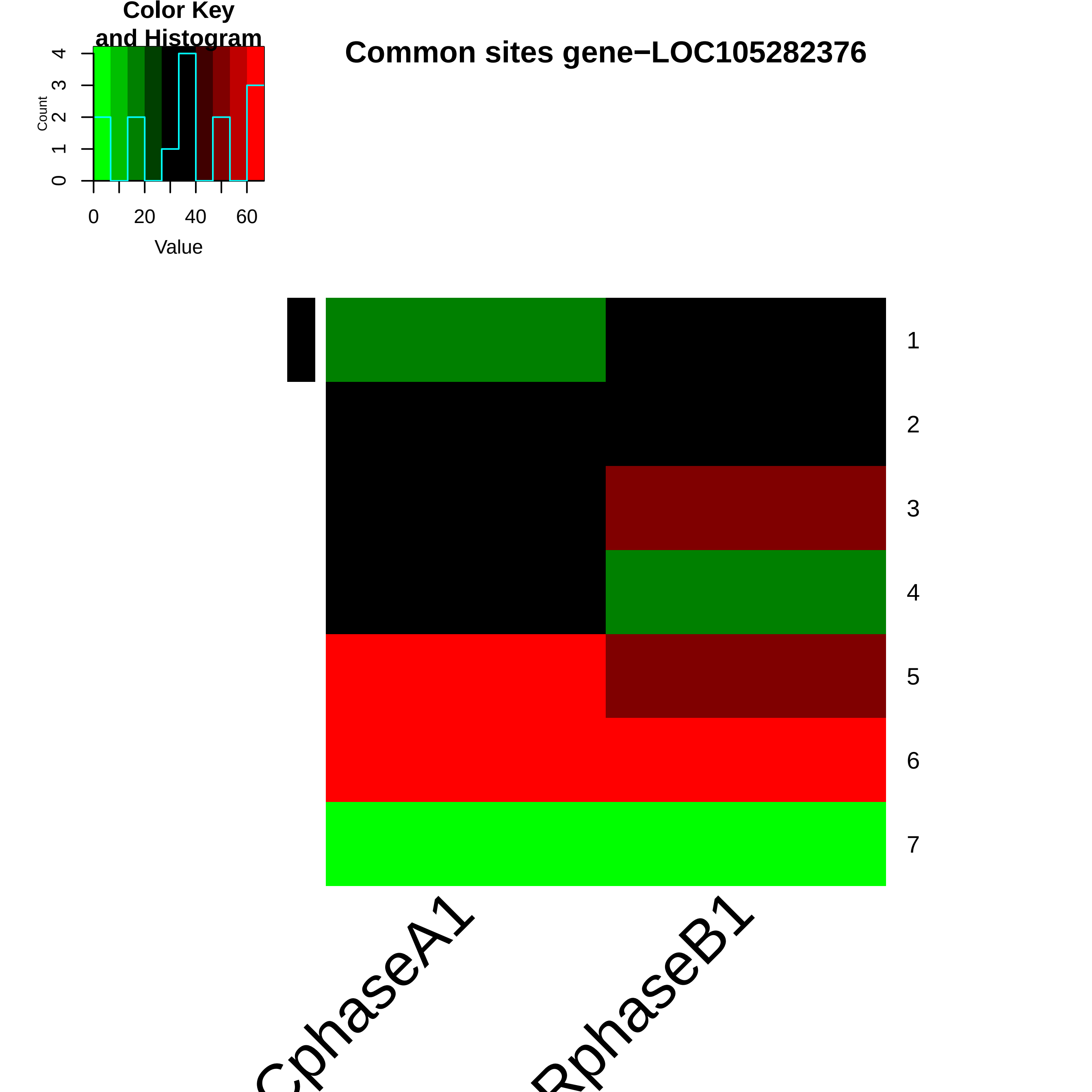
Color Key
and Histogram
Count
Value
0
20
40
60
0
1
2
3
4
Common sites gene−LOC105282376
1
2
3
4
5
6
7

### Common sites gene−LOC105282390

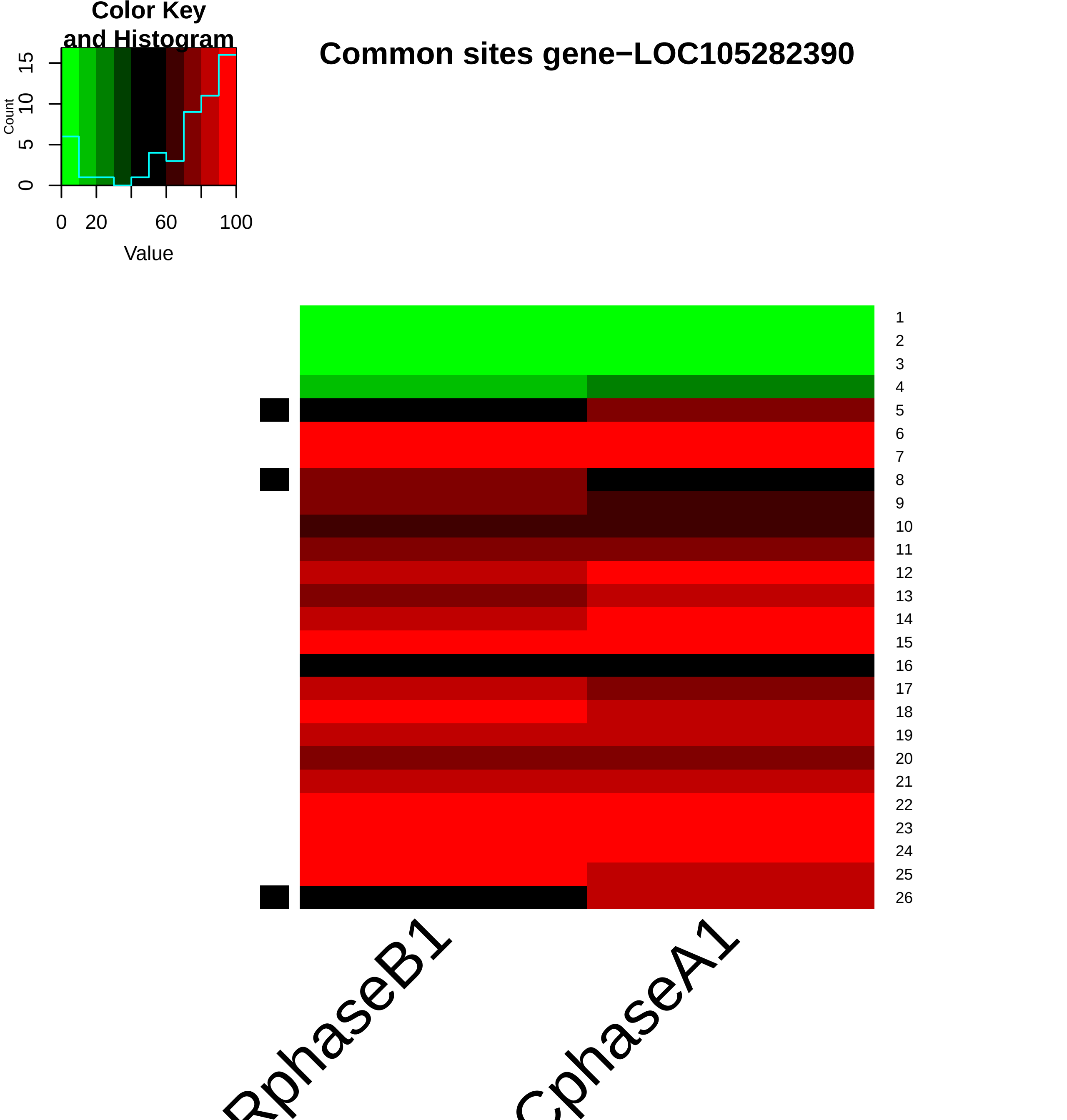

### Common sites gene−LOC105282481

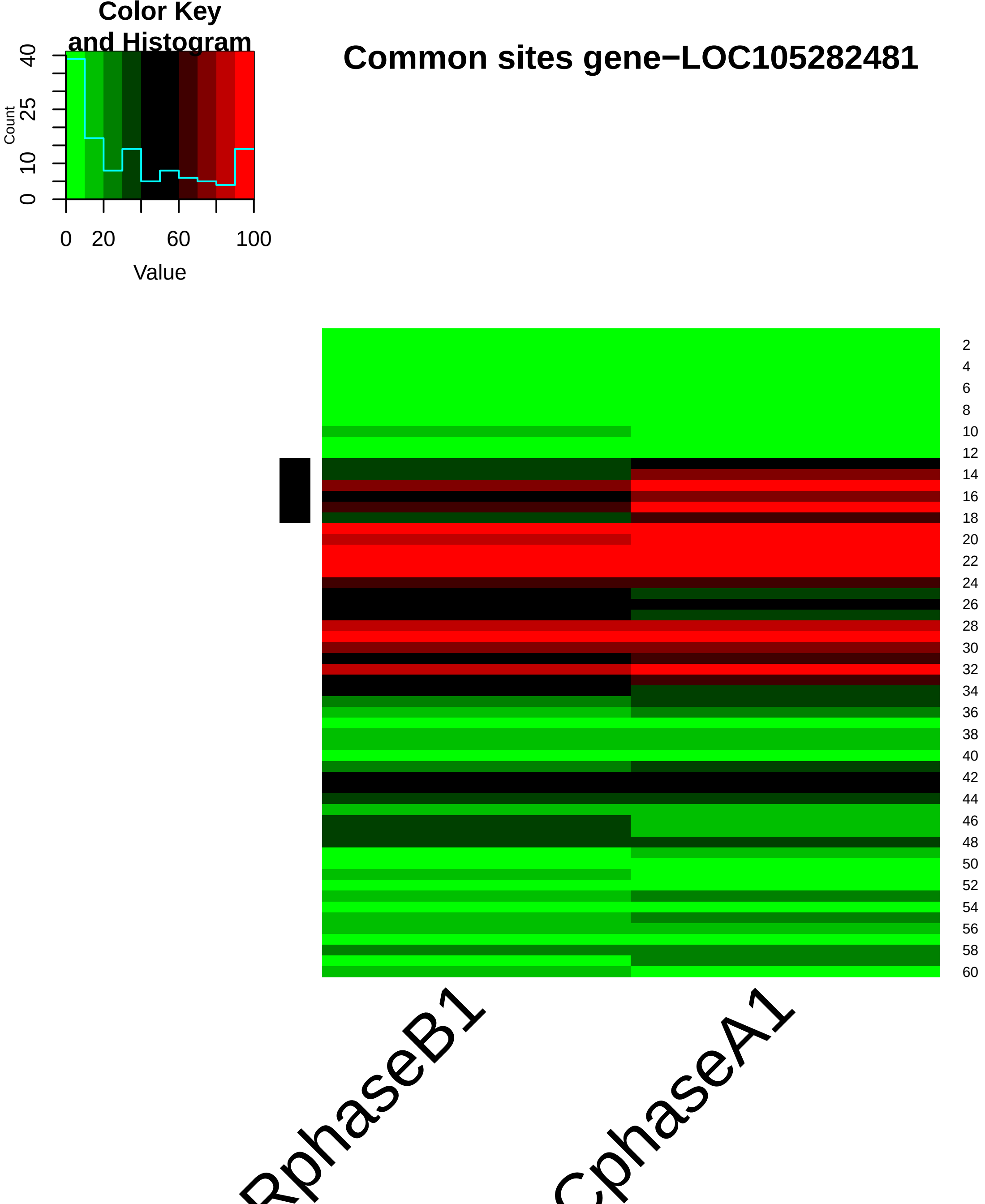

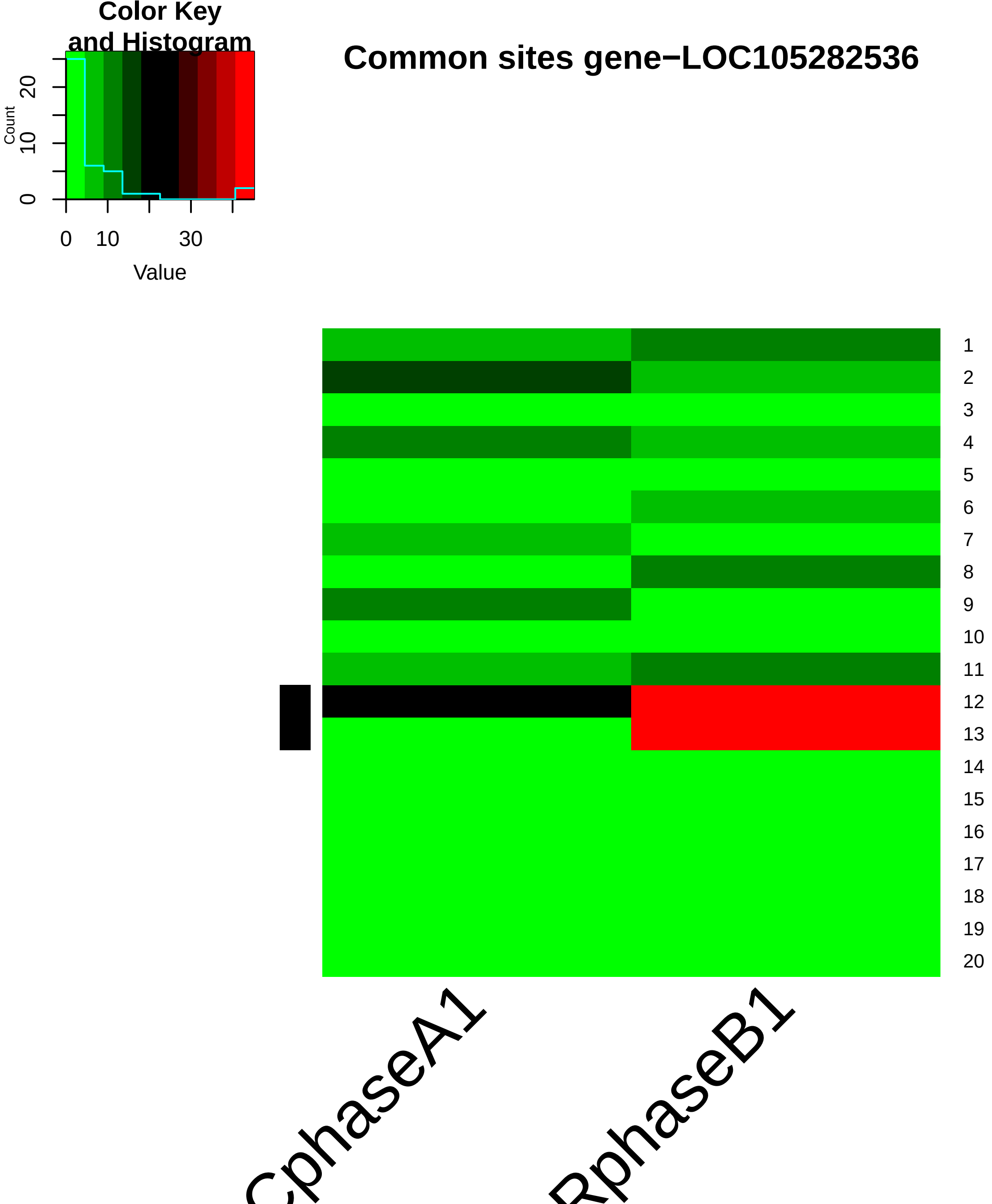
Color Key
and Histogram
Count
0
10
20
0
10
30
Value
Common sites gene−LOC105282536
CphaseA1
RphaseB1

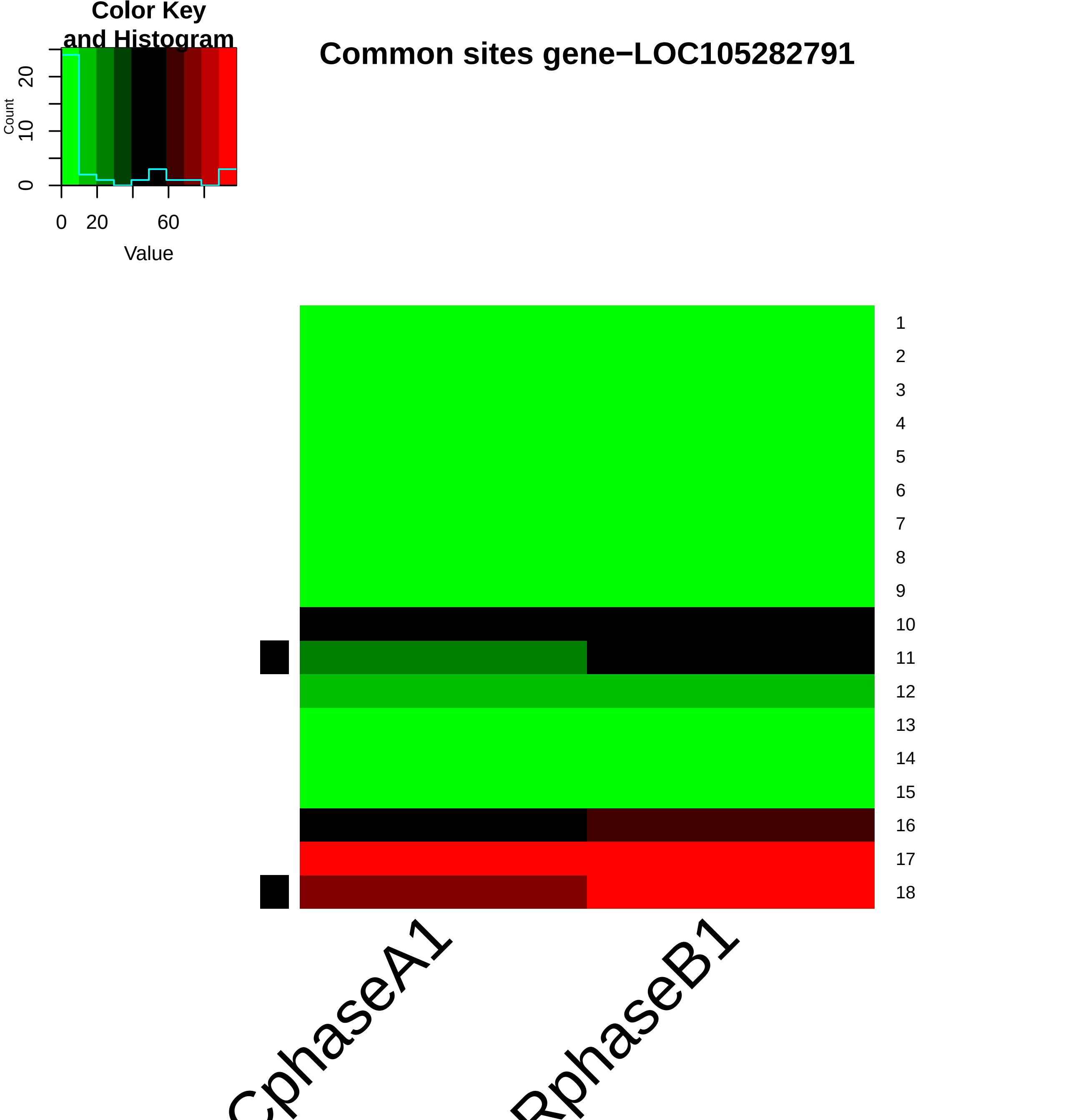

Color Key
and Histogram
Count
0
10
20
0
20
60
Value
Common sites gene−LOC105282791
CphaseA1
RphaseB1

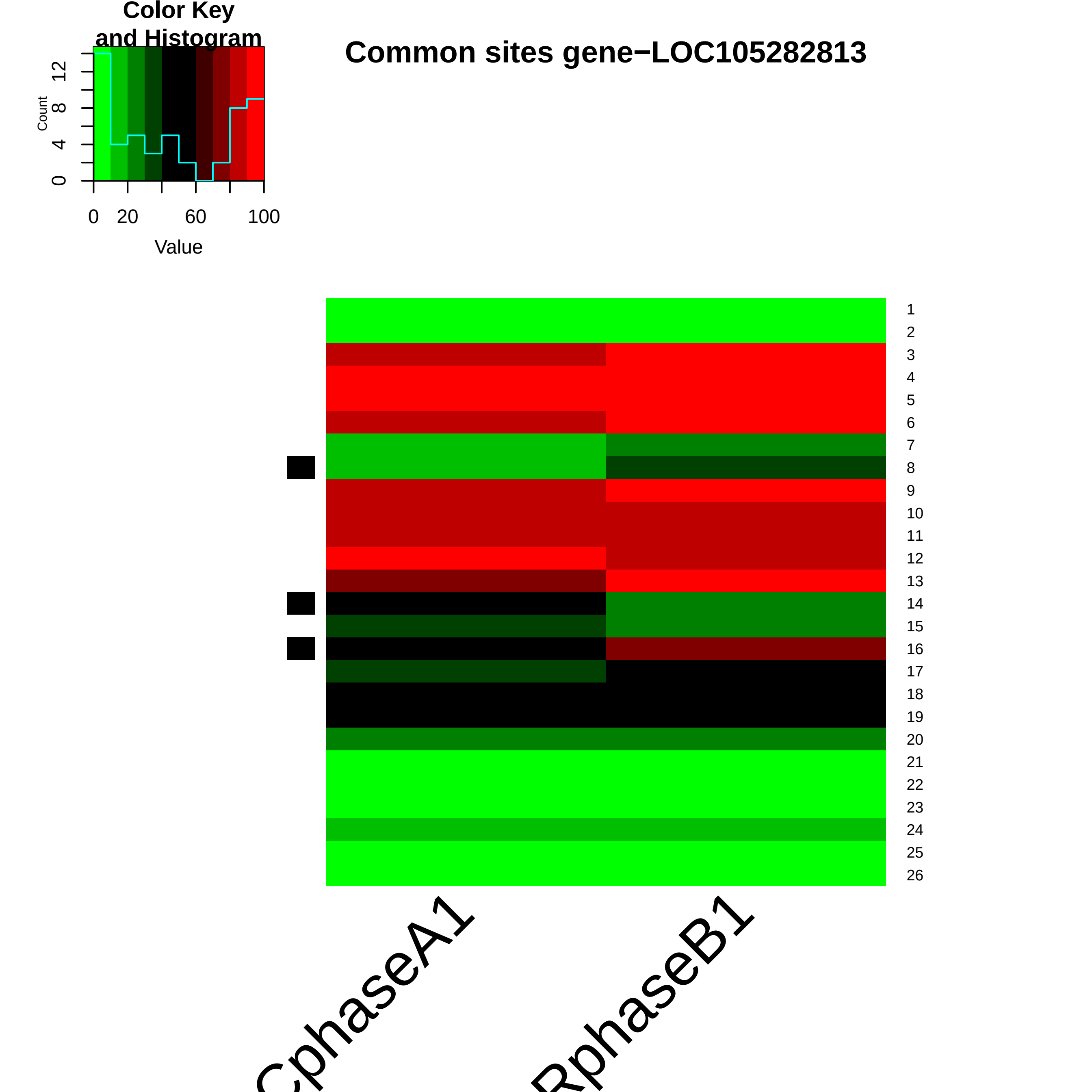
Color Key
and Histogram
Count
0
4
8
12
0
20
60
100
Value
Common sites gene−LOC105282813
CphaseA1
RphaseB1

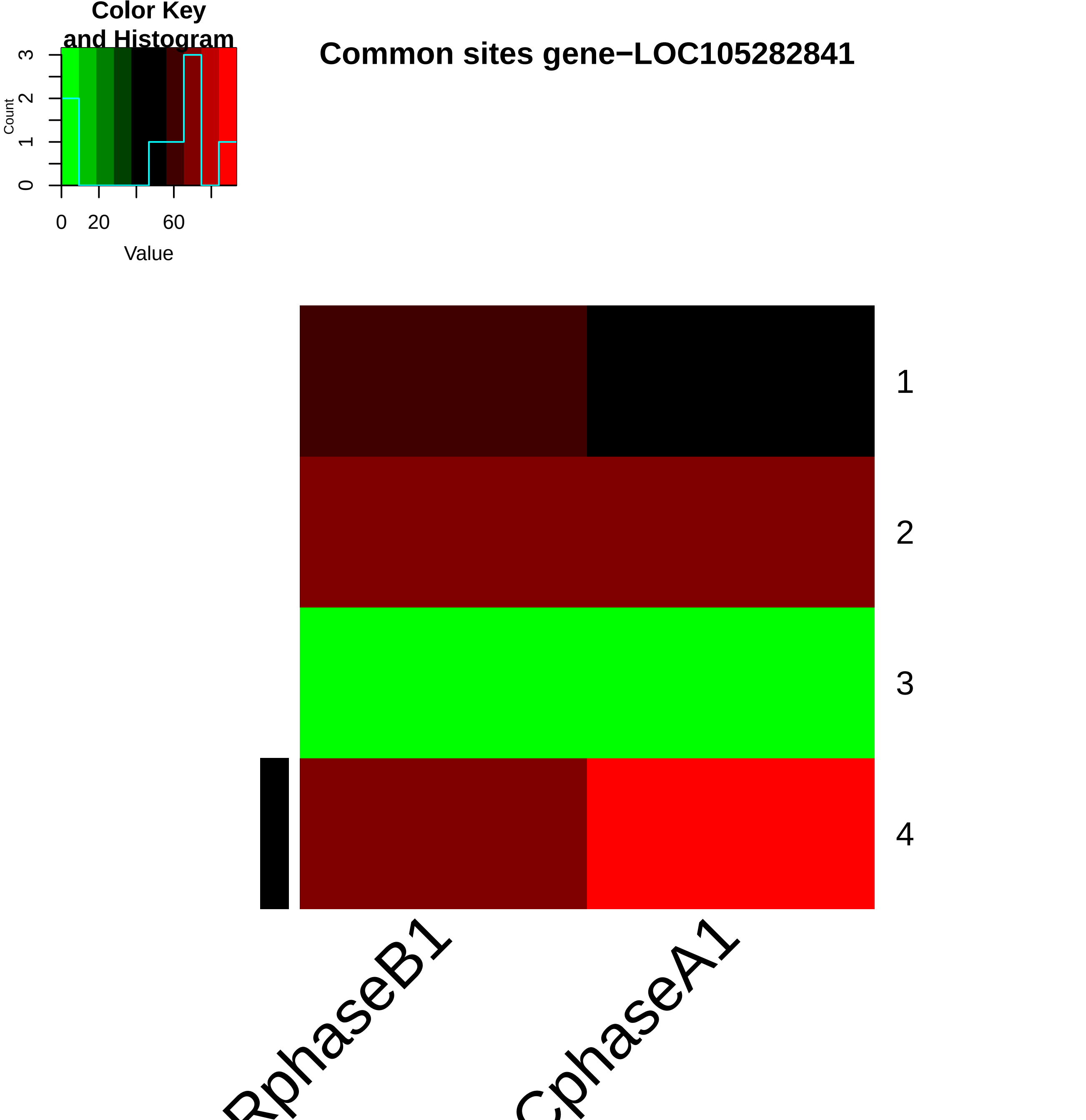

Color Key
and Histogram
Count
0
1
2
3
0
20
60
Value
Common sites gene−LOC105282841
1
2
3
4

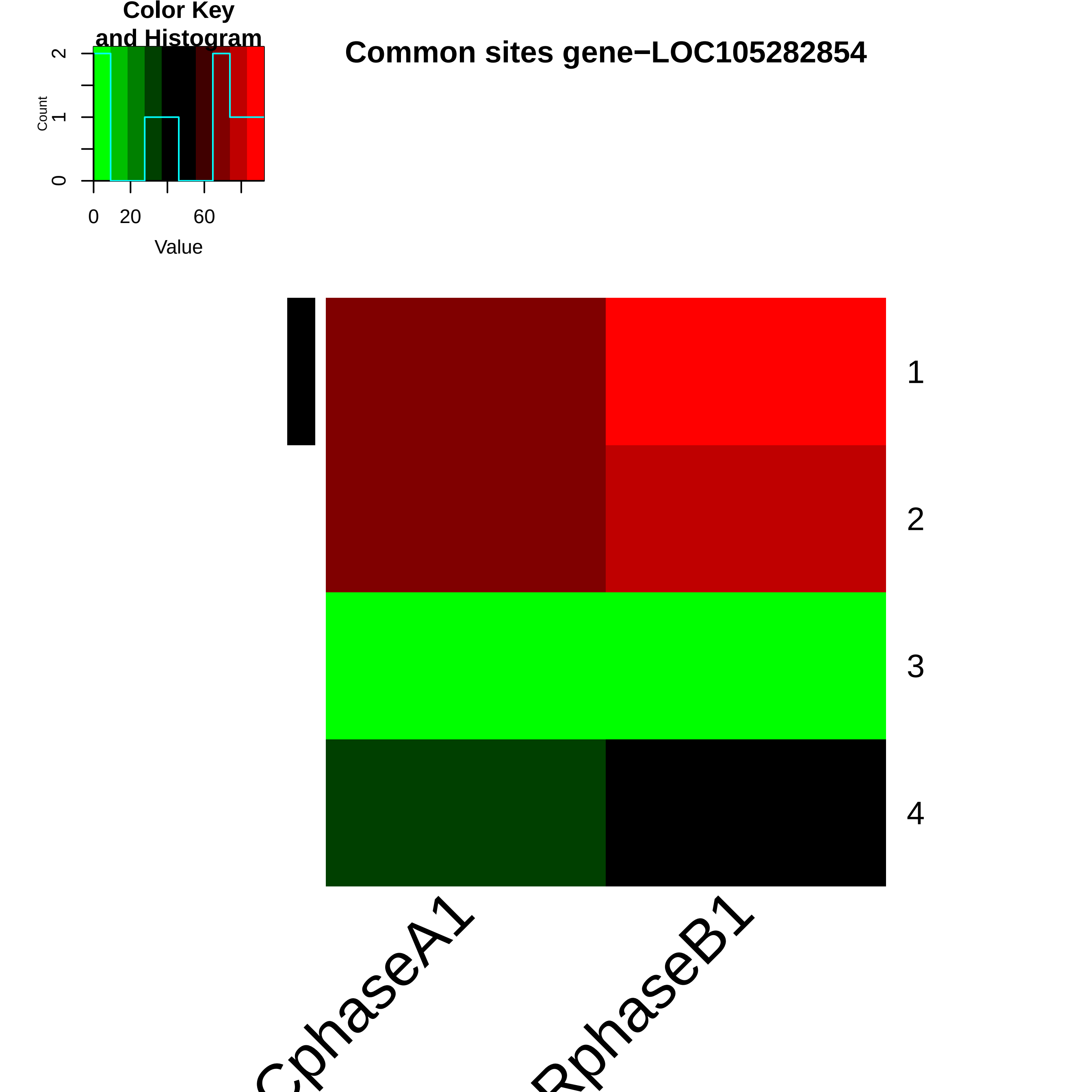
Color Key
and Histogram
Count
0
1
2
0
20
60
Value
Common sites gene−LOC105282854
1
2
3
4
CphaseA1
RphaseB1

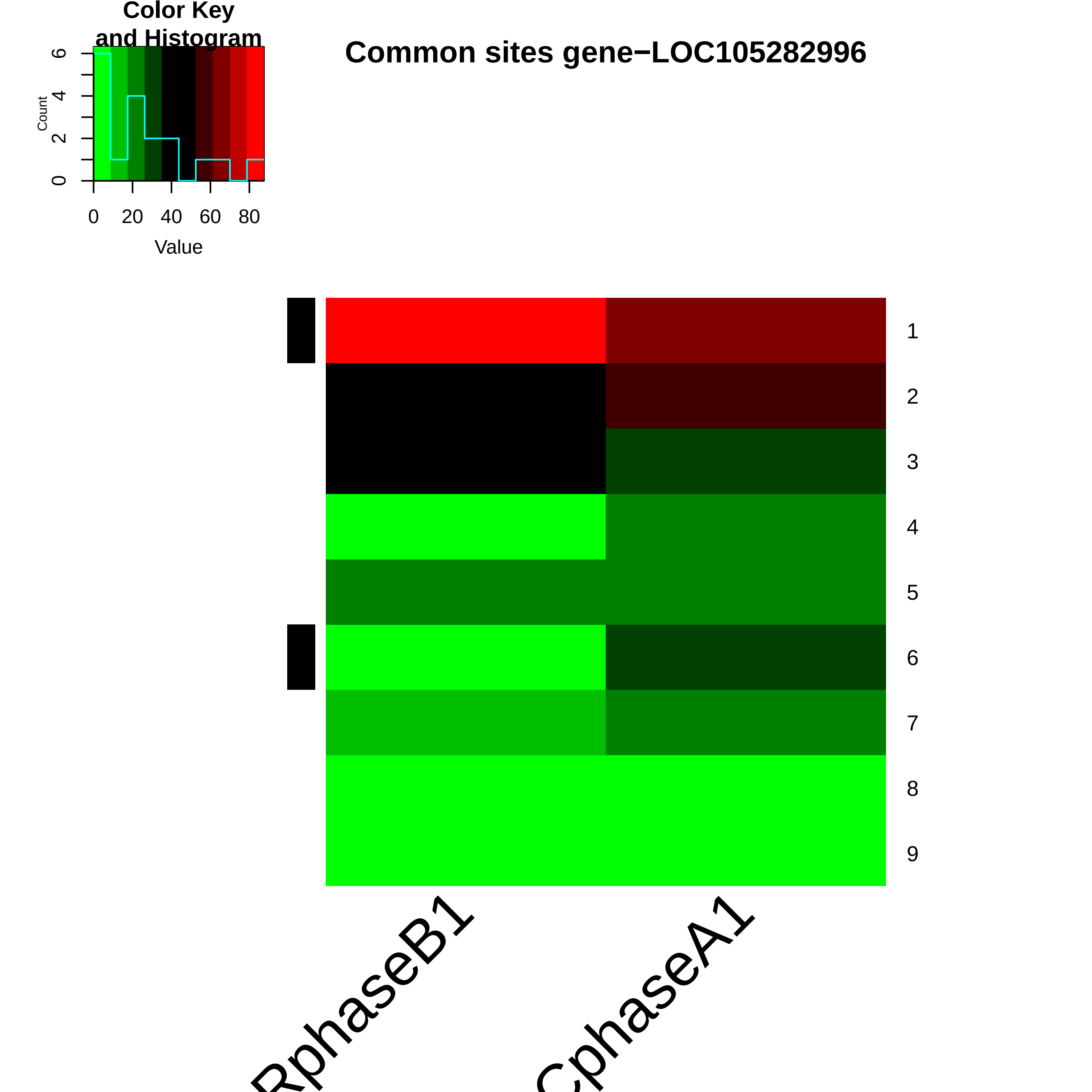

Color Key
and Histogram
Count
0
2
4
6
0
20
40
60
80
Value
Common sites gene−LOC105282996
1
2
3
4
5
6
7
8
9
RphaseB1
CphaseA1

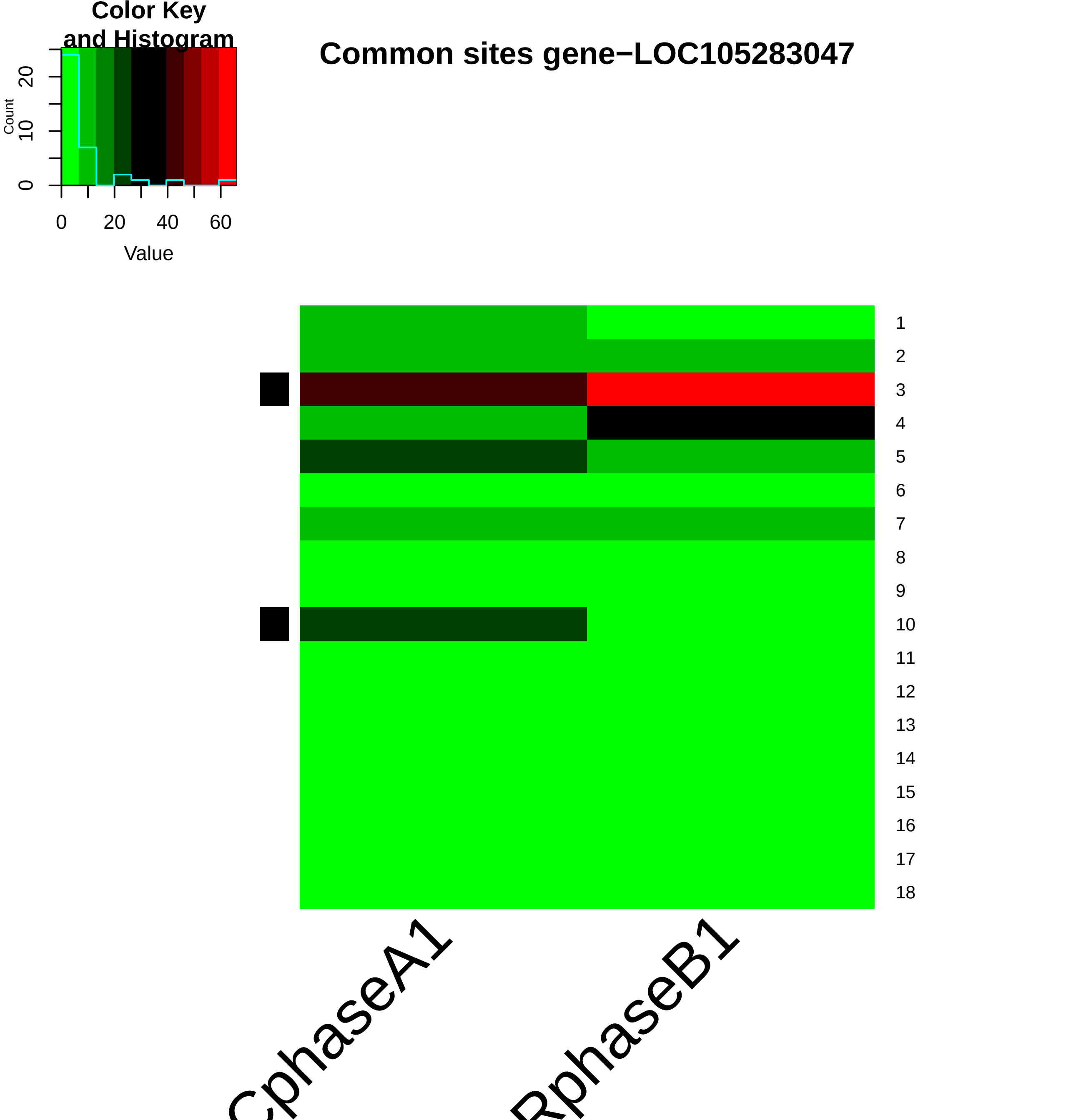
Color Key
and Histogram
Count
0
10
20
0
20
40
60
Value
Common sites gene−LOC105283047
CphaseA1
RphaseB1

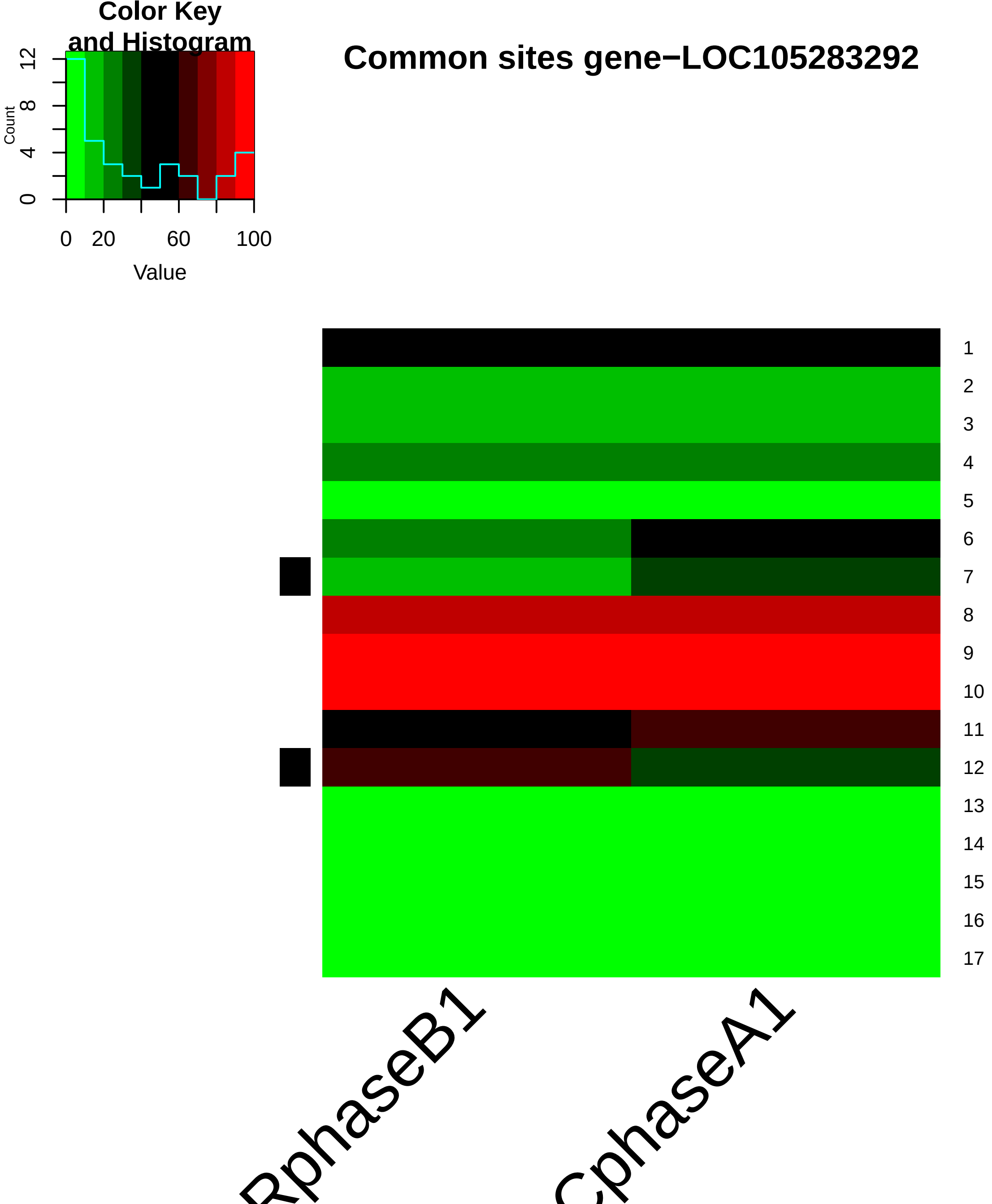
Color Key
and Histogram
Count
Value
0
4
8
12
0
20
60
100
Common sites gene−LOC105283292
RphaseB1
CphaseA1

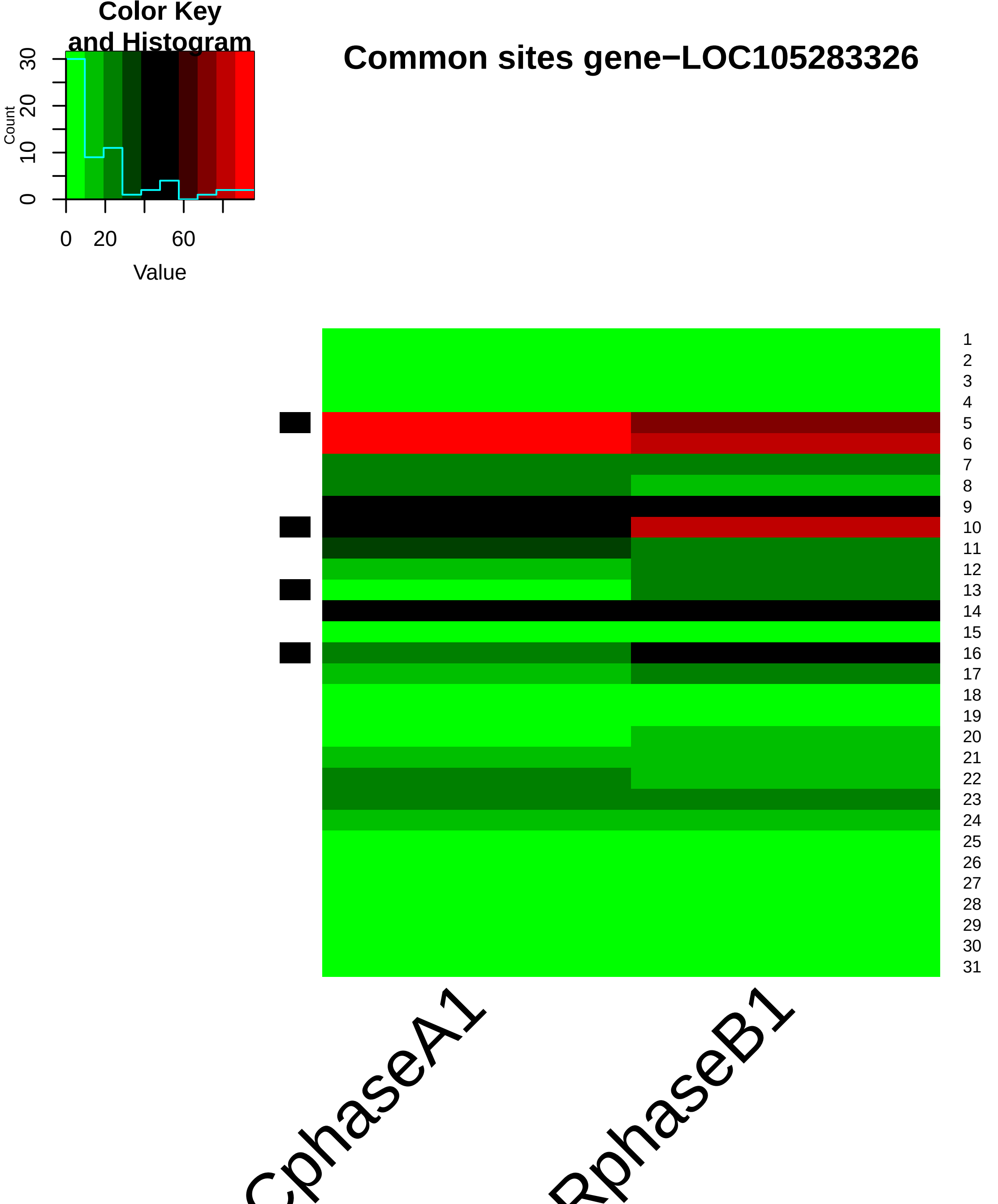
Color Key
and Histogram
Count
0
10
20
30
0
20
60
Value
Common sites gene−LOC105283326
CphaseA1
RphaseB1

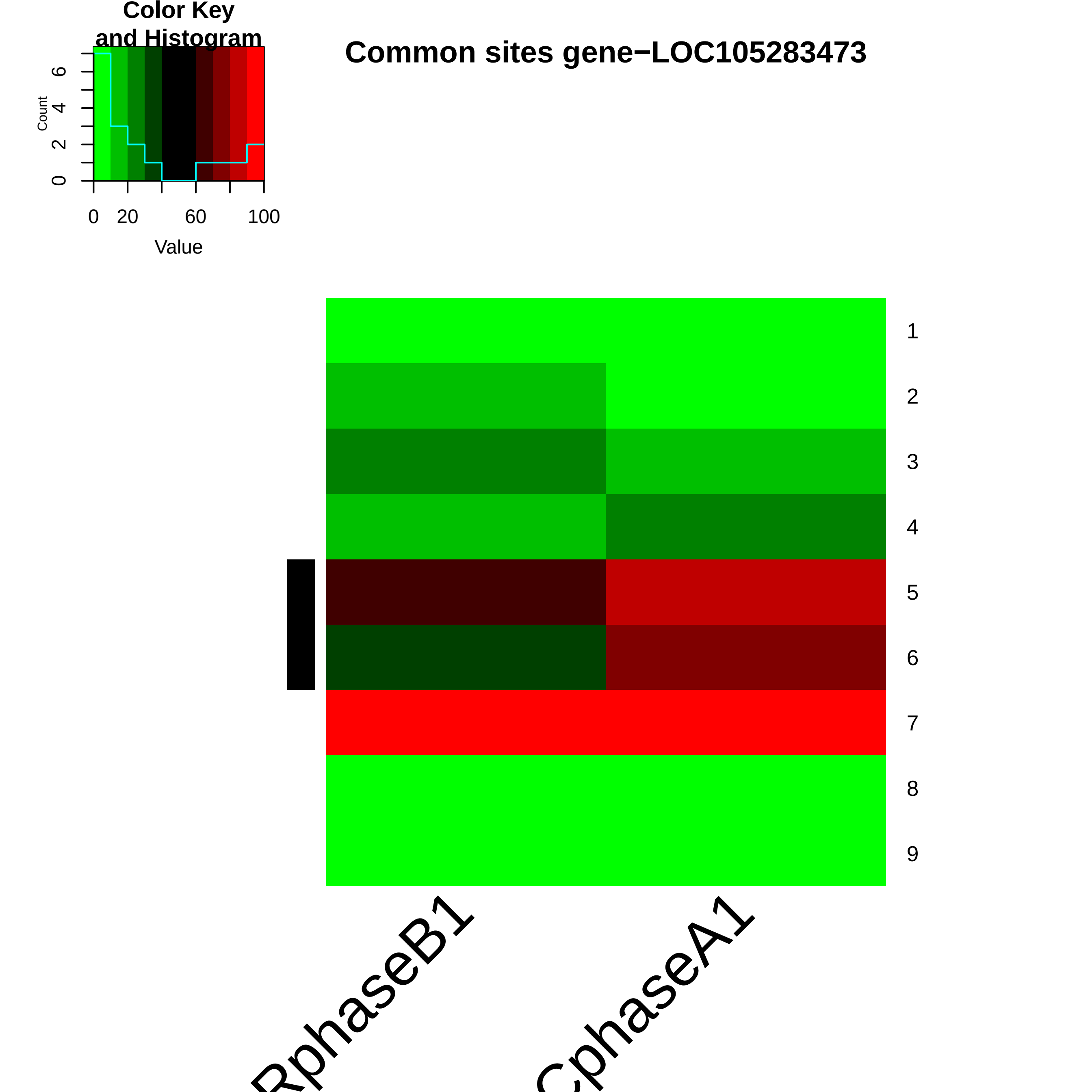
Color Key
and Histogram
Count
0
2
4
6
0
20
60
100
Value
Common sites gene−LOC105283473
1
2
3
4
5
6
7
8
9
RphaseB1
CphaseA1

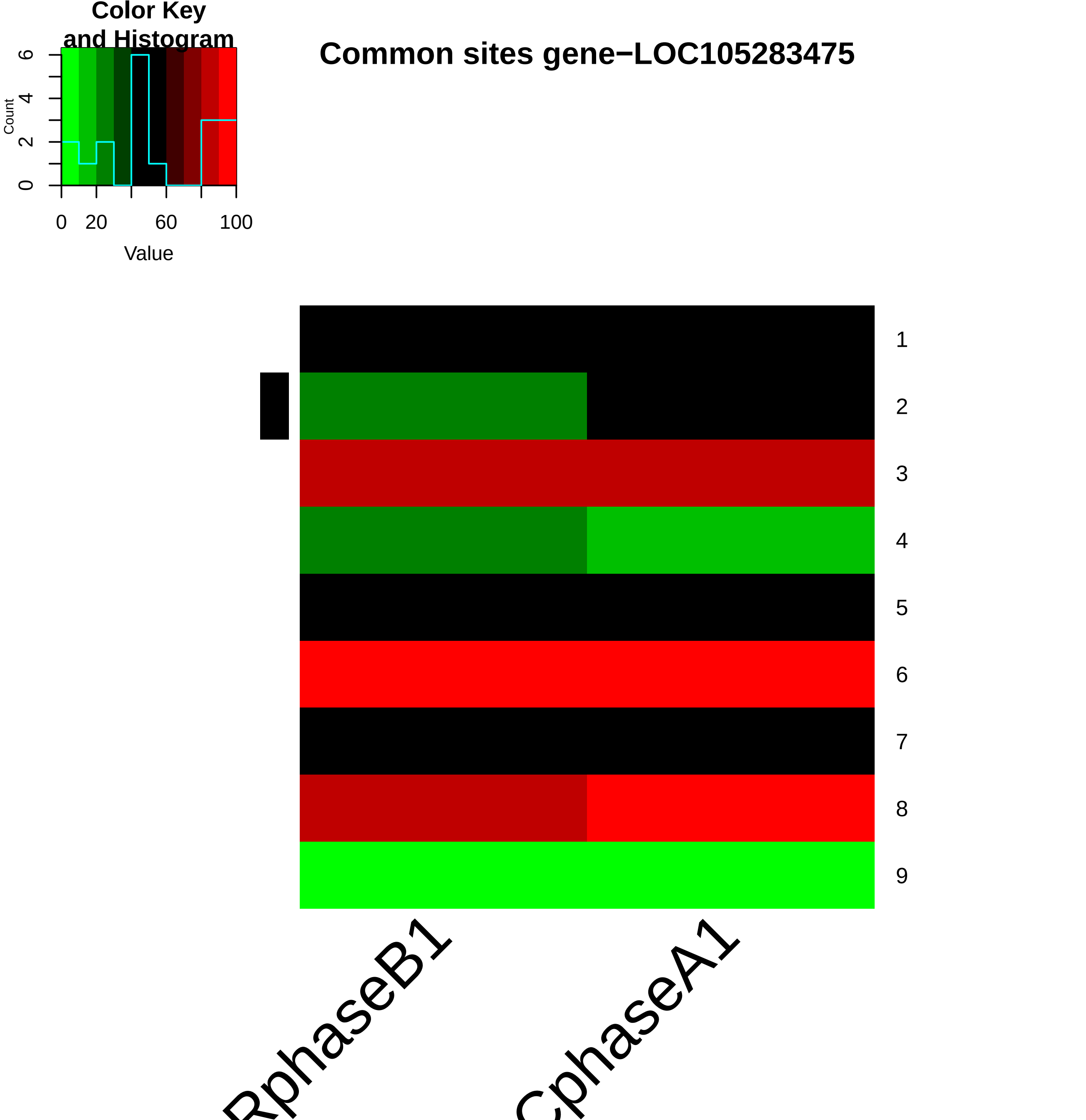

Color Key
and Histogram
Count
0
2
4
6
0
20
60
100
Value
Common sites gene−LOC105283475
1
2
3
4
5
6
7
8
9
RphaseB1
CphaseA1

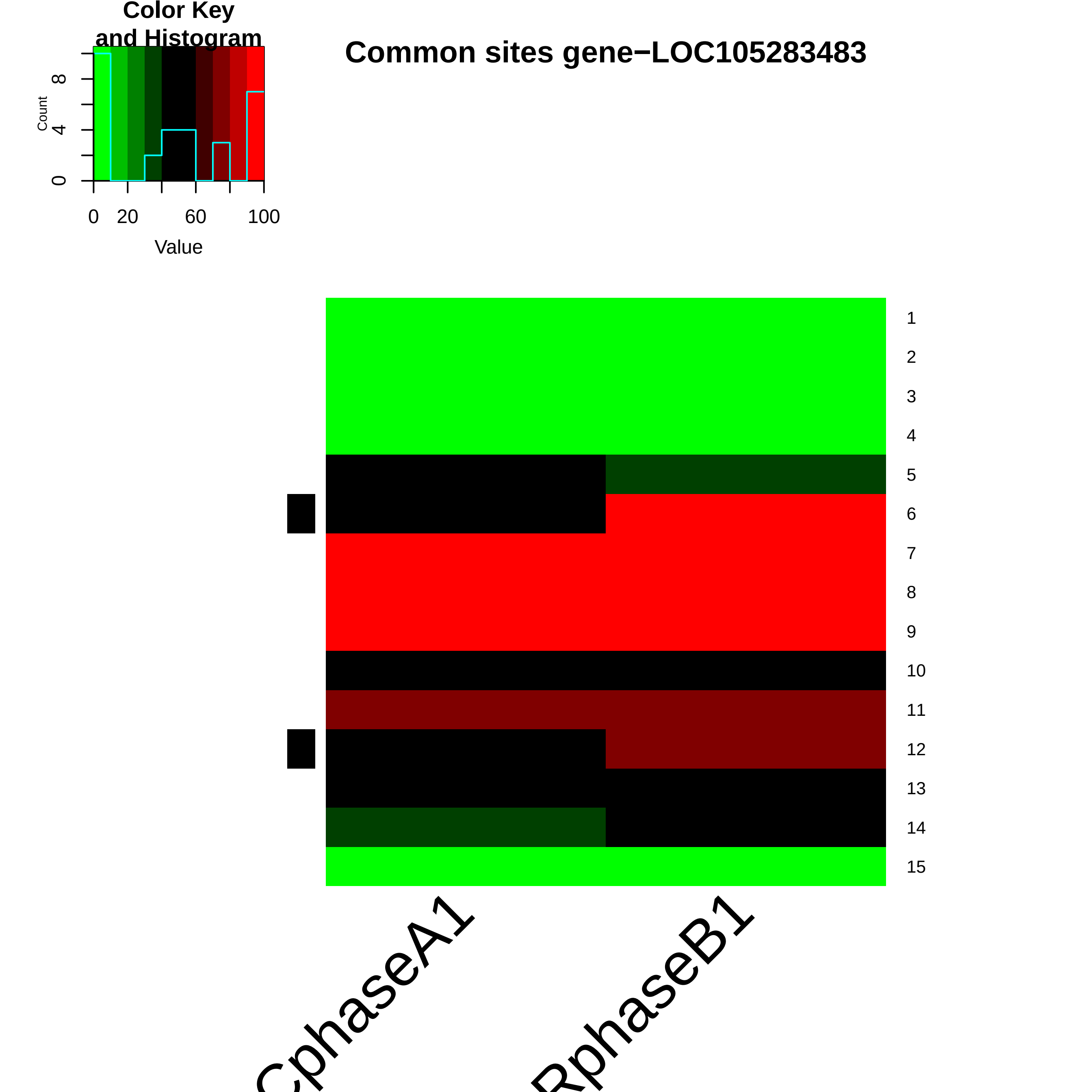

Color Key
and Histogram
Count
0
4
8
0
20
60
100
Value
Common sites gene−LOC105283483
CphaseA1
RphaseB1

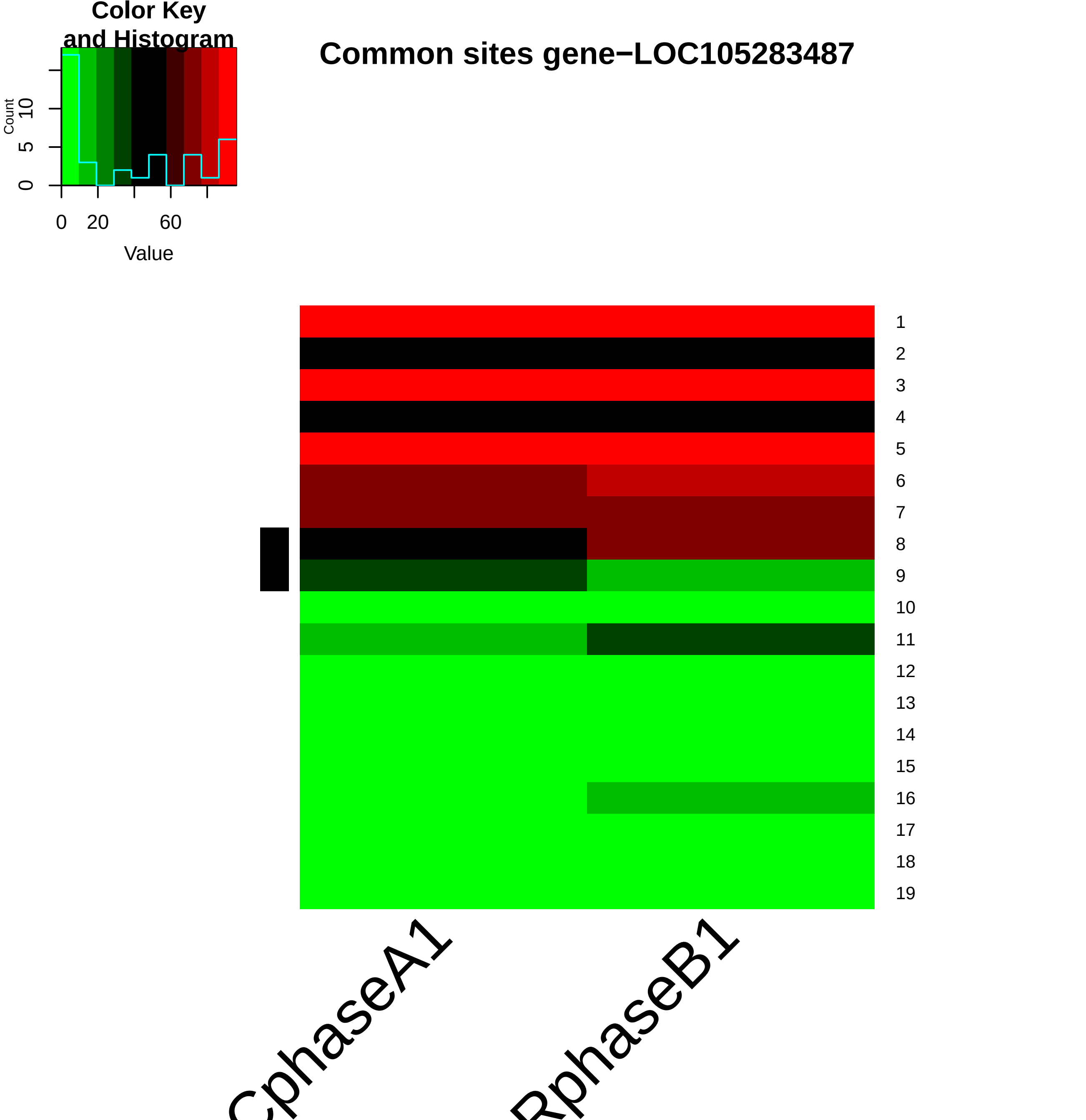
Color Key
and Histogram
Count
0
5
10
0
20
60
Value
Common sites gene−LOC105283487
CphaseA1
RphaseB1

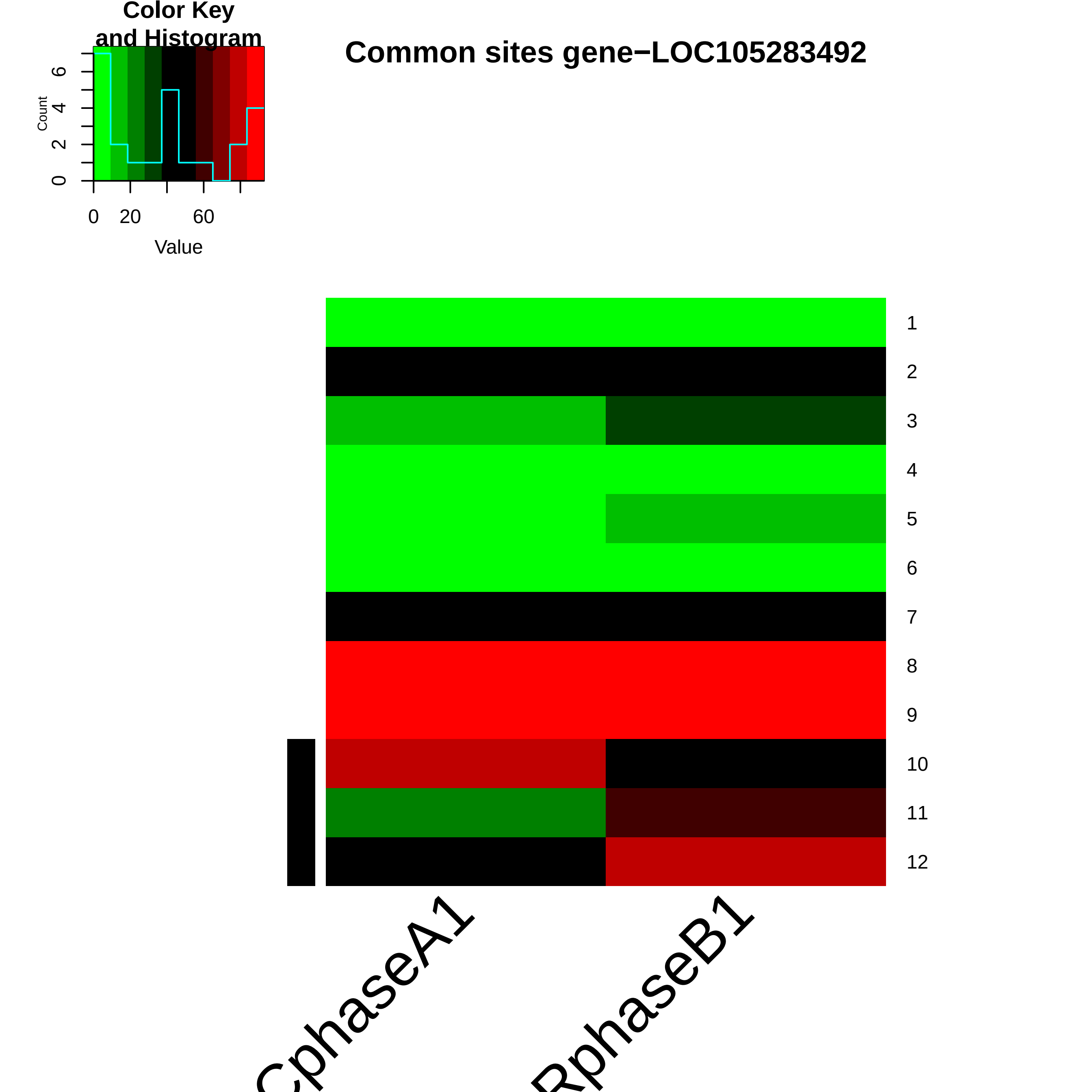
Color Key
and Histogram
Count
0
2
4
6
0
20
60
Value
Common sites gene−LOC105283492
1
2
3
4
5
6
7
8
9
10
11
12
CphaseA1
RphaseB1

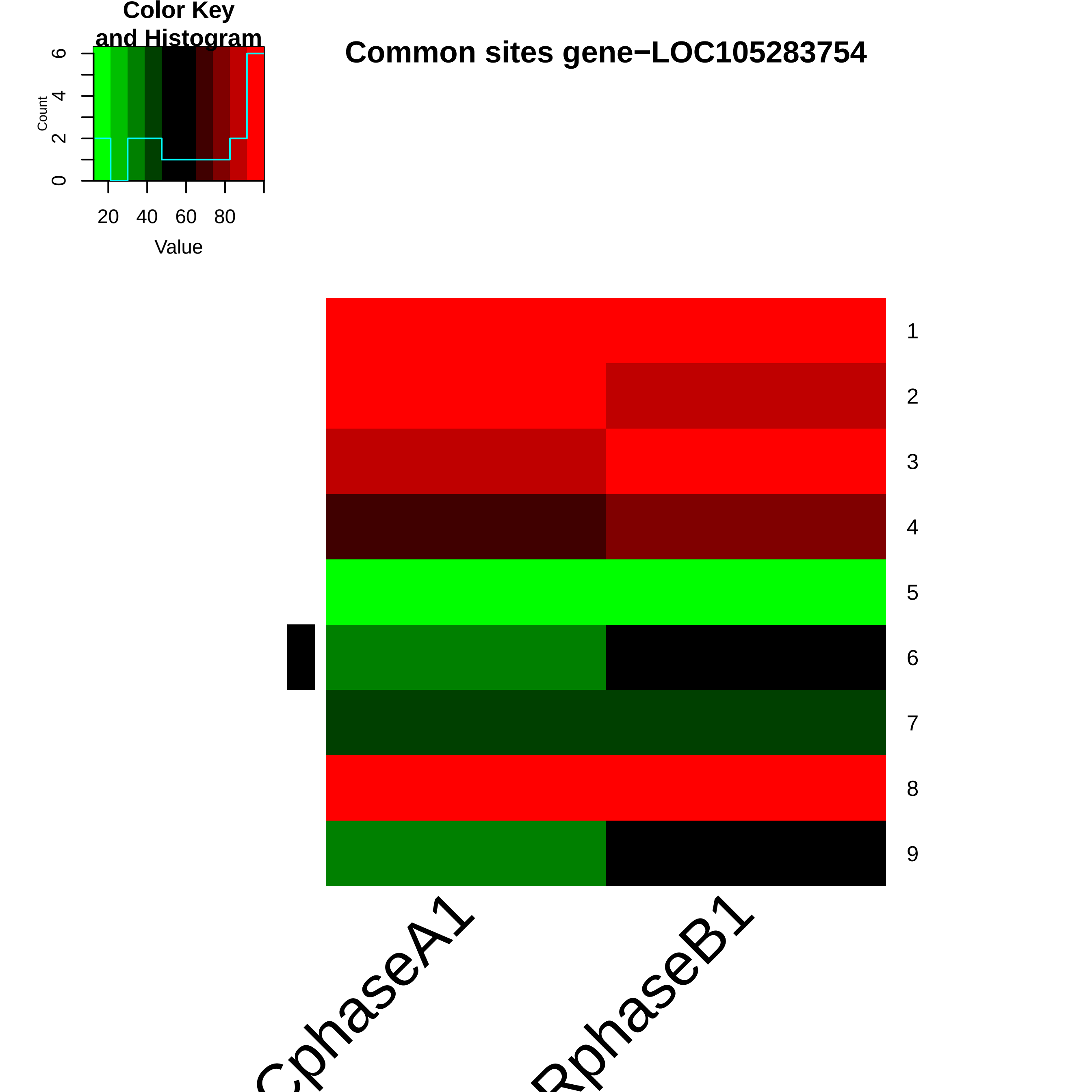

Color Key
and Histogram
Count
0
2
4
6
20
40
60
80
Value
Common sites gene−LOC105283754
1
2
3
4
5
6
7
8
9
CphaseA1
RphaseB1

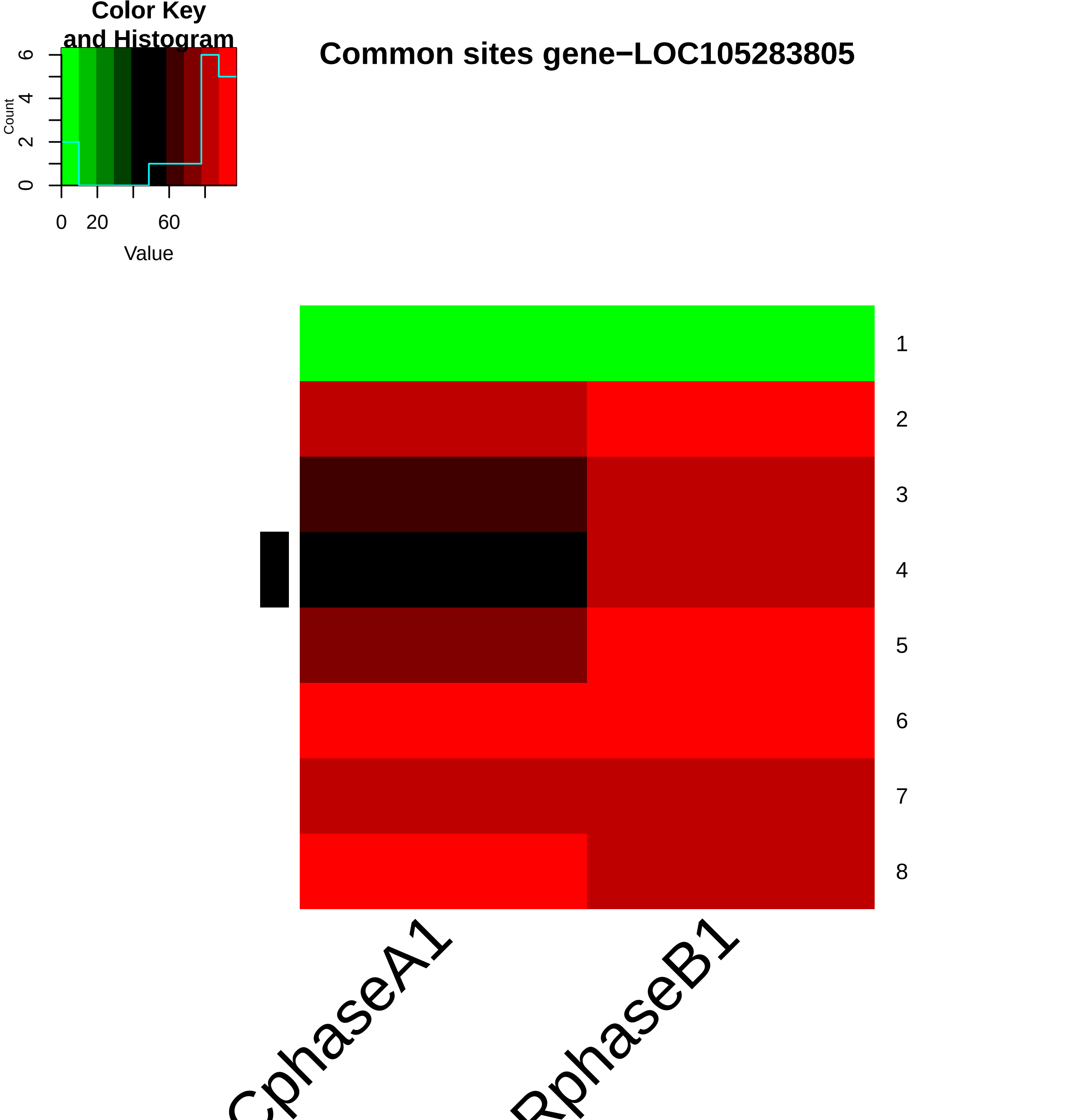

Color Key
and Histogram
Count
0
2
4
6
0
20
60
Value
Common sites gene−LOC105283805
1
2
3
4
5
6
7
8
CphaseA1
RphaseB1

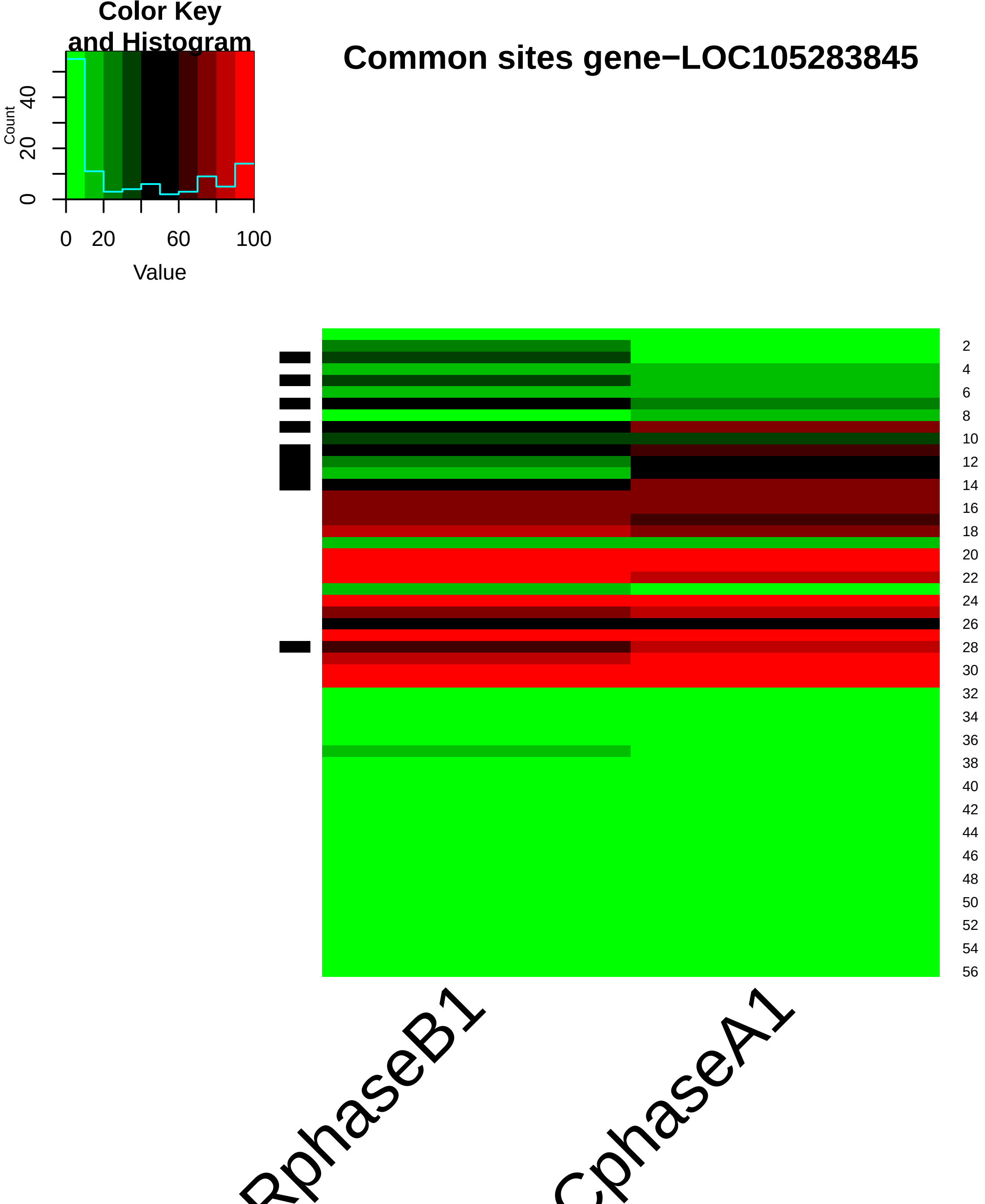

Color Key
and Histogram
Count
0
20
40
0
20
60
100
Value
Common sites gene−LOC105283845
2
4
6
8
10
12
14
16
18
20
22
24
26
28
30
32
34
36
38
40
42
44
46
48
50
52
54
56
RphaseB1
CphaseA1

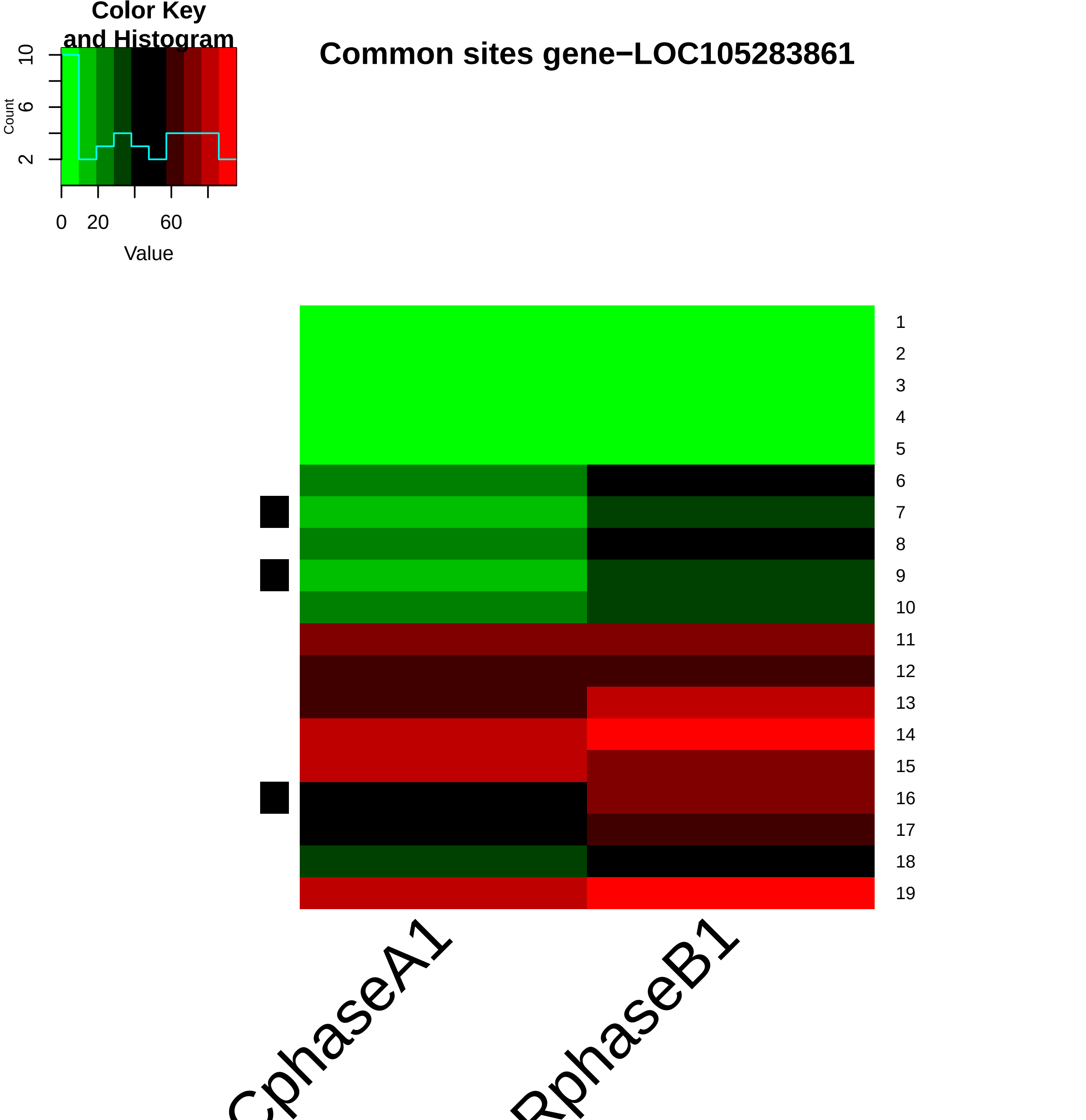

Color Key
and Histogram
Count
2
6
10
0
20
60
Value
Common sites gene−LOC105283861
CphaseA1
RphaseB1

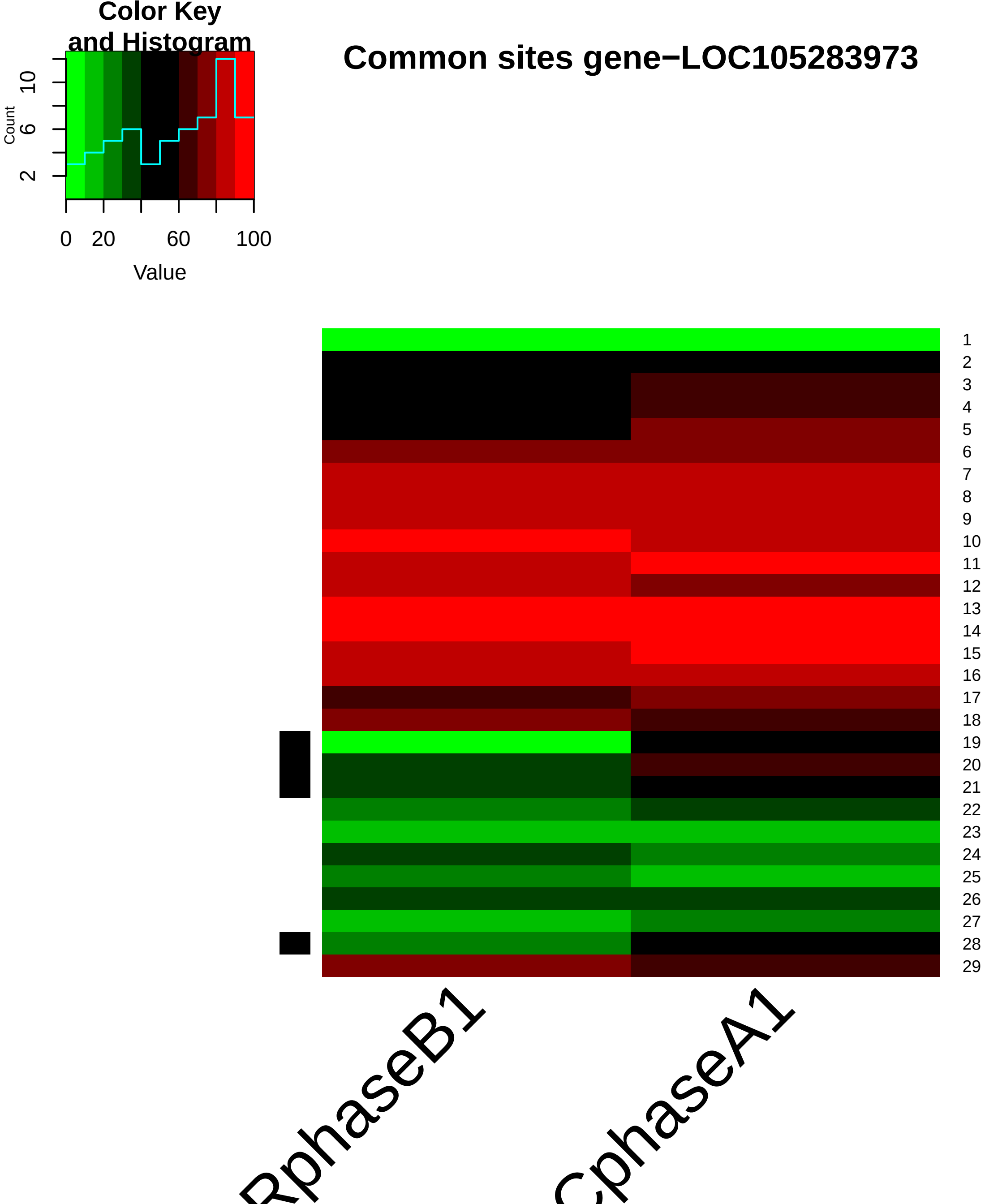
Color Key
and Histogram
Count
Value
0
20
60
100
2
6
10
Common sites gene−LOC105283973
RphaseB1
CphaseA1

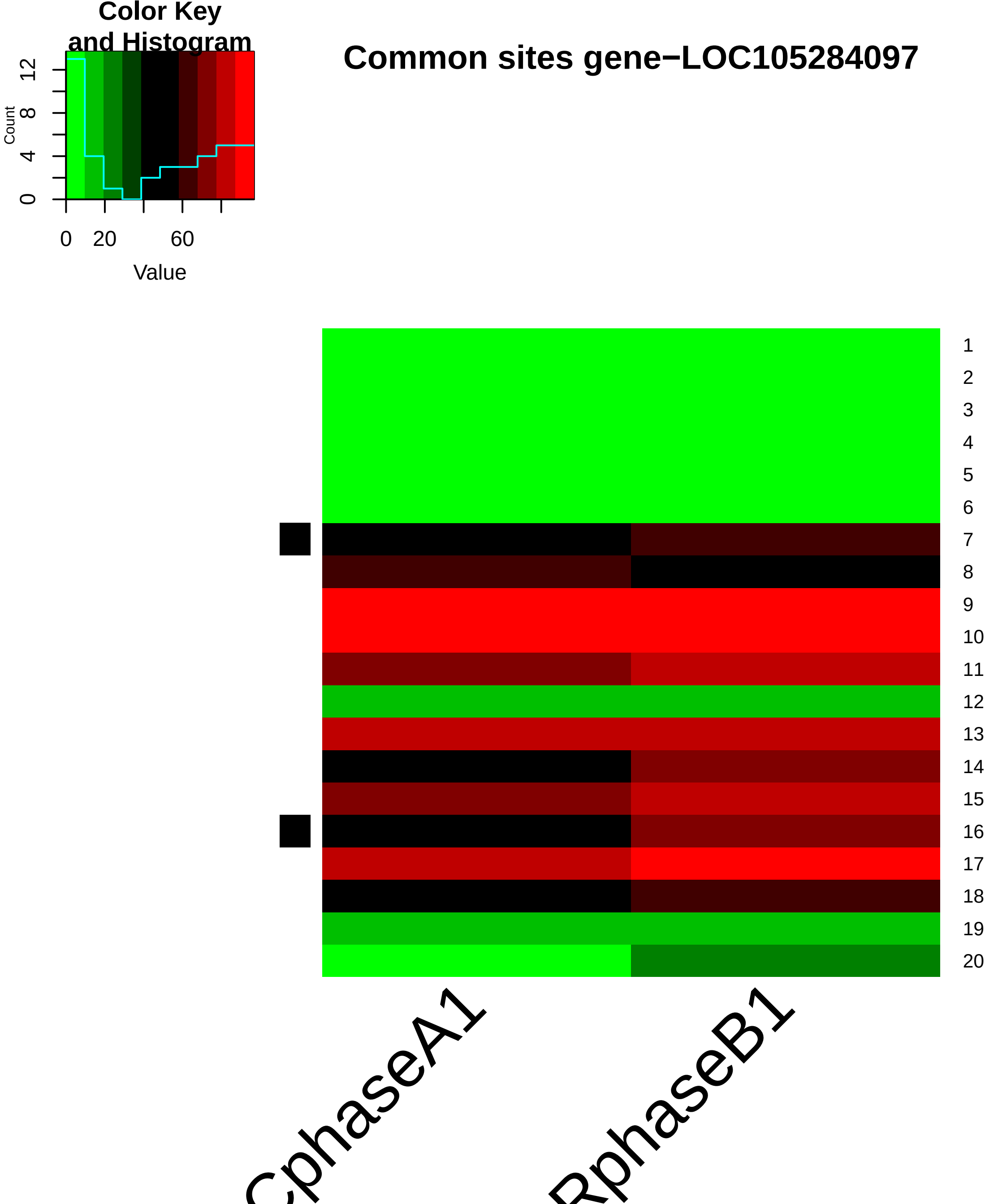

Color Key
and Histogram
Count
Value
0
20
60
0
4
8
12
Common sites gene−LOC105284097
CphaseA1
RphaseB1

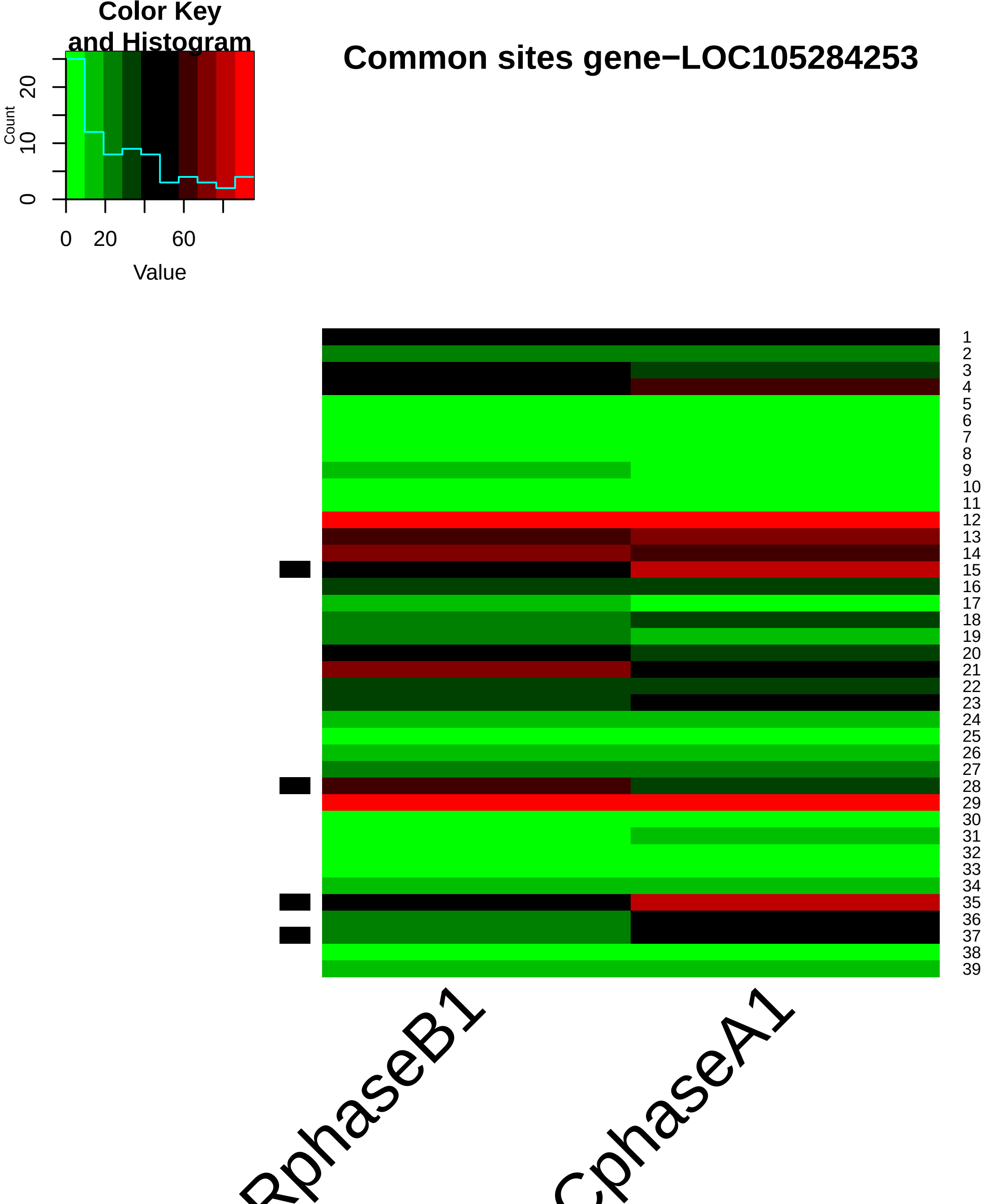

Color Key
and Histogram
Count
0
10
20
0
20
60
Value
Common sites gene−LOC105284253
RphaseB1
CphaseA1

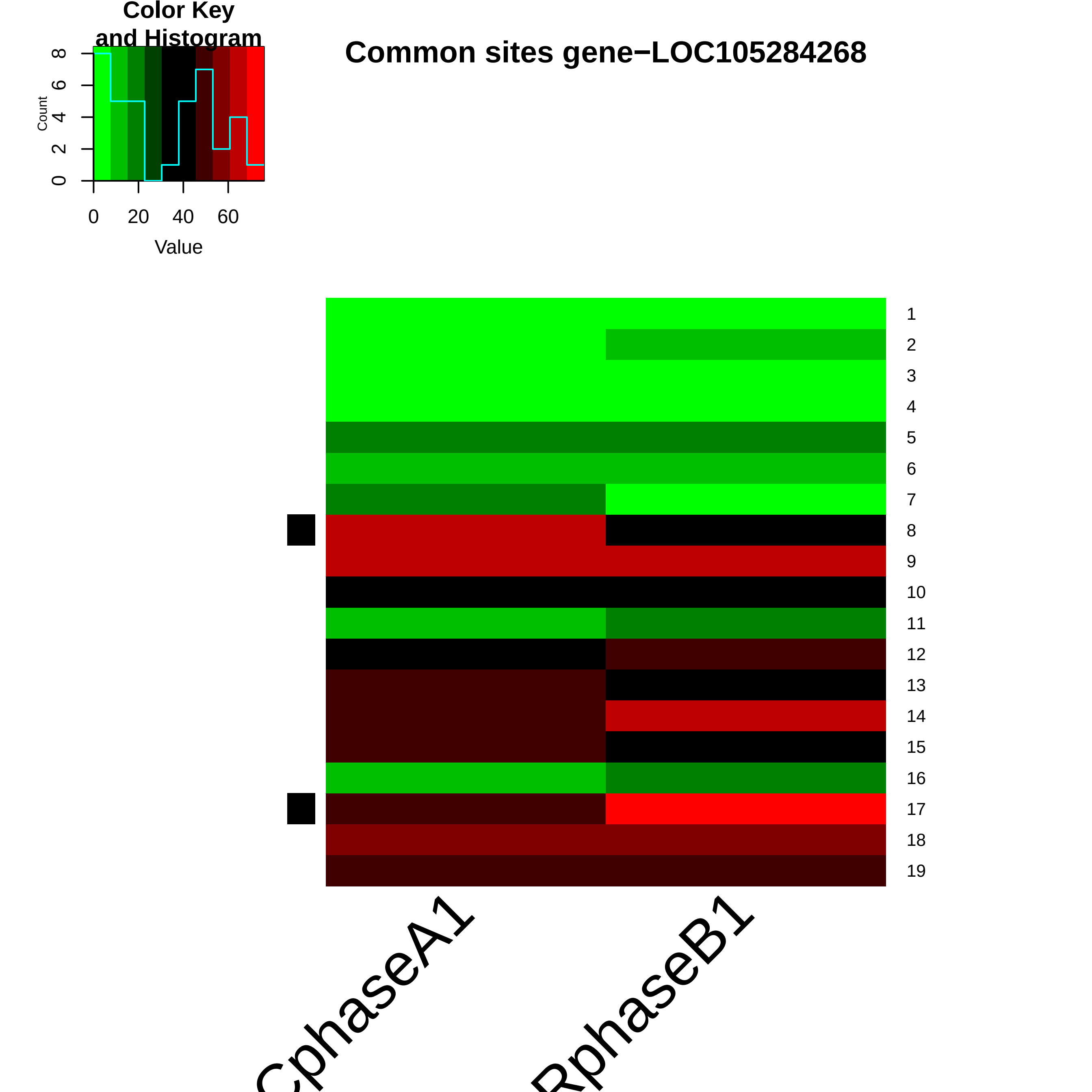

Color Key
and Histogram
Count
0
2
4
6
8
0
20
40
60
Value
Common sites gene−LOC105284268
CphaseA1
RphaseB1

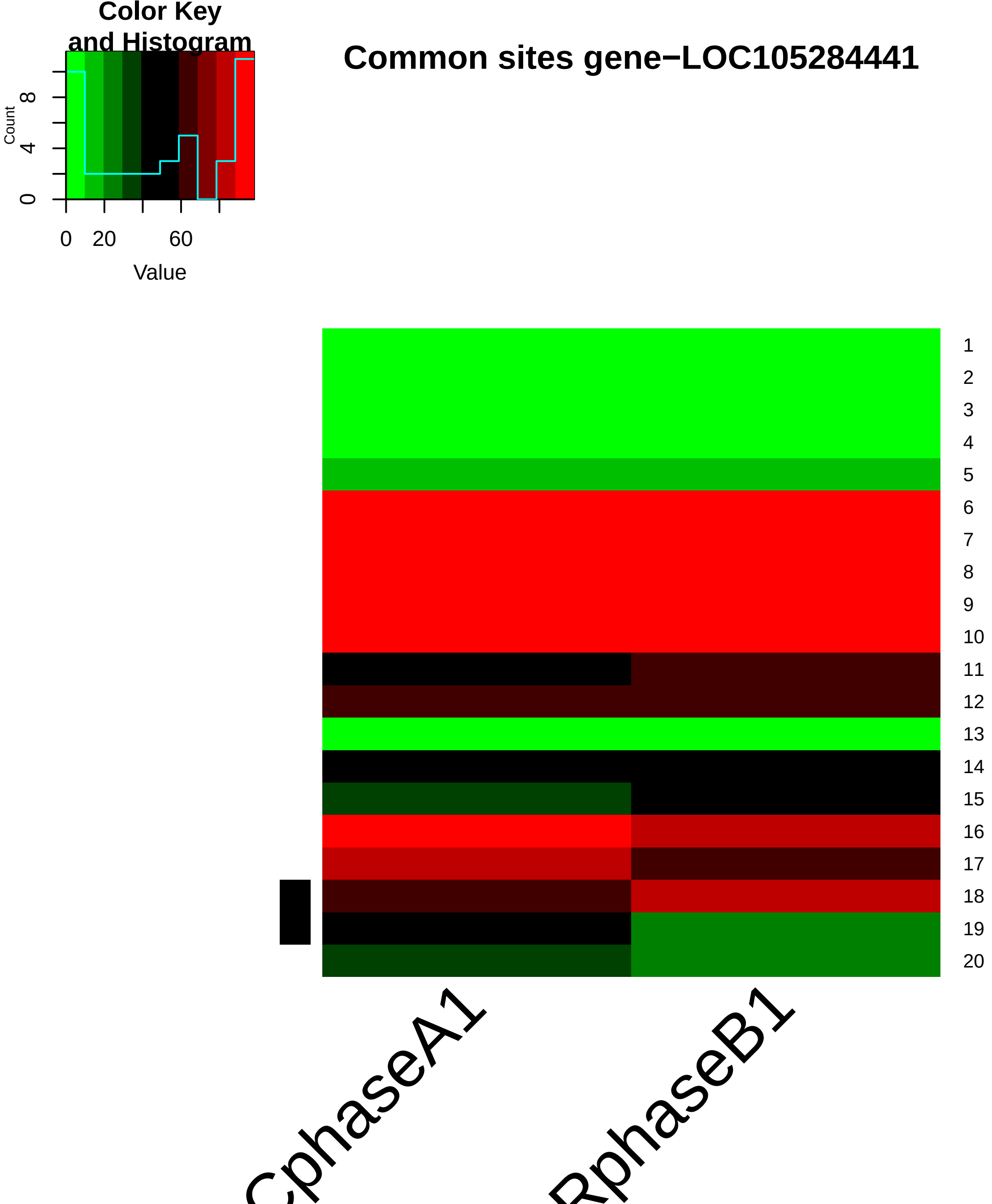
Color Key
and Histogram
Count
0
4
8
0
20
60
Value
Common sites gene−LOC105284441
CphaseA1
RphaseB1

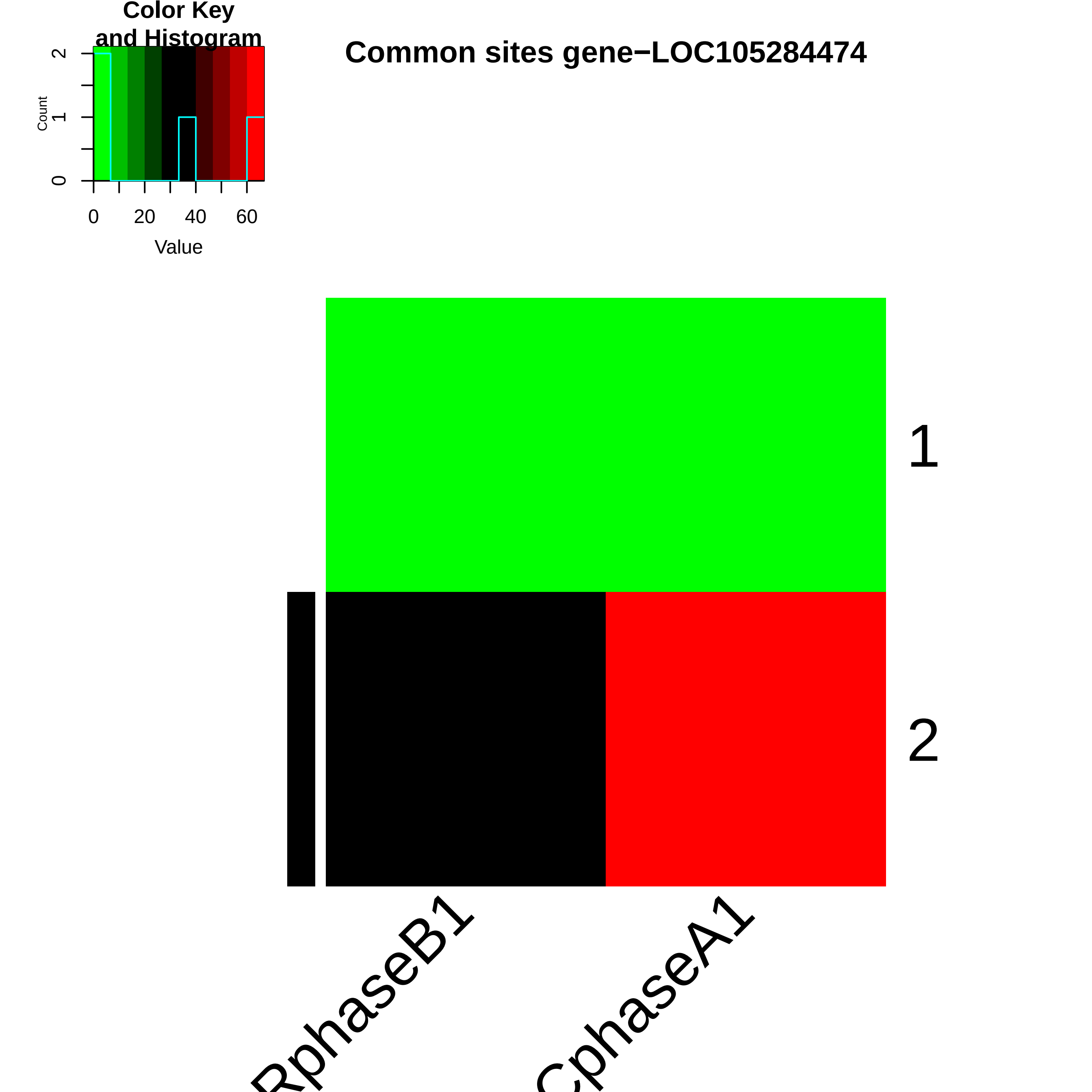

Color Key
and Histogram
Count
0
1
2
0
20
40
60
Value
Common sites gene−LOC105284474
1
2
RphaseB1
CphaseA1

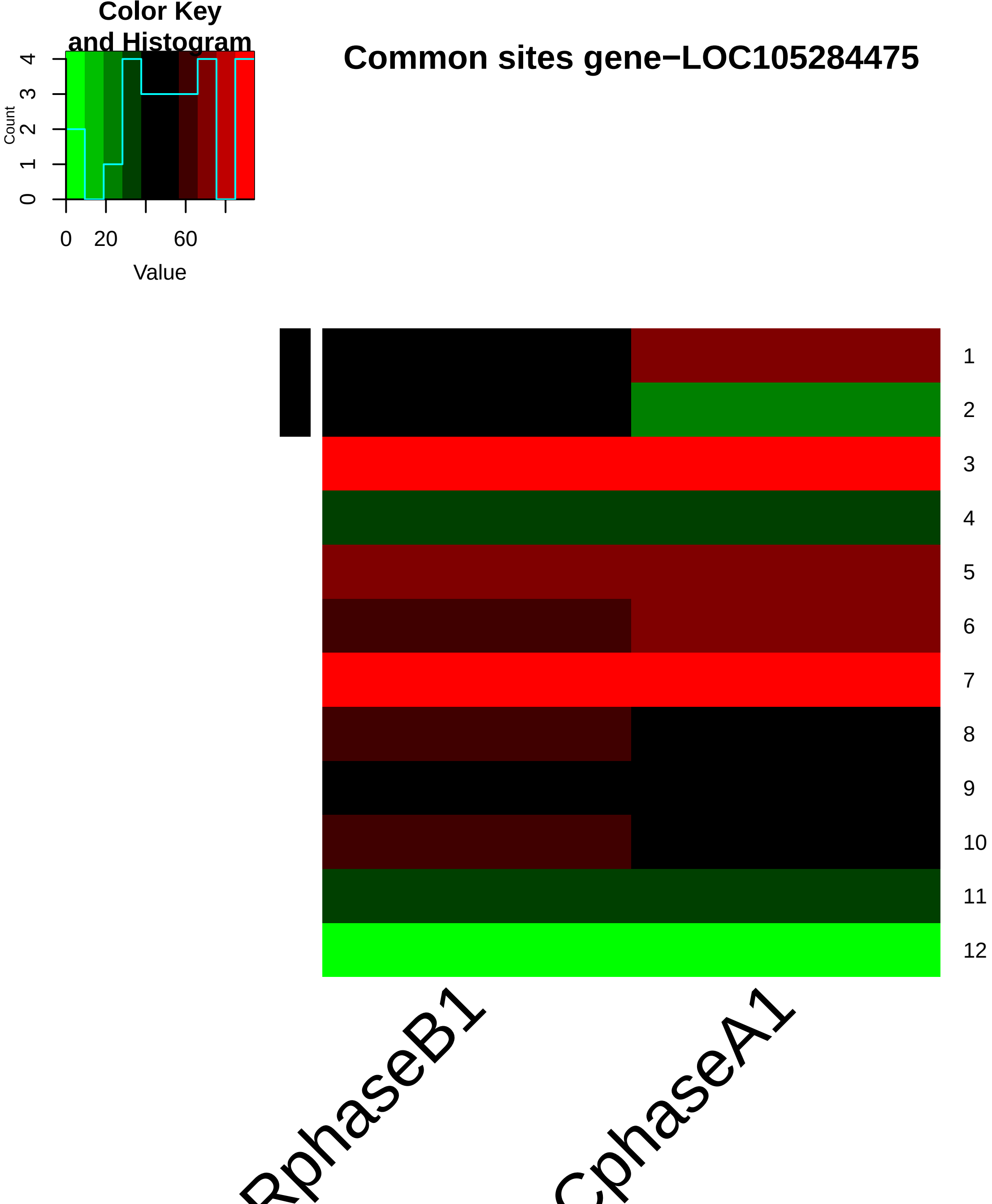
Color Key
and Histogram
Count
0
1
2
3
4
0
20
60
Value
Common sites gene−LOC105284475
1
2
3
4
5
6
7
8
9
10
11
12
RphaseB1
CphaseA1

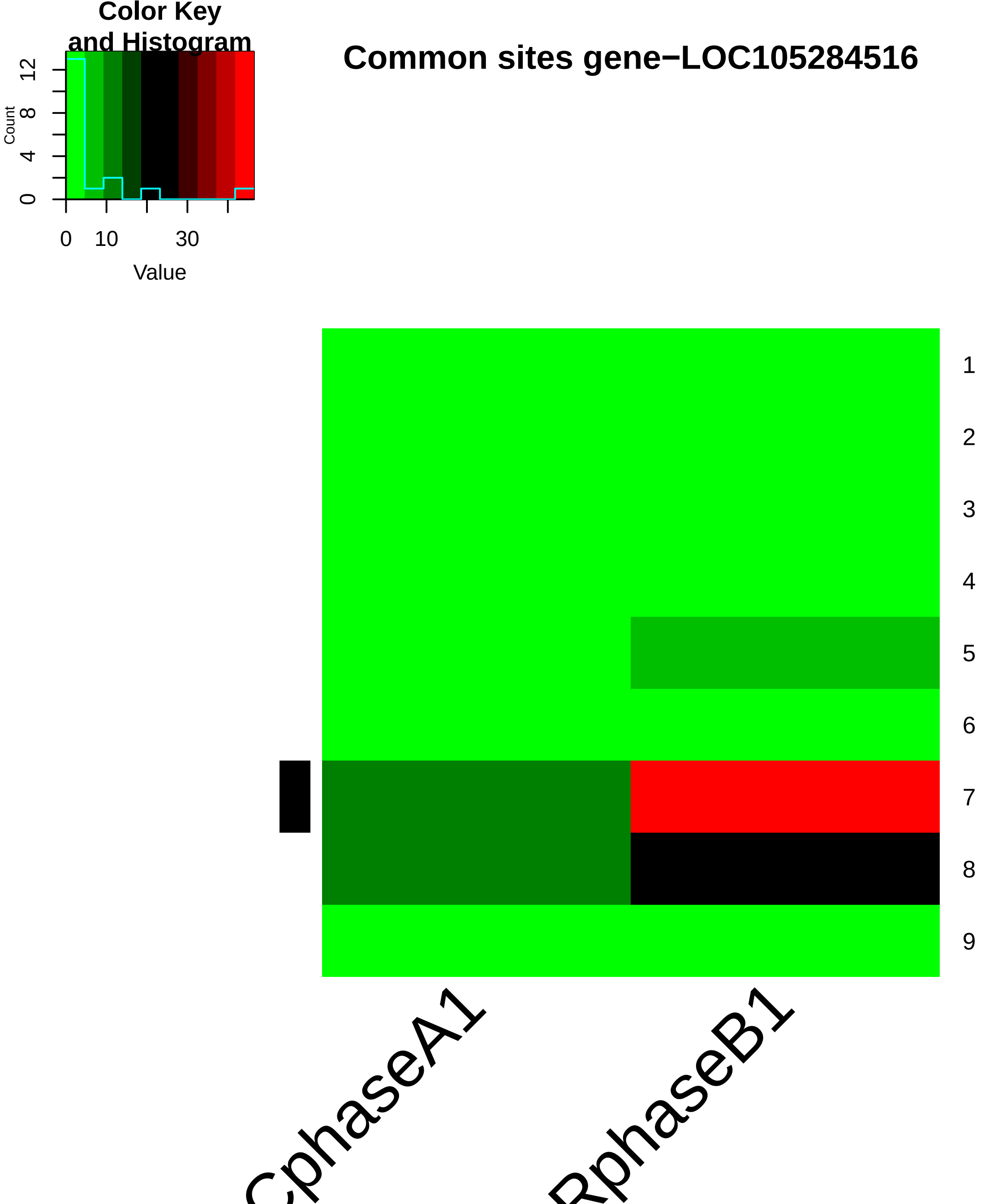

Color Key
and Histogram
Count
0
4
8
12
0
10
30
Value
Common sites gene−LOC105284516
1
2
3
4
5
6
7
8
9
CphaseA1
RphaseB1

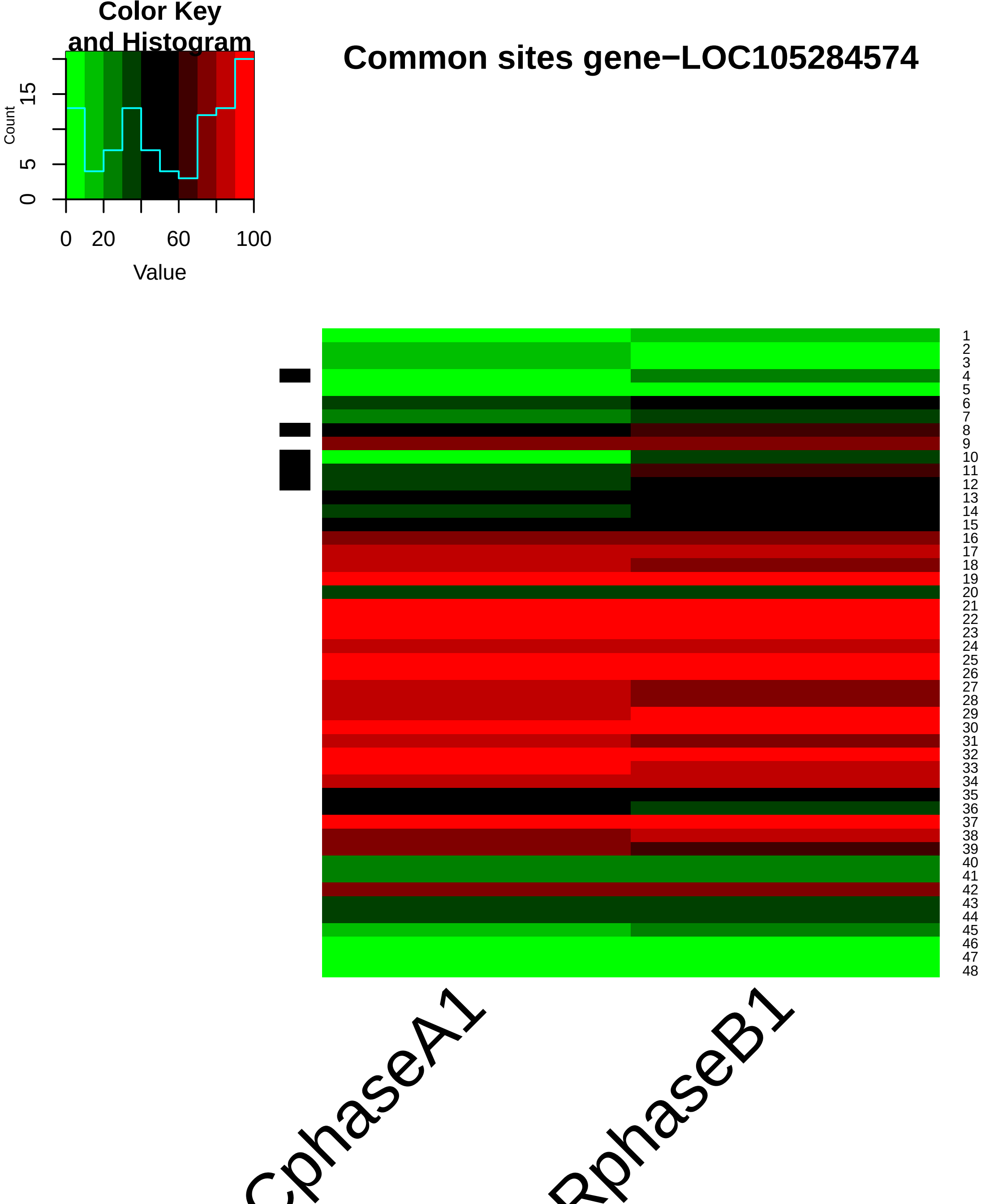
Color Key
and Histogram
Count
0
5
15
0
20
60
100
Value
Common sites gene−LOC105284574
CphaseA1
RphaseB1

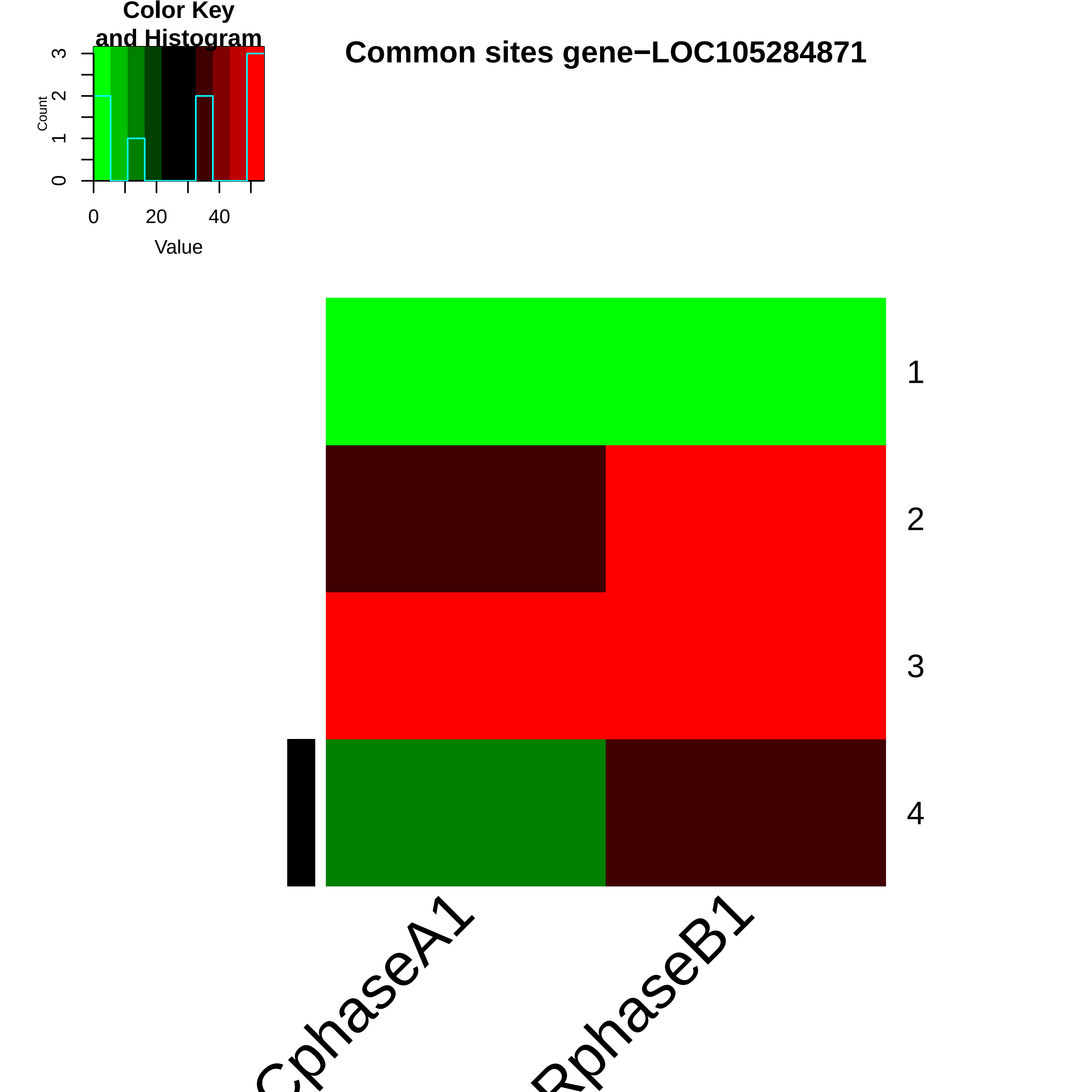
Color Key
and Histogram
Count
0
1
2
3
0
20
40
Value
Common sites gene−LOC105284871
1
2
3
4
CphaseA1
RphaseB1

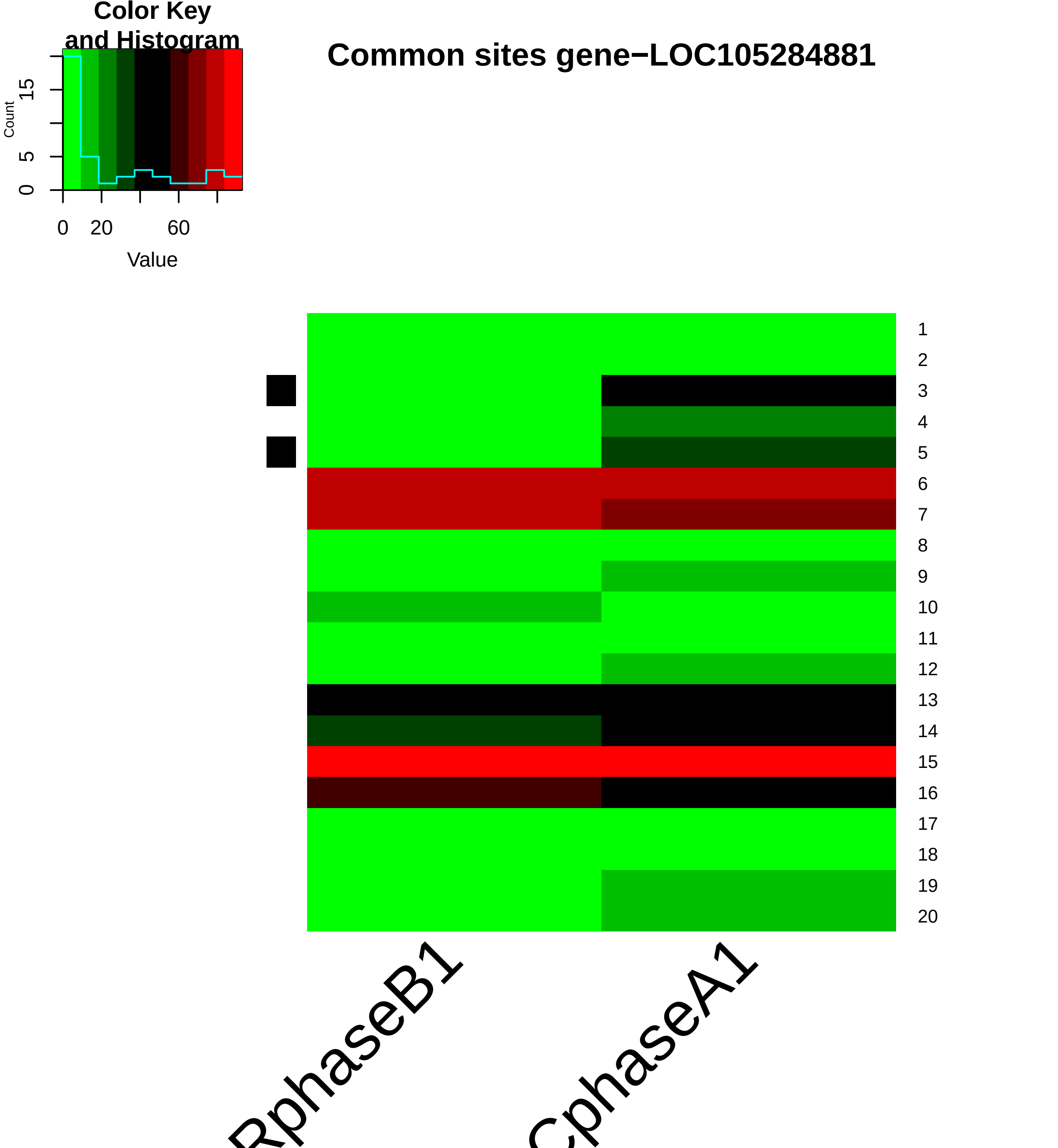
Color Key
and Histogram
Count
0
5
15
0
20
60
Value
Common sites gene−LOC105284881
RphaseB1
CphaseA1

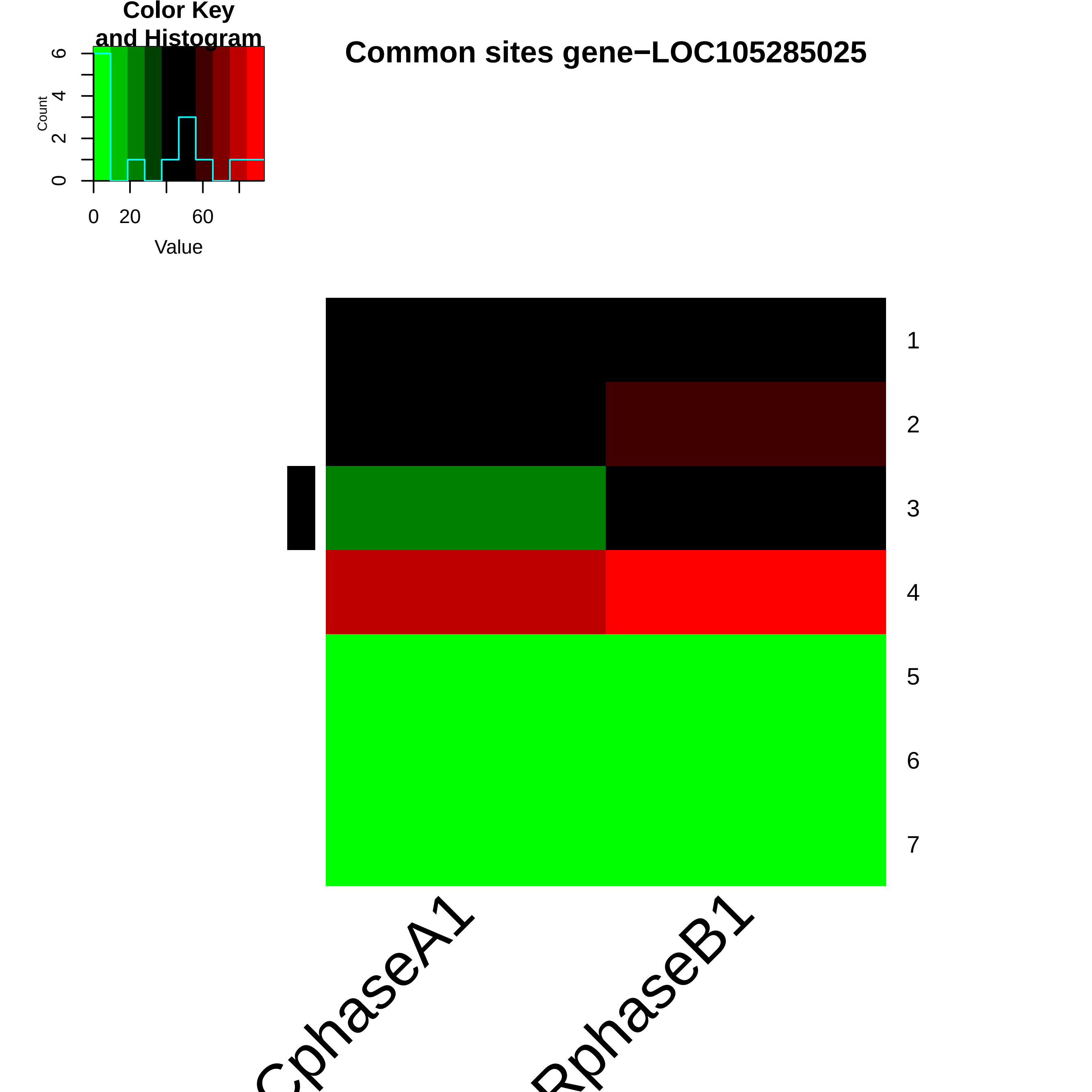

Color Key
and Histogram
Count
0
2
4
6
0
20
60
Value
Common sites gene−LOC105285025
1
2
3
4
5
6
7
CphaseA1
RphaseB1

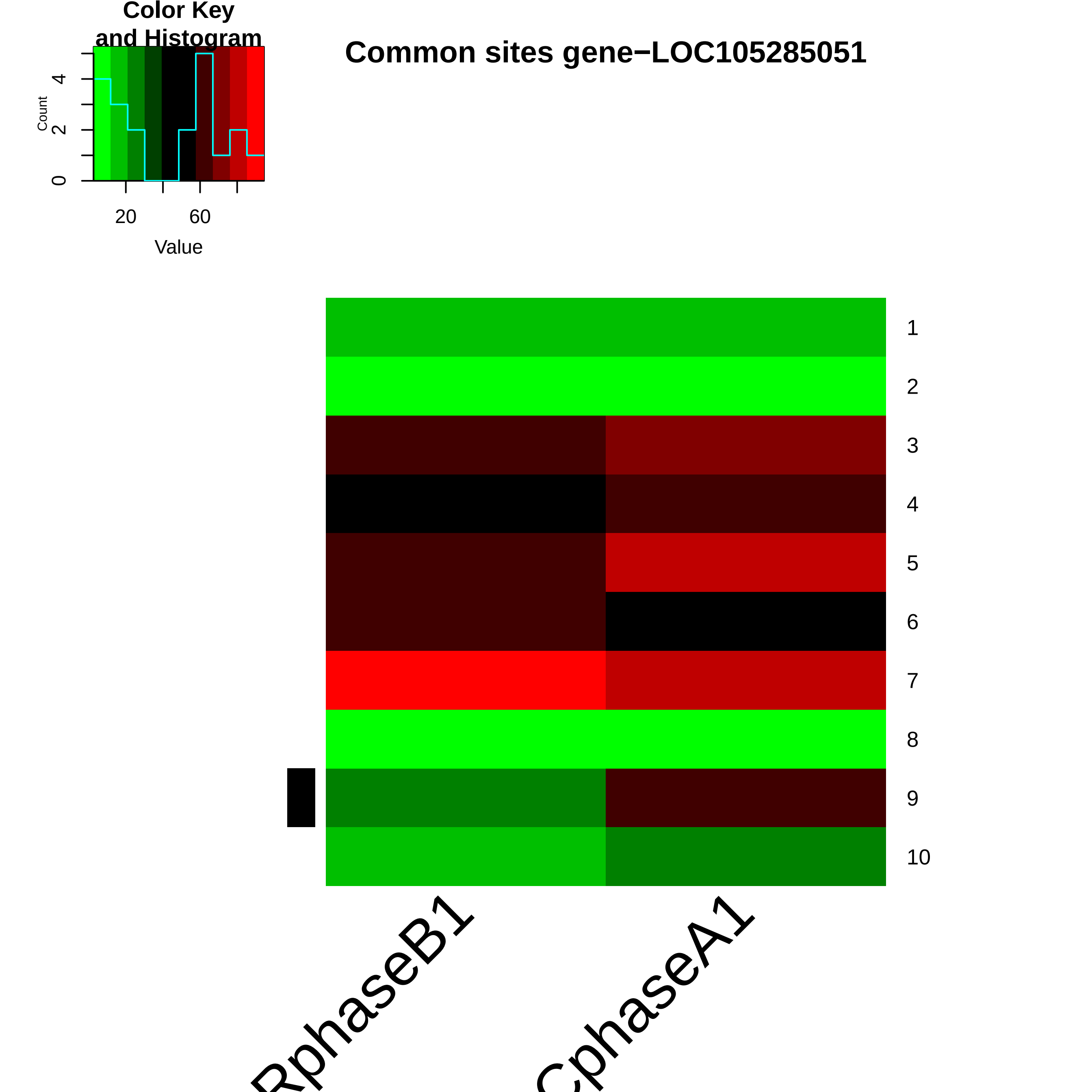
Color Key
and Histogram
Count
0
2
4
20
60
Value
Common sites gene−LOC105285051
1
2
3
4
5
6
7
8
9
10
RphaseB1
CphaseA1

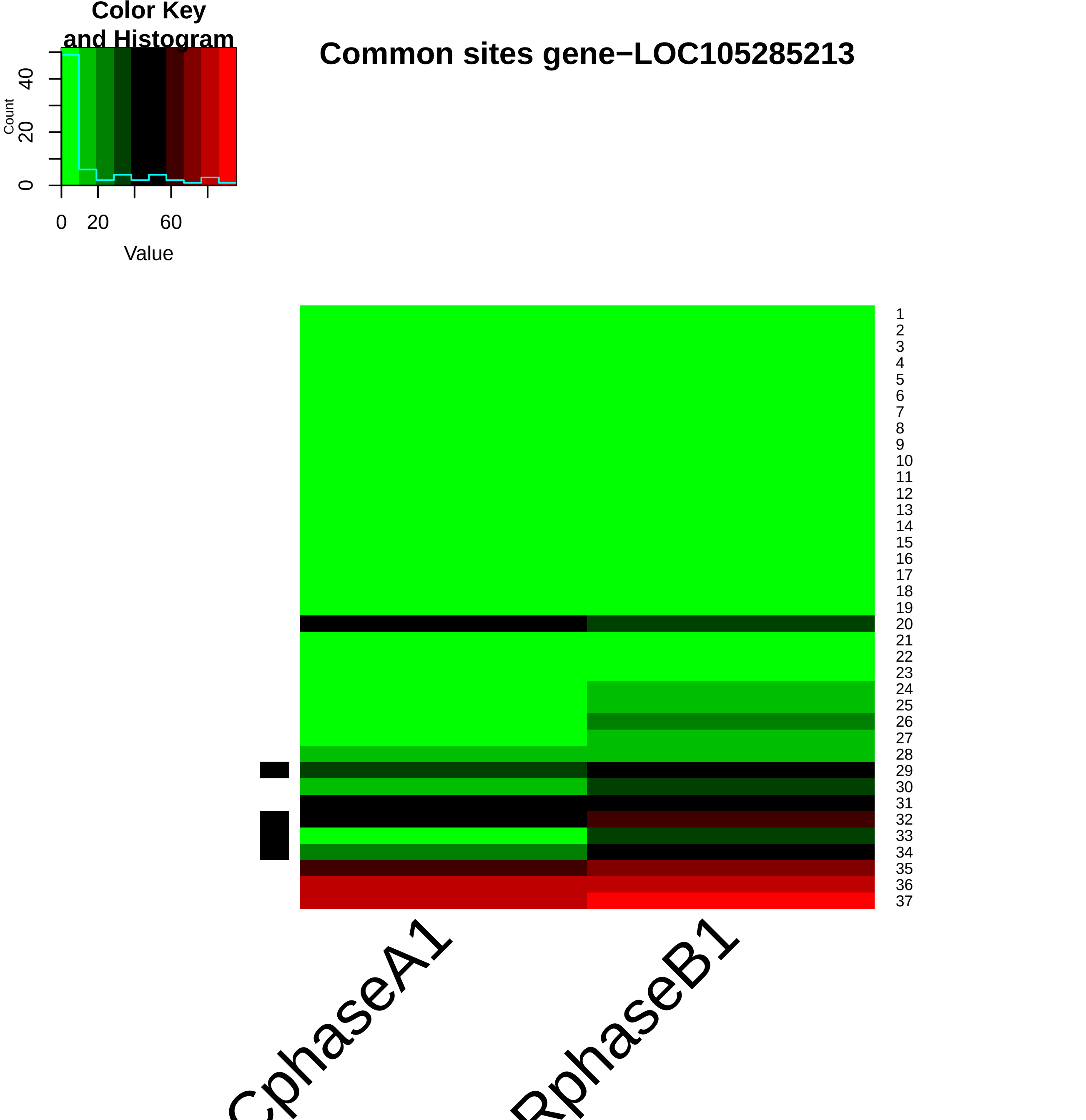
Color Key
and Histogram
Count
0
20
40
0
20
60
Value
Common sites gene−LOC105285213
CphaseA1
RphaseB1

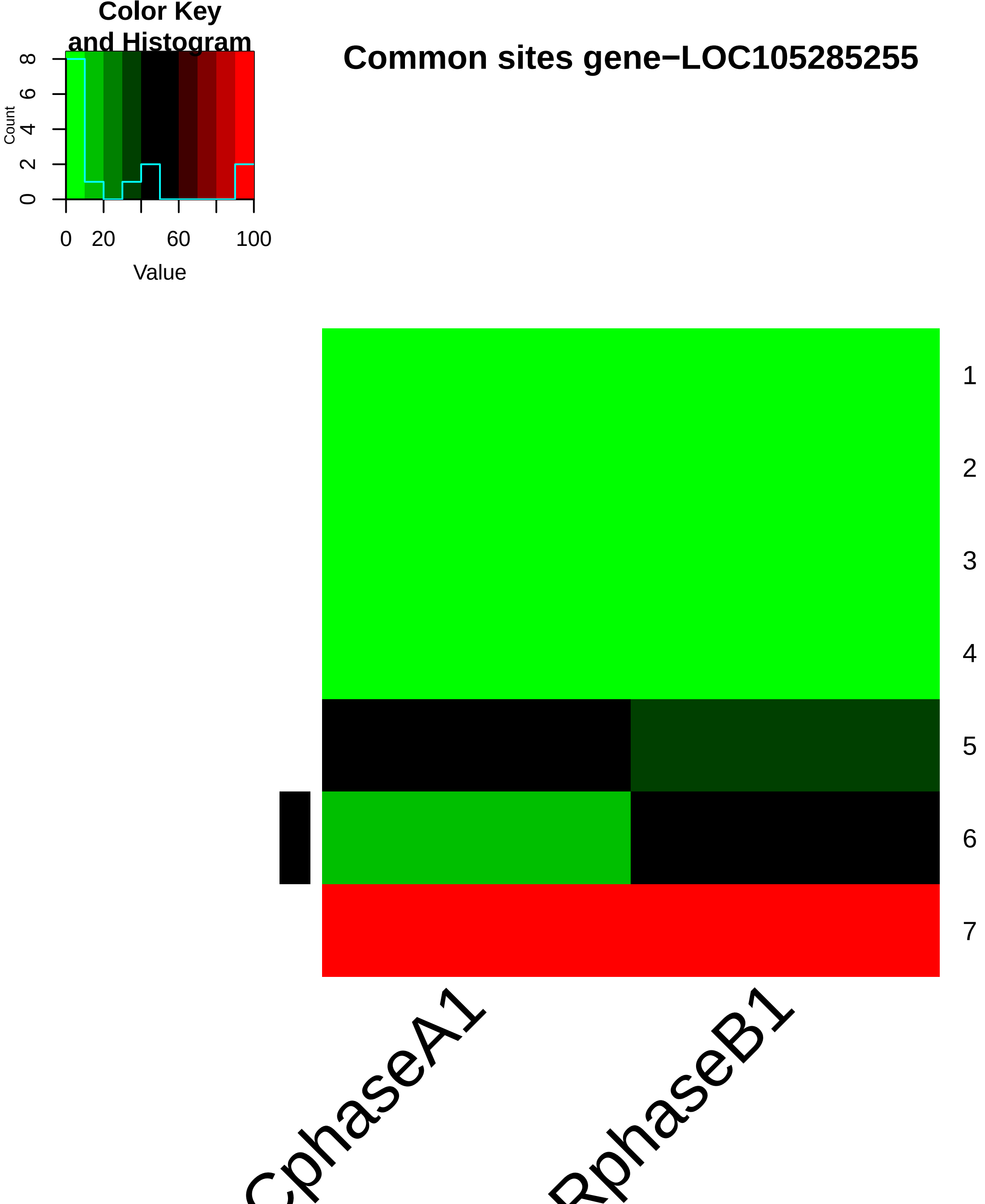

Color Key
and Histogram
Count
0
2
4
6
8
0
20
60
100
Value
Common sites gene−LOC105285255
1
2
3
4
5
6
7
CphaseA1
RphaseB1

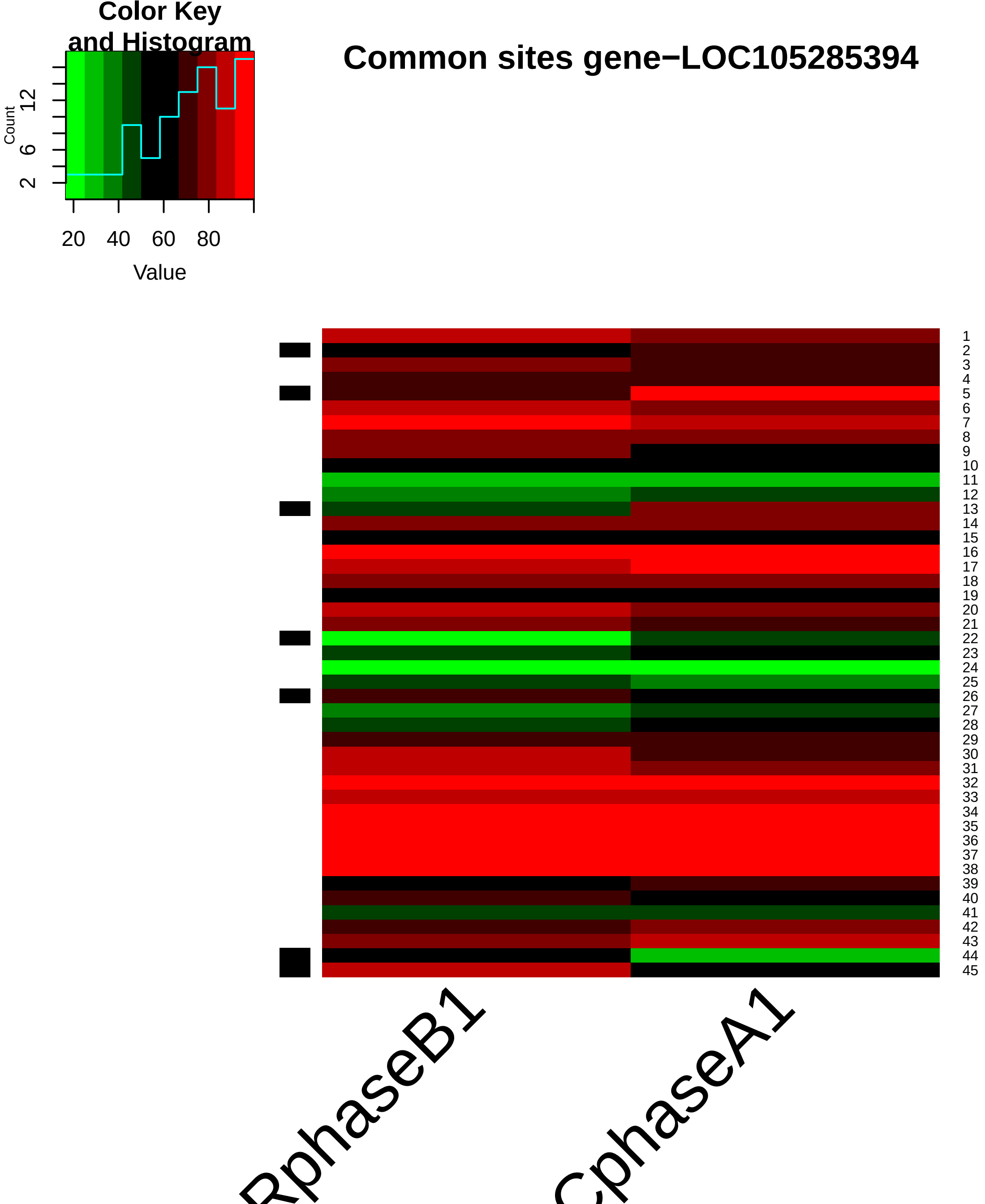

Color Key
and Histogram
Count
2
6
12
20
40
60
80
Value
Common sites gene−LOC105285394
RphaseB1
CphaseA1

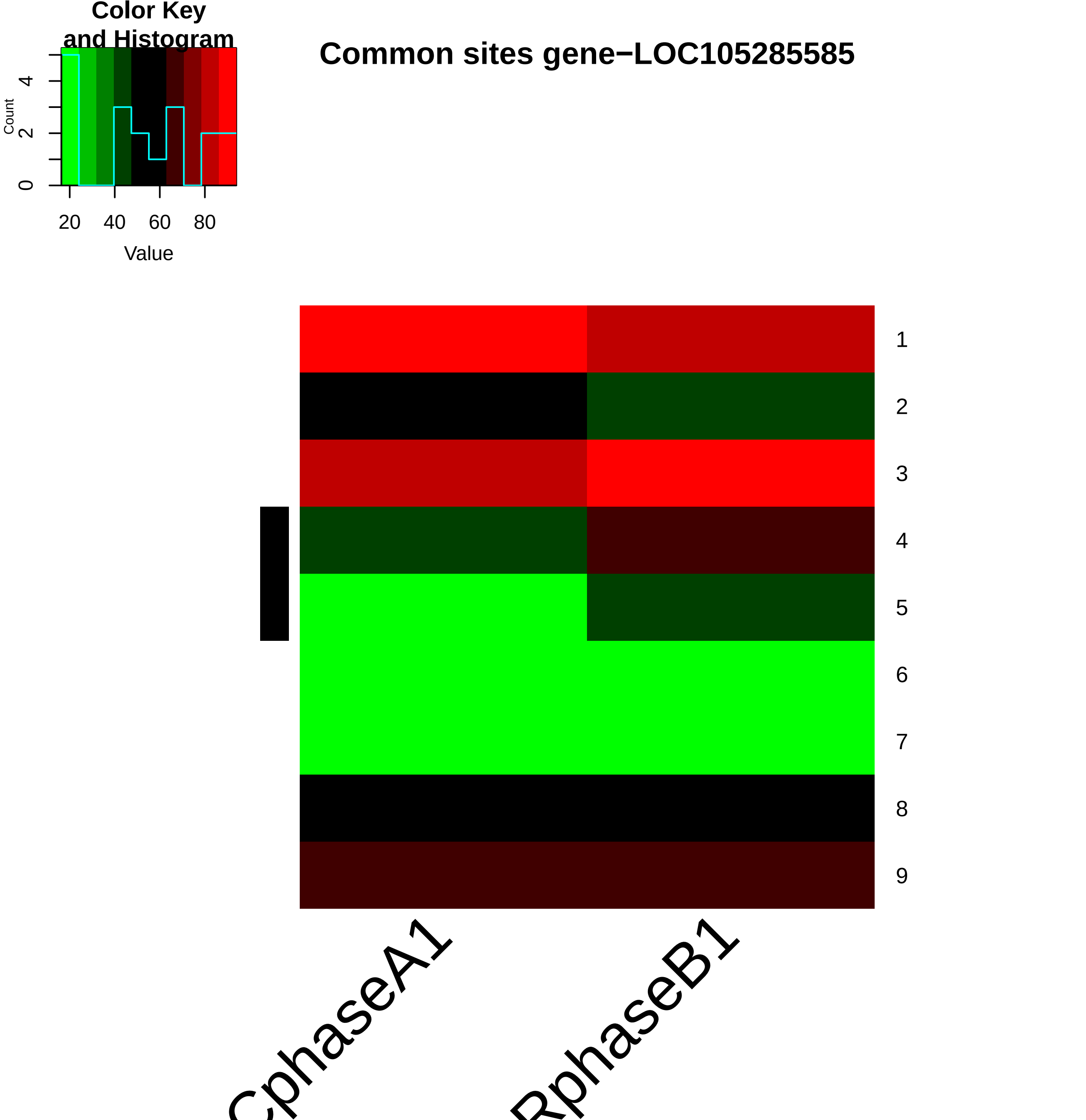

Color Key
and Histogram
Count
0
2
4
20
40
60
80
Value
Common sites gene−LOC105285585
1
2
3
4
5
6
7
8
9
CphaseA1
RphaseB1

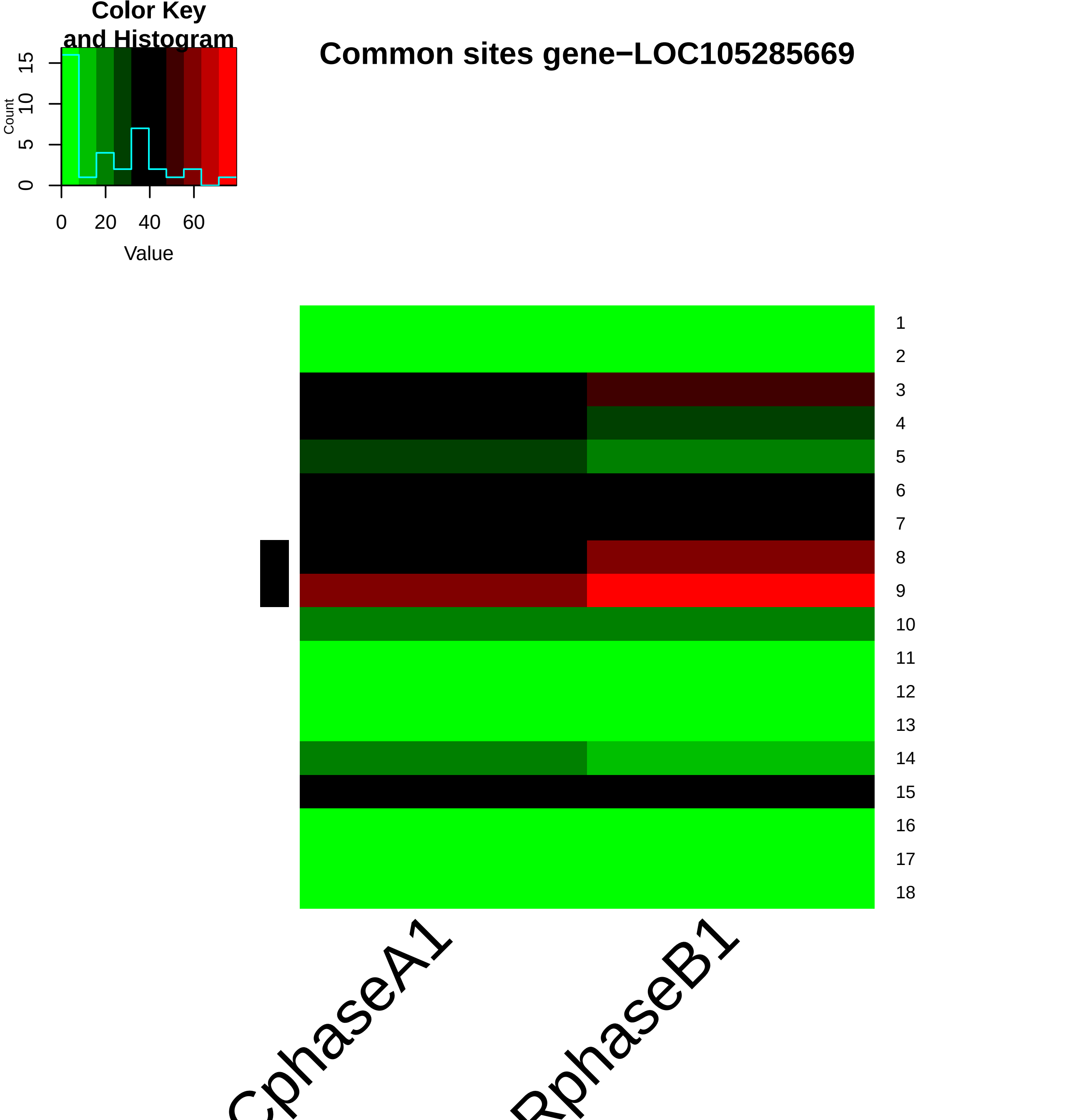
Color Key
and Histogram
Count
0
5
10
15
0
20
40
60
Value
Common sites gene−LOC105285669
CphaseA1
RphaseB1

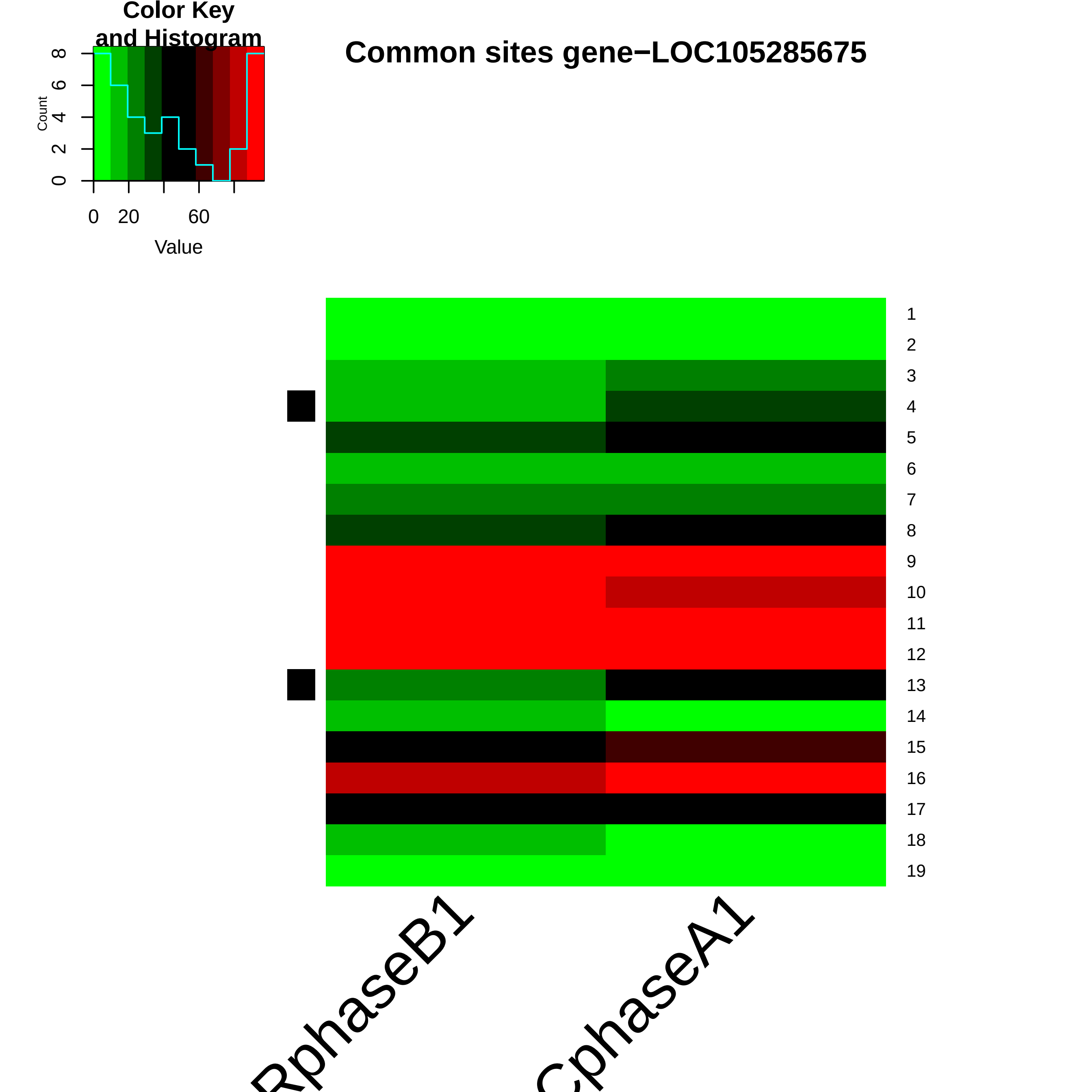
Color Key
and Histogram
Count
Value
Common sites gene−LOC105285675
RphaseB1
CphaseA1

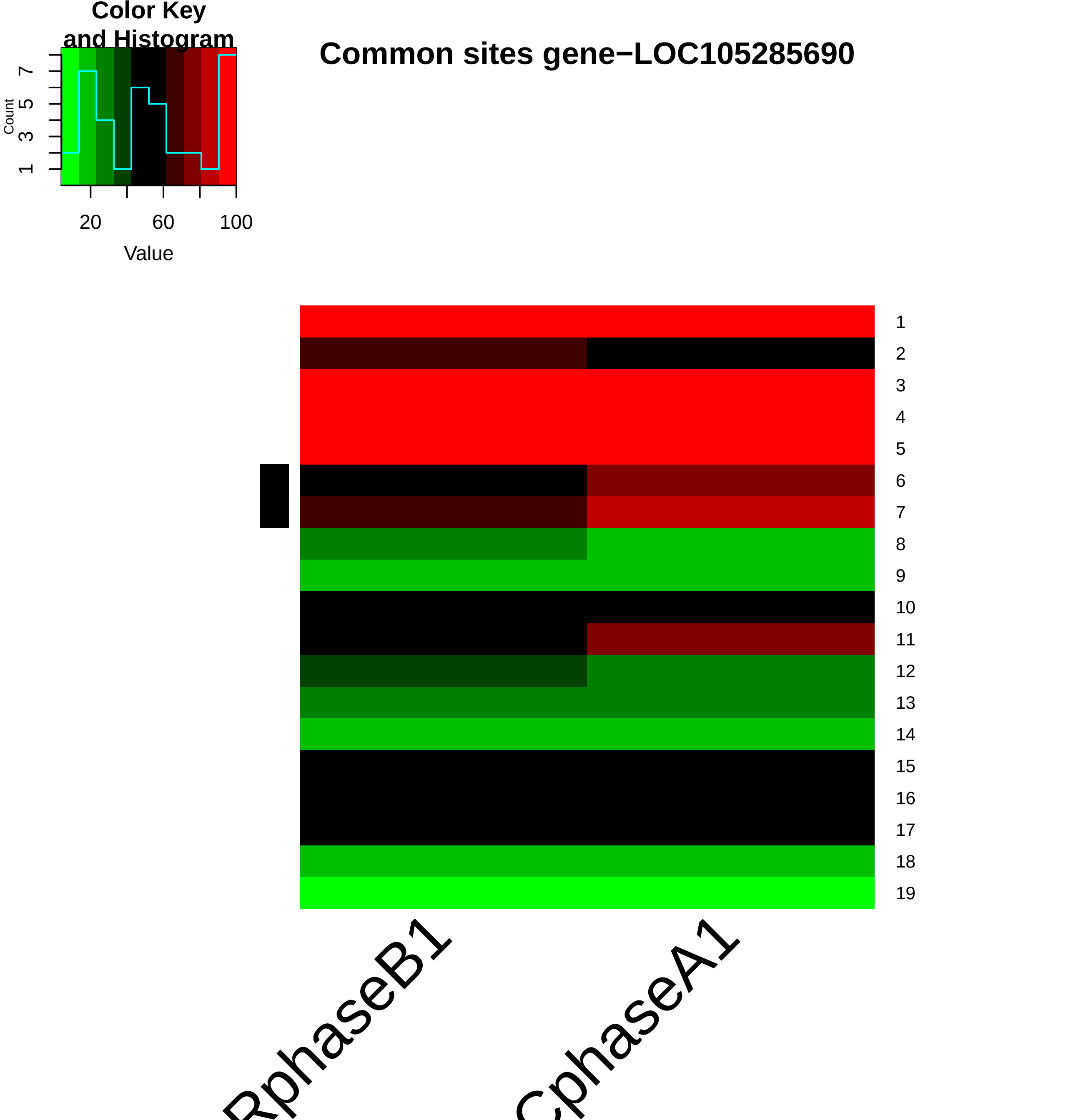
Color Key
and Histogram
Count
1
3
5
7
20
60
100
Value
Common sites gene−LOC105285690
RphaseB1
CphaseA1

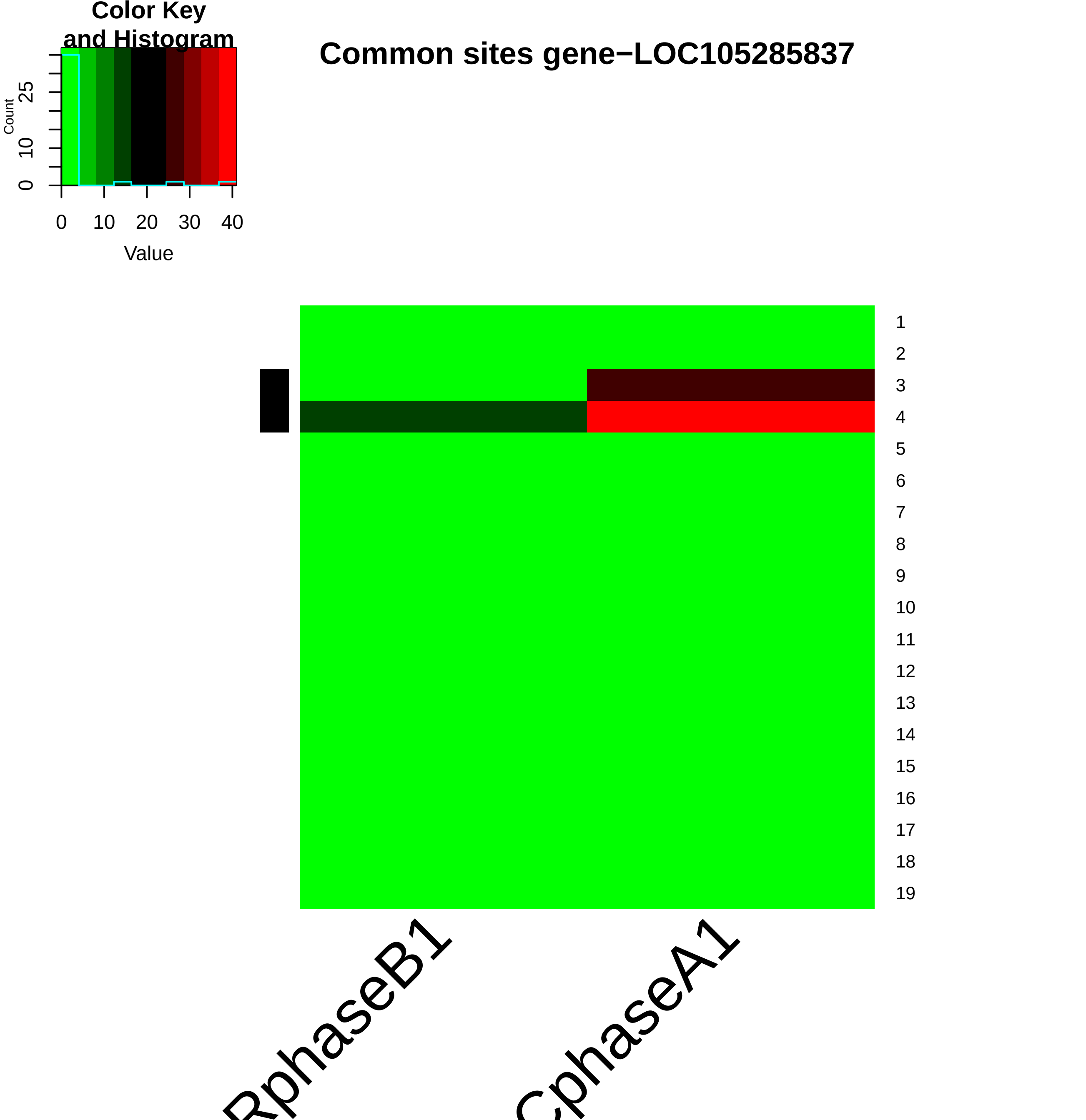

Color Key and Histogram
Count
0
10
25
0
10
20
30
40
Value
Common sites gene−LOC105285837
RphaseB1
CphaseA1

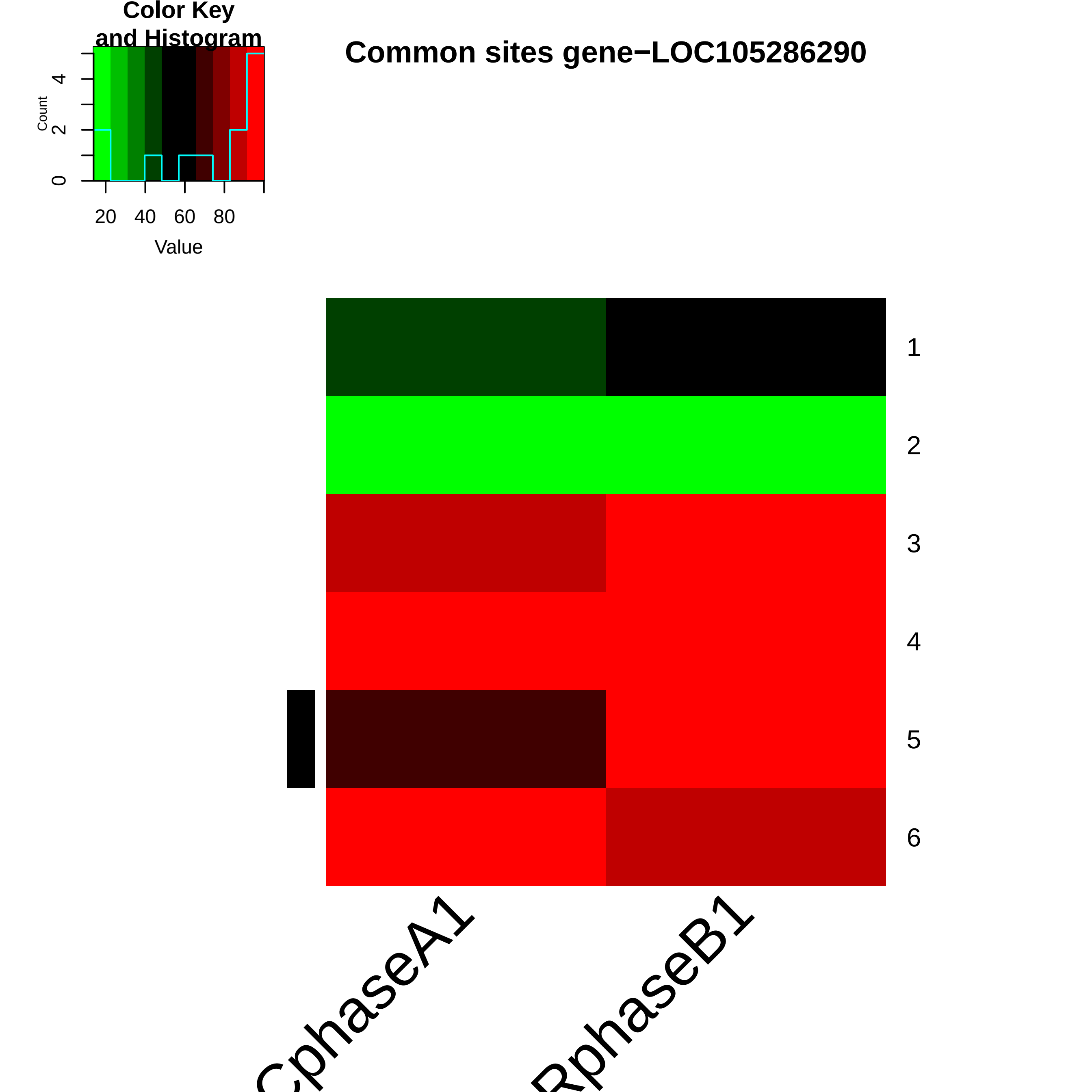

Color Key
and Histogram
Count
0
2
4
20
40
60
80
Value
Common sites gene−LOC105286290
1
2
3
4
5
6
CphaseA1
RphaseB1

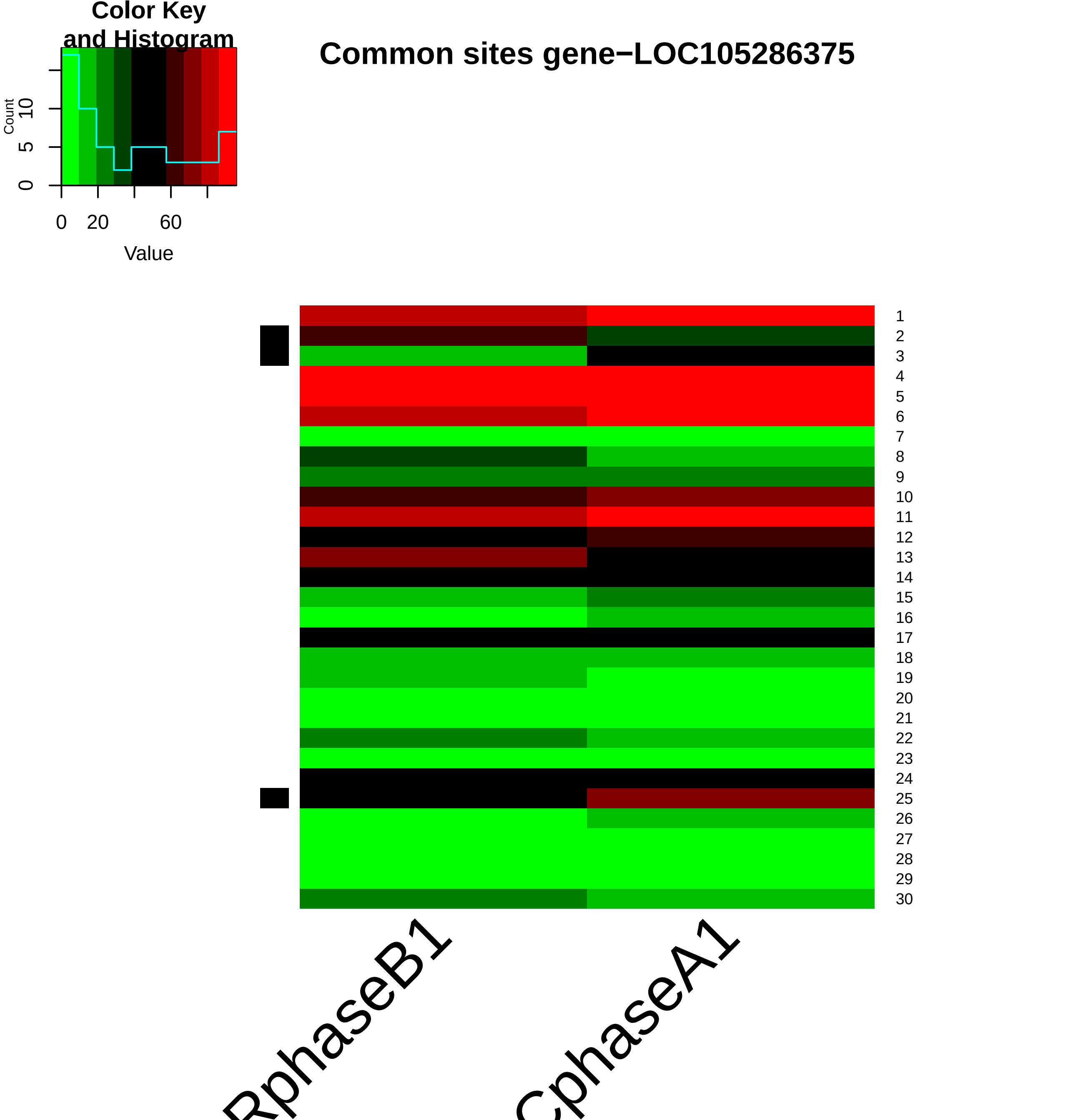
Color Key
and Histogram
Count
0
5
10
0
20
60
Value
Common sites gene−LOC105286375
RphaseB1
CphaseA1

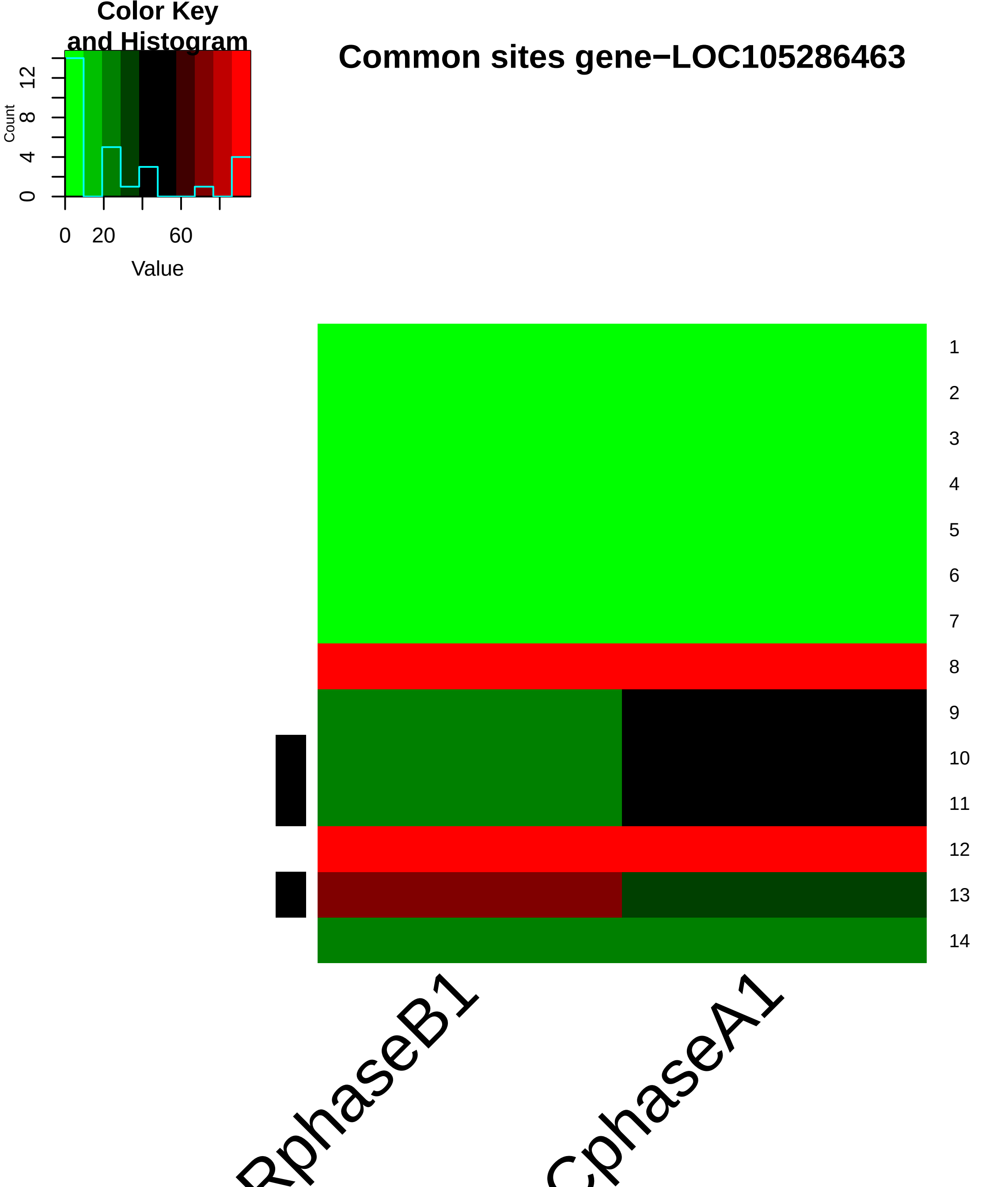

Color Key
and Histogram
Count
0
4
8
12
0
20
60
Value
Common sites gene−LOC105286463
1
2
3
4
5
6
7
8
9
10
11
12
13
14
RphaseB1
CphaseA1

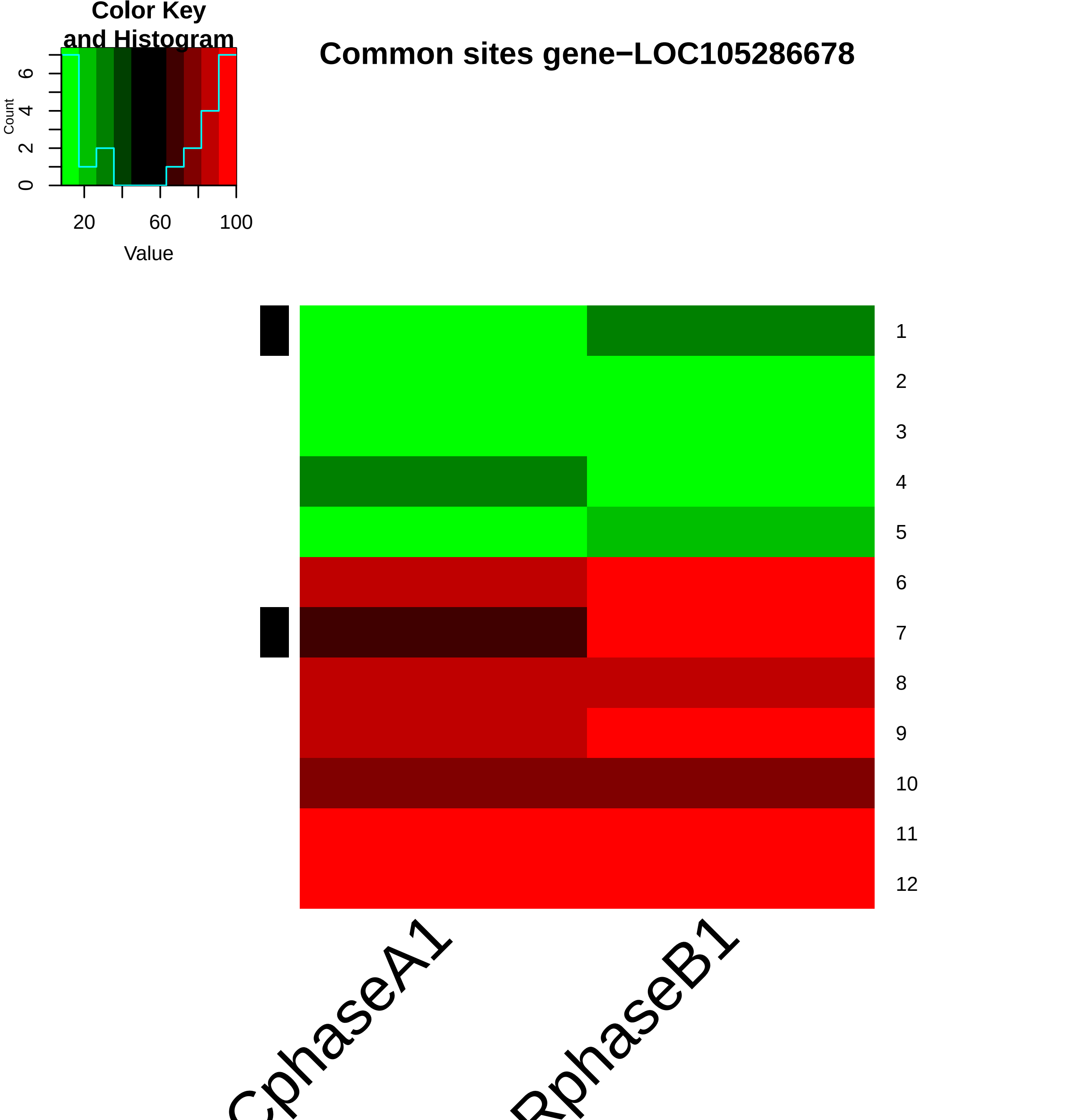
Color Key
and Histogram
Count
0
2
4
6
20
60
100
Value
Common sites gene−LOC105286678
1
2
3
4
5
6
7
8
9
10
11
12
CphaseA1
RphaseB1

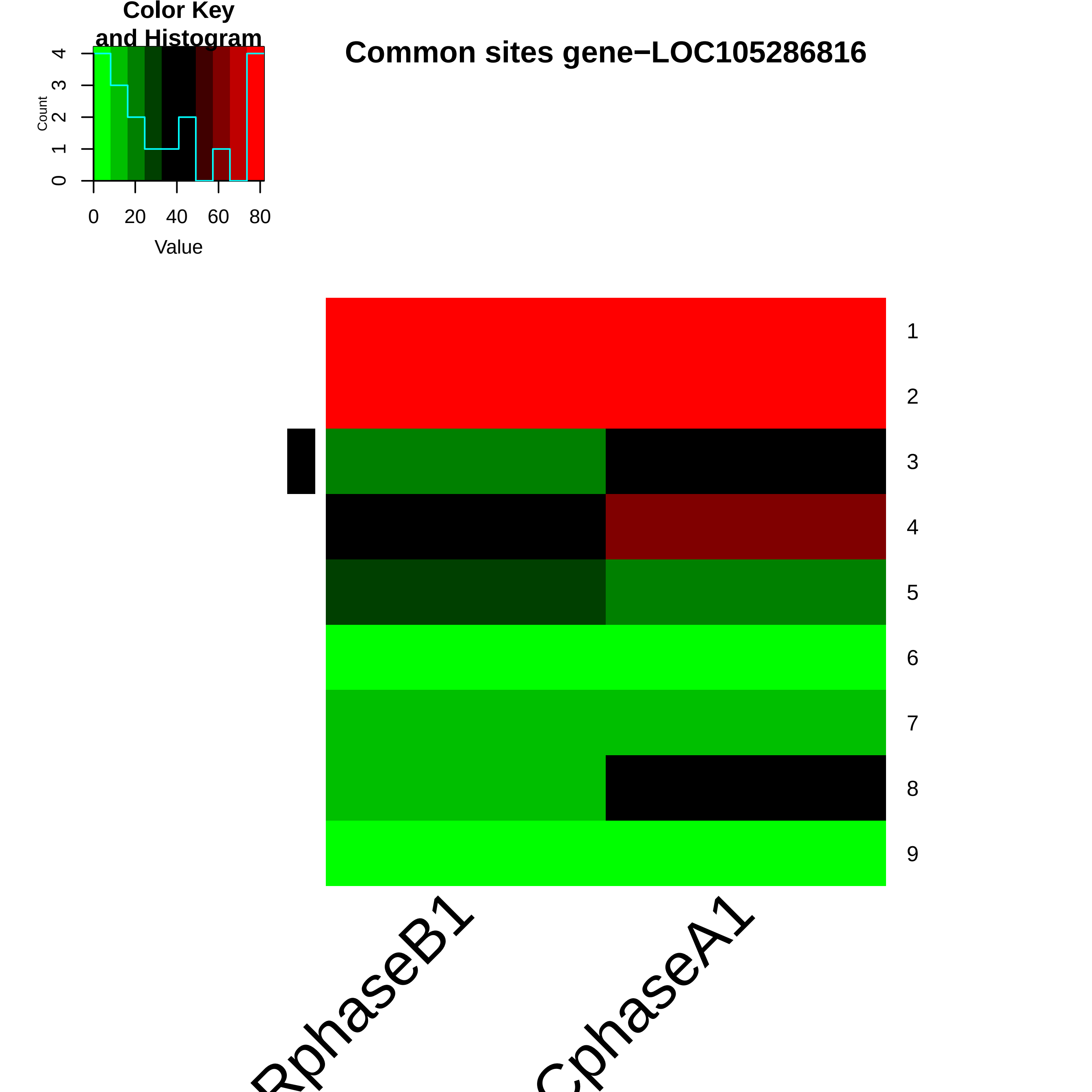
Color Key
and Histogram
Count
0
1
2
3
4
0
20
40
60
80
Value
Common sites gene−LOC105286816
1
2
3
4
5
6
7
8
9

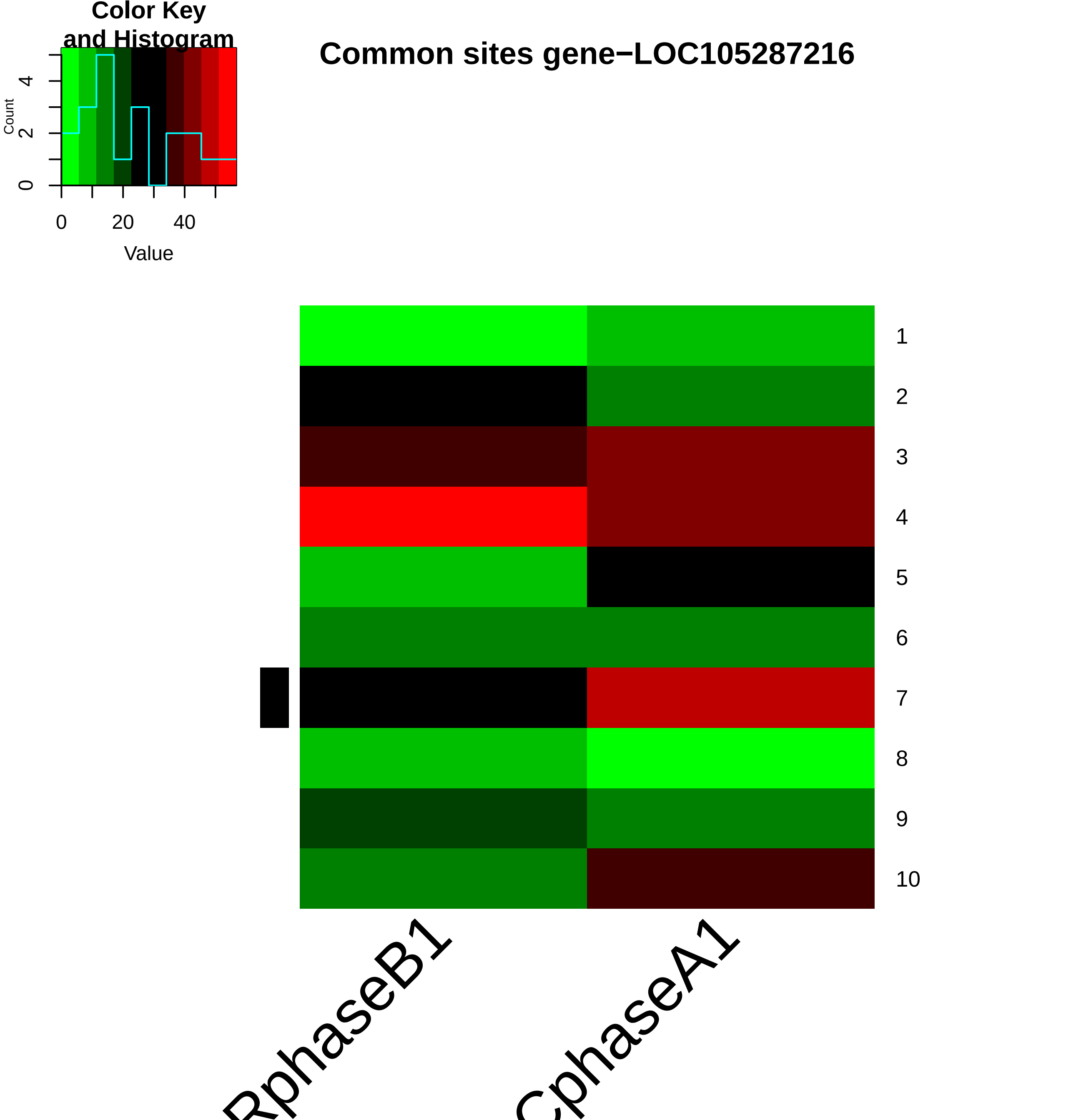
Color Key
and Histogram
Count
0
2
4
0
20
40
Value
Common sites gene−LOC105287216
1
2
3
4
5
6
7
8
9
10
RphaseB1
CphaseA1

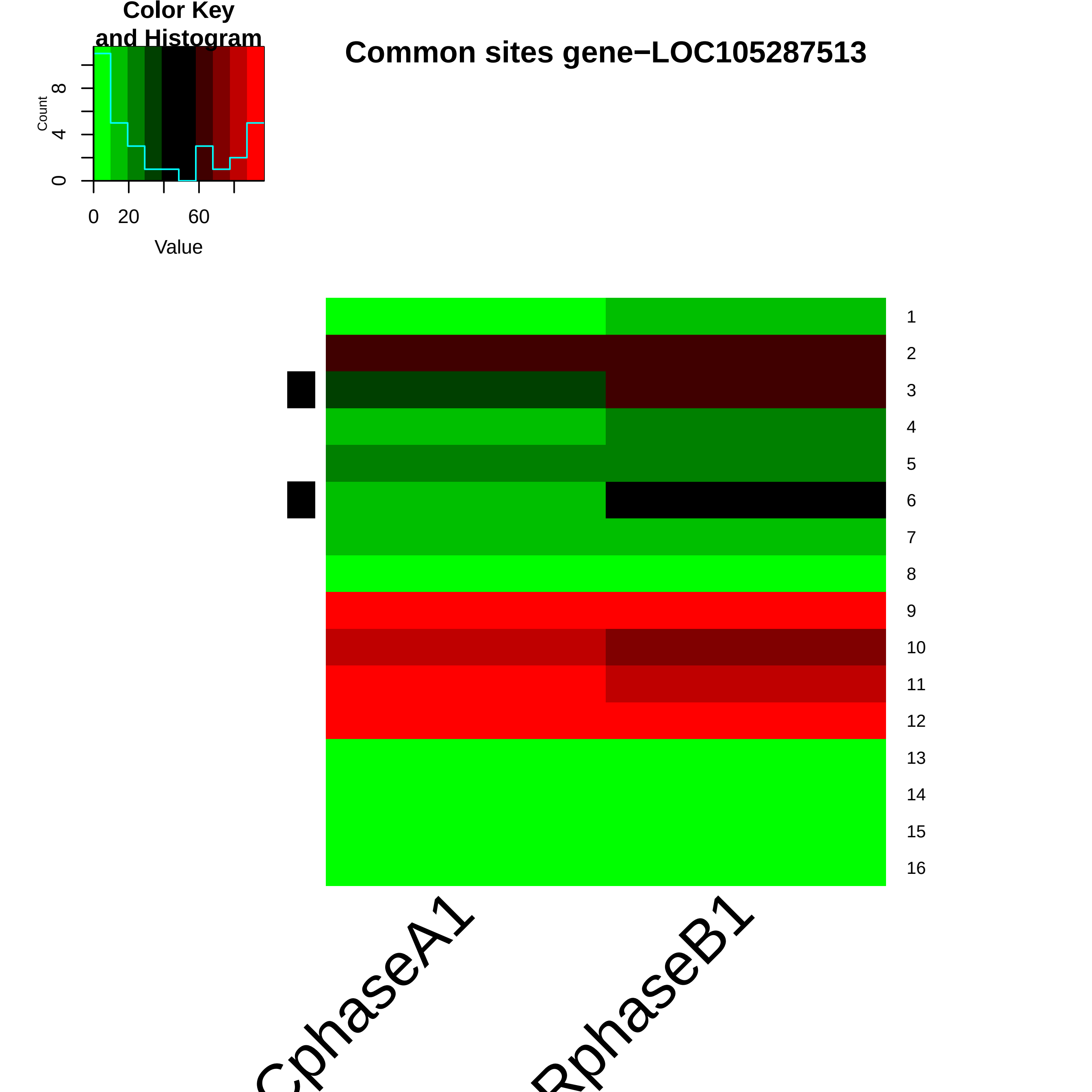

Color Key
and Histogram
Count
0
4
8
0
20
60
Value
Common sites gene−LOC105287513
CphaseA1
RphaseB1

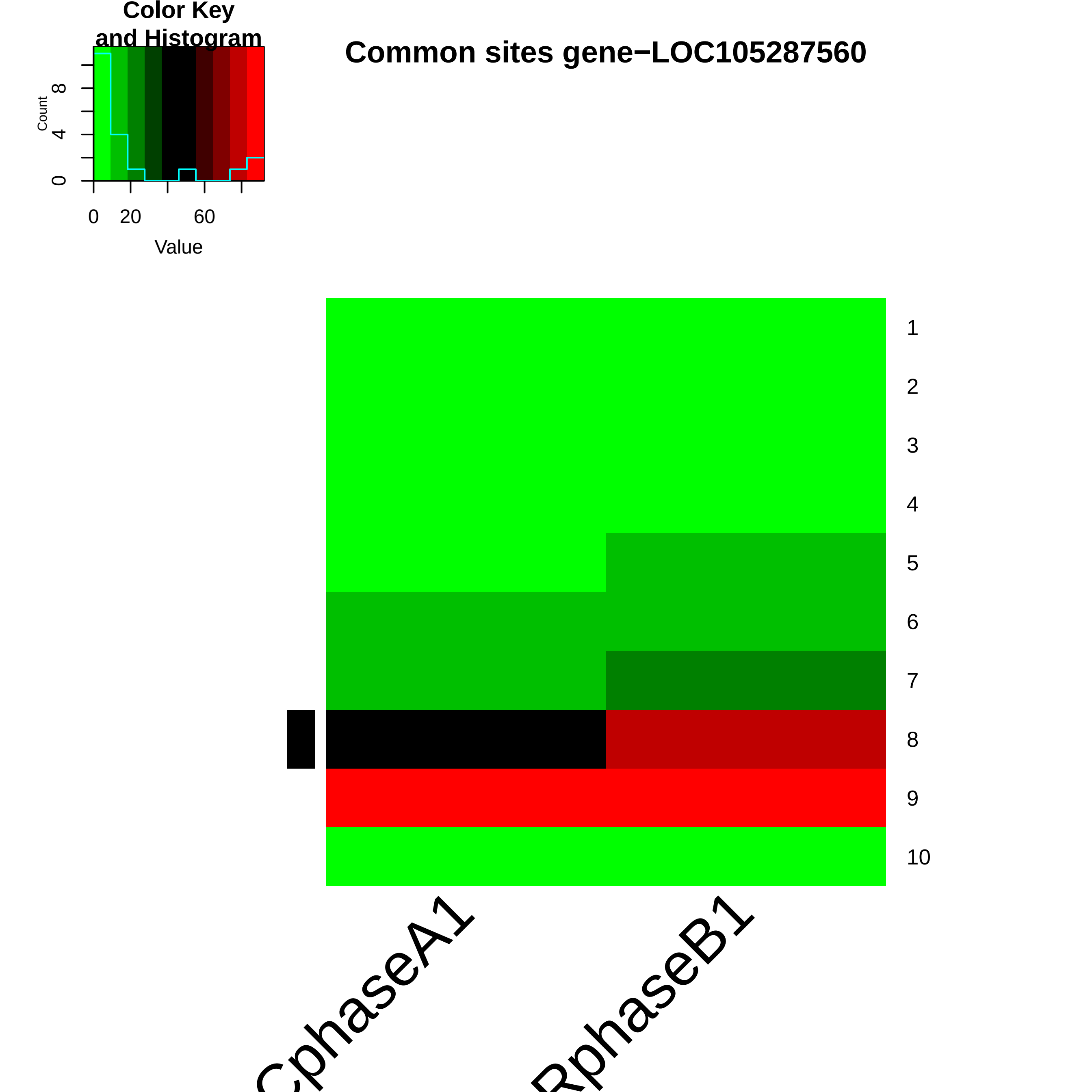

Color Key
and Histogram
Count
0
4
8
0
20
60
Value
Common sites gene−LOC105287560
1
2
3
4
5
6
7
8
9
10
CphaseA1
RphaseB1

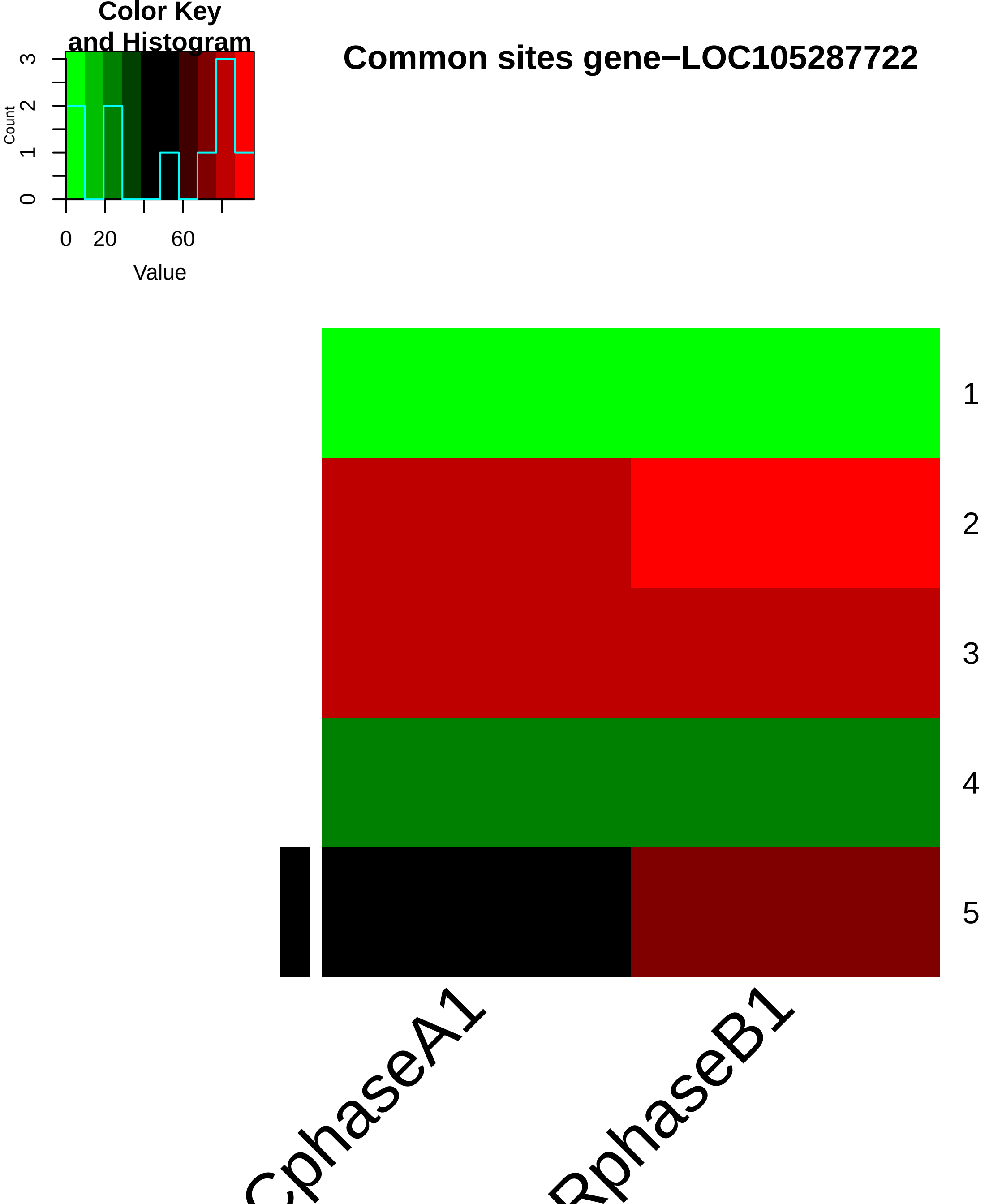

Color Key
and Histogram
Count
0
1
2
3
0
20
60
Value
Common sites gene−LOC105287722
1
2
3
4
5
CphaseA1
RphaseB1

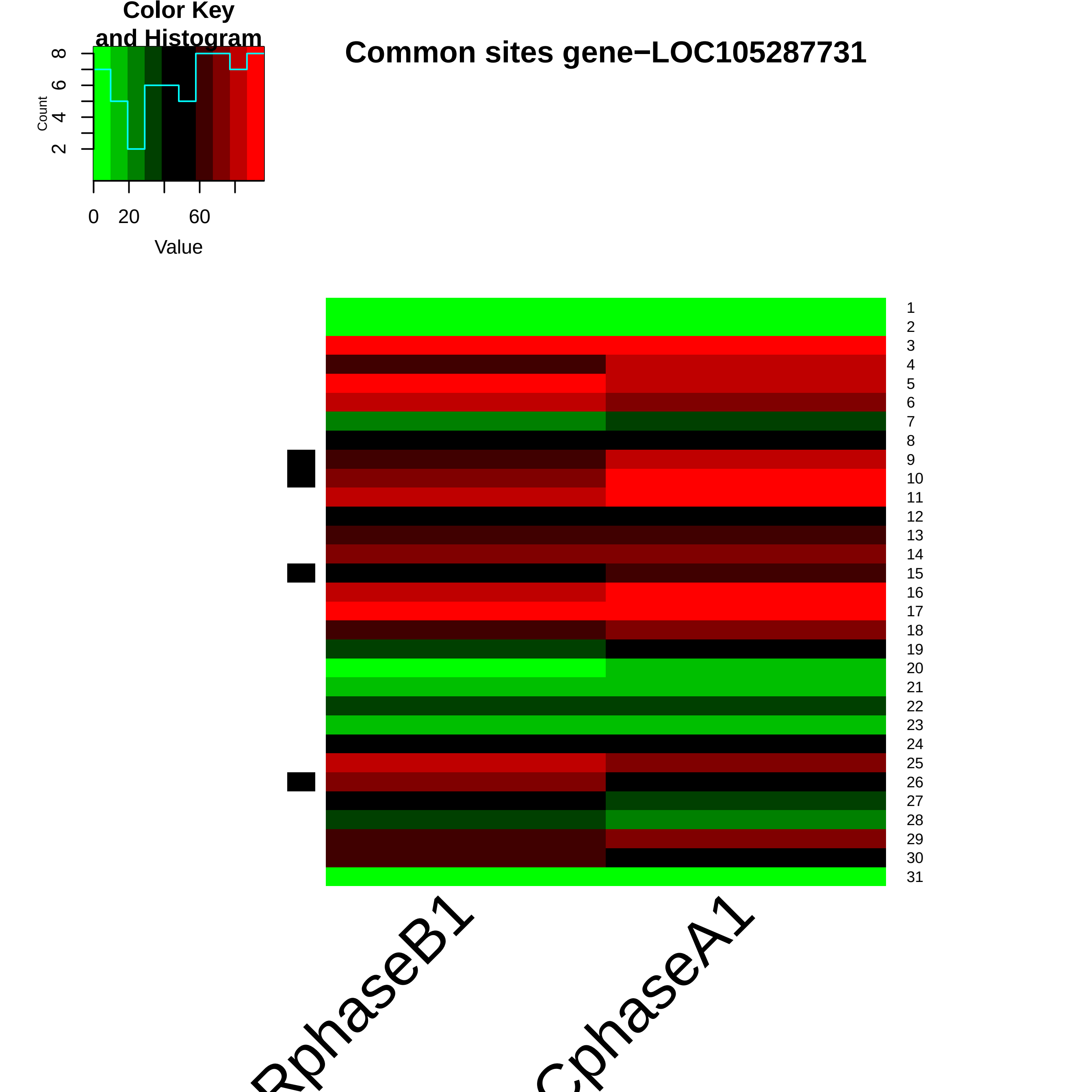

Color Key
and Histogram
Count
2
4
6
8
0
20
60
Value
Common sites gene−LOC105287731
RphaseB1
CphaseA1

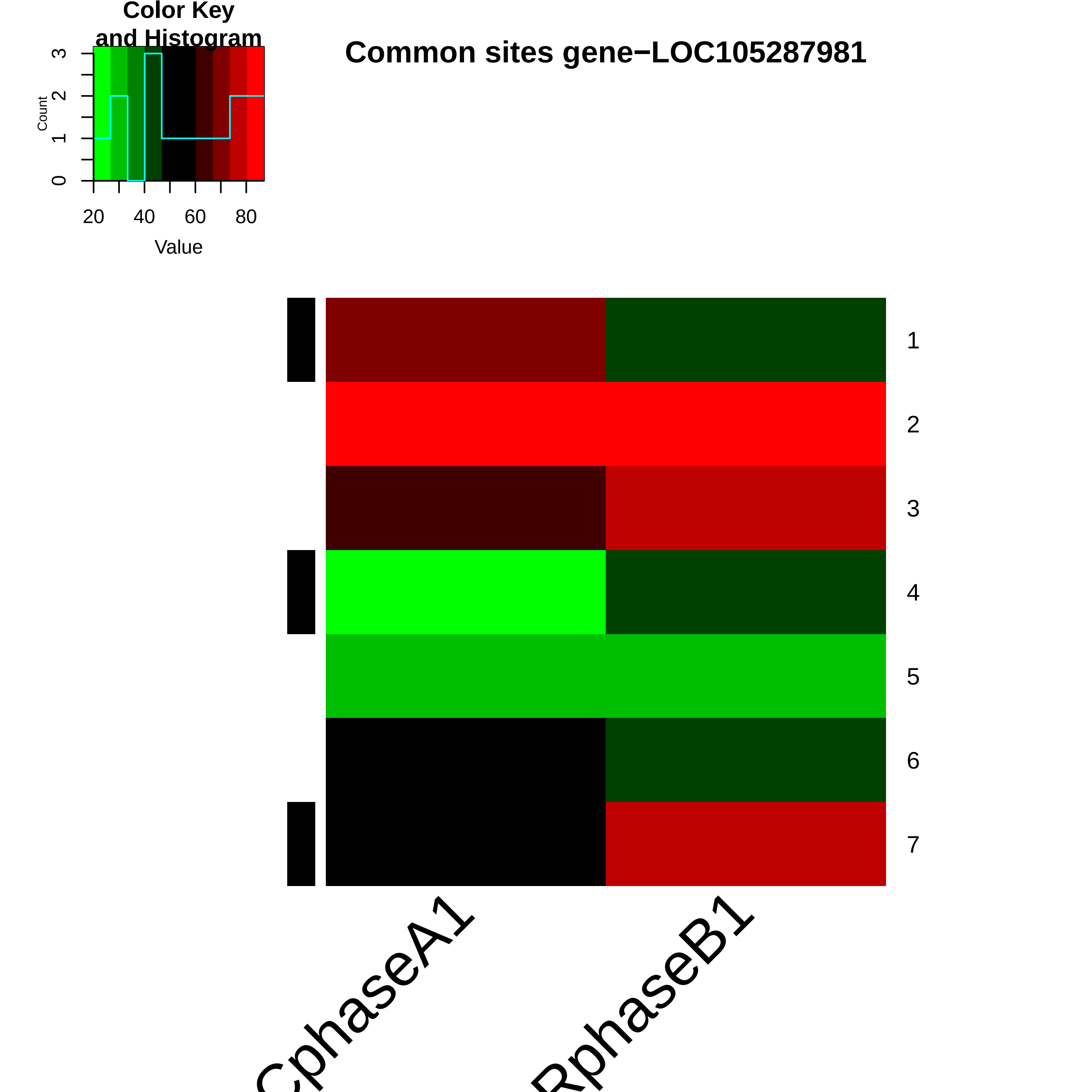
Color Key
and Histogram
Count
0
1
2
3
20
40
60
80
Value
Common sites gene−LOC105287981
1
2
3
4
5
6
7
CphaseA1
RphaseB1

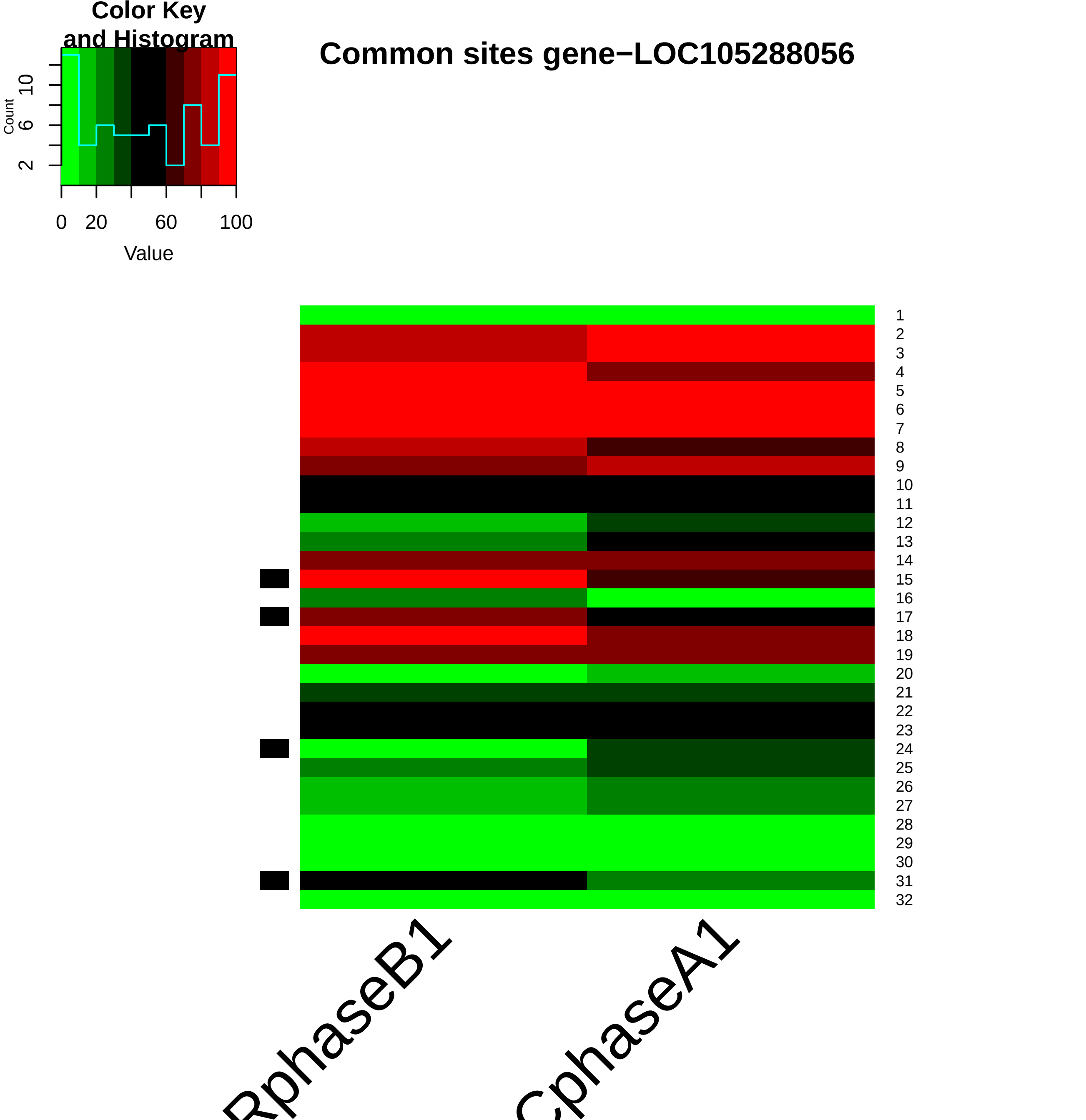
Color Key
and Histogram
Count
2
6
10
0
20
60
100
Value
Common sites gene−LOC105288056
RphaseB1
CphaseA1

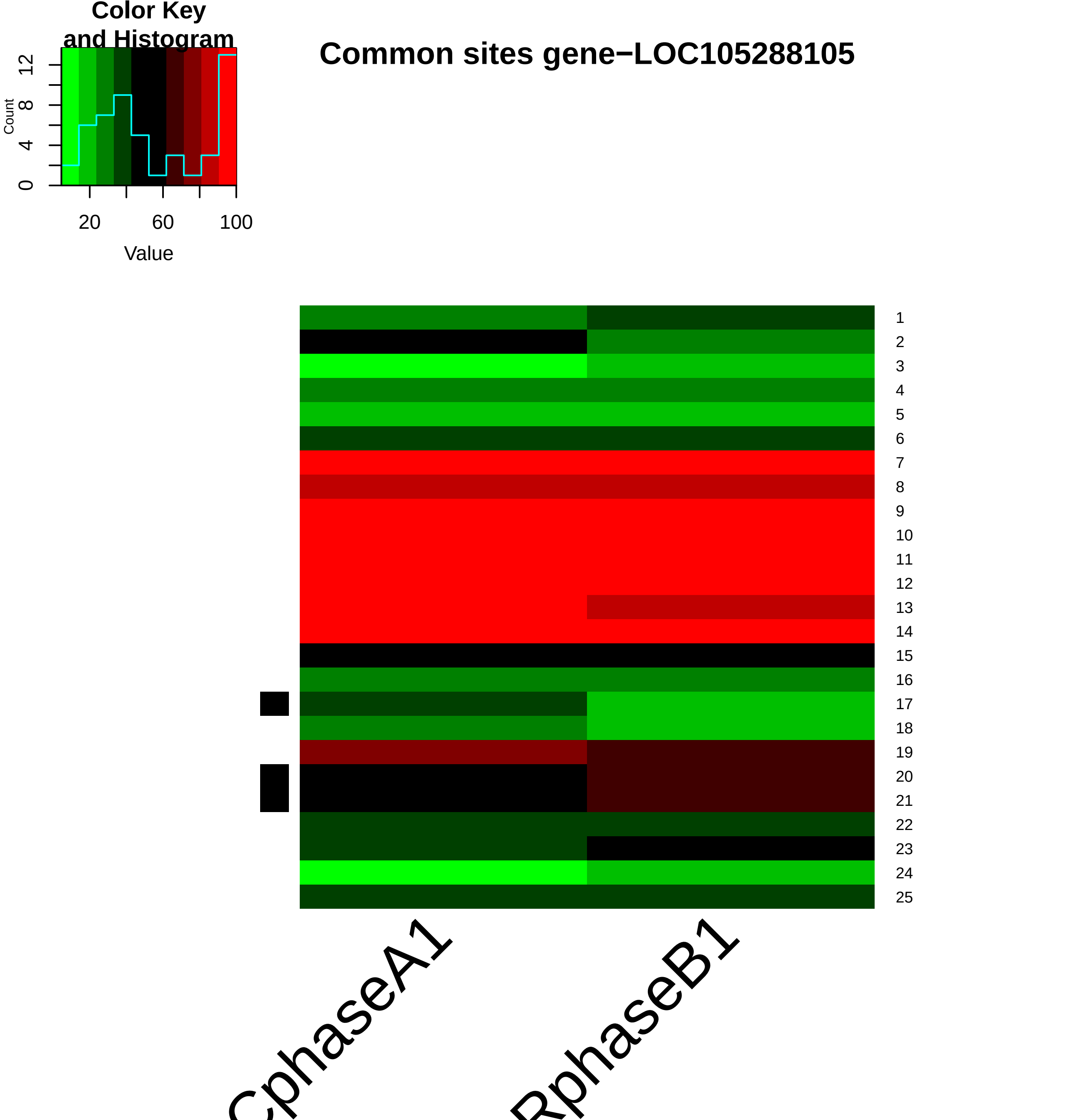
Color Key
and Histogram
Count
0
4
8
12
20
60
100
Value
Common sites gene−LOC105288105
CphaseA1
RphaseB1

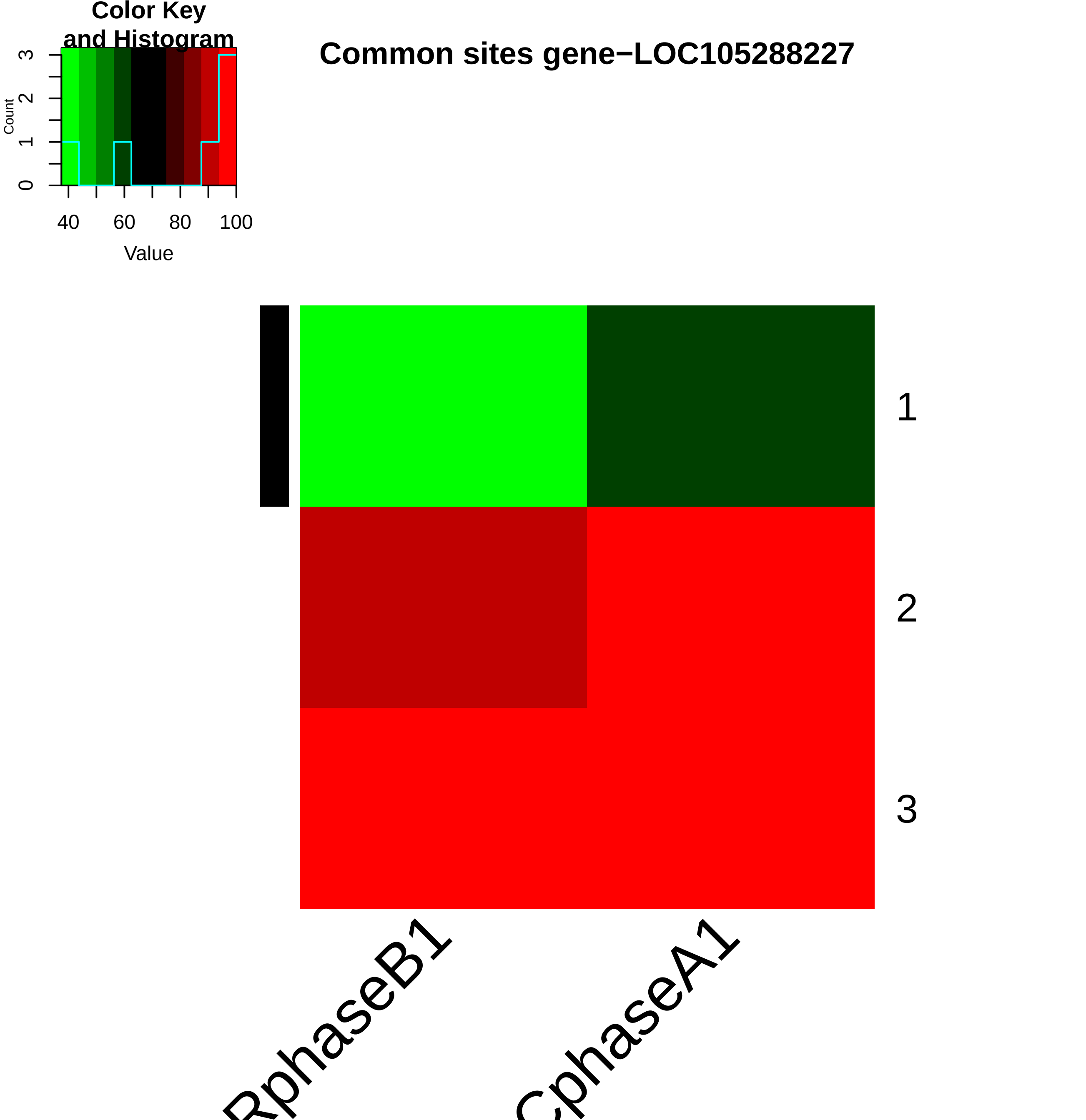

Color Key
and Histogram
Count
0
1
2
3
40
60
80
100
Value
Common sites gene−LOC105288227
1
2
3
RphaseB1
CphaseA1

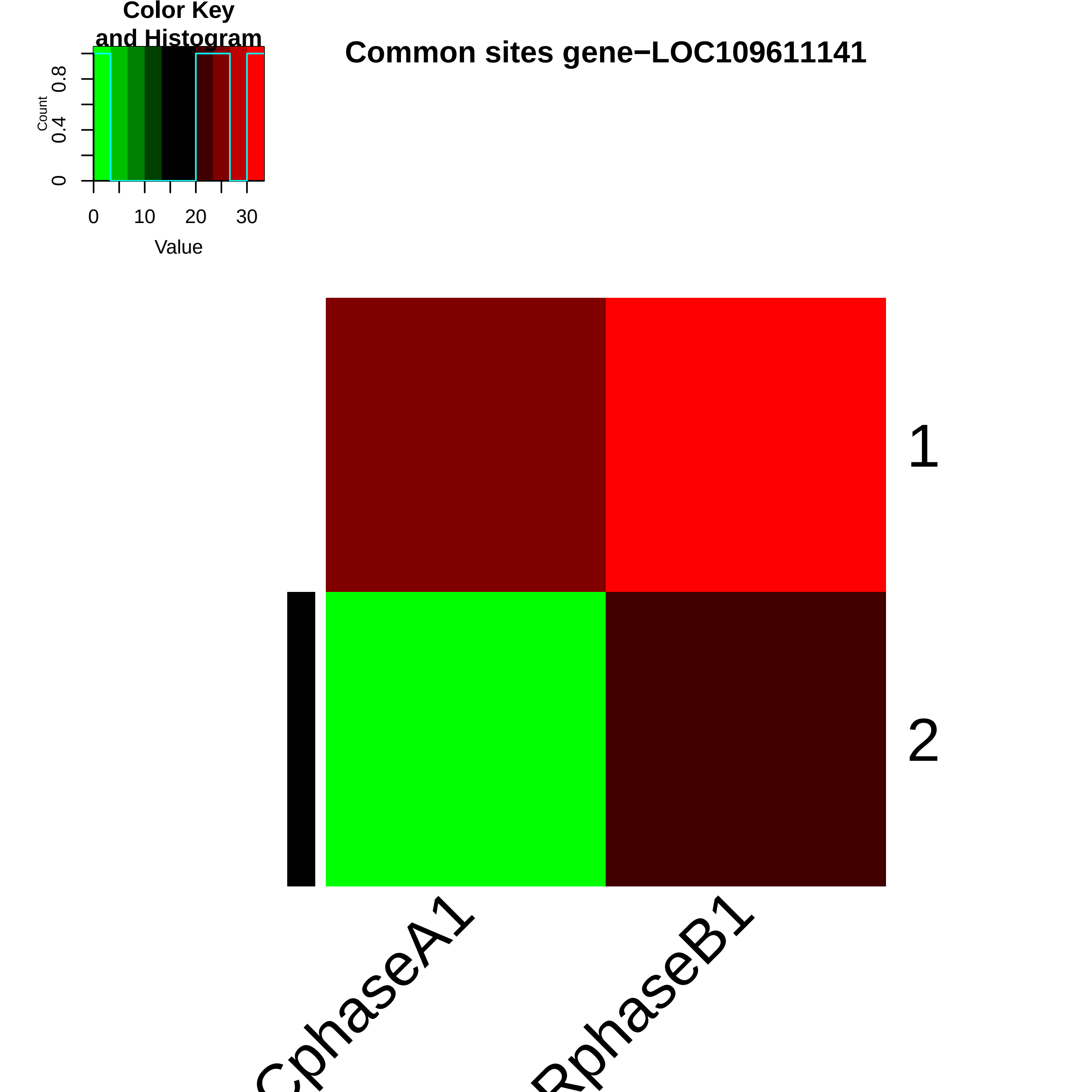
Color Key
and Histogram
Count
0
0.4
0.8
0
10
20
30
Value
Common sites gene−LOC109611141
1
2
CphaseA1
RphaseB1

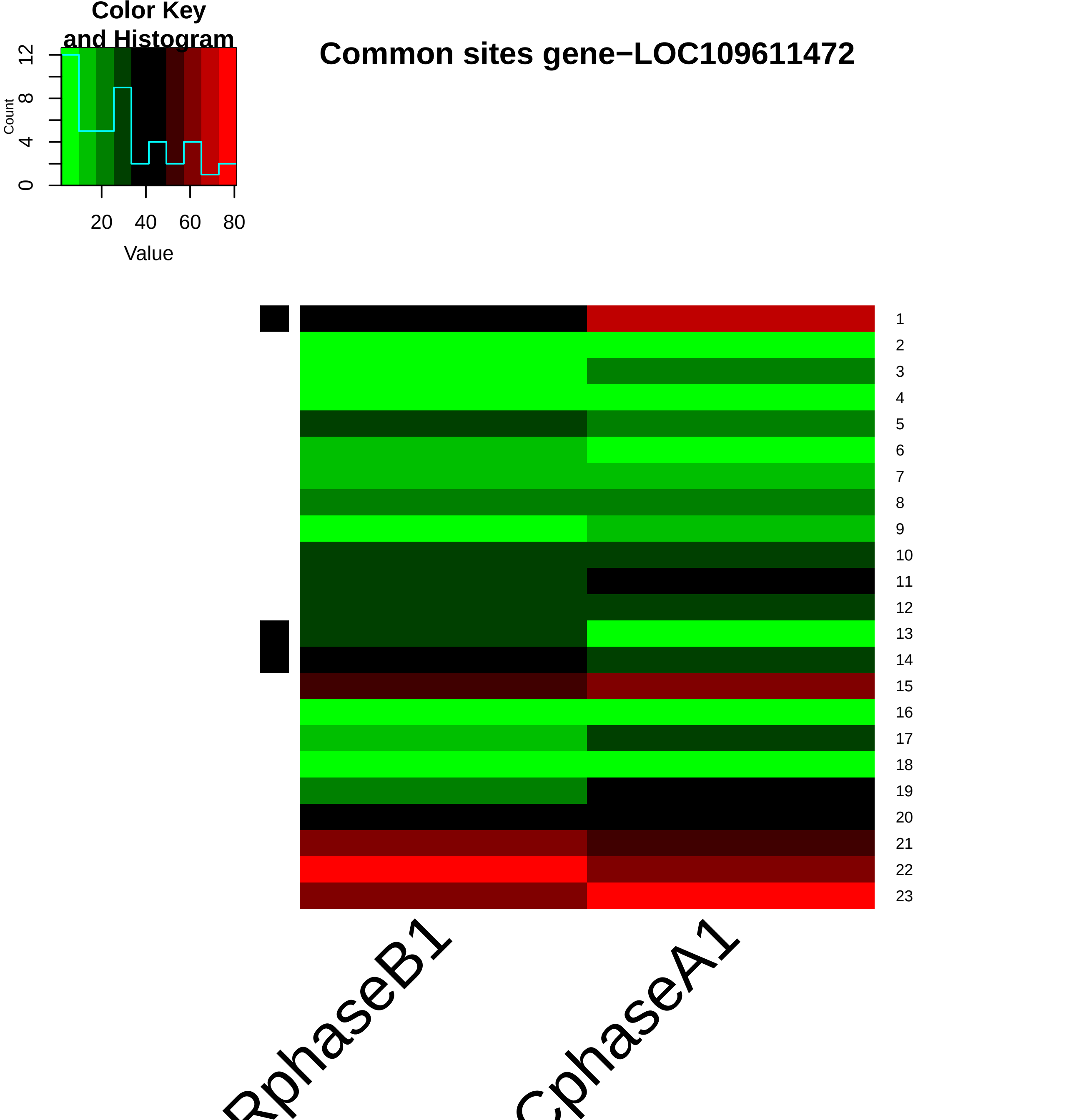
Color Key
and Histogram
Count
0
4
8
12
20
40
60
80
Value
Common sites gene−LOC109611472
RphaseB1
CphaseA1

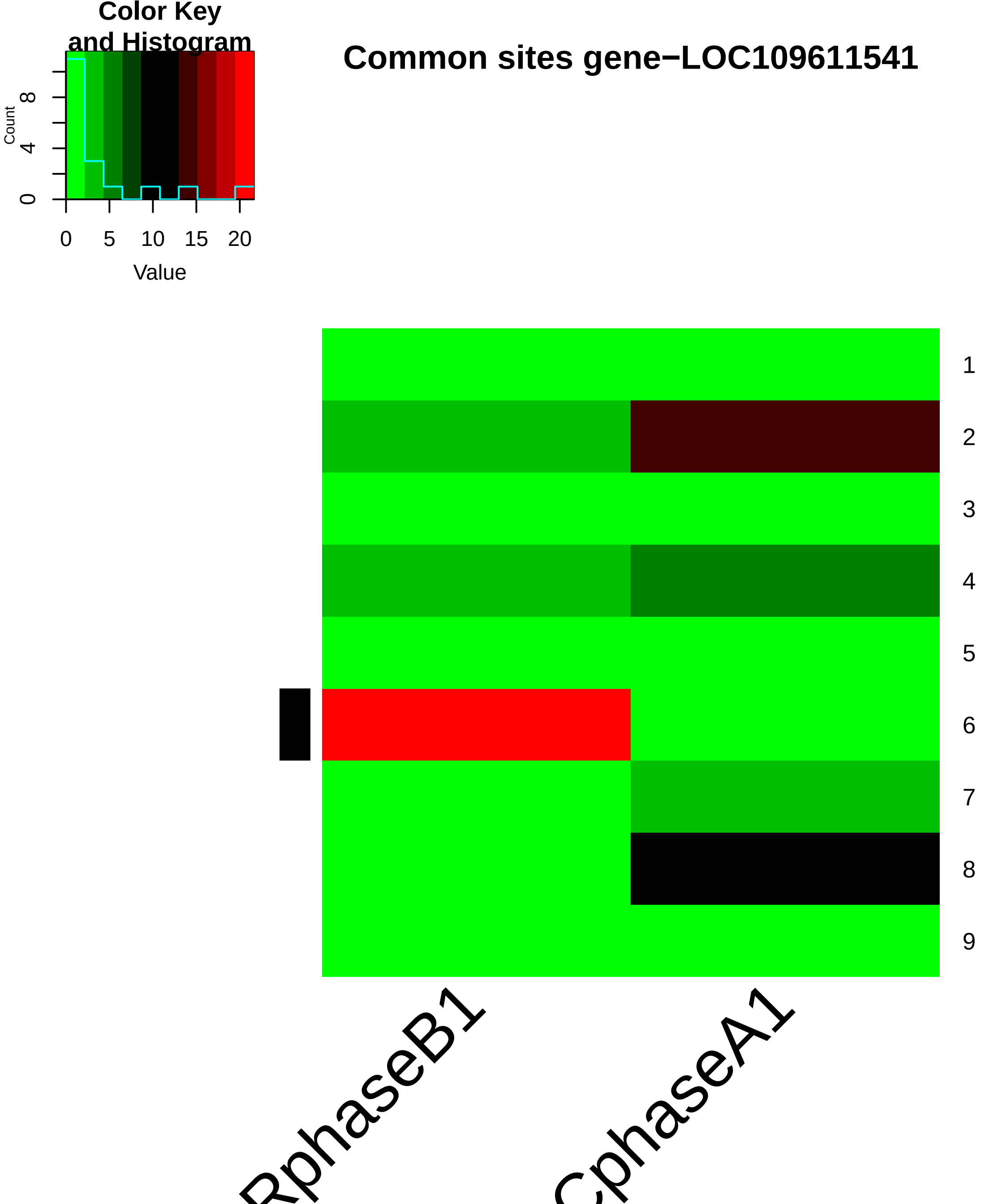

Color Key
and Histogram
Count
0
4
8
0
5
10
15
20
Value
Common sites gene−LOC109611541
1
2
3
4
5
6
7
8
9
RphaseB1
CphaseA1

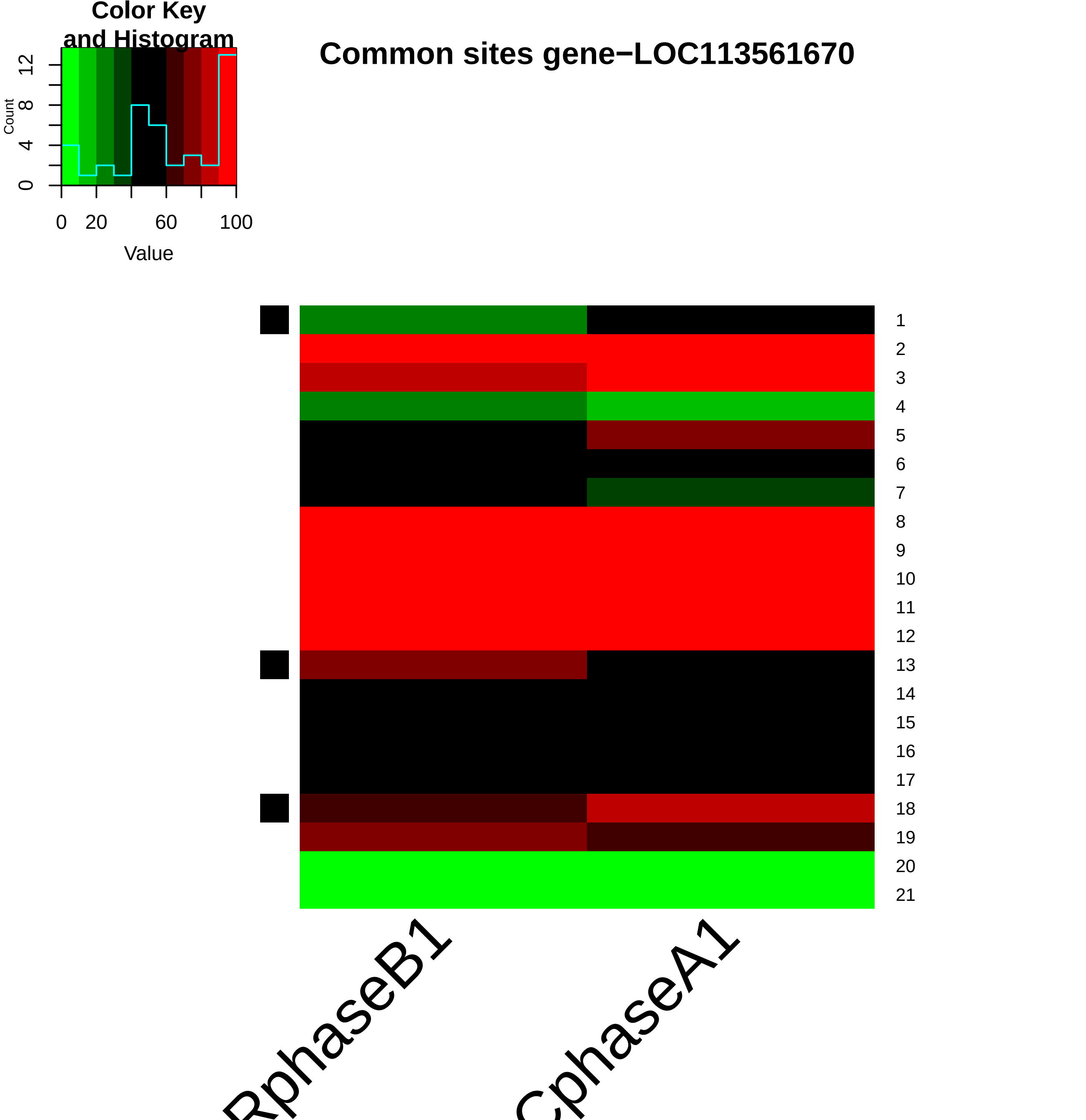
Color Key
and Histogram
Count
Value
0
4
8
12
0
20
60
100
Common sites gene−LOC113561670
RphaseB1
CphaseA1

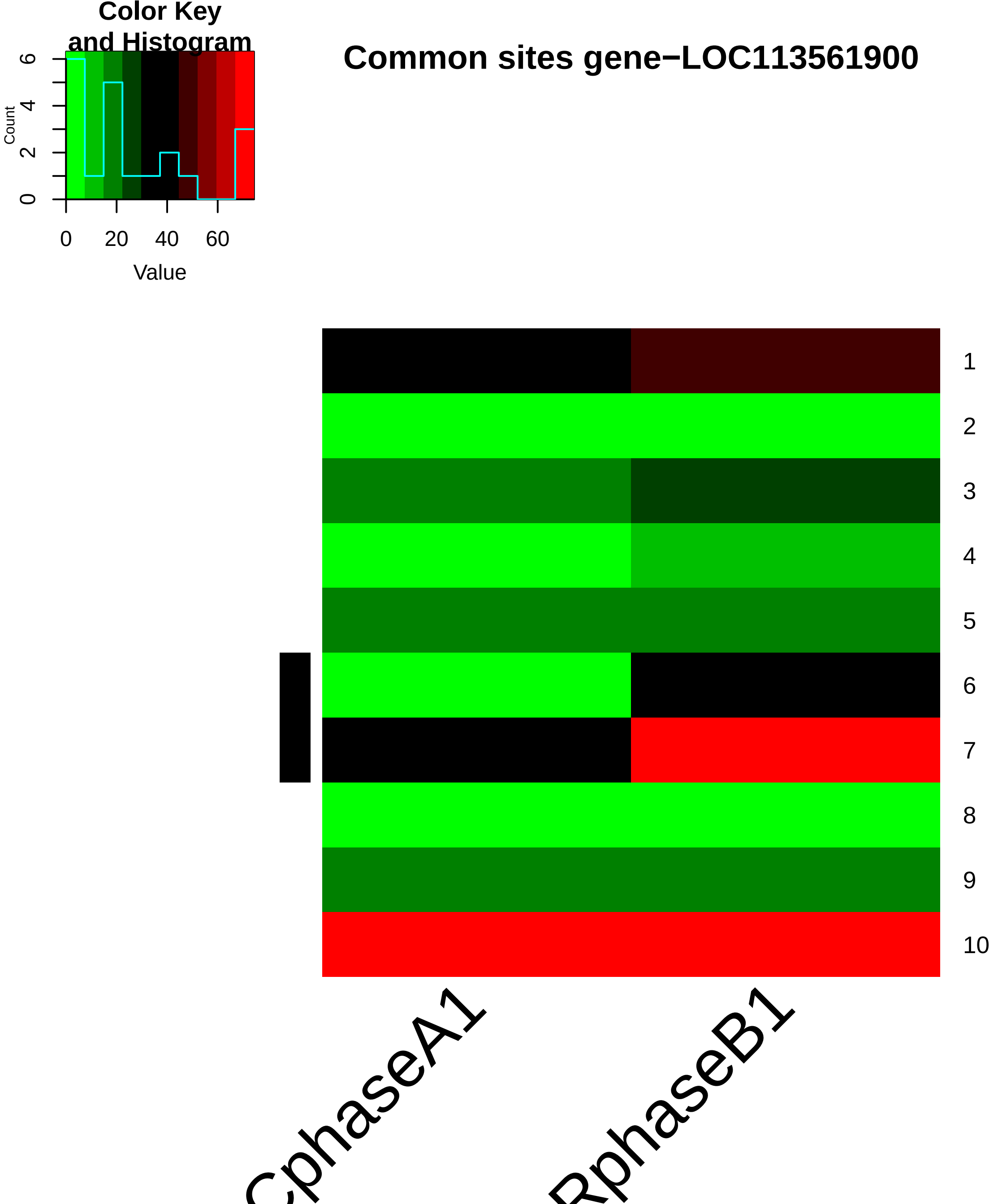

Color Key
and Histogram
Common sites gene−LOC113561900
Count
0
2
4
6
0
20
40
60
Value
1
2
3
4
5
6
7
8
9
10

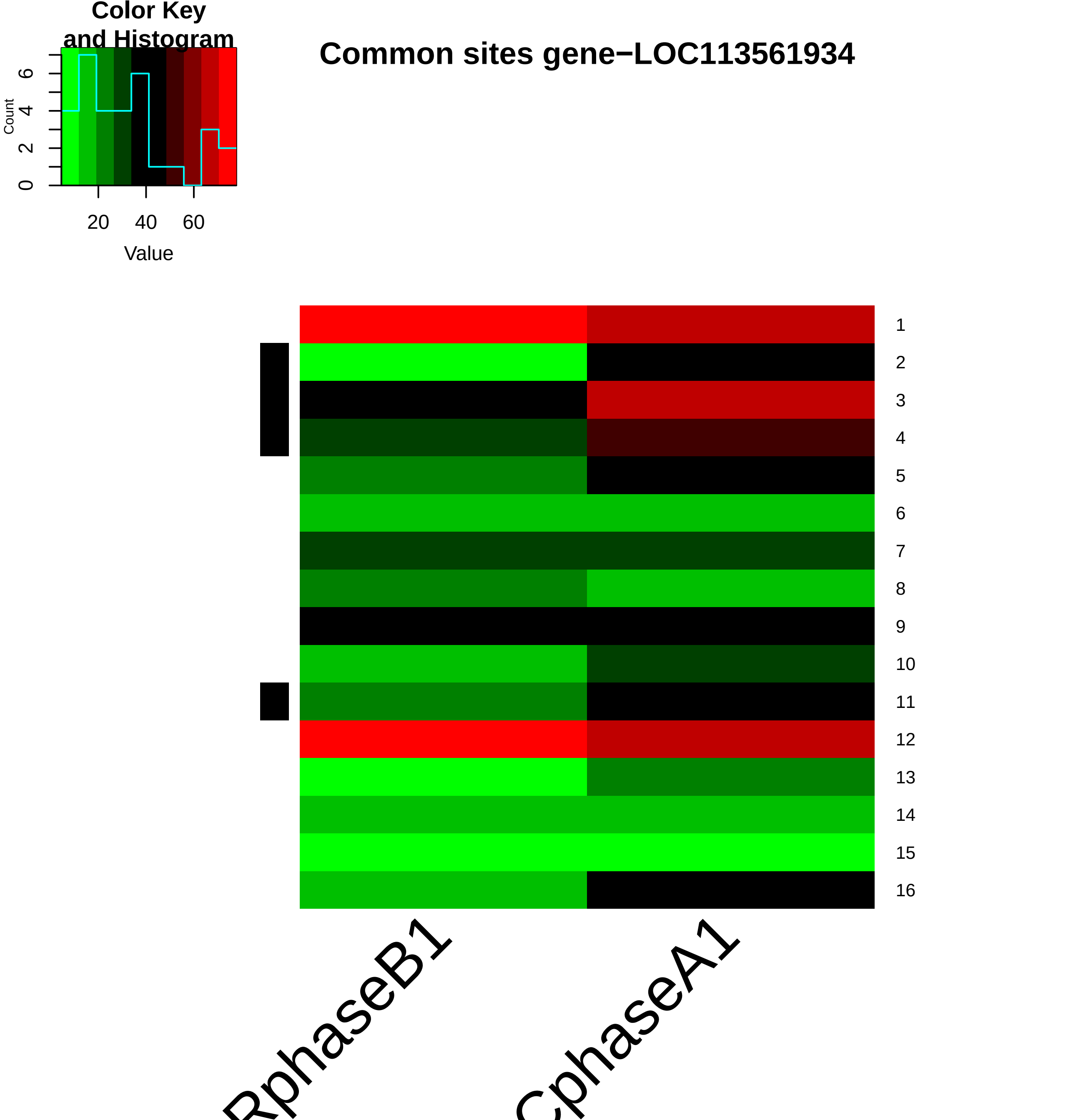

Color Key
and Histogram
Count
0
2
4
6
20
40
60
Value
Common sites gene−LOC113561934
RphaseB1
CphaseA1

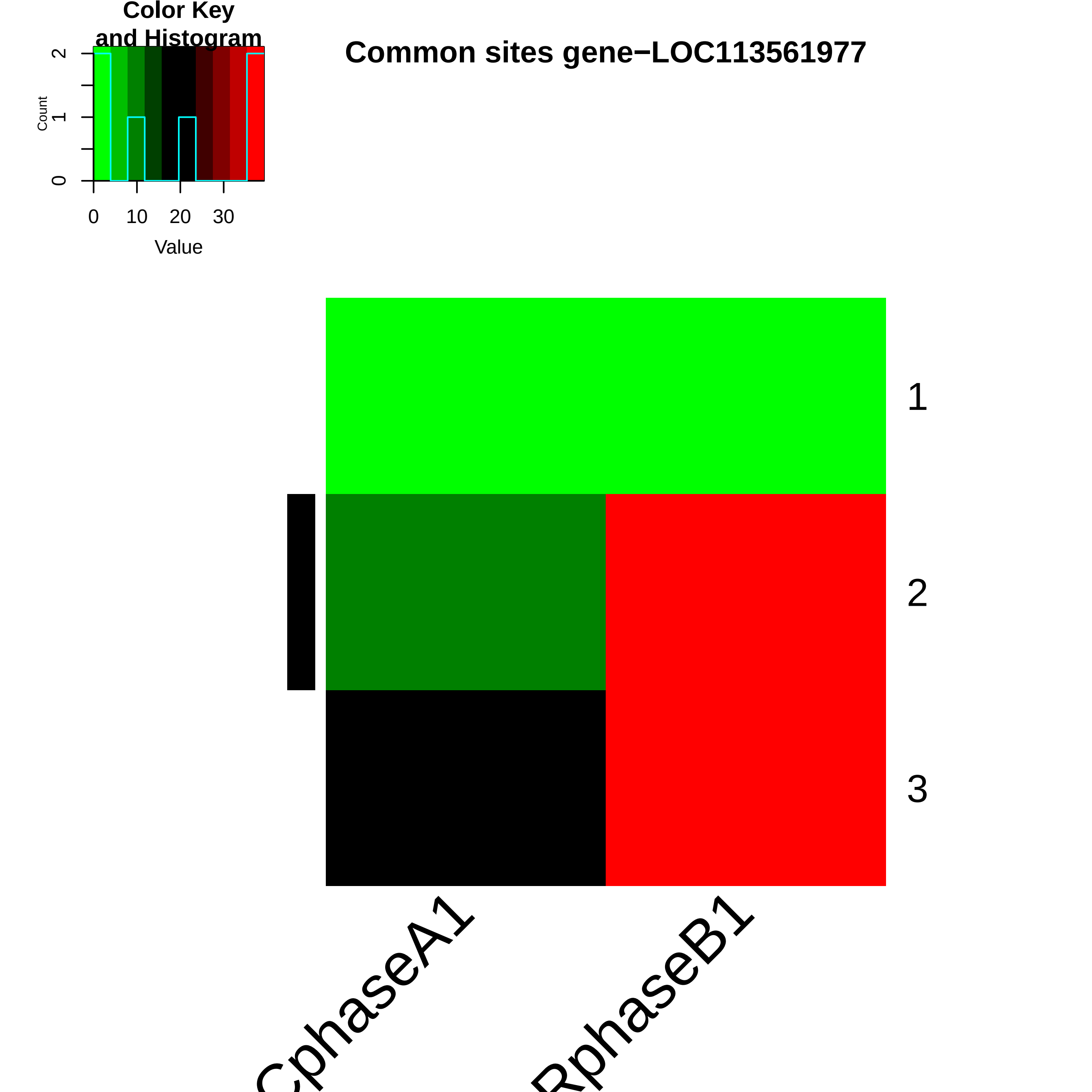
Color Key
and Histogram
Count
0
1
2
0
10
20
30
Value
Common sites gene−LOC113561977
1
2
3
CphaseA1
RphaseB1

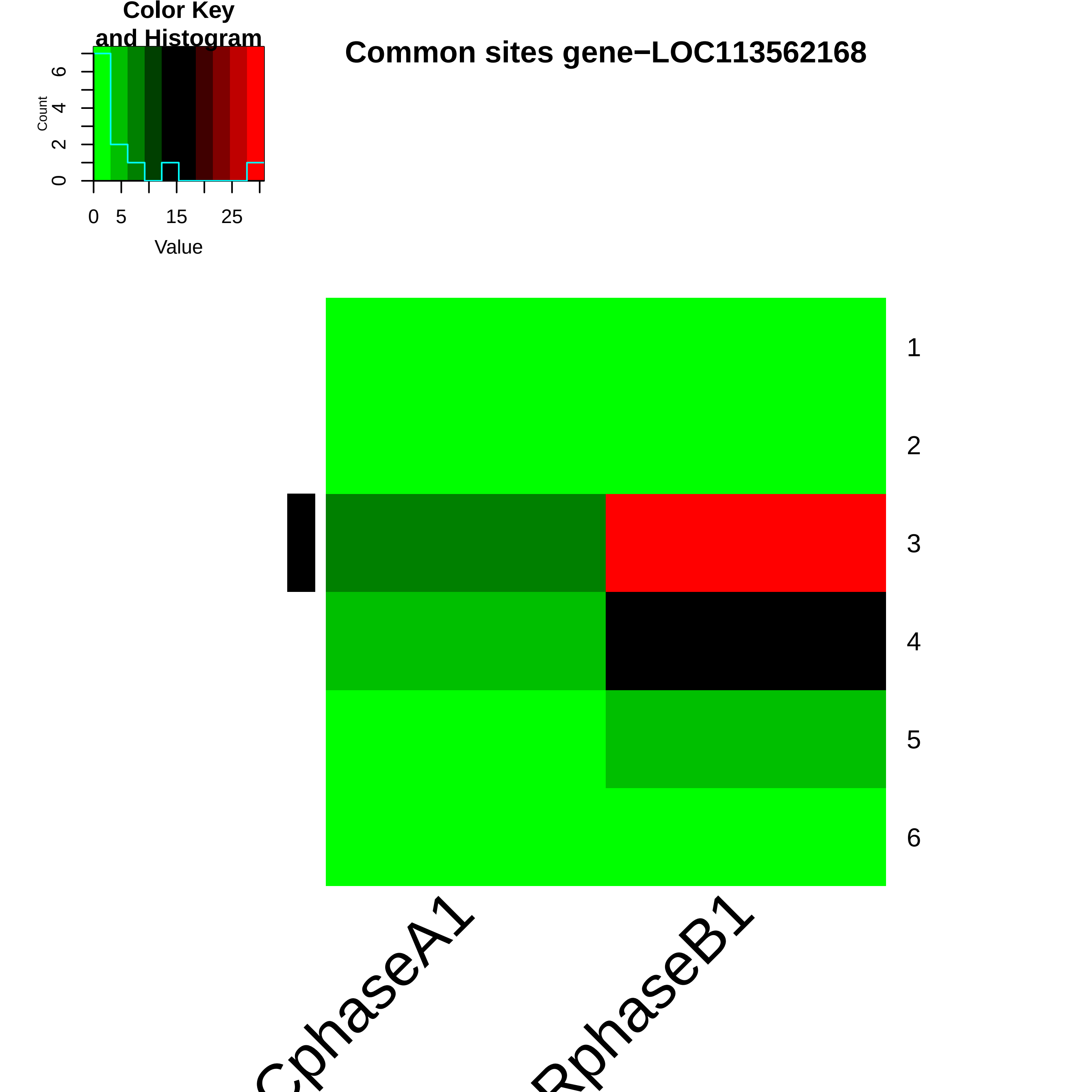

Color Key
and Histogram
Count
0
2
4
6
0
5
15
25
Value
Common sites gene−LOC113562168
1
2
3
4
5
6
CphaseA1
RphaseB1

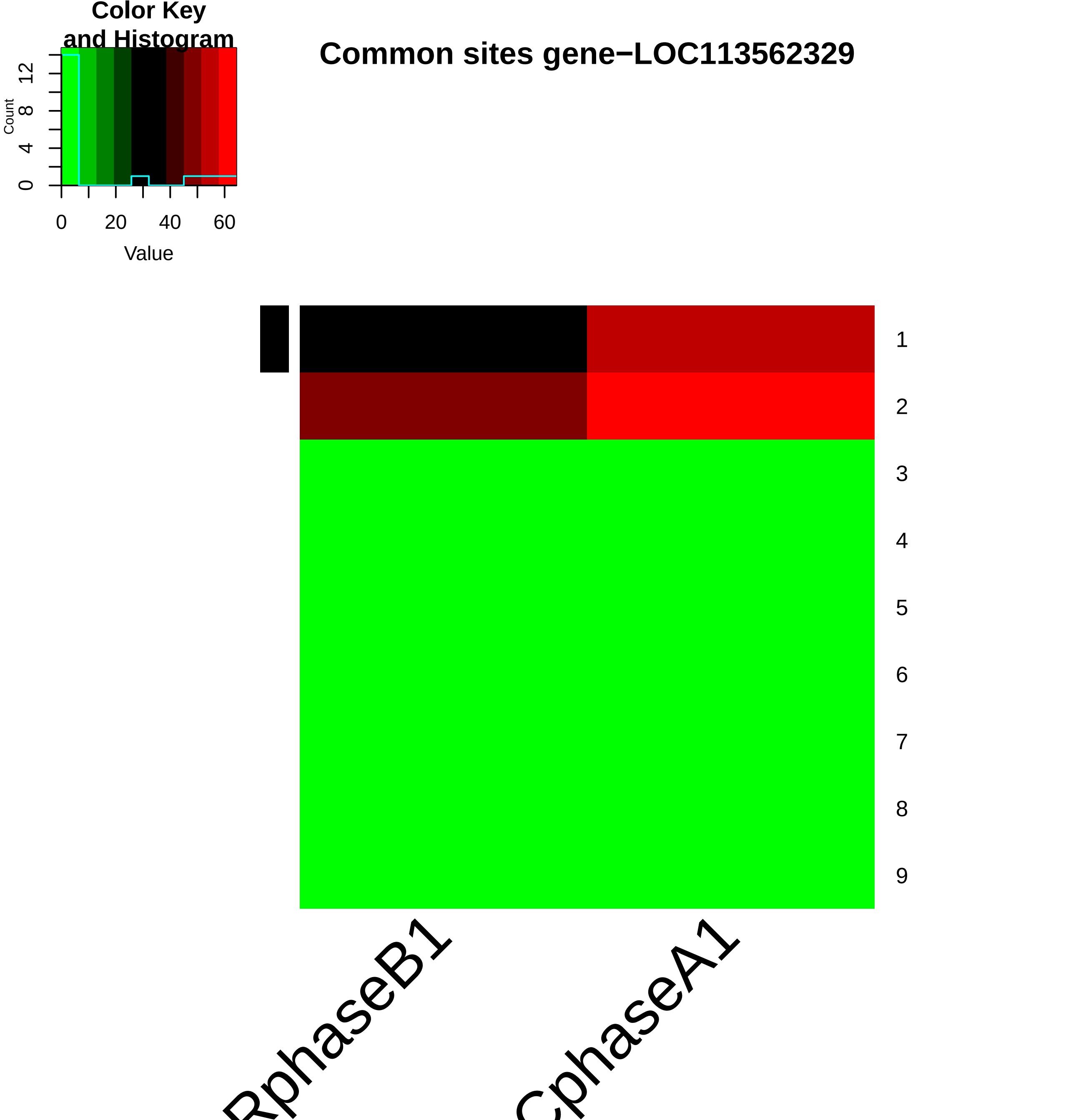
Color Key
and Histogram
Count
0
4
8
12
0
20
40
60
Value
Common sites gene−LOC113562329
1
2
3
4
5
6
7
8
9
RphaseB1
CphaseA1

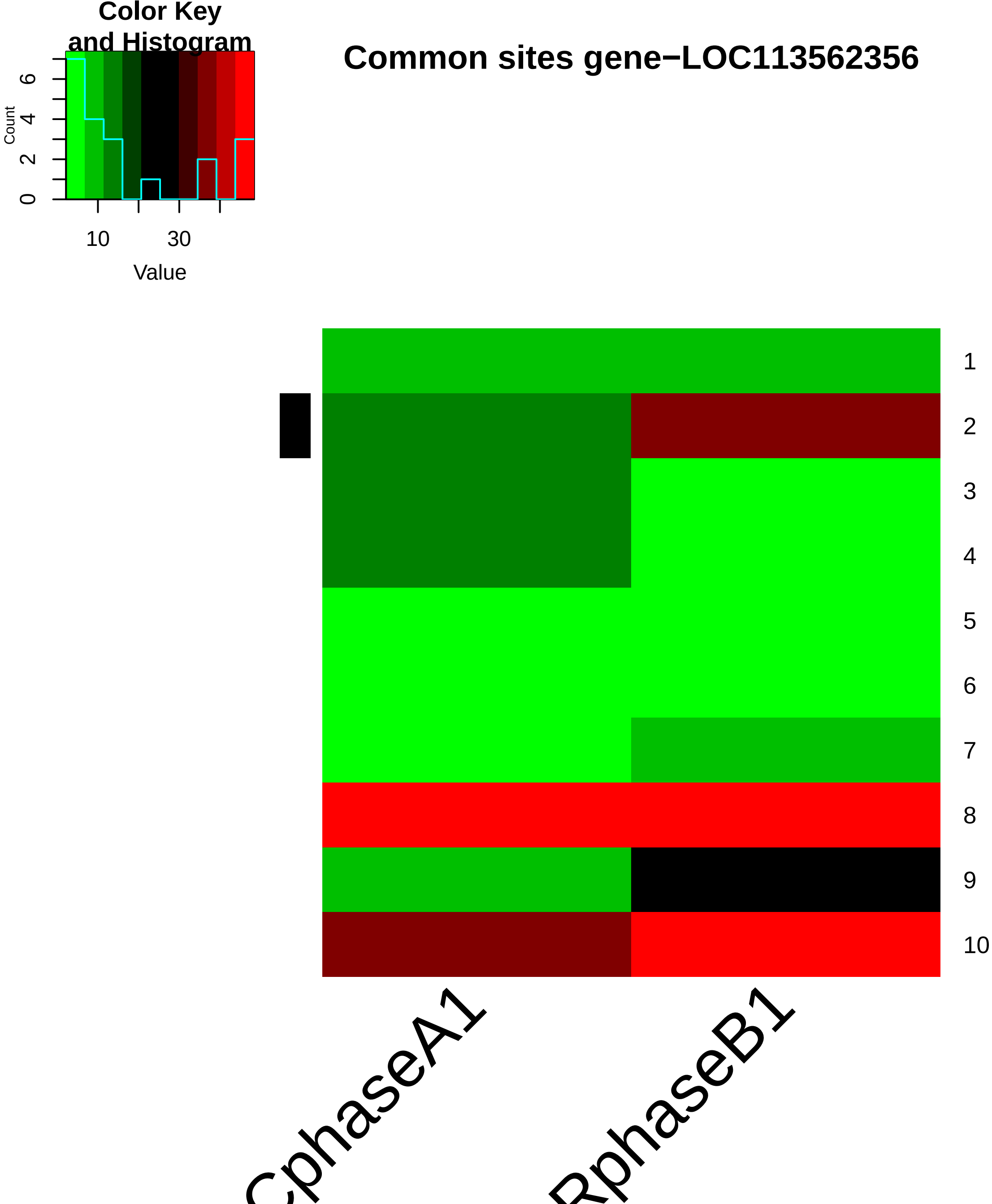

Color Key
and Histogram
Count
0
2
4
6
10
30
Value
Common sites gene−LOC113562356
1
2
3
4
5
6
7
8
9
10
CphaseA1
RphaseB1

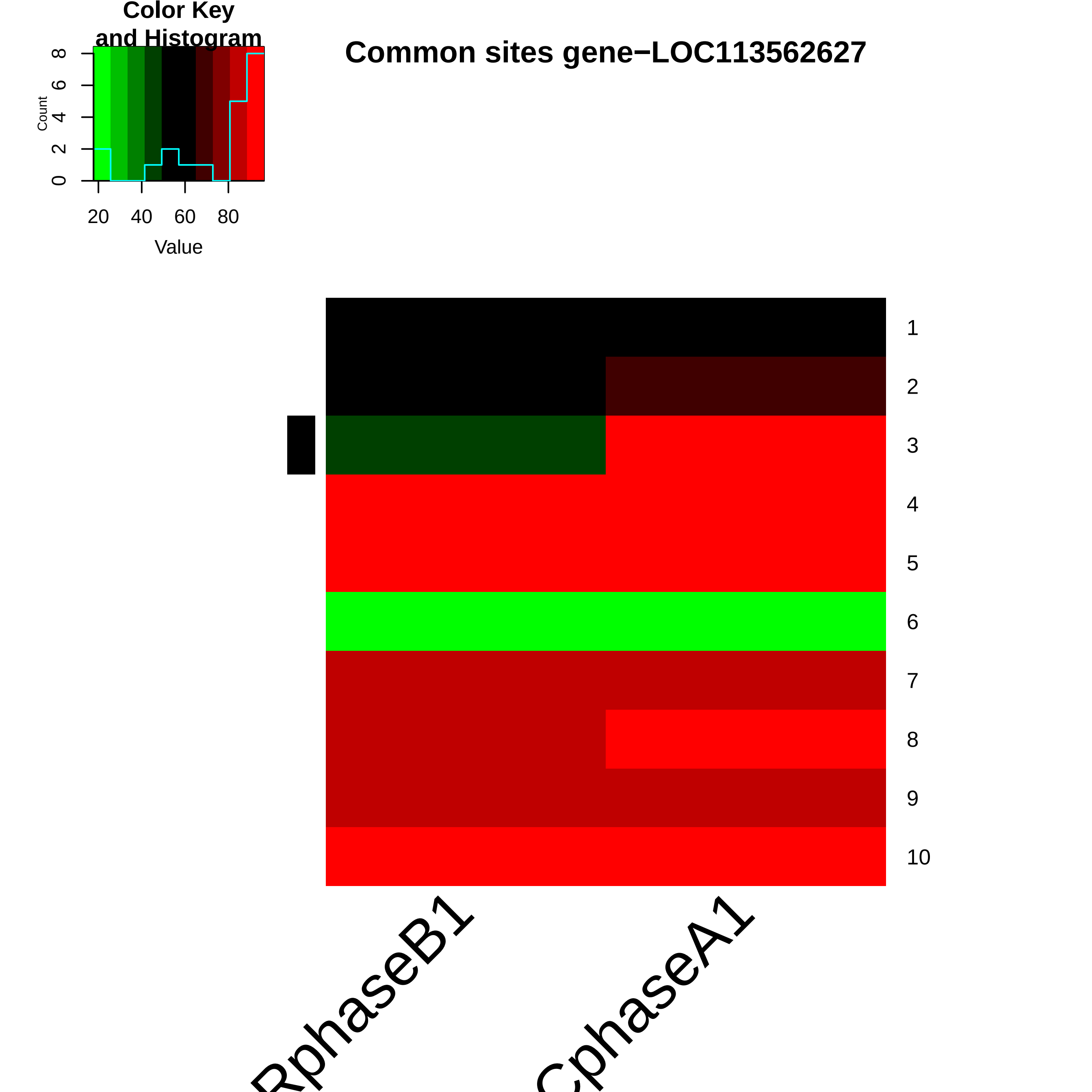

Color Key
and Histogram
Count
0
2
4
6
8
20
40
60
80
Value
Common sites gene−LOC113562627
1
2
3
4
5
6
7
8
9
10
RphaseB1
CphaseA1

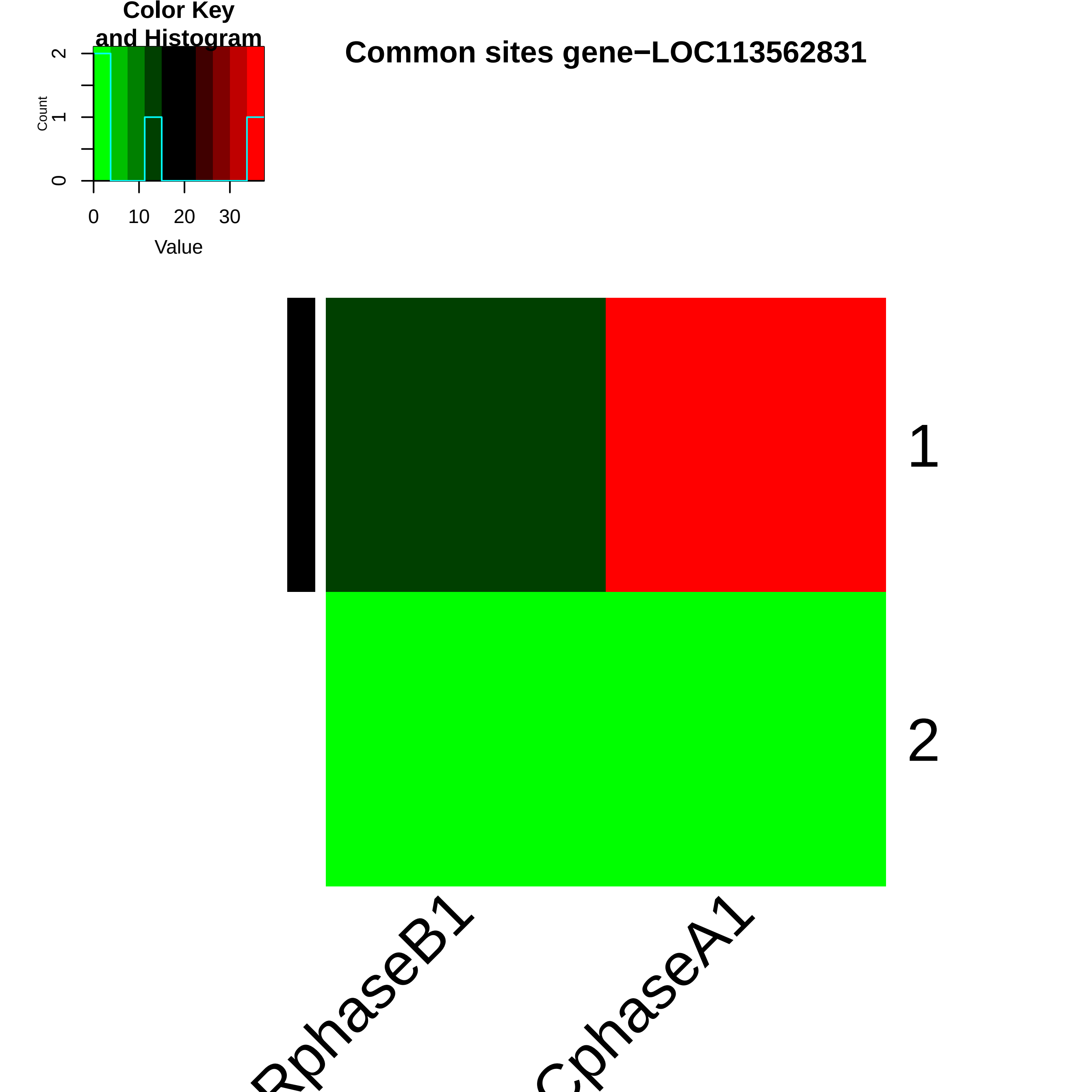
Color Key
and Histogram
Count
0
1
2
0
10
20
30
Value
Common sites gene−LOC113562831
1
2
RphaseB1
CphaseA1

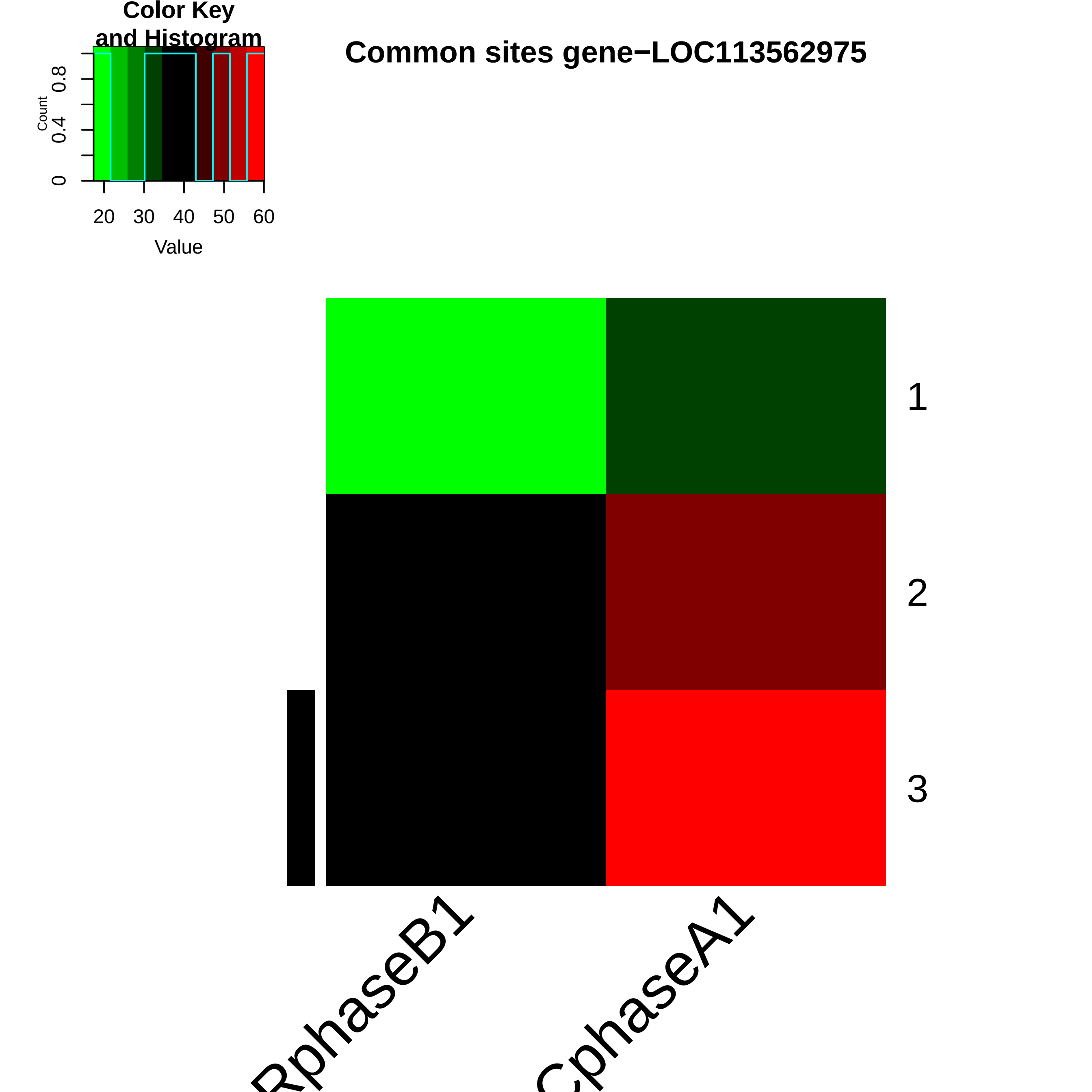

Color Key
and Histogram
Count
0
0.4
0.8
20
30
40
50
60
Value
Common sites gene−LOC113562975
1
2
3
RphaseB1
CphaseA1

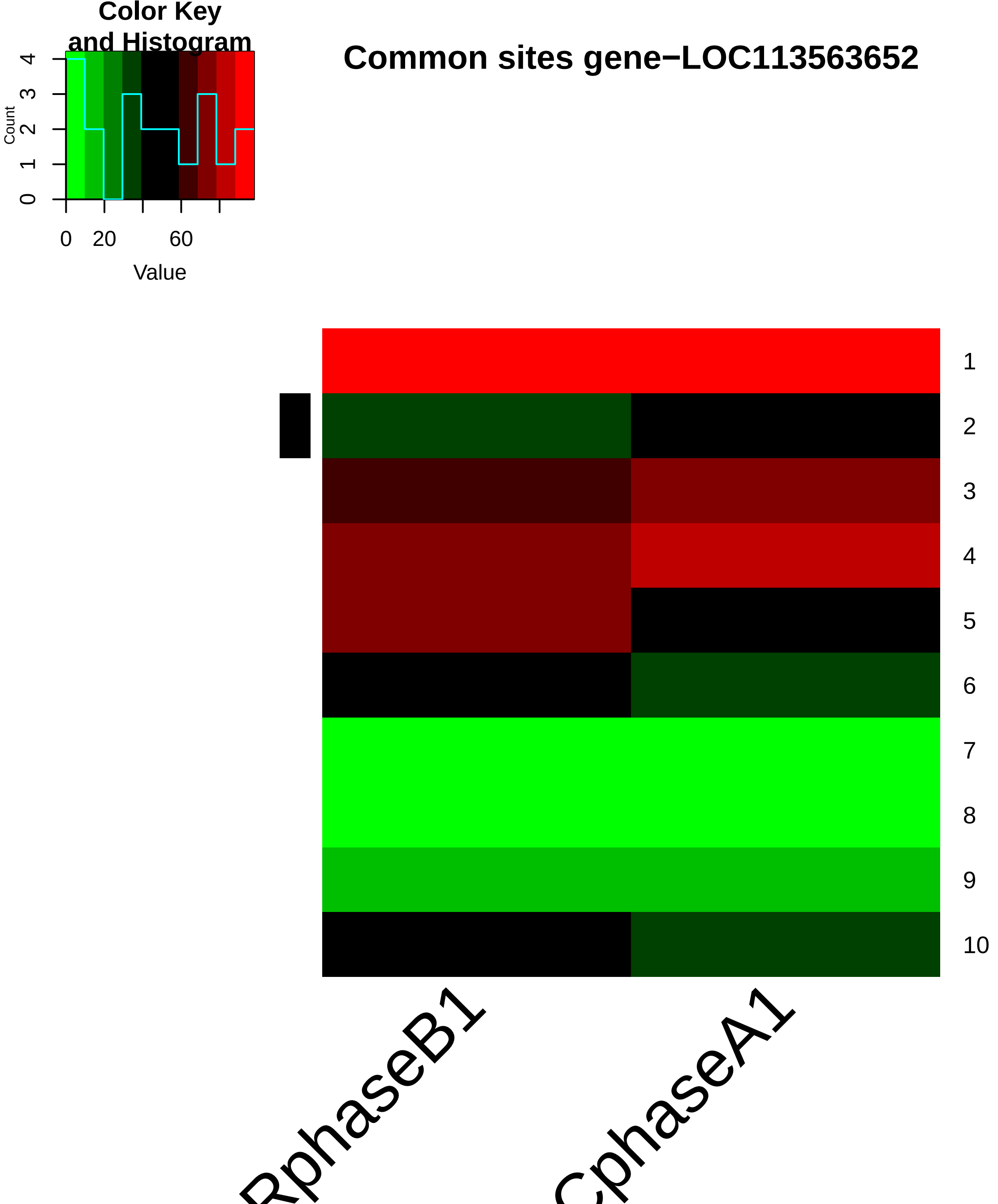

Color Key
and Histogram
Count
0
1
2
3
4
0
20
60
Value
Common sites gene−LOC113563652
1
2
3
4
5
6
7
8
9
10
RphaseB1
CphaseA1

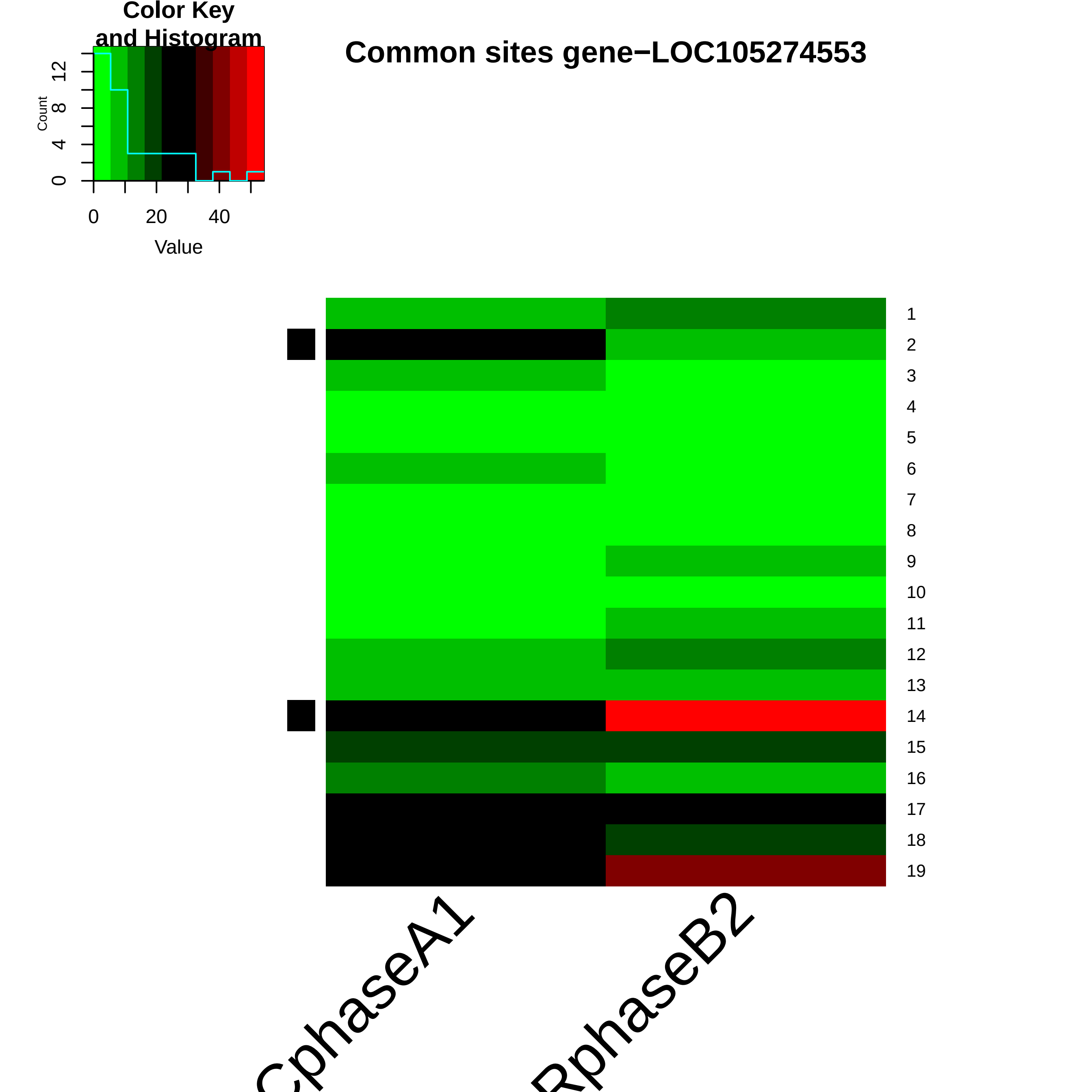

Color Key
and Histogram
Count
0
4
8
12
0
20
40
Value
Common sites gene−LOC105274553
CphaseA1
RphaseB2

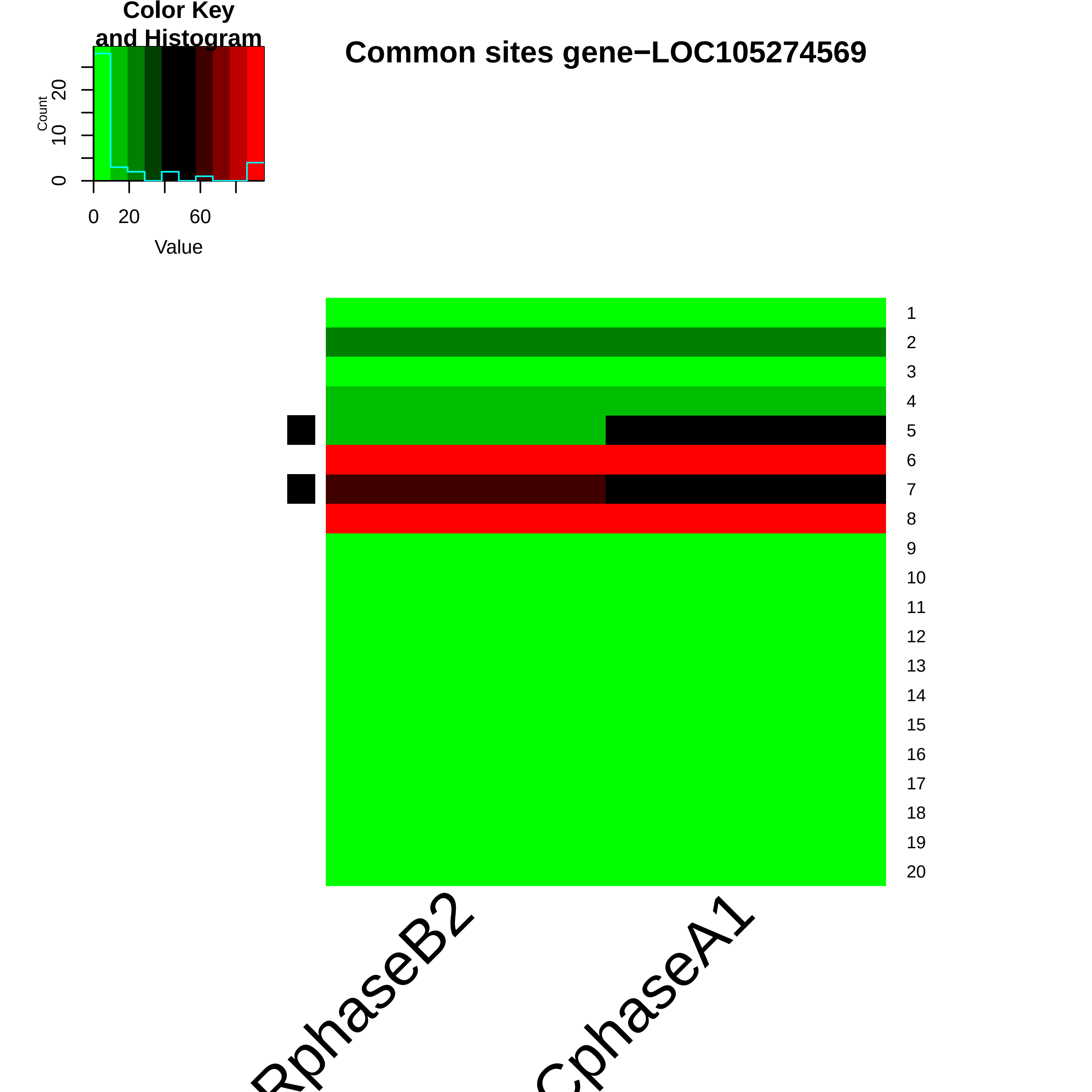

Color Key and Histogram
Count
0
10
20
0
20
60
Value
Common sites gene−LOC105274569
RphaseB2
CphaseA1

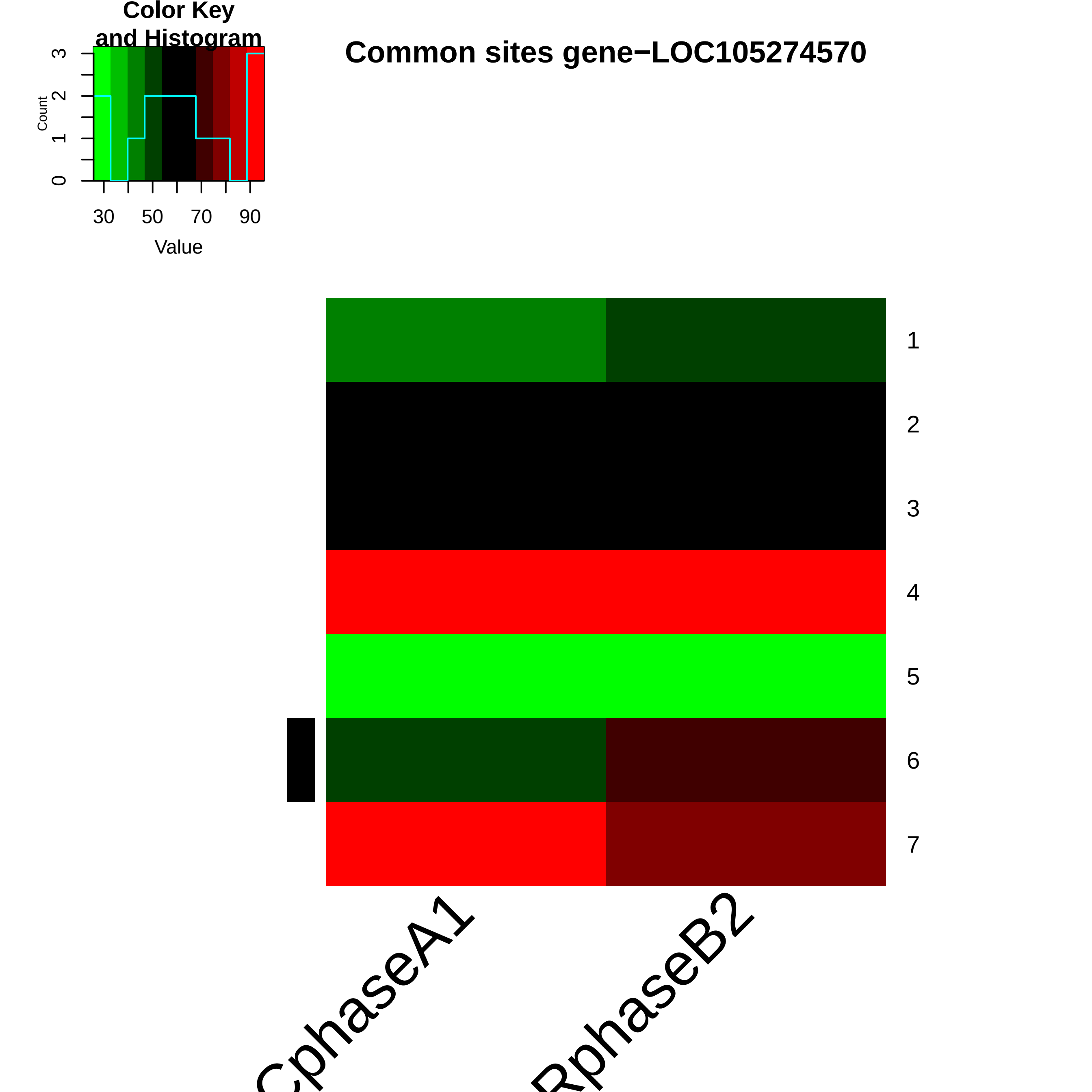

Color Key
and Histogram
Count
0
1
2
3
30
50
70
90
Value
Common sites gene−LOC105274570
1
2
3
4
5
6
7
CphaseA1
RphaseB2

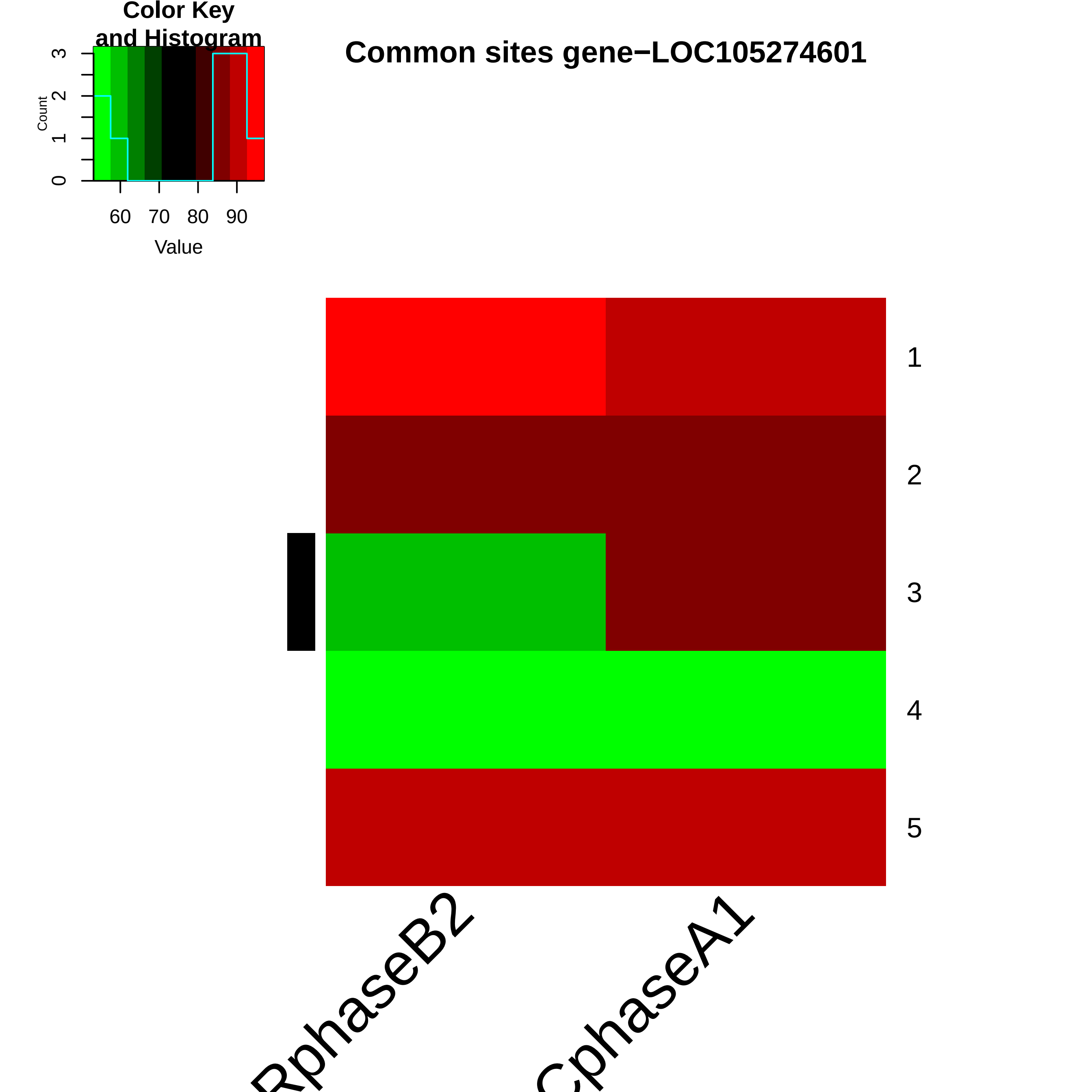
Color Key
and Histogram
Count
0
1
2
3
60
70
80
90
Value
Common sites gene−LOC105274601
1
2
3
4
5
RphaseB2
CphaseA1

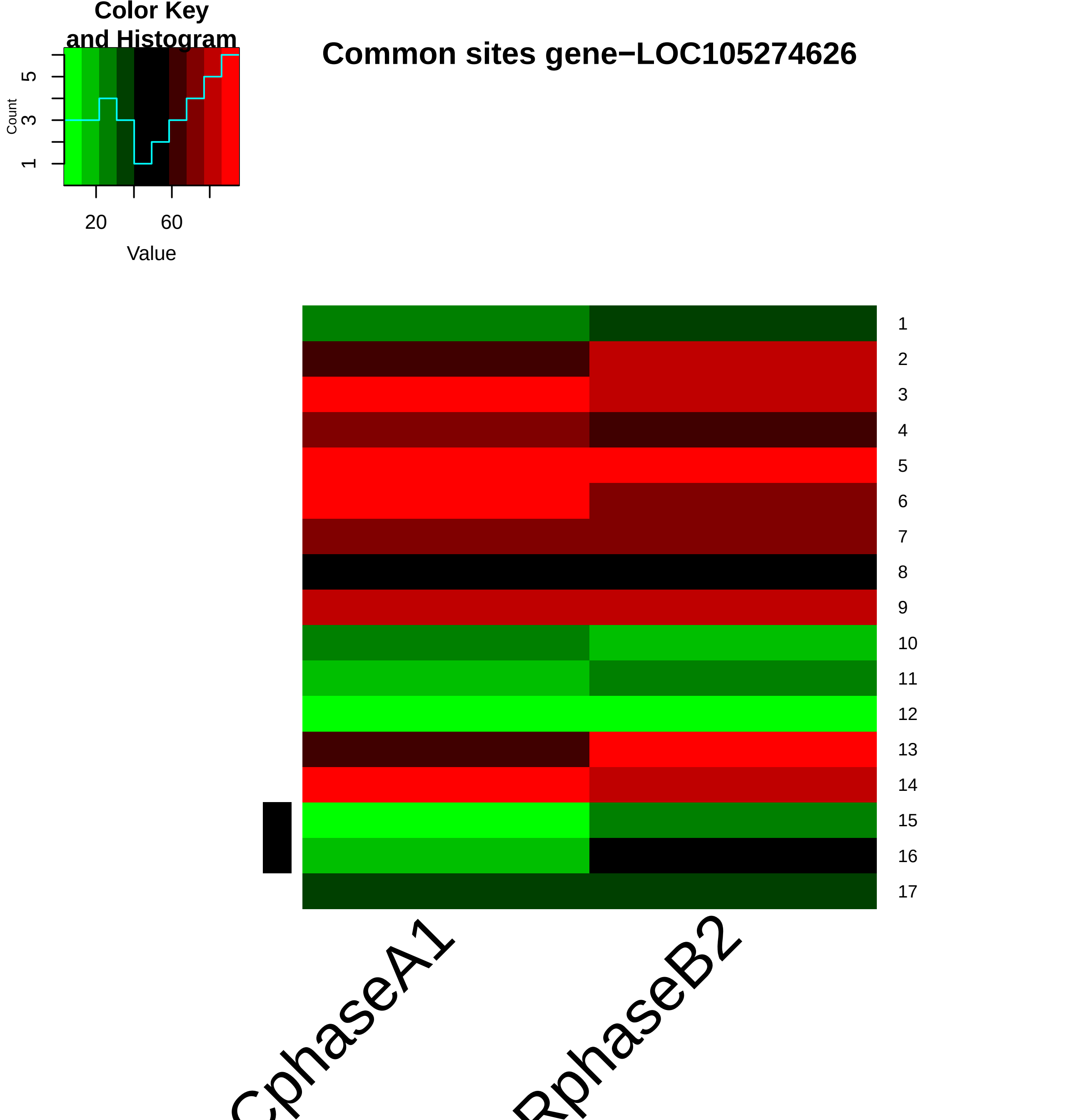

Color Key
and Histogram
Count
1
3
5
20
60
Value
Common sites gene−LOC105274626
CphaseA1
RphaseB2

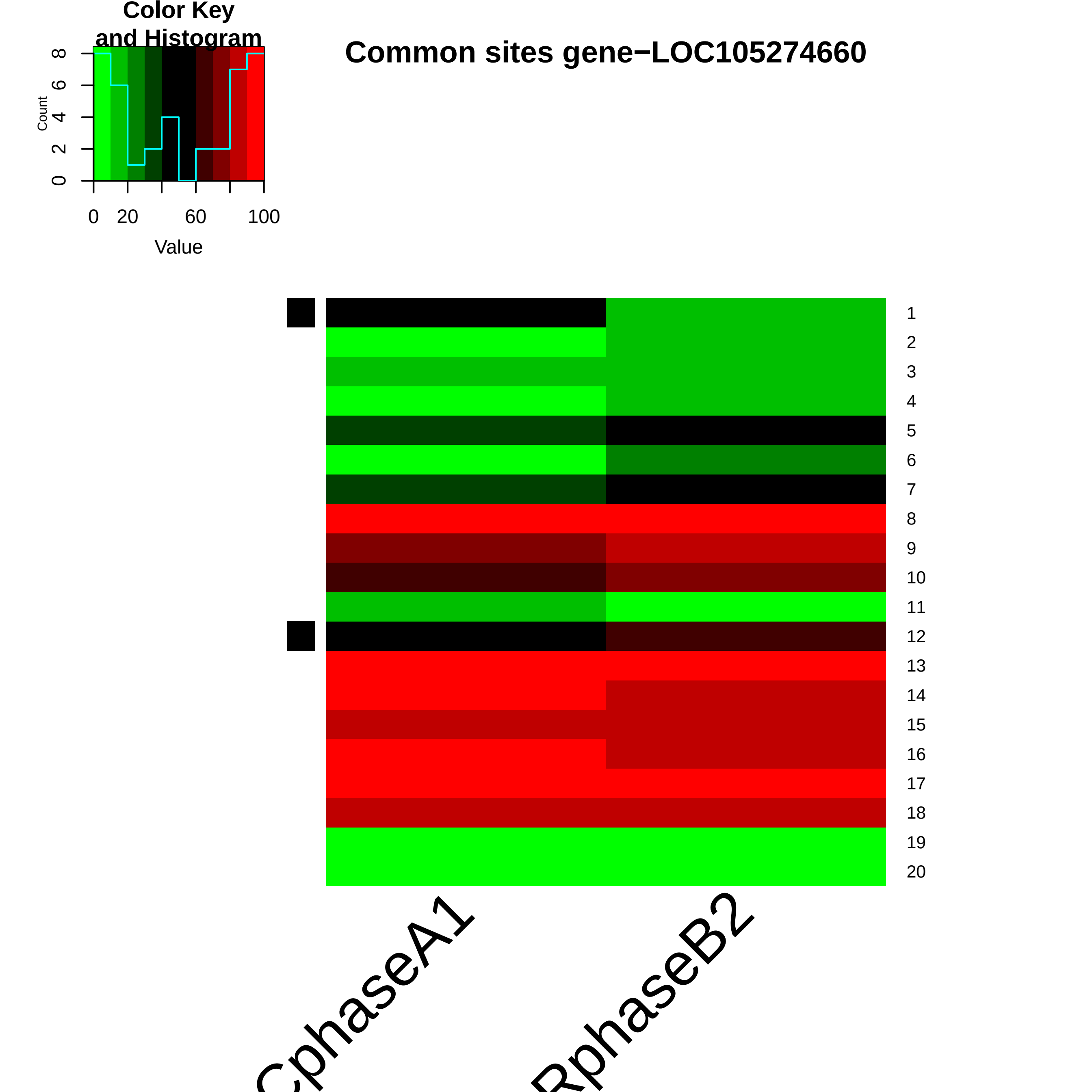

Color Key
and Histogram
Count
0
2
4
6
8
0
20
60
100
Value
Common sites gene−LOC105274660
CphaseA1
RphaseB2

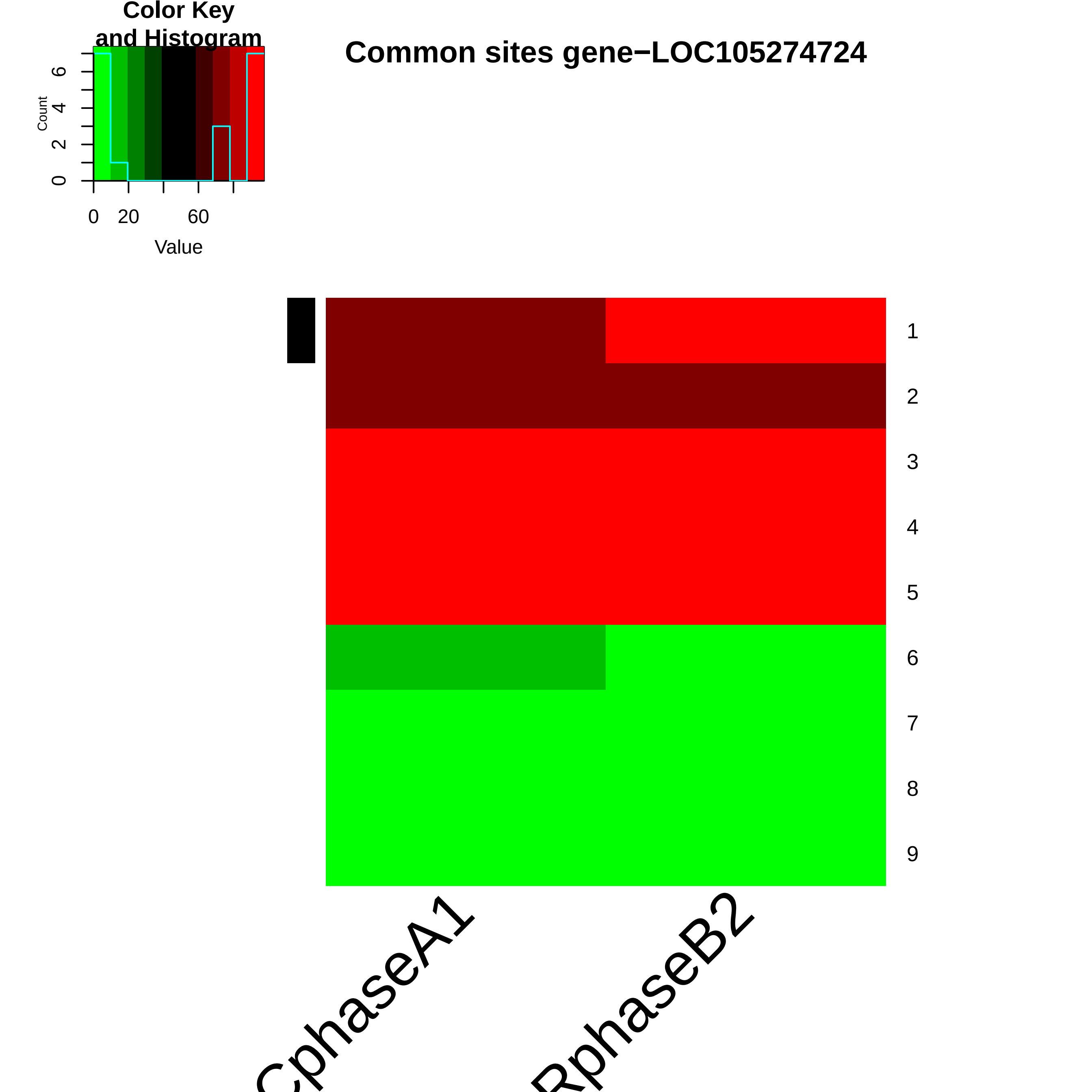
Color Key
and Histogram
Count
0
2
4
6
0
20
60
Value
Common sites gene−LOC105274724
1
2
3
4
5
6
7
8
9

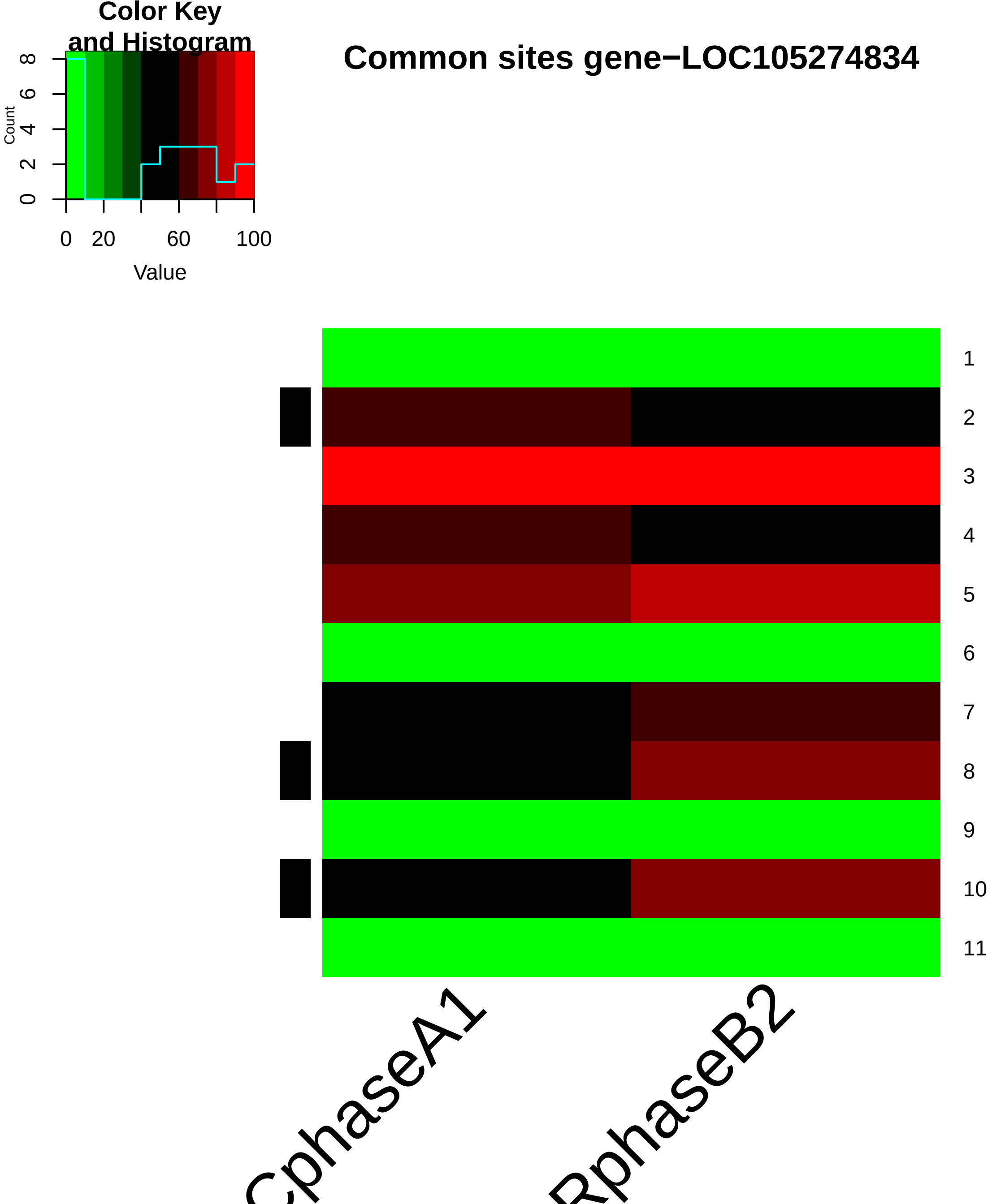

Color Key
and Histogram
Count
0
2
4
6
8
0
20
60
100
Value
Common sites gene−LOC105274834
1
2
3
4
5
6
7
8
9
10
11
CphaseA1
RphaseB2

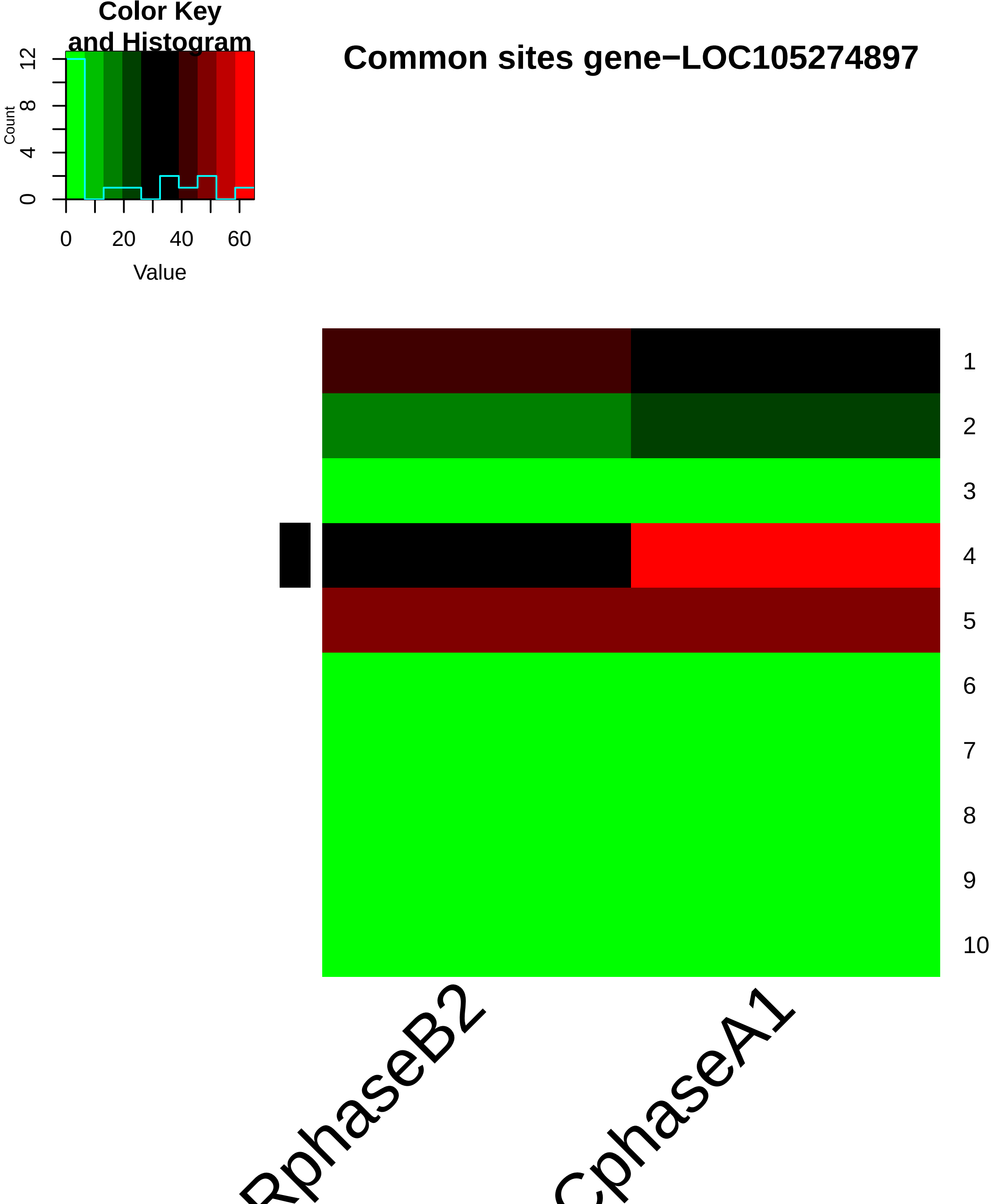
Color Key
and Histogram
Count
0
4
8
12
0
20
40
60
Value
Common sites gene−LOC105274897
1
2
3
4
5
6
7
8
9
10
RphaseB2
CphaseA1

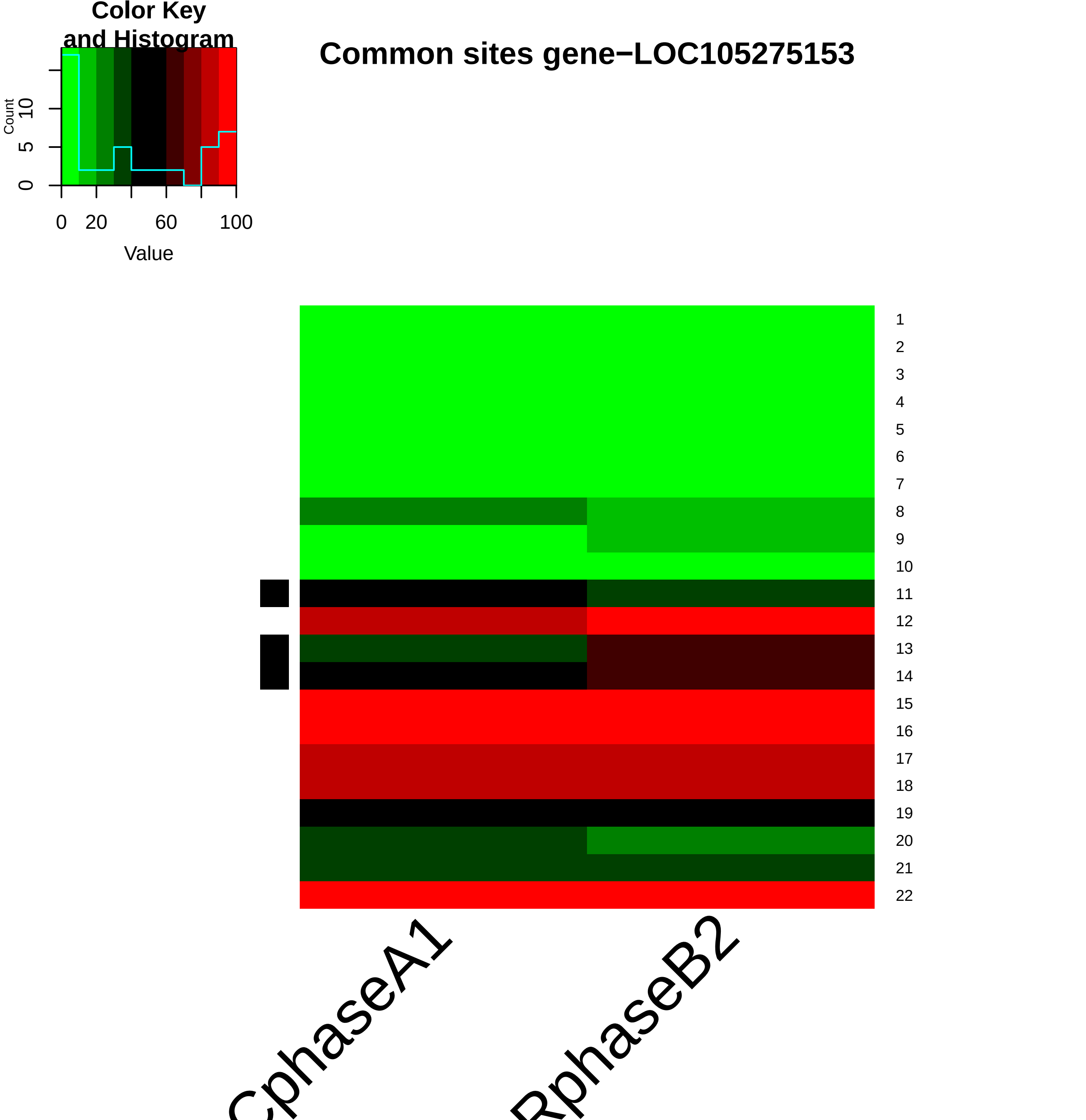
Color Key
and Histogram
Count
0
5
10
0
20
60
100
Value
Common sites gene−LOC105275153
CphaseA1
RphaseB2

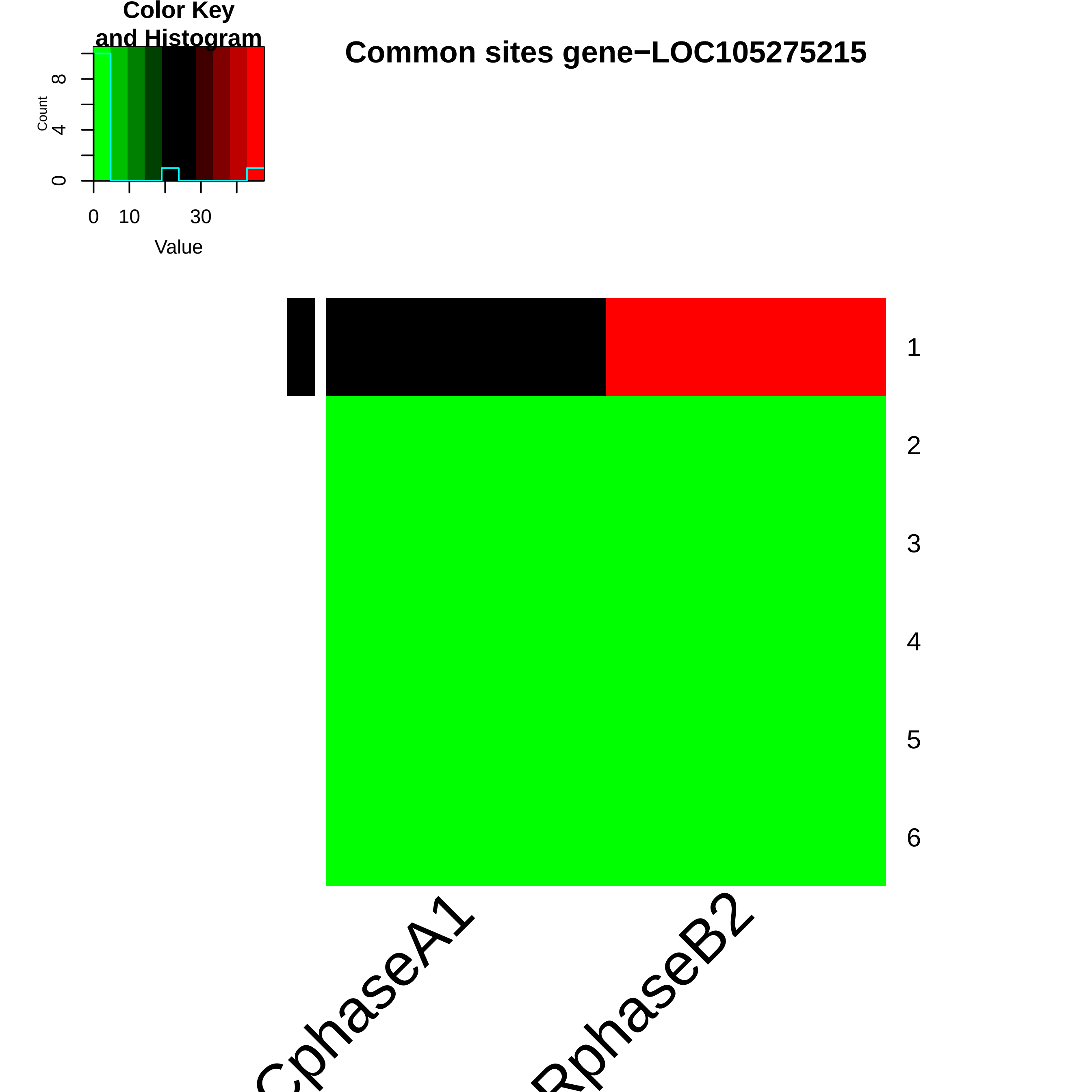

Color Key
and Histogram
Count
0
4
8
0
10
30
Value
Common sites gene−LOC105275215
1
2
3
4
5
6
CphaseA1
RphaseB2

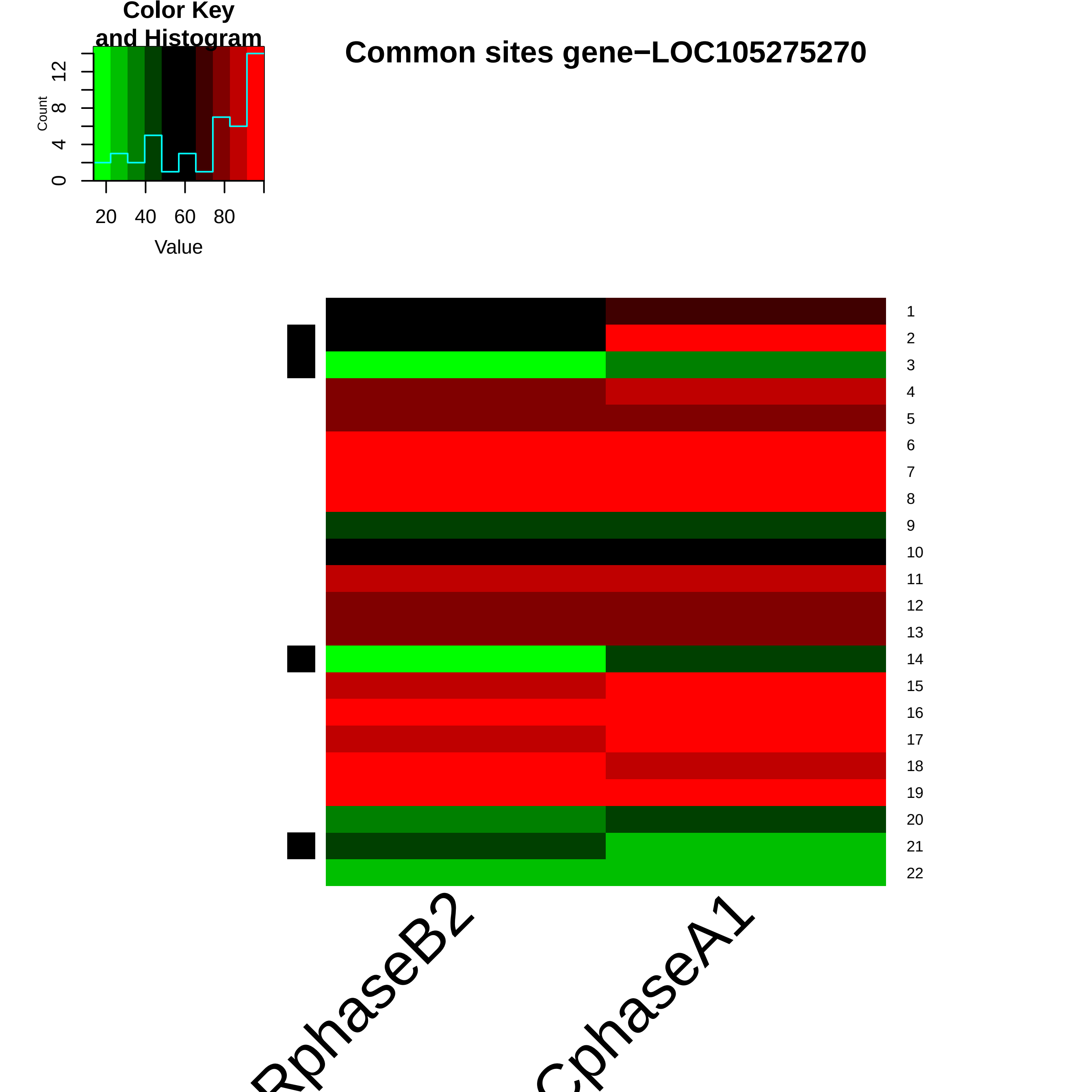
Color Key
and Histogram
Count
0
4
8
12
20
40
60
80
Value
Common sites gene−LOC105275270
RphaseB2
CphaseA1

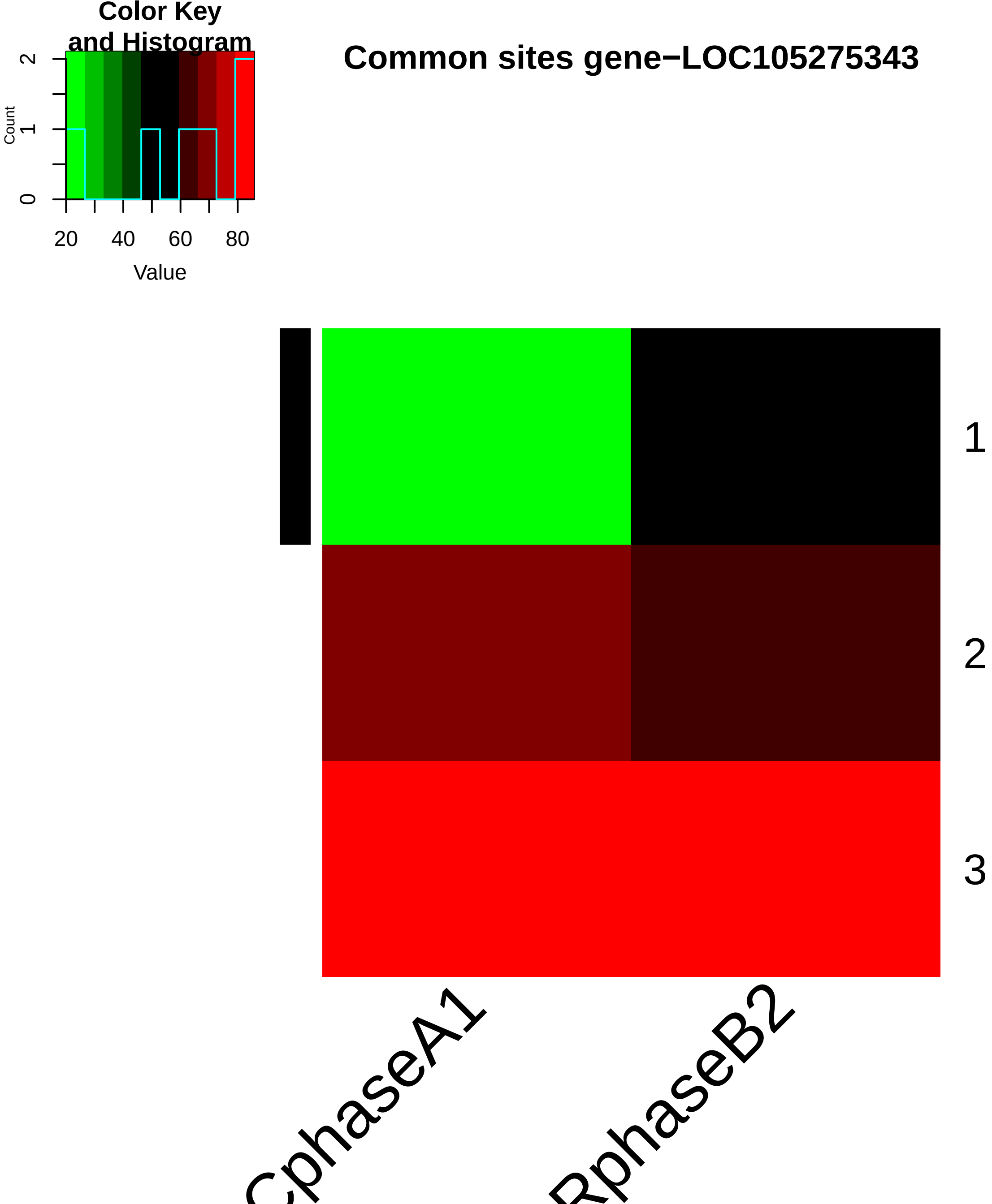
Color Key
and Histogram
Count
0
1
2
20
40
60
80
Value
Common sites gene−LOC105275343
1
2
3

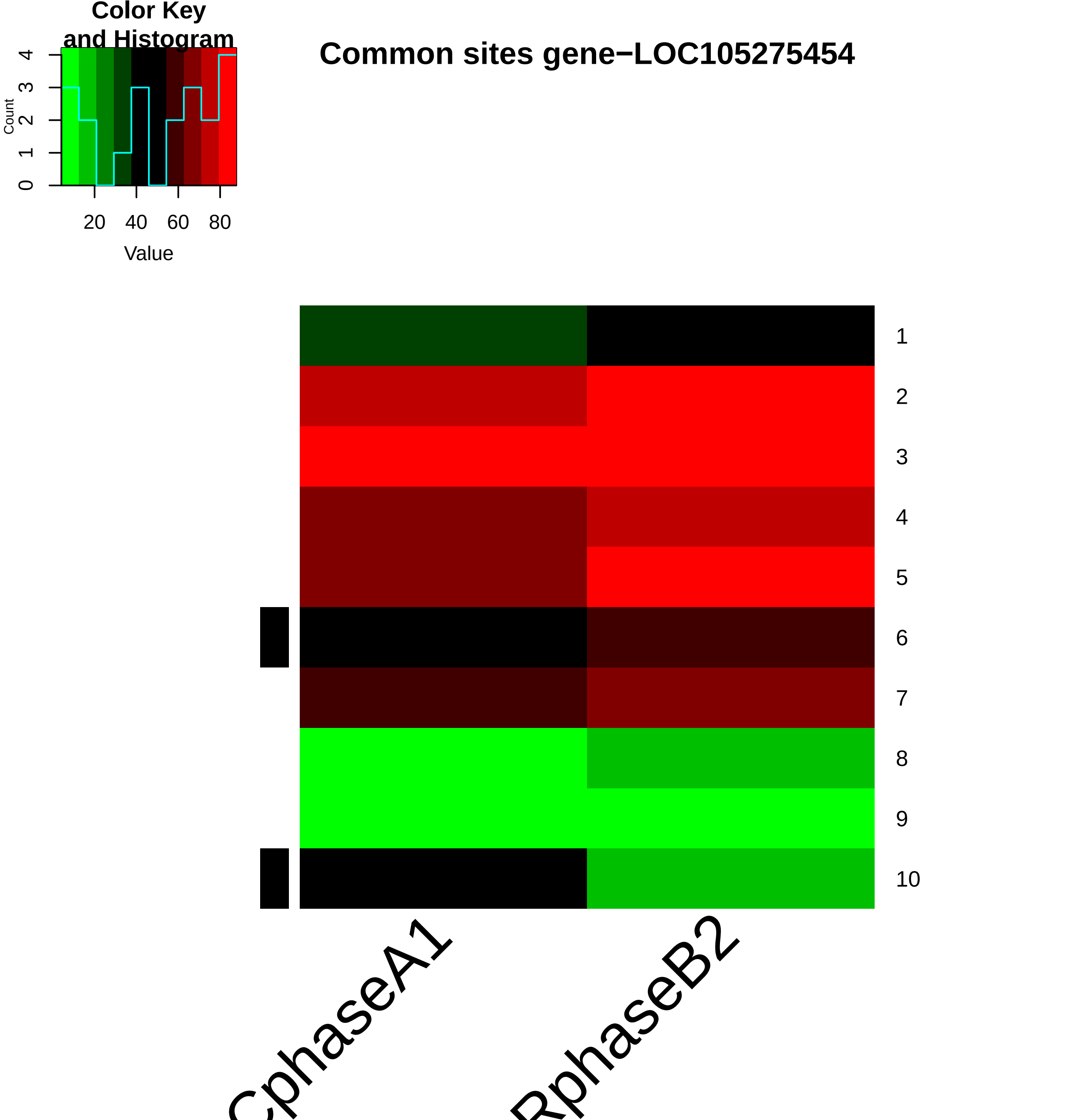

Color Key
and Histogram
Count
0
1
2
3
4
20
40
60
80
Value
Common sites gene−LOC105275454
1
2
3
4
5
6
7
8
9
10
CphaseA1
RphaseB2

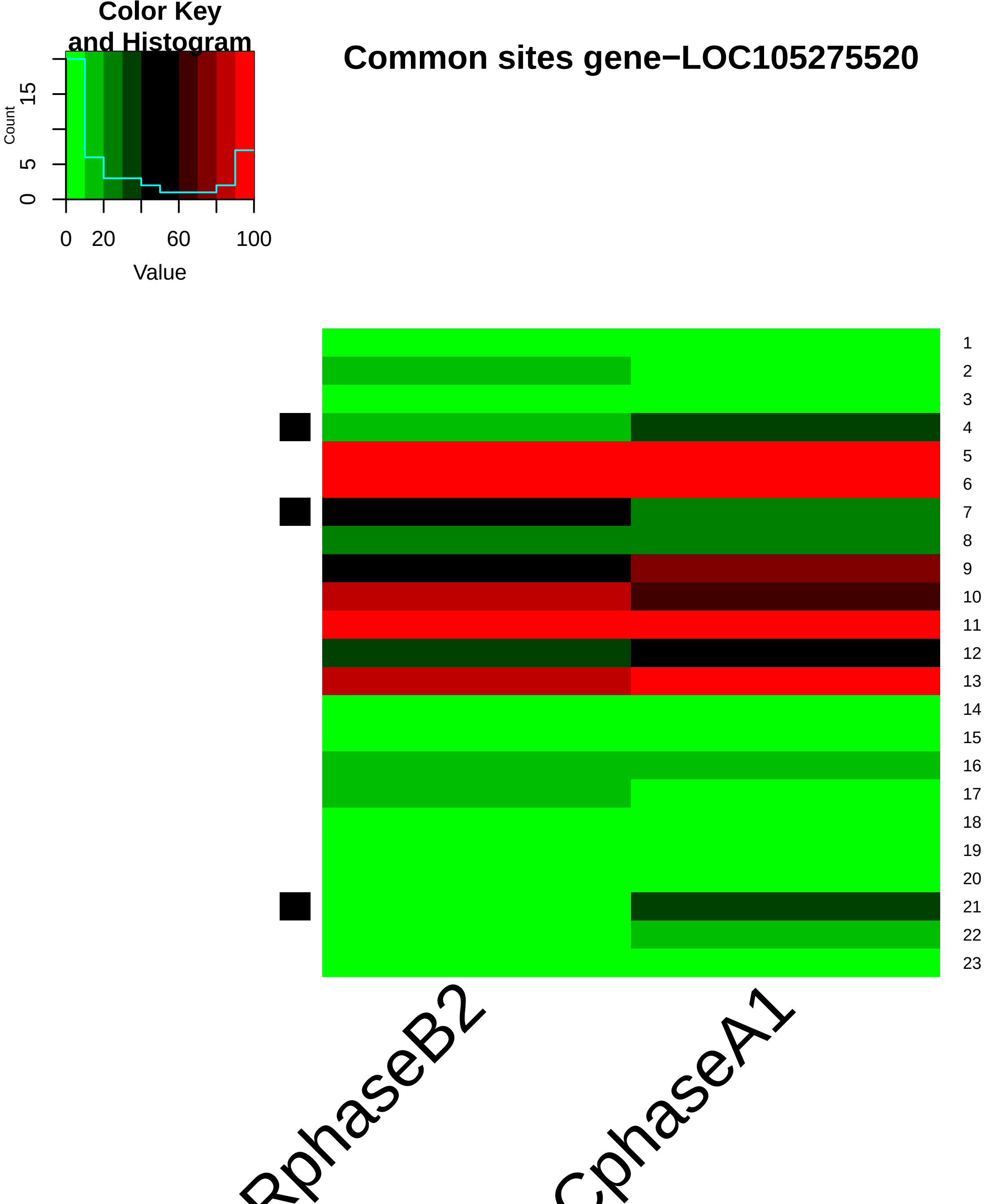
Color Key
and Histogram
Count
0
5
15
0
20
60
100
Value
Common sites gene−LOC105275520
RphaseB2
CphaseA1

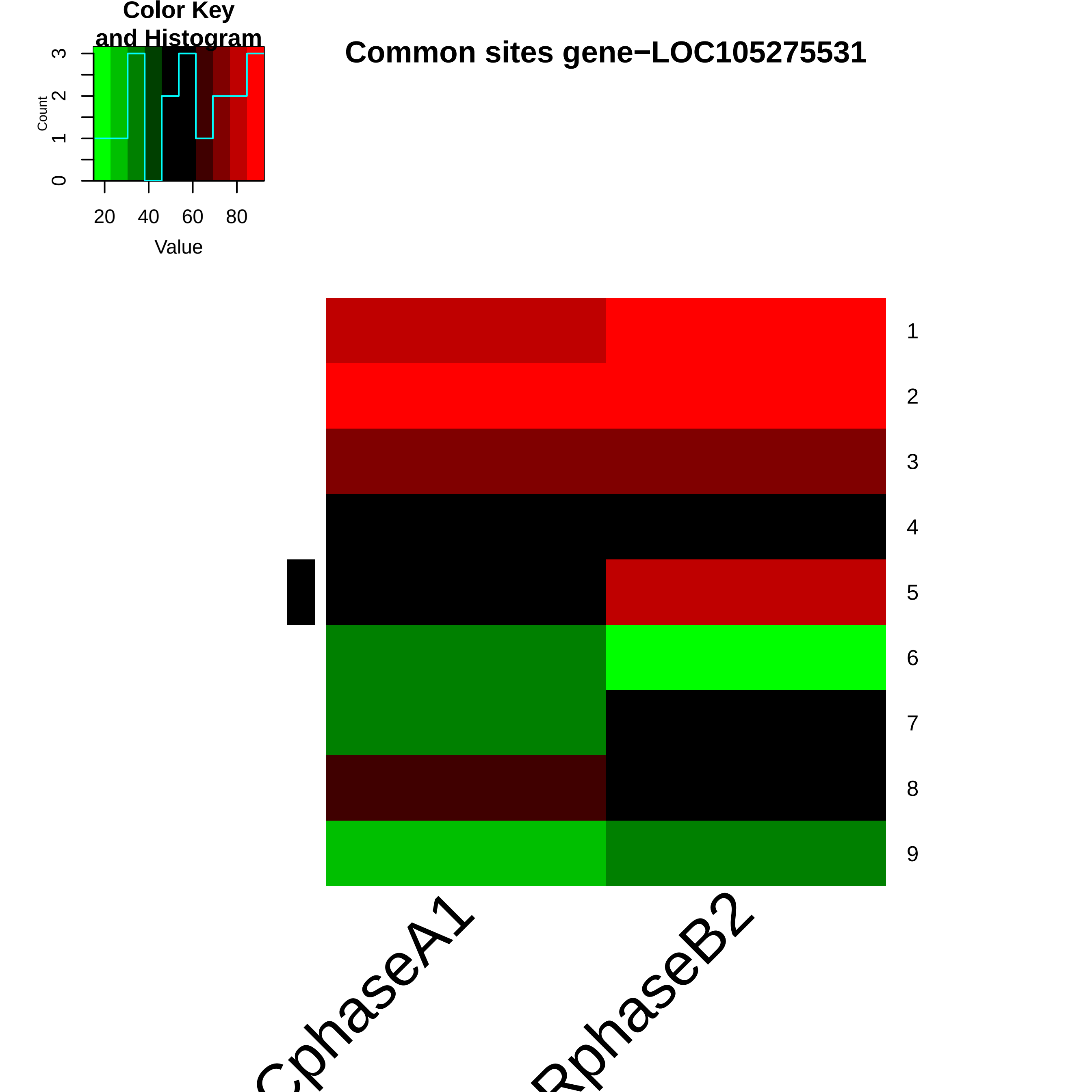

Color Key
and Histogram
Common sites gene−LOC105275531
Count
0
1
2
3
20
40
60
80
Value
1
2
3
4
5
6
7
8
9

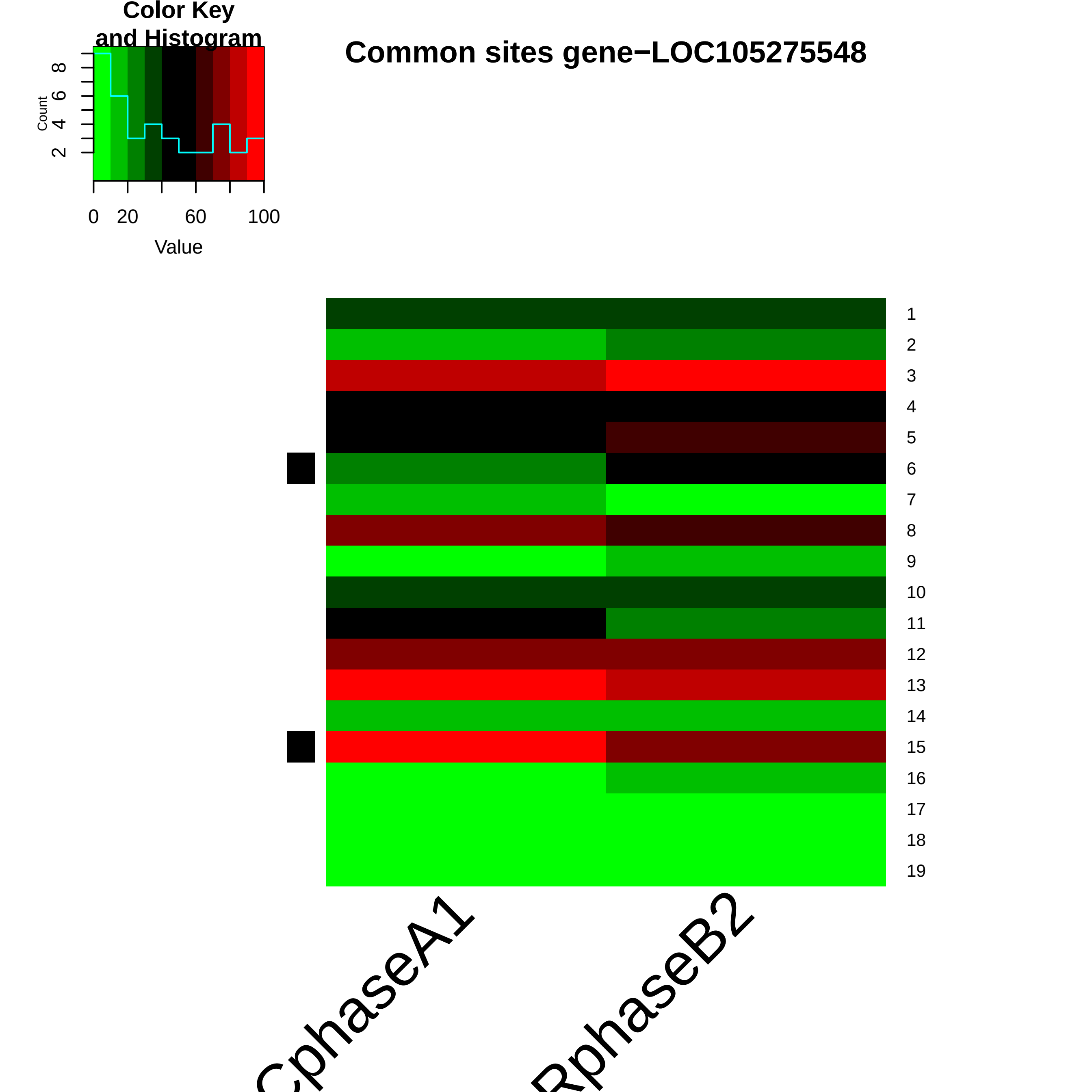

Color Key
and Histogram
Count
2
4
6
8
0
20
60
100
Value
Common sites gene−LOC105275548
CphaseA1
RphaseB2

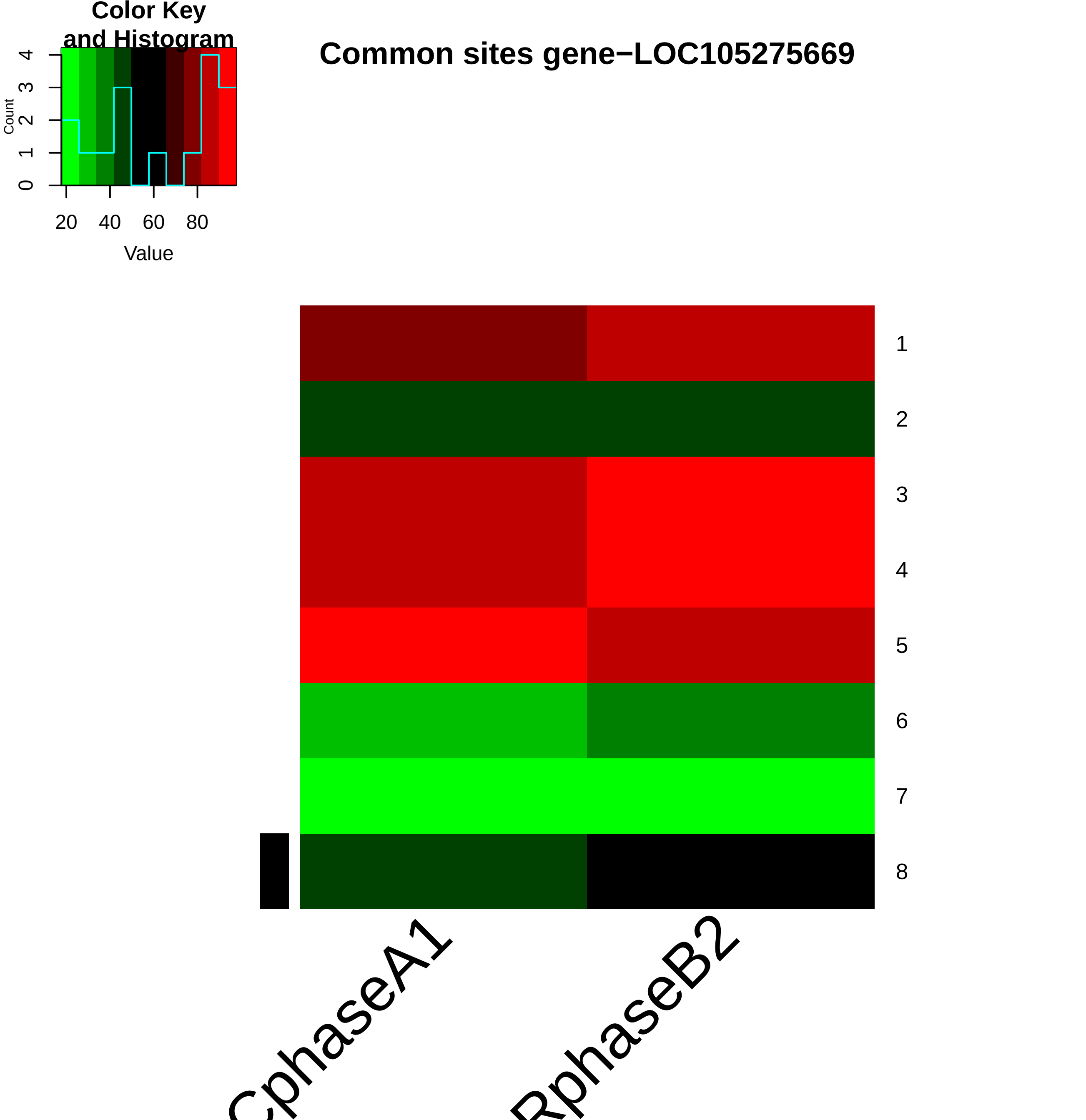
Color Key
and Histogram
Count
0
1
2
3
4
20
40
60
80
Value
Common sites gene−LOC105275669
1
2
3
4
5
6
7
8
CphaseA1
RphaseB2

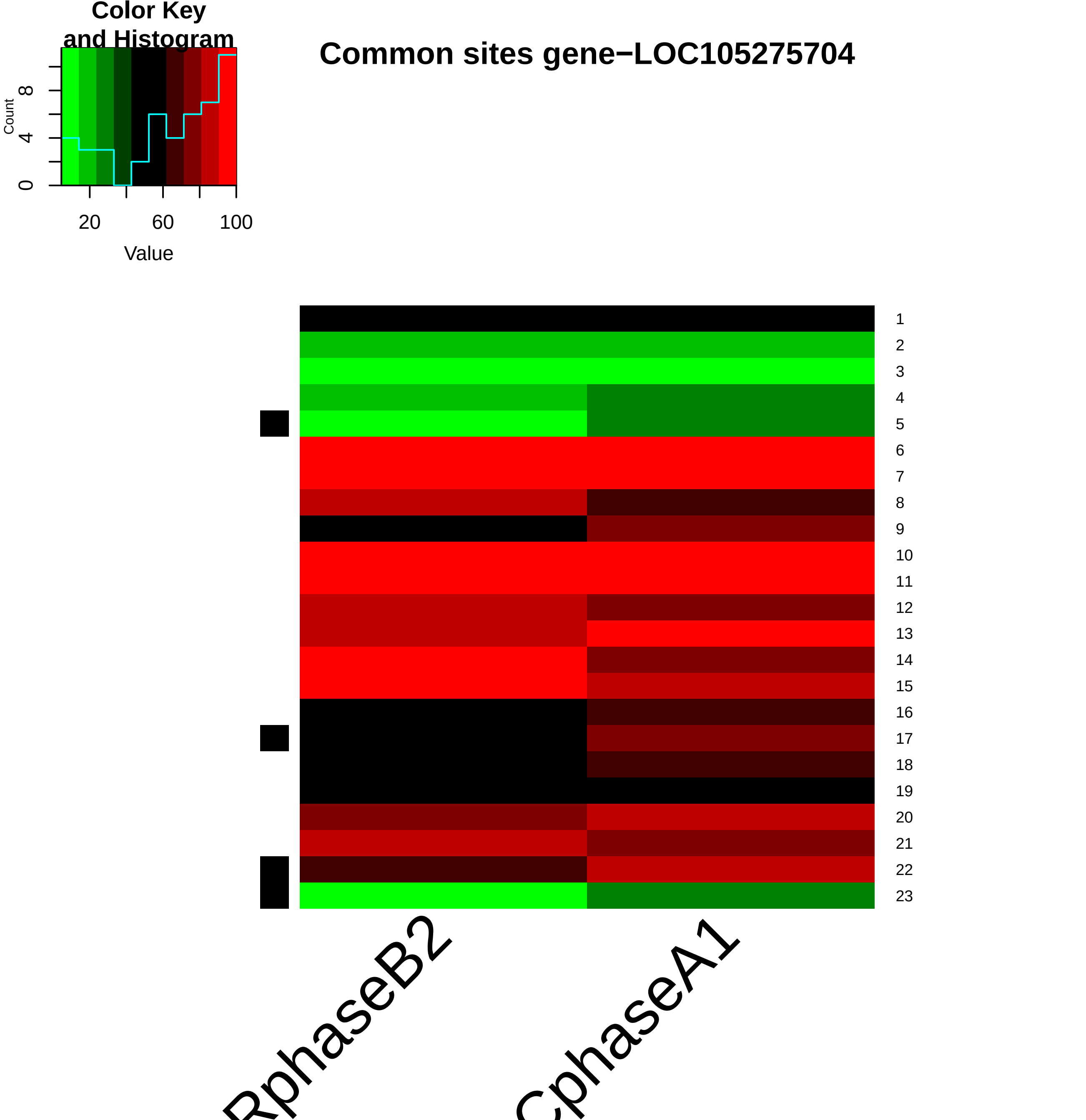

Color Key
and Histogram
Count
0
4
8
20
60
100
Value
Common sites gene−LOC105275704
RphaseB2
CphaseA1

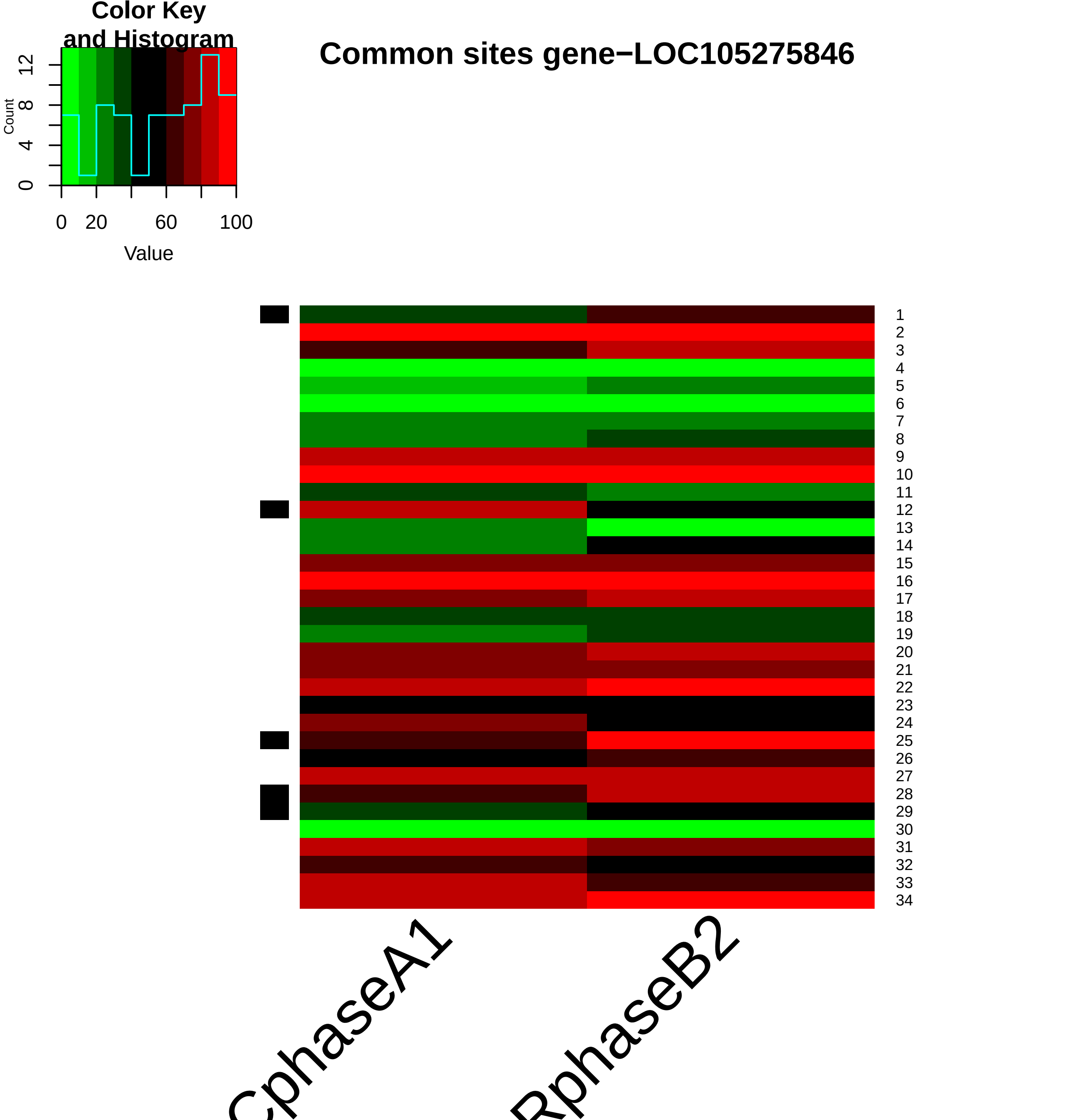
Color Key
and Histogram
Count
0
4
8
12
0
20
60
100
Value
Common sites gene−LOC105275846
CphaseA1
RphaseB2

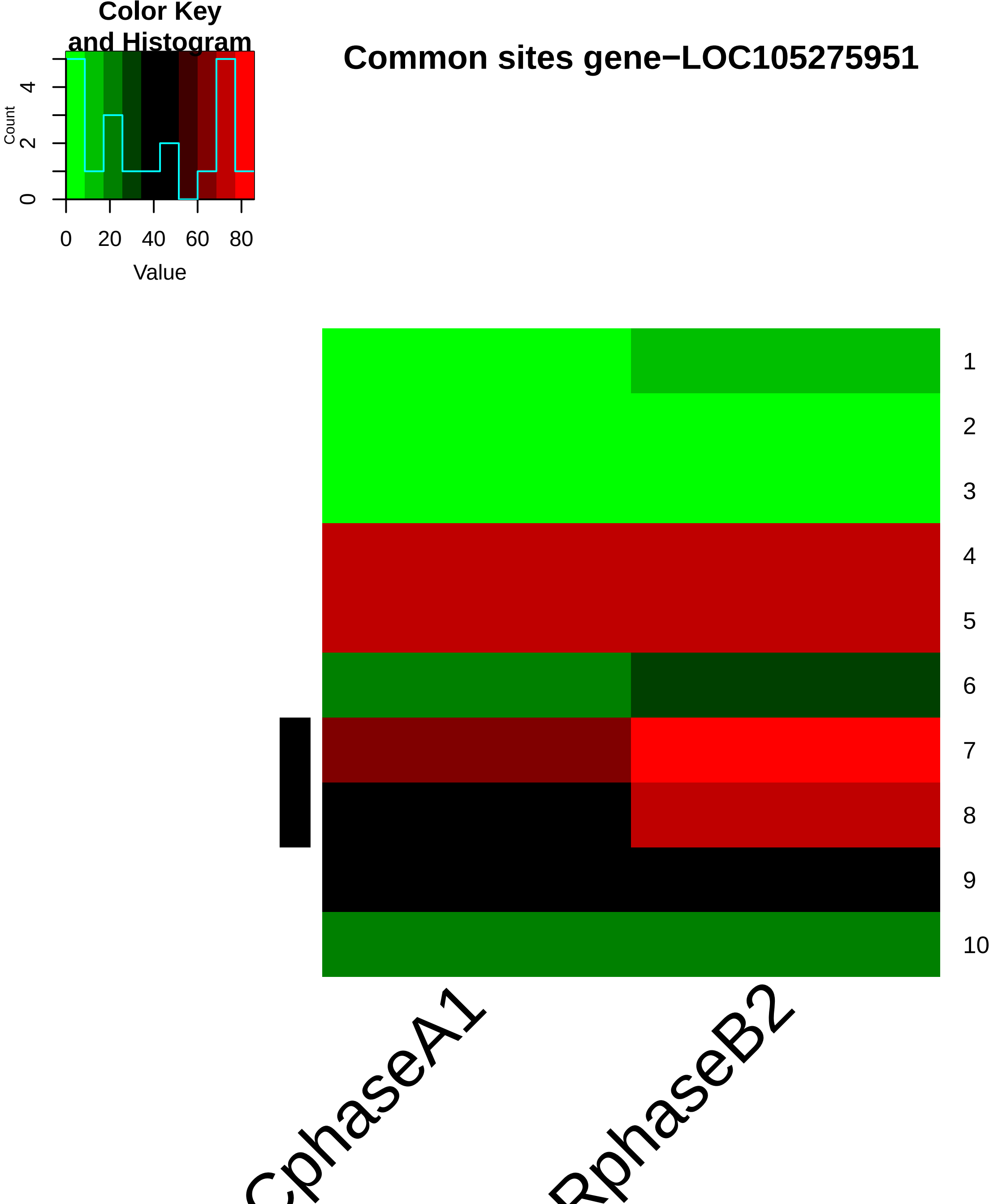

Color Key
and Histogram
Count
Value
0
20
40
60
80
0
2
4
Common sites gene−LOC105275951
1
2
3
4
5
6
7
8
9
10
CphaseA1
RphaseB2

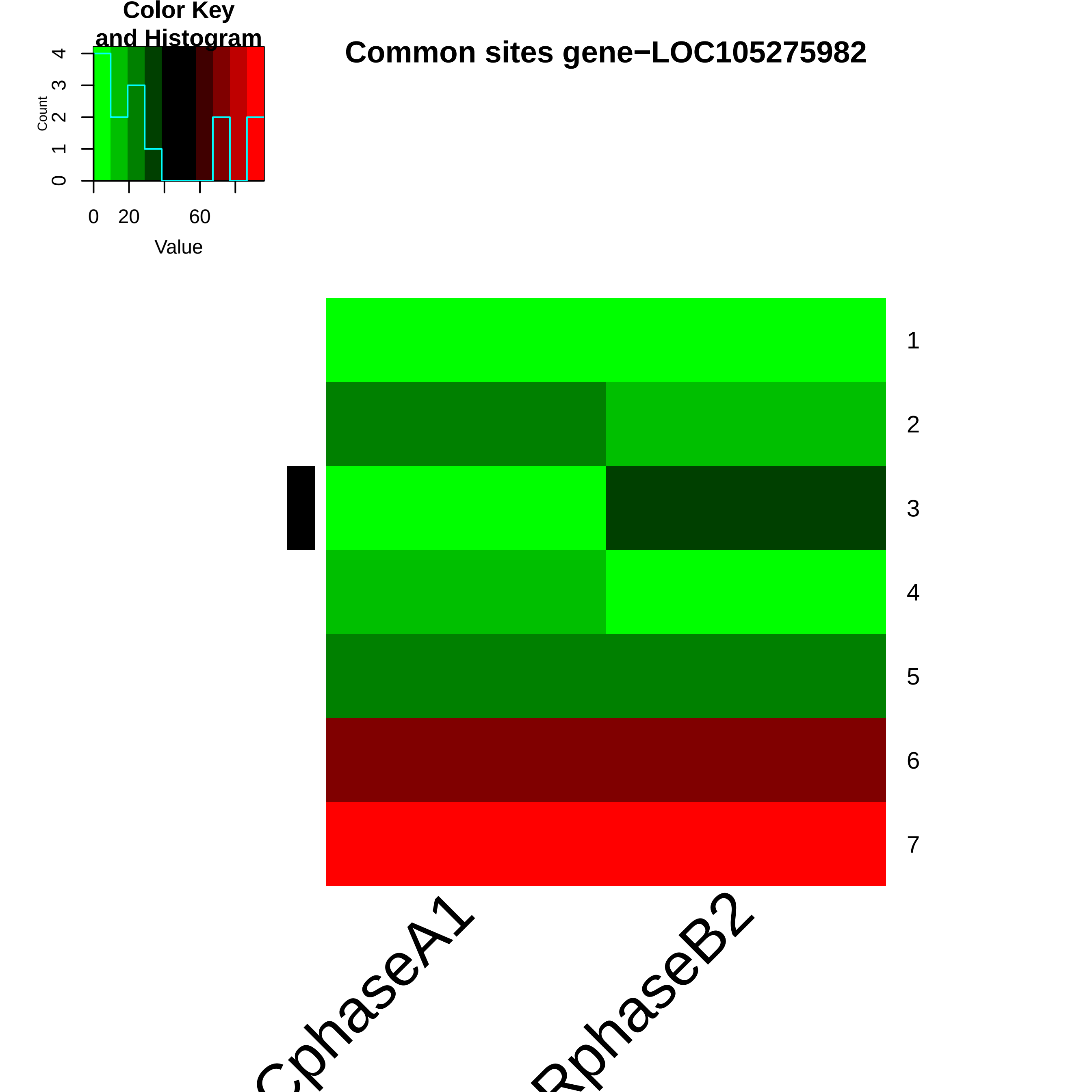
Color Key
and Histogram
Count
0
1
2
3
4
0
20
60
Value
Common sites gene−LOC105275982
1
2
3
4
5
6
7

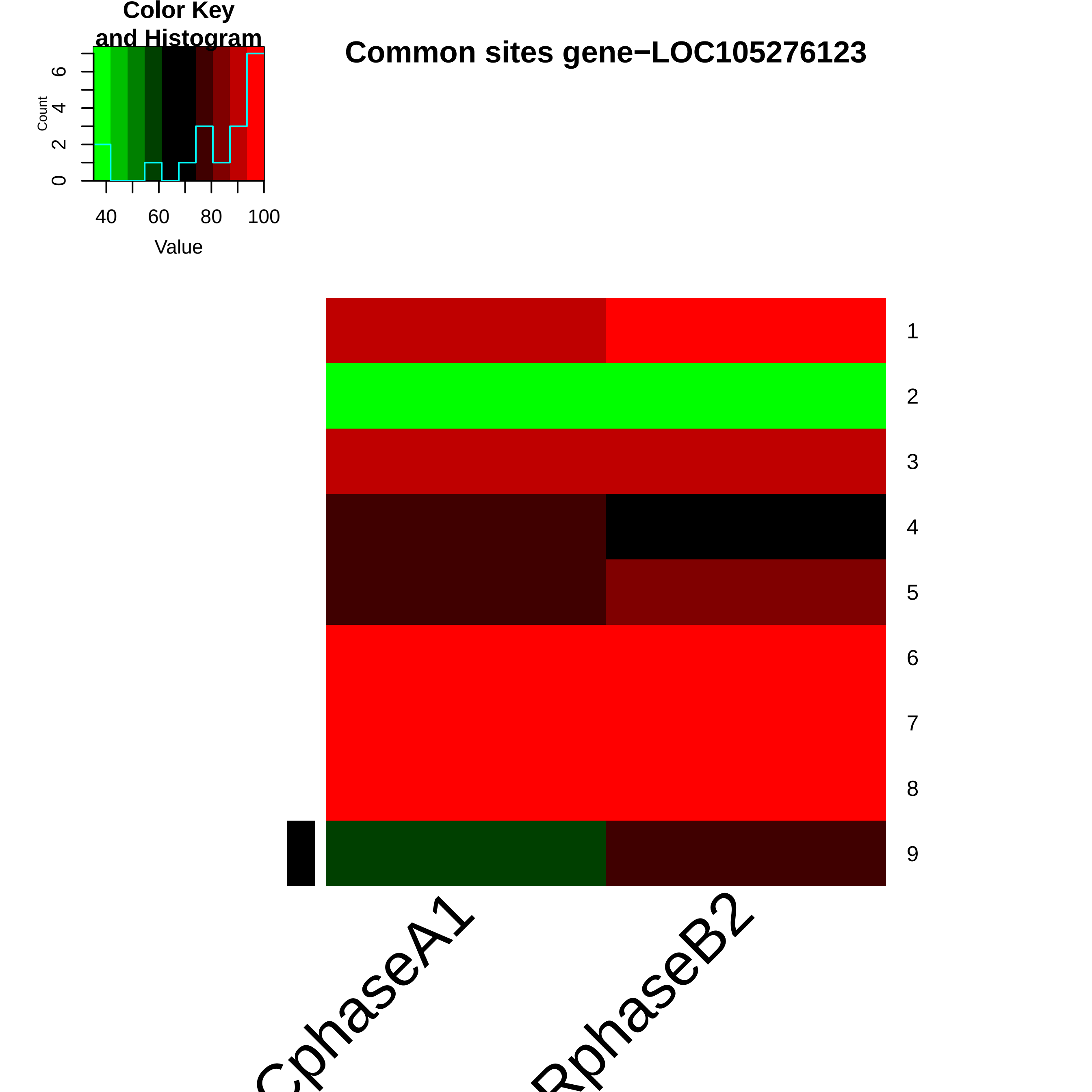

Color Key
and Histogram
Count
Value
Common sites gene−LOC105276123
1
2
3
4
5
6
7
8
9
CphaseA1
RphaseB2

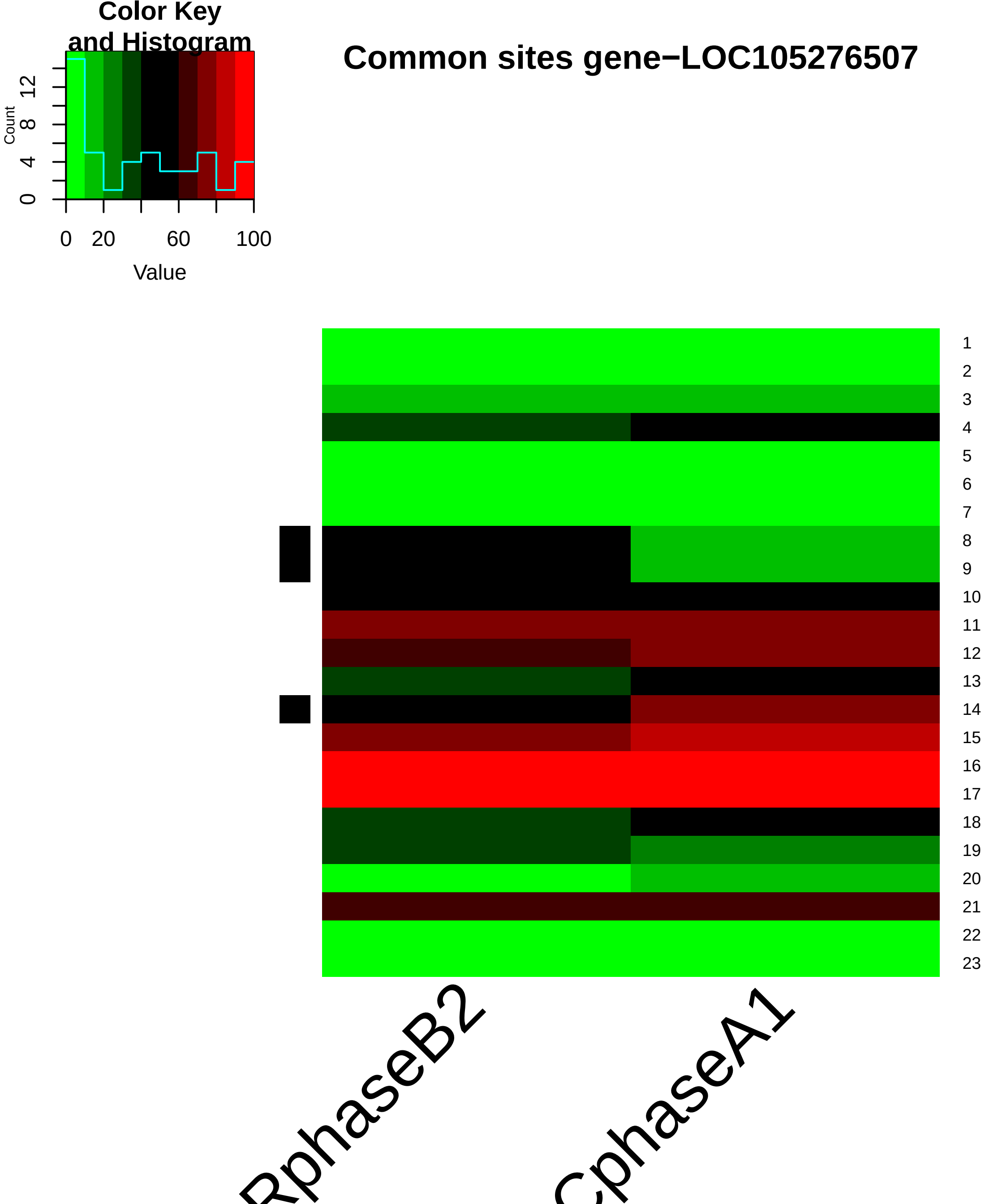
Color Key
and Histogram
Count
0
4
8
12
0
20
60
100
Value
Common sites gene−LOC105276507
RphaseB2
CphaseA1

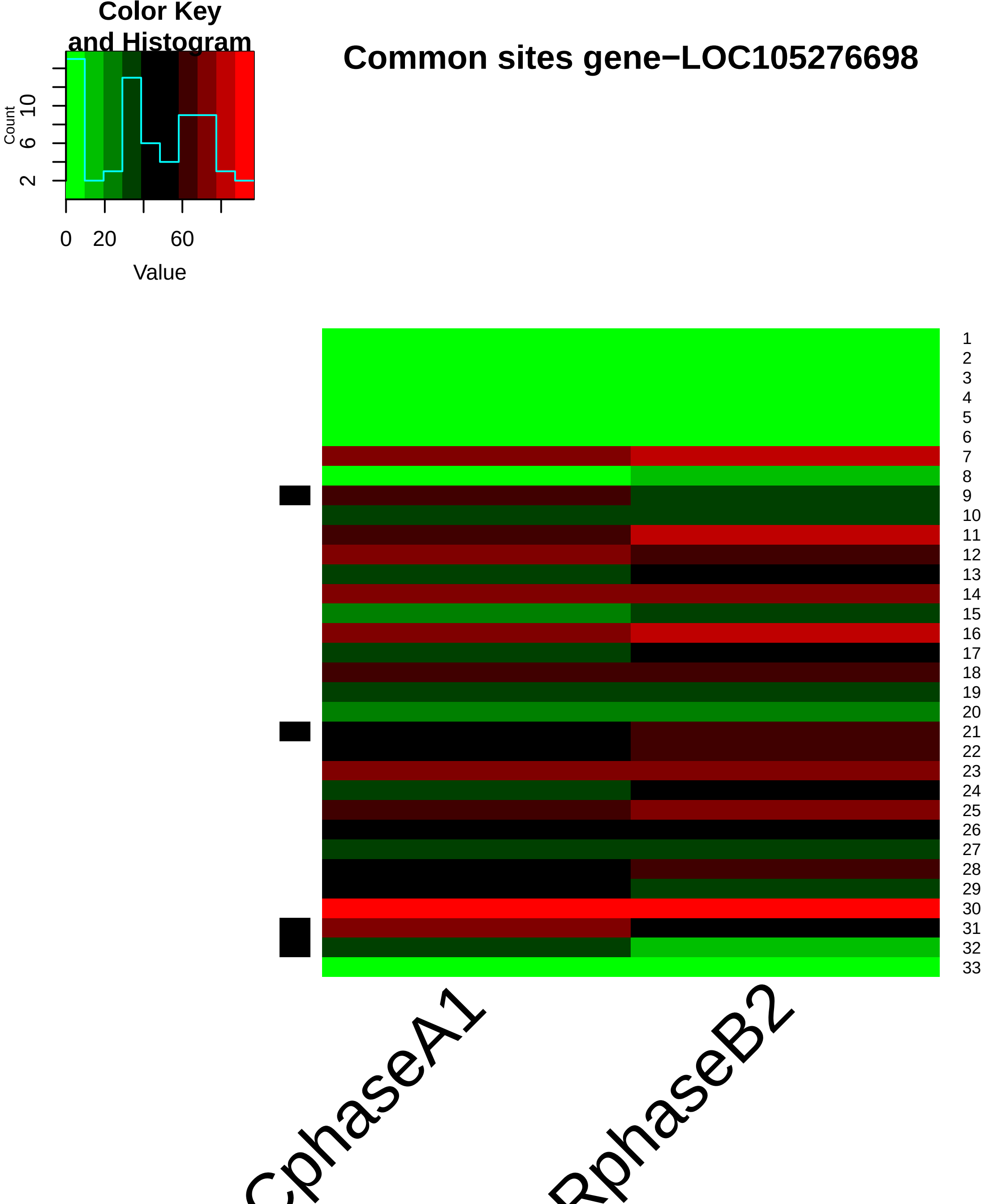
Color Key
and Histogram
Count
2
6
10
0
20
60
Value
Common sites gene−LOC105276698
CphaseA1
RphaseB2

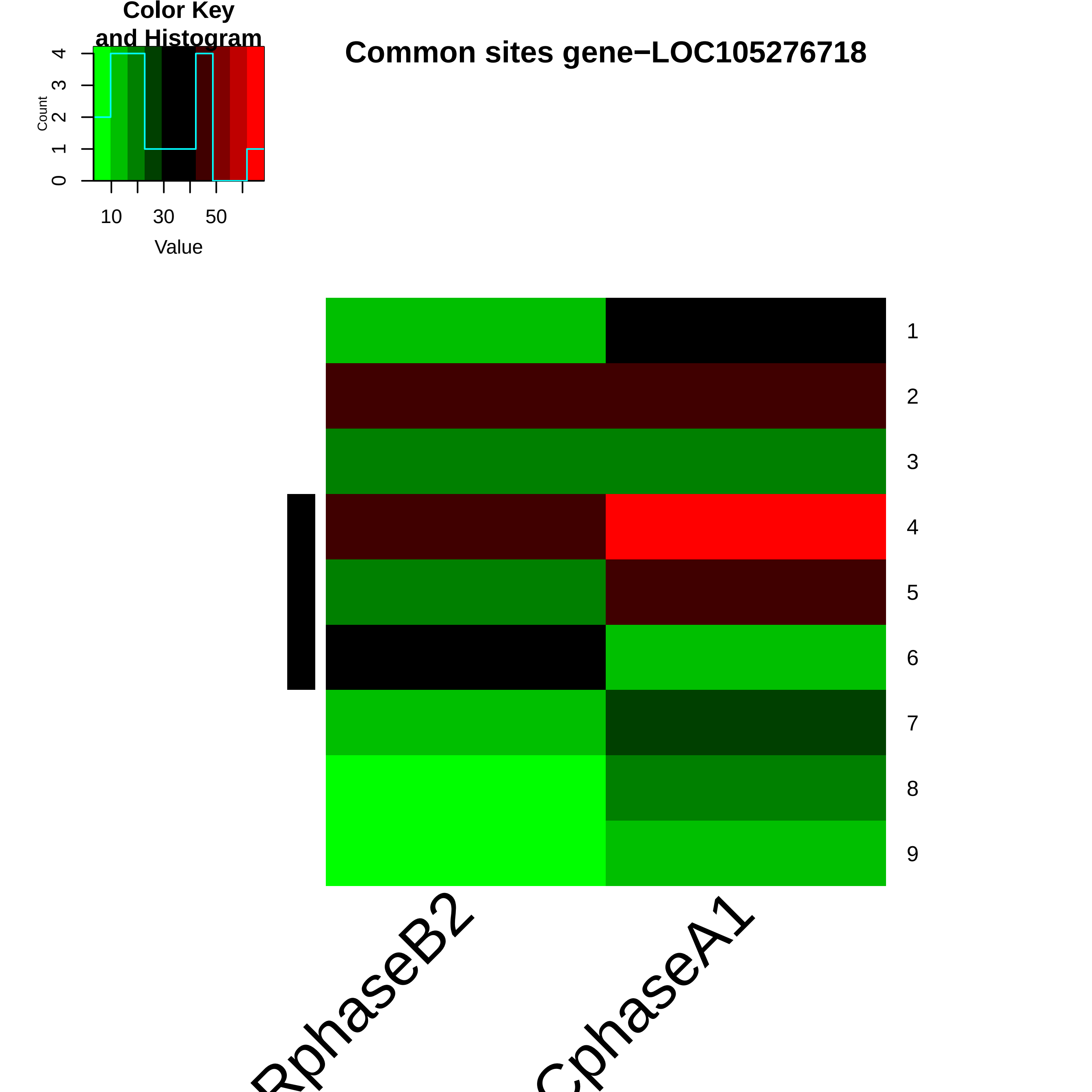
Color Key
and Histogram
Count
0
1
2
3
4
10
30
50
Value
Common sites gene−LOC105276718
1
2
3
4
5
6
7
8
9
RphaseB2
CphaseA1

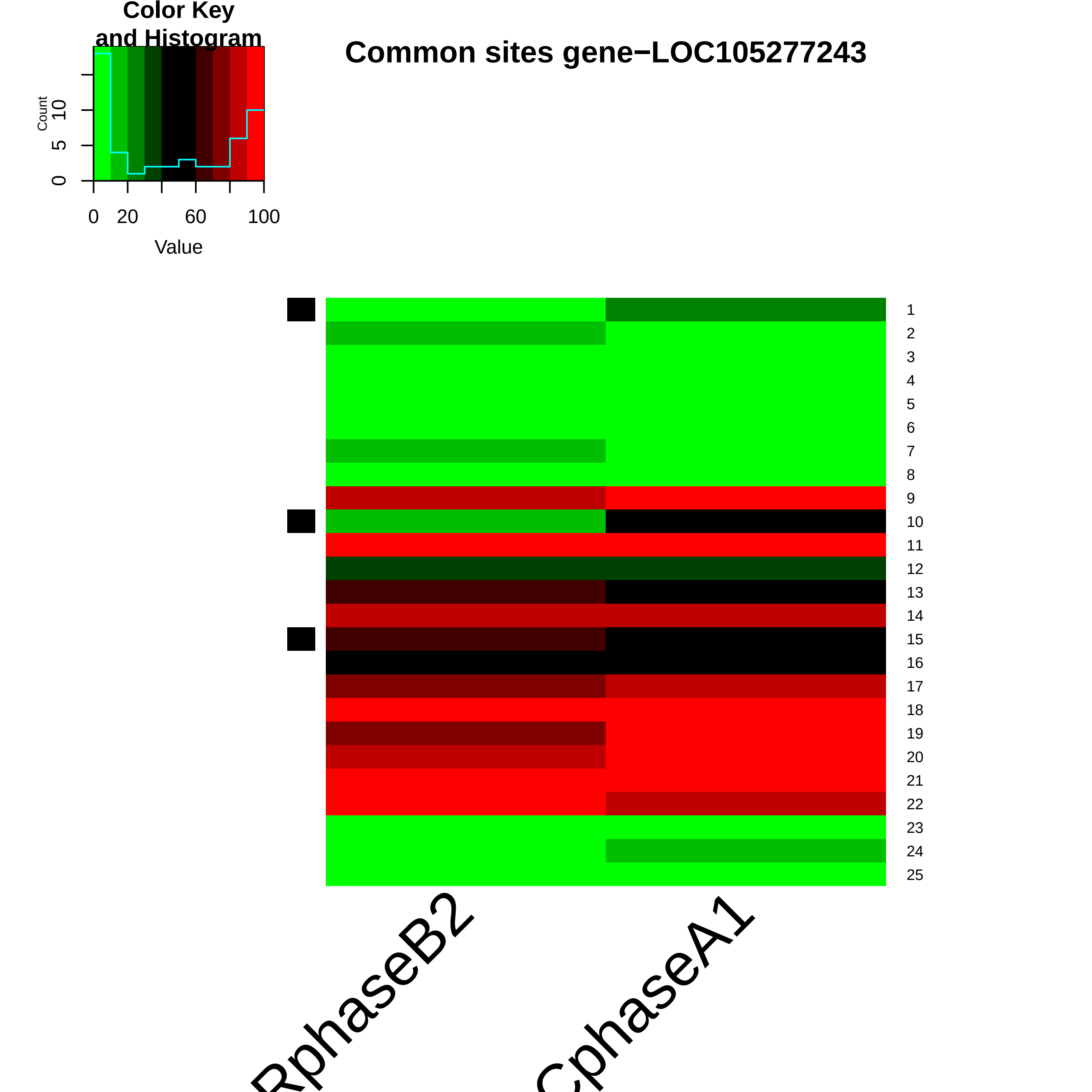
Color Key
and Histogram
Count
0
5
10
0
20
60
100
Value
Common sites gene−LOC105277243
RphaseB2
CphaseA1

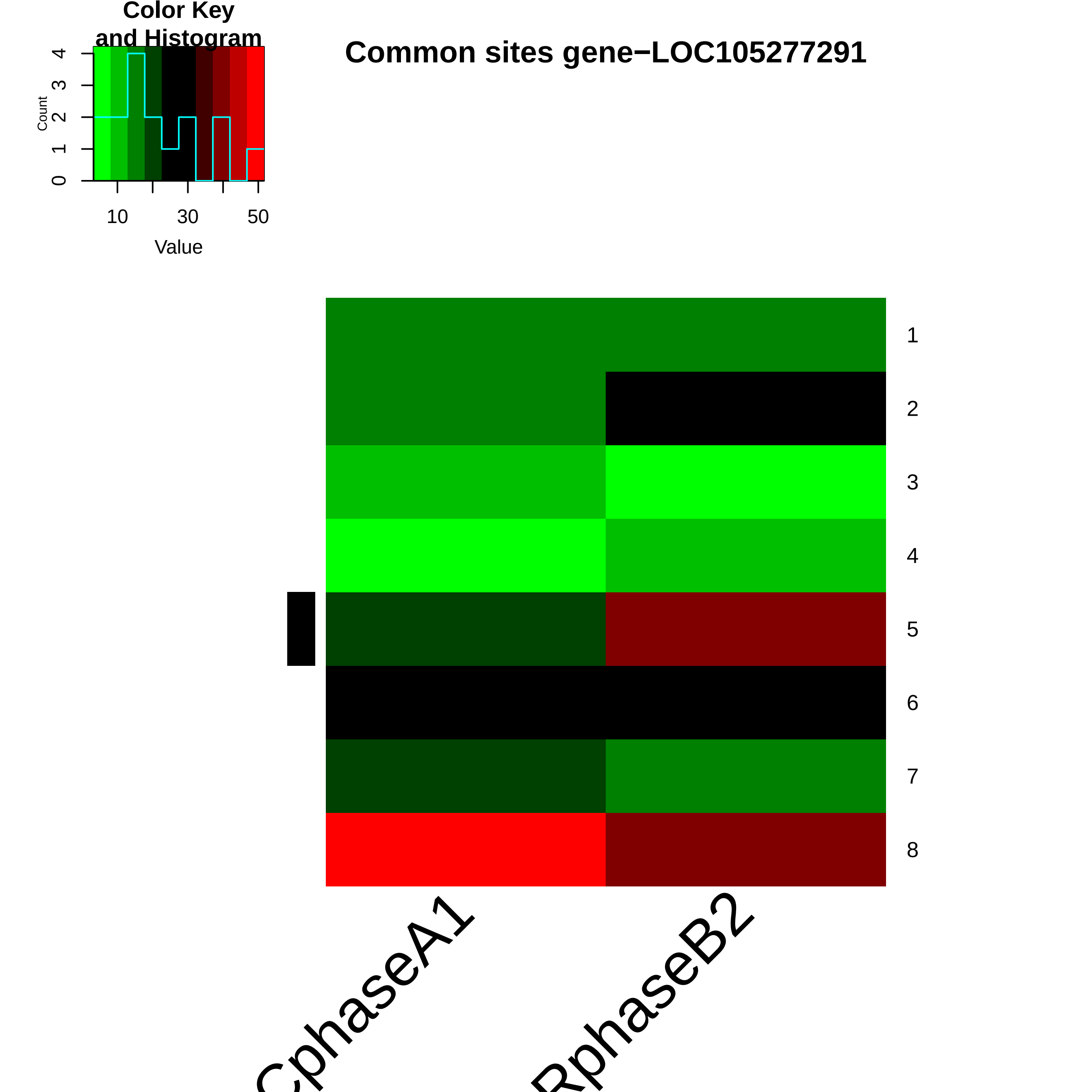
Color Key
and Histogram
Count
Value
Common sites gene−LOC105277291
1
2
3
4
5
6
7
8
CphaseA1
RphaseB2

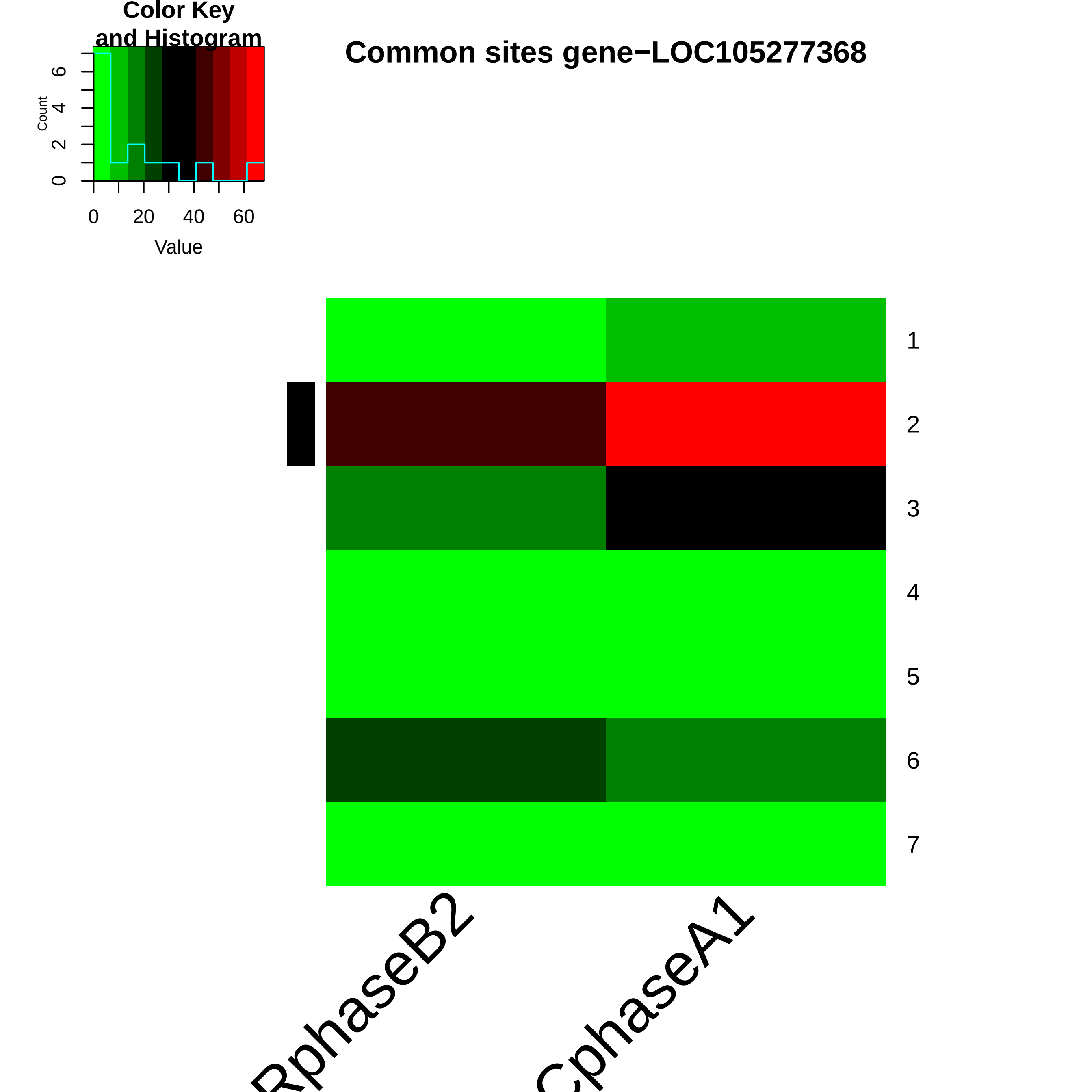
Color Key
and Histogram
Count
0
2
4
6
0
20
40
60
Value
Common sites gene−LOC105277368
1
2
3
4
5
6
7
RphaseB2
CphaseA1

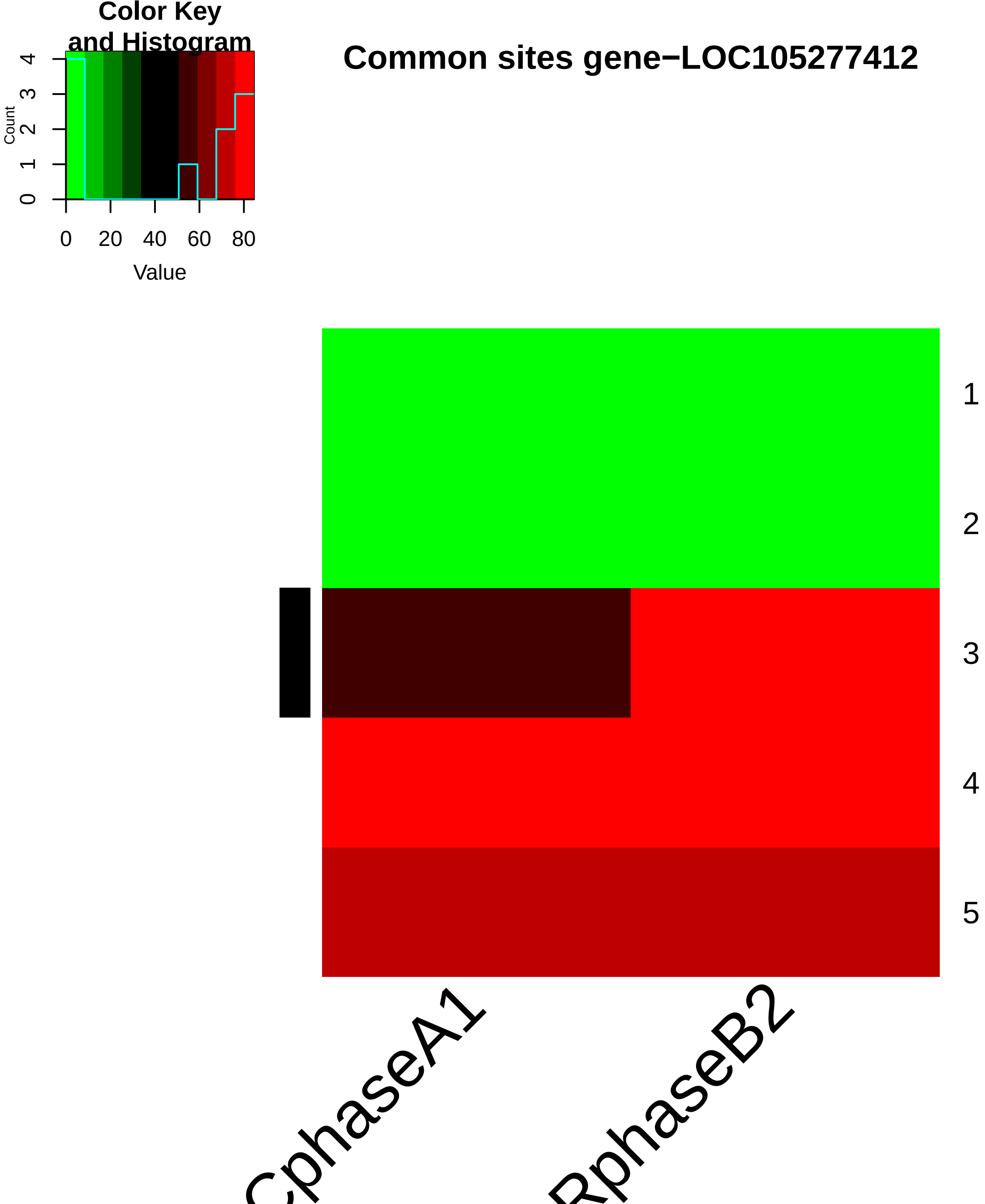

Color Key
and Histogram
Count
0
1
2
3
4
0
20
40
60
80
Value
Common sites gene−LOC105277412
1
2
3
4
5
CphaseA1
RphaseB2

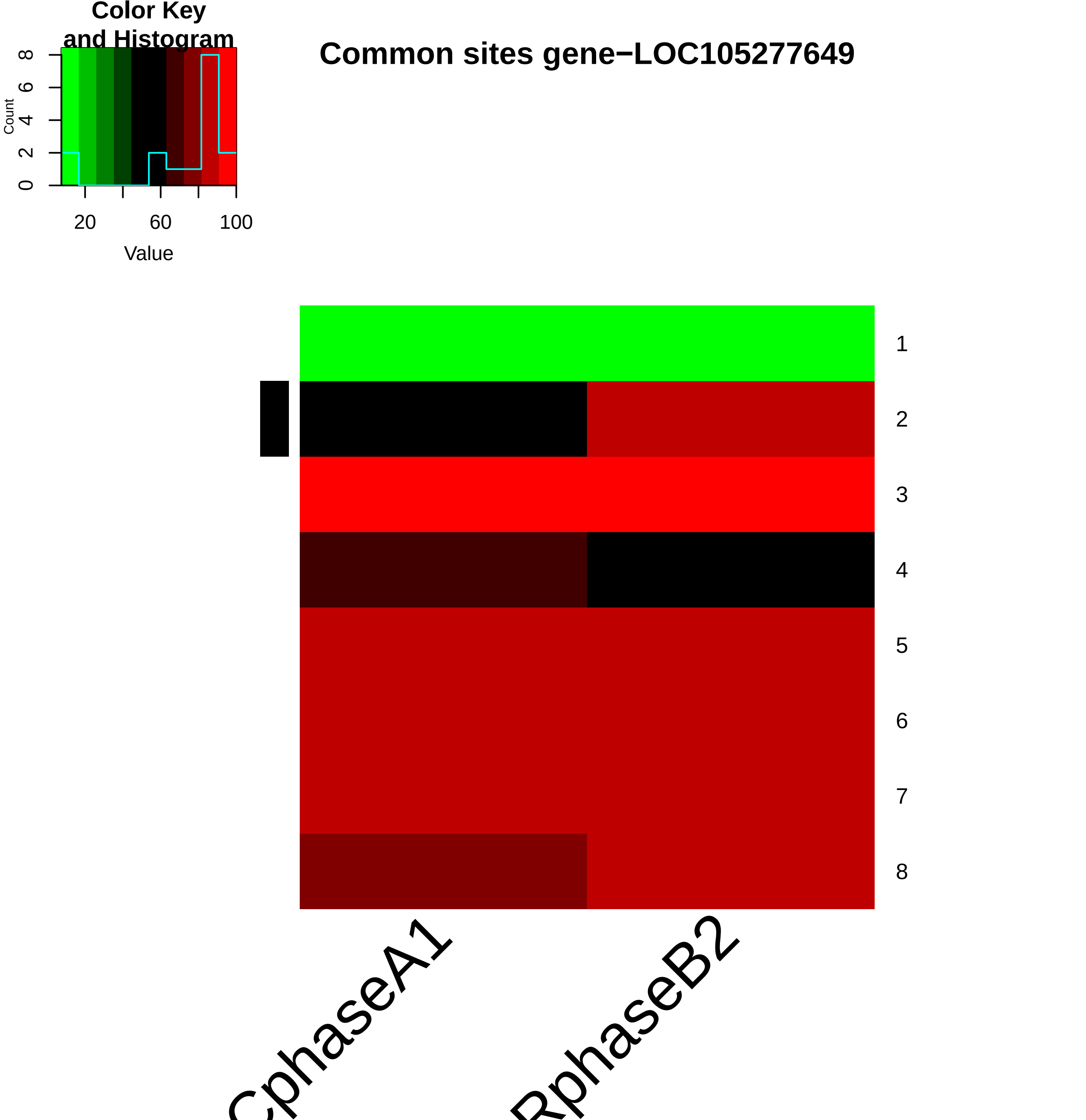

Color Key
and Histogram
Count
Value
20
60
100
0
2
4
6
8
Common sites gene−LOC105277649
1
2
3
4
5
6
7
8
CphaseA1
RphaseB2

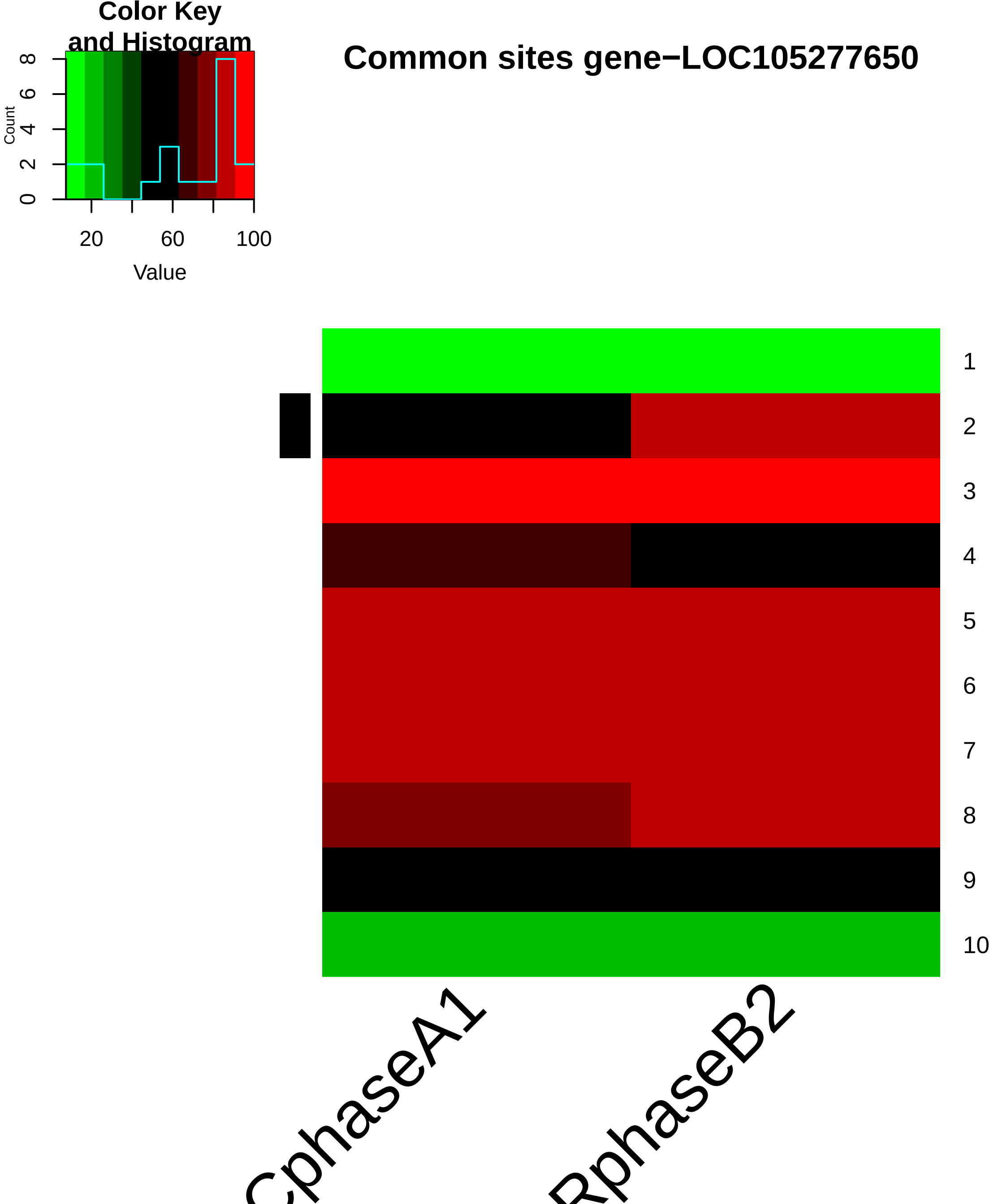
Color Key
and Histogram
Count
Value
0
2
4
6
8
20
60
100
Common sites gene−LOC105277650
1
2
3
4
5
6
7
8
9
10
CphaseA1
RphaseB2

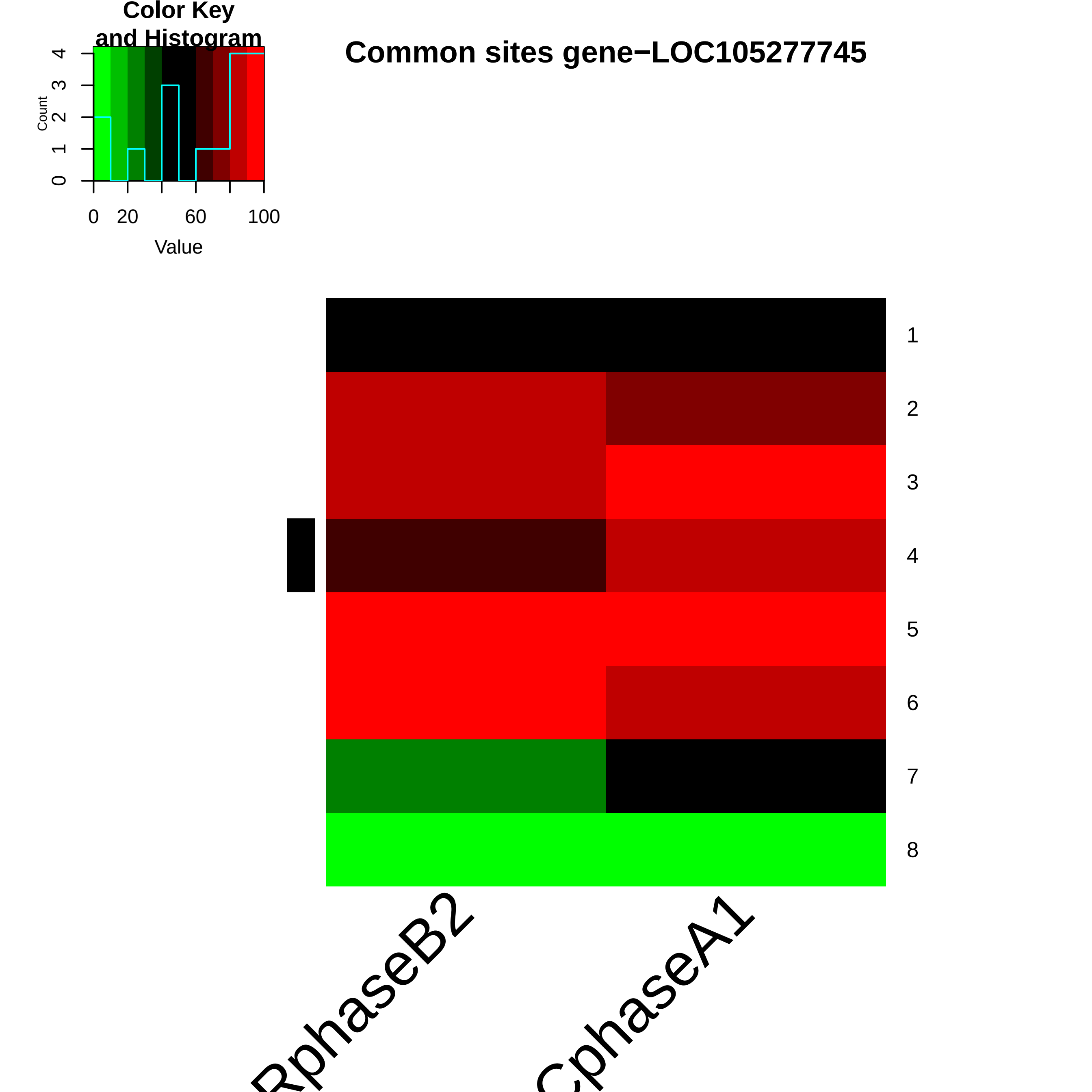
Color Key
and Histogram
Count
0
1
2
3
4
0
20
60
100
Value
Common sites gene−LOC105277745
1
2
3
4
5
6
7
8
RphaseB2
CphaseA1

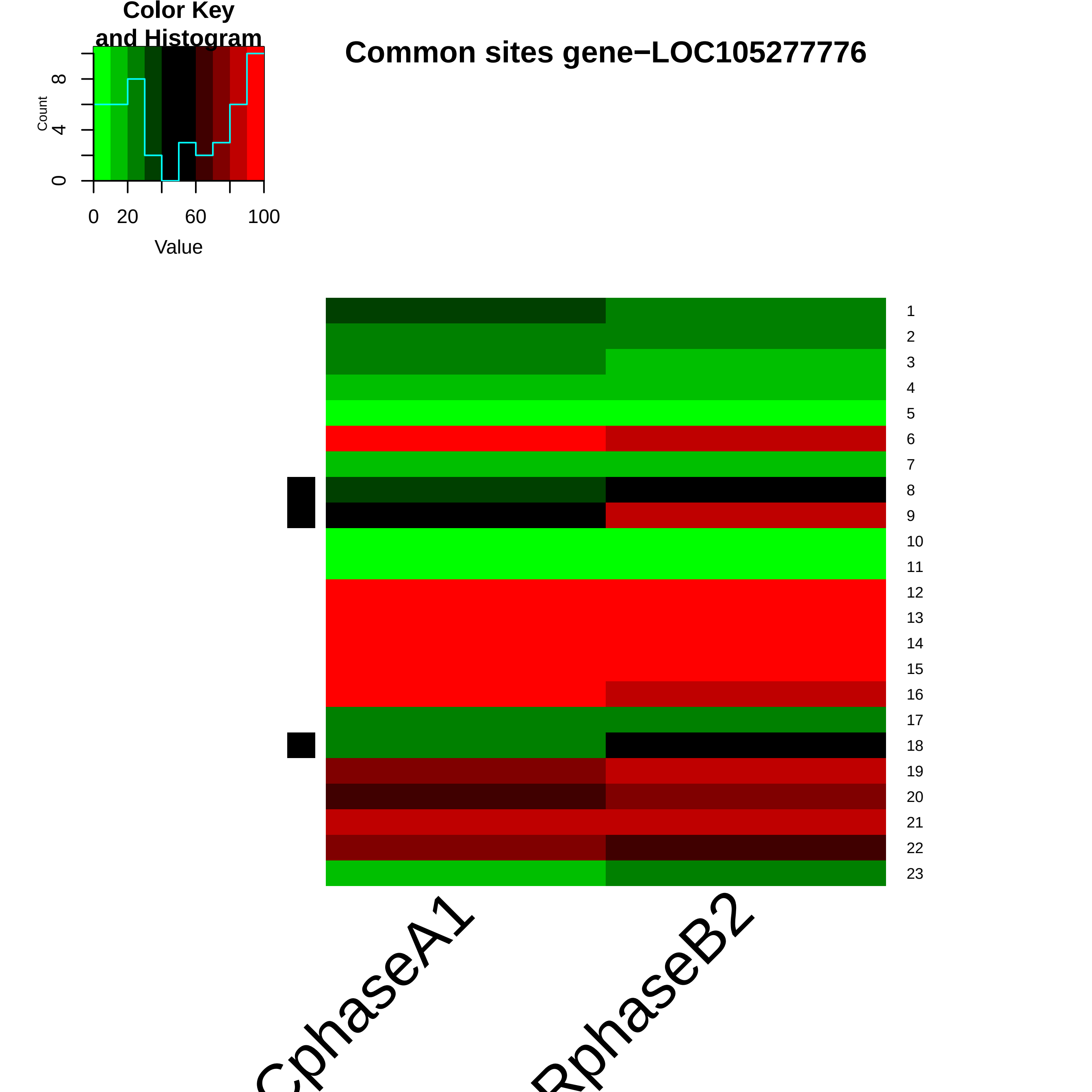

Color Key
and Histogram
Count
0
4
8
0
20
60
100
Value
Common sites gene−LOC105277776
CphaseA1
RphaseB2

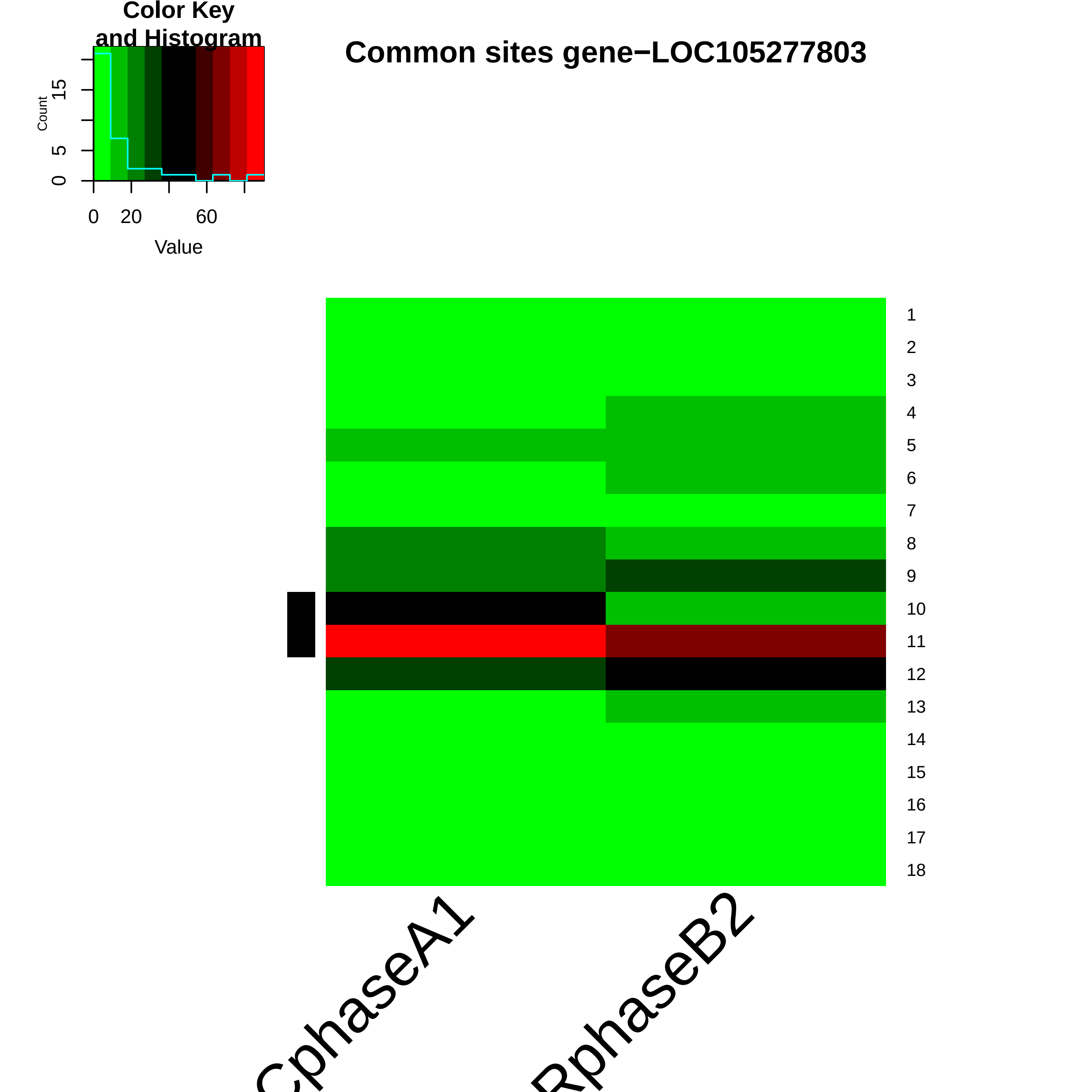
Color Key
and Histogram
Count
0
5
15
0
20
60
Value
Common sites gene−LOC105277803
CphaseA1
RphaseB2

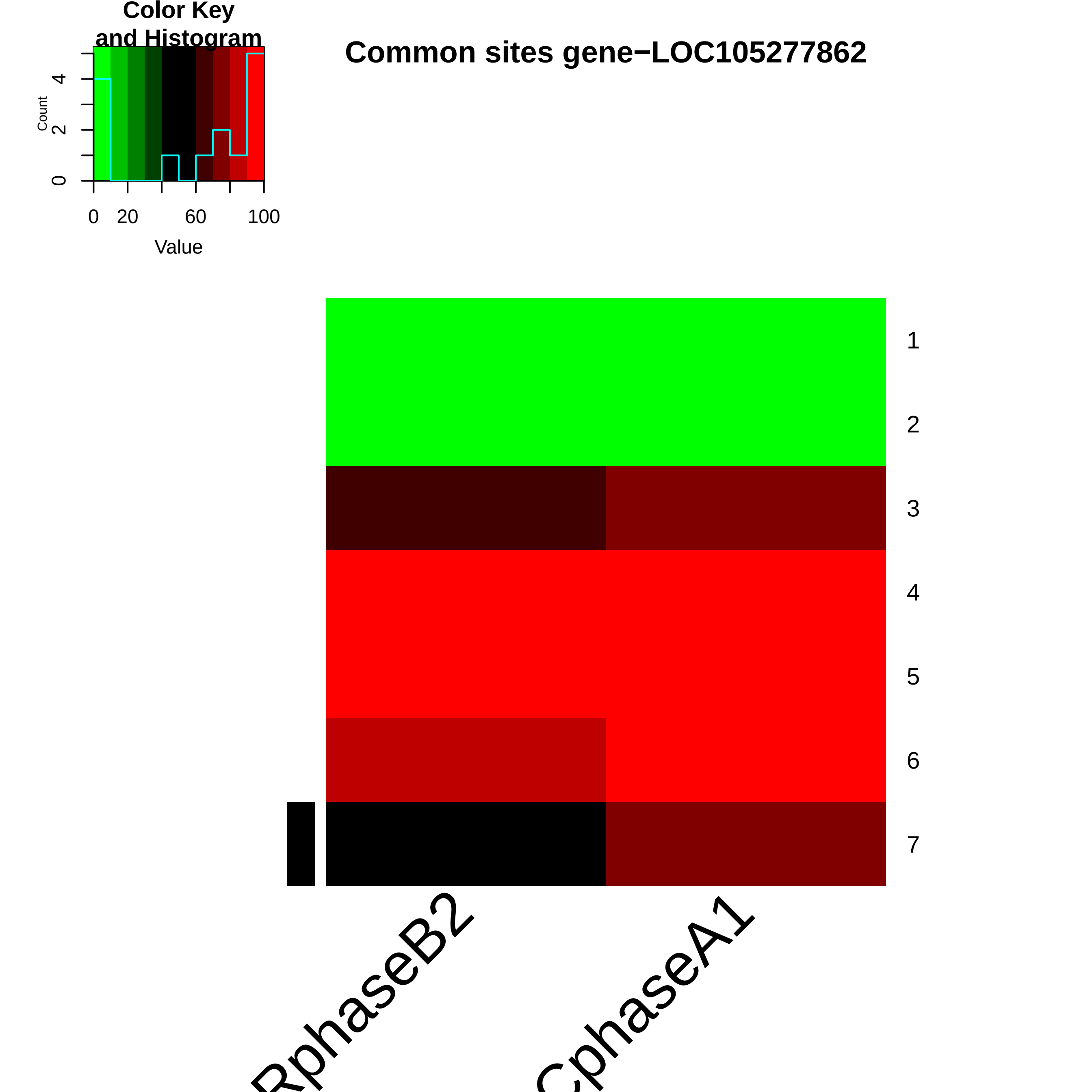

Color Key
and Histogram
Count
0
2
4
0
20
60
100
Value
Common sites gene−LOC105277862
1
2
3
4
5
6
7
RphaseB2
CphaseA1

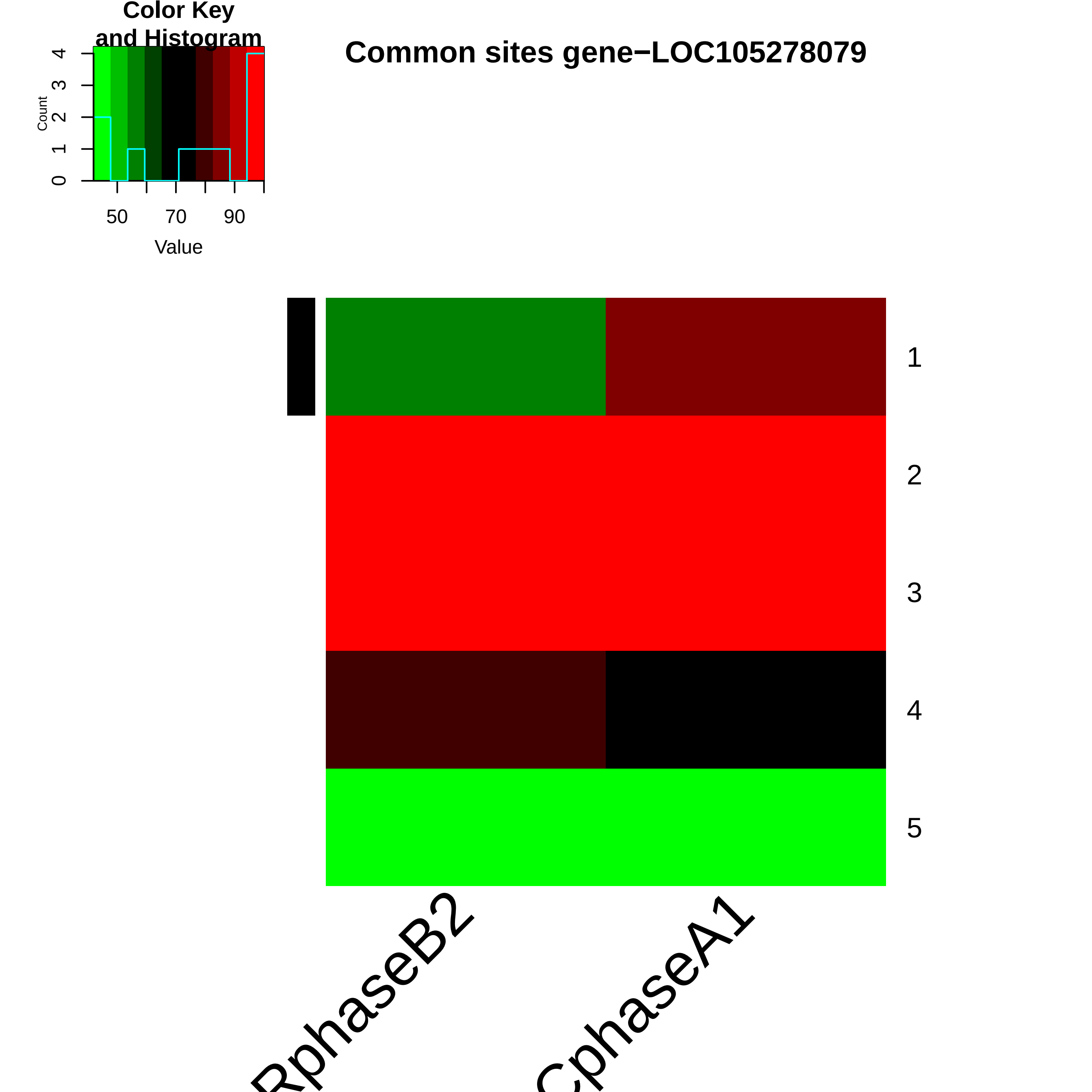

Color Key
and Histogram
Count
0
1
2
3
4
50
70
90
Value
Common sites gene−LOC105278079
1
2
3
4
5
RphaseB2
CphaseA1

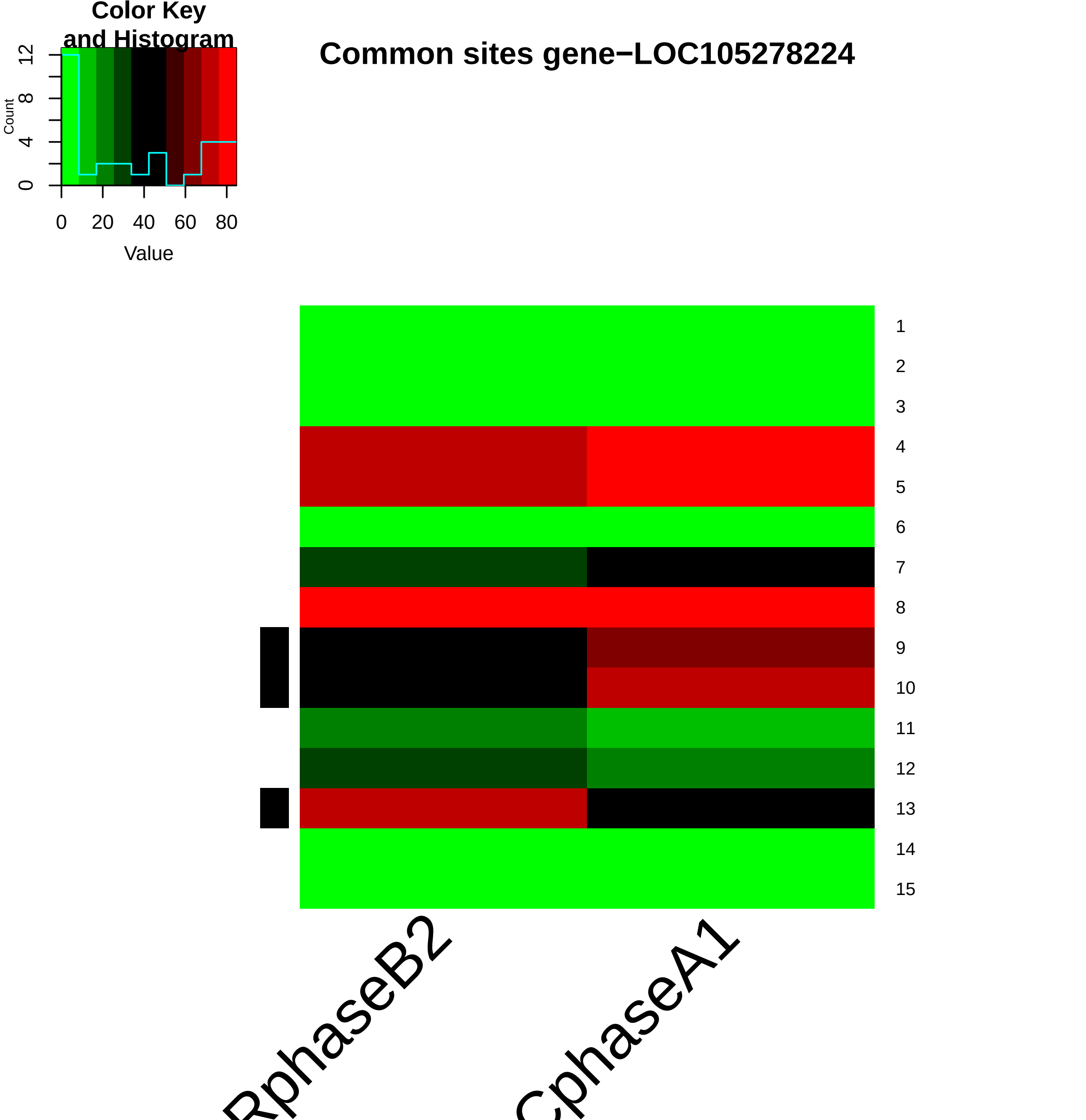
Color Key
and Histogram
Count
0
4
8
12
0
20
40
60
80
Value
Common sites gene−LOC105278224
RphaseB2
CphaseA1

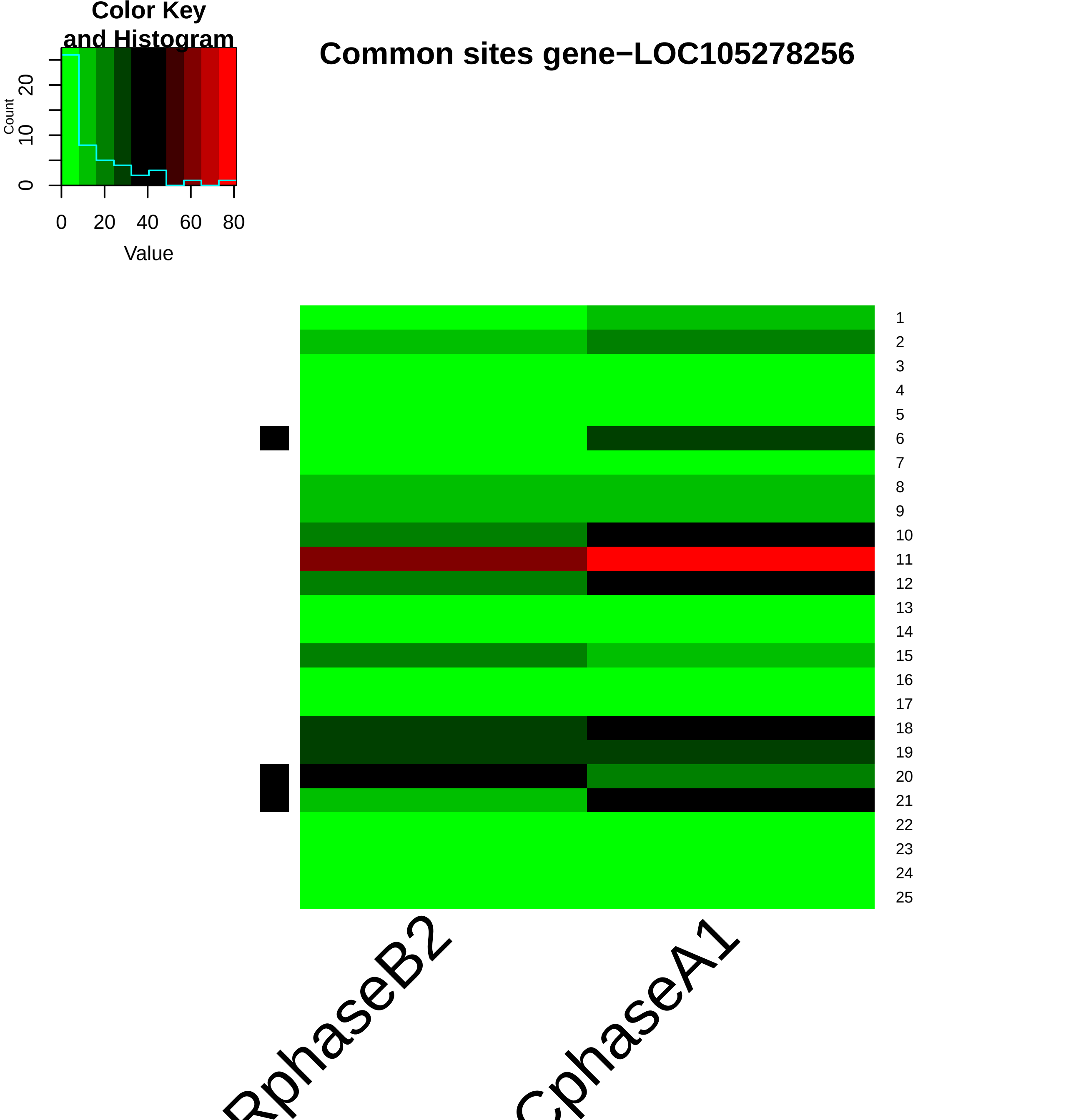

Color Key
and Histogram
Count
0
10
20
0
20
40
60
80
Value
Common sites gene−LOC105278256
RphaseB2
CphaseA1

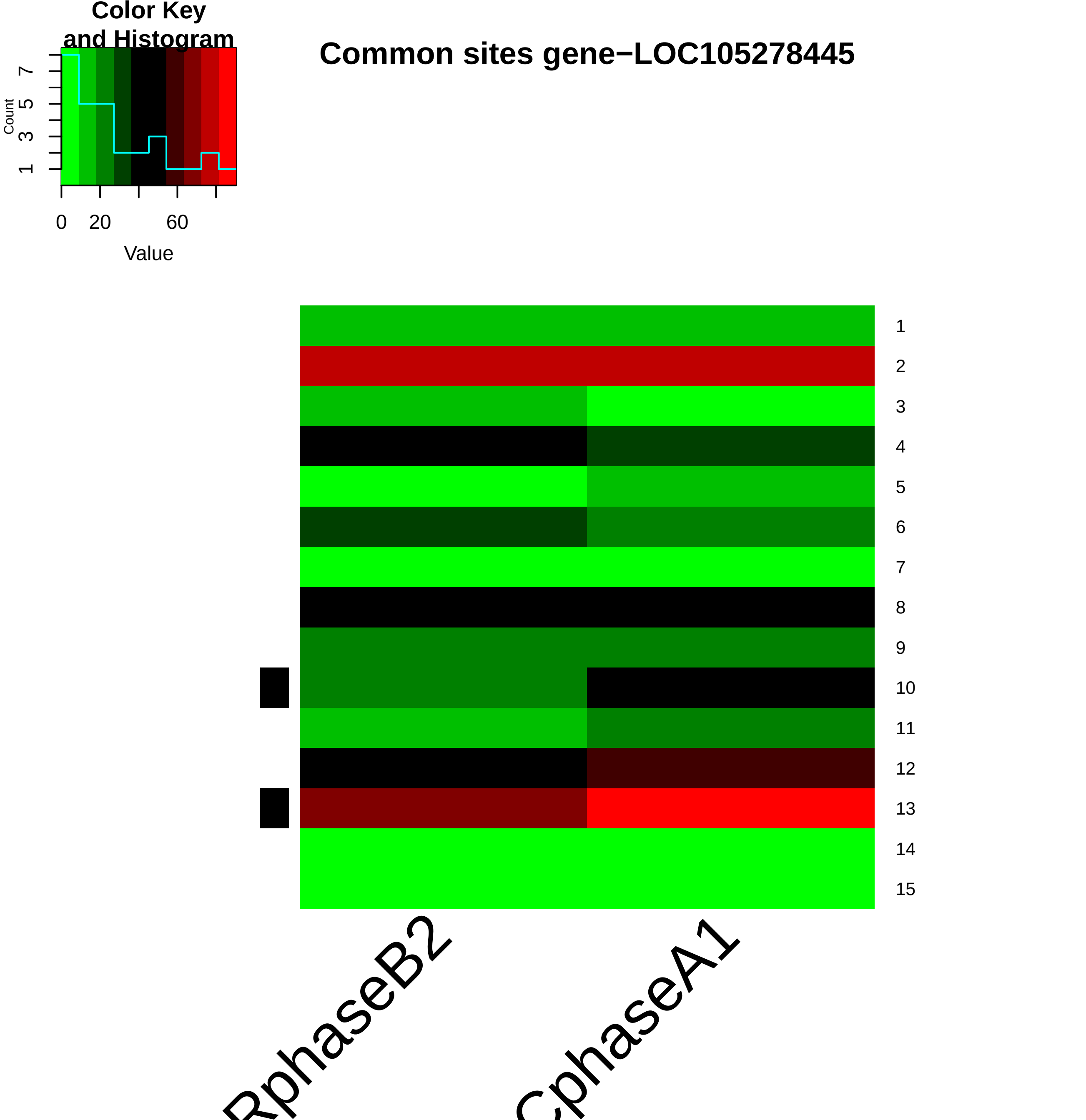

Color Key
and Histogram
Count
1
3
5
7
0
20
60
Value
Common sites gene−LOC105278445
RphaseB2
CphaseA1

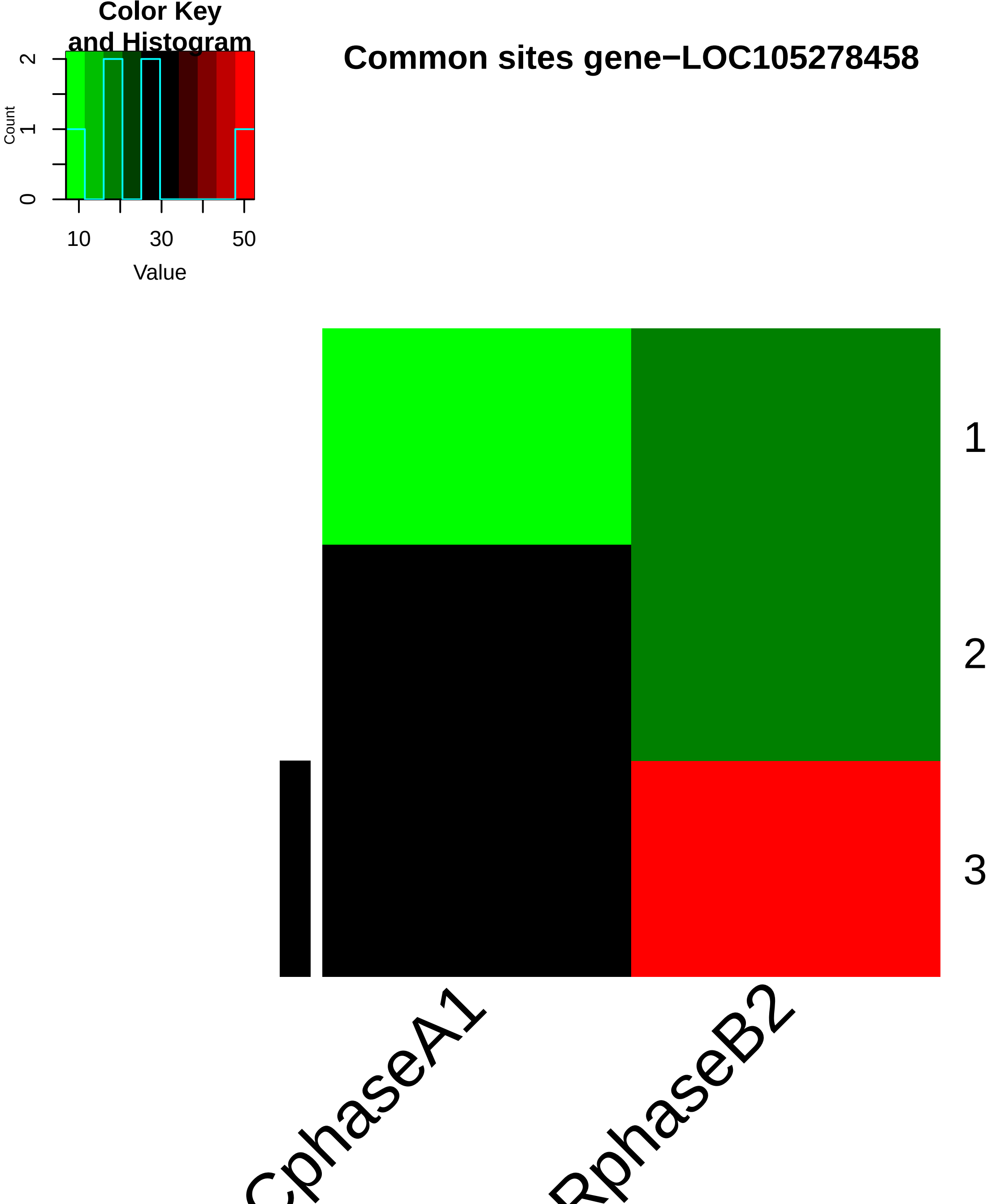
Color Key
and Histogram
Count
0
1
2
10
30
50
Value
Common sites gene−LOC105278458
1
2
3
CphaseA1
RphaseB2

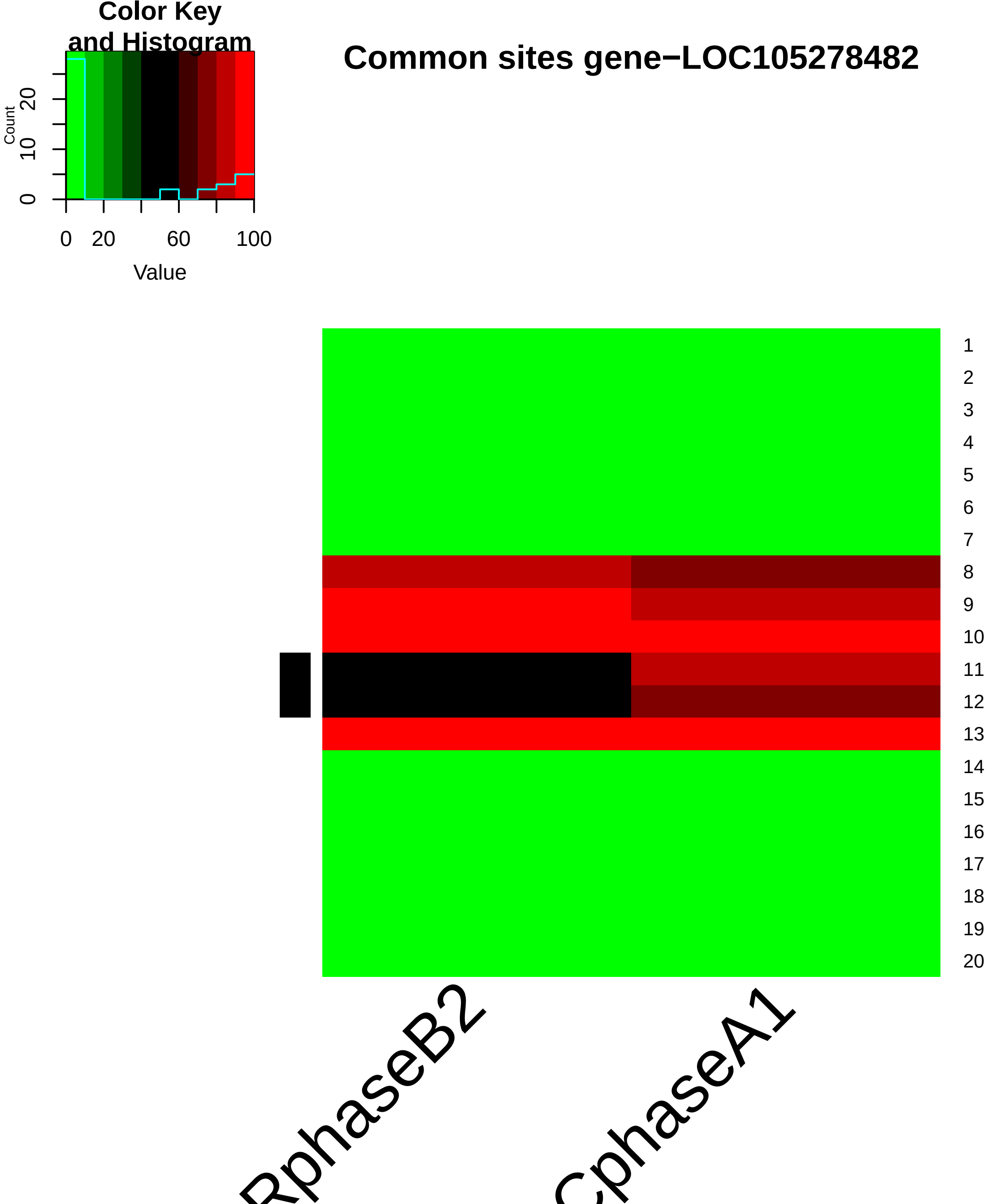
Color Key
and Histogram
Count
0
10
20
0
20
60
100
Value
Common sites gene−LOC105278482
RphaseB2
CphaseA1

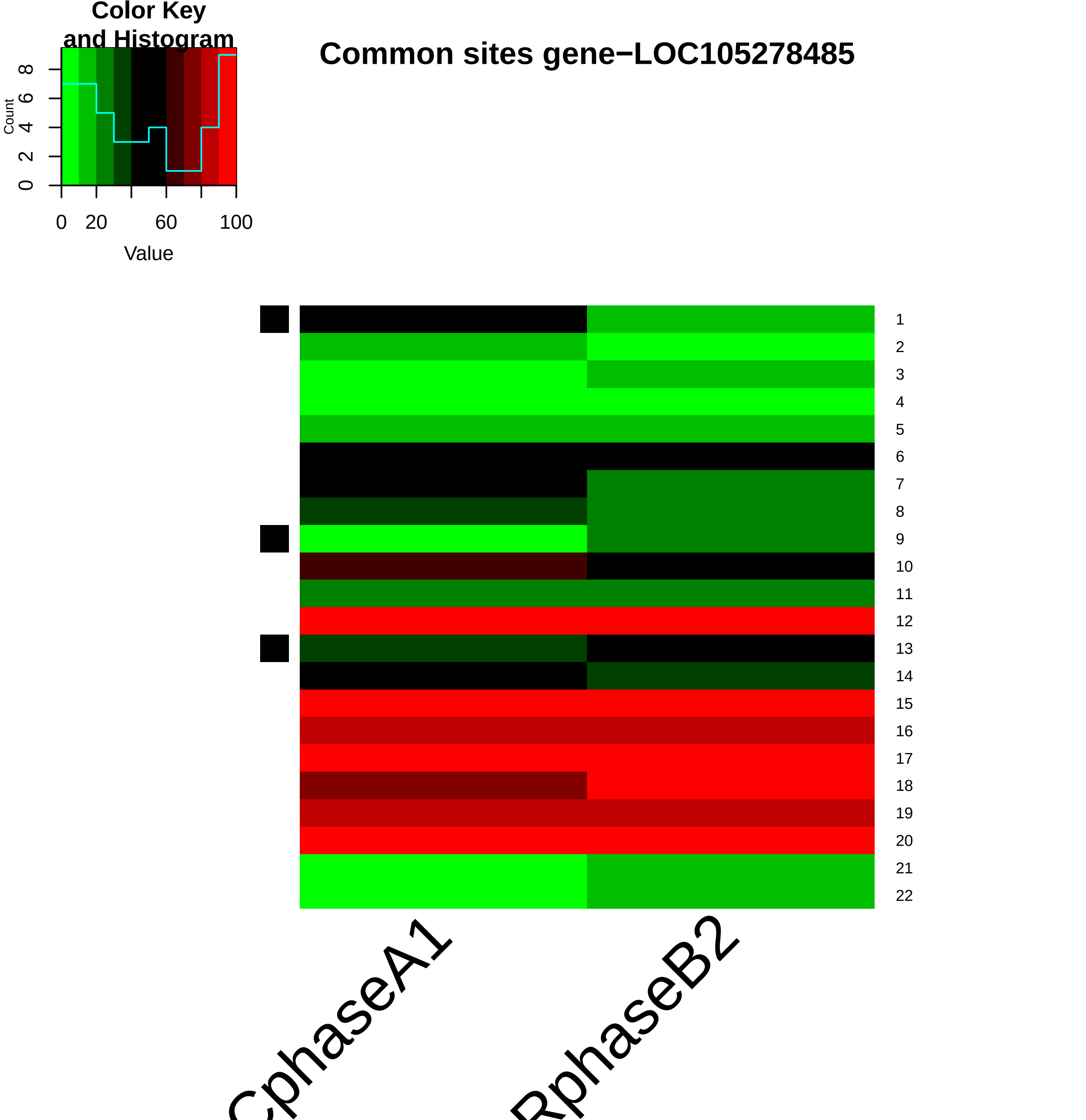

Color Key
and Histogram
Count
0
2
4
6
8
0
20
60
100
Value
Common sites gene−LOC105278485
CphaseA1
RphaseB2

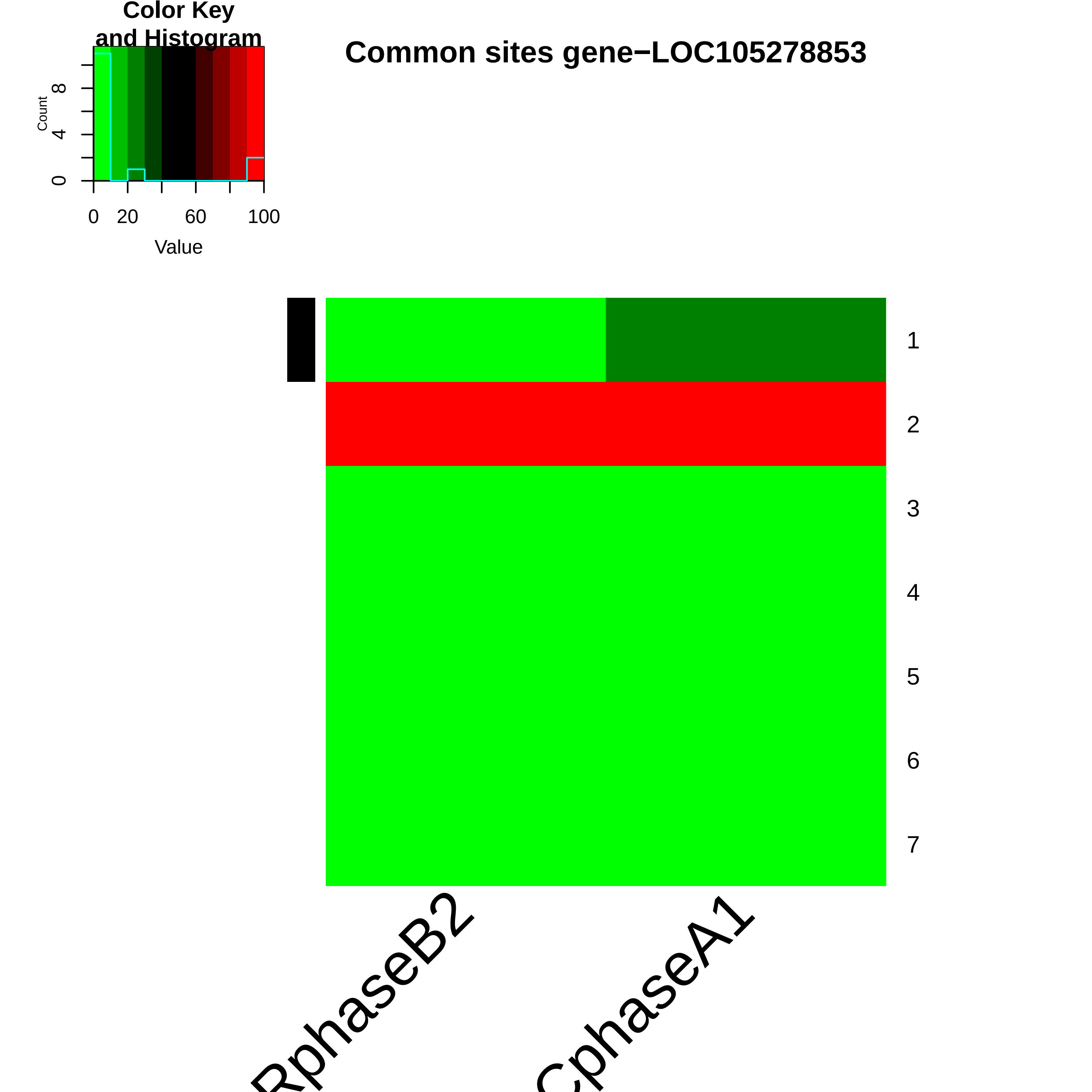

Color Key and Histogram
Count
0
4
8
0
20
60
100
Value
Common sites gene−LOC105278853
1
2
3
4
5
6
7
RphaseB2
CphaseA1

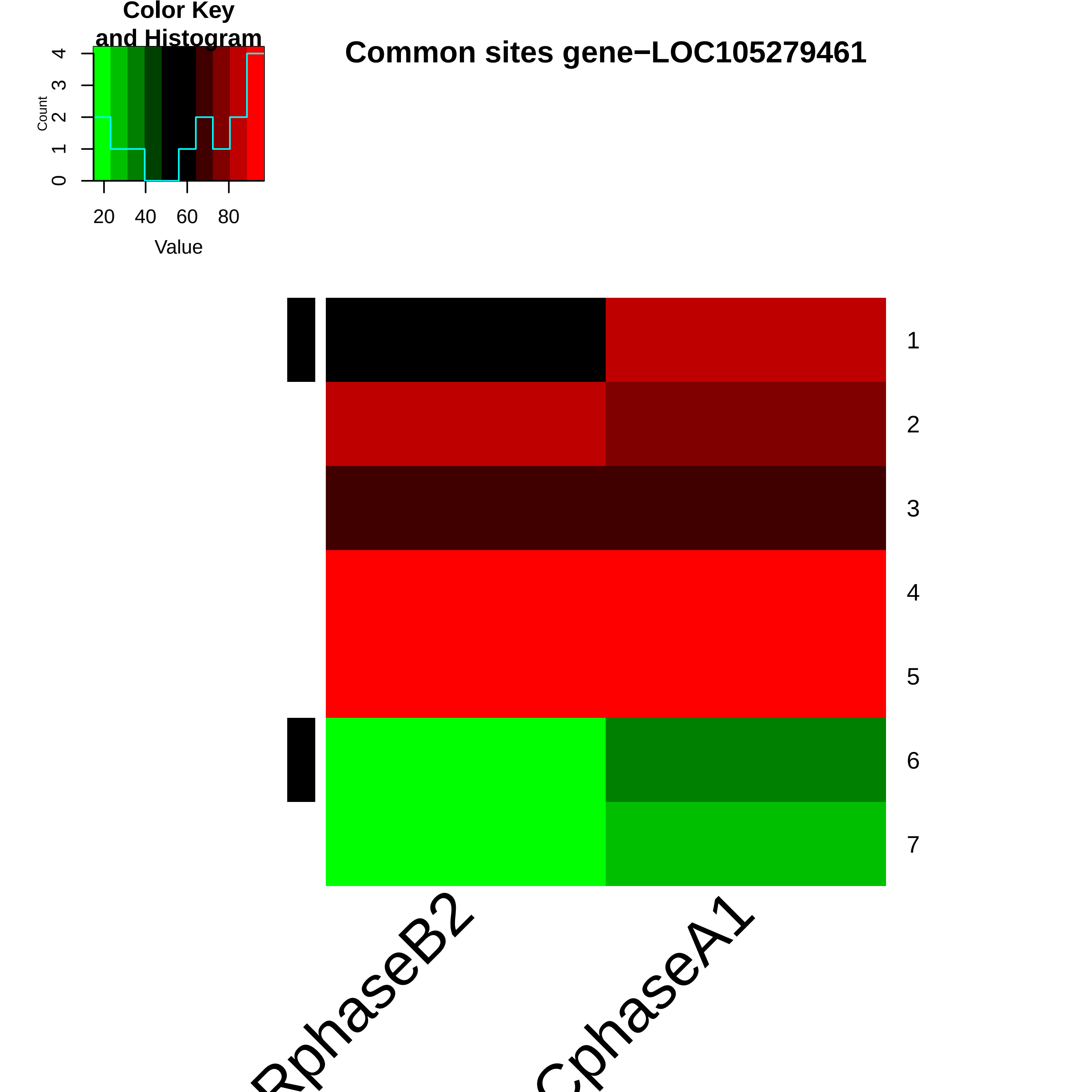

Color Key
and Histogram
Count
0
1
2
3
4
20
40
60
80
Value
Common sites gene−LOC105279461
1
2
3
4
5
6
7
RphaseB2
CphaseA1

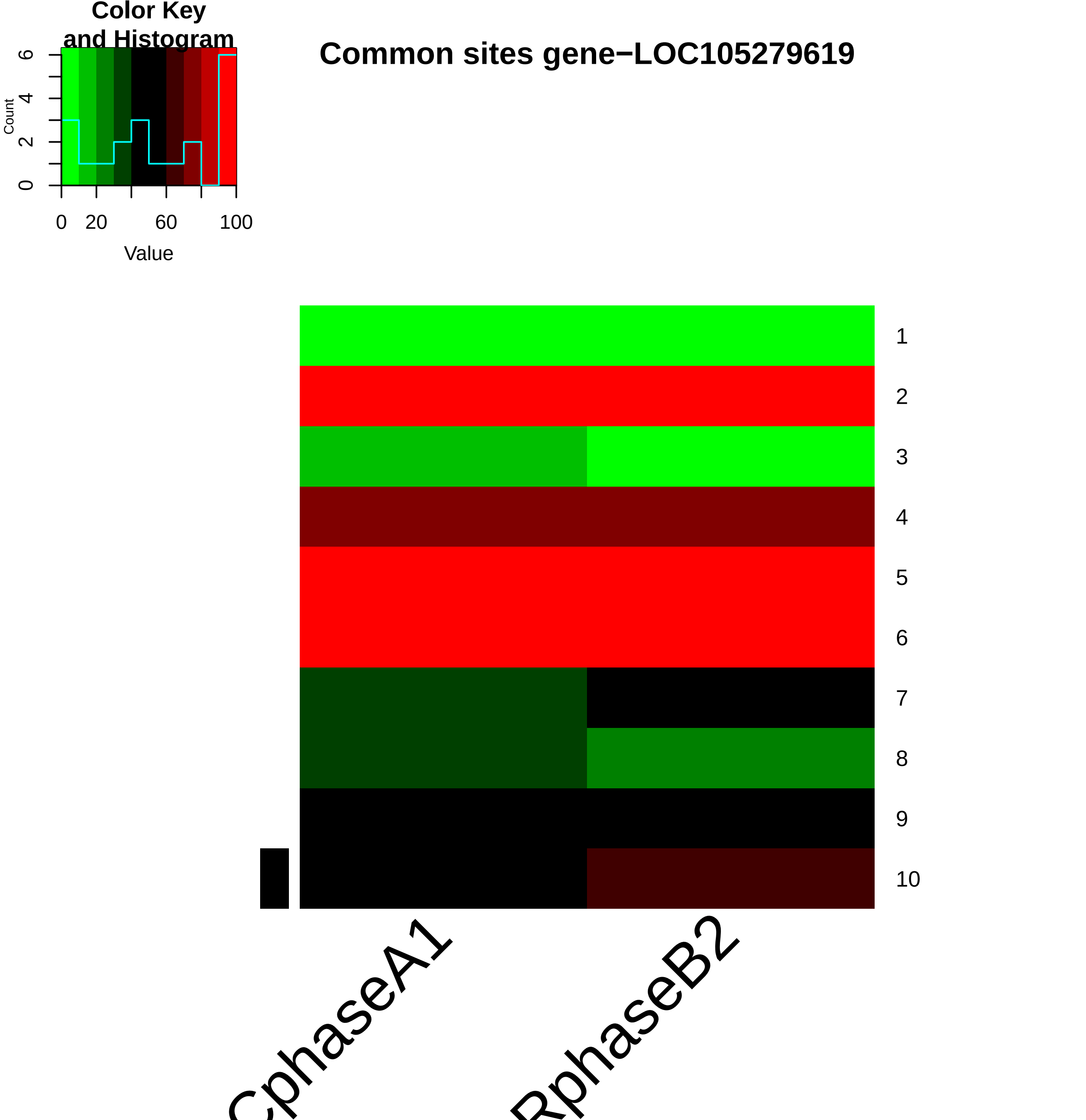

Color Key
and Histogram
Count
0
2
4
6
0
20
60
100
Value
Common sites gene−LOC105279619
1
2
3
4
5
6
7
8
9
10
CphaseA1
RphaseB2

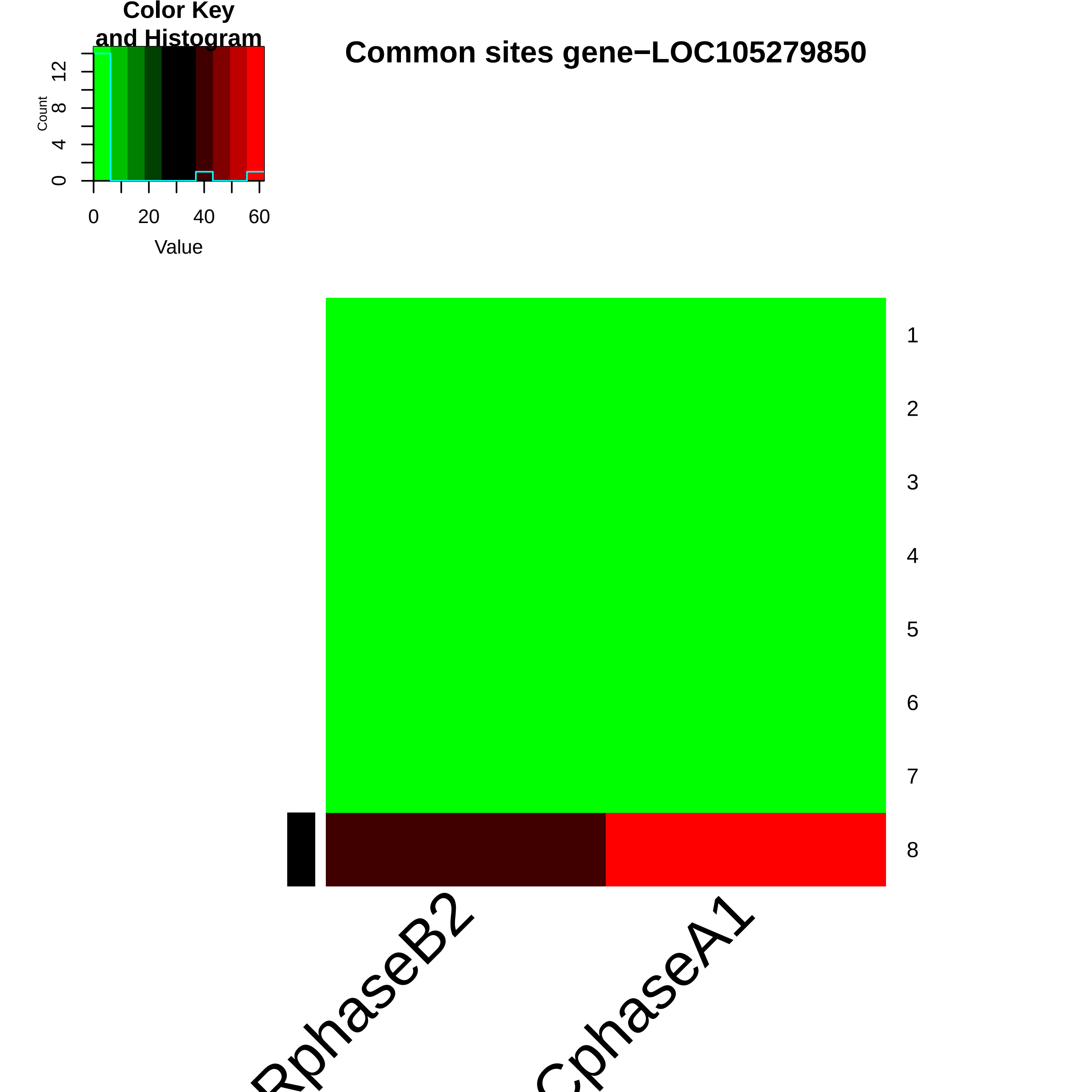

Color Key
and Histogram
Count
0
4
8
12
0
20
40
60
Value
Common sites gene−LOC105279850
1
2
3
4
5
6
7
8
RphaseB2
CphaseA1

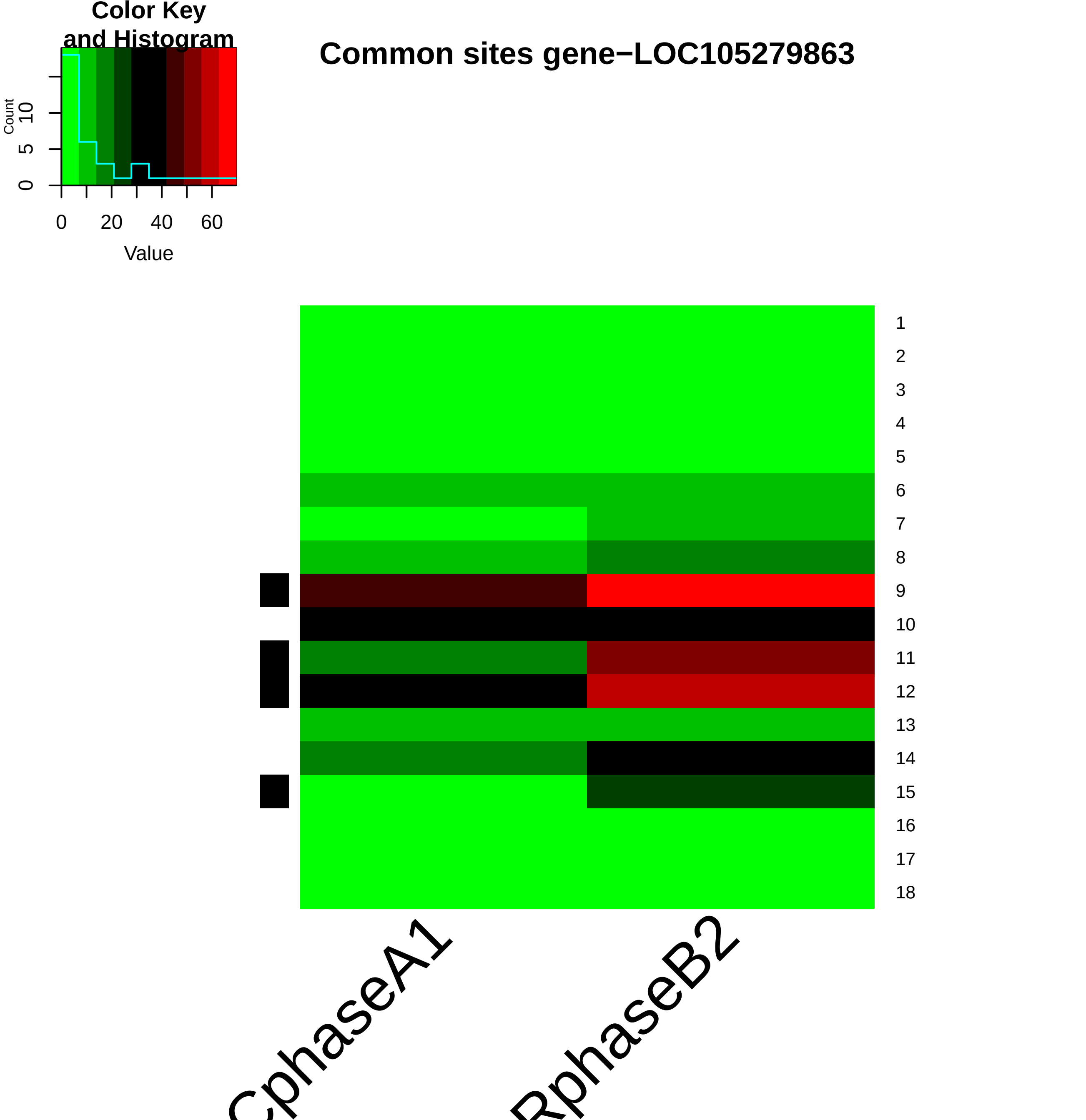

Color Key
and Histogram
Count
0
5
10
0
20
40
60
Value
Common sites gene−LOC105279863
CphaseA1
RphaseB2

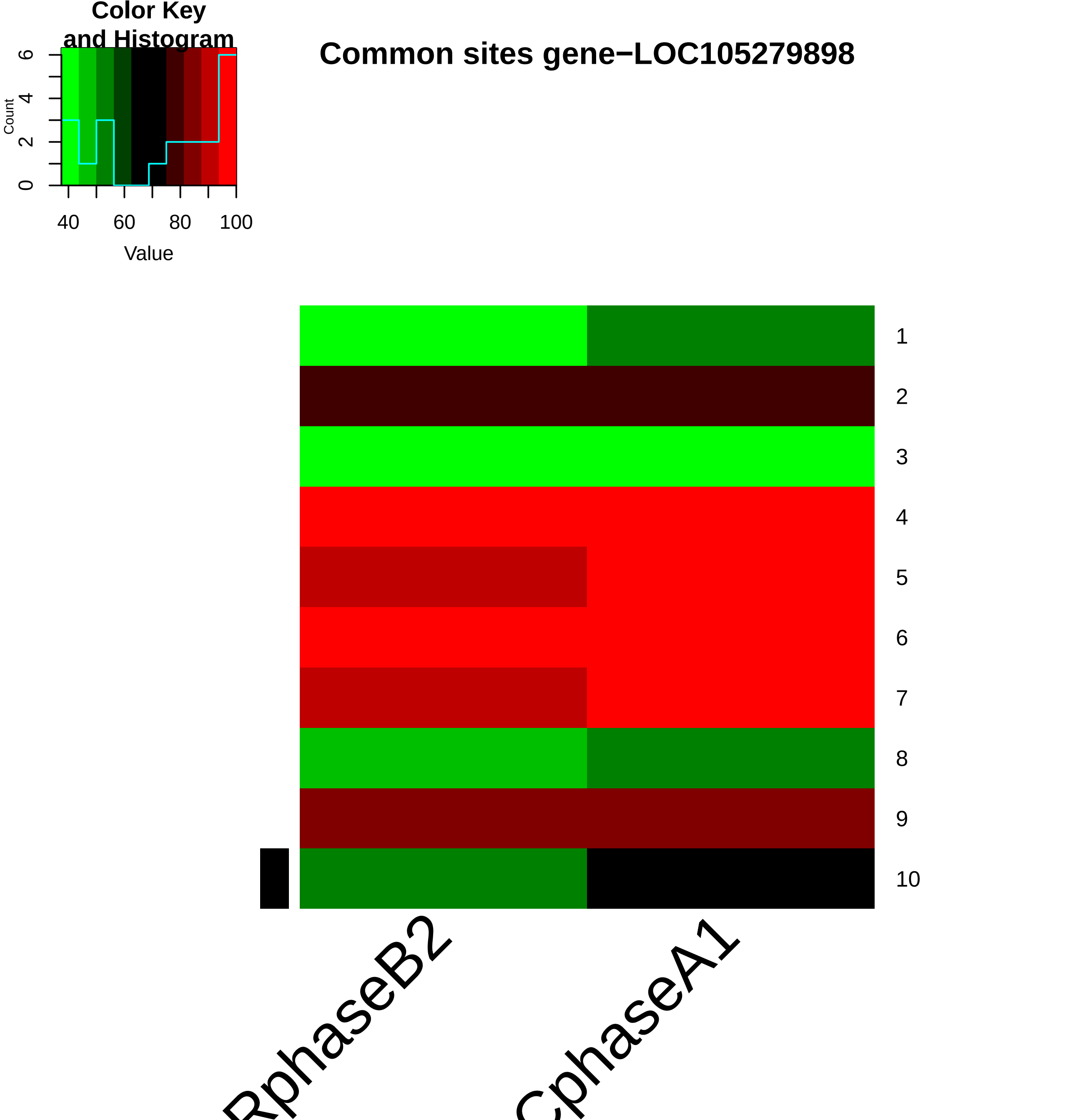
Color Key
and Histogram
Count
Value
0
2
4
6
40
60
80
100
Common sites gene−LOC105279898
1
2
3
4
5
6
7
8
9
10
RphaseB2
CphaseA1

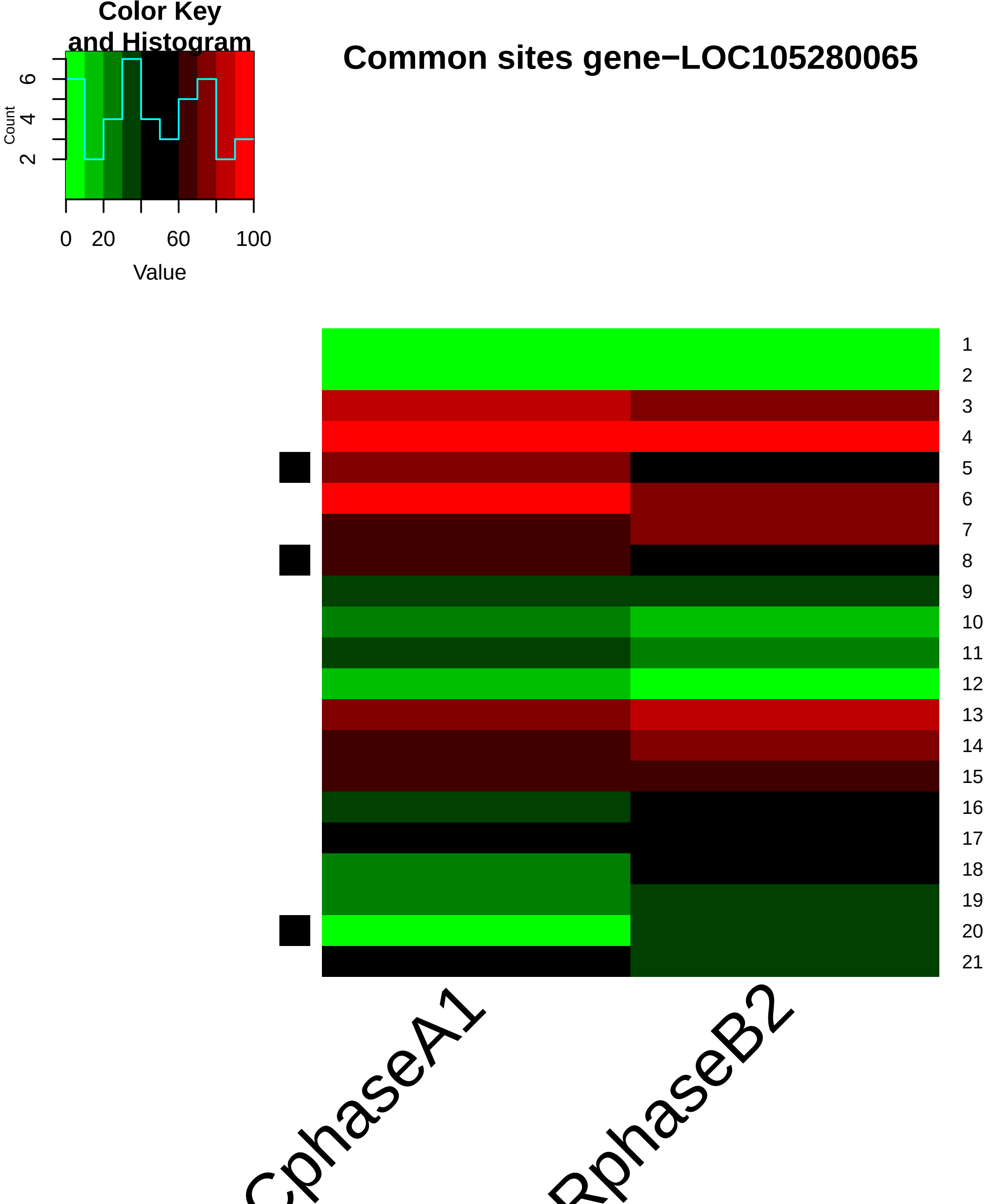

Color Key
and Histogram
Count
2
4
6
0
20
60
100
Value
Common sites gene−LOC105280065
CphaseA1
RphaseB2

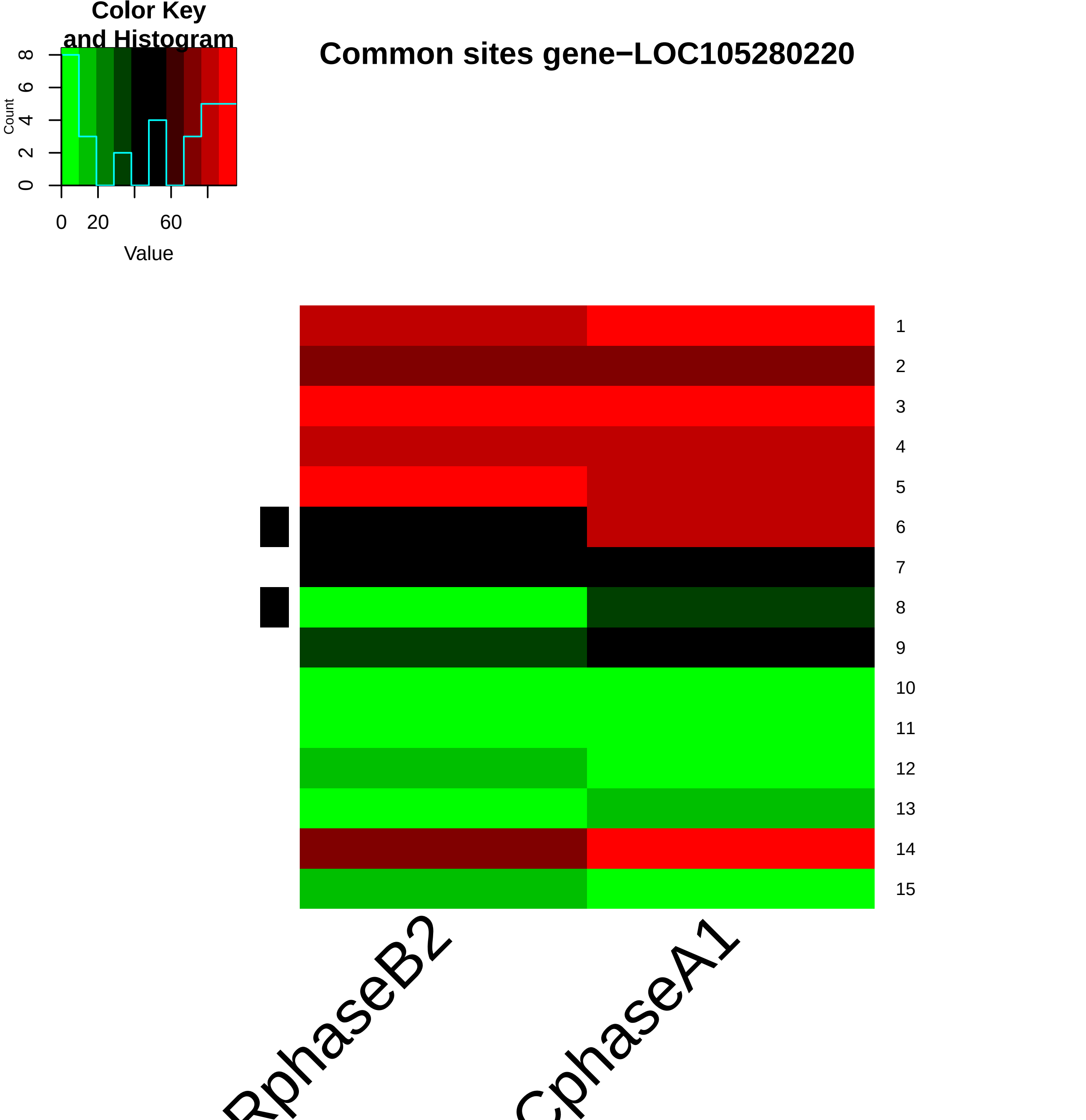

Color Key
and Histogram
Count
0
2
4
6
8
0
20
60
Value
Common sites gene−LOC105280220
RphaseB2
CphaseA1

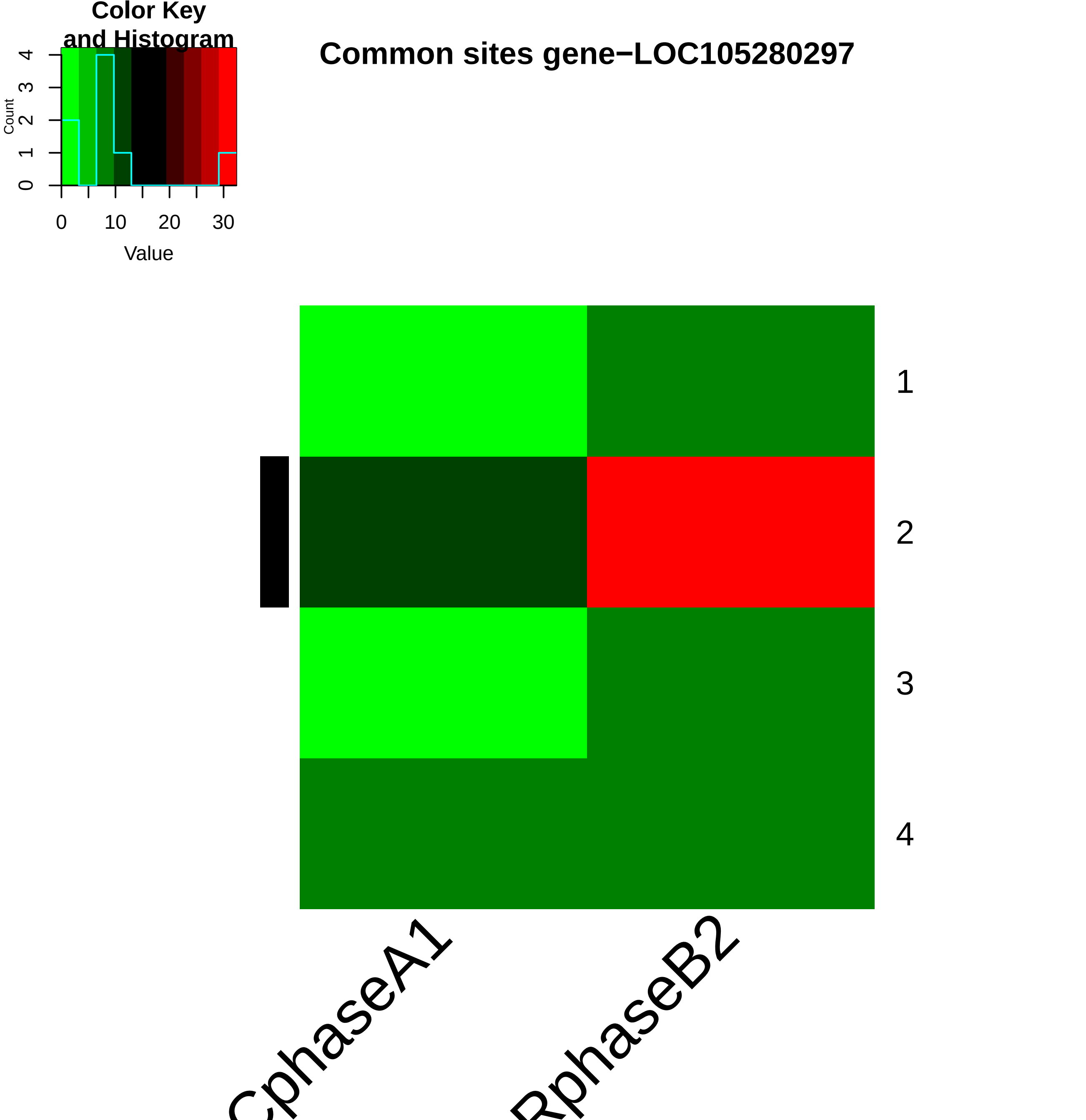
Color Key
and Histogram
Count
Value
0
10
20
30
0
1
2
3
4
Common sites gene−LOC105280297
1
2
3
4
CphaseA1
RphaseB2

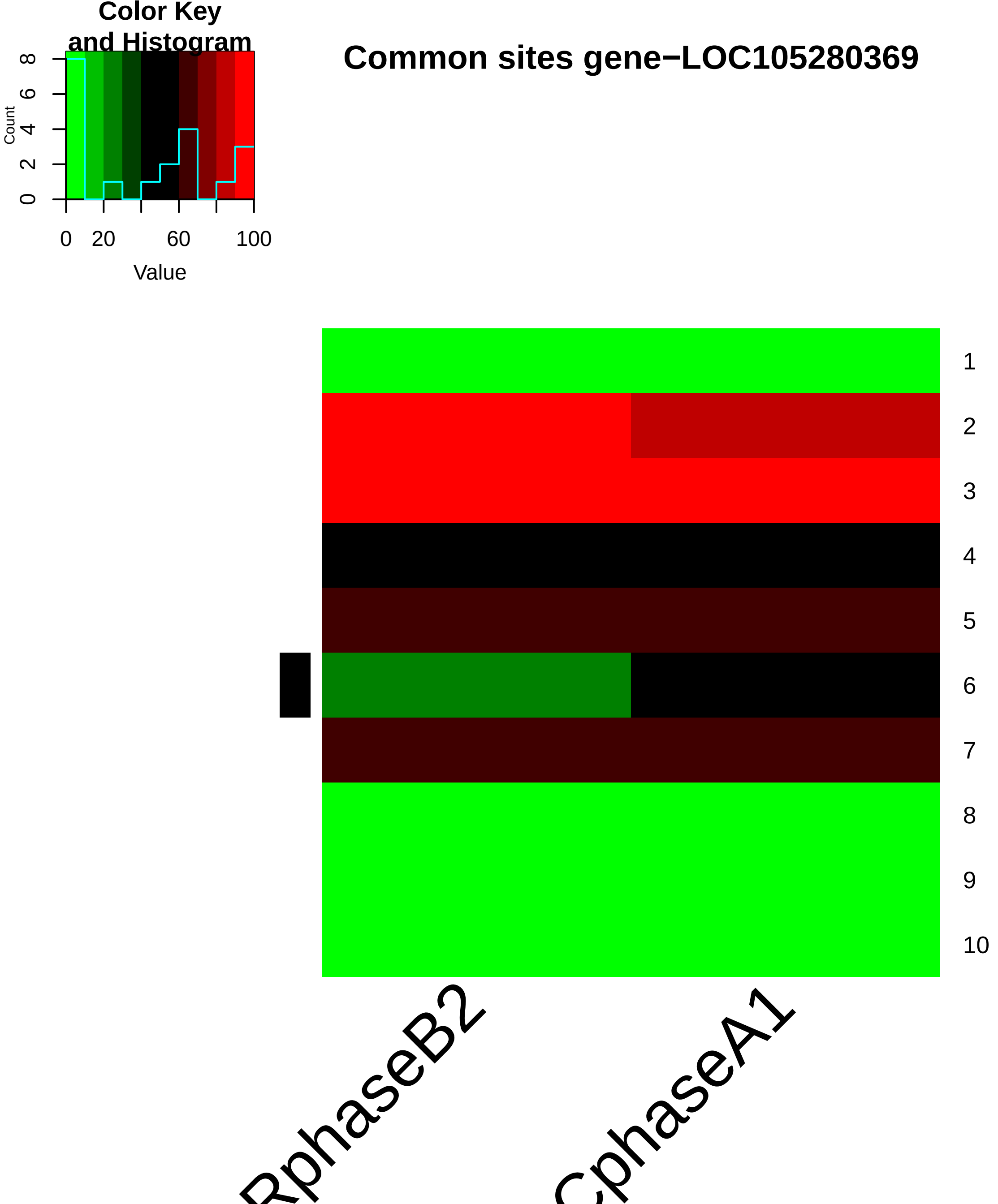

Color Key
and Histogram
Count
0
2
4
6
8
0
20
60
100
Value
Common sites gene−LOC105280369
1
2
3
4
5
6
7
8
9
10
RphaseB2
CphaseA1

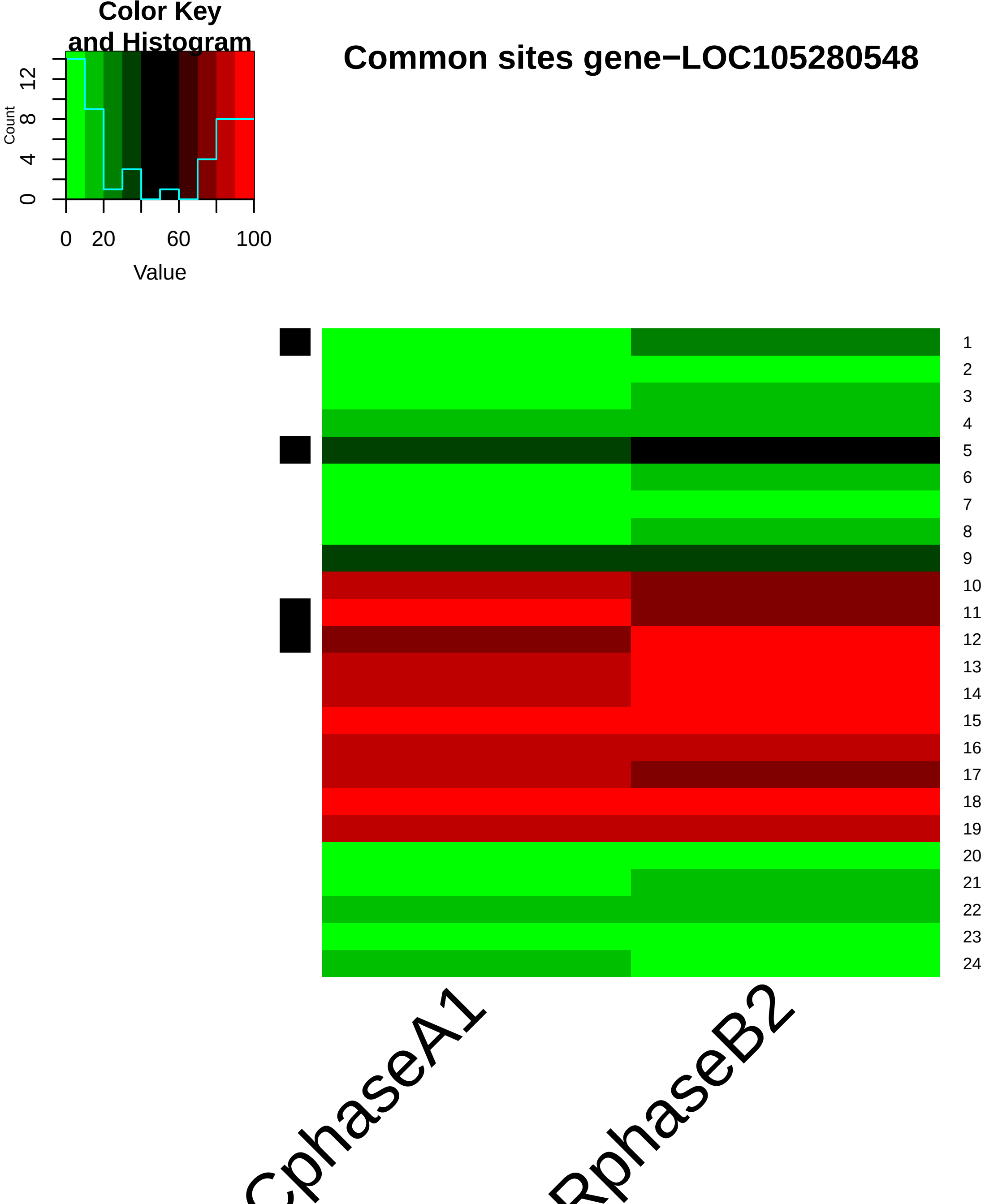

Color Key
and Histogram
Count
0
4
8
12
0
20
60
100
Value
Common sites gene−LOC105280548
CphaseA1
RphaseB2

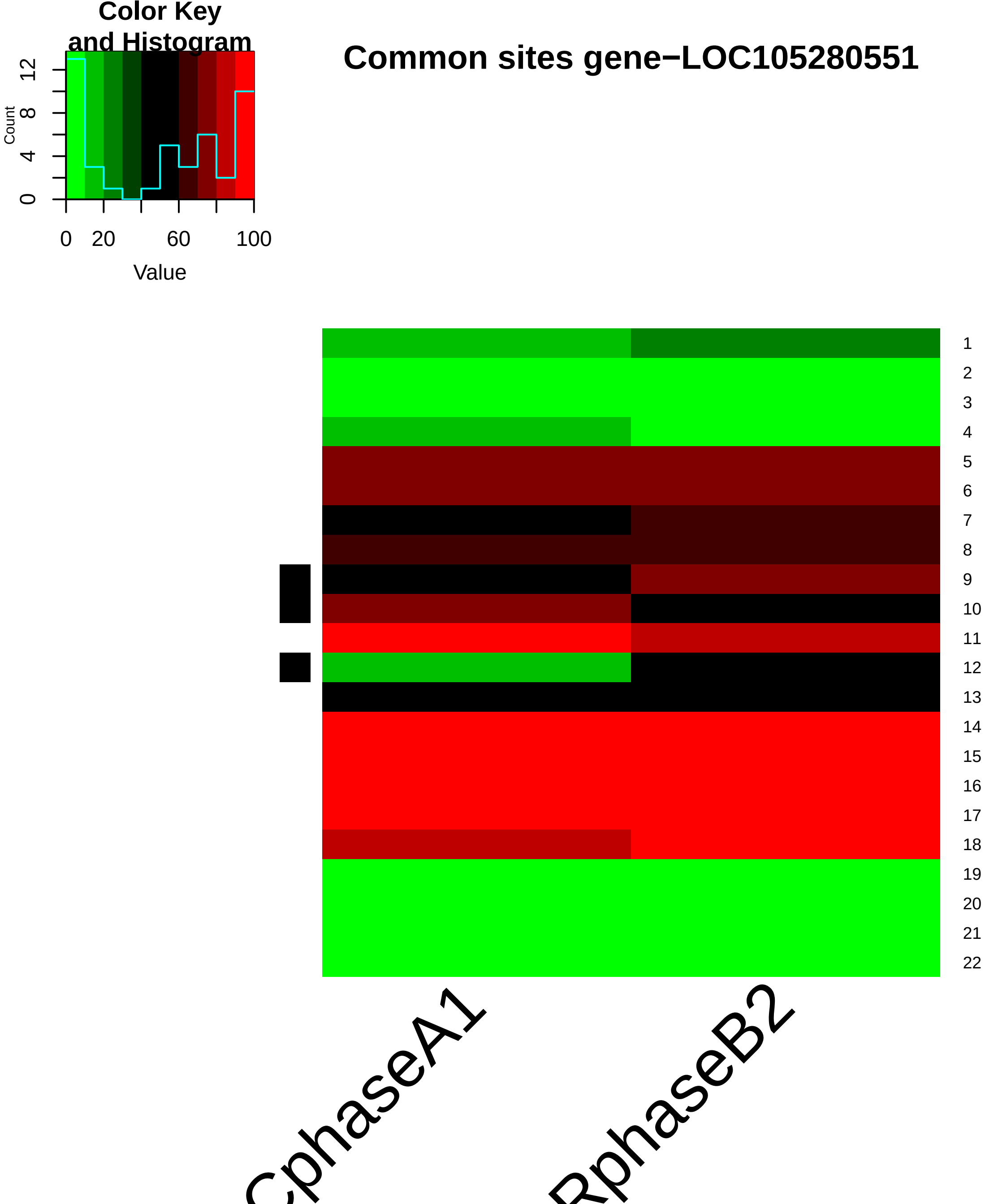
Color Key
and Histogram
Count
0
4
8
12
0
20
60
100
Value
Common sites gene−LOC105280551
CphaseA1
RphaseB2

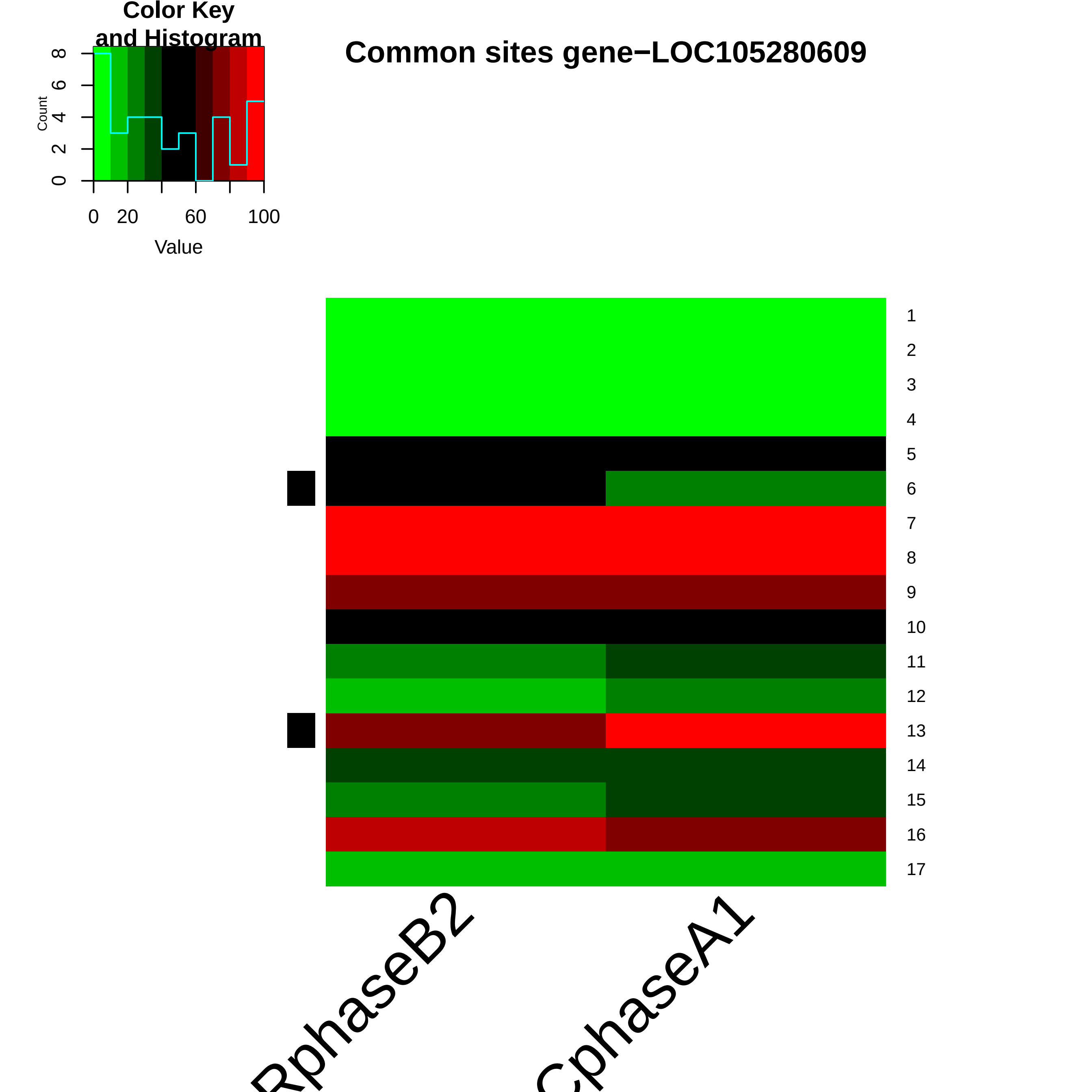

Color Key
and Histogram
Count
0
2
4
6
8
0
20
60
100
Value
Common sites gene−LOC105280609
RphaseB2
CphaseA1

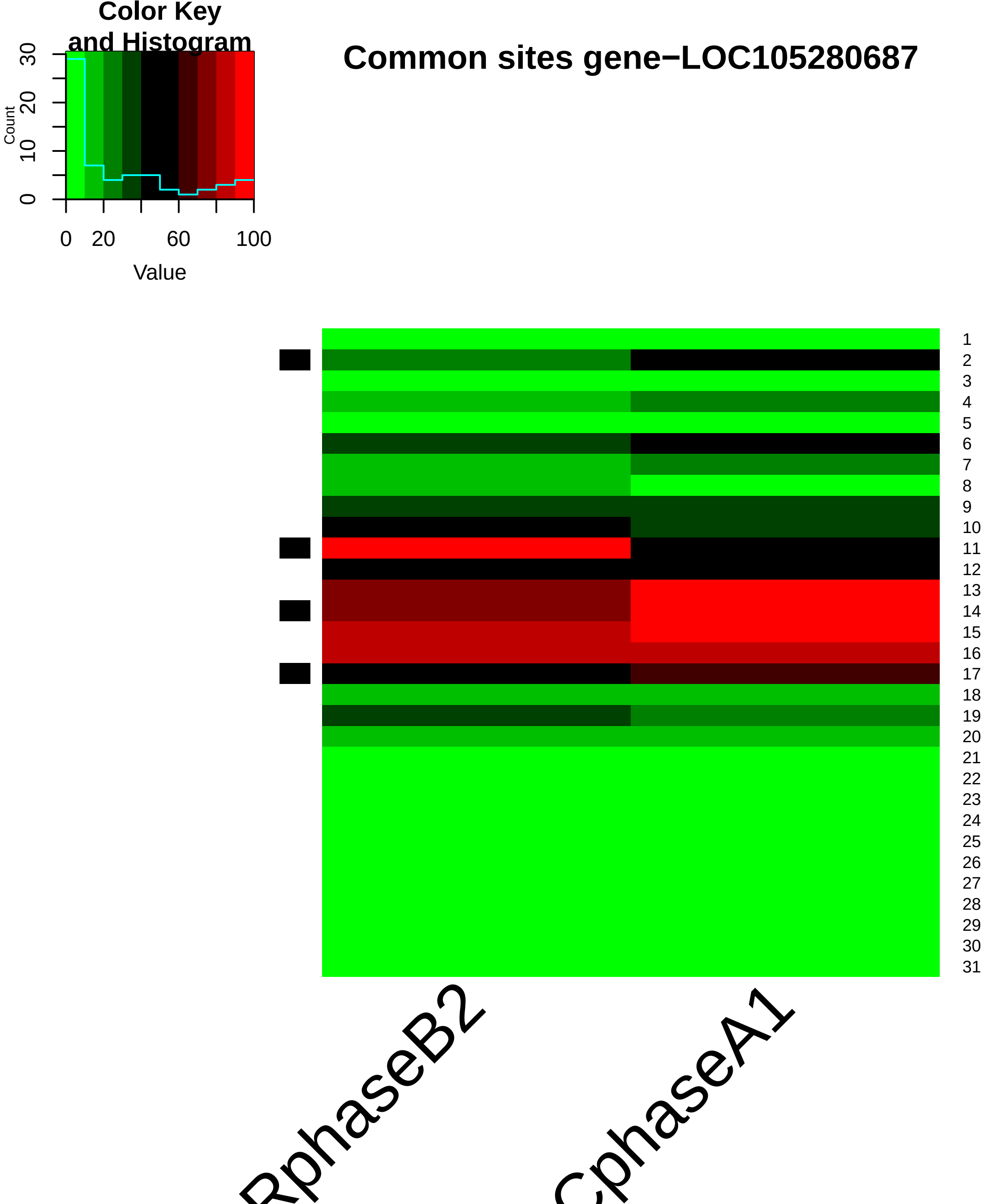

Color Key and Histogram
Count
0
10
20
30
0
20
60
100
Value
Common sites gene−LOC105280687
RphaseB2
CphaseA1

#### Common sites gene−LOC105280924

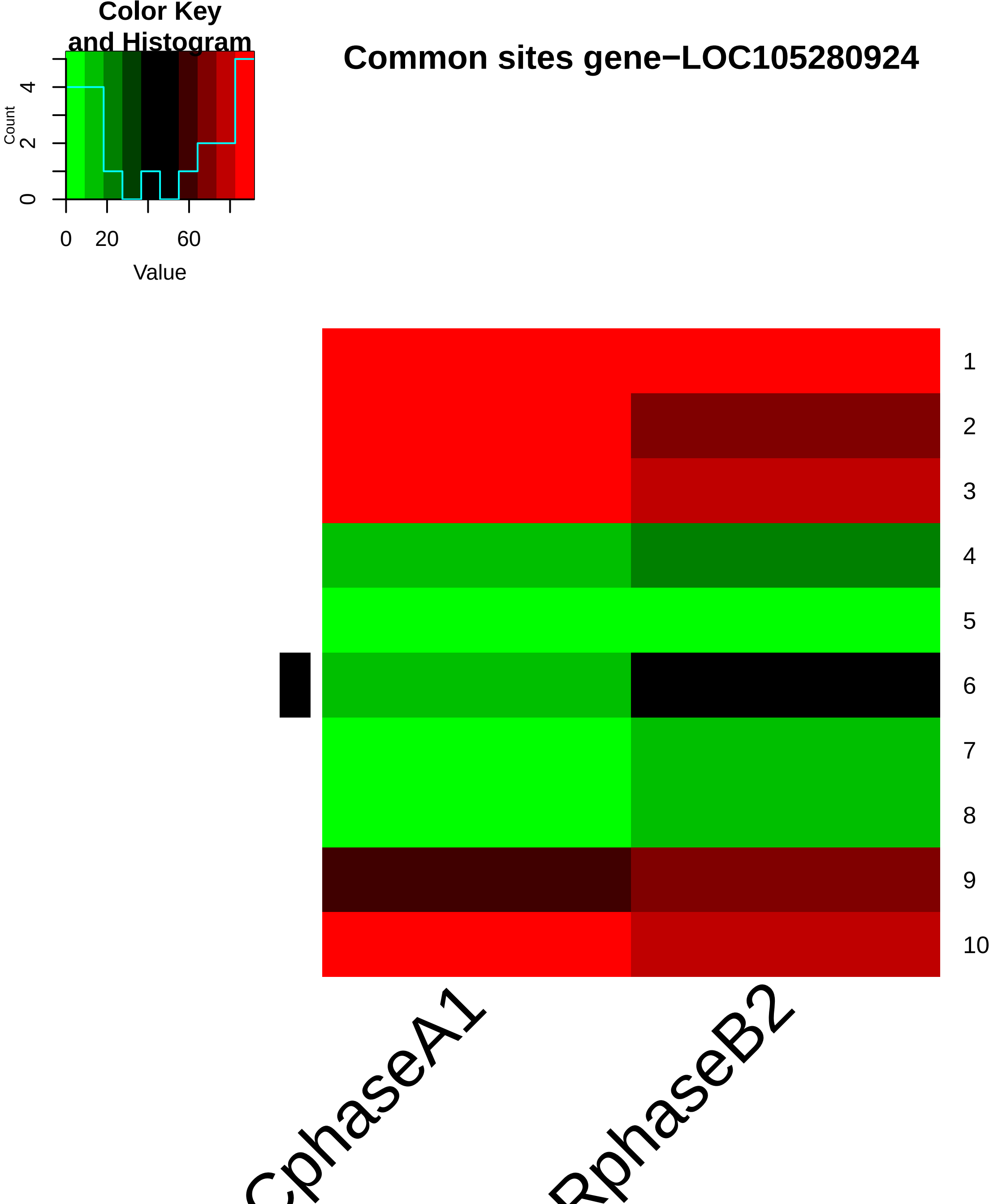

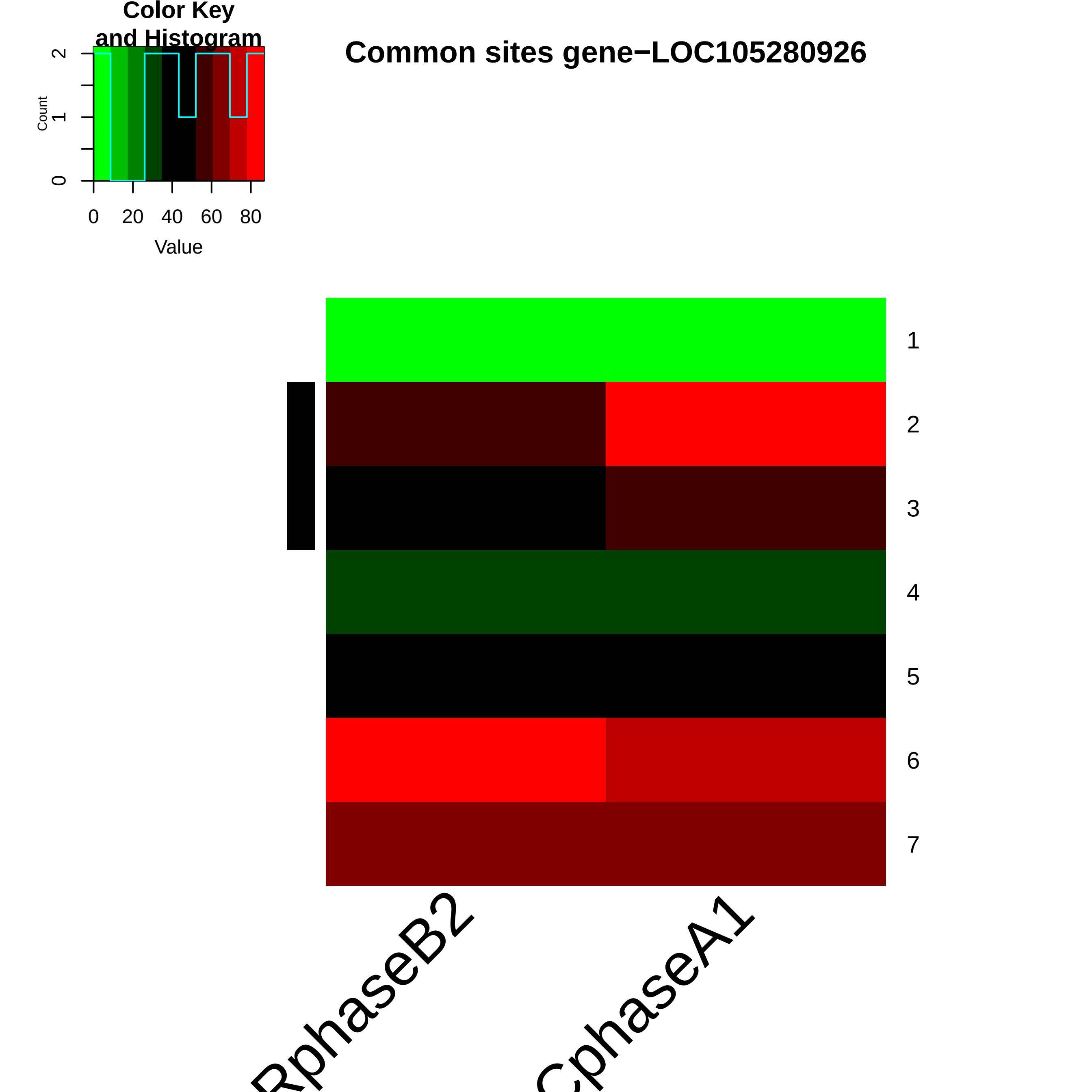

Color Key
and Histogram
Count
0
1
2
0
20
40
60
80
Value
Common sites gene−LOC105280926
1
2
3
4
5
6
7
RphaseB2
CphaseA1

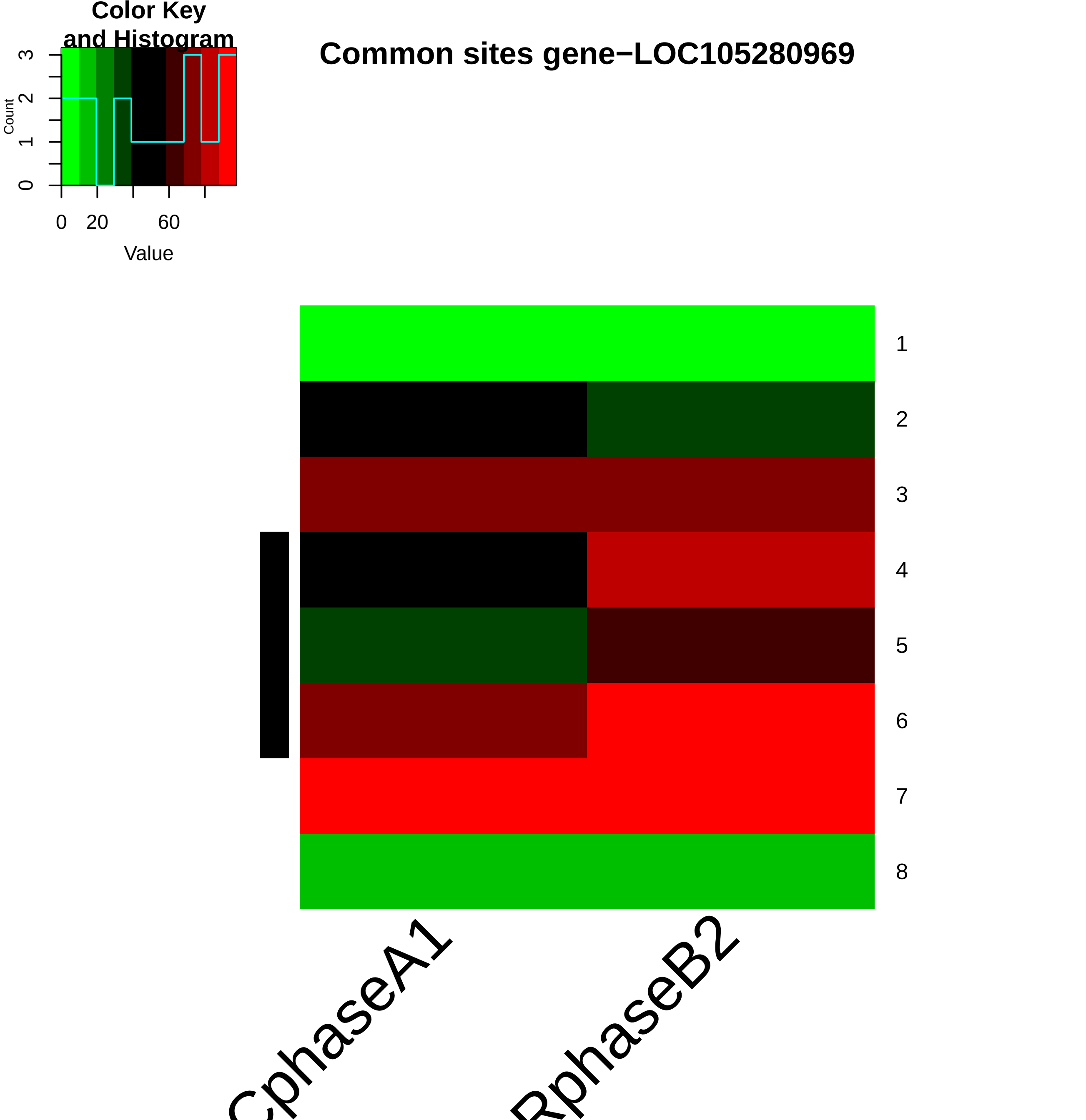
Color Key
and Histogram
Count
0
1
2
3
0
20
60
Value
Common sites gene−LOC105280969
1
2
3
4
5
6
7
8
CphaseA1
RphaseB2

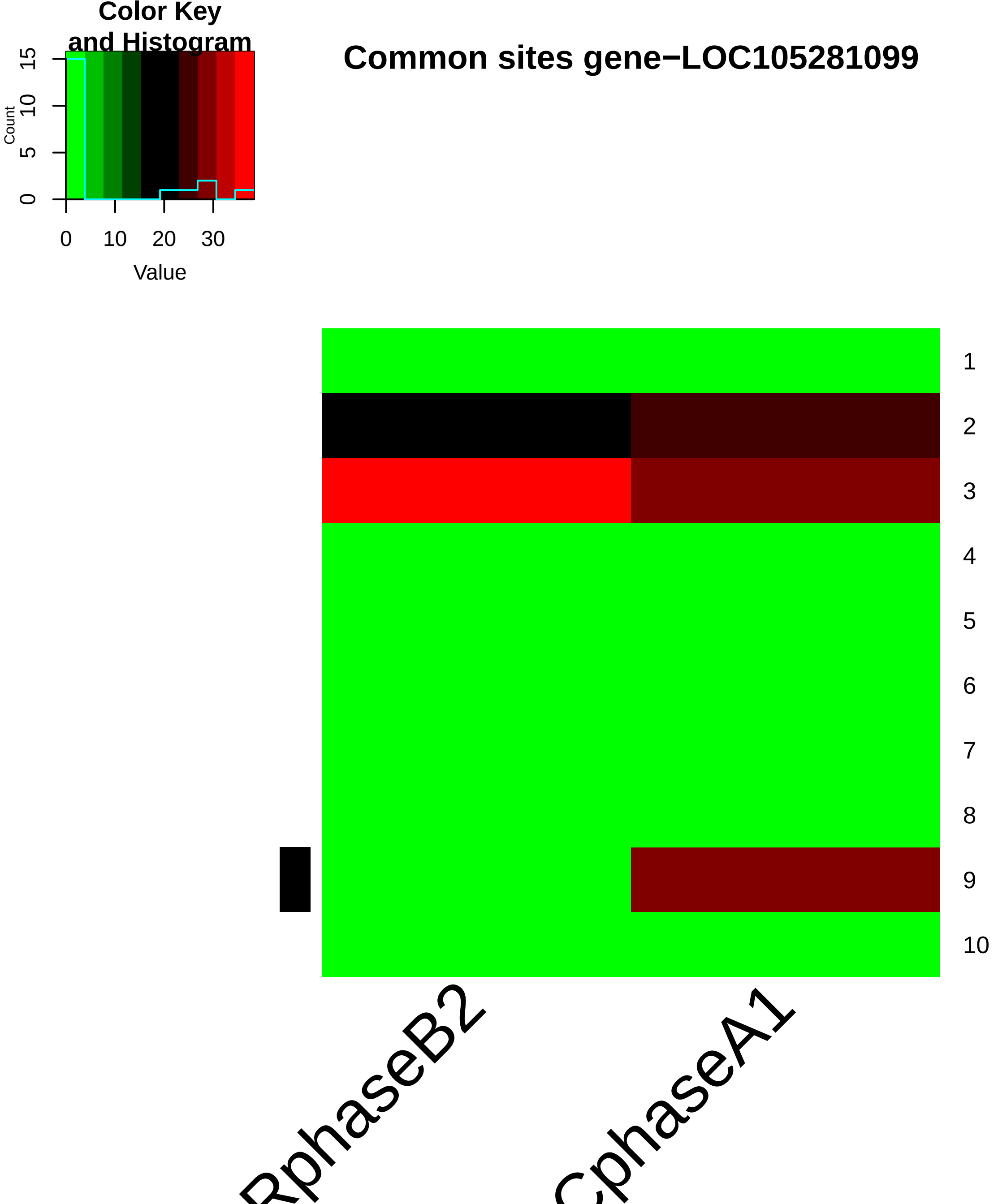
Color Key
and Histogram
Count
0
5
10
15
0
10
20
30
Value
Common sites gene−LOC105281099
1
2
3
4
5
6
7
8
9
10
RphaseB2
CphaseA1

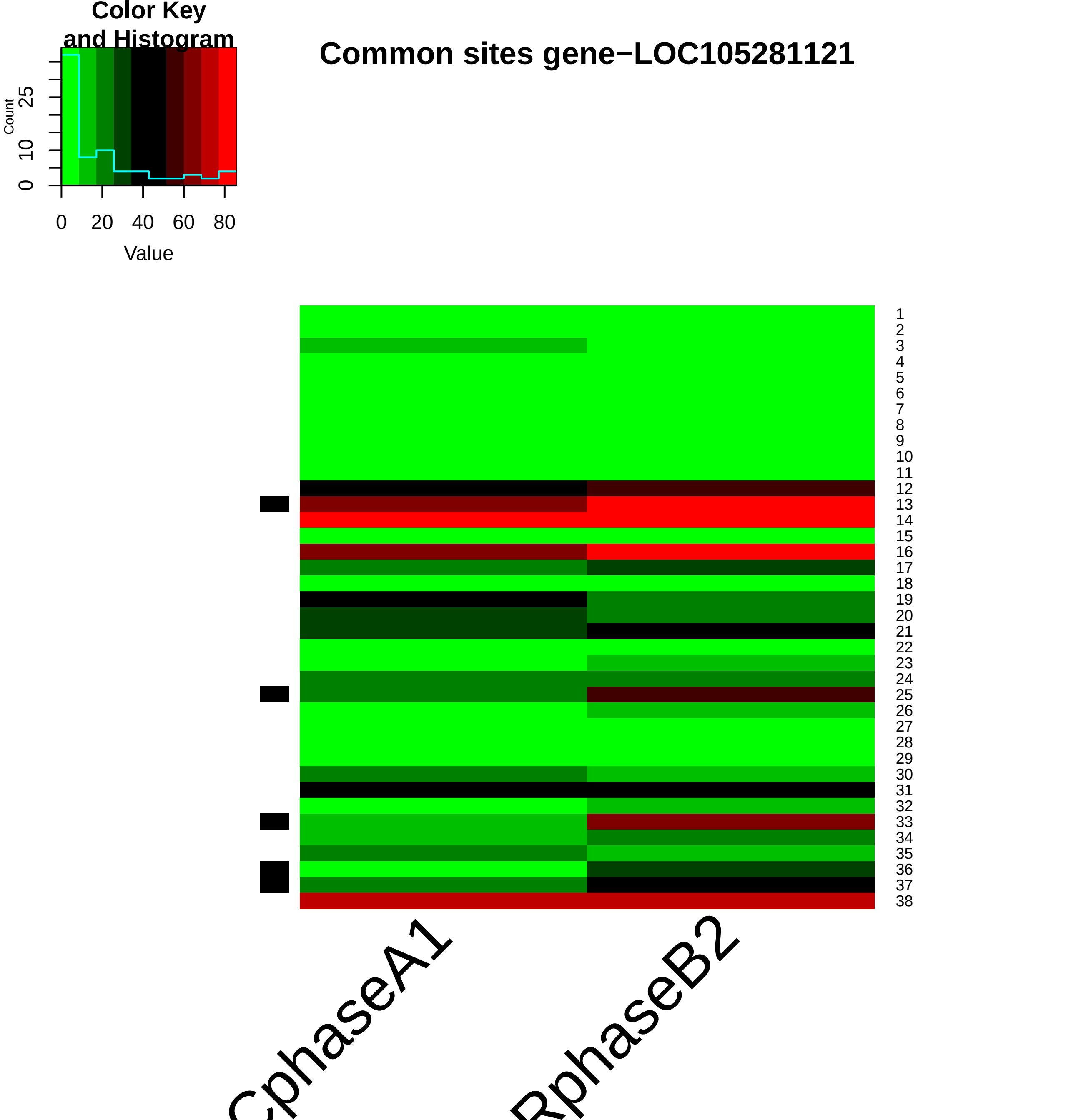

Color Key
and Histogram
Count
0
10
25
0
20
40
60
80
Value
Common sites gene−LOC105281121

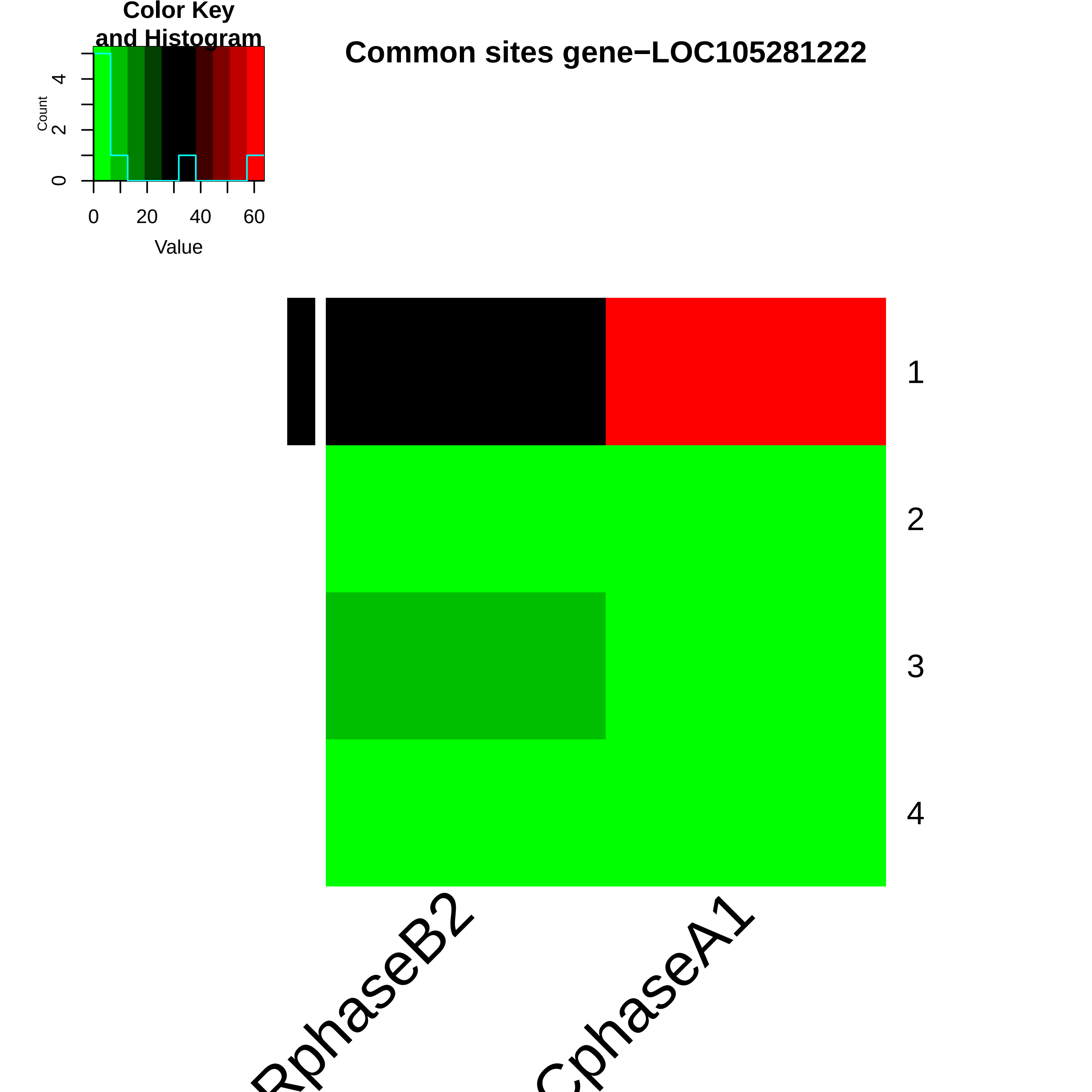
Color Key
and Histogram
Count
0
2
4
0
20
40
60
Value
Common sites gene−LOC105281222
1
2
3
4

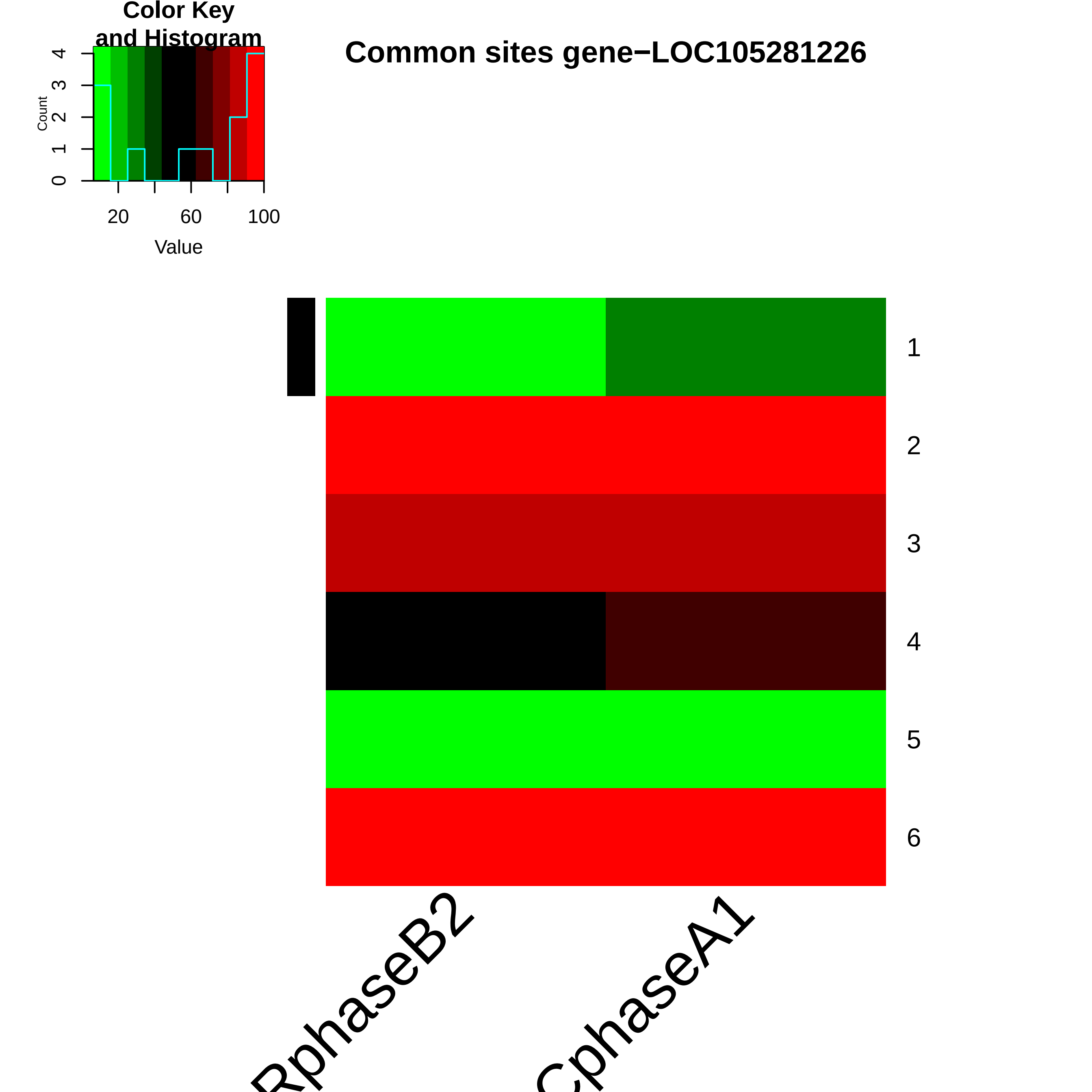

Color Key
and Histogram
Count
0
1
2
3
4
20
60
100
Value
Common sites gene−LOC105281226
1
2
3
4
5
6
RphaseB2
CphaseA1

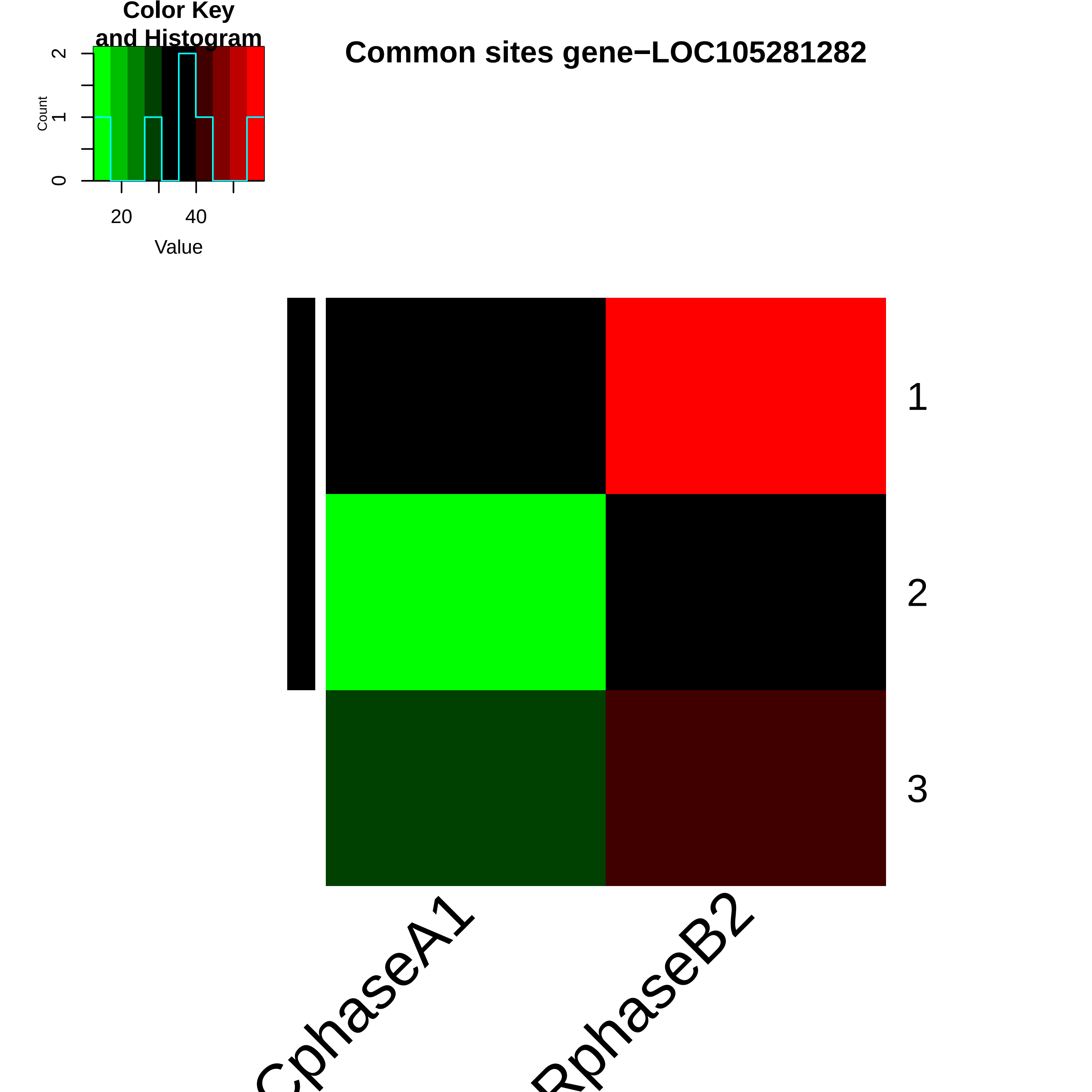
Color Key
and Histogram
Count
0
1
2
20
40
Value
Common sites gene−LOC105281282
1
2
3
CphaseA1
RphaseB2

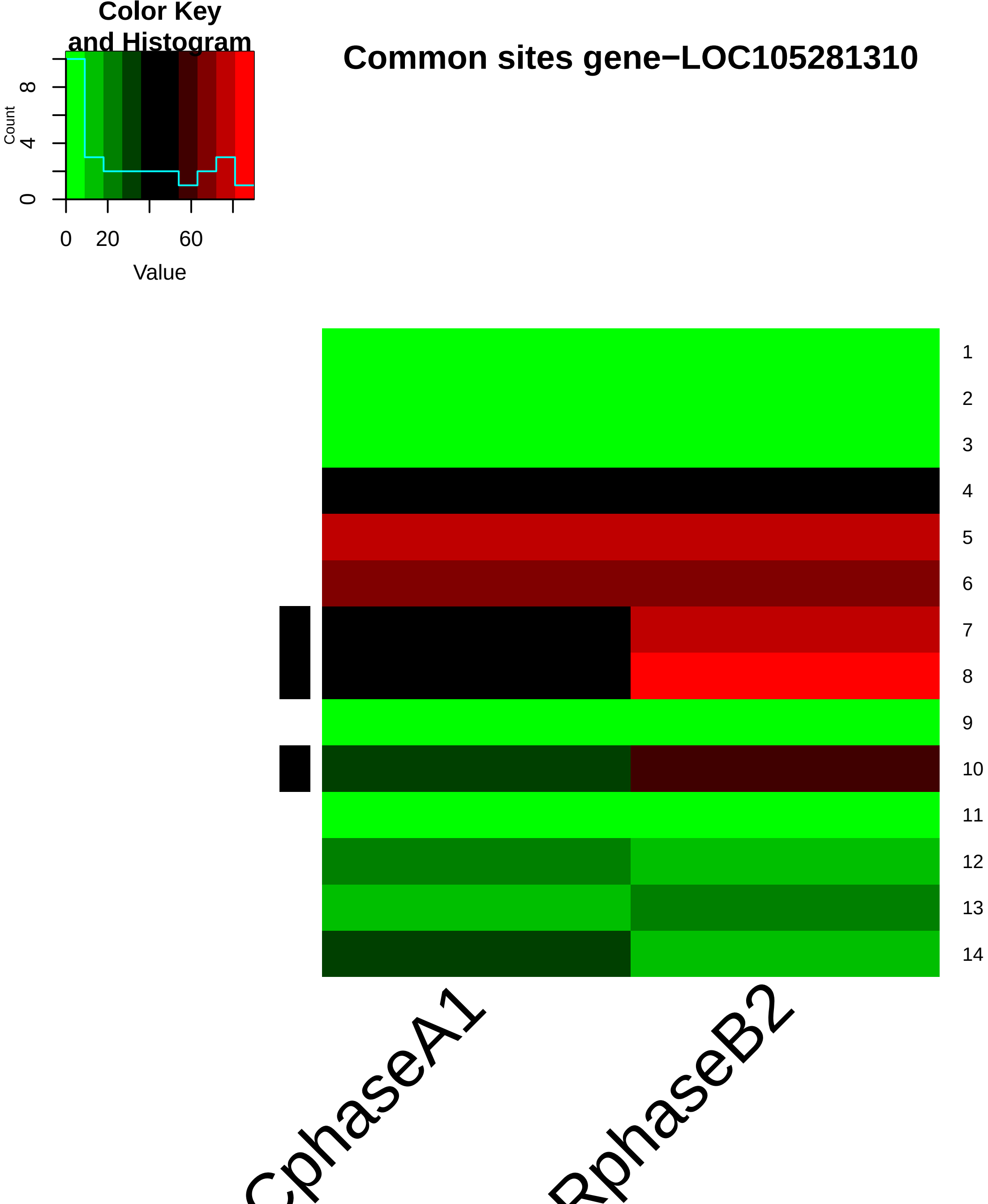
Color Key
and Histogram
Count
0
4
8
0
20
60
Value
Common sites gene−LOC105281310
1
2
3
4
5
6
7
8
9
10
11
12
13
14
CphaseA1
RphaseB2

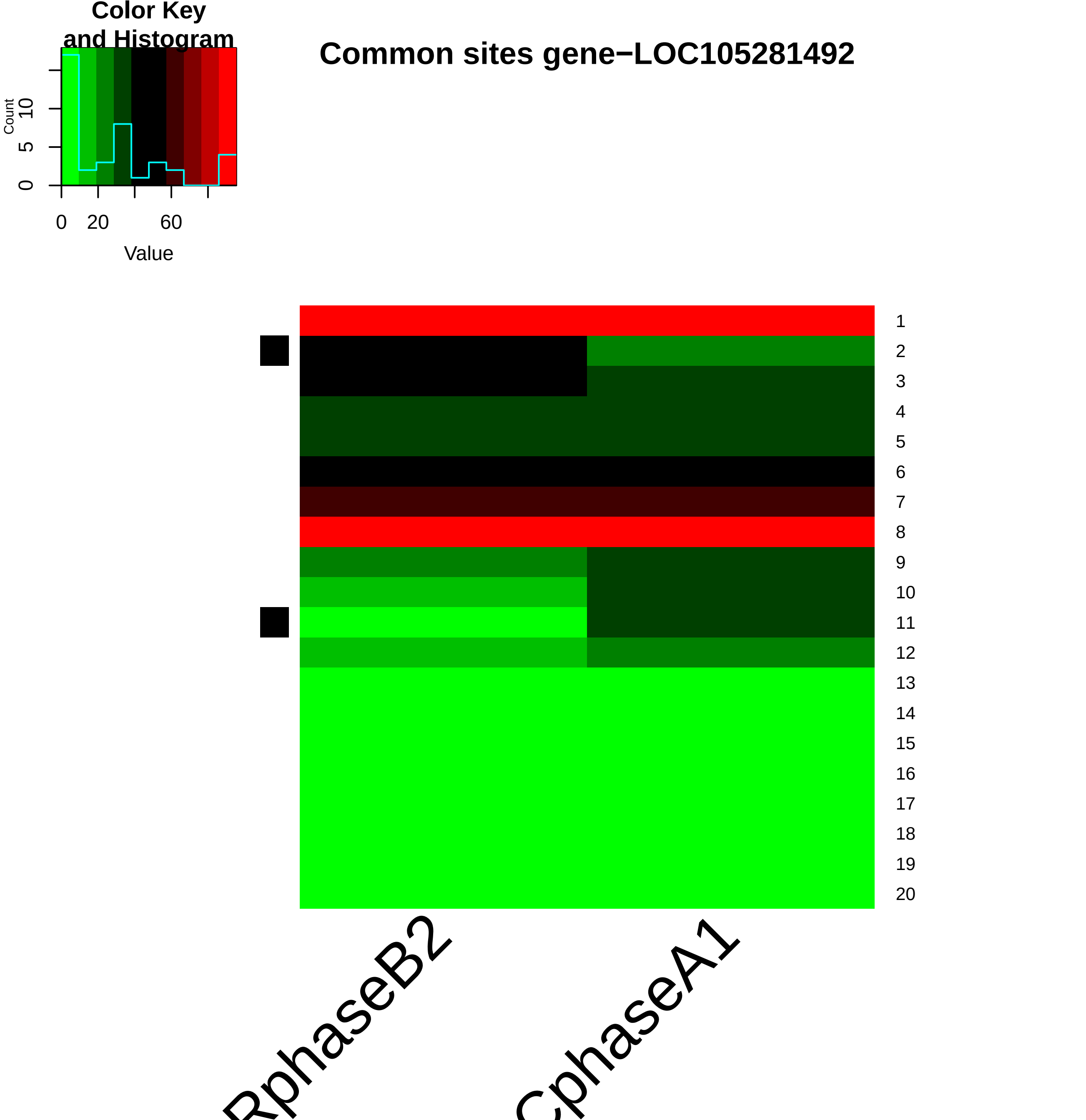

Color Key
and Histogram
Count
0
5
10
0
20
60
Value
Common sites gene−LOC105281492
RphaseB2
CphaseA1

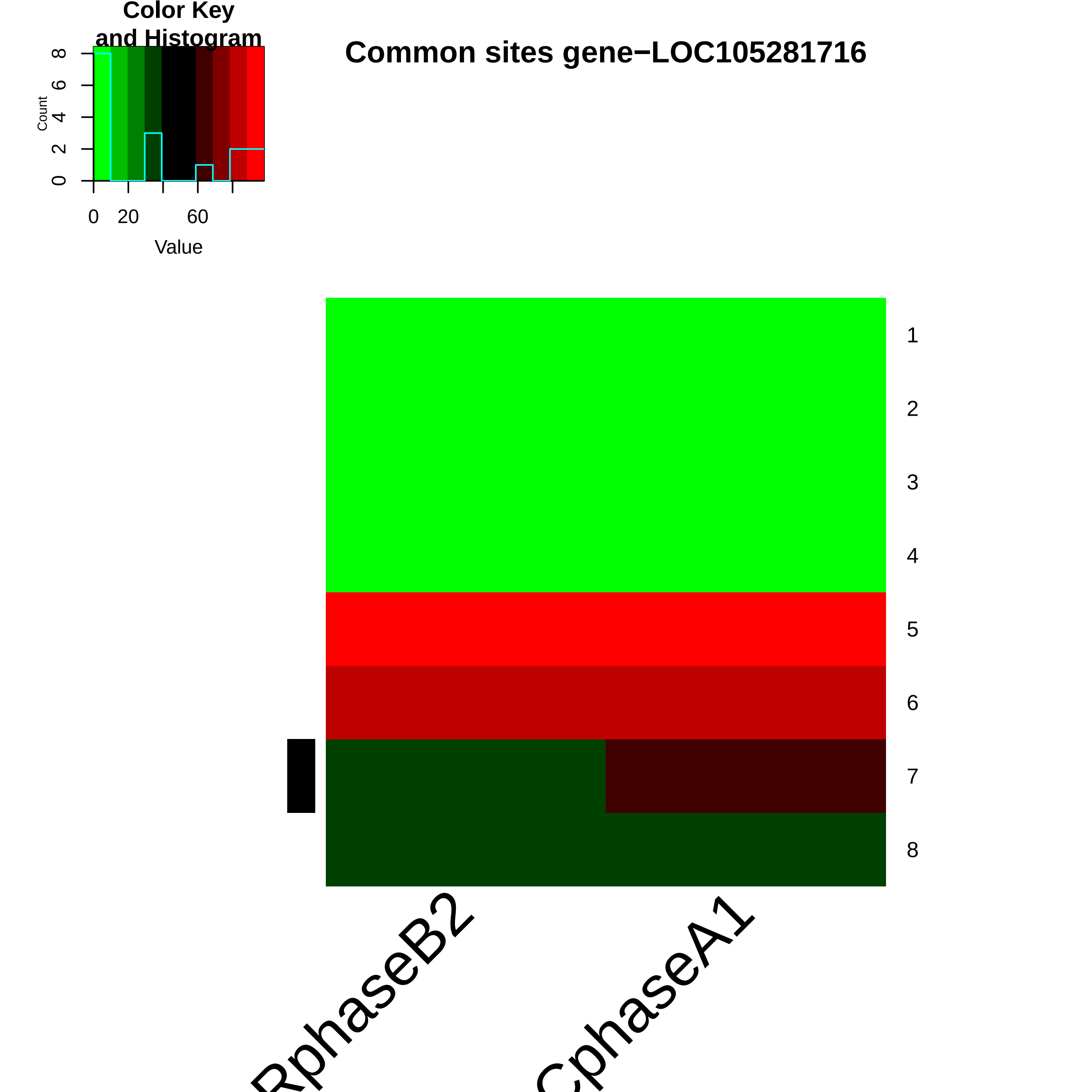

Color Key
and Histogram
Count
0
2
4
6
8
0
20
60
Value
Common sites gene−LOC105281716
1
2
3
4
5
6
7
8
RphaseB2
CphaseA1

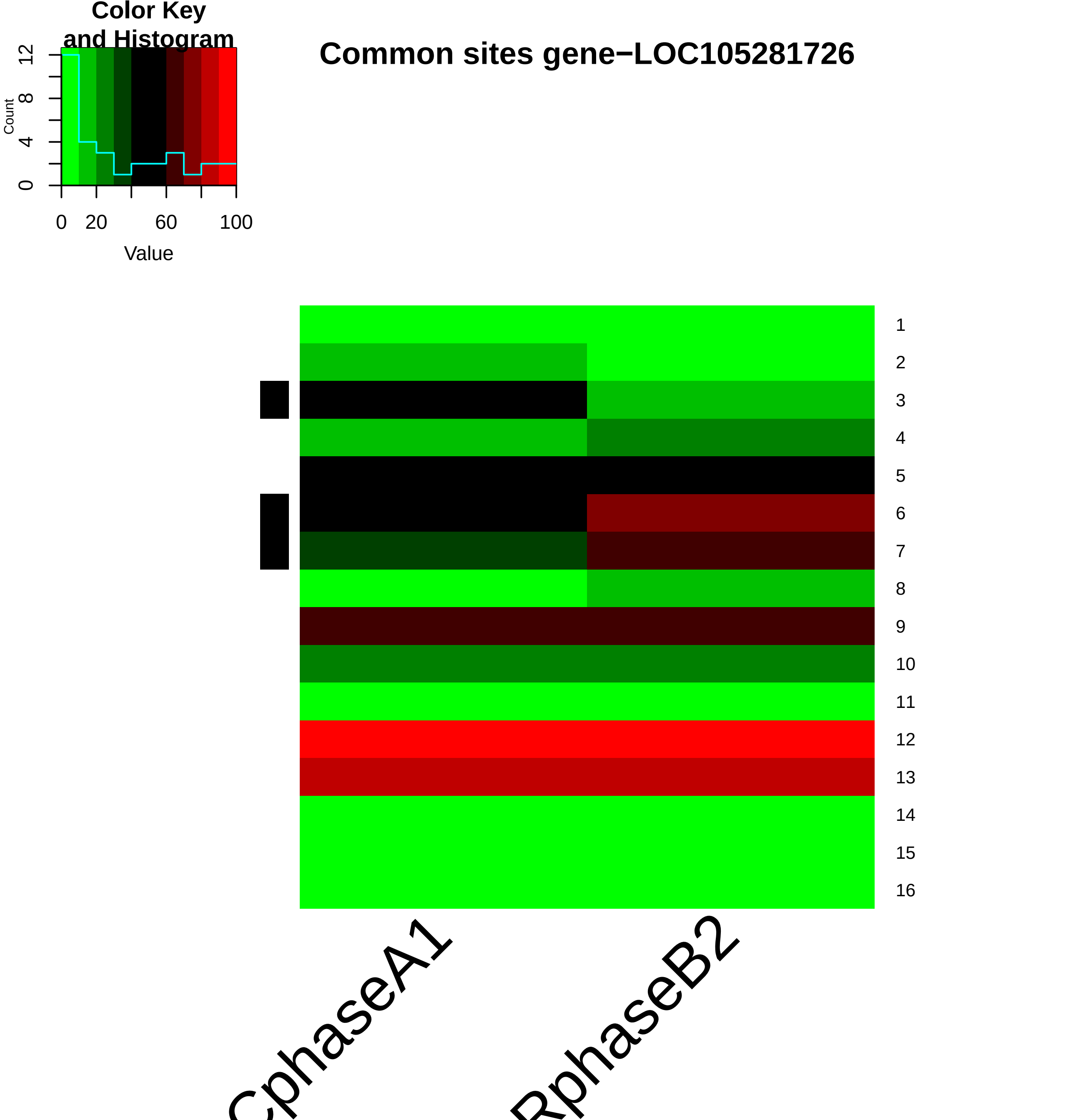
Color Key
and Histogram
Count
0
4
8
12
0
20
60
100
Value
Common sites gene−LOC105281726
CphaseA1
RphaseB2

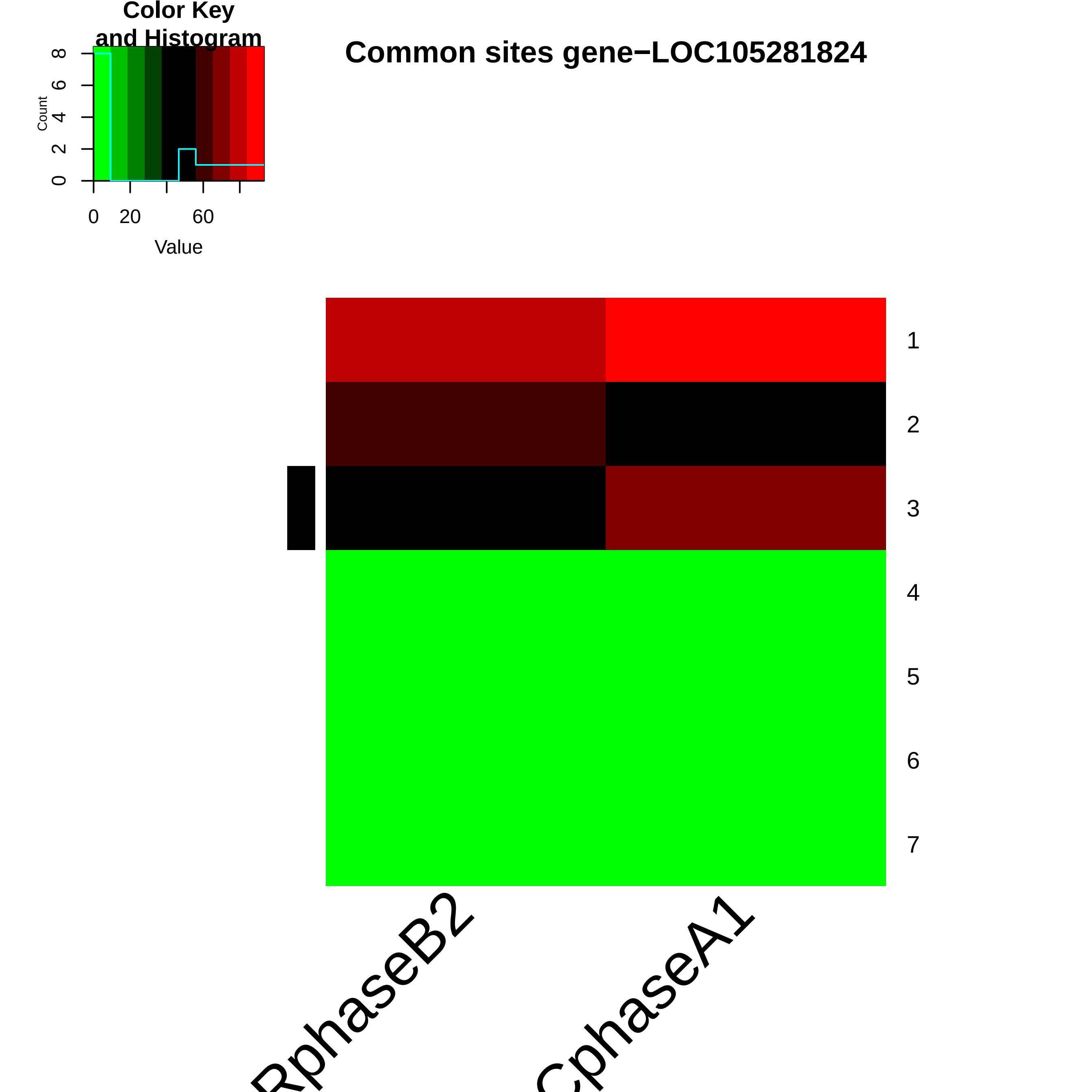
Color Key
and Histogram
Count
0
2
4
6
8
0
20
60
Value
Common sites gene−LOC105281824
1
2
3
4
5
6
7
RphaseB2
CphaseA1

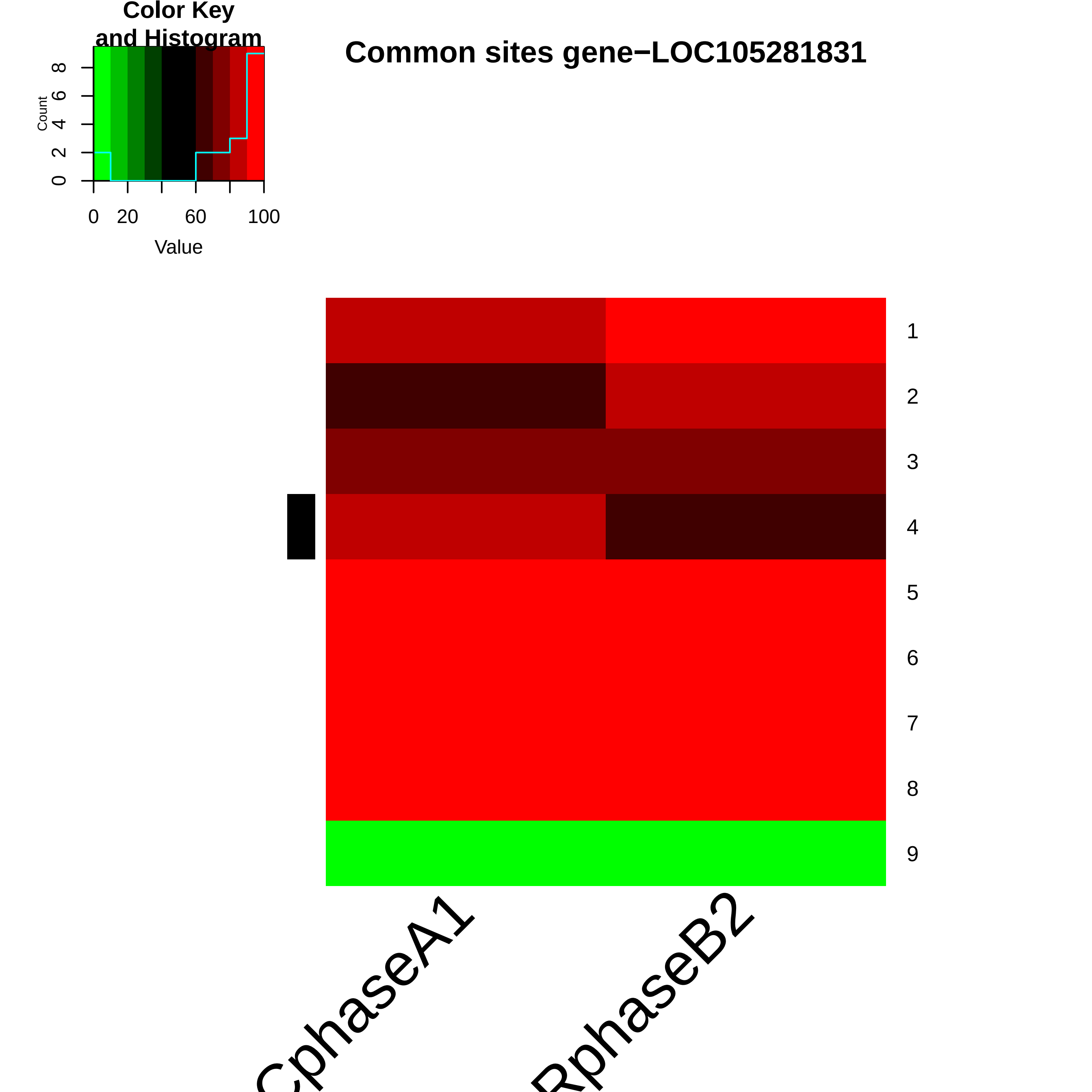
Color Key
and Histogram
Count
0
2
4
6
8
0
20
60
100
Value
Common sites gene−LOC105281831
1
2
3
4
5
6
7
8
9

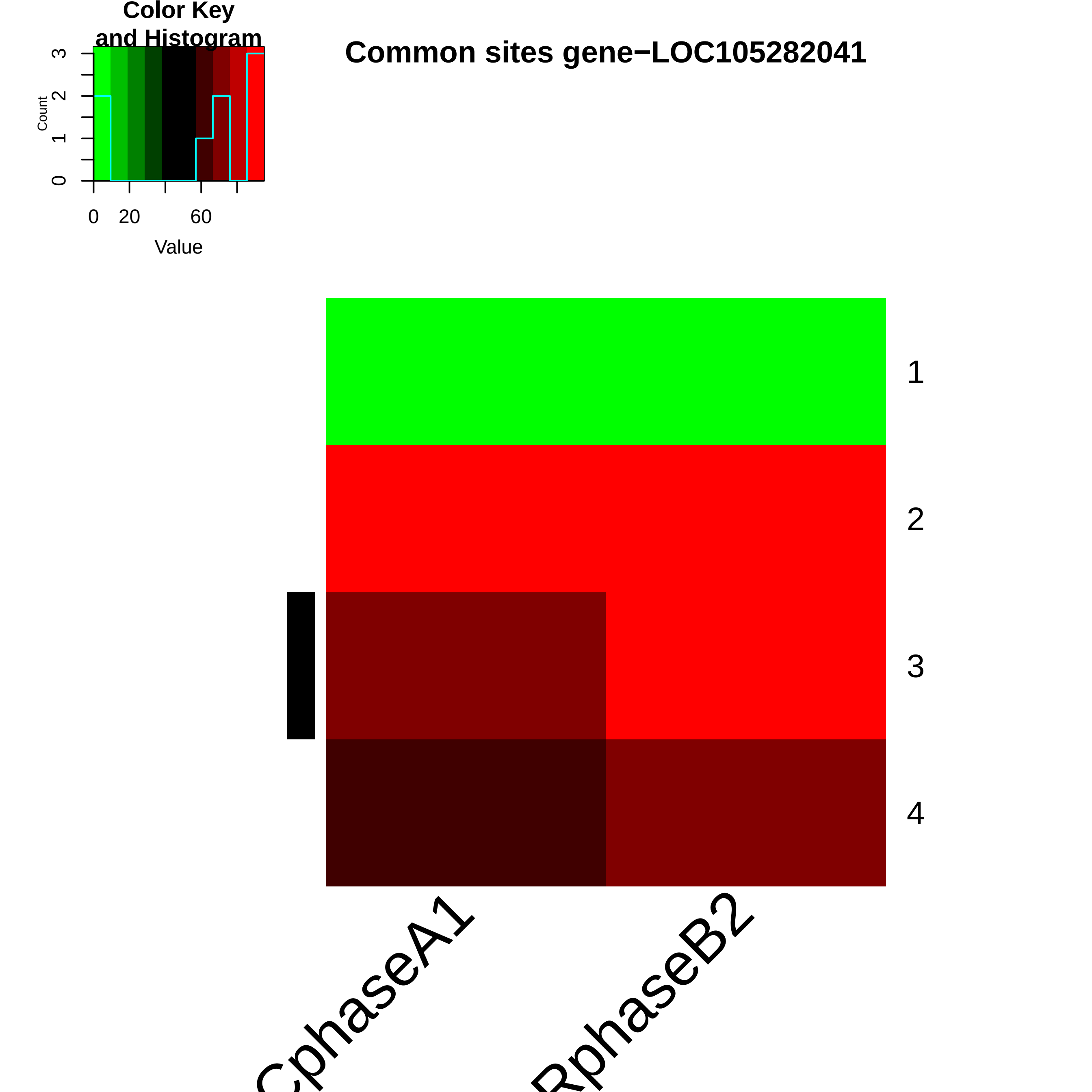

Color Key
and Histogram
Count
0
1
2
3
0
20
60
Value
Common sites gene−LOC105282041
1
2
3
4
CphaseA1
RphaseB2

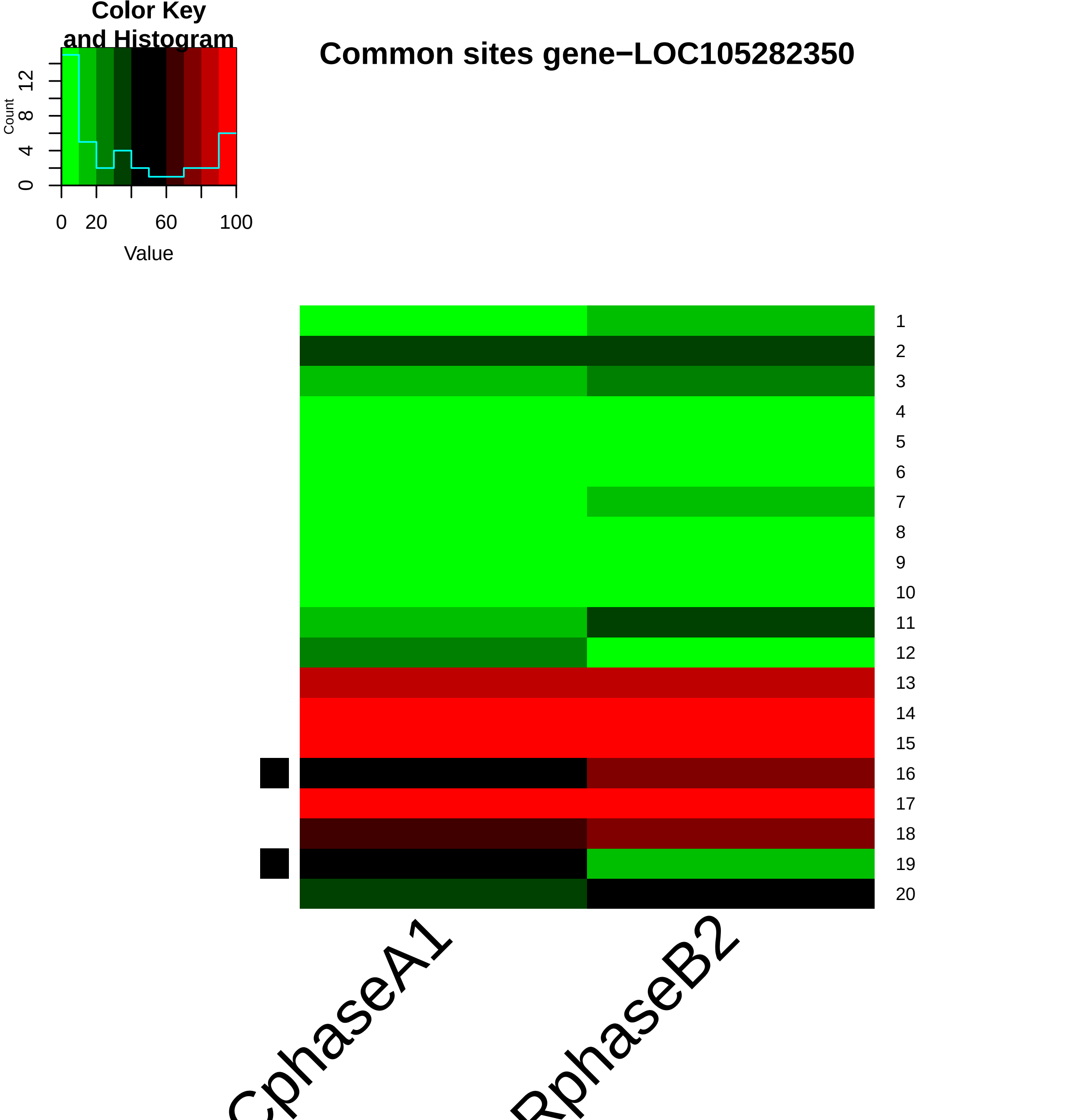
Color Key
and Histogram
Count
0
4
8
12
0
20
60
100
Value
Common sites gene−LOC105282350
CphaseA1
RphaseB2

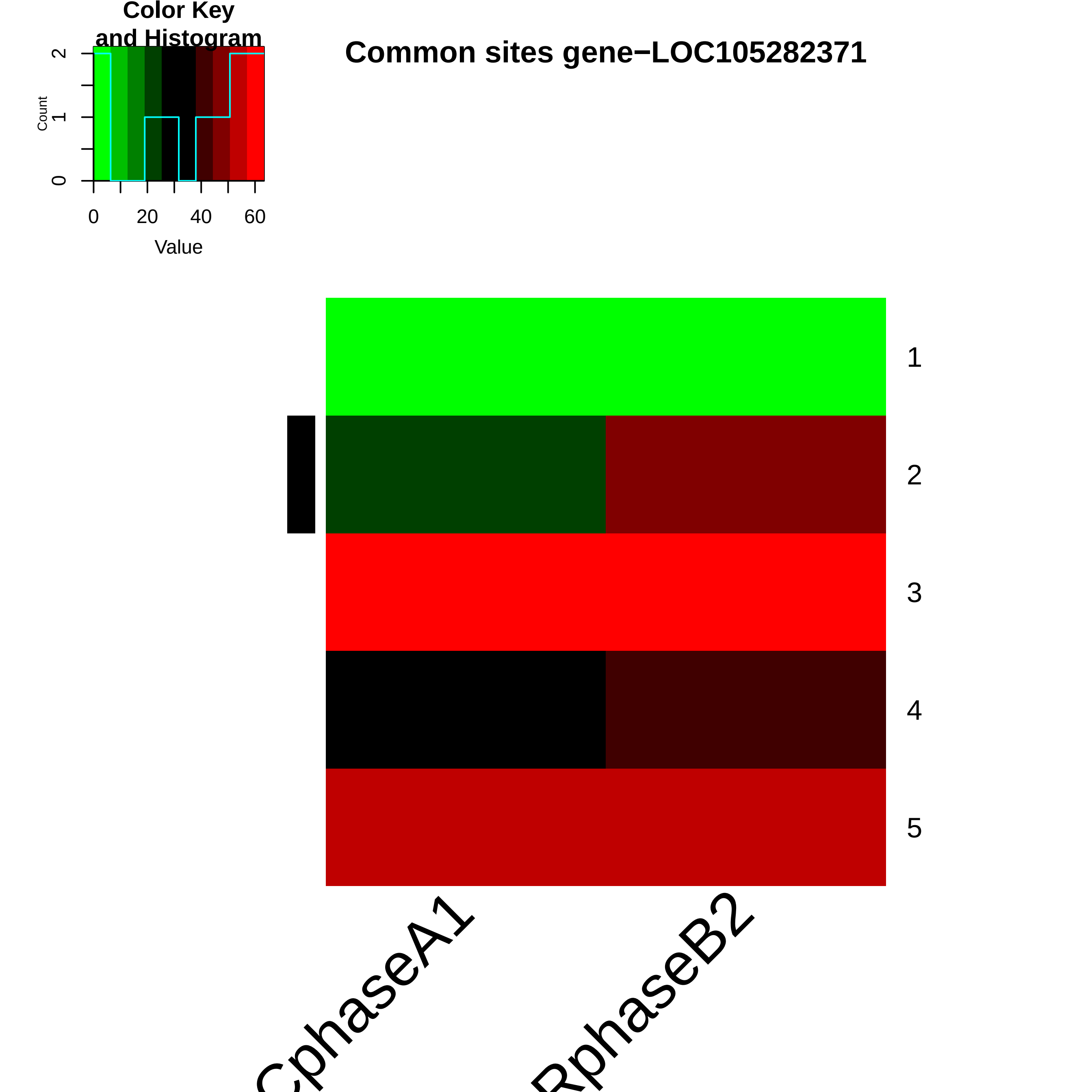

Color Key
and Histogram
Count
0
1
2
0
20
40
60
Value
Common sites gene−LOC105282371
1
2
3
4
5
CphaseA1
RphaseB2

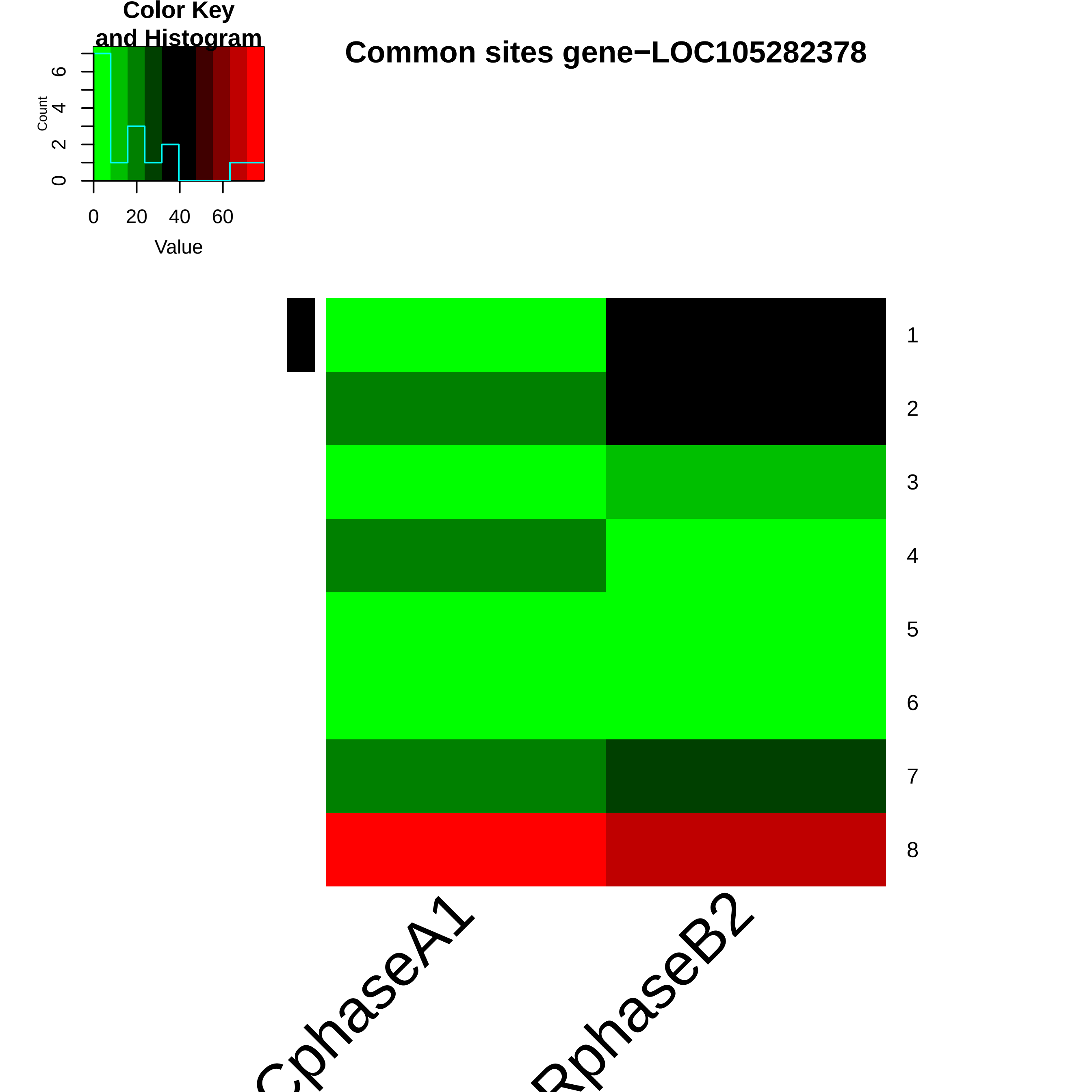

Color Key
and Histogram
Count
0
2
4
6
0
20
40
60
Value
Common sites gene−LOC105282378
1
2
3
4
5
6
7
8
CphaseA1
RphaseB2

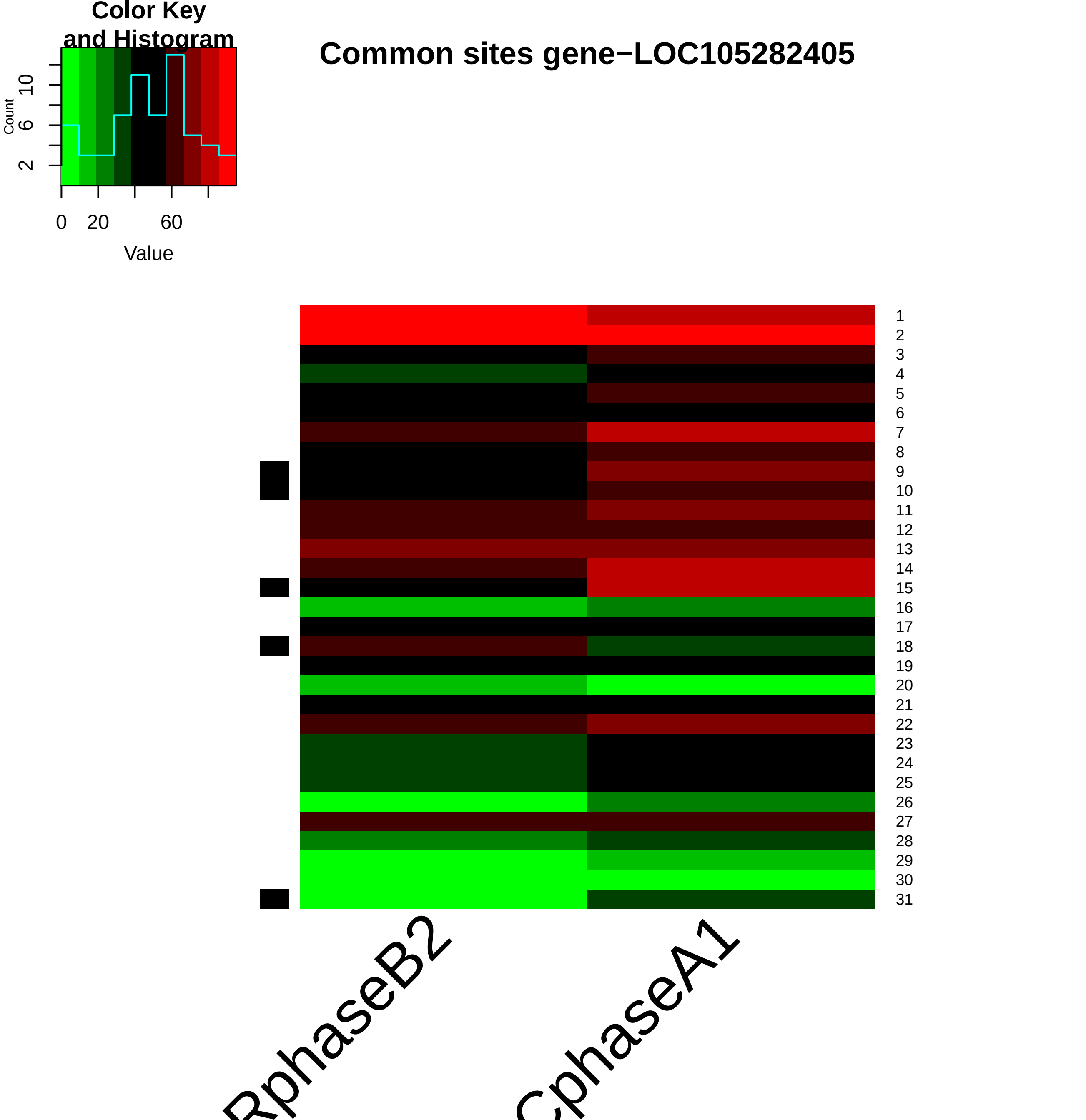

Color Key
and Histogram
Count
2
6
10
0
20
60
Value
Common sites gene−LOC105282405
RphaseB2
CphaseA1

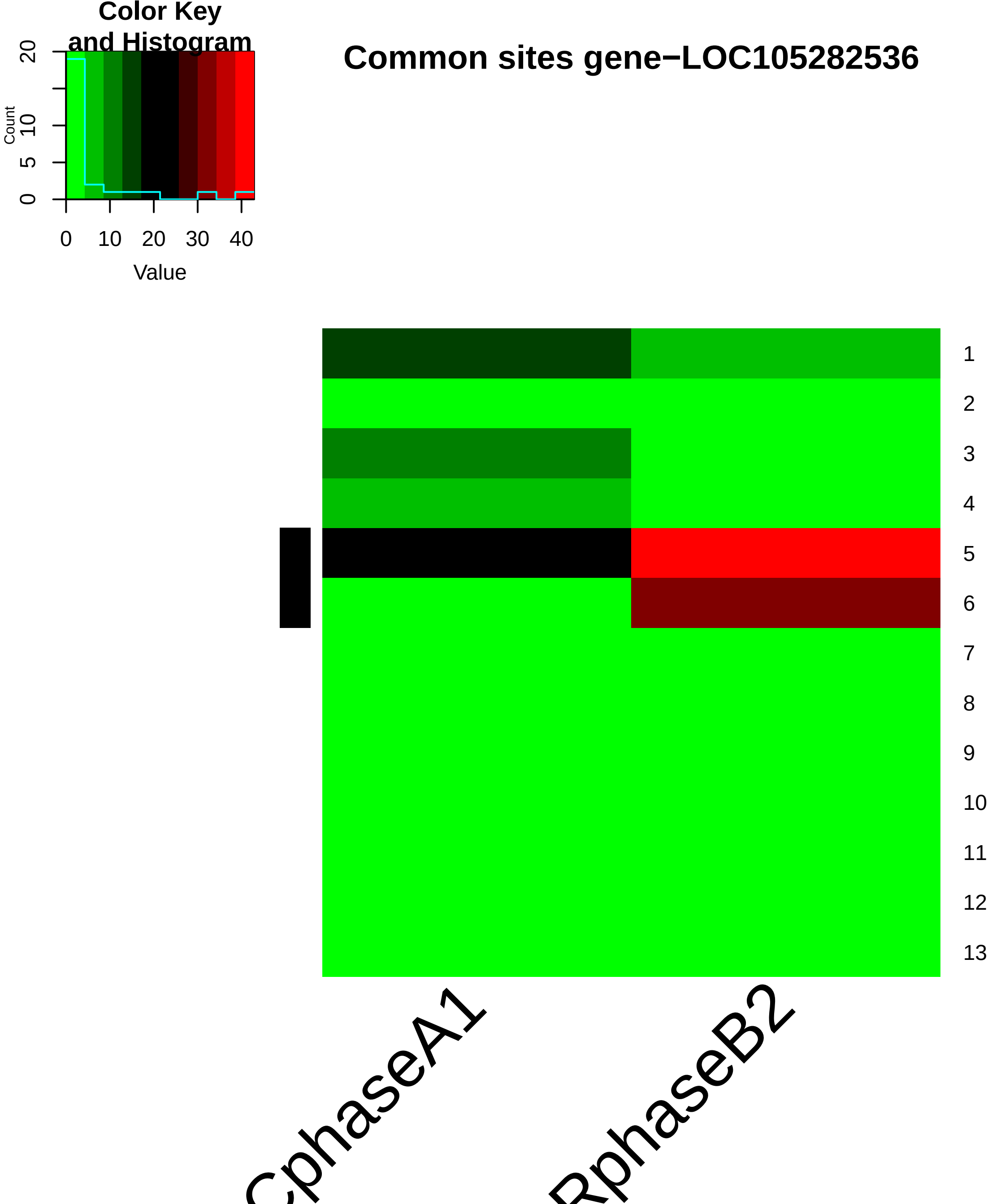

Color Key
and Histogram
Count
Value
Common sites gene−LOC105282536
1
2
3
4
5
6
7
8
9
10
11
12
13

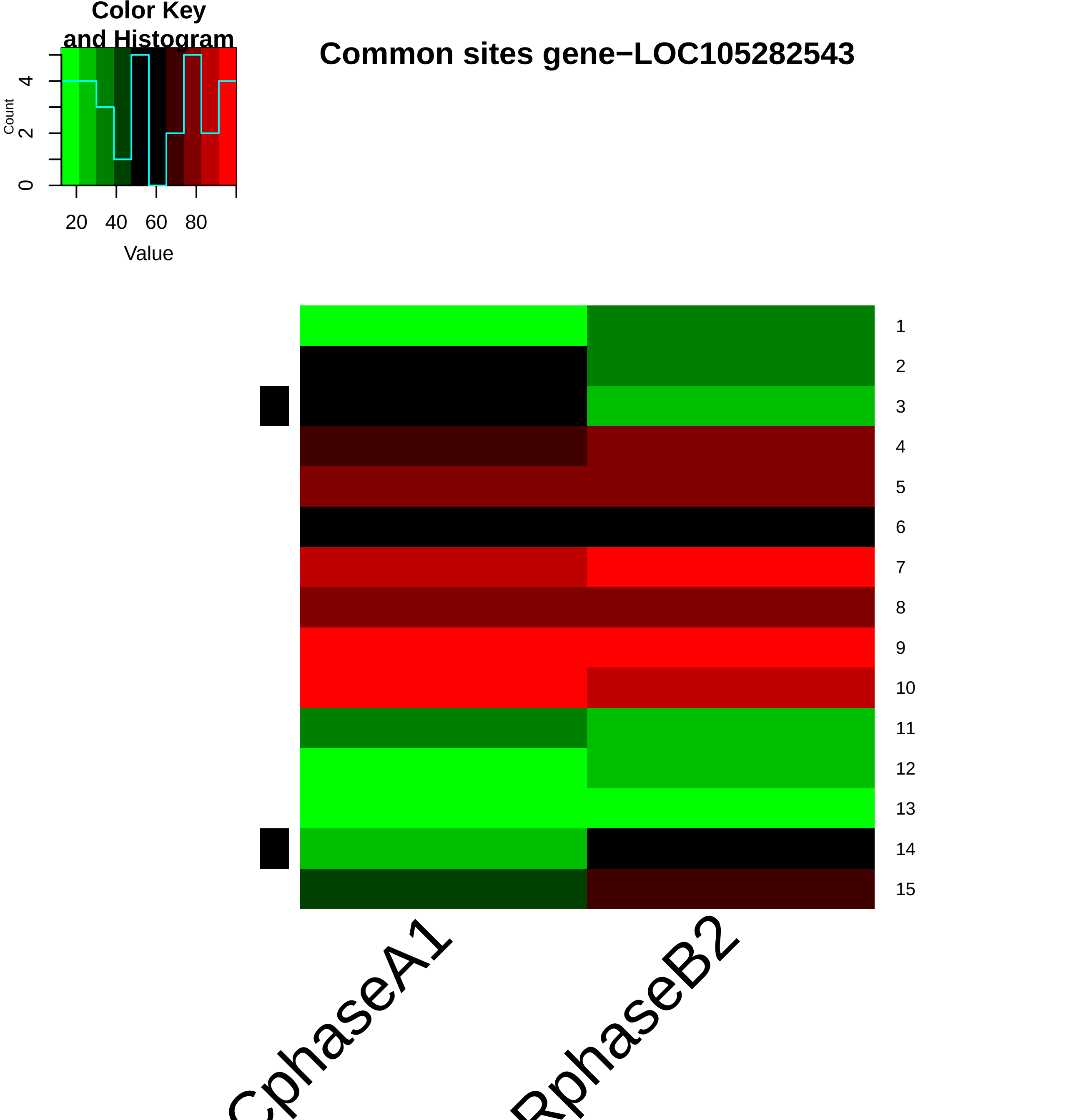

Color Key
and Histogram
Count
0
2
4
20
40
60
80
Value
Common sites gene−LOC105282543
CphaseA1
RphaseB2

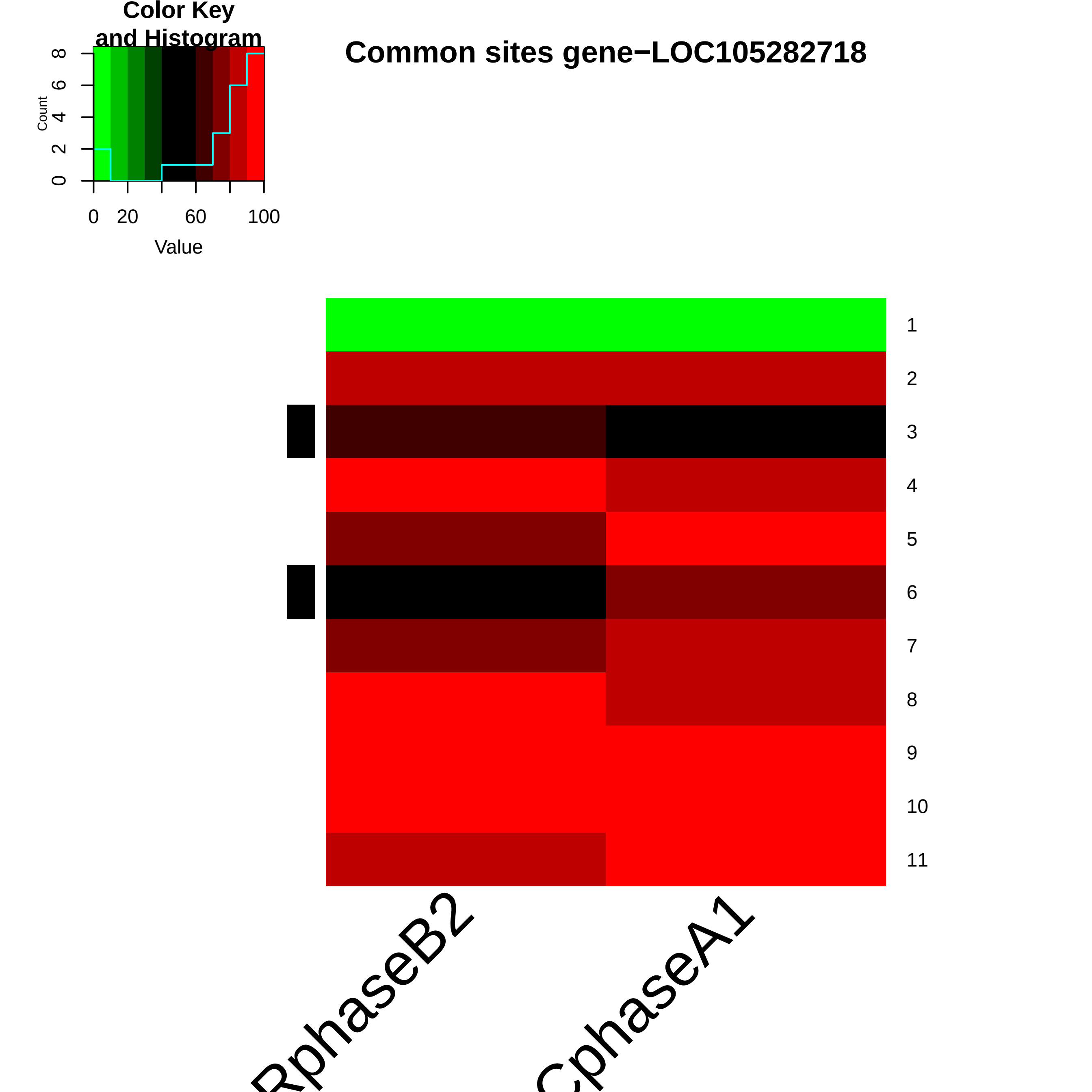

Color Key
and Histogram
Count
0
2
4
6
8
0
20
60
100
Value
Common sites gene−LOC105282718
1
2
3
4
5
6
7
8
9
10
11
RphaseB2
CphaseA1

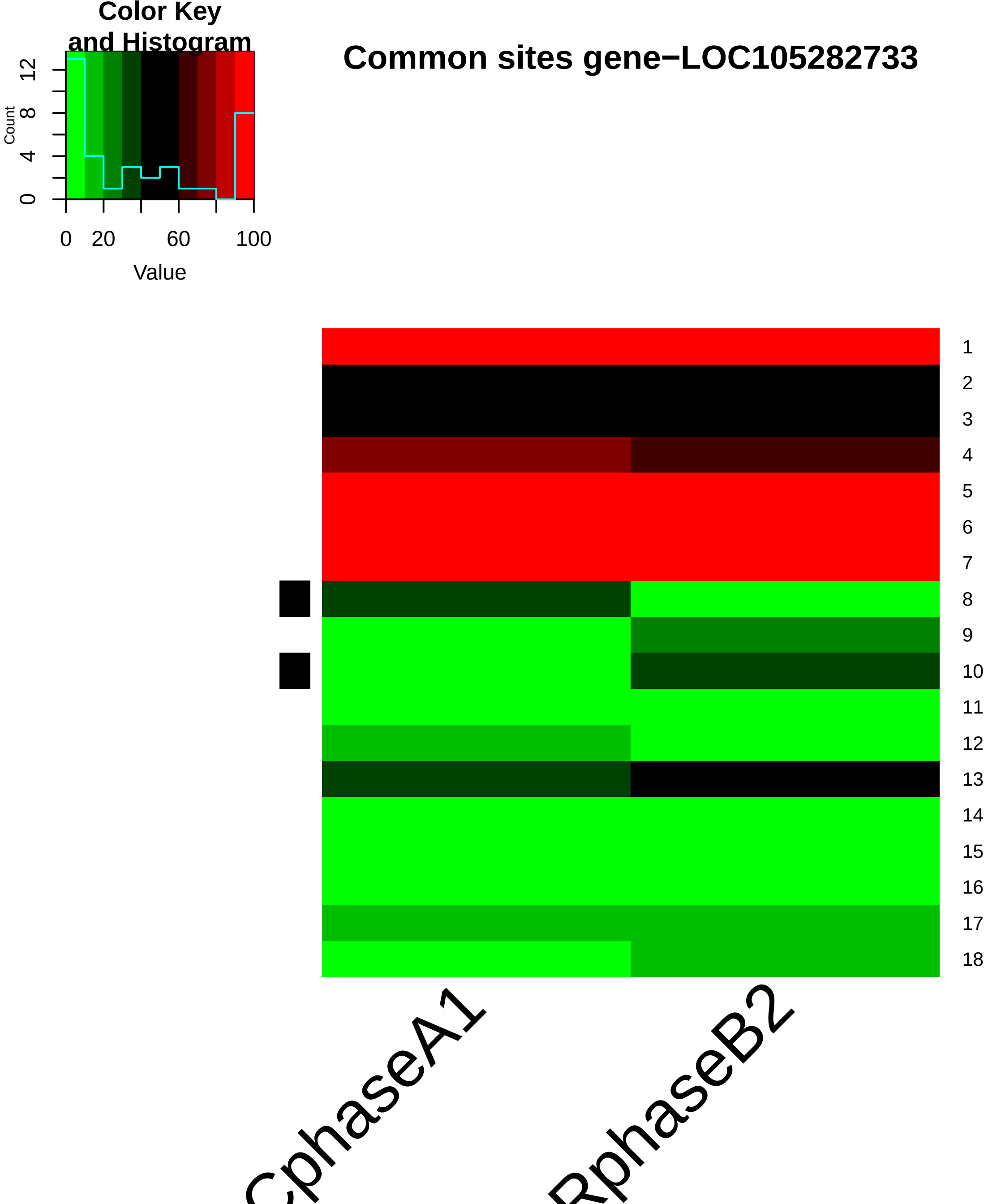

Color Key
and Histogram
Count
0
4
8
12
0
20
60
100
Value
Common sites gene−LOC105282733
CphaseA1
RphaseB2

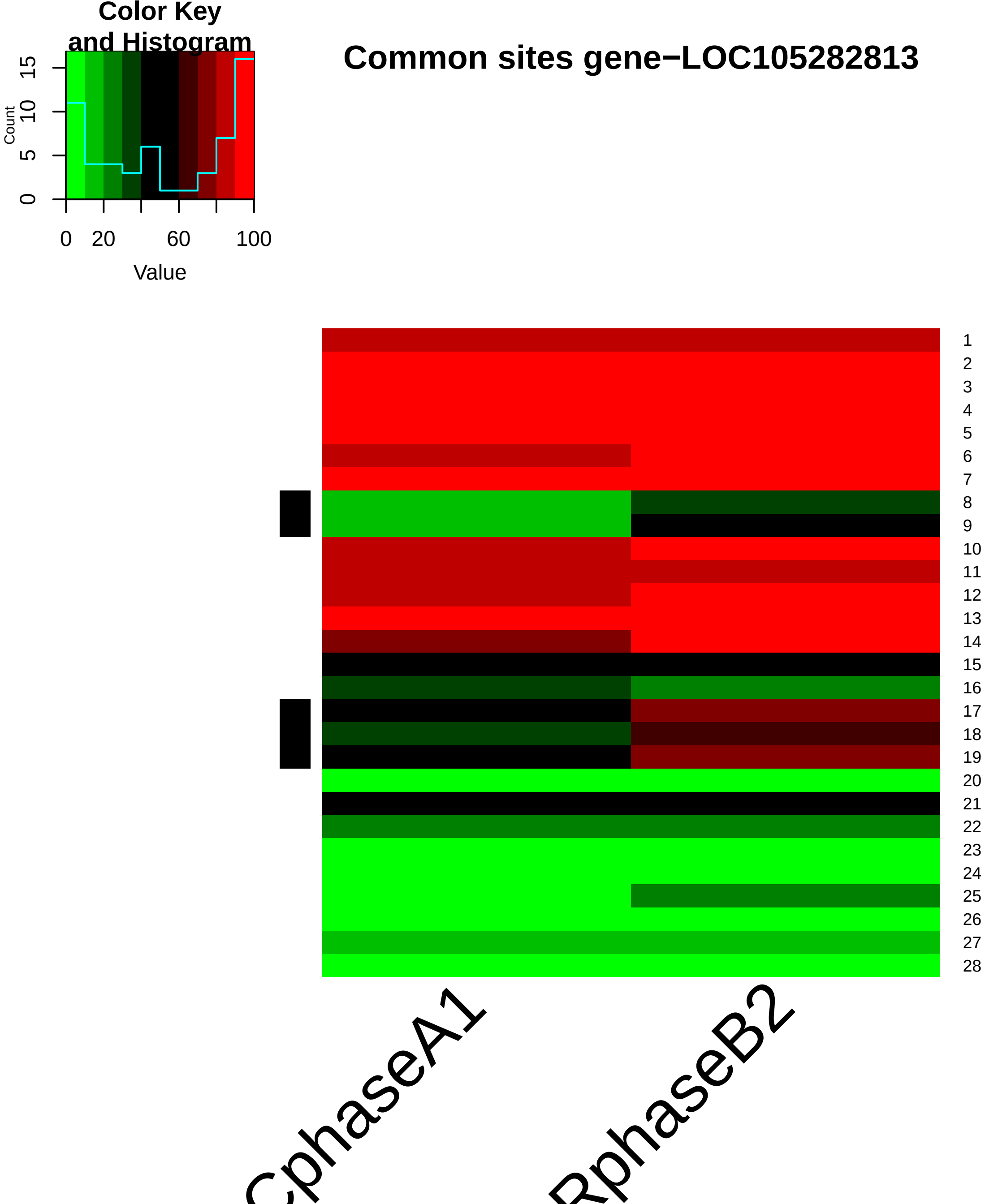
Color Key
and Histogram
Count
0
5
10
15
0
20
60
100
Value
Common sites gene−LOC105282813

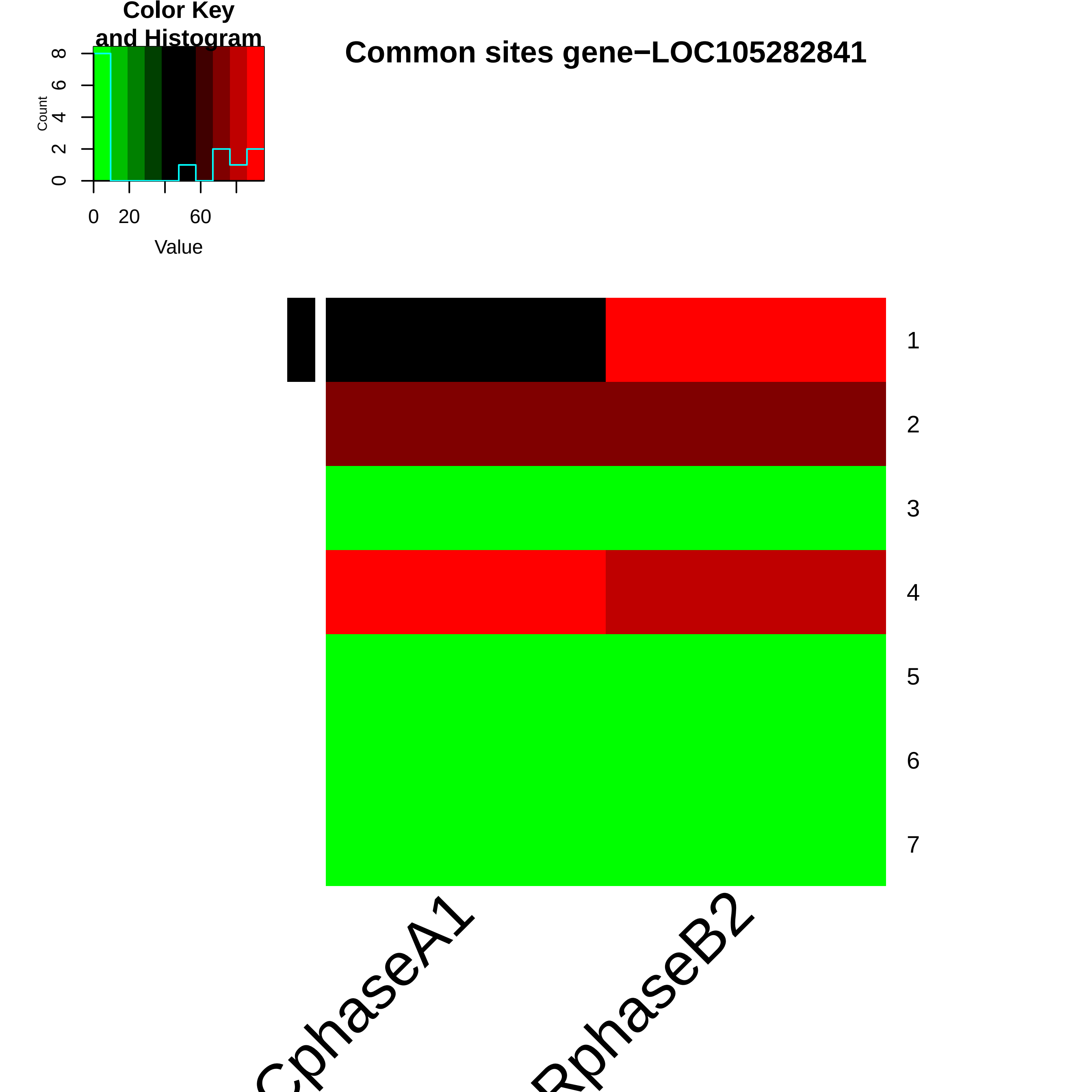
Color Key
and Histogram
Count
Value
0
20
60
0
2
4
6
8
Common sites gene−LOC105282841
1
2
3
4
5
6
7
CphaseA1
RphaseB2

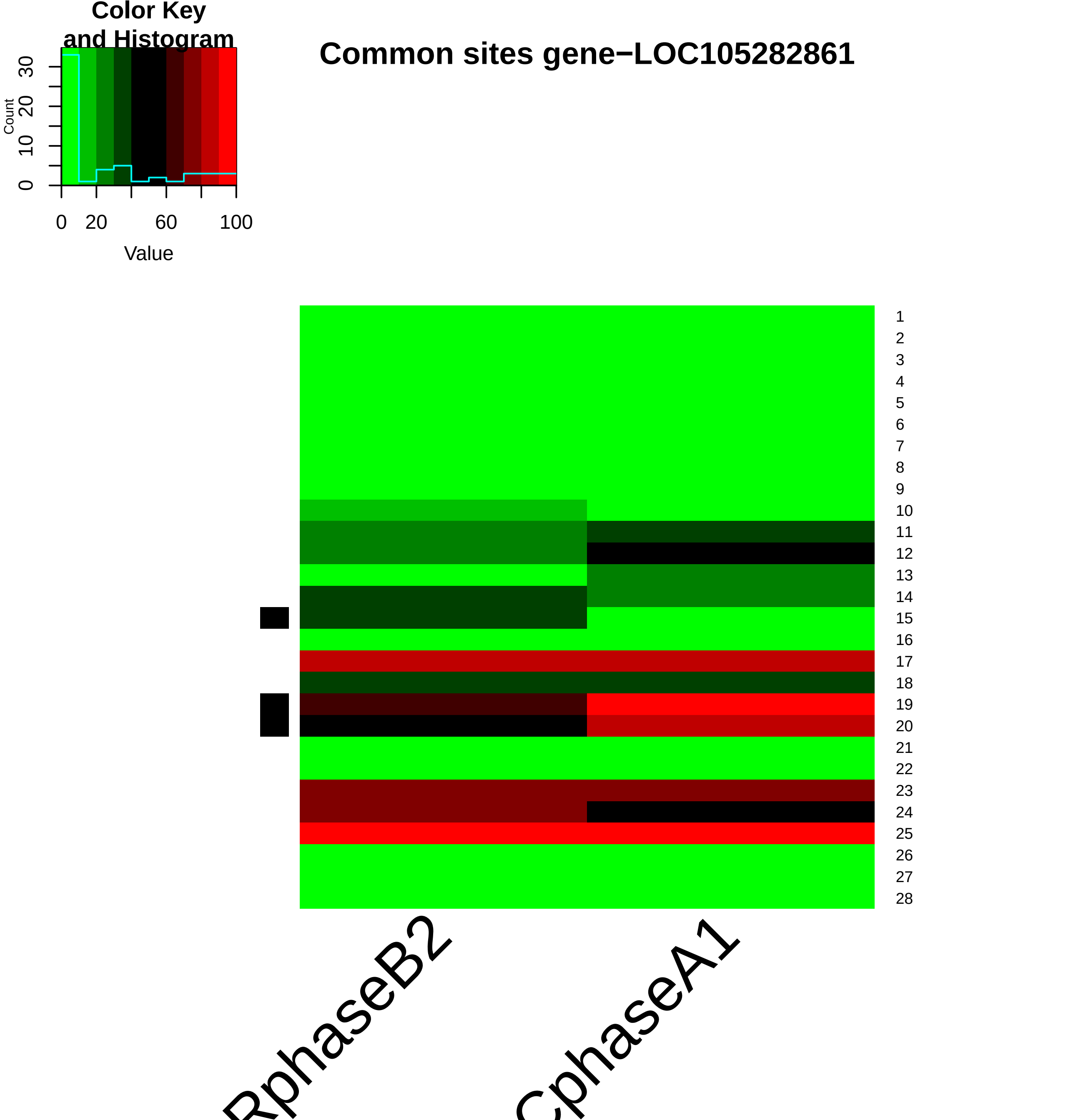

Color Key
and Histogram
Count
Value
0
10
20
30
0
20
60
100
Common sites gene−LOC105282861
RphaseB2
CphaseA1

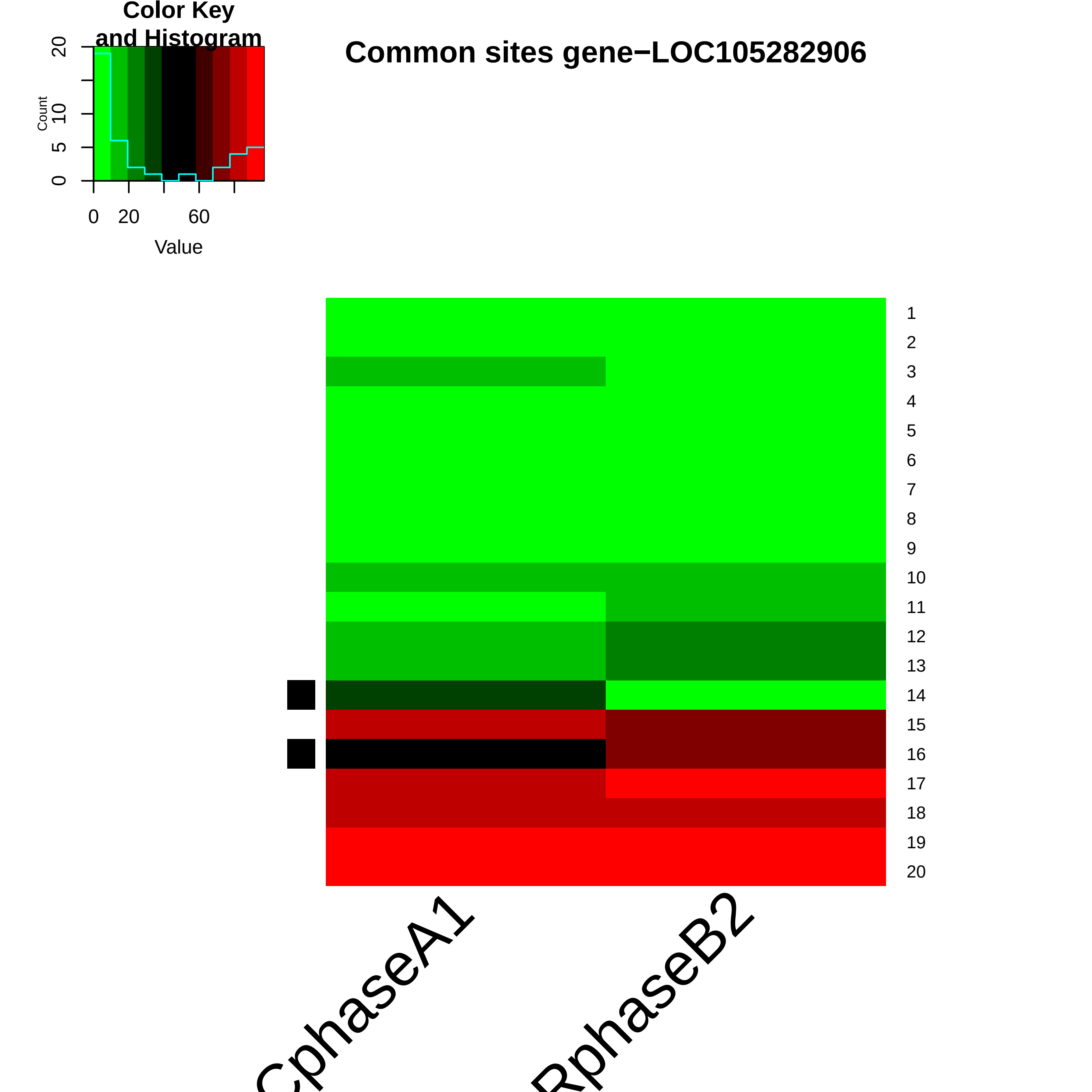

Color Key and Histogram
Count
0 5 10 20
0 20 60
Value
Common sites gene−LOC105282906
CphaseA1
RphaseB2

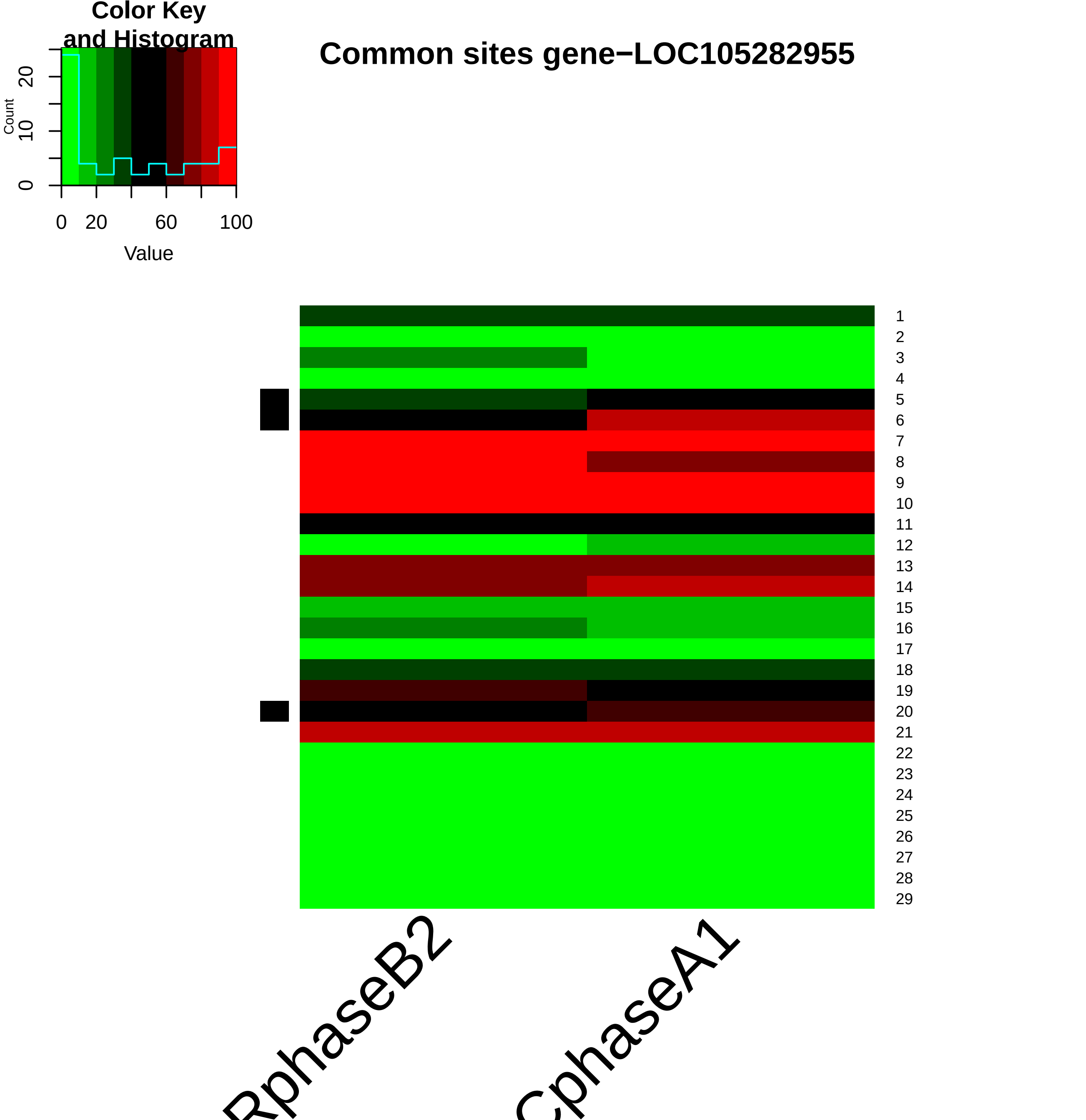

Color Key
and Histogram
Count
0
10
20
0
20
60
100
Value
Common sites gene−LOC105282955
RphaseB2
CphaseA1

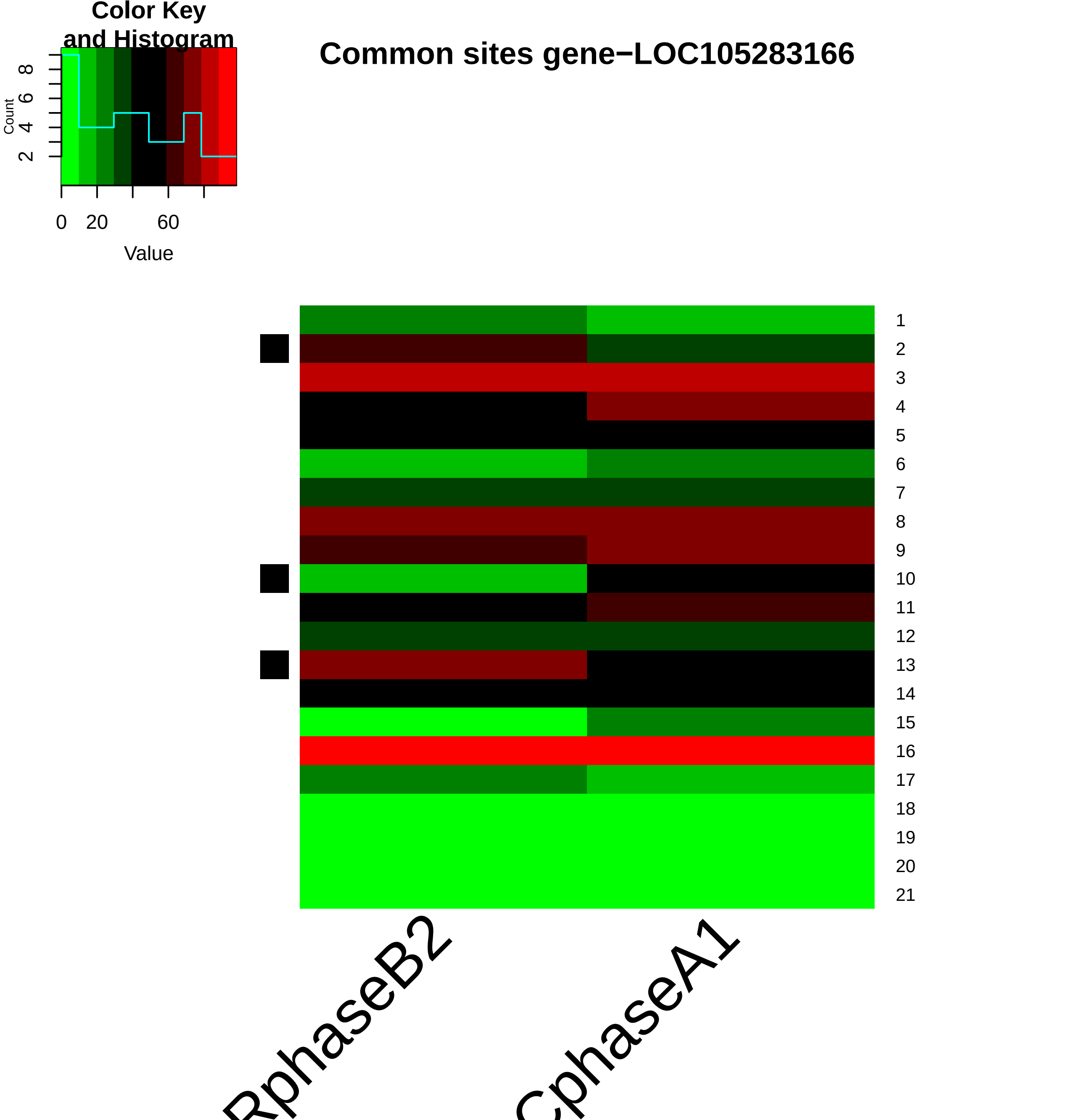
Color Key
and Histogram
Count
2
4
6
8
0
20
60
Value
Common sites gene−LOC105283166
RphaseB2
CphaseA1

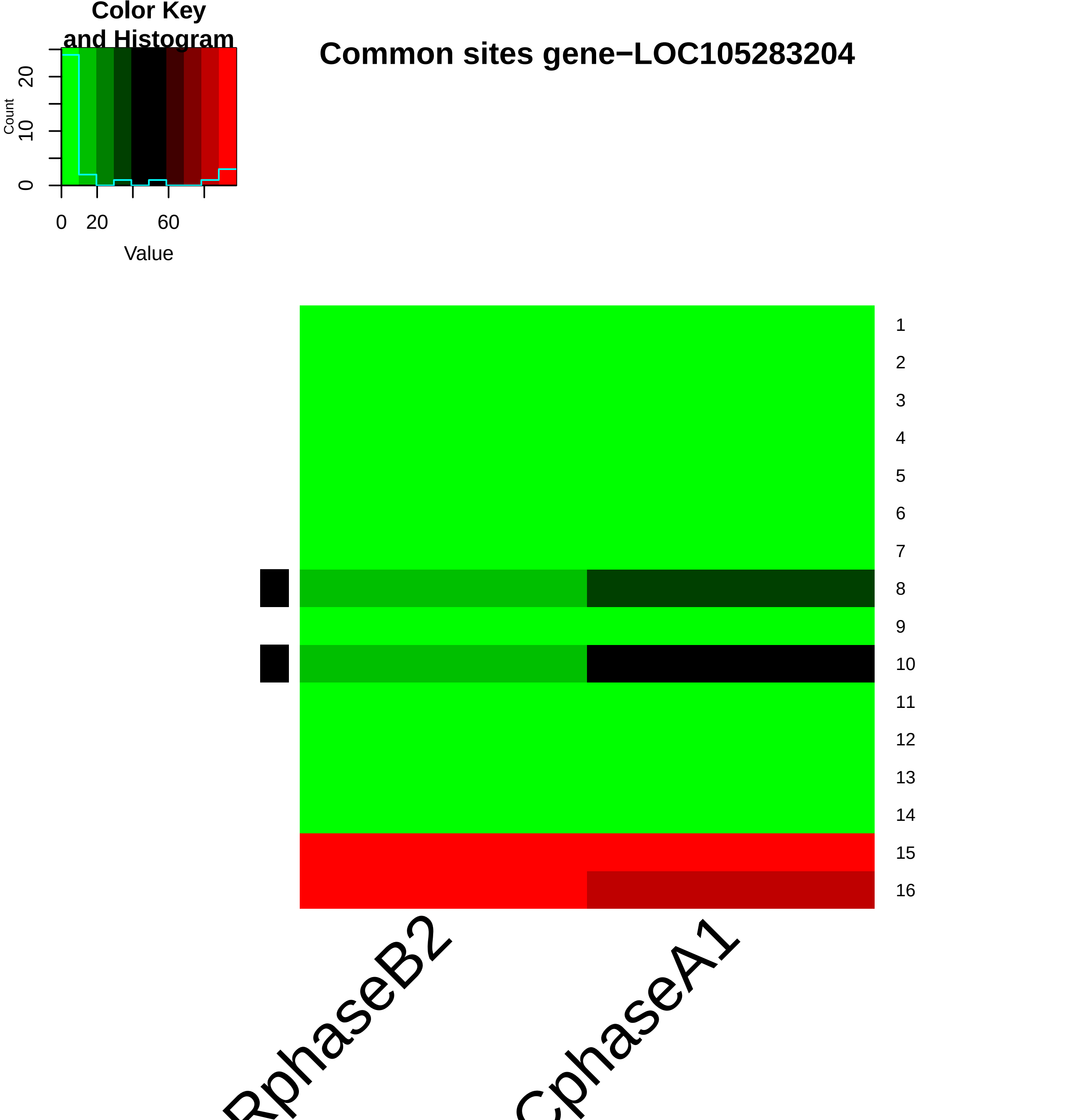

Color Key
and Histogram
Count
0
10
20
0
20
60
Value
Common sites gene−LOC105283204
RphaseB2
CphaseA1

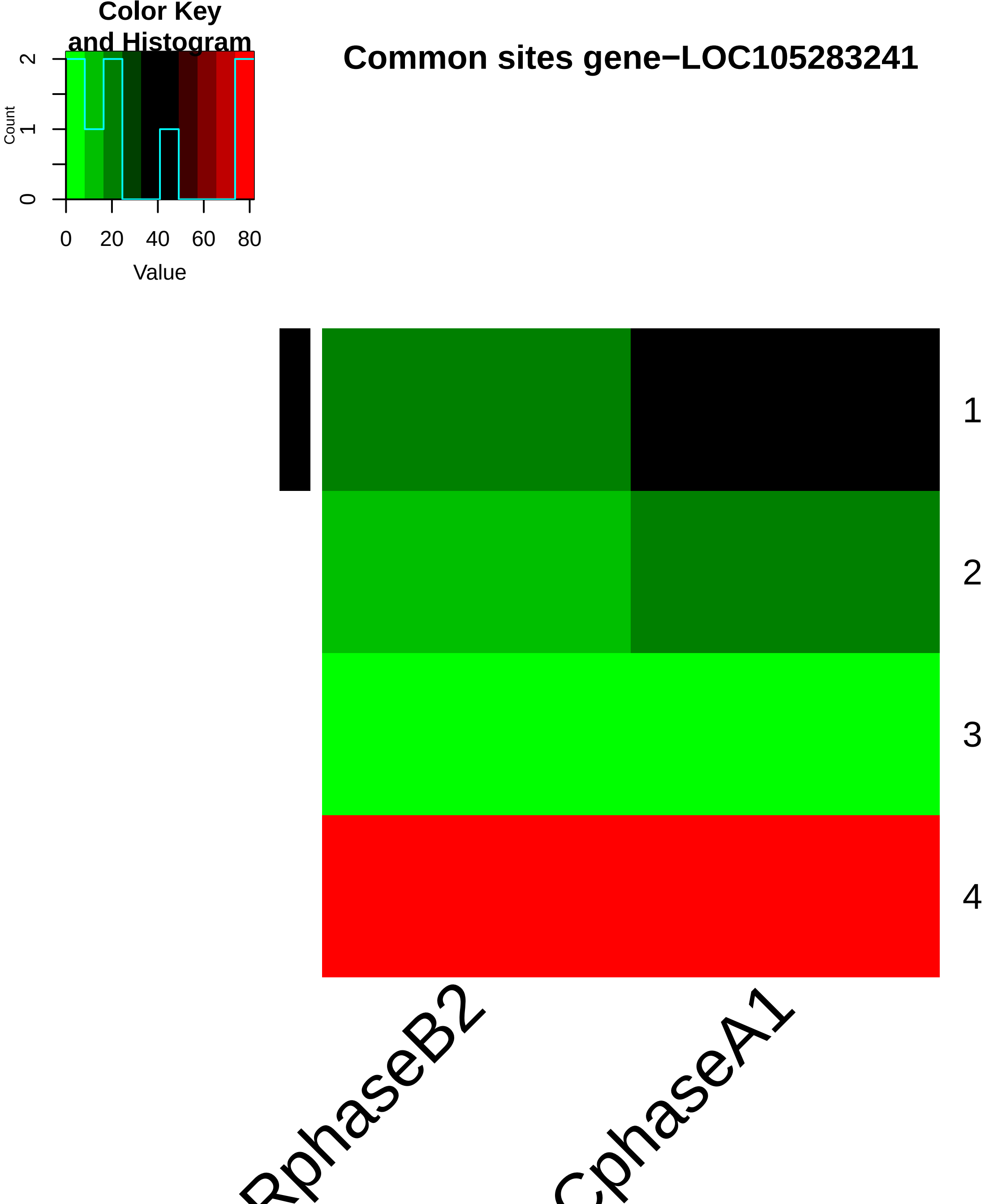
Color Key
and Histogram
Count
0
1
2
0
20
40
60
80
Value
Common sites gene−LOC105283241
1
2
3
4
RphaseB2
CphaseA1

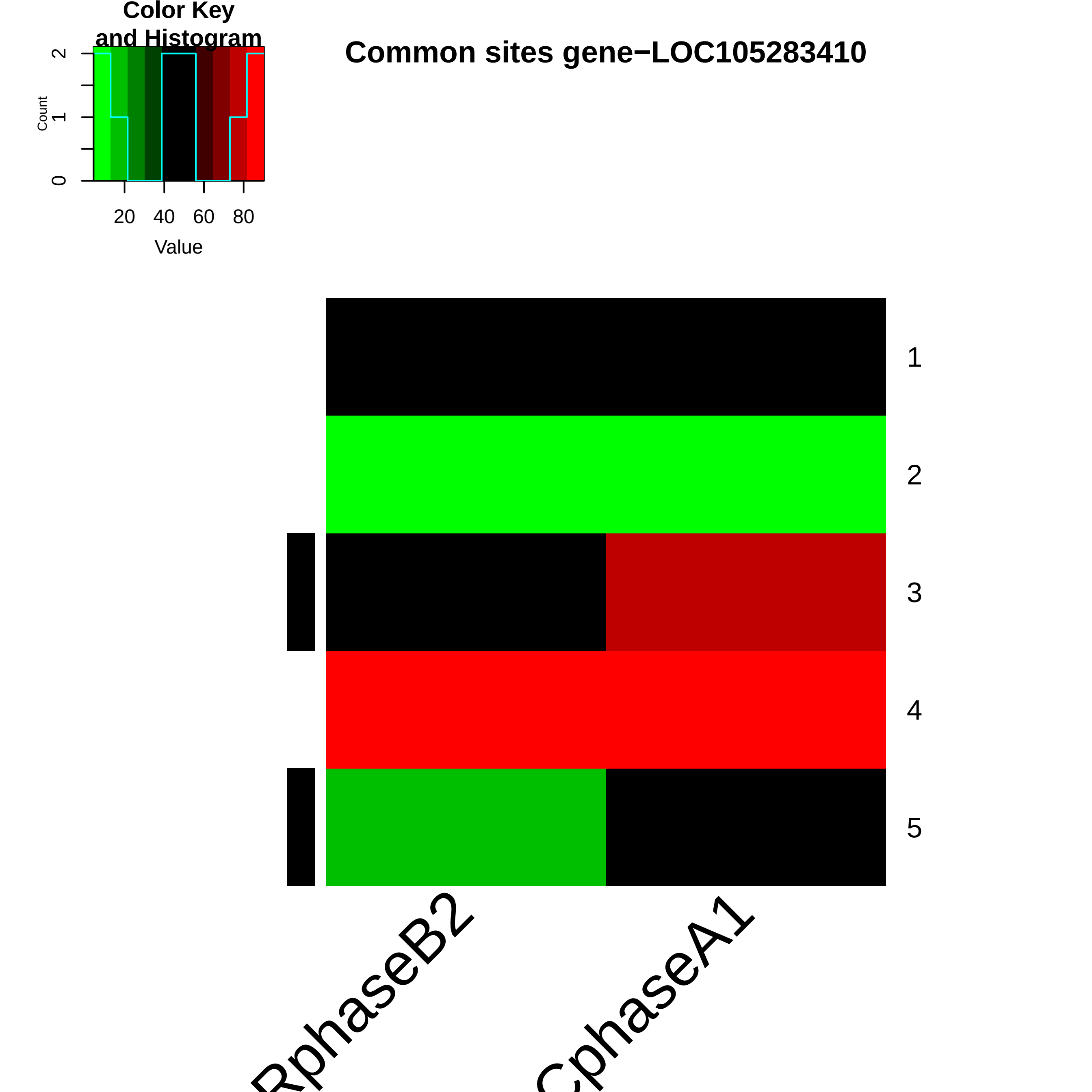
Color Key
and Histogram
Count
0
1
2
20
40
60
80
Value
Common sites gene−LOC105283410
1
2
3
4
5
RphaseB2
CphaseA1

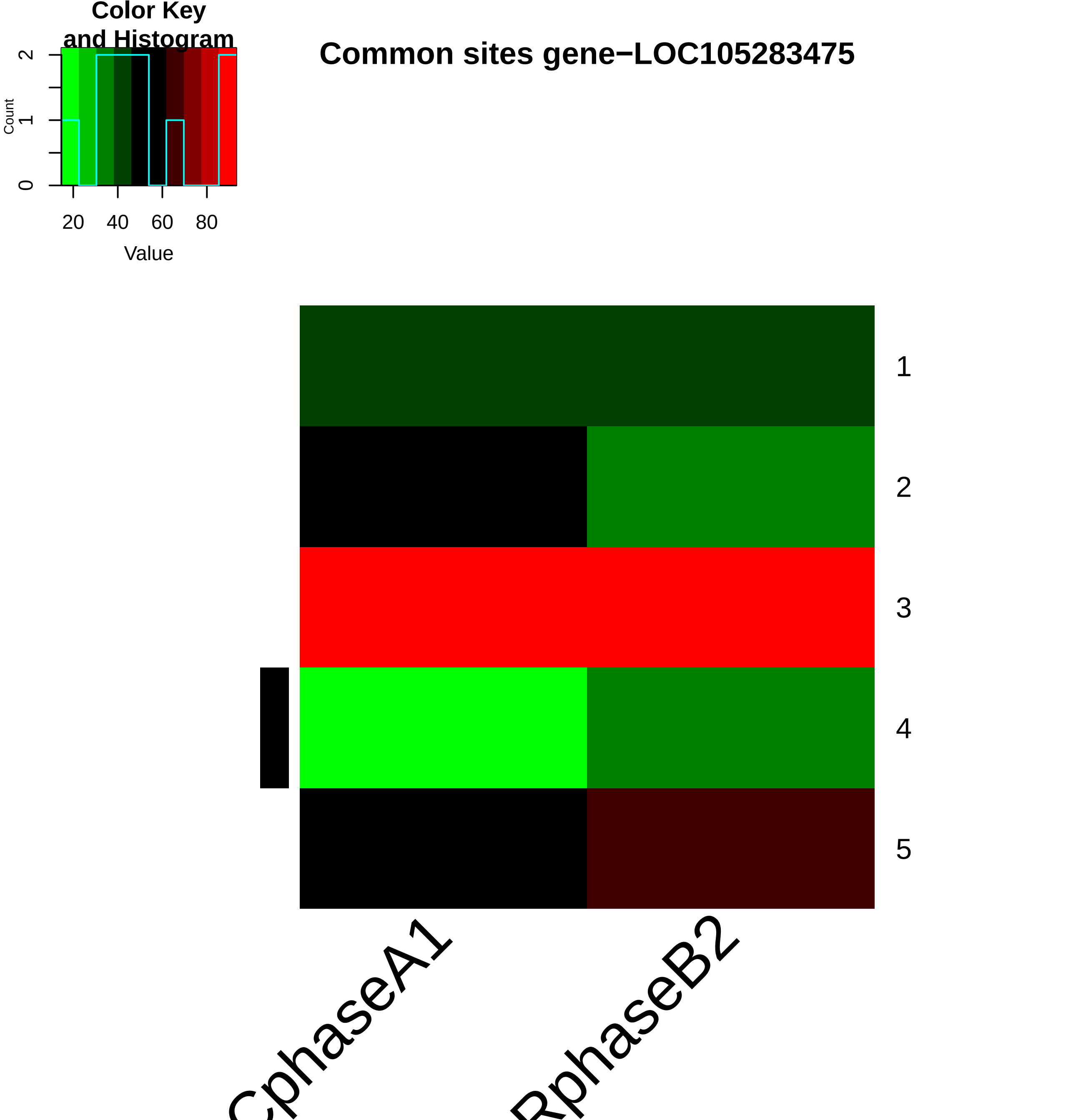
Color Key
and Histogram
Count
0
1
2
20
40
60
80
Value
Common sites gene−LOC105283475
1
2
3
4
5
CphaseA1
RphaseB2

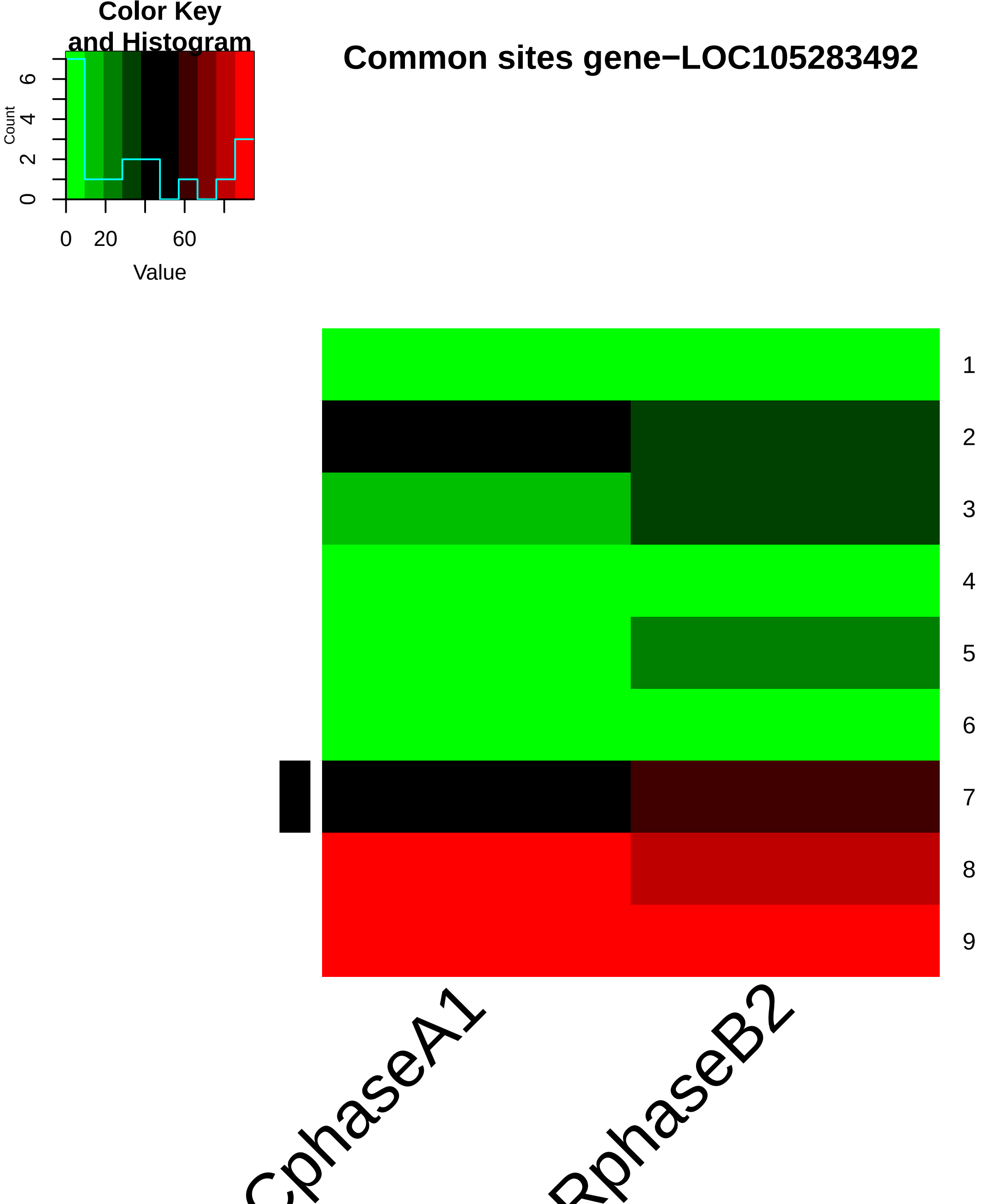
Color Key
and Histogram
Count
0
2
4
6
0
20
60
Value
Common sites gene−LOC105283492
1
2
3
4
5
6
7
8
9
CphaseA1
RphaseB2

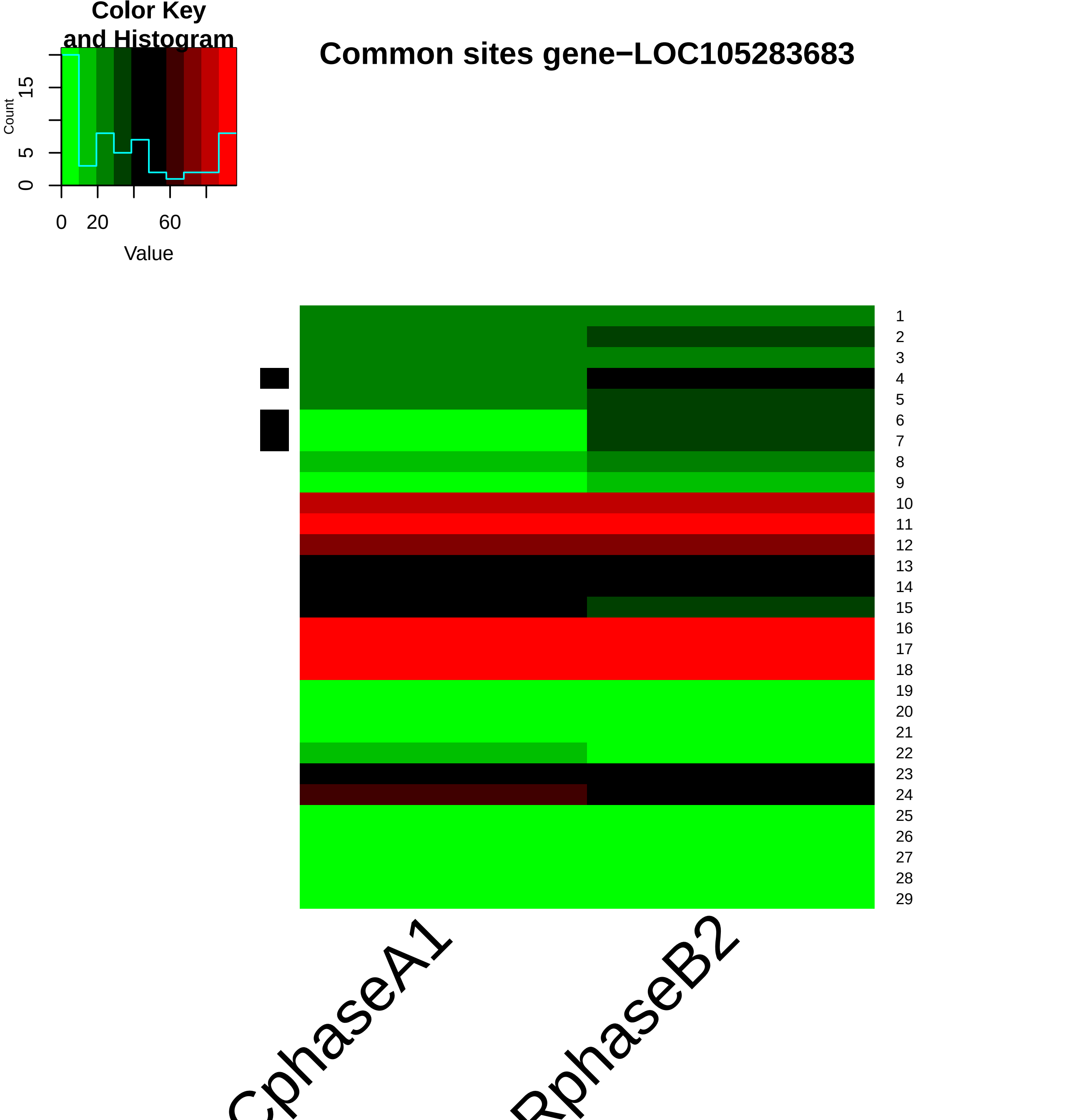
Color Key
and Histogram
Count
0
5
15
0
20
60
Value
Common sites gene−LOC105283683
CphaseA1
RphaseB2

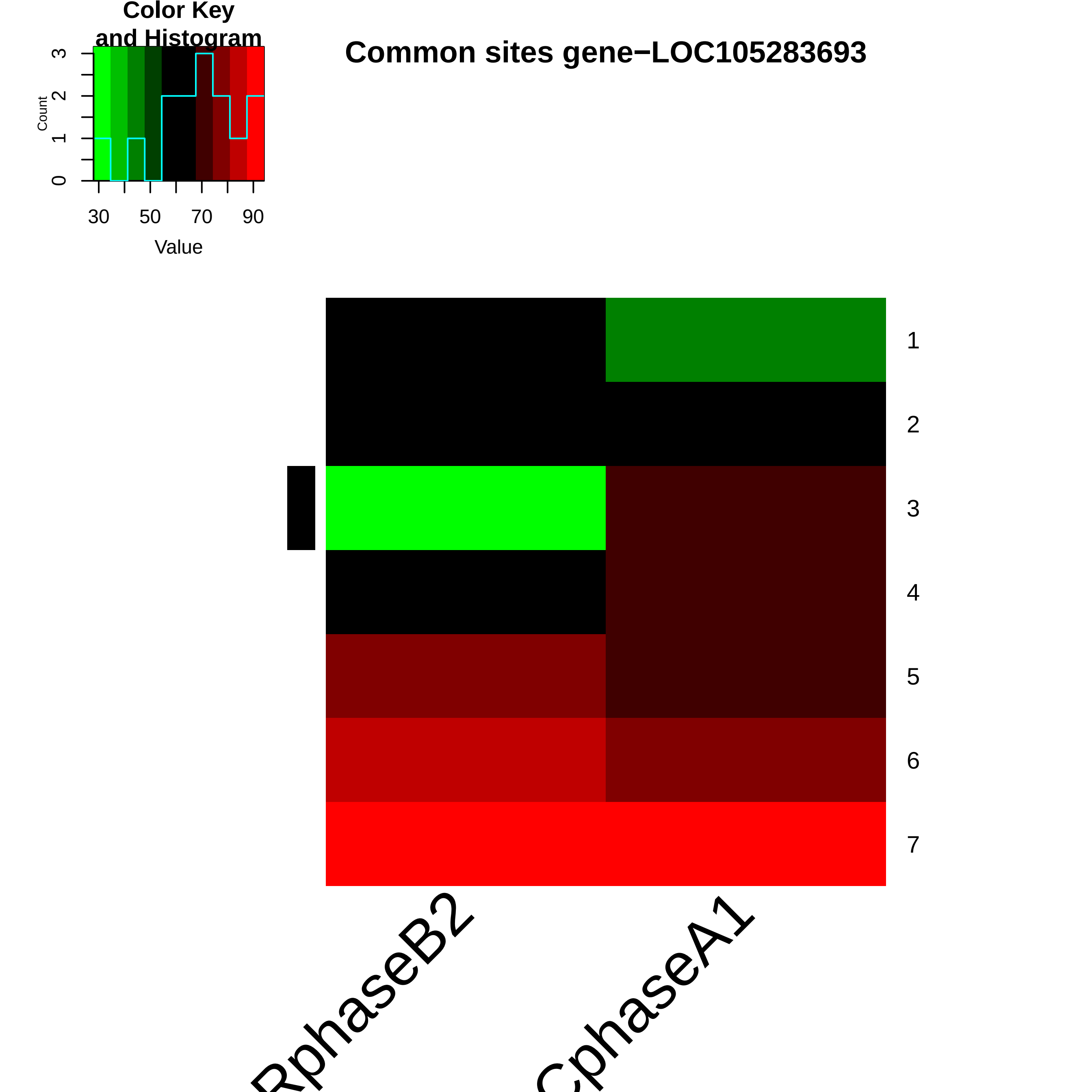

Color Key
and Histogram
Count
0
1
2
3
30
50
70
90
Value
Common sites gene−LOC105283693
1
2
3
4
5
6
7
RphaseB2
CphaseA1

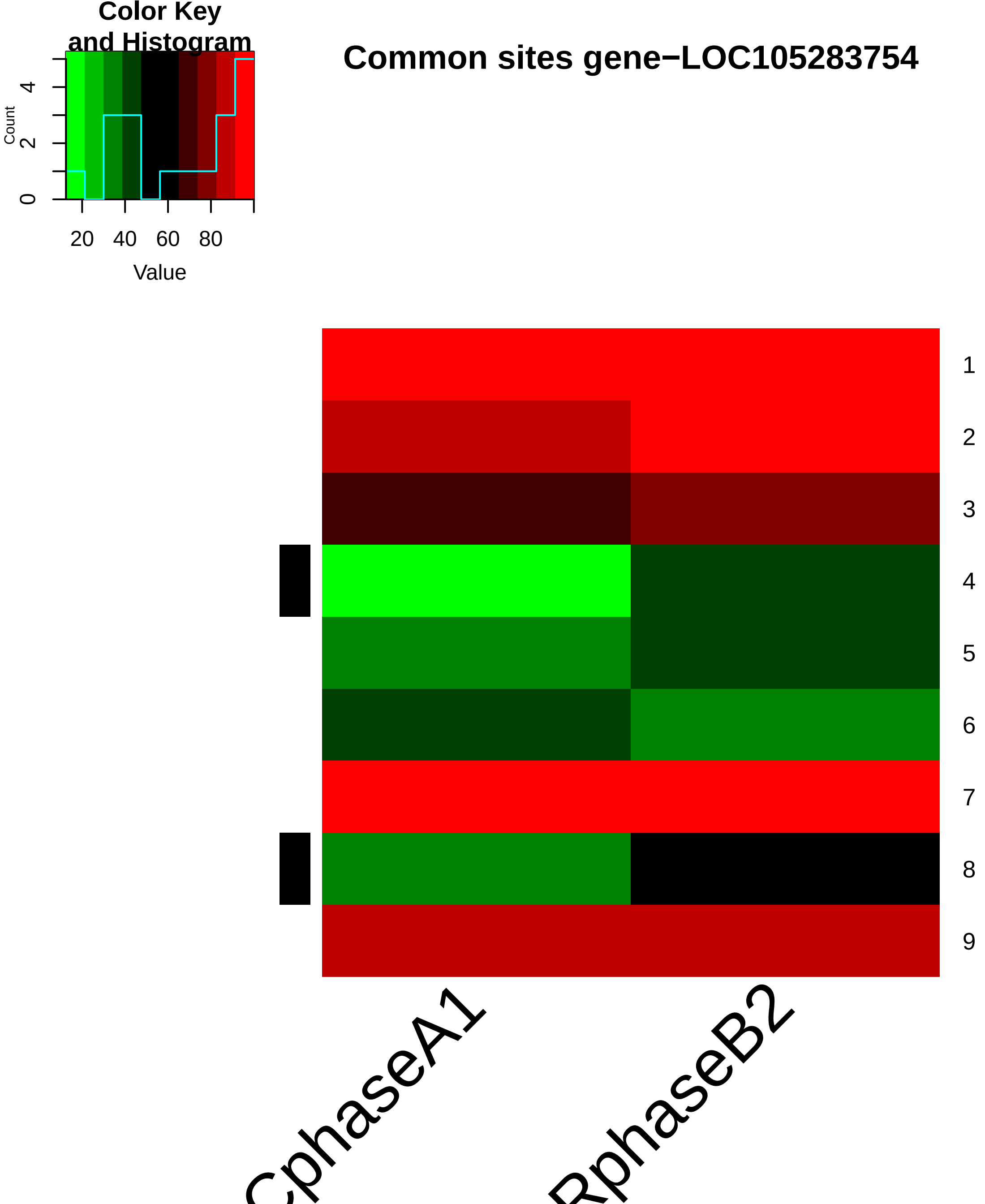
Color Key
and Histogram
Count
0
2
4
20
40
60
80
Value
Common sites gene−LOC105283754
1
2
3
4
5
6
7
8
9
CphaseA1
RphaseB2

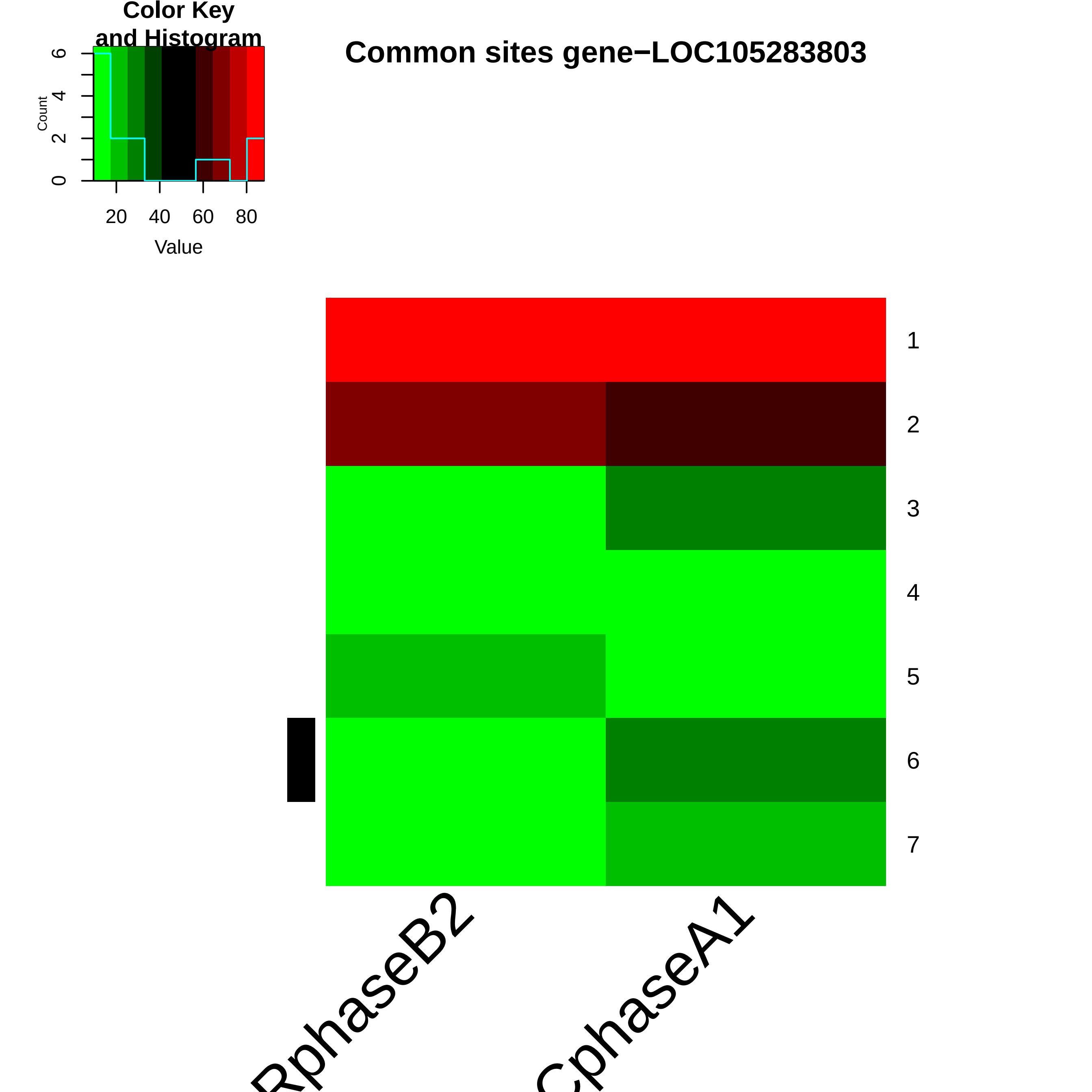

Color Key
and Histogram
Count
0
2
4
6
20
40
60
80
Value
Common sites gene−LOC105283803
1
2
3
4
5
6
7
RphaseB2
CphaseA1

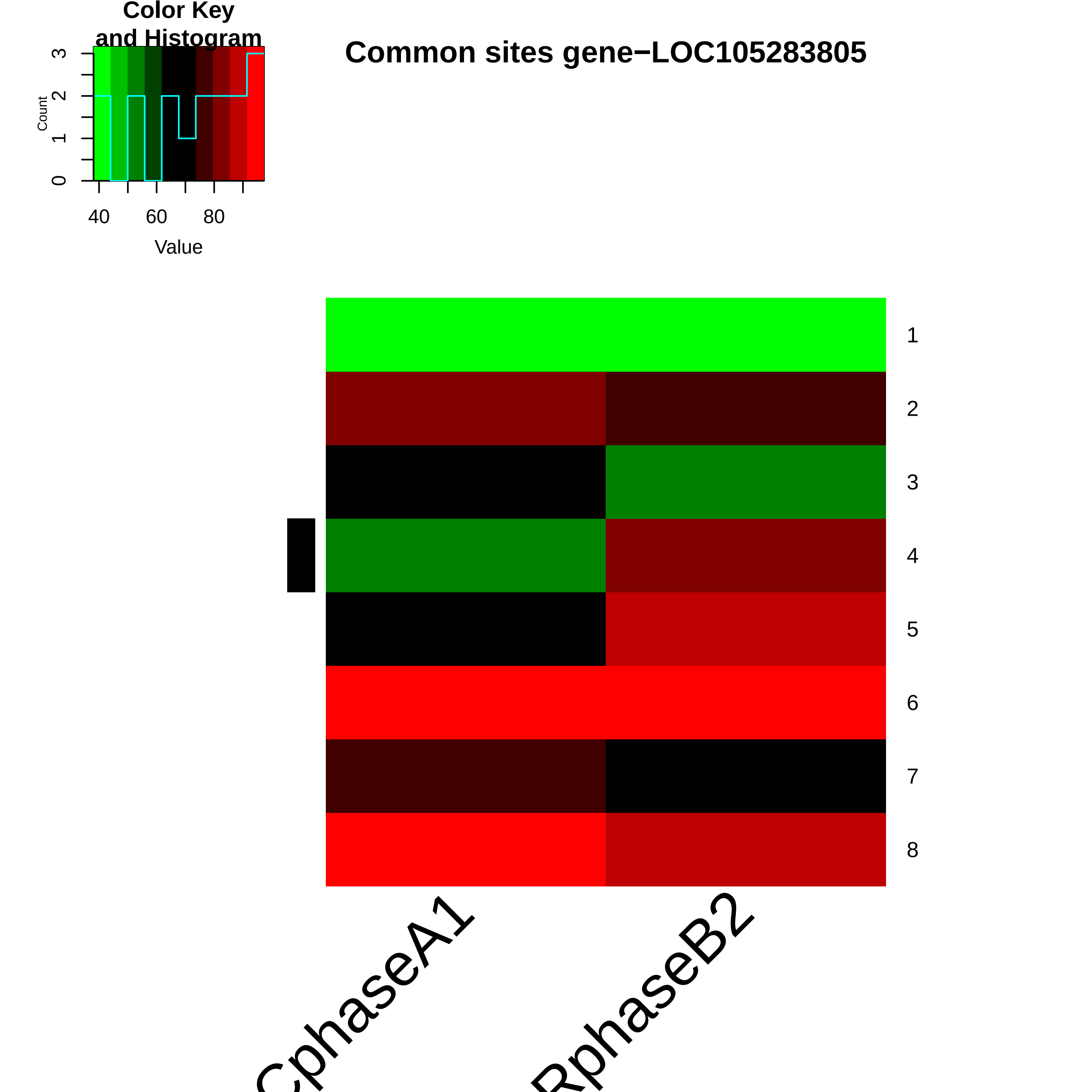

Color Key
and Histogram
Count
0
1
2
3
40
60
80
Value
Common sites gene−LOC105283805
1
2
3
4
5
6
7
8
CphaseA1
RphaseB2

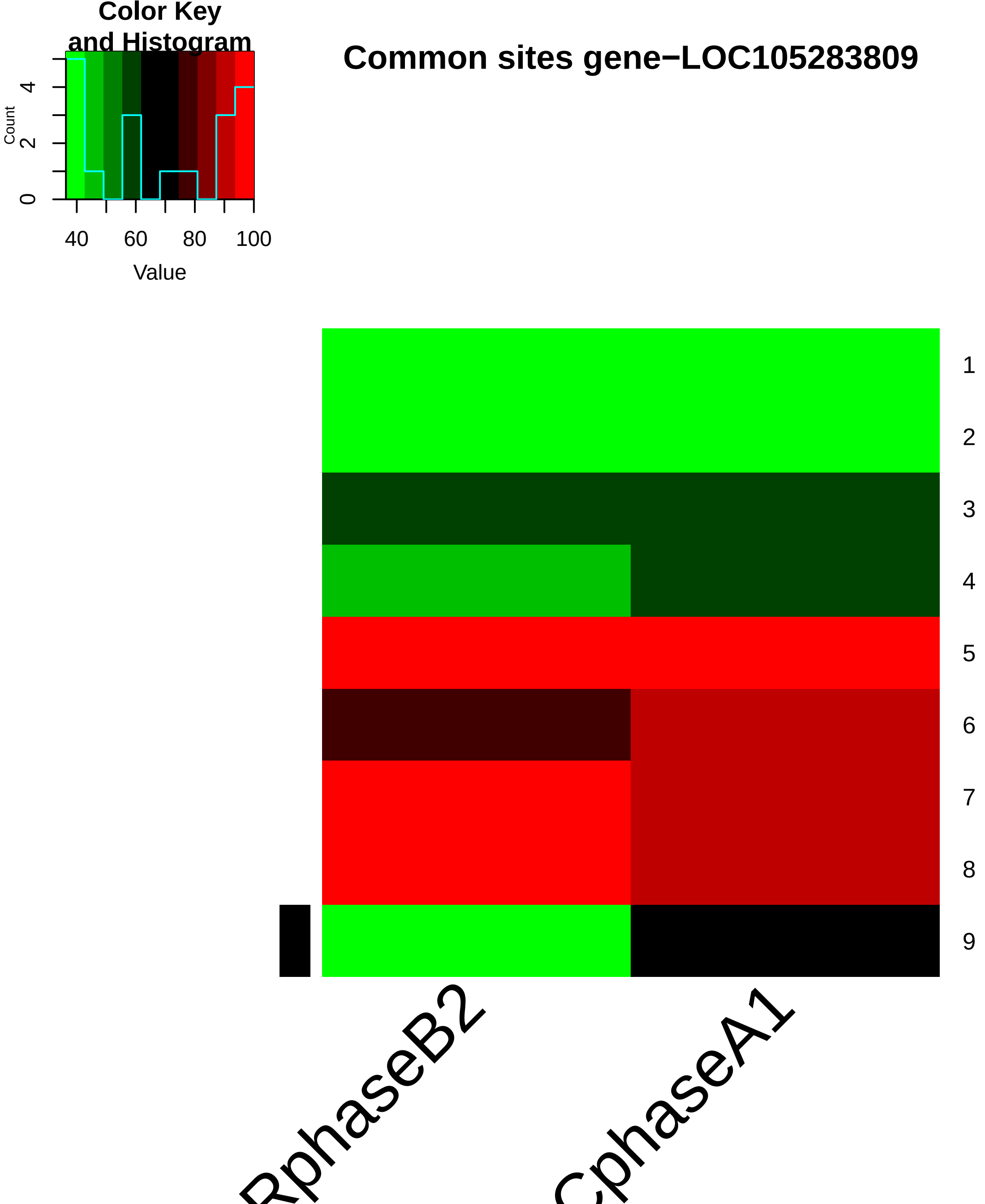
Color Key
and Histogram
Count
0
2
4
40
60
80
100
Value
Common sites gene−LOC105283809
1
2
3
4
5
6
7
8
9
RphaseB2
CphaseA1

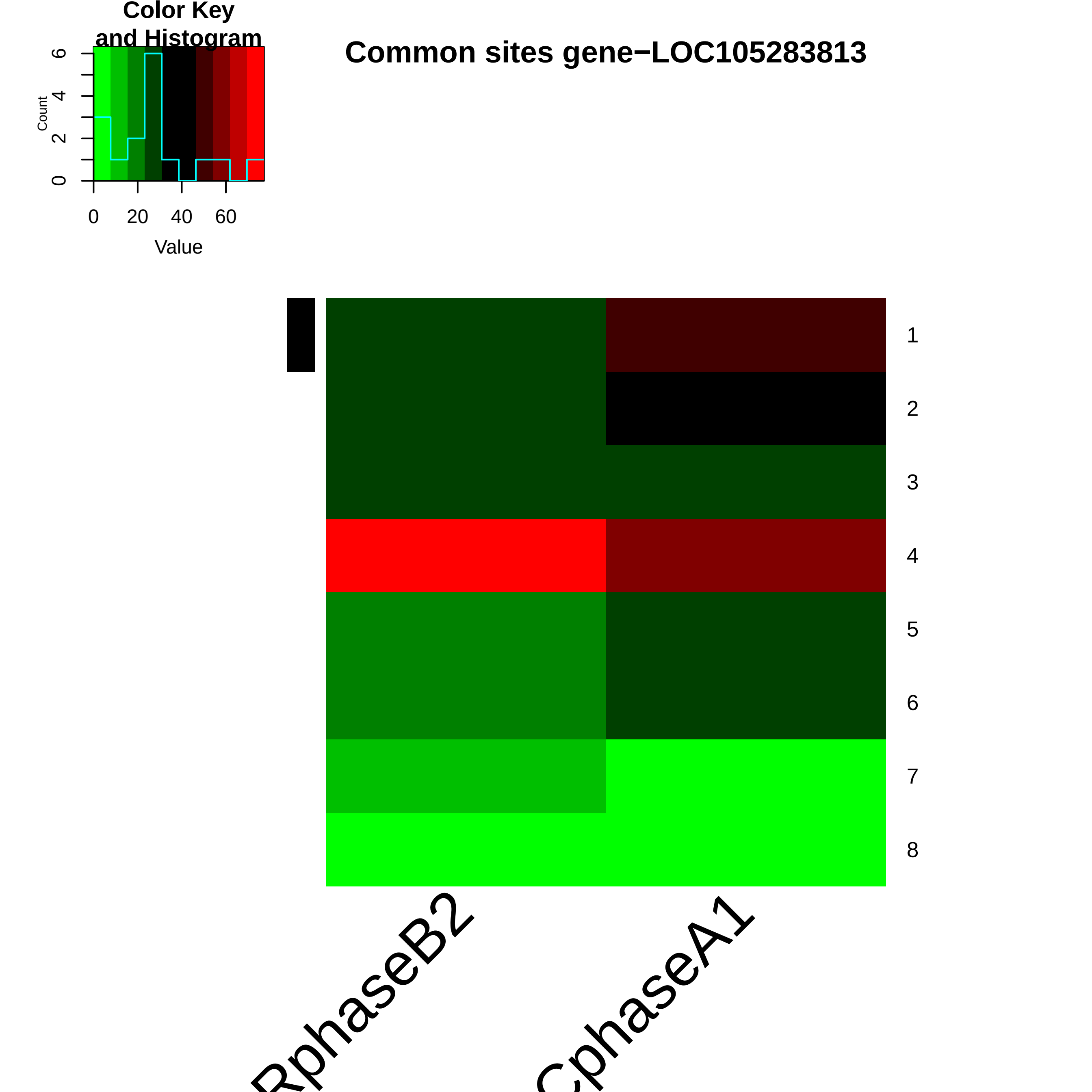
Color Key
and Histogram
Count
0
2
4
6
0
20
40
60
Value
Common sites gene−LOC105283813
1
2
3
4
5
6
7
8
RphaseB2
CphaseA1

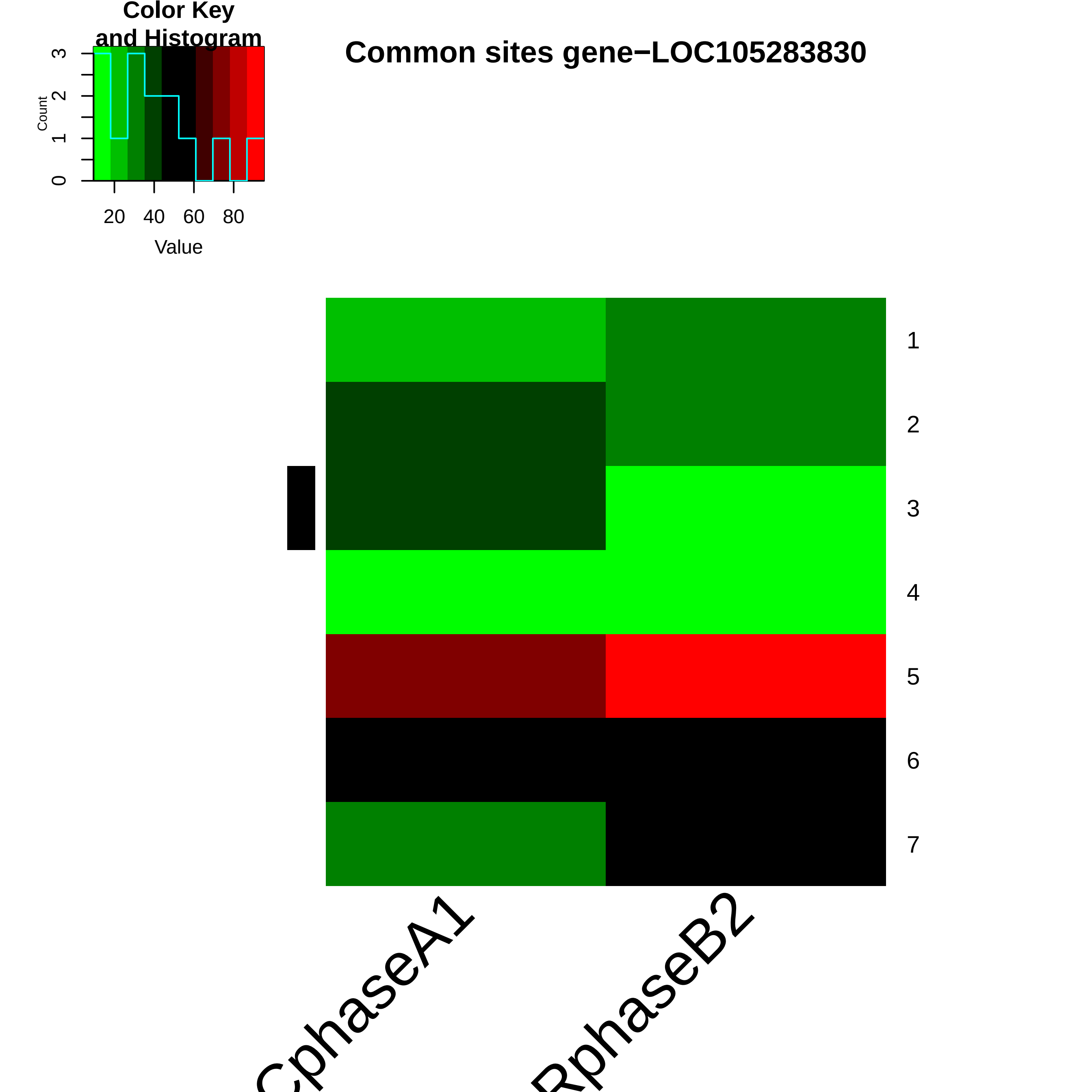

Color Key
and Histogram
Count
0
1
2
3
20
40
60
80
Value
Common sites gene−LOC105283830
1
2
3
4
5
6
7
CphaseA1
RphaseB2

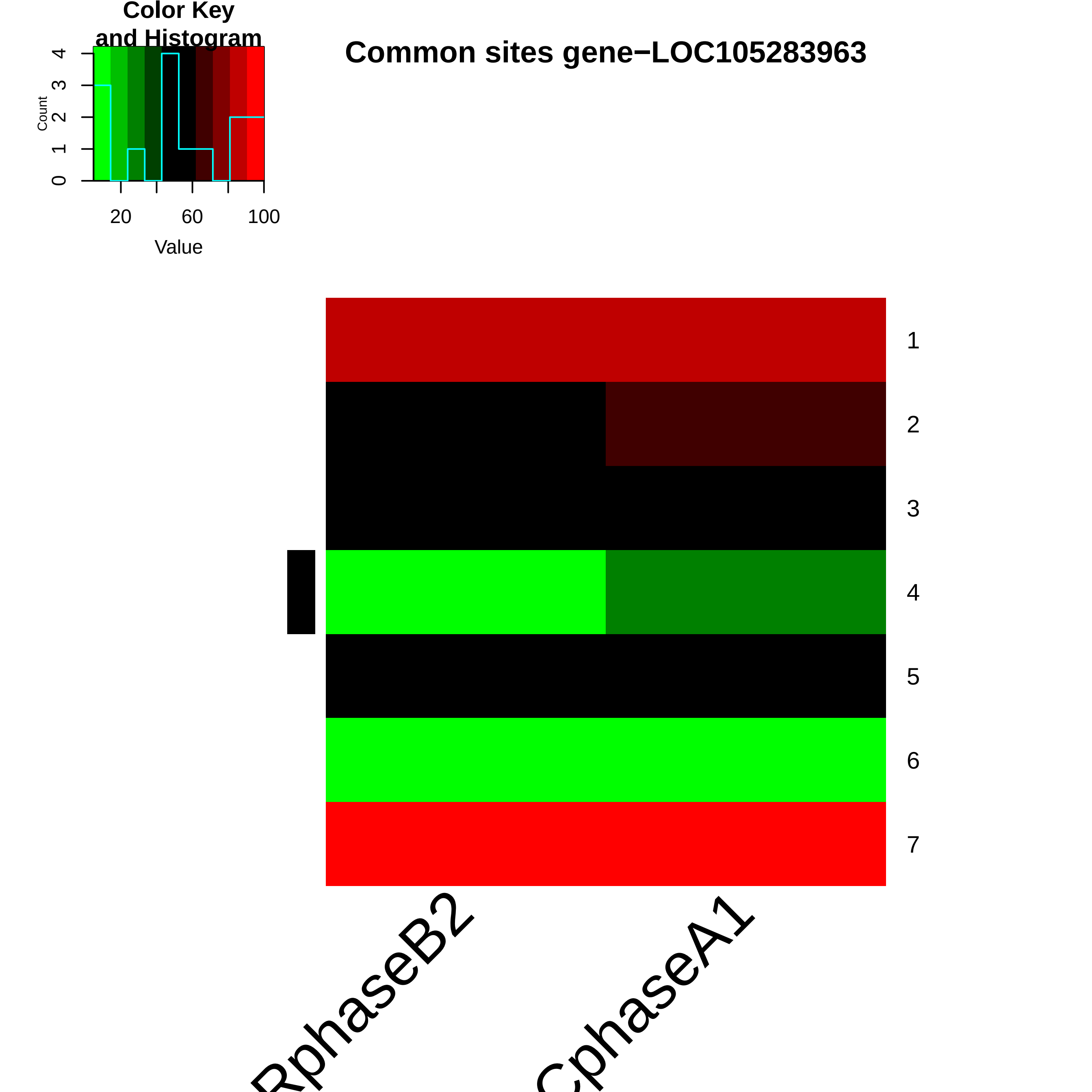

Color Key
and Histogram
Count
Value
20
60
100
0
1
2
3
4
Common sites gene−LOC105283963
1
2
3
4
5
6
7
RphaseB2
CphaseA1

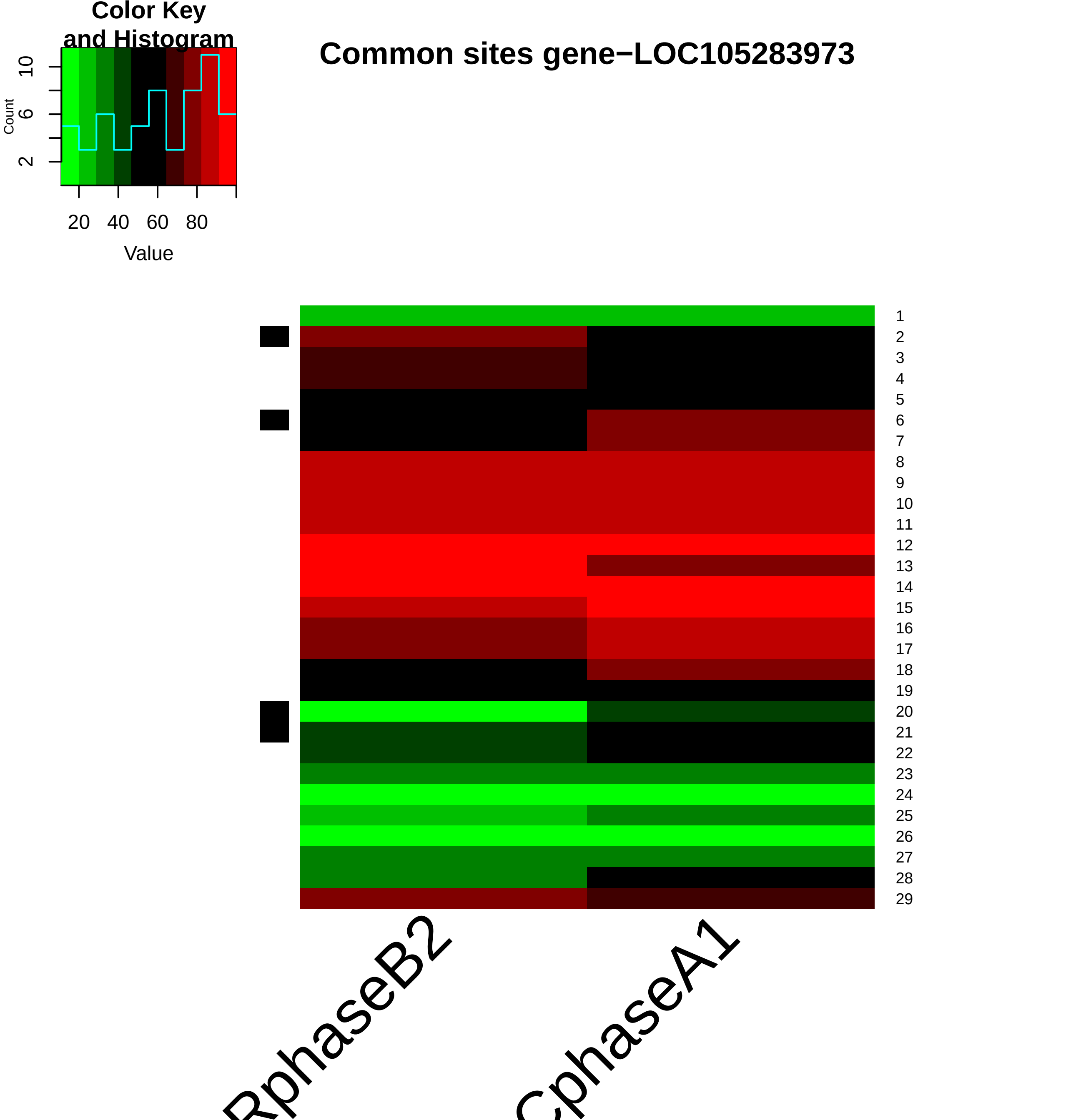

Color Key
and Histogram
Count
2
6
10
20
40
60
80
Value
Common sites gene−LOC105283973
RphaseB2
CphaseA1

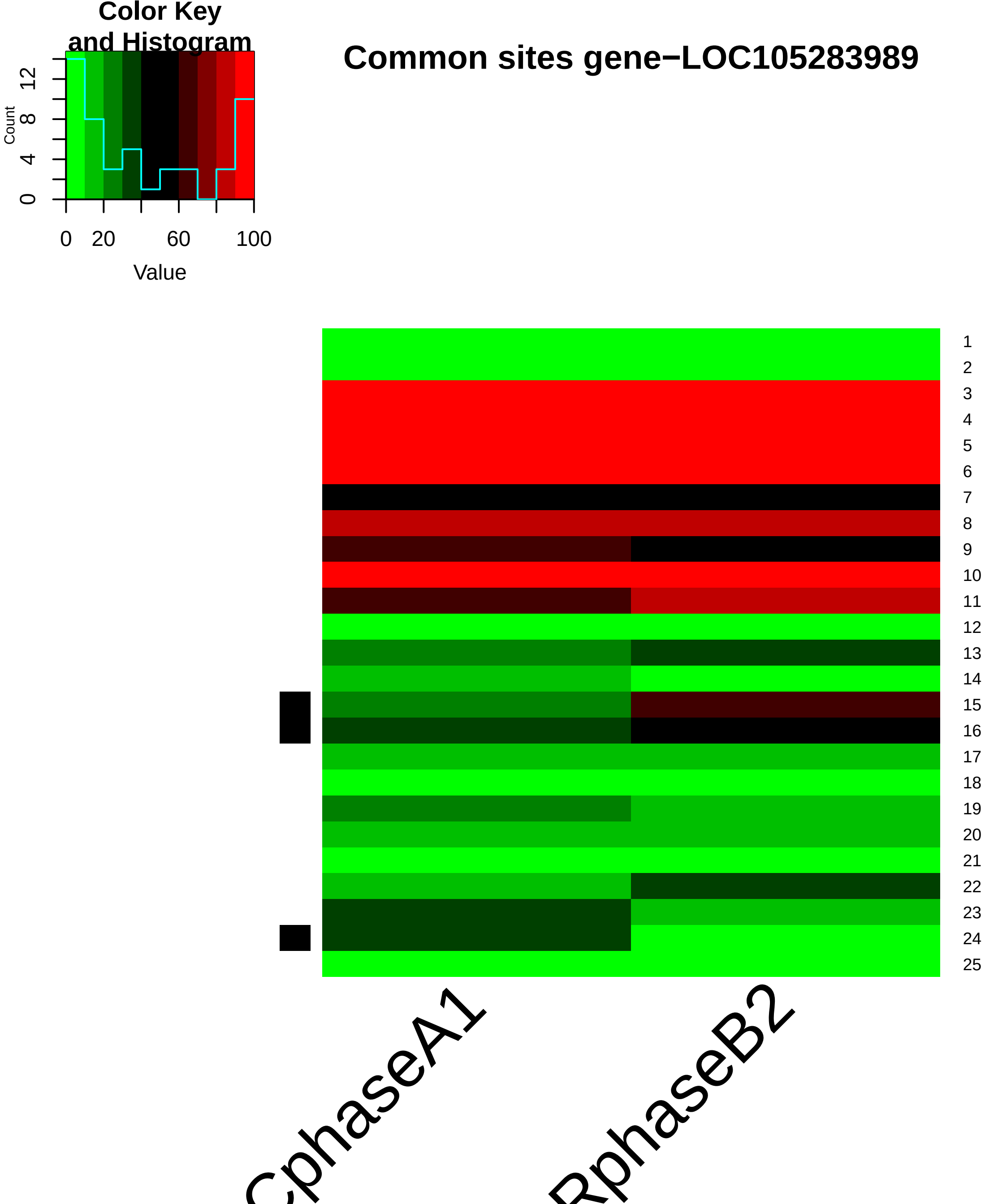
Color Key
and Histogram
Count
0
4
8
12
0
20
60
100
Value
Common sites gene−LOC105283989

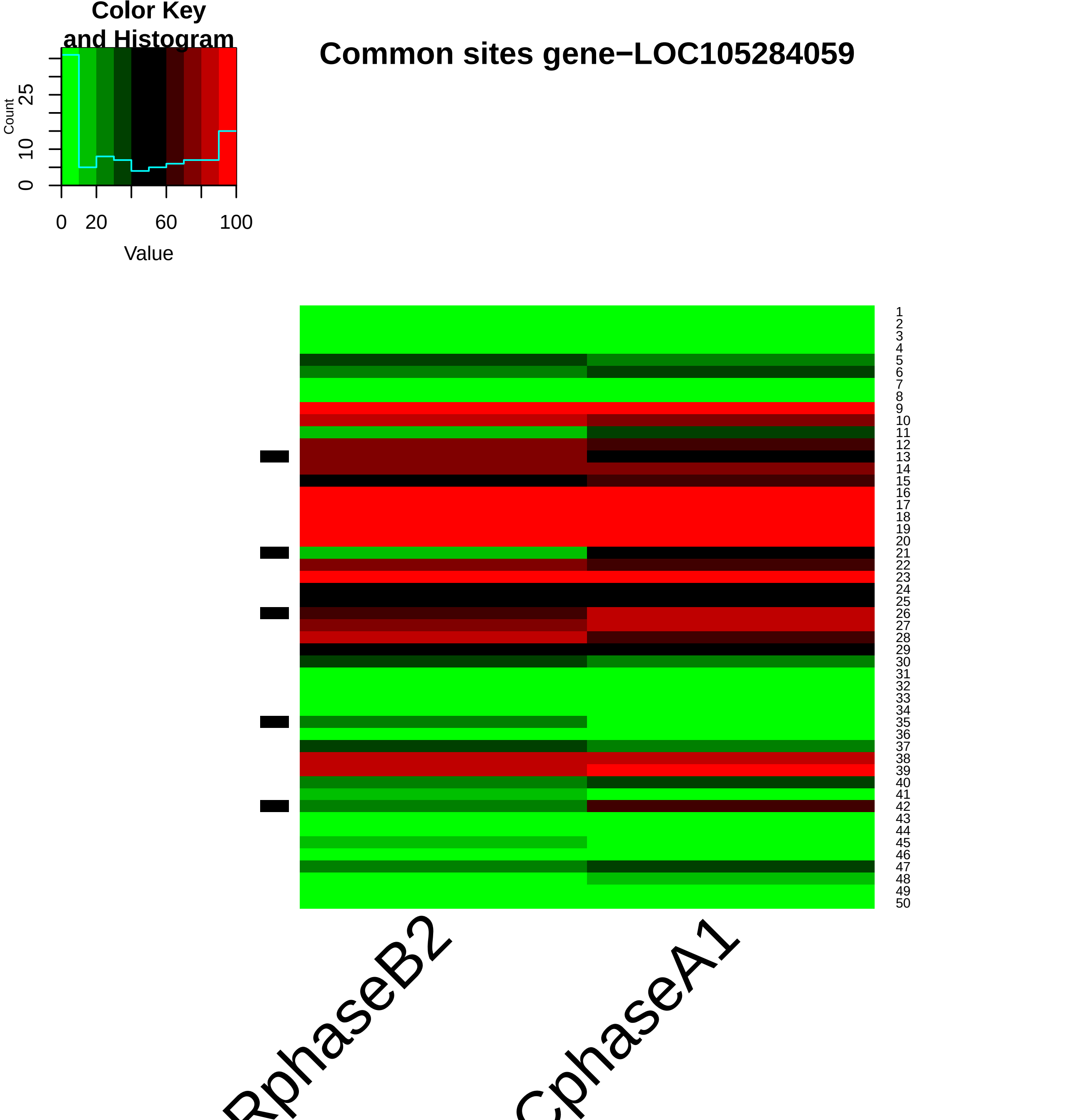
Color Key
and Histogram
Count
0
10
25
0
20
60
100
Value
Common sites gene−LOC105284059
RphaseB2
CphaseA1

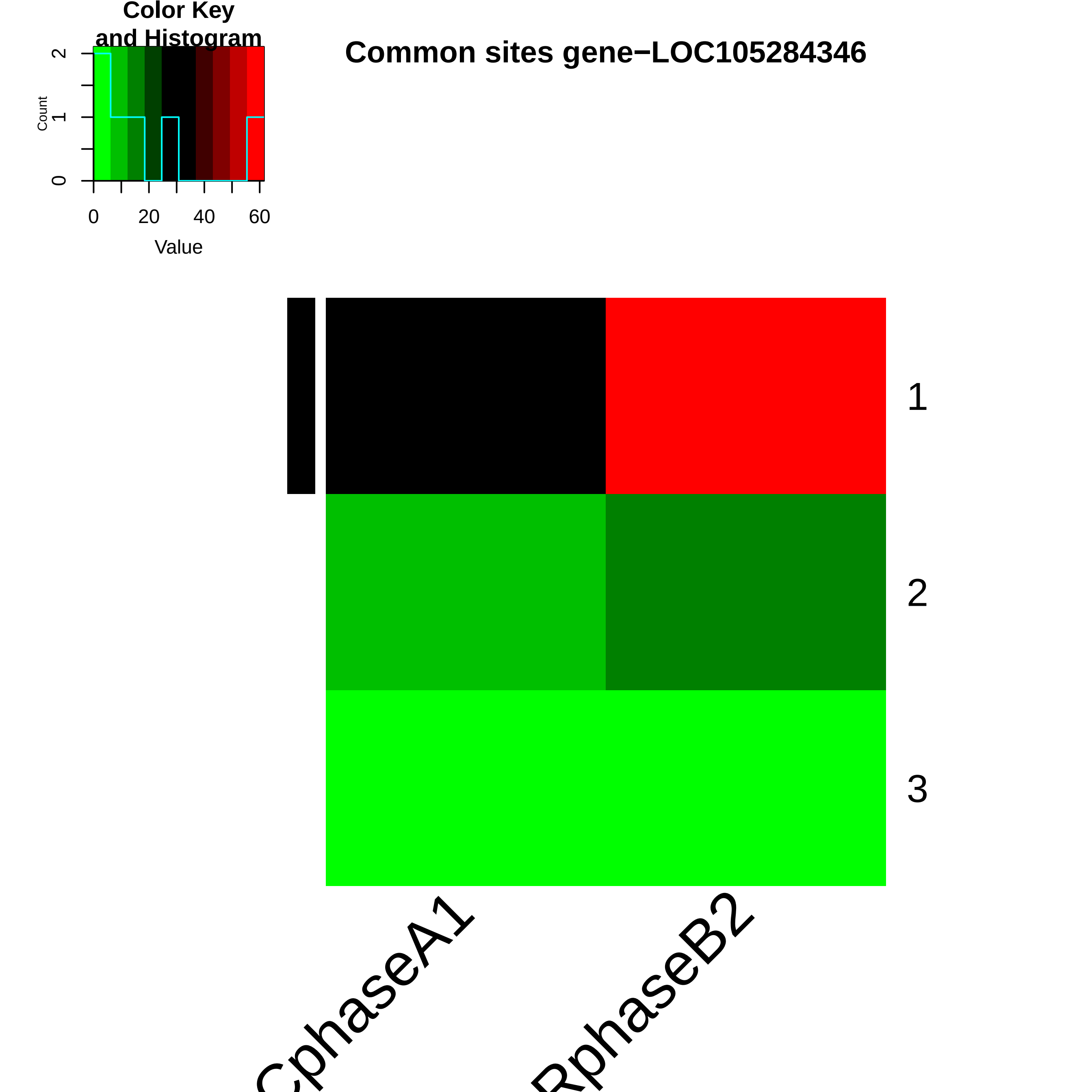
Color Key
and Histogram
Count
Value
0
20
40
60
0
1
2
Common sites gene−LOC105284346
1
2
3

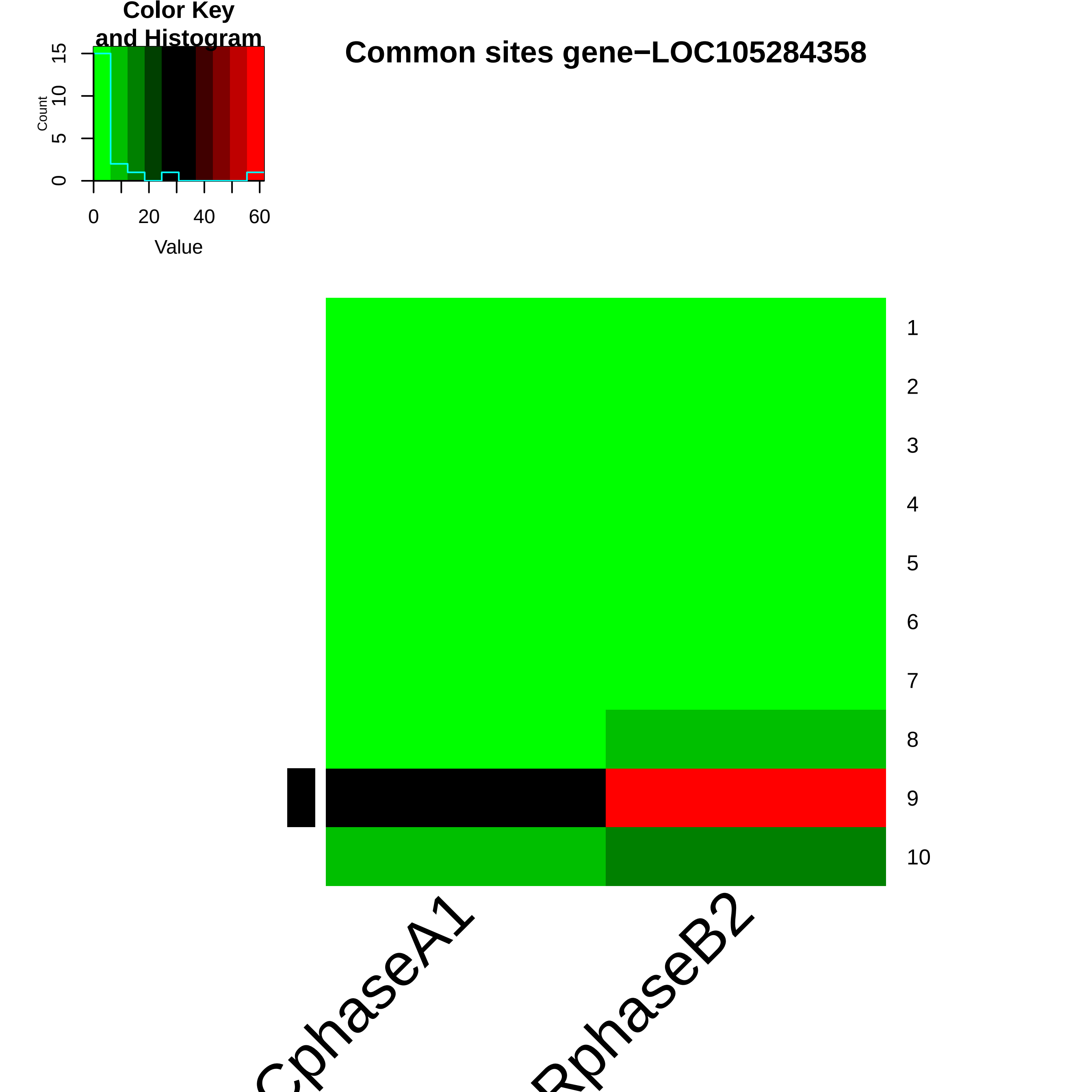
Color Key
and Histogram
Count
0
5
10
15
0
20
40
60
Value
Common sites gene−LOC105284358
1
2
3
4
5
6
7
8
9
10
CphaseA1
RphaseB2

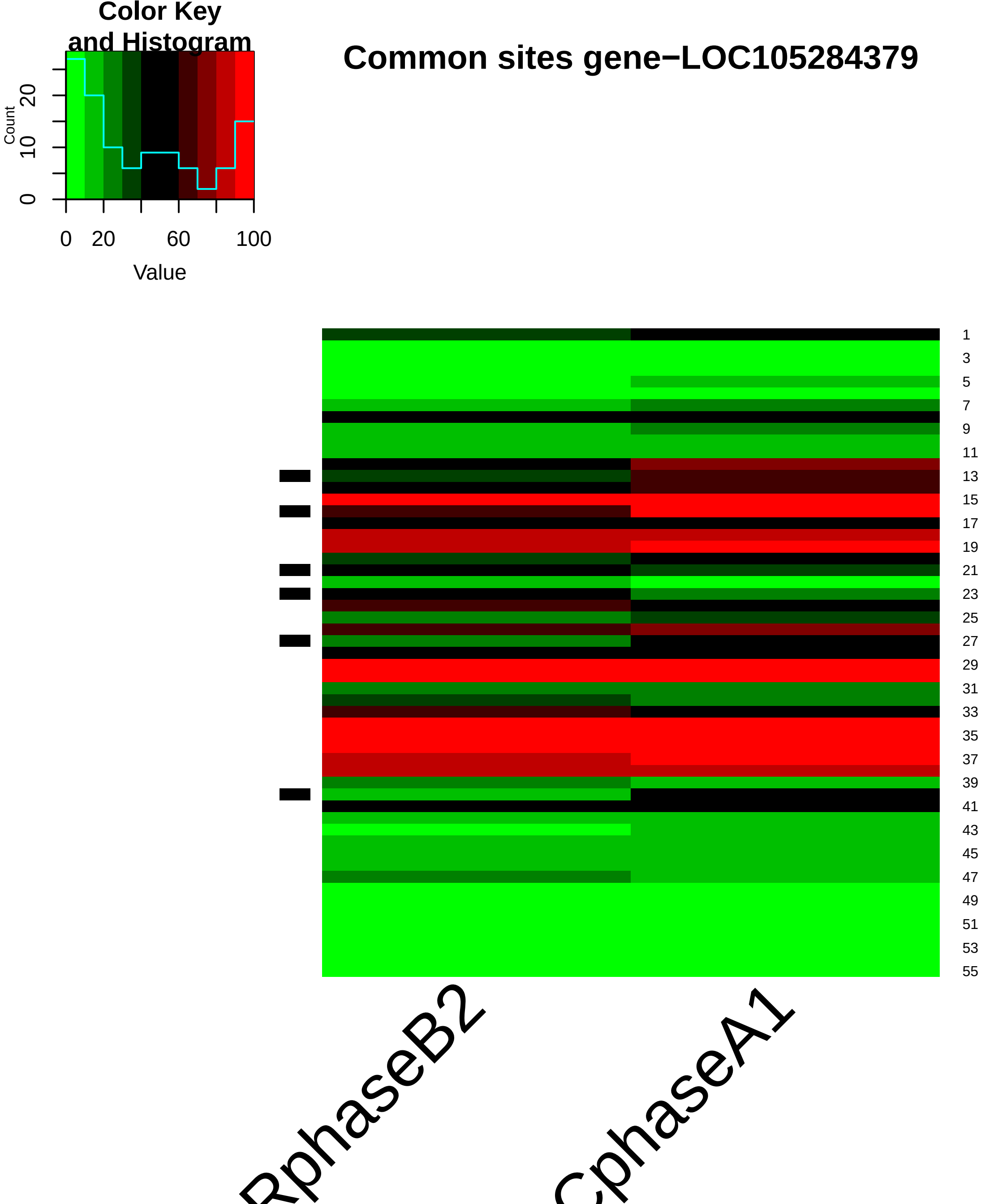

Color Key
and Histogram
Count
0
10
20
0
20
60
100
Value
Common sites gene−LOC105284379
1
3
5
7
9
11
13
15
17
19
21
23
25
27
29
31
33
35
37
39
41
43
45
47
49
51
53
55
RphaseB2
CphaseA1

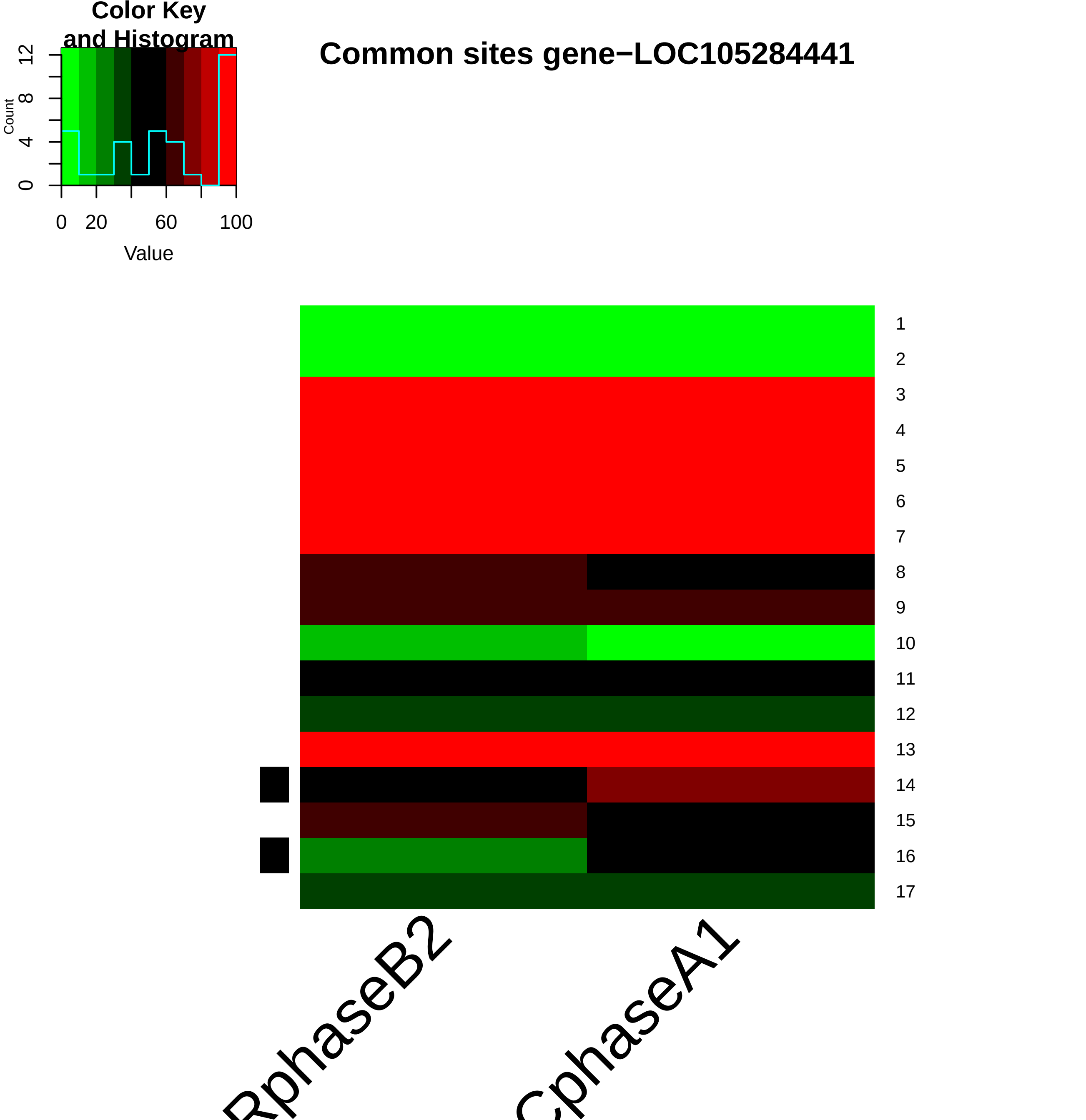
Color Key
and Histogram
Count
0
4
8
12
0
20
60
100
Value
Common sites gene−LOC105284441
RphaseB2
CphaseA1

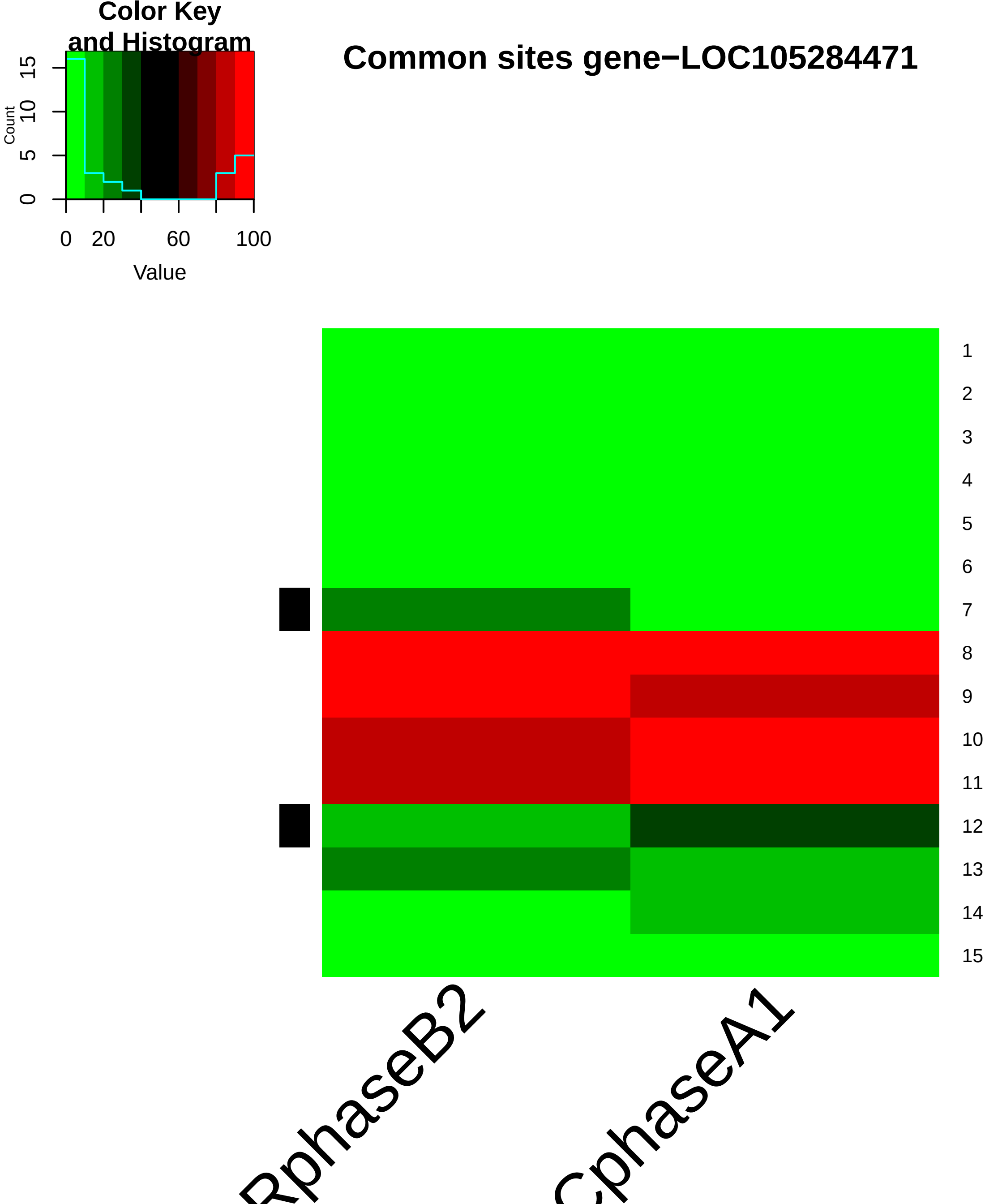

Color Key
and Histogram
Count
0
5
10
15
0
20
60
100
Value
Common sites gene−LOC105284471
RphaseB2
CphaseA1

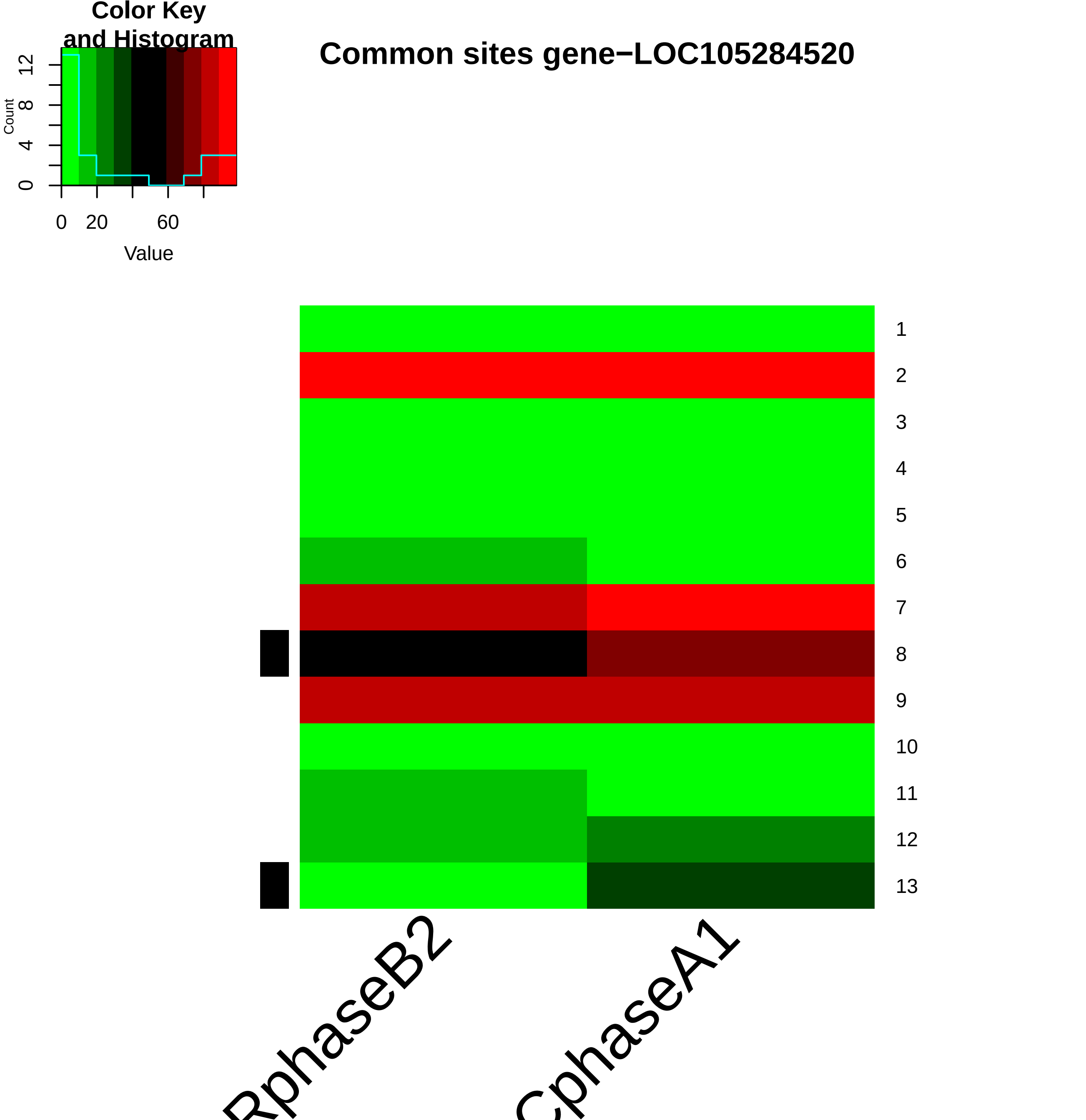
Color Key
and Histogram
Count
0
4
8
12
0
20
60
Value
Common sites gene−LOC105284520
1
2
3
4
5
6
7
8
9
10
11
12
13
RphaseB2
CphaseA1

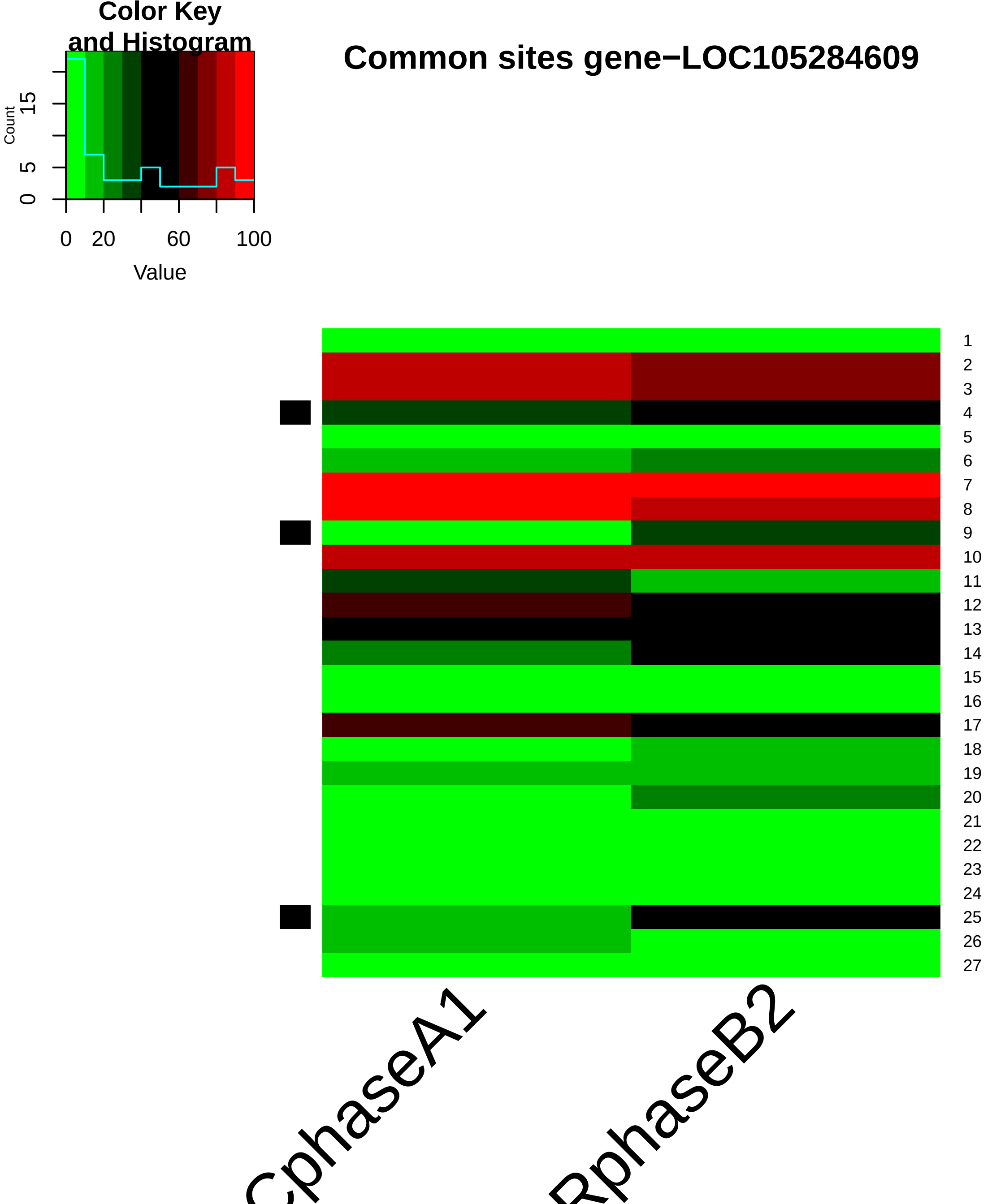

Color Key
and Histogram
Count
0
5
15
0
20
60
100
Value
Common sites gene−LOC105284609
CphaseA1
RphaseB2

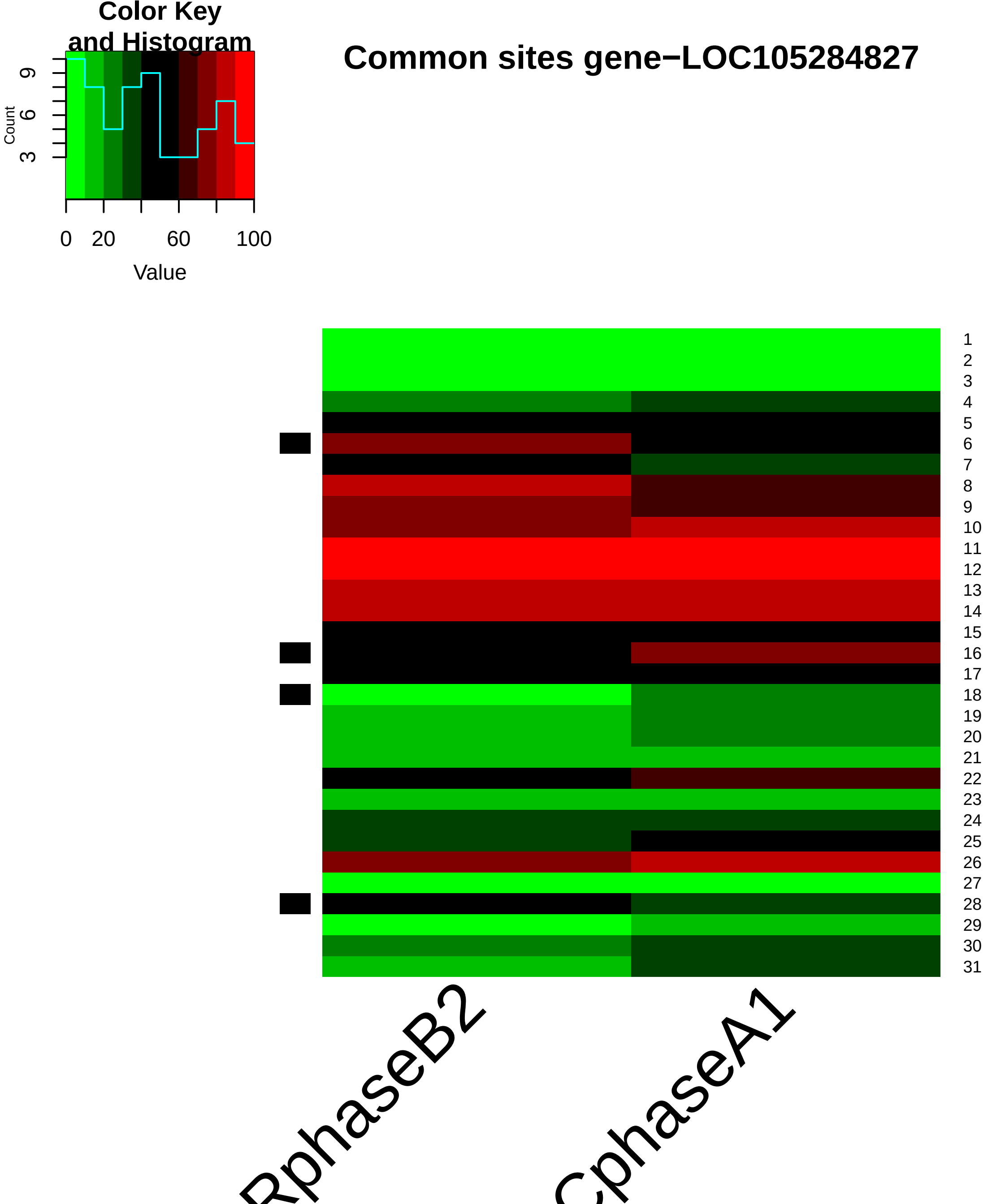
Color Key
and Histogram
Count
3
6
9
0
20
60
100
Value
Common sites gene−LOC105284827
RphaseB2
CphaseA1

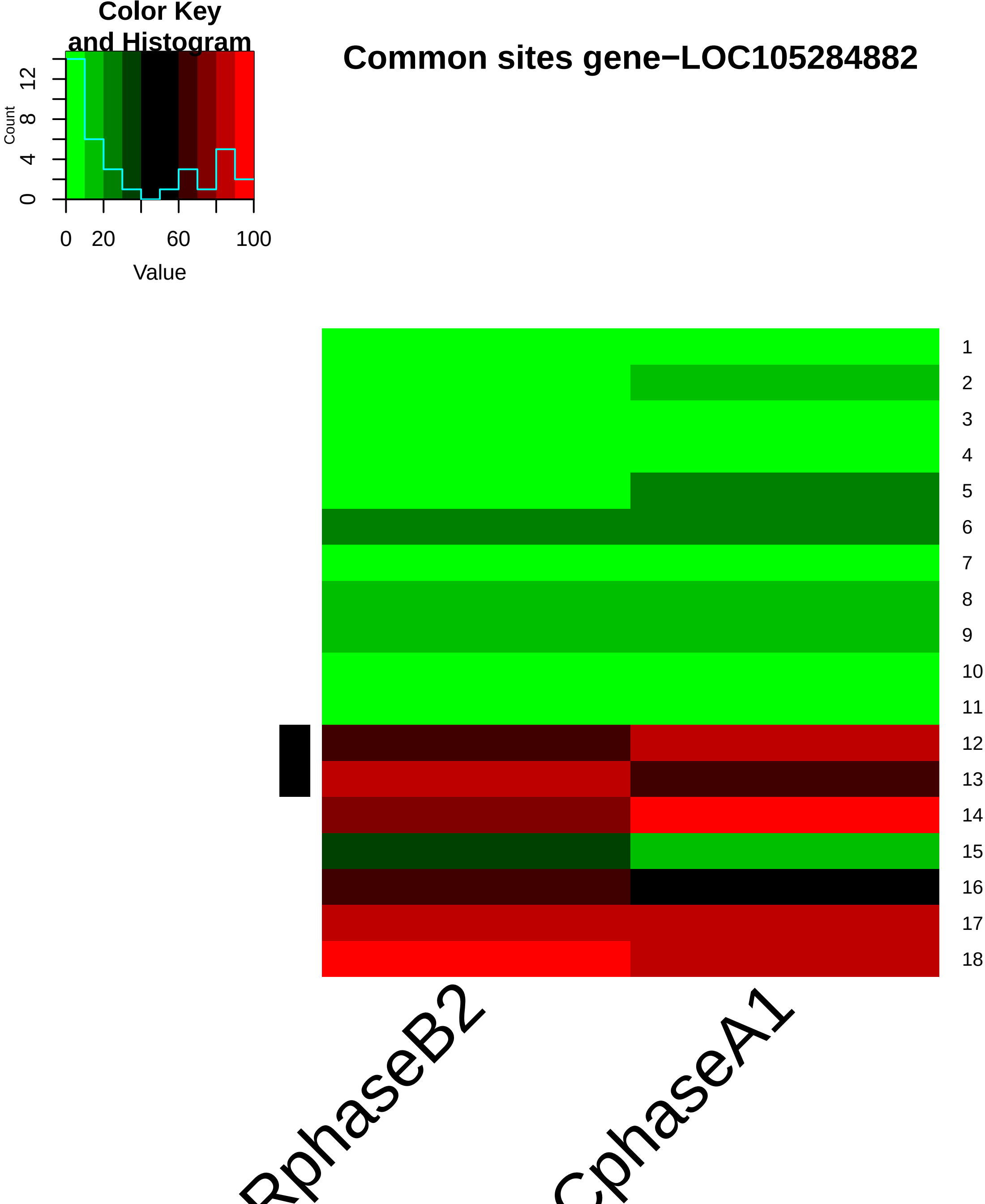
Color Key
and Histogram
Count
0
4
8
12
0
20
60
100
Value
Common sites gene−LOC105284882
RphaseB2
CphaseA1

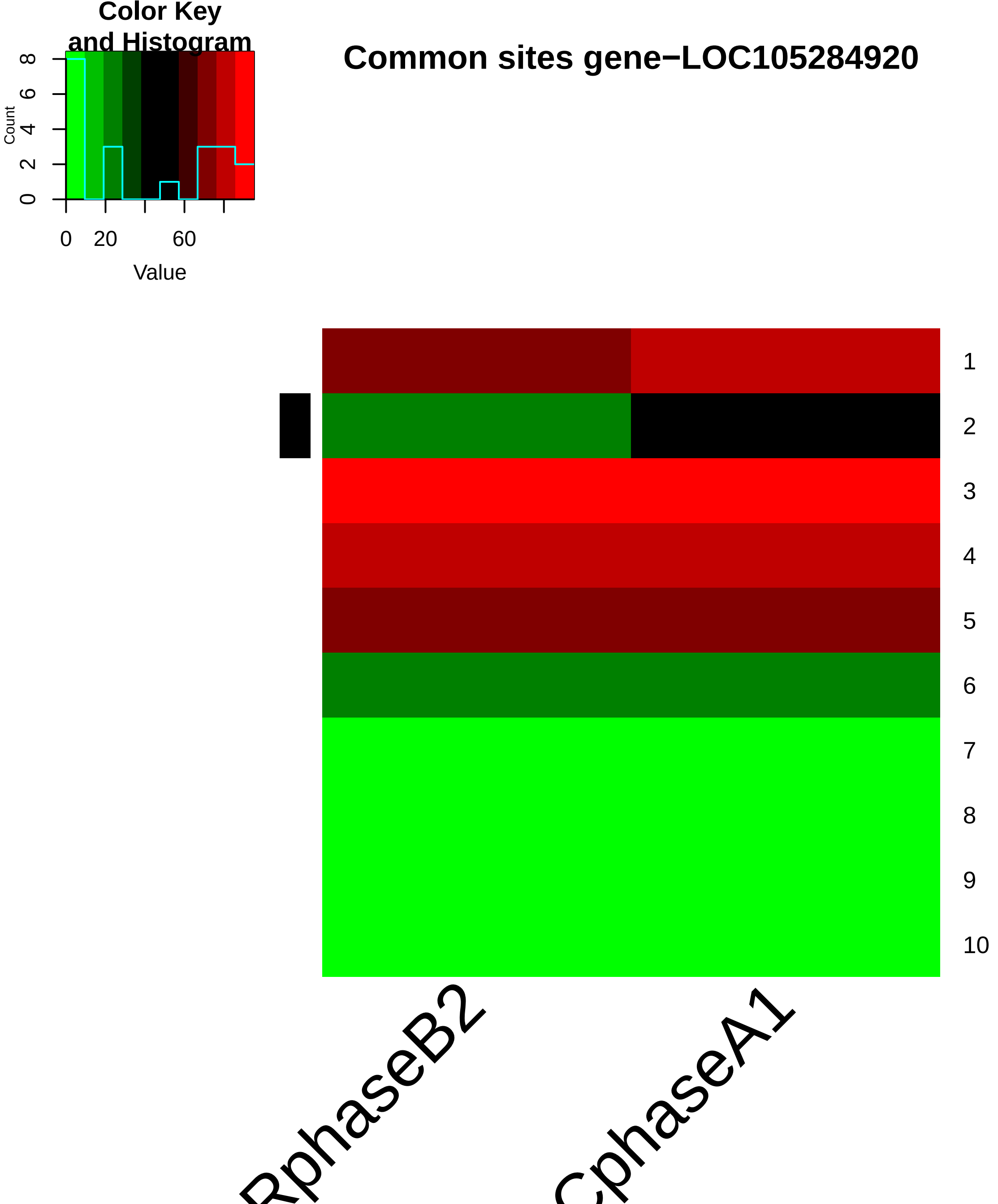
Color Key
and Histogram
Count
Value
0
20
60
0
2
4
6
8
Common sites gene−LOC105284920
1
2
3
4
5
6
7
8
9
10
RphaseB2
CphaseA1

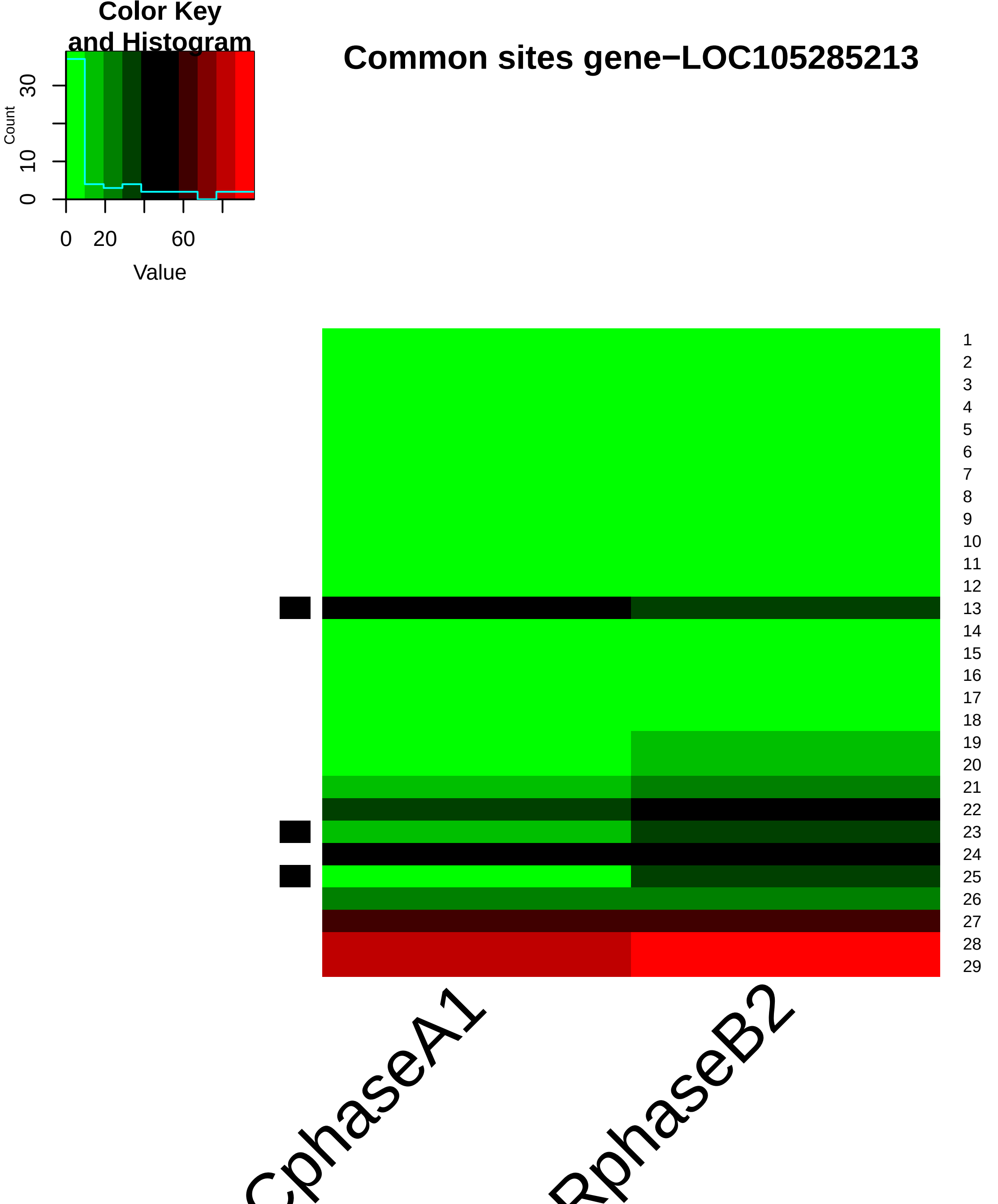
Color Key
and Histogram
Count
0
10
30
0
20
60
Value
Common sites gene−LOC105285213
CphaseA1
RphaseB2

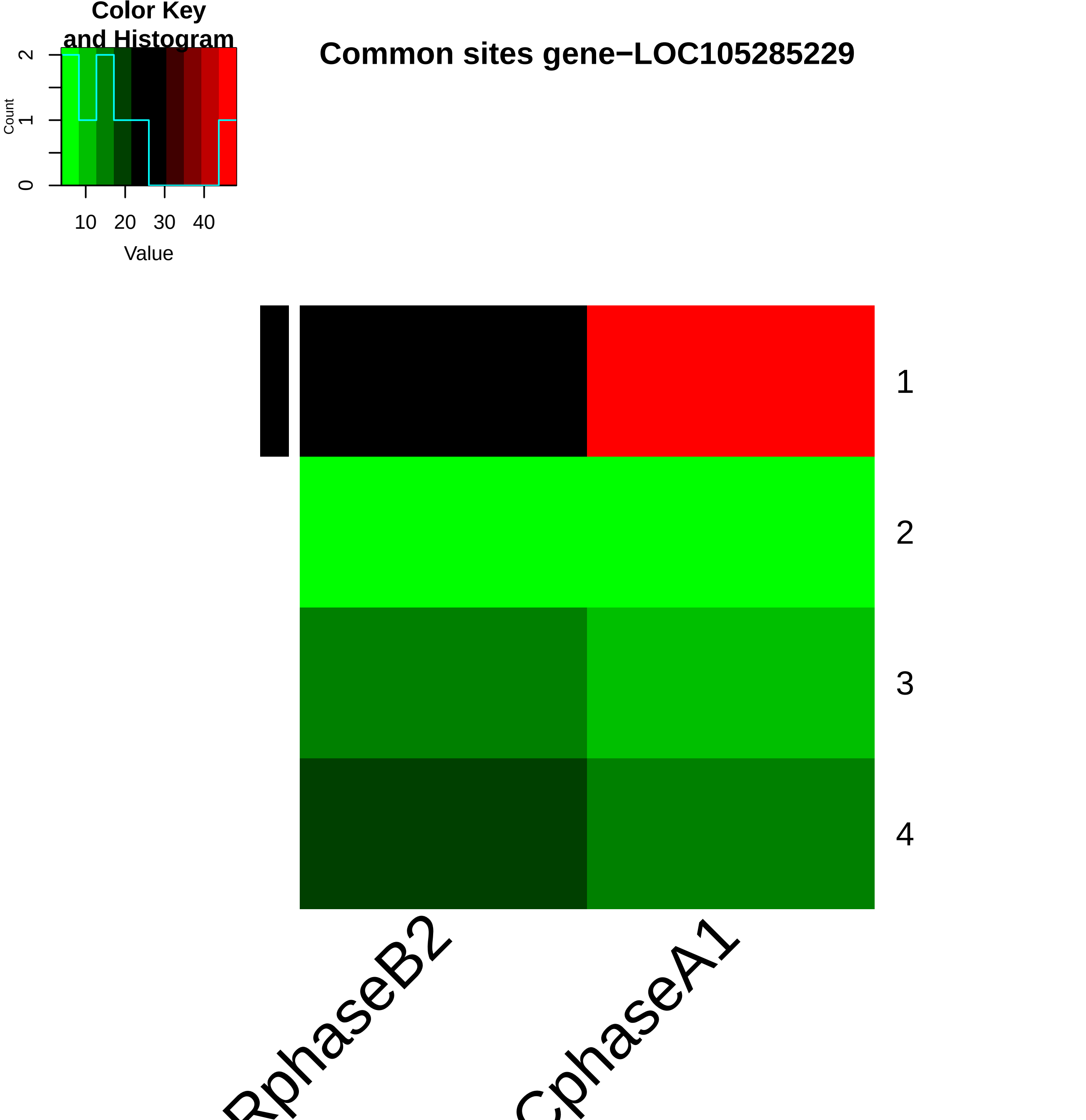

Color Key
and Histogram
Count
0
1
2
10
20
30
40
Value
Common sites gene−LOC105285229
1
2
3
4
RphaseB2
CphaseA1

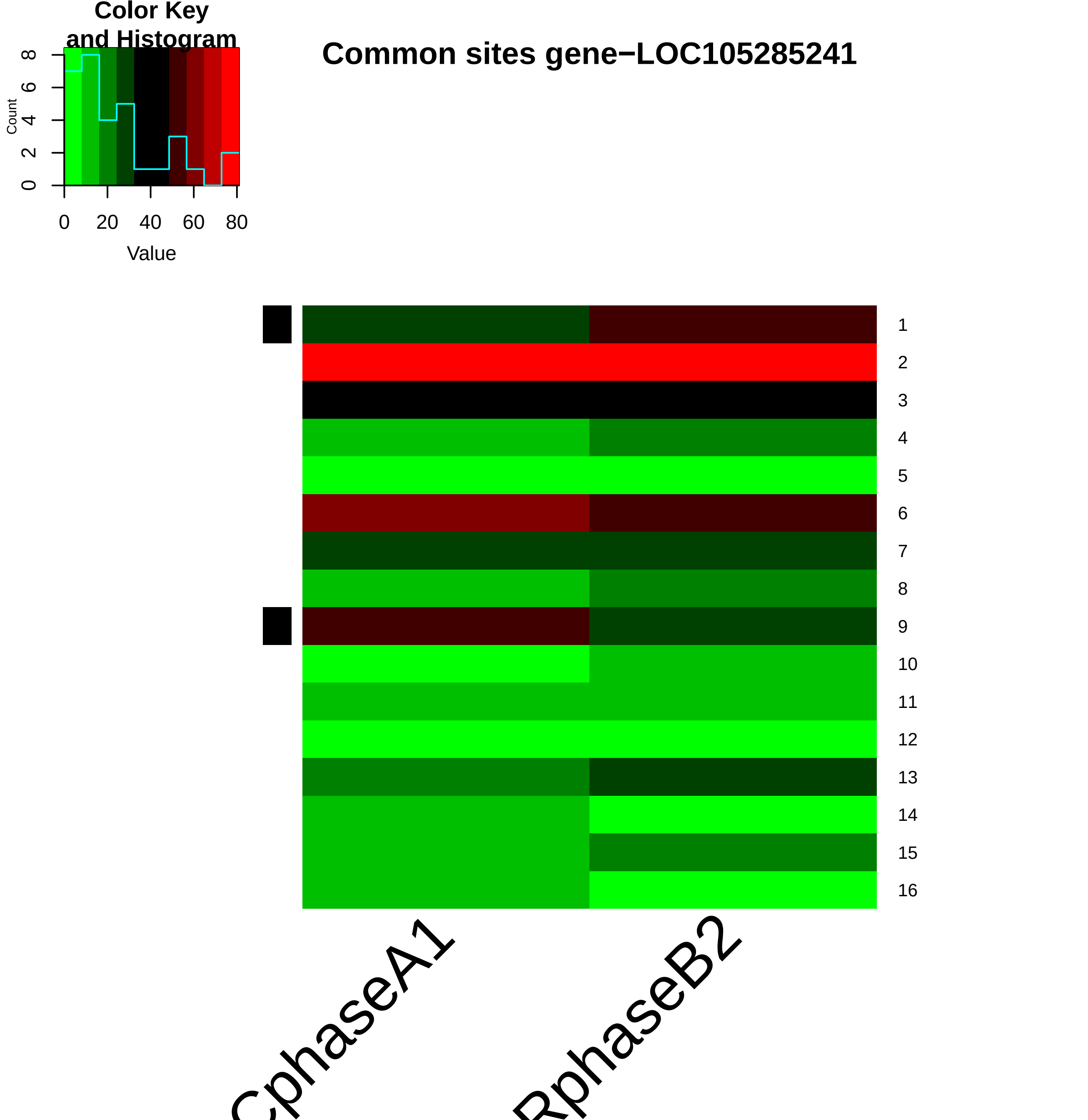
Color Key
and Histogram
Count
0
2
4
6
8
0
20
40
60
80
Value
Common sites gene−LOC105285241
CphaseA1
RphaseB2

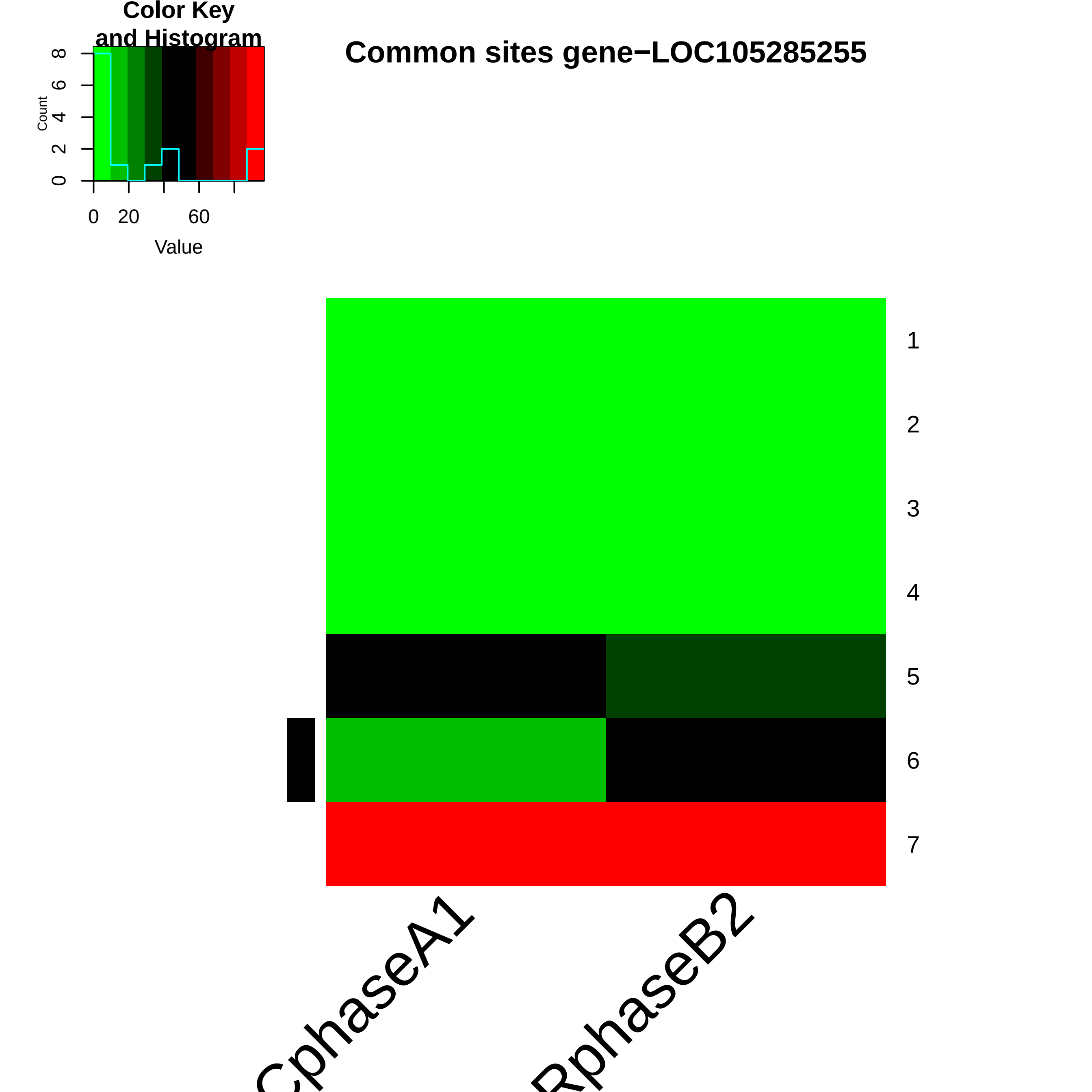
Color Key
and Histogram
Count
0
2
4
6
8
0
20
60
Value
Common sites gene−LOC105285255
1
2
3
4
5
6
7
CphaseA1
RphaseB2

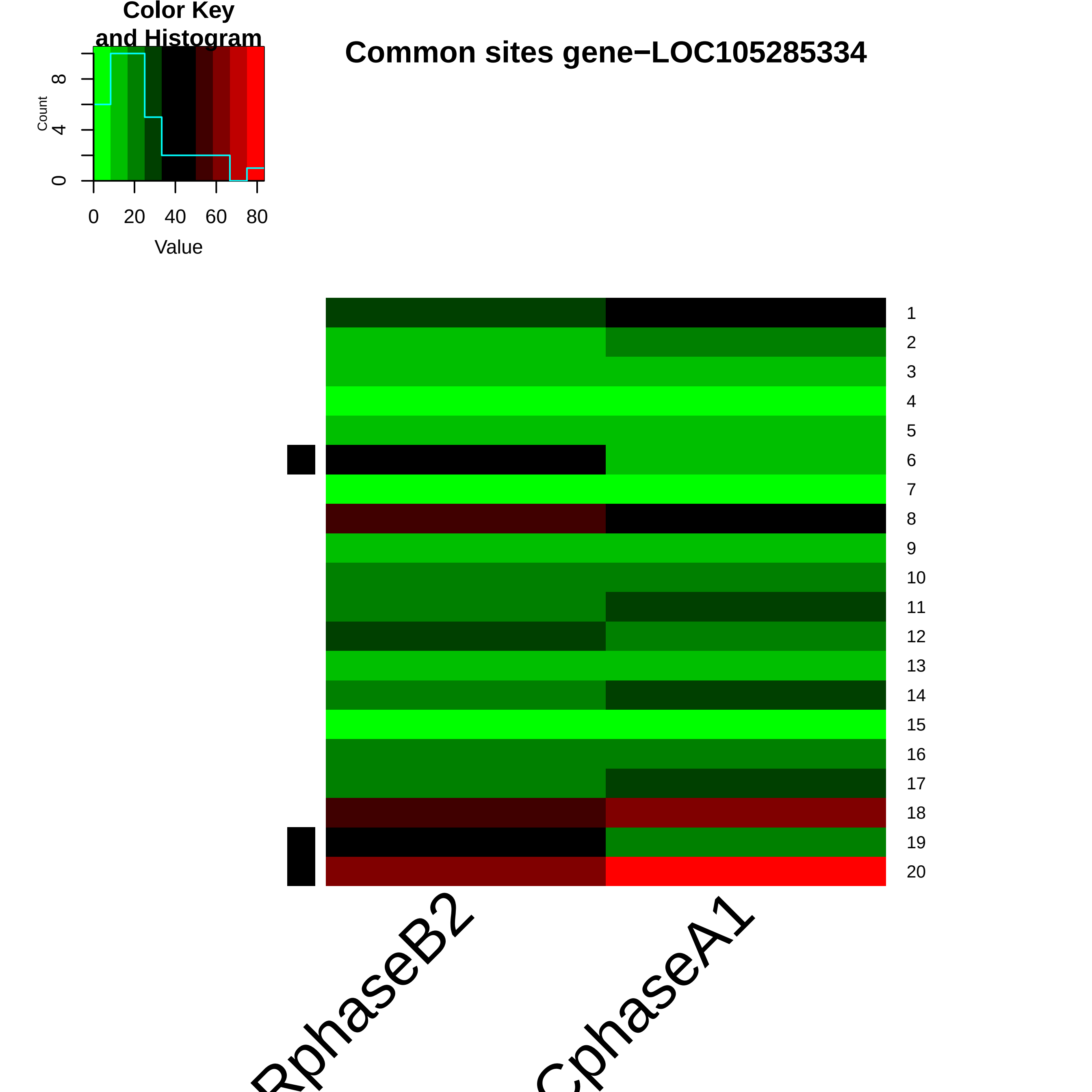

Color Key
and Histogram
Count
0
4
8
0
20
40
60
80
Value
Common sites gene−LOC105285334
RphaseB2
CphaseA1

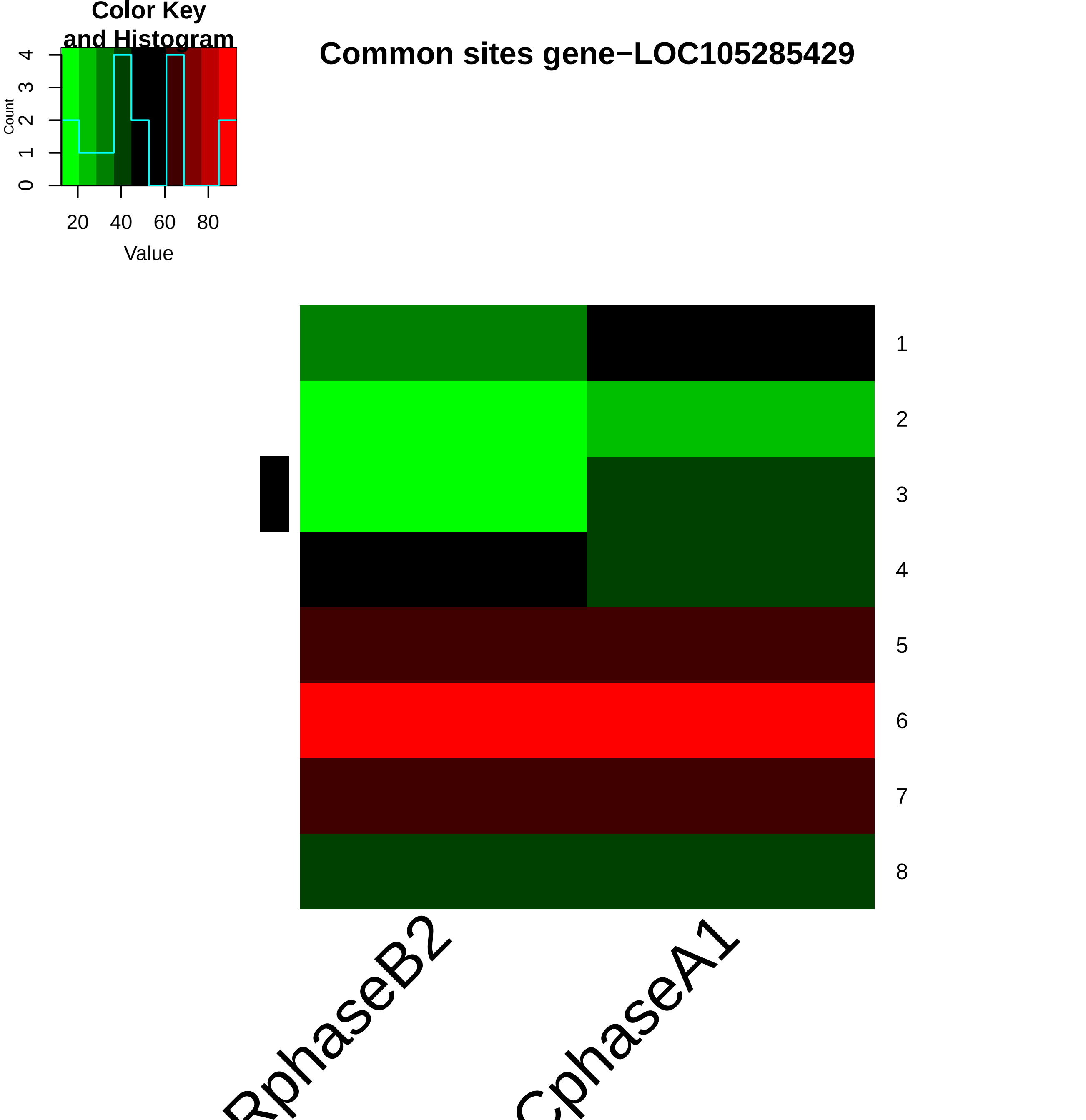
Color Key
and Histogram
Count
0
1
2
3
4
20
40
60
80
Value
Common sites gene−LOC105285429
1
2
3
4
5
6
7
8
RphaseB2
CphaseA1

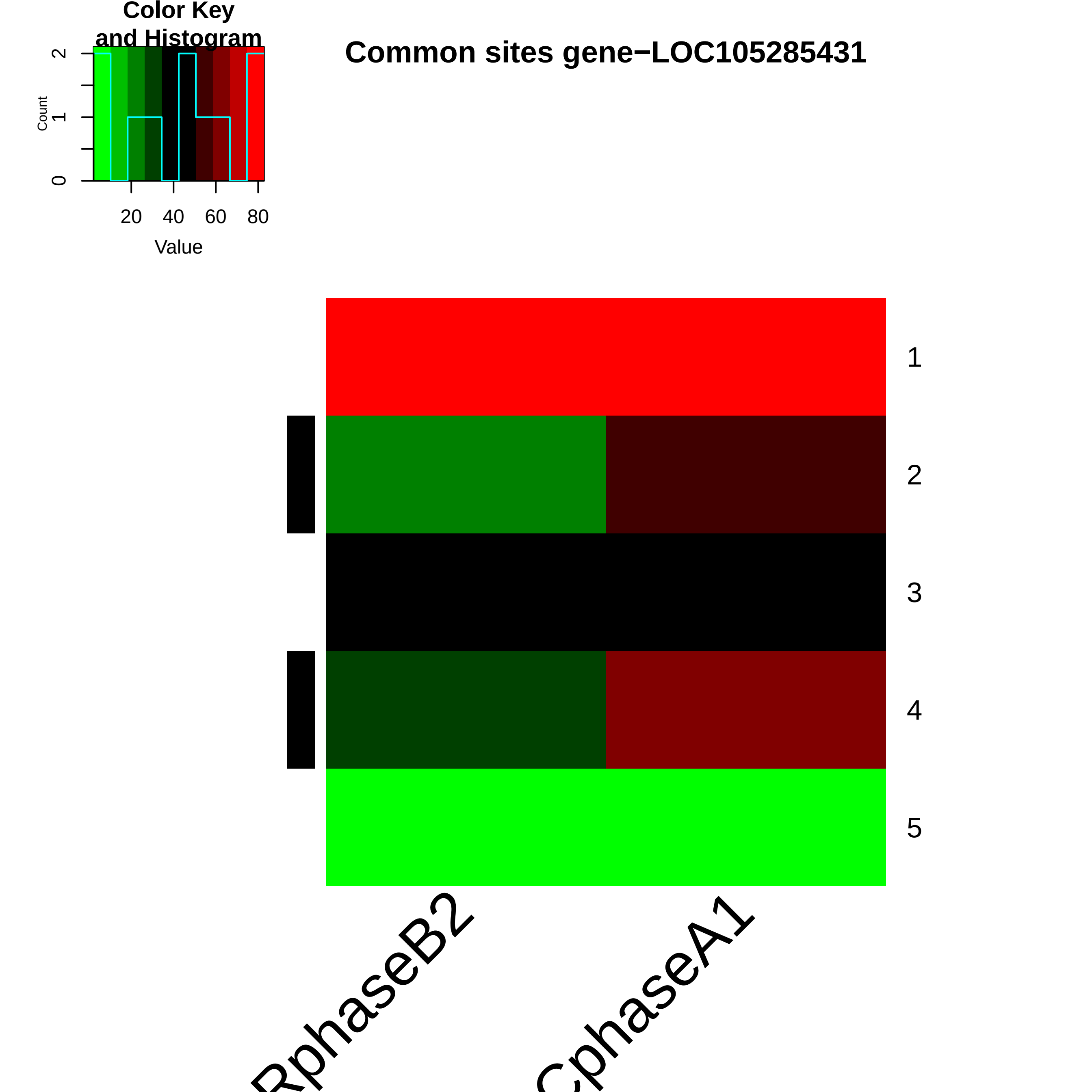

Color Key
and Histogram
Count
0
1
2
20
40
60
80
Value
Common sites gene−LOC105285431
1
2
3
4
5
RphaseB2
CphaseA1

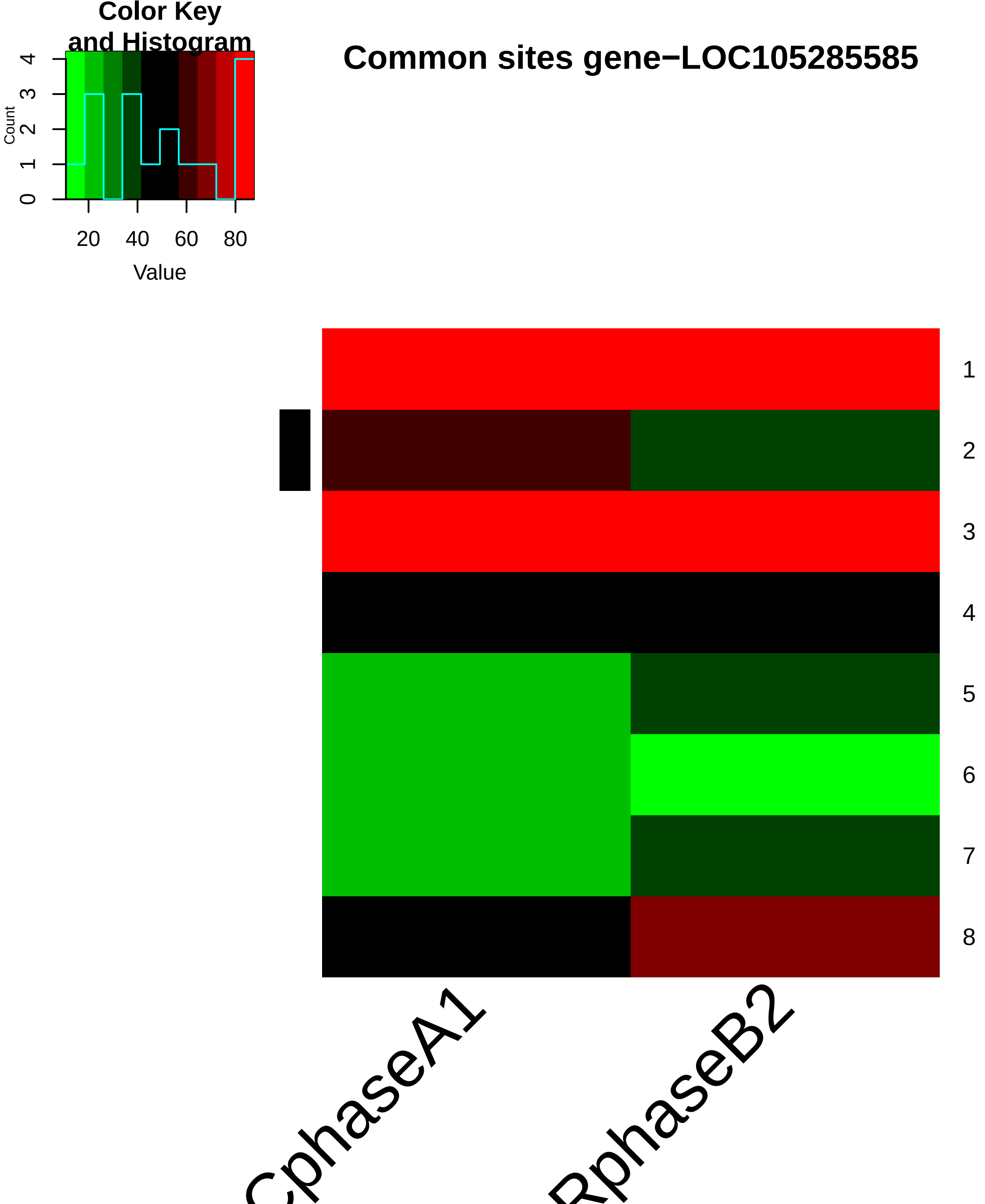

Color Key
and Histogram
Count
0
1
2
3
4
20
40
60
80
Value
Common sites gene−LOC105285585
1
2
3
4
5
6
7
8

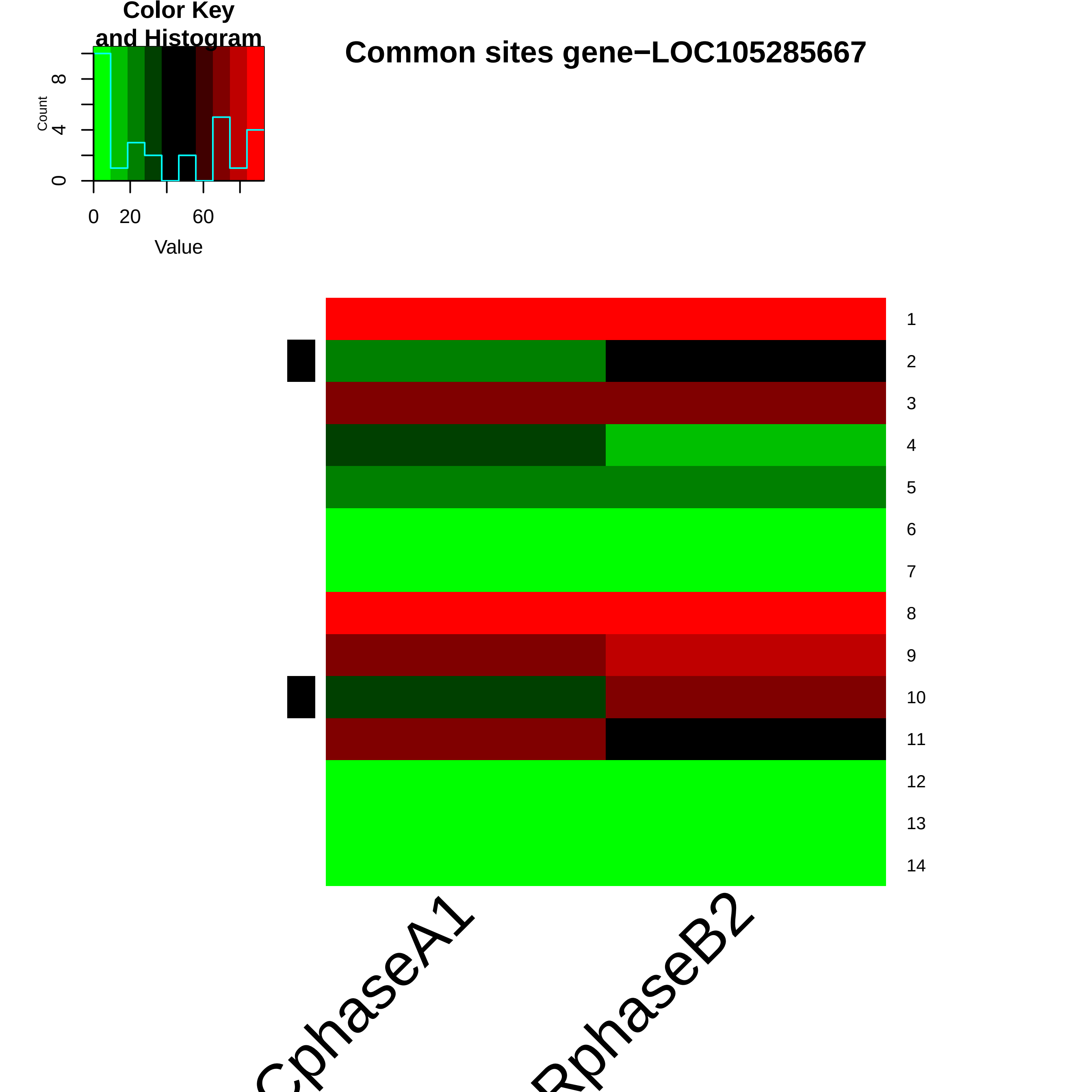

Color Key
and Histogram
Count
0
4
8
0
20
60
Value
Common sites gene−LOC105285667
1
2
3
4
5
6
7
8
9
10
11
12
13
14
CphaseA1
RphaseB2

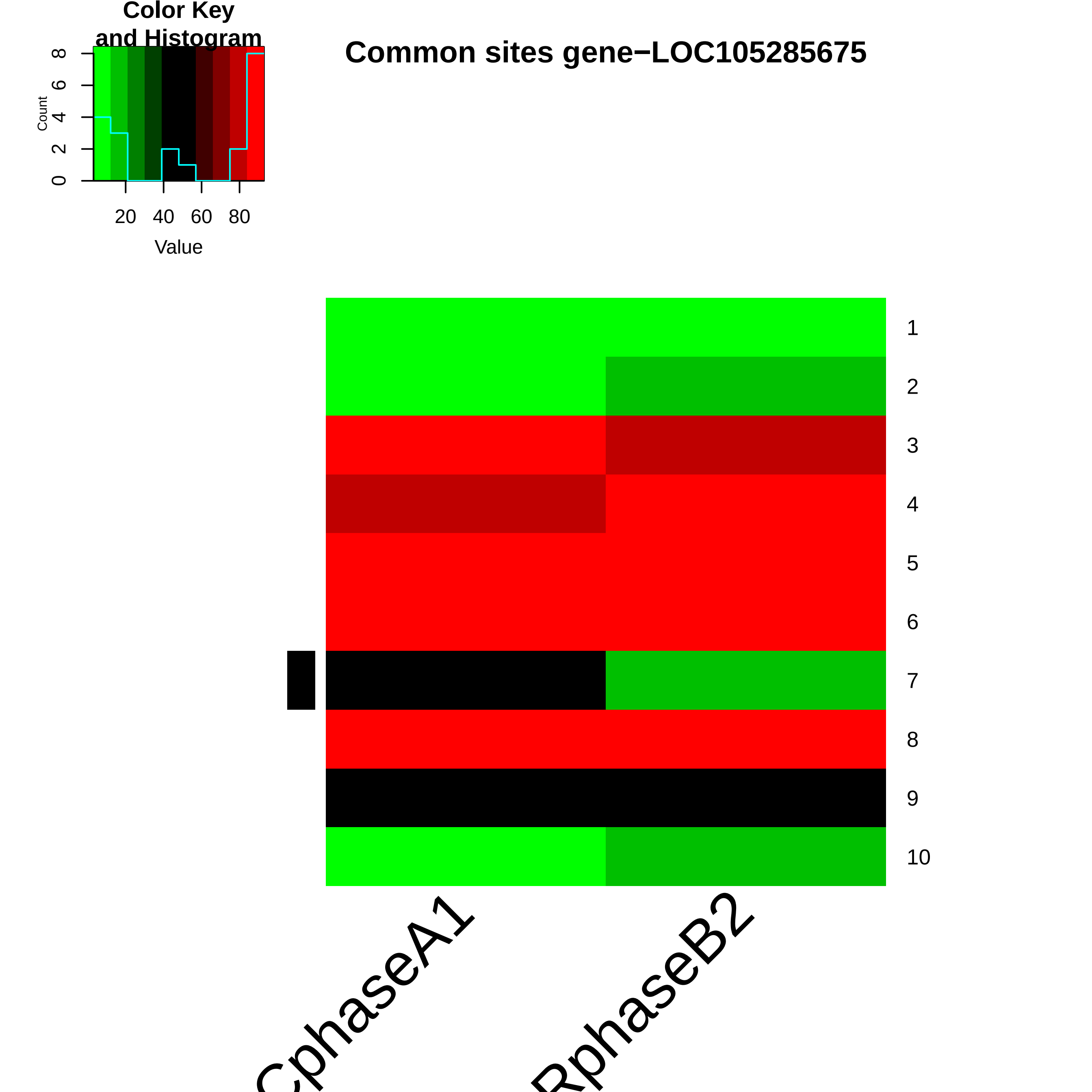

Color Key
and Histogram
Count
Value
Common sites gene−LOC105285675
1
2
3
4
5
6
7
8
9
10
CphaseA1
RphaseB2

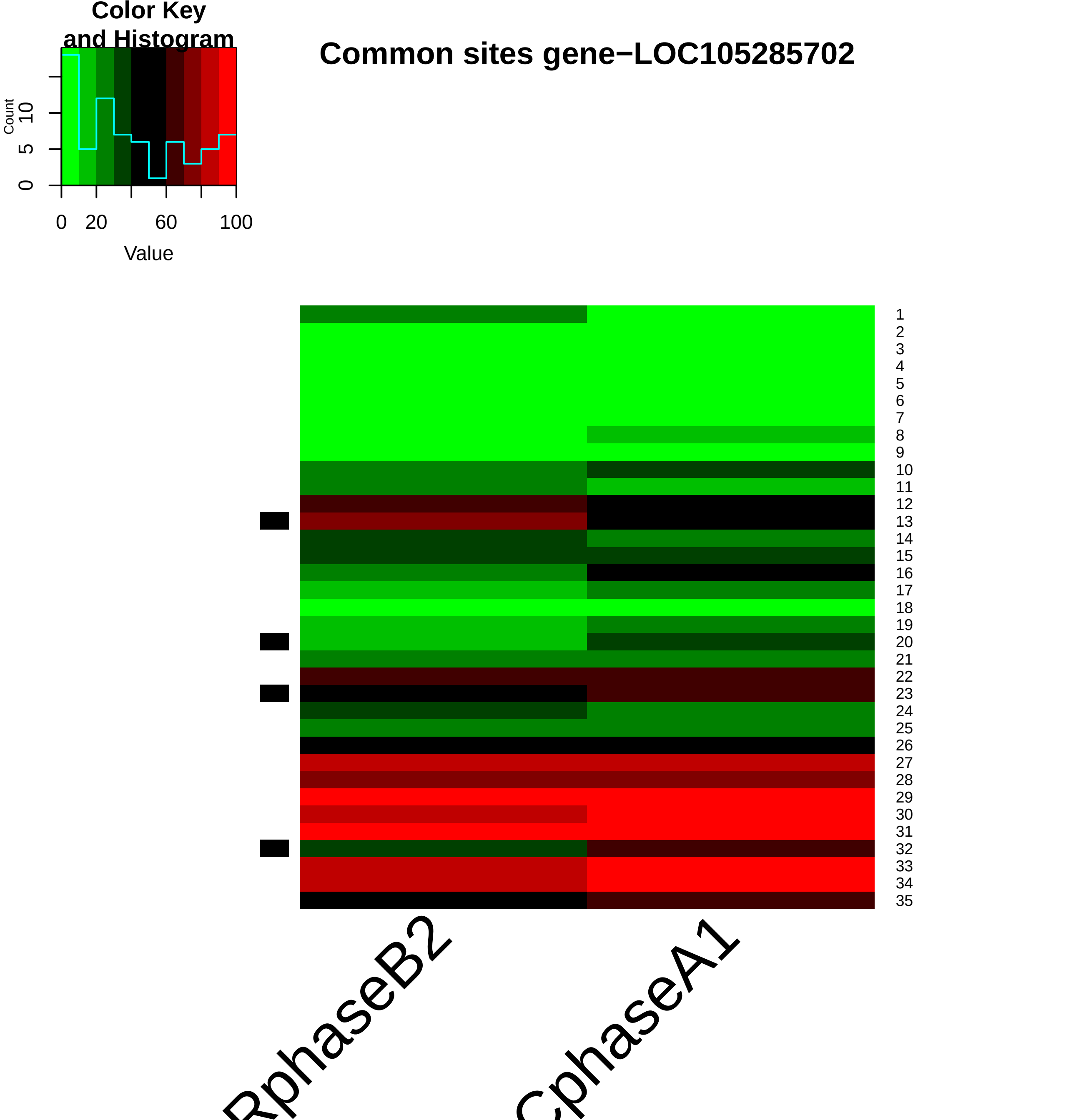

Color Key
and Histogram
Count
0
5
10
0
20
60
100
Value
Common sites gene−LOC105285702
RphaseB2
CphaseA1

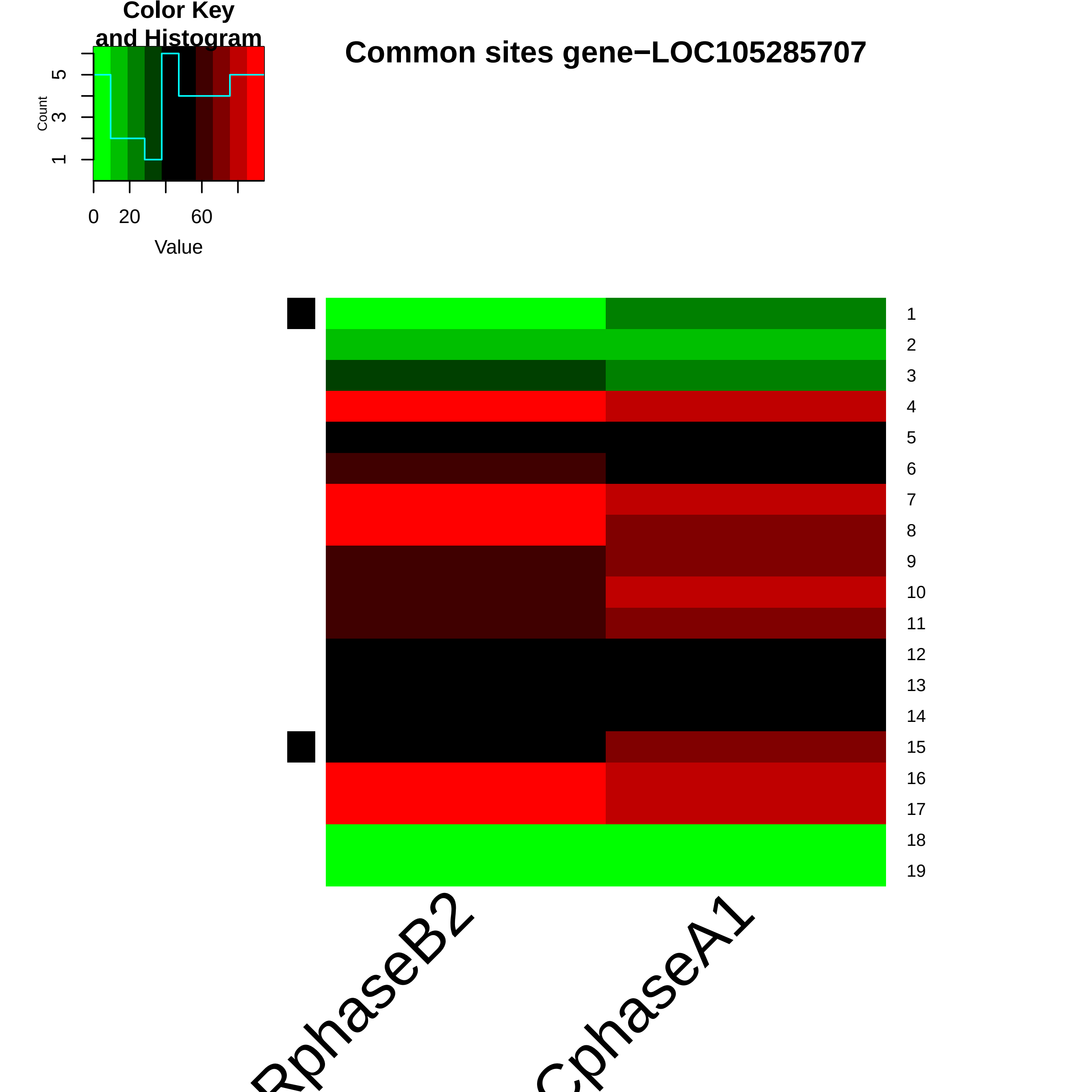

Color Key
and Histogram
Count
1
3
5
0
20
60
Value
Common sites gene−LOC105285707
RphaseB2
CphaseA1

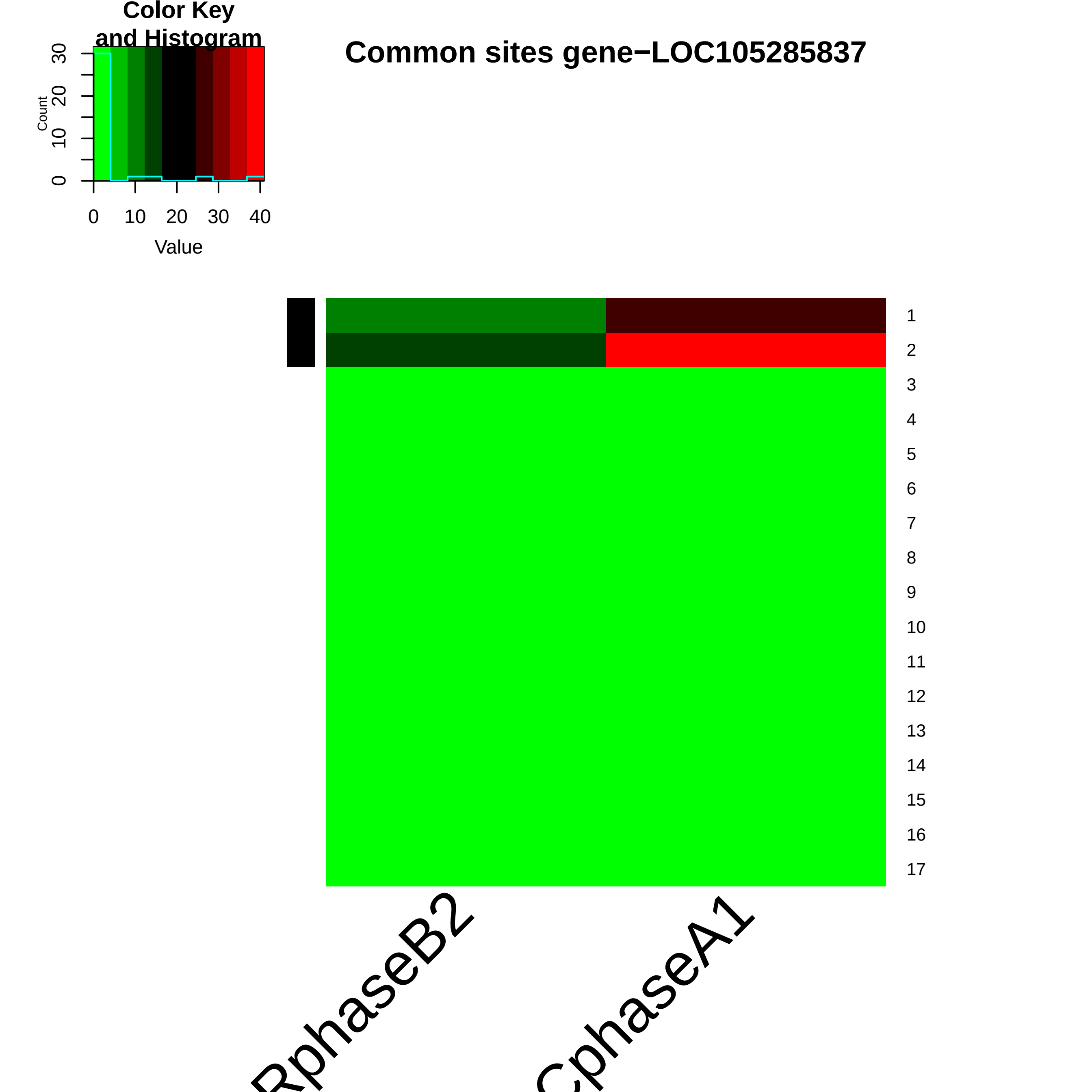

Color Key
and Histogram
Count
0
10
20
30
0
10
20
30
40
Value
Common sites gene−LOC105285837
RphaseB2
CphaseA1

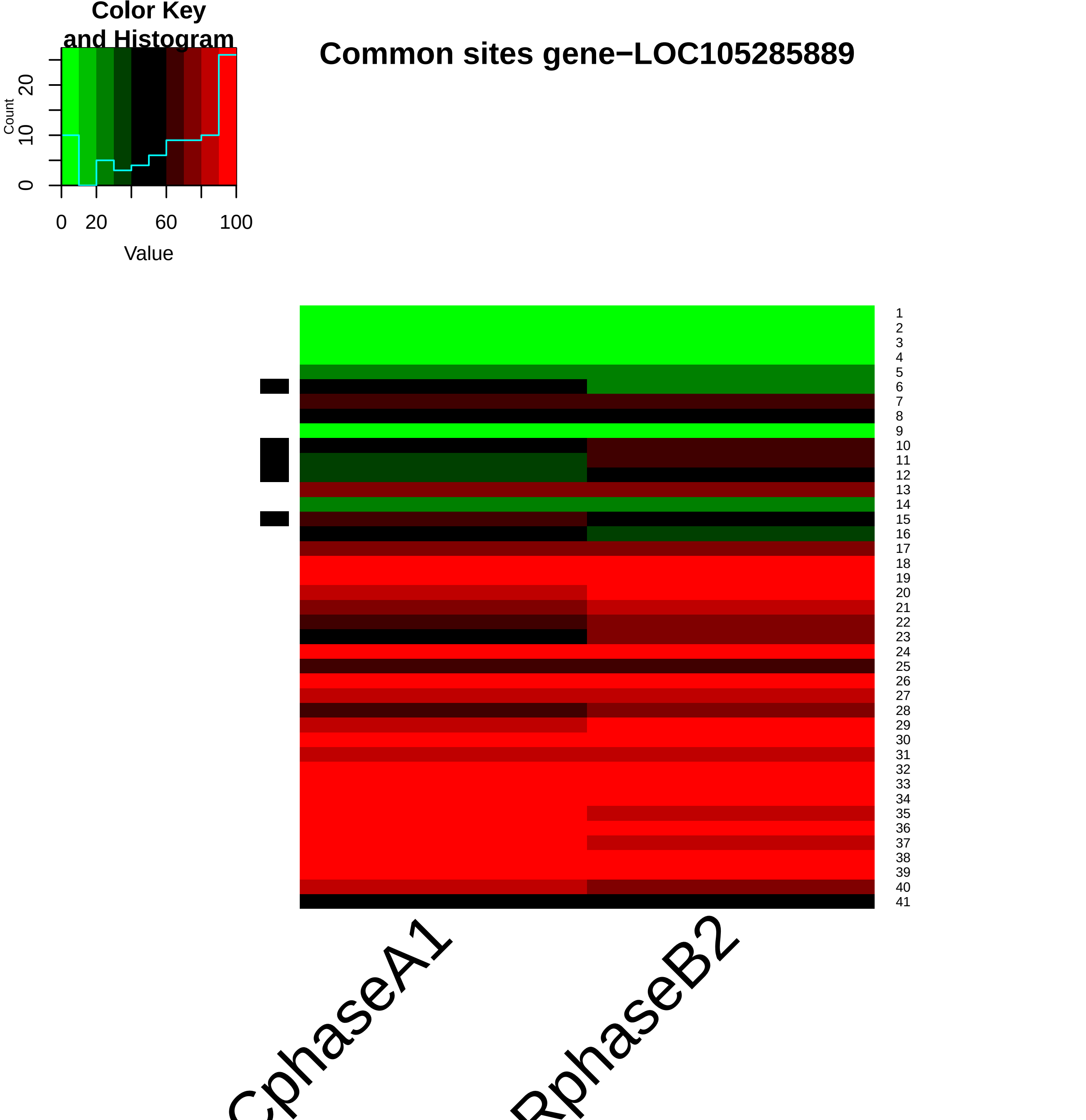

Common sites gene−LOC105285889
Color Key
and Histogram
Count
0
10
20
0
20
60
100
Value
CphaseA1
RphaseB2

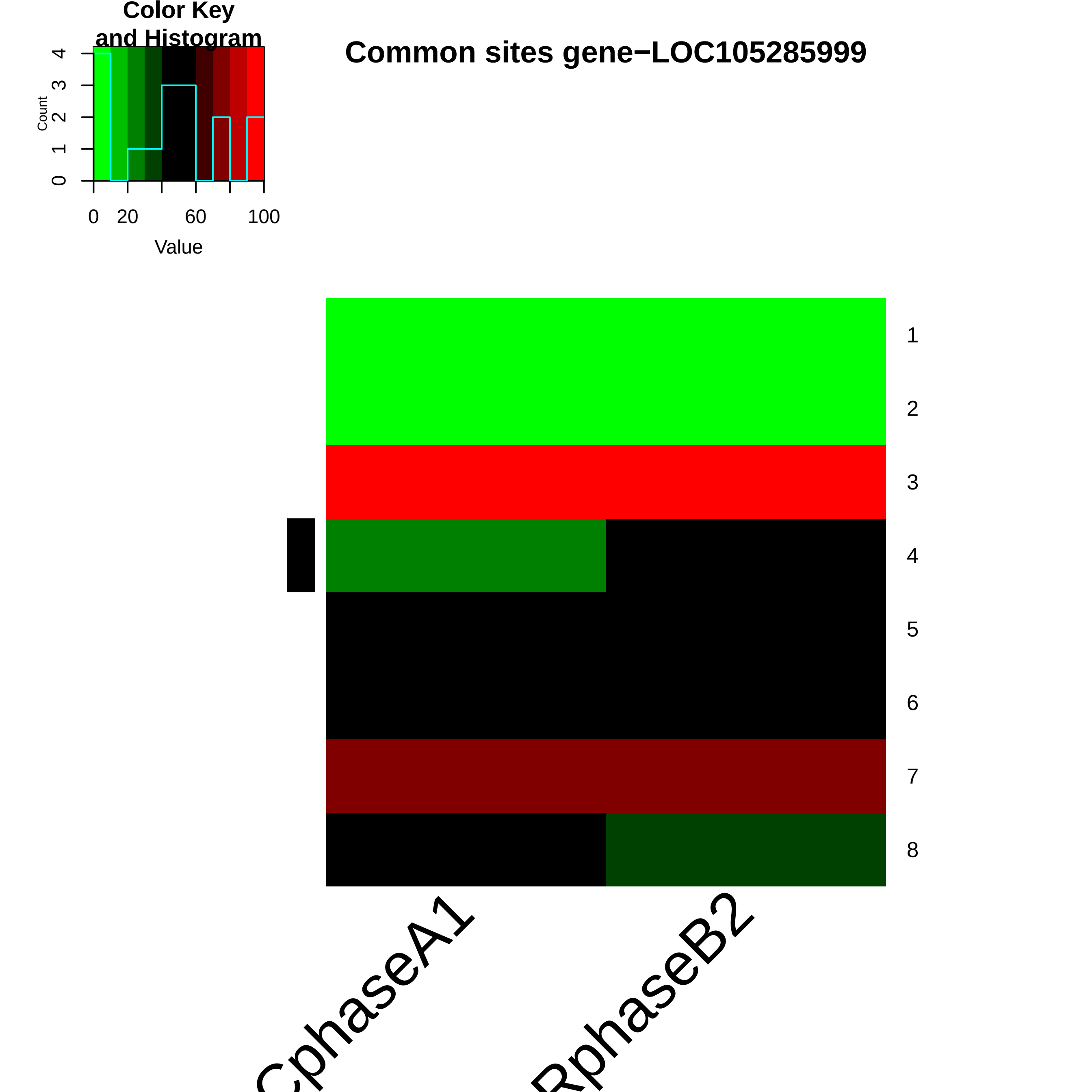

Color Key
and Histogram
Count
0
1
2
3
4
0
20
60
100
Value
Common sites gene−LOC105285999
1
2
3
4
5
6
7
8
CphaseA1
RphaseB2

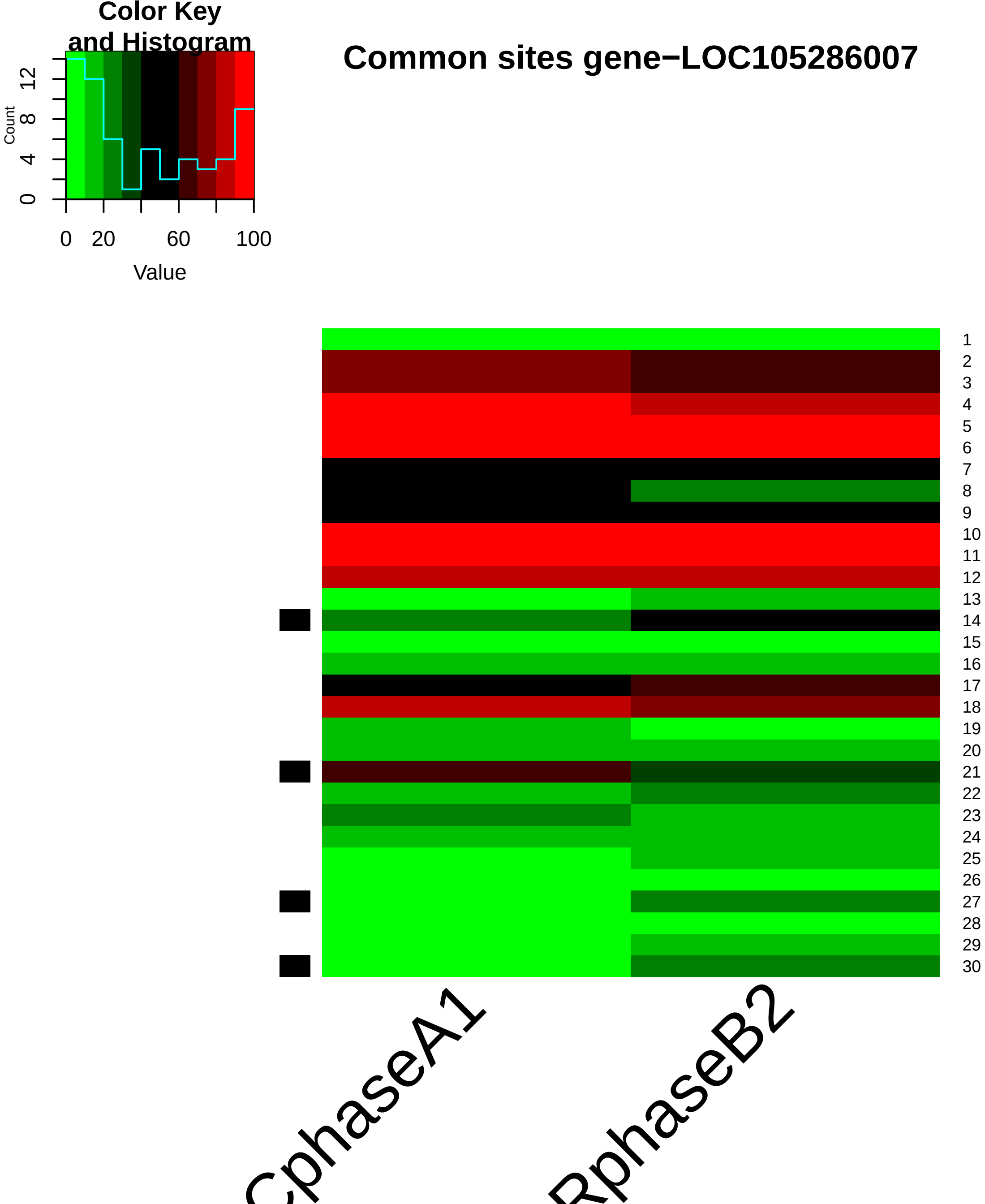

Color Key
and Histogram
Count
0
4
8
12
0
20
60
100
Value
Common sites gene−LOC105286007
CphaseA1
RphaseB2

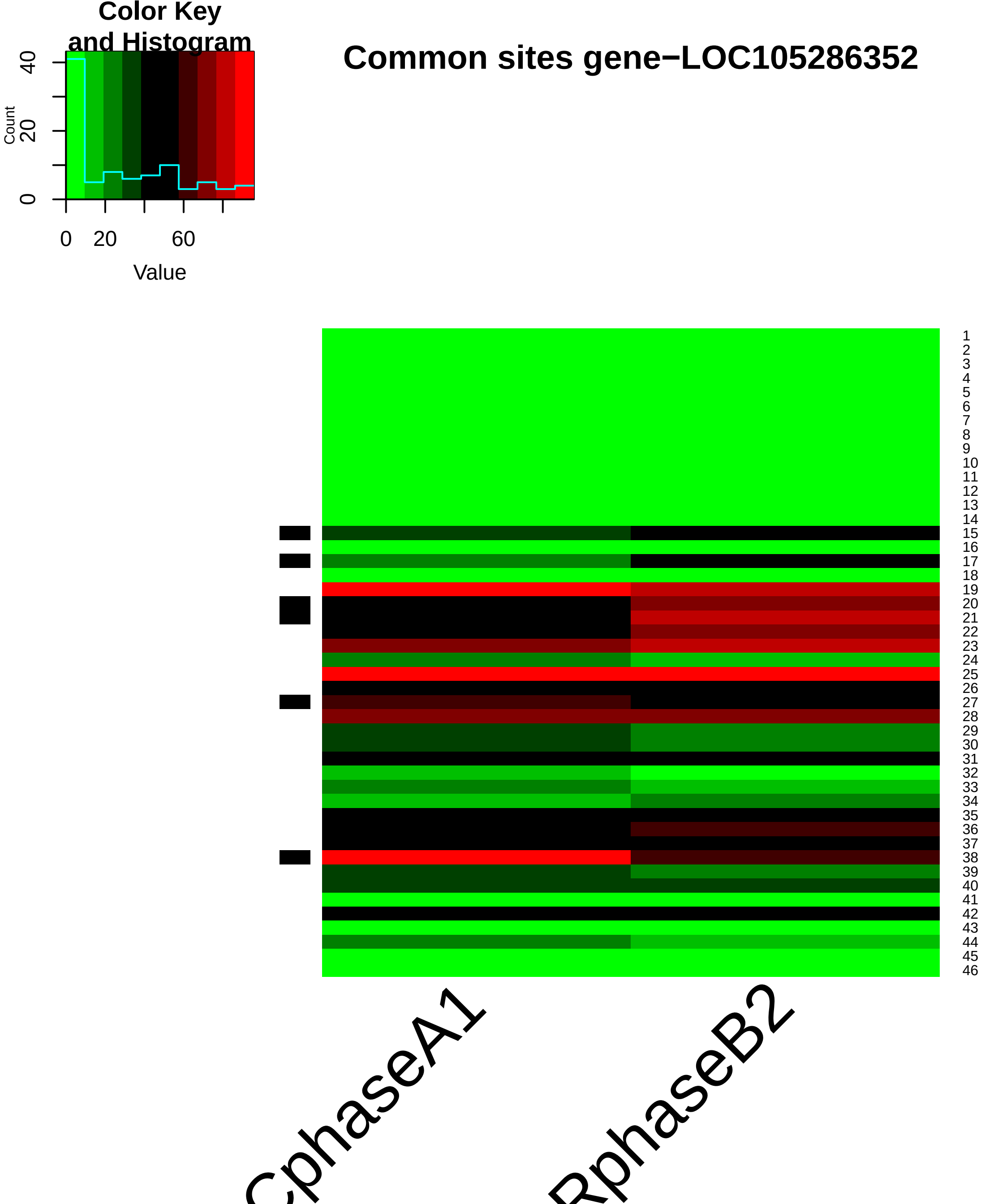
Color Key
and Histogram
Count
0
20
40
0
20
60
Value
Common sites gene−LOC105286352
CphaseA1
RphaseB2

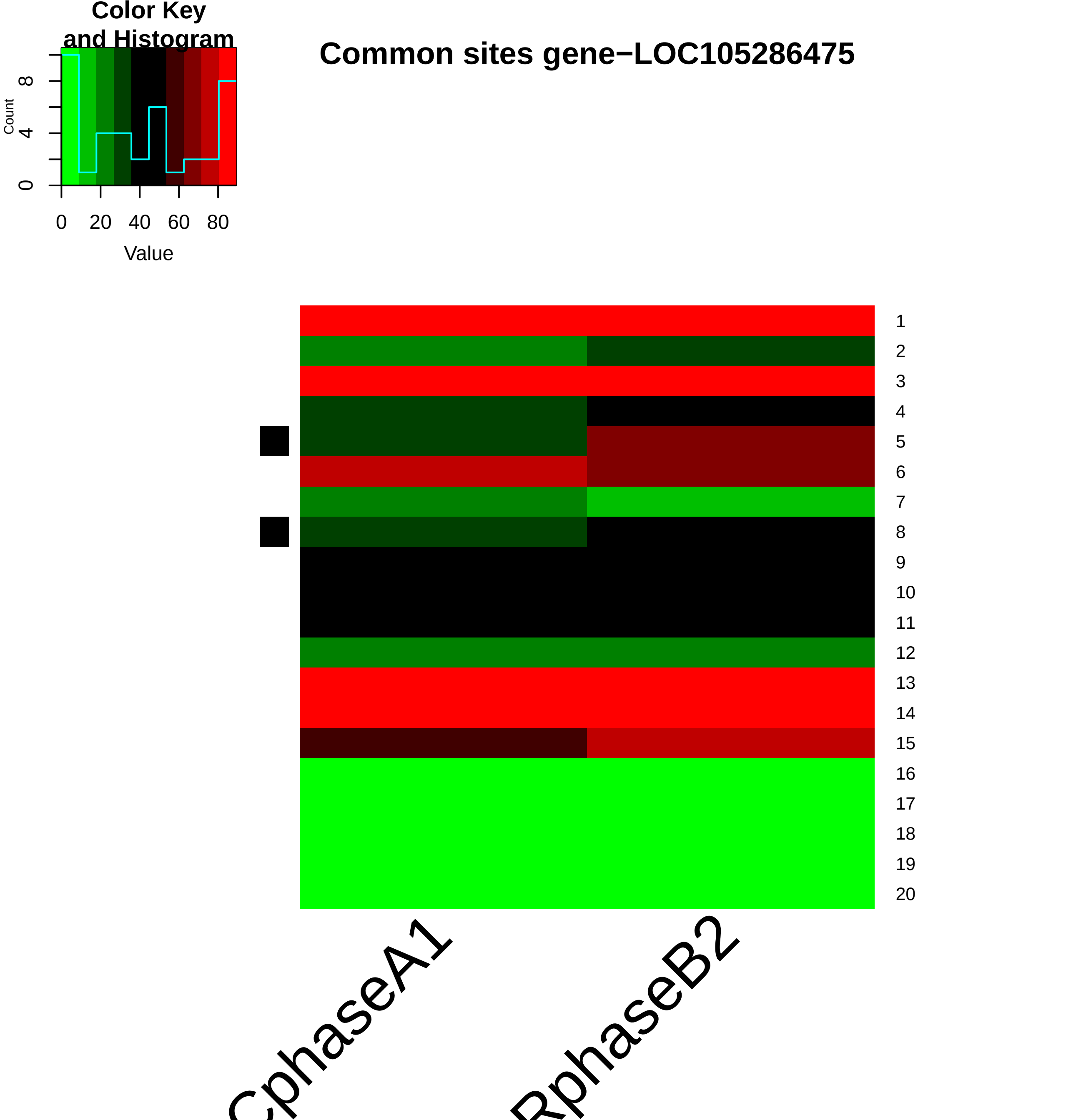
Color Key
and Histogram
Count
0
4
8
0
20
40
60
80
Value
Common sites gene−LOC105286475
CphaseA1
RphaseB2

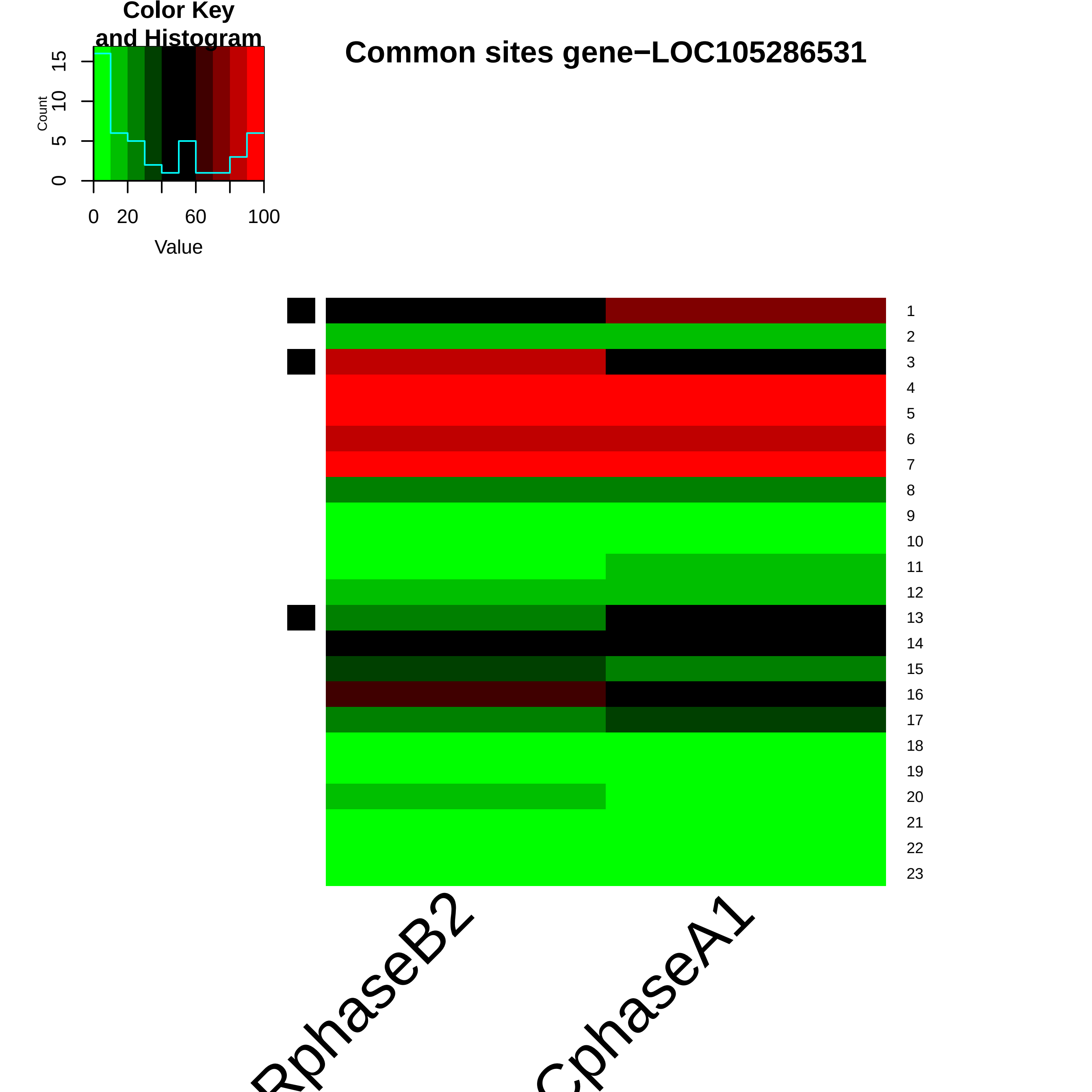

Color Key
and Histogram
Count
0
5
10
15
0
20
60
100
Value
Common sites gene−LOC105286531
RphaseB2
CphaseA1

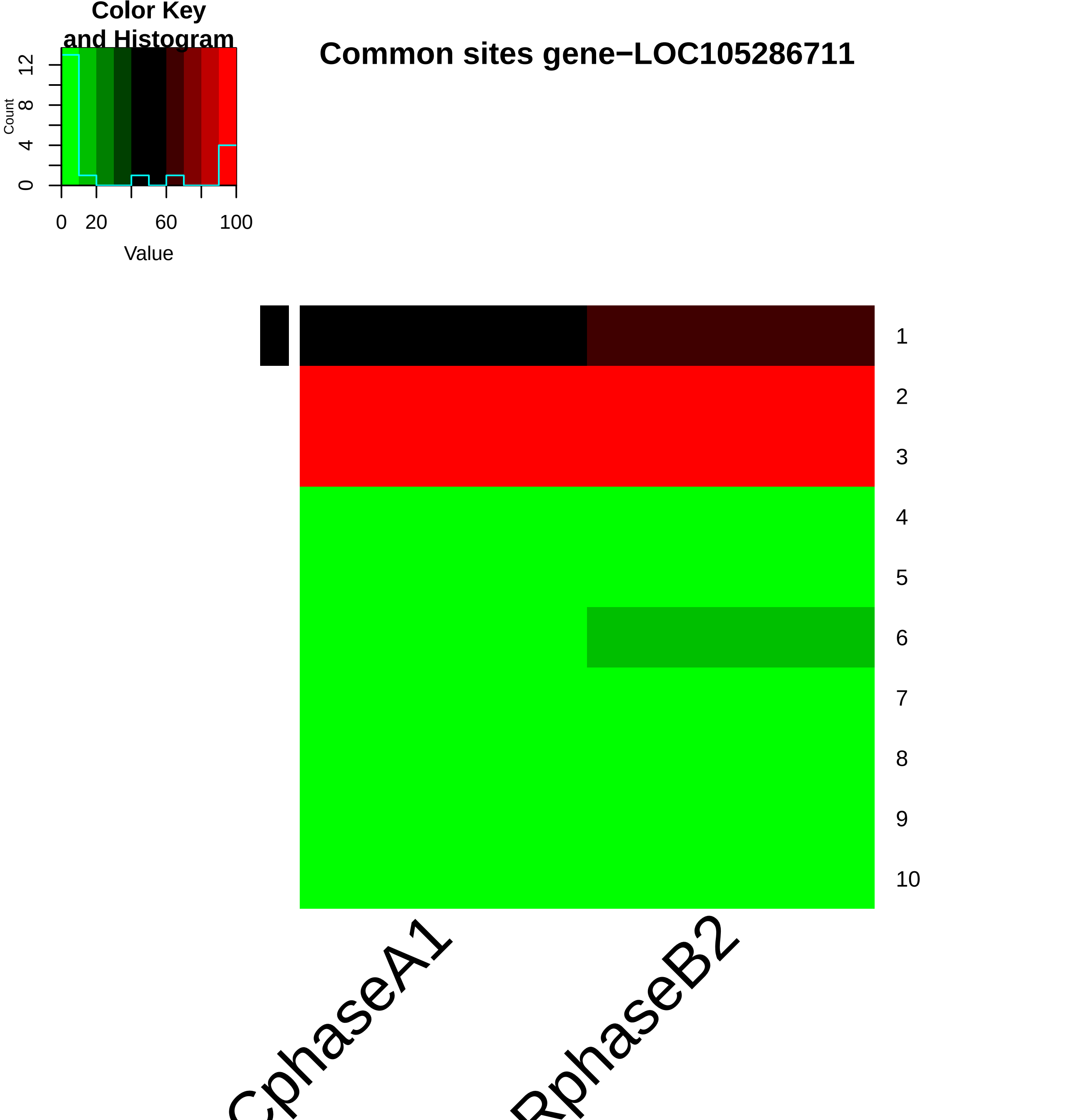

Color Key
and Histogram
Count
0
4
8
12
0
20
60
100
Value
Common sites gene−LOC105286711
1
2
3
4
5
6
7
8
9
10

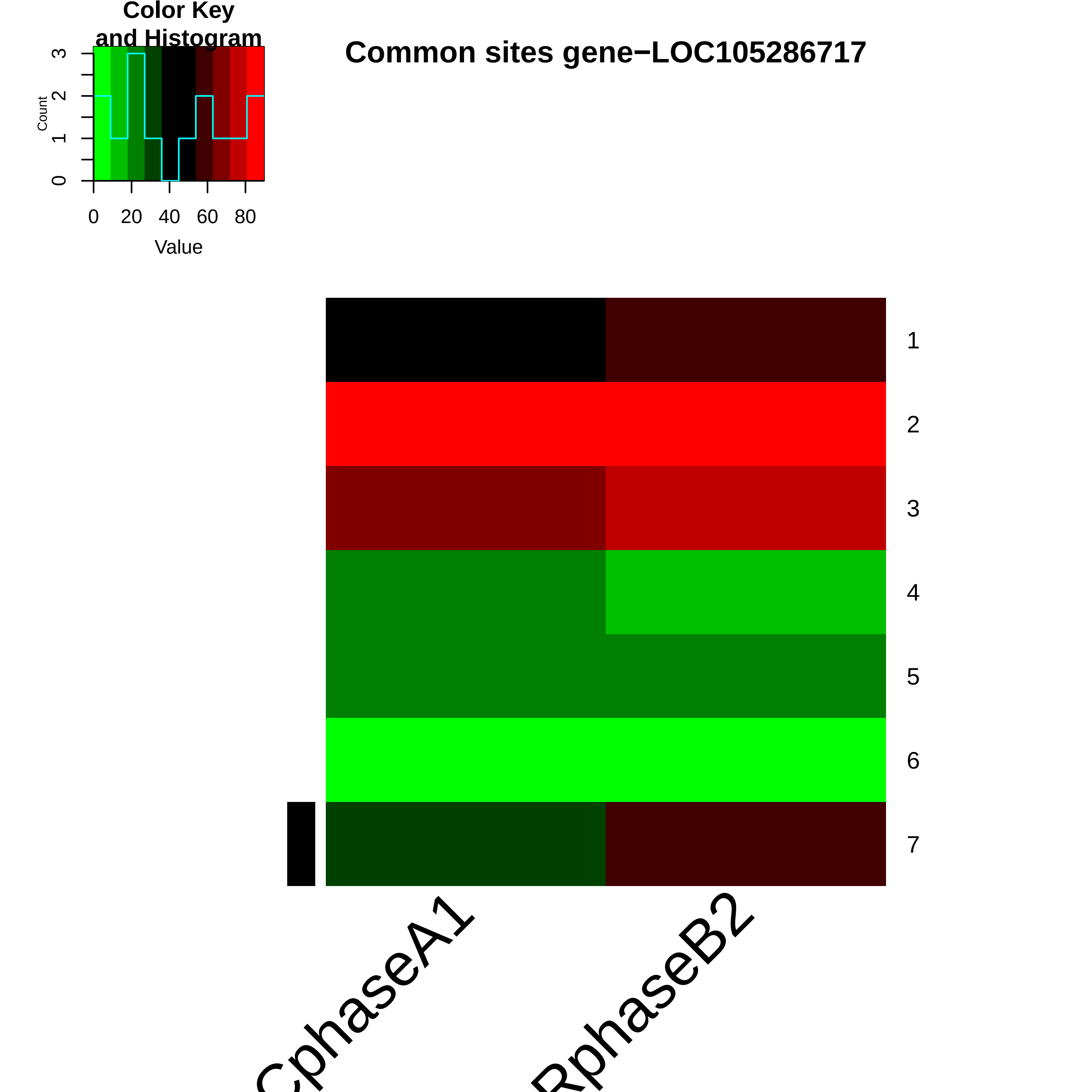

Color Key
and Histogram
Count
0
1
2
3
0
20
40
60
80
Value
Common sites gene−LOC105286717
1
2
3
4
5
6
7
CphaseA1
RphaseB2

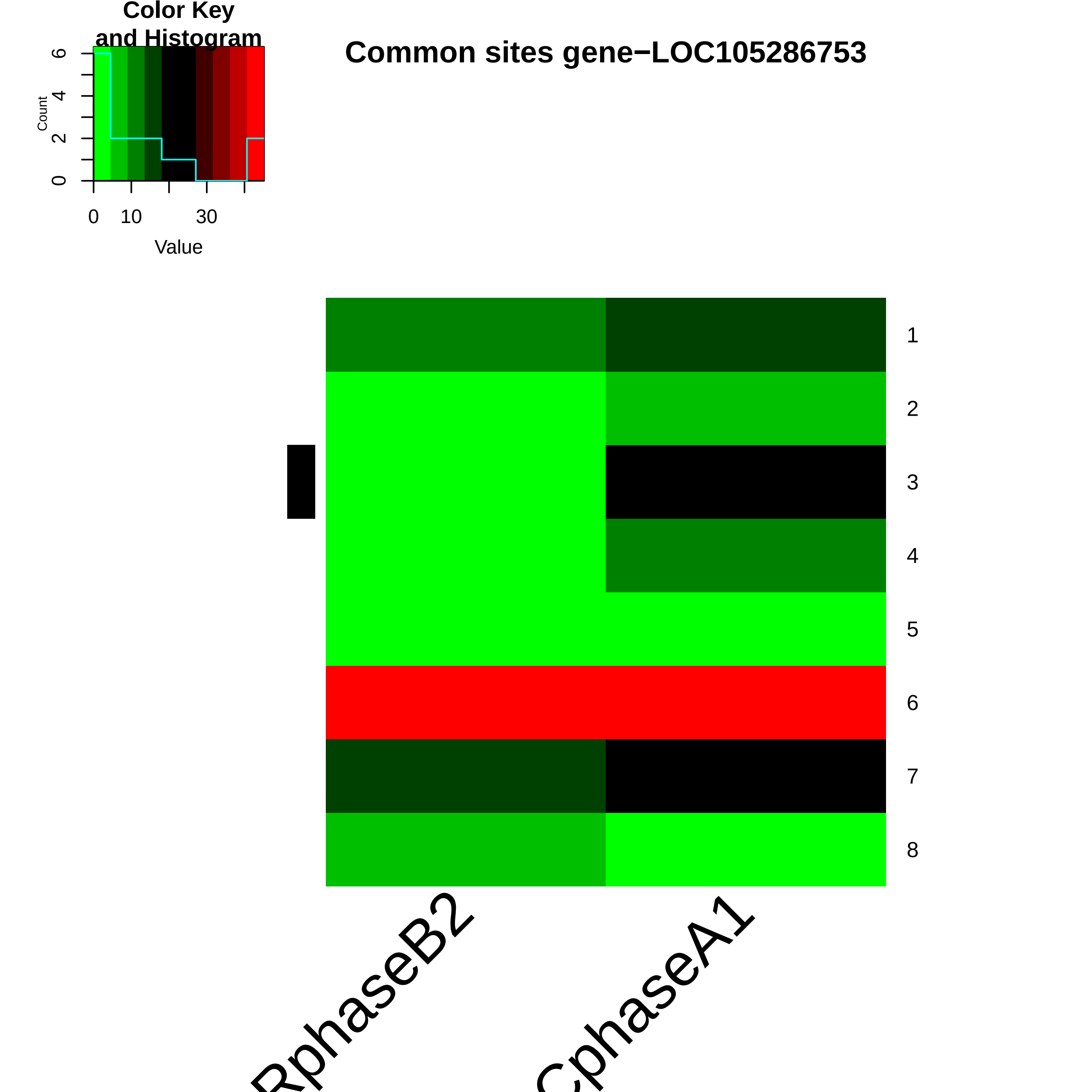

Color Key
and Histogram
Common sites gene−LOC105286753
Count
0
2
4
6
0
10
30
Value
1
2
3
4
5
6
7
8
RphaseB2
CphaseA1

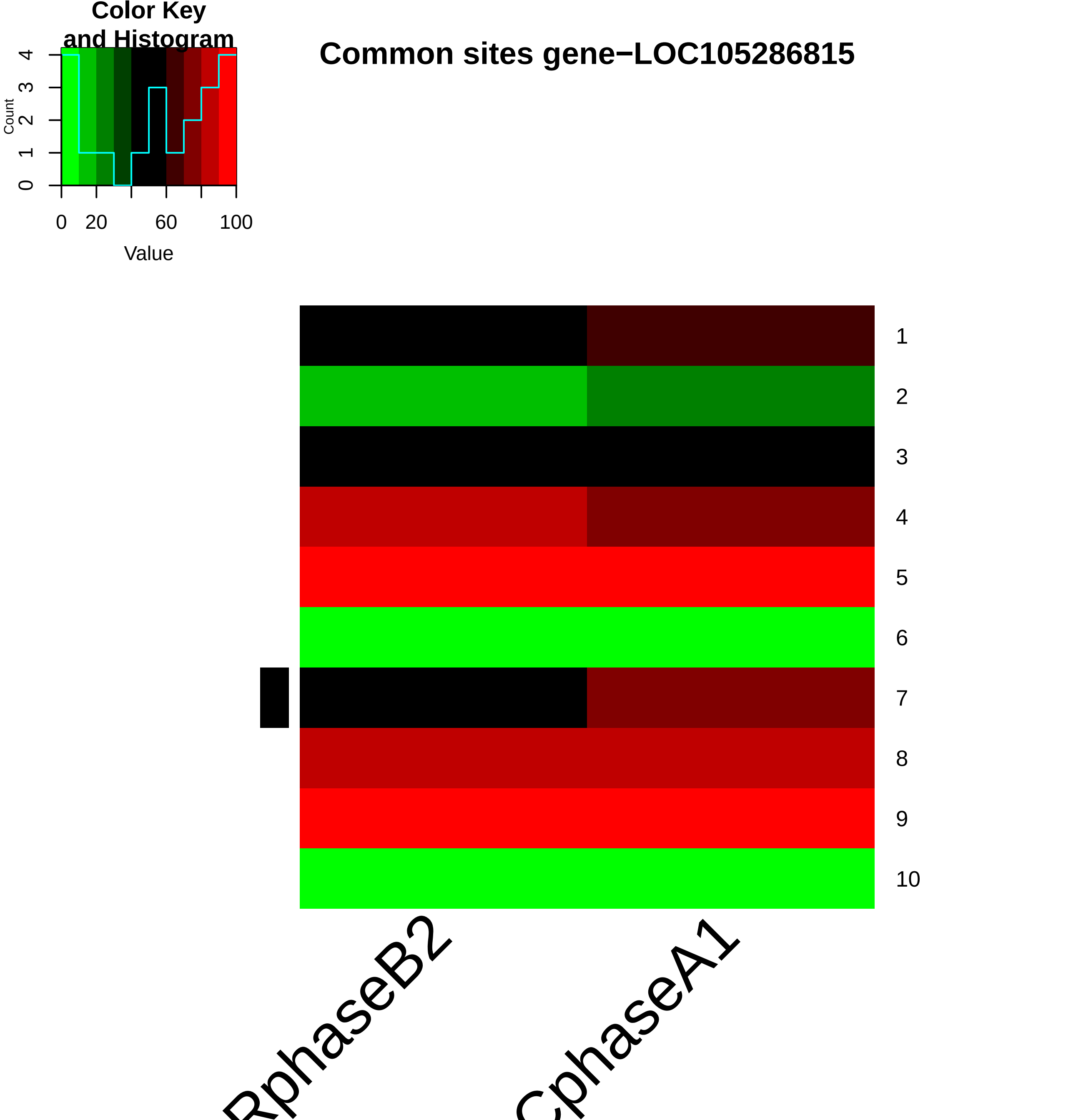

Color Key
and Histogram
Count
0
1
2
3
4
0
20
60
100
Value
Common sites gene−LOC105286815
1
2
3
4
5
6
7
8
9
10
RphaseB2
CphaseA1

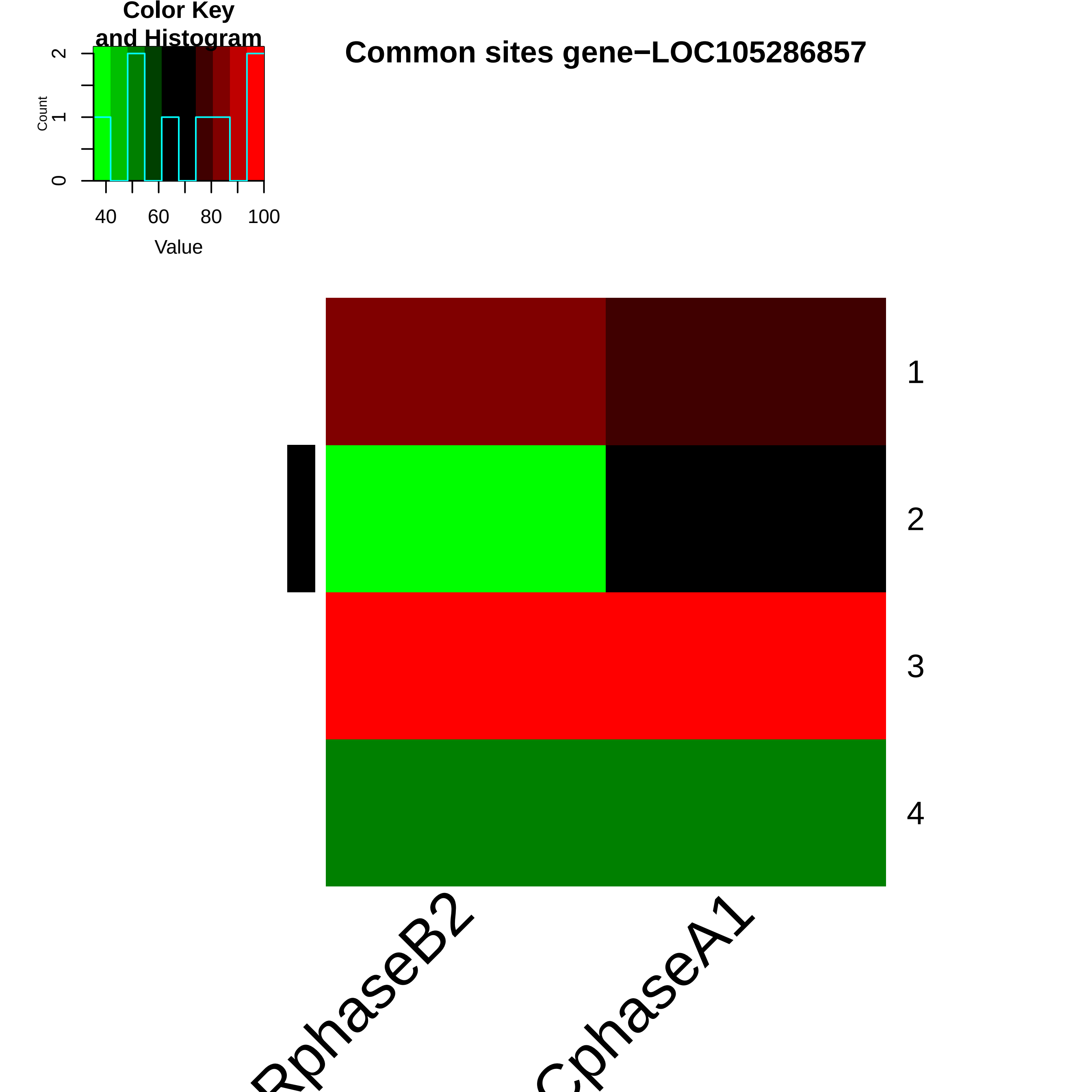

Color Key
and Histogram
Count
0
1
2
40
60
80
100
Value
Common sites gene−LOC105286857
1
2
3
4

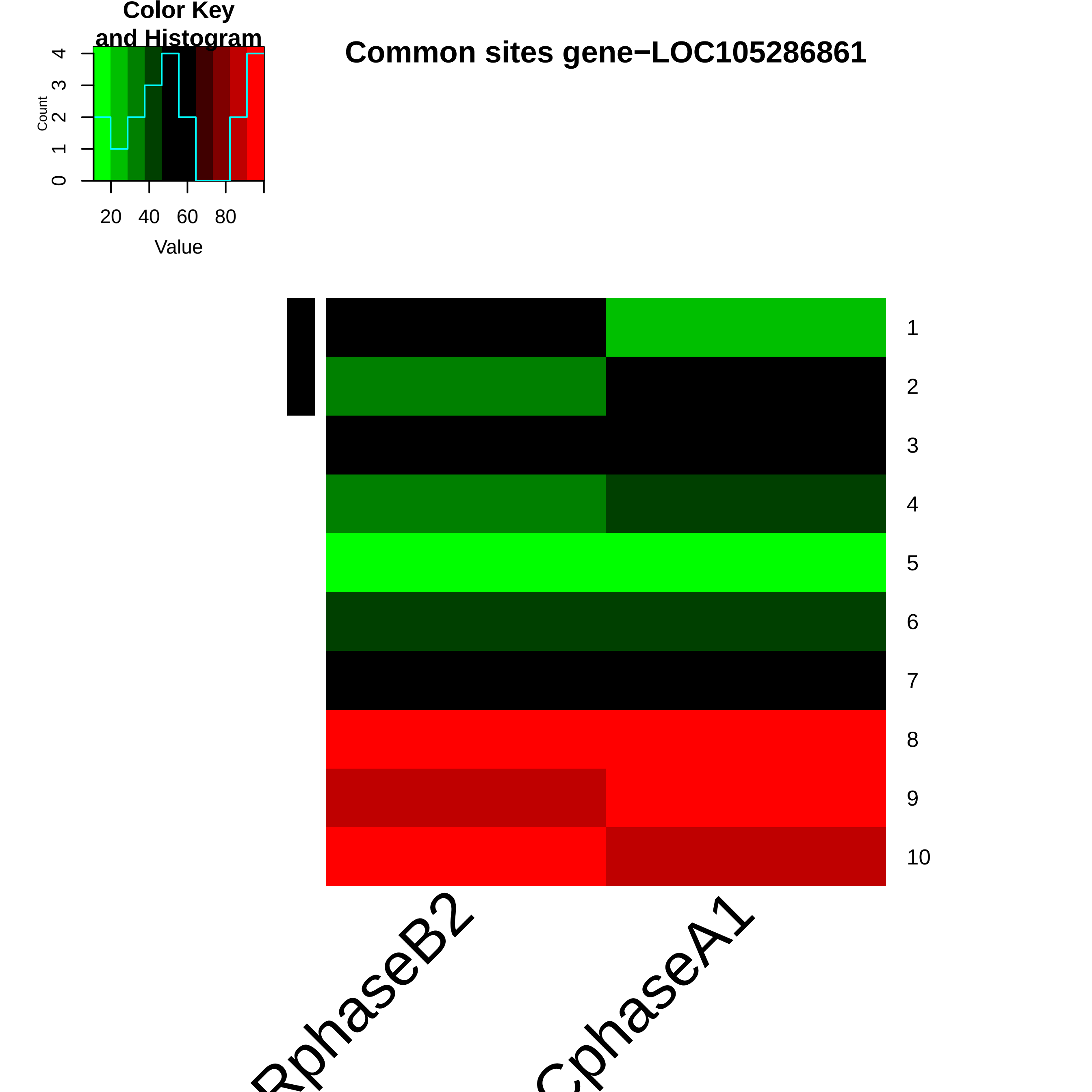

Color Key
and Histogram
Count
0
1
2
3
4
20
40
60
80
Value
Common sites gene−LOC105286861
1
2
3
4
5
6
7
8
9
10
RphaseB2
CphaseA1

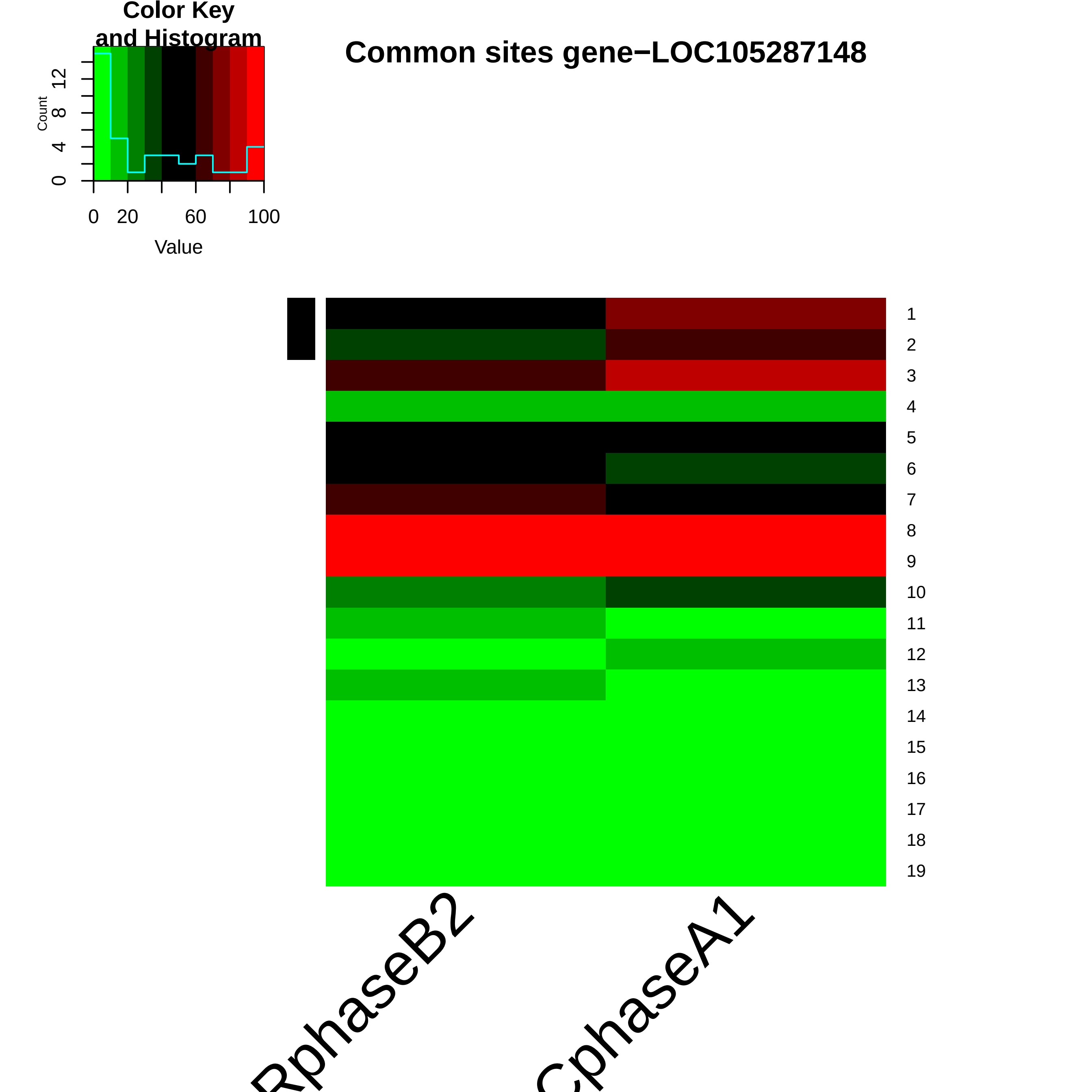

Color Key
and Histogram
Count
0
4
8
12
0
20
60
100
Value
Common sites gene−LOC105287148
RphaseB2
CphaseA1

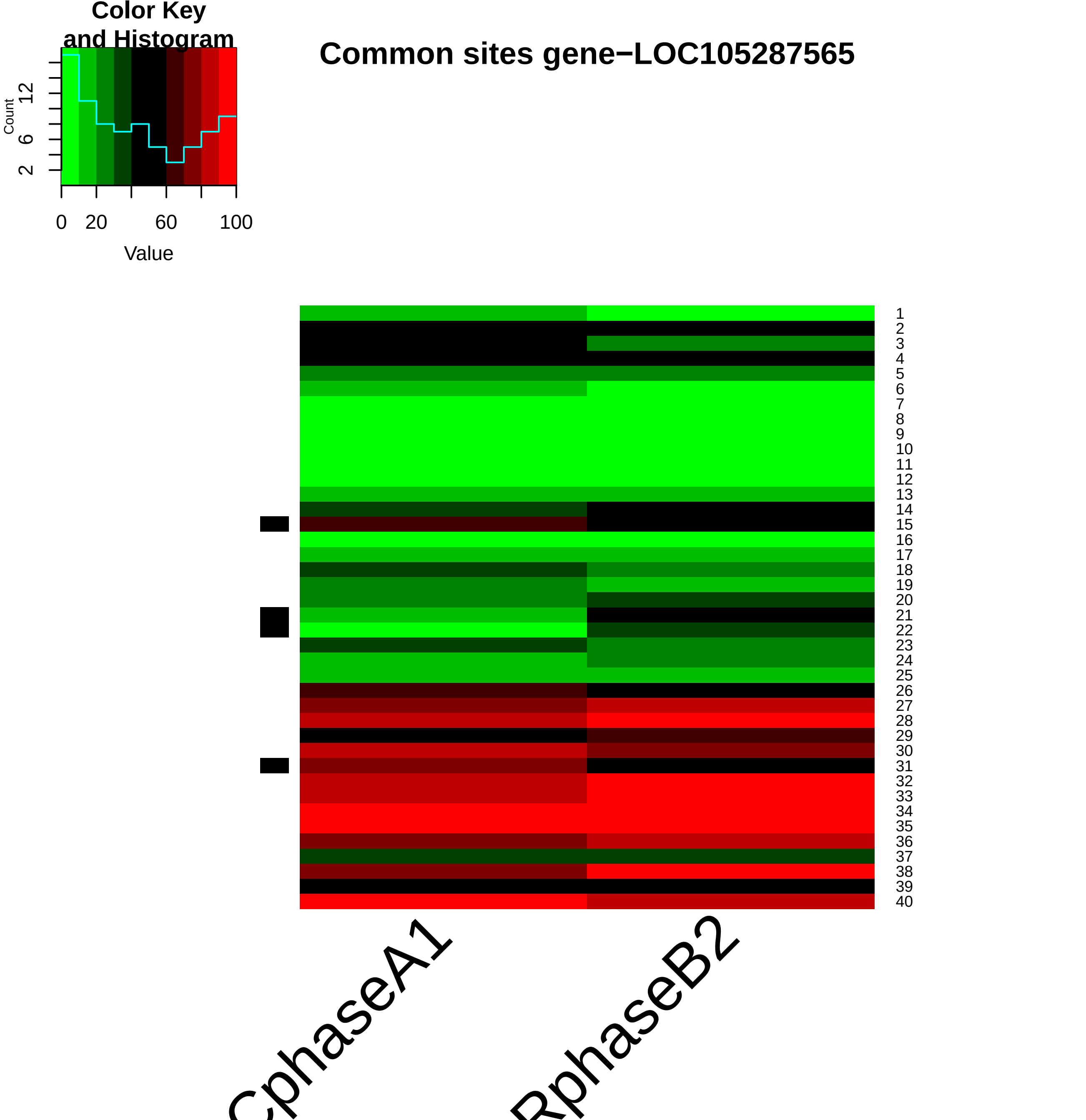
Color Key
and Histogram
Count
2
6
12
0
20
60
100
Value
Common sites gene−LOC105287565
CphaseA1
RphaseB2

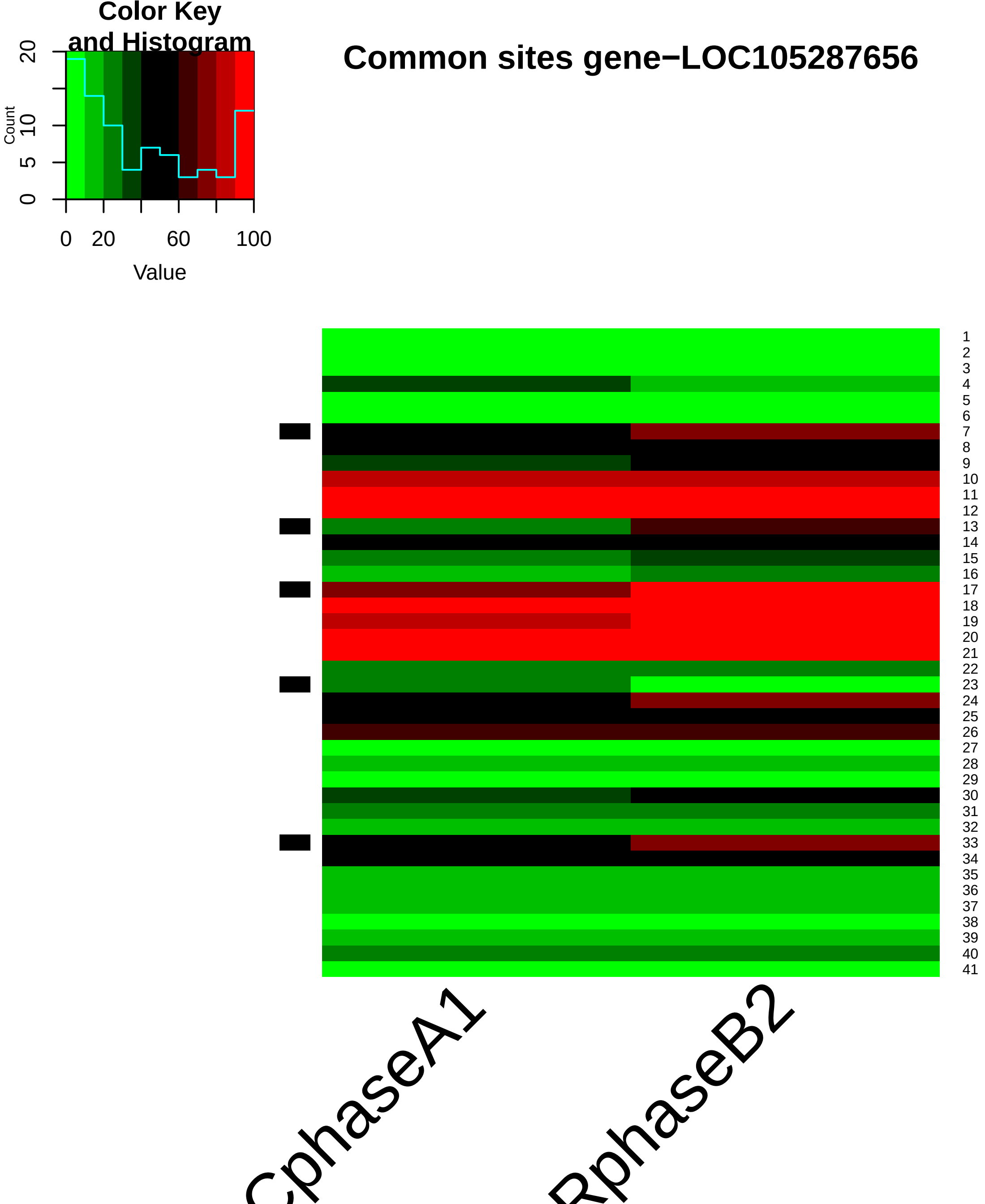

Color Key
and Histogram
Count
Value
0
20
60
100
0
5
10
20
Common sites gene−LOC105287656
CphaseA1
RphaseB2

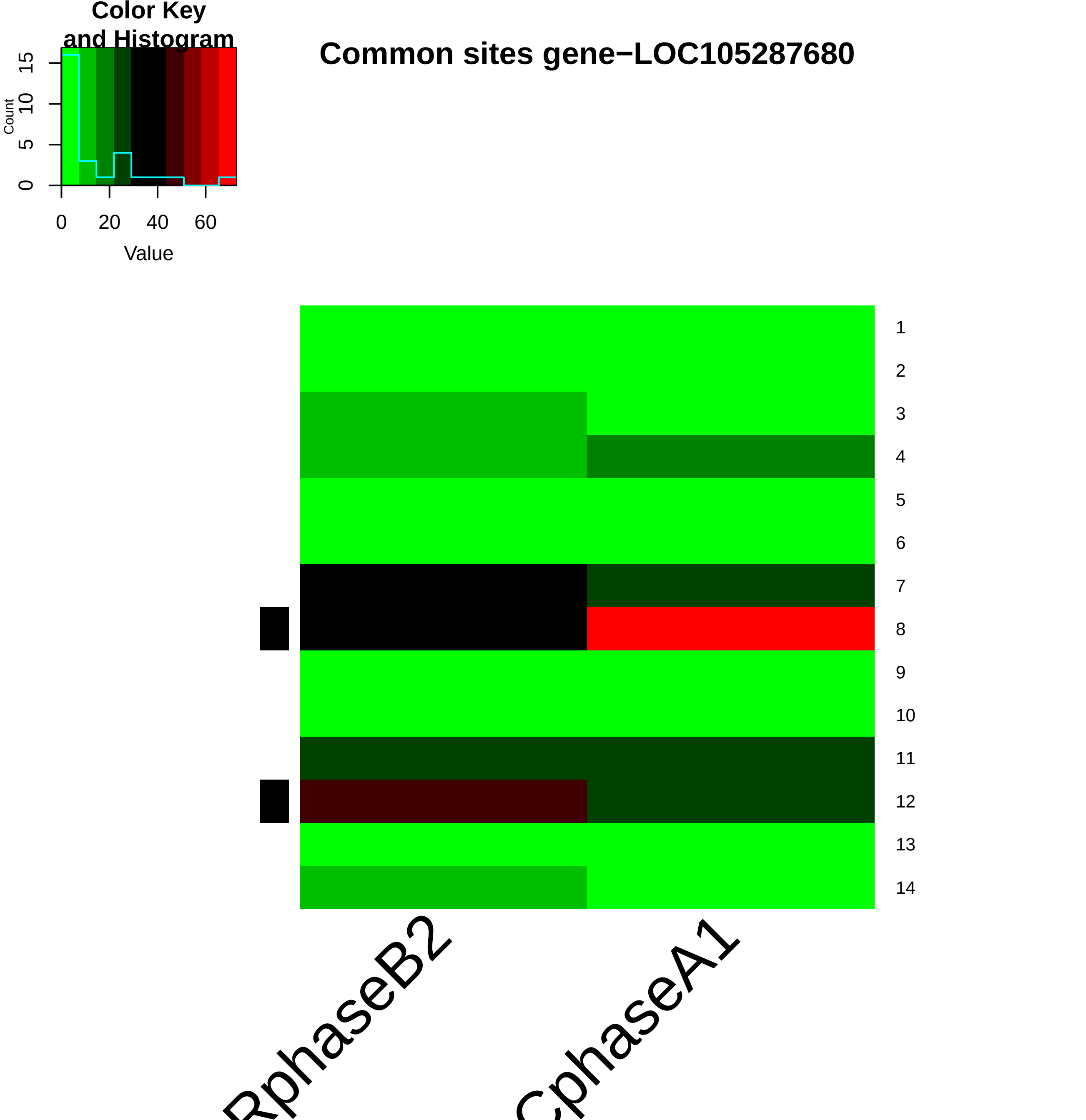
Color Key
and Histogram
Count
0
5
10
15
0
20
40
60
Value
Common sites gene−LOC105287680
1
2
3
4
5
6
7
8
9
10
11
12
13
14
RphaseB2
CphaseA1

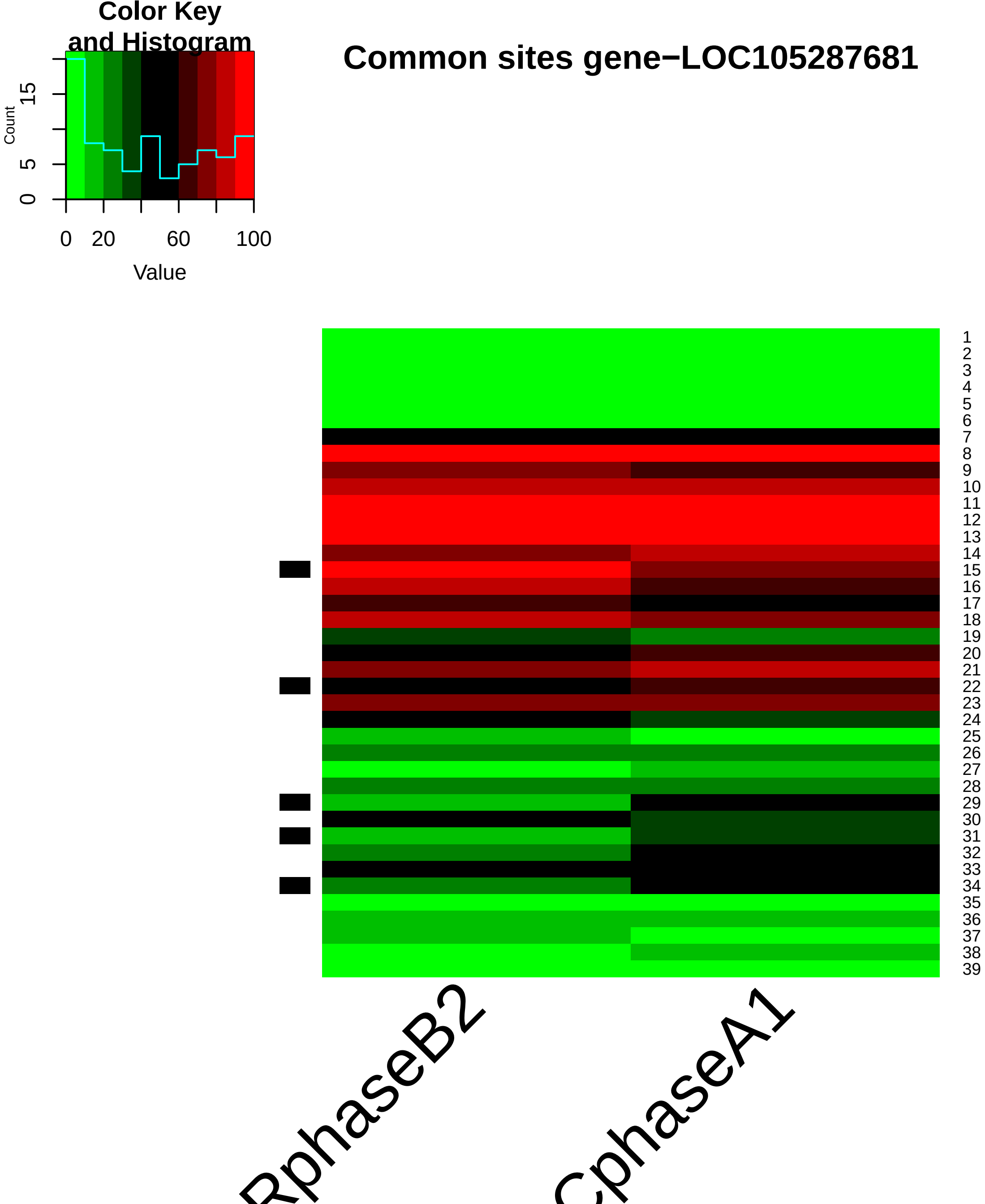
Color Key
and Histogram
Count
0
5
15
0
20
60
100
Value
Common sites gene−LOC105287681
RphaseB2
CphaseA1

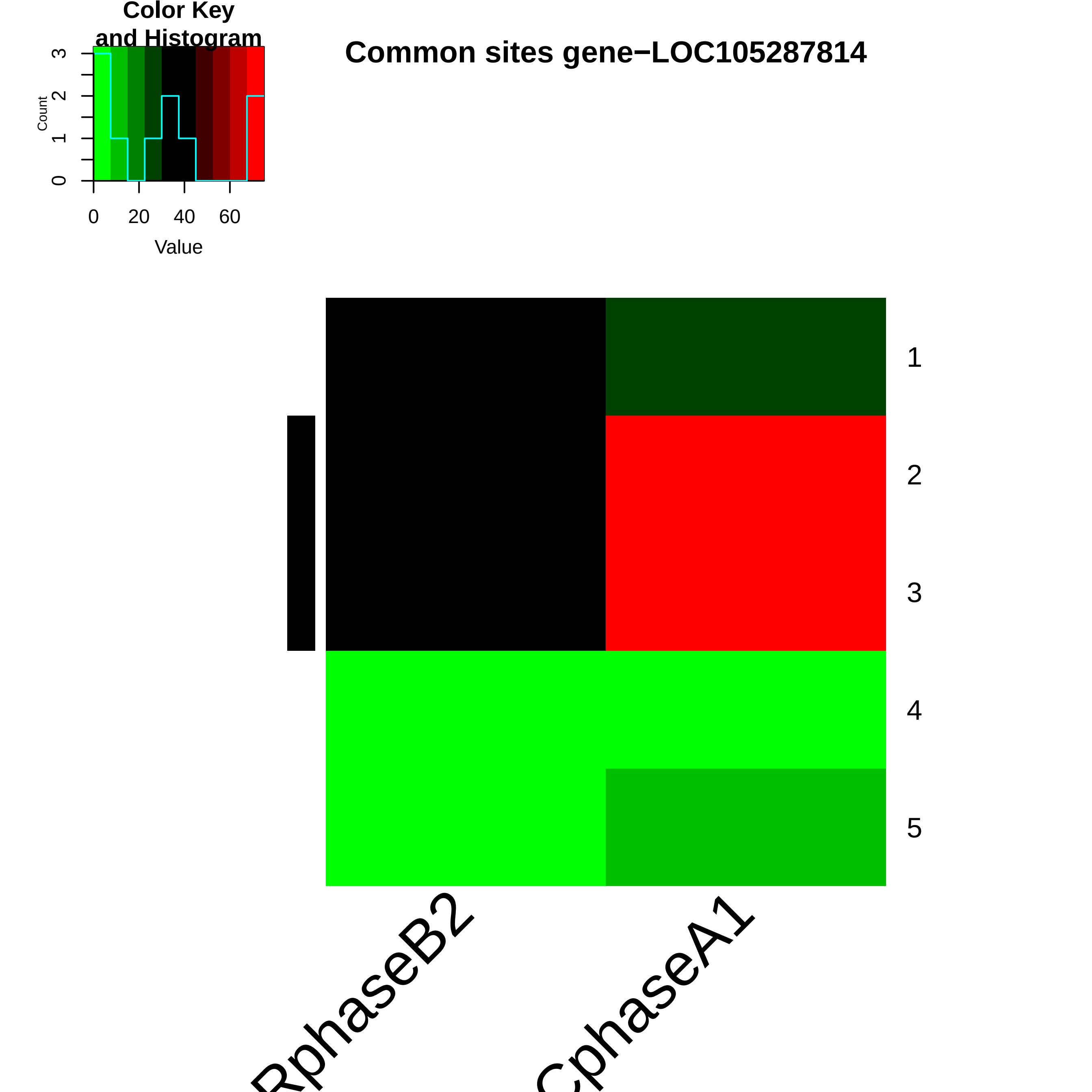

Color Key
and Histogram
Count
0
1
2
3
0
20
40
60
Value
Common sites gene−LOC105287814
1
2
3
4
5
RphaseB2
CphaseA1

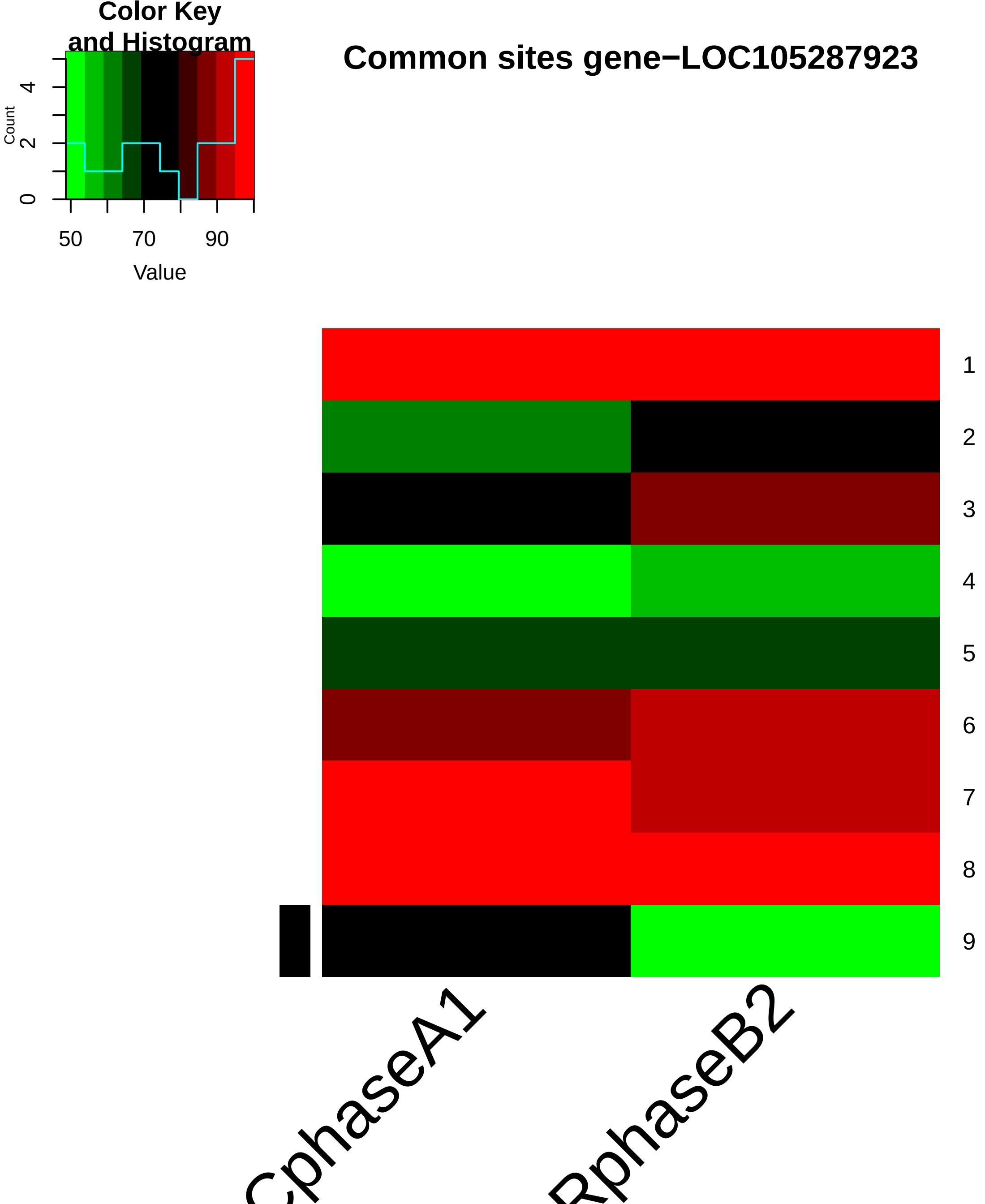
Color Key
and Histogram
Count
0
2
4
50
70
90
Value
Common sites gene−LOC105287923
1
2
3
4
5
6
7
8
9

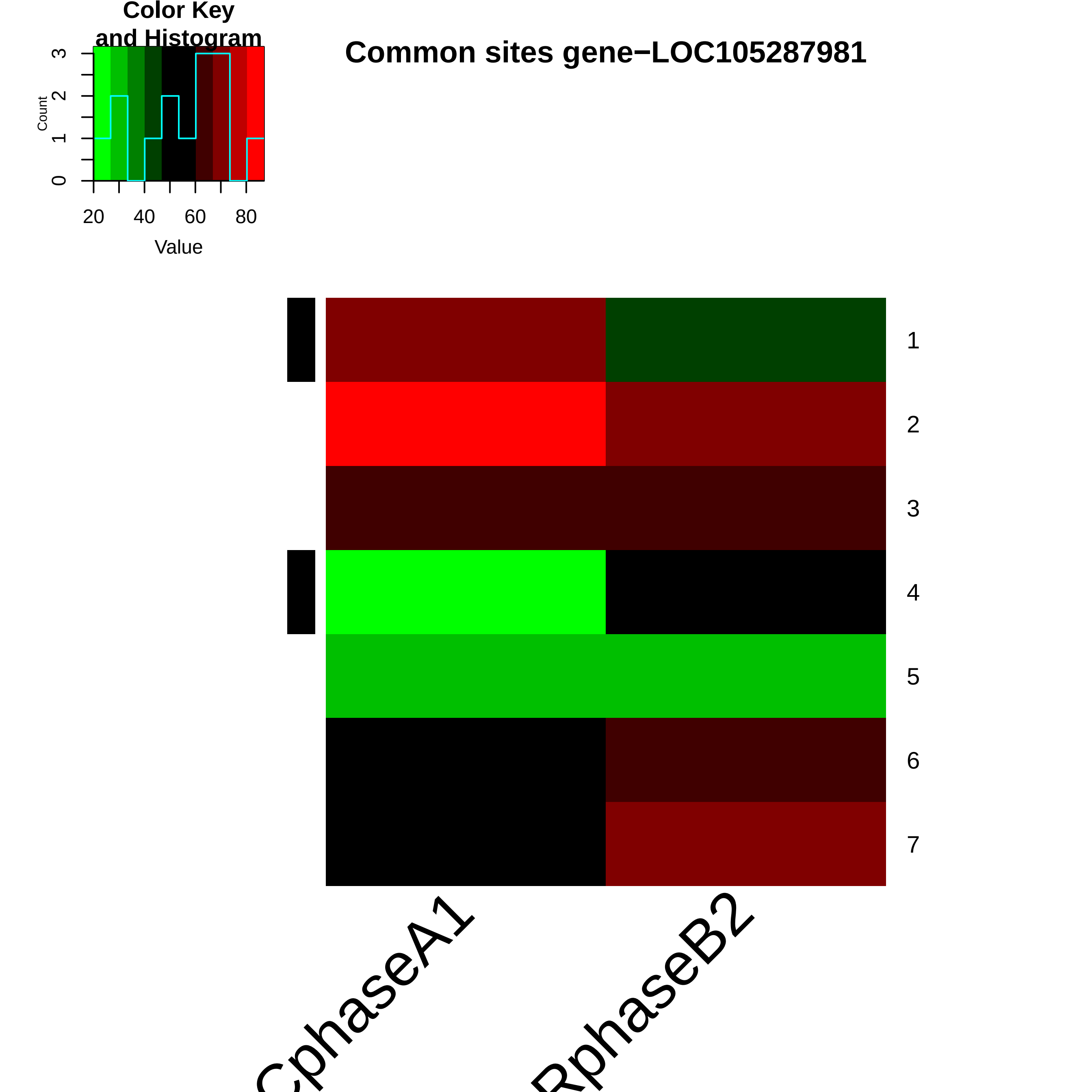
Color Key
and Histogram
Count
Value
Common sites gene−LOC105287981
1
2
3
4
5
6
7
CphaseA1
RphaseB2

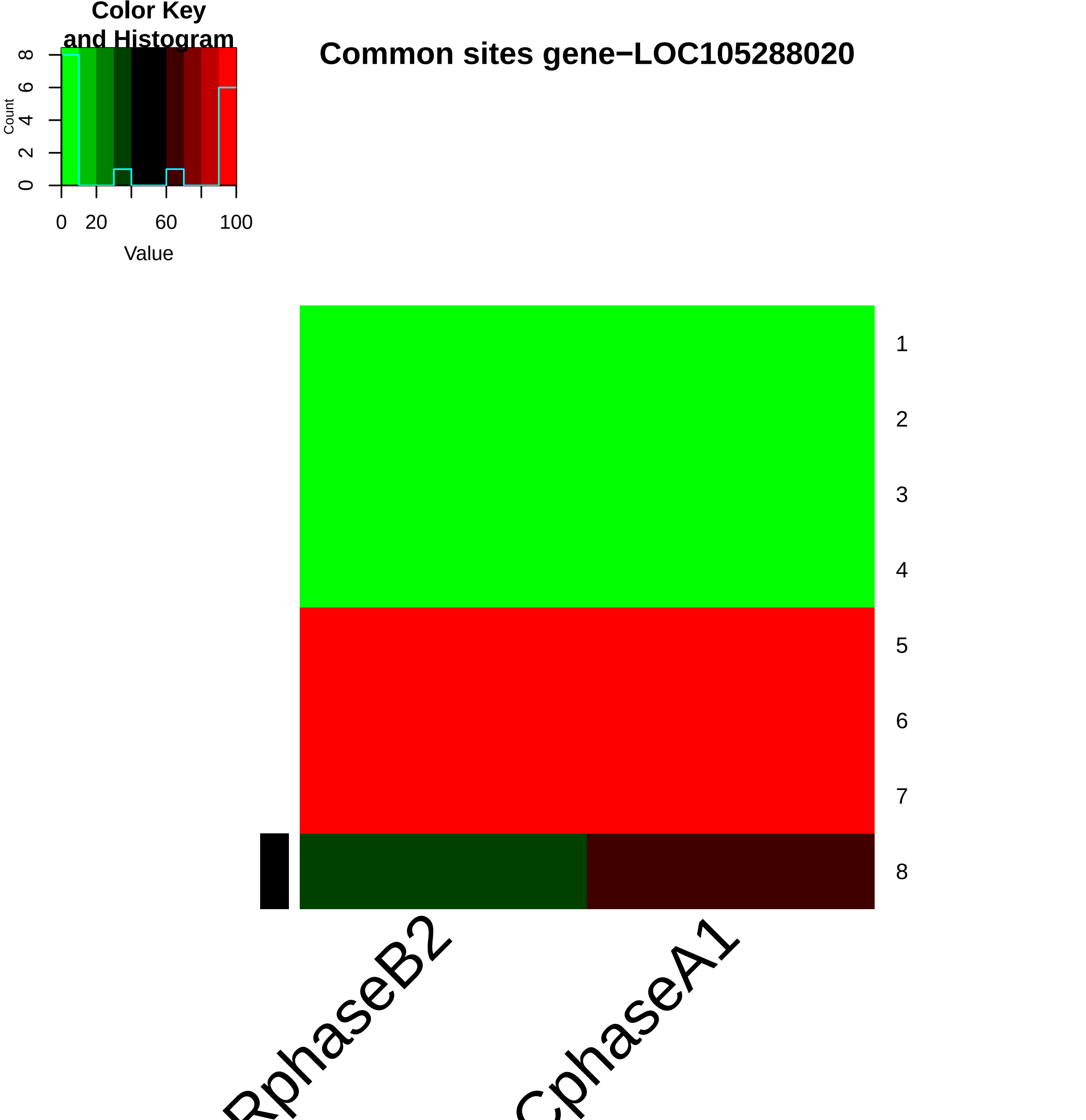

Color Key
and Histogram
Count
Value
0
20
60
100
0
2
4
6
8
Common sites gene−LOC105288020
1
2
3
4
5
6
7
8
RphaseB2
CphaseA1

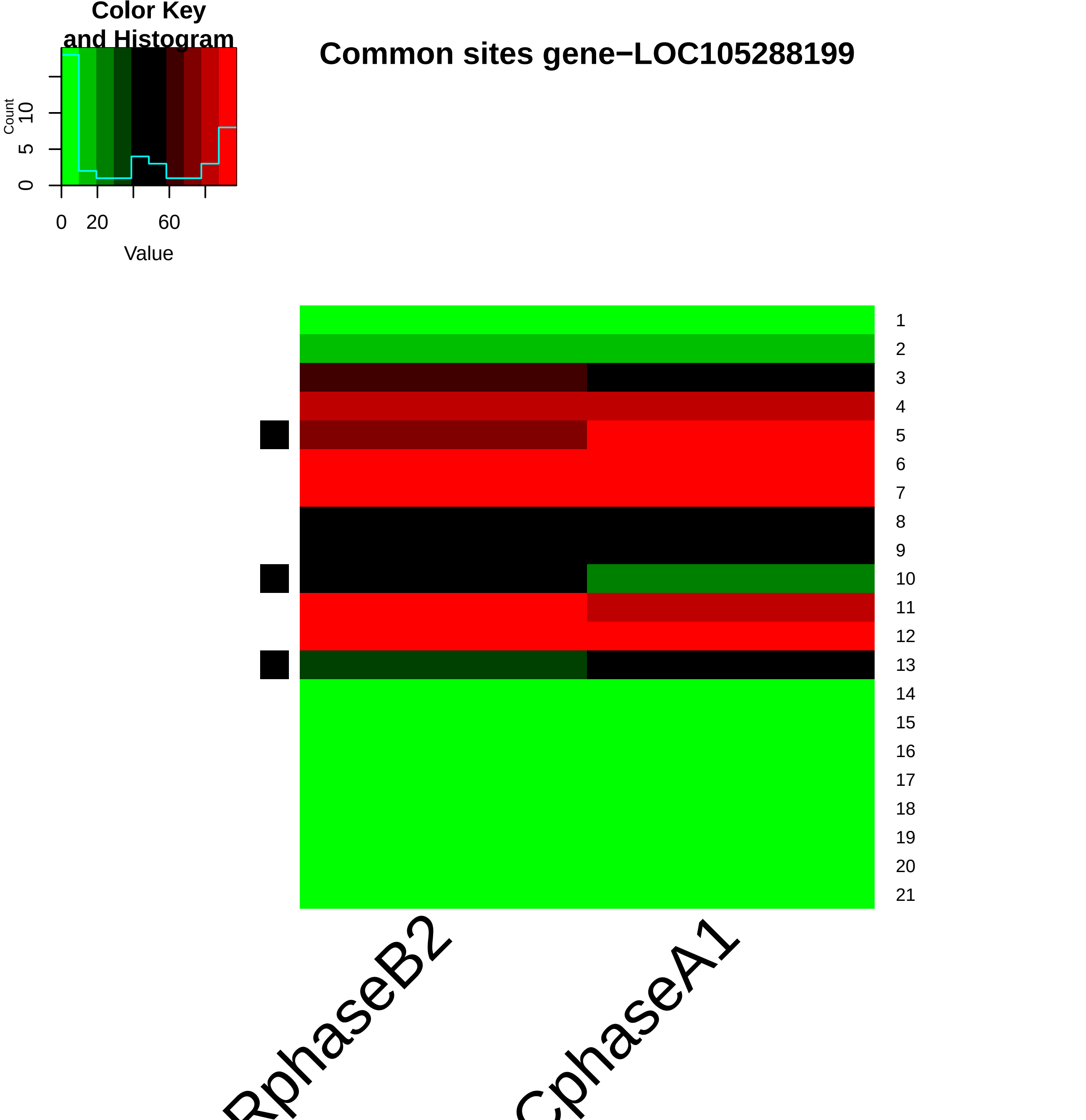
Color Key
and Histogram
Count
0
5
10
0
20
60
Value
Common sites gene−LOC105288199
RphaseB2
CphaseA1

#### Common sites gene−LOC105288225

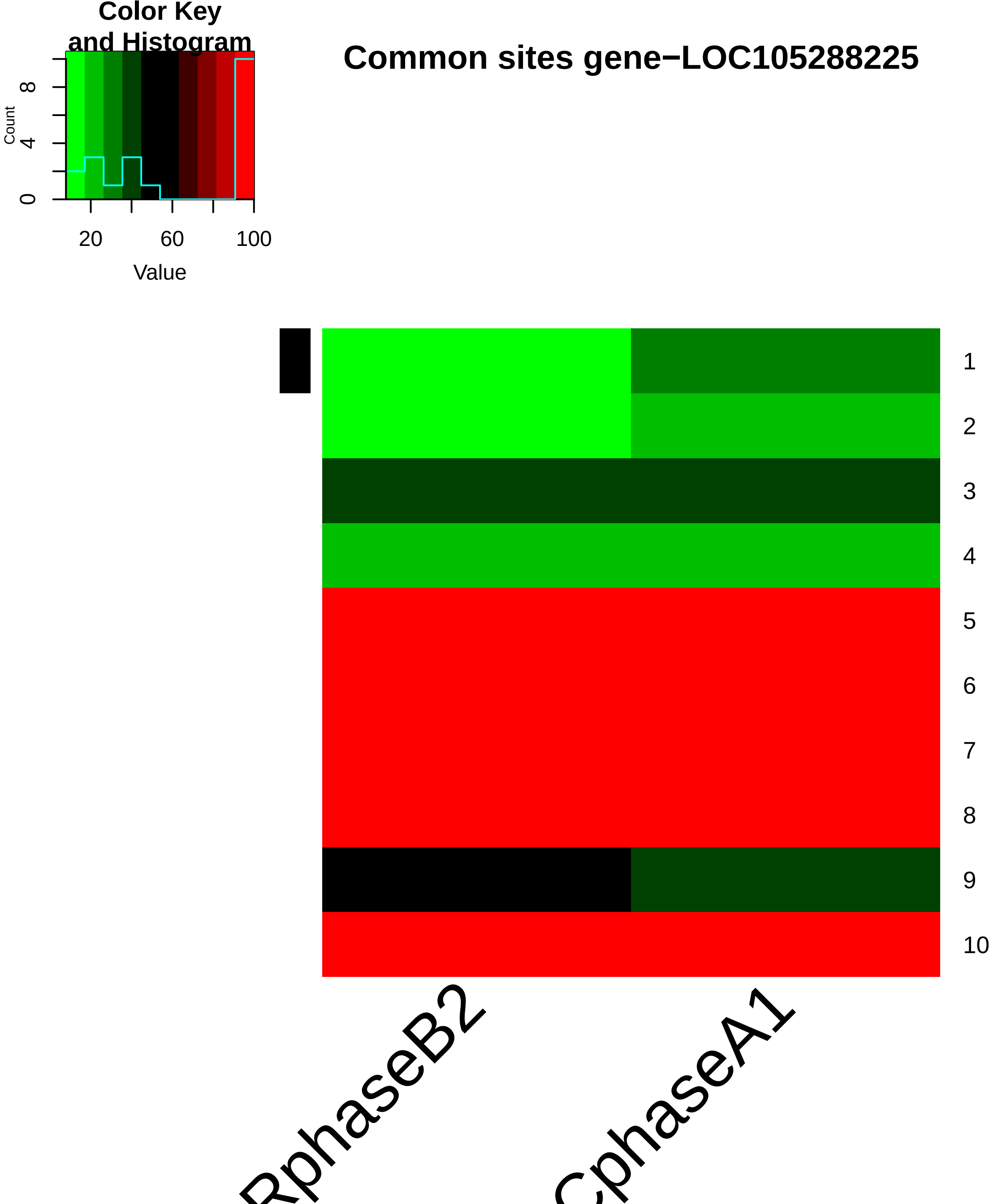

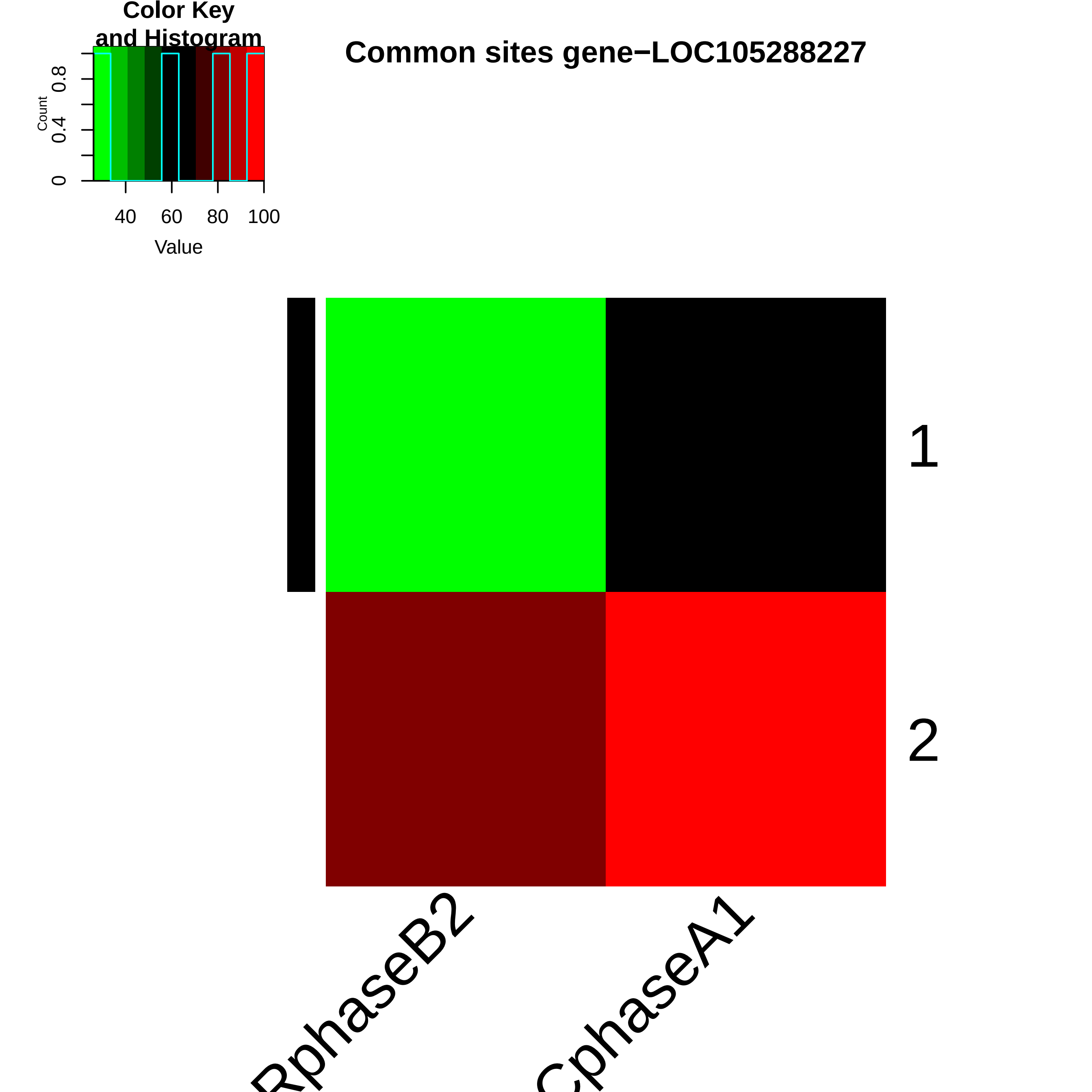

Color Key
and Histogram
Count
0
0.4
0.8
40
60
80
100
Value
Common sites gene−LOC105288227
1
2
RphaseB2
CphaseA1

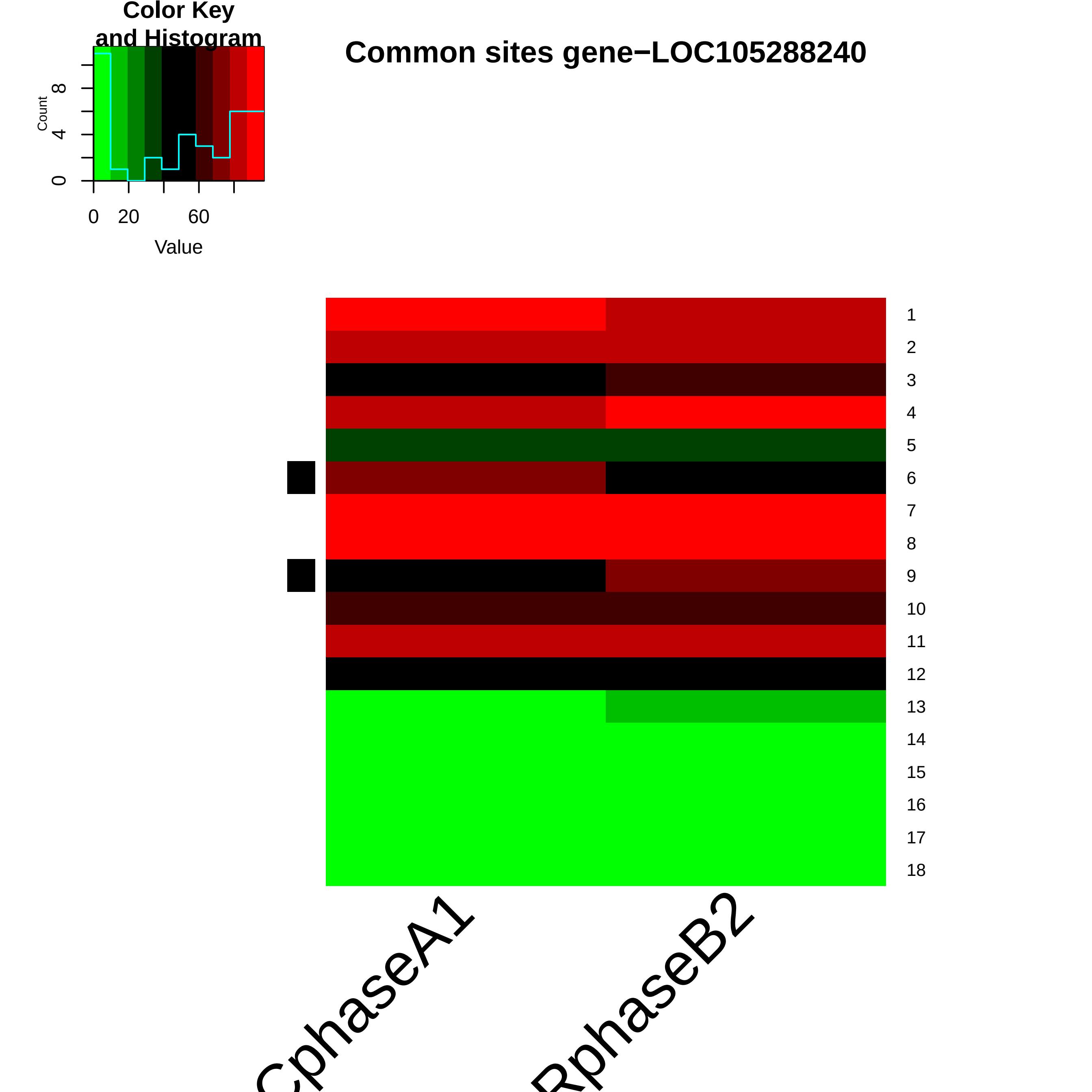

Color Key
and Histogram
Count
0
4
8
0
20
60
Value
Common sites gene−LOC105288240
CphaseA1
RphaseB2

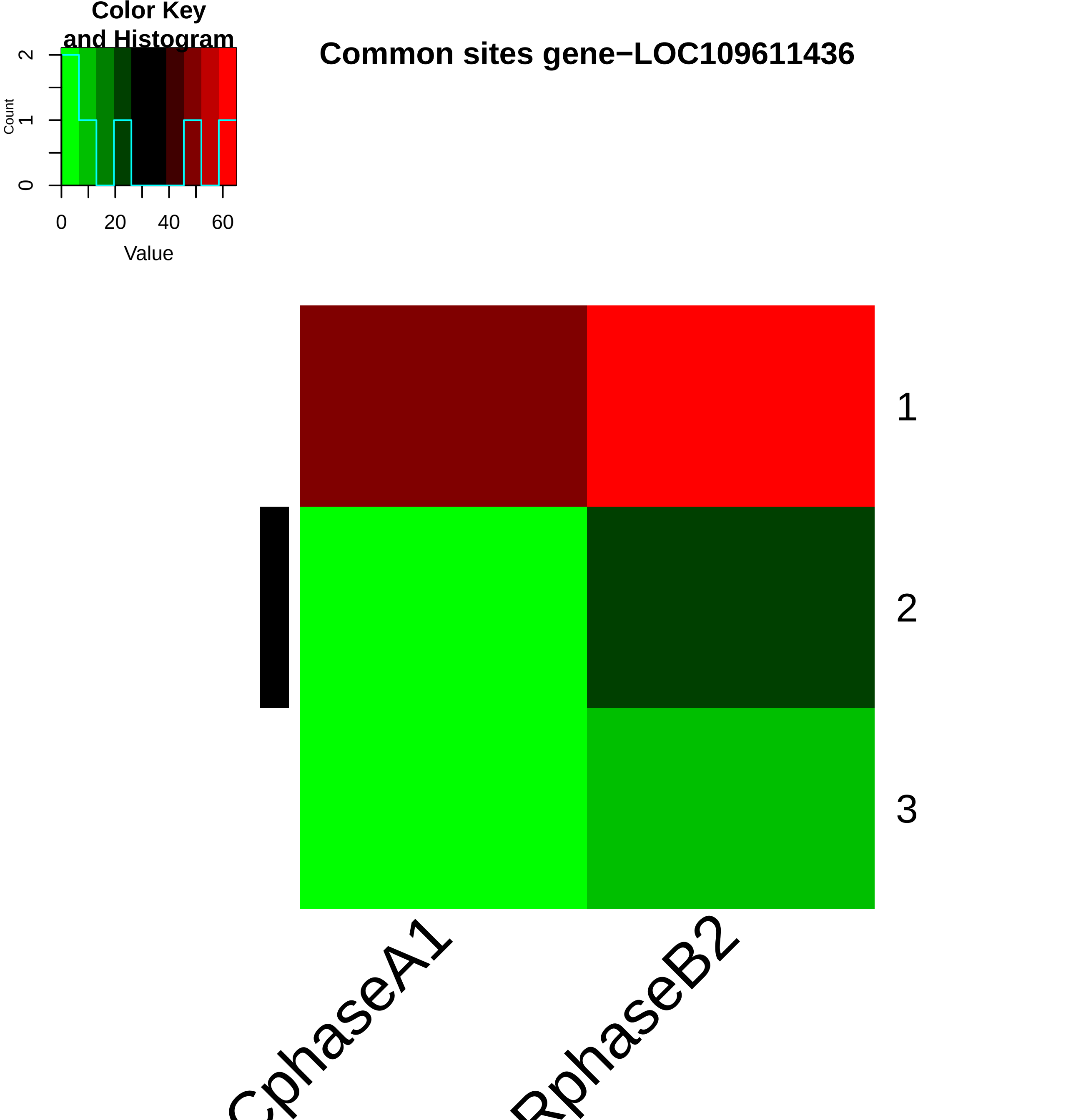
Color Key
and Histogram
Count
0
1
2
0
20
40
60
Value
Common sites gene−LOC109611436
1
2
3

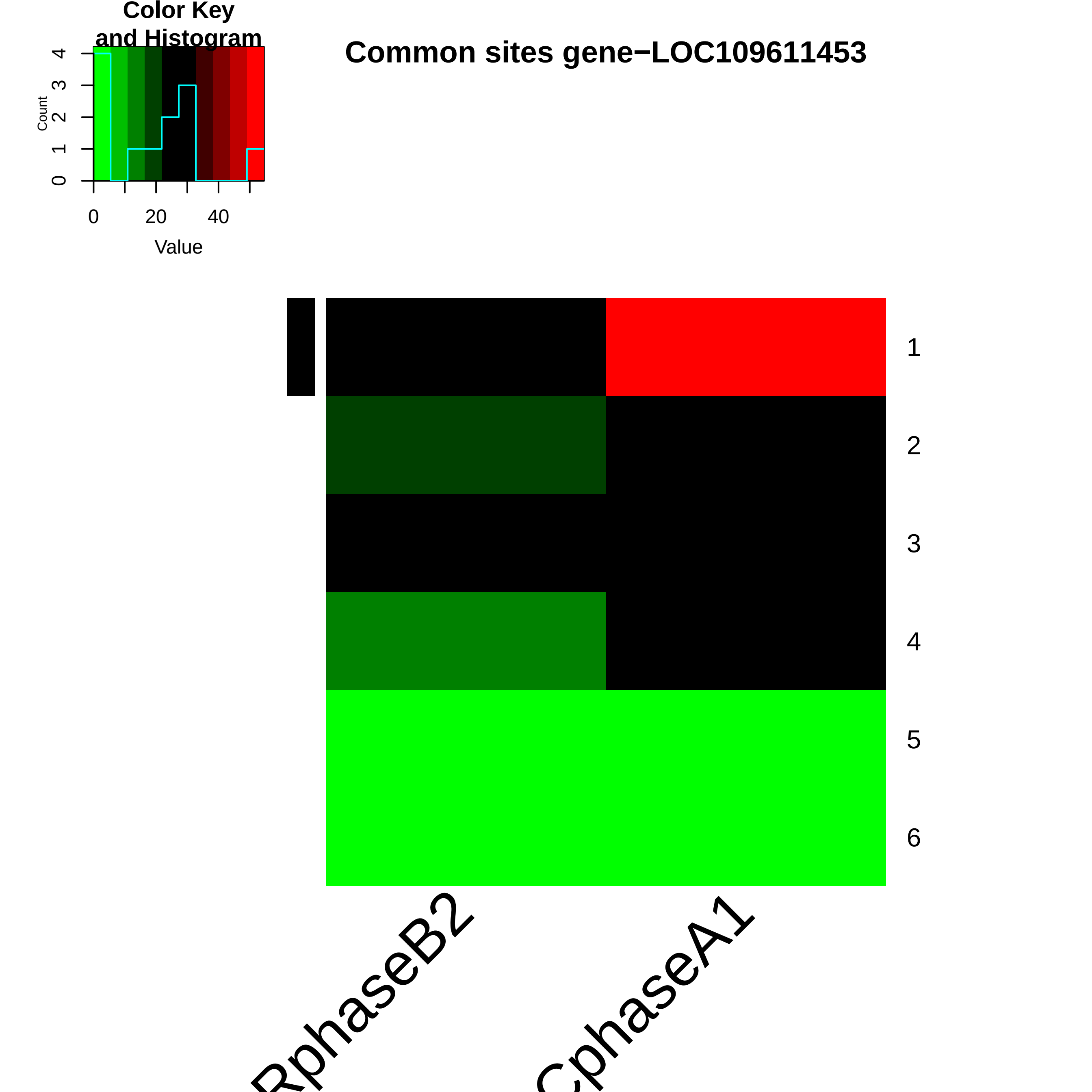
Color Key
and Histogram
Count
Value
0
20
40
0
1
2
3
4
Common sites gene−LOC109611453
1
2
3
4
5
6
RphaseB2
CphaseA1

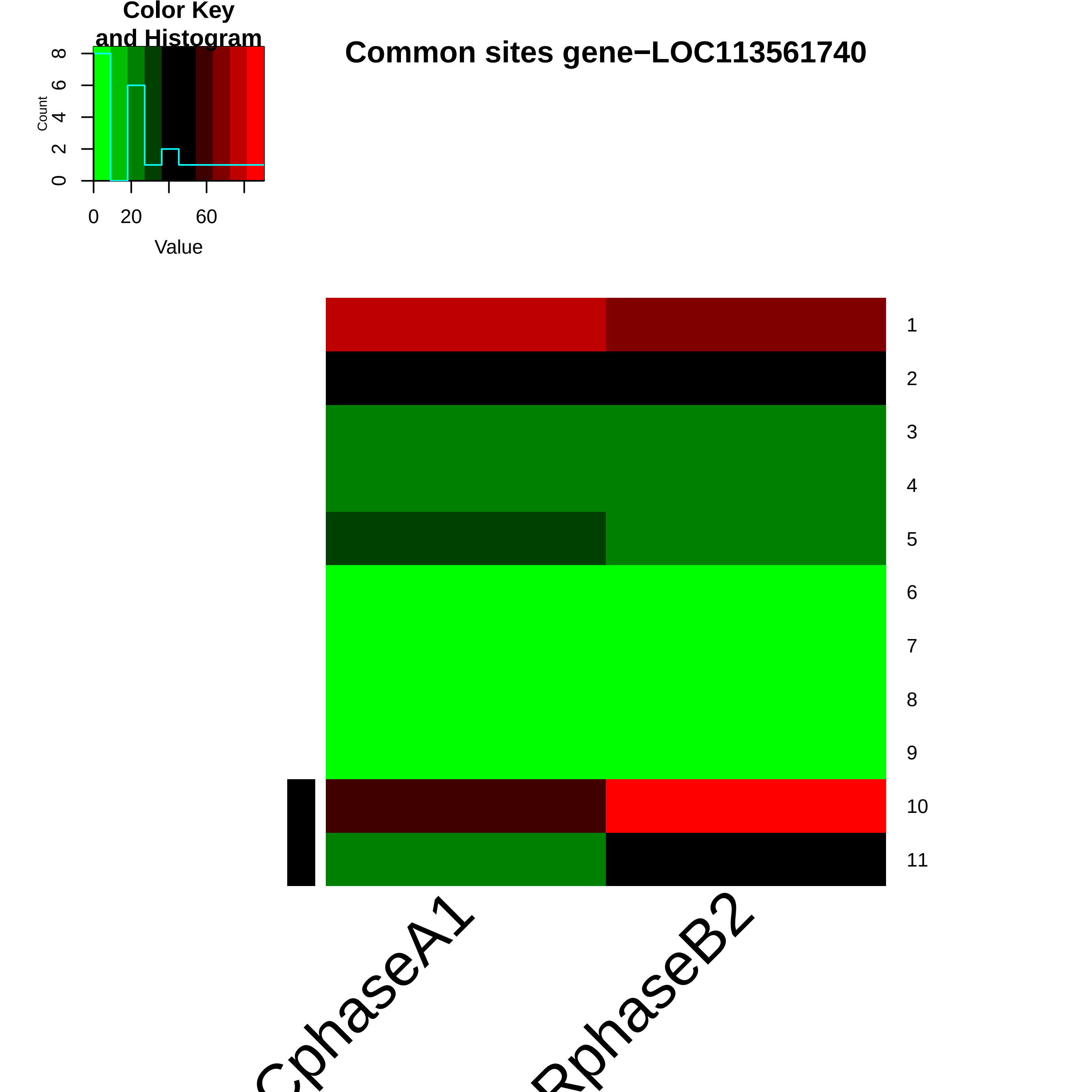
Color Key
and Histogram
Count
0
2
4
6
8
0
20
60
Value
Common sites gene−LOC113561740
1
2
3
4
5
6
7
8
9
10
11
CphaseA1
RphaseB2

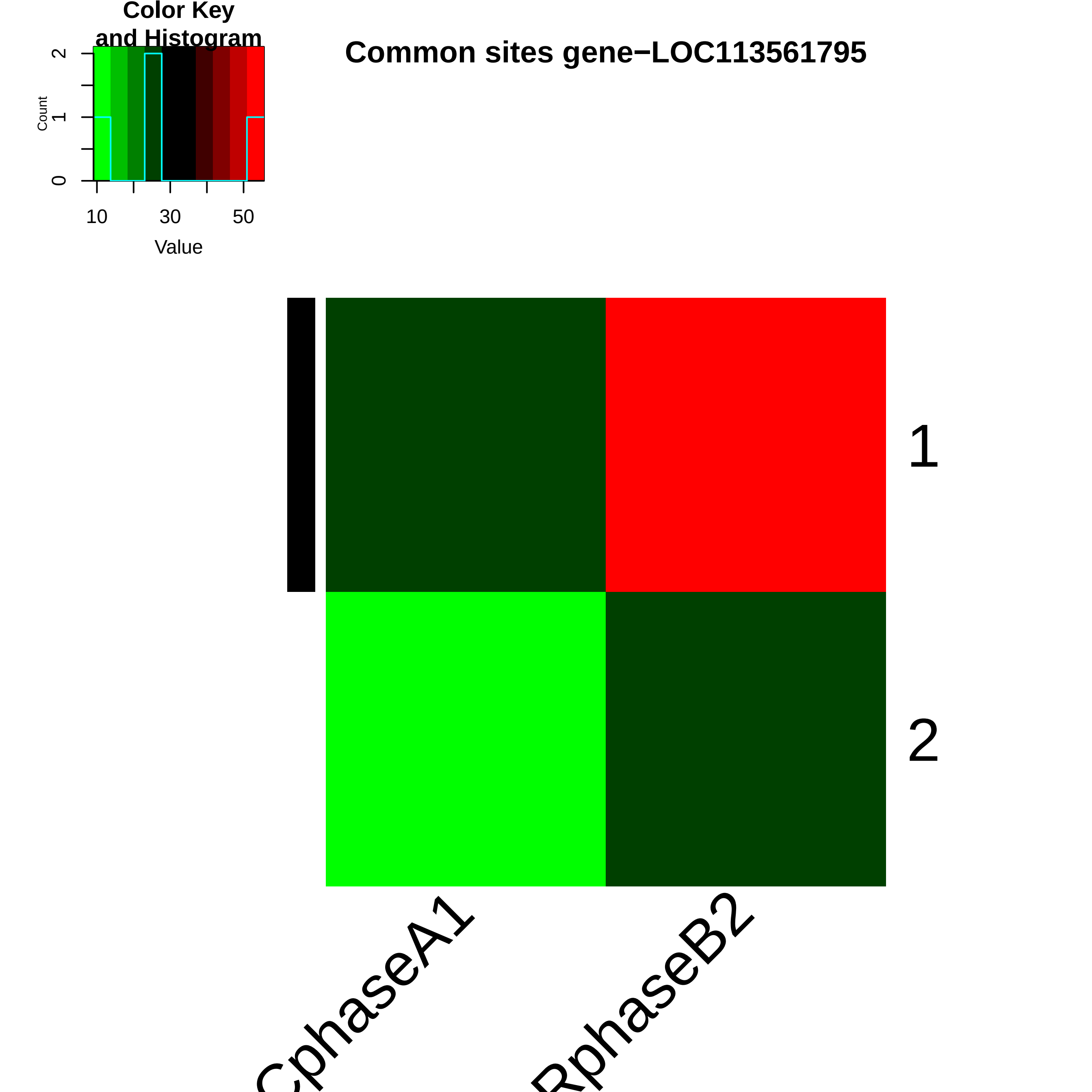
Color Key
and Histogram
Count
0
1
2
10
30
50
Value
Common sites gene−LOC113561795
1
2
CphaseA1
RphaseB2

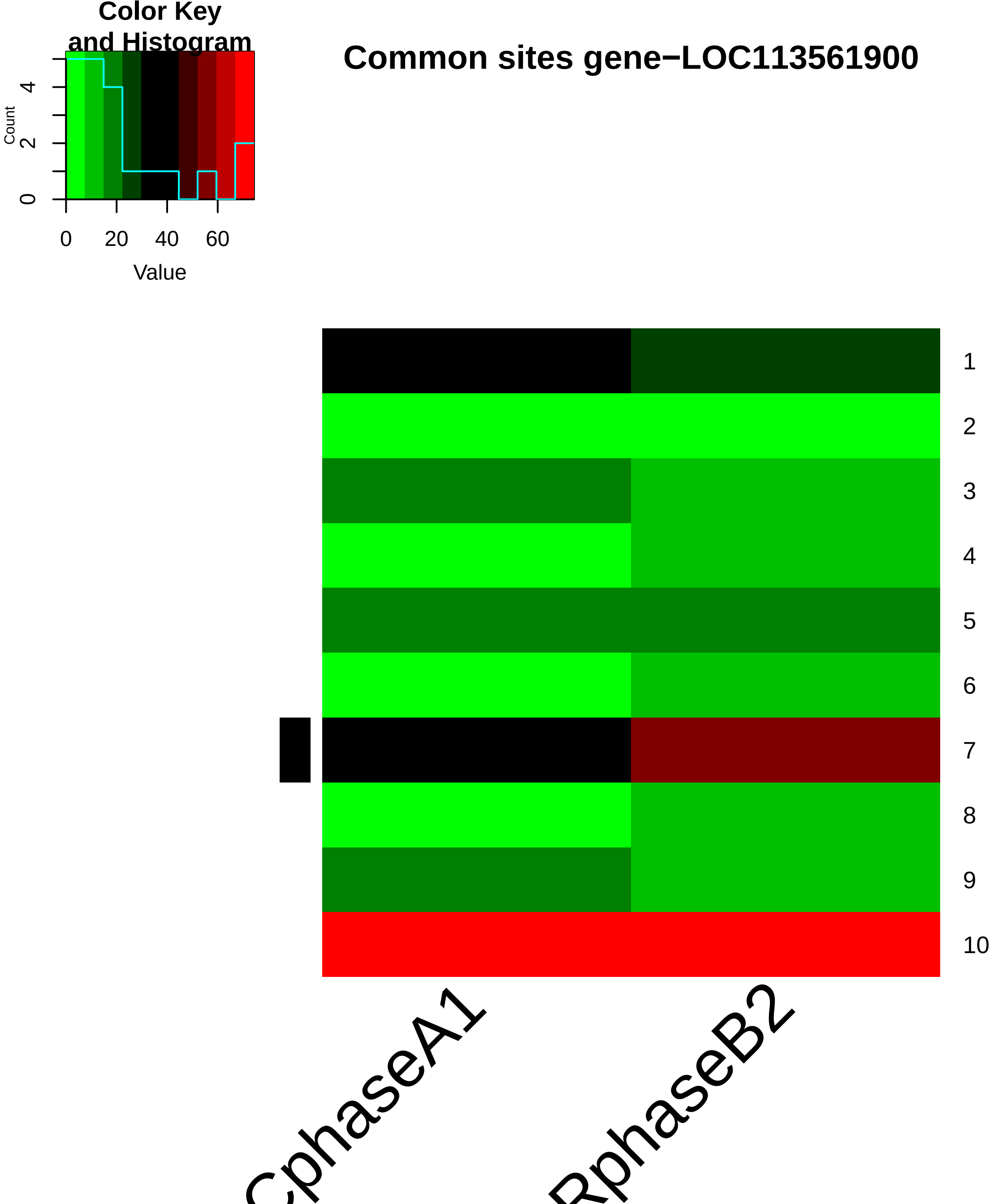

Color Key
and Histogram
Count
0
2
4
0
20
40
60
Value
Common sites gene−LOC113561900
1
2
3
4
5
6
7
8
9
10
CphaseA1
RphaseB2

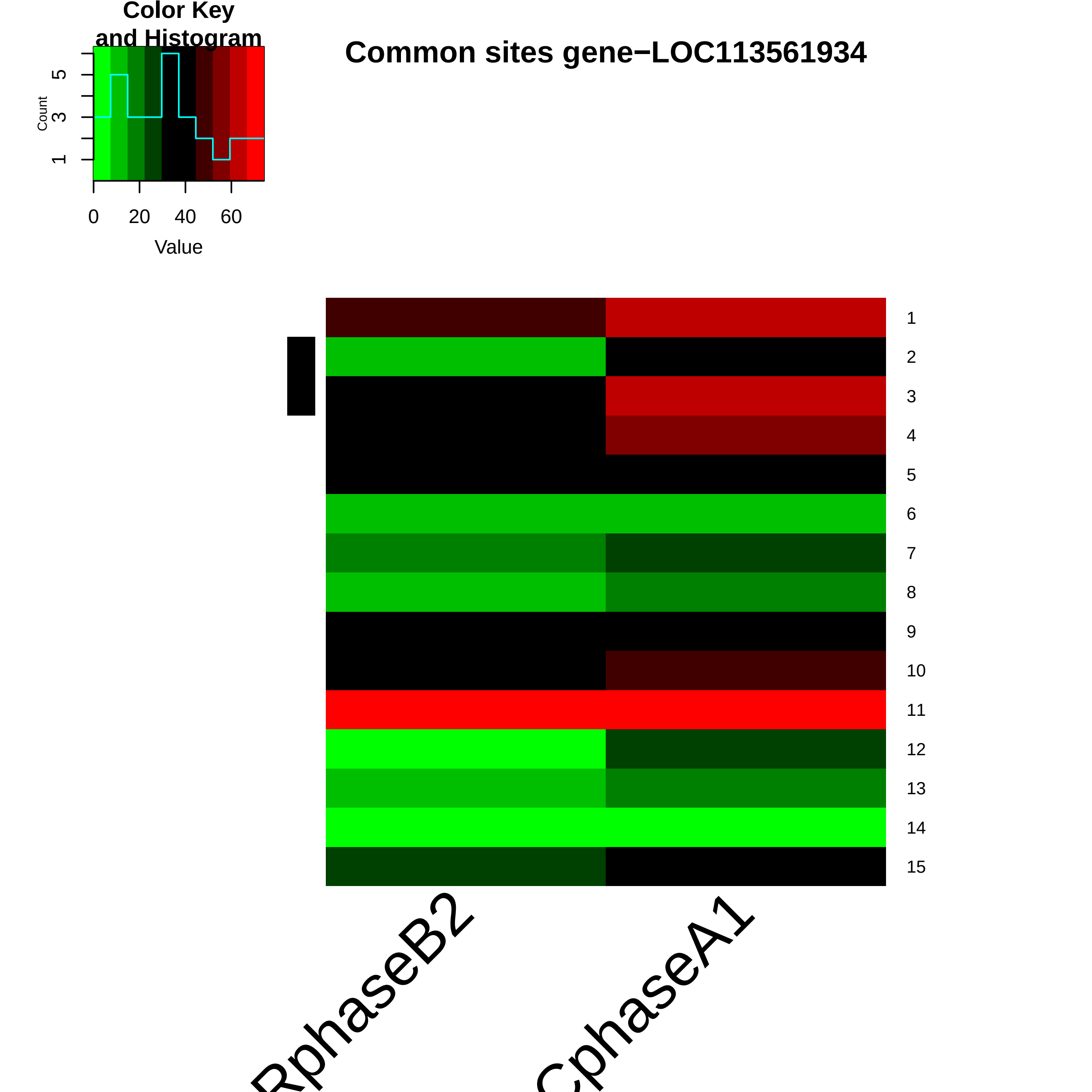

Color Key
and Histogram
Count
Value
0
20
40
60
1
3
5
Common sites gene−LOC113561934
RphaseB2
CphaseA1

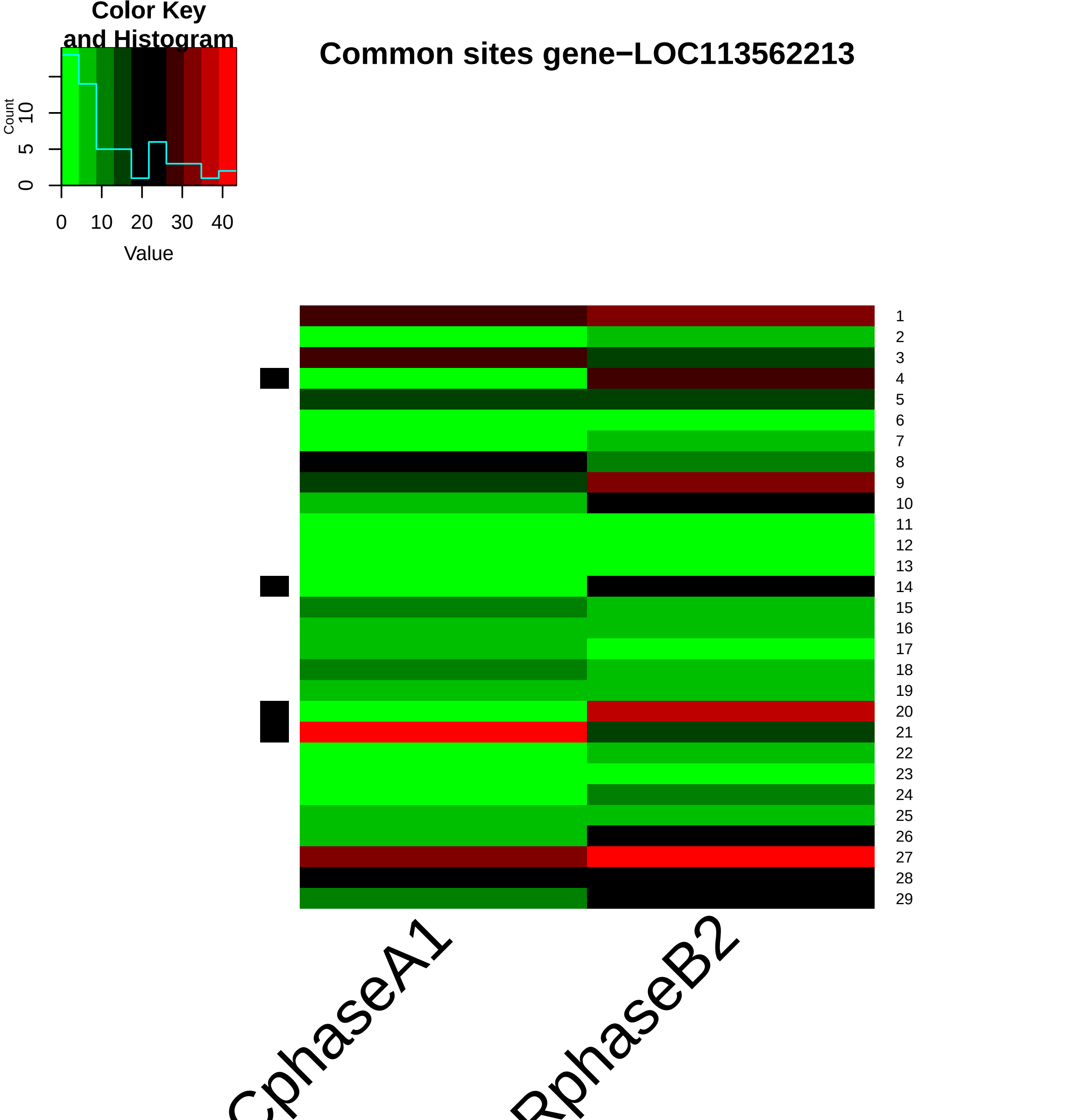

Color Key
and Histogram
Count
0
5
10
0
10
20
30
40
Value
Common sites gene−LOC113562213
CphaseA1
RphaseB2

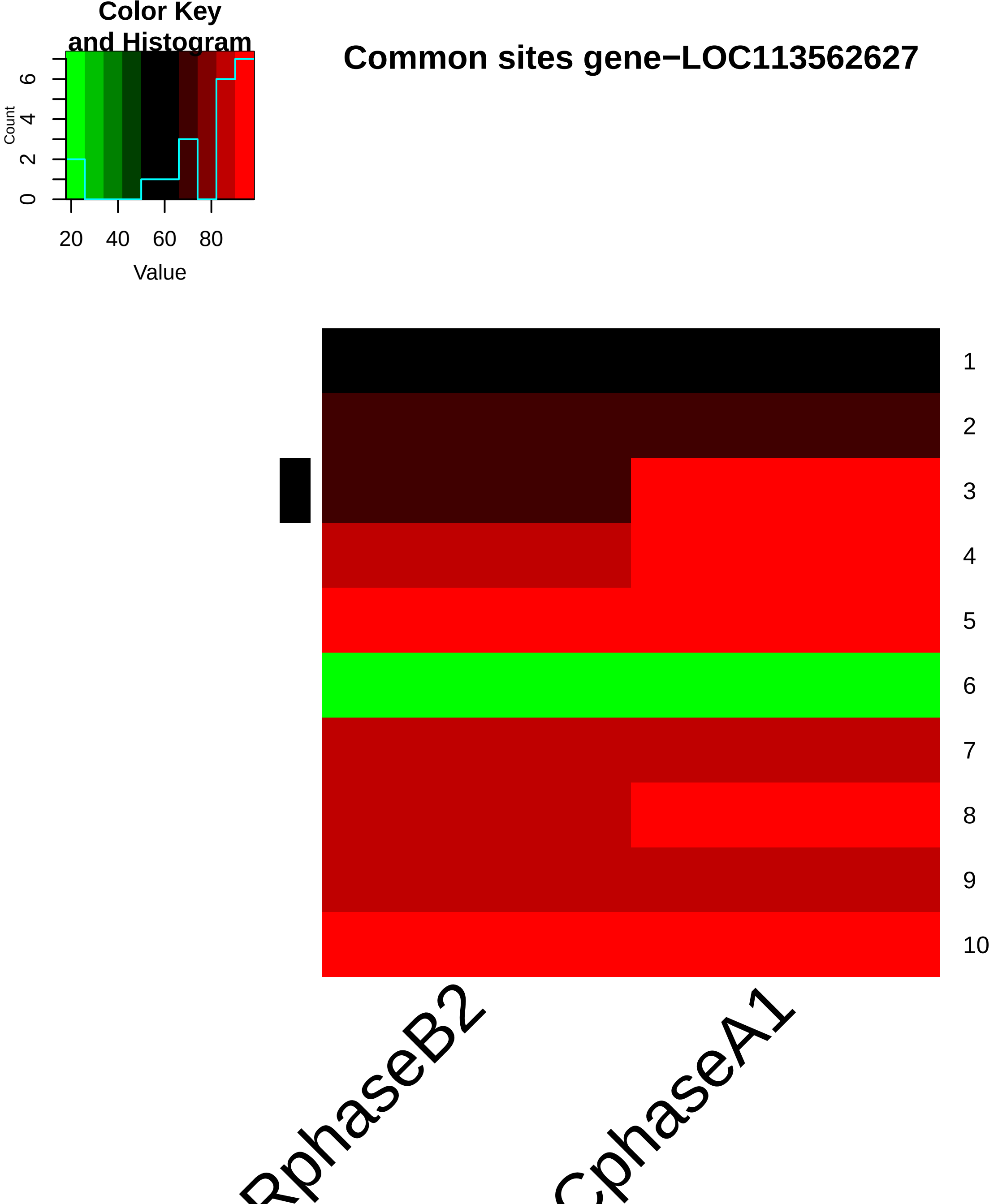
Color Key
and Histogram
Count
0
2
4
6
20
40
60
80
Value
Common sites gene−LOC113562627
1
2
3
4
5
6
7
8
9
10
RphaseB2
CphaseA1

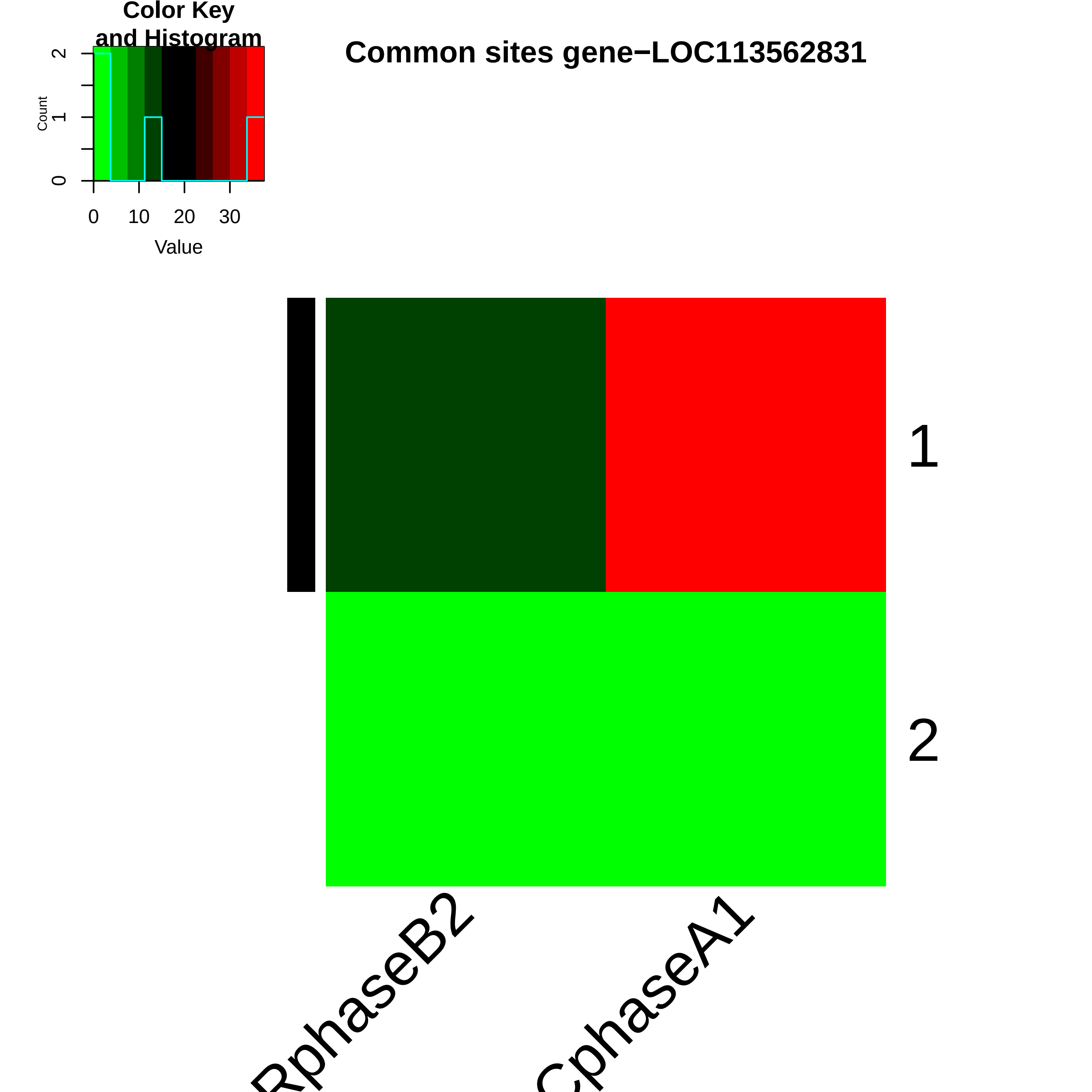
Color Key
and Histogram
Count
0
1
2
0
10
20
30
Value
Common sites gene−LOC113562831
1
2
RphaseB2
CphaseA1

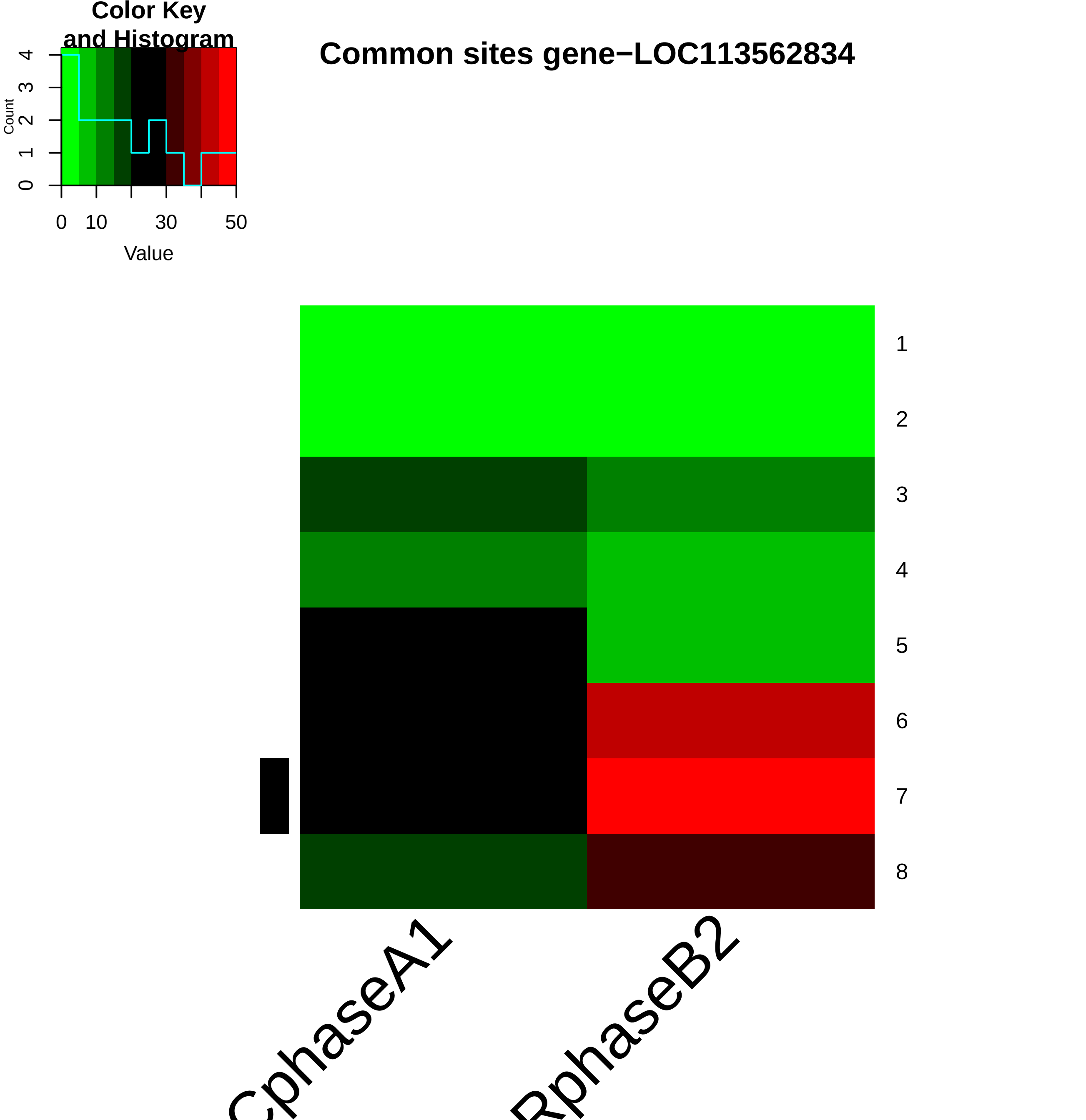

Color Key
and Histogram
Count
0
1
2
3
4
0
10
30
50
Value
Common sites gene−LOC113562834
1
2
3
4
5
6
7
8
CphaseA1
RphaseB2

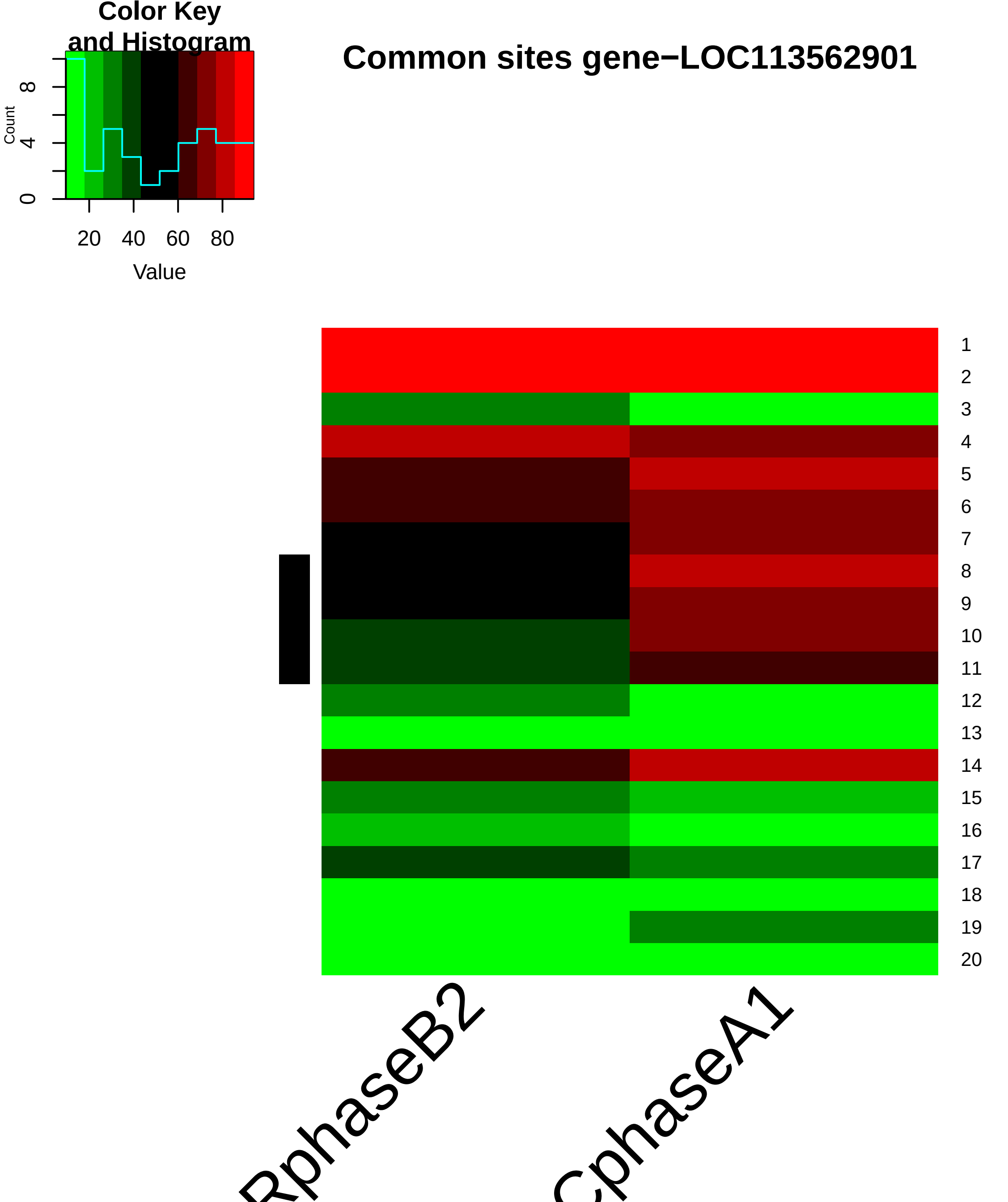

Color Key
and Histogram
Count
0
4
8
20
40
60
80
Value
Common sites gene−LOC113562901
RphaseB2
CphaseA1

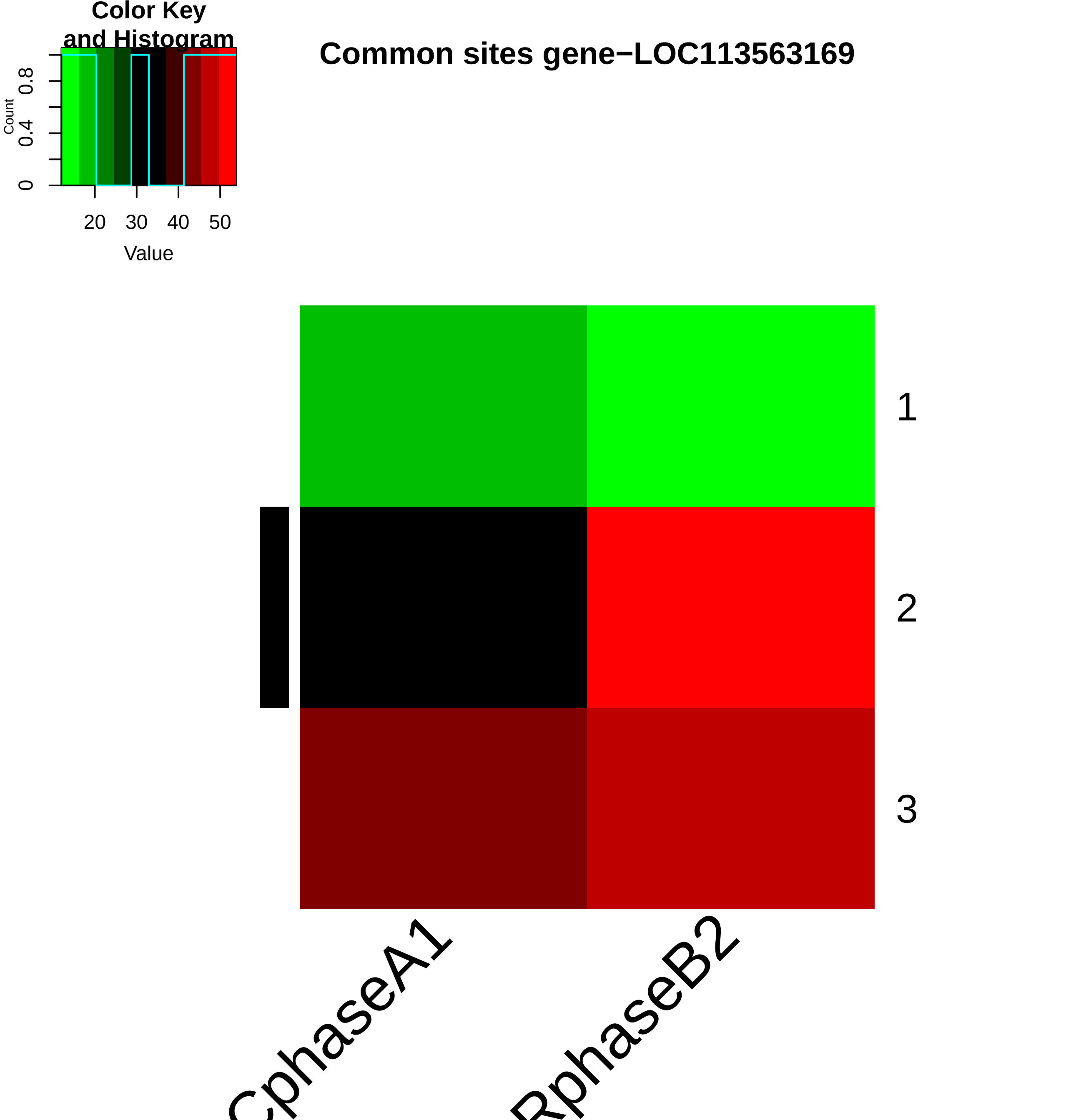

Color Key
and Histogram
Count
0
0.4
0.8
20
30
40
50
Value
Common sites gene−LOC113563169
1
2
3
CphaseA1
RphaseB2

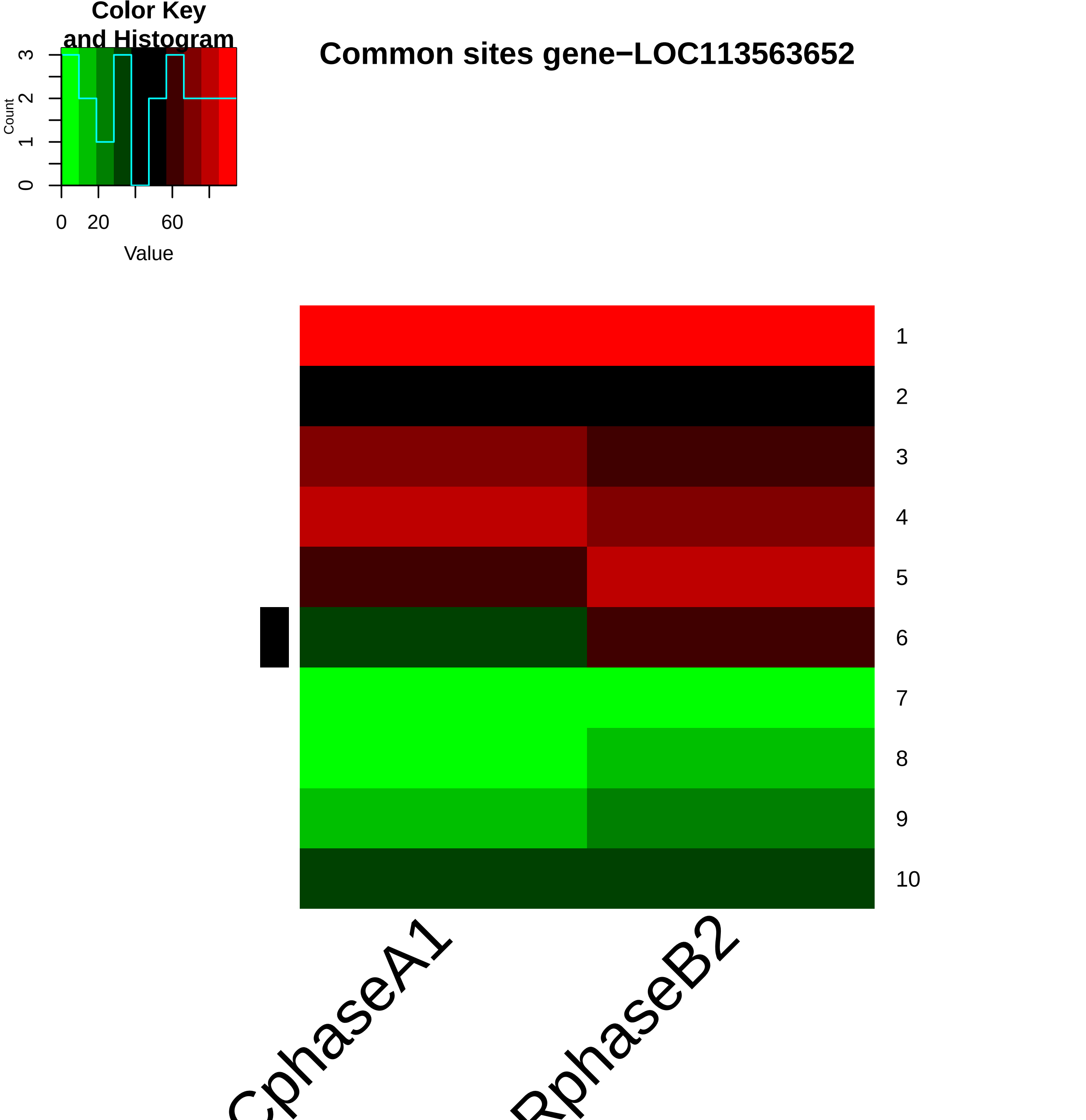

Color Key
and Histogram
Count
0
1
2
3
0
20
60
Value
Common sites gene−LOC113563652
1
2
3
4
5
6
7
8
9
10
CphaseA1
RphaseB2

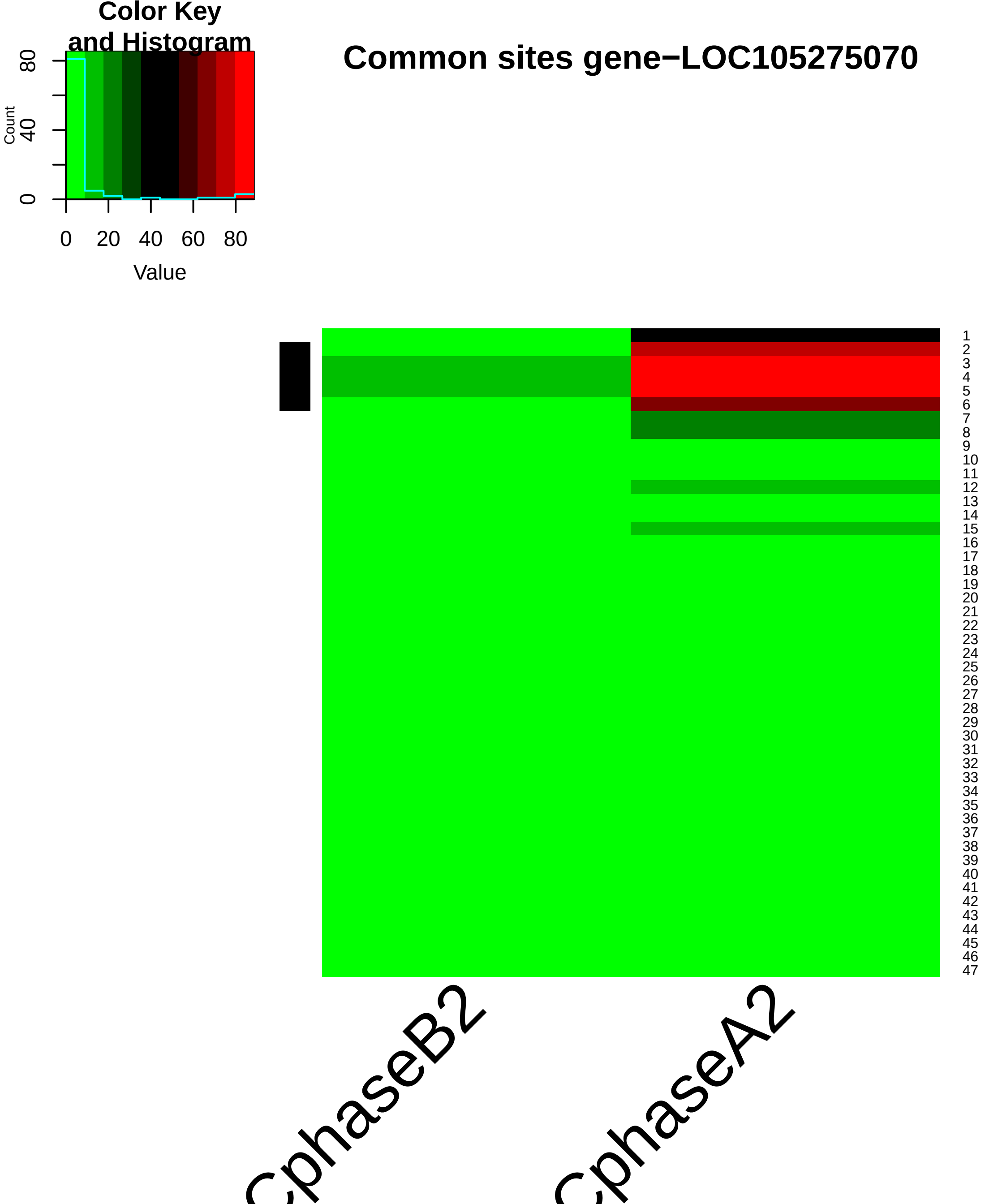

Color Key
and Histogram
Count
0
40
80
0
20
40
60
80
Value
Common sites gene−LOC105275070
CphaseB2
CphaseA2

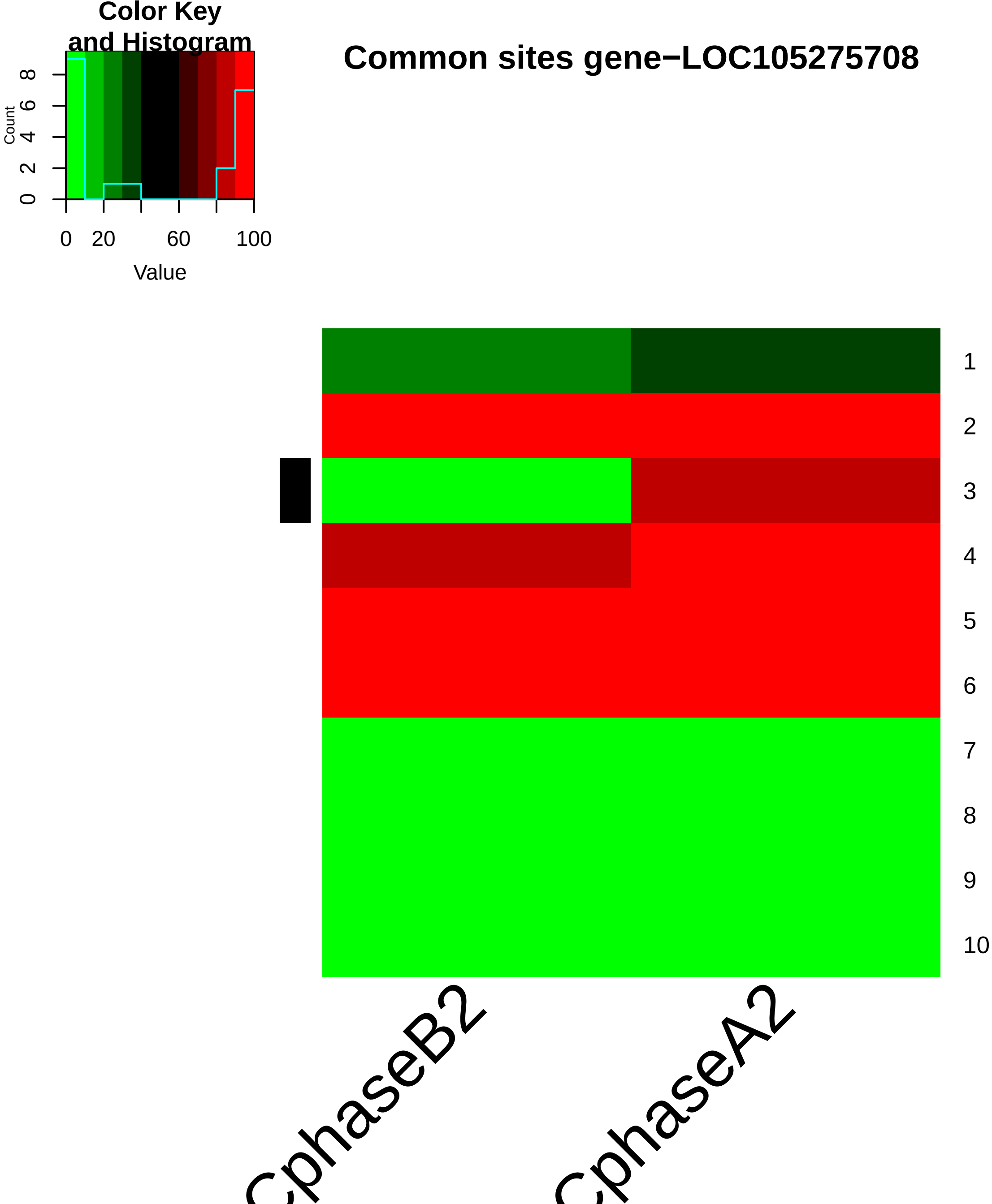
Color Key
and Histogram
Count
0
2
4
6
8
0
20
60
100
Value
Common sites gene−LOC105275708
1
2
3
4
5
6
7
8
9
10
CphaseB2
CphaseA2

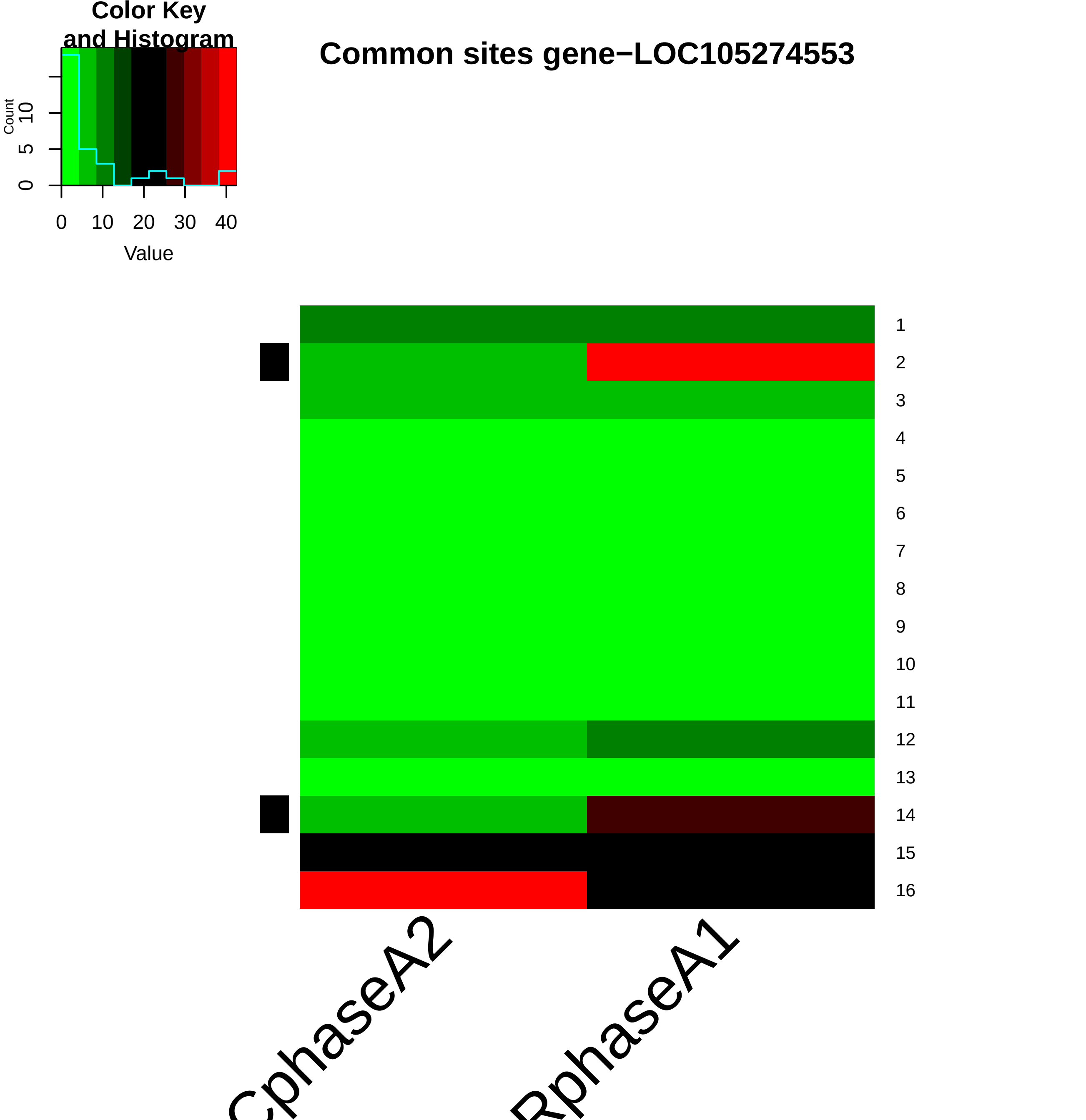

Color Key
and Histogram
Count
0
5
10
0
10
20
30
40
Value
Common sites gene−LOC105274553

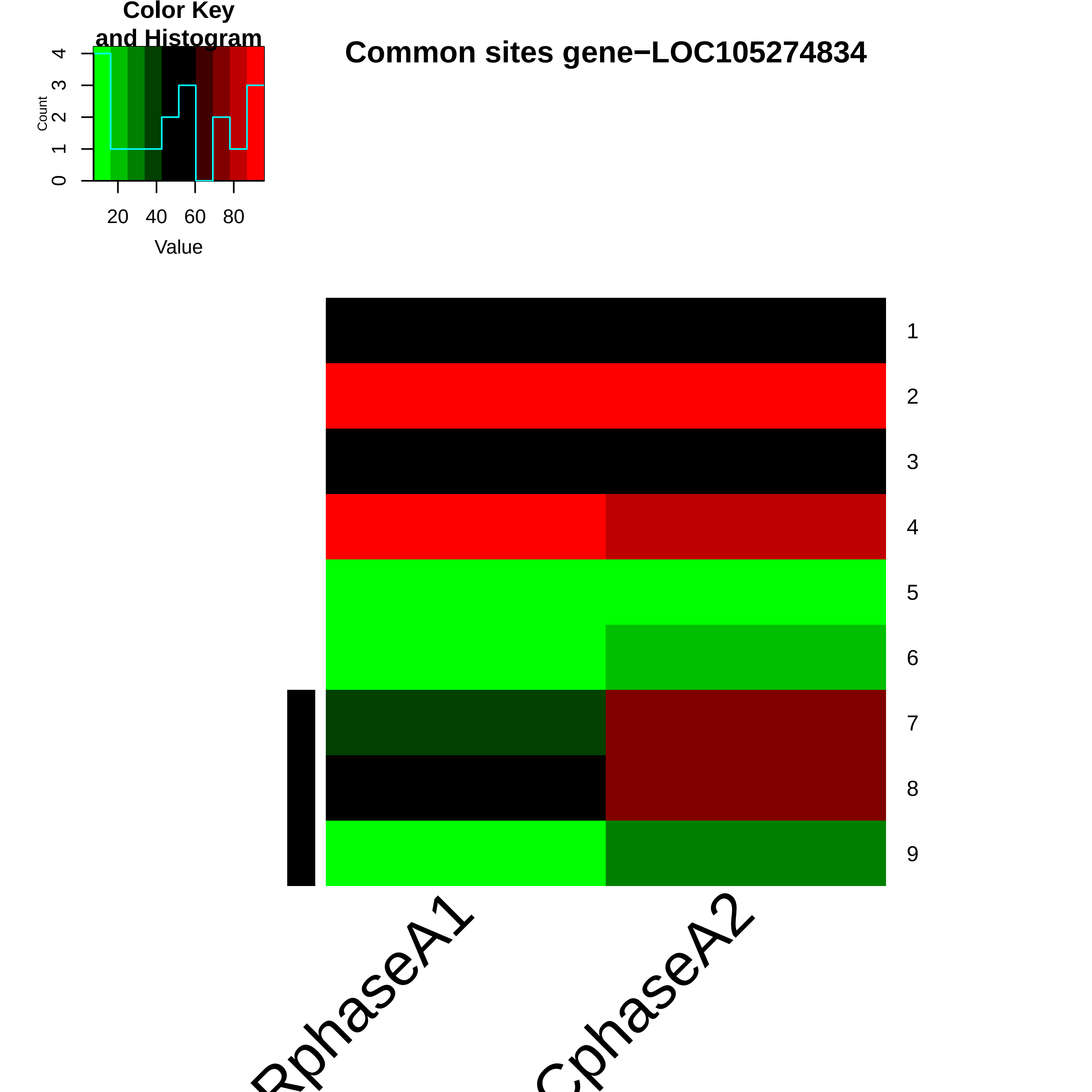

Color Key
and Histogram
Count
Value
0 1 2 3 4
20 40 60 80
Common sites gene−LOC105274834
1
2
3
4
5
6
7
8
9
RphaseA1
CphaseA2

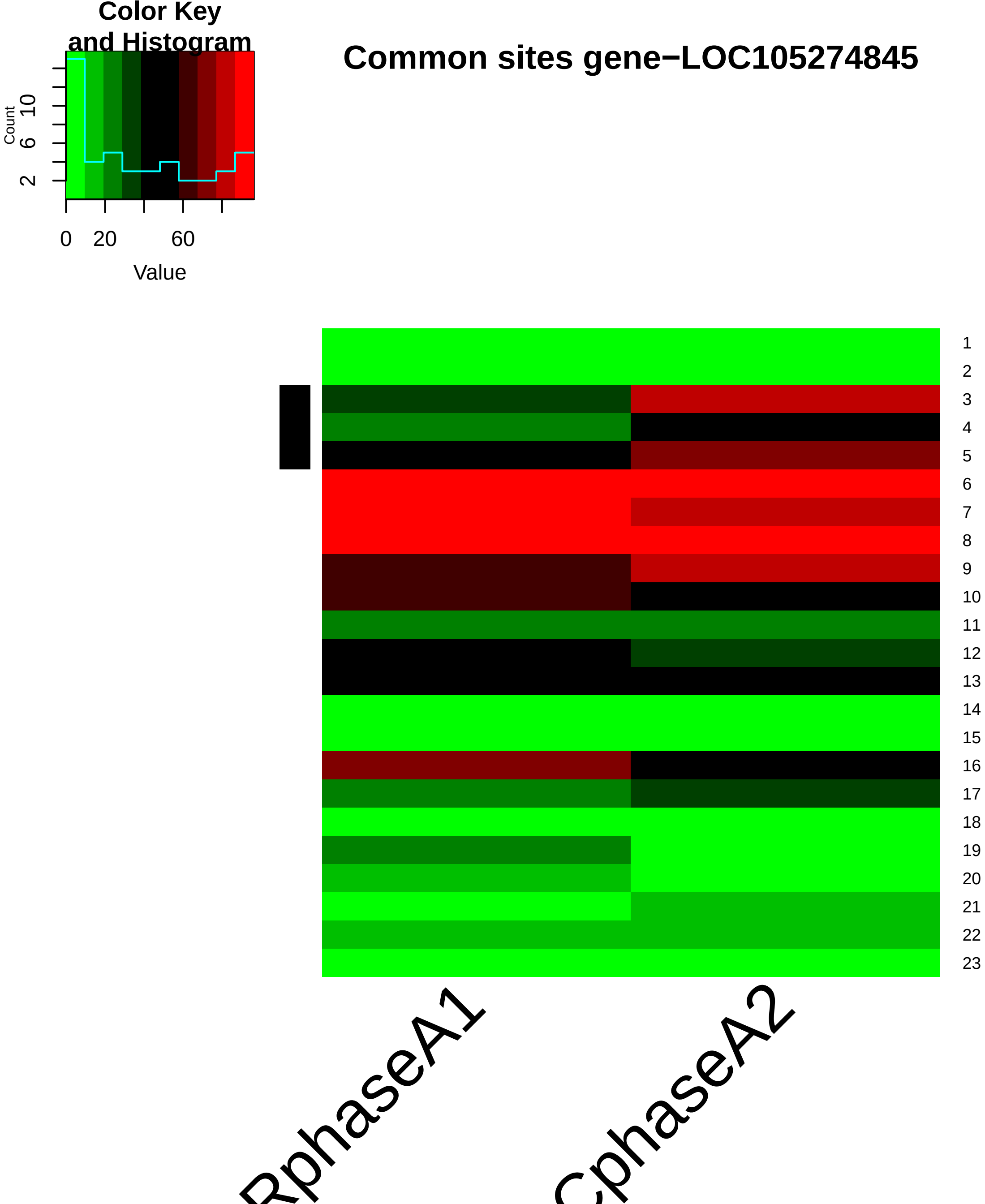

Color Key
and Histogram
Count
2
6
10
0
20
60
Value
Common sites gene−LOC105274845
RphaseA1
CphaseA2

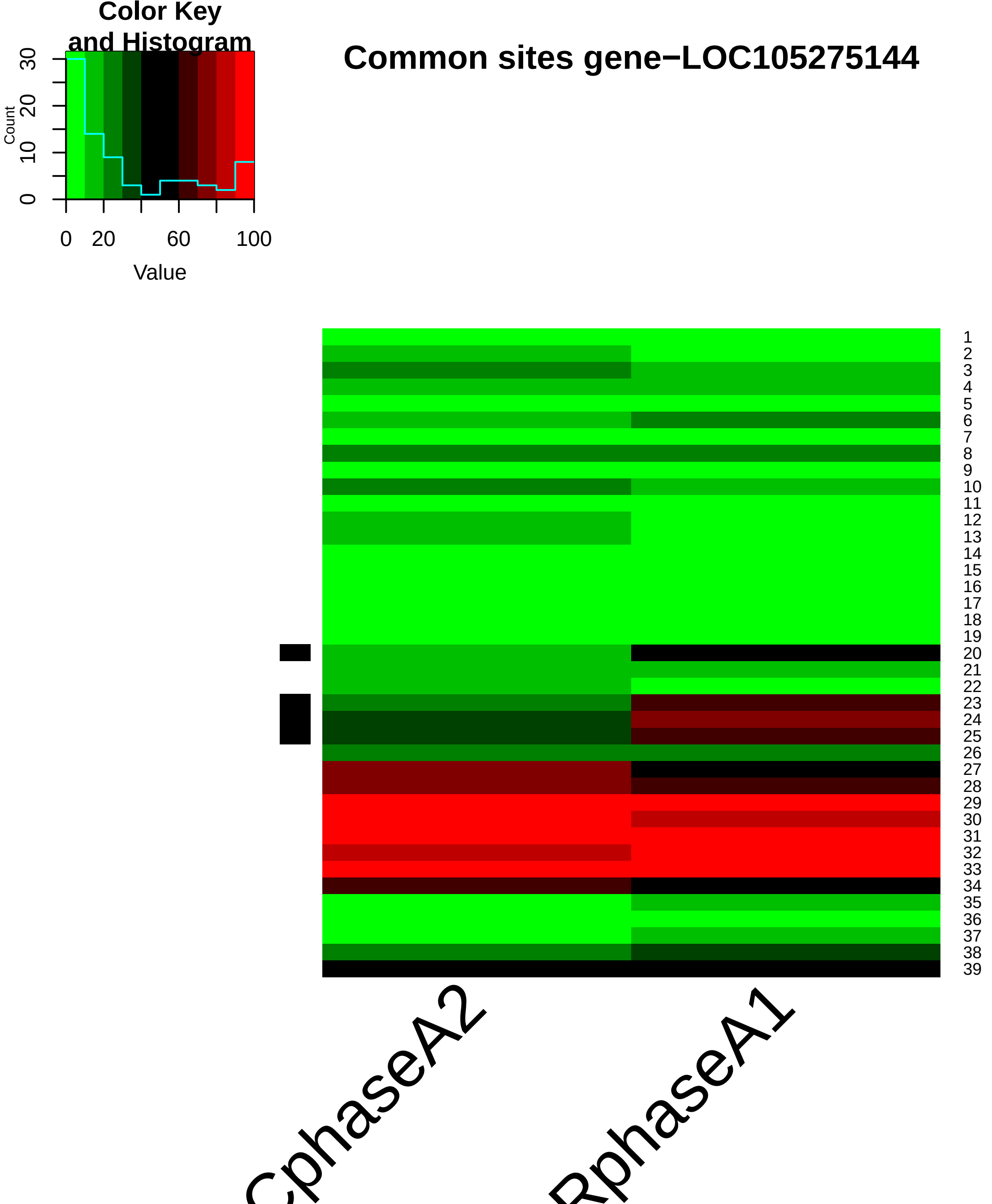

Color Key
and Histogram
Count
Value
0
10
20
30
0
20
60
100
Common sites gene−LOC105275144

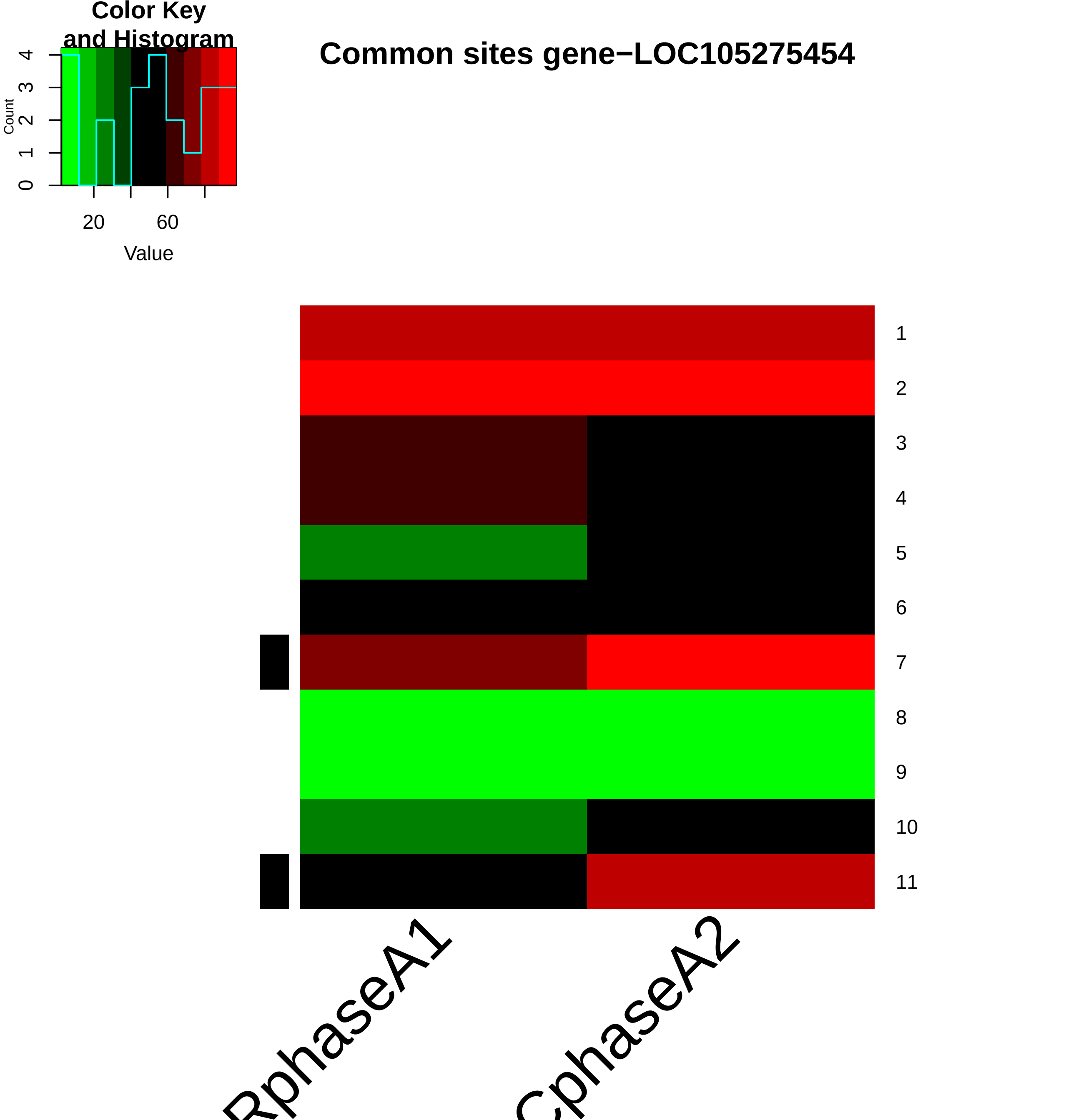
Color Key
and Histogram
Count
0
1
2
3
4
20
60
Value
Common sites gene−LOC105275454
1
2
3
4
5
6
7
8
9
10
11
RphaseA1
CphaseA2

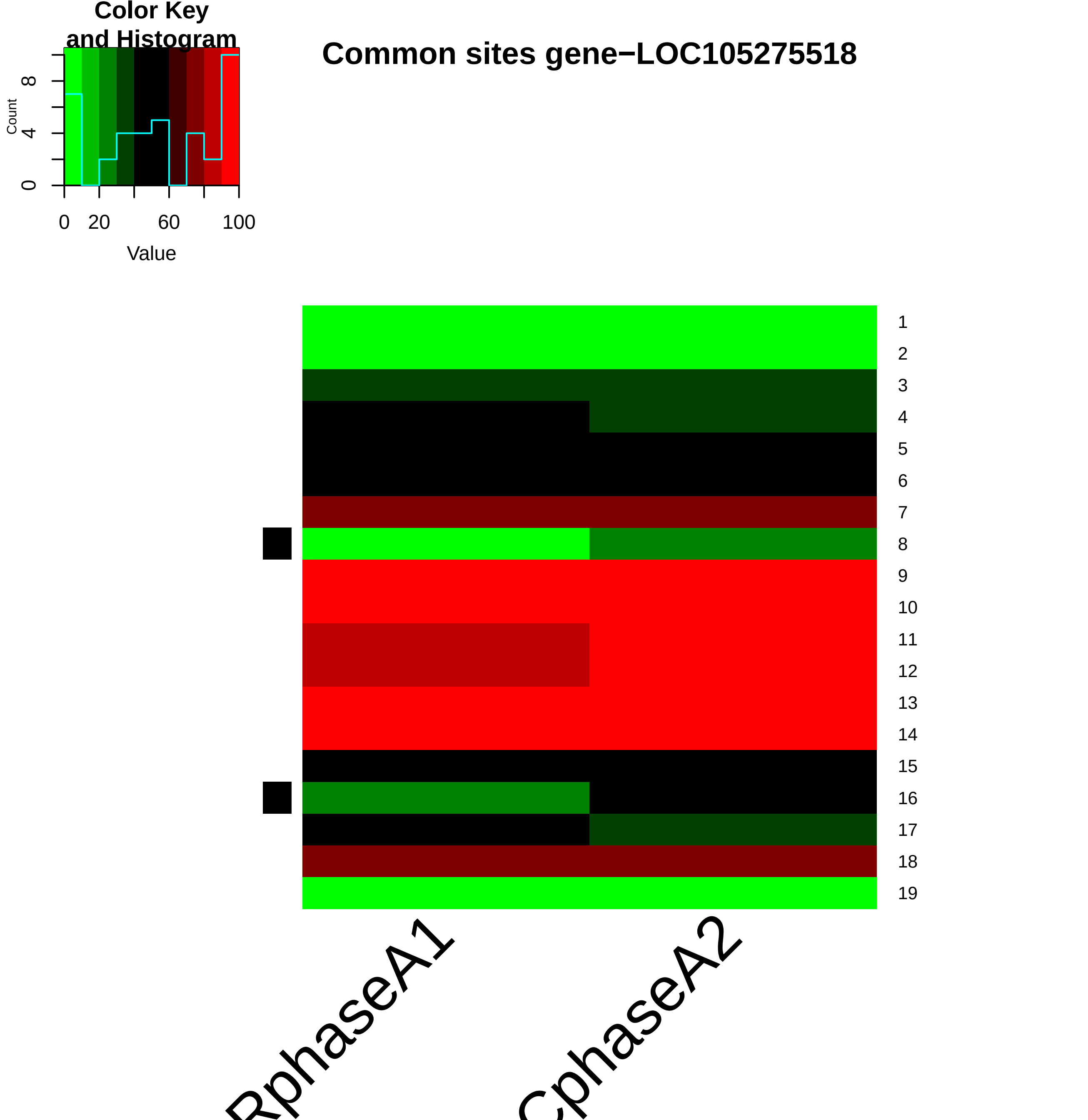

Color Key
and Histogram
Count
0
4
8
0
20
60
100
Value
Common sites gene−LOC105275518
RphaseA1
CphaseA2

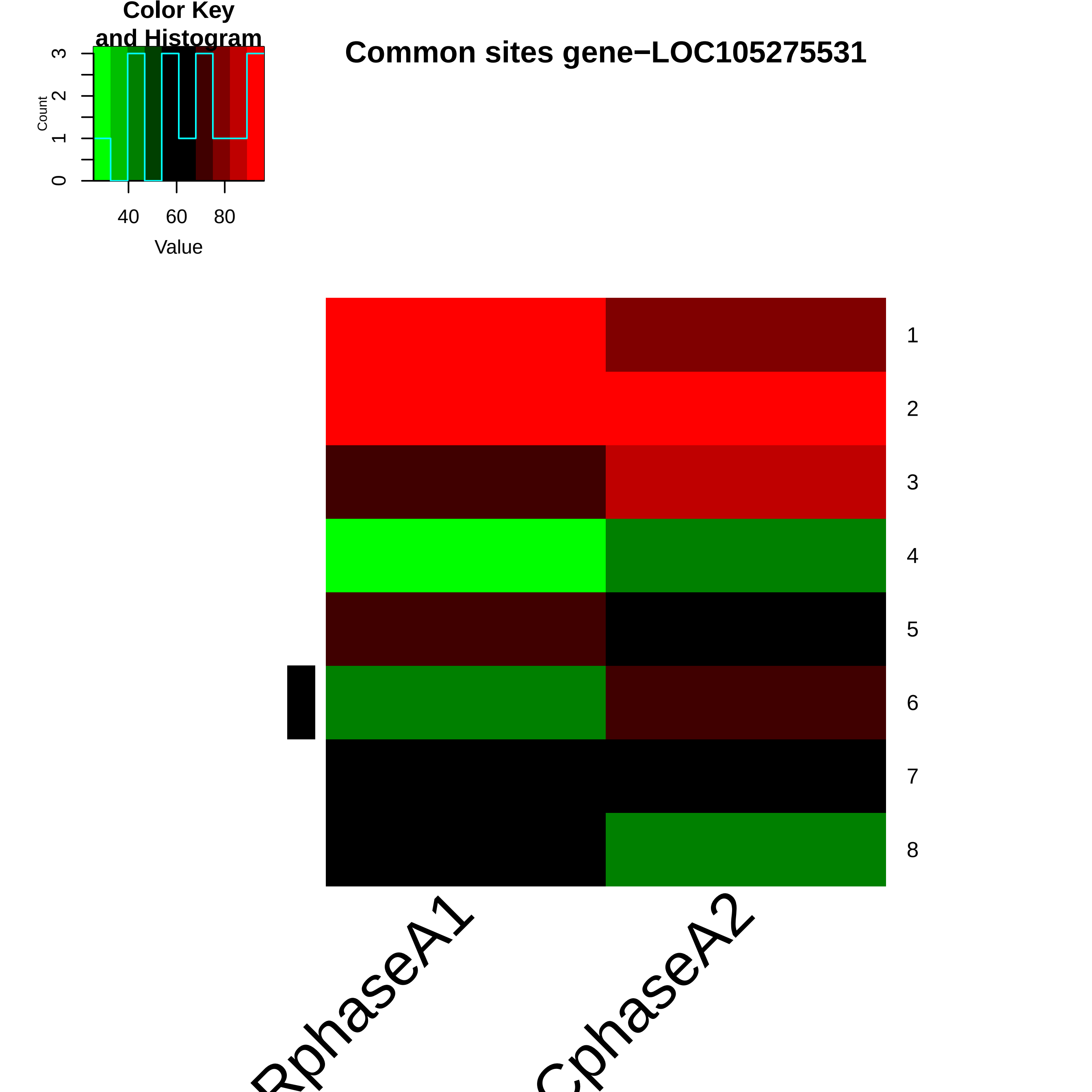
Color Key
and Histogram
Count
0
1
2
3
40
60
80
Value
Common sites gene−LOC105275531
1
2
3
4
5
6
7
8
RphaseA1
CphaseA2

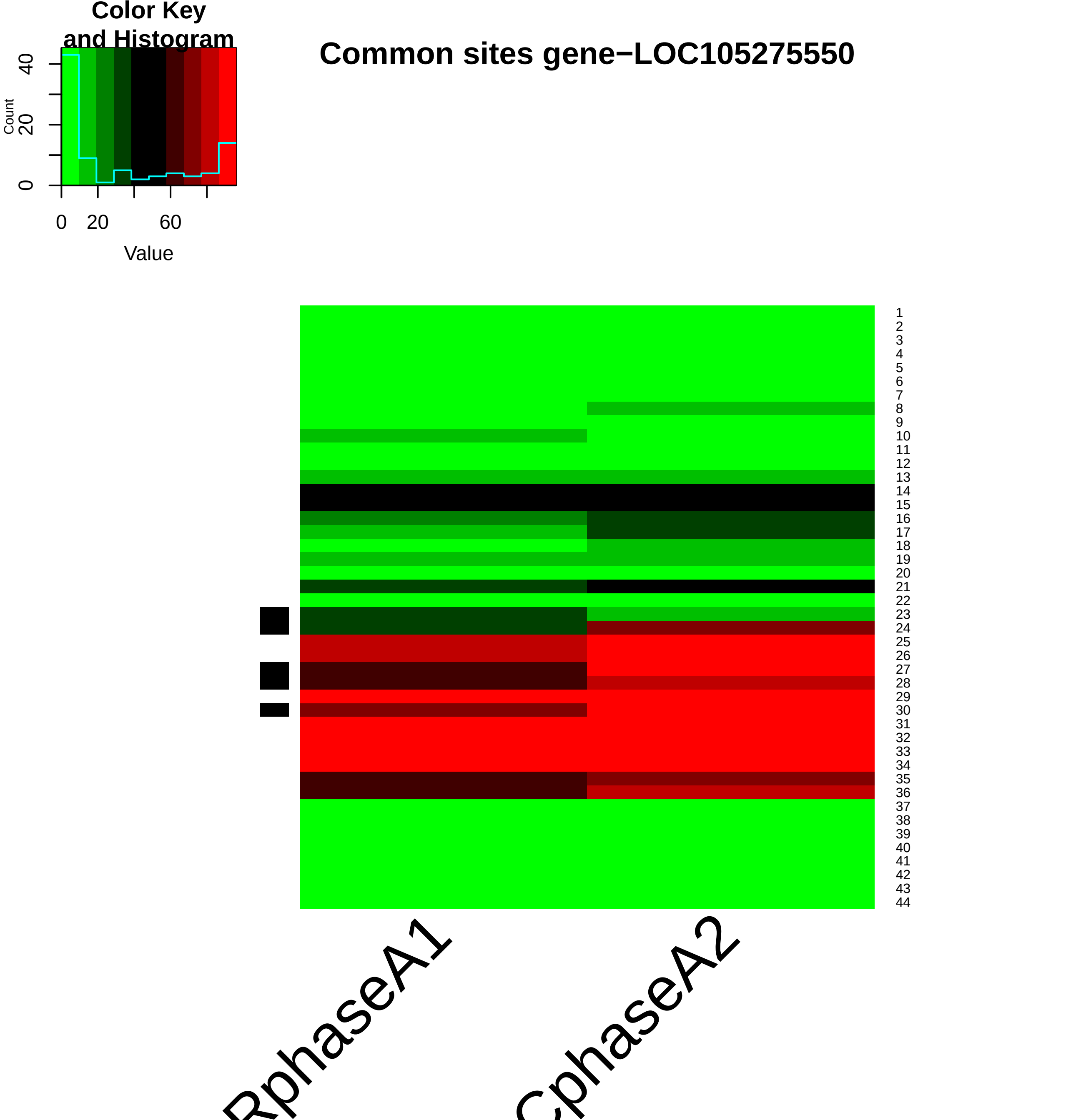
Color Key
and Histogram
Count
0
20
40
0
20
60
Value
Common sites gene−LOC105275550
RphaseA1
CphaseA2

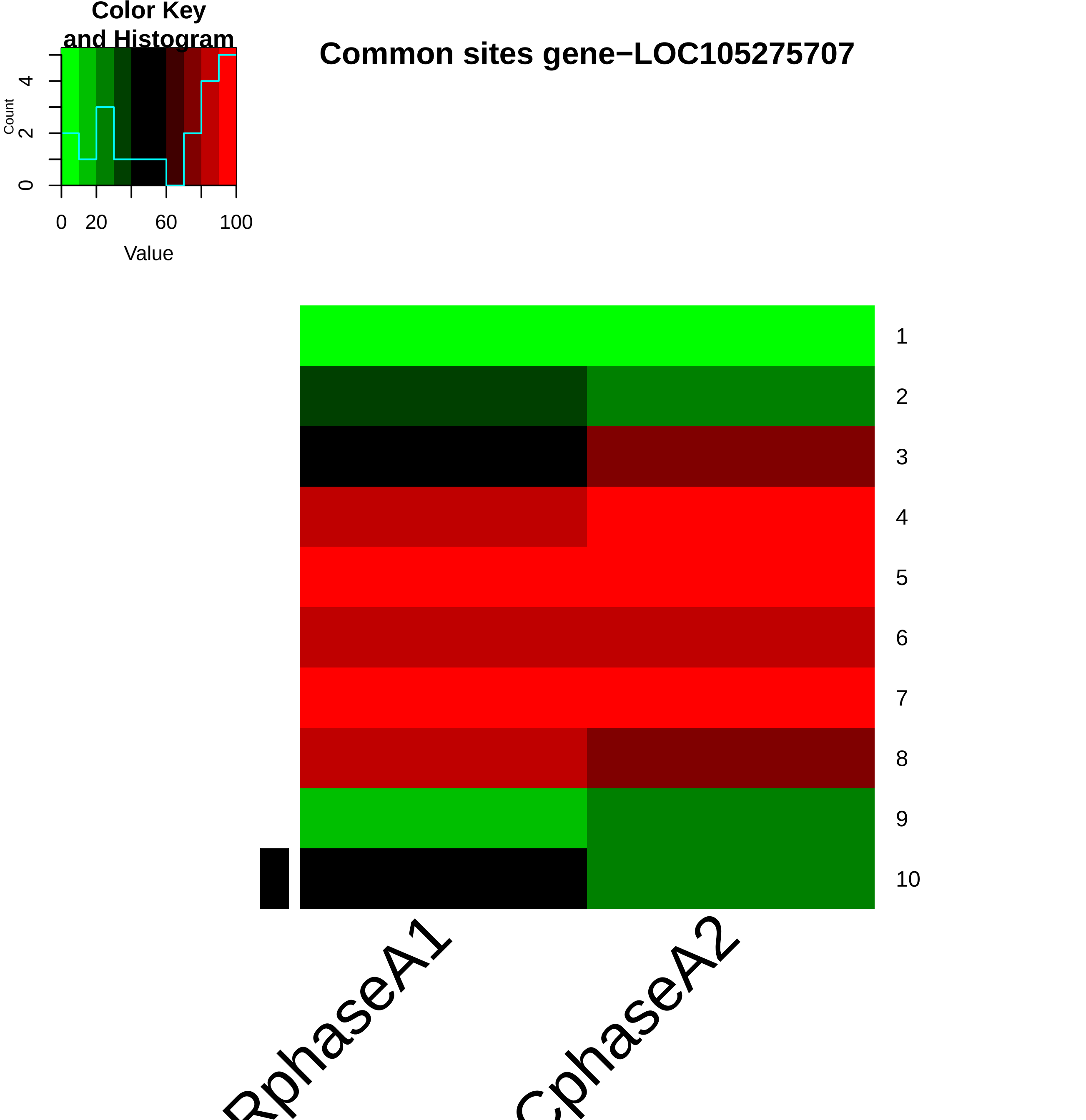
Color Key
and Histogram
Count
0
2
4
0
20
60
100
Value
Common sites gene−LOC105275707
1
2
3
4
5
6
7
8
9
10

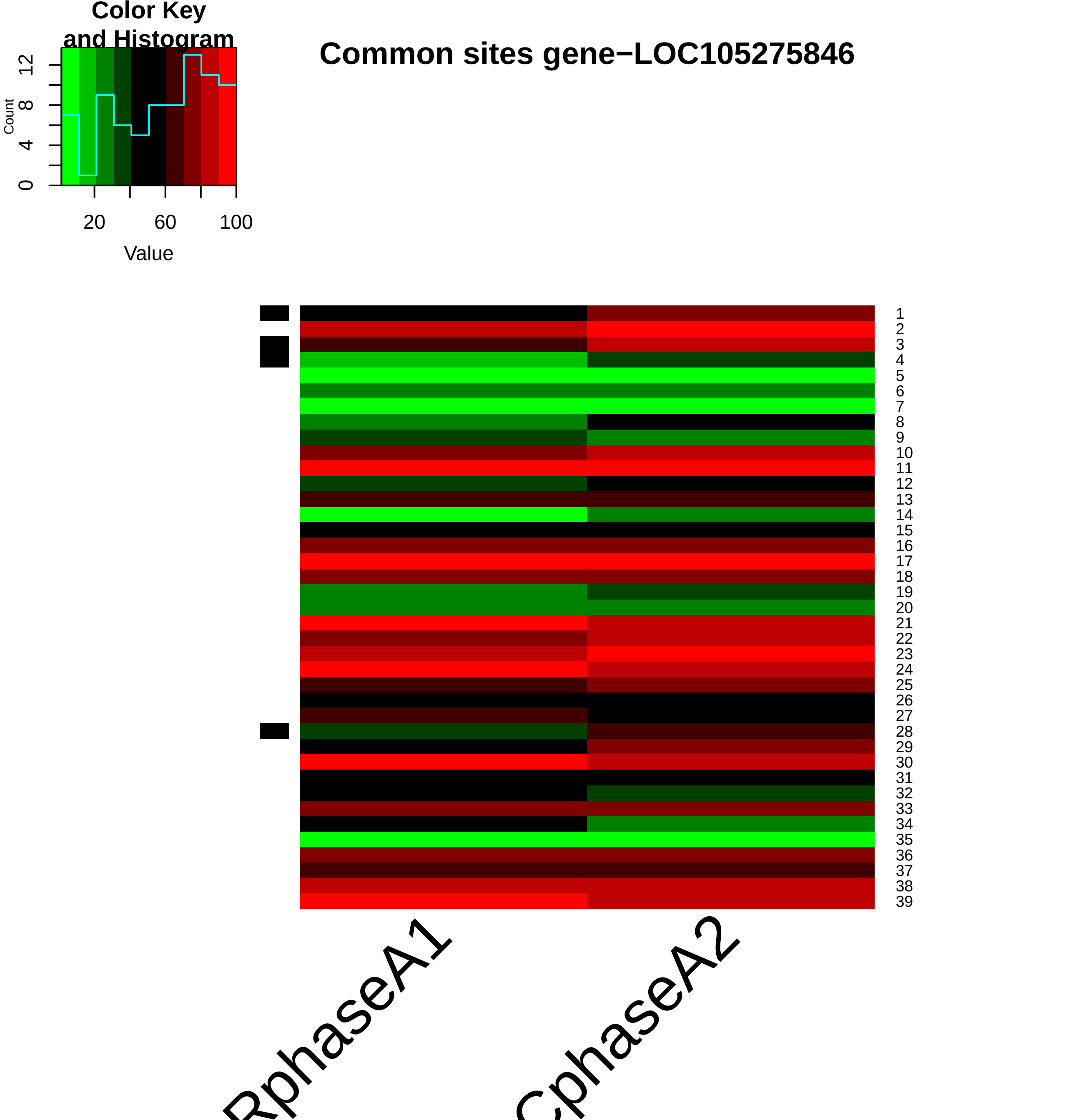
Color Key
and Histogram
Count
0
4
8
12
20
60
100
Value
Common sites gene−LOC105275846
RphaseA1
CphaseA2

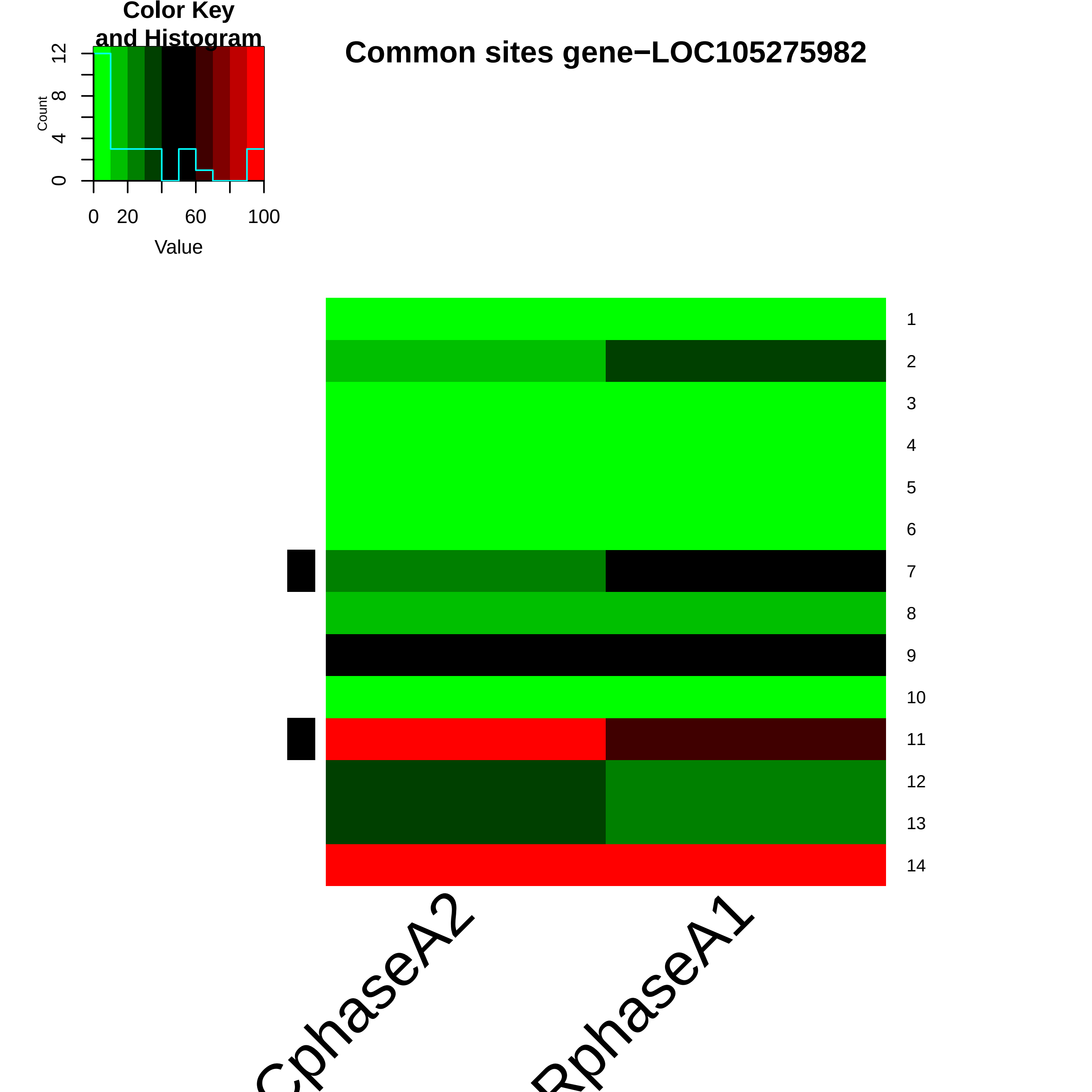

Color Key
and Histogram
Count
0
4
8
12
0
20
60
100
Value
Common sites gene−LOC105275982
1
2
3
4
5
6
7
8
9
10
11
12
13
14
CphaseA2
RphaseA1

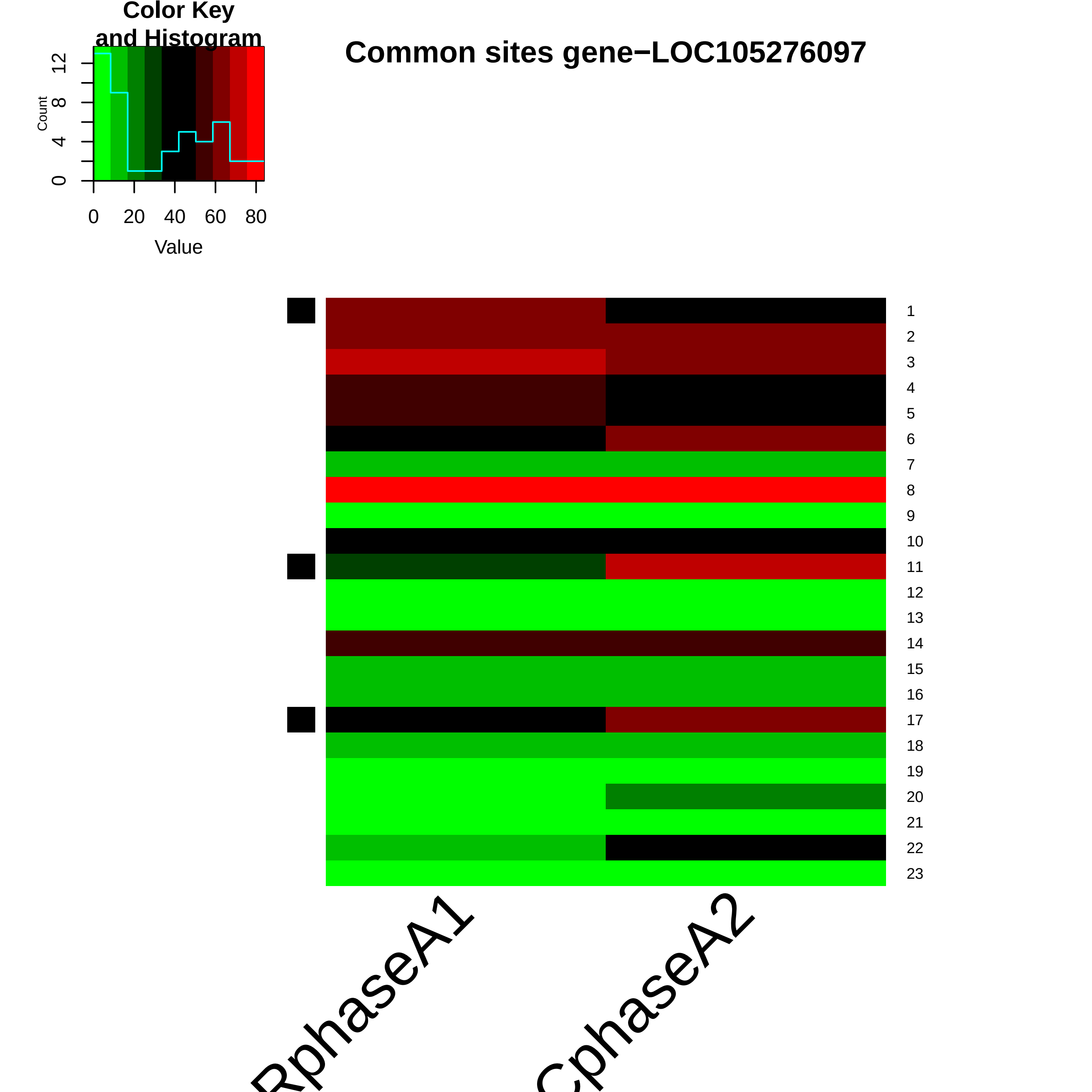

Color Key
and Histogram
Count
0
4
8
12
0
20
40
60
80
Value
Common sites gene−LOC105276097
RphaseA1
CphaseA2

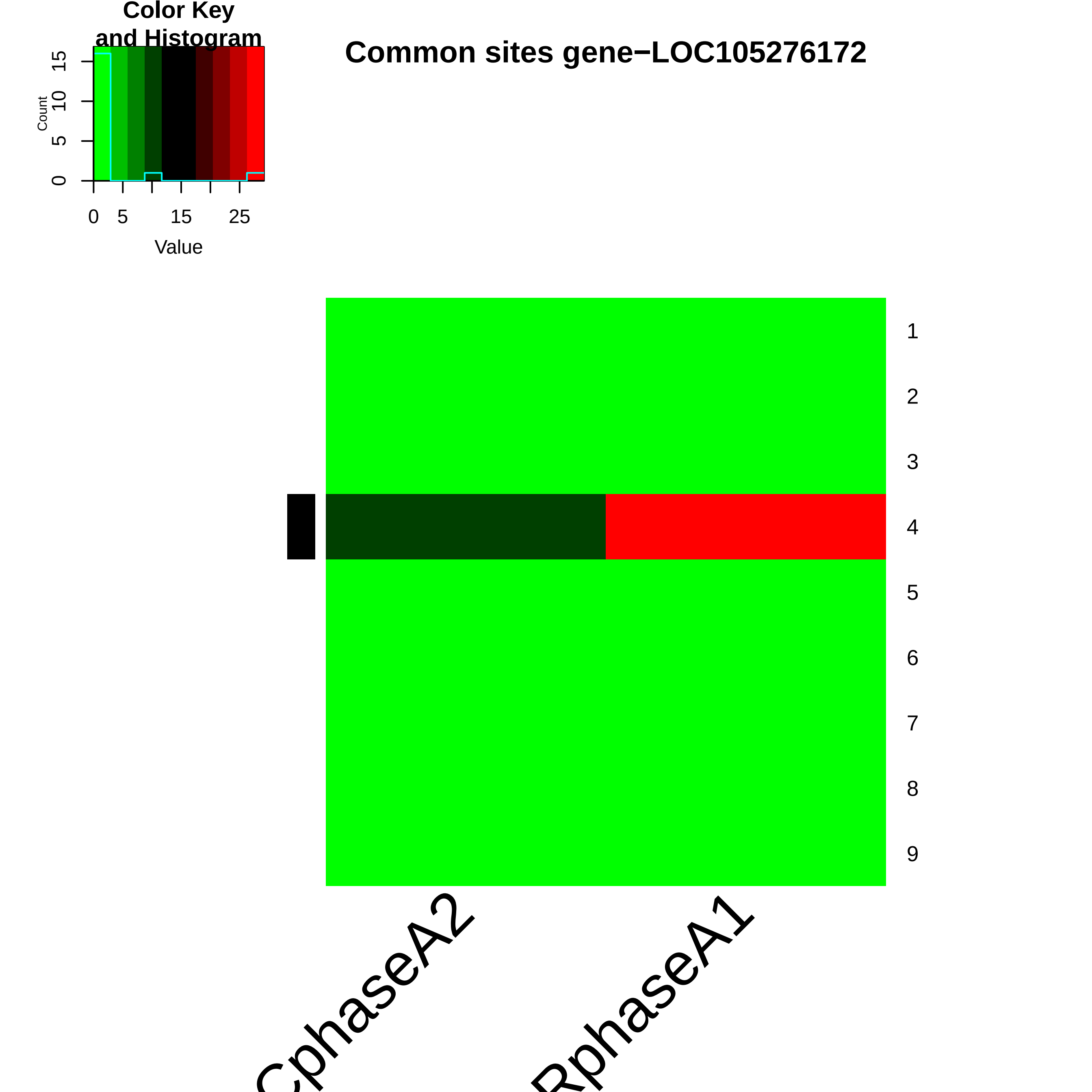

Color Key
and Histogram
Count
0
5
10
15
0
5
15
25
Value
Common sites gene−LOC105276172
1
2
3
4
5
6
7
8
9
CphaseA2
RphaseA1

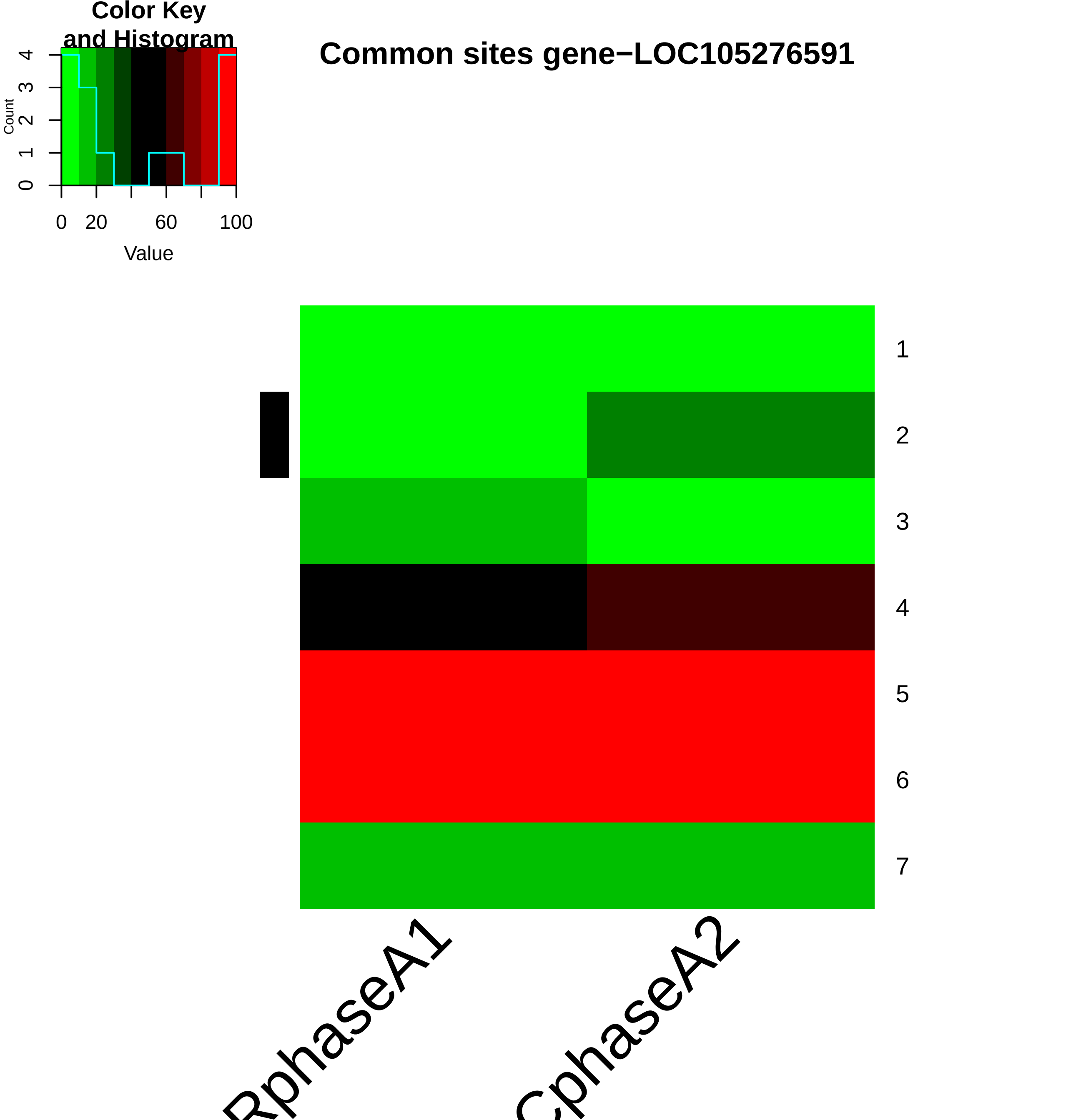
Color Key
and Histogram
Count
0
1
2
3
4
0
20
60
100
Value
Common sites gene−LOC105276591
1
2
3
4
5
6
7
RphaseA1
CphaseA2

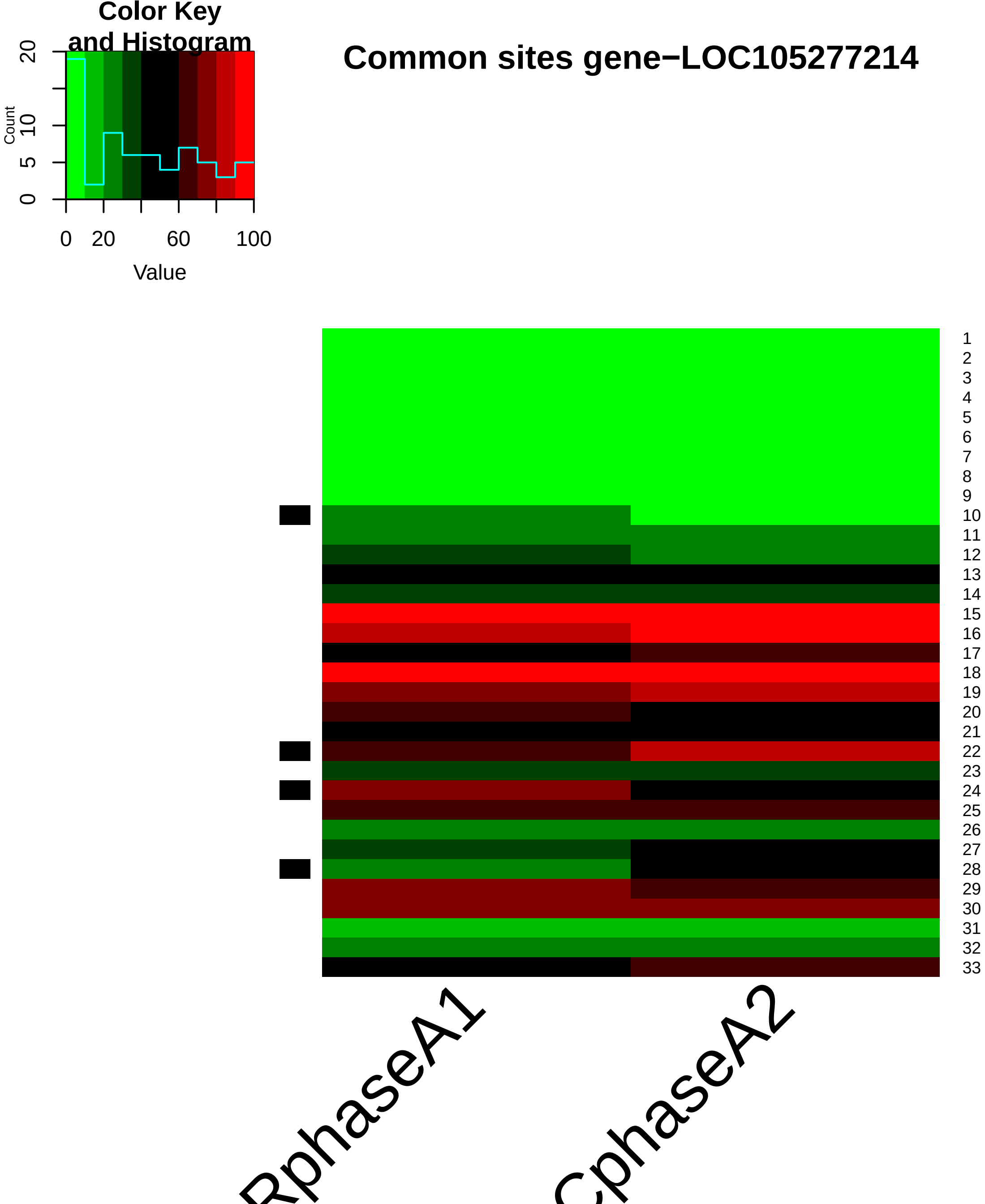
Color Key
and Histogram
Count
Value
0
5
10
20
0
20
60
100
Common sites gene−LOC105277214
RphaseA1
CphaseA2

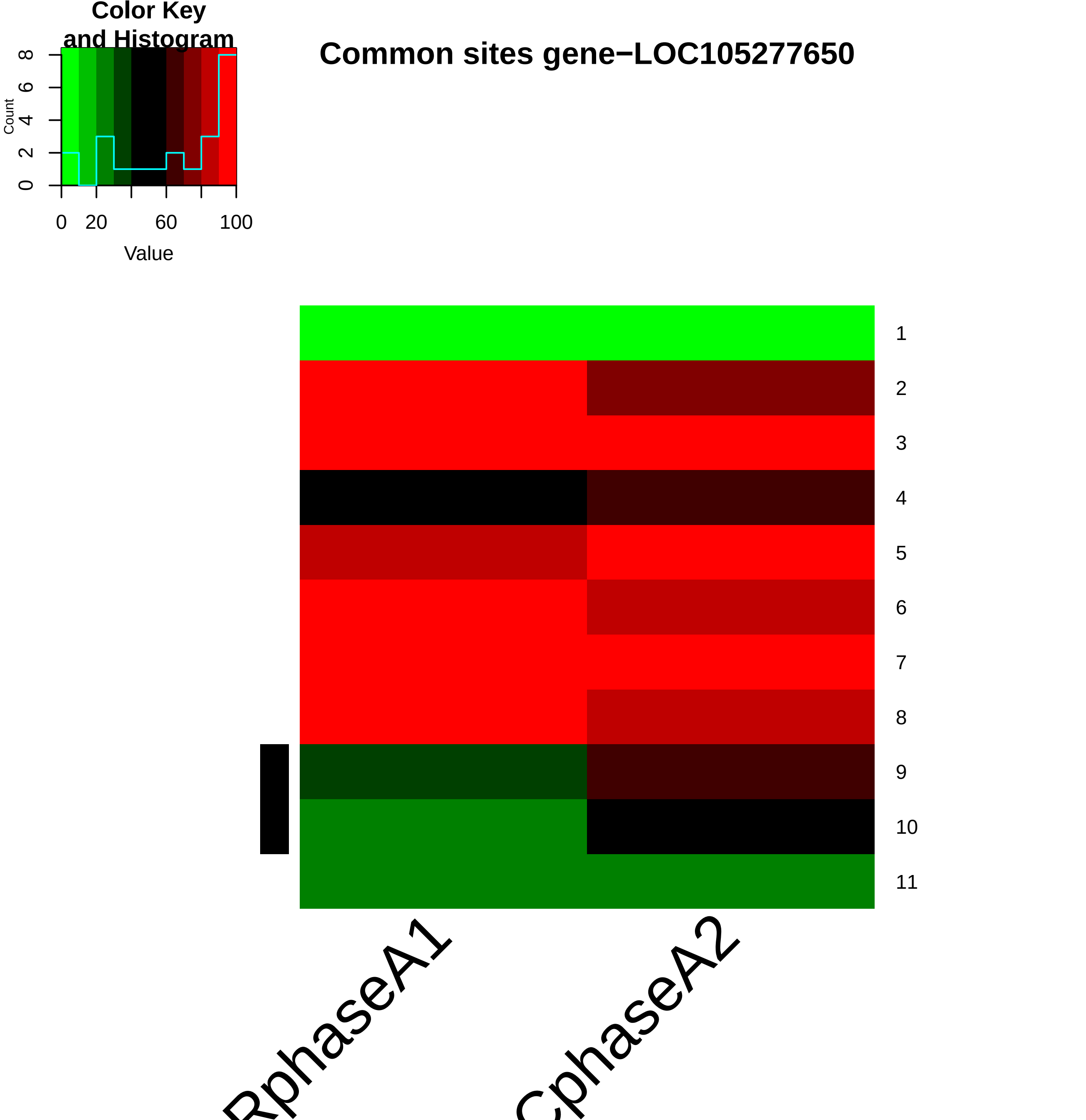

Color Key
and Histogram
Count
0
2
4
6
8
0
20
60
100
Value
Common sites gene−LOC105277650
1
2
3
4
5
6
7
8
9
10
11
RphaseA1
CphaseA2

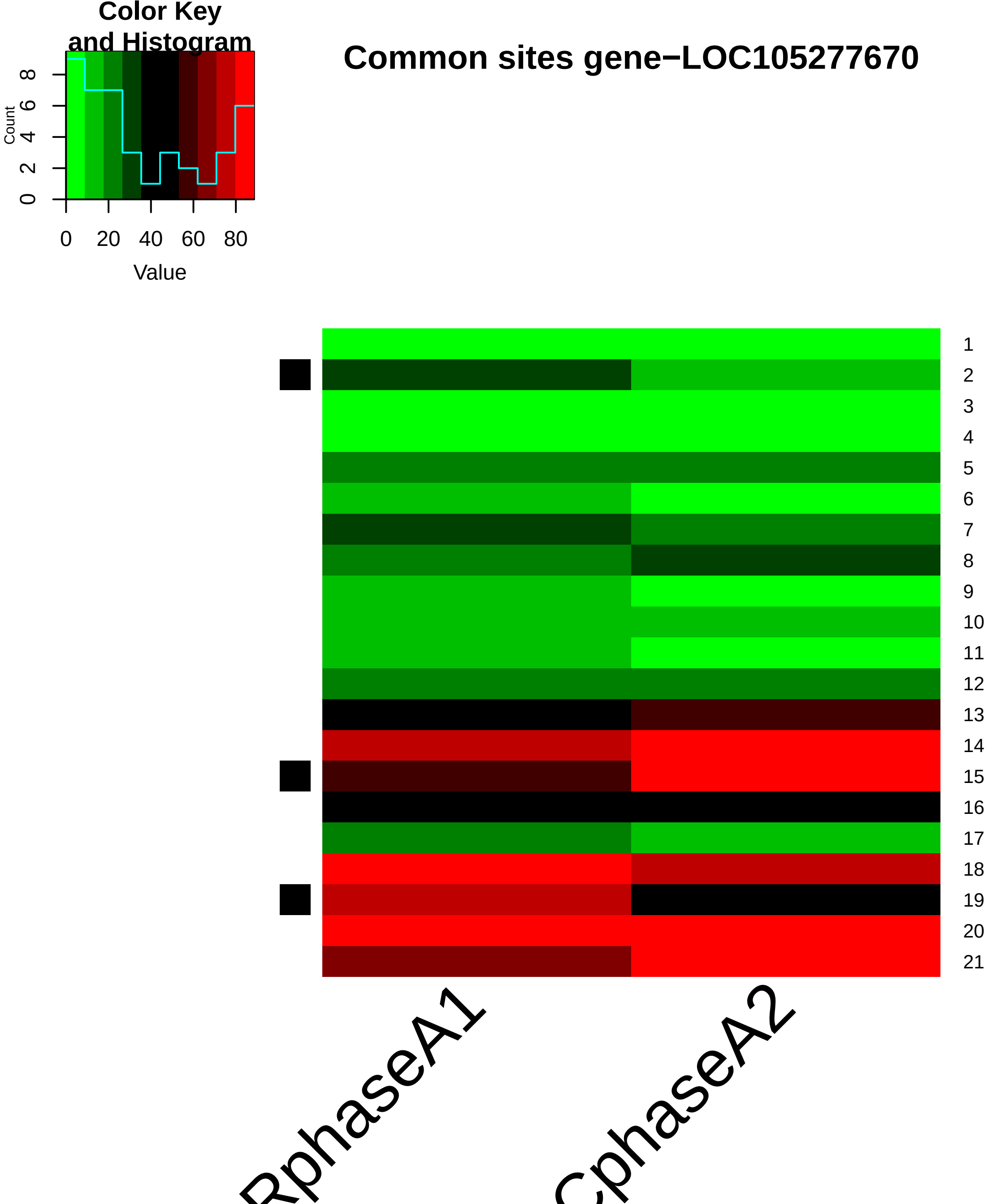

Color Key
and Histogram
Count
0
2
4
6
8
0
20
40
60
80
Value
Common sites gene−LOC105277670
RphaseA1
CphaseA2

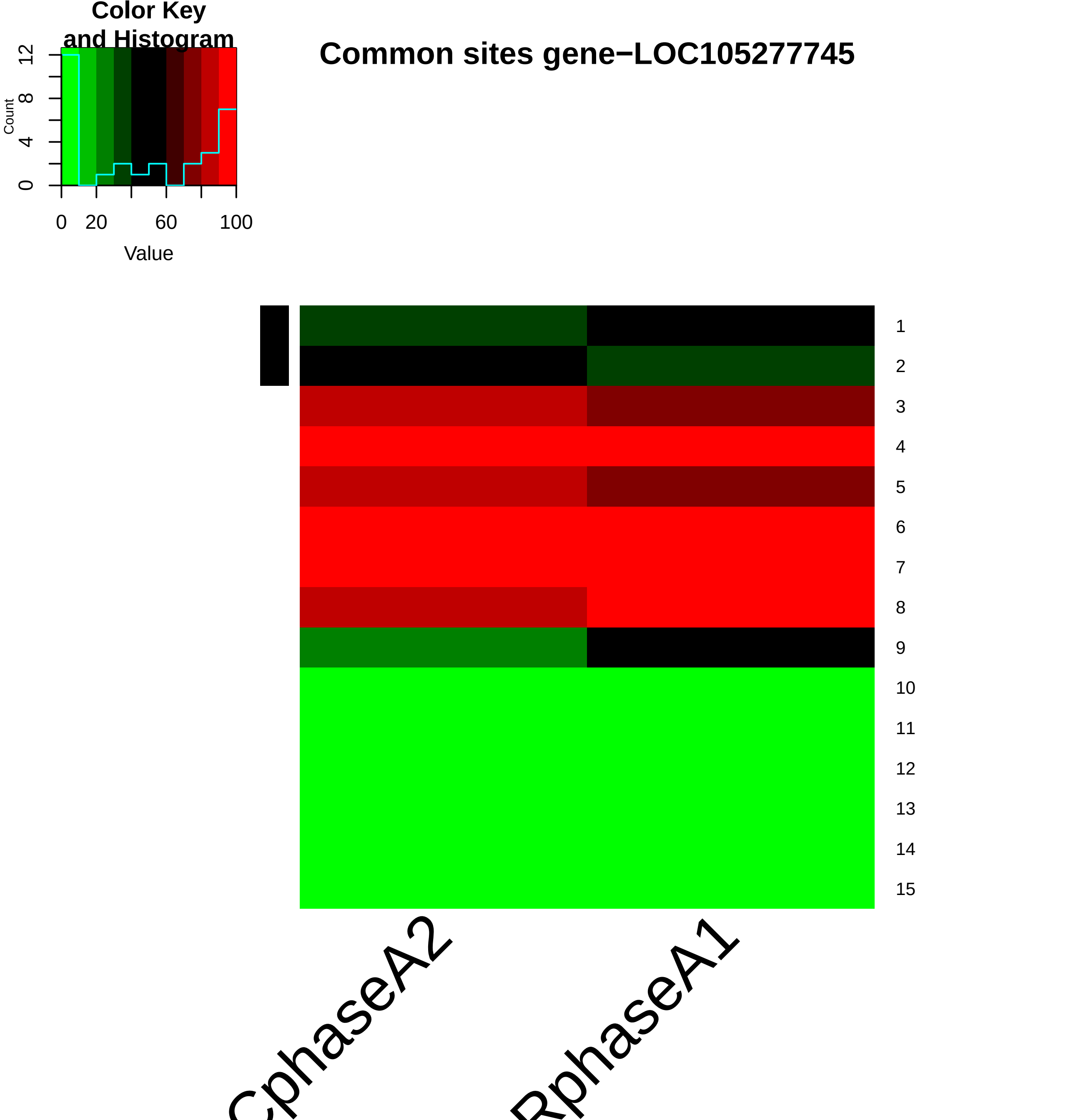

Color Key
and Histogram
Count
0
4
8
12
0
20
60
100
Value
Common sites gene−LOC105277745
CphaseA2
RphaseA1

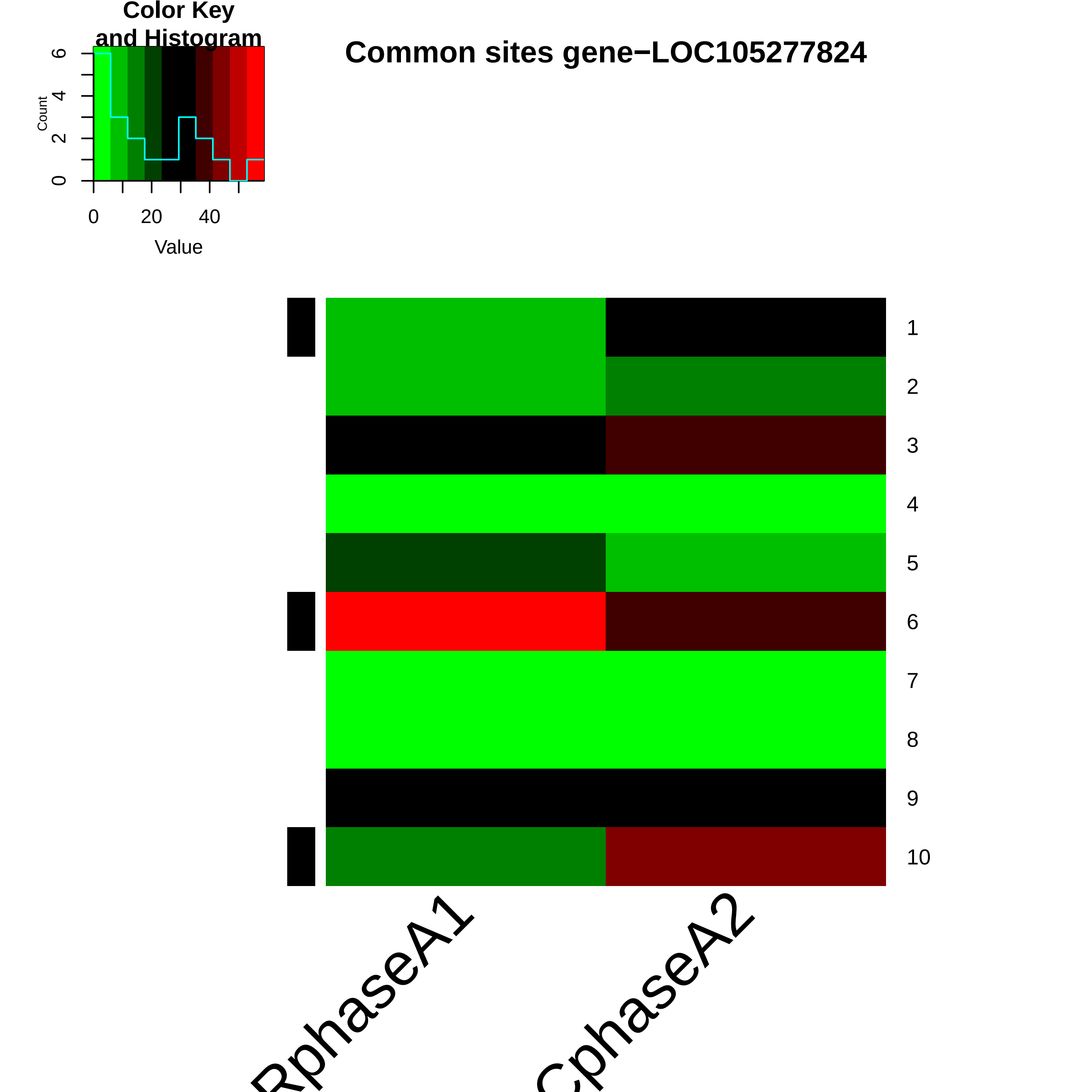

Color Key
and Histogram
Count
Value
0
20
40
0
2
4
6
Common sites gene−LOC105277824
1
2
3
4
5
6
7
8
9
10
RphaseA1
CphaseA2

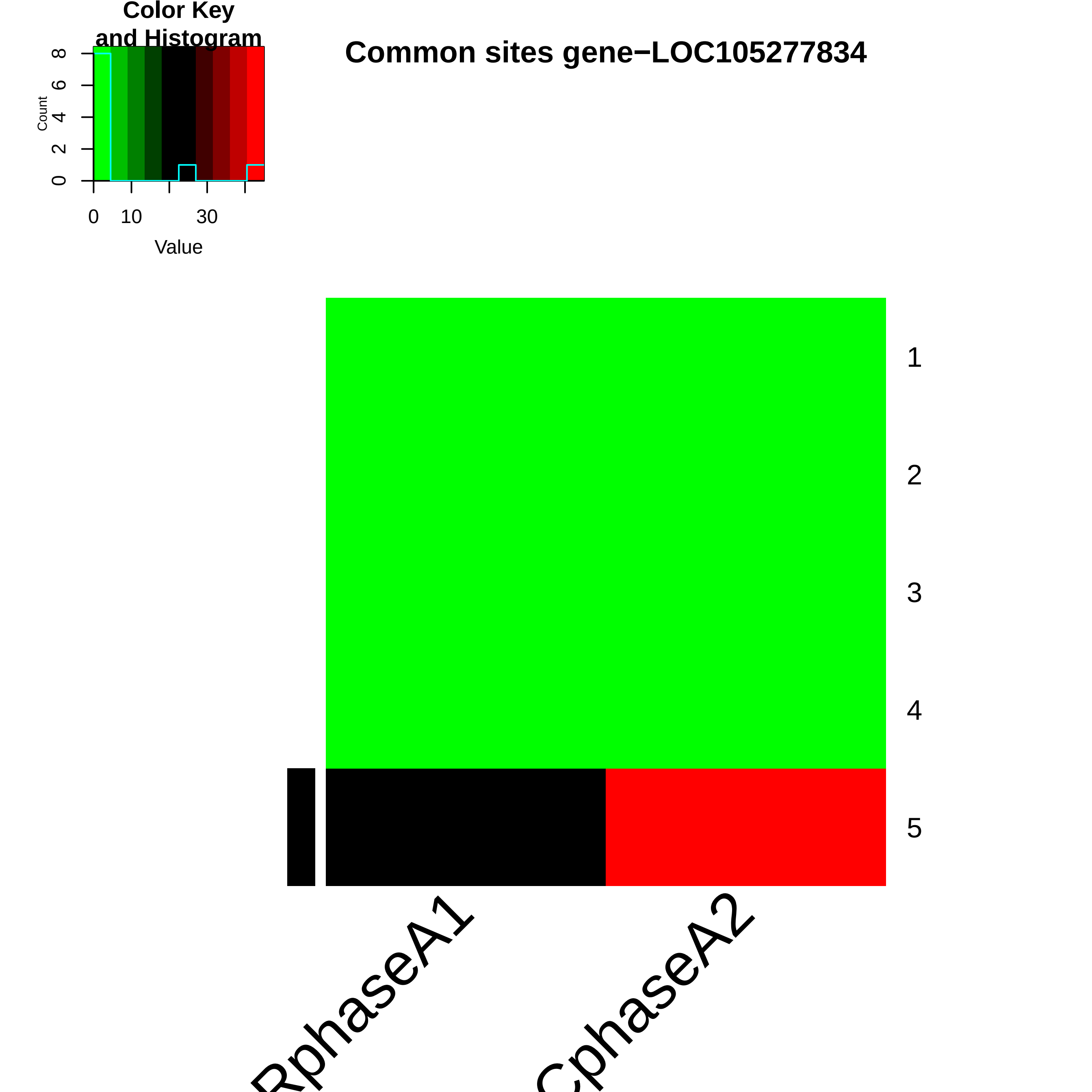

Color Key
and Histogram
Count
0
2
4
6
8
0
10
30
Value
Common sites gene−LOC105277834
1
2
3
4
5
RphaseA1
CphaseA2

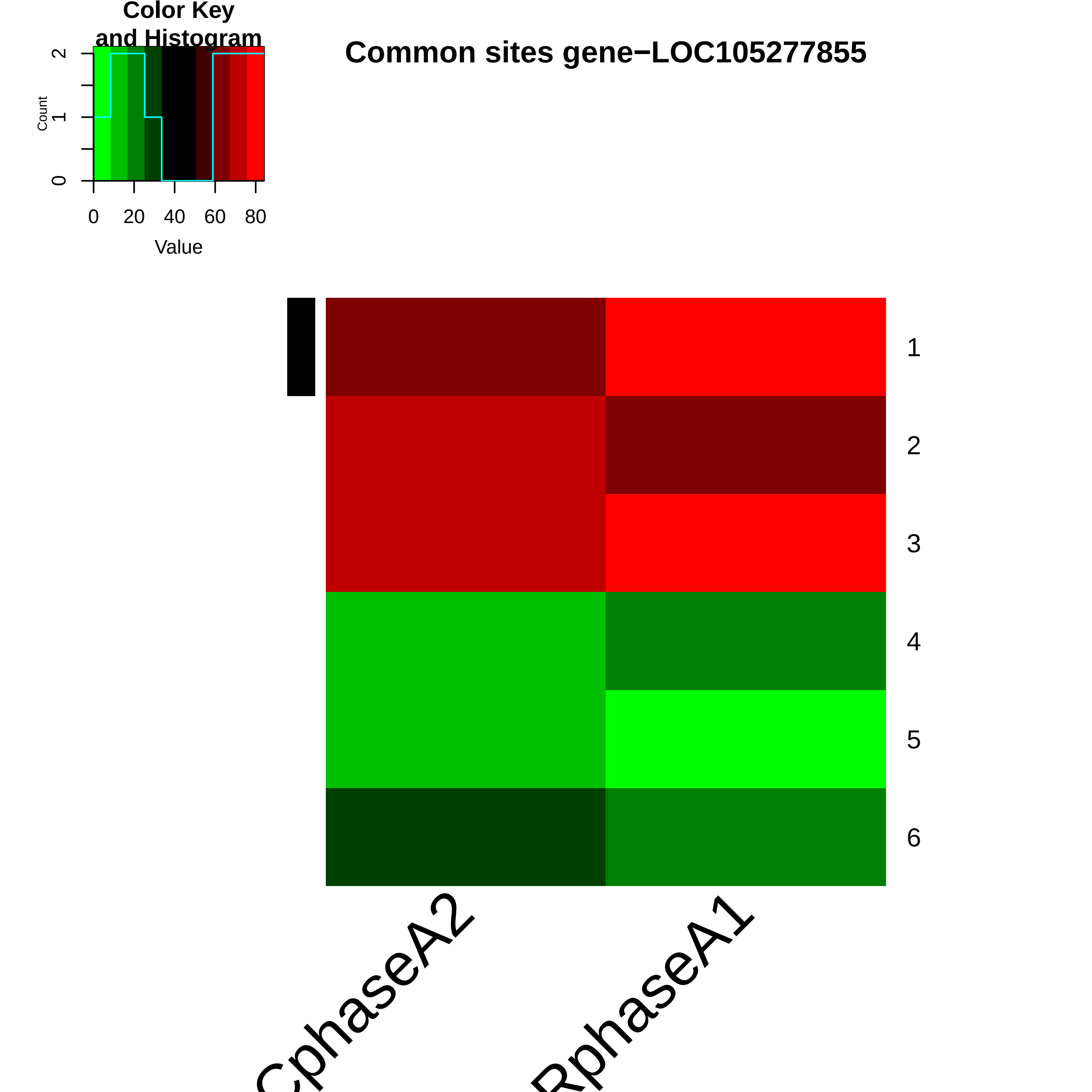
Color Key
and Histogram
Count
0
1
2
0
20
40
60
80
Value
Common sites gene−LOC105277855
1
2
3
4
5
6
CphaseA2
RphaseA1

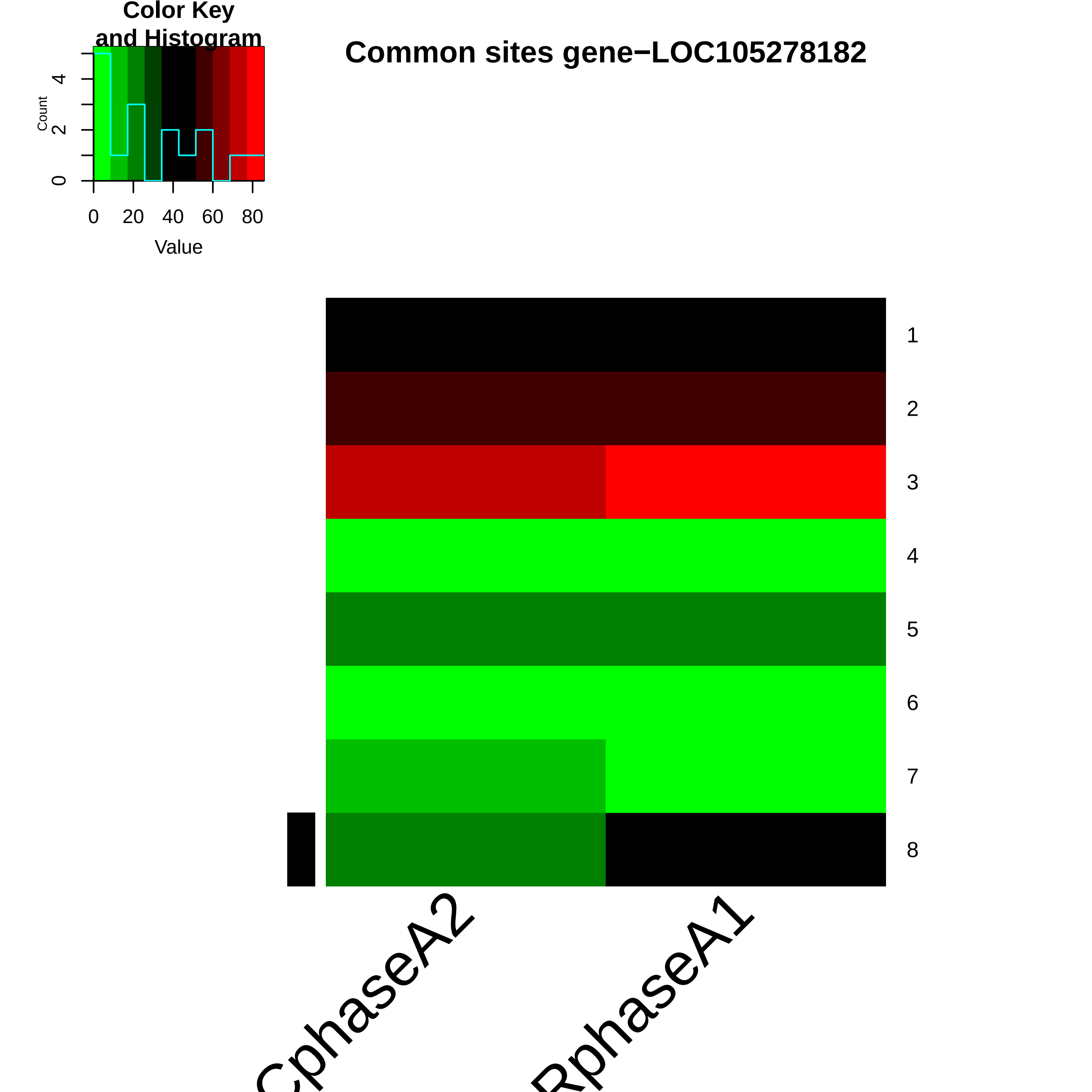

Color Key
and Histogram
Count
0
2
4
0
20
40
60
80
Value
Common sites gene−LOC105278182
1
2
3
4
5
6
7
8
CphaseA2
RphaseA1

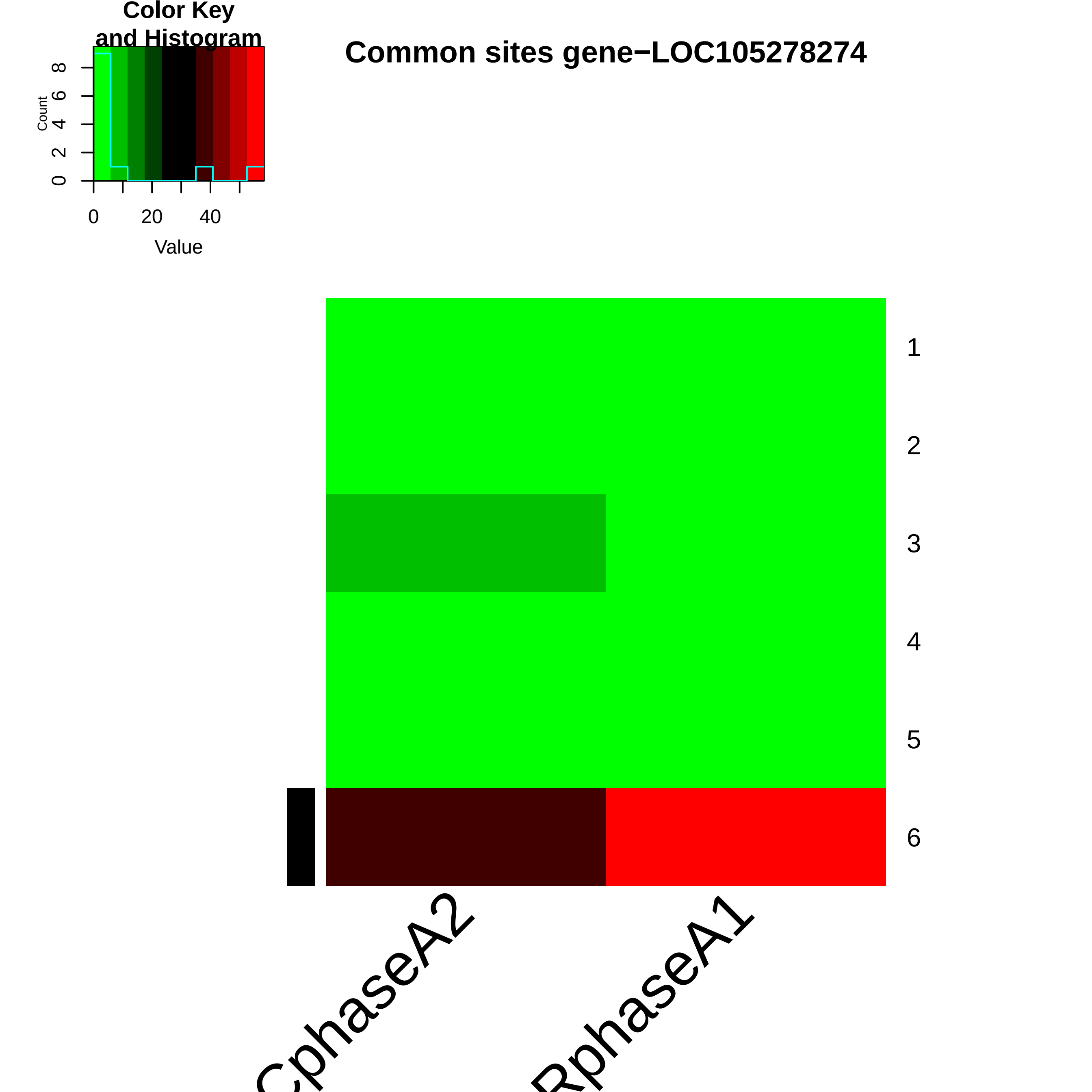
Color Key
and Histogram
Count
0
2
4
6
8
0
20
40
Value
Common sites gene−LOC105278274
1
2
3
4
5
6
CphaseA2
RphaseA1

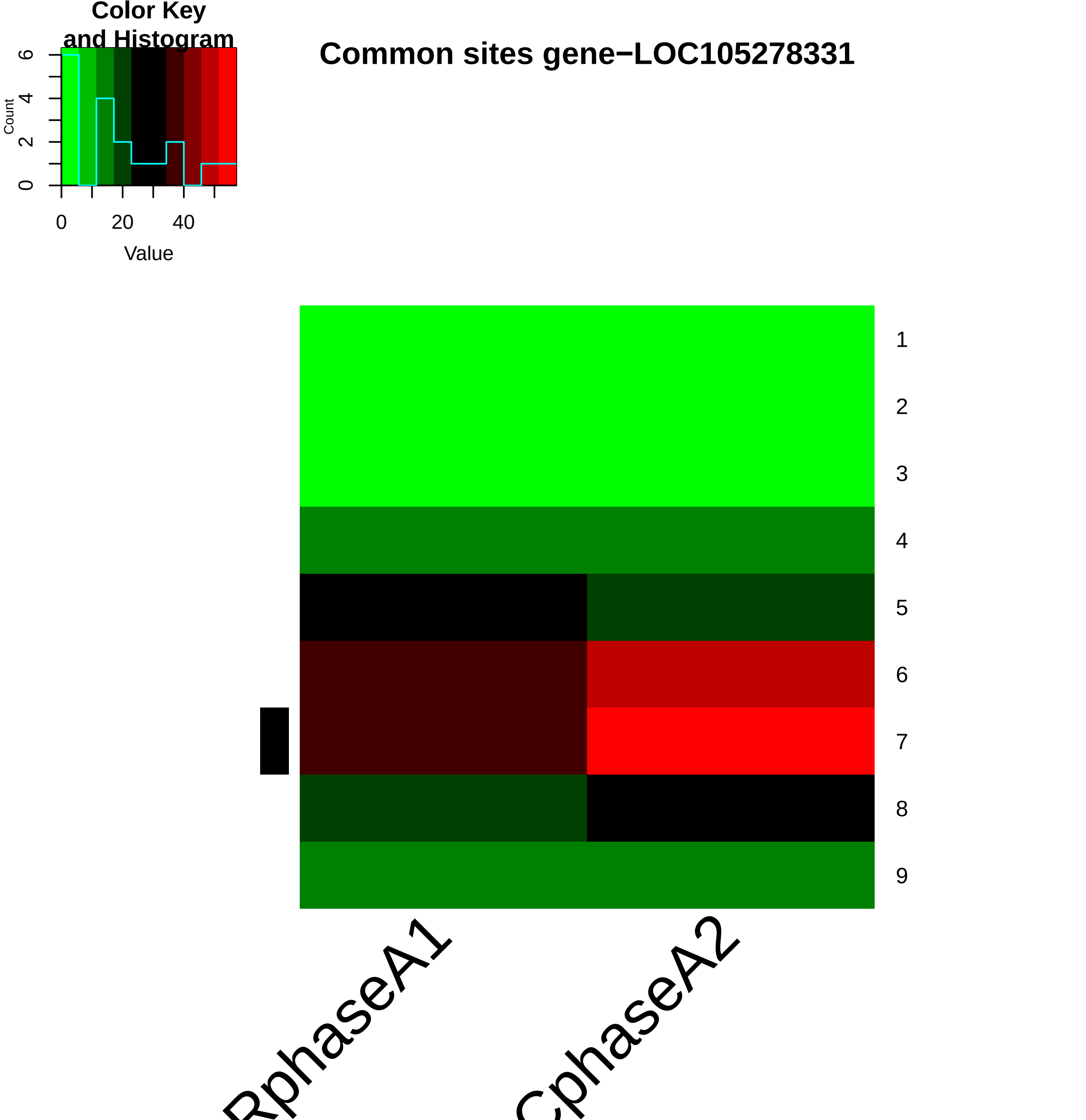
Color Key
and Histogram
Count
0
2
4
6
0
20
40
Value
Common sites gene−LOC105278331
1
2
3
4
5
6
7
8
9
RphaseA1
CphaseA2

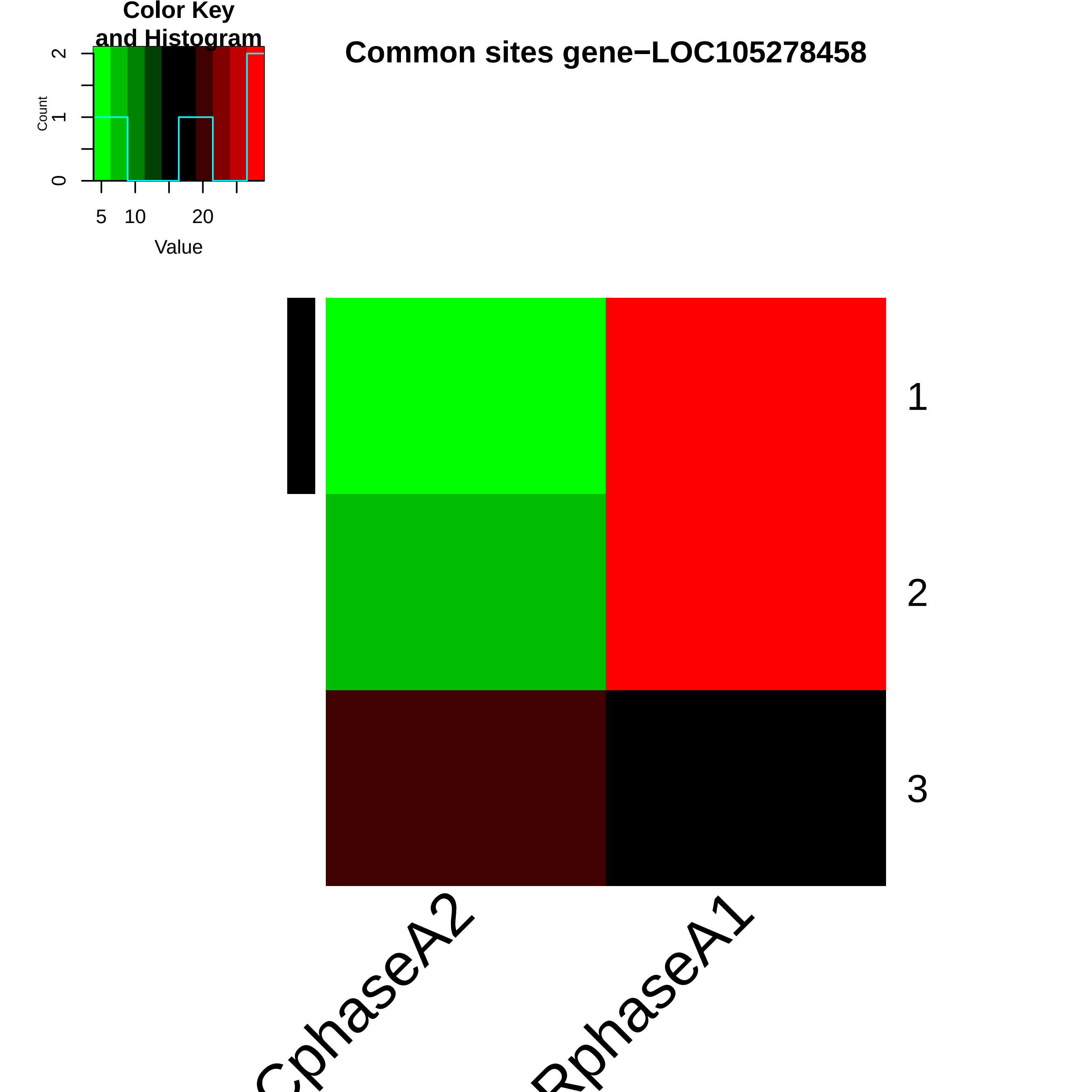

Color Key
and Histogram
Common sites gene−LOC105278458
Count
0
1
2
5
10
20
Value
1
2
3
CphaseA2
RphaseA1

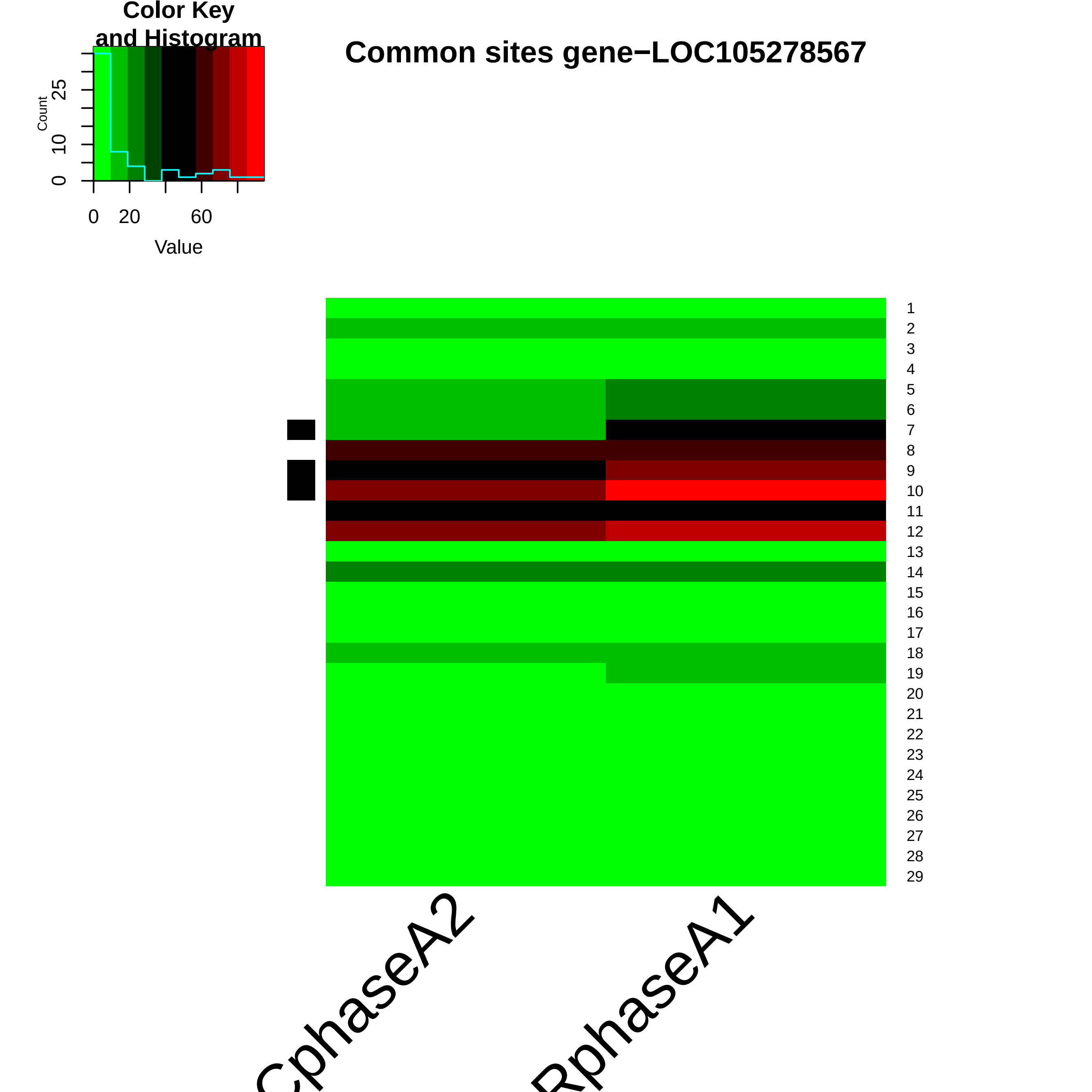

Color Key
and Histogram
Count
0
10
25
0
20
60
Value
Common sites gene−LOC105278567
CphaseA2
RphaseA1

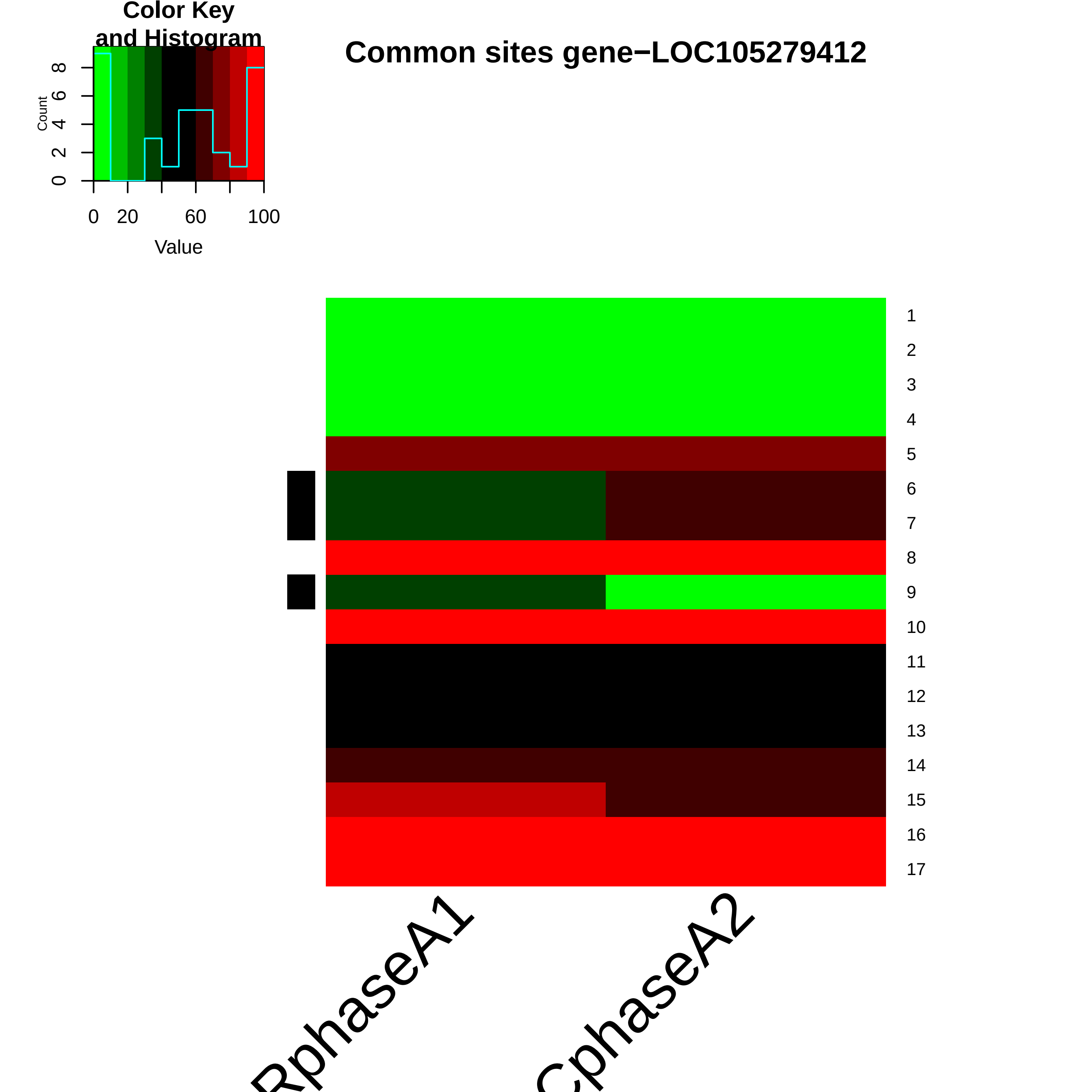

Color Key and Histogram
Count
0
2
4
6
8
0
20
60
100
Value
Common sites gene−LOC105279412
RphaseA1
CphaseA2

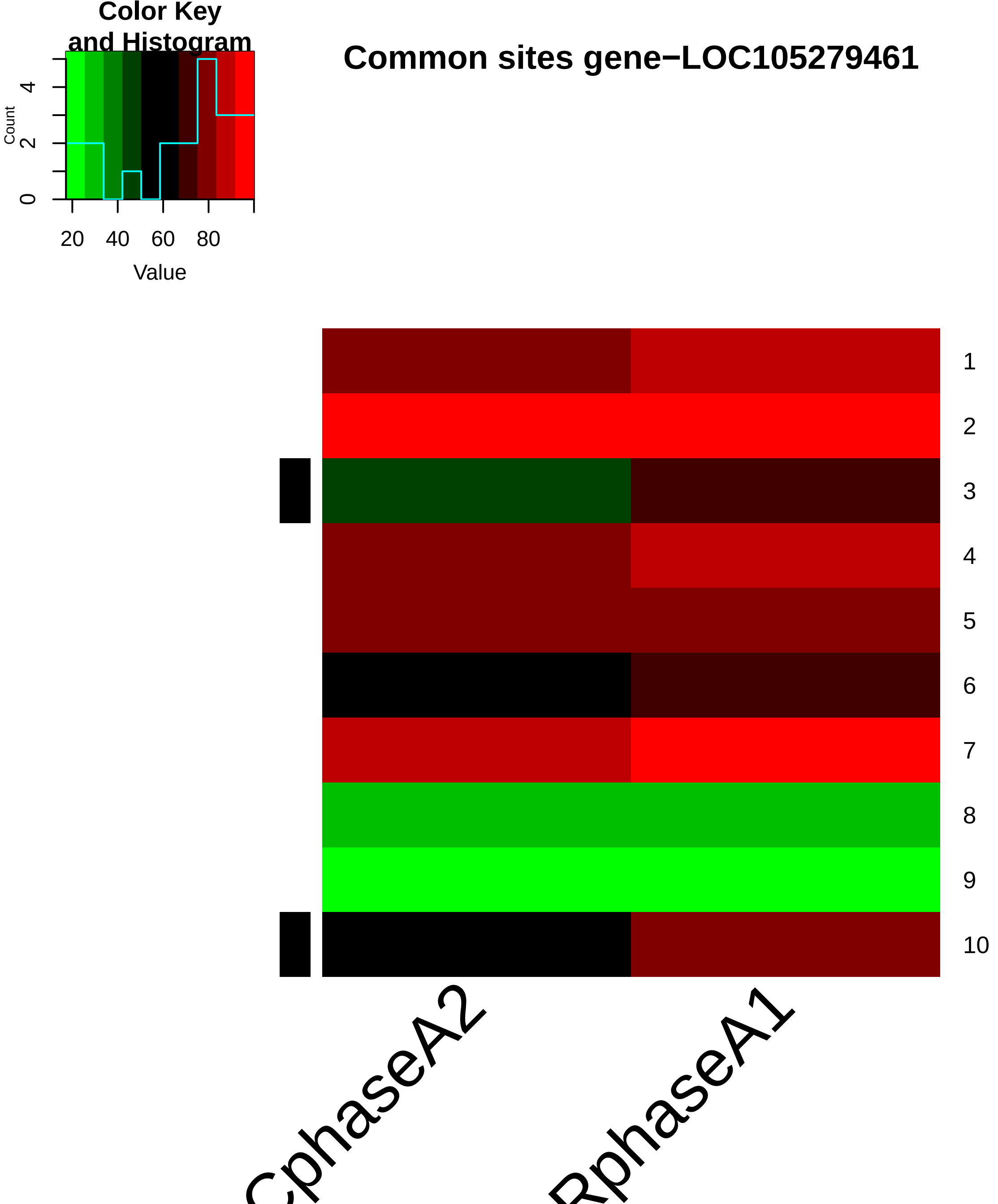
Color Key
and Histogram
Count
0
2
4
20
40
60
80
Value
Common sites gene−LOC105279461
1
2
3
4
5
6
7
8
9
10
CphaseA2
RphaseA1

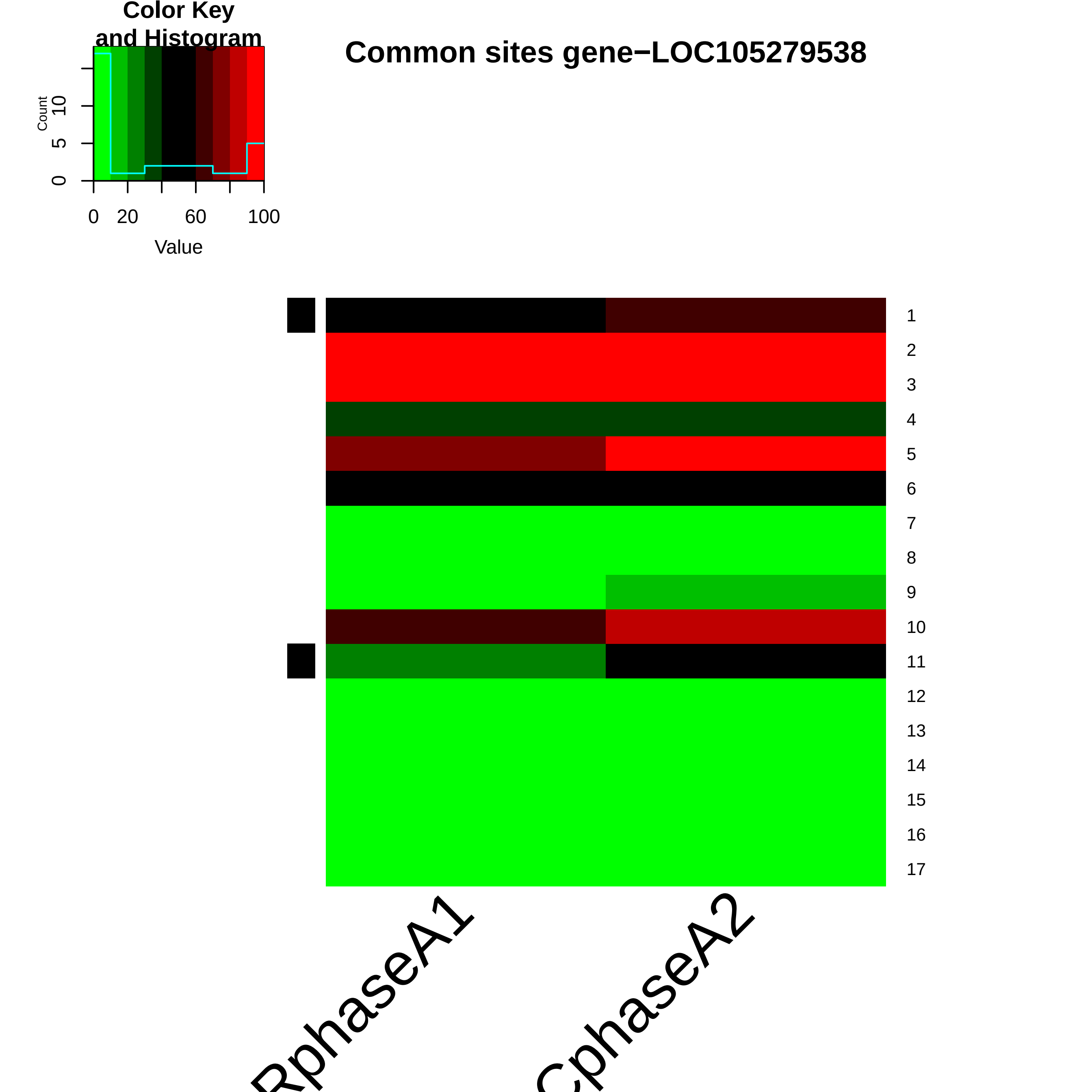

Color Key
and Histogram
Count
0
5
10
0
20
60
100
Value
Common sites gene−LOC105279538
RphaseA1
CphaseA2

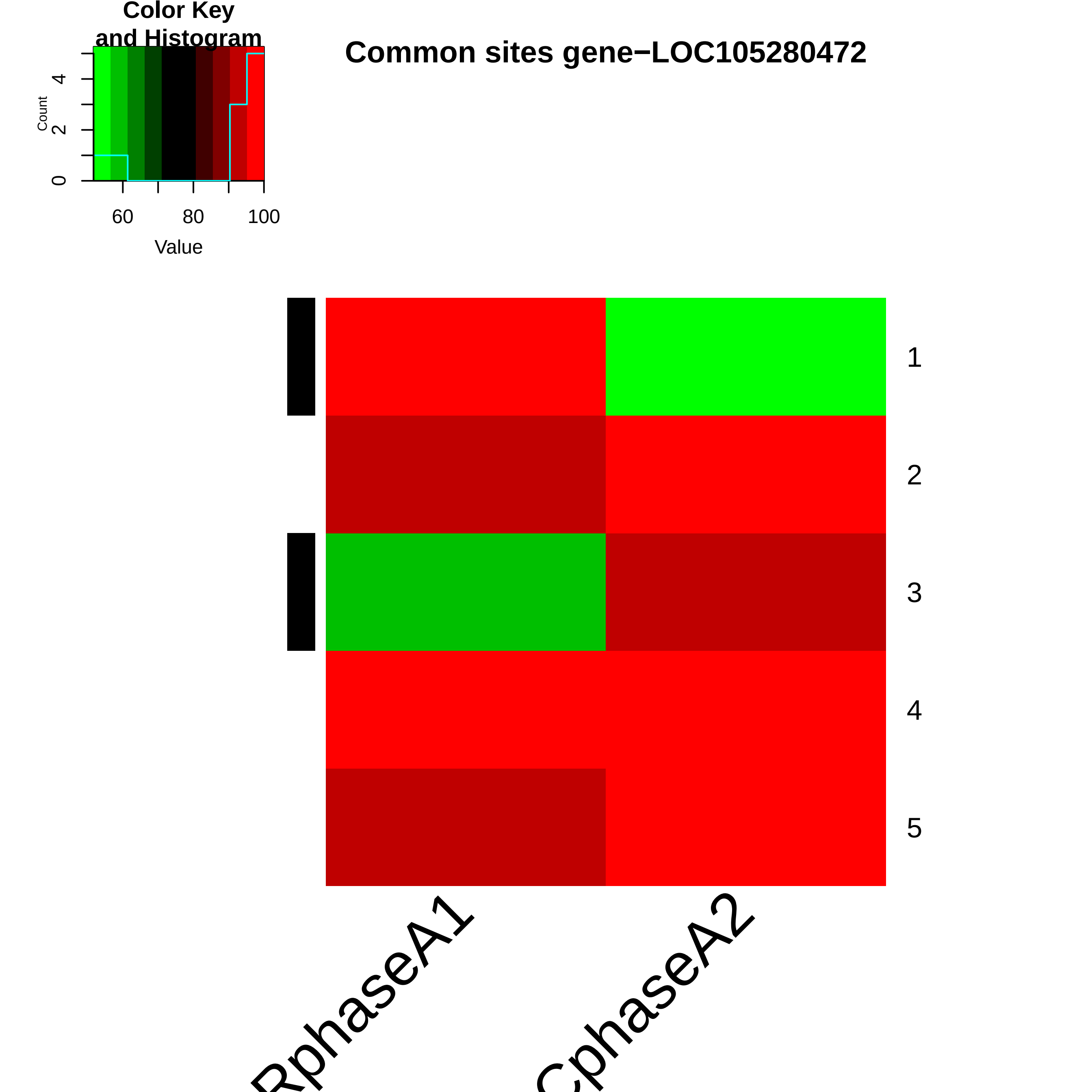
Color Key
and Histogram
Count
Value
Common sites gene−LOC105280472
1
2
3
4
5

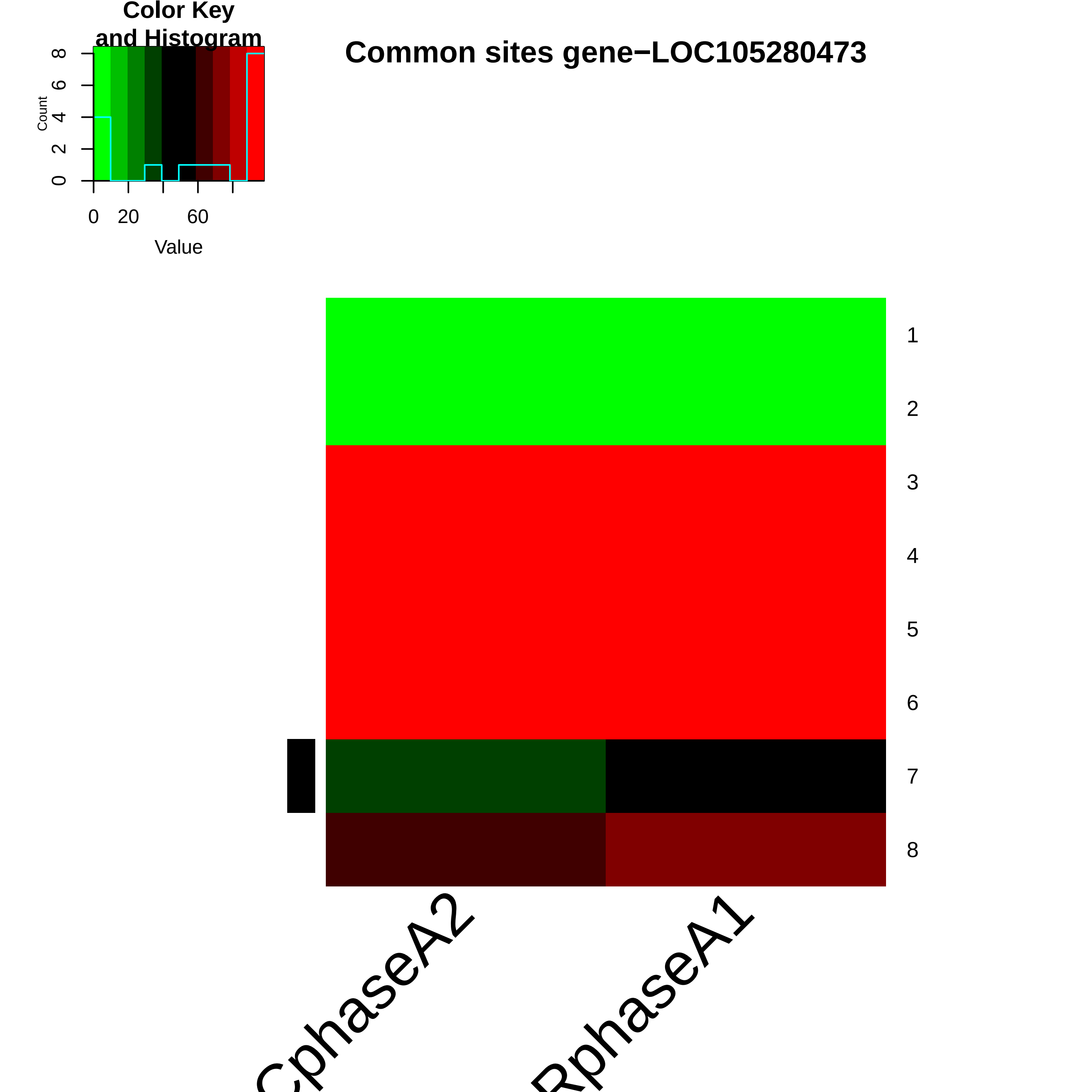

Color Key
and Histogram
Count
Value
0
20
60
0
2
4
6
8
Common sites gene−LOC105280473
1
2
3
4
5
6
7
8
CphaseA2
RphaseA1

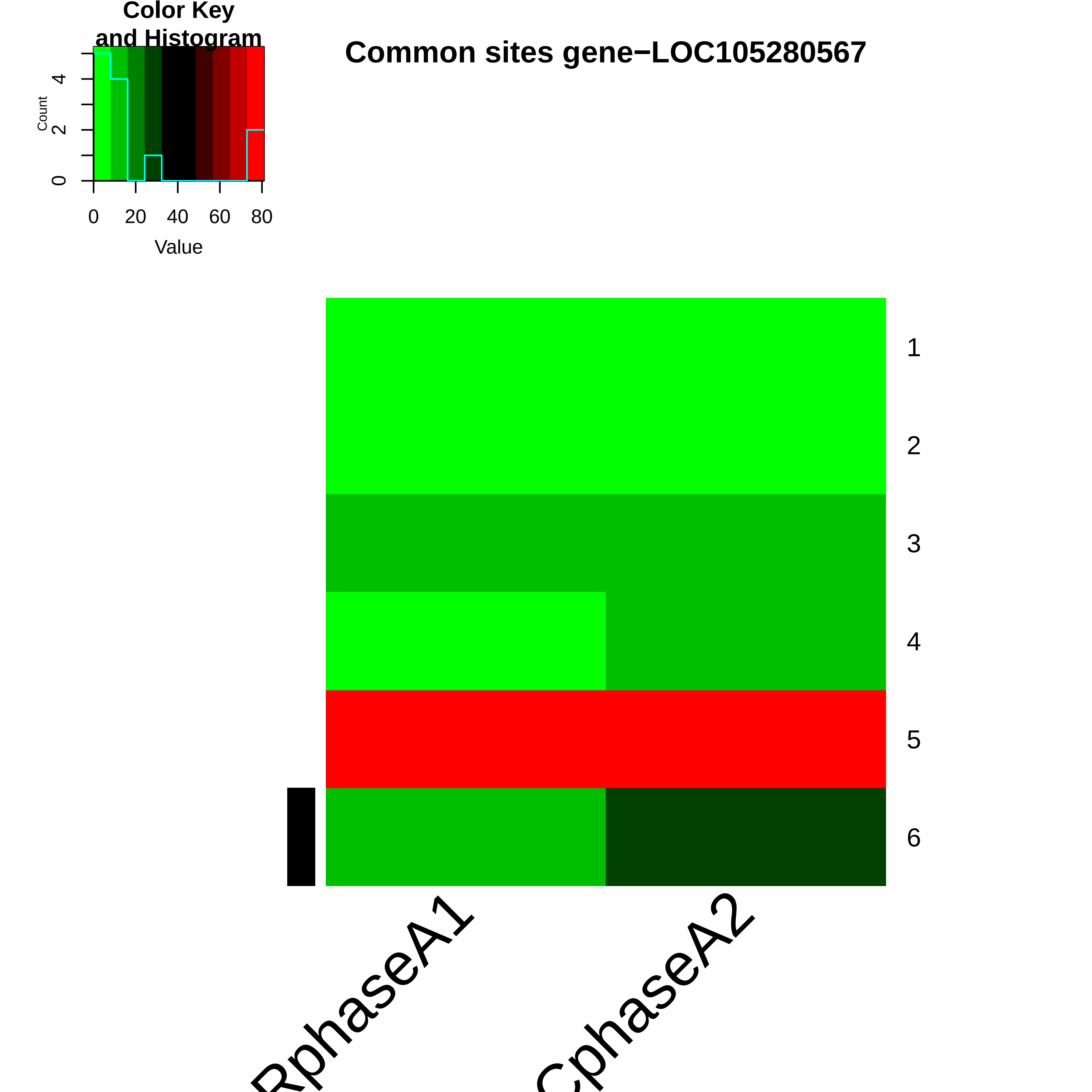

Color Key
and Histogram
Count
0
2
4
0
20
40
60
80
Value
Common sites gene−LOC105280567
1
2
3
4
5
6
RphaseA1
CphaseA2

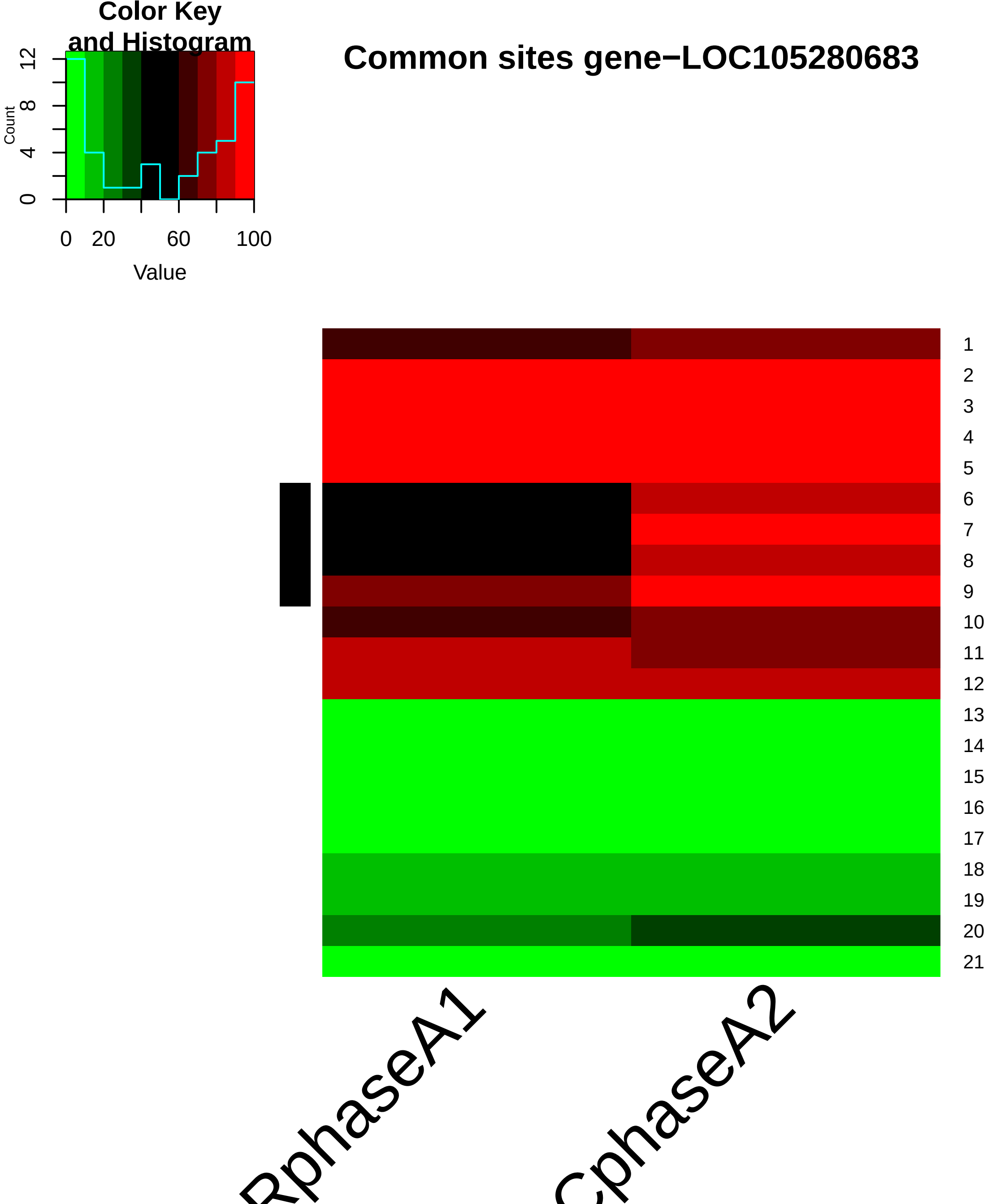

Color Key
and Histogram
Count
0
4
8
12
0
20
60
100
Value
Common sites gene−LOC105280683
RphaseA1
CphaseA2

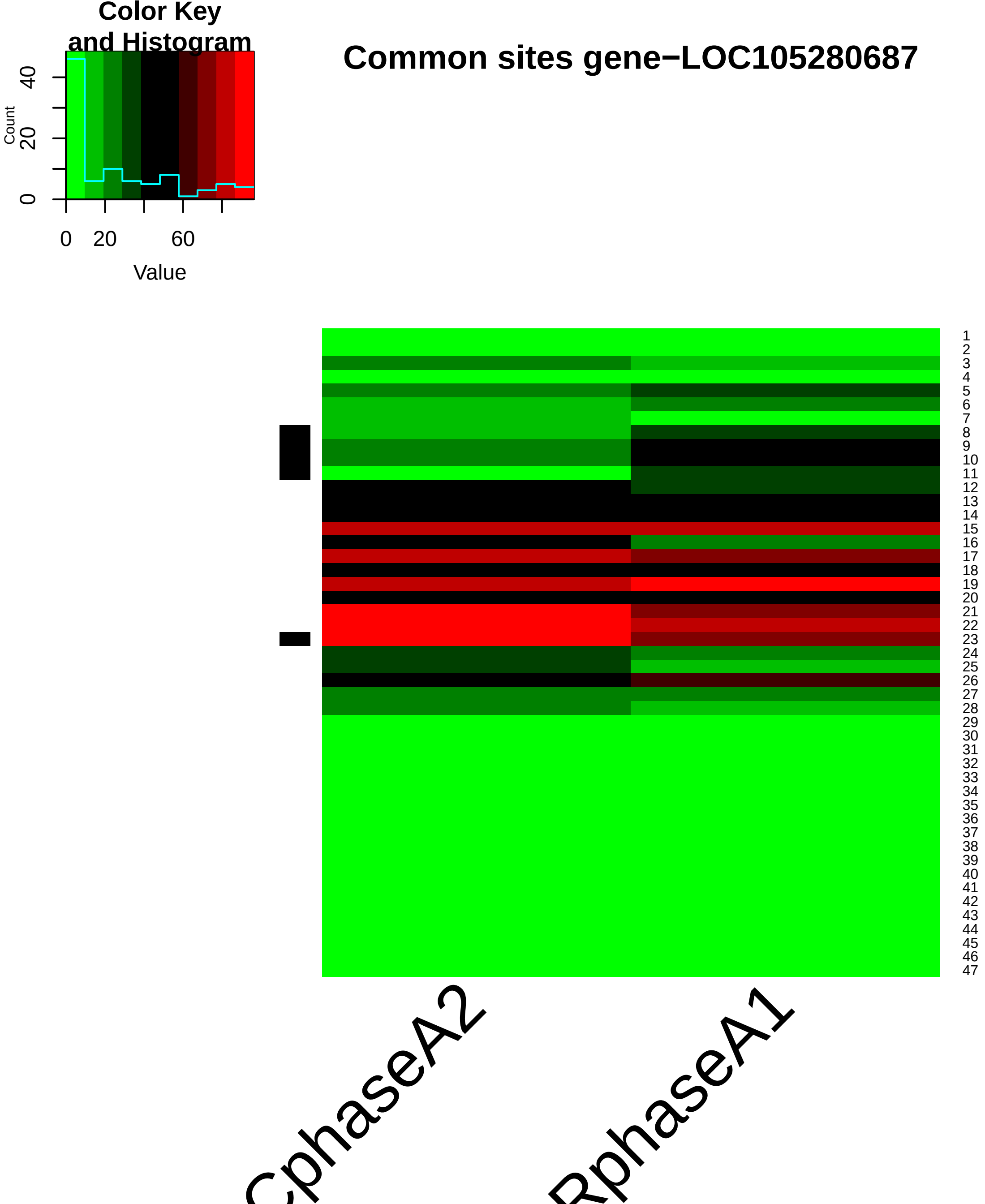
Color Key
and Histogram
Count
0
20
40
0
20
60
Value
Common sites gene−LOC105280687
CphaseA2
RphaseA1

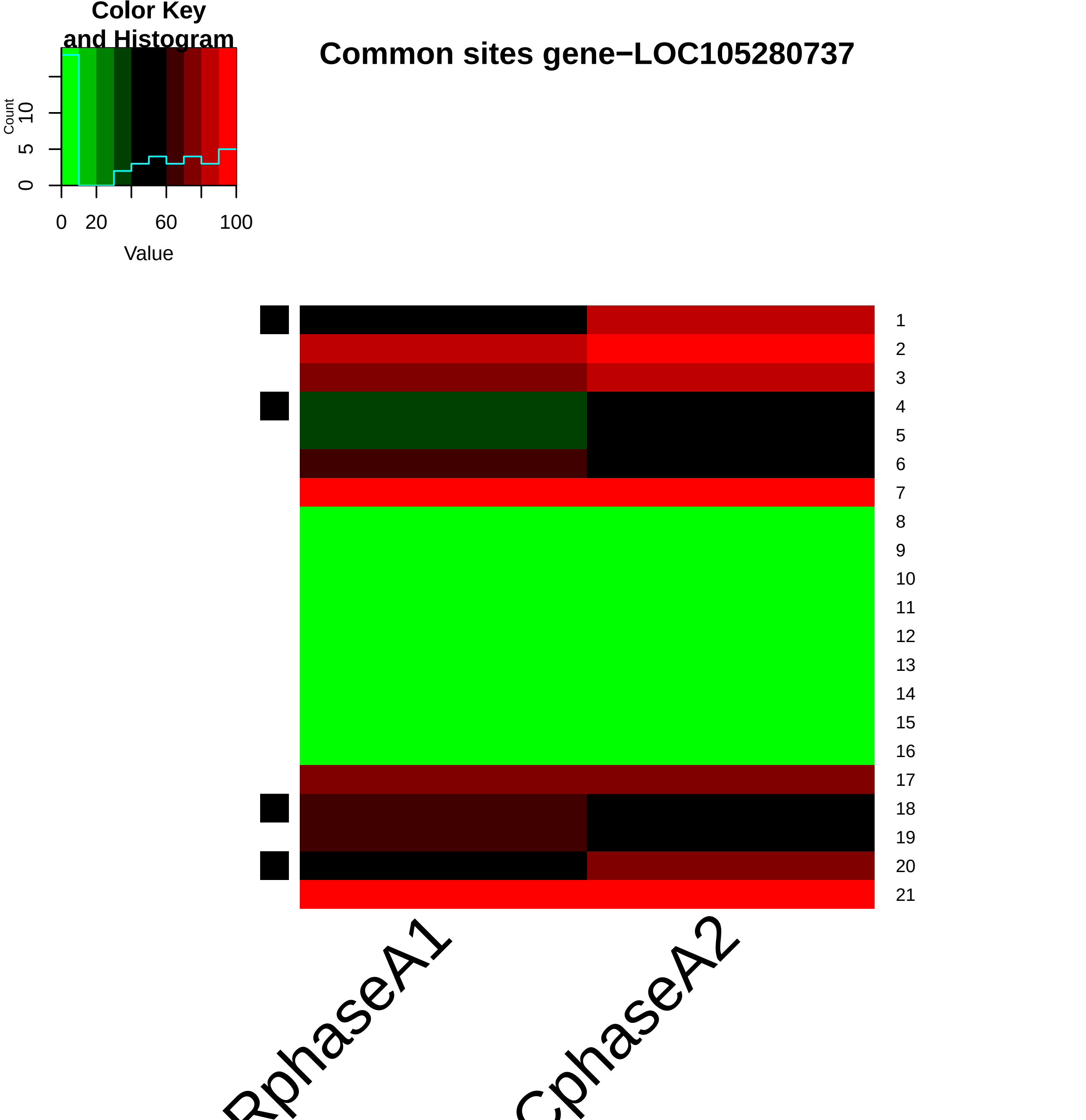

Color Key
and Histogram
Count
0
5
10
0
20
60
100
Value
Common sites gene−LOC105280737
RphaseA1
CphaseA2

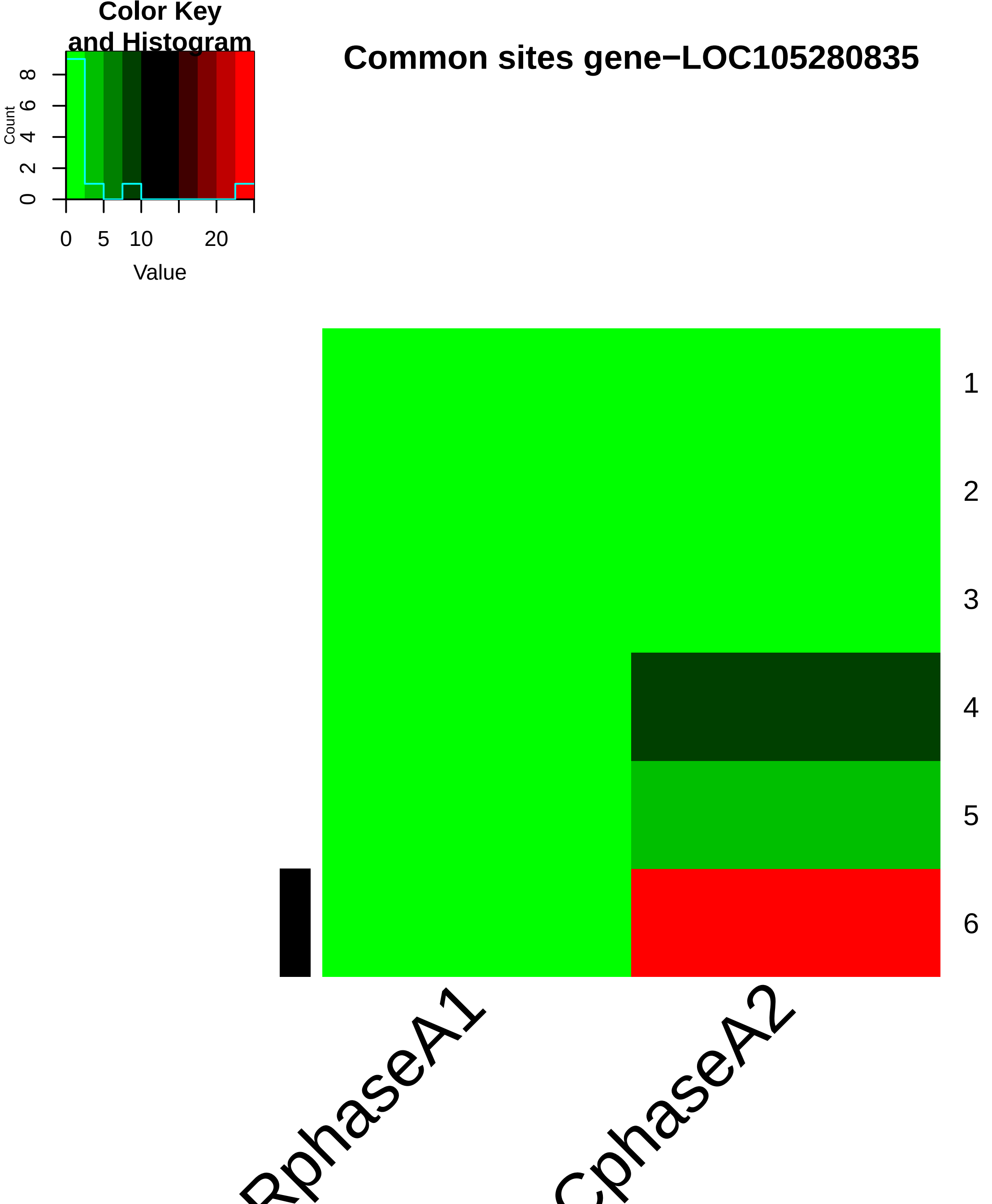
Color Key
and Histogram
Count
0
2
4
6
8
0
5
10
20
Value
Common sites gene−LOC105280835
1
2
3
4
5
6
RphaseA1
CphaseA2

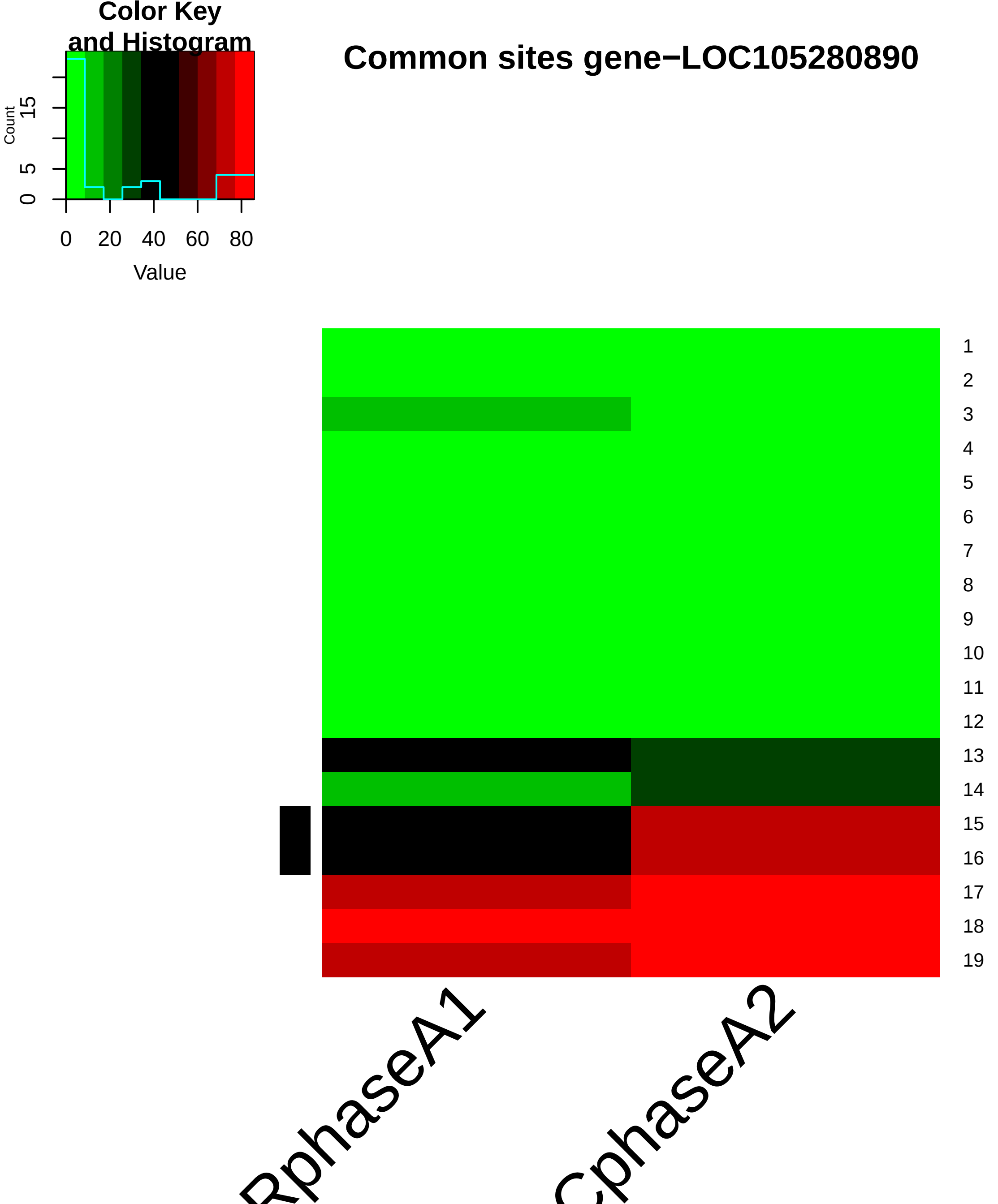

Color Key
and Histogram
Count
0
5
15
0
20
40
60
80
Value
Common sites gene−LOC105280890
RphaseA1
CphaseA2

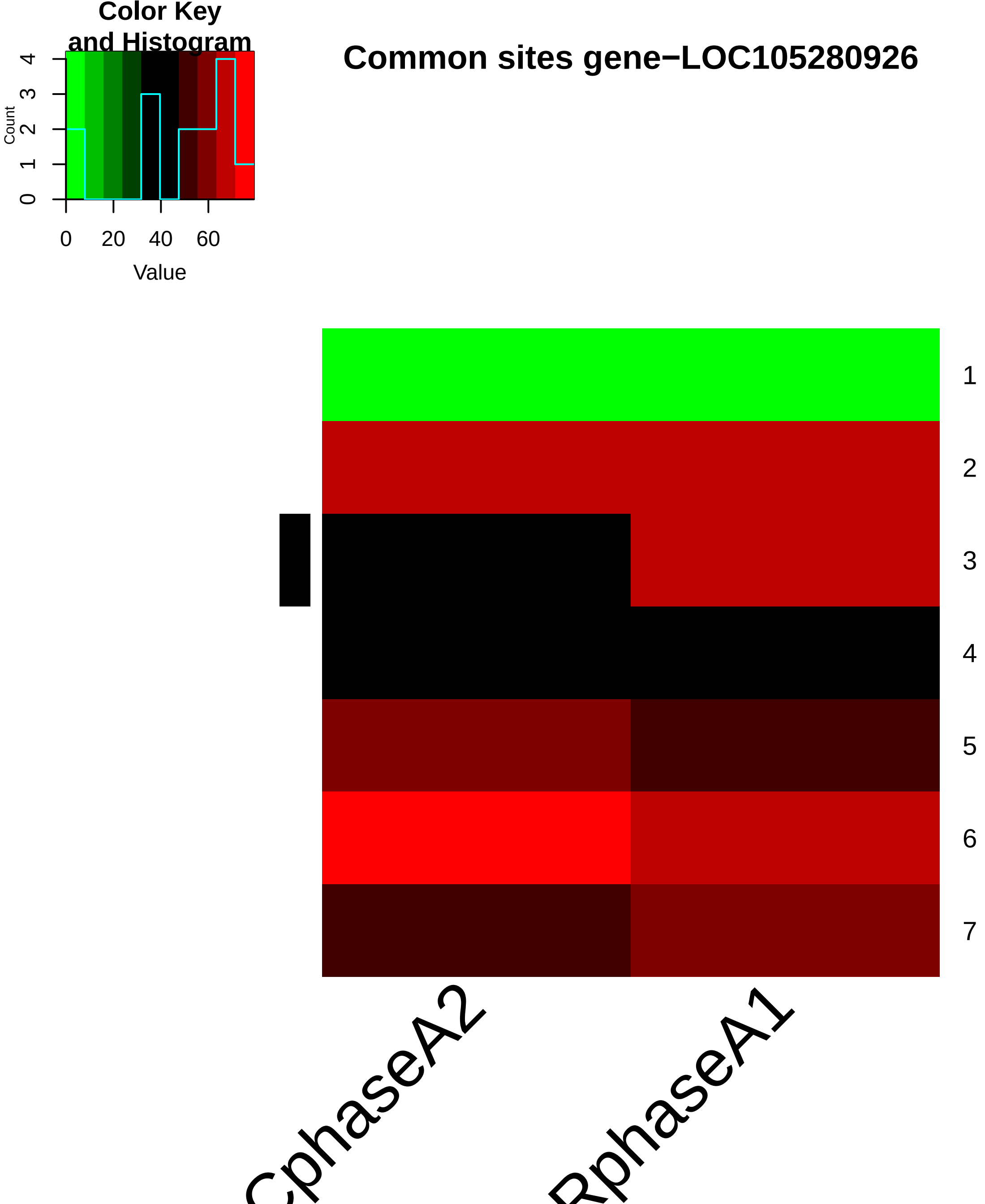
Color Key
and Histogram
Count
0
1
2
3
4
0
20
40
60
Value
Common sites gene−LOC105280926
1
2
3
4
5
6
7
CphaseA2
RphaseA1

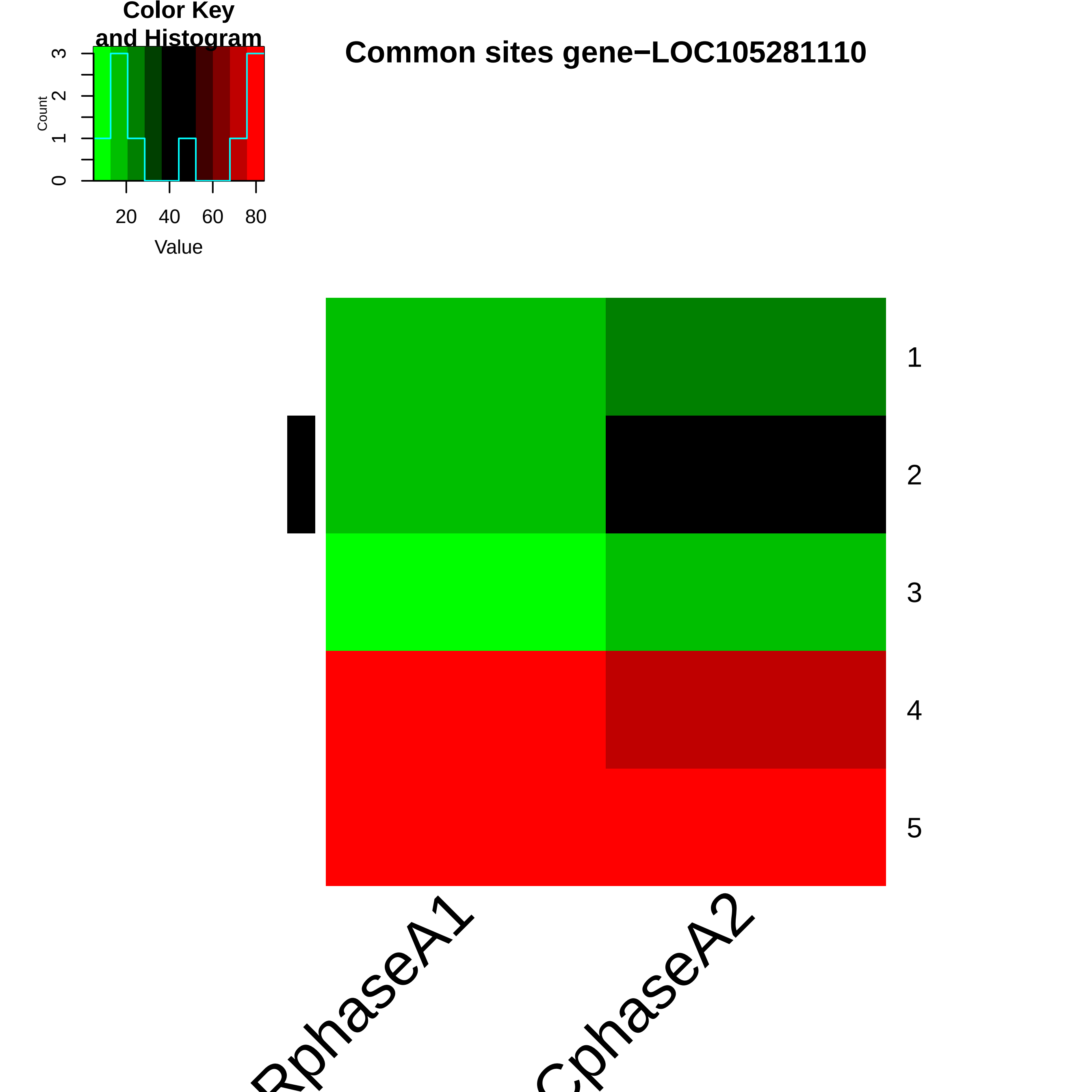
Color Key
and Histogram
Count
0
1
2
3
20
40
60
80
Value
Common sites gene−LOC105281110
1
2
3
4
5

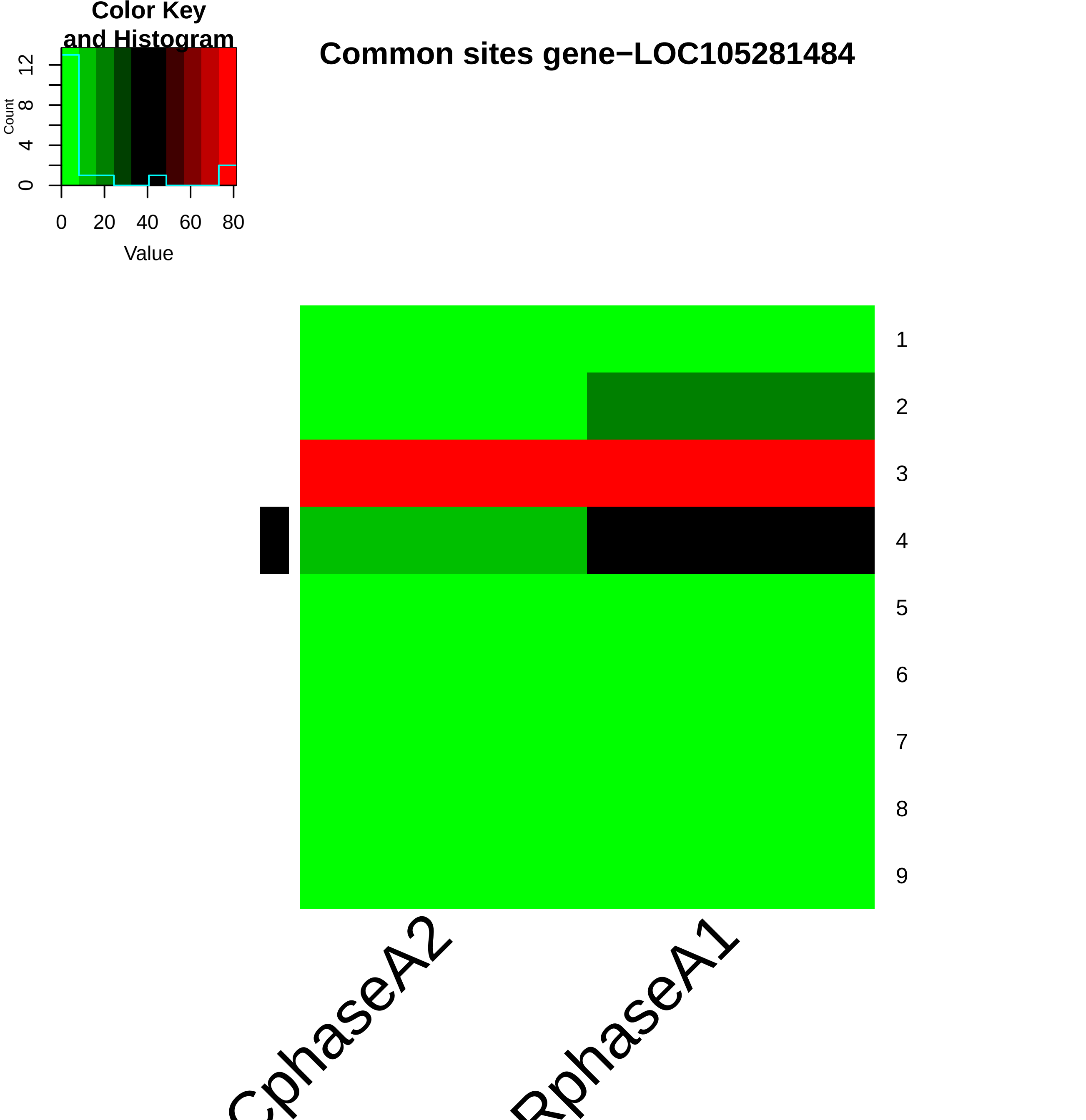

Color Key
and Histogram
Count
Value
Common sites gene−LOC105281484
1
2
3
4
5
6
7
8
9

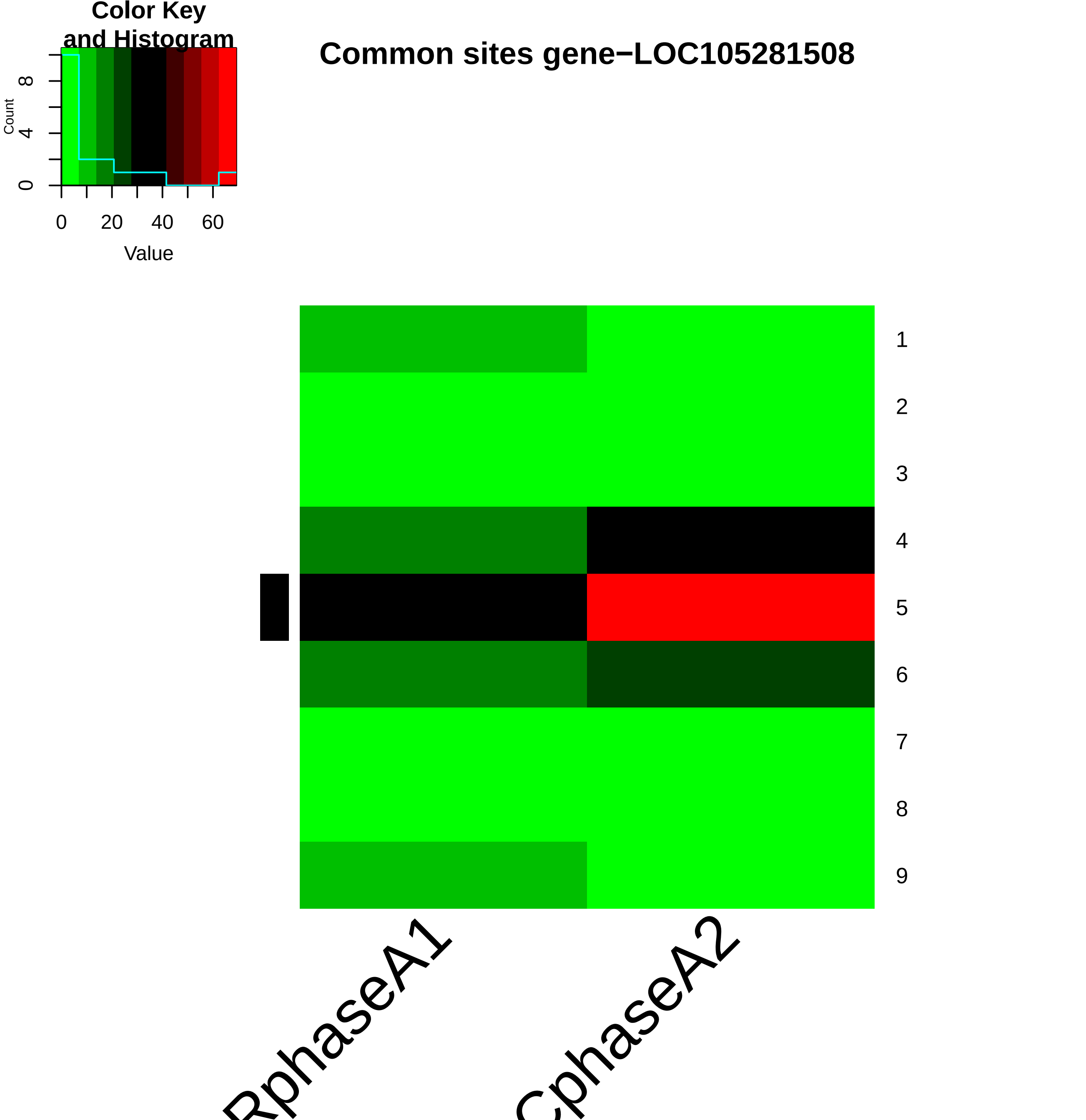

Color Key
and Histogram
Count
0
4
8
0
20
40
60
Value
Common sites gene−LOC105281508
1
2
3
4
5
6
7
8
9
RphaseA1
CphaseA2

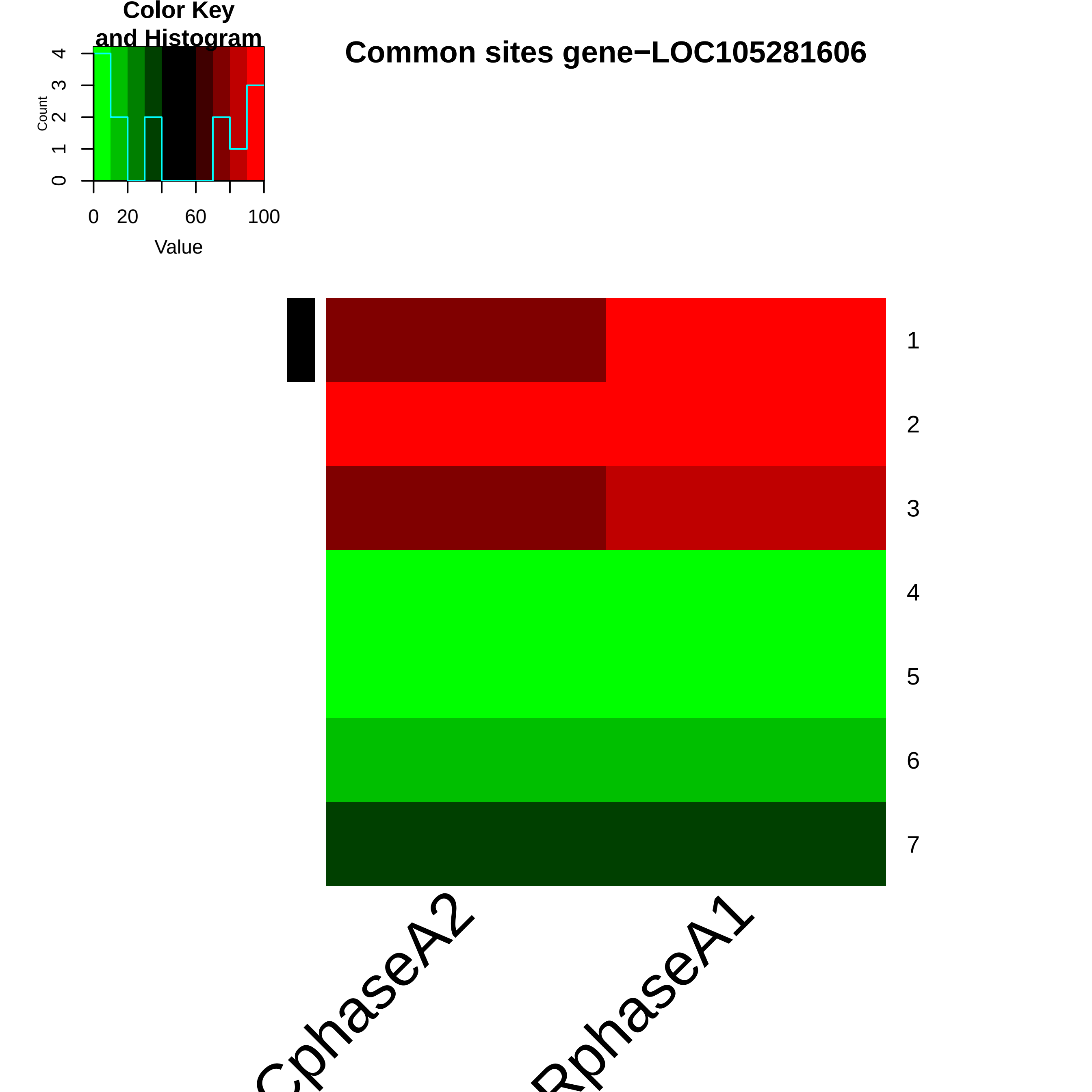

Color Key
and Histogram
Count
0
1
2
3
4
0
20
60
100
Value
Common sites gene−LOC105281606
1
2
3
4
5
6
7
CphaseA2
RphaseA1

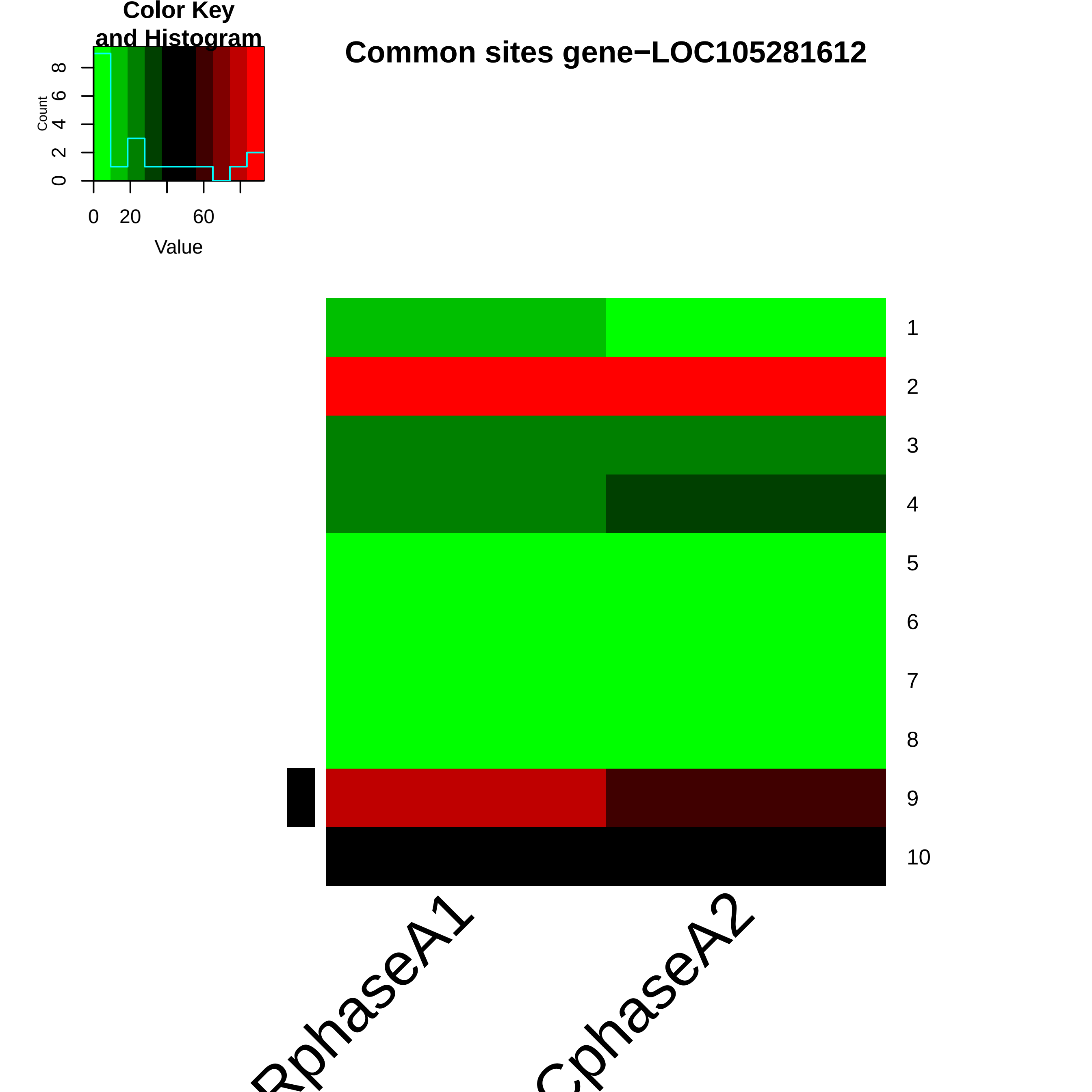

Color Key
and Histogram
Count
0 2 4 6 8
0 20 60
Value
Common sites gene−LOC105281612
1
2
3
4
5
6
7
8
9
10
RphaseA1
CphaseA2

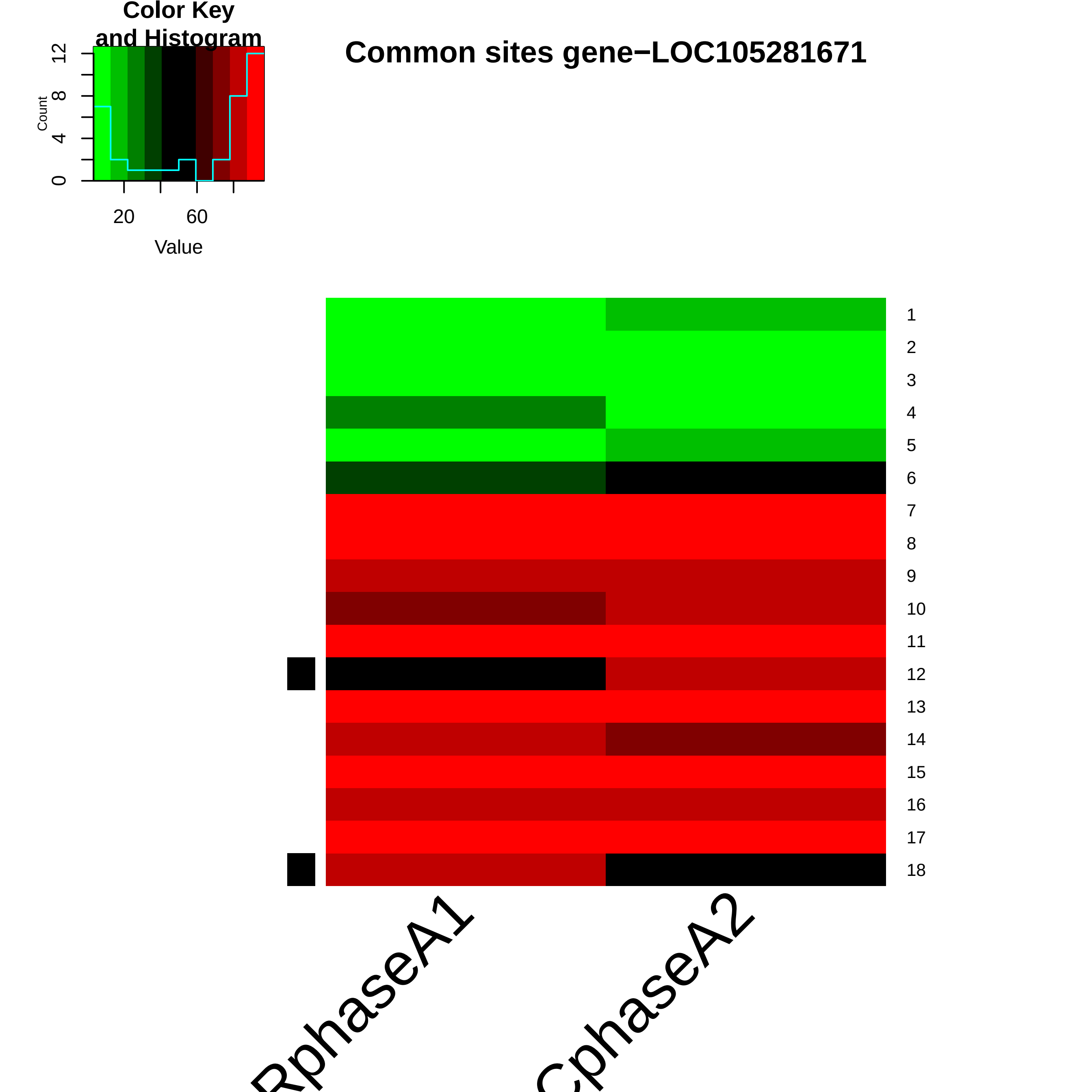

Color Key
and Histogram
Count
0
4
8
12
20
60
Value
Common sites gene−LOC105281671
RphaseA1
CphaseA2

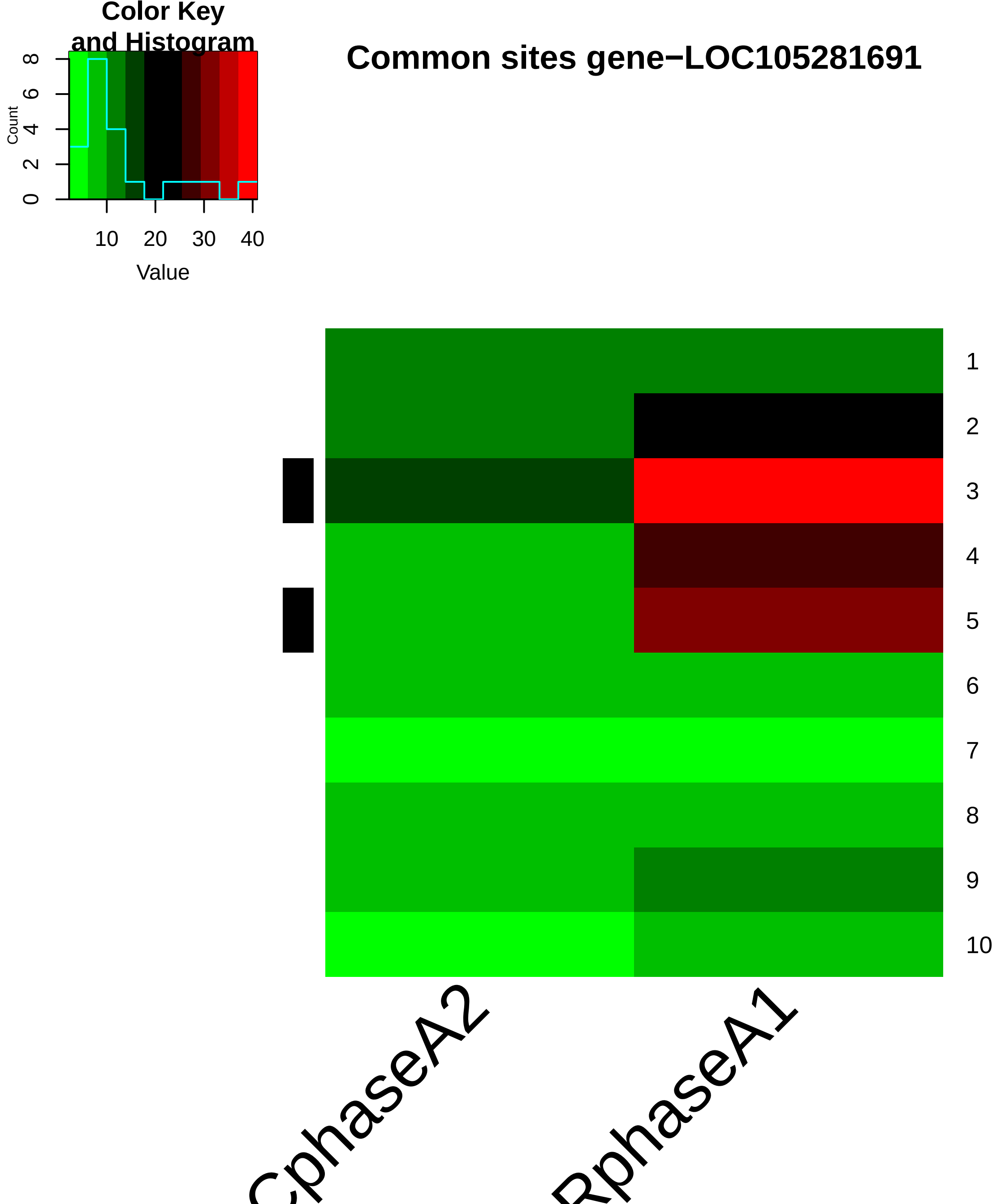
Color Key
and Histogram
Count
0
2
4
6
8
10
20
30
40
Value
Common sites gene−LOC105281691
1
2
3
4
5
6
7
8
9
10
CphaseA2
RphaseA1

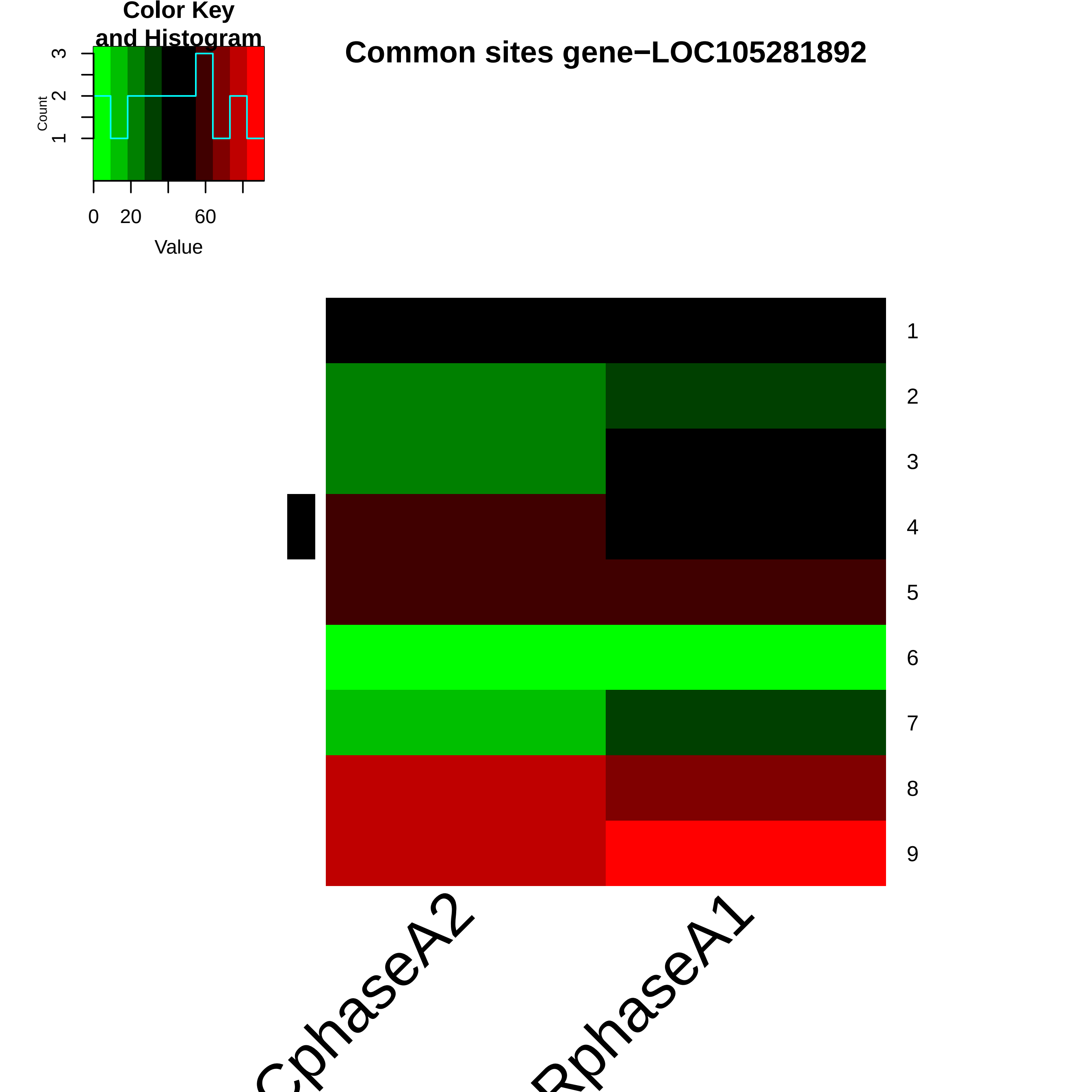

Color Key
and Histogram
Count
1
2
3
0
20
60
Value
Common sites gene−LOC105281892
1
2
3
4
5
6
7
8
9
CphaseA2
RphaseA1

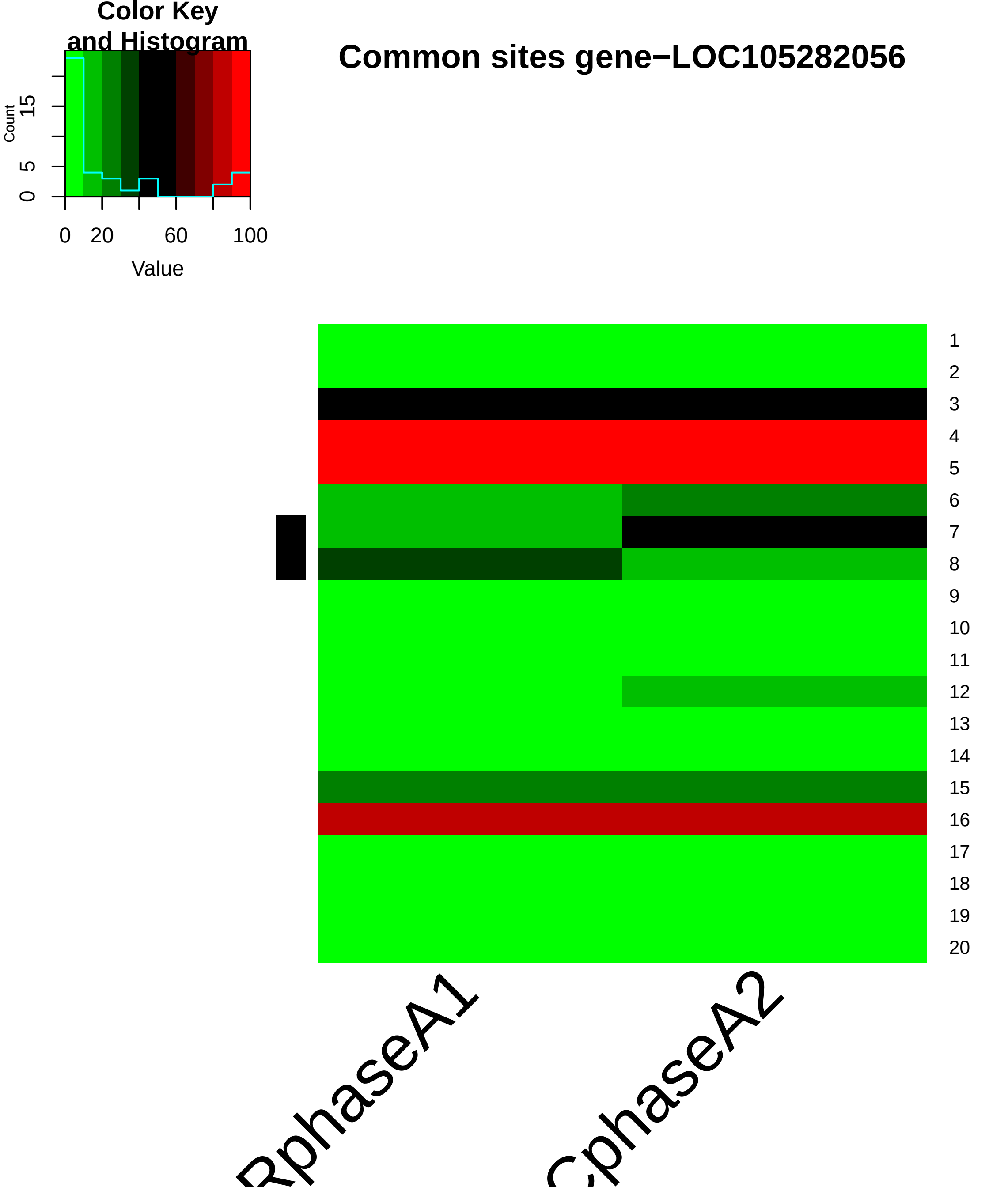
Color Key
and Histogram
Count
0
5
15
0
20
60
100
Value
Common sites gene−LOC105282056
RphaseA1
CphaseA2

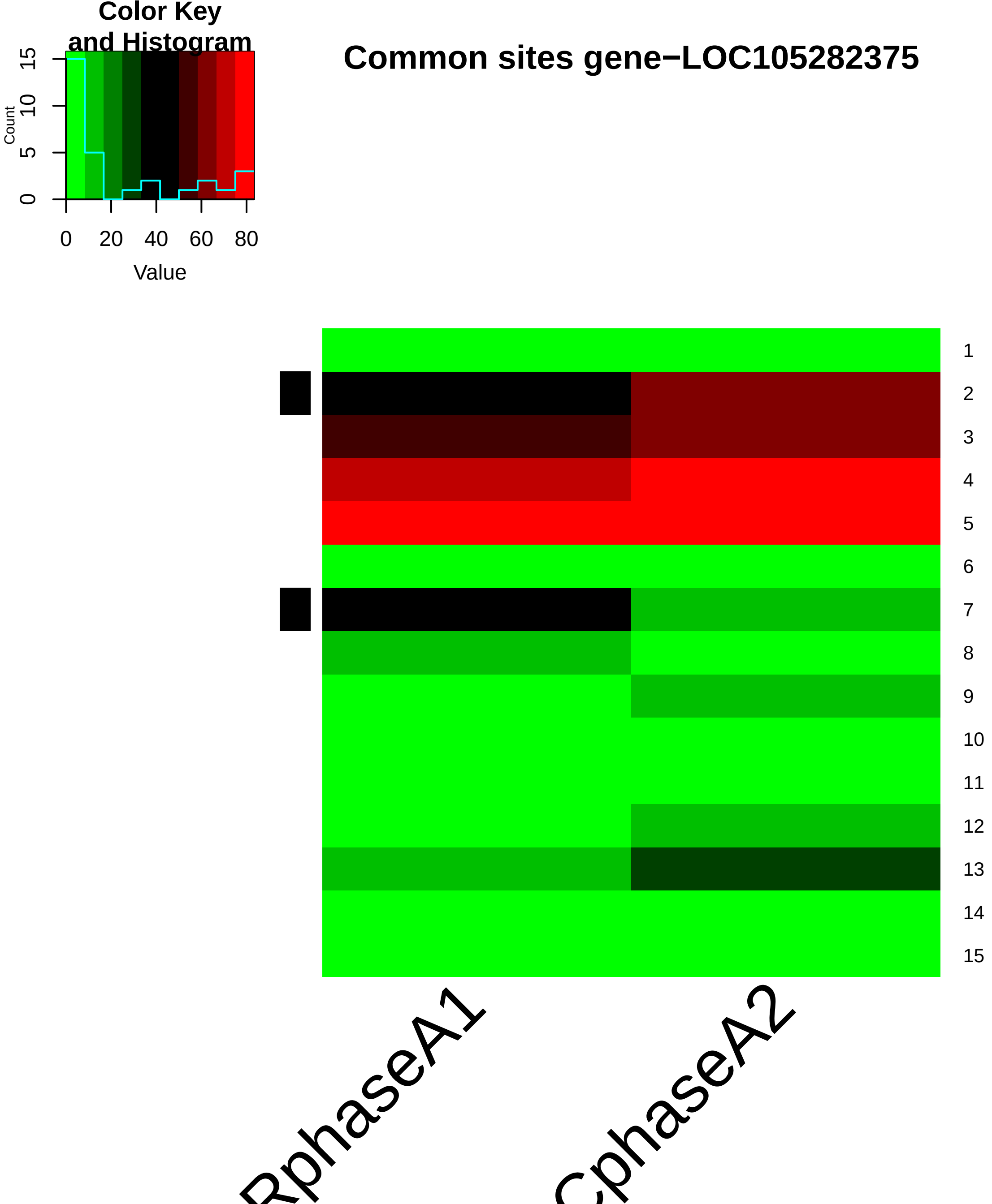

Color Key
and Histogram
Count
Value
0
5
10
15
0
20
40
60
80
Common sites gene−LOC105282375
RphaseA1
CphaseA2

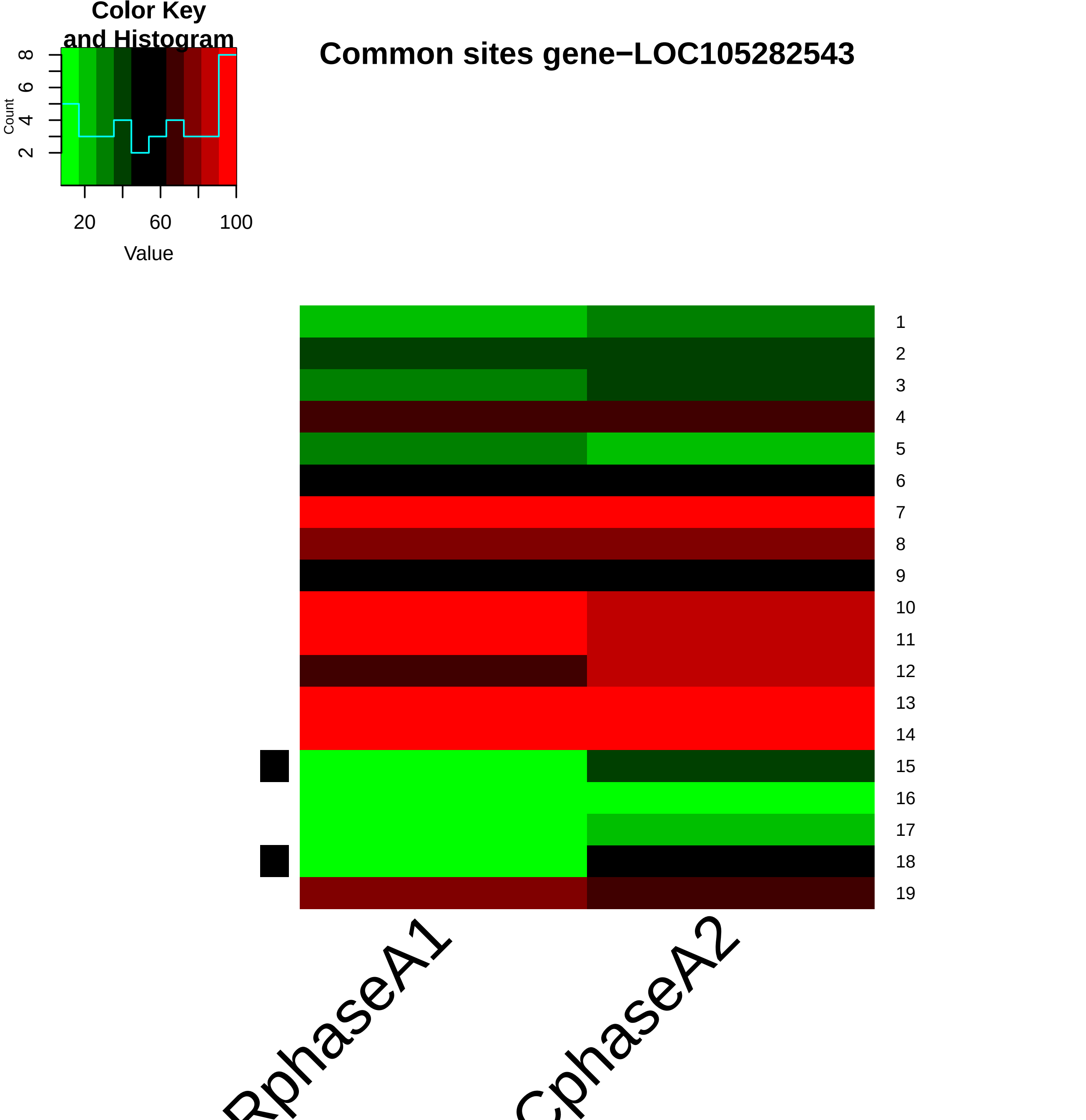
Color Key
and Histogram
Count
2
4
6
8
20
60
100
Value
Common sites gene−LOC105282543
RphaseA1
CphaseA2

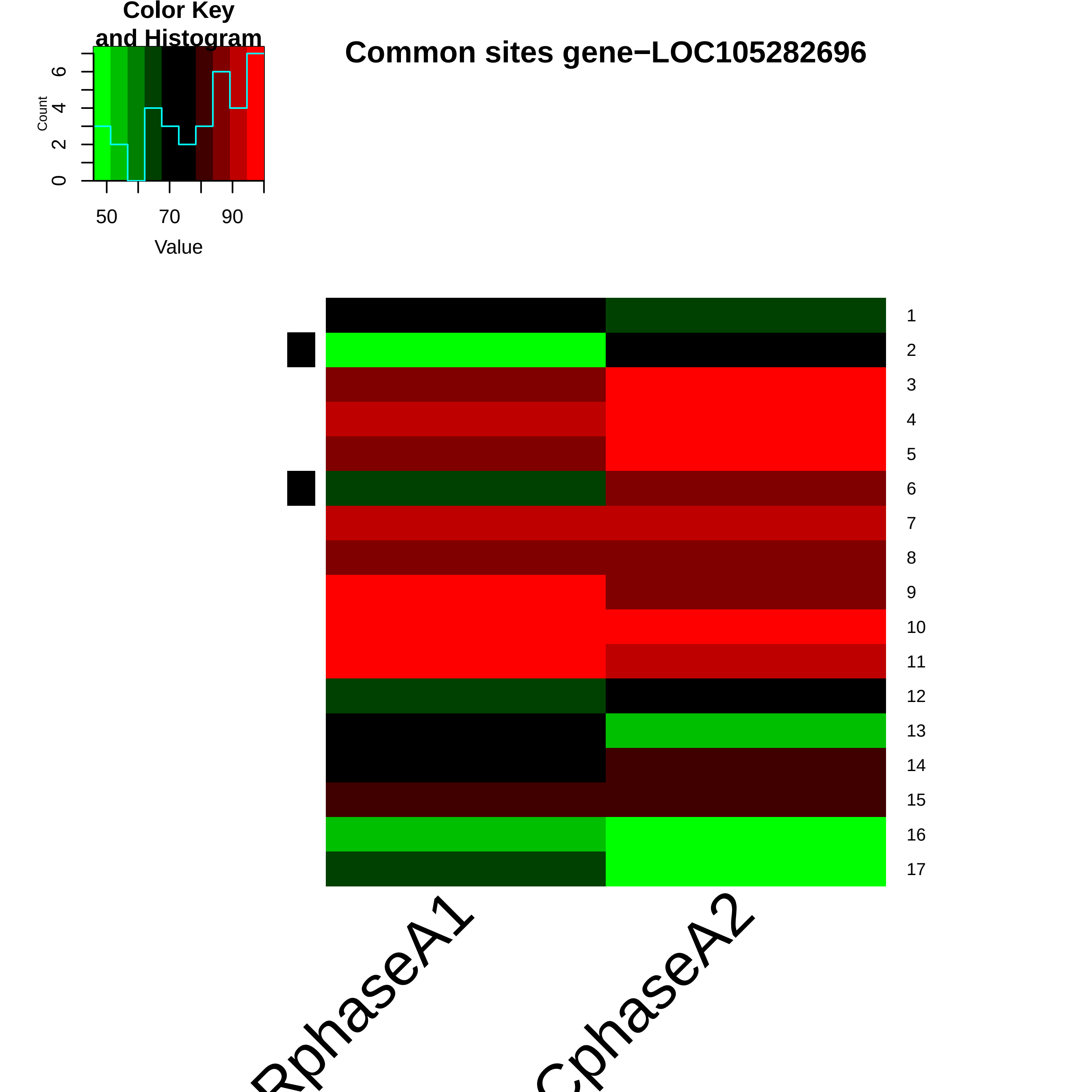
Color Key
and Histogram
Count
0
2
4
6
50
70
90
Value
Common sites gene−LOC105282696
RphaseA1
CphaseA2

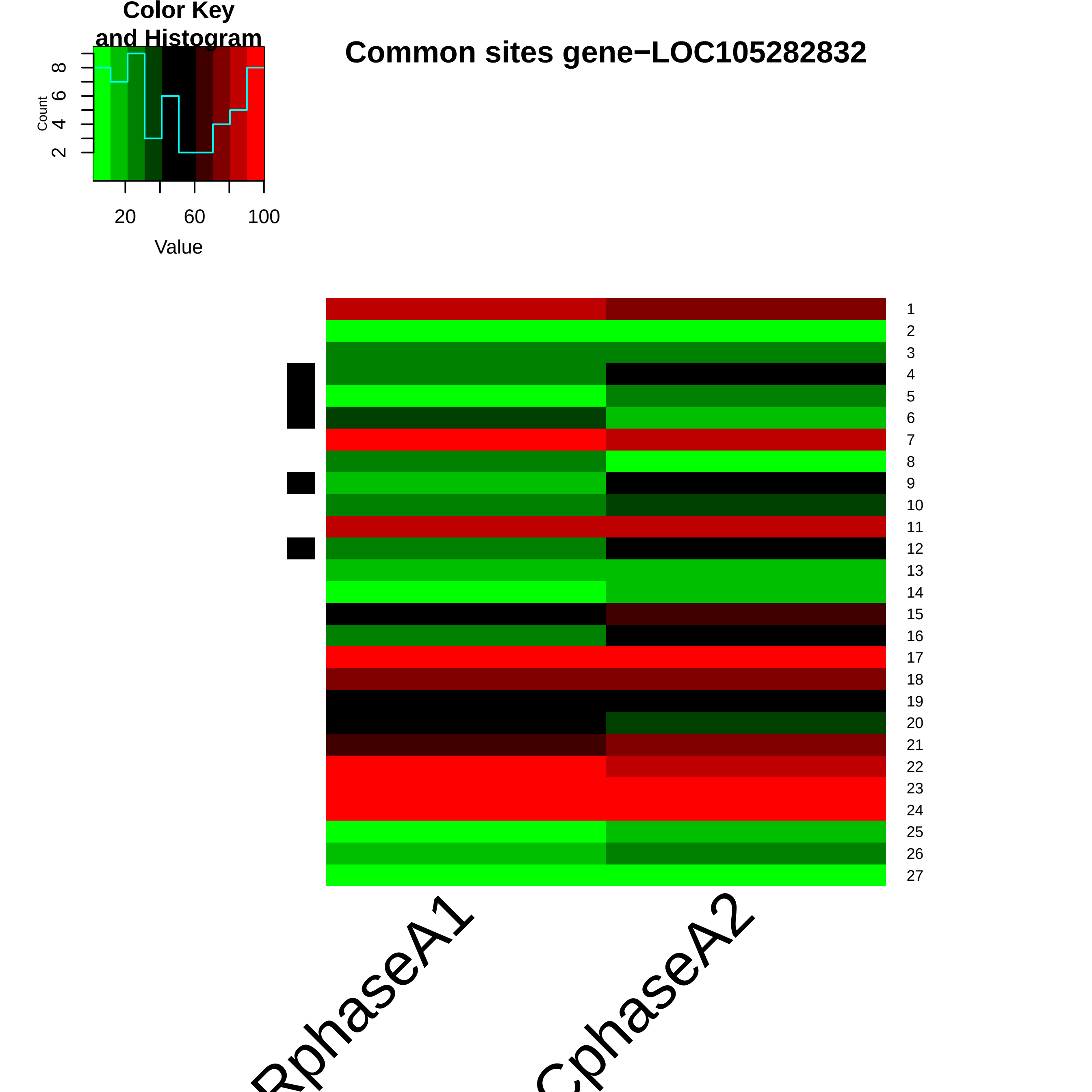

Color Key
and Histogram
Count
Value
20
60
100
2
4
6
8
Common sites gene−LOC105282832
RphaseA1
CphaseA2

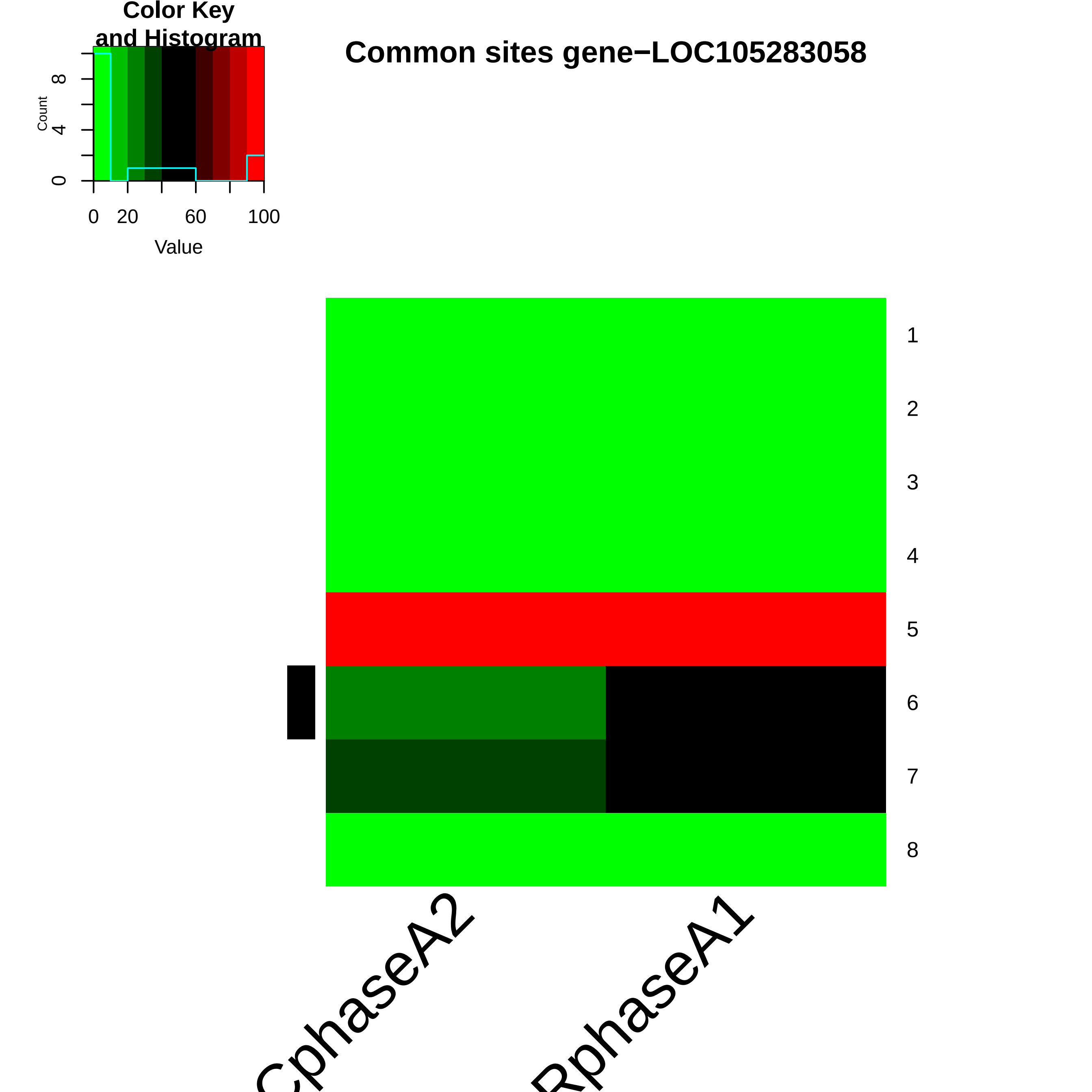

Color Key
and Histogram
Count
0
4
8
0
20
60
100
Value
Common sites gene−LOC105283058
1
2
3
4
5
6
7
8
CphaseA2
RphaseA1

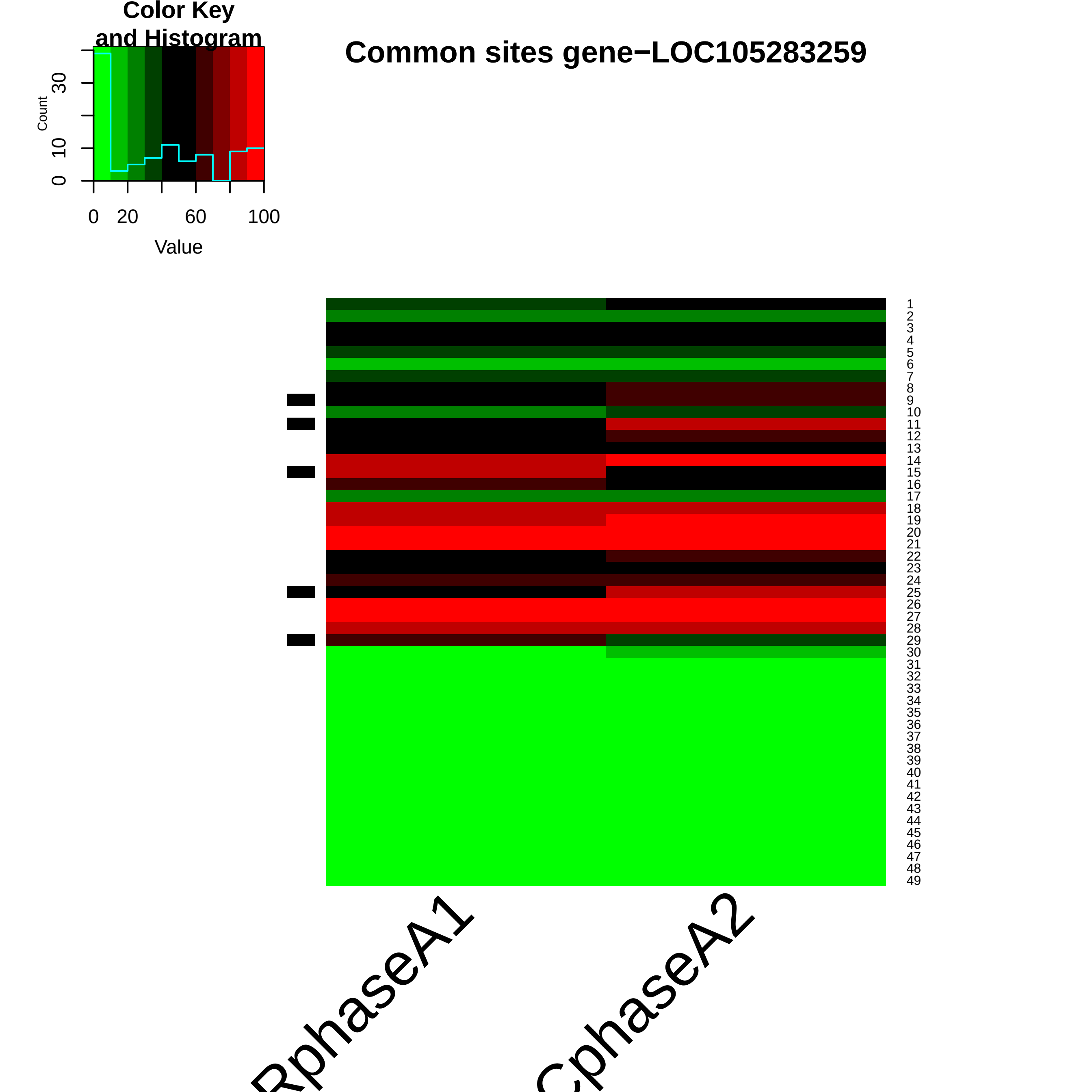
Color Key
and Histogram
Count
0
10
30
0
20
60
100
Value
Common sites gene−LOC105283259
RphaseA1
CphaseA2

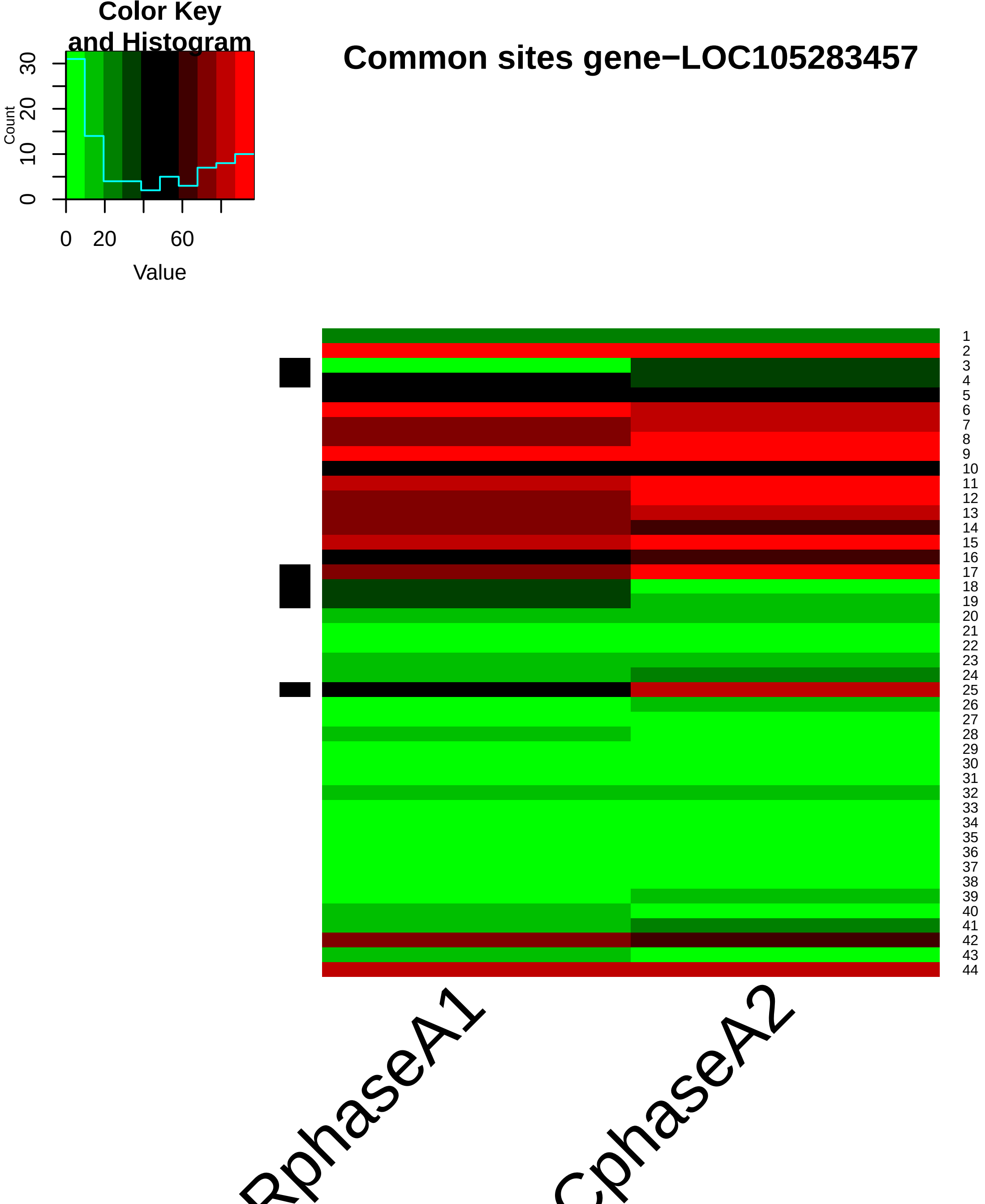
Color Key
and Histogram
Count
0
10
20
30
0
20
60
Value
Common sites gene−LOC105283457
RphaseA1
CphaseA2

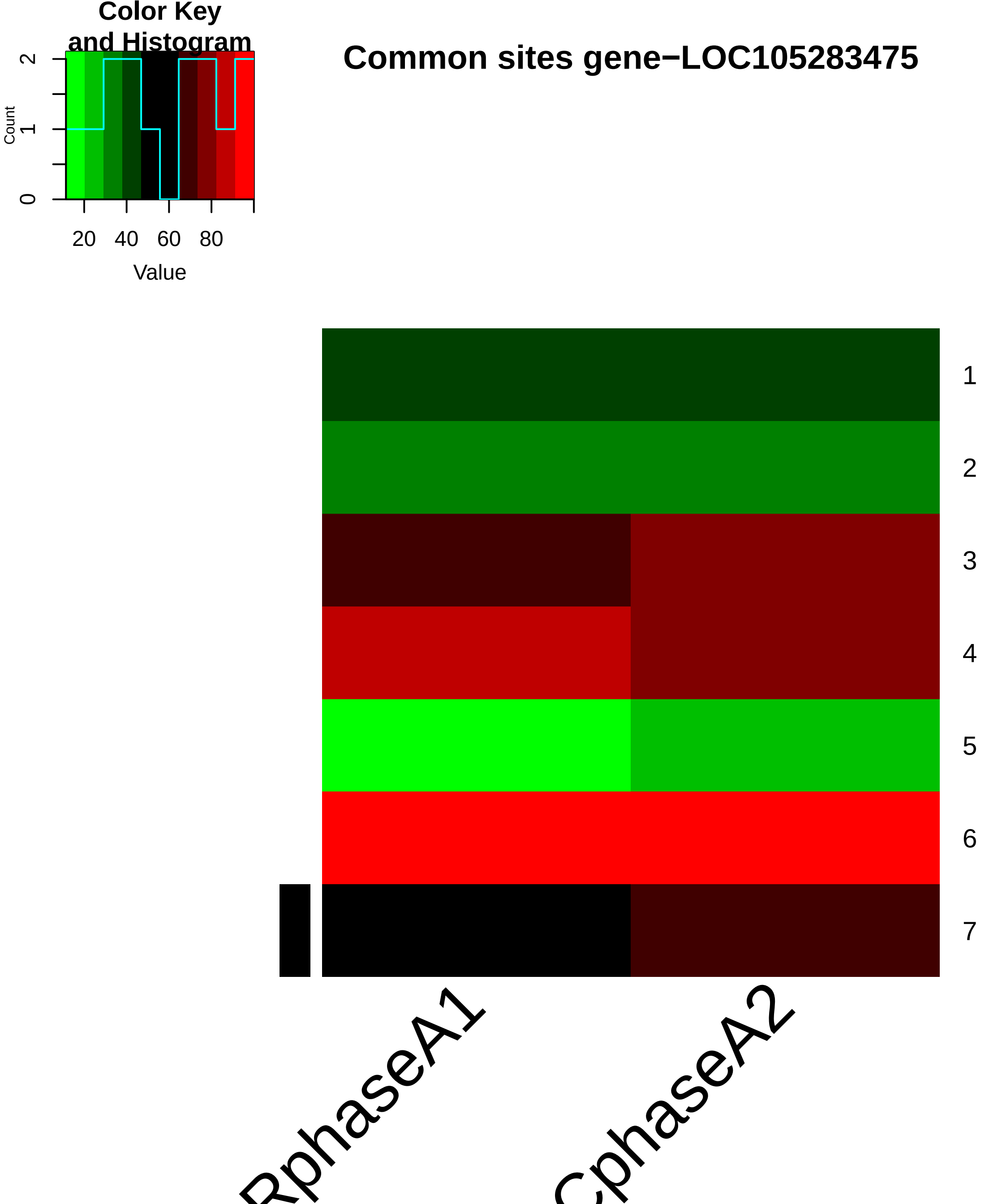

Color Key
and Histogram
Count
0
1
2
20
40
60
80
Value
Common sites gene−LOC105283475
1
2
3
4
5
6
7
RphaseA1
CphaseA2

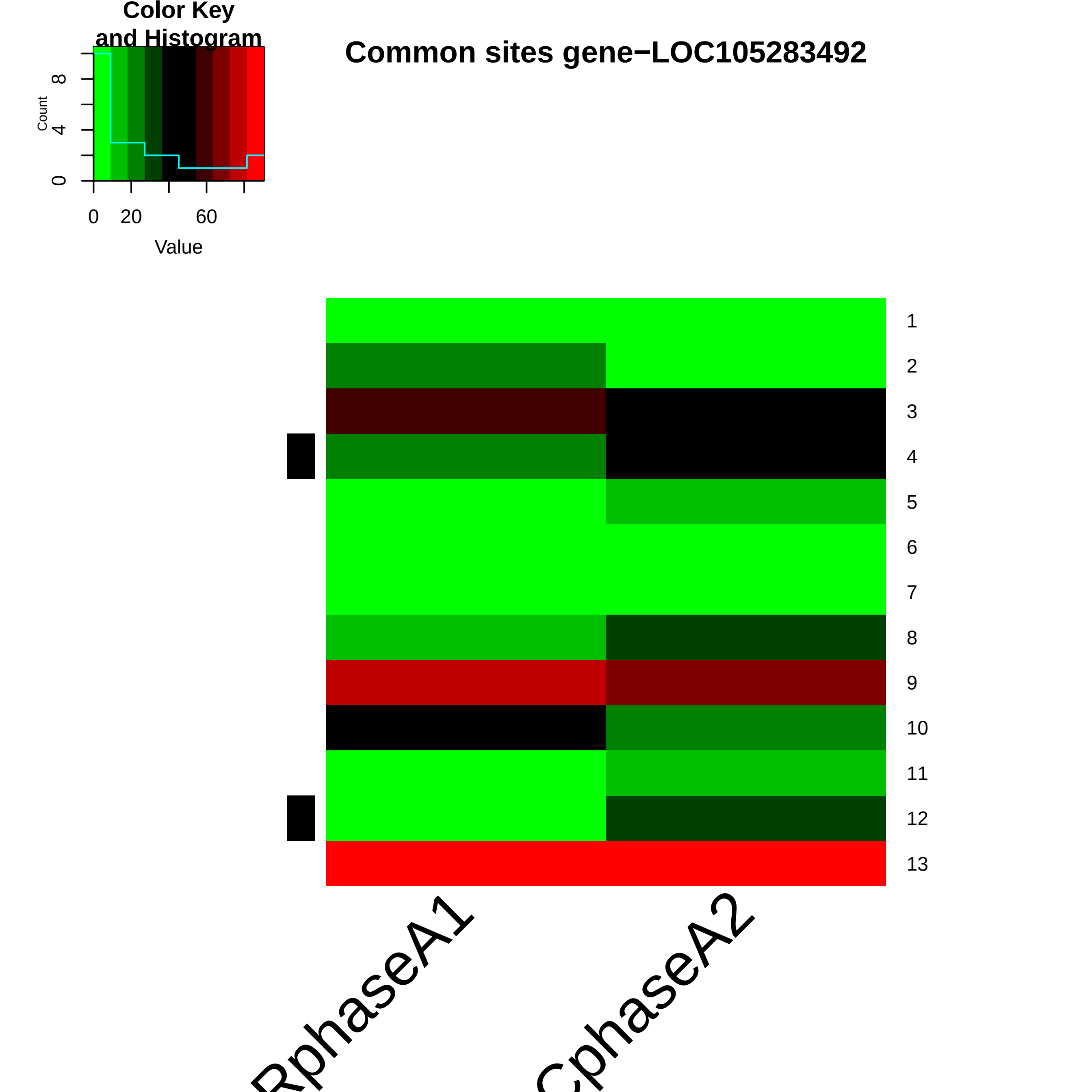
Color Key
and Histogram
Count
0
4
8
0
20
60
Value
Common sites gene−LOC105283492
1
2
3
4
5
6
7
8
9
10
11
12
13
RphaseA1
CphaseA2

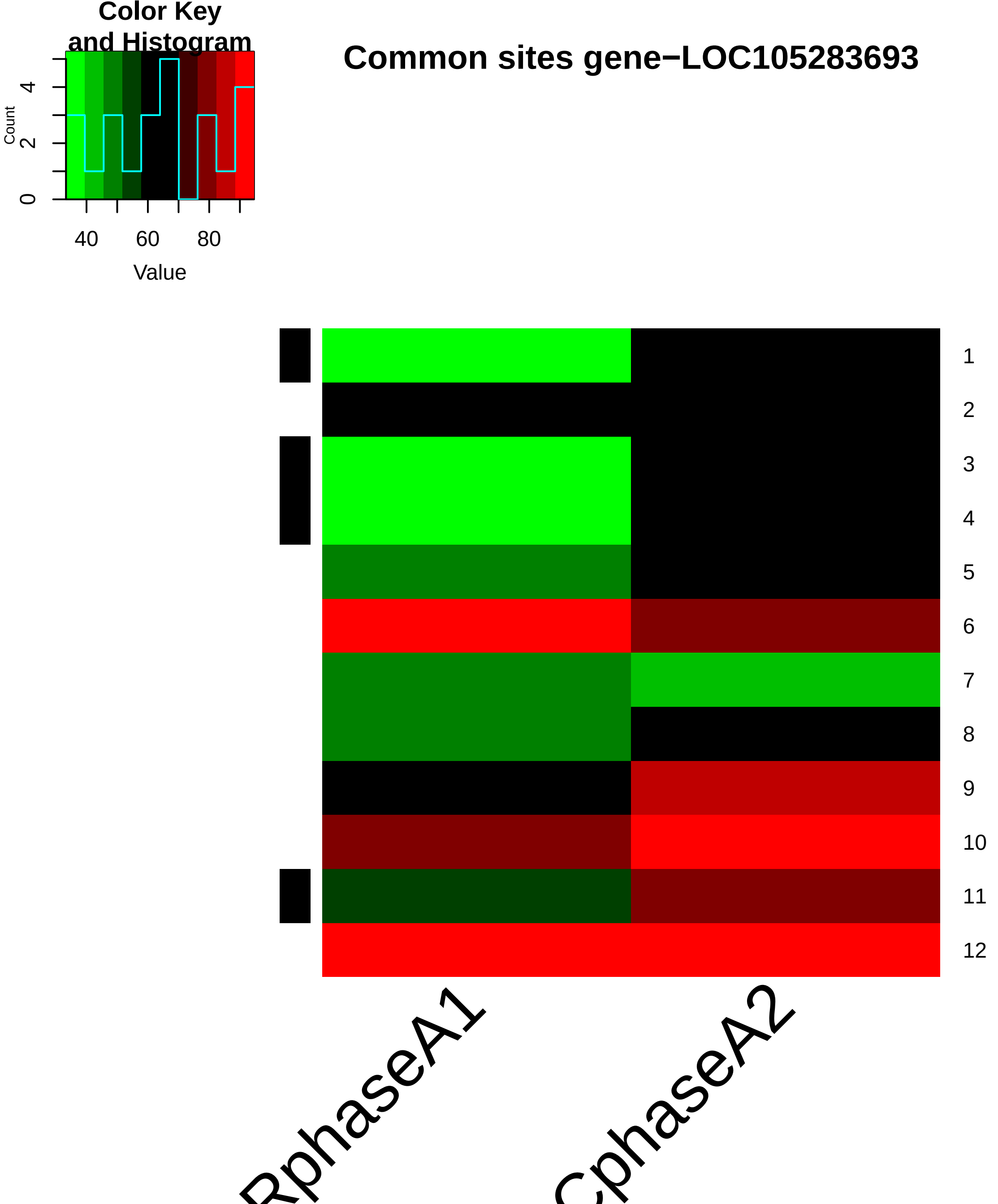

Color Key
and Histogram
Count
0
2
4
40
60
80
Value
Common sites gene−LOC105283693
1
2
3
4
5
6
7
8
9
10
11
12
RphaseA1
CphaseA2

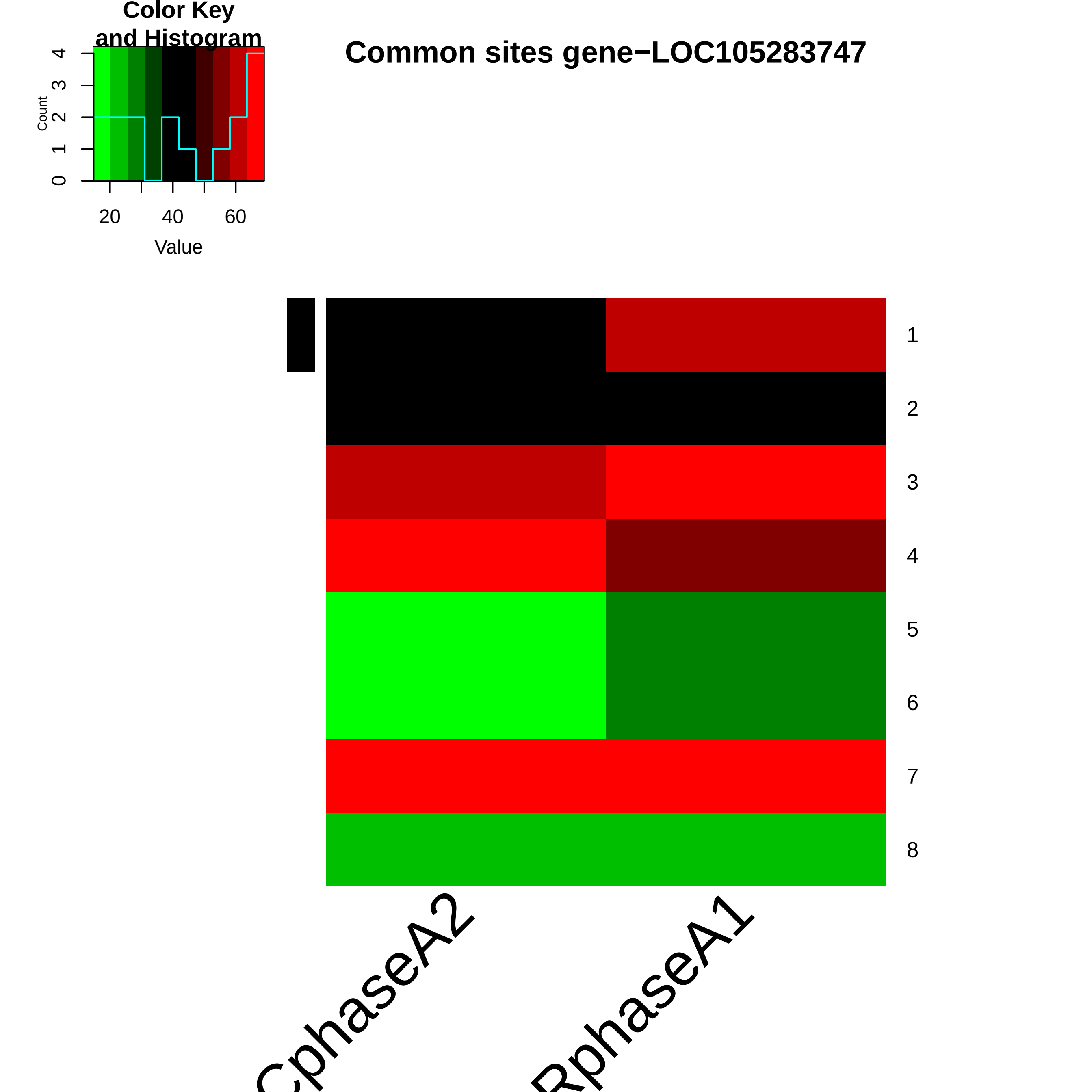
Color Key
and Histogram
Count
0
1
2
3
4
20
40
60
Value
Common sites gene−LOC105283747
1
2
3
4
5
6
7
8
CphaseA2
RphaseA1

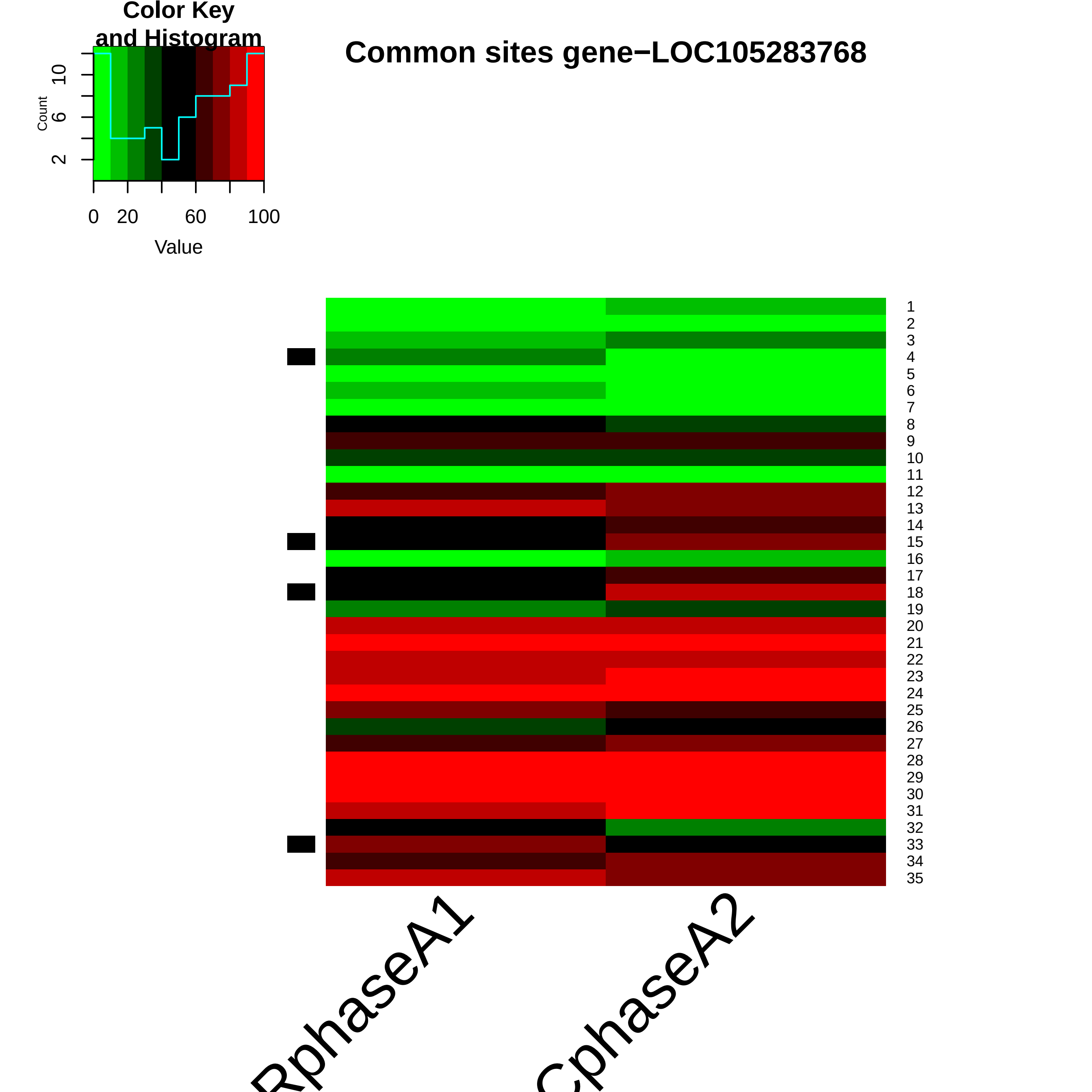

Color Key
and Histogram
Count
2
6
10
0
20
60
100
Value
Common sites gene−LOC105283768
RphaseA1
CphaseA2

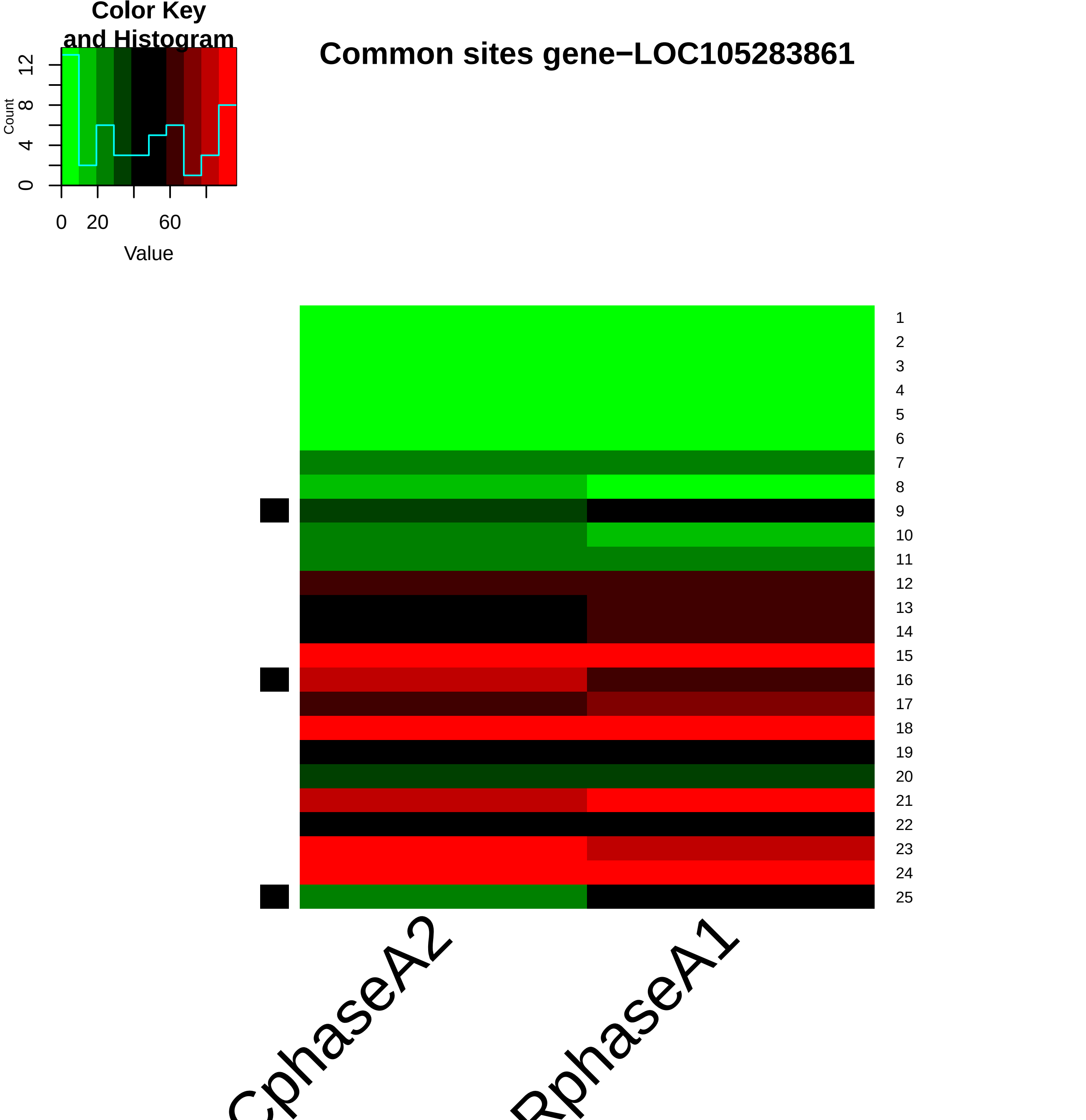

Color Key
and Histogram
Count
0
4
8
12
0
20
60
Value
Common sites gene−LOC105283861
CphaseA2
RphaseA1

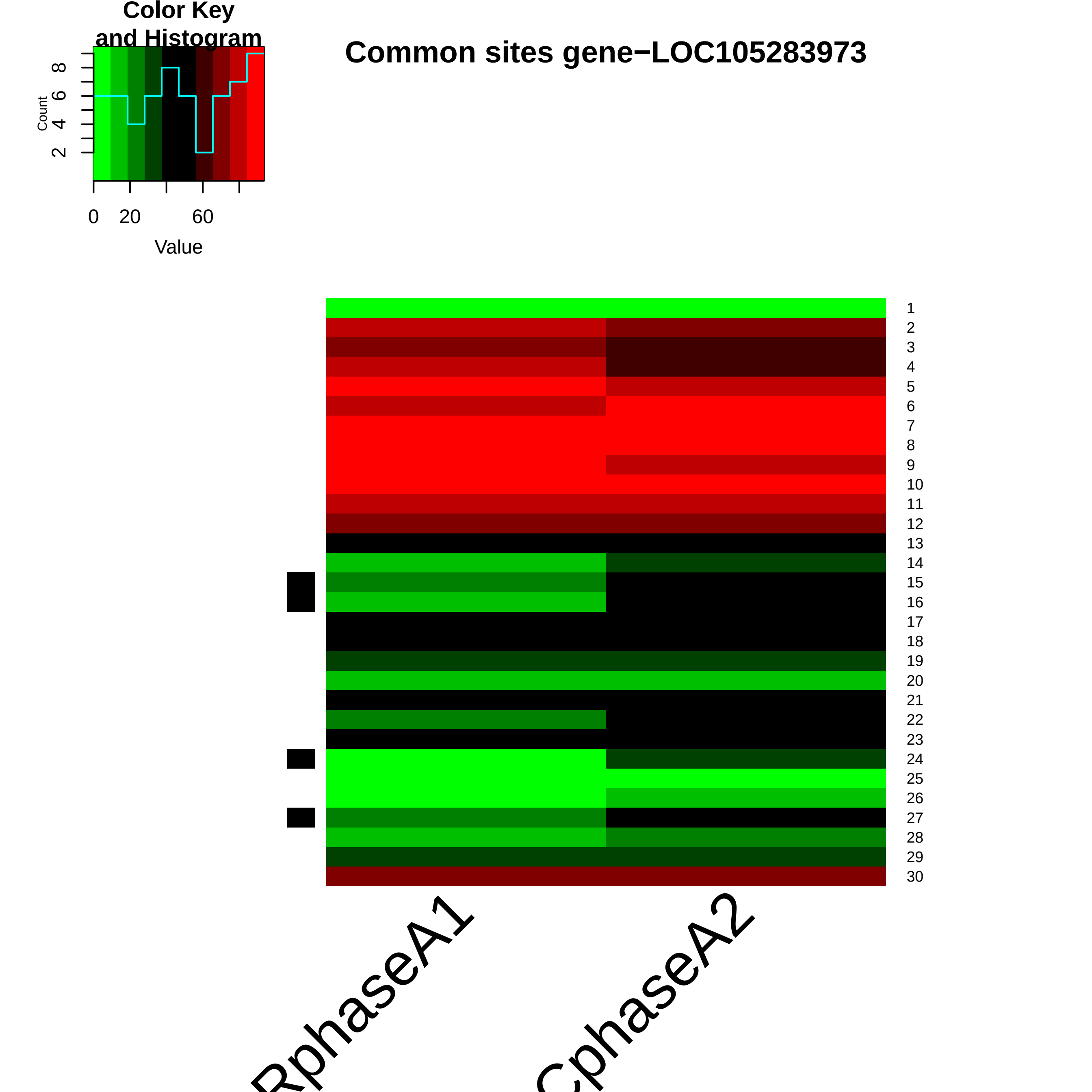

Color Key
and Histogram
Count
2
4
6
8
0
20
60
Value
Common sites gene−LOC105283973
RphaseA1
CphaseA2

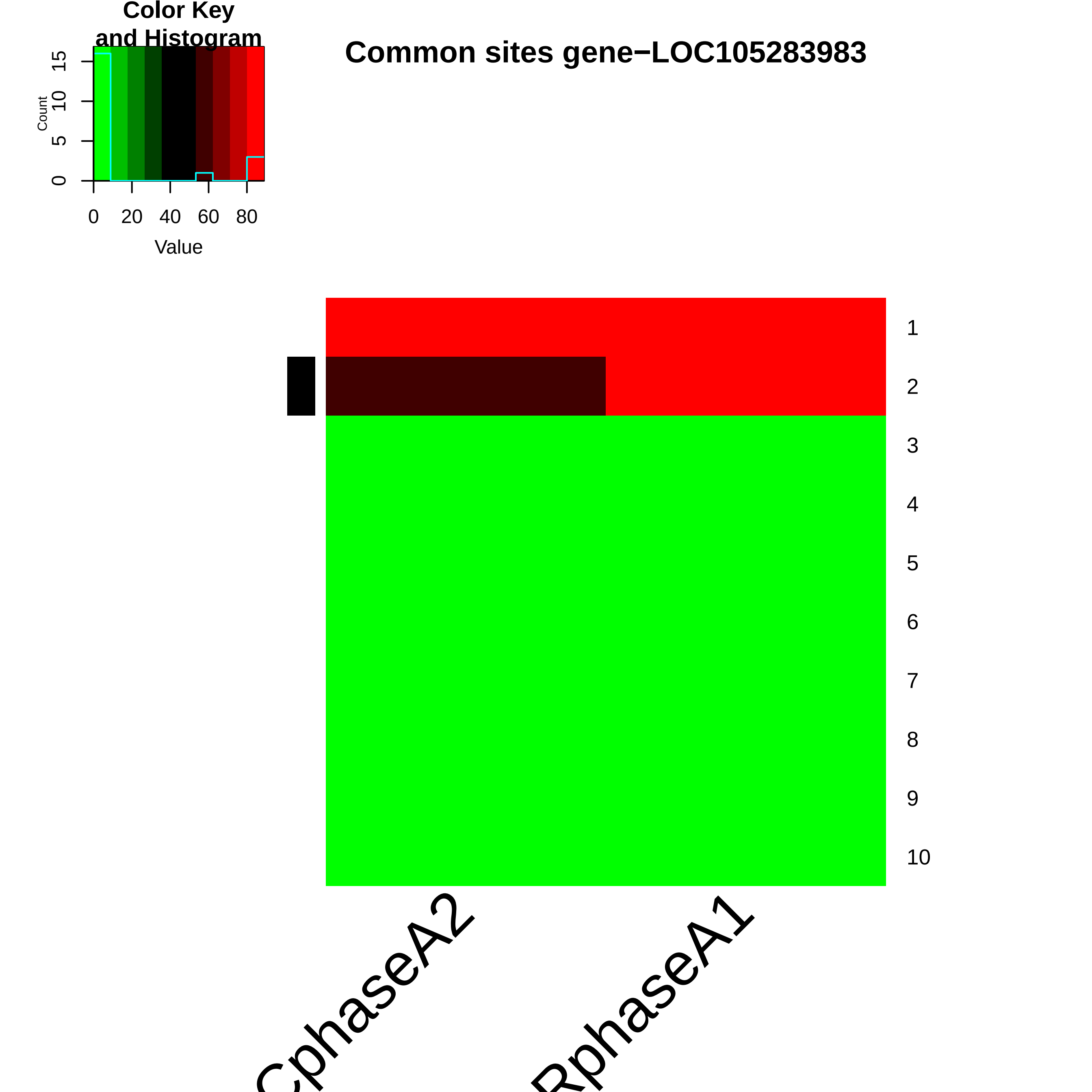
Color Key
and Histogram
Count
0
5
10
15
0
20
40
60
80
Value
Common sites gene−LOC105283983
1
2
3
4
5
6
7
8
9
10

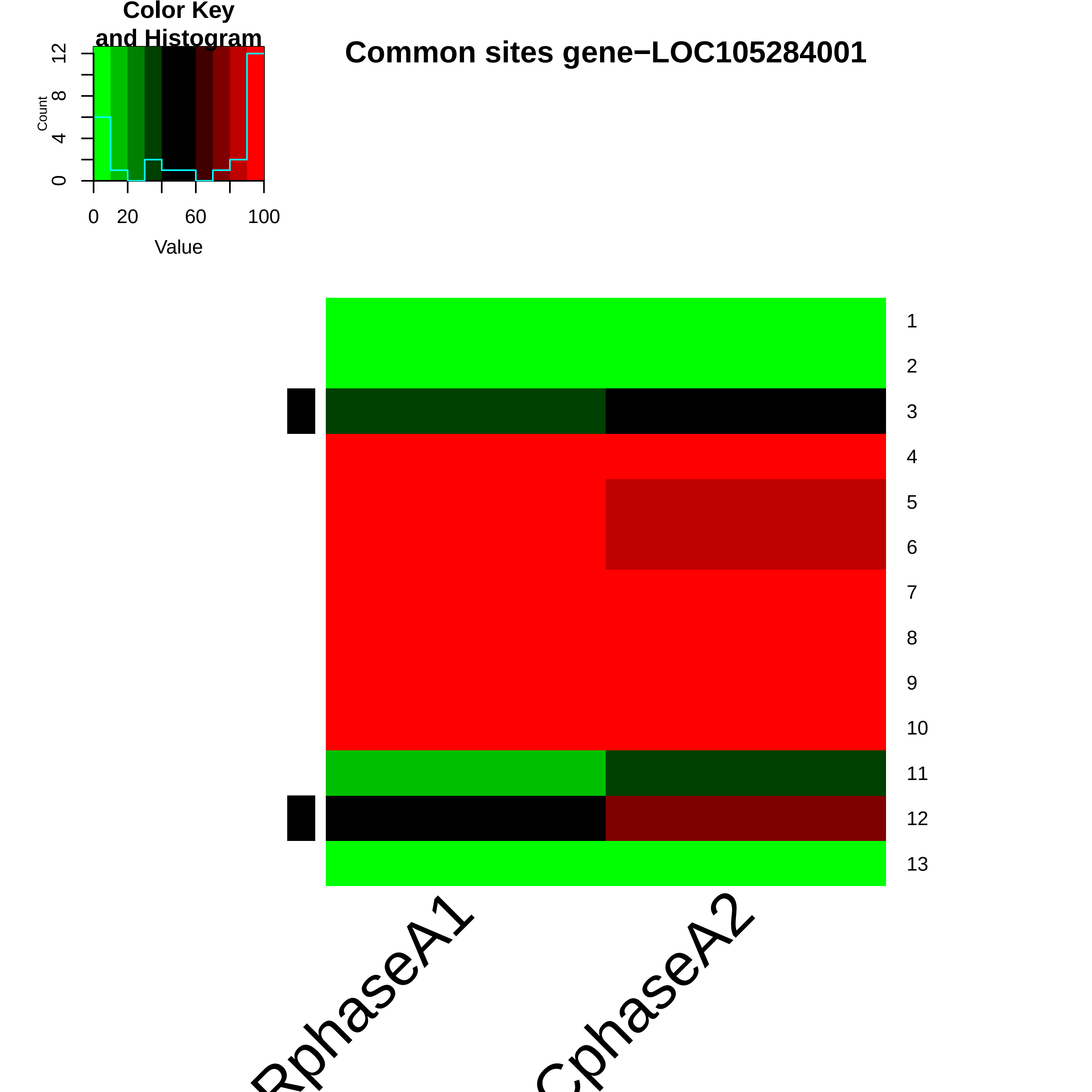
Color Key
and Histogram
Count
0
4
8
12
0
20
60
100
Value
Common sites gene−LOC105284001
1
2
3
4
5
6
7
8
9
10
11
12
13
RphaseA1
CphaseA2

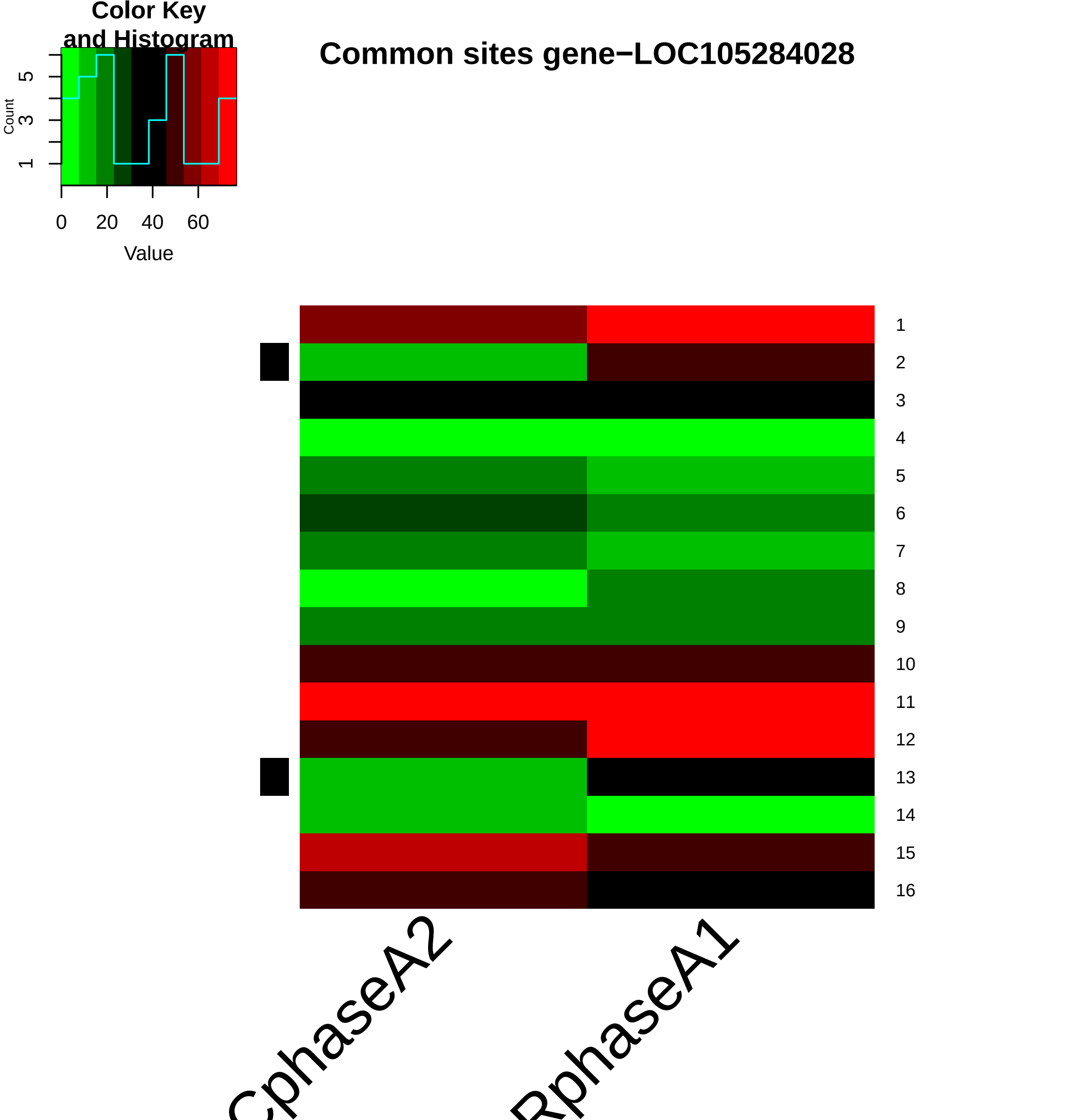
Color Key
and Histogram
Count
1
3
5
0
20
40
60
Value
Common sites gene−LOC105284028
CphaseA2
RphaseA1

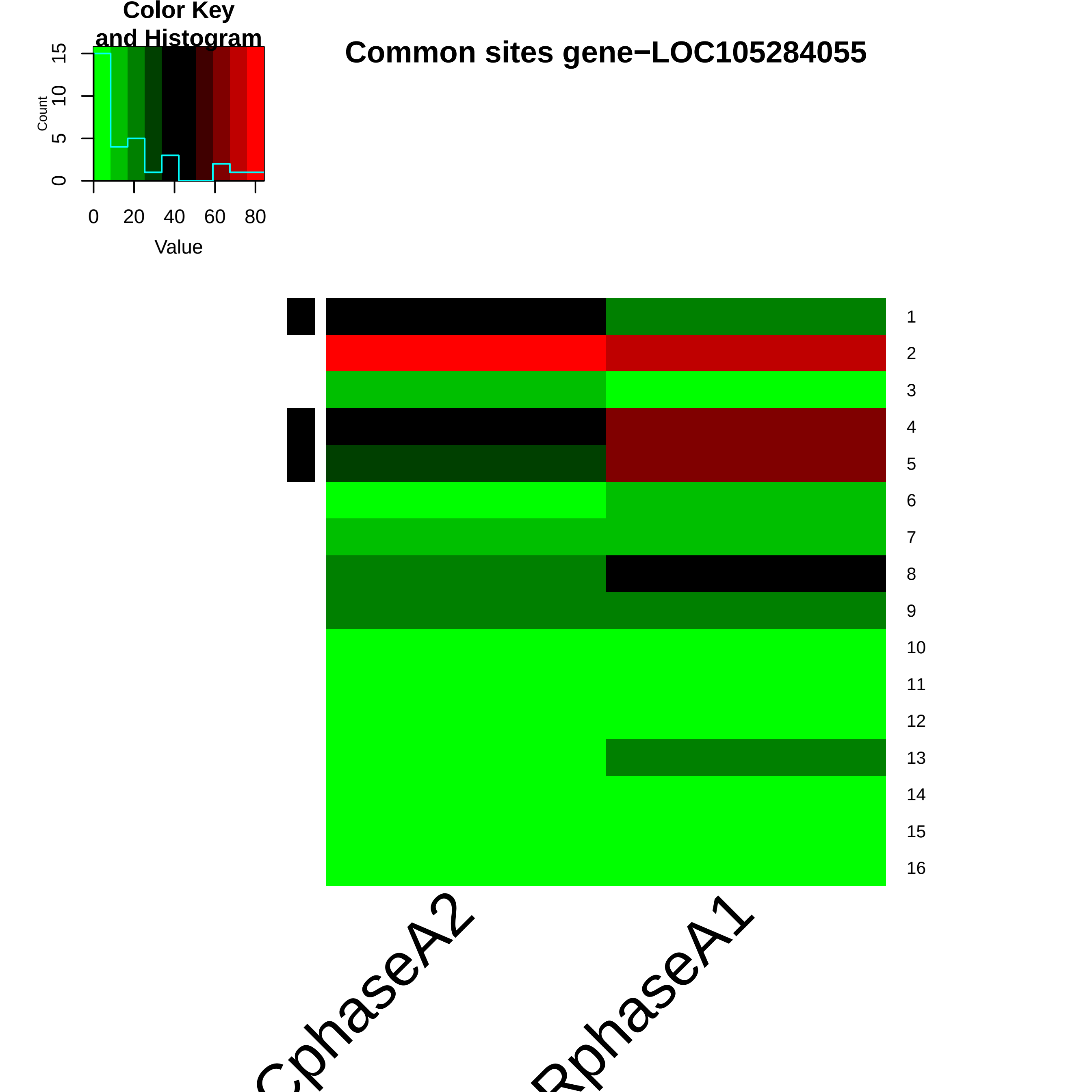

Color Key
and Histogram
Count
0
5
10
15
0
20
40
60
80
Value
Common sites gene−LOC105284055
CphaseA2
RphaseA1

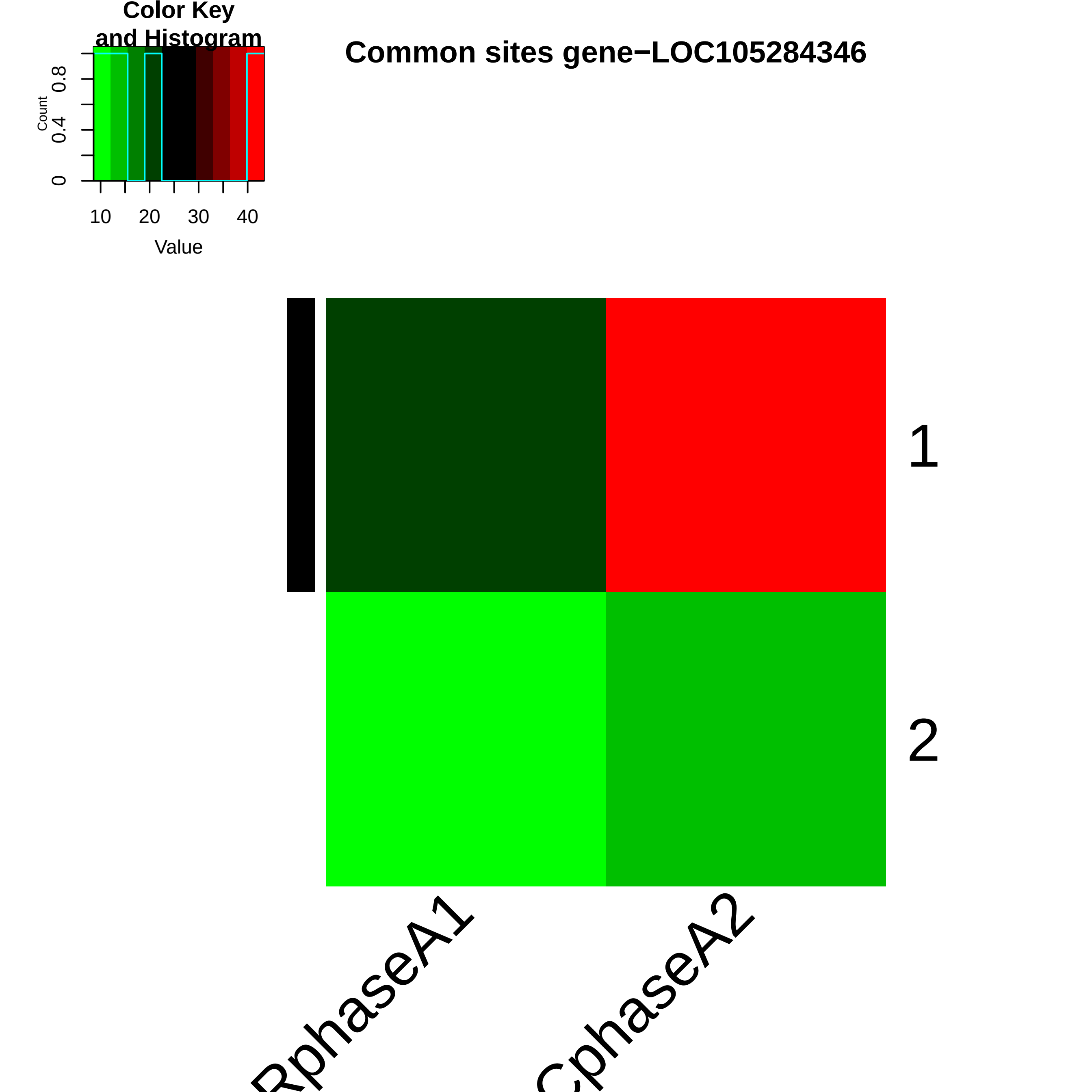

Color Key
and Histogram
Count
0
0.4
0.8
10
20
30
40
Value
Common sites gene−LOC105284346
1
2
RphaseA1
CphaseA2

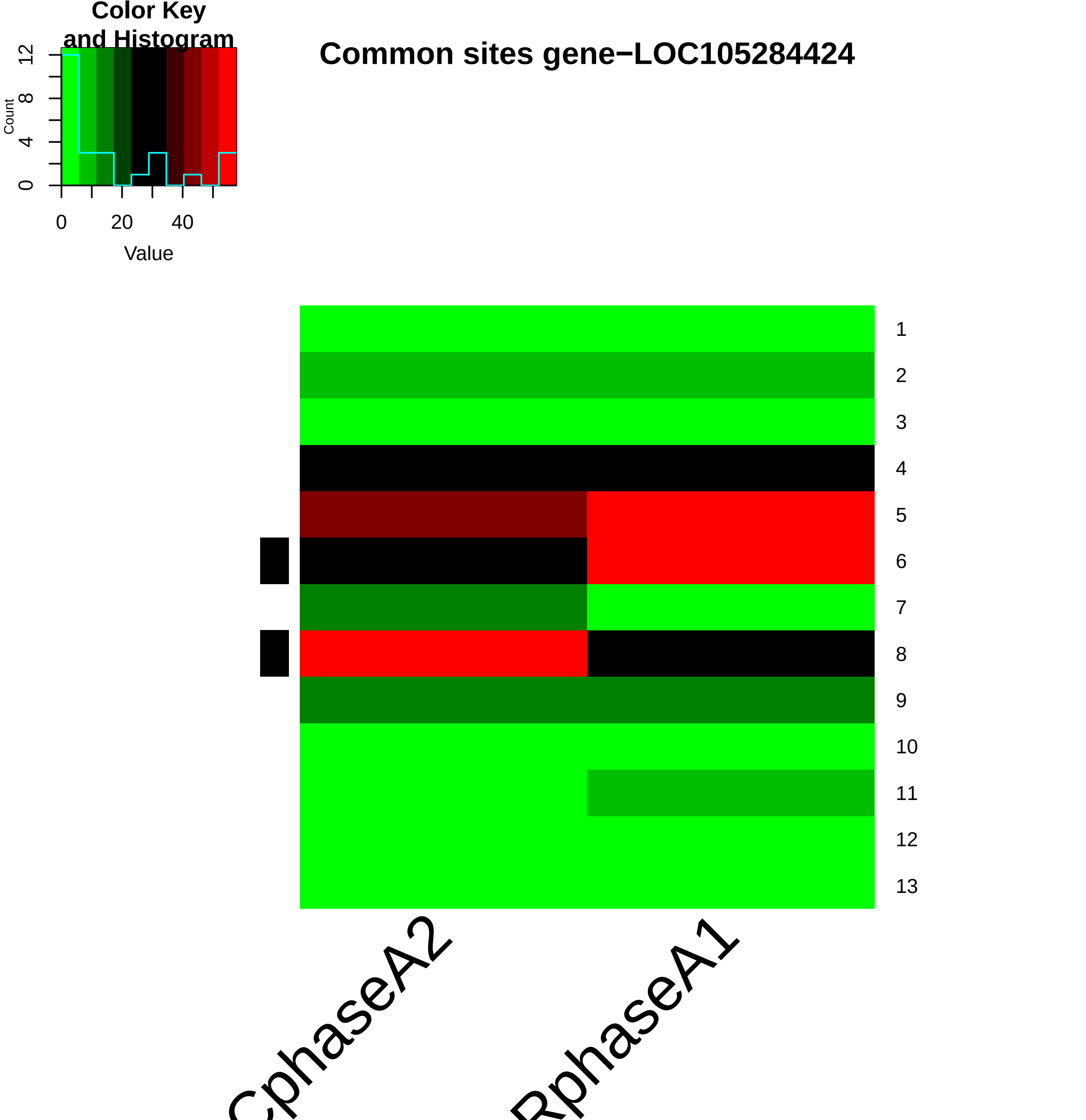
Color Key
and Histogram
Count
0
4
8
12
0
20
40
Value
Common sites gene−LOC105284424
1
2
3
4
5
6
7
8
9
10
11
12
13
CphaseA2
RphaseA1

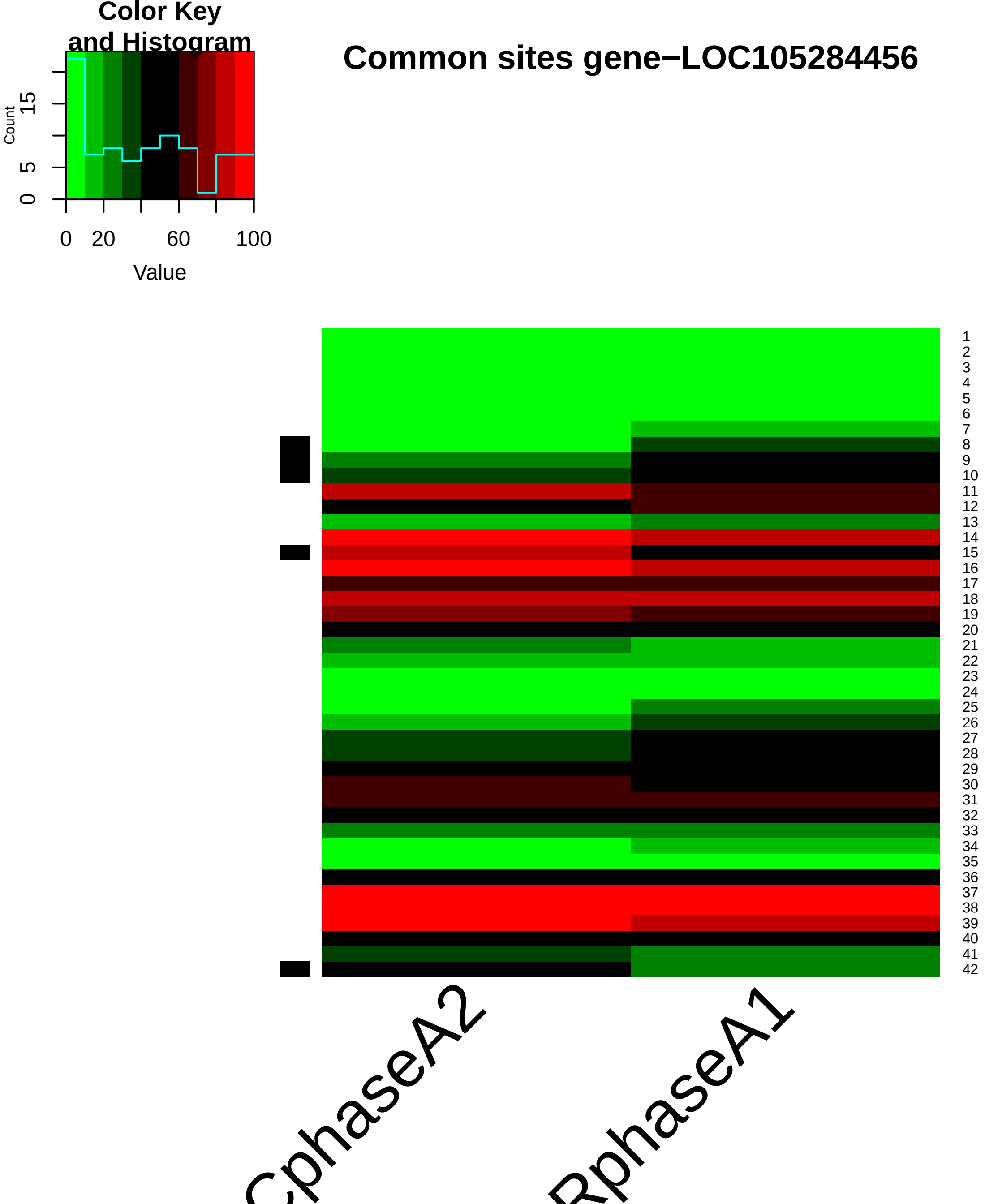
Color Key
and Histogram
Count
0
5
15
0
20
60
100
Value
Common sites gene−LOC105284456
CphaseA2
RphaseA1

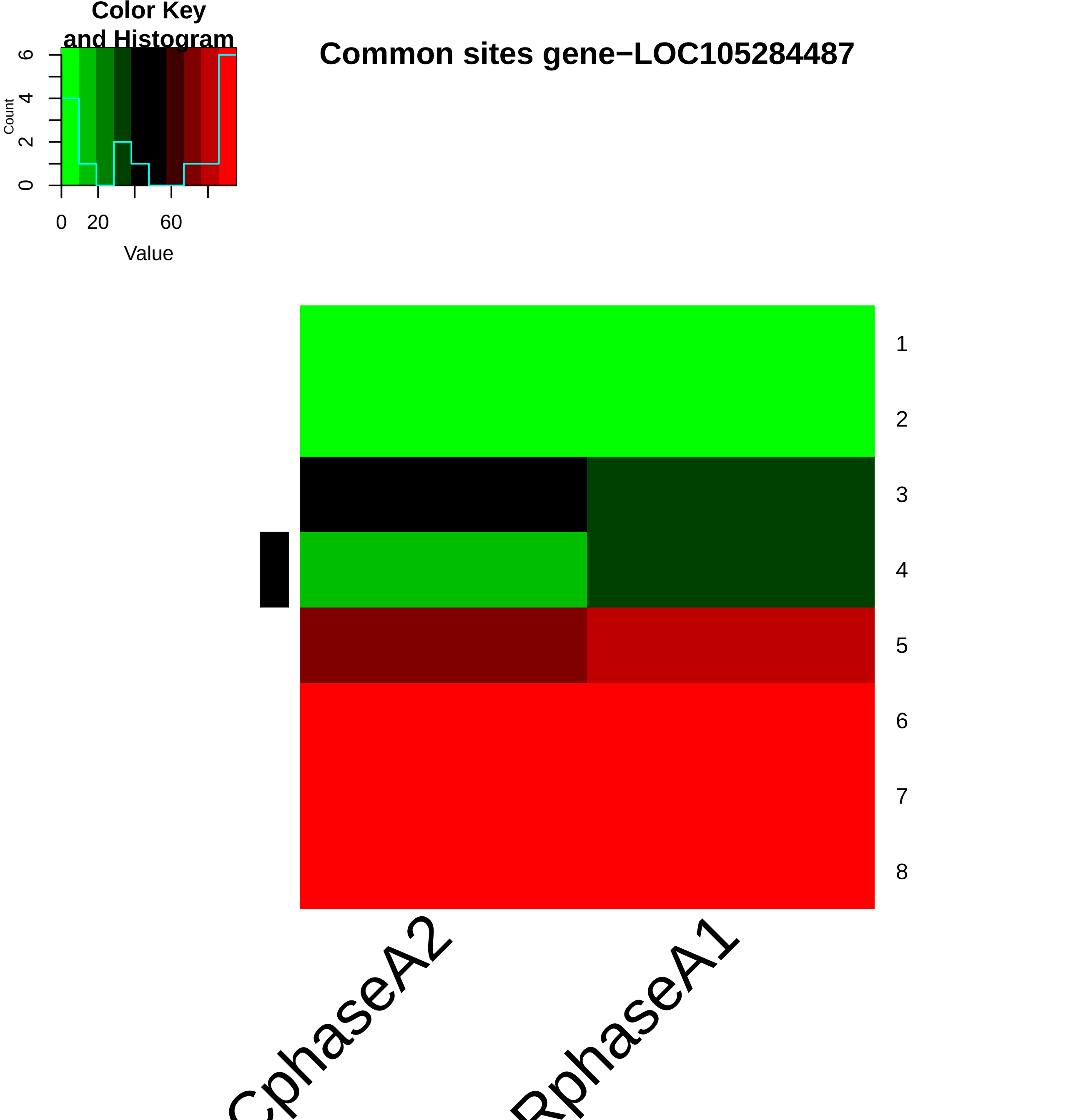
Color Key
and Histogram
Count
0
2
4
6
0
20
60
Value
Common sites gene−LOC105284487
1
2
3
4
5
6
7
8
CphaseA2
RphaseA1

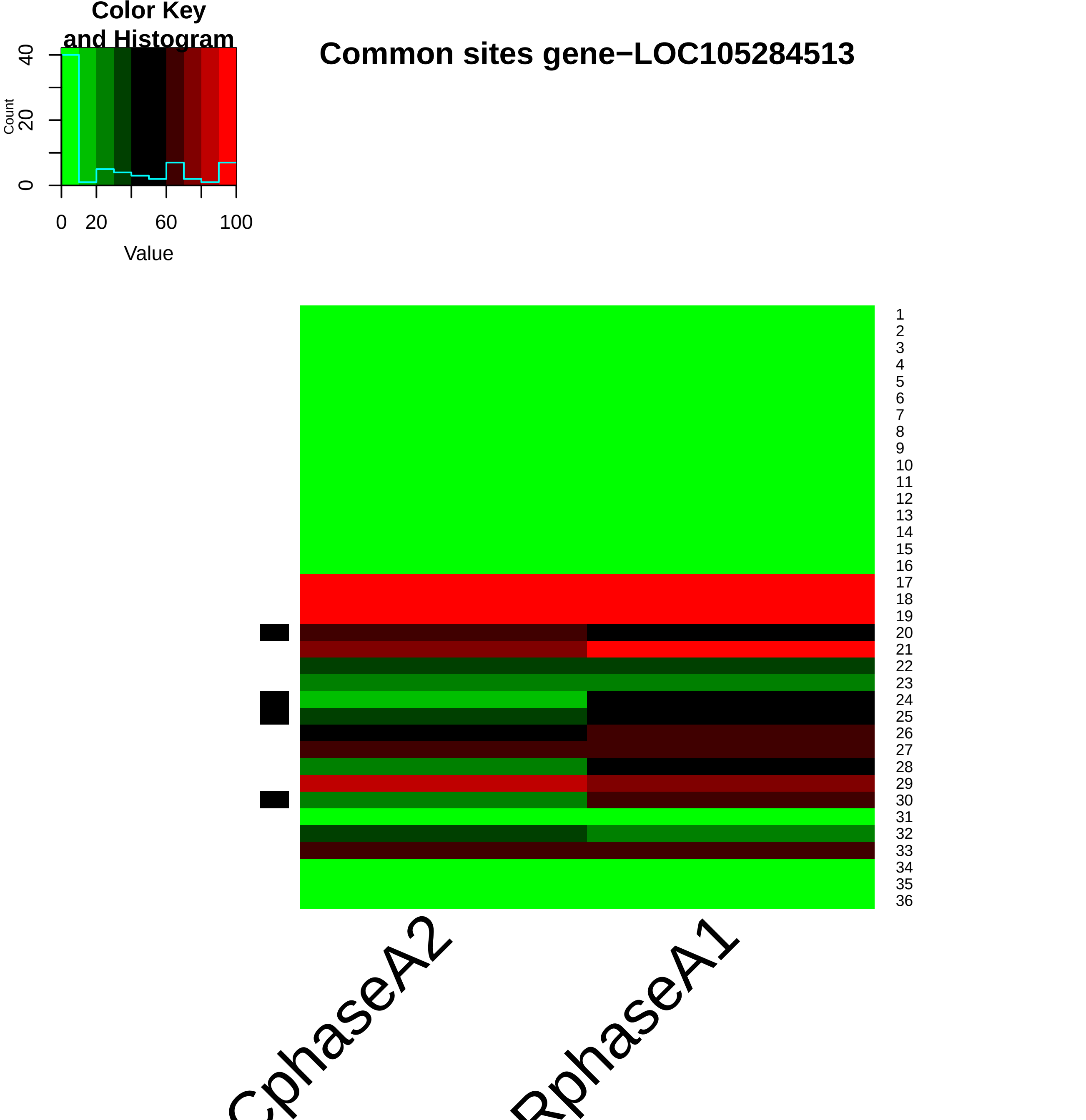
Color Key
and Histogram
Count
0
20
40
0
20
60
100
Value
Common sites gene−LOC105284513
CphaseA2
RphaseA1

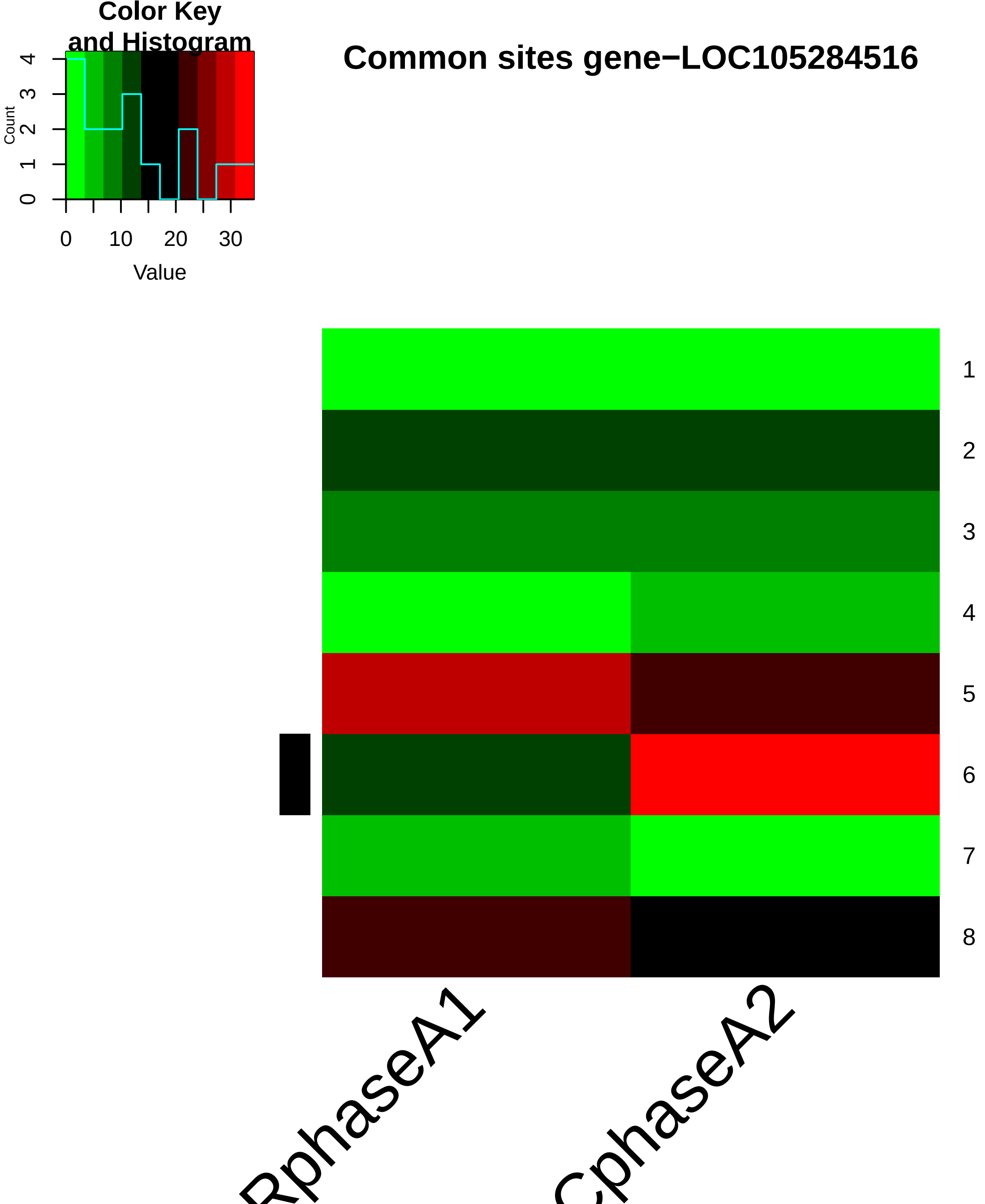
Color Key
and Histogram
Count
0
1
2
3
4
0
10
20
30
Value
Common sites gene−LOC105284516
1
2
3
4
5
6
7
8
RphaseA1
CphaseA2

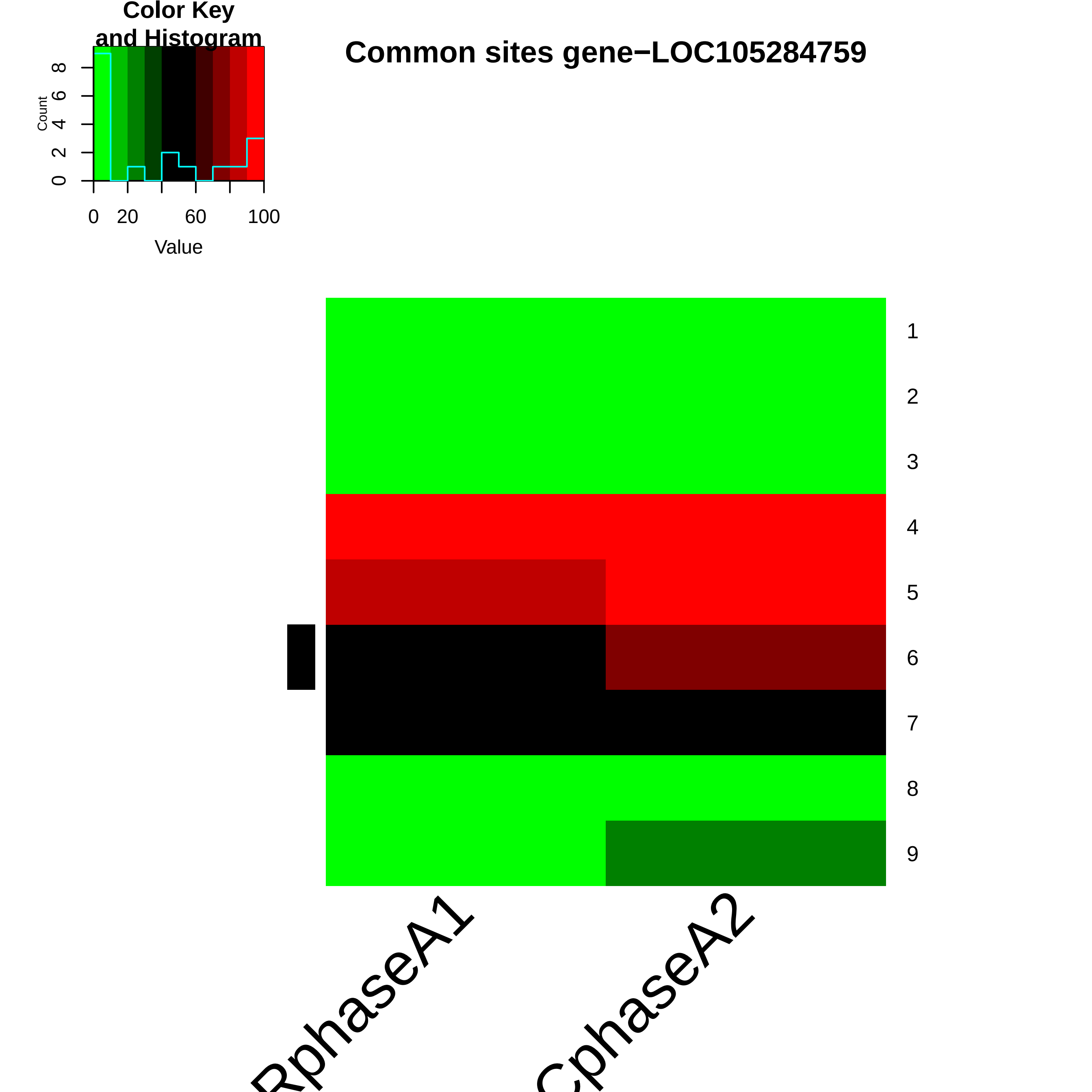

Color Key
and Histogram
Count
0
2
4
6
8
0
20
60
100
Value
Common sites gene−LOC105284759
1
2
3
4
5
6
7
8
9
RphaseA1
CphaseA2

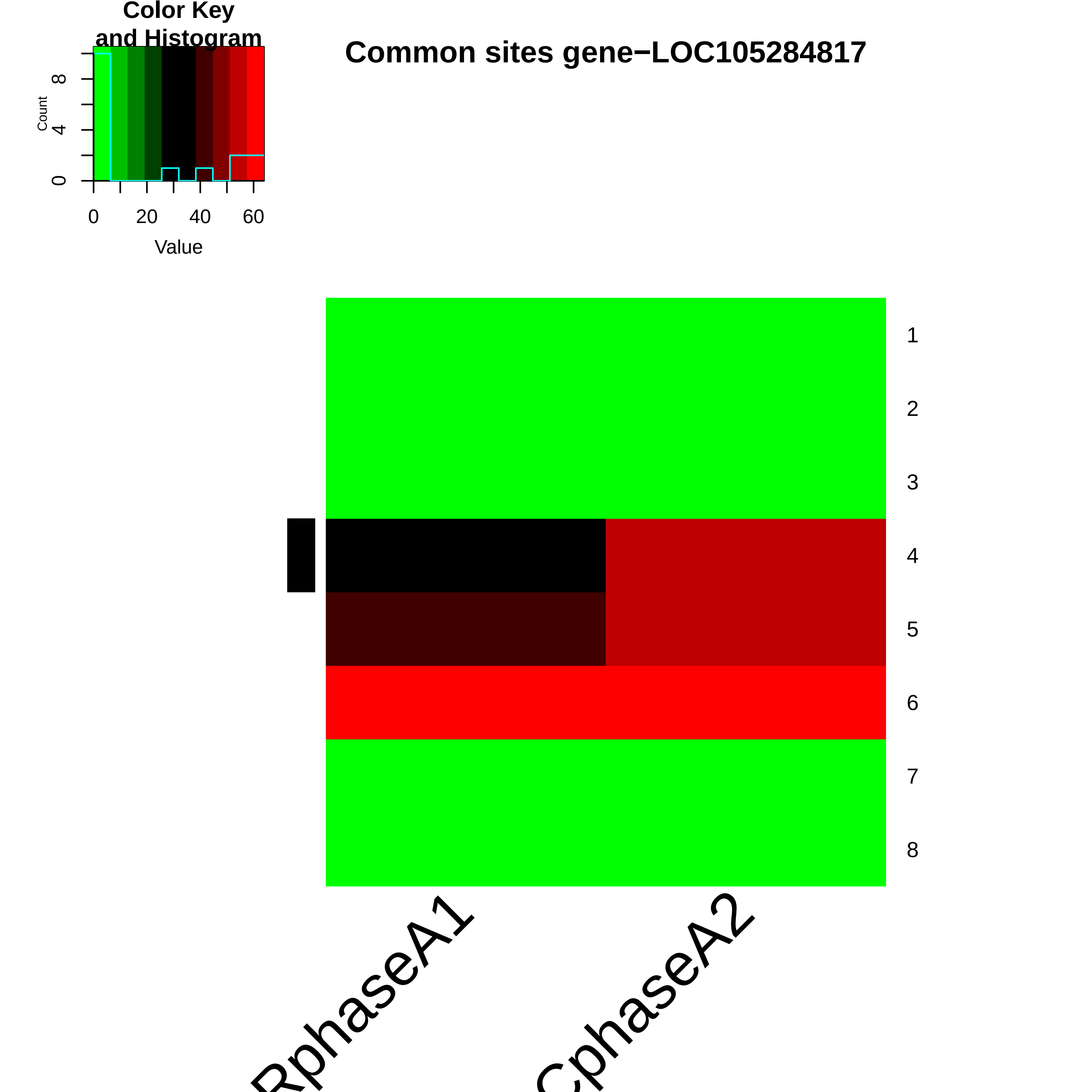
Color Key
and Histogram
Count
0
4
8
0
20
40
60
Value
Common sites gene−LOC105284817
1
2
3
4
5
6
7
8
RphaseA1
CphaseA2

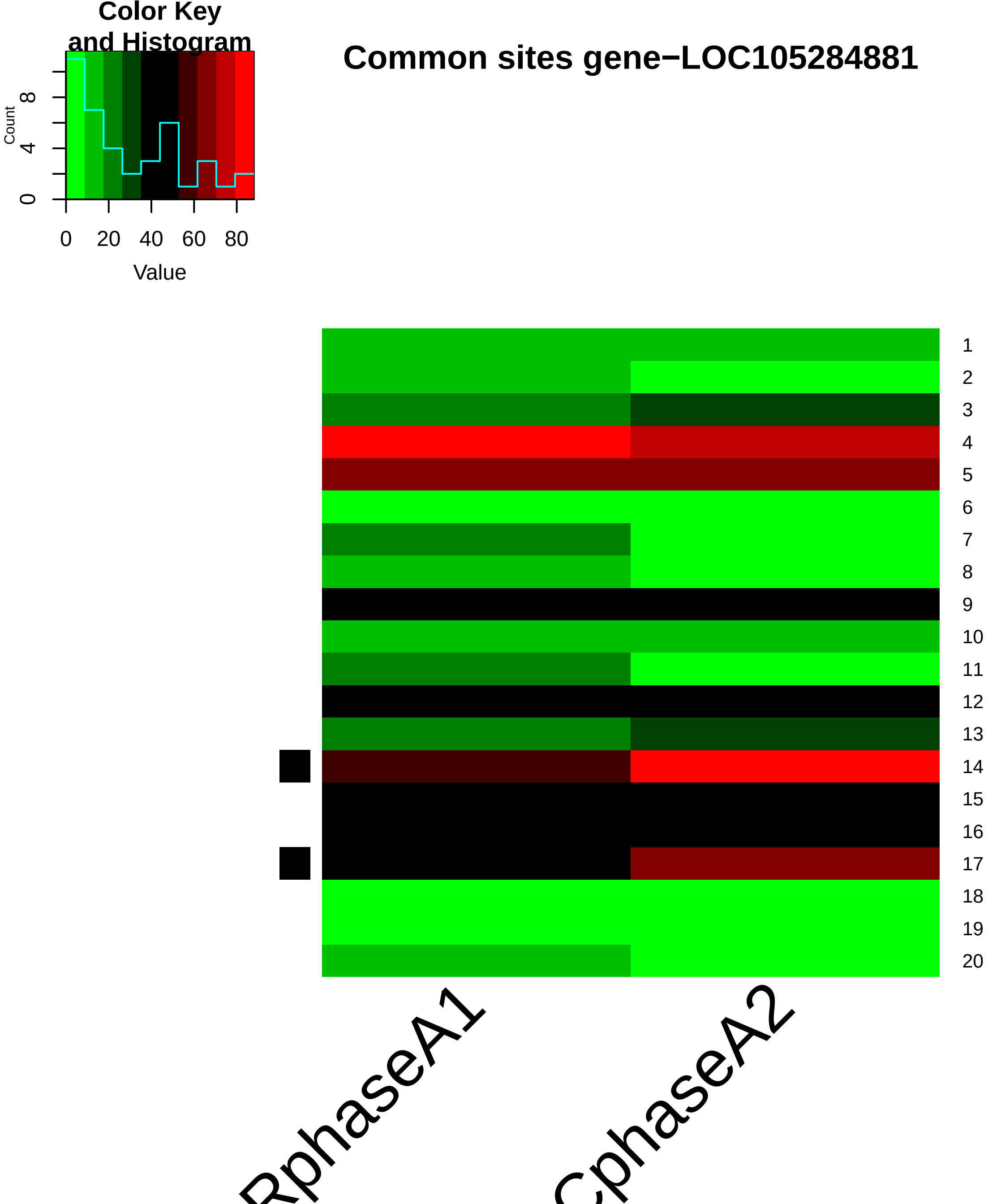
Color Key
and Histogram
Count
0
4
8
0
20
40
60
80
Value
Common sites gene−LOC105284881
RphaseA1
CphaseA2

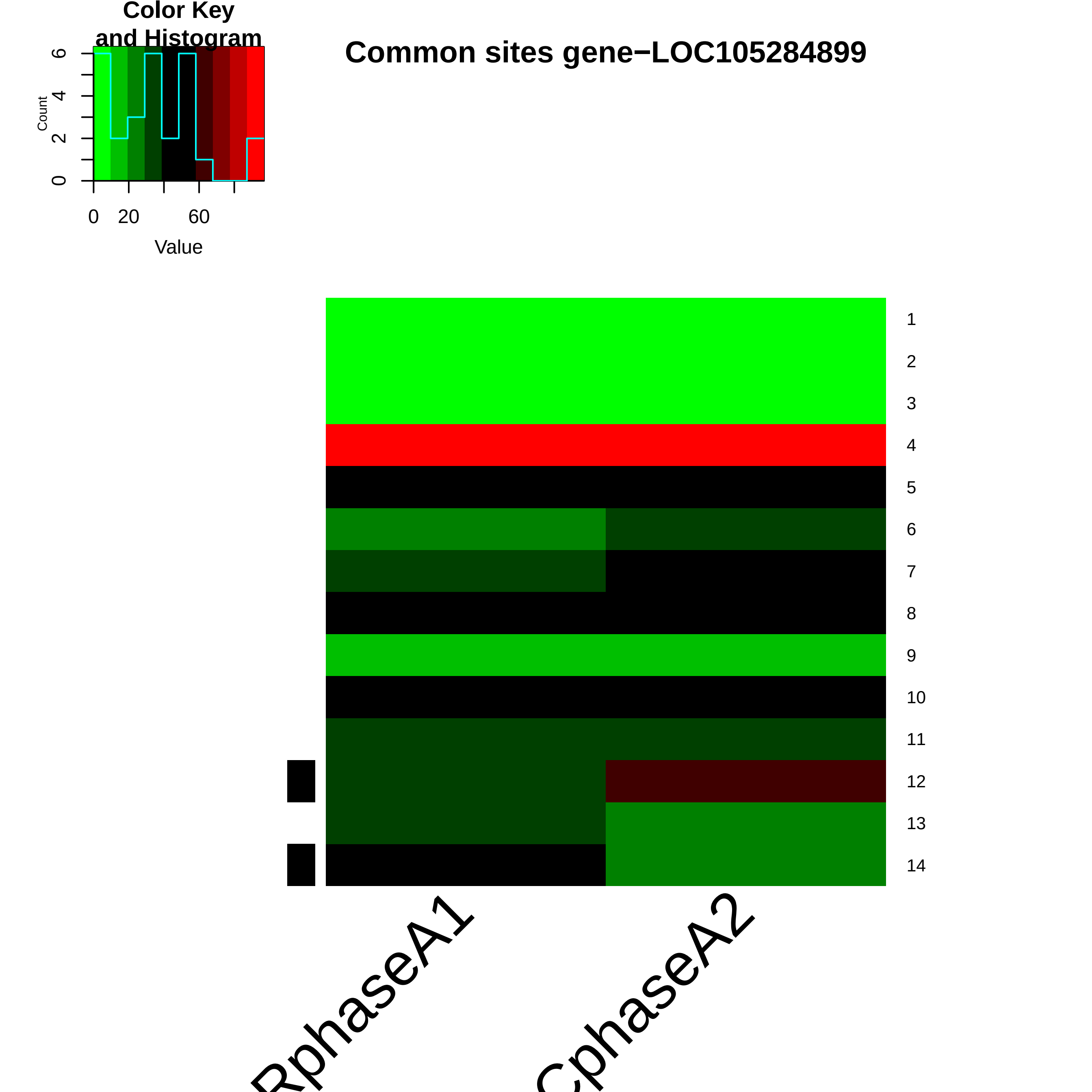
Color Key
and Histogram
Count
0
2
4
6
0
20
60
Value
Common sites gene−LOC105284899
1
2
3
4
5
6
7
8
9
10
11
12
13
14
RphaseA1
CphaseA2

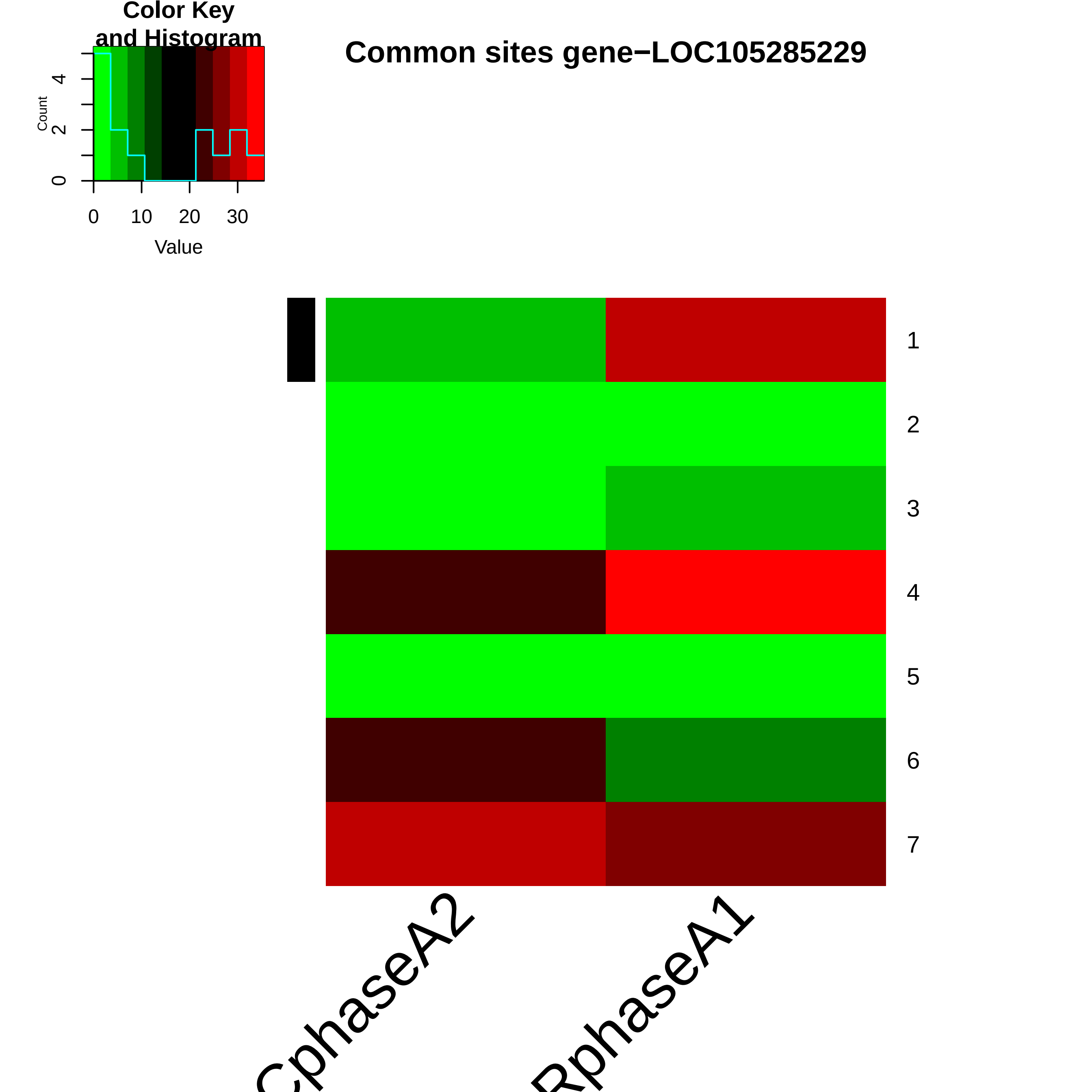
Color Key
and Histogram
Count
0
2
4
0
10
20
30
Value
Common sites gene−LOC105285229
1
2
3
4
5
6
7
CphaseA2
RphaseA1

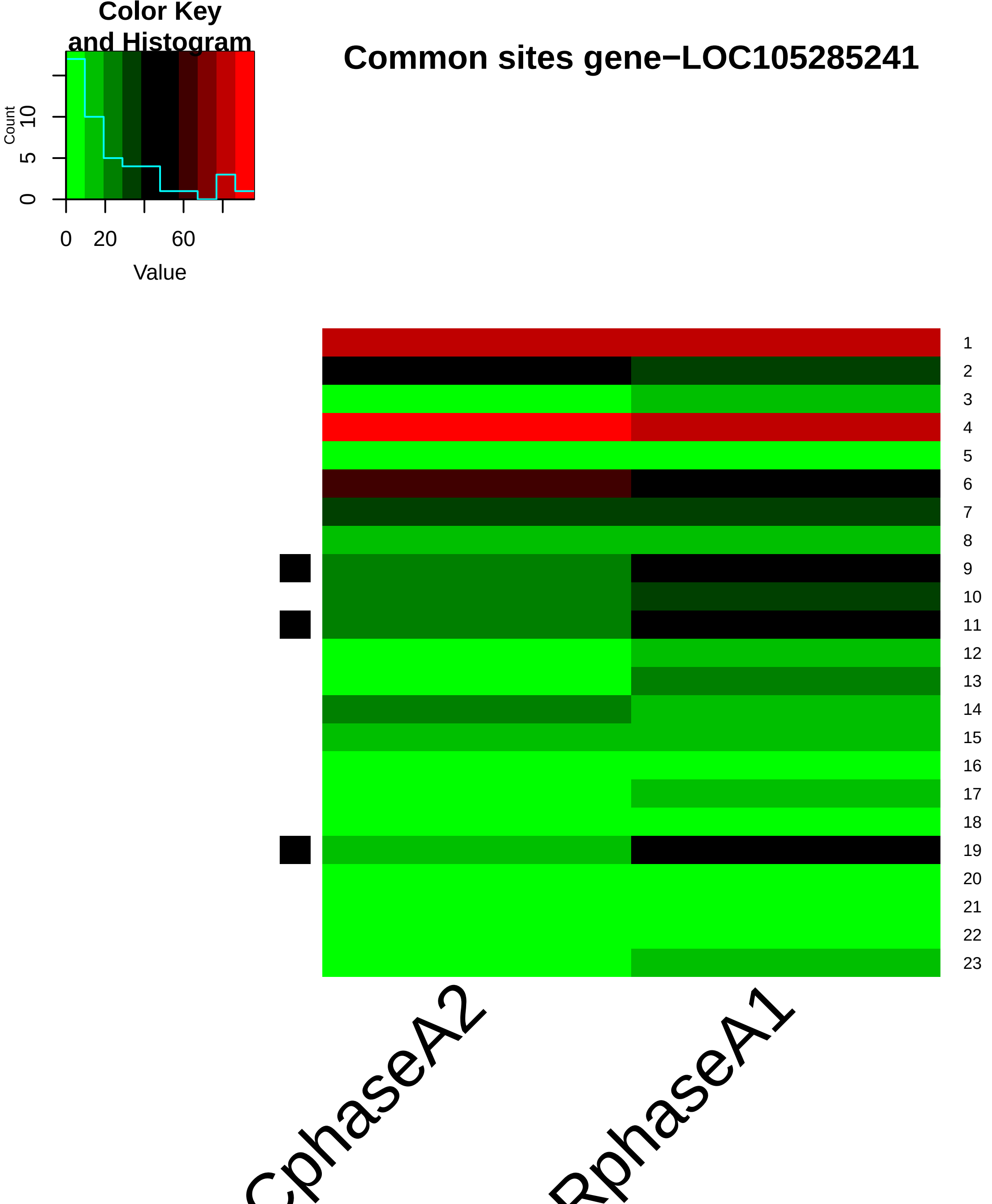
Color Key
and Histogram
Count
0
5
10
0
20
60
Value
Common sites gene−LOC105285241
CphaseA2
RphaseA1

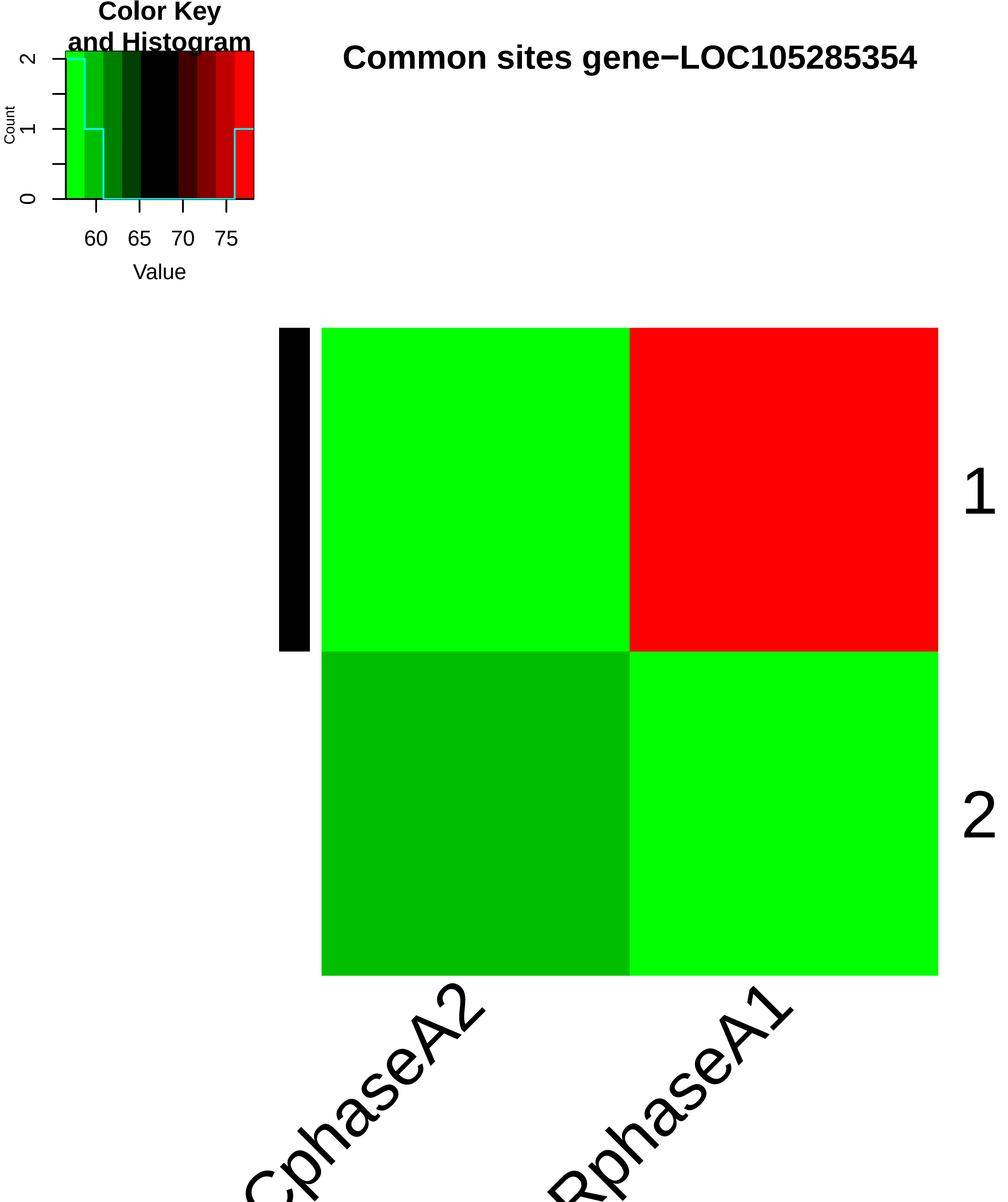
Color Key
and Histogram
Count
0
1
2
60
65
70
75
Value
Common sites gene−LOC105285354
1
2
CphaseA2
RphaseA1

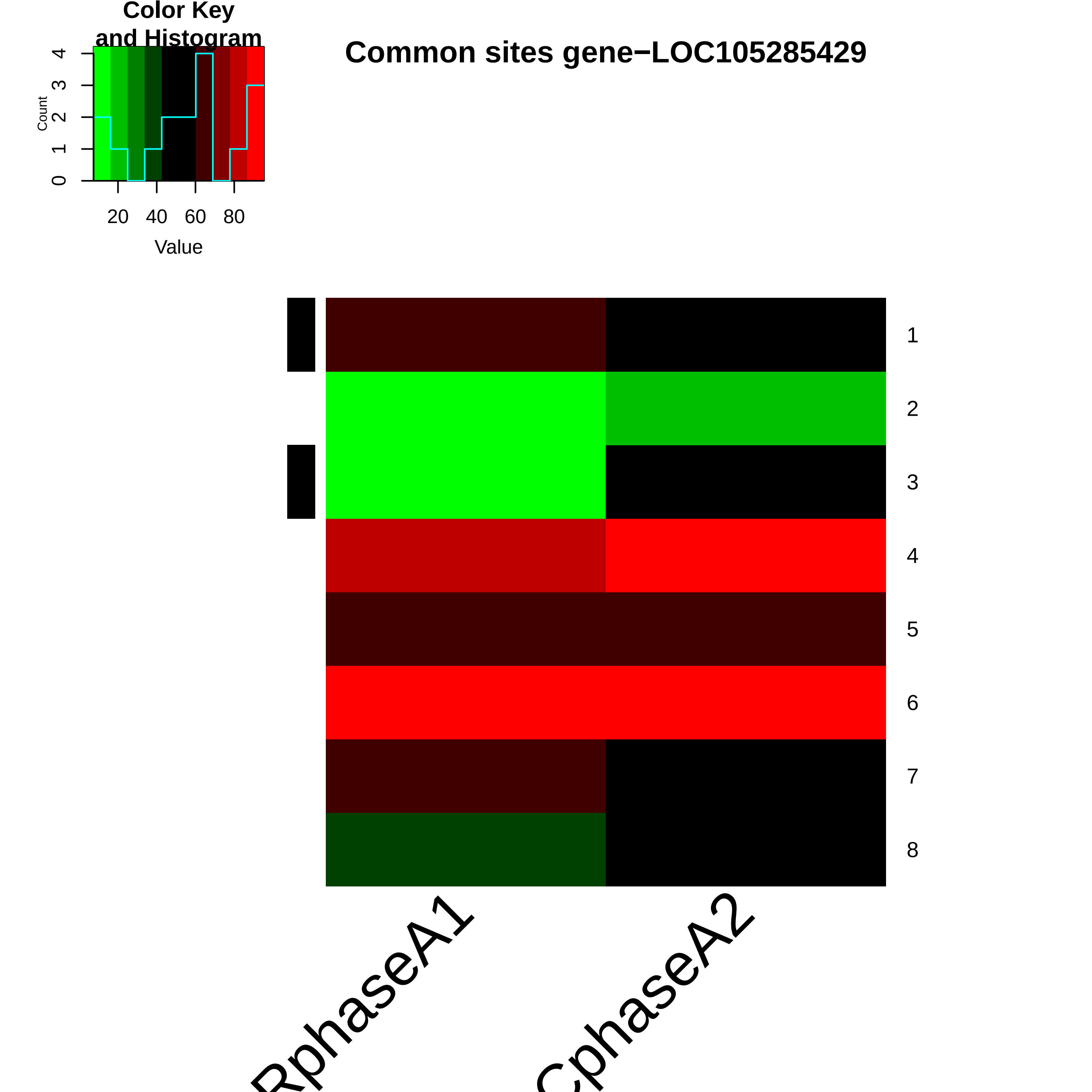
Color Key
and Histogram
Count
Value
Common sites gene−LOC105285429
1
2
3
4
5
6
7
8
RphaseA1
CphaseA2

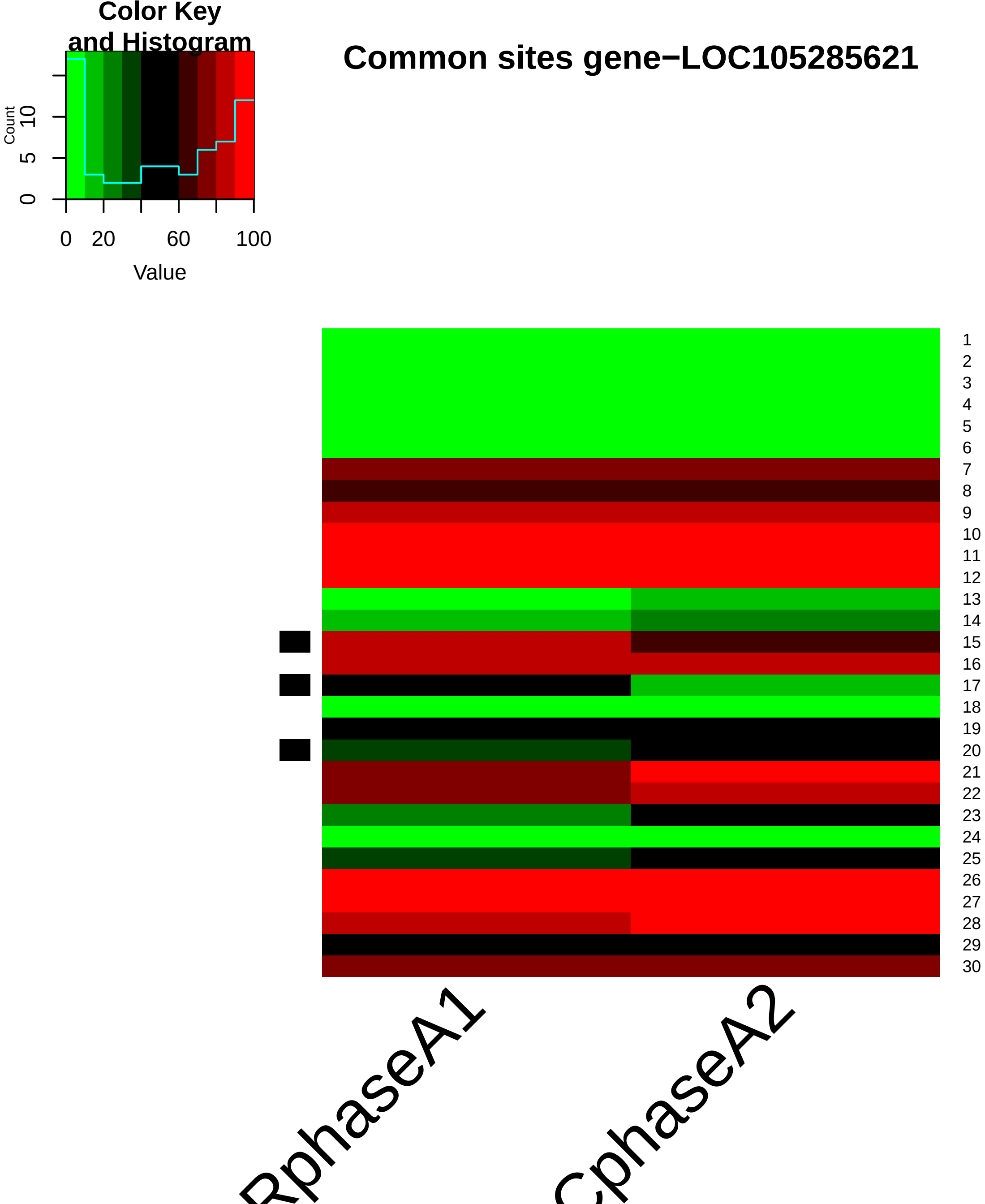
Color Key
and Histogram
Count
0
5
10
0
20
60
100
Value
Common sites gene−LOC105285621
RphaseA1
CphaseA2

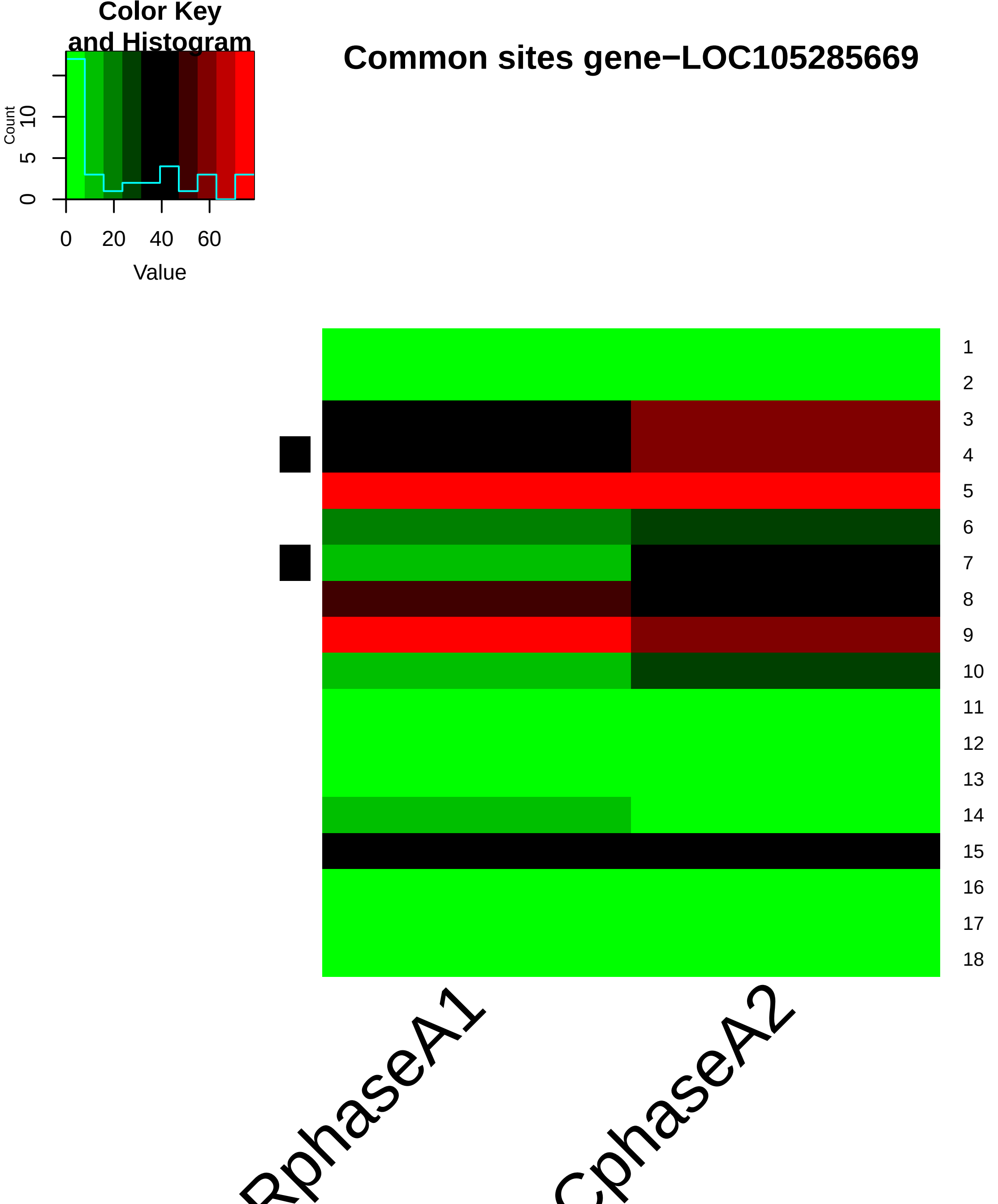
Color Key
and Histogram
Count
0
5
10
0
20
40
60
Value
Common sites gene−LOC105285669
RphaseA1
CphaseA2

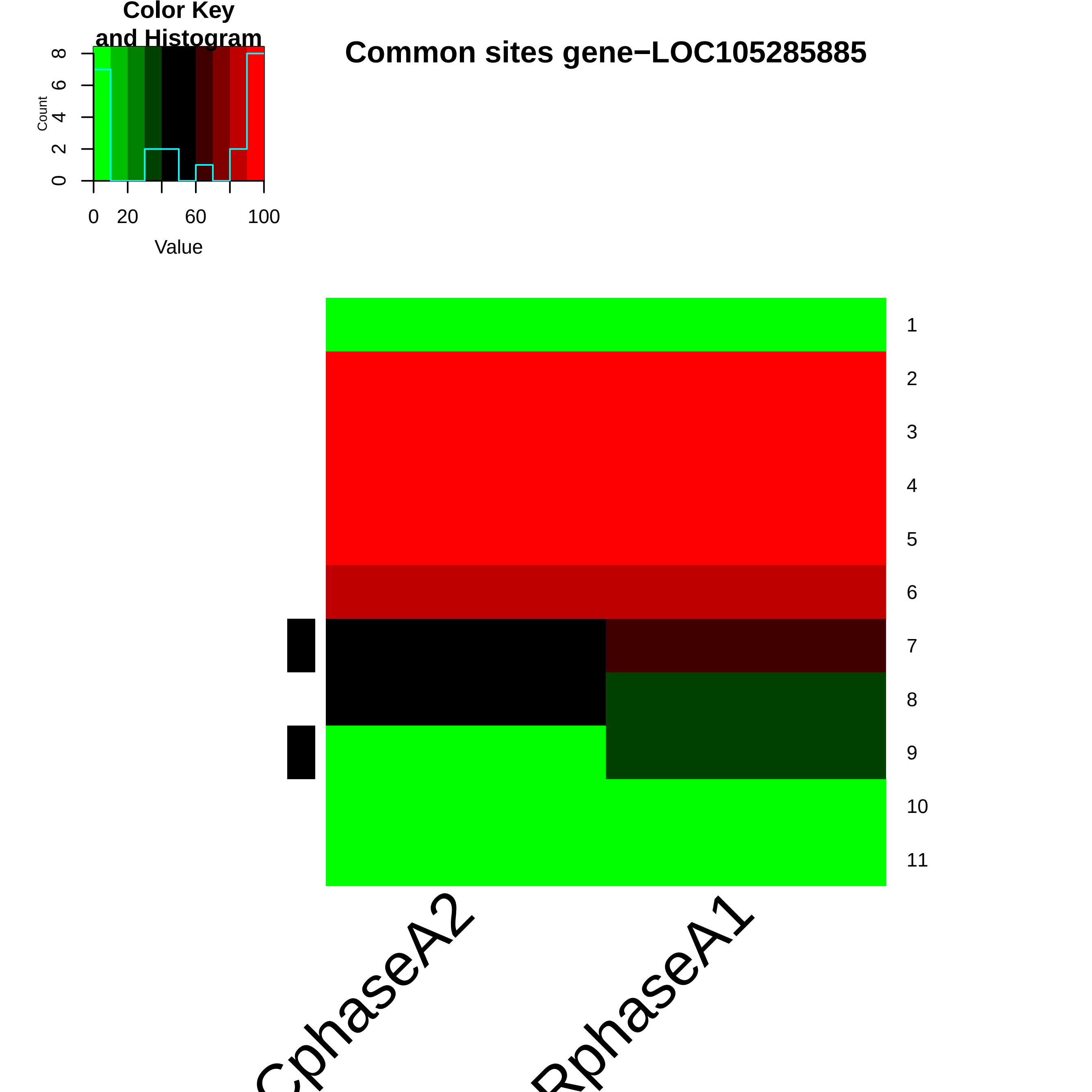
Color Key
and Histogram
Count
0
2
4
6
8
0
20
60
100
Value
Common sites gene−LOC105285885
1
2
3
4
5
6
7
8
9
10
11
CphaseA2
RphaseA1

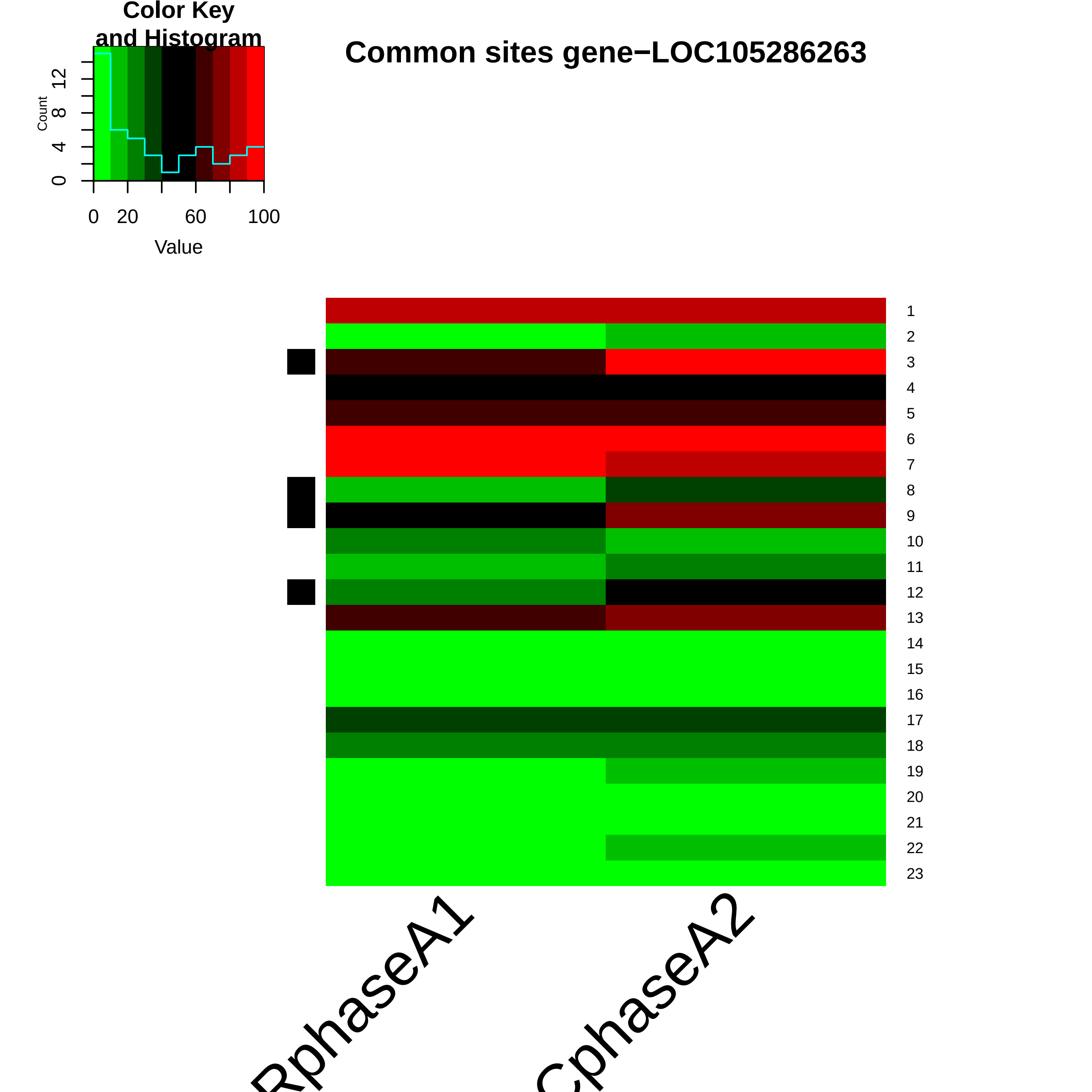

Color Key
and Histogram
Count
0
4
8
12
0
20
60
100
Value
Common sites gene−LOC105286263
RphaseA1
CphaseA2

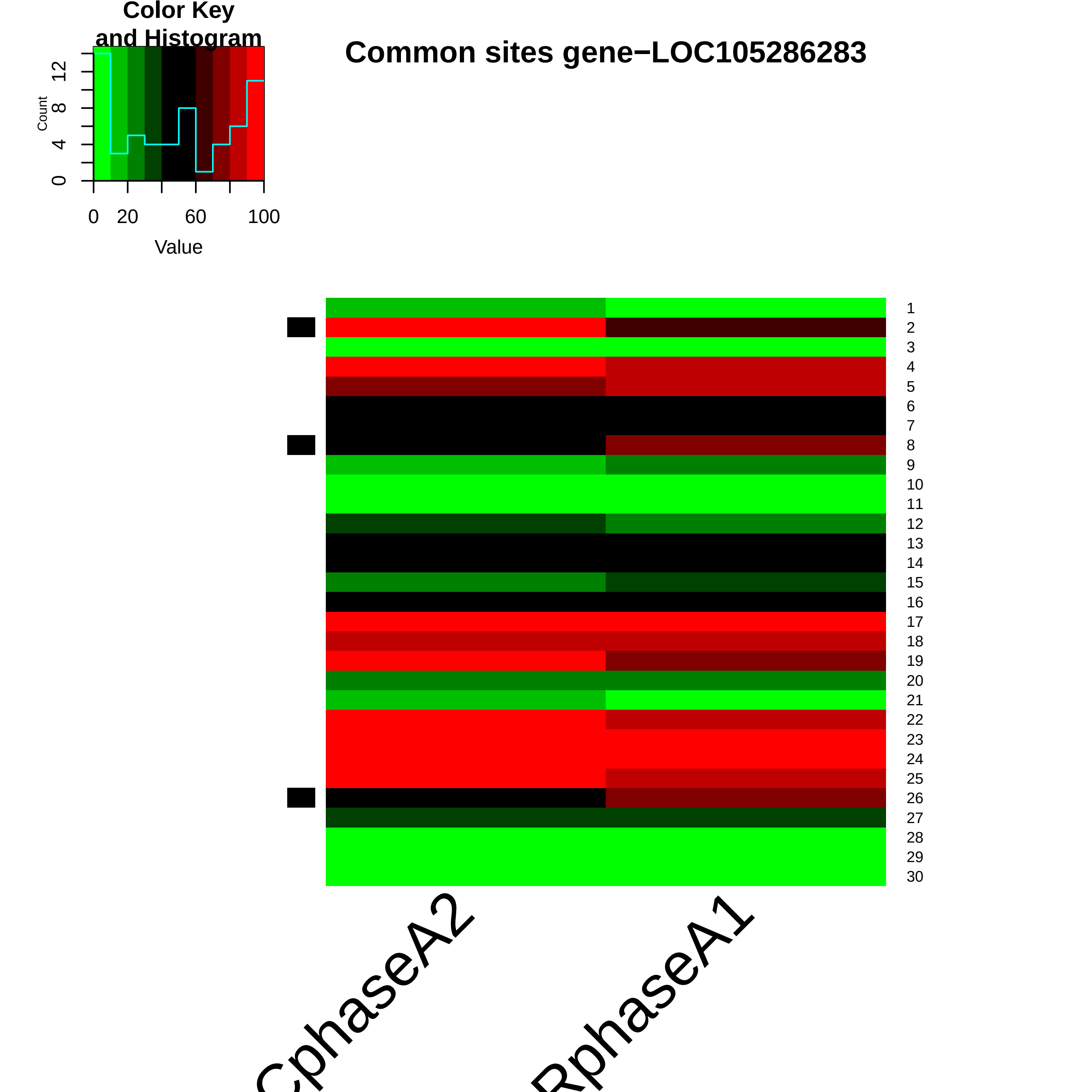

Color Key
and Histogram
Count
0
4
8
12
0
20
60
100
Value
Common sites gene−LOC105286283
CphaseA2
RphaseA1

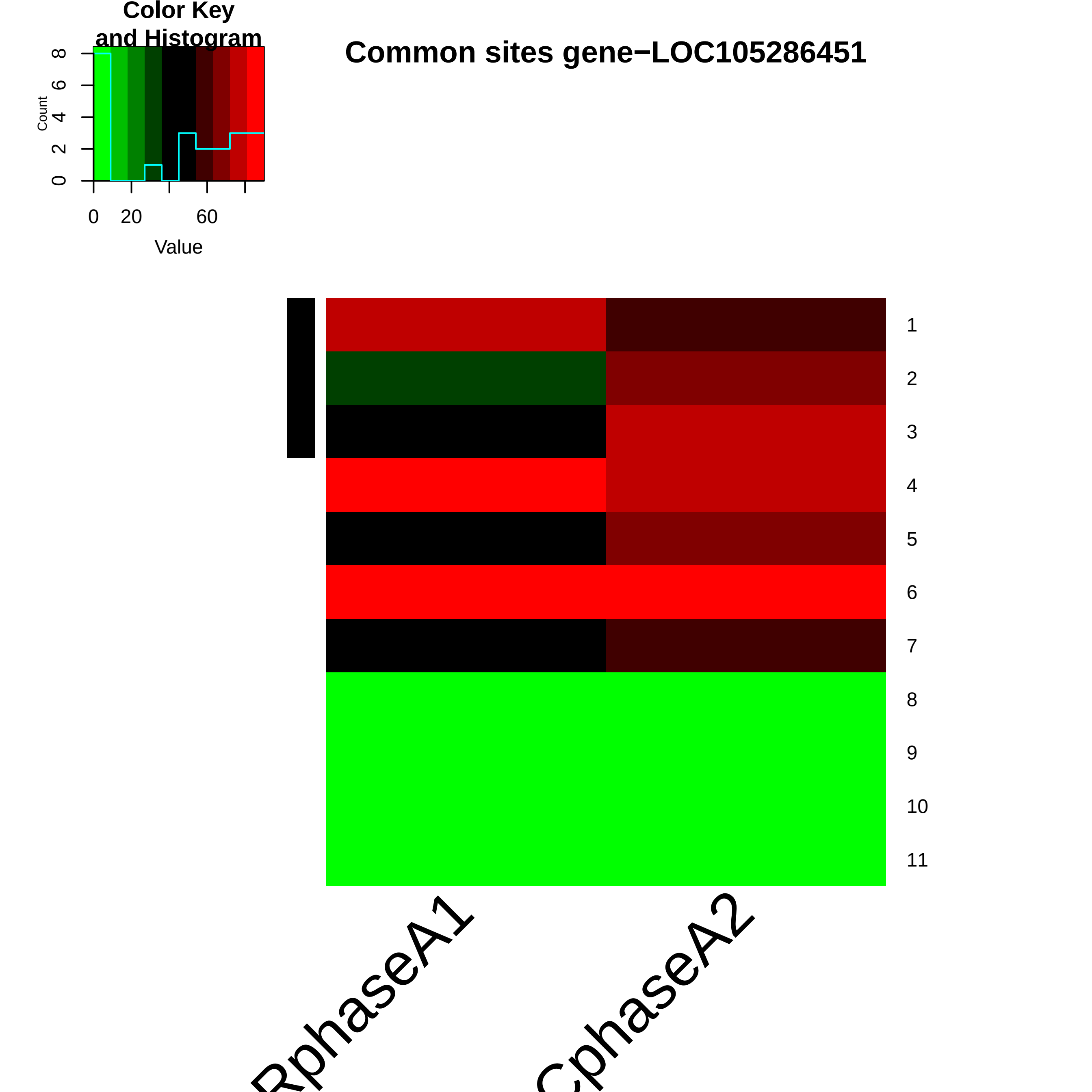

Color Key
and Histogram
Count
0
2
4
6
8
0
20
60
Value
Common sites gene−LOC105286451
1
2
3
4
5
6
7
8
9
10
11
RphaseA1
CphaseA2

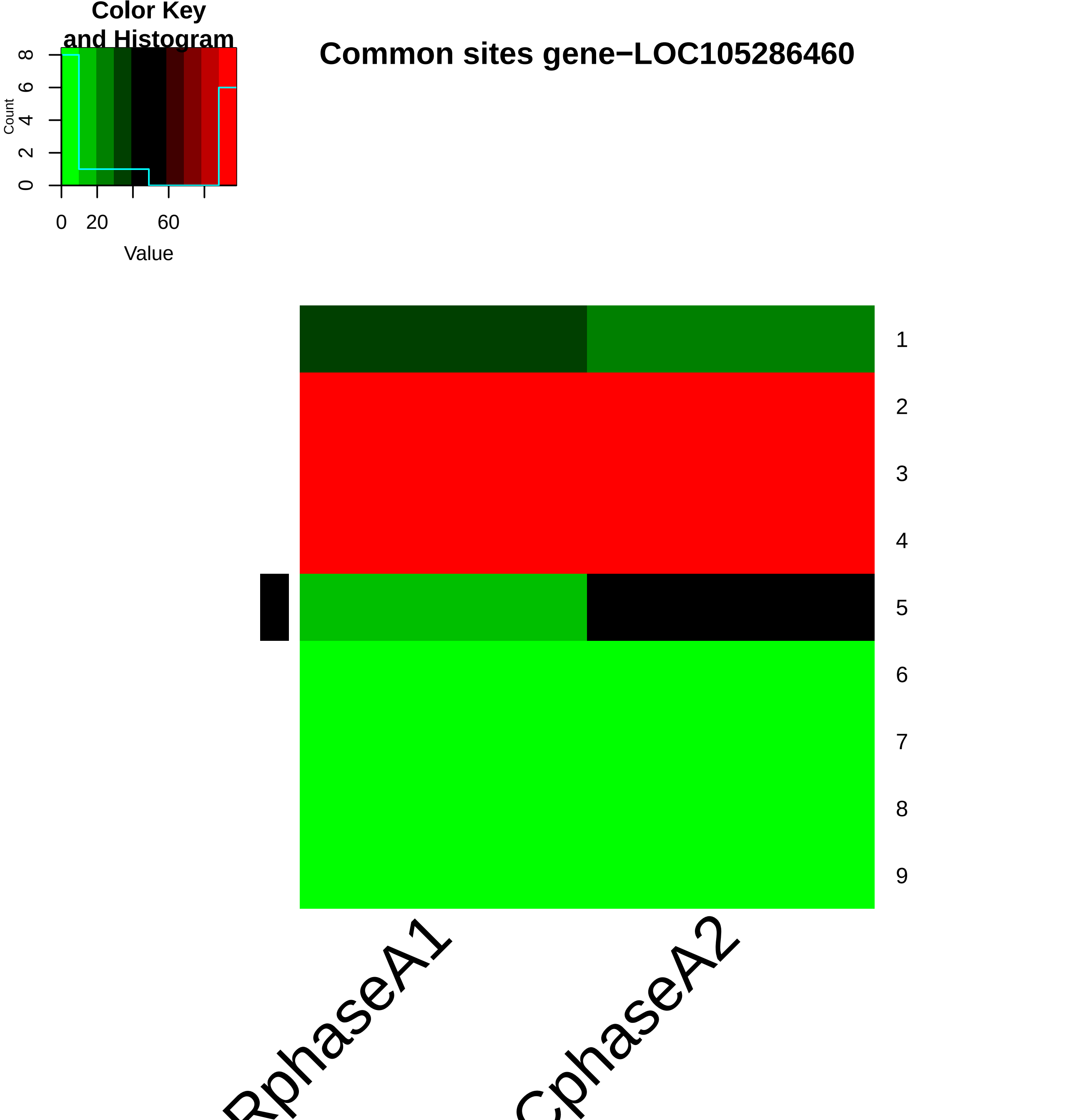

Color Key
and Histogram
Count
0
2
4
6
8
0
20
60
Value
Common sites gene−LOC105286460
1
2
3
4
5
6
7
8
9
RphaseA1
CphaseA2

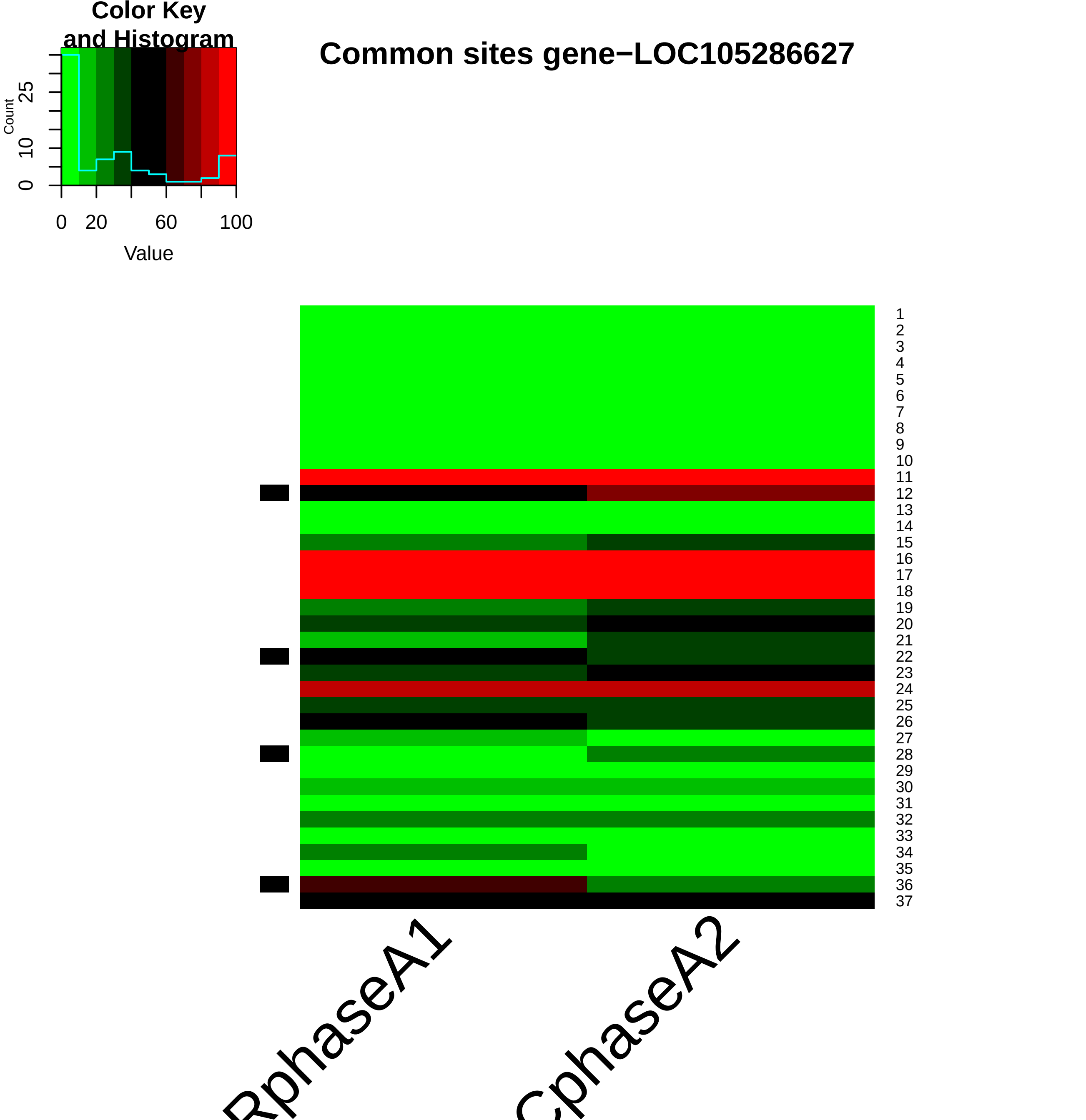
Color Key
and Histogram
Count
0
10
25
0
20
60
100
Value
Common sites gene−LOC105286627
RphaseA1
CphaseA2

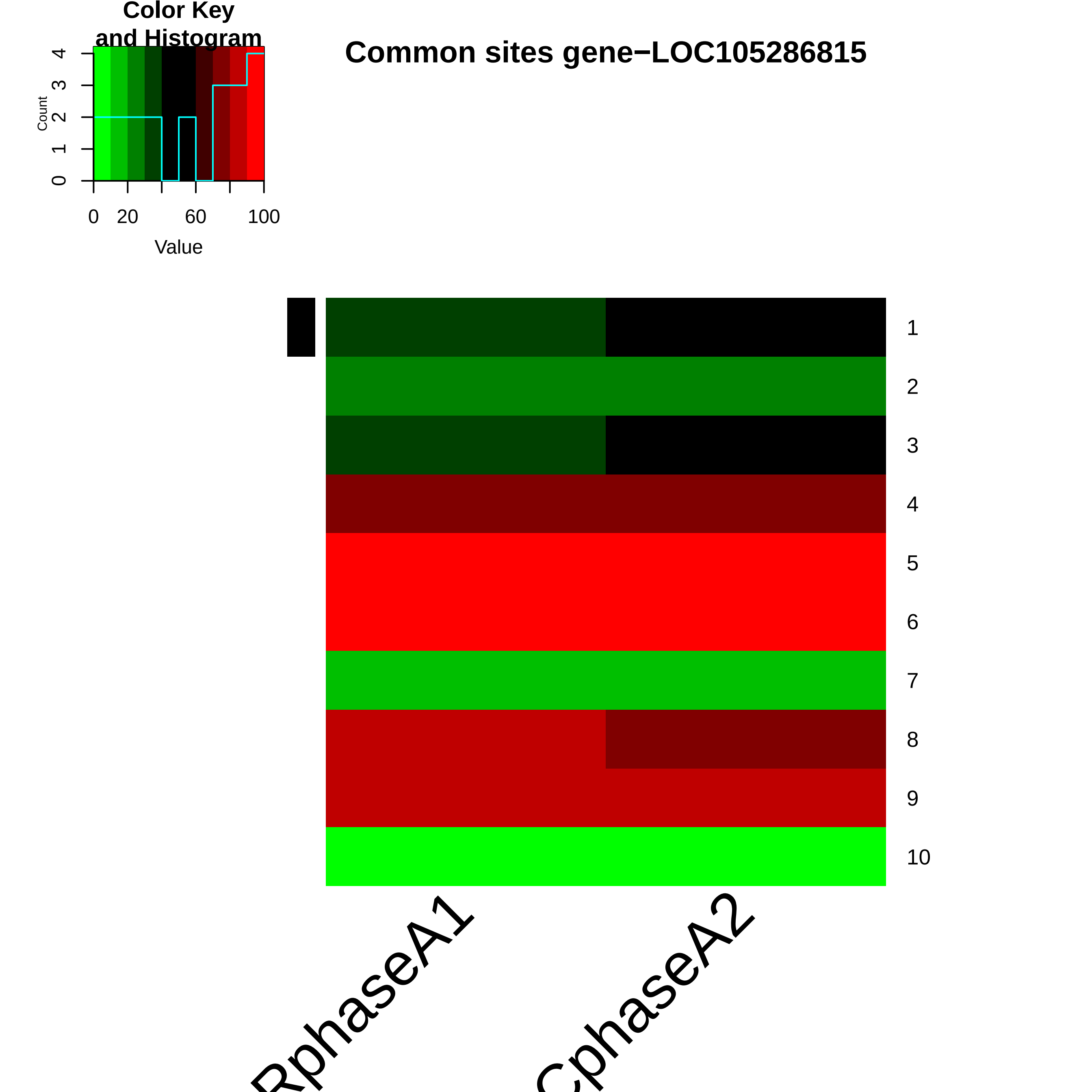

Color Key
and Histogram
Count
Value
Common sites gene−LOC105286815
0
20
60
100
0
1
2
3
4
1
2
3
4
5
6
7
8
9
10
RphaseA1
CphaseA2

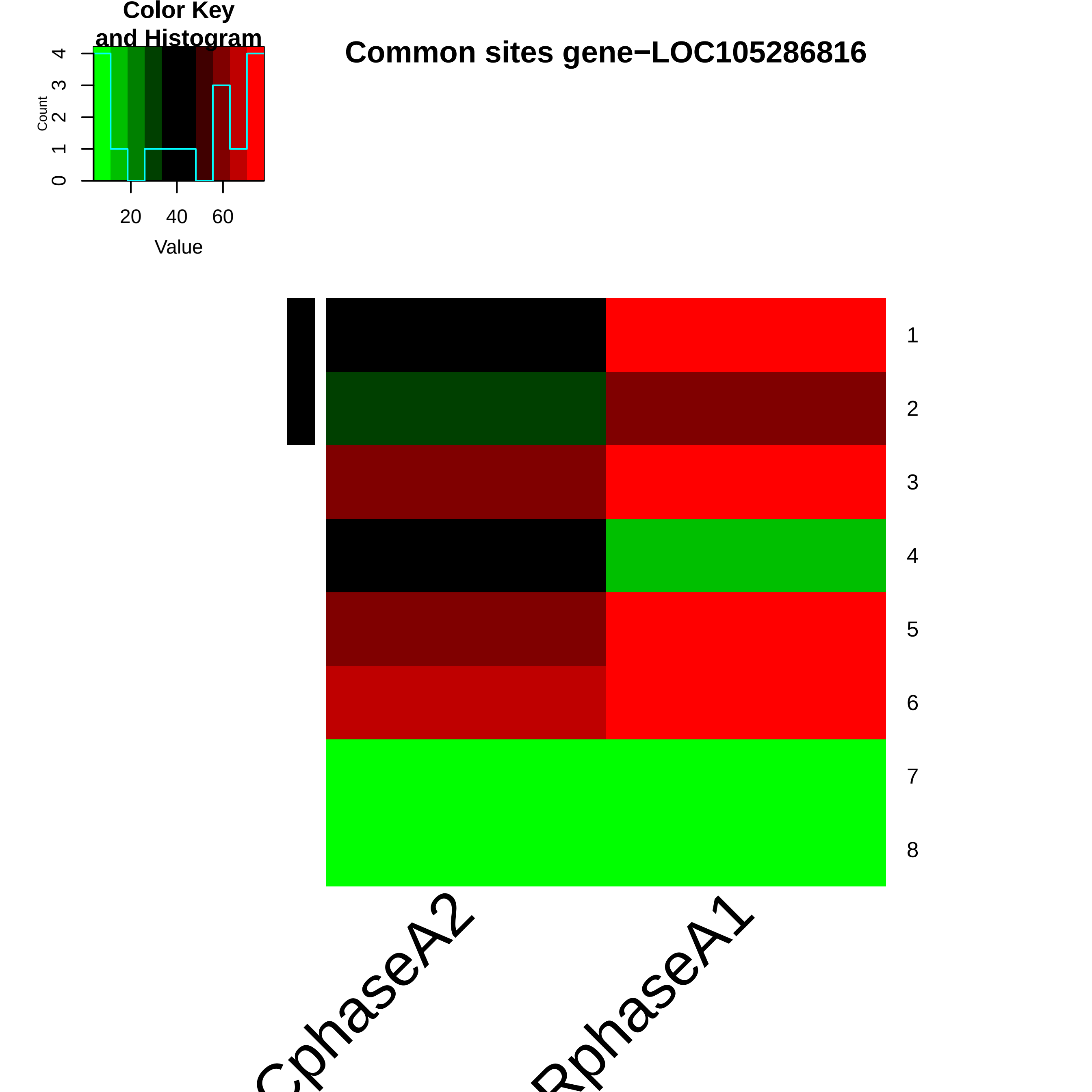

Color Key
and Histogram
Count
0
1
2
3
4
20
40
60
Value
Common sites gene−LOC105286816
1
2
3
4
5
6
7
8
CphaseA2
RphaseA1

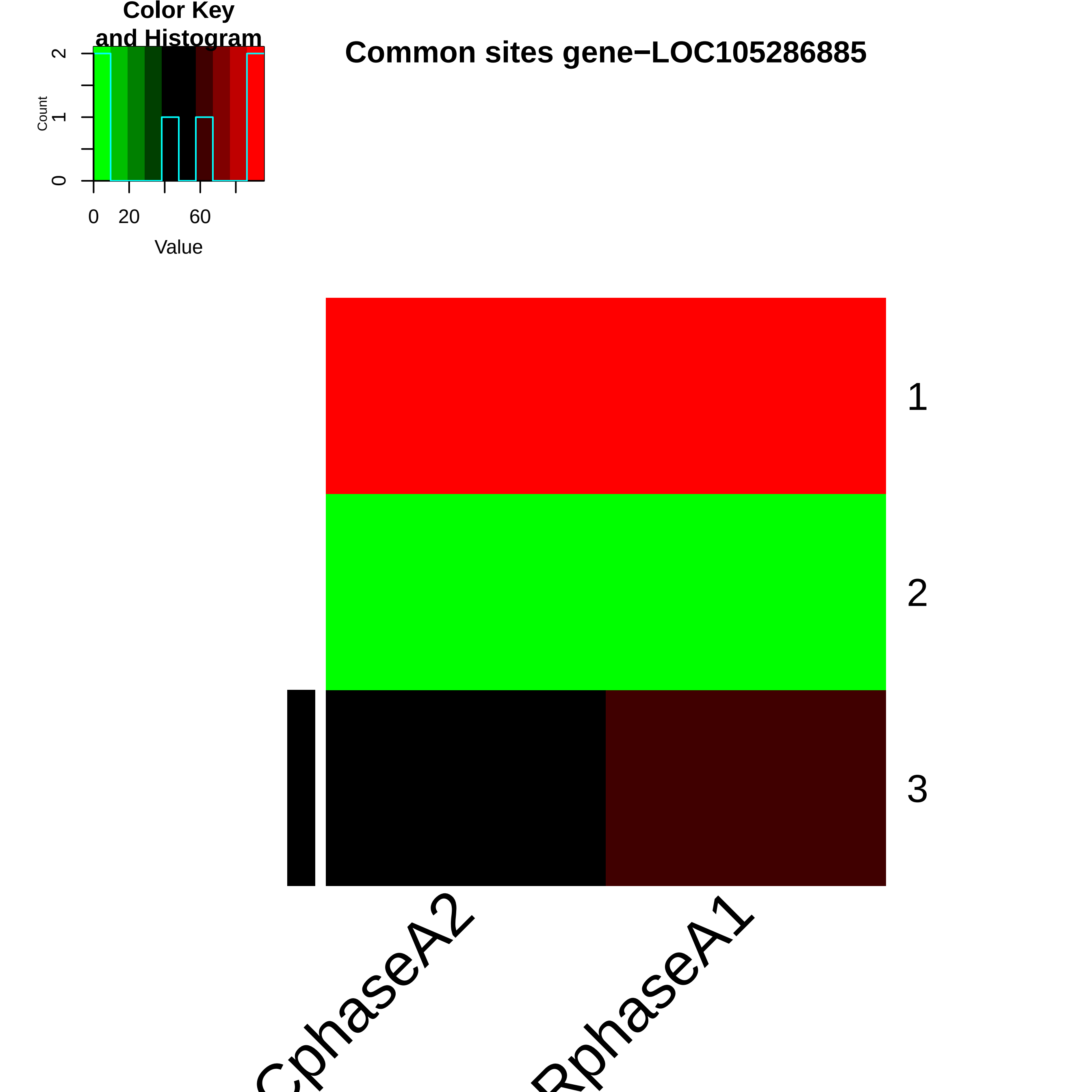
Color Key
and Histogram
Count
0
1
2
0
20
60
Value
Common sites gene−LOC105286885
1
2
3
CphaseA2
RphaseA1

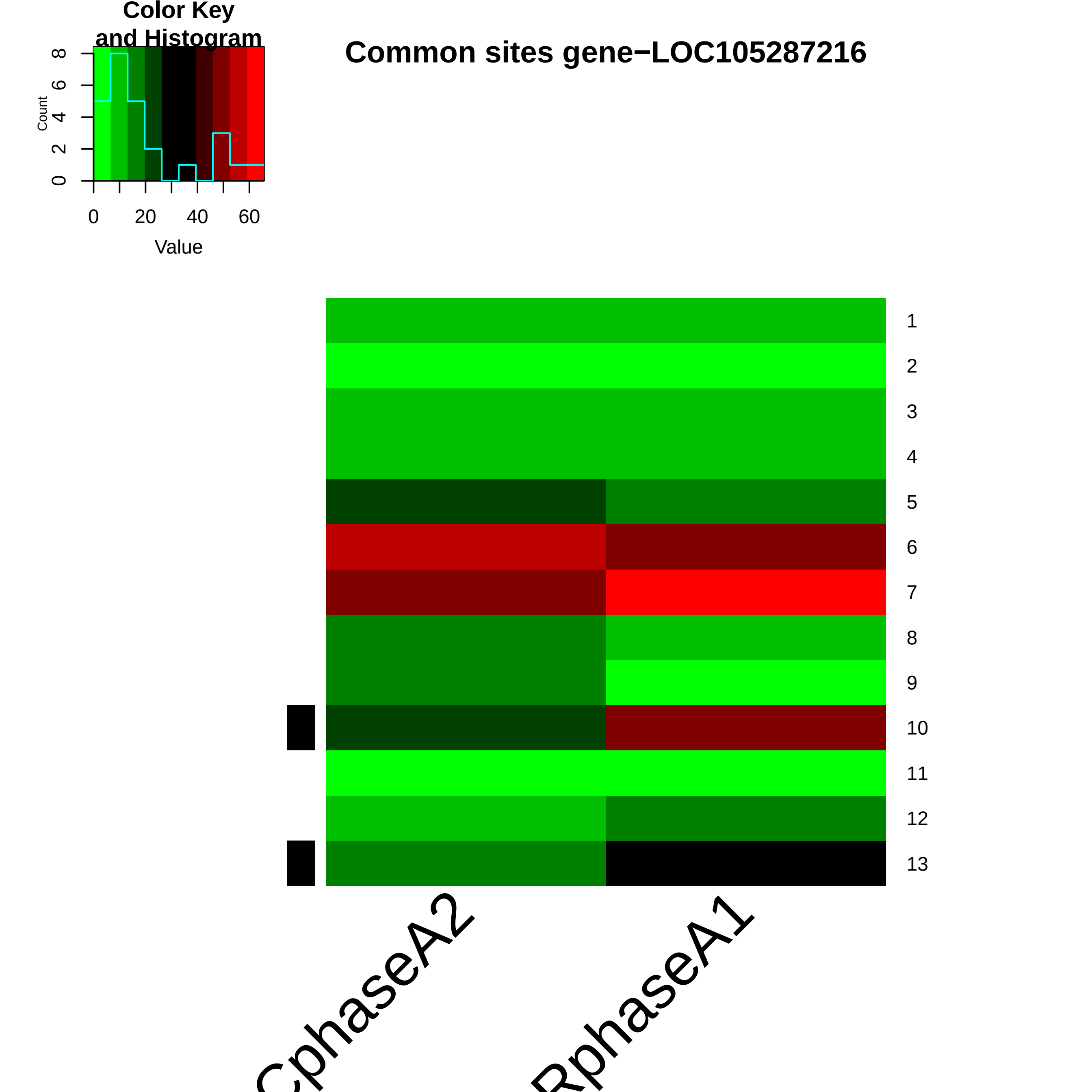

Color Key
and Histogram
Count
Value
Common sites gene−LOC105287216
1
2
3
4
5
6
7
8
9
10
11
12
13
CphaseA2
RphaseA1

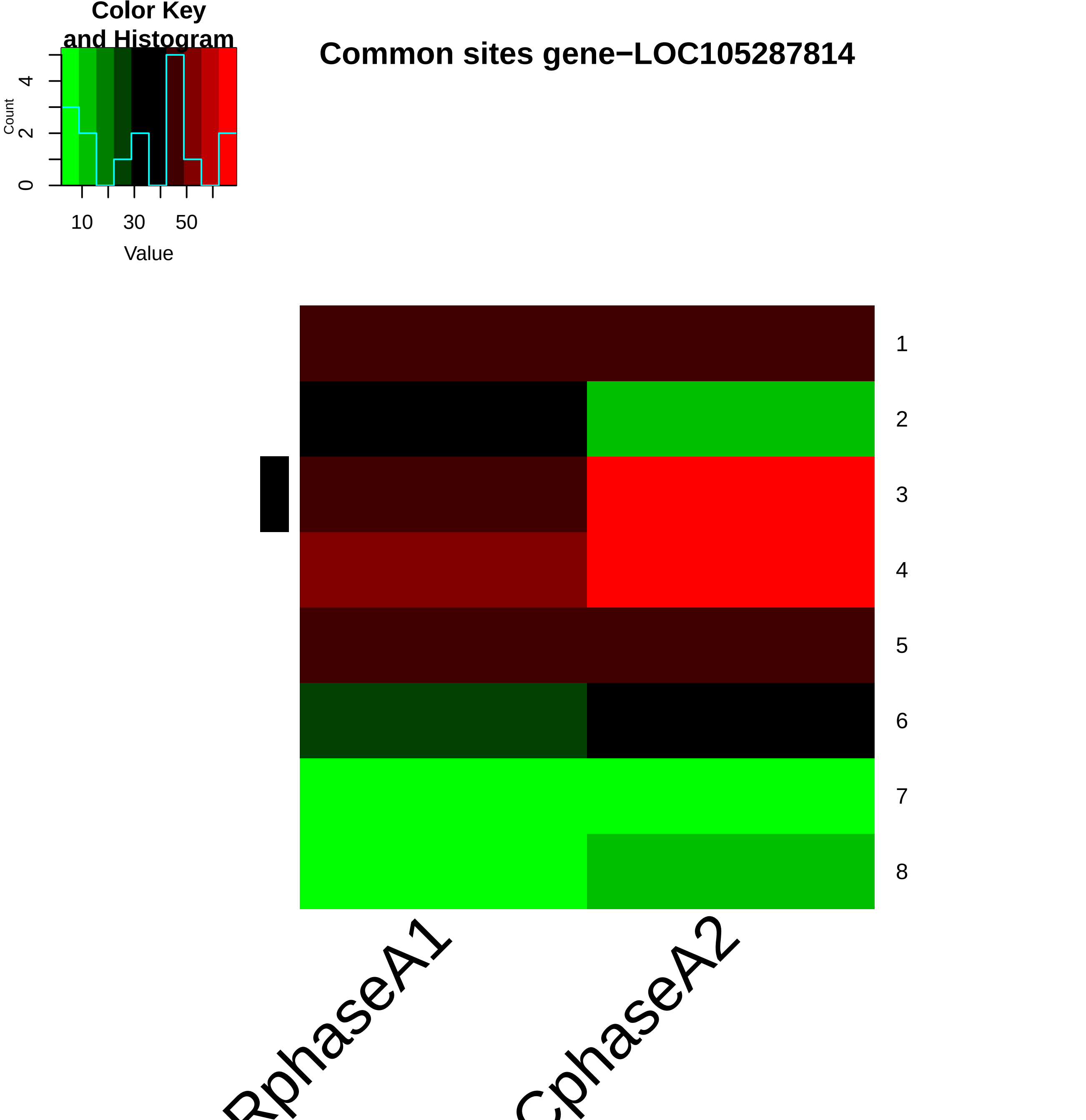
Color Key
and Histogram
Count
0
2
4
10
30
50
Value
Common sites gene−LOC105287814
1
2
3
4
5
6
7
8
RphaseA1
CphaseA2

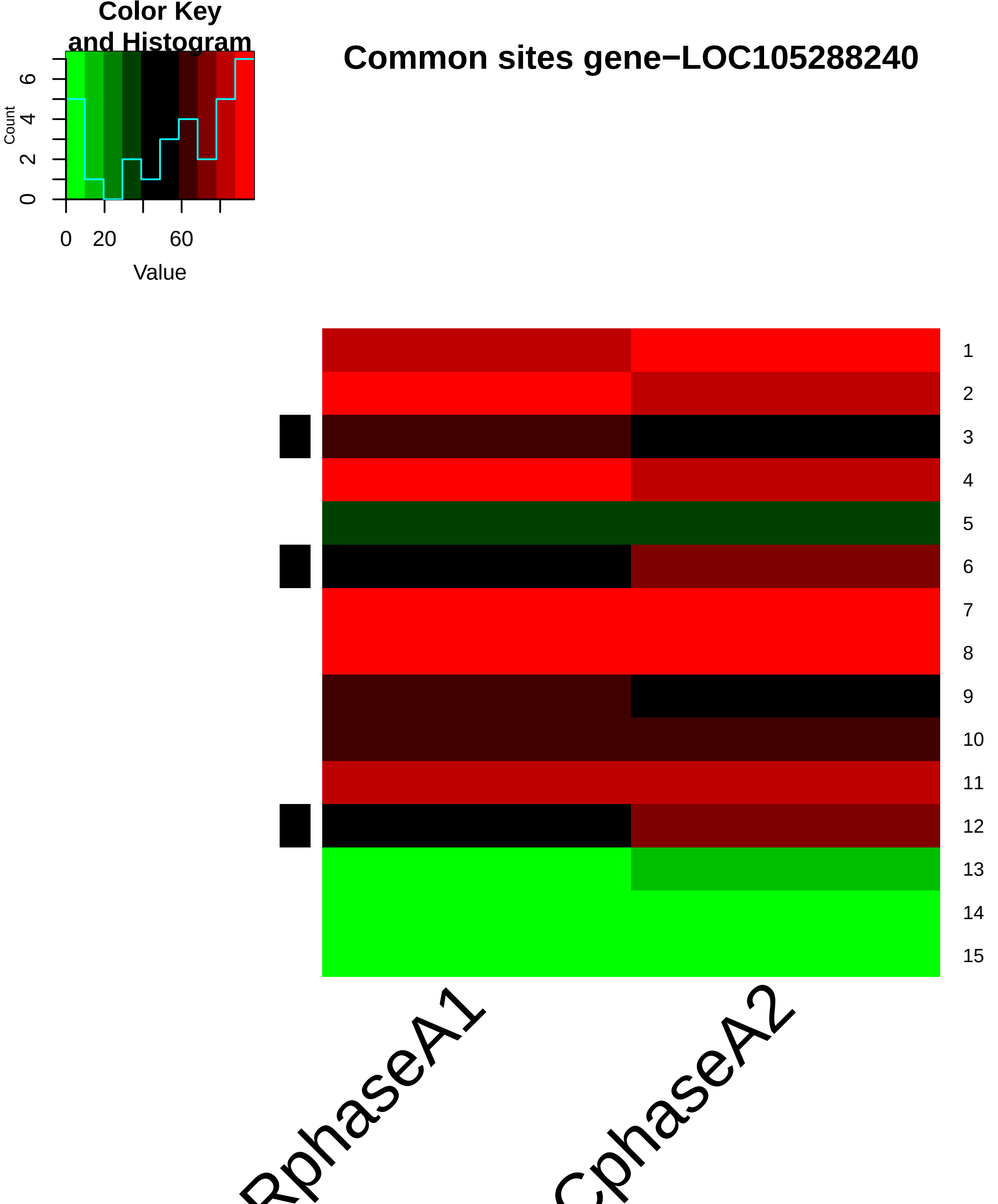

Color Key
and Histogram
Count
0
2
4
6
0
20
60
Value
Common sites gene−LOC105288240
RphaseA1
CphaseA2

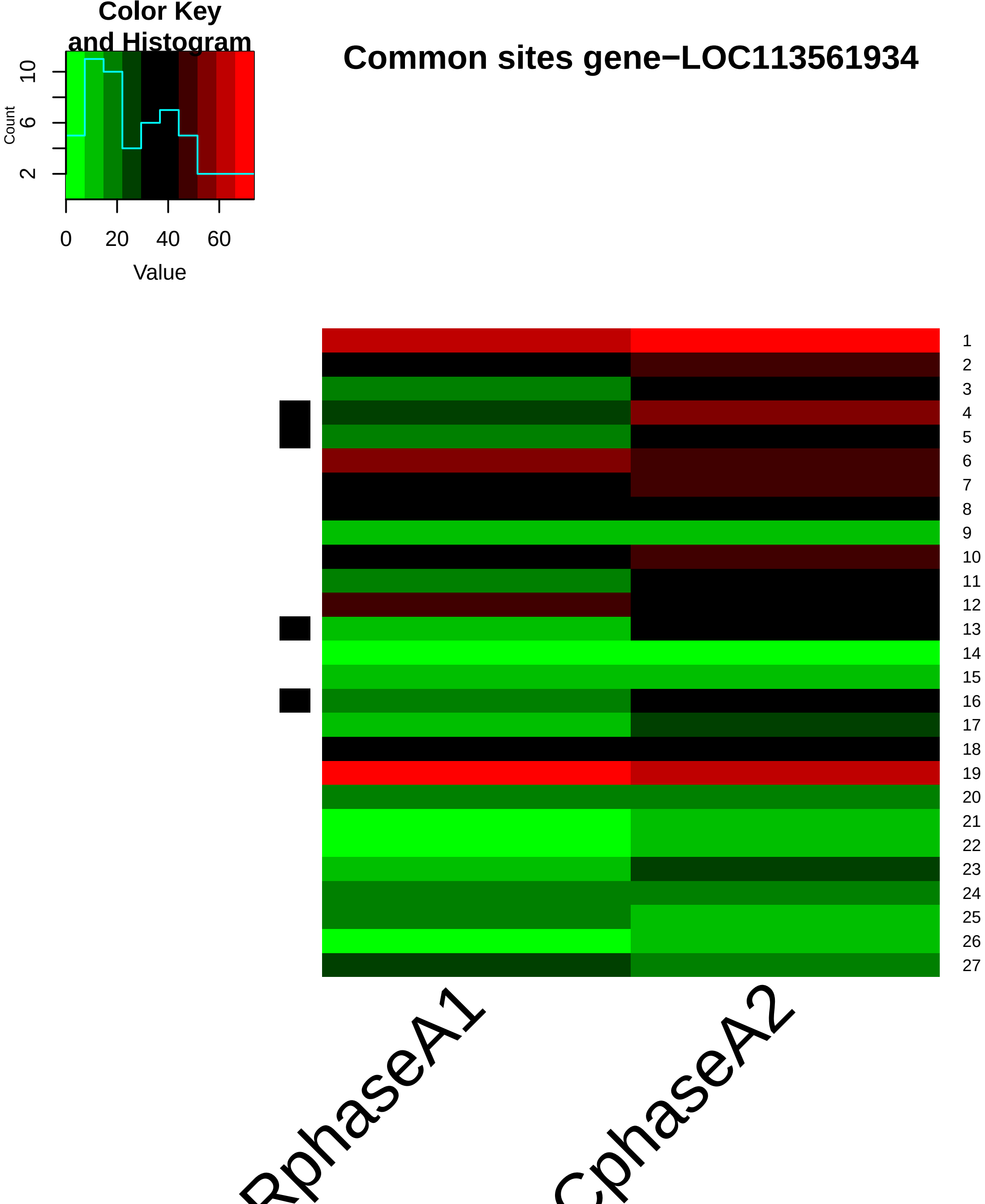

Color Key
and Histogram
Count
10
6
2
0
20
40
60
Value
Common sites gene−LOC113561934
RphaseA1
CphaseA2

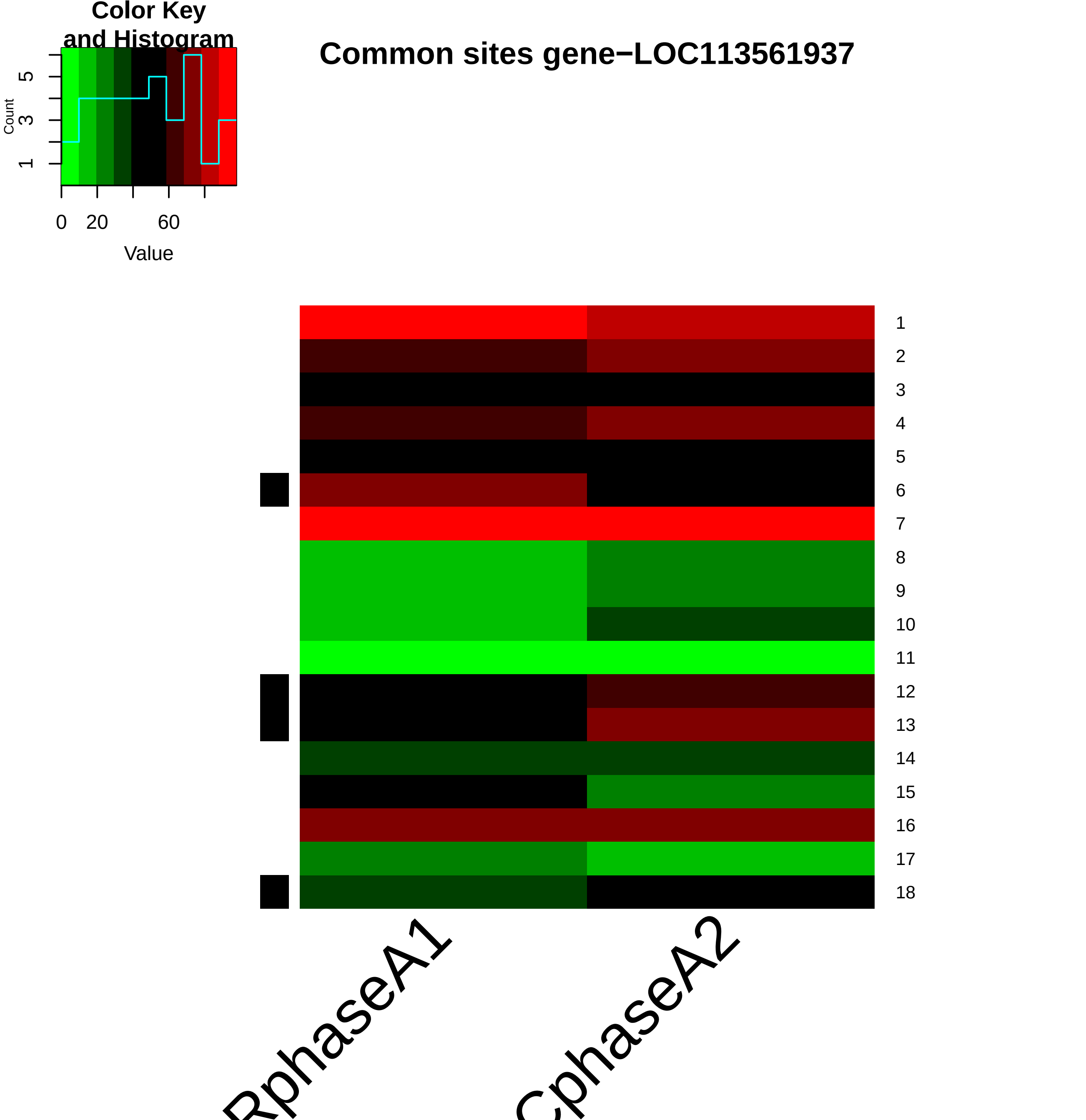

Color Key
and Histogram
Count
1
3
5
0
20
60
Value
Common sites gene−LOC113561937
RphaseA1
CphaseA2

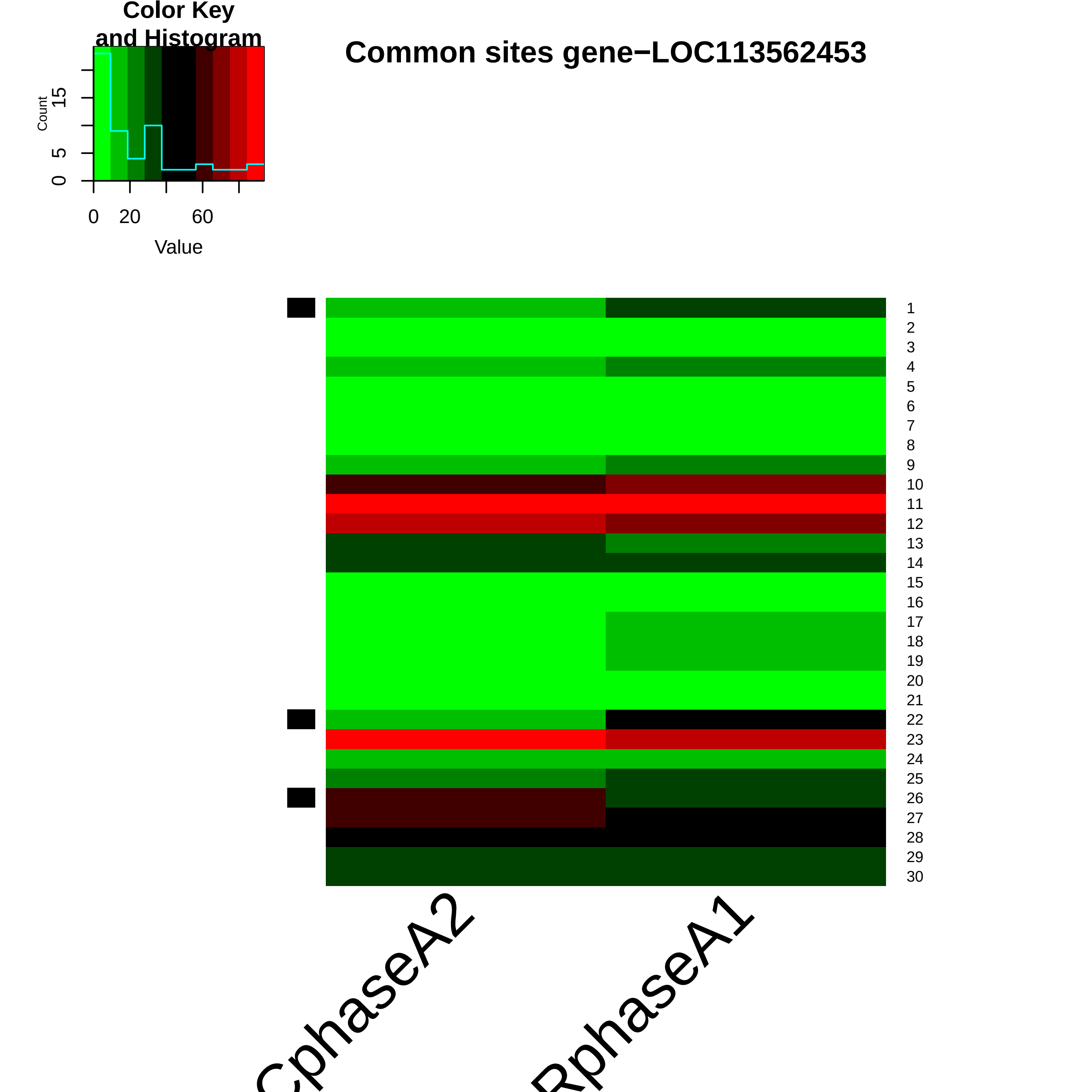

Color Key
and Histogram
Count
0
5
15
0
20
60
Value
Common sites gene−LOC113562453
CphaseA2
RphaseA1

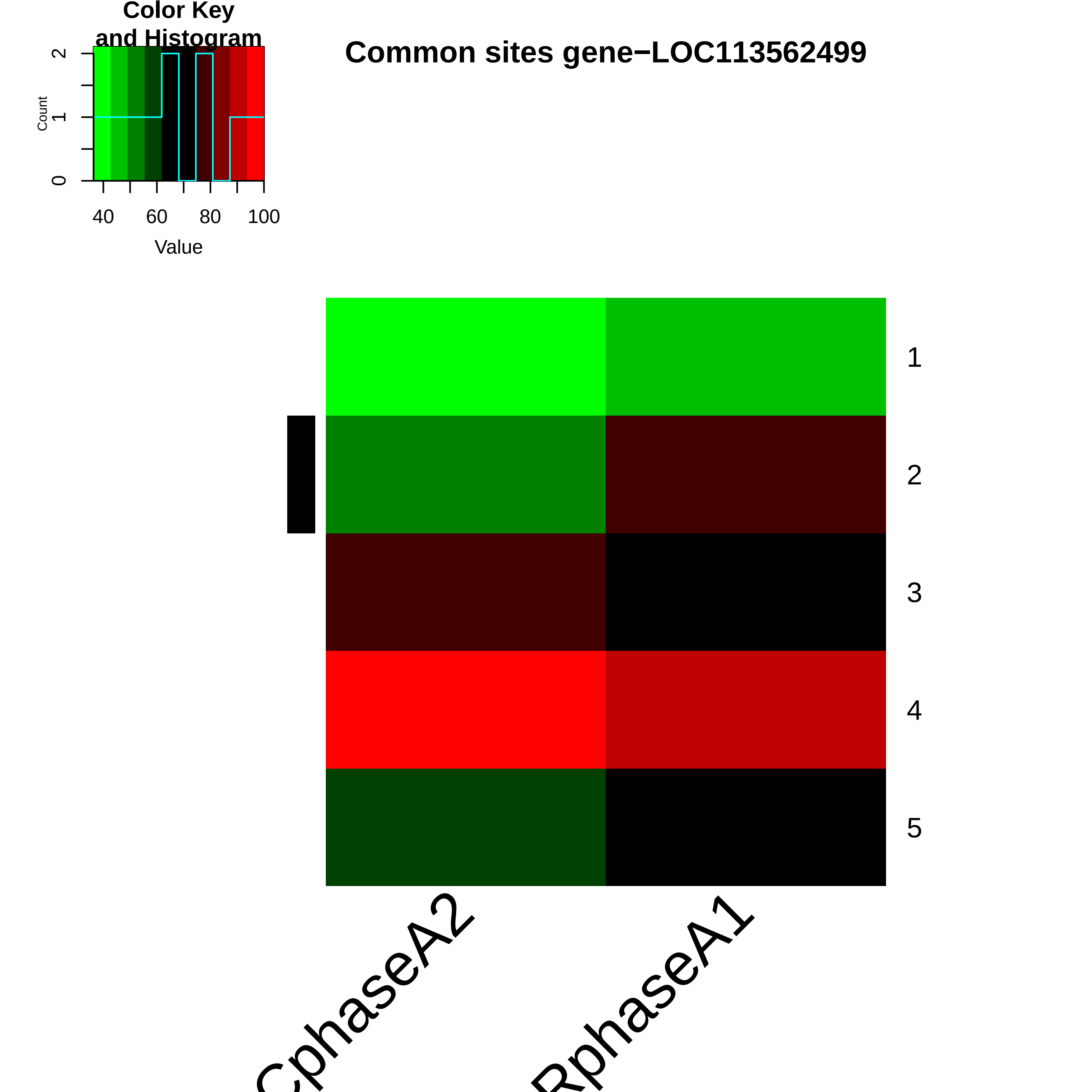

Color Key
and Histogram
Count
0
1
2
40
60
80
100
Value
Common sites gene−LOC113562499
1
2
3
4
5
CphaseA2
RphaseA1

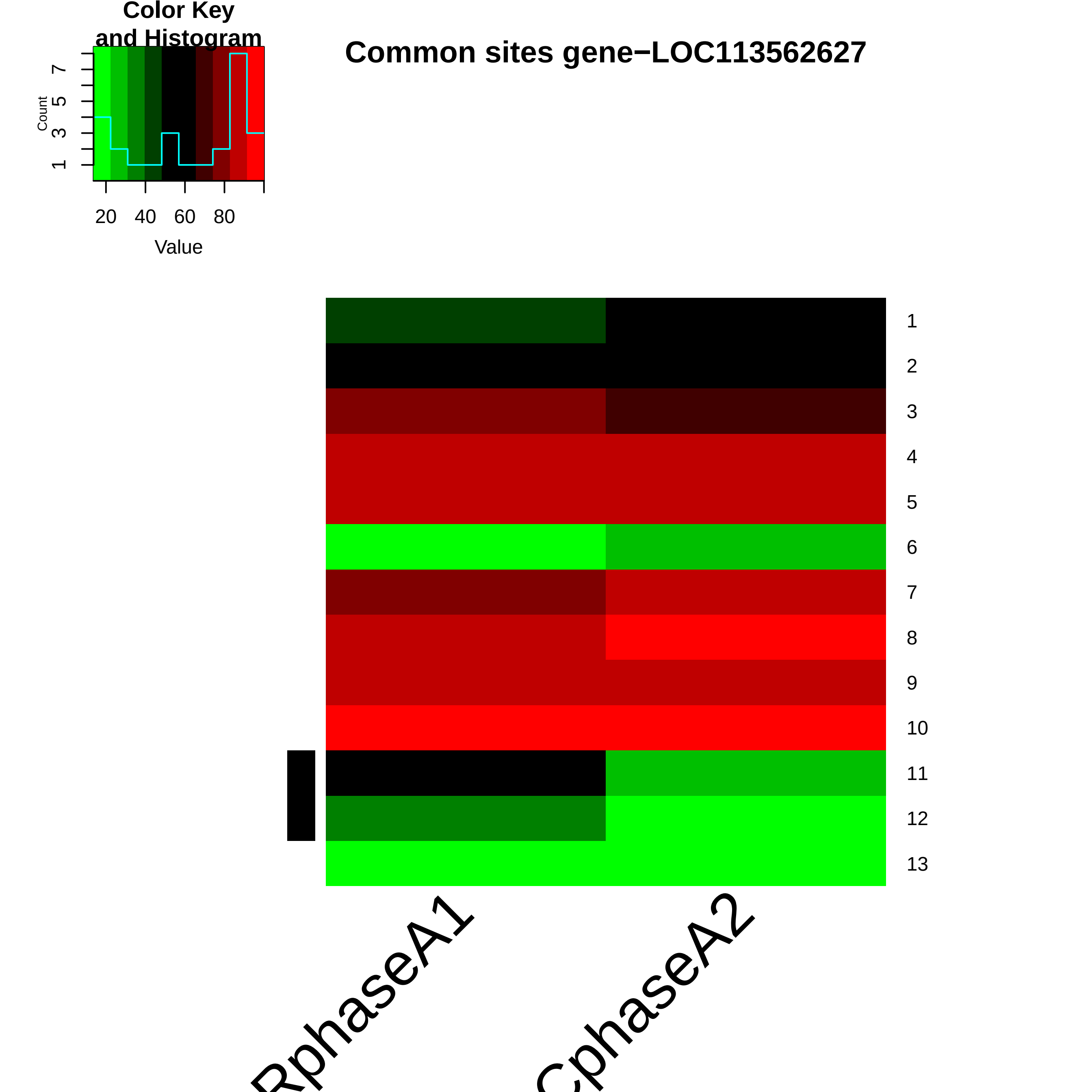

Color Key
and Histogram
Count
1
3
5
7
20
40
60
80
Value
Common sites gene−LOC113562627
1
2
3
4
5
6
7
8
9
10
11
12
13
RphaseA1
CphaseA2

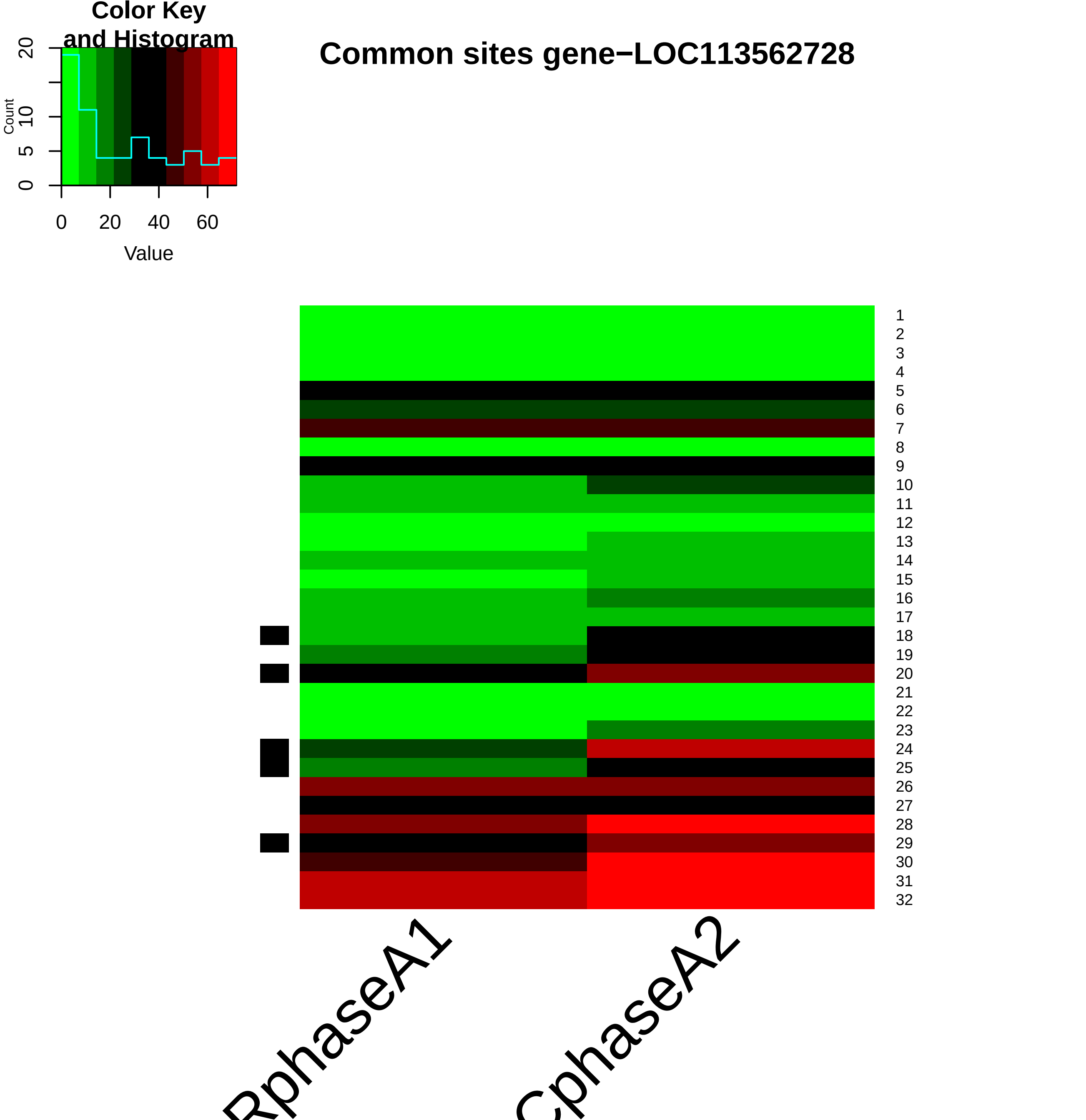

Color Key
and Histogram
Count
0
5
10
20
0
20
40
60
Value
Common sites gene−LOC113562728
RphaseA1
CphaseA2

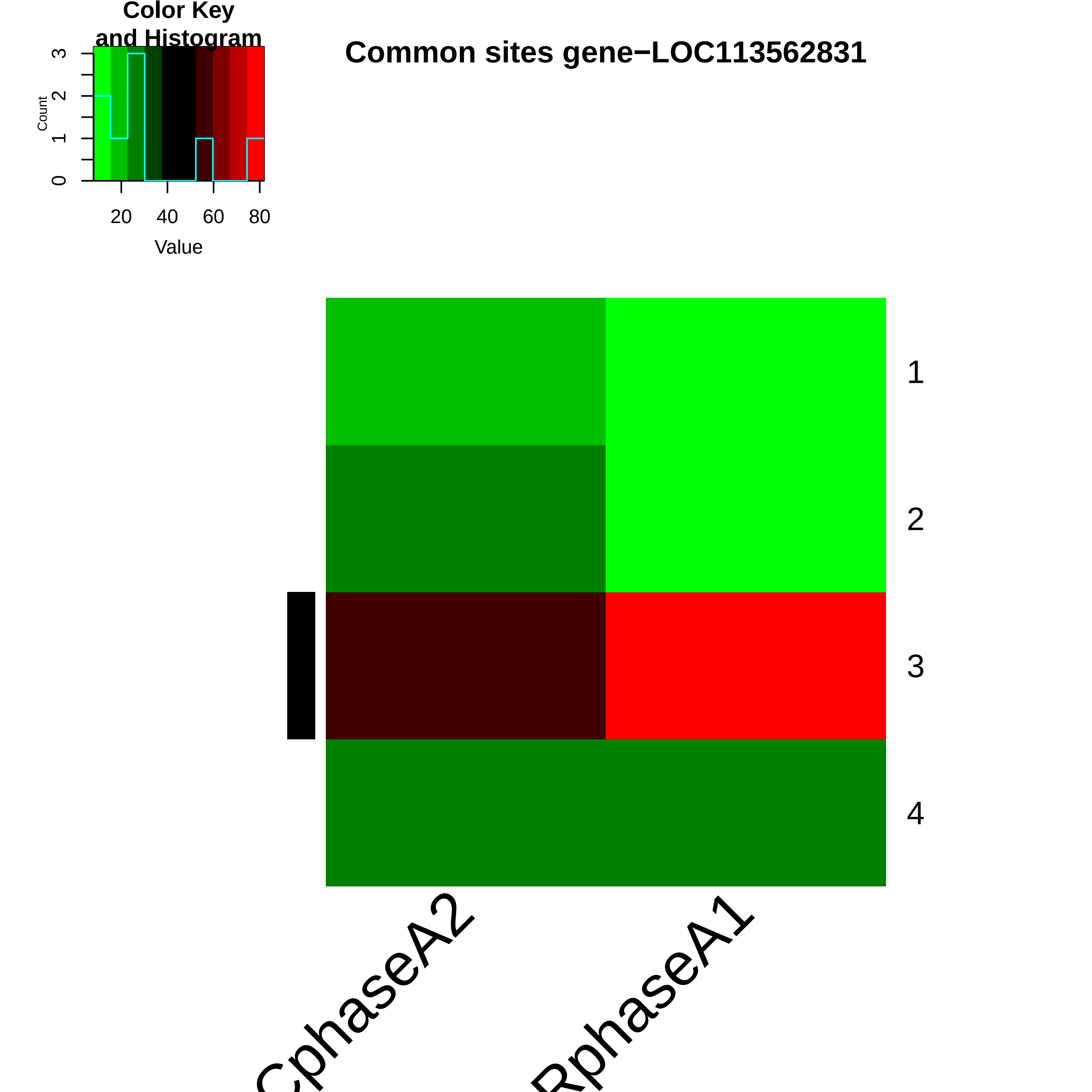

Color Key
and Histogram
Count
Value
Common sites gene−LOC113562831
1
2
3
4

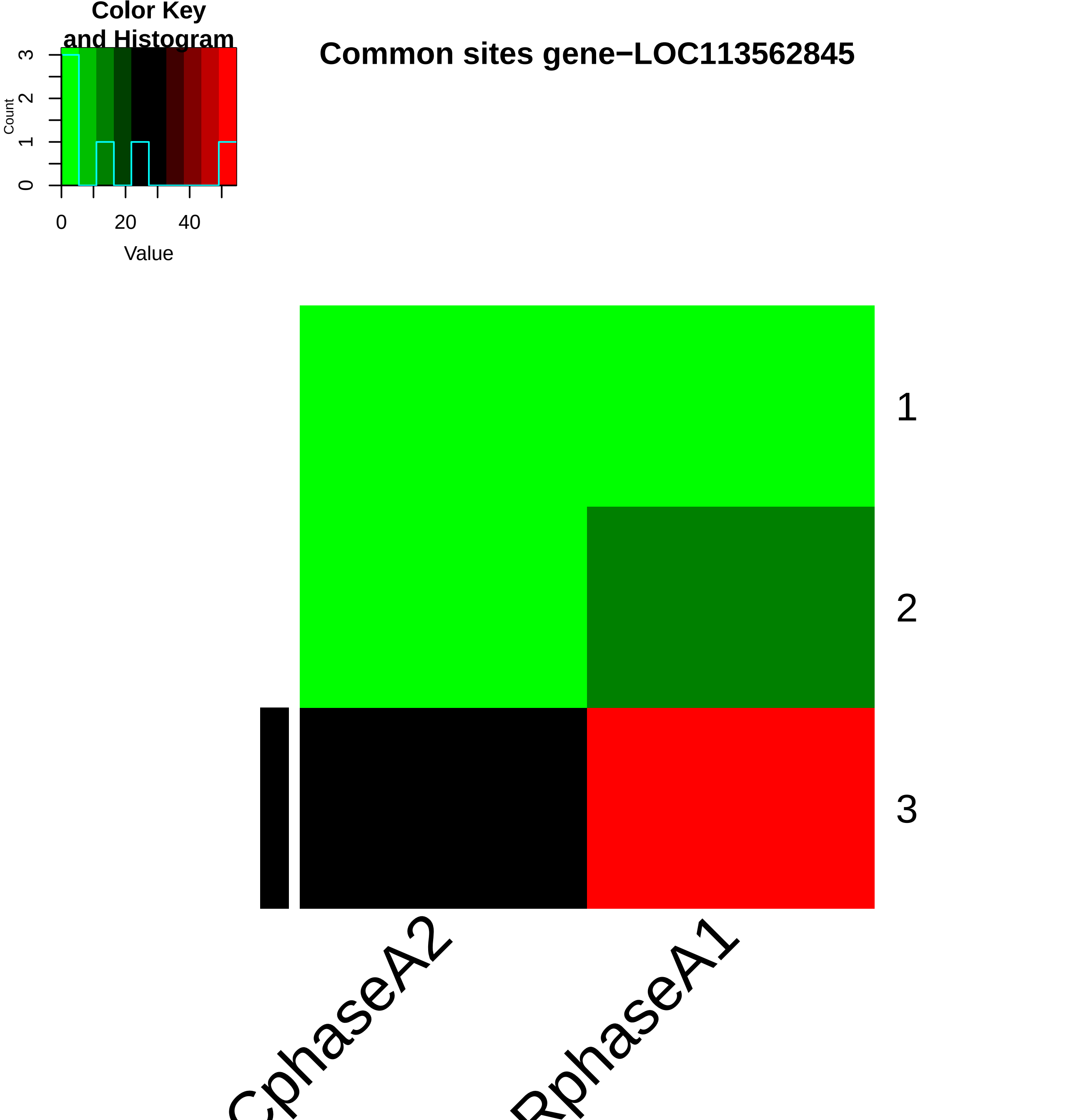
Color Key
and Histogram
Count
0
1
2
3
0
20
40
Value
Common sites gene−LOC113562845
1
2
3

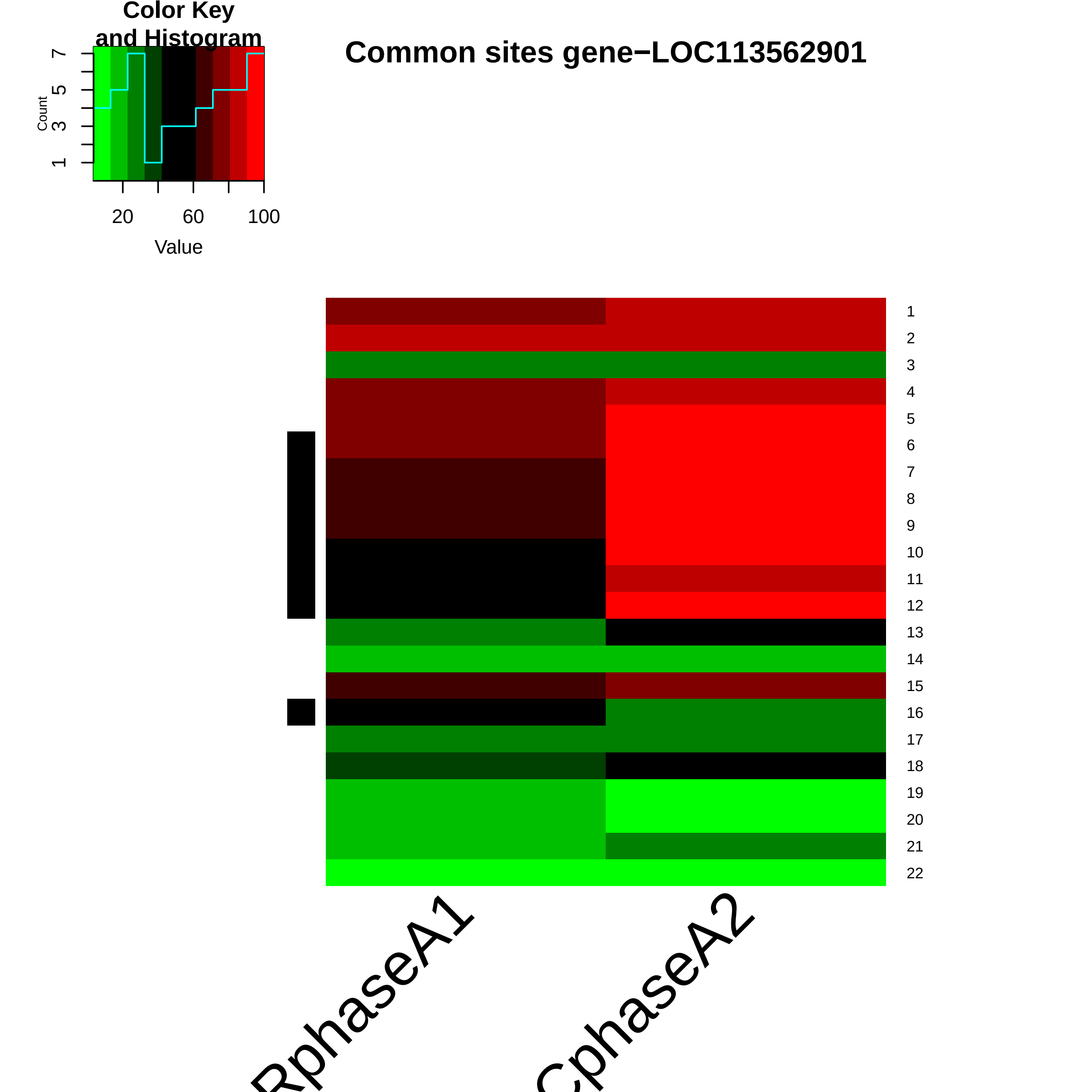
Color Key
and Histogram
Count
1
3
5
7
20
60
100
Value
Common sites gene−LOC113562901
RphaseA1
CphaseA2

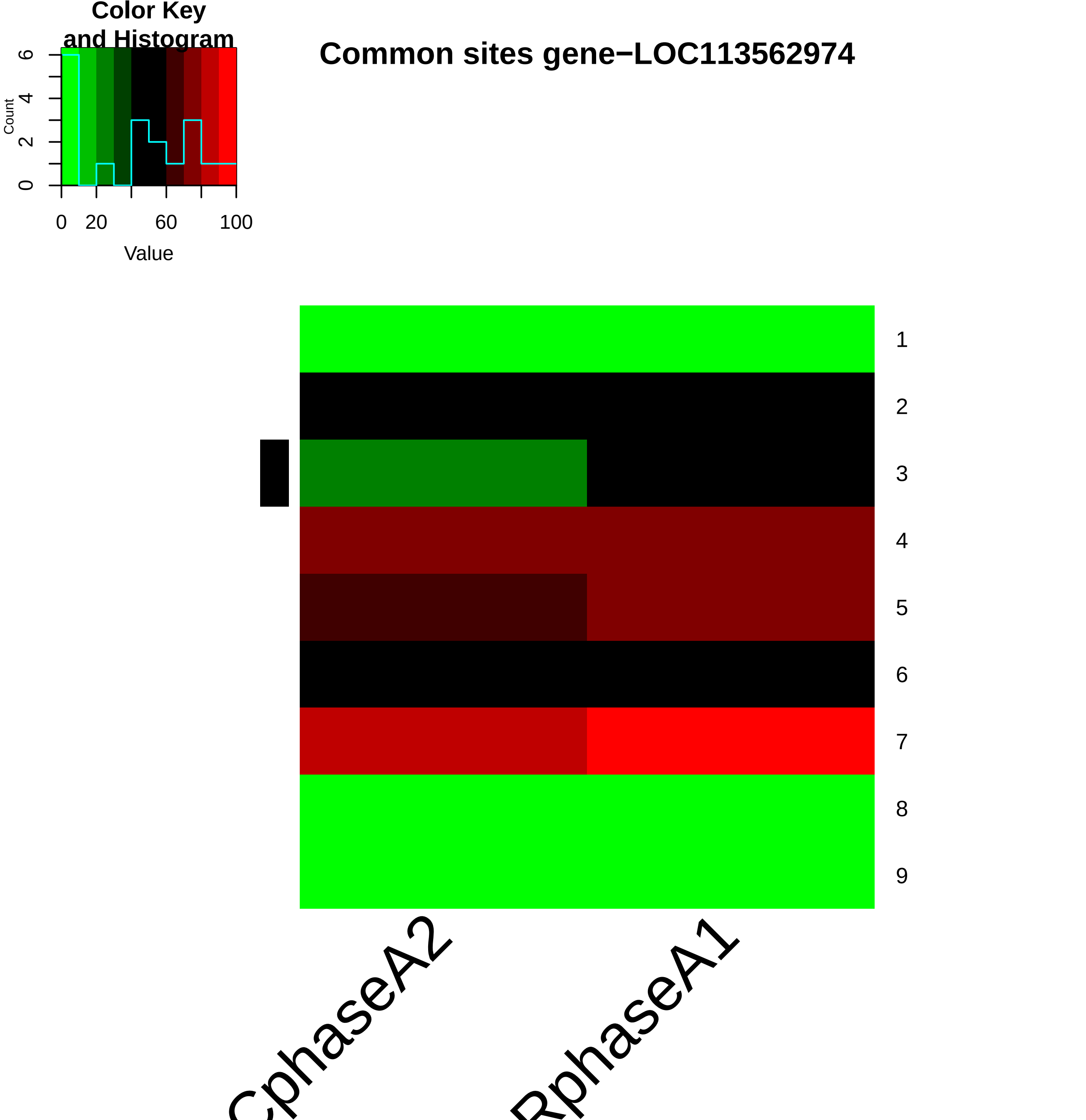
Color Key
and Histogram
Count
0
2
4
6
0
20
60
100
Value
Common sites gene−LOC113562974
1
2
3
4
5
6
7
8
9
CphaseA2
RphaseA1

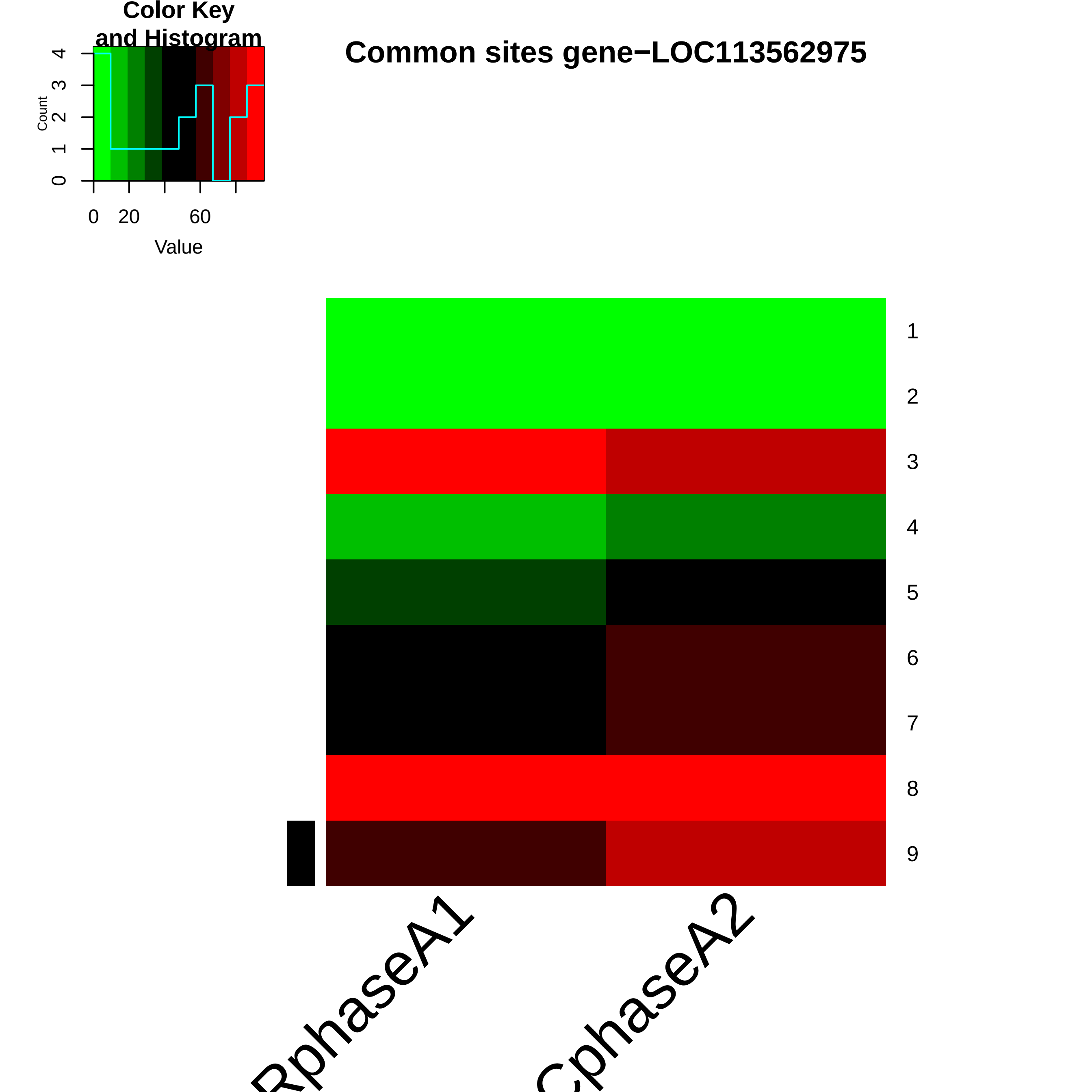
Color Key
and Histogram
Count
Value
0
20
60
0
1
2
3
4
Common sites gene−LOC113562975
1
2
3
4
5
6
7
8
9
RphaseA1
CphaseA2

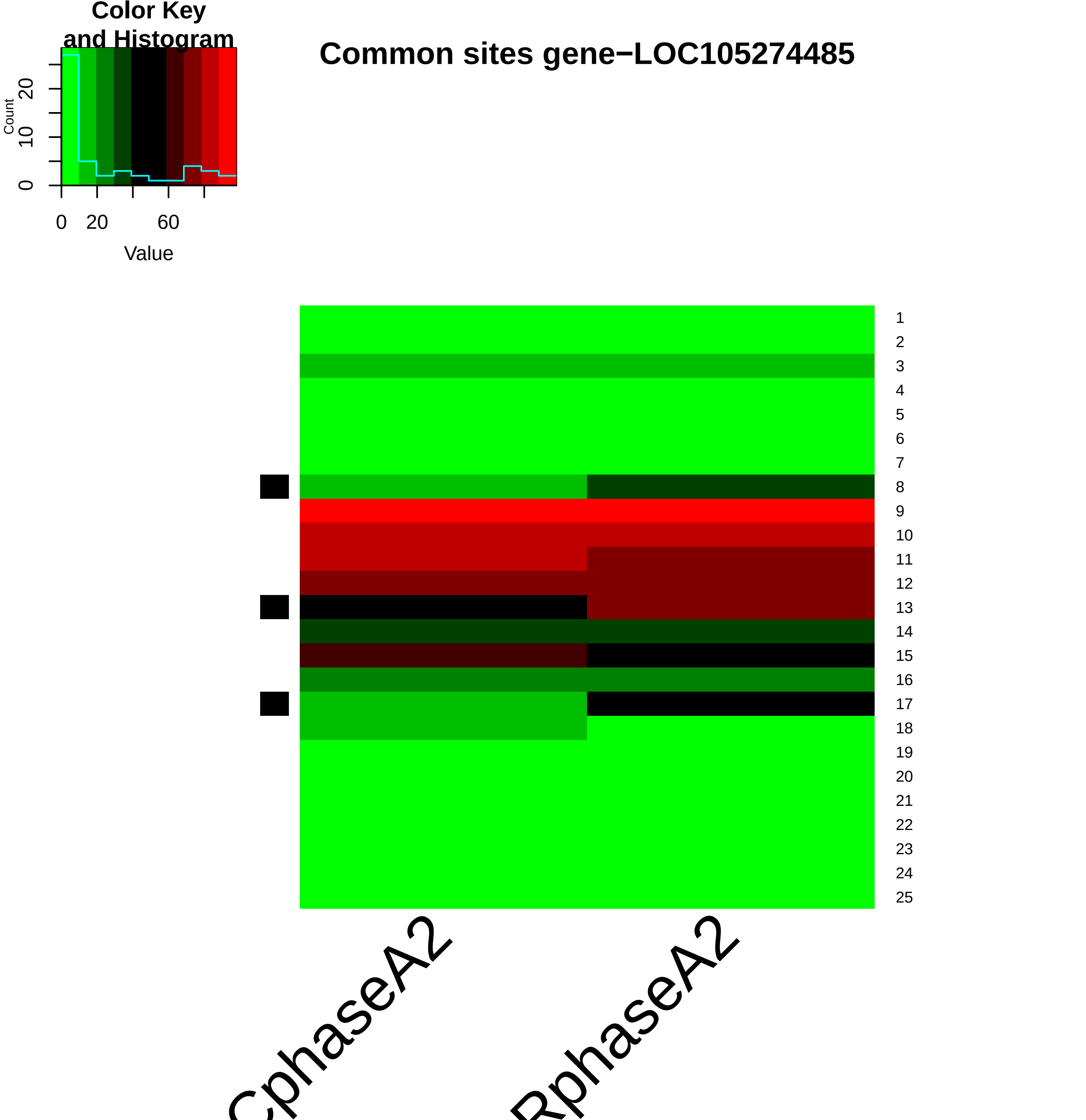
Color Key
and Histogram
Count
0
10
20
0
20
60
Value
Common sites gene−LOC105274485
CphaseA2
RphaseA2

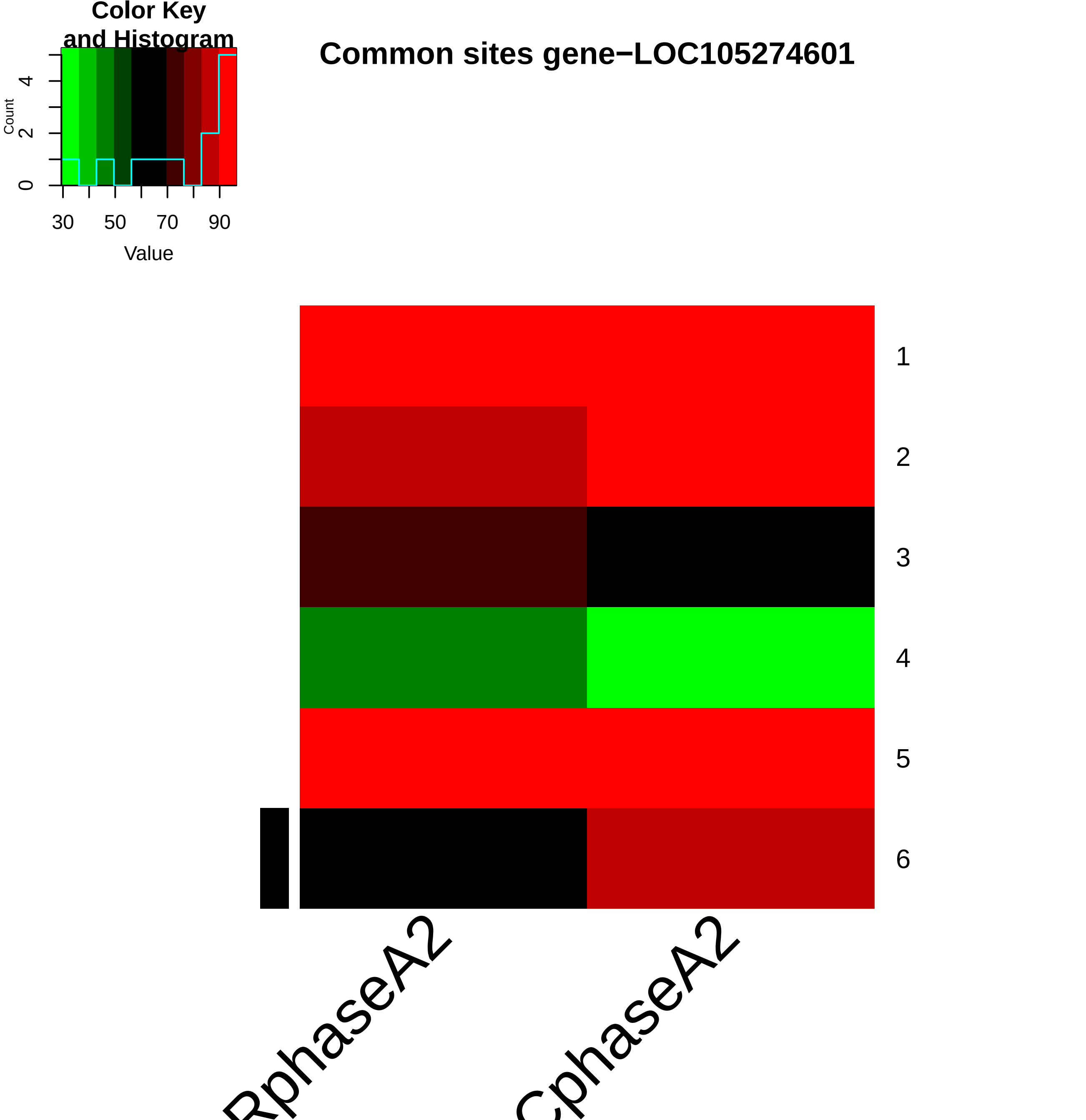

Color Key
and Histogram
Count
0
2
4
30
50
70
90
Value
Common sites gene−LOC105274601
1
2
3
4
5
6
RphaseA2
CphaseA2

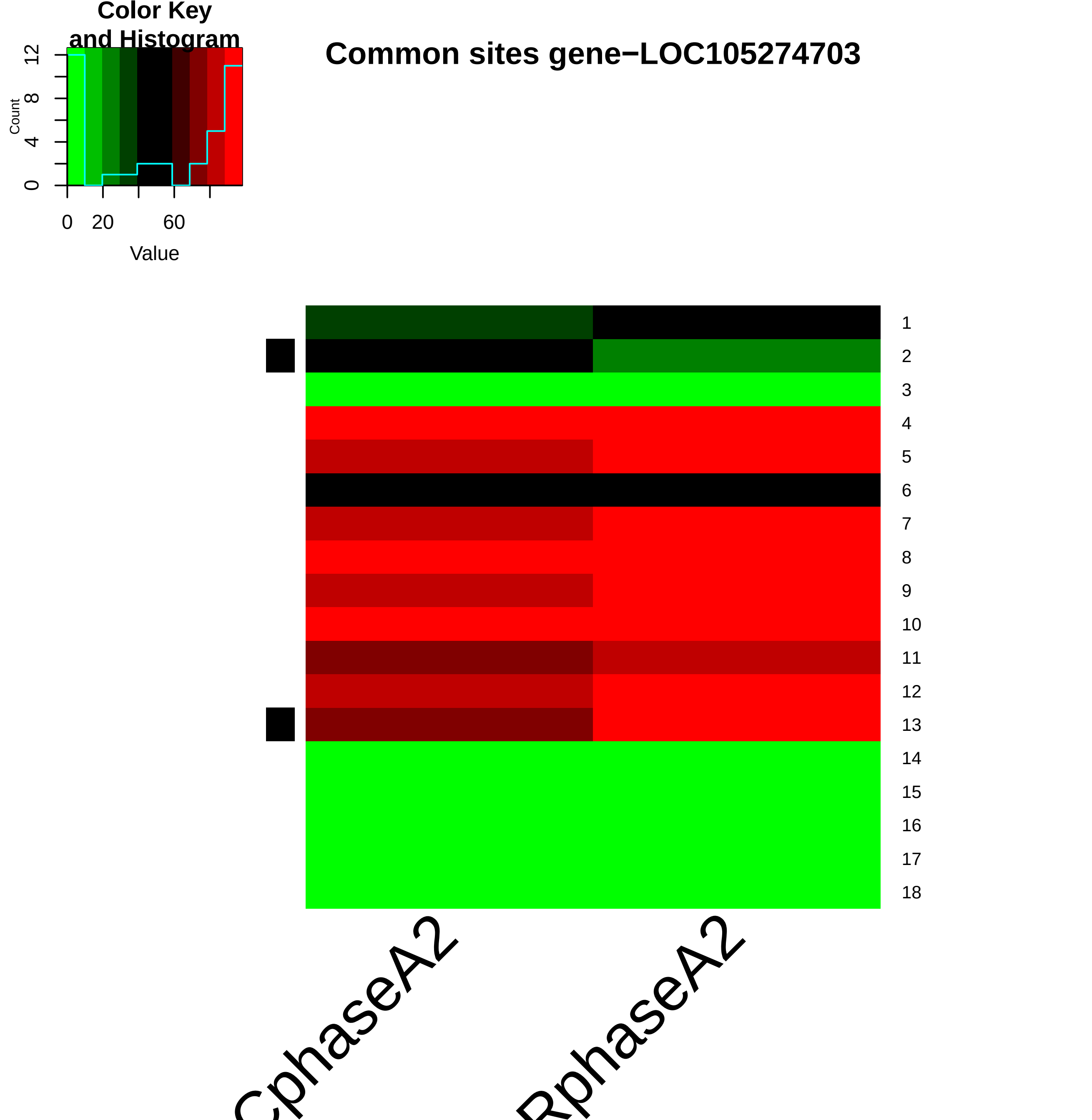

Color Key
and Histogram
Count
0
4
8
12
0
20
60
Value
Common sites gene−LOC105274703
CphaseA2
RphaseA2

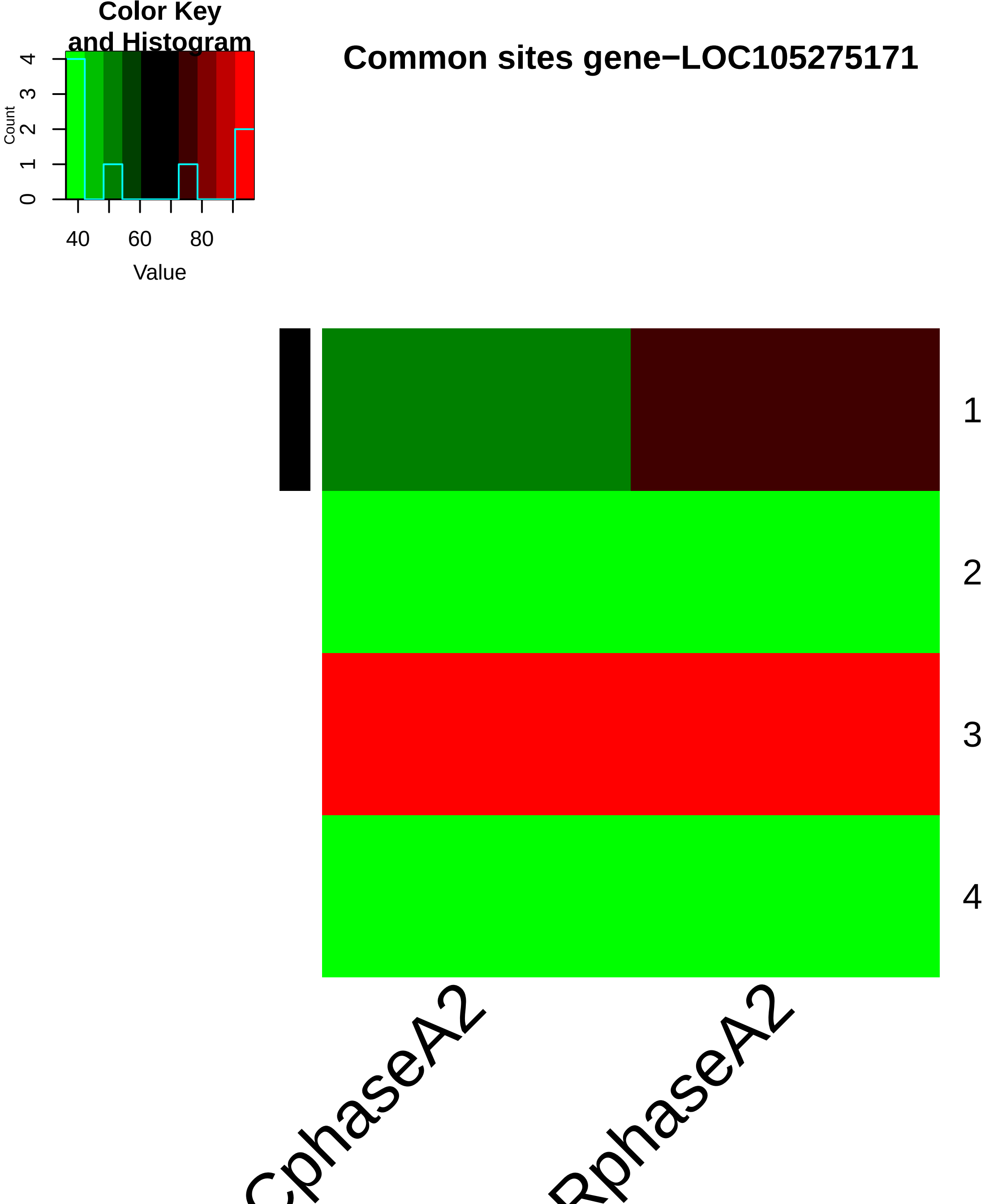

Color Key
and Histogram
Count
0
1
2
3
4
40
60
80
Value
Common sites gene−LOC105275171
1
2
3
4

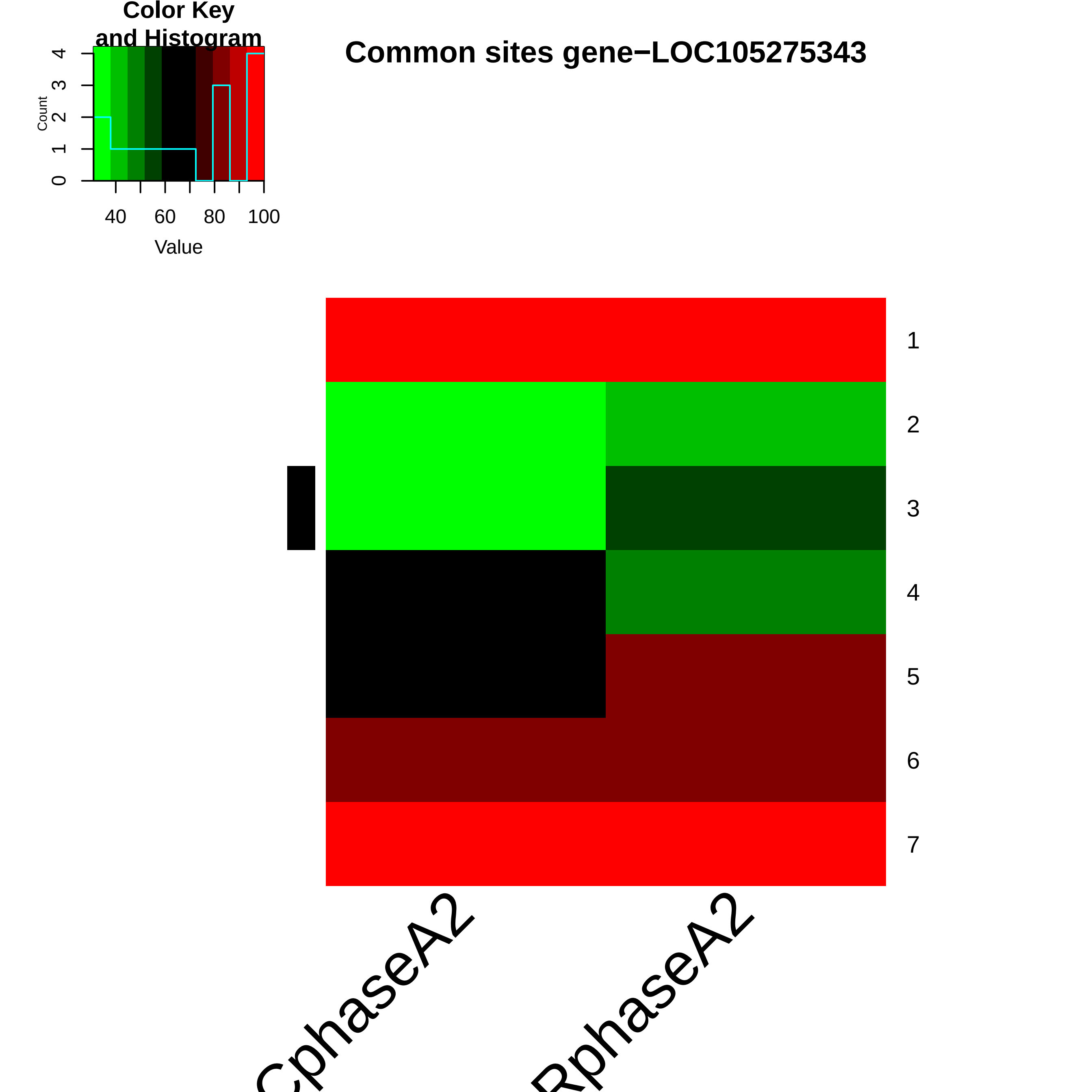
Color Key
and Histogram
Count
0
1
2
3
4
40
60
80
100
Value
Common sites gene−LOC105275343
1
2
3
4
5
6
7

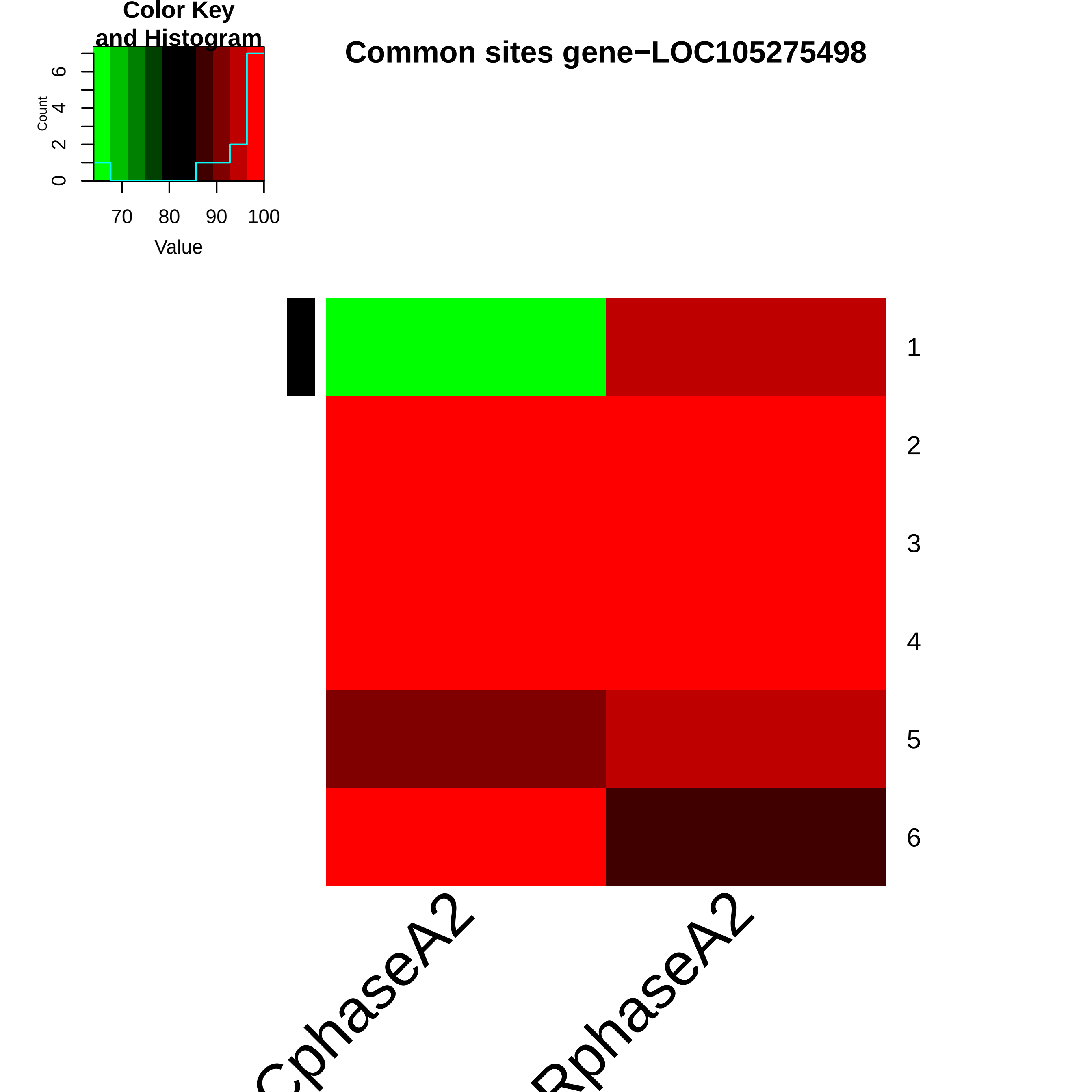

Color Key
and Histogram
Count
0
2
4
6
70
80
90
100
Value
Common sites gene−LOC105275498
1
2
3
4
5
6
CphaseA2
RphaseA2

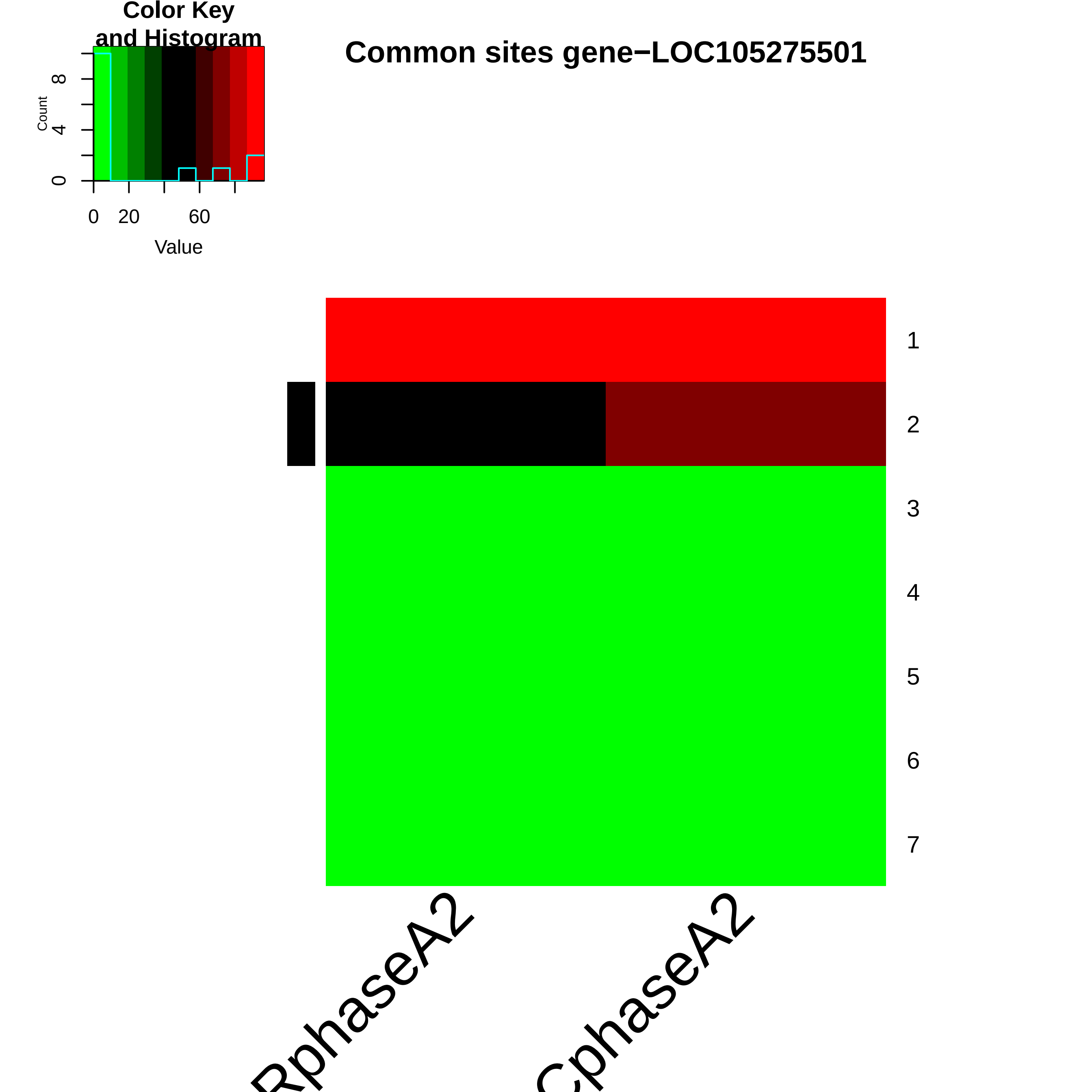

Color Key and Histogram
Count
0
4
8
0
20
60
Value
Common sites gene−LOC105275501
1
2
3
4
5
6
7
RphaseA2
CphaseA2

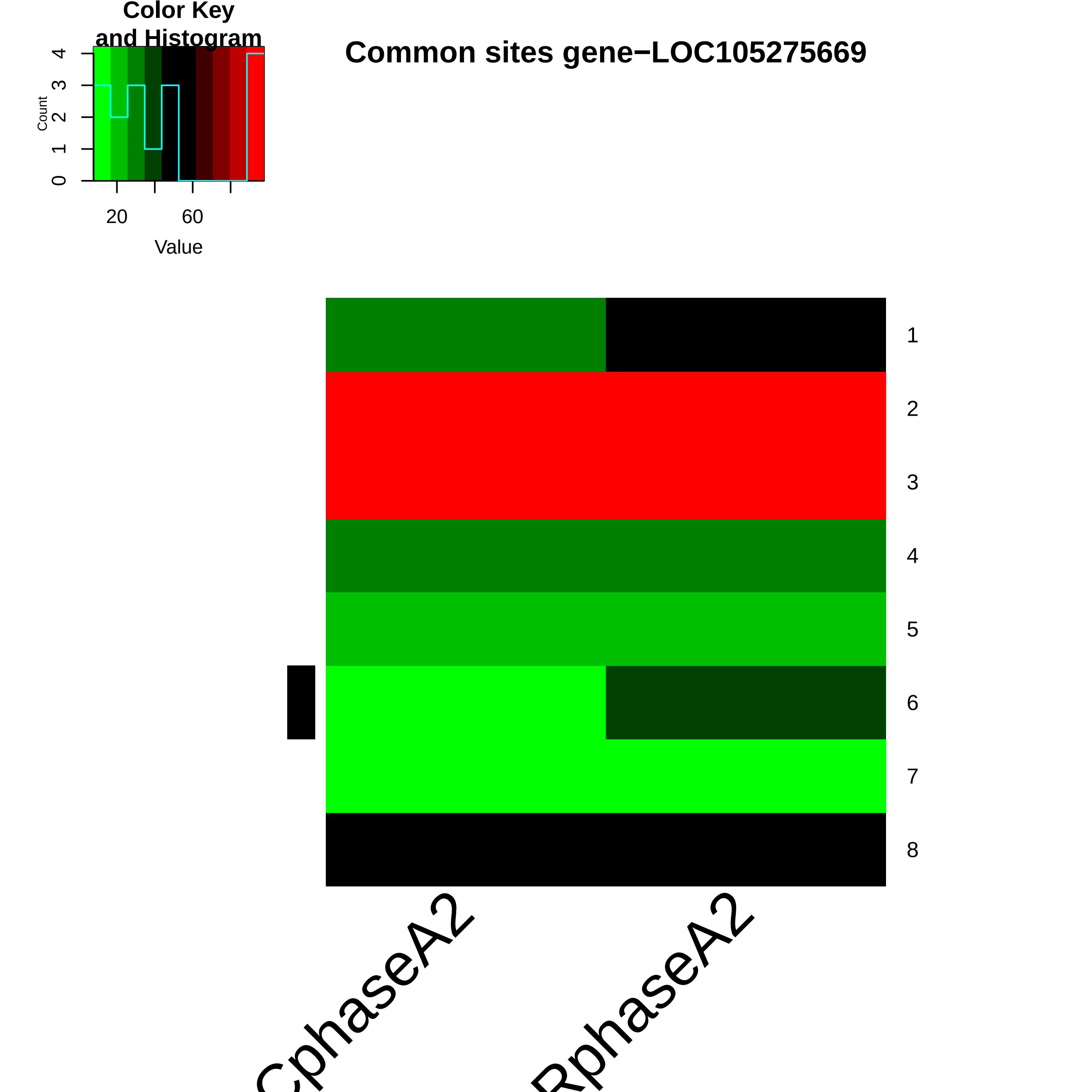
Color Key
and Histogram
Count
0
1
2
3
4
20
60
Value
Common sites gene−LOC105275669
1
2
3
4
5
6
7
8
CphaseA2
RphaseA2

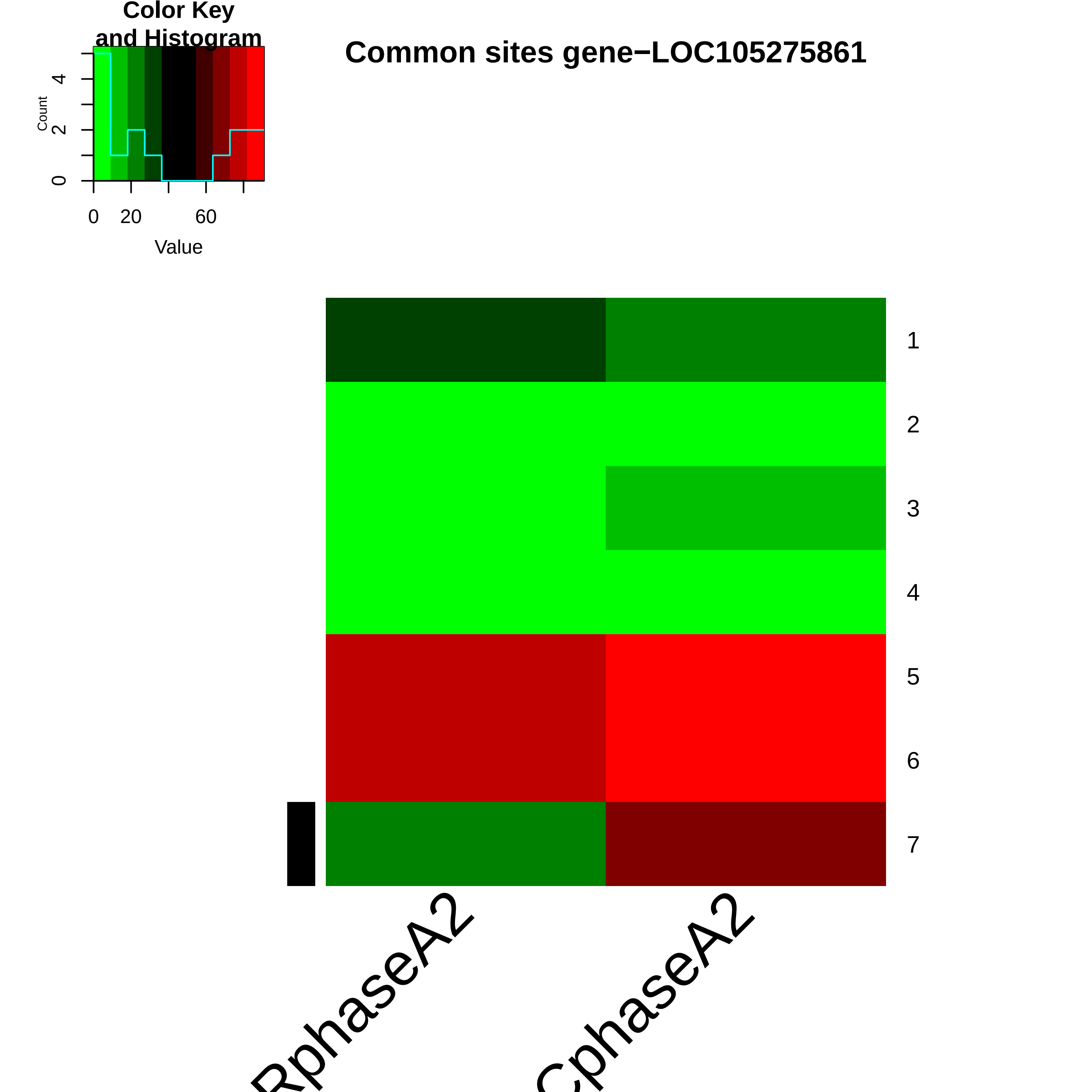

Color Key
and Histogram
Count
0
2
4
0
20
60
Value
Common sites gene−LOC105275861
1
2
3
4
5
6
7
RphaseA2
CphaseA2

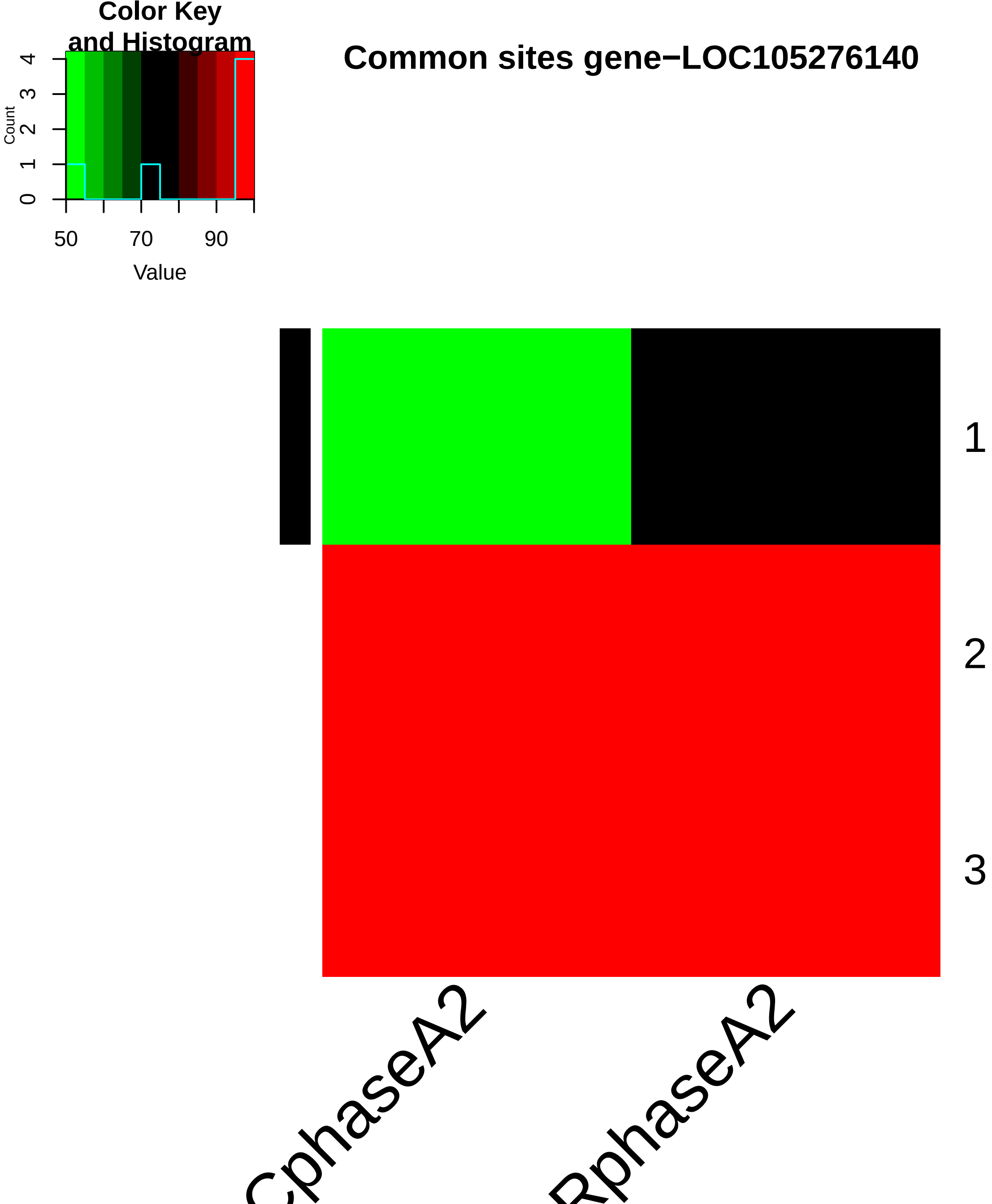
Color Key
and Histogram
Count
0
1
2
3
4
50
70
90
Value
Common sites gene−LOC105276140
1
2
3

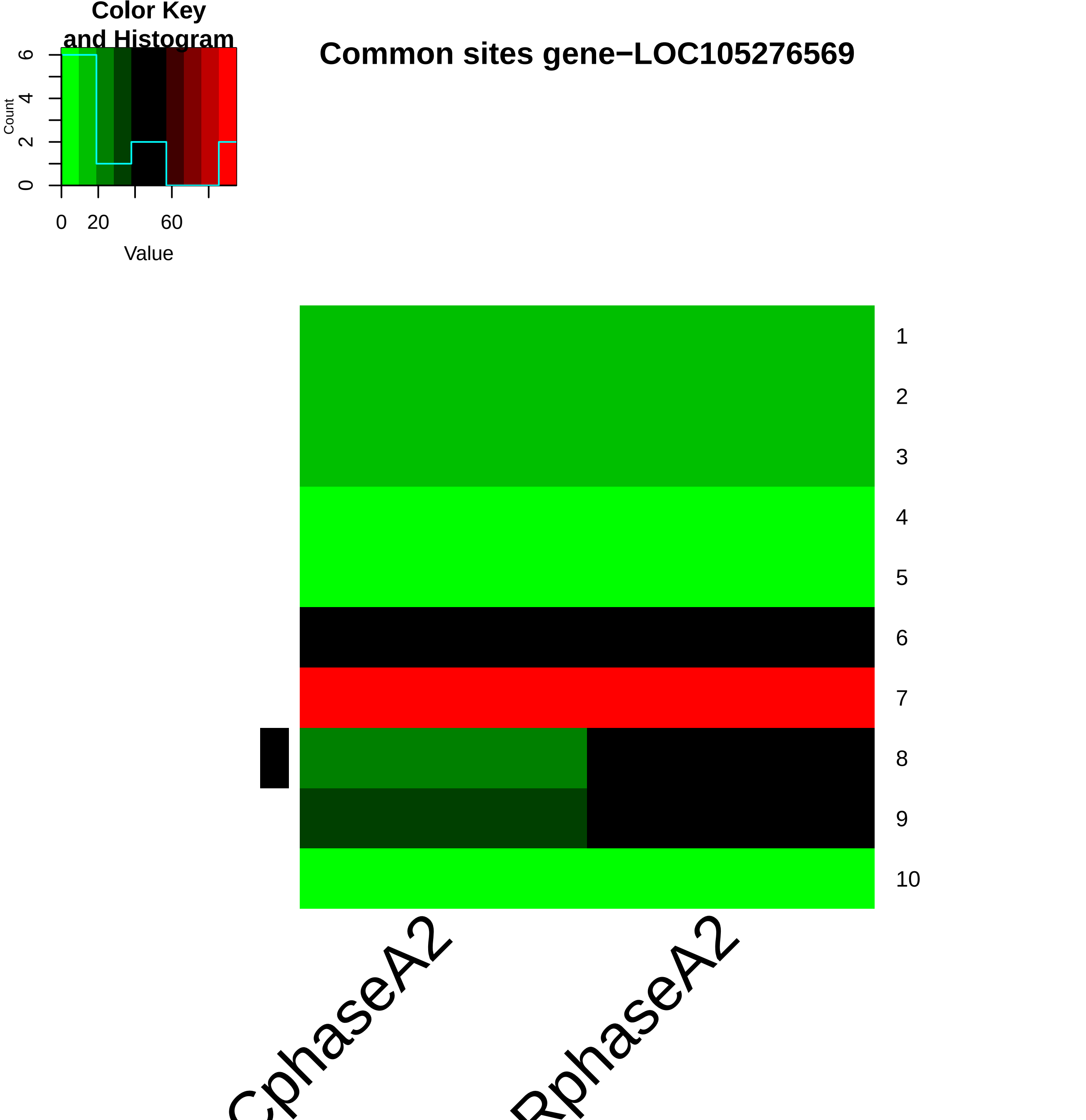
Color Key
and Histogram
Count
Value
0
20
60
0
2
4
6
Common sites gene−LOC105276569
1
2
3
4
5
6
7
8
9
10
CphaseA2
RphaseA2

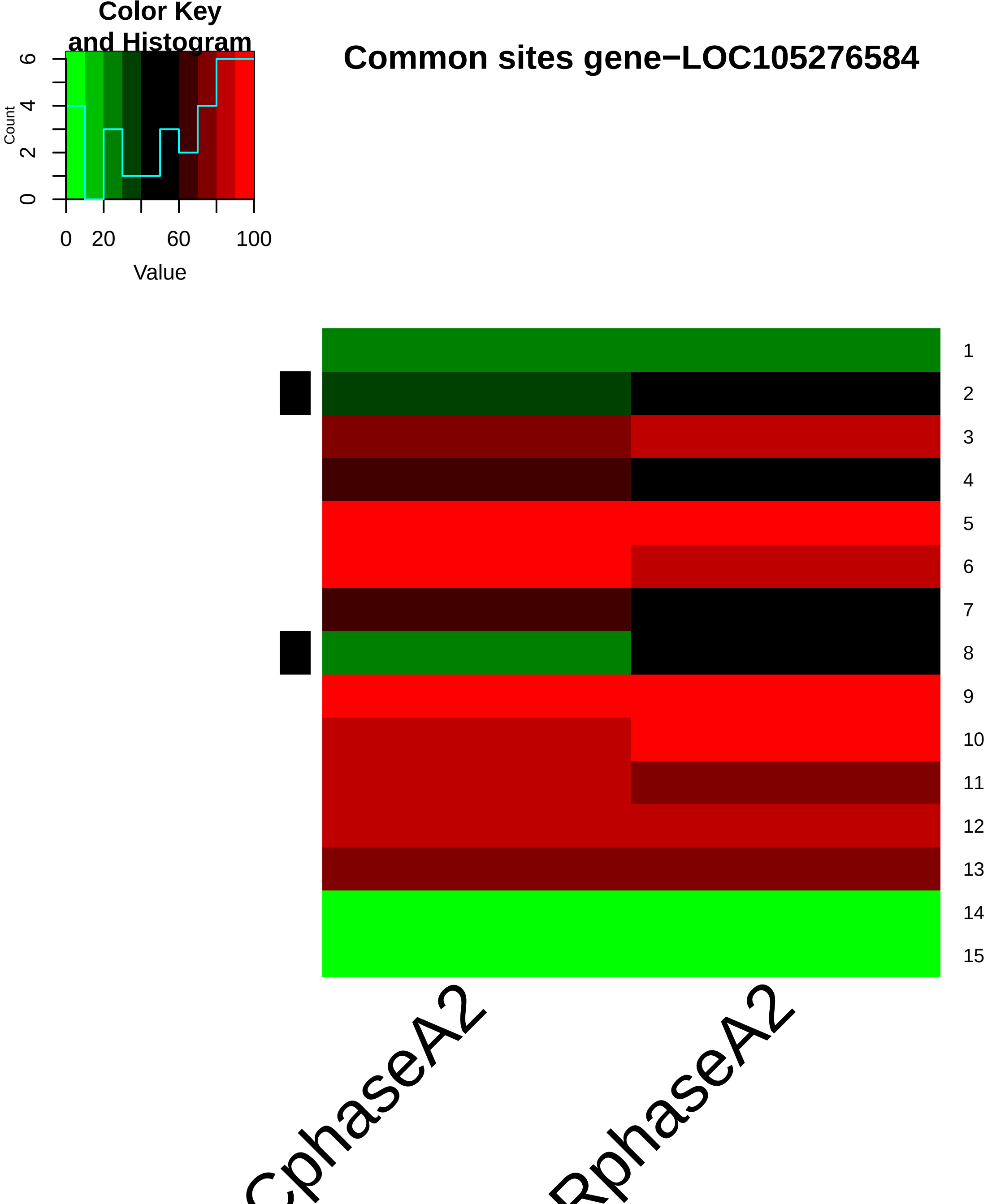
Color Key
and Histogram
Count
0
2
4
6
0
20
60
100
Value
Common sites gene−LOC105276584
CphaseA2
RphaseA2

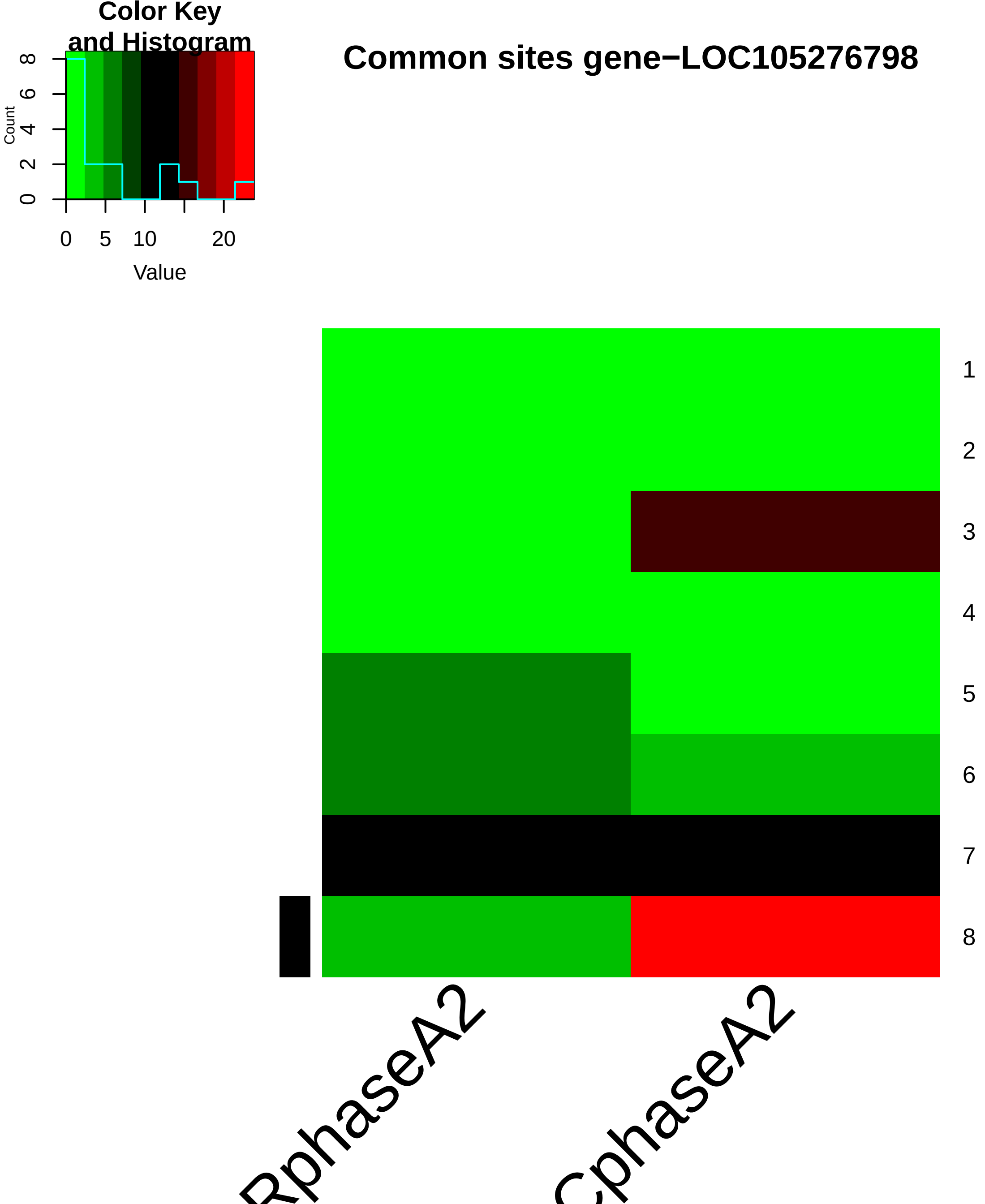

Color Key
and Histogram
Count
0
2
4
6
8
0
5
10
20
Value
Common sites gene−LOC105276798
1
2
3
4
5
6
7
8
RphaseA2
CphaseA2

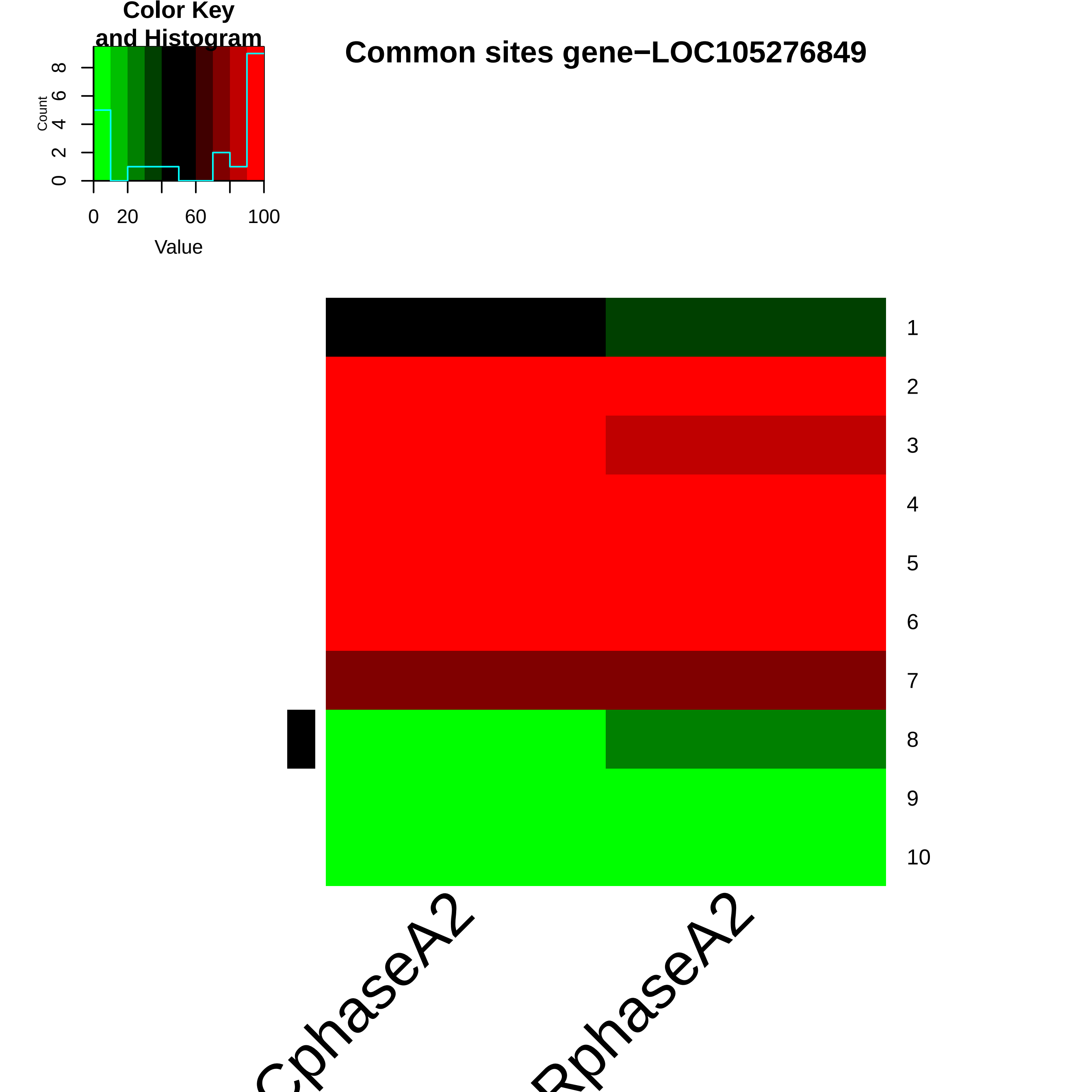
Color Key
and Histogram
Count
0 2 4 6 8
0 20 60 100
Value
Common sites gene−LOC105276849
1
2
3
4
5
6
7
8
9
10
CphaseA2
RphaseA2

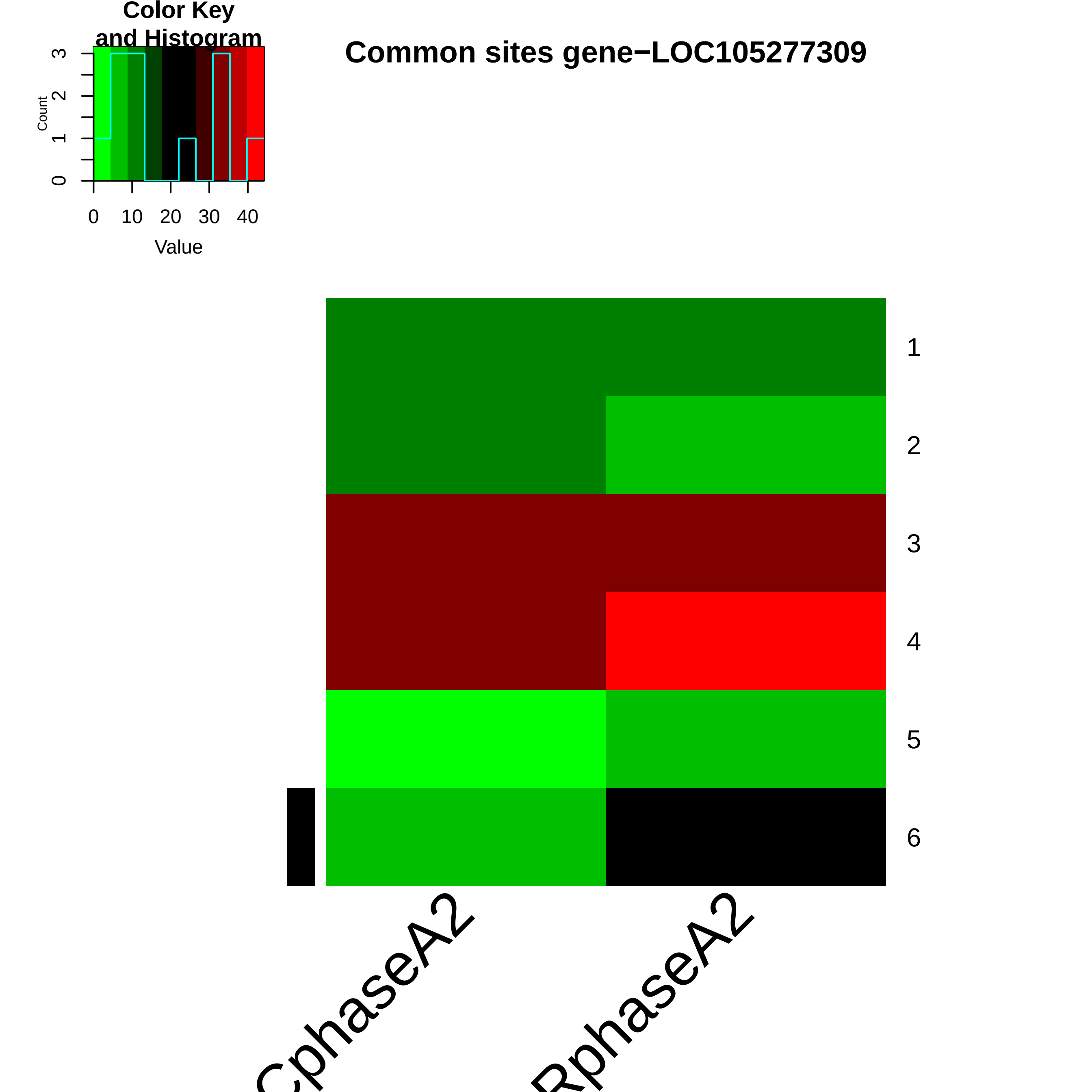

Color Key
and Histogram
Count
Value
0
1
2
3
0
10
20
30
40
Common sites gene−LOC105277309
1
2
3
4
5
6

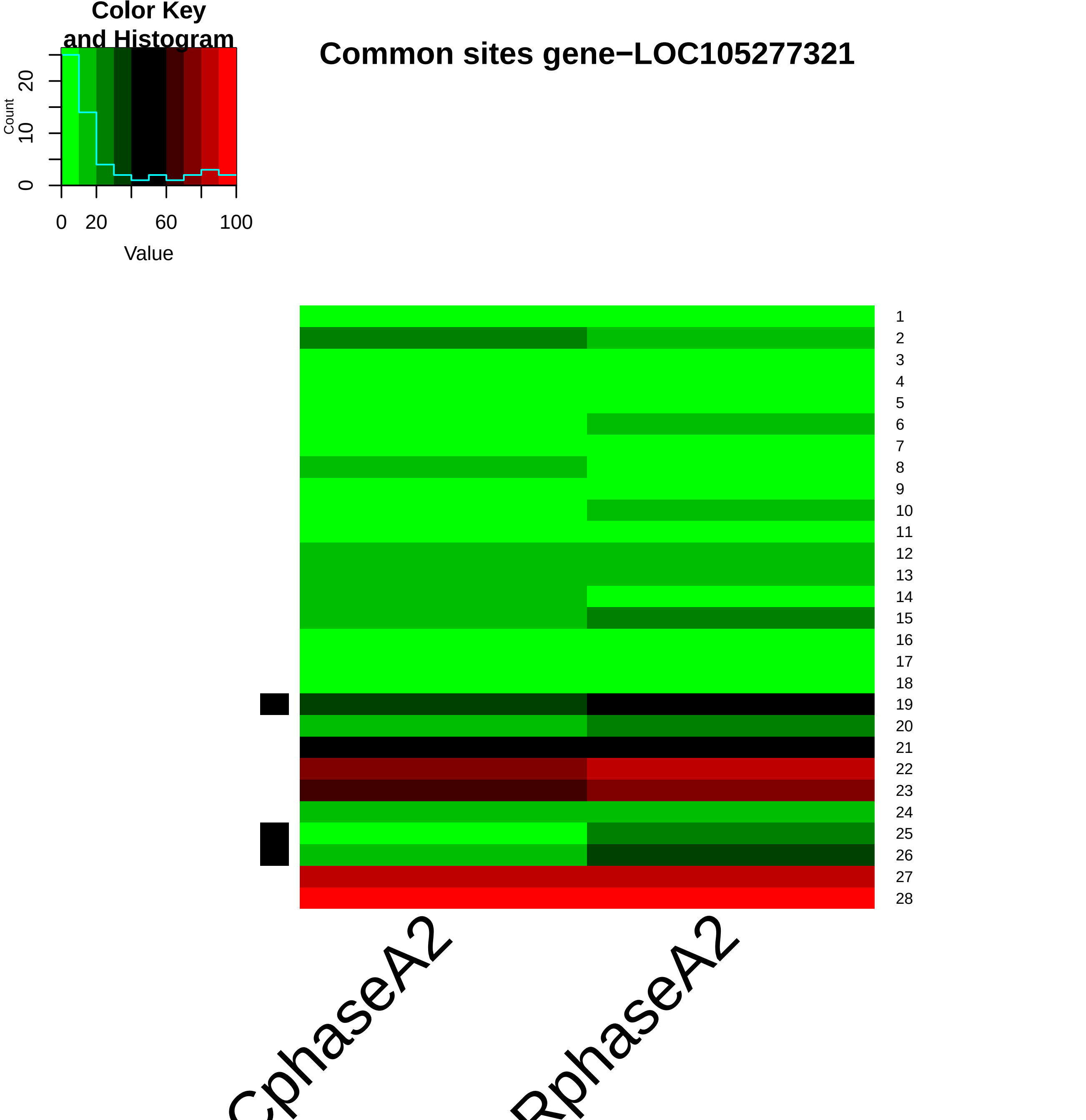

Color Key
and Histogram
Count
0
10
20
0
20
60
100
Value
Common sites gene−LOC105277321

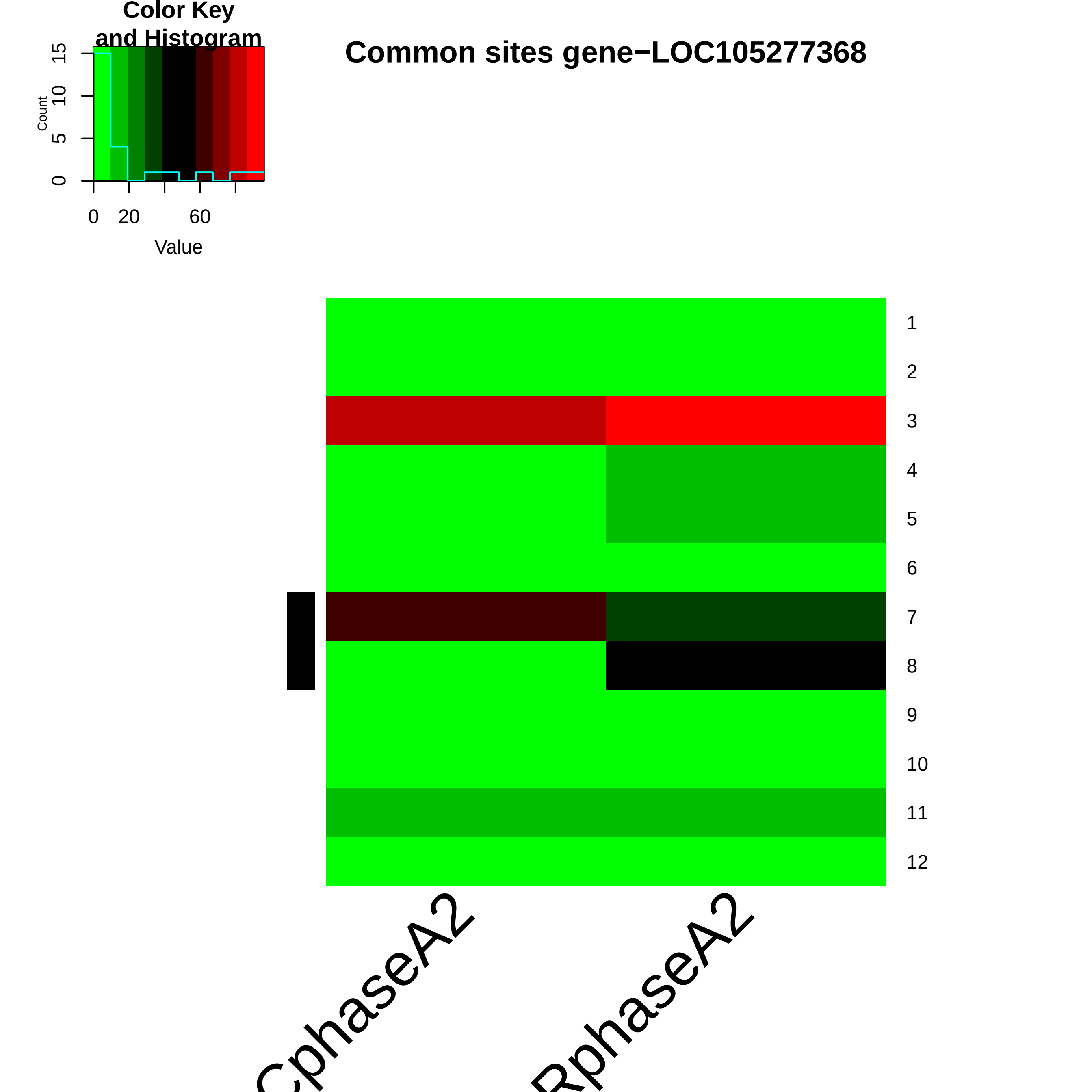

Color Key
and Histogram
Count
0
5
10
15
0
20
60
Value
Common sites gene−LOC105277368
1
2
3
4
5
6
7
8
9
10
11
12

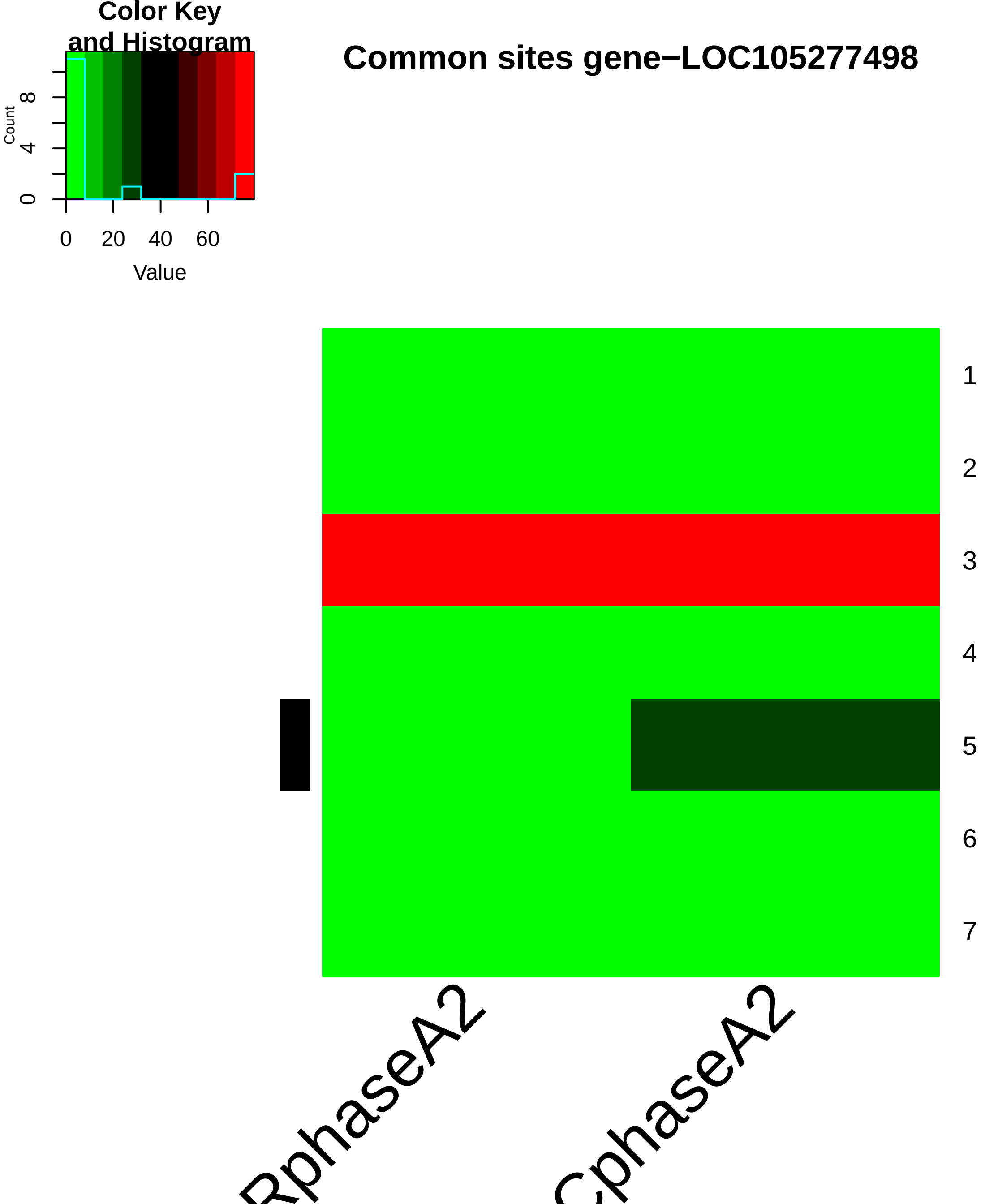

Color Key
and Histogram
Count
0
4
8
0
20
40
60
Value
Common sites gene−LOC105277498
1
2
3
4
5
6
7
RphaseA2
CphaseA2

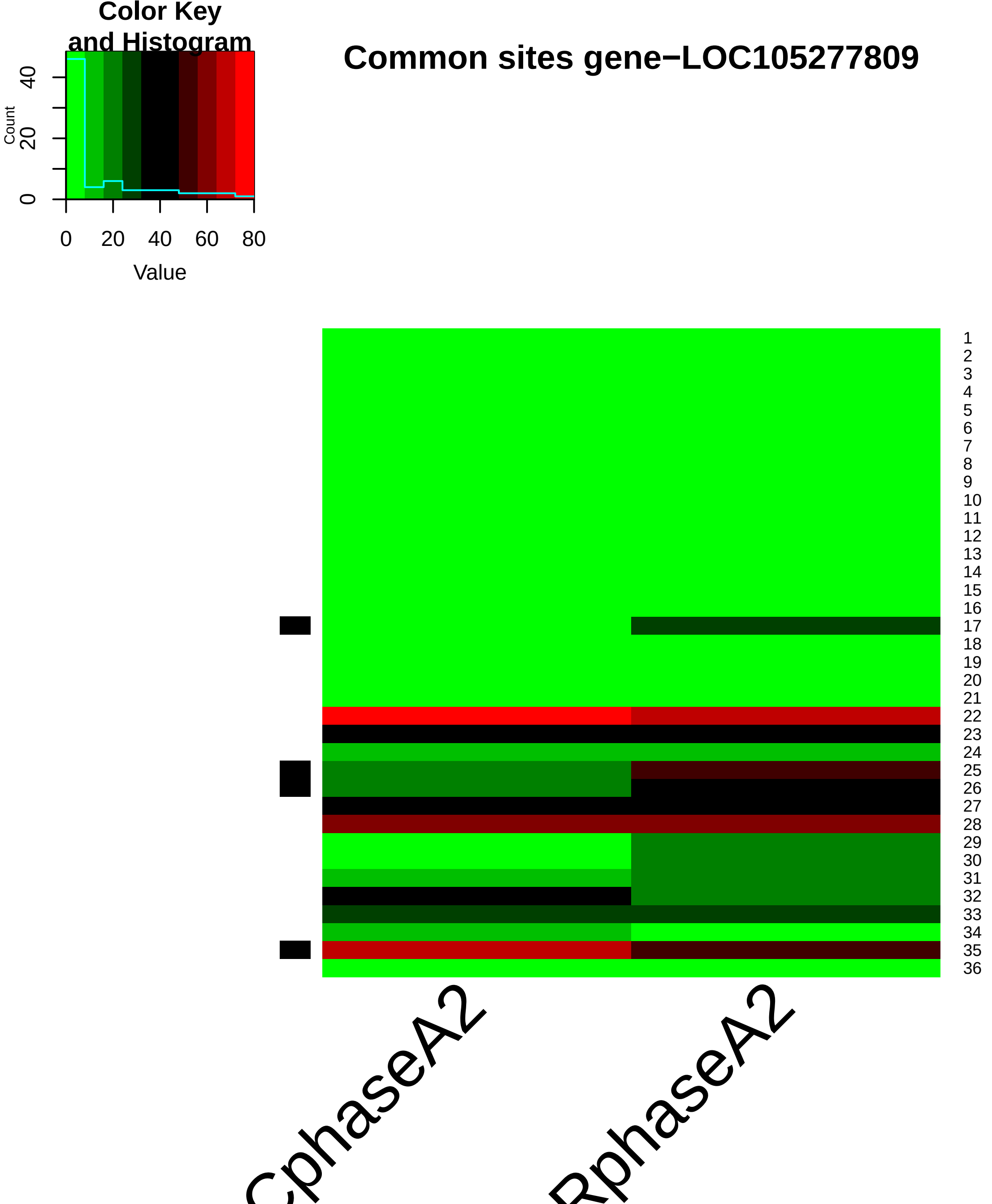
Color Key
and Histogram
Count
0
20
40
0
20
40
60
80
Value
Common sites gene−LOC105277809
CphaseA2
RphaseA2

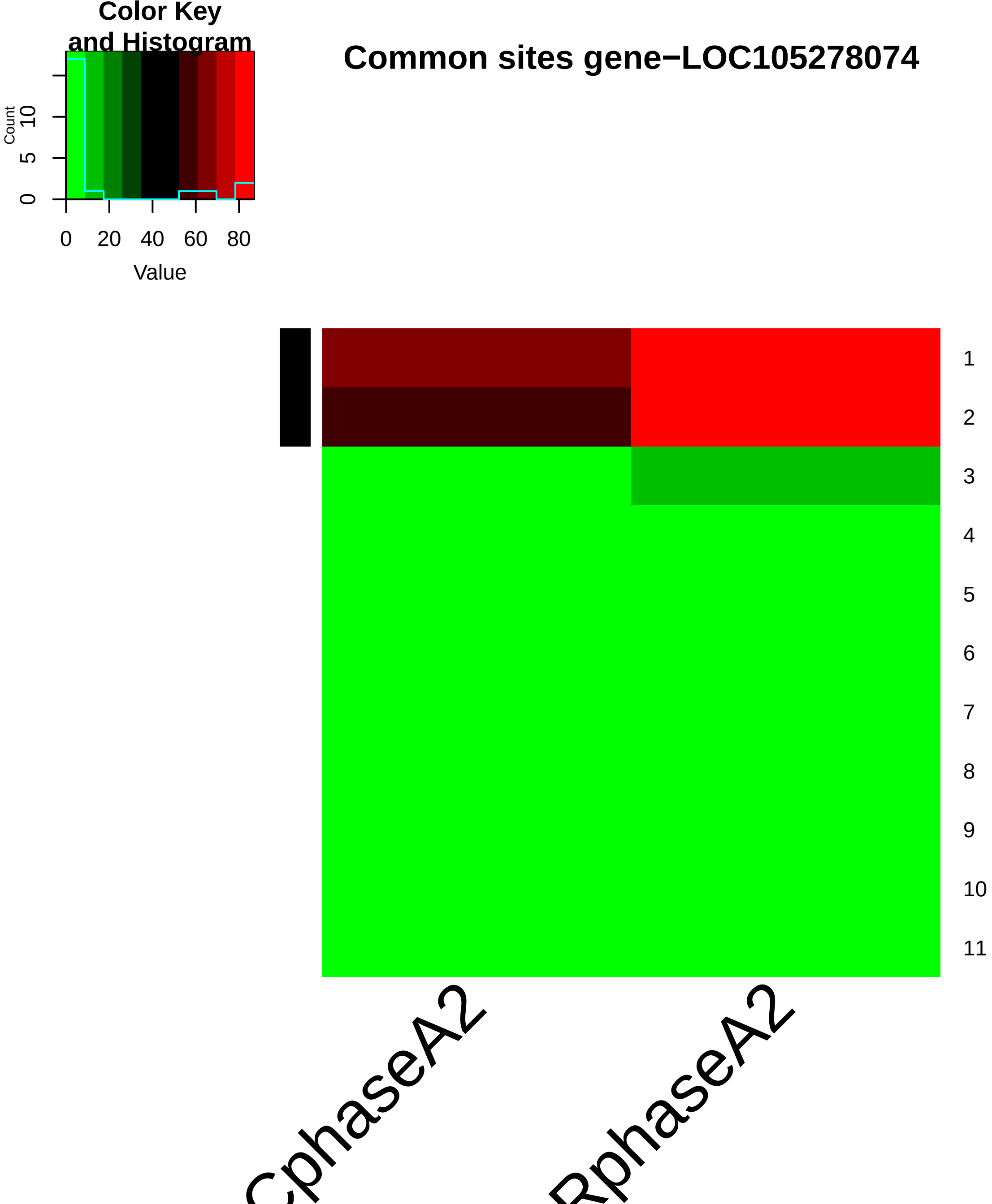

Color Key
and Histogram
Count
0
5
10
0
20
40
60
80
Value
Common sites gene−LOC105278074
1
2
3
4
5
6
7
8
9
10
11
CphaseA2
RphaseA2

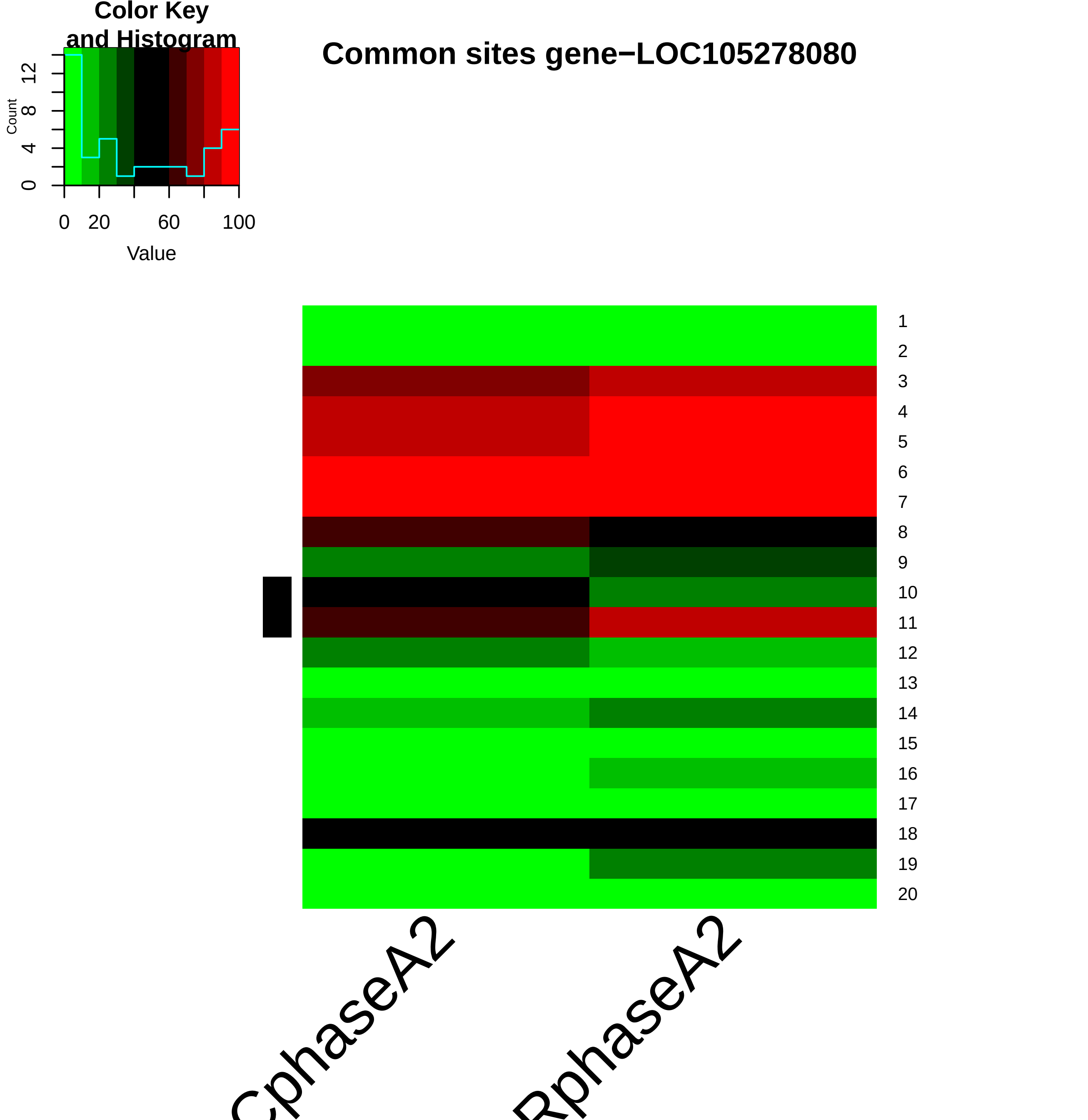
Color Key
and Histogram
Count
0
4
8
12
0
20
60
100
Value
Common sites gene−LOC105278080
CphaseA2
RphaseA2

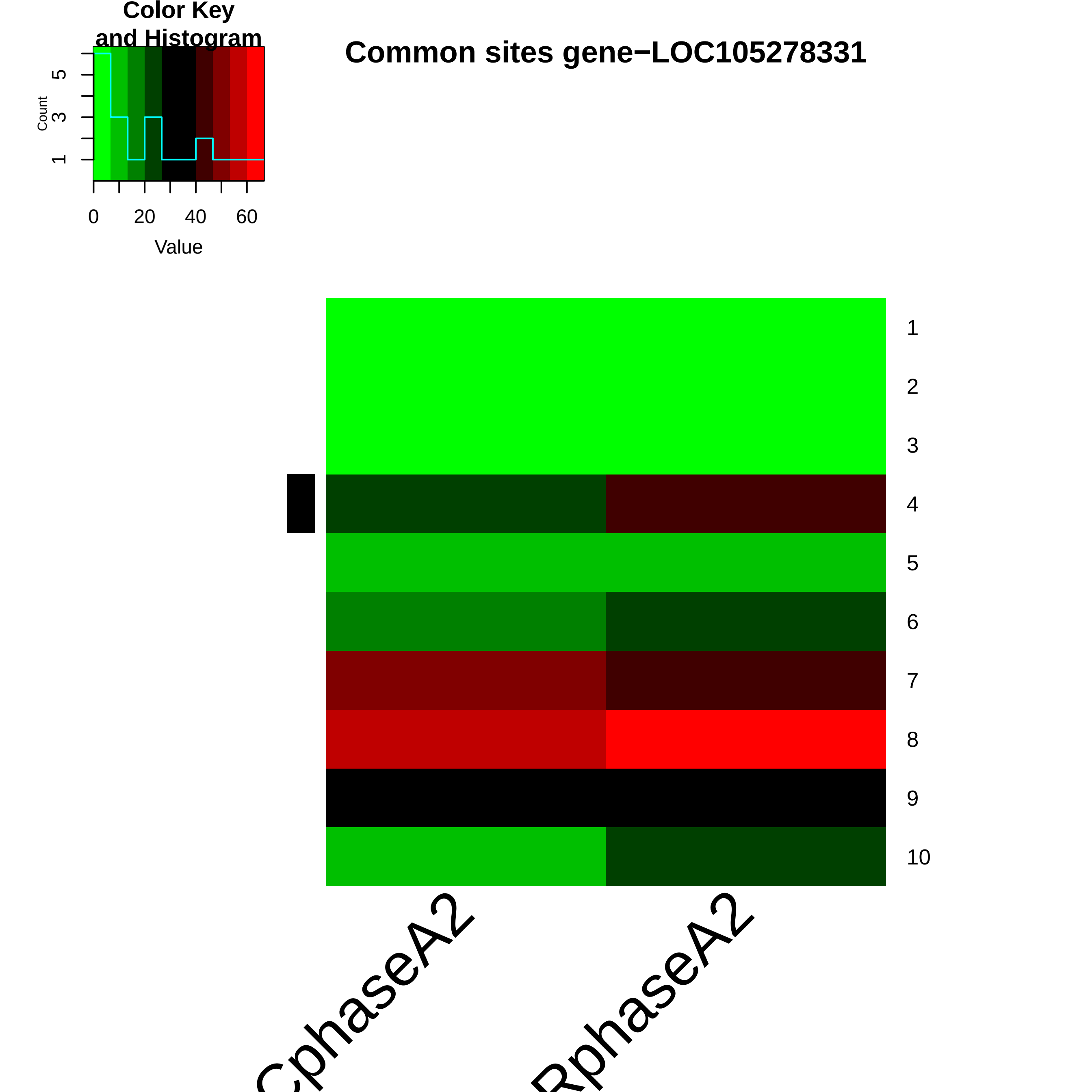
Color Key
and Histogram
Count
1
3
5
0
20
40
60
Value
Common sites gene−LOC105278331
1
2
3
4
5
6
7
8
9
10
CphaseA2
RphaseA2

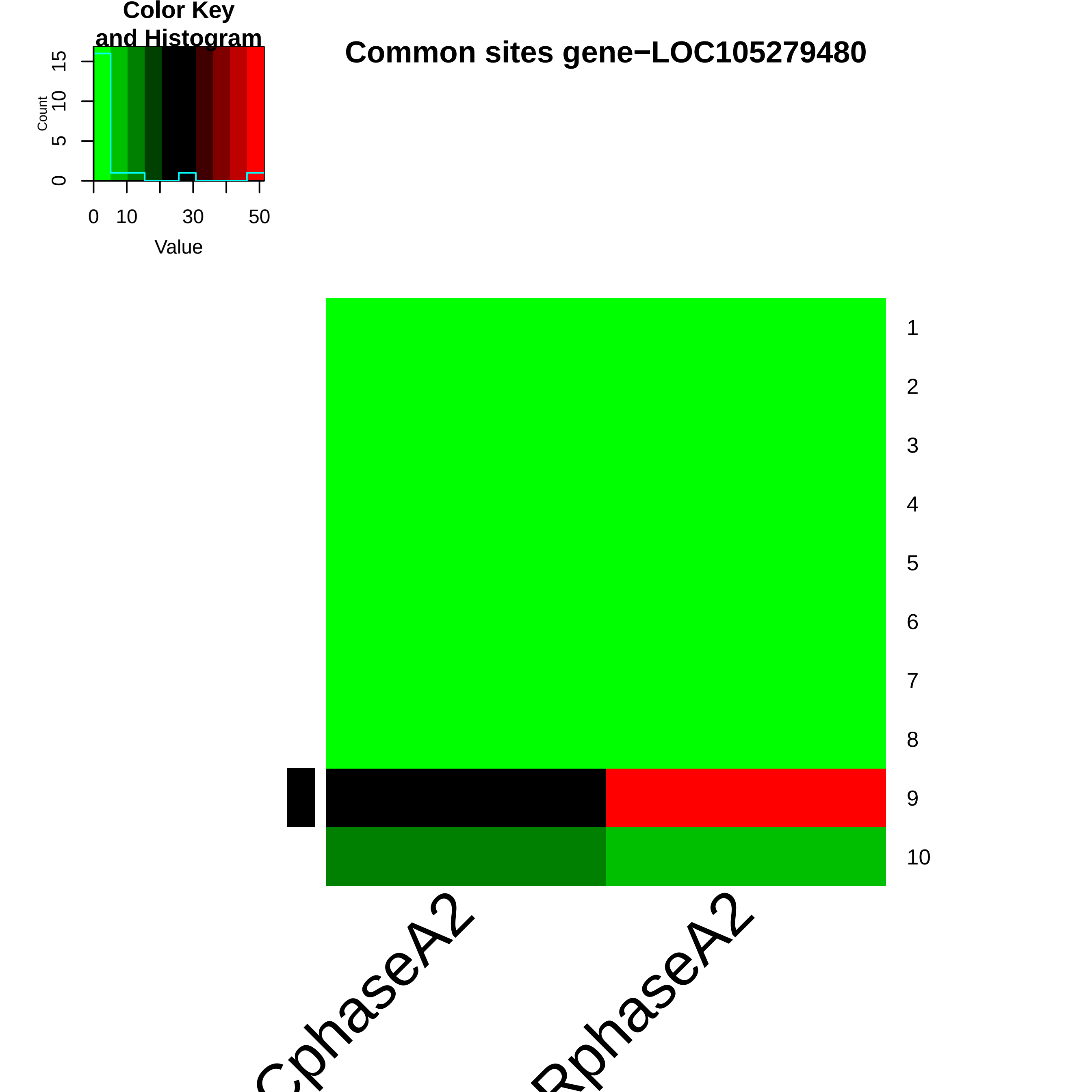
Color Key
and Histogram
Count
0
5
10
15
0
10
30
50
Value
Common sites gene−LOC105279480
1
2
3
4
5
6
7
8
9
10
CphaseA2
RphaseA2

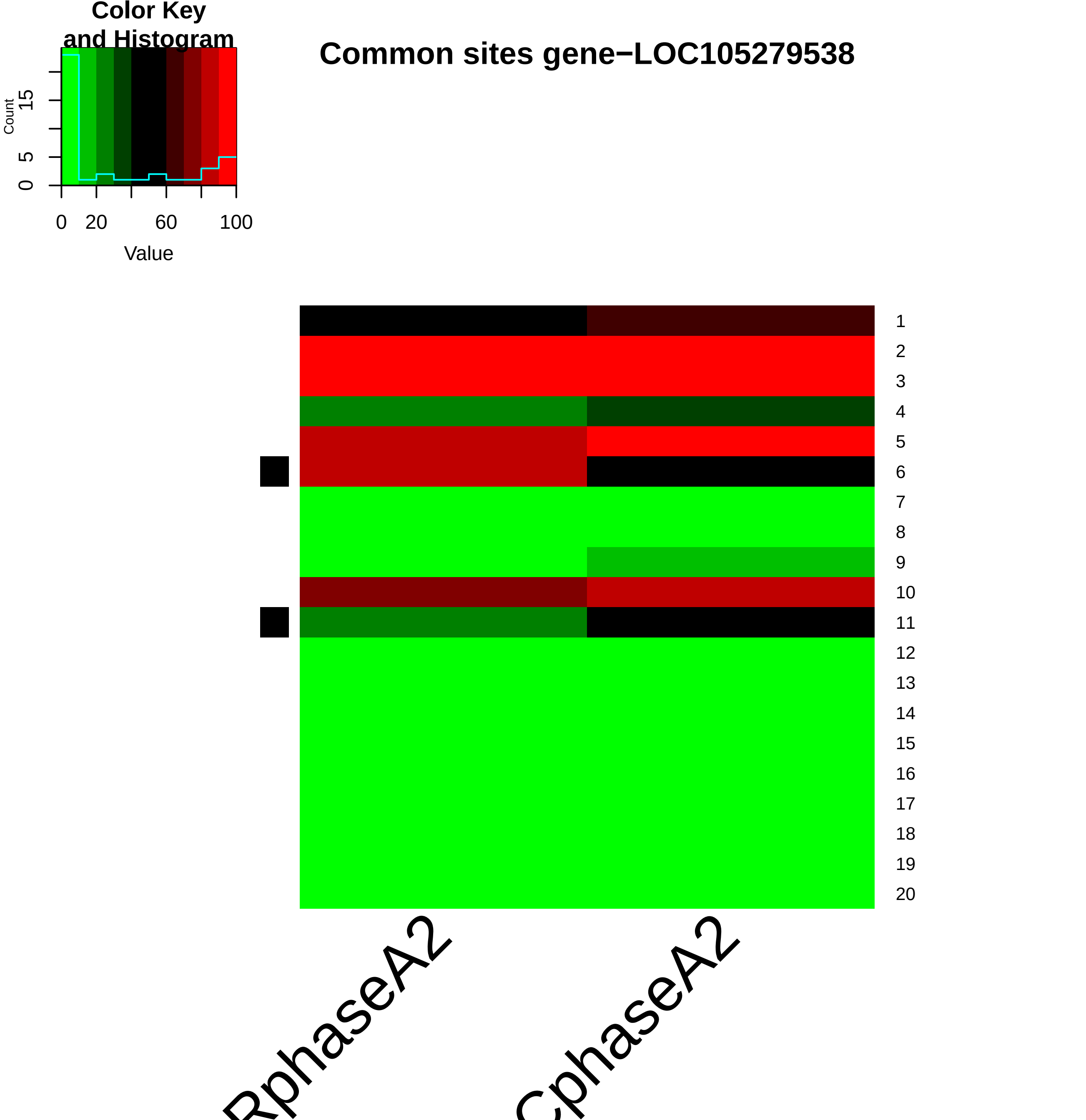
Color Key
and Histogram
Count
0
5
15
0
20
60
100
Value
Common sites gene−LOC105279538
RphaseA2
CphaseA2

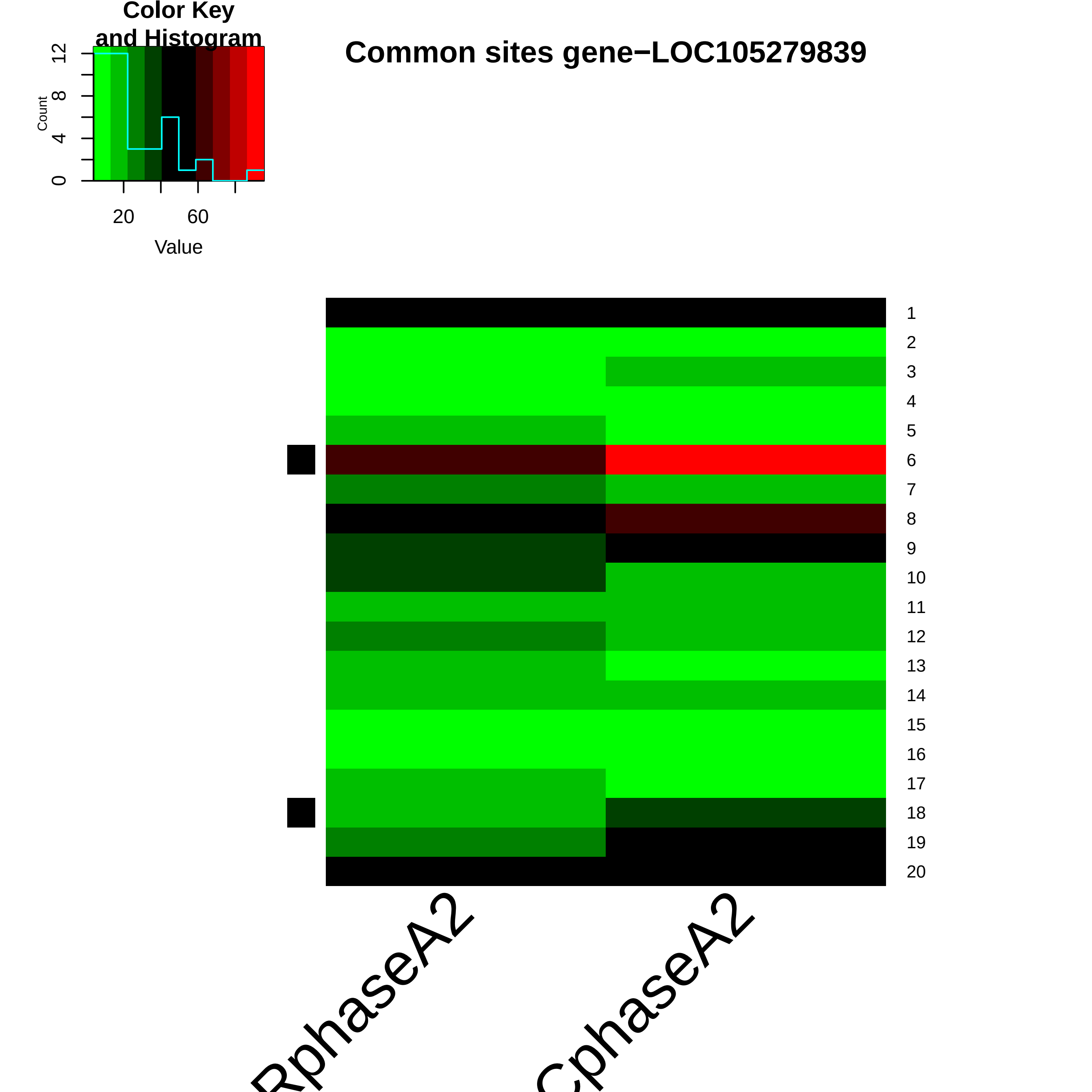
Color Key
and Histogram
Count
0
4
8
12
20
60
Value
Common sites gene−LOC105279839
RphaseA2
CphaseA2

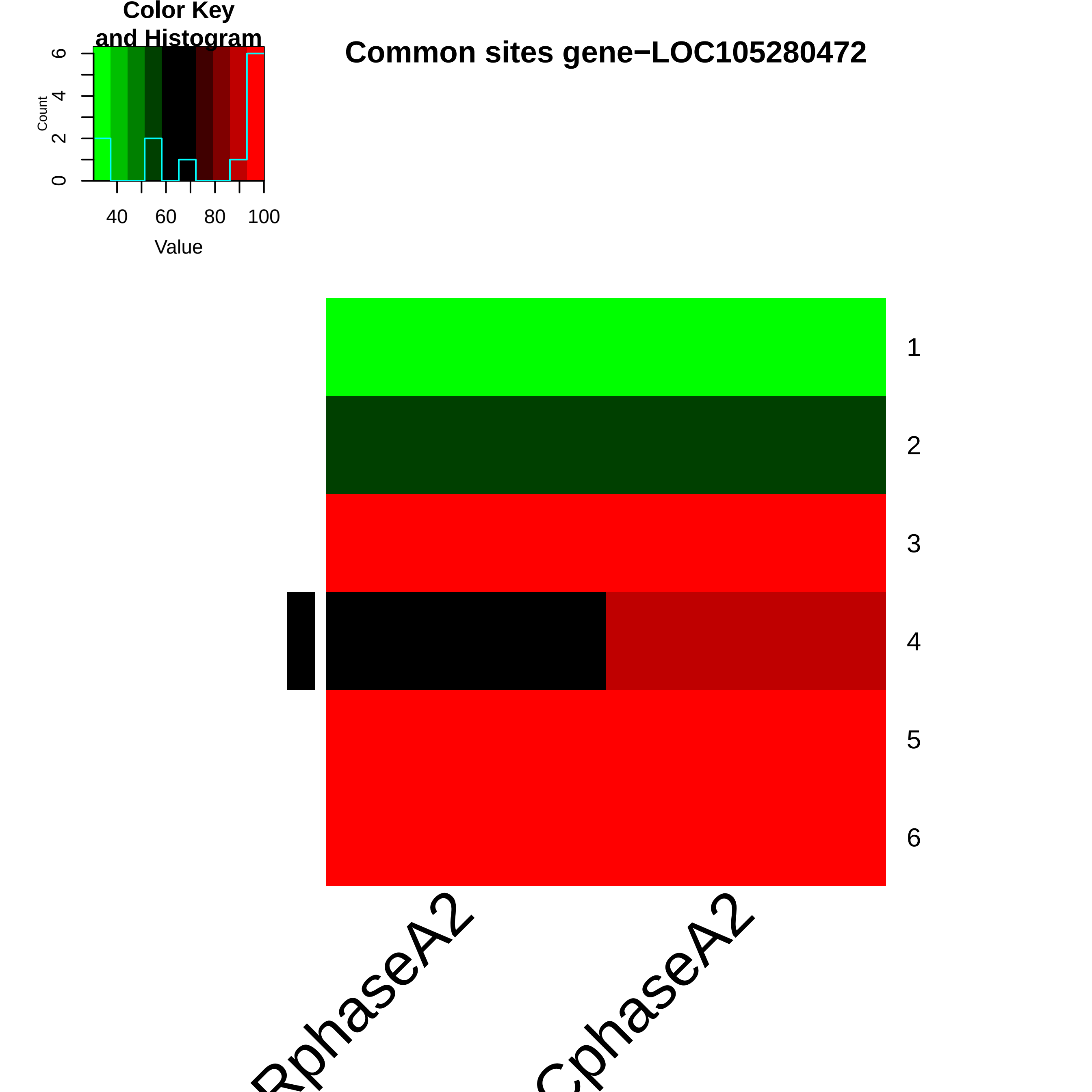

Color Key
and Histogram
Count
0
2
4
6
40
60
80
100
Value
Common sites gene−LOC105280472
1
2
3
4
5
6
RphaseA2
CphaseA2

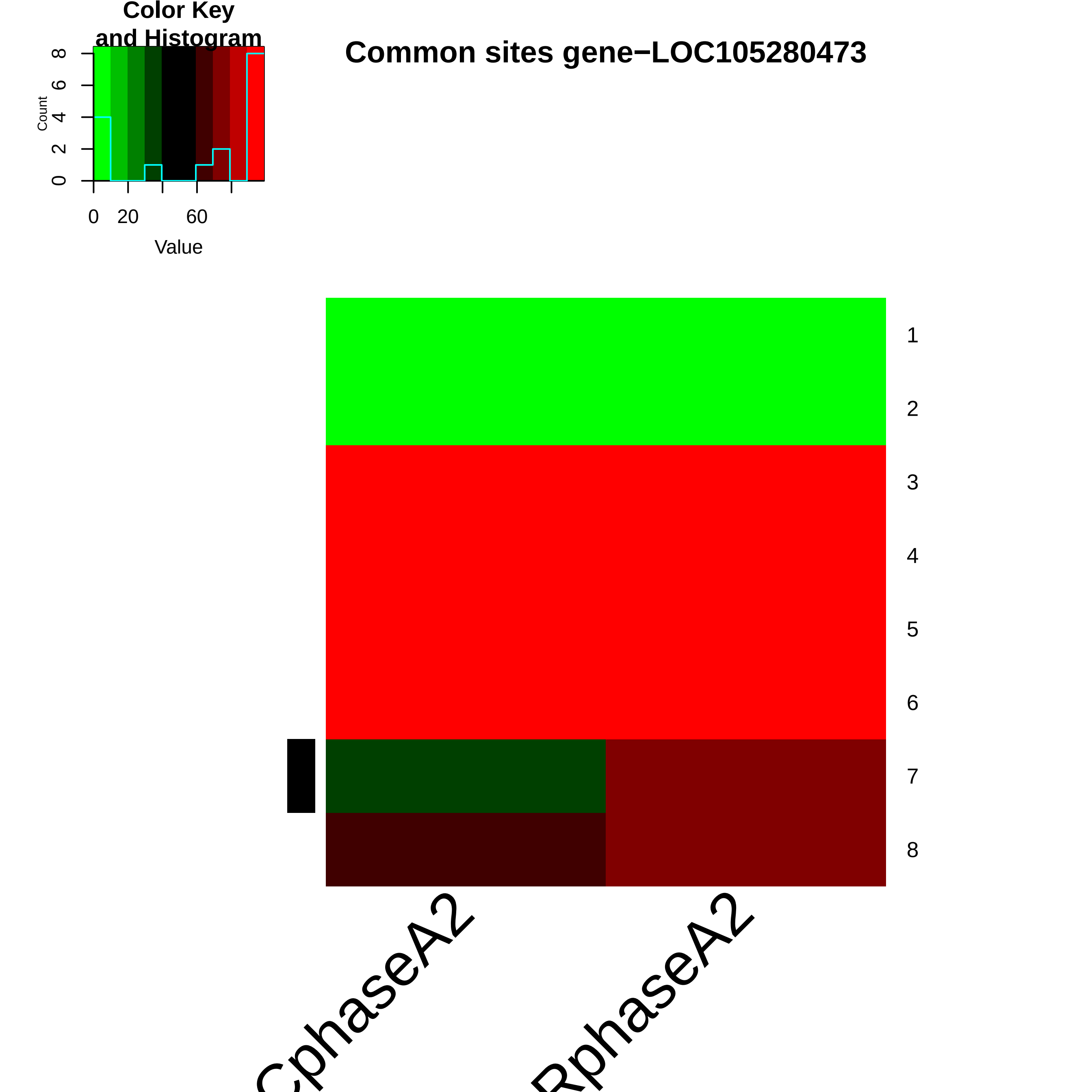
Color Key
and Histogram
Count
Value
0
20
60
0
2
4
6
8
Common sites gene−LOC105280473
1
2
3
4
5
6
7
8
CphaseA2
RphaseA2

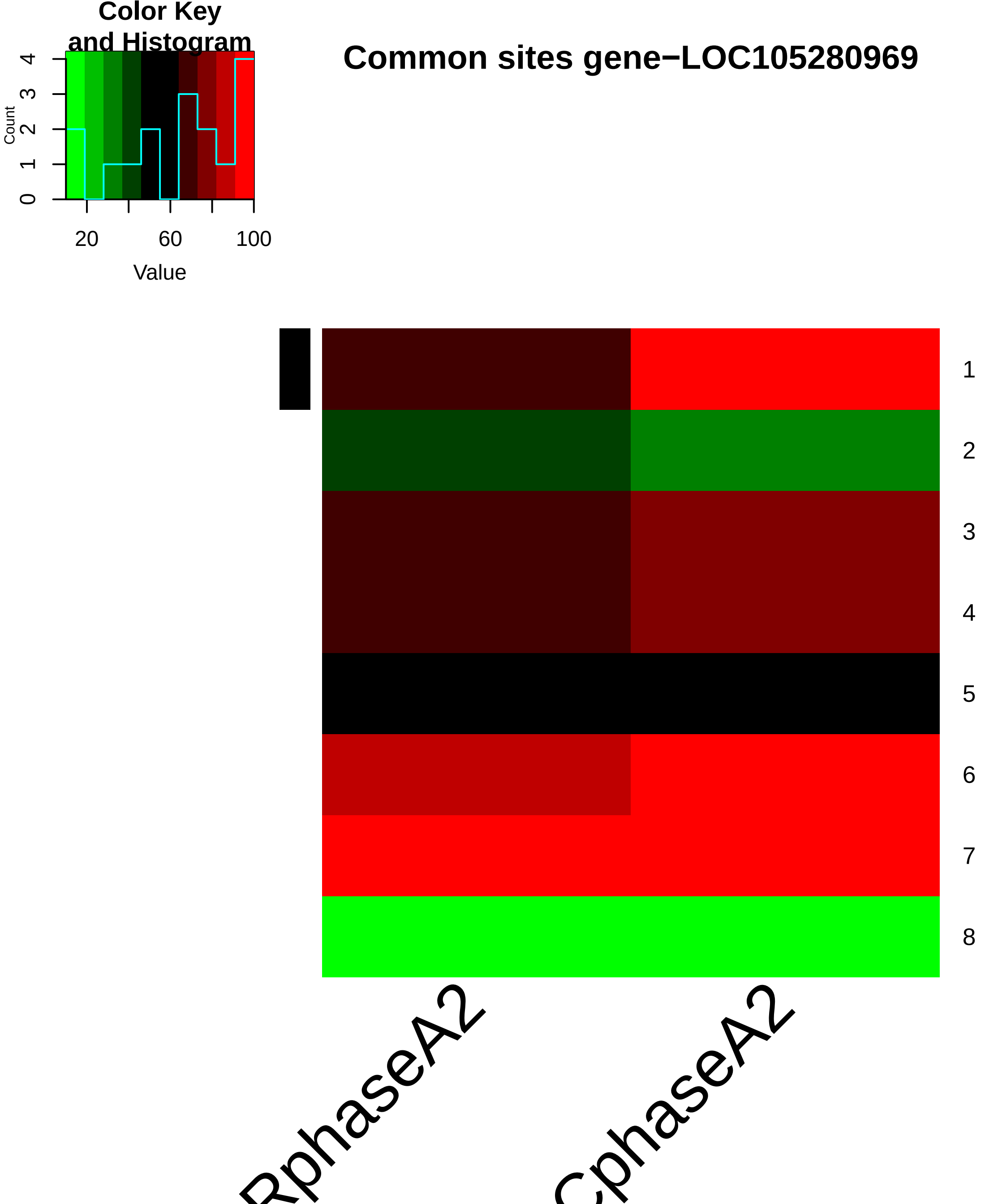

Color Key
and Histogram
Count
0
1
2
3
4
20
60
100
Value
Common sites gene−LOC105280969
1
2
3
4
5
6
7
8
RphaseA2
CphaseA2

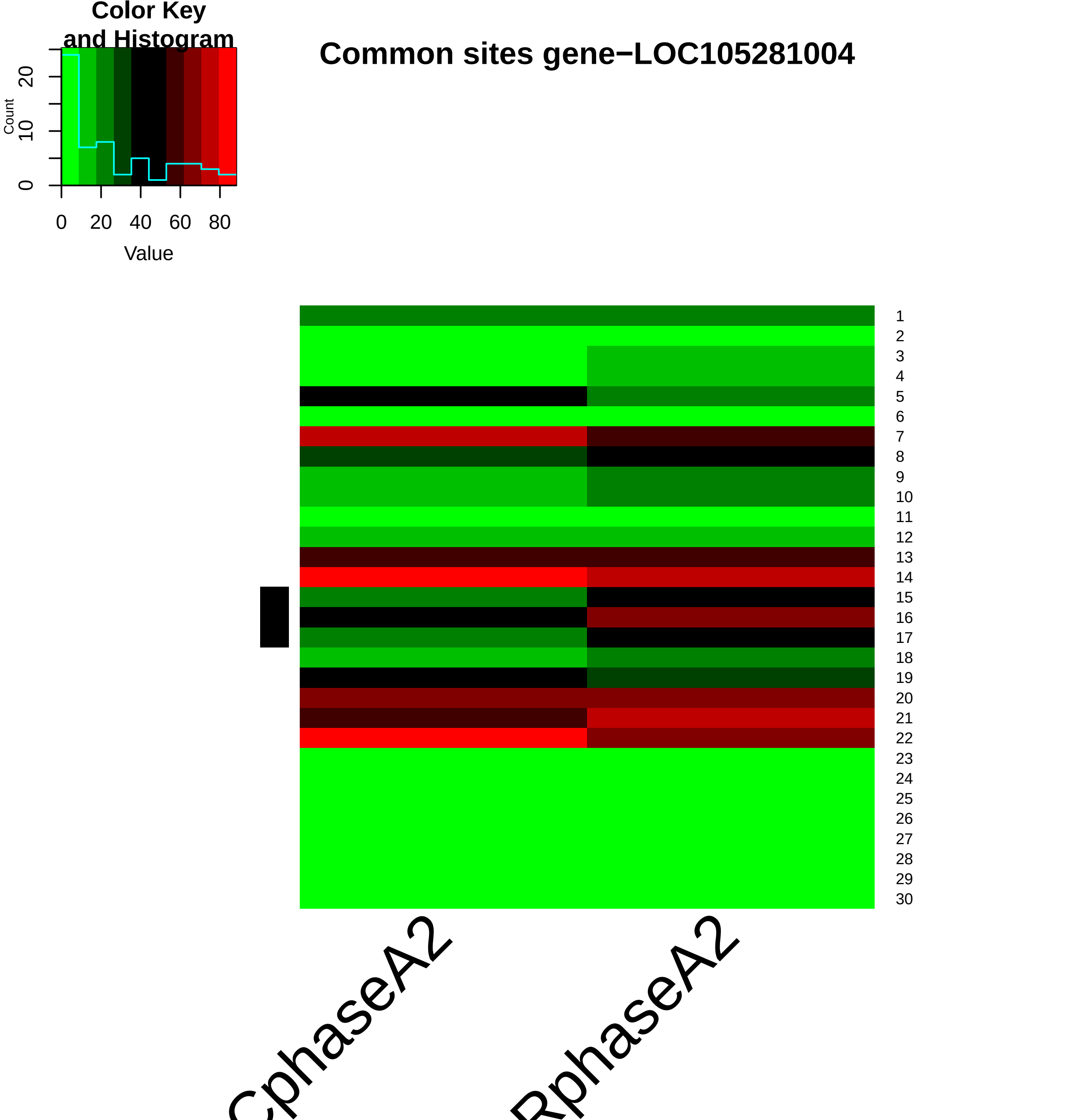

Color Key
and Histogram
Count
0
10
20
0
20
40
60
80
Value
Common sites gene−LOC105281004
CphaseA2
RphaseA2

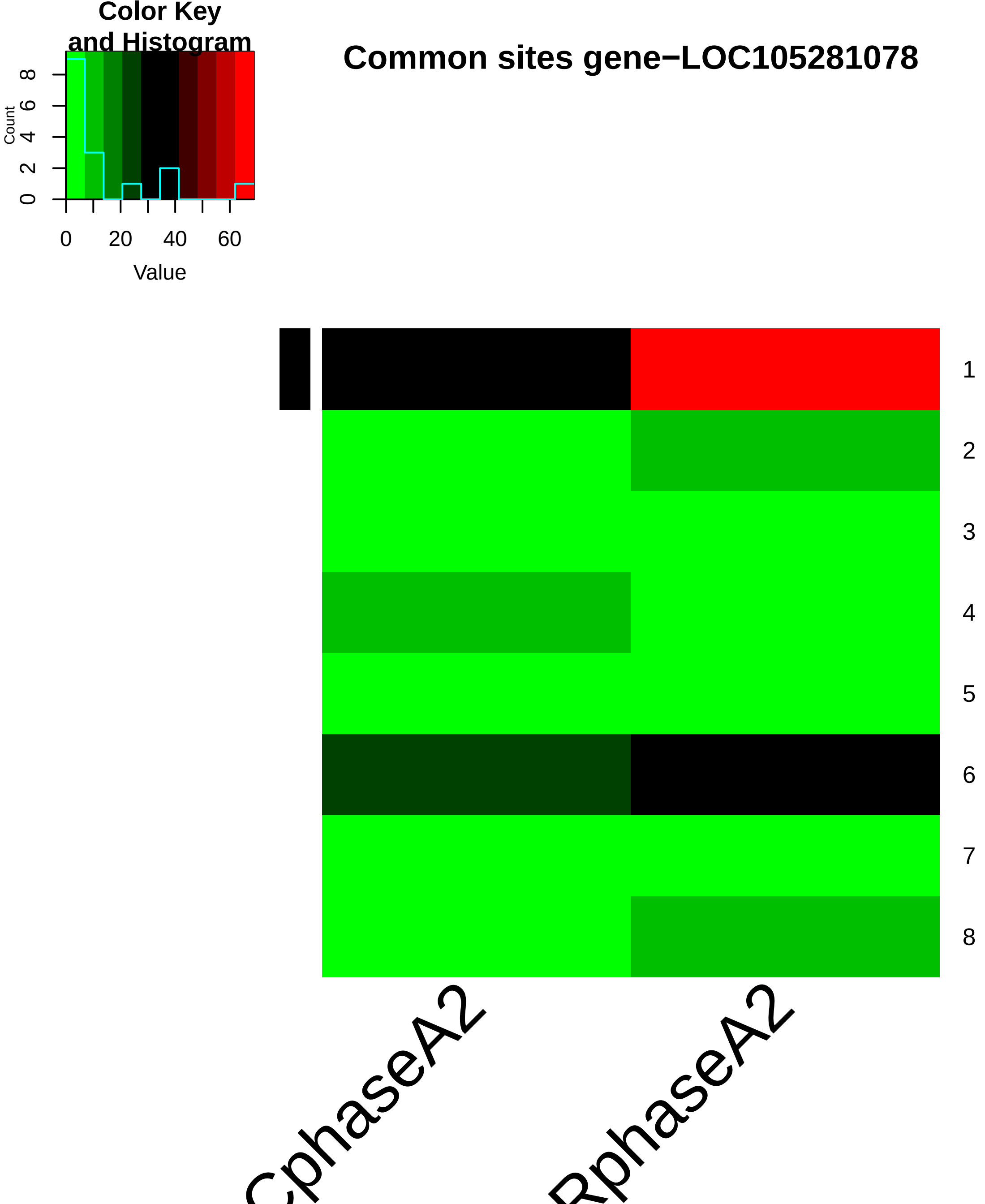
Color Key
and Histogram
Count
0
2
4
6
8
0
20
40
60
Value
Common sites gene−LOC105281078
1
2
3
4
5
6
7
8
CphaseA2
RphaseA2

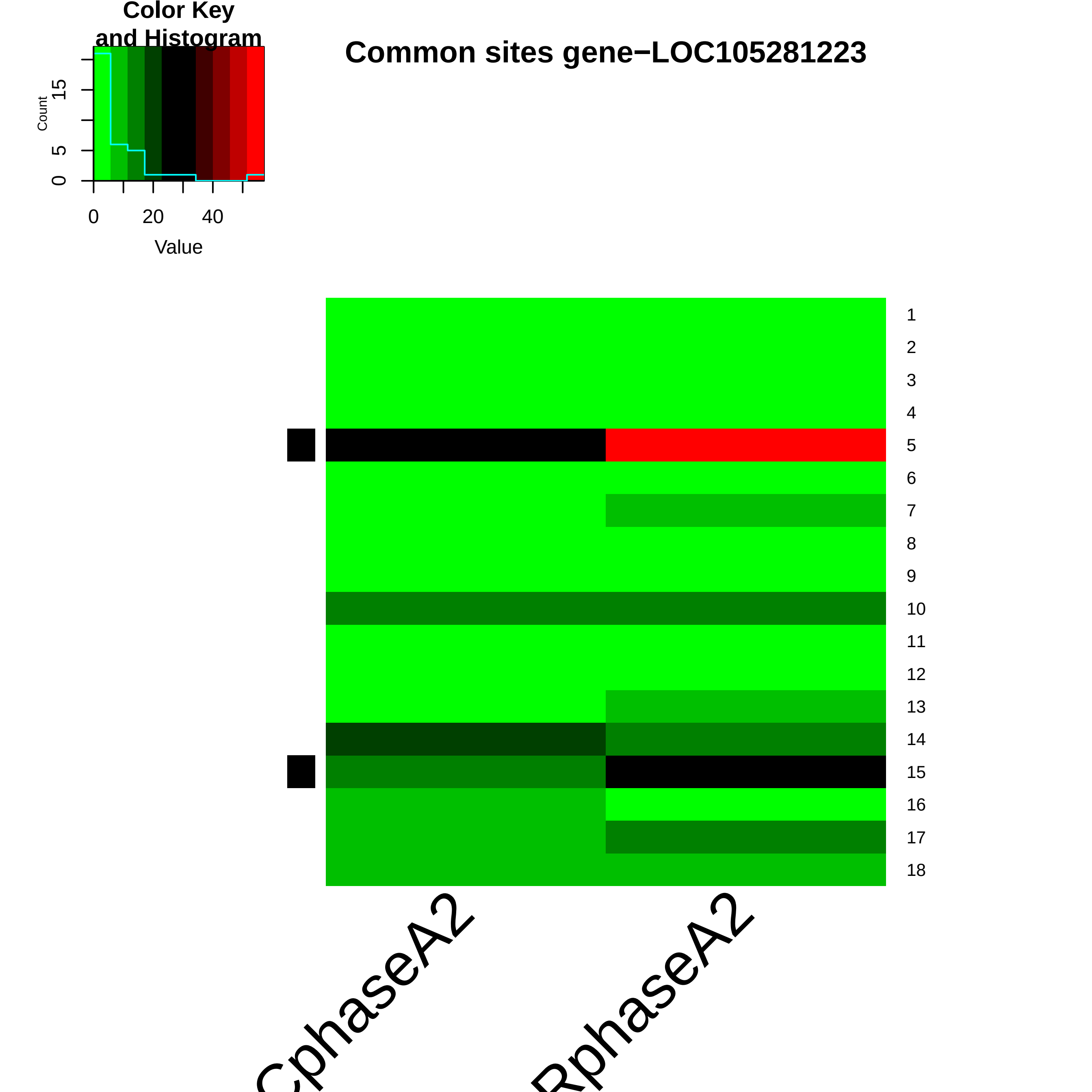

Color Key
and Histogram
Count
0
5
15
0
20
40
Value
Common sites gene−LOC105281223
CphaseA2
RphaseA2

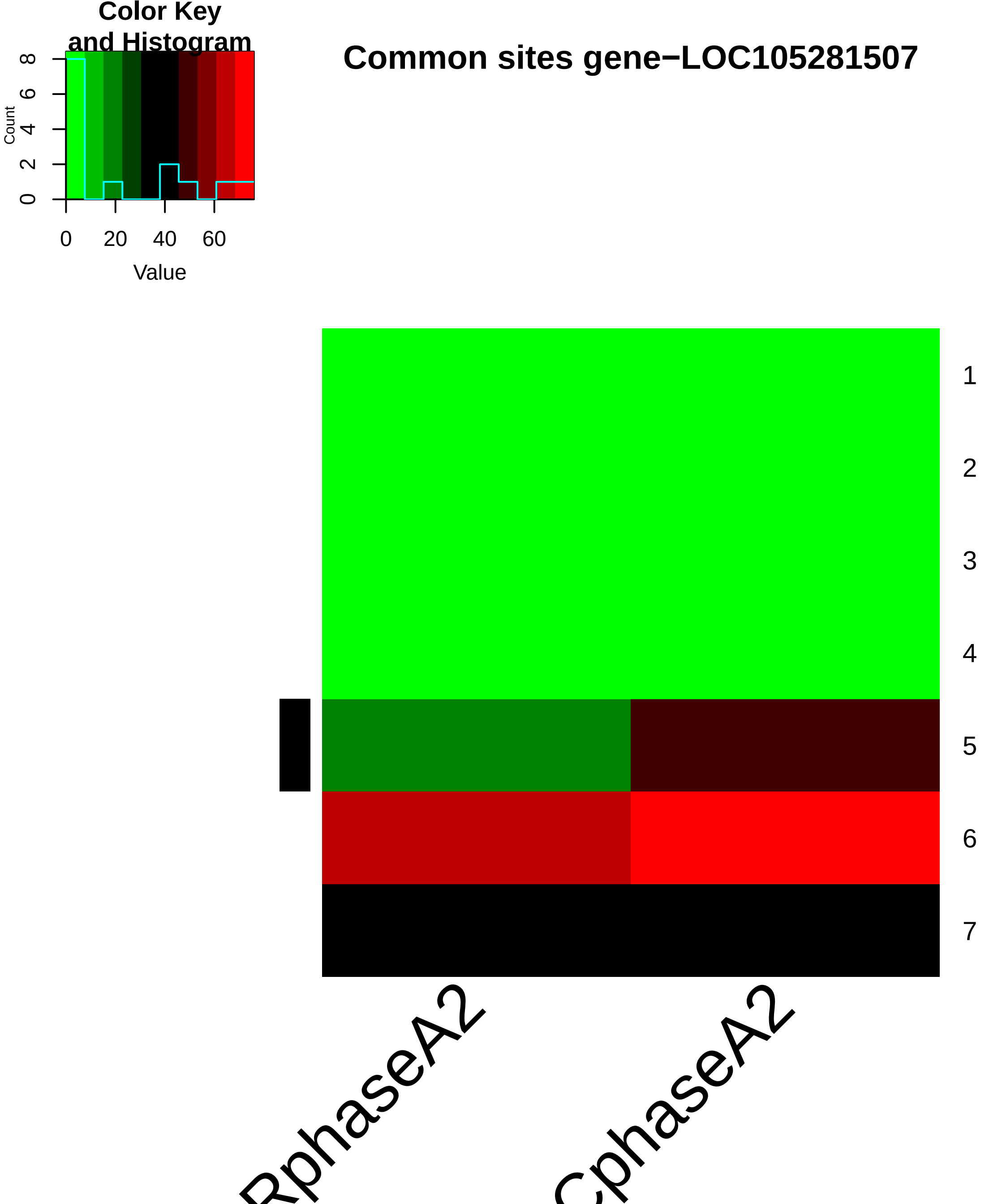
Color Key
and Histogram
Count
0 2 4 6 8
0 20 40 60
Value
Common sites gene−LOC105281507
1
2
3
4
5
6
7
RphaseA2
CphaseA2

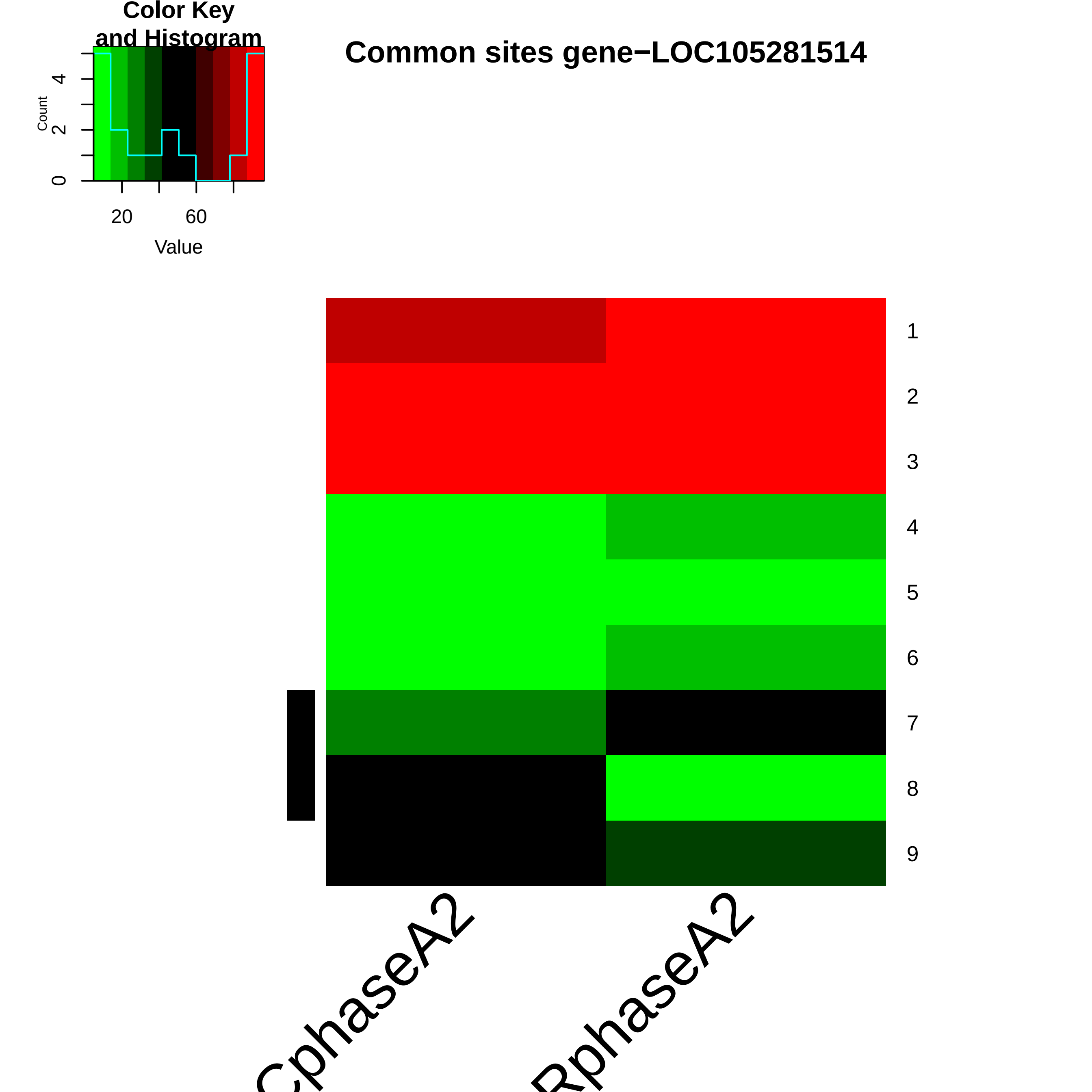

Color Key
and Histogram
Count
0
2
4
20
60
Value
Common sites gene−LOC105281514
1
2
3
4
5
6
7
8
9

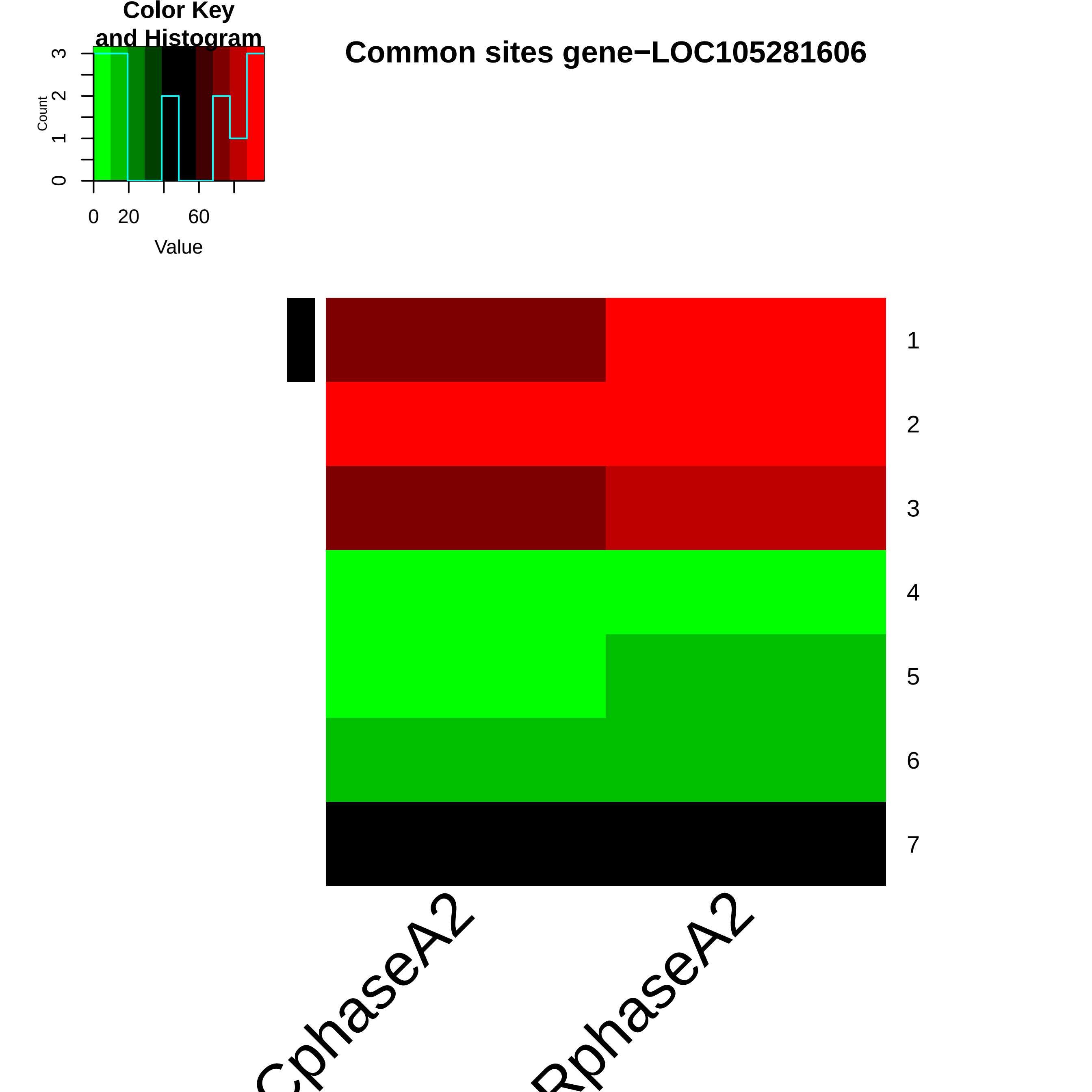

Color Key
and Histogram
Count
0
1
2
3
0
20
60
Value
Common sites gene−LOC105281606
1
2
3
4
5
6
7
CphaseA2
RphaseA2

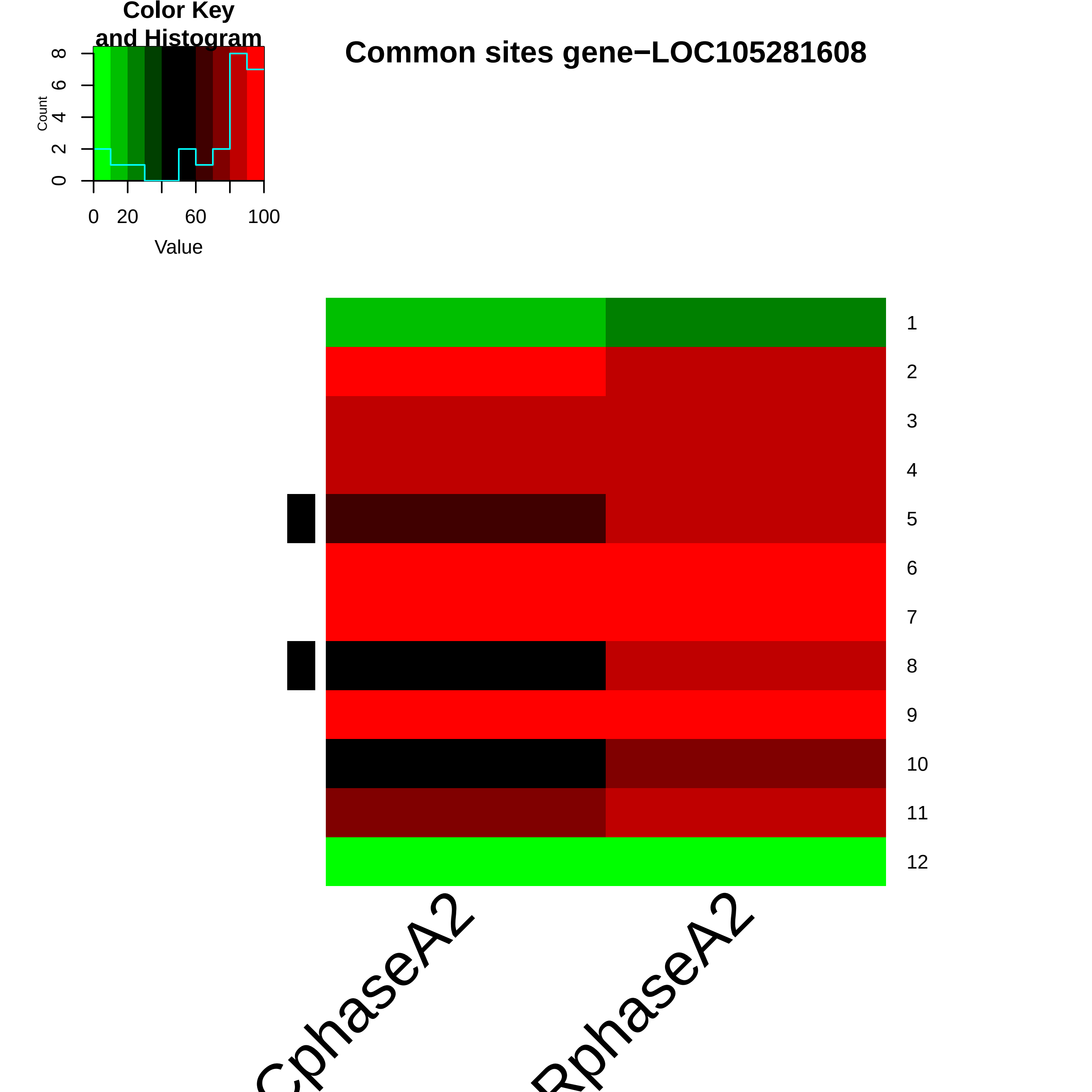

Color Key
and Histogram
Count
0
2
4
6
8
0
20
60
100
Value
Common sites gene−LOC105281608
1
2
3
4
5
6
7
8
9
10
11
12
CphaseA2
RphaseA2

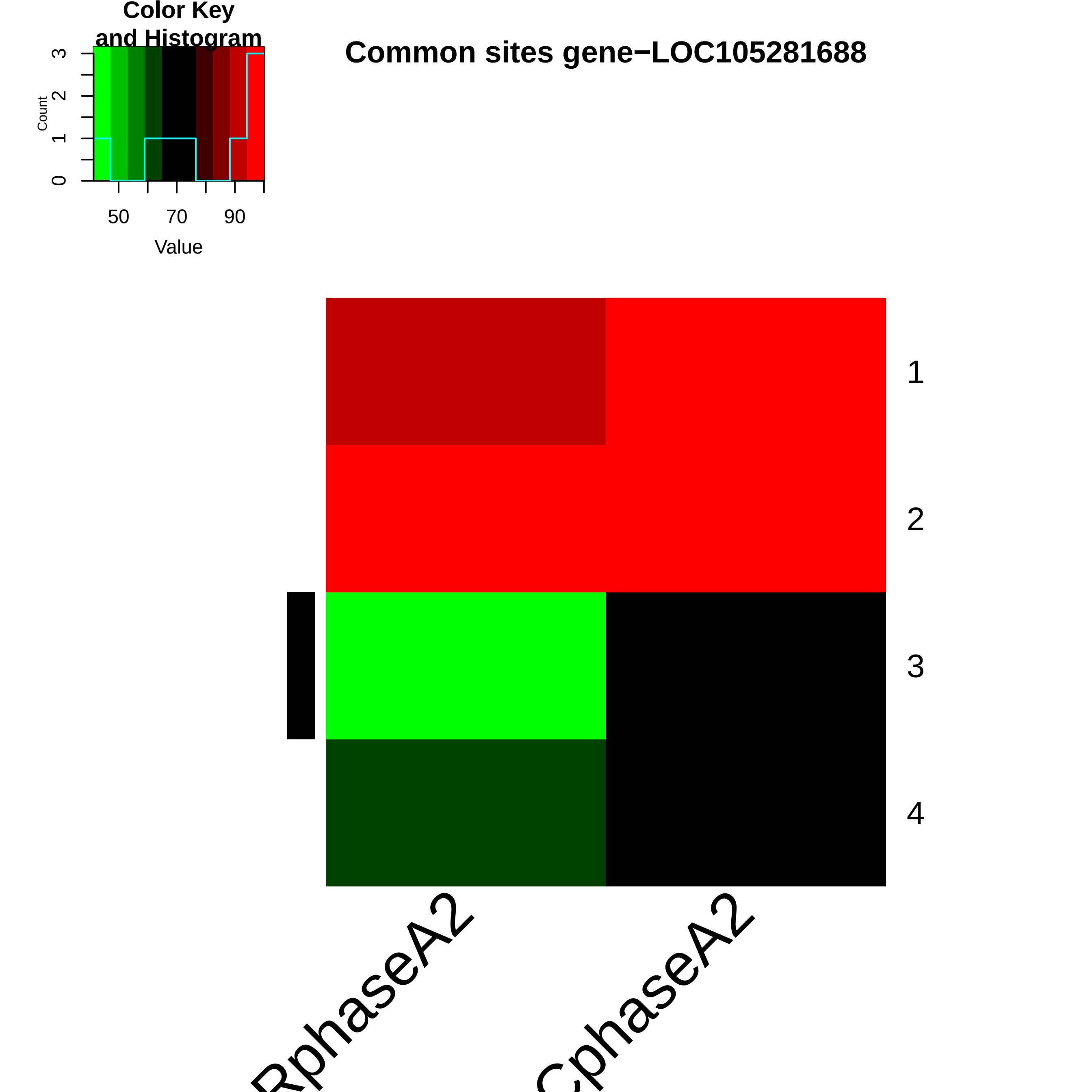

Color Key
and Histogram
Count
0
1
2
3
50
70
90
Value
Common sites gene−LOC105281688
1
2
3
4
RphaseA2
CphaseA2

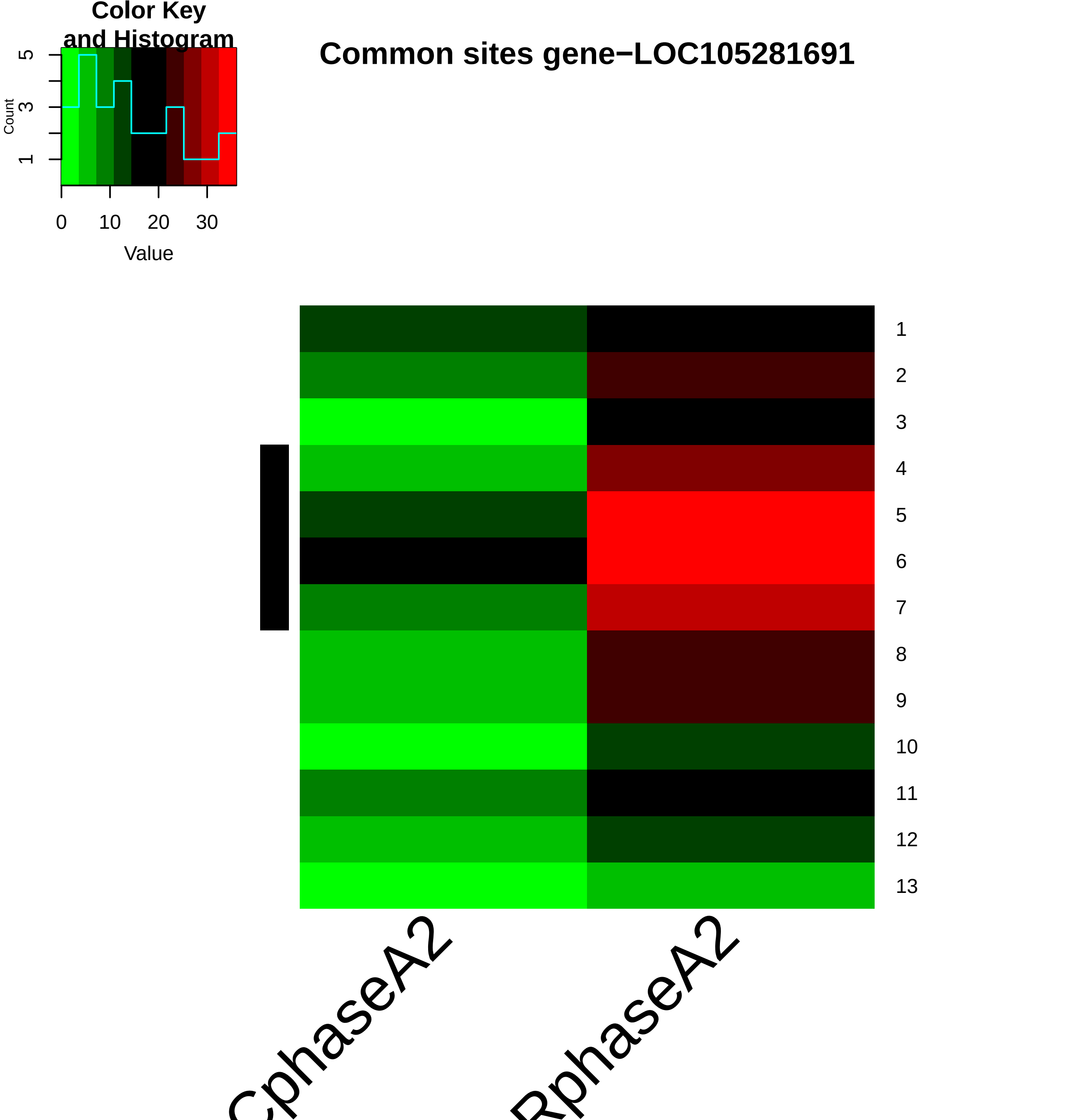
Color Key
and Histogram
Count
Value
Common sites gene−LOC105281691
1
2
3
4
5
6
7
8
9
10
11
12
13
CphaseA2
RphaseA2

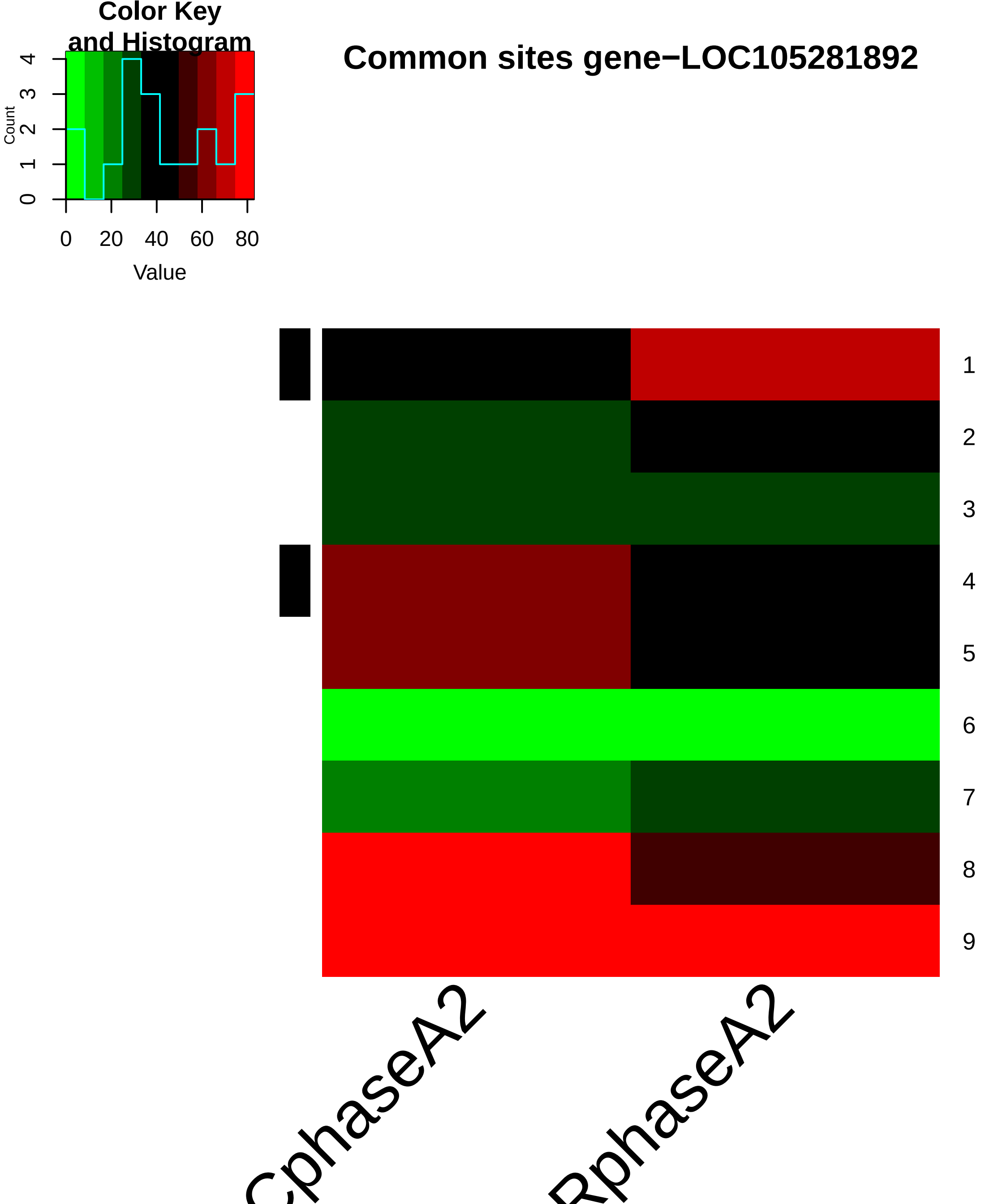

Color Key
and Histogram
Count
0
1
2
3
4
0
20
40
60
80
Value
Common sites gene−LOC105281892
1
2
3
4
5
6
7
8
9
CphaseA2
RphaseA2

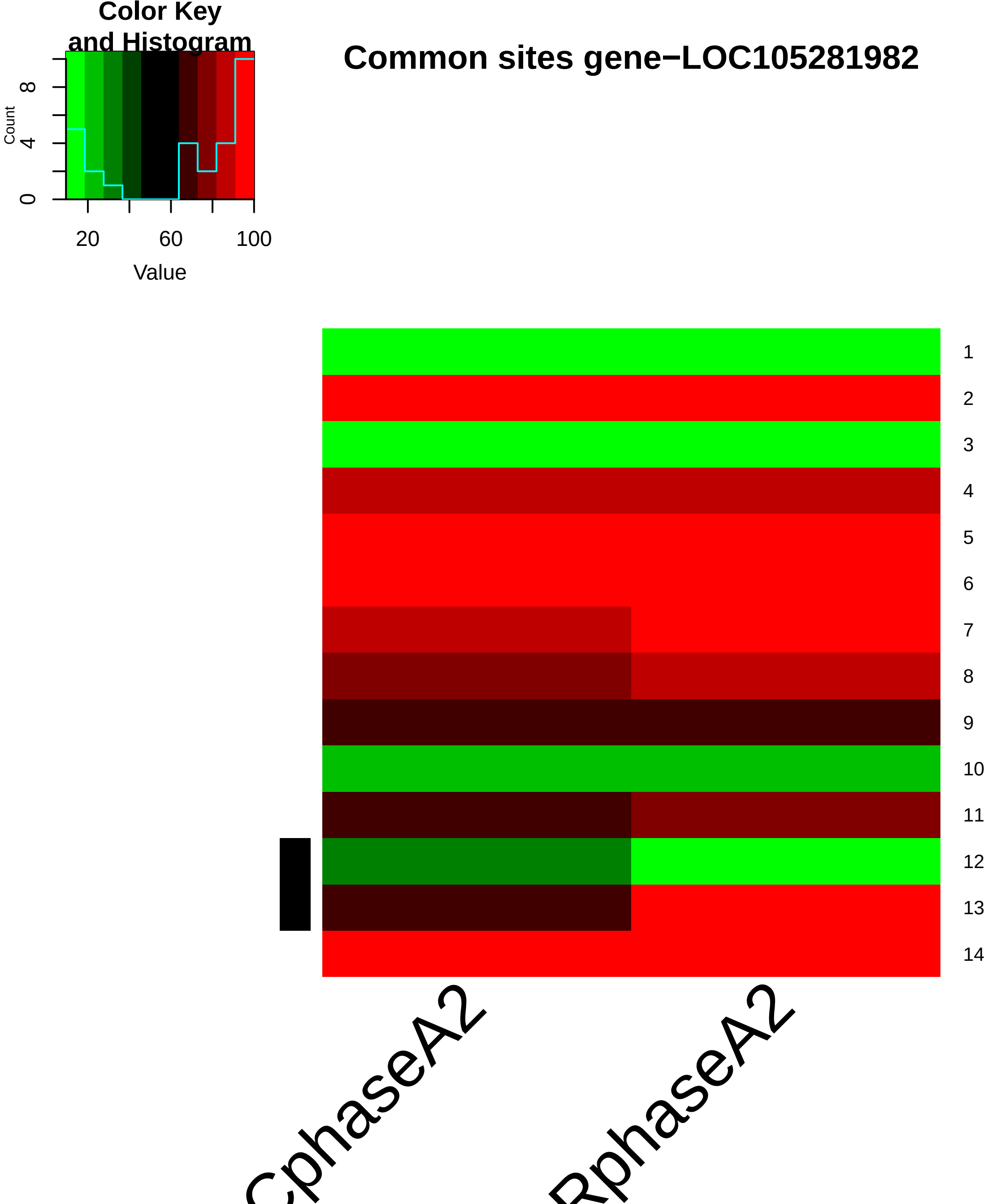
Color Key
and Histogram
Count
0
4
8
20
60
100
Value
Common sites gene−LOC105281982
1
2
3
4
5
6
7
8
9
10
11
12
13
14
CphaseA2
RphaseA2

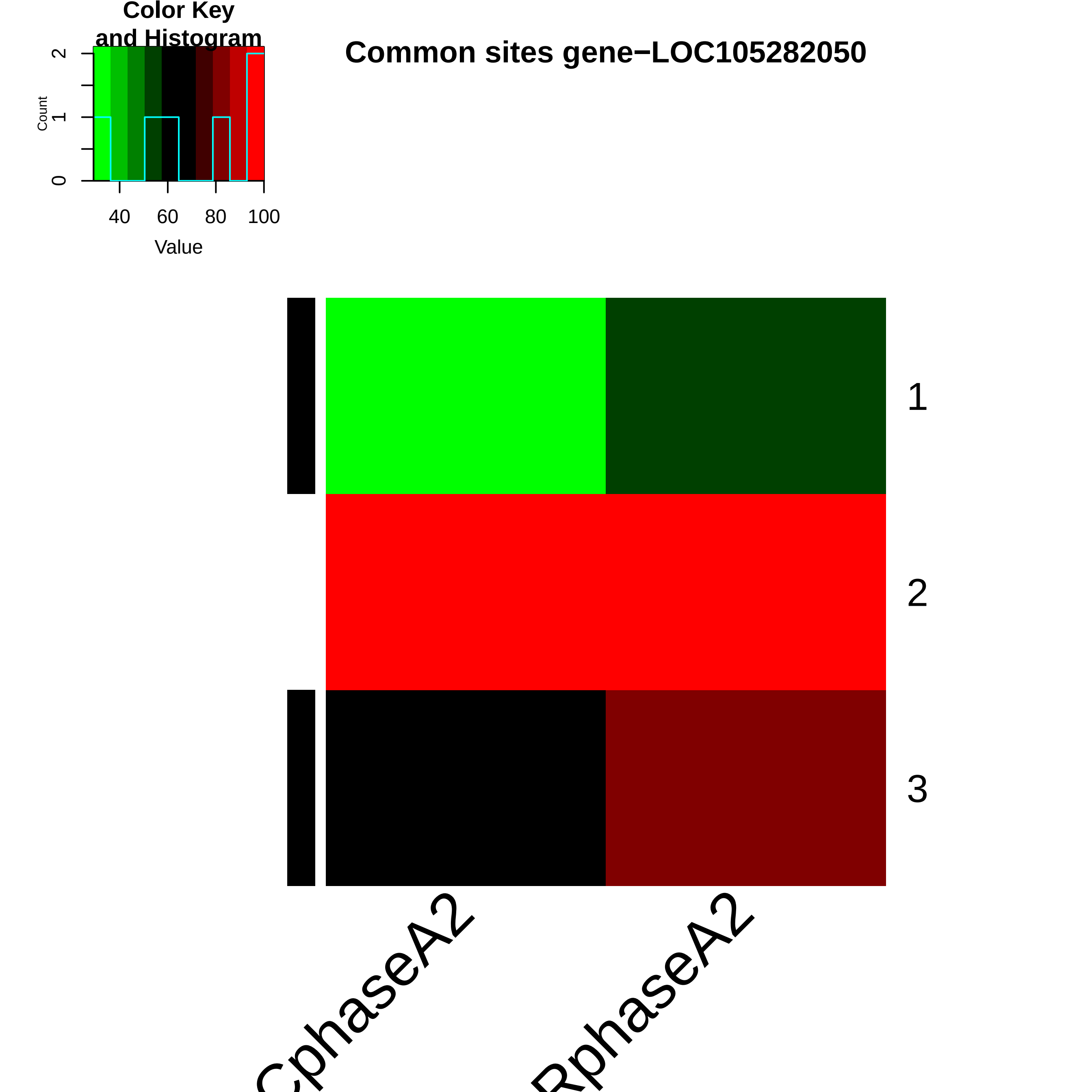

Color Key
and Histogram
Count
0
1
2
40
60
80
100
Value
Common sites gene−LOC105282050
1
2
3
CphaseA2
RphaseA2

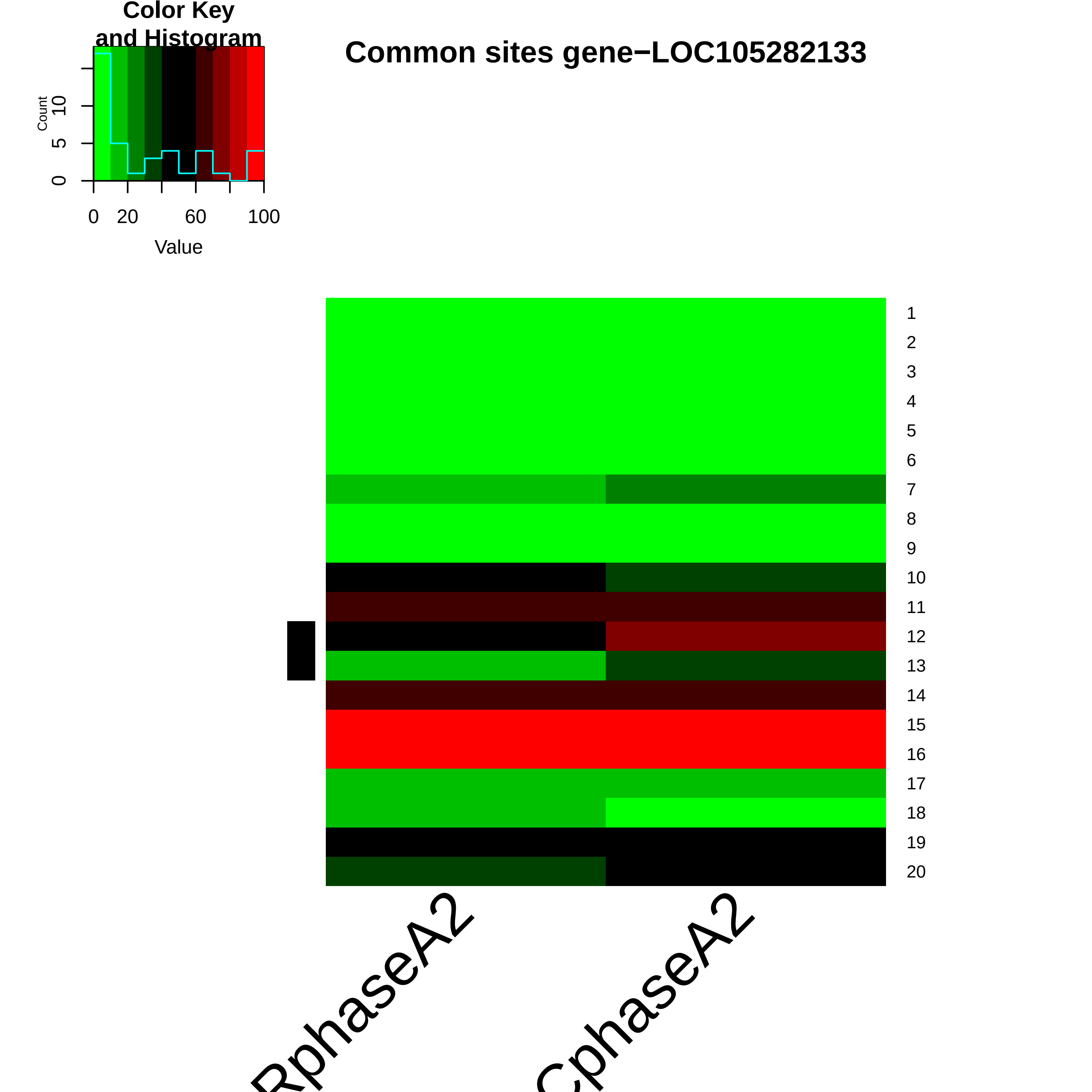
Color Key
and Histogram
Count
0
5
10
0
20
60
100
Value
Common sites gene−LOC105282133
RphaseA2
CphaseA2

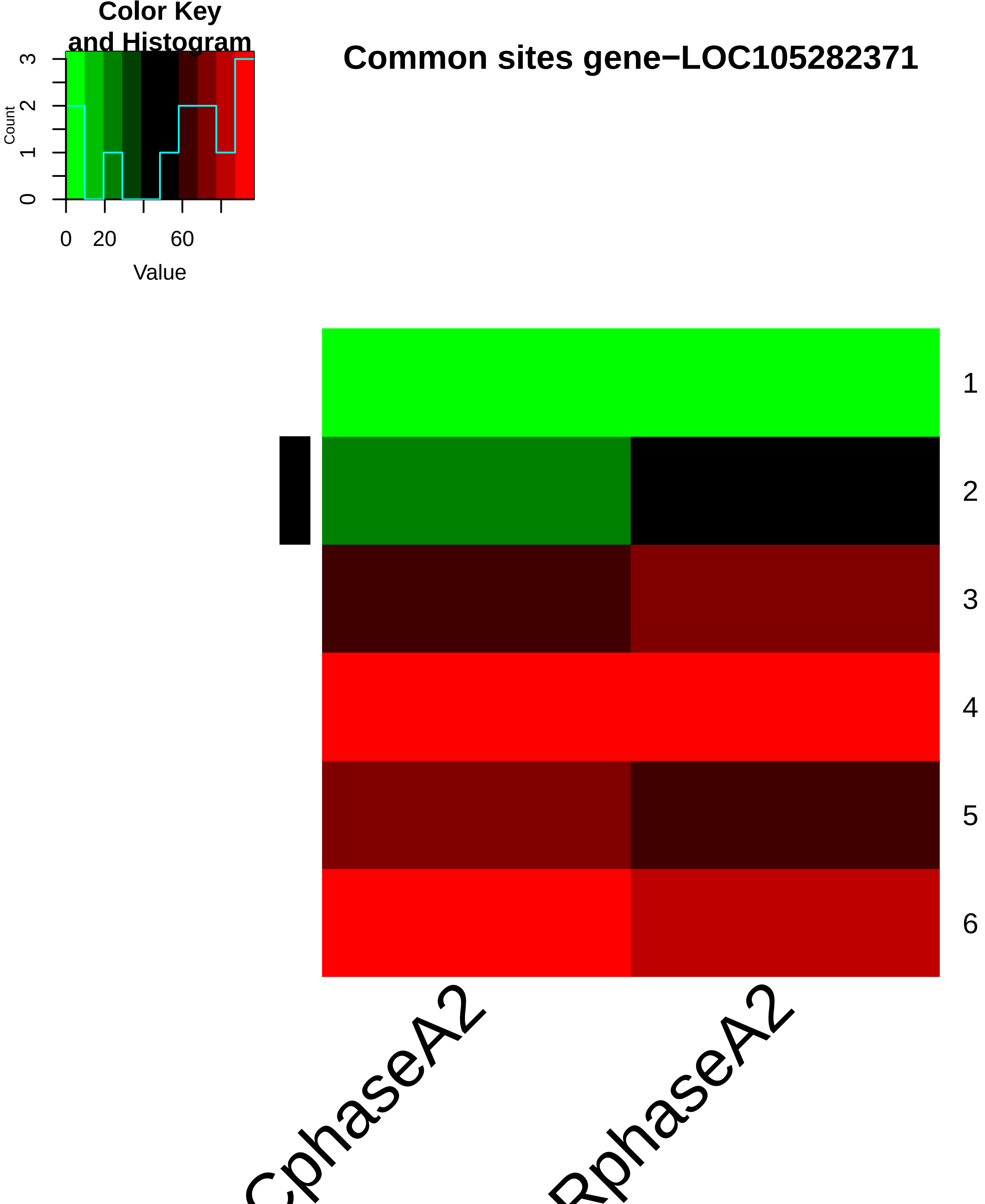
Color Key
and Histogram
Count
0
1
2
3
0
20
60
Value
Common sites gene−LOC105282371
1
2
3
4
5
6
CphaseA2
RphaseA2

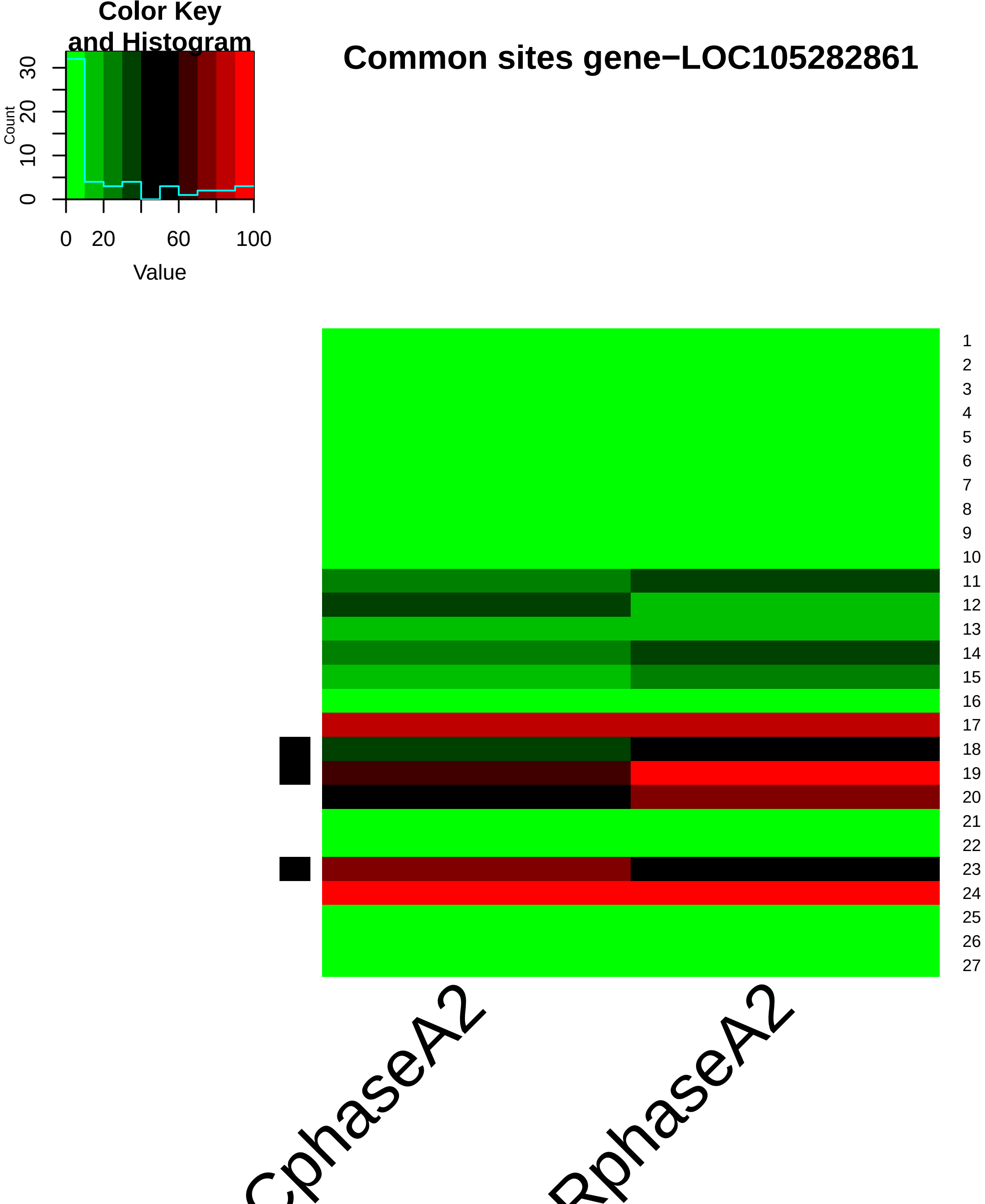

Color Key
and Histogram
Count
0
10
20
30
0
20
60
100
Value
Common sites gene−LOC105282861
CphaseA2
RphaseA2

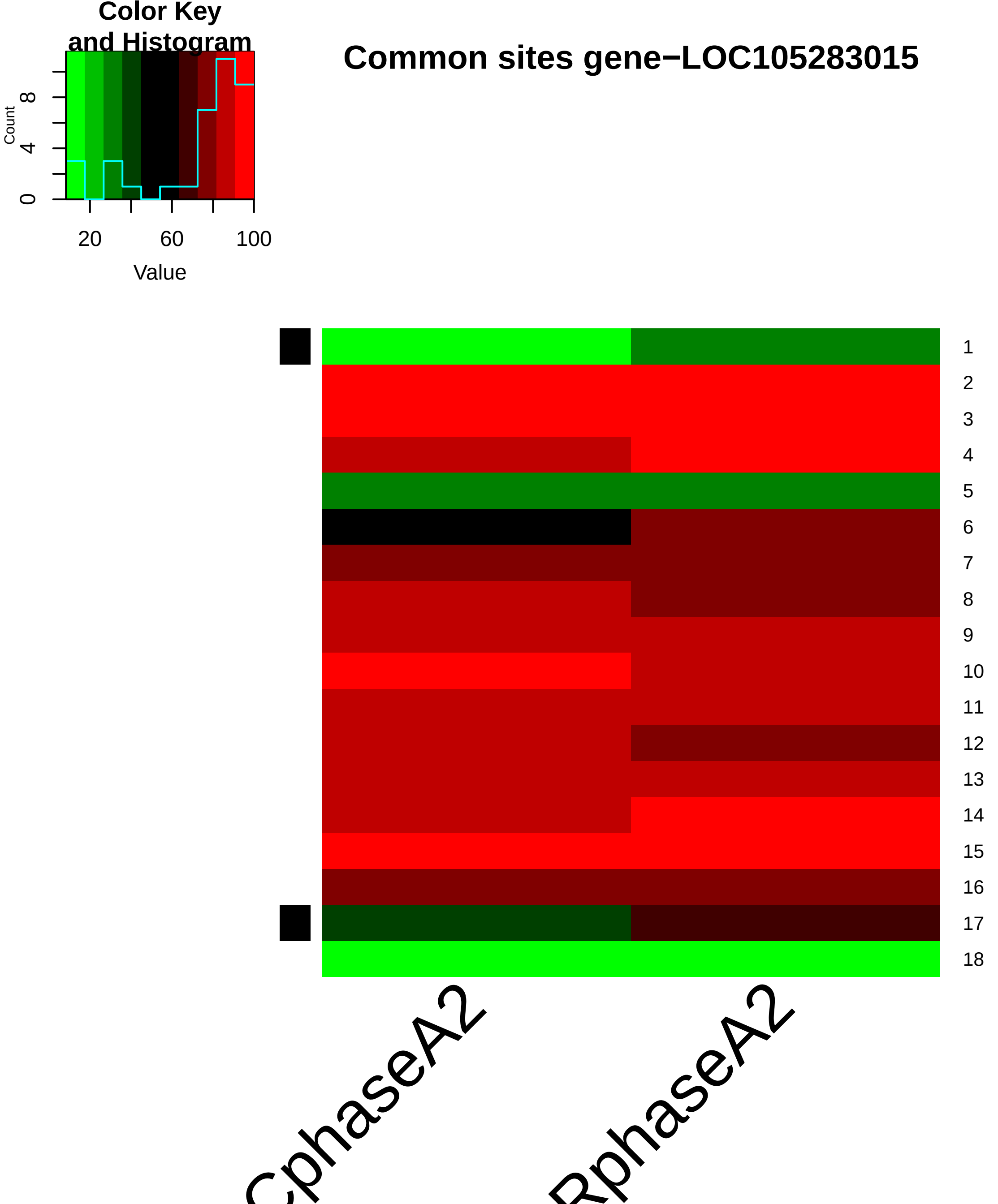
Color Key
and Histogram
Count
0
4
8
20
60
100
Value
Common sites gene−LOC105283015
CphaseA2
RphaseA2

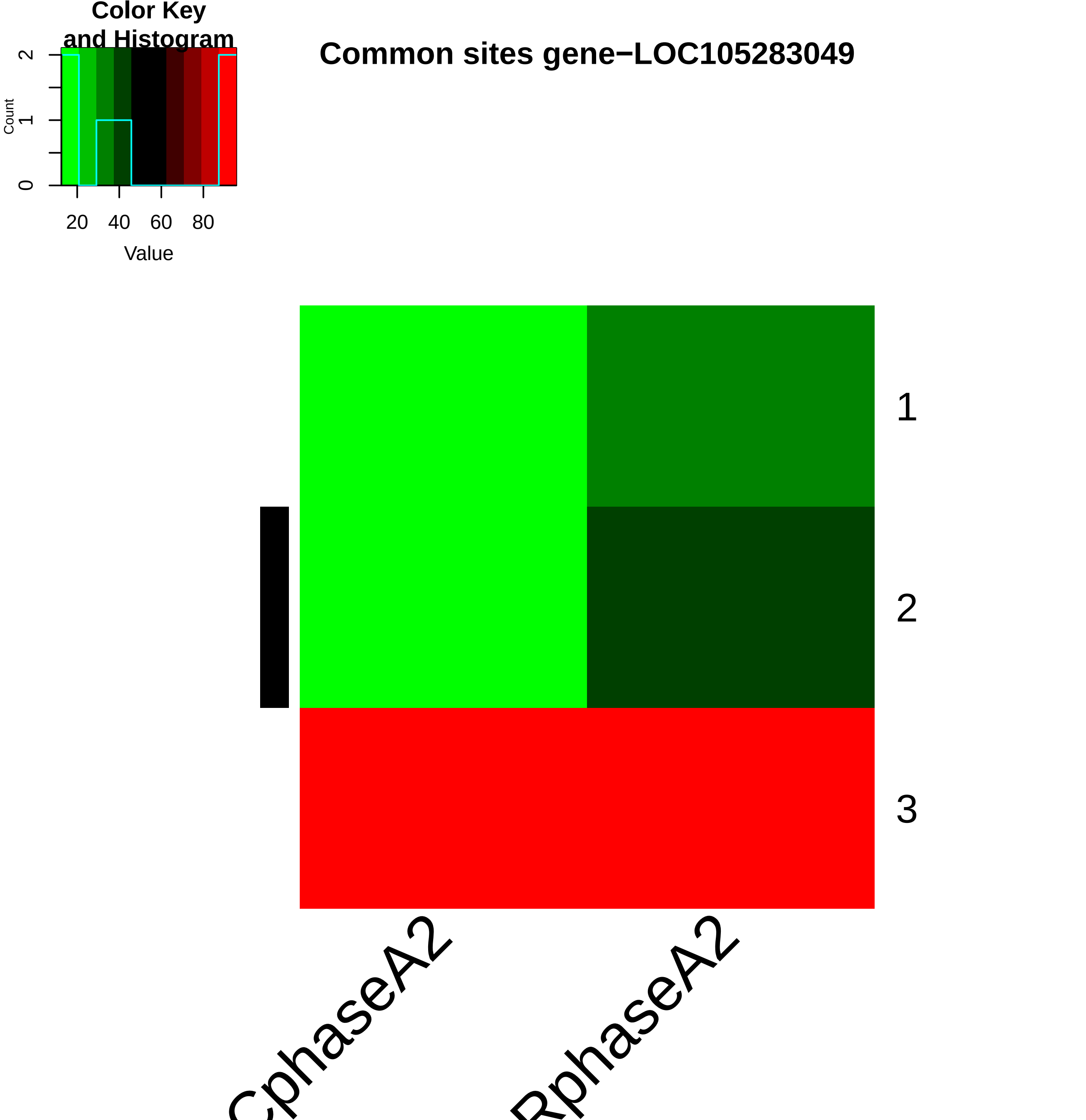

Color Key
and Histogram
Count
0
1
2
20
40
60
80
Value
Common sites gene−LOC105283049
1
2
3

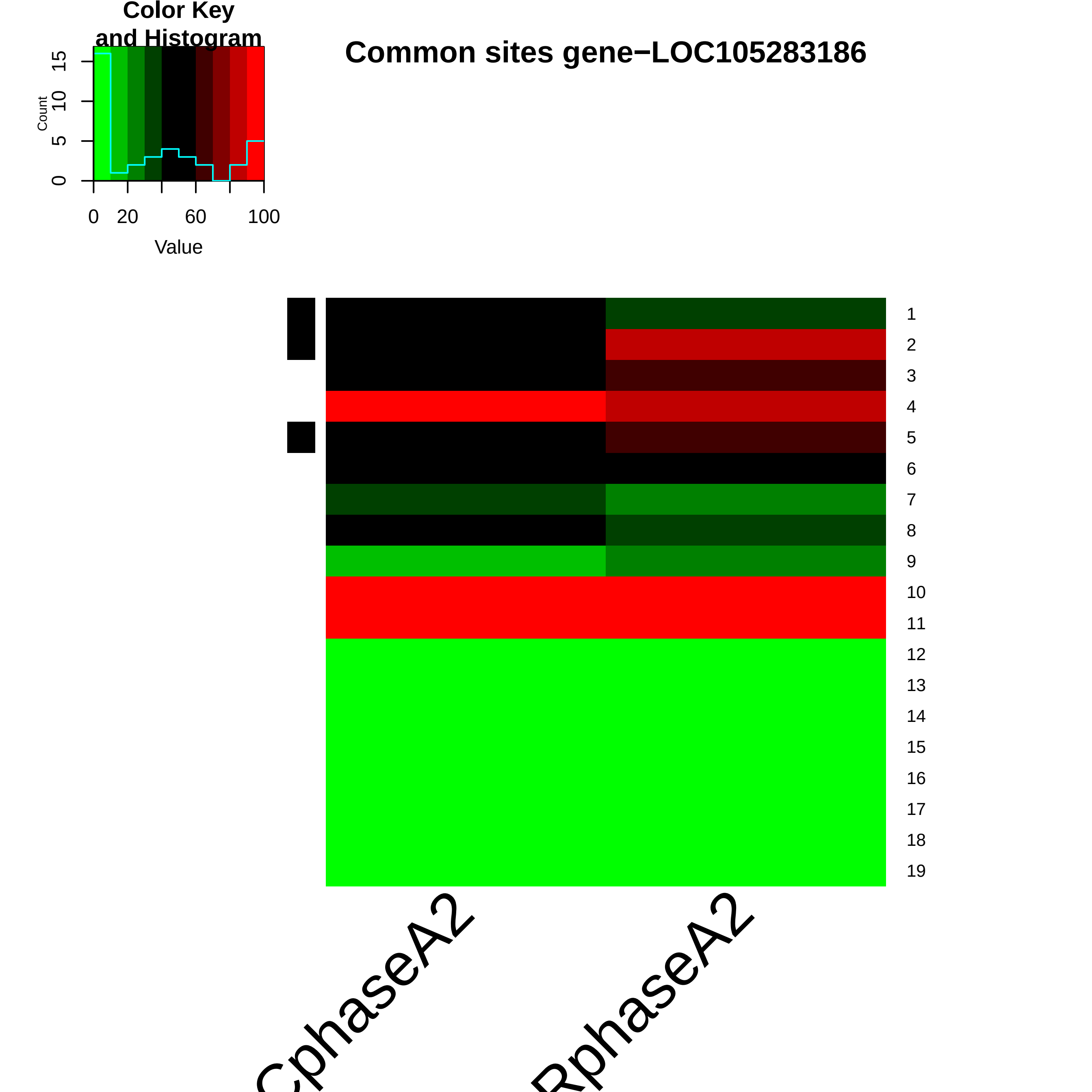
Color Key
and Histogram
Count
0
5
10
15
0
20
60
100
Value
Common sites gene−LOC105283186
CphaseA2
RphaseA2

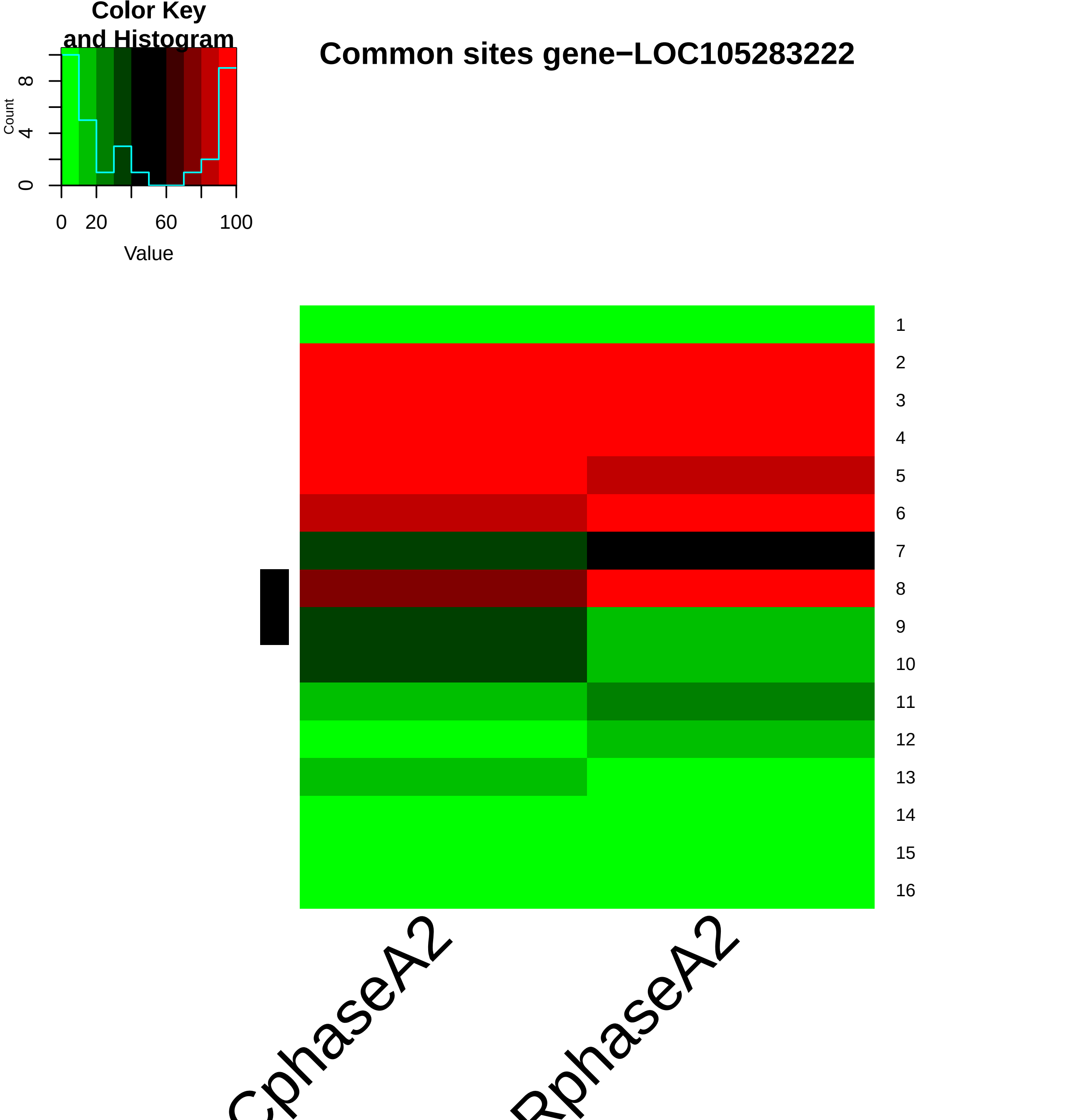

Color Key
and Histogram
Count
0
4
8
0
20
60
100
Value
Common sites gene−LOC105283222
CphaseA2
RphaseA2

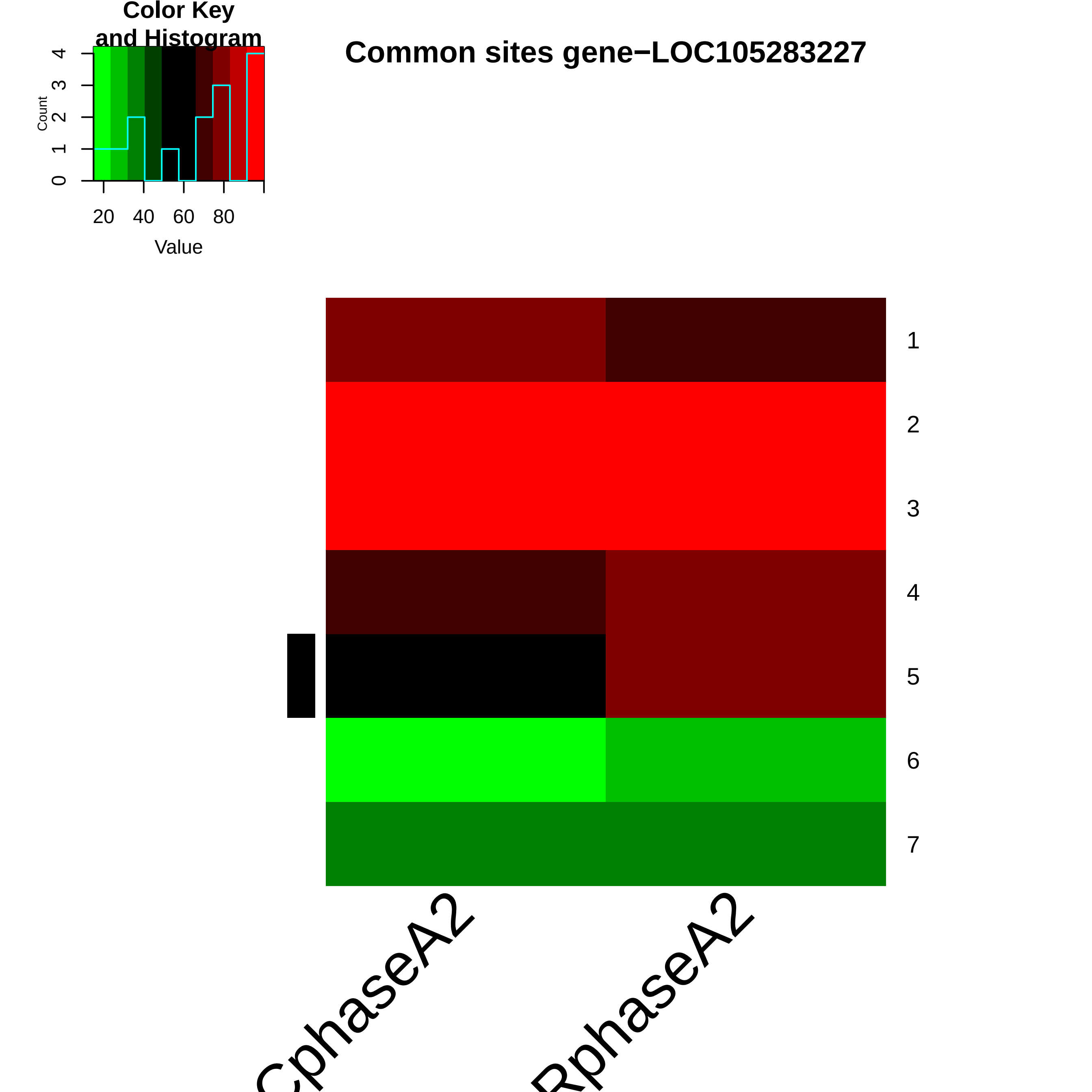

Color Key
and Histogram
Count
Value
20
40
60
80
0
1
2
3
4
Common sites gene−LOC105283227
1
2
3
4
5
6
7
CphaseA2
RphaseA2

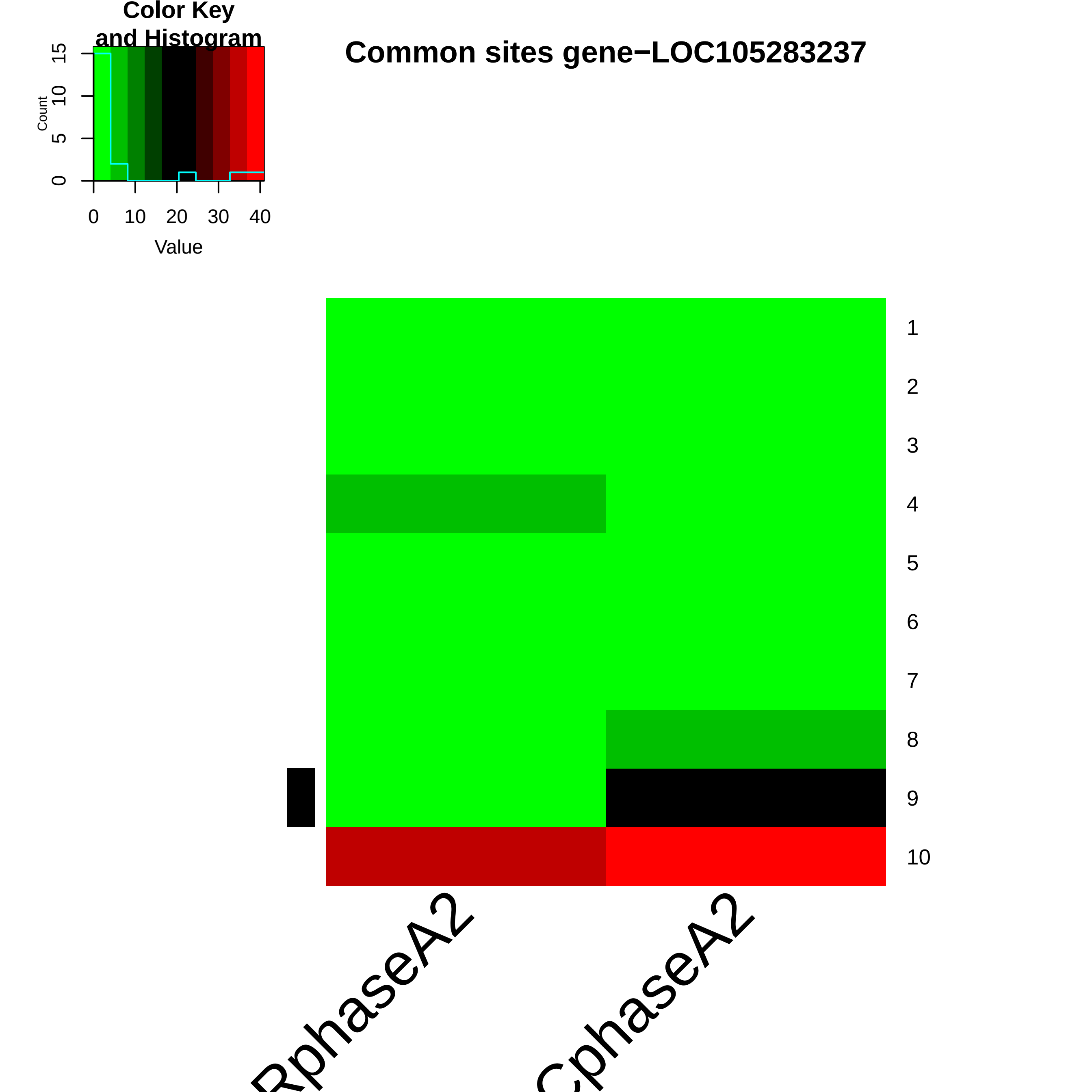

Color Key
and Histogram
Count
0
5
10
15
0
10
20
30
40
Value
Common sites gene−LOC105283237
1
2
3
4
5
6
7
8
9
10
RphaseA2
CphaseA2

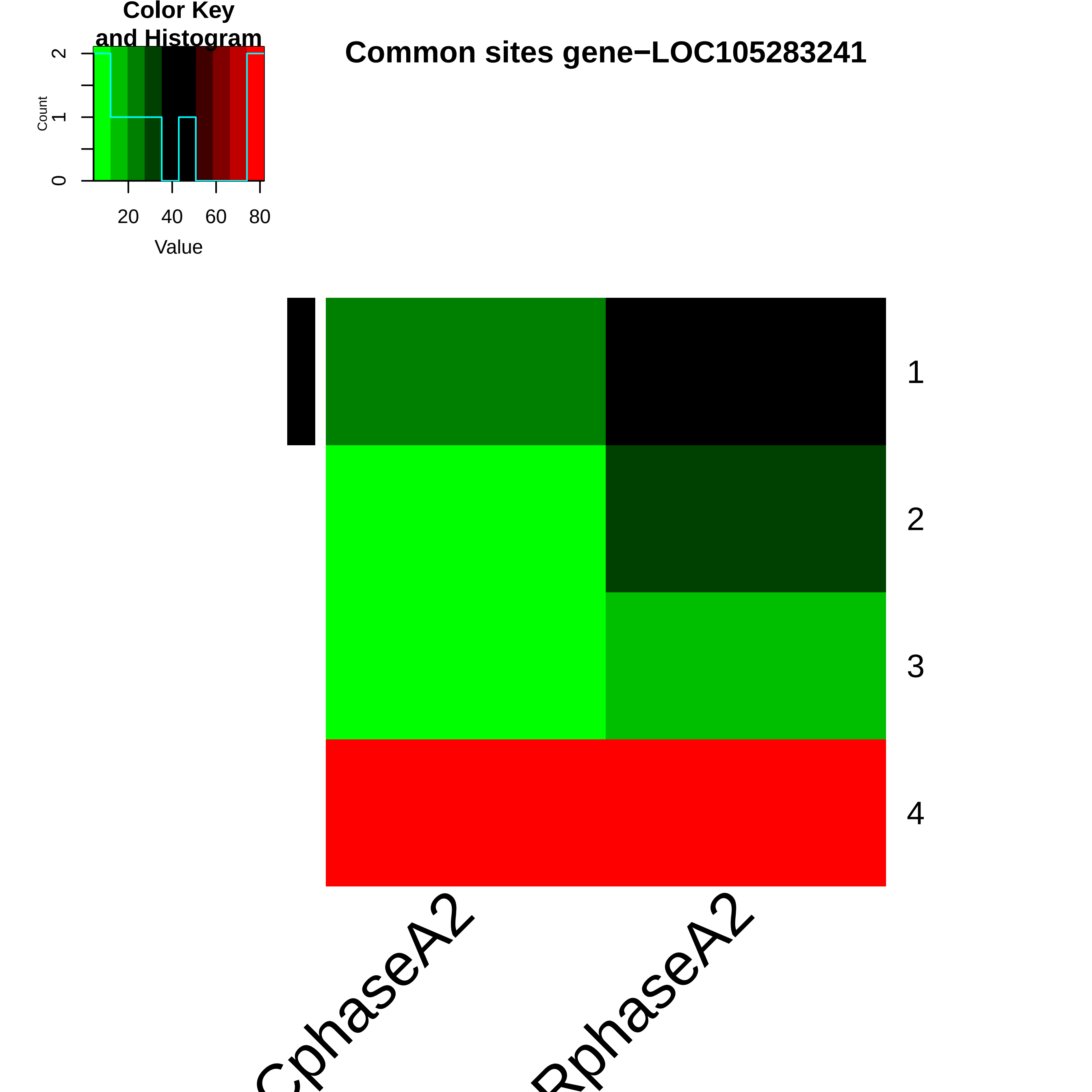

Color Key
and Histogram
Count
0
1
2
20
40
60
80
Value
Common sites gene−LOC105283241
1
2
3
4
CphaseA2
RphaseA2

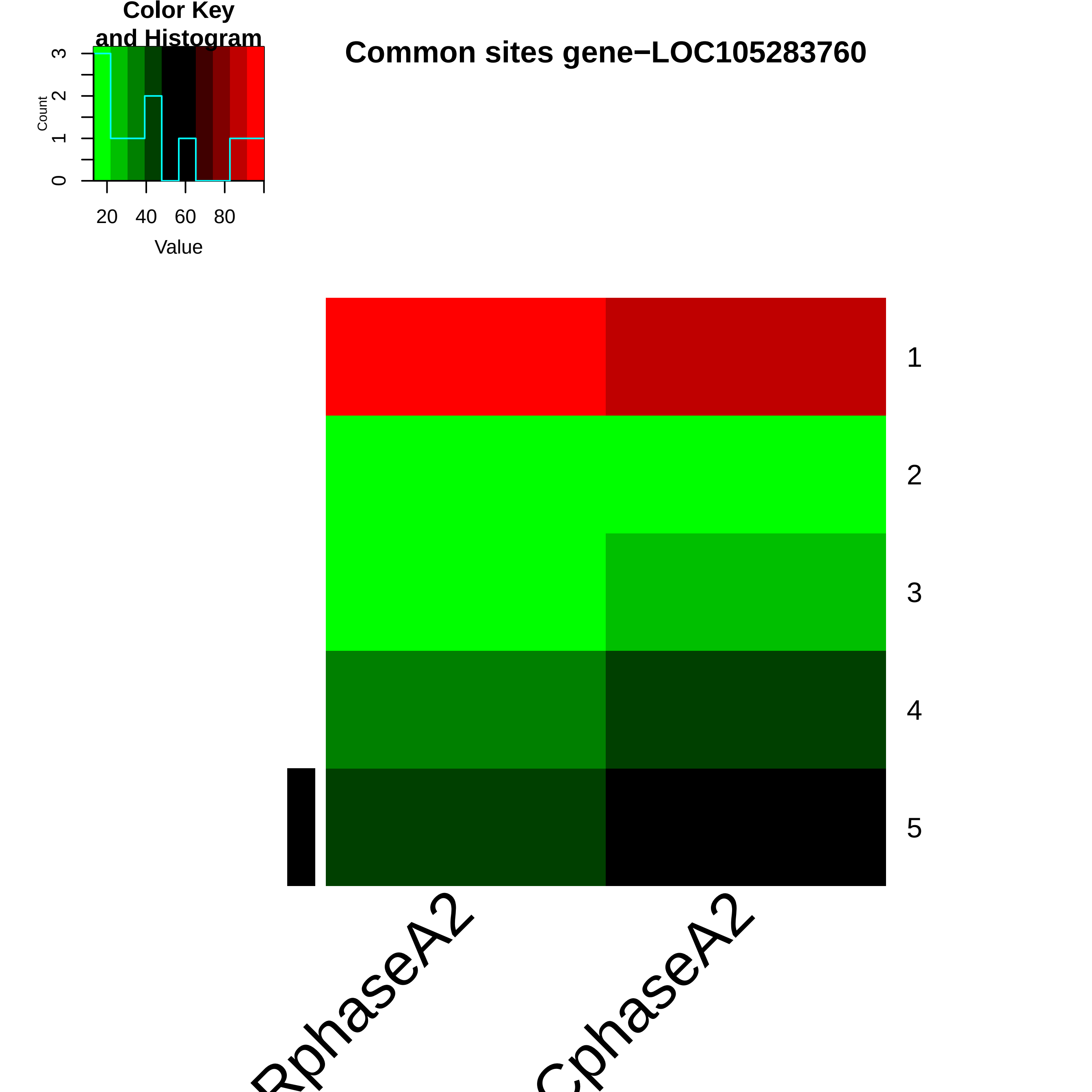
Color Key
and Histogram
Count
Value
20
40
60
80
0
1
2
3
Common sites gene−LOC105283760
1
2
3
4
5
RphaseA2
CphaseA2

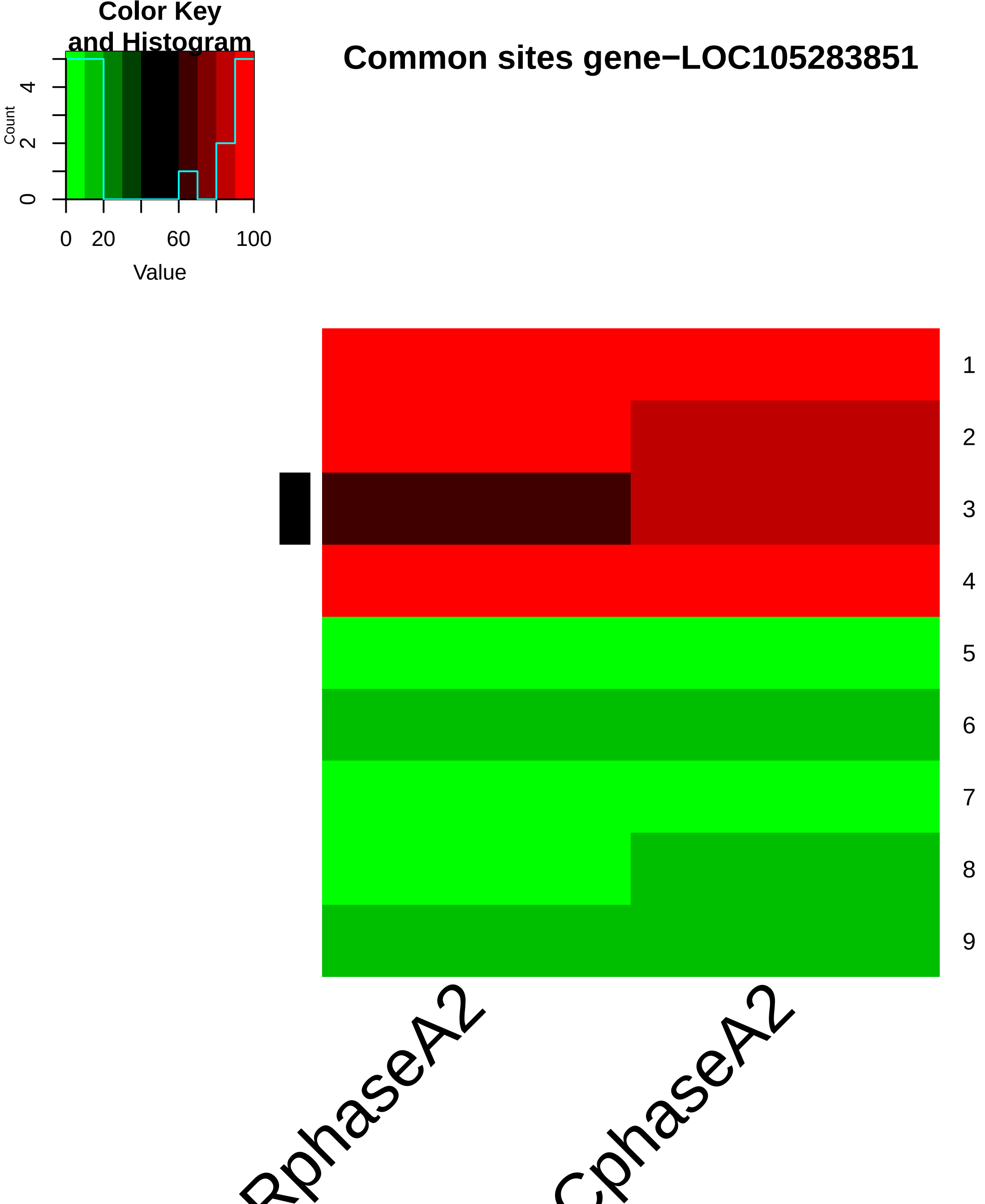
Color Key
and Histogram
Count
0
2
4
0
20
60
100
Value
Common sites gene−LOC105283851
1
2
3
4
5
6
7
8
9
RphaseA2
CphaseA2

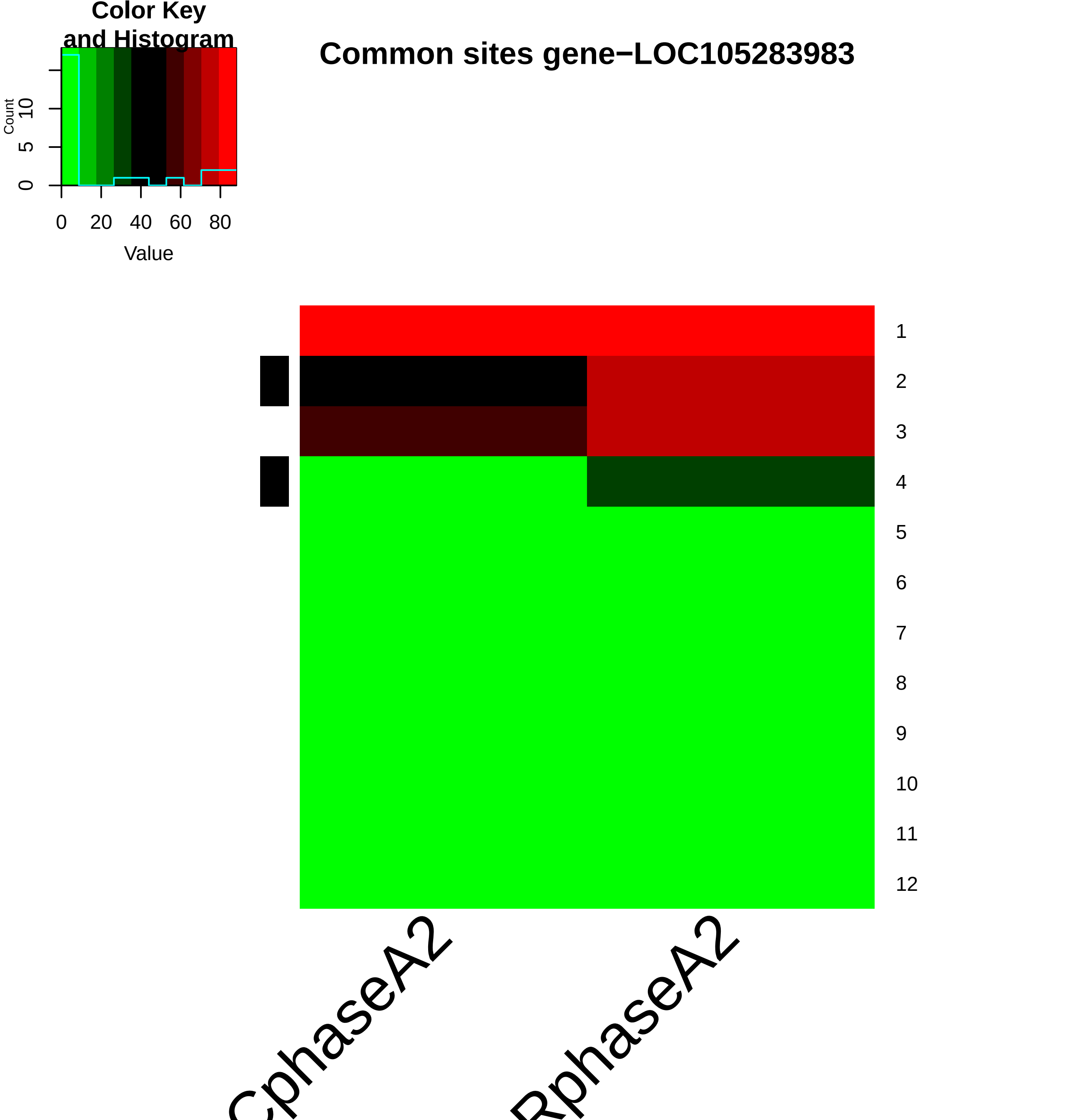
Color Key
and Histogram
Count
0
5
10
0
20
40
60
80
Value
Common sites gene−LOC105283983
1
2
3
4
5
6
7
8
9
10
11
12
CphaseA2
RphaseA2

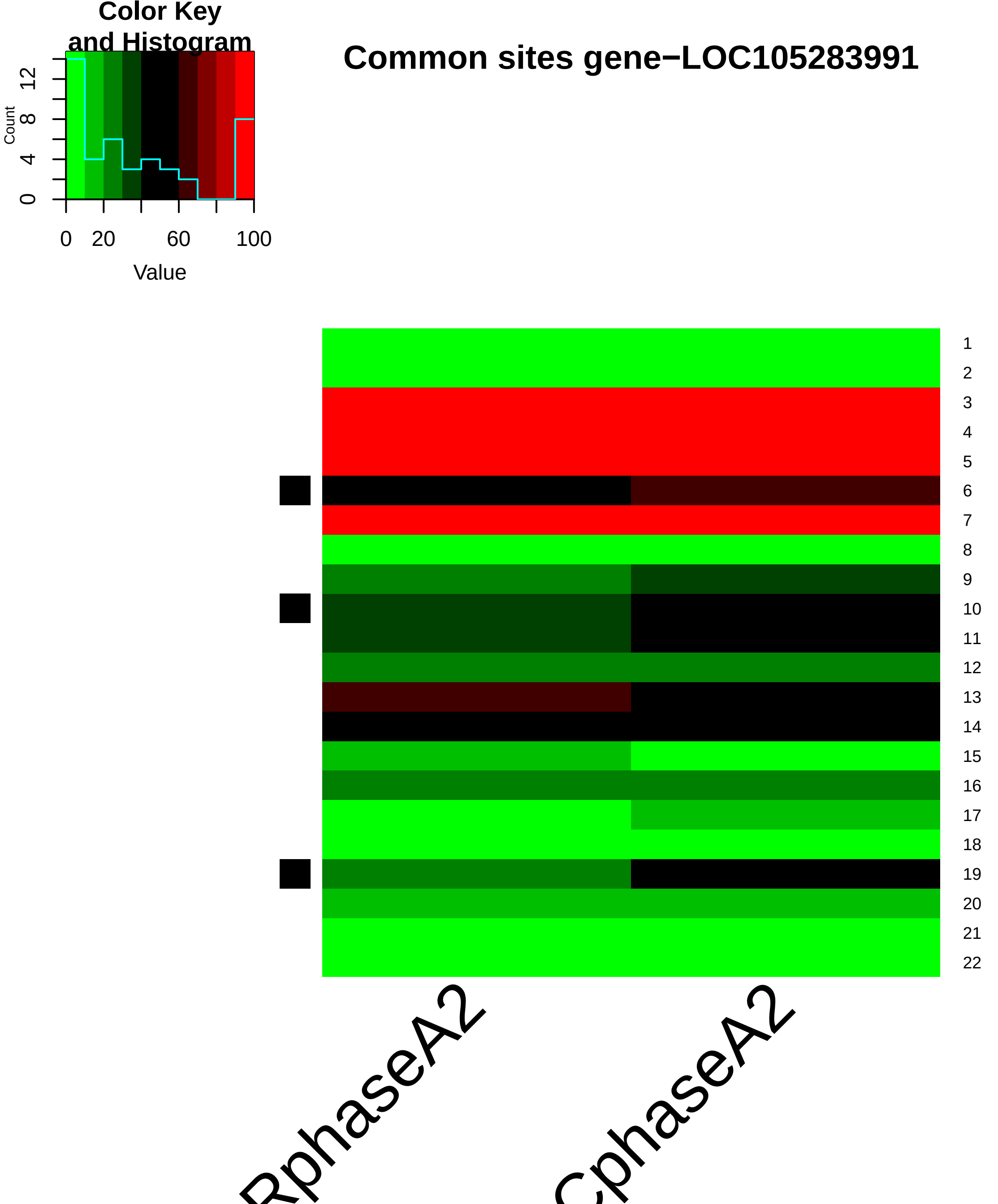

Color Key
and Histogram
Count
0
4
8
12
0
20
60
100
Value
Common sites gene−LOC105283991
RphaseA2
CphaseA2

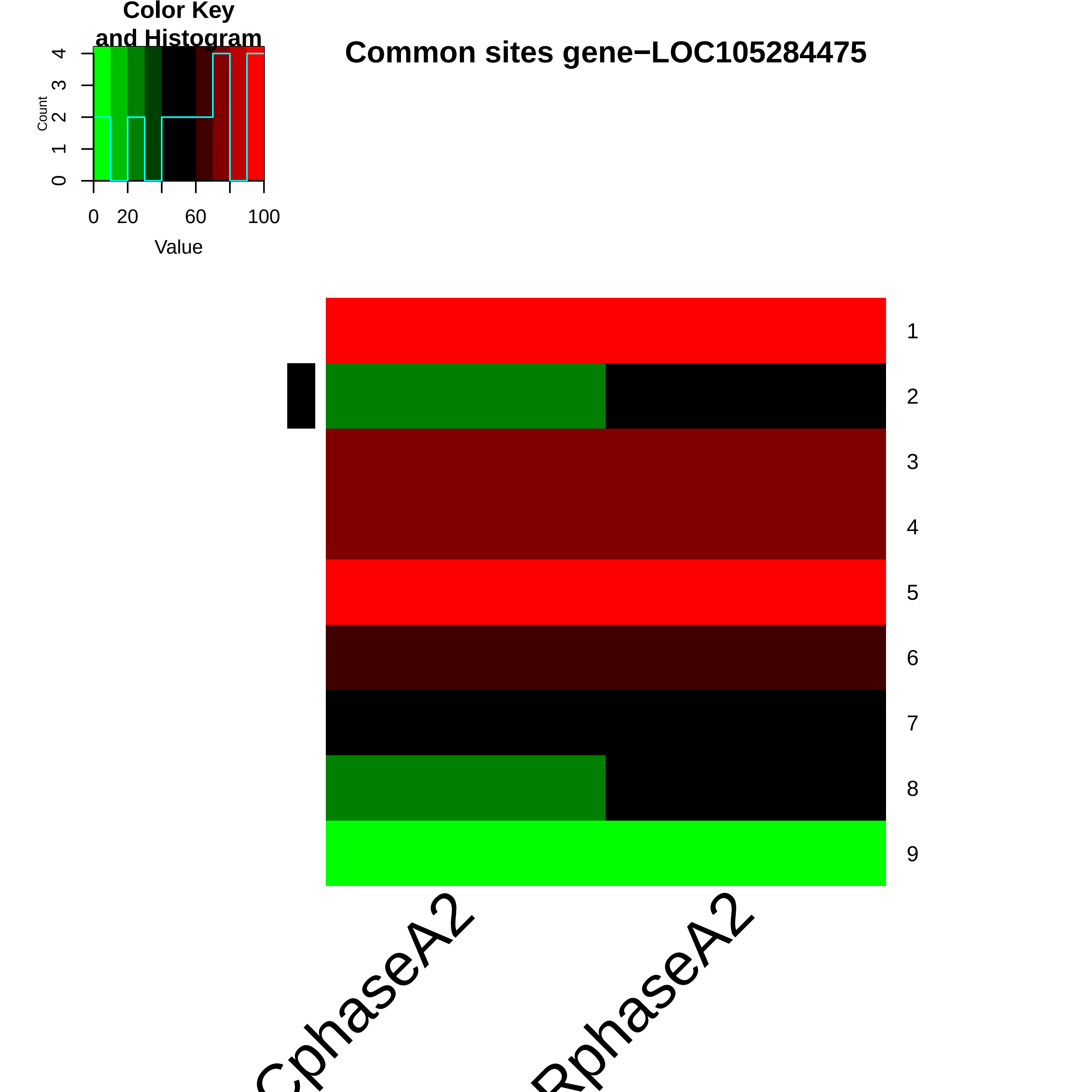
Color Key
and Histogram
Count
0
1
2
3
4
0
20
60
100
Value
Common sites gene−LOC105284475
1
2
3
4
5
6
7
8
9

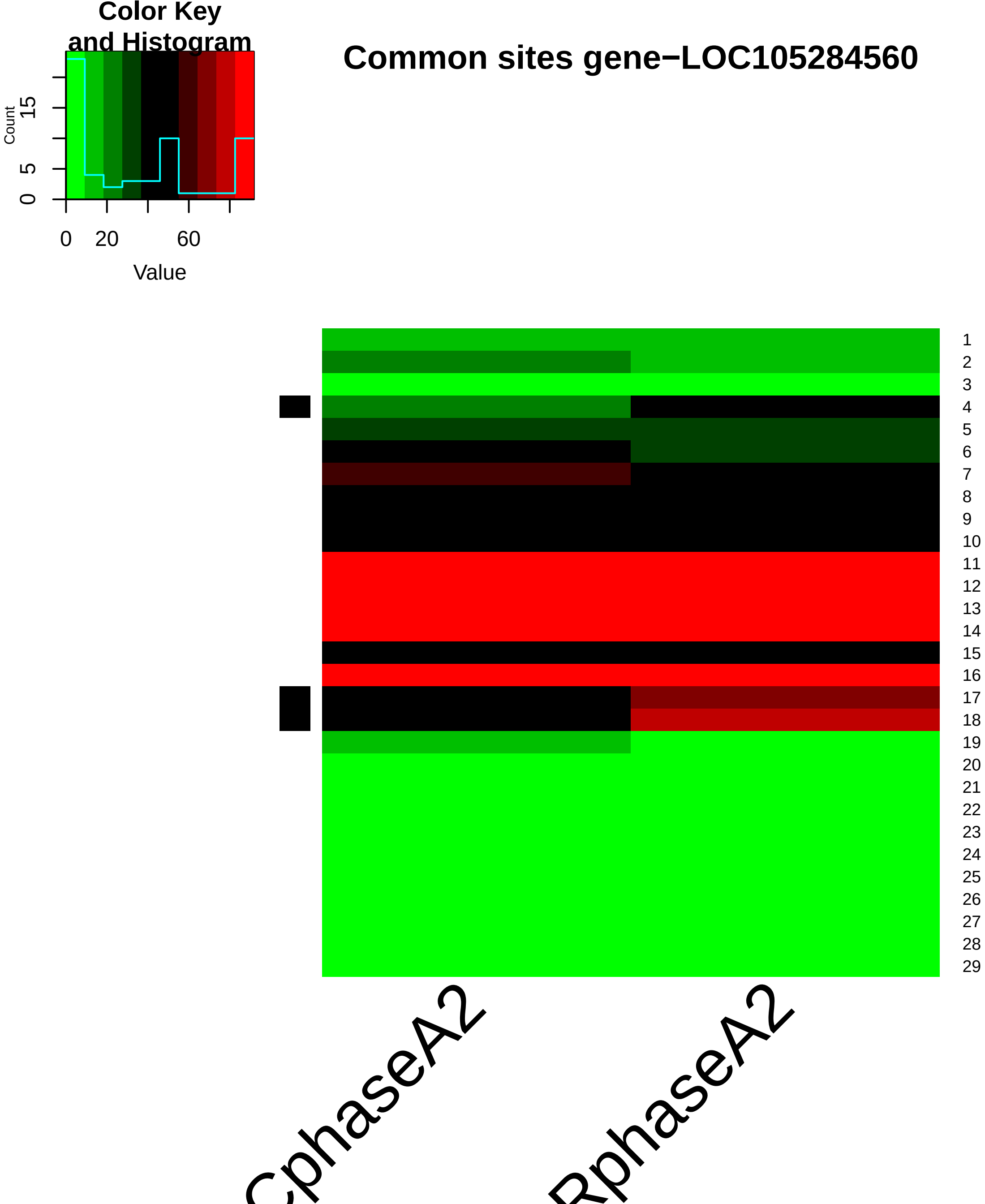

Color Key
and Histogram
Count
0
5
15
0
20
60
Value
Common sites gene−LOC105284560
CphaseA2
RphaseA2

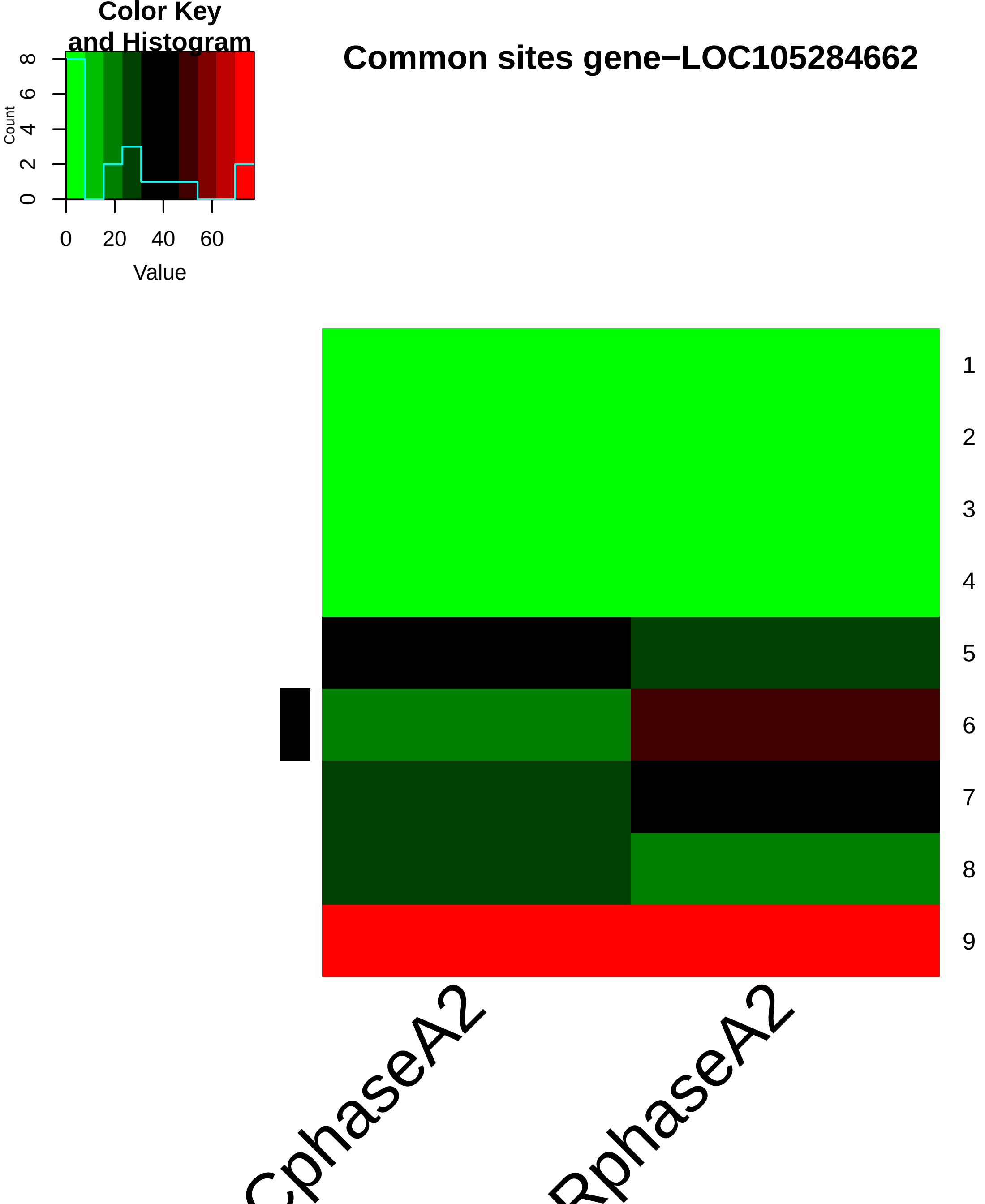

Color Key
and Histogram
Count
0
2
4
6
8
0
20
40
60
Value
Common sites gene−LOC105284662
1
2
3
4
5
6
7
8
9
CphaseA2
RphaseA2

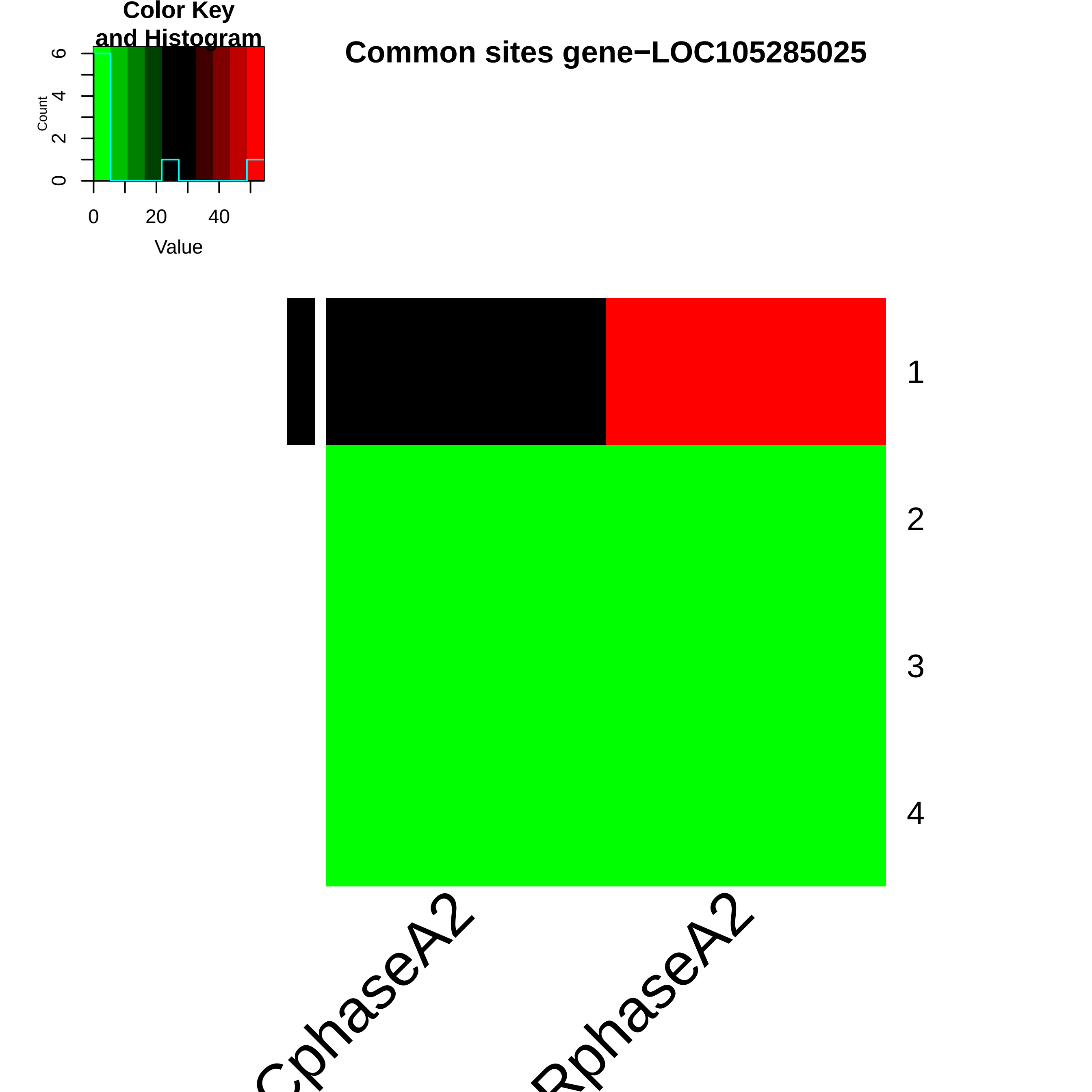
Color Key
and Histogram
Count
Value
0
2
4
6
0
20
40
Common sites gene−LOC105285025
1
2
3
4
CphaseA2
RphaseA2

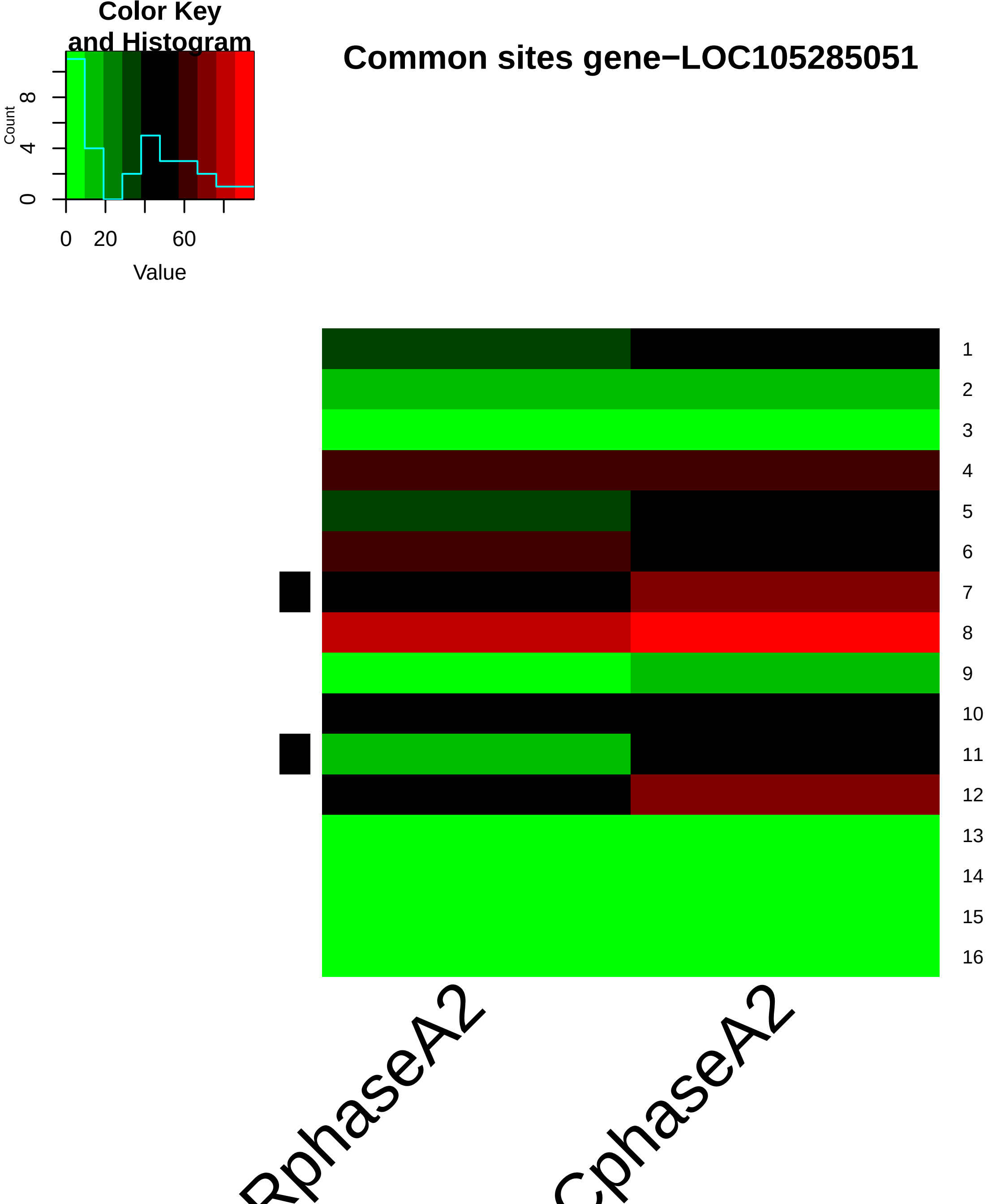
Color Key
and Histogram
Common sites gene−LOC105285051
Count
0
4
8
0
20
60
Value
RphaseA2
CphaseA2

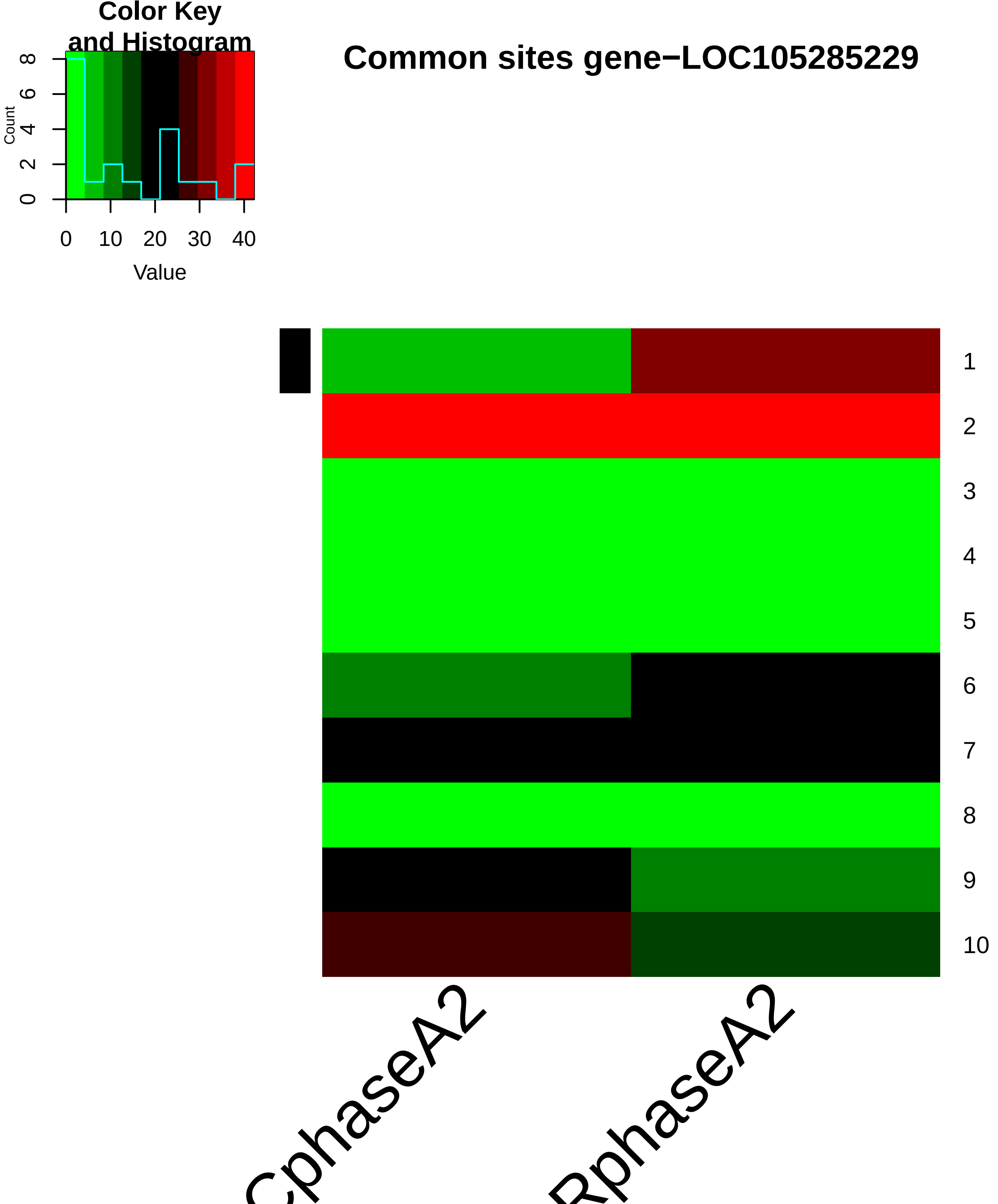
Color Key
and Histogram
Count
Value
0
10
20
30
40
0
2
4
6
8
Common sites gene−LOC105285229
1
2
3
4
5
6
7
8
9
10
CphaseA2
RphaseA2

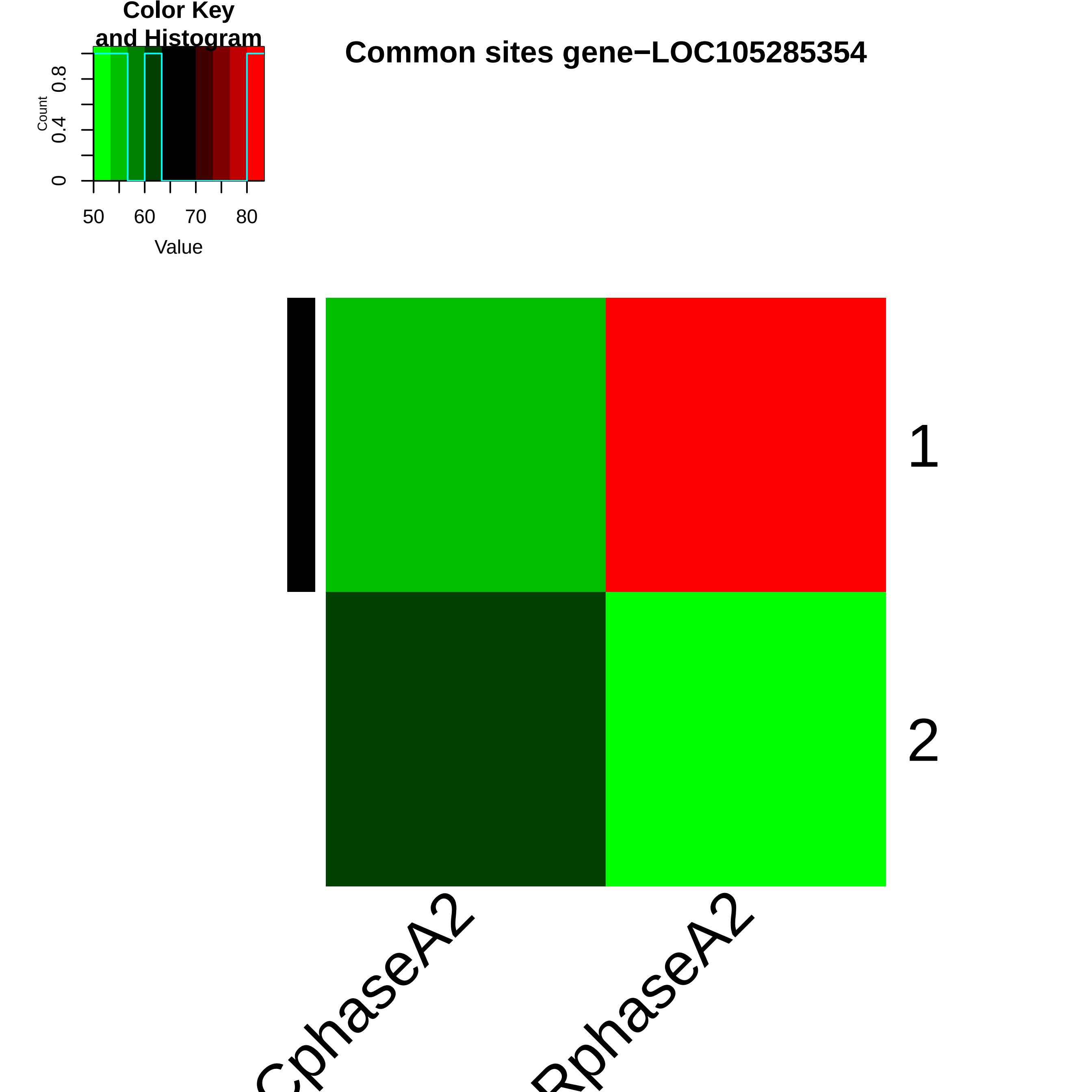
Color Key
and Histogram
Count
0
0.4
0.8
50
60
70
80
Value
Common sites gene−LOC105285354
1
2
CphaseA2
RphaseA2

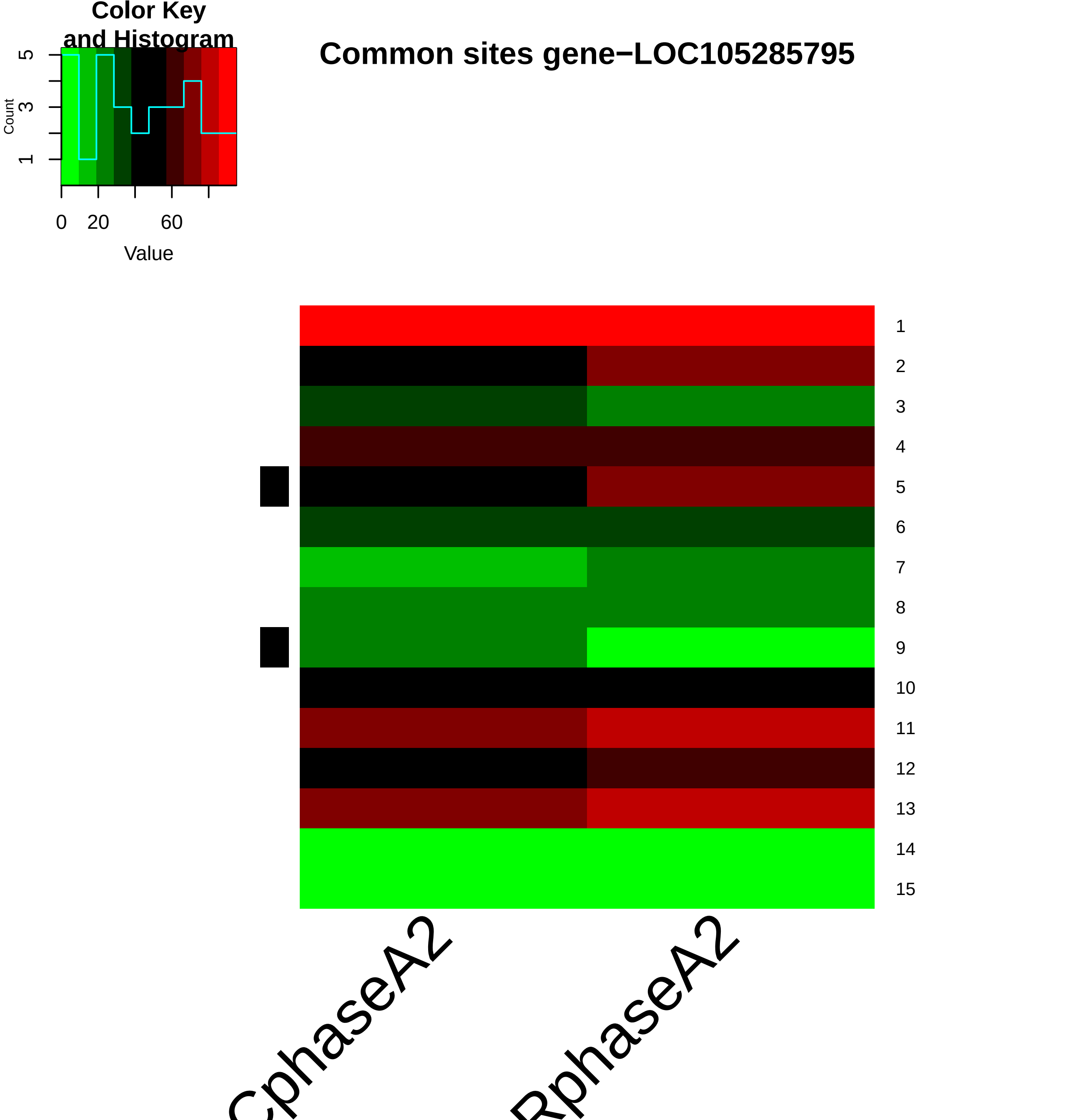
Color Key
and Histogram
Count
1
3
5
0
20
60
Value
Common sites gene−LOC105285795
CphaseA2
RphaseA2

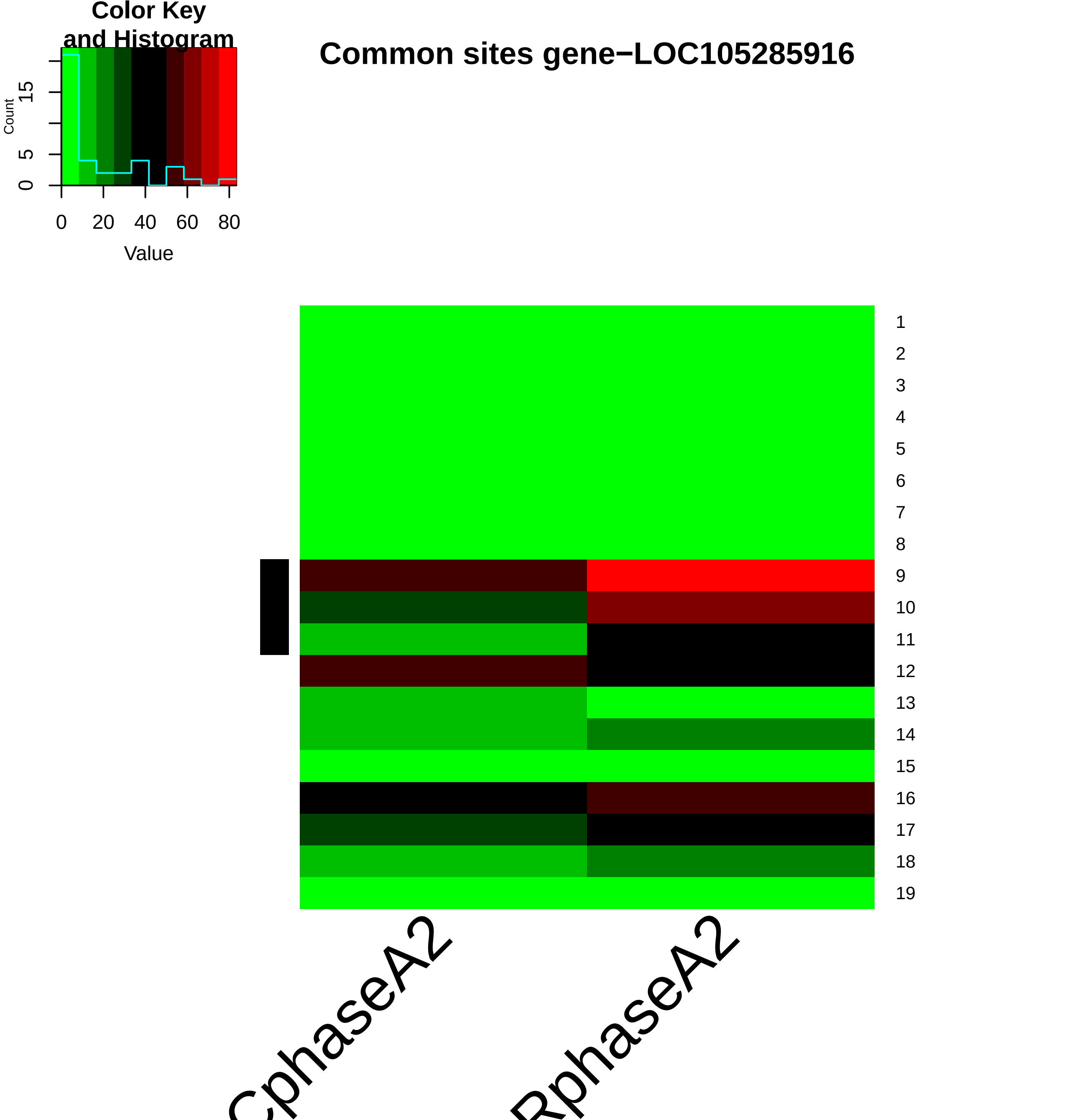

Color Key
and Histogram
Count
0
5
15
0
20
40
60
80
Value
Common sites gene−LOC105285916
CphaseA2
RphaseA2

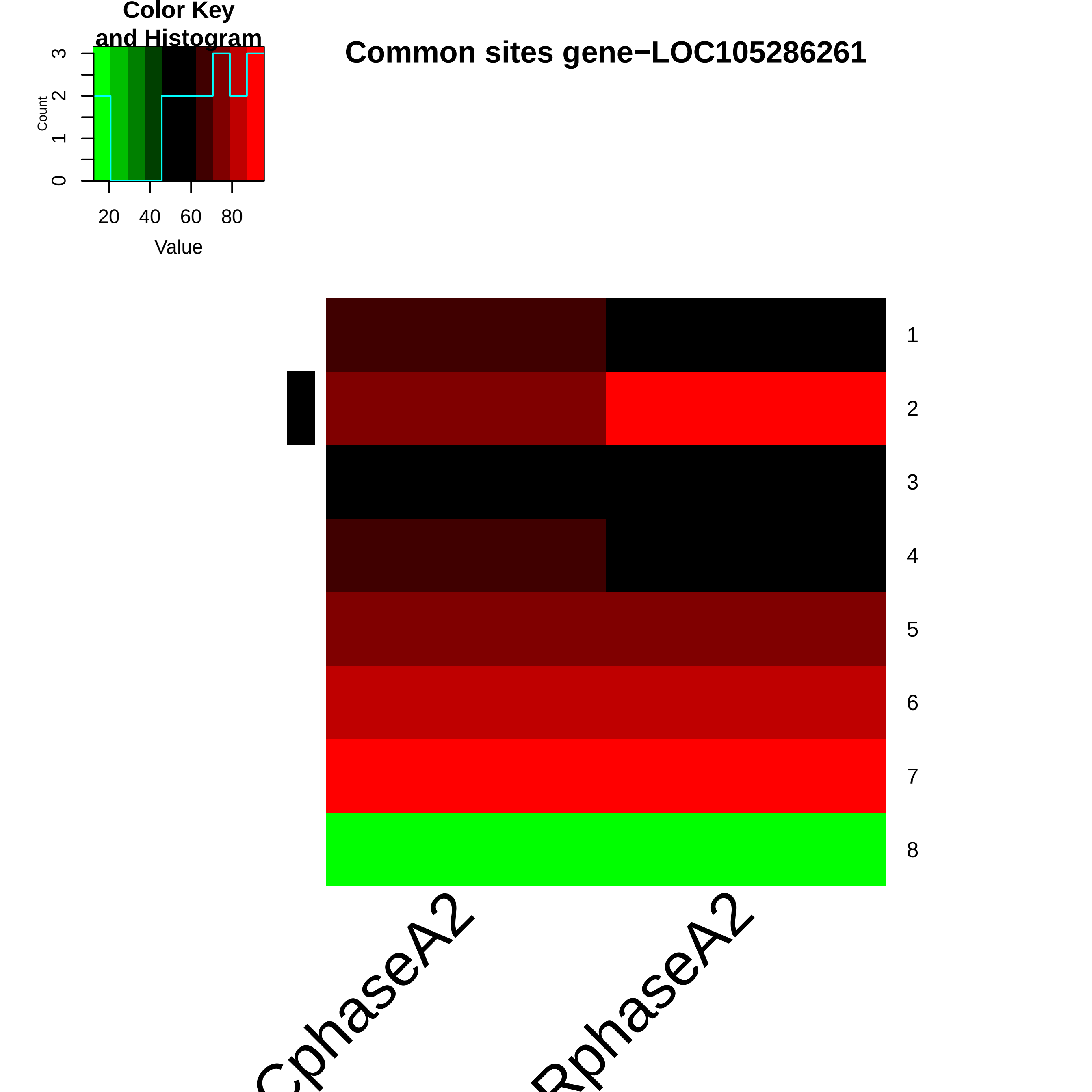
Color Key
and Histogram
Count
0
1
2
3
20
40
60
80
Value
Common sites gene−LOC105286261
1
2
3
4
5
6
7
8
CphaseA2
RphaseA2

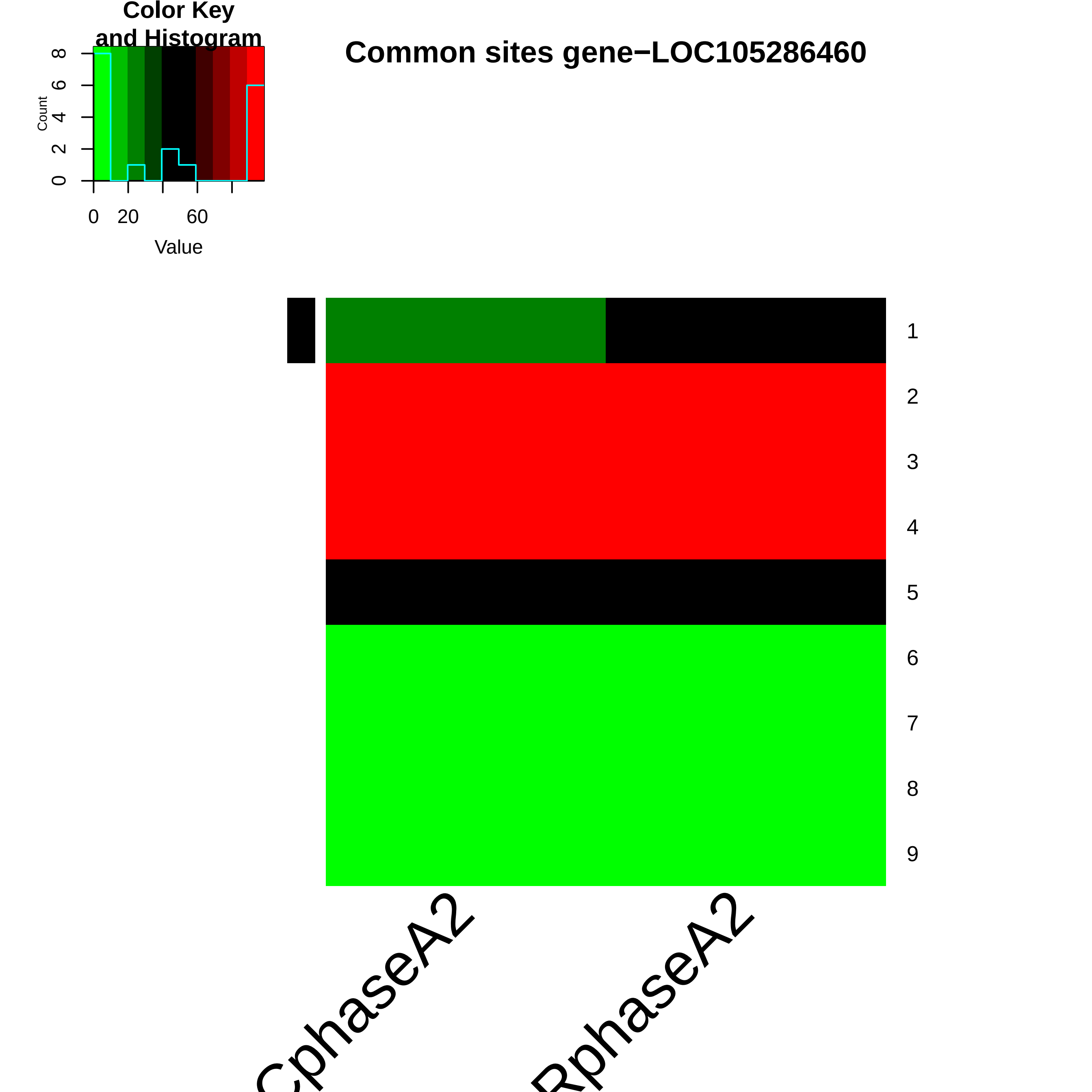
Color Key
and Histogram
Count
0
2
4
6
8
0
20
60
Value
Common sites gene−LOC105286460
1
2
3
4
5
6
7
8
9
CphaseA2
RphaseA2

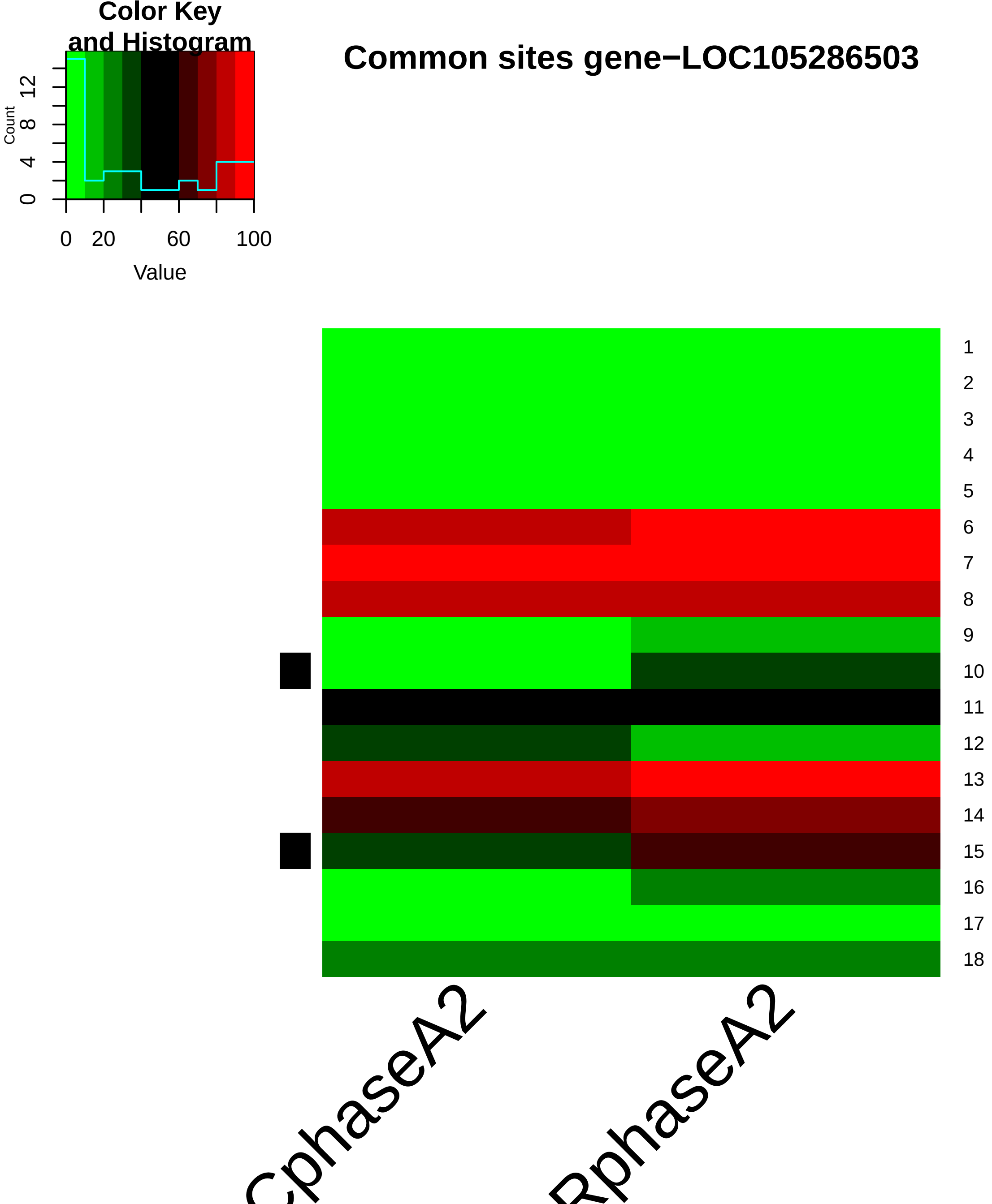
Color Key
and Histogram
Count
0
4
8
12
0
20
60
100
Value
Common sites gene−LOC105286503
CphaseA2
RphaseA2

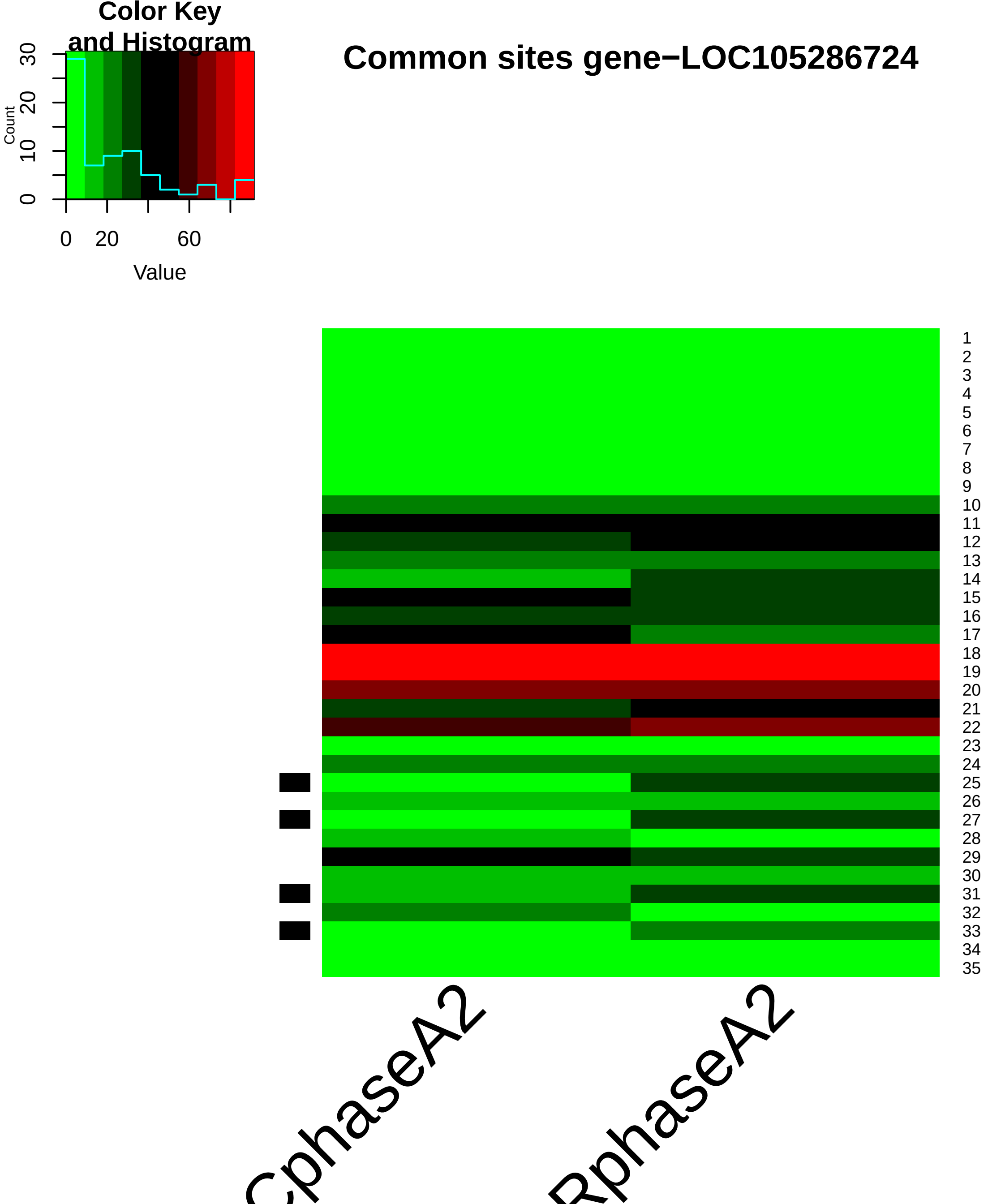
Color Key
and Histogram
Count
0
10
20
30
0
20
60
Value
Common sites gene−LOC105286724
CphaseA2
RphaseA2

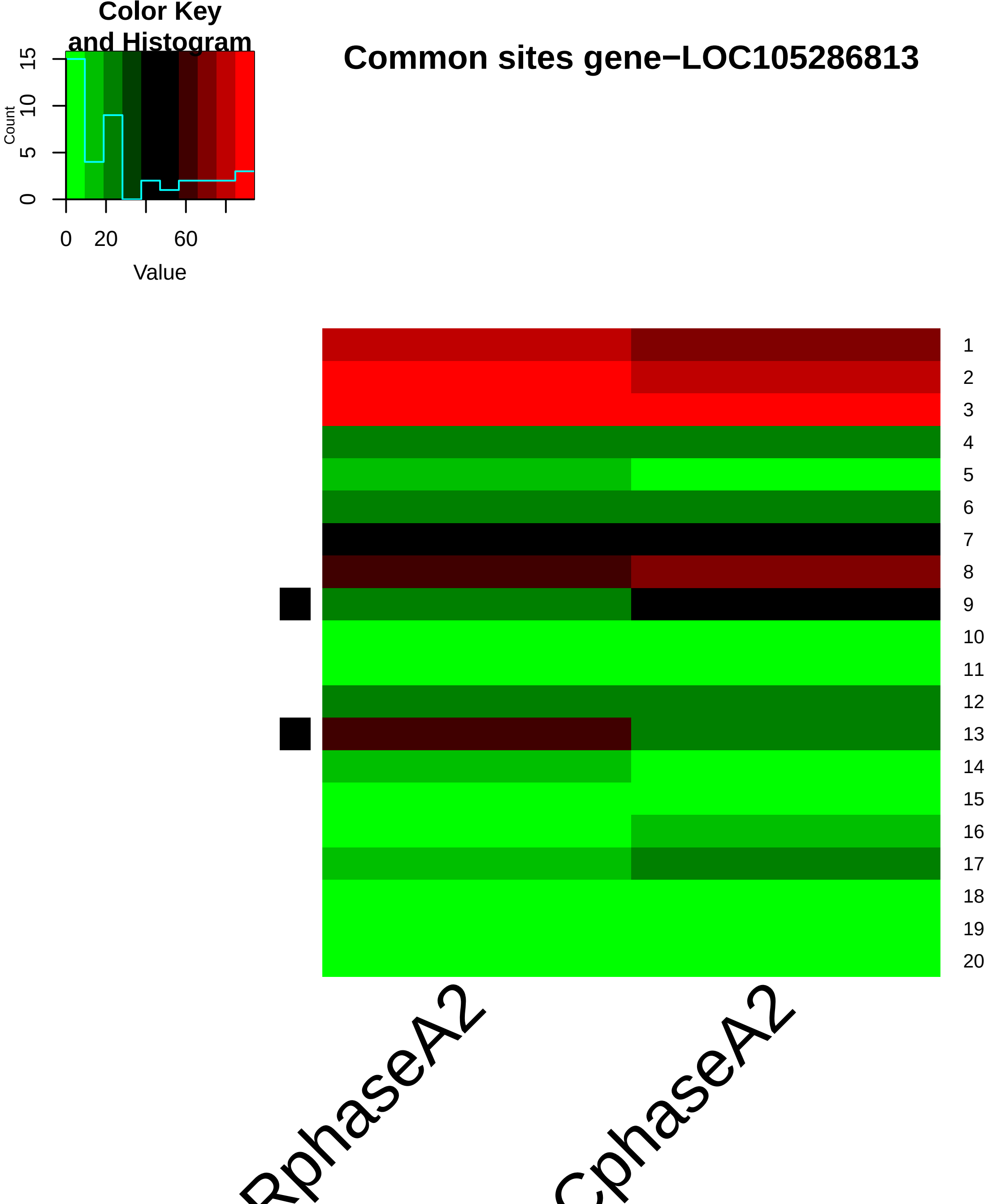
Color Key
and Histogram
Count
0
5
10
15
0
20
60
Value
Common sites gene−LOC105286813
RphaseA2
CphaseA2

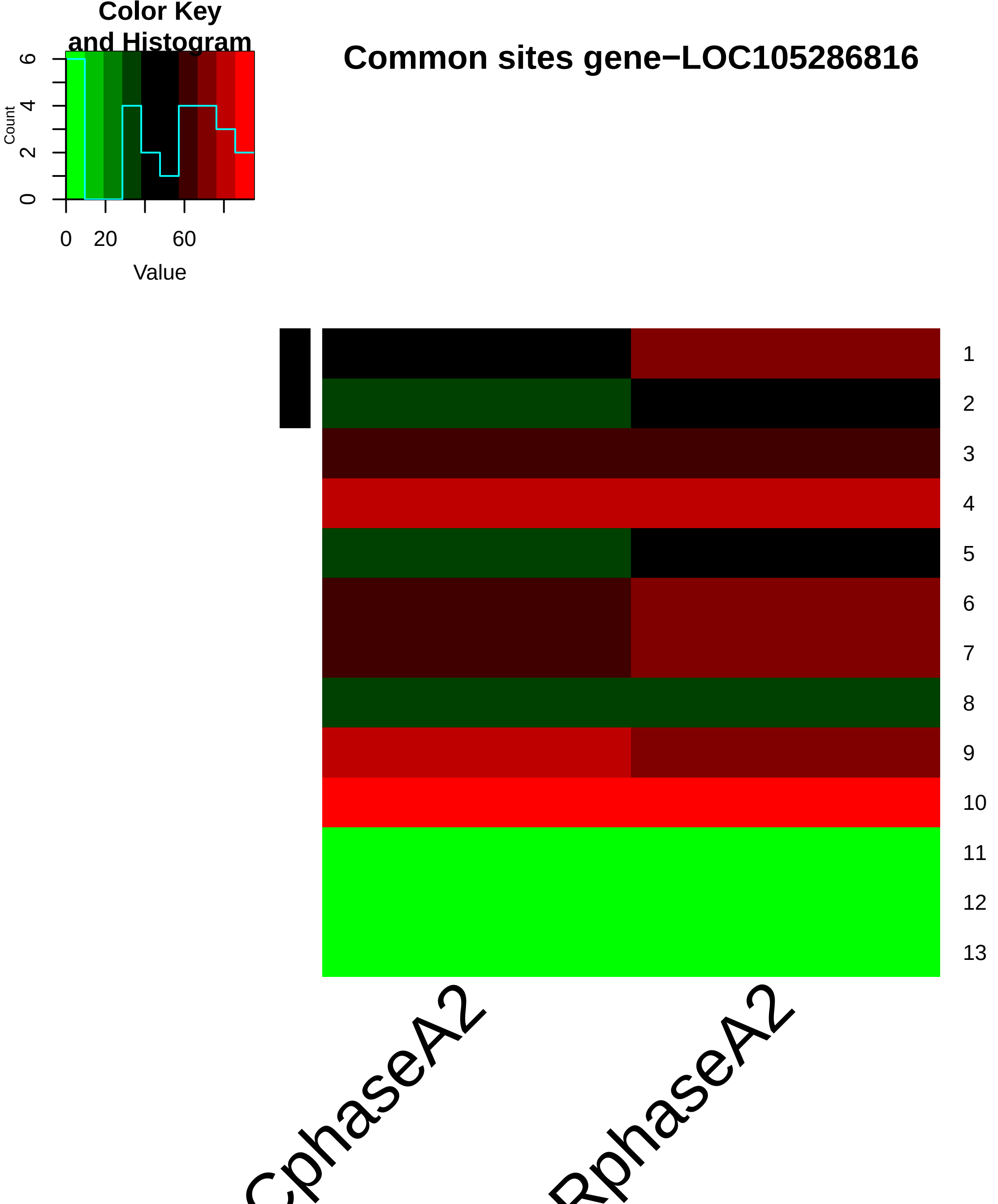

Color Key
and Histogram
Count
Value
0
20
60
0
2
4
6
Common sites gene−LOC105286816
1
2
3
4
5
6
7
8
9
10
11
12
13
CphaseA2
RphaseA2

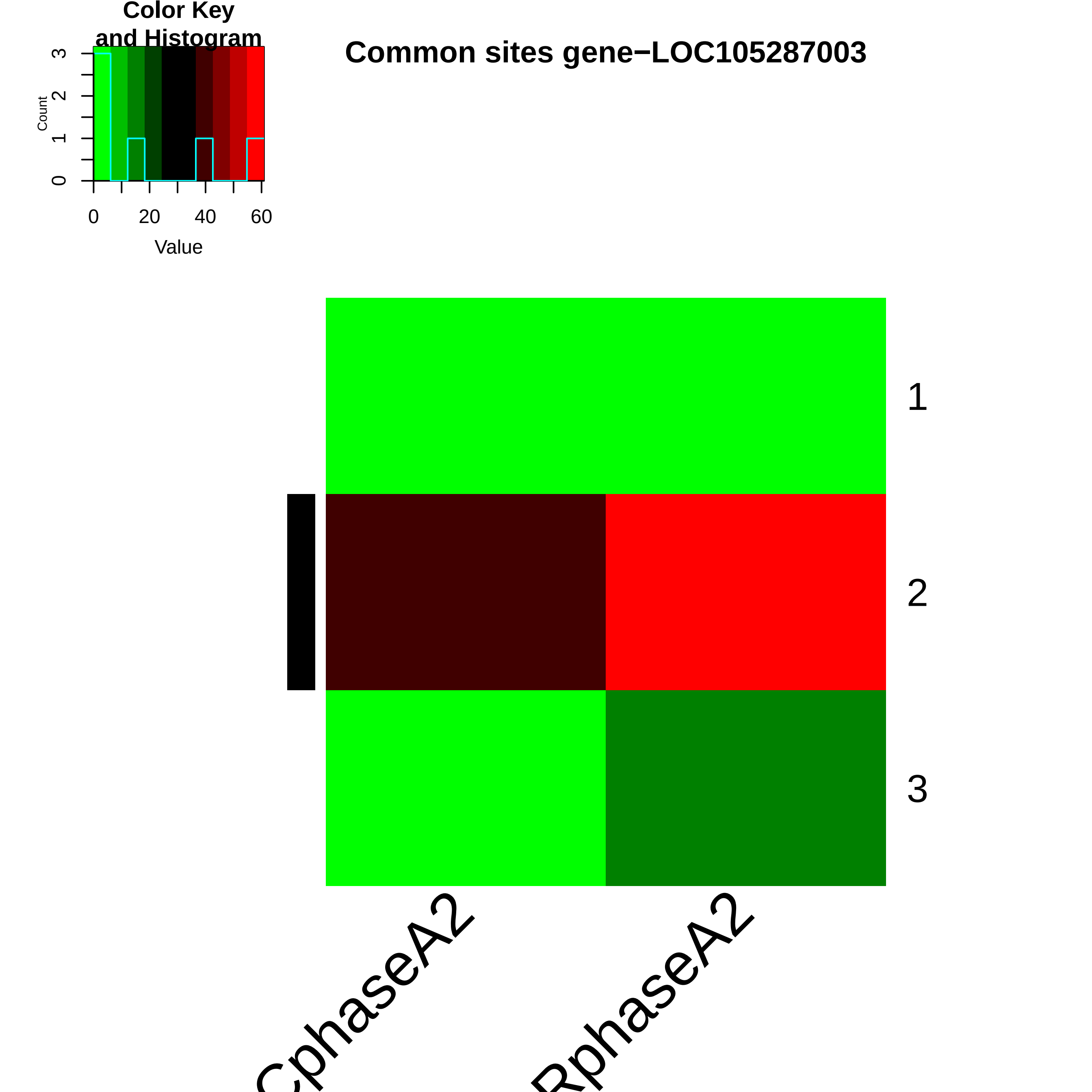

Color Key
and Histogram
Count
Value
0
20
40
60
0
1
2
3
Common sites gene−LOC105287003
1
2
3
CphaseA2
RphaseA2

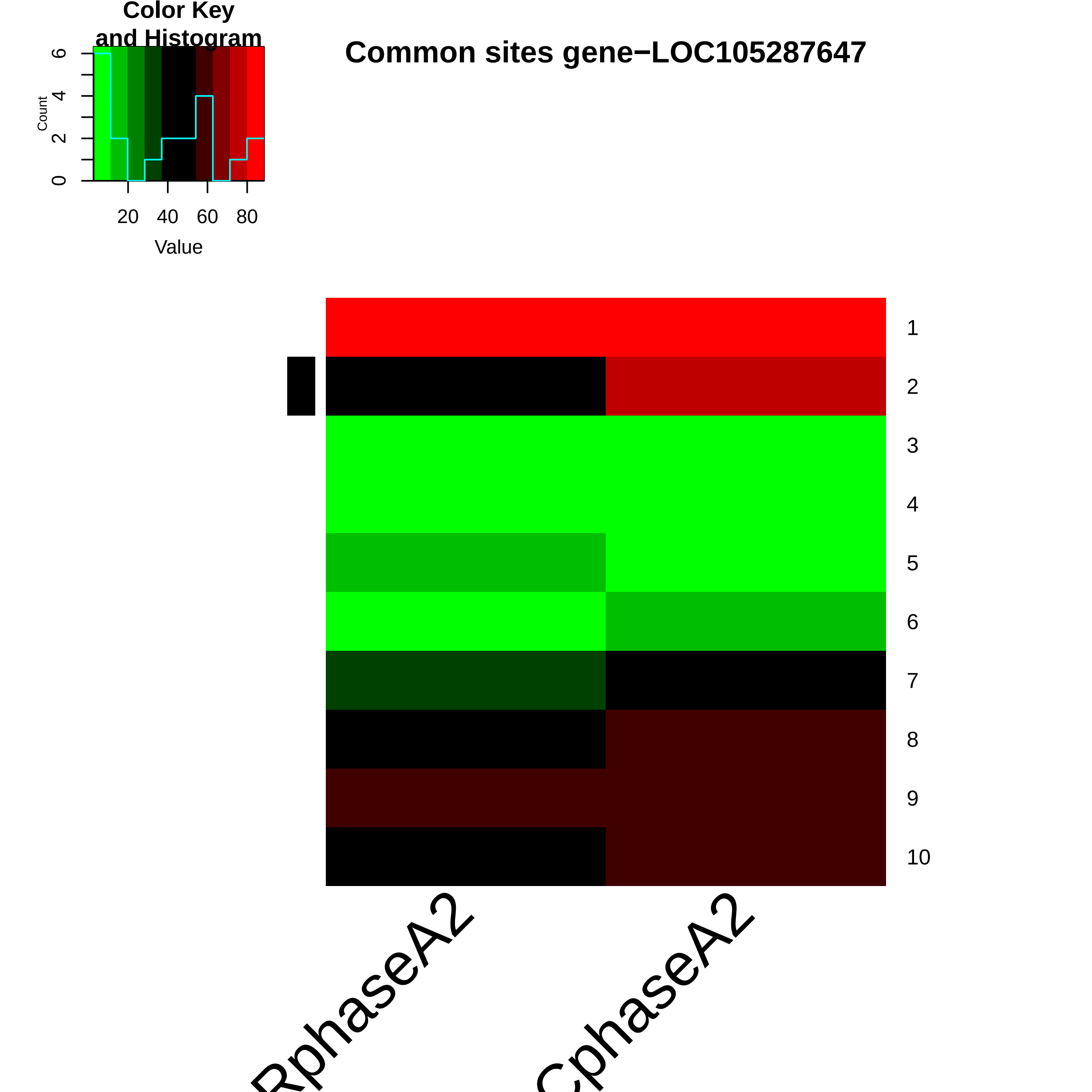

Color Key
and Histogram
Count
Value
0
2
4
6
20
40
60
80
Common sites gene−LOC105287647
1
2
3
4
5
6
7
8
9
10
RphaseA2
CphaseA2

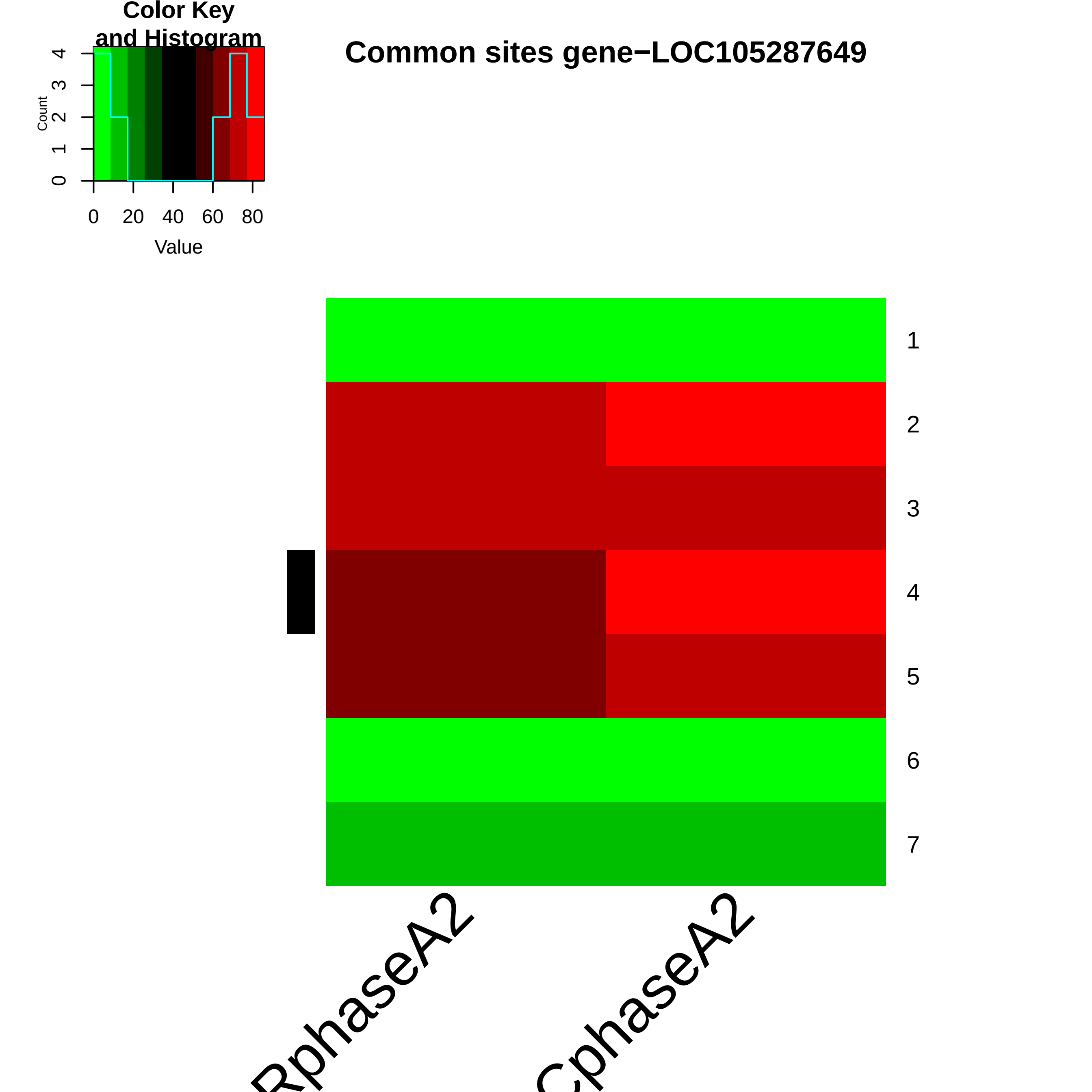
Color Key
and Histogram
Count
0
1
2
3
4
0
20
40
60
80
Value
Common sites gene−LOC105287649
1
2
3
4
5
6
7
RphaseA2
CphaseA2

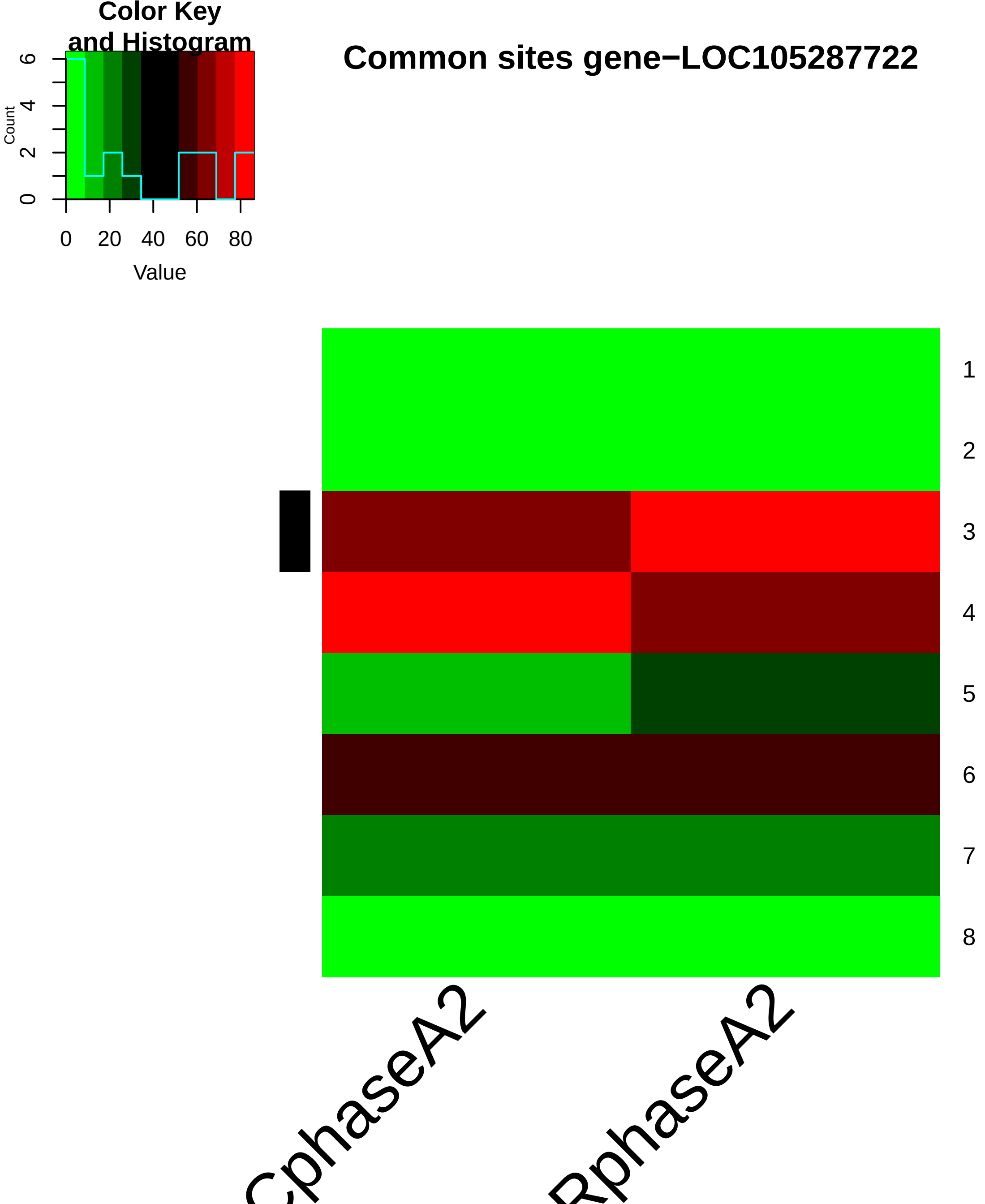
Color Key
and Histogram
Count
0
2
4
6
0
20
40
60
80
Value
Common sites gene−LOC105287722
1
2
3
4
5
6
7
8
CphaseA2
RphaseA2

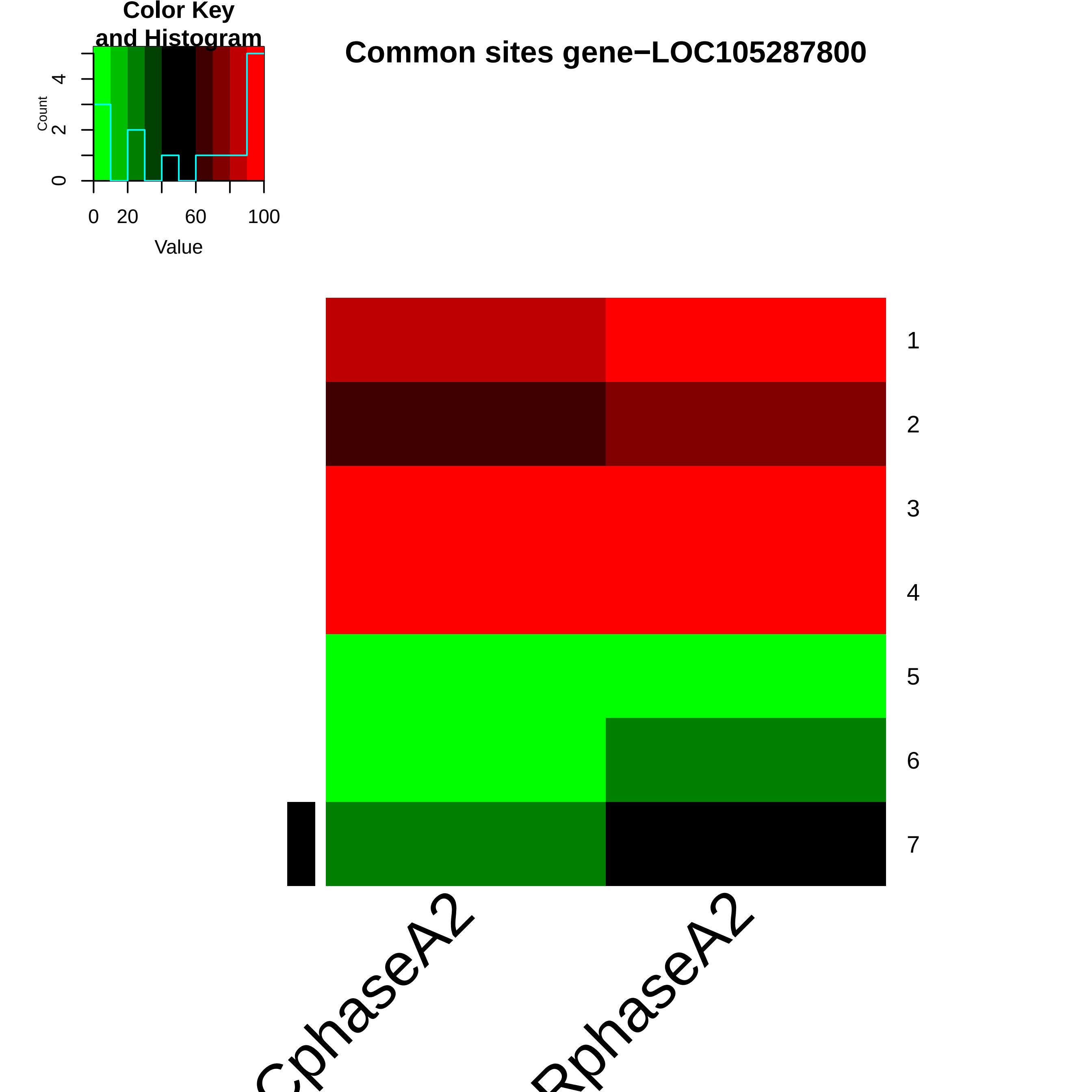

Color Key
and Histogram
Count
0
2
4
0
20
60
100
Value
Common sites gene−LOC105287800
1
2
3
4
5
6
7
CphaseA2
RphaseA2

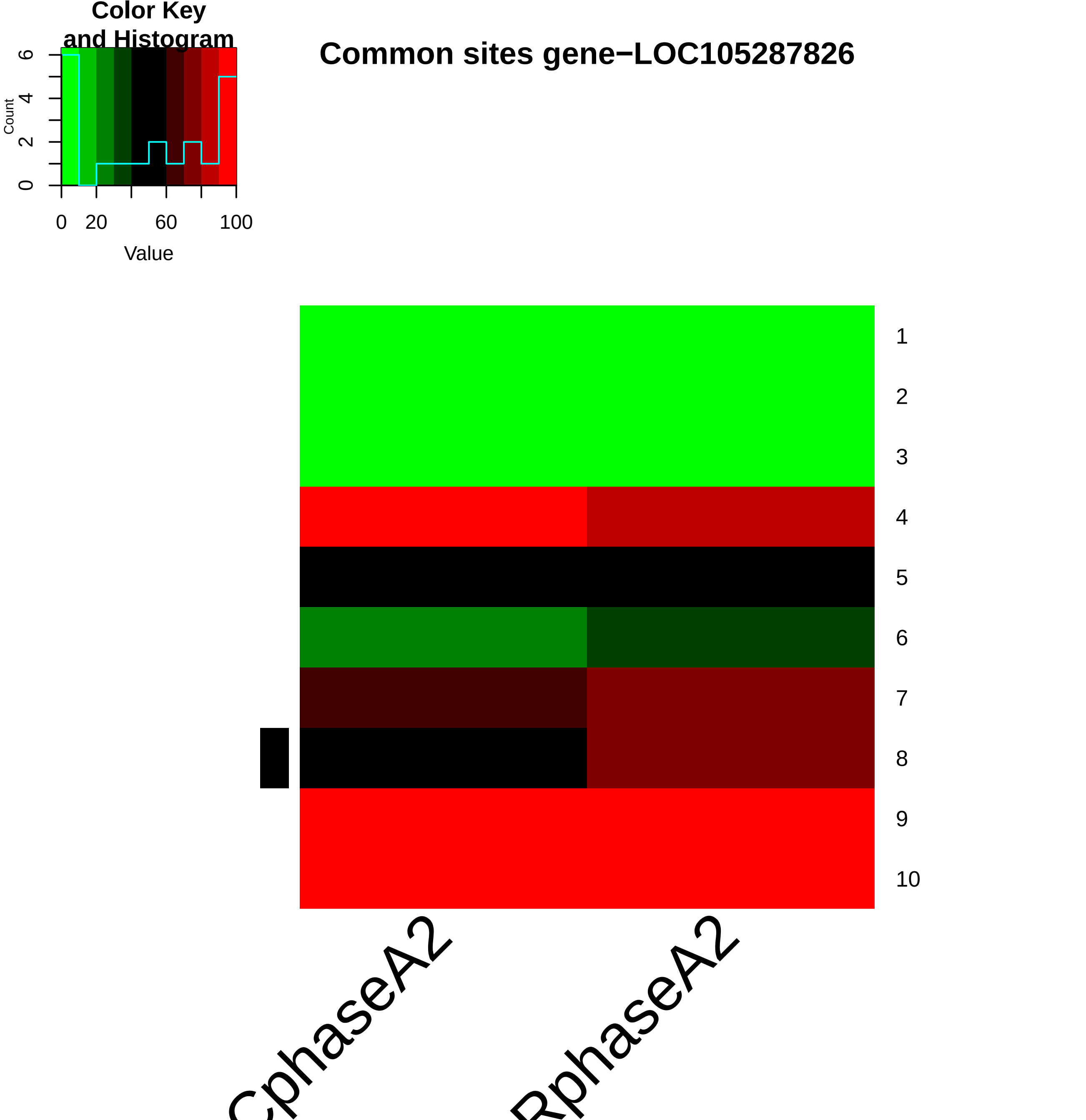

Color Key
and Histogram
Count
Value
0
20
60
100
0
2
4
6
Common sites gene−LOC105287826
1
2
3
4
5
6
7
8
9
10

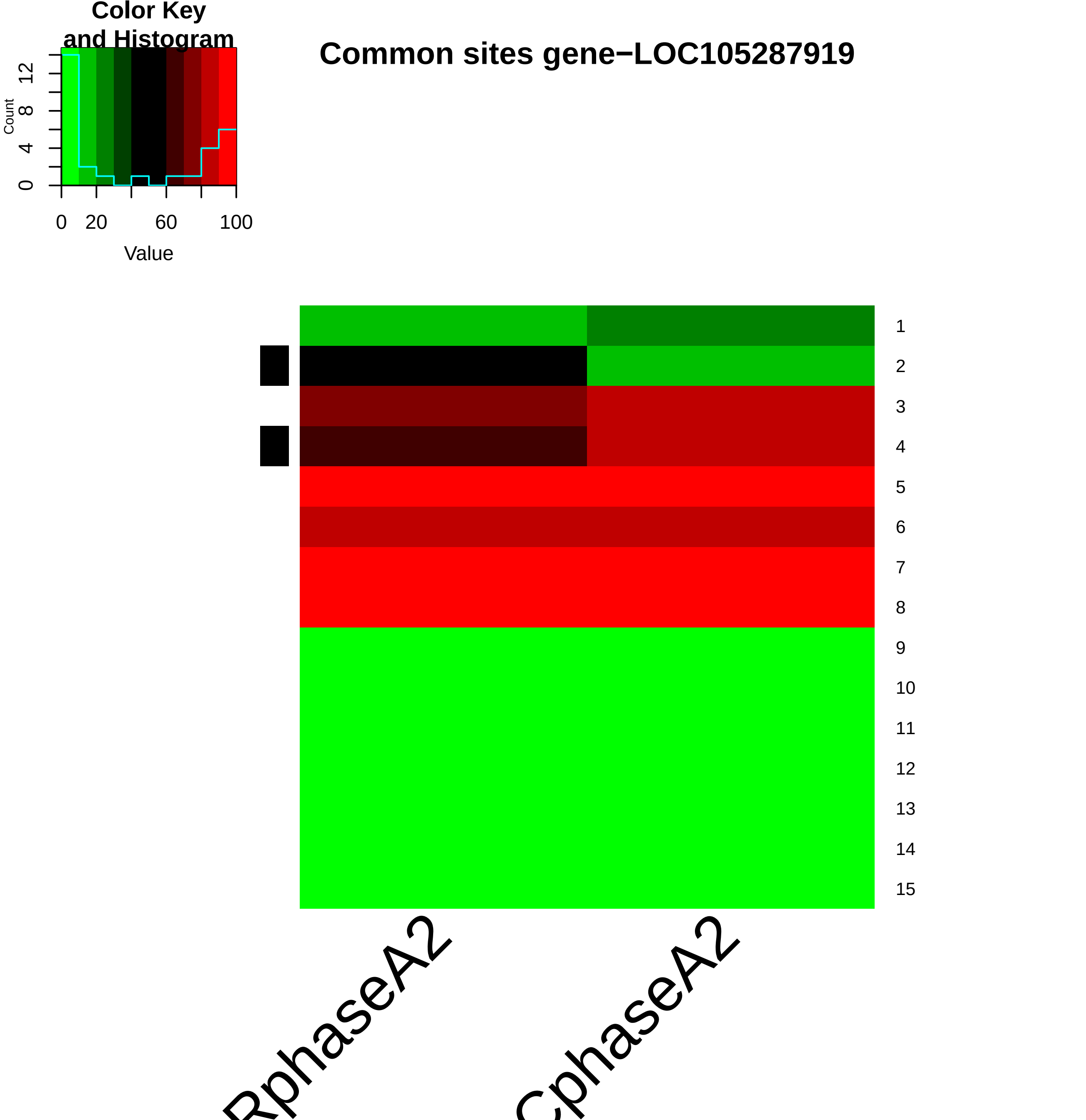
Color Key
and Histogram
Count
0
4
8
12
0
20
60
100
Value
Common sites gene−LOC105287919
RphaseA2
CphaseA2

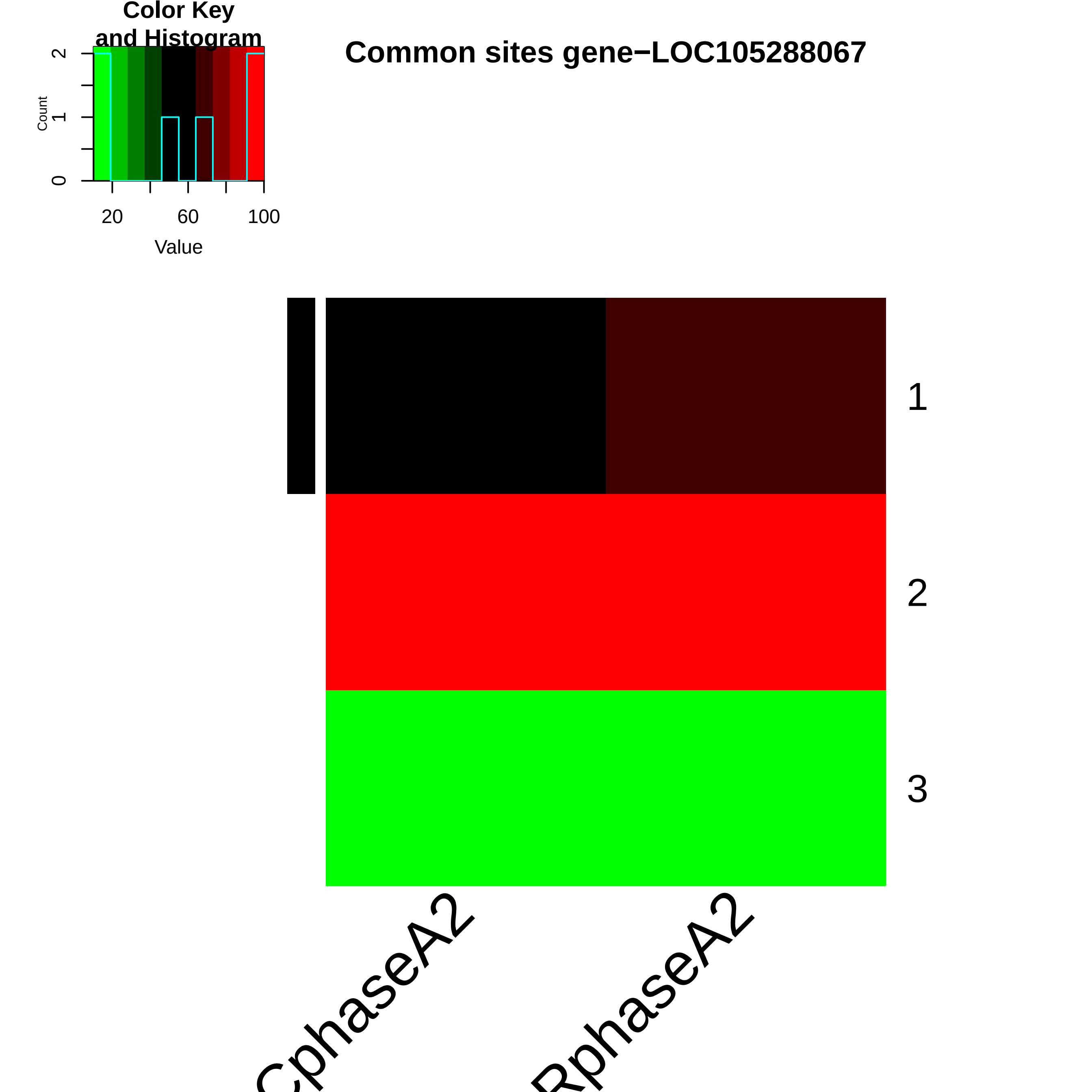

Color Key
and Histogram
Count
0
1
2
20
60
100
Value
Common sites gene−LOC105288067
1
2
3
CphaseA2
RphaseA2

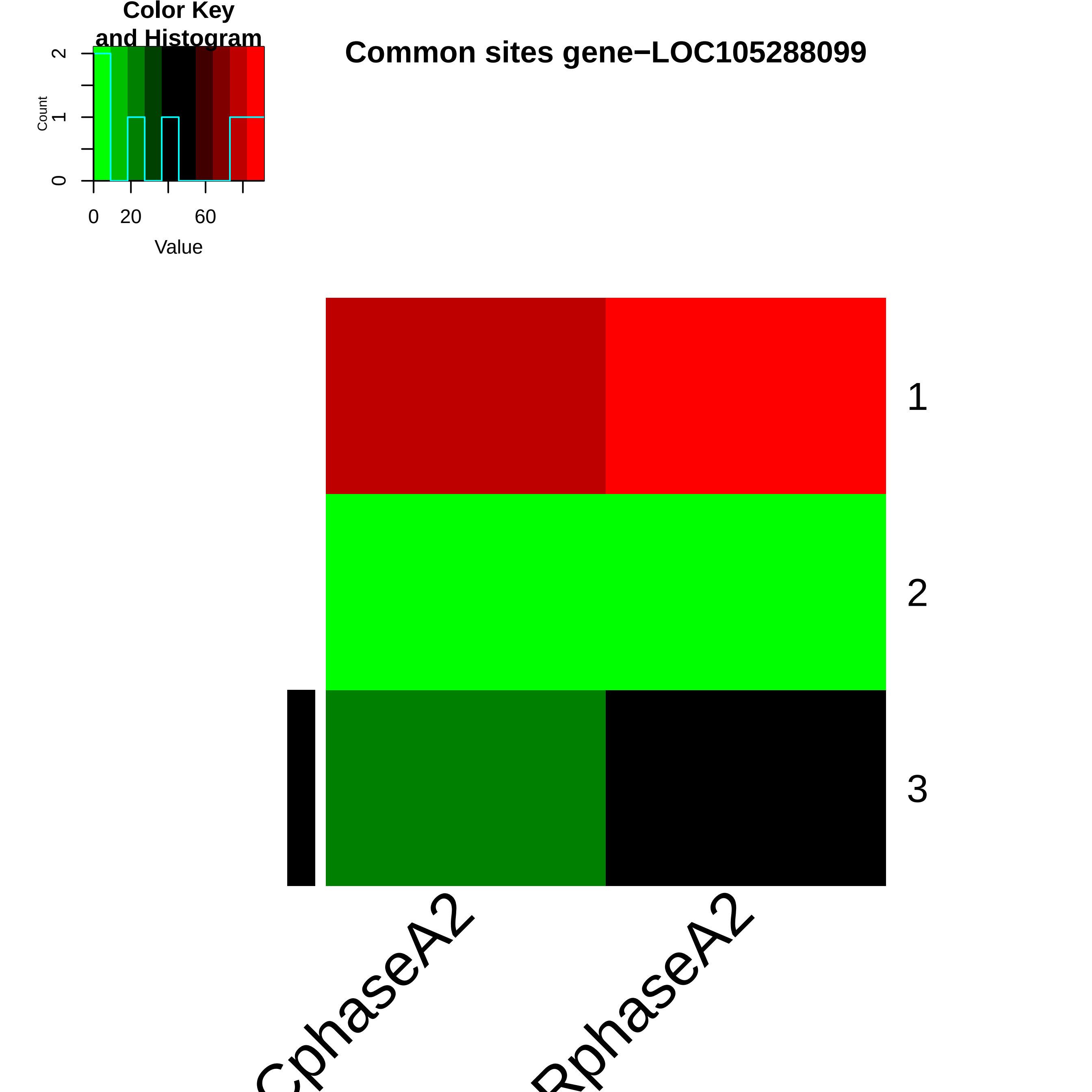

Color Key
and Histogram
Count
Value
Common sites gene−LOC105288099
1
2
3

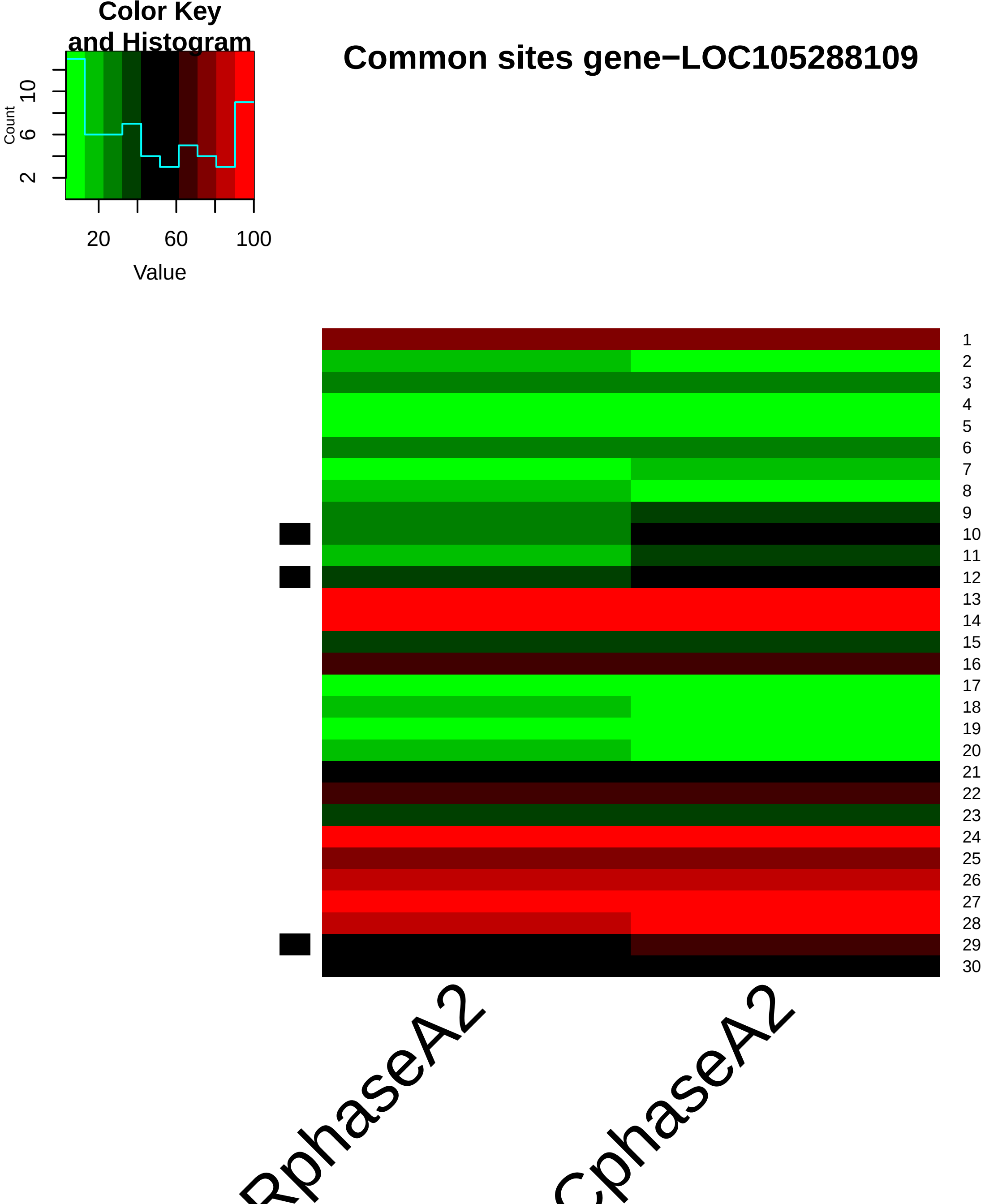
Color Key
and Histogram
Count
2
6
10
20
60
100
Value
Common sites gene−LOC105288109
RphaseA2
CphaseA2

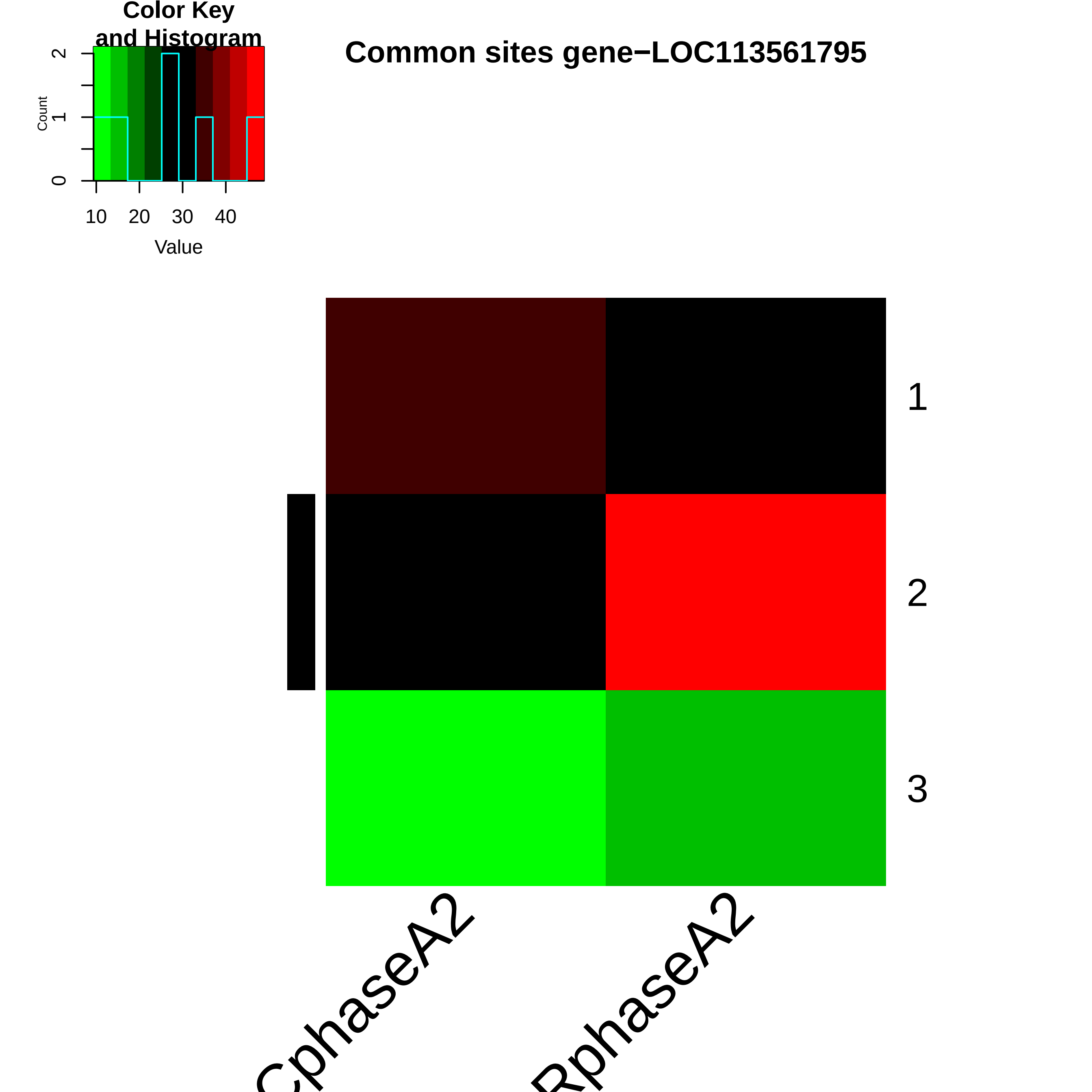

Color Key
and Histogram
Count
0
1
2
10
20
30
40
Value
Common sites gene−LOC113561795
1
2
3

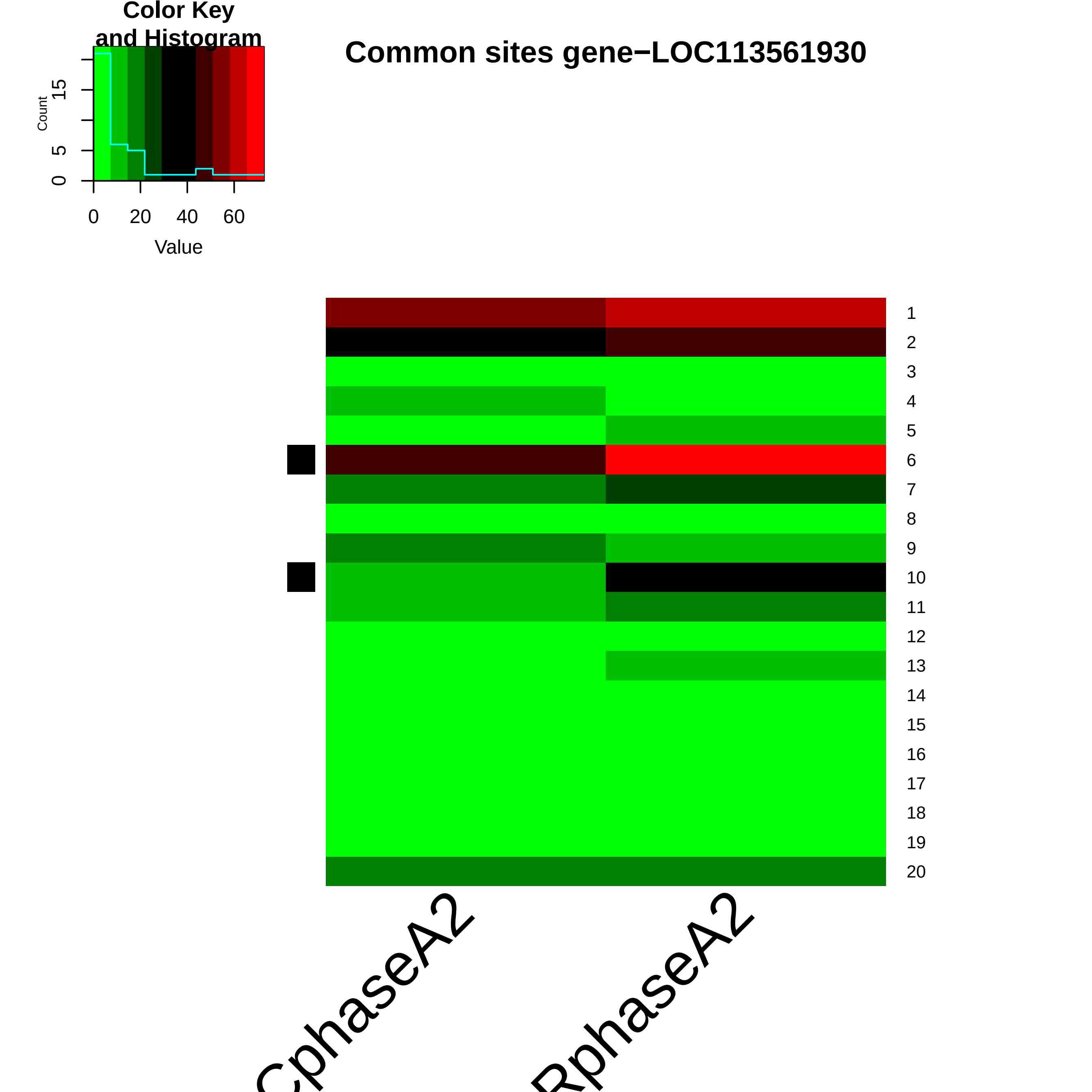
Color Key
and Histogram
Count
0
5
15
0
20
40
60
Value
Common sites gene−LOC113561930
CphaseA2
RphaseA2

#### Common sites gene−LOC113562627

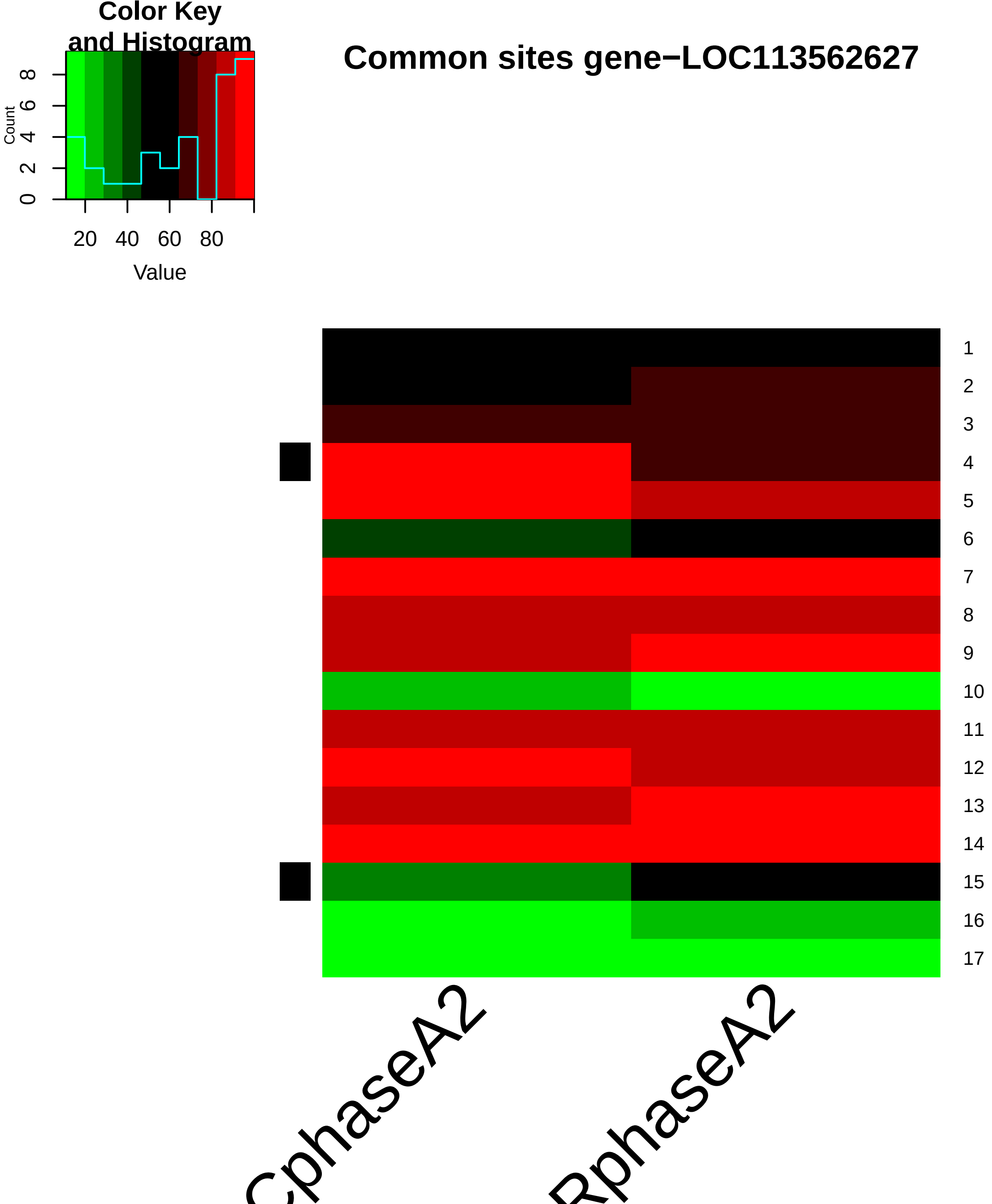

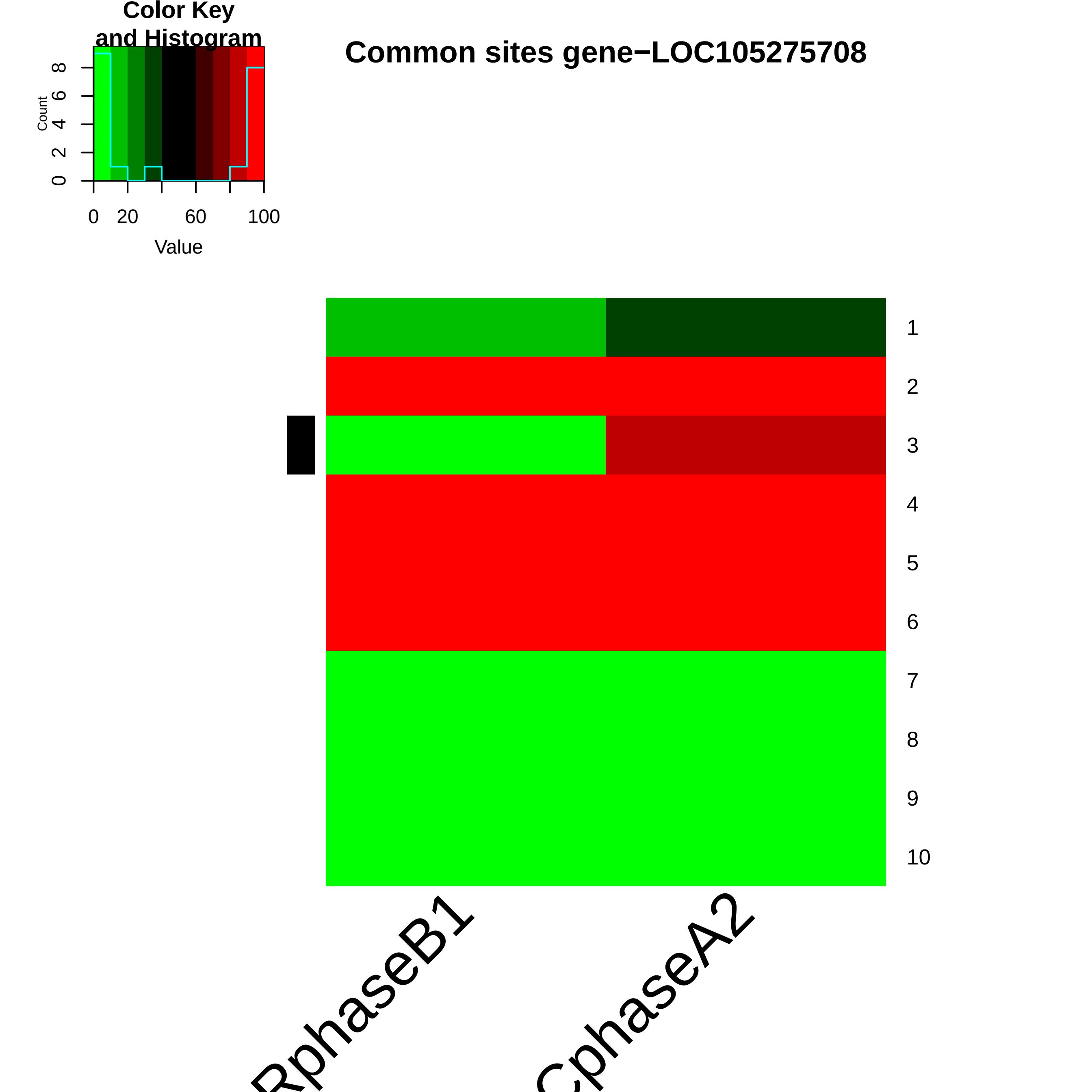

Color Key
and Histogram
Count
0
2
4
6
8
0
20
60
100
Value
Common sites gene−LOC105275708
1
2
3
4
5
6
7
8
9
10
RphaseB1
CphaseA2

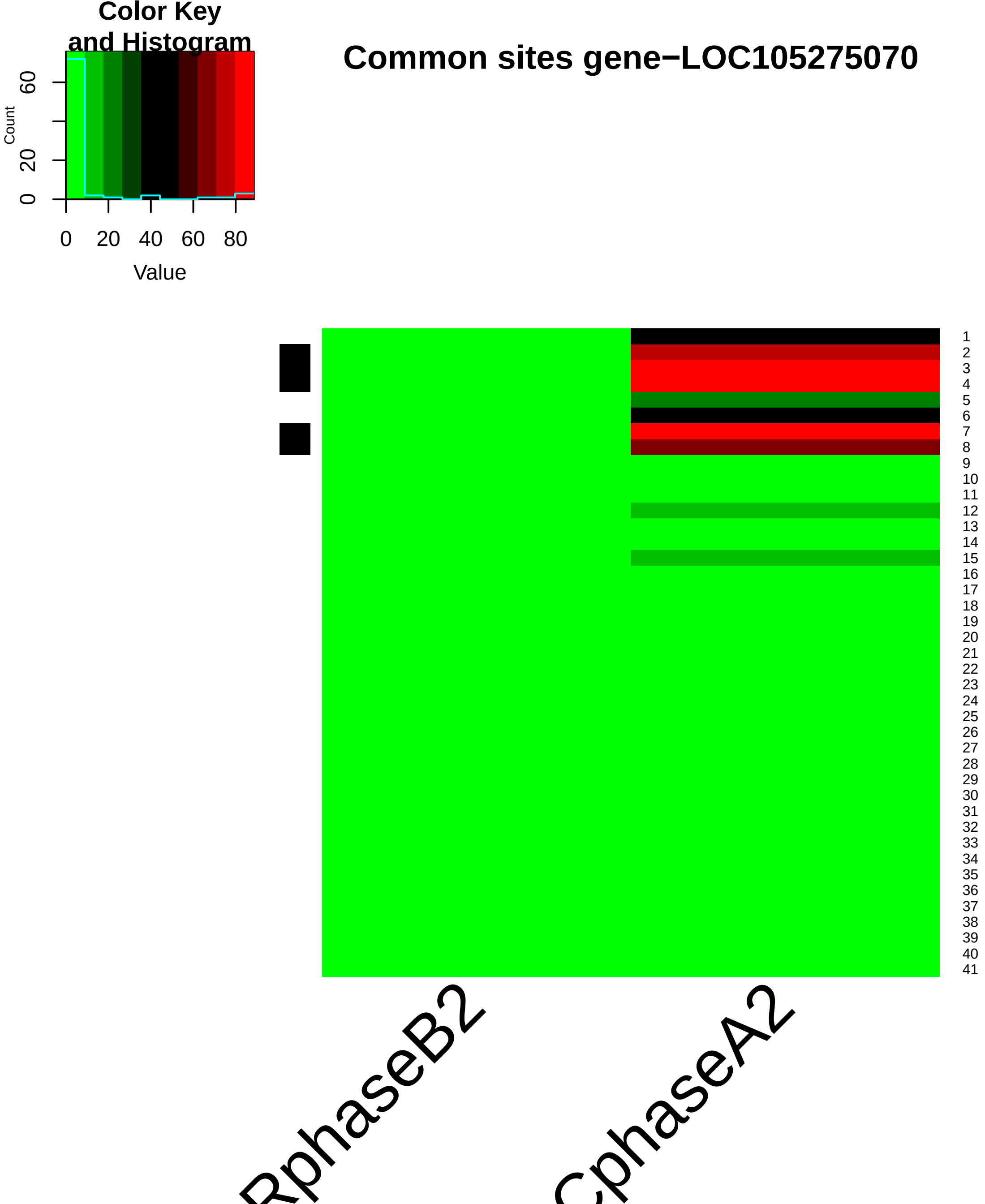

Color Key
and Histogram
Common sites gene−LOC105275070
Count
0
20
60
0
20
40
60
80
Value
RphaseB2
CphaseA2

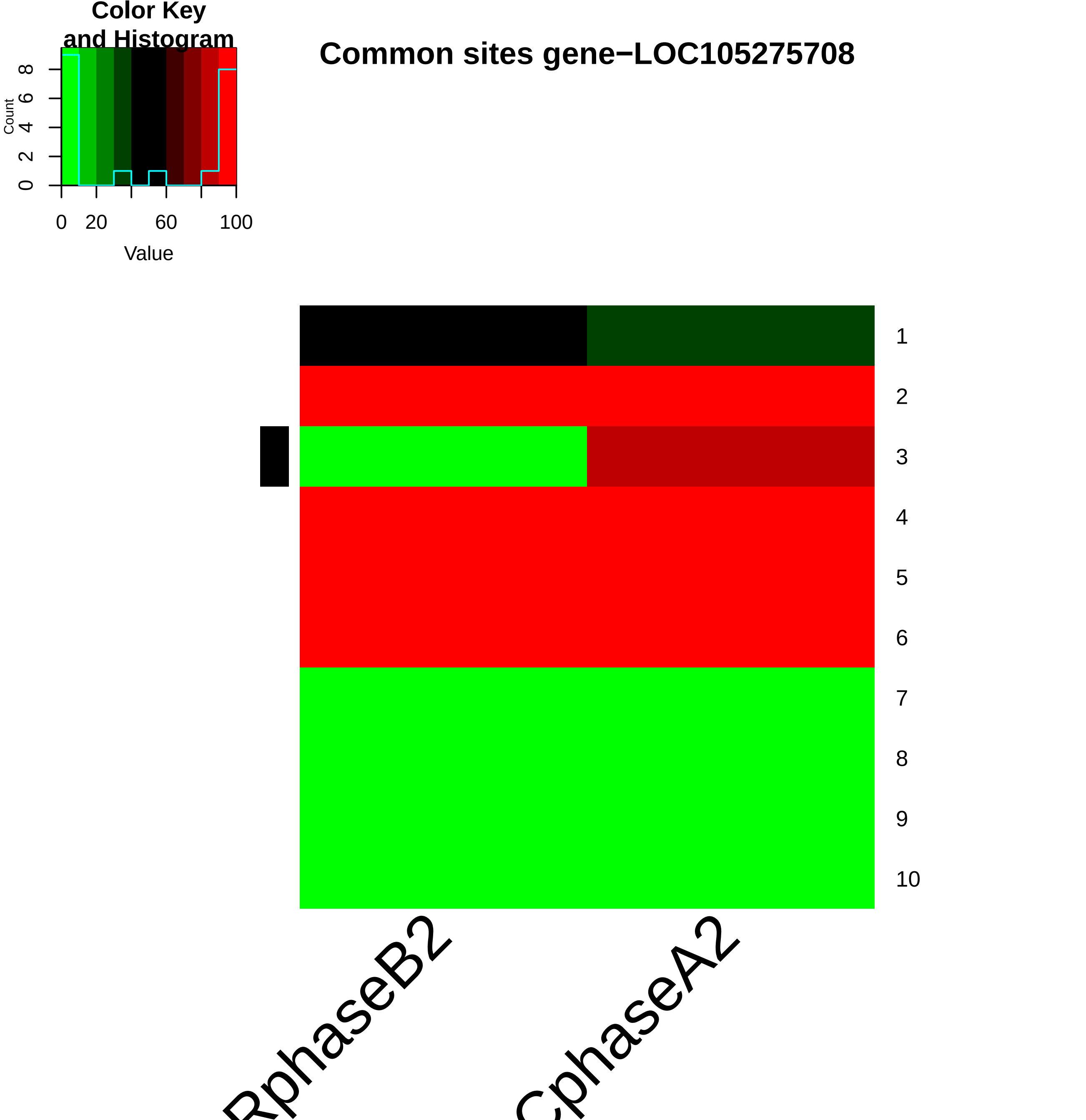

Color Key
and Histogram
Count
0 2 4 6 8
0 20 60 100
Value
Common sites gene−LOC105275708
1
2
3
4
5
6
7
8
9
10
RphaseB2
CphaseA2

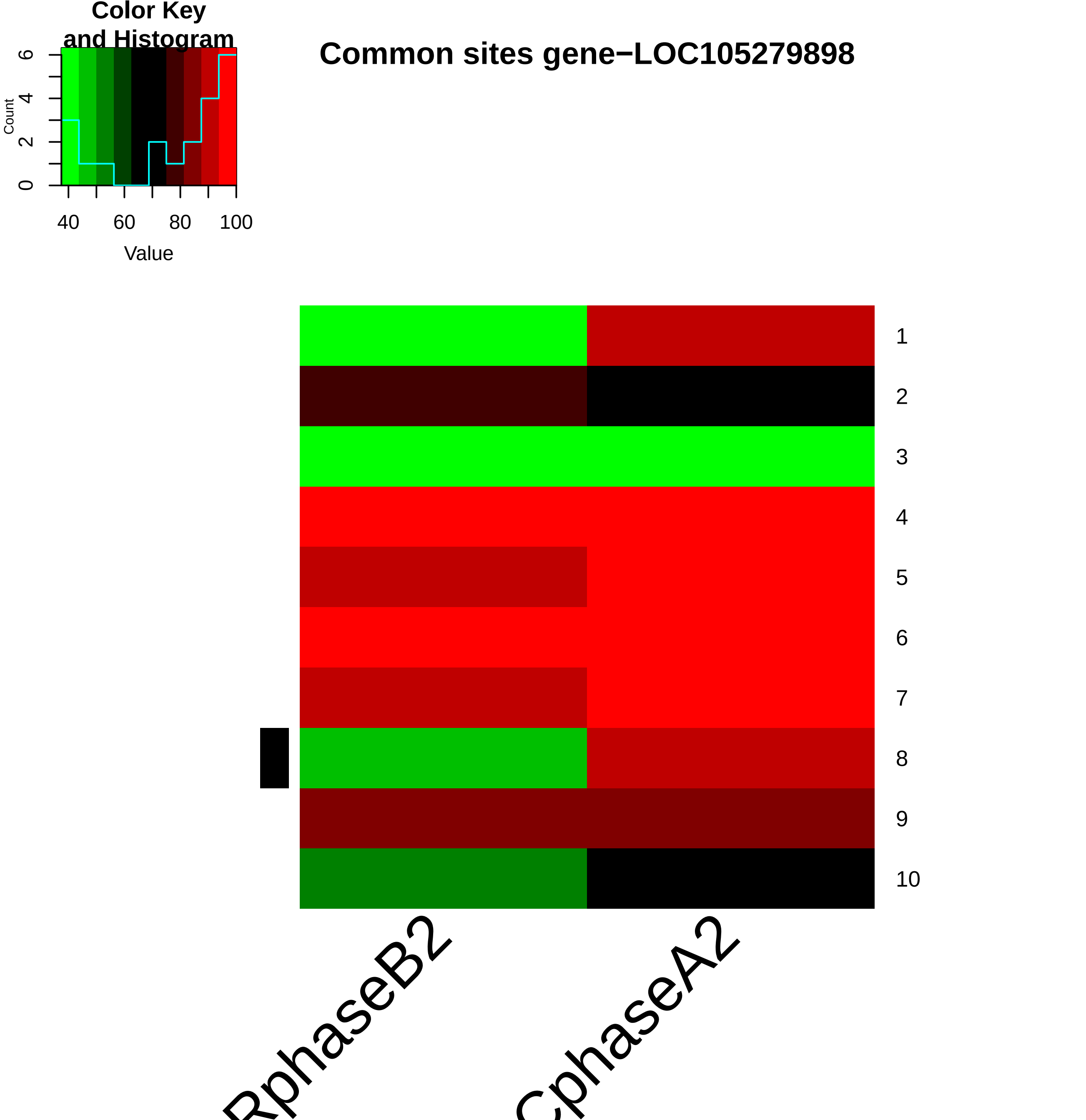

Color Key
and Histogram
Count
Value
0
2
4
6
40
60
80
100
Common sites gene−LOC105279898
1
2
3
4
5
6
7
8
9
10
RphaseB2
CphaseA2

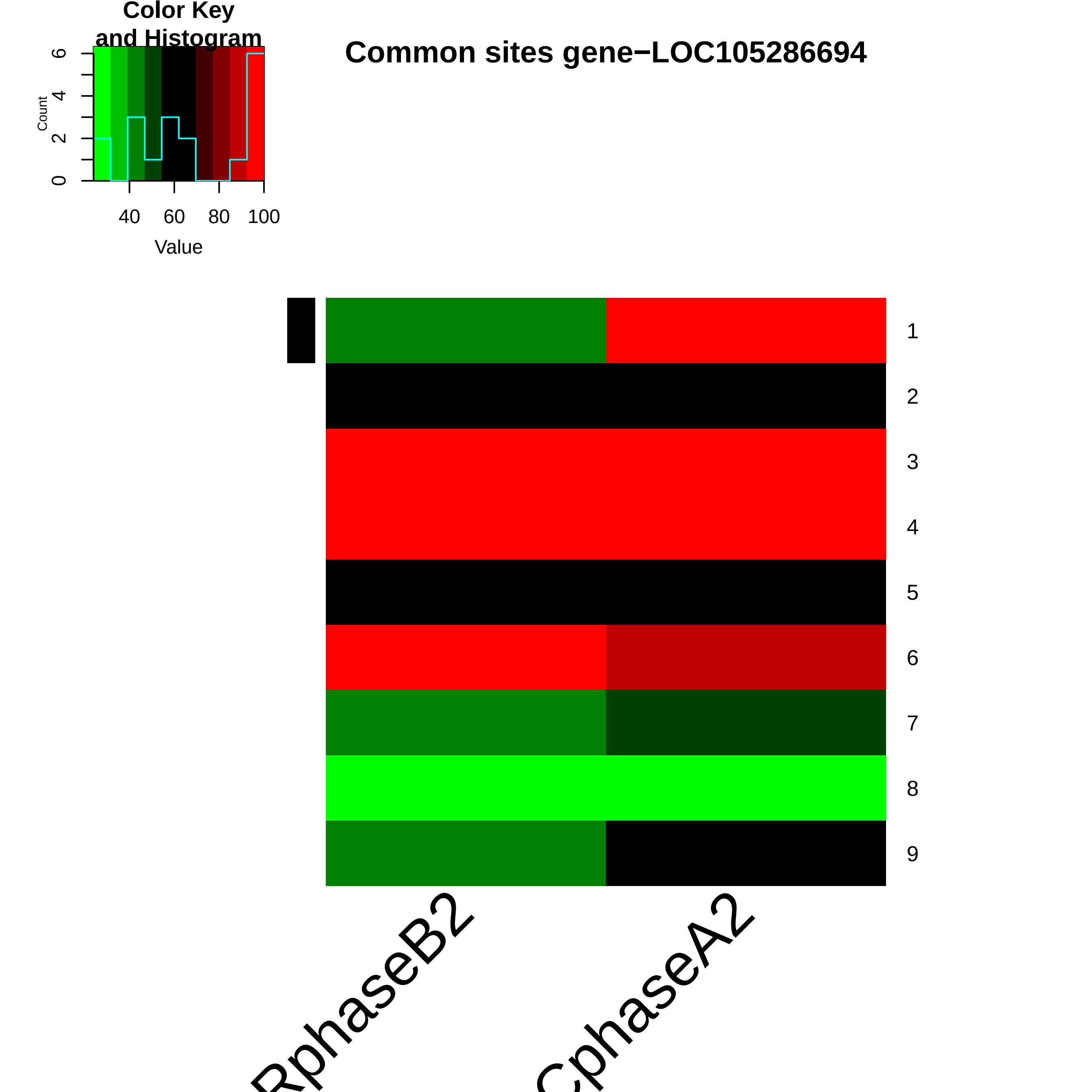

Color Key
and Histogram
Count
0
2
4
6
40
60
80
100
Value
Common sites gene−LOC105286694
1
2
3
4
5
6
7
8
9

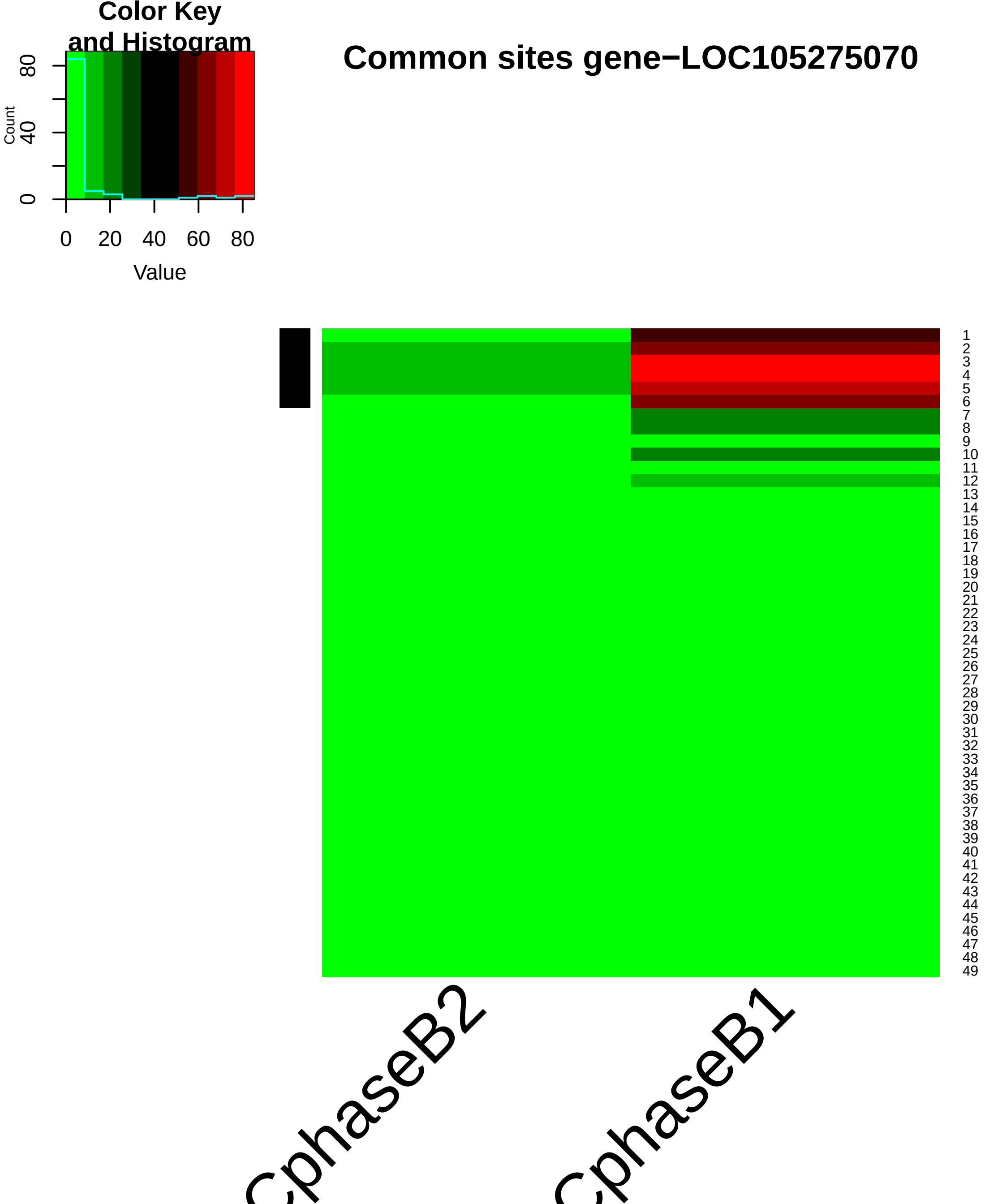

Color Key
and Histogram
Count
0
40
80
0
20
40
60
80
Value
Common sites gene−LOC105275070
CphaseB2
CphaseB1

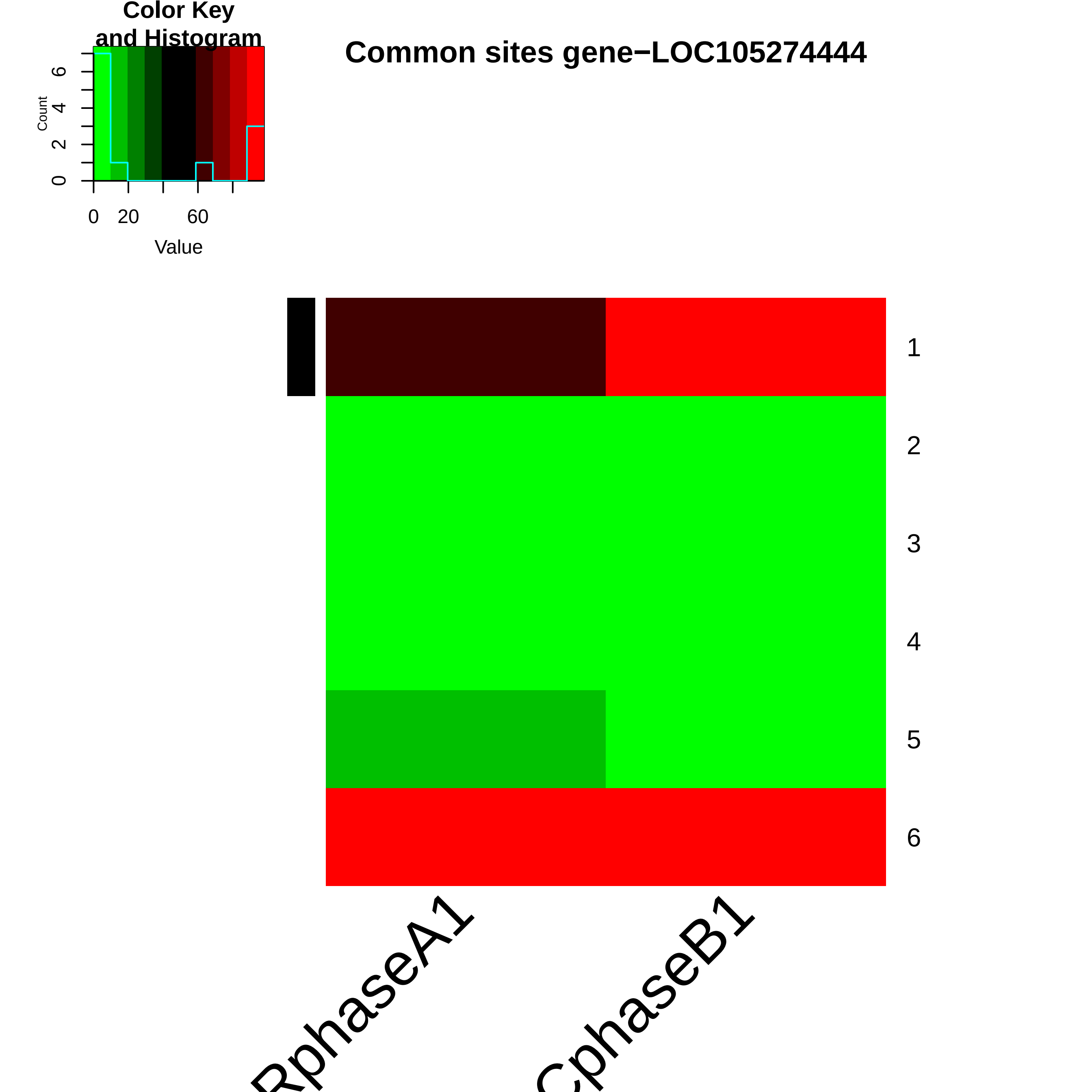

Color Key
and Histogram
Count
0
2
4
6
0
20
60
Value
Common sites gene−LOC105274444
1
2
3
4
5
6
RphaseA1
CphaseB1

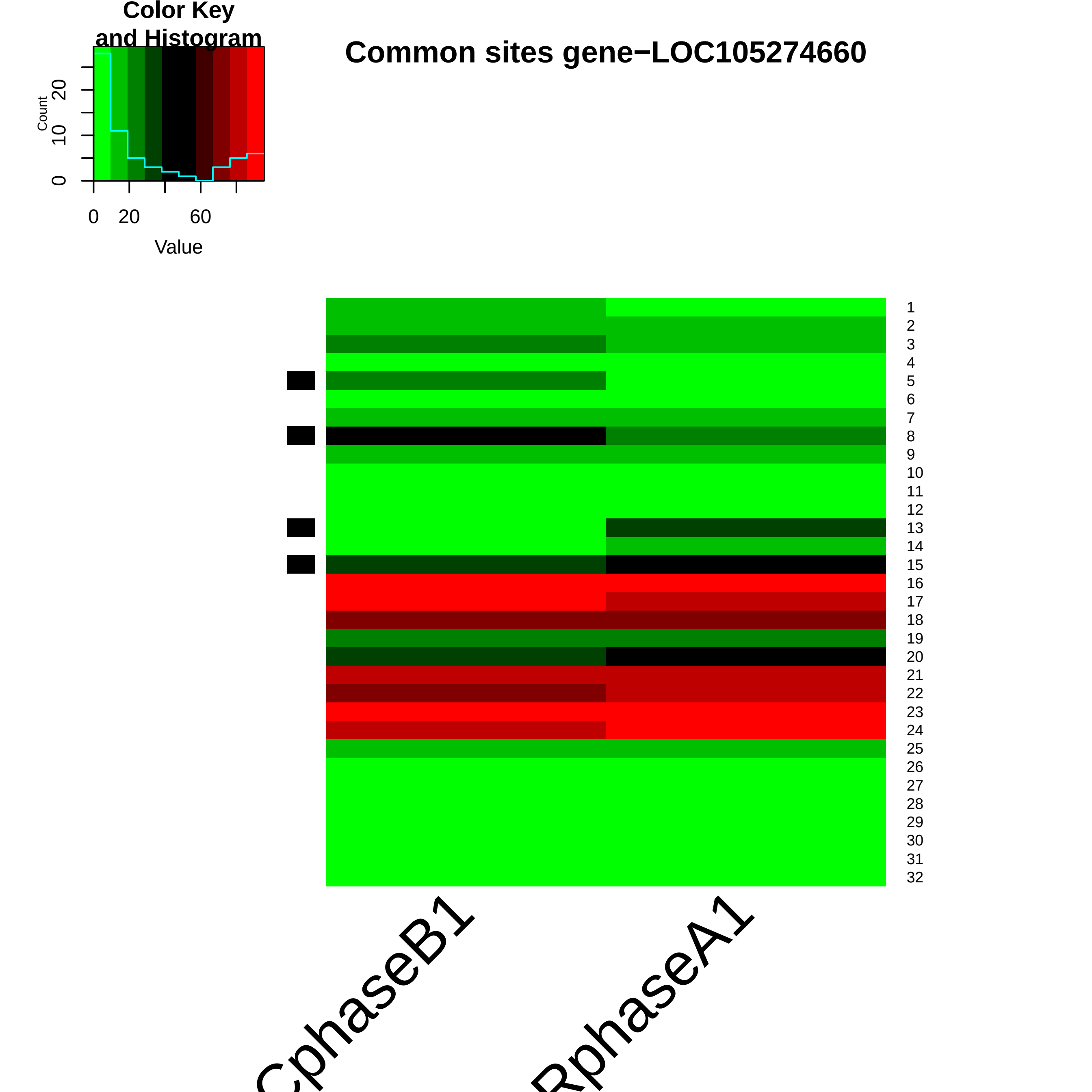
Color Key
and Histogram
Count
0
10
20
0
20
60
Value
Common sites gene−LOC105274660
CphaseB1
RphaseA1

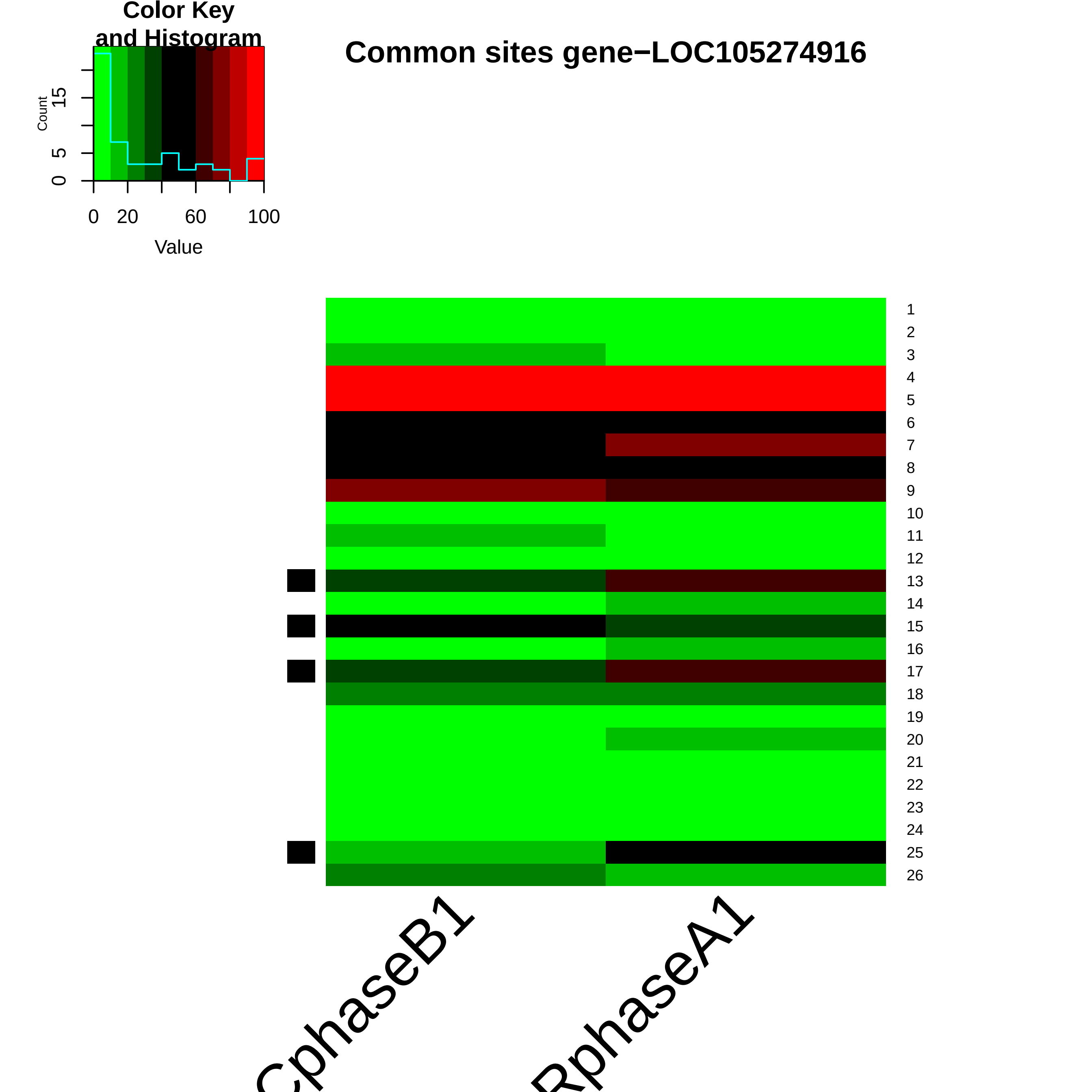

Color Key
and Histogram
Count
0
5
15
0
20
60
100
Value
Common sites gene−LOC105274916
CphaseB1
RphaseA1

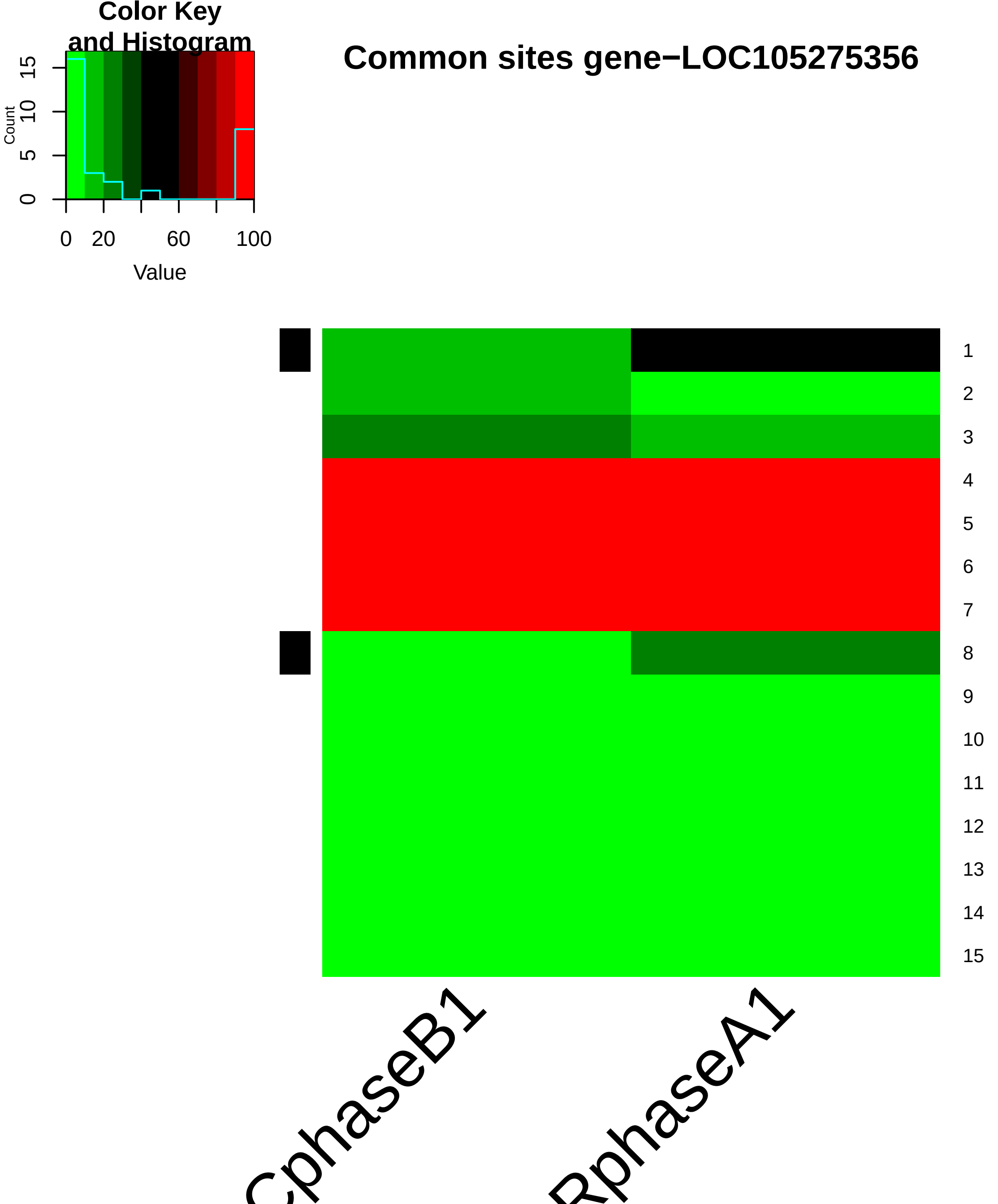
Color Key
and Histogram
Count
0
5
10
15
0
20
60
100
Value
Common sites gene−LOC105275356
CphaseB1
RphaseA1

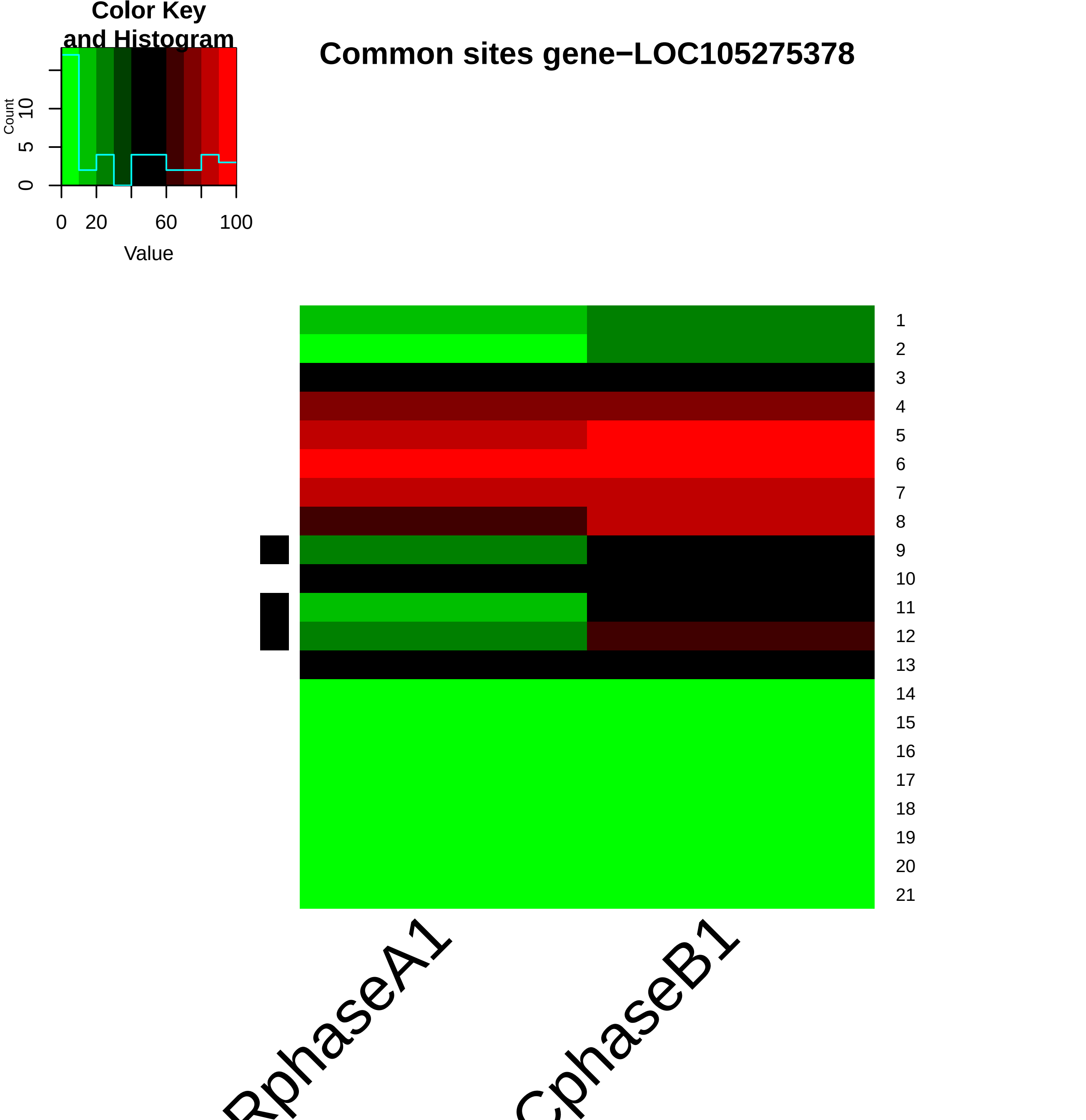

Color Key
and Histogram
Count
0
5
10
0
20
60
100
Value
Common sites gene−LOC105275378
RphaseA1
CphaseB1

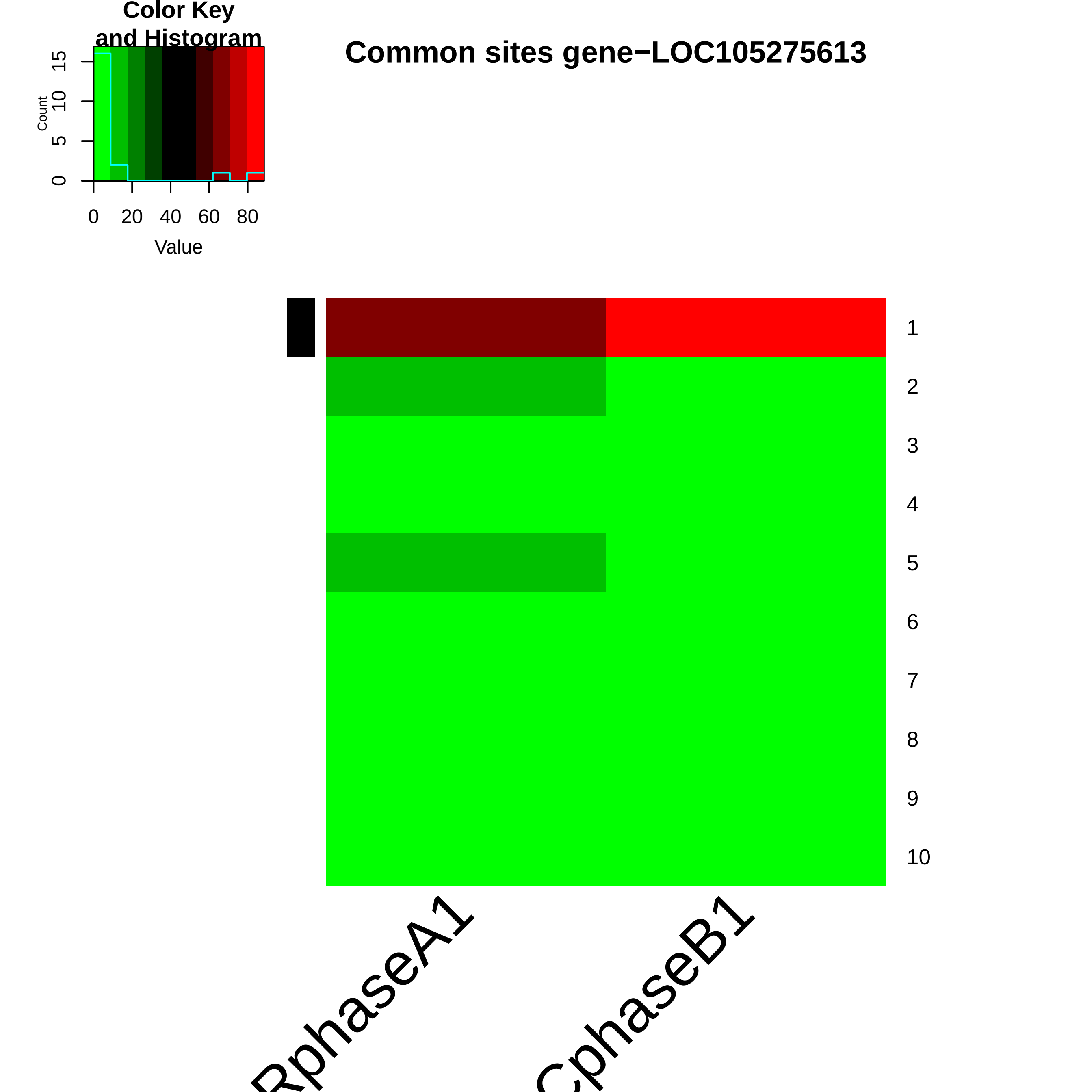
Color Key
and Histogram
Count
0
5
10
15
0
20
40
60
80
Value
Common sites gene−LOC105275613
1
2
3
4
5
6
7
8
9
10

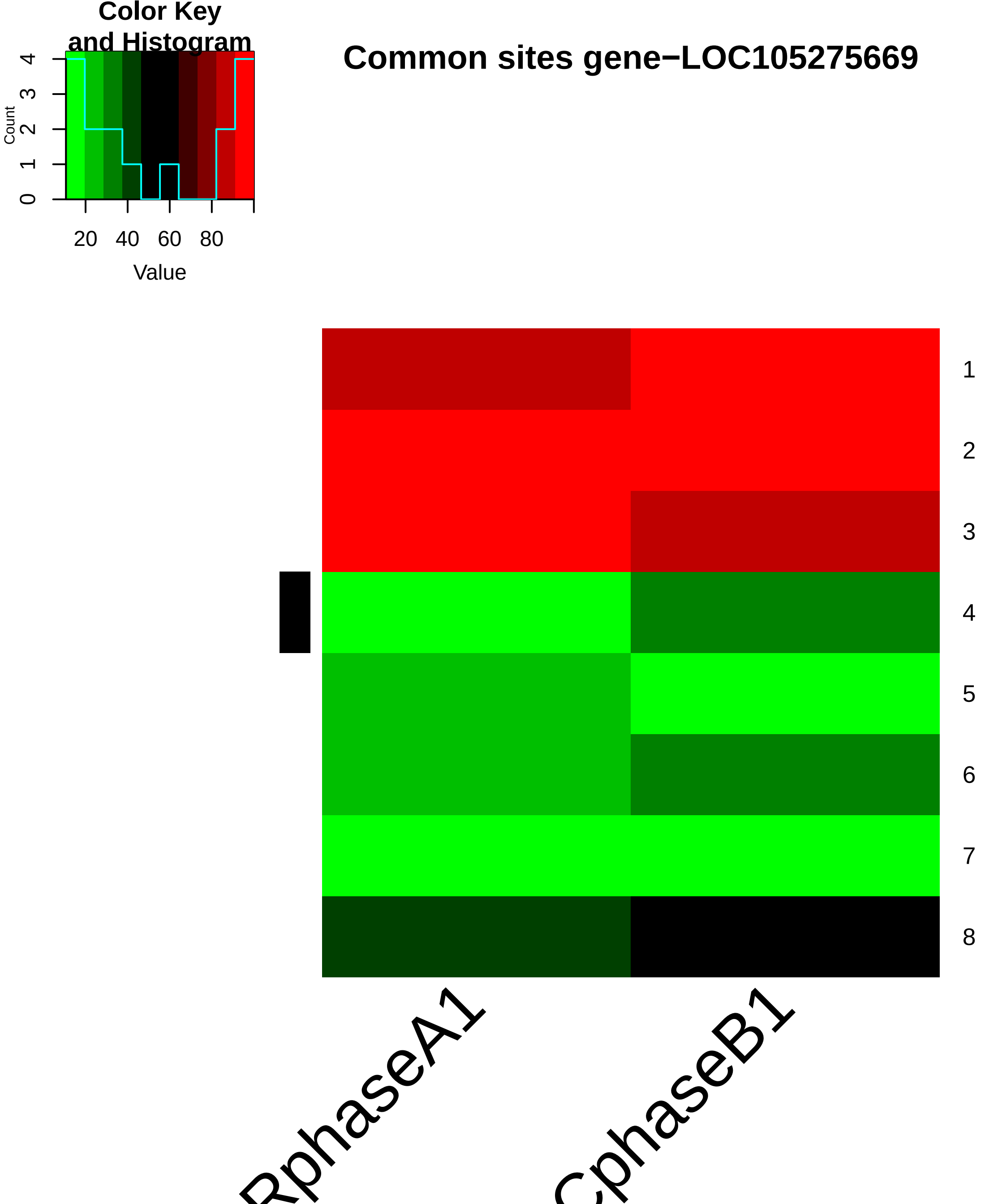
Color Key
and Histogram
Count
0
1
2
3
4
20
40
60
80
Value
Common sites gene−LOC105275669
1
2
3
4
5
6
7
8
RphaseA1
CphaseB1

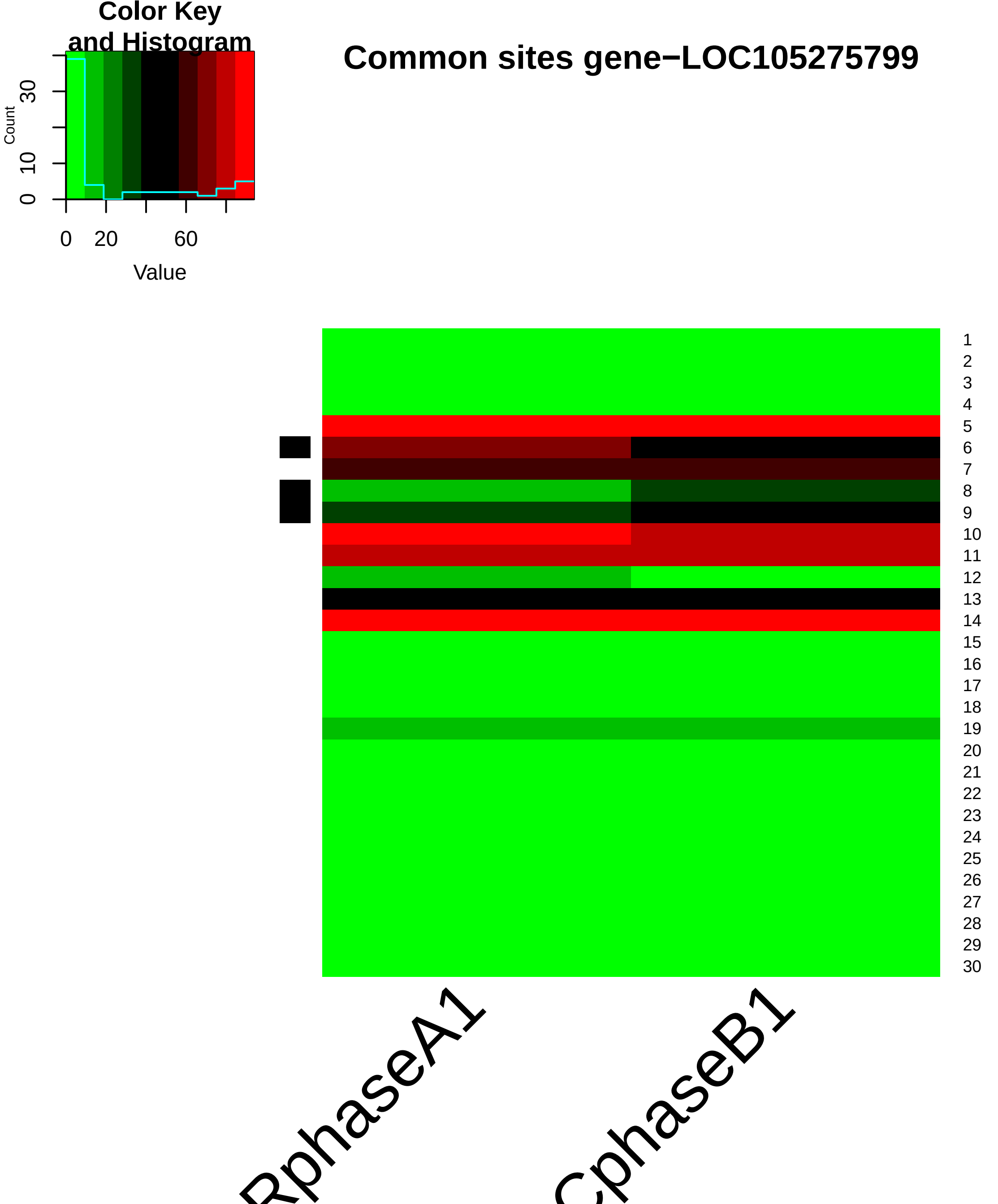
Color Key
and Histogram
Count
0
10
30
0
20
60
Value
Common sites gene−LOC105275799
RphaseA1
CphaseB1

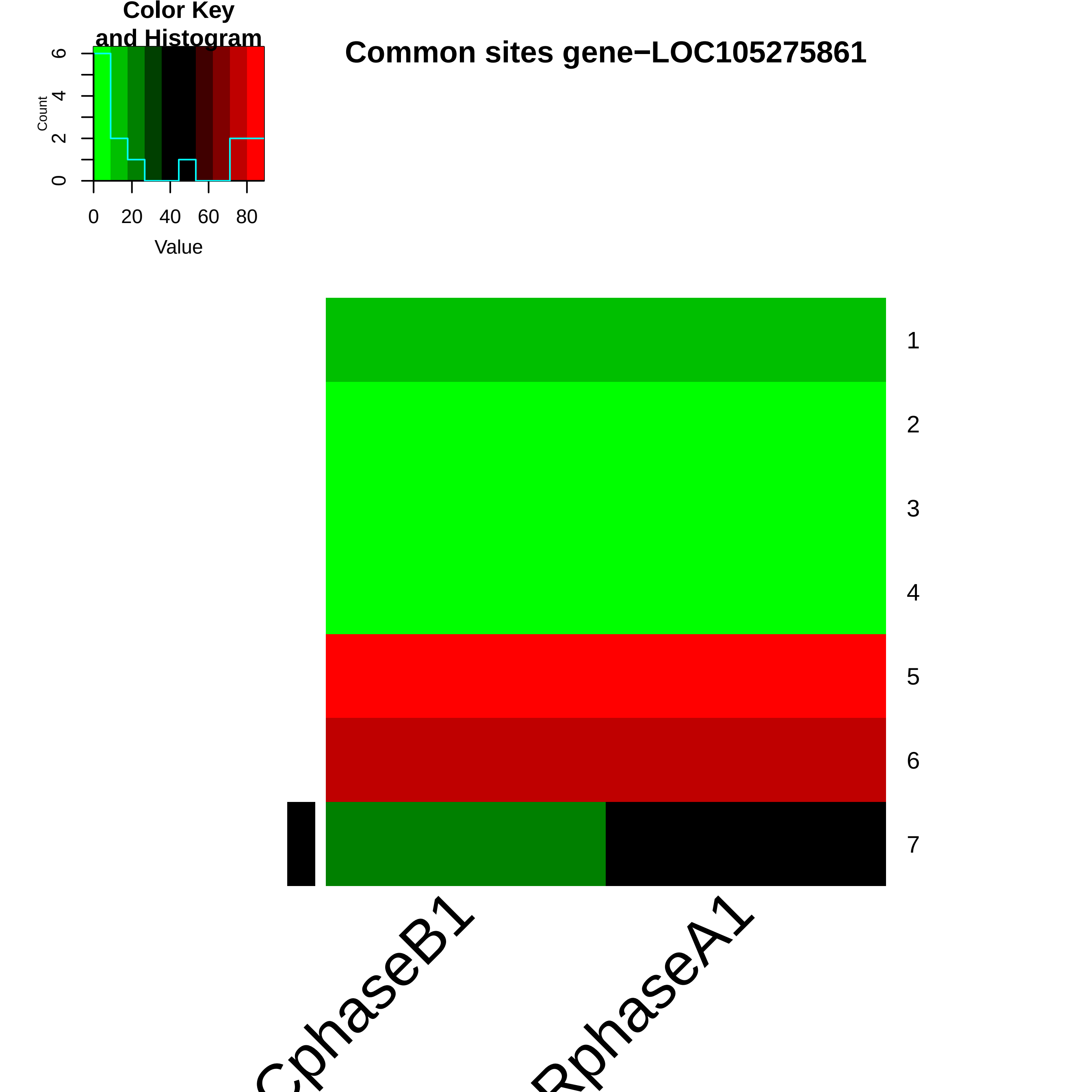
Color Key and Histogram
Count
0
2
4
6
0
20
40
60
80
Value
Common sites gene−LOC105275861
1
2
3
4
5
6
7

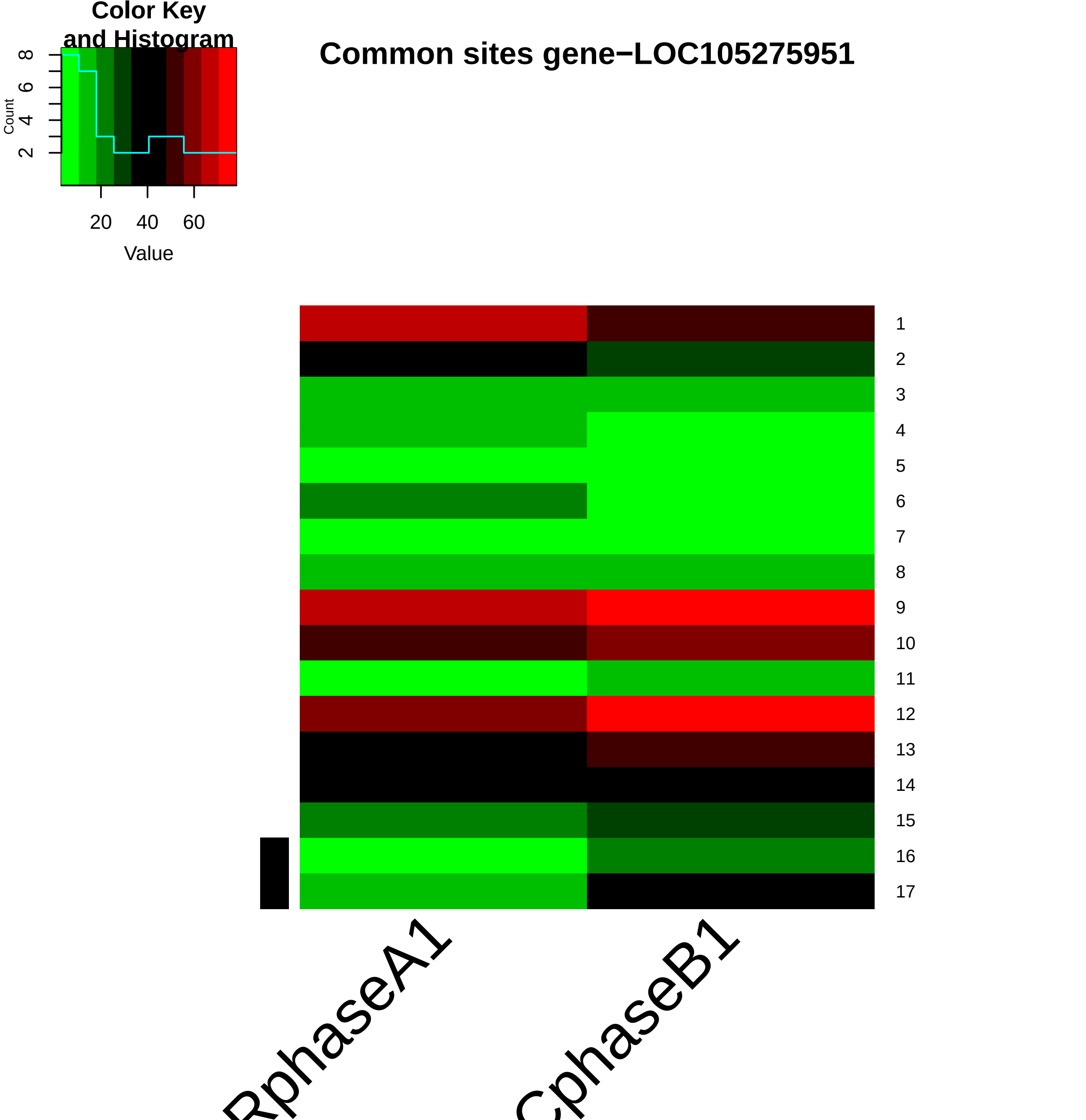

Color Key
and Histogram
Count
Value
Common sites gene−LOC105275951
RphaseA1
CphaseB1

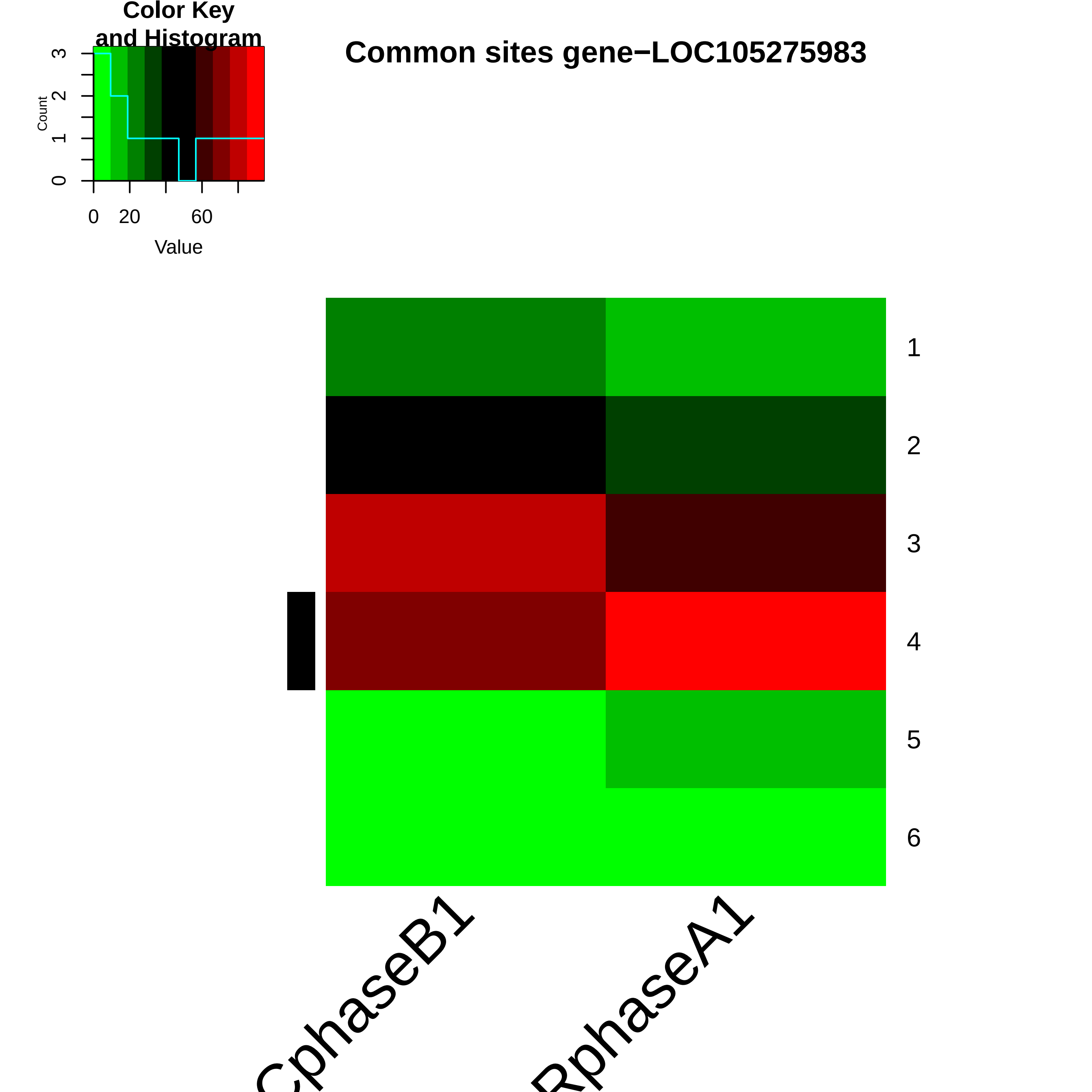

Color Key
and Histogram
Count
Value
0
20
60
0
1
2
3
Common sites gene−LOC105275983
1
2
3
4
5
6
CphaseB1
RphaseA1

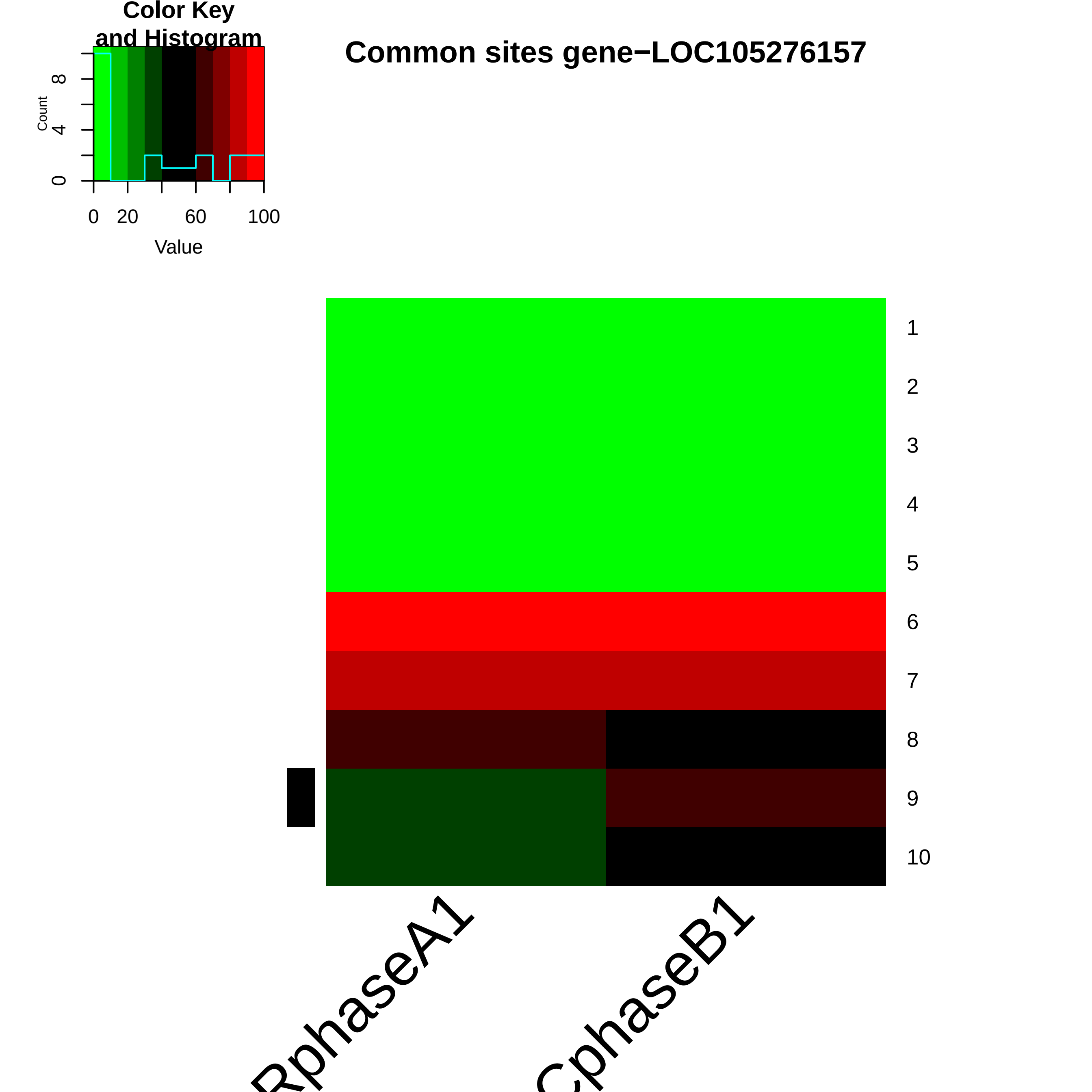

Color Key
and Histogram
Count
0
4
8
0
20
60
100
Value
Common sites gene−LOC105276157
1
2
3
4
5
6
7
8
9
10
RphaseA1
CphaseB1

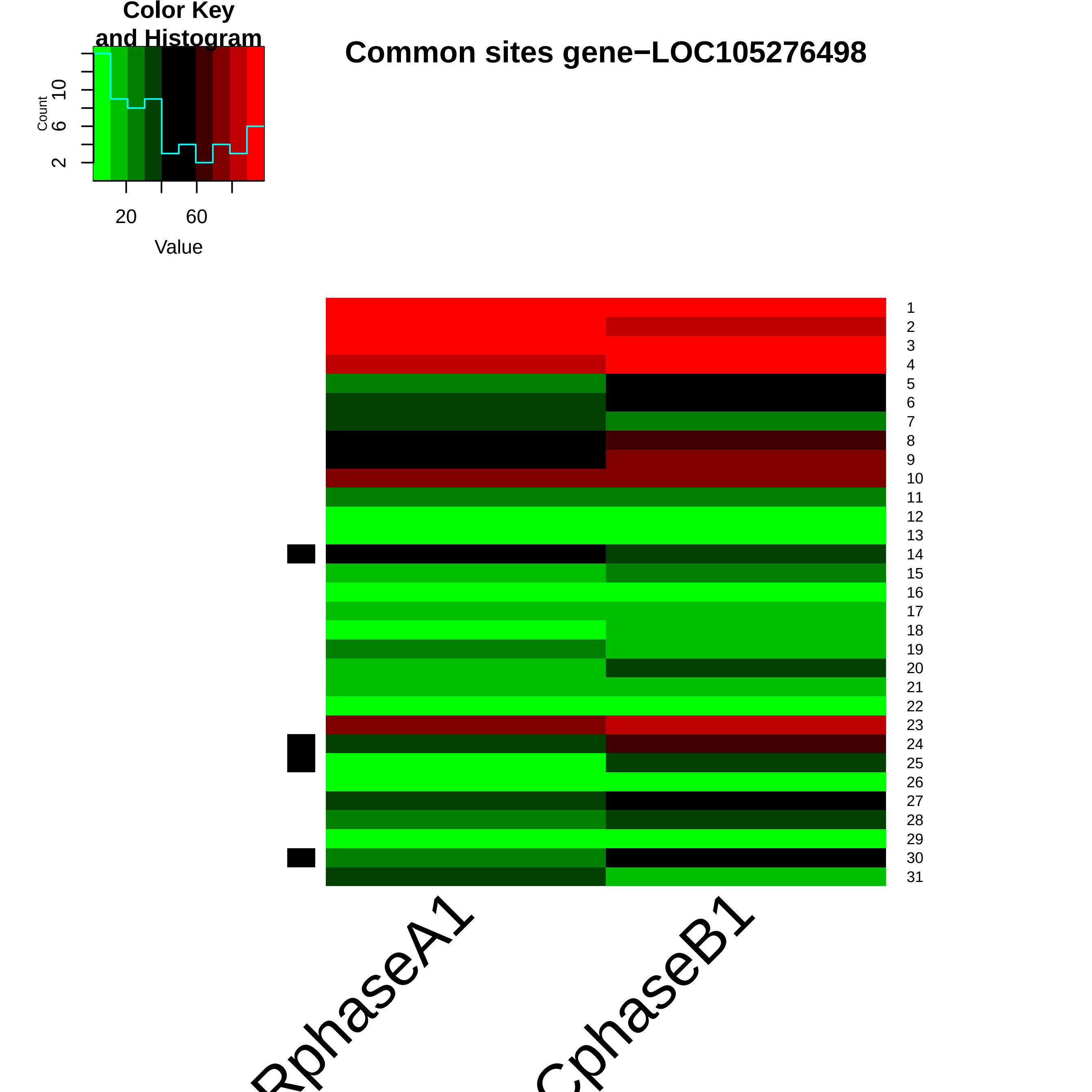

Color Key
and Histogram
Count
2
6
10
20
60
Value
Common sites gene−LOC105276498
RphaseA1
CphaseB1

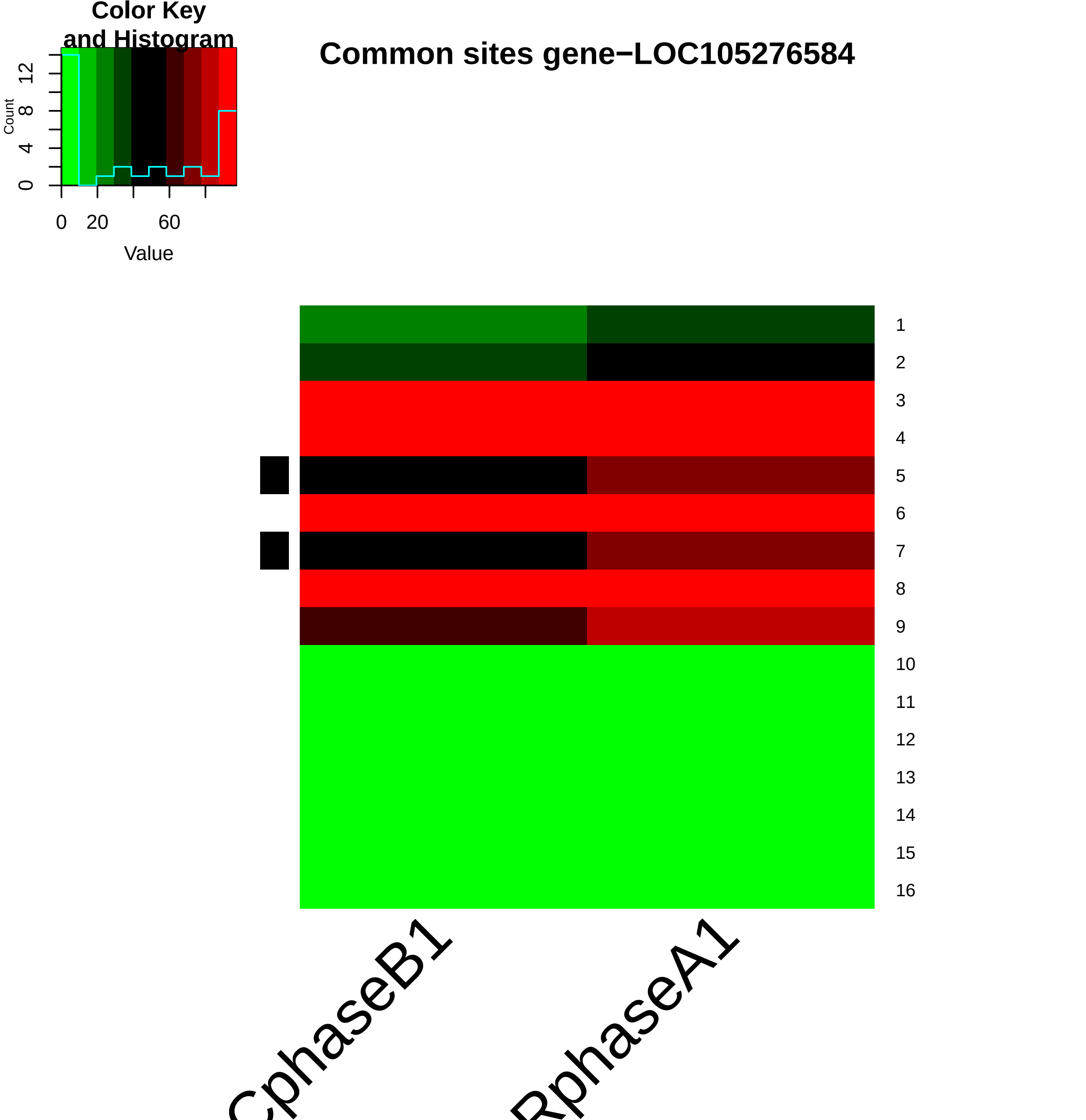
Color Key
and Histogram
Count
0
4
8
12
0
20
60
Value
Common sites gene−LOC105276584
CphaseB1
RphaseA1

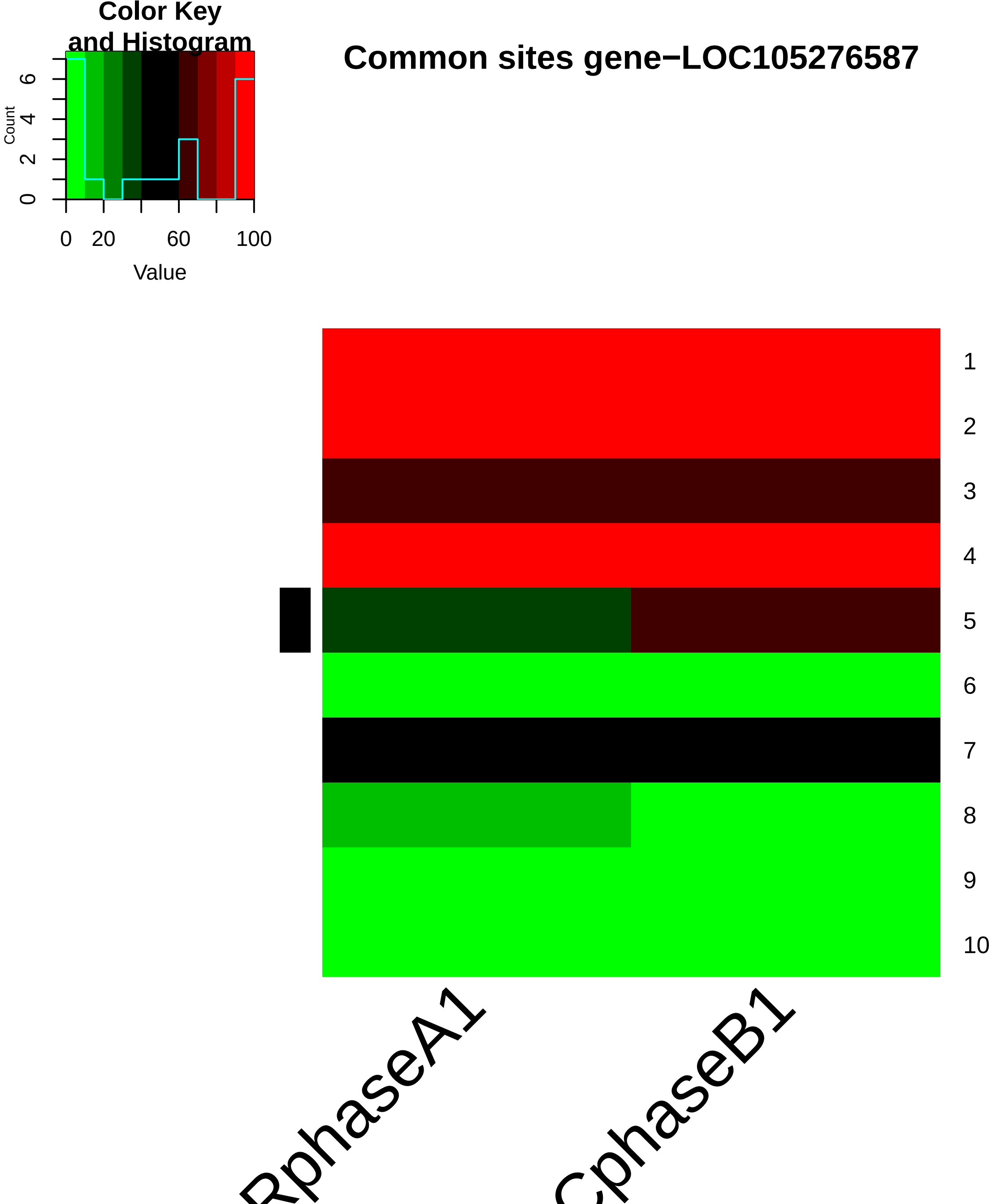

Color Key
and Histogram
Count
0
2
4
6
0
20
60
100
Value
Common sites gene−LOC105276587
1
2
3
4
5
6
7
8
9
10
RphaseA1
CphaseB1

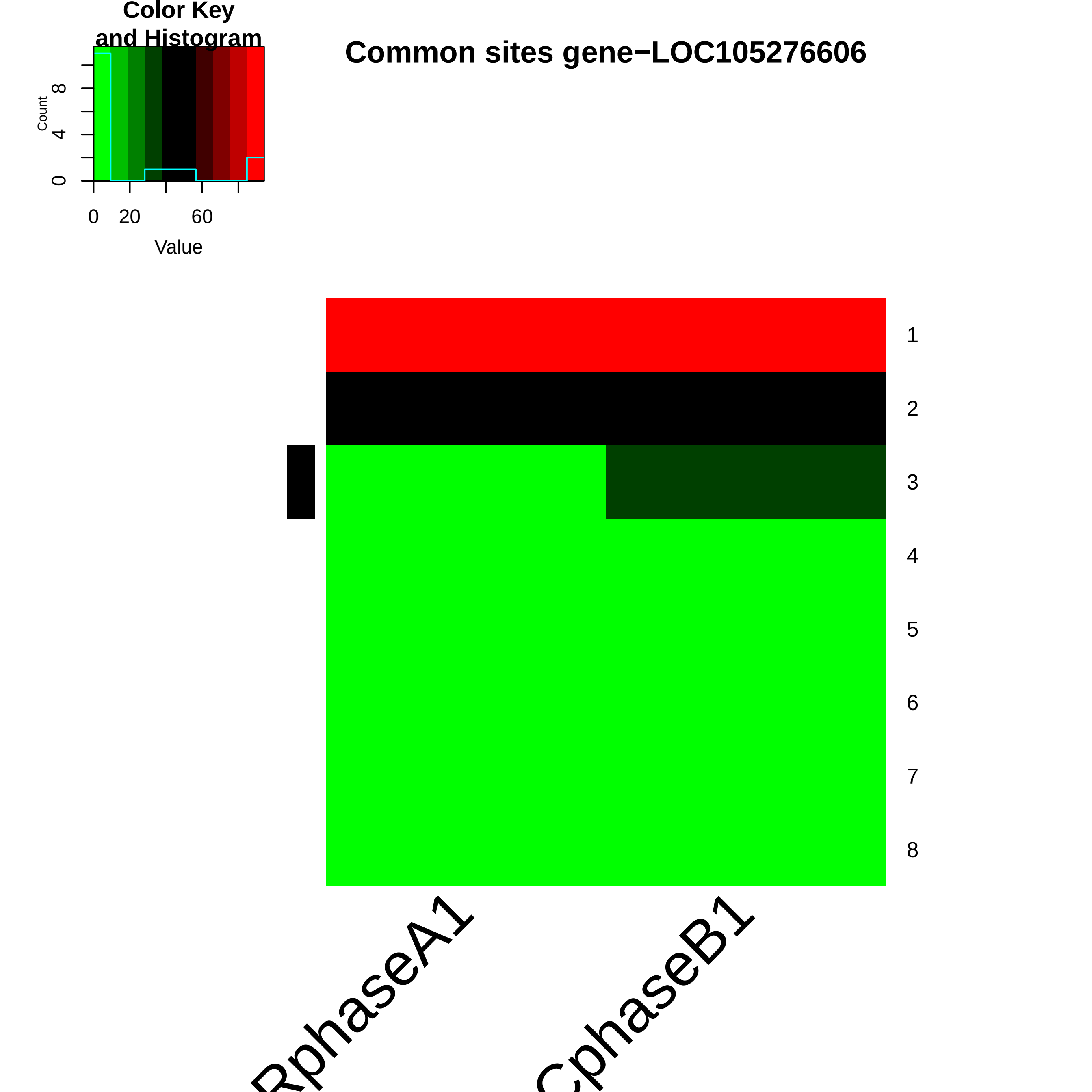

Color Key
and Histogram
Count
0
4
8
0
20
60
Value
Common sites gene−LOC105276606
1
2
3
4
5
6
7
8
RphaseA1
CphaseB1

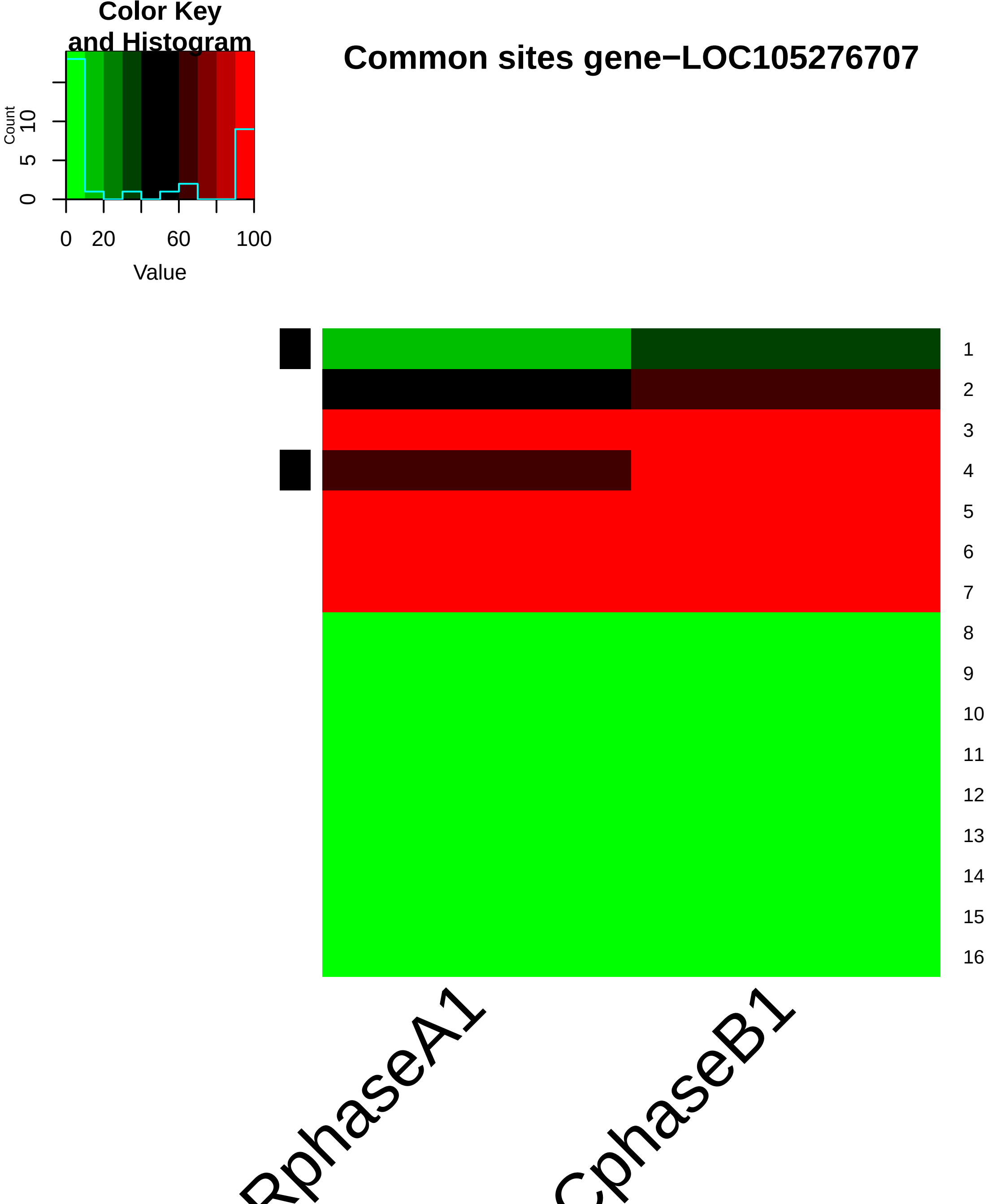
Color Key
and Histogram
Count
0
5
10
0
20
60
100
Value
Common sites gene−LOC105276707
RphaseA1
CphaseB1

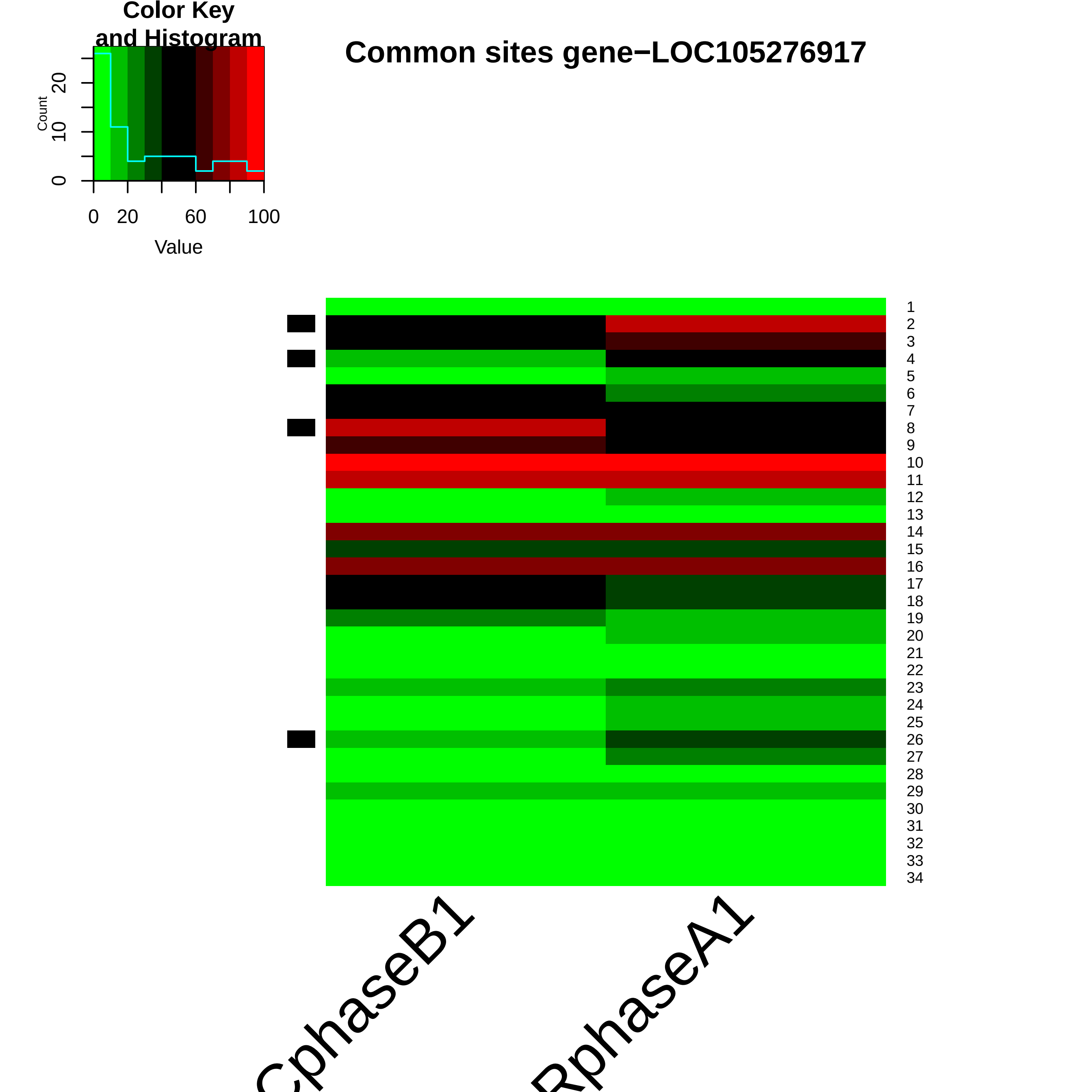

Color Key
and Histogram
Count
0
10
20
0
20
60
100
Value
Common sites gene−LOC105276917
CphaseB1
RphaseA1

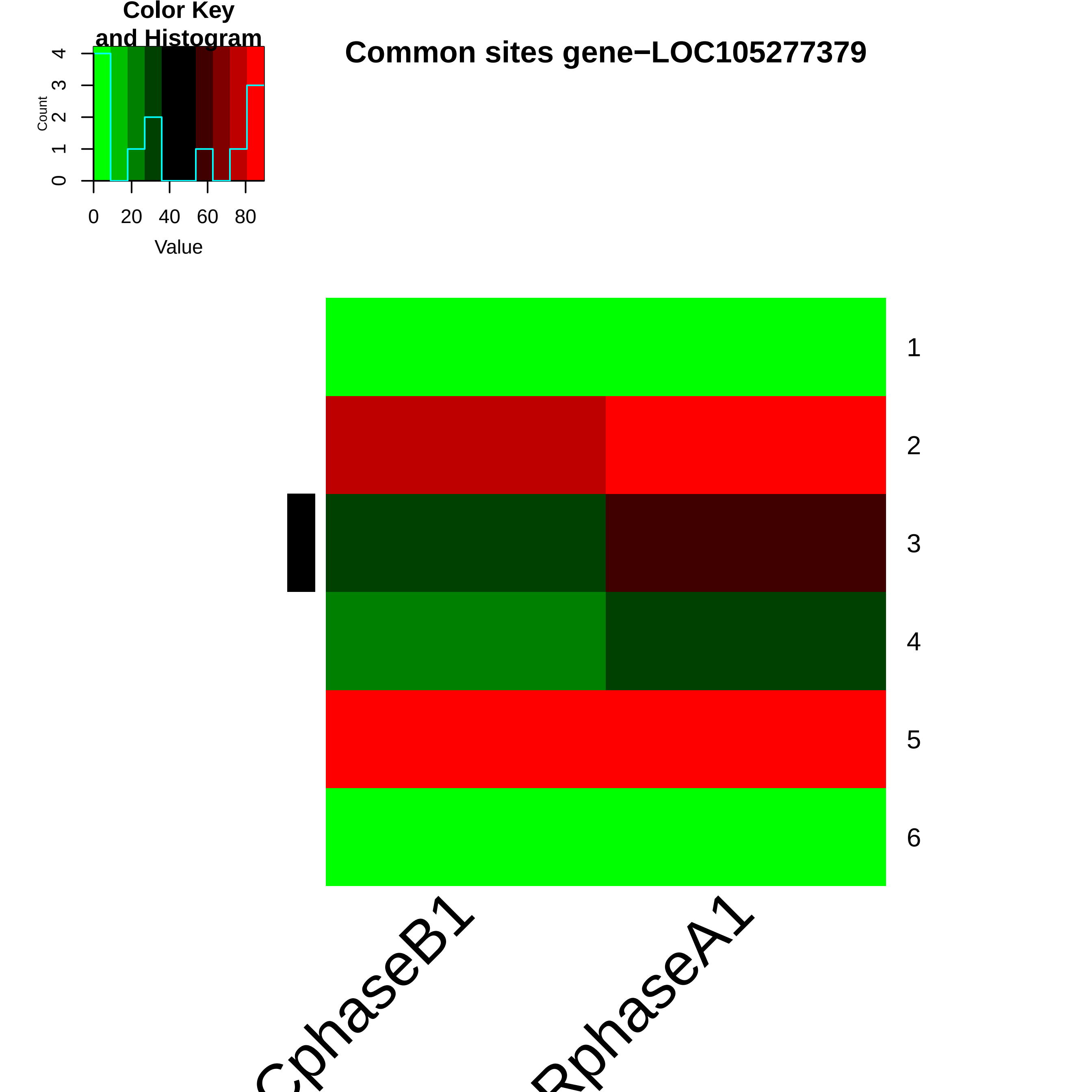

Color Key
and Histogram
Count
0
1
2
3
4
0
20
40
60
80
Value
Common sites gene−LOC105277379
1
2
3
4
5
6
CphaseB1
RphaseA1

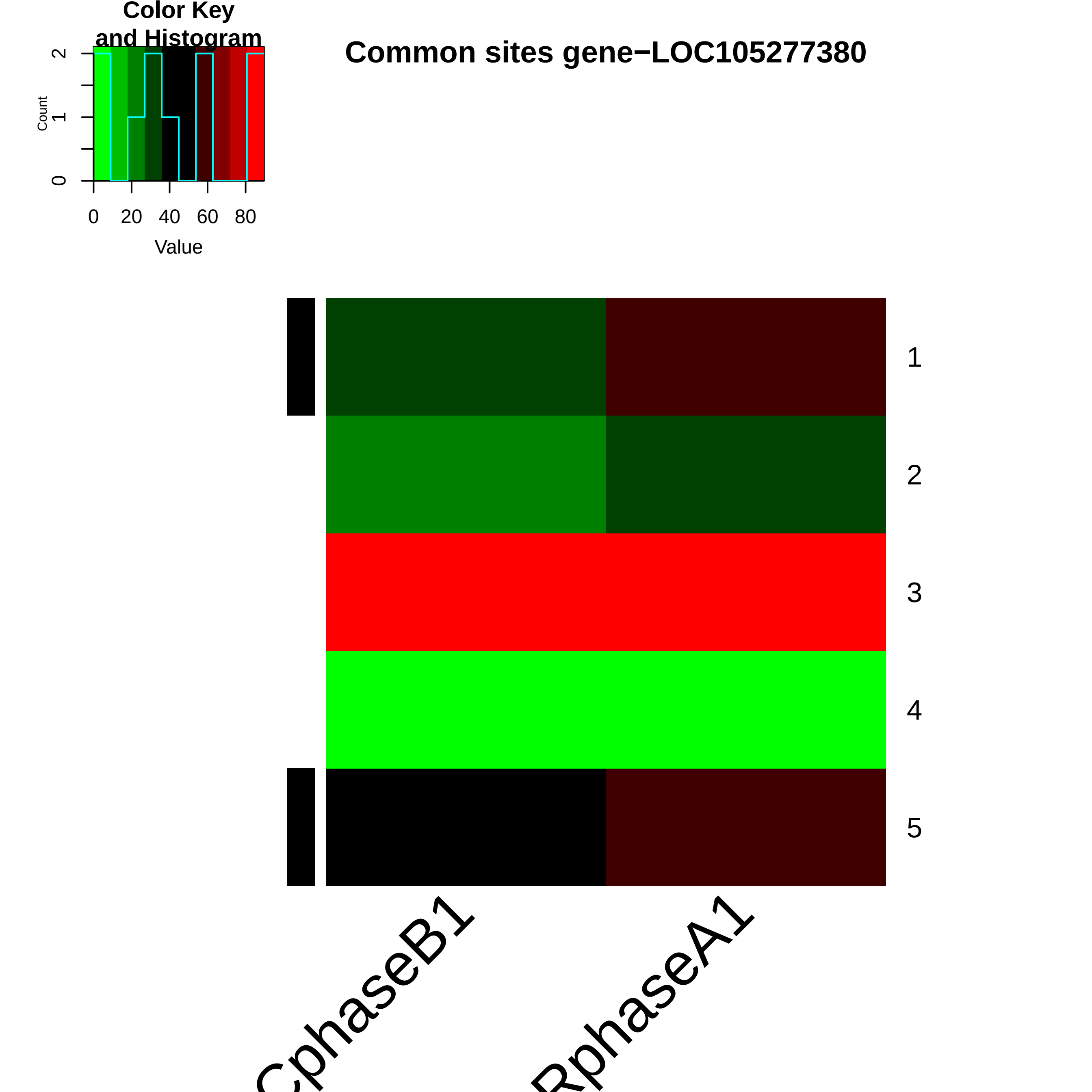

Color Key and Histogram
Count
0
1
2
0
20
40
60
80
Value
Common sites gene−LOC105277380
1
2
3
4
5
CphaseB1
RphaseA1

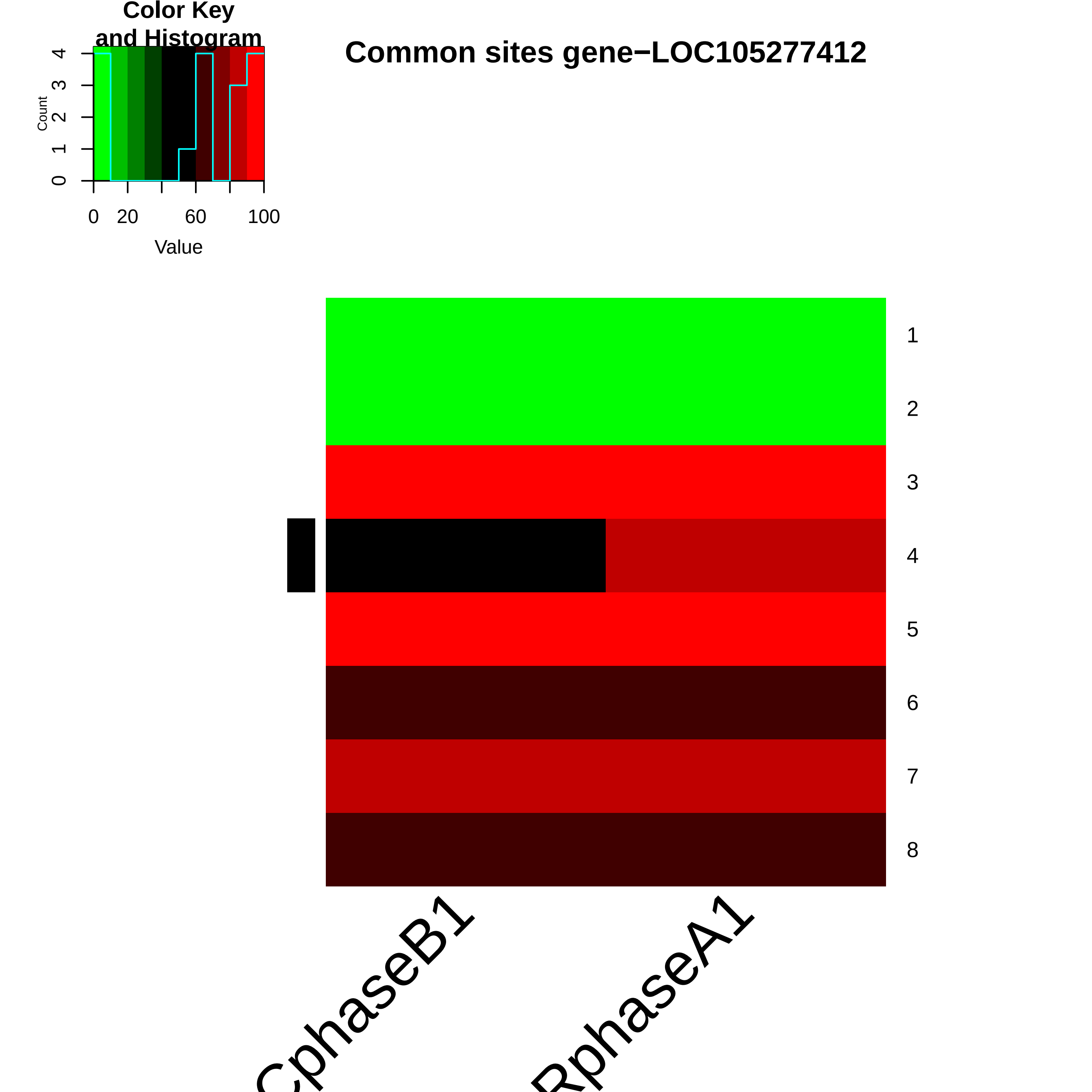
Color Key
and Histogram
Count
0
1
2
3
4
0
20
60
100
Value
Common sites gene−LOC105277412
1
2
3
4
5
6
7
8
CphaseB1
RphaseA1

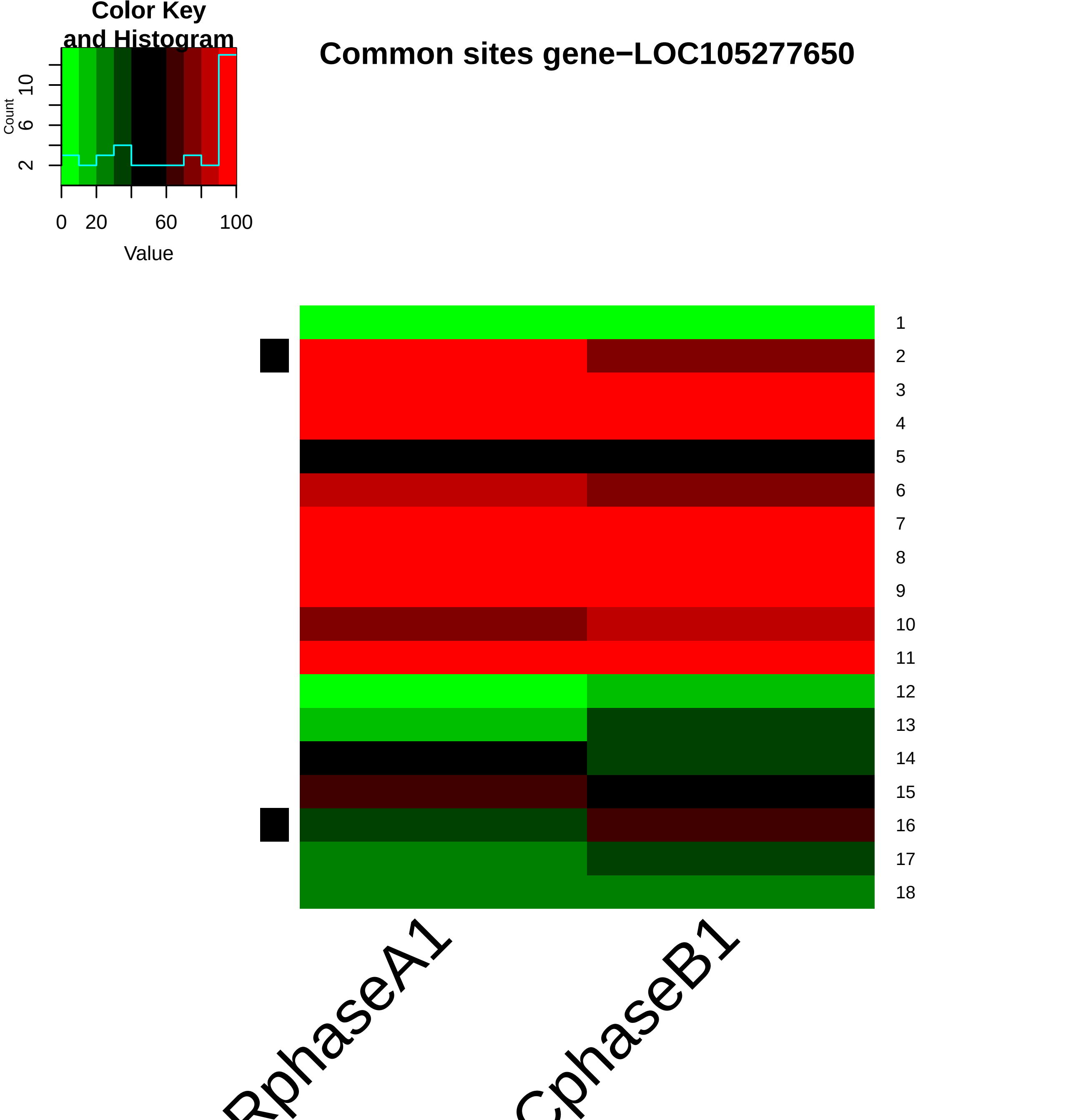
Color Key
and Histogram
Count
2
6
10
0
20
60
100
Value
Common sites gene−LOC105277650
RphaseA1
CphaseB1

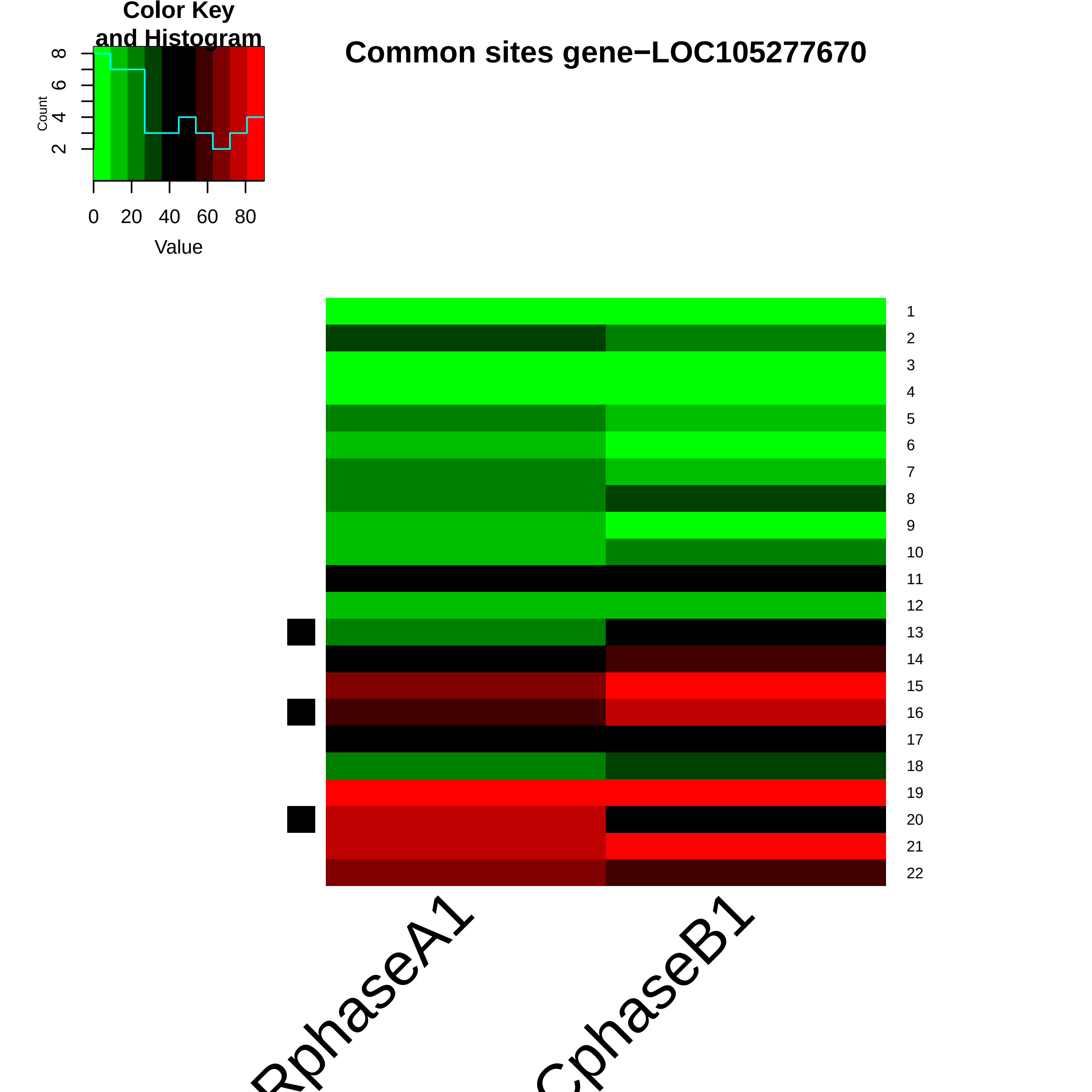
Color Key
and Histogram
Count
2
4
6
8
0
20
40
60
80
Value
Common sites gene−LOC105277670
RphaseA1
CphaseB1

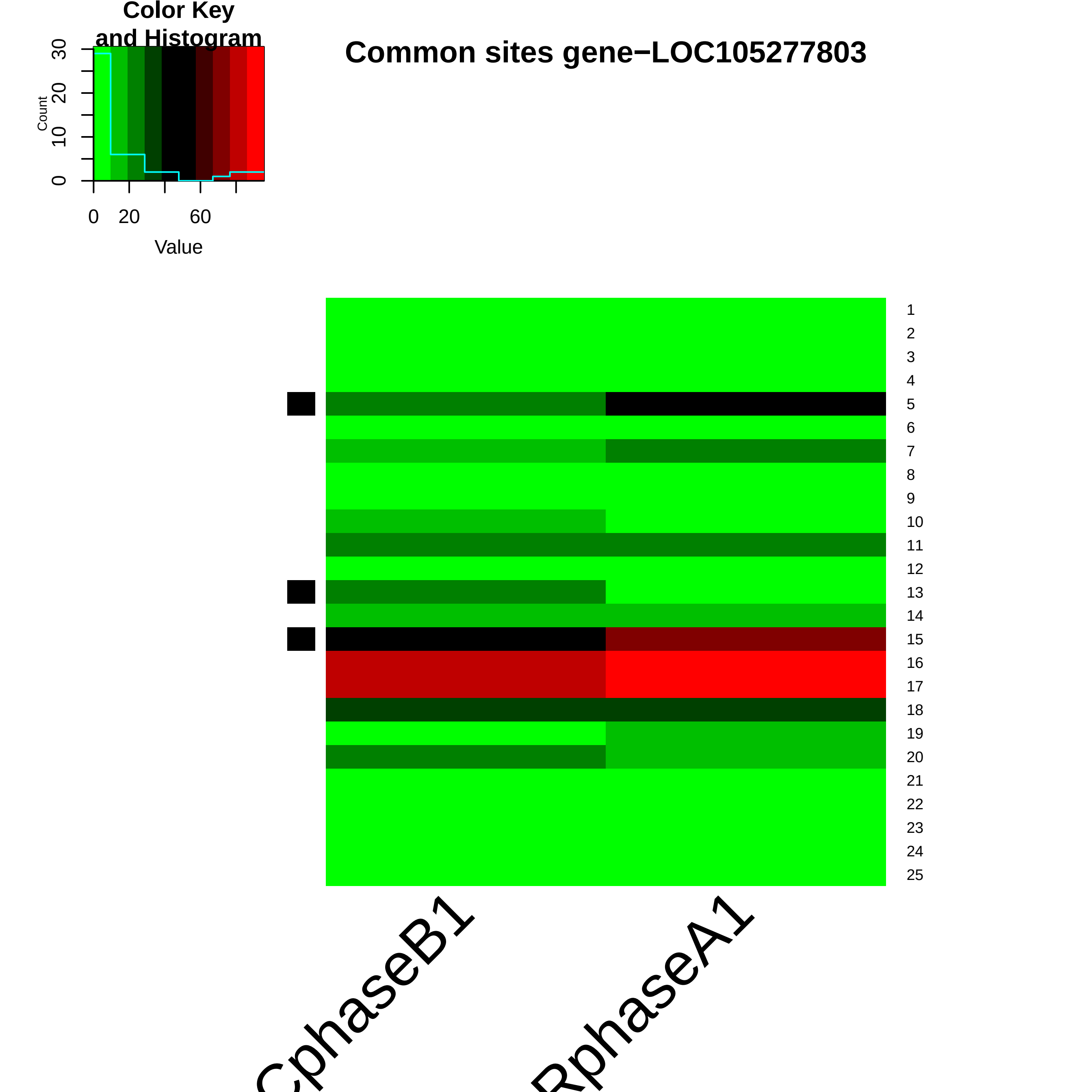
Color Key
and Histogram
Count
0
10
20
30
0
20
60
Value
Common sites gene−LOC105277803
CphaseB1
RphaseA1

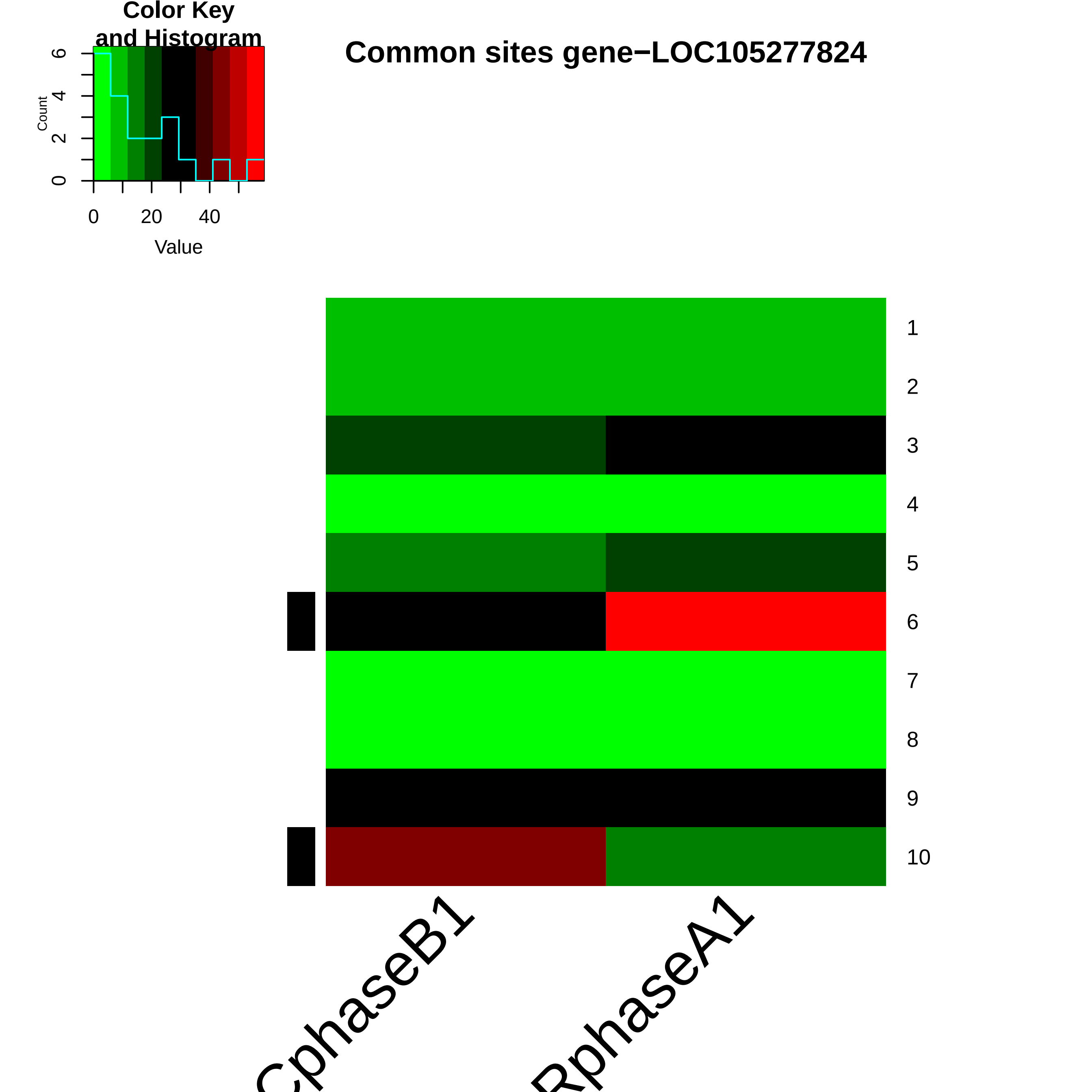
Color Key
and Histogram
Count
0
2
4
6
0
20
40
Value
Common sites gene−LOC105277824
1
2
3
4
5
6
7
8
9
10
CphaseB1
RphaseA1

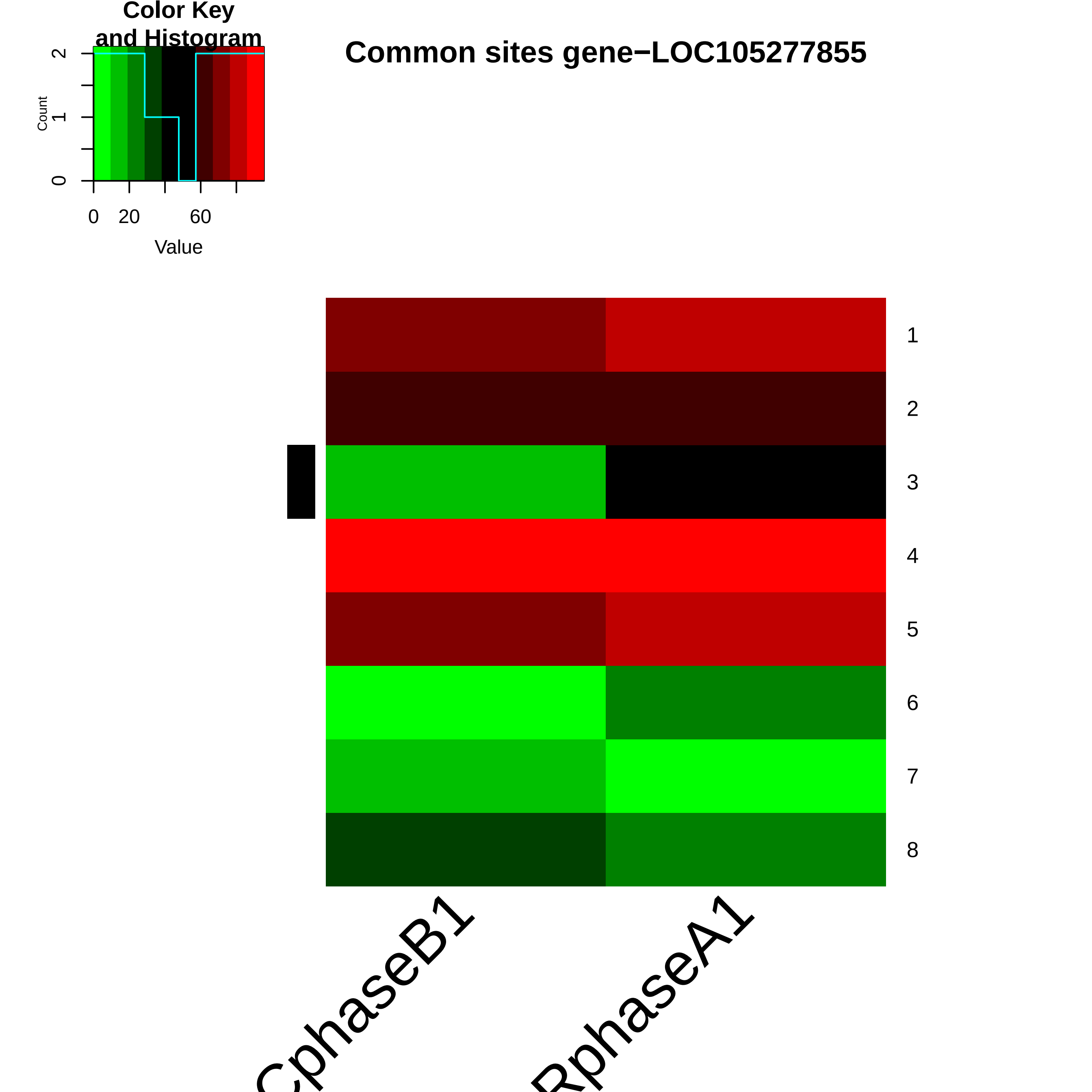
Color Key
and Histogram
Count
0
1
2
0
20
60
Value
Common sites gene−LOC105277855
1
2
3
4
5
6
7
8
CphaseB1
RphaseA1

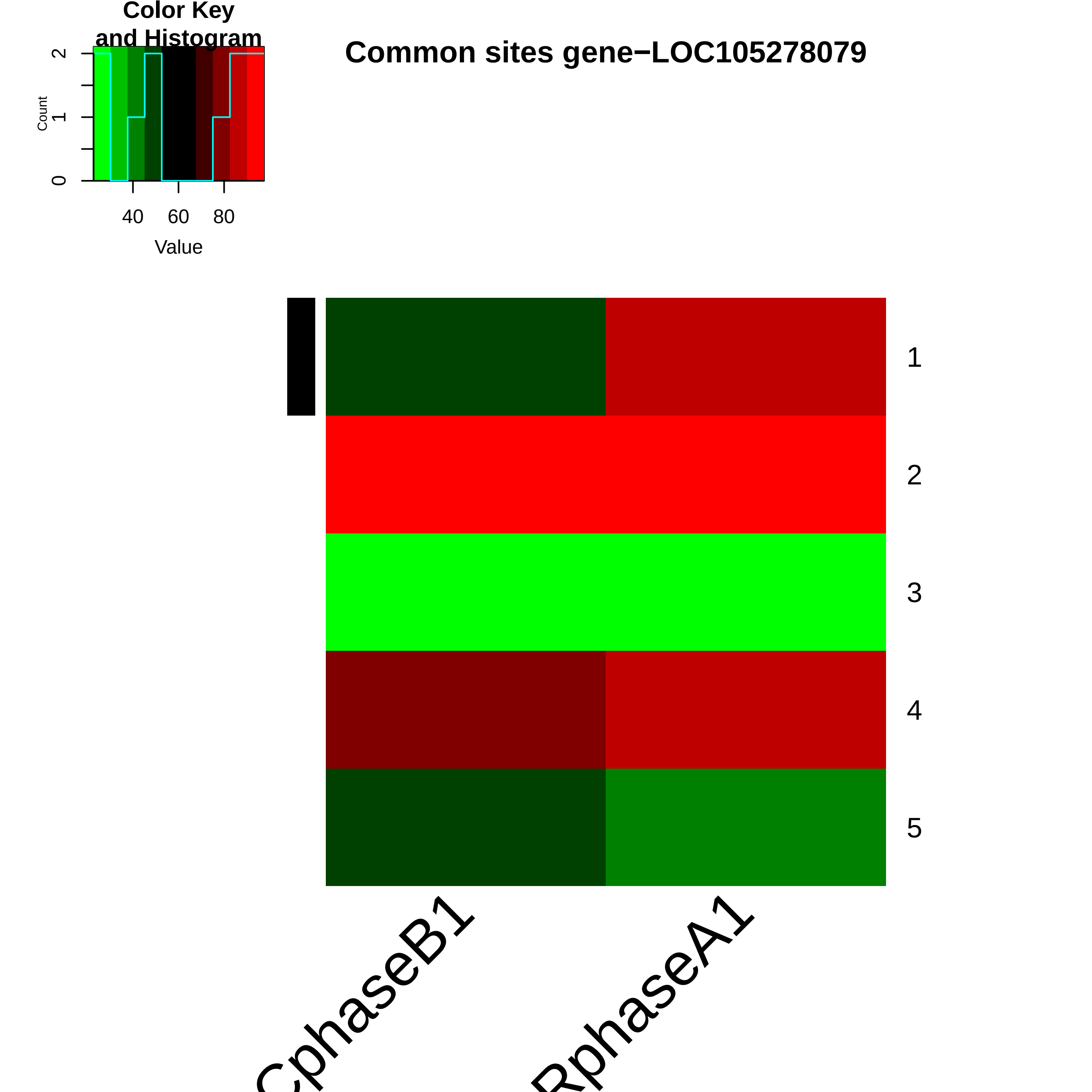
Color Key
and Histogram
Count
0
1
2
40
60
80
Value
Common sites gene−LOC105278079
1
2
3
4
5

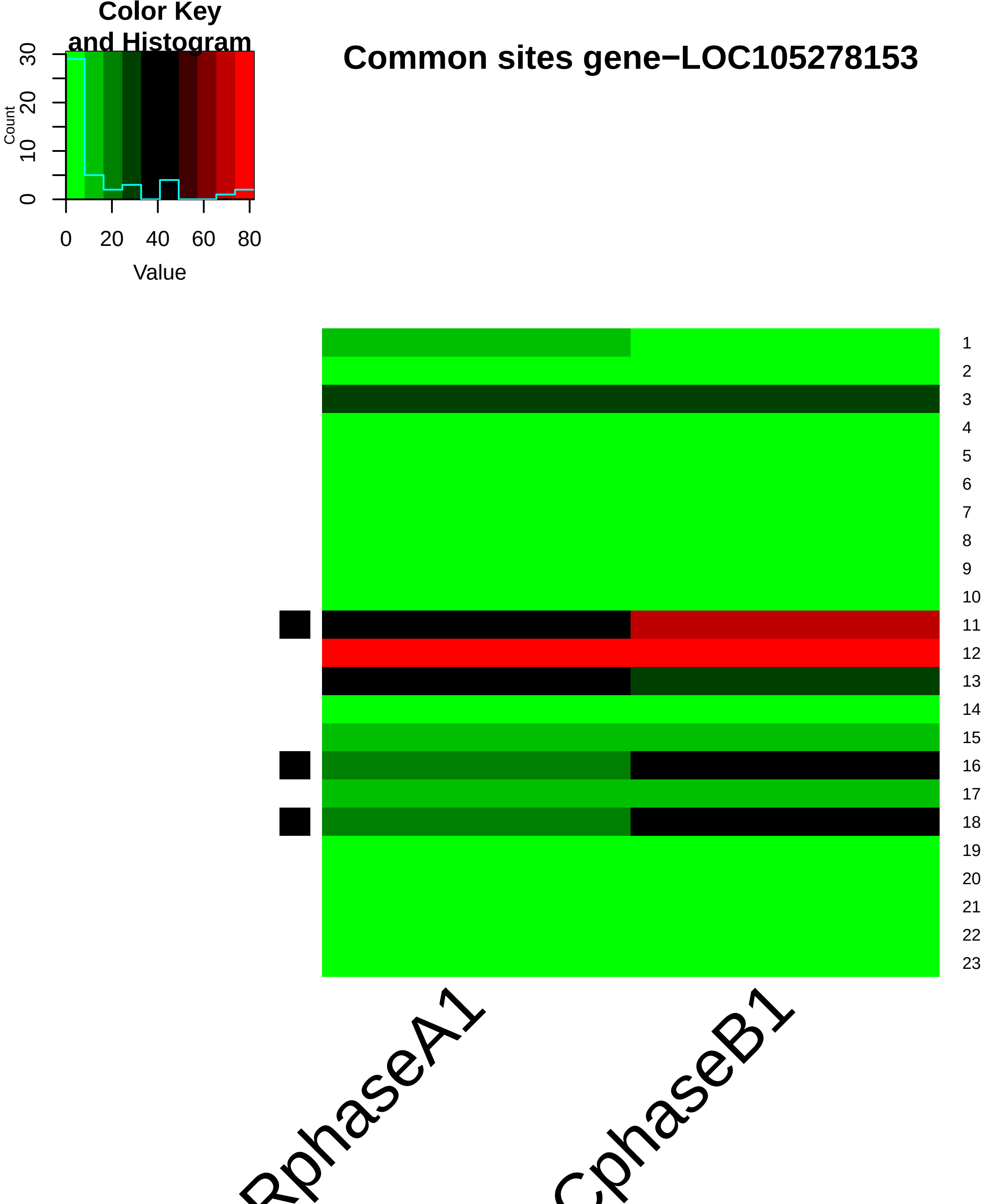
Color Key
and Histogram
Count
0
10
20
30
0
20
40
60
80
Value
Common sites gene−LOC105278153
RphaseA1
CphaseB1

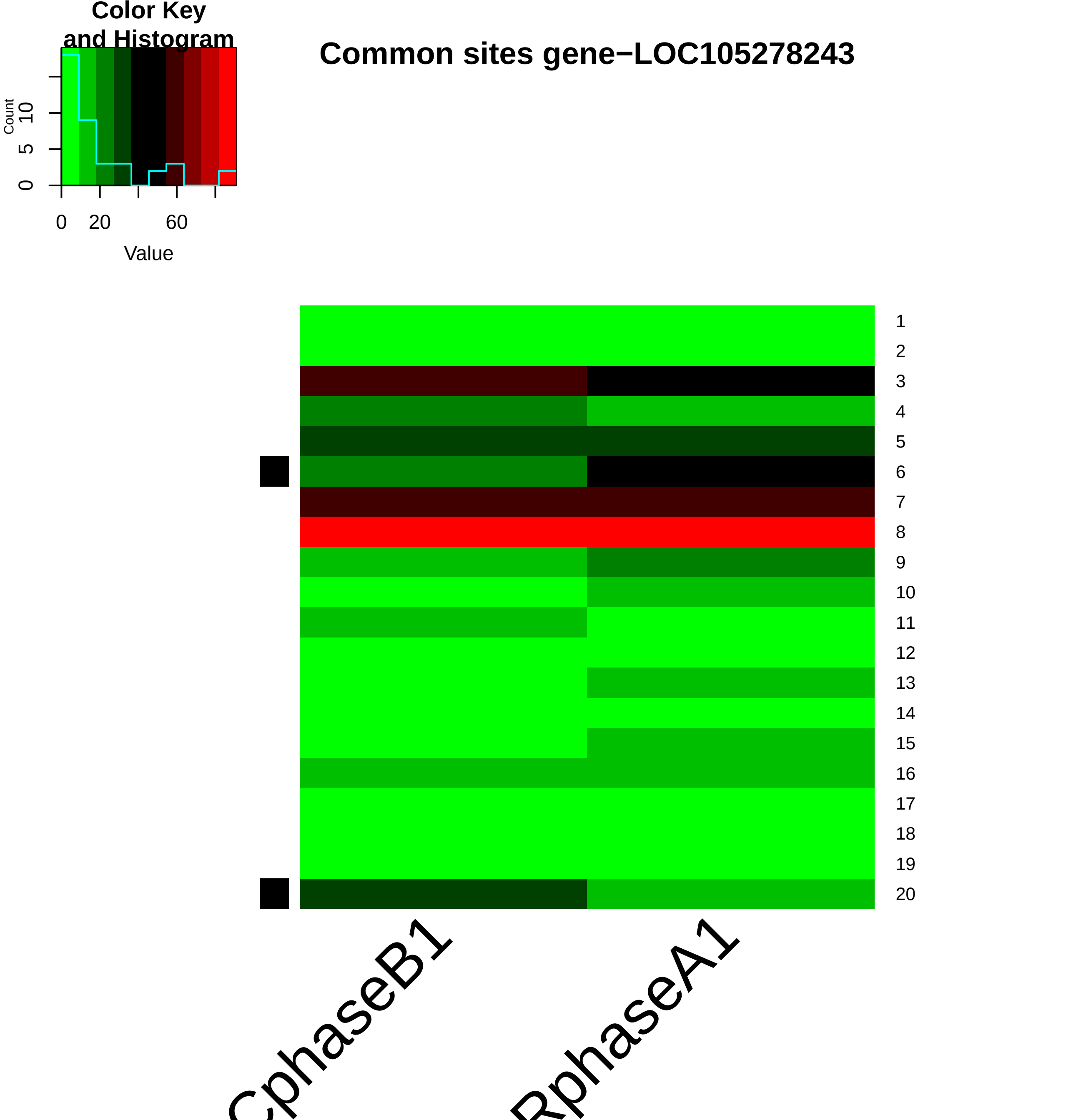
Color Key
and Histogram
Count
0
5
10
0
20
60
Value
Common sites gene−LOC105278243
CphaseB1
RphaseA1

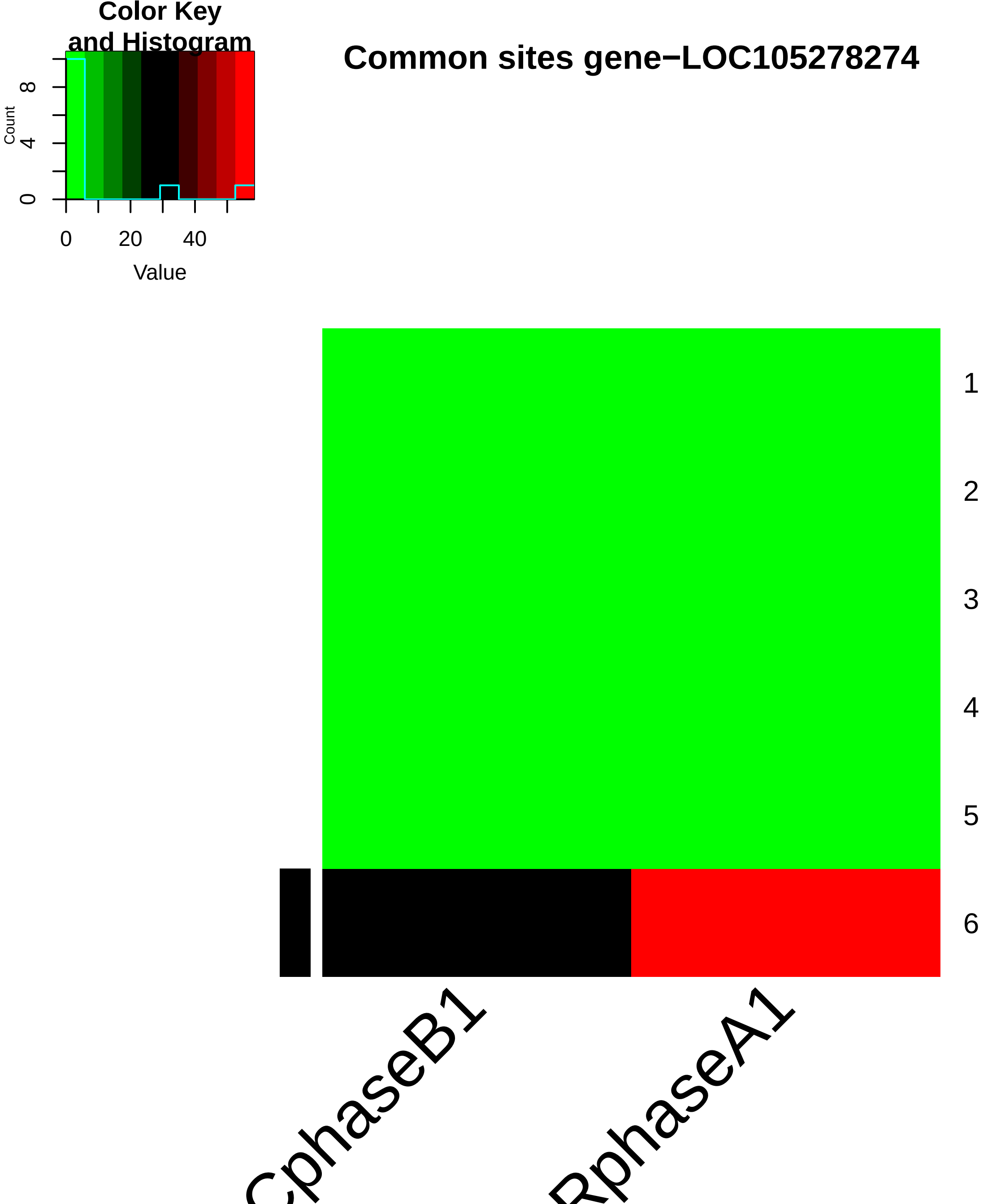

Color Key
and Histogram
Count
0
4
8
0
20
40
Value
Common sites gene−LOC105278274
1
2
3
4
5
6

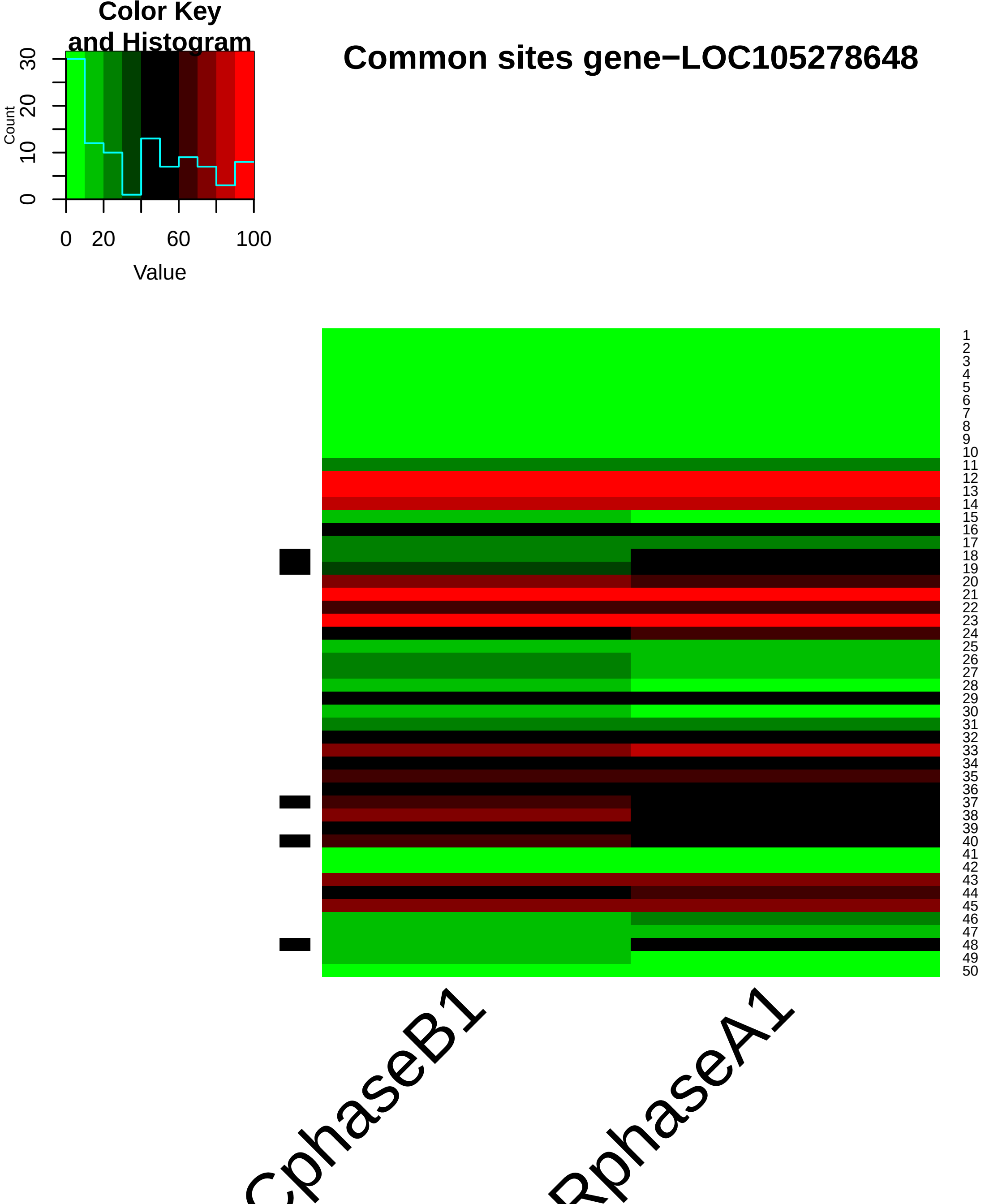

Color Key
and Histogram
Count
0
10
20
30
0
20
60
100
Value
Common sites gene−LOC105278648
CphaseB1
RphaseA1

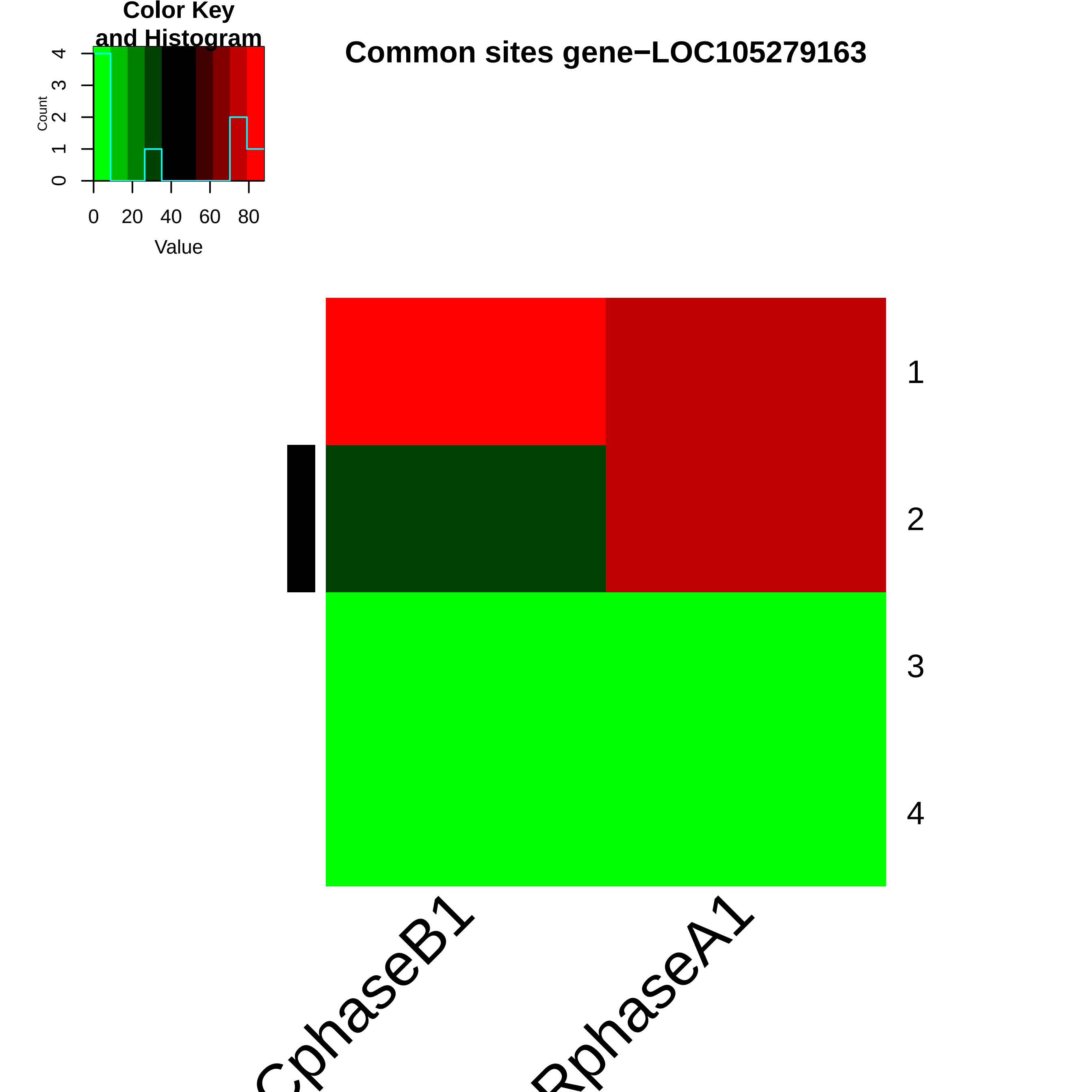

Color Key
and Histogram
Count
0
1
2
3
4
0
20
40
60
80
Value
Common sites gene−LOC105279163
1
2
3
4
CphaseB1
RphaseA1

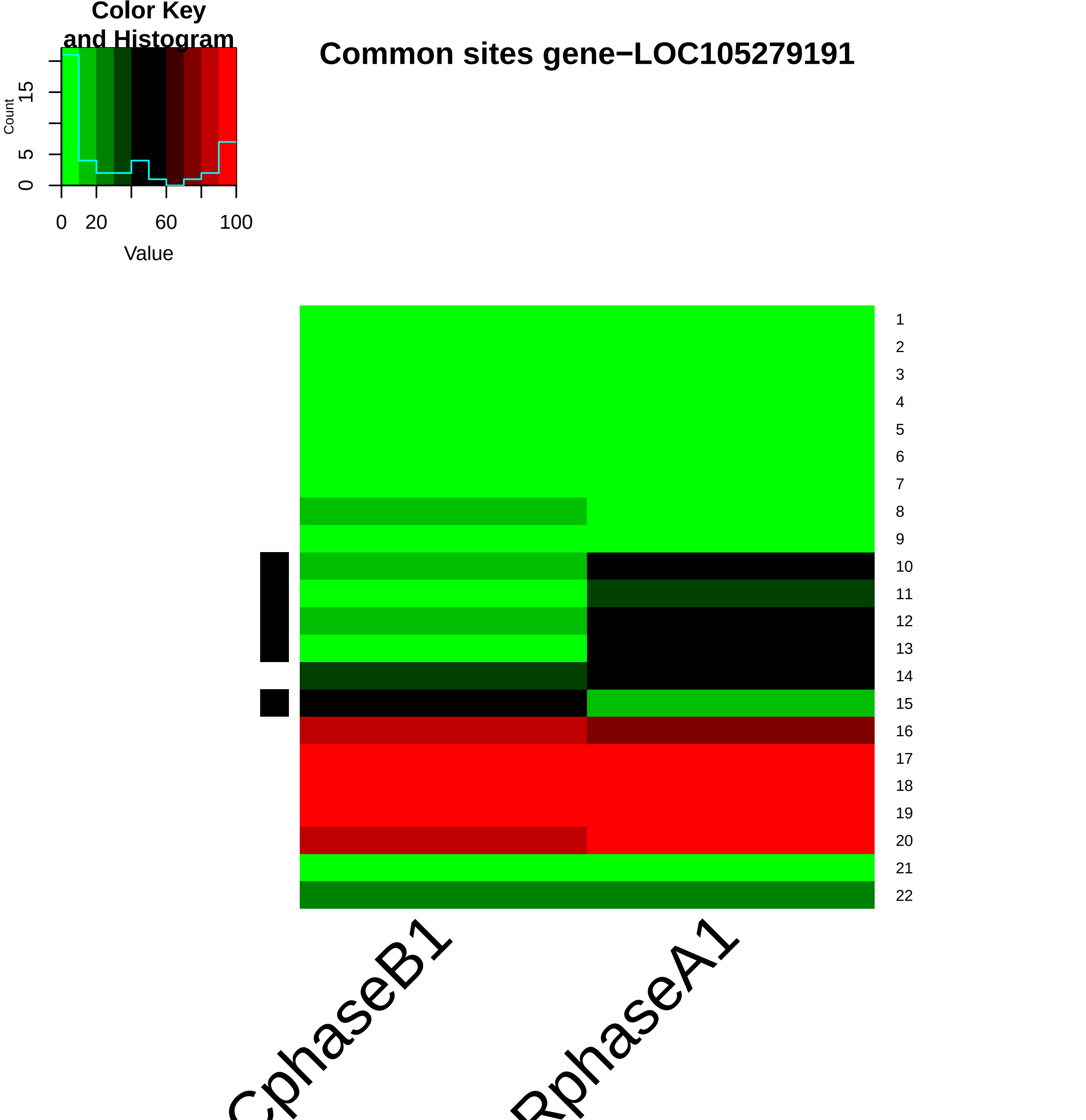
Color Key
and Histogram
Count
0
5
15
0
20
60
100
Value
Common sites gene−LOC105279191
CphaseB1
RphaseA1

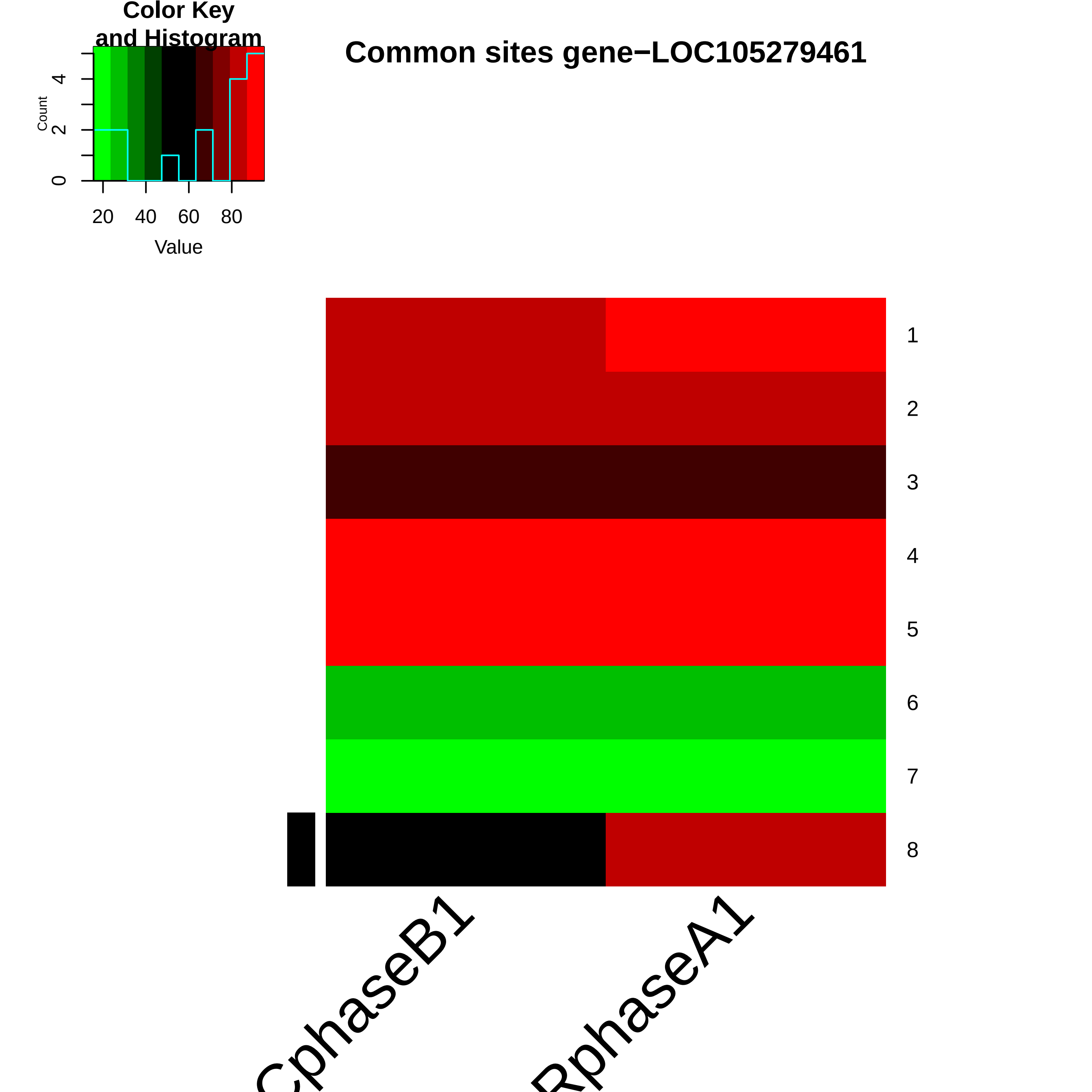

Color Key
and Histogram
Count
0
2
4
20
40
60
80
Value
Common sites gene−LOC105279461
1
2
3
4
5
6
7
8
CphaseB1
RphaseA1

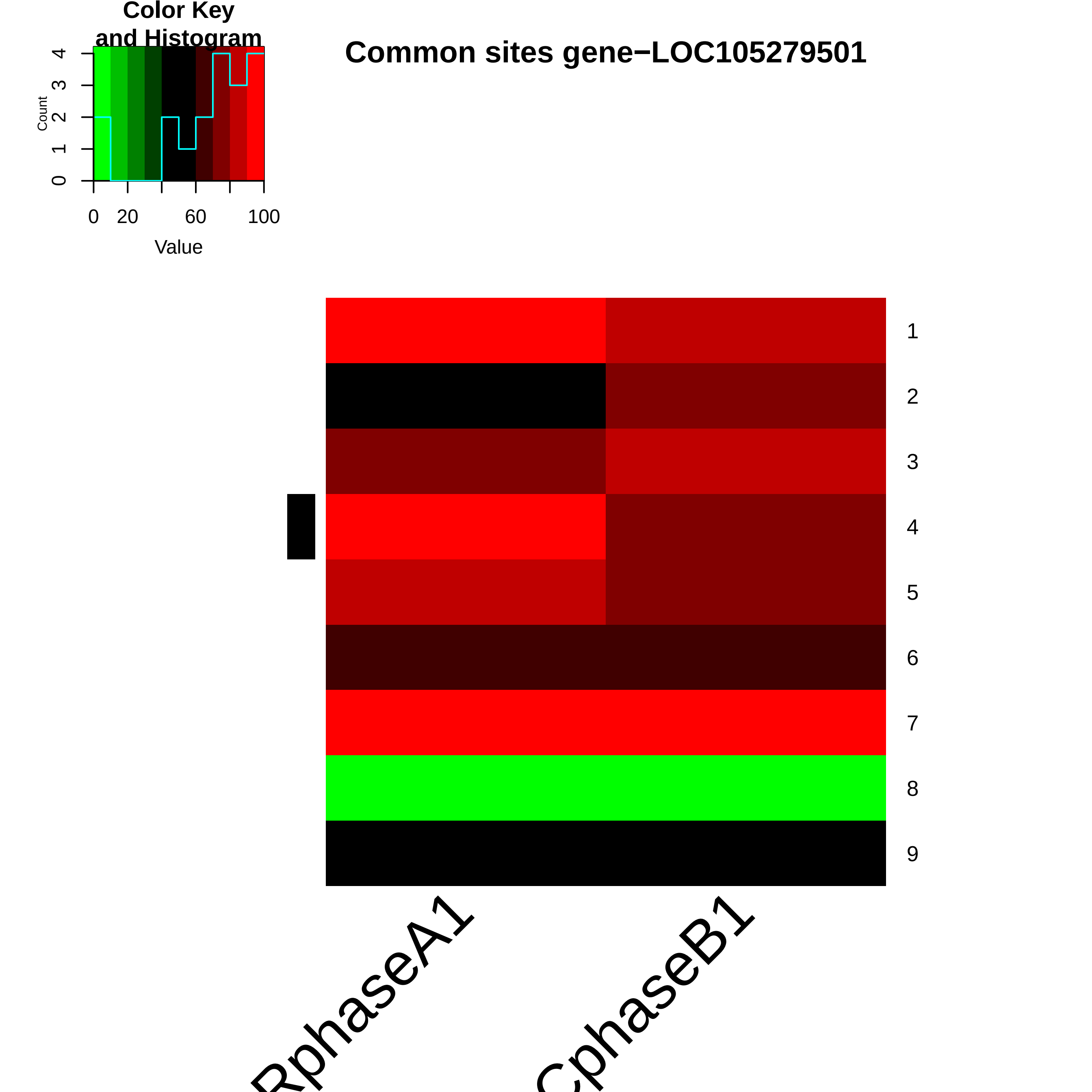
Color Key
and Histogram
Count
0
1
2
3
4
0
20
60
100
Value
Common sites gene−LOC105279501
1
2
3
4
5
6
7
8
9
RphaseA1
CphaseB1

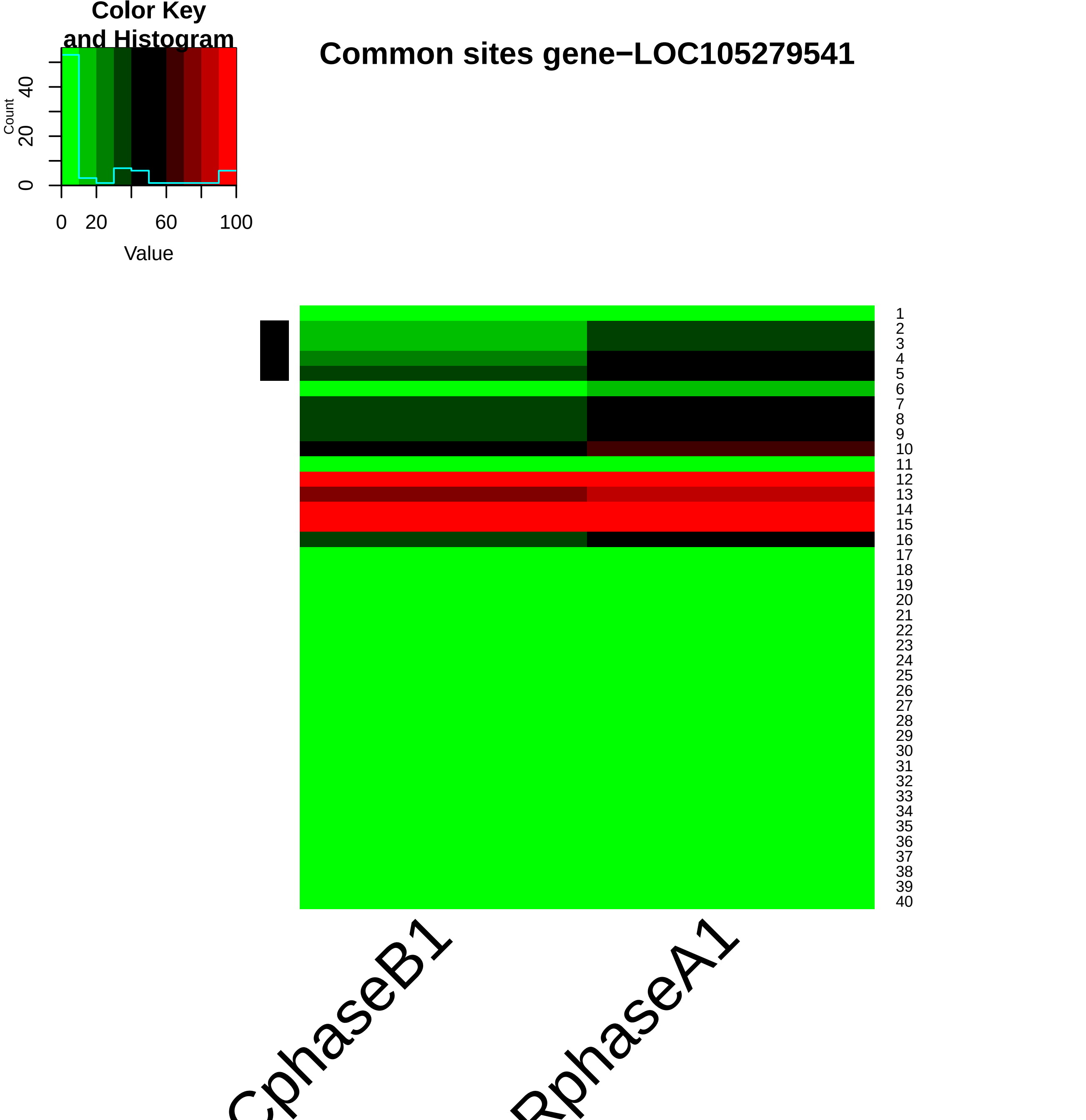

Color Key
and Histogram
Count
0
20
40
0
20
60
100
Value
Common sites gene−LOC105279541
CphaseB1
RphaseA1

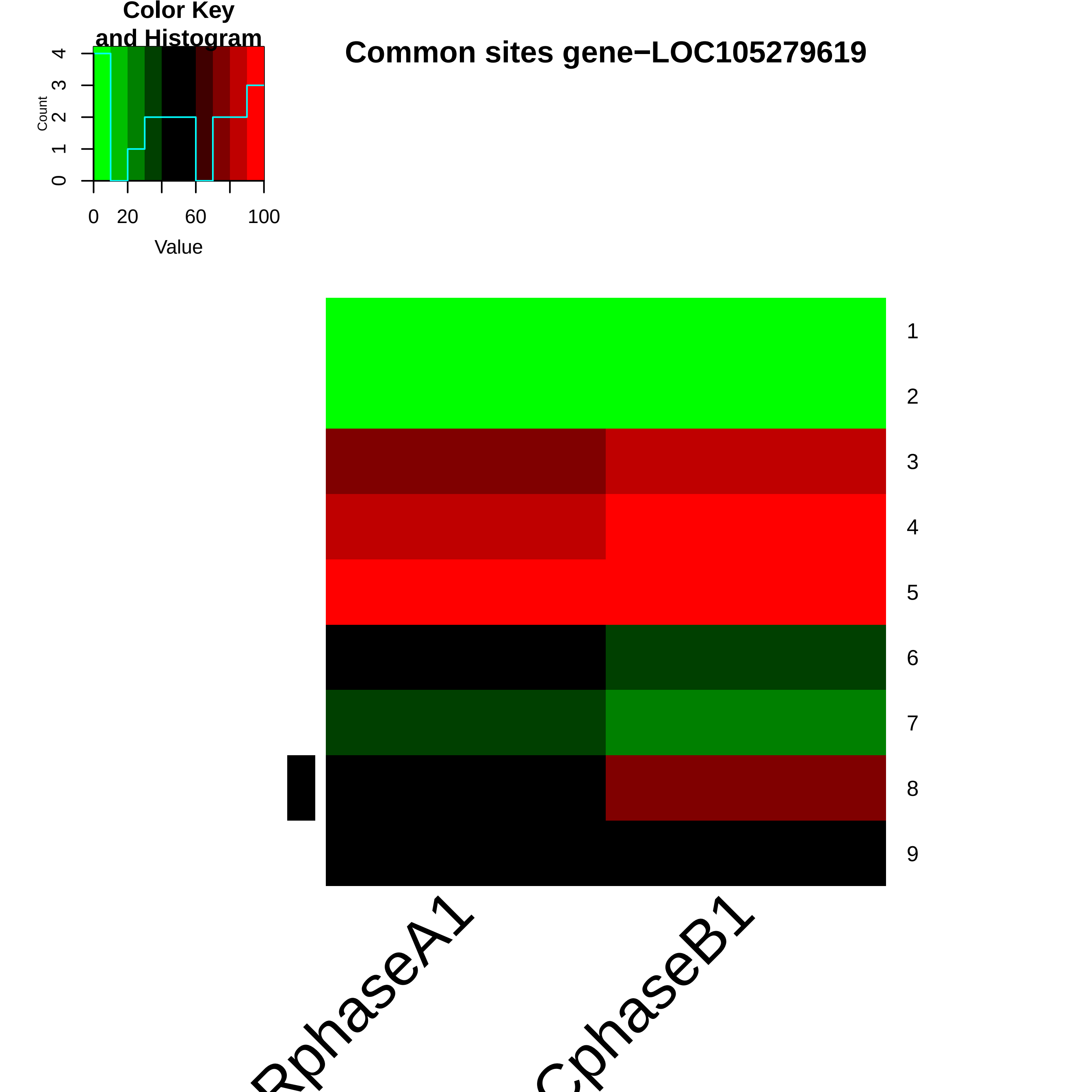

Color Key
and Histogram
Count
0
1
2
3
4
0
20
60
100
Value
Common sites gene−LOC105279619
1
2
3
4
5
6
7
8
9
RphaseA1
CphaseB1

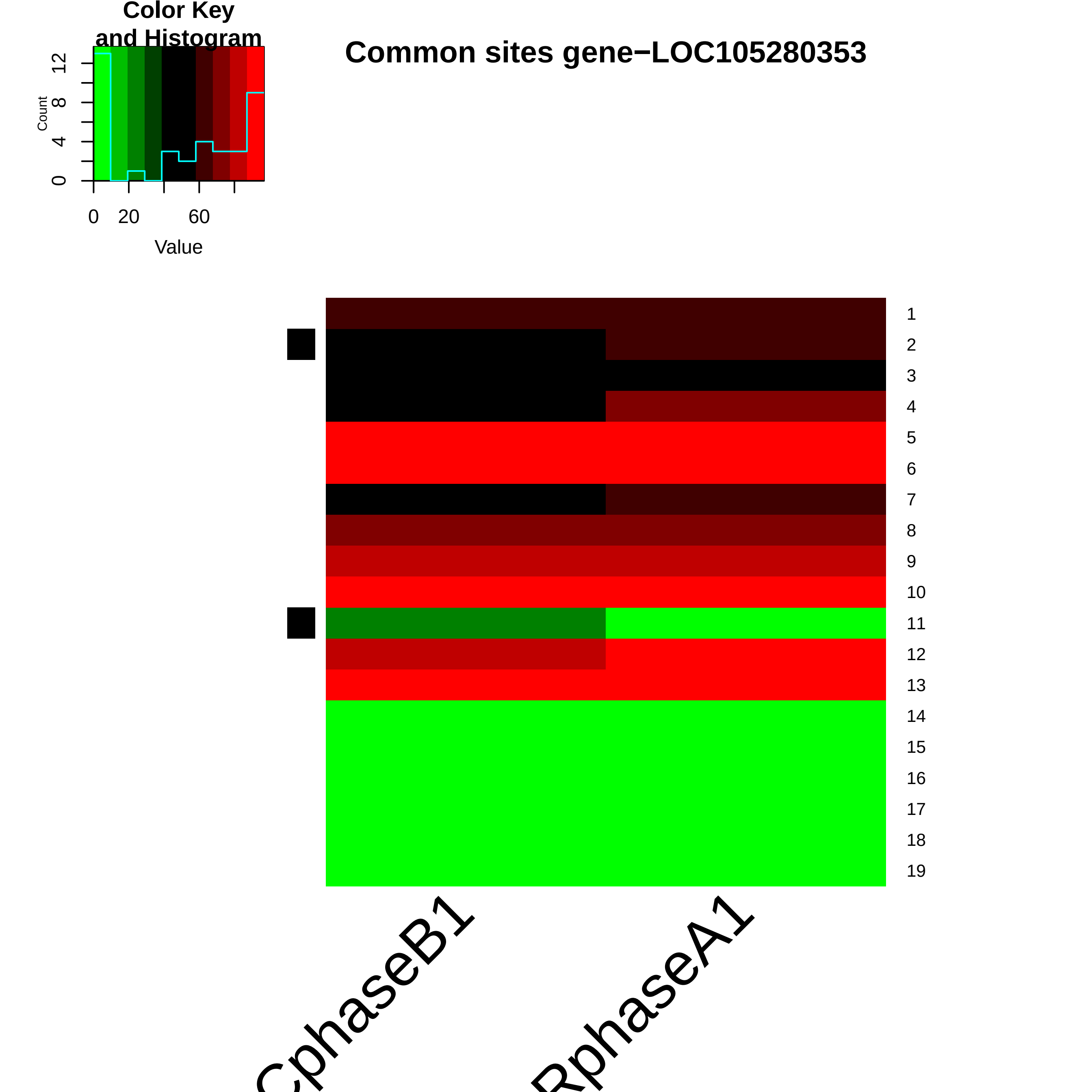
Color Key
and Histogram
Count
0
4
8
12
0
20
60
Value
Common sites gene−LOC105280353
CphaseB1
RphaseA1

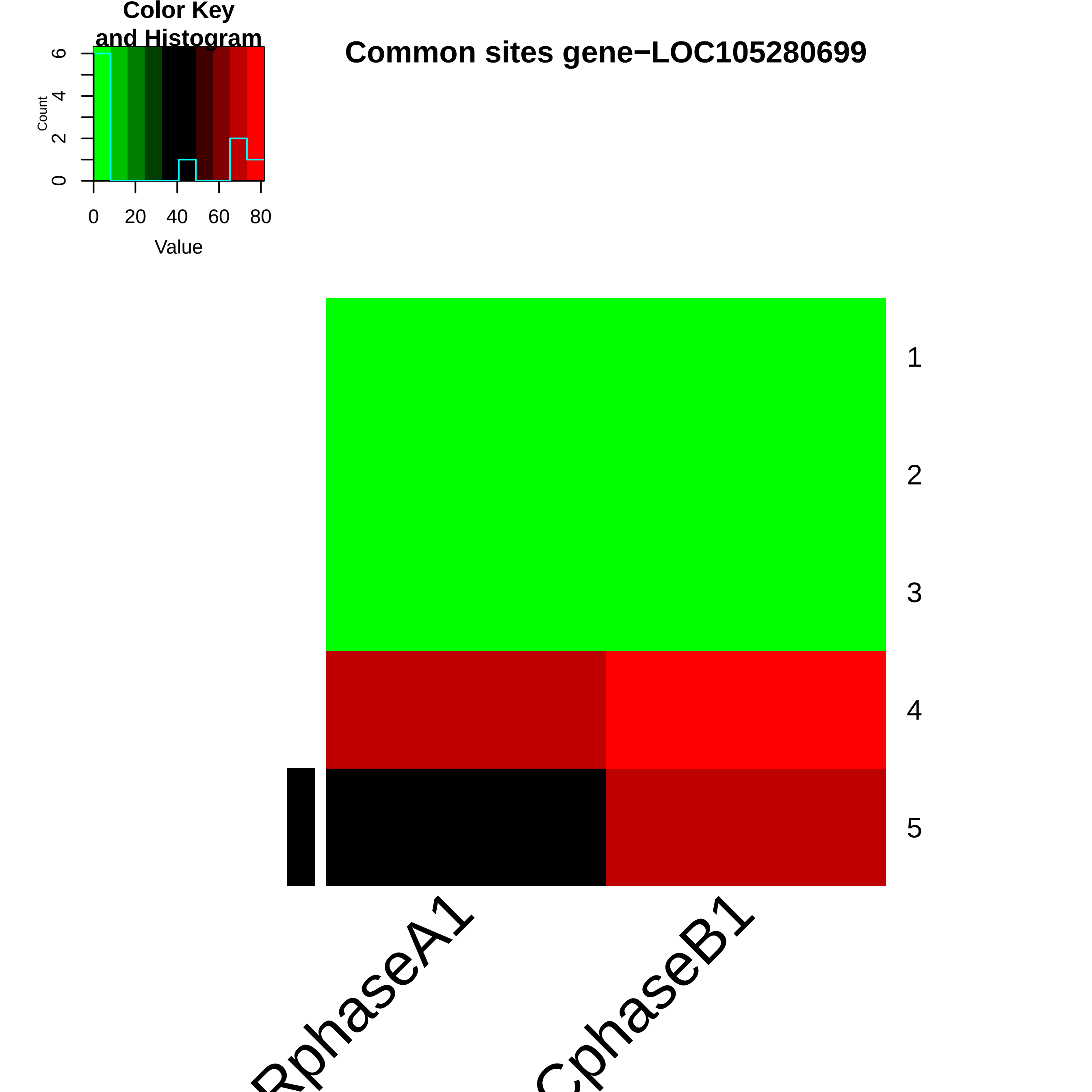

Color Key
and Histogram
Count
Value
0
20
40
60
80
0
2
4
6
Common sites gene−LOC105280699
1
2
3
4
5
RphaseA1
CphaseB1

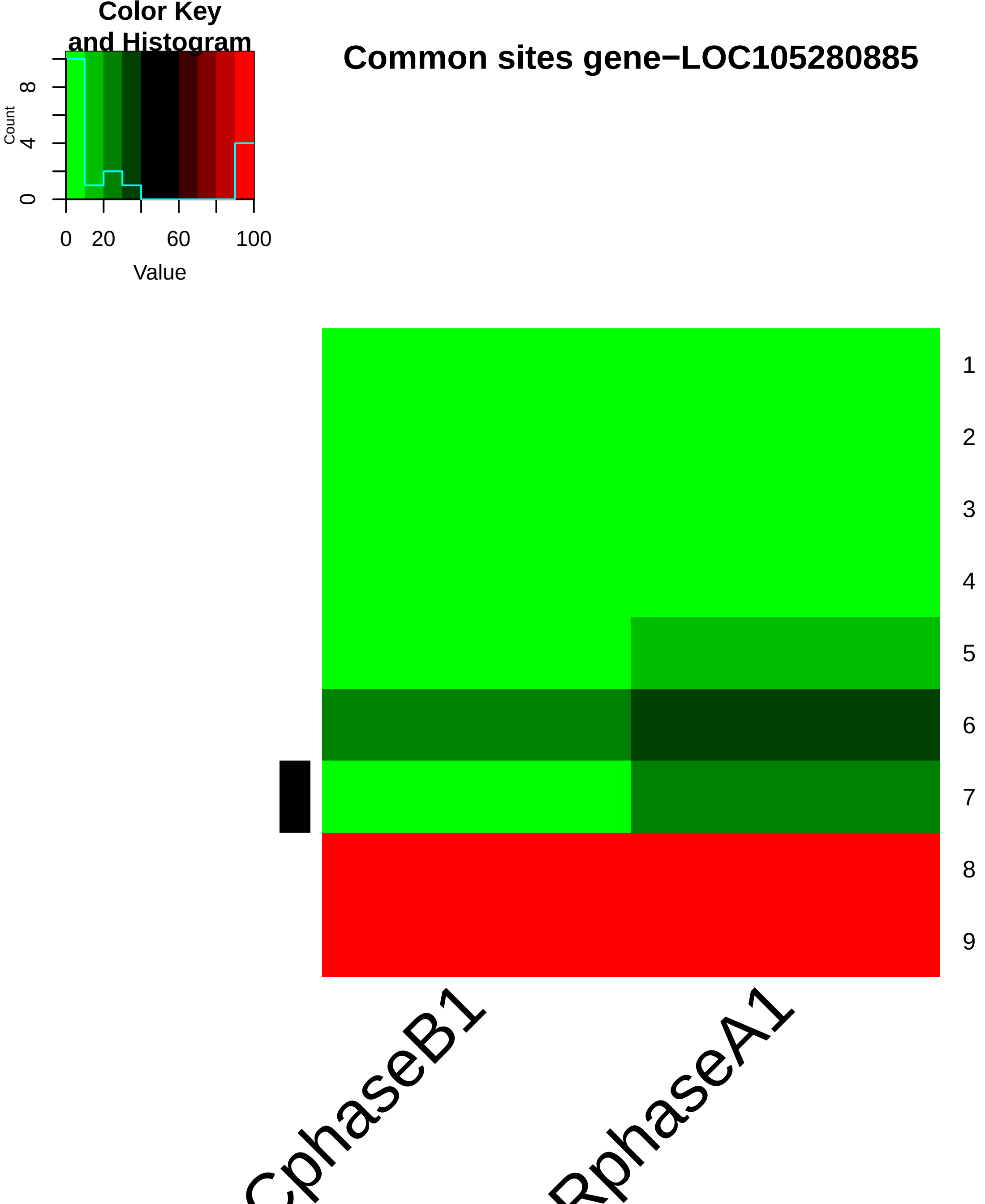

Color Key
and Histogram
Count
0
4
8
0
20
60
100
Value
Common sites gene−LOC105280885
1
2
3
4
5
6
7
8
9
phaseB1
RphaseA1

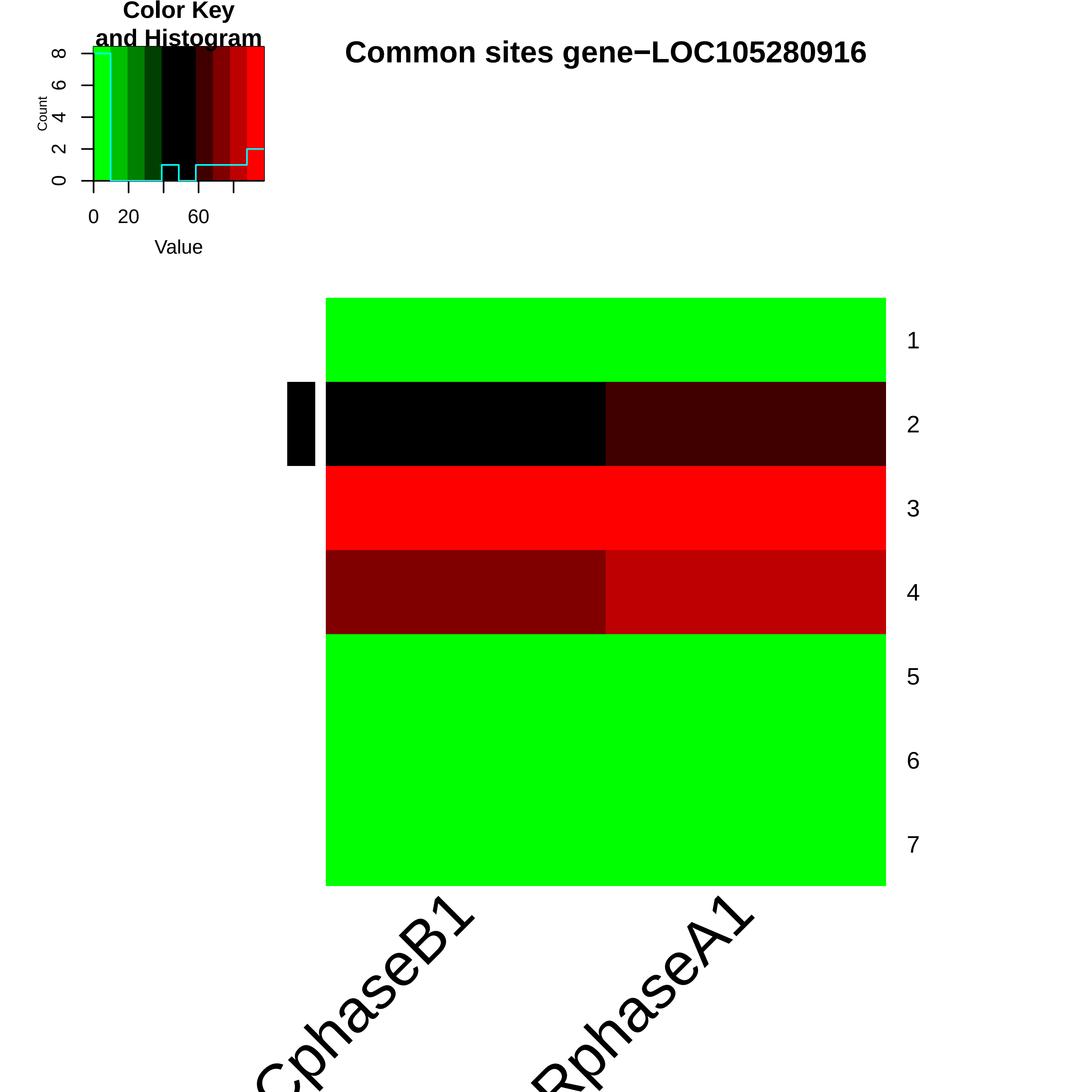
Color Key
and Histogram
Count
0
2
4
6
8
0
20
60
Value
Common sites gene−LOC105280916
1
2
3
4
5
6
7
CphaseB1
RphaseA1

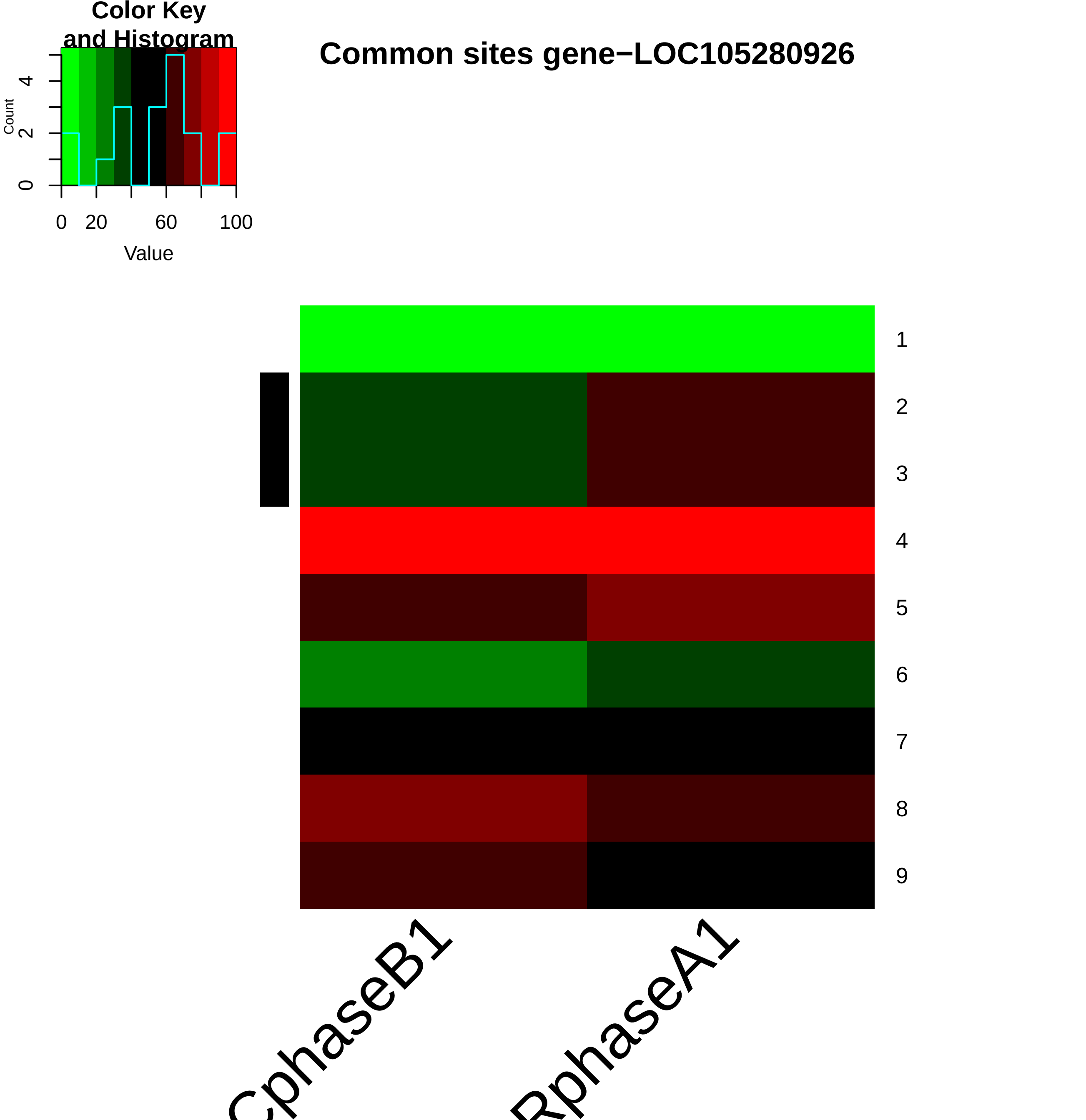
Color Key
and Histogram
Count
0
2
4
0
20
60
100
Value
Common sites gene−LOC105280926
1
2
3
4
5
6
7
8
9
CphaseB1
RphaseA1

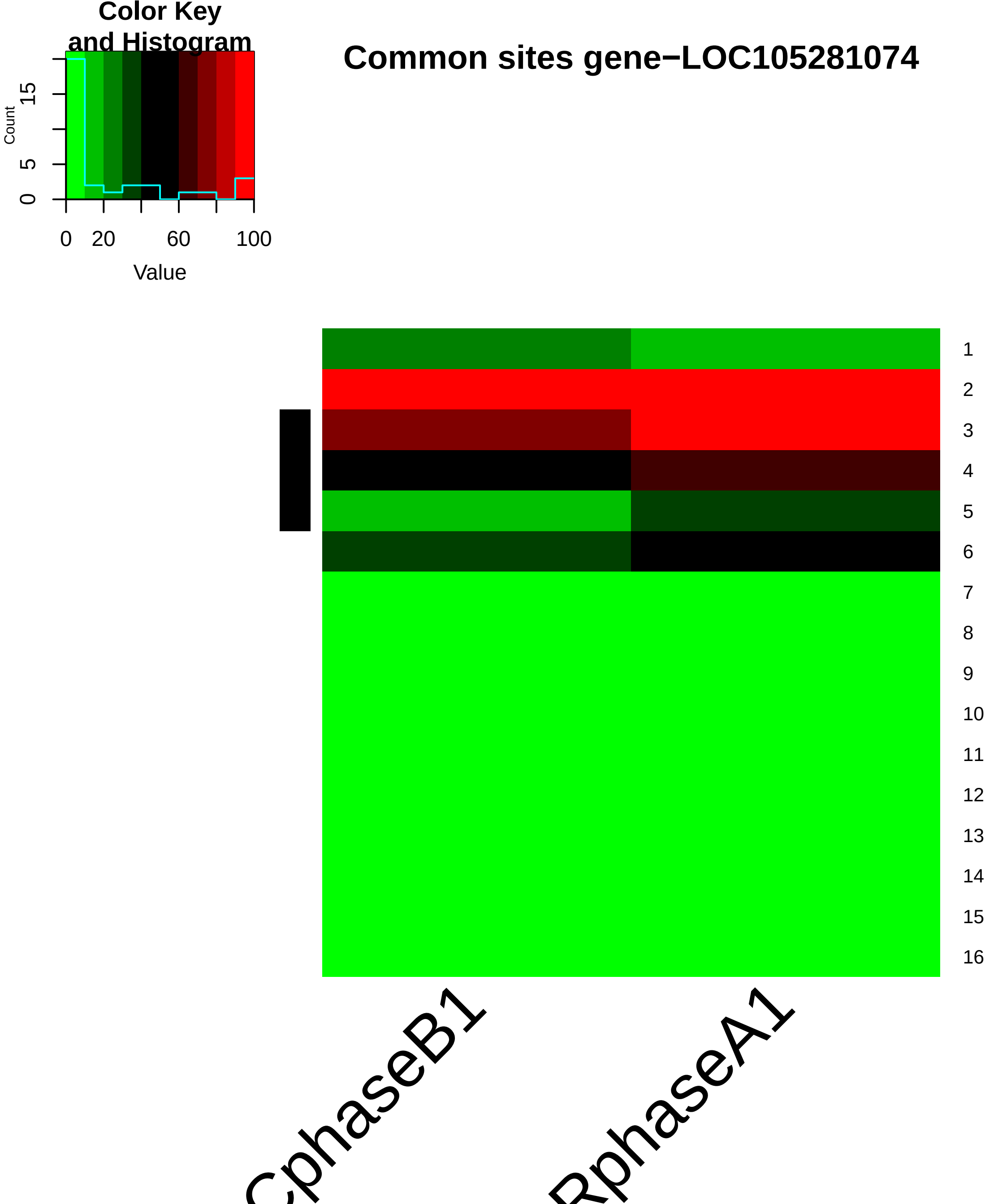
Color Key
and Histogram
Count
0
5
15
0
20
60
100
Value
Common sites gene−LOC105281074
CphaseB1
RphaseA1

### Common sites gene−LOC105281375

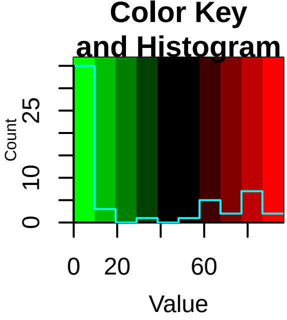

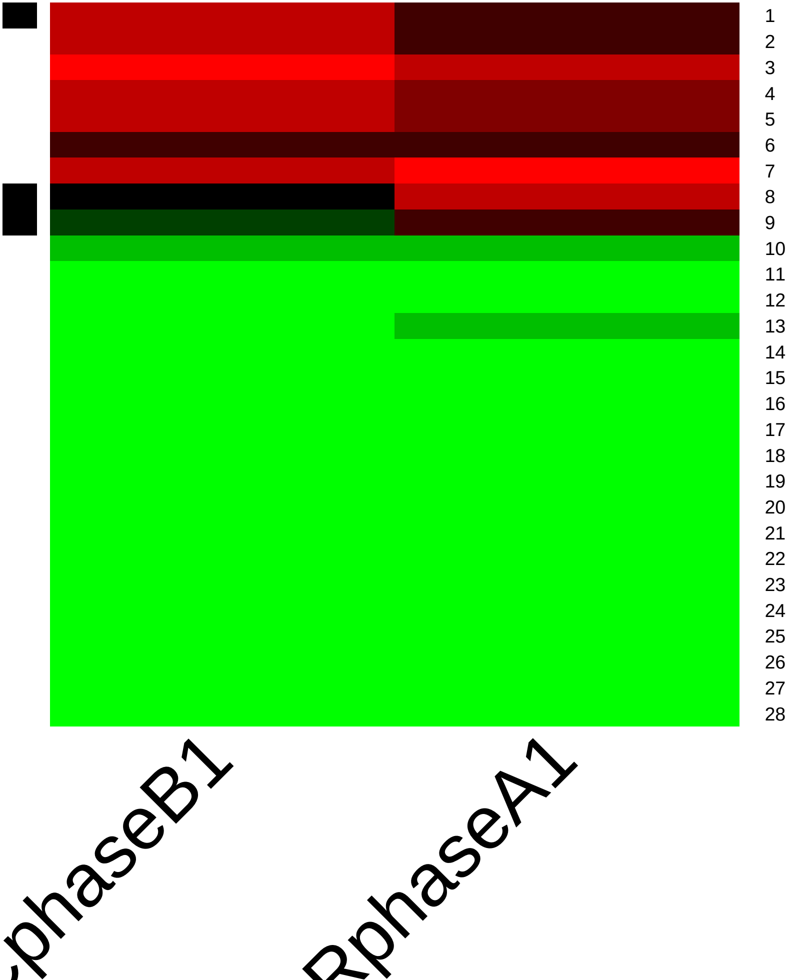

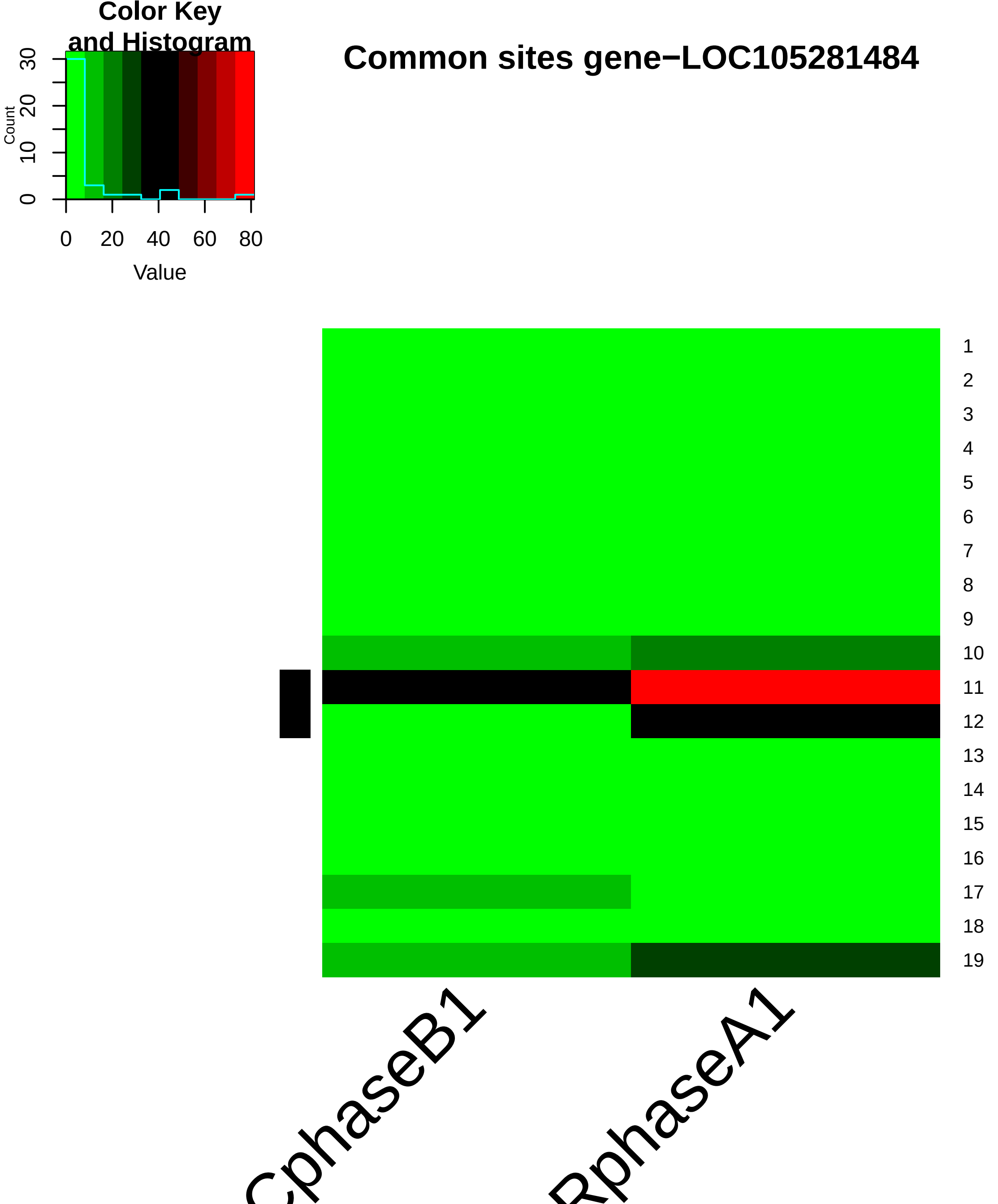
Color Key
and Histogram
Count
0
10
20
30
0
20
40
60
80
Value
Common sites gene−LOC105281484
CphaseB1
RphaseA1

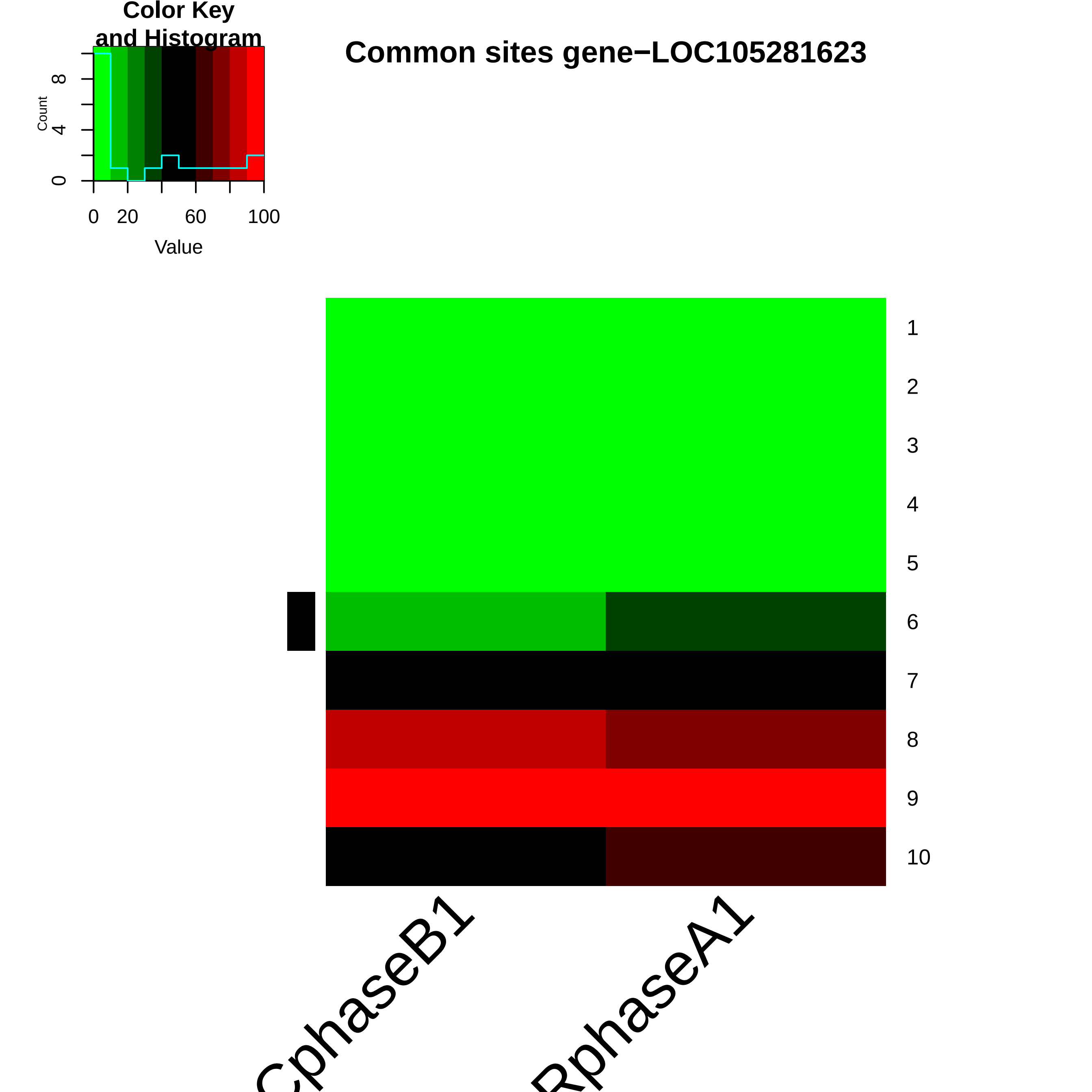
Color Key
and Histogram
Count
0
4
8
0
20
60
100
Value
Common sites gene−LOC105281623
1
2
3
4
5
6
7
8
9
10
CphaseB1
RphaseA1

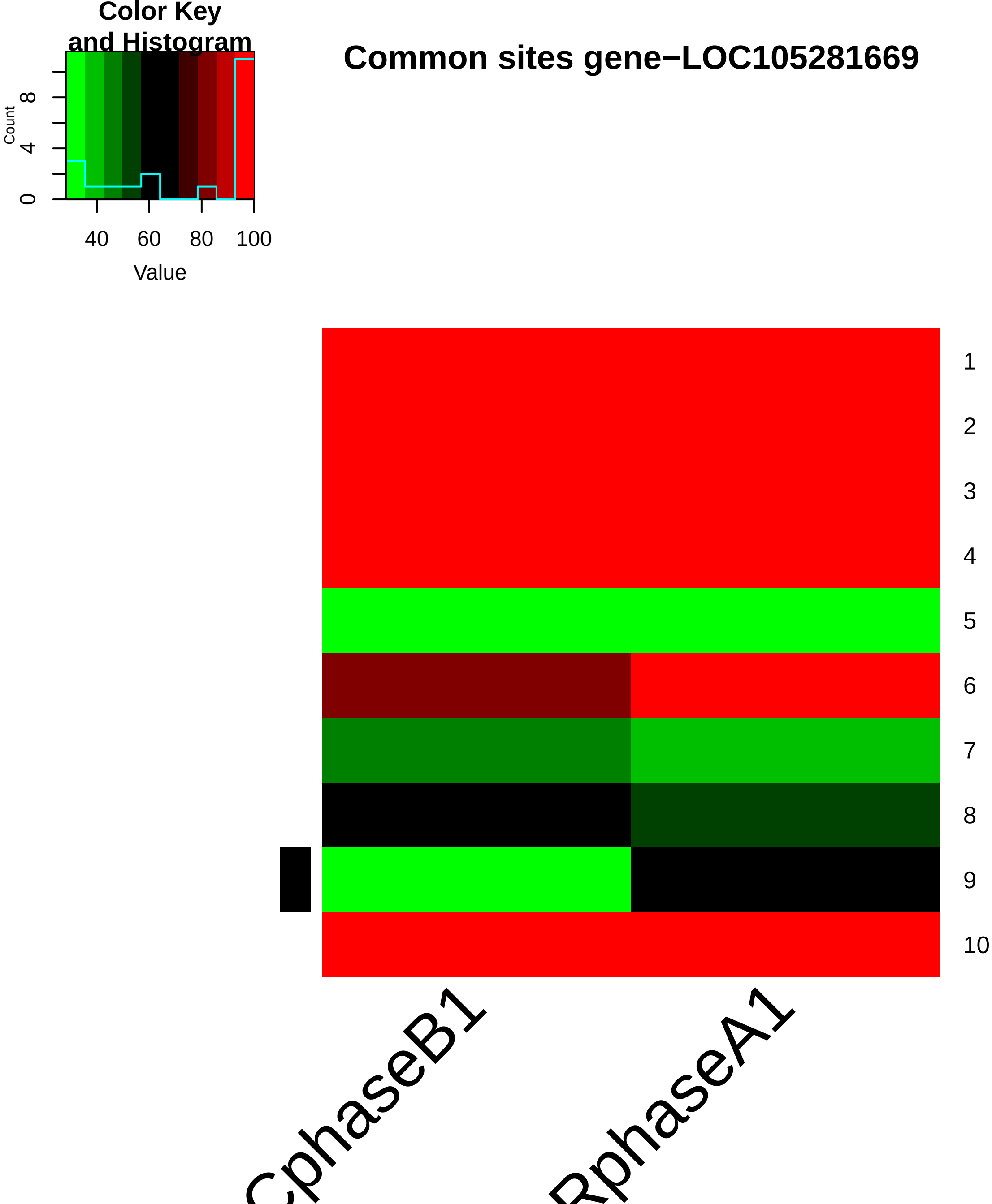

Color Key
and Histogram
Count
0
4
8
40
60
80
100
Value
Common sites gene−LOC105281669
1
2
3
4
5
6
7
8
9
10
CphaseB1
RphaseA1

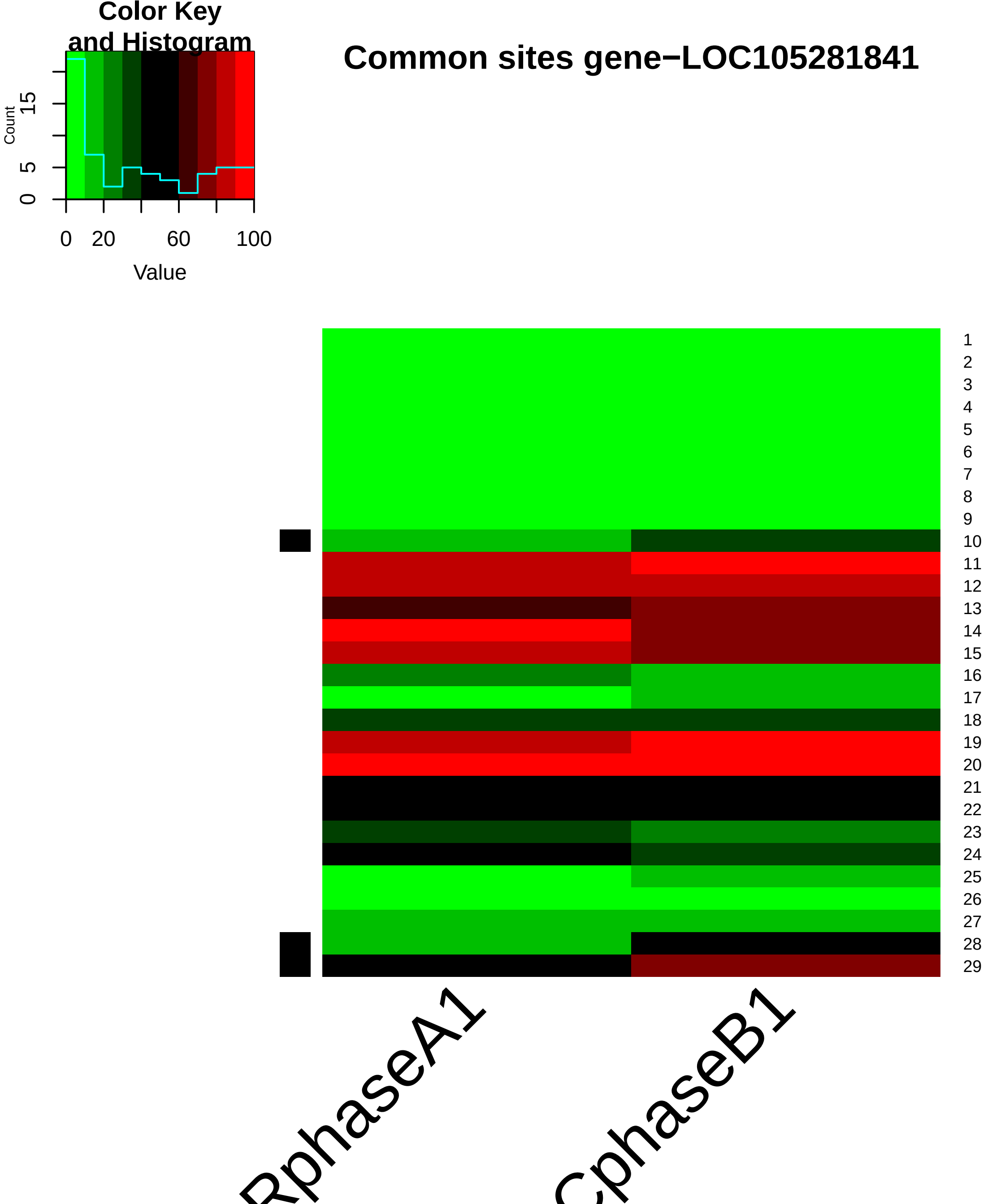

Color Key
and Histogram
Count
0
5
15
0
20
60
100
Value
Common sites gene−LOC105281841
RphaseA1
CphaseB1

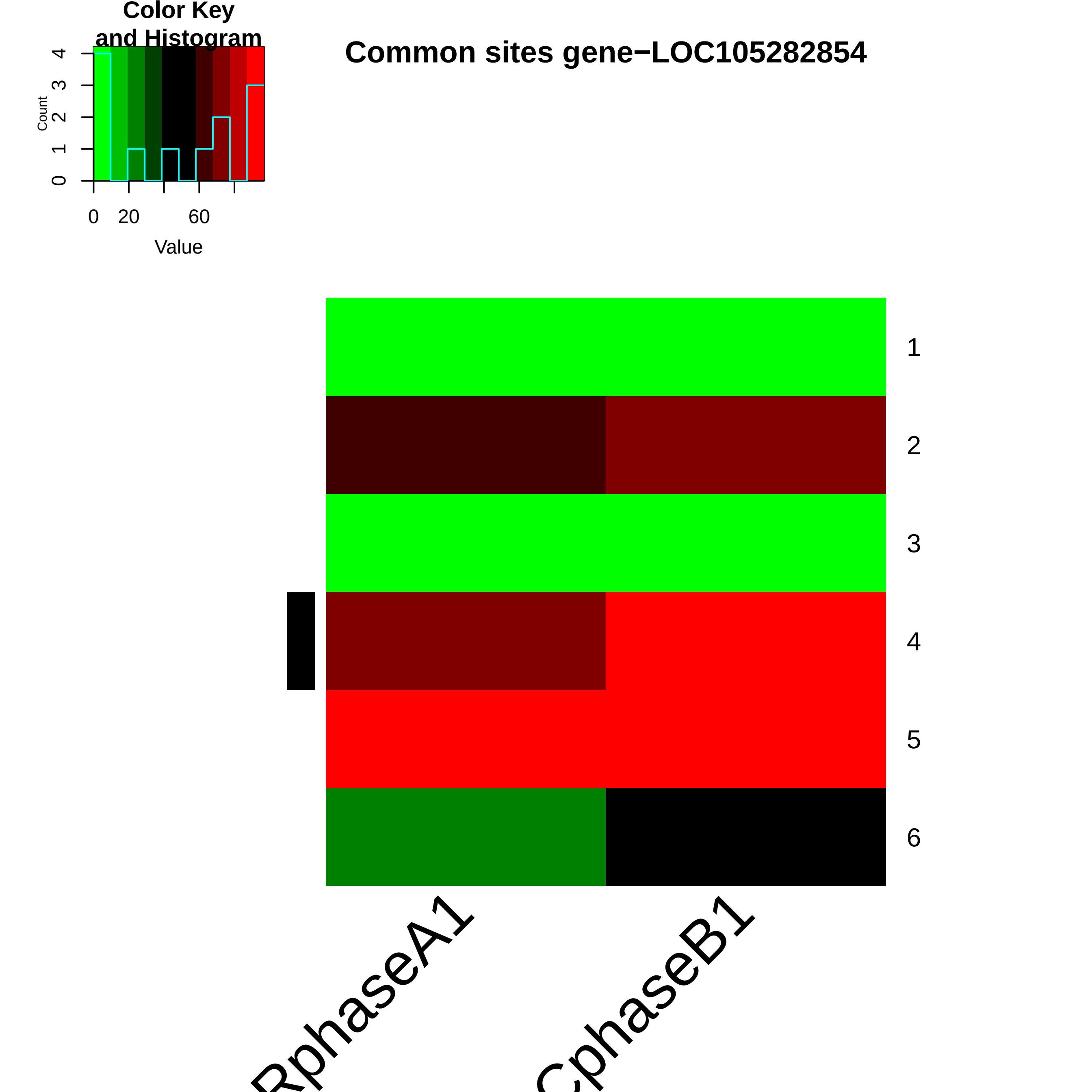

Color Key
and Histogram
Count
Value
0
20
60
Common sites gene−LOC105282854
1
2
3
4
5
6
RphaseA1
CphaseB1

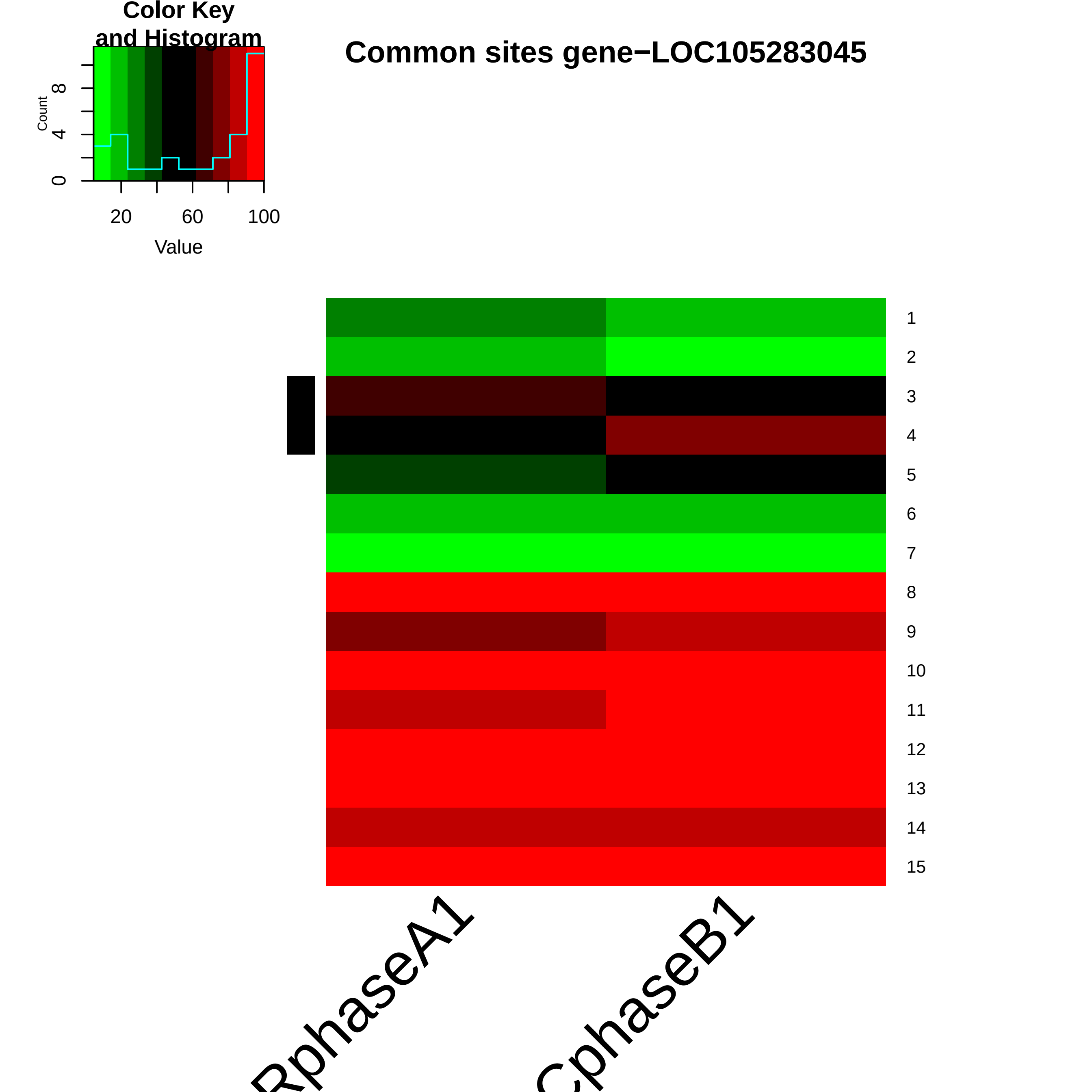
Color Key
and Histogram
Count
0
4
8
20
60
100
Value
Common sites gene−LOC105283045
RphaseA1
CphaseB1

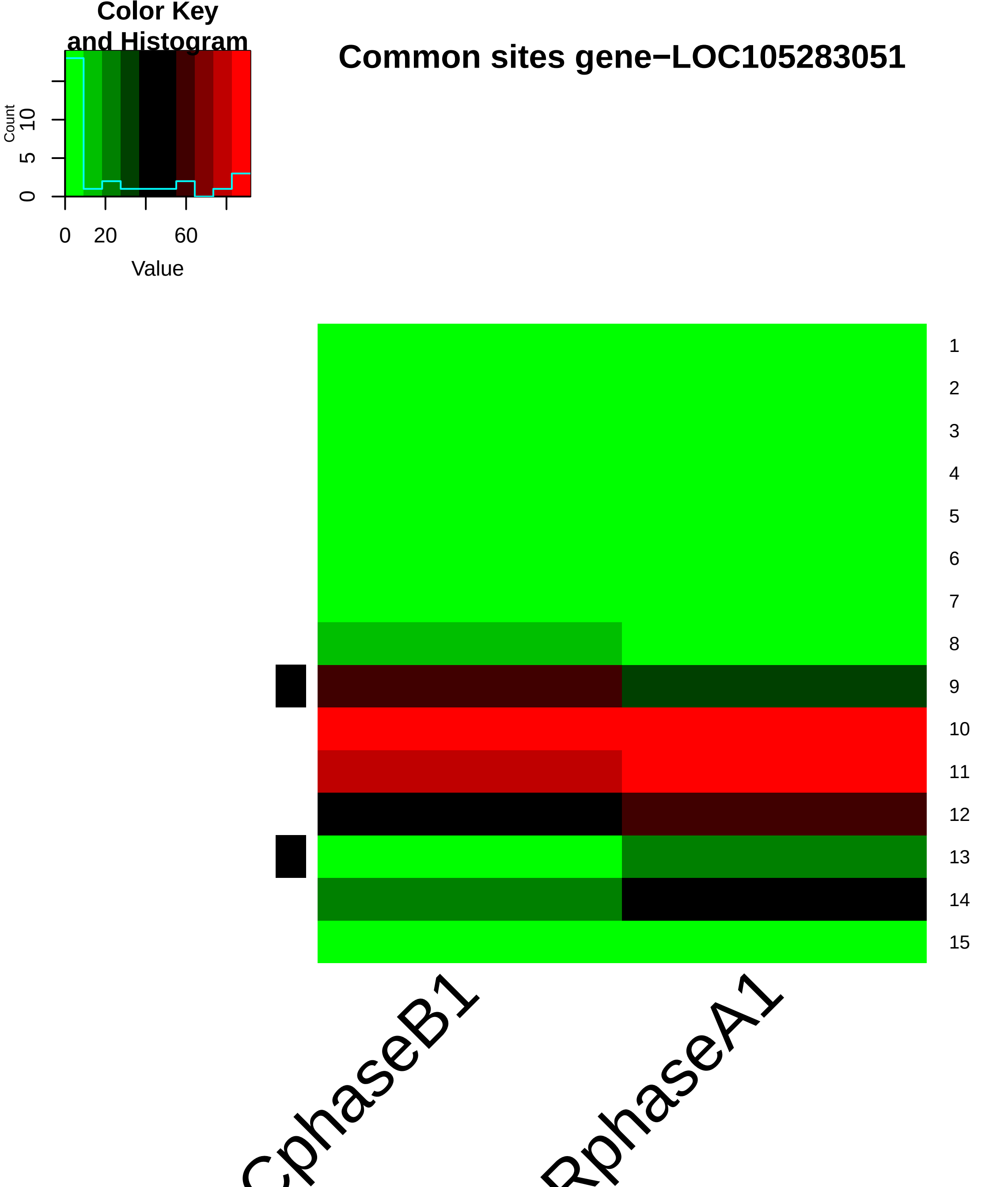

Color Key
and Histogram
Count
0
5
10
0
20
60
Value
Common sites gene−LOC105283051
CphaseB1
RphaseA1

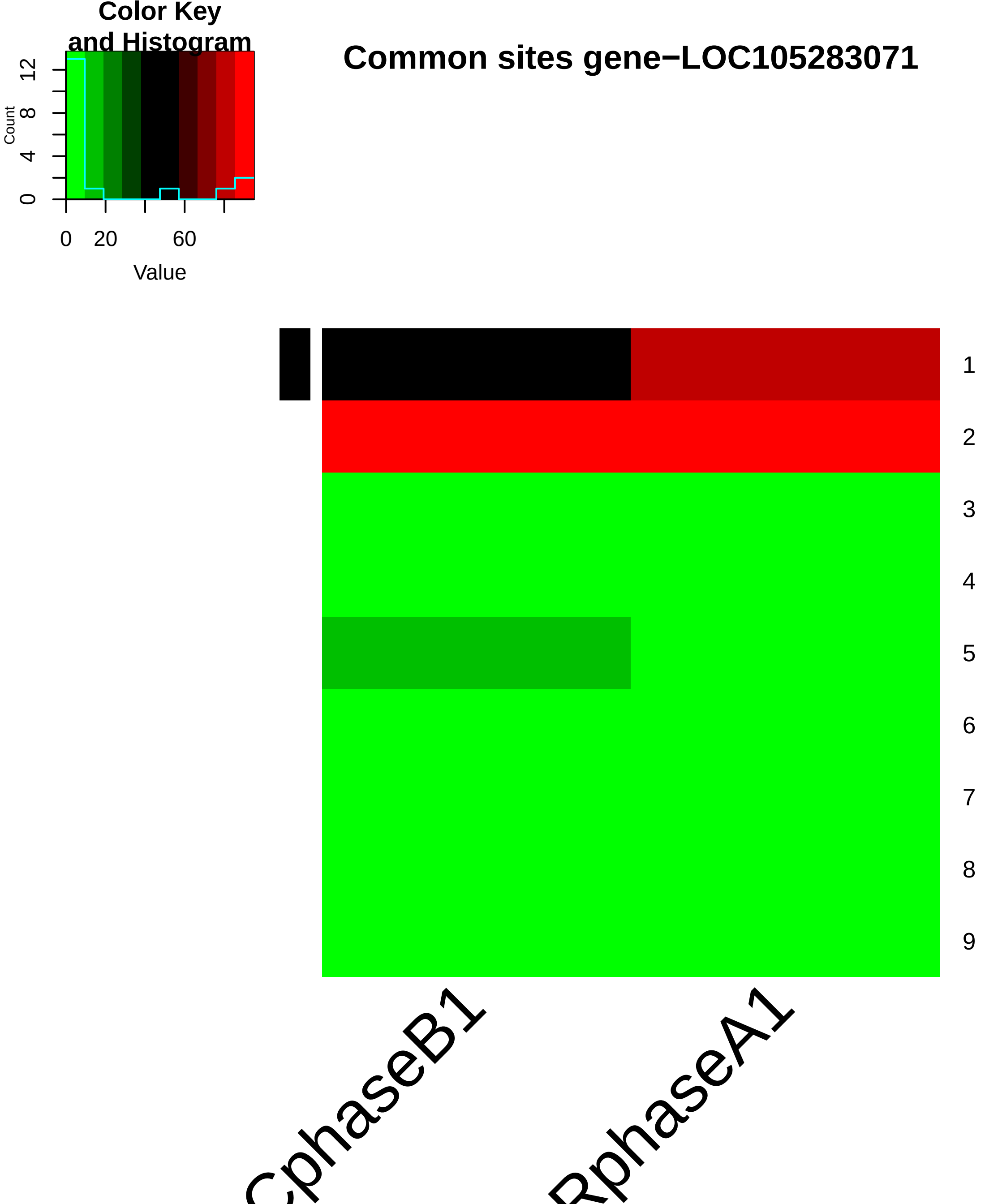

Color Key
and Histogram
Count
0
4
8
12
0
20
60
Value
Common sites gene−LOC105283071
1
2
3
4
5
6
7
8
9
CphaseB1
RphaseA1

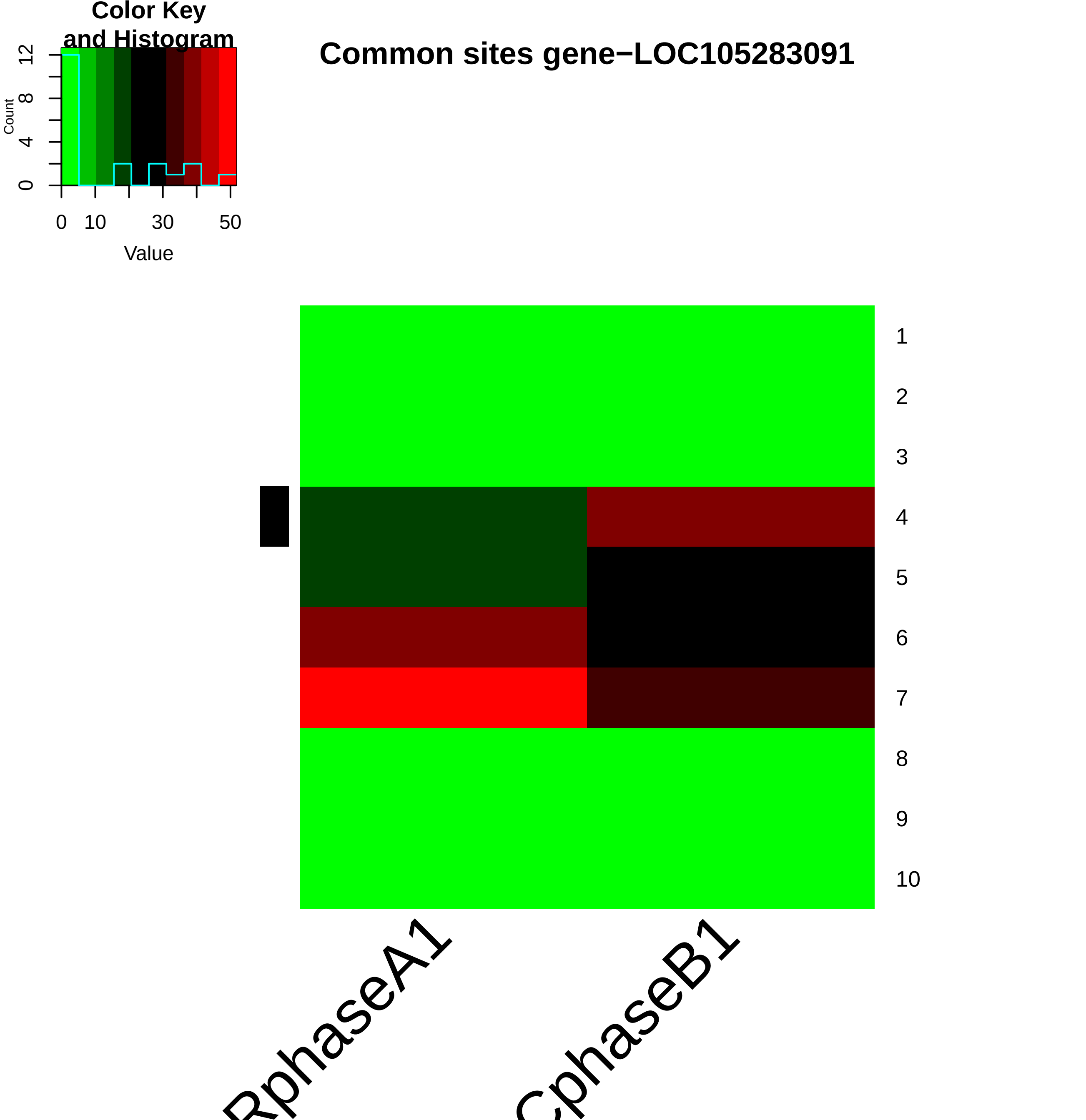
Color Key
and Histogram
Count
0
4
8
12
0
10
30
50
Value
Common sites gene−LOC105283091
1
2
3
4
5
6
7
8
9
10
RphaseA1
CphaseB1

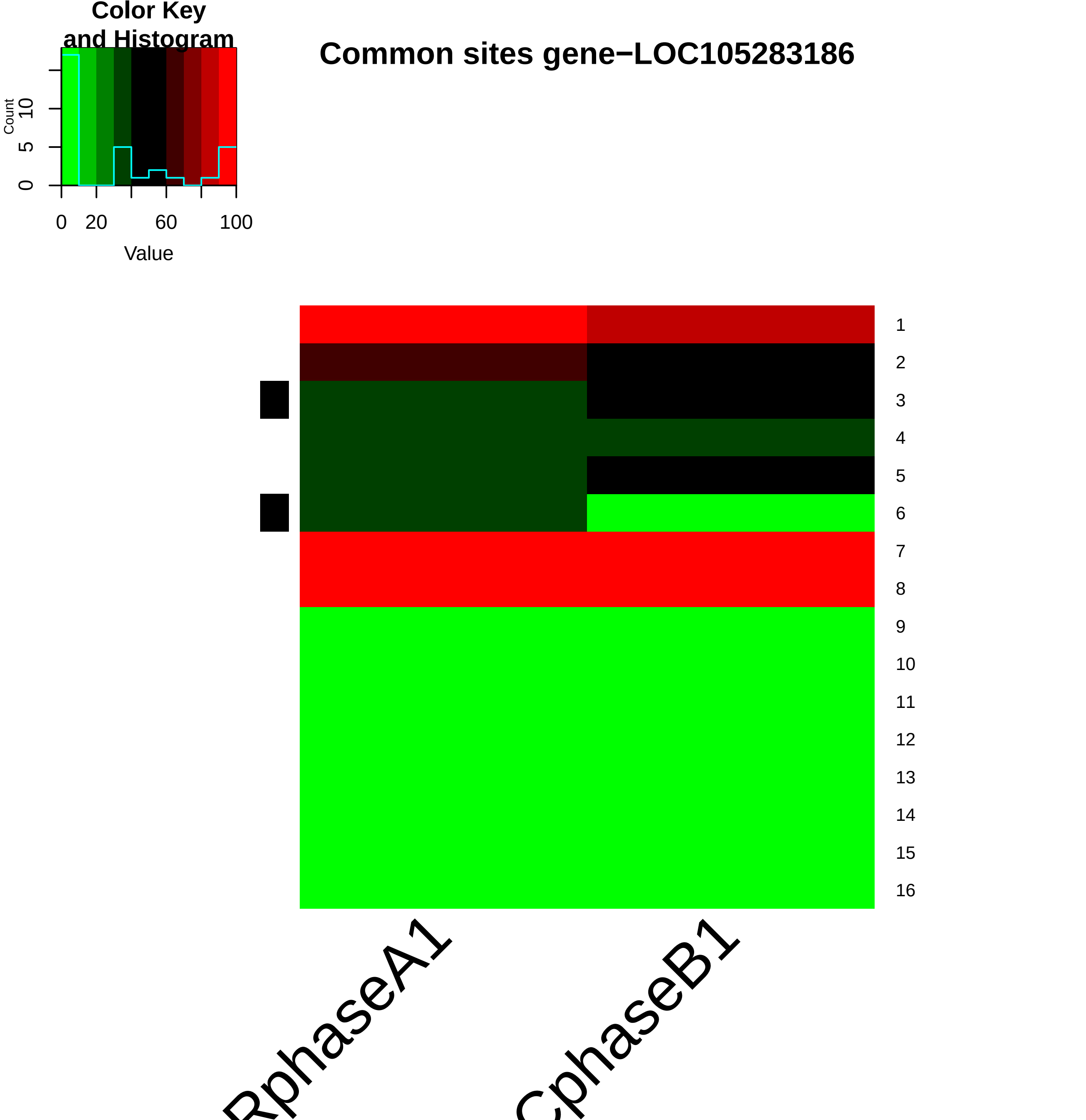
Color Key
and Histogram
Count
0
5
10
0
20
60
100
Value
Common sites gene−LOC105283186
RphaseA1
CphaseB1

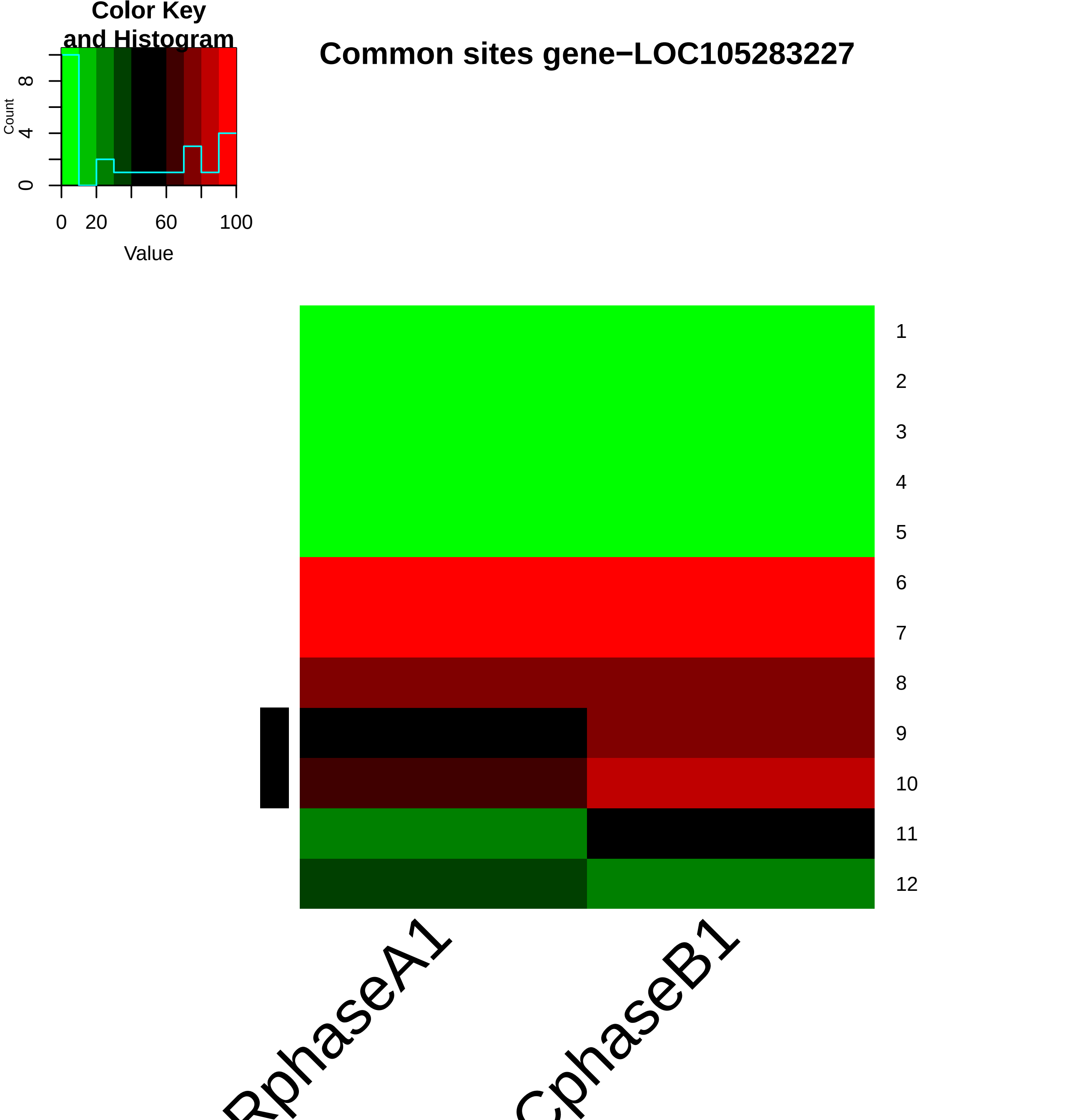

Color Key
and Histogram
Count
0
4
8
0
20
60
100
Value
Common sites gene−LOC105283227
1
2
3
4
5
6
7
8
9
10
11
12
RphaseA1
CphaseB1

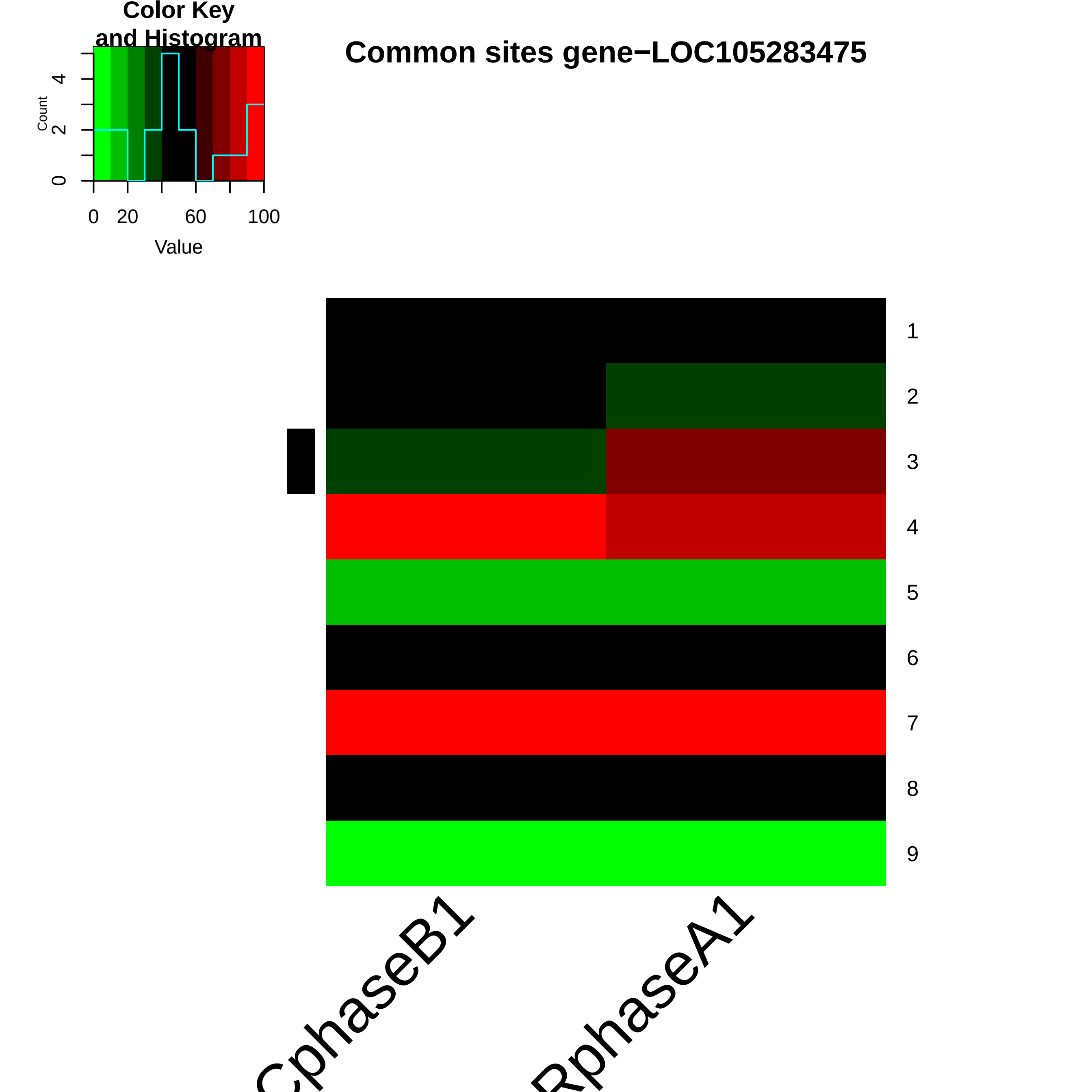

Color Key
and Histogram
Count
0
2
4
0
20
60
100
Value
Common sites gene−LOC105283475
1
2
3
4
5
6
7
8
9
CphaseB1
RphaseA1

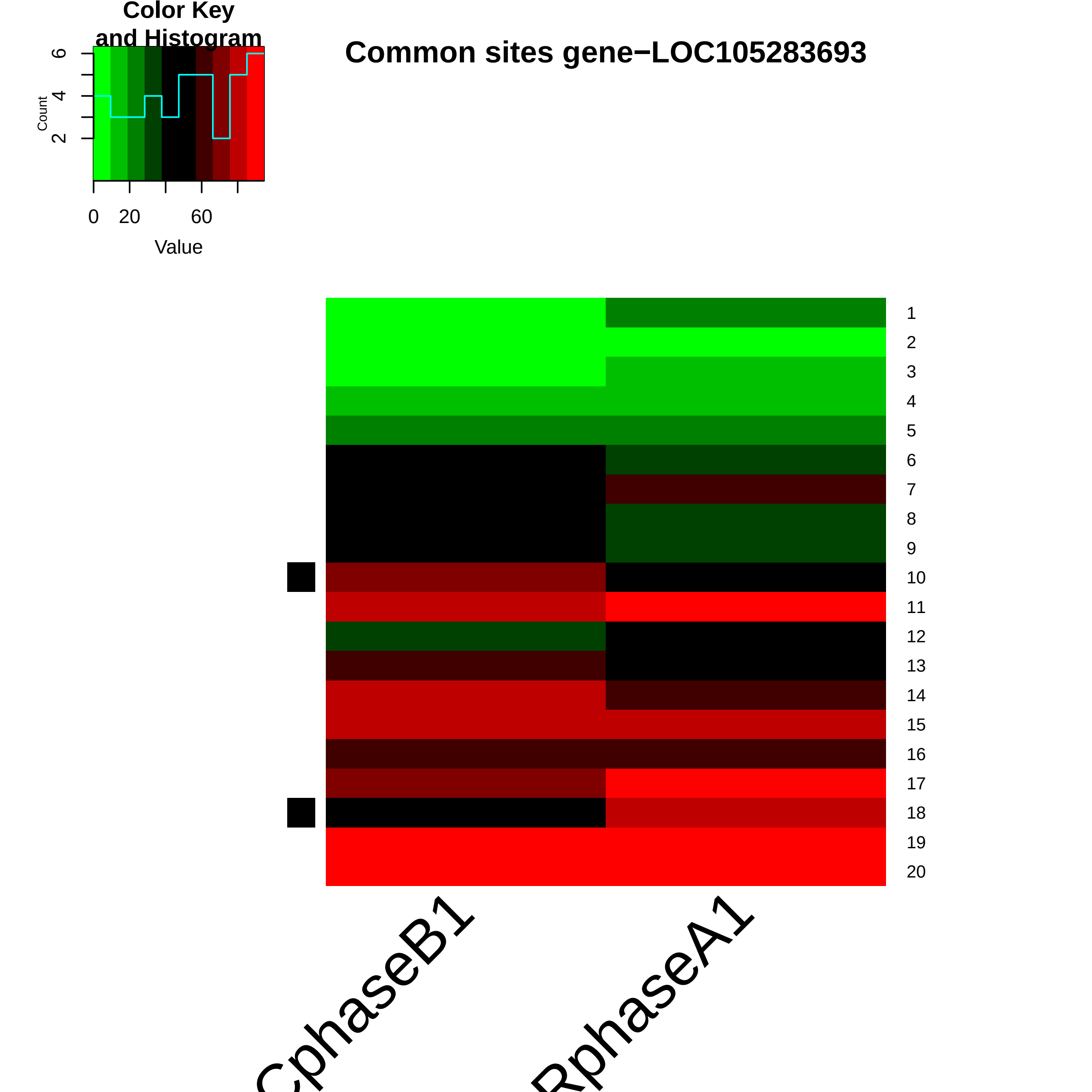
Color Key
and Histogram
Count
Value
0
20
60
2
4
6
Common sites gene−LOC105283693
CphaseB1
RphaseA1

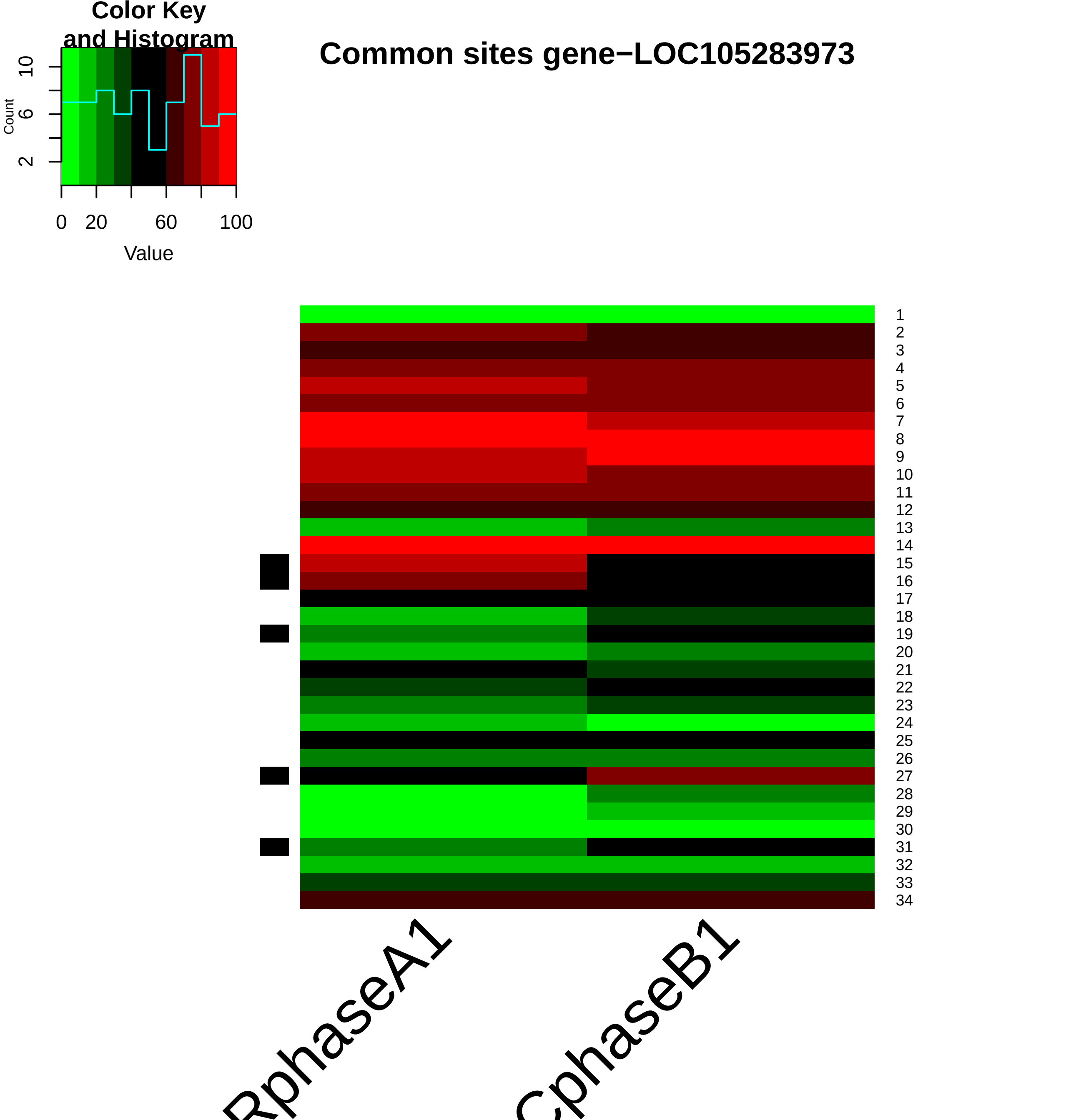

Color Key
and Histogram
Count
2
6
10
0
20
60
100
Value
Common sites gene−LOC105283973
RphaseA1
CphaseB1

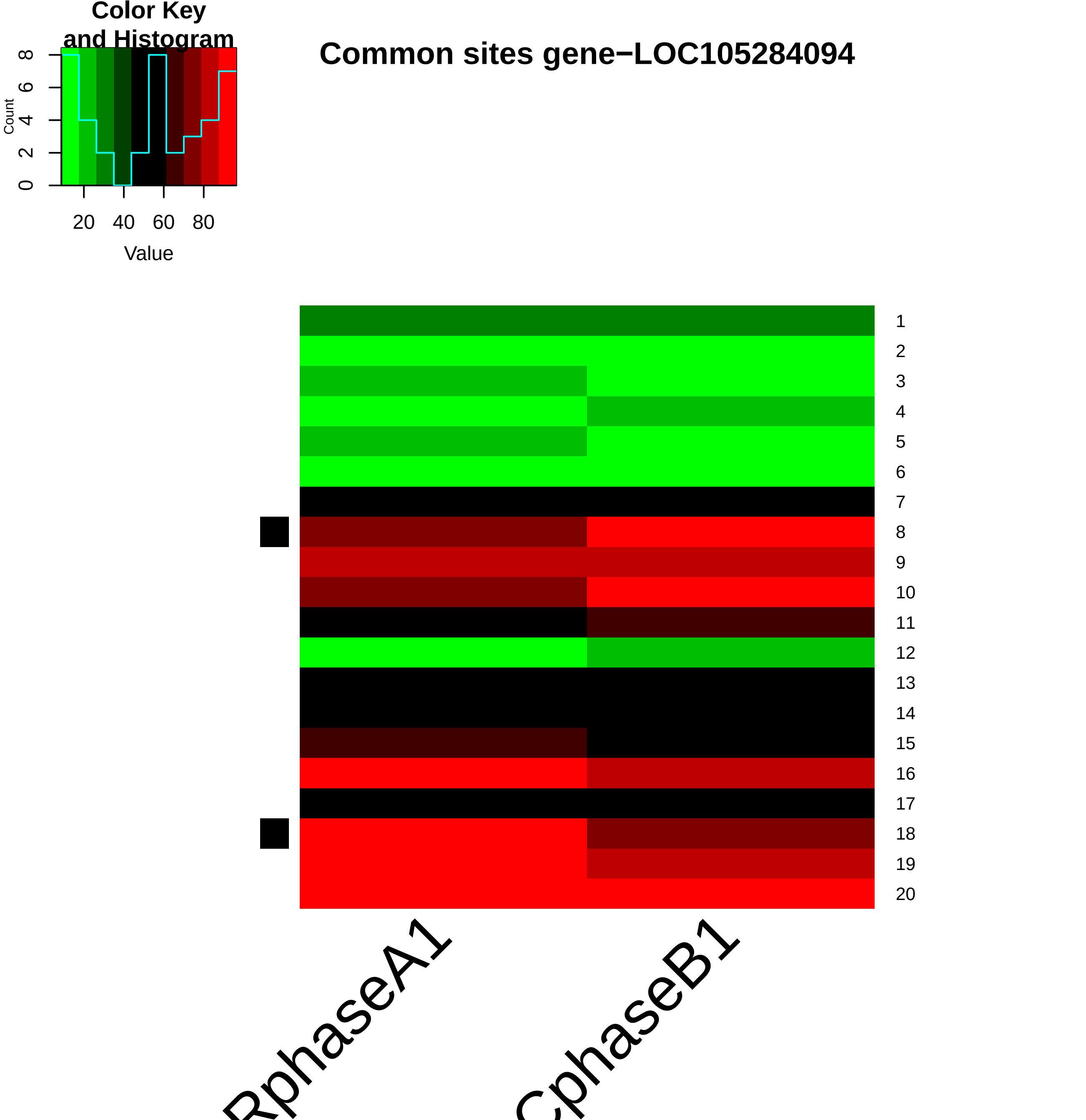

Color Key
and Histogram
Count
0
2
4
6
8
20
40
60
80
Value
Common sites gene−LOC105284094
RphaseA1
CphaseB1

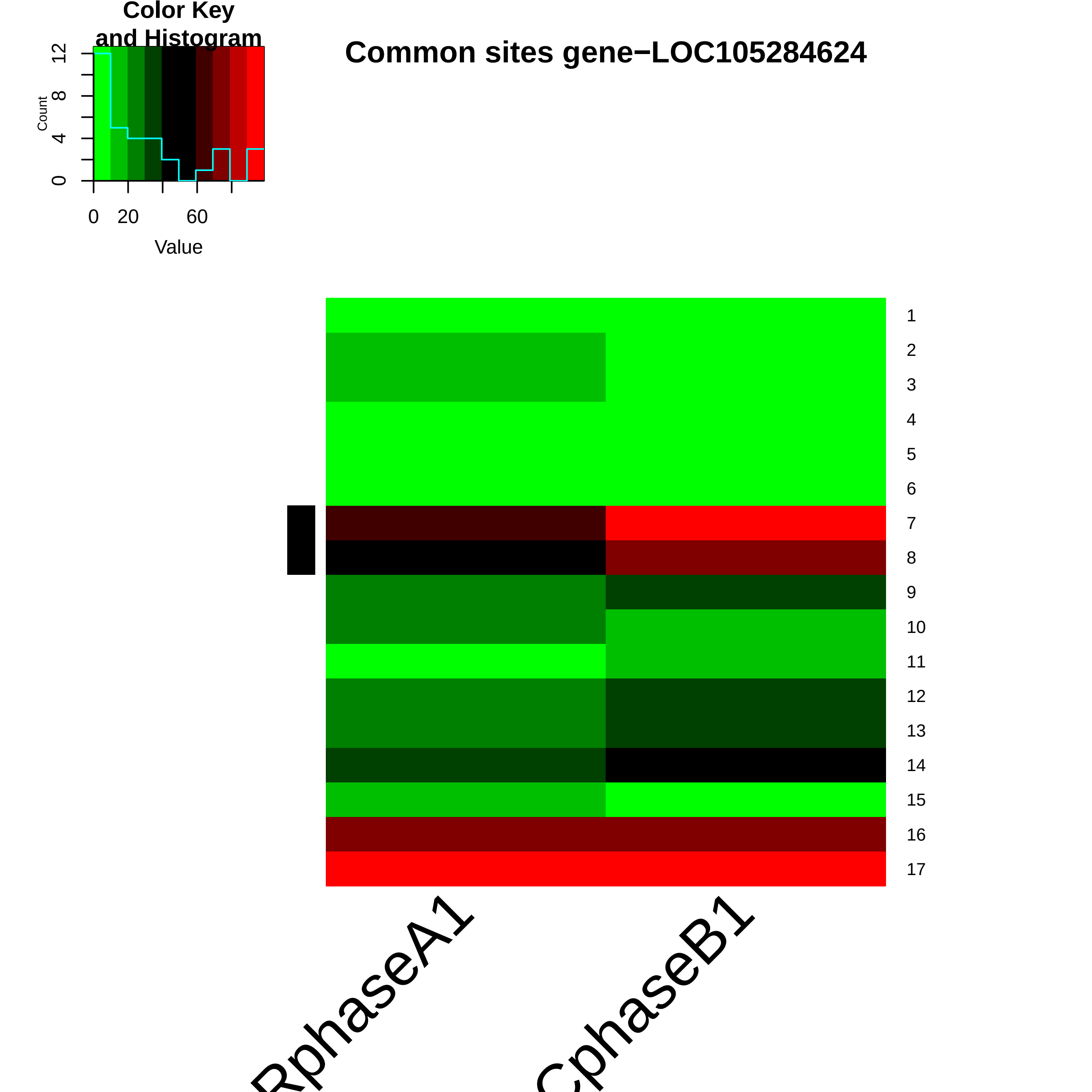
Color Key
and Histogram
Count
0
4
8
12
0
20
60
Value
Common sites gene−LOC105284624
RphaseA1
CphaseB1

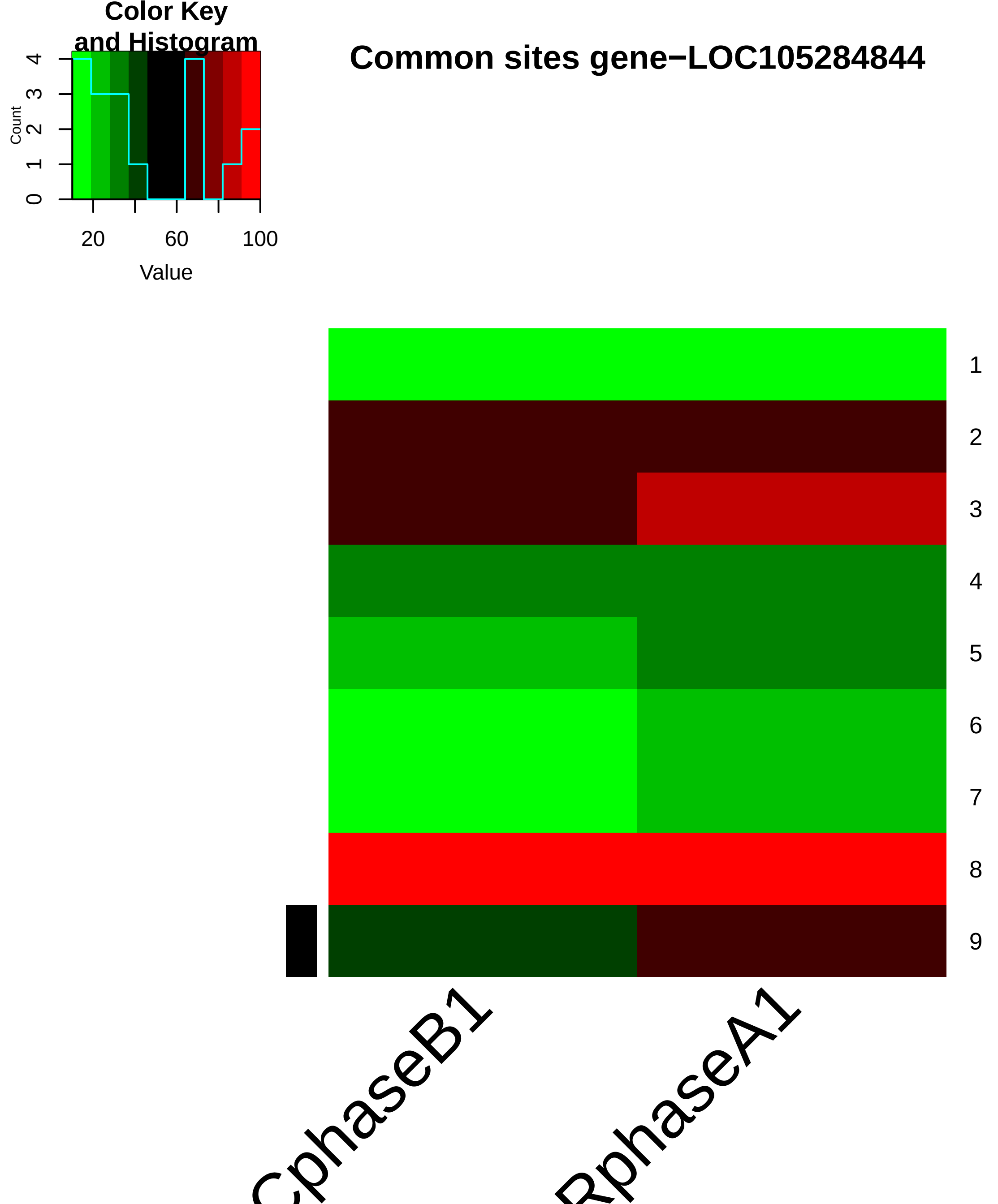
Color Key
and Histogram
Count
0
1
2
3
4
20
60
100
Value
Common sites gene−LOC105284844
1
2
3
4
5
6
7
8
9
CphaseB1
RphaseA1

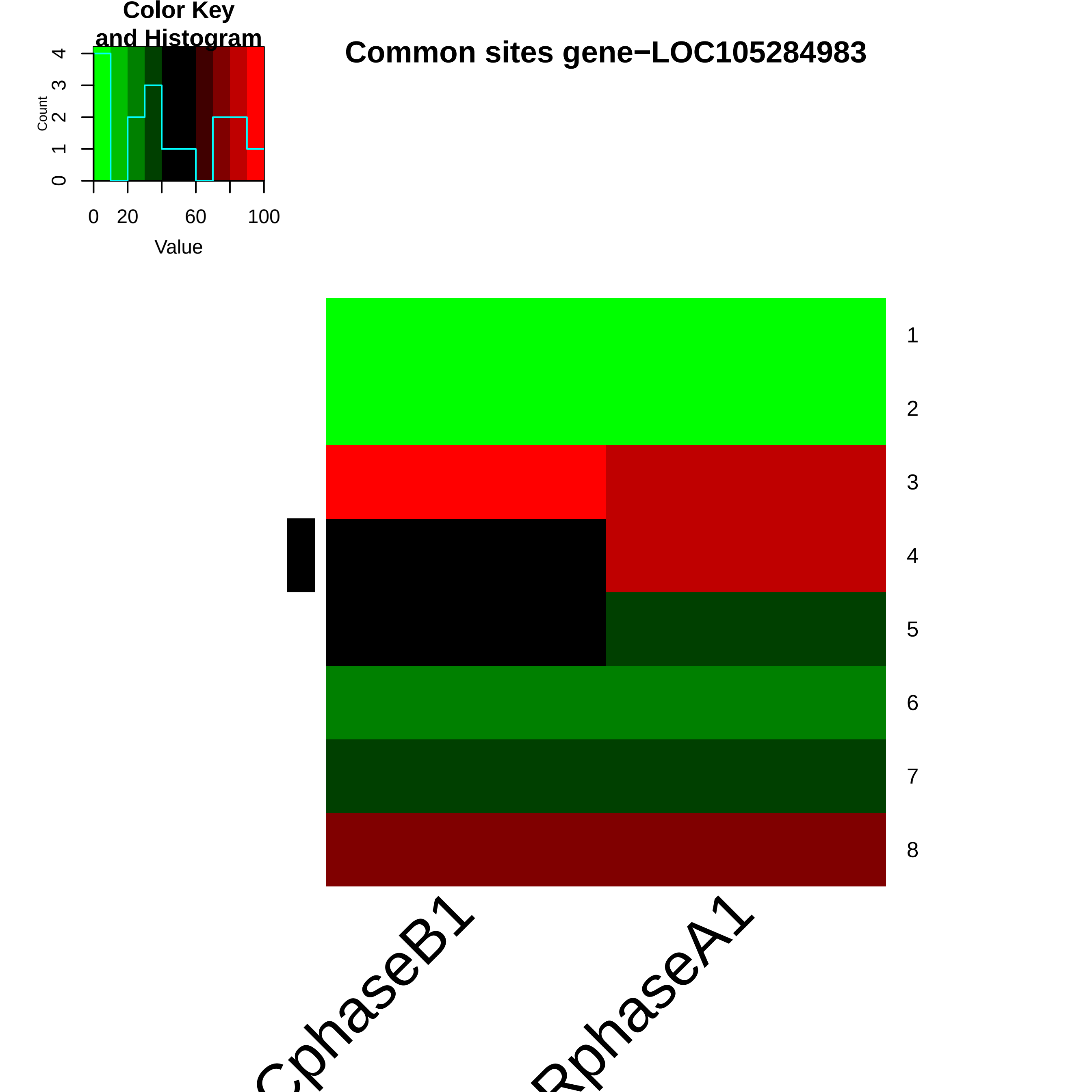

Color Key
and Histogram
Count
Value
0
20
60
100
Common sites gene−LOC105284983
1
2
3
4
5
6
7
8

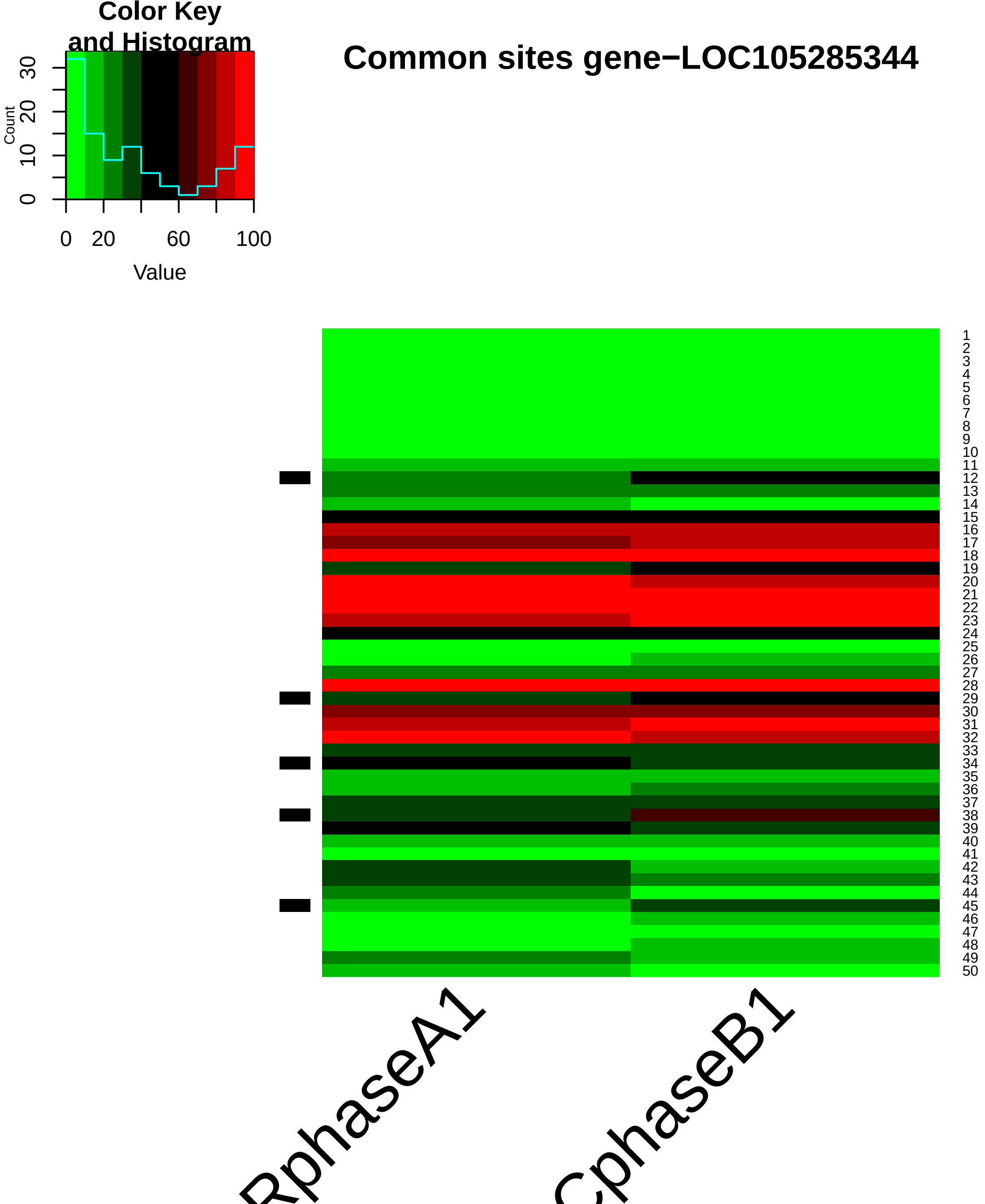
Color Key
and Histogram
Count
0
10
20
30
0
20
60
100
Value
Common sites gene−LOC105285344
RphaseA1
CphaseB1

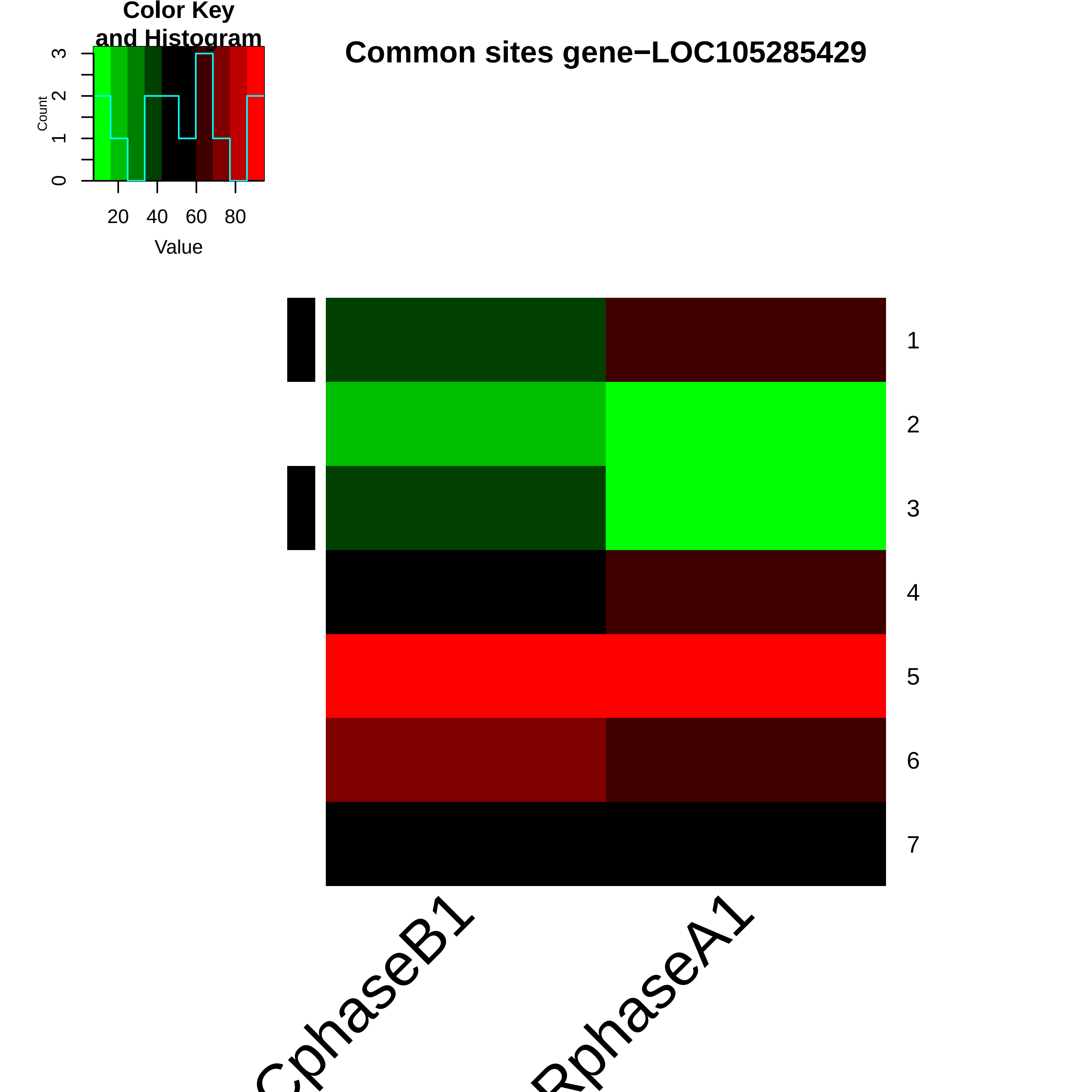

Color Key
and Histogram
Count
Value
0
1
2
3
20
40
60
80
Common sites gene−LOC105285429
1
2
3
4
5
6
7
CphaseB1
RphaseA1

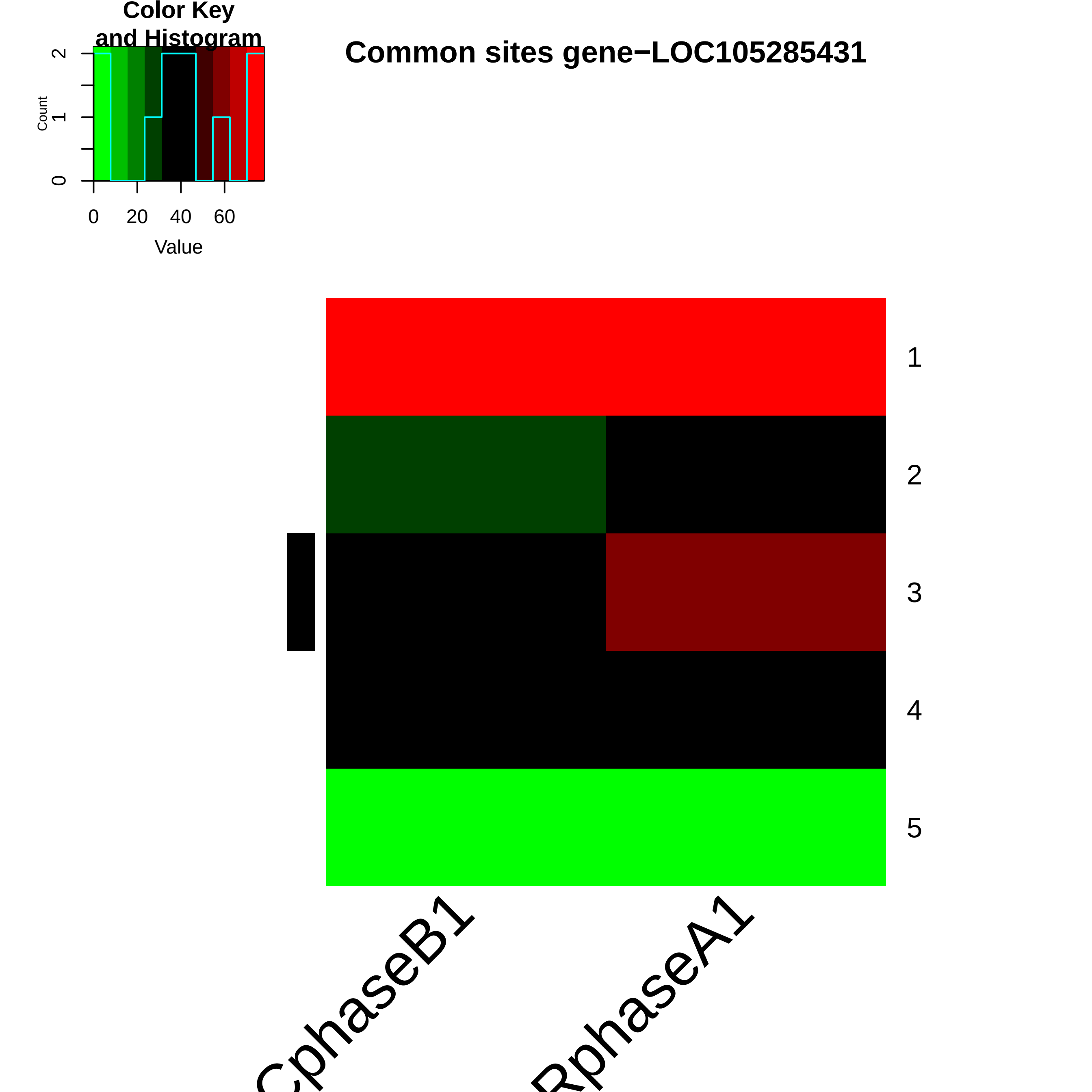
Color Key
and Histogram
Count
0
1
2
0
20
40
60
Value
Common sites gene−LOC105285431
1
2
3
4
5
CphaseB1
RphaseA1

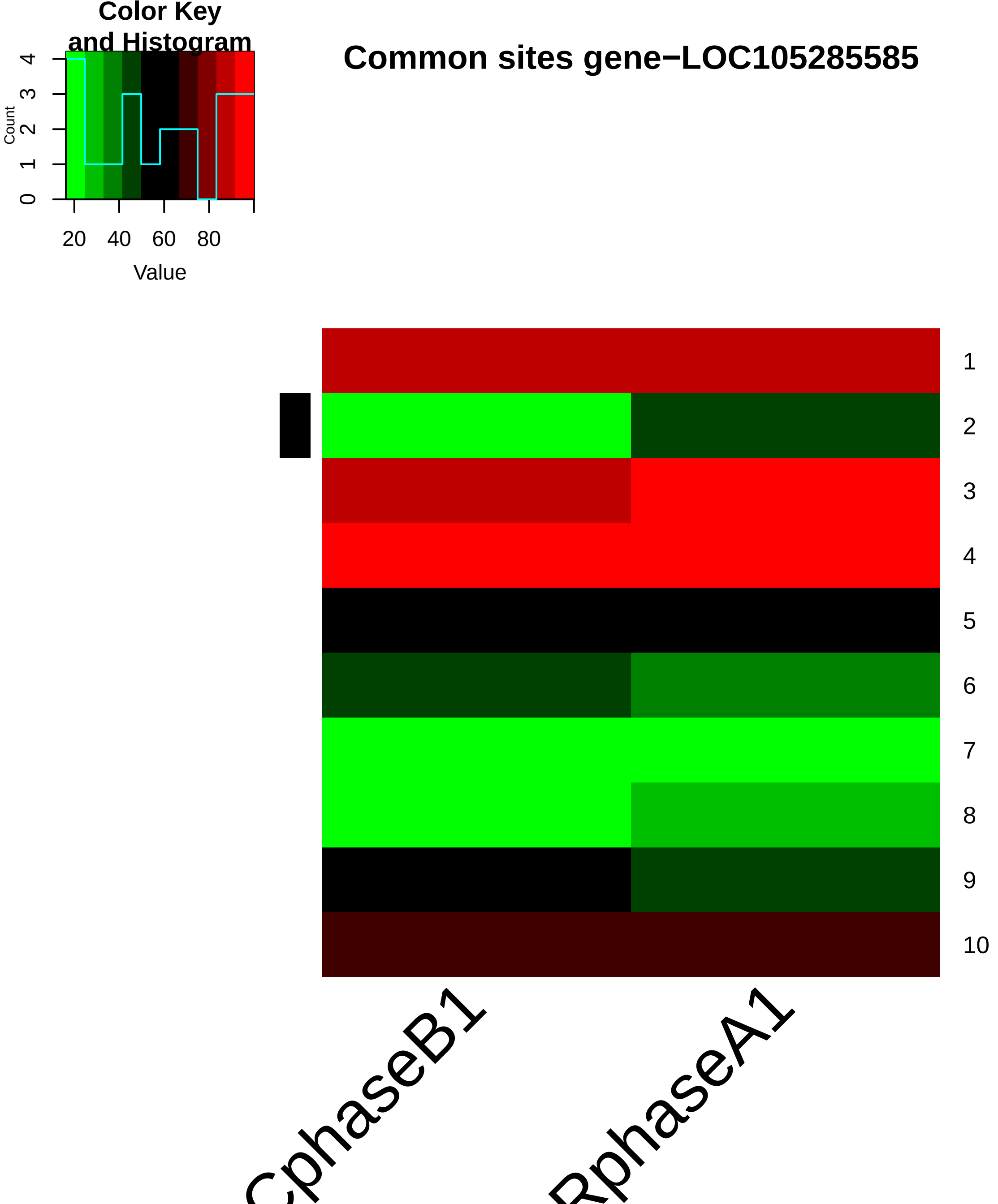

Color Key
and Histogram
Count
0
1
2
3
4
20
40
60
80
Value
Common sites gene−LOC105285585
1
2
3
4
5
6
7
8
9
10
CphaseB1
RphaseA1

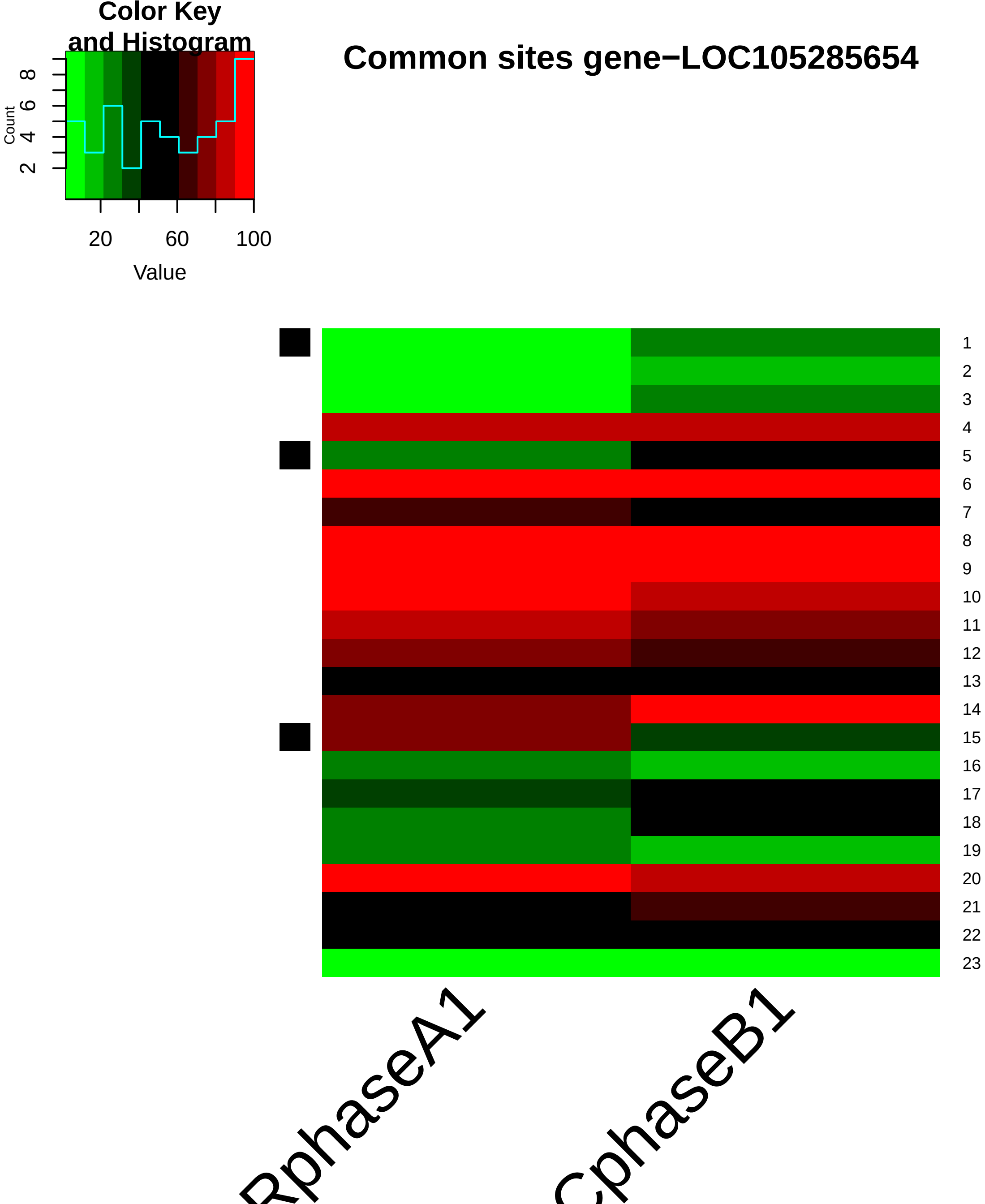
Color Key
and Histogram
Count
2
4
6
8
20
60
100
Value
Common sites gene−LOC105285654
RphaseA1
CphaseB1

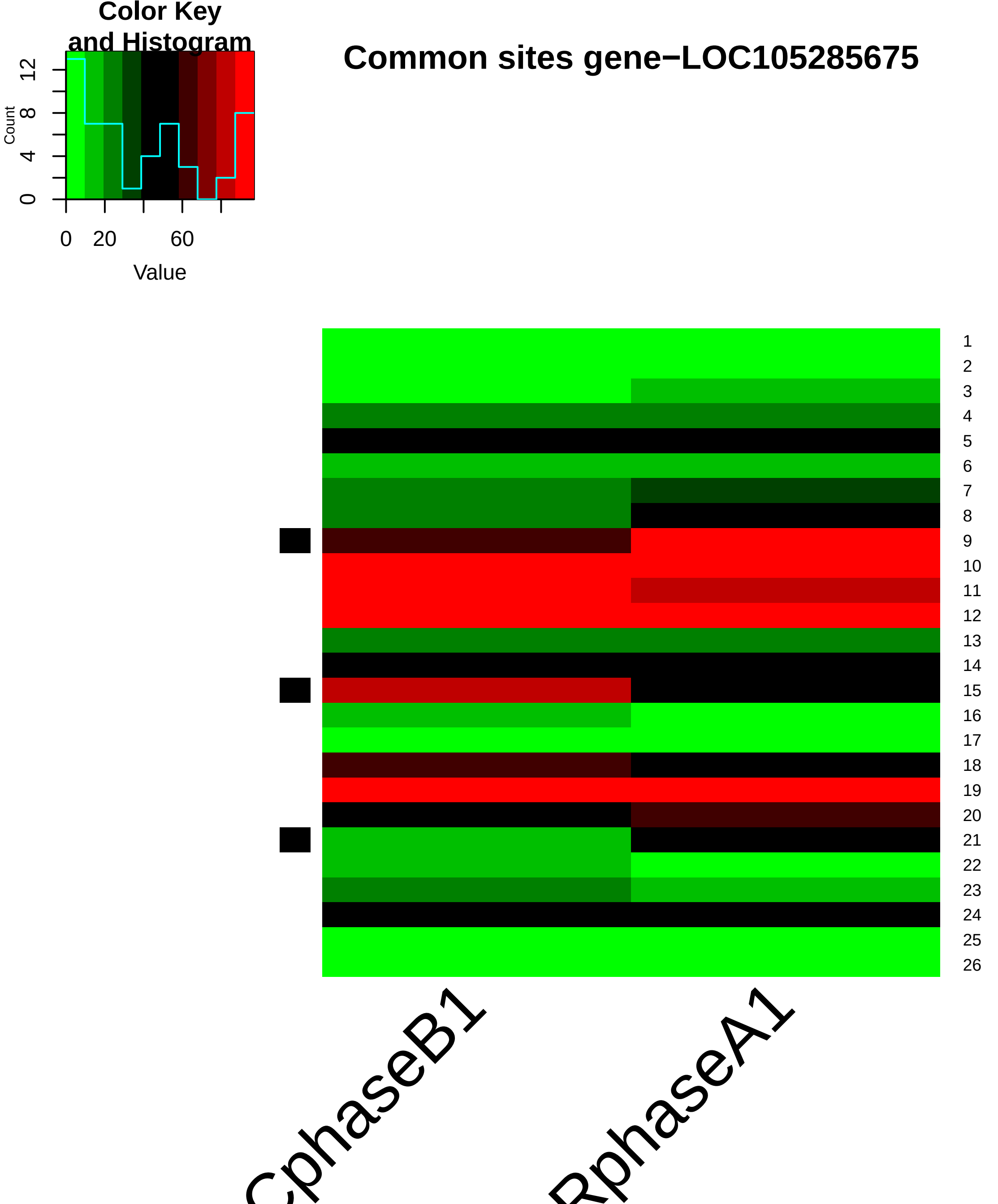
Color Key
and Histogram
Count
0
4
8
12
0
20
60
Value
Common sites gene−LOC105285675
CphaseB1
RphaseA1

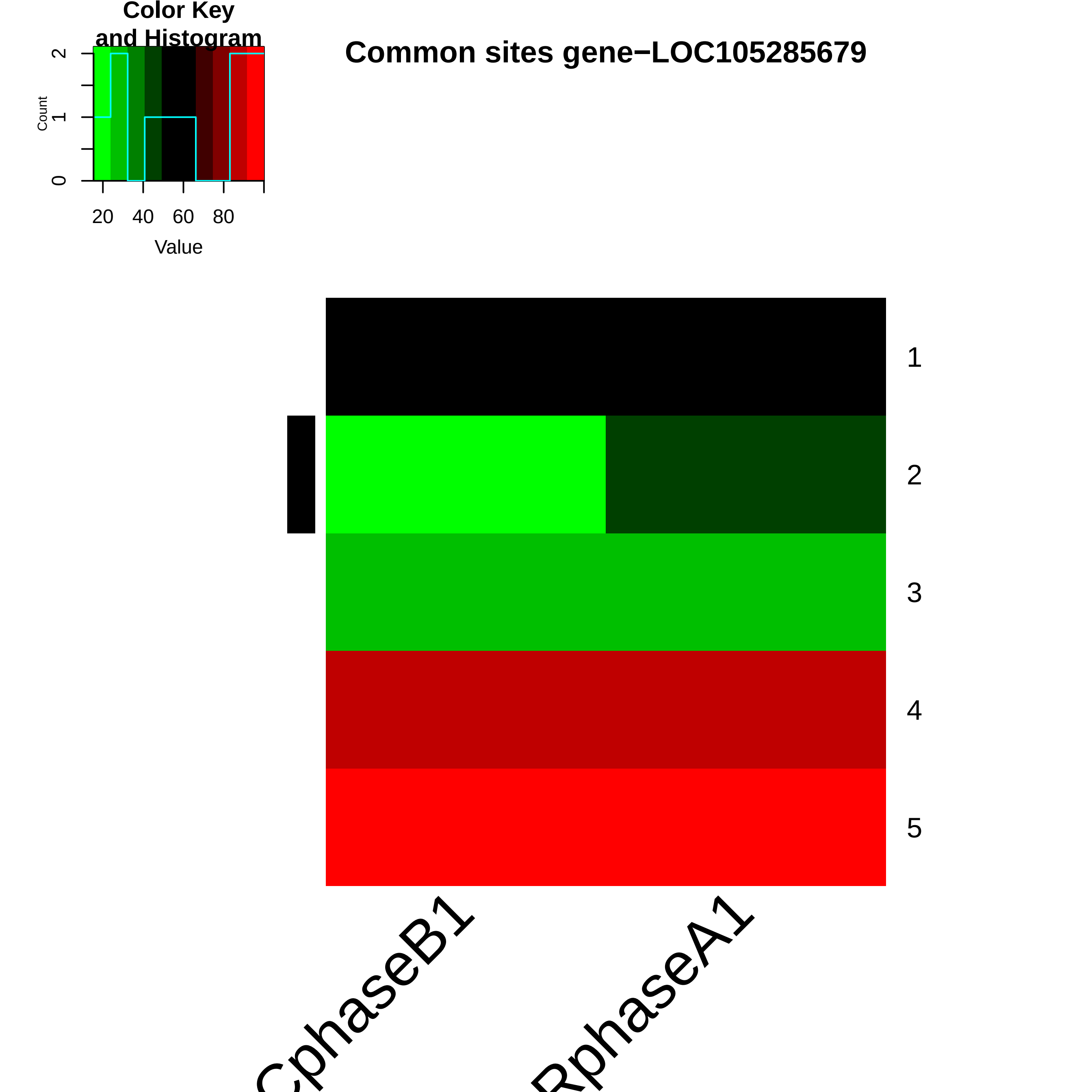
Color Key
and Histogram
Count
0
1
2
20
40
60
80
Value
Common sites gene−LOC105285679
1
2
3
4
5
CphaseB1
RphaseA1

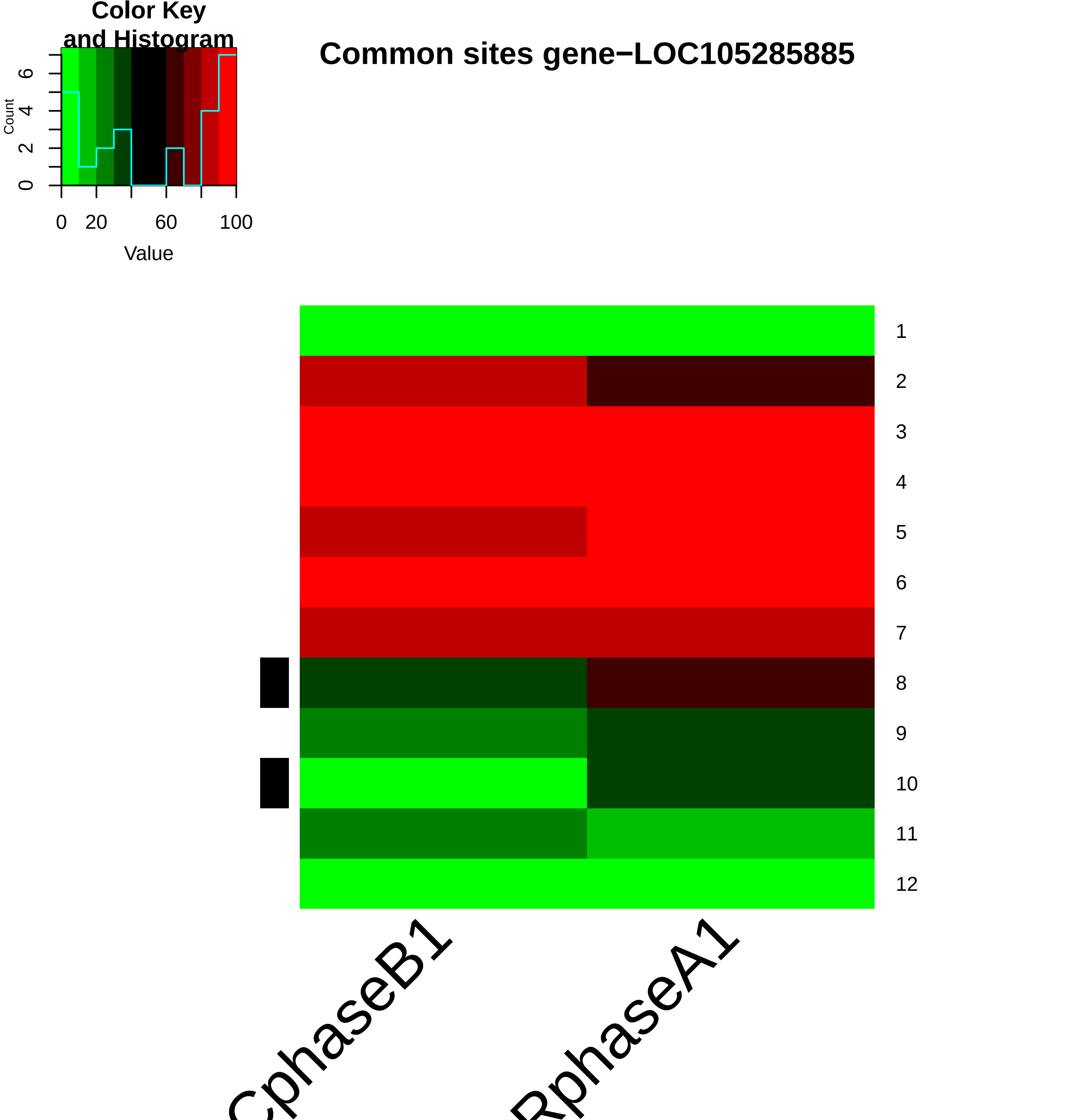

Color Key
and Histogram
Count
0
2
4
6
0
20
60
100
Value
Common sites gene−LOC105285885
1
2
3
4
5
6
7
8
9
10
11
12
CphaseB1
RphaseA1

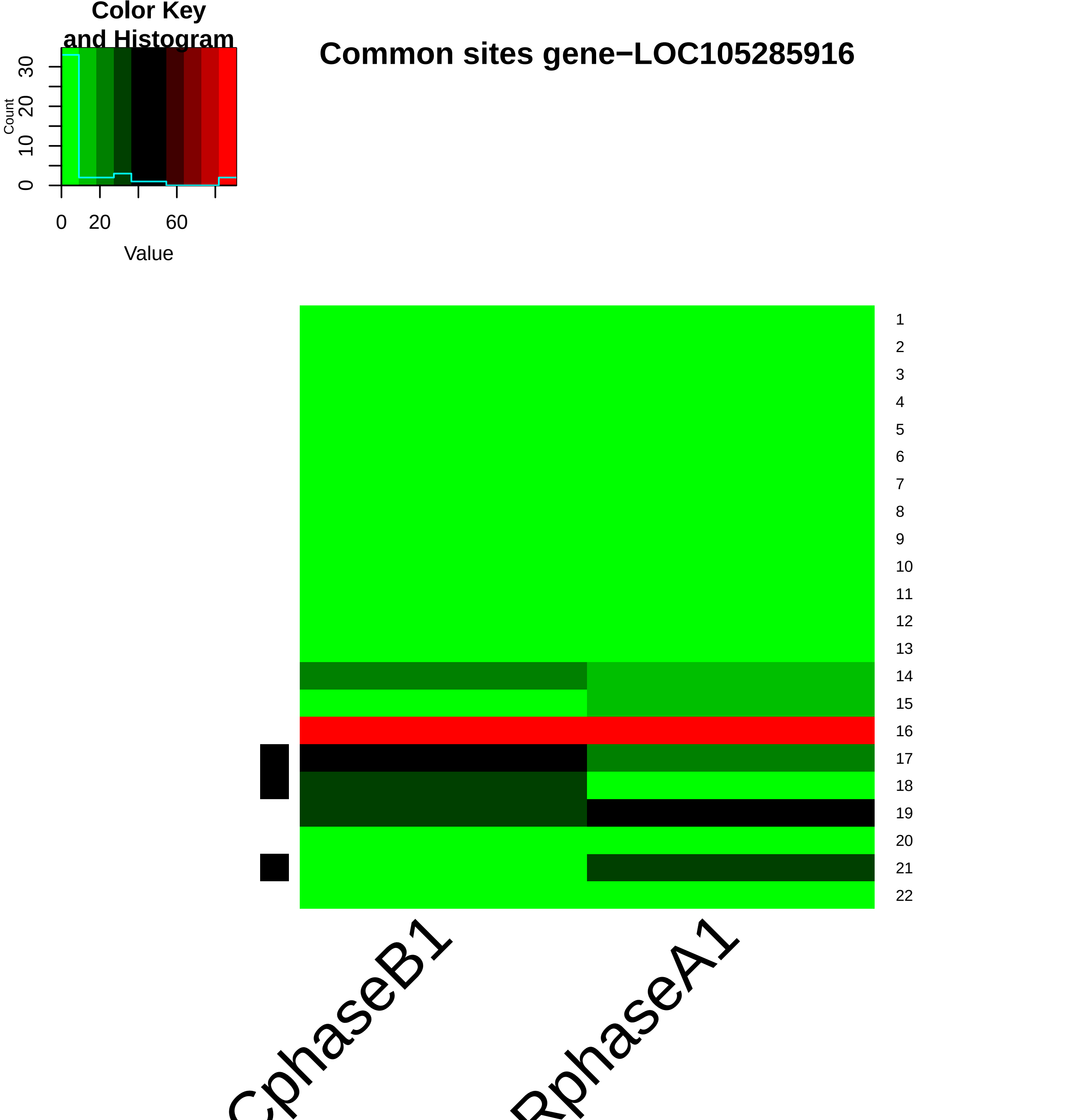

Color Key
and Histogram
Count
0
10
20
30
0
20
60
Value
Common sites gene−LOC105285916
CphaseB1
RphaseA1

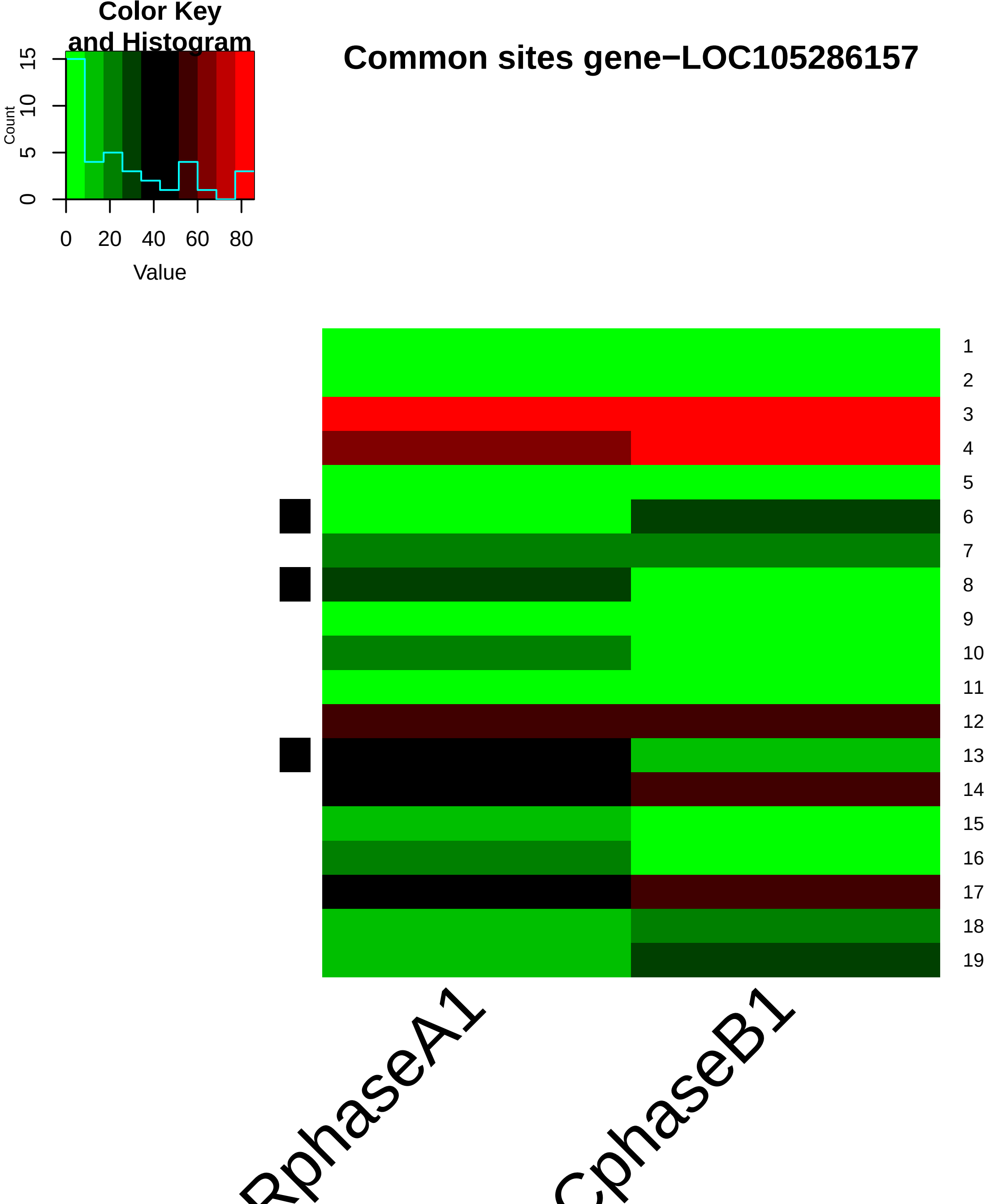
Color Key
and Histogram
Count
0
5
10
15
0
20
40
60
80
Value
Common sites gene−LOC105286157
RphaseA1
CphaseB1

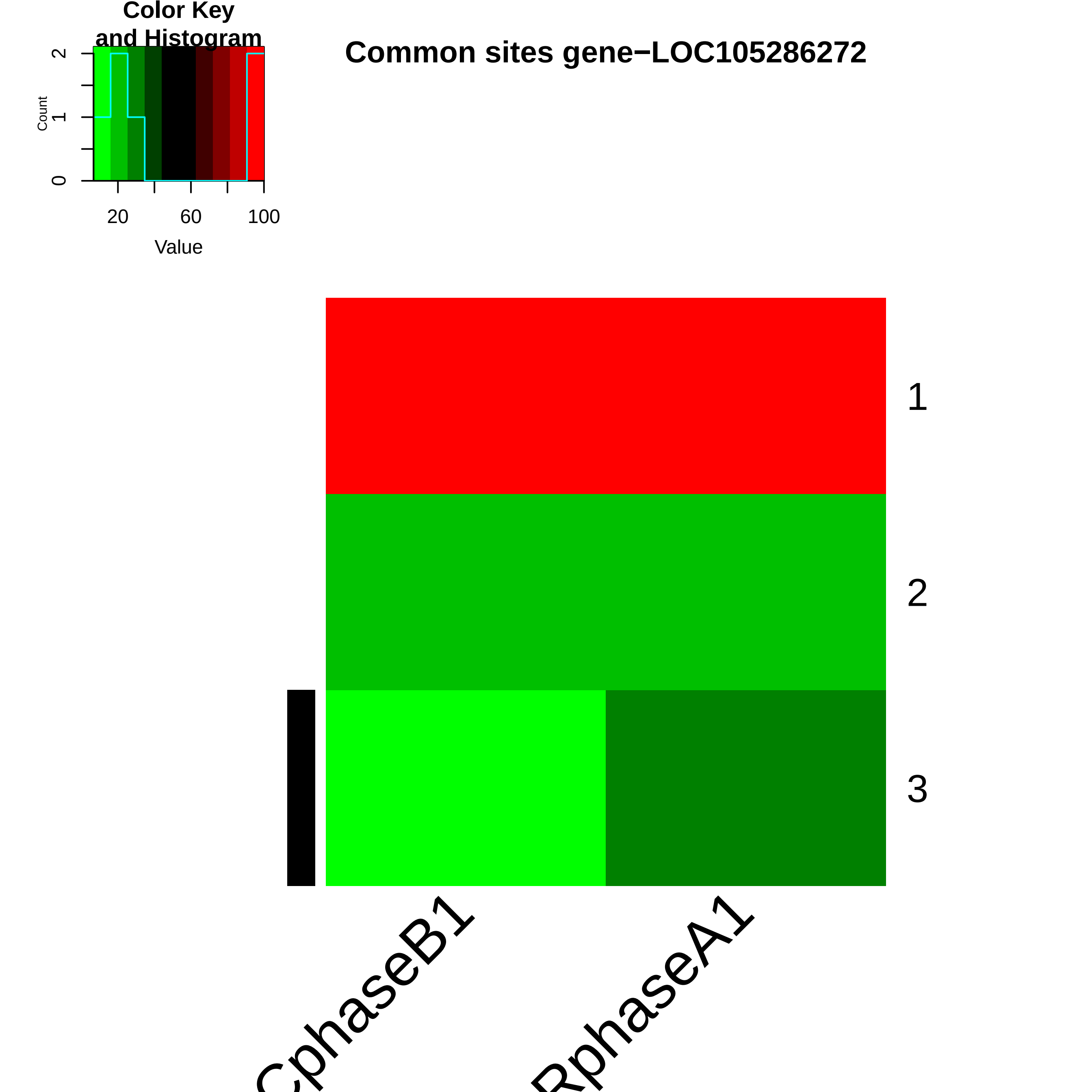

Color Key
and Histogram
Count
0
1
2
20
60
100
Value
Common sites gene−LOC105286272
1
2
3
CphaseB1
RphaseA1

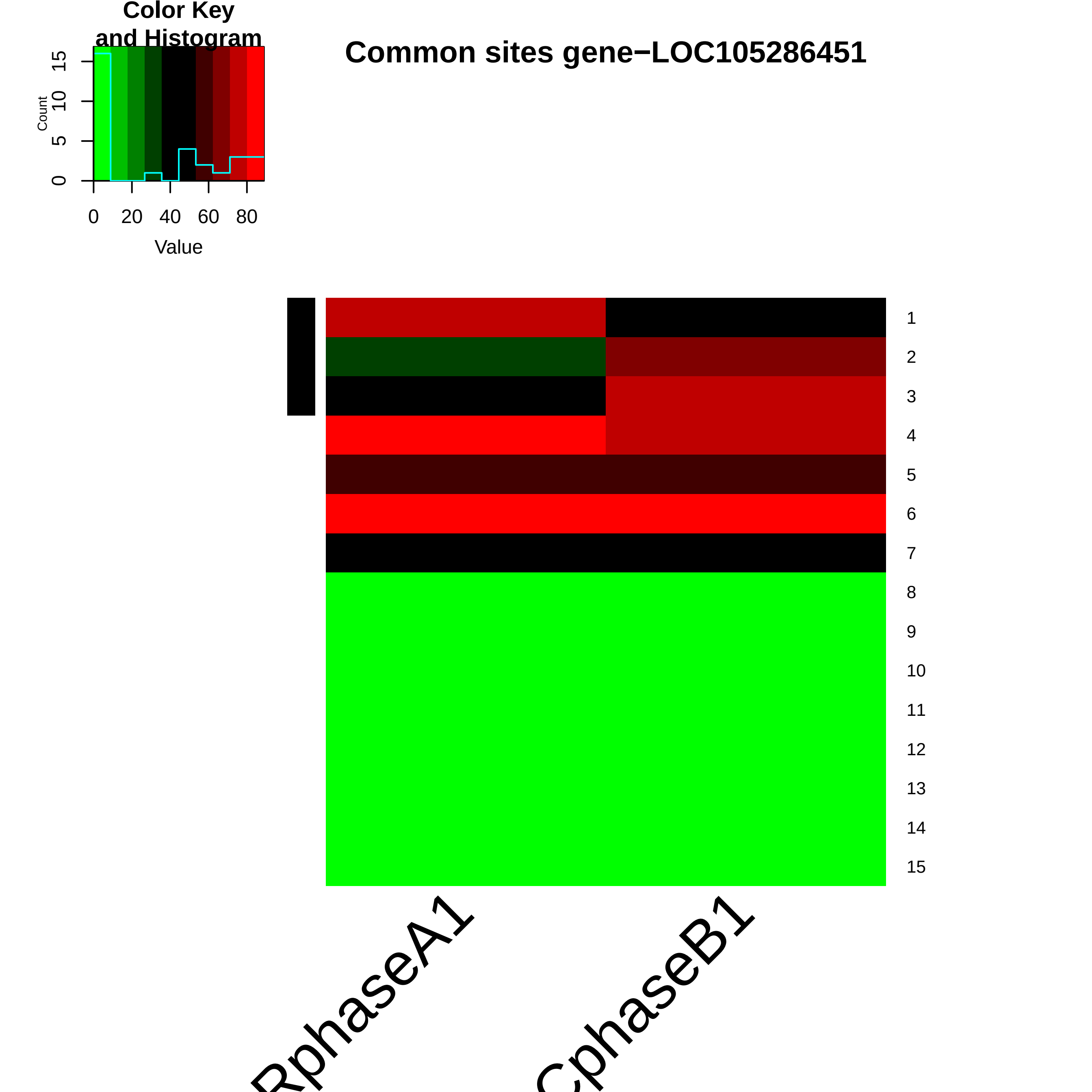
Color Key
and Histogram
Count
0
5
10
15
0
20
40
60
80
Value
Common sites gene−LOC105286451
RphaseA1
CphaseB1

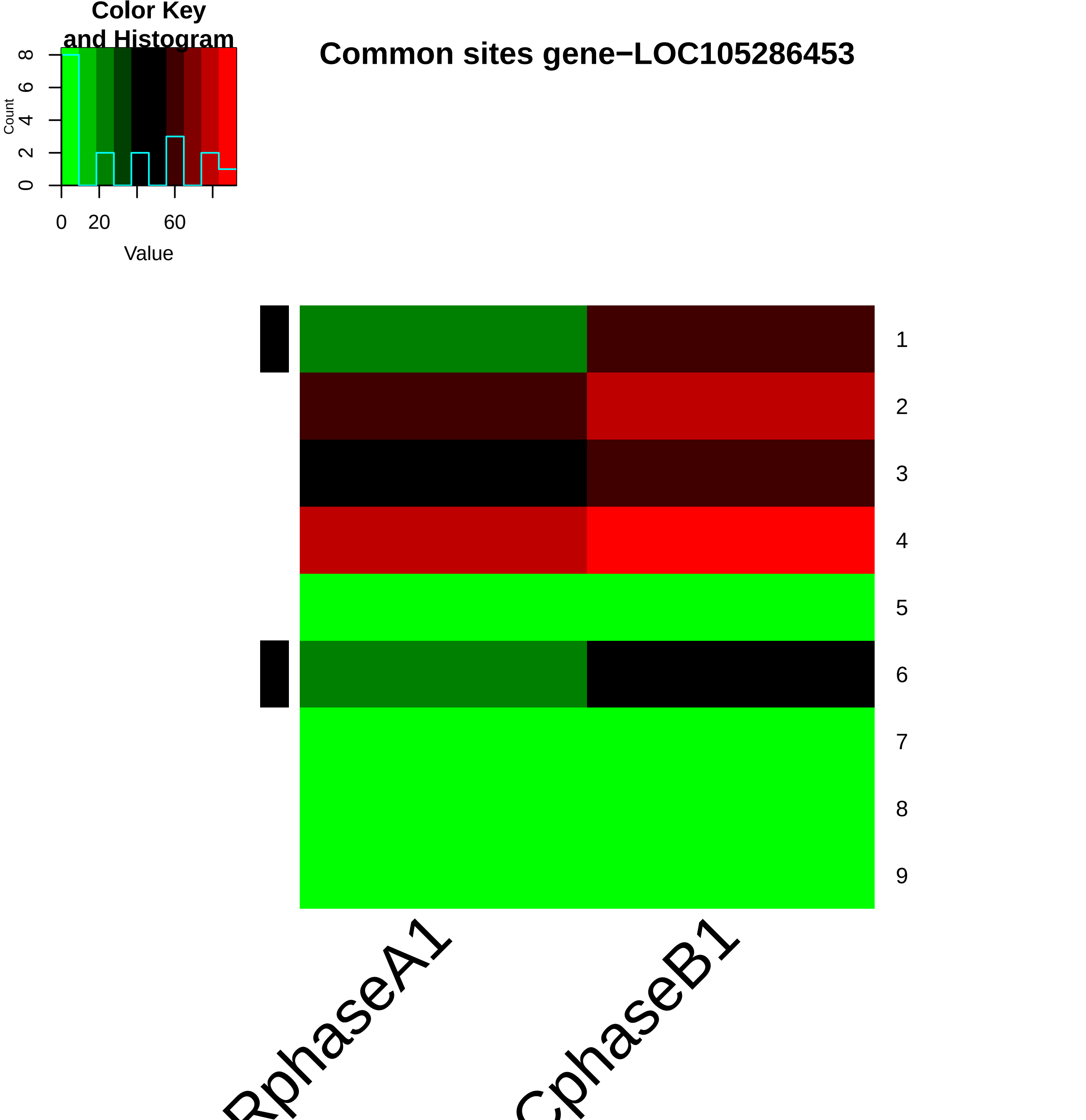

Color Key
and Histogram
Count
0
2
4
6
8
0
20
60
Value
Common sites gene−LOC105286453
1
2
3
4
5
6
7
8
9
RphaseA1
CphaseB1

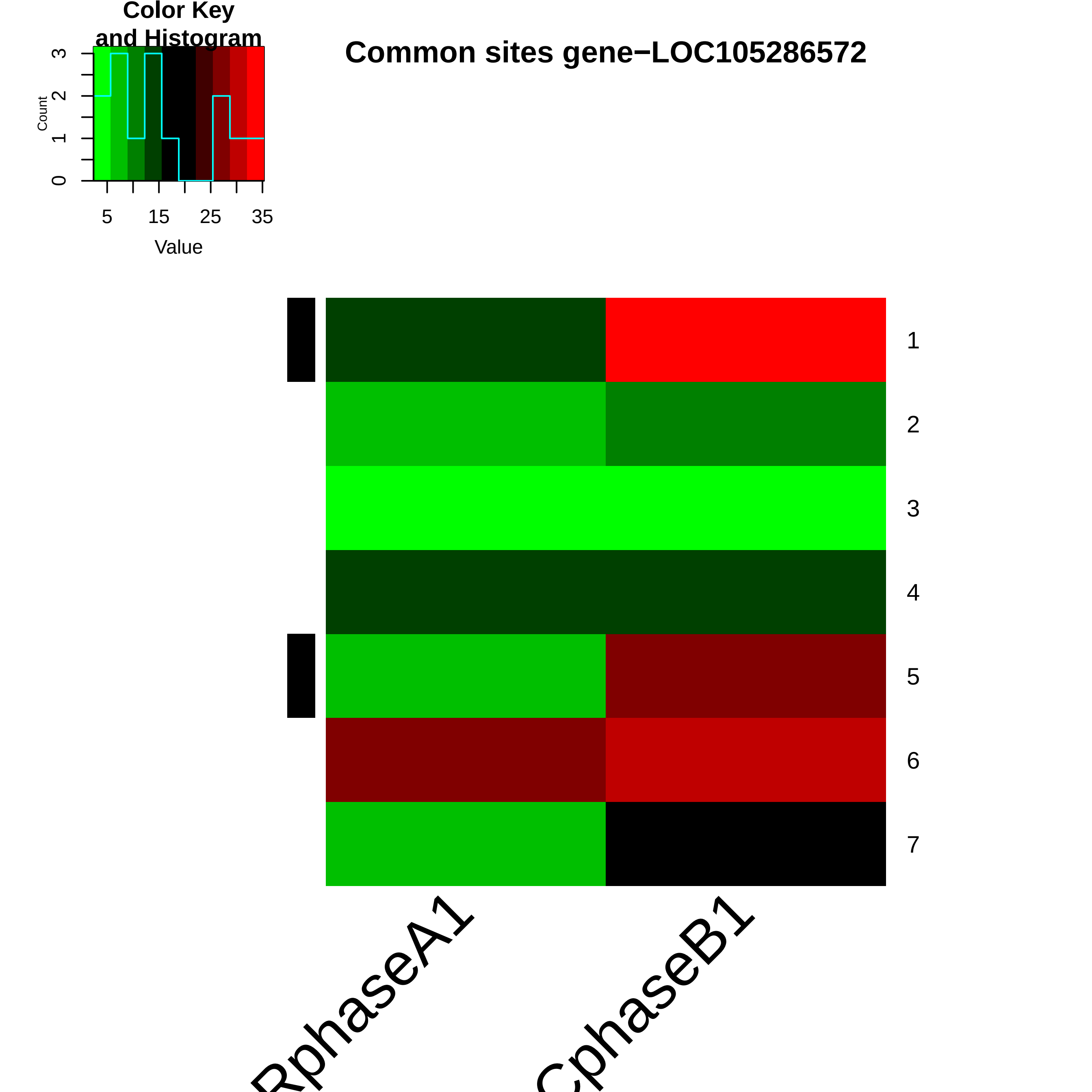
Color Key
and Histogram
Count
Value
Common sites gene−LOC105286572
1
2
3
4
5
6
7
RphaseA1
CphaseB1

#### Common sites gene−LOC105286627

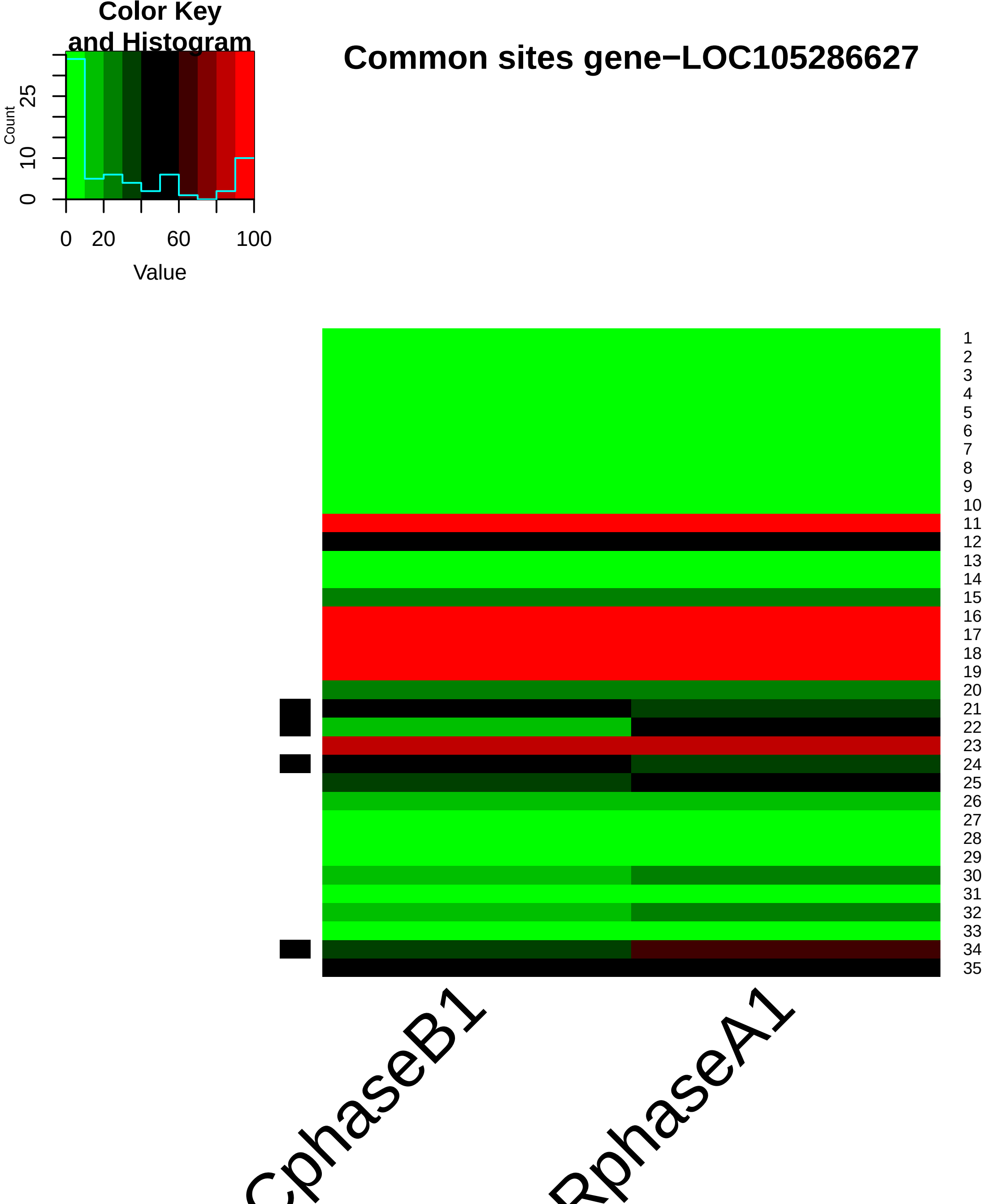

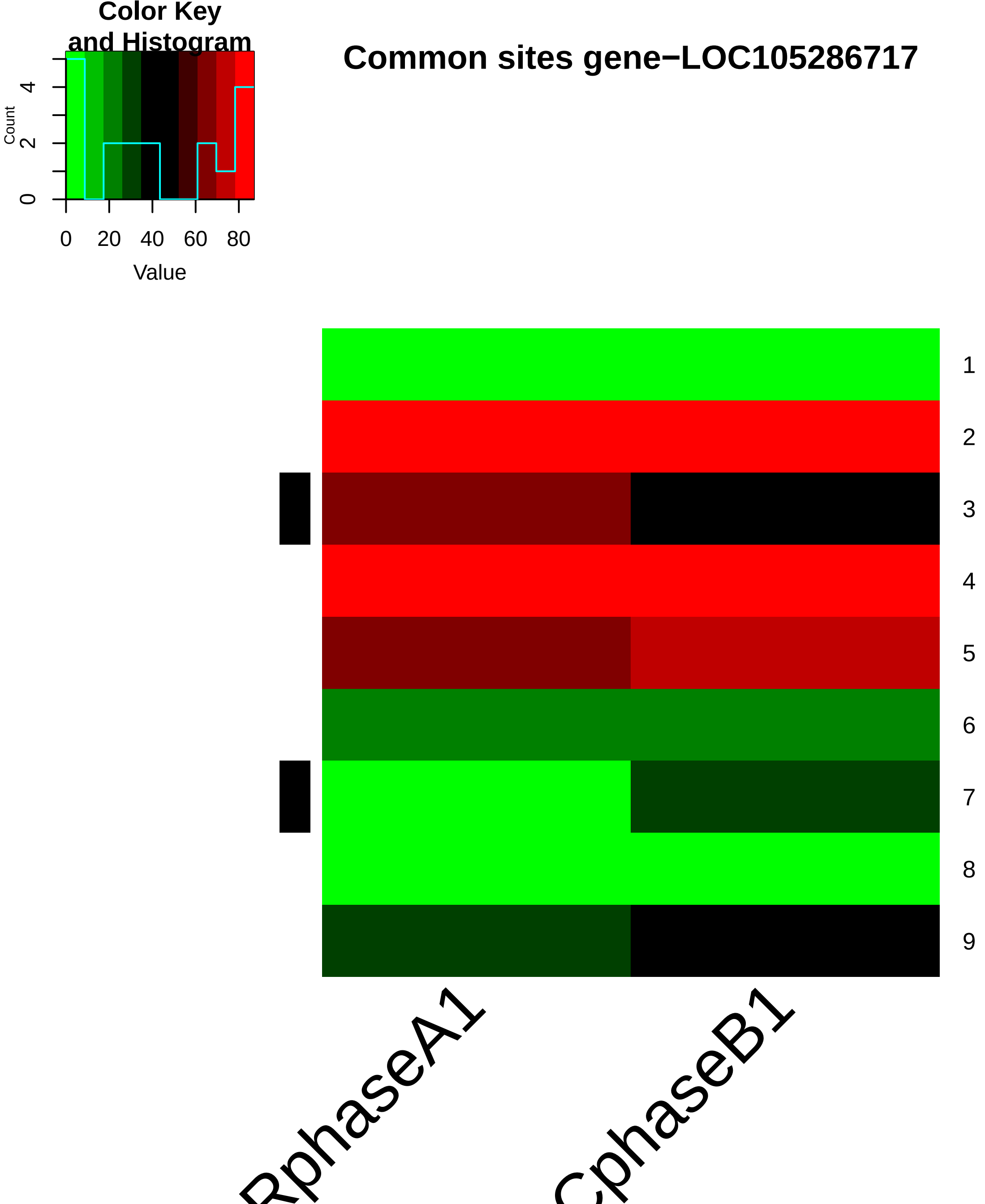
Color Key
and Histogram
Count
0
2
4
0
20
40
60
80
Value
Common sites gene−LOC105286717
1
2
3
4
5
6
7
8
9
RphaseA1
CphaseB1

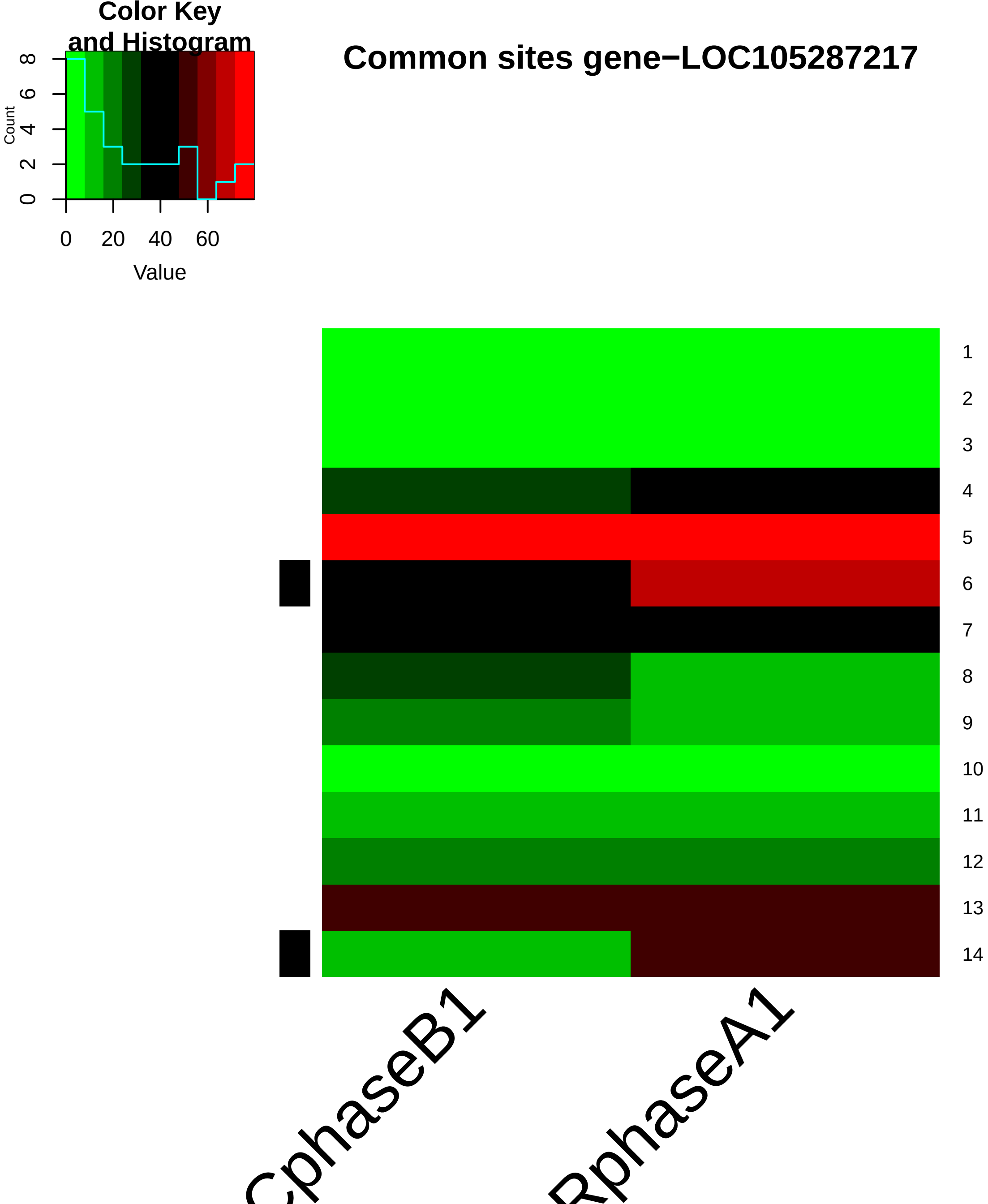

Color Key
and Histogram
Count
0
2
4
6
8
0
20
40
60
Value
Common sites gene−LOC105287217
1
2
3
4
5
6
7
8
9
10
11
12
13
14
CphaseB1
RphaseA1

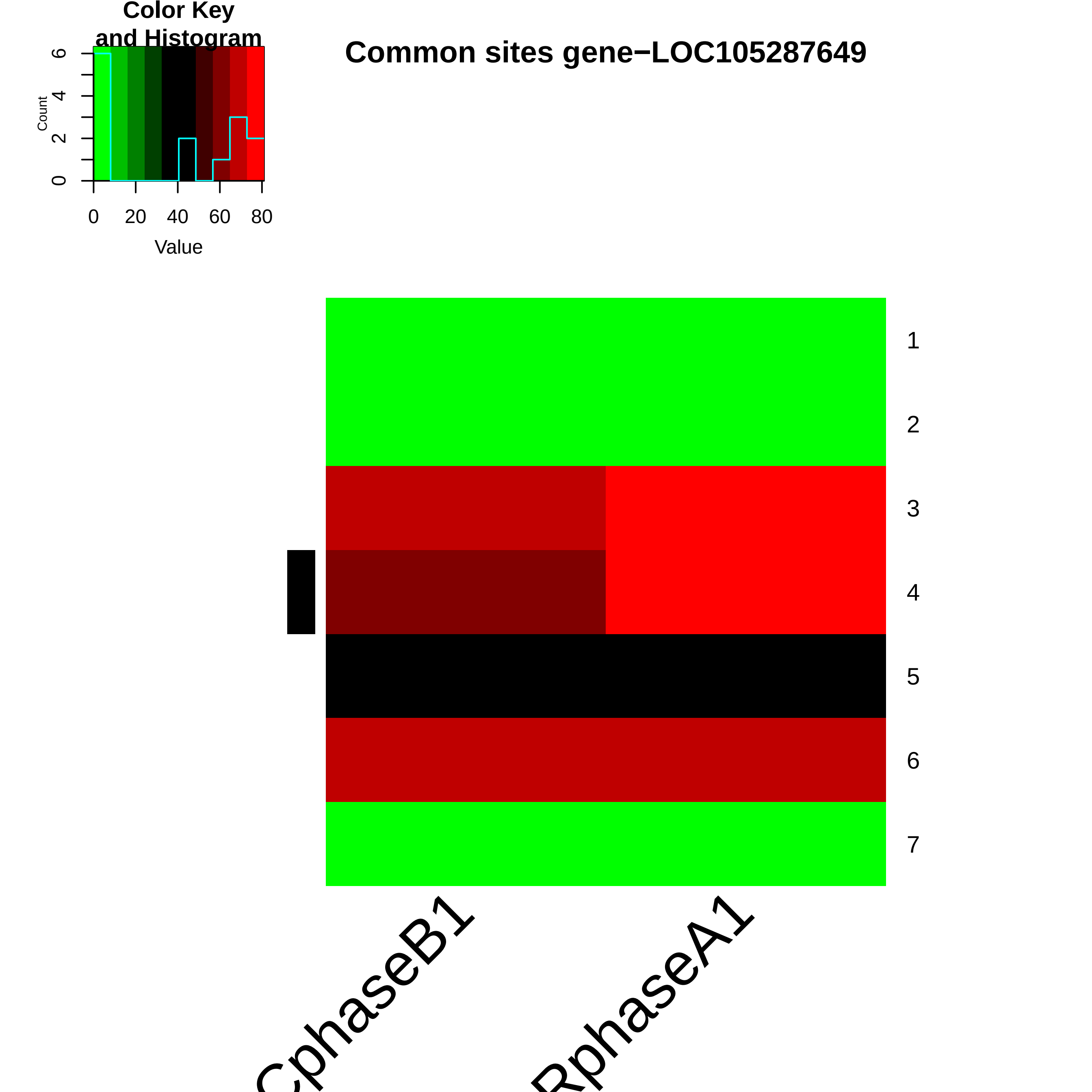

Color Key
and Histogram
Count
0
2
4
6
0
20
40
60
80
Value
Common sites gene−LOC105287649
1
2
3
4
5
6
7
CphaseB1
RphaseA1

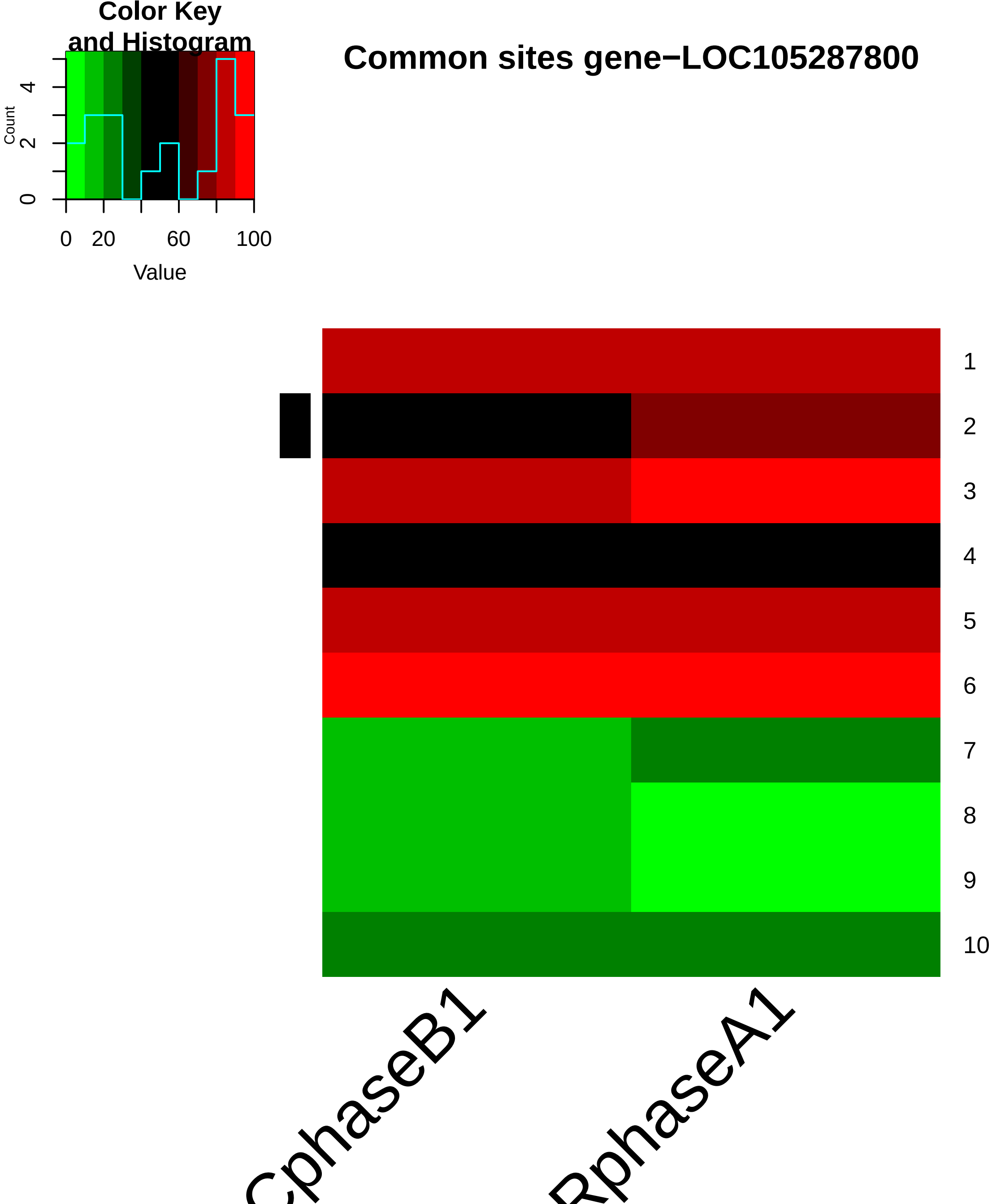

Color Key and Histogram
Count
0
2
4
0
20
60
100
Value
Common sites gene−LOC105287800
1
2
3
4
5
6
7
8
9
10

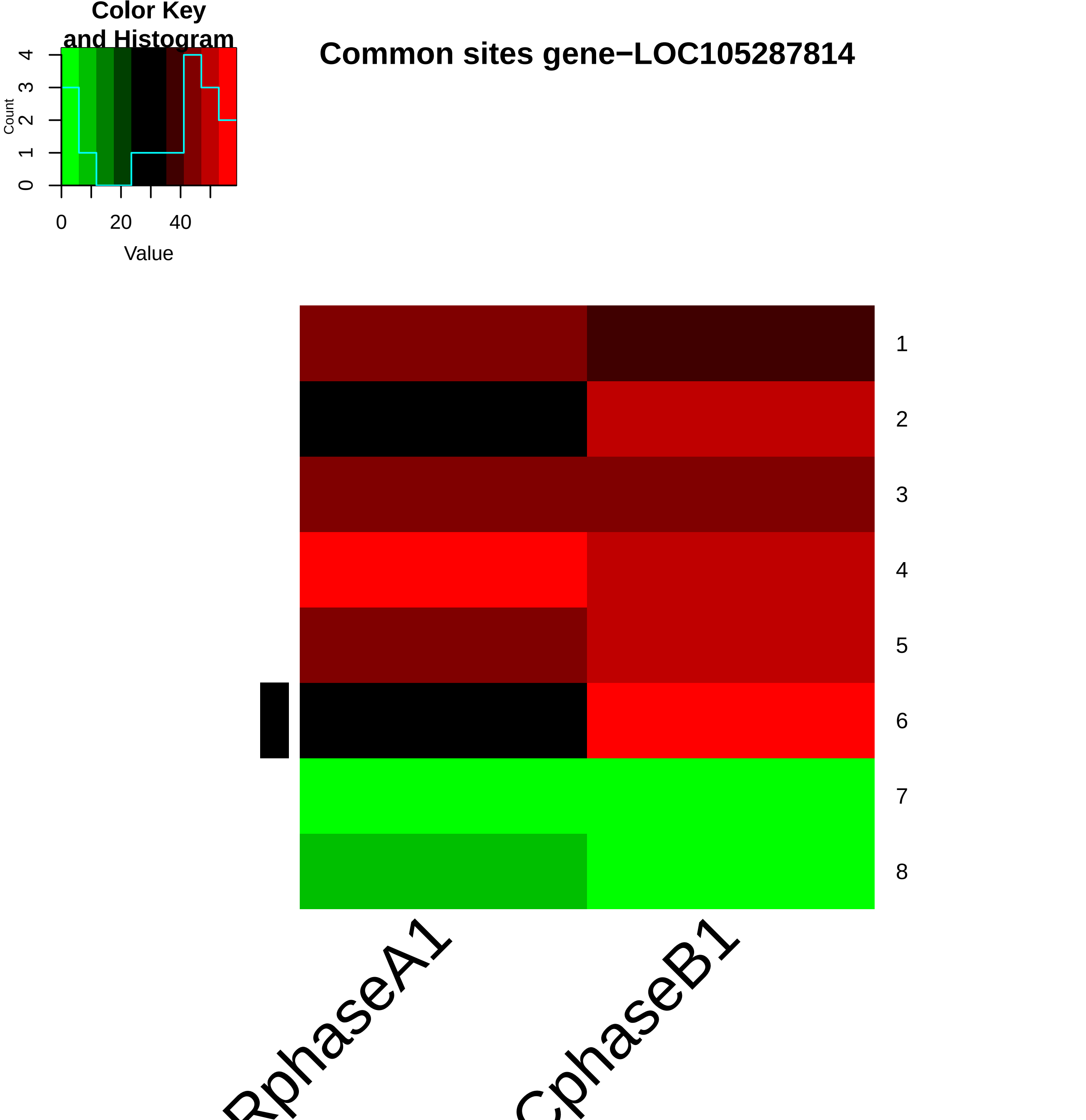
Color Key
and Histogram
Count
0
1
2
3
4
0
20
40
Value
Common sites gene−LOC105287814
1
2
3
4
5
6
7
8
RphaseA1
CphaseB1

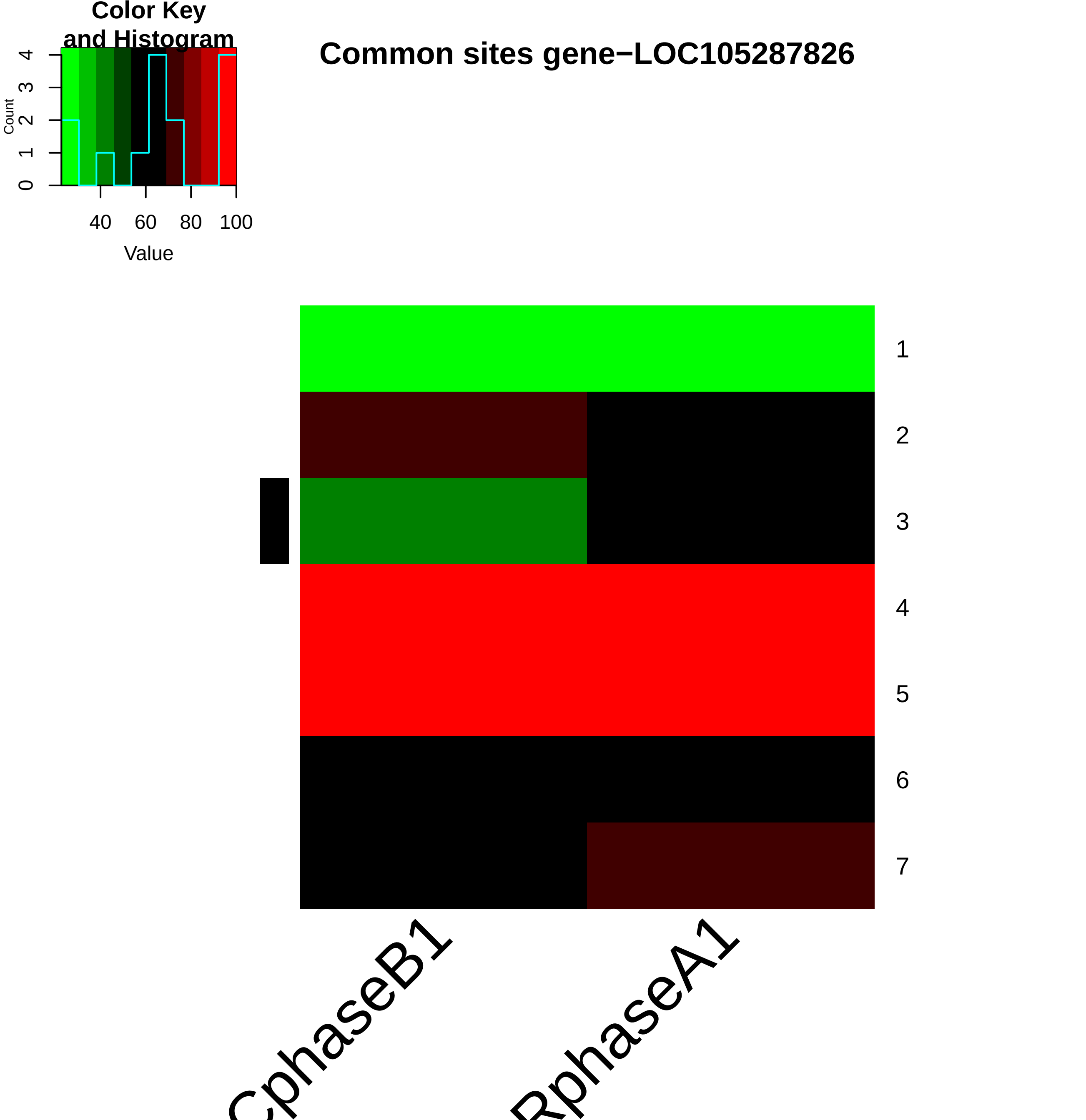
Color Key
and Histogram
Count
0
1
2
3
4
40
60
80
100
Value
Common sites gene−LOC105287826
1
2
3
4
5
6
7
CphaseB1
RphaseA1

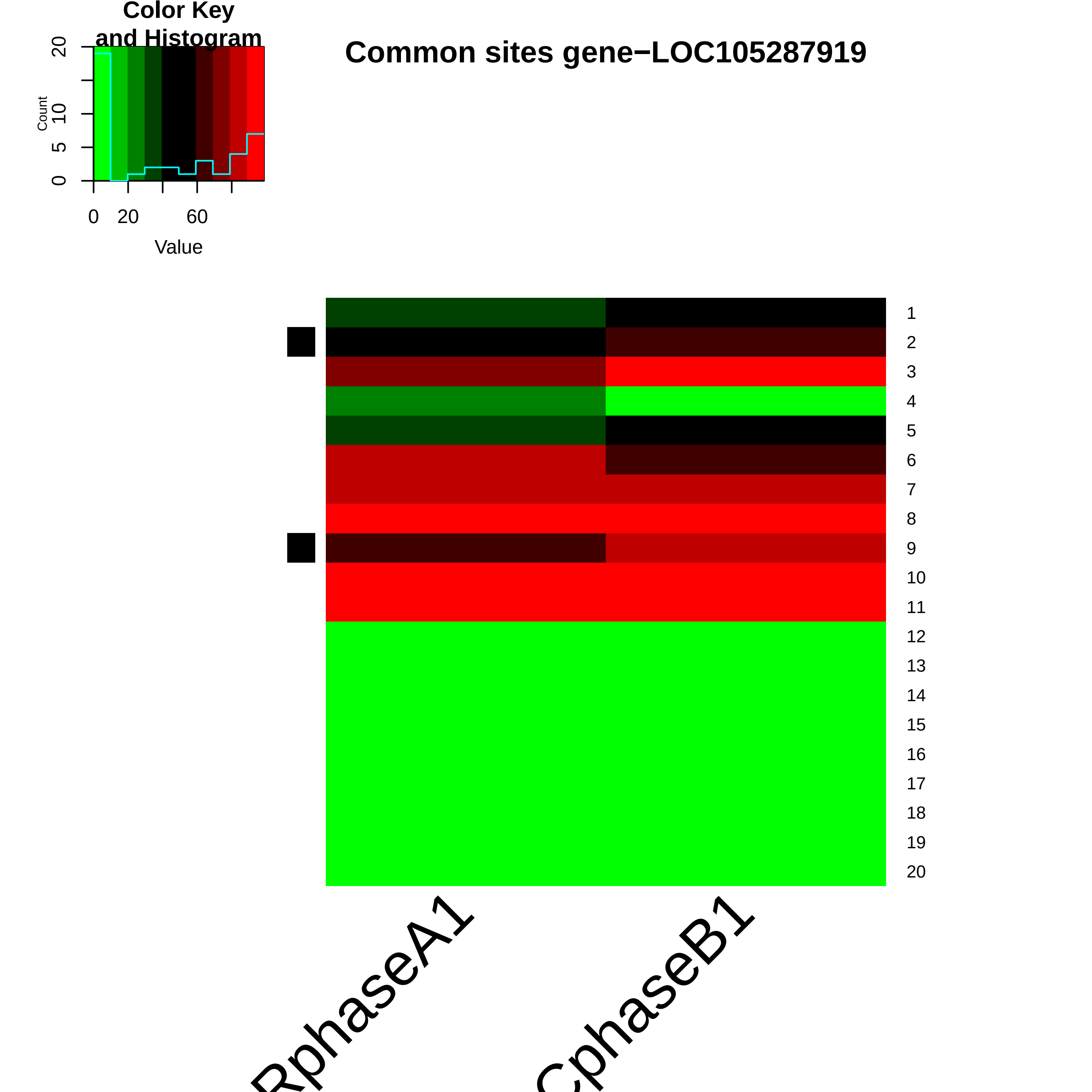
Color Key
and Histogram
Count
Value
0
20
60
0
5
10
20
Common sites gene−LOC105287919
RphaseA1
CphaseB1

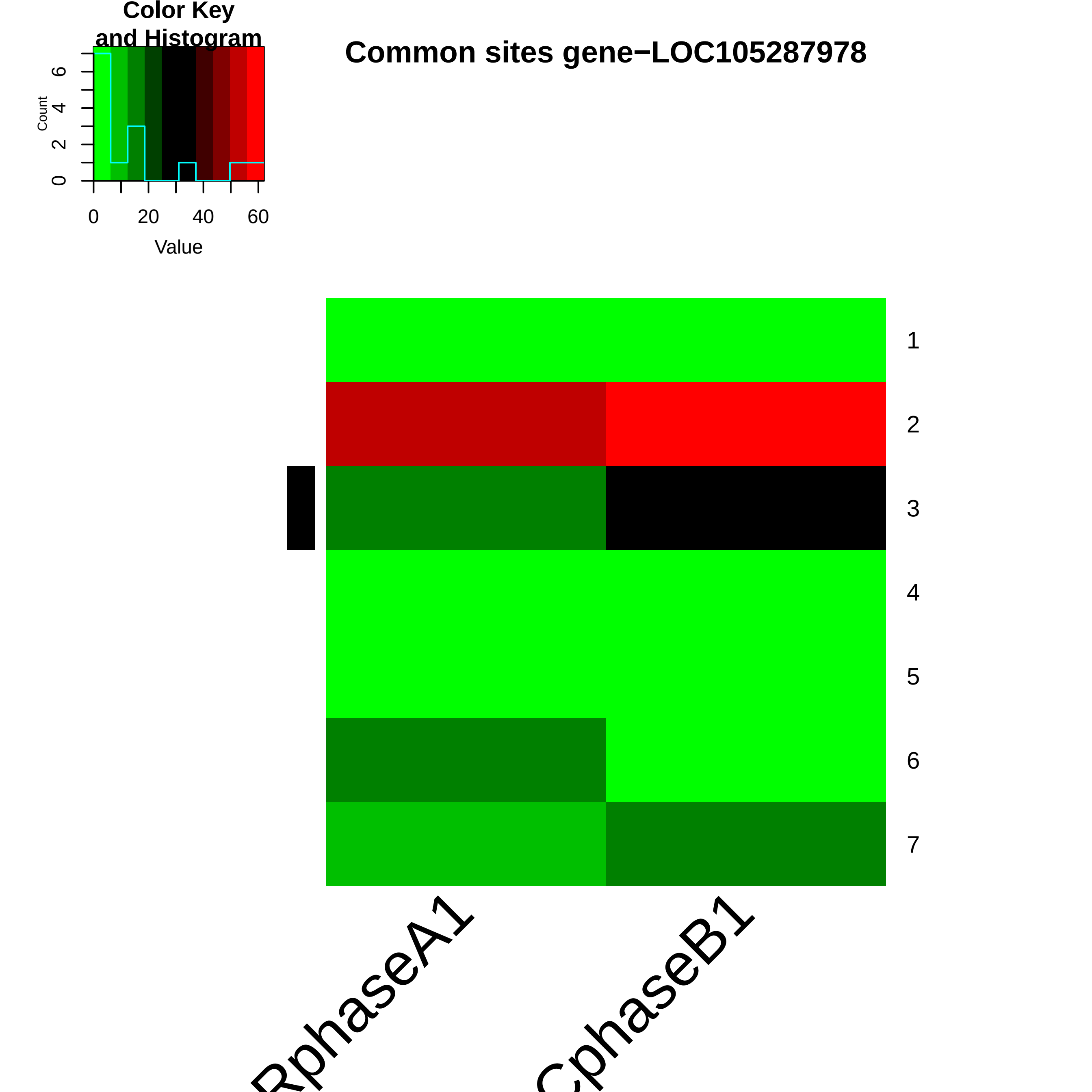
Color Key
and Histogram
Count
0
2
4
6
0
20
40
60
Value
Common sites gene−LOC105287978
1
2
3
4
5
6
7
RphaseA1
CphaseB1

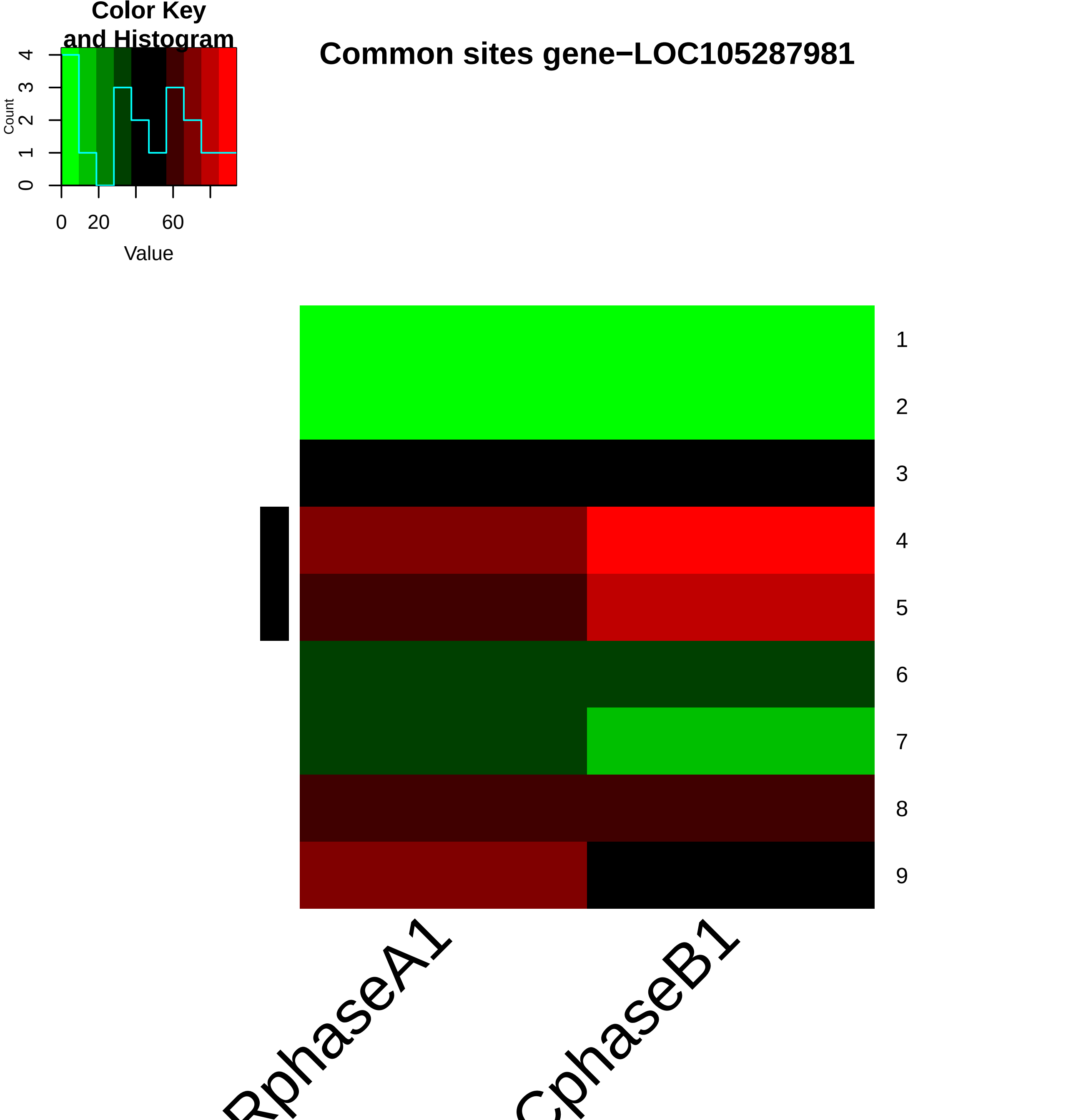

Color Key
and Histogram
Count
0
1
2
3
4
0
20
60
Value
Common sites gene−LOC105287981
1
2
3
4
5
6
7
8
9
RphaseA1
CphaseB1

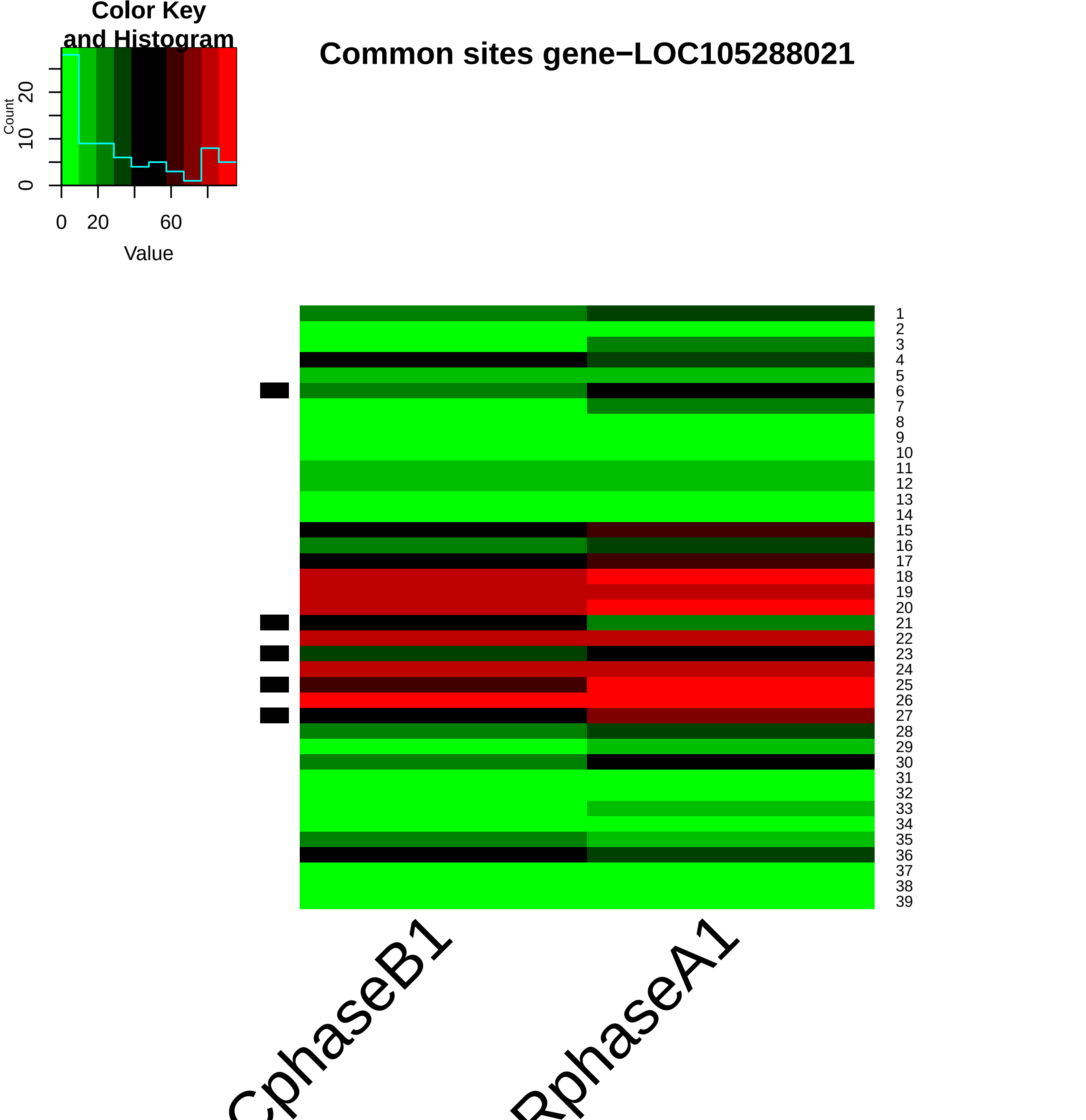

Color Key
and Histogram
Count
0
10
20
0
20
60
Value
Common sites gene−LOC105288021
CphaseB1
RphaseA1

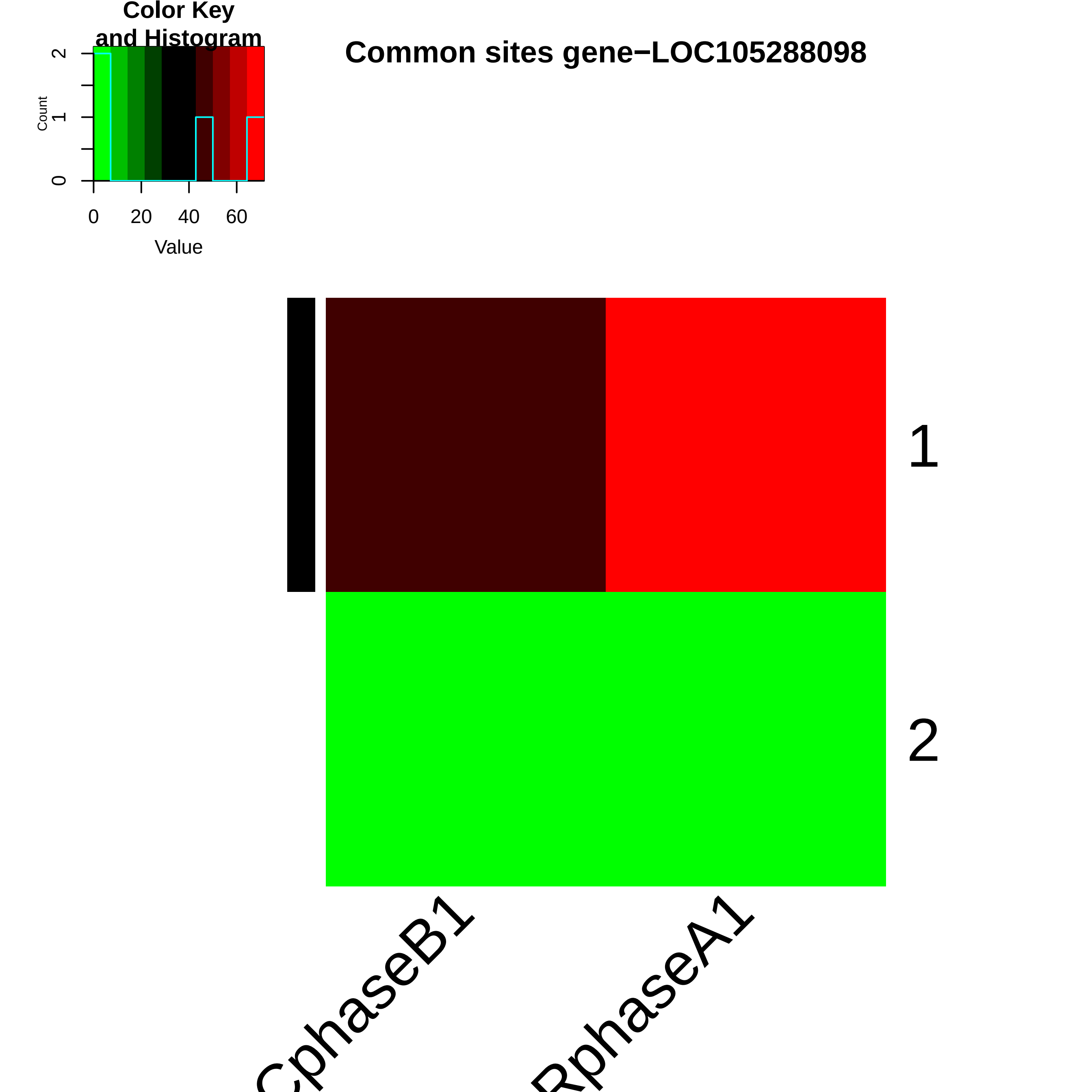

Color Key
and Histogram
Count
0
1
2
0
20
40
60
Value
Common sites gene−LOC105288098
1
2
CphaseB1
RphaseA1

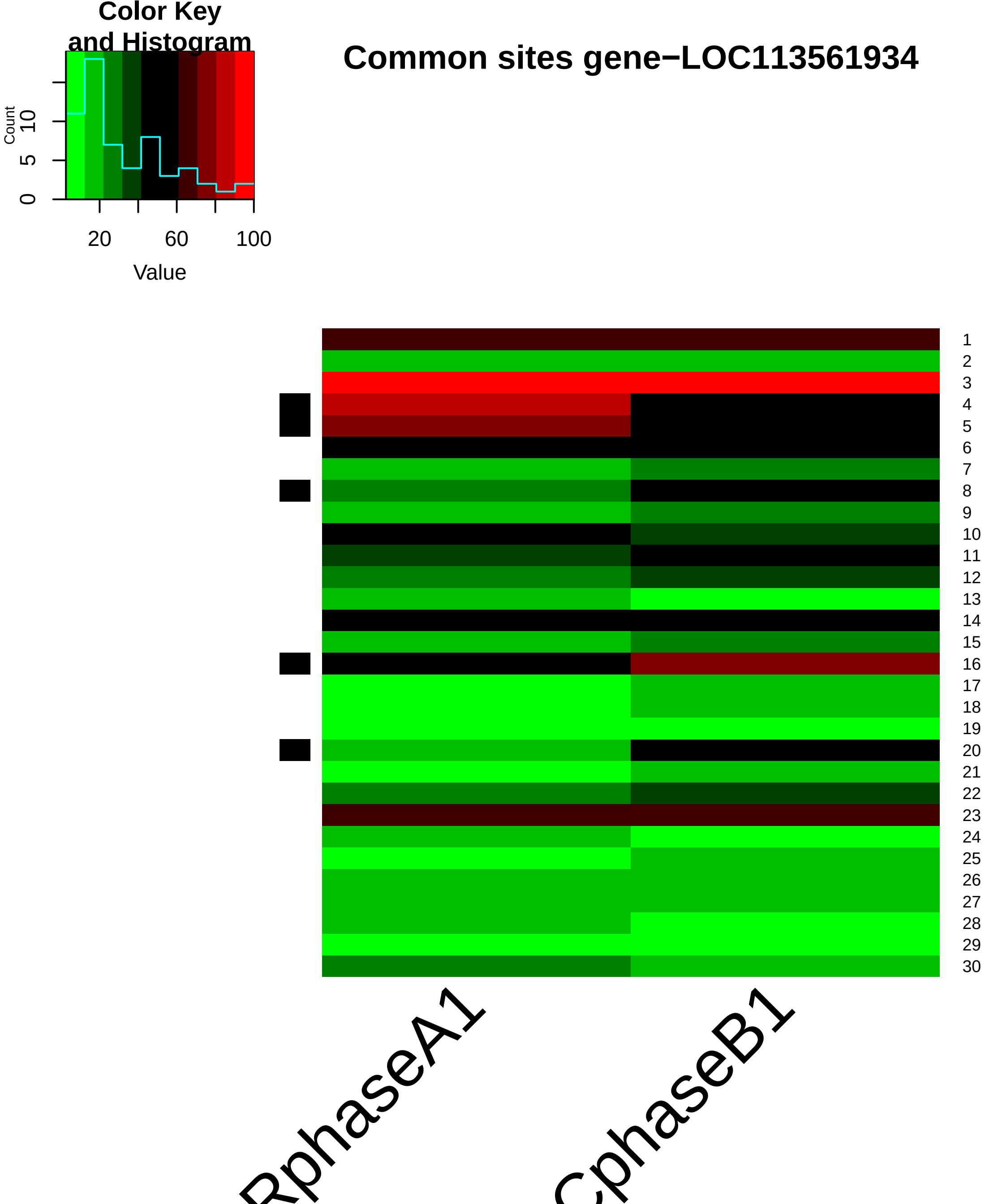

Color Key
and Histogram
Count
0
5
10
20
60
100
Value
Common sites gene−LOC113561934
RphaseA1
CphaseB1

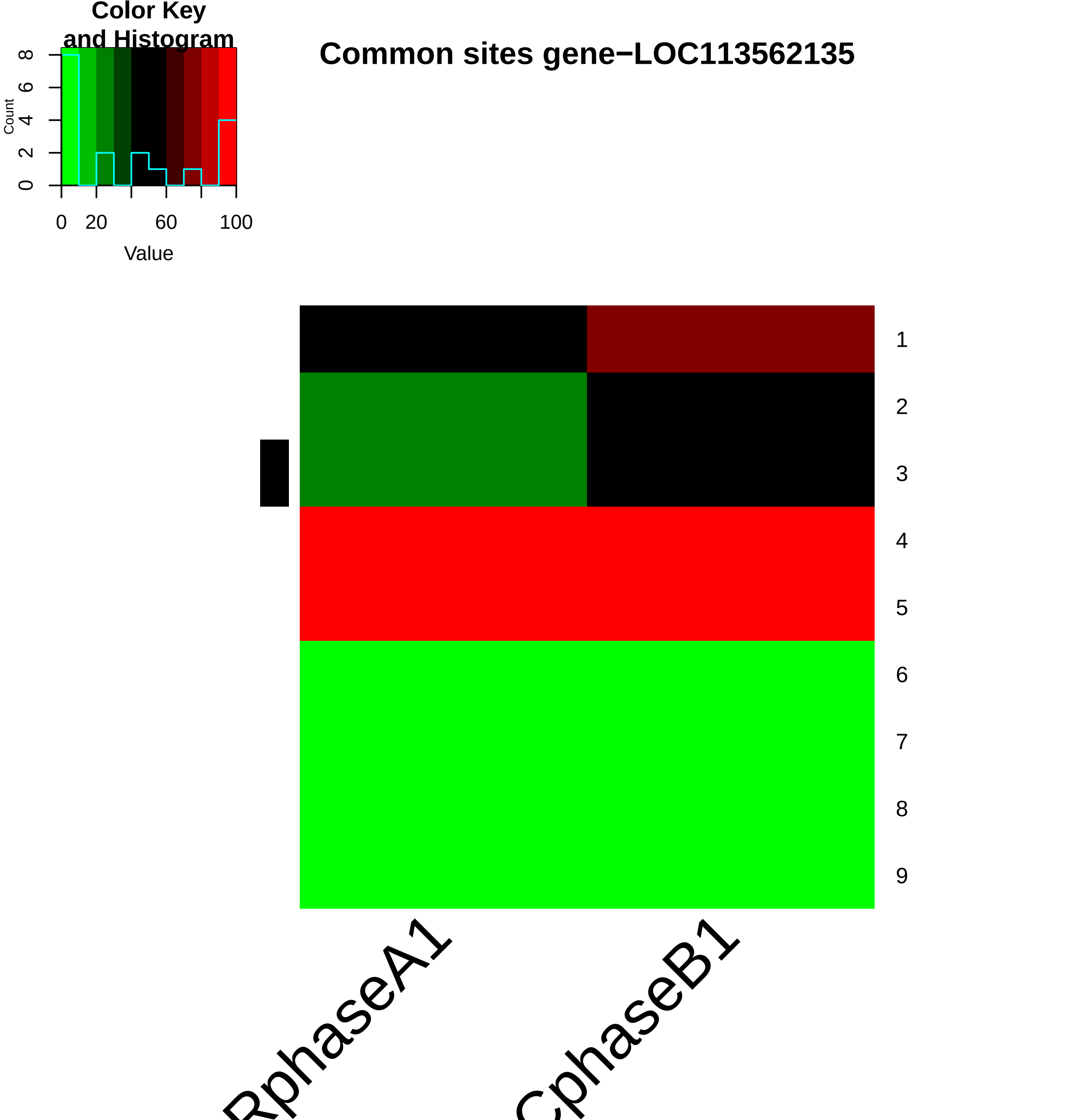

Color Key
and Histogram
Count
0
2
4
6
8
0
20
60
100
Value
Common sites gene−LOC113562135
1
2
3
4
5
6
7
8
9
RphaseA1
CphaseB1

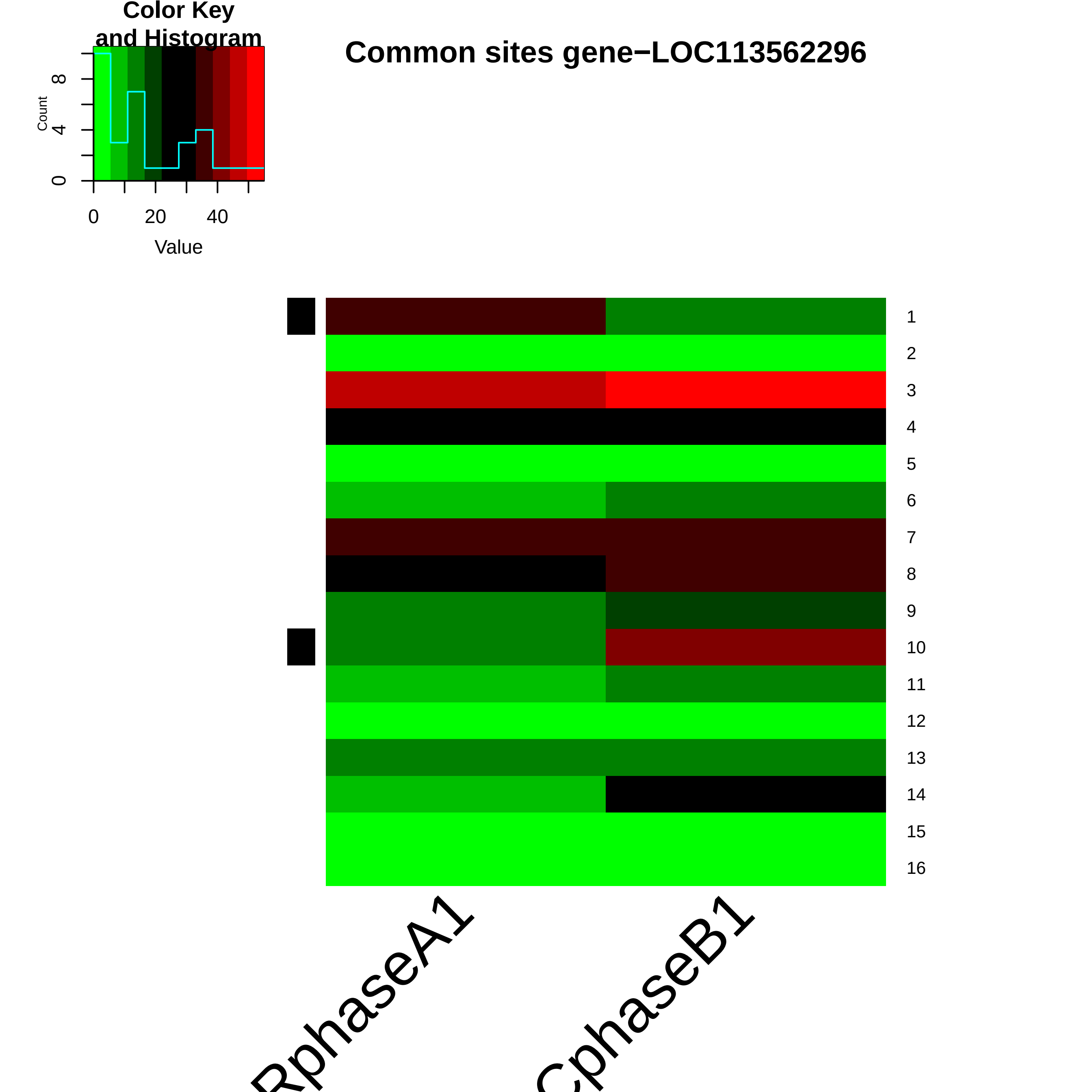
Color Key
and Histogram
Count
0
4
8
0
20
40
Value
Common sites gene−LOC113562296
RphaseA1
CphaseB1

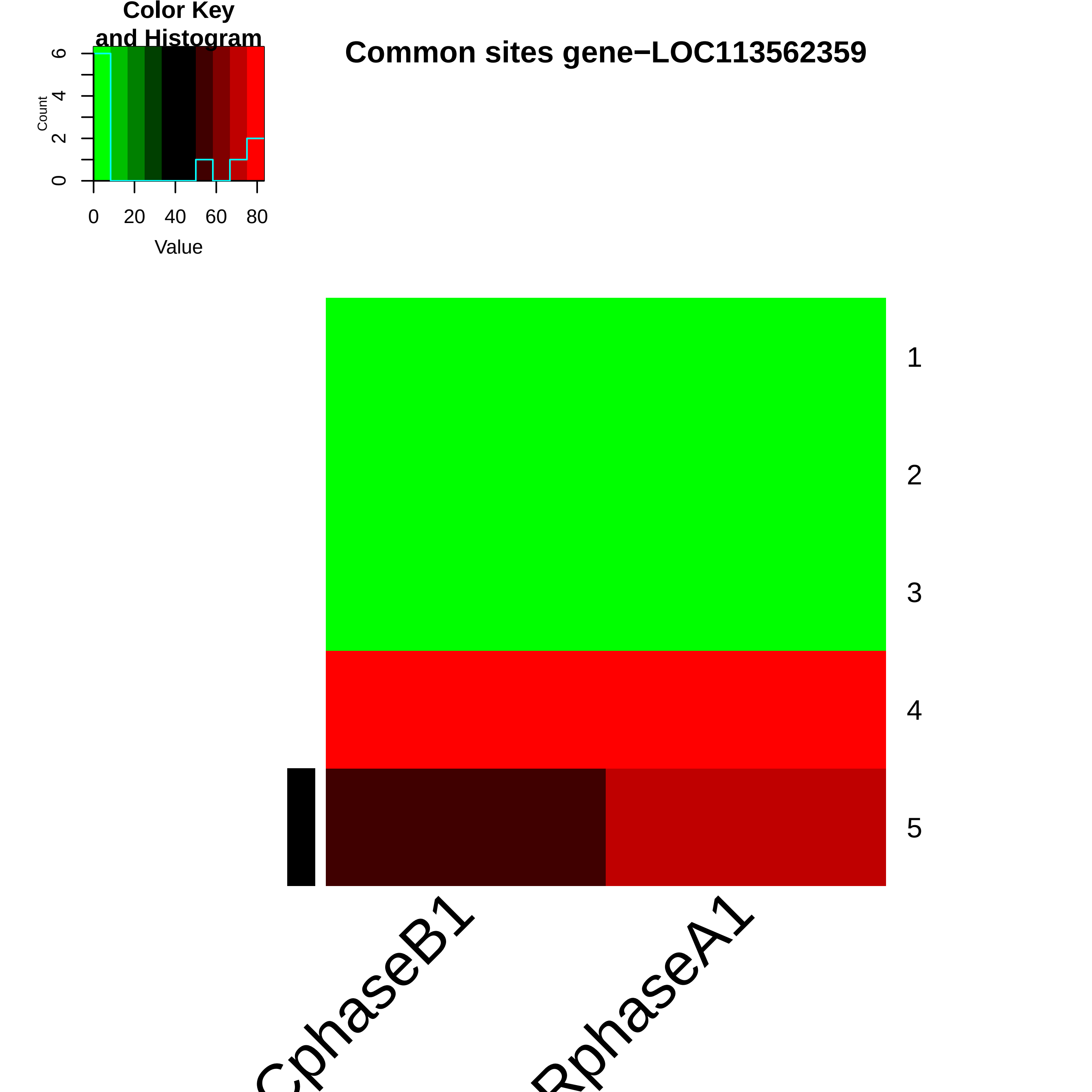

Color Key
and Histogram
Count
Value
0
20
40
60
80
0
2
4
6
Common sites gene−LOC113562359
1
2
3
4
5
CphaseB1
RphaseA1

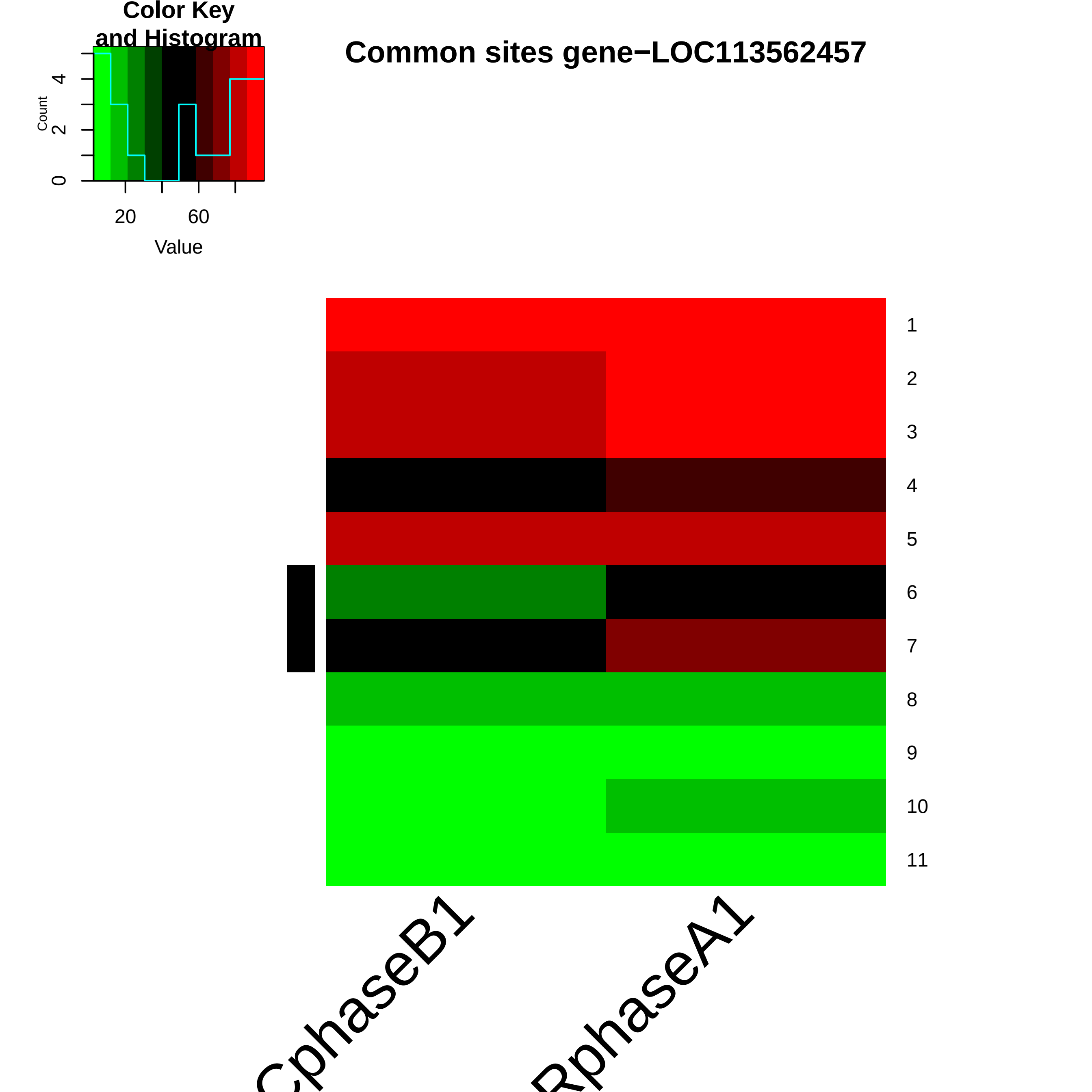

Color Key
and Histogram
Count
0
2
4
20
60
Value
Common sites gene−LOC113562457
1
2
3
4
5
6
7
8
9
10
11
CphaseB1
RphaseA1

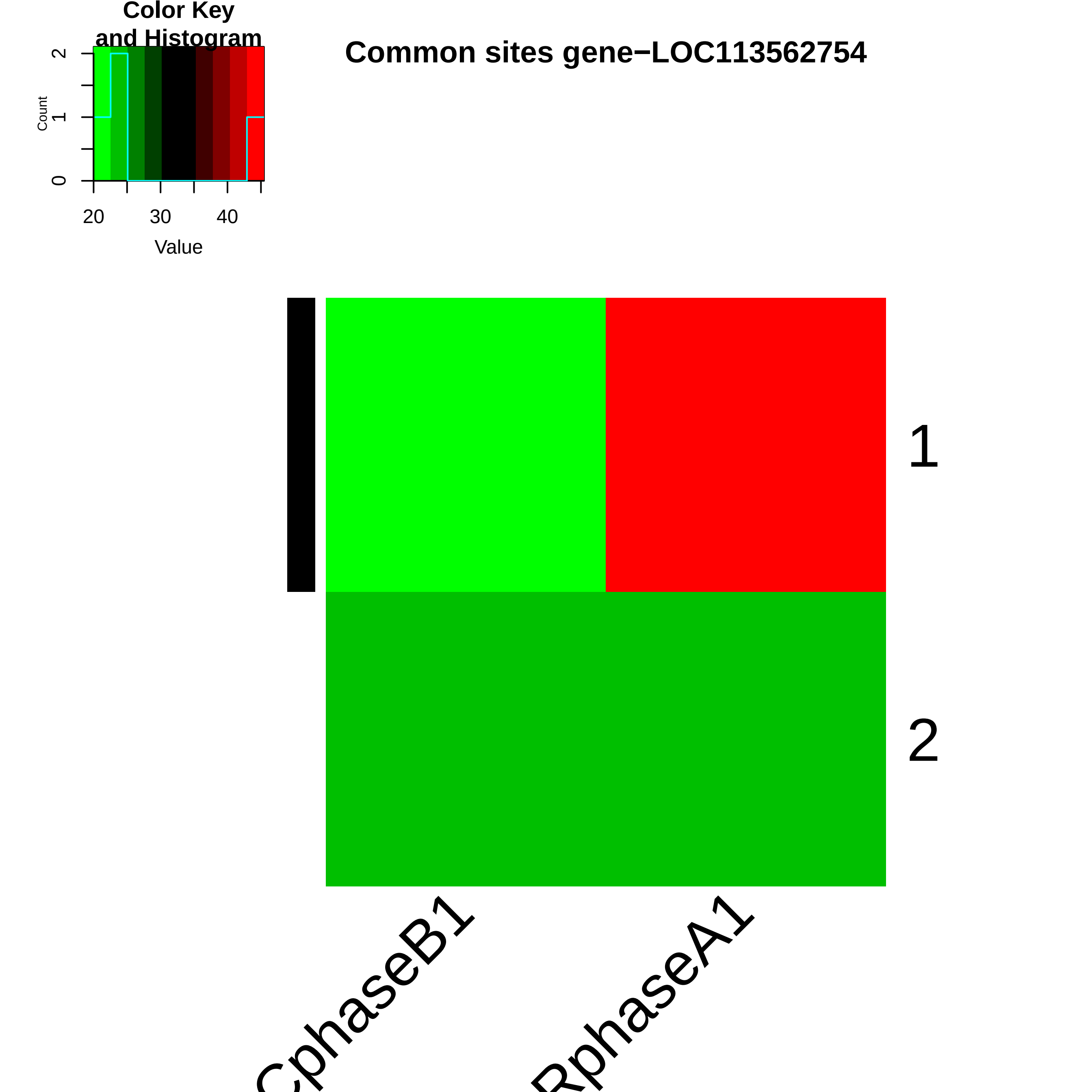

Color Key
and Histogram
Count
0
1
2
20
30
40
Value
Common sites gene−LOC113562754
1
2
CphaseB1
RphaseA1

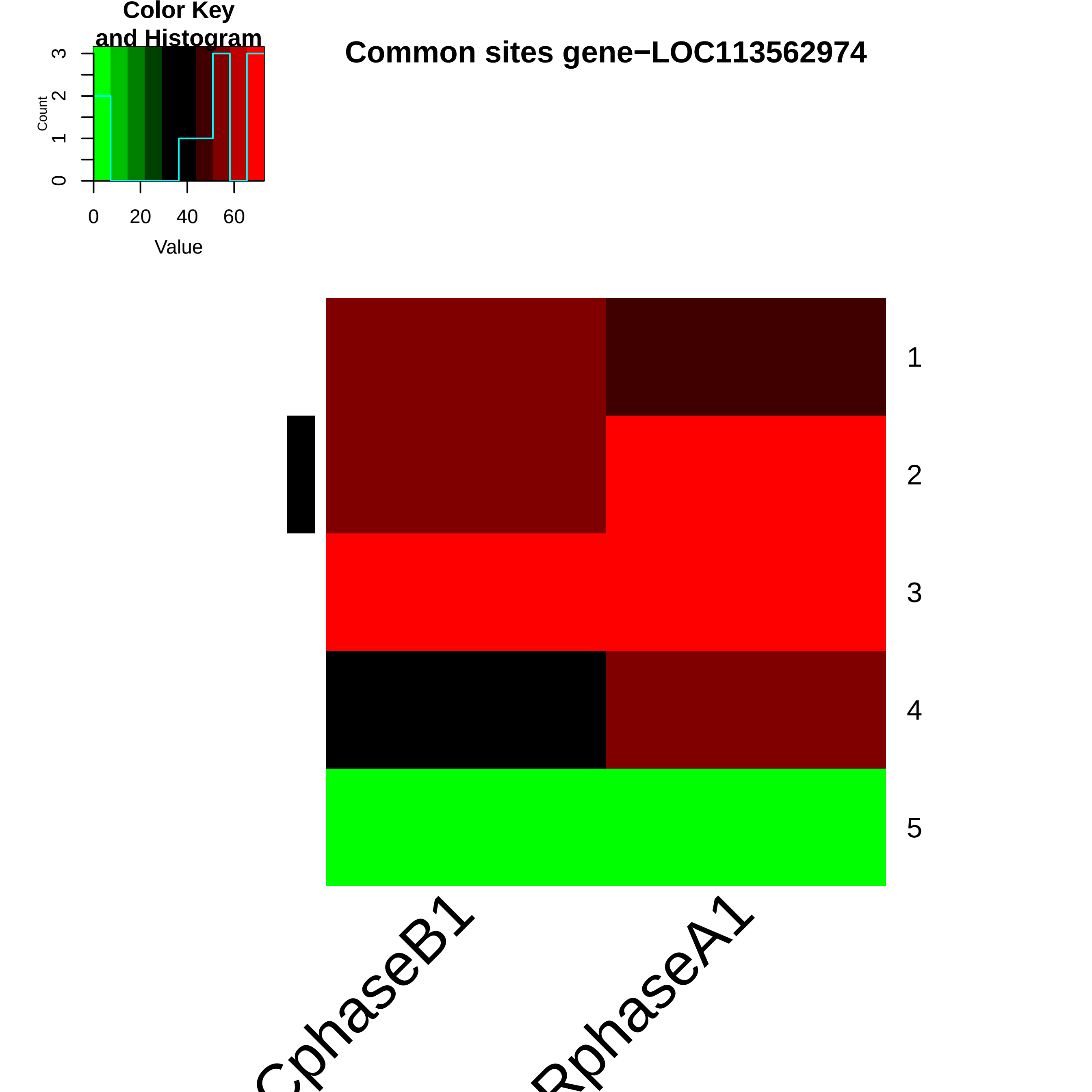

Color Key
and Histogram
Count
0
1
2
3
0
20
40
60
Value
Common sites gene−LOC113562974
1
2
3
4
5
CphaseB1
RphaseA1

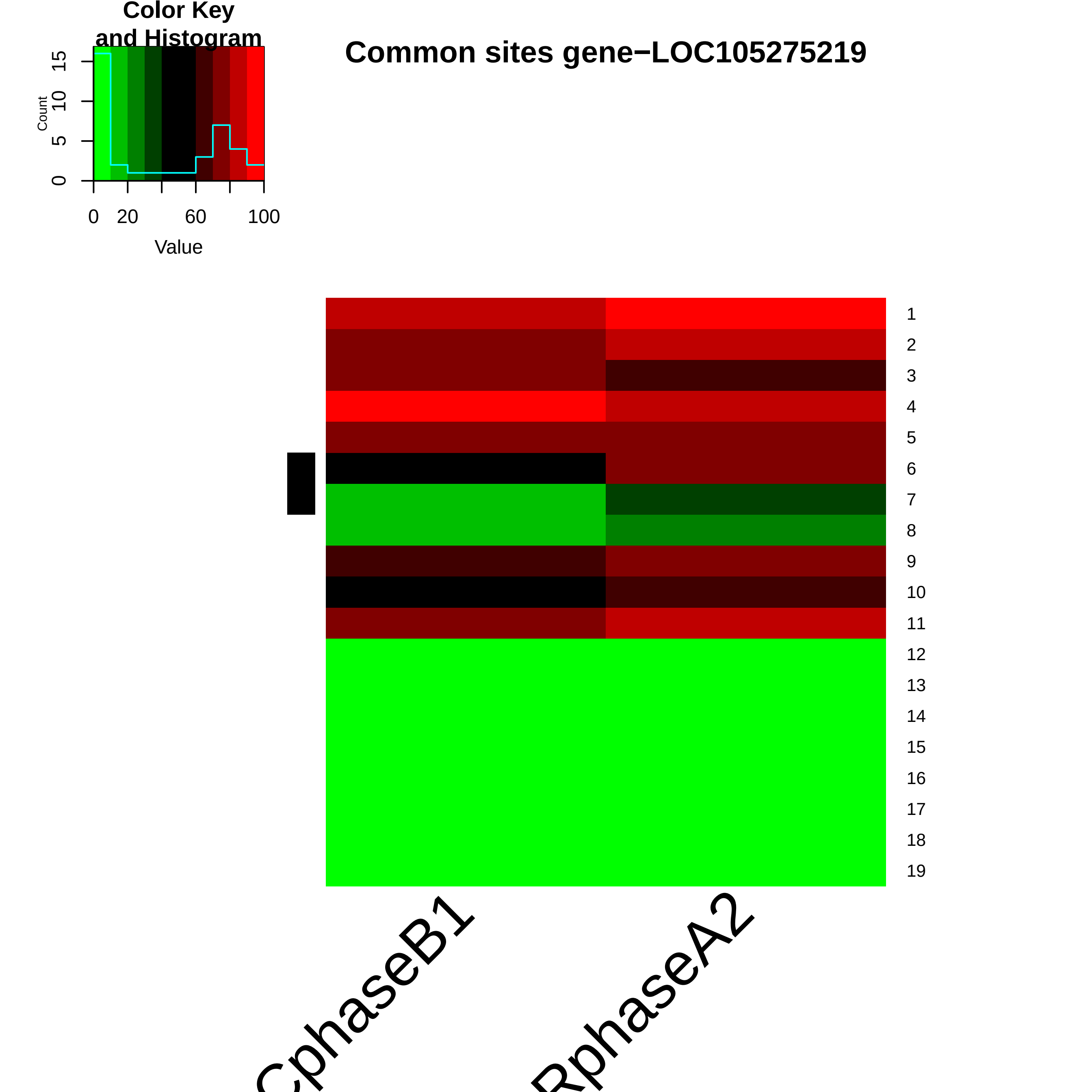

Color Key
and Histogram
Count
0
5
10
15
0
20
60
100
Value
Common sites gene−LOC105275219
CphaseB1
RphaseA2

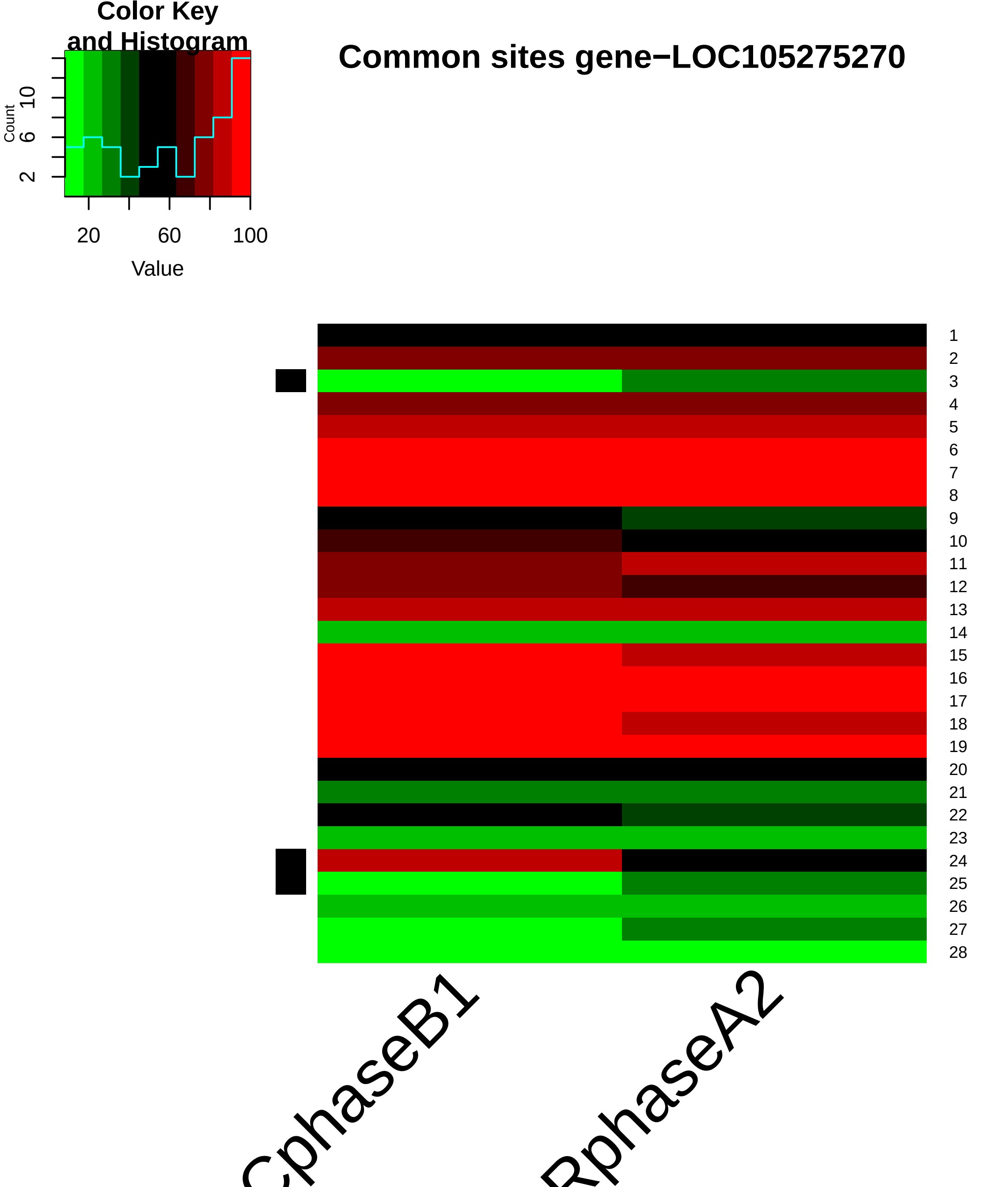
Color Key
and Histogram
Count
2
6
10
20
60
100
Value
Common sites gene−LOC105275270
CphaseB1
RphaseA2

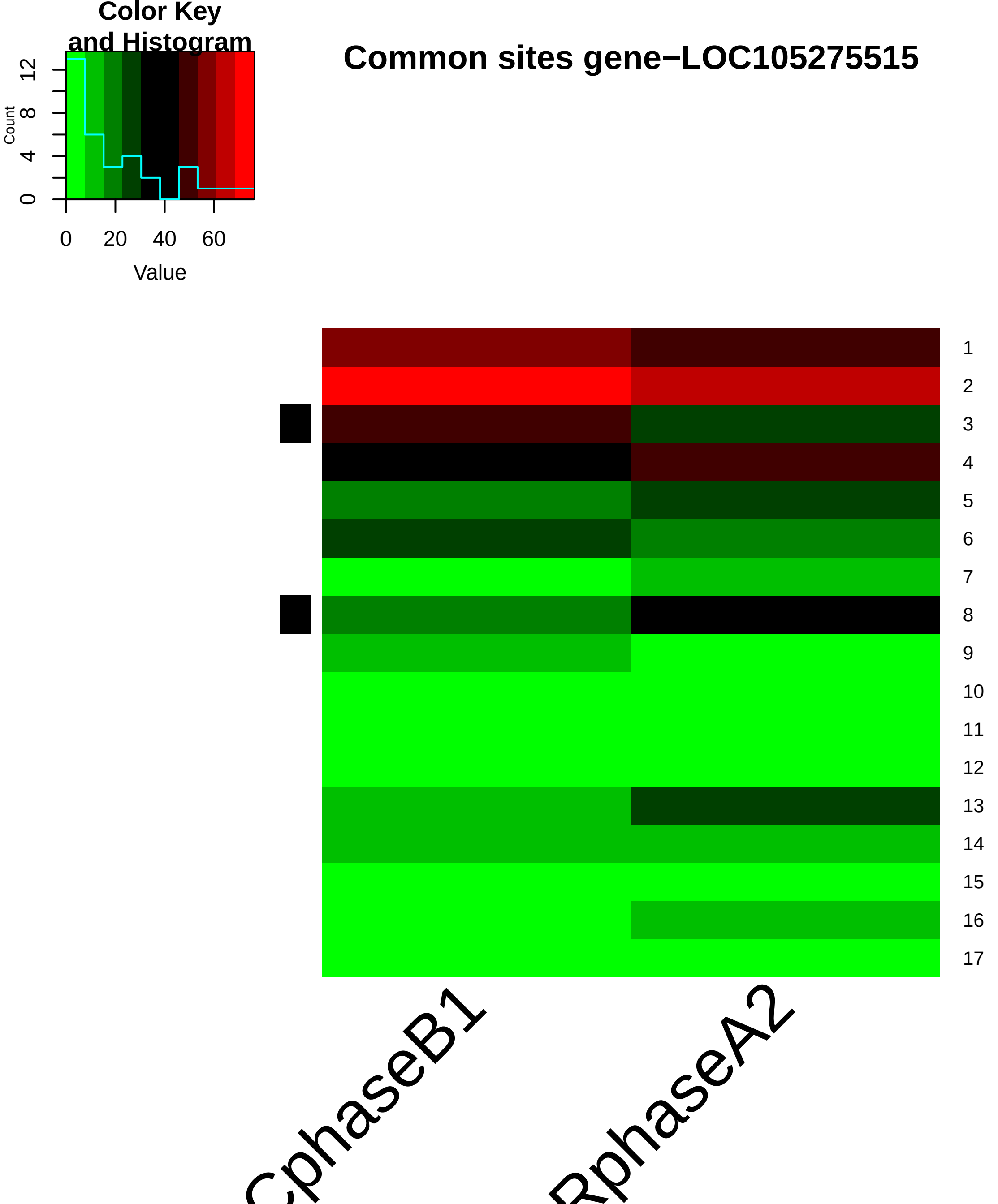
Color Key
and Histogram
Count
0
4
8
12
0
20
40
60
Value
Common sites gene−LOC105275515
CphaseB1
RphaseA2

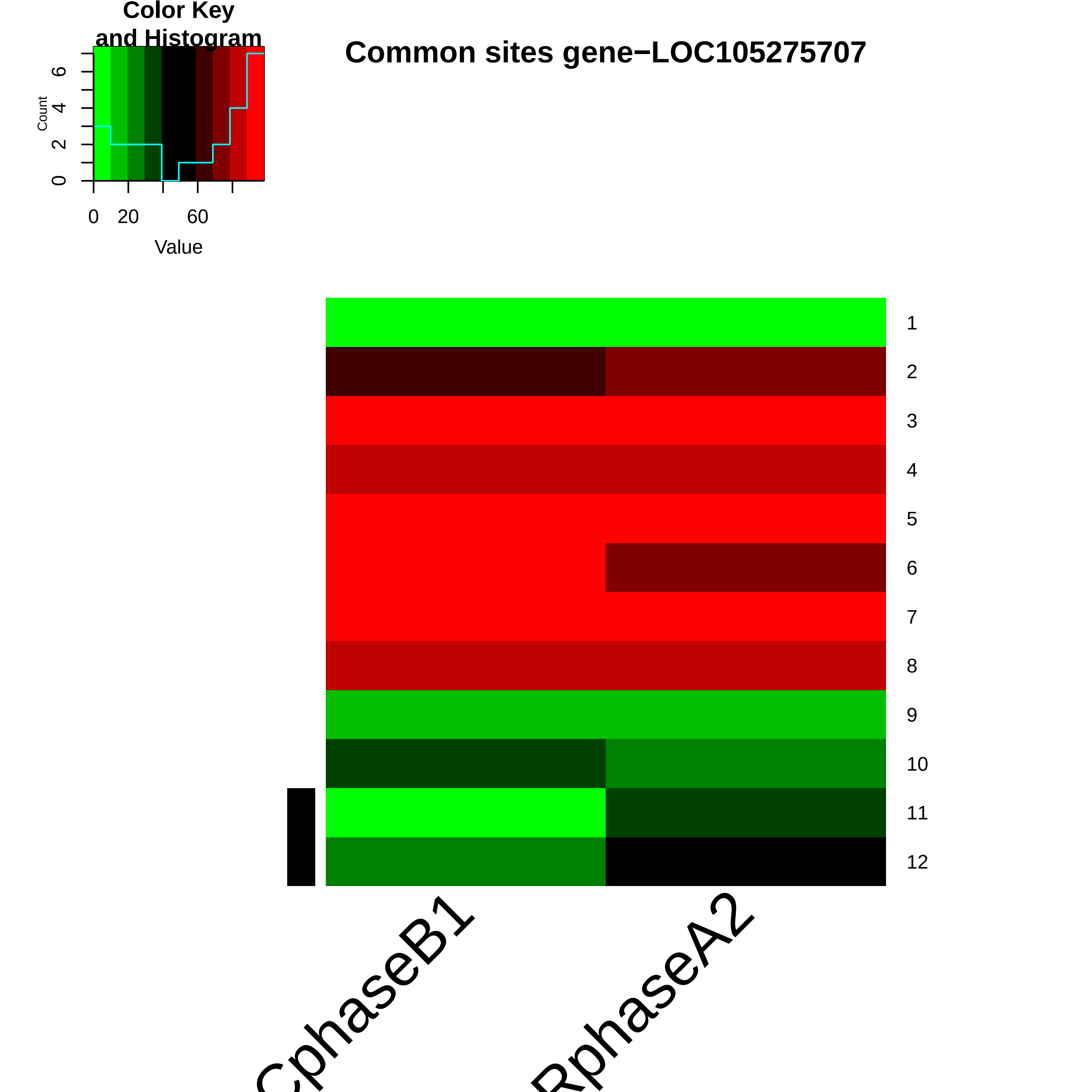
Color Key
and Histogram
Count
0
2
4
6
0
20
60
Value
Common sites gene−LOC105275707
1
2
3
4
5
6
7
8
9
10
11
12

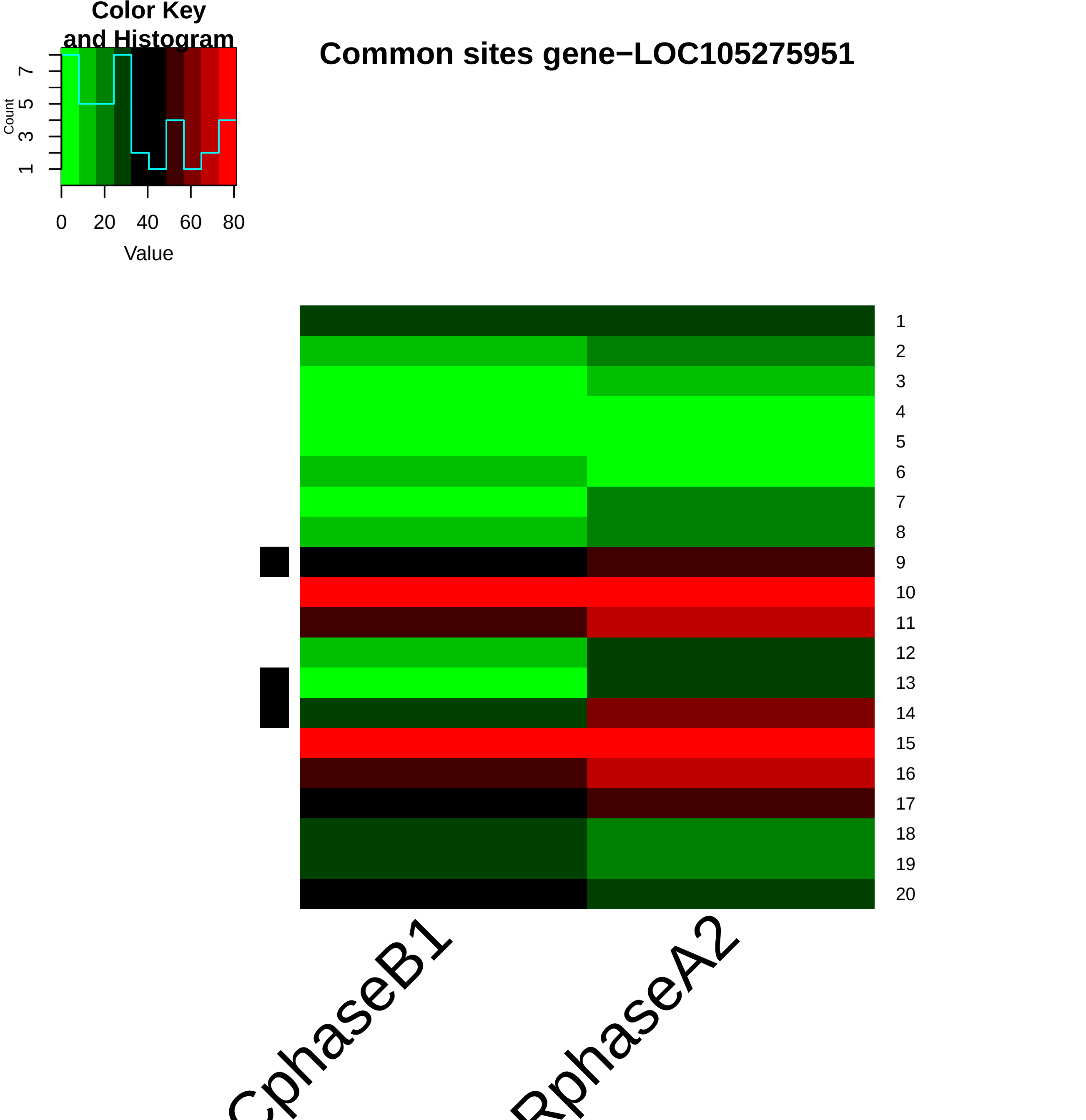
Color Key
and Histogram
Count
1
3
5
7
0
20
40
60
80
Value
Common sites gene−LOC105275951
CphaseB1
RphaseA2

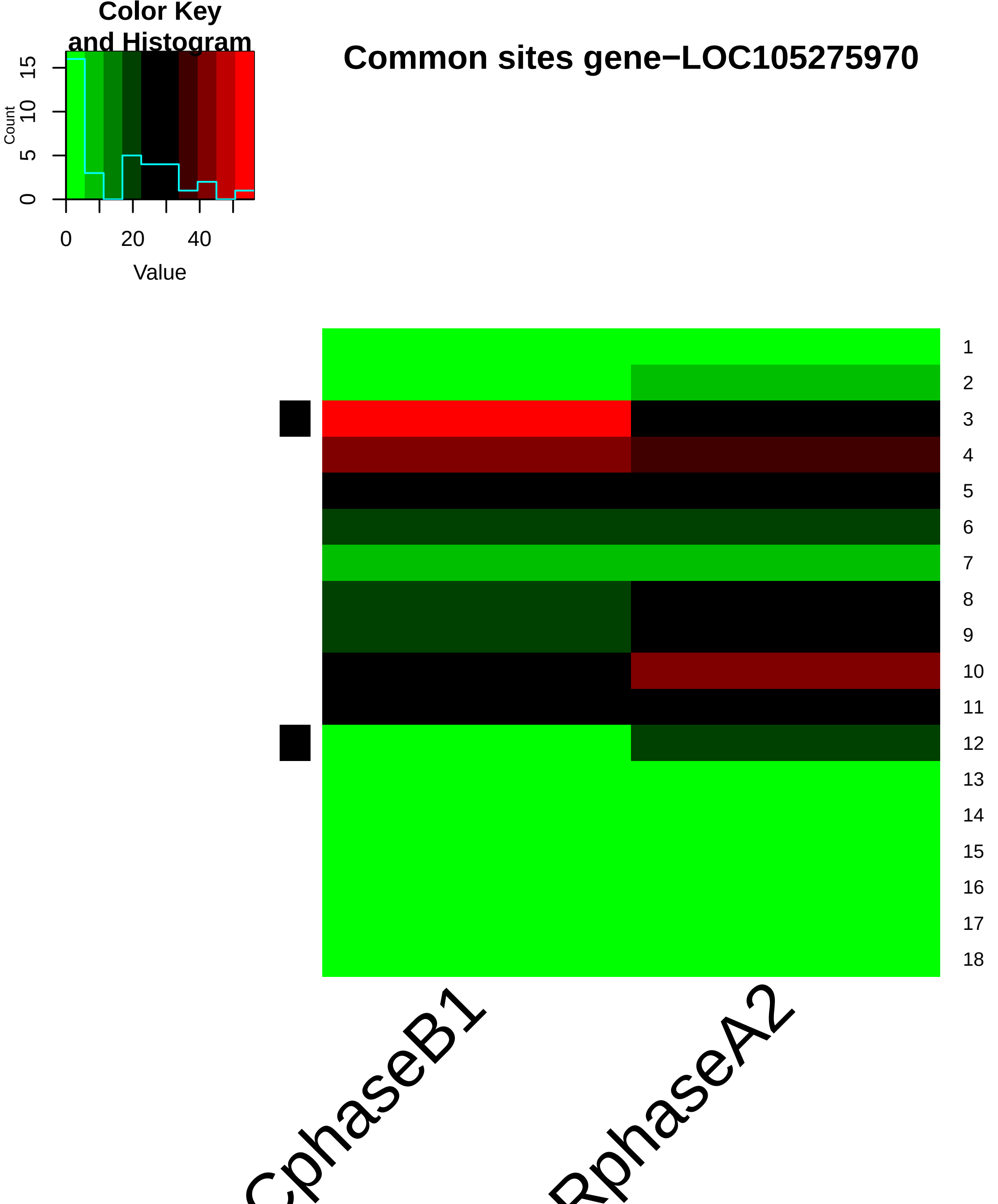

Color Key
and Histogram
Count
0
5
10
15
0
20
40
Value
Common sites gene−LOC105275970
CphaseB1
RphaseA2

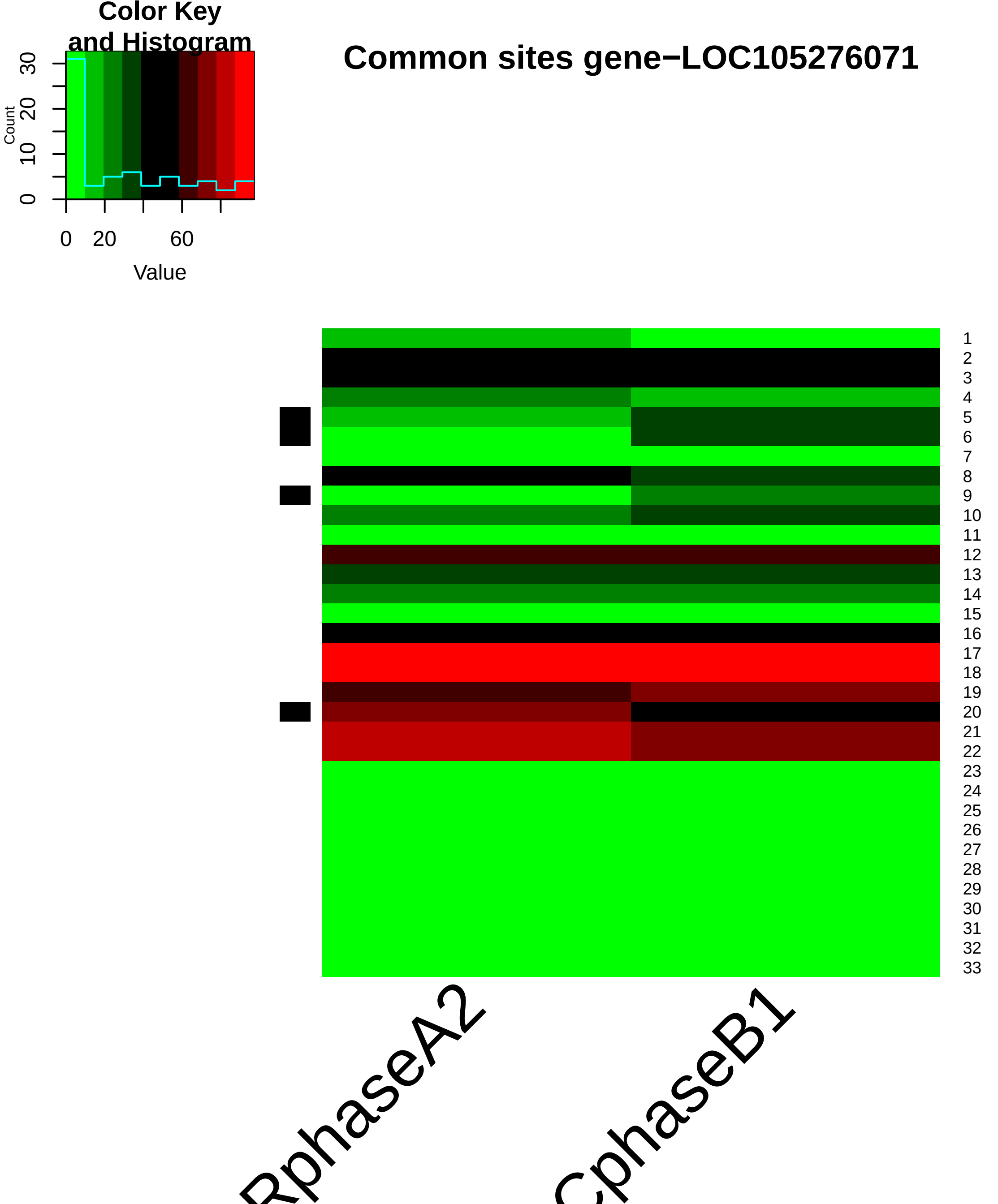

Color Key
and Histogram
Count
0
10
20
30
0
20
60
Value
Common sites gene−LOC105276071
RphaseA2
CphaseB1

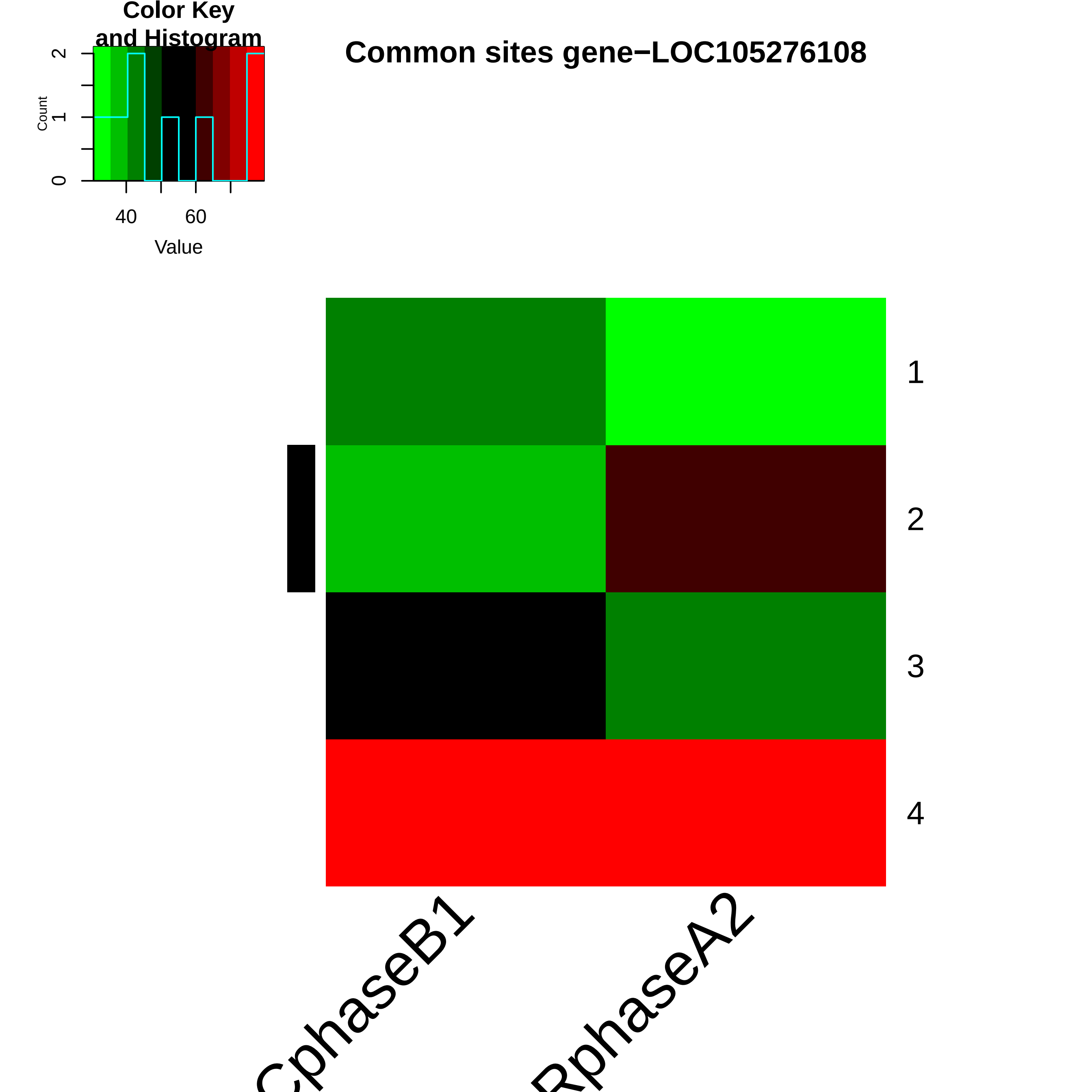

Color Key
and Histogram
Count
0
1
2
40
60
Value
Common sites gene−LOC105276108
1
2
3
4
CphaseB1
RphaseA2

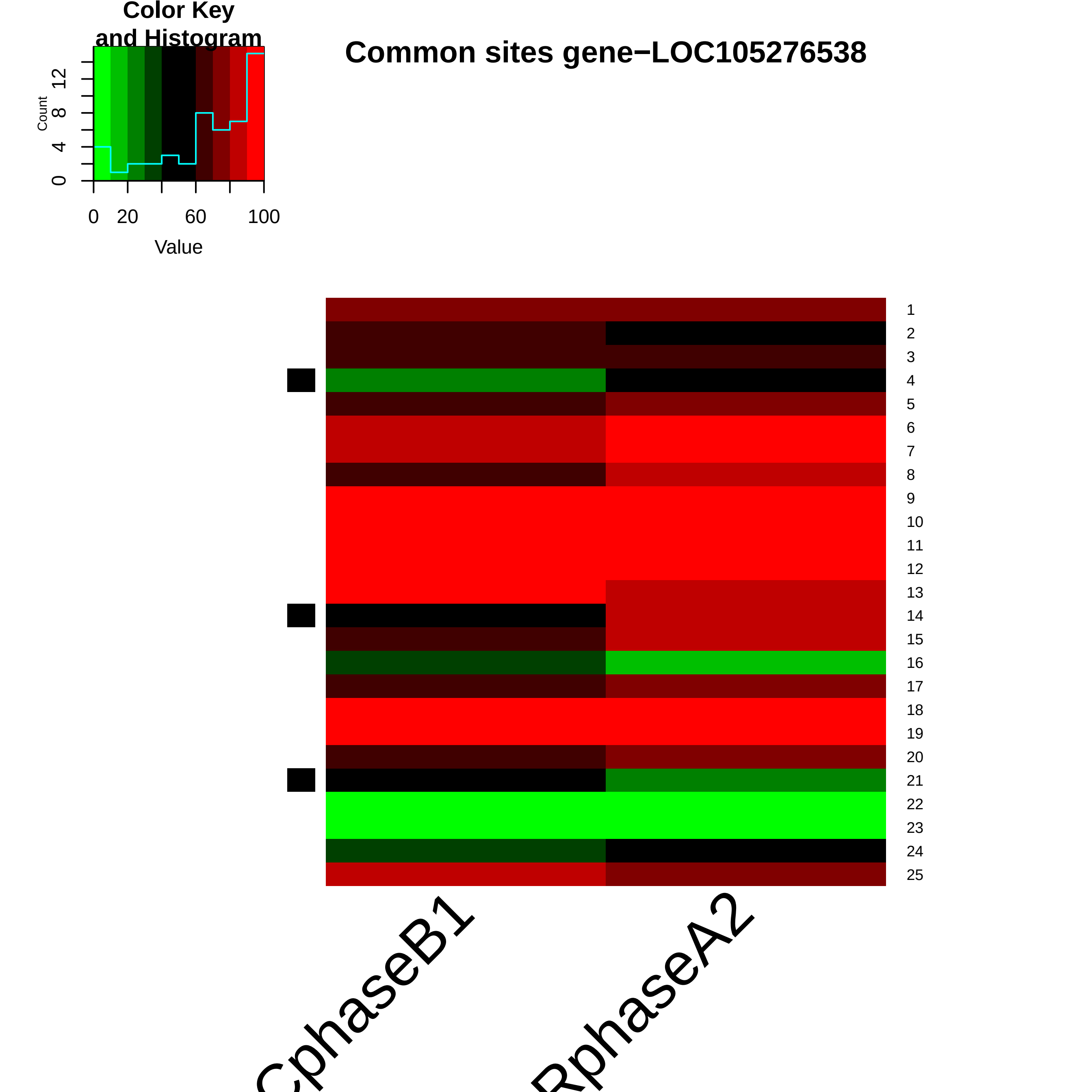

Color Key
and Histogram
Count
0
4
8
12
0
20
60
100
Value
Common sites gene−LOC105276538
CphaseB1
RphaseA2

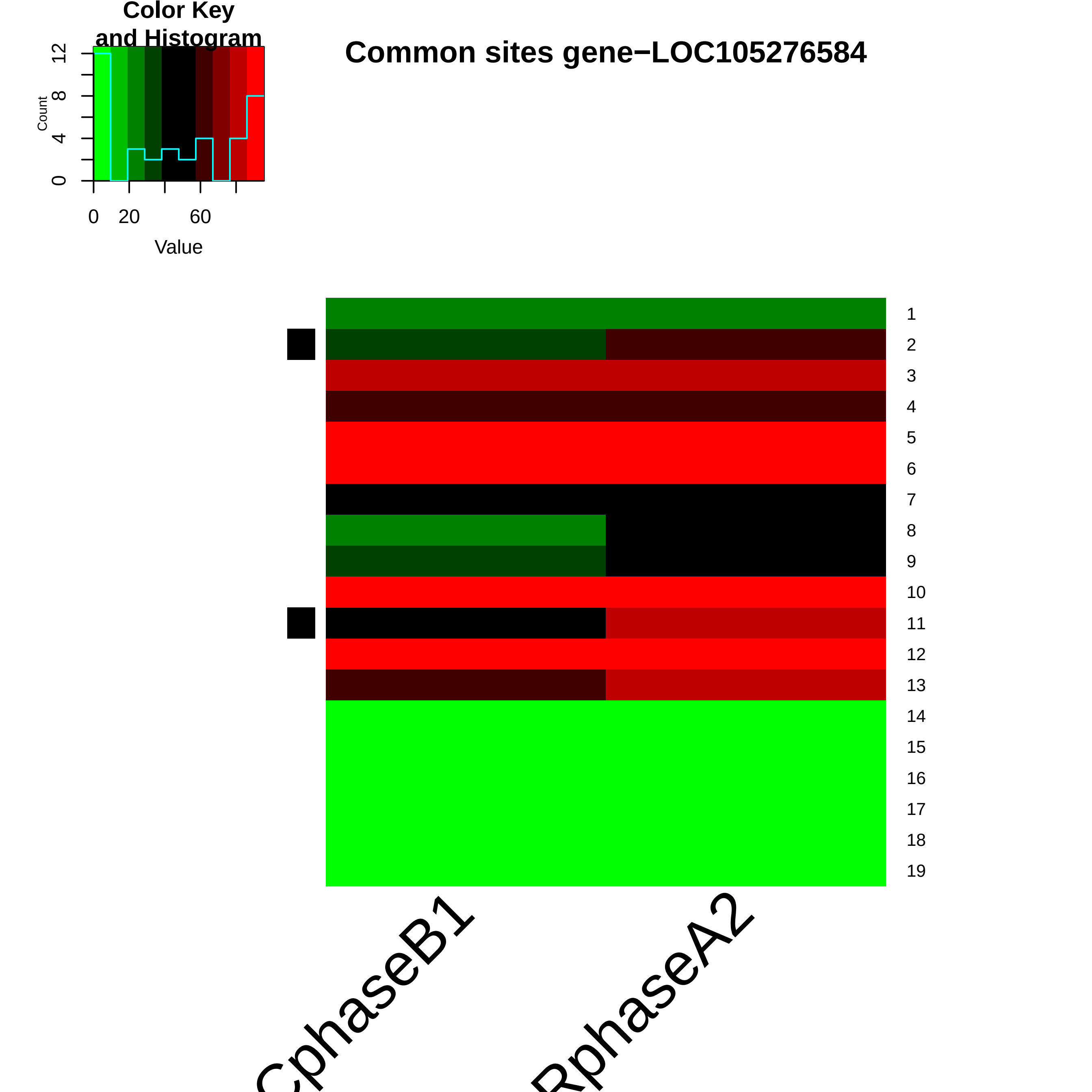
Color Key
and Histogram
Count
0
4
8
12
0
20
60
Value
Common sites gene−LOC105276584
CphaseB1
RphaseA2

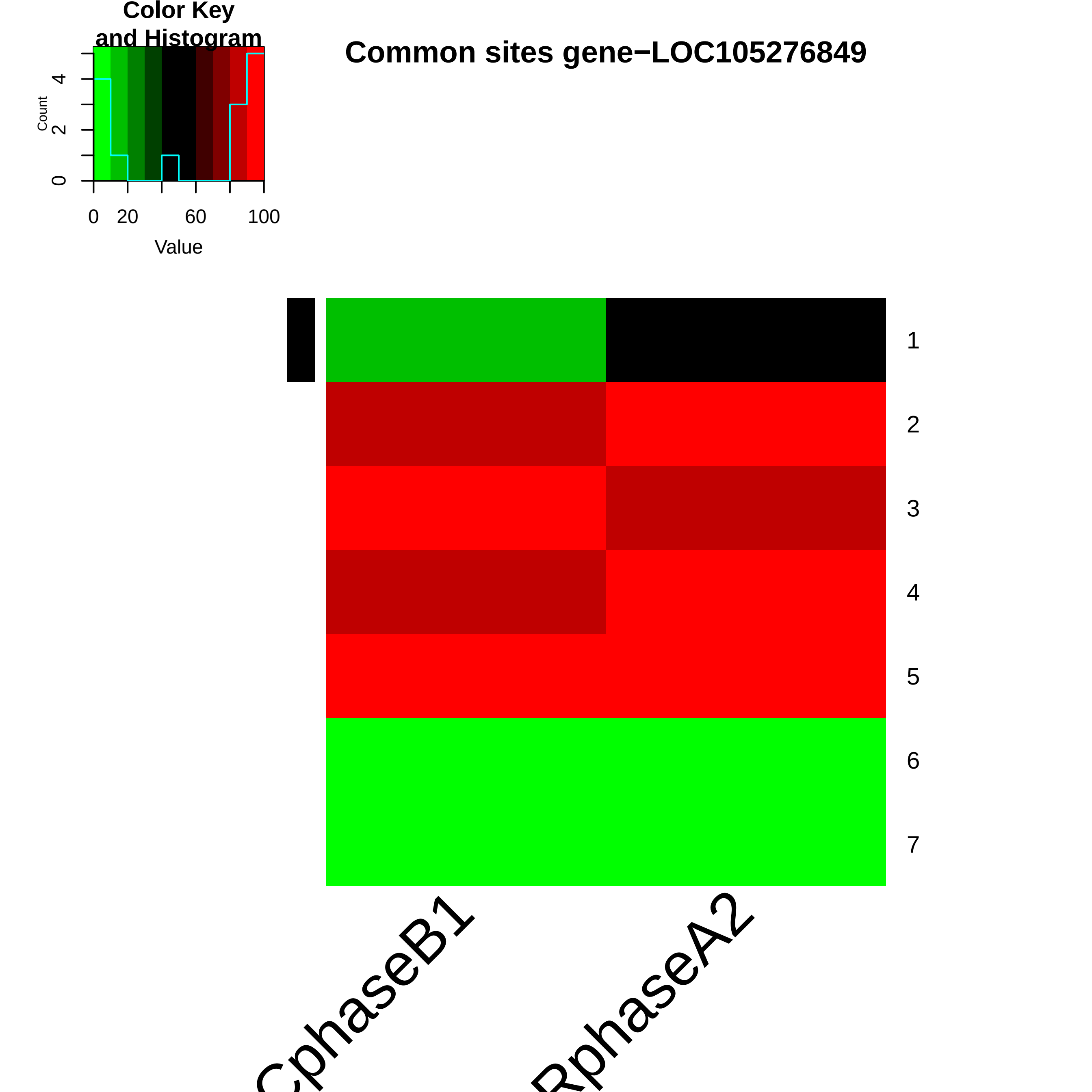
Color Key
and Histogram
Count
0
2
4
0
20
60
100
Value
Common sites gene−LOC105276849
1
2
3
4
5
6
7
CphaseB1
RphaseA2

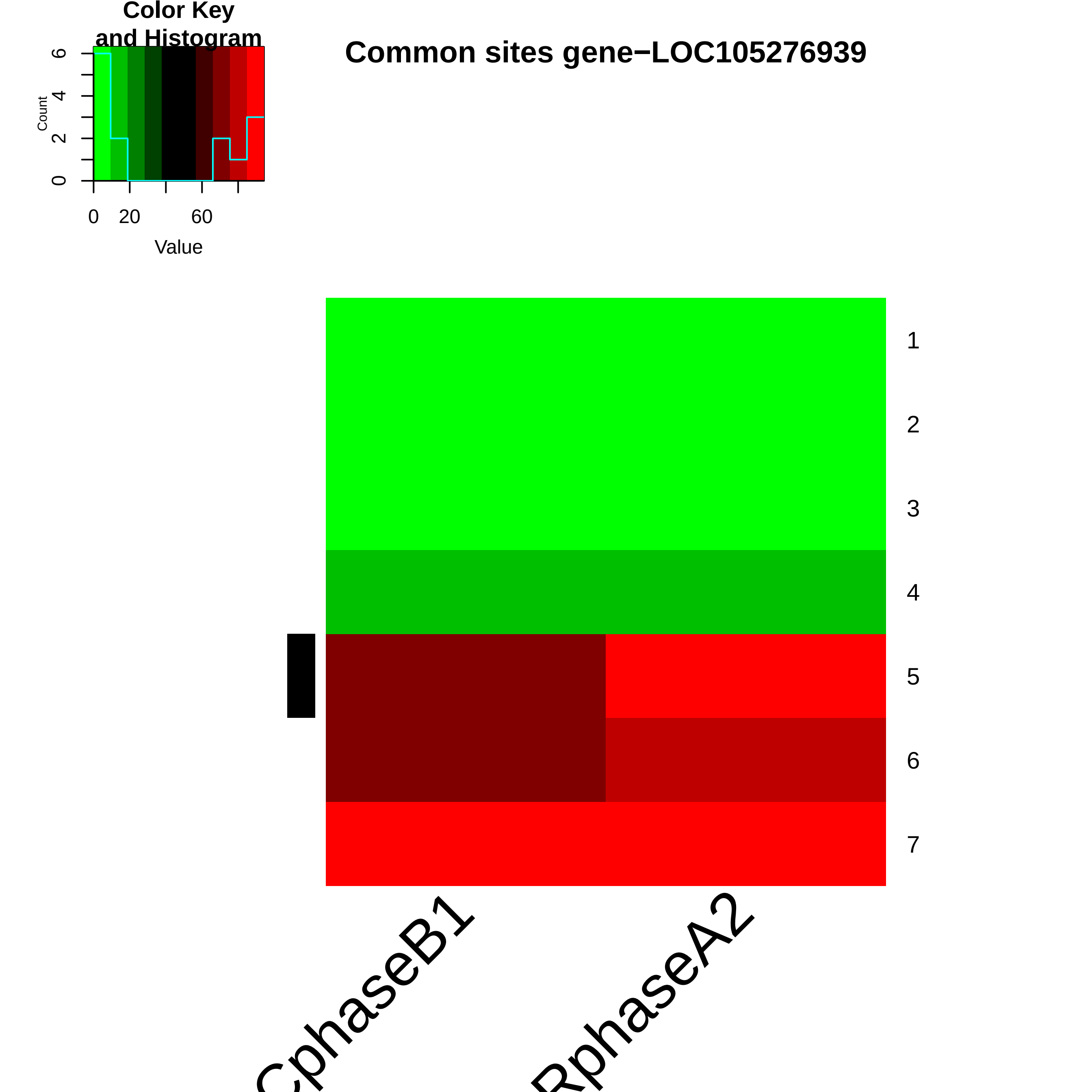
Color Key
and Histogram
Count
0
2
4
6
0
20
60
Value
Common sites gene−LOC105276939
1
2
3
4
5
6
7
CphaseB1
RphaseA2

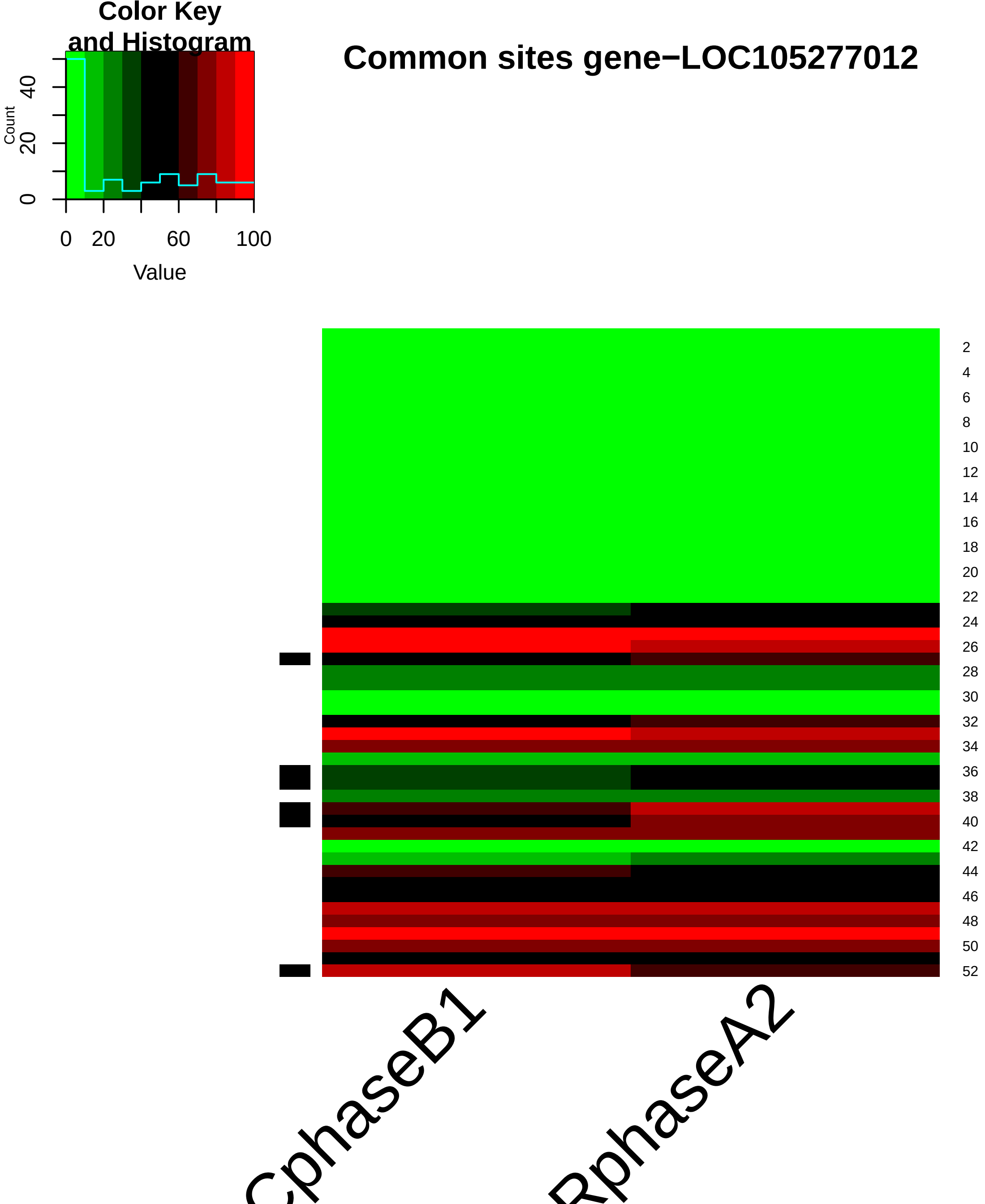

Color Key
and Histogram
Count
0
20
40
0
20
60
100
Value
Common sites gene−LOC105277012
2
4
6
8
10
12
14
16
18
20
22
24
26
28
30
32
34
36
38
40
42
44
46
48
50
52
CphaseB1
RphaseA2

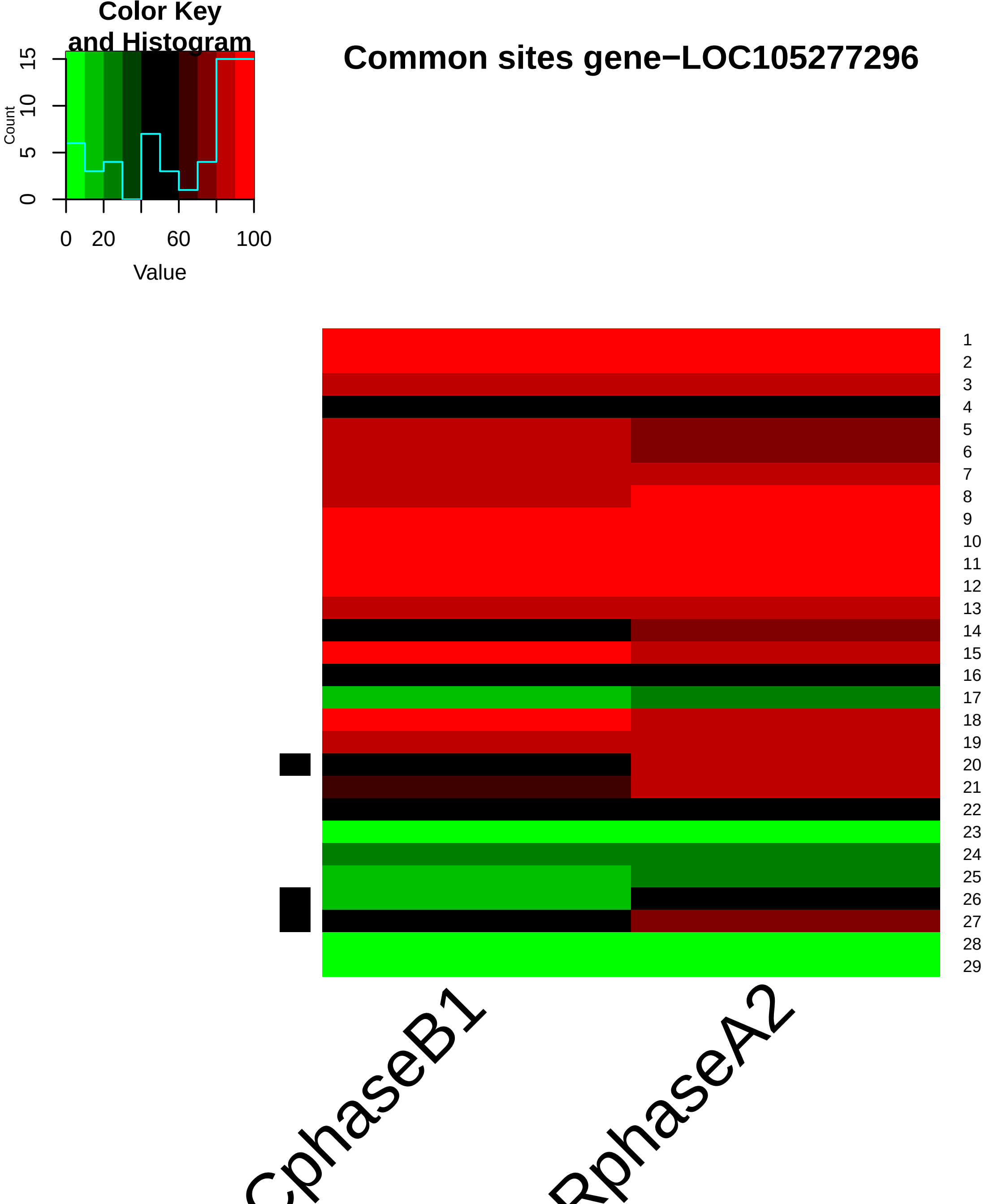
Color Key
and Histogram
Count
0
5
10
15
0
20
60
100
Value
Common sites gene−LOC105277296
CphaseB1
RphaseA2

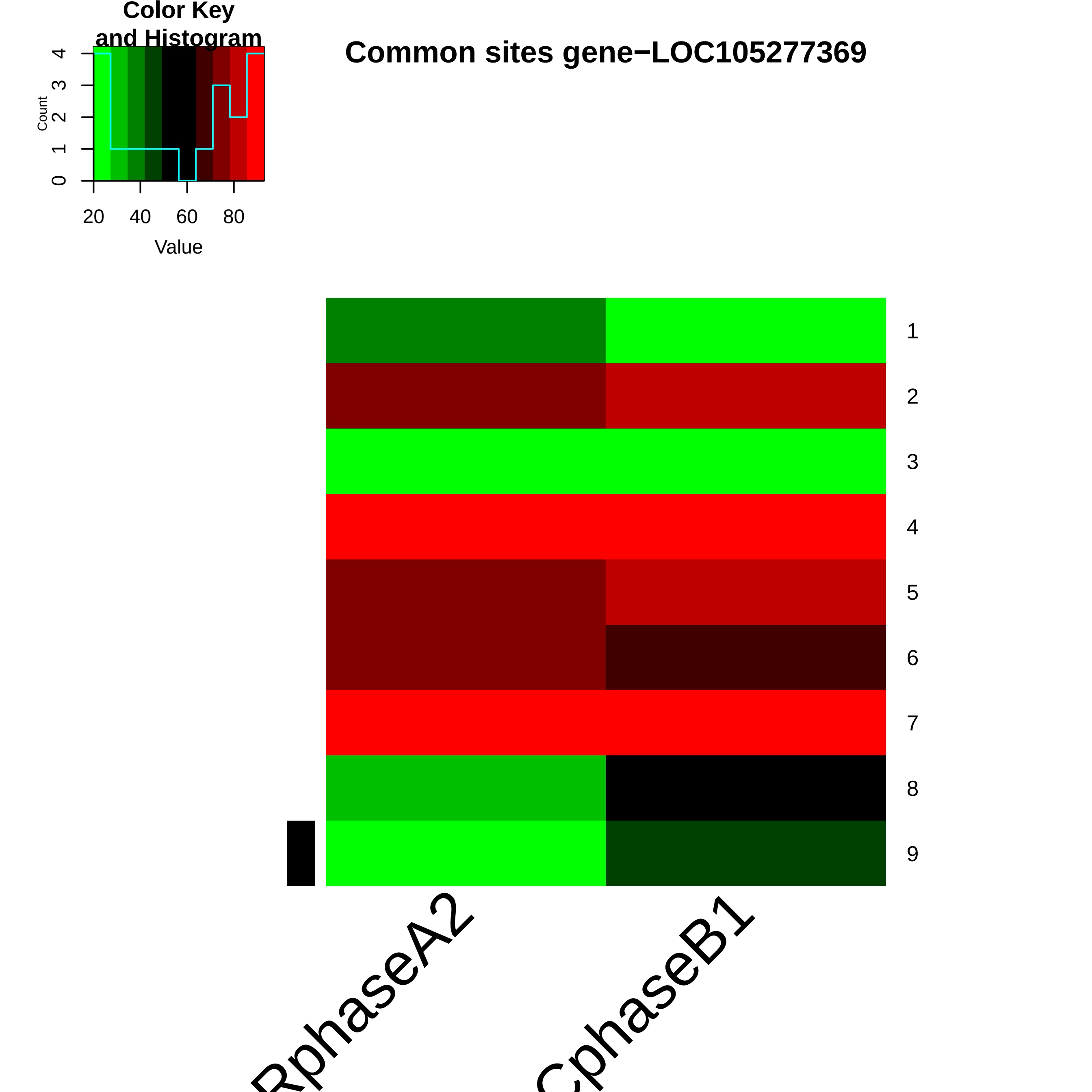
Color Key
and Histogram
Count
0
1
2
3
4
20
40
60
80
Value
Common sites gene−LOC105277369
1
2
3
4
5
6
7
8
9
RphaseA2
CphaseB1

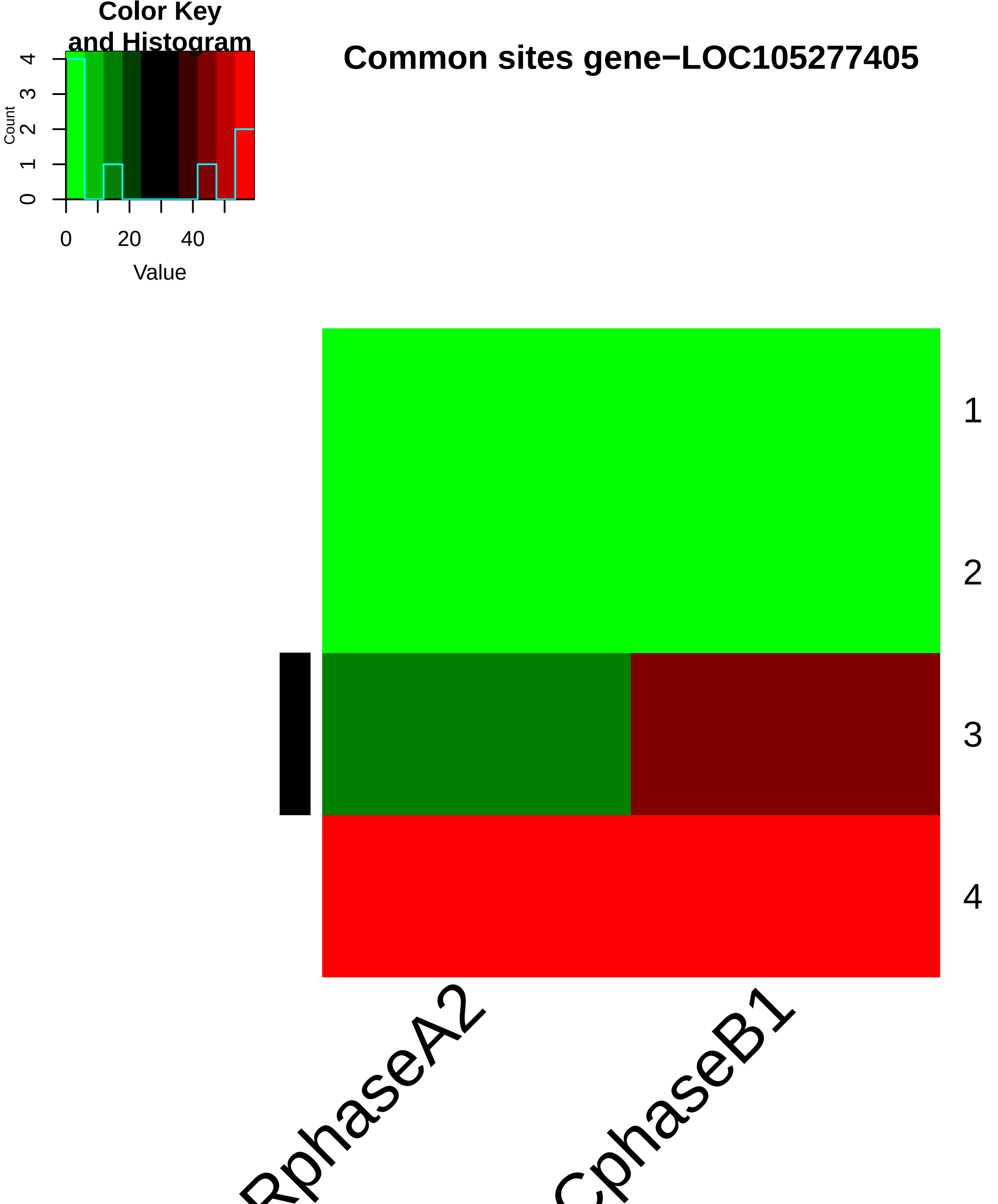
Color Key
and Histogram
Count
Value
Common sites gene−LOC105277405
1
2
3
4
RphaseA2
CphaseB1

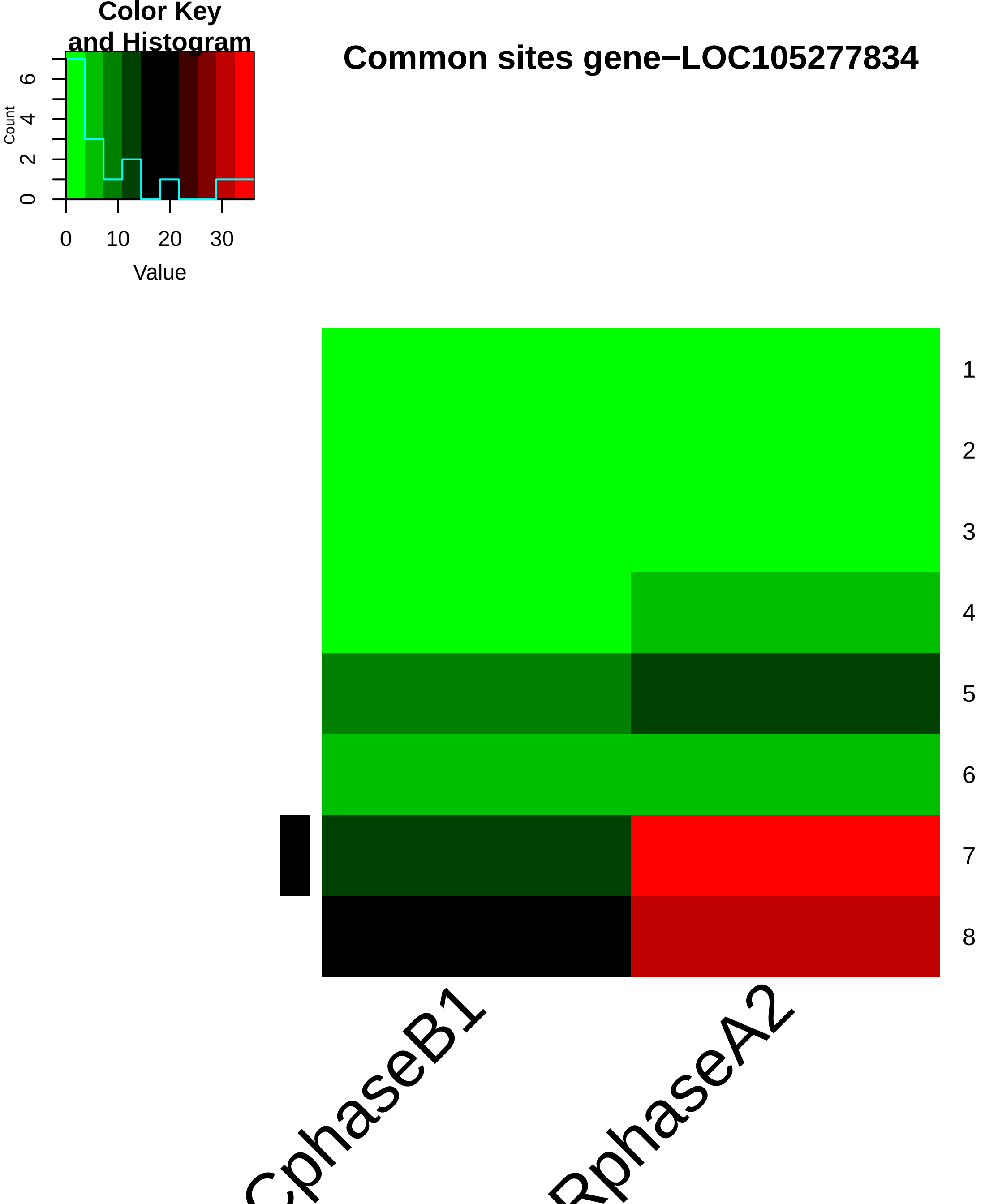

Color Key
and Histogram
Count
0
2
4
6
0
10
20
30
Value
Common sites gene−LOC105277834
1
2
3
4
5
6
7
8
CphaseB1
RphaseA2

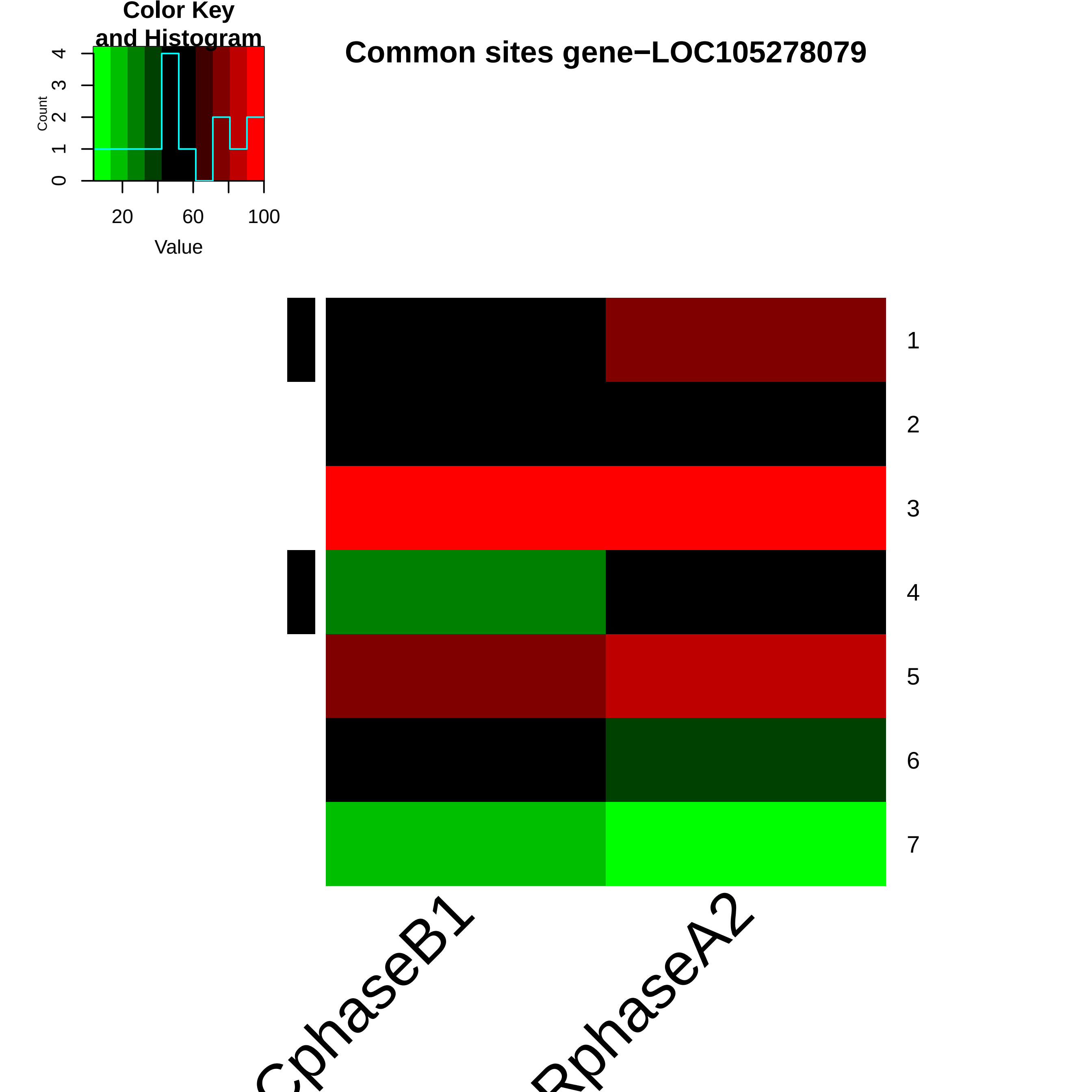

Color Key
and Histogram
Count
0
1
2
3
4
20
60
100
Value
Common sites gene−LOC105278079
1
2
3
4
5
6
7
CphaseB1
RphaseA2

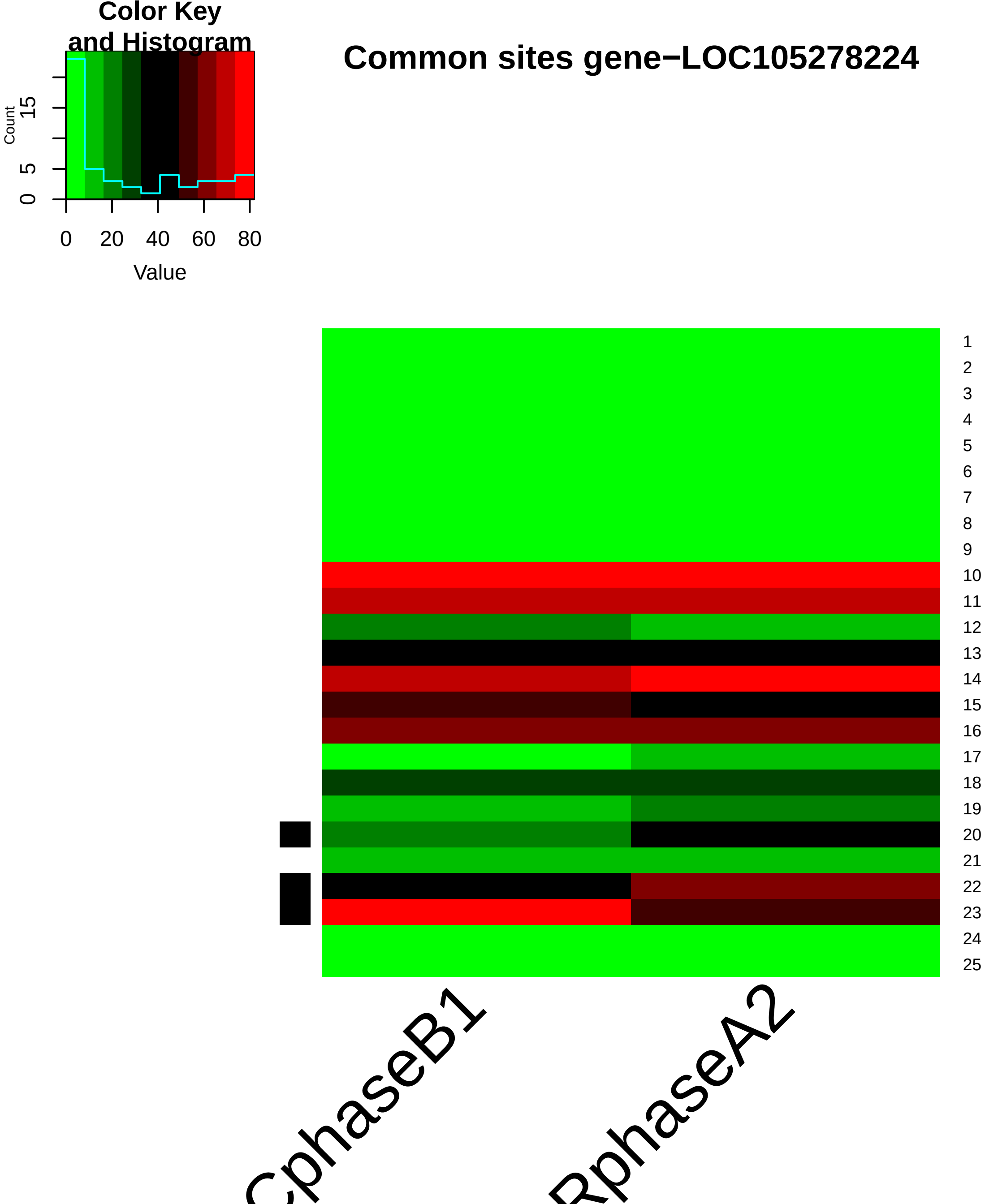

Color Key
and Histogram
Count
0
5
15
0
20
40
60
80
Value
Common sites gene−LOC105278224
CphaseB1
RphaseA2

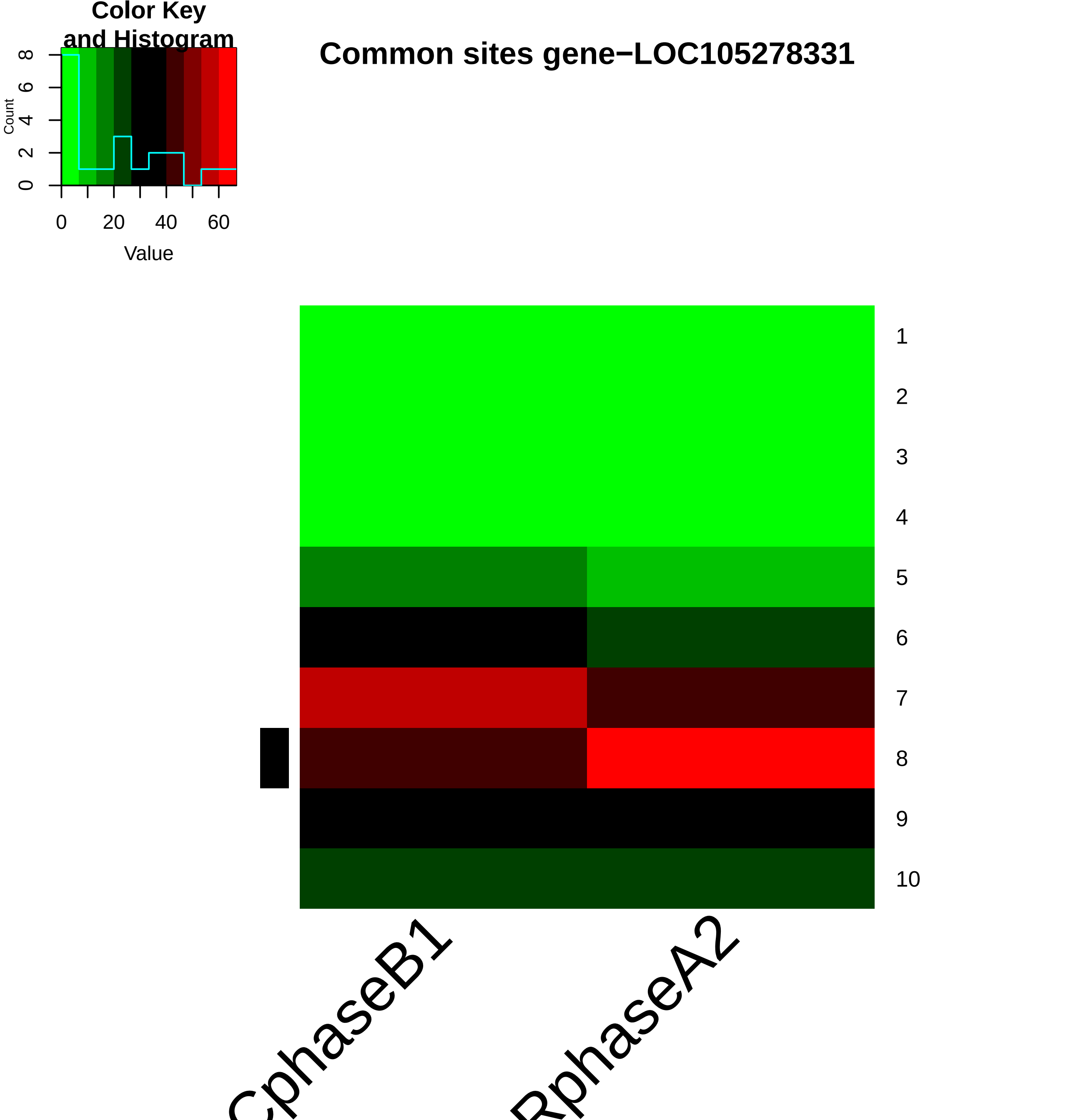
Color Key
and Histogram
Count
Value
0
20
40
60
0
2
4
6
8
Common sites gene−LOC105278331
1
2
3
4
5
6
7
8
9
10
CphaseB1
RphaseA2

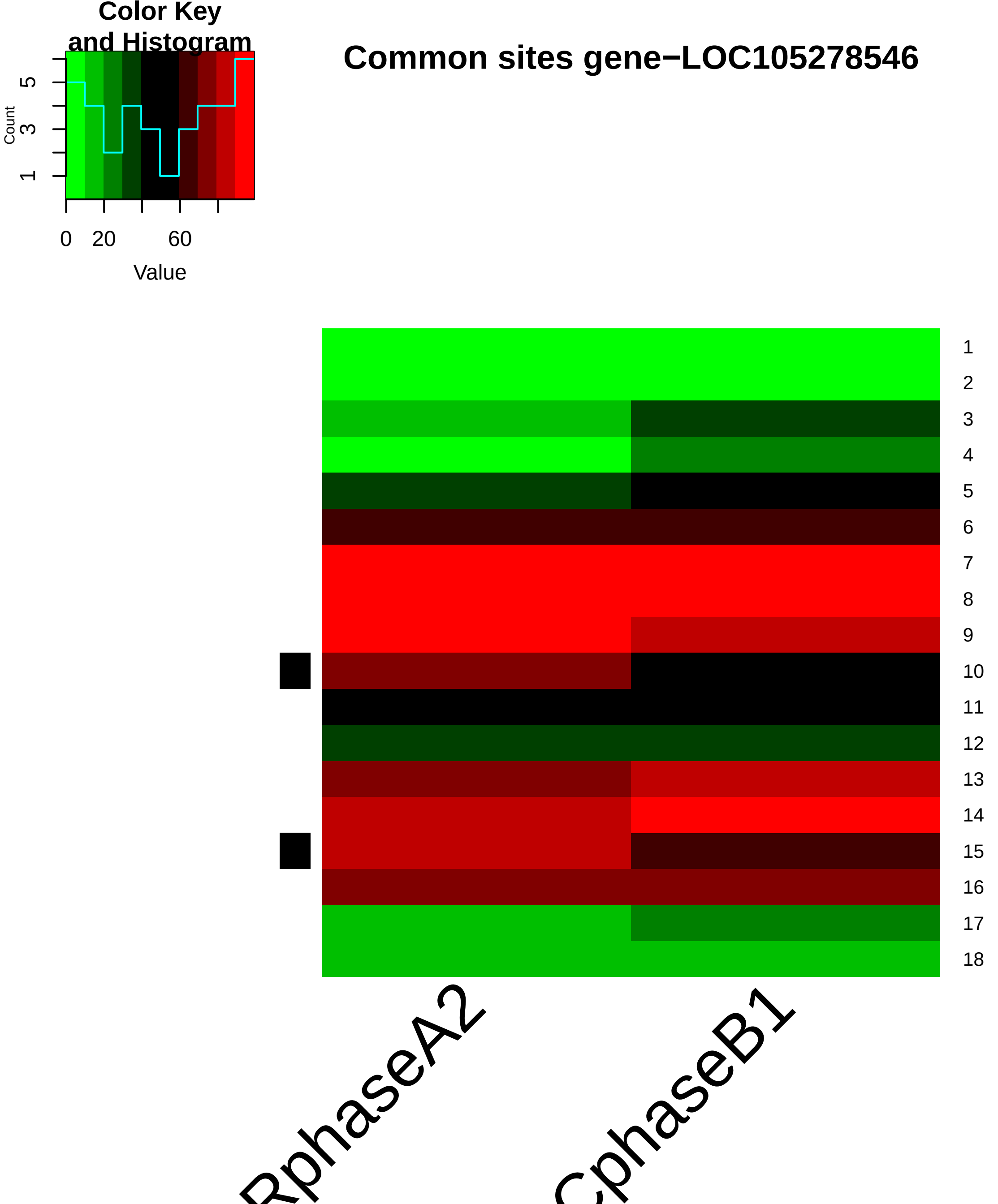
Color Key
and Histogram
Count
1
3
5
0
20
60
Value
Common sites gene−LOC105278546
RphaseA2
CphaseB1

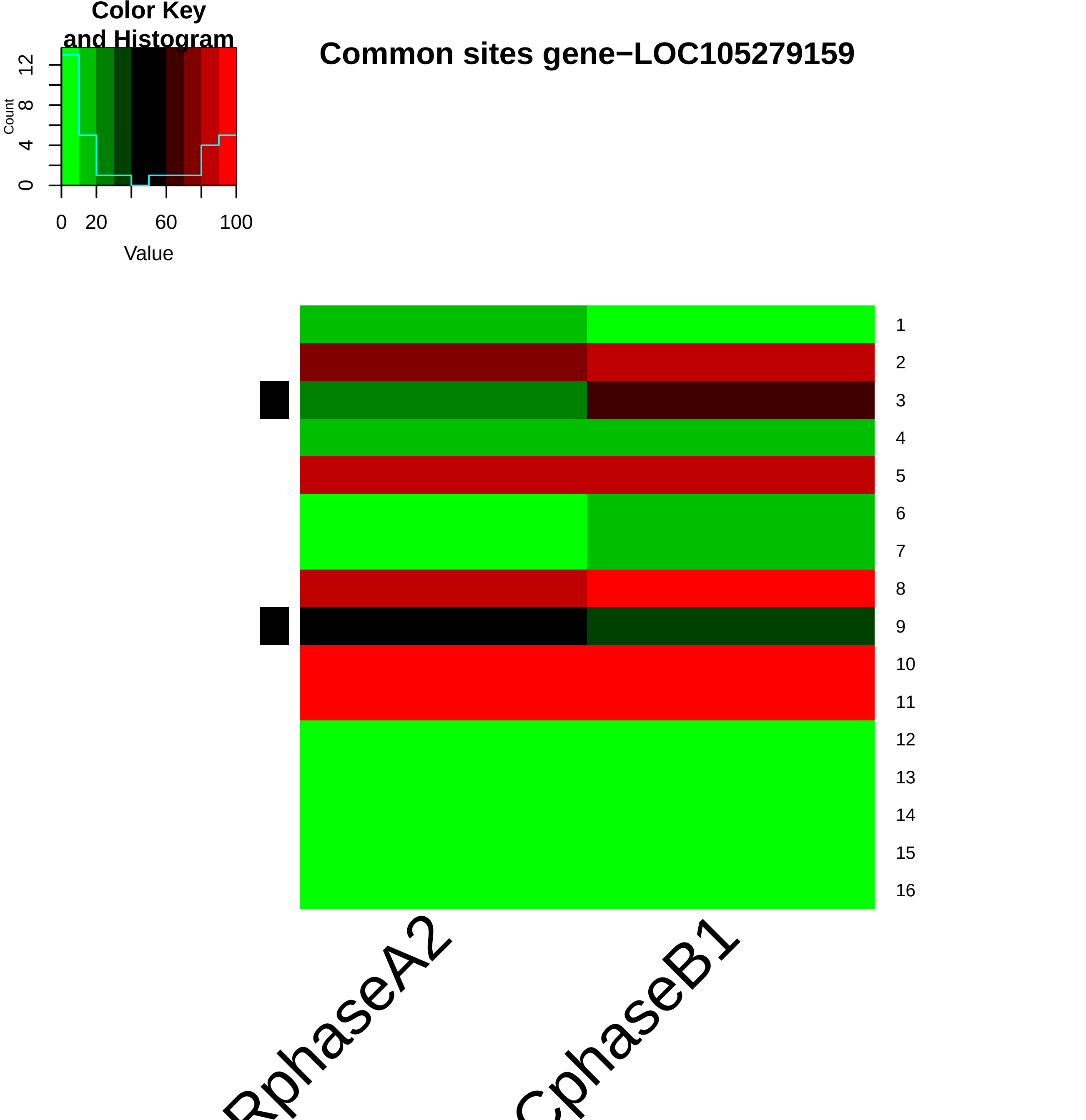
Color Key
and Histogram
Count
0
4
8
12
0
20
60
100
Value
Common sites gene−LOC105279159
RphaseA2
CphaseB1

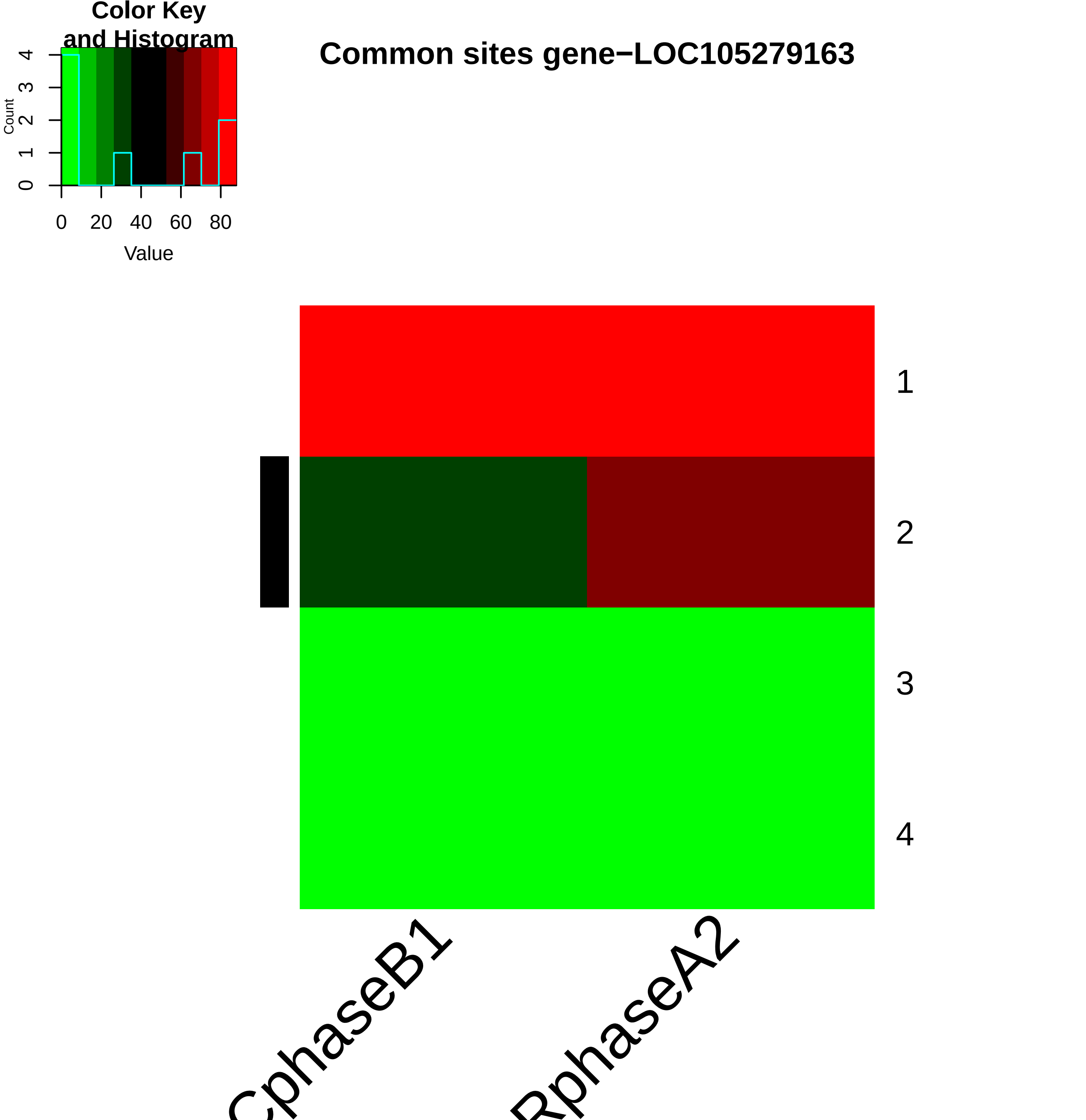

Color Key
and Histogram
Count
Value
0
20
40
60
80
0
1
2
3
4
Common sites gene−LOC105279163
1
2
3
4
CphaseB1
RphaseA2

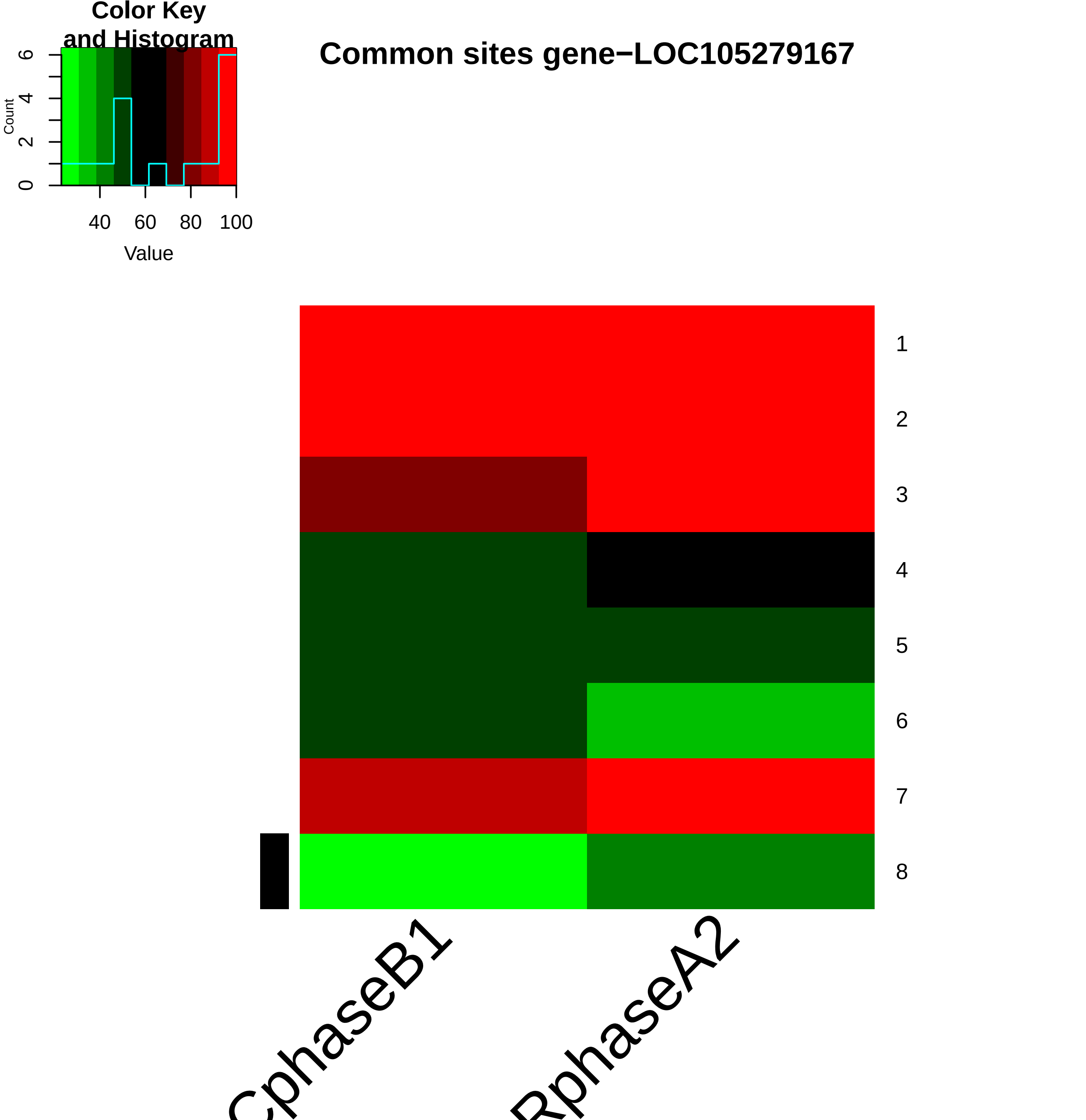

Color Key
and Histogram
Count
Value
40
60
80
100
0
2
4
6
Common sites gene−LOC105279167
1
2
3
4
5
6
7
8
CphaseB1
RphaseA2

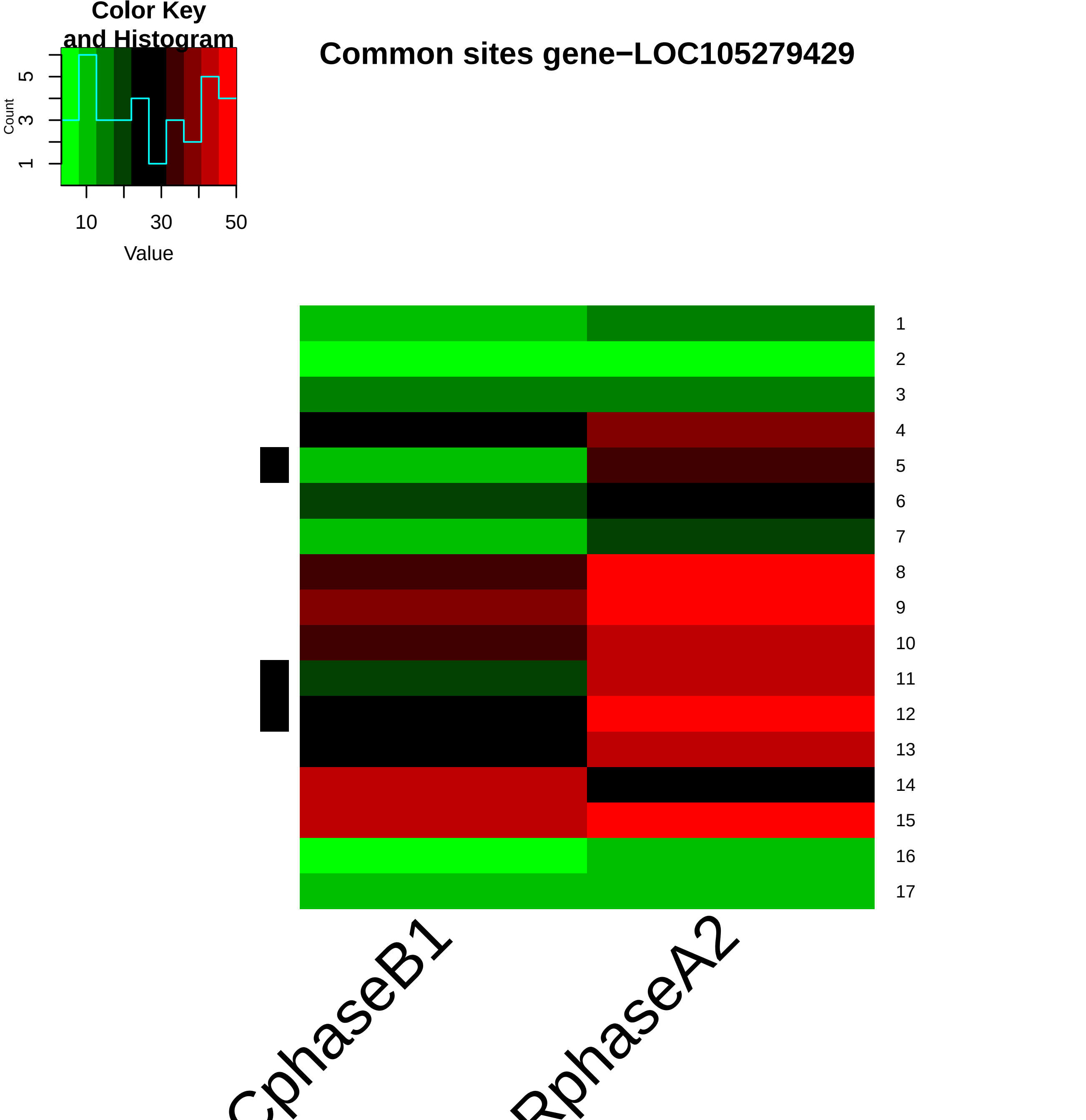
Color Key
and Histogram
Count
1
3
5
10
30
50
Value
Common sites gene−LOC105279429
CphaseB1
RphaseA2

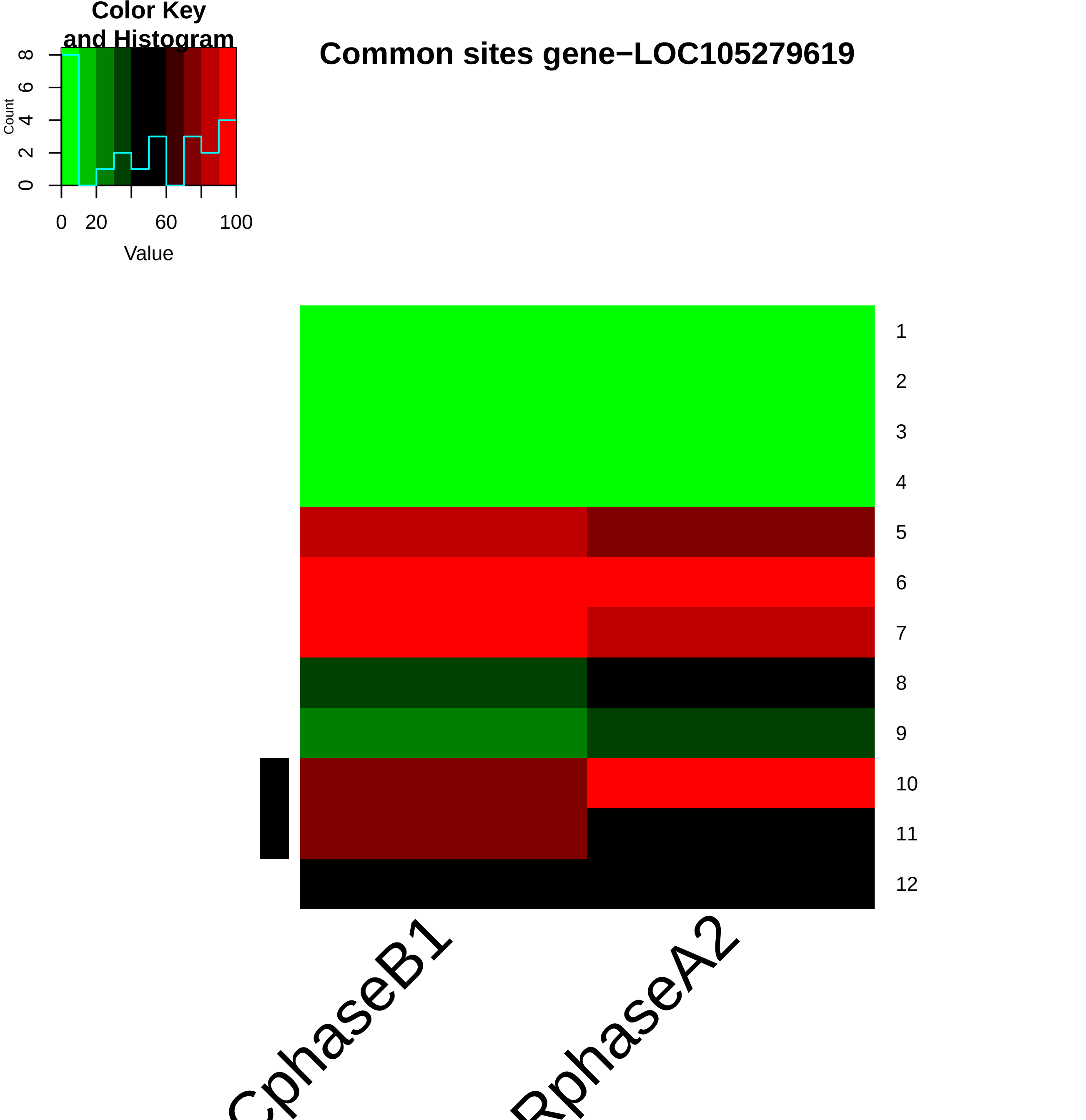
Color Key
and Histogram
Count
Value
0
20
60
100
0
2
4
6
8
Common sites gene−LOC105279619
1
2
3
4
5
6
7
8
9
10
11
12
CphaseB1
RphaseA2

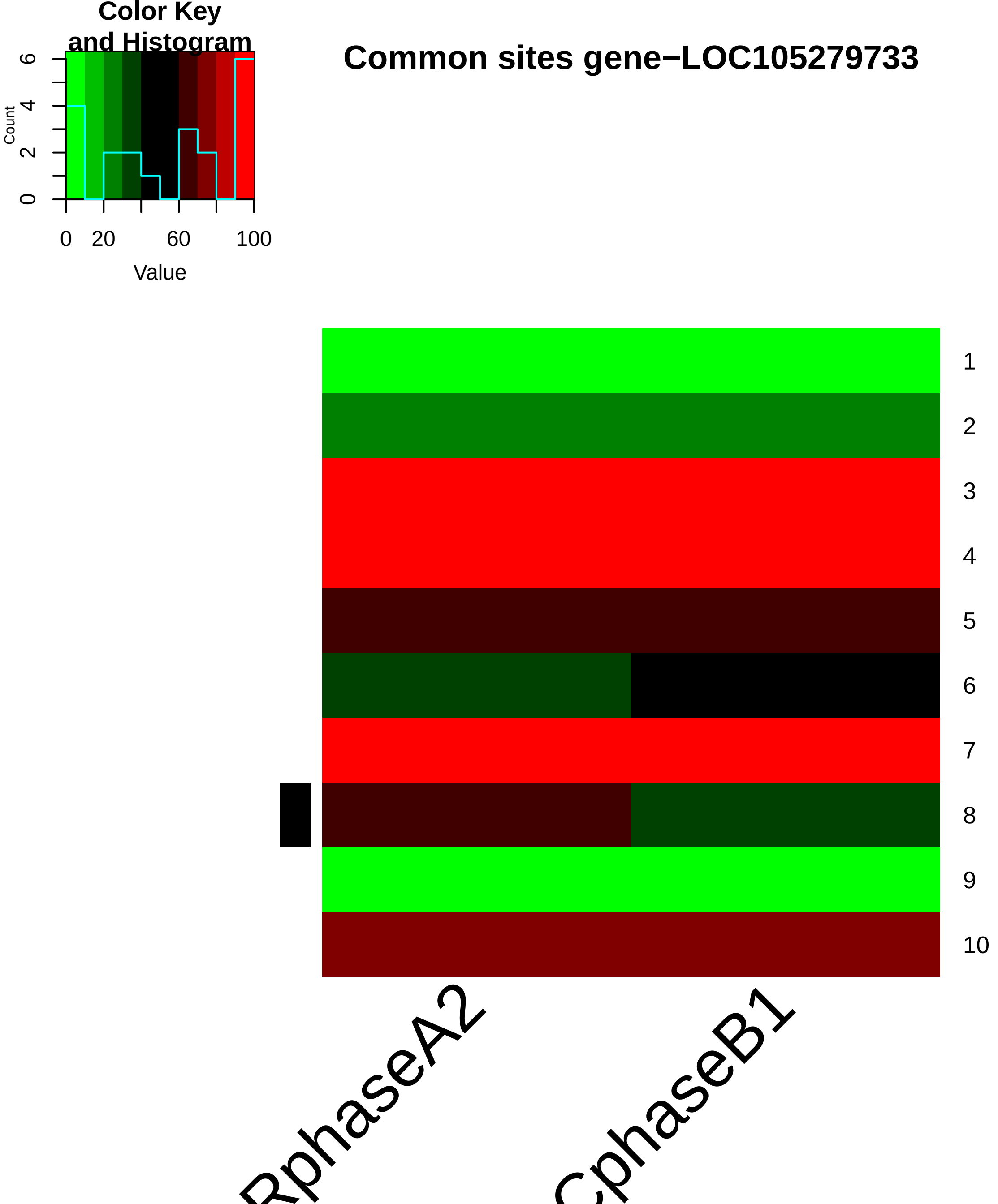

Color Key
and Histogram
Count
0
2
4
6
0
20
60
100
Value
Common sites gene−LOC105279733
1
2
3
4
5
6
7
8
9
10
RphaseA2
CphaseB1

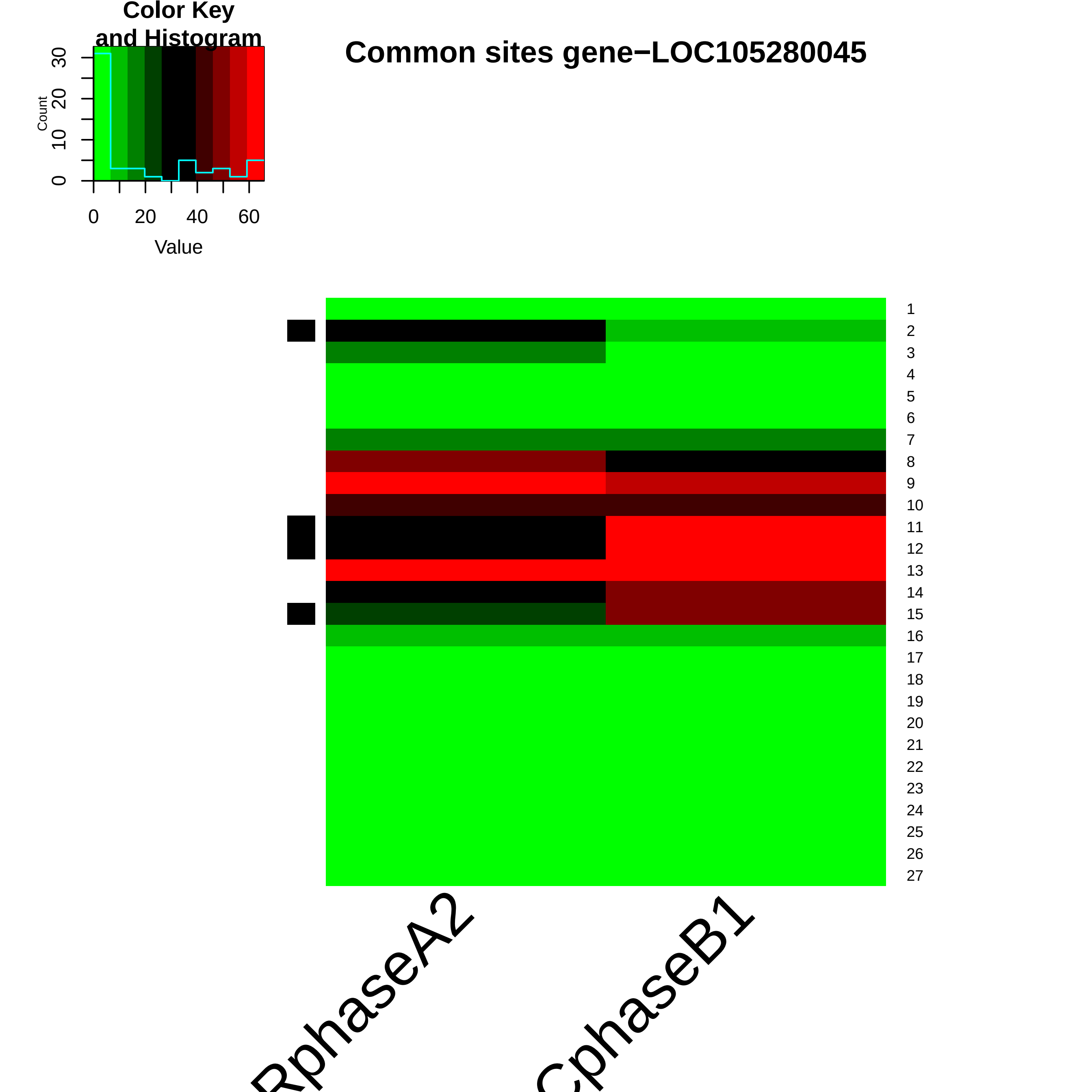

Color Key
and Histogram
Count
0
10
20
30
0
20
40
60
Value
Common sites gene−LOC105280045
RphaseA2
CphaseB1

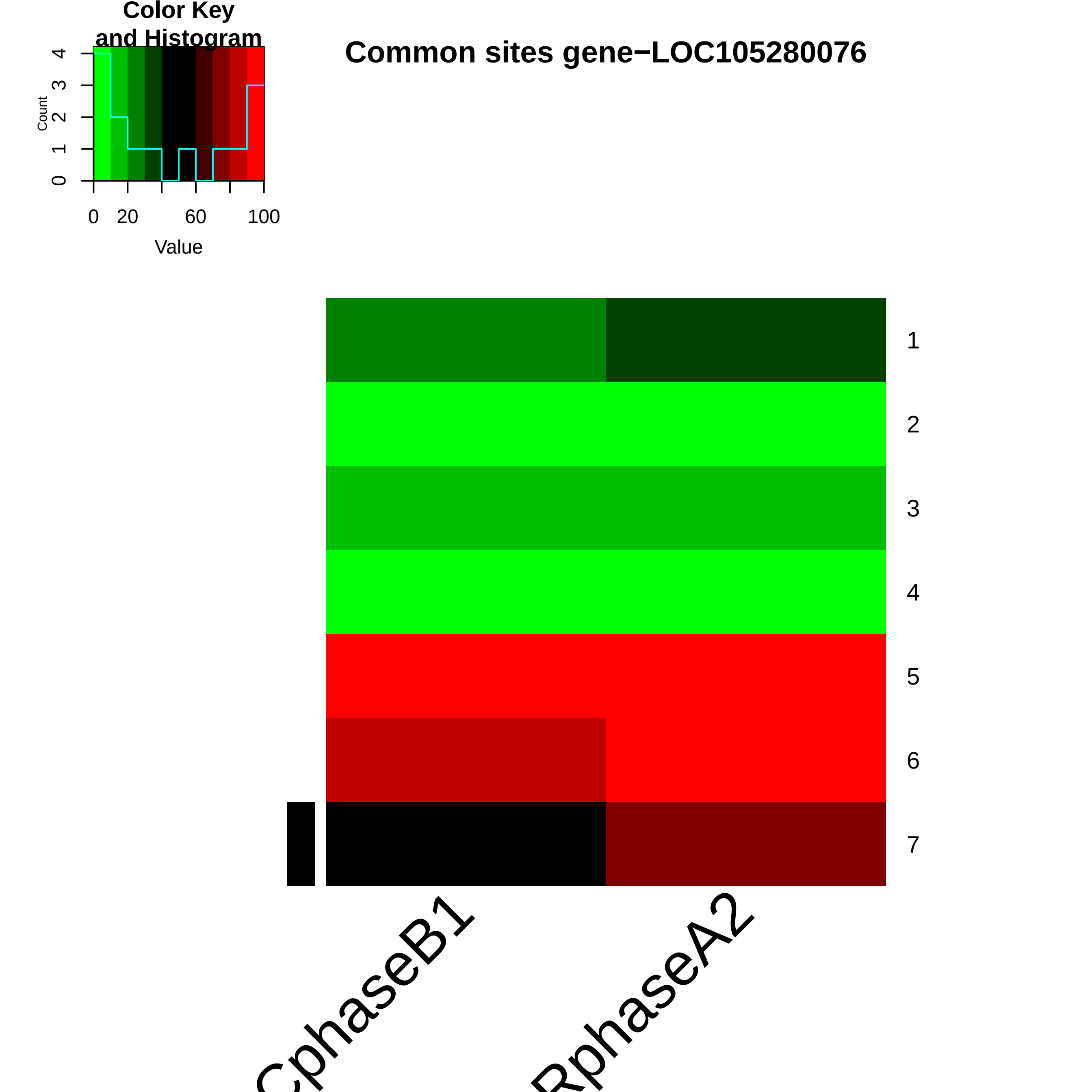

Color Key
and Histogram
Count
0
1
2
3
4
0
20
60
100
Value
Common sites gene−LOC105280076
1
2
3
4
5
6
7
CphaseB1
RphaseA2

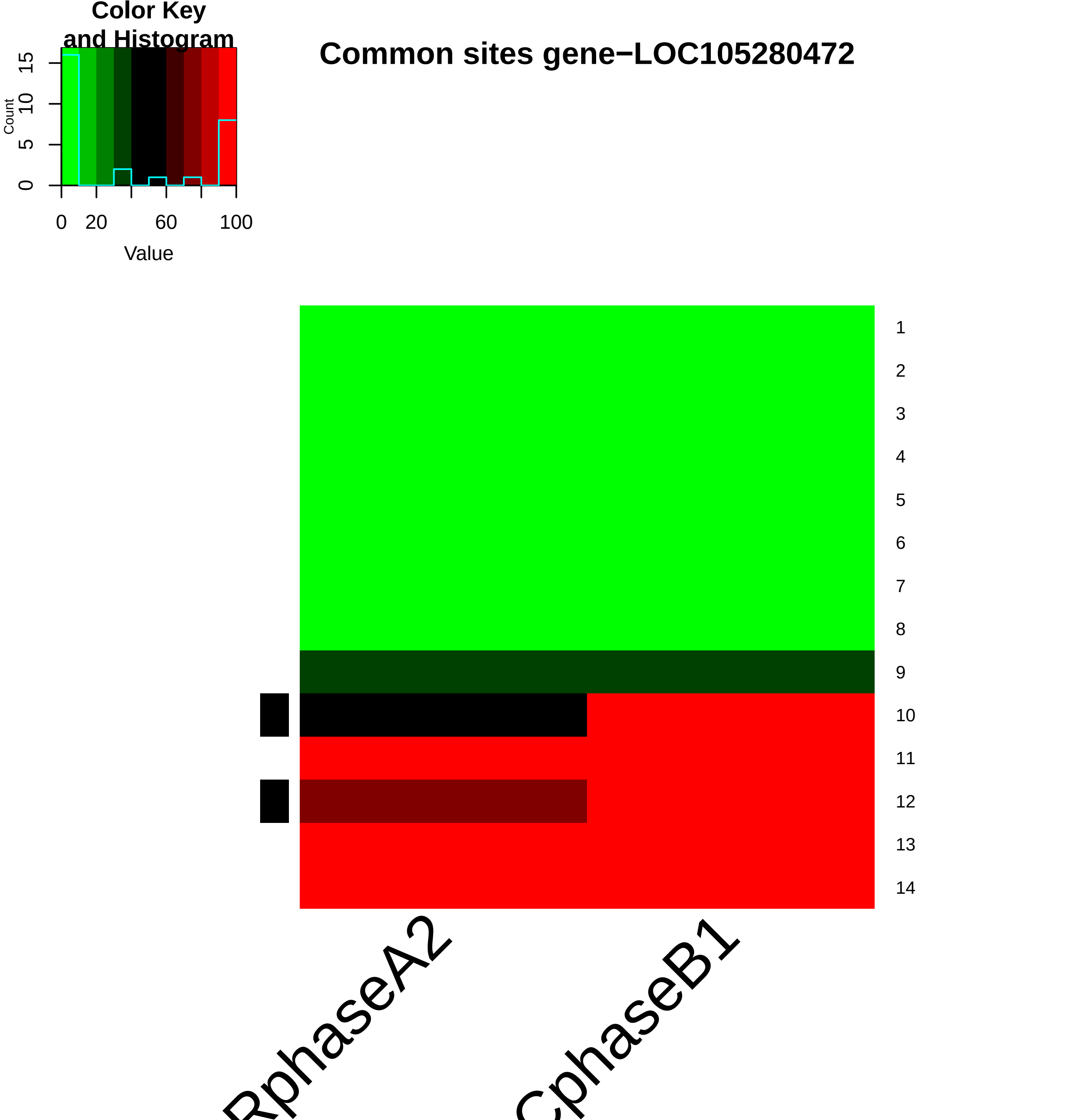

Color Key
and Histogram
Count
0
5
10
15
0
20
60
100
Value
Common sites gene−LOC105280472
1
2
3
4
5
6
7
8
9
10
11
12
13
14
RphaseA2
CphaseB1

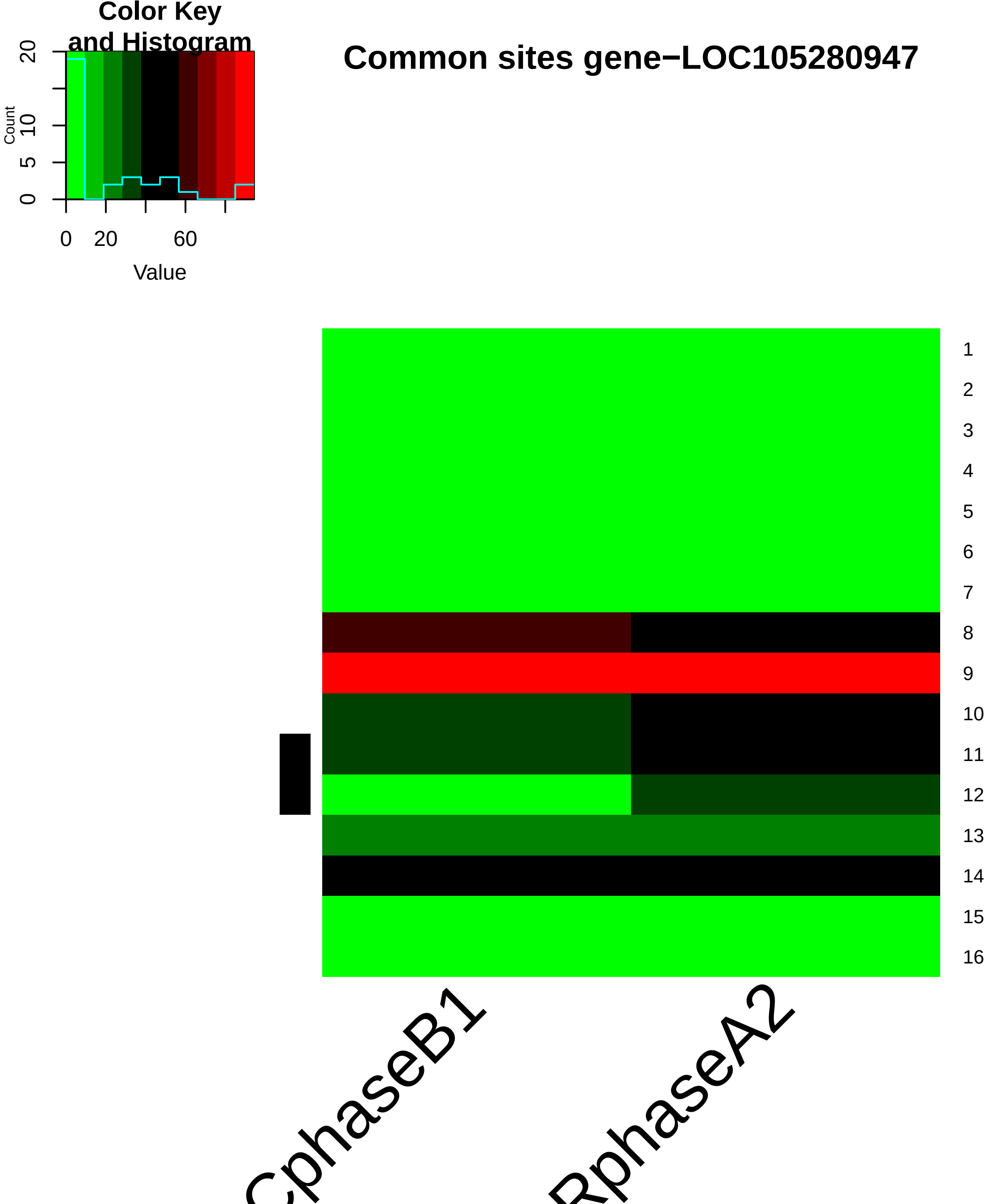
Color Key
and Histogram
Count
Value
0
20
60
0
5
10
20
Common sites gene−LOC105280947
CphaseB1
RphaseA2

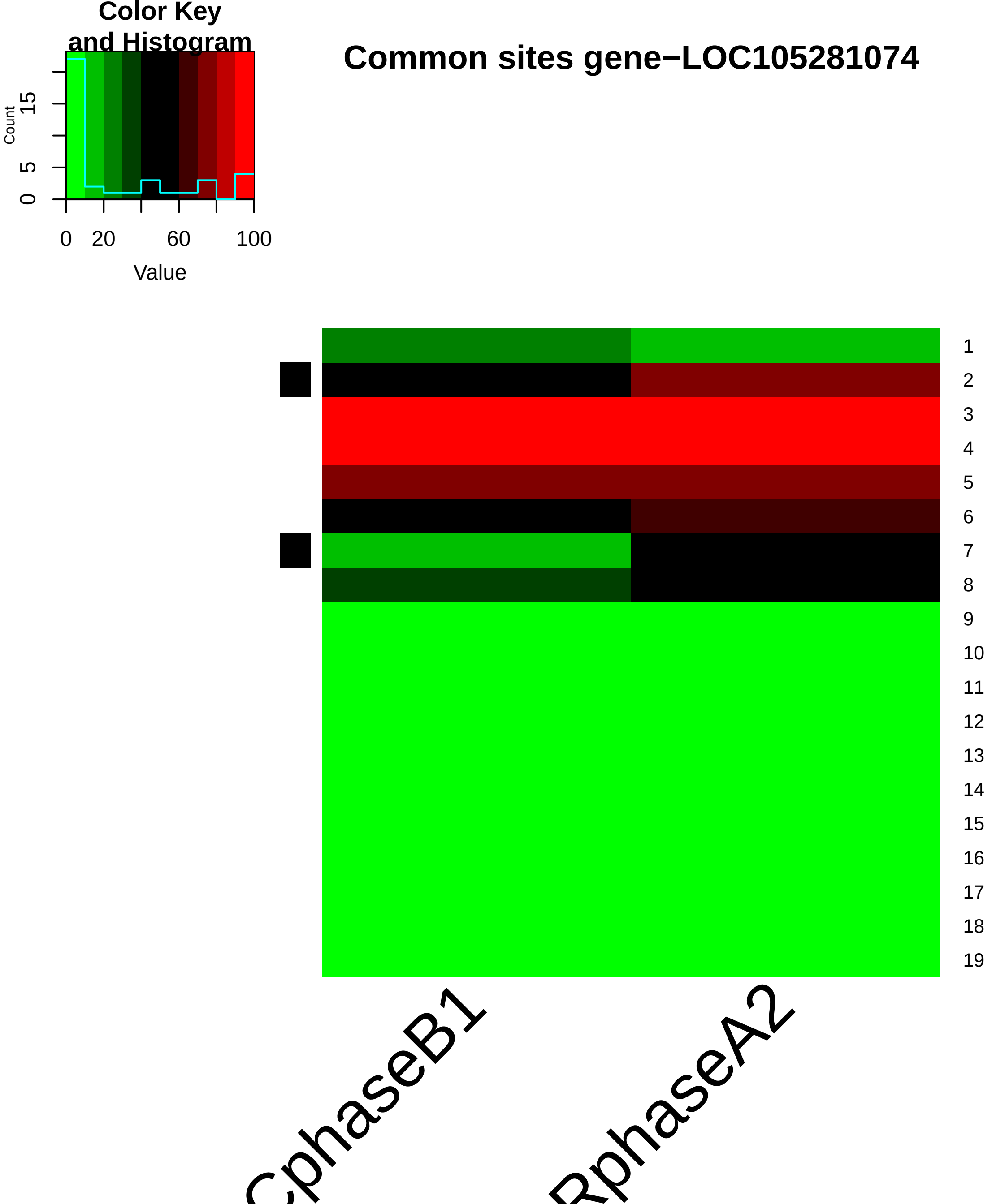

Color Key
and Histogram
Count
0
5
15
0
20
60
100
Value
Common sites gene−LOC105281074
CphaseB1
RphaseA2

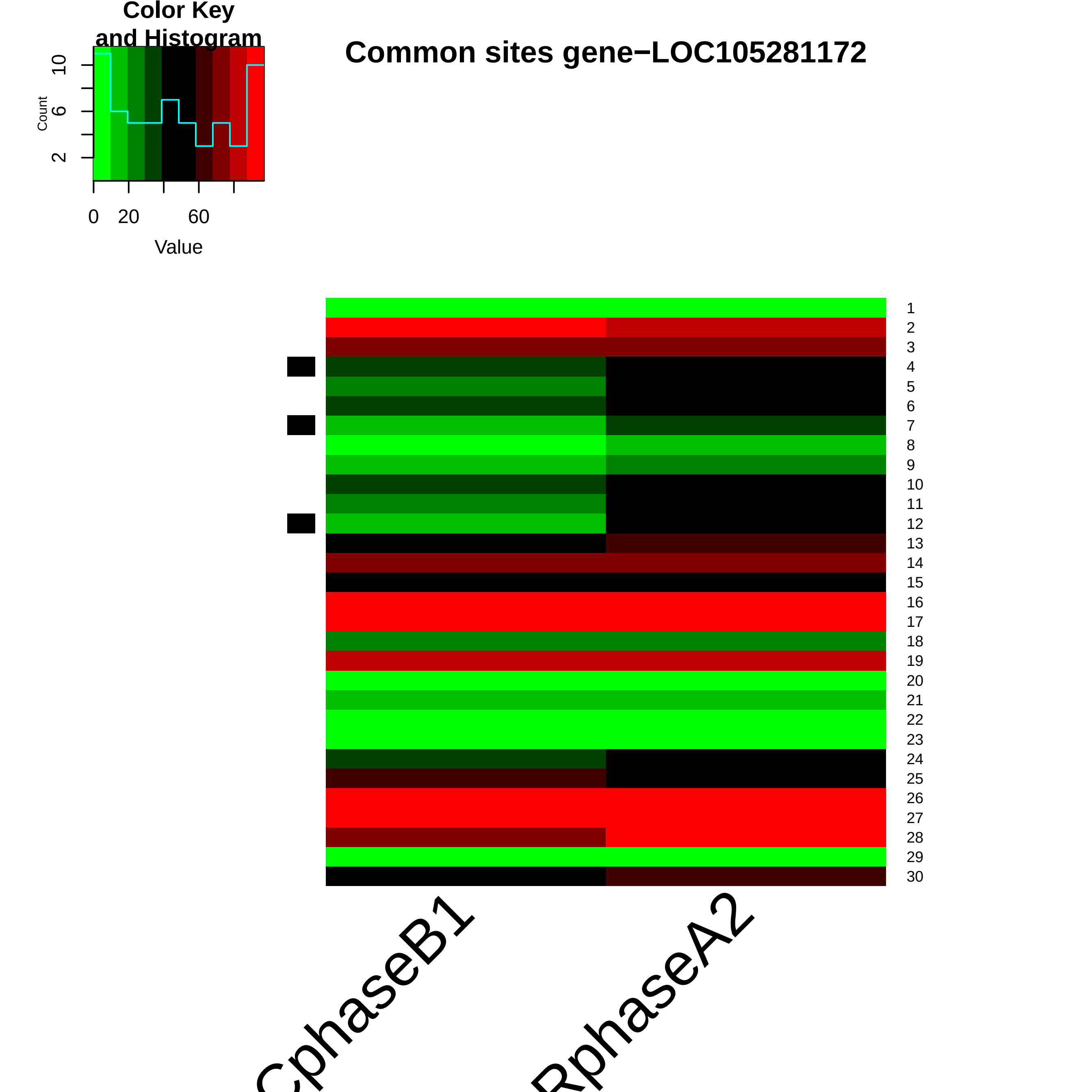
Color Key
and Histogram
Count
10
6
2
0
20
60
Value
Common sites gene−LOC105281172
CphaseB1
RphaseA2

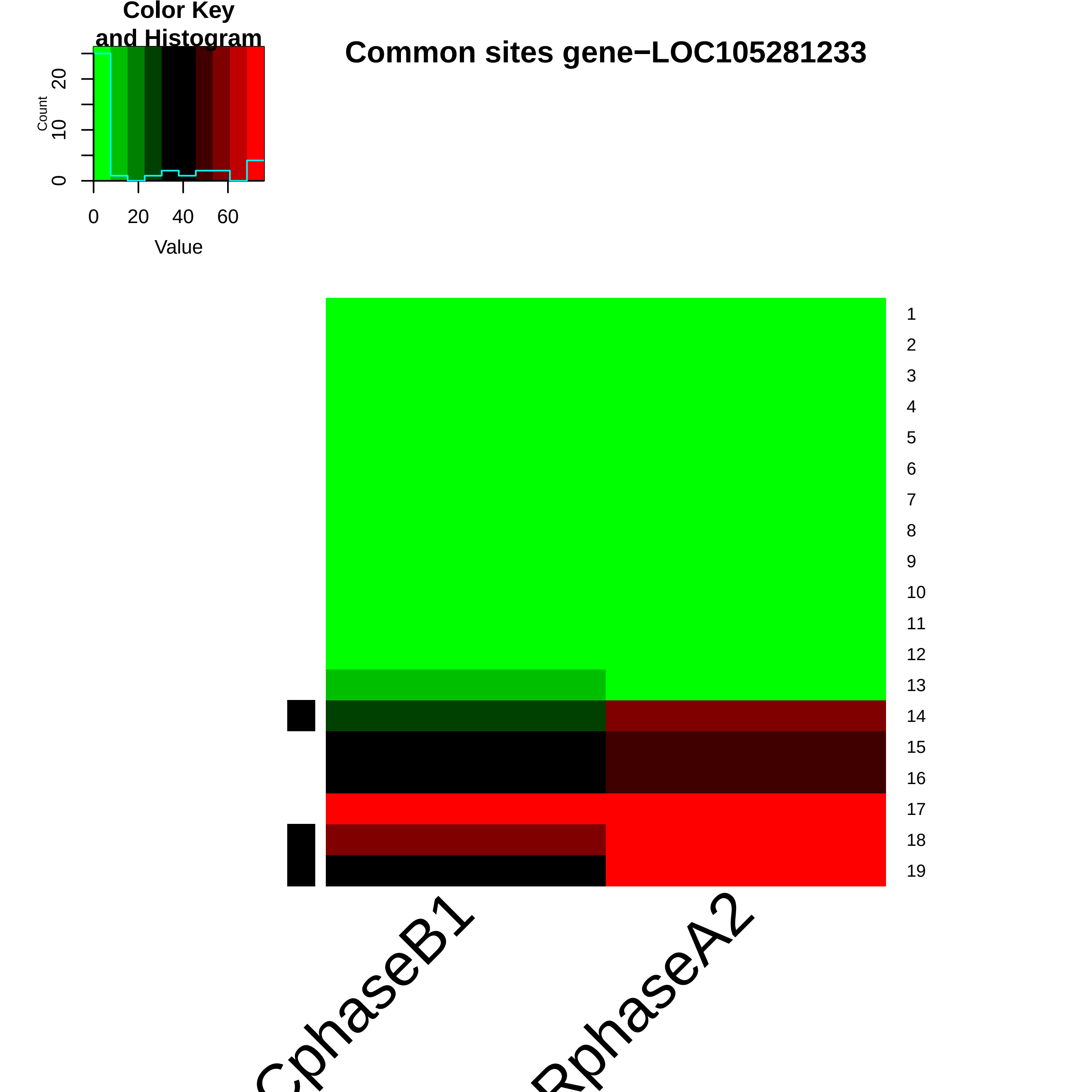

Color Key
and Histogram
Count
0
10
20
0
20
40
60
Value
Common sites gene−LOC105281233
CphaseB1
RphaseA2

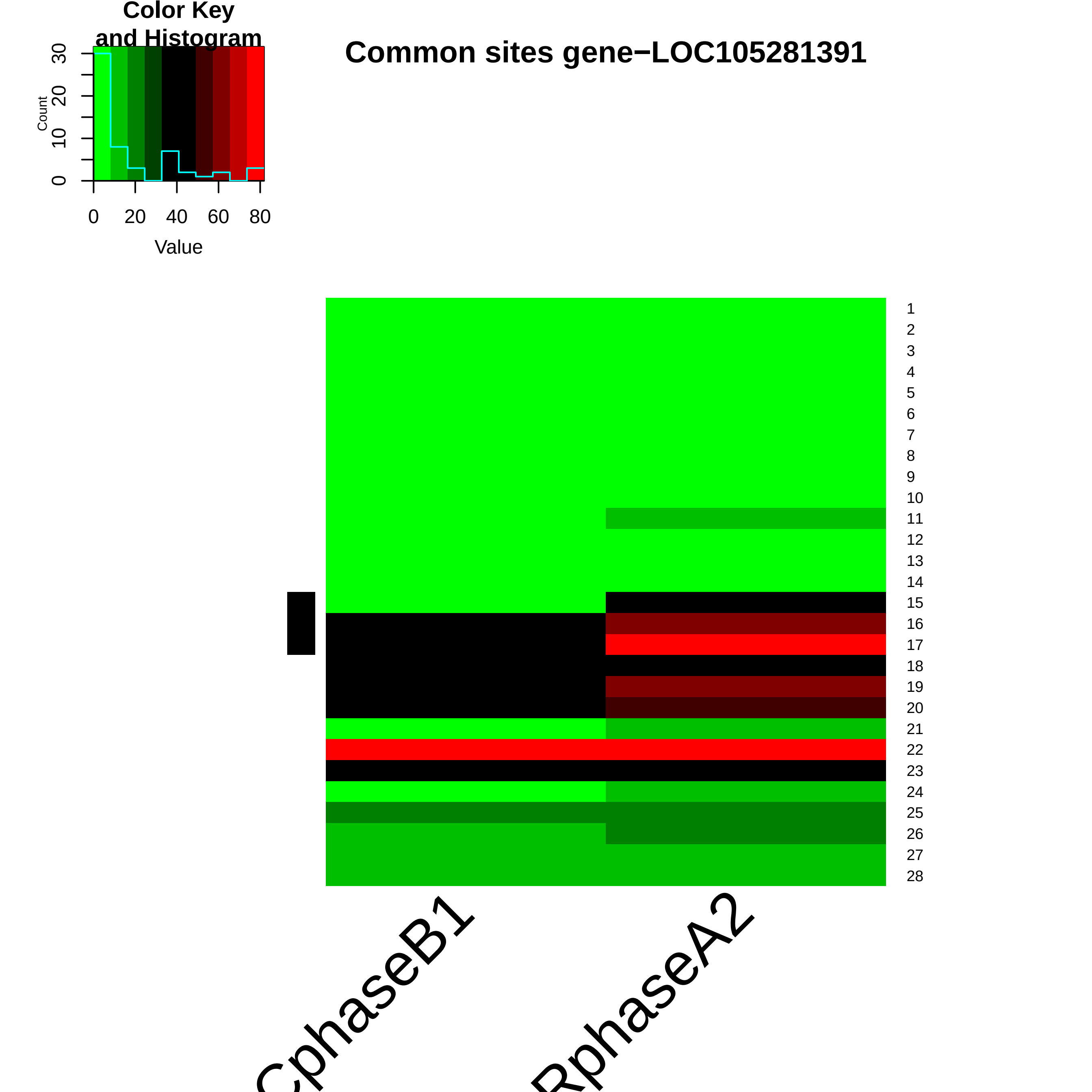

Color Key
and Histogram
Count
0
10
20
30
0
20
40
60
80
Value
Common sites gene−LOC105281391
CphaseB1
RphaseA2

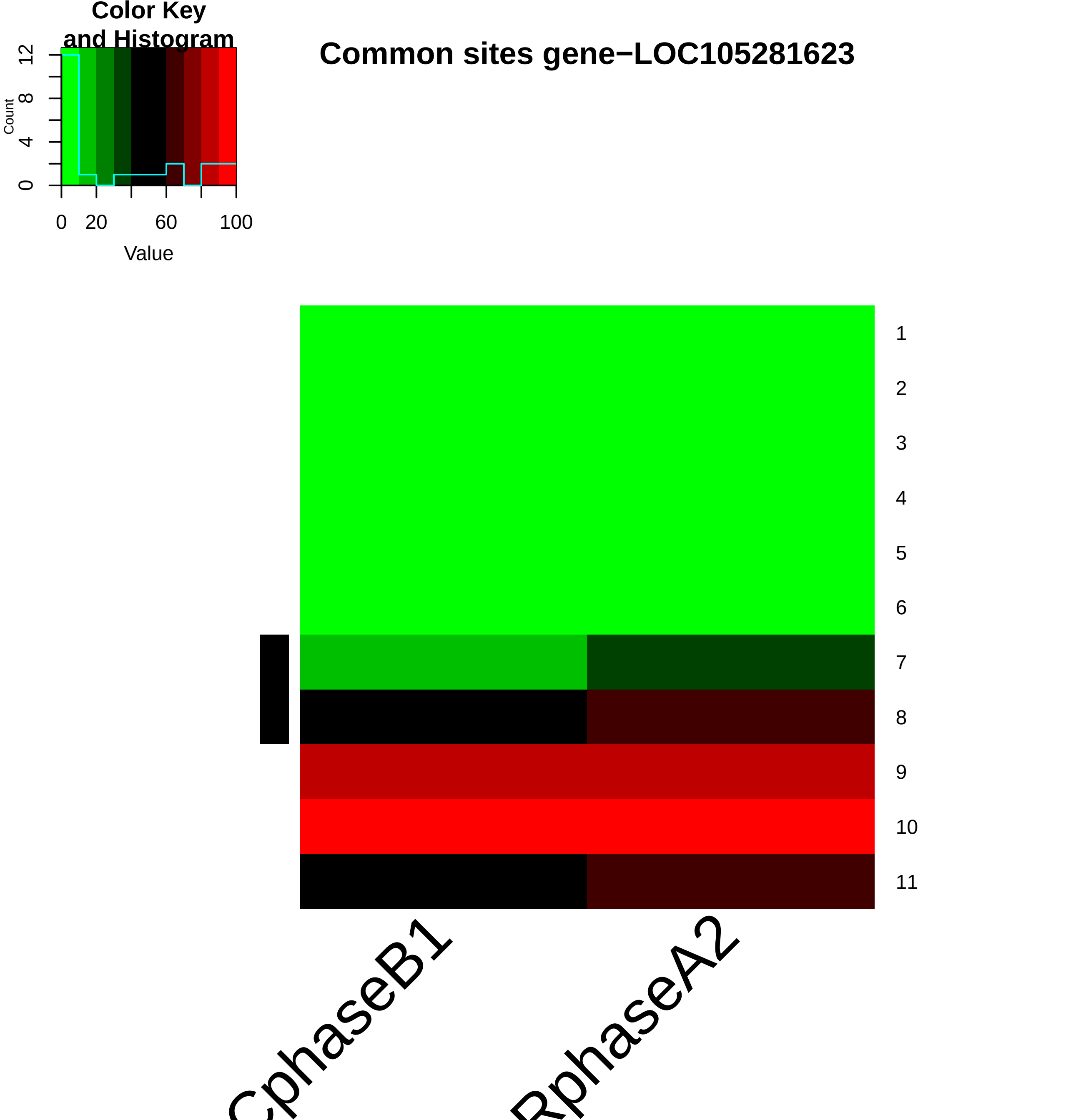

Color Key
and Histogram
Count
0
4
8
12
0
20
60
100
Value
Common sites gene−LOC105281623
1
2
3
4
5
6
7
8
9
10
11
CphaseB1
RphaseA2

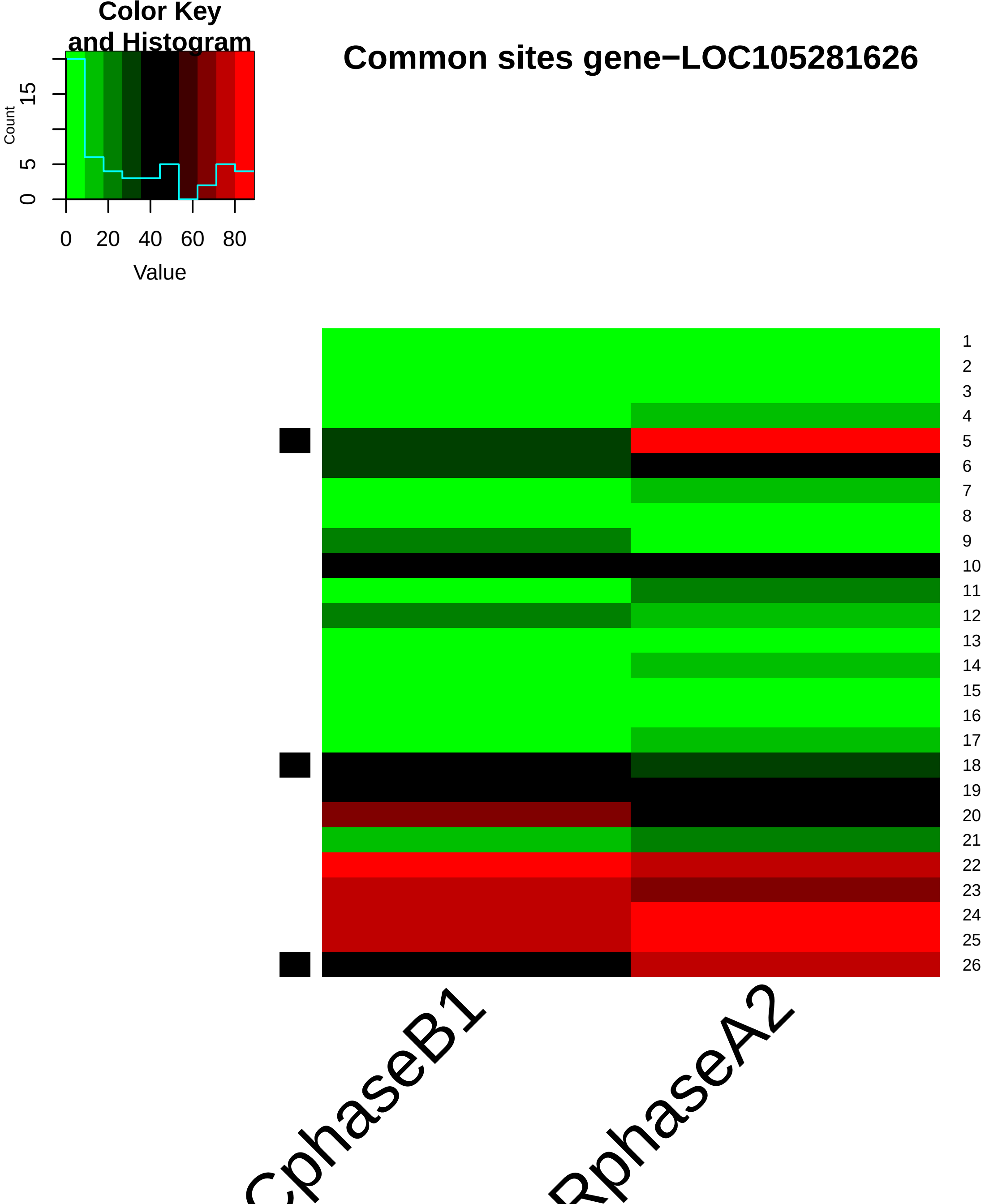

Color Key
and Histogram
Count
0
5
15
0
20
40
60
80
Value
Common sites gene−LOC105281626
CphaseB1
RphaseA2

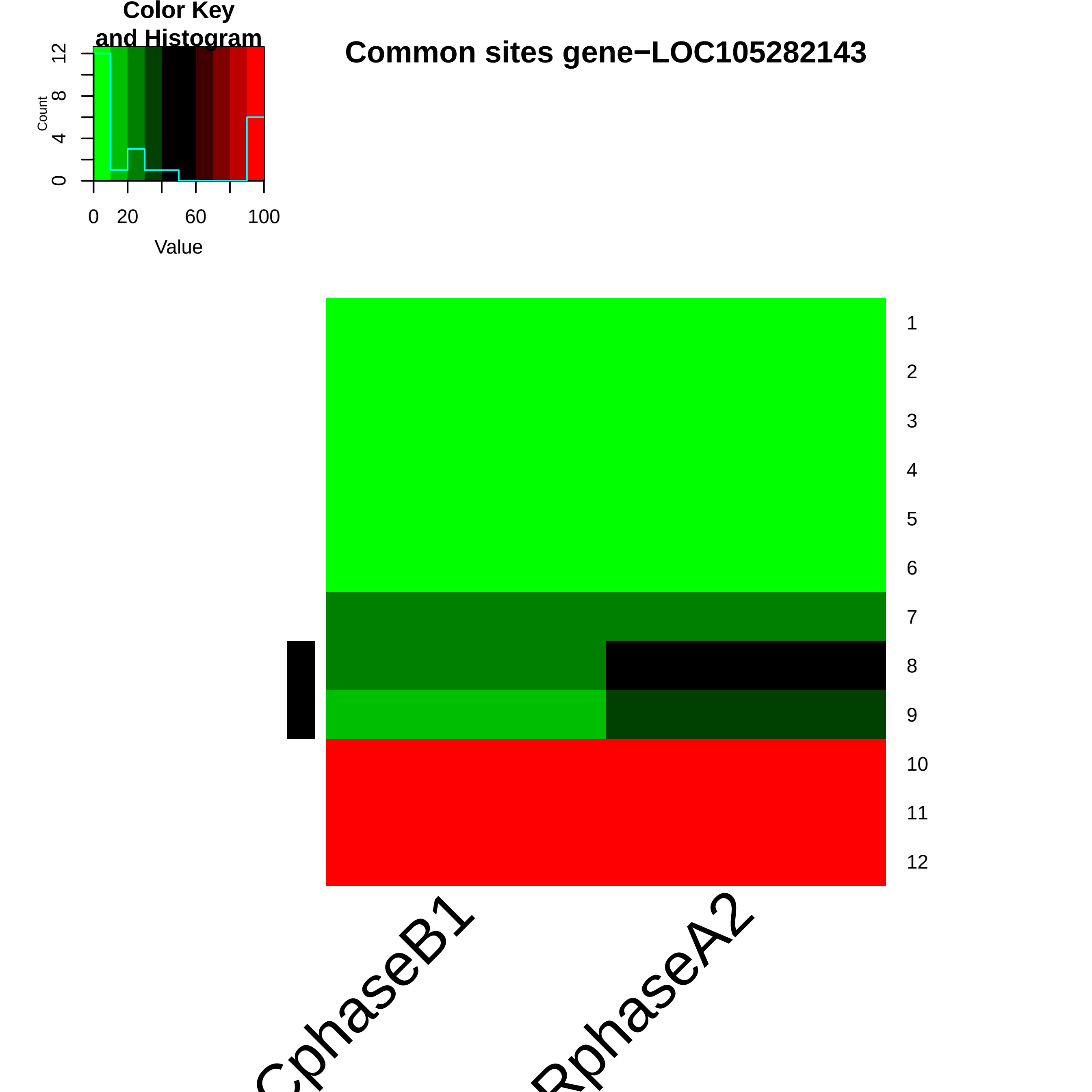
Color Key
and Histogram
Count
0
4
8
12
0
20
60
100
Value
Common sites gene−LOC105282143
1
2
3
4
5
6
7
8
9
10
11
12
CphaseB1
RphaseA2

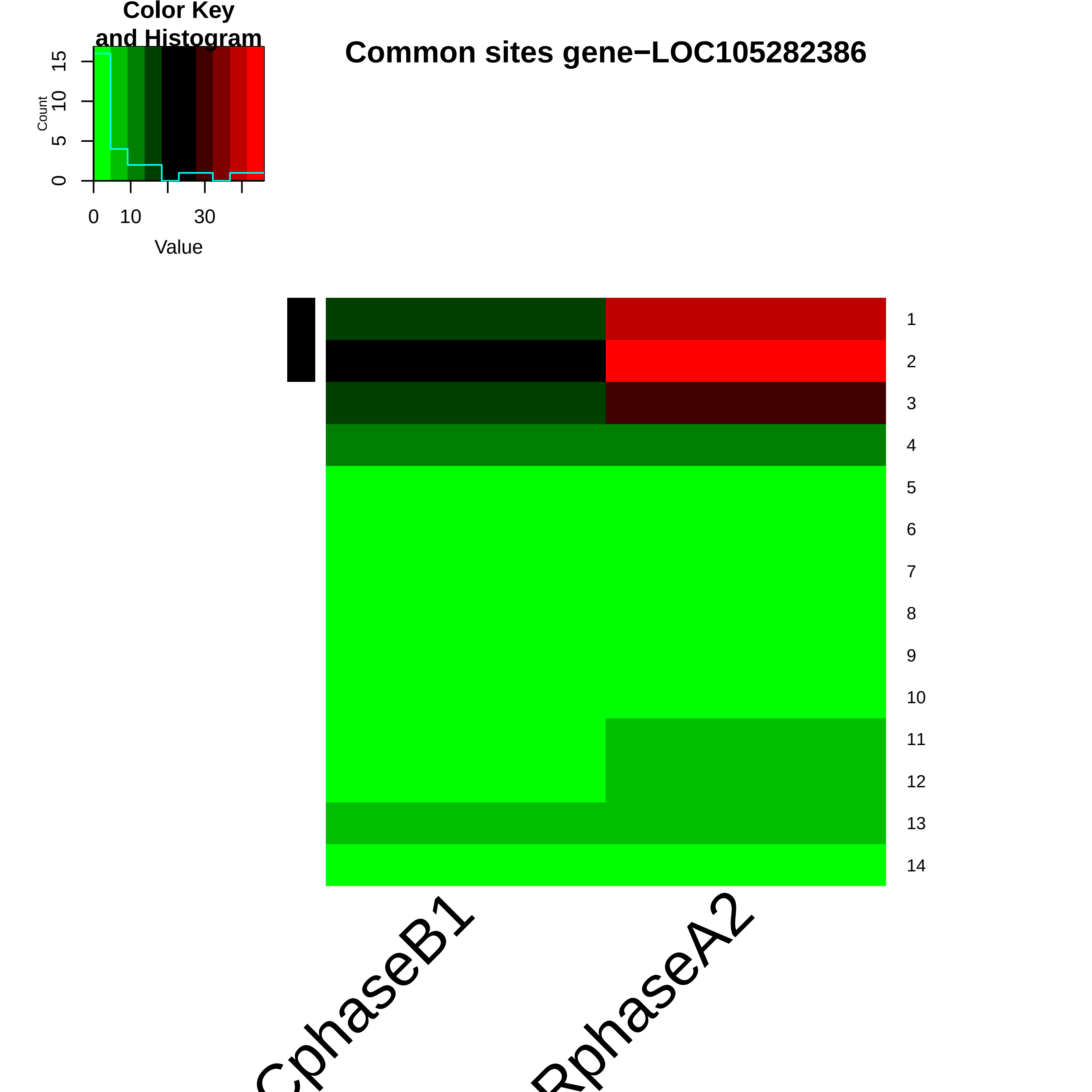
Color Key
and Histogram
Count
0
5
10
15
0
10
30
Value
Common sites gene−LOC105282386
1
2
3
4
5
6
7
8
9
10
11
12
13
14
CphaseB1
RphaseA2

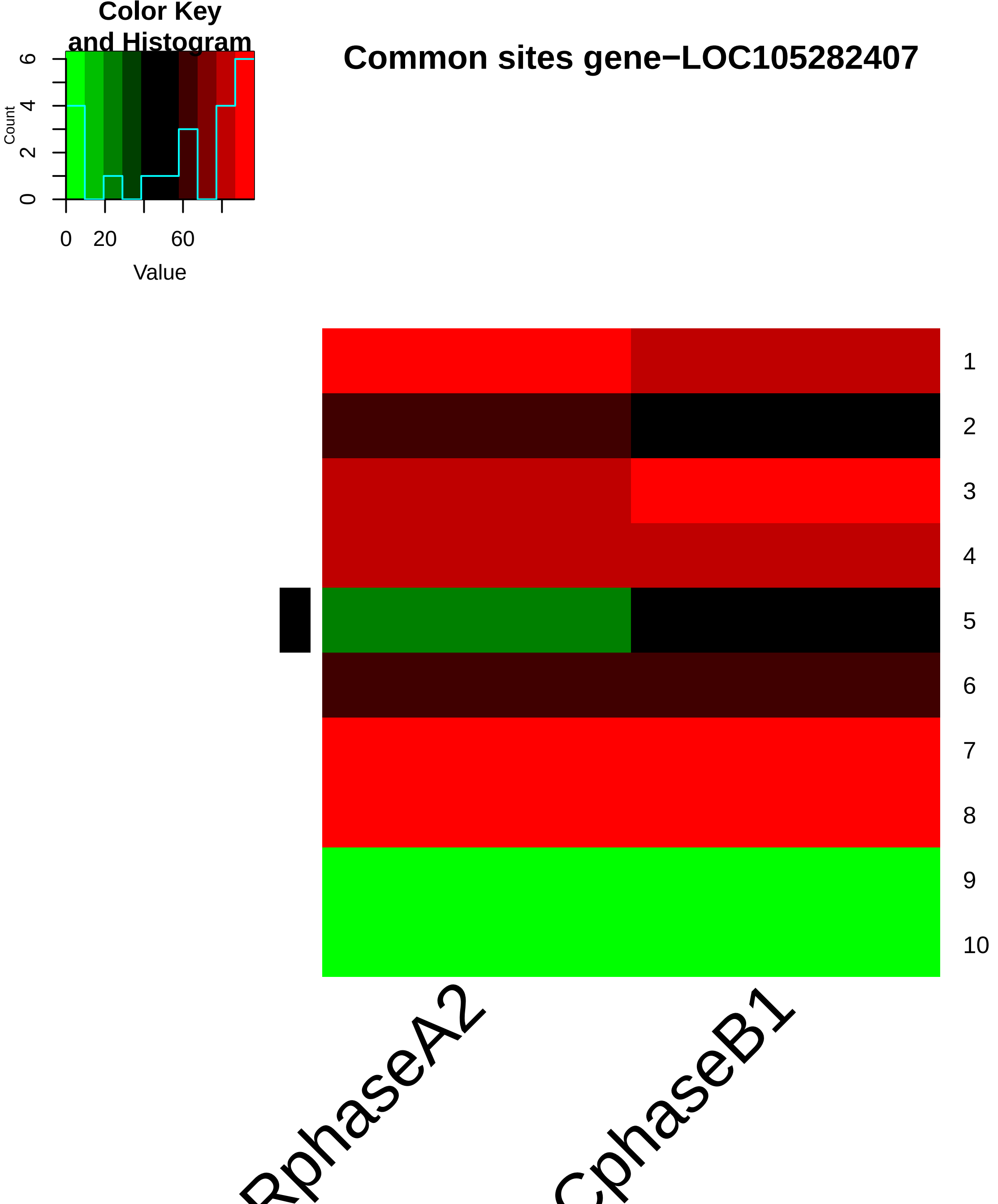
Color Key
and Histogram
Count
0
2
4
6
0
20
60
Value
Common sites gene−LOC105282407
1
2
3
4
5
6
7
8
9
10
RphaseA2
CphaseB1

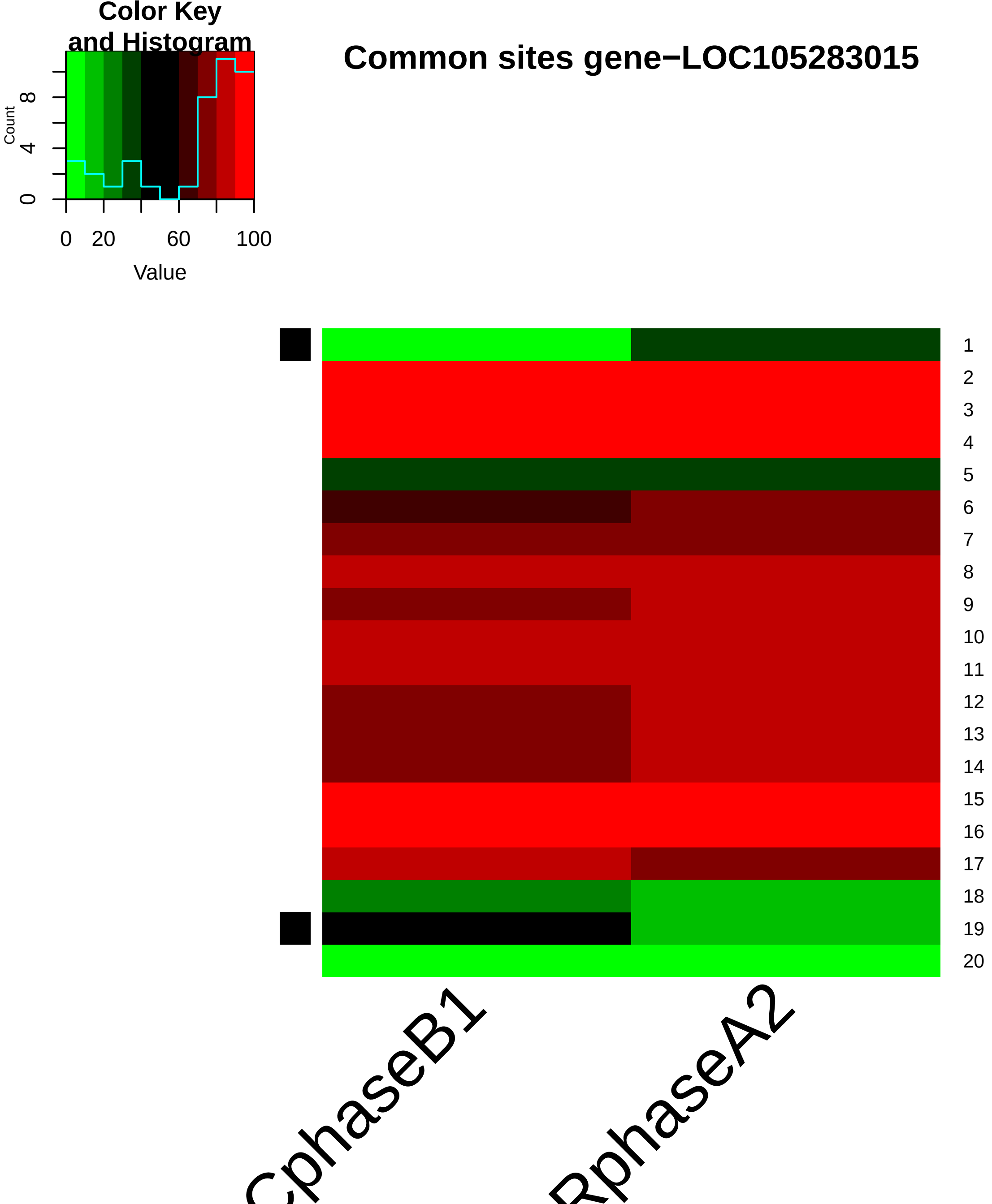
Color Key
and Histogram
Count
0
4
8
0
20
60
100
Value
Common sites gene−LOC105283015

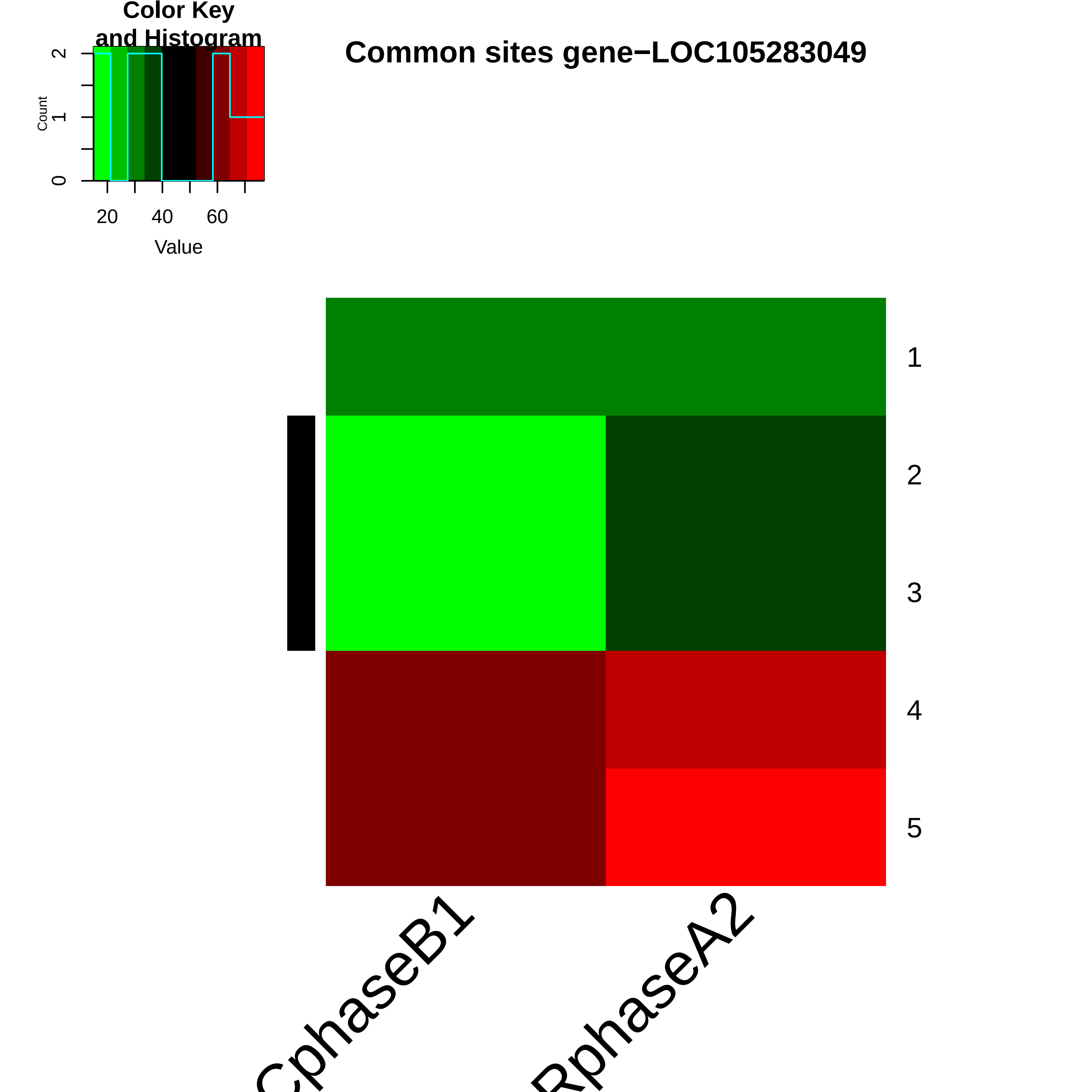
Color Key
and Histogram
Count
0
1
2
20
40
60
Value
Common sites gene−LOC105283049
1
2
3
4
5
CphaseB1
RphaseA2

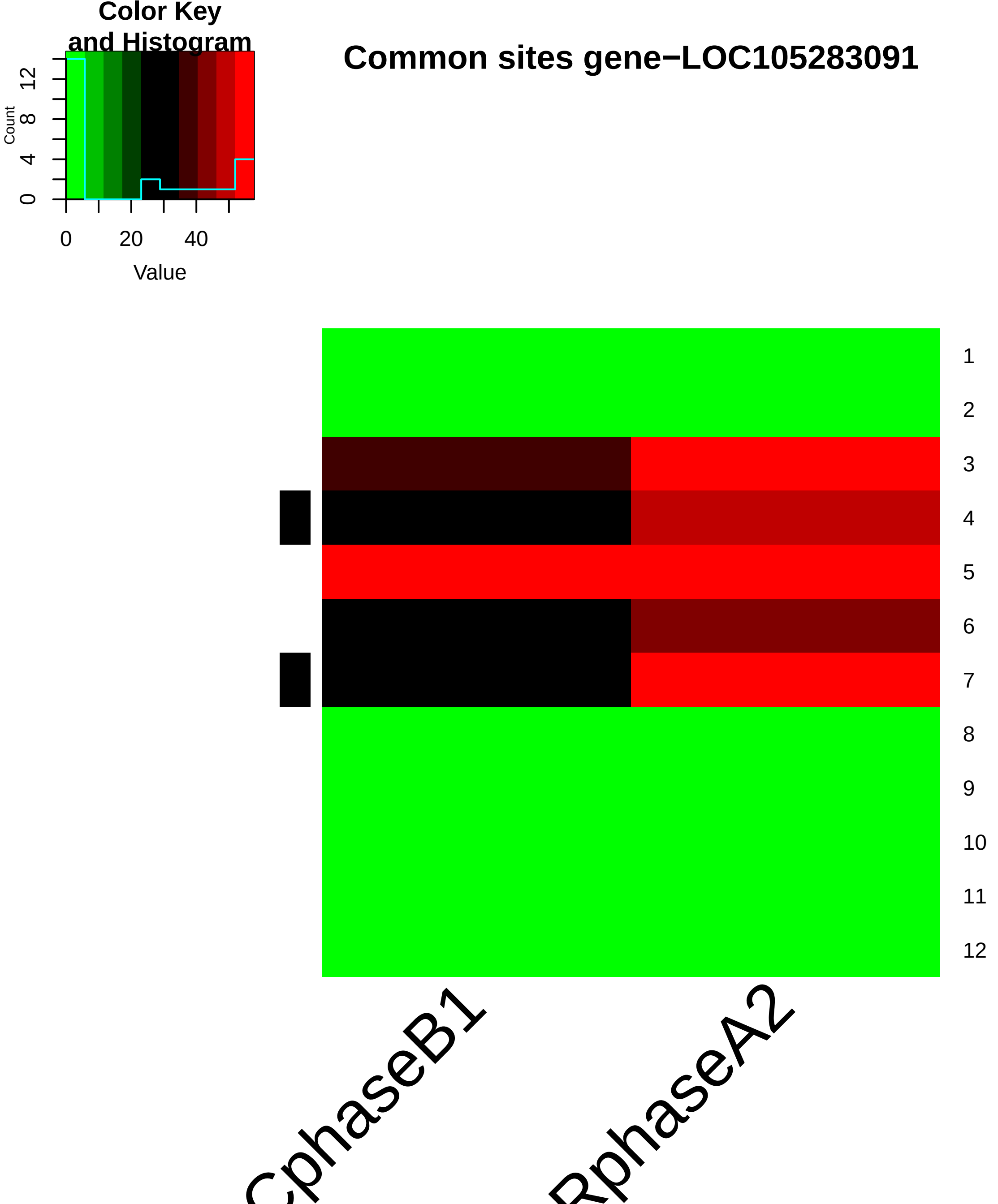

Color Key
and Histogram
Count
0
4
8
12
0
20
40
Value
Common sites gene−LOC105283091
1
2
3
4
5
6
7
8
9
10
11
12
CphaseB1
RphaseA2

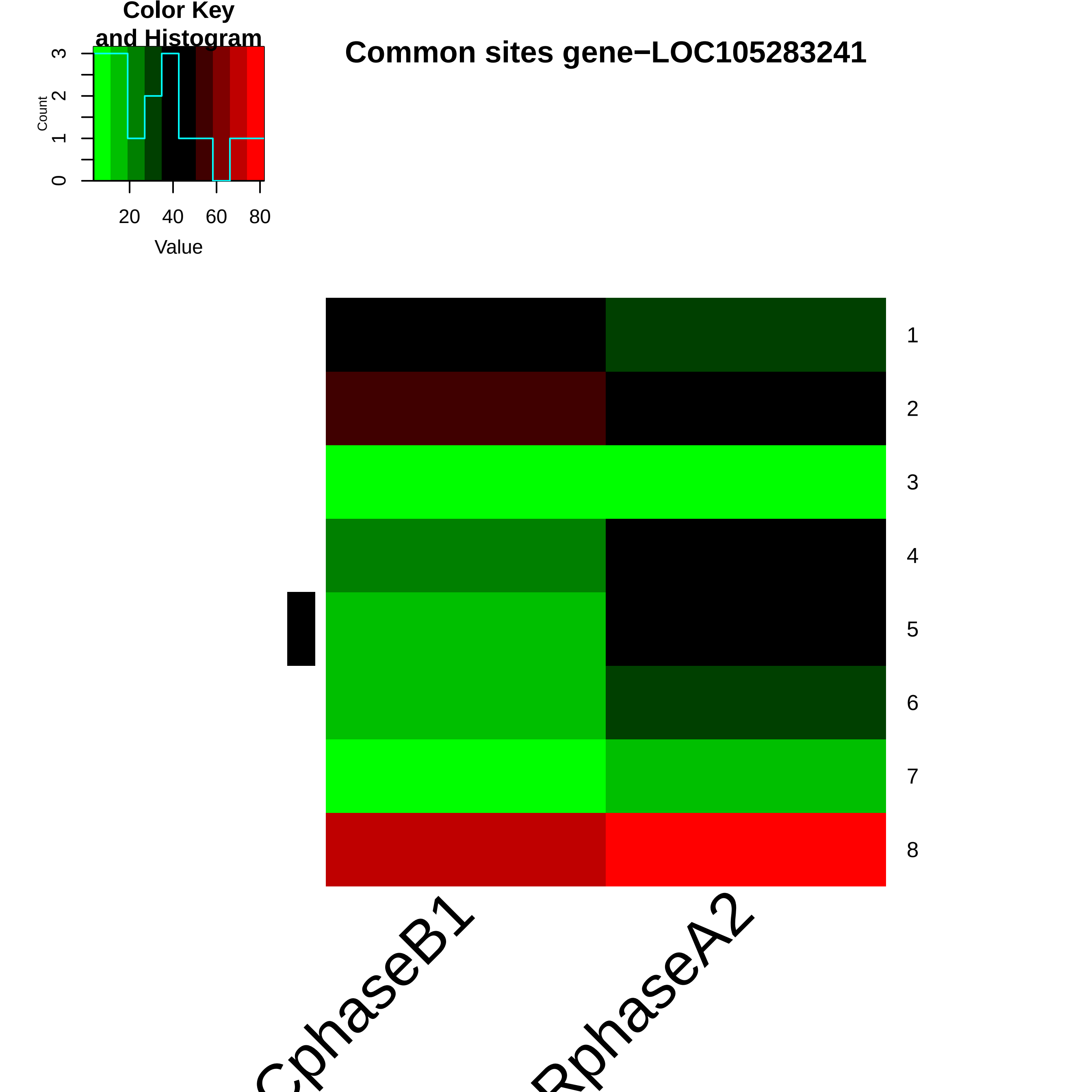

Color Key
and Histogram
Count
0
1
2
3
20
40
60
80
Value
Common sites gene−LOC105283241
1
2
3
4
5
6
7
8
CphaseB1
RphaseA2

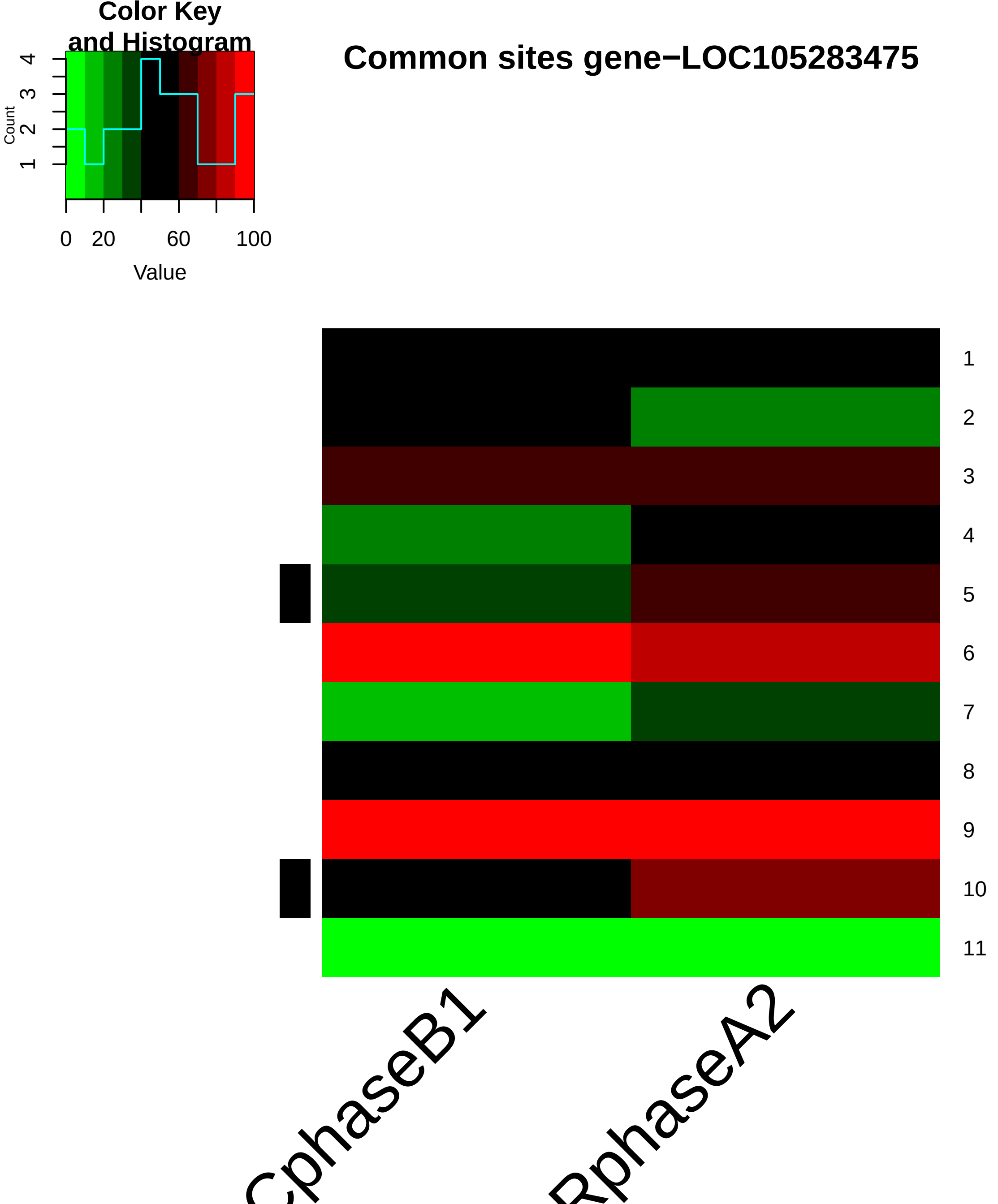
Color Key
and Histogram
Count
1
2
3
4
0
20
60
100
Value
Common sites gene−LOC105283475
1
2
3
4
5
6
7
8
9
10
11
CphaseB1
RphaseA2

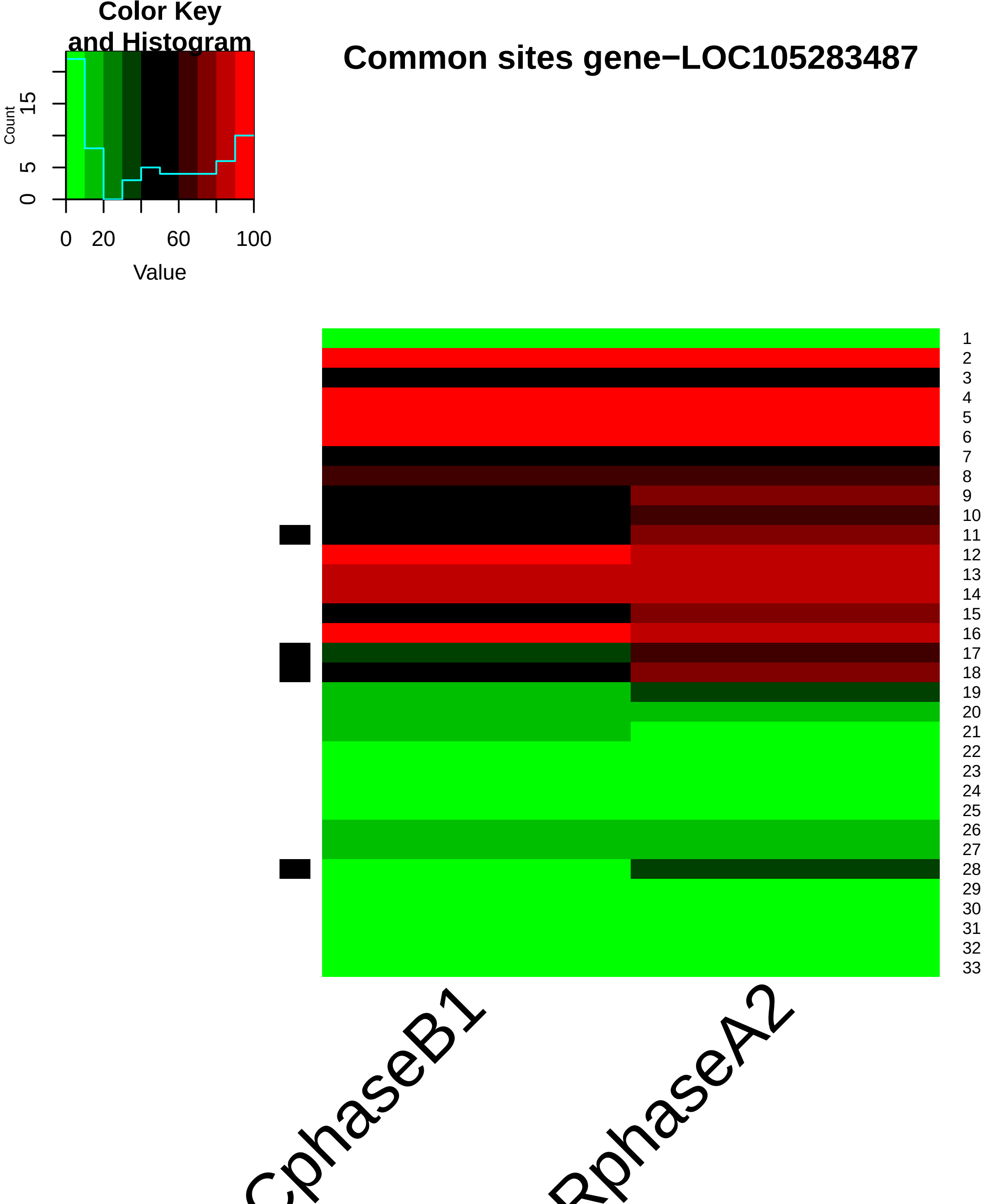

Color Key and Histogram
Count
0
5
15
0
20
60
100
Value
Common sites gene−LOC105283487
CphaseB1
RphaseA2

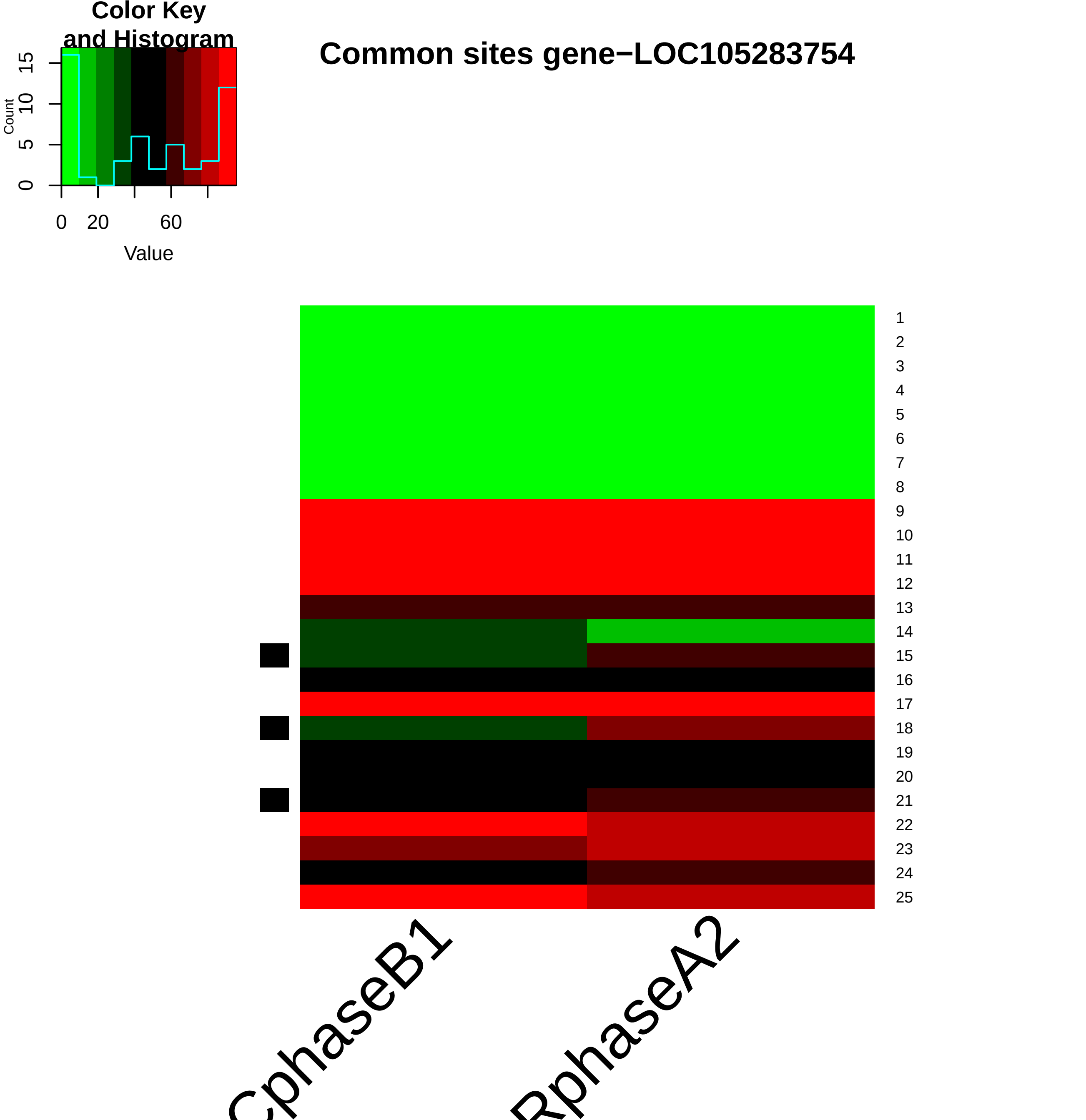
Color Key
and Histogram
Count
0
5
10
15
0
20
60
Value
Common sites gene−LOC105283754
CphaseB1
RphaseA2

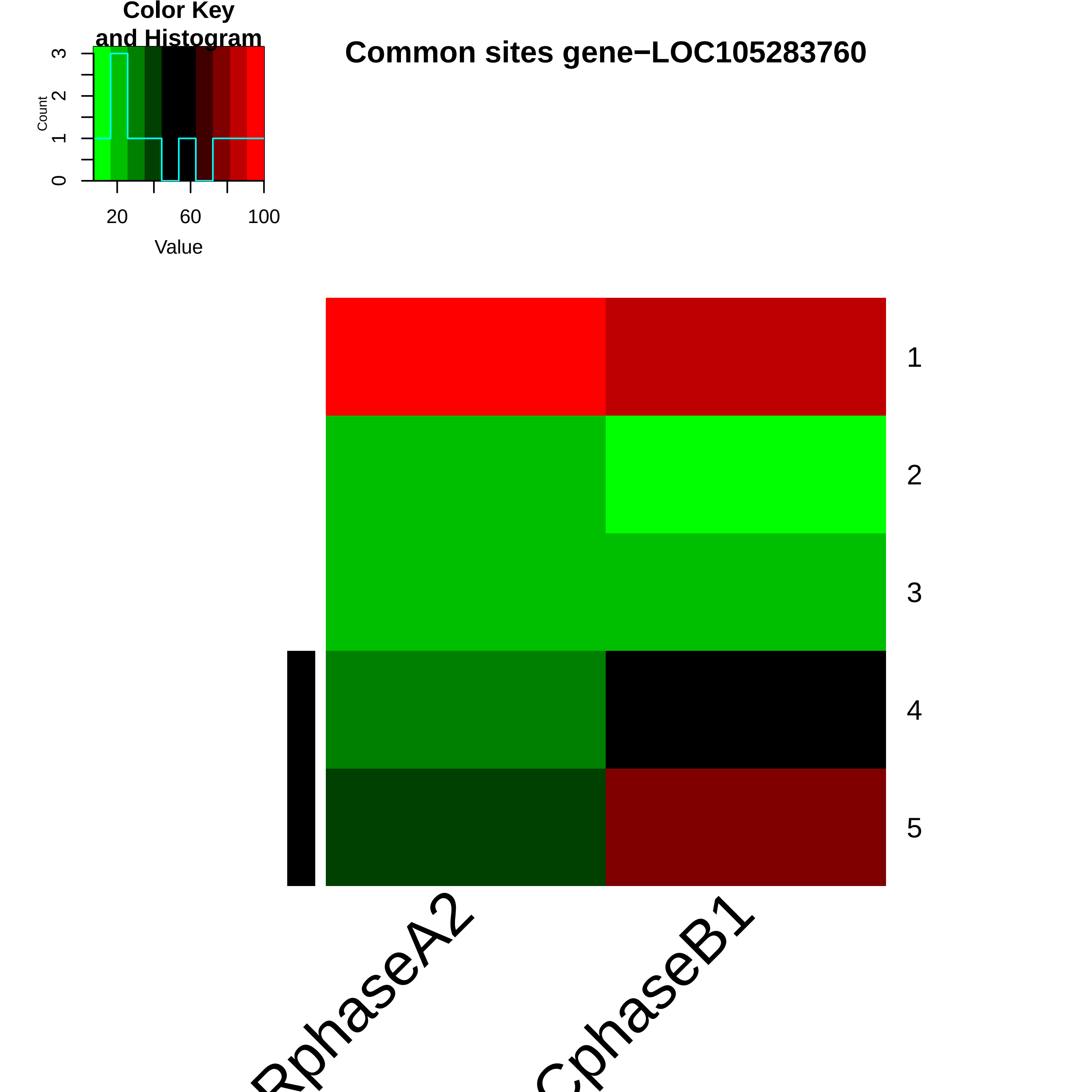
Color Key
and Histogram
Count
0
1
2
3
20
60
100
Value
Common sites gene−LOC105283760
1
2
3
4
5
RphaseA2
CphaseB1

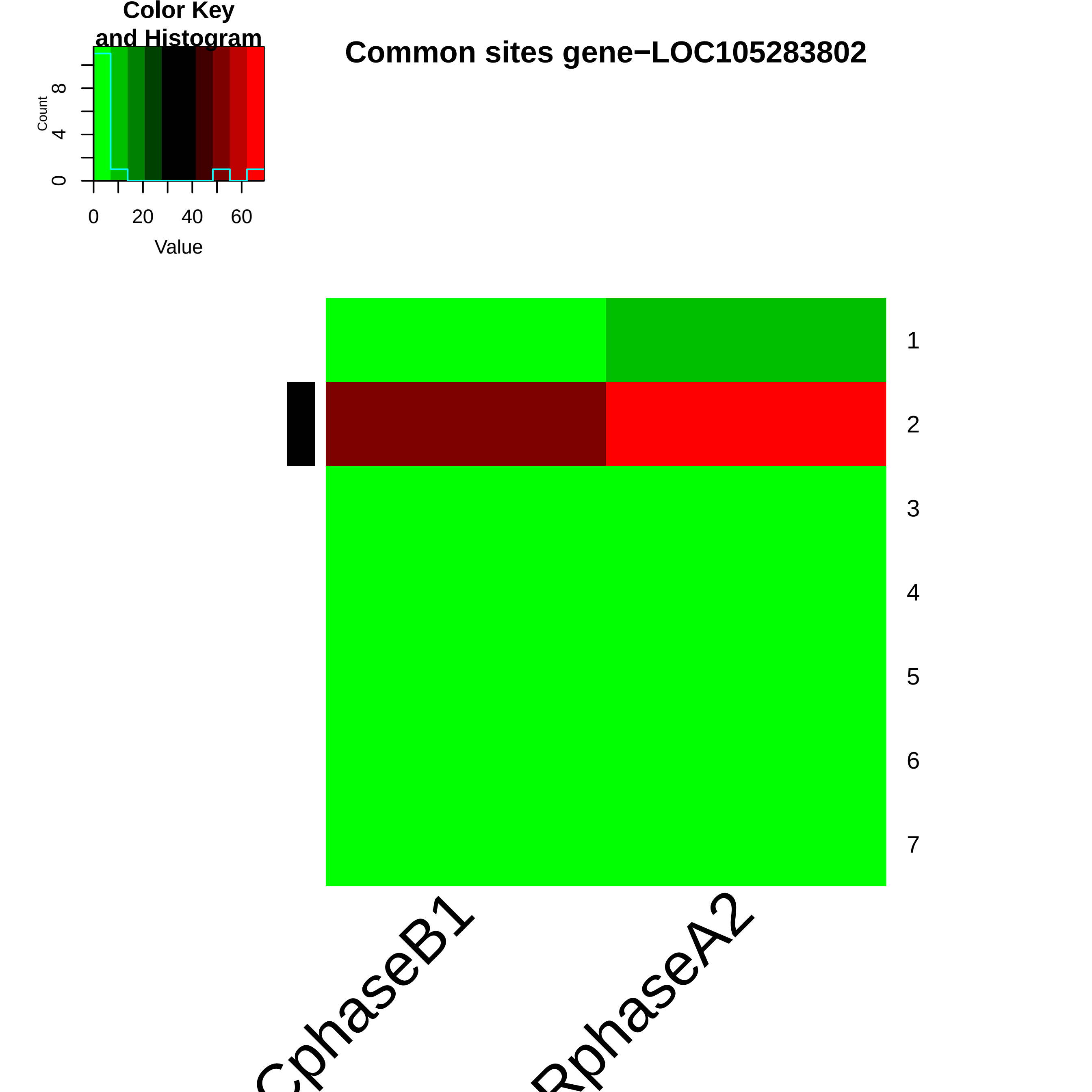
Color Key
and Histogram
Count
0
4
8
0
20
40
60
Value
Common sites gene−LOC105283802
1
2
3
4
5
6
7
CphaseB1
RphaseA2

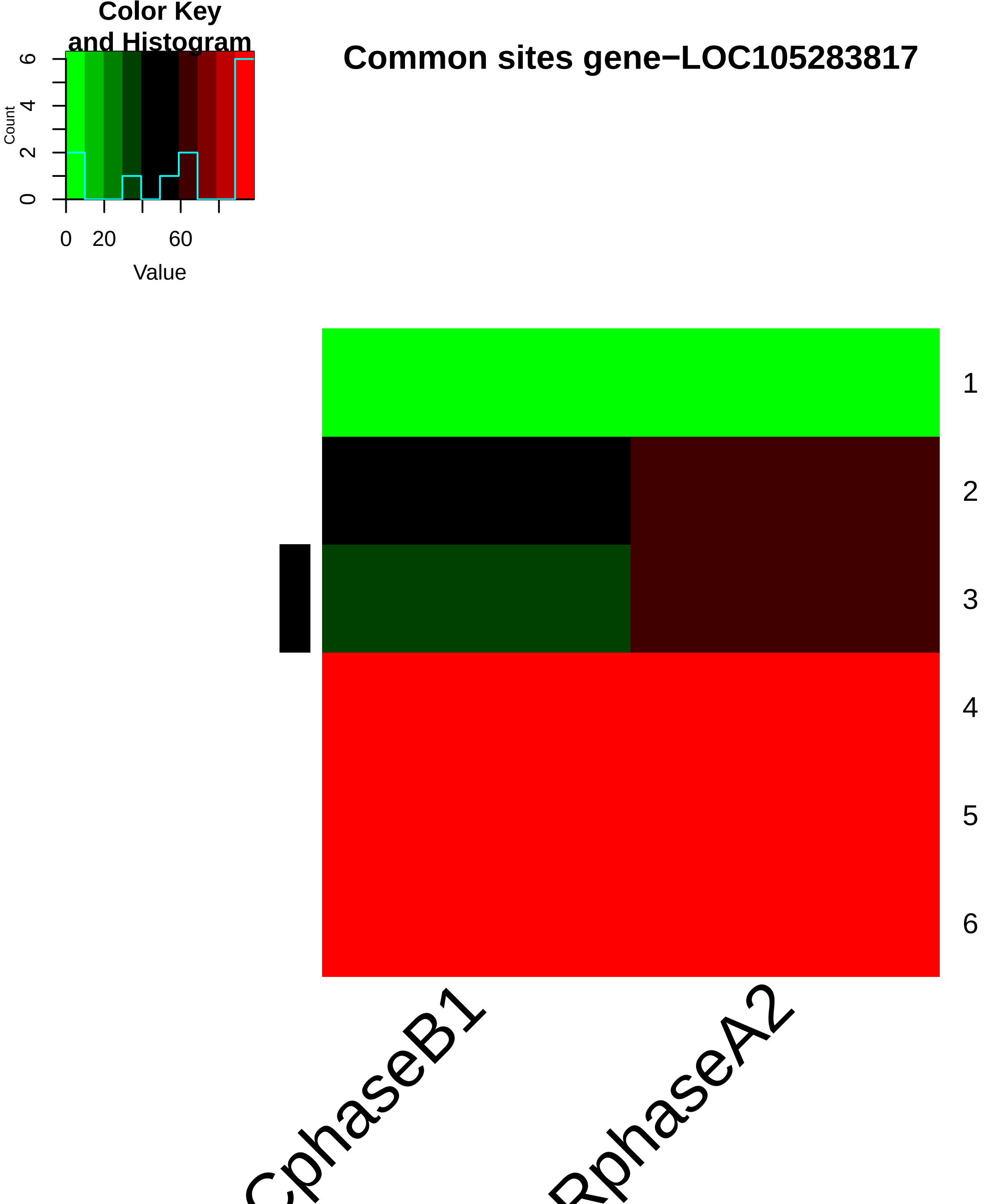
Color Key
and Histogram
Count
0
2
4
6
0
20
60
Value
Common sites gene−LOC105283817
1
2
3
4
5
6
CphaseB1
RphaseA2

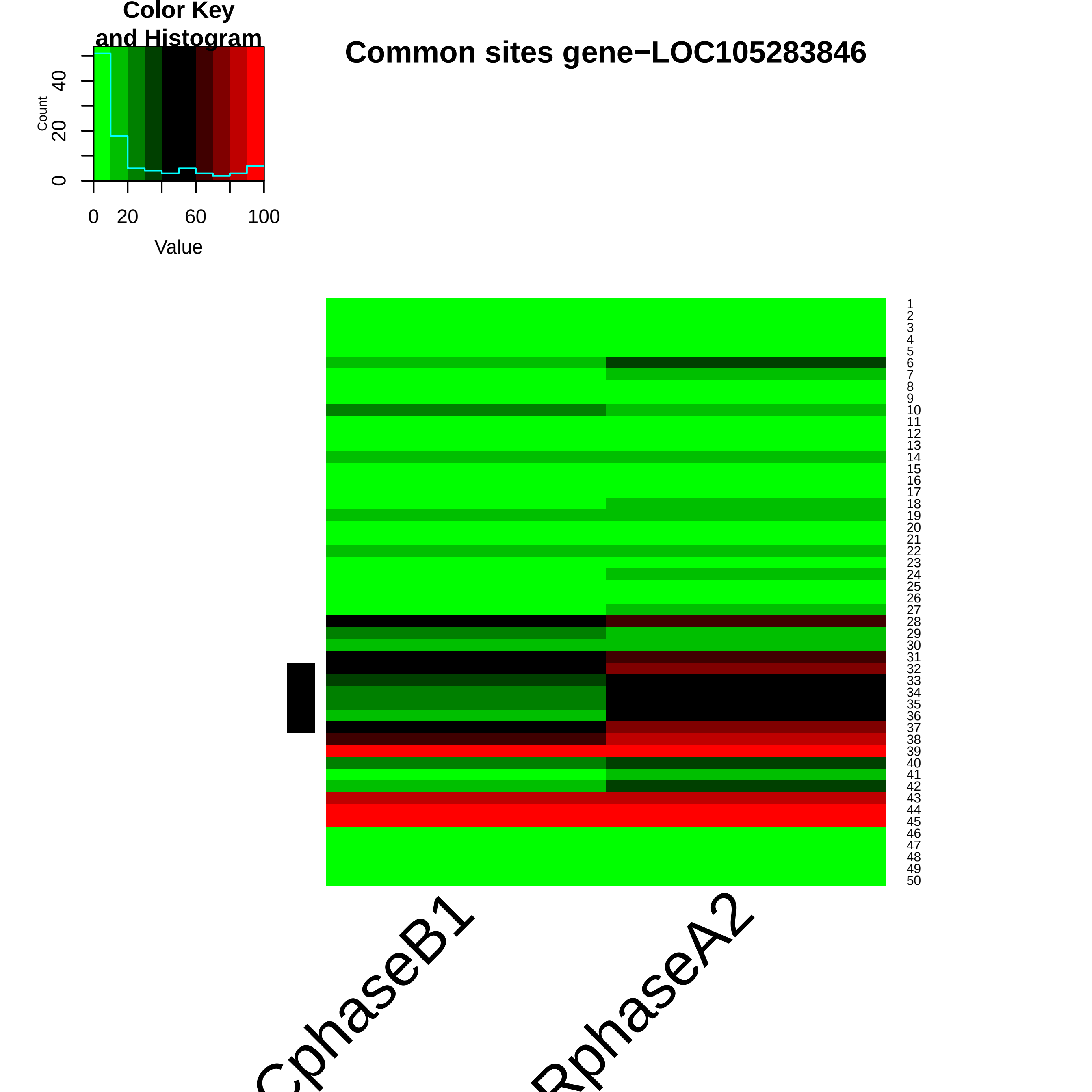
Color Key
and Histogram
Count
0
20
40
0
20
60
100
Value
Common sites gene−LOC105283846
1 2 3 4 5 6 7 8 9 10 11 12 13 14 15 16 17 18 19 20 21 22 23 24 25 26 27 28 29 30 31 32 33 34 35 36 37 38 39 40 41 42 43 44 45 46 47 48 49 50
CphaseB1
RphaseA2

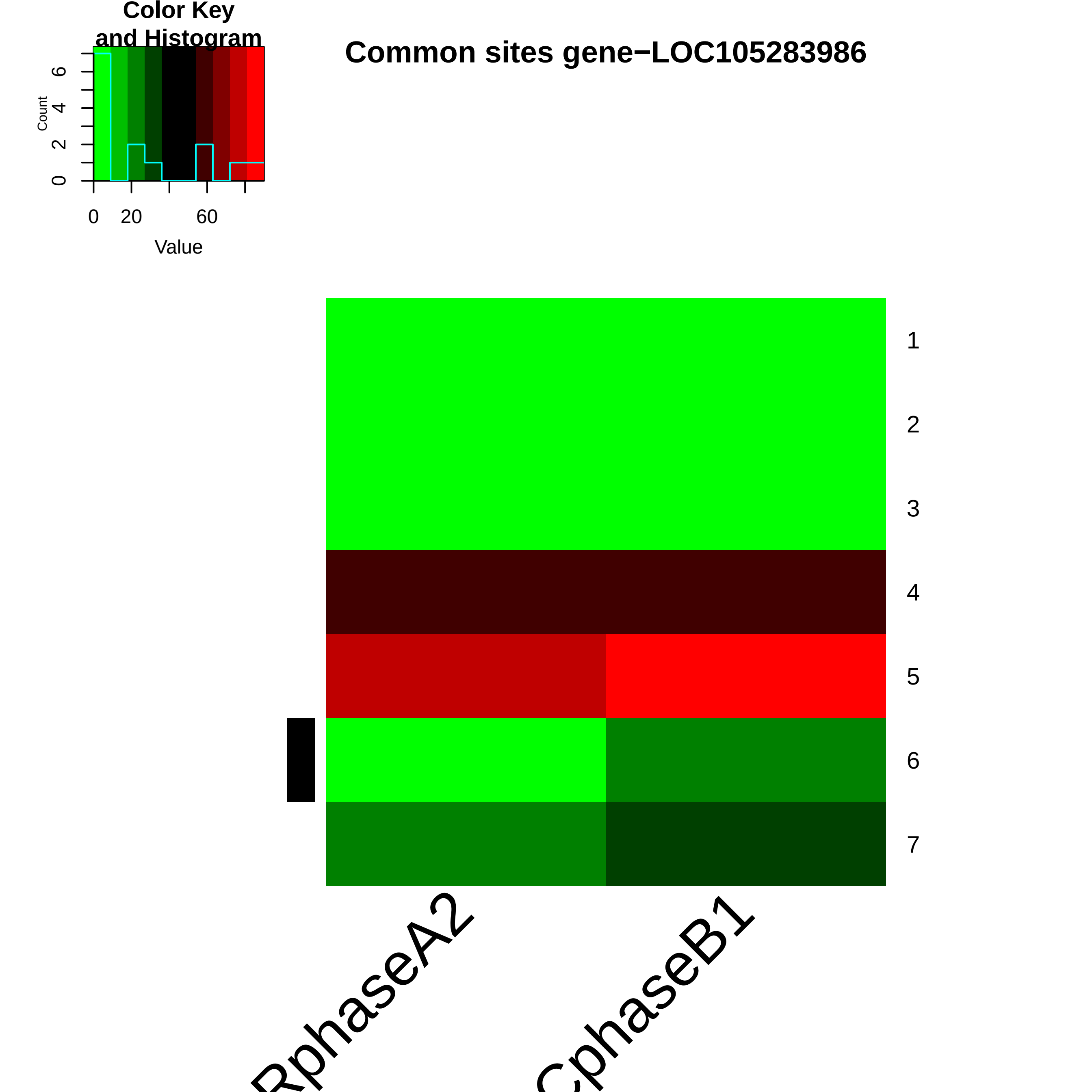
Color Key
and Histogram
Count
0
2
4
6
0
20
60
Value
Common sites gene−LOC105283986
1
2
3
4
5
6
7
RphaseA2
CphaseB1

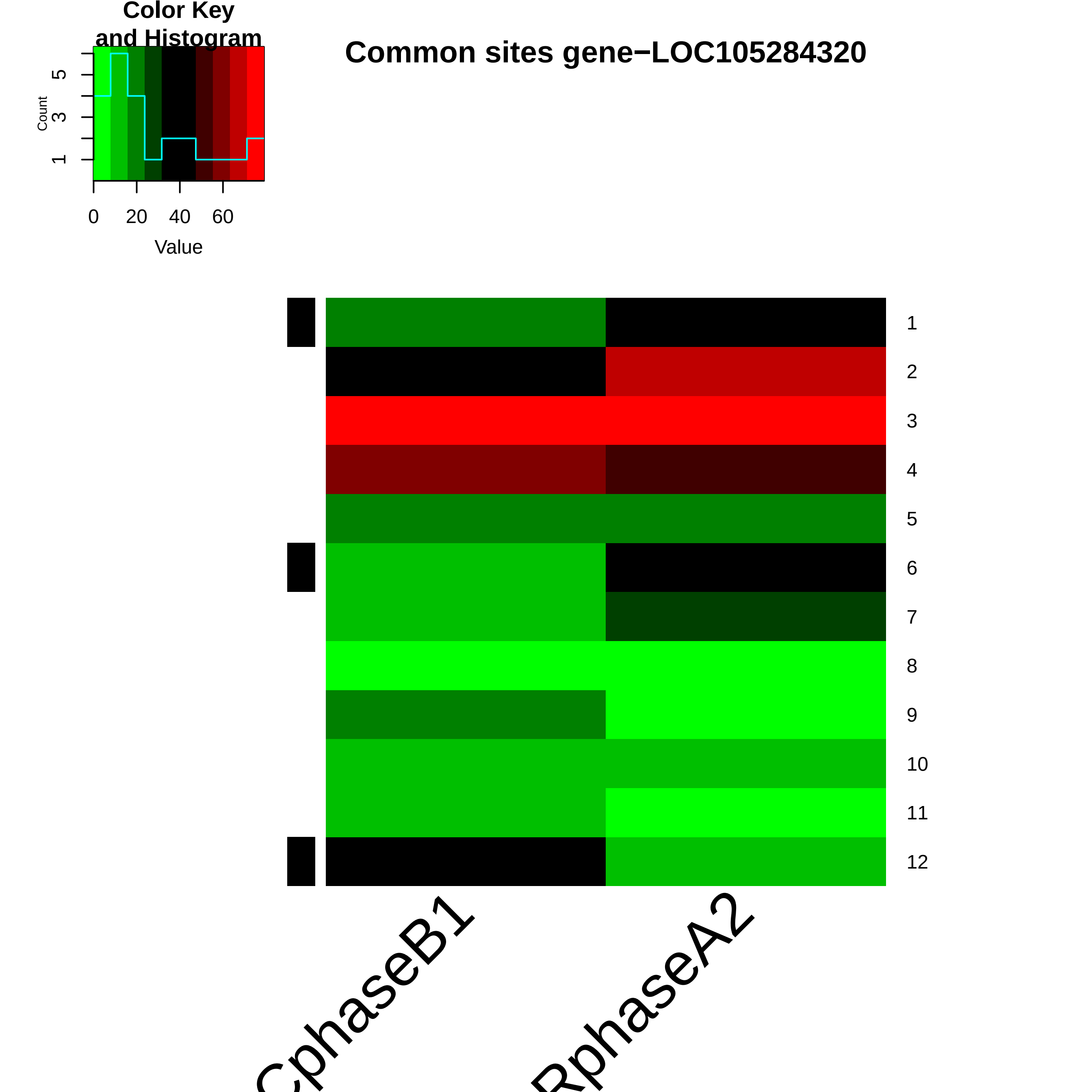

Color Key
and Histogram
Count
1
3
5
0
20
40
60
Value
Common sites gene−LOC105284320
1
2
3
4
5
6
7
8
9
10
11
12
CphaseB1
RphaseA2

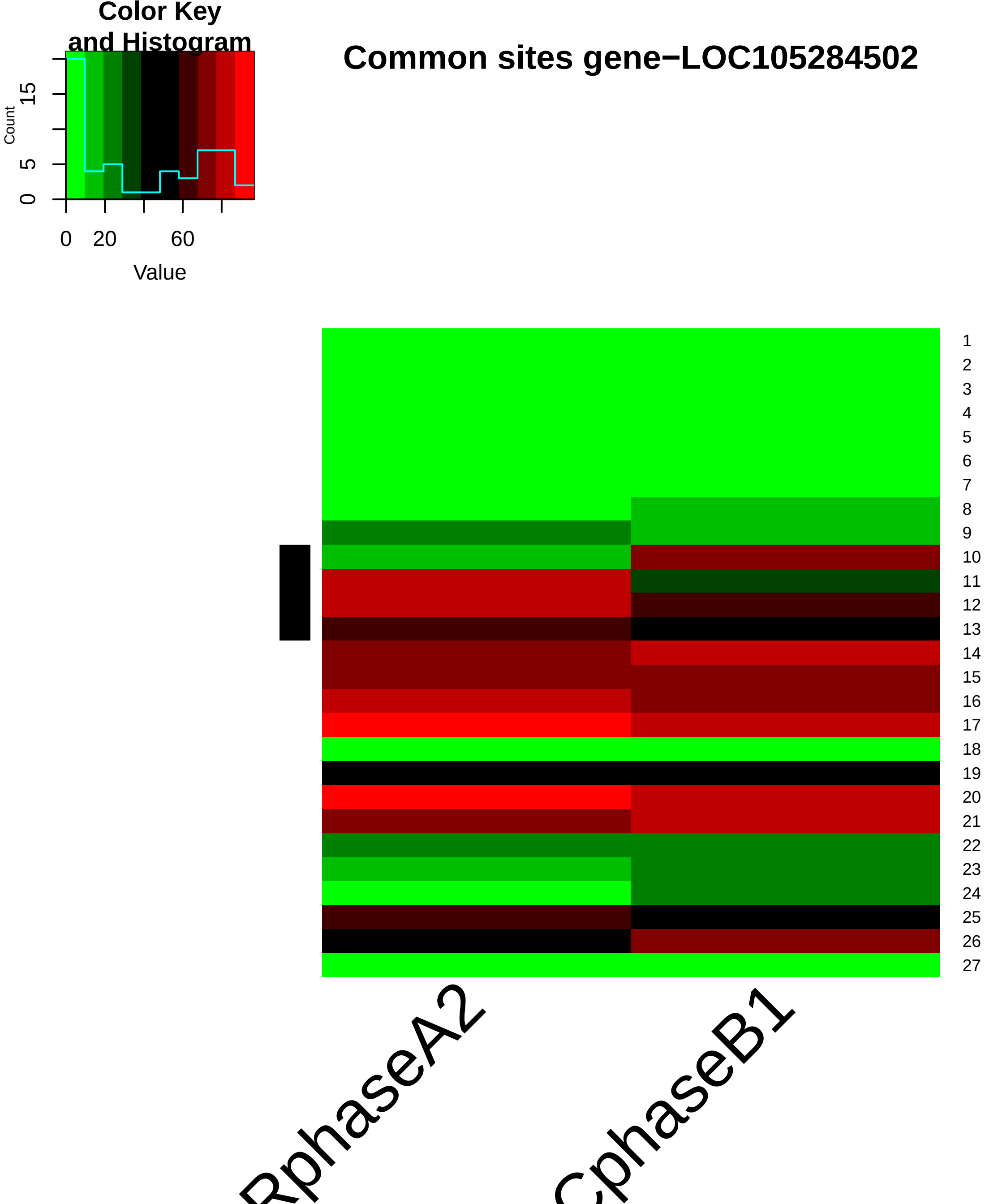

Color Key
and Histogram
Count
0
5
15
0
20
60
Value
Common sites gene−LOC105284502
RphaseA2
CphaseB1

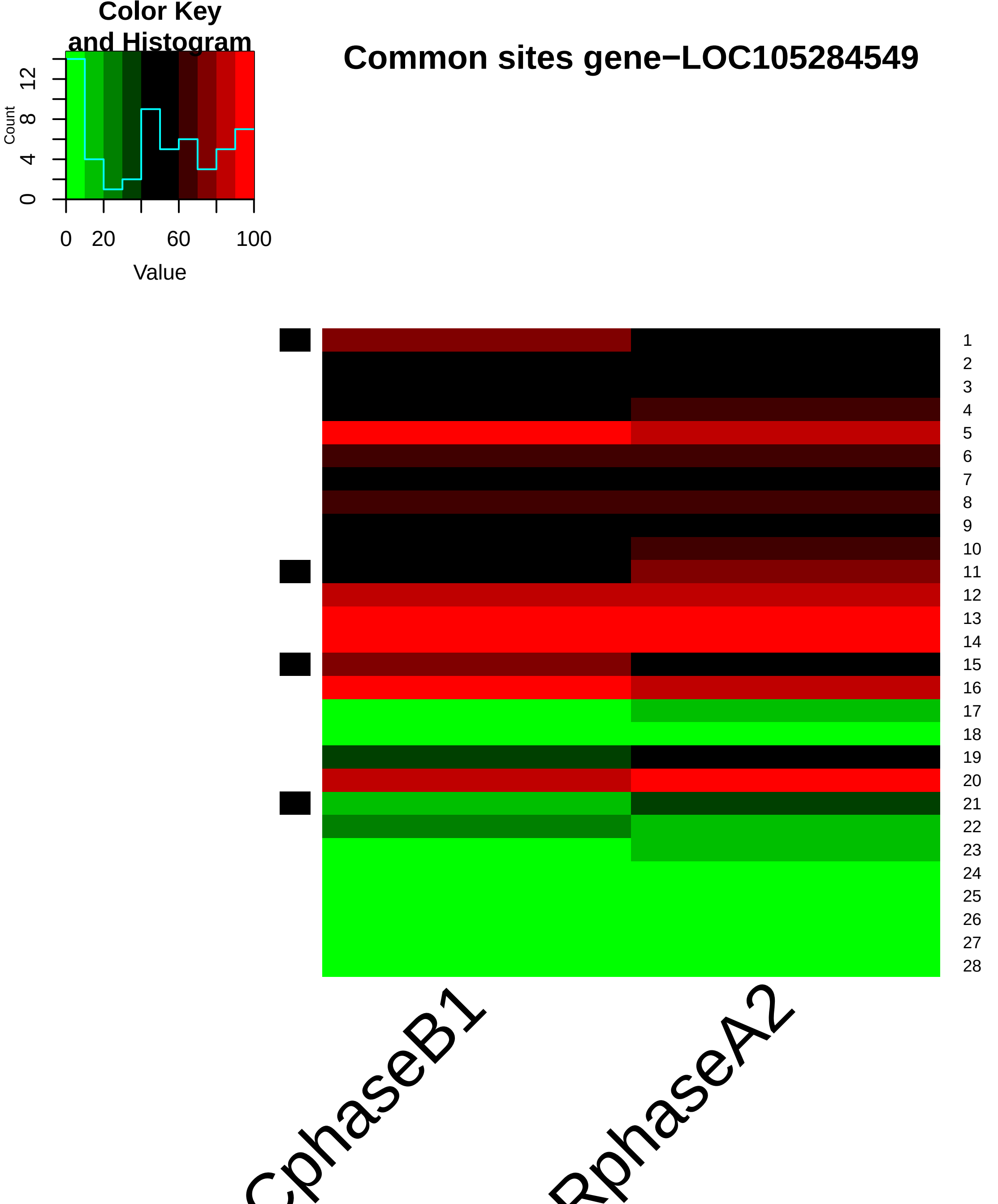
Color Key
and Histogram
Count
0
4
8
12
0
20
60
100
Value
Common sites gene−LOC105284549
CphaseB1
RphaseA2

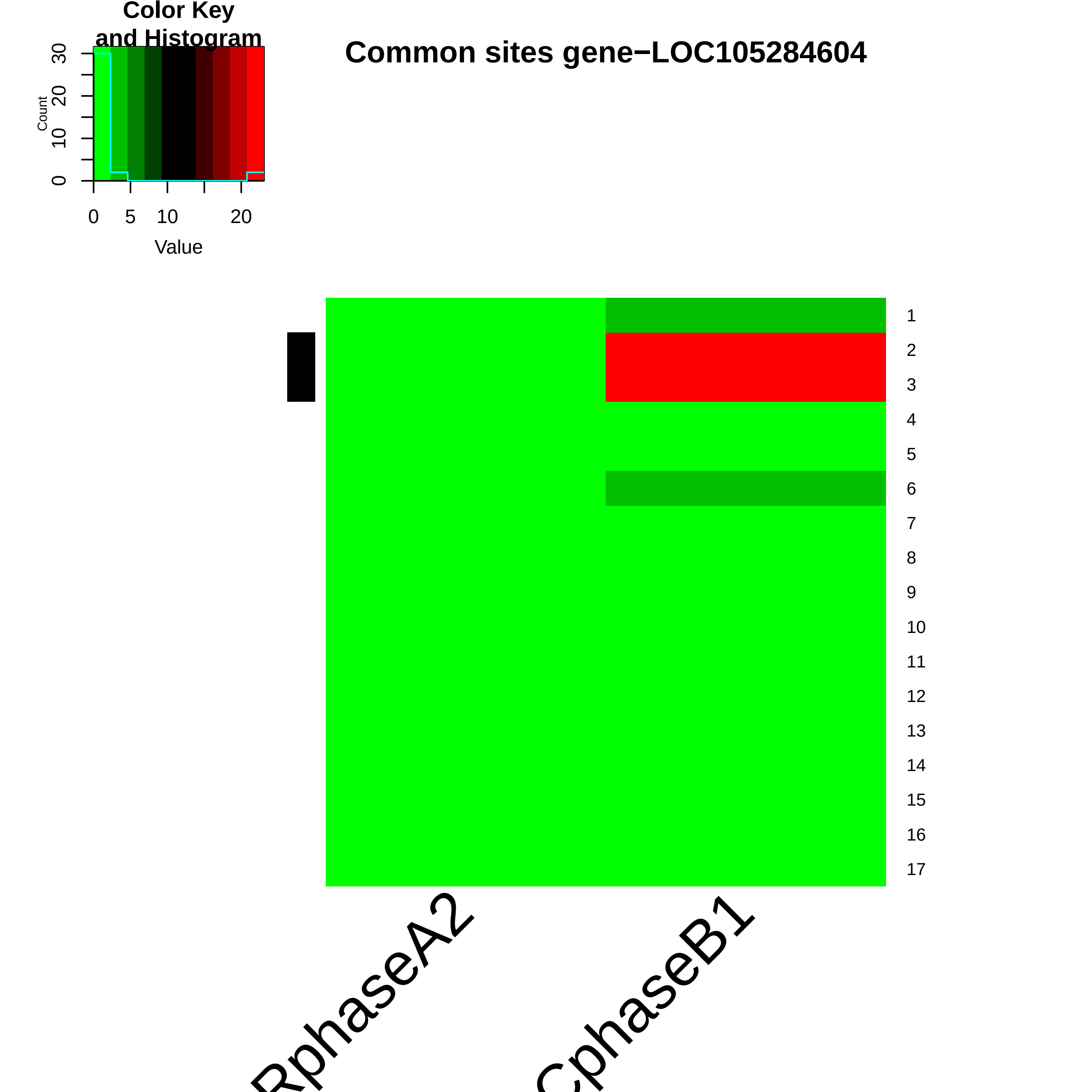

Color Key
and Histogram
Count
0
10
20
30
0
5
10
20
Value
Common sites gene−LOC105284604
RphaseA2
CphaseB1

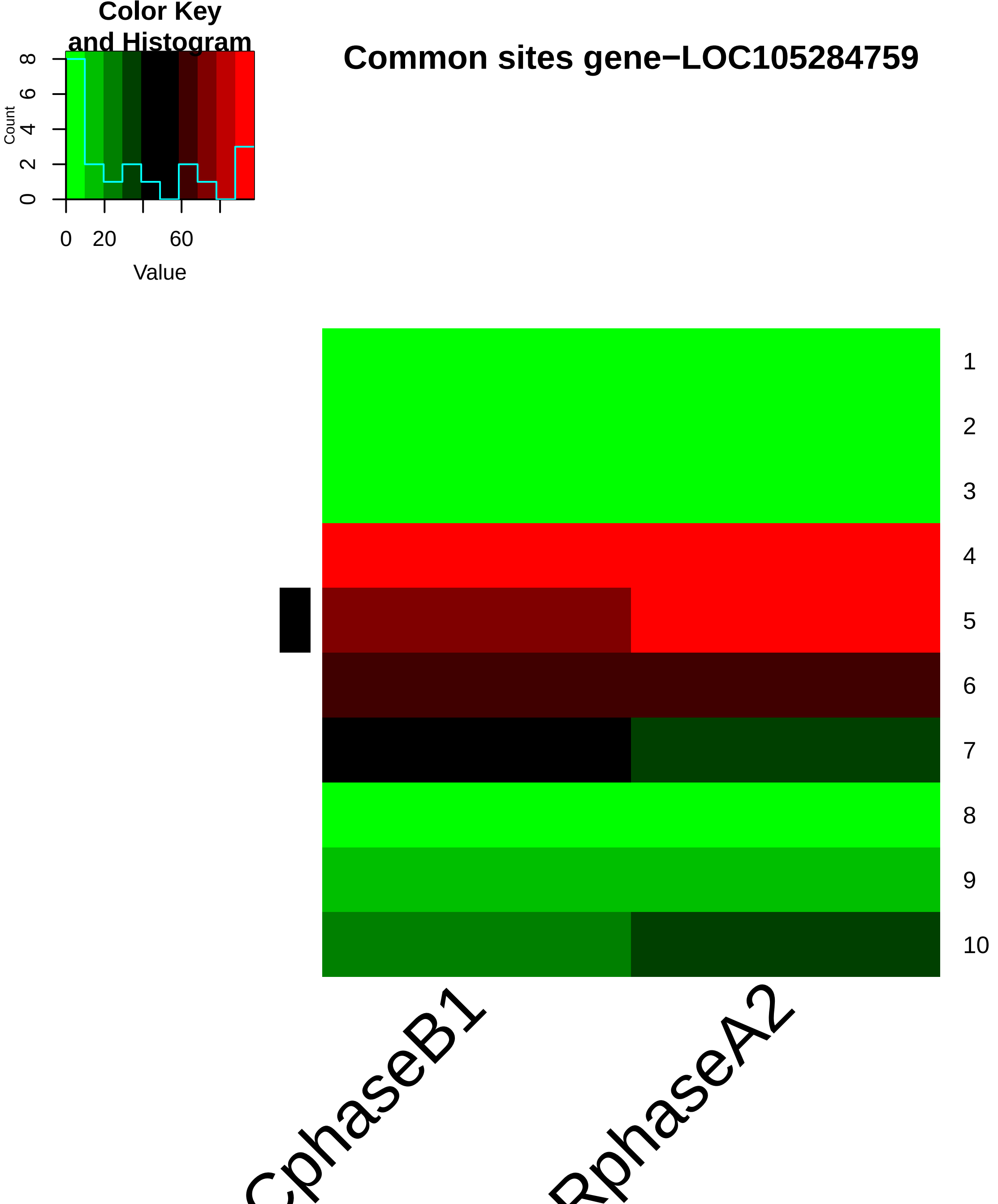
Color Key
and Histogram
Common sites gene−LOC105284759
Count
0
2
4
6
8
0
20
60
Value
1
2
3
4
5
6
7
8
9
10
CphaseB1
RphaseA2

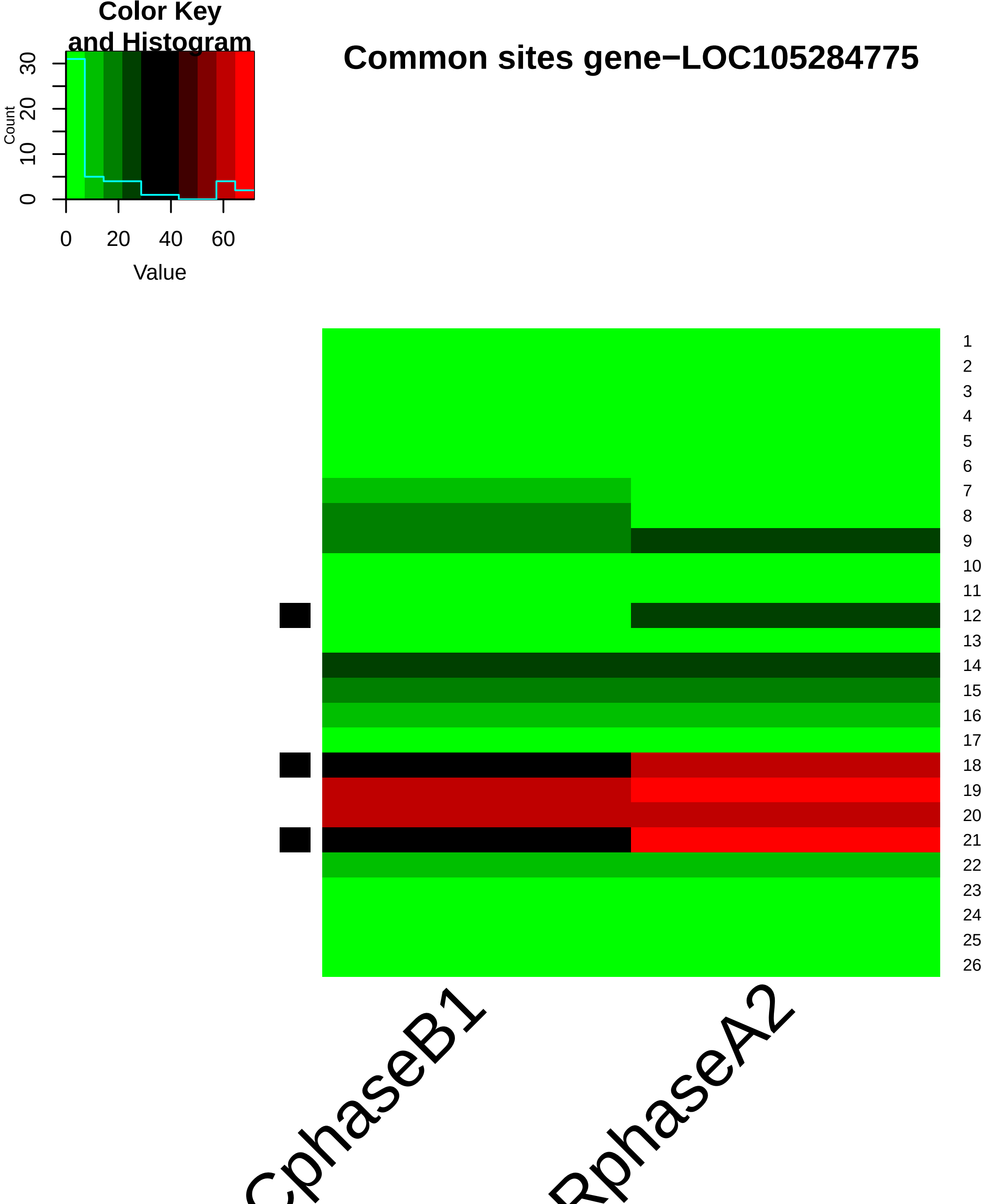

Color Key
and Histogram
Count
0
10
20
30
0
20
40
60
Value
Common sites gene−LOC105284775
CphaseB1
RphaseA2

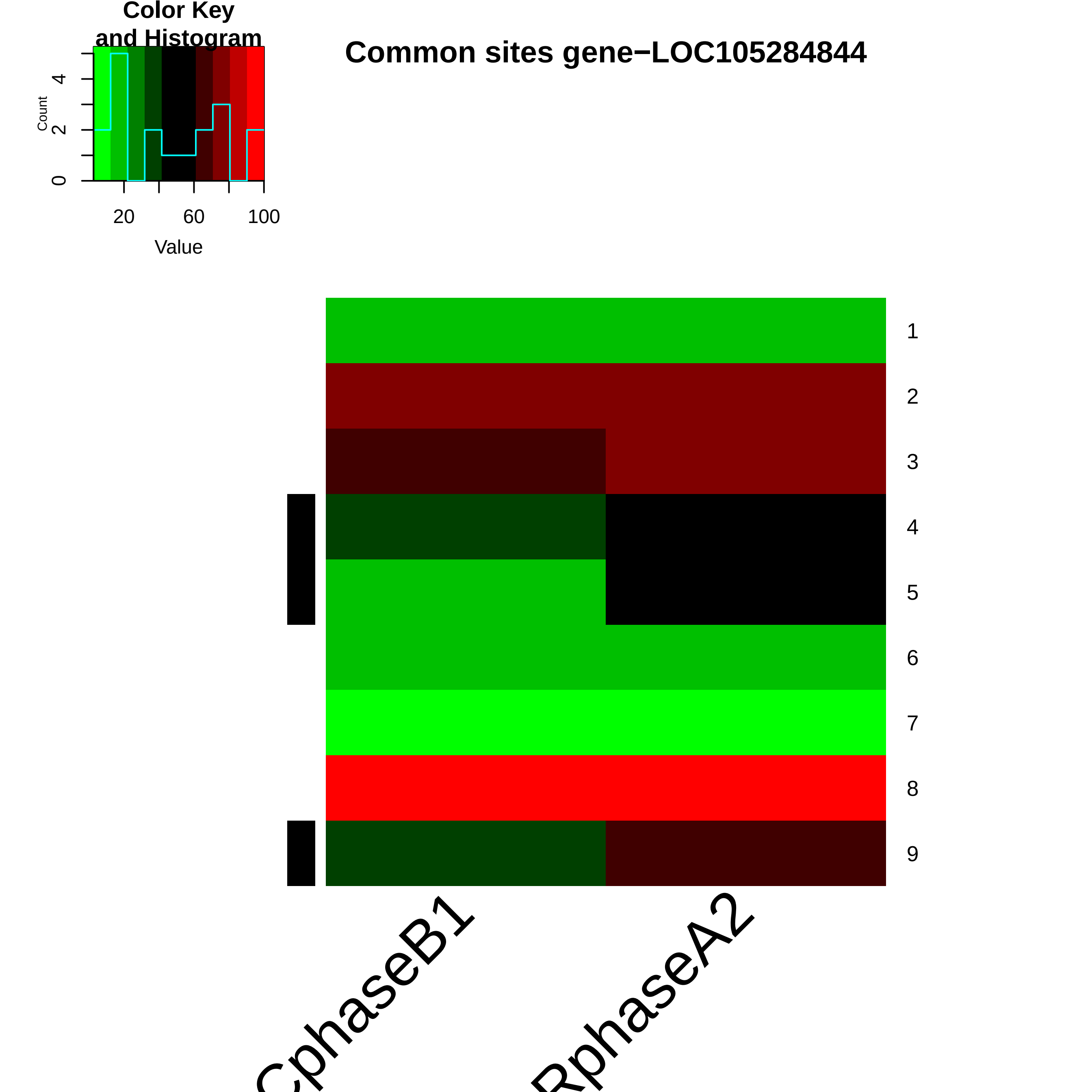

Color Key
and Histogram
Common sites gene−LOC105284844
Count
0
2
4
20
60
100
Value
1
2
3
4
5
6
7
8
9
CphaseB1
RphaseA2

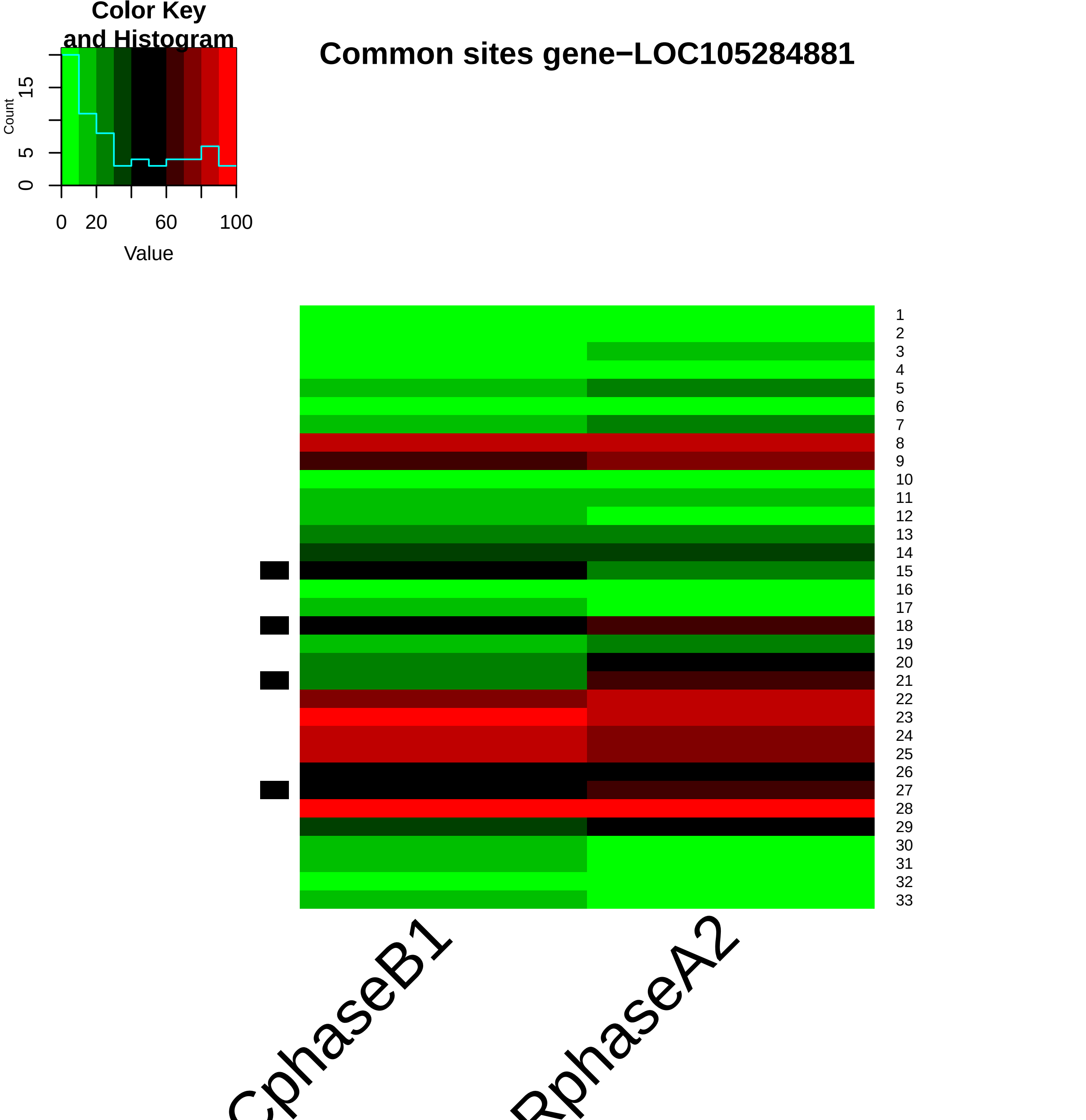

Color Key
and Histogram
Count
0
5
15
0
20
60
100
Value
Common sites gene−LOC105284881
CphaseB1
RphaseA2

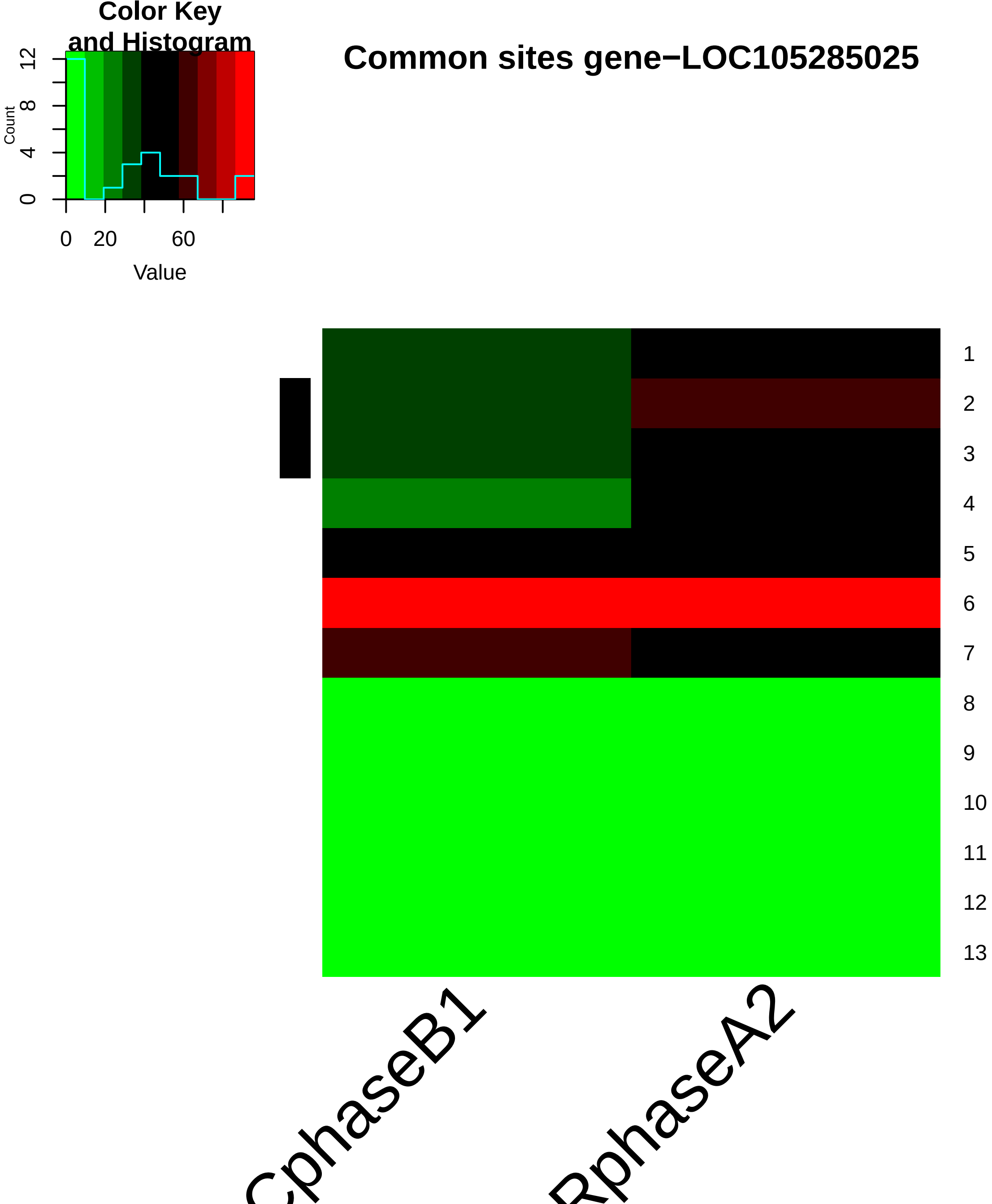
Color Key
and Histogram
Count
0
4
8
12
0
20
60
Value
Common sites gene−LOC105285025
1
2
3
4
5
6
7
8
9
10
11
12
13
CphaseB1
RphaseA2

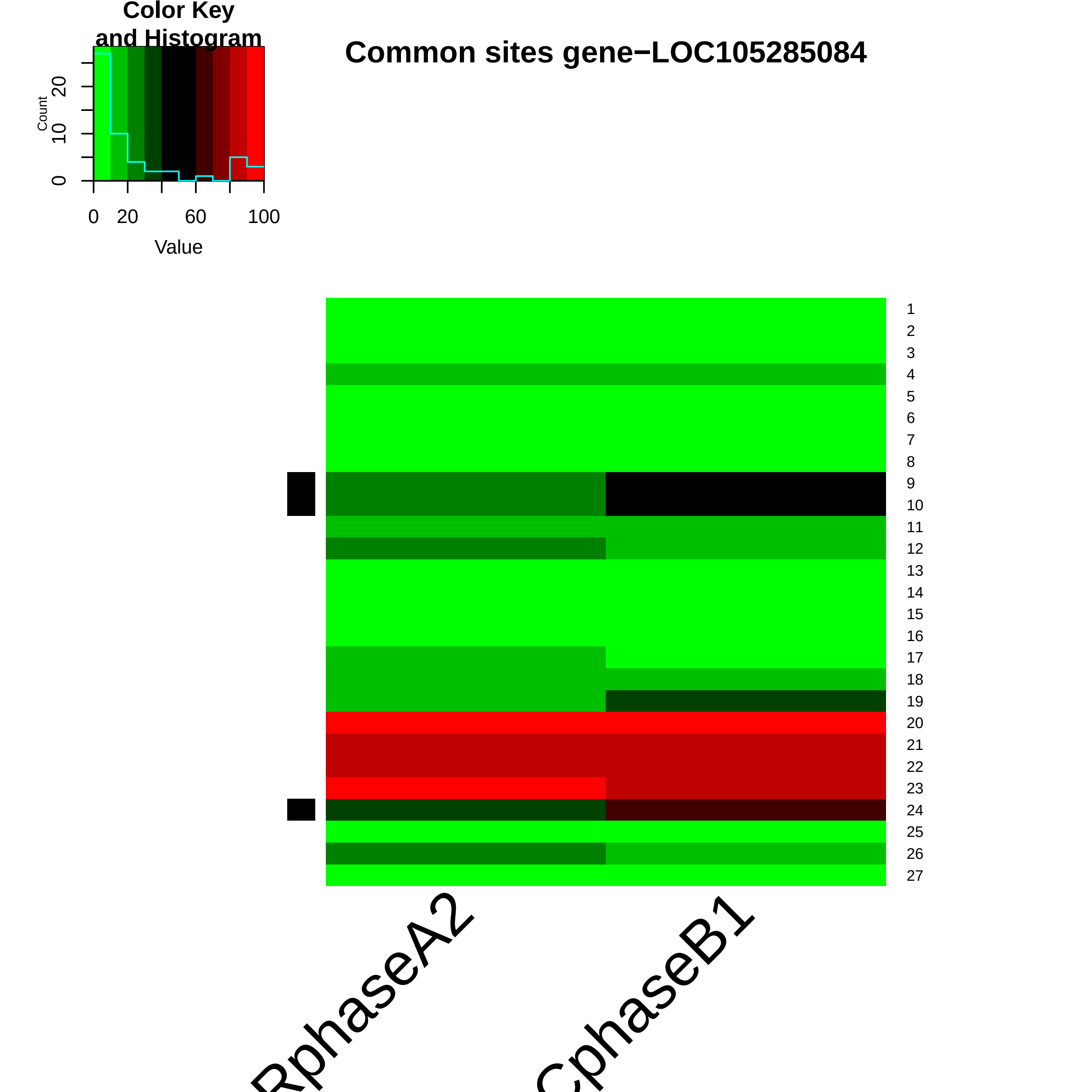

Color Key
and Histogram
Count
0
10
20
0
20
60
100
Value
Common sites gene−LOC105285084
RphaseA2
CphaseB1

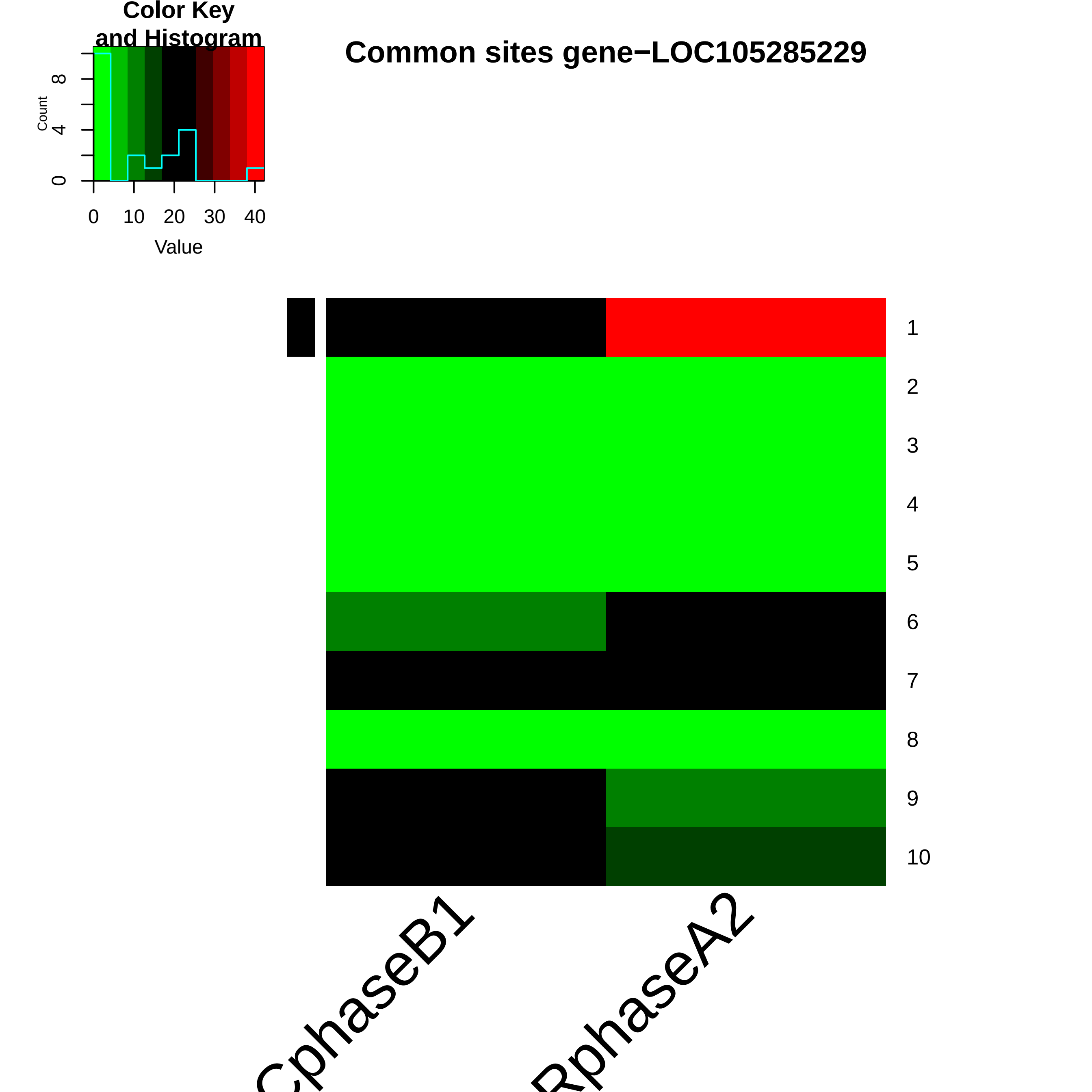
Color Key
and Histogram
Count
0
4
8
0
10
20
30
40
Value
Common sites gene−LOC105285229
1
2
3
4
5
6
7
8
9
10
CphaseB1
RphaseA2

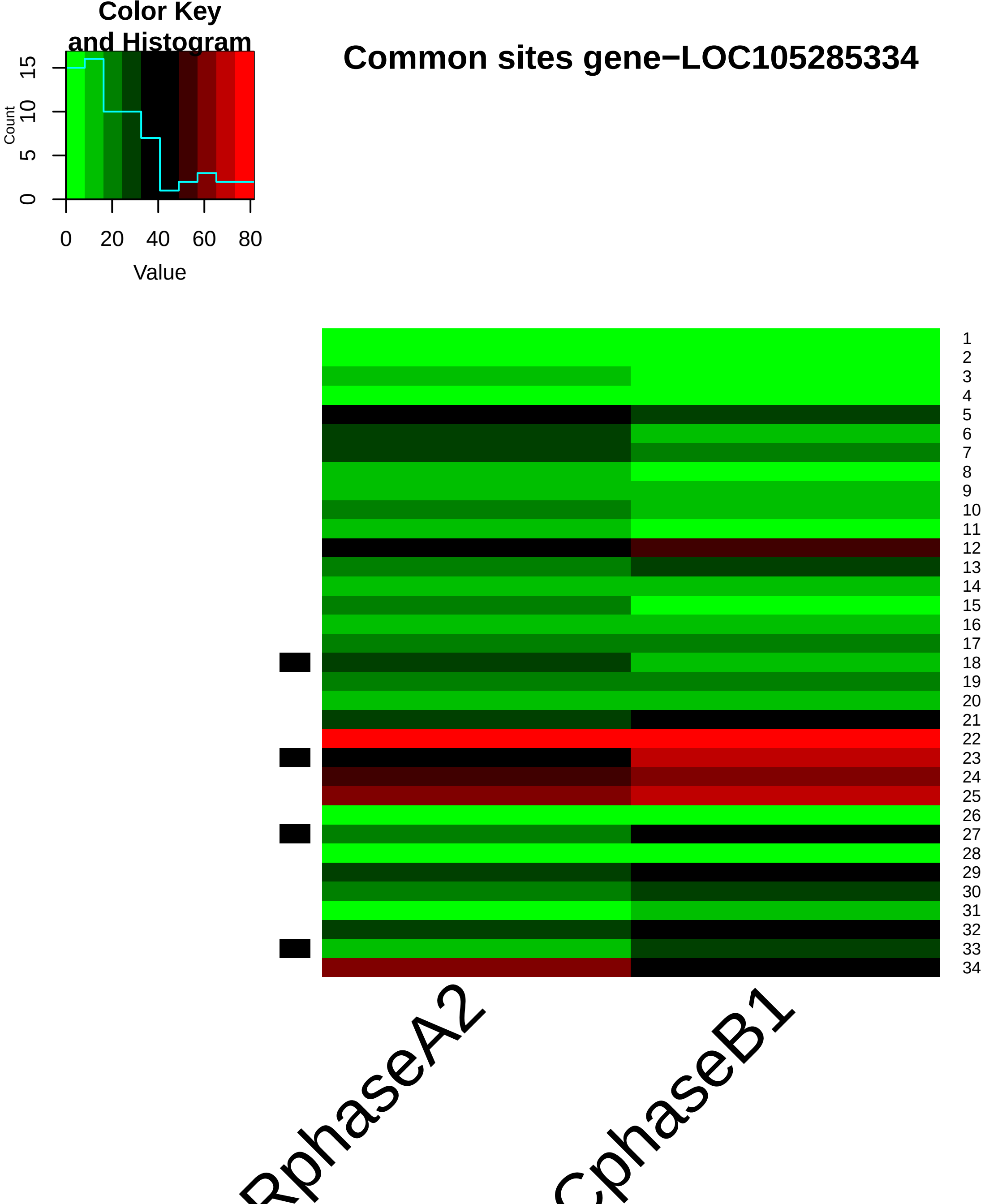
Color Key
and Histogram
Count
0
5
10
15
0
20
40
60
80
Value
Common sites gene−LOC105285334
RphaseA2
CphaseB1

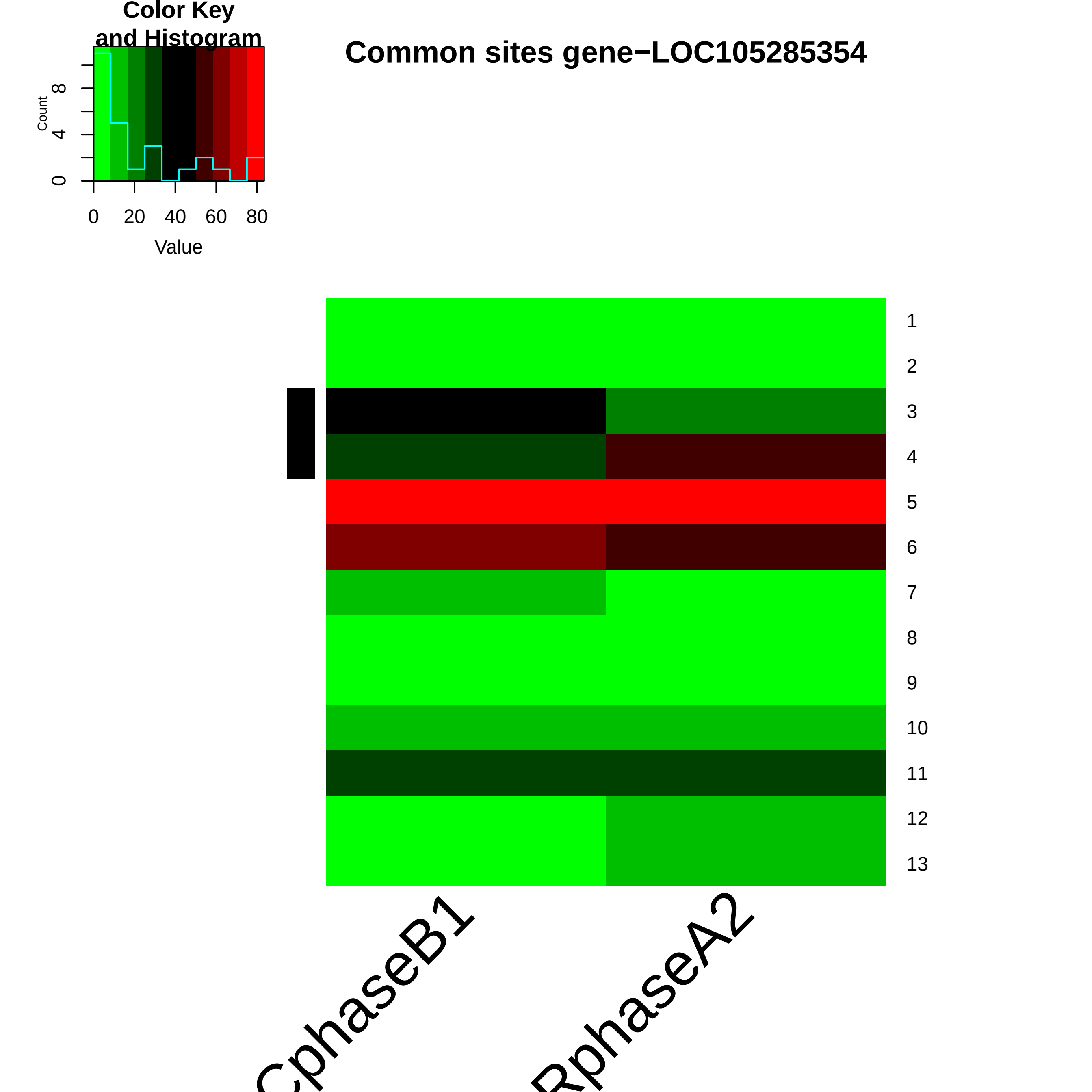
Color Key
and Histogram
Count
0
4
8
0
20
40
60
80
Value
Common sites gene−LOC105285354
1
2
3
4
5
6
7
8
9
10
11
12
13
CphaseB1
RphaseA2

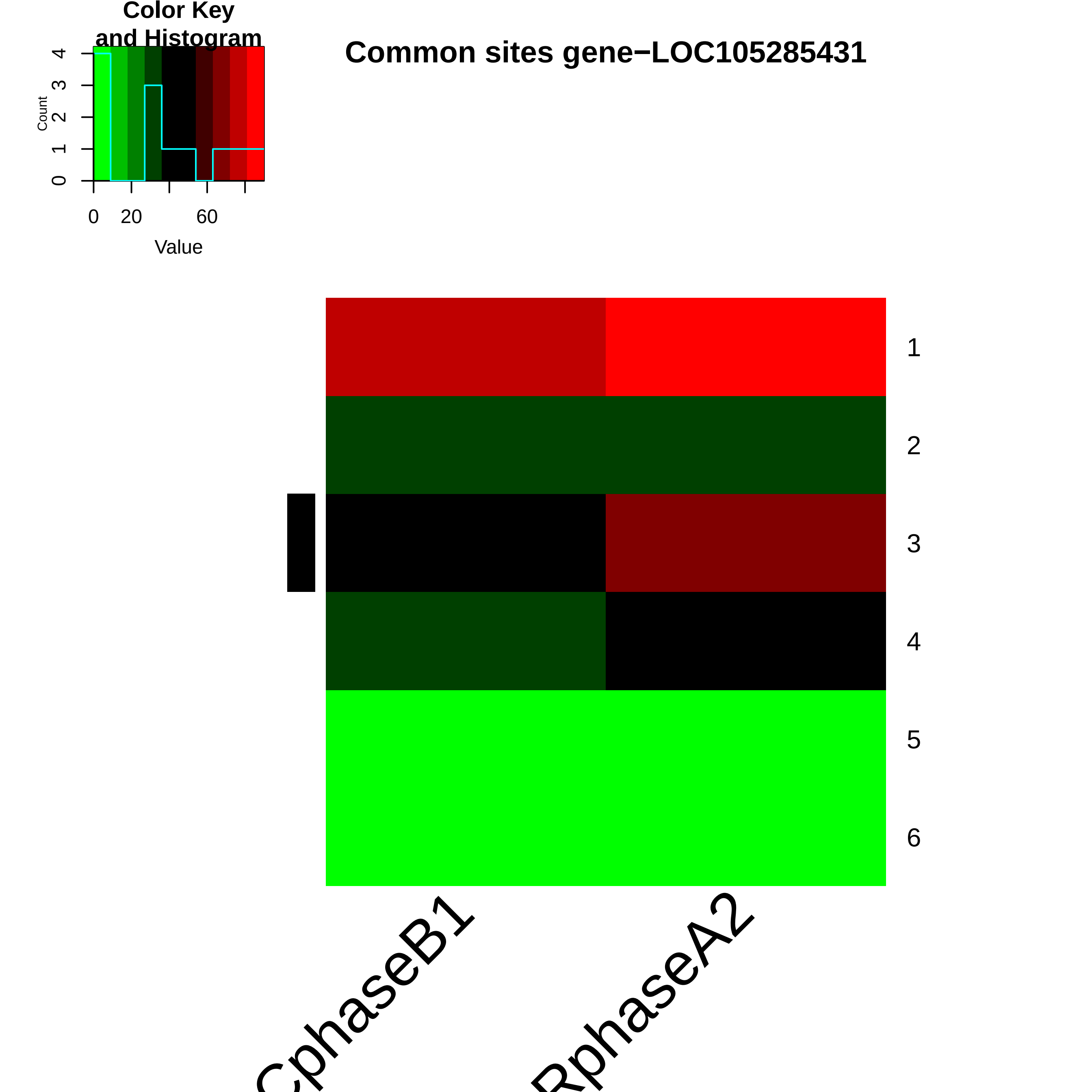

Color Key
and Histogram
Count
0
1
2
3
4
0
20
60
Value
Common sites gene−LOC105285431
1
2
3
4
5
6
CphaseB1
RphaseA2

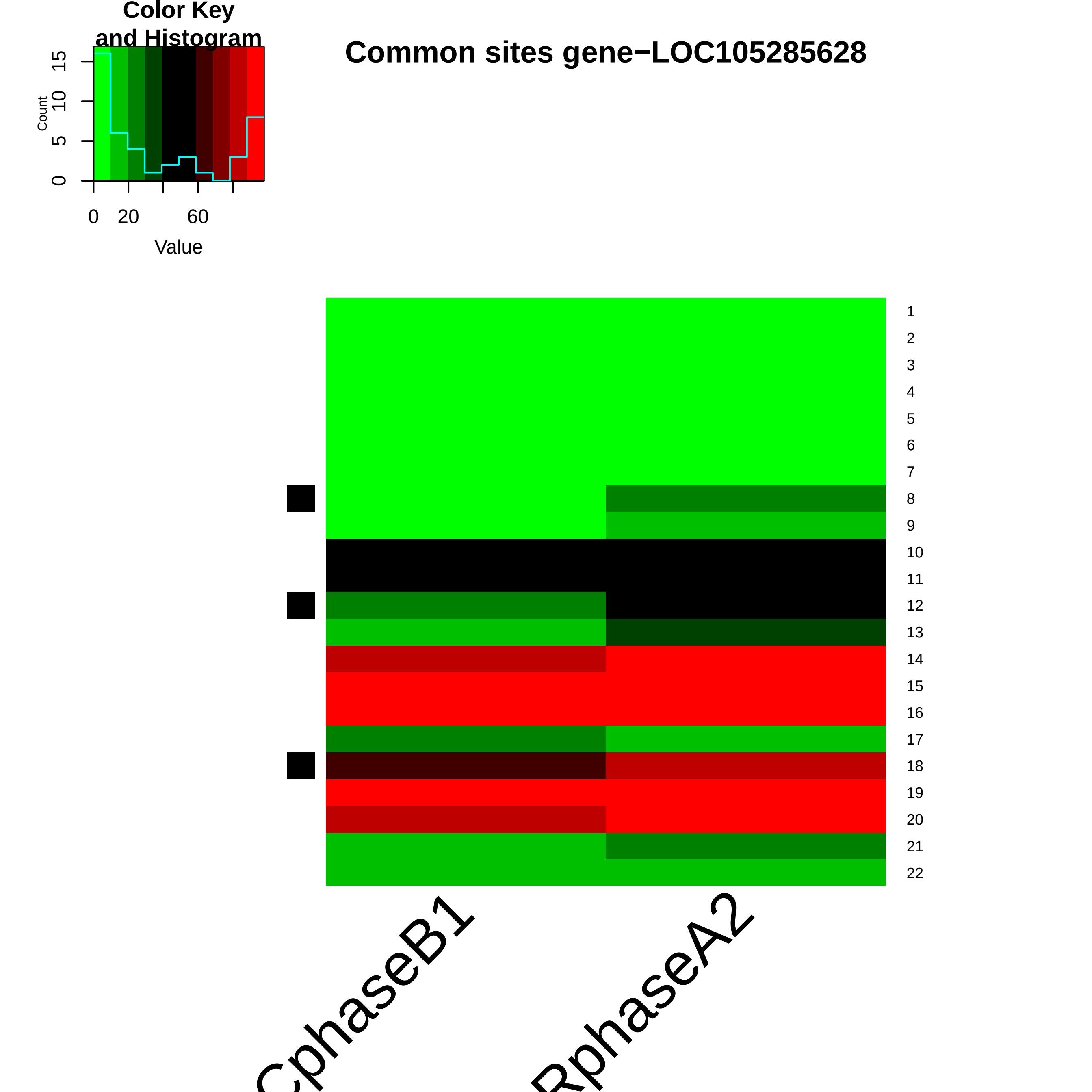
Color Key
and Histogram
Count
0
5
10
15
0
20
60
Value
Common sites gene−LOC105285628
CphaseB1
RphaseA2

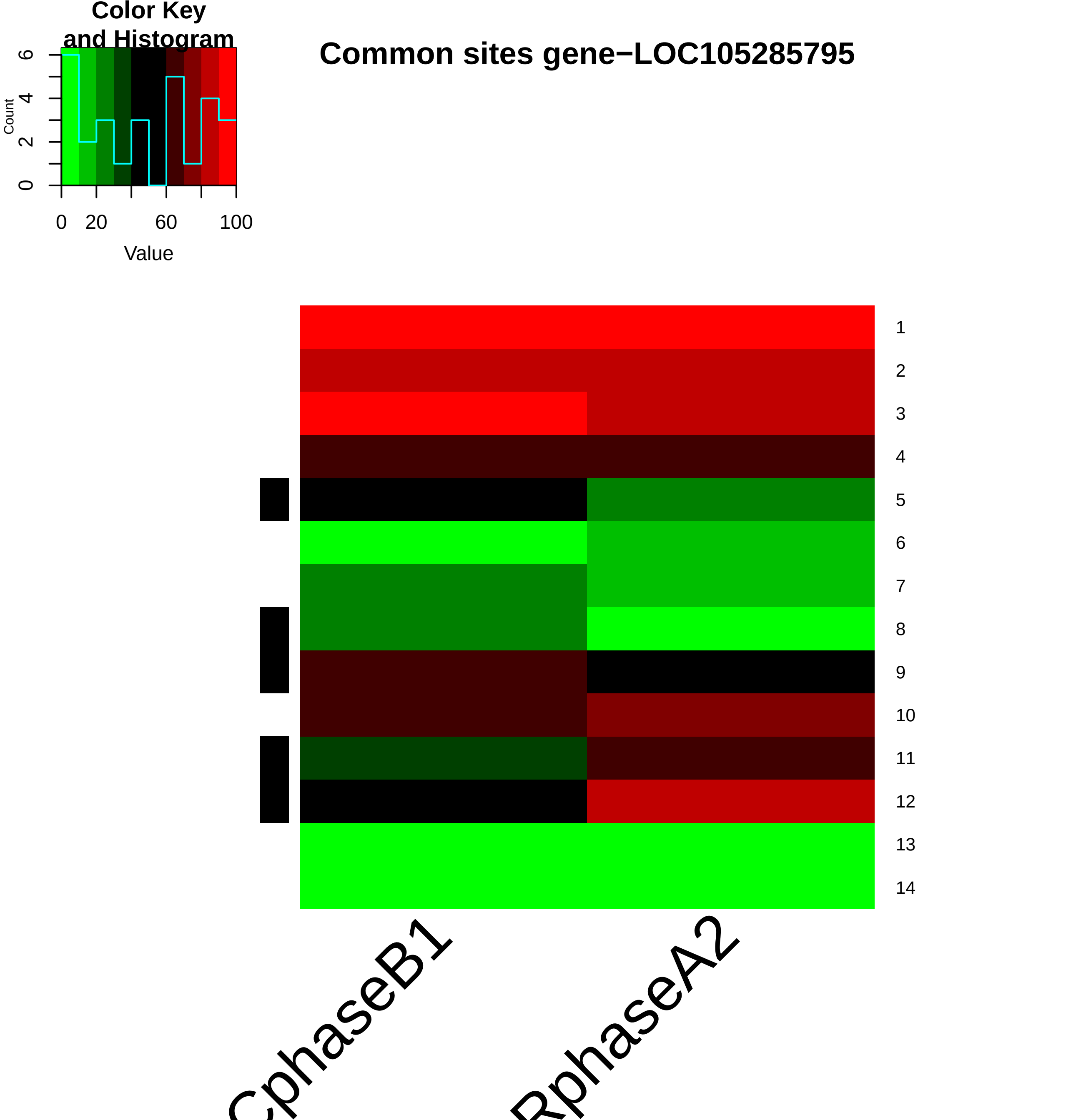

Color Key
and Histogram
Count
Value
0
2
4
6
0
20
60
100
Common sites gene−LOC105285795
1
2
3
4
5
6
7
8
9
10
11
12
13
14
CphaseB1
RphaseA2

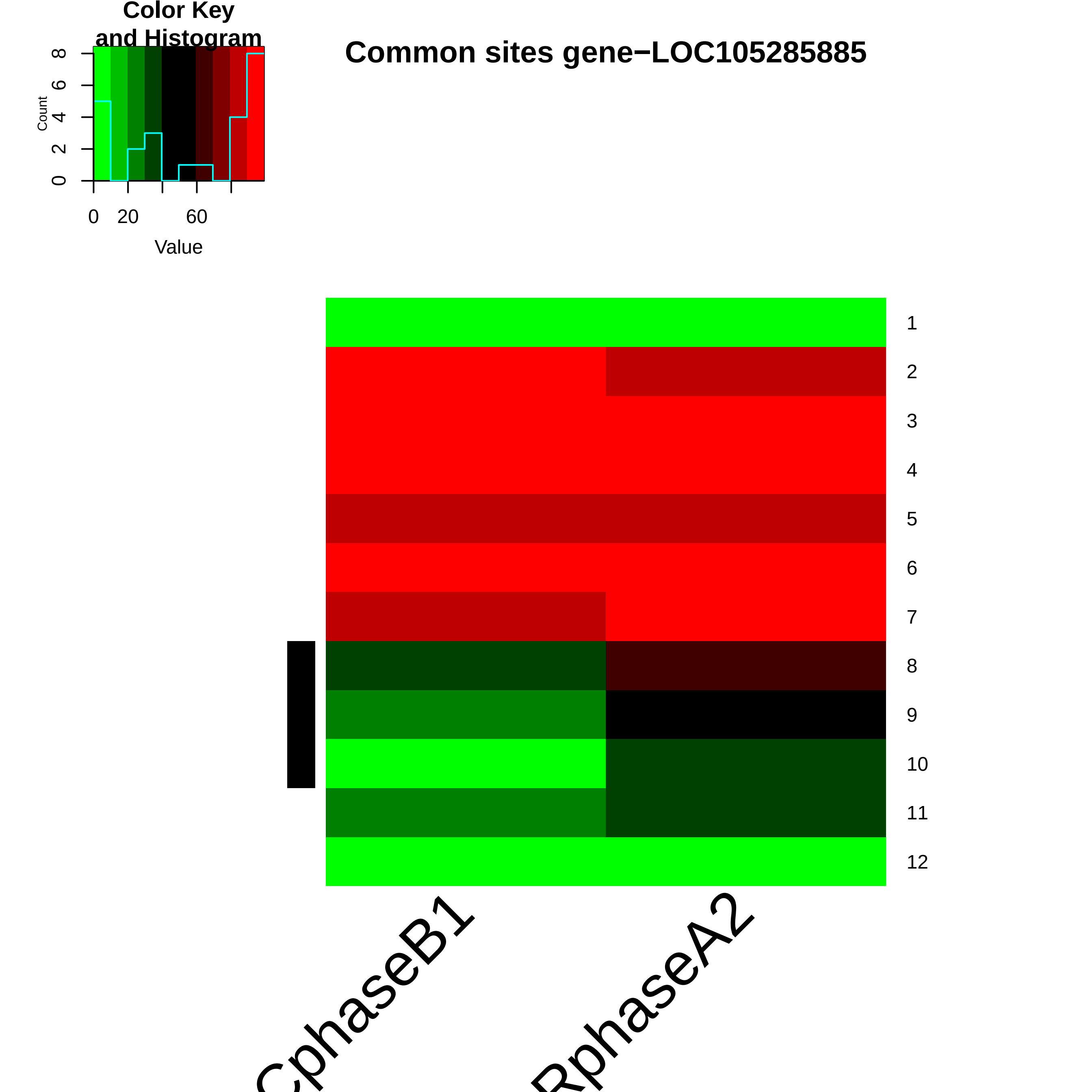

Color Key
and Histogram
Count
0
2
4
6
8
0
20
60
Value
Common sites gene−LOC105285885
1
2
3
4
5
6
7
8
9
10
11
12
CphaseB1
RphaseA2

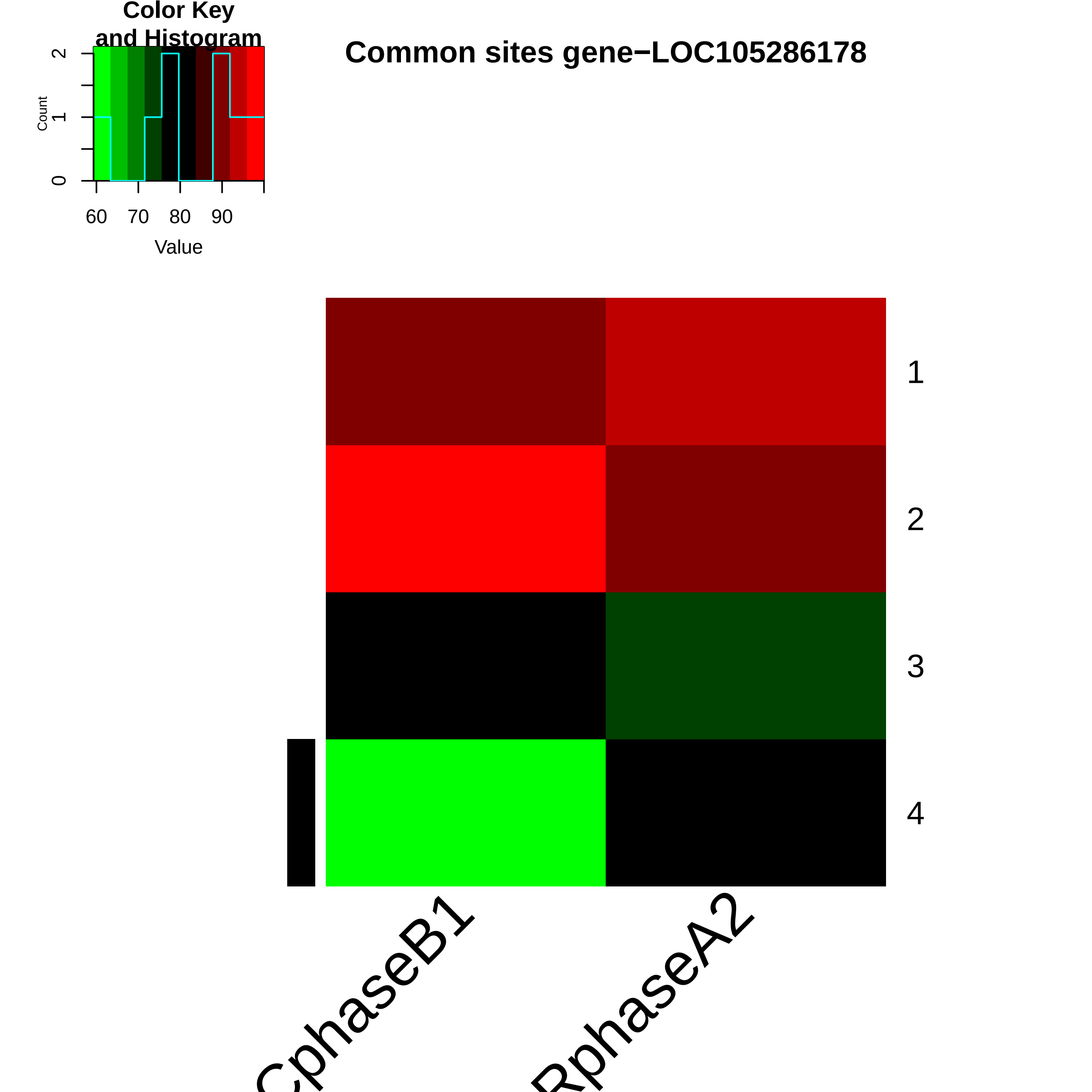

Color Key
and Histogram
Count
0
1
2
60
70
80
90
Value
Common sites gene−LOC105286178
1
2
3
4
CphaseB1
RphaseA2

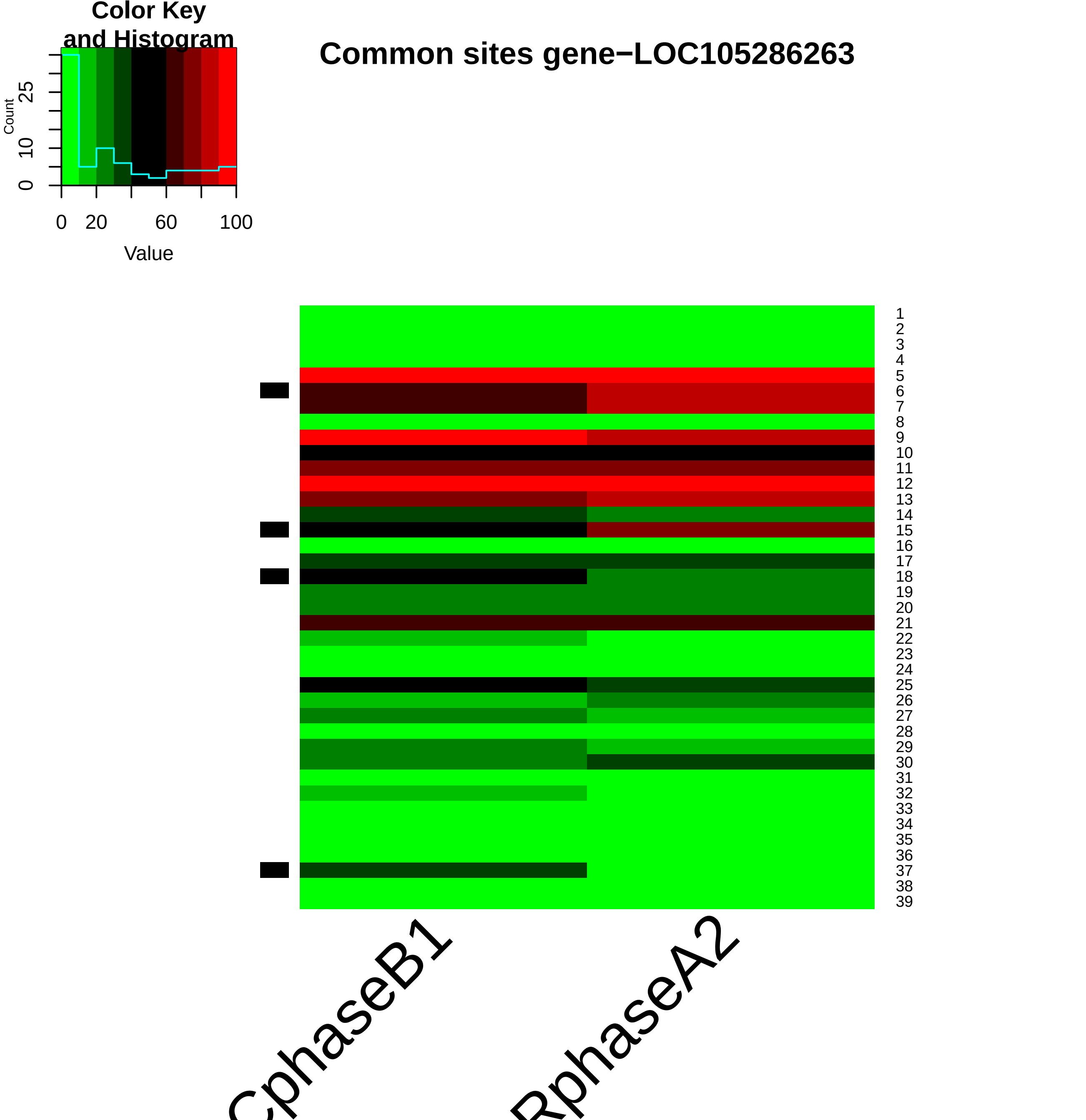
Color Key
and Histogram
Count
0
10
25
0
20
60
100
Value
Common sites gene−LOC105286263
CphaseB1
RphaseA2

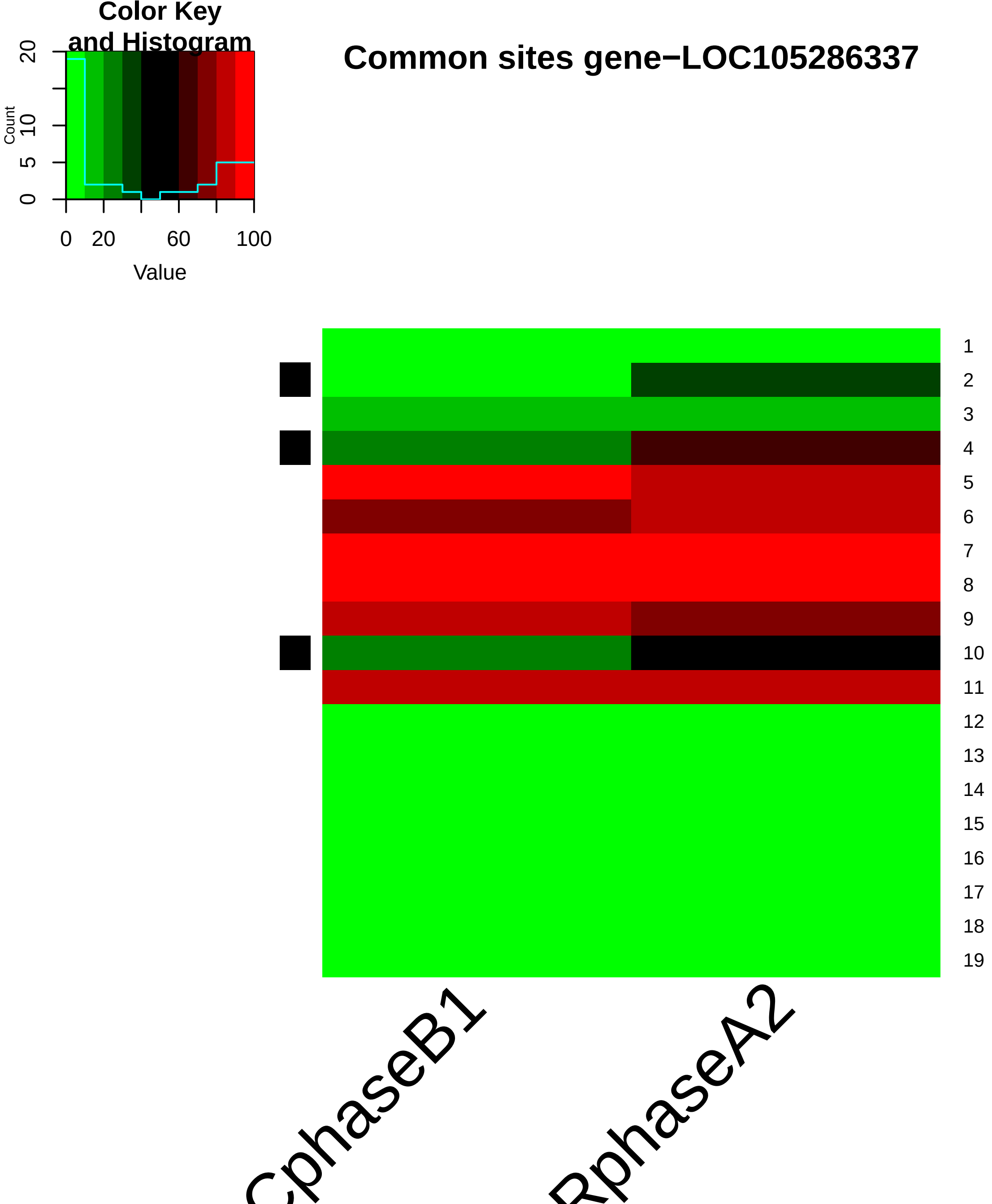
Color Key
and Histogram
Count
0
5
10
20
0
20
60
100
Value
Common sites gene−LOC105286337
CphaseB1
RphaseA2

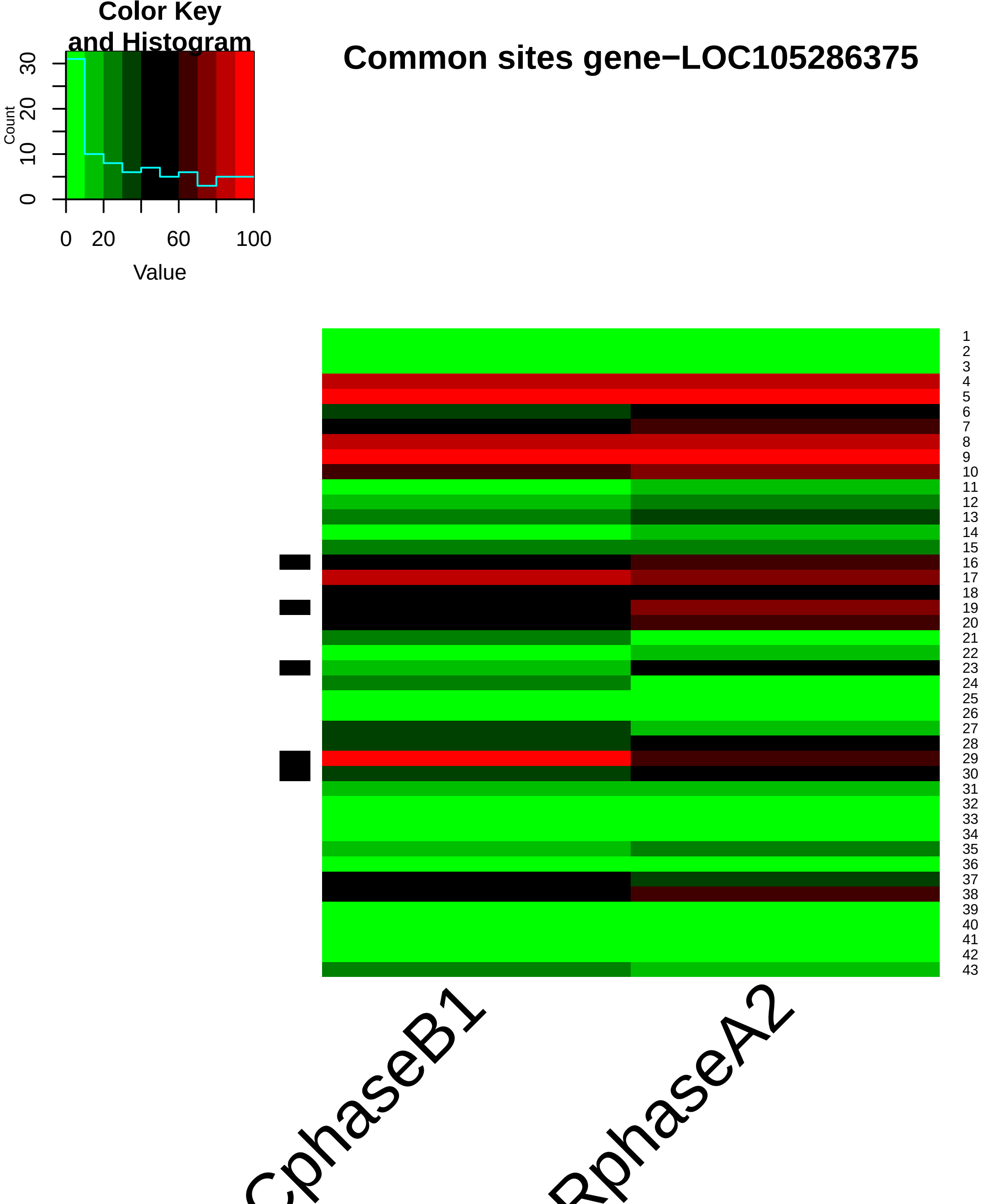

Color Key
and Histogram
Count
0
10
20
30
0
20
60
100
Value
Common sites gene−LOC105286375
CphaseB1
RphaseA2

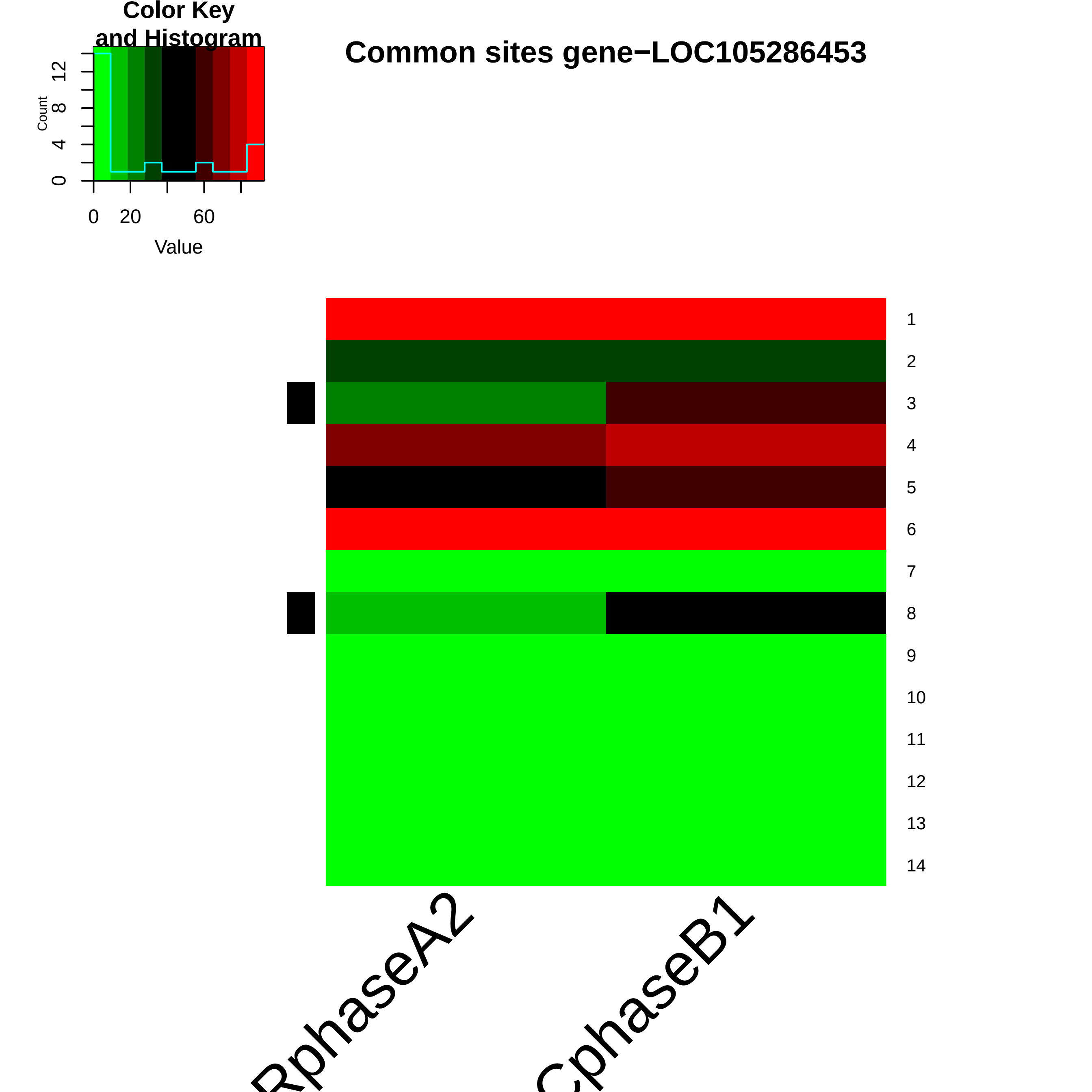
Color Key
and Histogram
Count
0
4
8
12
0
20
60
Value
Common sites gene−LOC105286453
1
2
3
4
5
6
7
8
9
10
11
12
13
14
RphaseA2
CphaseB1

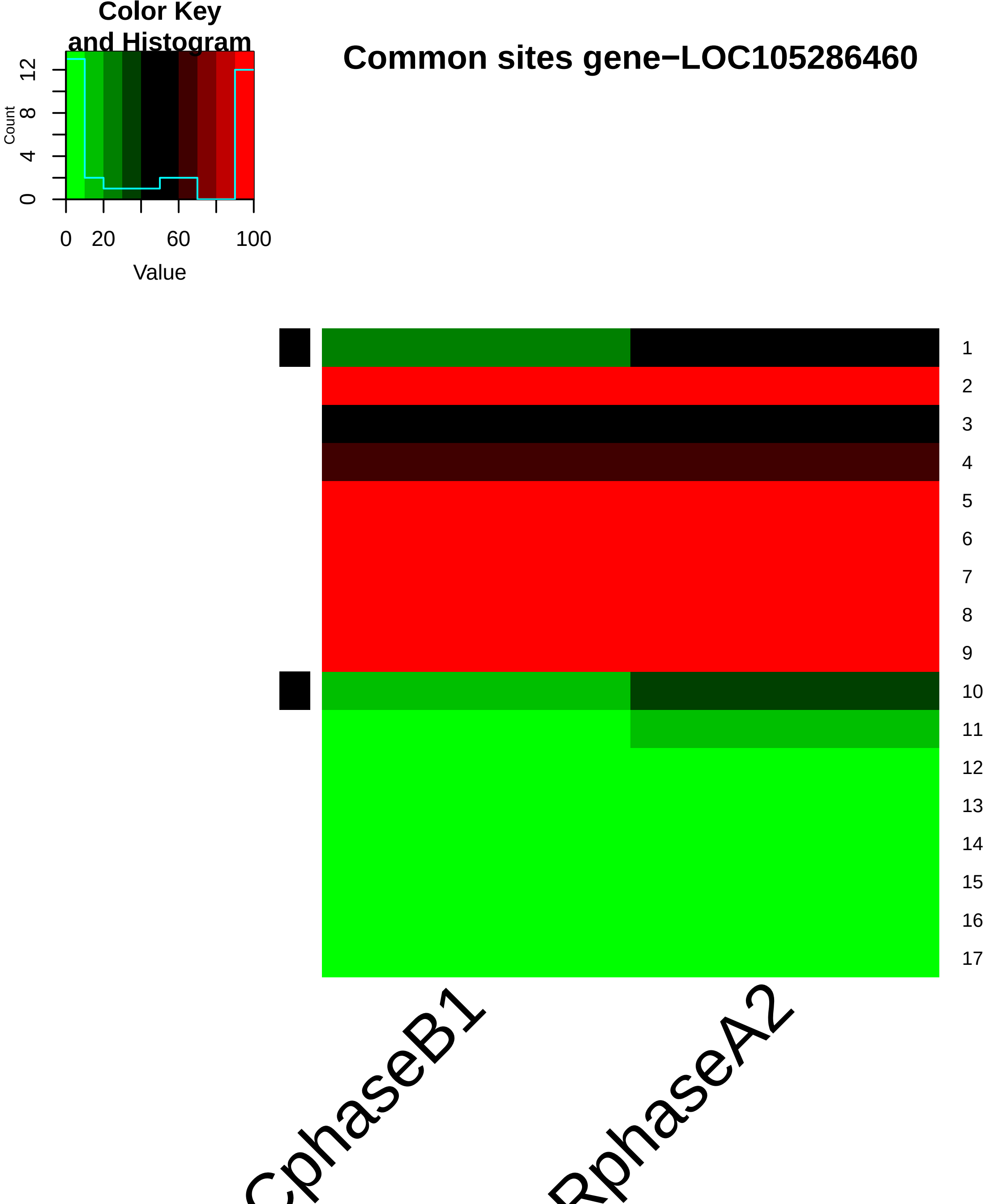
Color Key
and Histogram
Count
0
4
8
12
0
20
60
100
Value
Common sites gene−LOC105286460
CphaseB1
RphaseA2

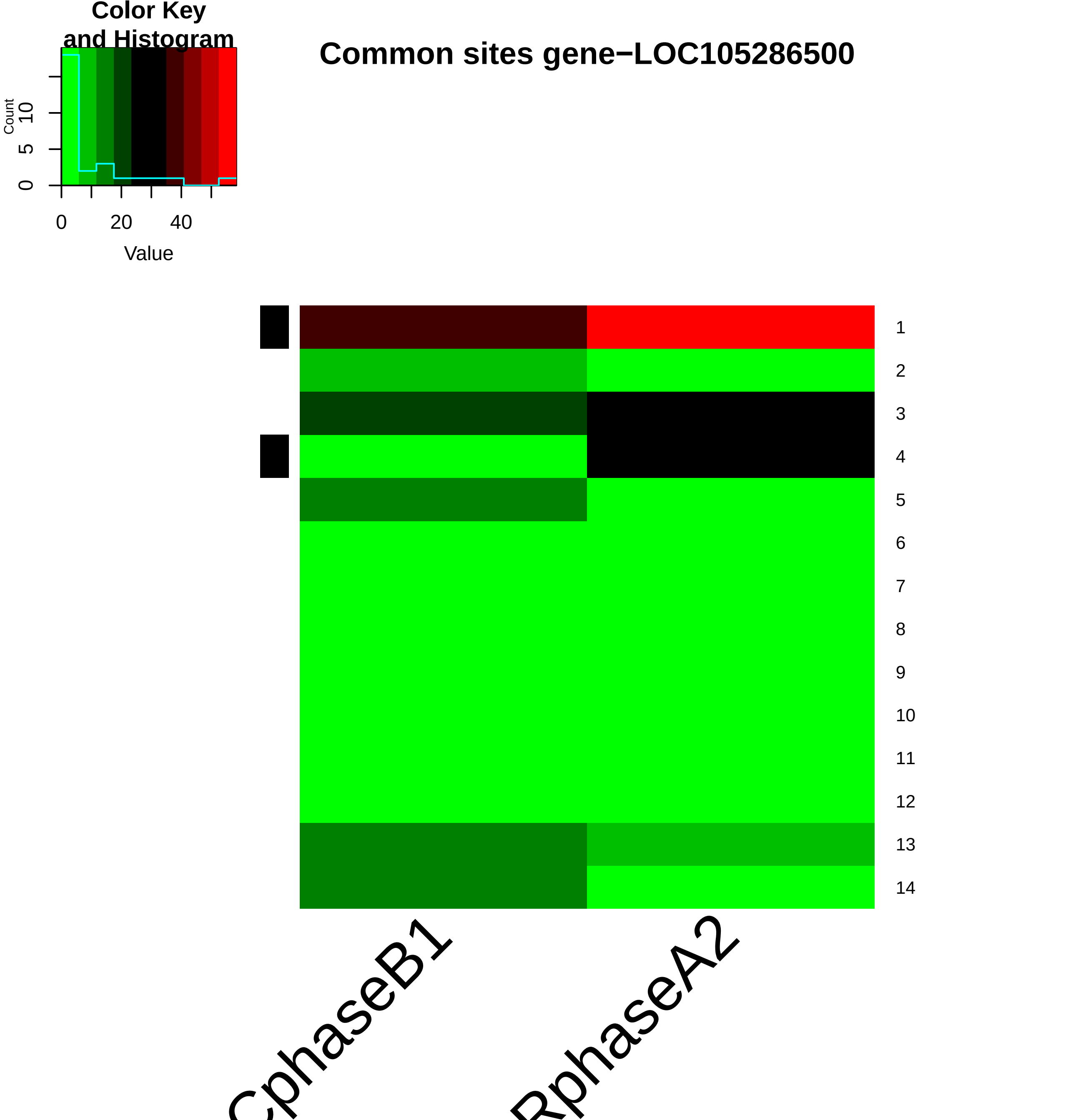

Color Key
and Histogram
Count
0
5
10
0
20
40
Value
Common sites gene−LOC105286500
1
2
3
4
5
6
7
8
9
10
11
12
13
14
CphaseB1
RphaseA2

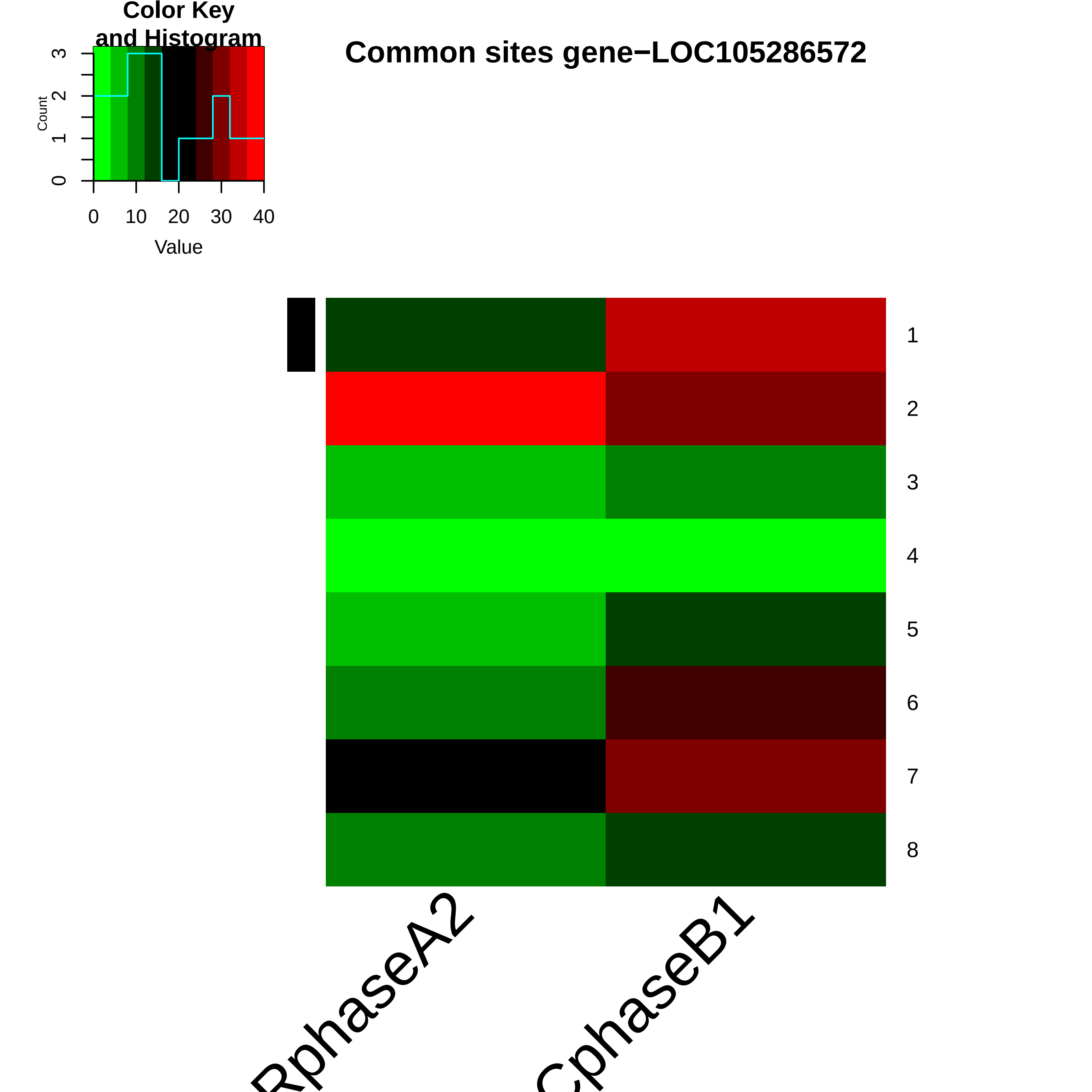

Color Key
and Histogram
Common sites gene−LOC105286572
Count
0
1
2
3
0
10
20
30
40
Value
1
2
3
4
5
6
7
8
RphaseA2
CphaseB1

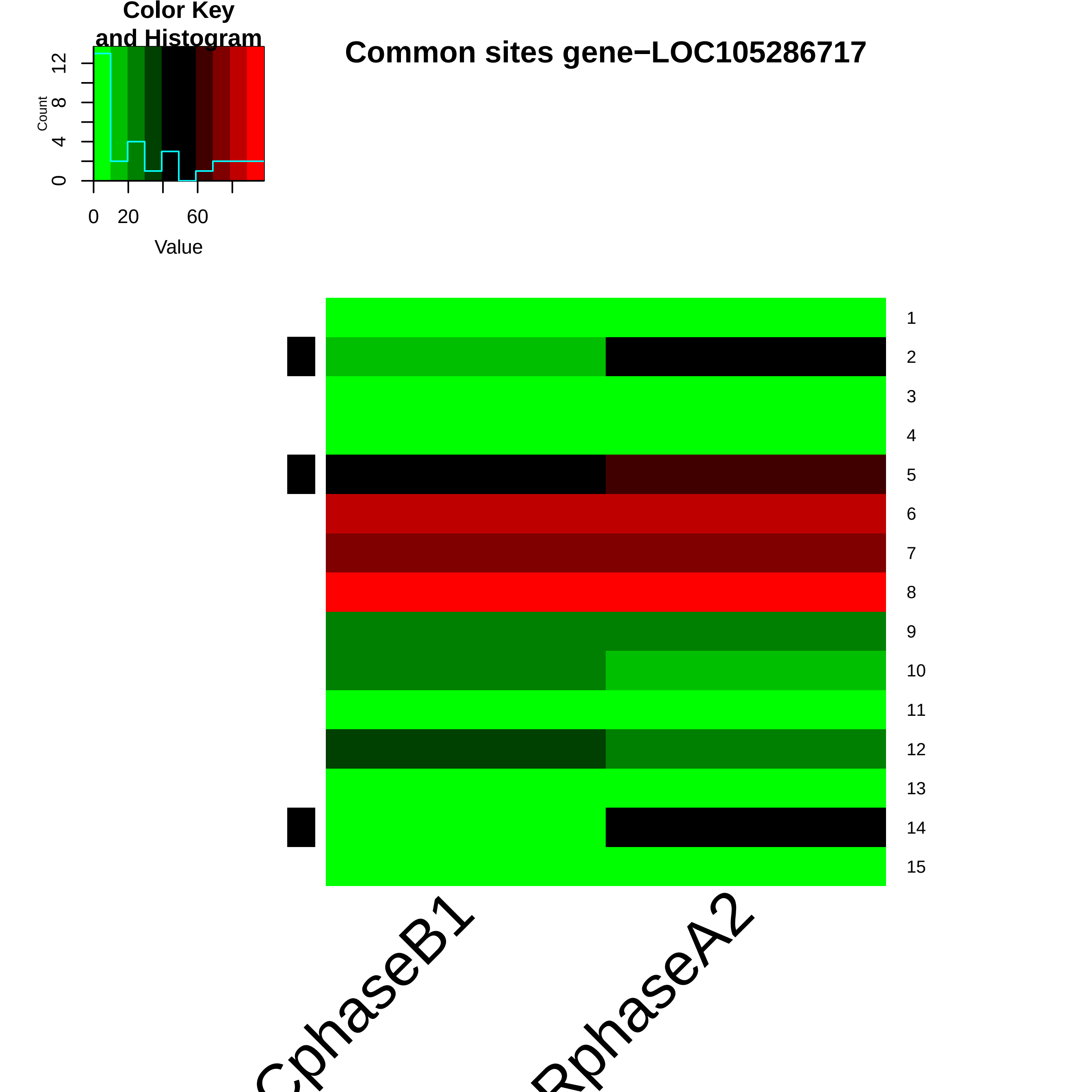

Color Key
and Histogram
Count
0
4
8
12
0
20
60
Value
Common sites gene−LOC105286717
CphaseB1
RphaseA2

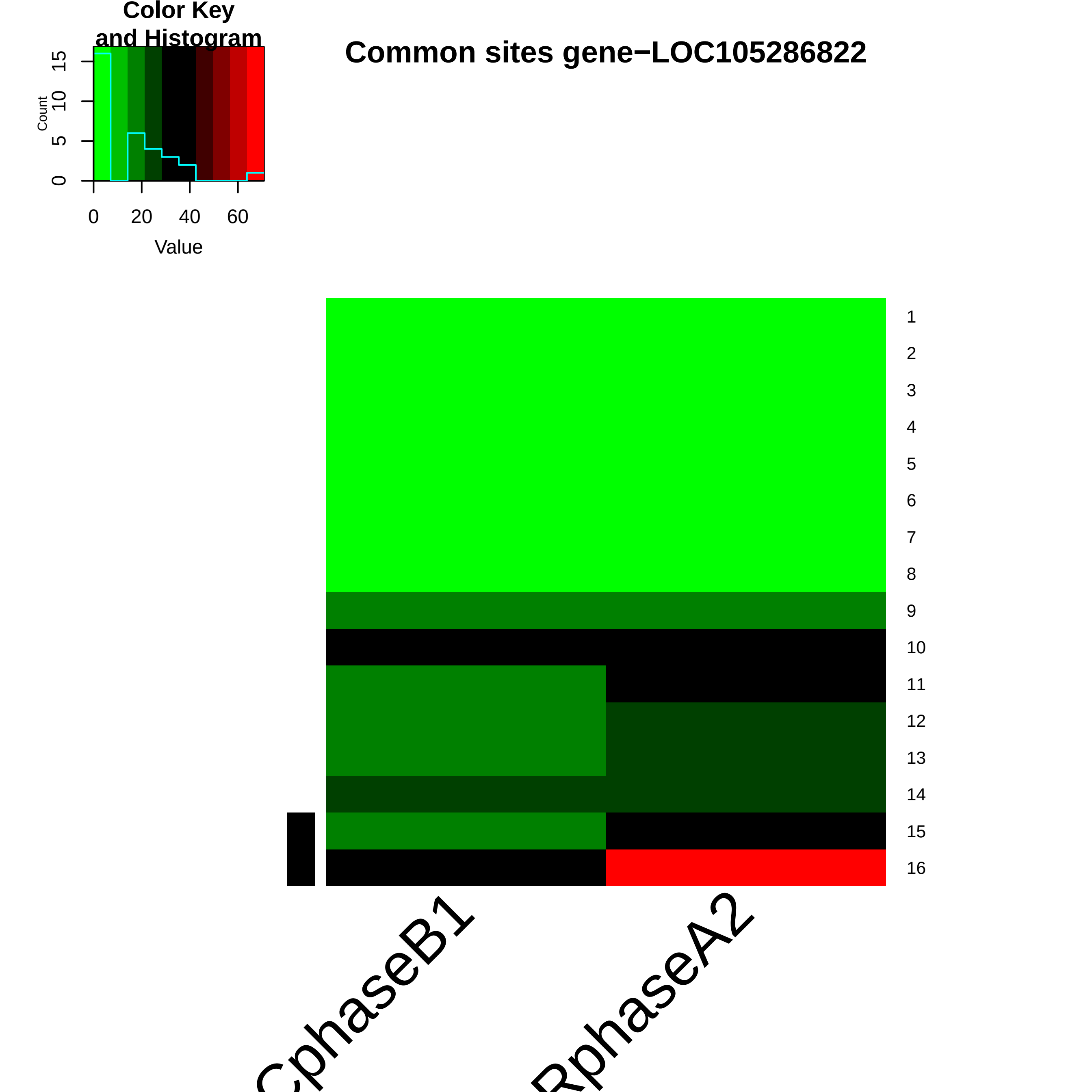

Color Key
and Histogram
Count
0
5
10
15
0
20
40
60
Value
Common sites gene−LOC105286822
CphaseB1
RphaseA2

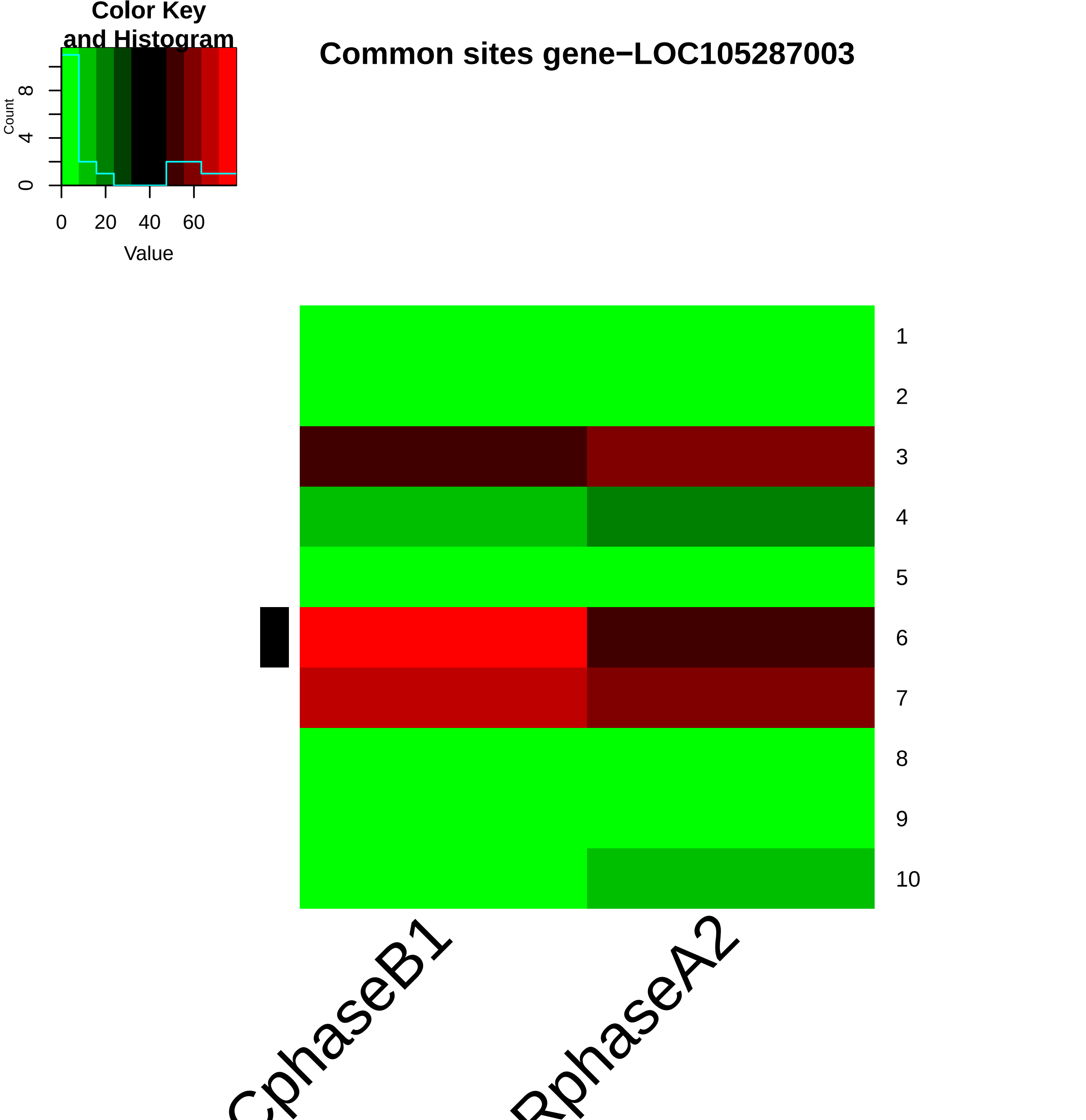

Color Key
and Histogram
Count
0
4
8
0
20
40
60
Value
Common sites gene−LOC105287003
1
2
3
4
5
6
7
8
9
10
CphaseB1
RphaseA2

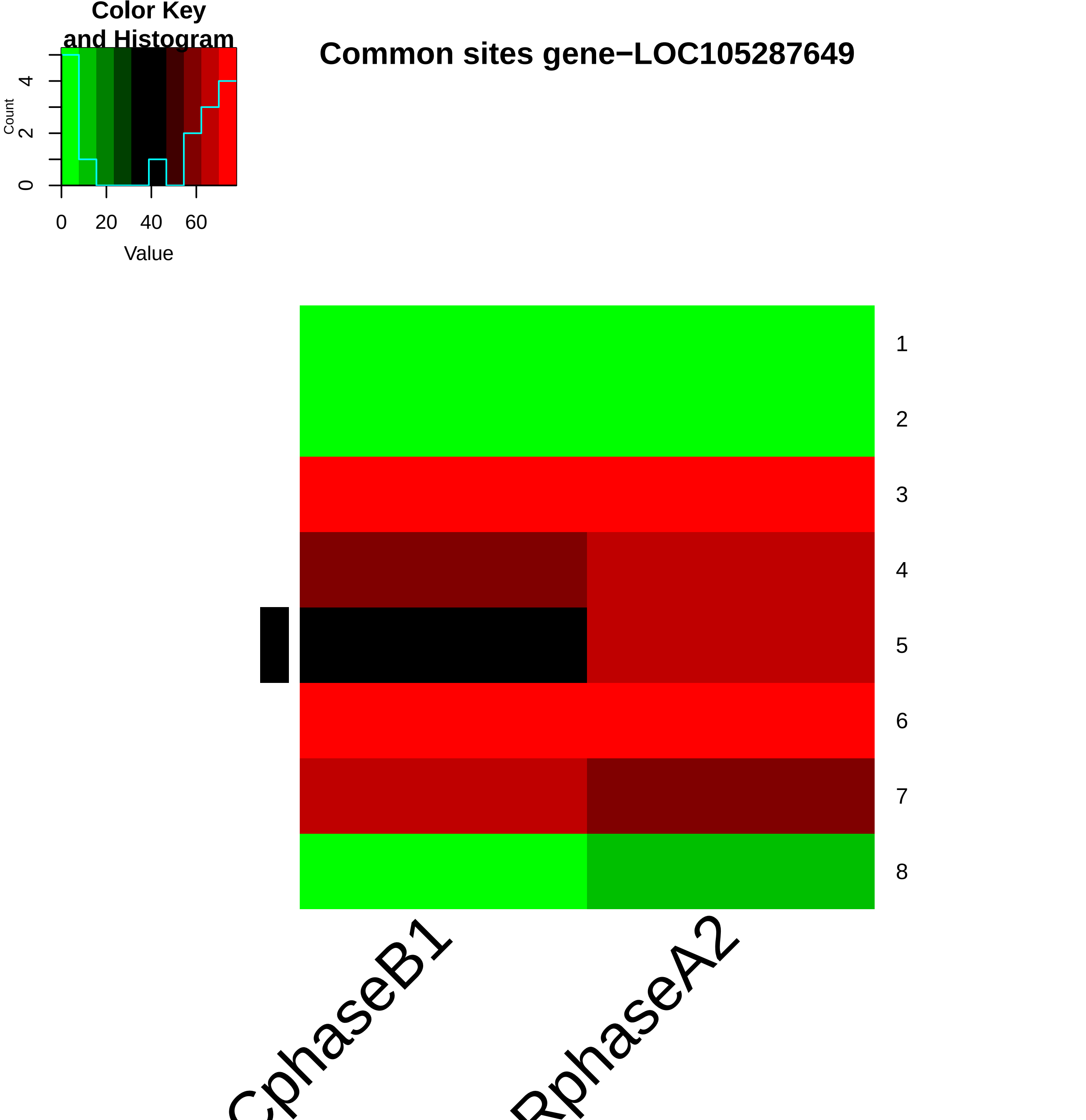
Color Key
and Histogram
Count
Value
0
20
40
60
0
2
4
Common sites gene−LOC105287649
1
2
3
4
5
6
7
8
CphaseB1
RphaseA2

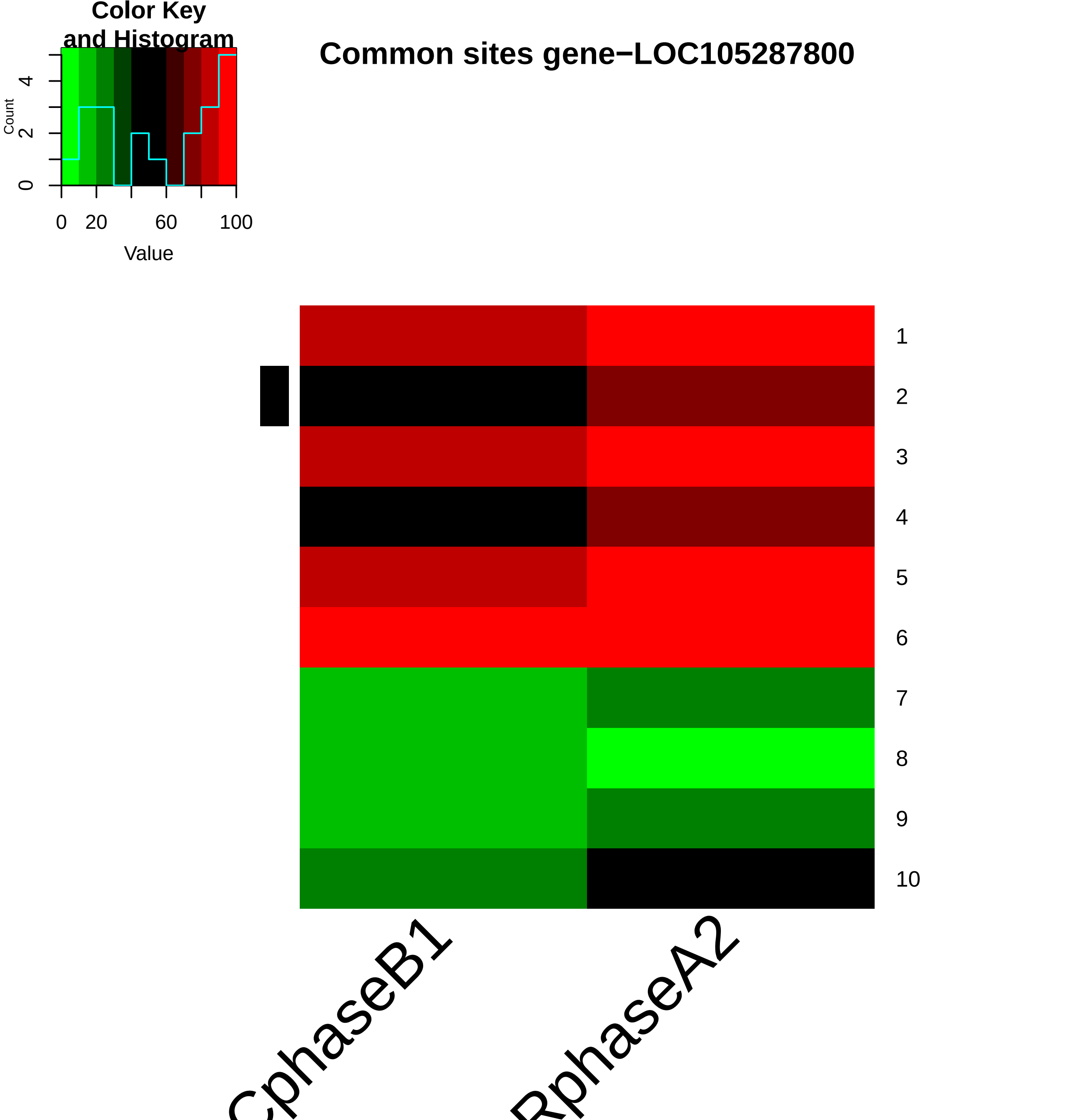
Color Key
and Histogram
Count
0
2
4
0
20
60
100
Value
Common sites gene−LOC105287800
1
2
3
4
5
6
7
8
9
10
CphaseB1
RphaseA2

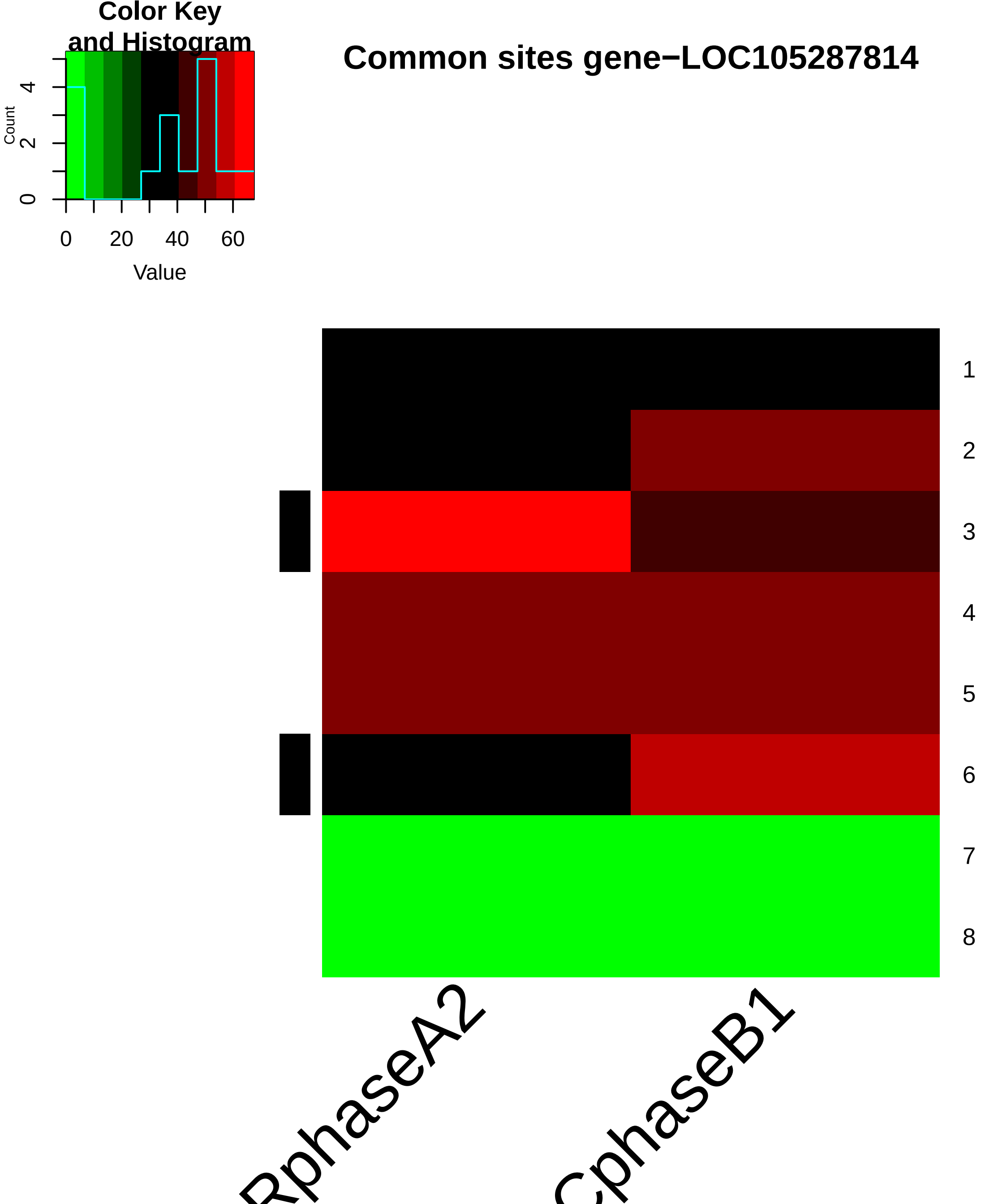
Color Key
and Histogram
Count
0
2
4
0
20
40
60
Value
Common sites gene−LOC105287814
1
2
3
4
5
6
7
8

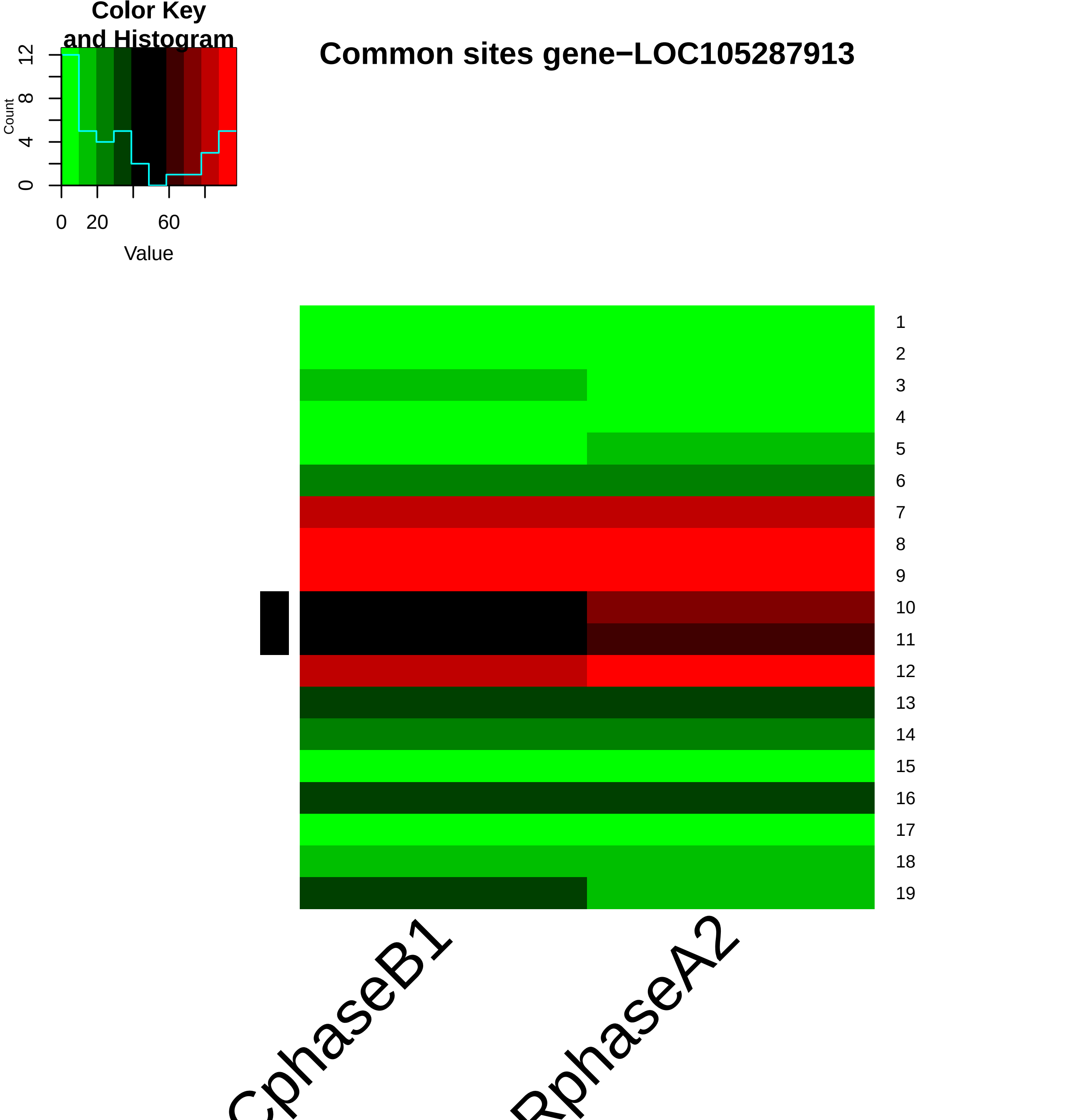

Color Key
and Histogram
Count
0
4
8
12
0
20
60
Value
Common sites gene−LOC105287913
CphaseB1
RphaseA2

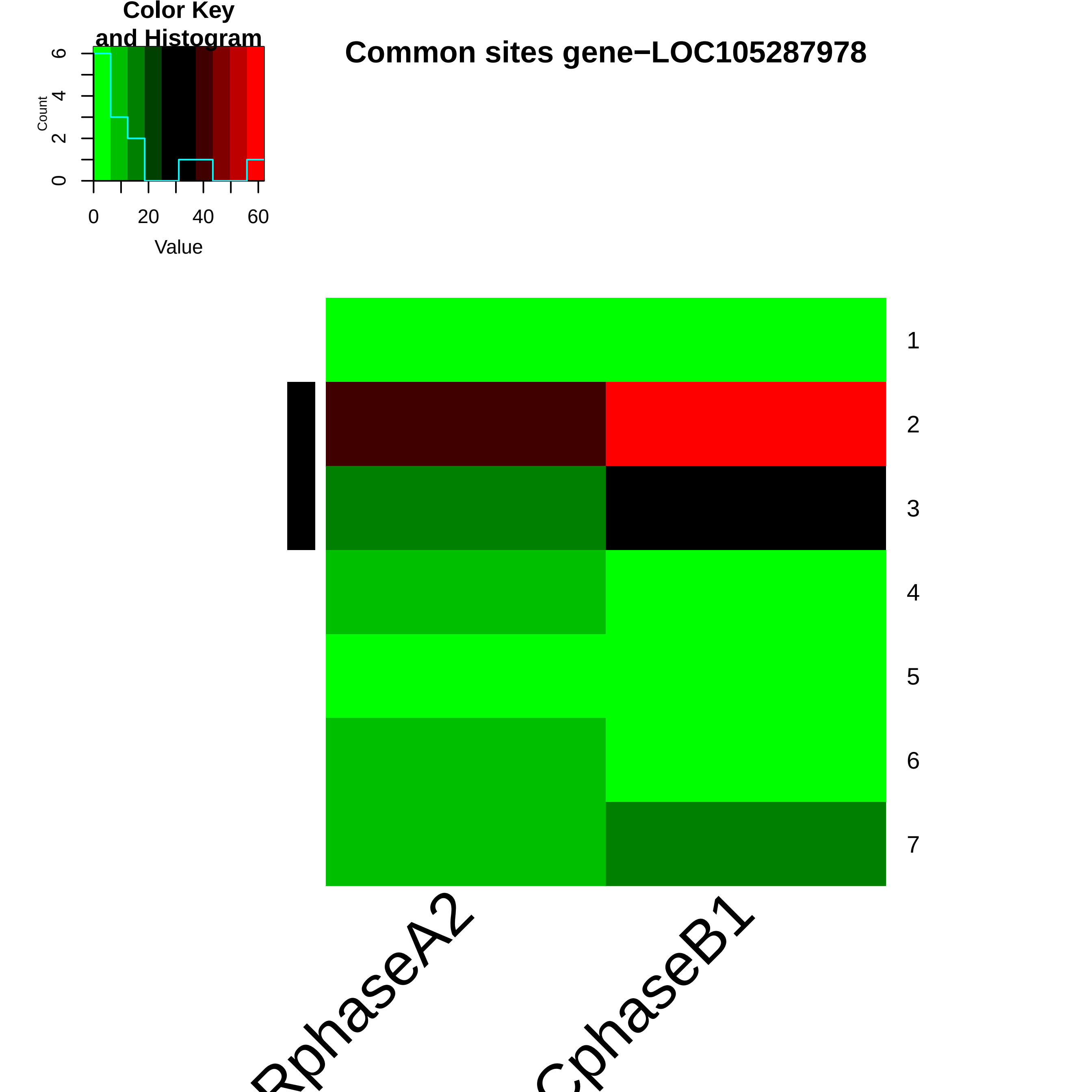

Color Key
and Histogram
Count
0
2
4
6
0
20
40
60
Value
Common sites gene−LOC105287978
1
2
3
4
5
6
7
RphaseA2
CphaseB1

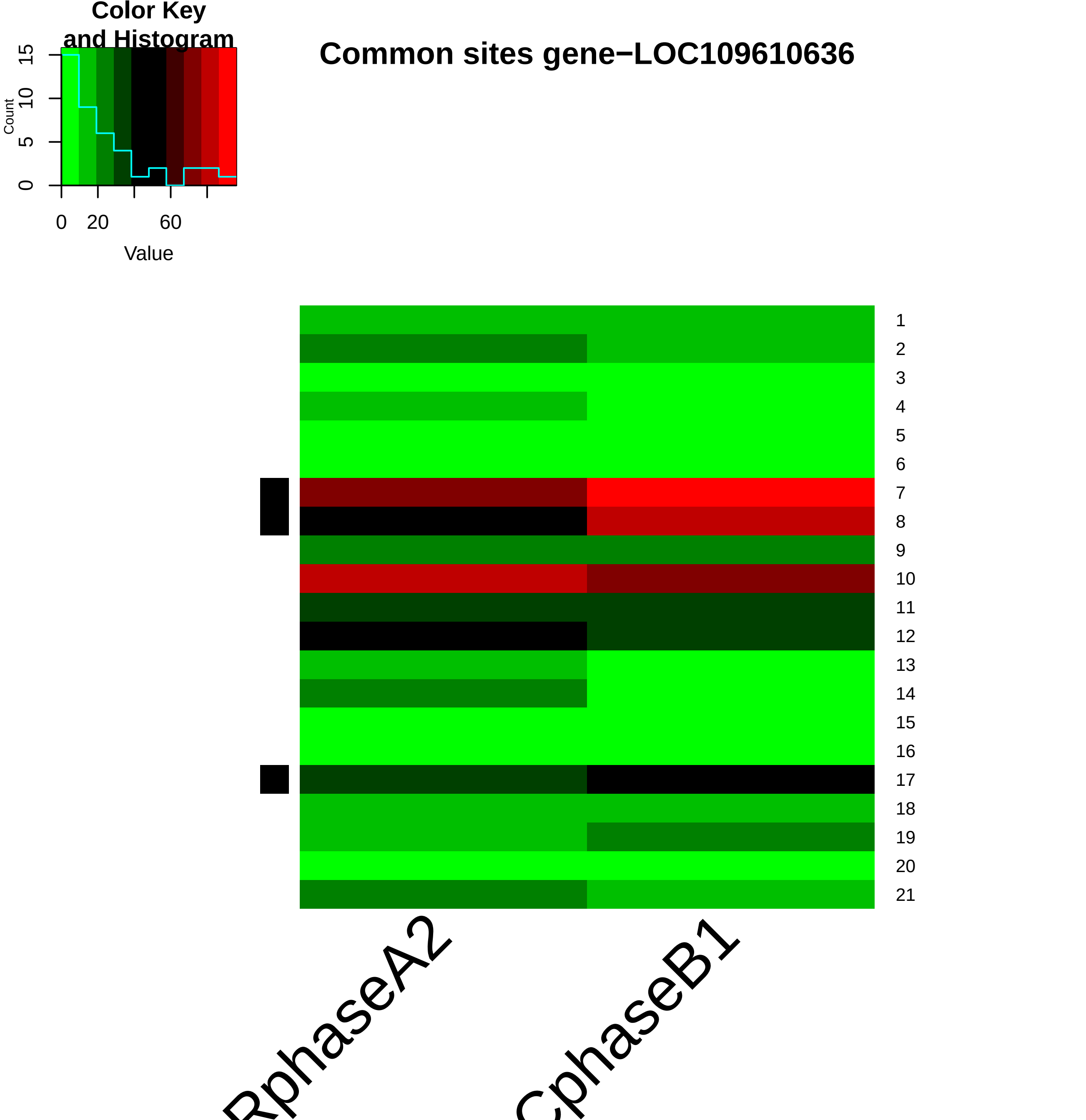

Color Key
and Histogram
Count
Value
0
20
60
0
5
10
15
Common sites gene−LOC109610636
RphaseA2
CphaseB1

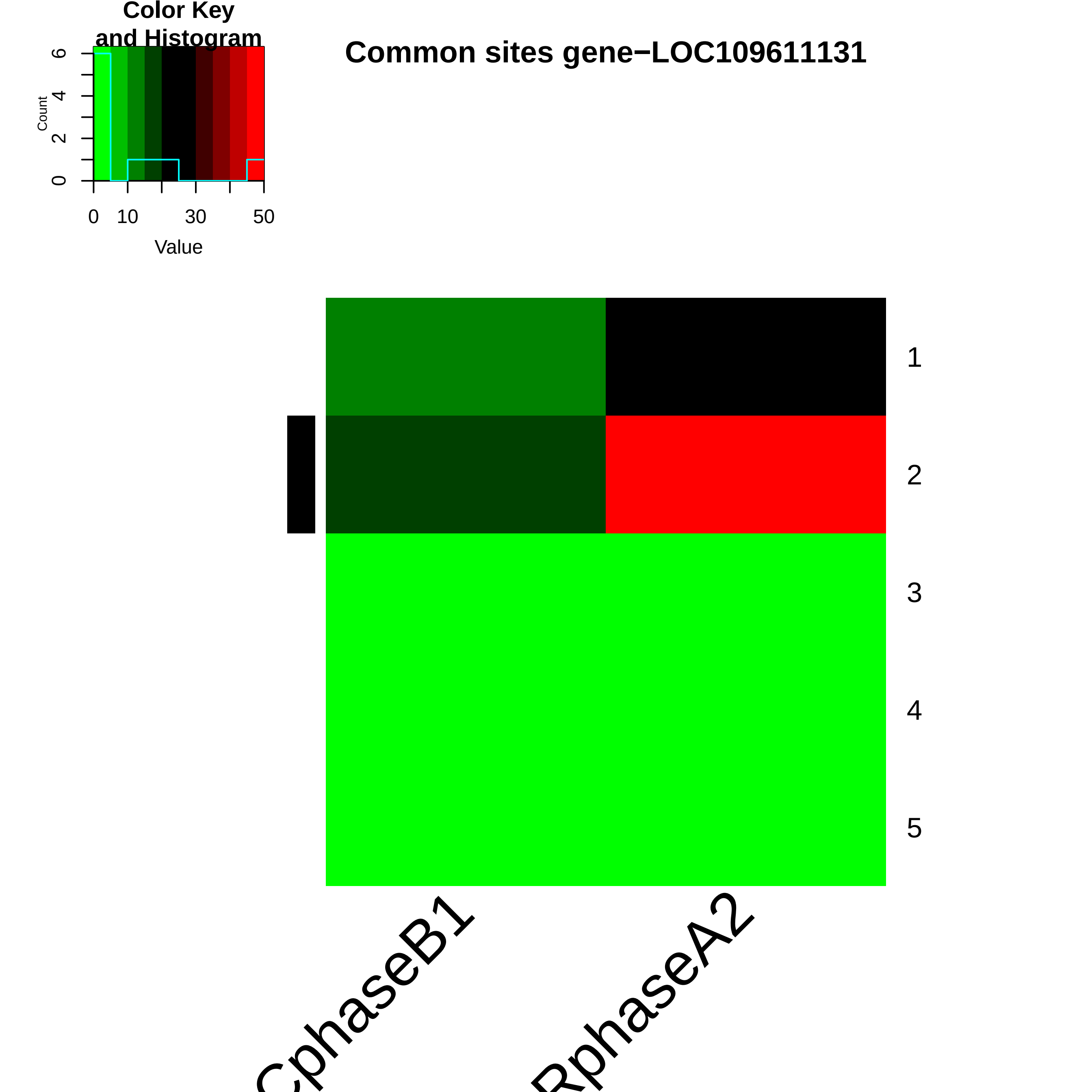
Color Key
and Histogram
Count
0
2
4
6
0
10
30
50
Value
Common sites gene−LOC109611131
1
2
3
4
5
CphaseB1
RphaseA2

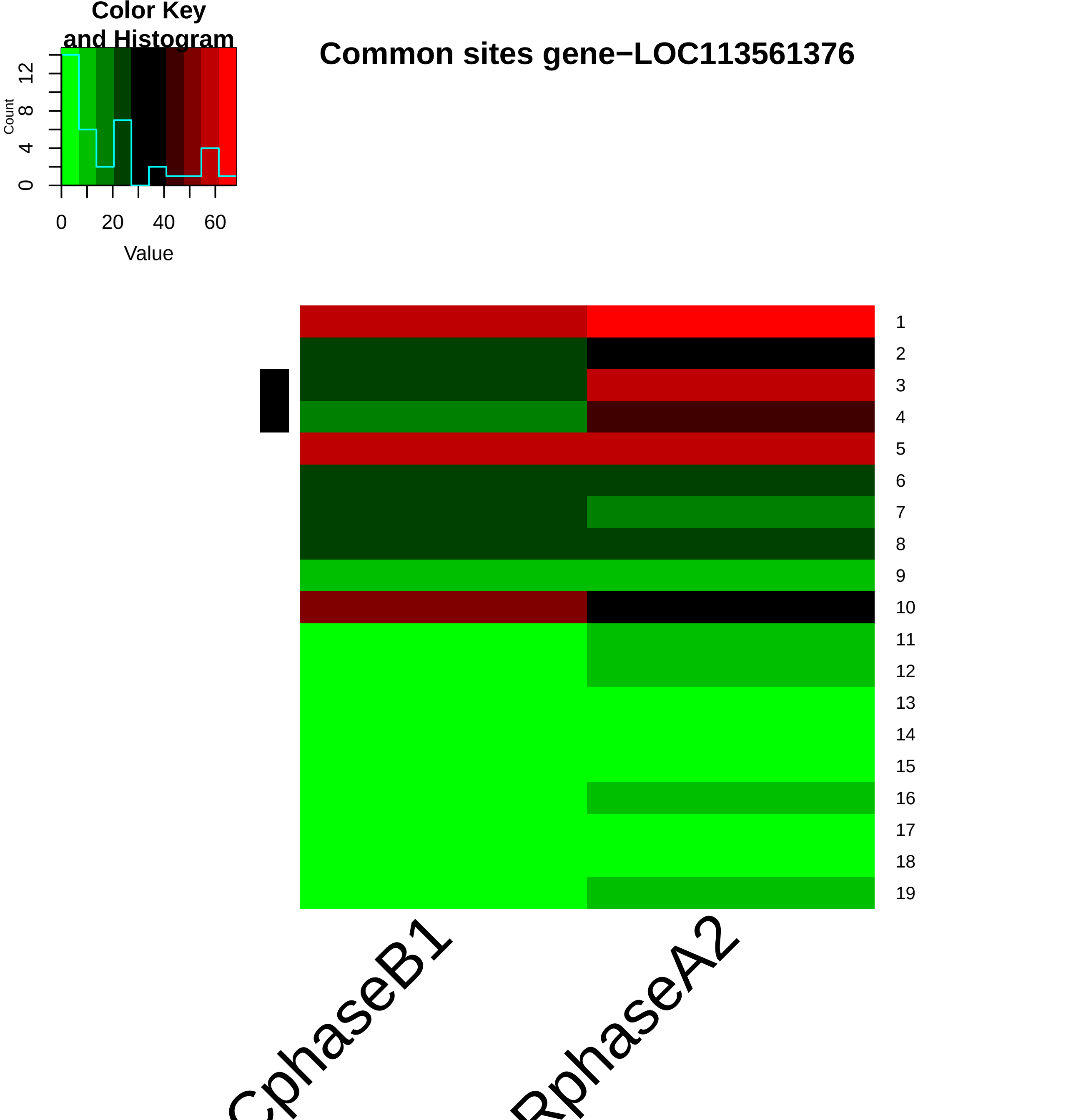
Color Key
and Histogram
Count
0
4
8
12
0
20
40
60
Value
Common sites gene−LOC113561376
CphaseB1
RphaseA2

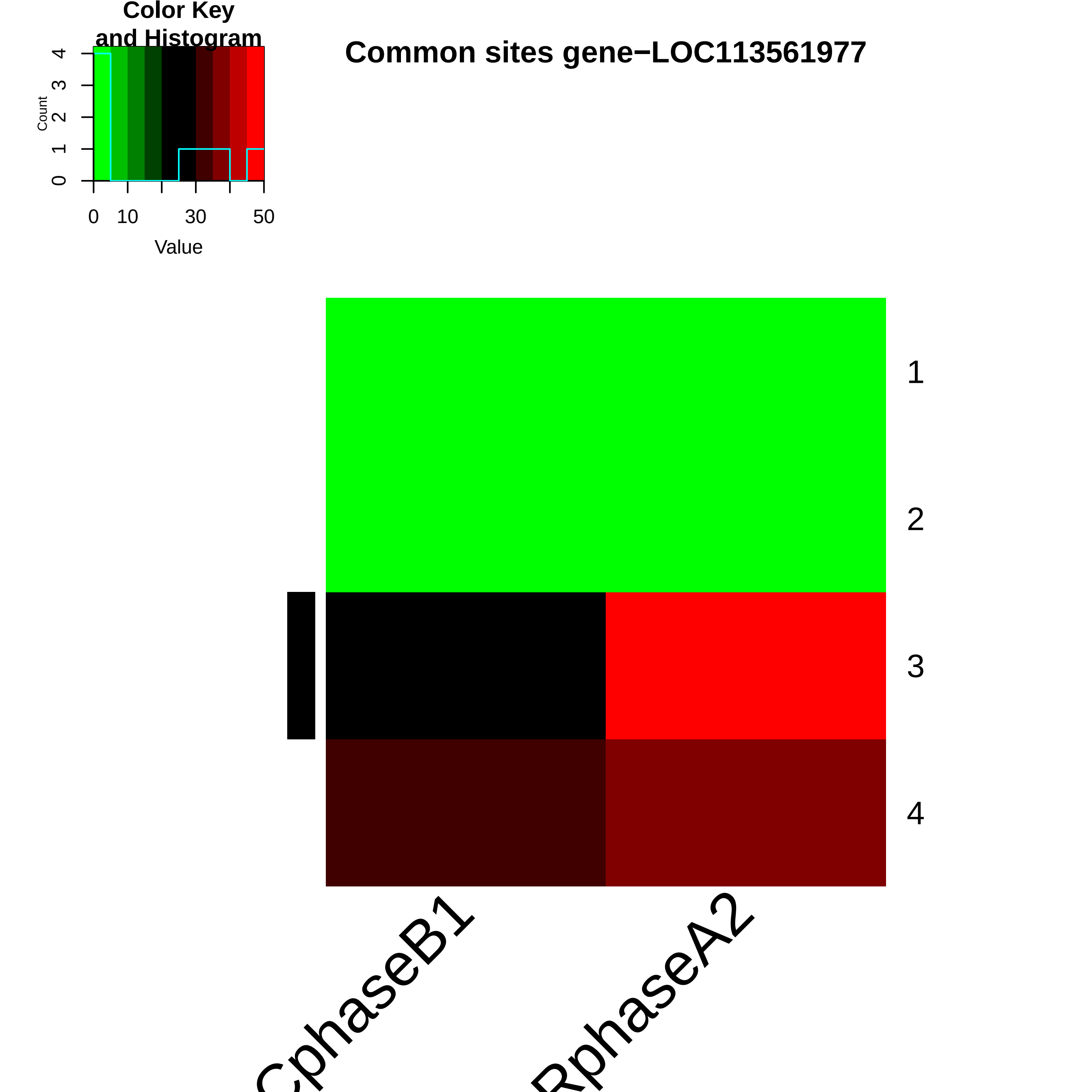

Color Key and Histogram
Count
Value
0
10
30
50
0
1
2
3
4
Common sites gene−LOC113561977
1
2
3
4
CphaseB1
RphaseA2

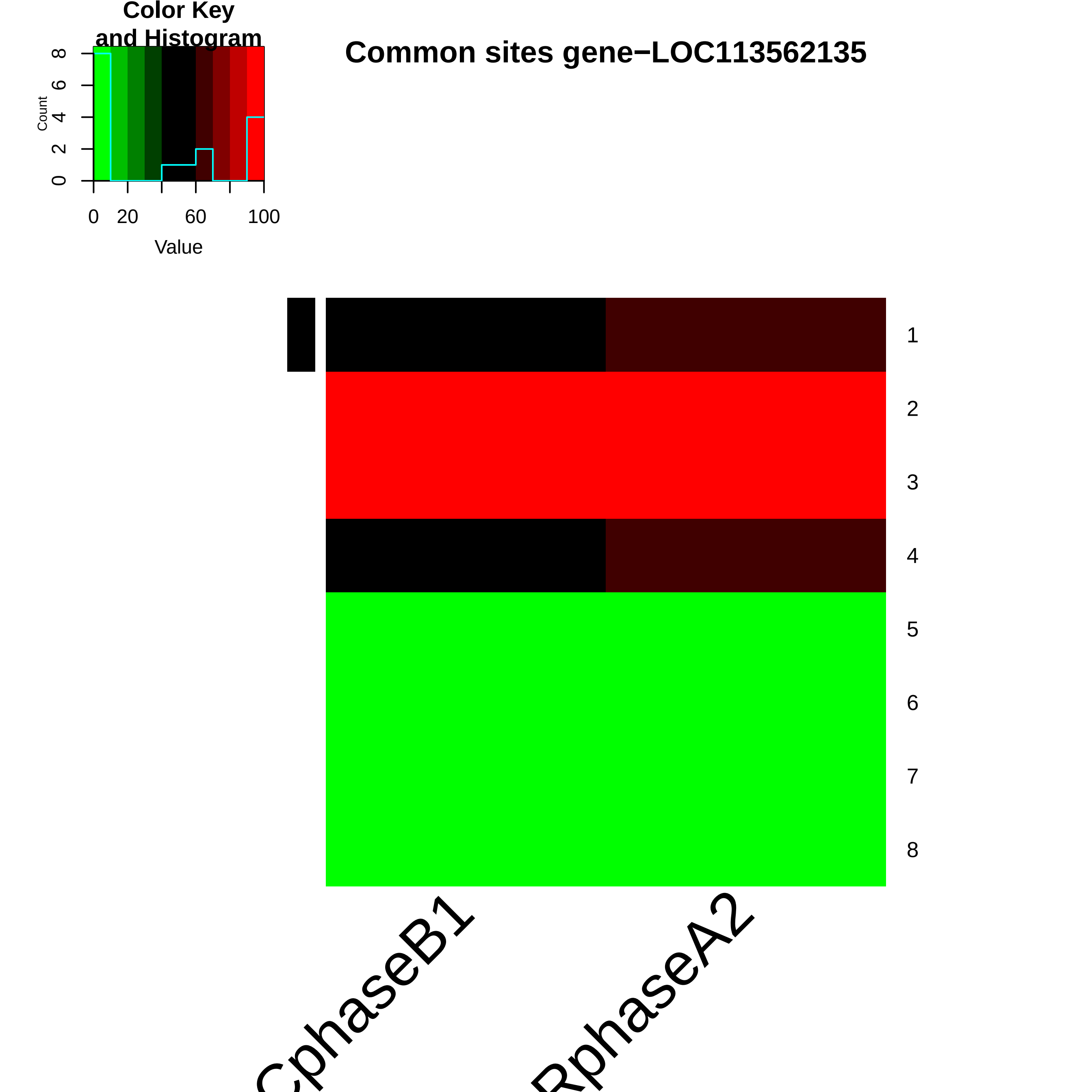
Color Key
and Histogram
Count
0
2
4
6
8
0
20
60
100
Value
Common sites gene−LOC113562135
1
2
3
4
5
6
7
8
CphaseB1
RphaseA2

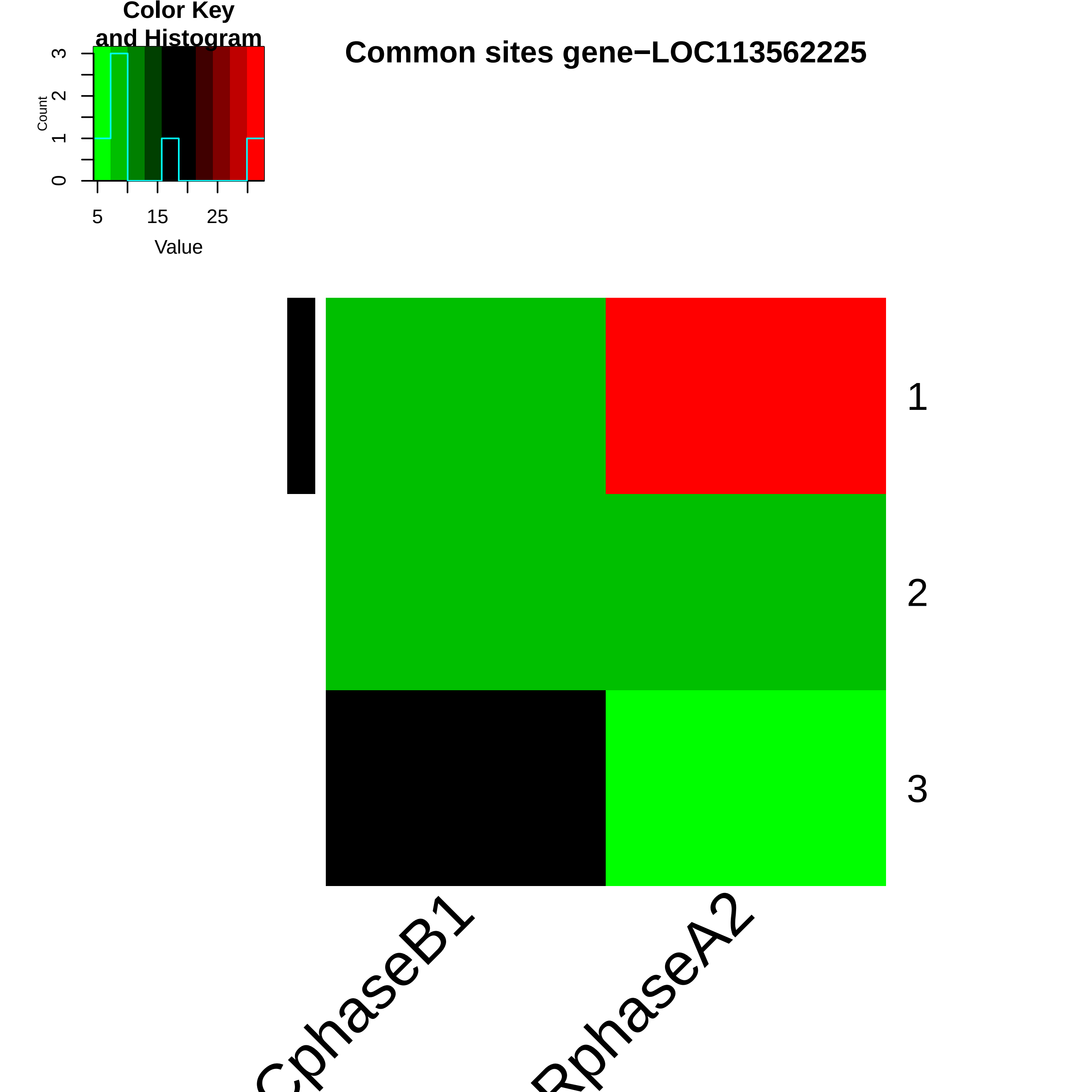
Color Key
and Histogram
Count
0
1
2
3
5
15
25
Value
Common sites gene−LOC113562225
1
2
3
CphaseB1
RphaseA2

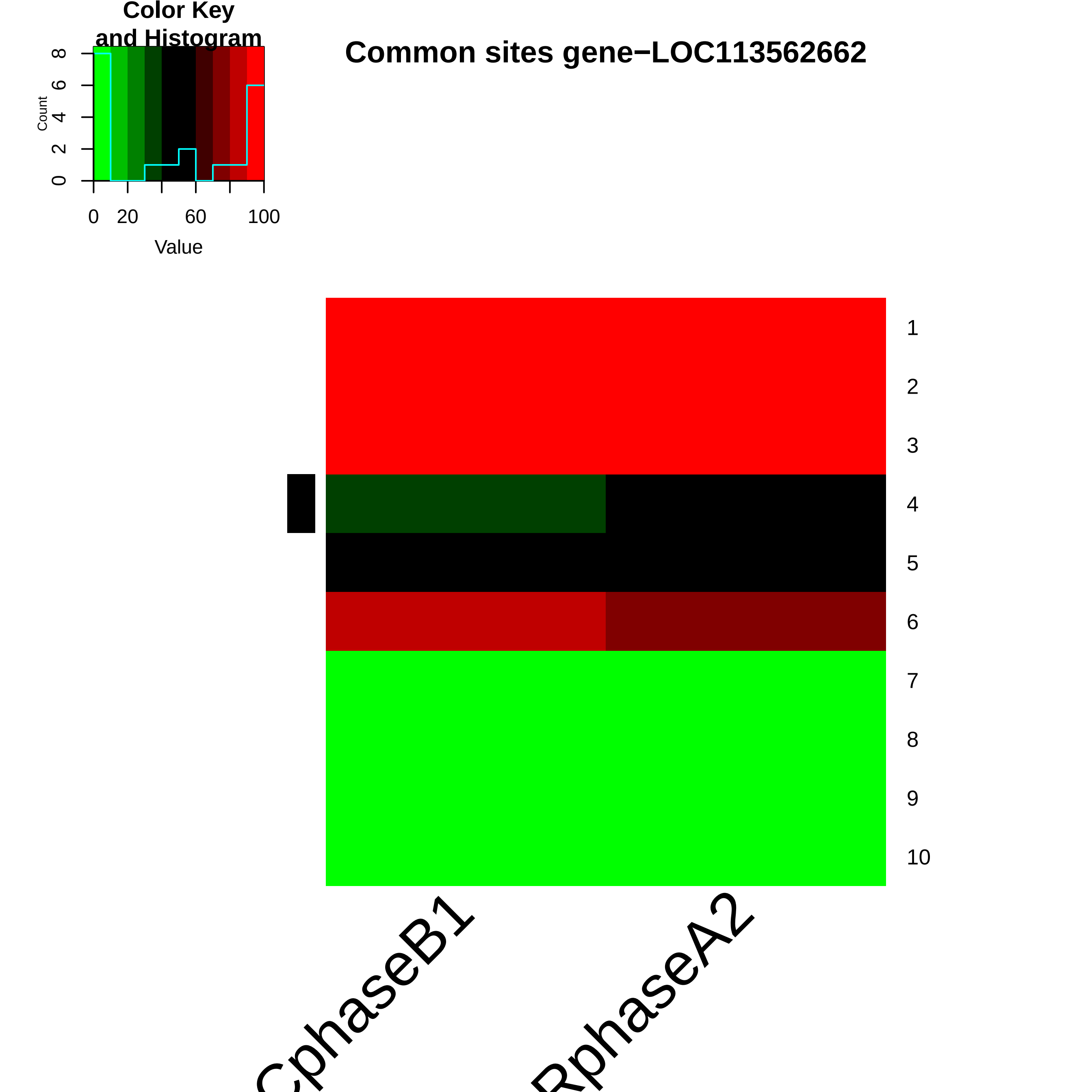

Color Key
and Histogram
Count
0
2
4
6
8
0
20
60
100
Value
Common sites gene−LOC113562662
1
2
3
4
5
6
7
8
9
10
CphaseB1
RphaseA2

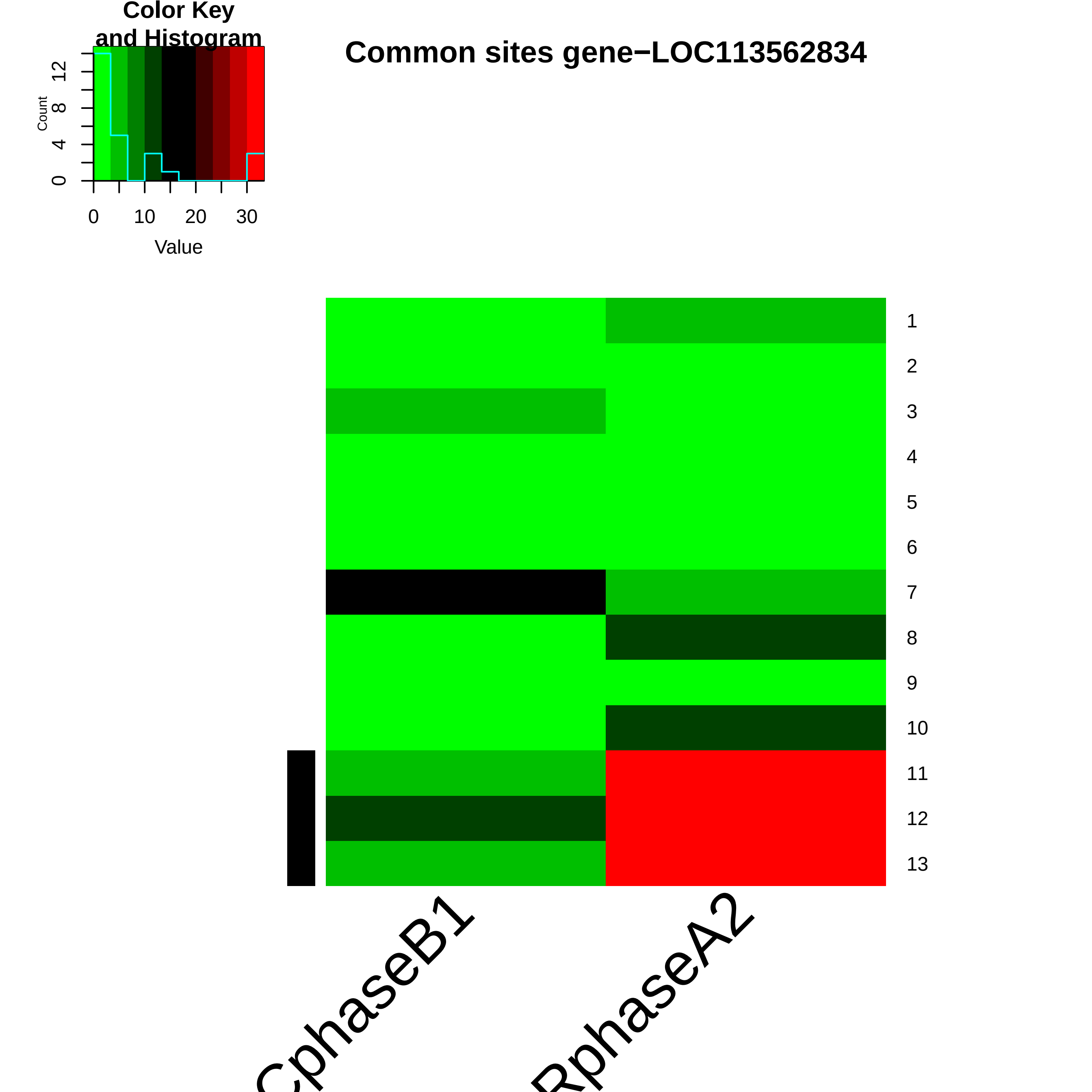

Color Key
and Histogram
Count
0
4
8
12
0
10
20
30
Value
Common sites gene−LOC113562834
1
2
3
4
5
6
7
8
9
10
11
12
13
CphaseB1
RphaseA2

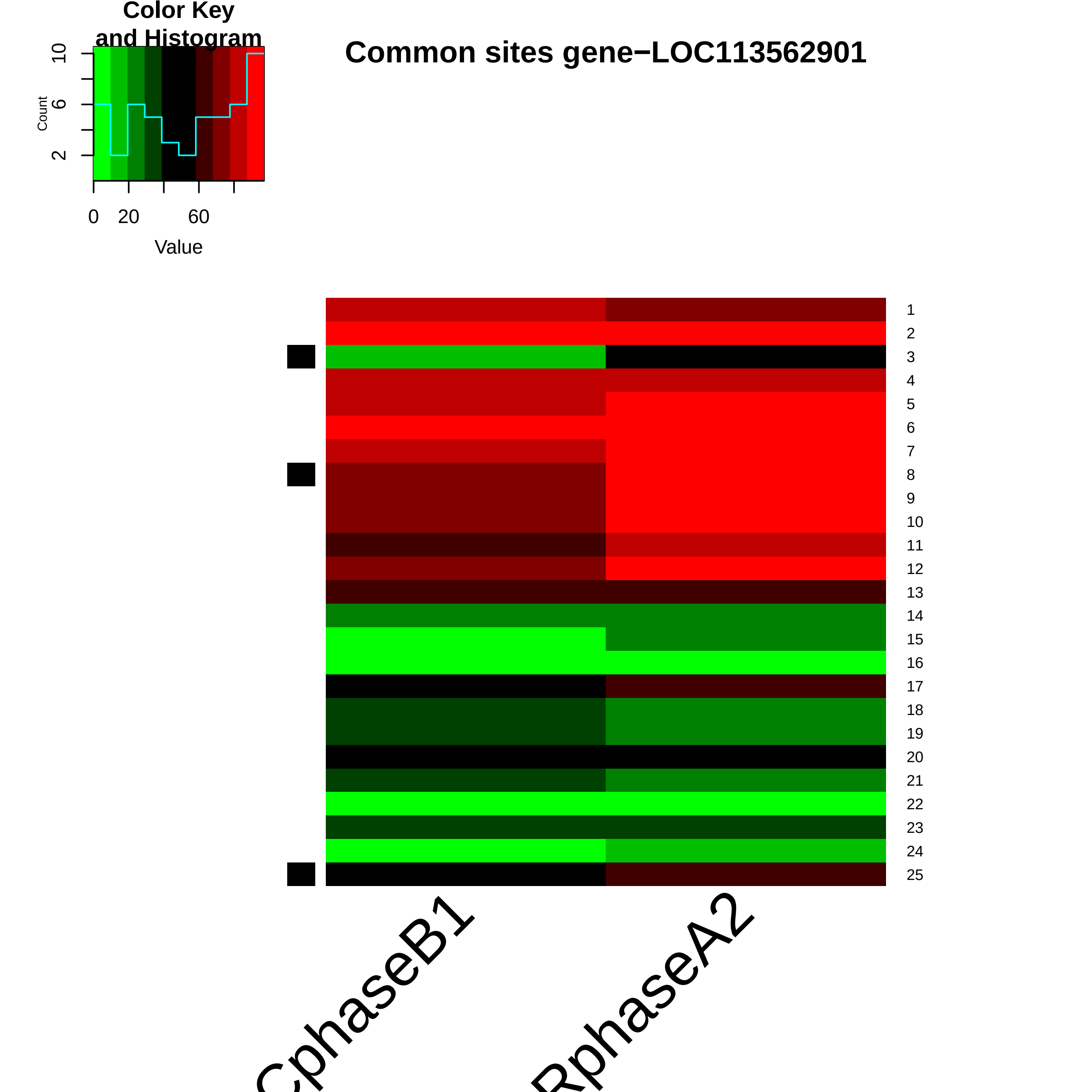
Color Key
and Histogram
Count
2
6
10
0
20
60
Value
Common sites gene−LOC113562901
CphaseB1
RphaseA2

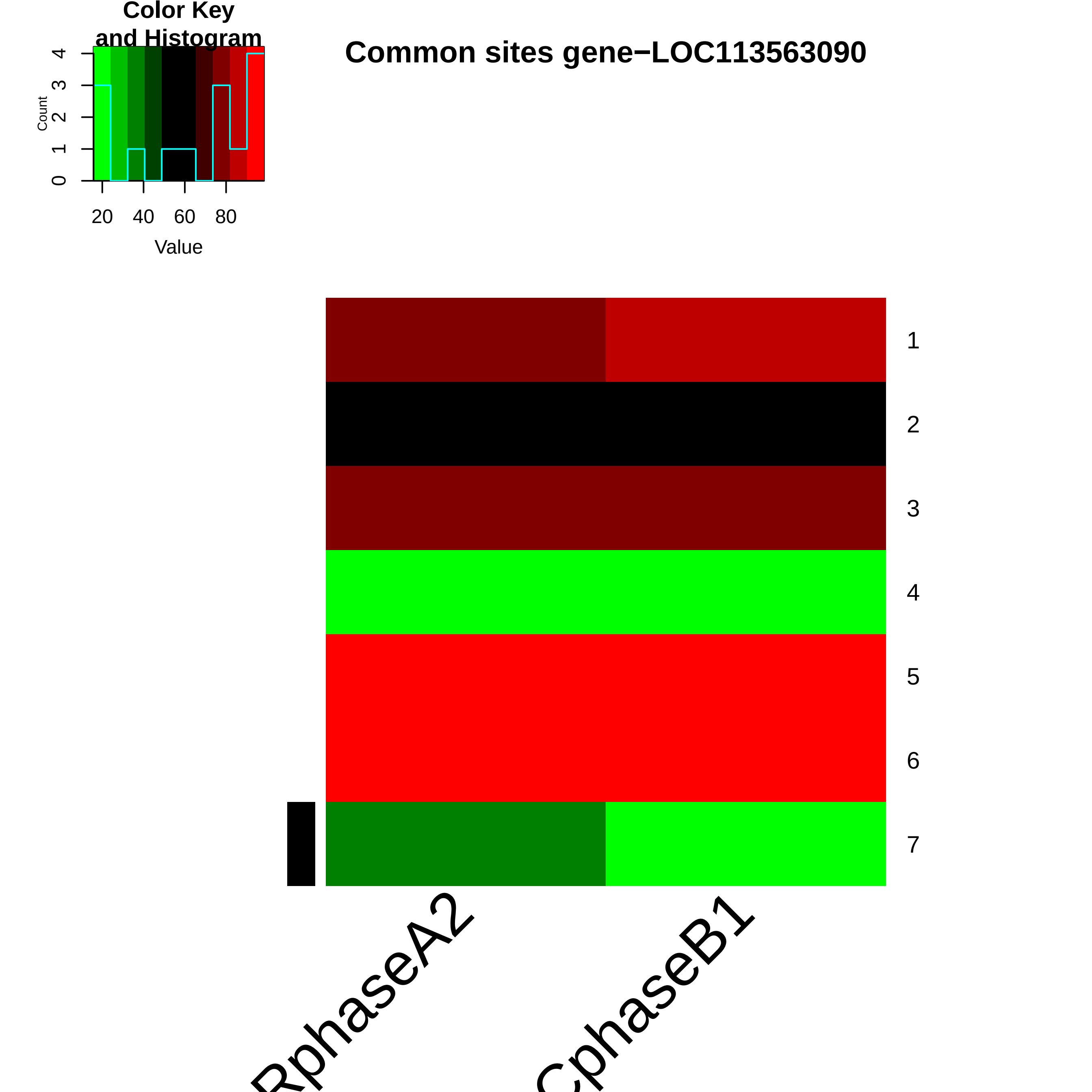

Color Key
and Histogram
Count
0
1
2
3
4
20
40
60
80
Value
Common sites gene−LOC113563090
1
2
3
4
5
6
7
RphaseA2
CphaseB1

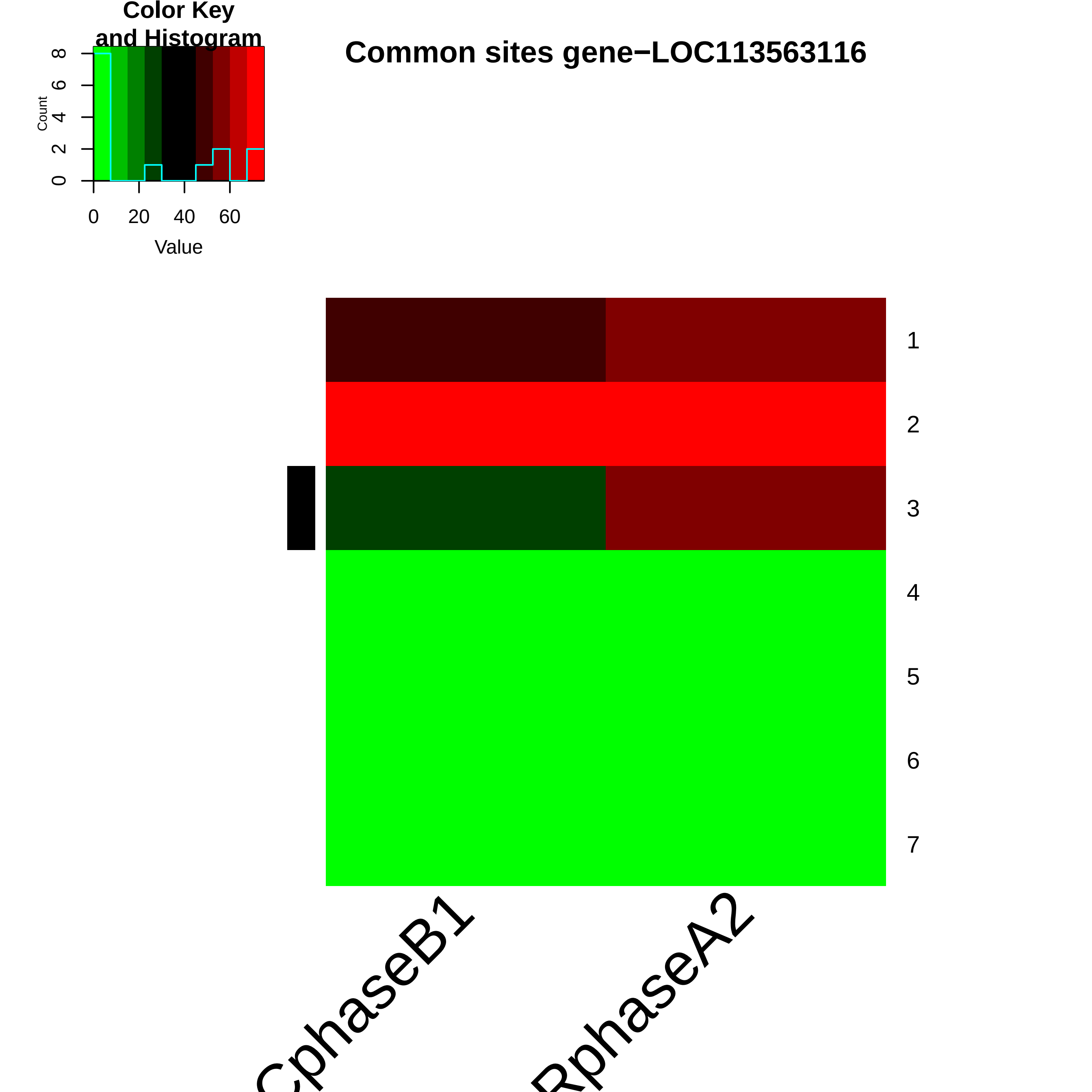
Color Key
and Histogram
Count
0
2
4
6
8
0
20
40
60
Value
Common sites gene−LOC113563116
1
2
3
4
5
6
7

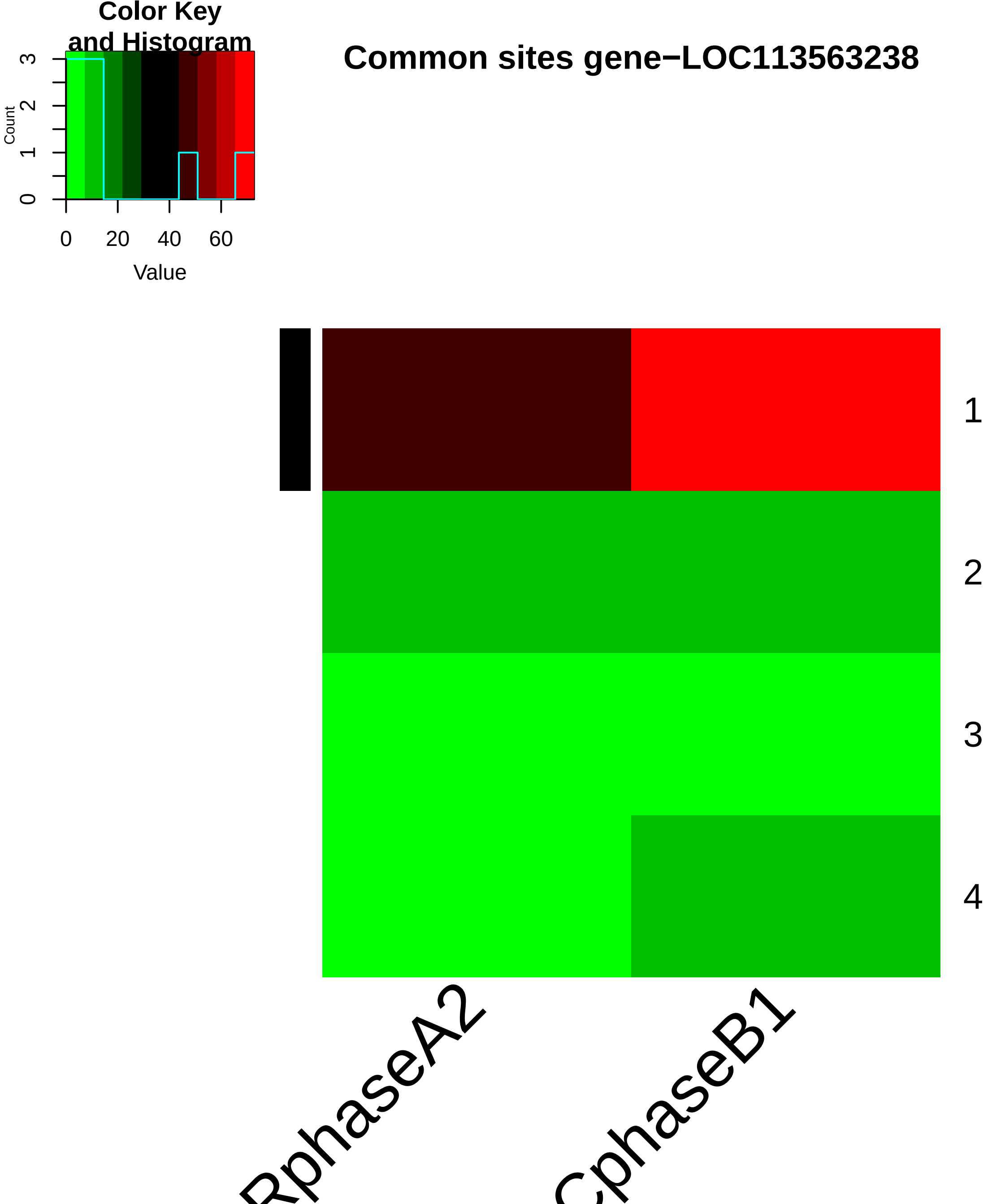
Color Key
and Histogram
Count
0
1
2
3
0
20
40
60
Value
Common sites gene−LOC113563238
1
2
3
4
RphaseA2
CphaseB1

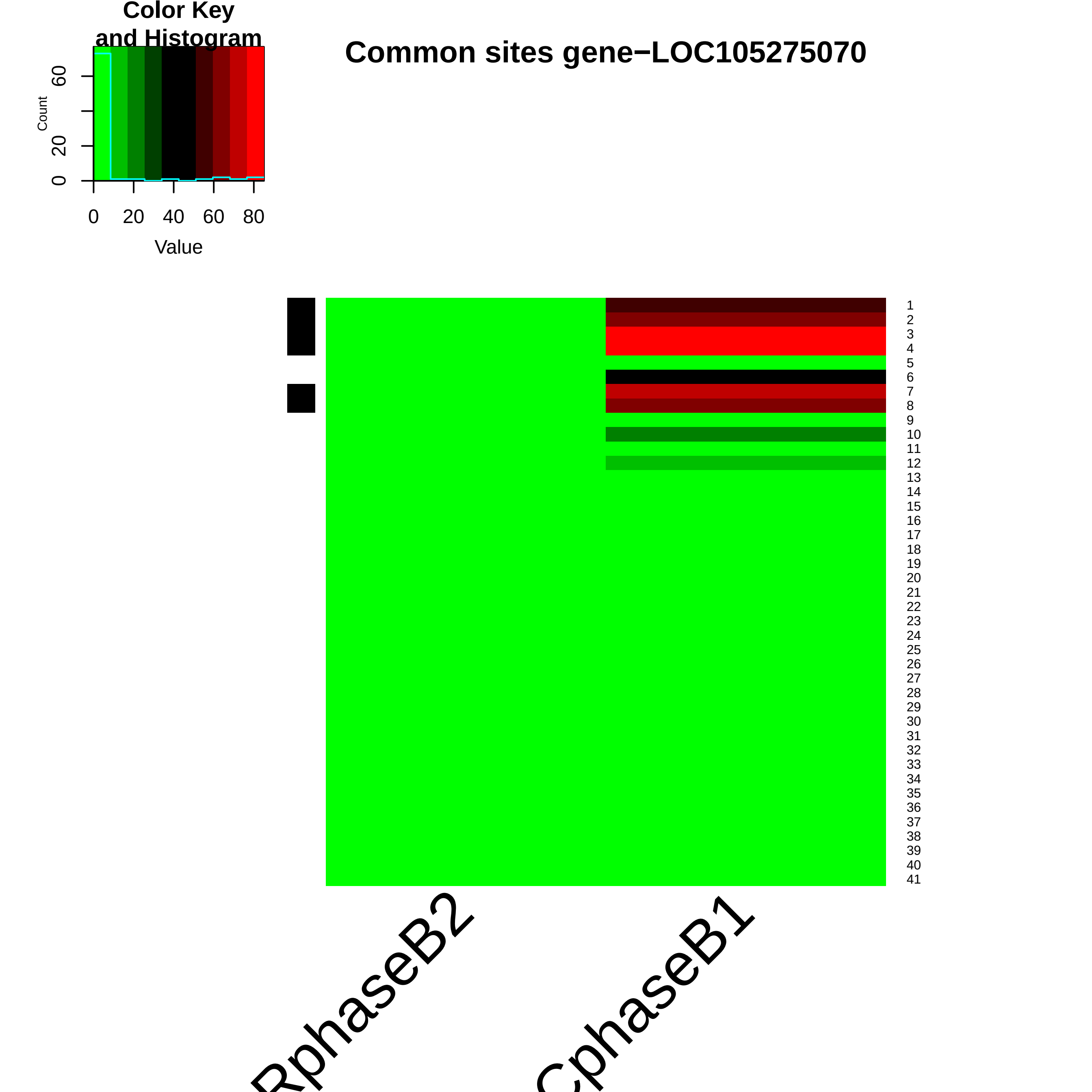

Color Key
and Histogram
Count
Value
Common sites gene−LOC105275070
RphaseB2
CphaseB1

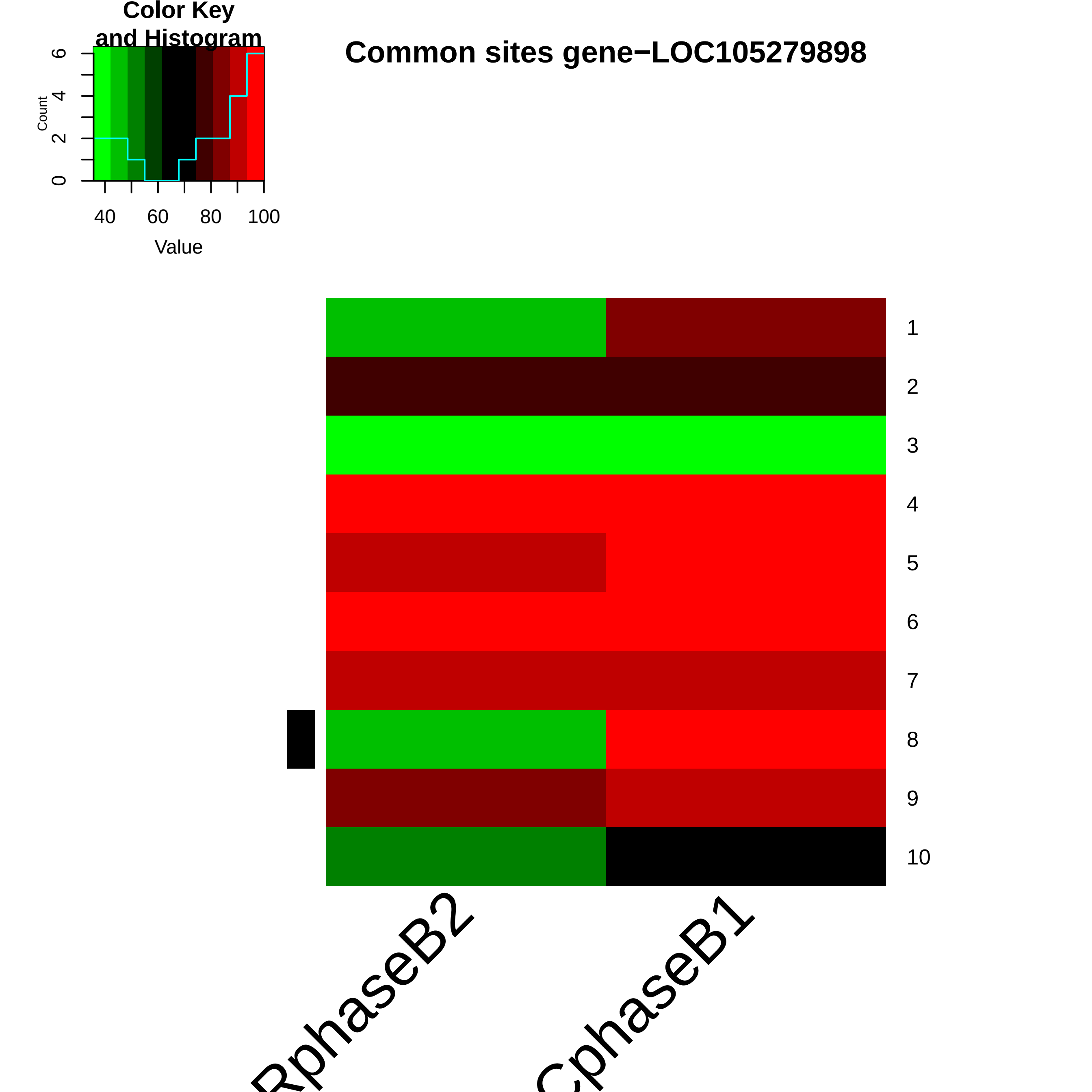
Color Key
and Histogram
Count
0
2
4
6
40
60
80
100
Value
Common sites gene−LOC105279898
1
2
3
4
5
6
7
8
9
10
RphaseB2
CphaseB1

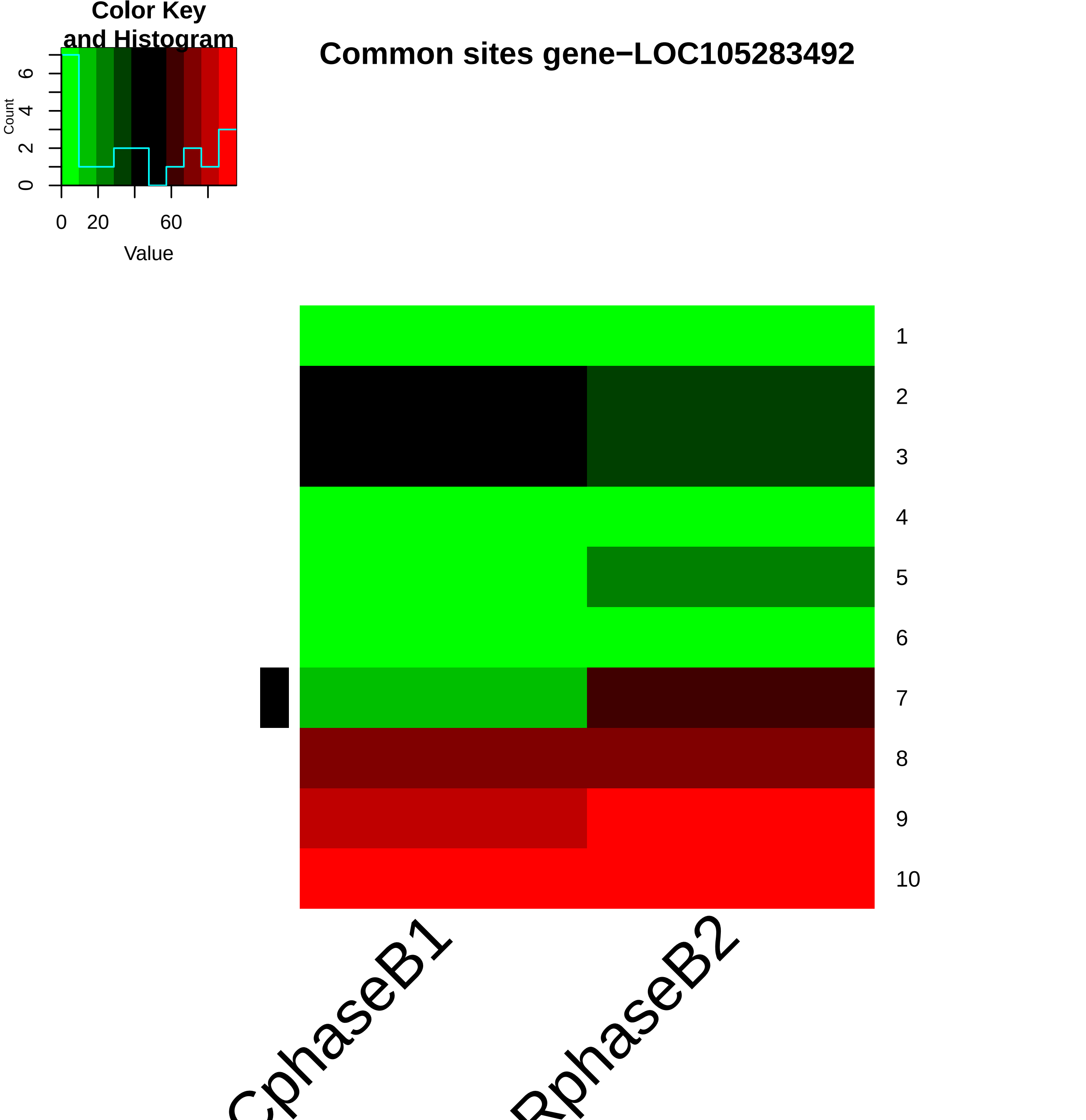

Color Key
and Histogram
Count
0
2
4
6
0
20
60
Value
Common sites gene−LOC105283492
1
2
3
4
5
6
7
8
9
10
CphaseB1
RphaseB2

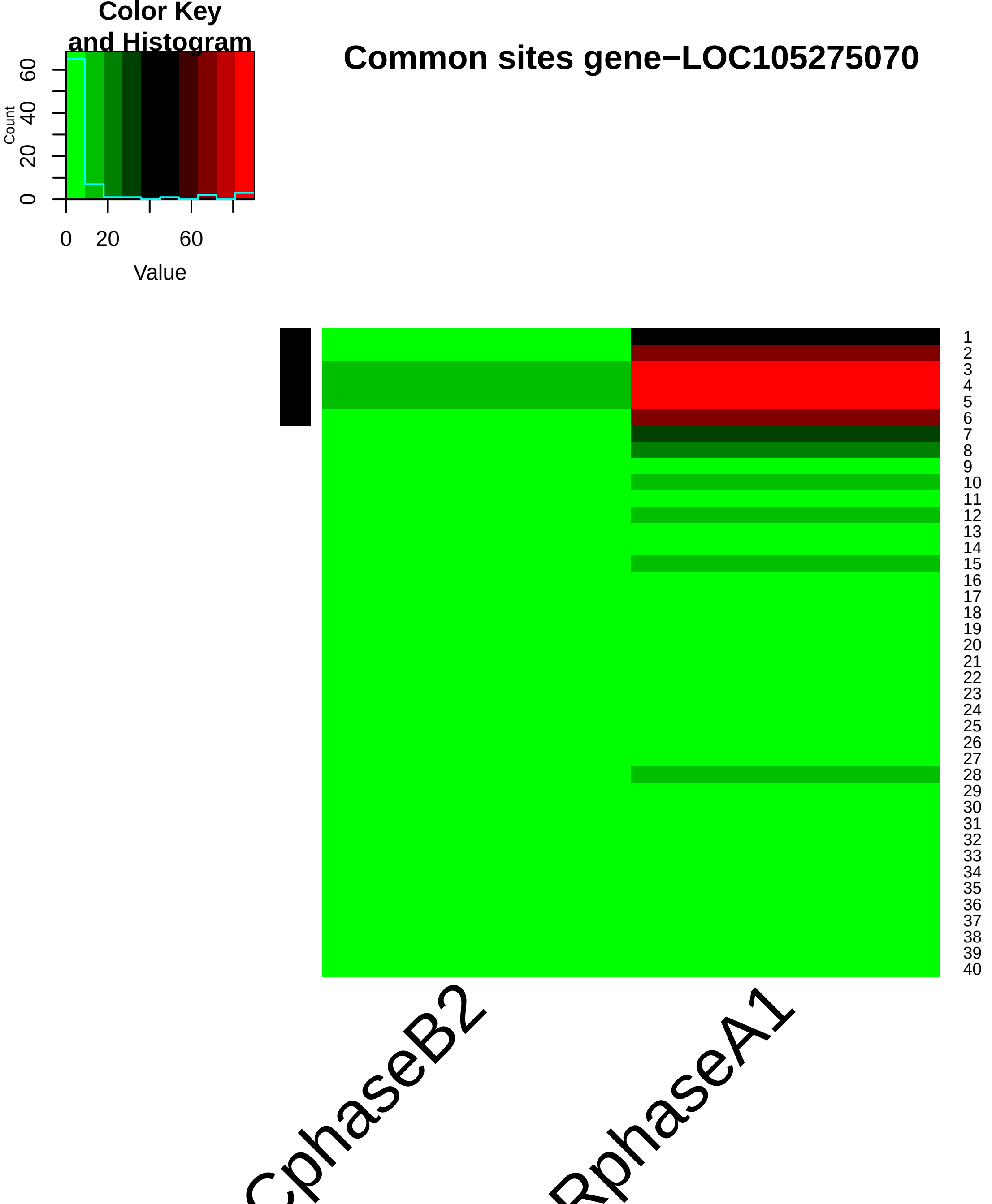

Color Key
and Histogram
Count
0
20
40
60
0
20
60
Value
Common sites gene−LOC105275070
CphaseB2
RphaseA1

#### Common sites gene−LOC105275070

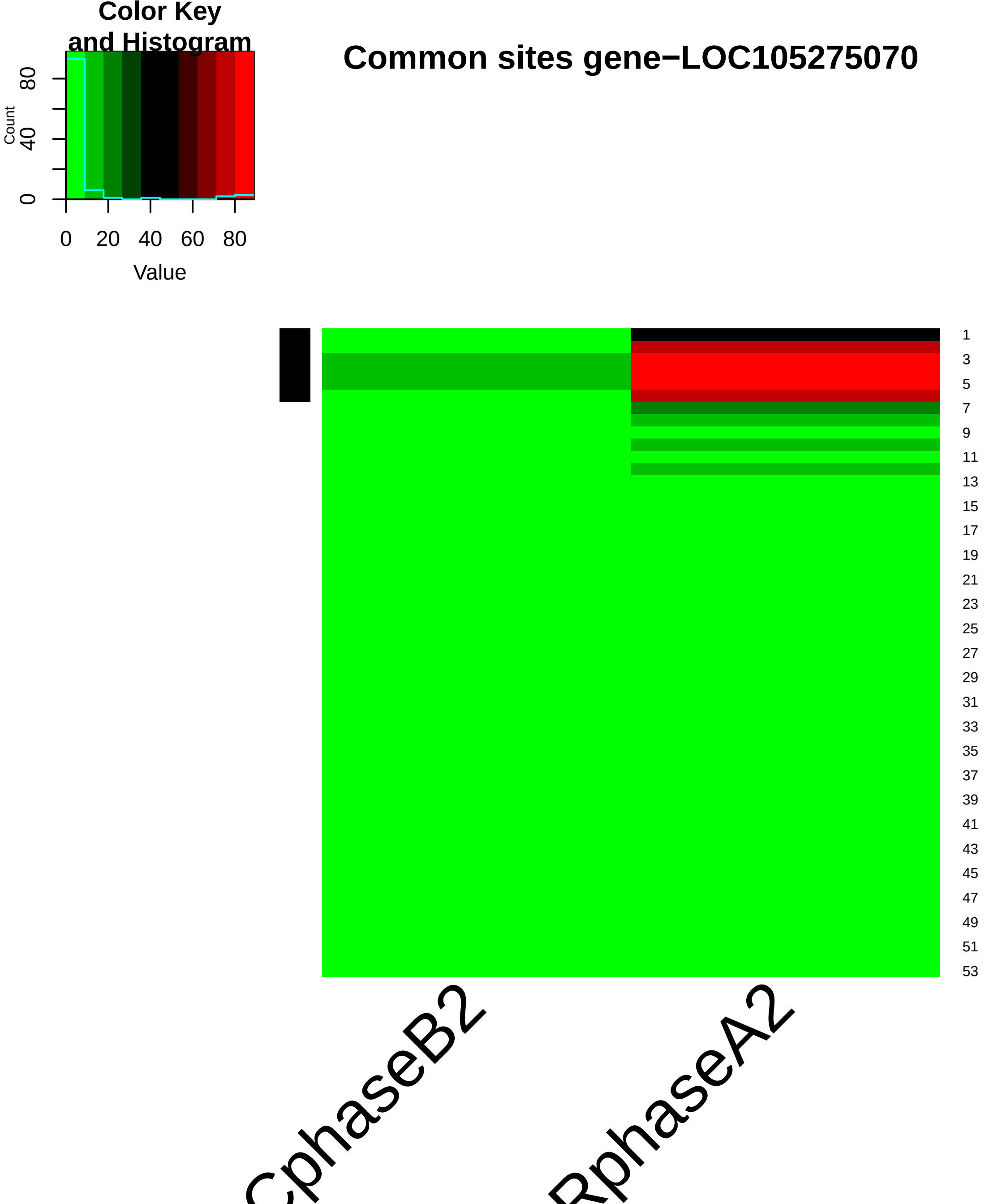

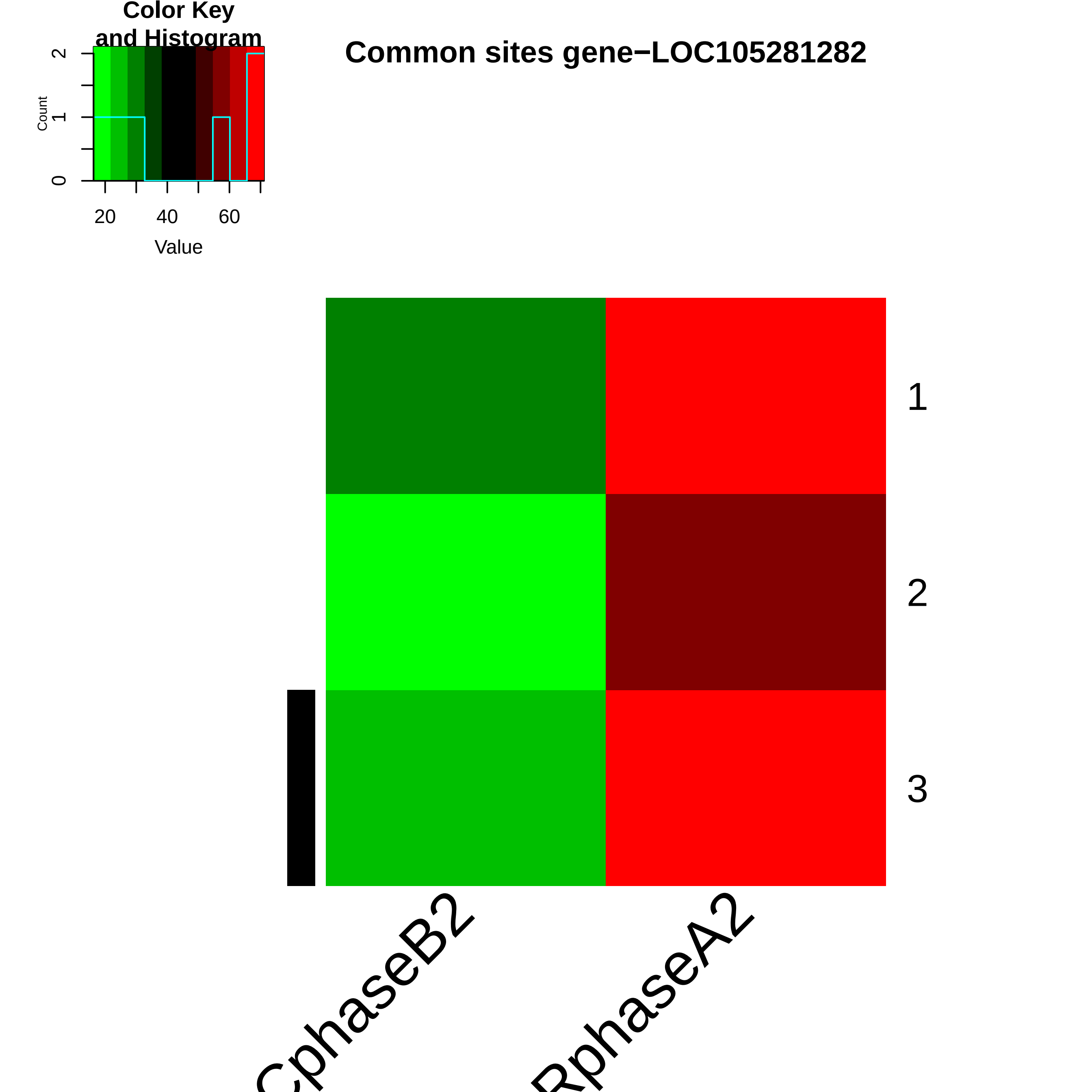
Color Key
and Histogram
Count
0
1
2
20
40
60
Value
Common sites gene−LOC105281282
1
2
3
CphaseB2
RphaseA2

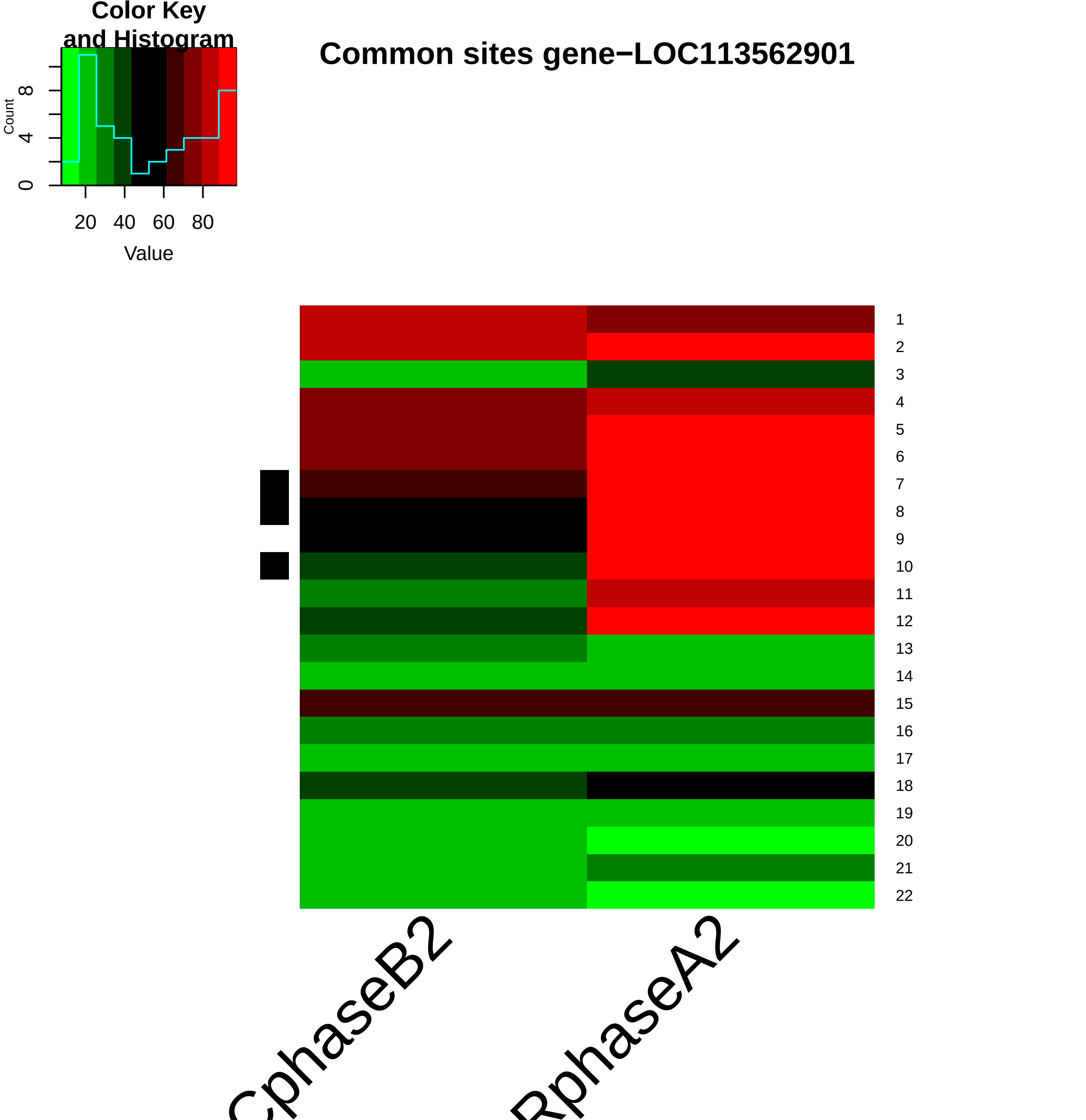

Color Key
and Histogram
Common sites gene−LOC113562901
Count
0
4
8
20
40
60
80
Value
CphaseB2
RphaseA2

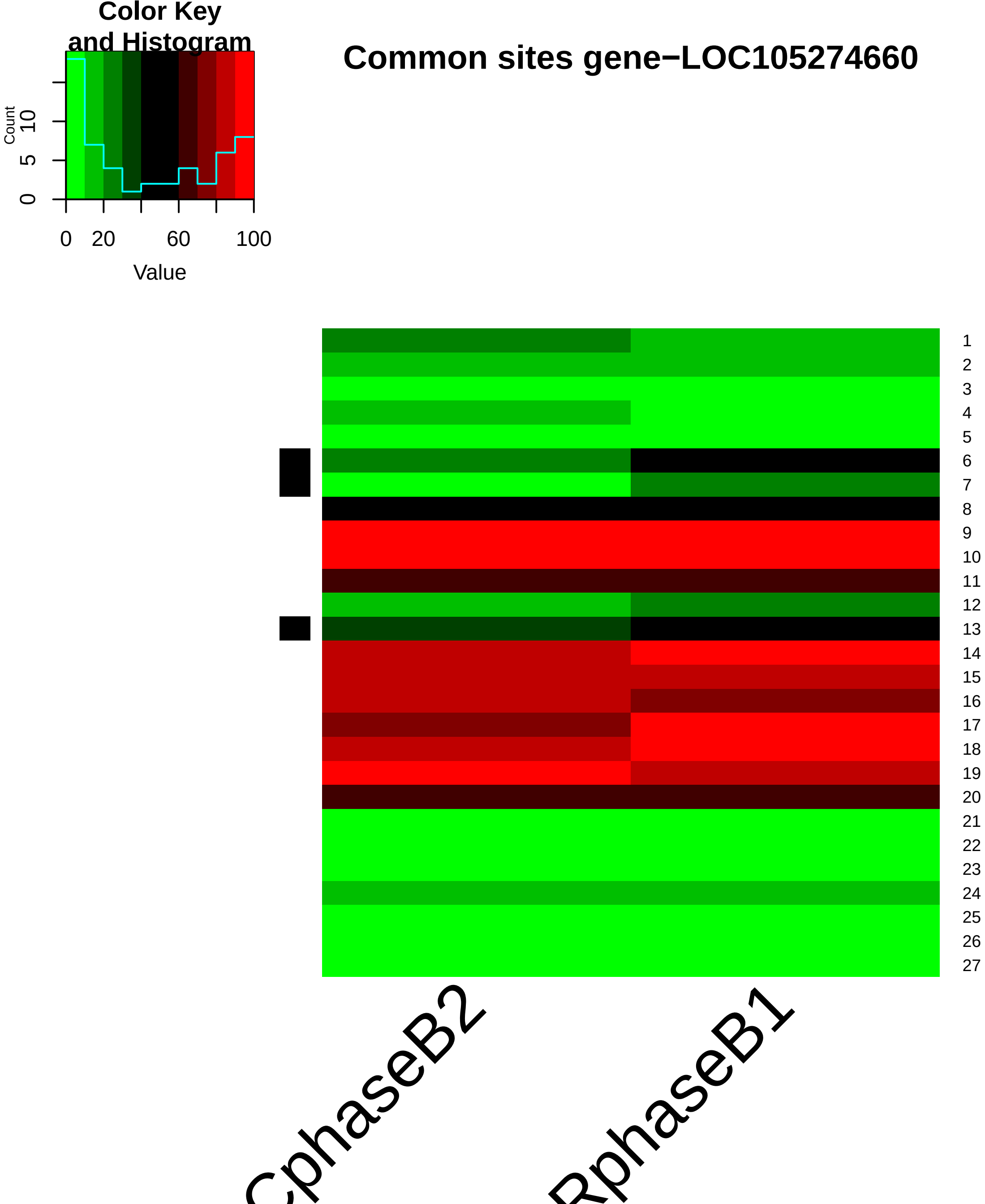

Color Key
and Histogram
Count
0
5
10
0
20
60
100
Value
Common sites gene−LOC105274660
CphaseB2
RphaseB1

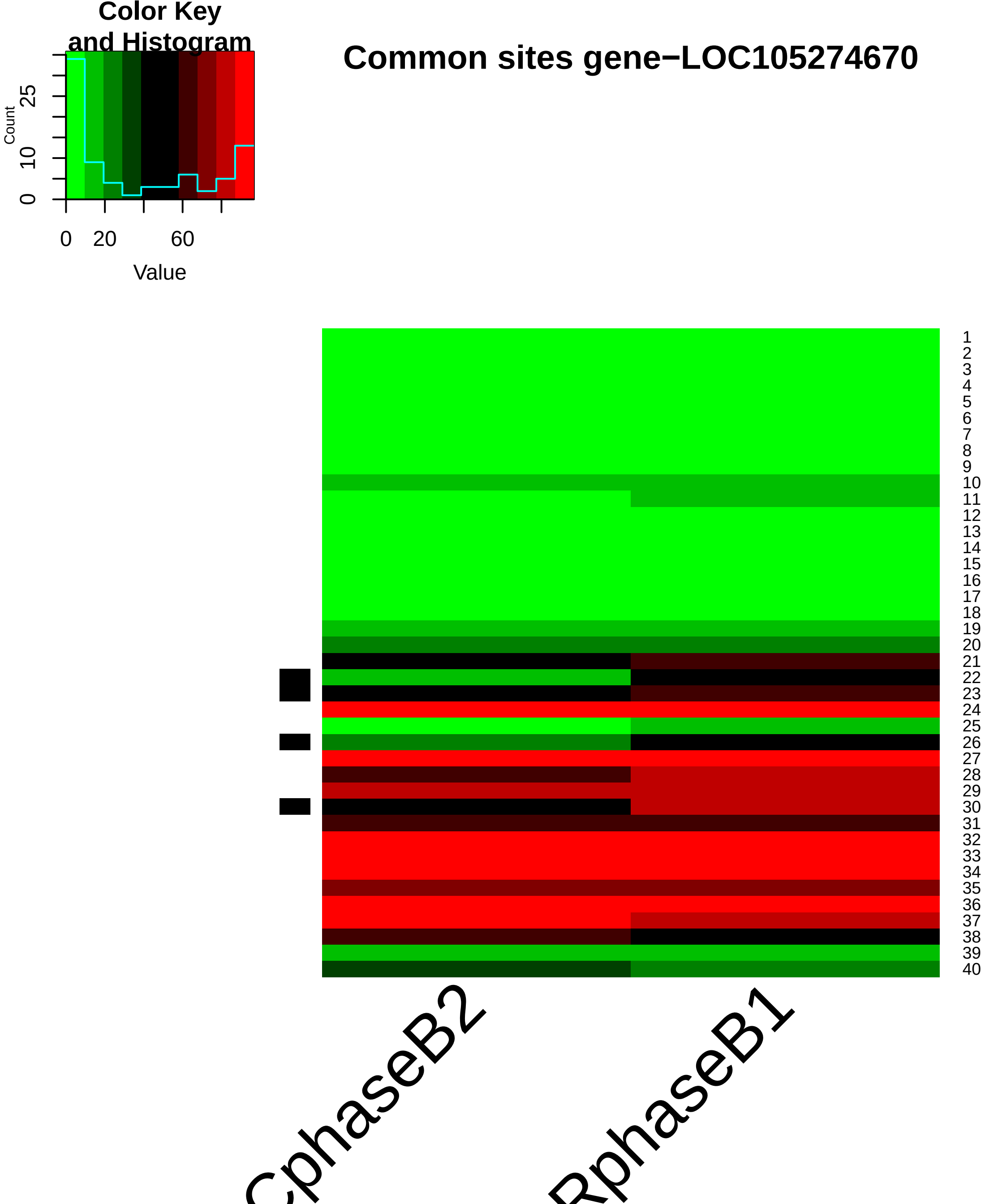

Color Key and Histogram
Count
0
10
25
0
20
60
Value
Common sites gene−LOC105274670
CphaseB2
RphaseB1

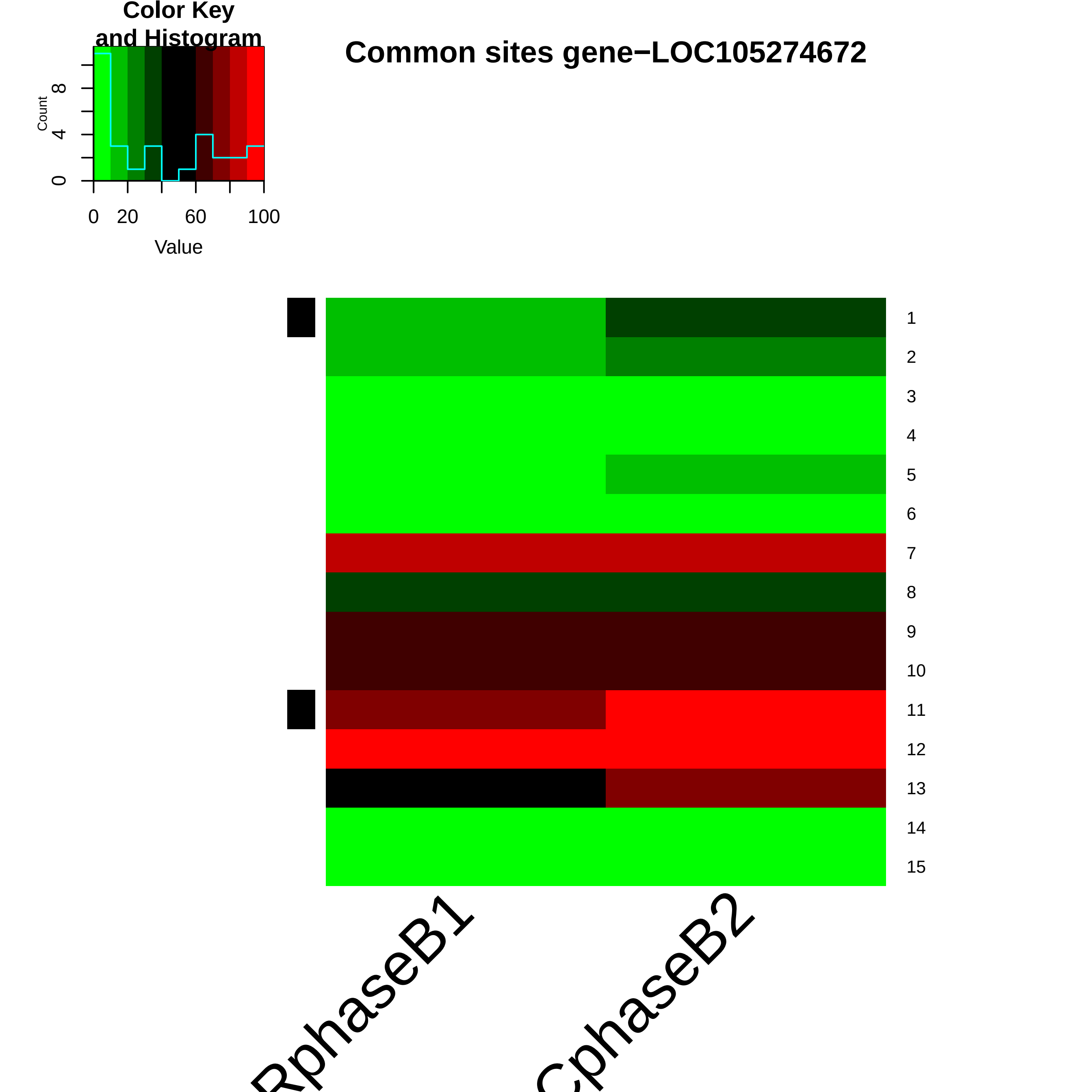
Color Key
and Histogram
Count
0
4
8
0
20
60
100
Value
Common sites gene−LOC105274672
RphaseB1
CphaseB2

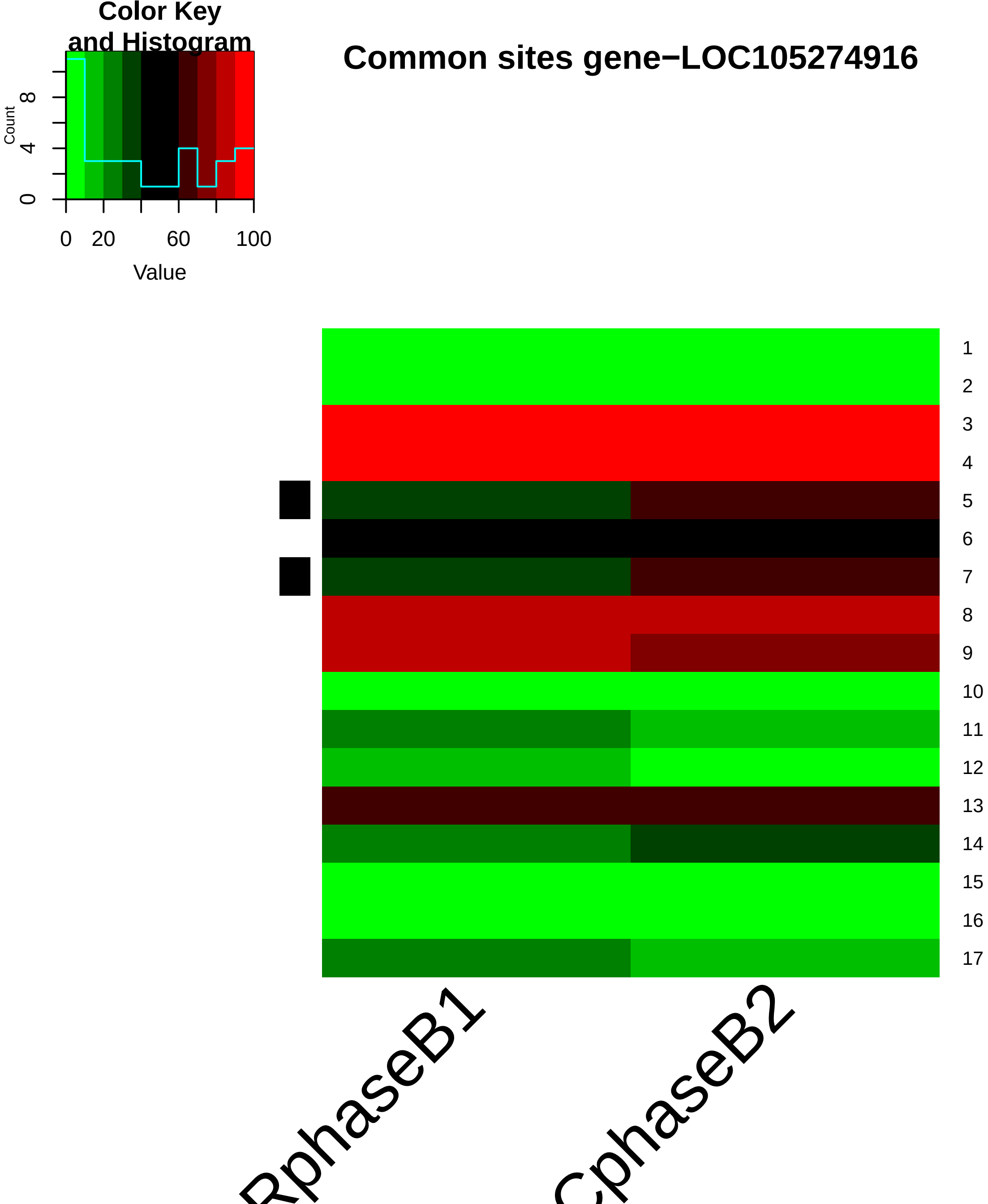
Color Key
and Histogram
Count
0
4
8
0
20
60
100
Value
Common sites gene−LOC105274916
RphaseB1
CphaseB2

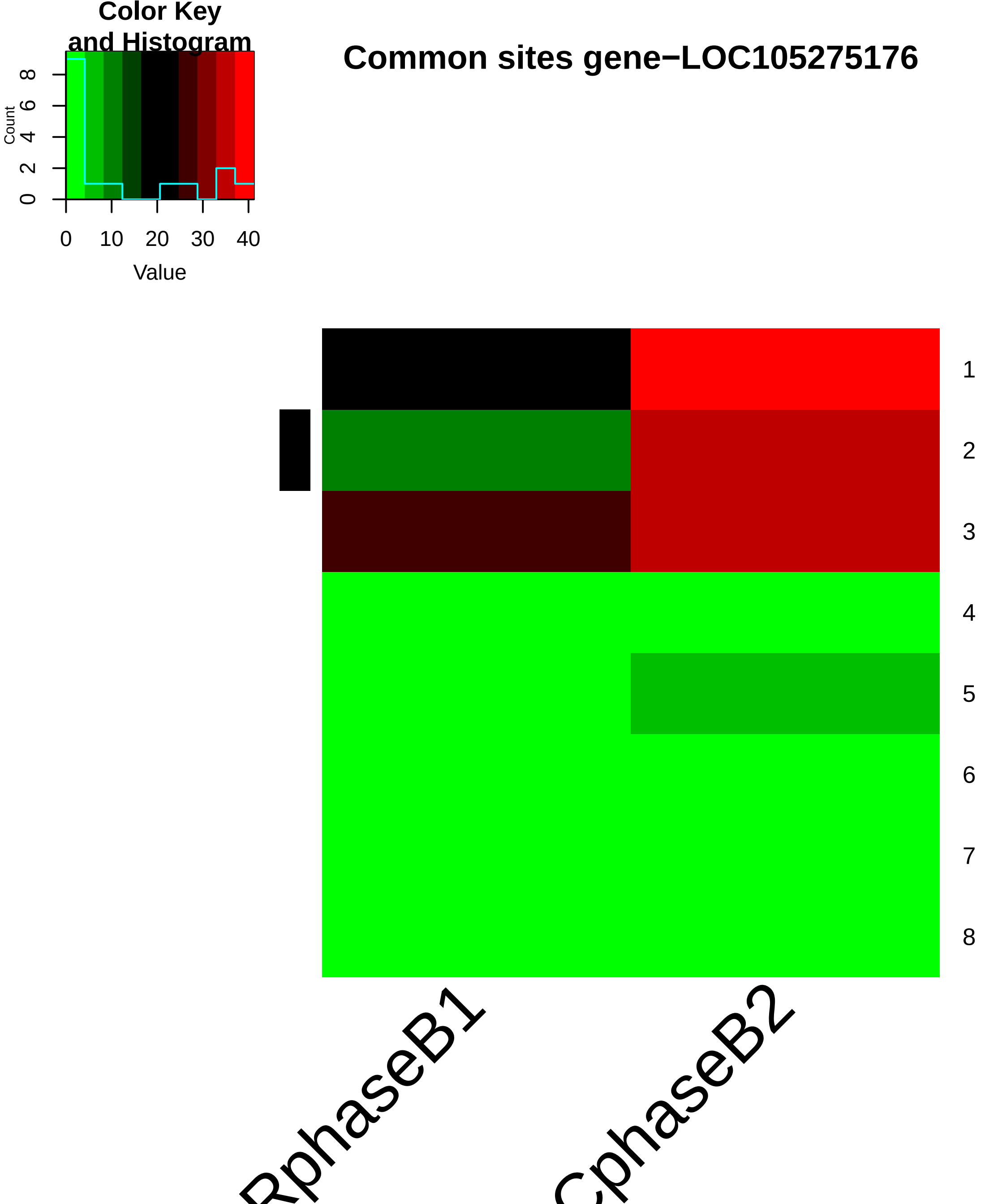
Color Key
and Histogram
Count
0 2 4 6 8
0 10 20 30 40
Value
Common sites gene−LOC105275176
1
2
3
4
5
6
7
8
RphaseB1
CphaseB2

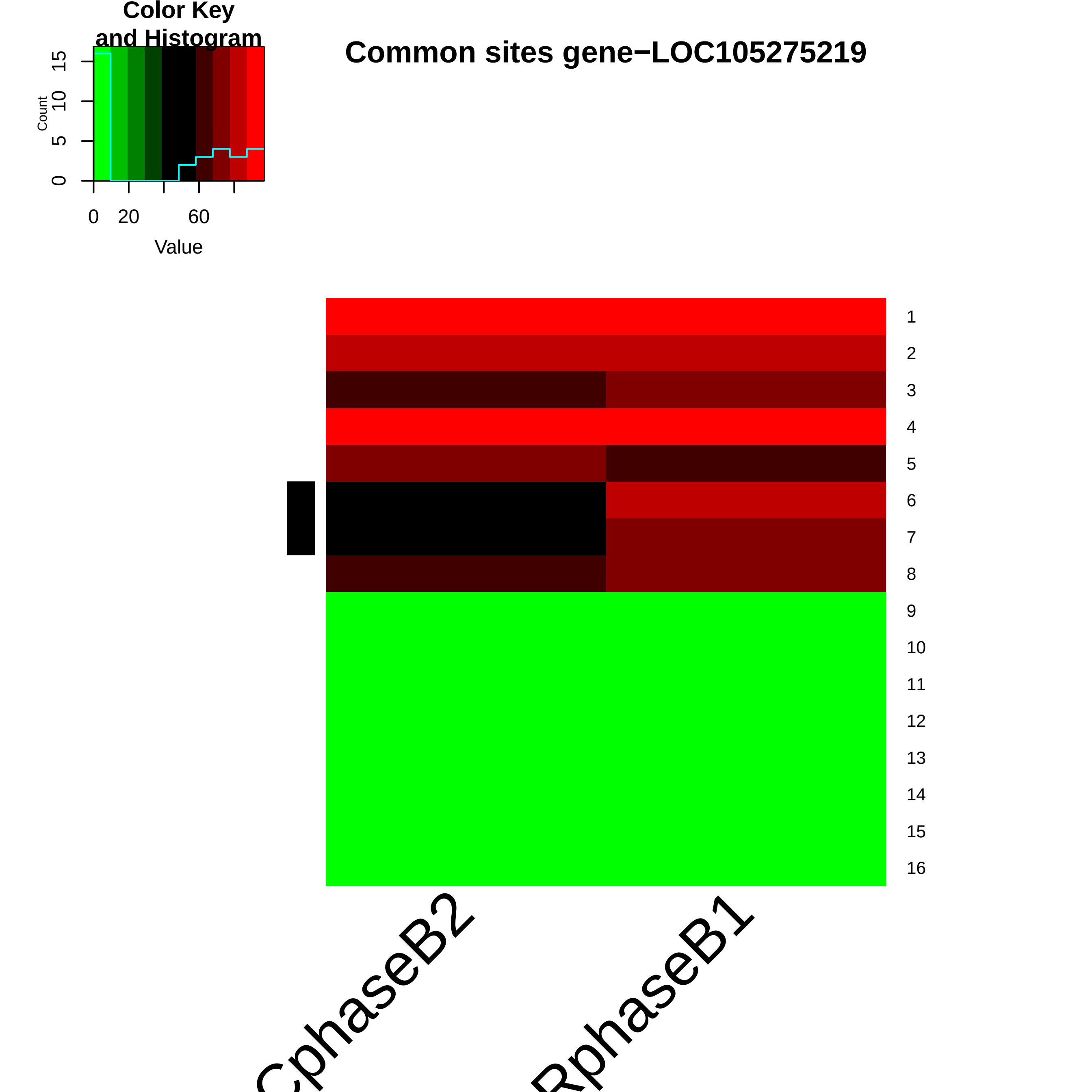

Color Key
and Histogram
Count
0
5
10
15
0
20
60
Value
Common sites gene−LOC105275219
CphaseB2
RphaseB1

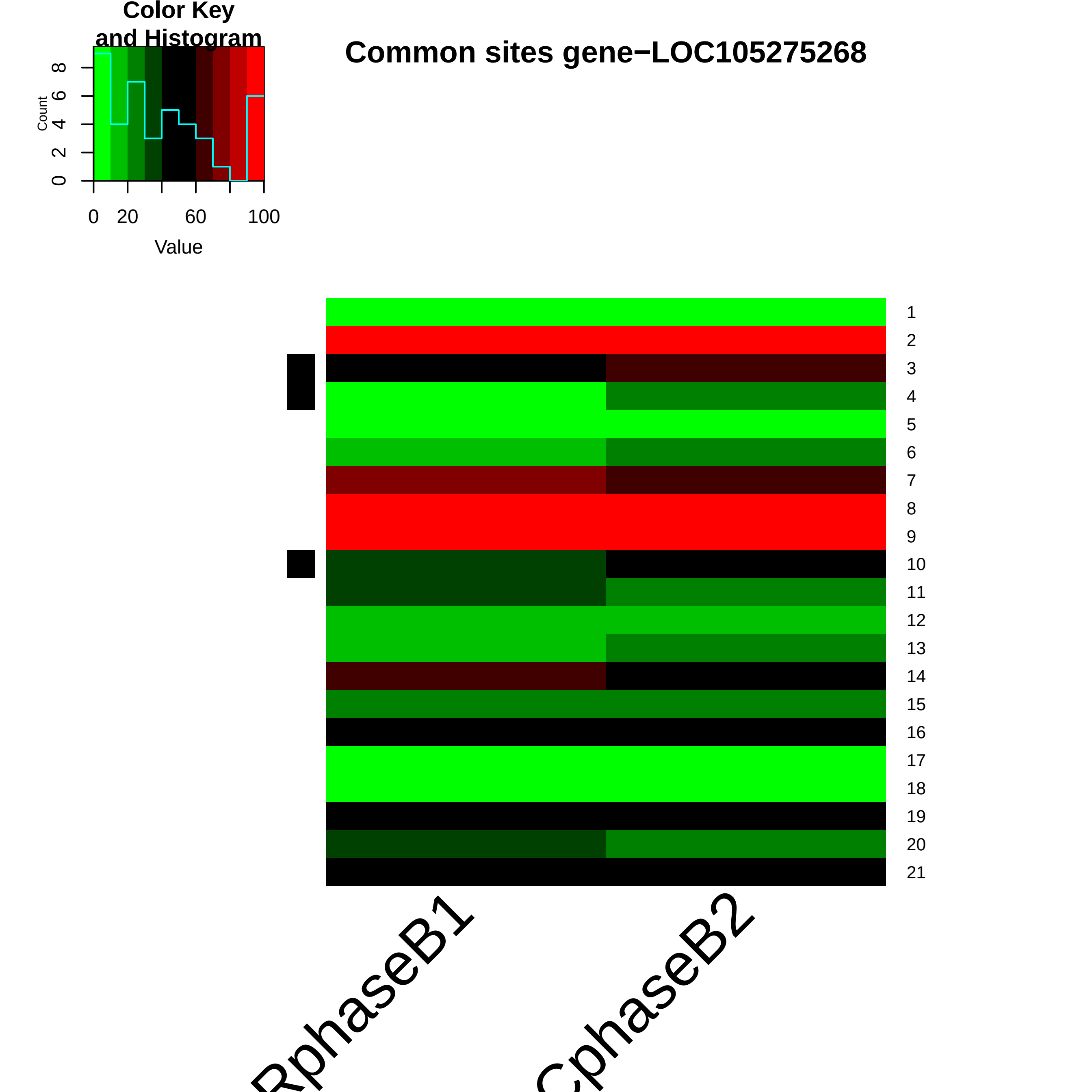

Color Key
and Histogram
Count
0
2
4
6
8
0
20
60
100
Value
Common sites gene−LOC105275268
RphaseB1
CphaseB2

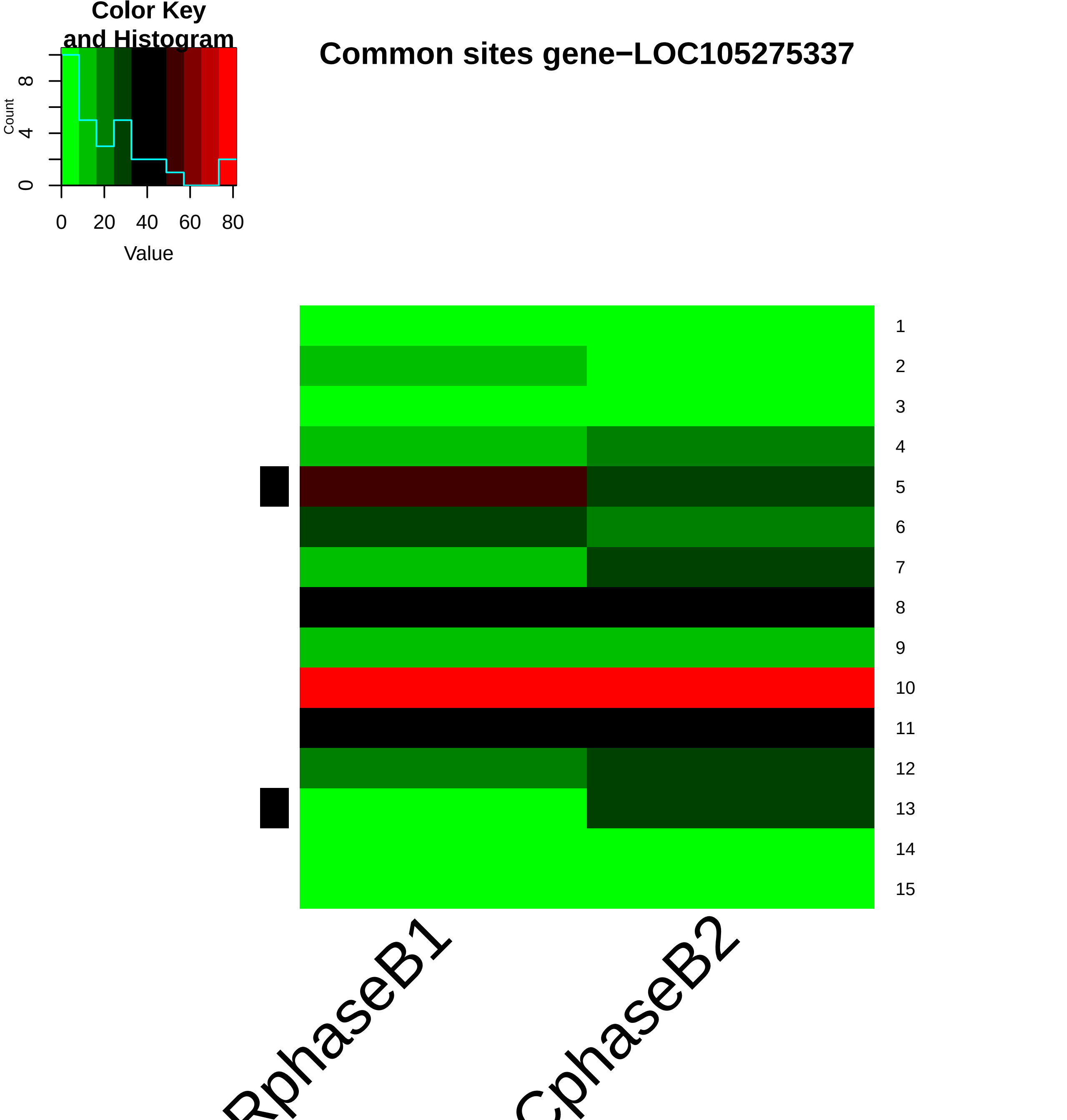
Color Key
and Histogram
Count
0
4
8
0
20
40
60
80
Value
Common sites gene−LOC105275337
RphaseB1
CphaseB2

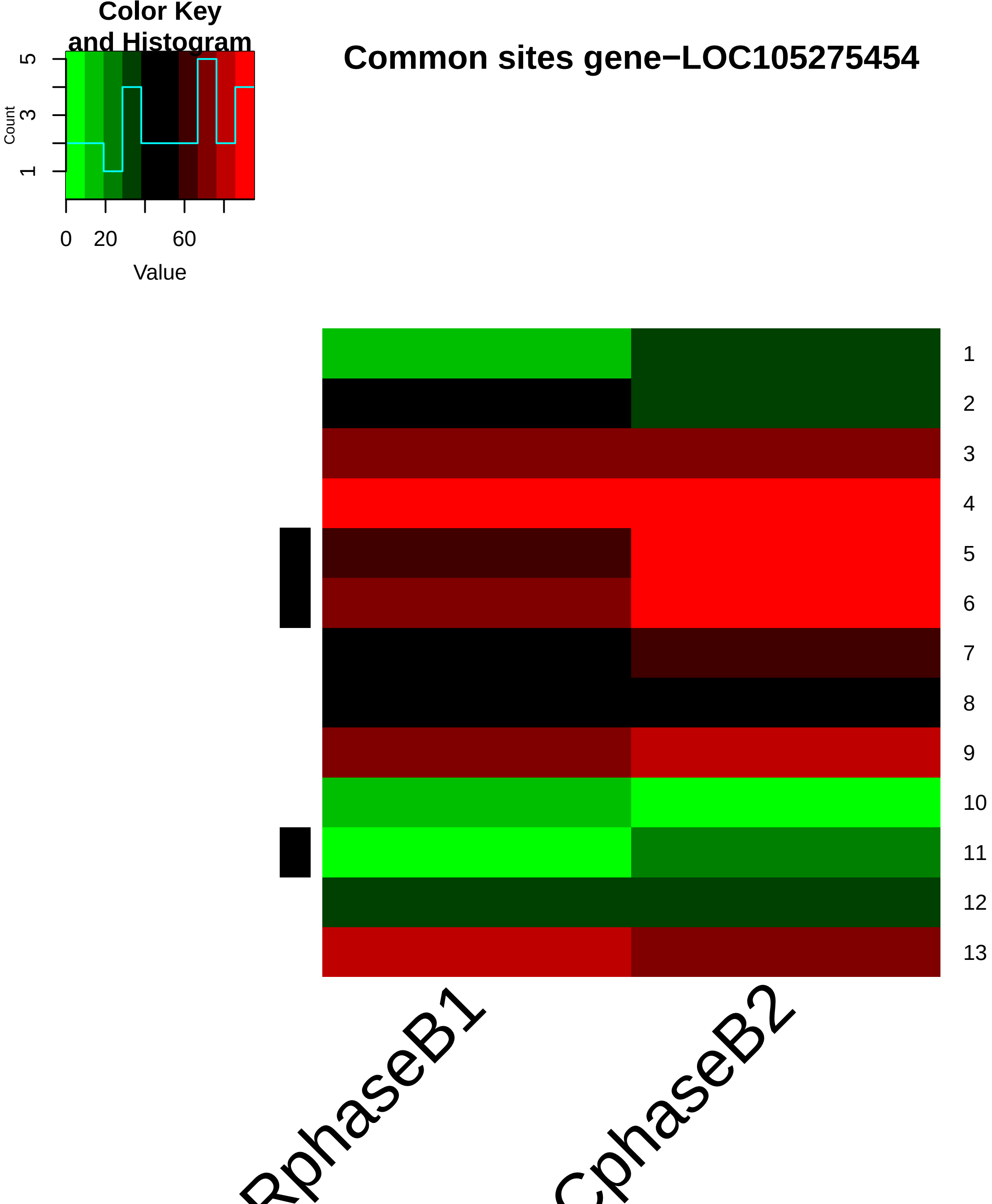

Color Key
and Histogram
Count
5
3
1
0
20
60
Value
Common sites gene−LOC105275454
1
2
3
4
5
6
7
8
9
10
11
12
13
RphaseB1
CphaseB2

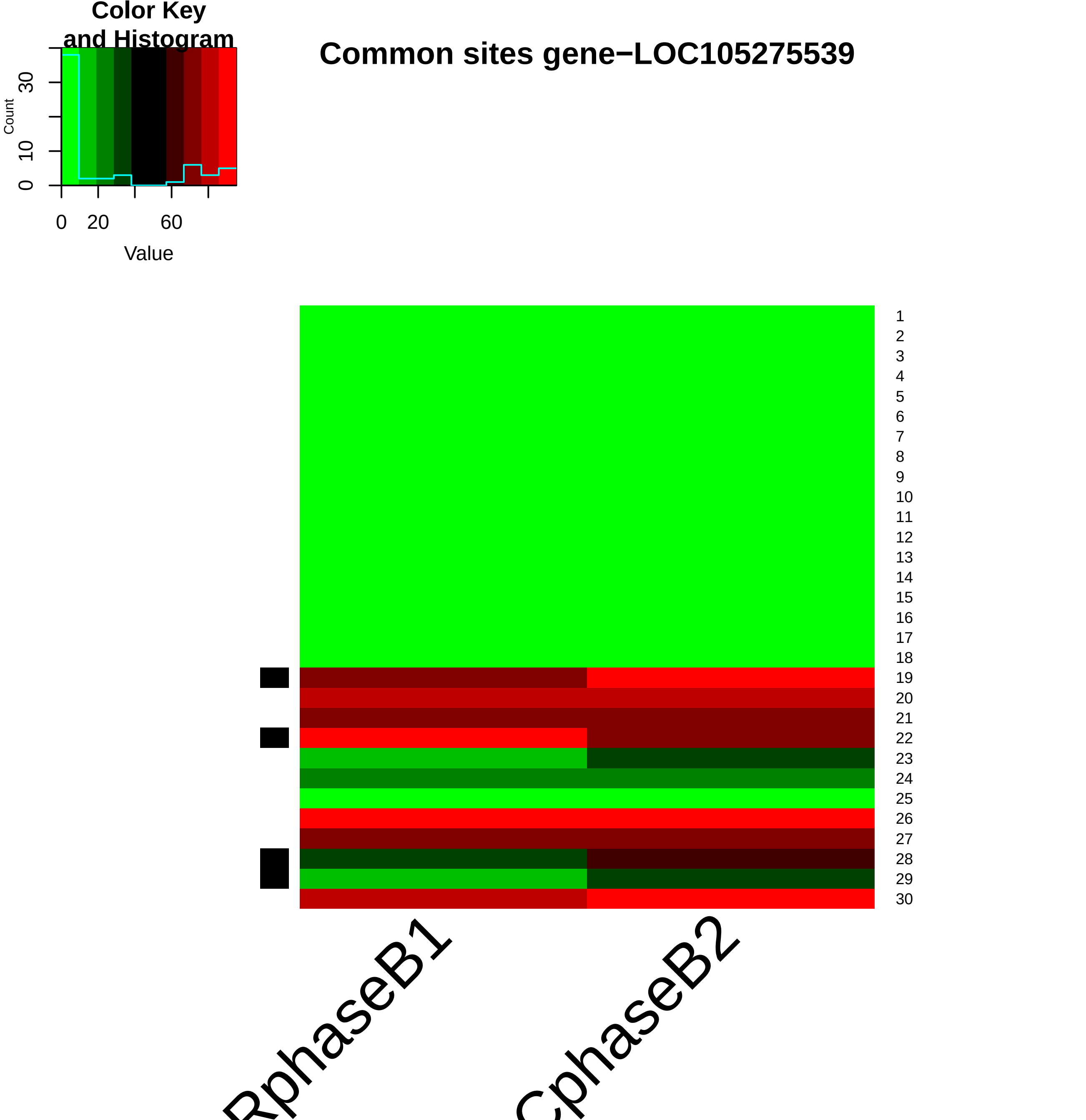

Color Key
and Histogram
Count
0
10
30
0
20
60
Value
Common sites gene−LOC105275539
RphaseB1
CphaseB2

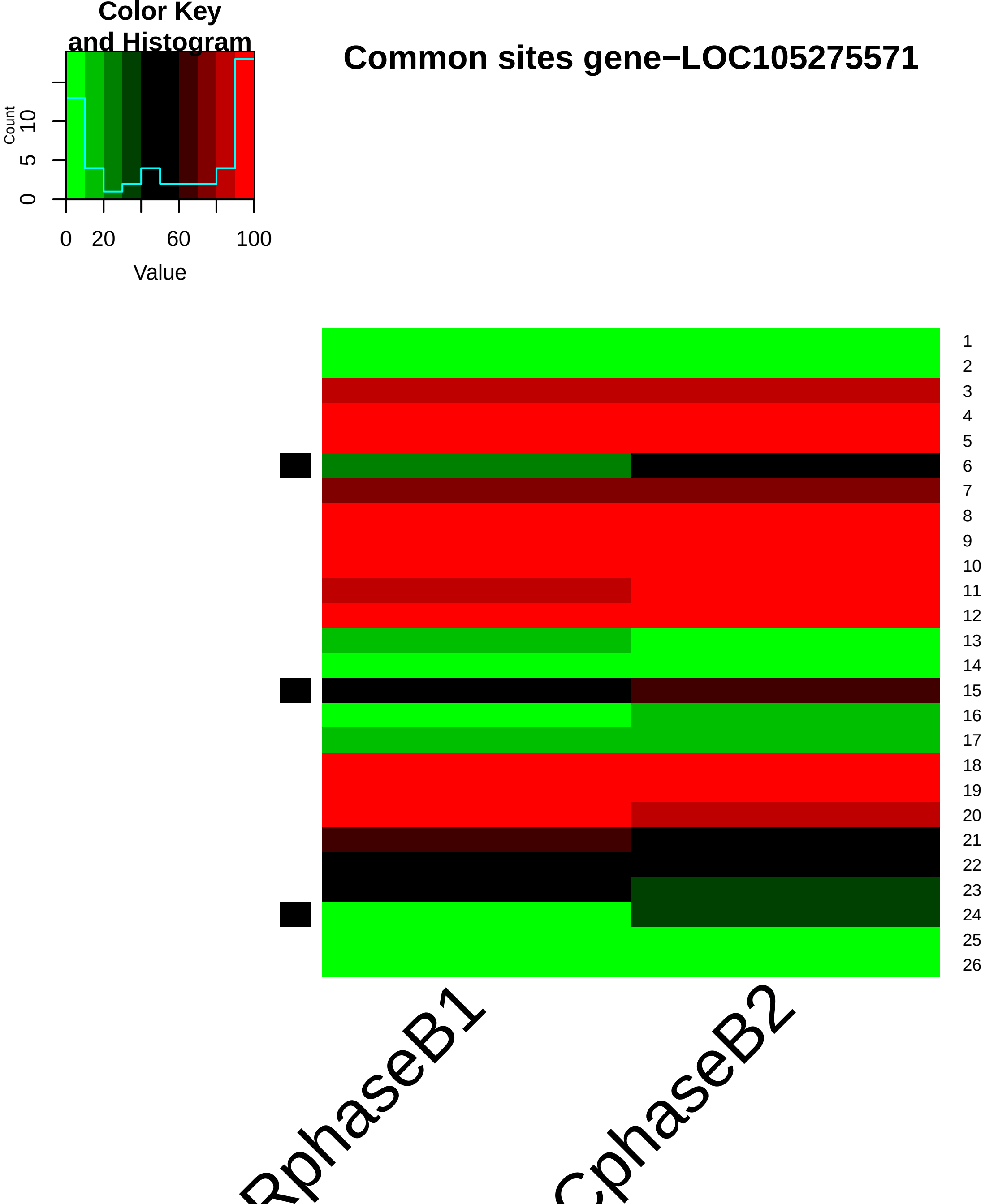

Color Key
and Histogram
Count
0
5
10
0
20
60
100
Value
Common sites gene−LOC105275571
RphaseB1
CphaseB2

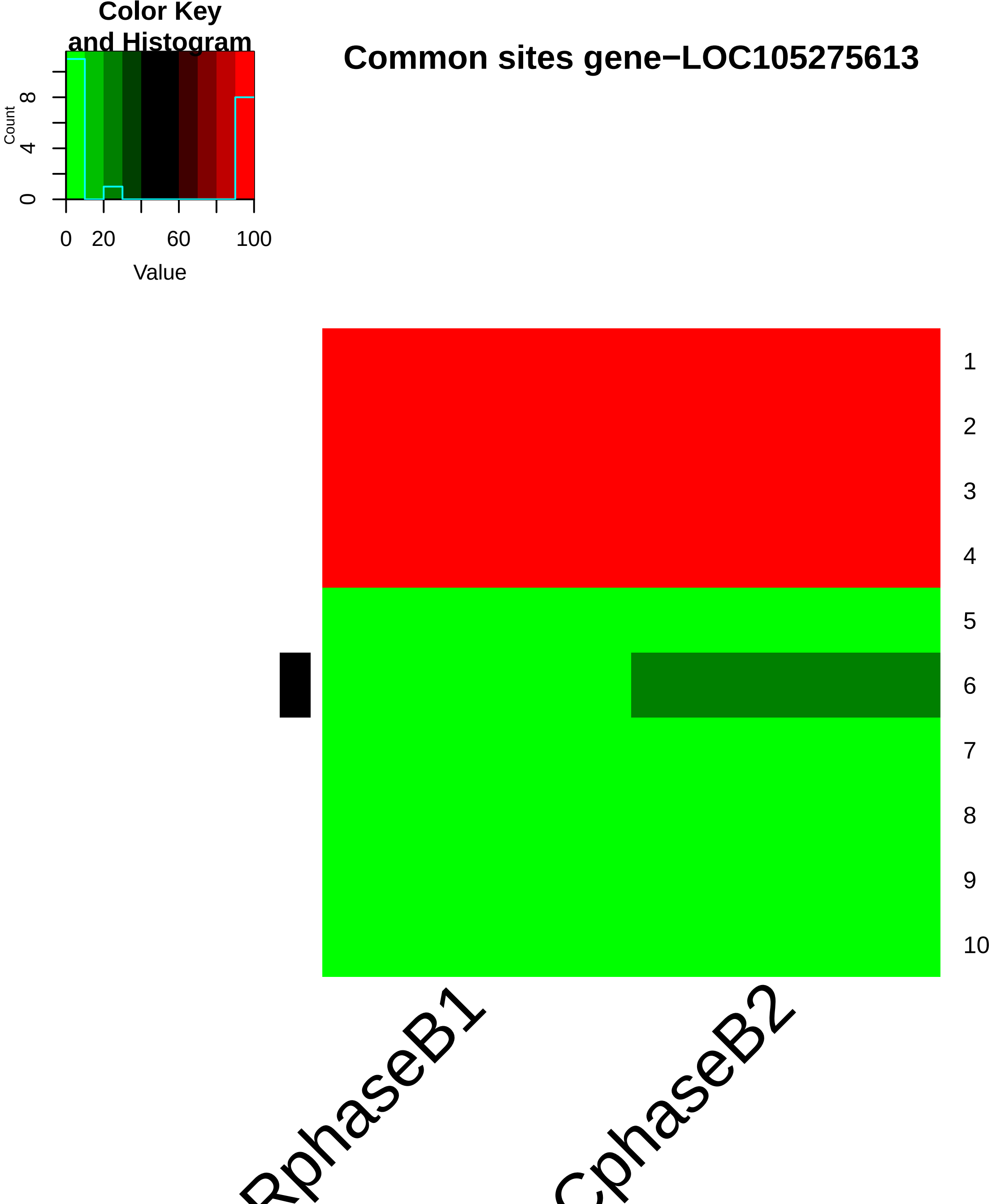

Color Key
and Histogram
Count
0
4
8
0
20
60
100
Value
Common sites gene−LOC105275613
1
2
3
4
5
6
7
8
9
10
RphaseB1
CphaseB2

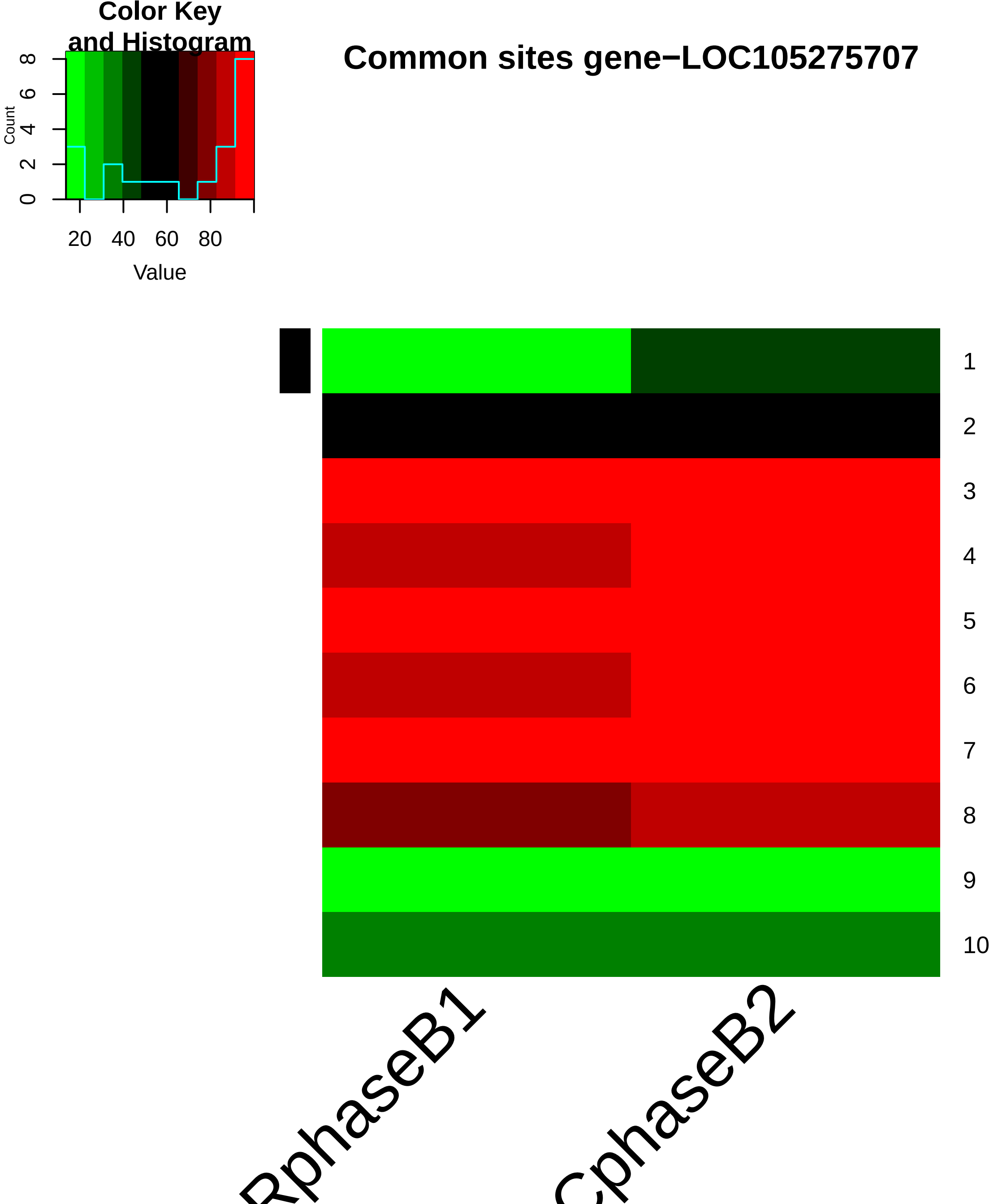

Color Key
and Histogram
Count
0
2
4
6
8
20
40
60
80
Value
Common sites gene−LOC105275707
1
2
3
4
5
6
7
8
9
10
RphaseB1
CphaseB2

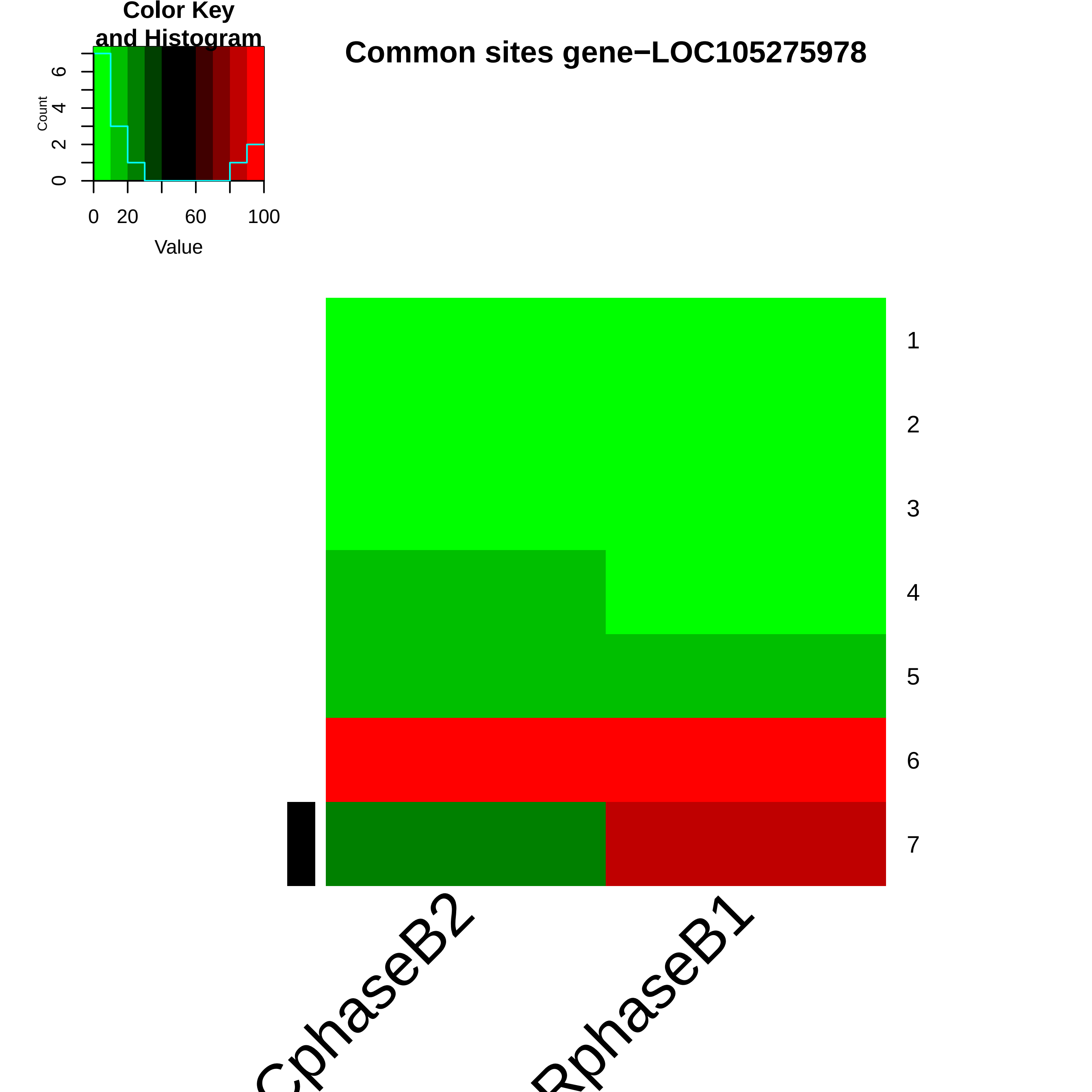
Color Key
and Histogram
Count
0
2
4
6
0
20
60
100
Value
Common sites gene−LOC105275978
1
2
3
4
5
6
7
CphaseB2
RphaseB1

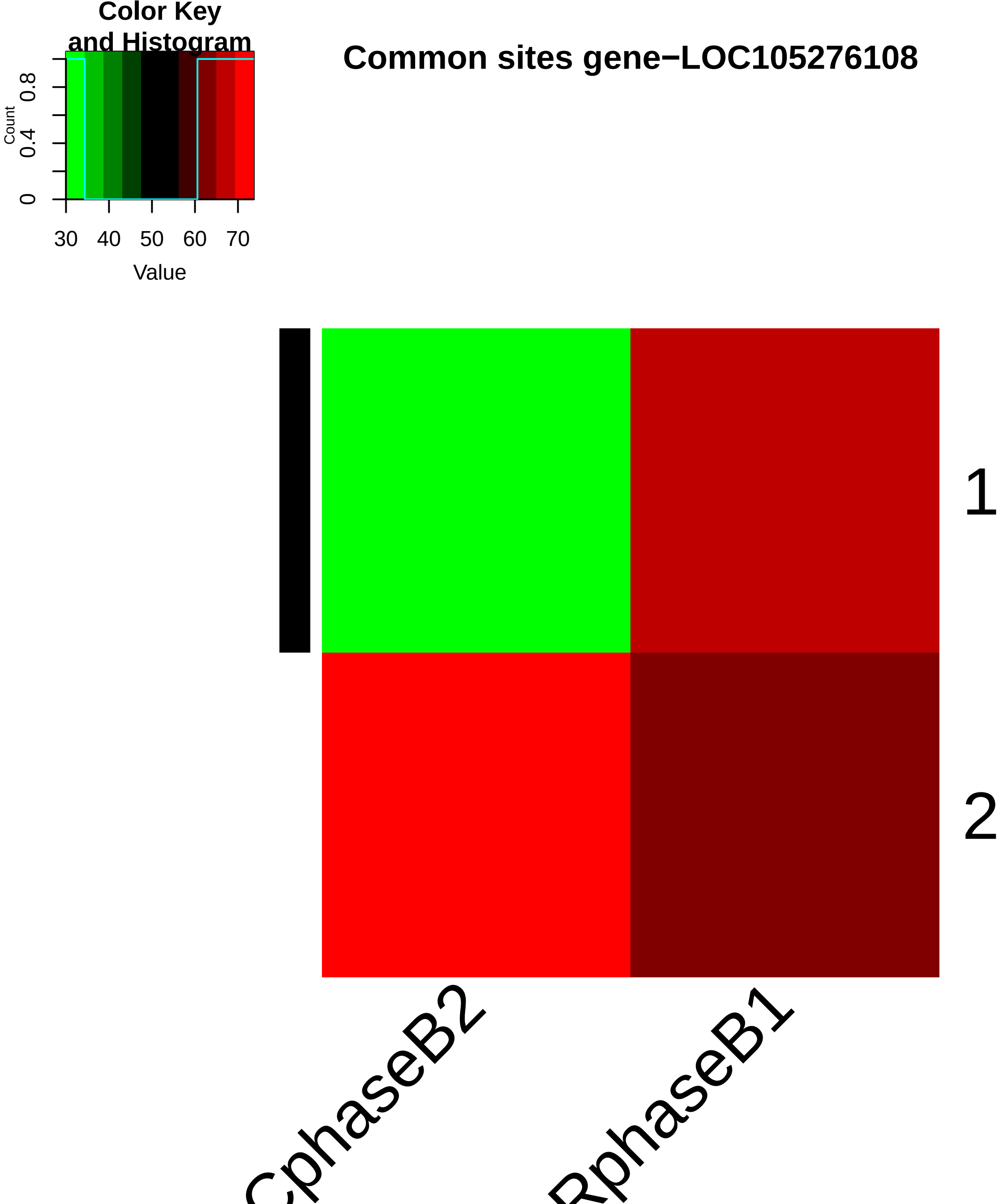

Color Key
and Histogram
Count
0
0.4
0.8
30
40
50
60
70
Value
Common sites gene−LOC105276108
1
2
CphaseB2
RphaseB1

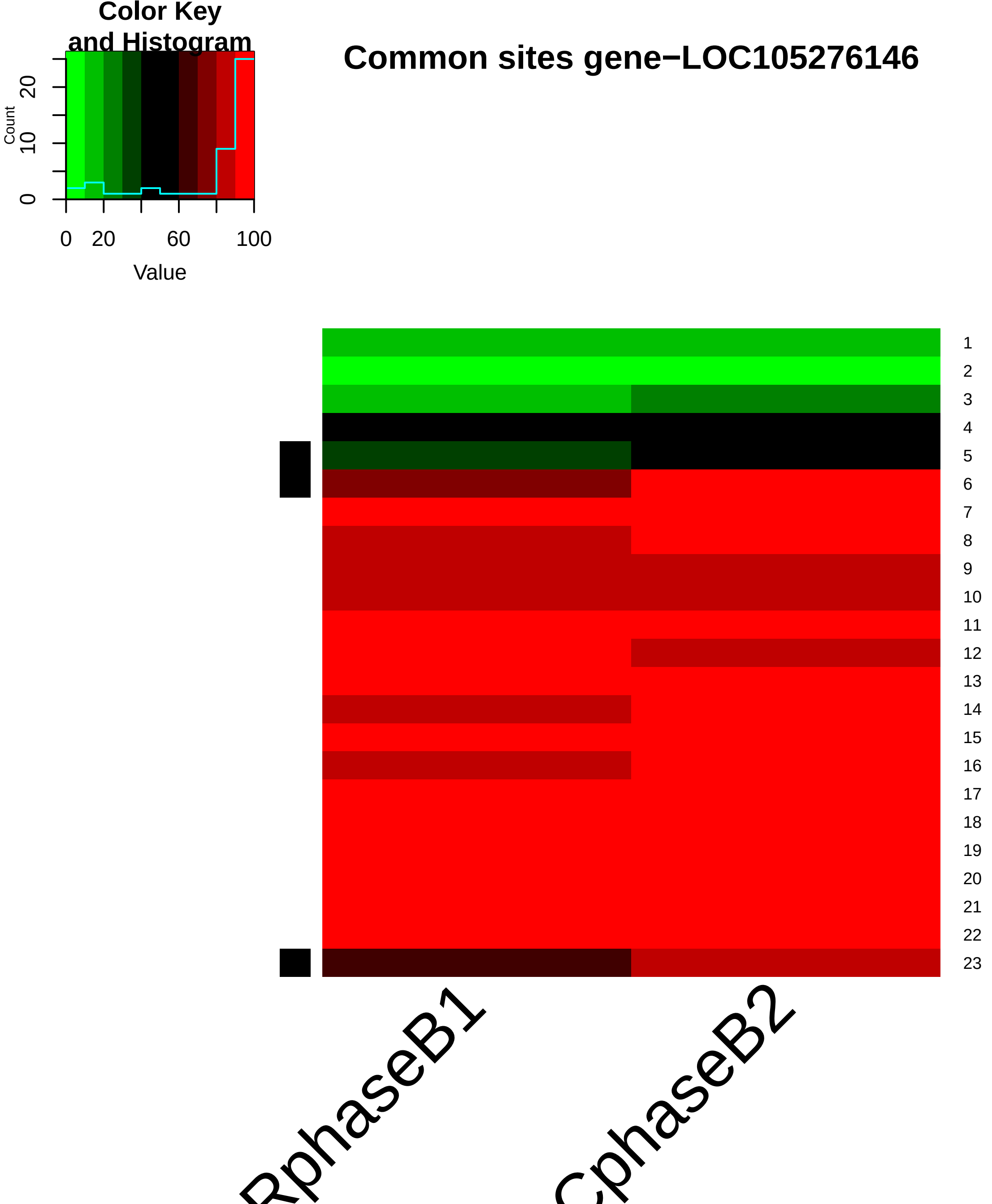
Color Key
and Histogram
Count
0
10
20
0
20
60
100
Value
Common sites gene−LOC105276146
RphaseB1
CphaseB2

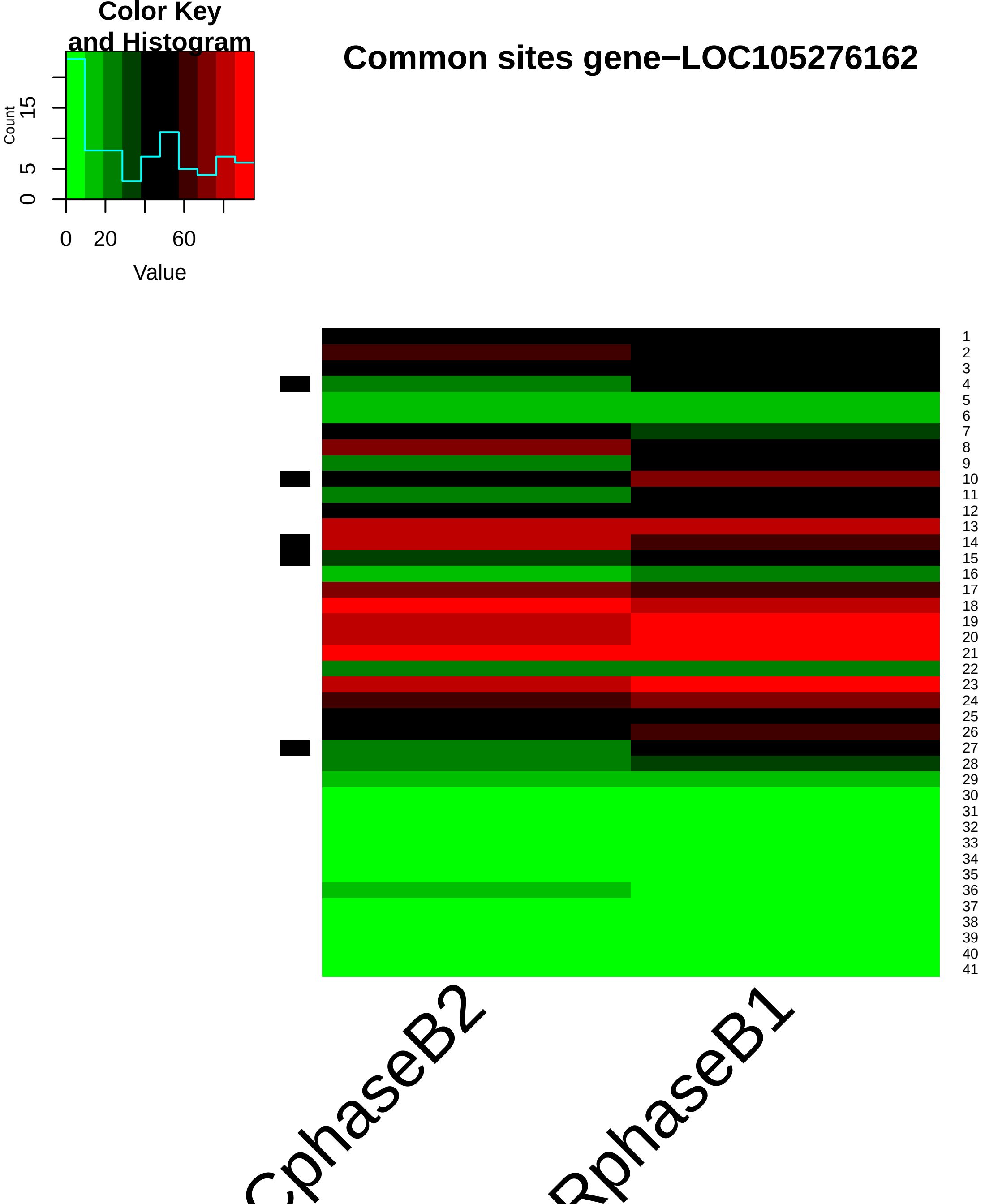
Color Key
and Histogram
Count
0
5
15
0
20
60
Value
Common sites gene−LOC105276162
CphaseB2
RphaseB1

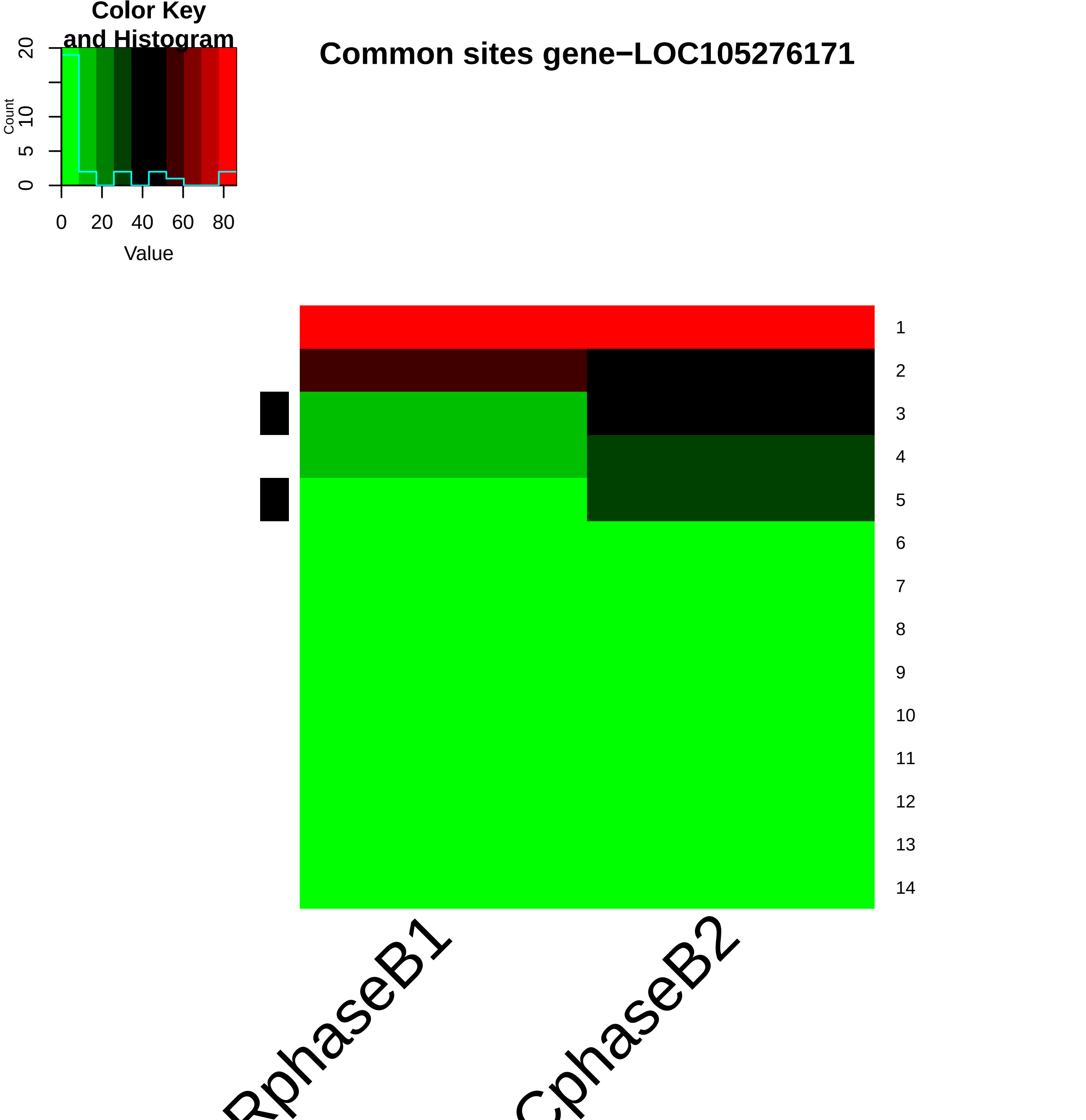
Color Key
and Histogram
Count
Value
0
5
10
20
0
20
40
60
80
Common sites gene−LOC105276171
1
2
3
4
5
6
7
8
9
10
11
12
13
14
RphaseB1
CphaseB2

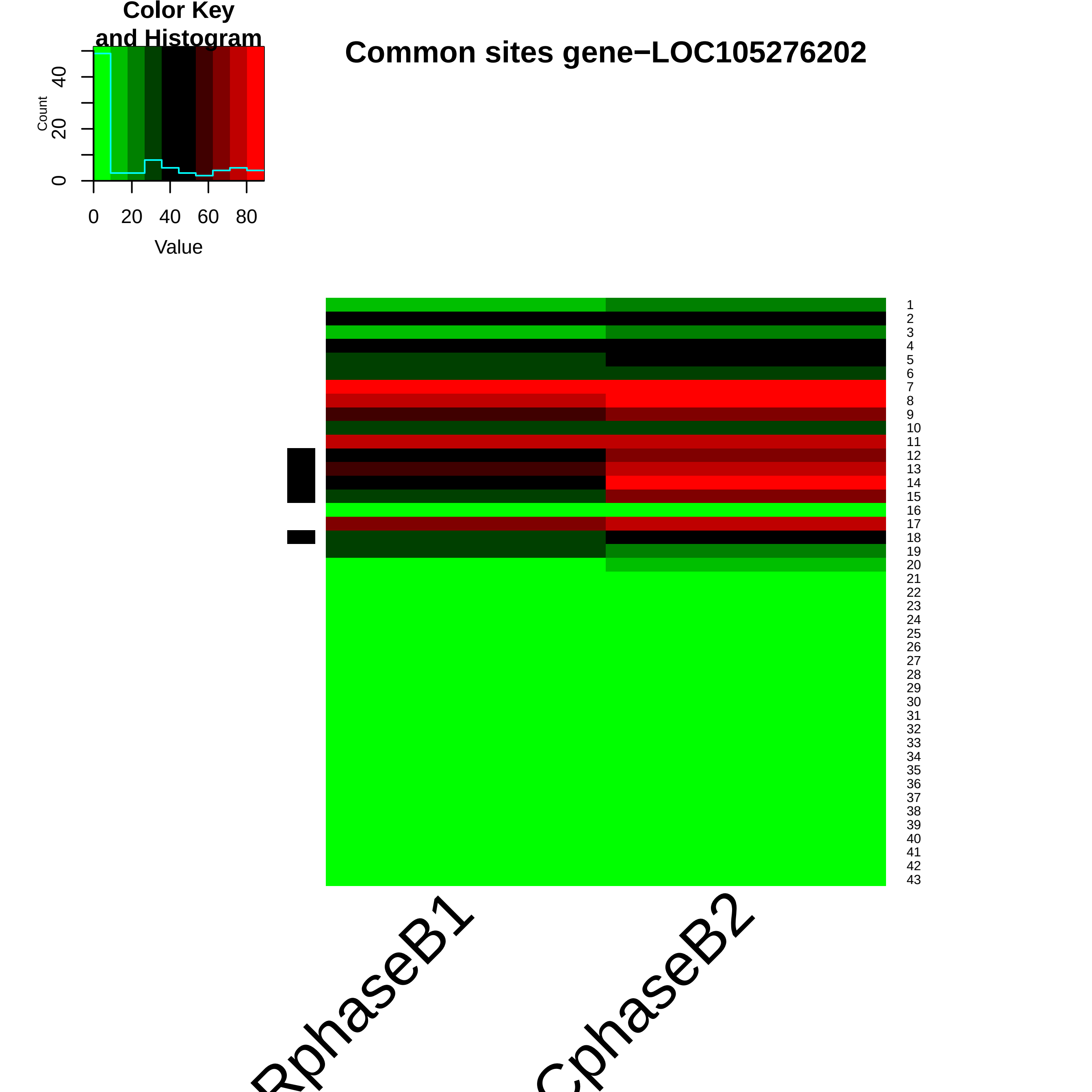
Color Key
and Histogram
Count
0
20
40
0
20
40
60
80
Value
Common sites gene−LOC105276202
RphaseB1
CphaseB2

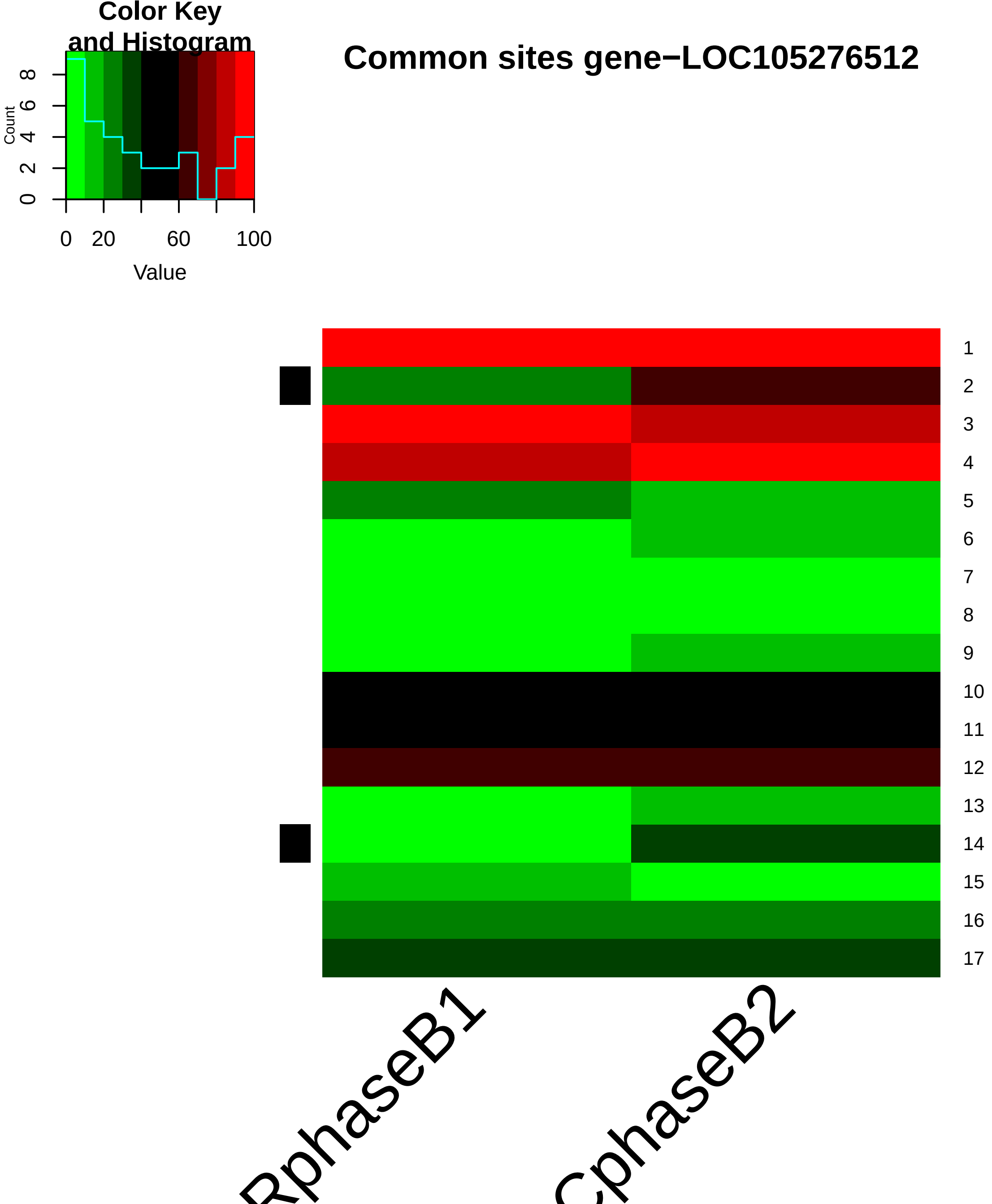

Color Key
and Histogram
Count
0
2
4
6
8
0
20
60
100
Value
Common sites gene−LOC105276512
RphaseB1
CphaseB2

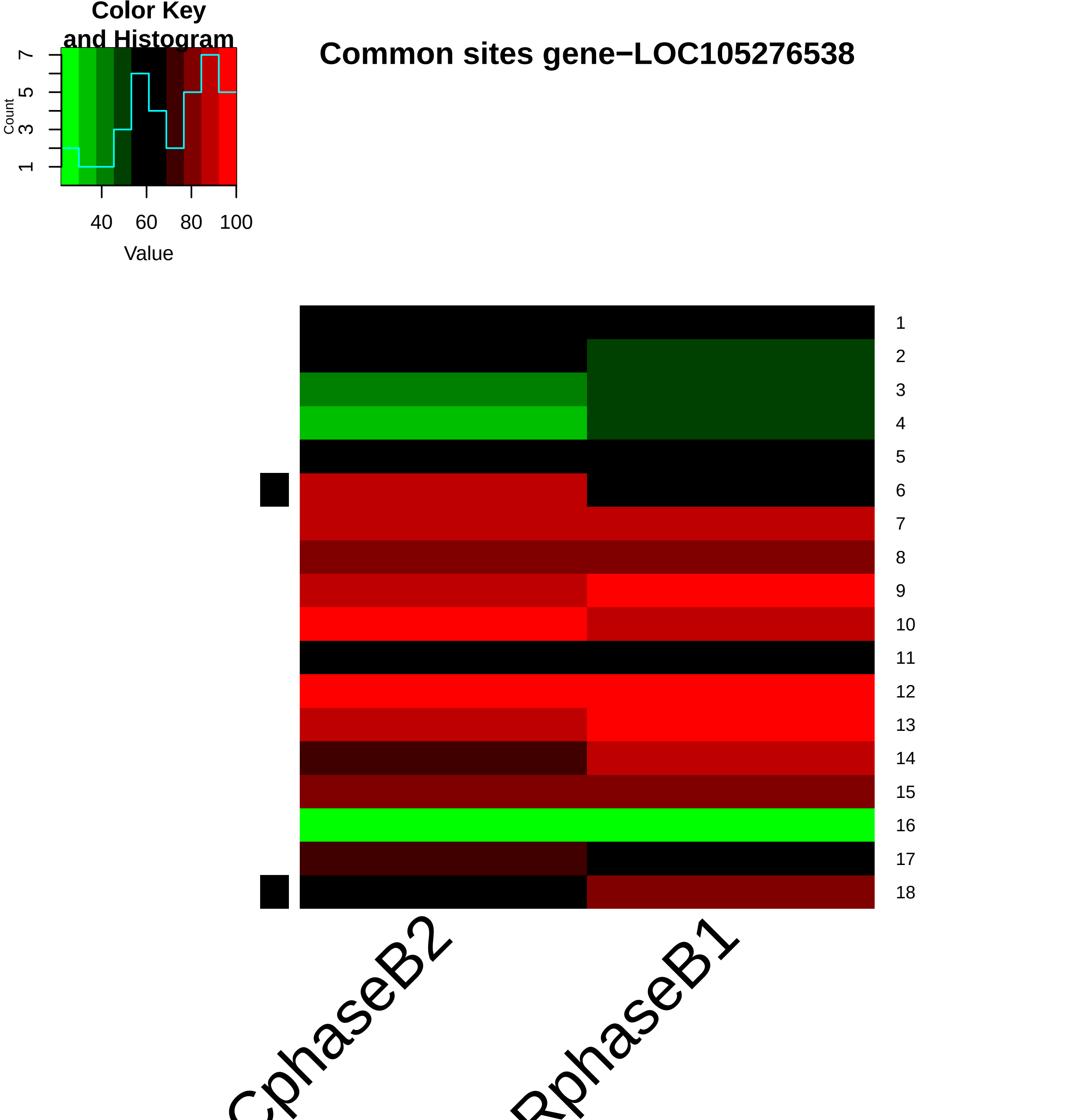

Color Key
and Histogram
Count
1
3
5
7
40
60
80
100
Value
Common sites gene−LOC105276538
CphaseB2
RphaseB1

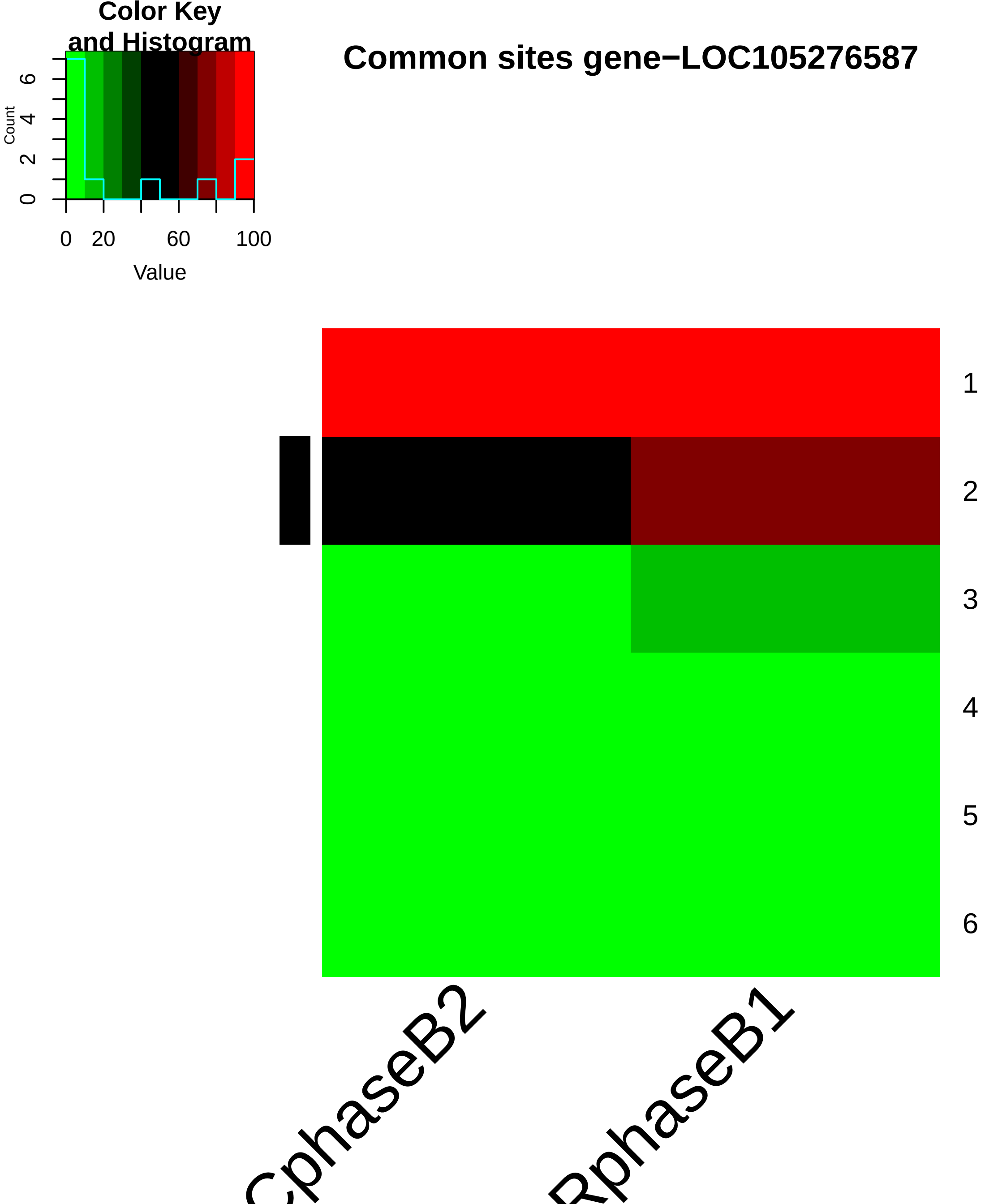

Color Key
and Histogram
Count
0
2
4
6
0
20
60
100
Value
Common sites gene−LOC105276587
1
2
3
4
5
6
CphaseB2
RphaseB1

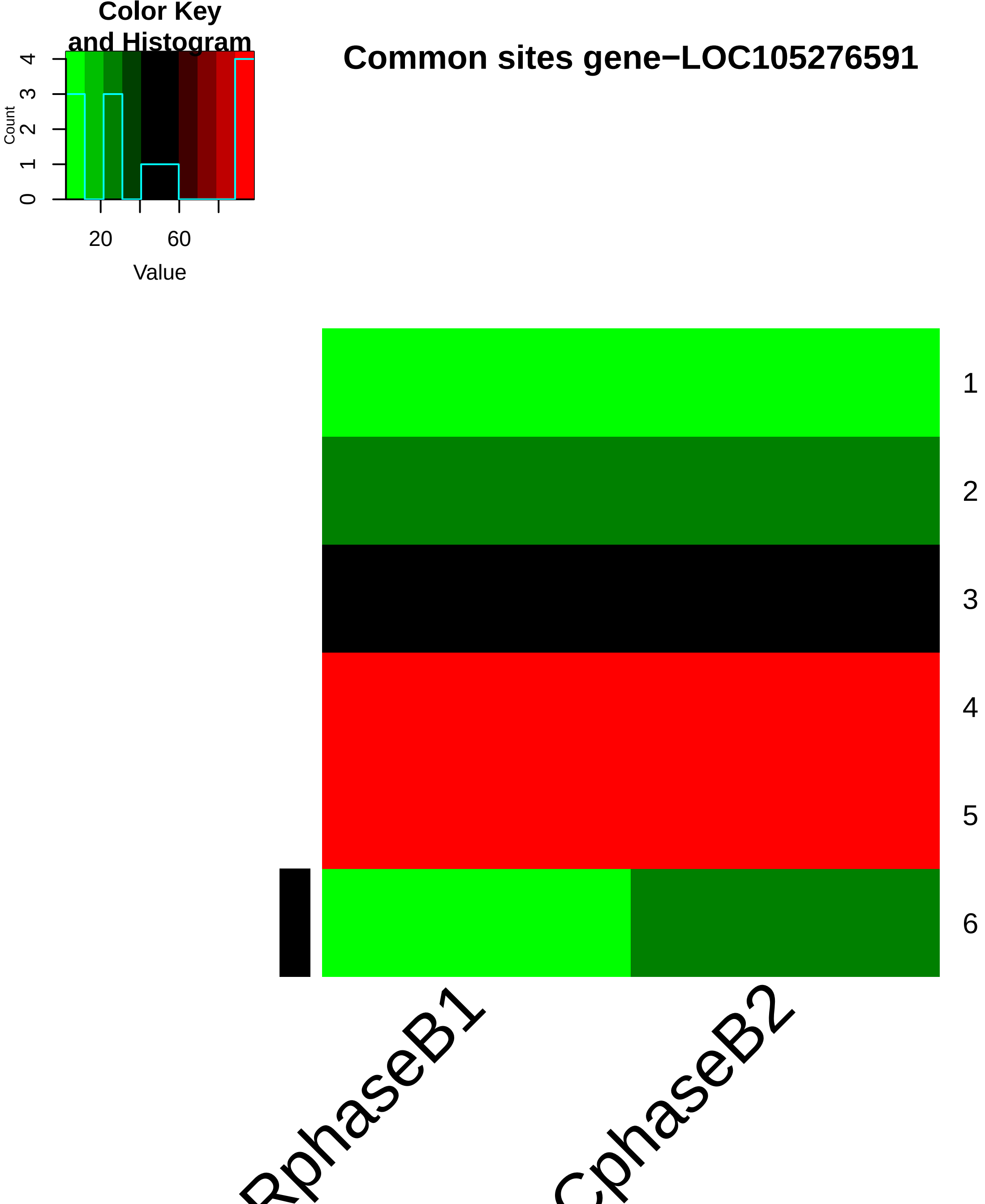

Color Key
and Histogram
Count
0
1
2
3
4
20
60
Value
Common sites gene−LOC105276591
1
2
3
4
5
6
RphaseB1
CphaseB2

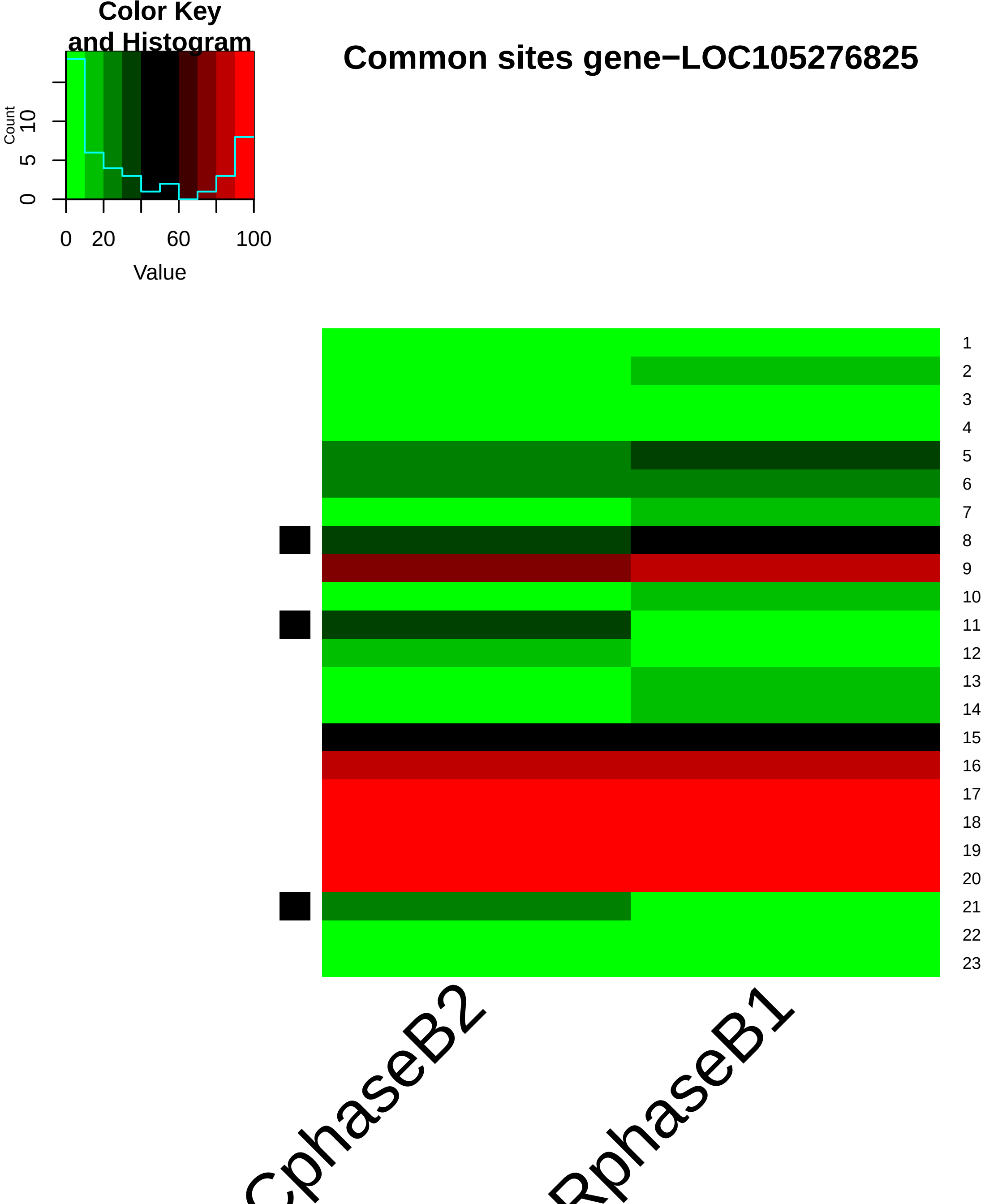
Color Key
and Histogram
Count
0
5
10
0
20
60
100
Value
Common sites gene−LOC105276825
CphaseB2
RphaseB1

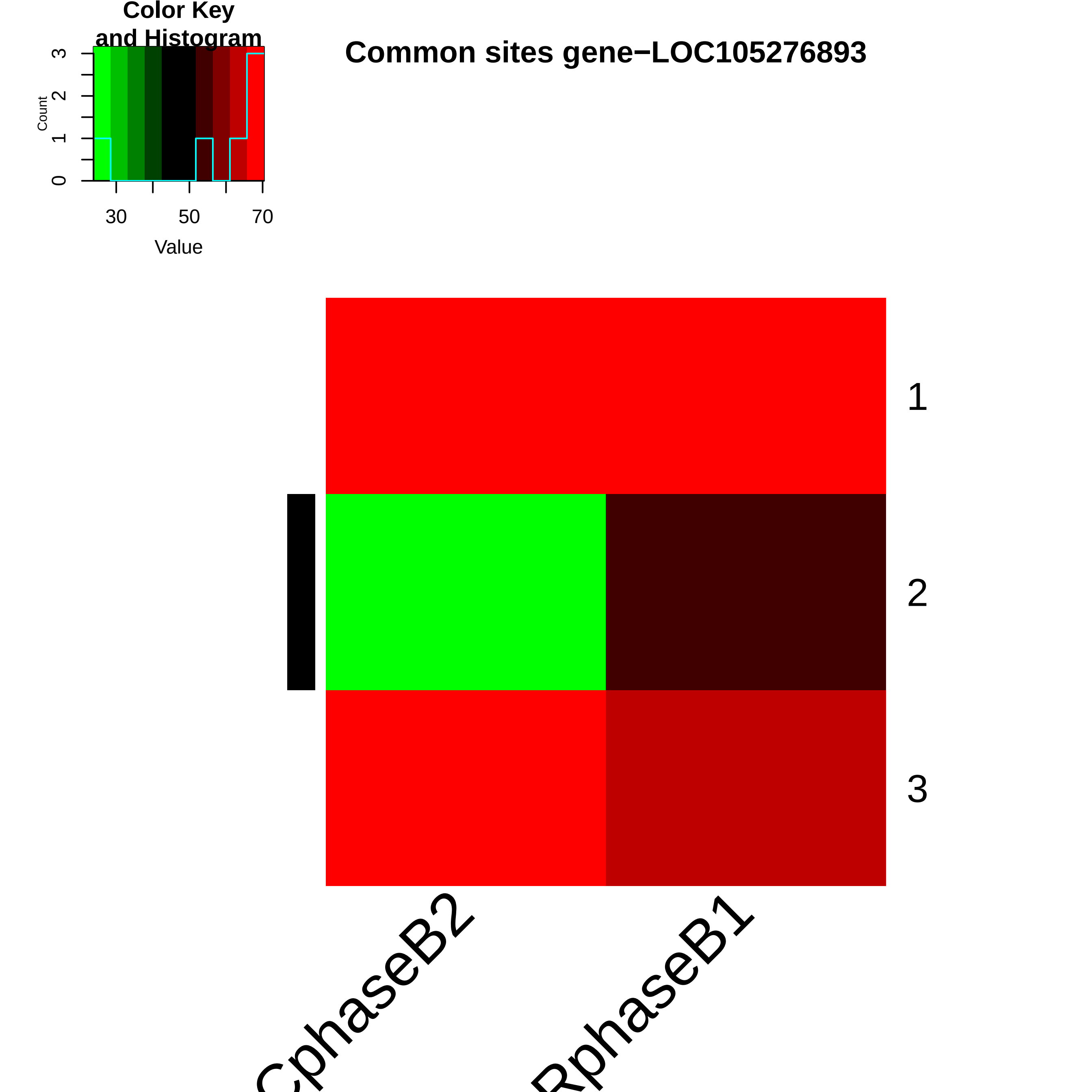
Color Key
and Histogram
Count
0
1
2
3
30
50
70
Value
Common sites gene−LOC105276893
1
2
3
CphaseB2
RphaseB1

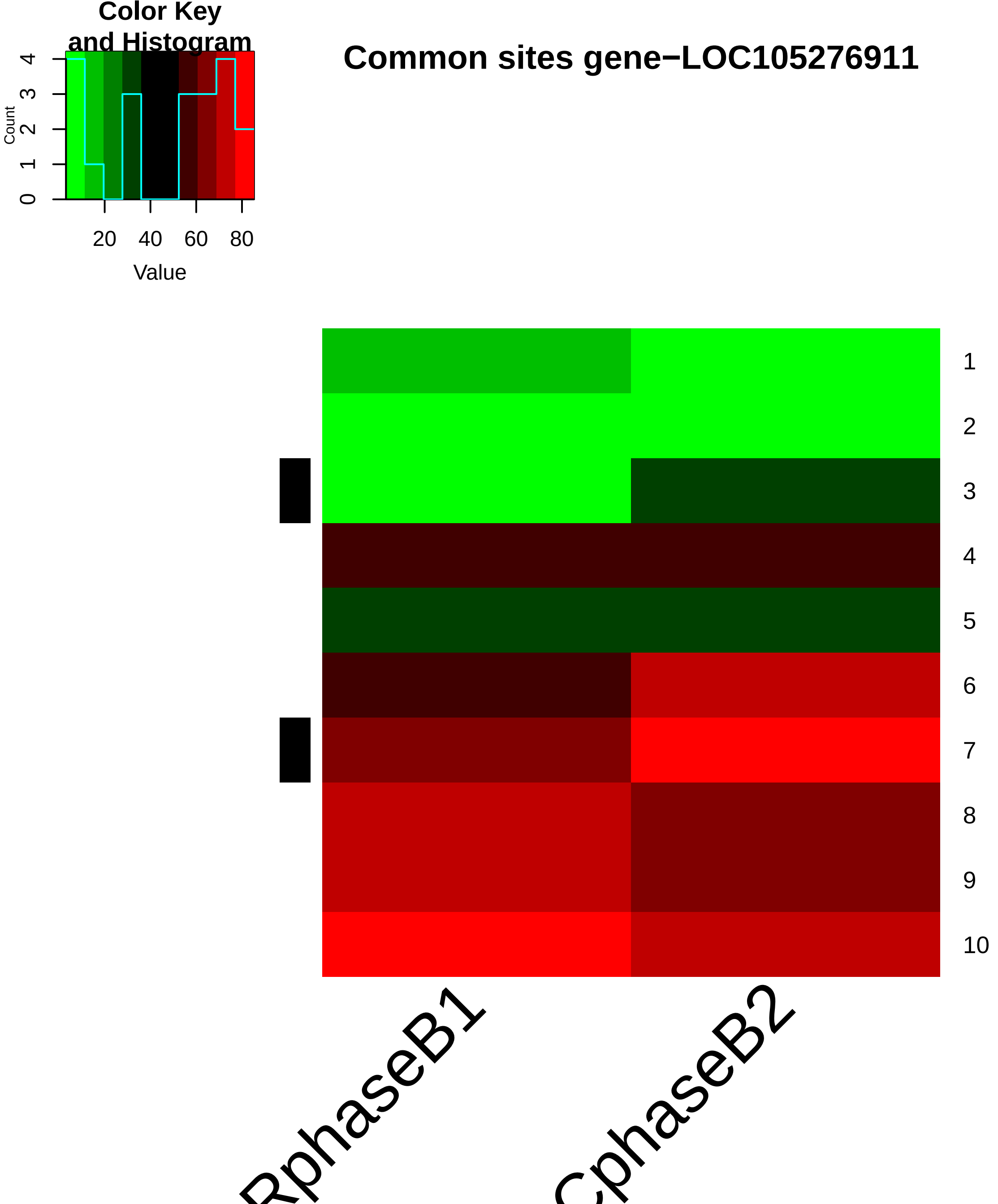
Color Key
and Histogram
Count
0
1
2
3
4
20
40
60
80
Value
Common sites gene−LOC105276911
1
2
3
4
5
6
7
8
9
10
RphaseB1
CphaseB2

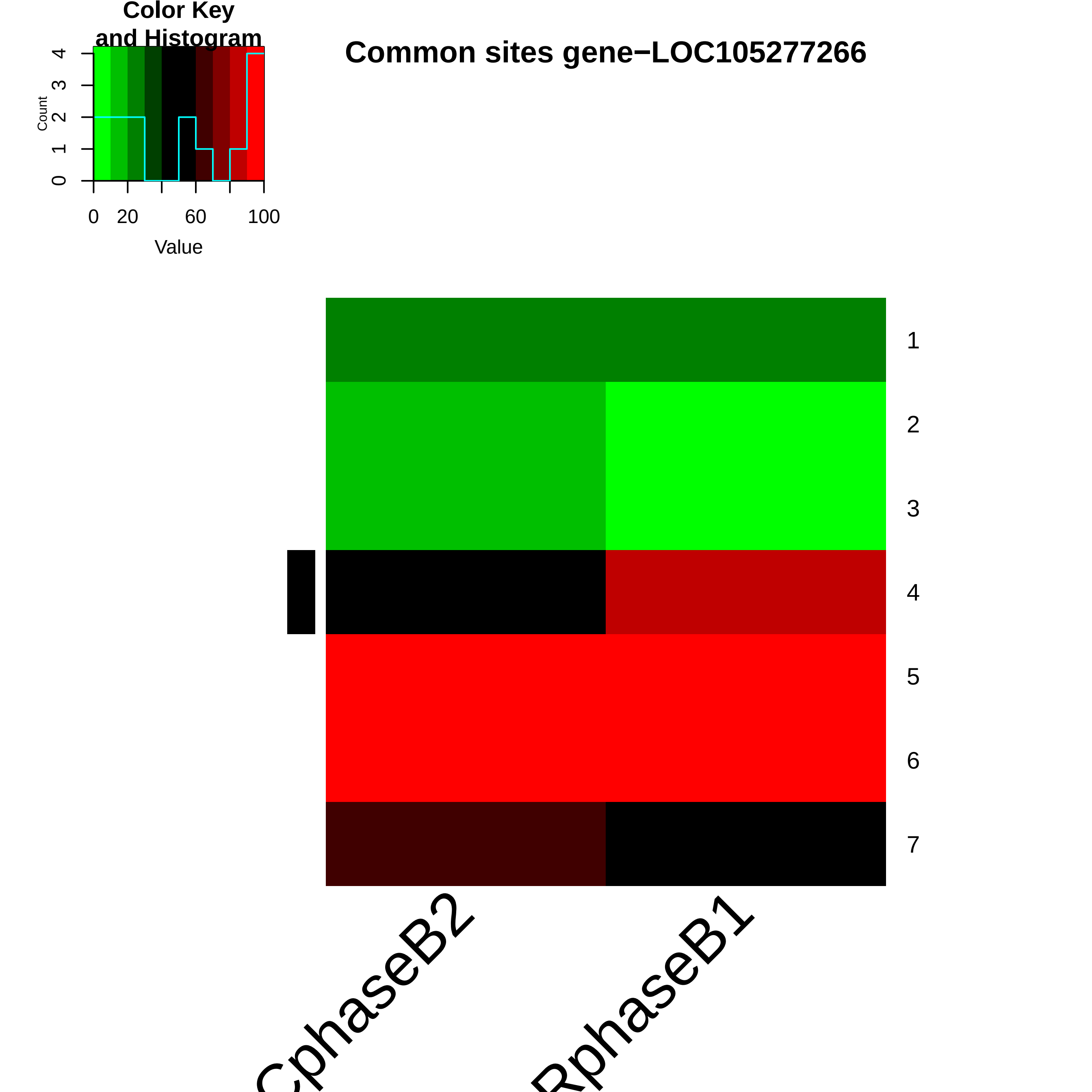

Color Key
and Histogram
Count
0
1
2
3
4
0
20
60
100
Value
Common sites gene−LOC105277266
1
2
3
4
5
6
7
CphaseB2
RphaseB1

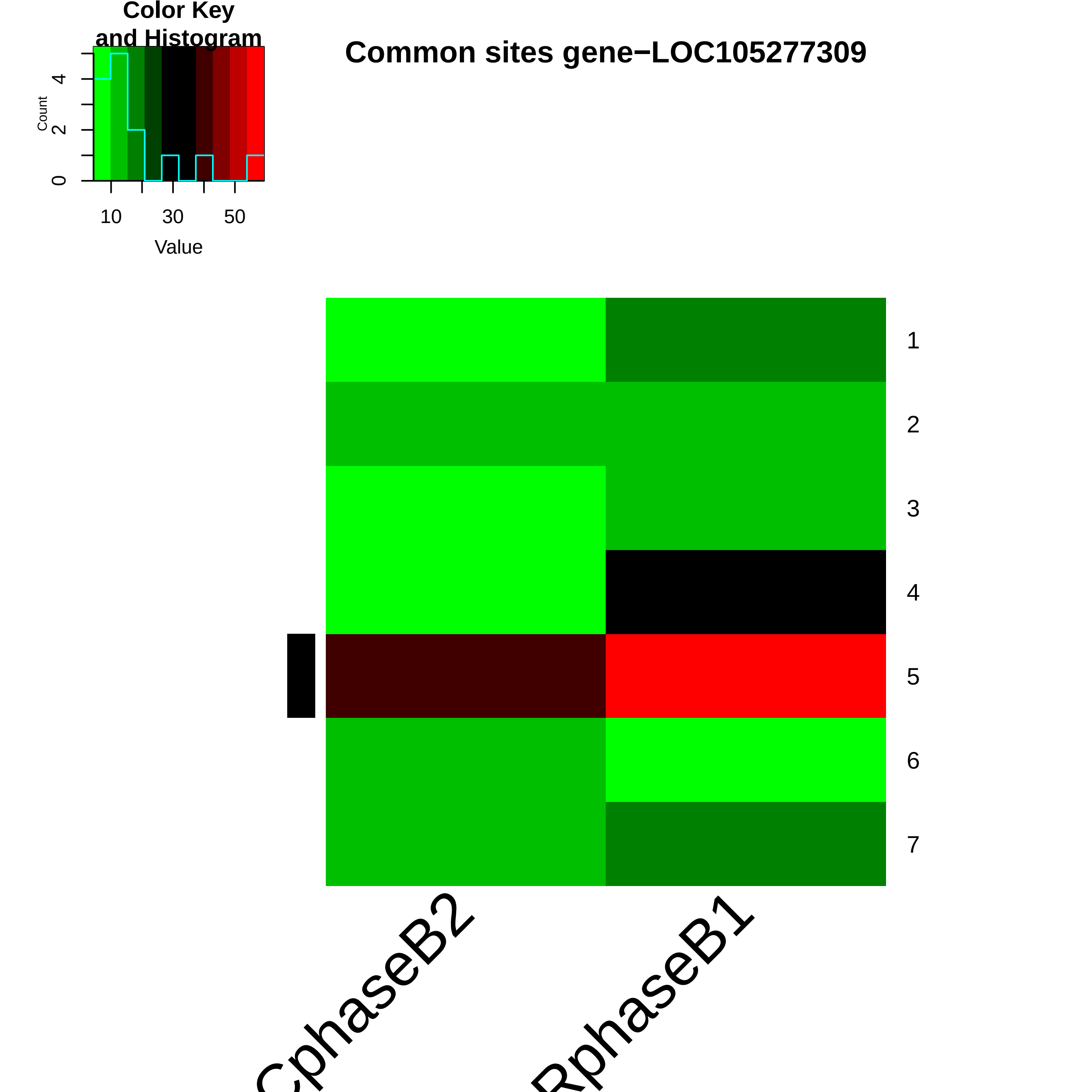

Color Key
and Histogram
Count
0
2
4
10
30
50
Value
Common sites gene−LOC105277309
1
2
3
4
5
6
7
CphaseB2
RphaseB1

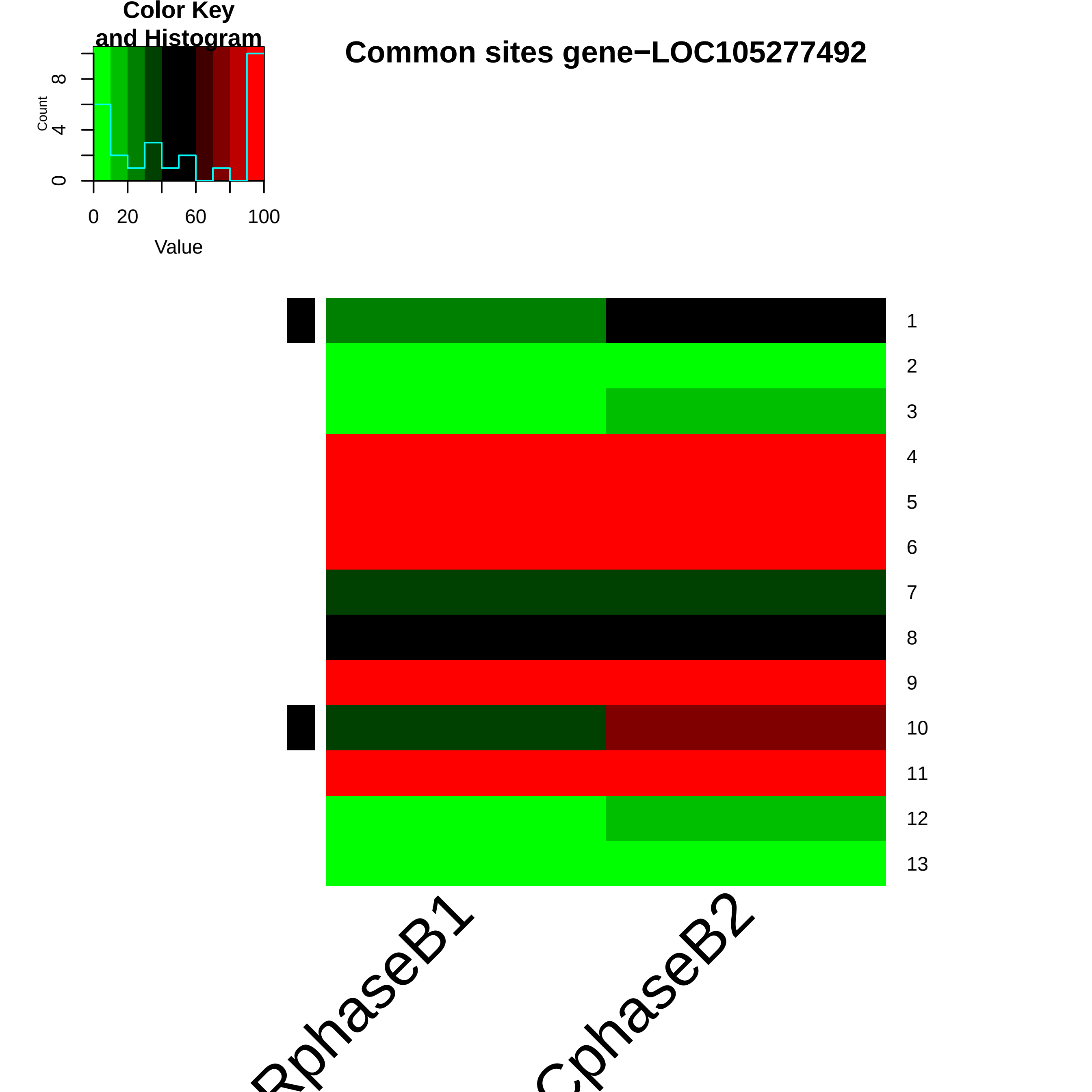

Color Key
and Histogram
Count
0
4
8
0
20
60
100
Value
Common sites gene−LOC105277492
1
2
3
4
5
6
7
8
9
10
11
12
13
RphaseB1
CphaseB2

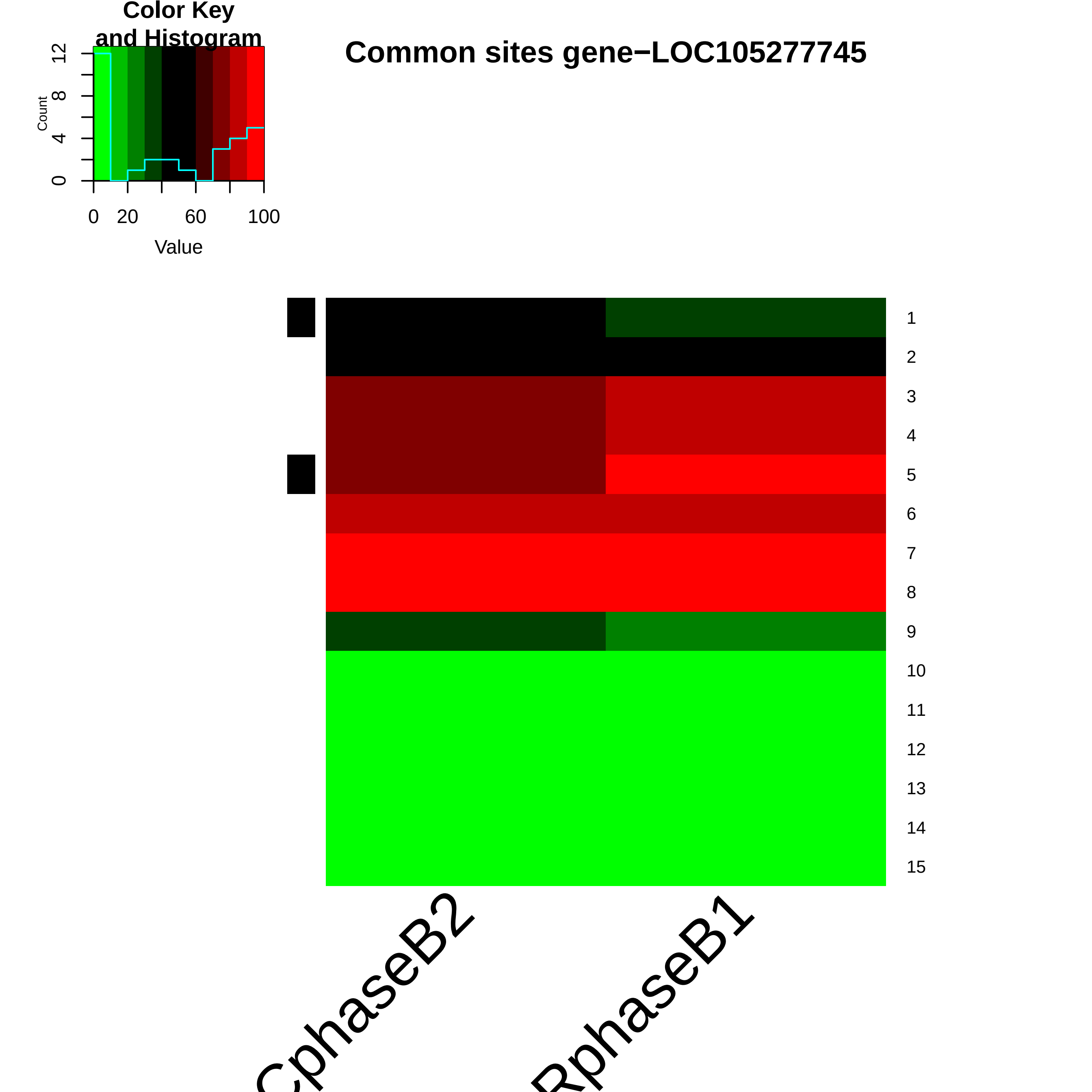
Color Key
and Histogram
Count
0
4
8
12
0
20
60
100
Value
Common sites gene−LOC105277745
CphaseB2
RphaseB1

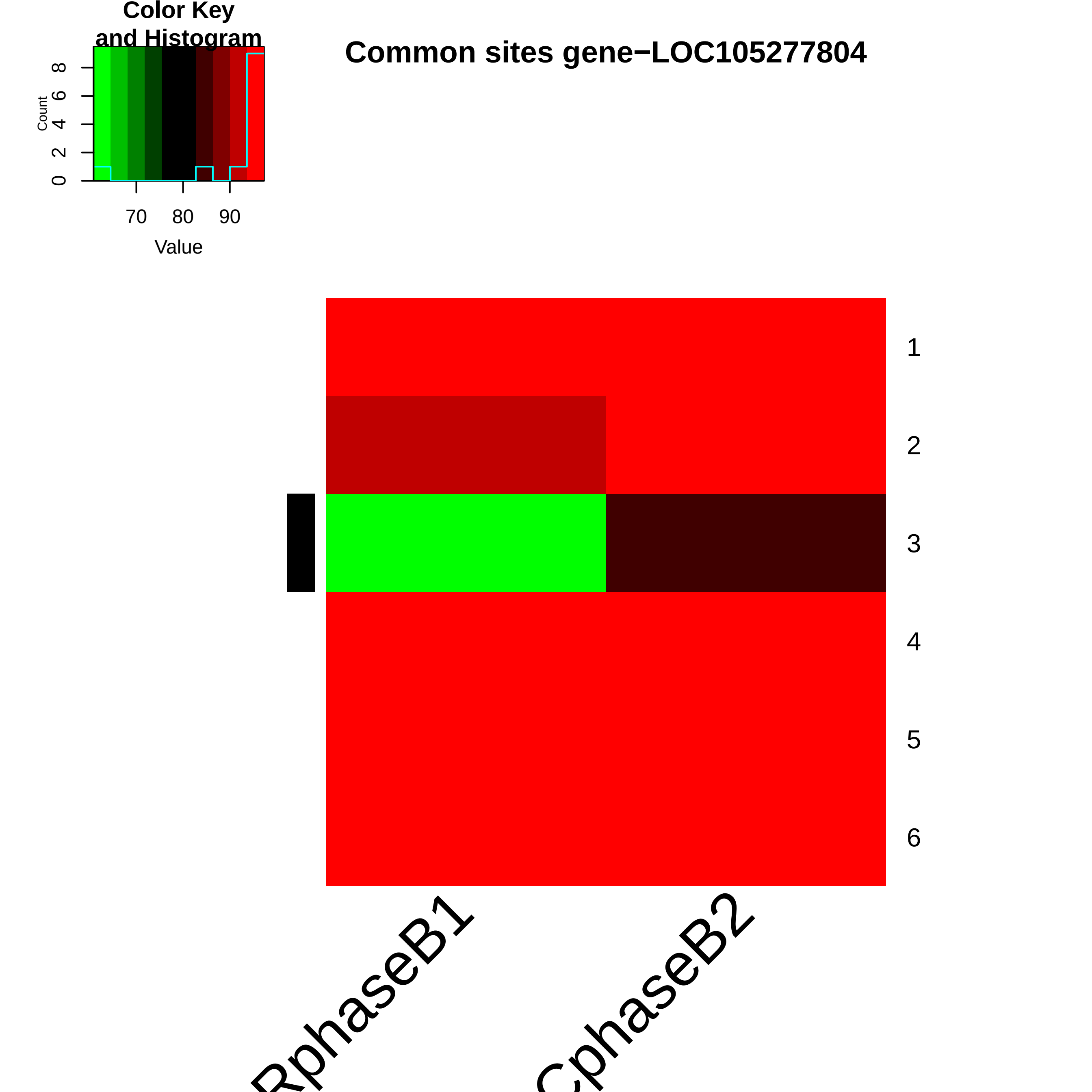

Color Key
and Histogram
Count
0 2 4 6 8
70 80 90
Value
Common sites gene−LOC105277804
1
2
3
4
5
6
RphaseB1
CphaseB2

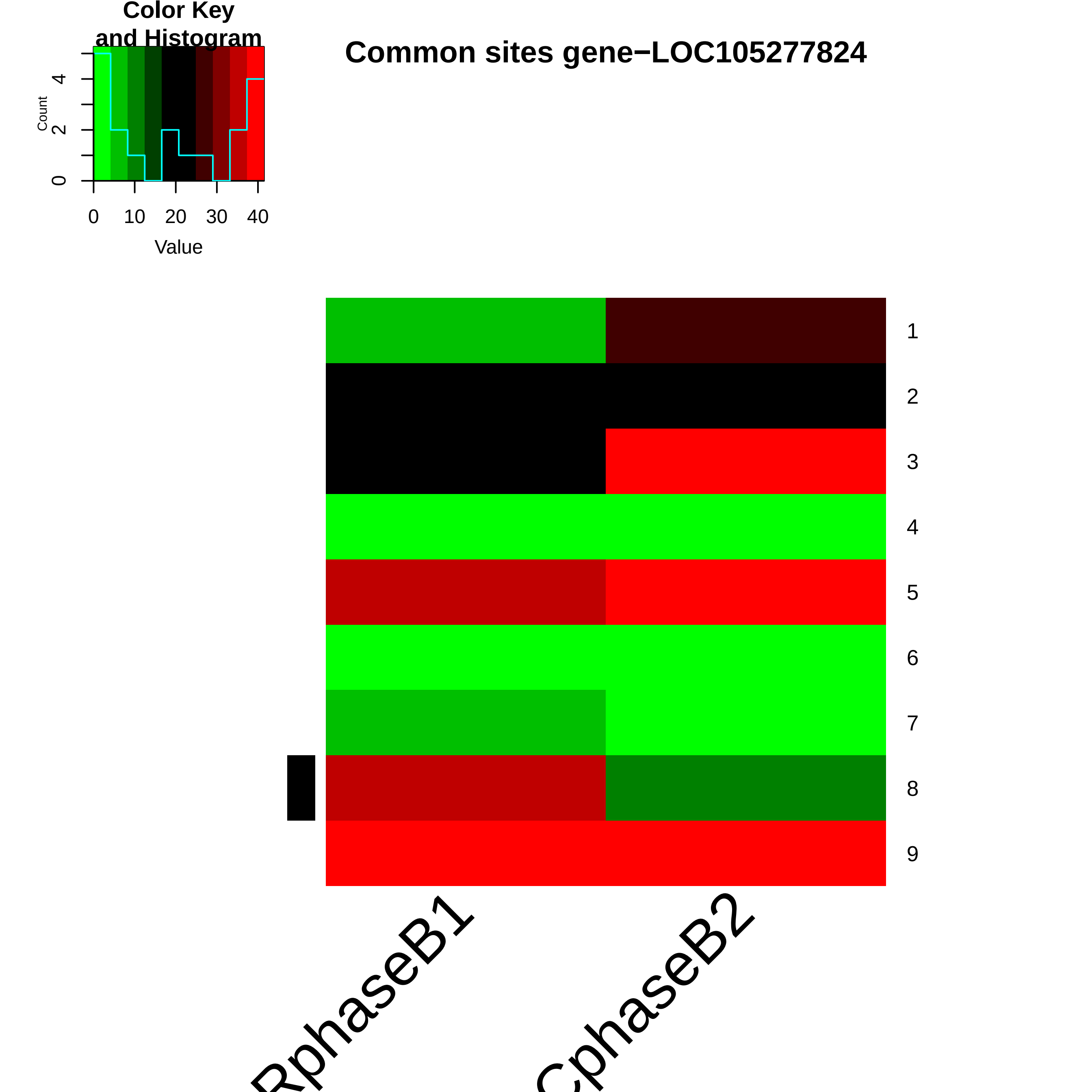

Color Key
and Histogram
Count
0
2
4
0
10
20
30
40
Value
Common sites gene−LOC105277824
1
2
3
4
5
6
7
8
9
RphaseB1
CphaseB2

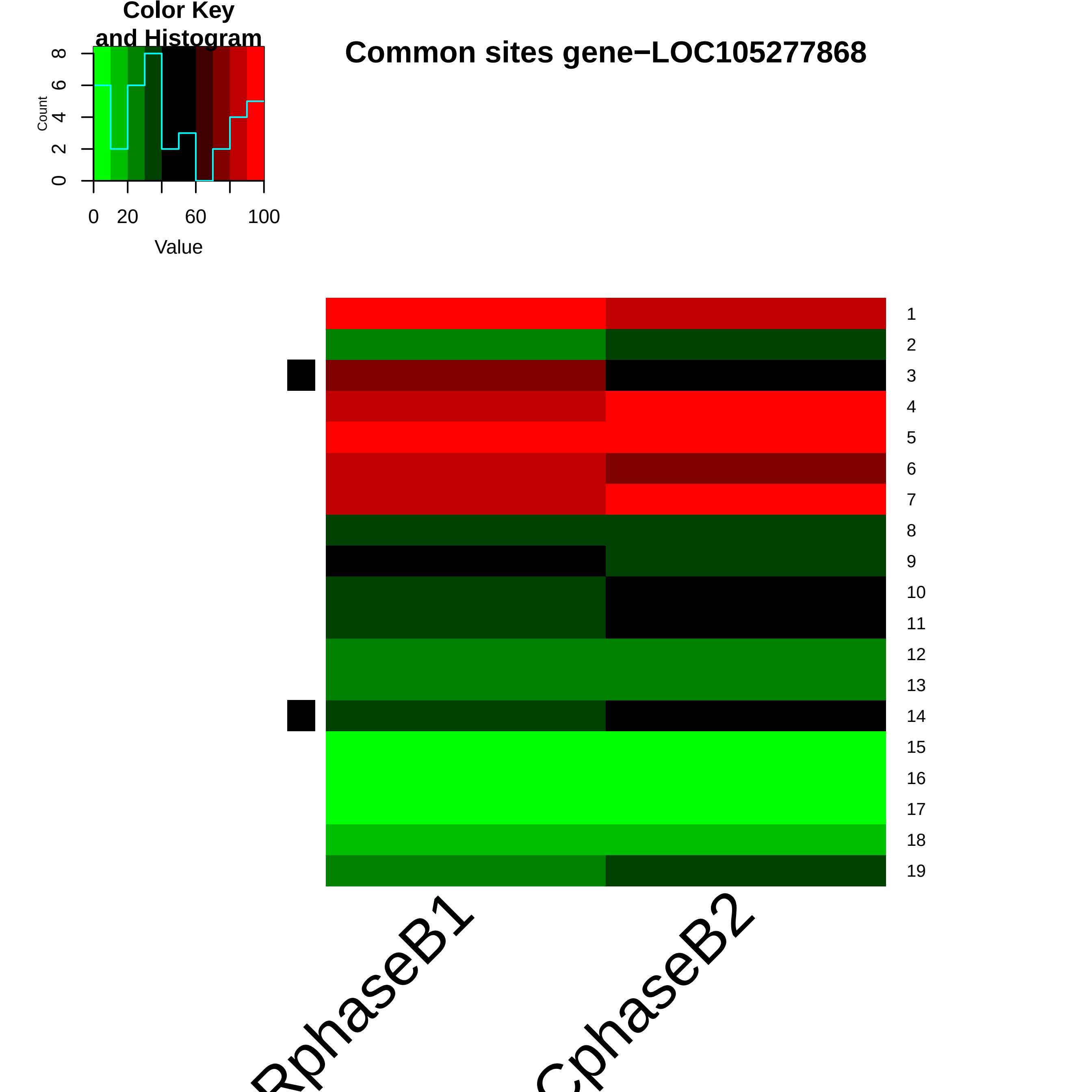
Color Key
and Histogram
Count
0
2
4
6
8
0
20
60
100
Value
Common sites gene−LOC105277868
RphaseB1
CphaseB2

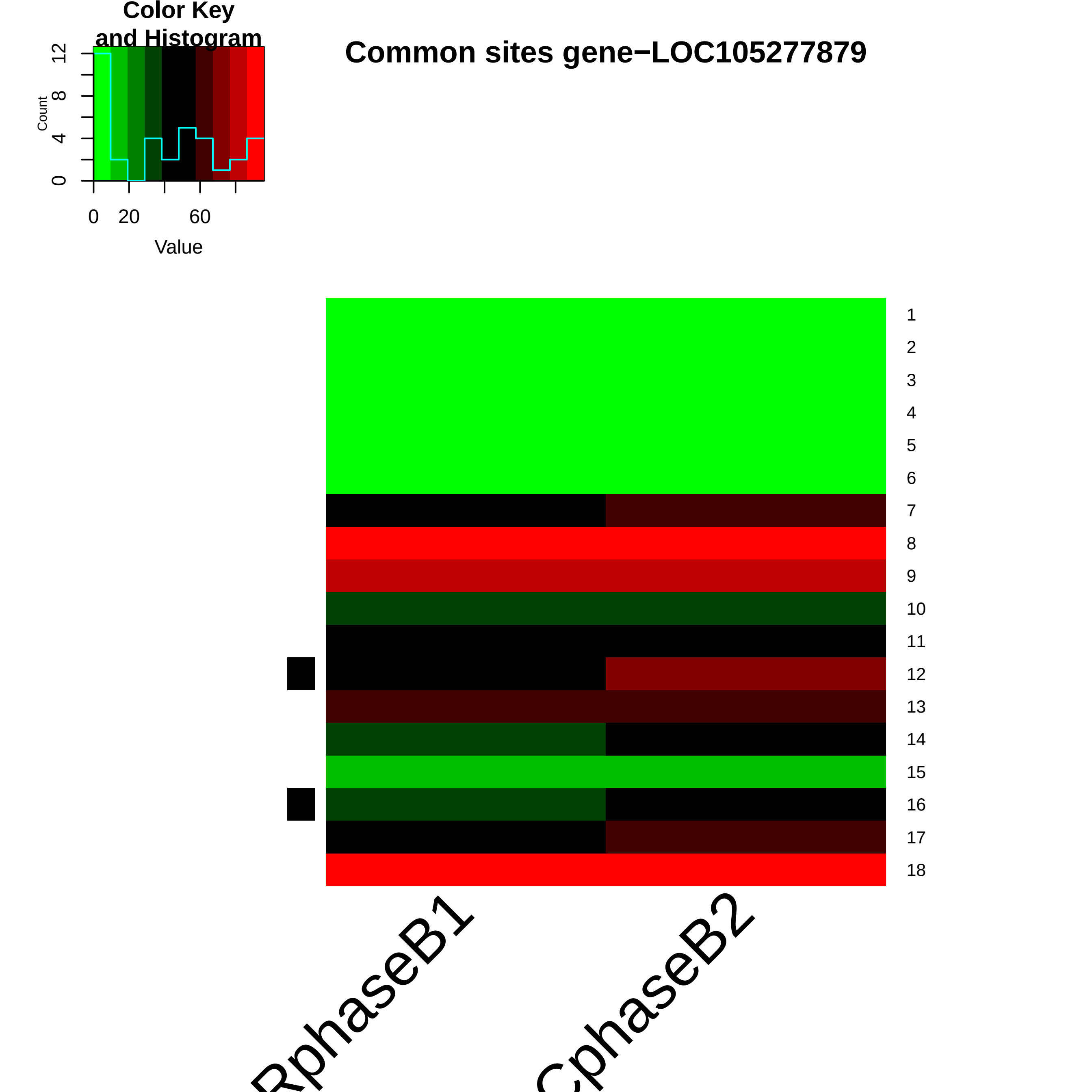

Color Key
and Histogram
Count
0
4
8
12
0
20
60
Value
Common sites gene−LOC105277879
RphaseB1
CphaseB2

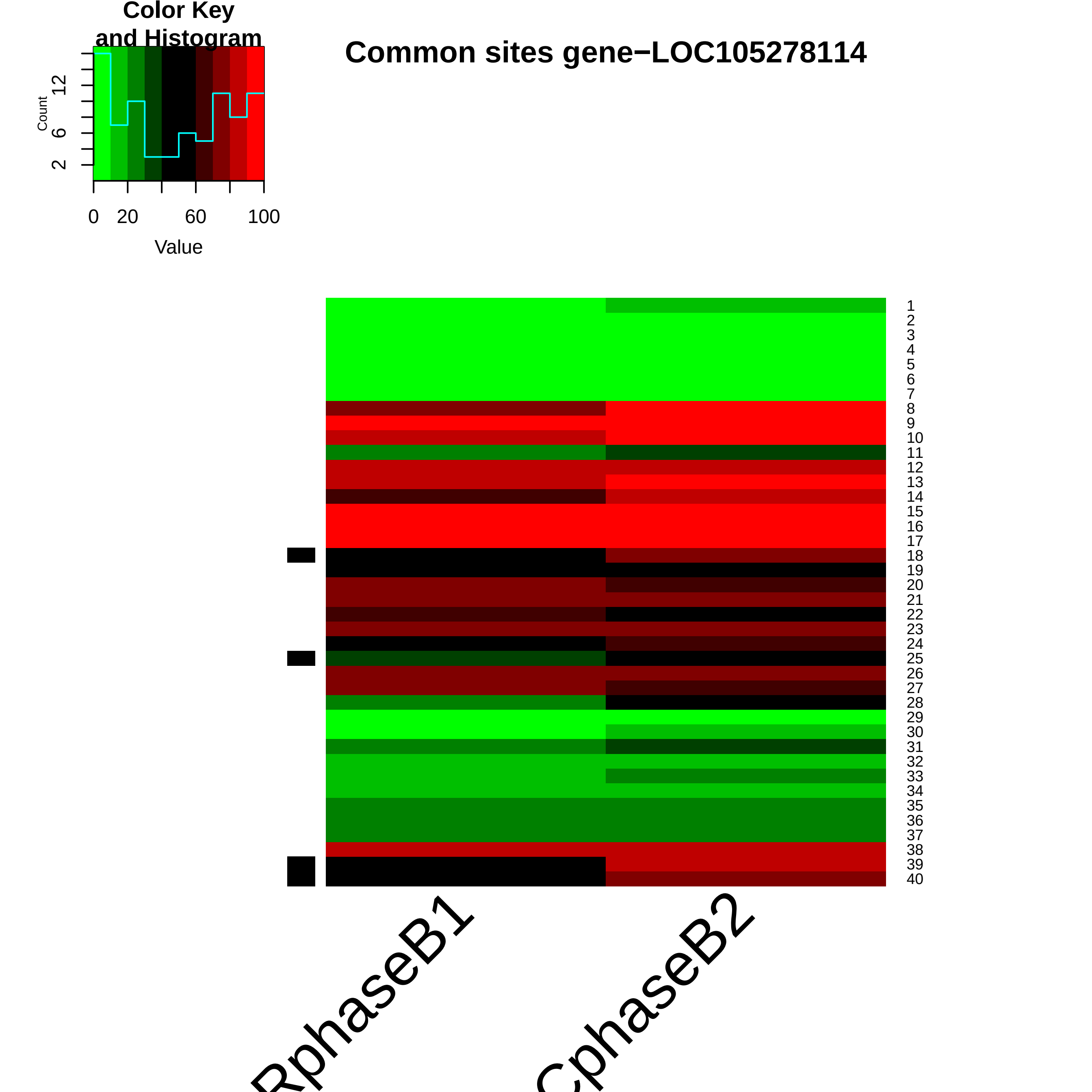
Color Key
and Histogram
Common sites gene−LOC105278114
Count
2
6
12
0
20
60
100
Value
RphaseB1
CphaseB2

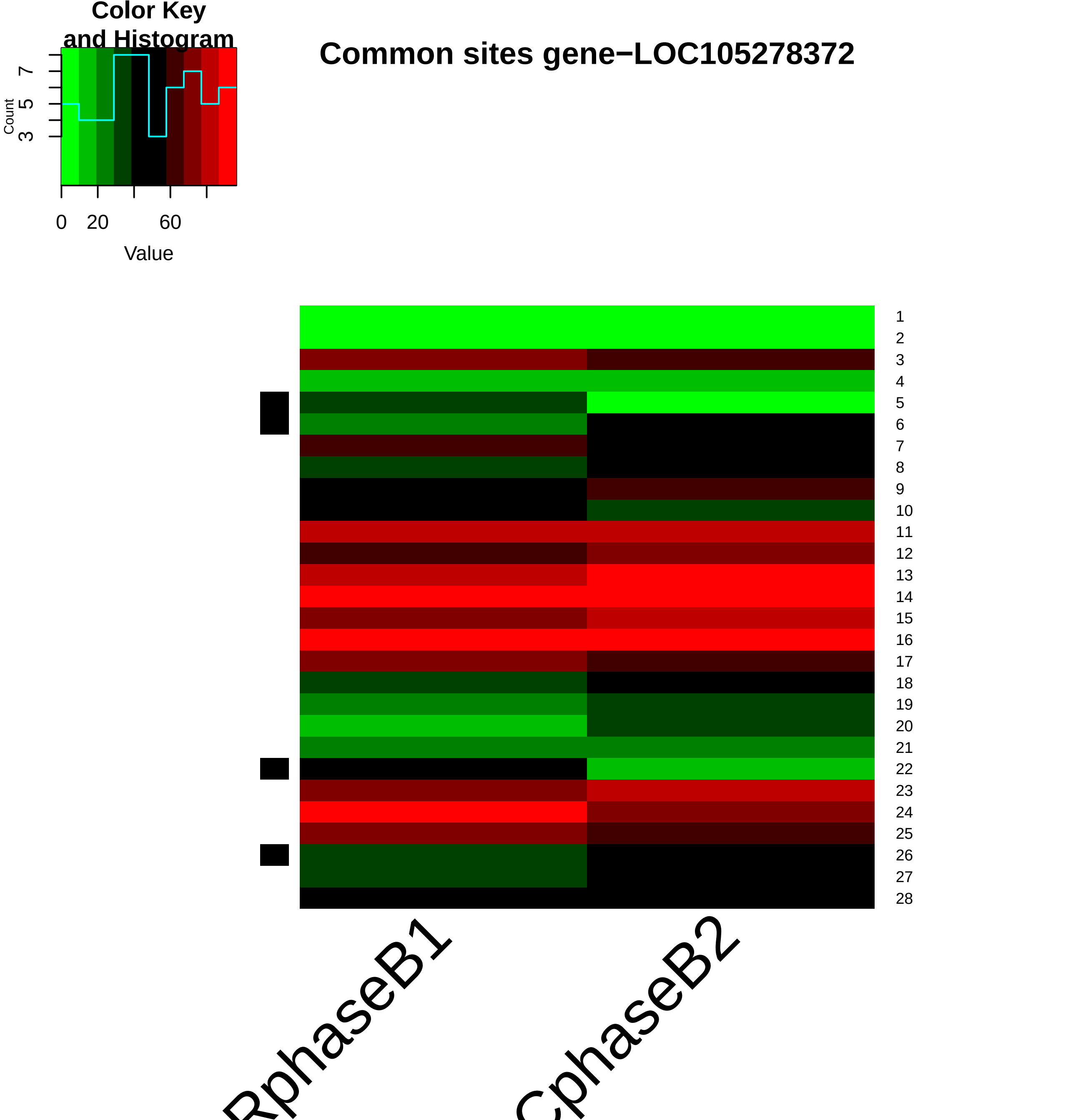
Color Key
and Histogram
Count
3
5
7
0
20
60
Value
Common sites gene−LOC105278372
RphaseB1
CphaseB2

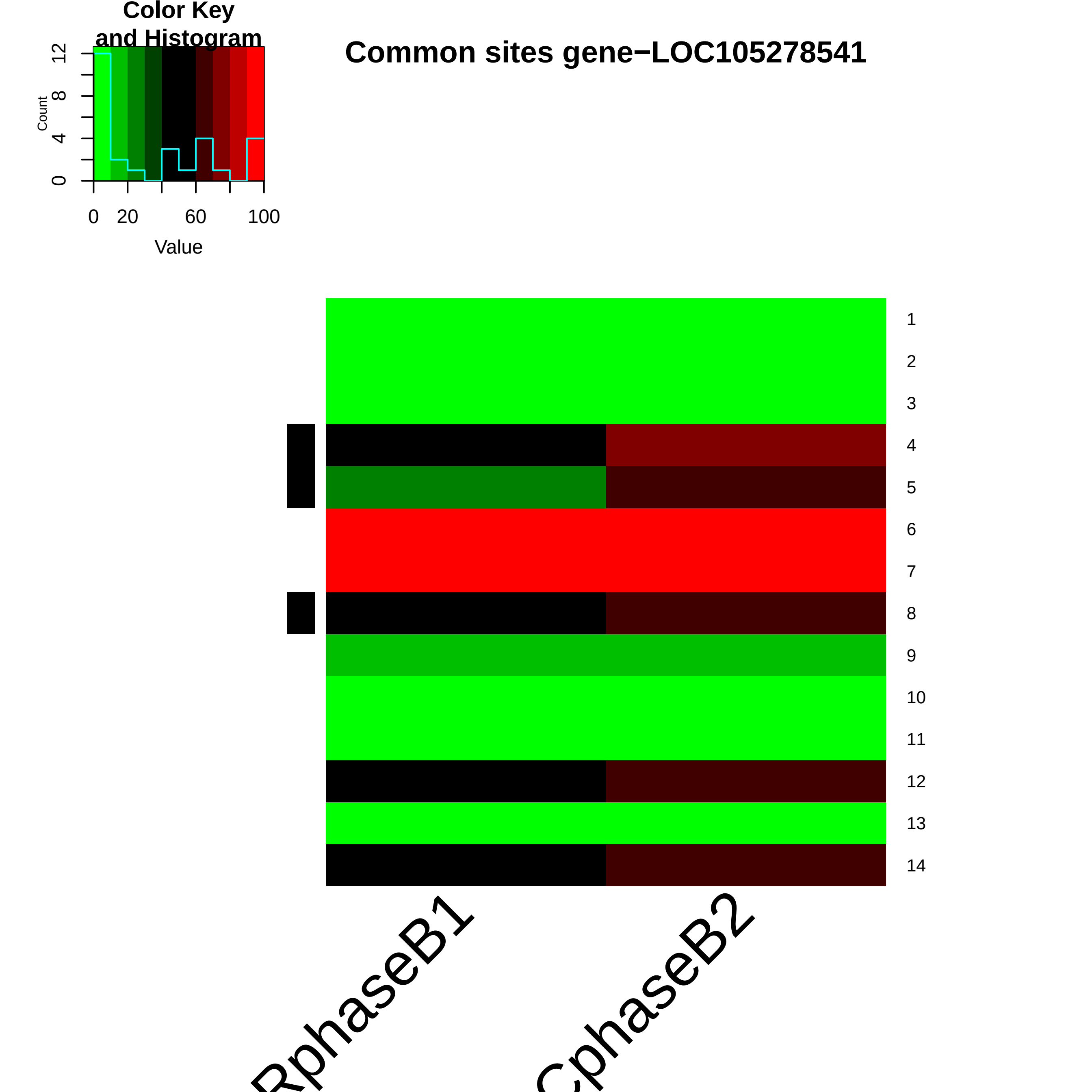

Color Key
and Histogram
Count
0
4
8
12
0
20
60
100
Value
Common sites gene−LOC105278541
1
2
3
4
5
6
7
8
9
10
11
12
13
14
RphaseB1
CphaseB2

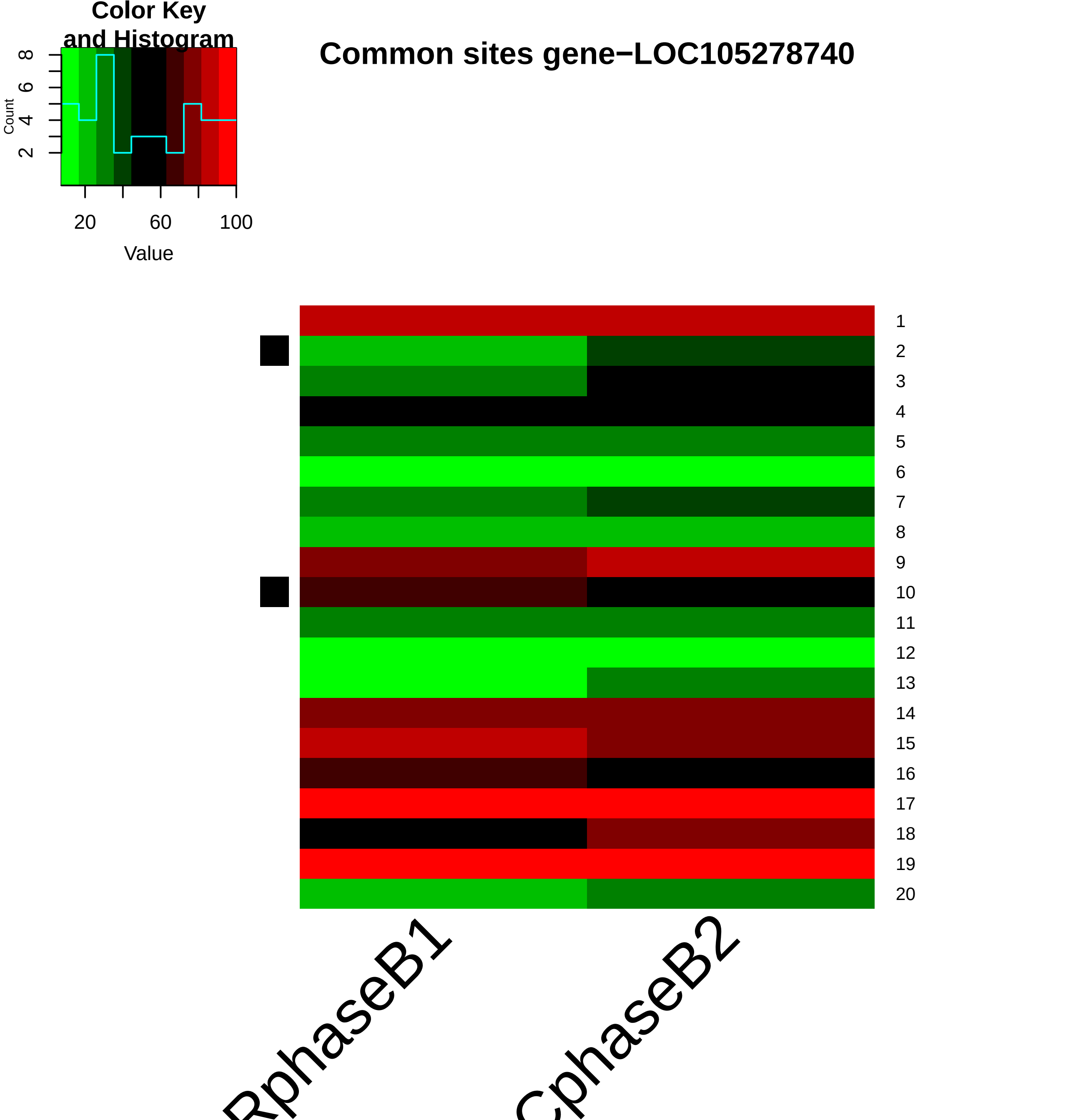
Color Key
and Histogram
Count
2
4
6
8
20
60
100
Value
Common sites gene−LOC105278740
RphaseB1
CphaseB2

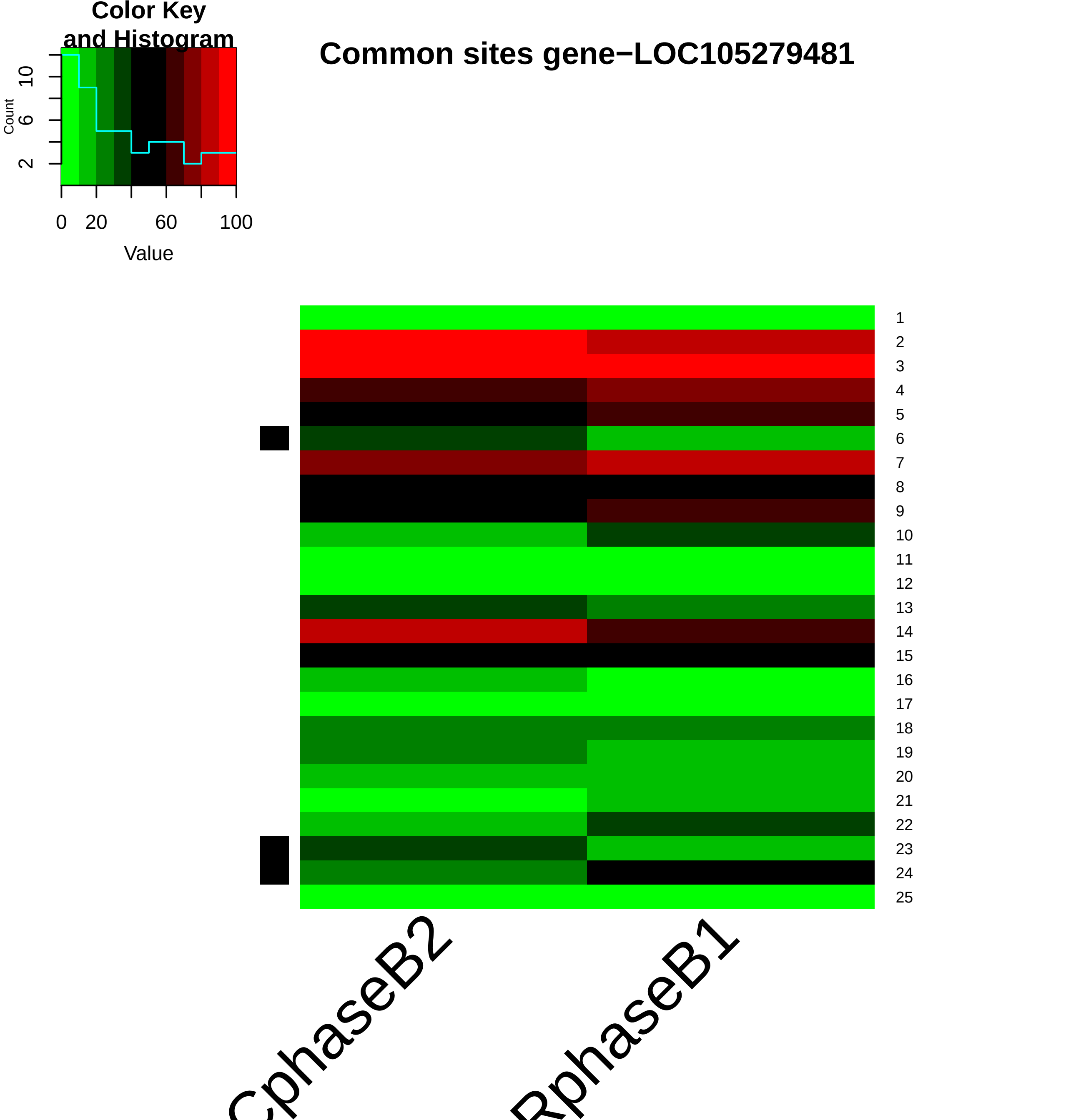

Color Key and Histogram
Count
2
6
10
0
20
60
100
Value
Common sites gene−LOC105279481
CphaseB2
RphaseB1

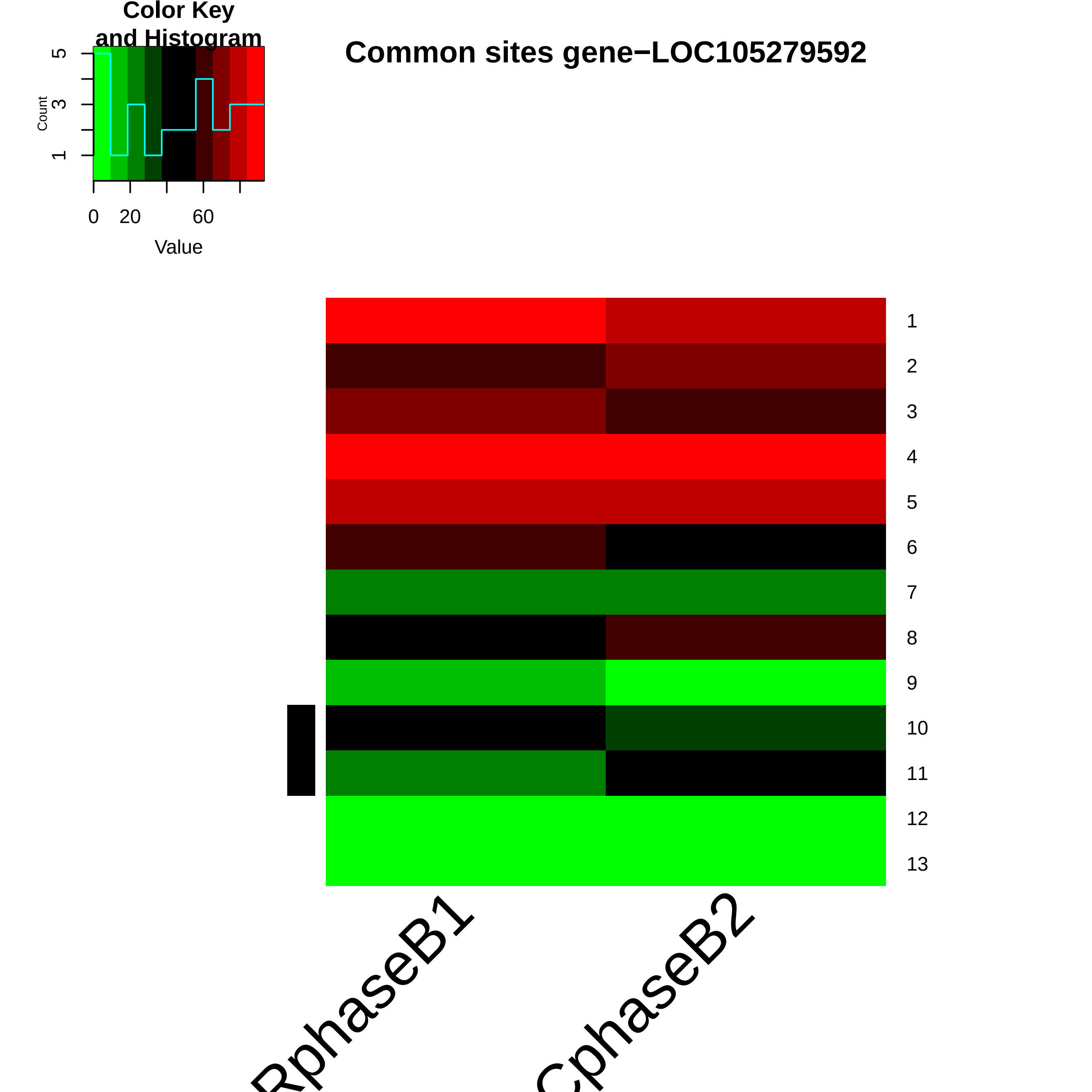

Color Key
and Histogram
Count
1
3
5
0
20
60
Value
Common sites gene−LOC105279592
1
2
3
4
5
6
7
8
9
10
11
12
13
RphaseB1
CphaseB2

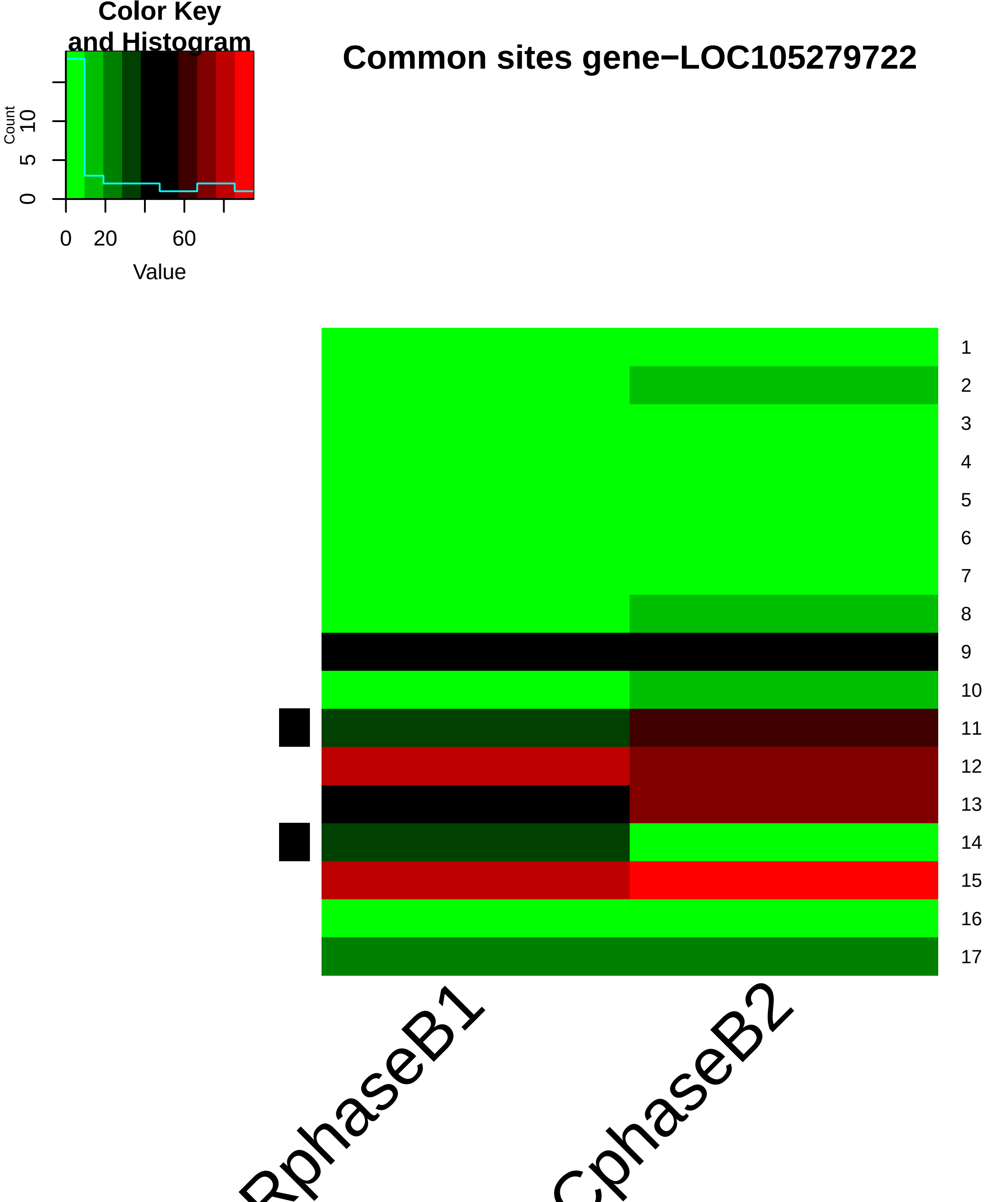
Color Key
and Histogram
Count
0
5
10
0
20
60
Value
Common sites gene−LOC105279722
RphaseB1
CphaseB2

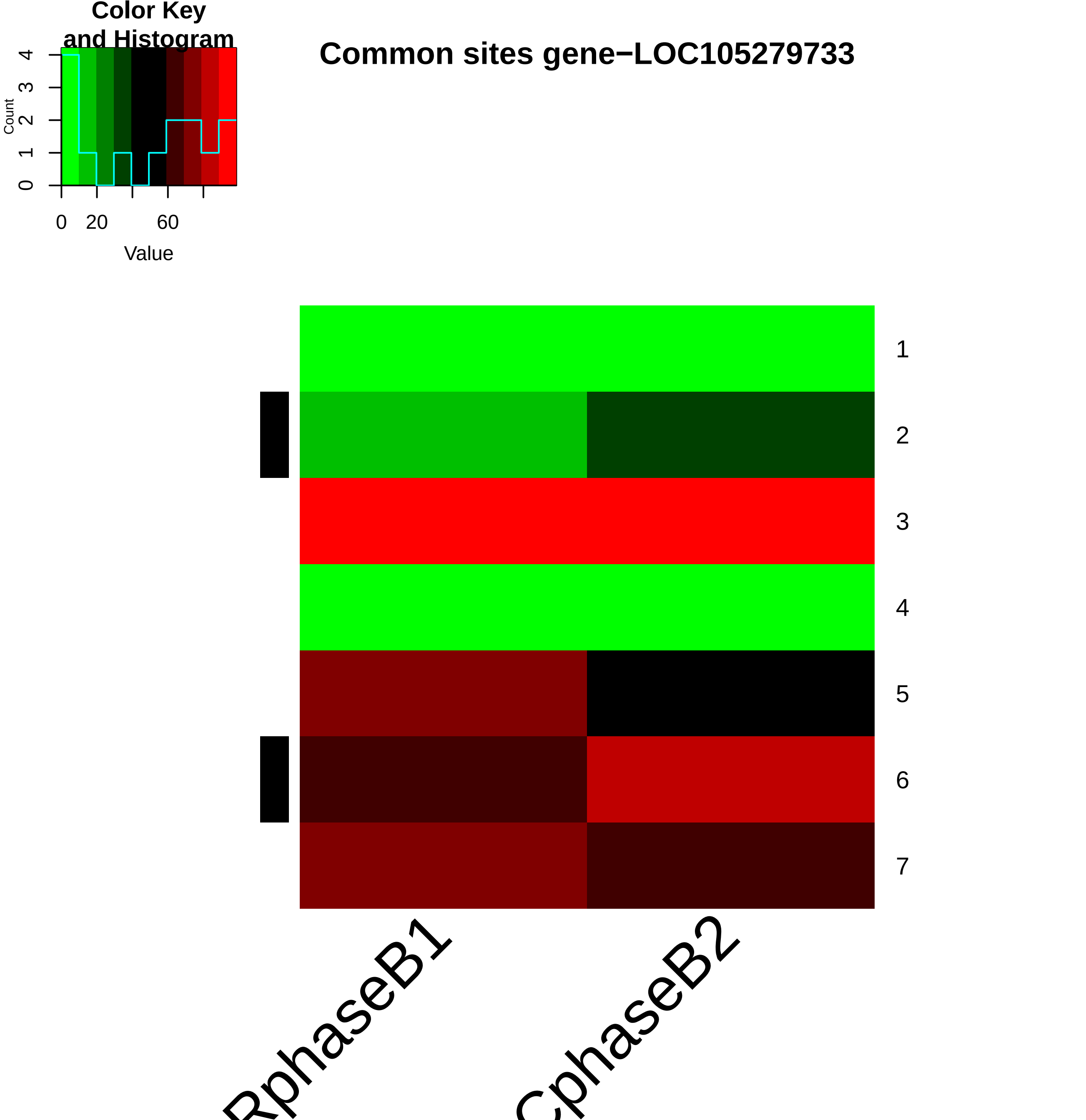

Color Key
and Histogram
Count
Value
0
20
60
0
1
2
3
4
Common sites gene−LOC105279733
1
2
3
4
5
6
7
RphaseB1
CphaseB2

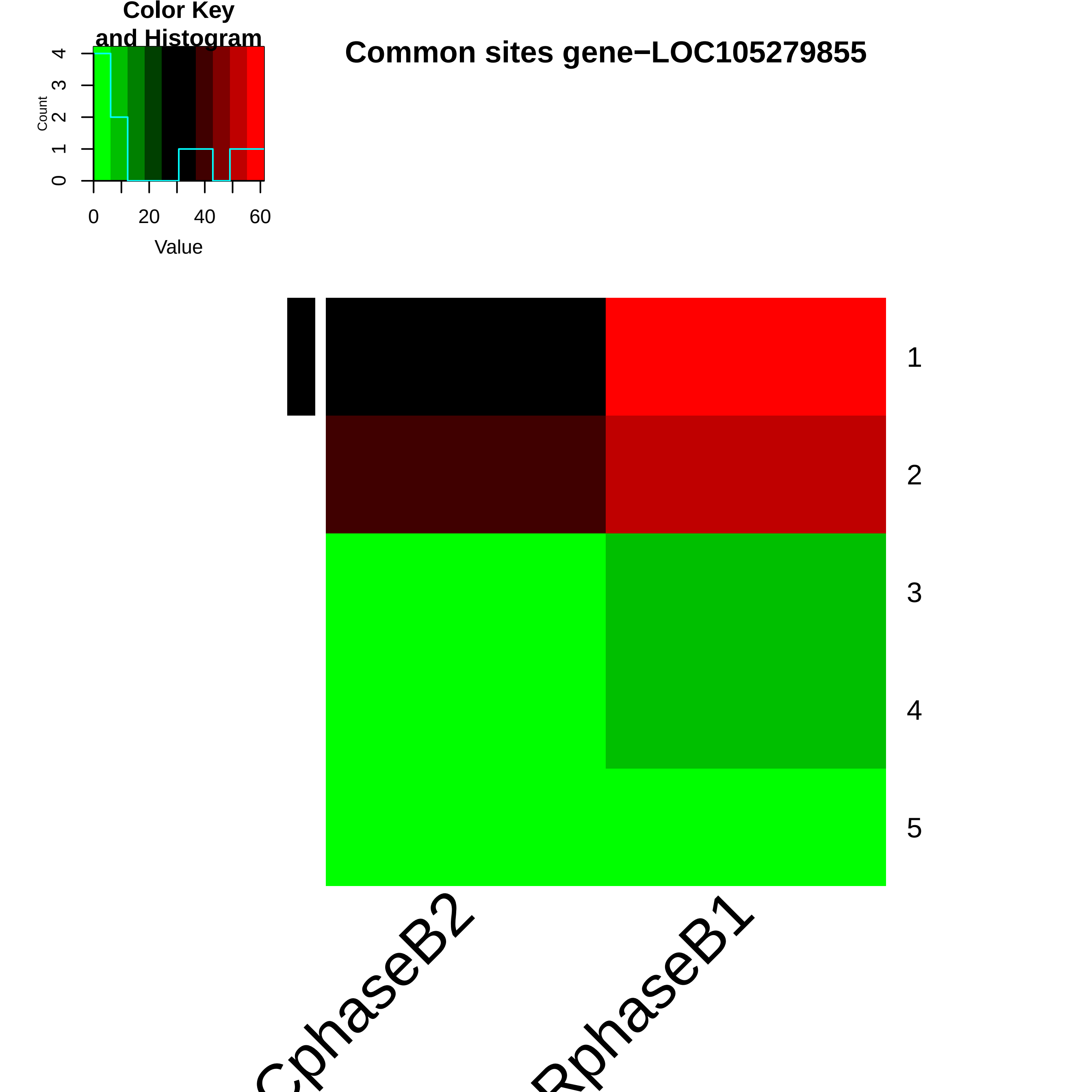
Color Key
and Histogram
Count
0
1
2
3
4
0
20
40
60
Value
Common sites gene−LOC105279855
1
2
3
4
5
CphaseB2
RphaseB1

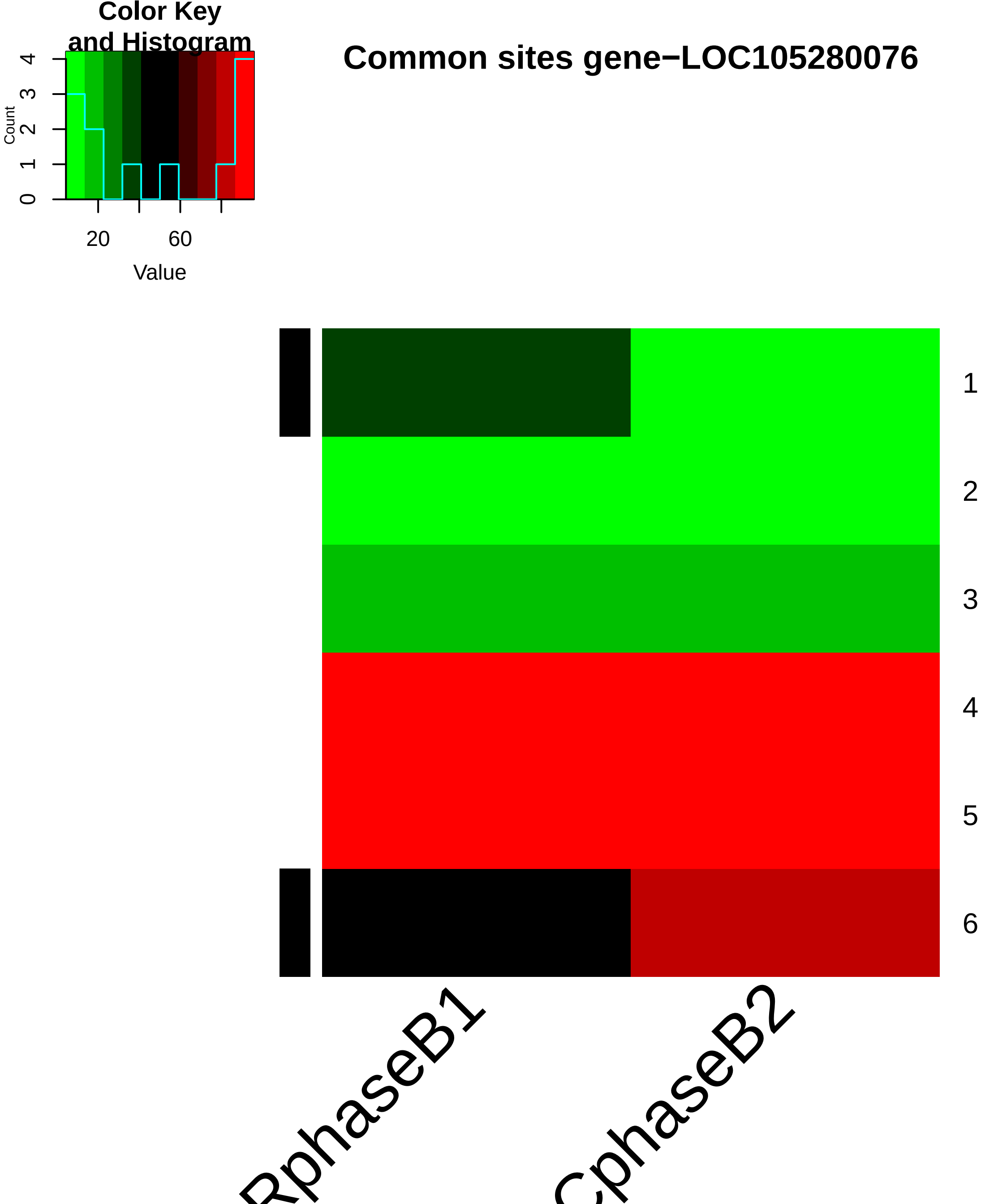
Color Key
and Histogram
Count
0
1
2
3
4
20
60
Value
Common sites gene−LOC105280076
1
2
3
4
5
6
RphaseB1
CphaseB2

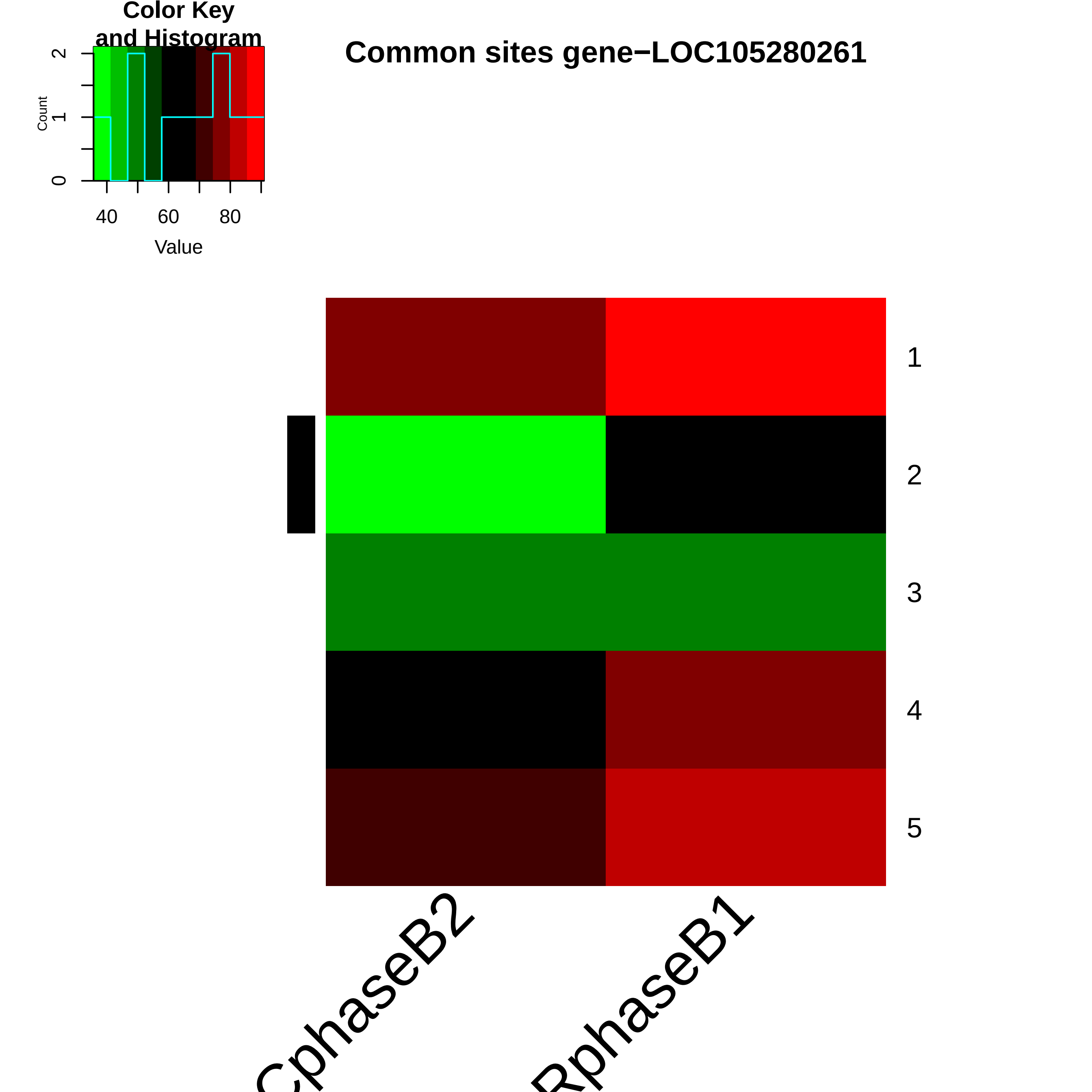

Color Key
and Histogram
Count
0
1
2
40
60
80
Value
Common sites gene−LOC105280261
1
2
3
4
5
CphaseB2
RphaseB1

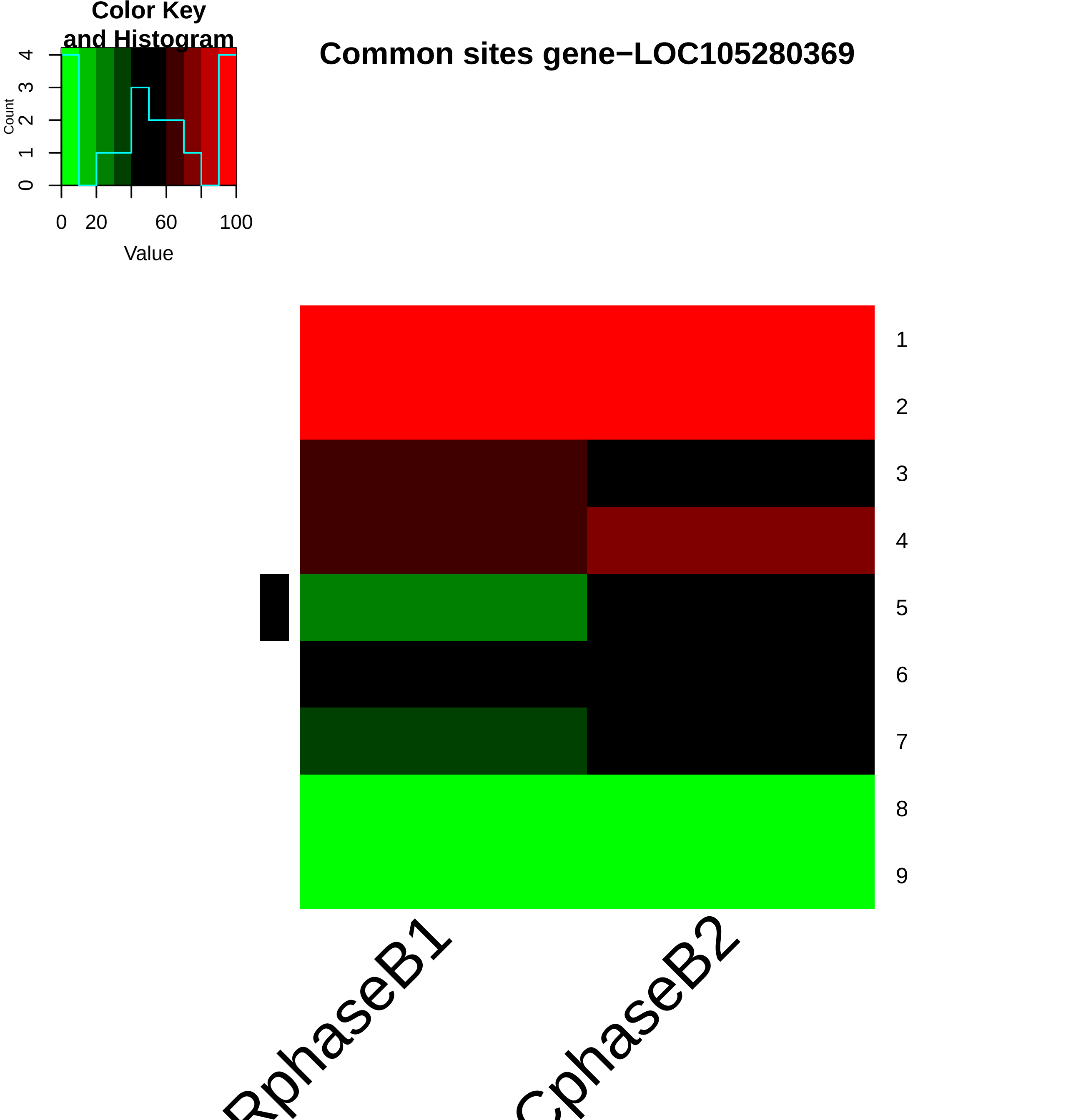
Color Key
and Histogram
Count
0 1 2 3 4
0 20 60 100
Value
Common sites gene−LOC105280369
1
2
3
4
5
6
7
8
9
RphaseB1
CphaseB2

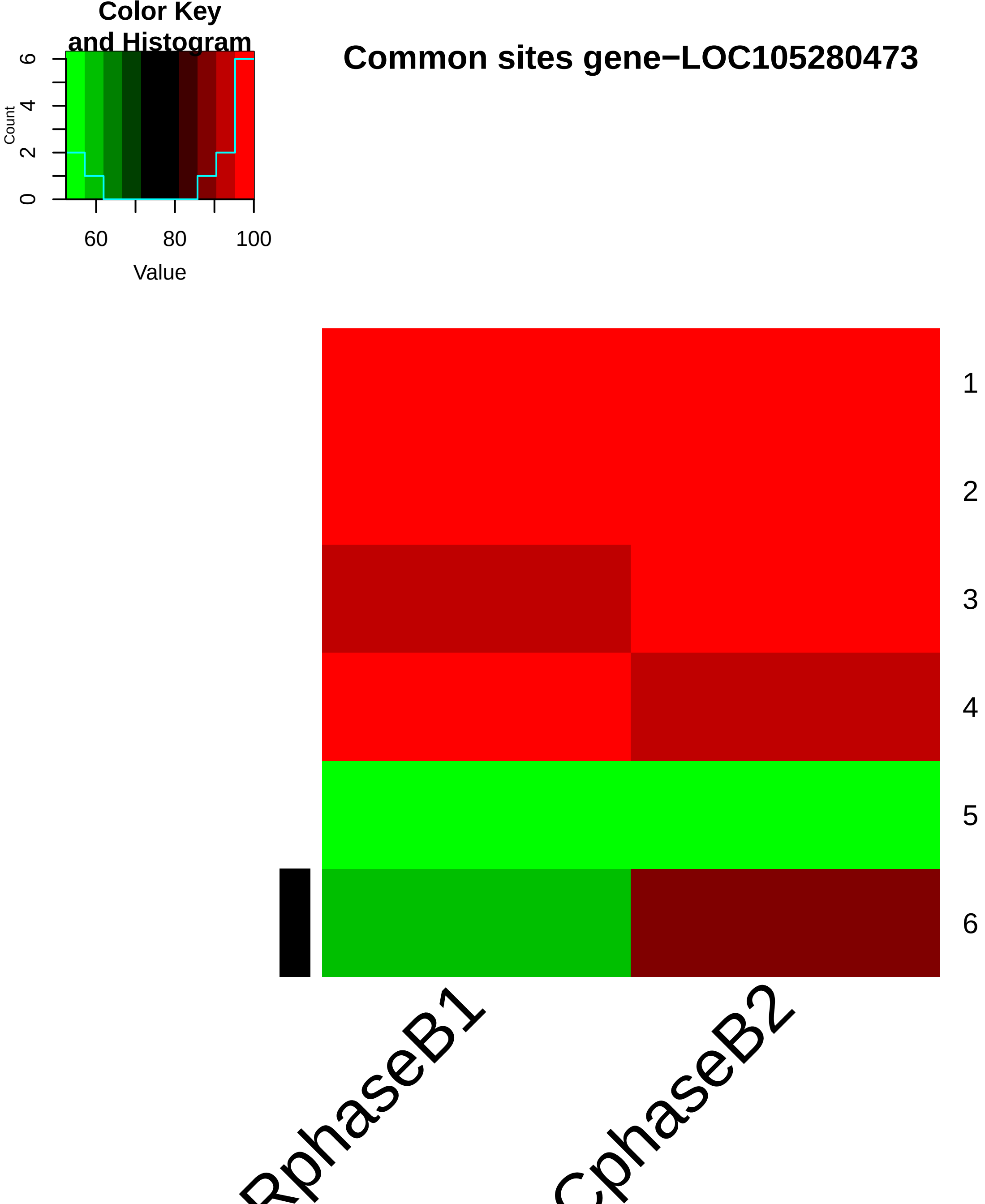

Color Key
and Histogram
Count
0
2
4
6
60
80
100
Value
Common sites gene−LOC105280473
1
2
3
4
5
6
RphaseB1
CphaseB2

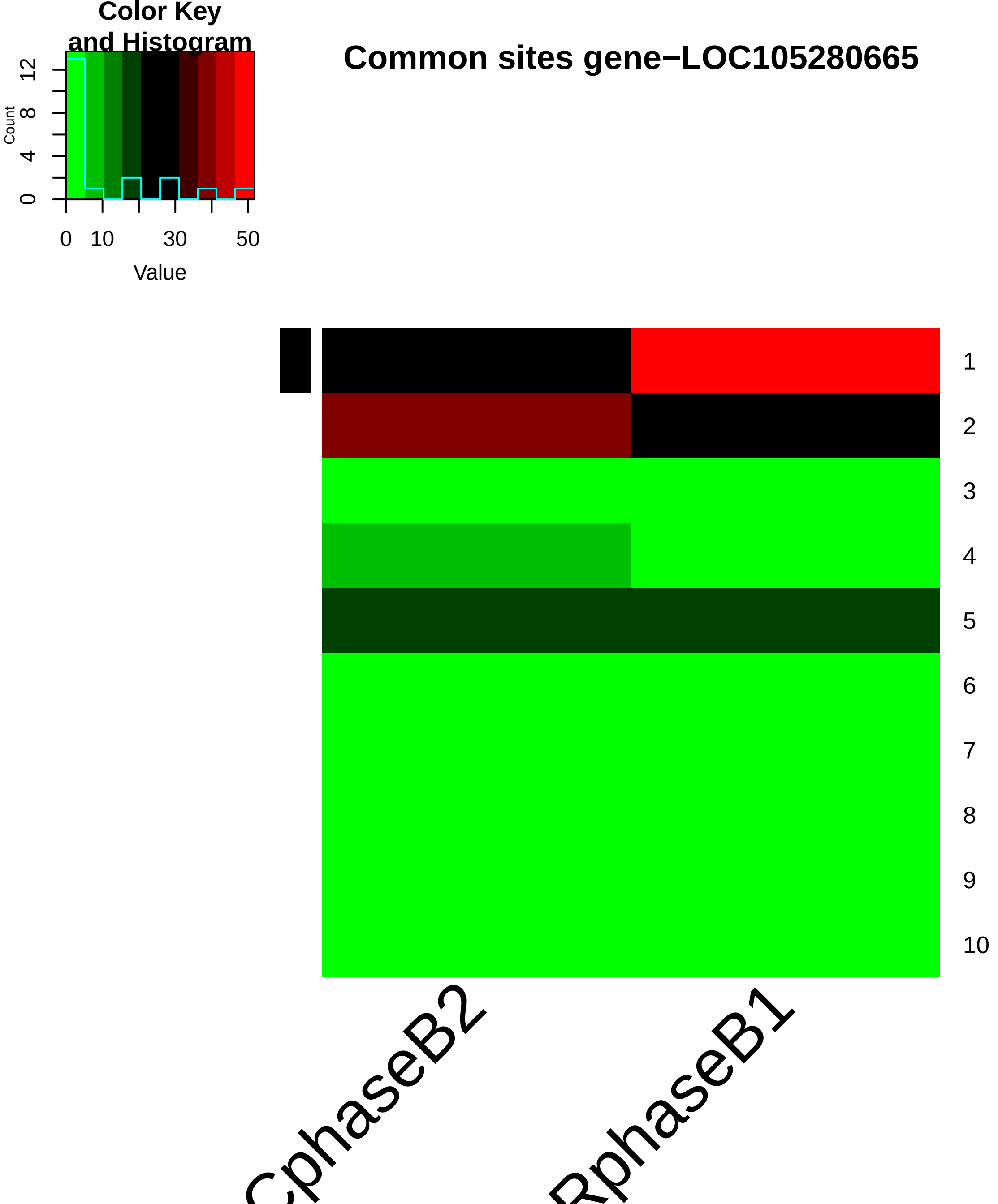

Color Key
and Histogram
Count
0
4
8
12
0
10
30
50
Value
Common sites gene−LOC105280665
1
2
3
4
5
6
7
8
9
10
CphaseB2
RphaseB1

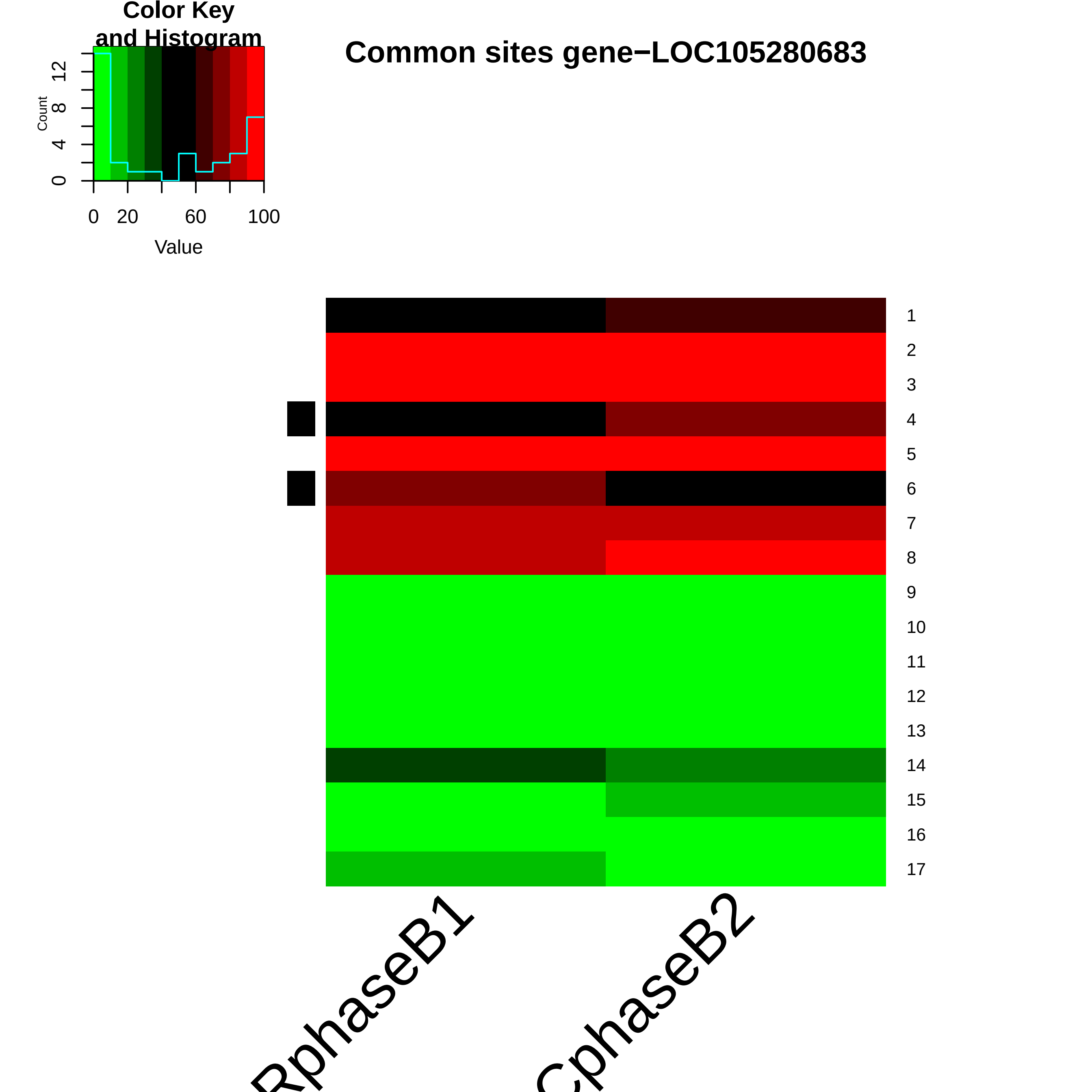

Color Key
and Histogram
Count
0
4
8
12
0
20
60
100
Value
Common sites gene−LOC105280683
RphaseB1
CphaseB2

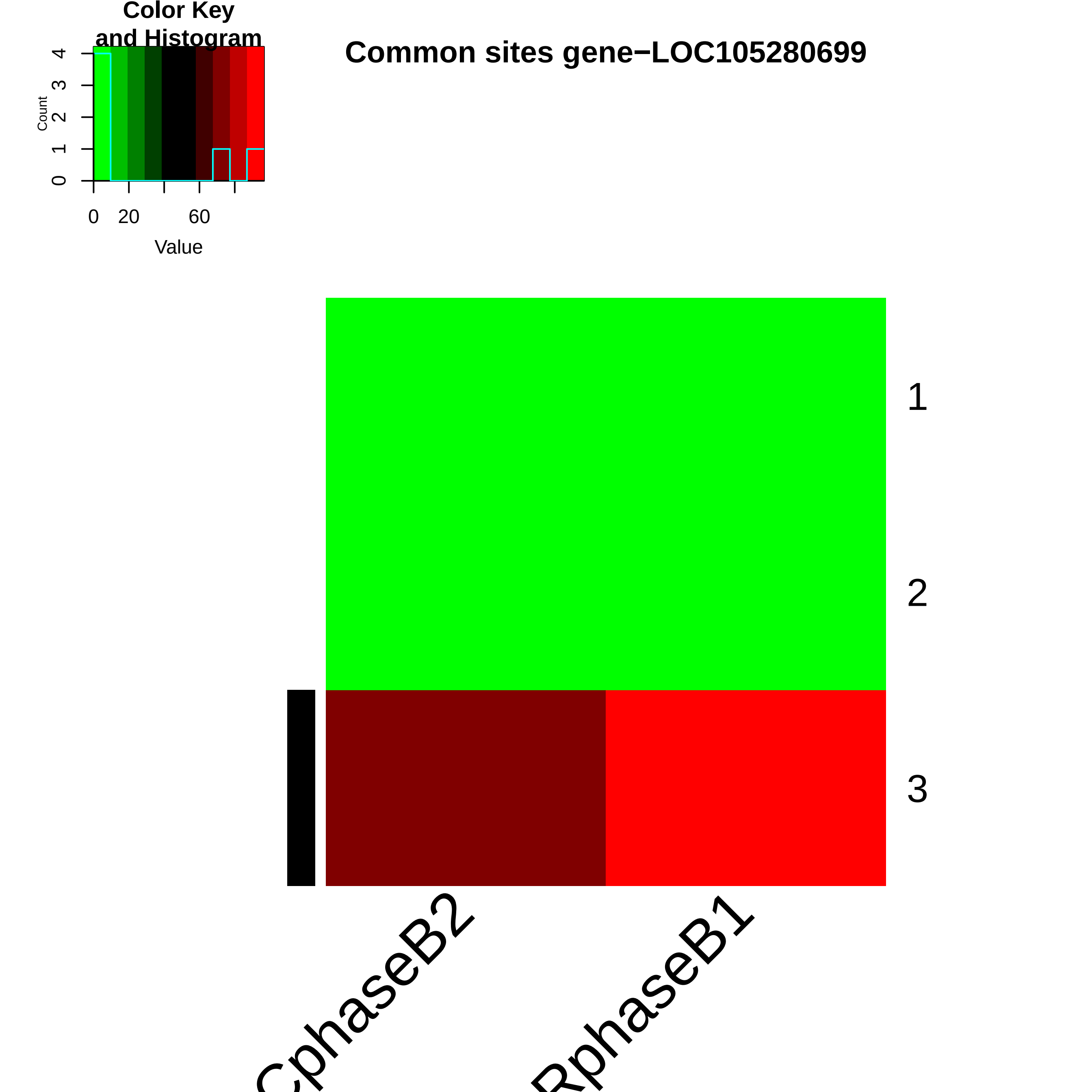

Color Key
and Histogram
Count
0
1
2
3
4
0
20
60
Value
Common sites gene−LOC105280699
1
2
3
CphaseB2
RphaseB1

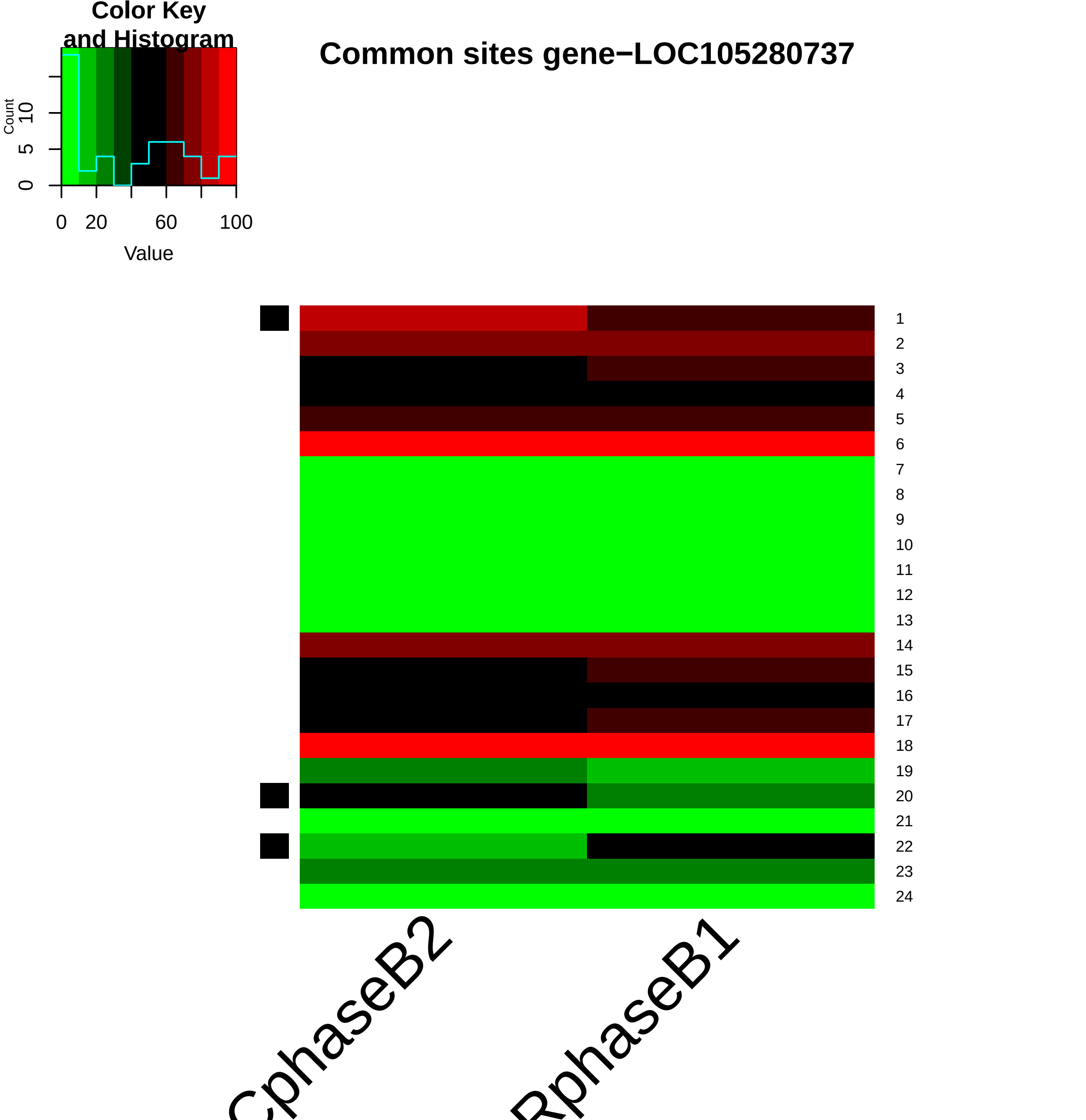
Color Key
and Histogram
Count
0
5
10
0
20
60
100
Value
Common sites gene−LOC105280737
CphaseB2
RphaseB1

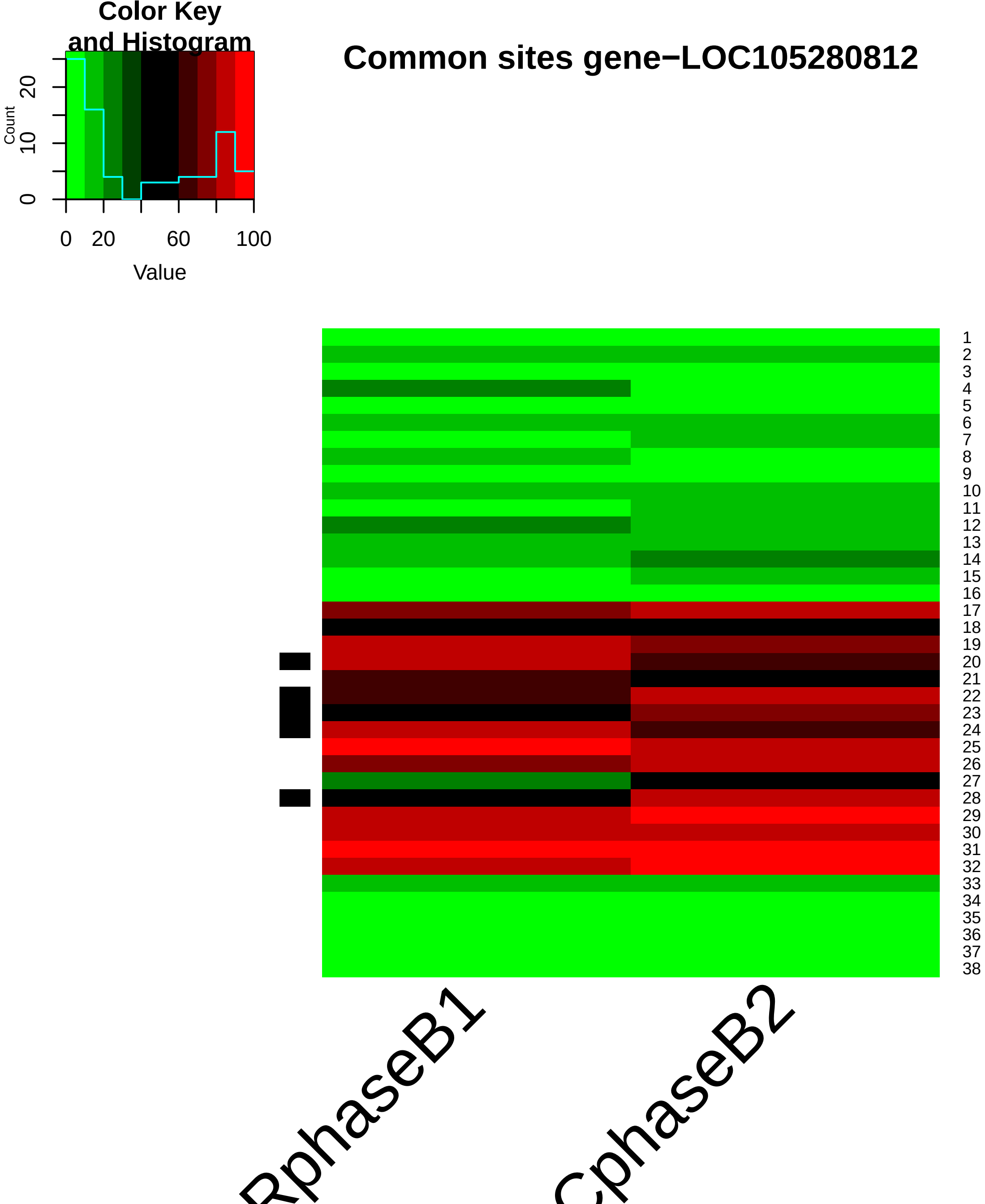
Color Key
and Histogram
Count
0
10
20
0
20
60
100
Value
Common sites gene−LOC105280812
RphaseB1
CphaseB2

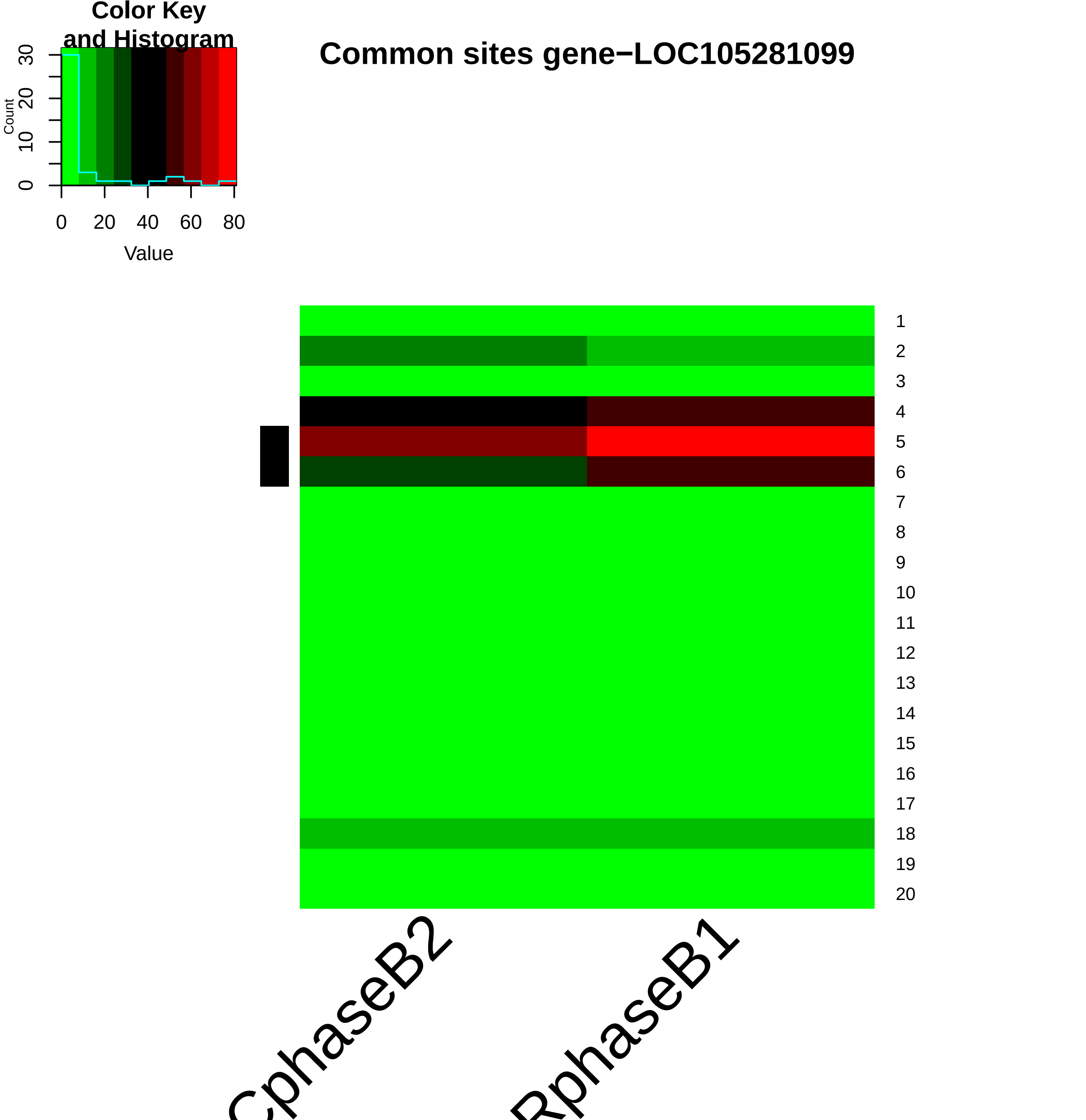
Color Key
and Histogram
Count
0
10
20
30
0
20
40
60
80
Value
Common sites gene−LOC105281099
CphaseB2
RphaseB1

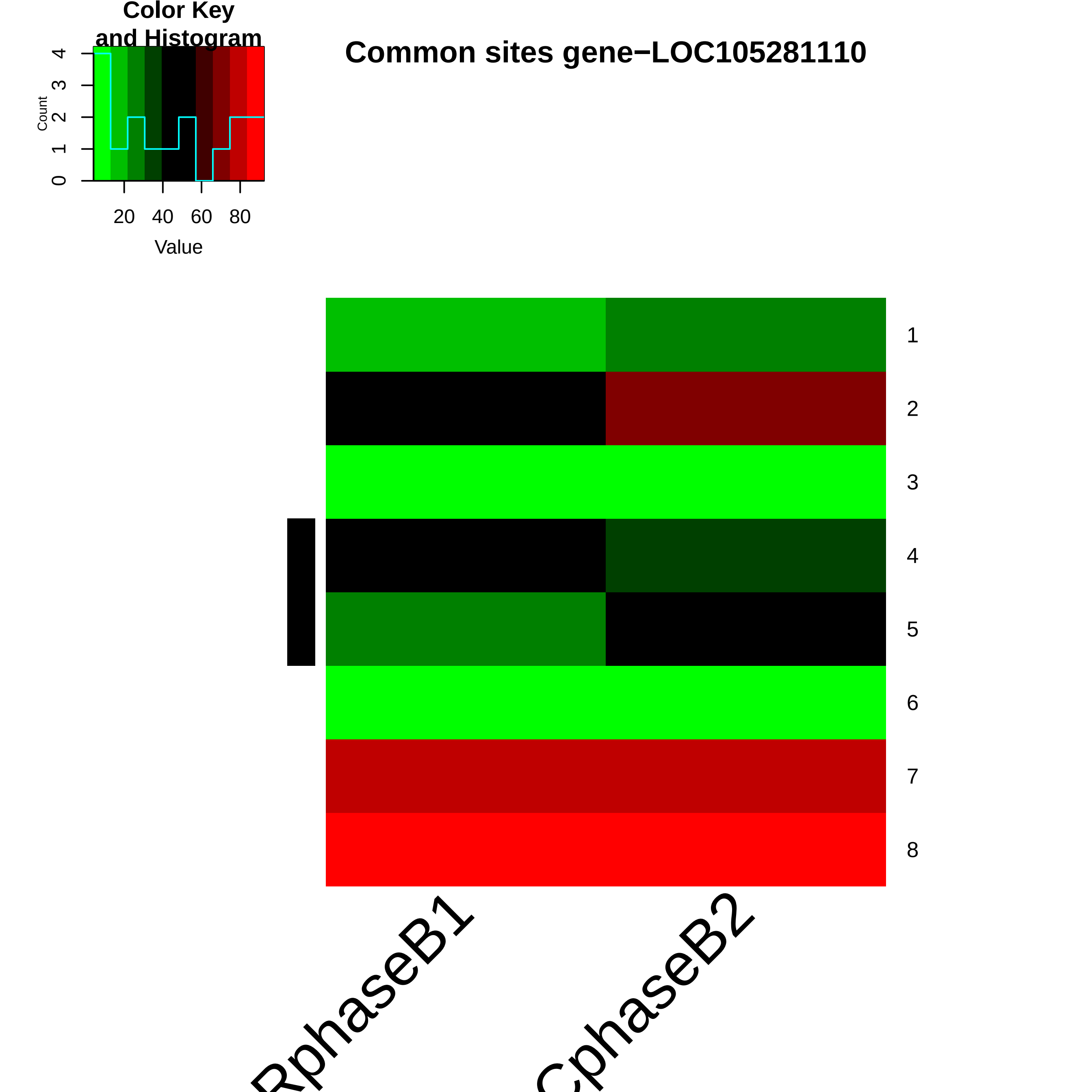

Color Key
and Histogram
Count
Value
20
40
60
80
0
1
2
3
4
Common sites gene−LOC105281110
1
2
3
4
5
6
7
8
RphaseB1
CphaseB2

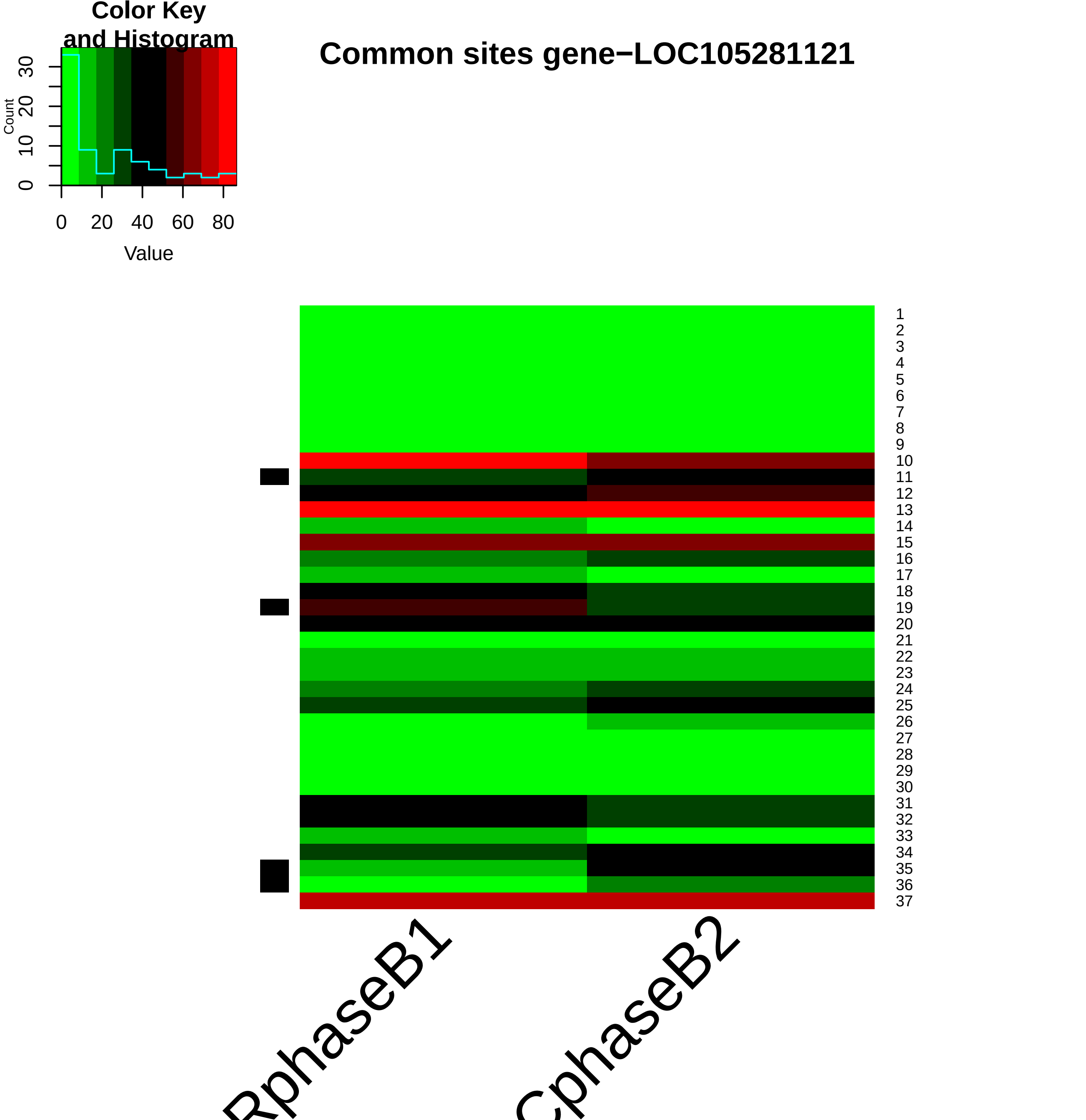
Color Key
and Histogram
Count
0
10
20
30
0
20
40
60
80
Value
Common sites gene−LOC105281121
RphaseB1
CphaseB2

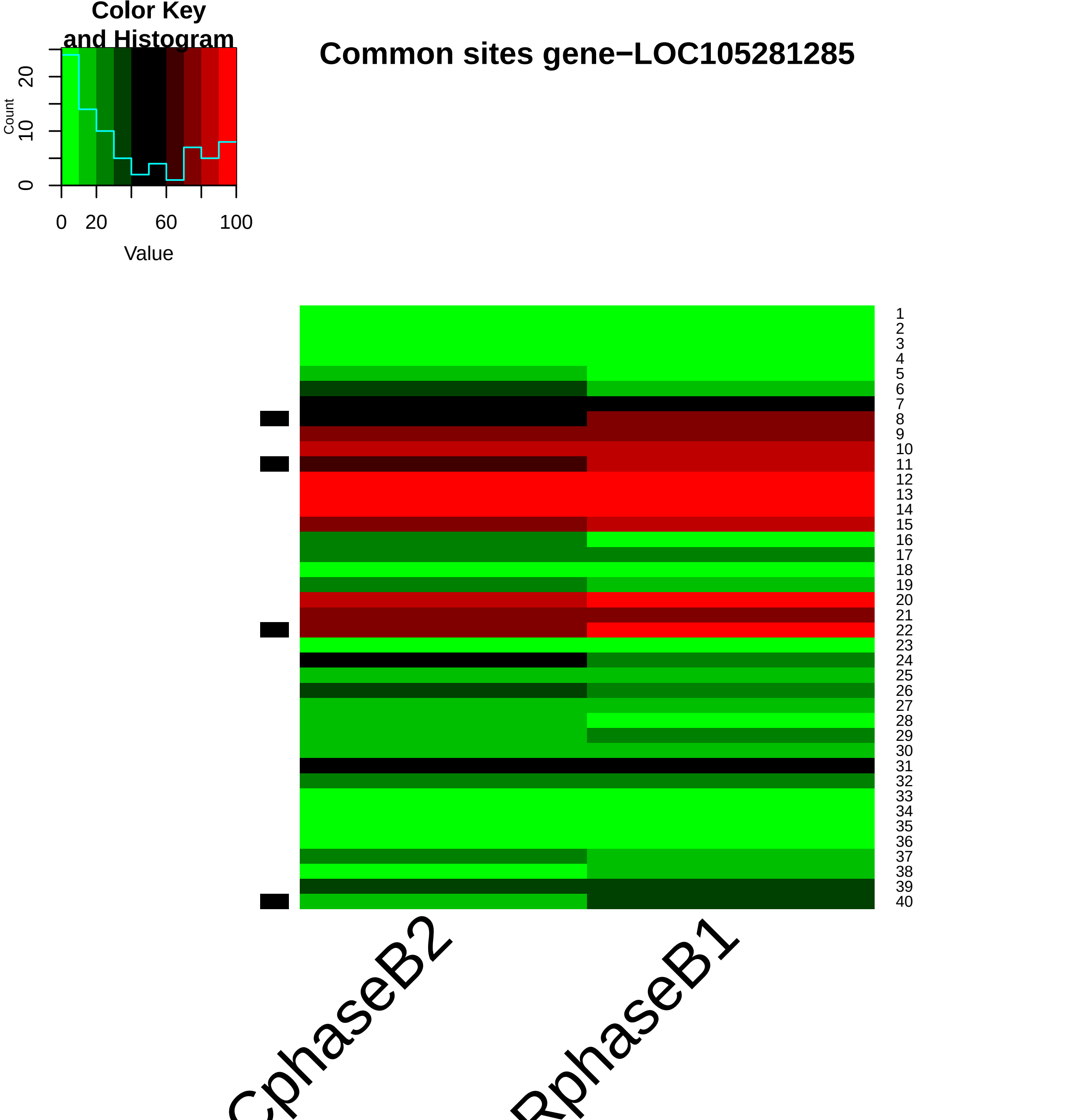
Color Key
and Histogram
Count
0
10
20
0
20
60
100
Value
Common sites gene−LOC105281285
CphaseB2
RphaseB1

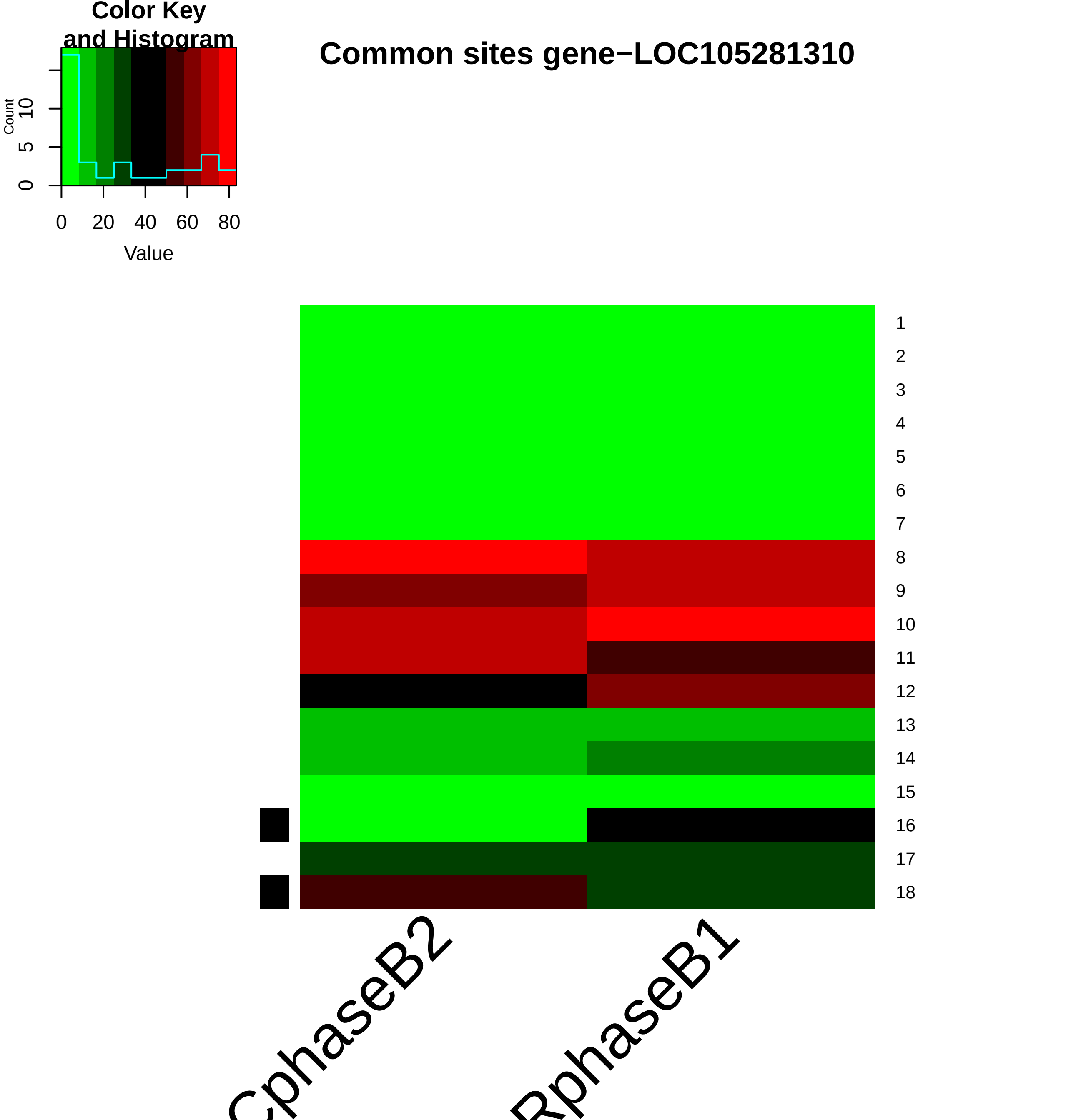
Color Key
and Histogram
Count
0
5
10
0
20
40
60
80
Value
Common sites gene−LOC105281310
CphaseB2
RphaseB1

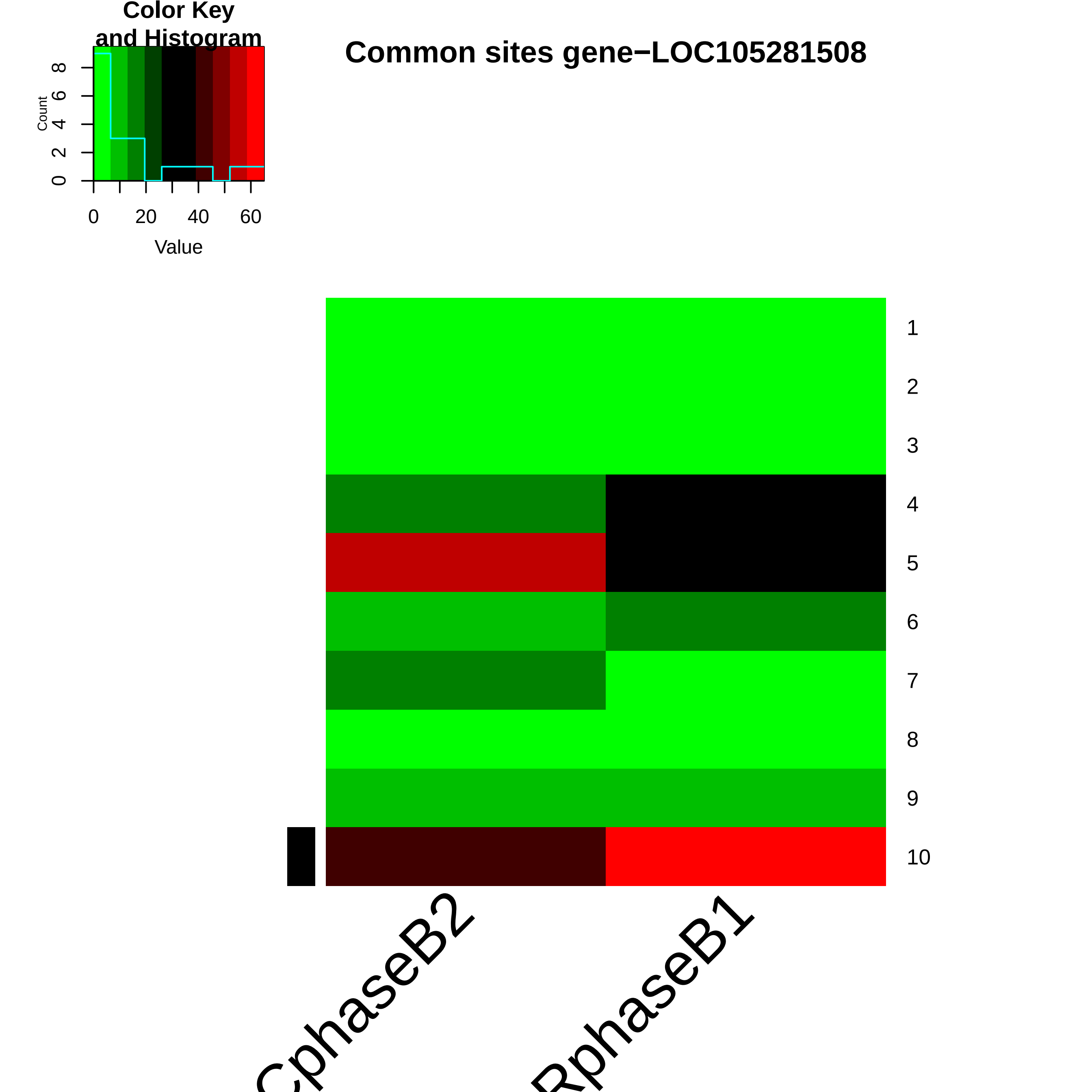

Color Key
and Histogram
Count
0 2 4 6 8
0 20 40 60
Value
Common sites gene−LOC105281508
1
2
3
4
5
6
7
8
9
10
CphaseB2
RphaseB1

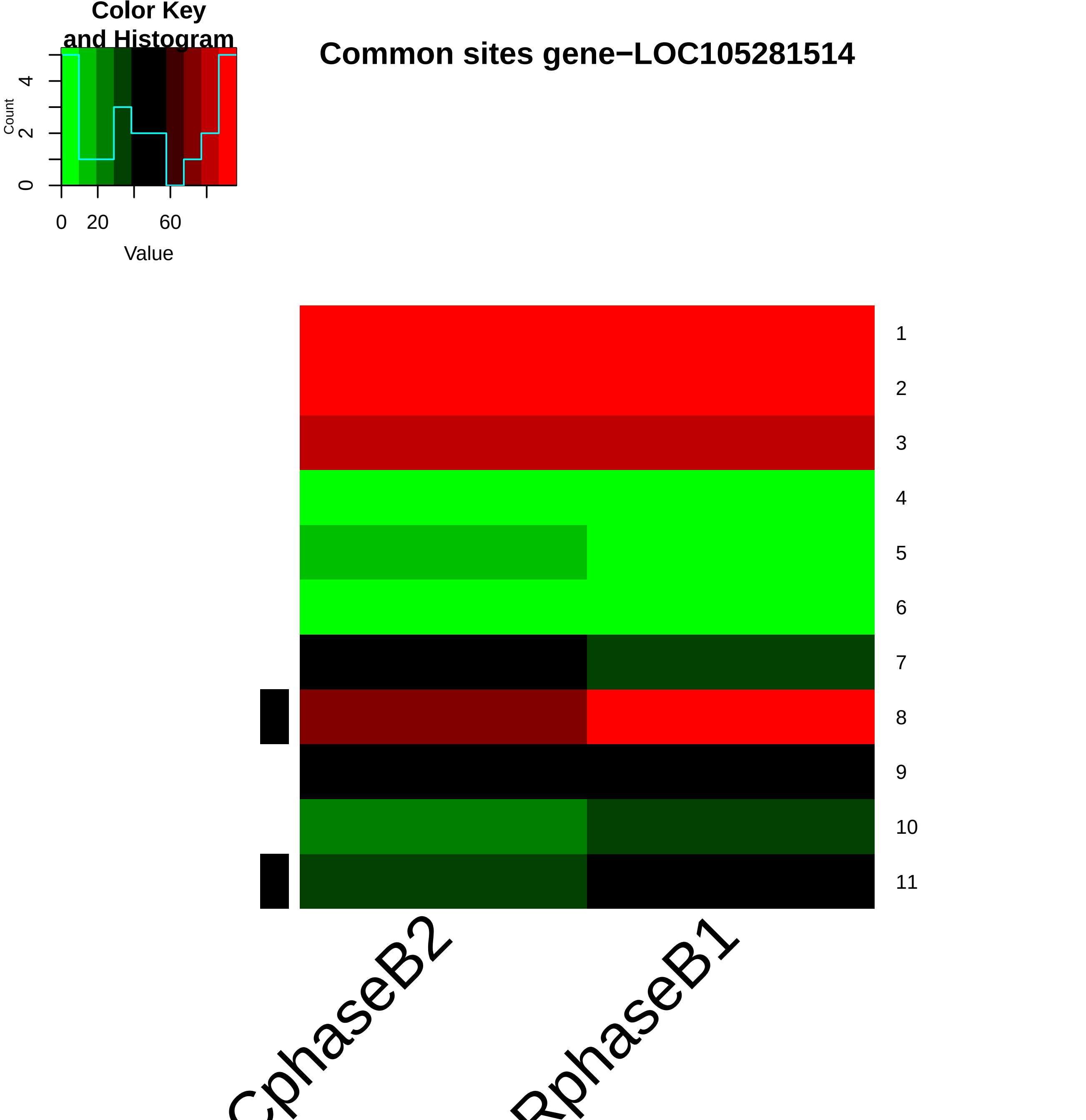

Color Key
and Histogram
Count
0
2
4
0
20
60
Value
Common sites gene−LOC105281514
1
2
3
4
5
6
7
8
9
10
11
CphaseB2
RphaseB1

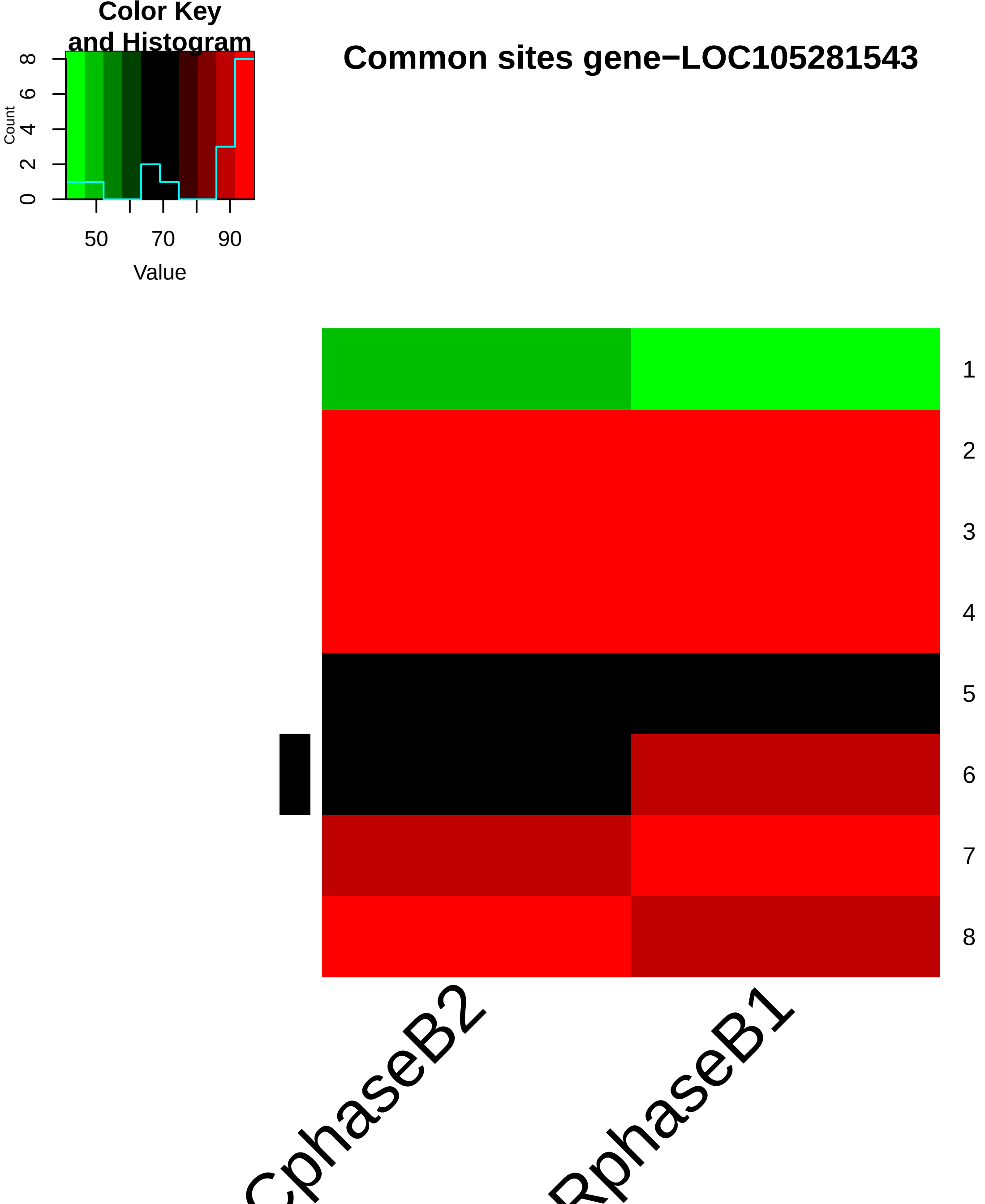
Color Key
and Histogram
Count
Value
0
2
4
6
8
50
70
90
Common sites gene−LOC105281543
1
2
3
4
5
6
7
8
CphaseB2
RphaseB1

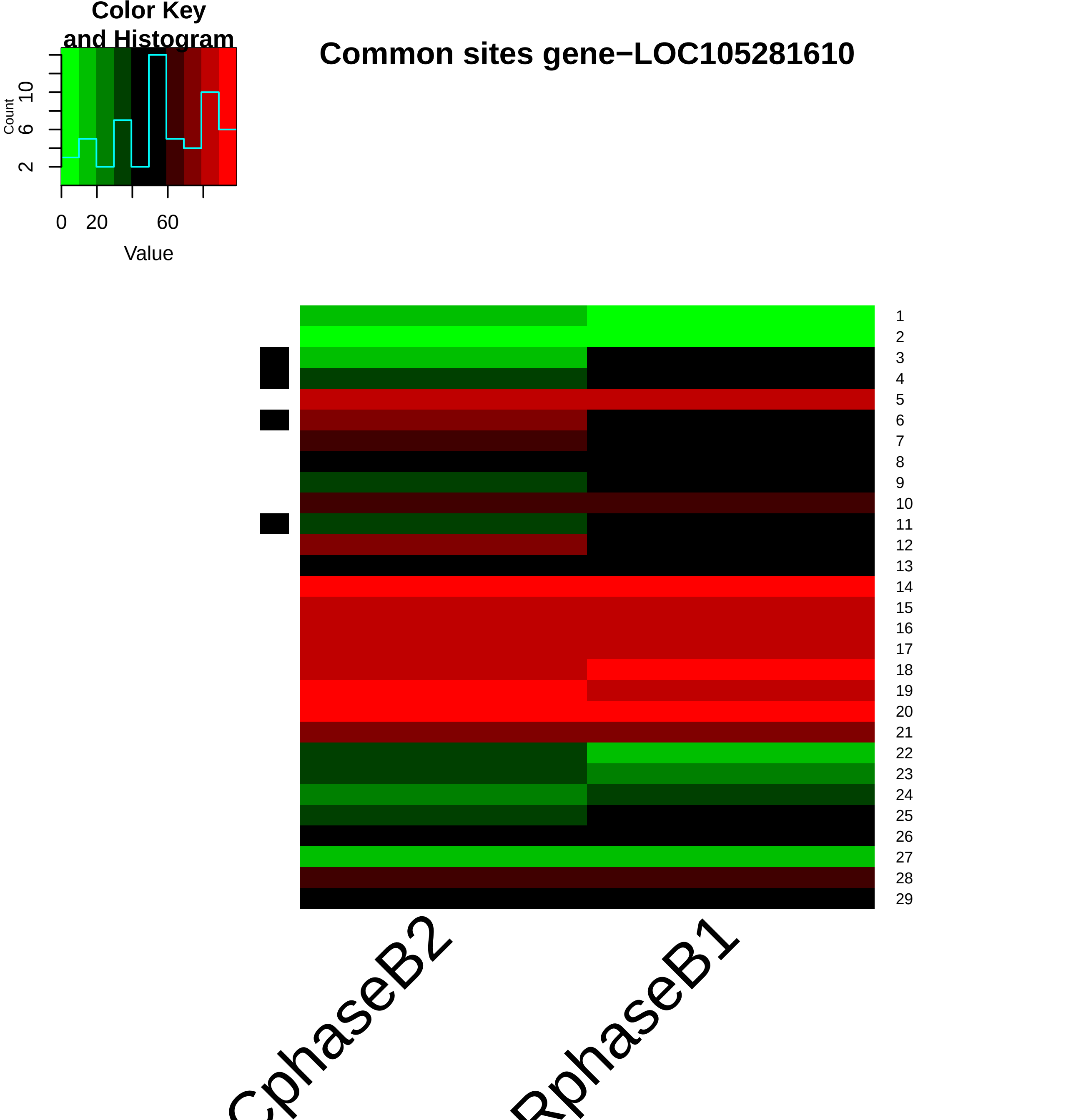

Color Key
and Histogram
Count
2
6
10
0
20
60
Value
Common sites gene−LOC105281610

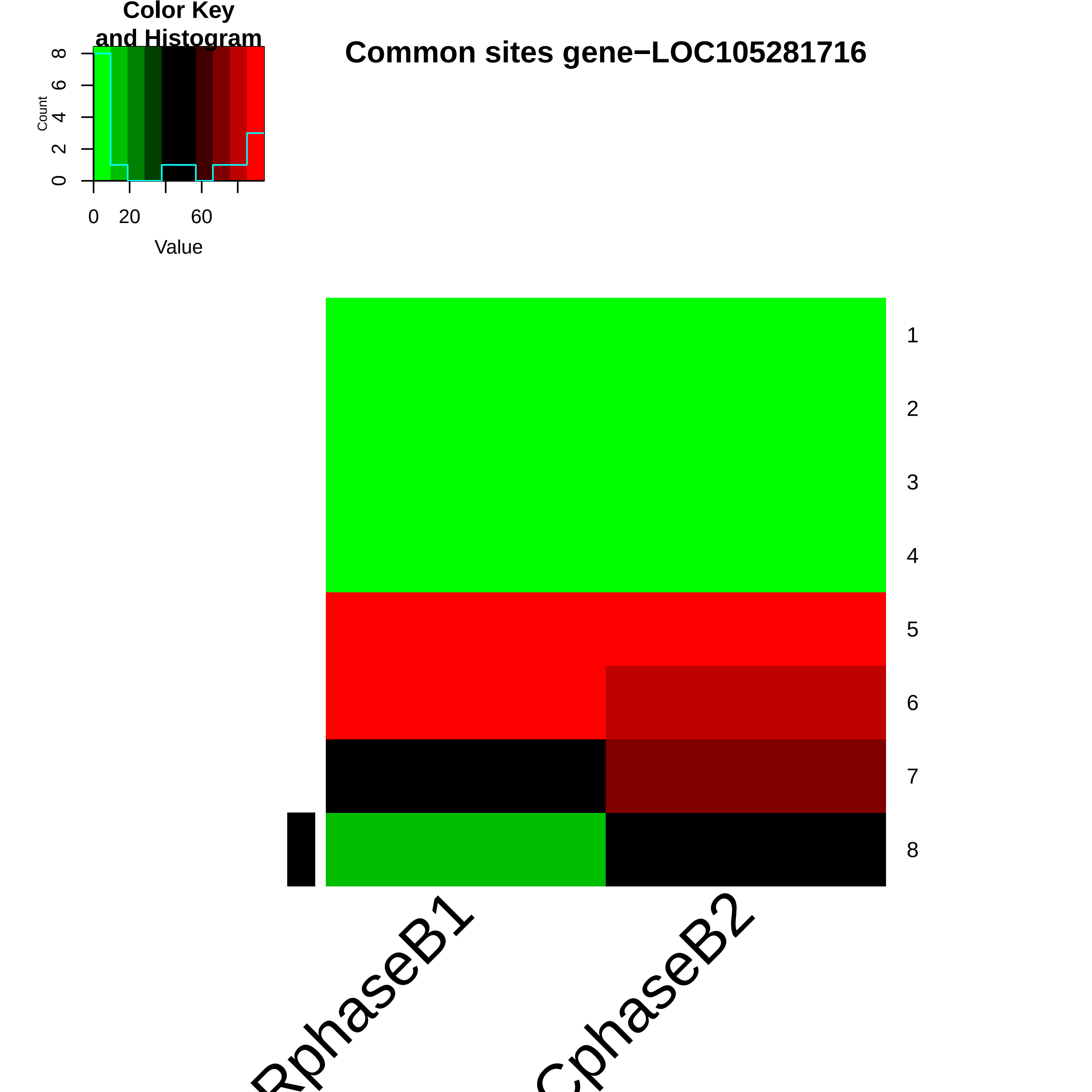
Color Key
and Histogram
Count
0
2
4
6
8
0
20
60
Value
Common sites gene−LOC105281716
1
2
3
4
5
6
7
8
RphaseB1
CphaseB2

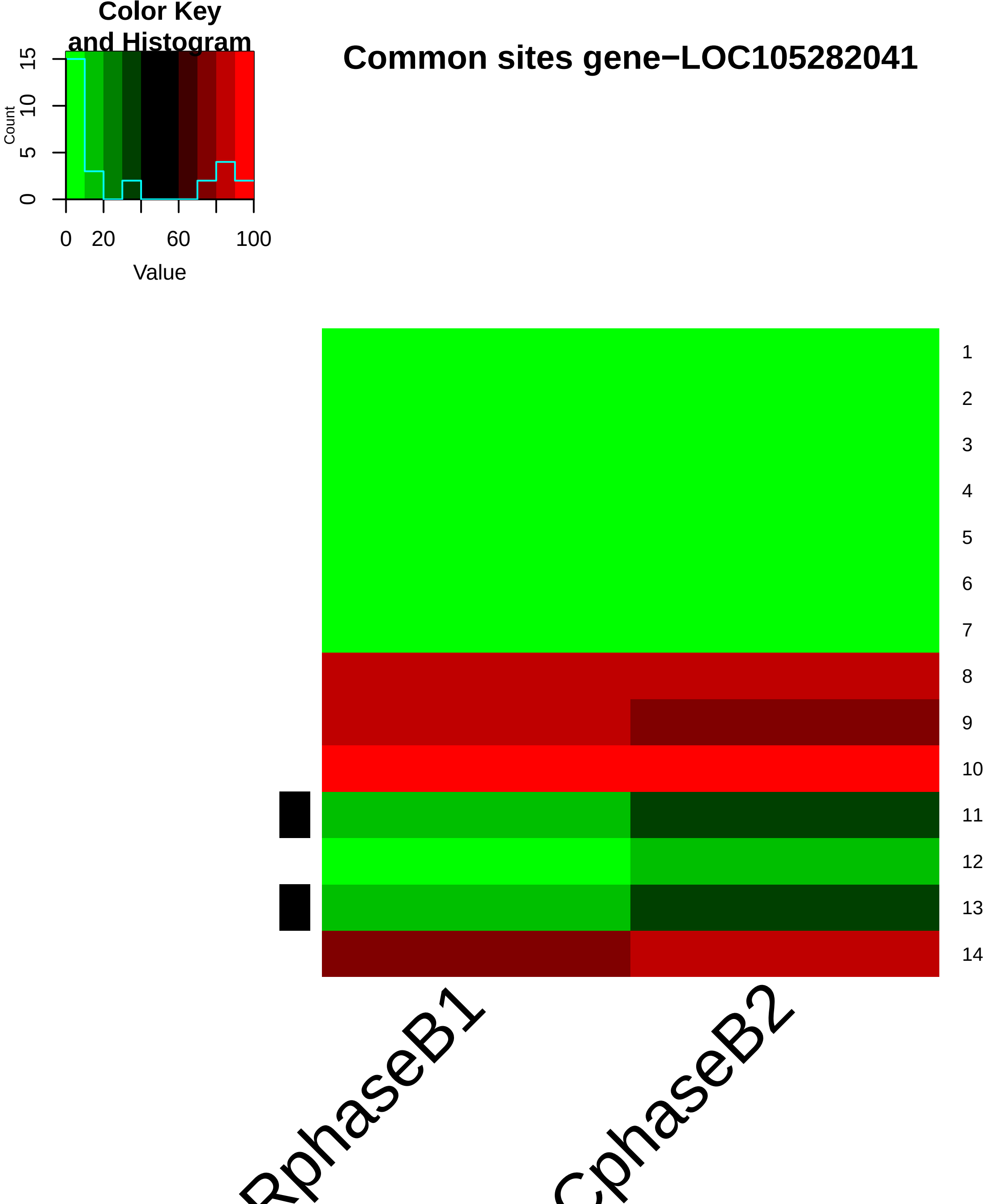

Color Key
and Histogram
Count
0
5
10
15
0
20
60
100
Value
Common sites gene−LOC105282041
1
2
3
4
5
6
7
8
9
10
11
12
13
14
RphaseB1
CphaseB2

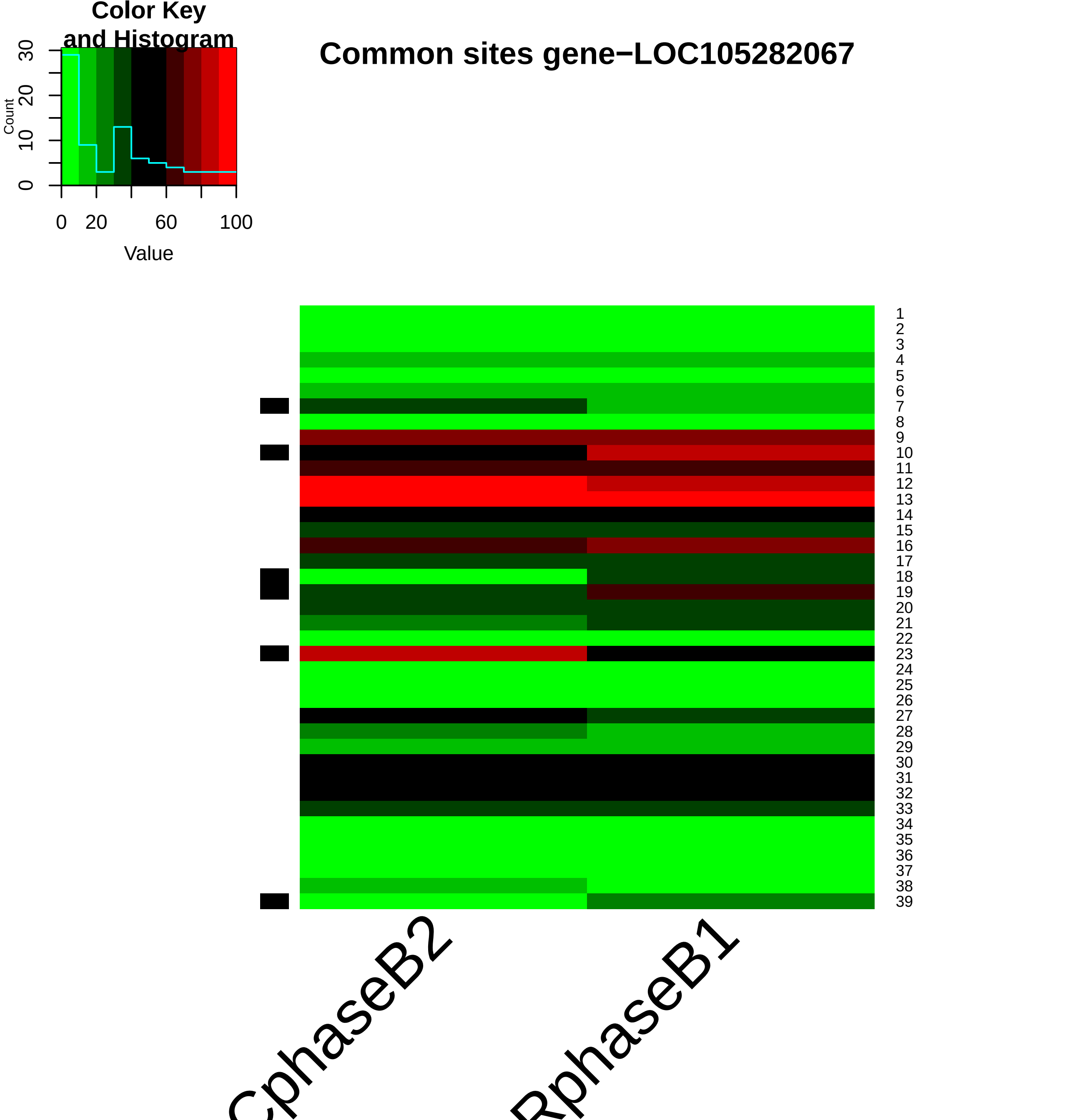
Color Key
and Histogram
Count
0
10
20
30
0
20
60
100
Value
Common sites gene−LOC105282067
1 2 3 4 5 6 7 8 9 10 11 12 13 14 15 16 17 18 19 20 21 22 23 24 25 26 27 28 29 30 31 32 33 34 35 36 37 38 39
CphaseB2
RphaseB1

### Common sites gene−LOC105282155

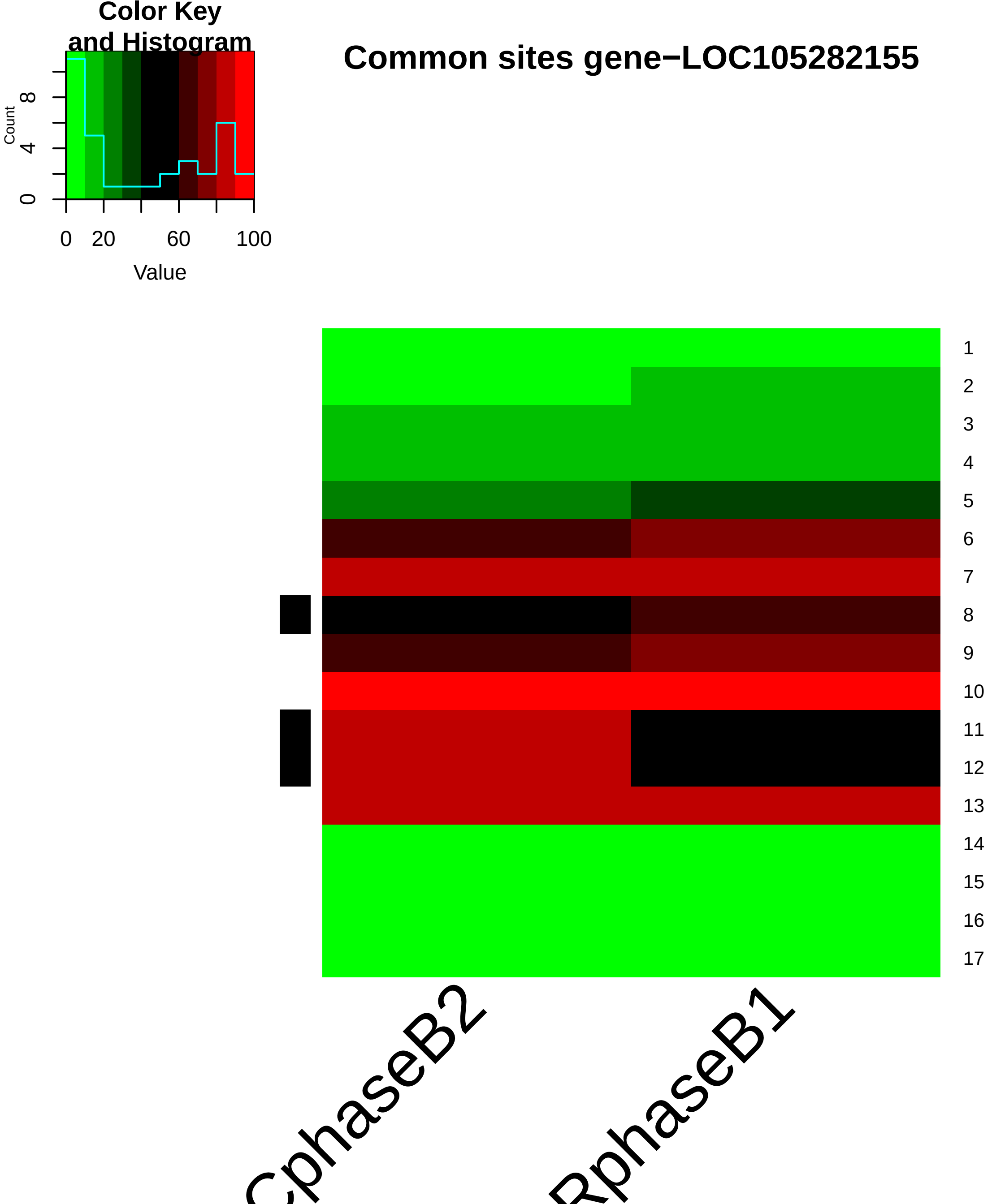

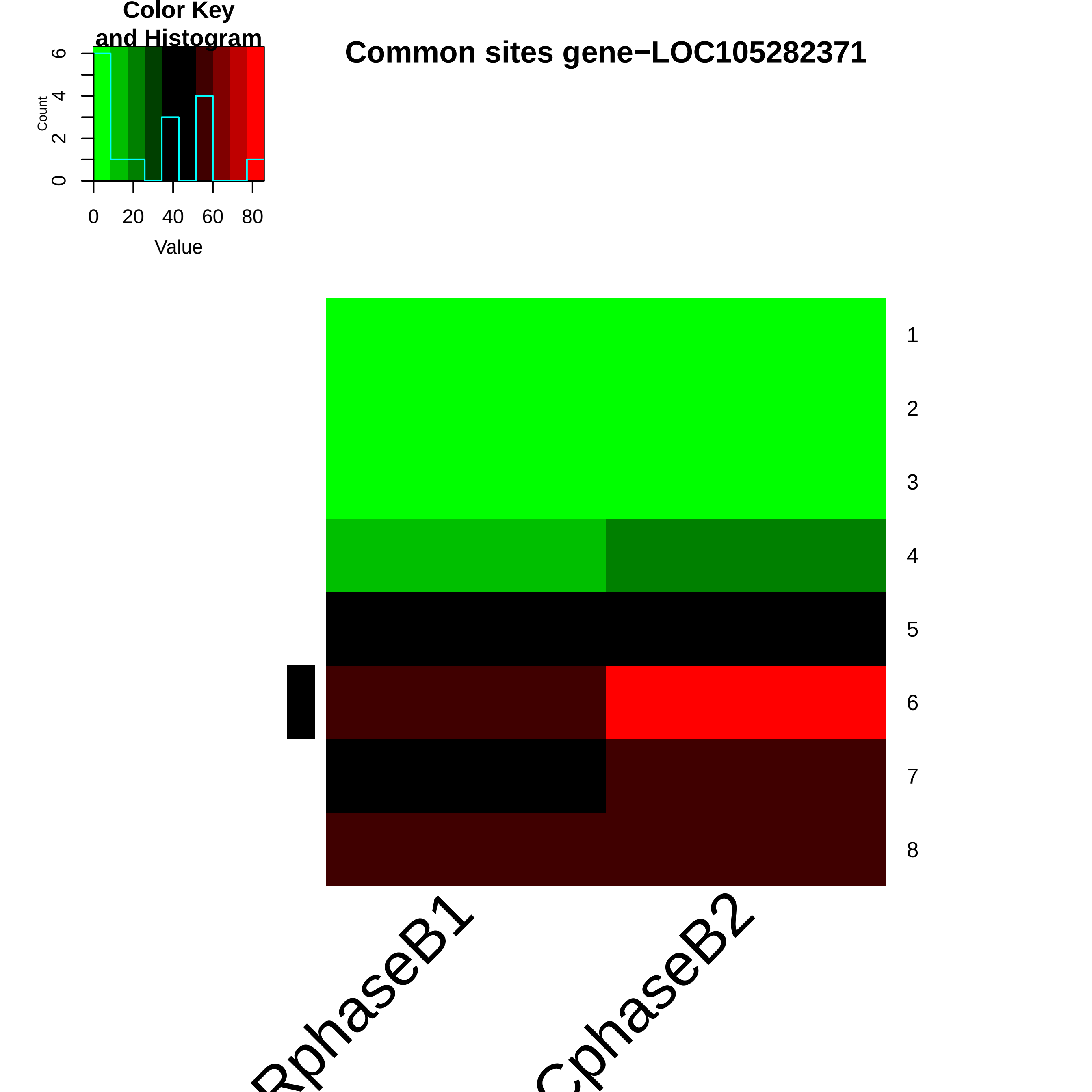
Color Key
and Histogram
Count
0
2
4
6
0
20
40
60
80
Value
Common sites gene−LOC105282371
1
2
3
4
5
6
7
8
RphaseB1
CphaseB2

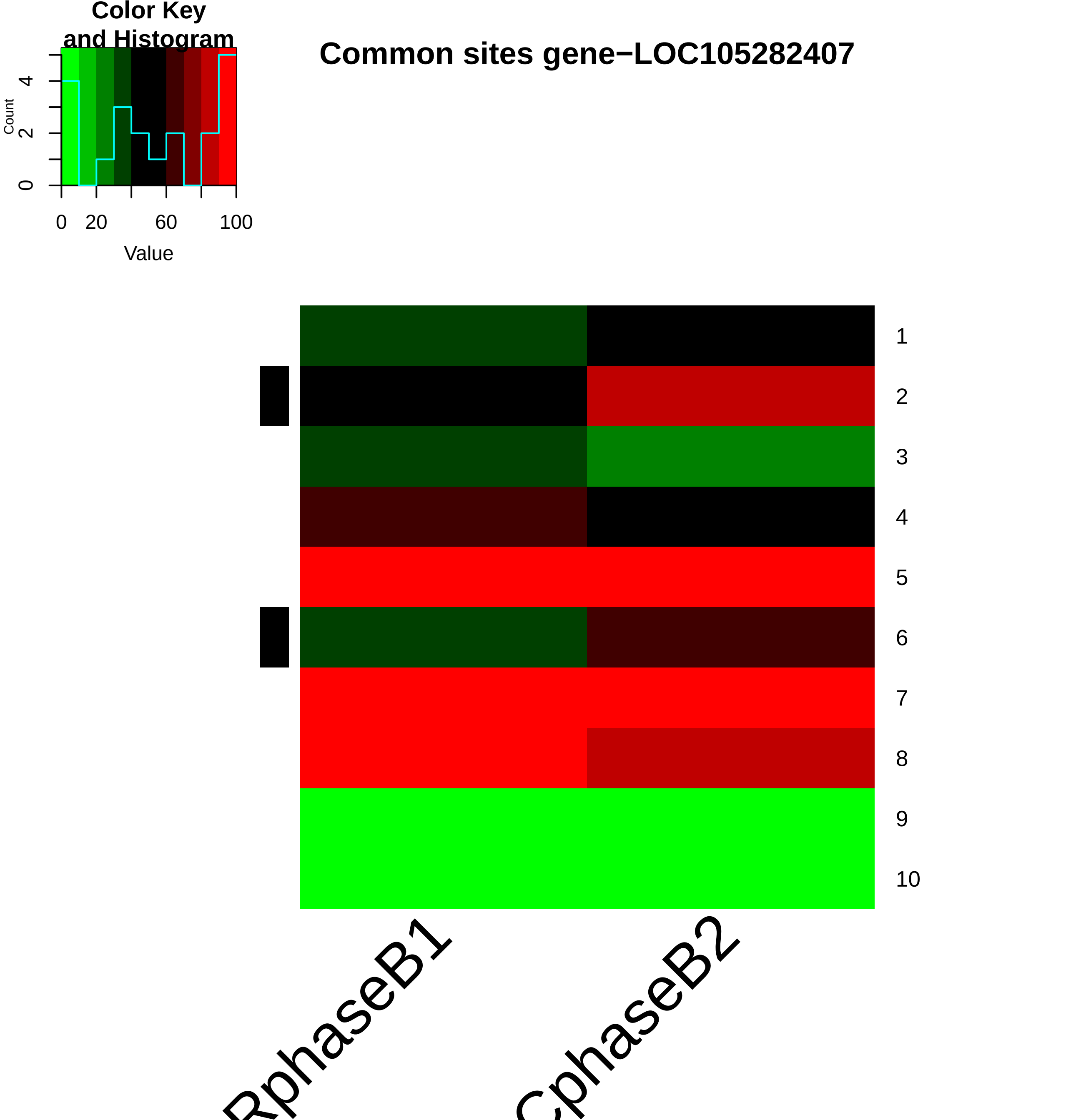

Color Key
and Histogram
Count
0
2
4
0
20
60
100
Value
Common sites gene−LOC105282407
1
2
3
4
5
6
7
8
9
10
RphaseB1
CphaseB2

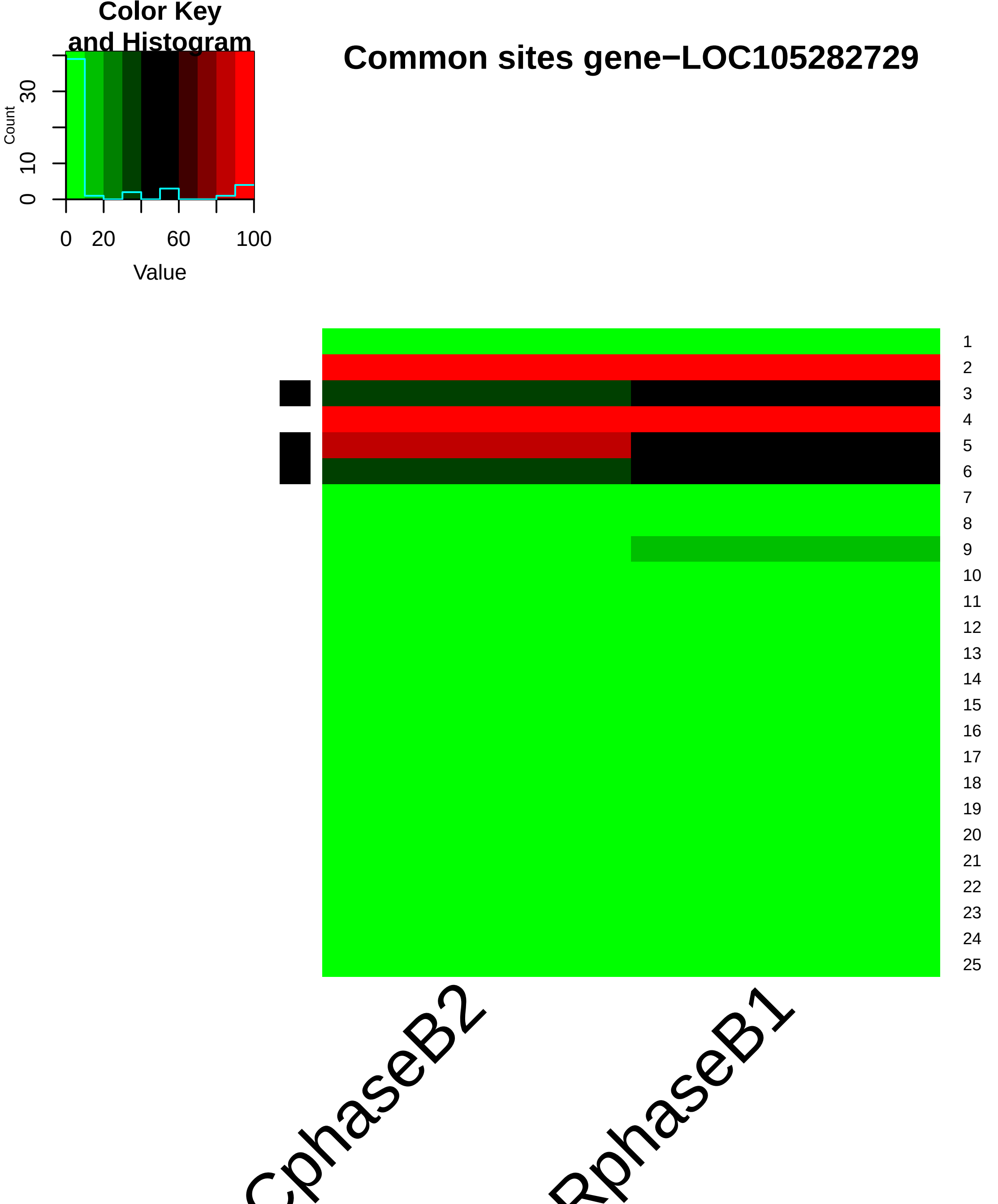

Color Key and Histogram
Count
0
10
30
0
20
60
100
Value
Common sites gene−LOC105282729
CphaseB2
RphaseB1

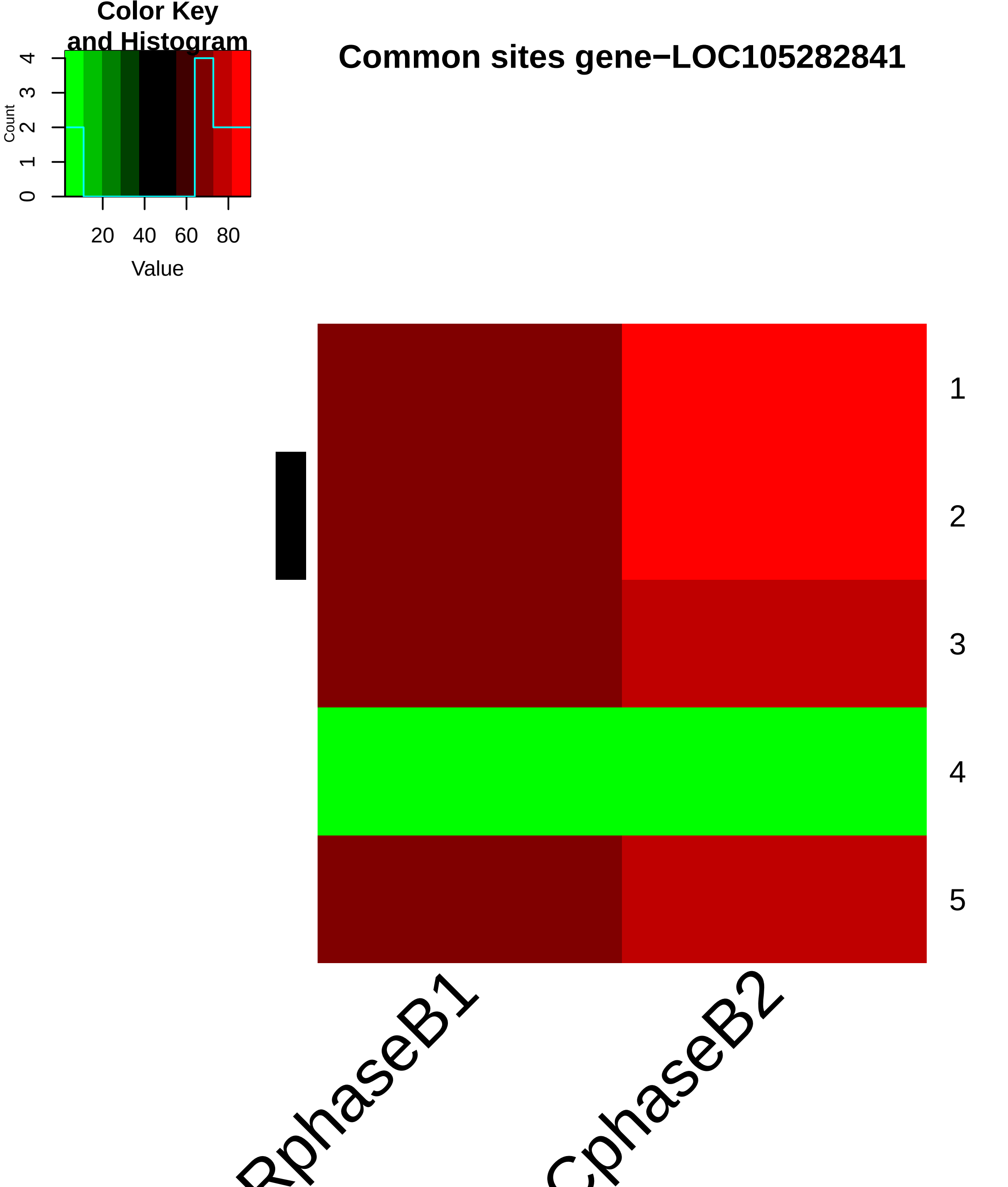

Color Key
and Histogram
Count
0
1
2
3
4
20
40
60
80
Value
Common sites gene−LOC105282841
1
2
3
4
5
RphaseB1
CphaseB2

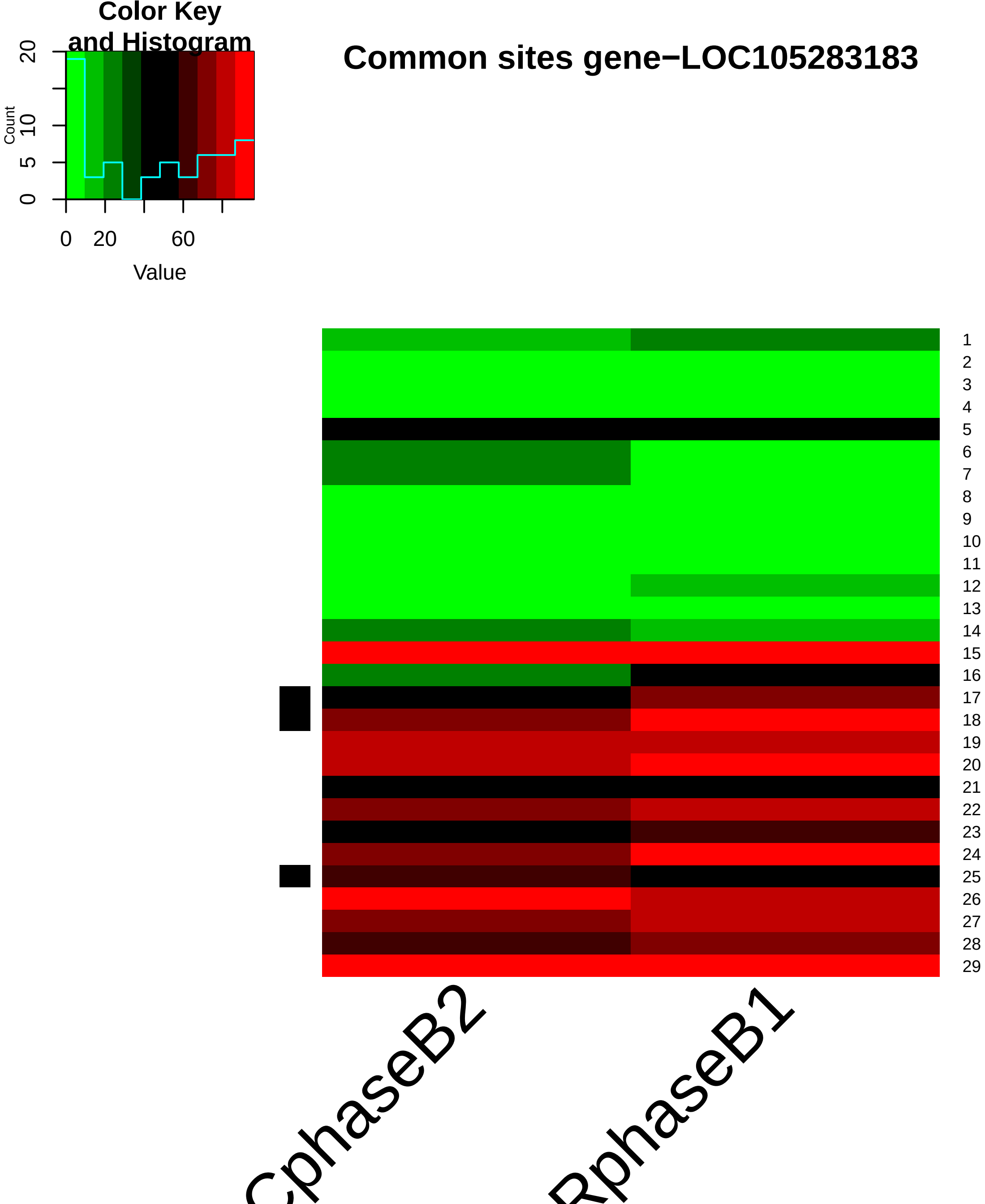
Color Key
and Histogram
Count
0
5
10
20
0
20
60
Value
Common sites gene−LOC105283183
CphaseB2
RphaseB1

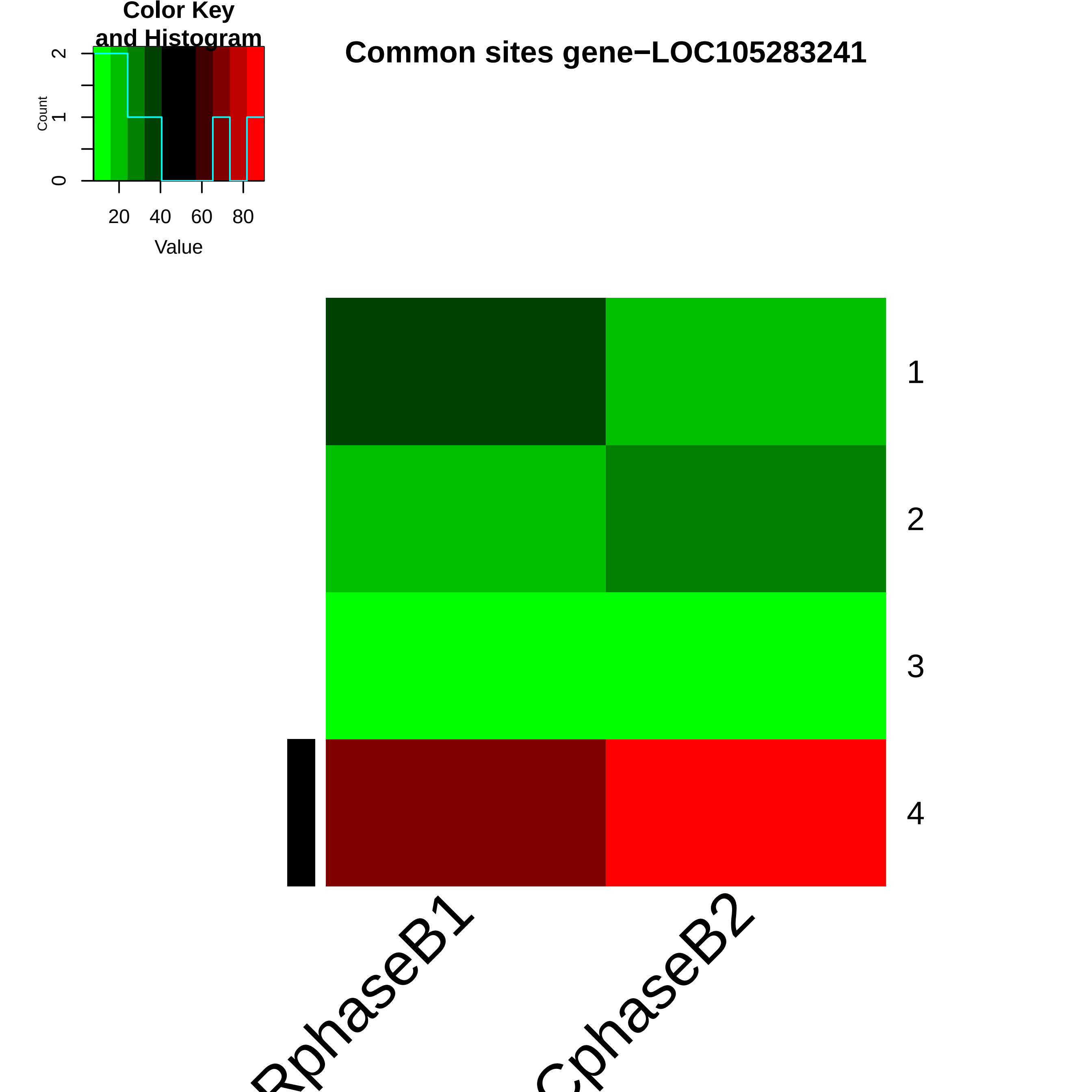

Color Key
and Histogram
Count
0
1
2
20
40
60
80
Value
Common sites gene−LOC105283241
1
2
3
4
RphaseB1
CphaseB2

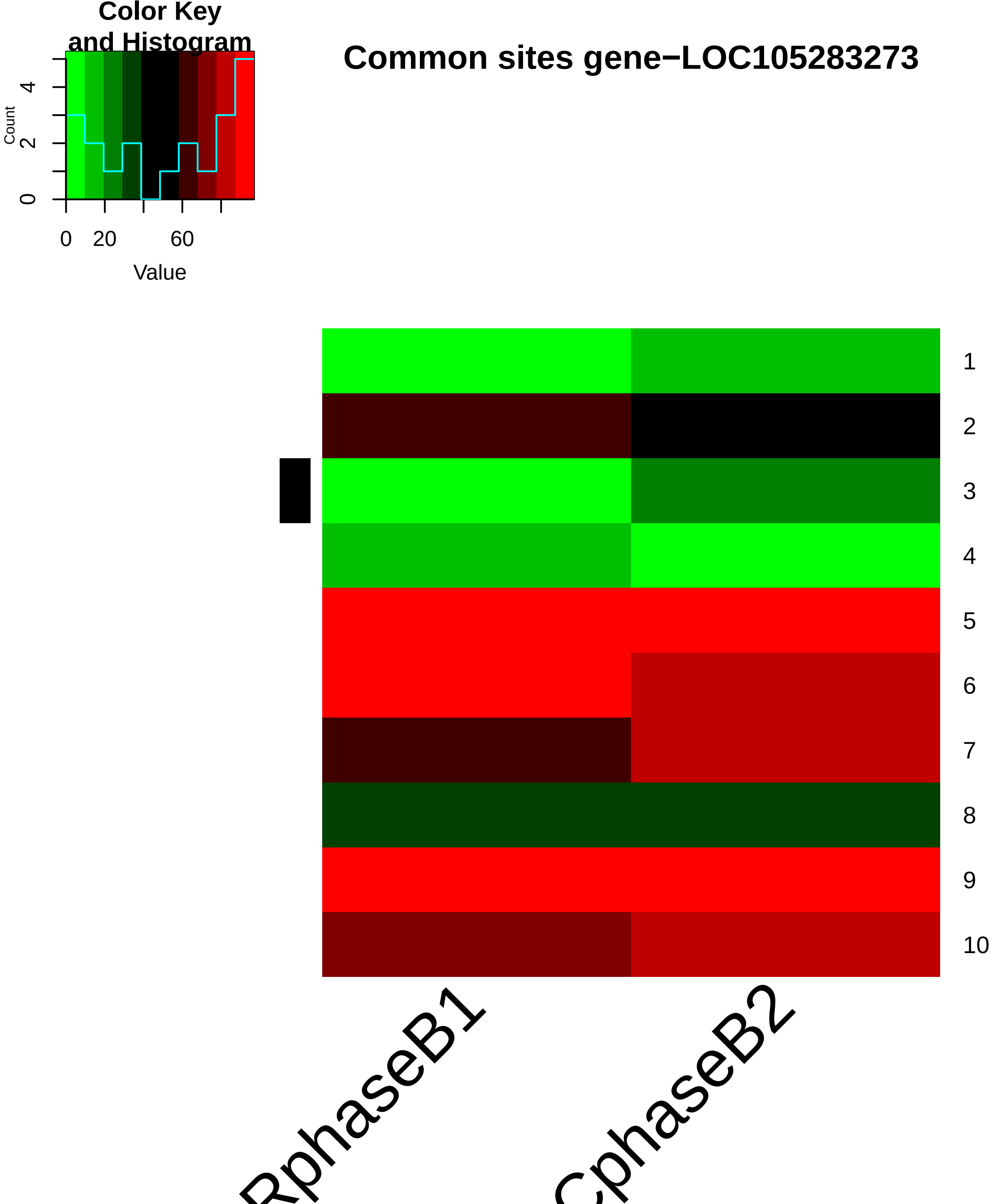
Color Key
and Histogram
Count
0
2
4
0
20
60
Value
Common sites gene−LOC105283273
1
2
3
4
5
6
7
8
9
10
RphaseB1
CphaseB2

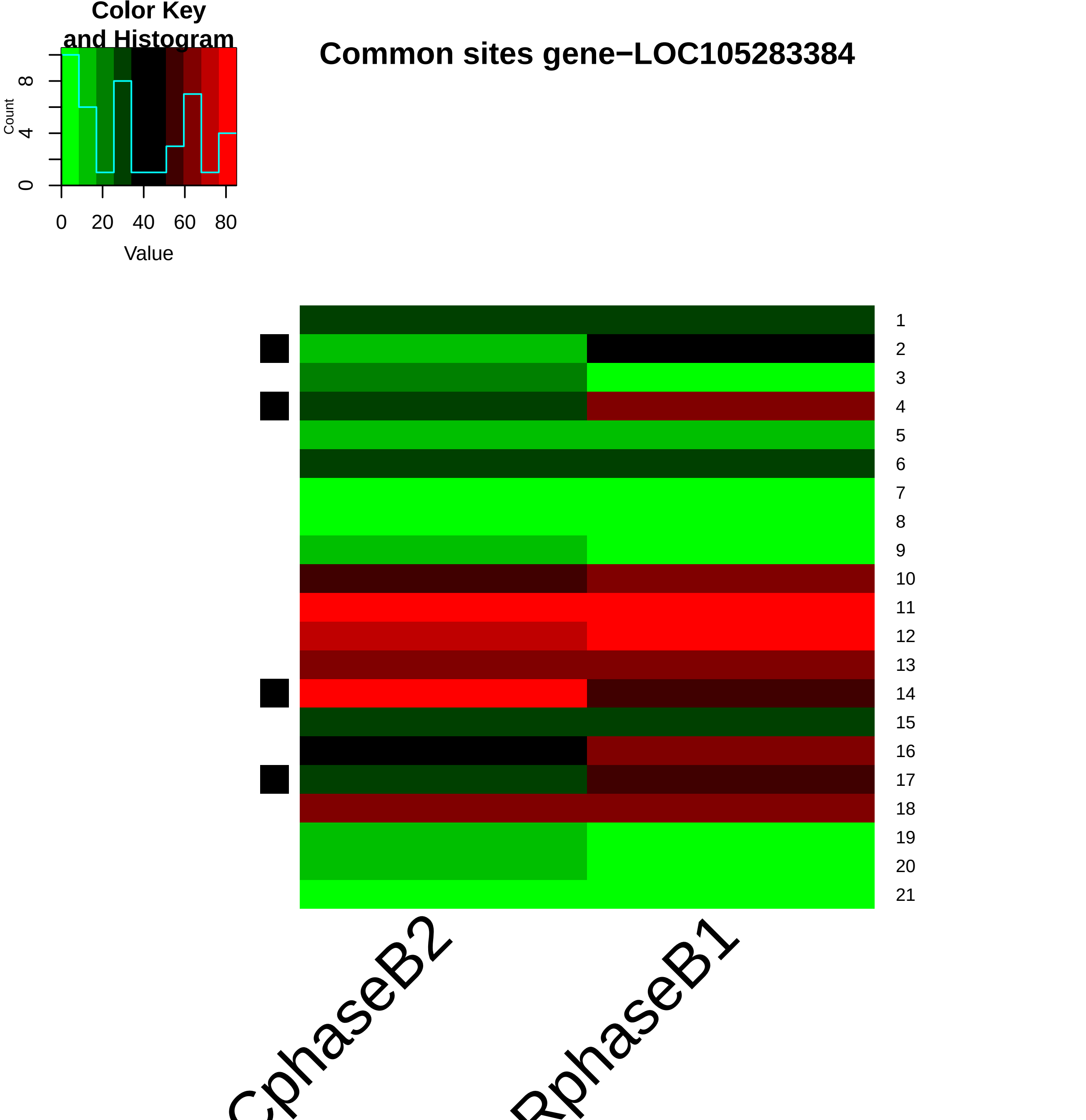
Color Key
and Histogram
Count
0
4
8
0
20
40
60
80
Value
Common sites gene−LOC105283384
CphaseB2
RphaseB1

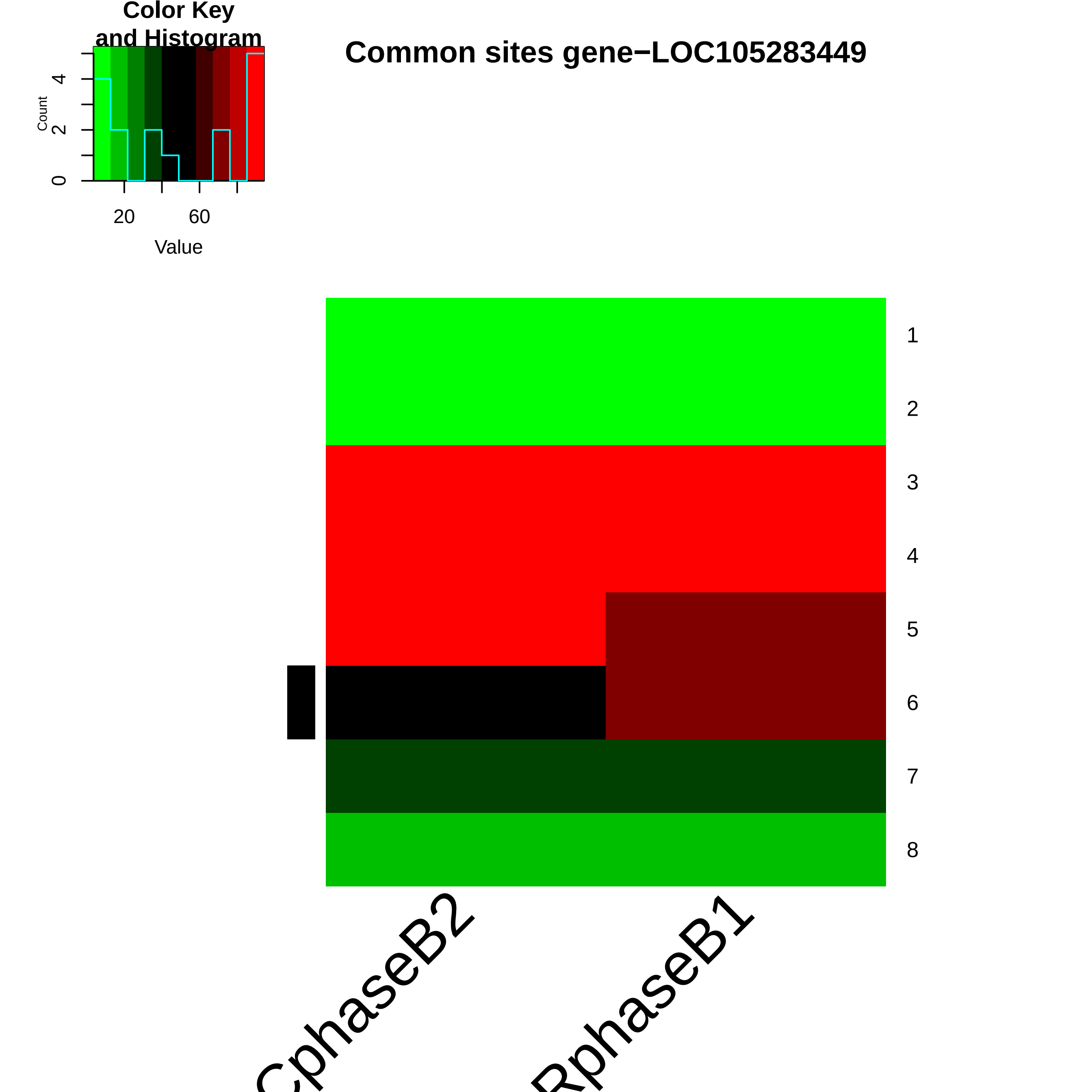
Color Key
and Histogram
Count
0
2
4
20
60
Value
Common sites gene−LOC105283449
1
2
3
4
5
6
7
8
CphaseB2
RphaseB1

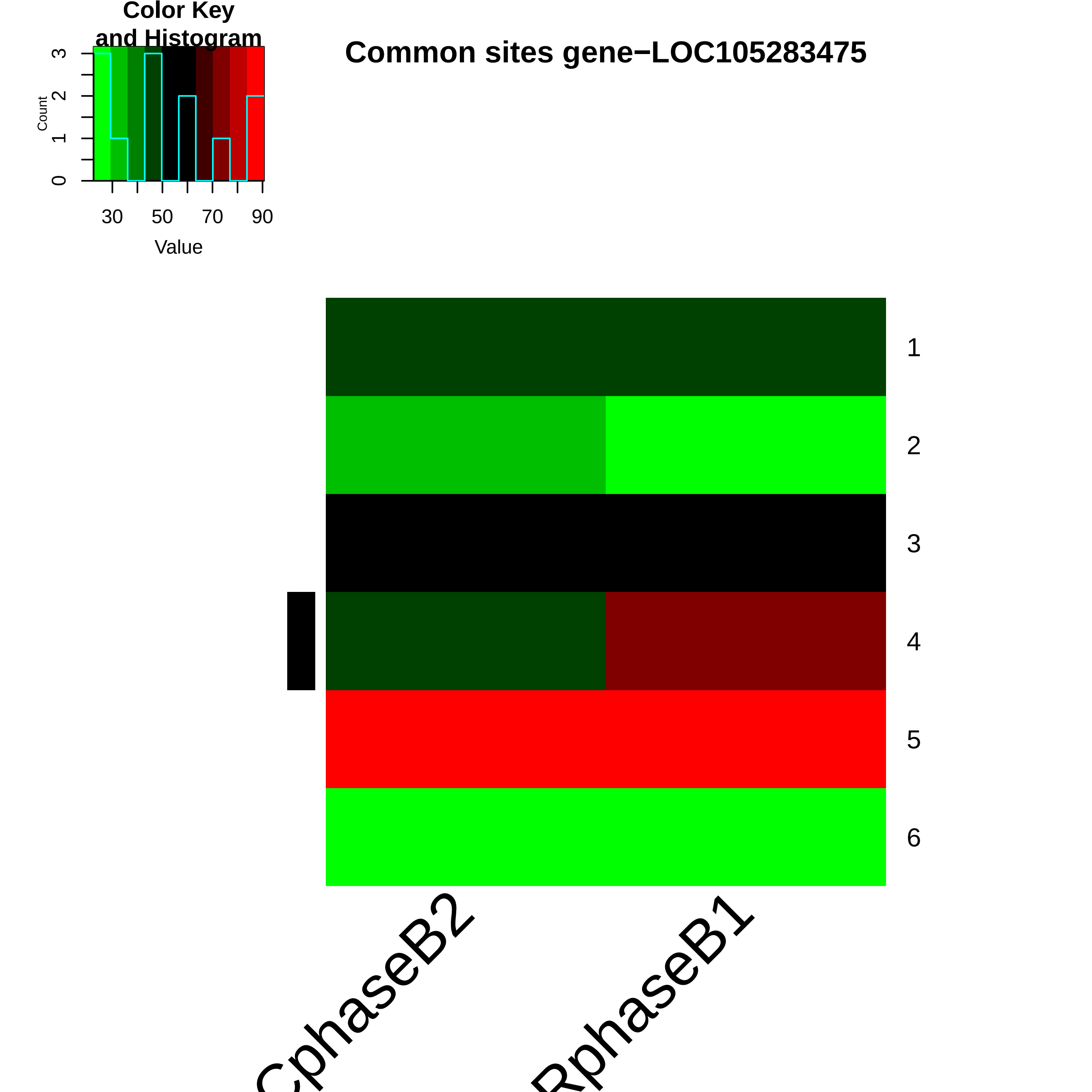
Color Key
and Histogram
Count
0
1
2
3
30
50
70
90
Value
Common sites gene−LOC105283475
1
2
3
4
5
6
CphaseB2
RphaseB1

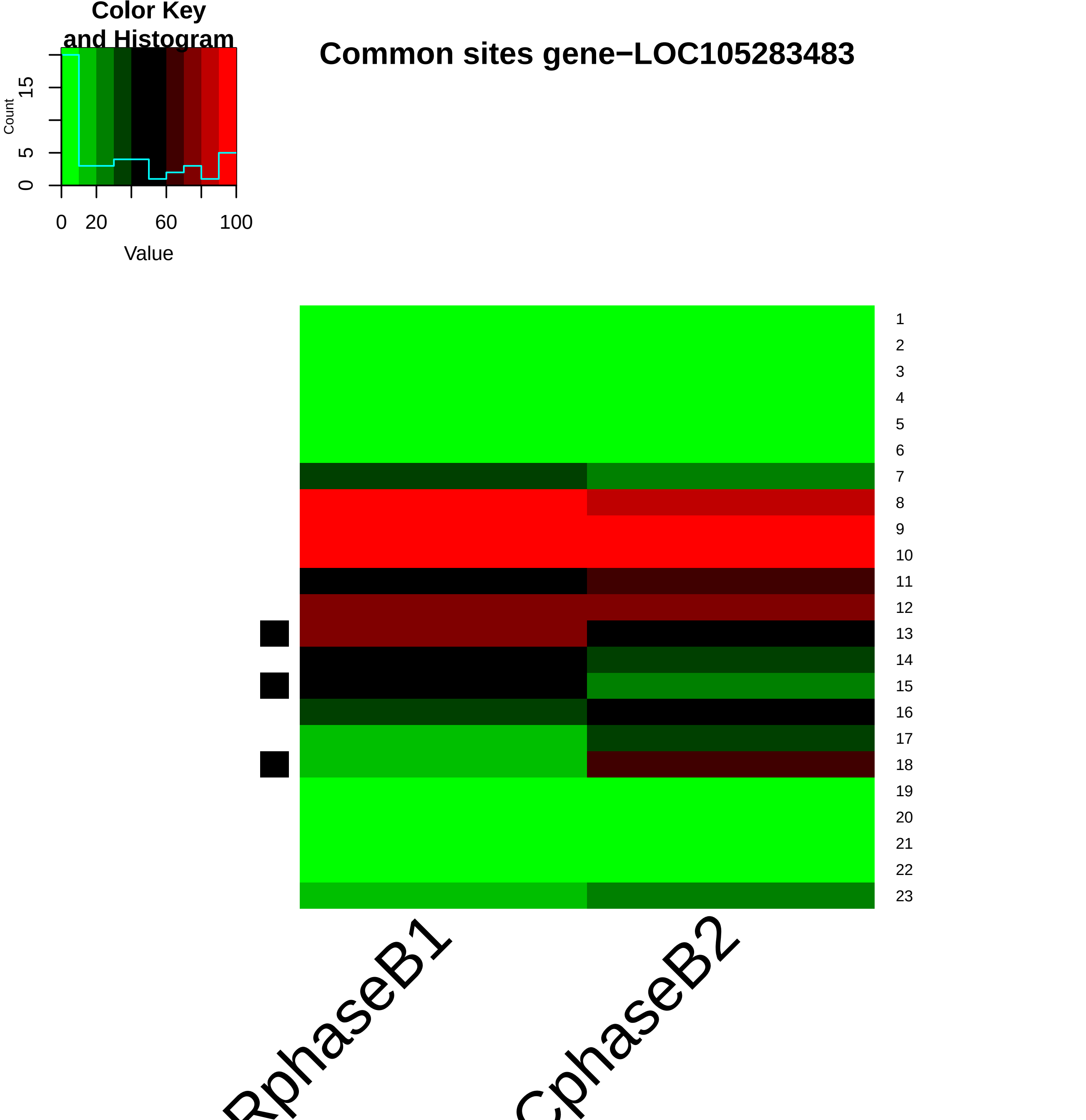
Common sites gene−LOC105283483
Color Key
and Histogram
Count
0
5
15
0
20
60
100
Value
RphaseB1
CphaseB2

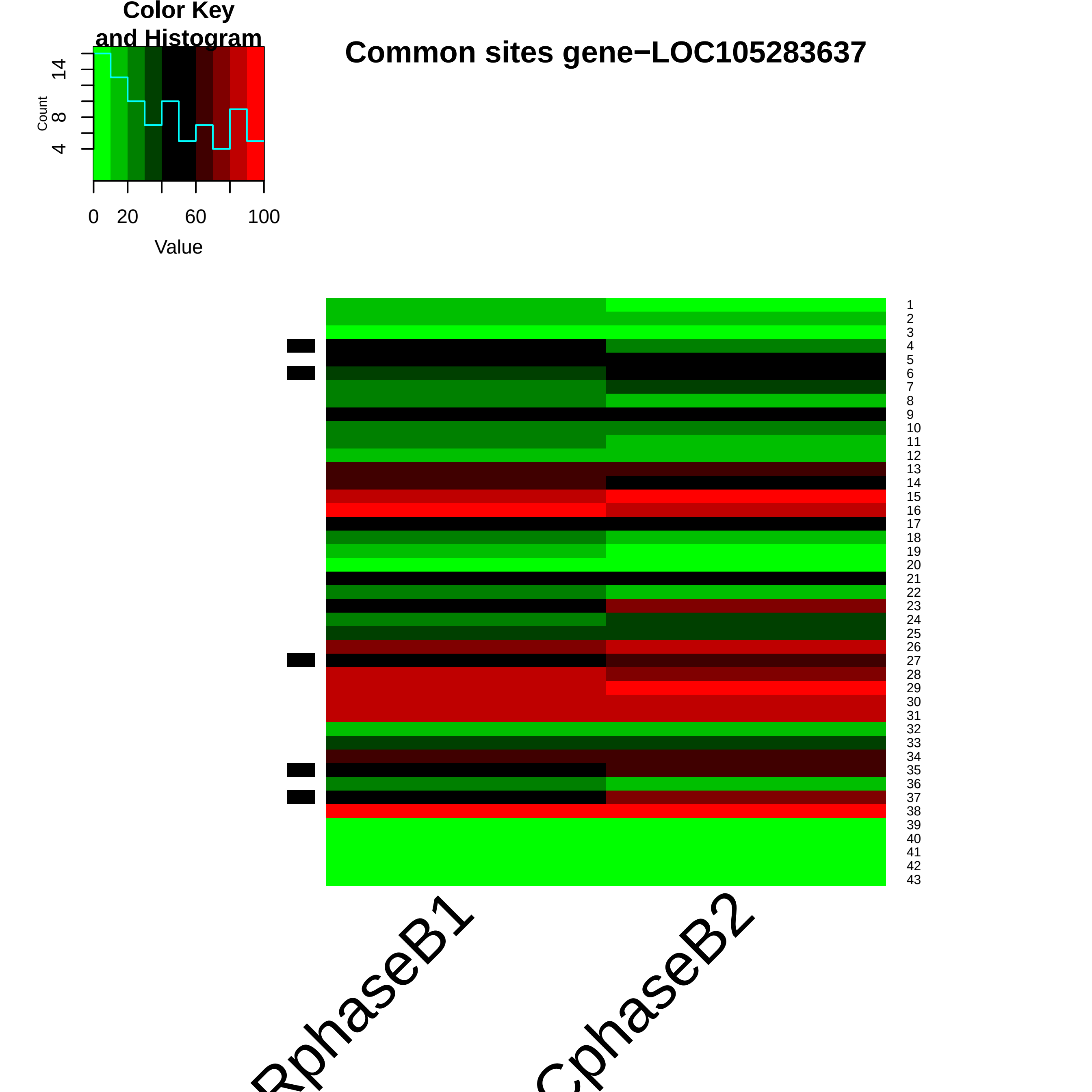
Color Key
and Histogram
Count
4
8
14
0
20
60
100
Value
Common sites gene−LOC105283637
RphaseB1
CphaseB2

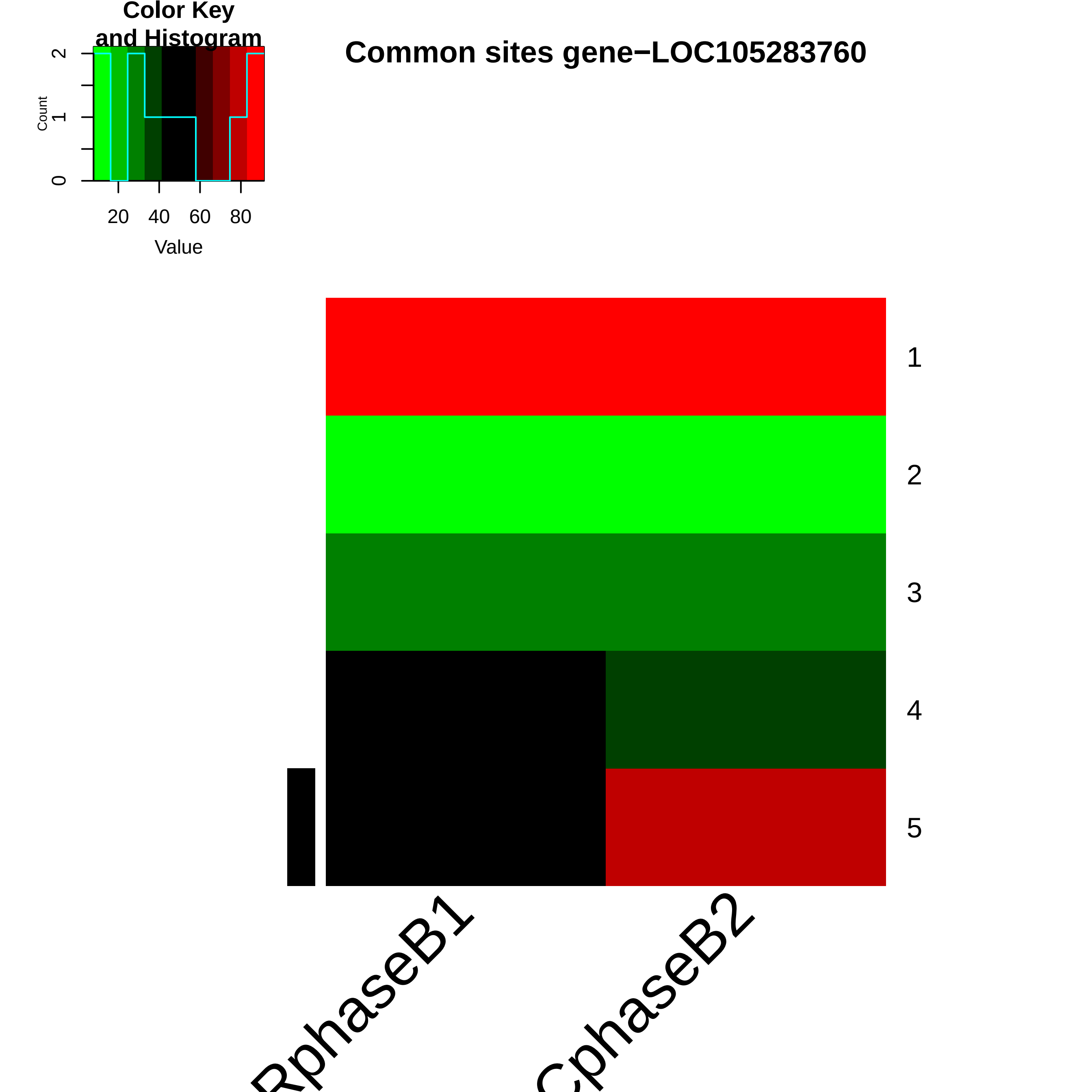
Color Key
and Histogram
Count
0
1
2
20
40
60
80
Value
Common sites gene−LOC105283760
1
2
3
4
5
RphaseB1
CphaseB2

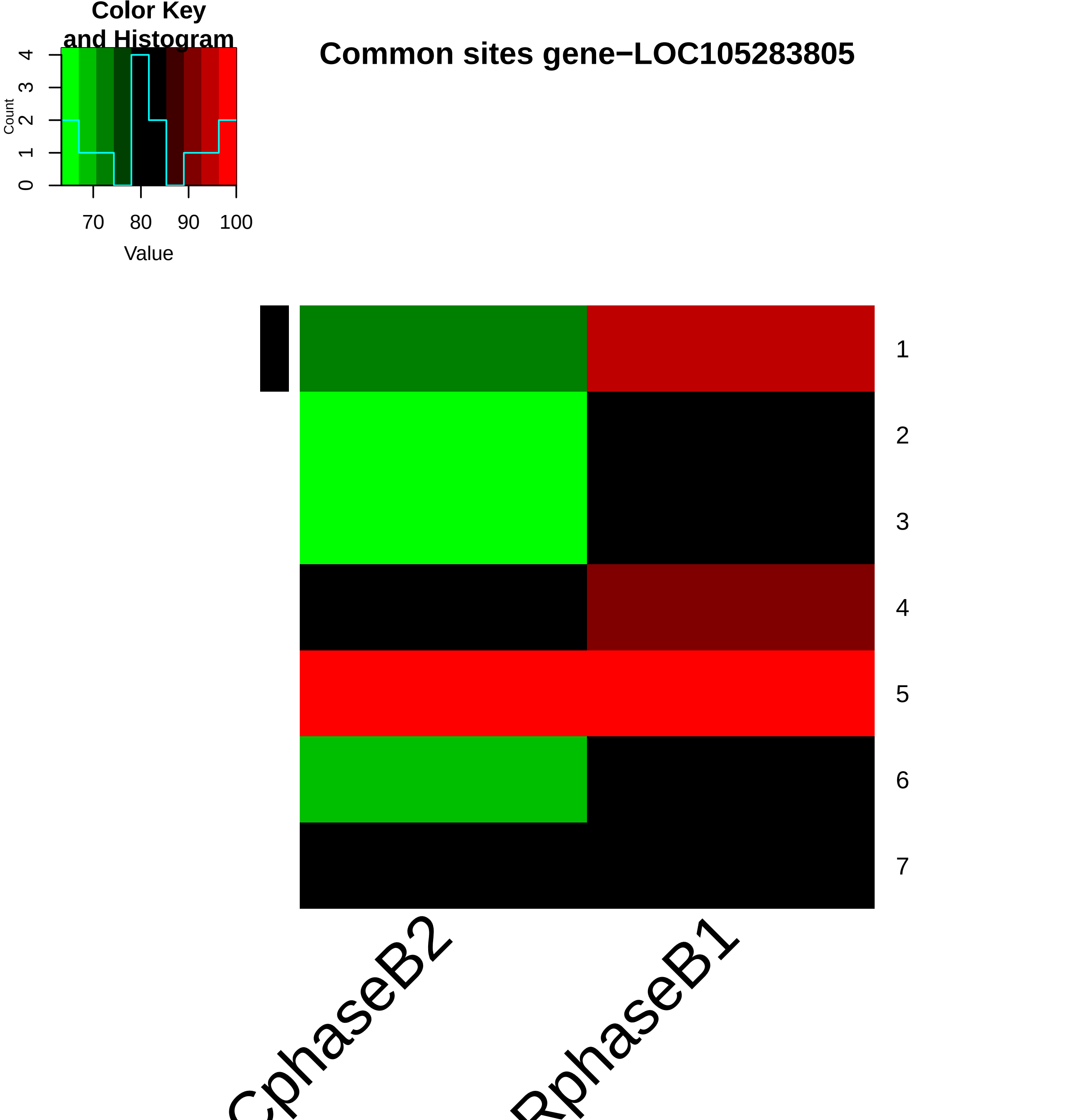
Color Key
and Histogram
Count
Value
70
80
90
100
0
1
2
3
4
Common sites gene−LOC105283805
1
2
3
4
5
6
7
CphaseB2
RphaseB1

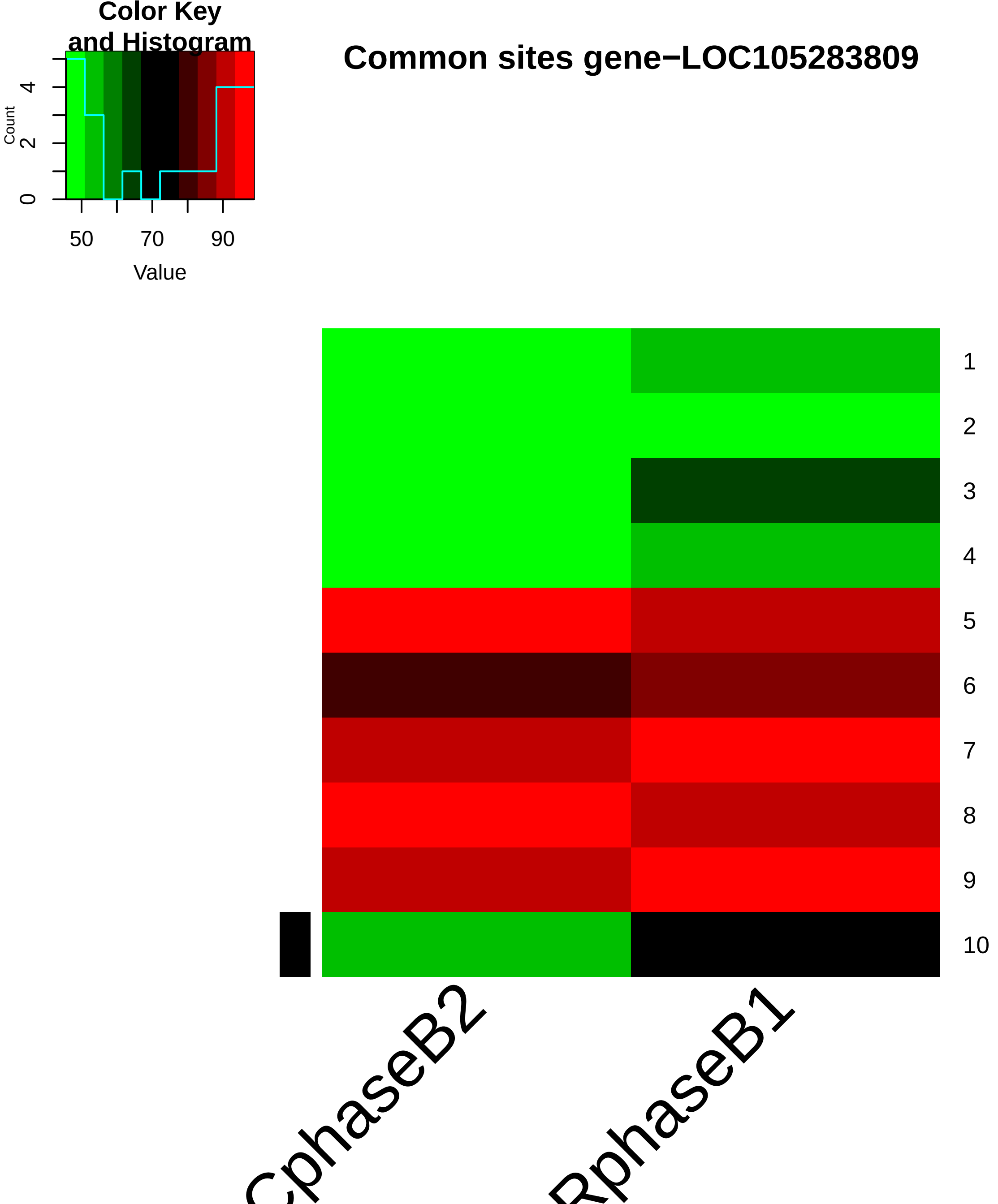

Color Key
and Histogram
Count
0
2
4
50
70
90
Value
Common sites gene−LOC105283809
1
2
3
4
5
6
7
8
9
10
CphaseB2
RphaseB1

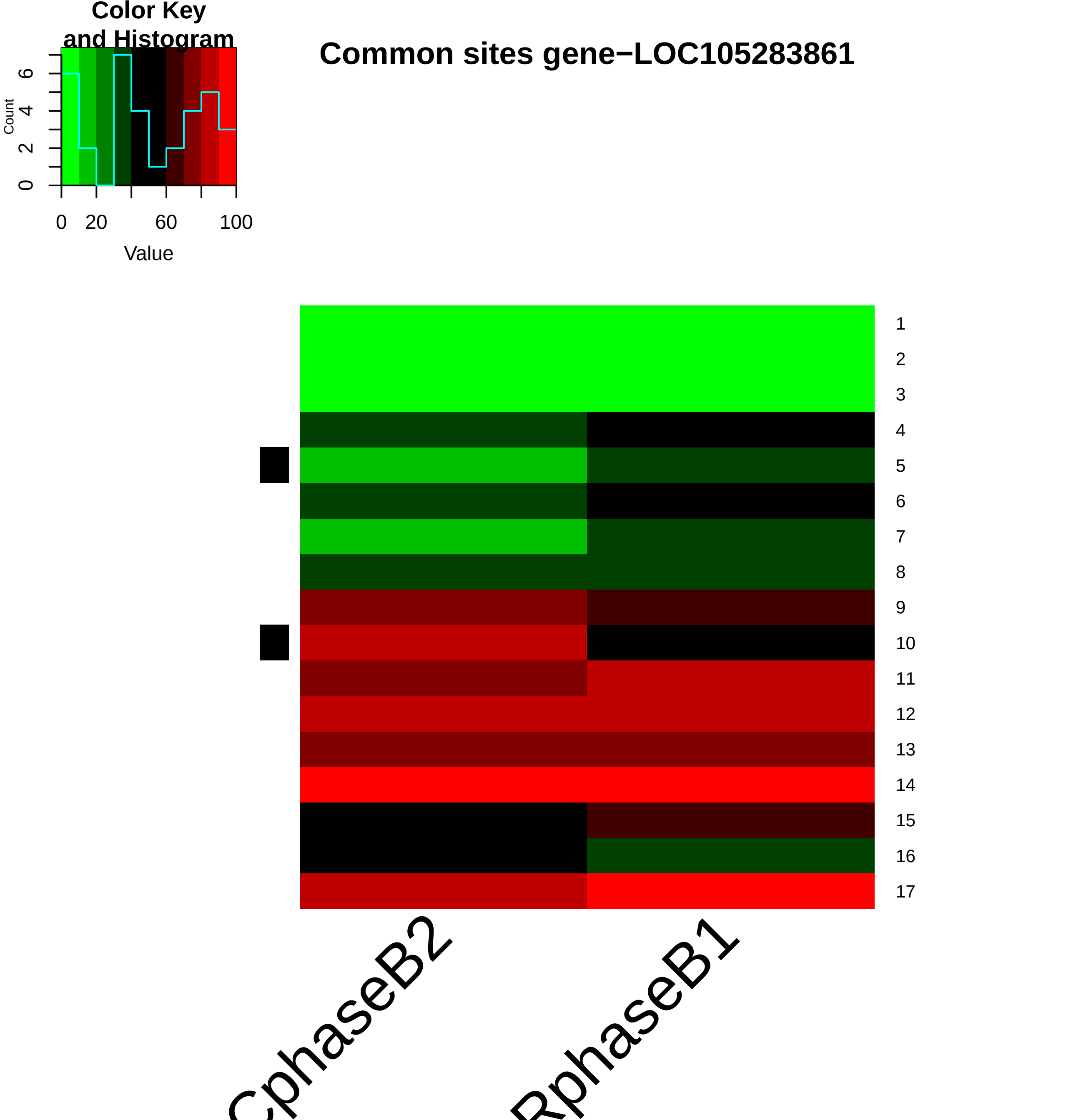
Color Key
and Histogram
Count
0
2
4
6
0
20
60
100
Value
Common sites gene−LOC105283861
CphaseB2
RphaseB1

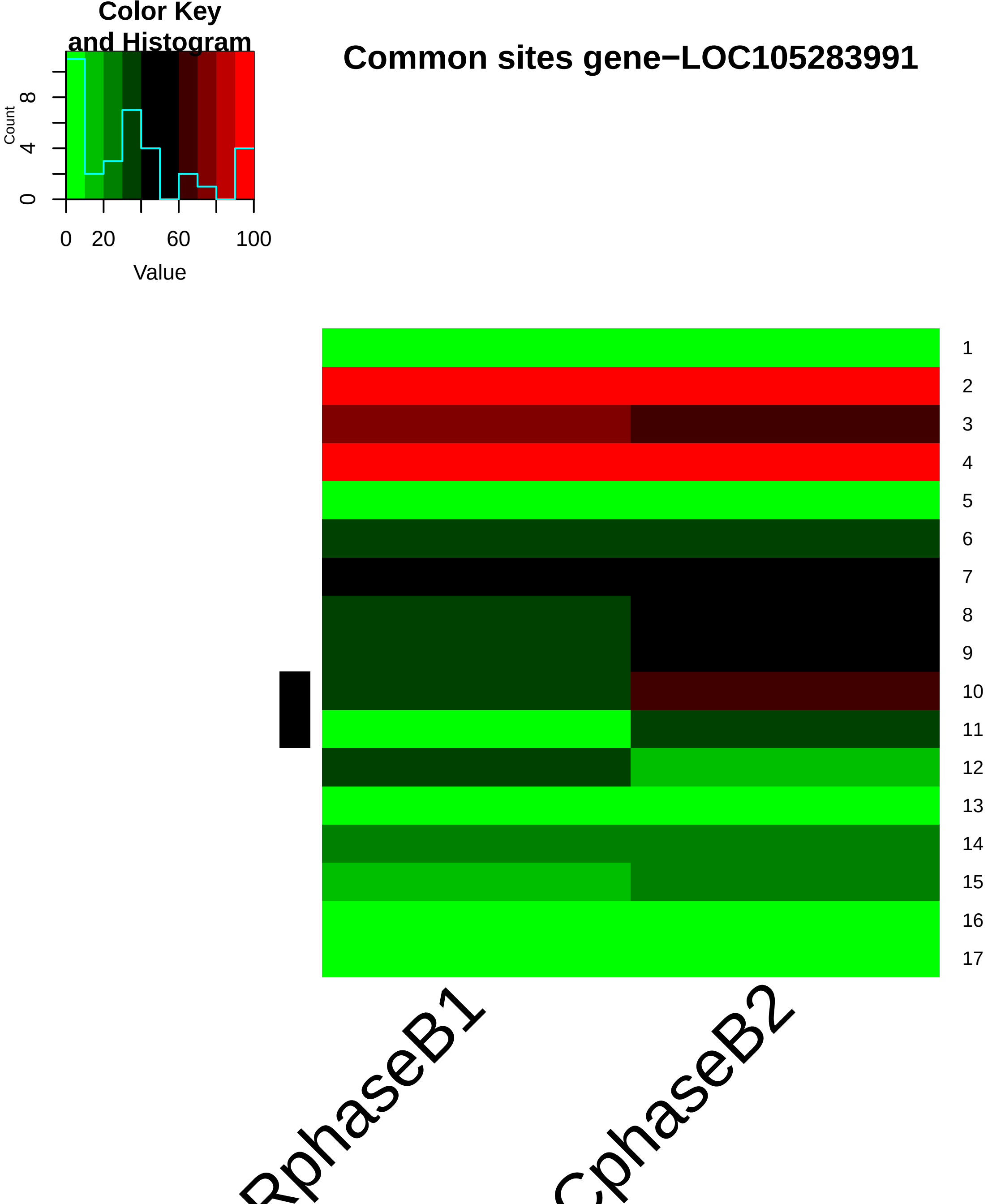
Color Key
and Histogram
Count
0
4
8
0
20
60
100
Value
Common sites gene−LOC105283991
RphaseB1
CphaseB2

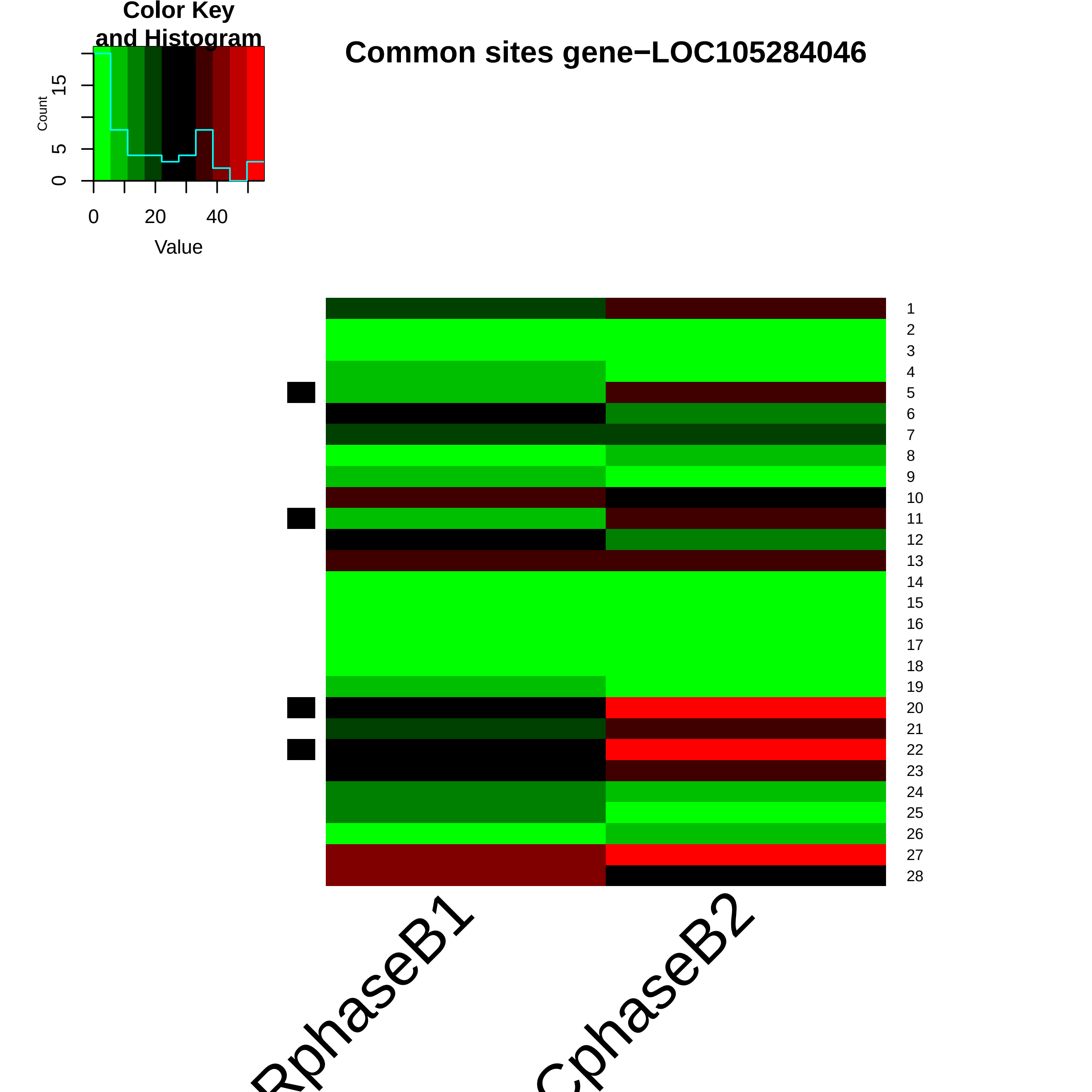
Color Key
and Histogram
Count
0
5
15
0
20
40
Value
Common sites gene−LOC105284046
RphaseB1
CphaseB2

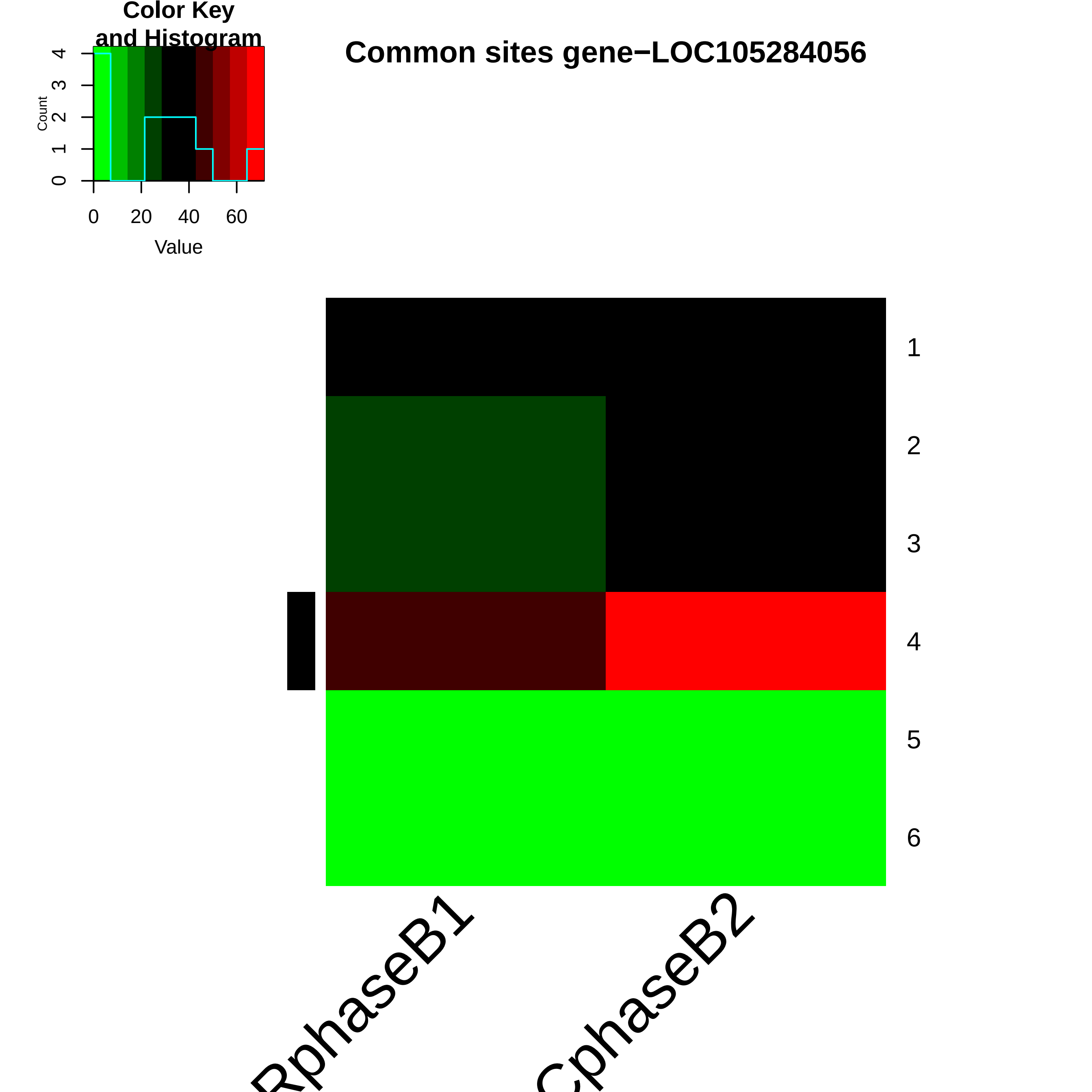
Color Key
and Histogram
Count
0
1
2
3
4
0
20
40
60
Value
Common sites gene−LOC105284056
1
2
3
4
5
6
RphaseB1
CphaseB2

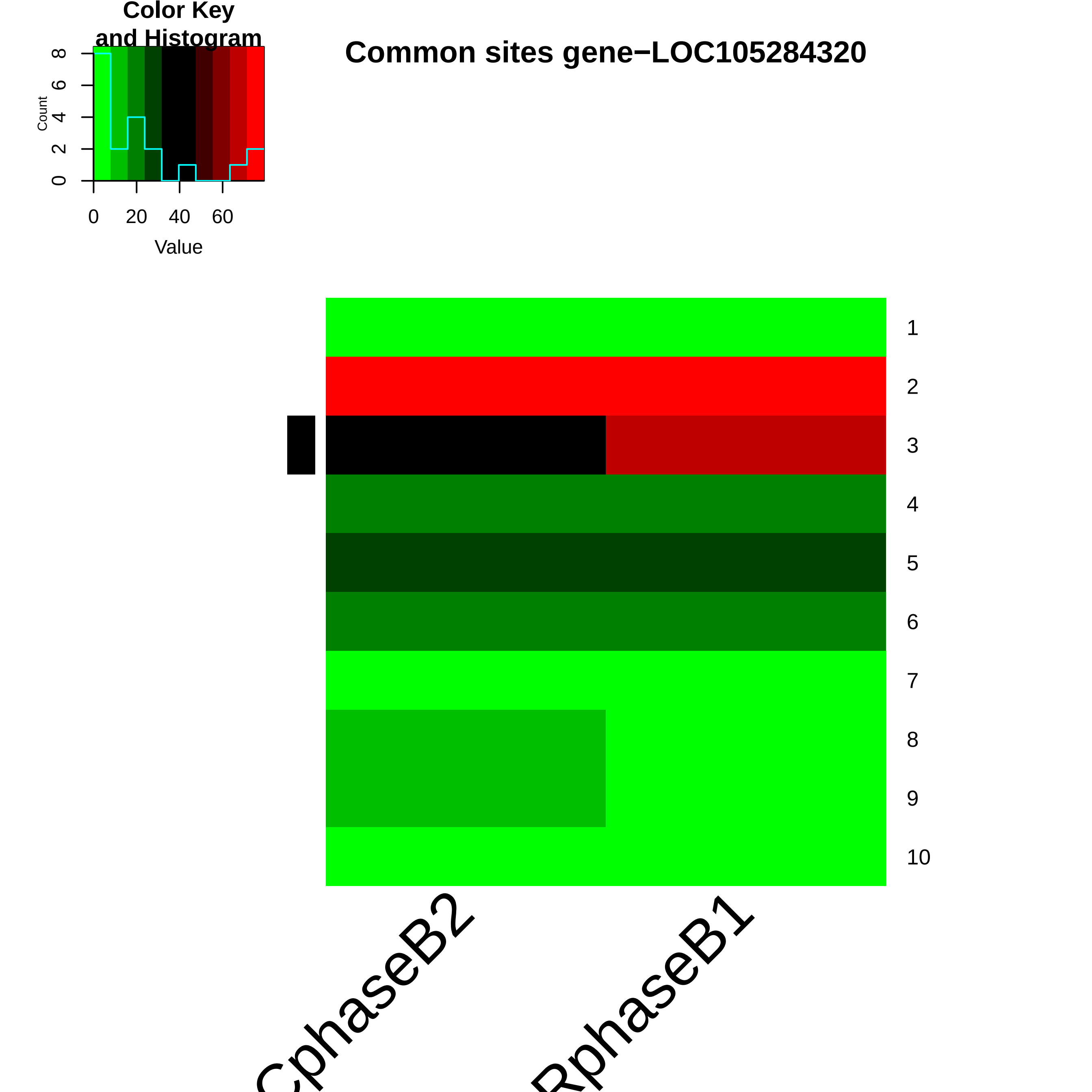

Color Key
and Histogram
Count
Value
0
2
4
6
8
0
20
40
60
Common sites gene−LOC105284320
1
2
3
4
5
6
7
8
9
10
CphaseB2
RphaseB1

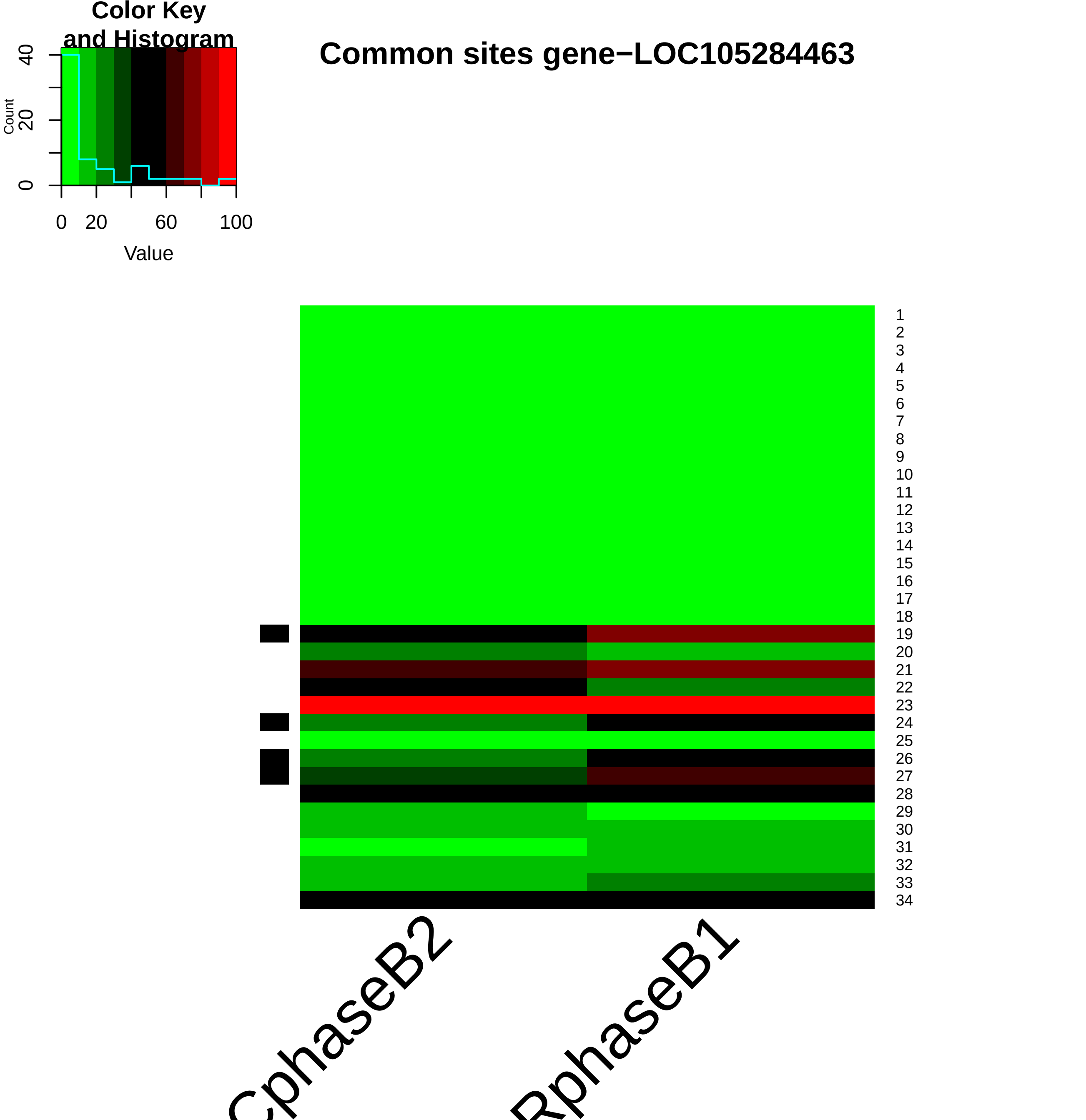

Color Key
and Histogram
Count
0
20
40
0
20
60
100
Value
Common sites gene−LOC105284463
CphaseB2
RphaseB1

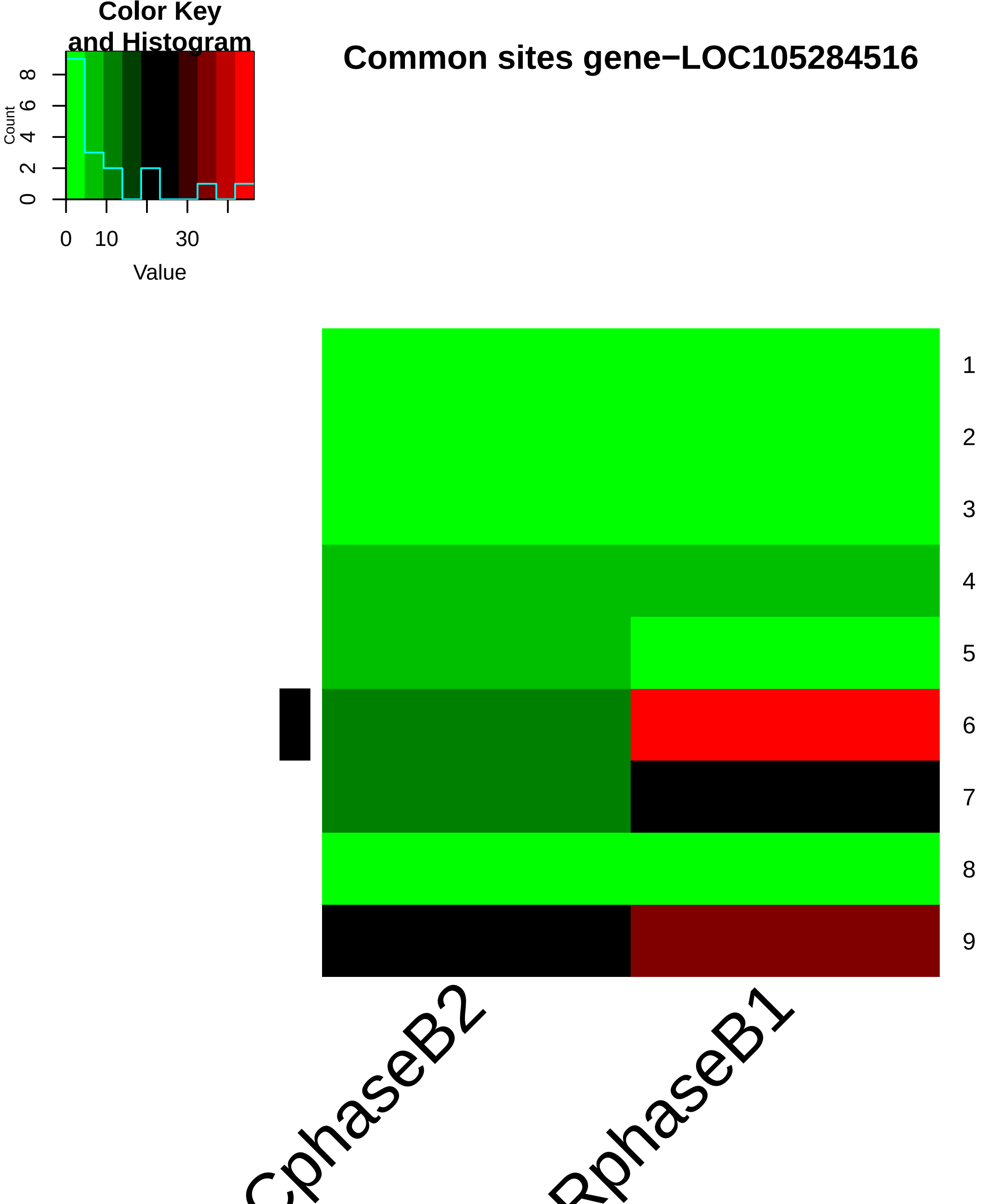
Color Key
and Histogram
Count
0
2
4
6
8
0
10
30
Value
Common sites gene−LOC105284516
1
2
3
4
5
6
7
8
9
CphaseB2
RphaseB1

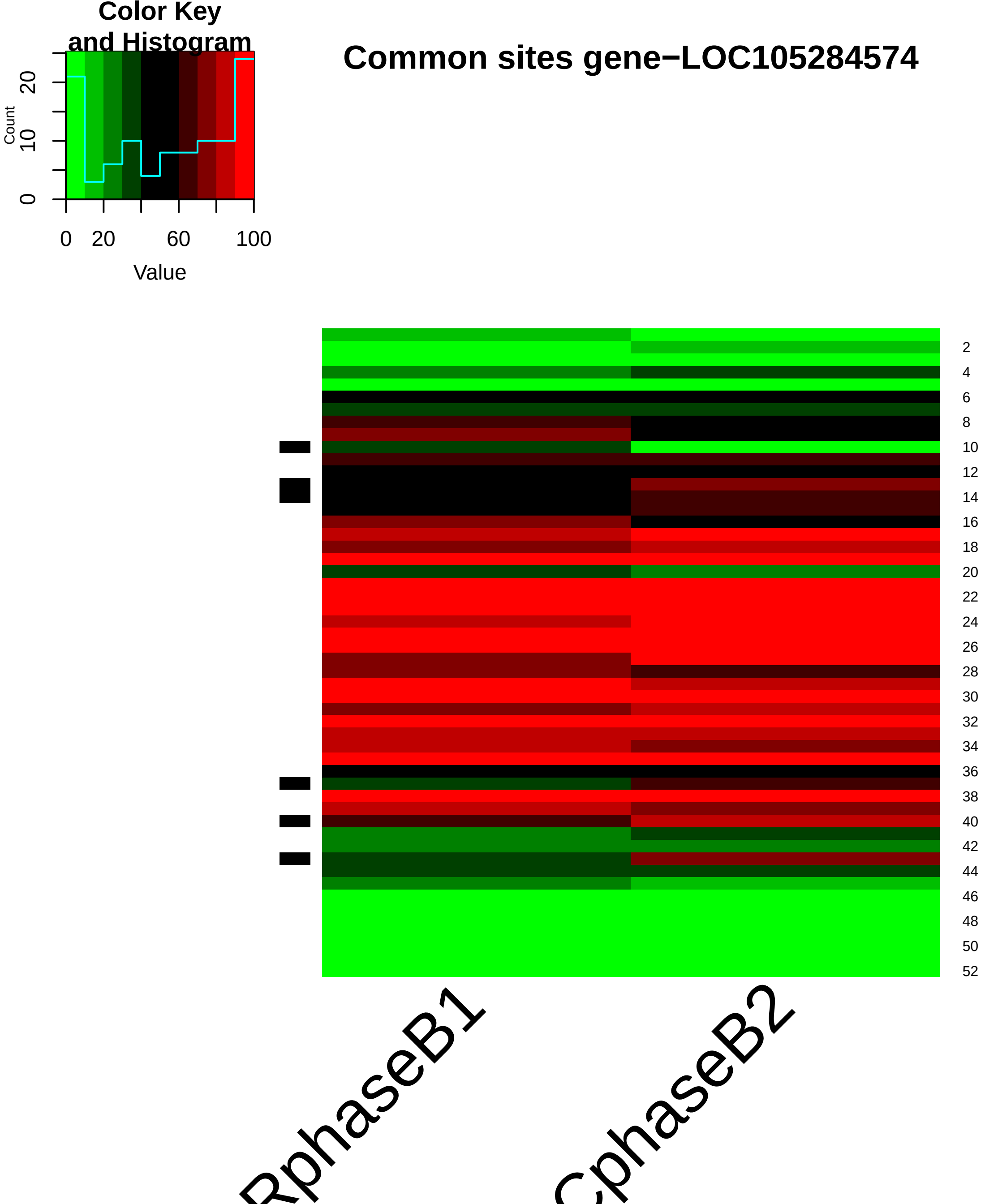

Color Key
and Histogram
Count
0
10
20
0
20
60
100
Value
Common sites gene−LOC105284574
2
4
6
8
10
12
14
16
18
20
22
24
26
28
30
32
34
36
38
40
42
44
46
48
50
52
RphaseB1
CphaseB2

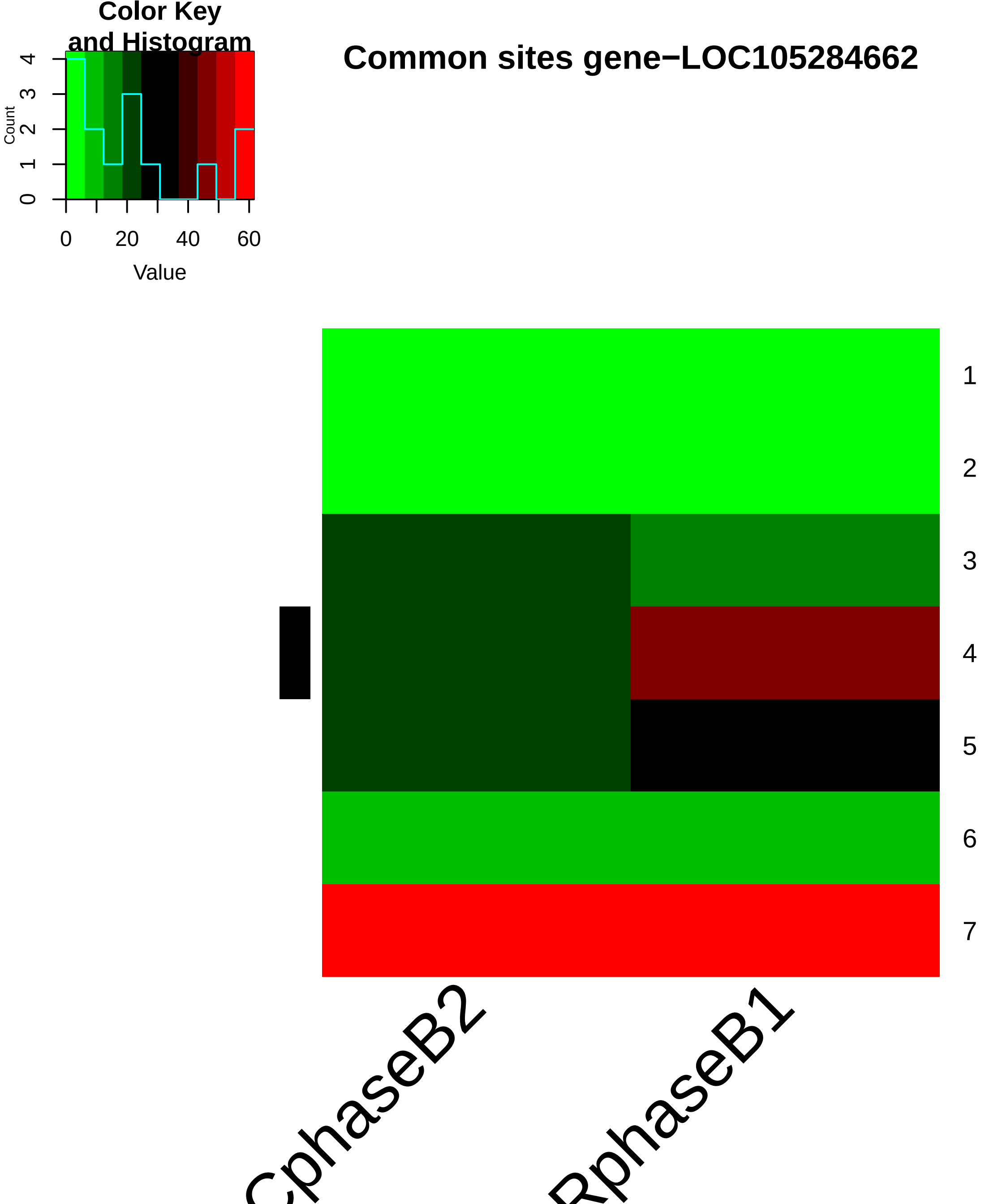
Color Key
and Histogram
Count
0
1
2
3
4
0
20
40
60
Value
Common sites gene−LOC105284662
1
2
3
4
5
6
7
CphaseB2
RphaseB1

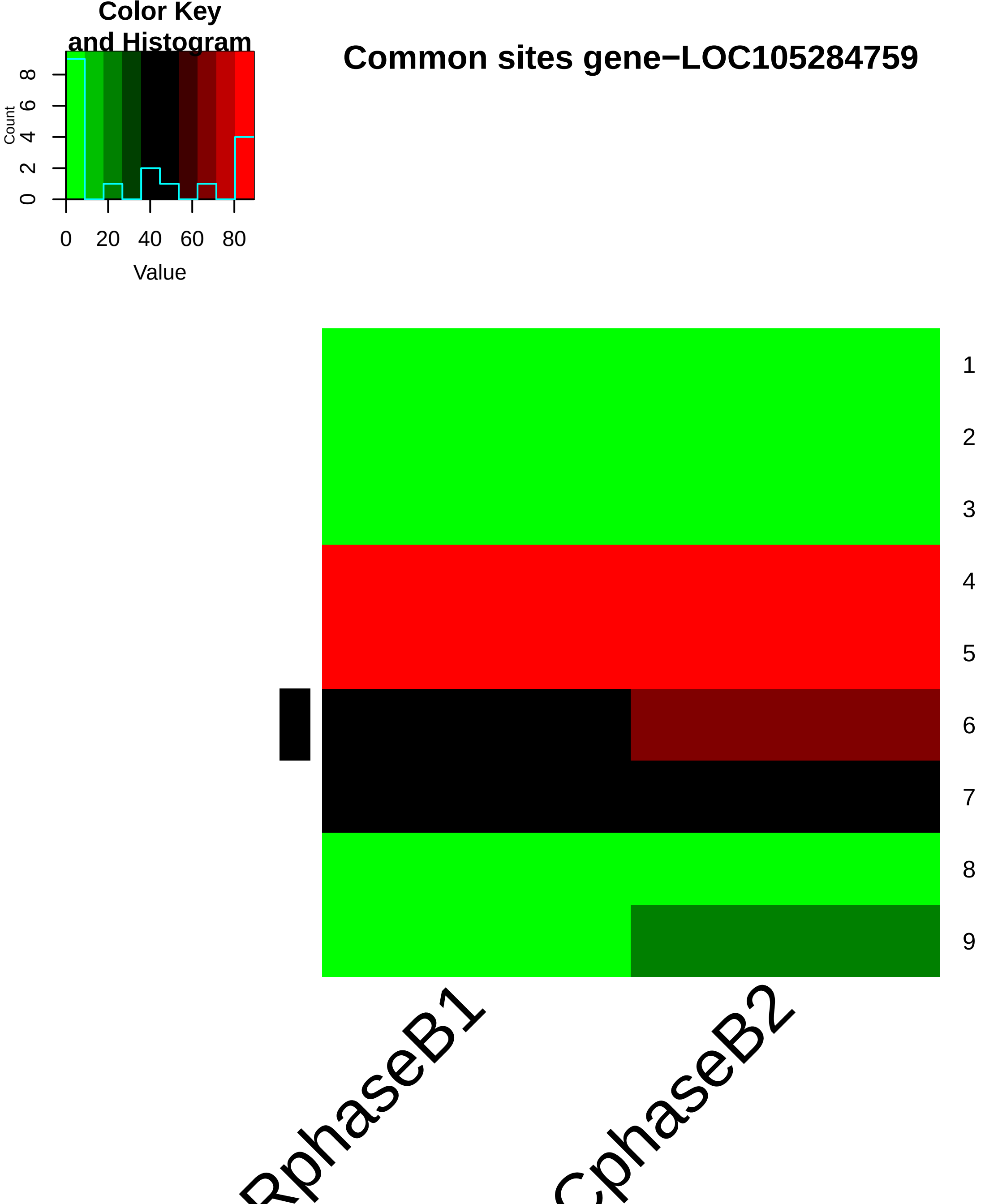
Color Key
and Histogram
Count
0
2
4
6
8
0
20
40
60
80
Value
Common sites gene−LOC105284759
1
2
3
4
5
6
7
8
9
RphaseB1
CphaseB2

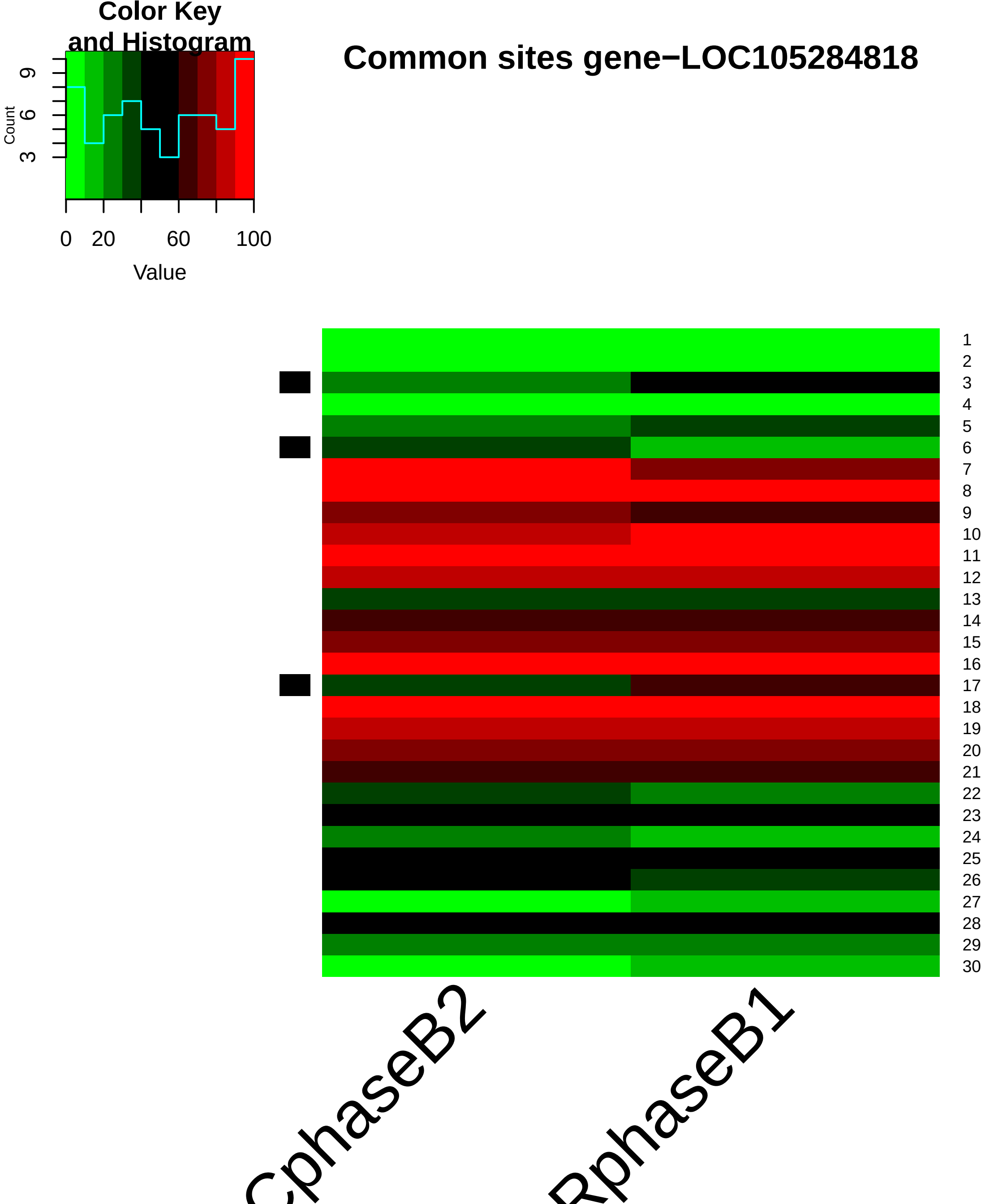
Color Key
and Histogram
Count
3
6
9
0
20
60
100
Value
Common sites gene−LOC105284818

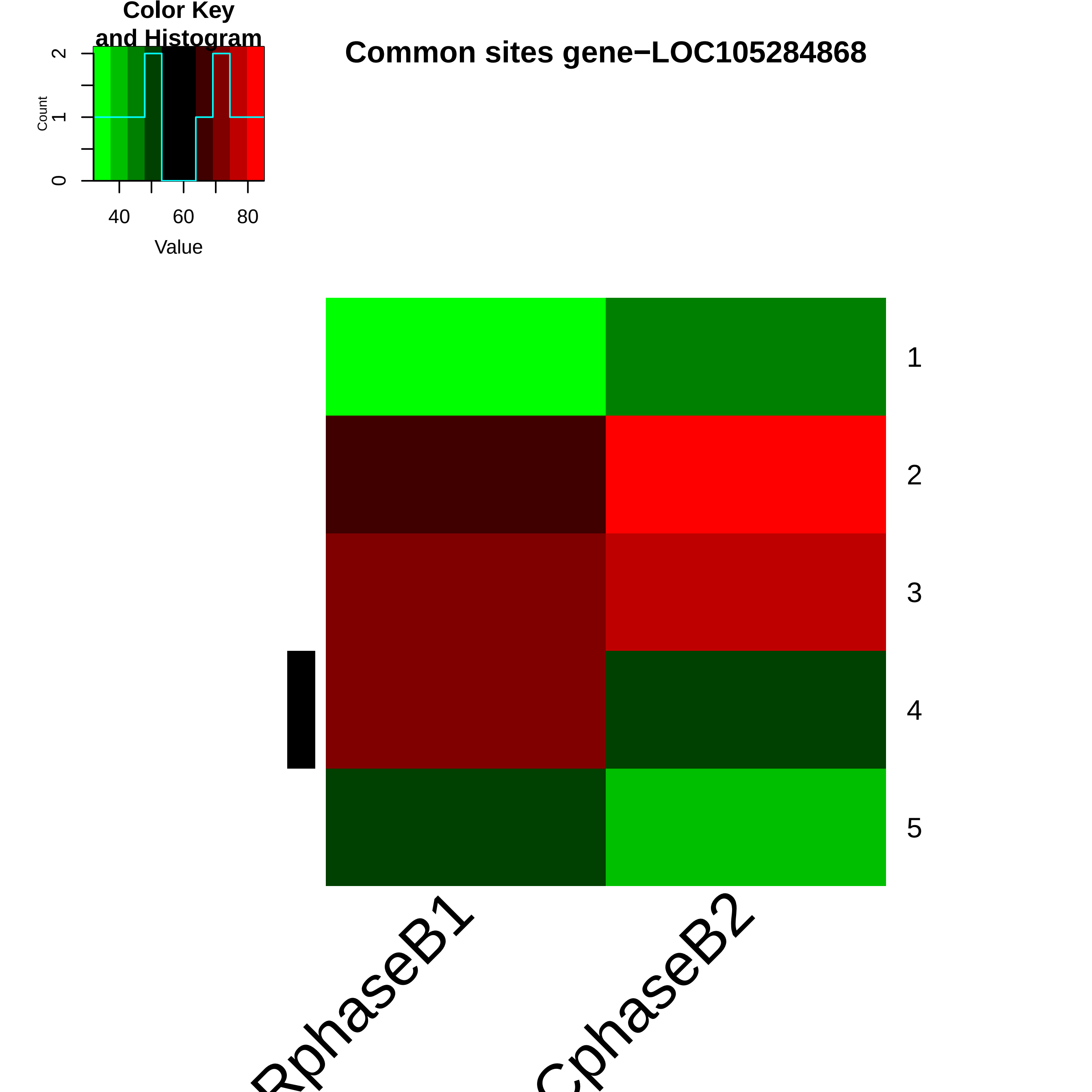

Color Key
and Histogram
Count
0
1
2
40
60
80
Value
Common sites gene−LOC105284868
1
2
3
4
5
RphaseB1
CphaseB2

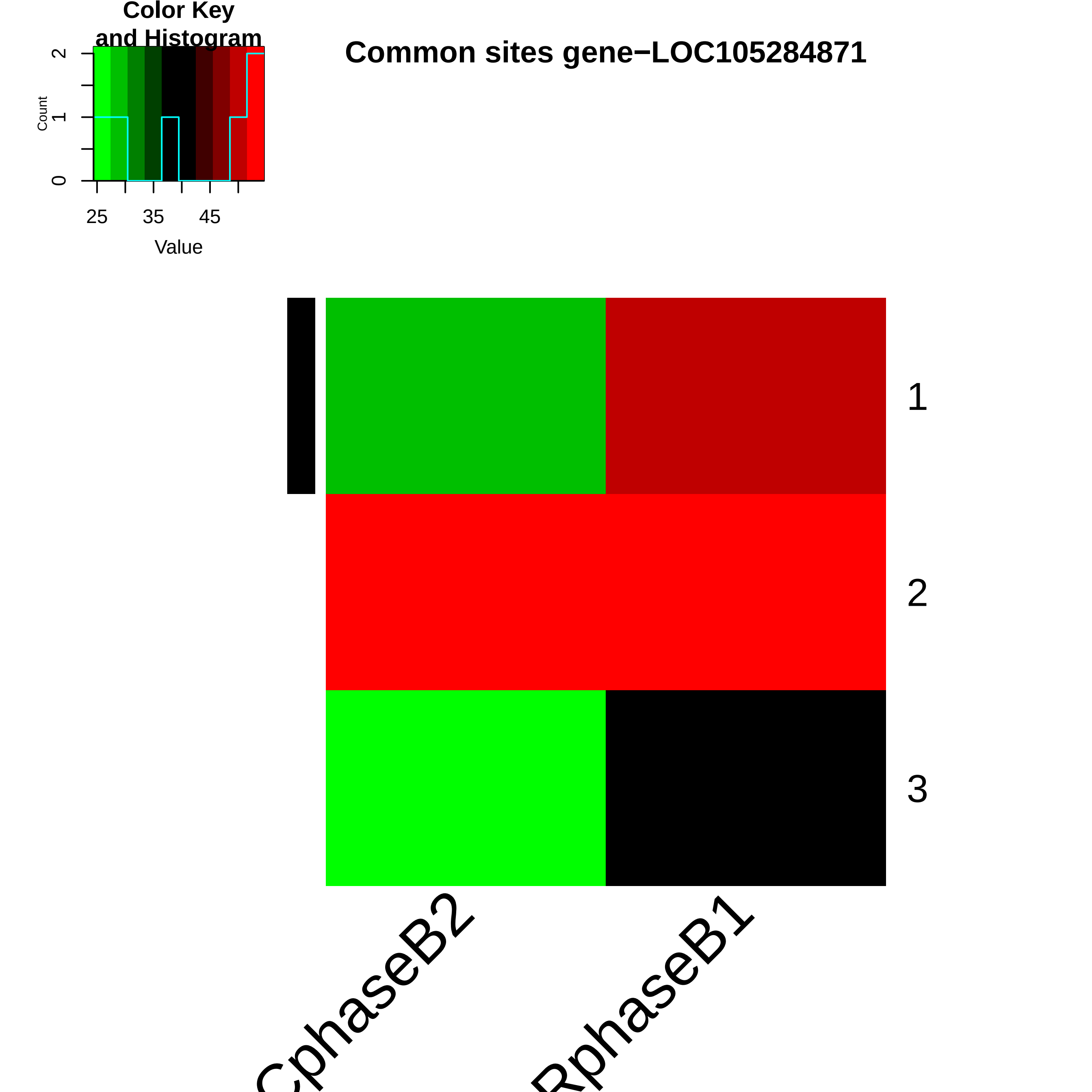
Color Key
and Histogram
Count
0
1
2
25
35
45
Value
Common sites gene−LOC105284871
1
2
3
CphaseB2
RphaseB1

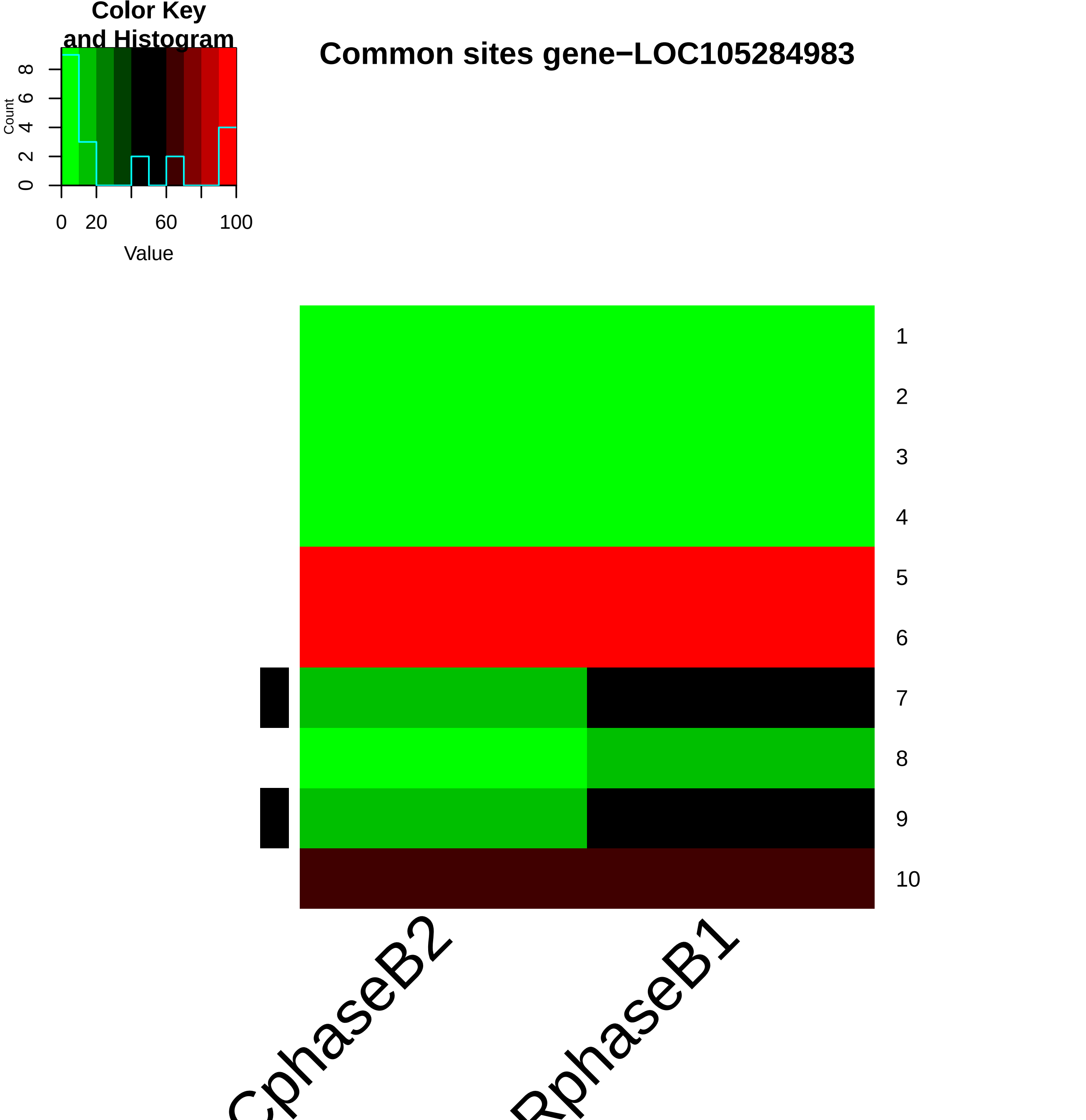

Color Key
and Histogram
Count
0 2 4 6 8
0 20 60 100
Value
Common sites gene−LOC105284983
1
2
3
4
5
6
7
8
9
10
CphaseB2
RphaseB1

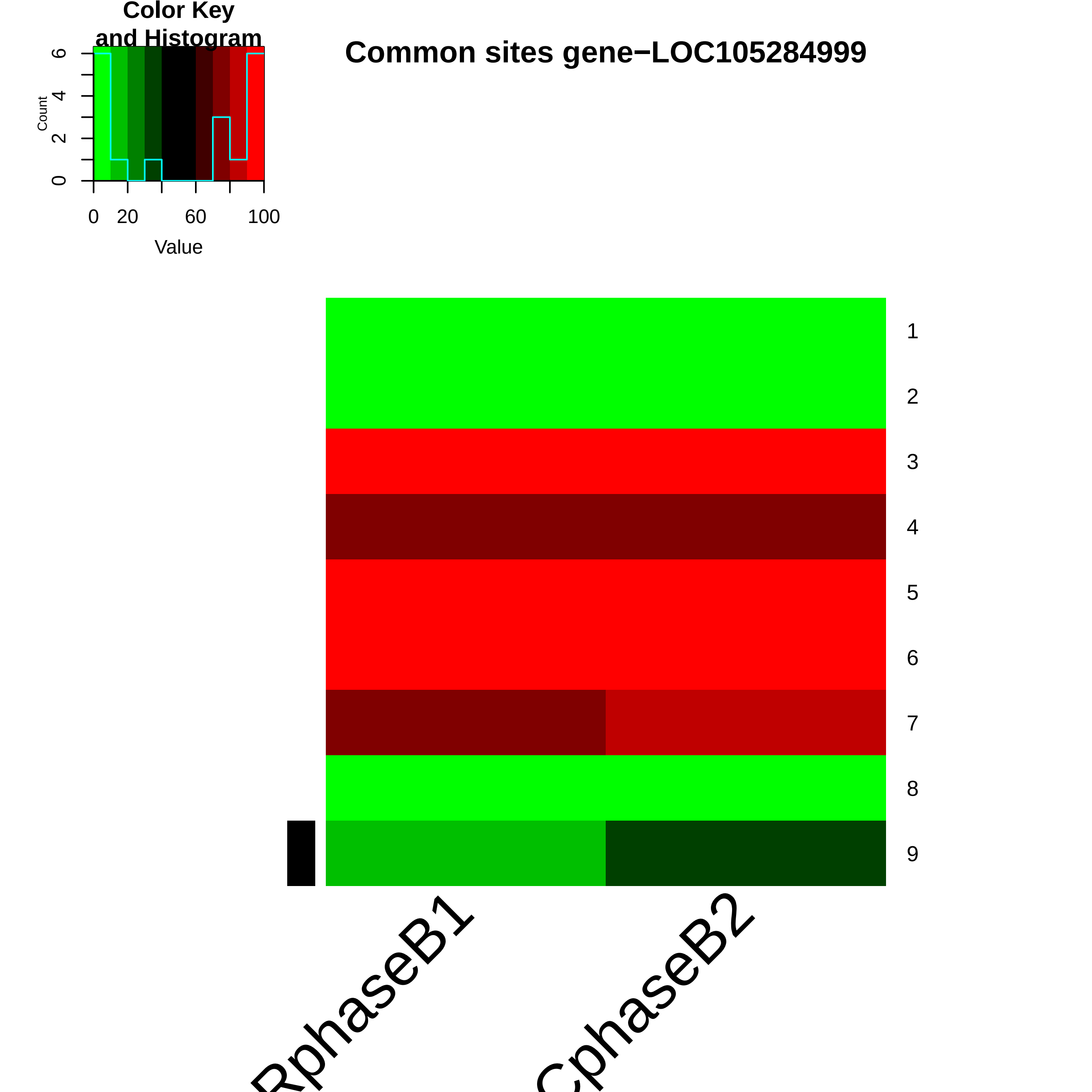

Color Key
and Histogram
Count
0
2
4
6
0
20
60
100
Value
Common sites gene−LOC105284999
1
2
3
4
5
6
7
8
9
RphaseB1
CphaseB2

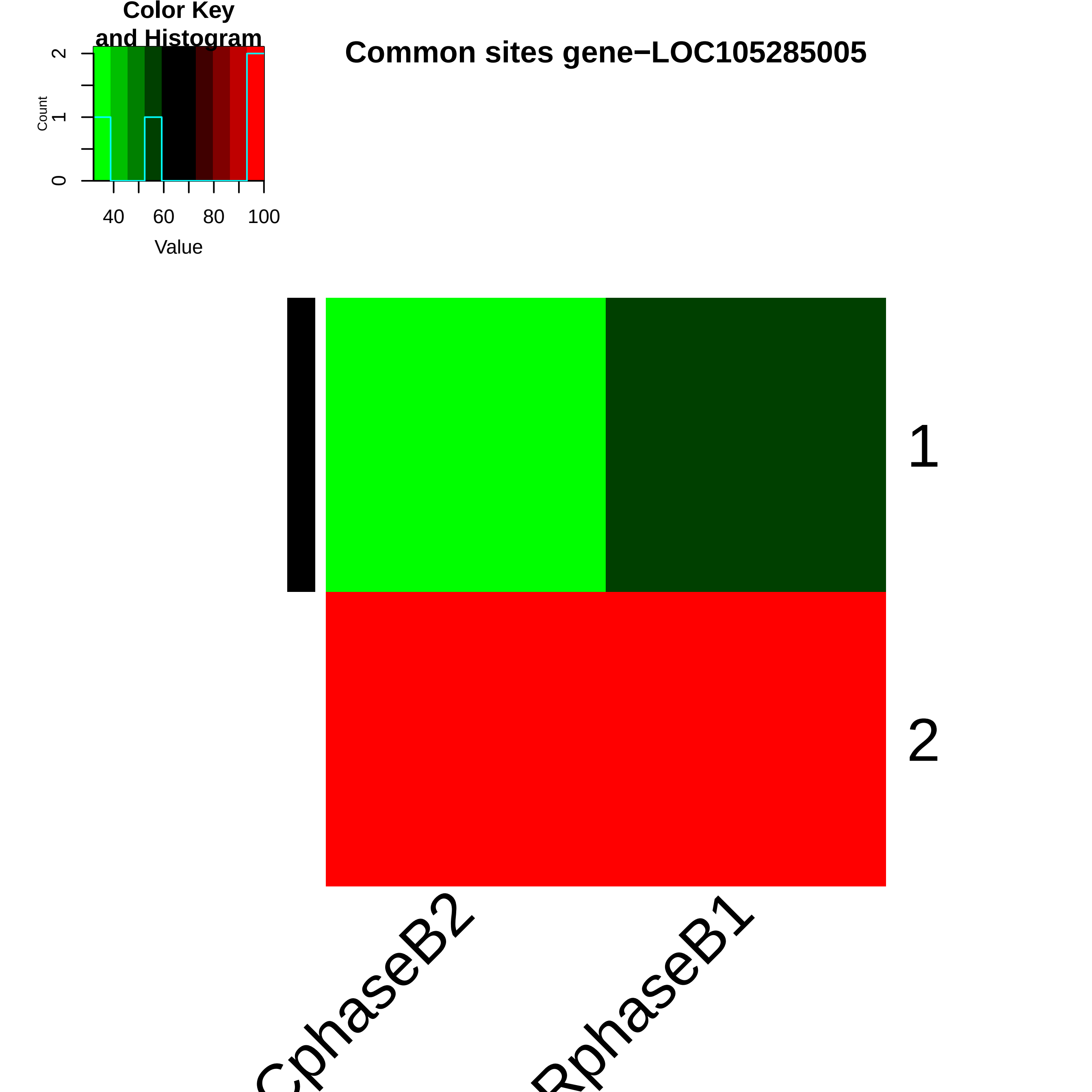

Color Key
and Histogram
Count
0
1
2
40
60
80
100
Value
Common sites gene−LOC105285005
1
2

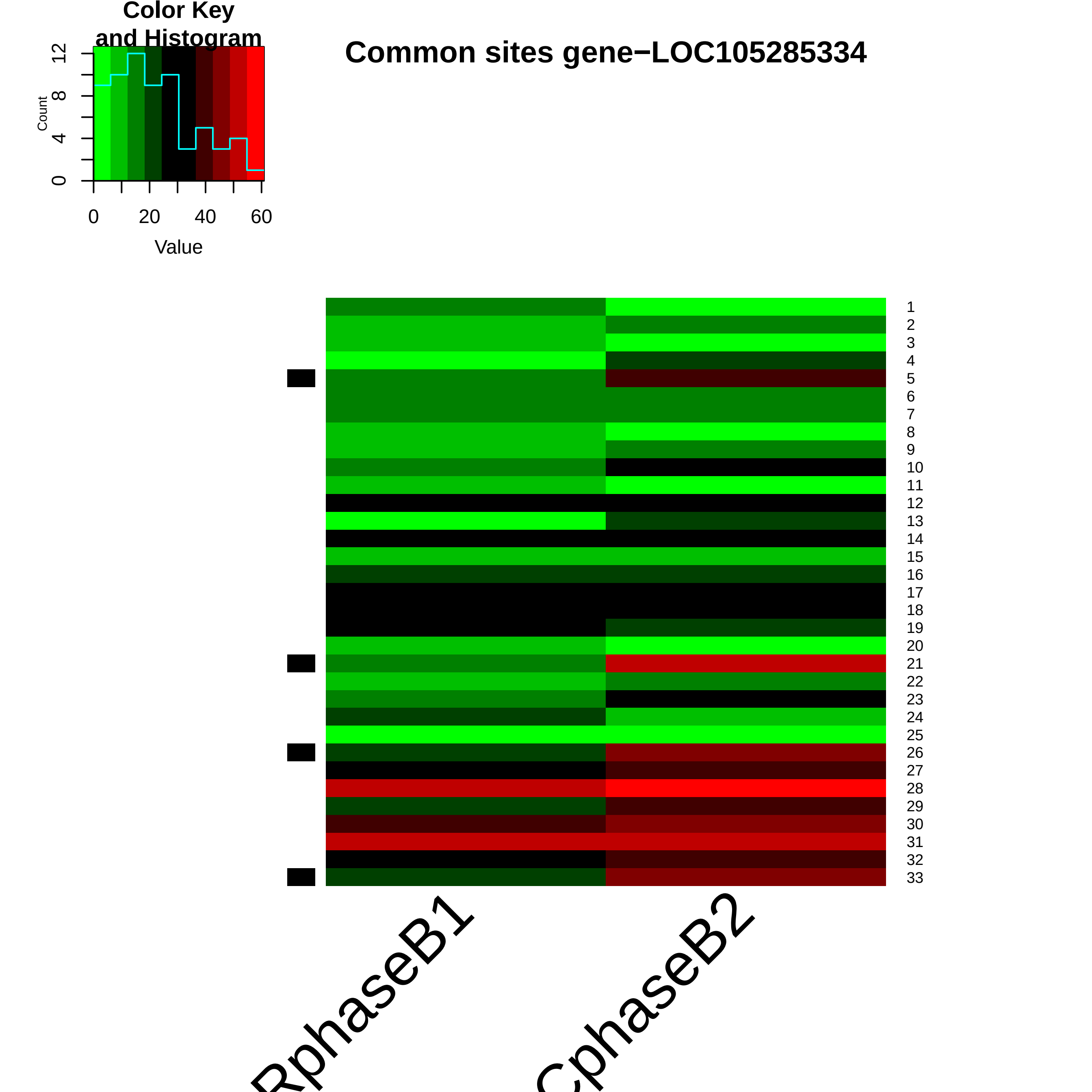
Color Key
and Histogram
Count
Value
Common sites gene−LOC105285334
RphaseB1
CphaseB2

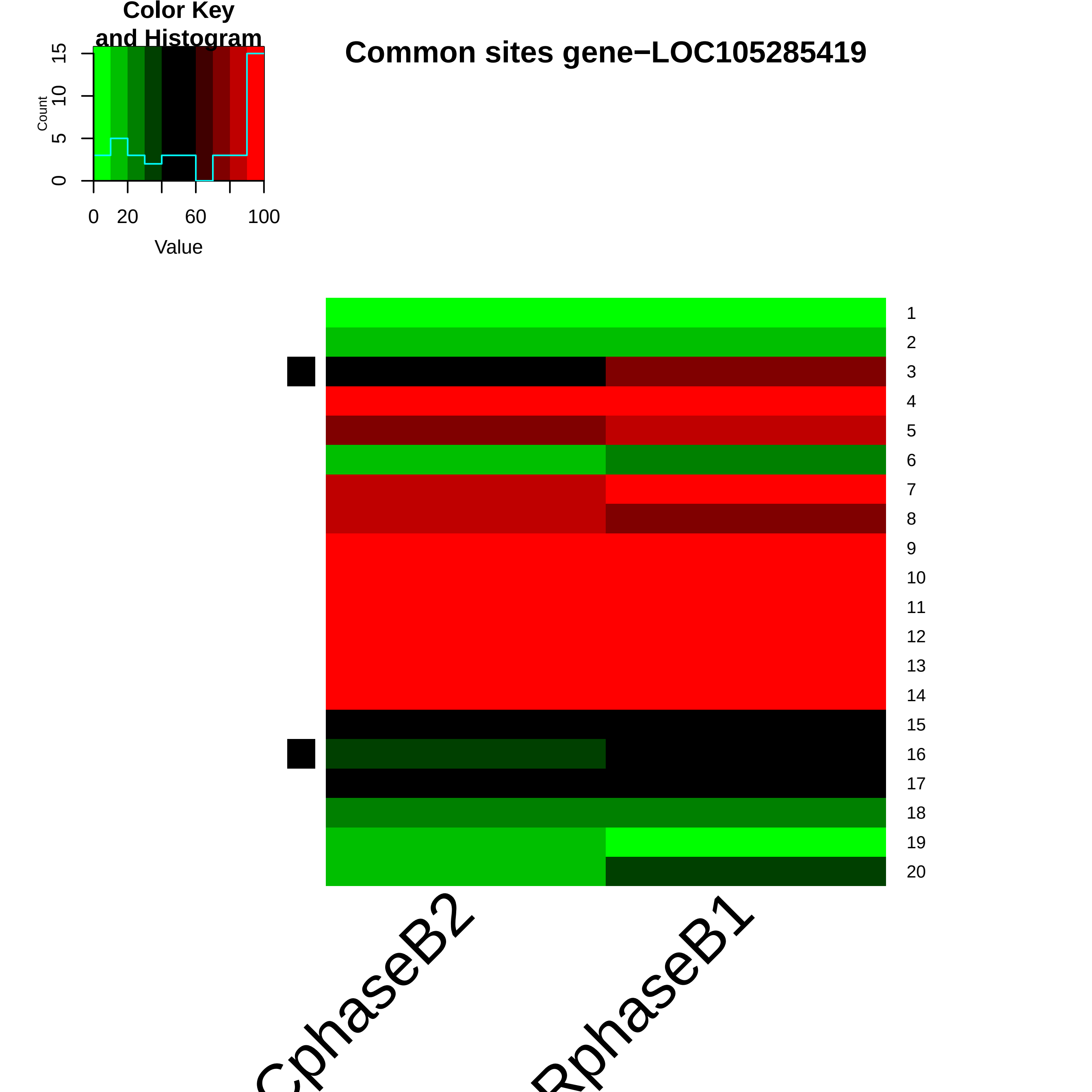

Color Key
and Histogram
Count
Value
0
5
10
15
0
20
60
100
Common sites gene−LOC105285419
CphaseB2
RphaseB1

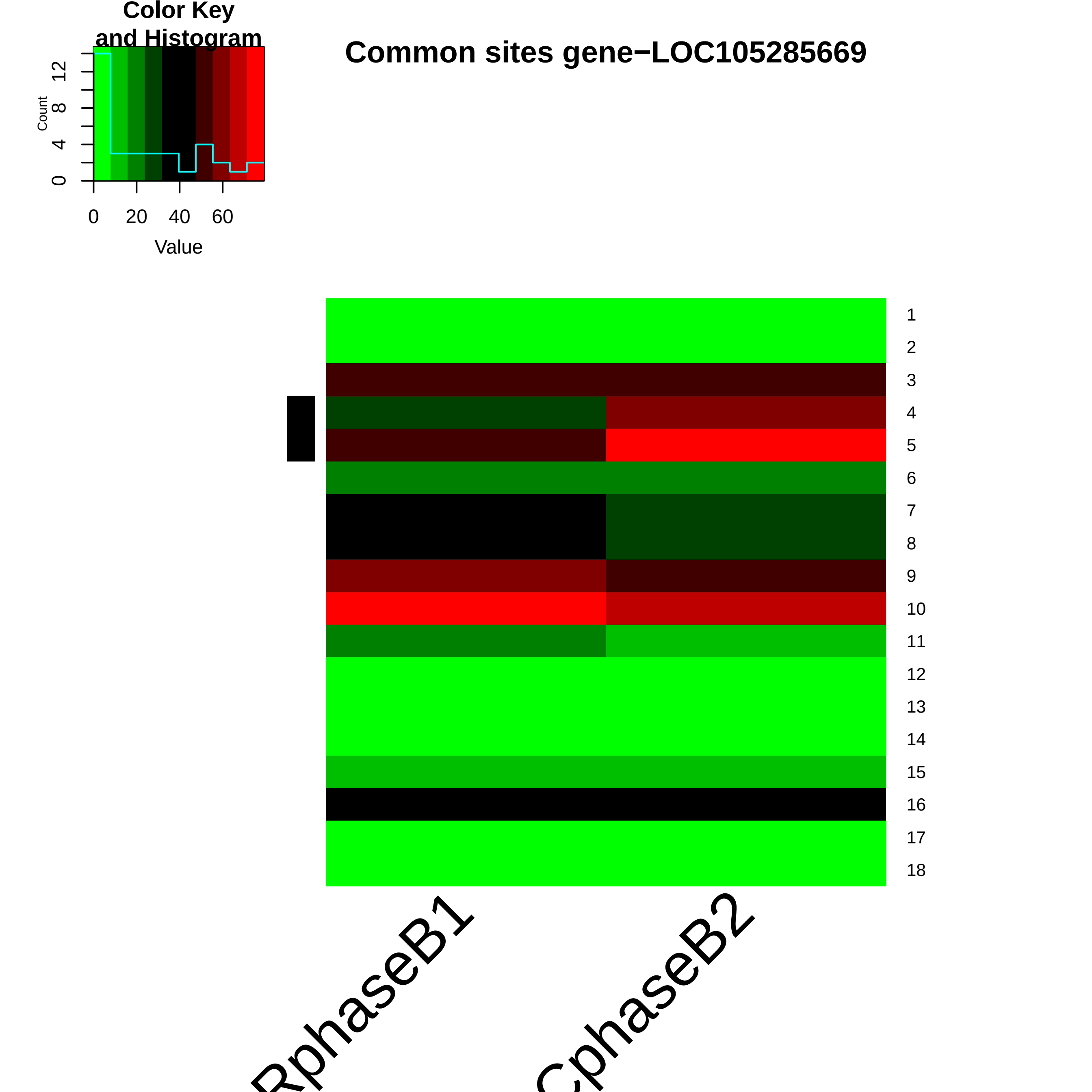
Color Key
and Histogram
Count
0
4
8
12
0
20
40
60
Value
Common sites gene−LOC105285669
RphaseB1
CphaseB2

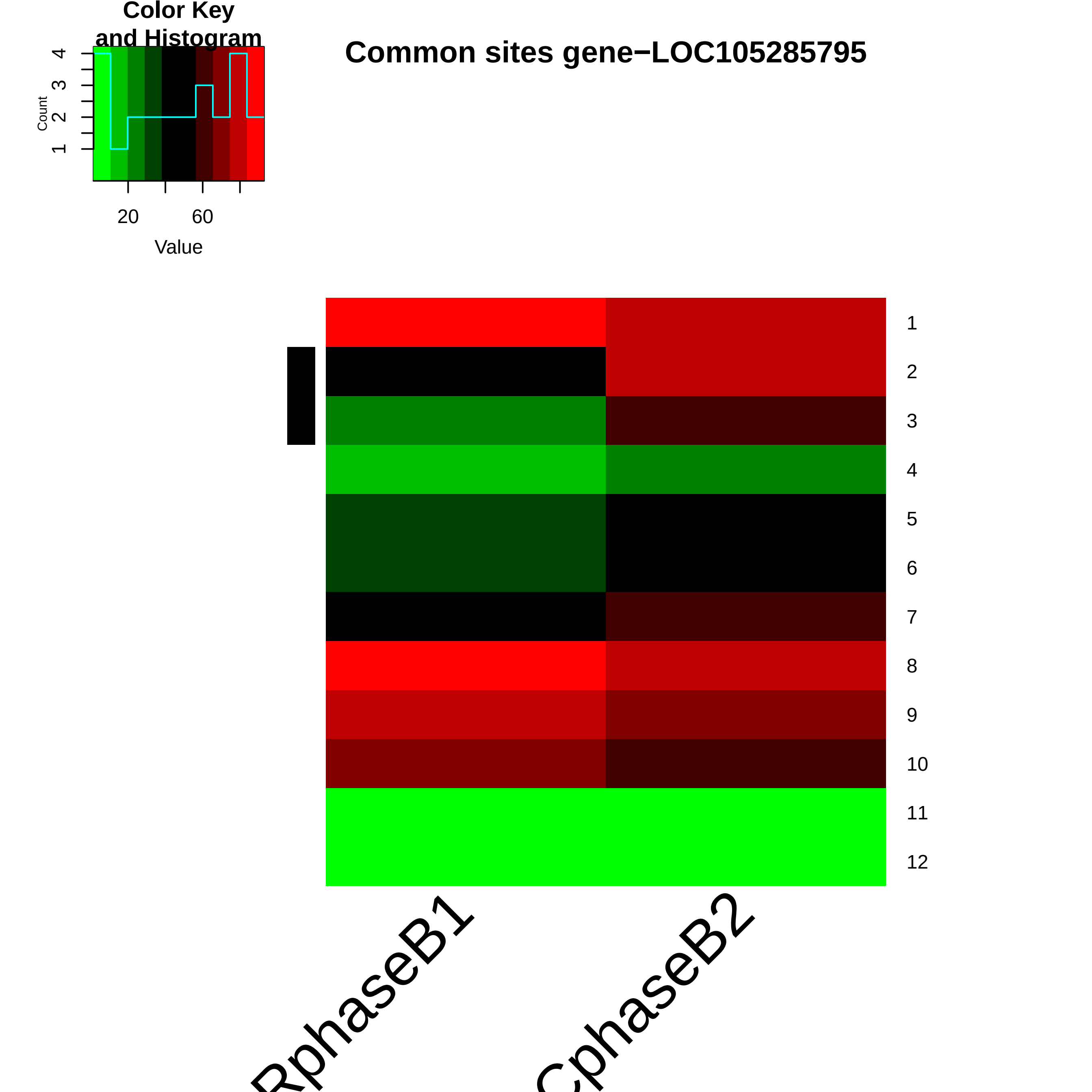

Color Key
and Histogram
Count
Value
Common sites gene−LOC105285795
RphaseB1
CphaseB2

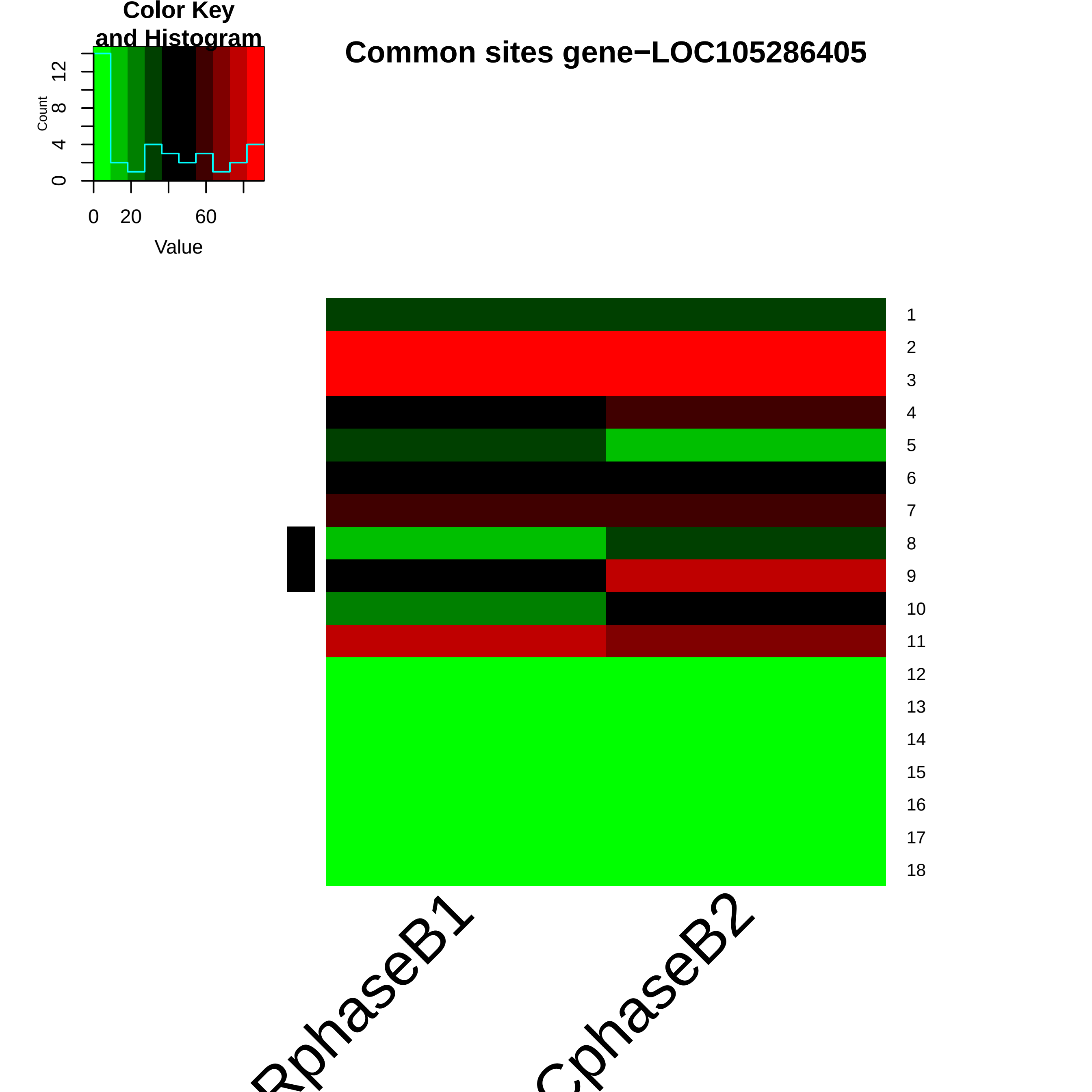

Color Key
and Histogram
Count
0
4
8
12
0
20
60
Value
Common sites gene−LOC105286405
RphaseB1
CphaseB2

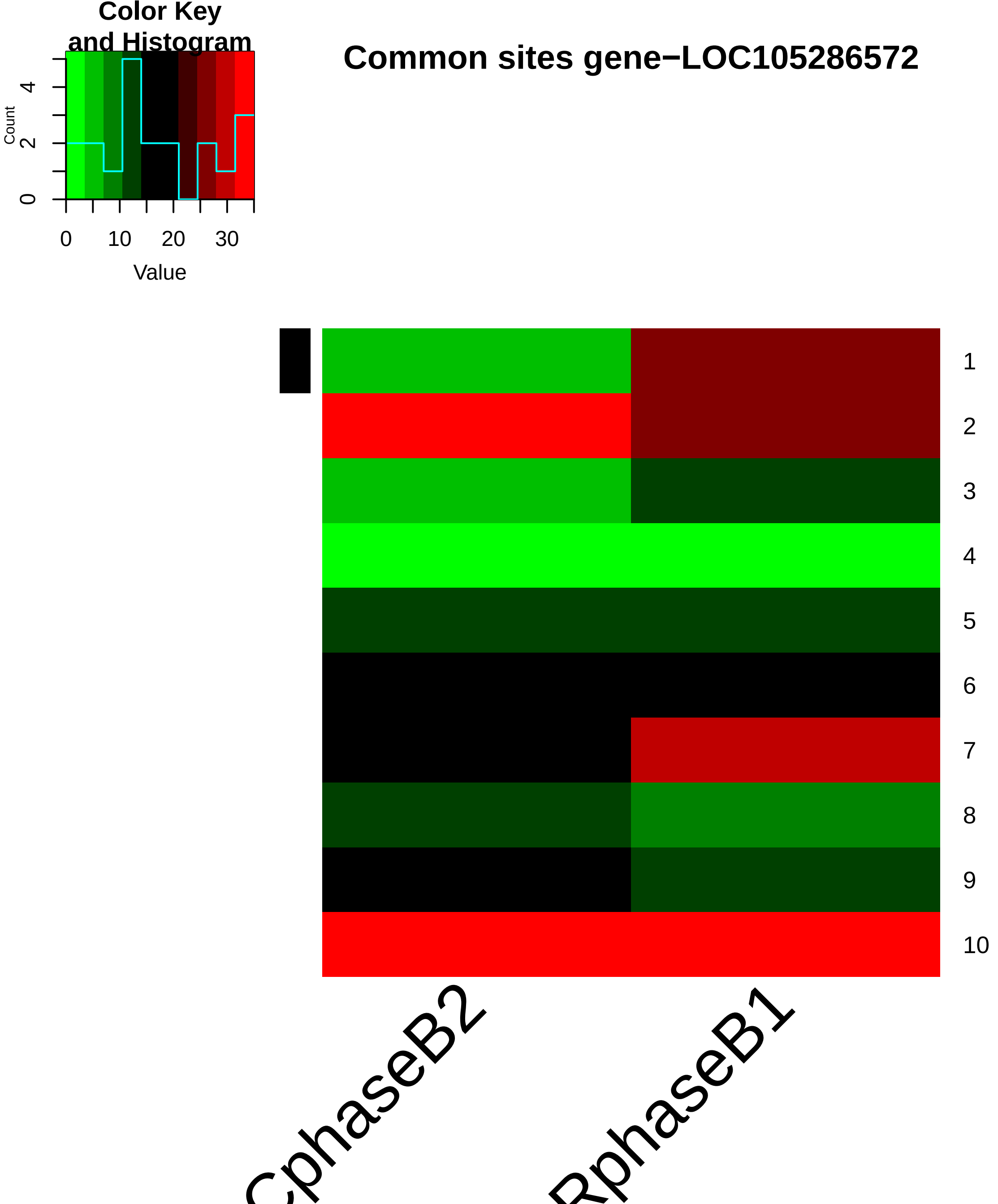

Color Key
and Histogram
Count
0
2
4
0
10
20
30
Value
Common sites gene−LOC105286572
1
2
3
4
5
6
7
8
9
10
CphaseB2
RphaseB1

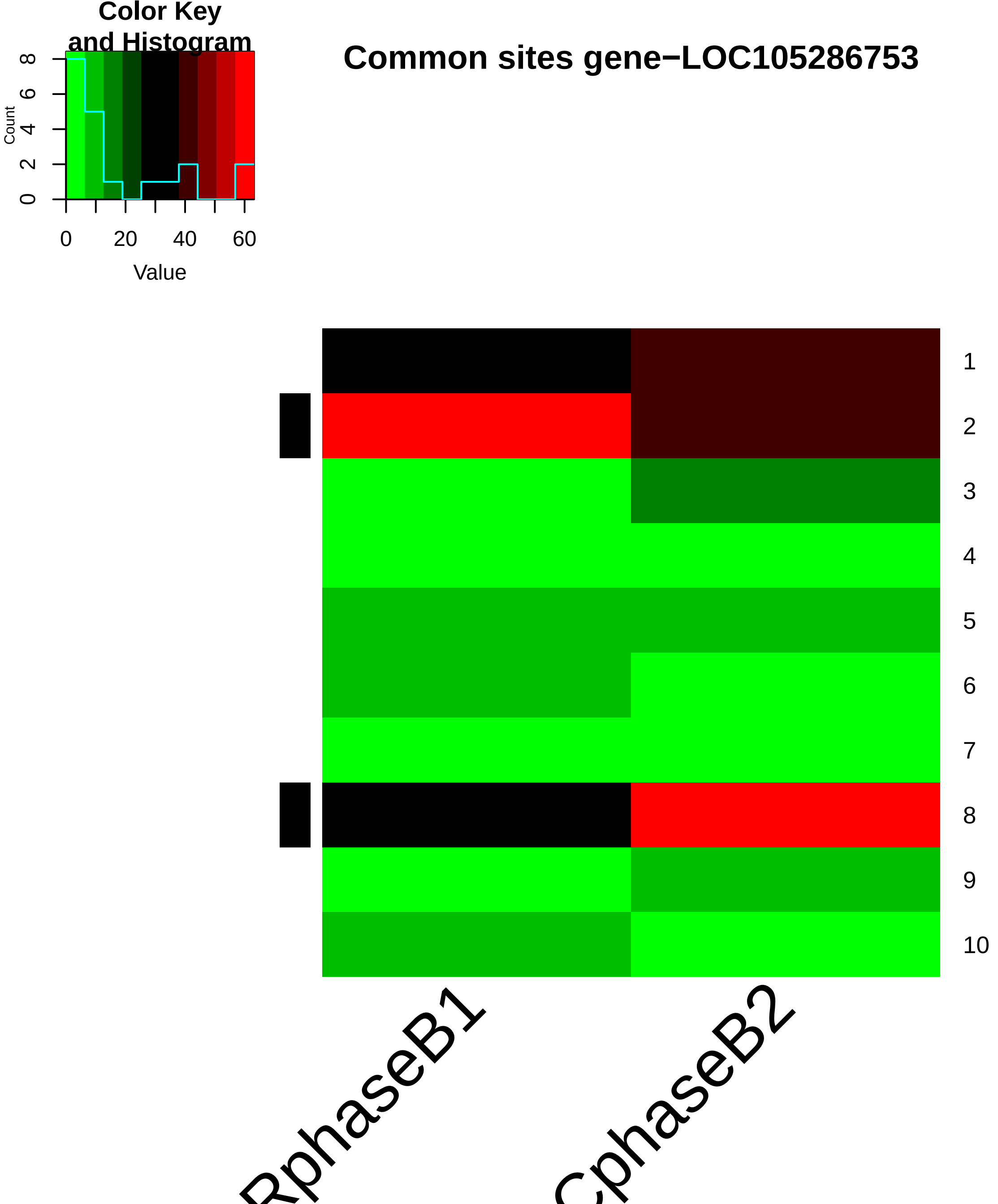

Color Key
and Histogram
Count
0
2
4
6
8
0
20
40
60
Value
Common sites gene−LOC105286753
1
2
3
4
5
6
7
8
9
10
RphaseB1
CphaseB2

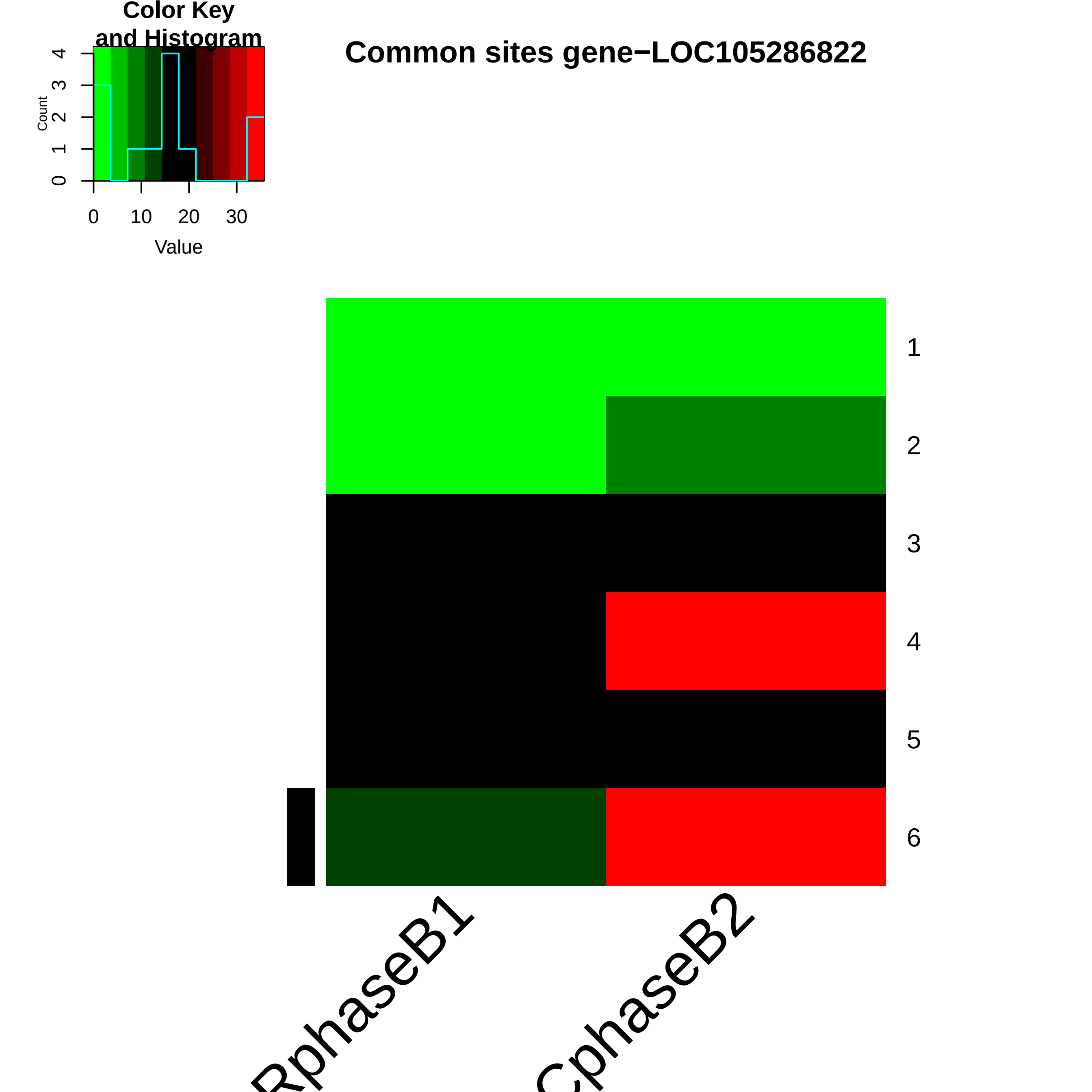
Color Key
and Histogram
Count
0
1
2
3
4
0
10
20
30
Value
Common sites gene−LOC105286822
1
2
3
4
5
6
RphaseB1
CphaseB2

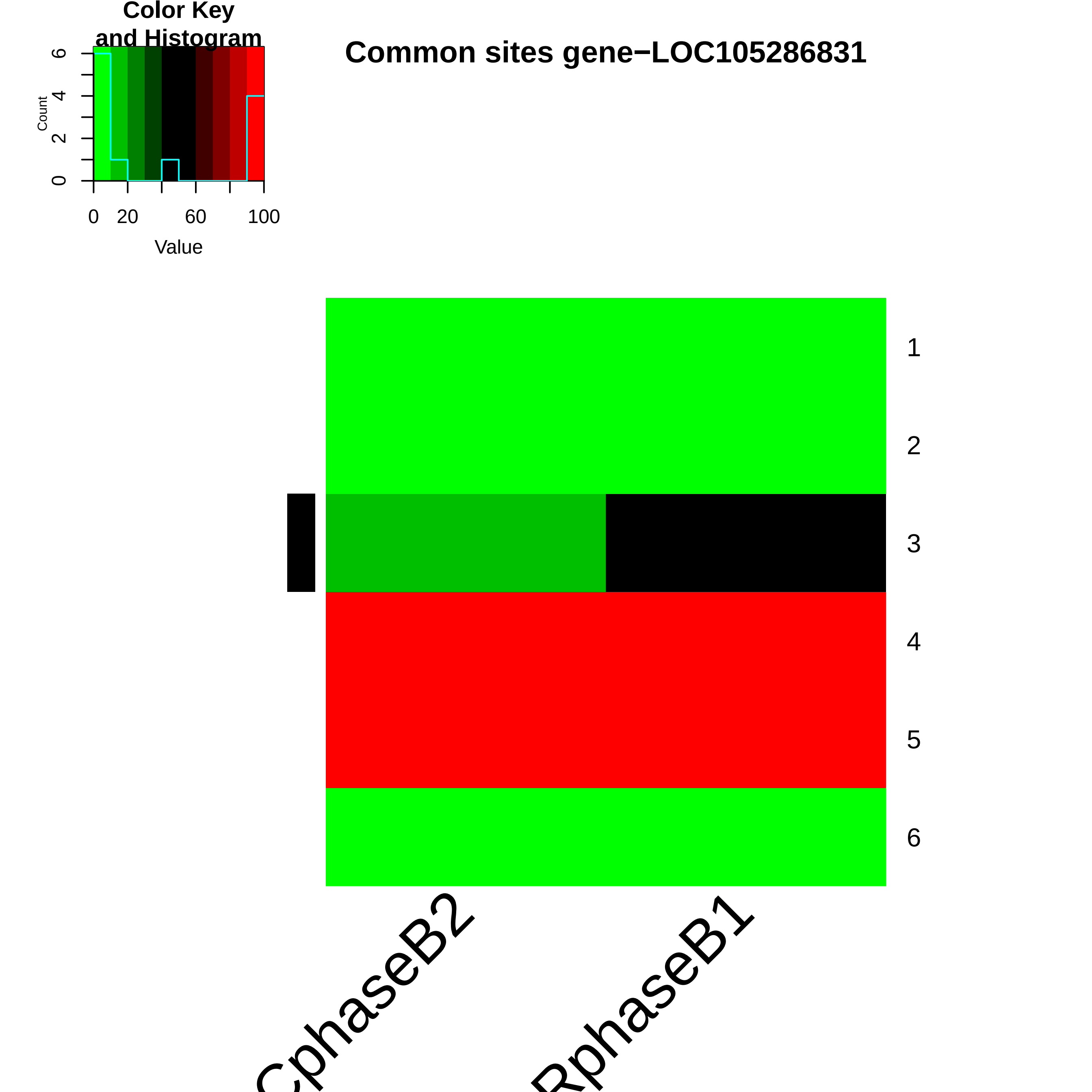

Color Key and Histogram
Count
0
2
4
6
0
20
60
100
Value
Common sites gene−LOC105286831
1
2
3
4
5
6
CphaseB2
RphaseB1

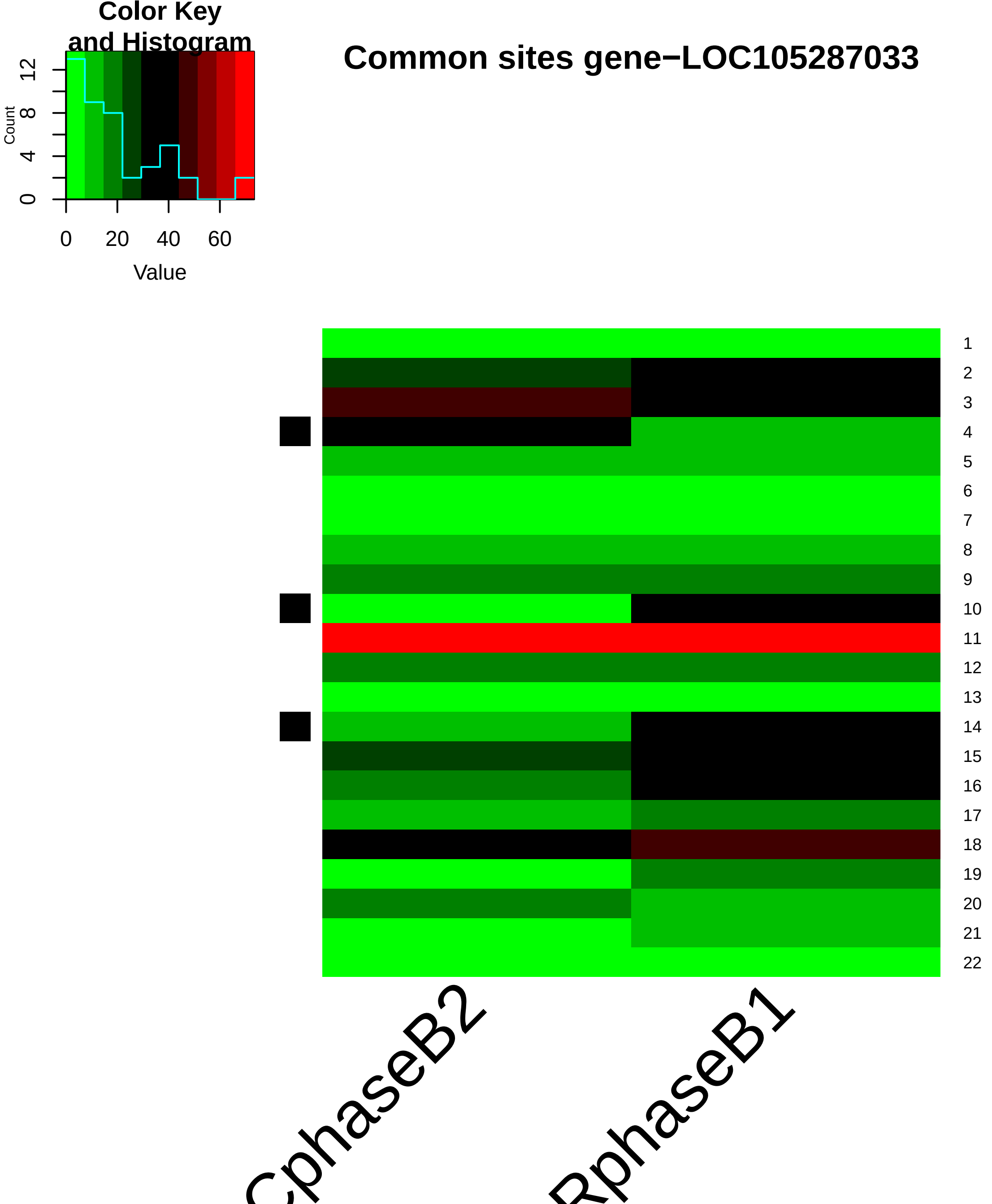

Color Key
and Histogram
Count
0
4
8
12
0
20
40
60
Value
Common sites gene−LOC105287033
CphaseB2
RphaseB1

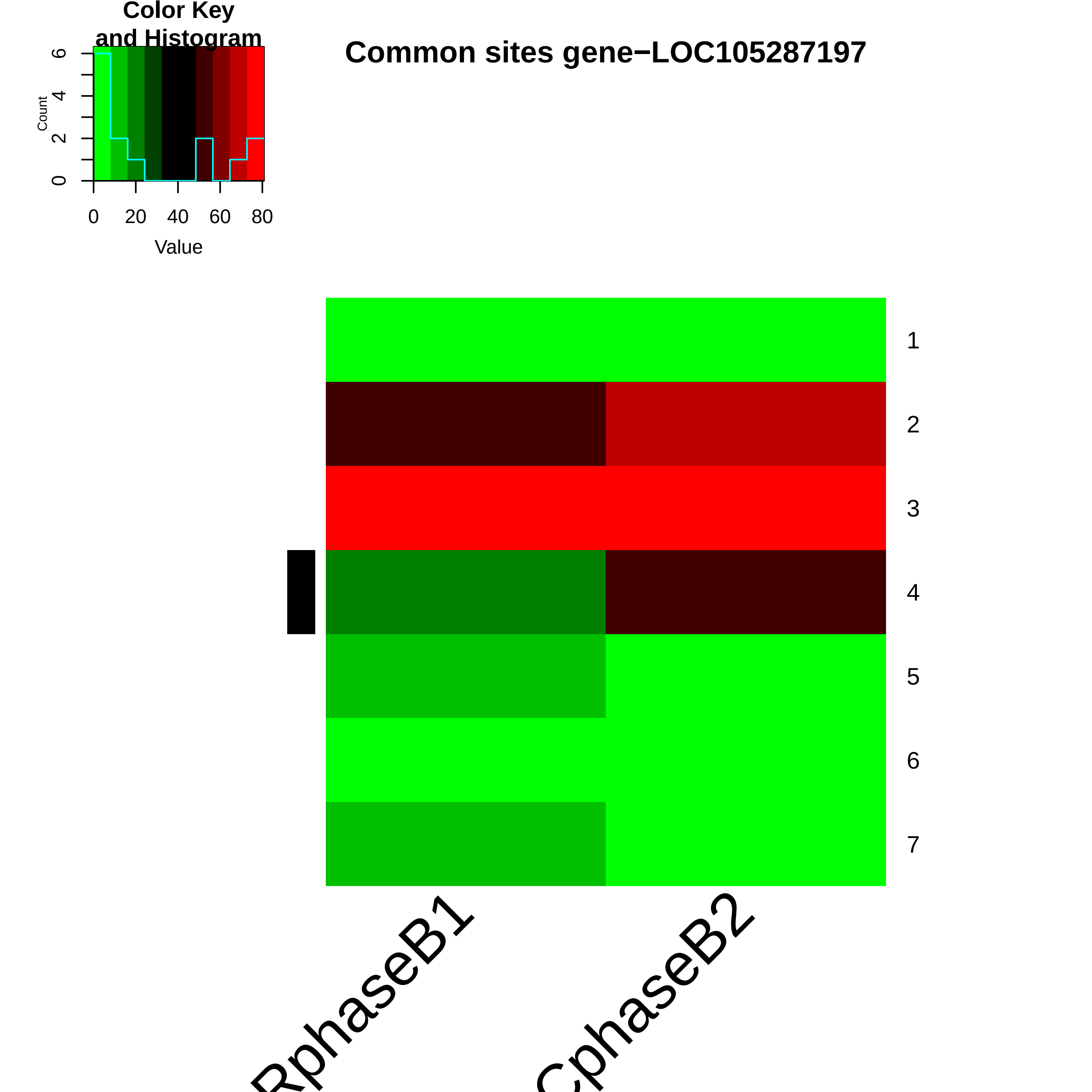

Color Key
and Histogram
Count
Value
Common sites gene−LOC105287197
1
2
3
4
5
6
7
RphaseB1
CphaseB2

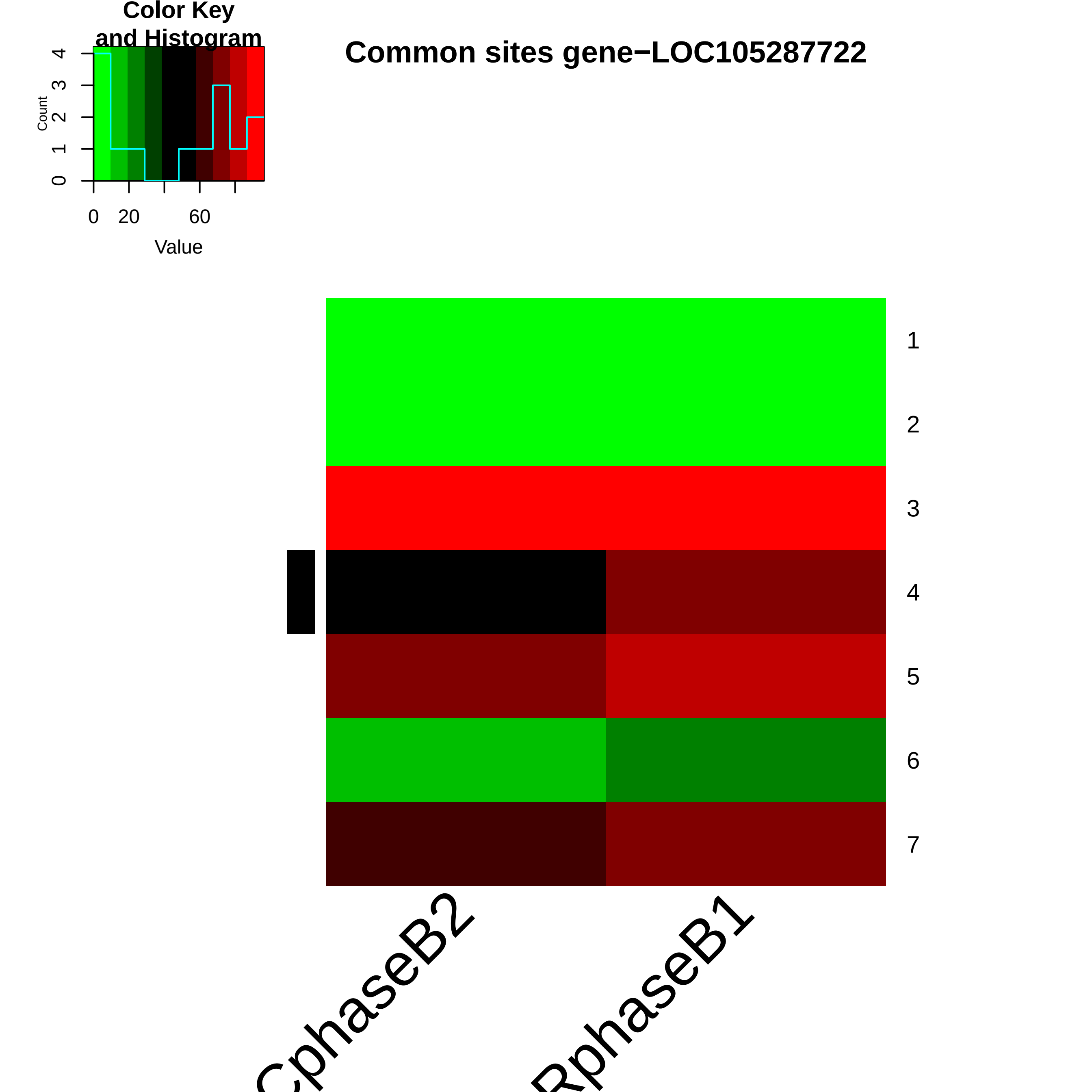
Color Key
and Histogram
Count
0
1
2
3
4
0
20
60
Value
Common sites gene−LOC105287722
1
2
3
4
5
6
7
CphaseB2
RphaseB1

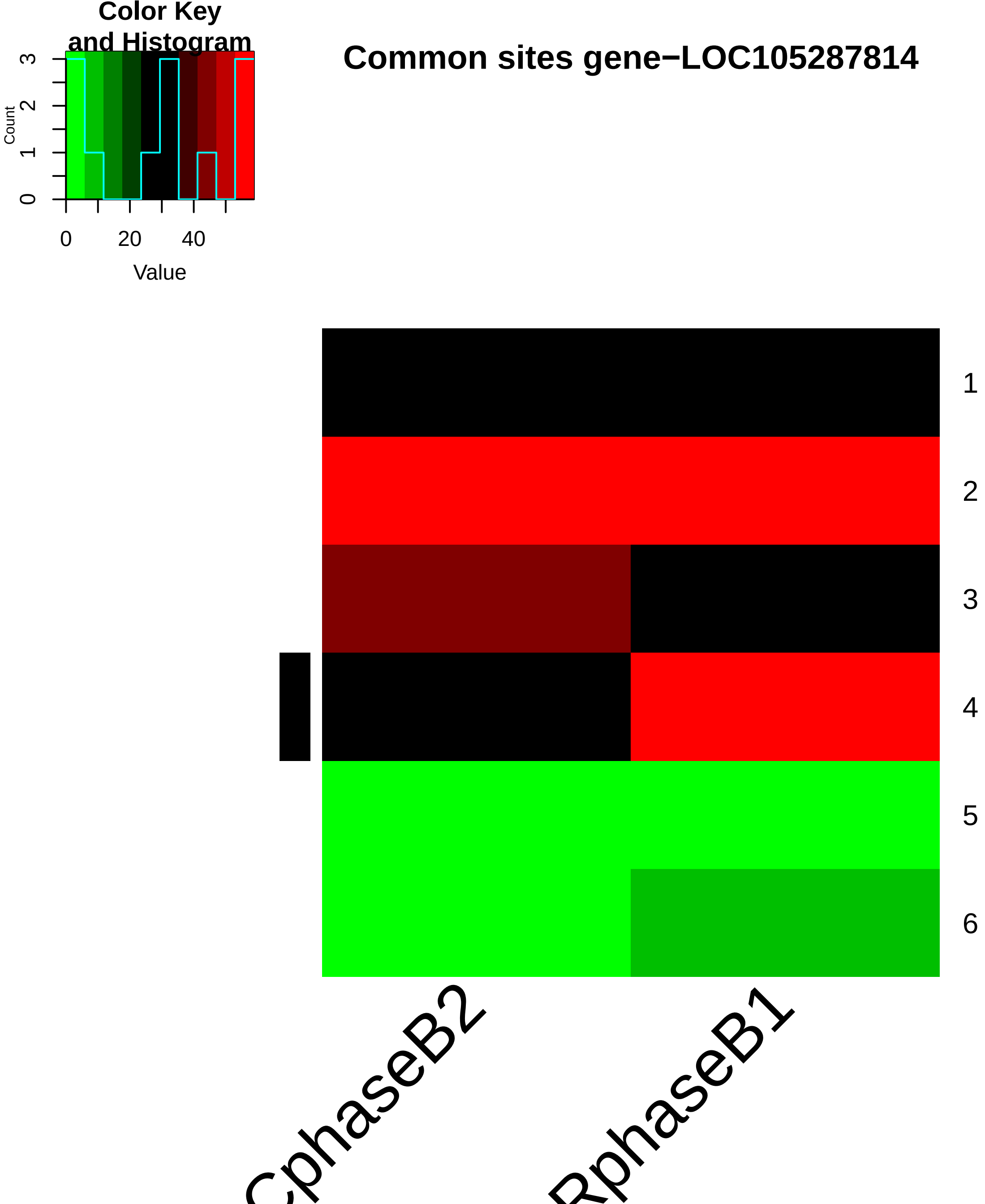
Color Key
and Histogram
Count
0
1
2
3
0
20
40
Value
Common sites gene−LOC105287814
1
2
3
4
5
6
CphaseB2
RphaseB1

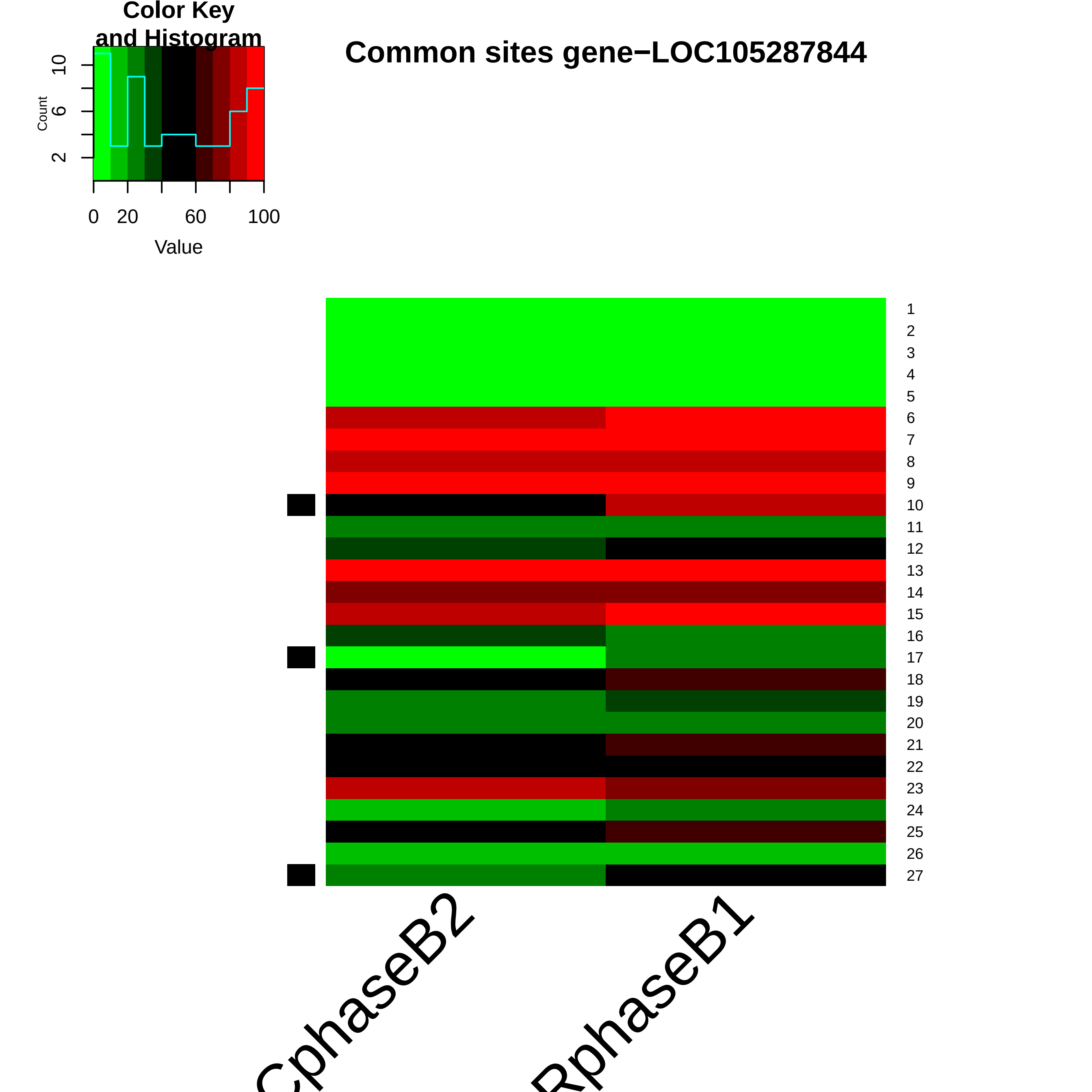

Color Key
and Histogram
Count
Value
Common sites gene−LOC105287844
CphaseB2
RphaseB1

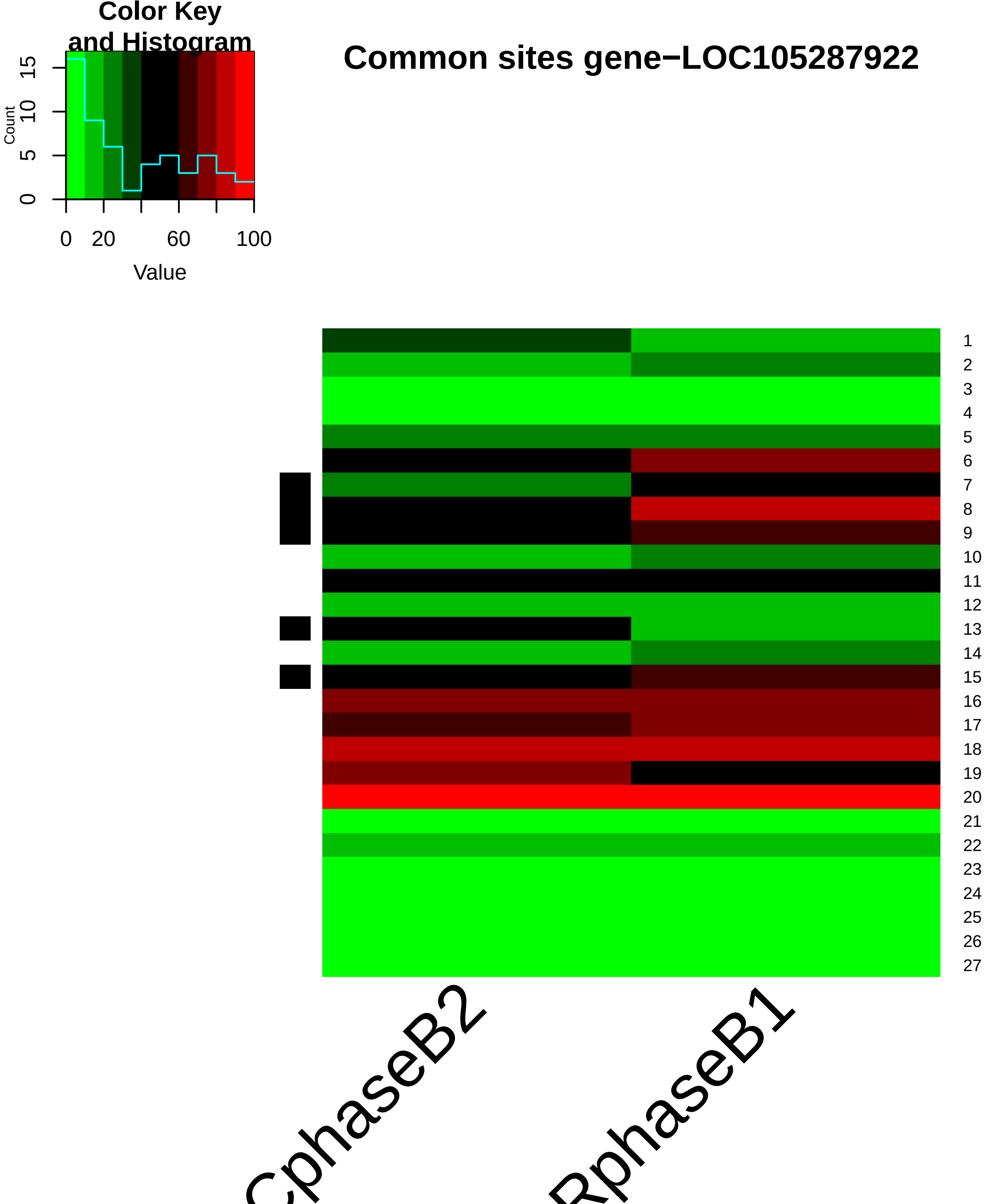
Color Key
and Histogram
Count
0
5
10
15
0
20
60
100
Value
Common sites gene−LOC105287922
CphaseB2
RphaseB1

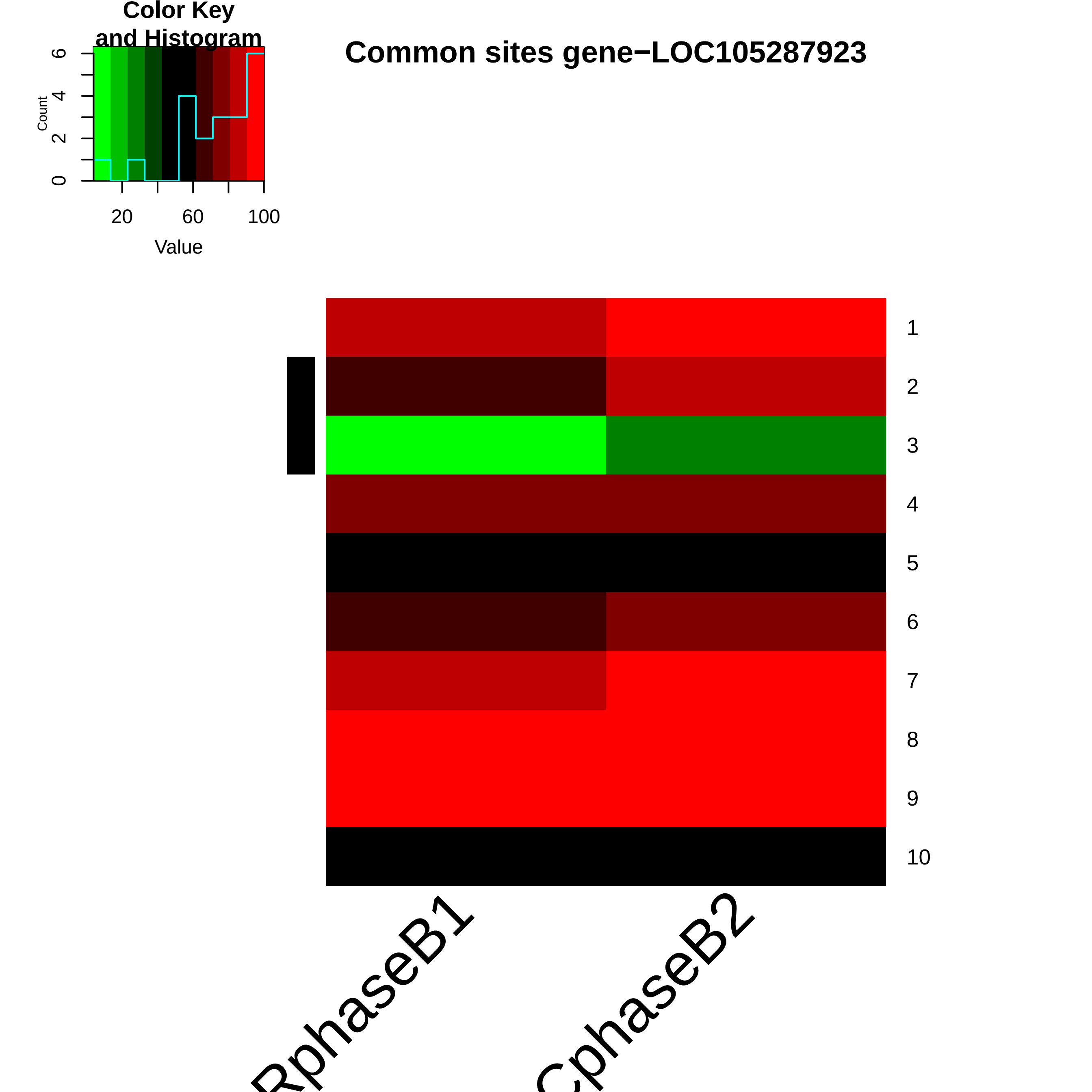

Color Key
and Histogram
Count
0
2
4
6
20
60
100
Value
Common sites gene−LOC105287923
1
2
3
4
5
6
7
8
9
10
RphaseB1
CphaseB2

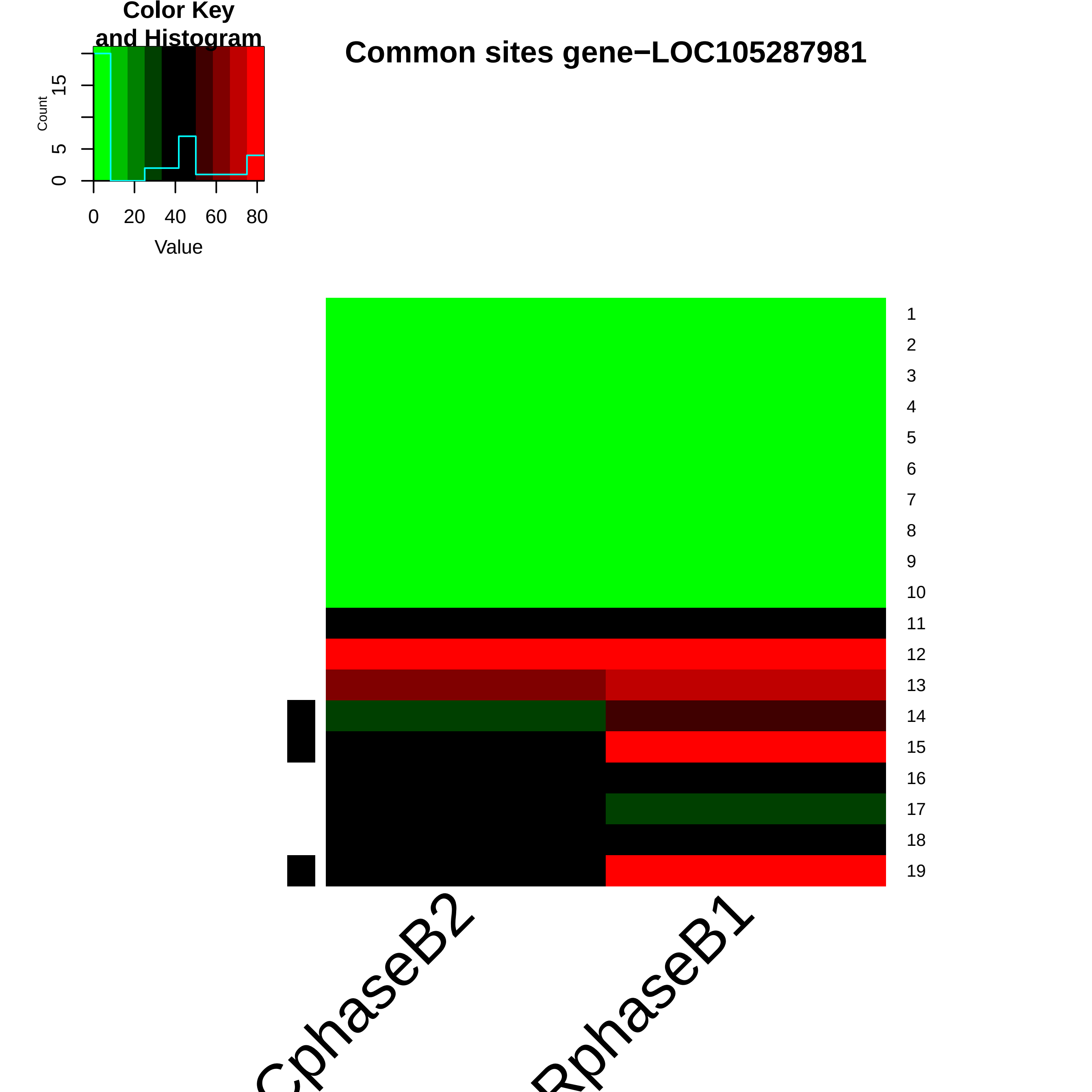

Color Key
and Histogram
Count
0
5
15
0
20
40
60
80
Value
Common sites gene−LOC105287981
CphaseB2
RphaseB1

### Common sites gene−LOC113561616

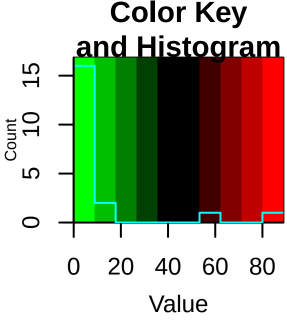

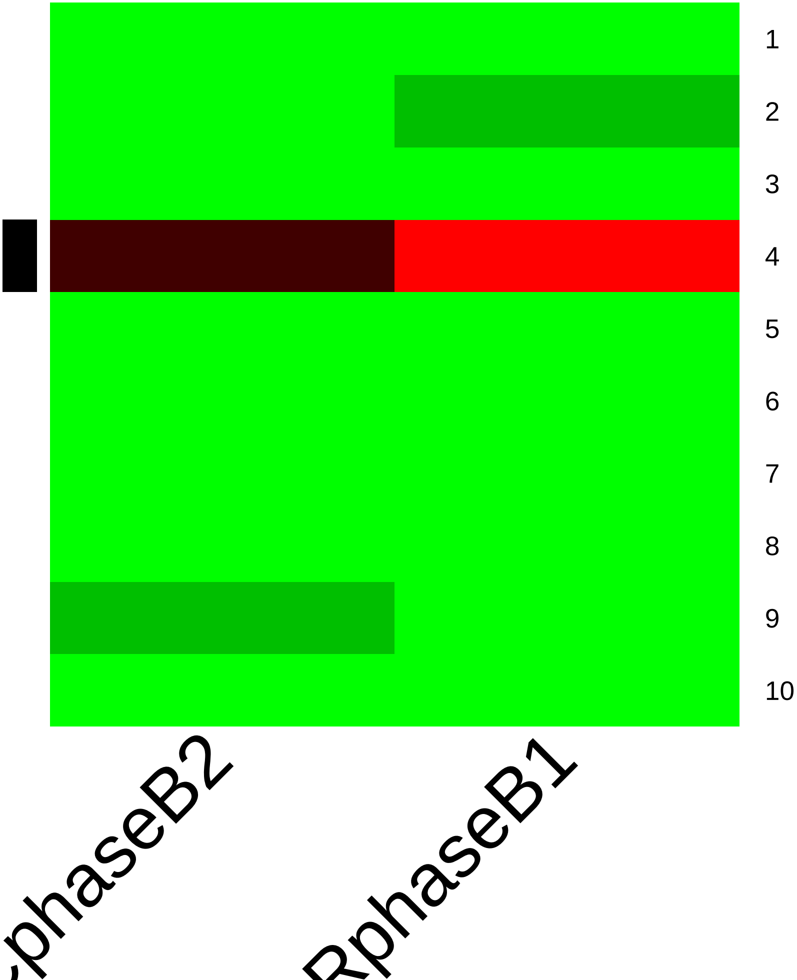

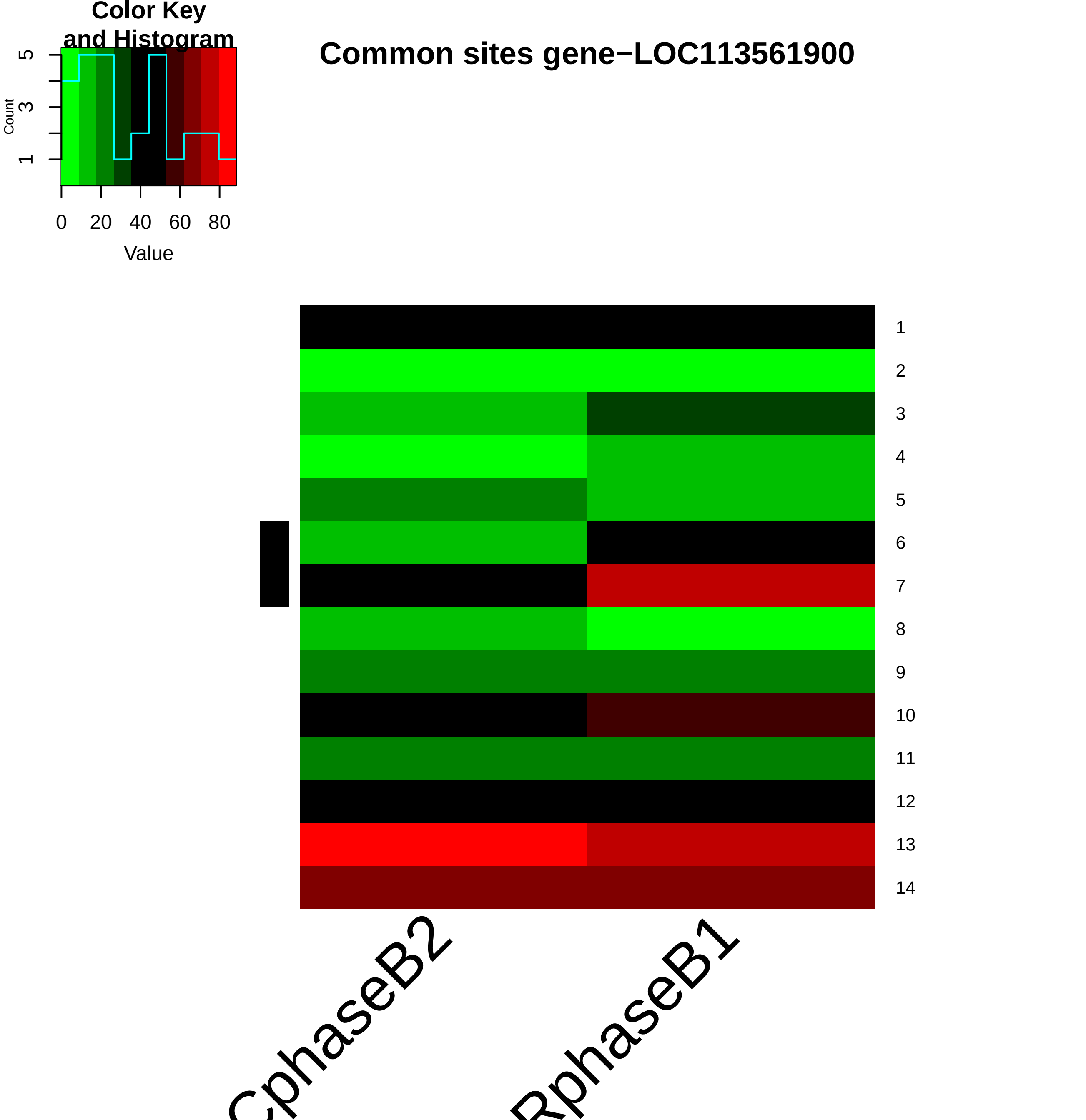
Color Key
and Histogram
Count
5
3
1
0
20
40
60
80
Value
Common sites gene−LOC113561900
1
2
3
4
5
6
7
8
9
10
11
12
13
14
CphaseB2
RphaseB1

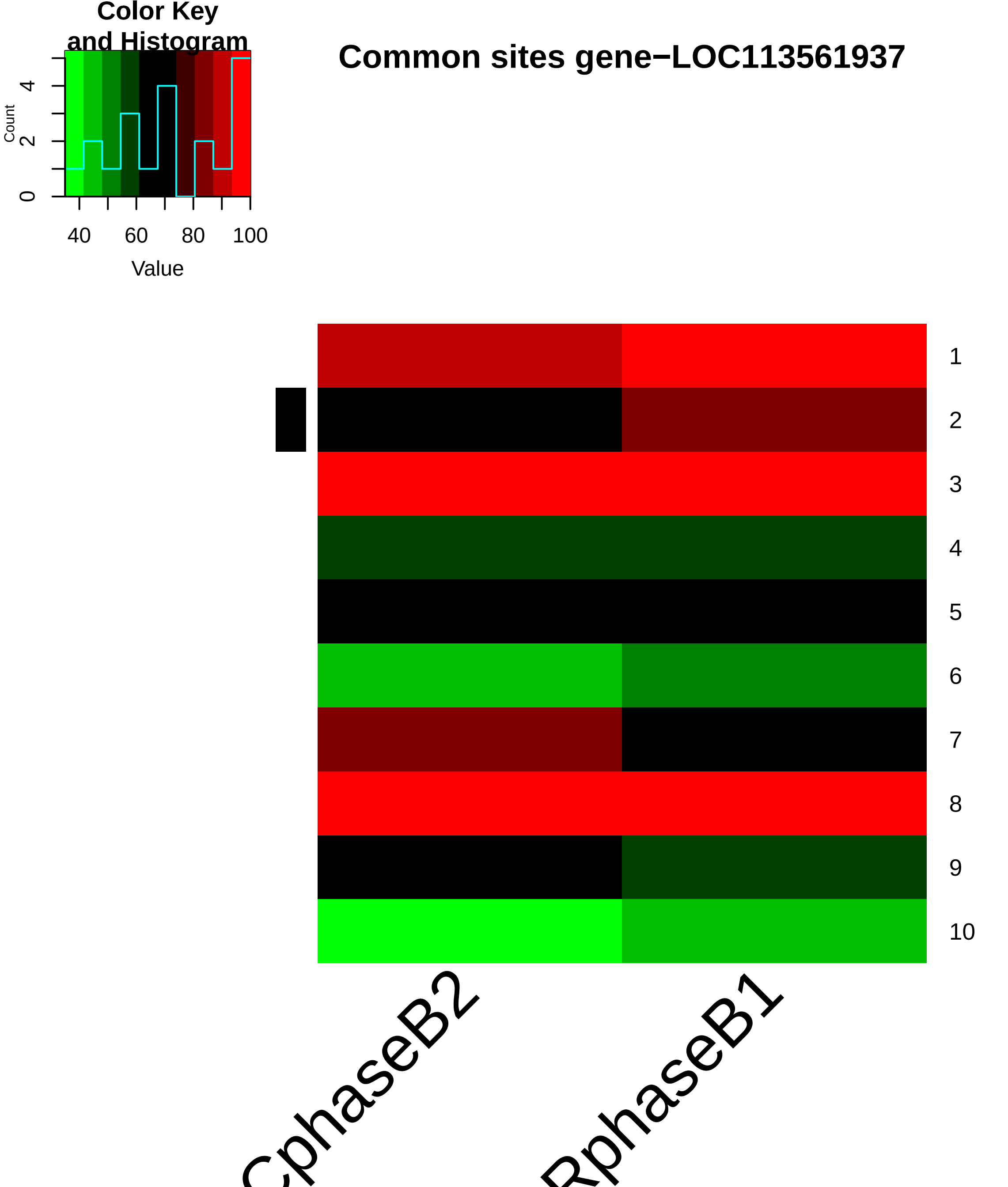
Color Key
and Histogram
Count
0
2
4
40
60
80
100
Value
Common sites gene−LOC113561937
1
2
3
4
5
6
7
8
9
10
CphaseB2
RphaseB1

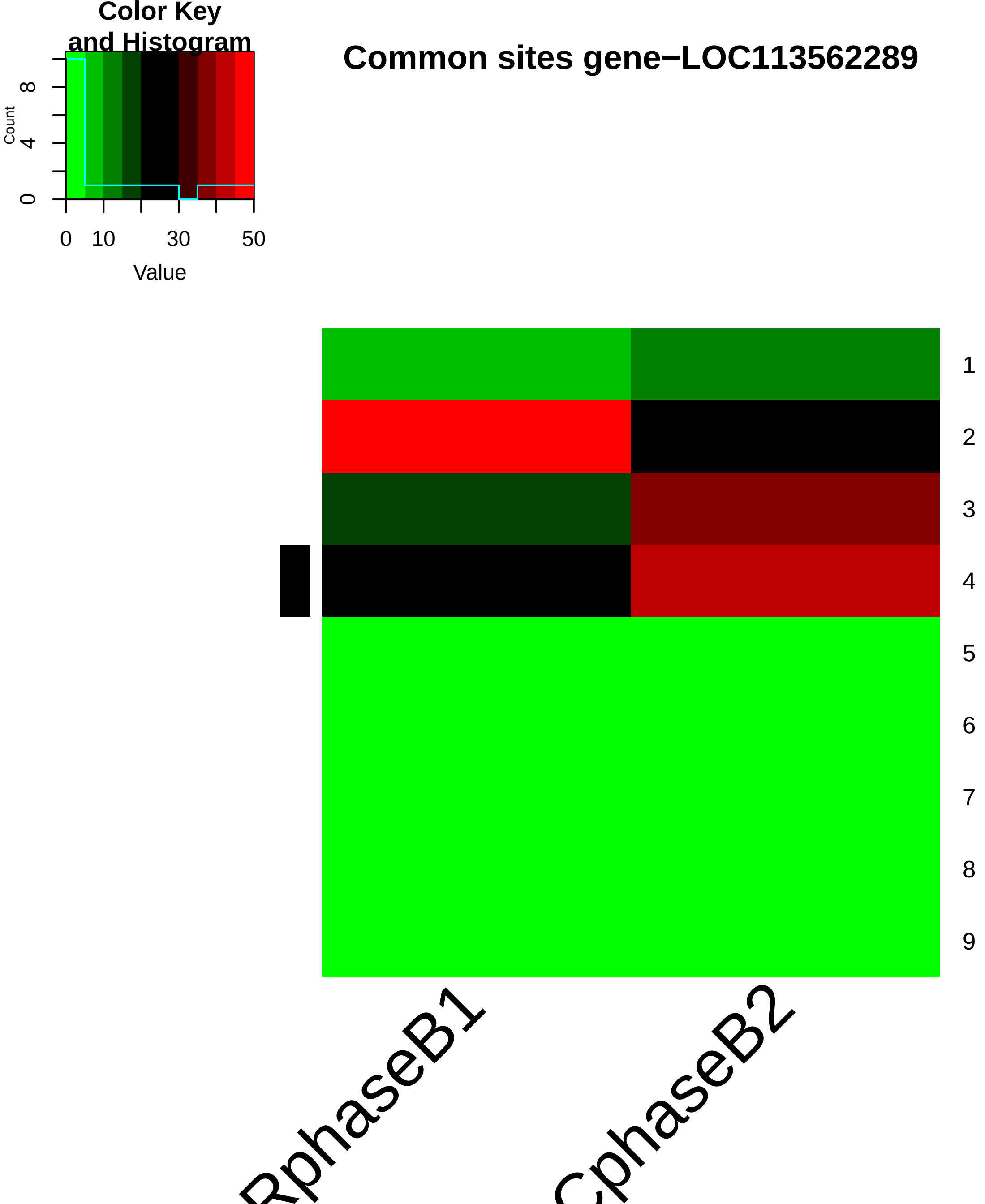

Color Key
and Histogram
Count
0
4
8
0
10
30
50
Value
Common sites gene−LOC113562289
1
2
3
4
5
6
7
8
9
RphaseB1
CphaseB2

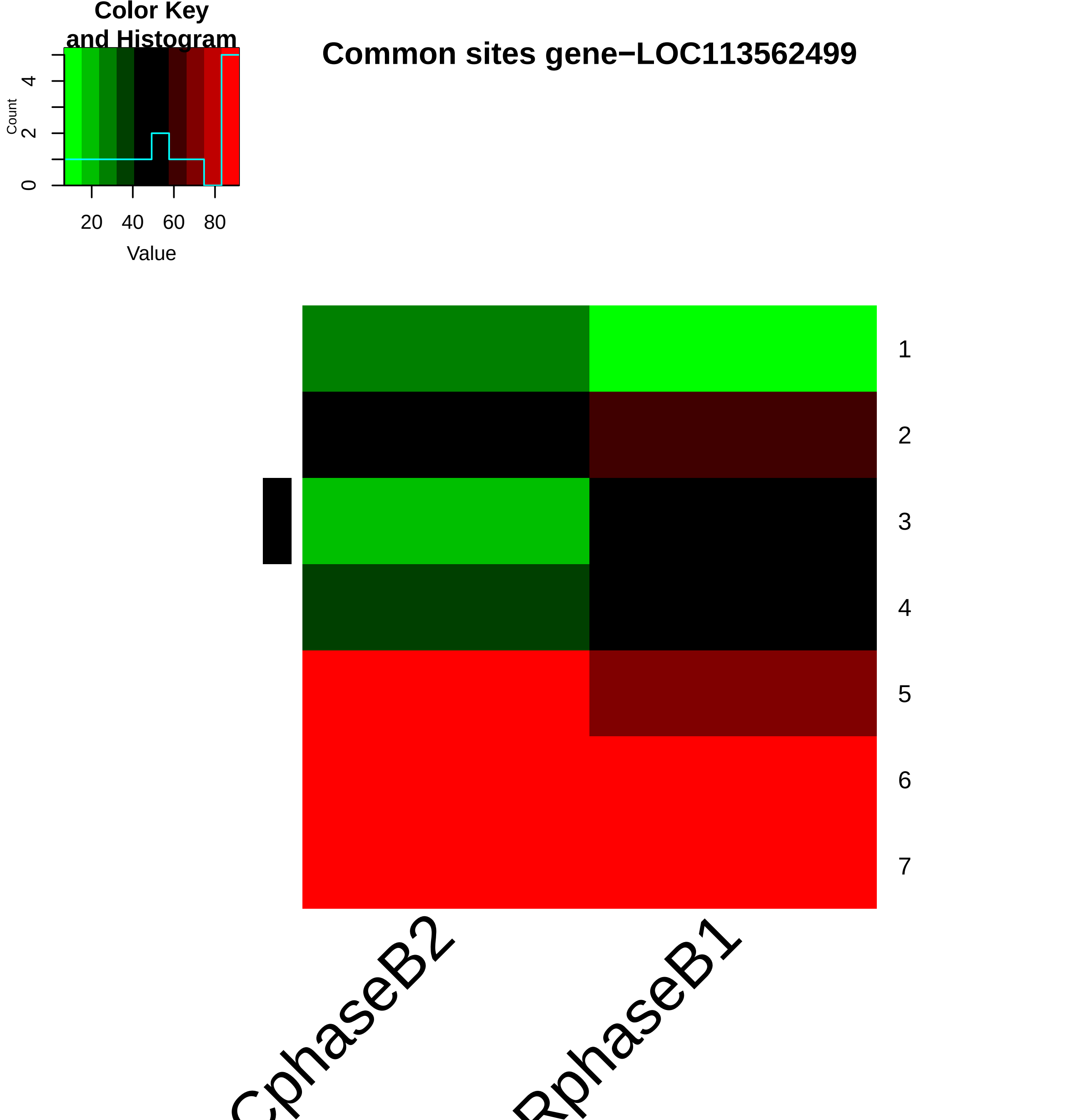

Color Key
and Histogram
Count
0
2
4
20
40
60
80
Value
Common sites gene−LOC113562499
1
2
3
4
5
6
7
CphaseB2
RphaseB1

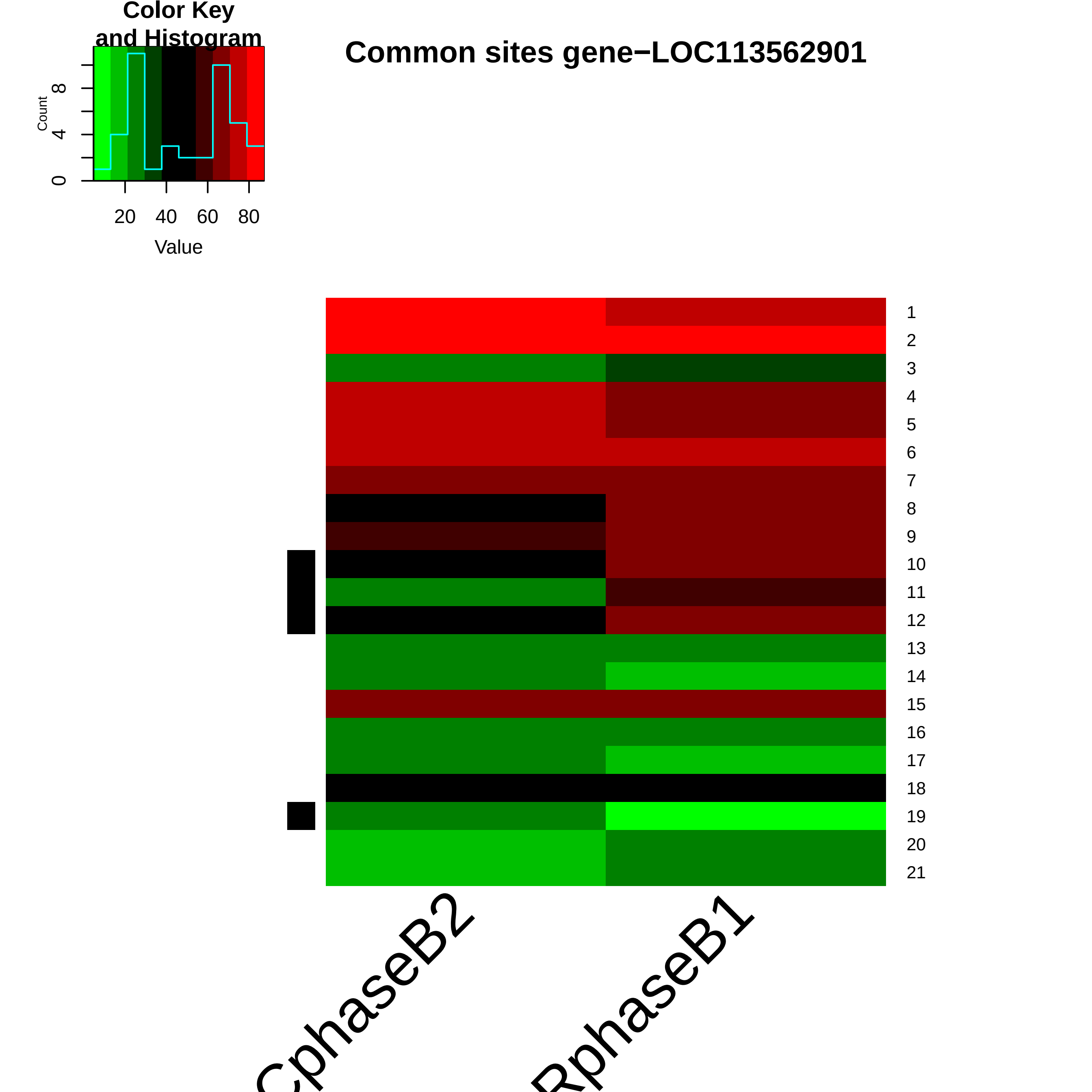

Color Key
and Histogram
Count
0
4
8
20
40
60
80
Value
Common sites gene−LOC113562901
CphaseB2
RphaseB1

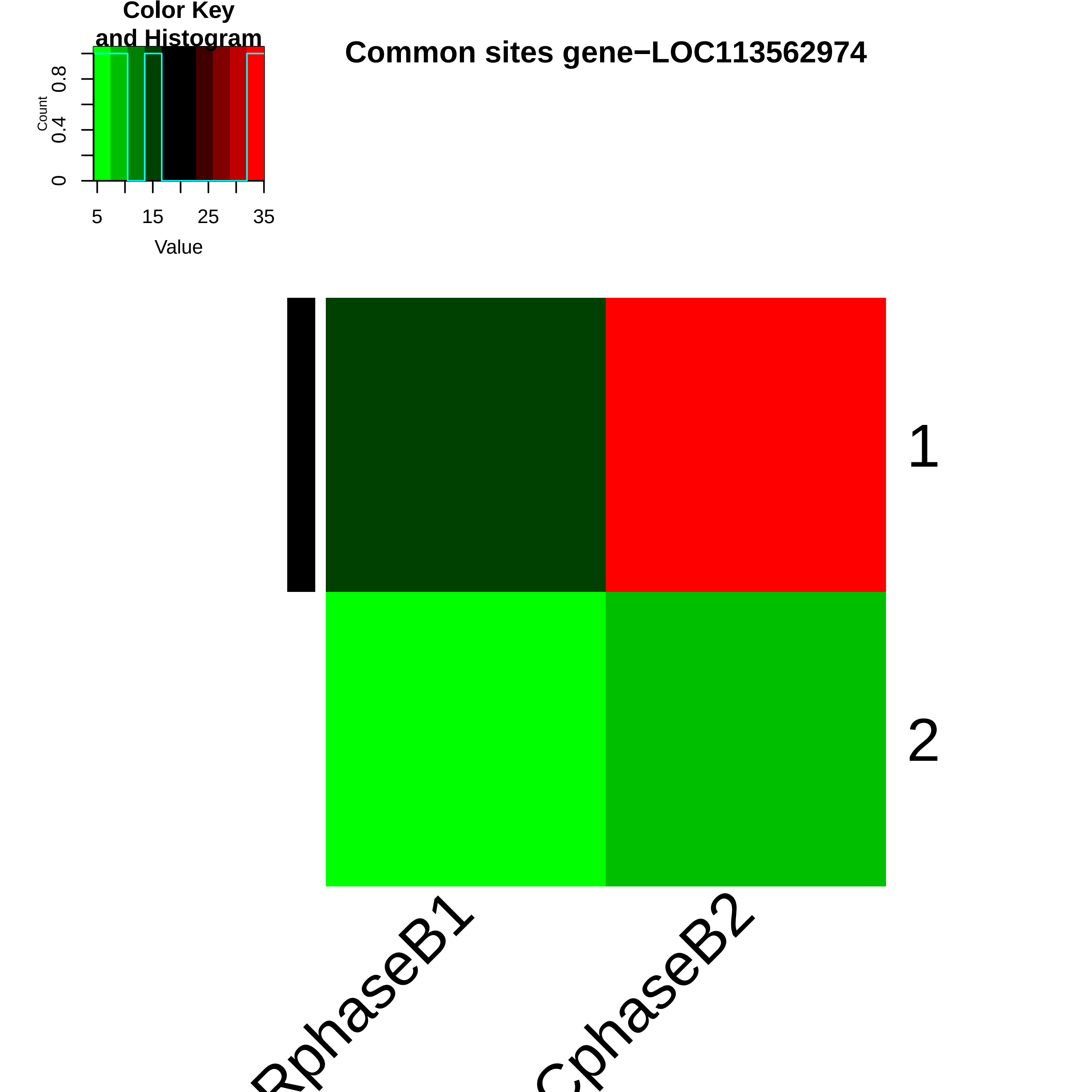
Color Key
and Histogram
Count
0
0.4
0.8
5
15
25
35
Value
Common sites gene−LOC113562974
1
2
RphaseB1
CphaseB2

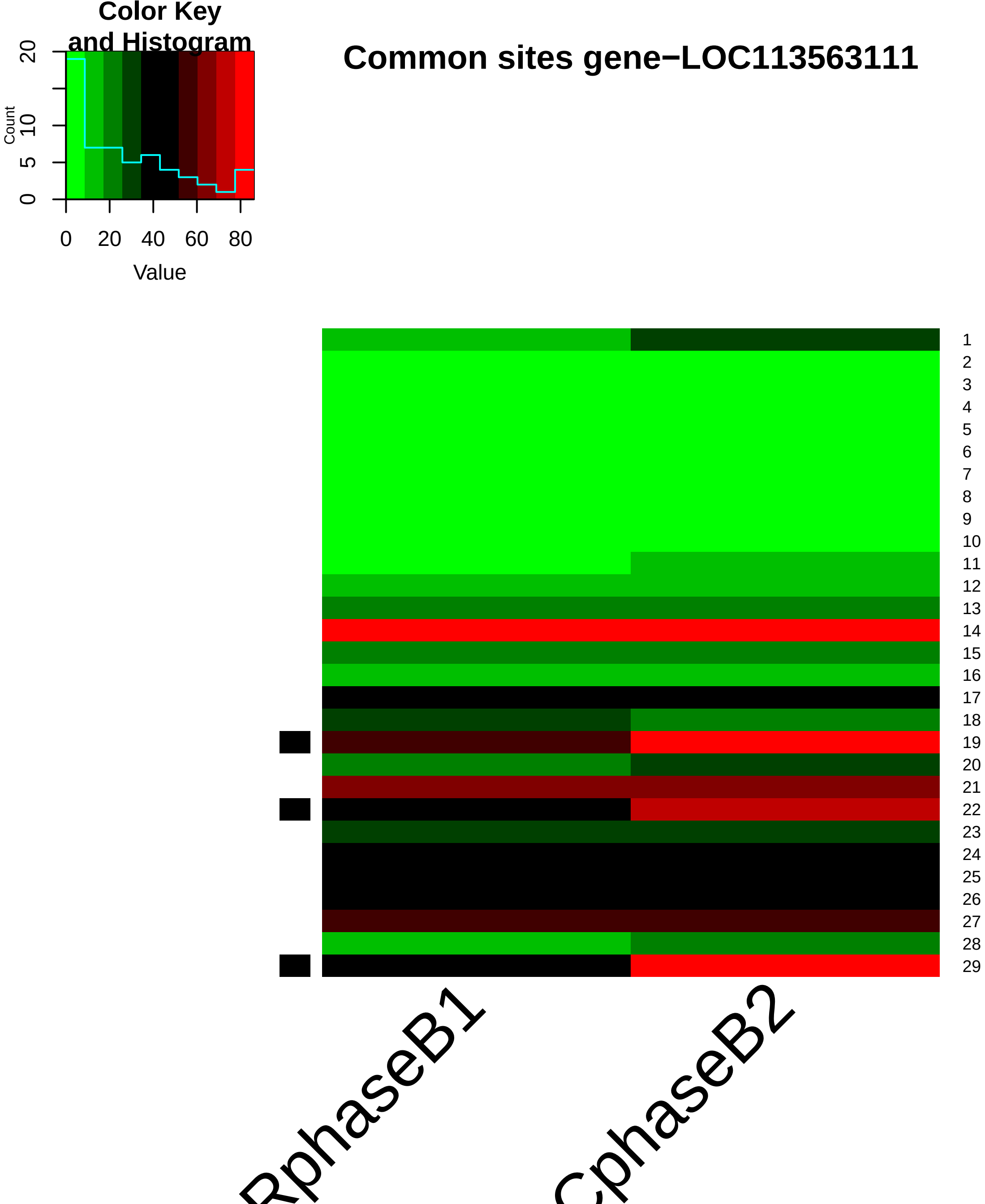

Color Key
and Histogram
Count
0
5
10
20
0
20
40
60
80
Value
Common sites gene−LOC113563111
RphaseB1
CphaseB2

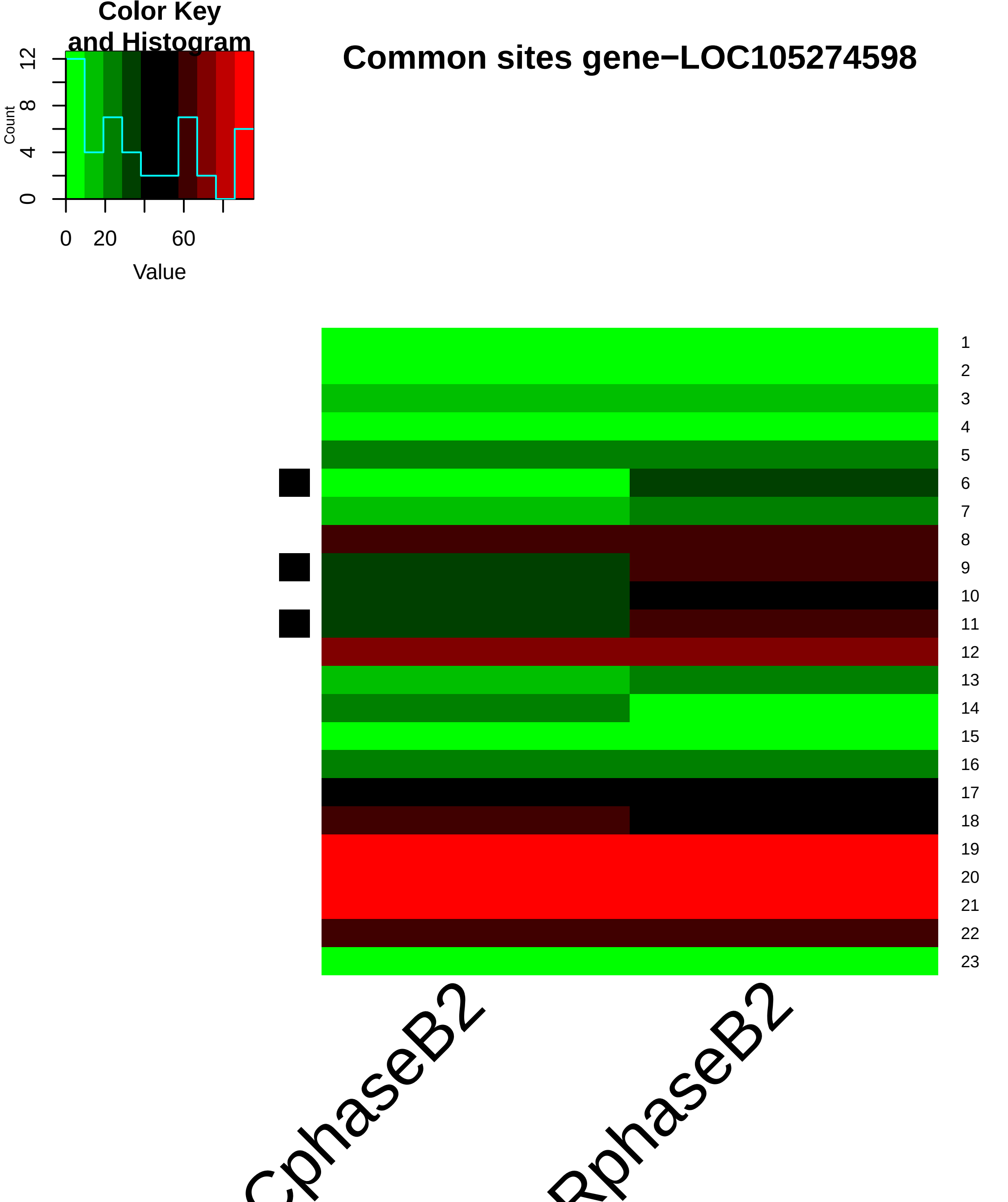
Color Key
and Histogram
Count
0
4
8
12
0
20
60
Value
Common sites gene−LOC105274598
CphaseB2
RphaseB2

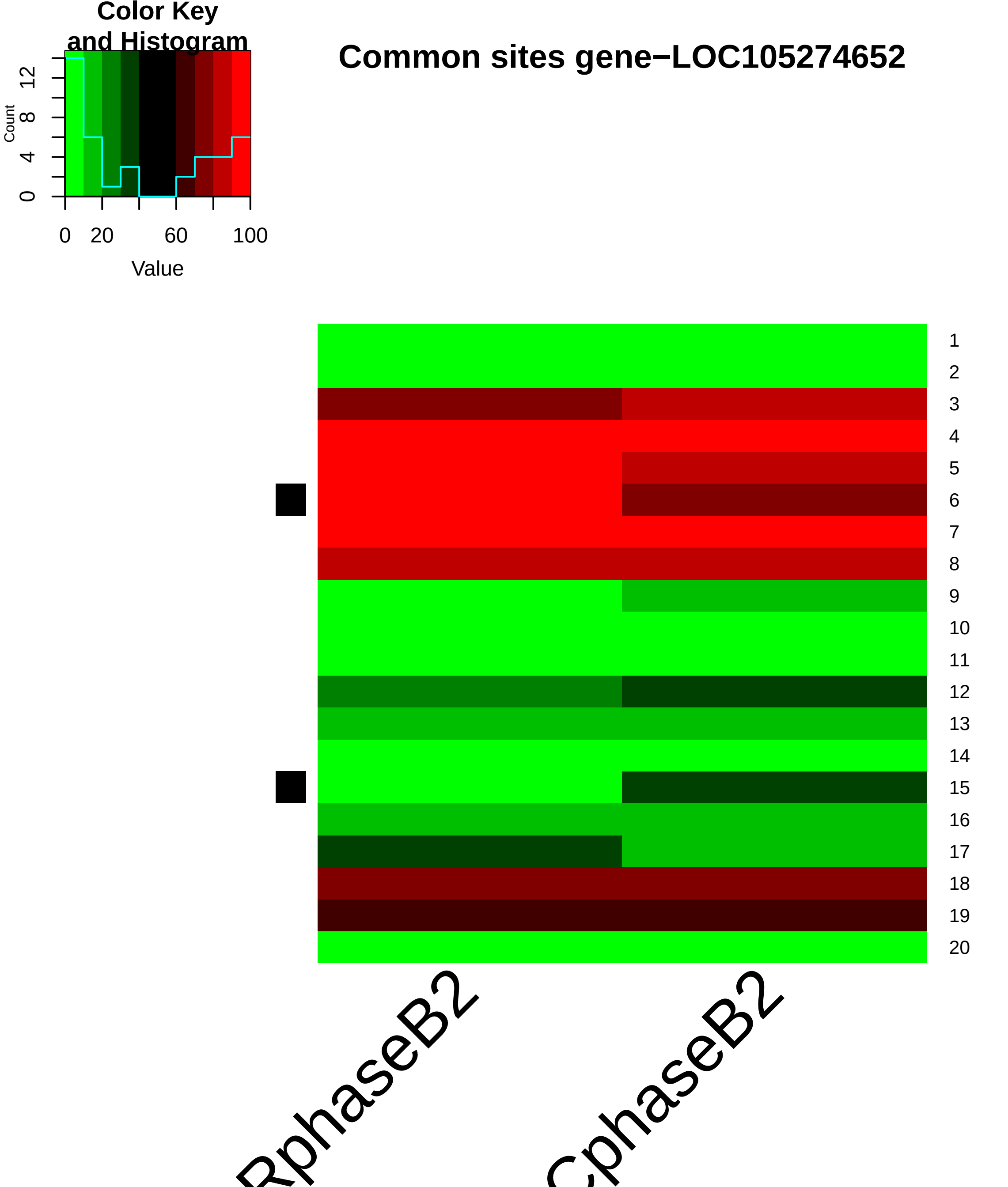
Color Key
and Histogram
Count
0
4
8
12
0
20
60
100
Value
Common sites gene−LOC105274652
RphaseB2
CphaseB2

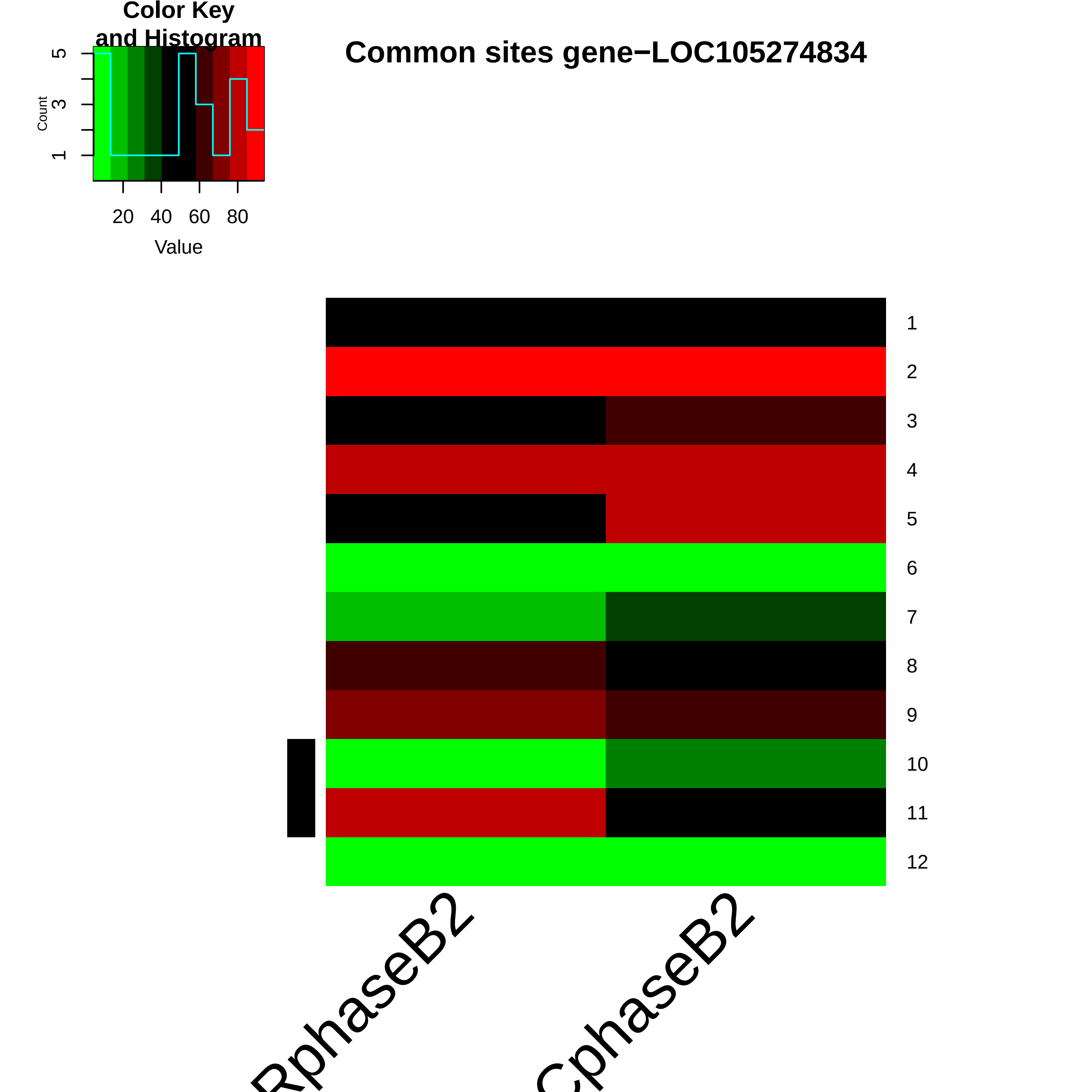
Color Key
and Histogram
Count
1
3
5
20
40
60
80
Value
Common sites gene−LOC105274834
1
2
3
4
5
6
7
8
9
10
11
12
RphaseB2
CphaseB2

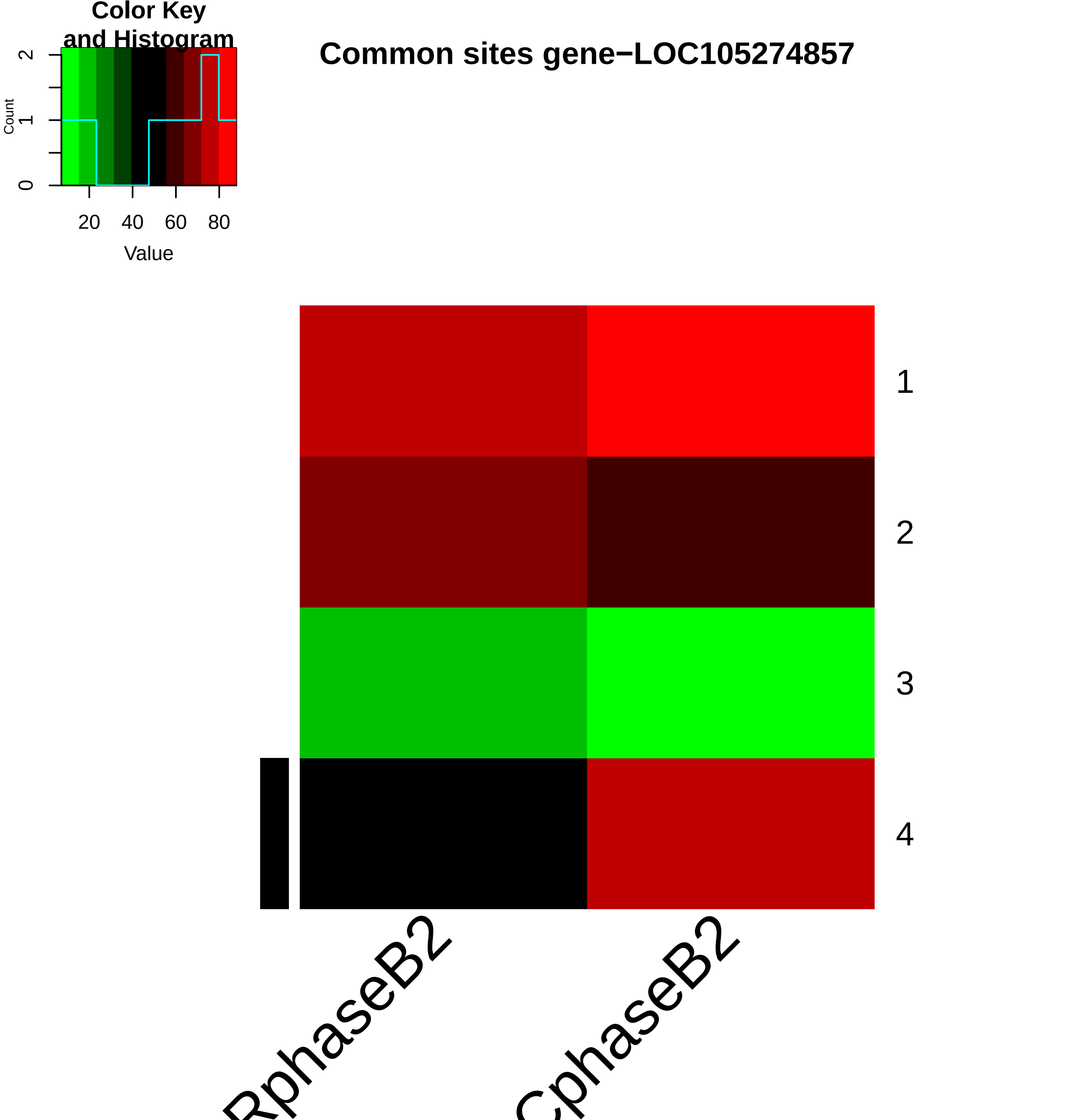
Color Key
and Histogram
Count
0
1
2
20
40
60
80
Value
Common sites gene−LOC105274857
1
2
3
4
RphaseB2
CphaseB2

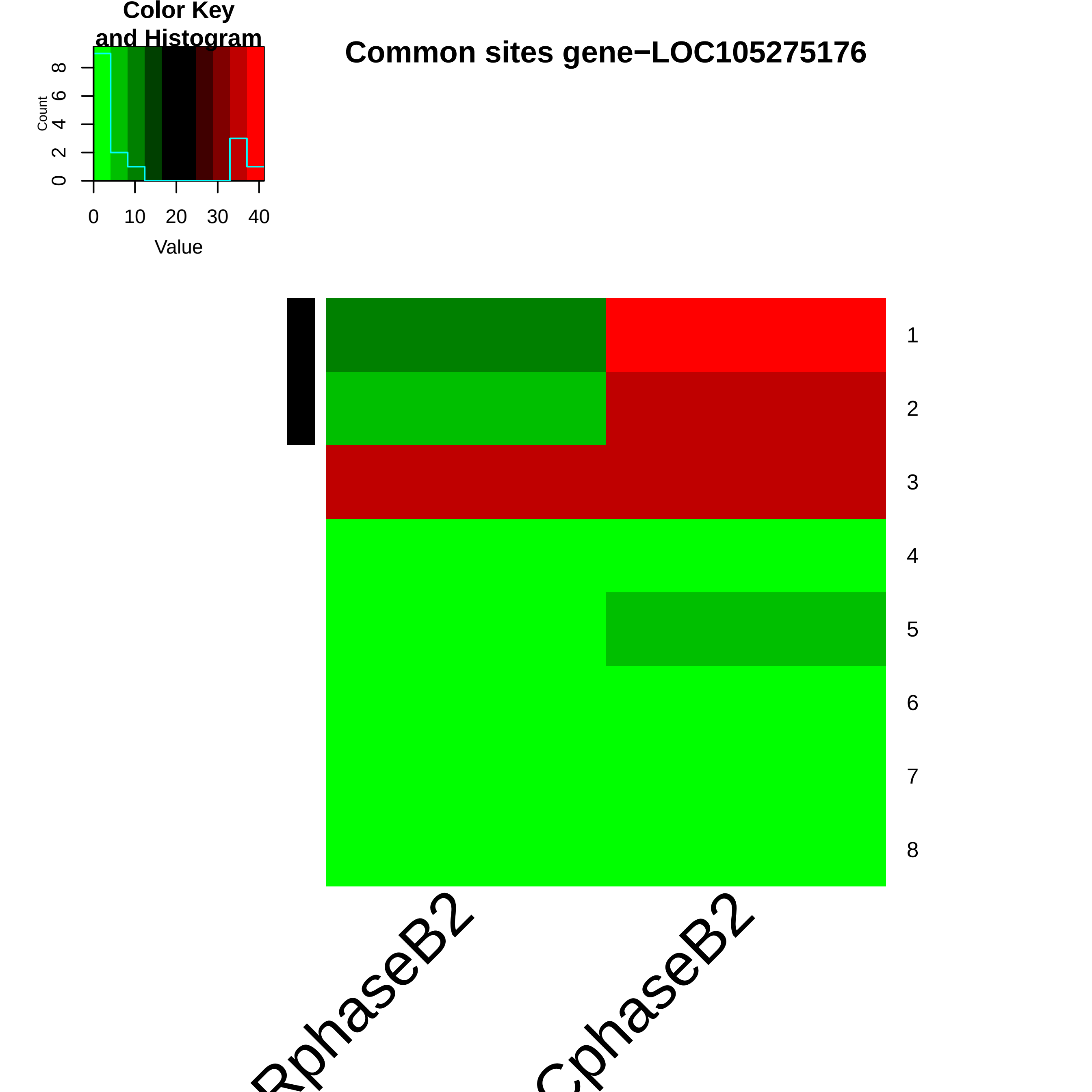

Color Key
and Histogram
Count
0
2
4
6
8
0
10
20
30
40
Value
Common sites gene−LOC105275176
1
2
3
4
5
6
7
8
RphaseB2
CphaseB2

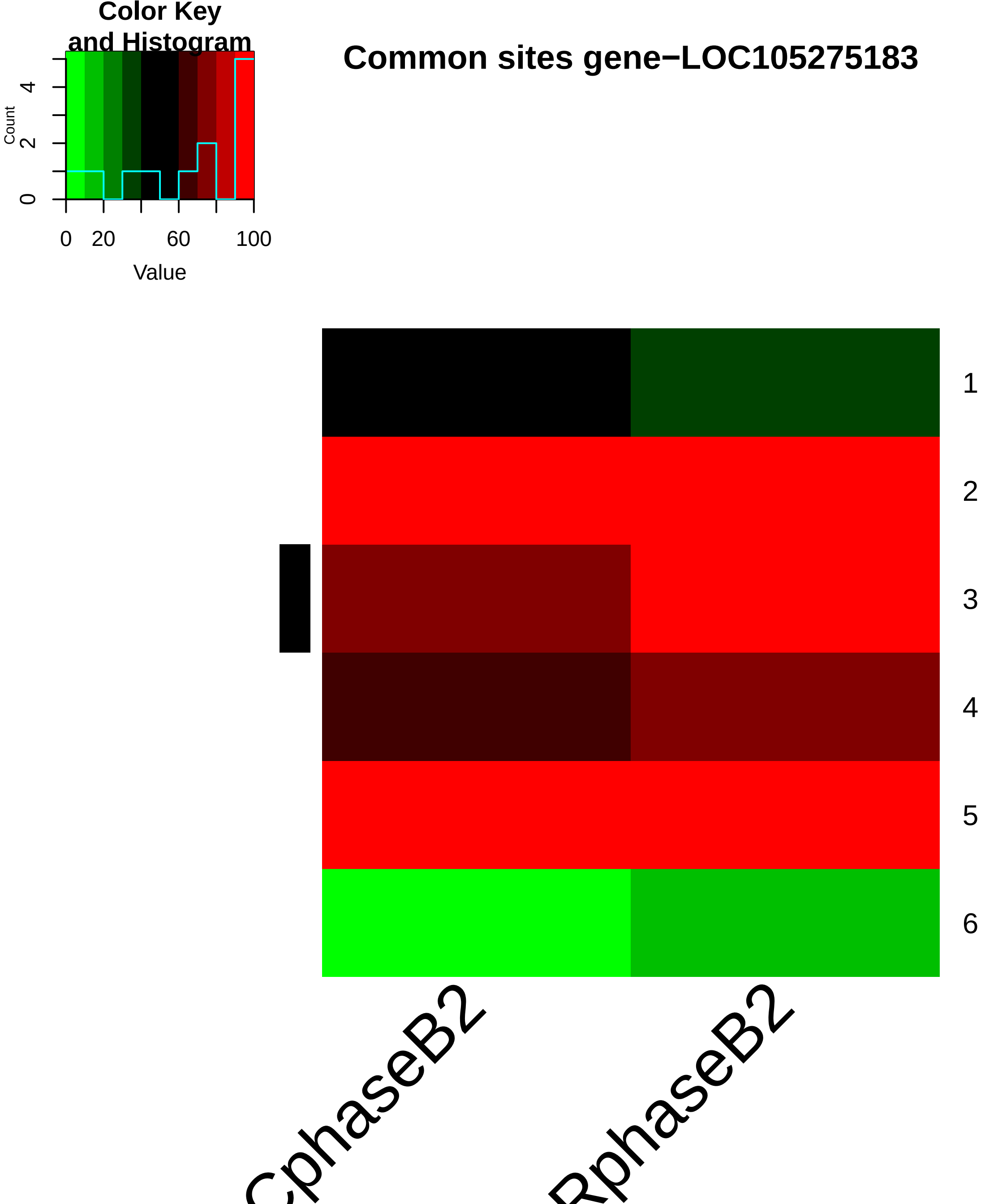
Color Key
and Histogram
Count
0
2
4
0
20
60
100
Value
Common sites gene−LOC105275183
1
2
3
4
5
6

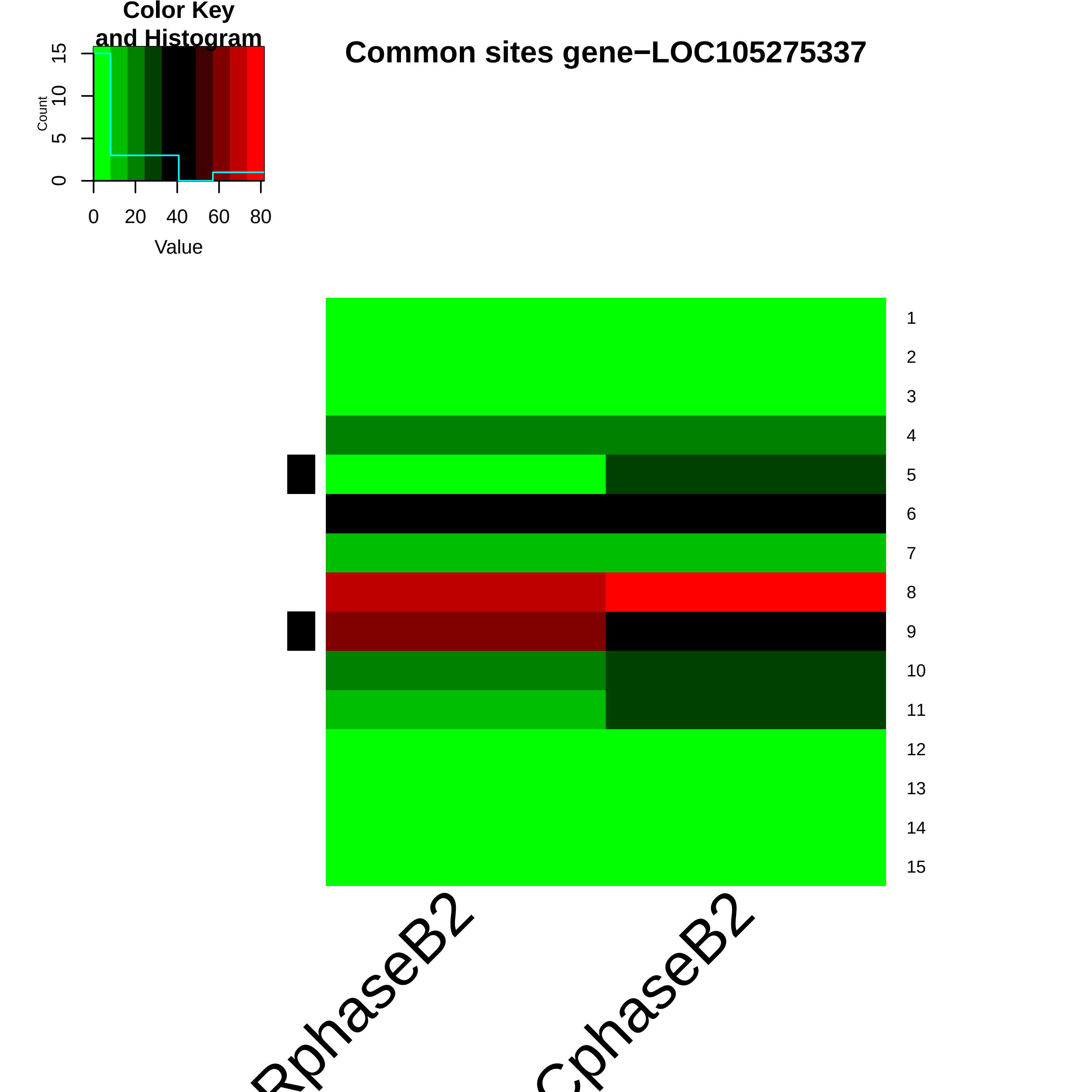
Color Key
and Histogram
Count
Value
Common sites gene−LOC105275337
RphaseB2
CphaseB2

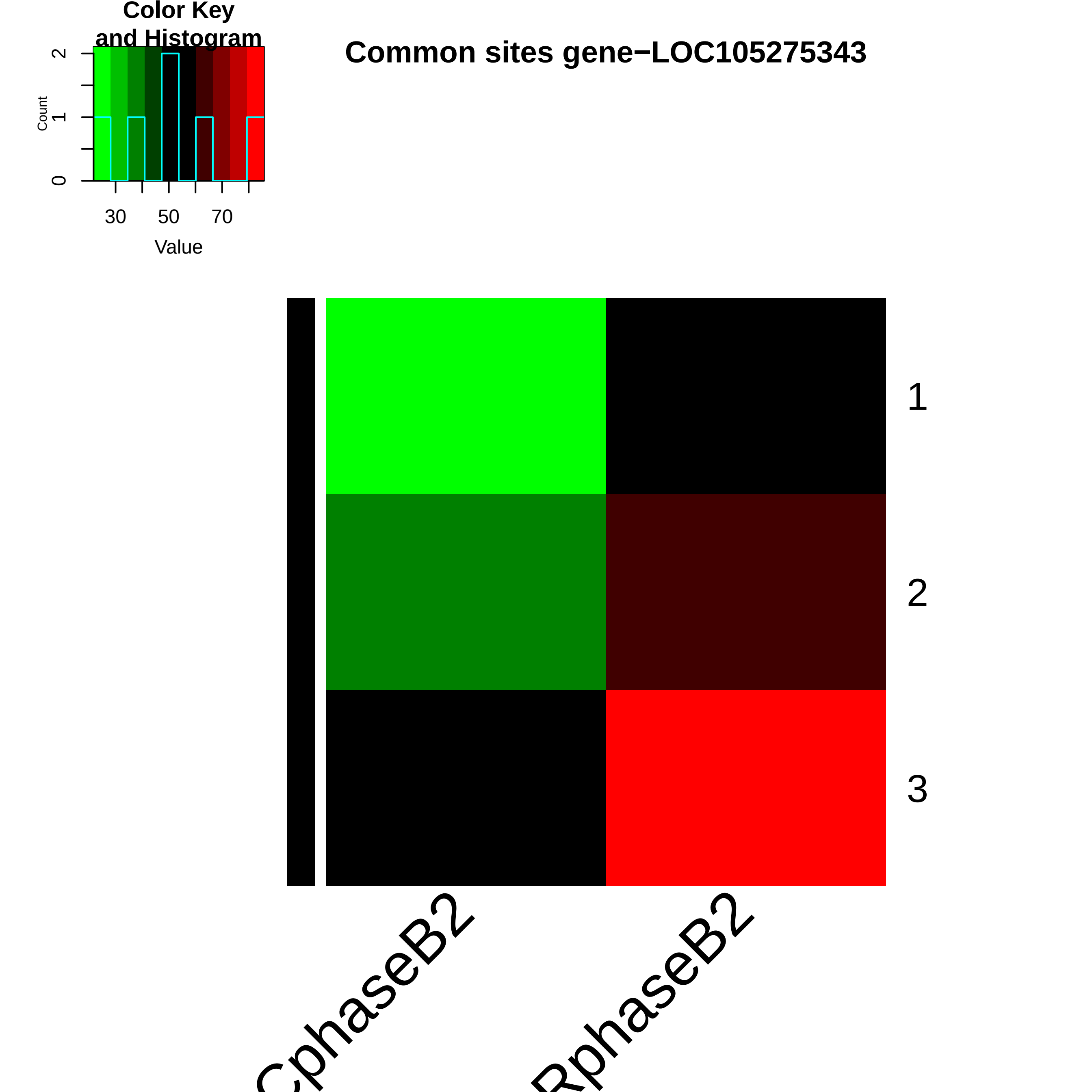

Color Key
and Histogram
Count
0
1
2
30
50
70
Value
Common sites gene−LOC105275343
1
2
3
CphaseB2
RphaseB2

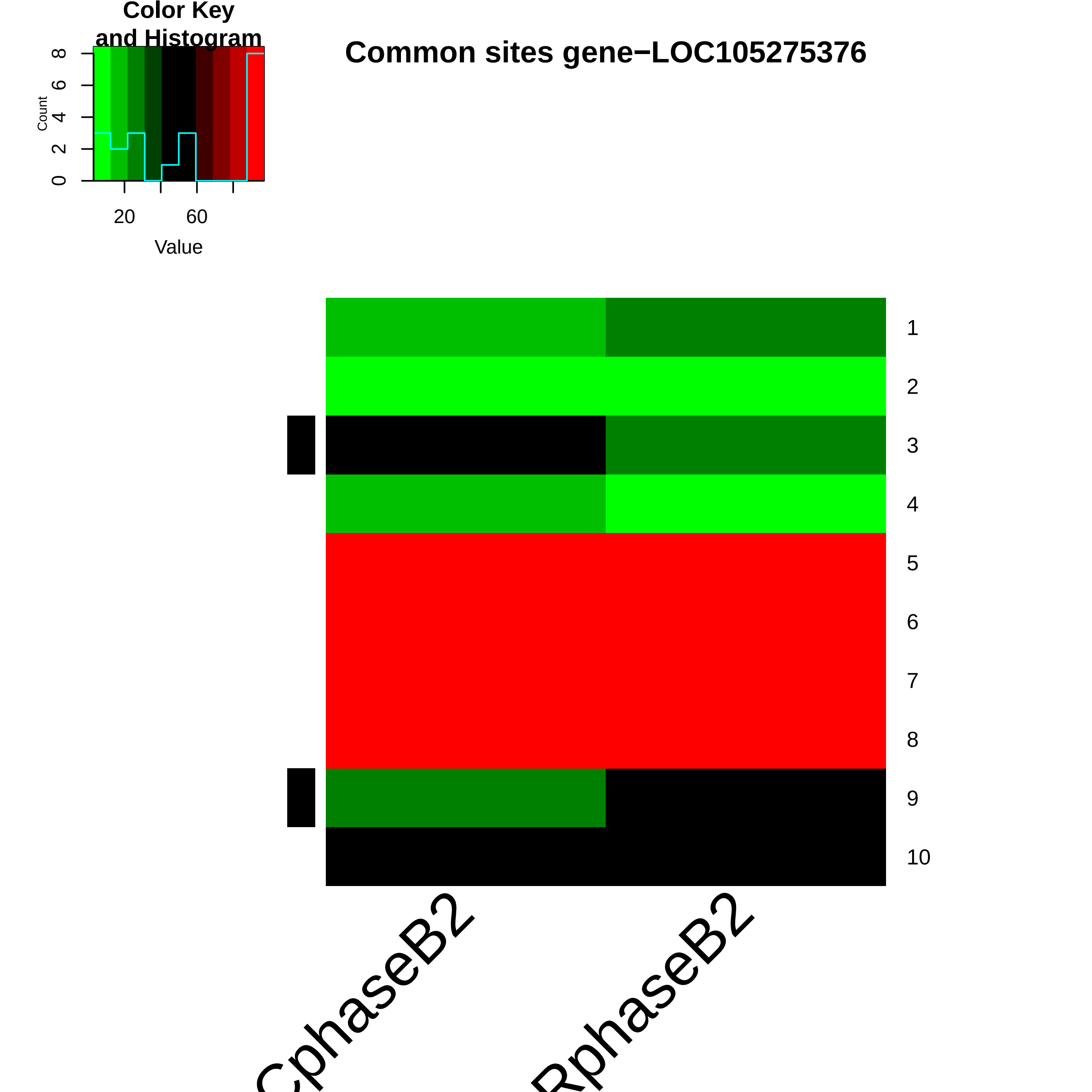
Color Key
and Histogram
Count
0
2
4
6
8
20
60
Value
Common sites gene−LOC105275376
1
2
3
4
5
6
7
8
9
10

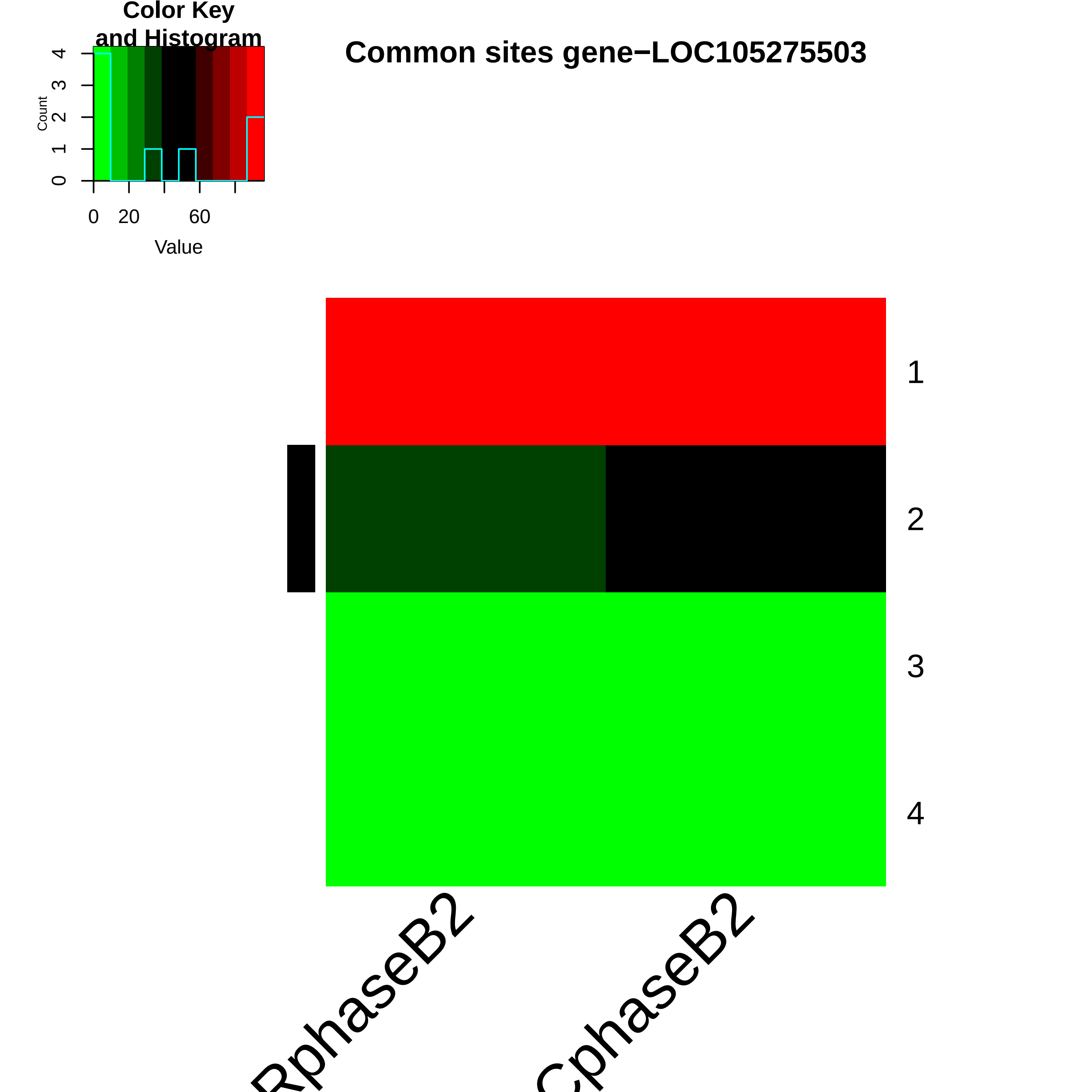
Color Key
and Histogram
Count
Value
0
20
60
0
1
2
3
4
Common sites gene−LOC105275503
1
2
3
4
RphaseB2
CphaseB2

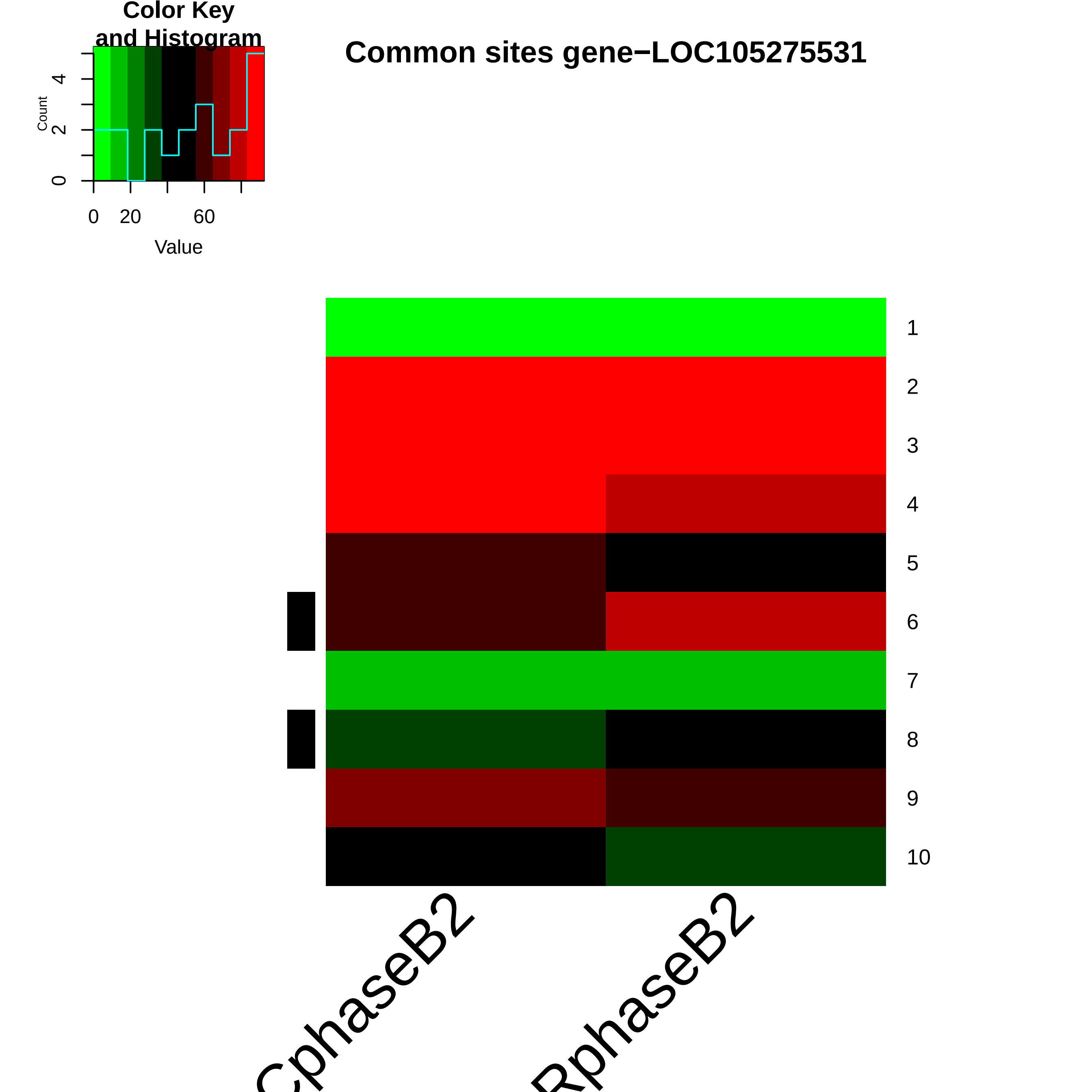
Color Key
and Histogram
Count
0
2
4
0
20
60
Value
Common sites gene−LOC105275531
1
2
3
4
5
6
7
8
9
10
CphaseB2
RphaseB2

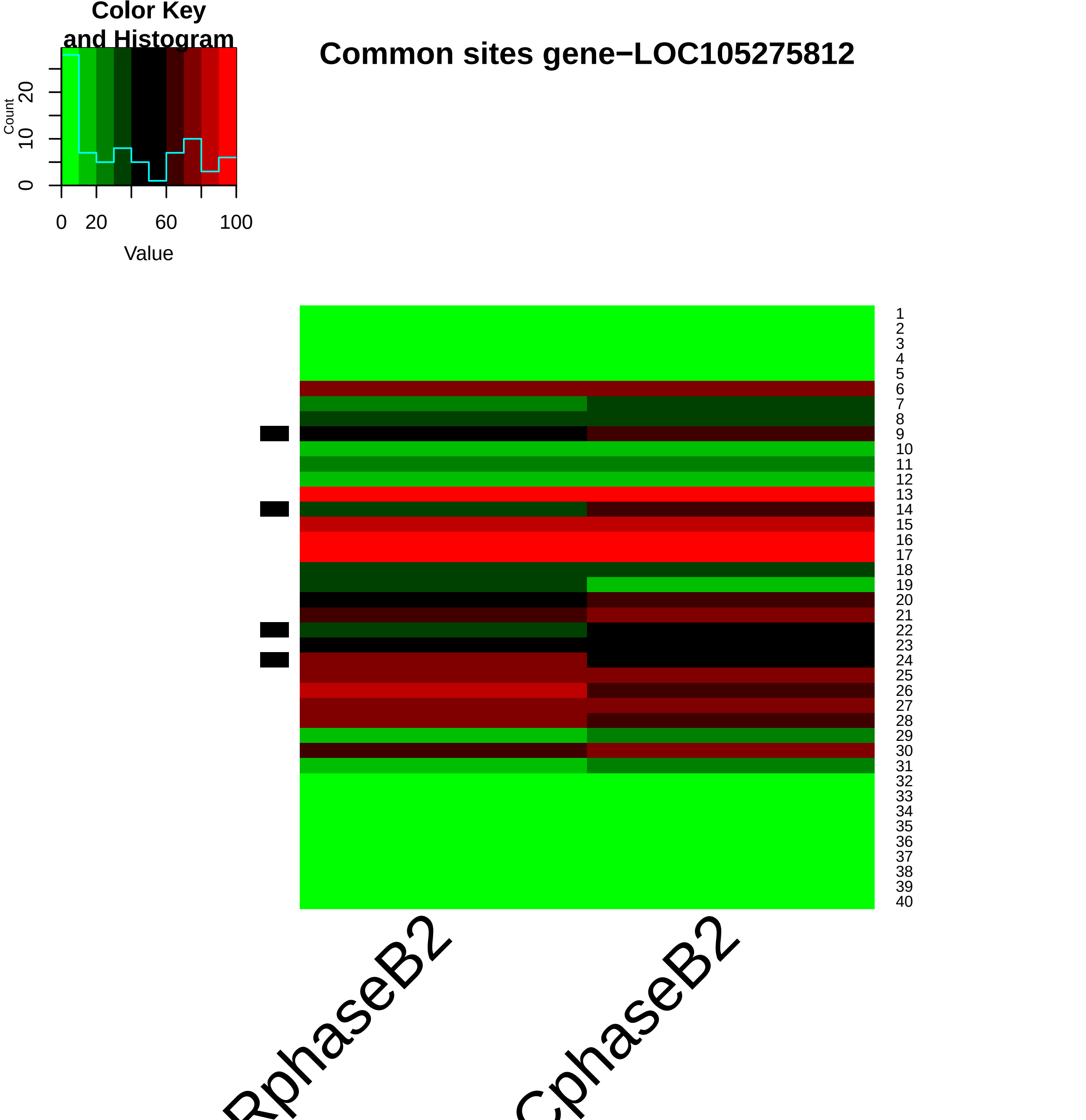
Color Key
and Histogram
Count
0
10
20
0
20
60
100
Value
Common sites gene−LOC105275812
RphaseB2
CphaseB2

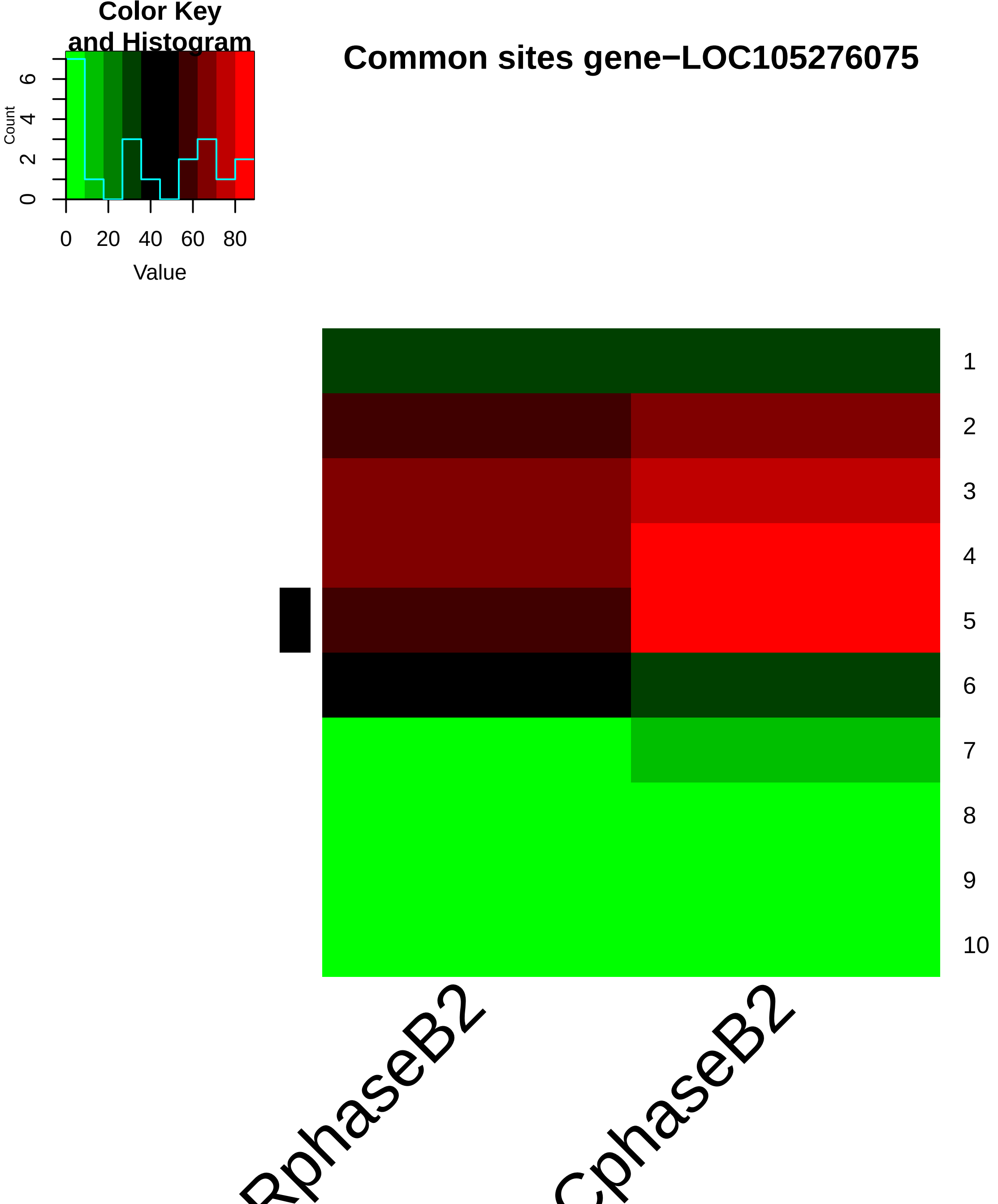

Color Key and Histogram
Count
0
2
4
6
0
20
40
60
80
Value
Common sites gene−LOC105276075
1
2
3
4
5
6
7
8
9
10
RphaseB2
CphaseB2

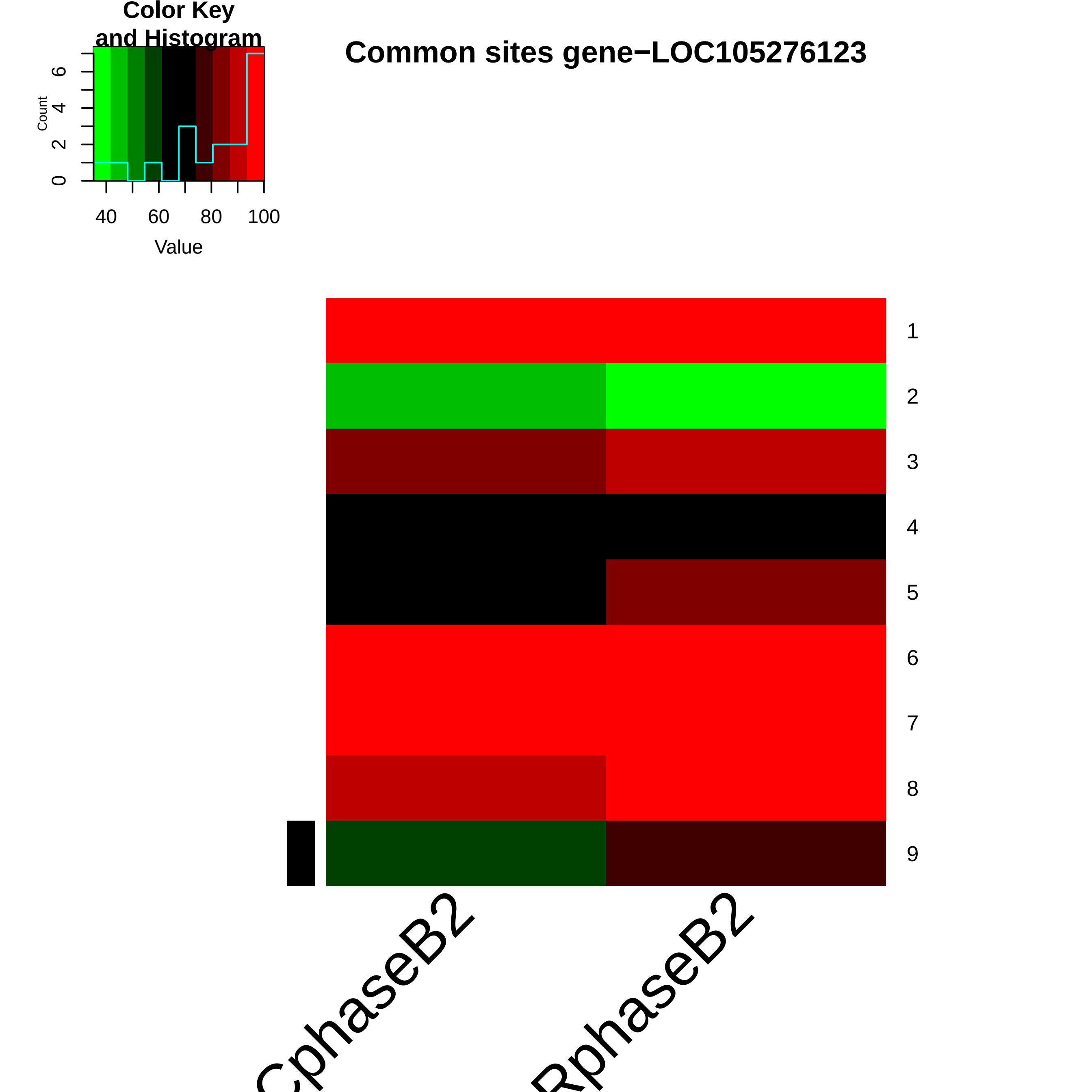
Color Key
and Histogram
Count
0
2
4
6
40
60
80
100
Value
Common sites gene−LOC105276123
1
2
3
4
5
6
7
8
9
CphaseB2
RphaseB2

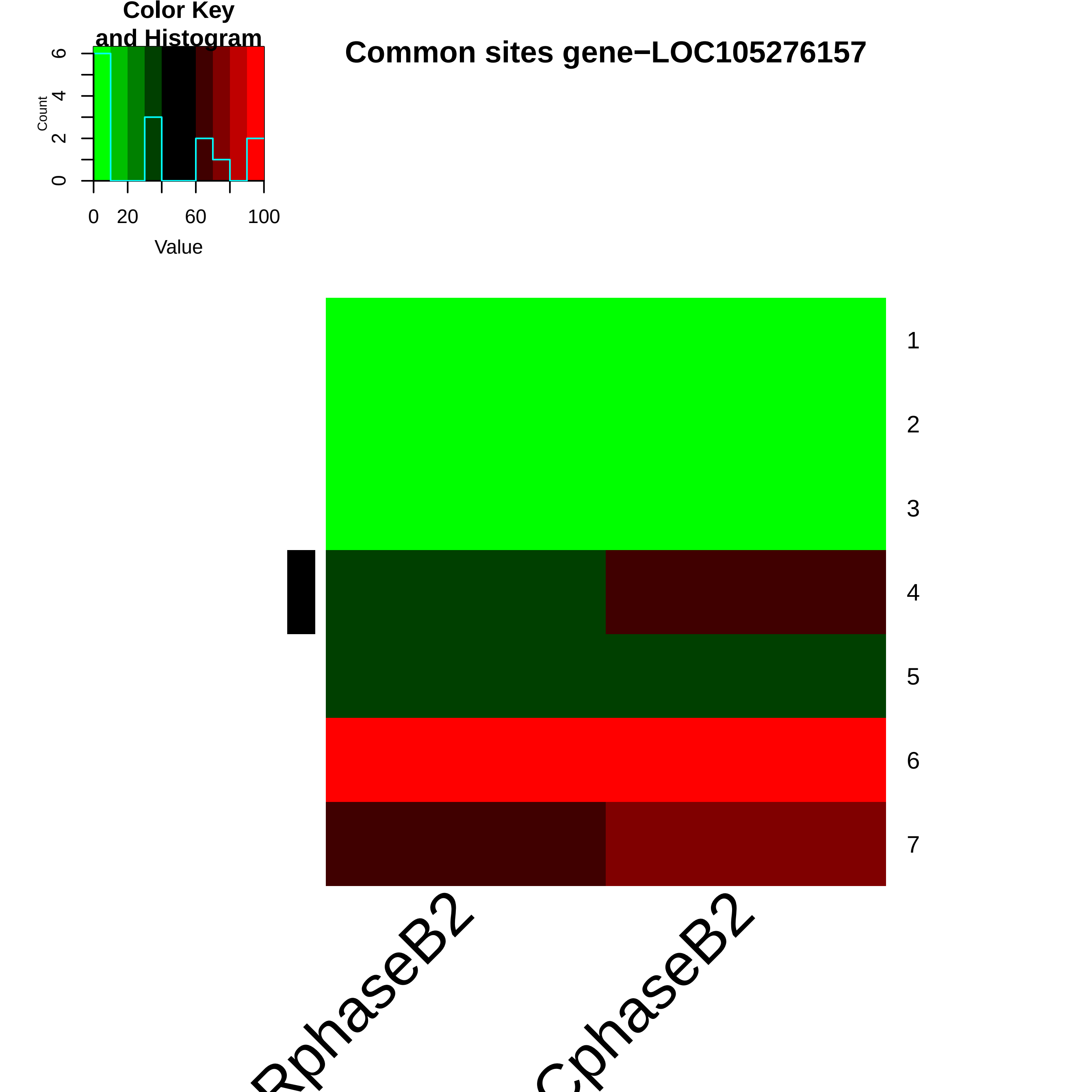
Color Key
and Histogram
Count
0
2
4
6
0
20
60
100
Value
Common sites gene−LOC105276157
1
2
3
4
5
6
7
RphaseB2
CphaseB2

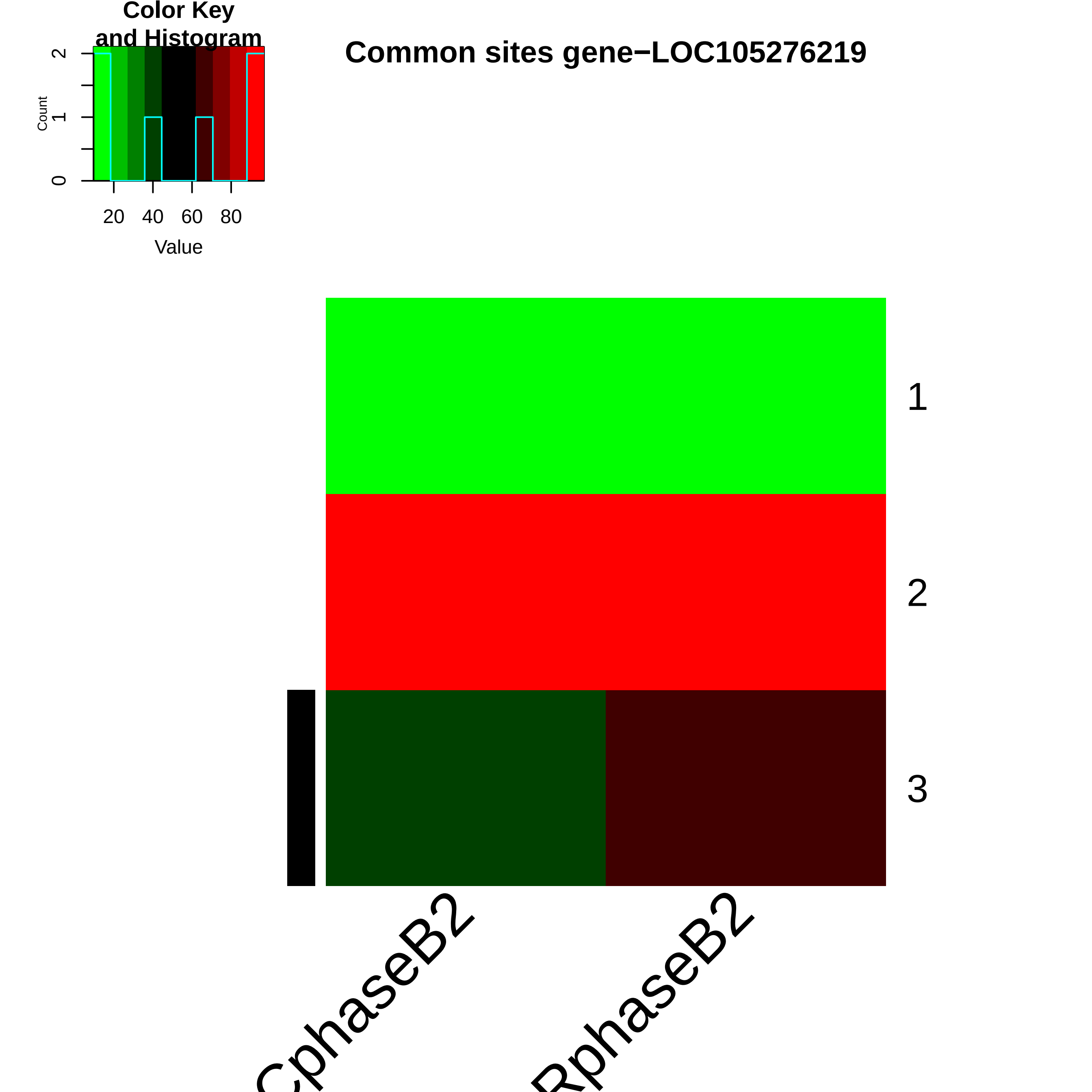

Color Key
and Histogram
Count
0
1
2
20
40
60
80
Value
Common sites gene−LOC105276219
1
2
3
CphaseB2
RphaseB2

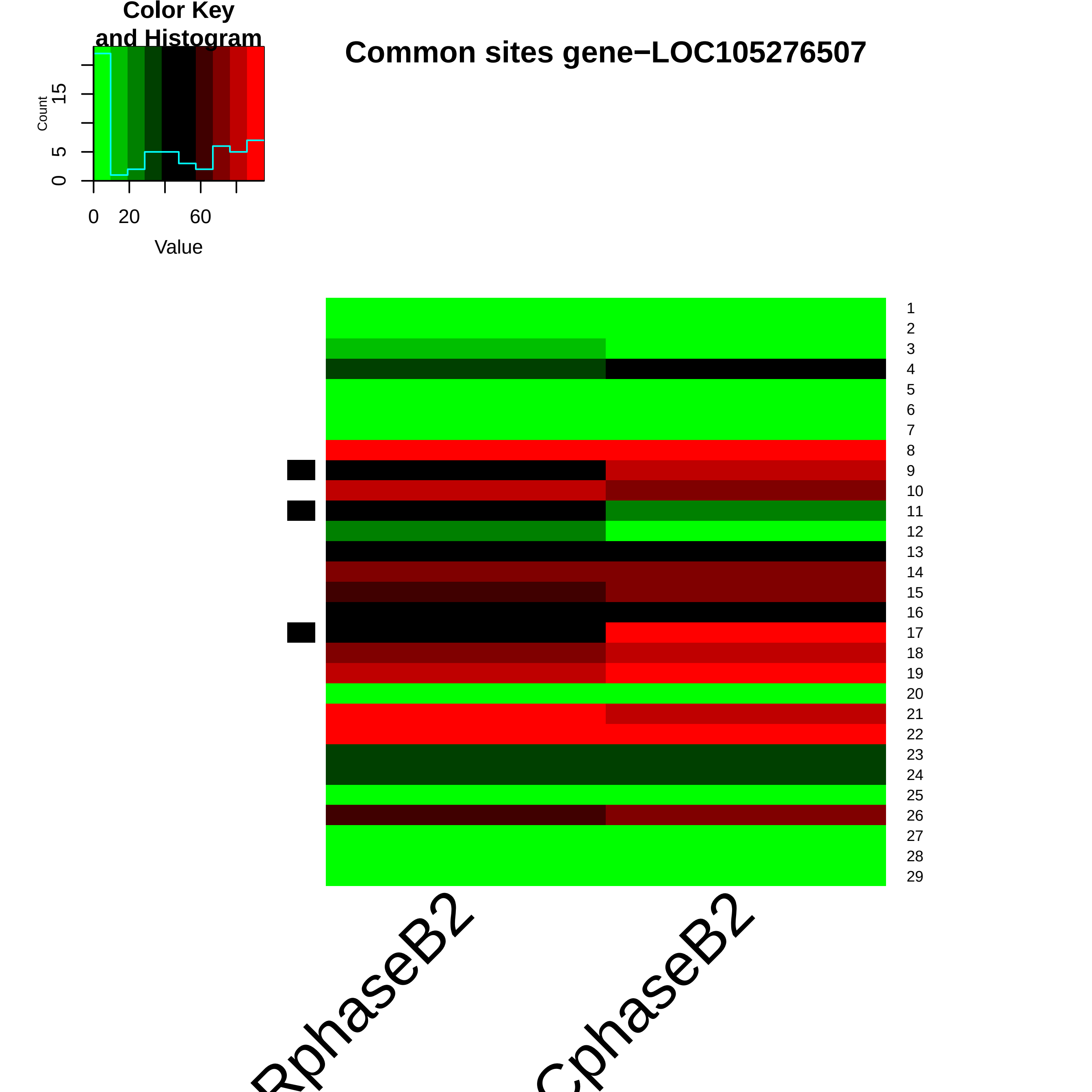
Color Key
and Histogram
Count
0
5
15
0
20
60
Value
Common sites gene−LOC105276507
RphaseB2
CphaseB2

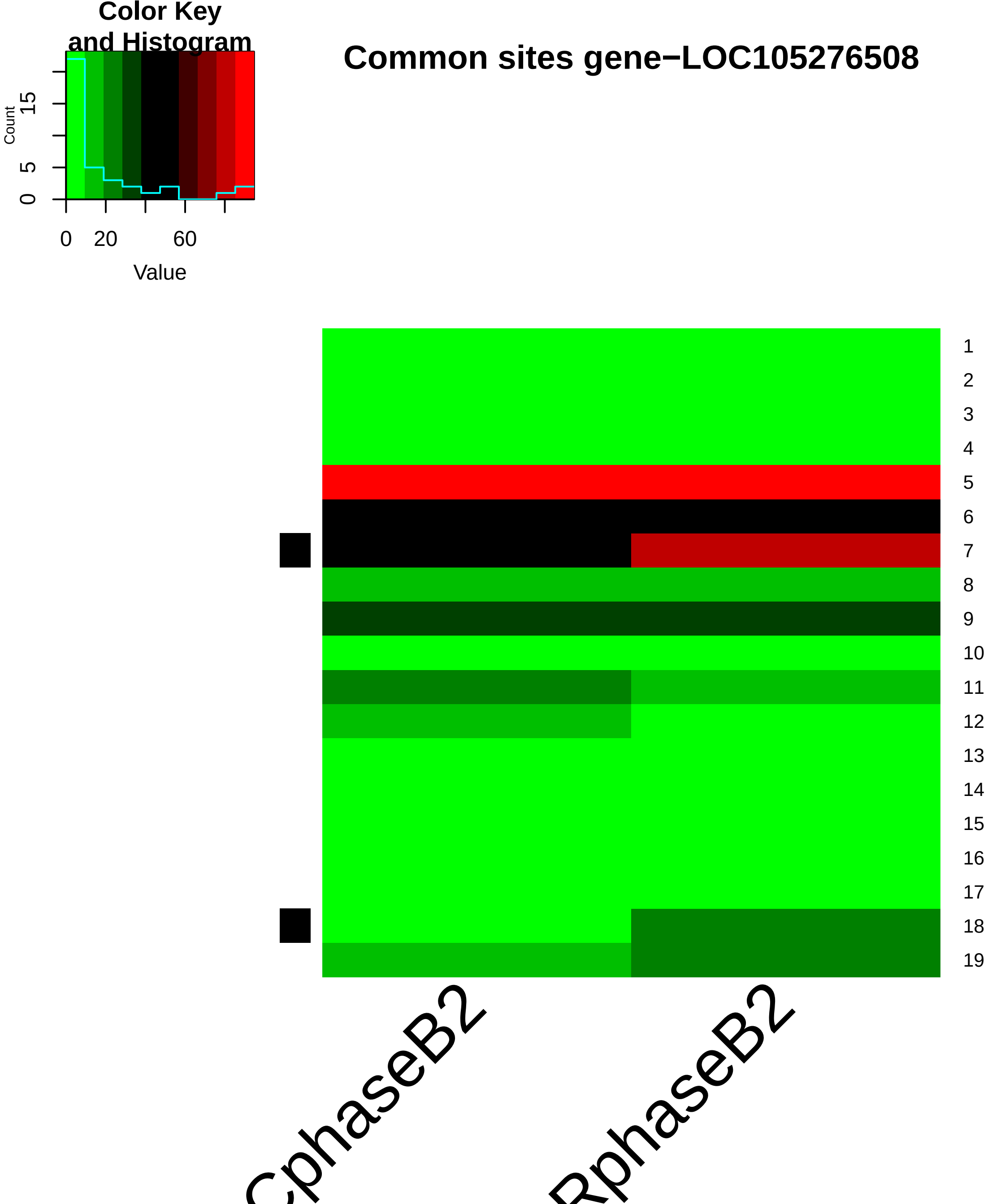
Color Key
and Histogram
Count
0
5
15
0
20
60
Value
Common sites gene−LOC105276508
CphaseB2
RphaseB2

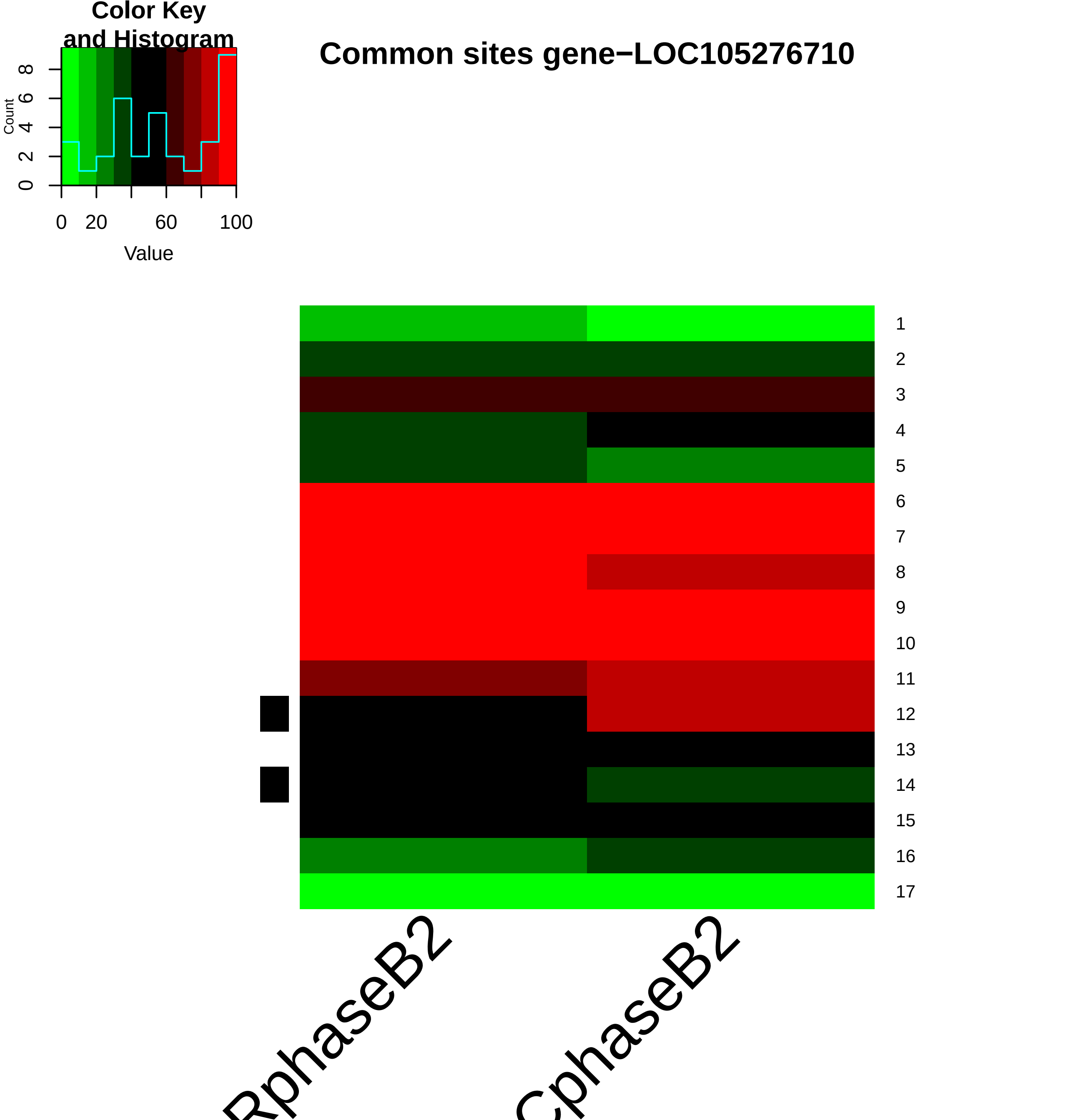
Color Key
and Histogram
Count
0
2
4
6
8
0
20
60
100
Value
Common sites gene−LOC105276710
RphaseB2
CphaseB2

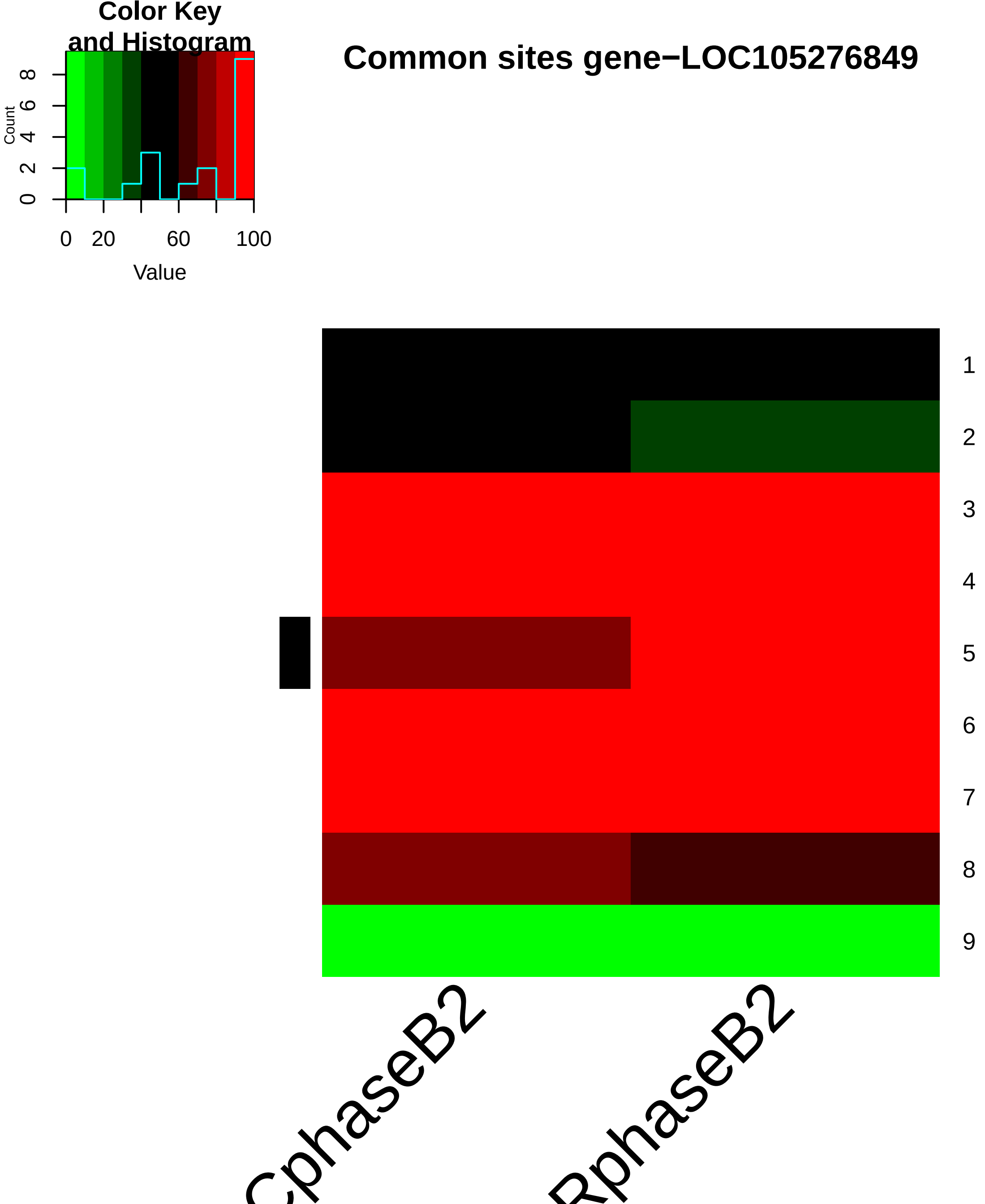
Color Key
and Histogram
Count
0 2 4 6 8
0 20 60 100
Value
Common sites gene−LOC105276849
1
2
3
4
5
6
7
8
9

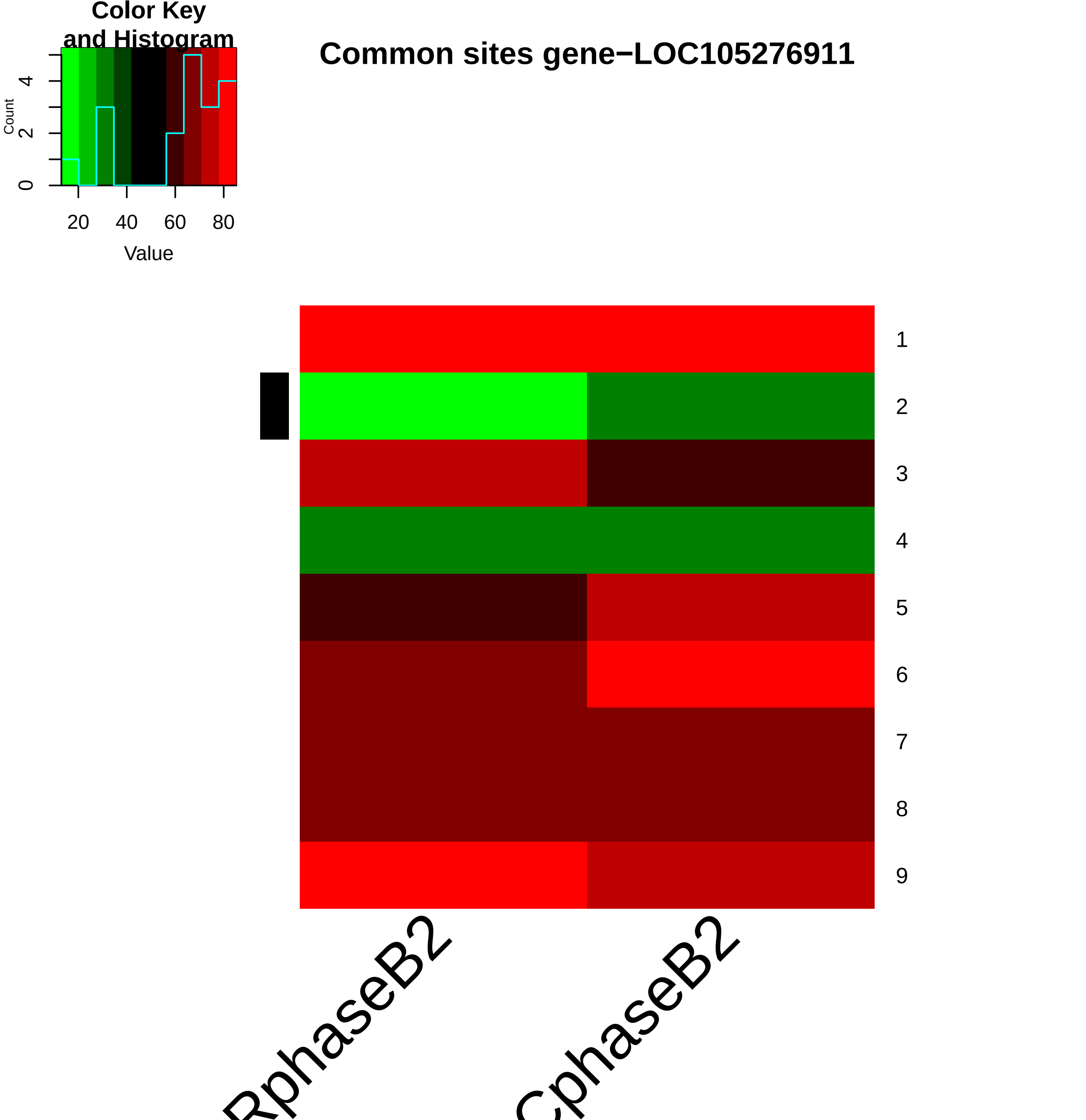
Color Key
and Histogram
Count
0
2
4
20
40
60
80
Value
Common sites gene−LOC105276911
1
2
3
4
5
6
7
8
9
RphaseB2
CphaseB2

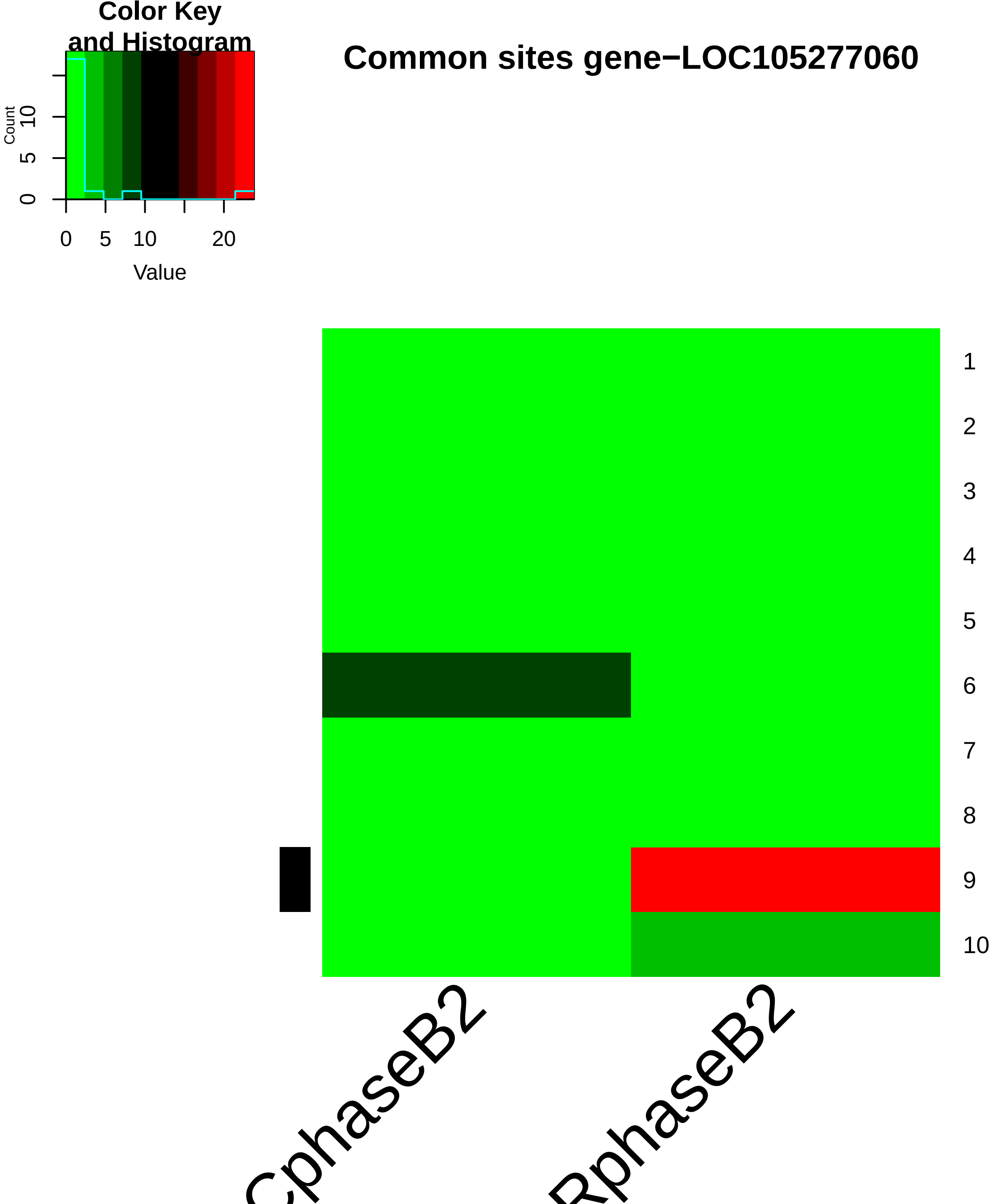

Color Key
and Histogram
Count
0
5
10
0
5
10
20
Value
Common sites gene−LOC105277060
1
2
3
4
5
6
7
8
9
10
CphaseB2
RphaseB2

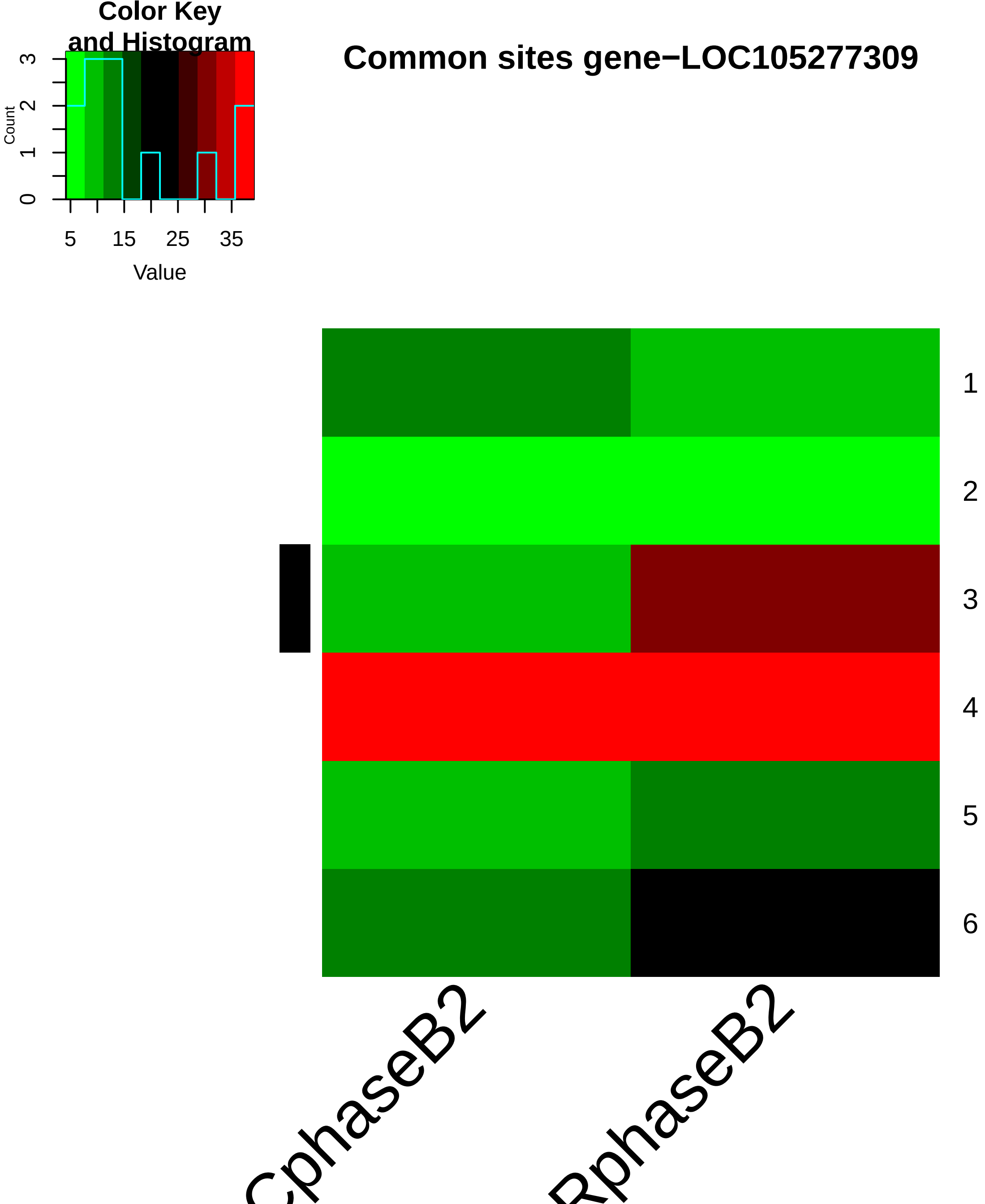
Color Key
and Histogram
Count
0
1
2
3
5
15
25
35
Value
Common sites gene−LOC105277309
1
2
3
4
5
6
CphaseB2
RphaseB2

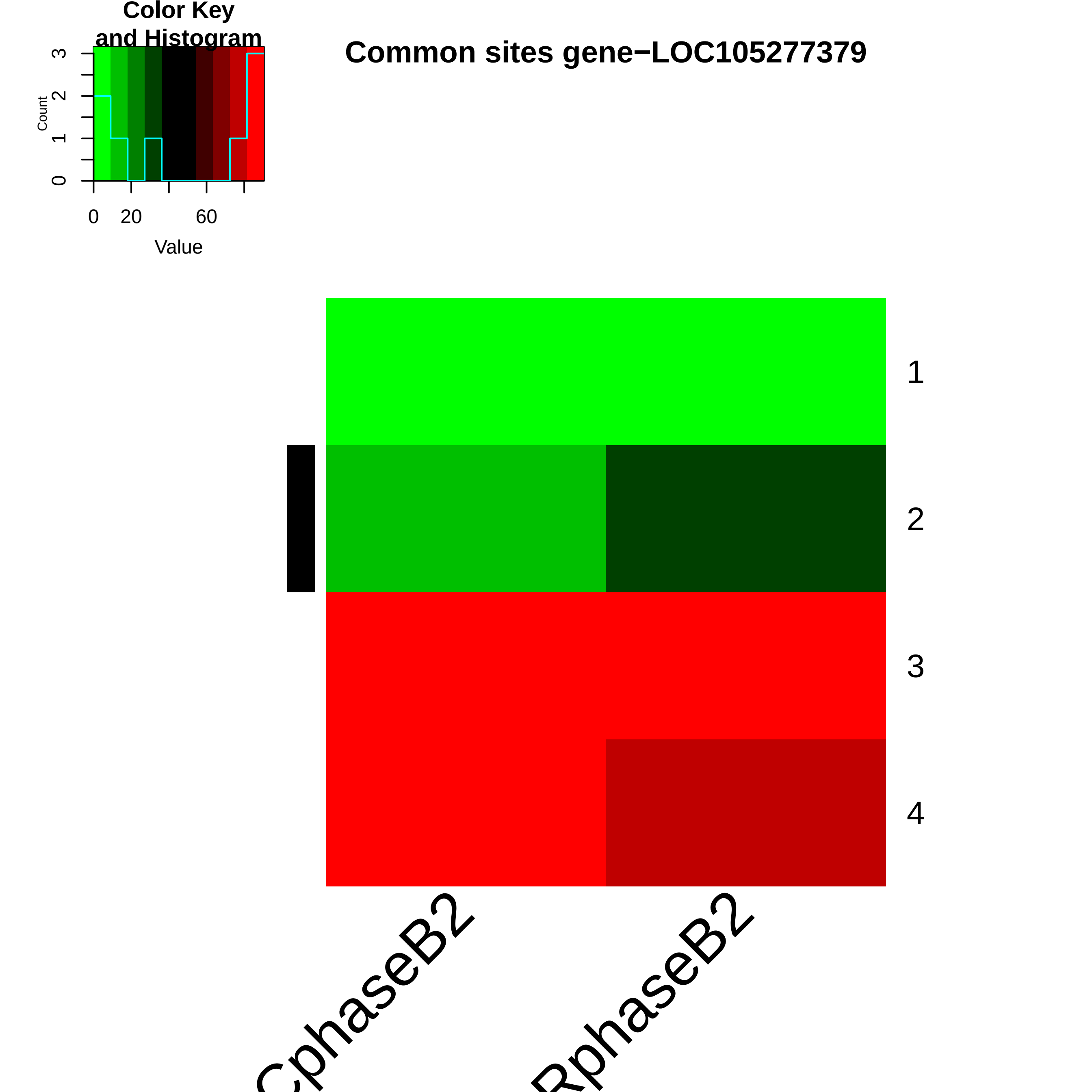
Color Key
and Histogram
Count
0
1
2
3
0
20
60
Value
Common sites gene−LOC105277379
1
2
3
4

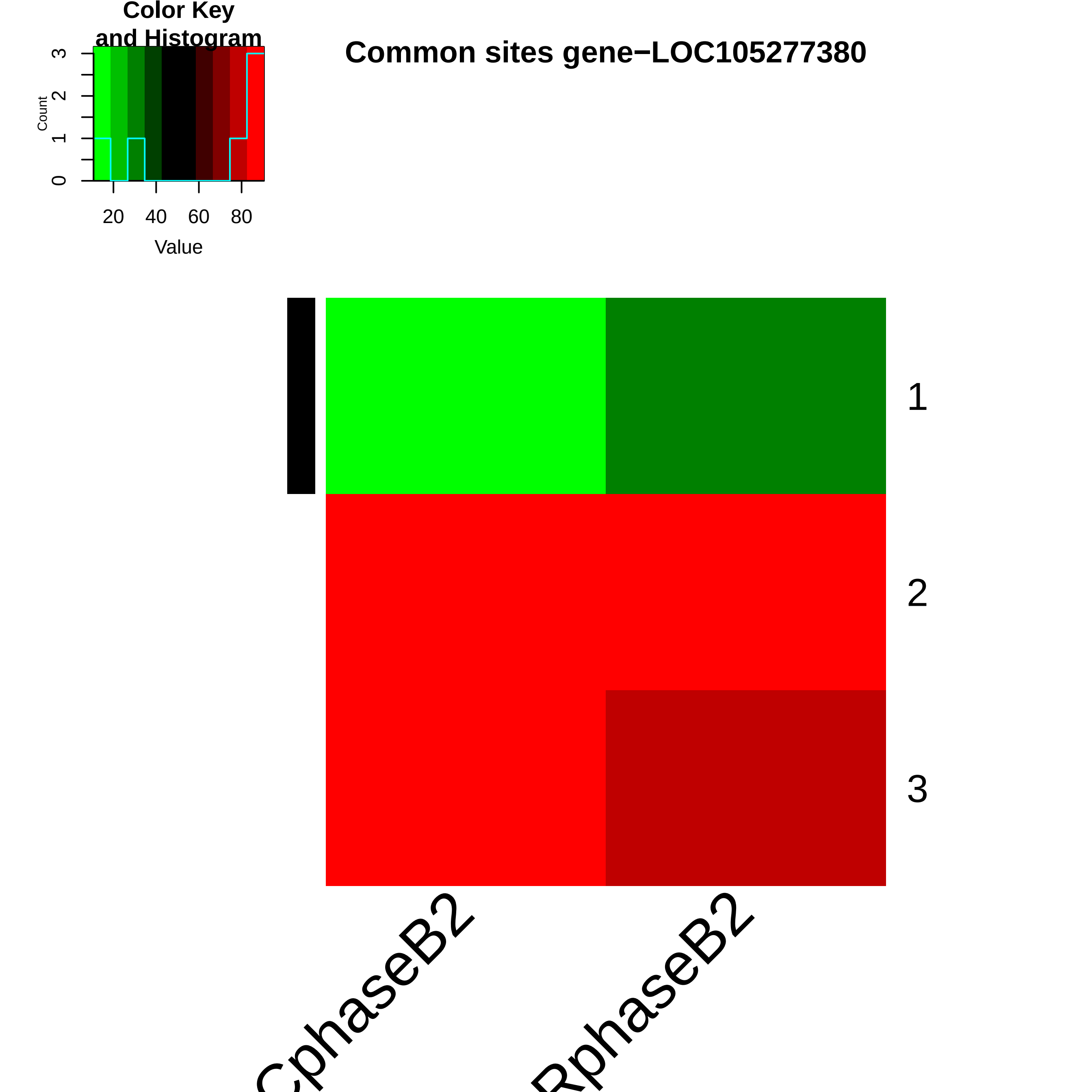
Color Key
and Histogram
Count
0
1
2
3
20
40
60
80
Value
Common sites gene−LOC105277380
1
2
3

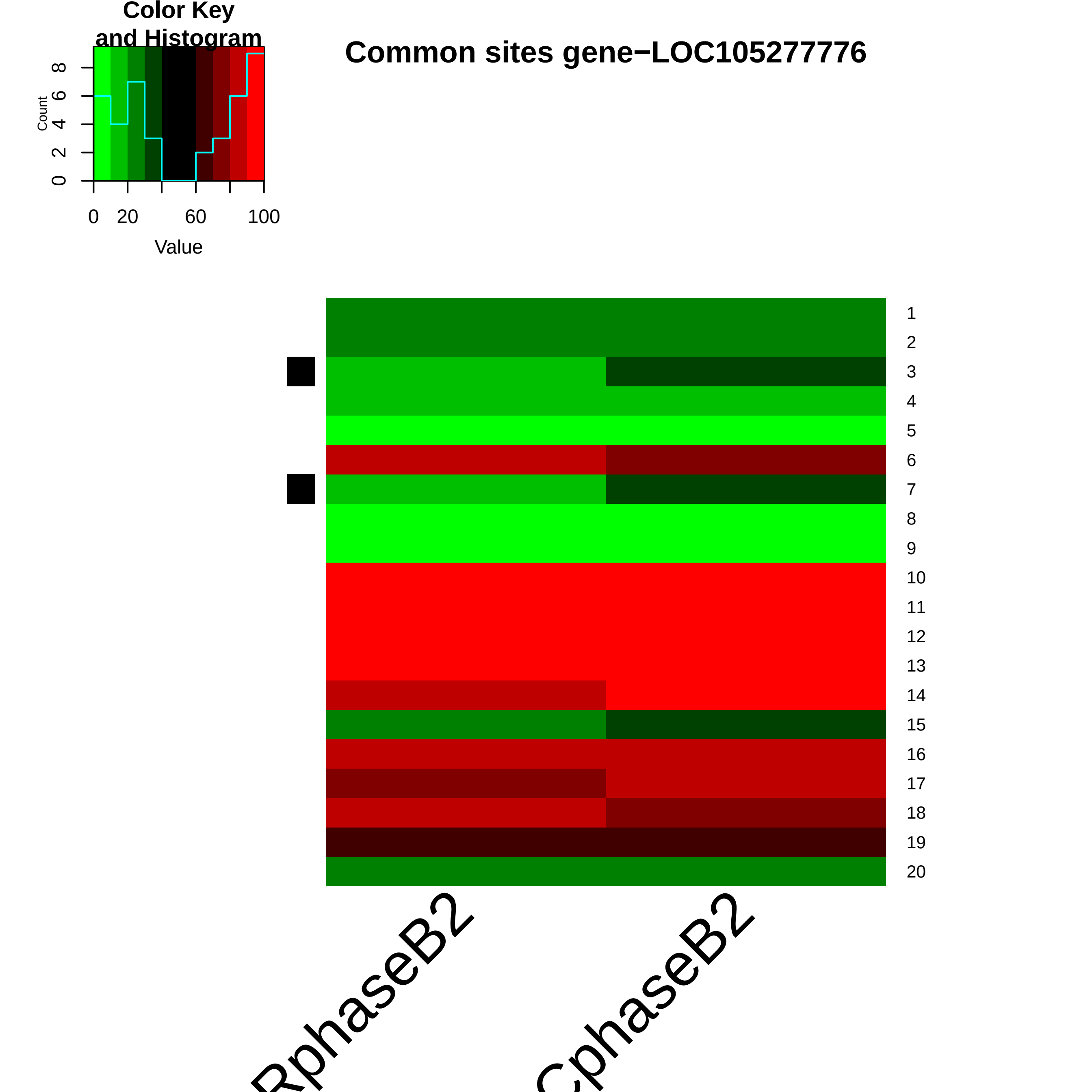

Color Key
and Histogram
Count
0 2 4 6 8
0 20 60 100
Value
Common sites gene−LOC105277776
RphaseB2
CphaseB2

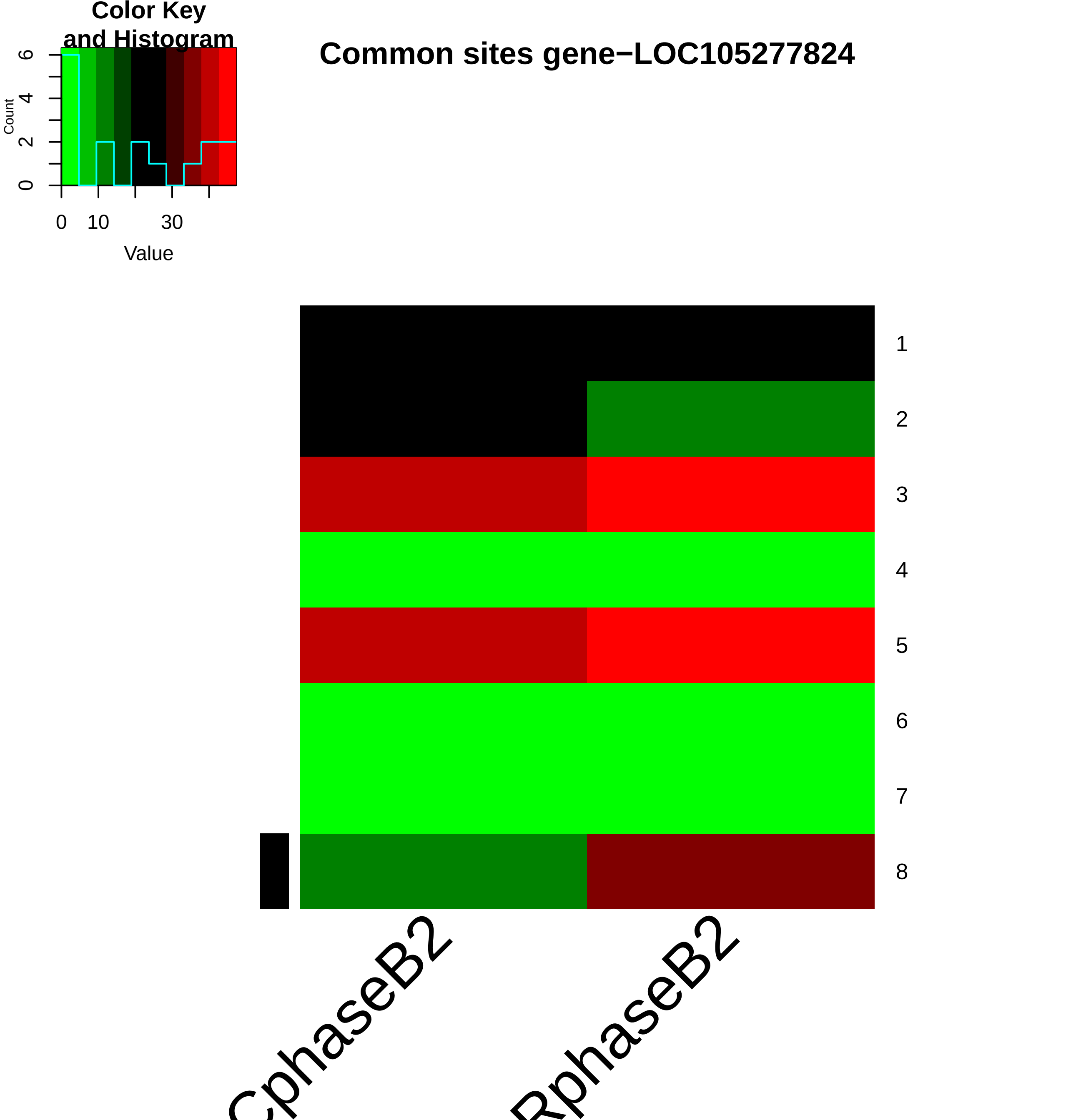

Color Key
and Histogram
Count
0
2
4
6
0
10
30
Value
Common sites gene−LOC105277824
1
2
3
4
5
6
7
8
CphaseB2
RphaseB2

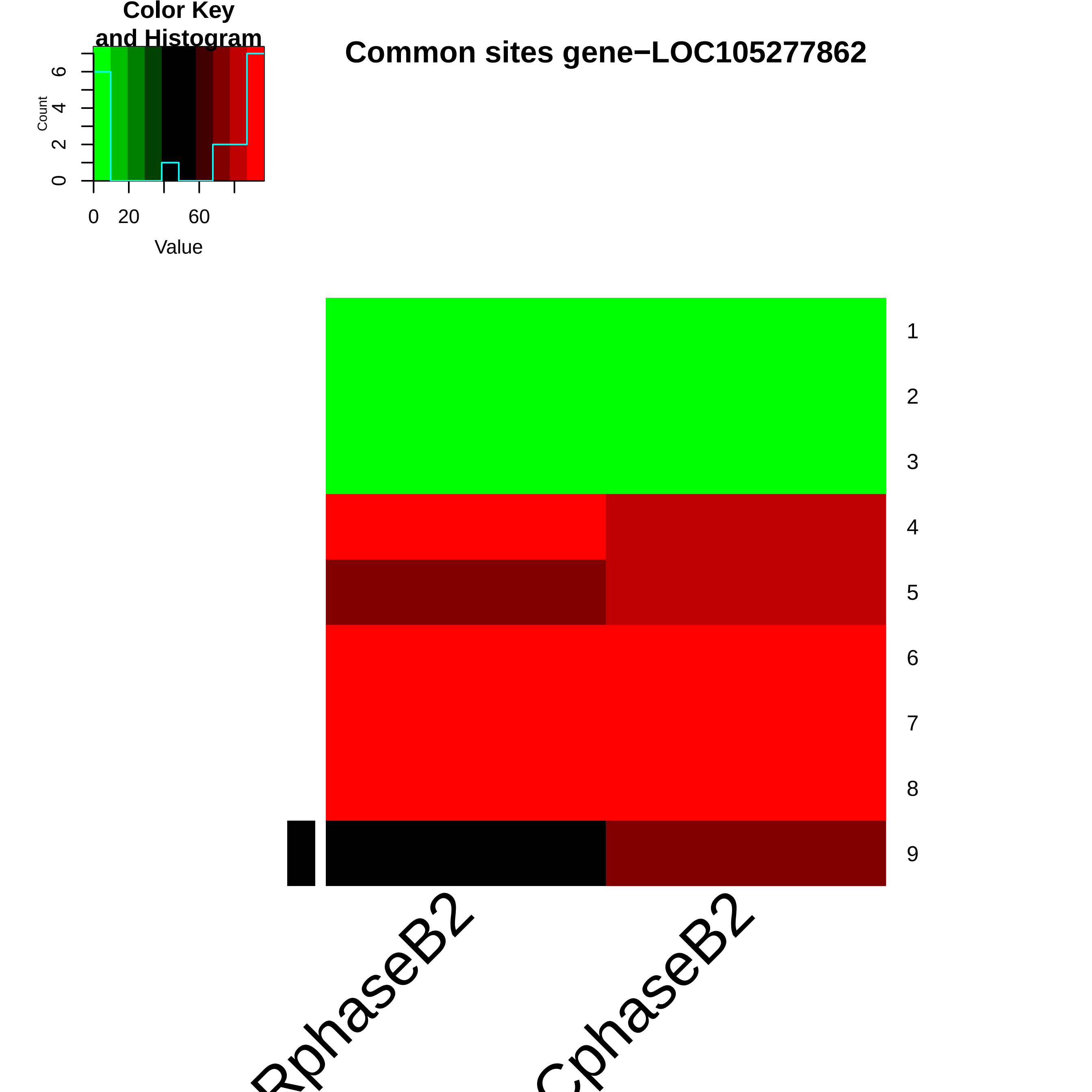

Color Key
and Histogram
Count
0
2
4
6
0
20
60
Value
Common sites gene−LOC105277862
1
2
3
4
5
6
7
8
9
RphaseB2
CphaseB2

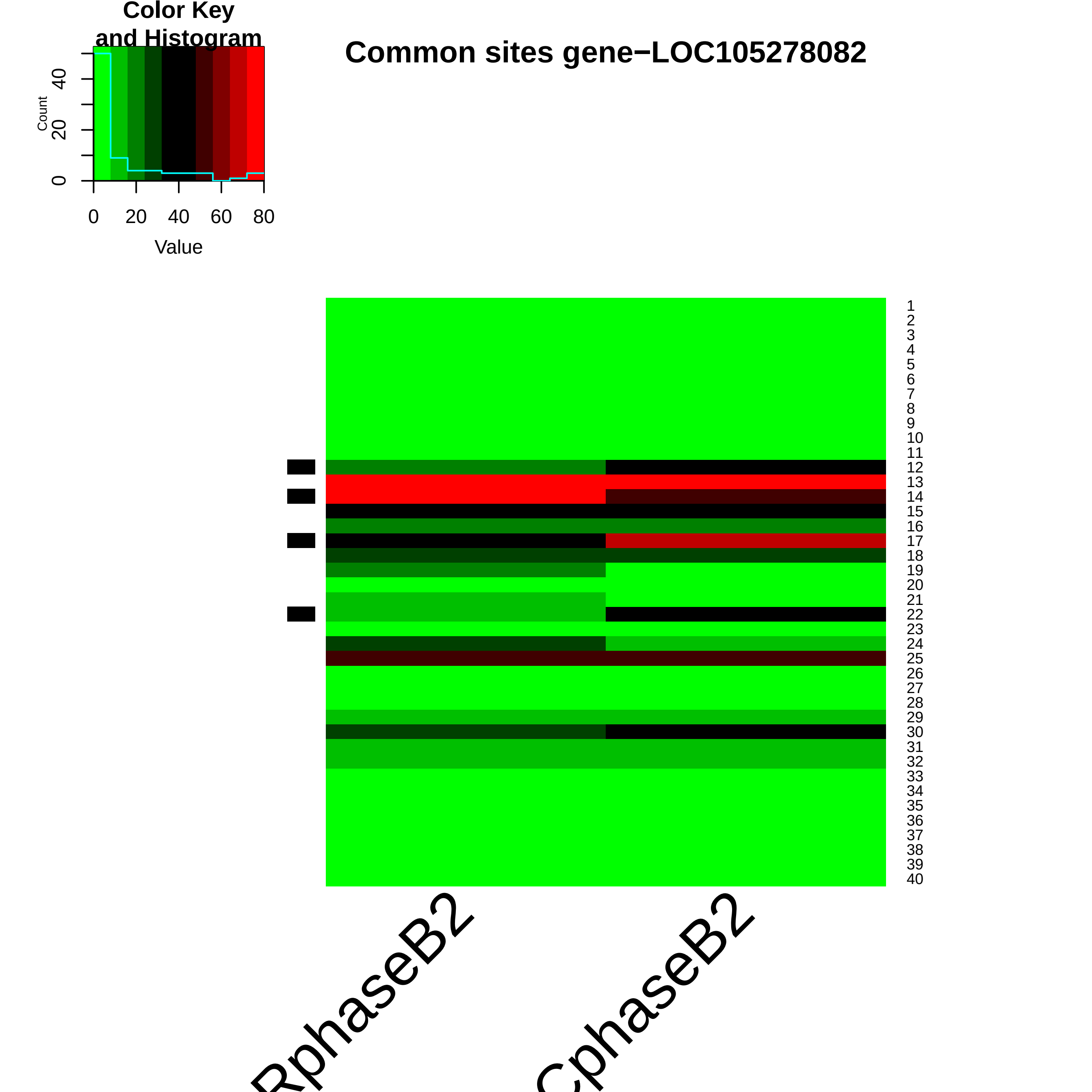
Color Key
and Histogram
Count
Value
0
20
40
60
80
Common sites gene−LOC105278082
RphaseB2
CphaseB2

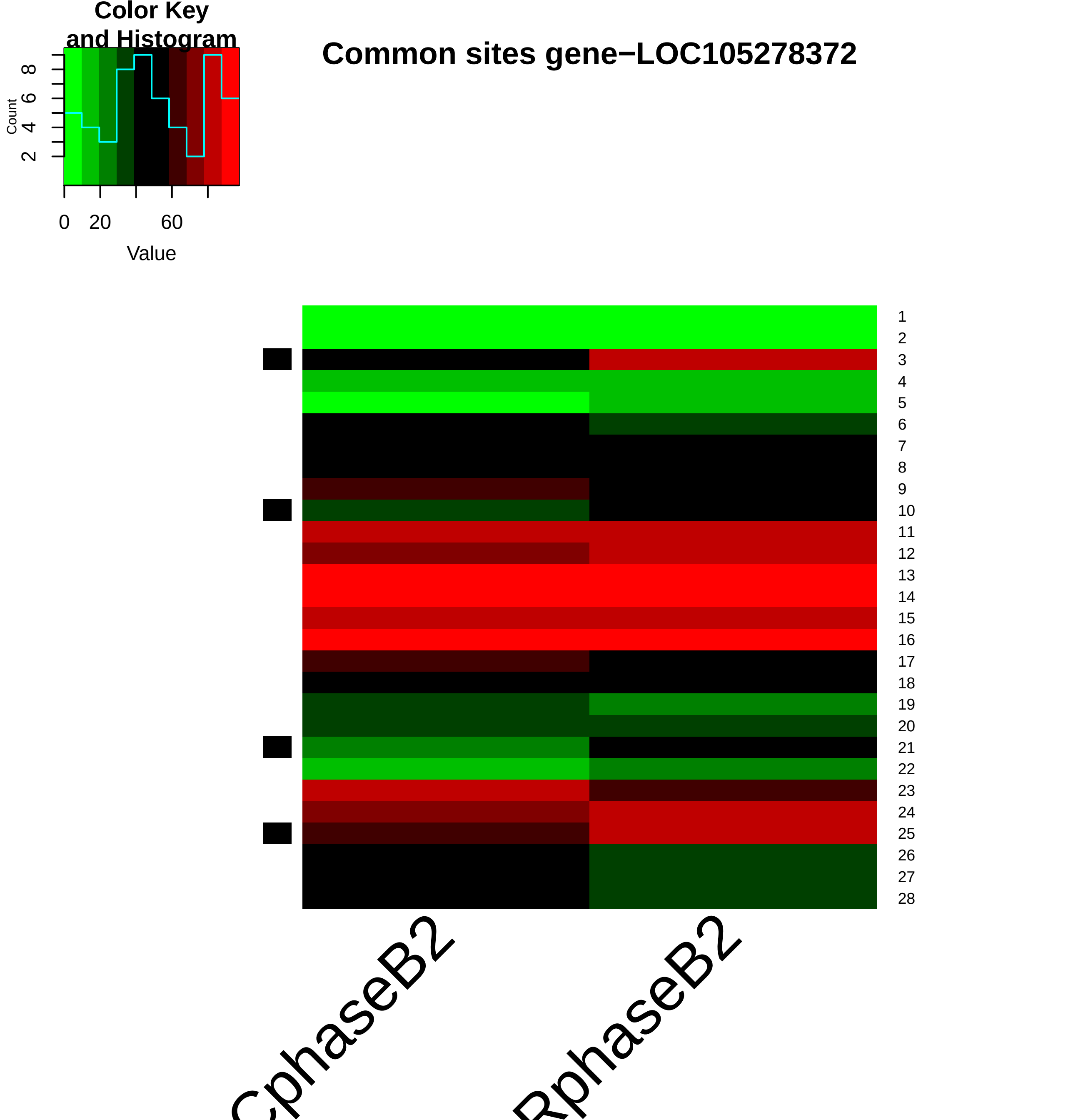
Color Key
and Histogram
Count
2
4
6
8
0
20
60
Value
Common sites gene−LOC105278372
CphaseB2
RphaseB2

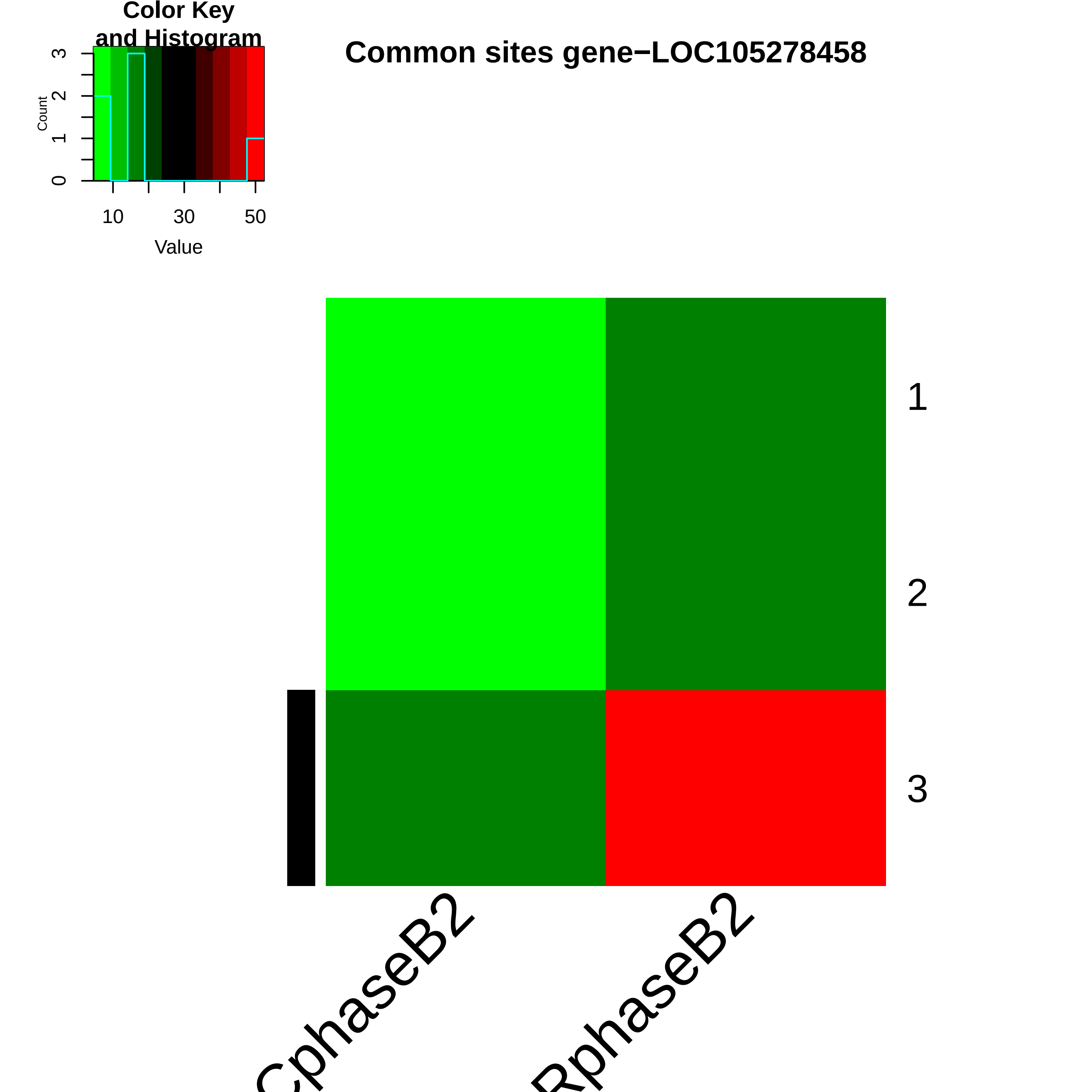

Color Key
and Histogram
Count
0
1
2
3
10
30
50
Value
Common sites gene−LOC105278458
1
2
3
CphaseB2
RphaseB2

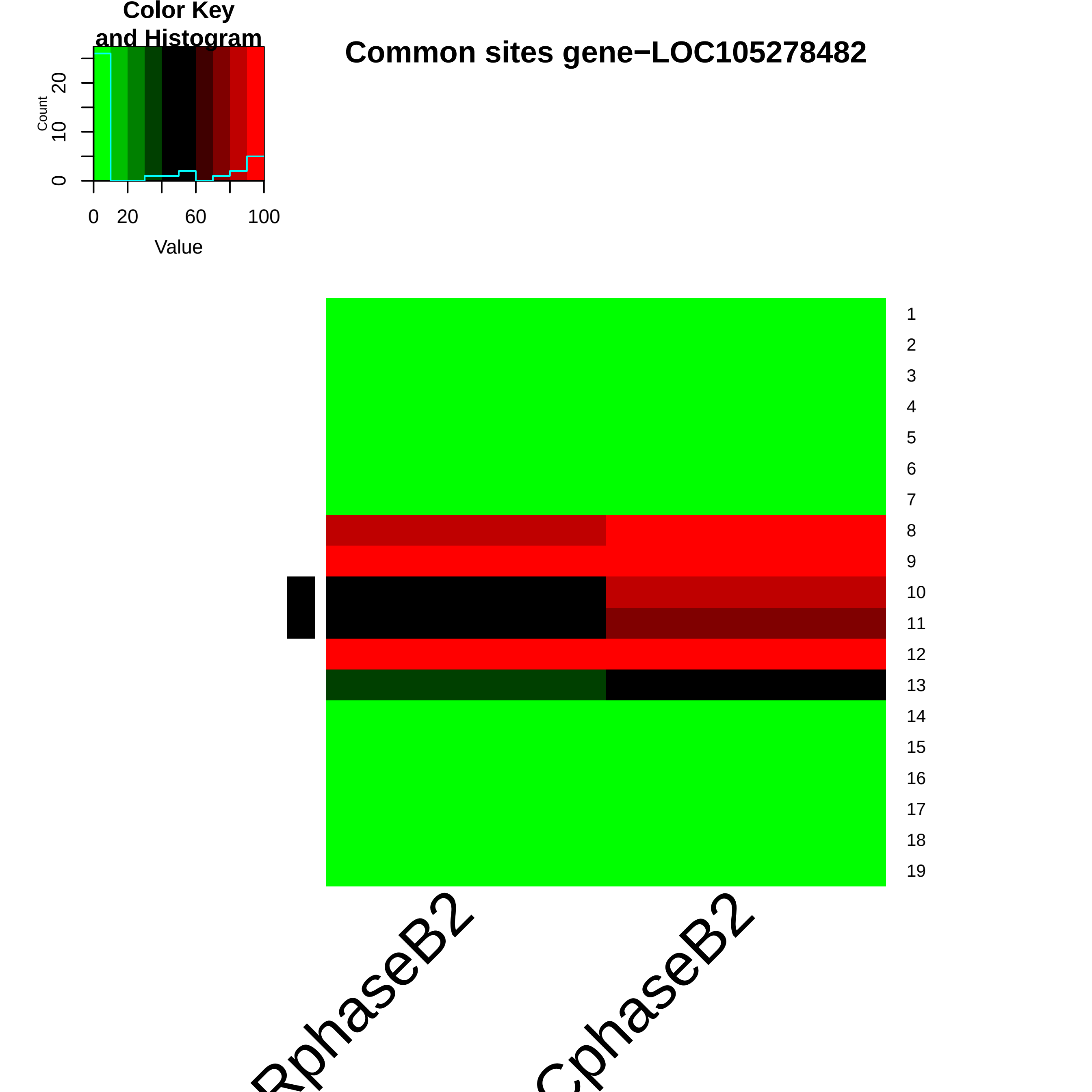
Color Key
and Histogram
Count
0
10
20
0
20
60
100
Value
Common sites gene−LOC105278482
RphaseB2
CphaseB2

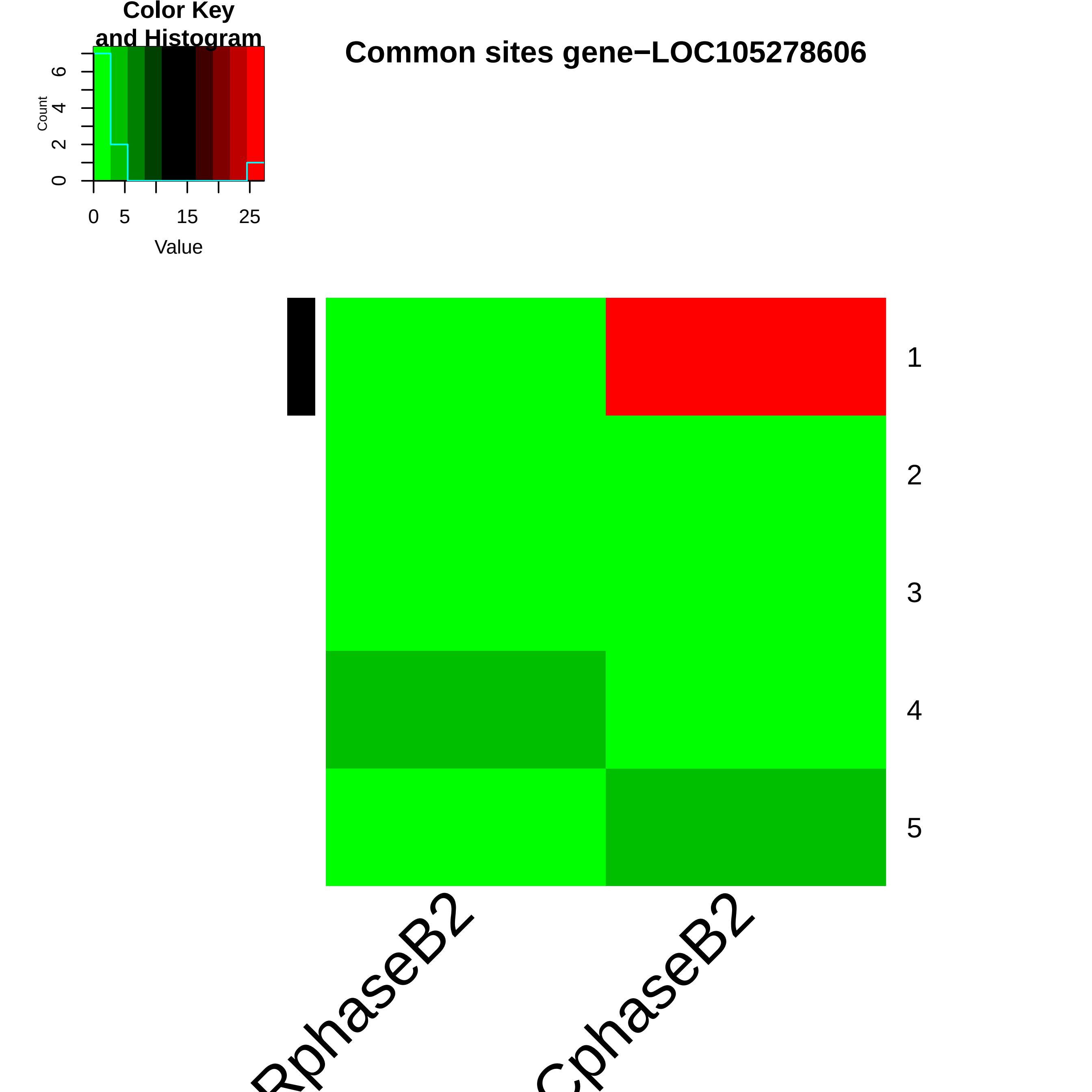
Color Key
and Histogram
Count
0
2
4
6
0
5
15
25
Value
Common sites gene−LOC105278606
1
2
3
4
5
RphaseB2
CphaseB2

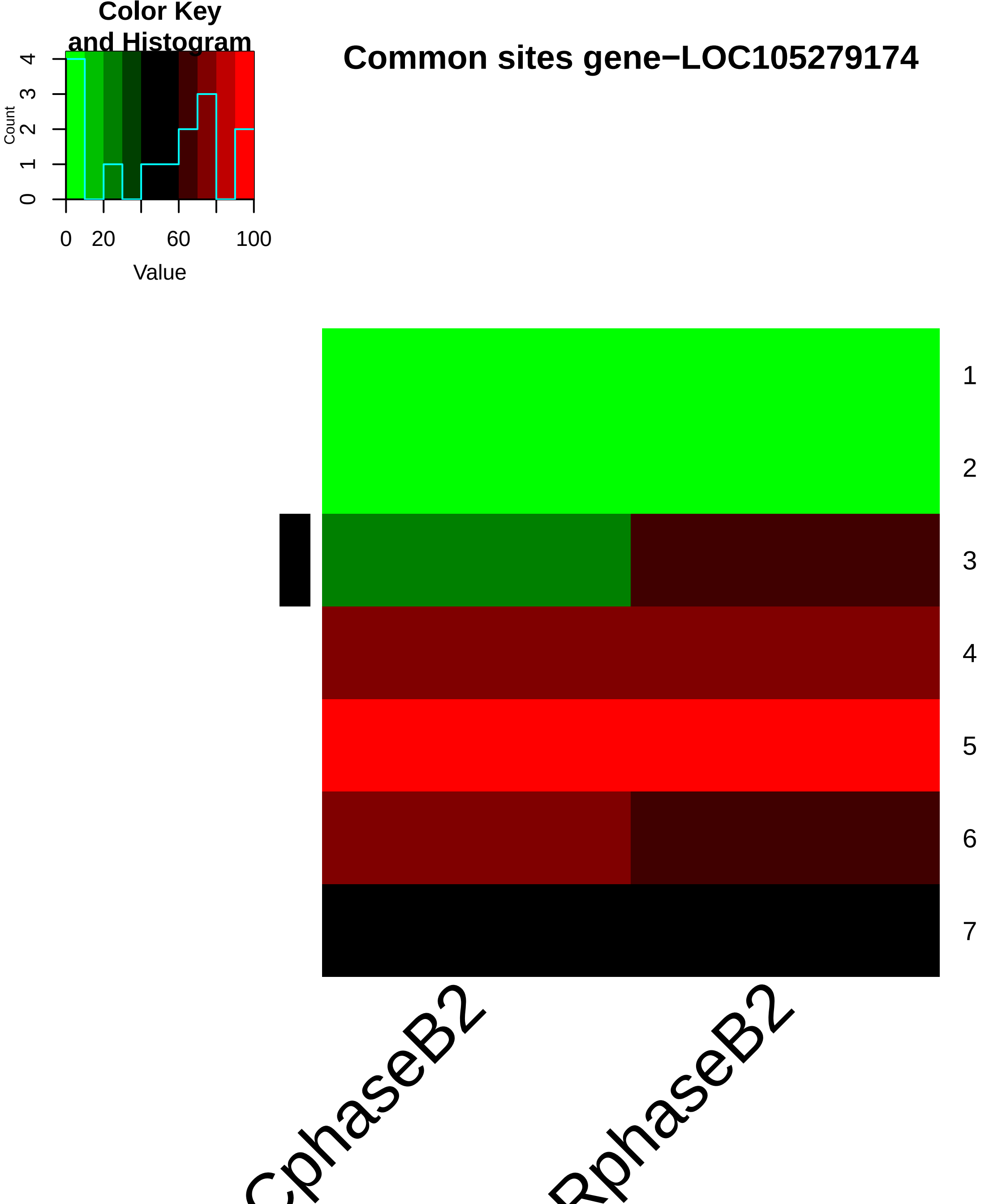
Color Key
and Histogram
Count
0
1
2
3
4
0
20
60
100
Value
Common sites gene−LOC105279174
1
2
3
4
5
6
7
CphaseB2
RphaseB2

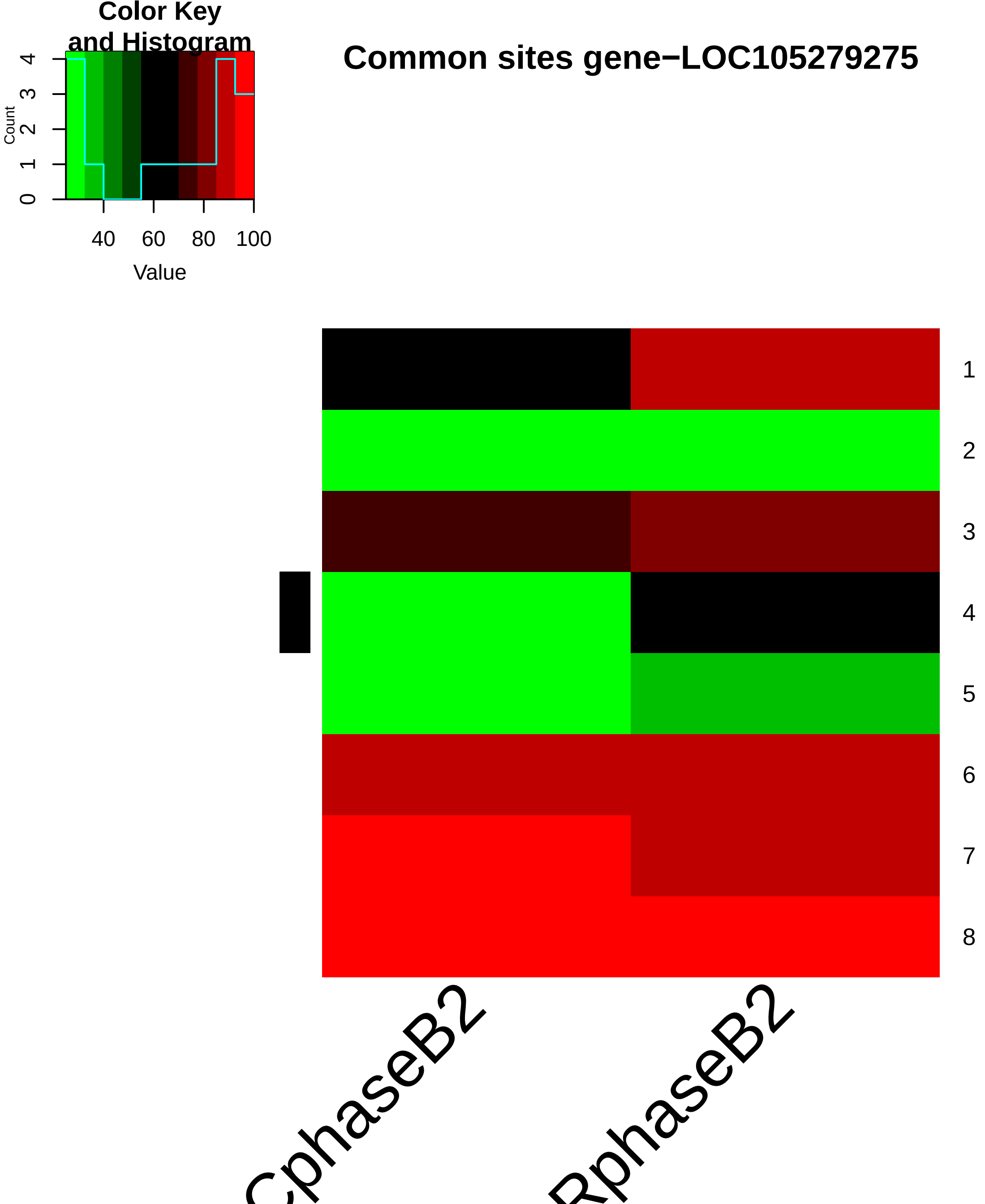
Color Key
and Histogram
Count
0
1
2
3
4
40
60
80
100
Value
Common sites gene−LOC105279275
1
2
3
4
5
6
7
8
CphaseB2
RphaseB2

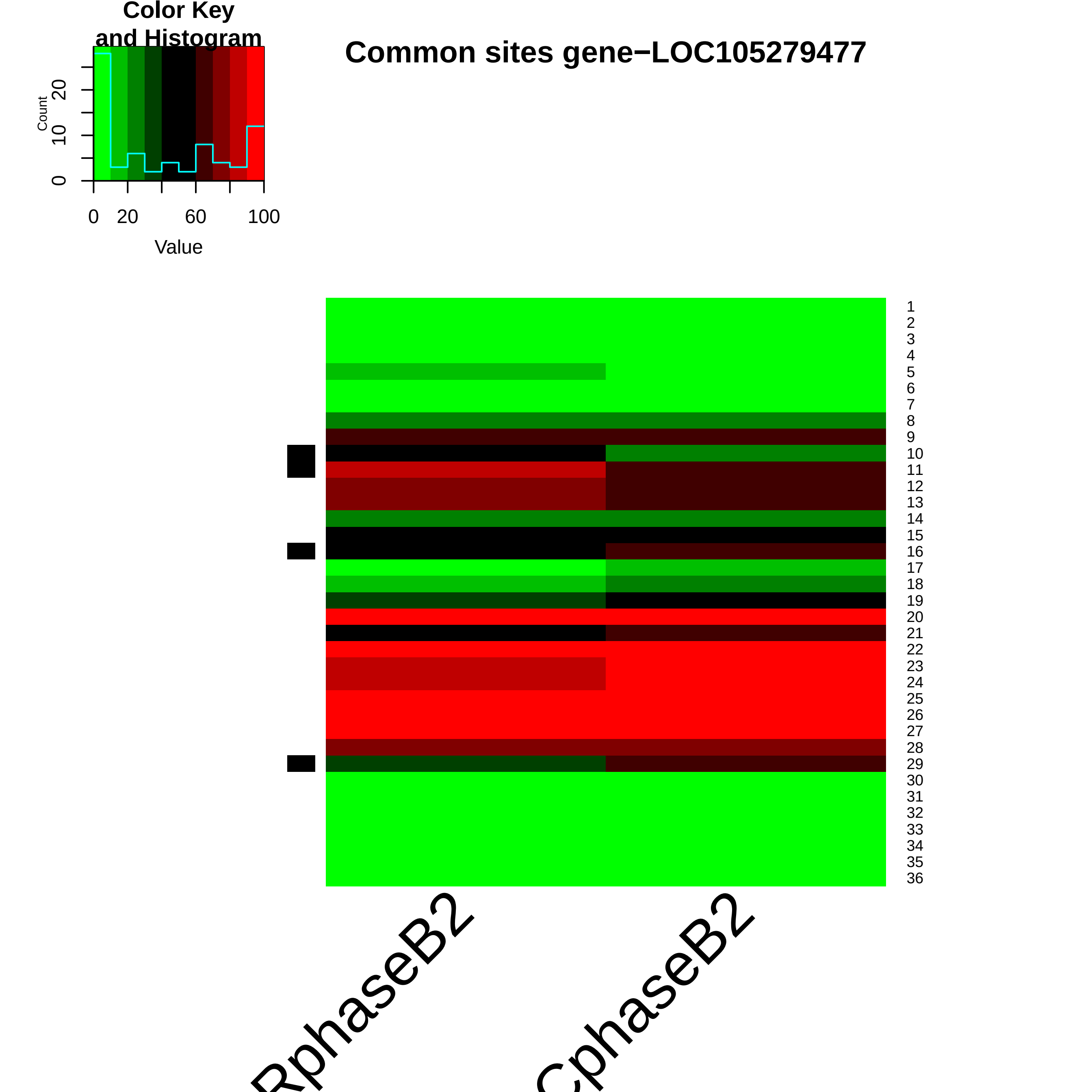
Color Key
and Histogram
Count
0
10
20
0
20
60
100
Value
Common sites gene−LOC105279477
RphaseB2
CphaseB2

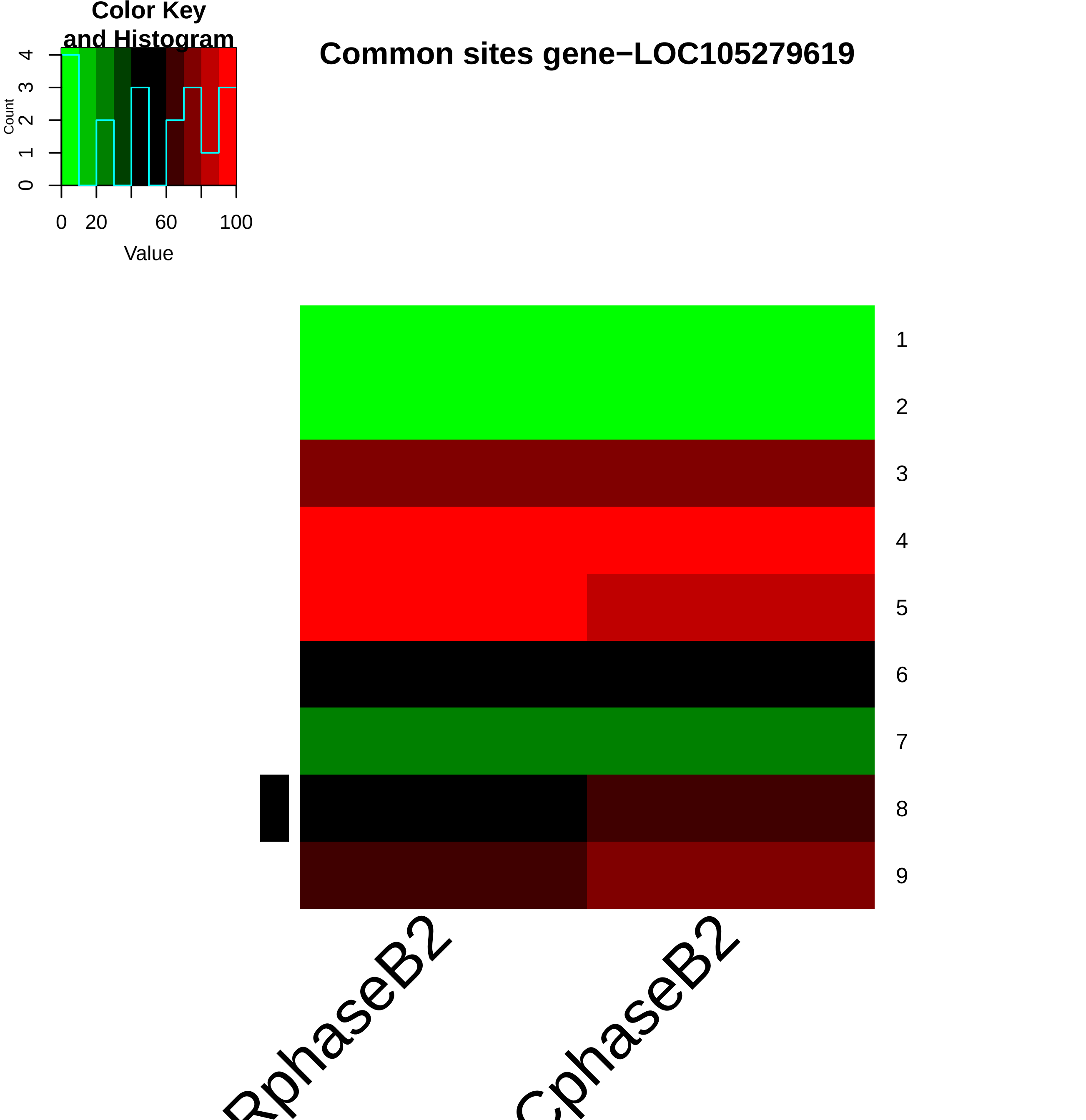
Color Key
and Histogram
Count
0
1
2
3
4
0
20
60
100
Value
Common sites gene−LOC105279619
1
2
3
4
5
6
7
8
9
RphaseB2
CphaseB2

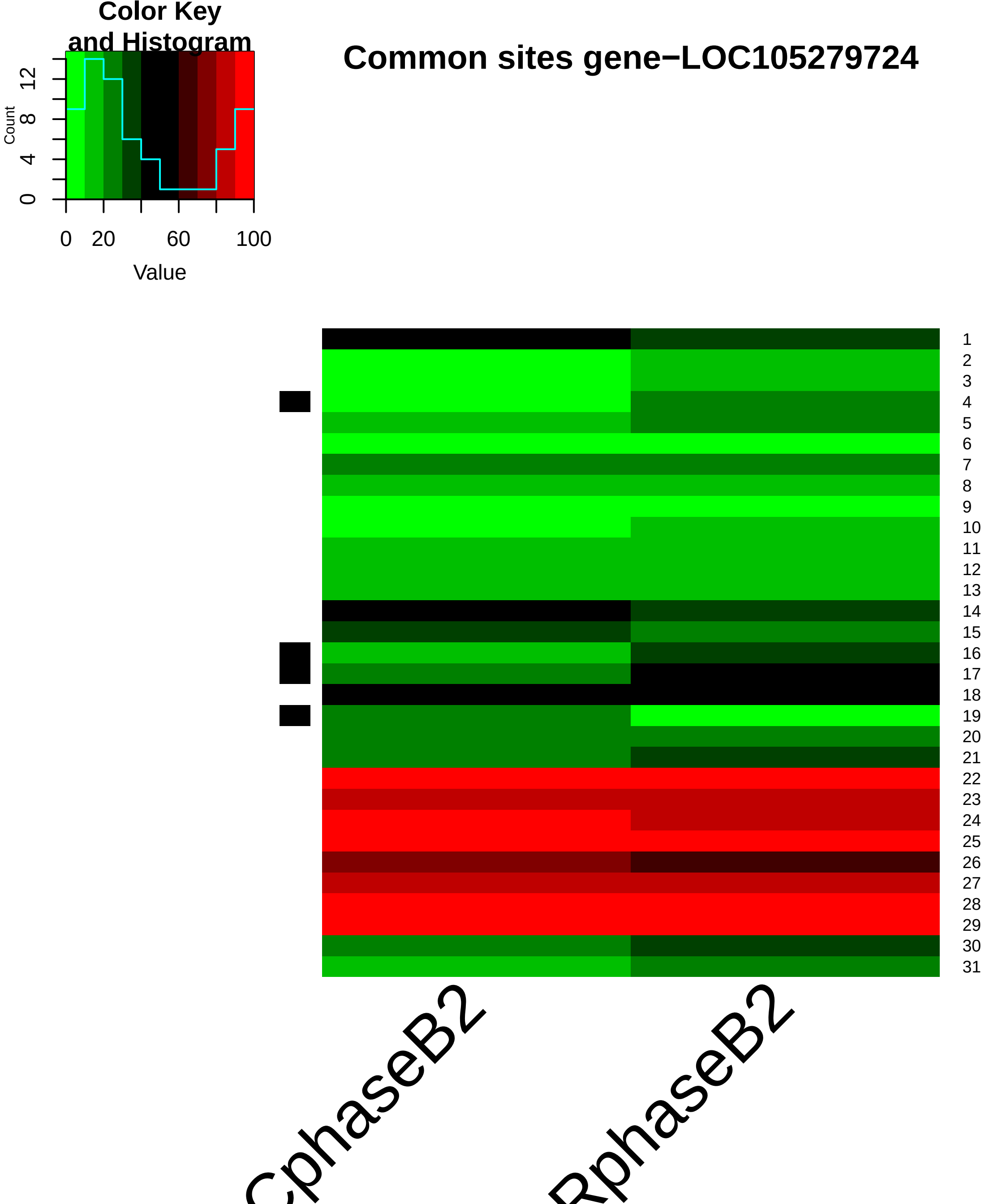

Color Key
and Histogram
Count
0
4
8
12
0
20
60
100
Value
Common sites gene−LOC105279724
CphaseB2
RphaseB2

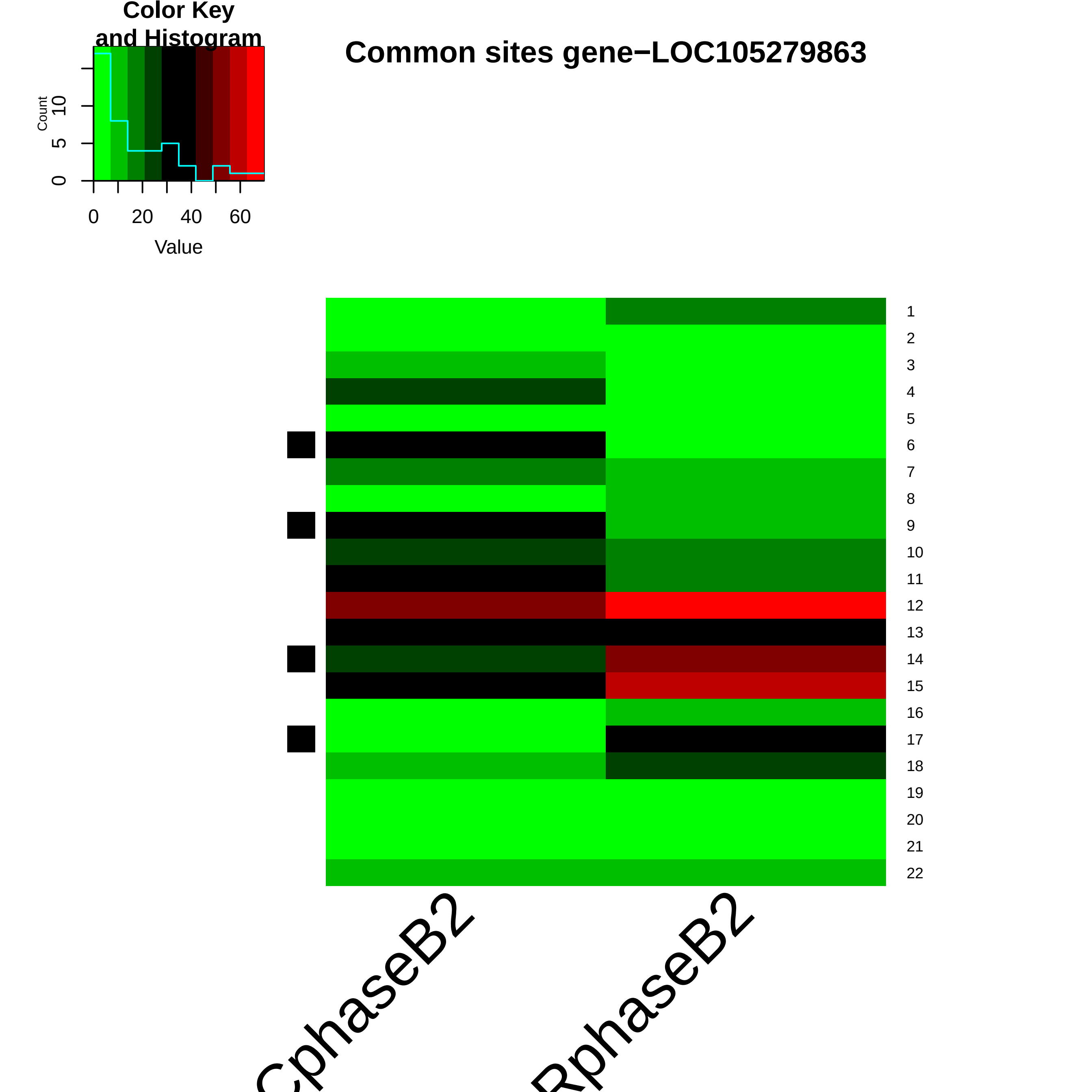

Color Key
and Histogram
Count
0
5
10
0
20
40
60
Value
Common sites gene−LOC105279863

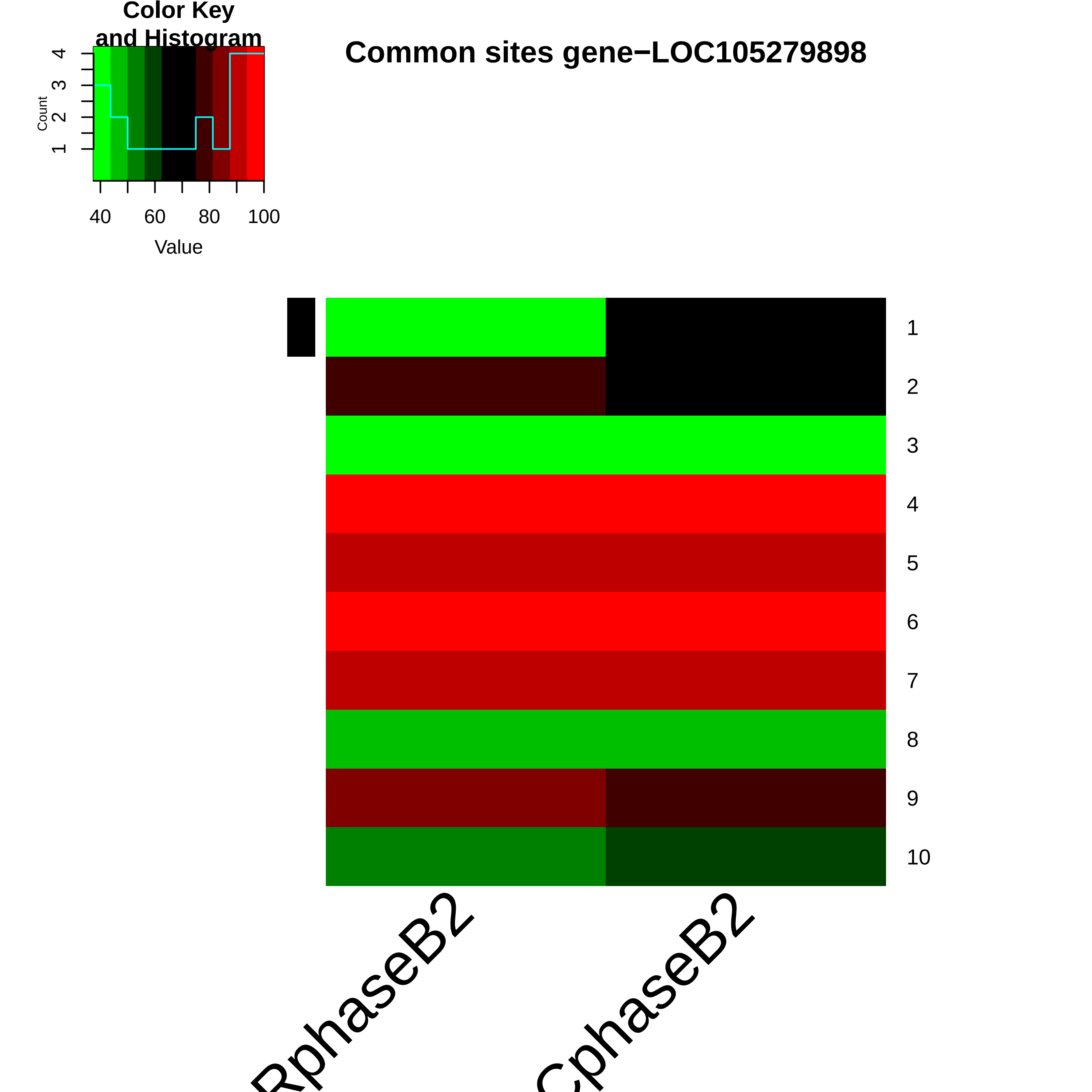
Color Key
and Histogram
Count
1
2
3
4
40
60
80
100
Value
Common sites gene−LOC105279898
1
2
3
4
5
6
7
8
9
10
RphaseB2
CphaseB2

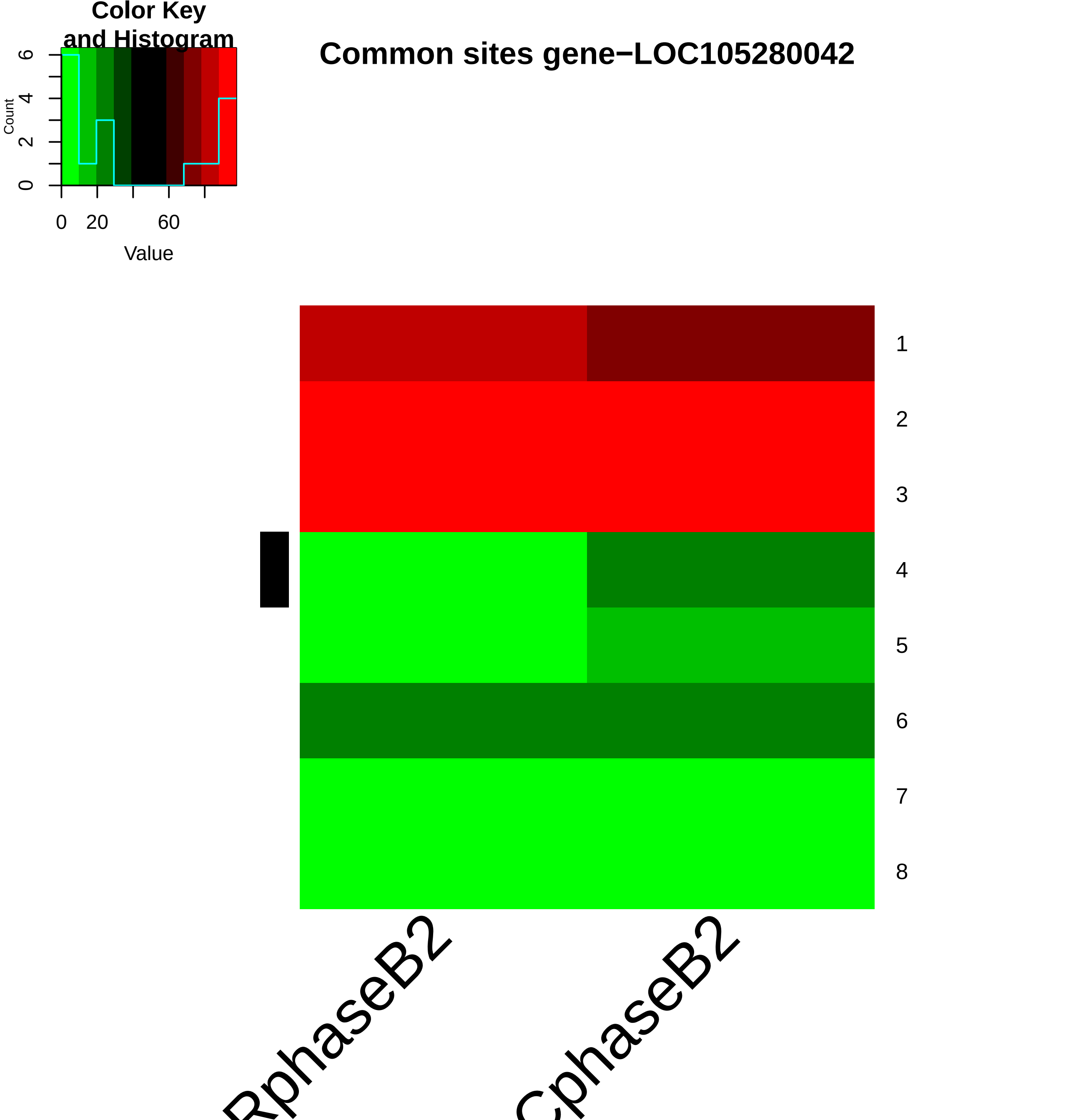
Color Key
and Histogram
Count
0
2
4
6
0
20
60
Value
Common sites gene−LOC105280042
1
2
3
4
5
6
7
8
RphaseB2
CphaseB2

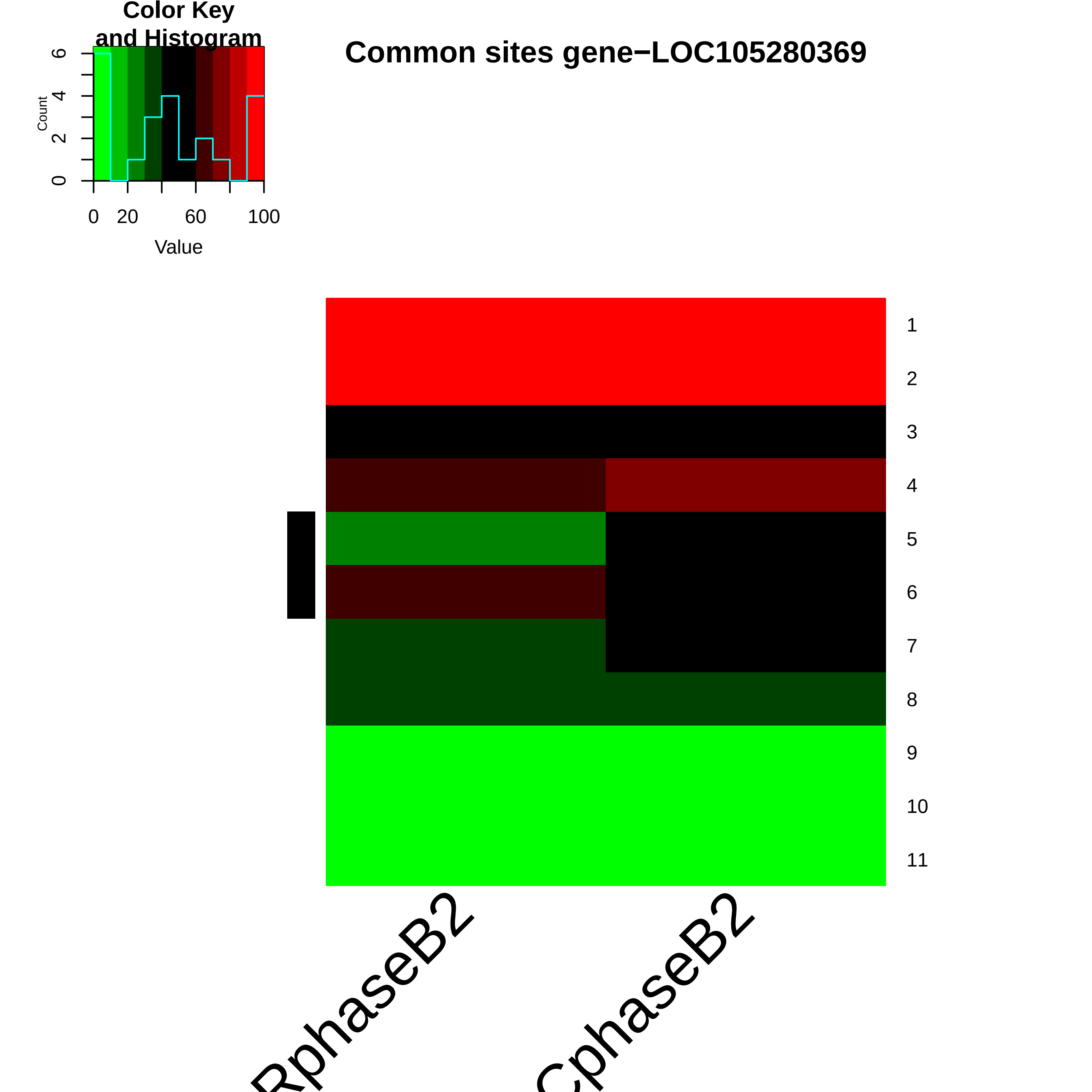

Color Key
and Histogram
Count
Value
0
20
60
100
0
2
4
6
Common sites gene−LOC105280369
1
2
3
4
5
6
7
8
9
10
11
RphaseB2
CphaseB2

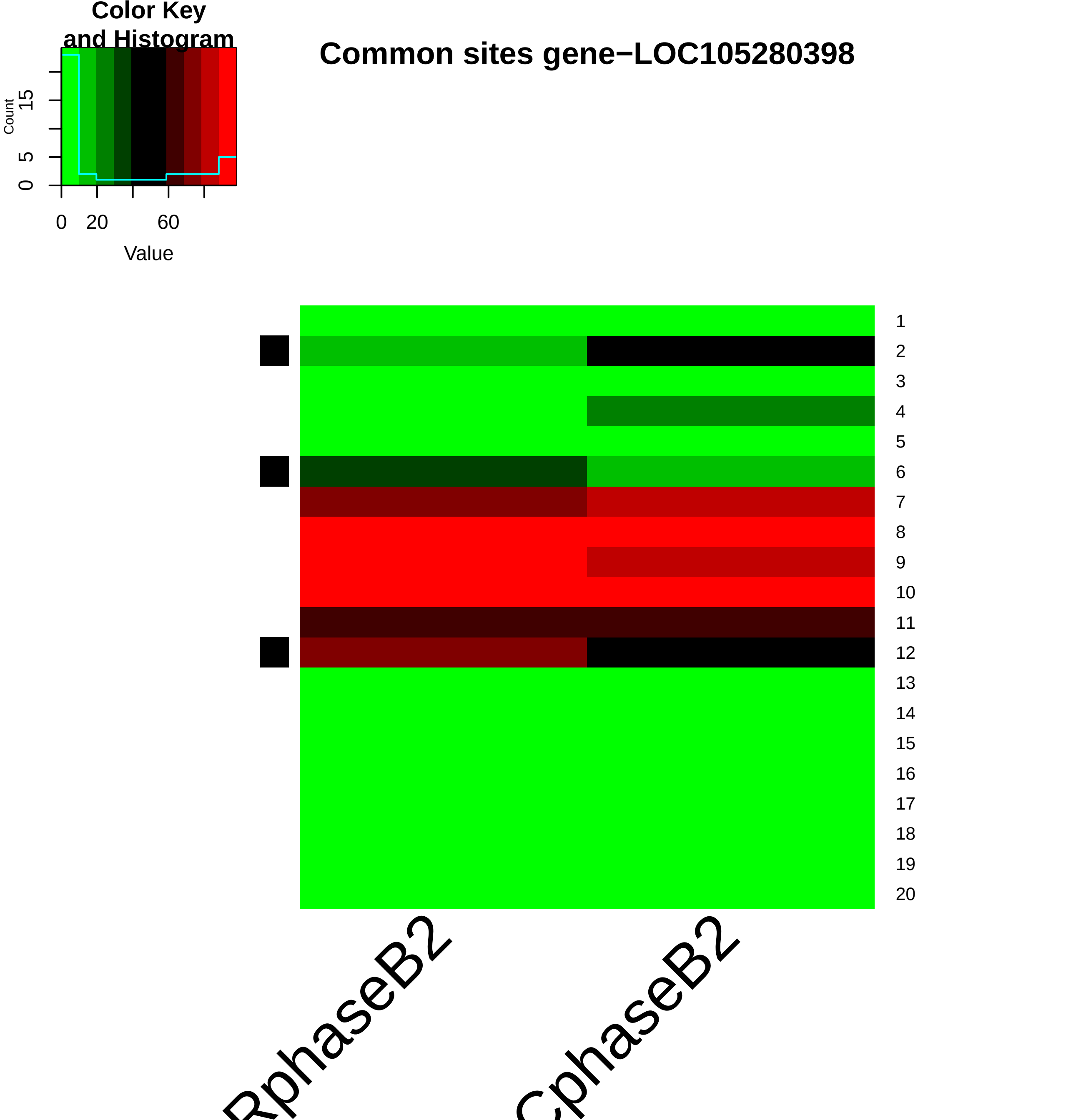
Color Key
and Histogram
Count
0
5
15
0
20
60
Value
Common sites gene−LOC105280398
RphaseB2
CphaseB2

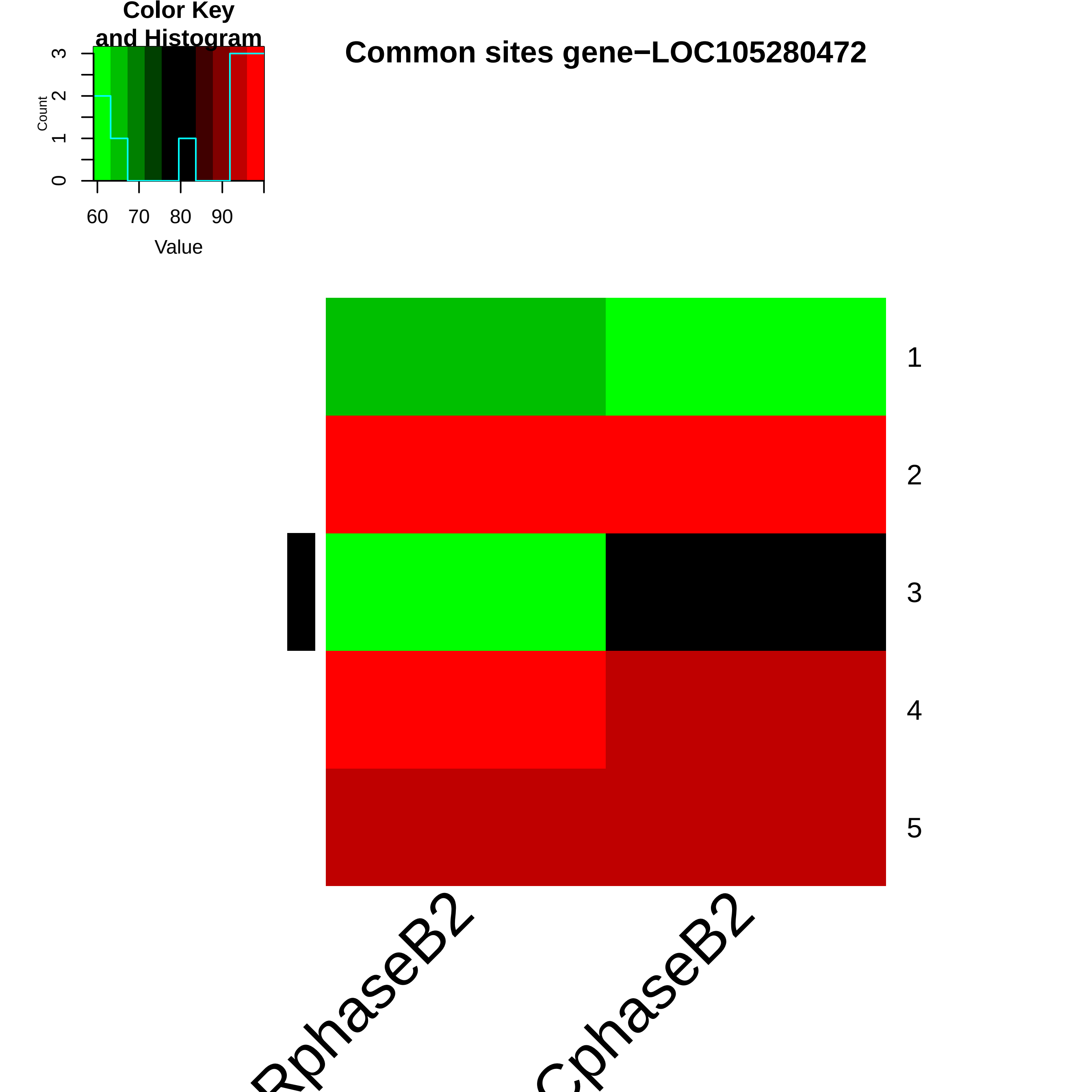

Color Key
and Histogram
Count
0
1
2
3
60
70
80
90
Value
Common sites gene−LOC105280472
1
2
3
4
5
RphaseB2
CphaseB2

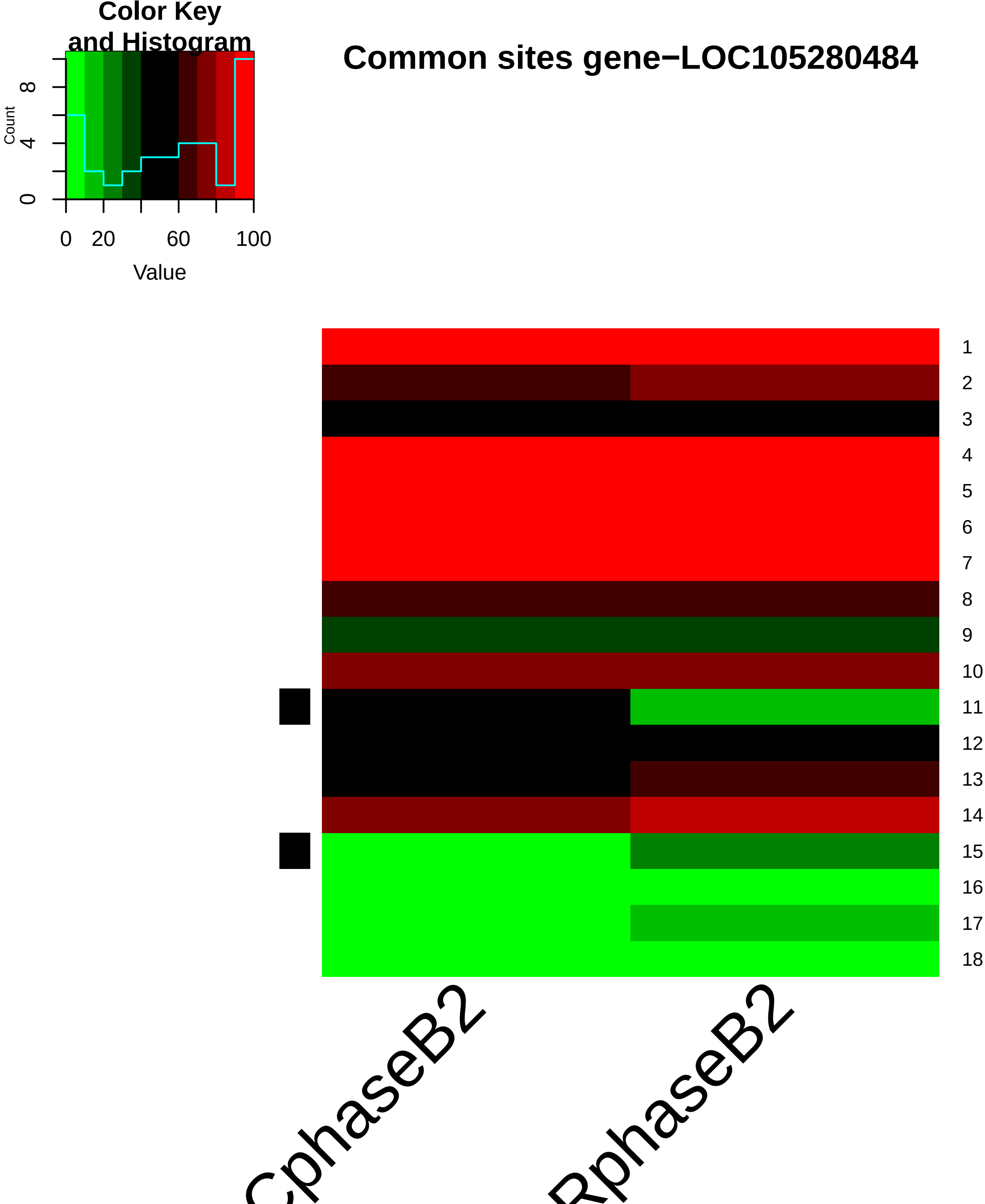

Color Key
and Histogram
Count
0
4
8
0
20
60
100
Value
Common sites gene−LOC105280484
CphaseB2
RphaseB2

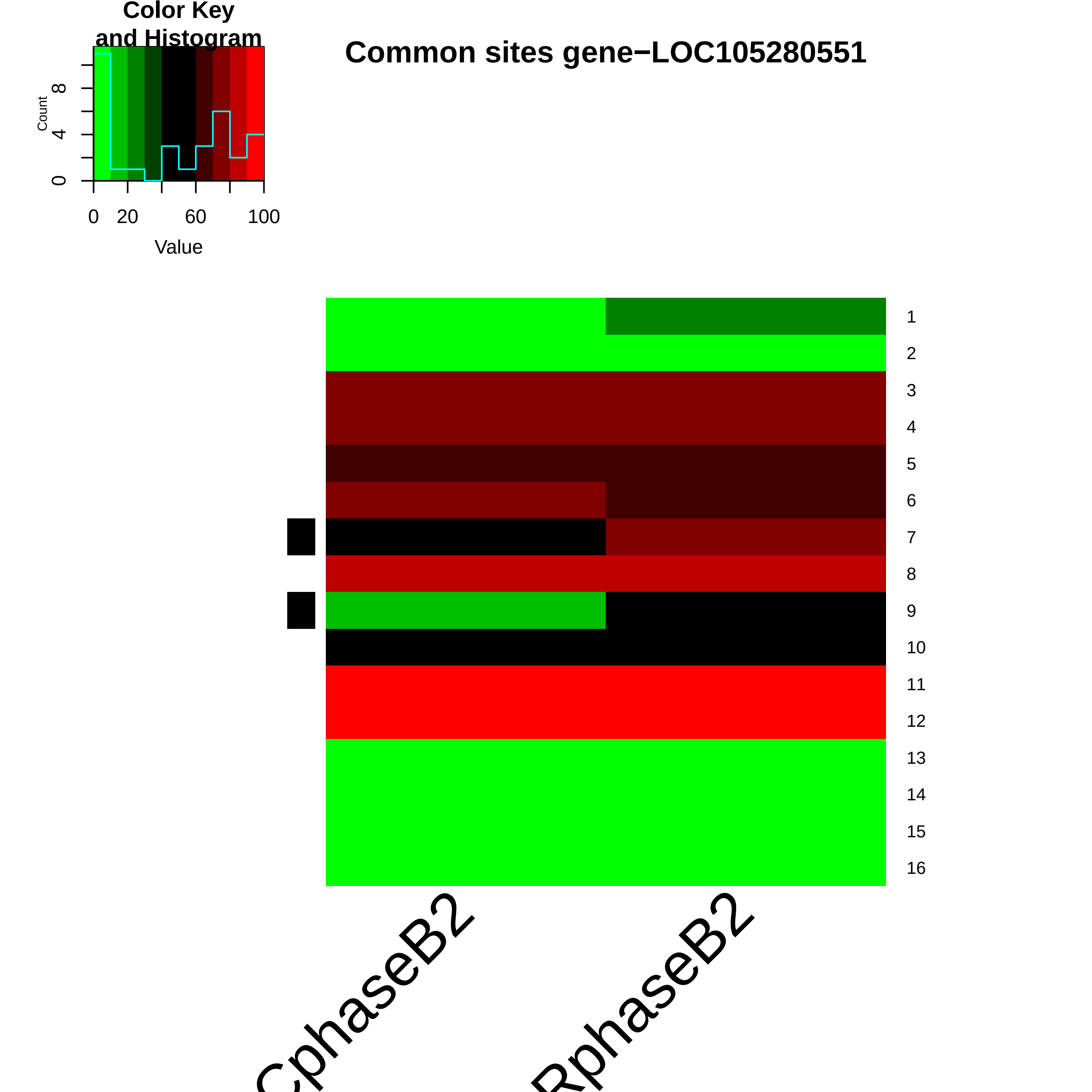
Color Key
and Histogram
Count
0
4
8
0
20
60
100
Value
Common sites gene−LOC105280551
CphaseB2
RphaseB2

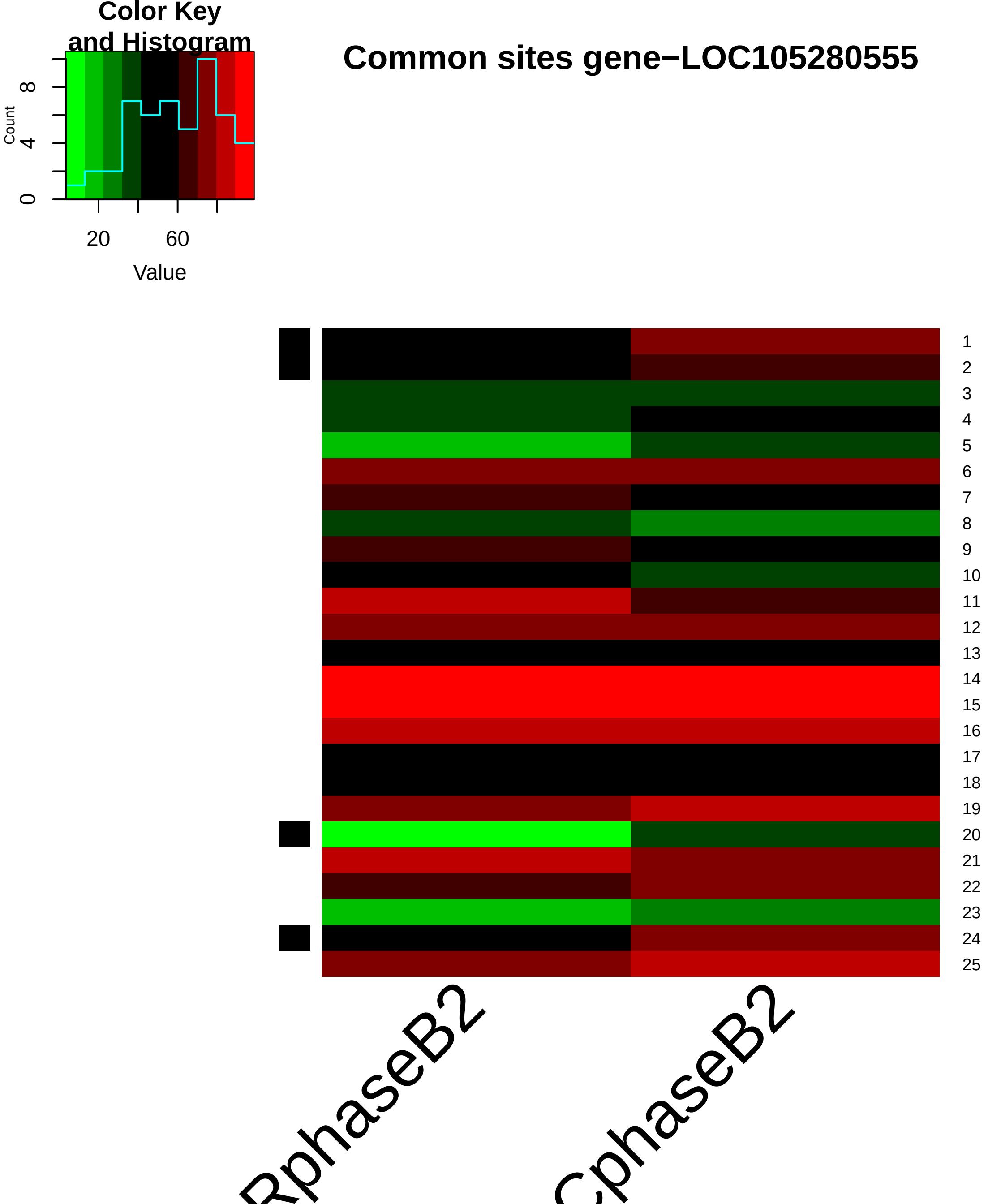

Color Key
and Histogram
Count
Value
0
4
8
20
60
Common sites gene−LOC105280555
RphaseB2
CphaseB2

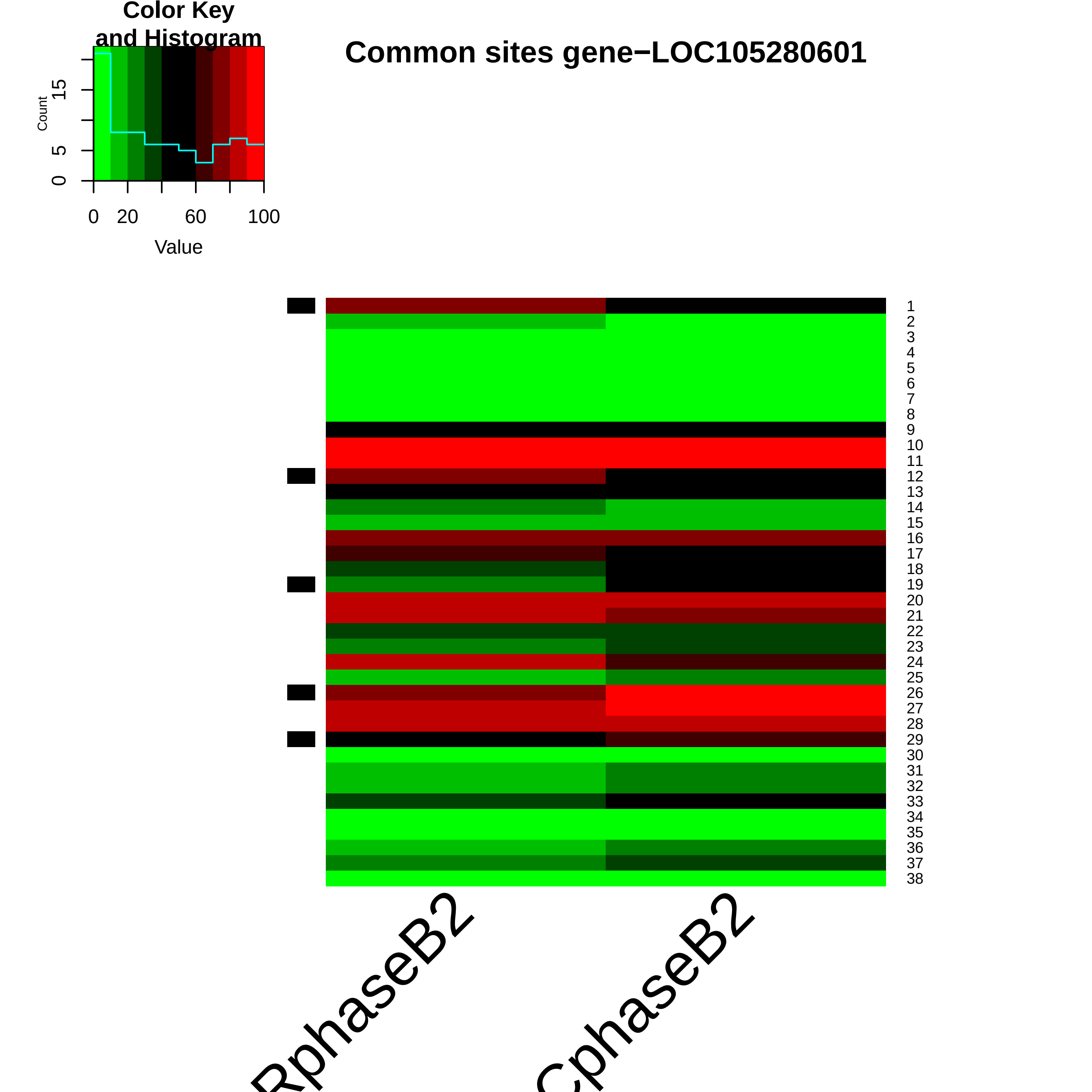
Color Key
and Histogram
Count
0
5
15
0
20
60
100
Value
Common sites gene−LOC105280601
RphaseB2
CphaseB2

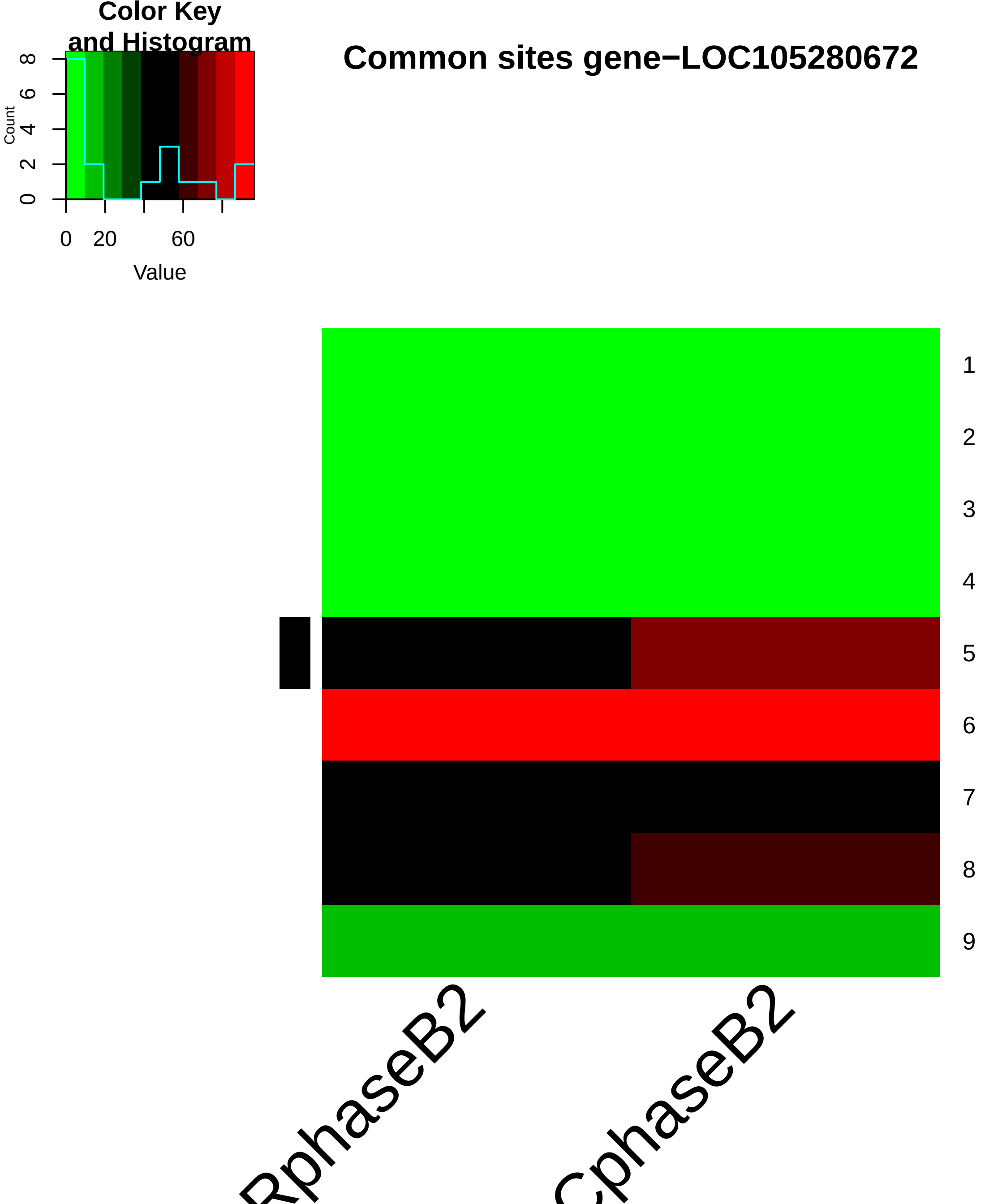

Color Key and Histogram
Count
0
2
4
6
8
0
20
60
Value
Common sites gene−LOC105280672
1
2
3
4
5
6
7
8
9
RphaseB2
CphaseB2

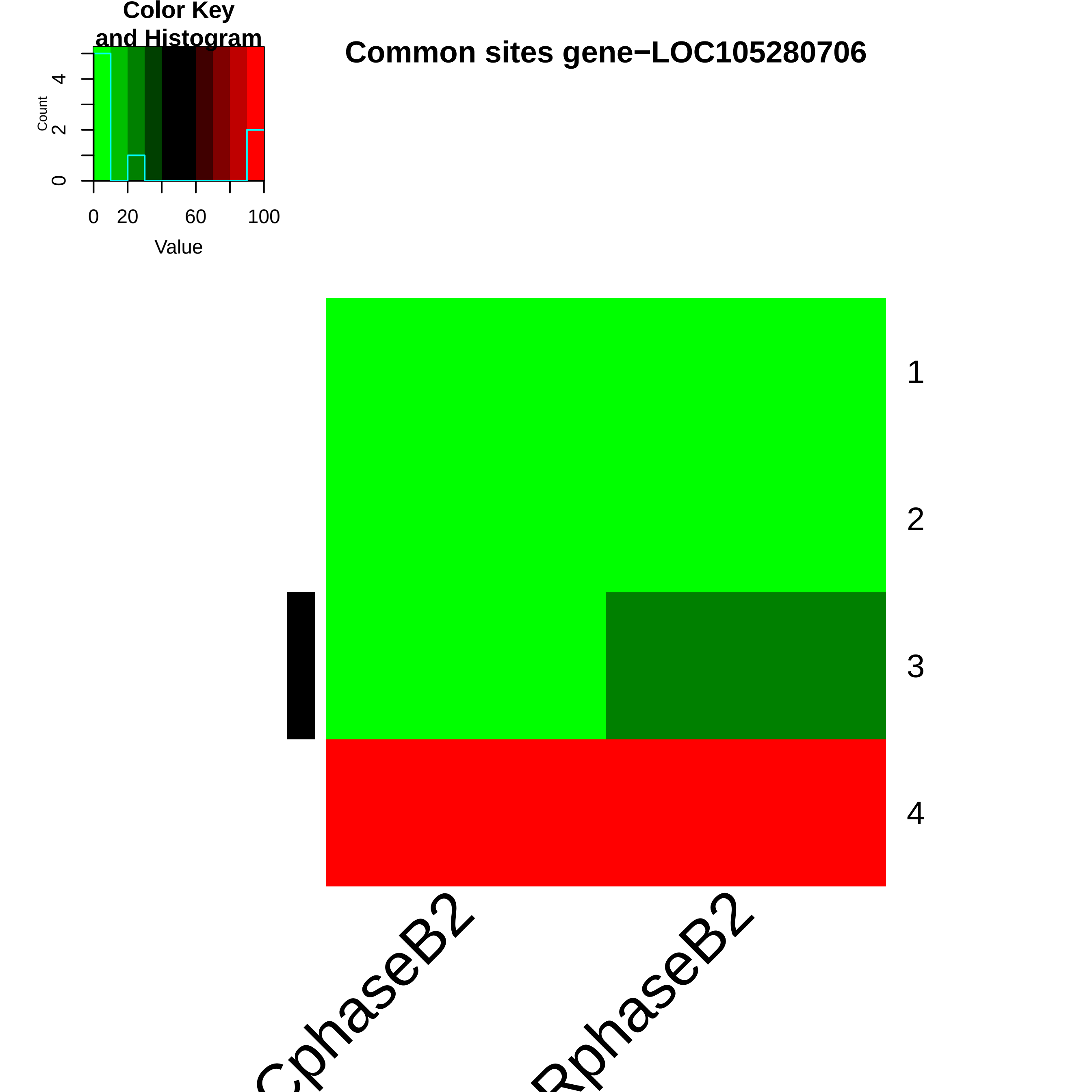

Color Key
and Histogram
Count
0
2
4
0
20
60
100
Value
Common sites gene−LOC105280706
1
2
3
4
CphaseB2
RphaseB2

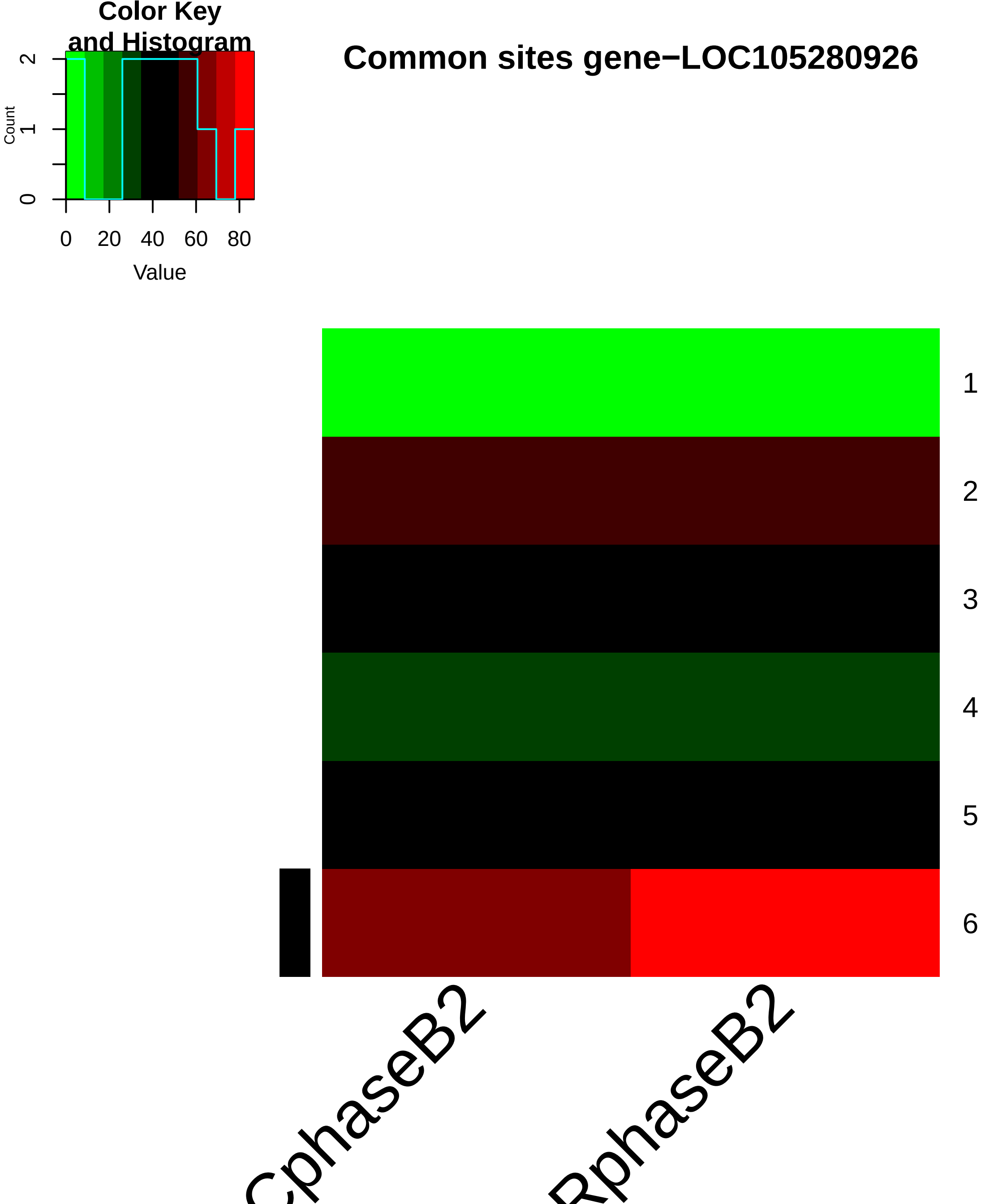
Color Key
and Histogram
Count
0
1
2
0
20
40
60
80
Value
Common sites gene−LOC105280926
1
2
3
4
5
6
CphaseB2
RphaseB2

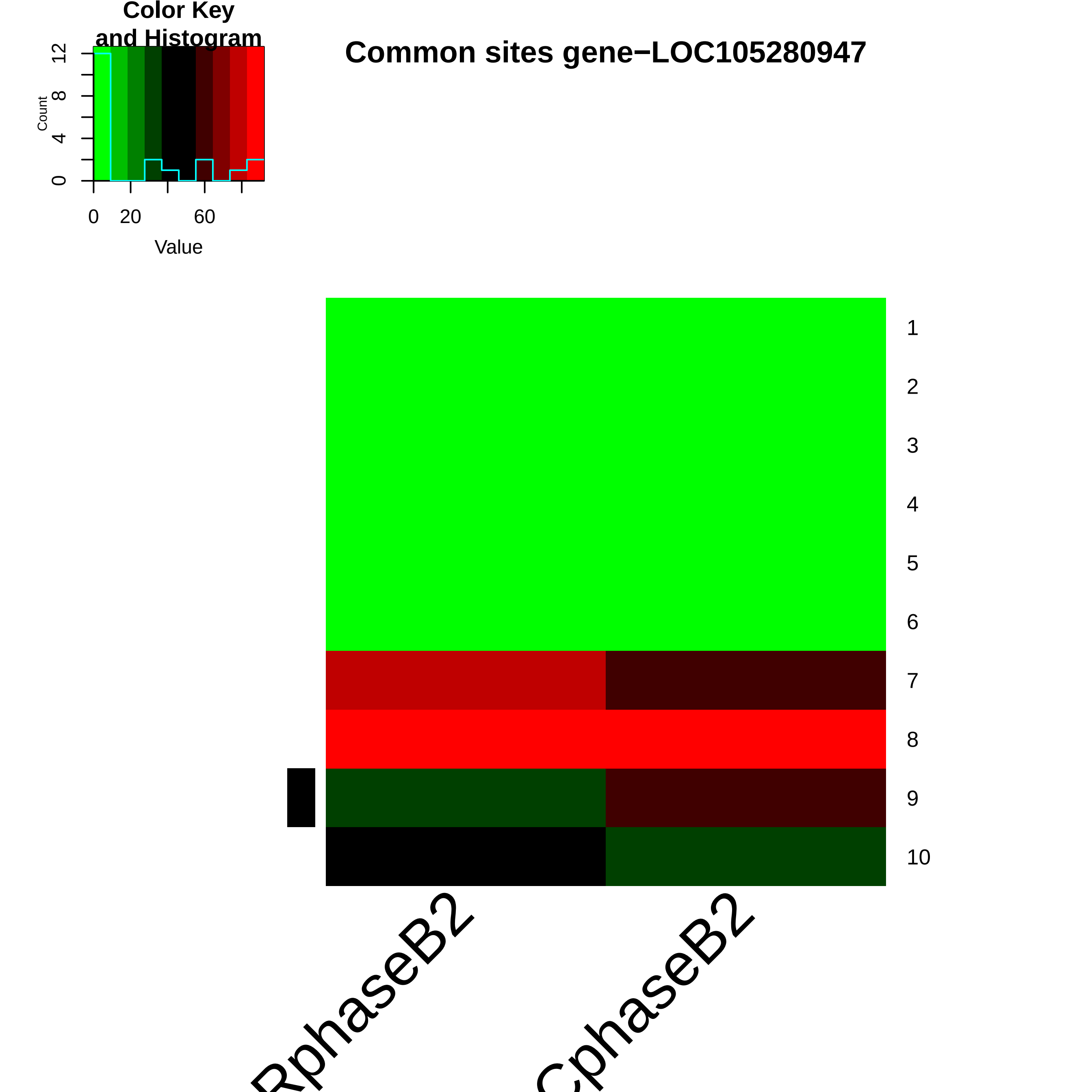

Color Key
and Histogram
Count
Value
0
4
8
12
0
20
60
Common sites gene−LOC105280947
1
2
3
4
5
6
7
8
9
10
RphaseB2
CphaseB2

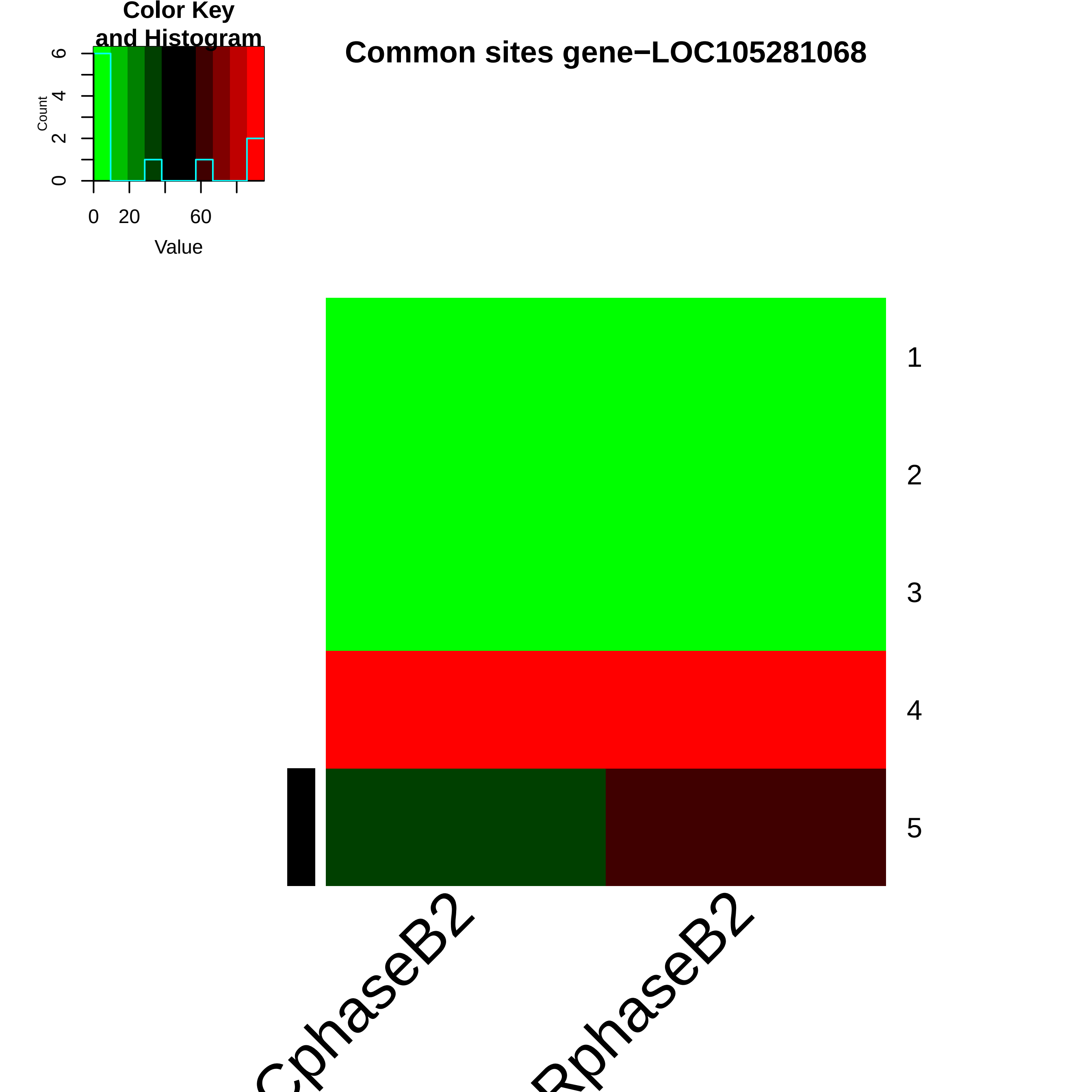

Color Key
and Histogram
Count
0
2
4
6
0
20
60
Value
Common sites gene−LOC105281068
1
2
3
4
5
CphaseB2
RphaseB2

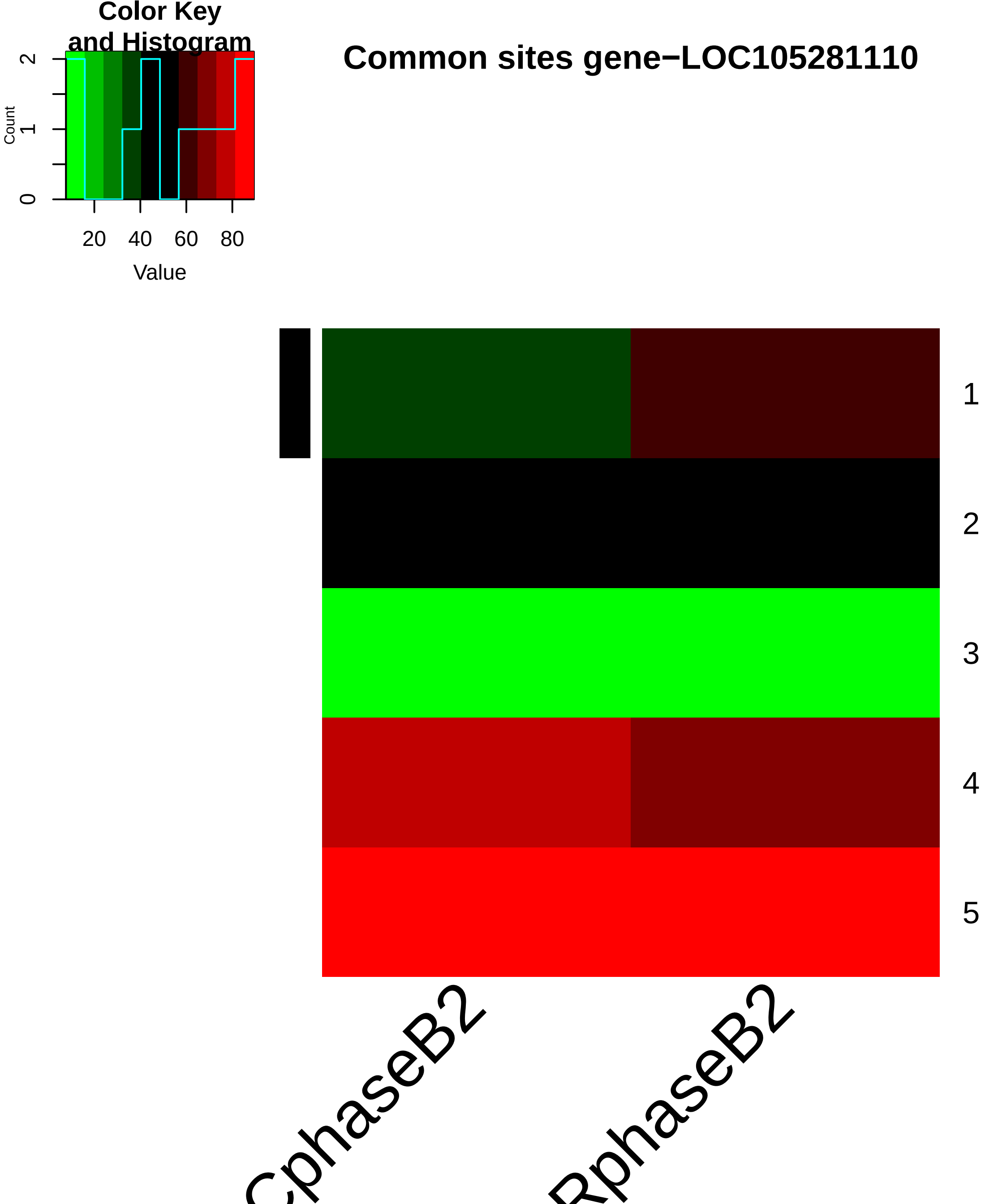

Color Key and Histogram
Count
0
1
2
20
40
60
80
Value
Common sites gene−LOC105281110
1
2
3
4
5
CphaseB2
RphaseB2

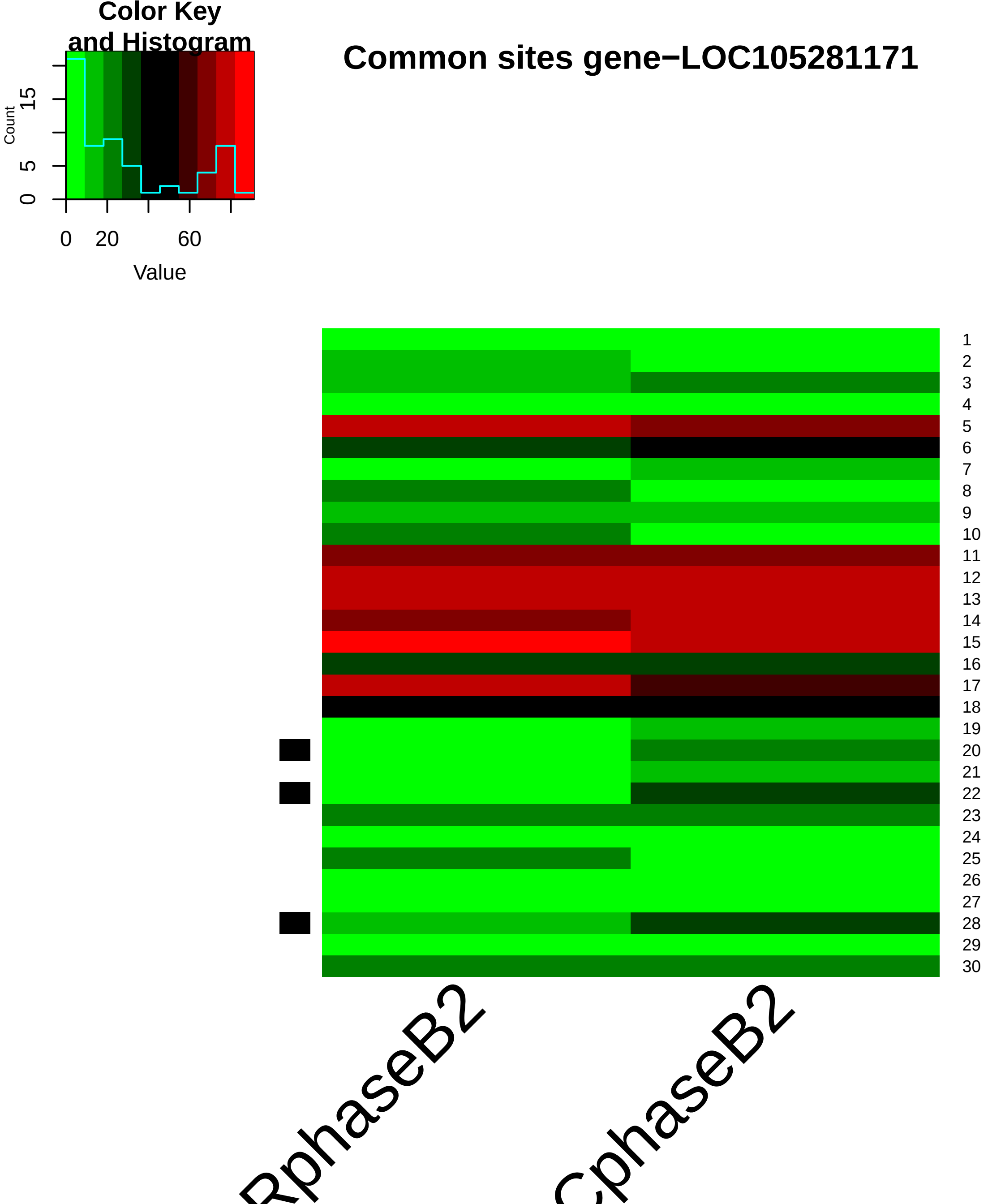

Color Key
and Histogram
Count
0
5
15
0
20
60
Value
Common sites gene−LOC105281171
RphaseB2
CphaseB2

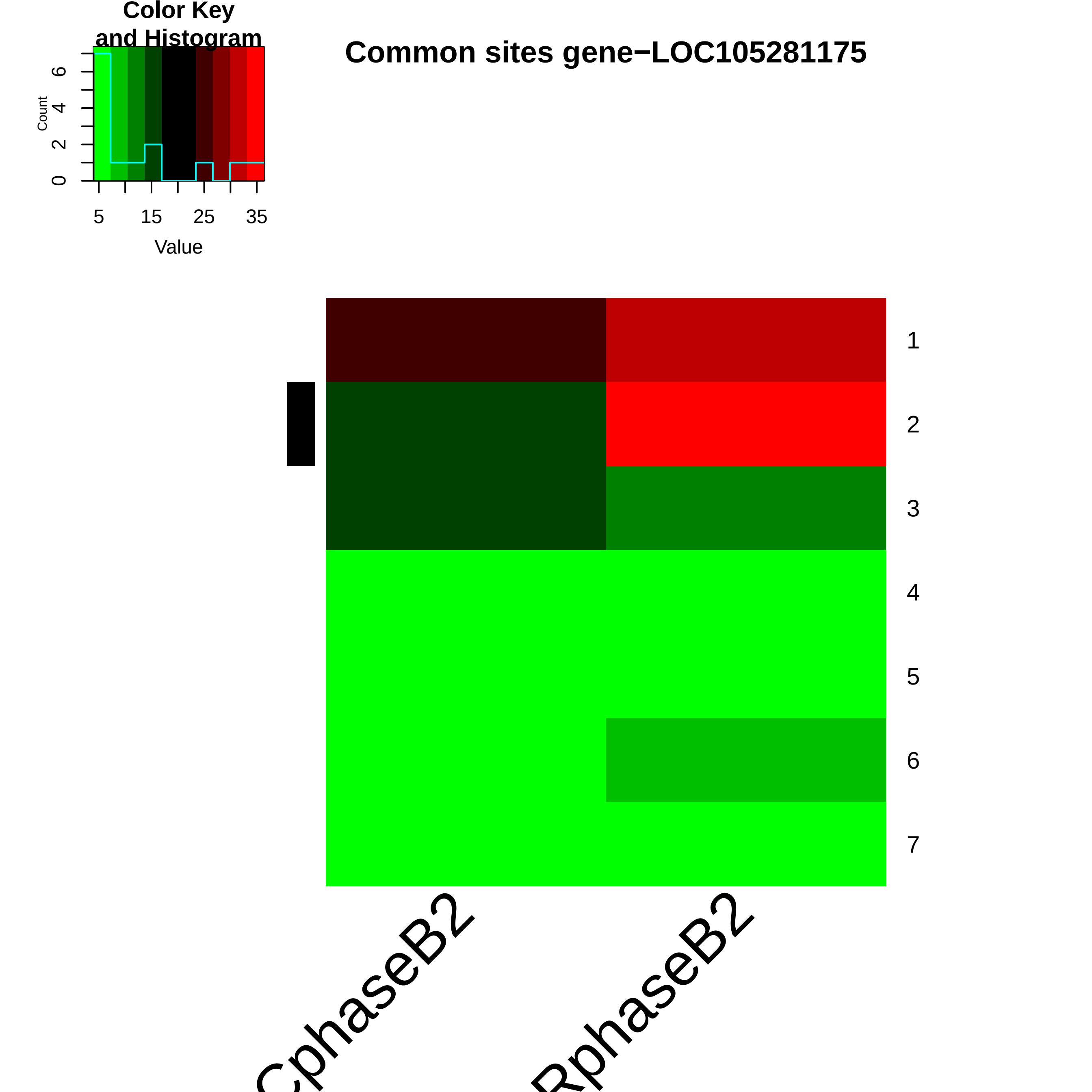
Color Key
and Histogram
Count
0
2
4
6
5
15
25
35
Value
Common sites gene−LOC105281175
1
2
3
4
5
6
7

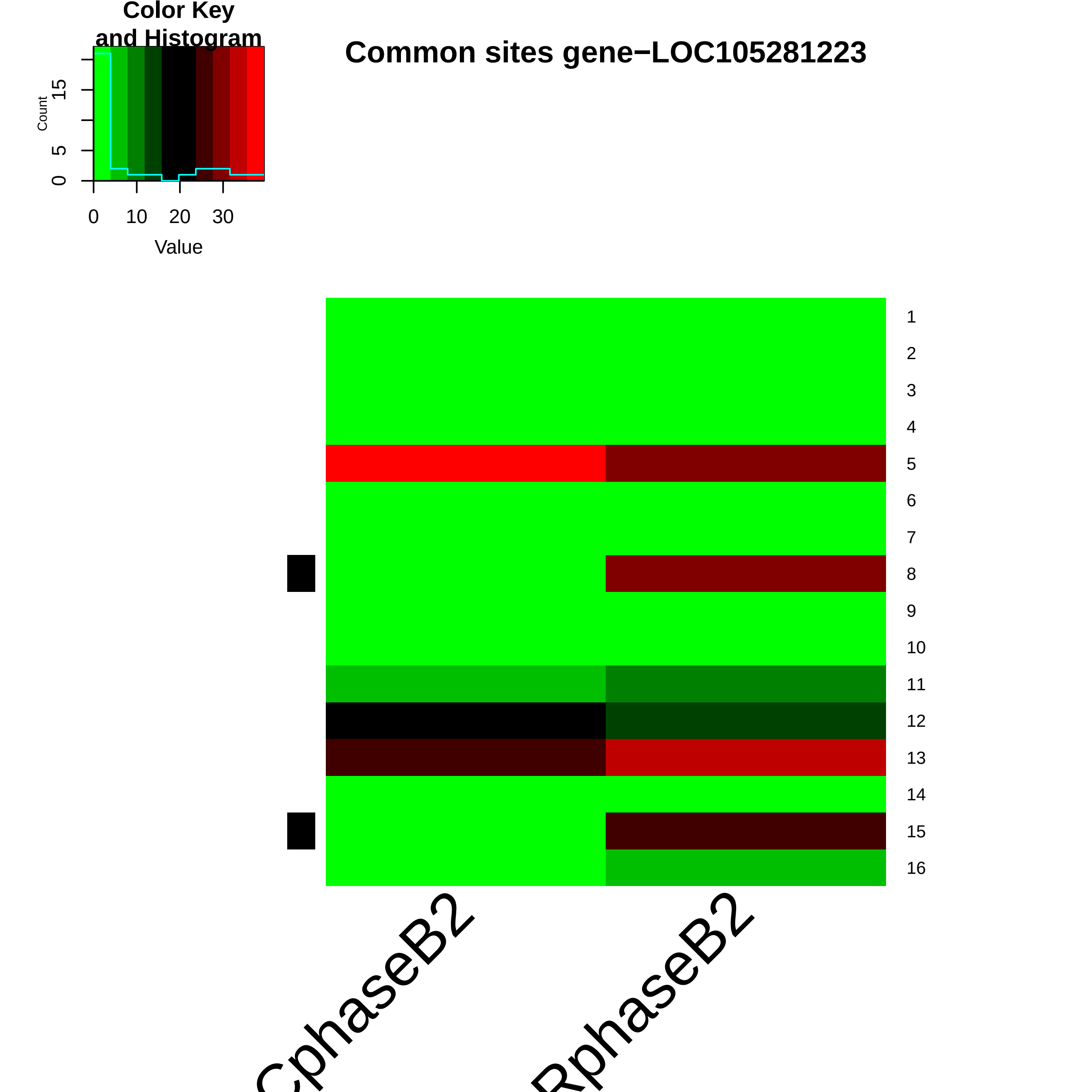

Color Key
and Histogram
Count
0
5
15
0
10
20
30
Value
Common sites gene−LOC105281223
CphaseB2
RphaseB2

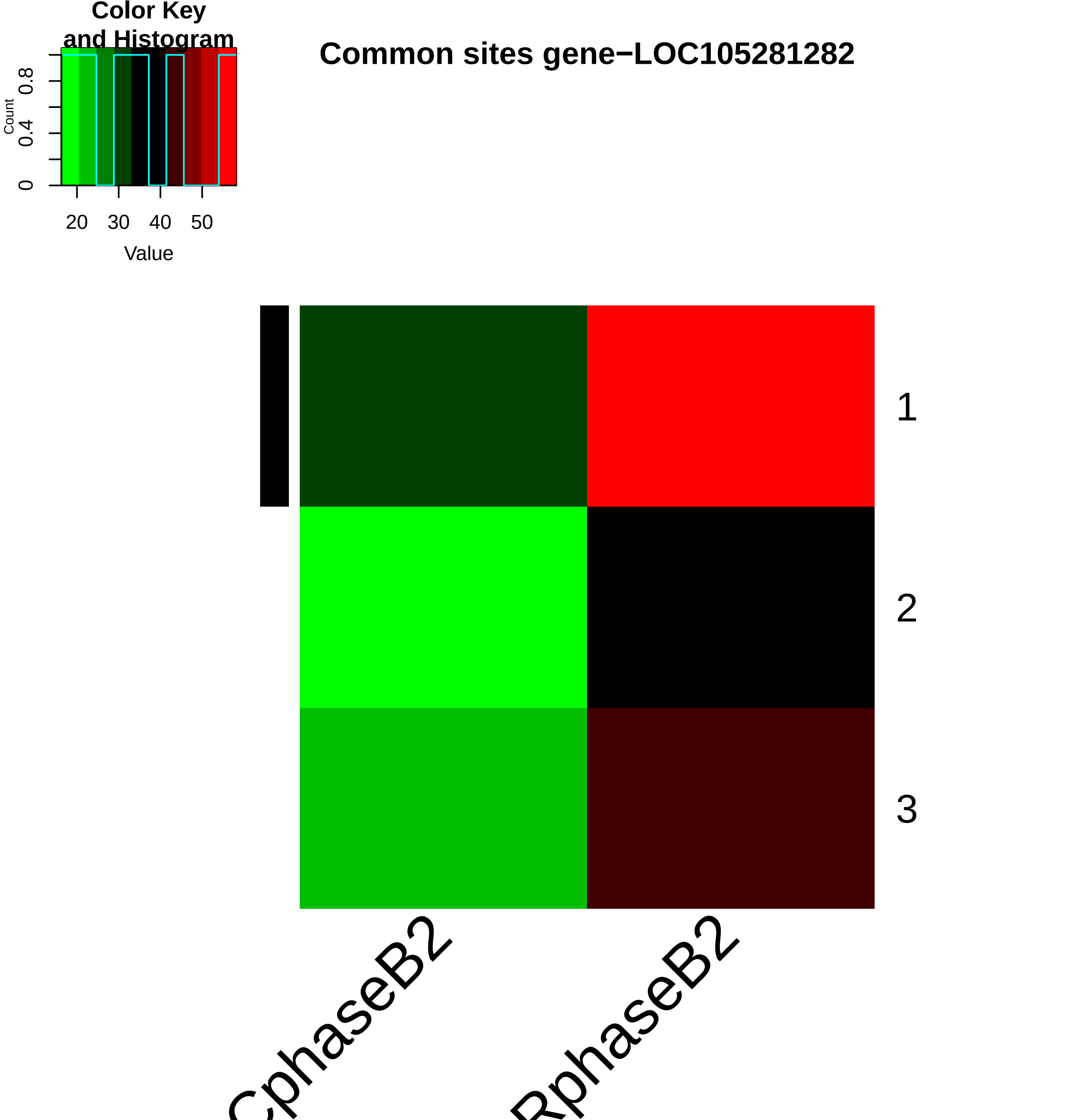

Color Key
and Histogram
Count
0
0.4
0.8
20
30
40
50
Value
Common sites gene−LOC105281282
1
2
3
CphaseB2
RphaseB2

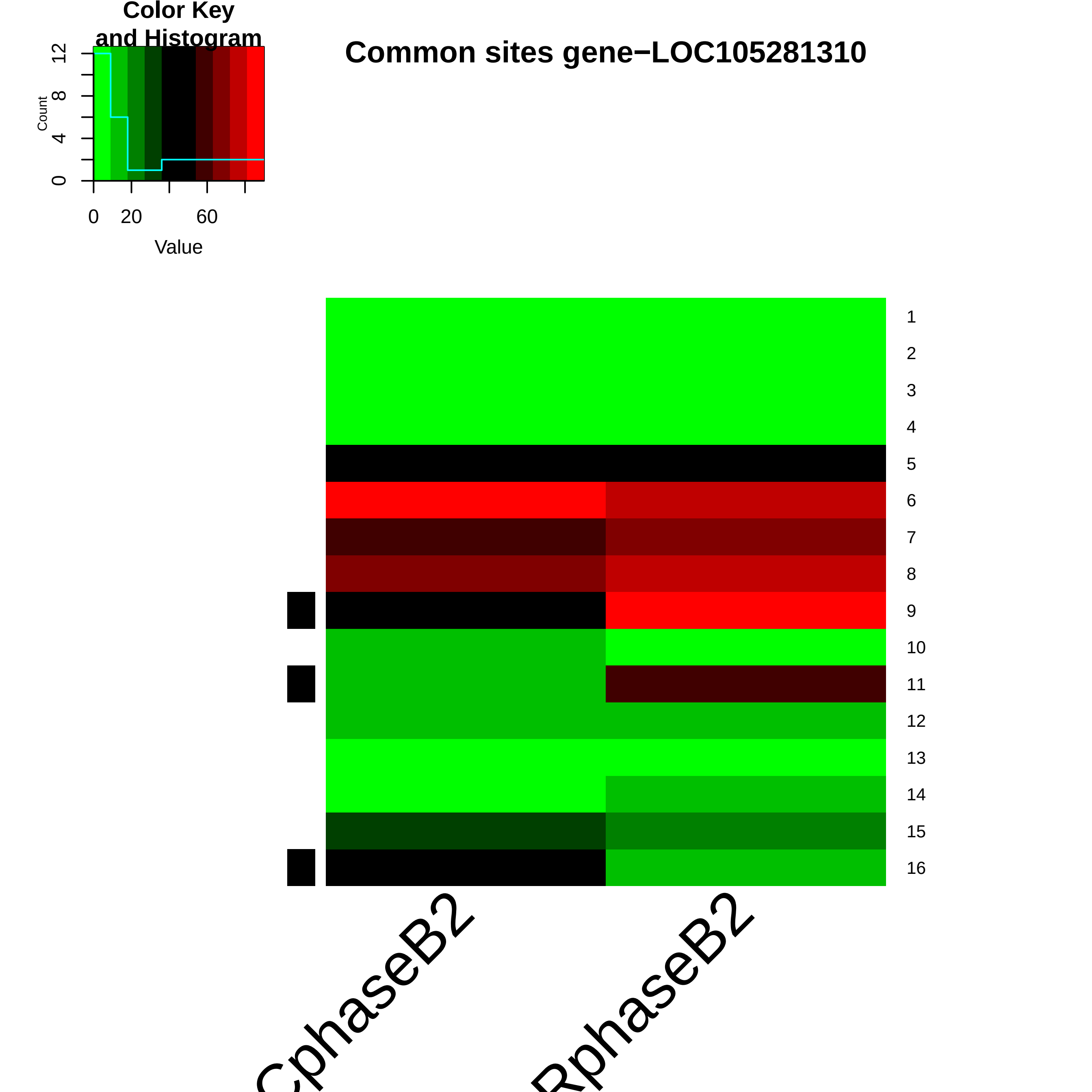
Color Key
and Histogram
Count
0
4
8
12
0
20
60
Value
Common sites gene−LOC105281310
CphaseB2
RphaseB2

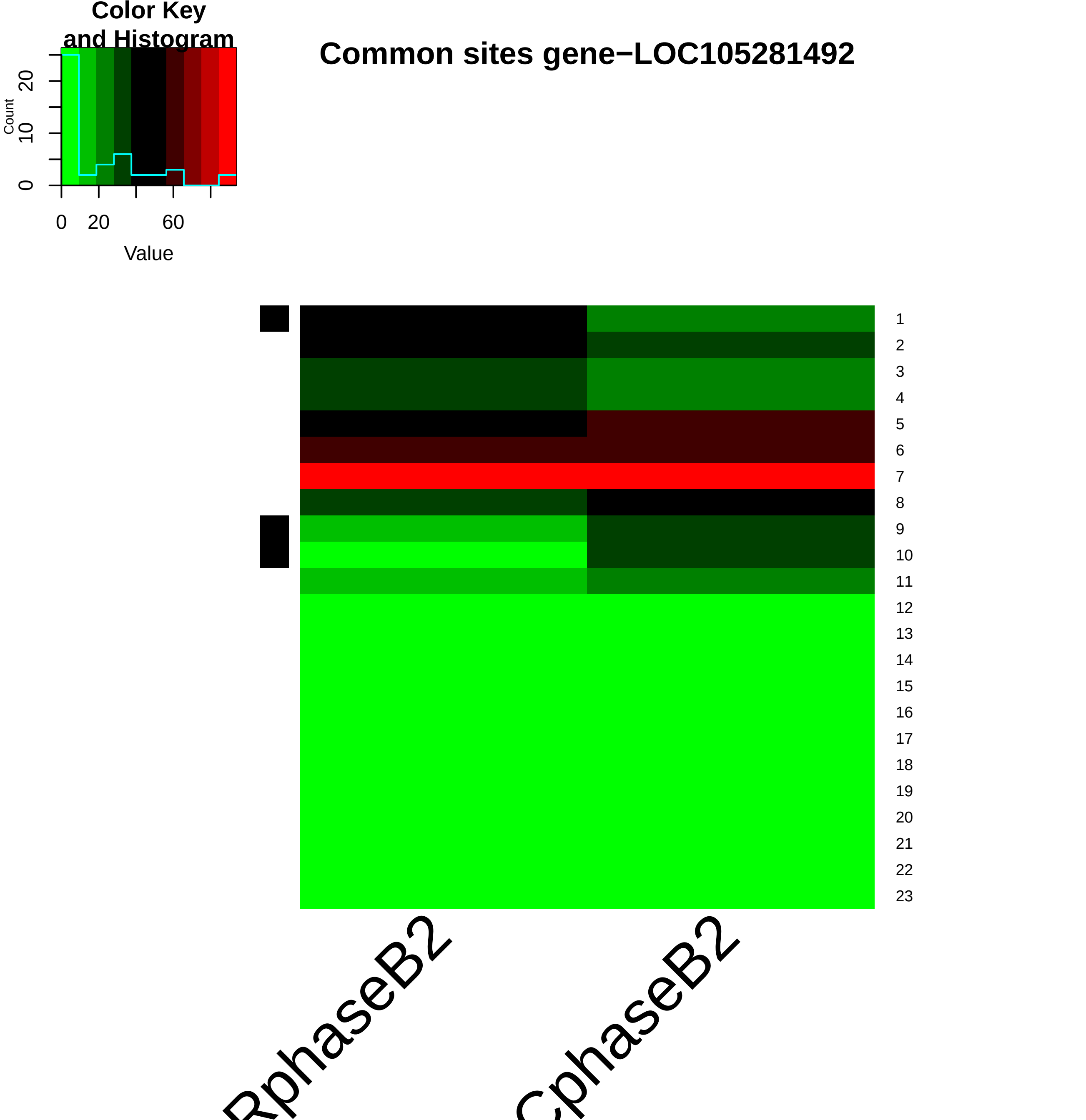
Color Key
and Histogram
Count
0
10
20
0
20
60
Value
Common sites gene−LOC105281492
RphaseB2
CphaseB2

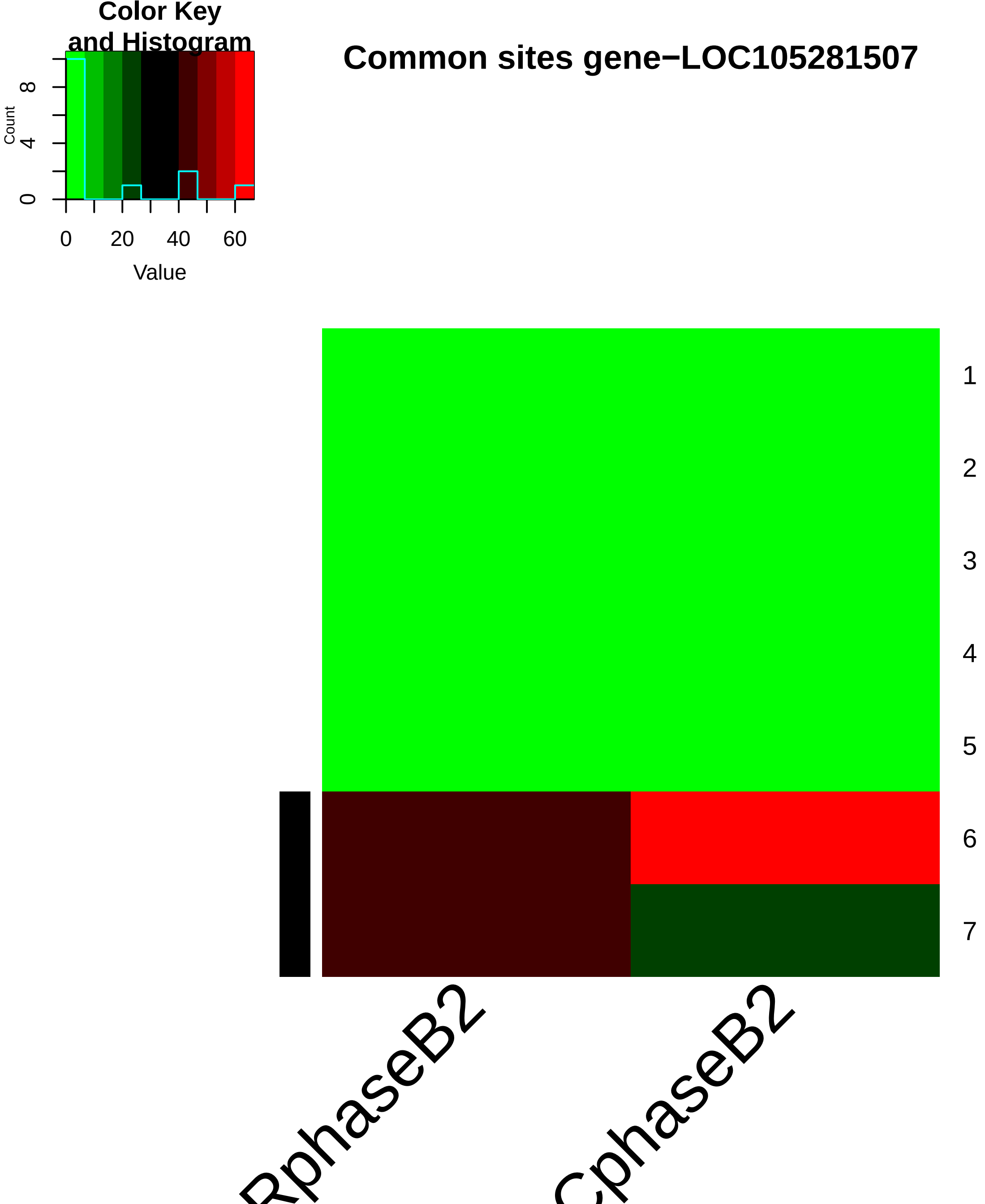
Color Key
and Histogram
Count
0
4
8
0
20
40
60
Value
Common sites gene−LOC105281507
1
2
3
4
5
6
7
RphaseB2
CphaseB2

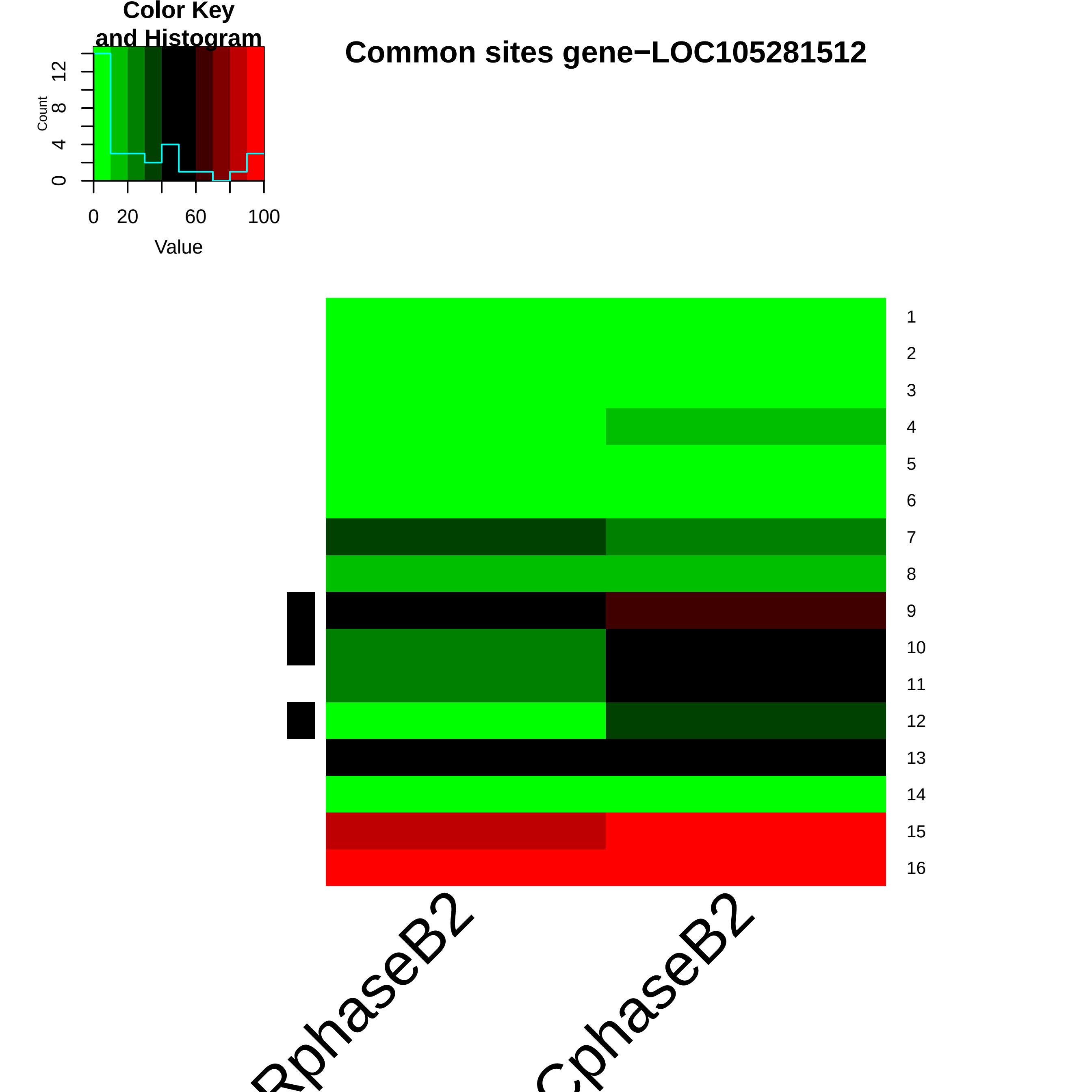

Color Key
and Histogram
Count
0
4
8
12
0
20
60
100
Value
Common sites gene−LOC105281512
RphaseB2
CphaseB2

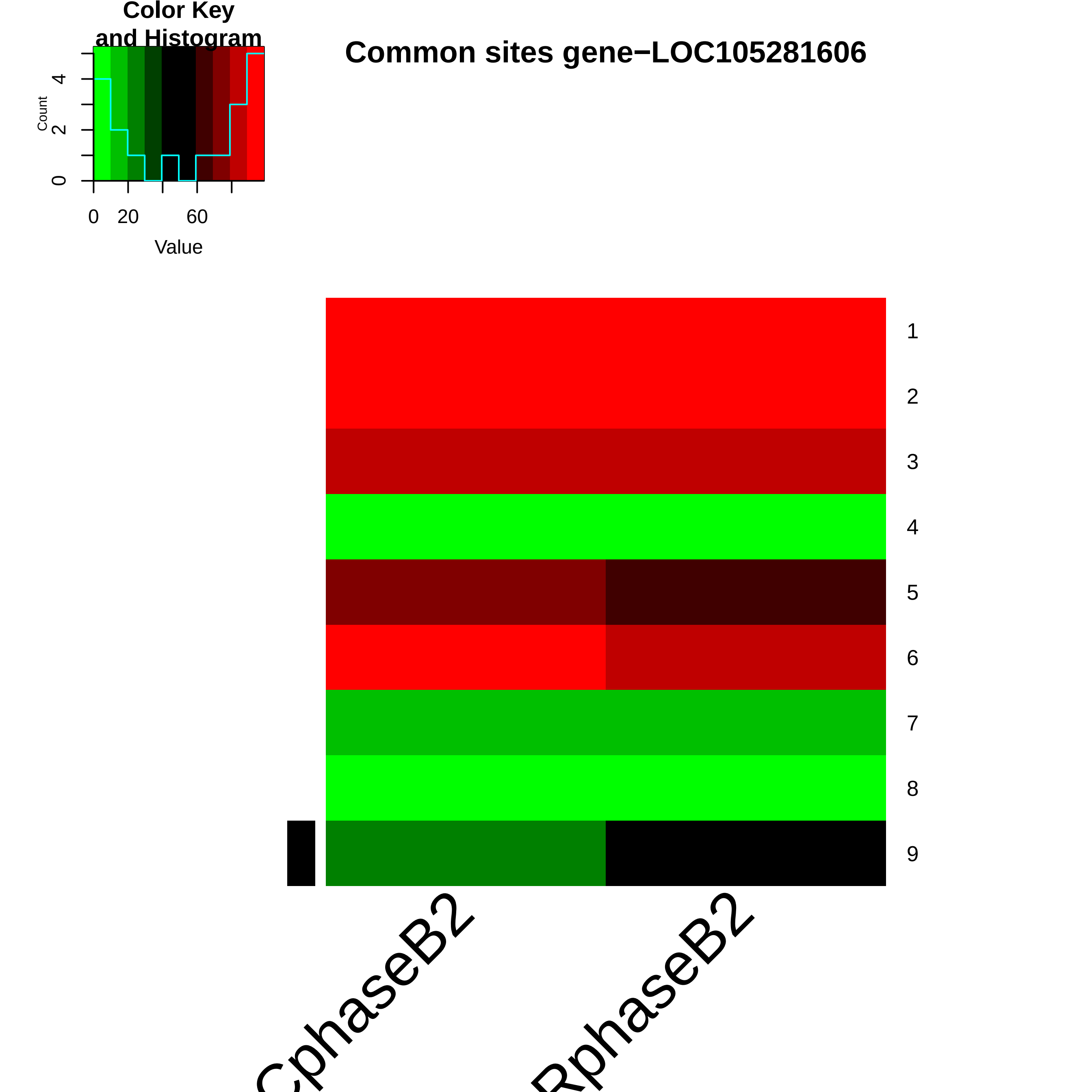
Color Key
and Histogram
Count
0
2
4
0
20
60
Value
Common sites gene−LOC105281606
1
2
3
4
5
6
7
8
9
CphaseB2
RphaseB2

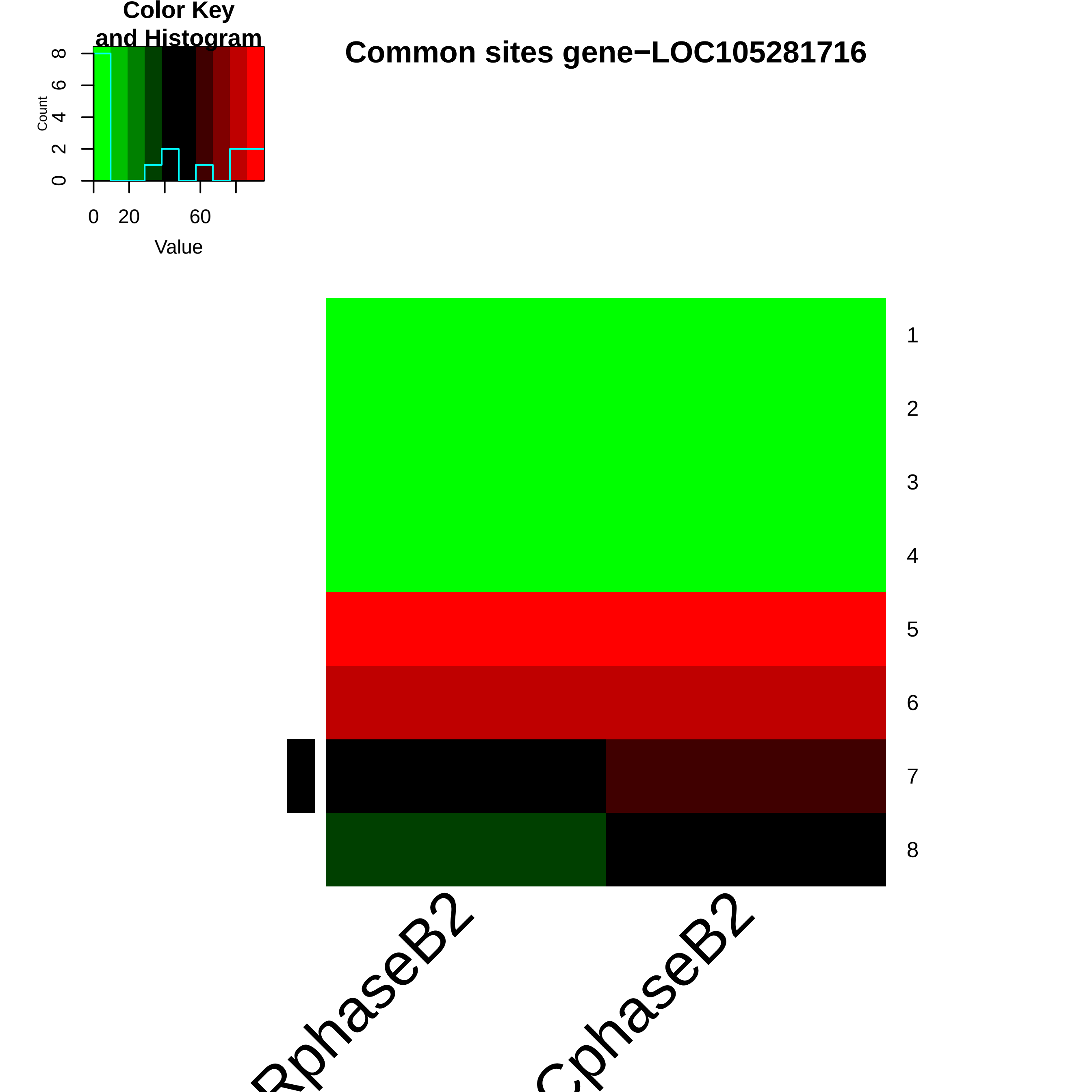

Color Key
and Histogram
Count
0
2
4
6
8
0
20
60
Value
Common sites gene−LOC105281716
1
2
3
4
5
6
7
8
RphaseB2
CphaseB2

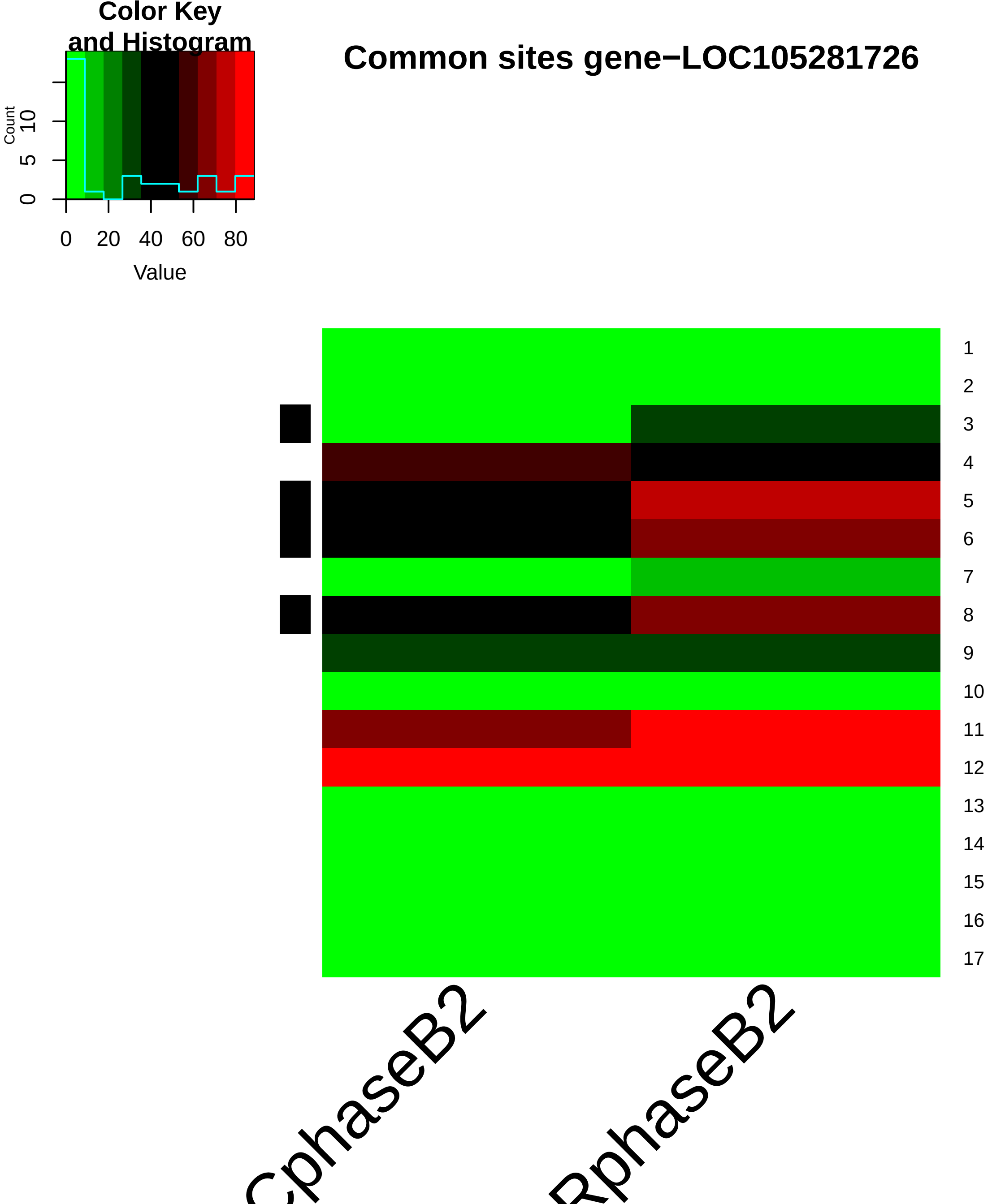
Color Key
and Histogram
Count
0
5
10
0
20
40
60
80
Value
Common sites gene−LOC105281726
CphaseB2
RphaseB2

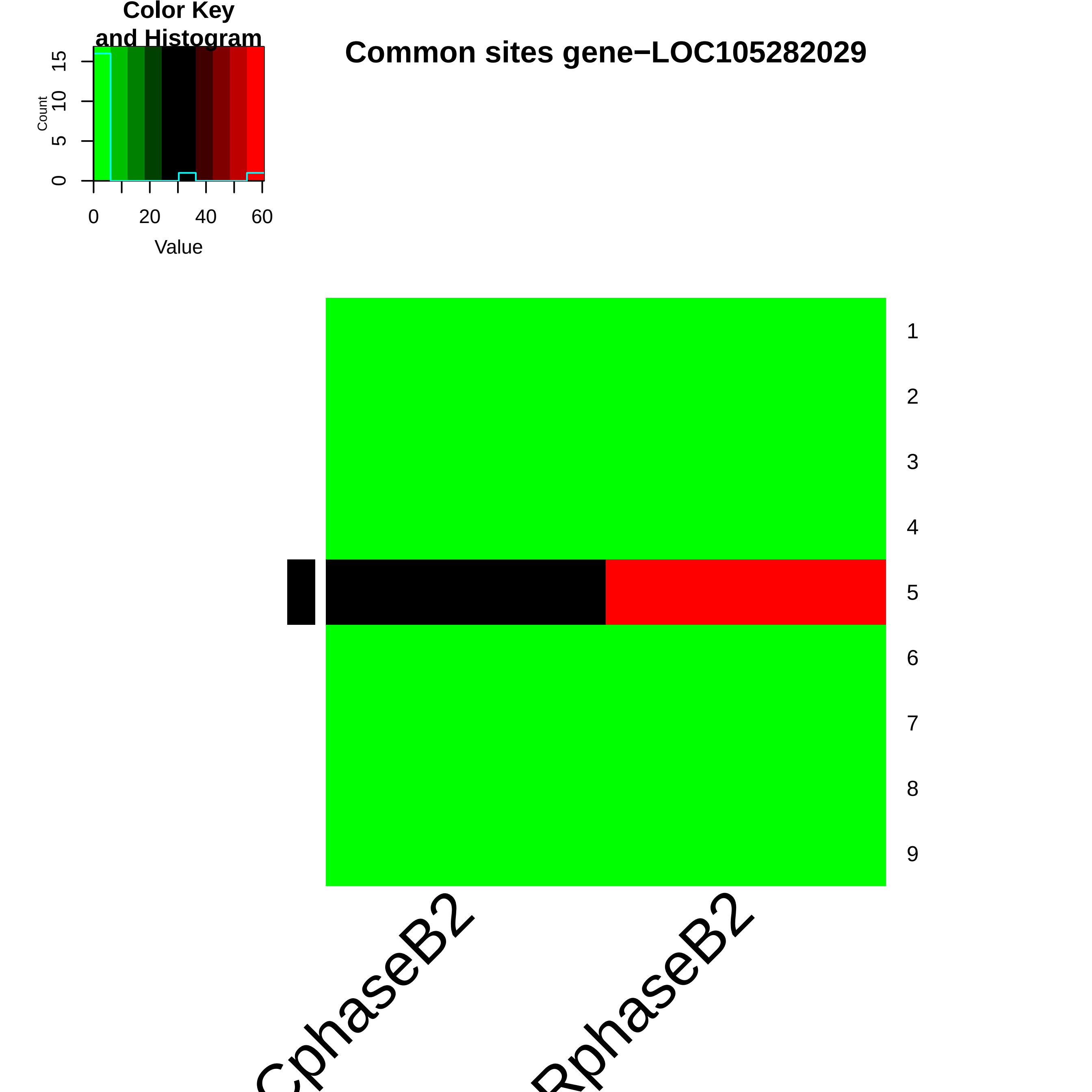

Color Key
and Histogram
Count
0
5
10
15
0
20
40
60
Value
Common sites gene−LOC105282029
1
2
3
4
5
6
7
8
9
CphaseB2
RphaseB2

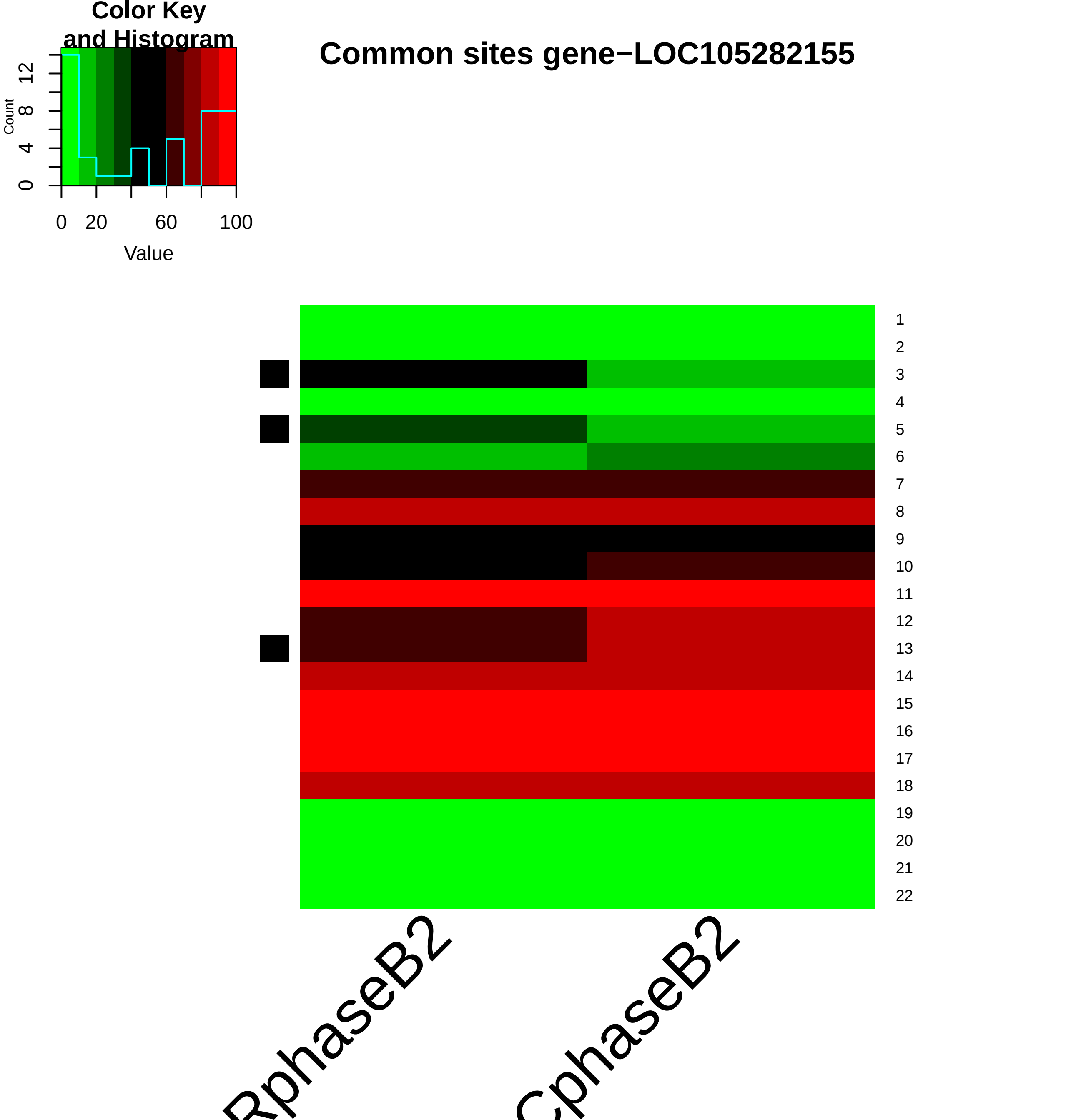
Color Key
and Histogram
Count
0
4
8
12
0
20
60
100
Value
Common sites gene−LOC105282155
RphaseB2
CphaseB2

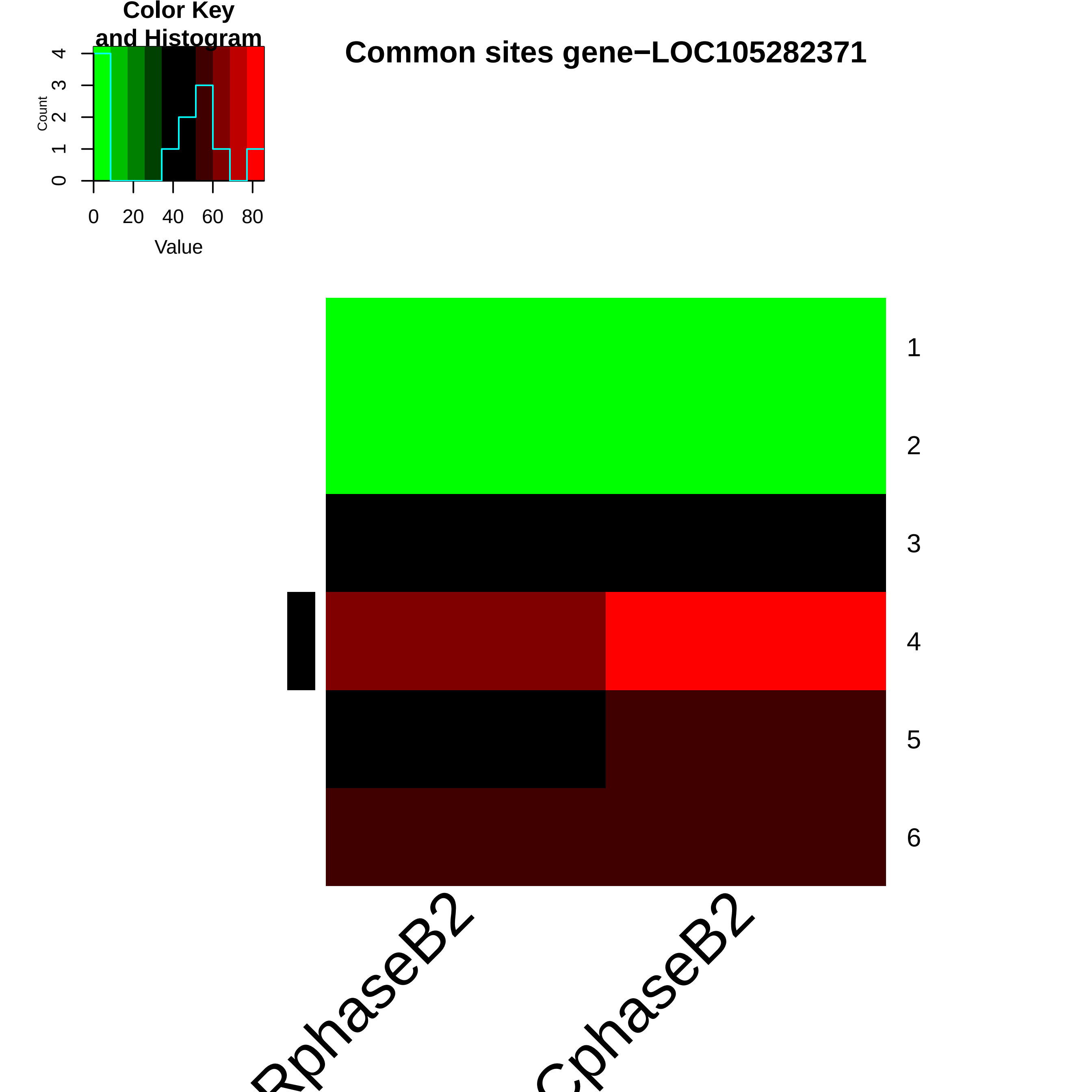

Color Key
and Histogram
Count
0
1
2
3
4
0
20
40
60
80
Value
Common sites gene−LOC105282371
1
2
3
4
5
6
RphaseB2
CphaseB2

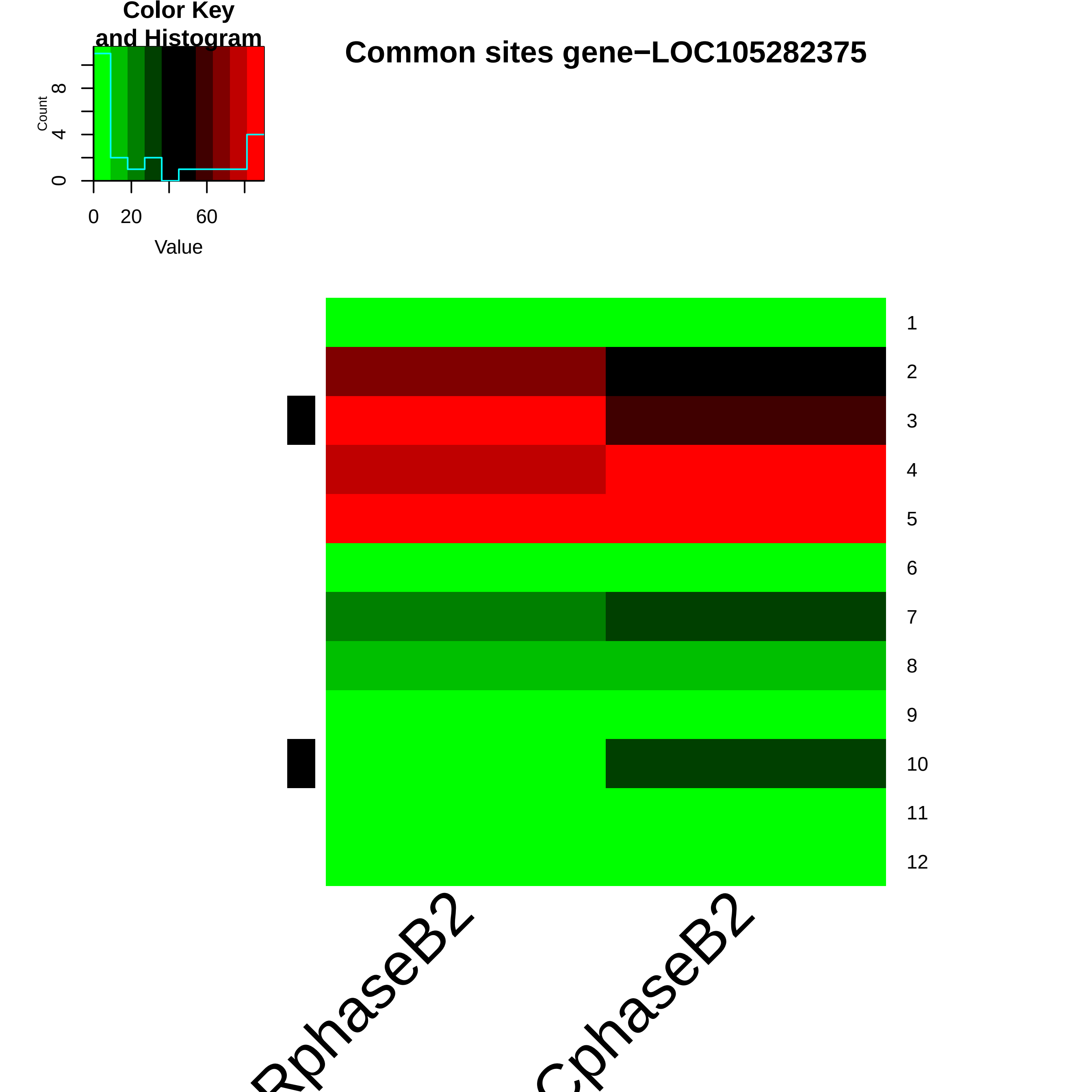

Color Key
and Histogram
Count
0
4
8
0
20
60
Value
Common sites gene−LOC105282375
1
2
3
4
5
6
7
8
9
10
11
12
RphaseB2
CphaseB2

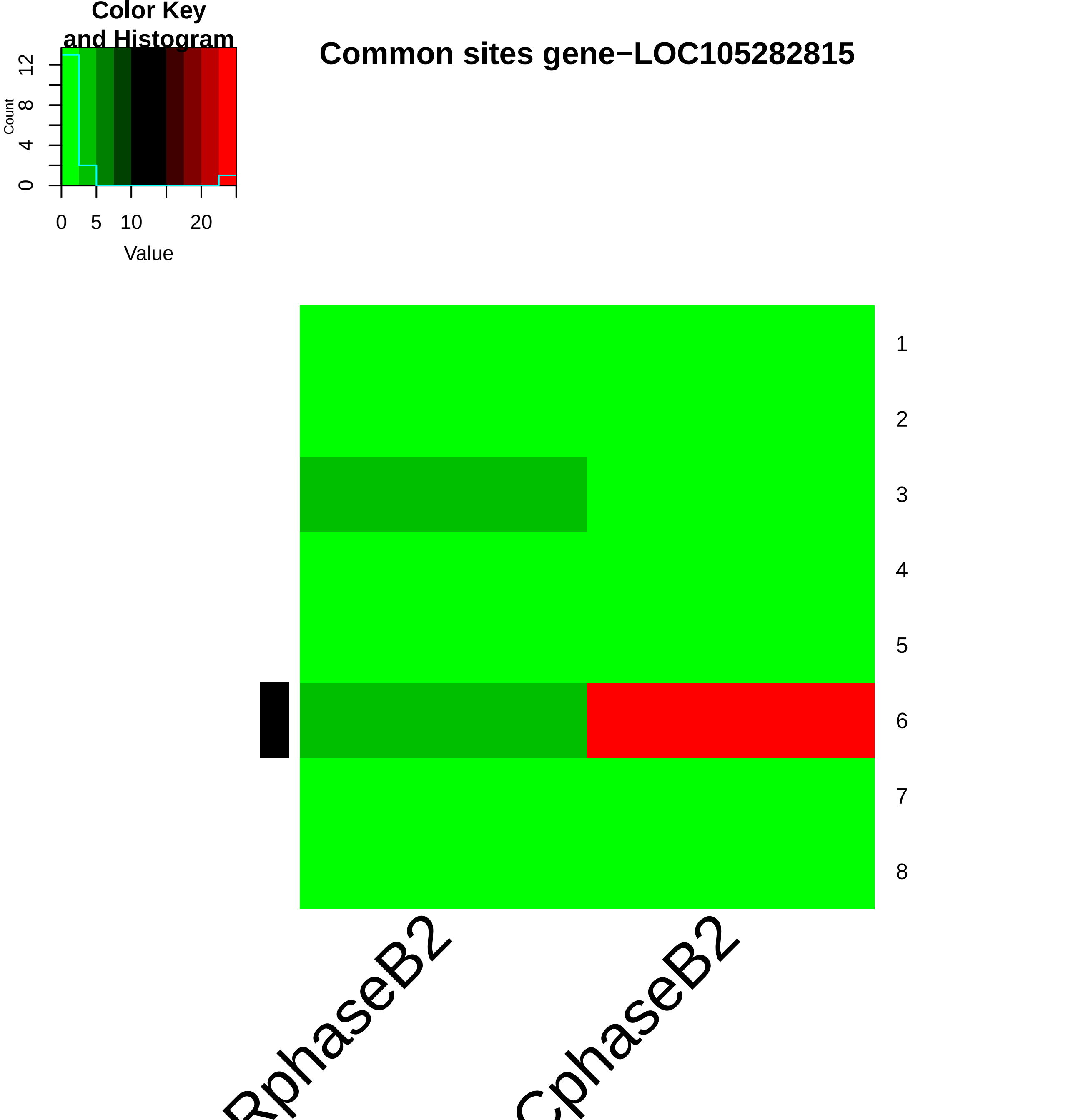
Color Key
and Histogram
Count
0
4
8
12
0
5
10
20
Value
Common sites gene−LOC105282815
1
2
3
4
5
6
7
8
RphaseB2
CphaseB2

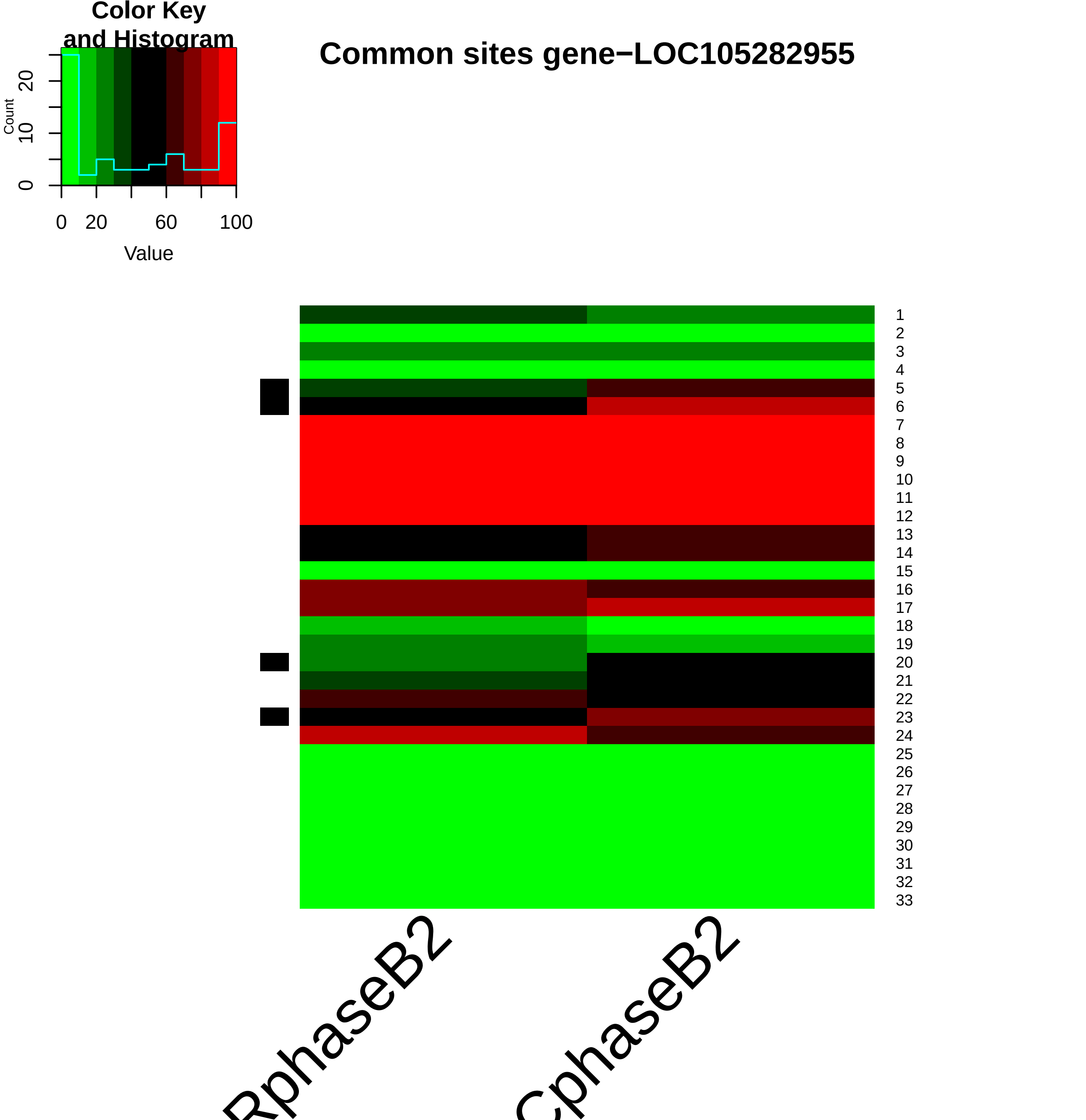
Color Key
and Histogram
Count
0
10
20
0
20
60
100
Value
Common sites gene−LOC105282955
RphaseB2
CphaseB2

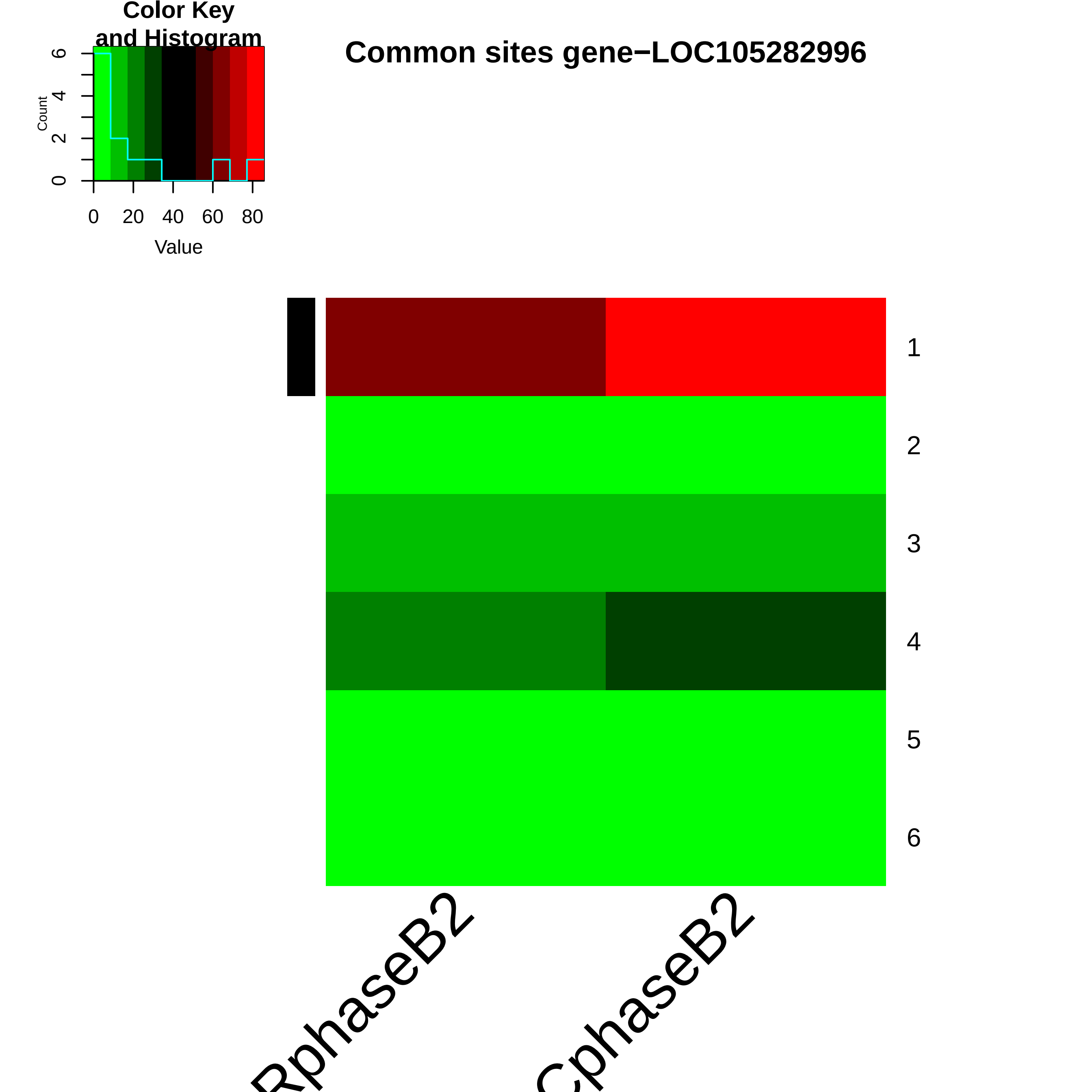

Color Key
and Histogram
Count
0
2
4
6
0
20
40
60
80
Value
Common sites gene−LOC105282996
1
2
3
4
5
6
RphaseB2
CphaseB2

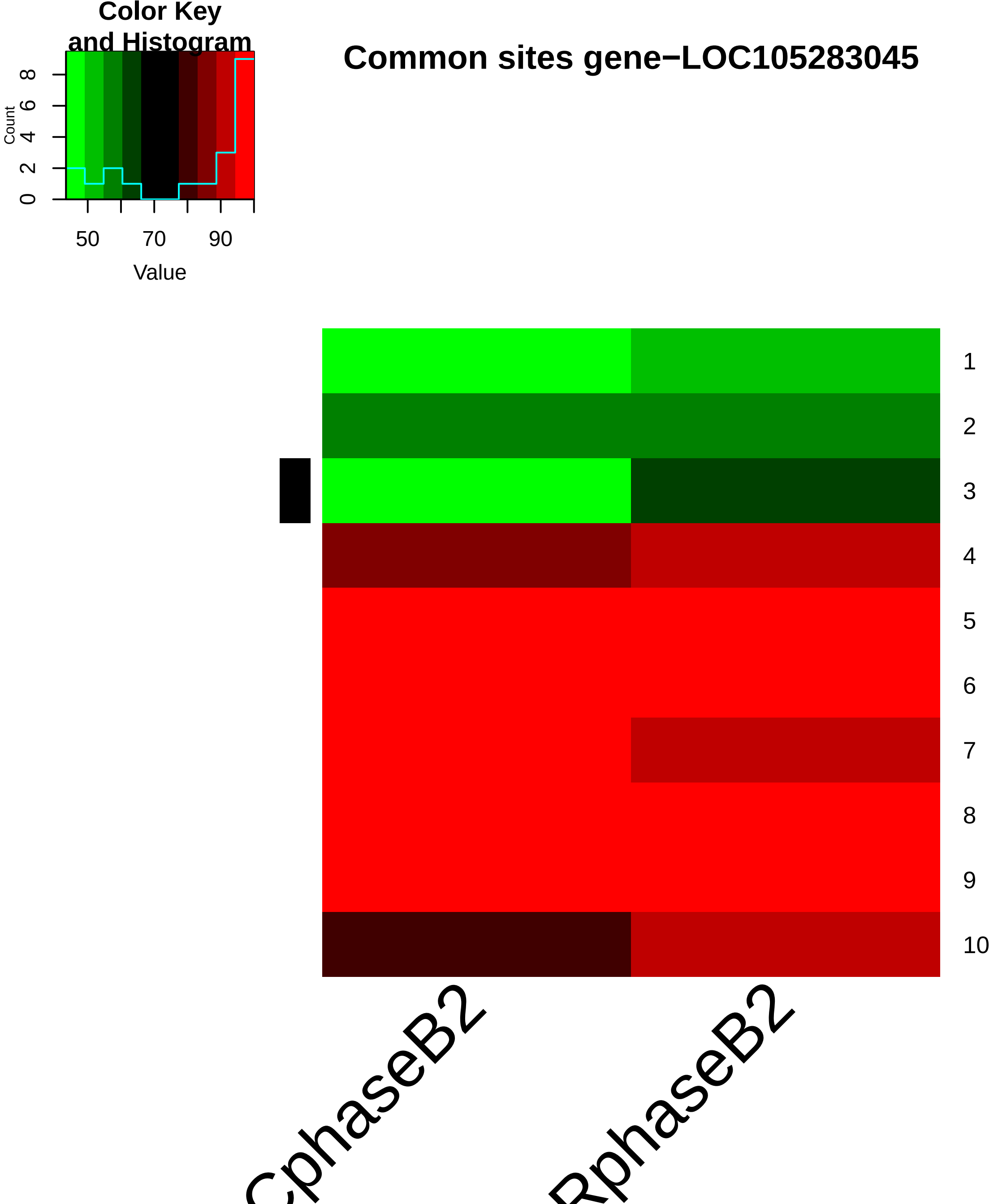
Color Key
and Histogram
Count
0 2 4 6 8
50 70 90
Value
Common sites gene−LOC105283045
1
2
3
4
5
6
7
8
9
10
CphaseB2
RphaseB2

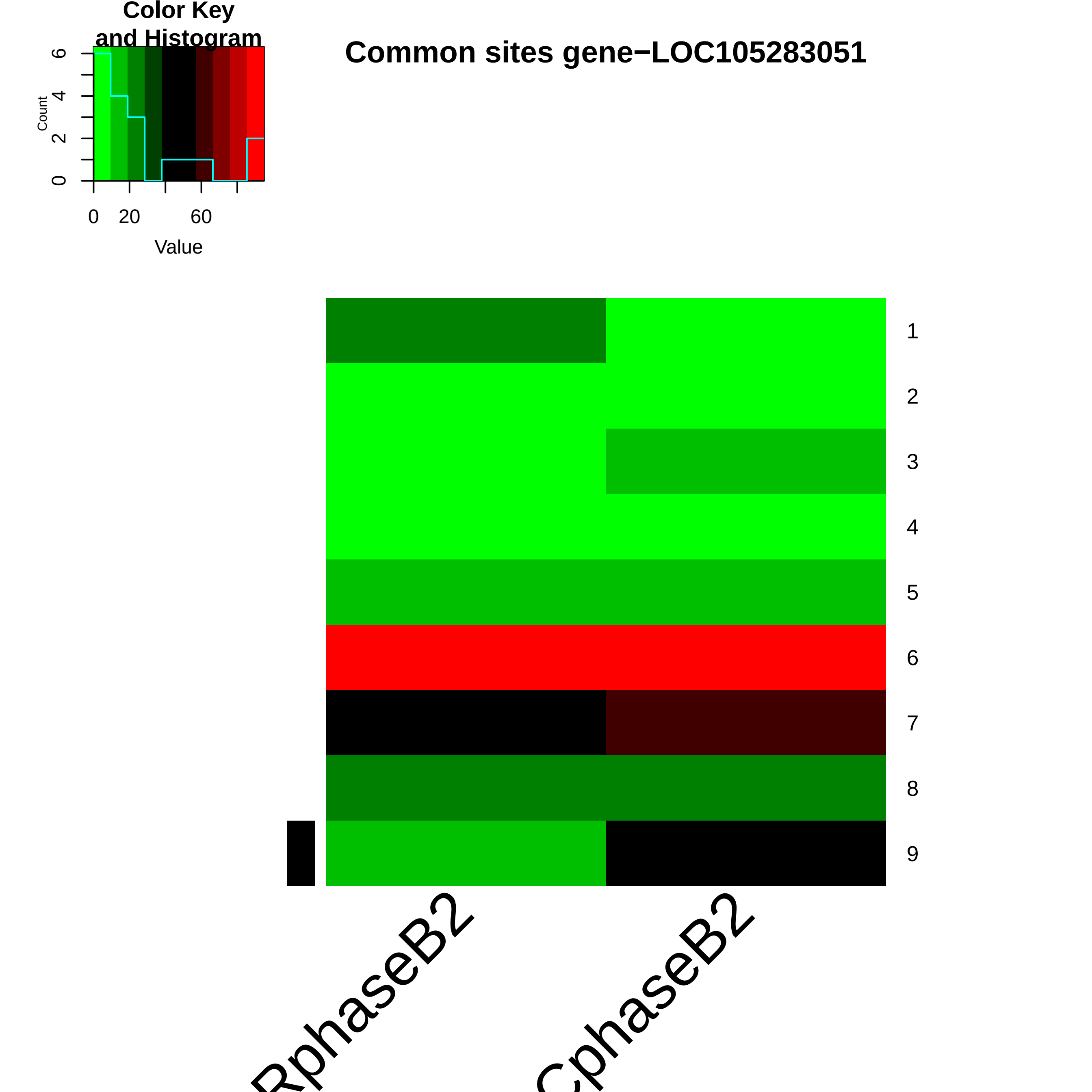

Color Key
and Histogram
Count
0
2
4
6
0
20
60
Value
Common sites gene−LOC105283051
1
2
3
4
5
6
7
8
9
RphaseB2
CphaseB2

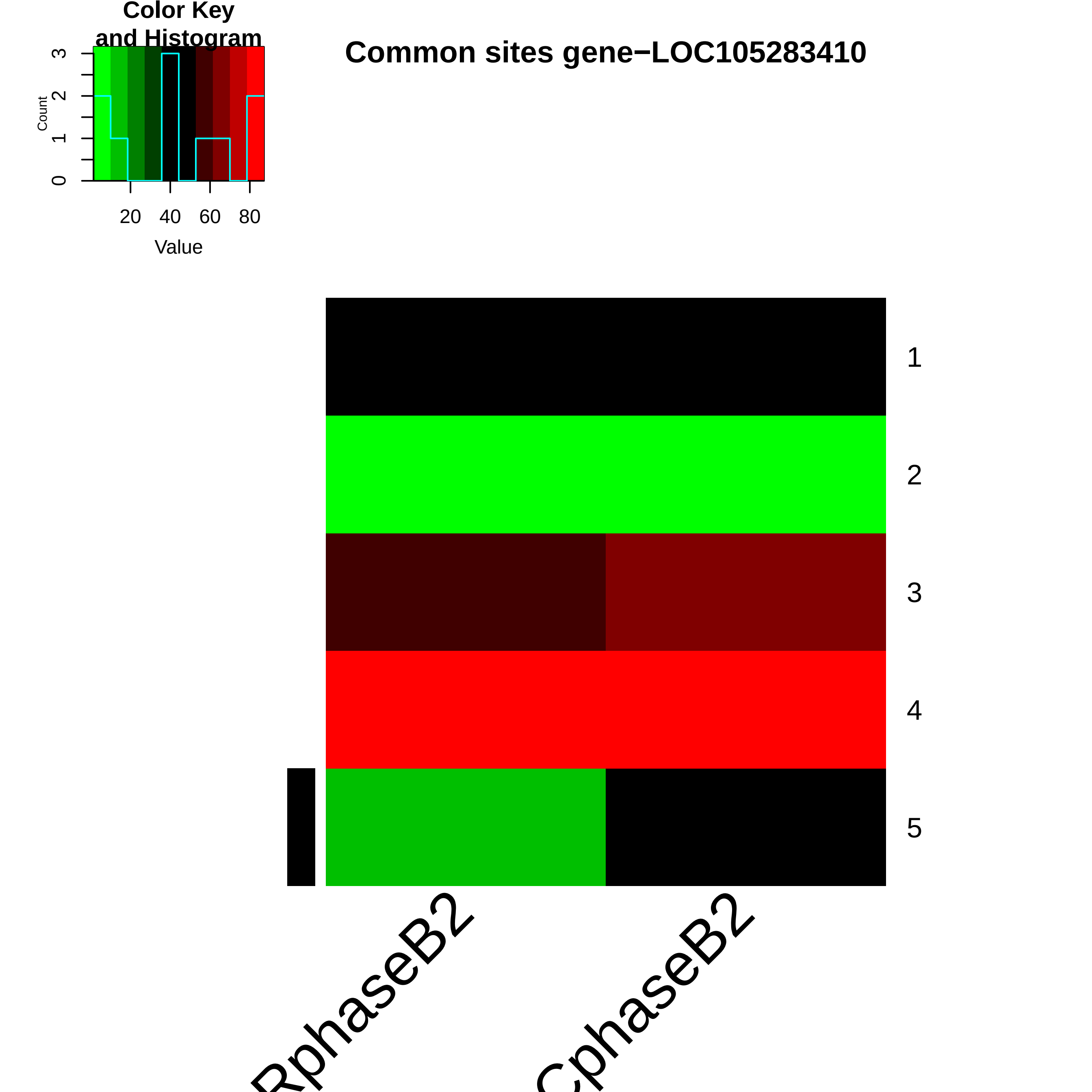

Color Key and Histogram
Count
0
1
2
3
20
40
60
80
Value
Common sites gene−LOC105283410
1
2
3
4
5
RphaseB2
CphaseB2

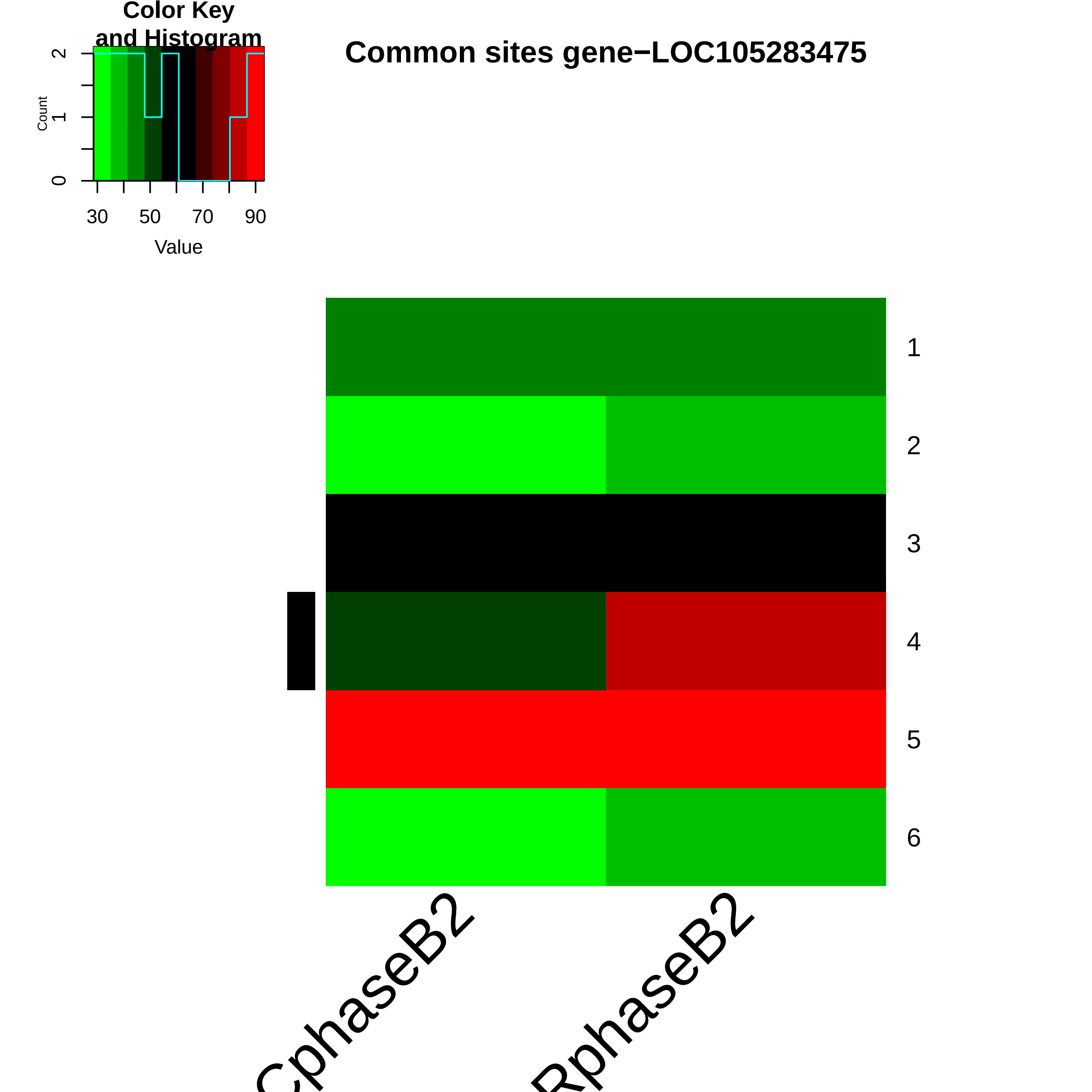
Color Key
and Histogram
Count
0
1
2
30
50
70
90
Value
Common sites gene−LOC105283475
1
2
3
4
5
6
CphaseB2
RphaseB2

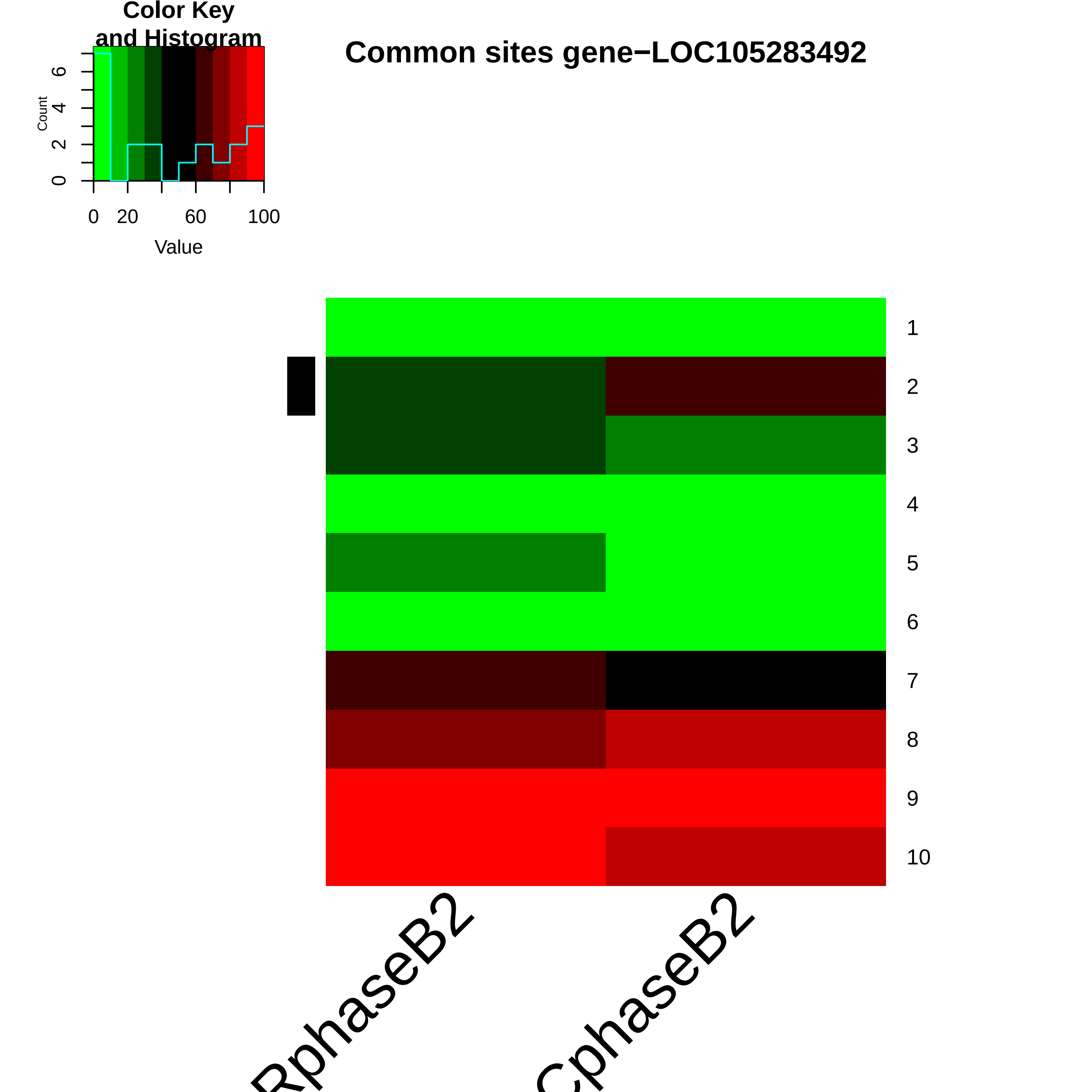
Color Key
and Histogram
Count
0
2
4
6
0
20
60
100
Value
Common sites gene−LOC105283492
1
2
3
4
5
6
7
8
9
10
RphaseB2
CphaseB2

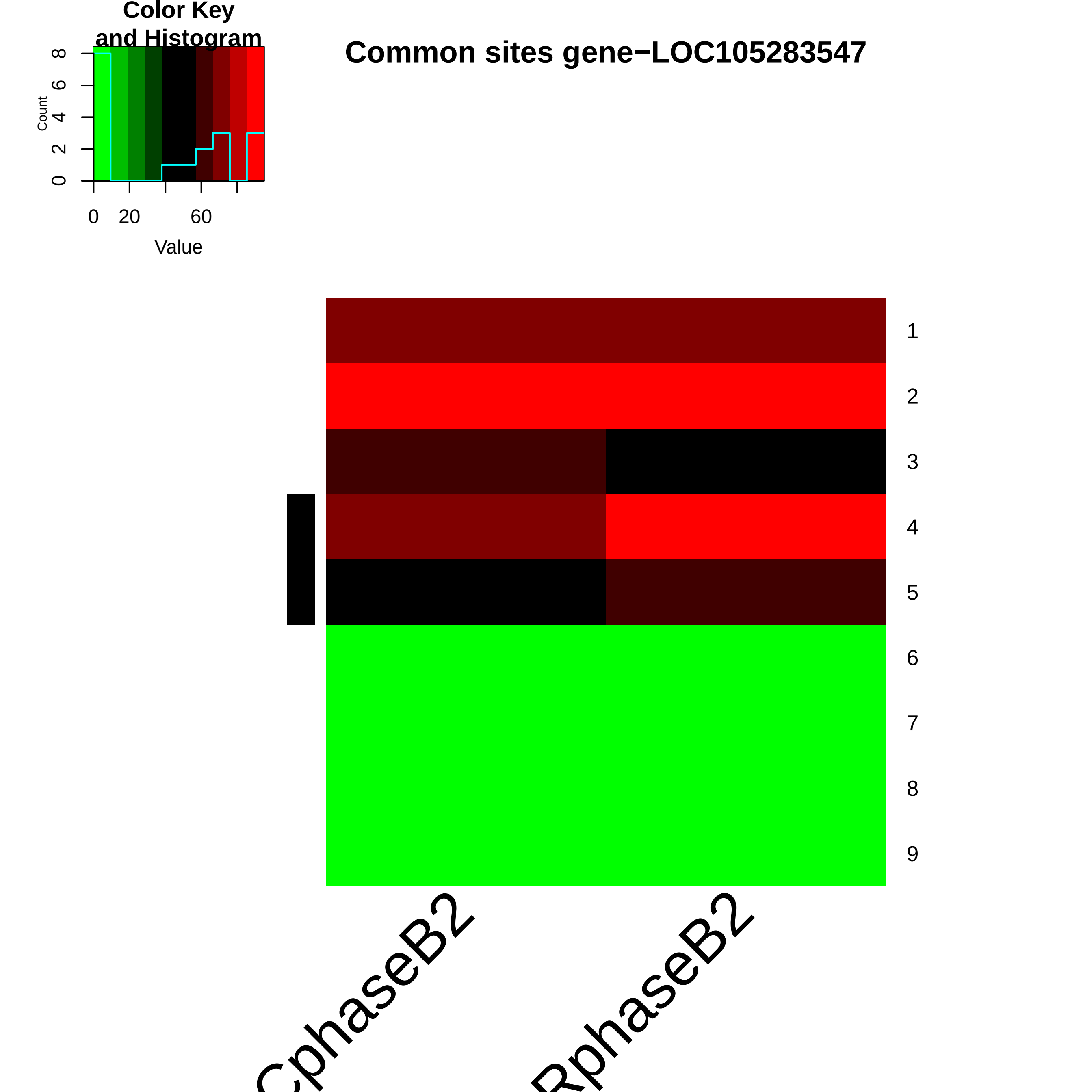
Color Key
and Histogram
Count
0
2
4
6
8
0
20
60
Value
Common sites gene−LOC105283547
1
2
3
4
5
6
7
8
9
CphaseB2
RphaseB2

### Common sites gene−LOC105283693

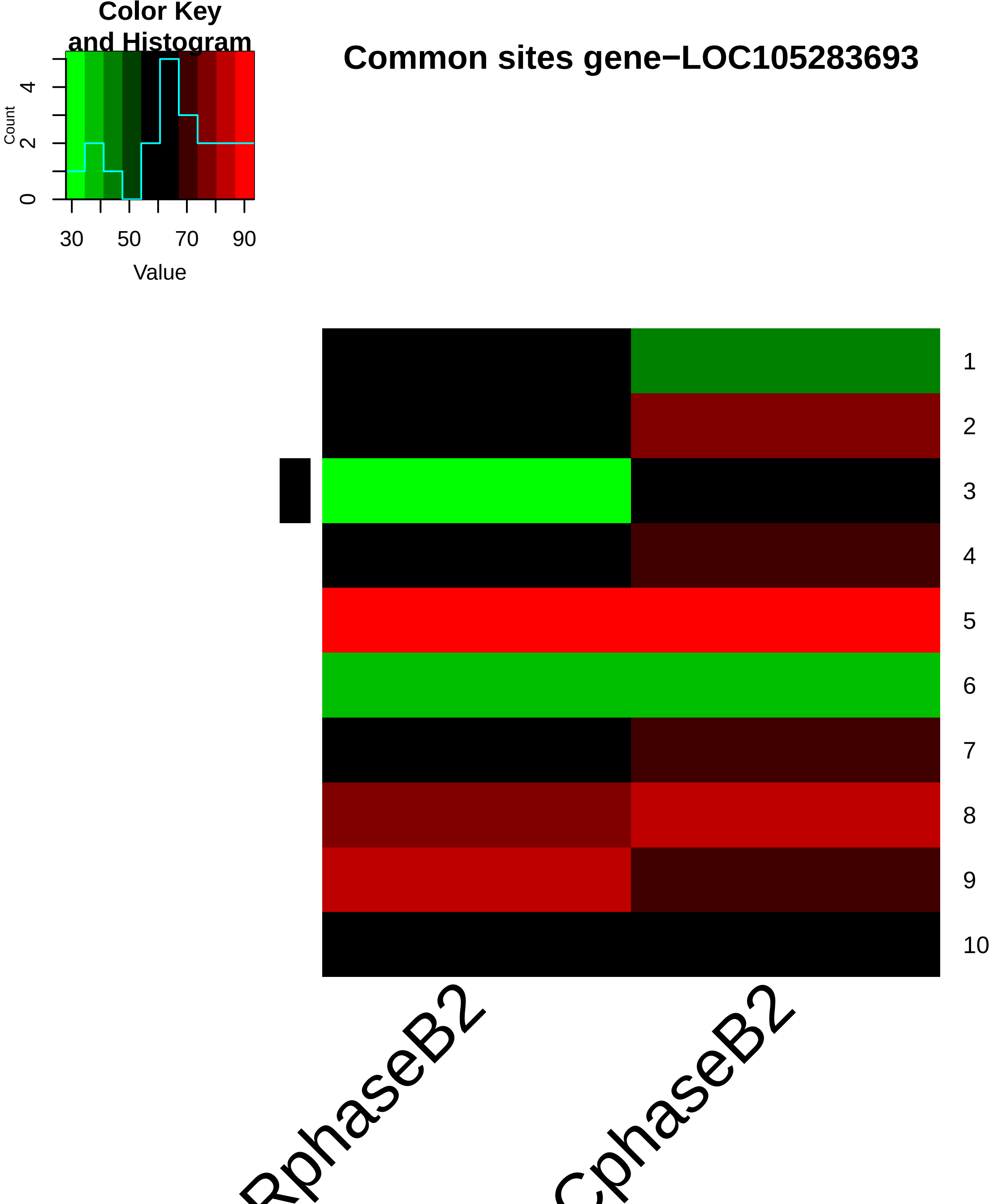

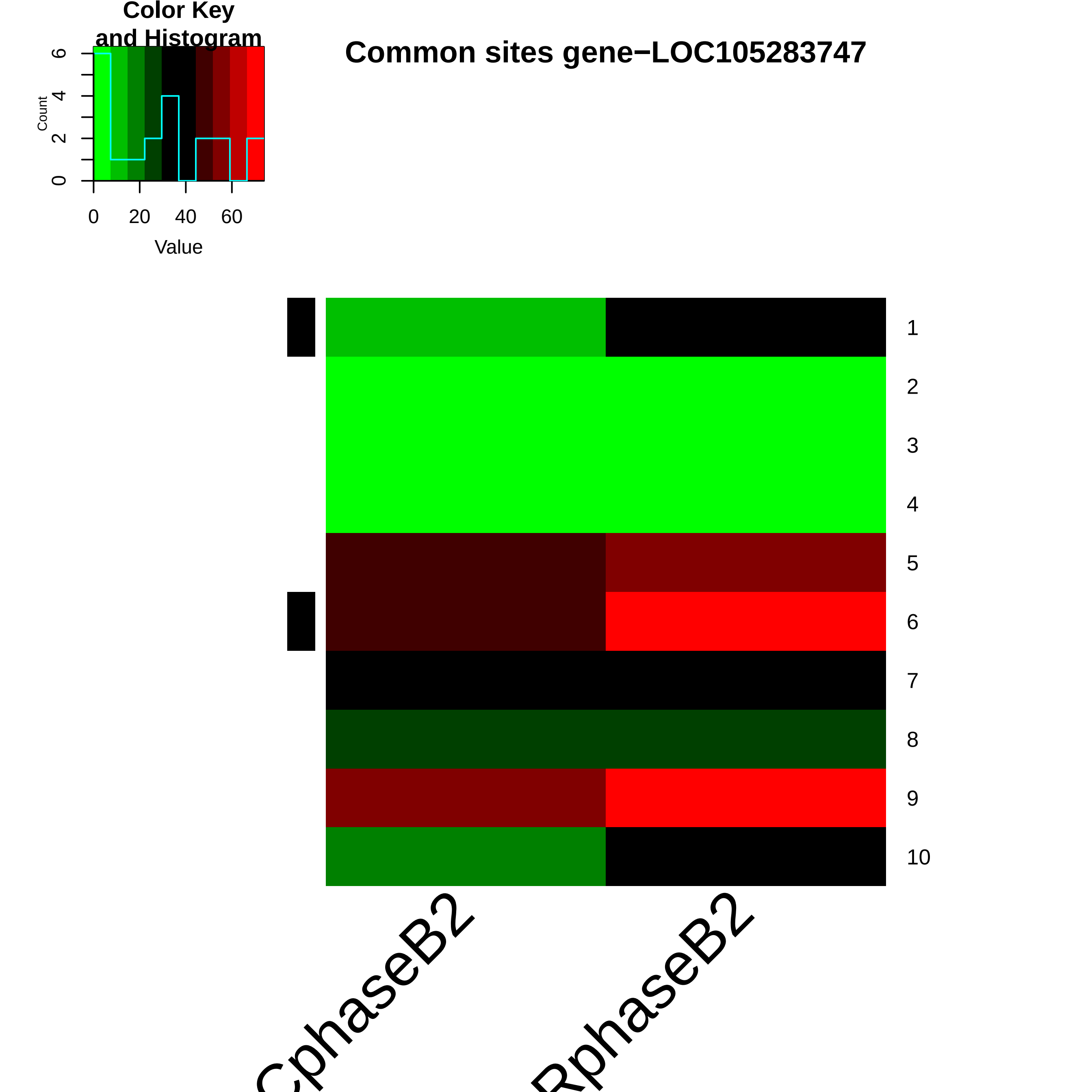
Color Key
and Histogram
Count
Value
0
2
4
6
0
20
40
60
Common sites gene−LOC105283747
1
2
3
4
5
6
7
8
9
10
CphaseB2
RphaseB2

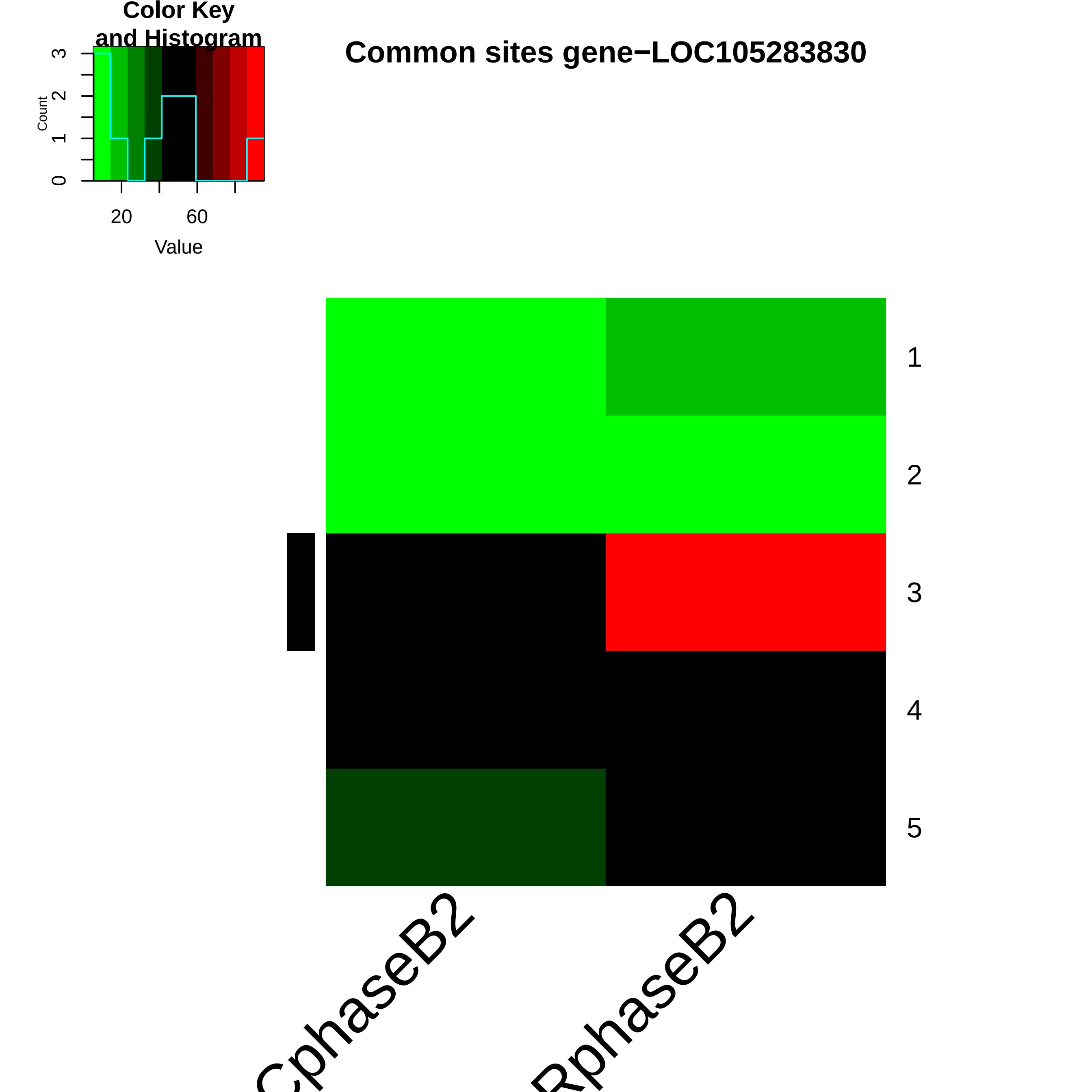

Color Key
and Histogram
Count
0
1
2
3
20
60
Value
Common sites gene−LOC105283830
1
2
3
4
5
CphaseB2
RphaseB2

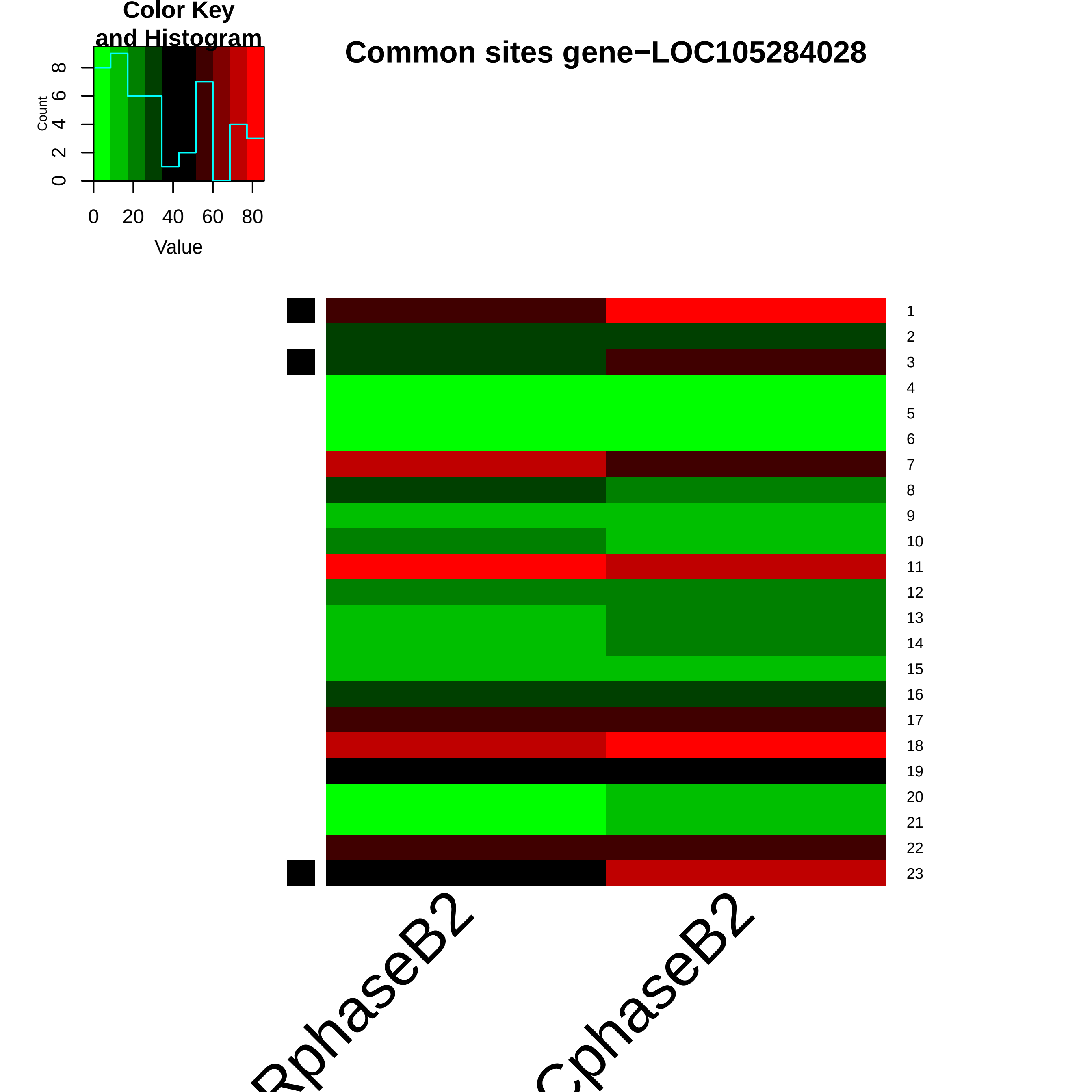

Color Key
and Histogram
Count
Value
0
2
4
6
8
0
20
40
60
80
Common sites gene−LOC105284028
RphaseB2
CphaseB2

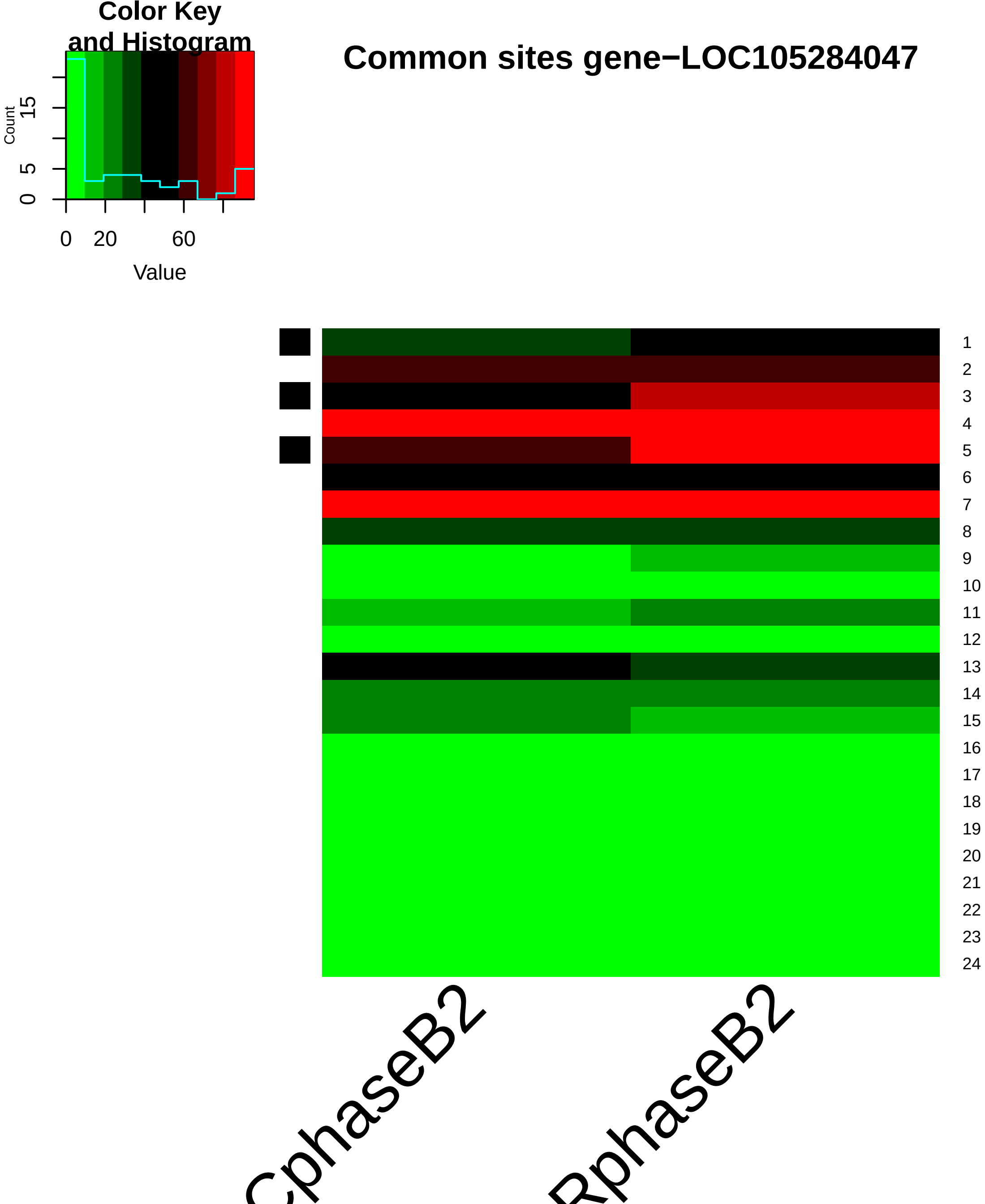

Color Key
and Histogram
Count
0
5
15
0
20
60
Value
Common sites gene−LOC105284047
CphaseB2
RphaseB2

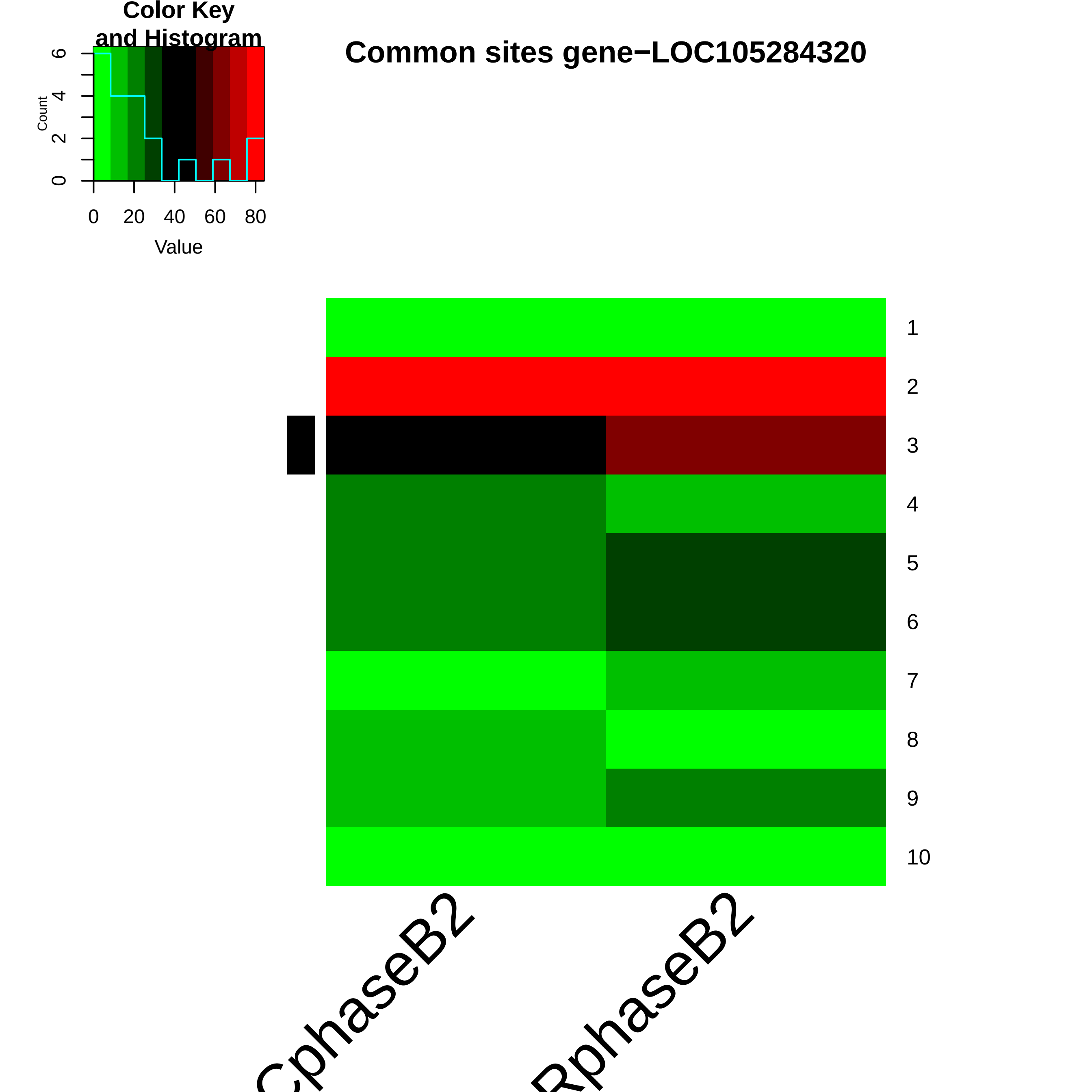
Color Key
and Histogram
Count
0
2
4
6
0
20
40
60
80
Value
Common sites gene−LOC105284320
1
2
3
4
5
6
7
8
9
10
CphaseB2
RphaseB2

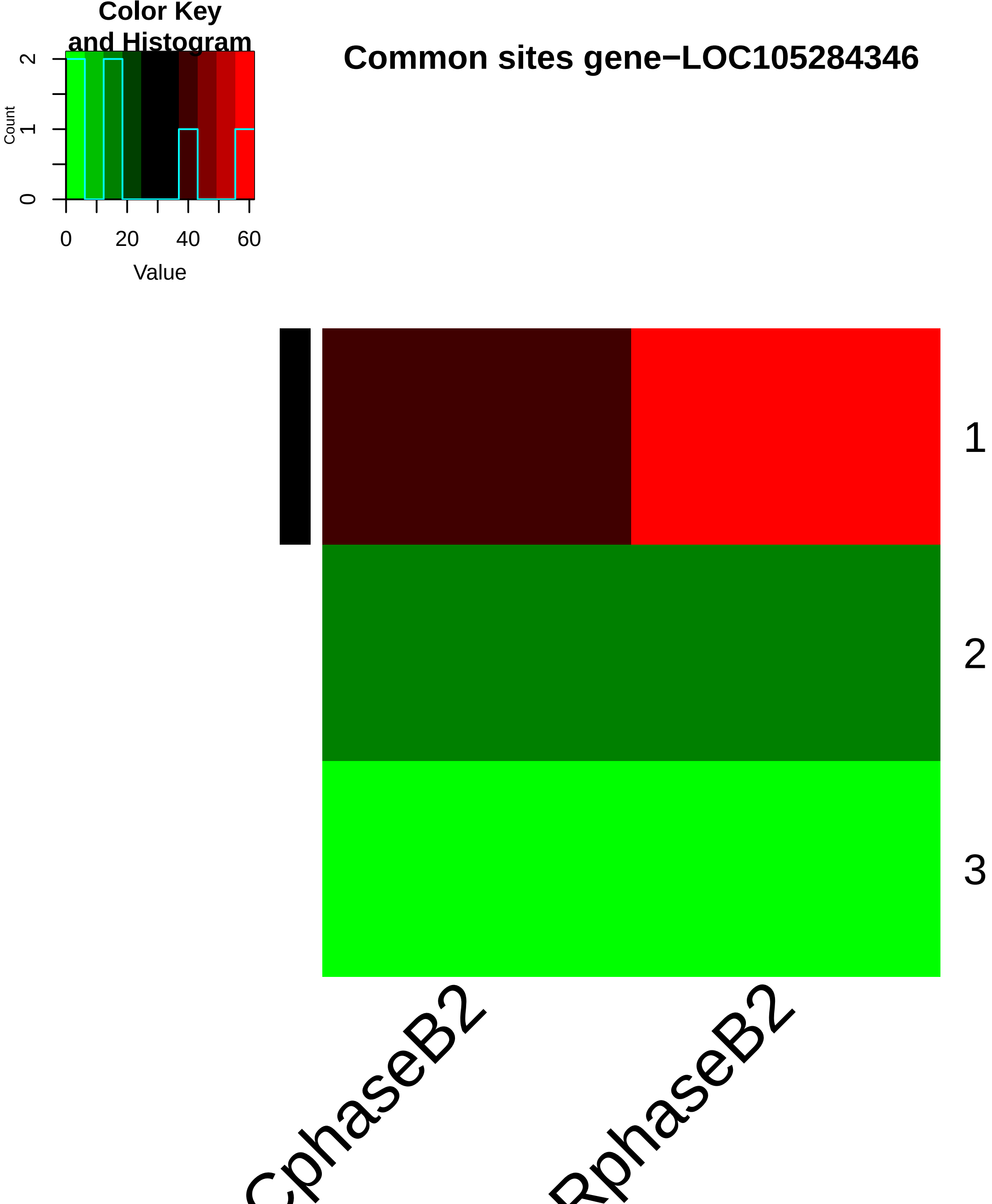
Color Key
and Histogram
Count
0
1
2
0
20
40
60
Value
Common sites gene−LOC105284346
1
2
3
CphaseB2
RphaseB2

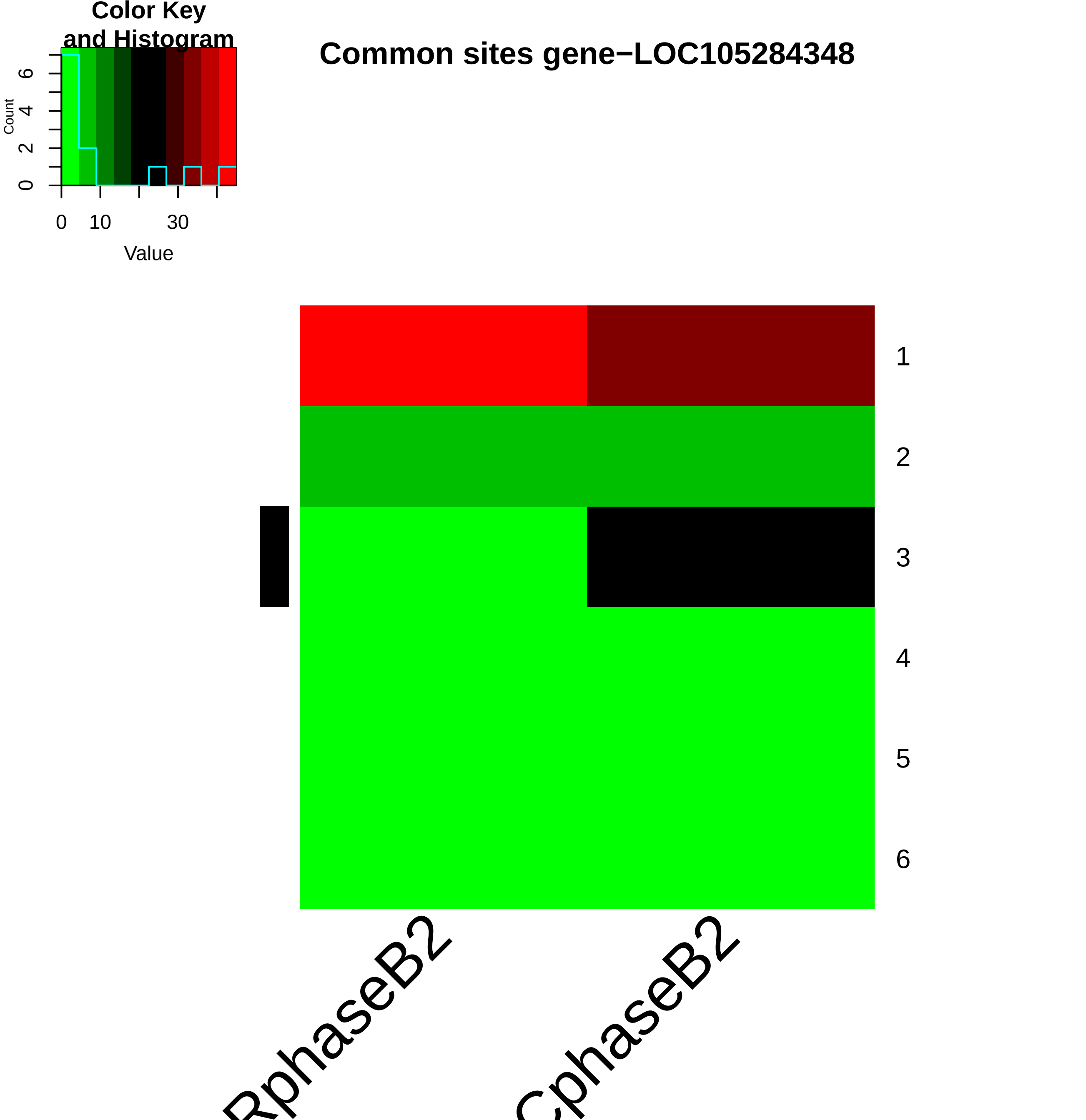
Color Key
and Histogram
Count
0
2
4
6
0
10
30
Value
Common sites gene−LOC105284348
1
2
3
4
5
6
RphaseB2
CphaseB2

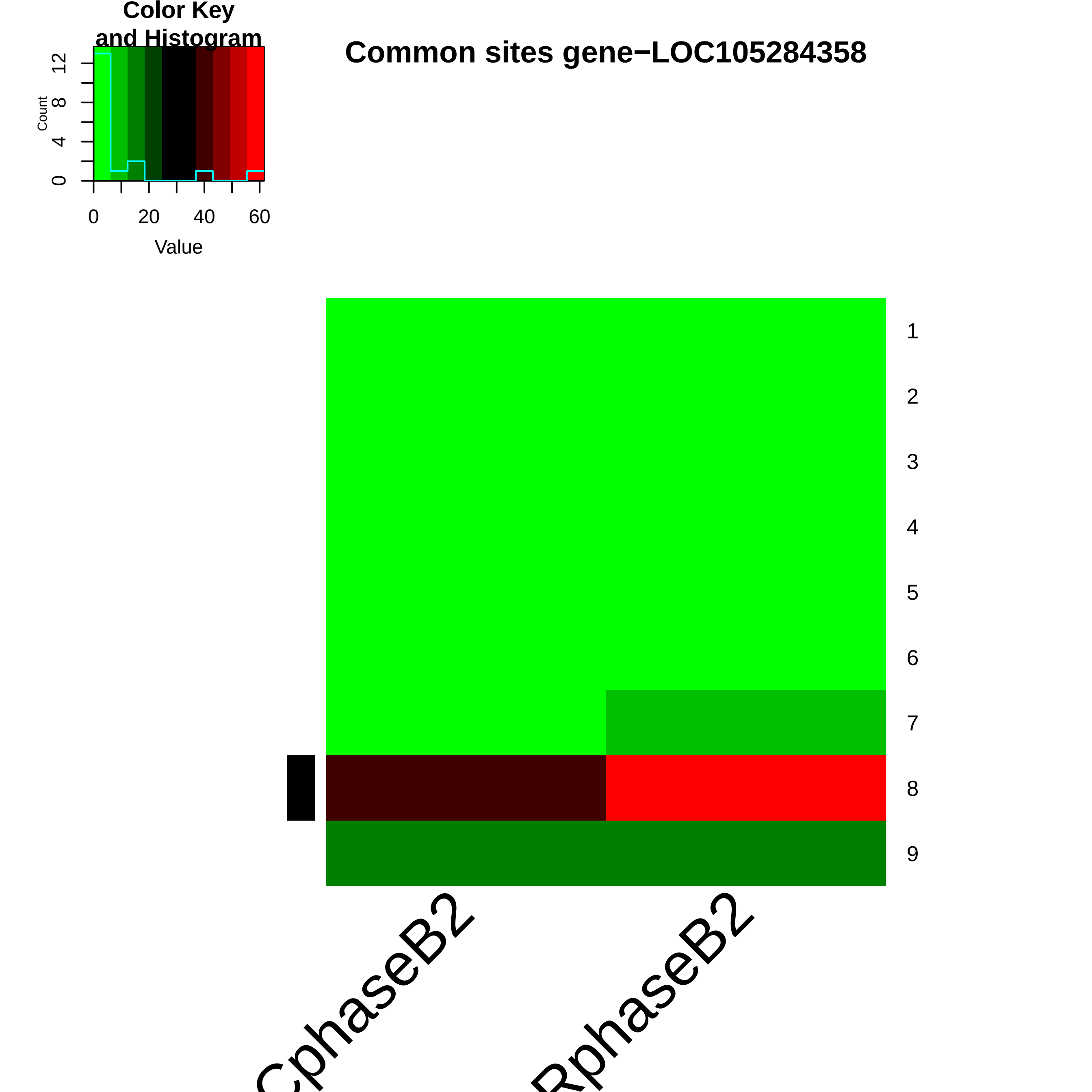

Color Key
and Histogram
Count
0
4
8
12
0
20
40
60
Value
Common sites gene−LOC105284358
1
2
3
4
5
6
7
8
9
CphaseB2
RphaseB2

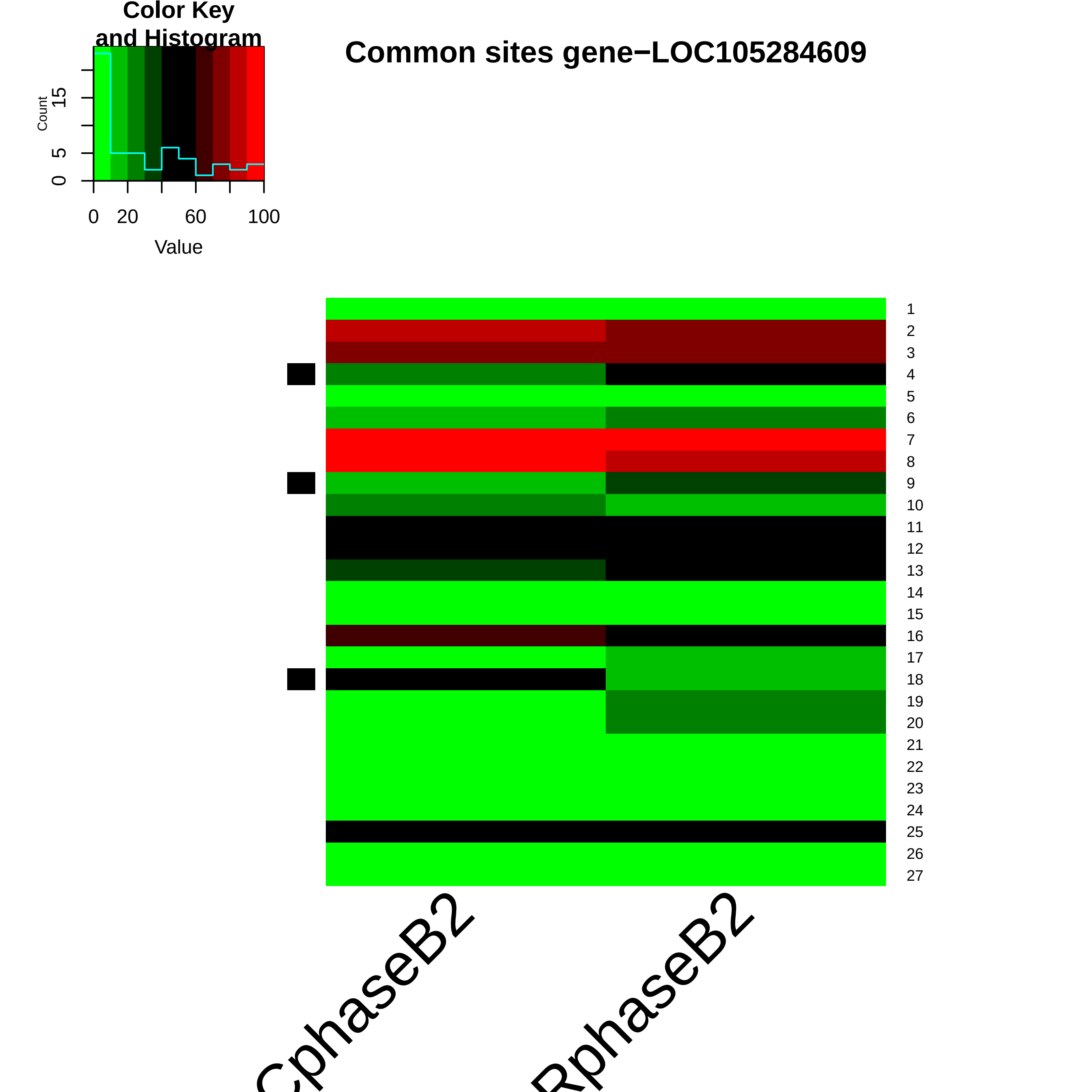
Color Key
and Histogram
Count
0
5
15
0
20
60
100
Value
Common sites gene−LOC105284609
CphaseB2
RphaseB2

### Common sites gene−LOC105284876

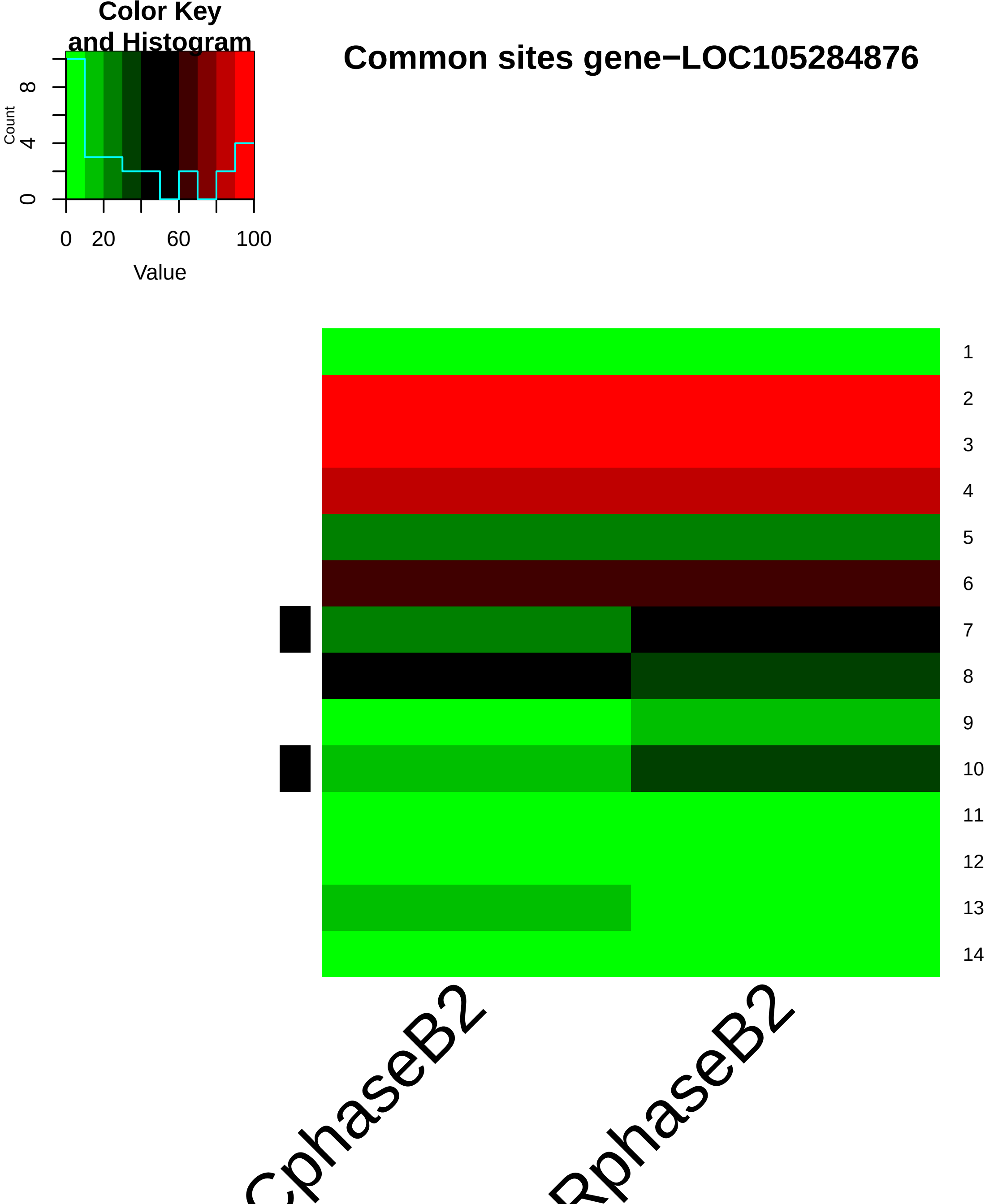

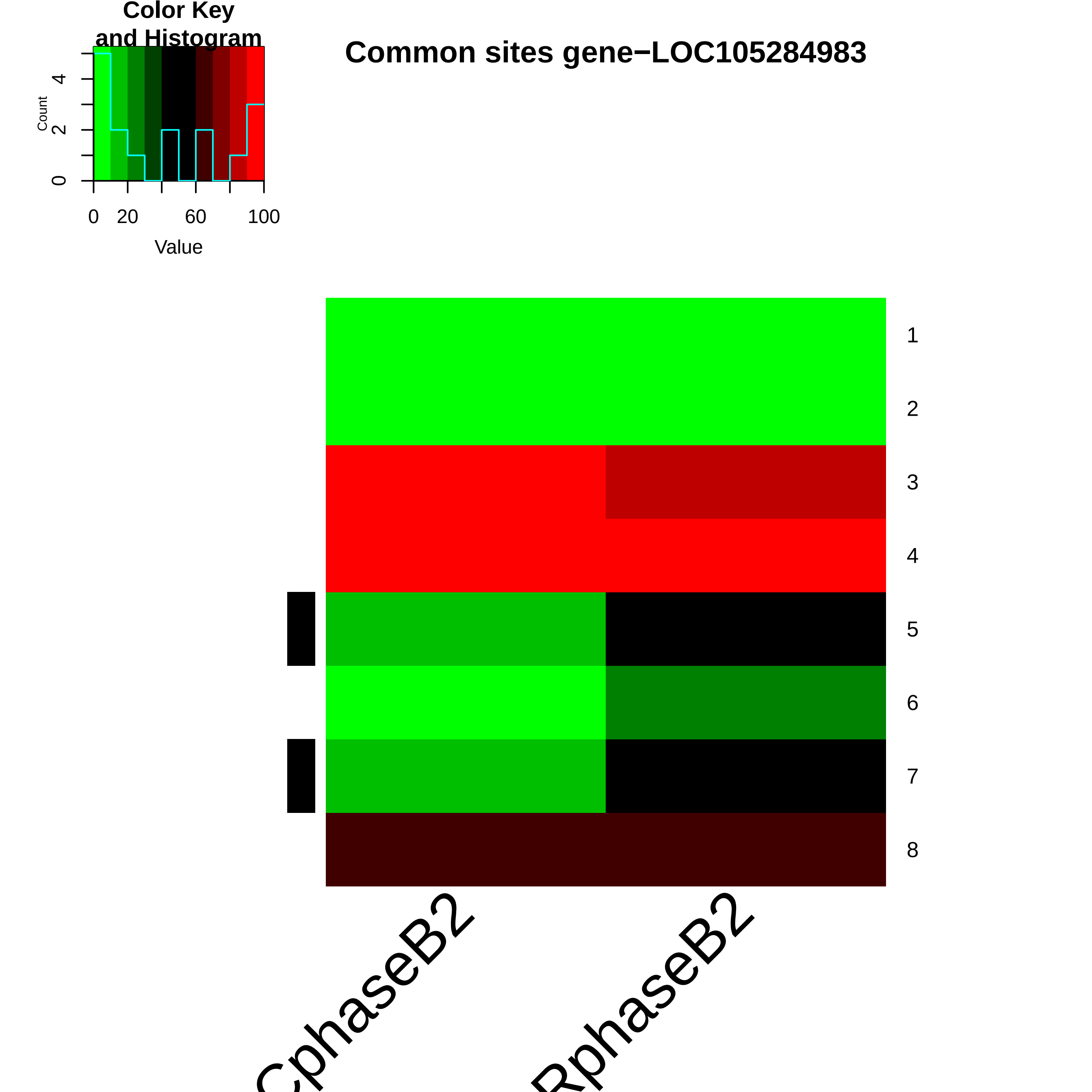

Color Key
and Histogram
Count
0
2
4
0
20
60
100
Value
Common sites gene−LOC105284983
1
2
3
4
5
6
7
8
CphaseB2
RphaseB2

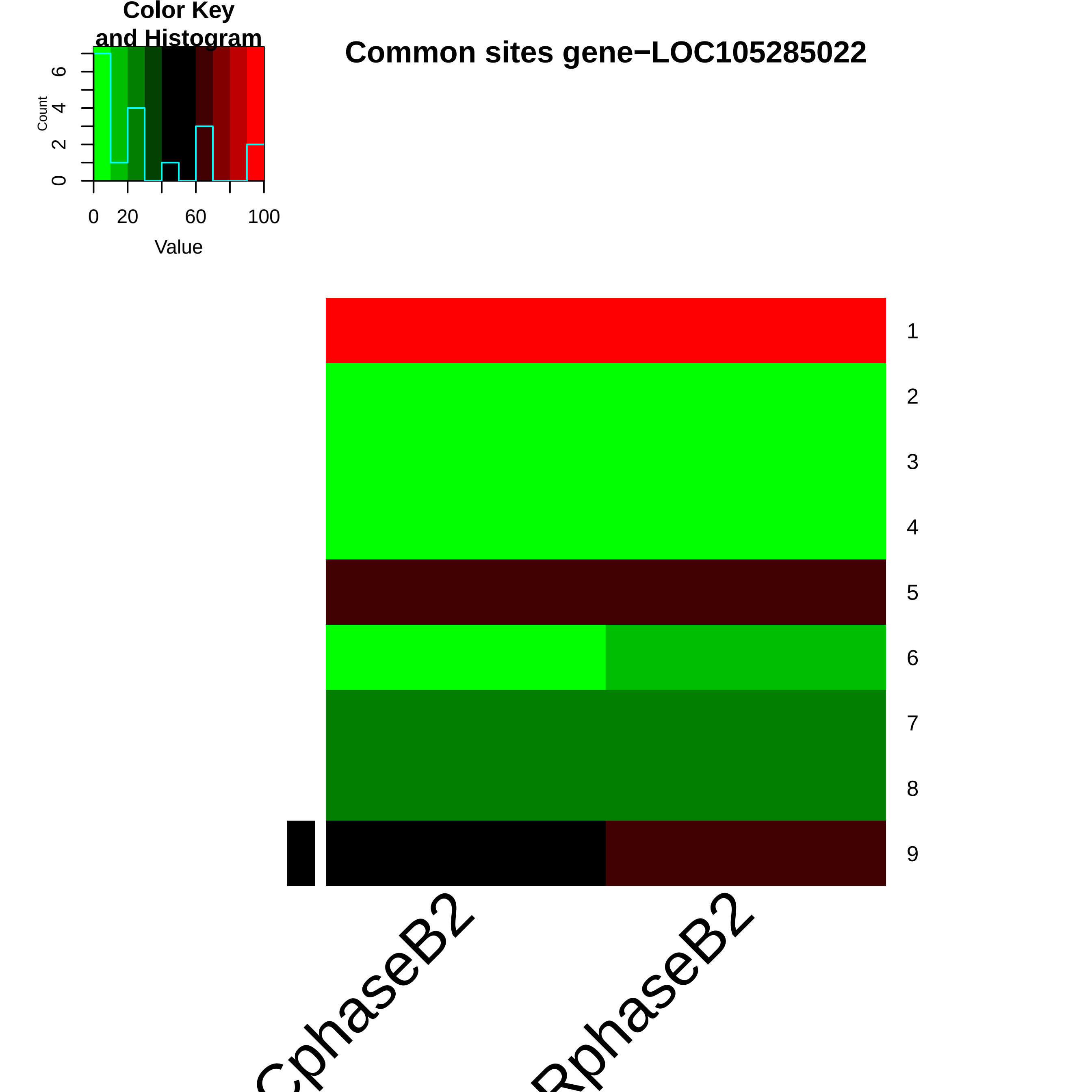

Color Key
and Histogram
Count
0
2
4
6
0
20
60
100
Value
Common sites gene−LOC105285022
1
2
3
4
5
6
7
8
9
CphaseB2
RphaseB2

#### Common sites gene−LOC105285060

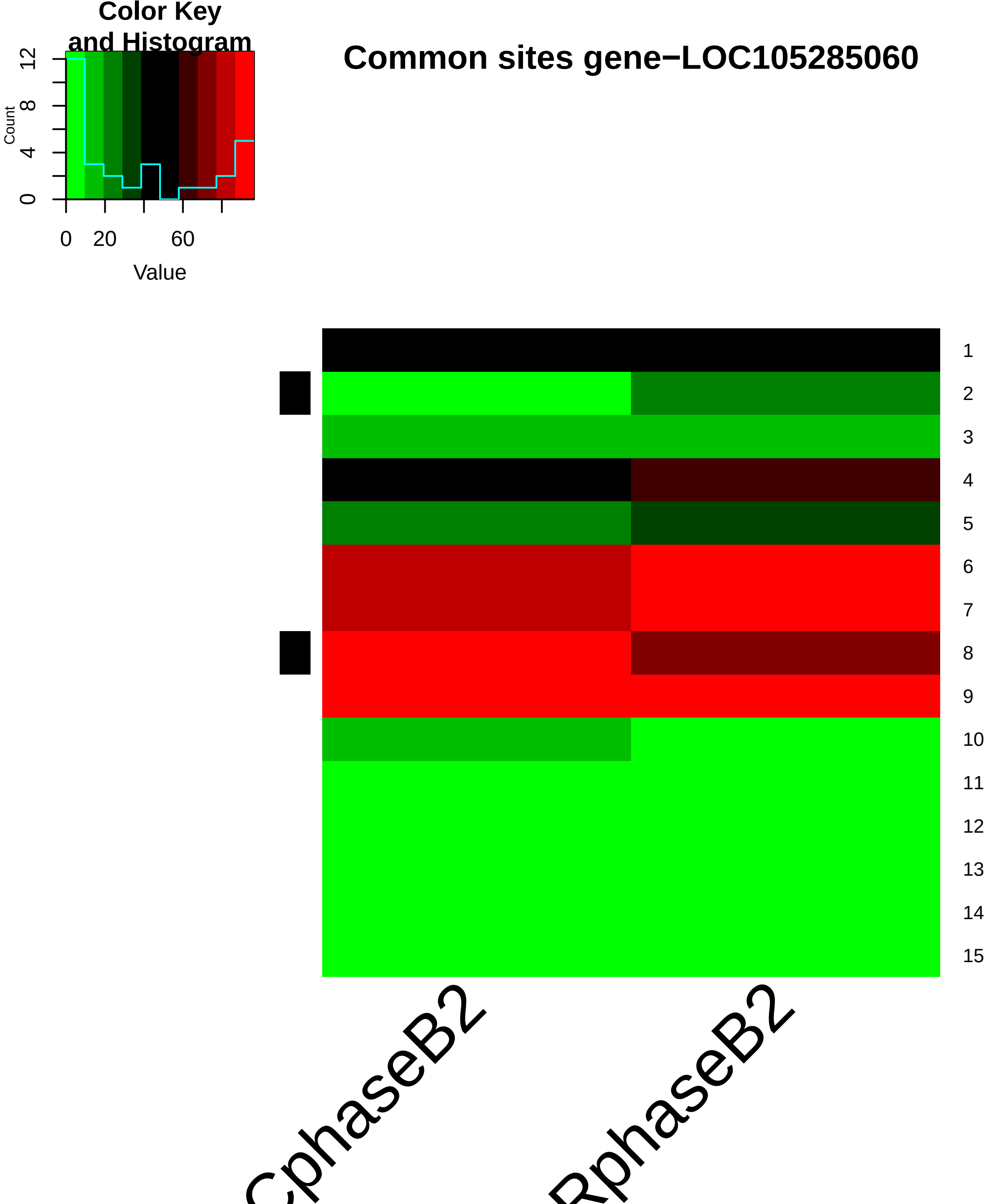

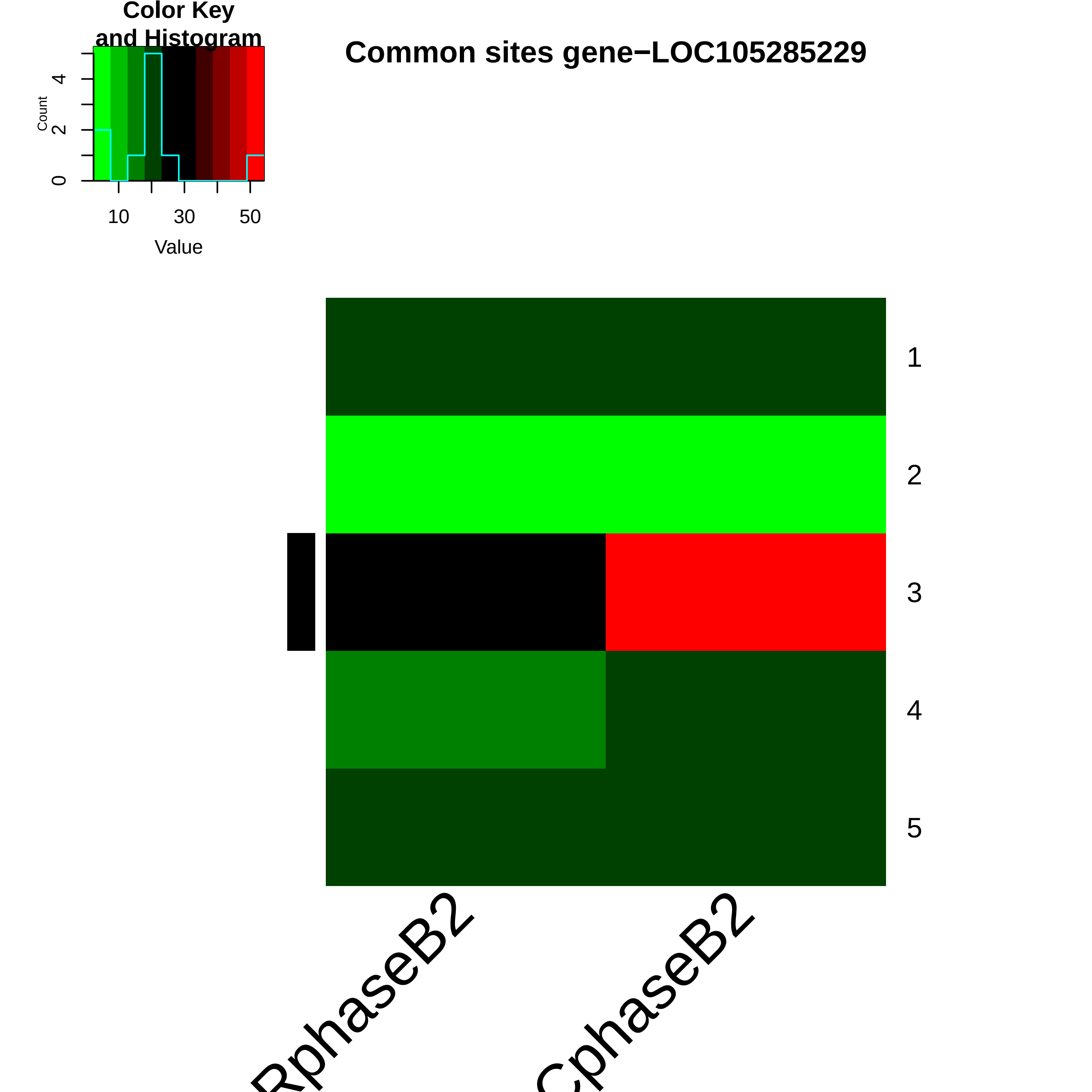
Color Key
and Histogram
Count
0
2
4
10
30
50
Value
Common sites gene−LOC105285229
1
2
3
4
5
RphaseB2
CphaseB2

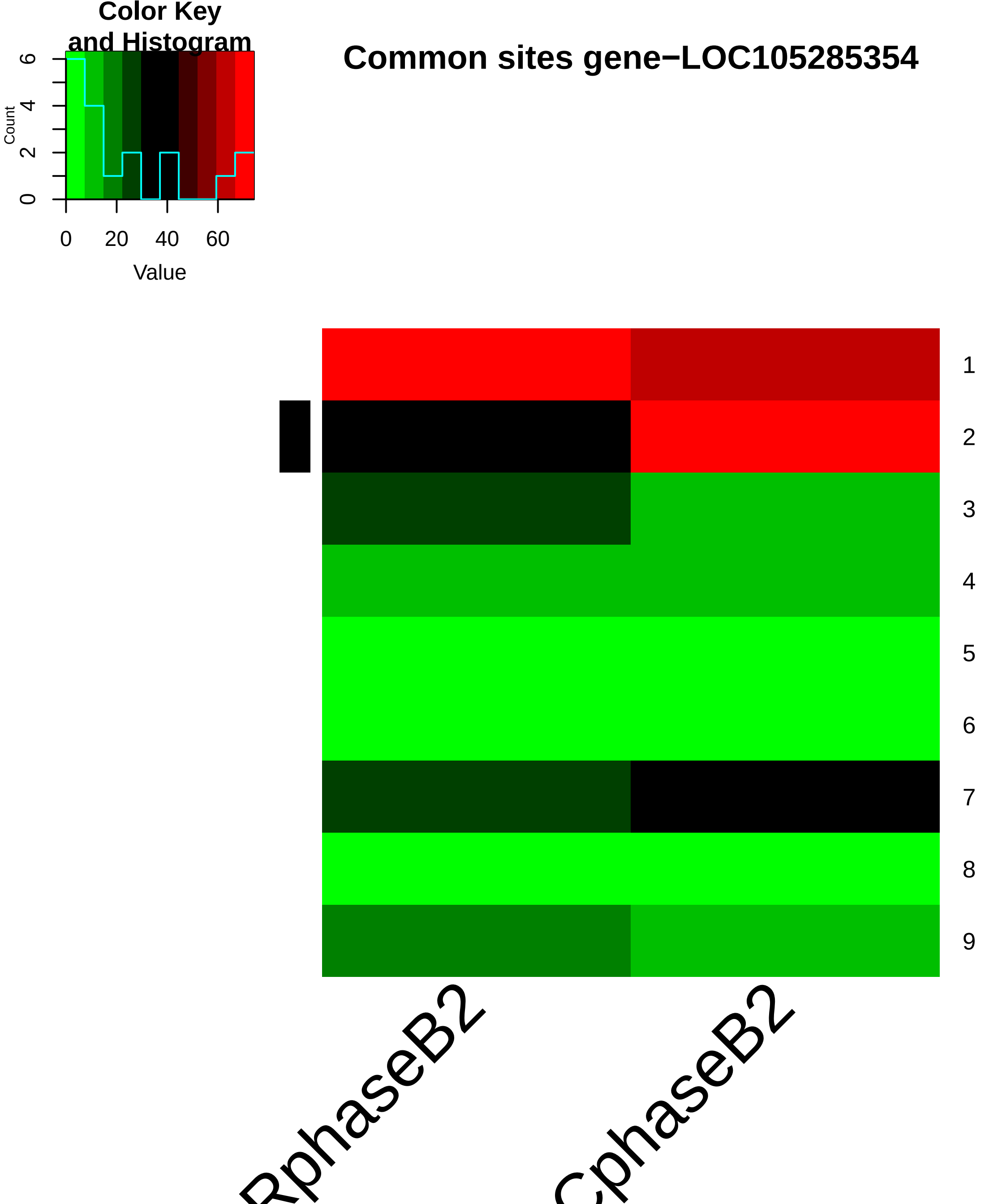

Color Key
and Histogram
Count
0
2
4
6
0
20
40
60
Value
Common sites gene−LOC105285354
1
2
3
4
5
6
7
8
9
RphaseB2
CphaseB2

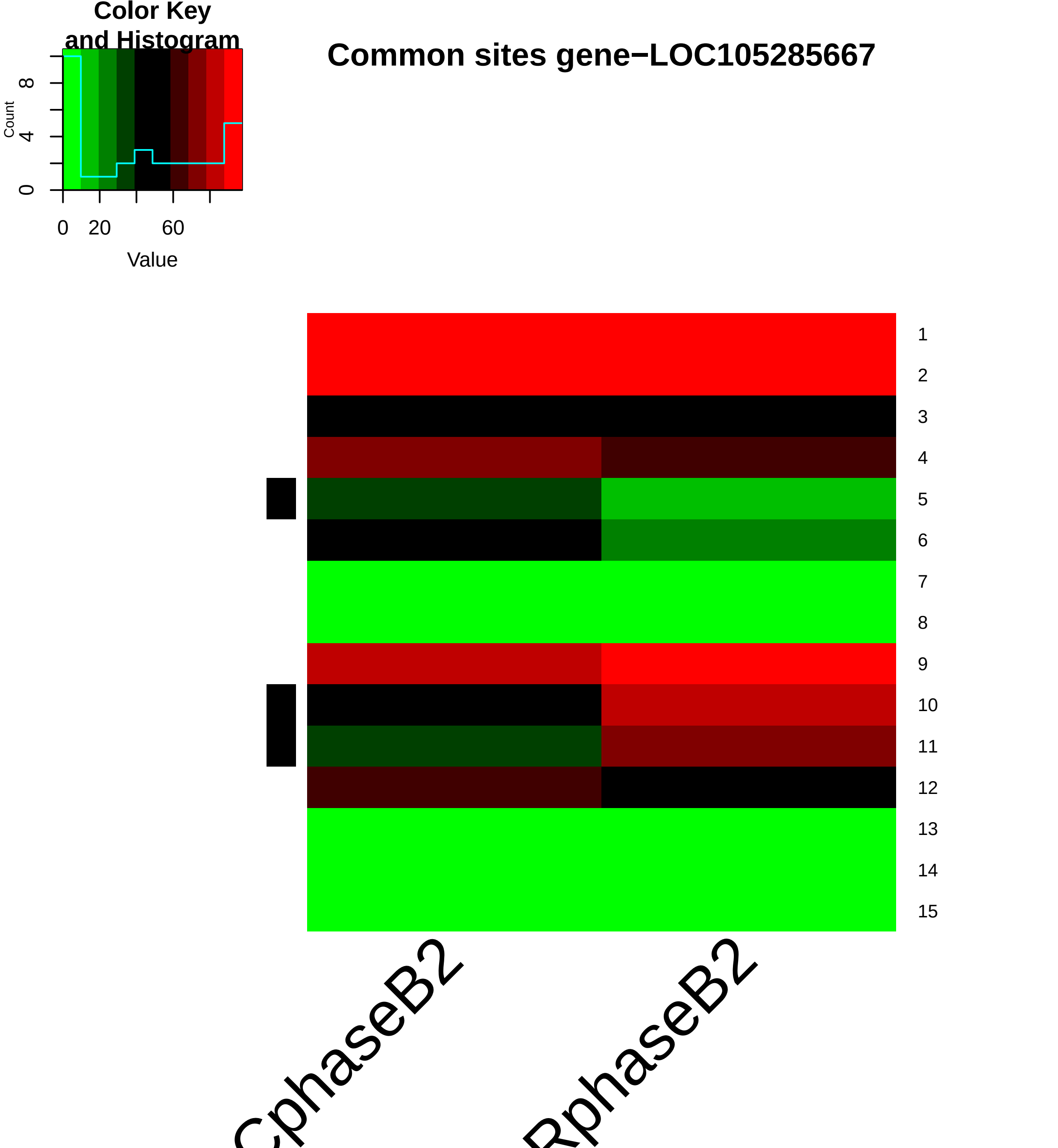

Color Key
and Histogram
Count
0
4
8
0
20
60
Value
Common sites gene−LOC105285667
CphaseB2
RphaseB2

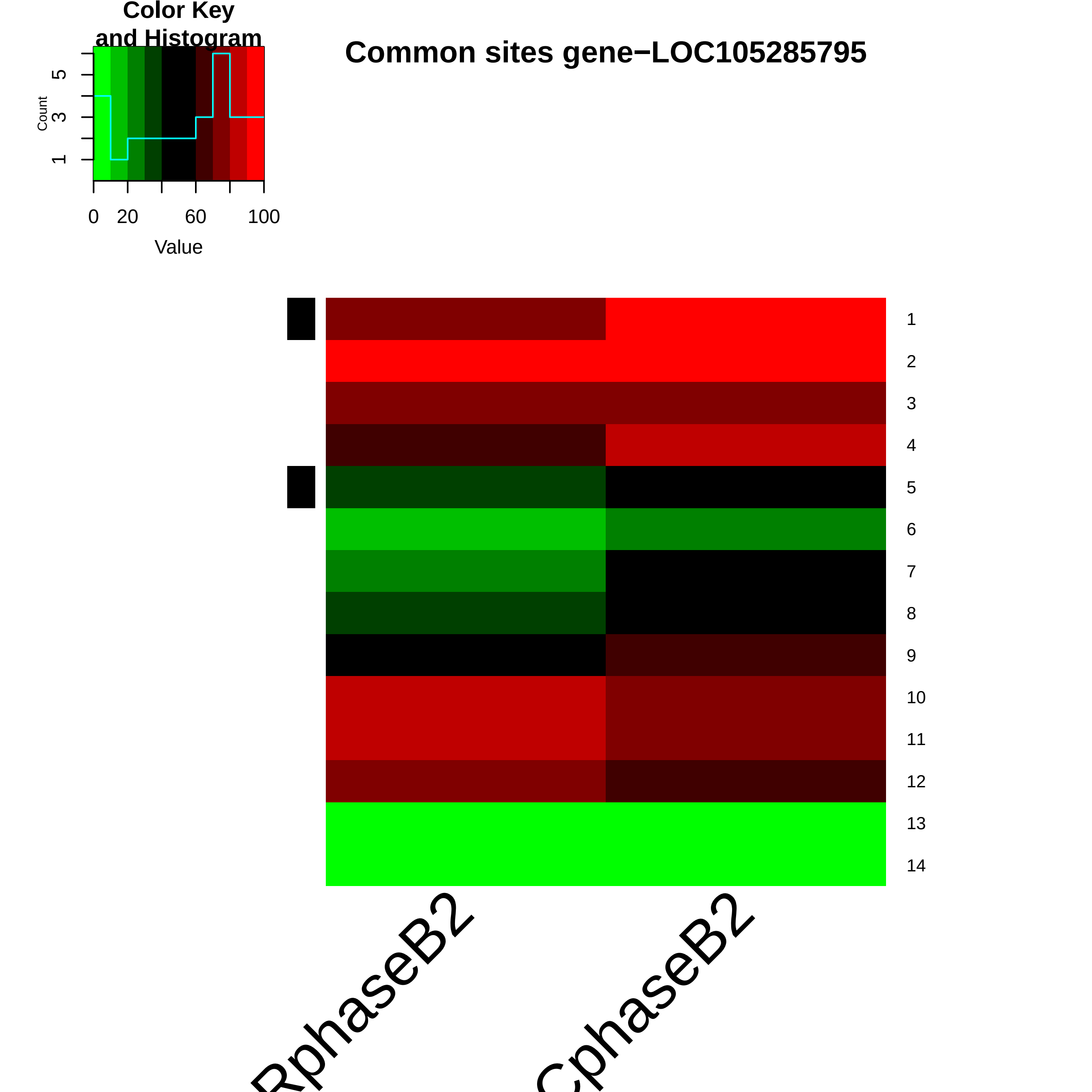

Color Key and Histogram
Count
1
3
5
0
20
60
100
Value
Common sites gene−LOC105285795
1
2
3
4
5
6
7
8
9
10
11
12
13
14
RphaseB2
CphaseB2

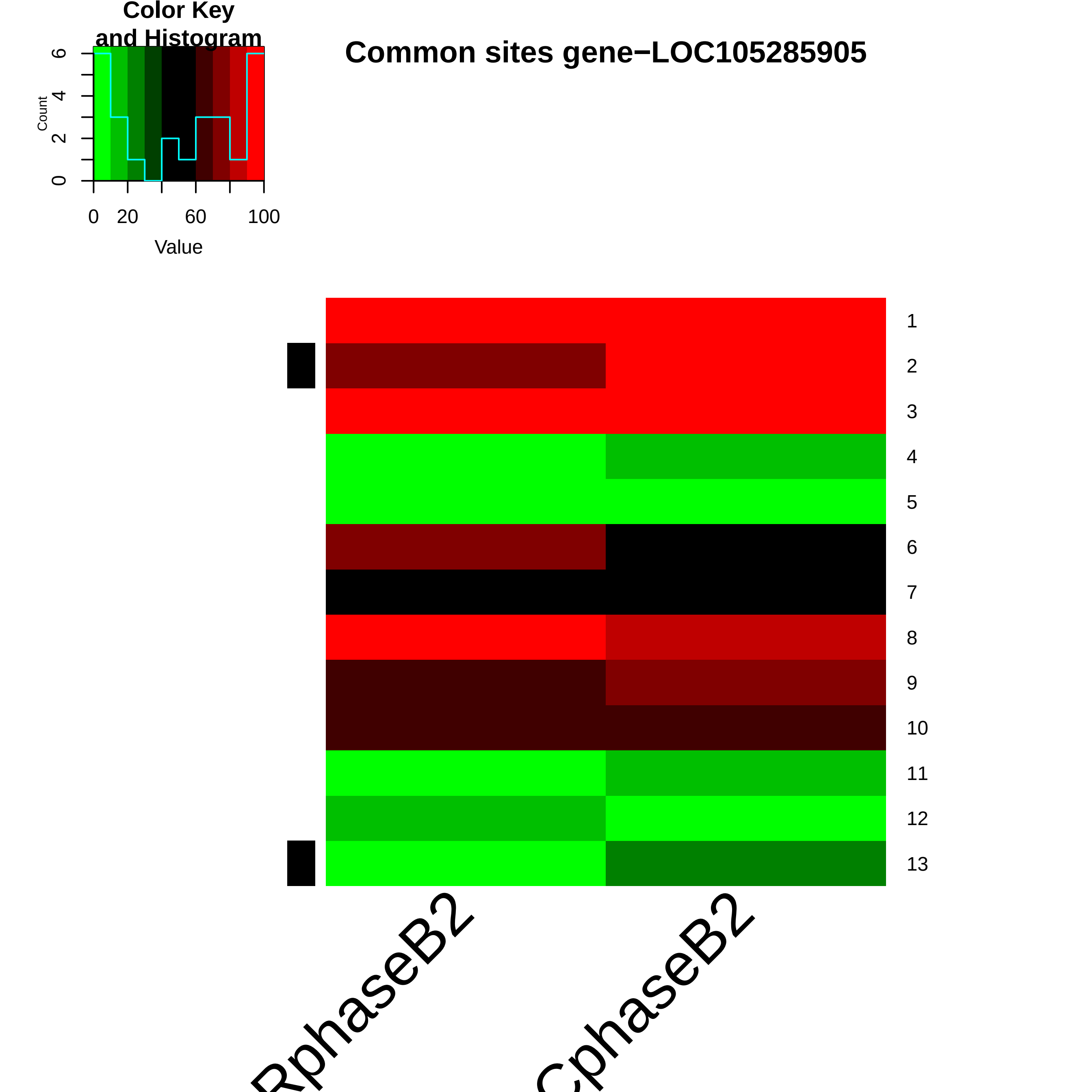
Color Key
and Histogram
Count
0
2
4
6
0
20
60
100
Value
Common sites gene−LOC105285905
1
2
3
4
5
6
7
8
9
10
11
12
13
RphaseB2
CphaseB2

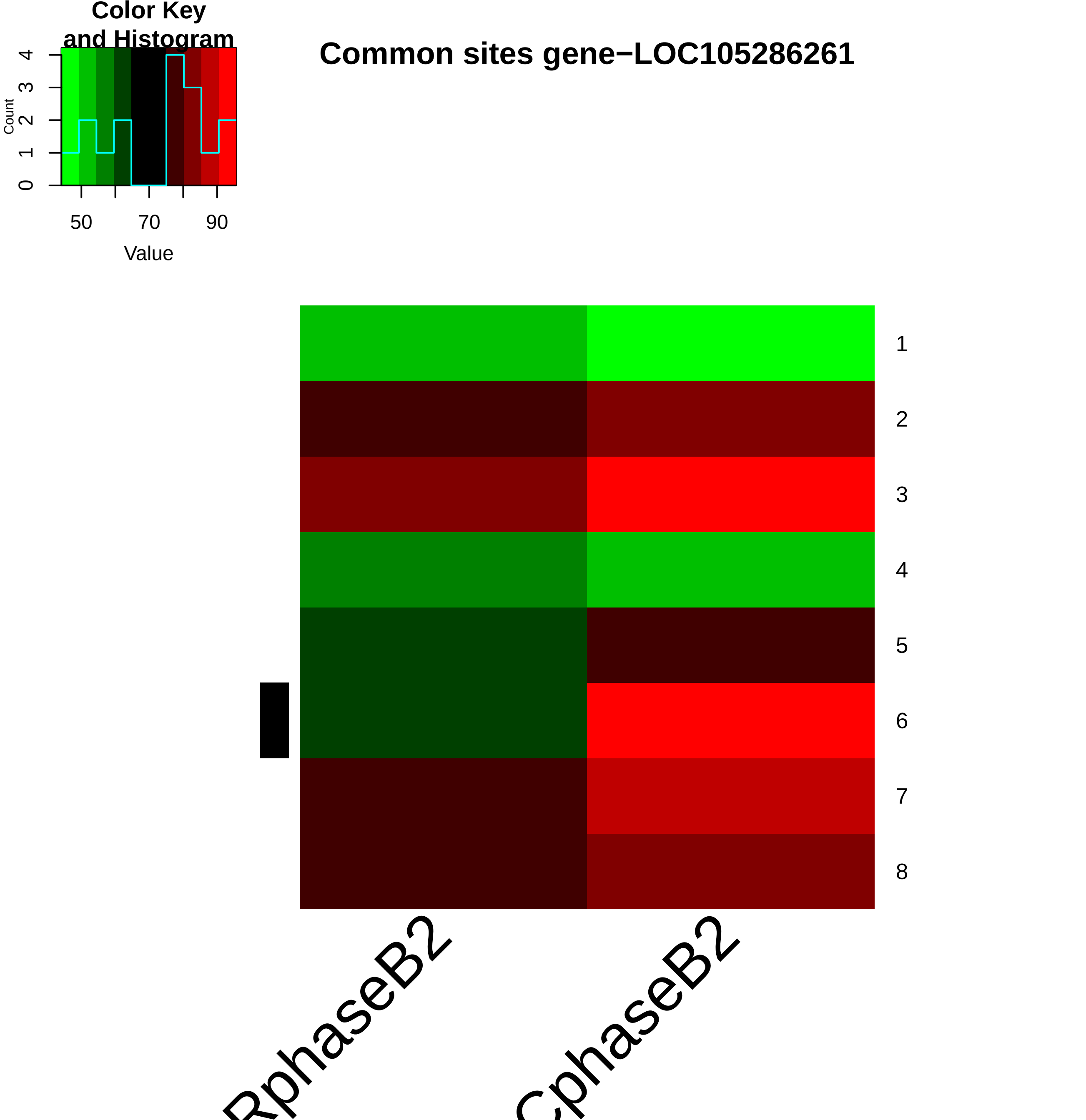
Color Key
and Histogram
Count
0
1
2
3
4
50
70
90
Value
Common sites gene−LOC105286261
1
2
3
4
5
6
7
8
RphaseB2
CphaseB2

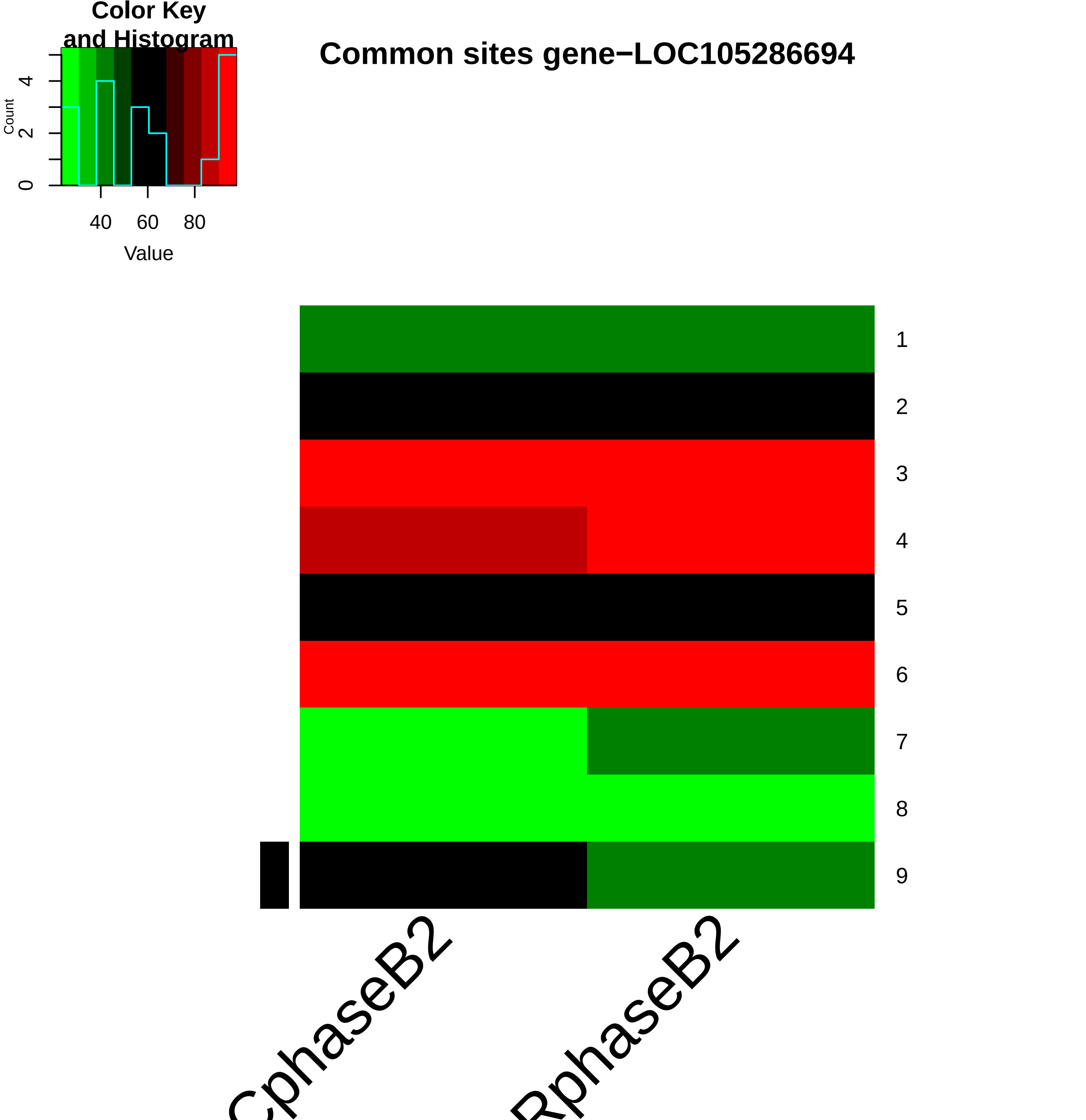

Color Key
and Histogram
Count
0
2
4
40
60
80
Value
Common sites gene−LOC105286694
1
2
3
4
5
6
7
8
9
CphaseB2
RphaseB2

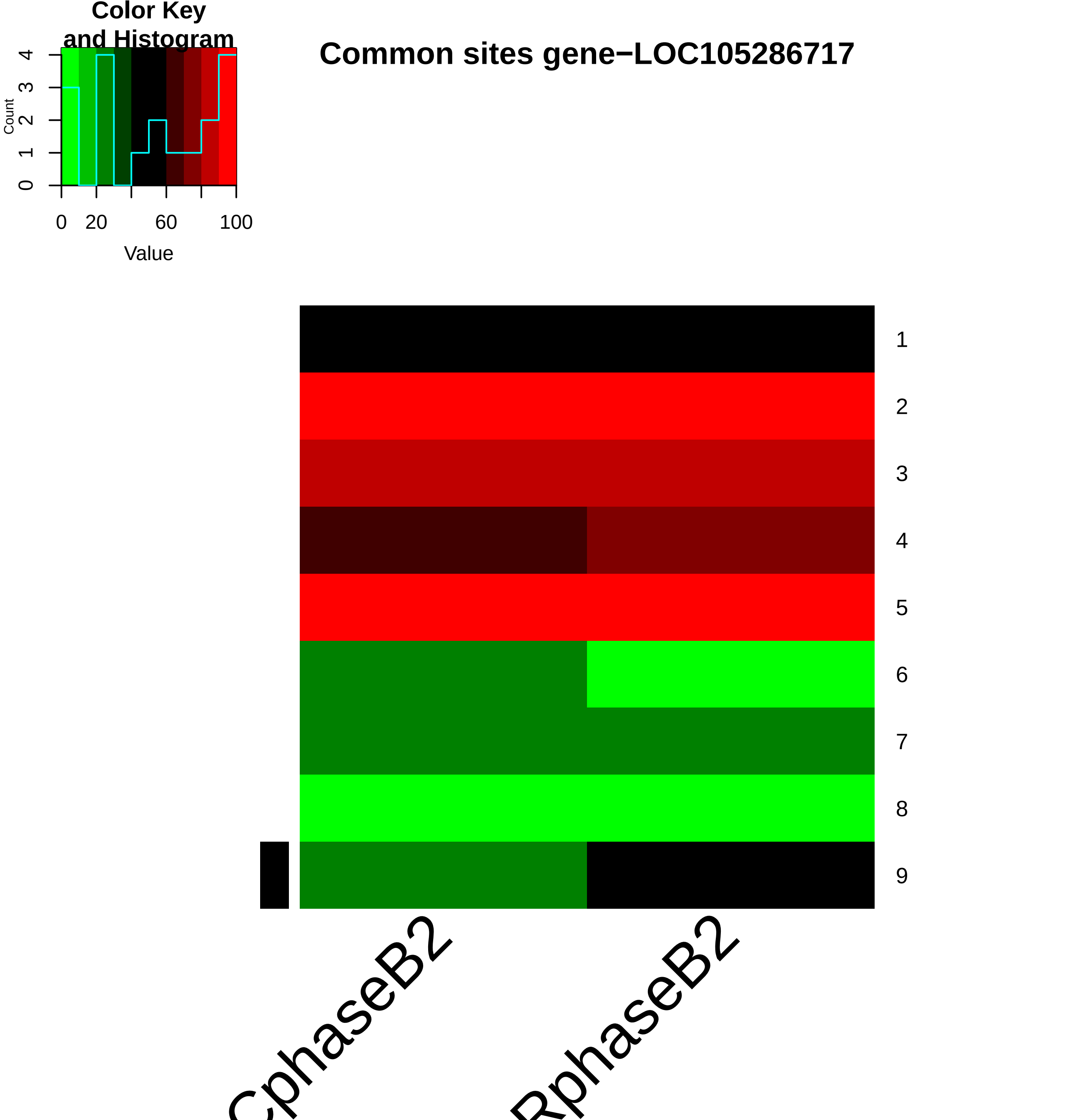

Color Key
and Histogram
Count
0
1
2
3
4
0
20
60
100
Value
Common sites gene−LOC105286717
1
2
3
4
5
6
7
8
9
CphaseB2
RphaseB2

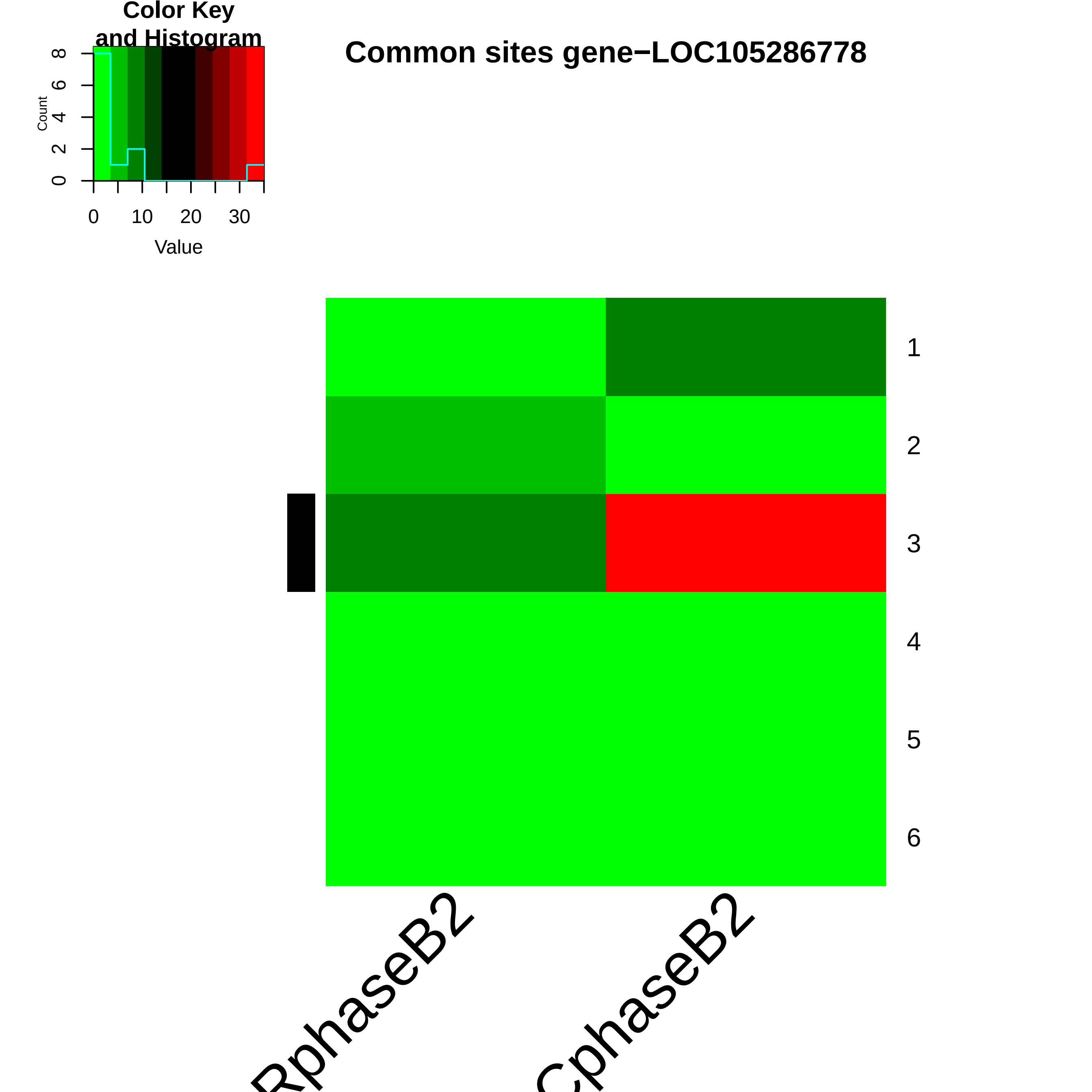

Color Key
and Histogram
Count
Value
0
2
4
6
8
0
10
20
30
Common sites gene−LOC105286778
1
2
3
4
5
6
RphaseB2
CphaseB2

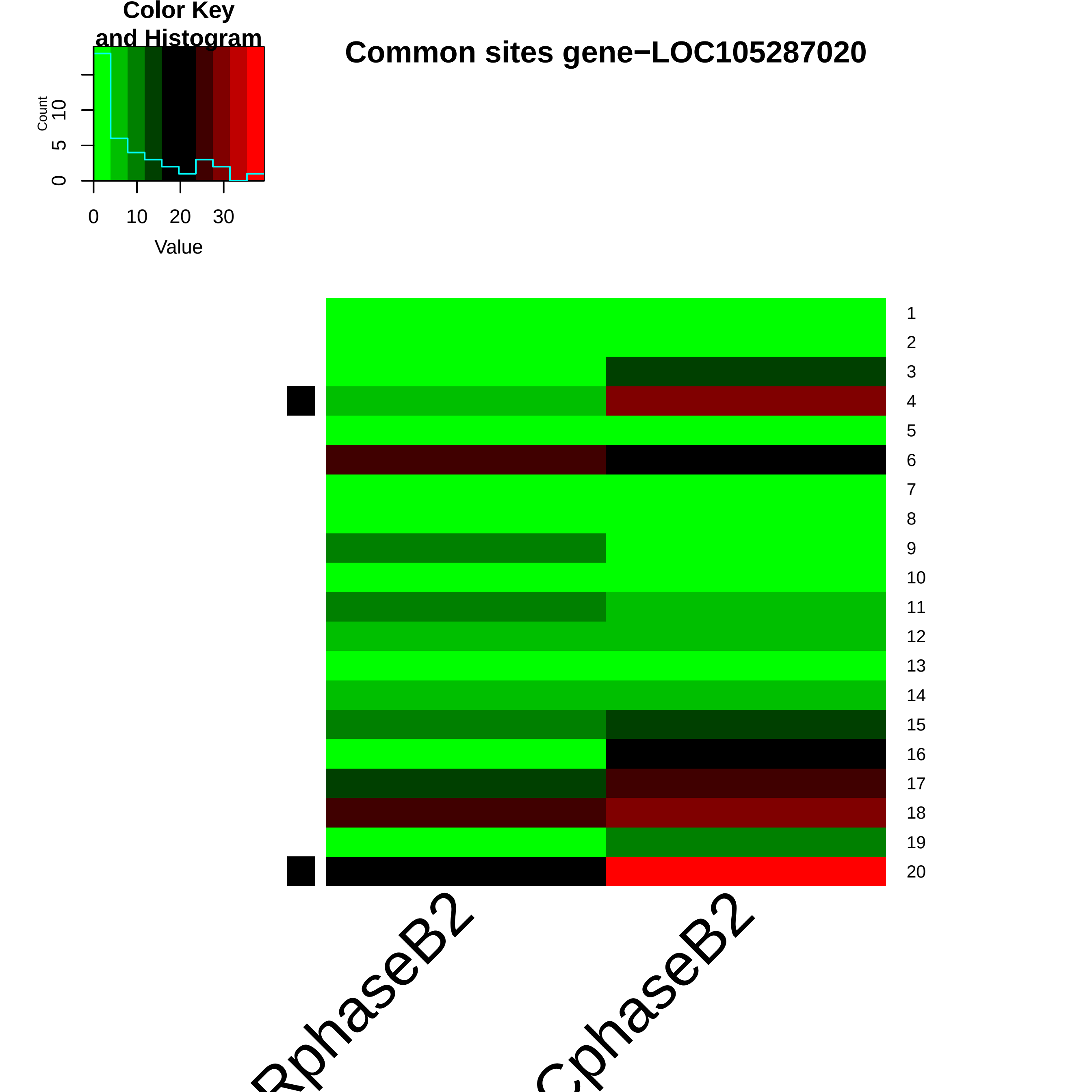
Color Key
and Histogram
Count
0
5
10
0
10
20
30
Value
Common sites gene−LOC105287020
RphaseB2
CphaseB2

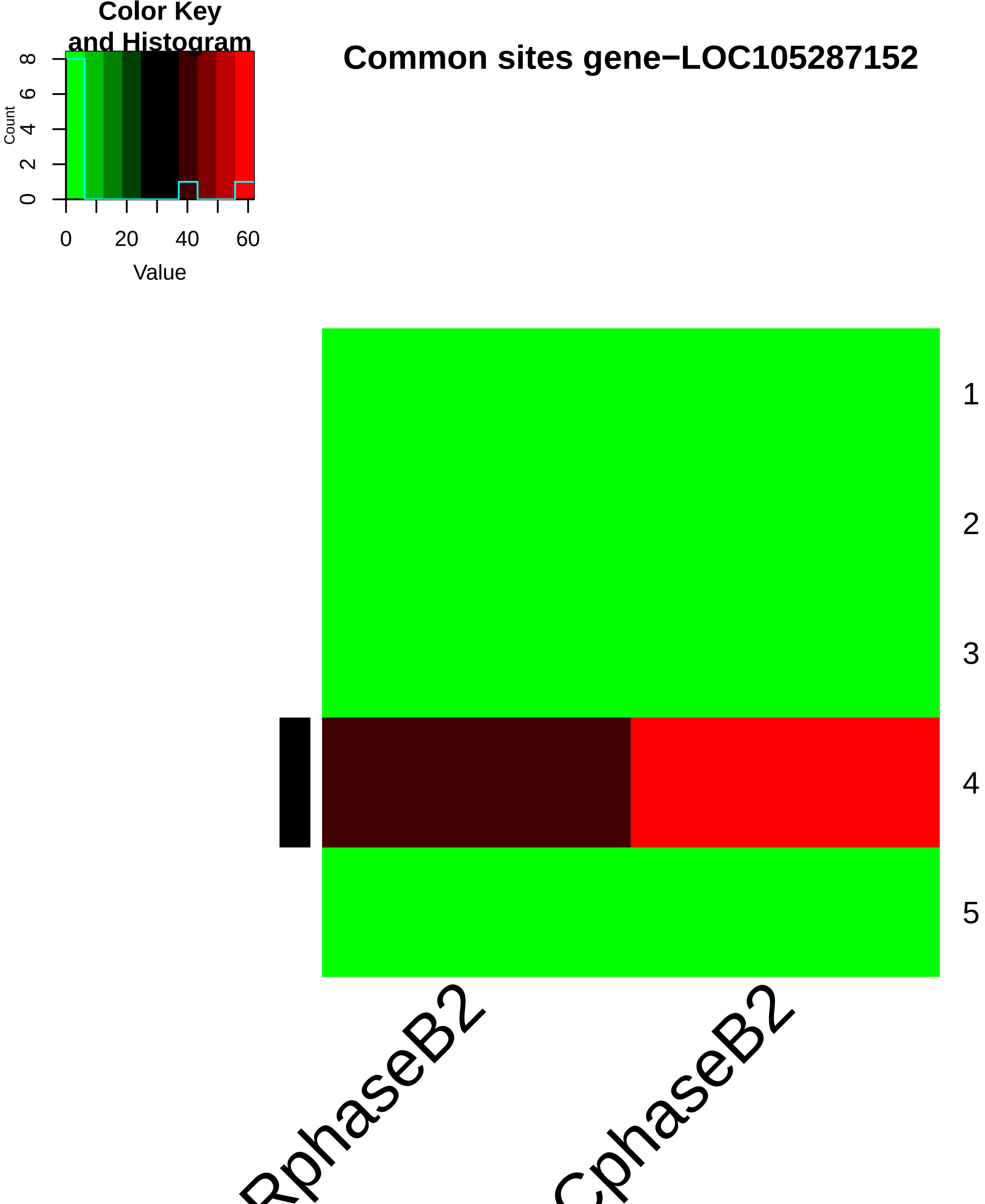
Color Key
and Histogram
Count
0
2
4
6
8
0
20
40
60
Value
Common sites gene−LOC105287152
1
2
3
4
5
RphaseB2
CphaseB2

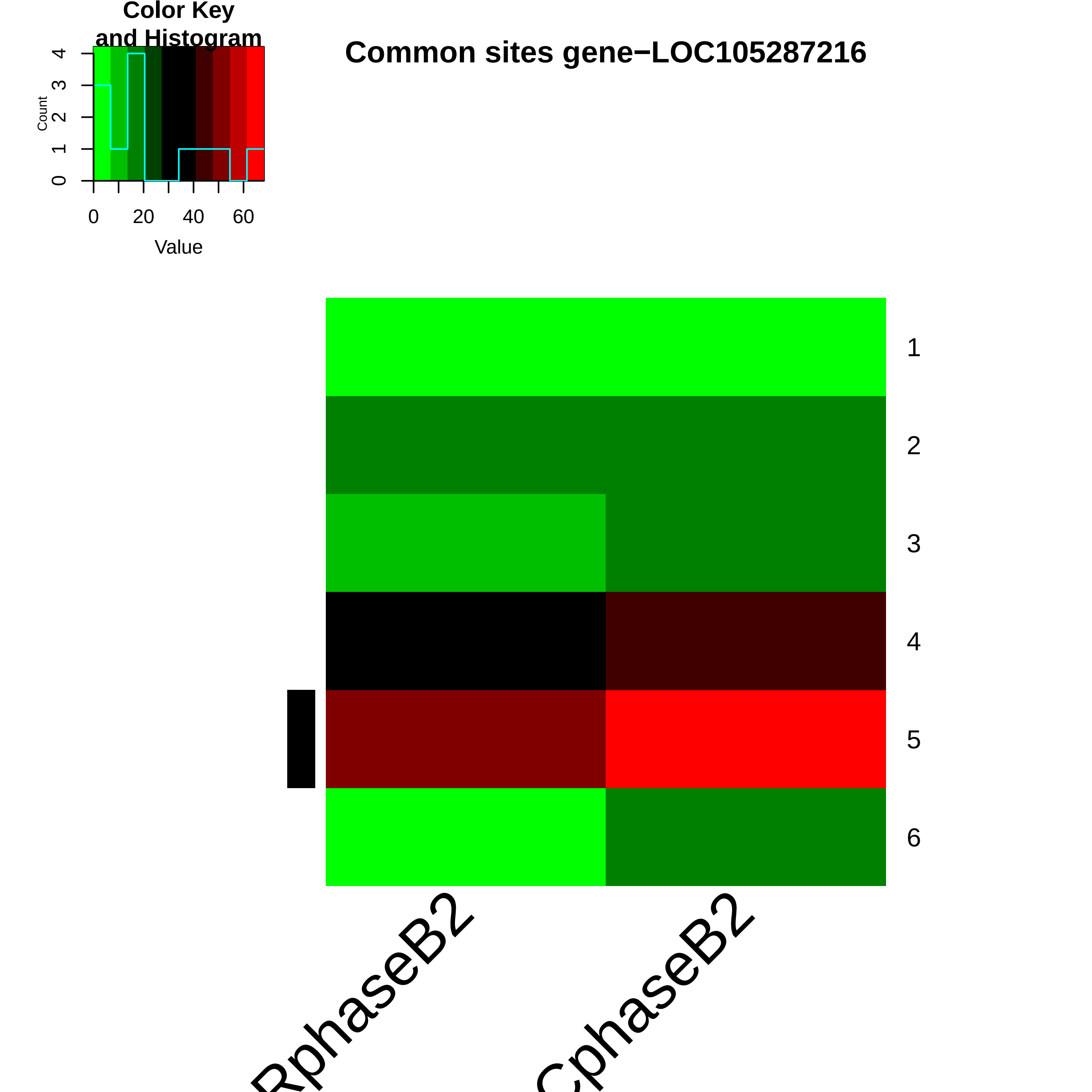

Color Key
and Histogram
Count
0
1
2
3
4
0
20
40
60
Value
Common sites gene−LOC105287216
1
2
3
4
5
6
RphaseB2
CphaseB2

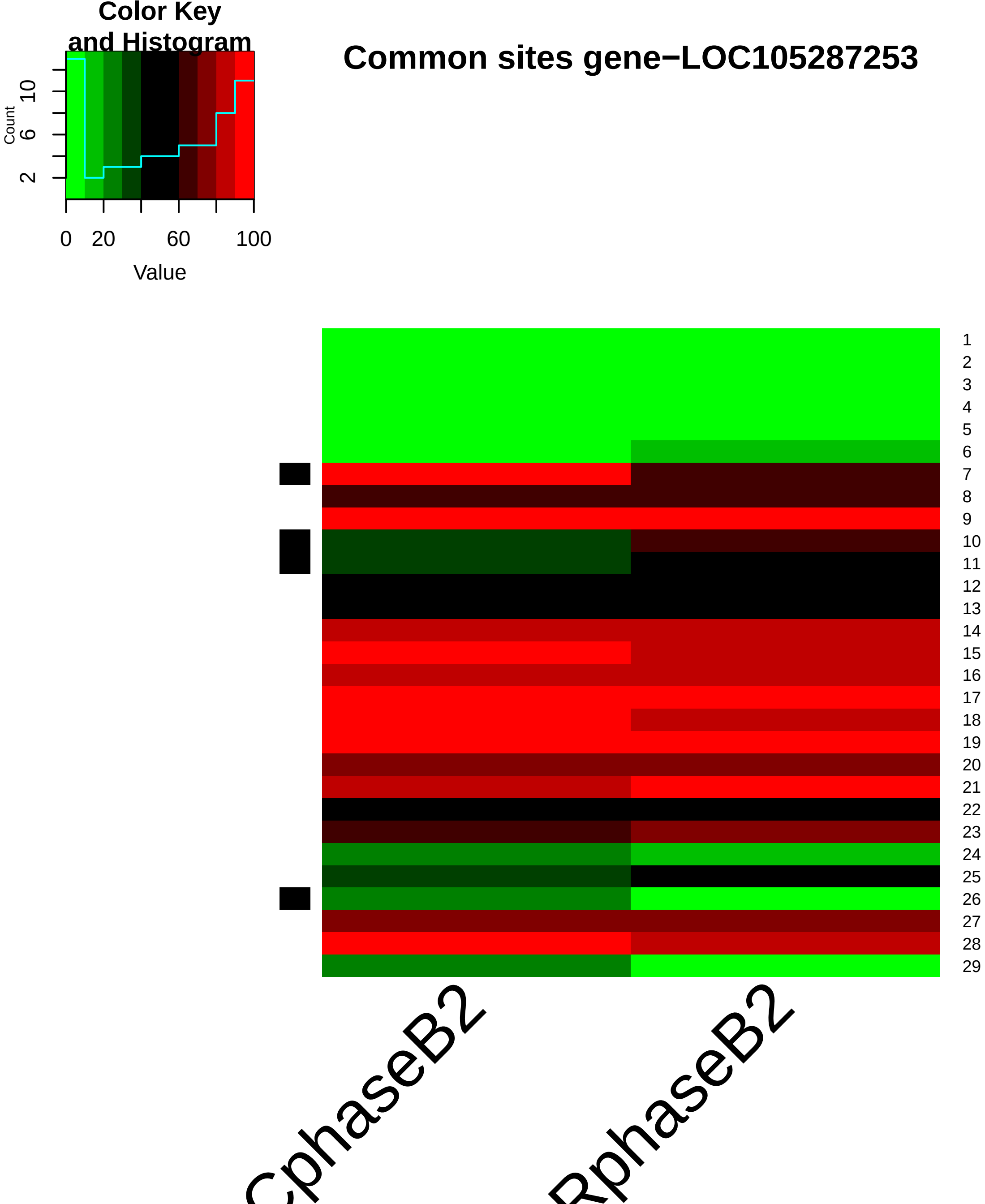
Color Key
and Histogram
Count
2
6
10
0
20
60
100
Value
Common sites gene−LOC105287253
CphaseB2
RphaseB2

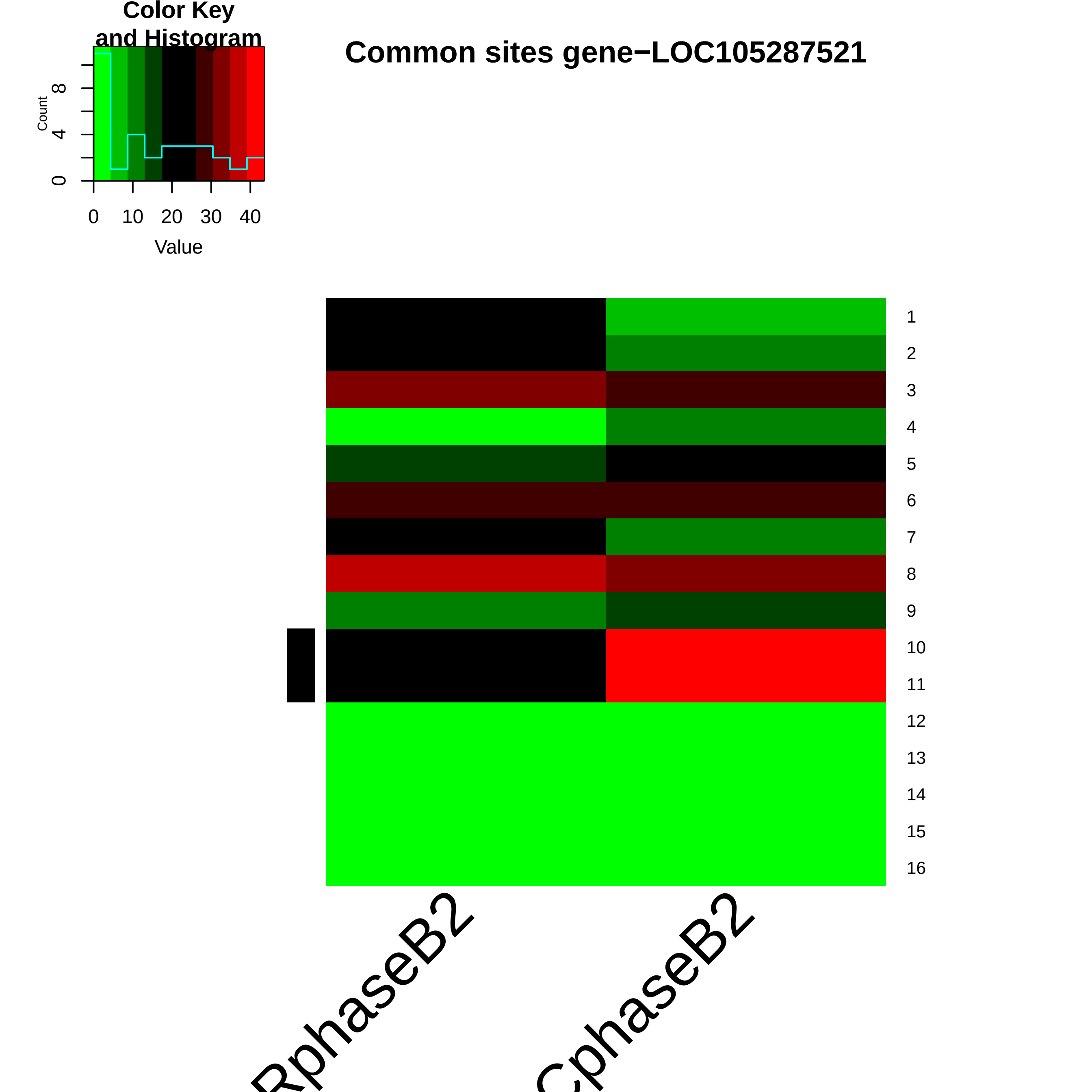
Color Key
and Histogram
Count
0
4
8
0
10
20
30
40
Value
Common sites gene−LOC105287521
RphaseB2
CphaseB2

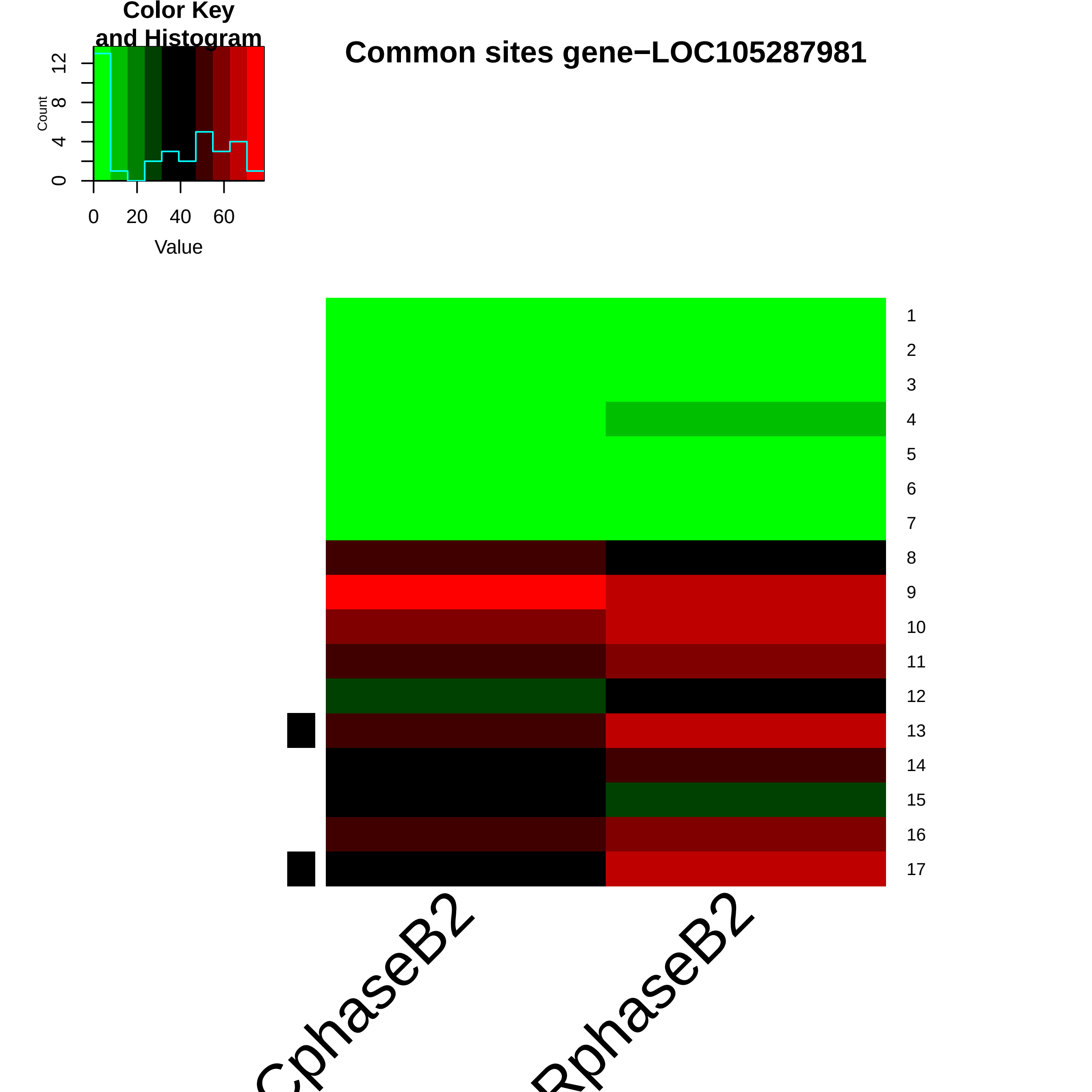
Color Key
and Histogram
Count
0
4
8
12
0
20
40
60
Value
Common sites gene−LOC105287981
CphaseB2
RphaseB2

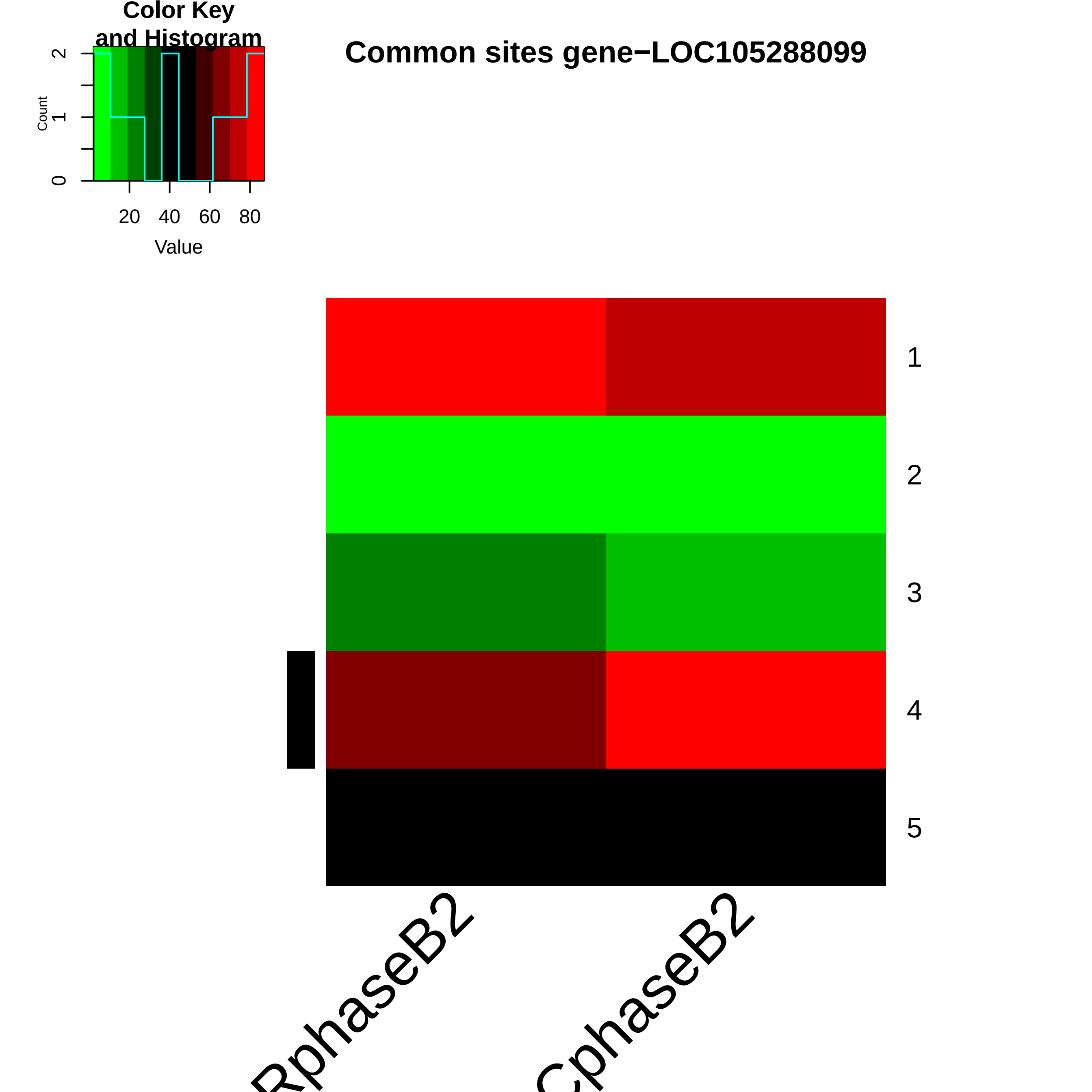

Color Key
and Histogram
Count
0
1
2
20
40
60
80
Value
Common sites gene−LOC105288099
1
2
3
4
5
RphaseB2
CphaseB2

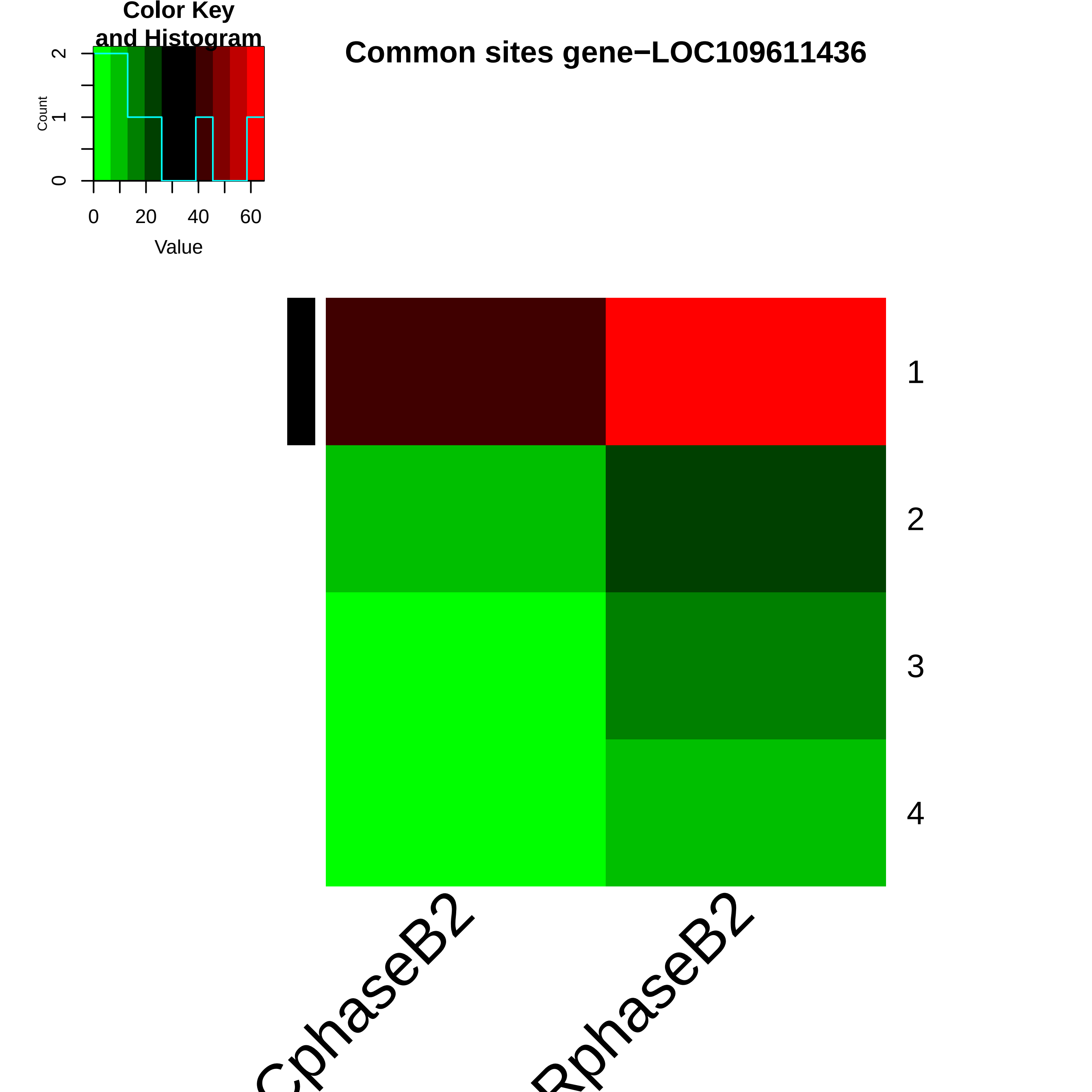

Color Key
and Histogram
Count
0
1
2
0
20
40
60
Value
Common sites gene−LOC109611436
1
2
3
4
CphaseB2
RphaseB2

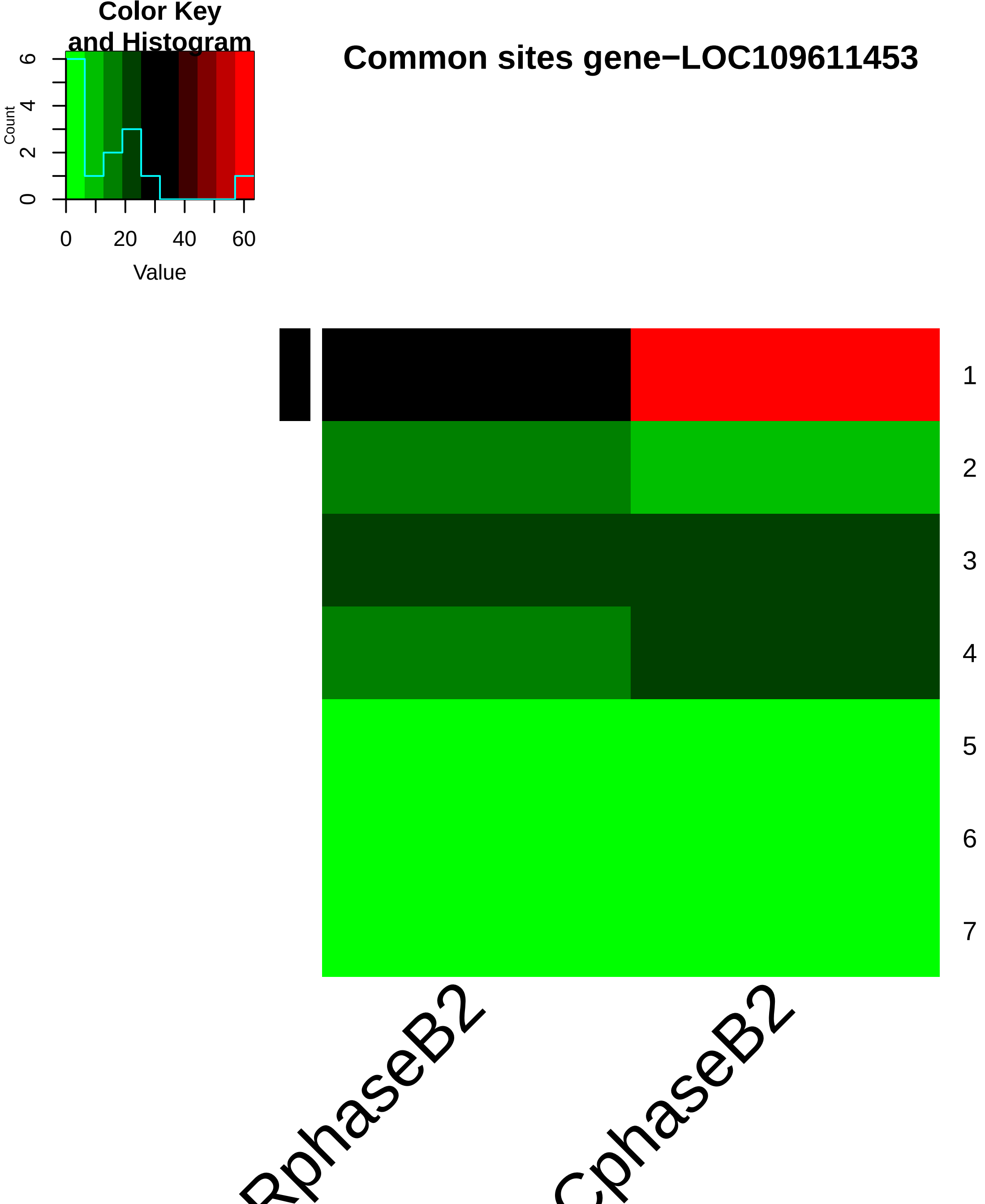
Color Key
and Histogram
Count
Value
0
2
4
6
0
20
40
60
Common sites gene−LOC109611453
1
2
3
4
5
6
7
RphaseB2
CphaseB2

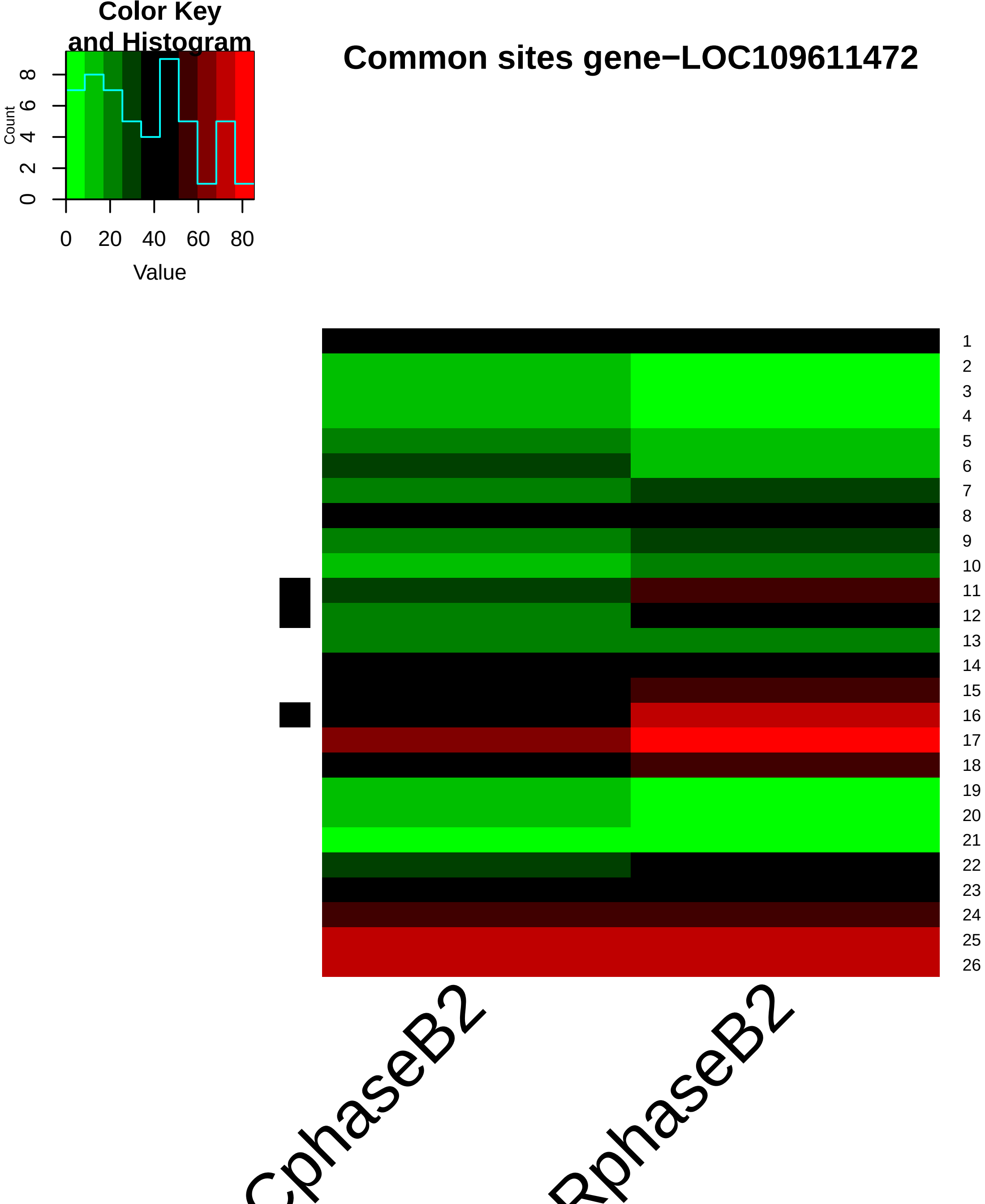

Color Key
and Histogram
Count
0 2 4 6 8
0 20 40 60 80
Value
Common sites gene−LOC109611472
CphaseB2
RphaseB2

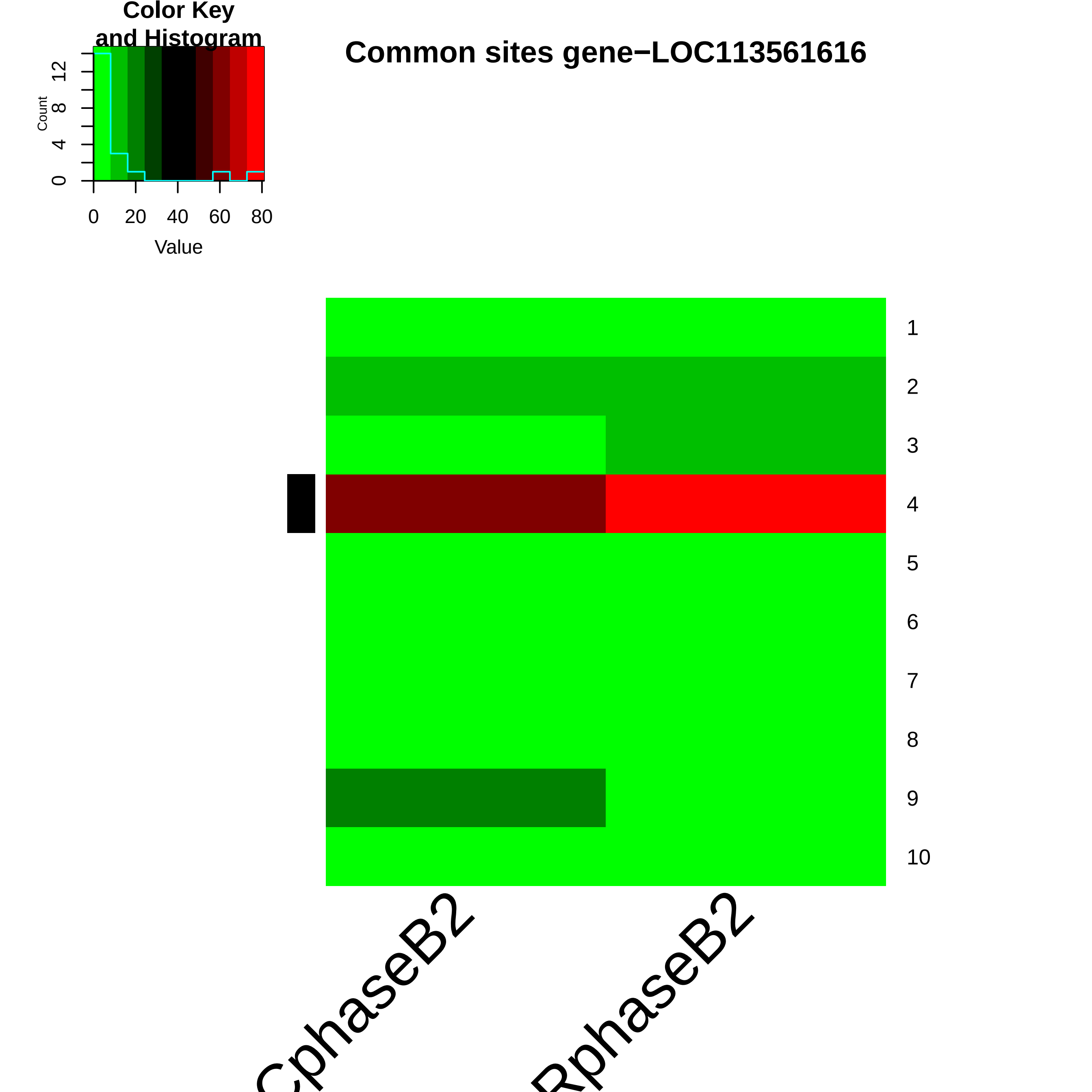

Color Key
and Histogram
Count
0
4
8
12
0
20
40
60
80
Value
Common sites gene−LOC113561616
1
2
3
4
5
6
7
8
9
10
CphaseB2
RphaseB2

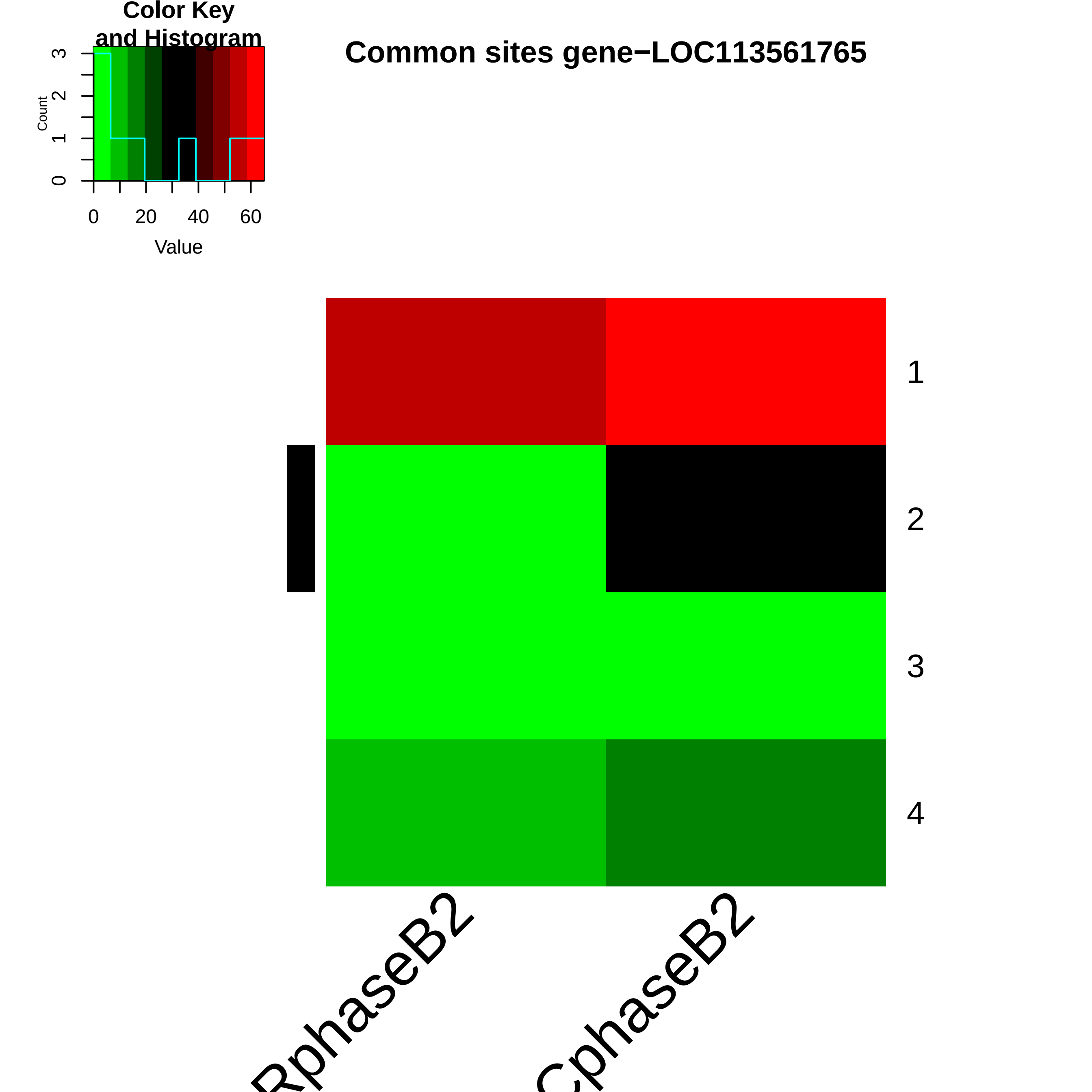

Color Key
and Histogram
Count
3
2
1
0
0
20
40
60
Value
Common sites gene−LOC113561765
1
2
3
4

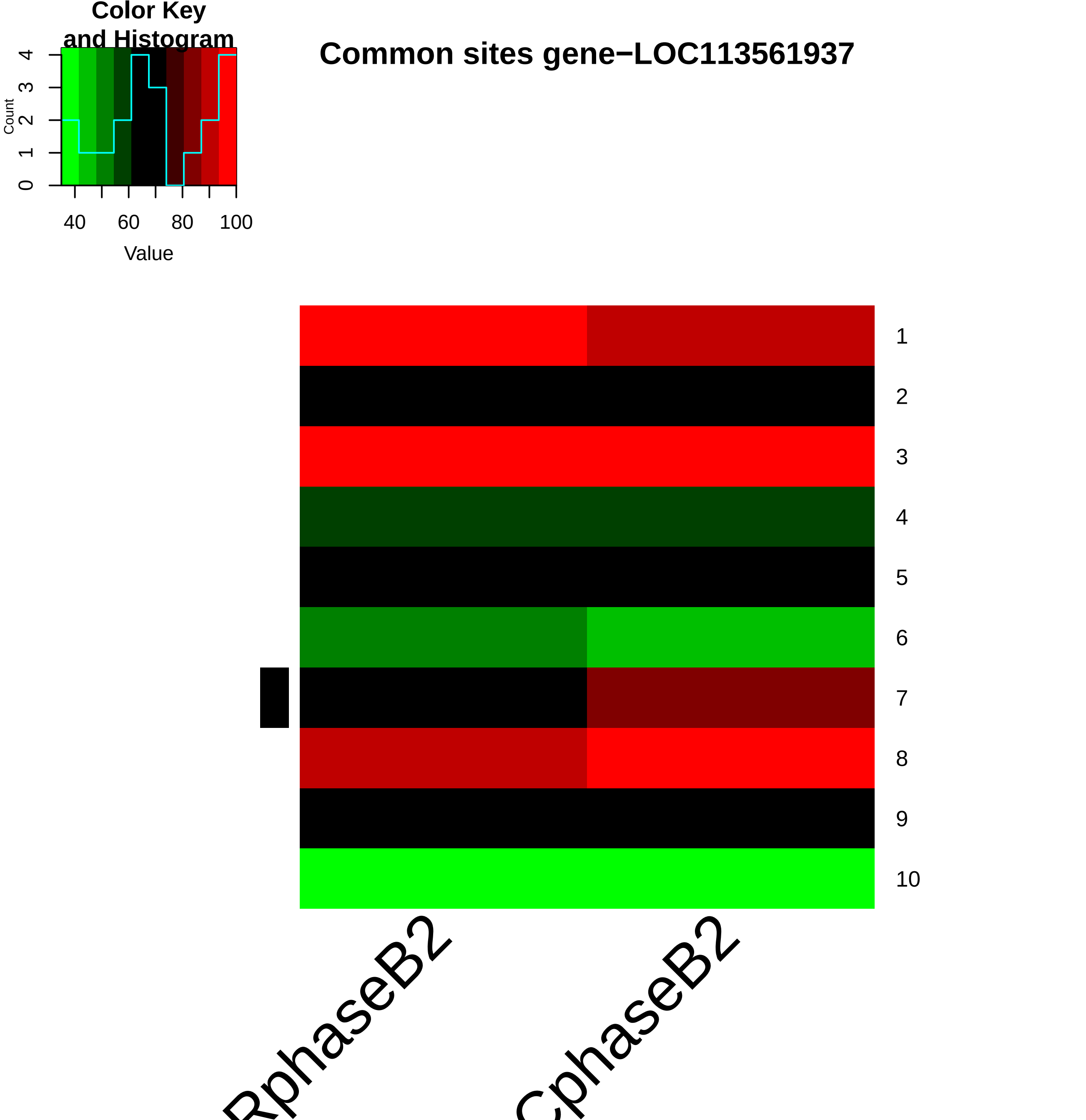

Color Key
and Histogram
Count
0
1
2
3
4
40
60
80
100
Value
Common sites gene−LOC113561937
1
2
3
4
5
6
7
8
9
10
RphaseB2
CphaseB2

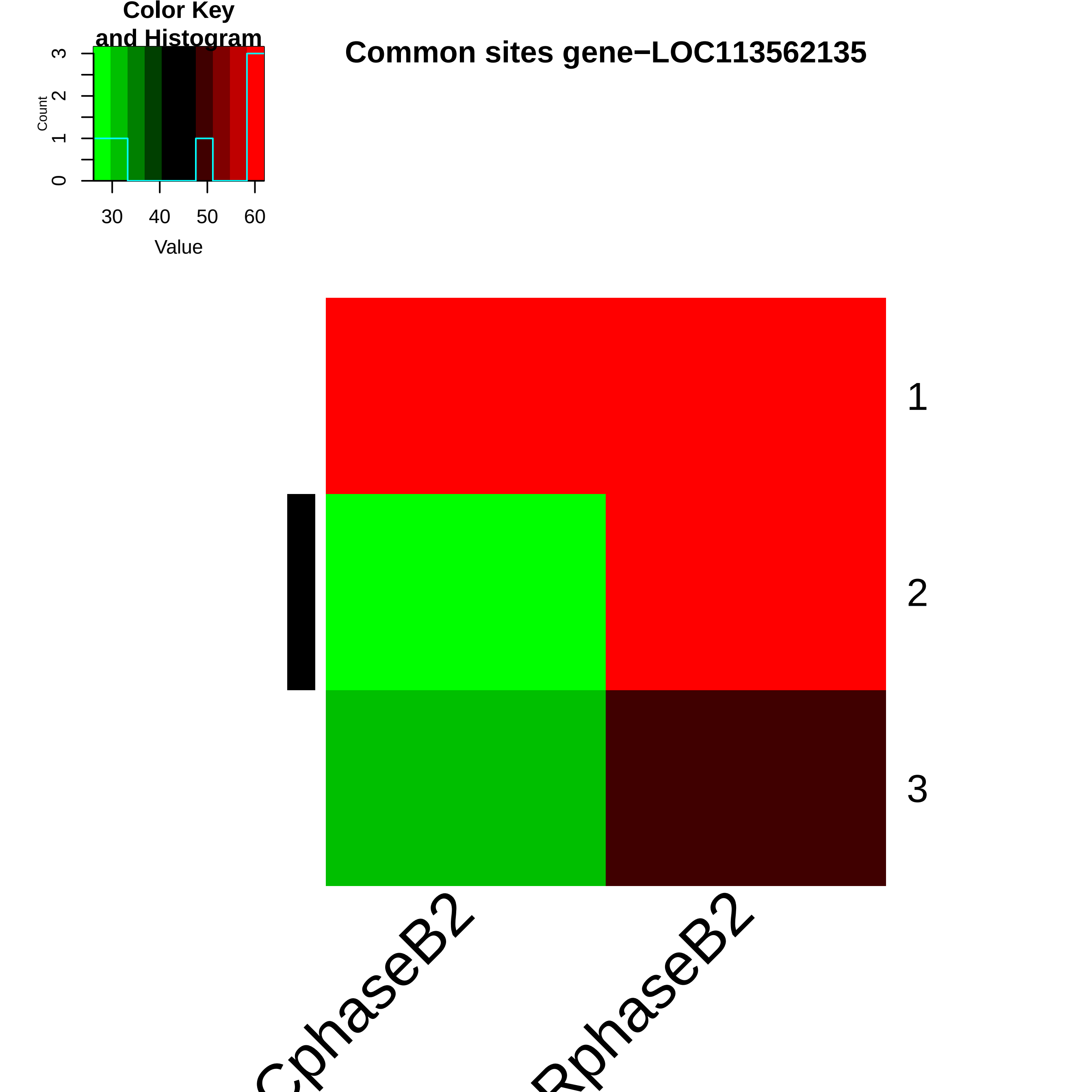

Color Key and Histogram
Count
0
1
2
3
30
40
50
60
Value
Common sites gene−LOC113562135
1
2
3
CphaseB2
RphaseB2

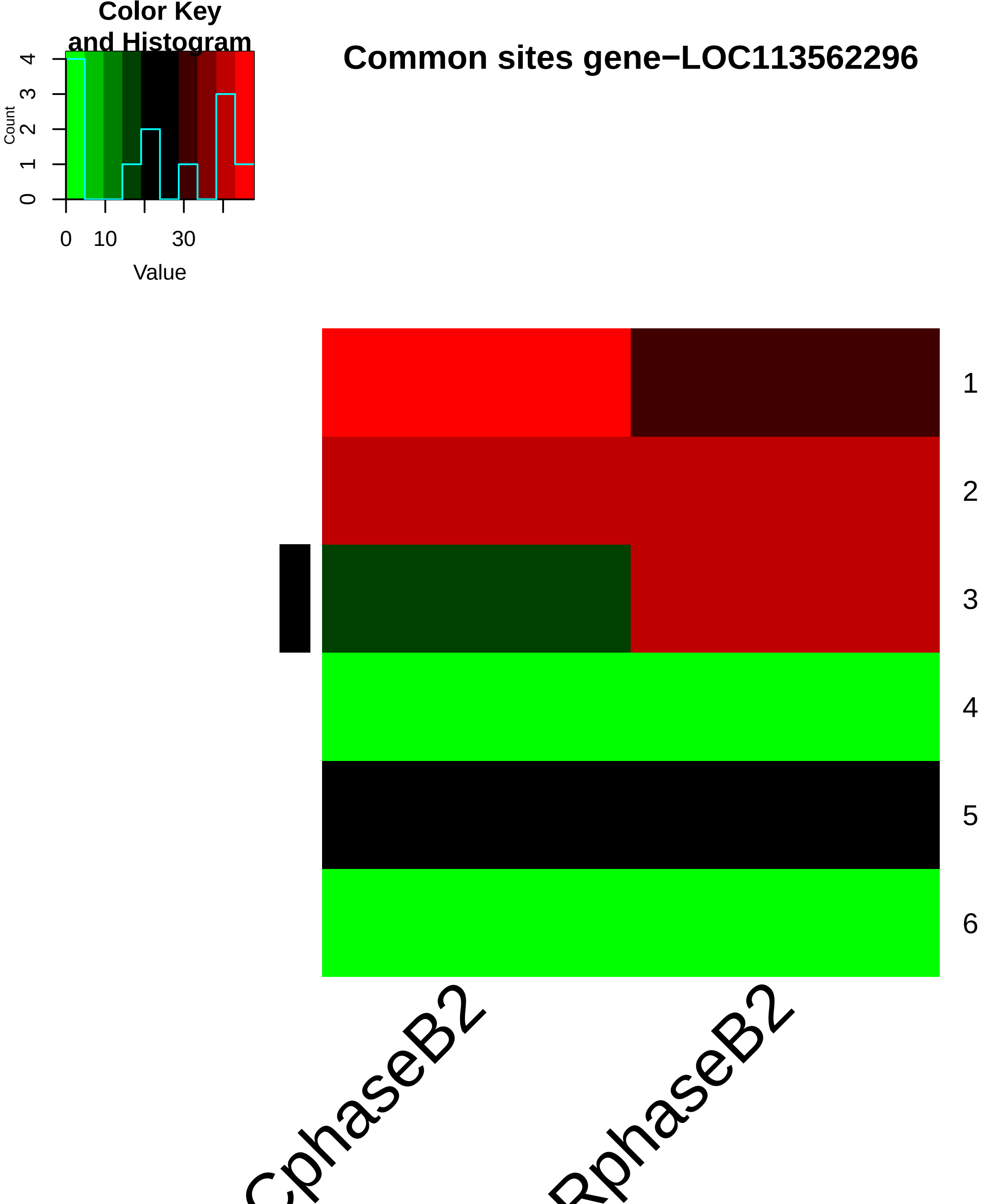
Color Key
and Histogram
Count
0
1
2
3
4
0
10
30
Value
Common sites gene−LOC113562296
1
2
3
4
5
6
CphaseB2
RphaseB2

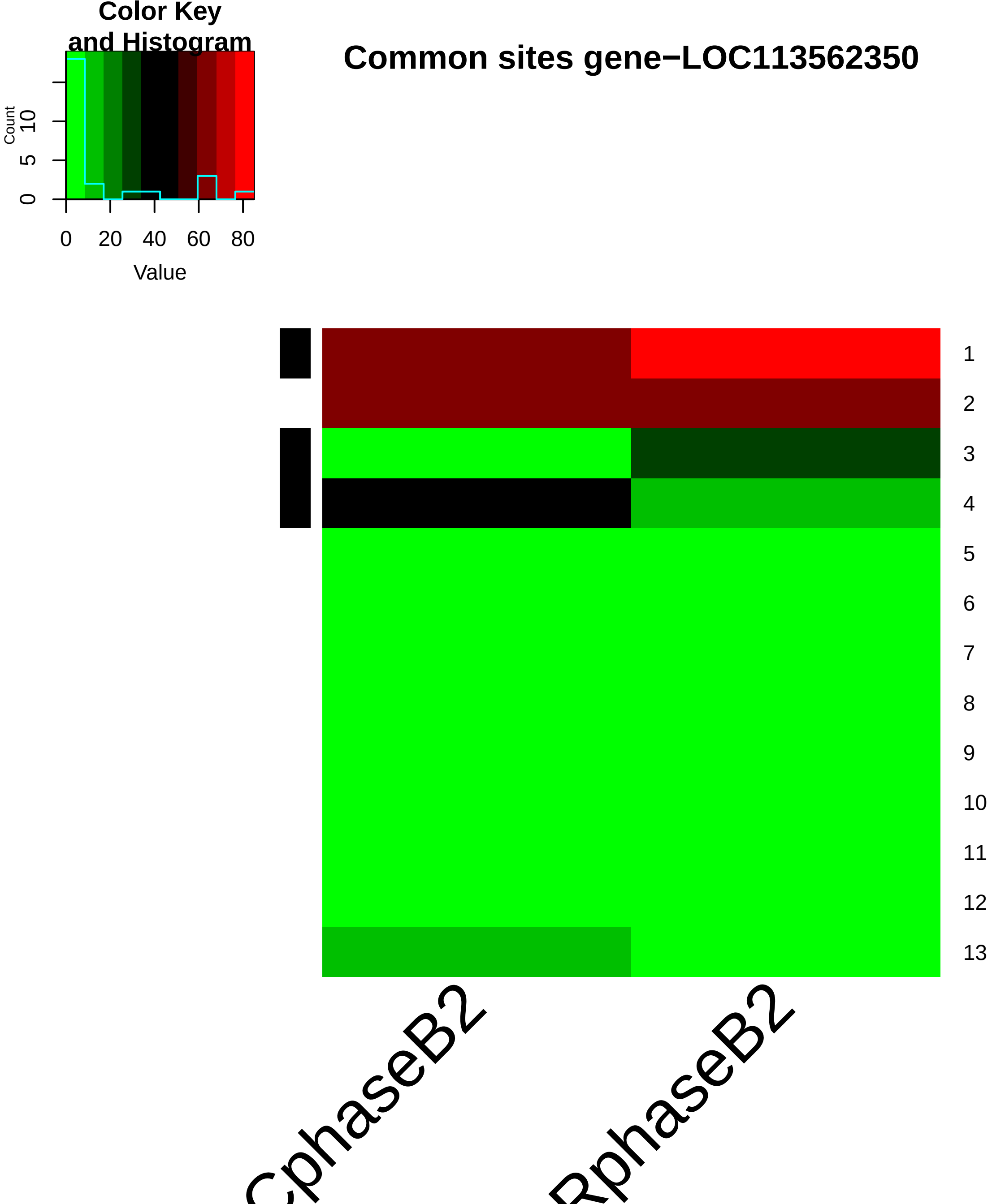
Color Key
and Histogram
Count
0
5
10
0
20
40
60
80
Value
Common sites gene−LOC113562350
1
2
3
4
5
6
7
8
9
10
11
12
13
CphaseB2
RphaseB2

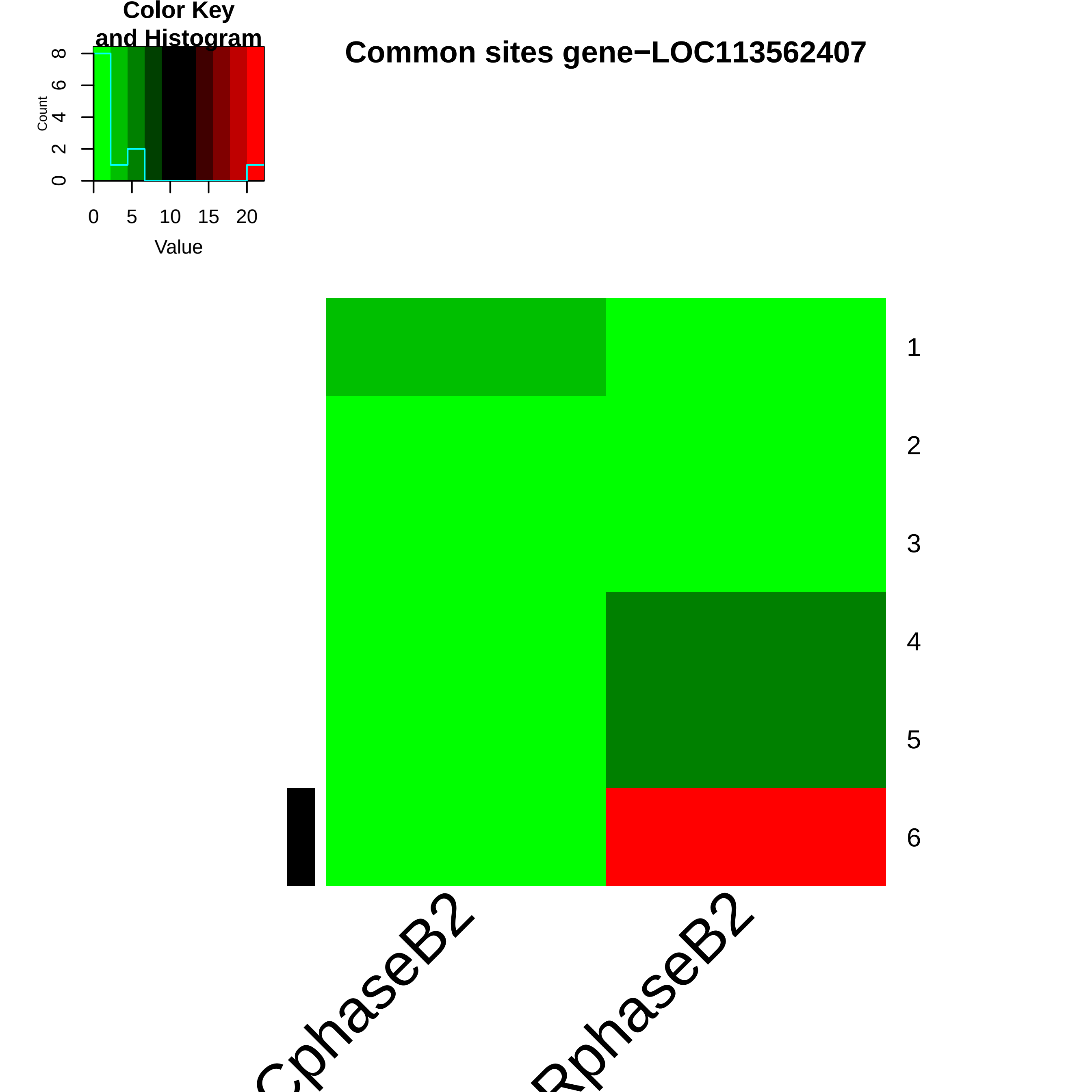

Color Key
and Histogram
Count
0
2
4
6
8
0
5
10
15
20
Value
Common sites gene−LOC113562407
1
2
3
4
5
6
CphaseB2
RphaseB2

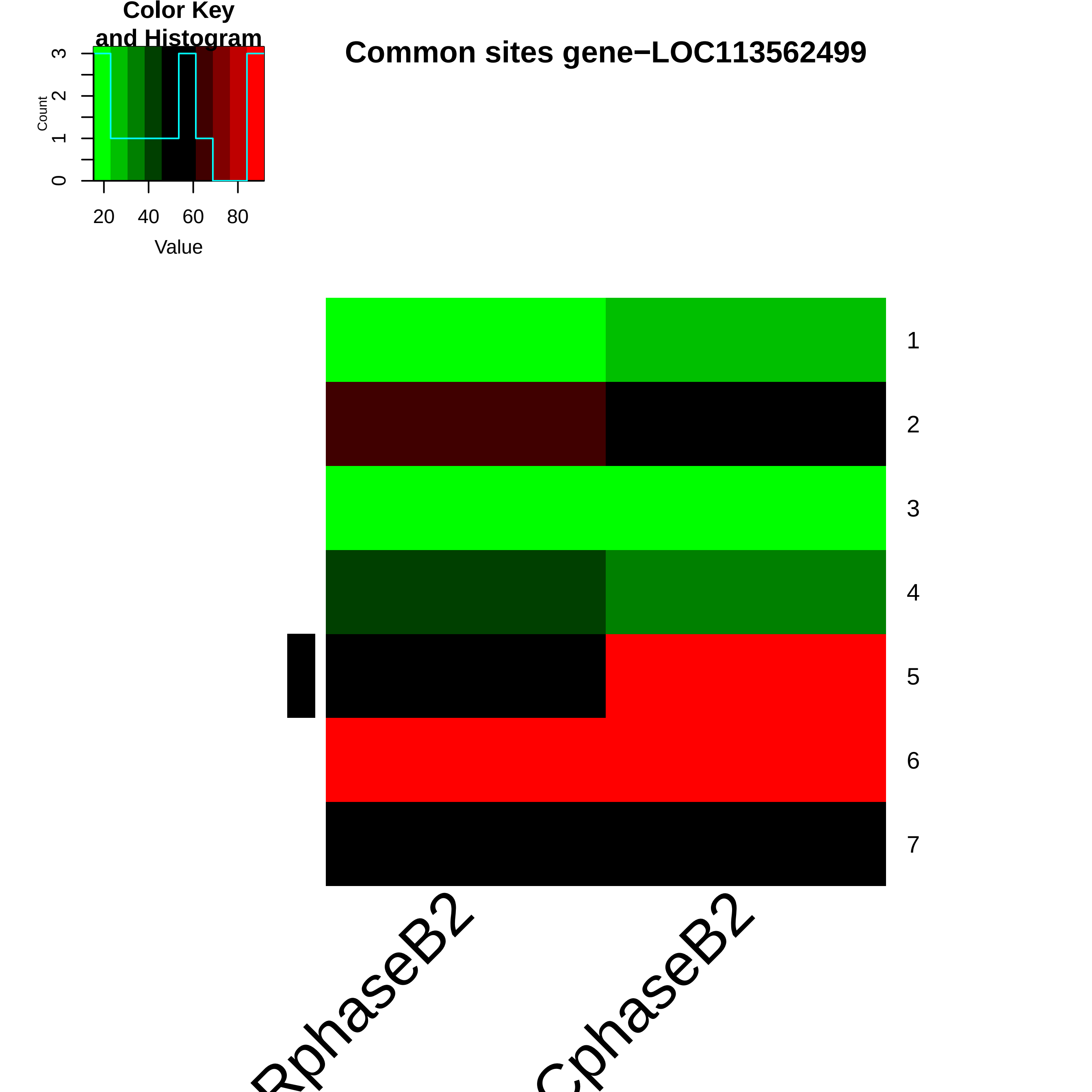
Color Key
and Histogram
Count
0
1
2
3
20
40
60
80
Value
Common sites gene−LOC113562499
1
2
3
4
5
6
7
RphaseB2
CphaseB2

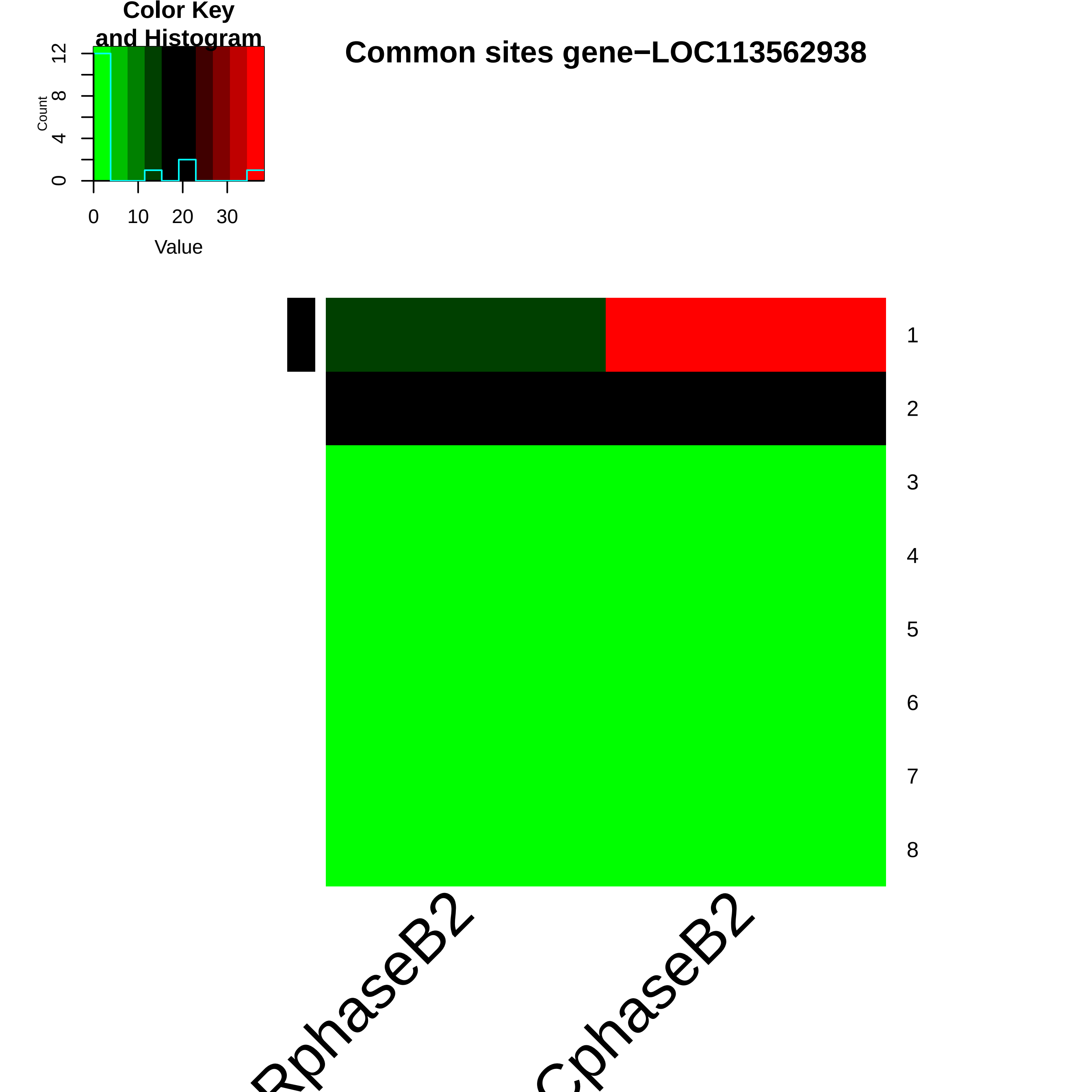
Color Key
and Histogram
Count
0
4
8
12
0
10
20
30
Value
Common sites gene−LOC113562938
1
2
3
4
5
6
7
8
RphaseB2
CphaseB2

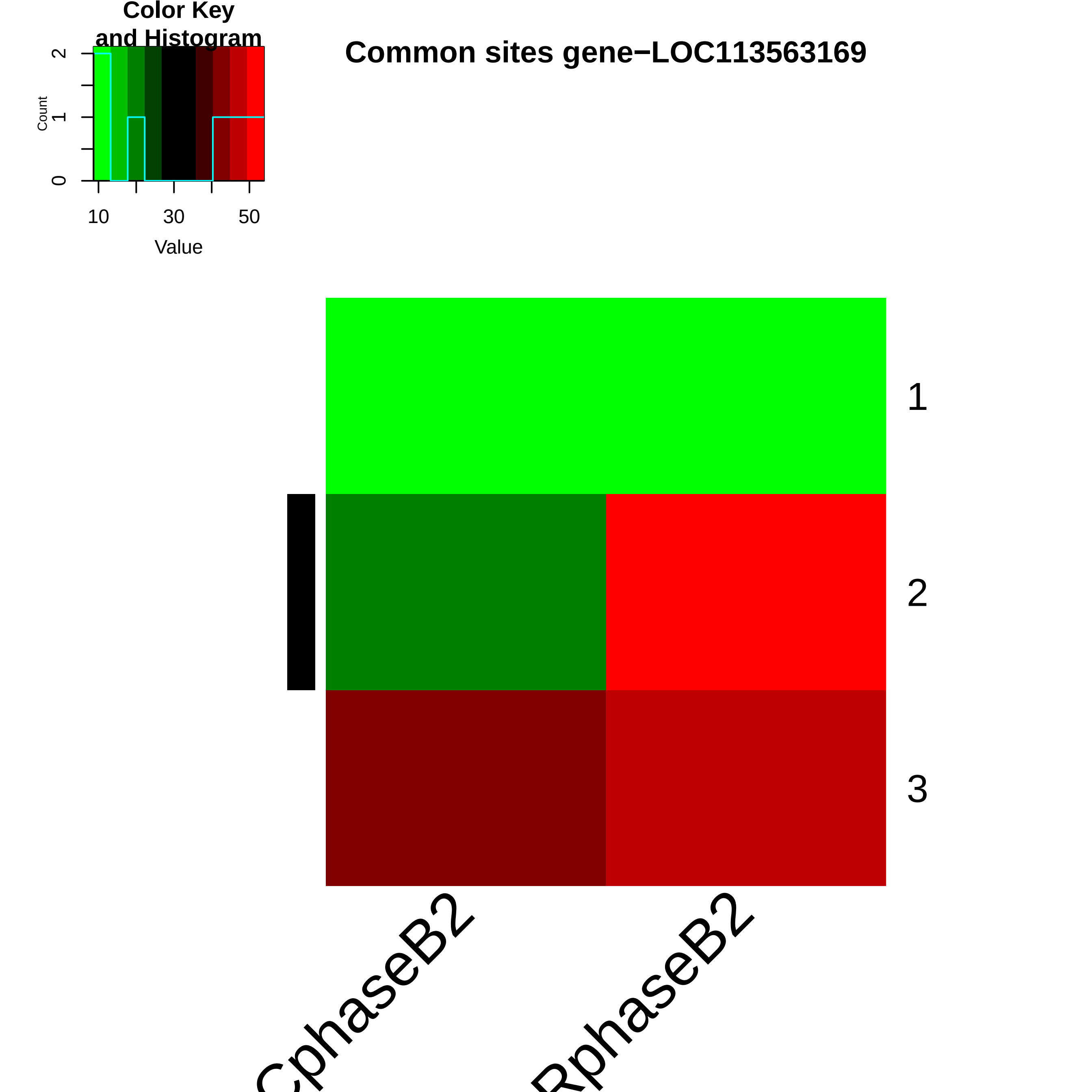

Color Key and Histogram
Count
0
1
2
10
30
50
Value
Common sites gene−LOC113563169
1
2
3
CphaseB2
RphaseB2

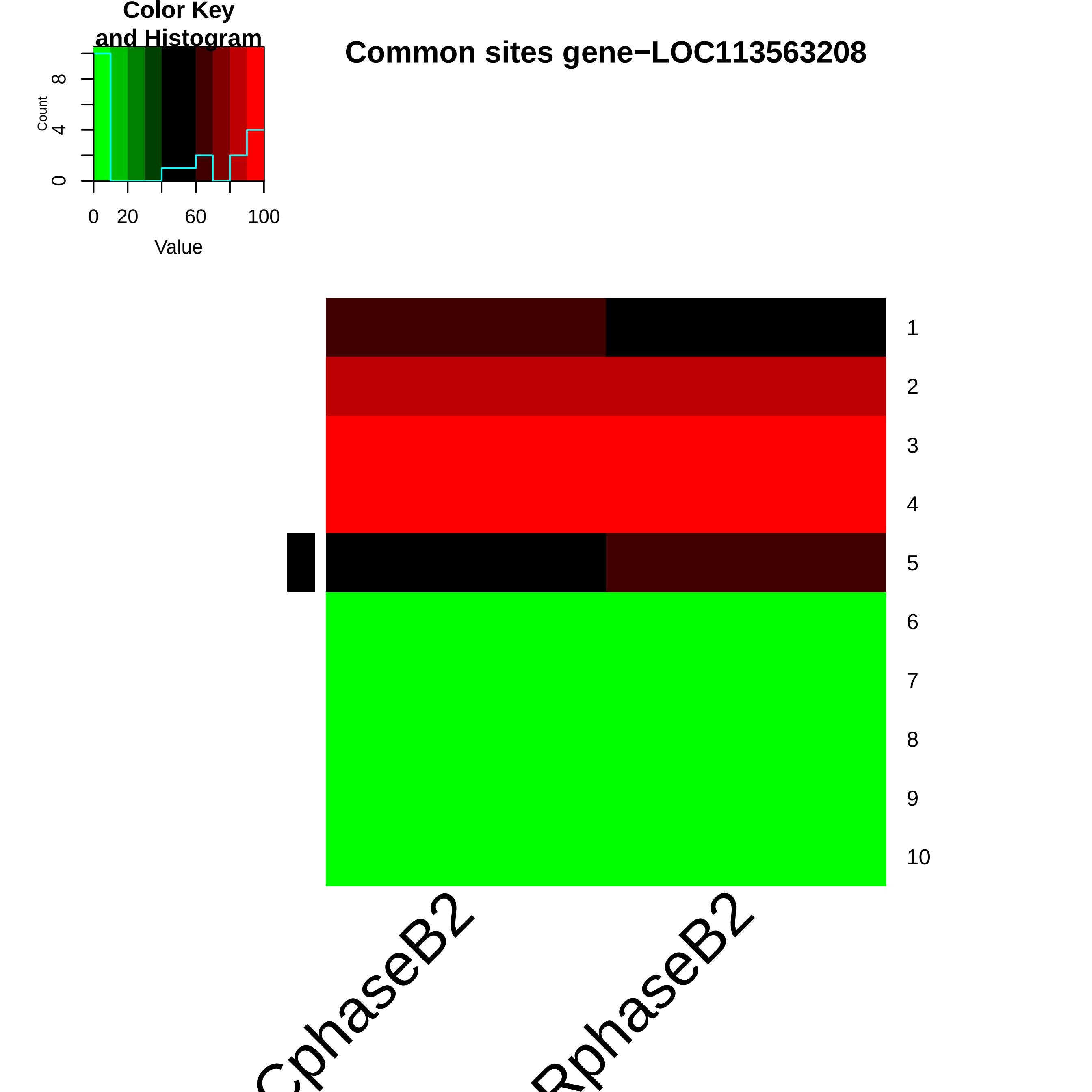

Color Key
and Histogram
Count
0
4
8
0
20
60
100
Value
Common sites gene−LOC113563208
1
2
3
4
5
6
7
8
9
10
CphaseB2
RphaseB2

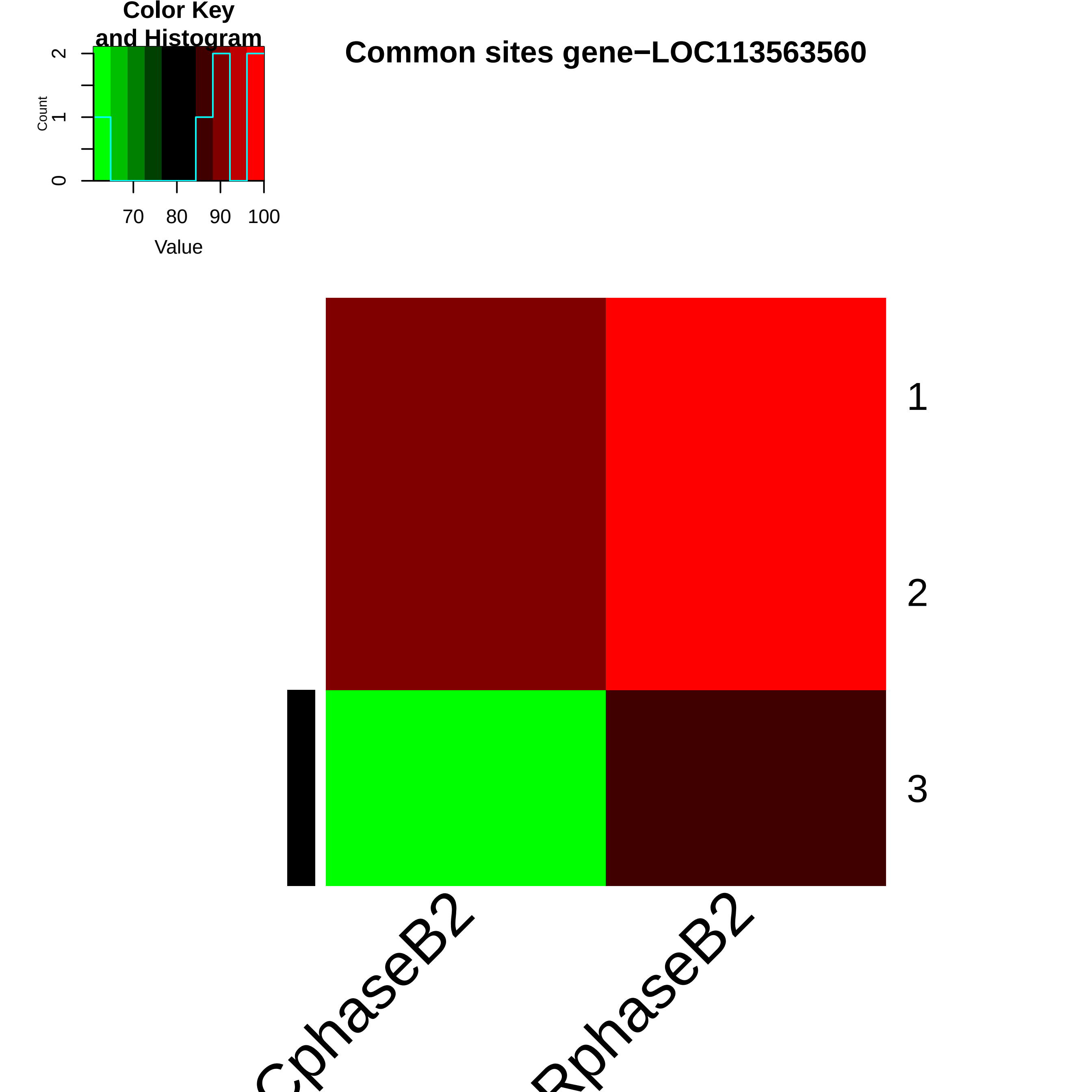
Color Key
and Histogram
Count
0
1
2
70
80
90
100
Value
Common sites gene−LOC113563560
1
2
3
CphaseB2
RphaseB2

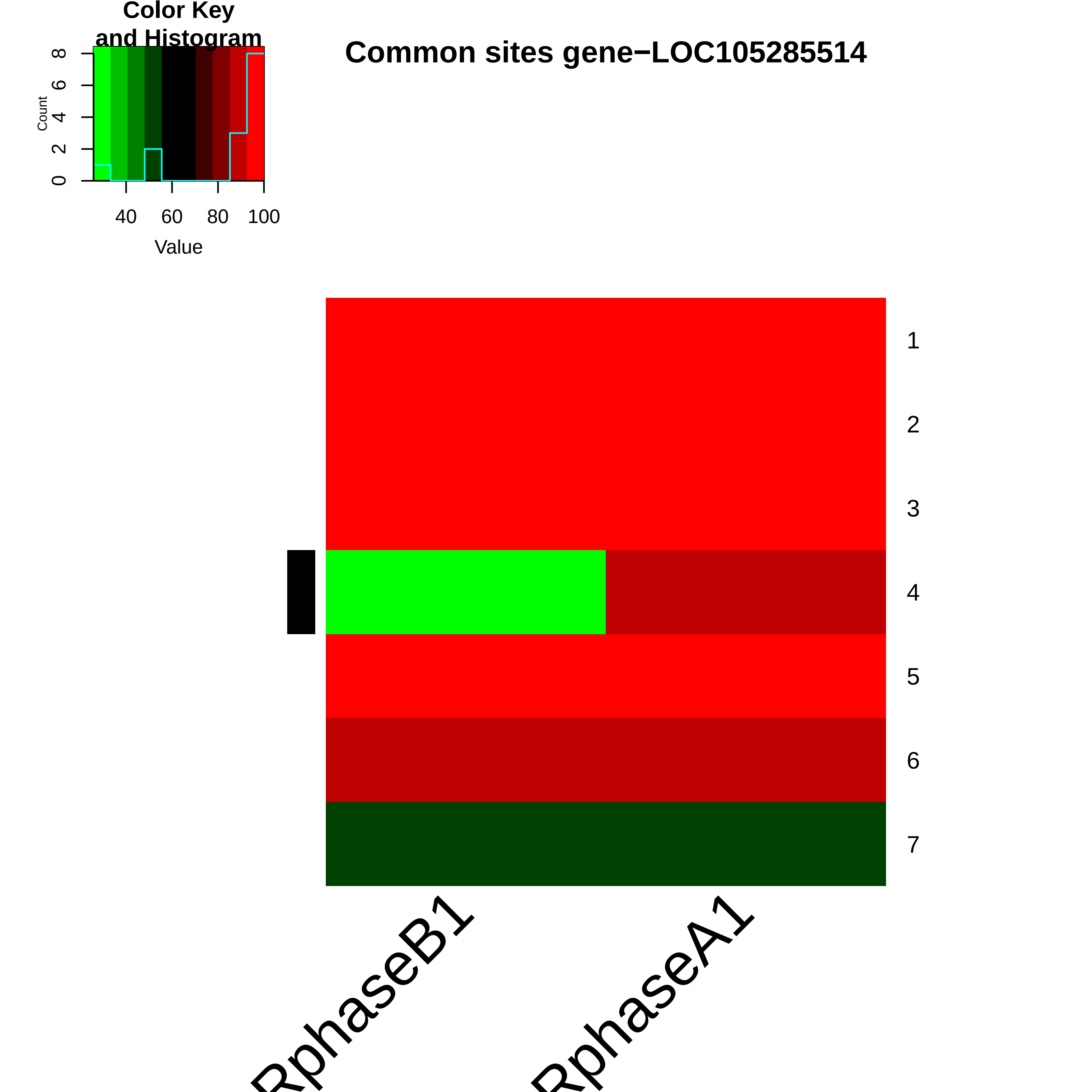

Color Key
and Histogram
Count
Value
0
2
4
6
8
40
60
80
100
Common sites gene−LOC105285514
1
2
3
4
5
6
7
RphaseB1
RphaseA1

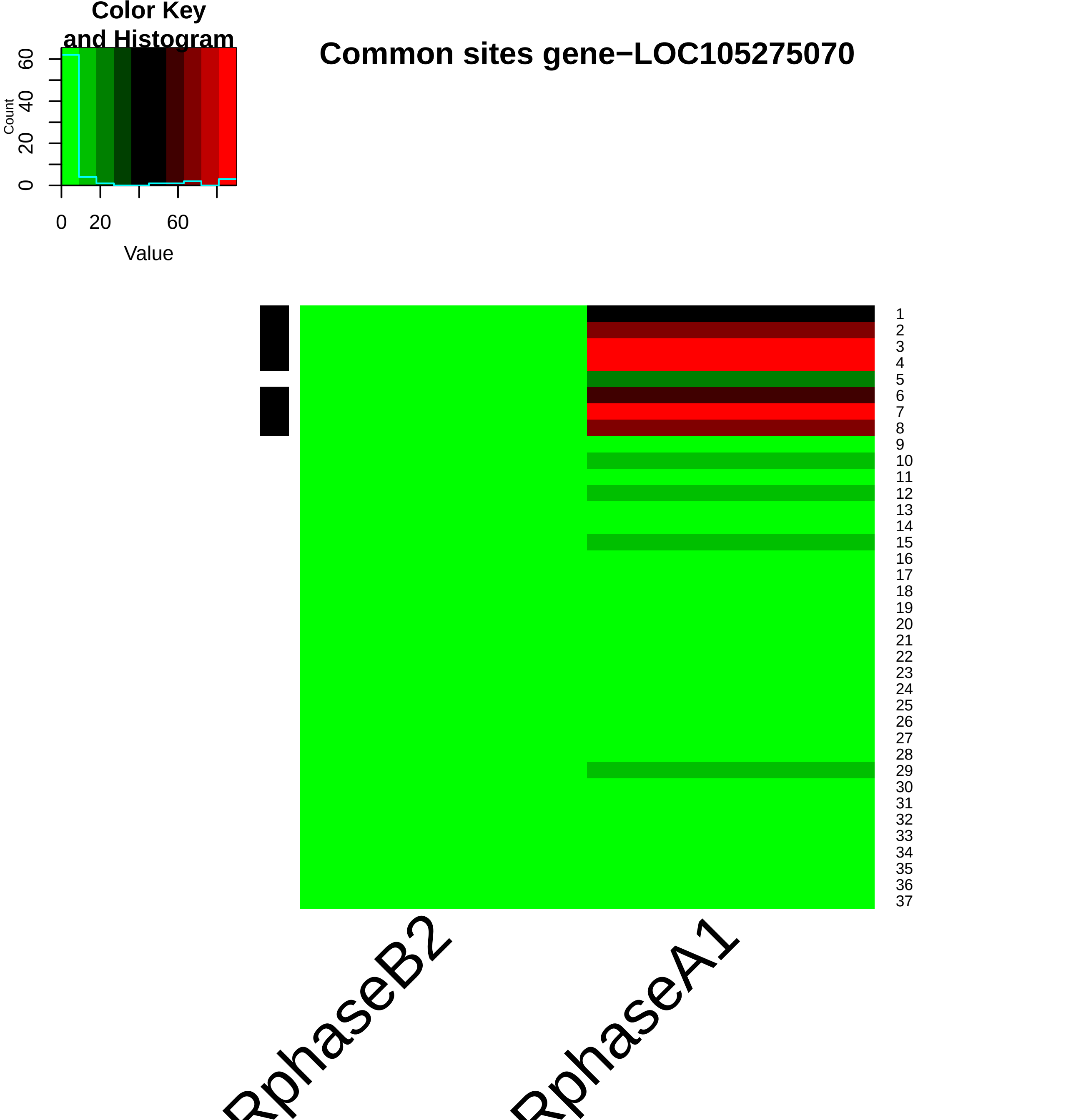

Color Key
and Histogram
Count
0
20
40
60
0
20
60
Value
Common sites gene−LOC105275070
RphaseB2
RphaseA1

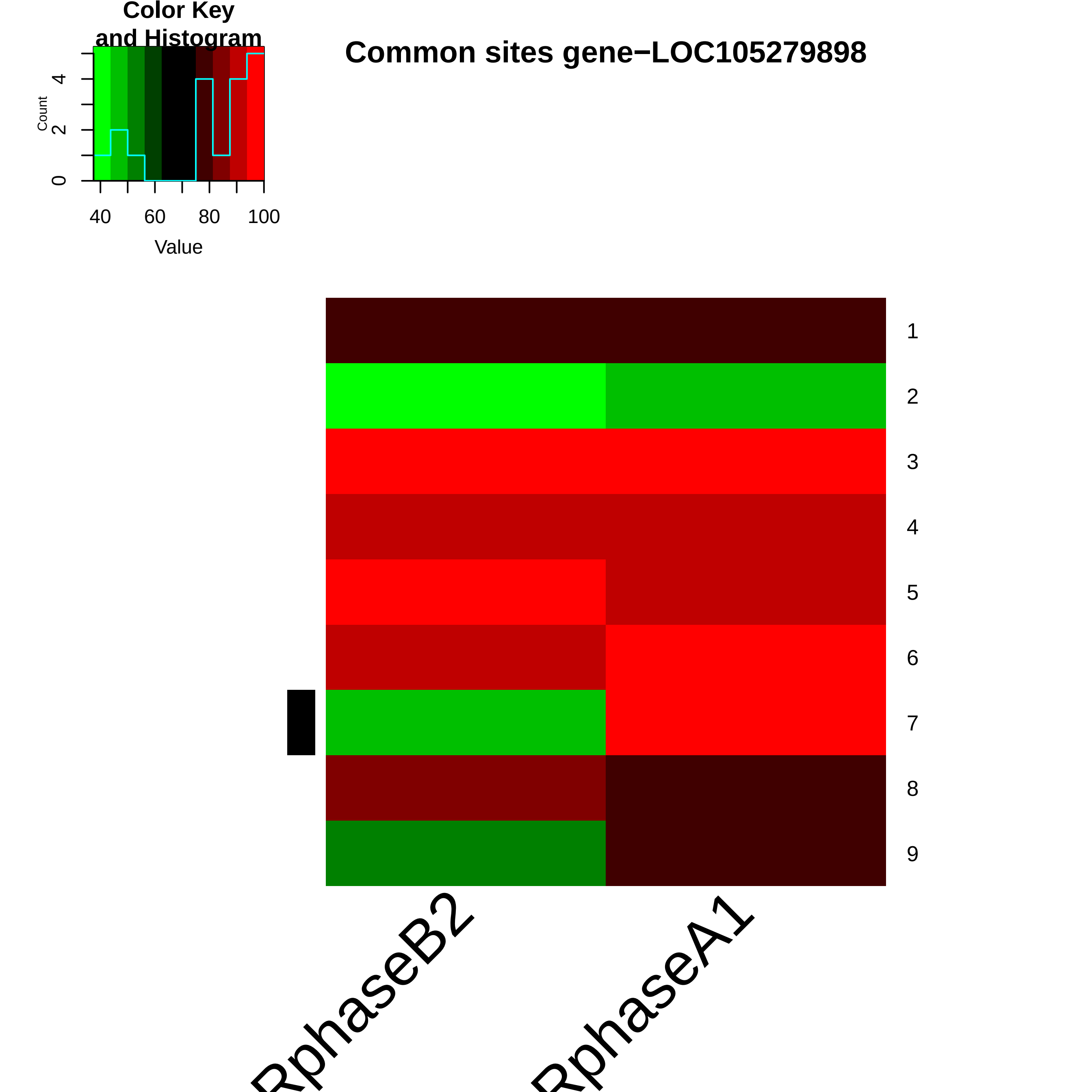
Color Key
and Histogram
Count
0
2
4
40
60
80
100
Value
Common sites gene−LOC105279898
1
2
3
4
5
6
7
8
9
RphaseB2
RphaseA1

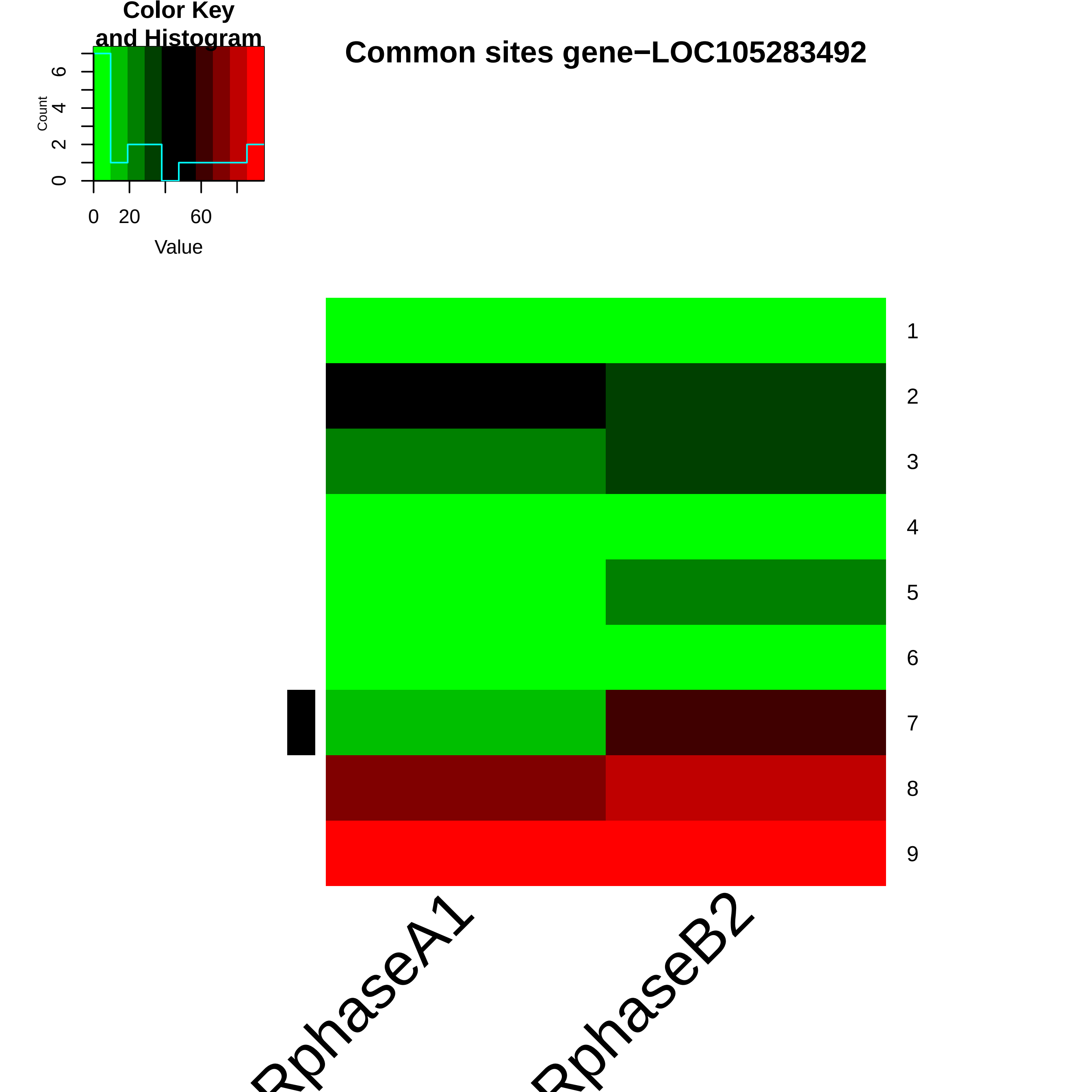
Color Key
and Histogram
Count
0
2
4
6
0
20
60
Value
Common sites gene−LOC105283492
1
2
3
4
5
6
7
8
9
RphaseA1
RphaseB2

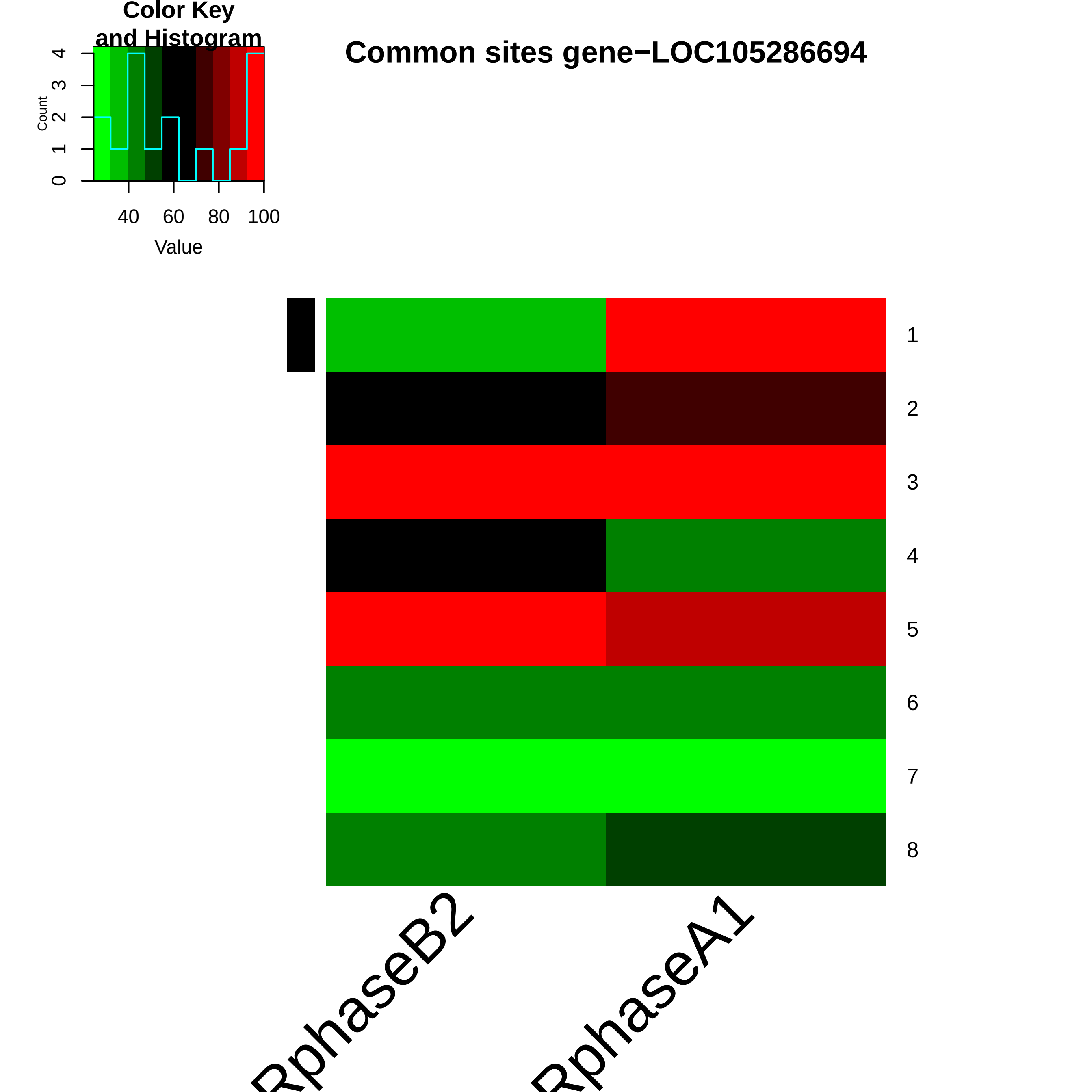

Color Key
and Histogram
Count
0
1
2
3
4
40
60
80
100
Value
Common sites gene−LOC105286694
1
2
3
4
5
6
7
8
RphaseB2
RphaseA1

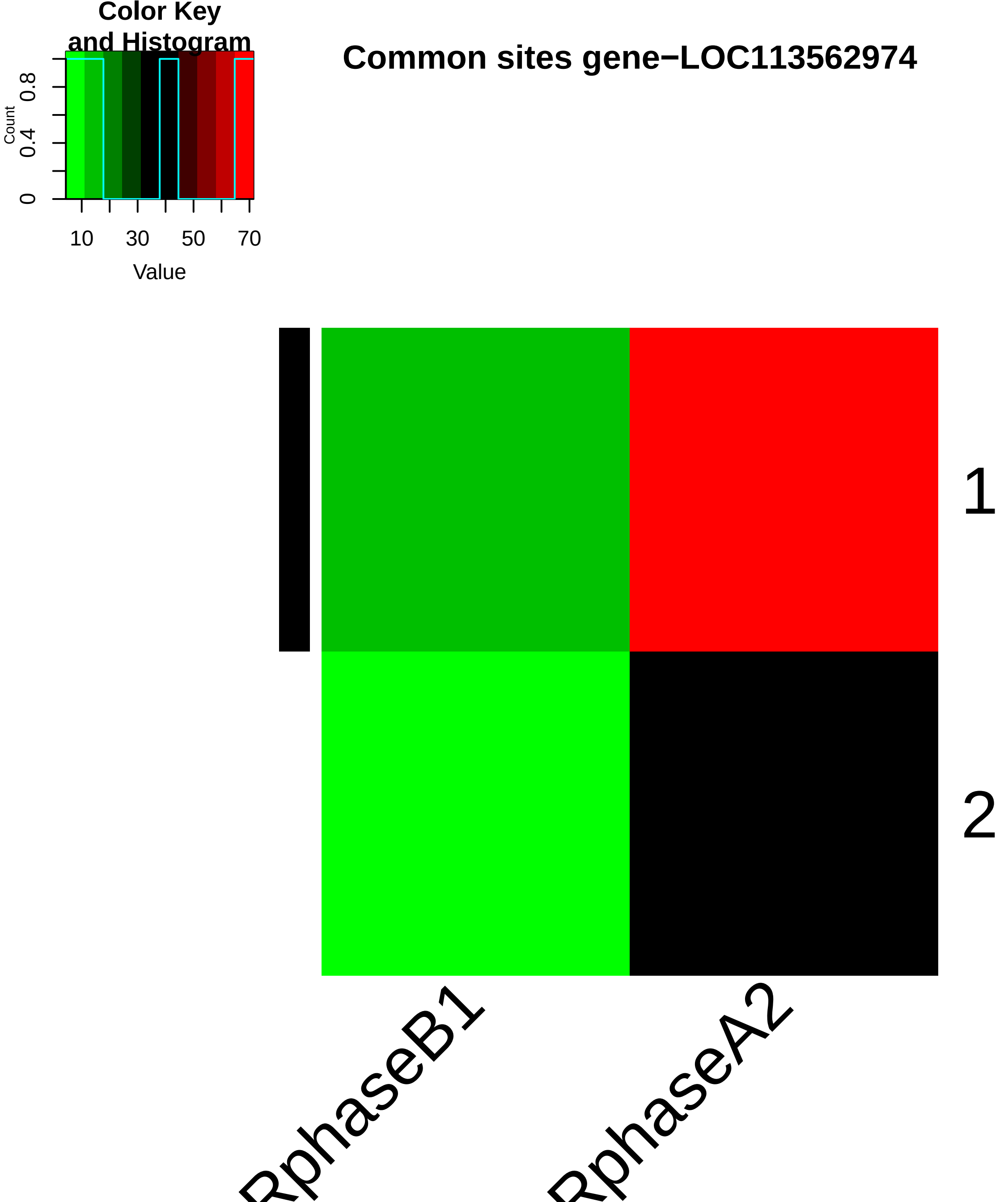

Color Key
and Histogram
Count
0
0.4
0.8
10
30
50
70
Value
Common sites gene−LOC113562974
1
2
RphaseB1
RphaseA2

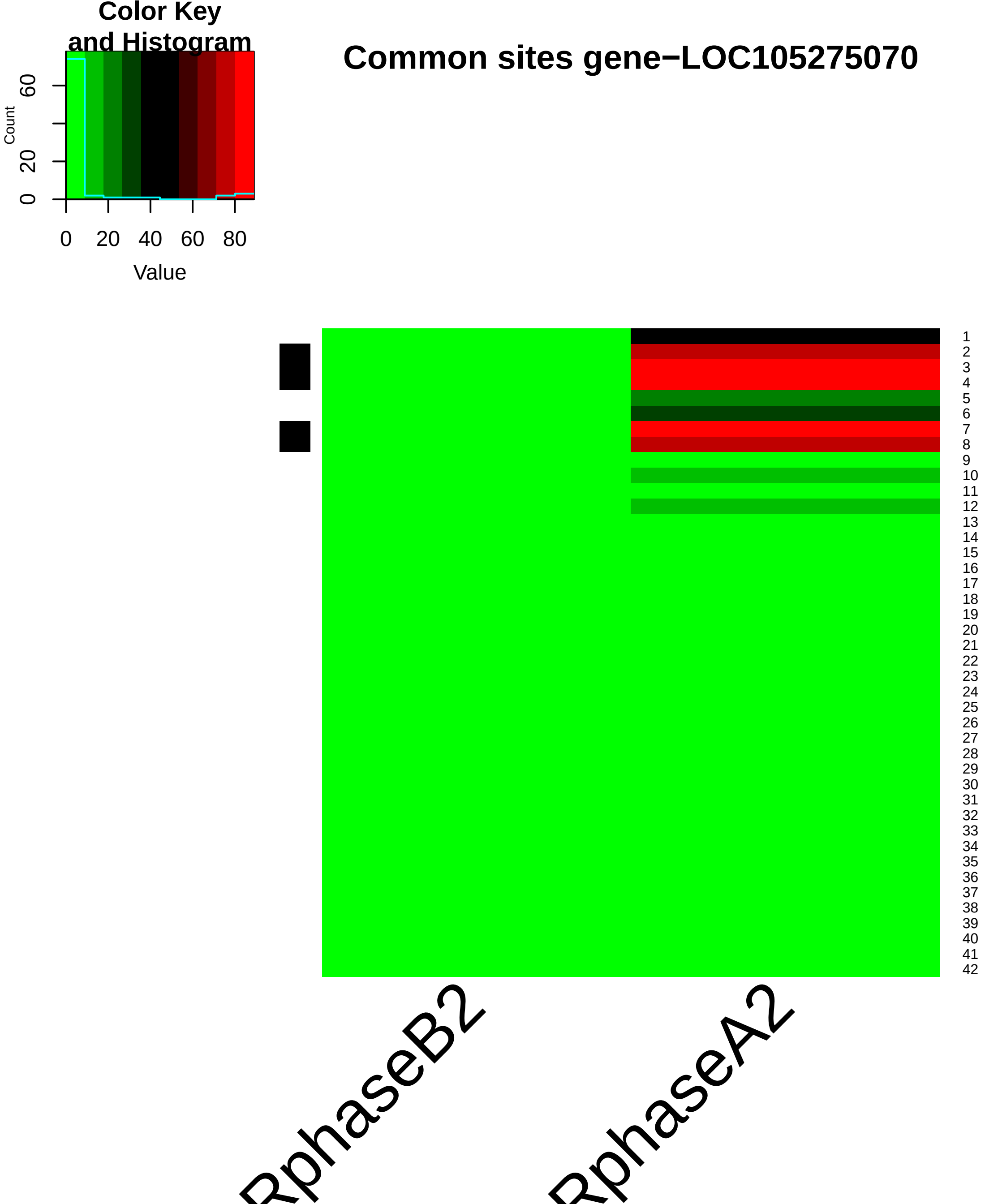
Color Key
and Histogram
Common sites gene−LOC105275070
Count
0
20
60
0
20
40
60
80
Value
RphaseB2
RphaseA2

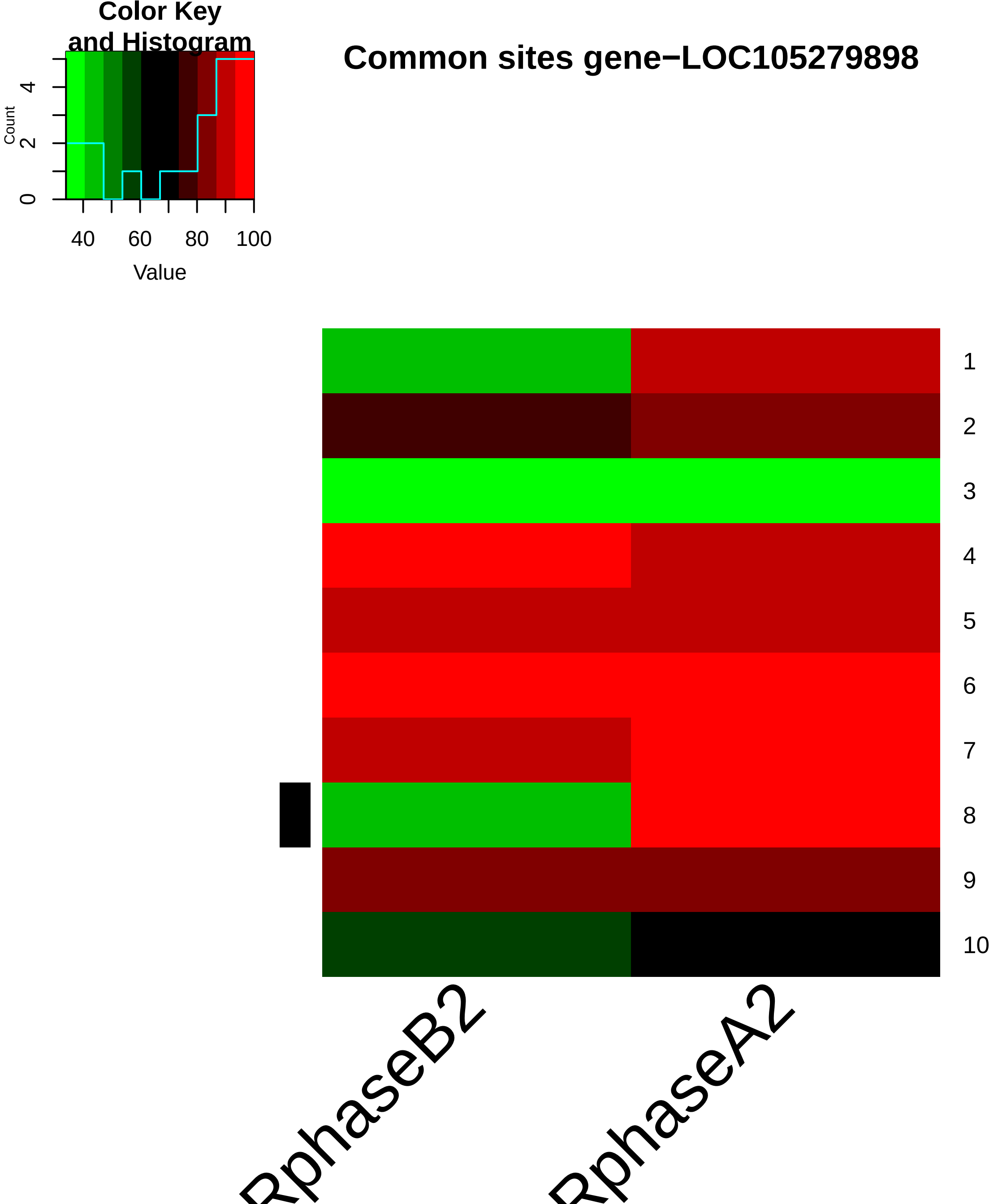
Color Key
and Histogram
Count
0
2
4
40
60
80
100
Value
Common sites gene−LOC105279898
1
2
3
4
5
6
7
8
9
10
RphaseB2
RphaseA2

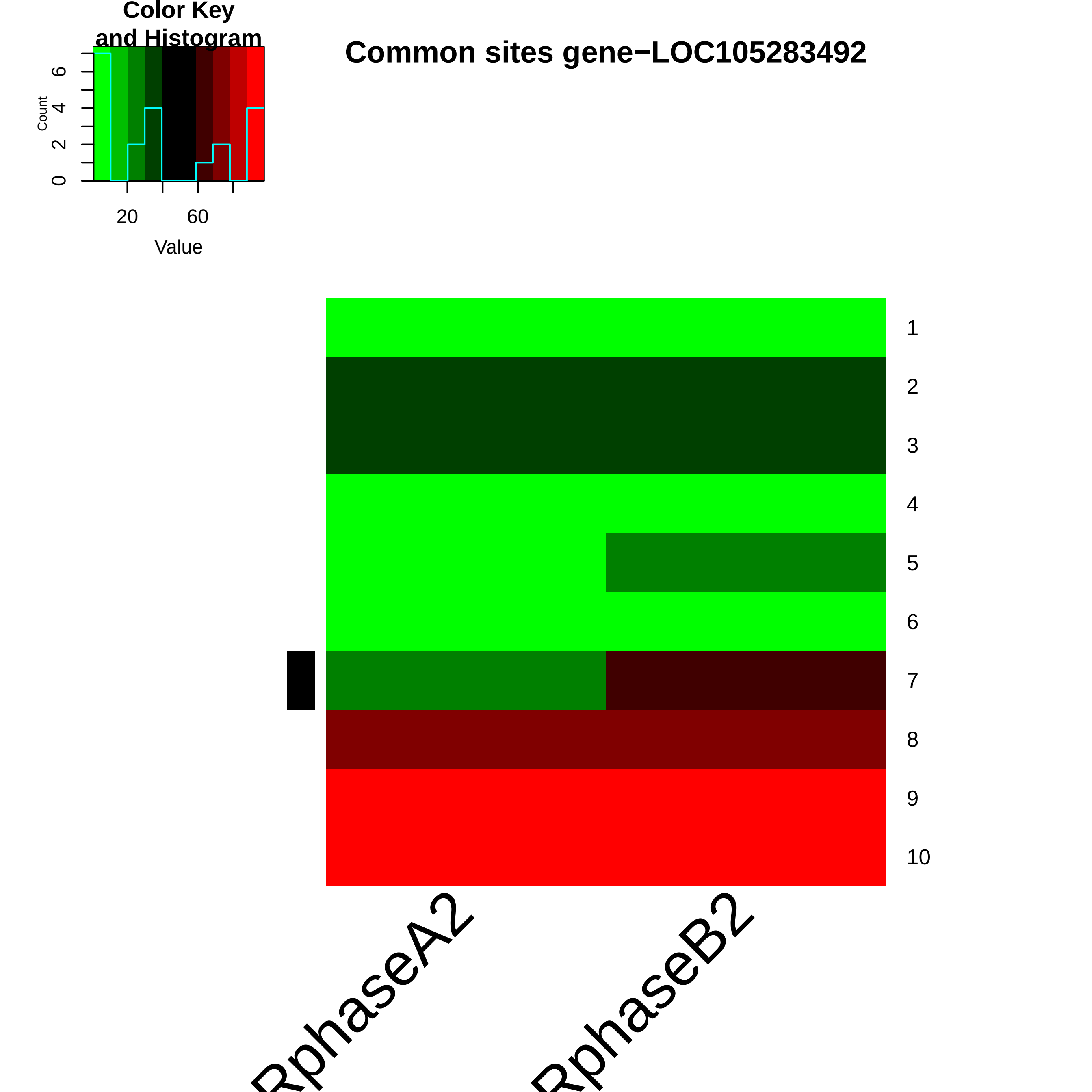
Color Key
and Histogram
Common sites gene−LOC105283492
Count
0
2
4
6
20
60
Value
1
2
3
4
5
6
7
8
9
10
RphaseA2
RphaseB2

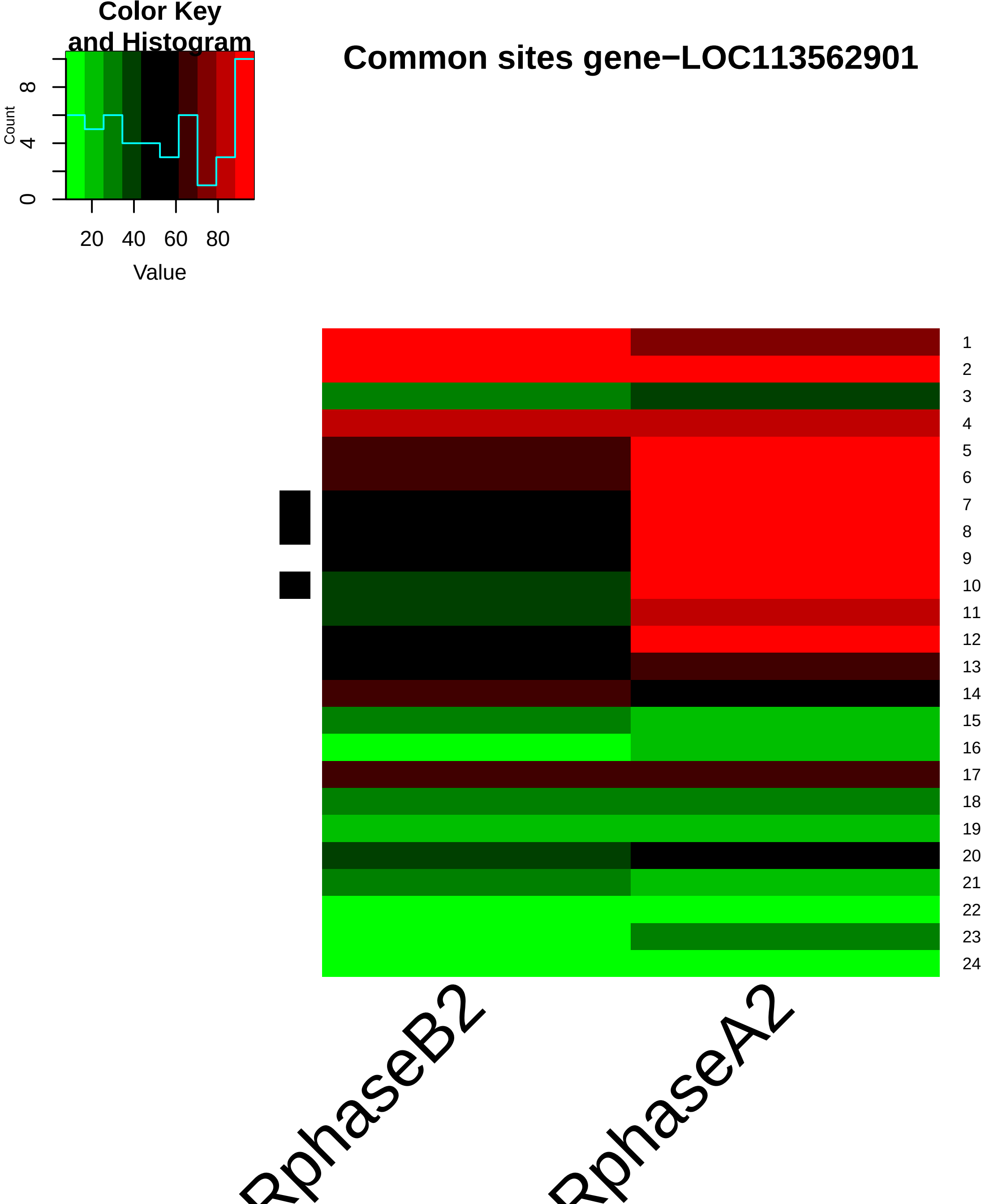
Color Key
and Histogram
Count
0
4
8
20
40
60
80
Value
Common sites gene−LOC113562901
RphaseB2
RphaseA2

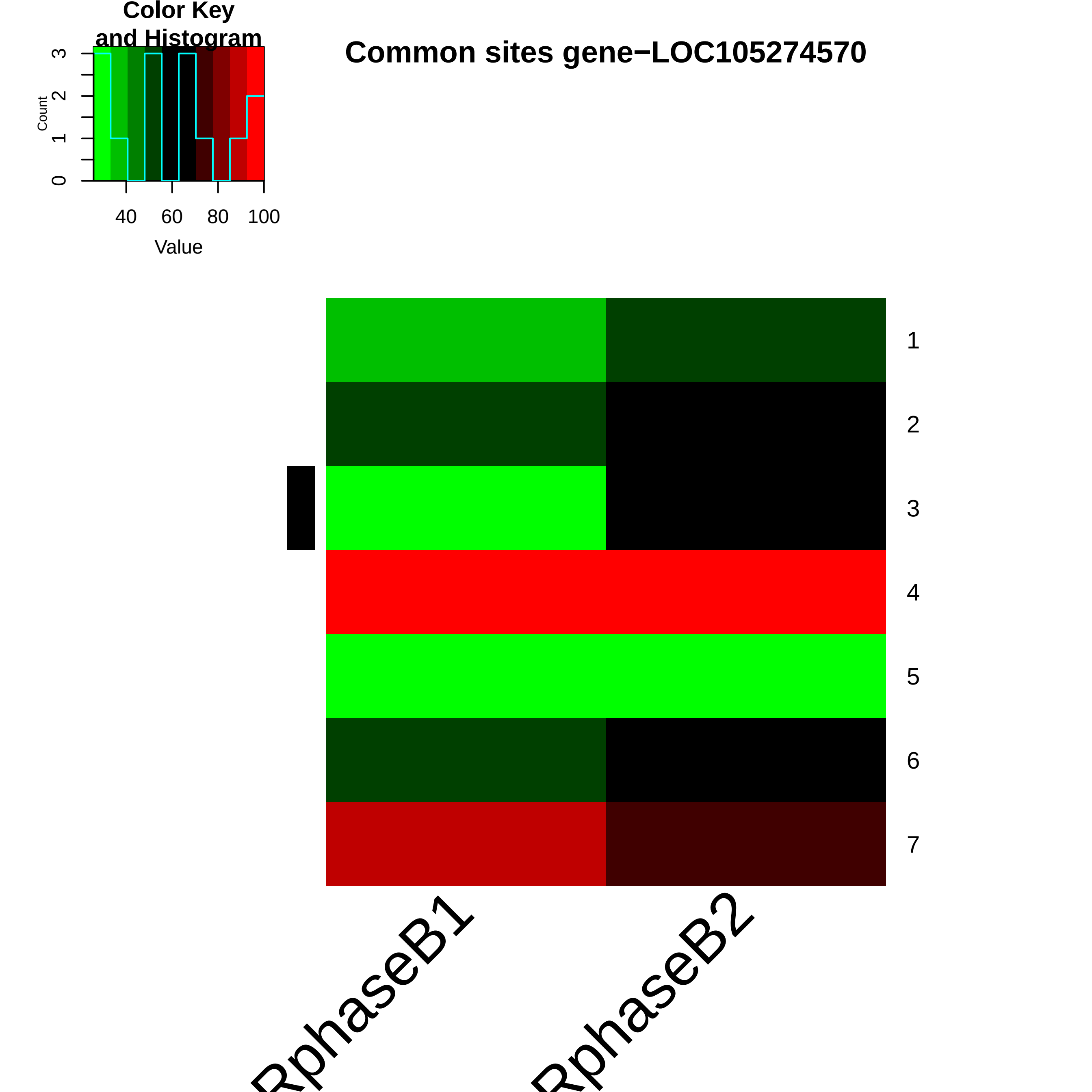

Color Key
and Histogram
Count
0
1
2
3
40
60
80
100
Value
Common sites gene−LOC105274570
1
2
3
4
5
6
7
RphaseB1
RphaseB2

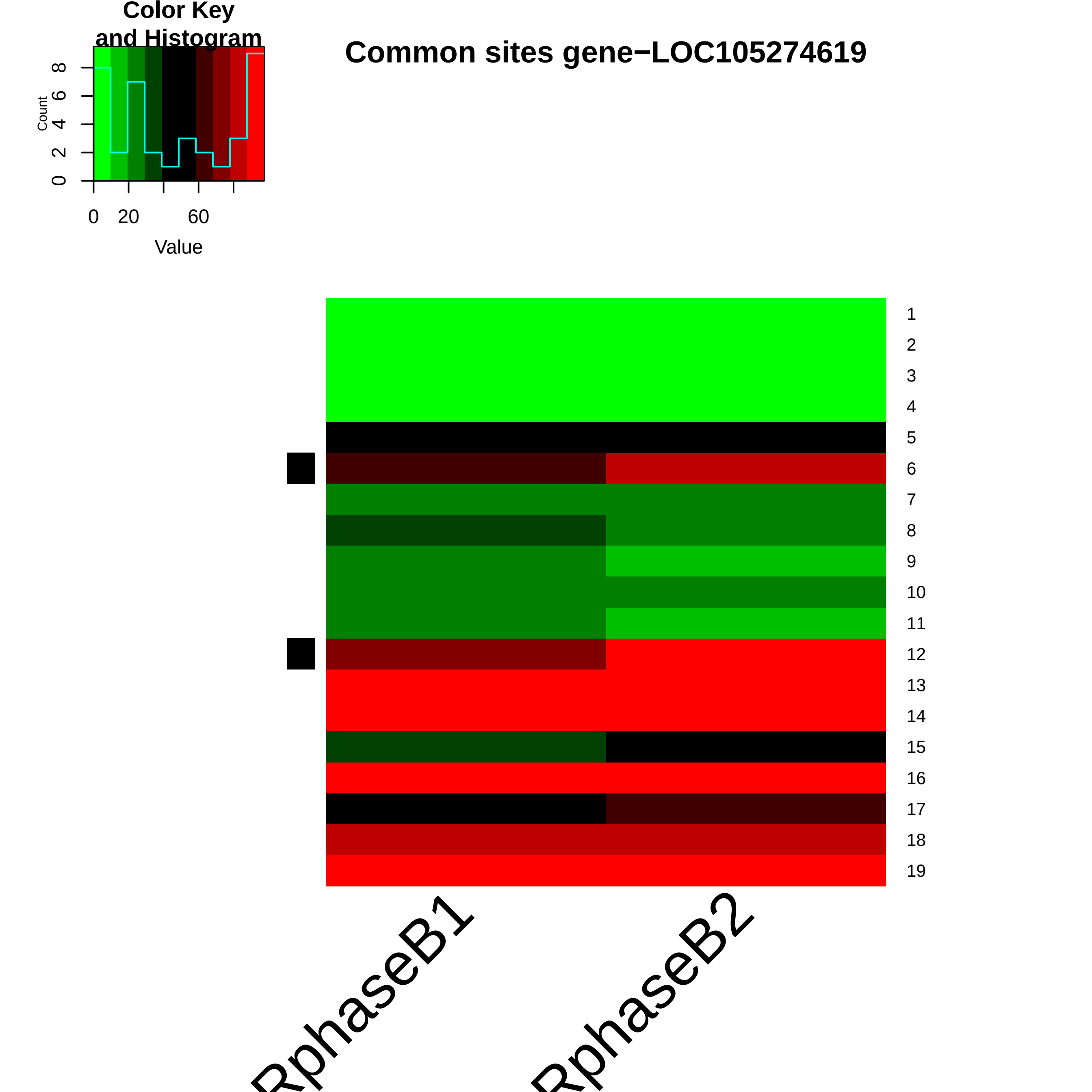

Color Key
and Histogram
Count
0 2 4 6 8
0 20 60
Value
Common sites gene−LOC105274619
RphaseB1
RphaseB2

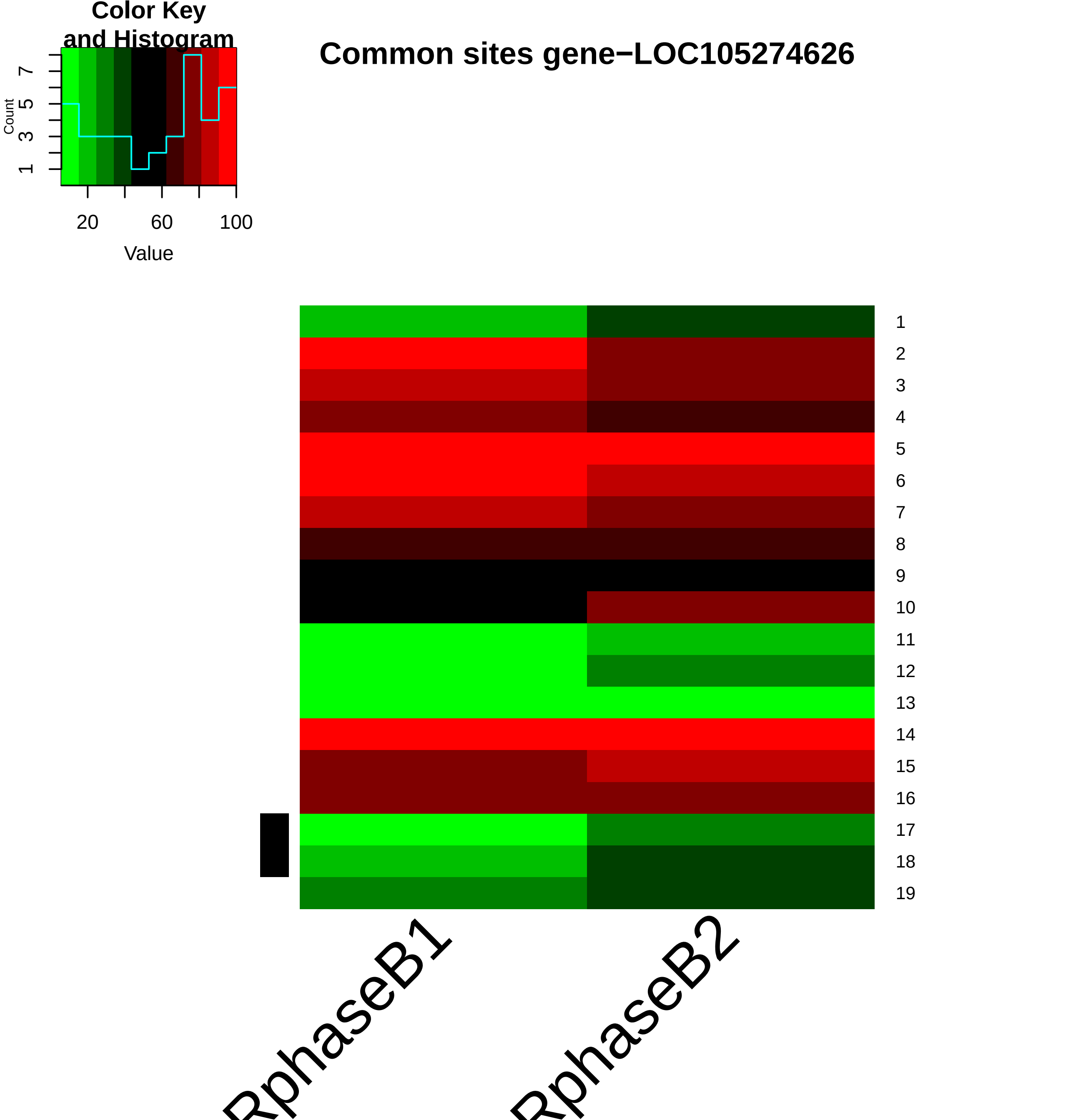
Color Key
and Histogram
Count
1
3
5
7
20
60
100
Value
Common sites gene−LOC105274626
RphaseB1
RphaseB2

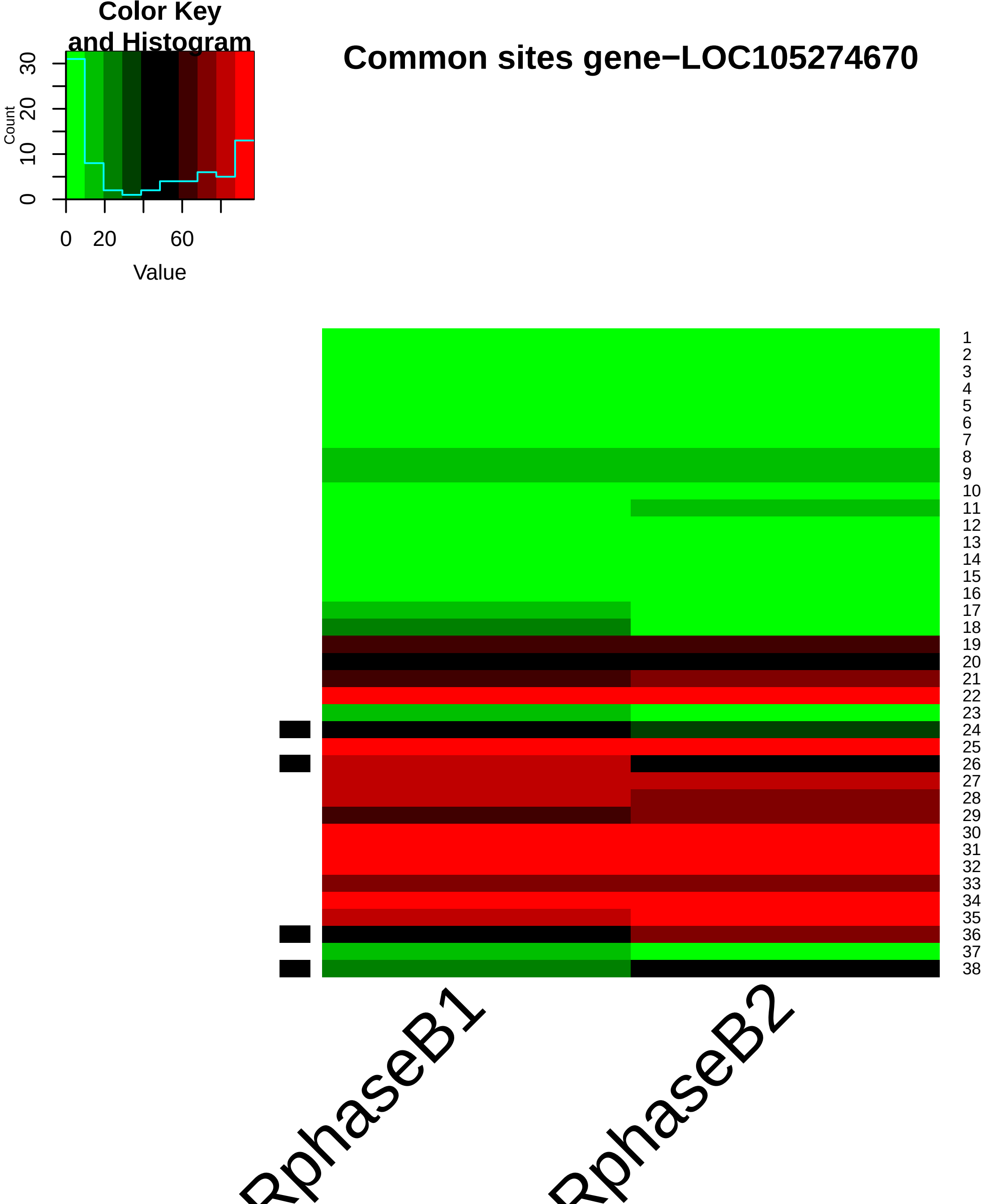
Color Key
and Histogram
Common sites gene−LOC105274670
Count
Value
0
10
20
30
0
20
60
RphaseB1
RphaseB2

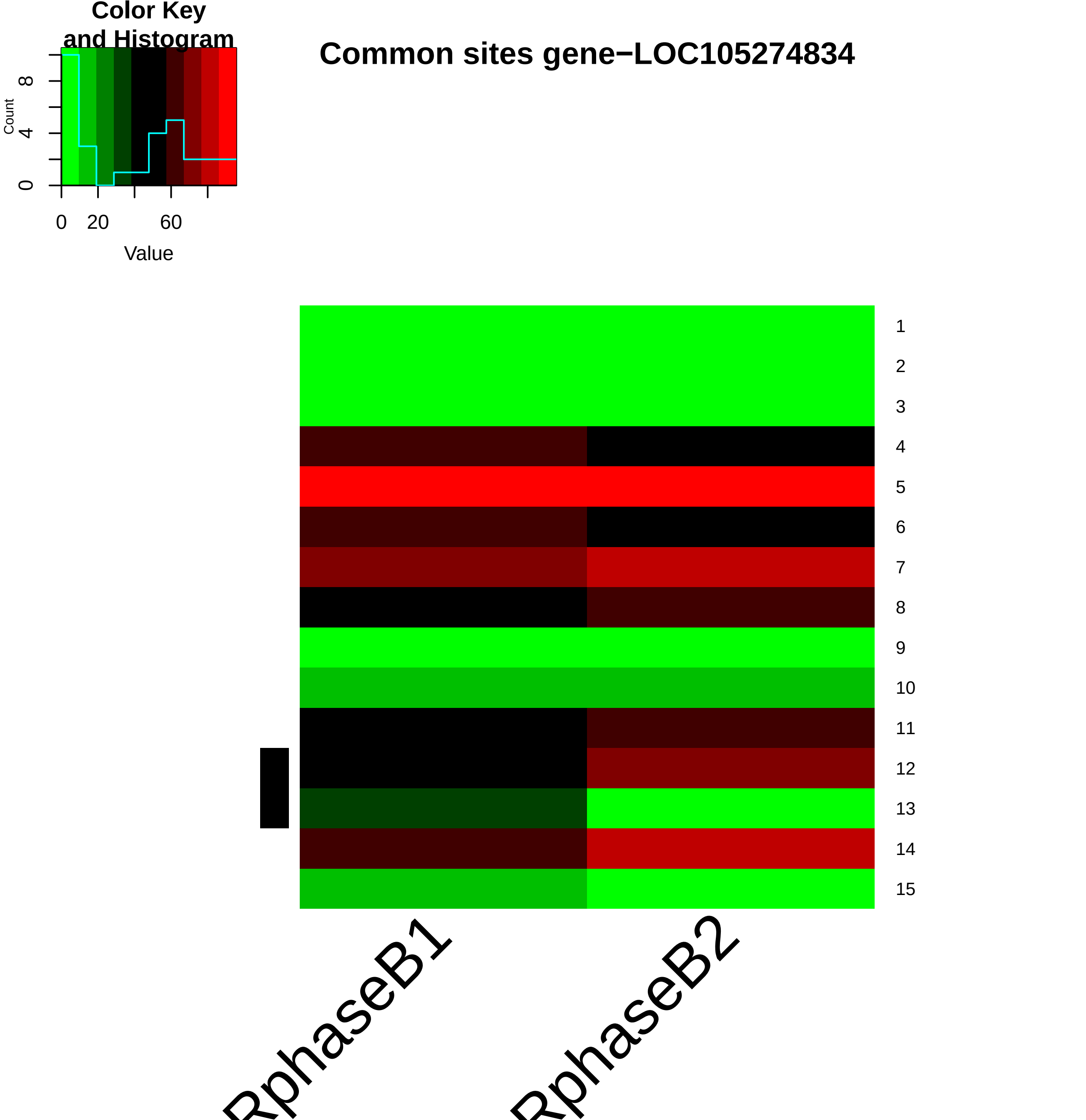

Color Key
and Histogram
Count
0
4
8
0
20
60
Value
Common sites gene−LOC105274834
RphaseB1
RphaseB2

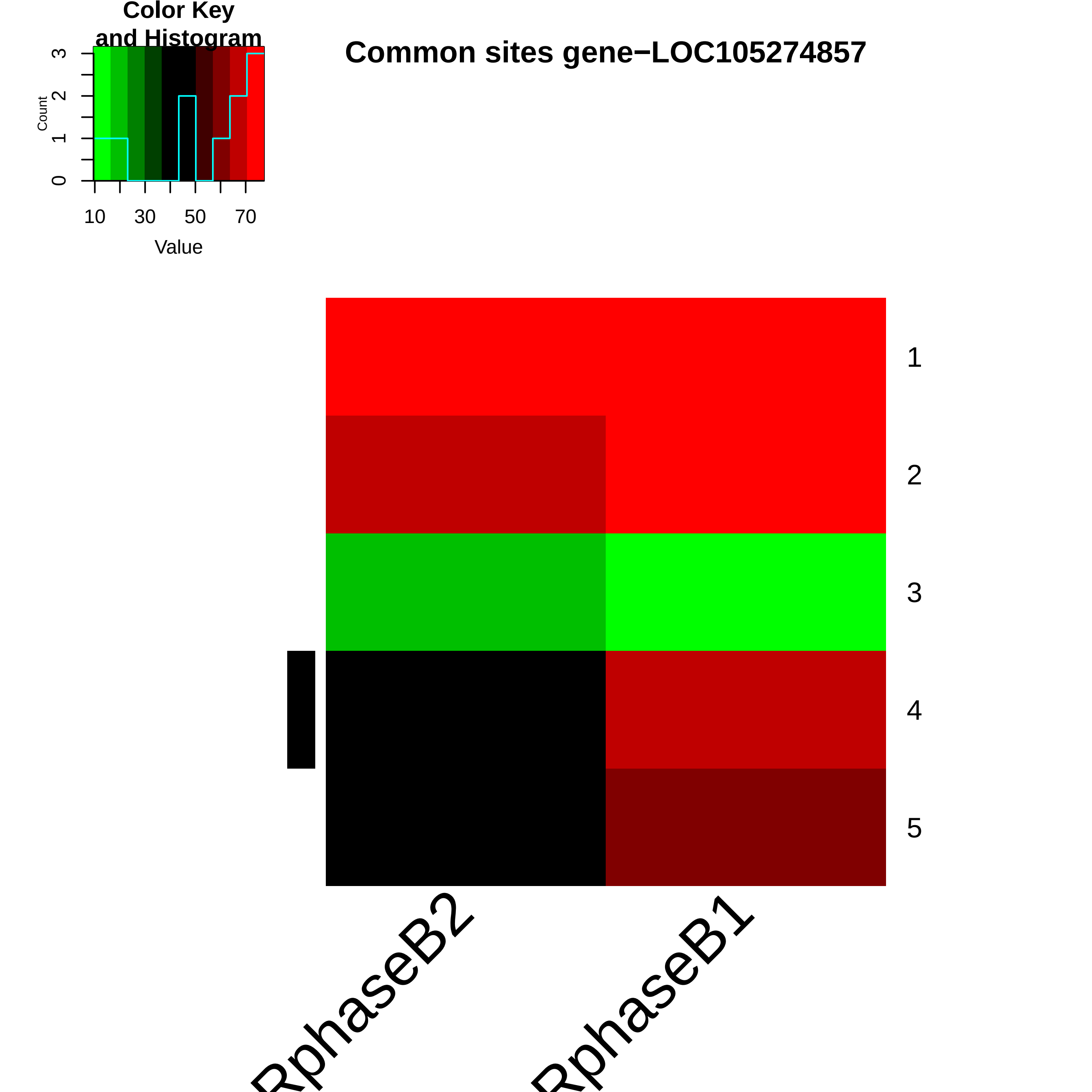

Color Key
and Histogram
Count
0
1
2
3
10
30
50
70
Value
Common sites gene−LOC105274857
1
2
3
4
5
RphaseB2
RphaseB1

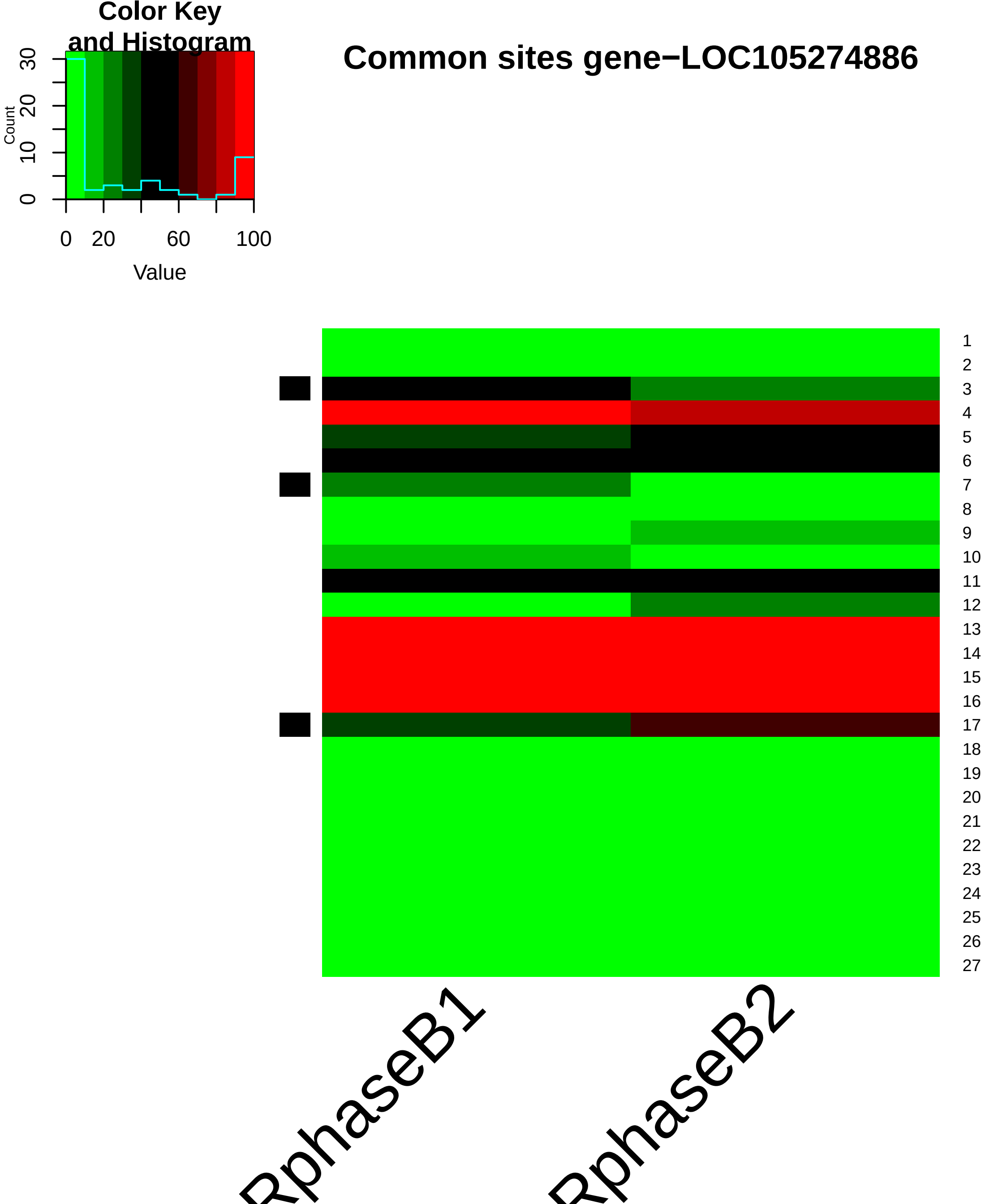
Color Key
and Histogram
Count
0
10
20
30
0
20
60
100
Value
Common sites gene−LOC105274886
RphaseB1
RphaseB2

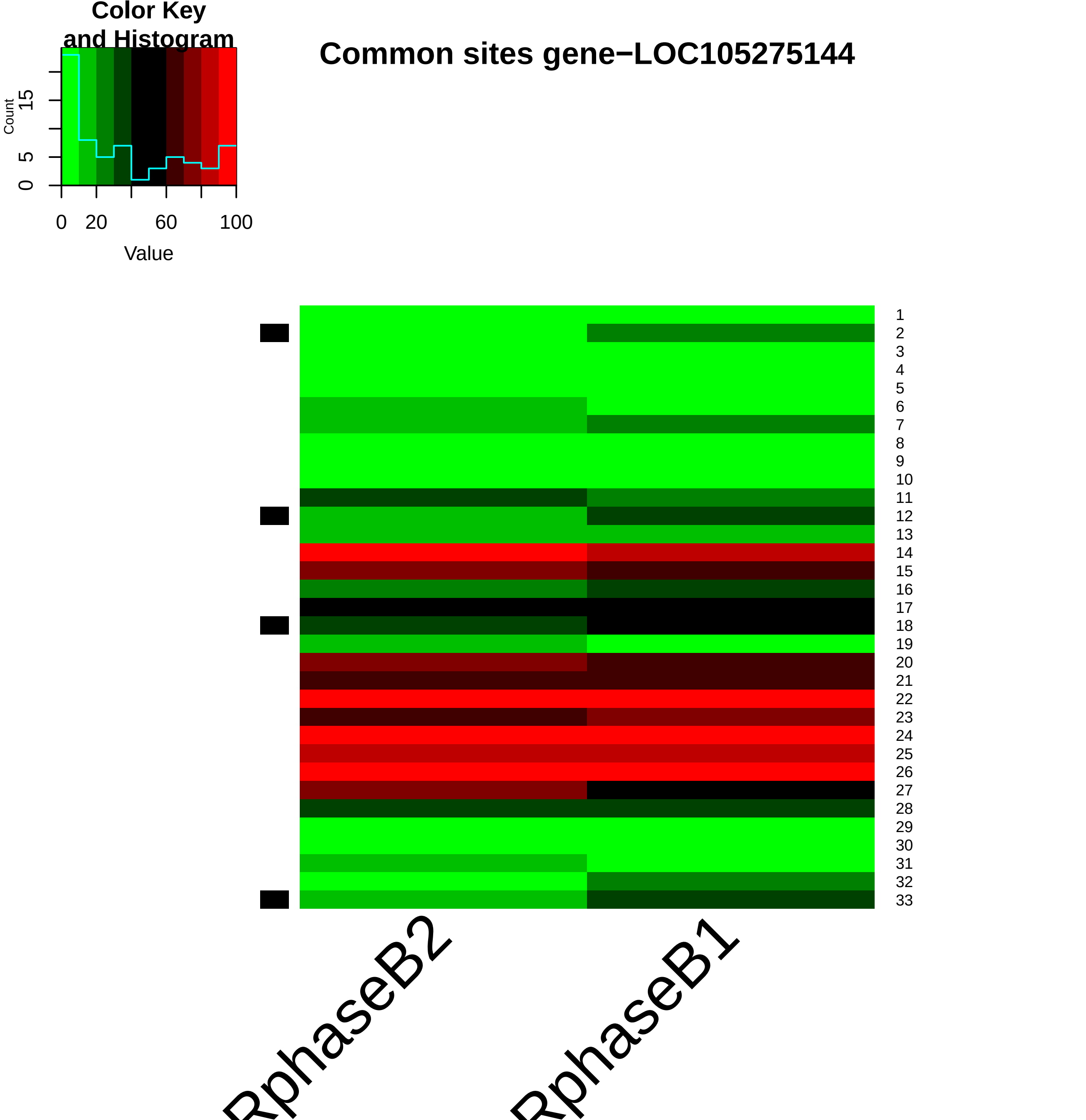

Color Key
and Histogram
Count
0
5
15
0
20
60
100
Value
Common sites gene−LOC105275144
RphaseB2
RphaseB1

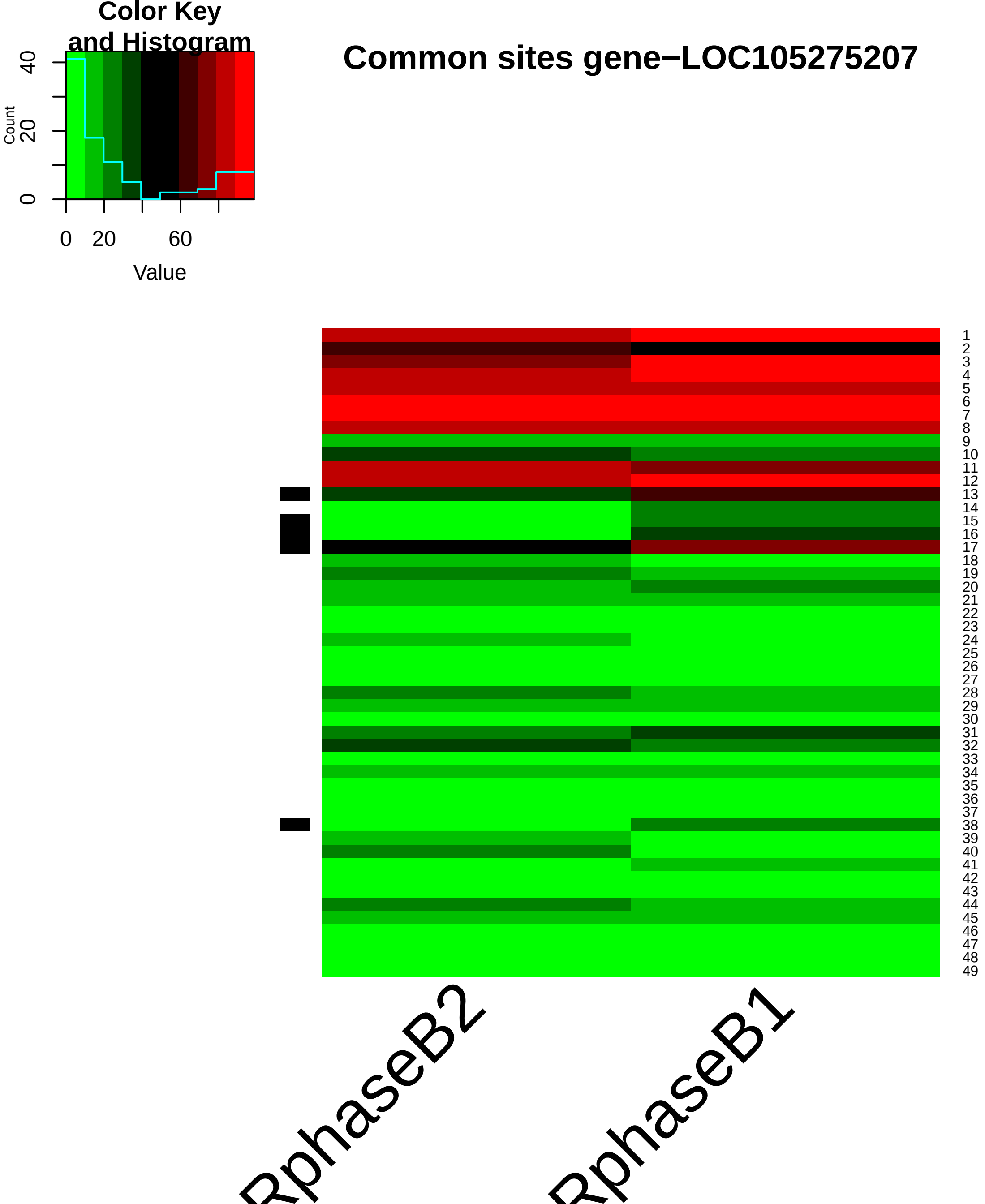
Color Key
and Histogram
Count
0
20
40
0
20
60
Value
Common sites gene−LOC105275207
RphaseB2
RphaseB1

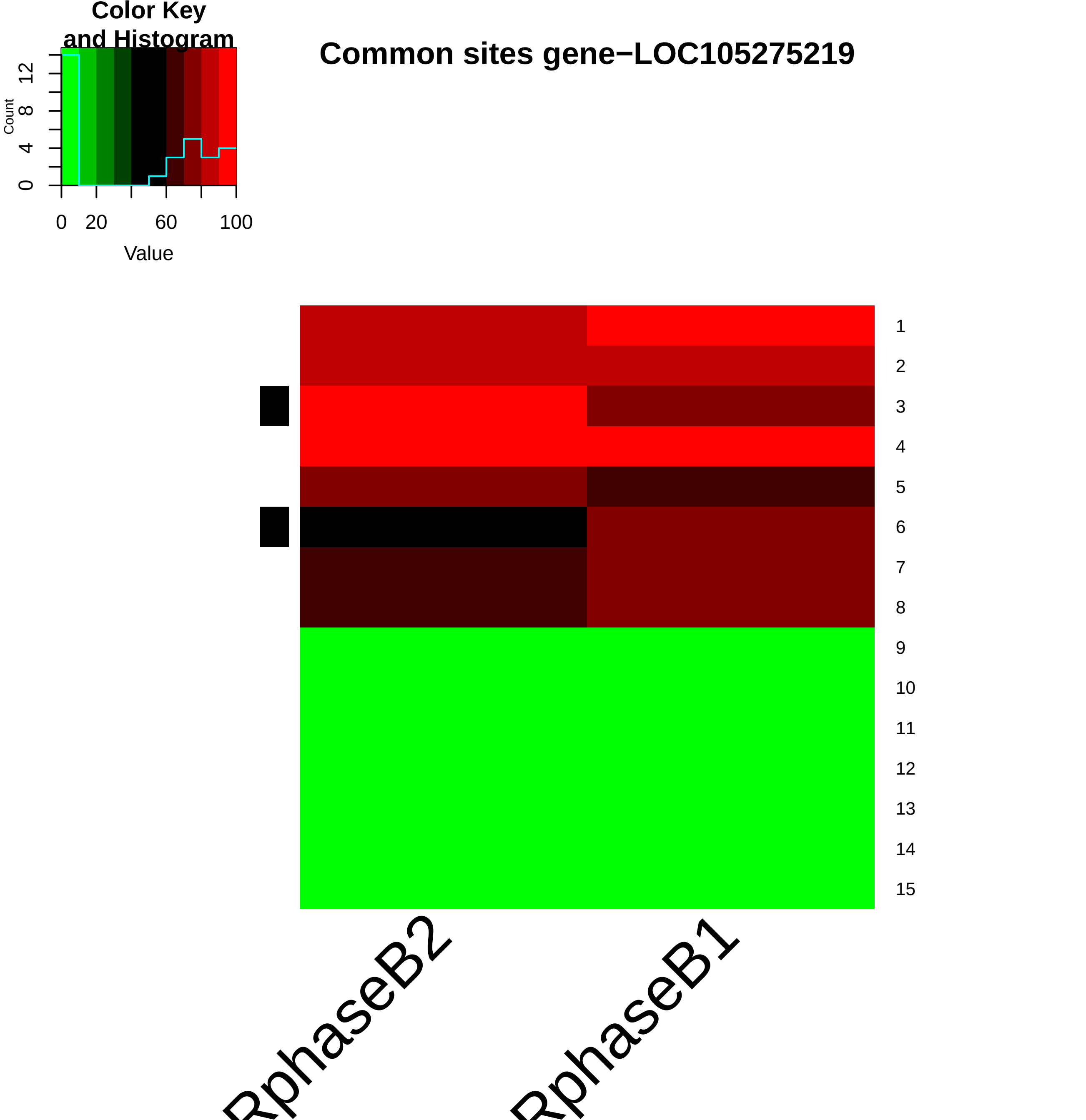

Color Key
and Histogram
Count
0
4
8
12
0
20
60
100
Value
Common sites gene−LOC105275219
RphaseB2
RphaseB1

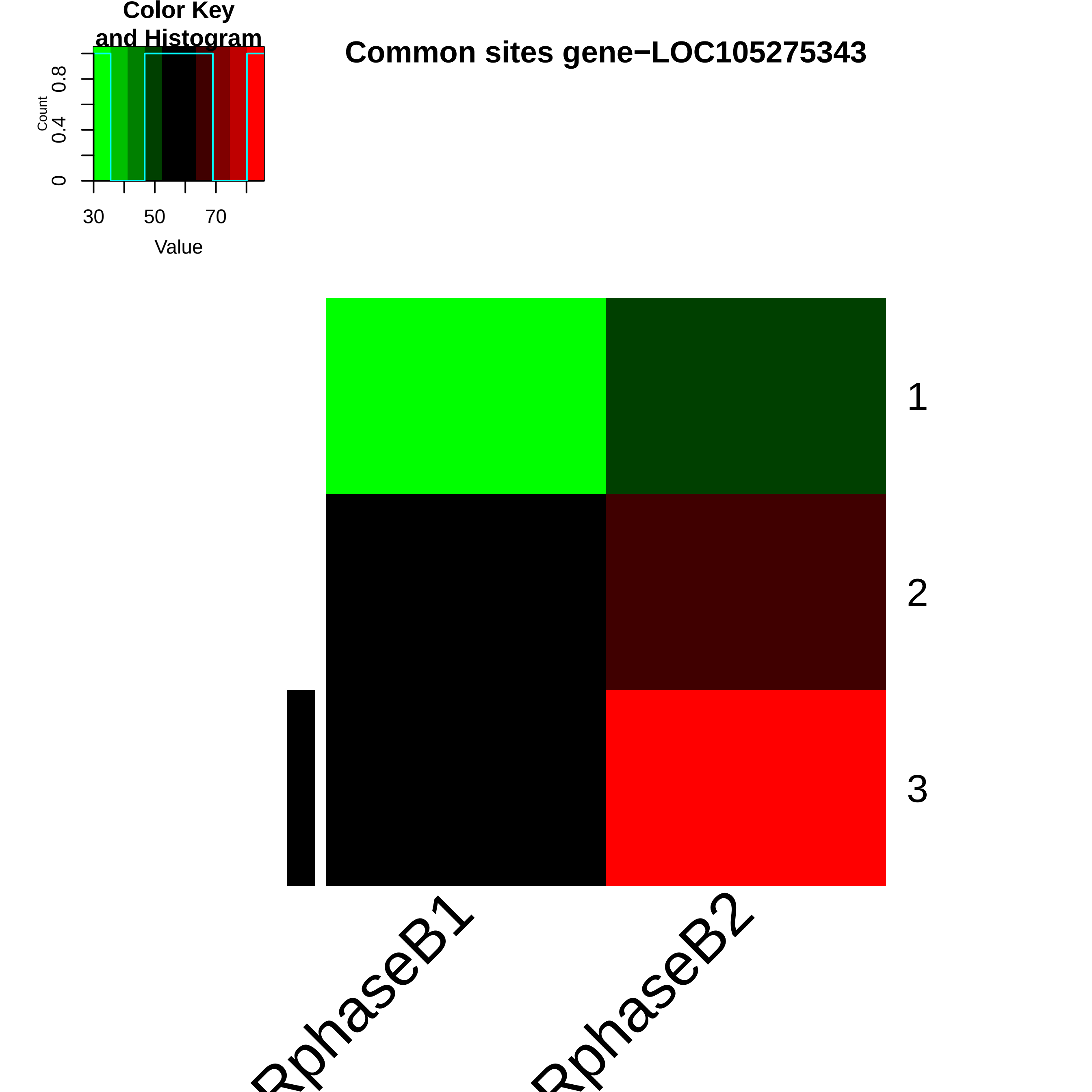

Color Key
and Histogram
Count
0
0.4
0.8
30
50
70
Value
Common sites gene−LOC105275343
1
2
3
RphaseB1
RphaseB2

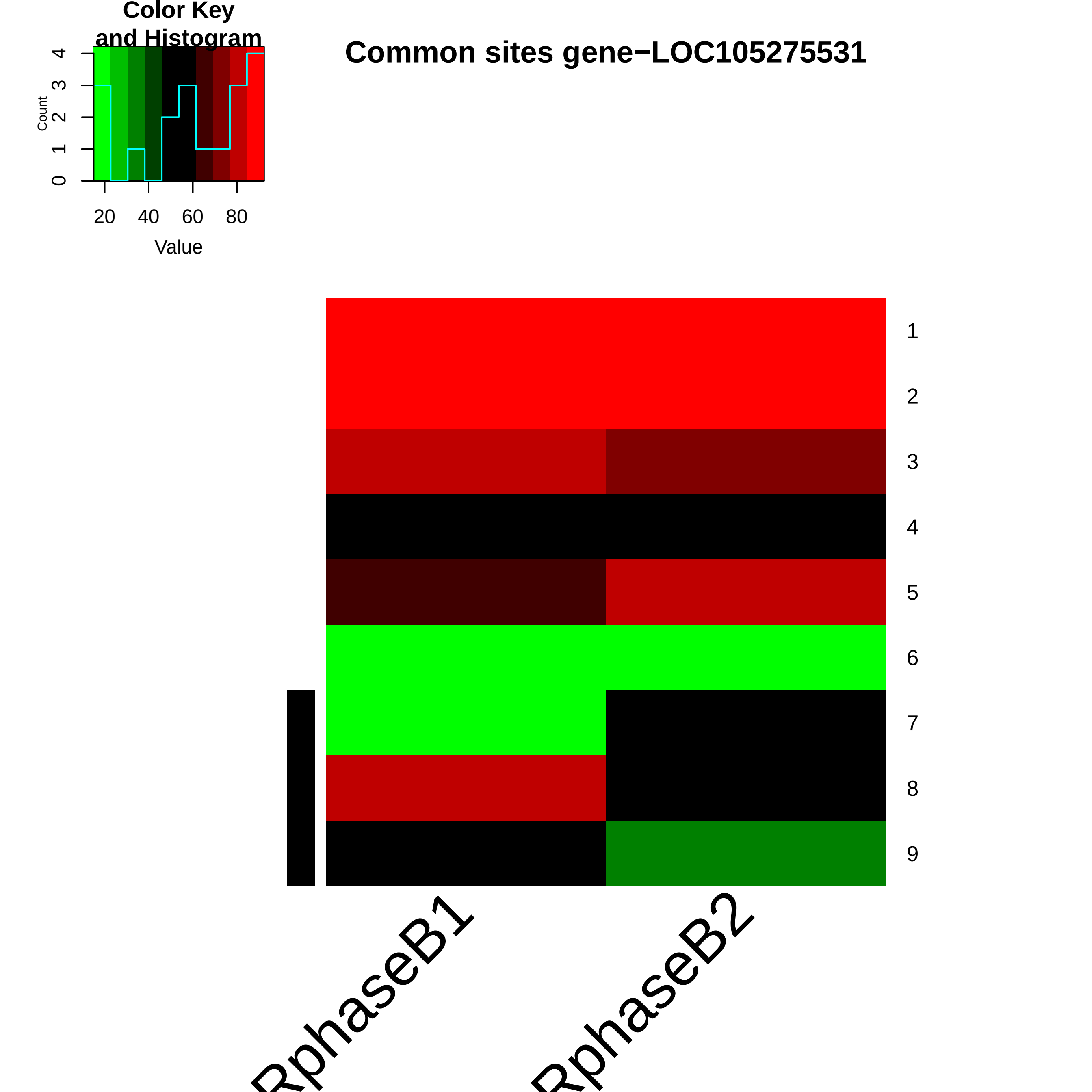

Color Key
and Histogram
Count
Value
0
1
2
3
4
20
40
60
80
Common sites gene−LOC105275531
1
2
3
4
5
6
7
8
9
RphaseB1
RphaseB2

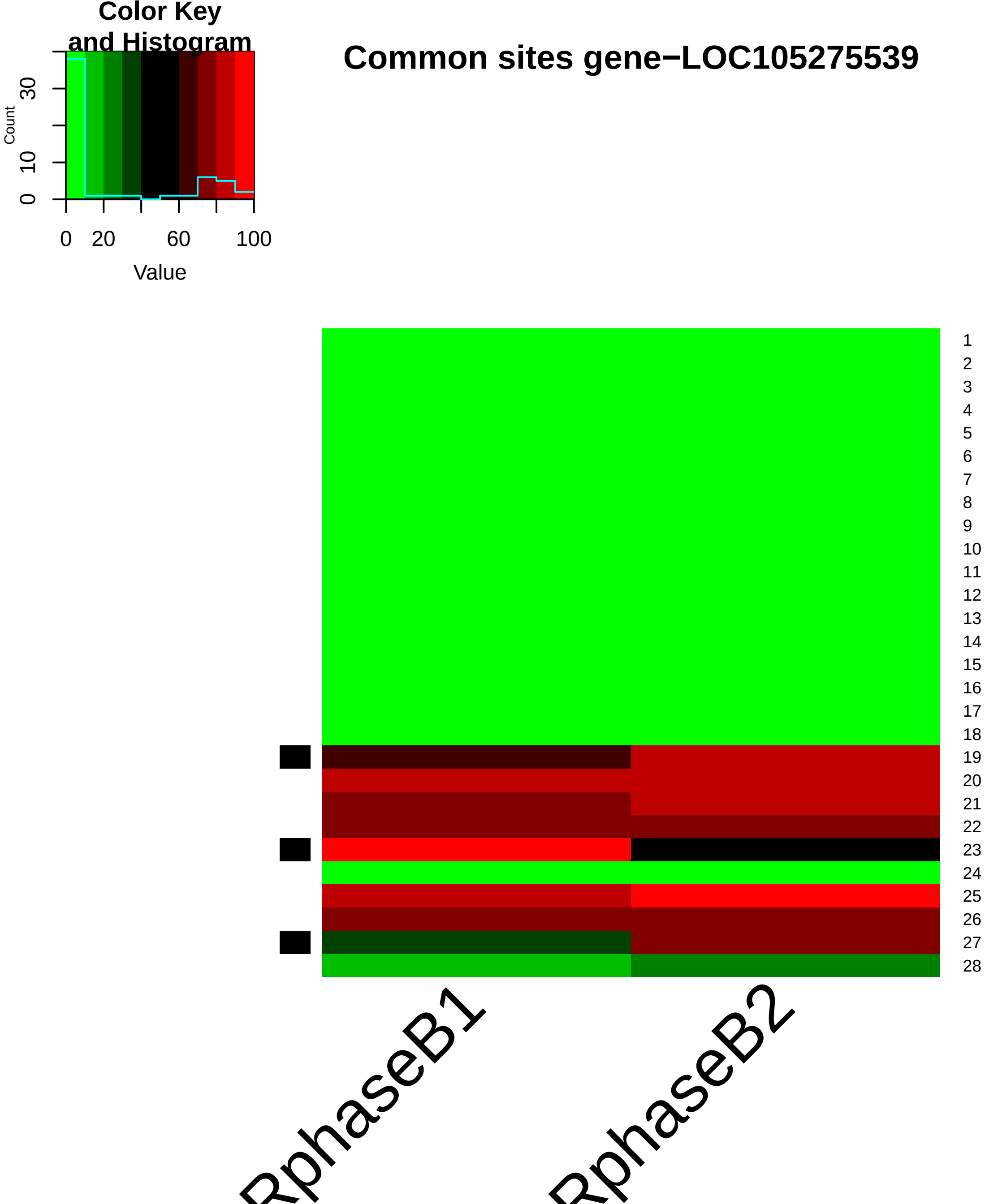

Color Key and Histogram
Count
0
10
30
0
20
60
100
Value
Common sites gene−LOC105275539
RphaseB1
RphaseB2

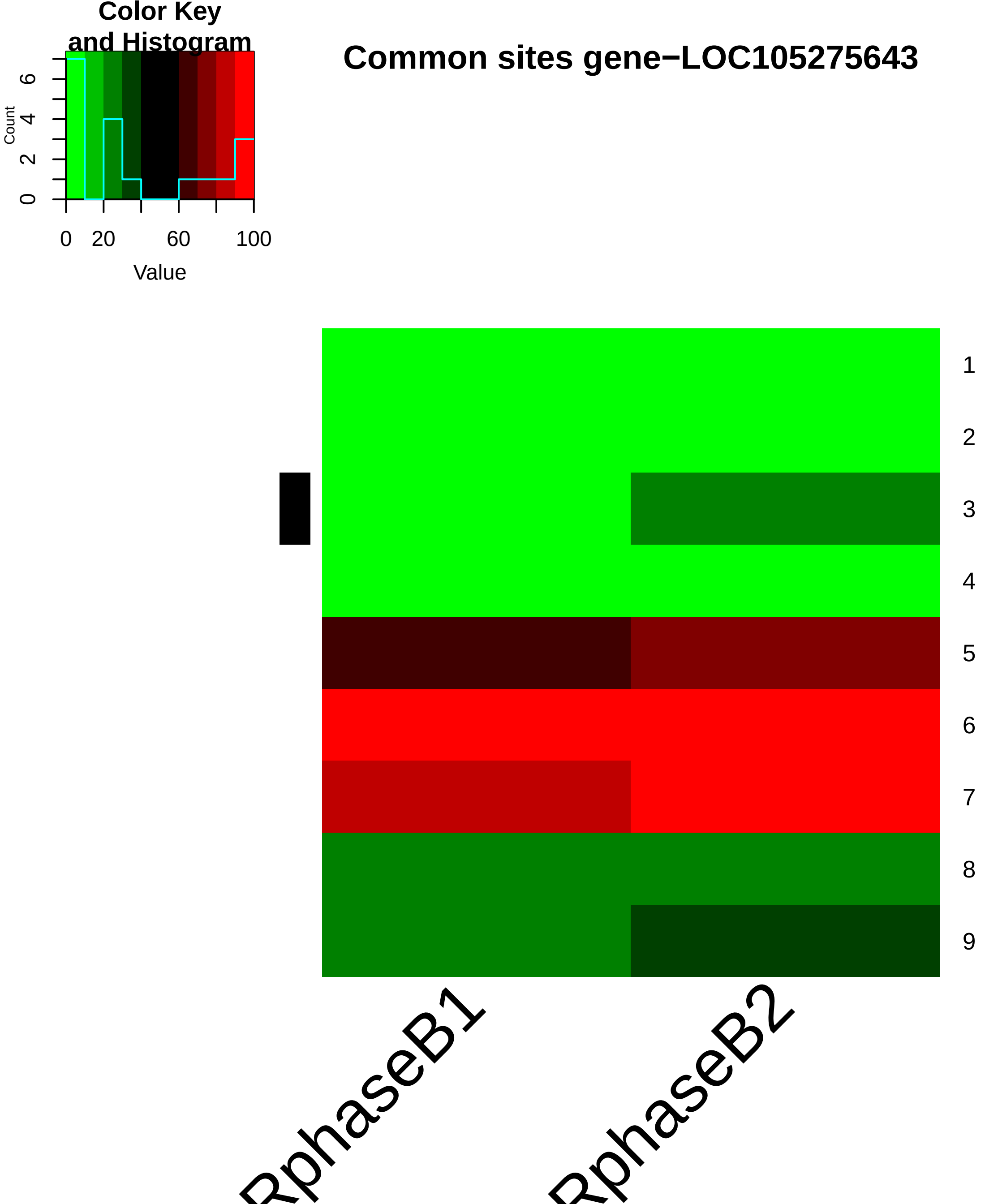
Color Key
and Histogram
Count
0
2
4
6
0
20
60
100
Value
Common sites gene−LOC105275643
1
2
3
4
5
6
7
8
9
RphaseB1
RphaseB2

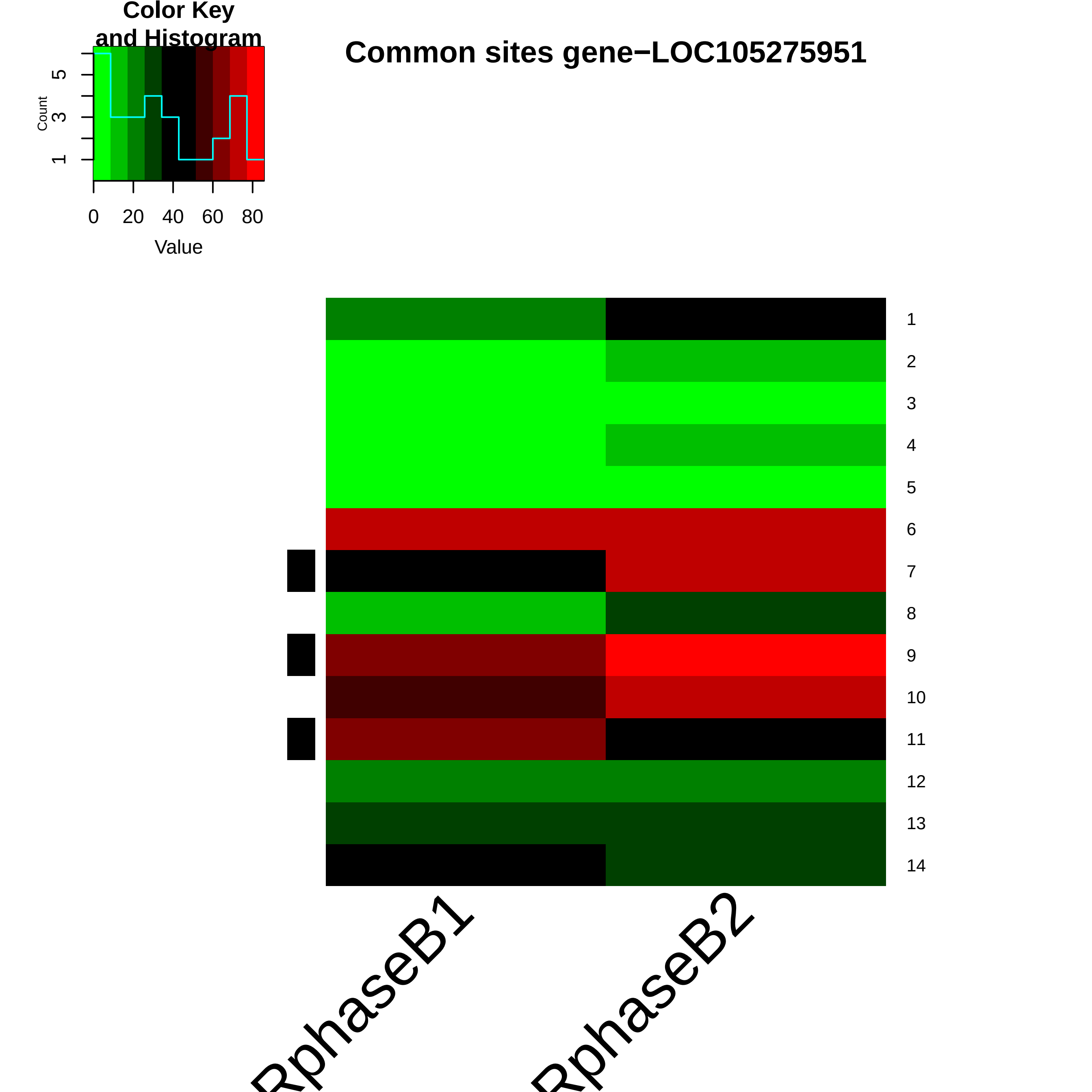

Color Key
and Histogram
Count
1
3
5
0
20
40
60
80
Value
Common sites gene−LOC105275951
1
2
3
4
5
6
7
8
9
10
11
12
13
14
RphaseB1
RphaseB2

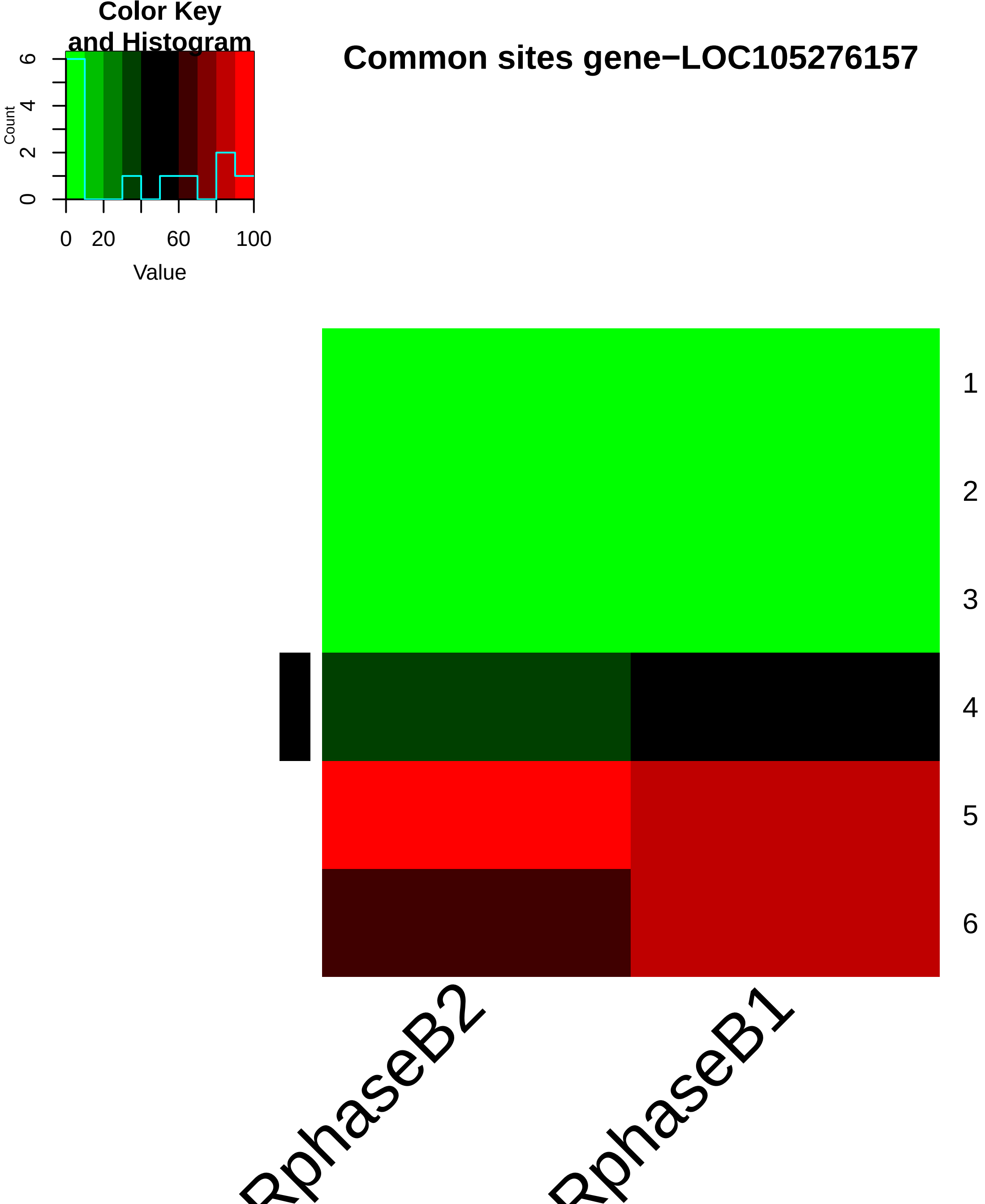
Color Key
and Histogram
Count
0
2
4
6
0
20
60
100
Value
Common sites gene−LOC105276157
1
2
3
4
5
6
RphaseB2
RphaseB1

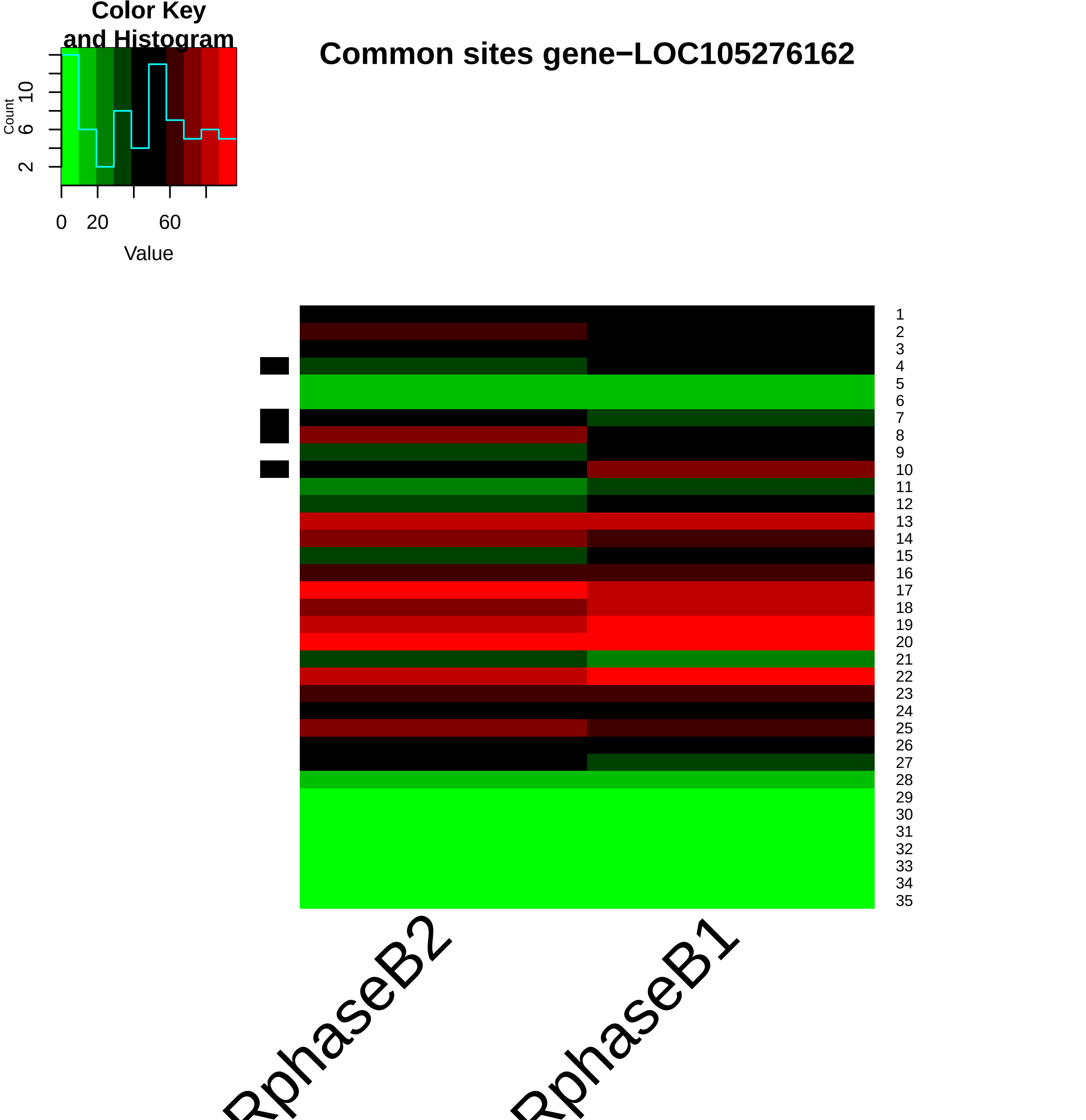

Color Key
and Histogram
Count
2
6
10
0
20
60
Value
Common sites gene−LOC105276162
RphaseB2
RphaseB1

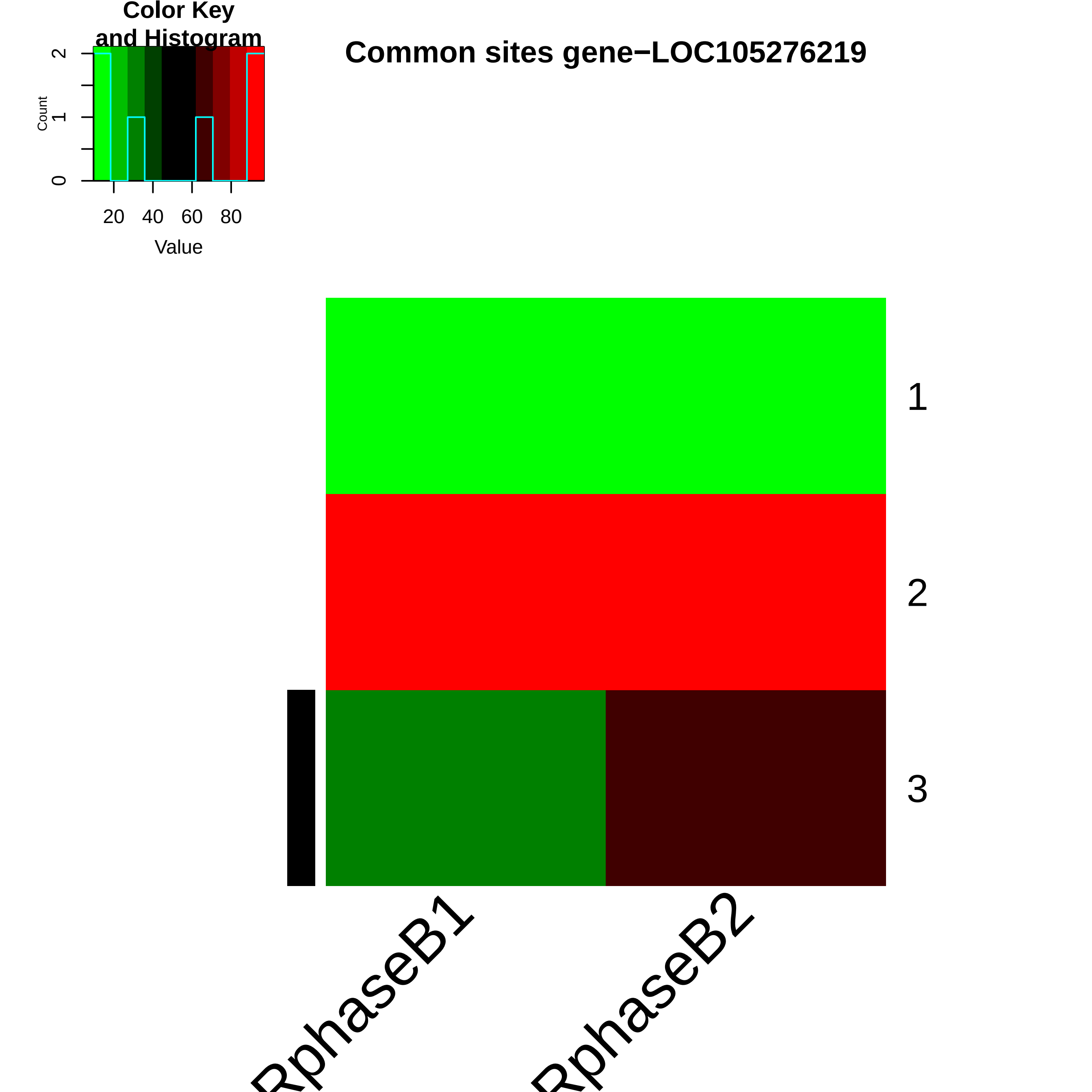

Color Key
and Histogram
Count
0
1
2
20
40
60
80
Value
Common sites gene−LOC105276219
1
2
3
RphaseB1
RphaseB2

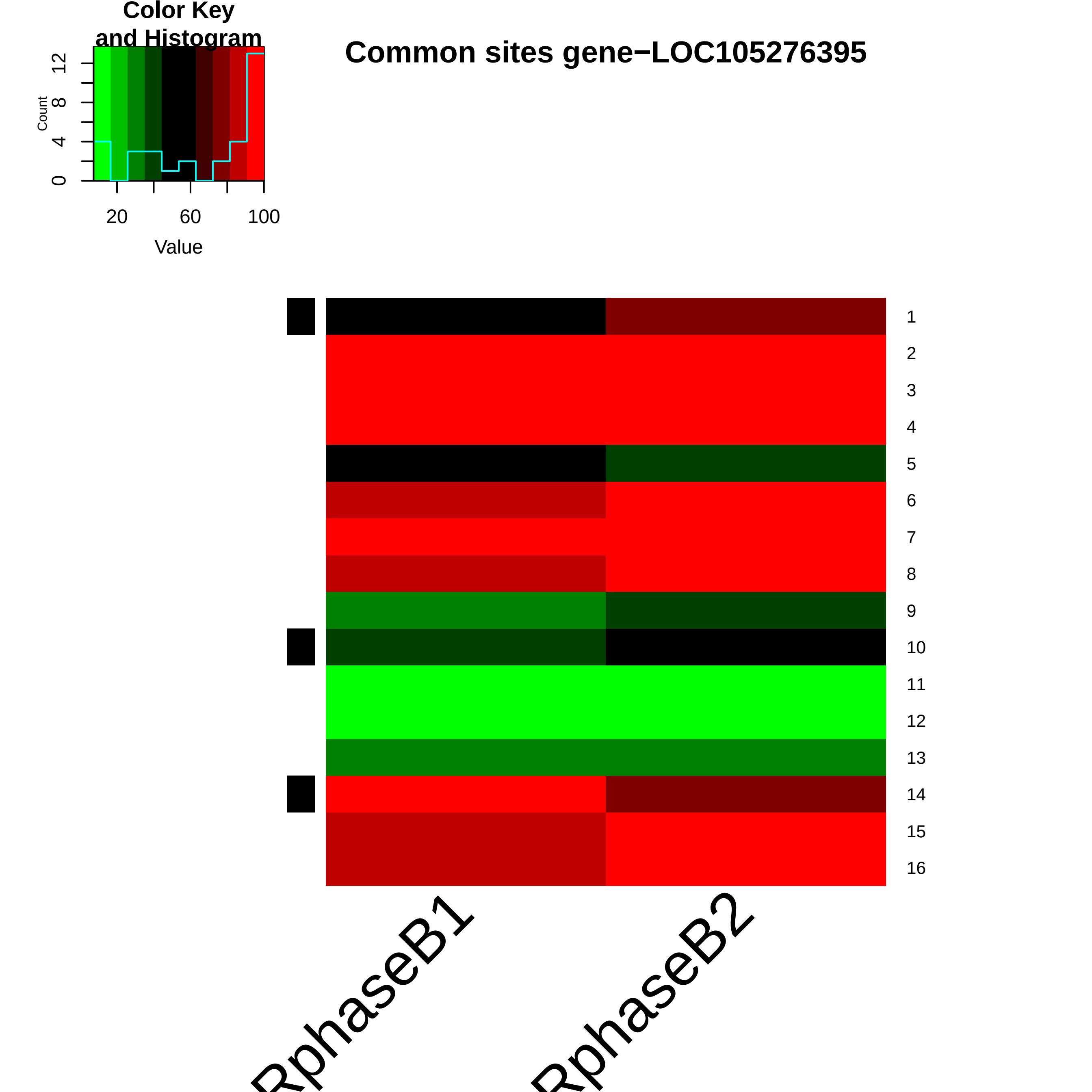

Color Key
and Histogram
Count
0
4
8
12
20
60
100
Value
Common sites gene−LOC105276395
RphaseB1
RphaseB2

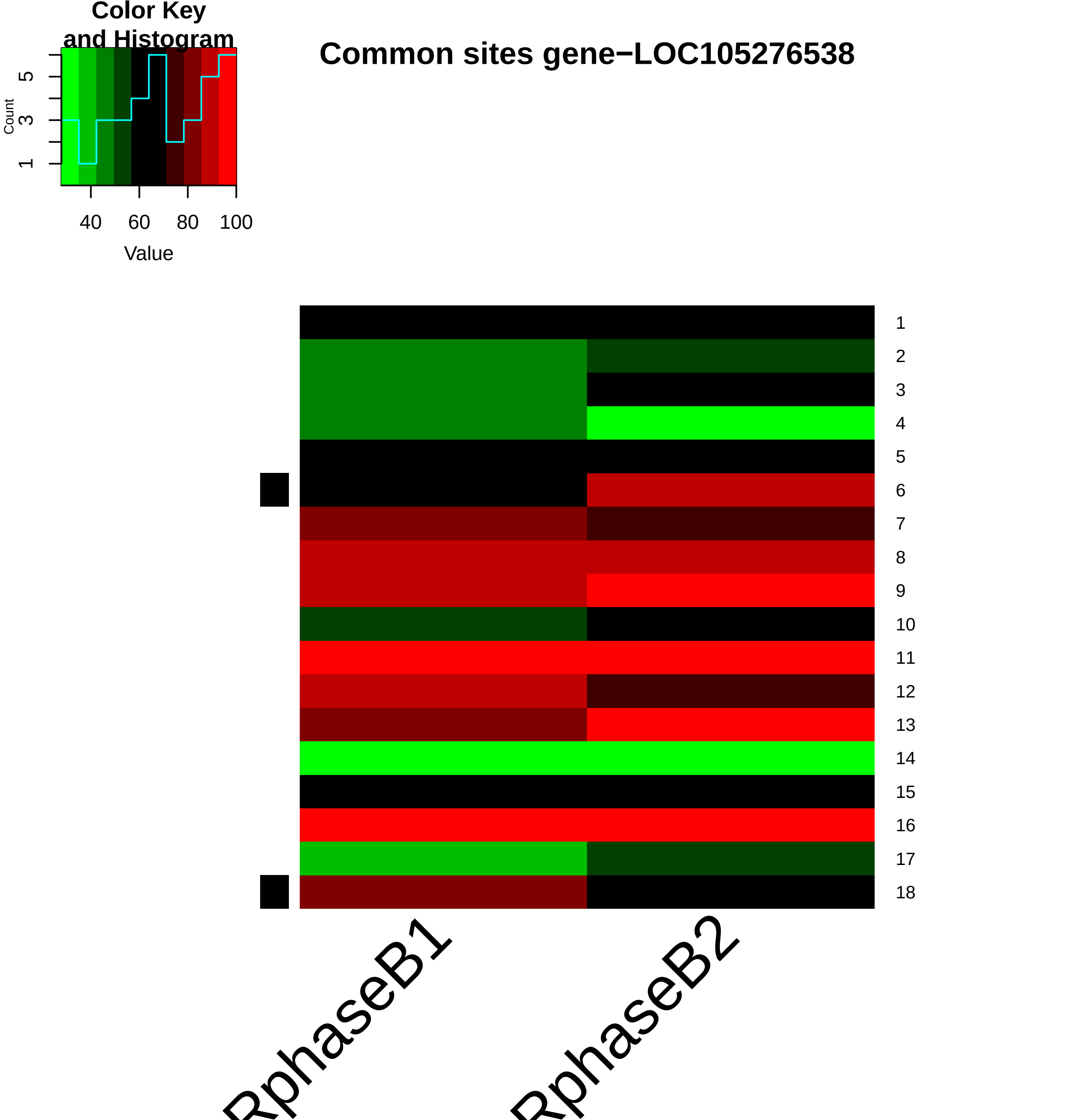
Color Key
and Histogram
Count
1
3
5
40
60
80
100
Value
Common sites gene−LOC105276538
RphaseB1
RphaseB2

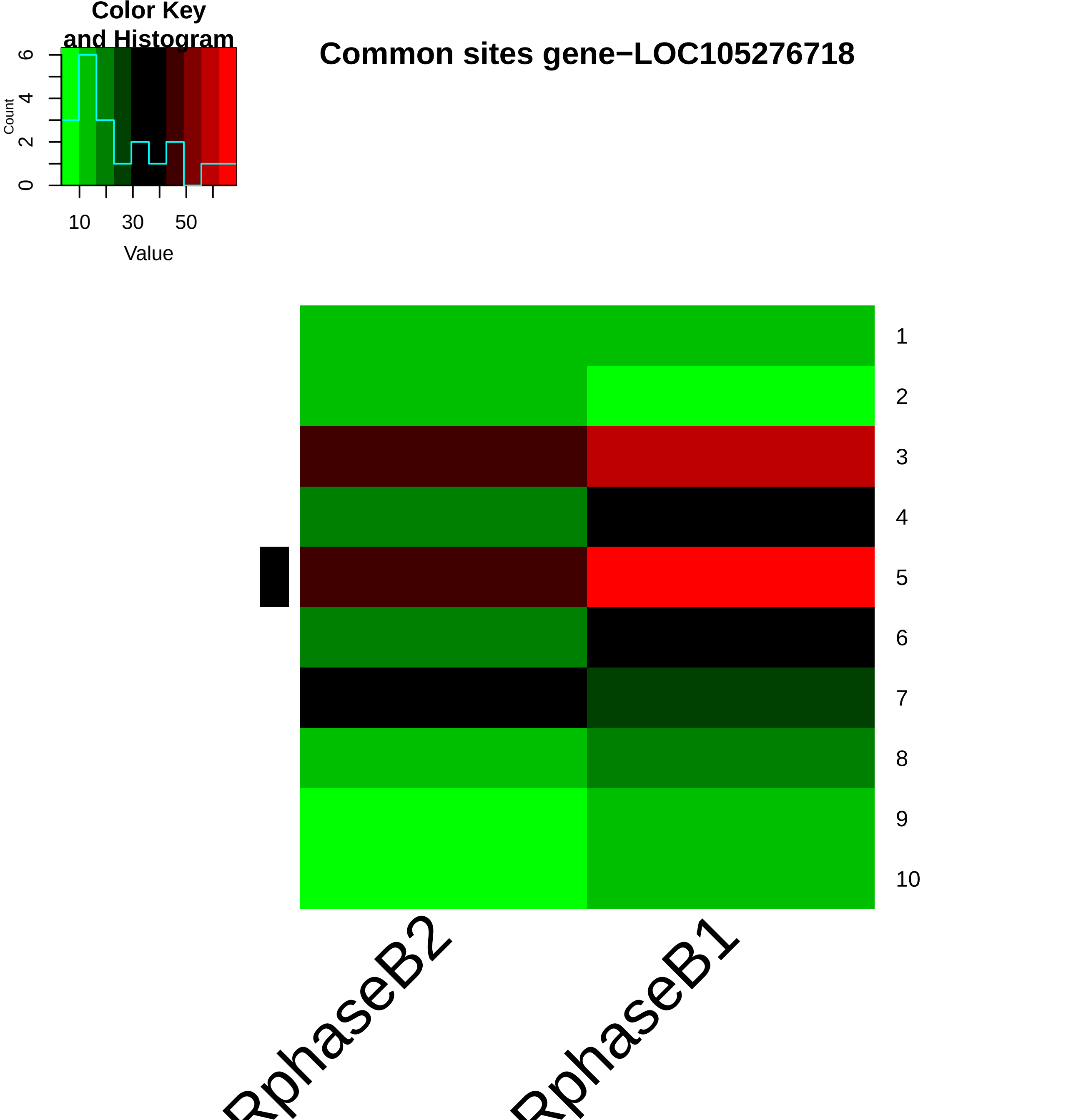
Color Key
and Histogram
Count
0
2
4
6
10
30
50
Value
Common sites gene−LOC105276718
1
2
3
4
5
6
7
8
9
10
RphaseB2
RphaseB1

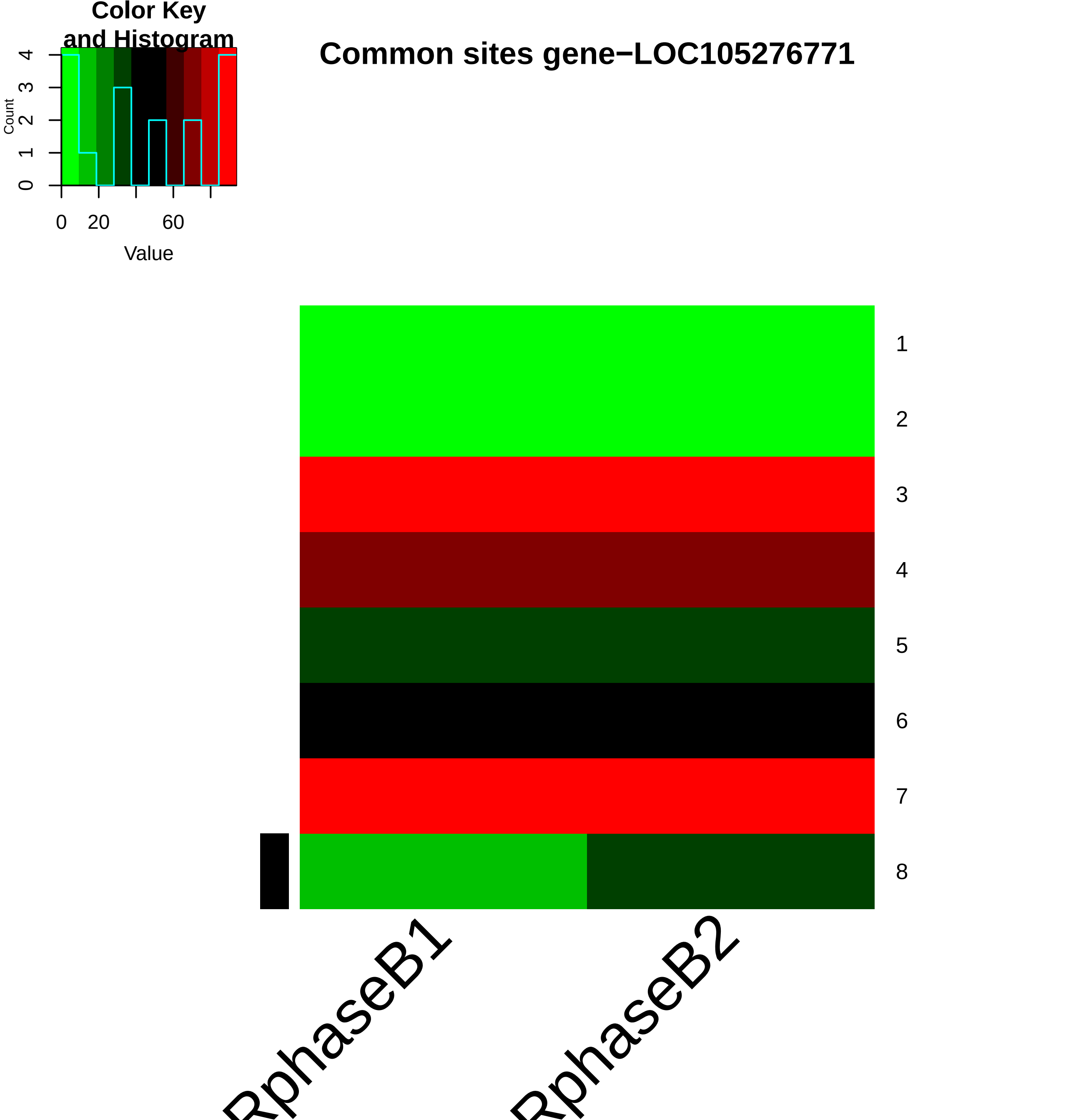

Color Key
and Histogram
Count
0
1
2
3
4
0
20
60
Value
Common sites gene−LOC105276771
1
2
3
4
5
6
7
8
RphaseB1
RphaseB2

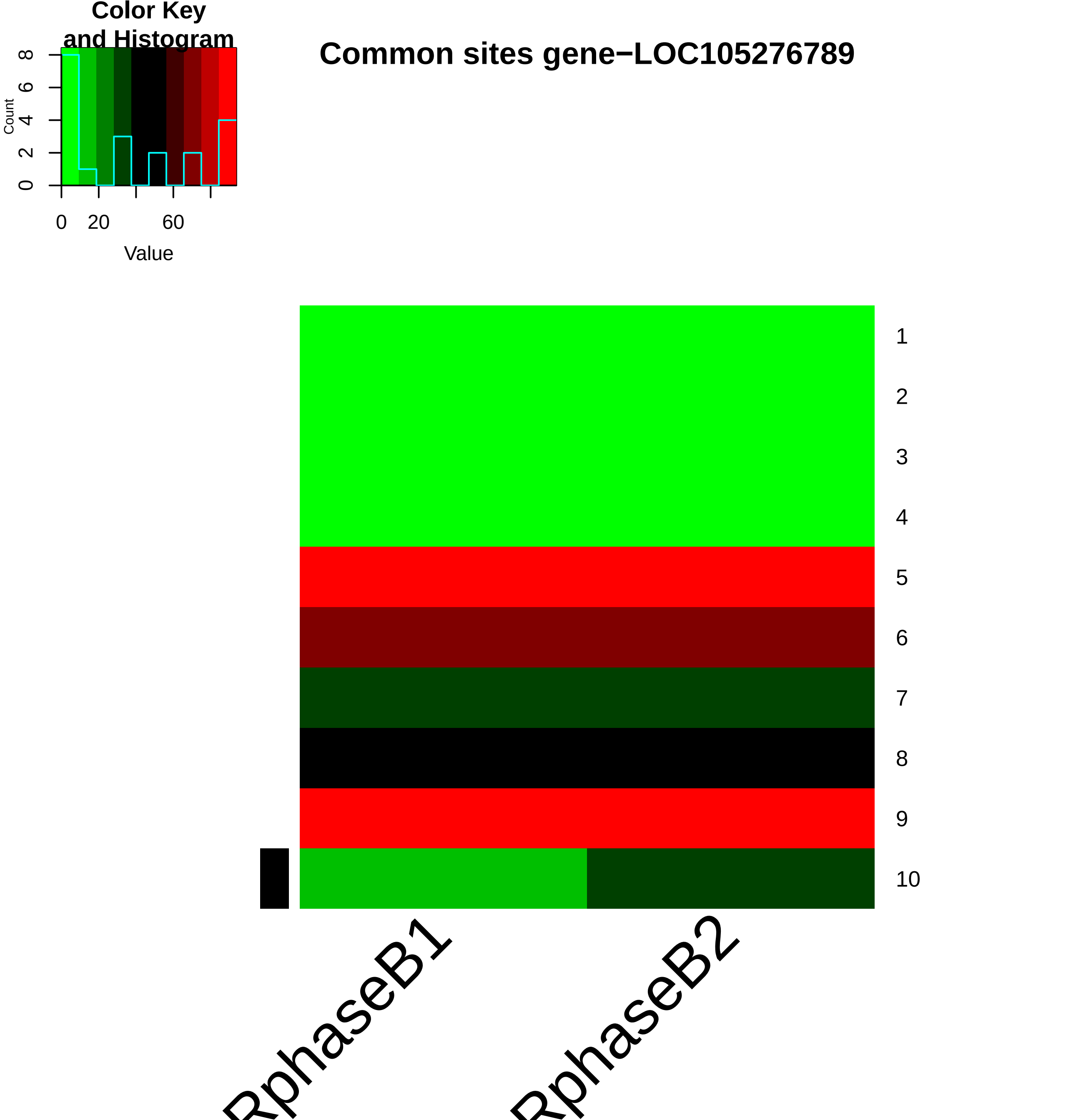
Color Key
and Histogram
Count
0
2
4
6
8
0
20
60
Value
Common sites gene−LOC105276789
1
2
3
4
5
6
7
8
9
10
RphaseB1
RphaseB2

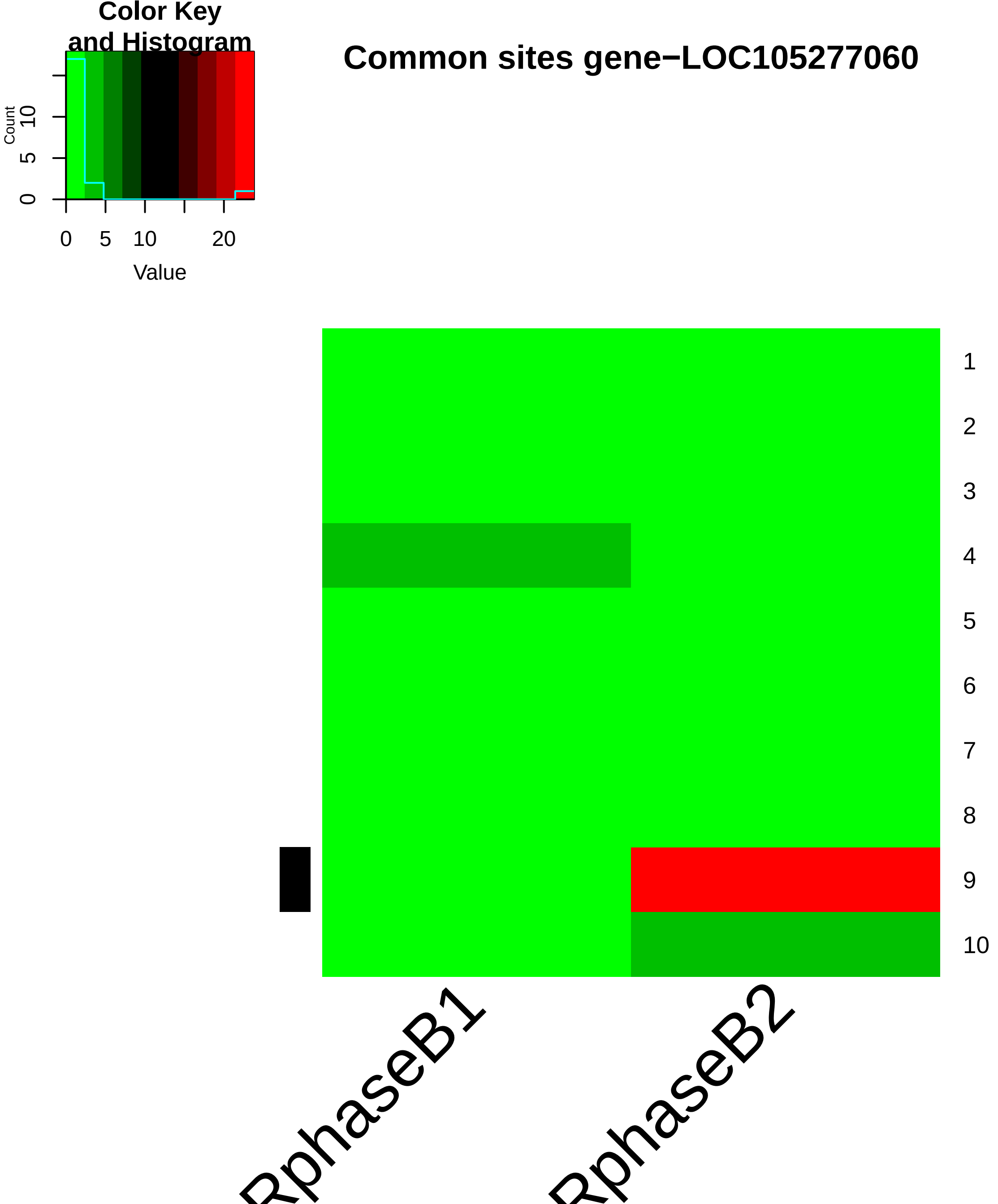
Color Key
and Histogram
Count
0
5
10
0
5
10
20
Value
Common sites gene−LOC105277060
1
2
3
4
5
6
7
8
9
10
RphaseB1
RphaseB2

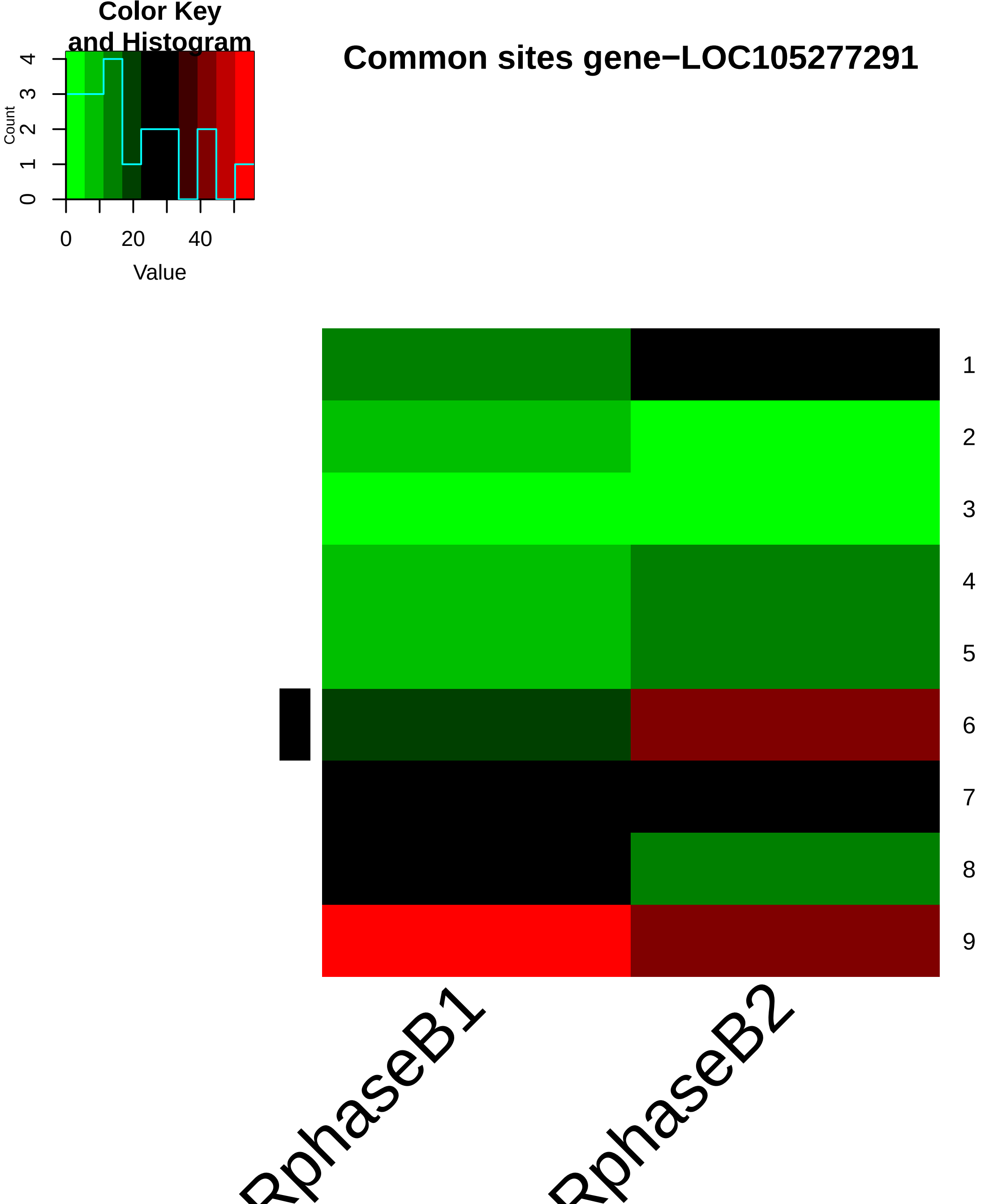

Color Key
and Histogram
Count
0
1
2
3
4
0
20
40
Value
Common sites gene−LOC105277291
1
2
3
4
5
6
7
8
9
RphaseB1
RphaseB2

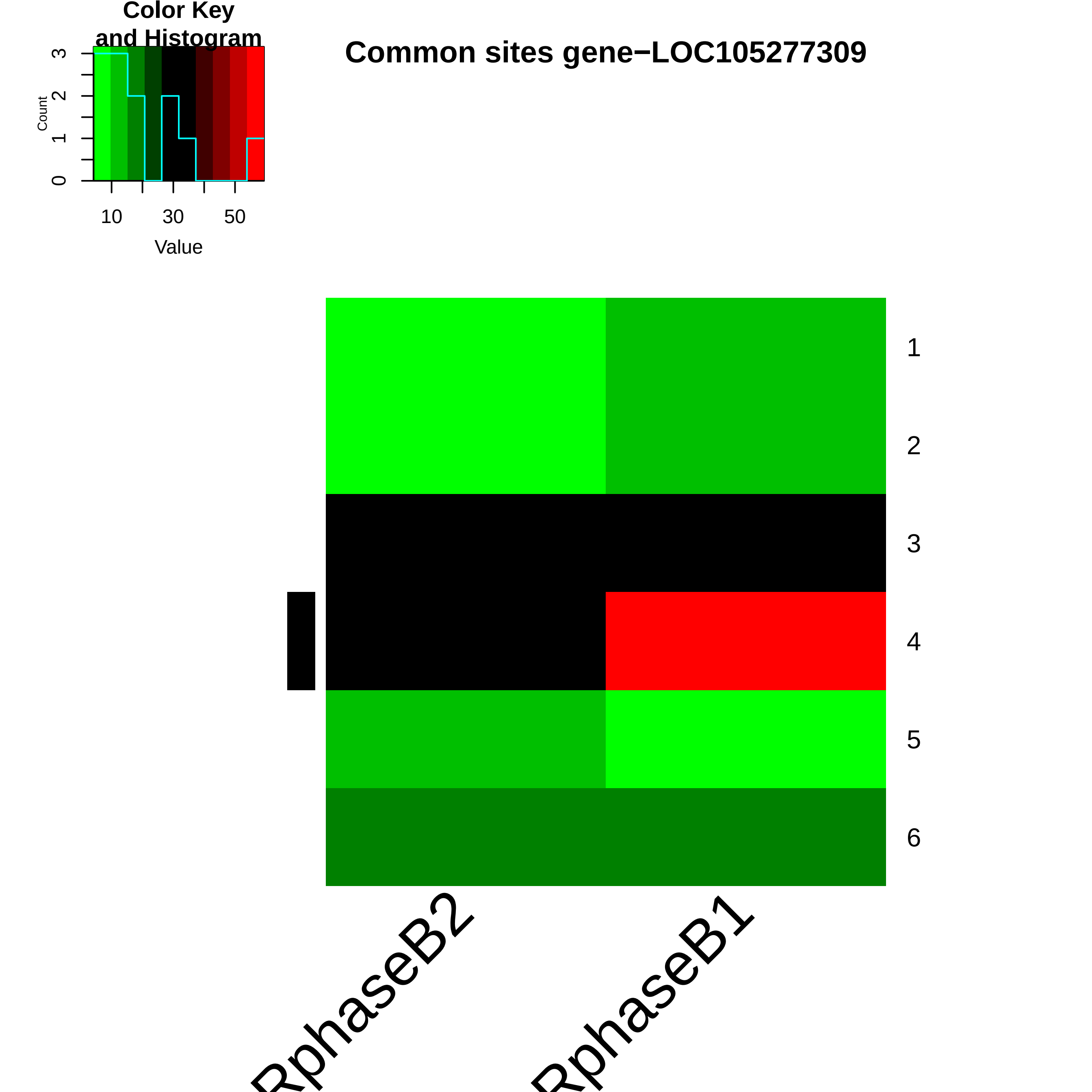

Color Key
and Histogram
Count
Value
10
30
50
0
1
2
3
Common sites gene−LOC105277309
1
2
3
4
5
6
RphaseB2
RphaseB1

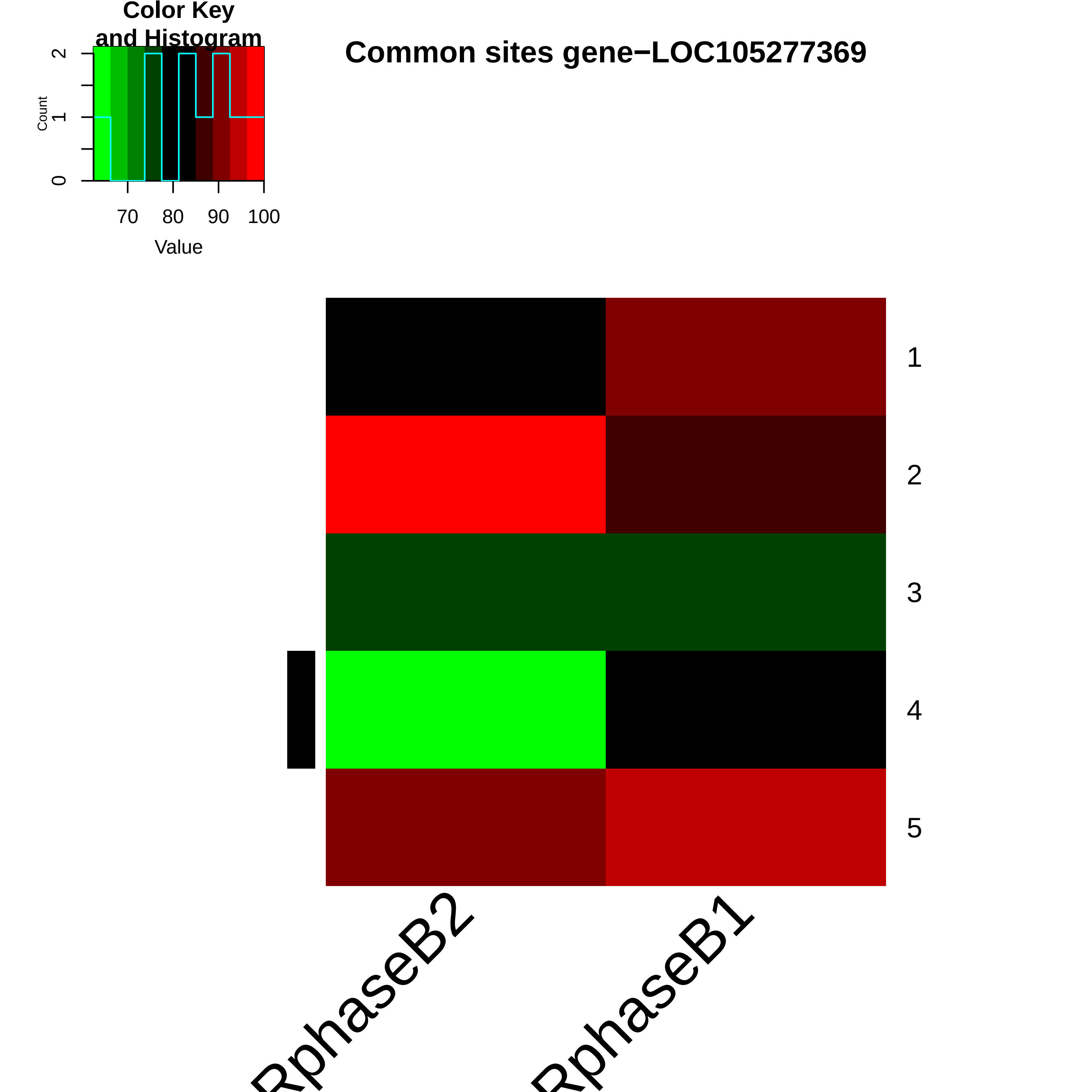
Color Key
and Histogram
Count
0
1
2
70
80
90
100
Value
Common sites gene−LOC105277369
1
2
3
4
5
RphaseB2
RphaseB1

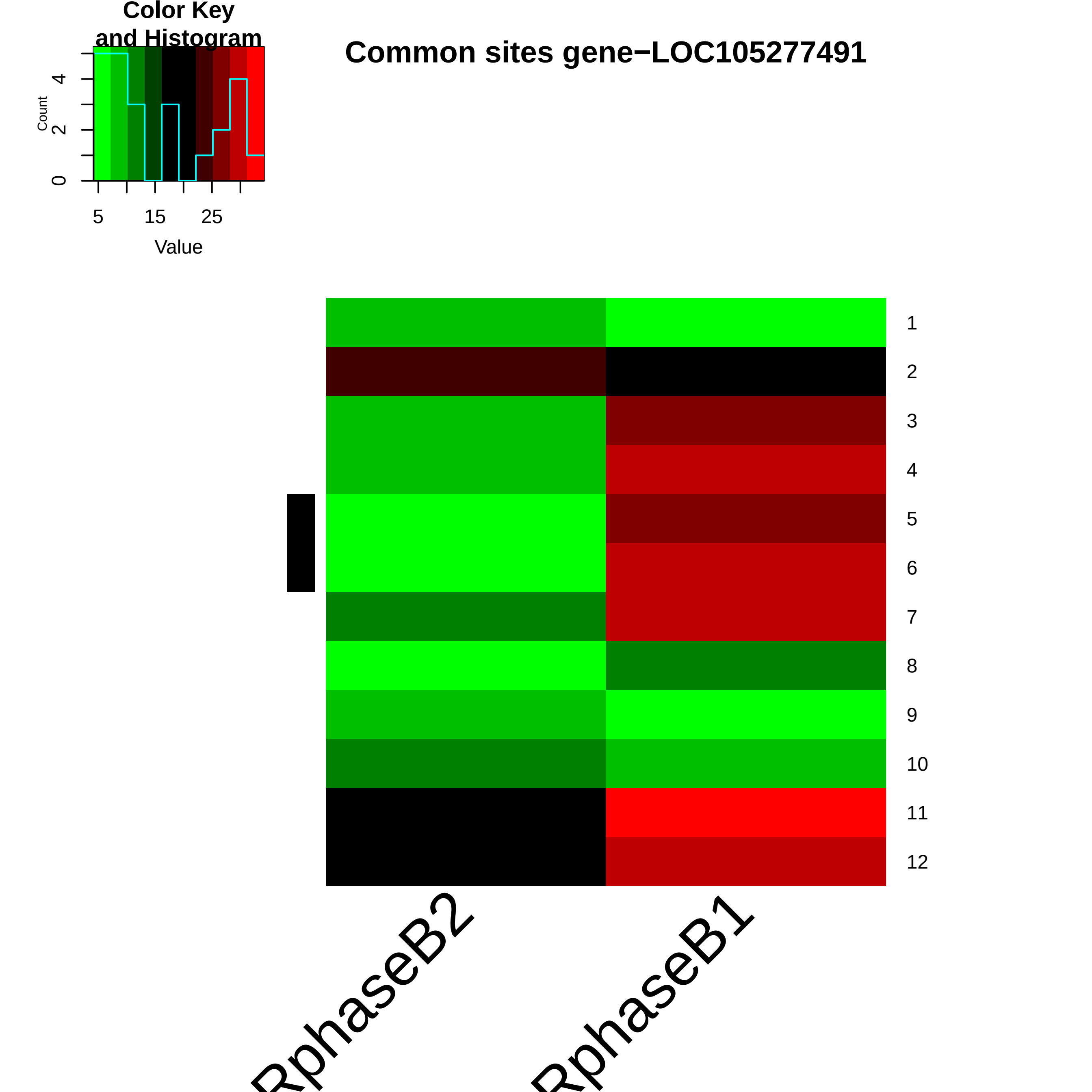

Color Key
and Histogram
Count
Value
Common sites gene−LOC105277491
1
2
3
4
5
6
7
8
9
10
11
12
RphaseB2
RphaseB1

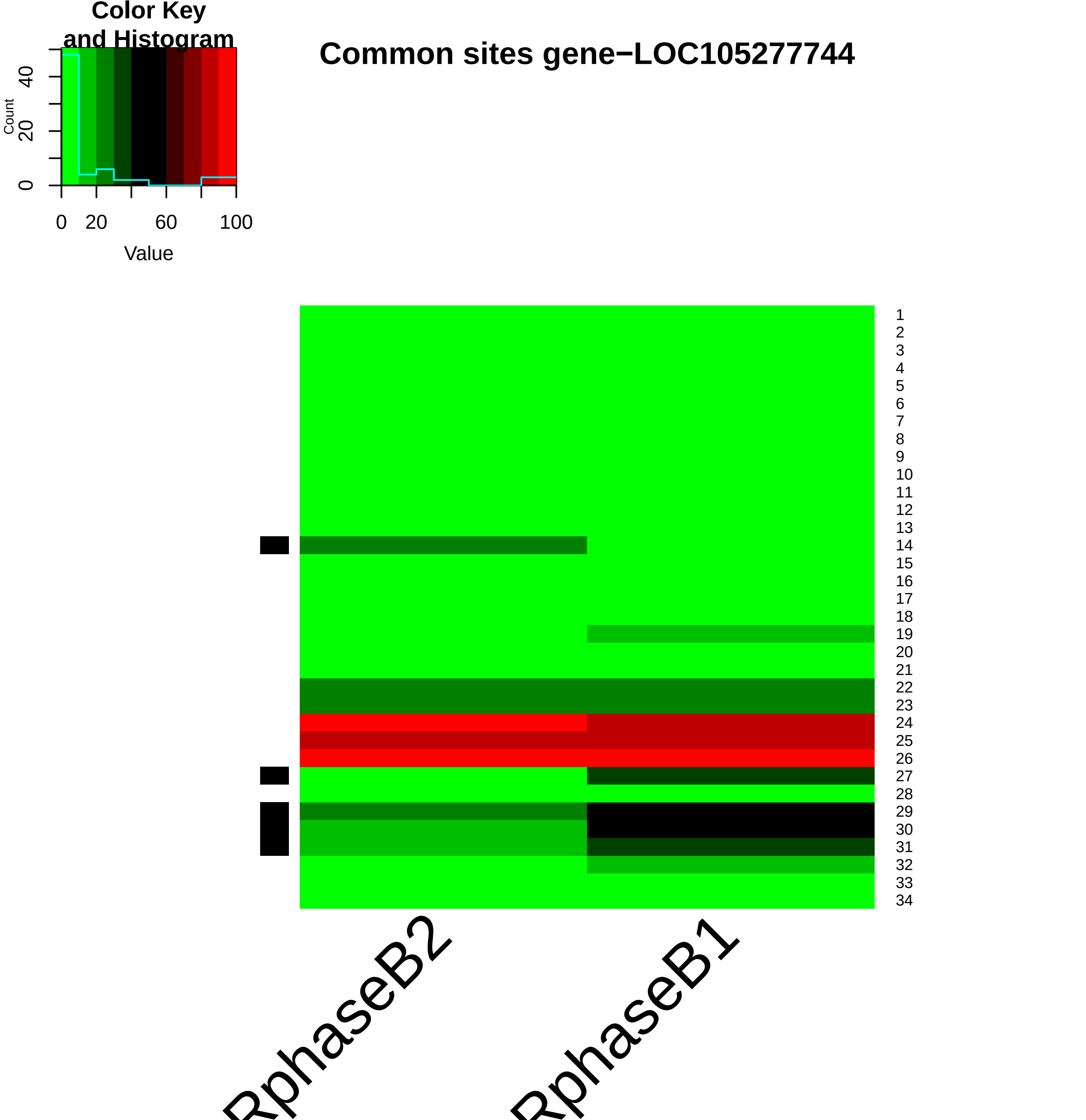
Color Key
and Histogram
Count
0
20
40
0
20
60
100
Value
Common sites gene−LOC105277744
RphaseB2
RphaseB1

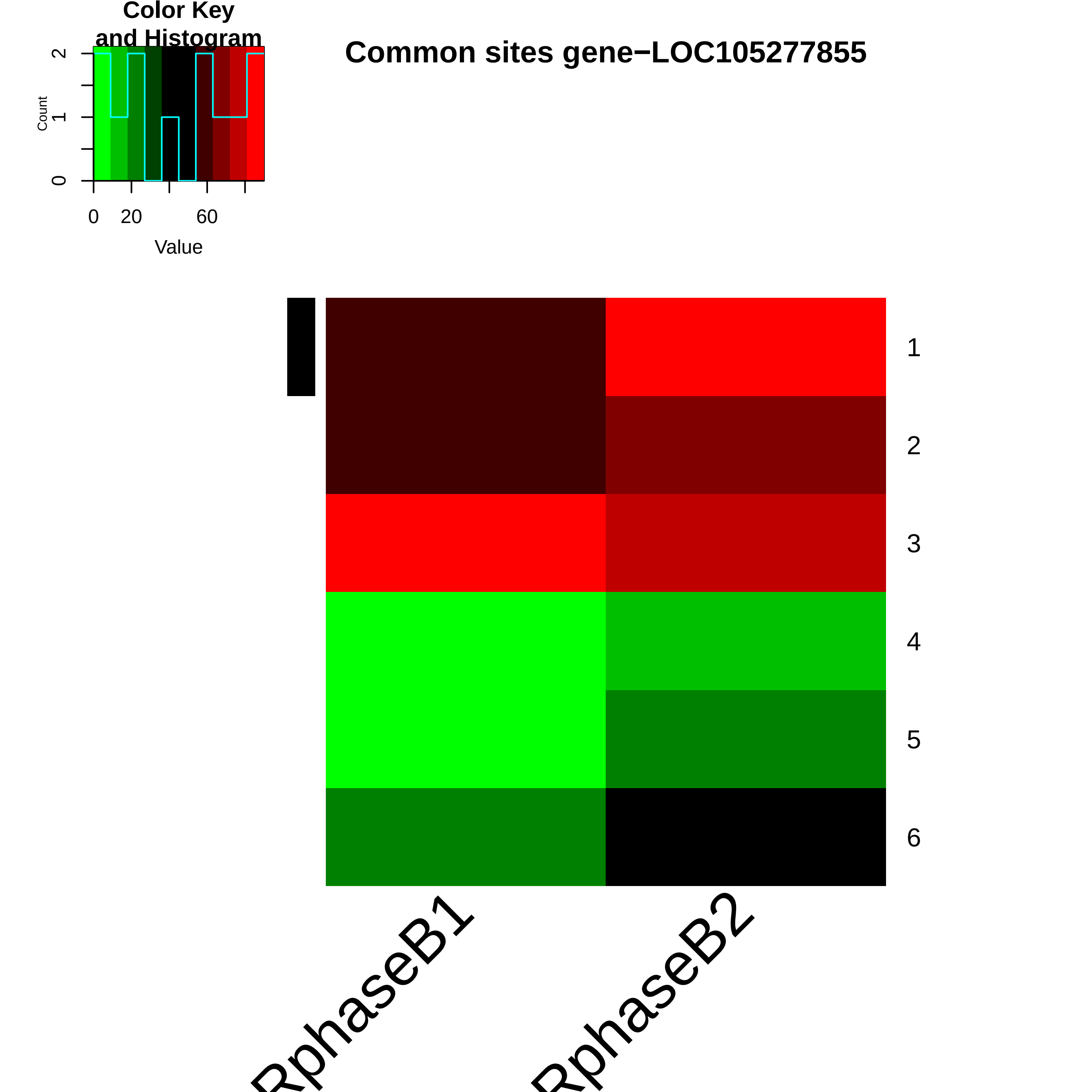

Color Key
and Histogram
Count
0
1
2
0
20
60
Value
Common sites gene−LOC105277855
1
2
3
4
5
6
RphaseB1
RphaseB2

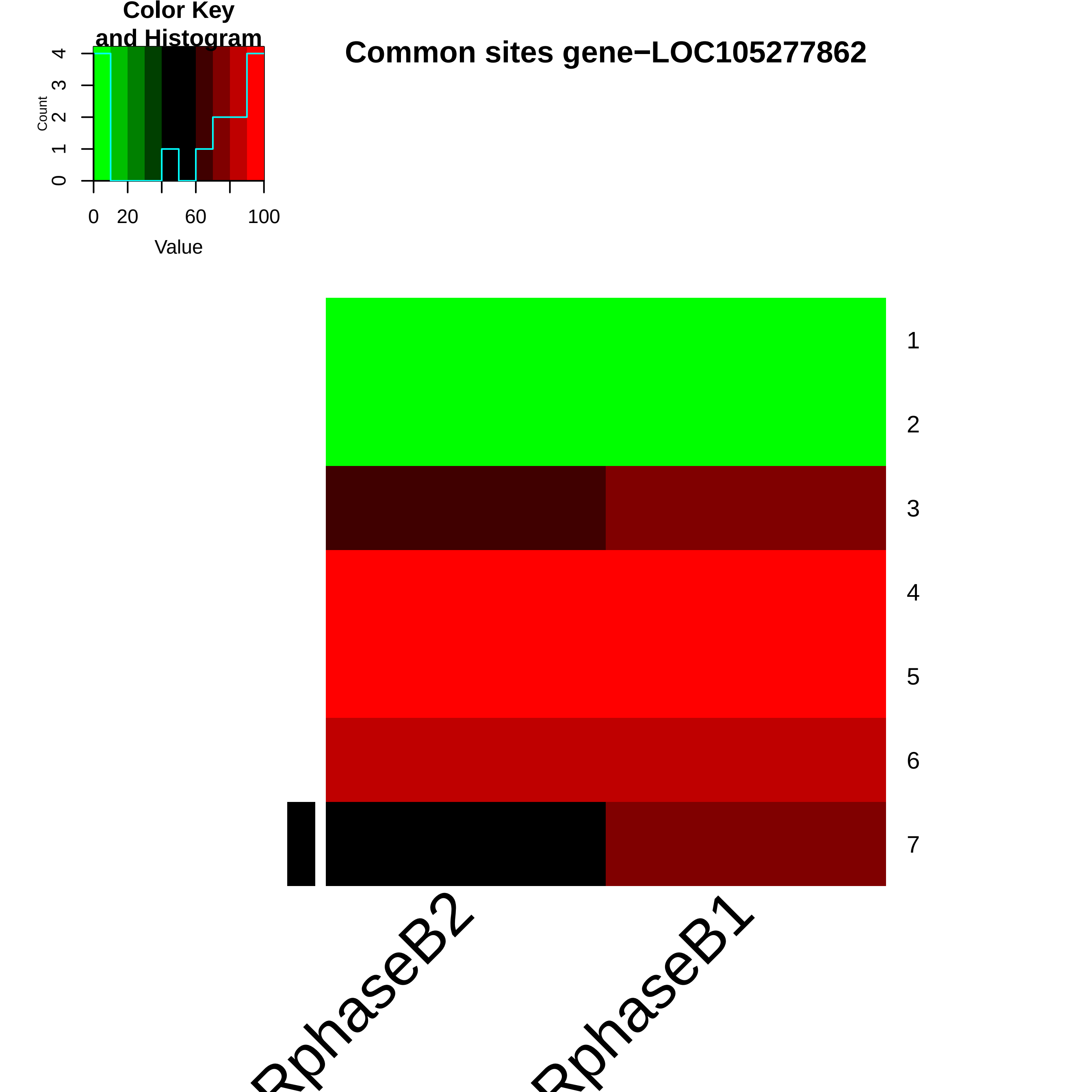

Color Key
and Histogram
Count
0
1
2
3
4
0
20
60
100
Value
Common sites gene−LOC105277862
1
2
3
4
5
6
7
RphaseB2
RphaseB1

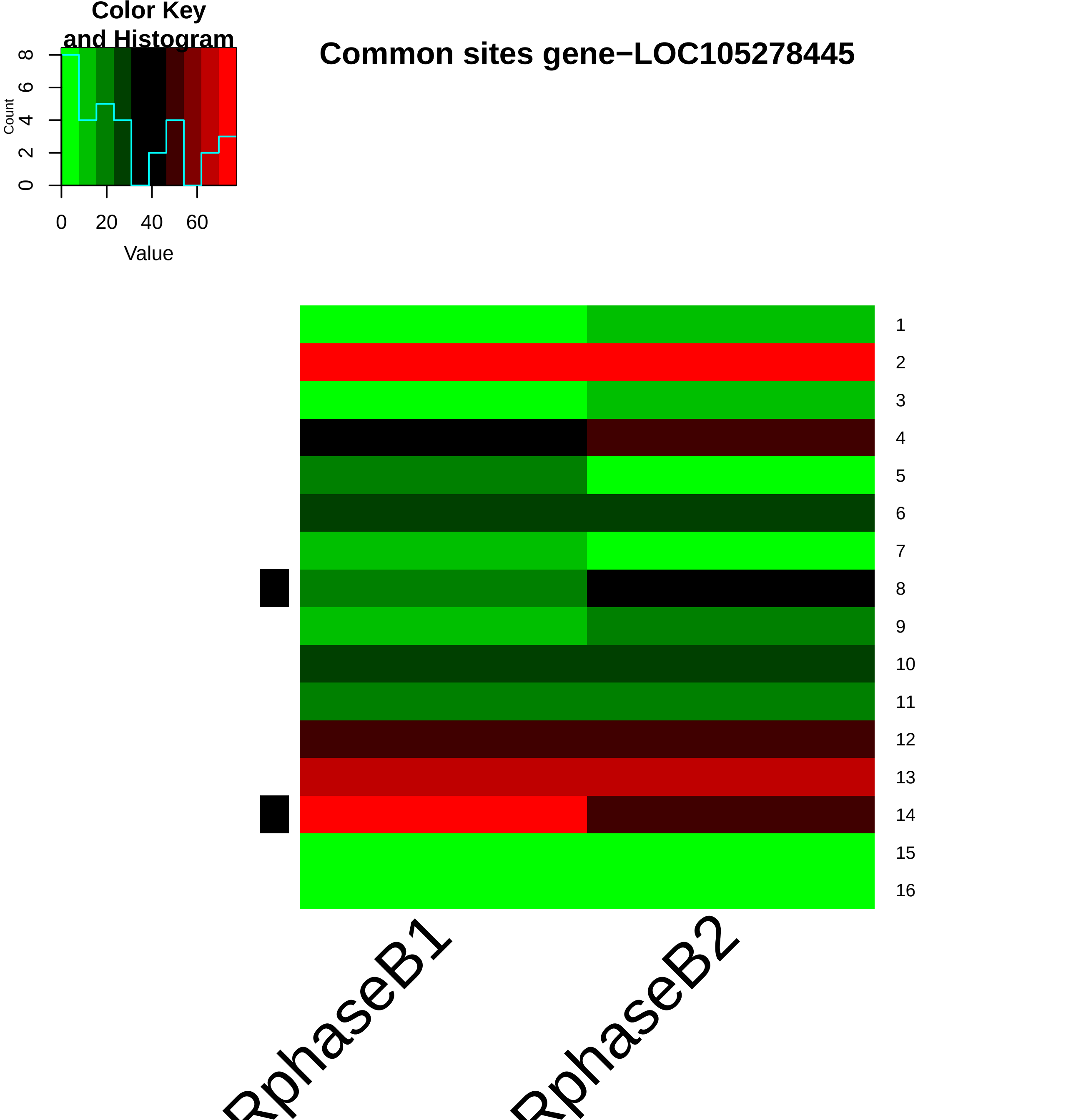
Color Key
and Histogram
Count
0
2
4
6
8
0
20
40
60
Value
Common sites gene−LOC105278445
RphaseB1
RphaseB2

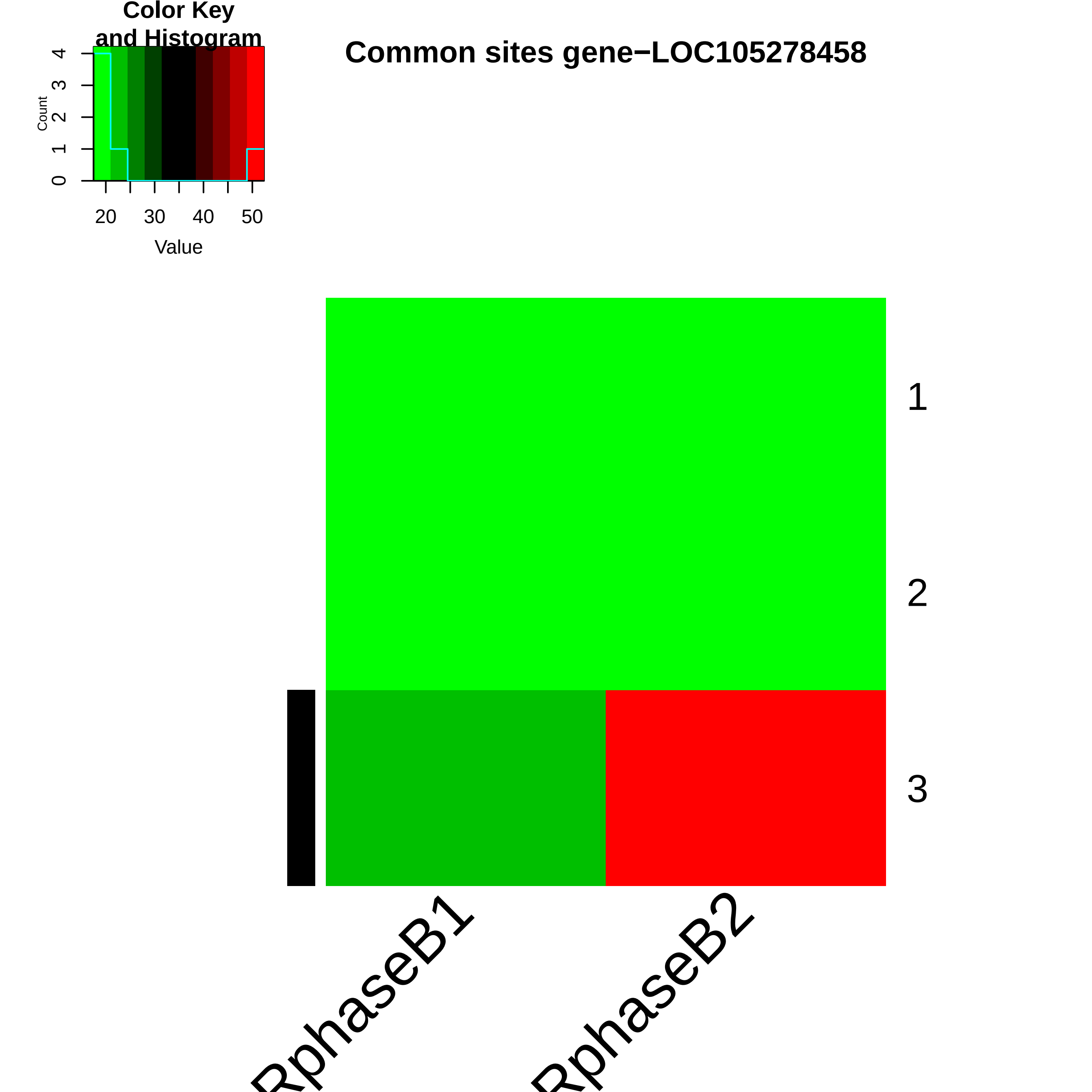

Color Key
and Histogram
Count
0
1
2
3
4
20
30
40
50
Value
Common sites gene−LOC105278458
1
2
3
RphaseB1
RphaseB2

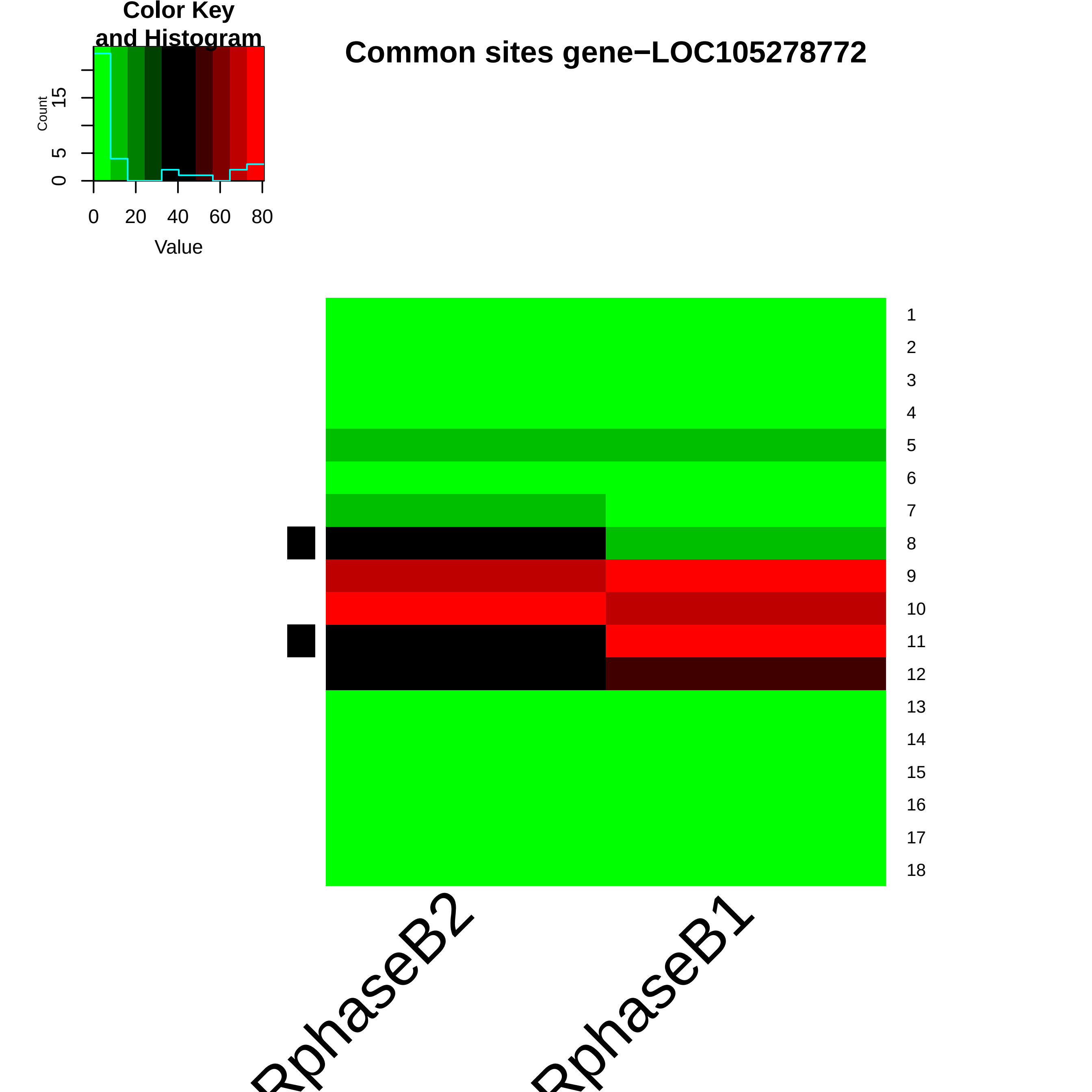
Color Key
and Histogram
Count
0
5
15
0
20
40
60
80
Value
Common sites gene−LOC105278772
RphaseB2
RphaseB1

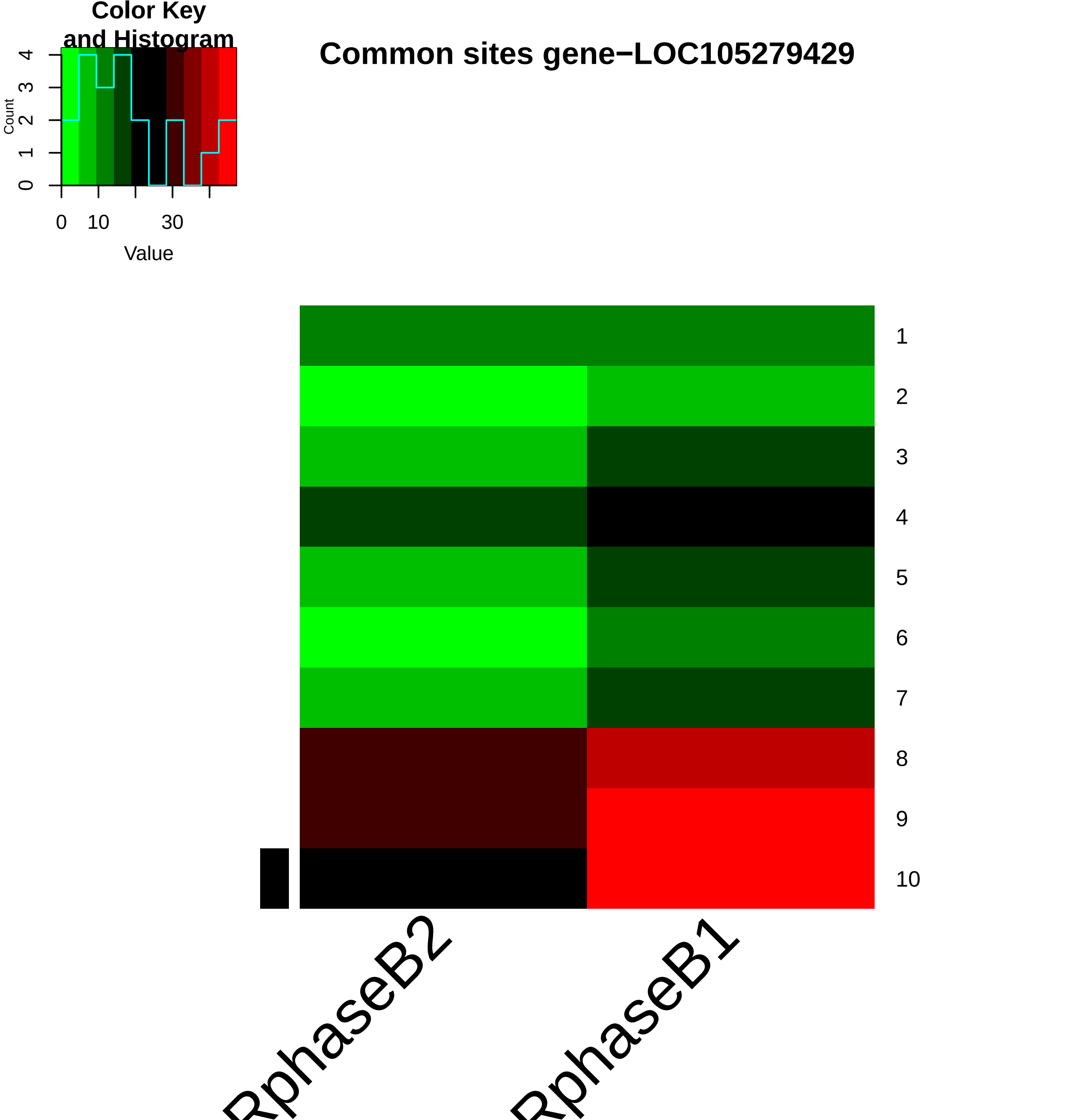
Color Key
and Histogram
Count
0
1
2
3
4
0
10
30
Value
Common sites gene−LOC105279429
1
2
3
4
5
6
7
8
9
10
RphaseB2
RphaseB1

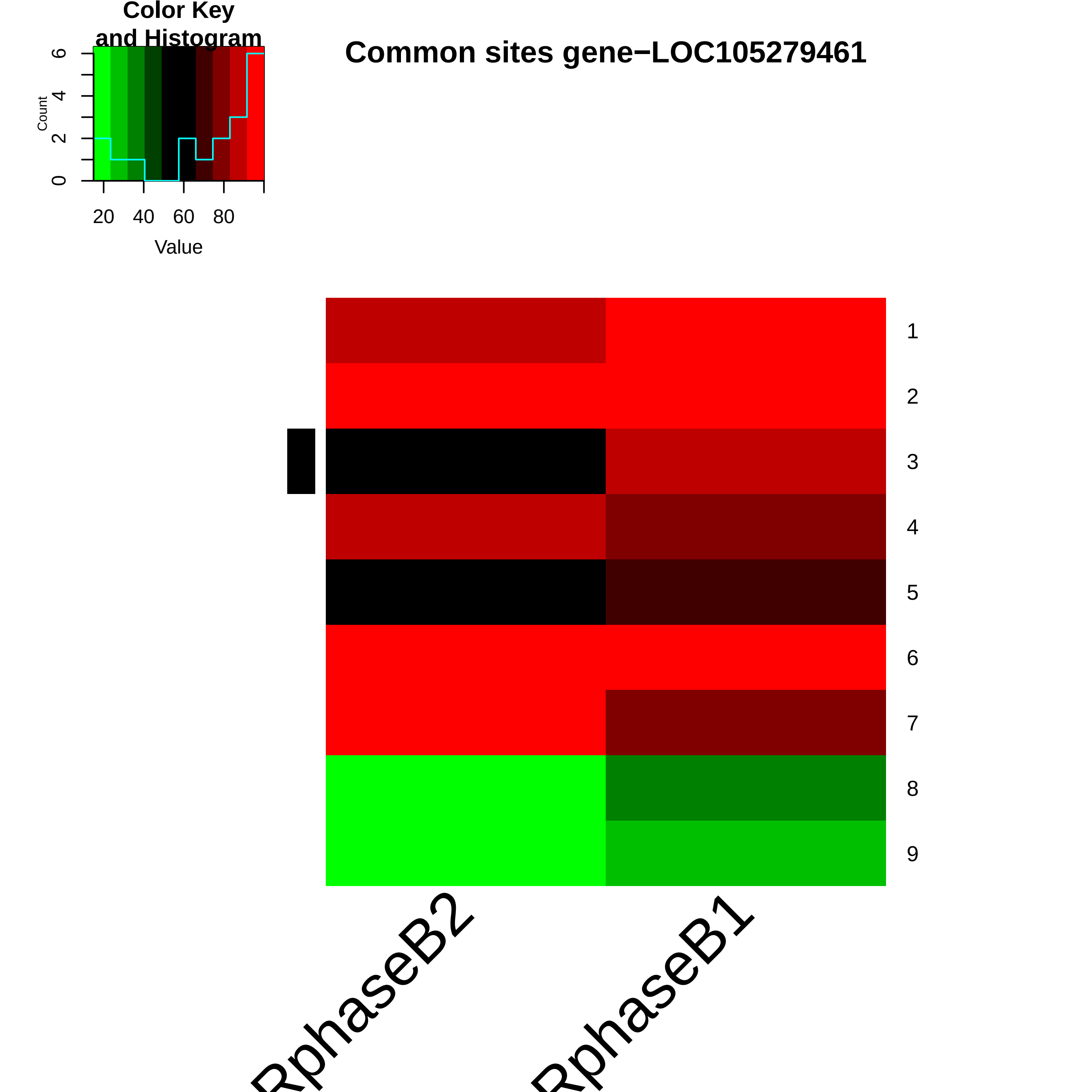
Color Key
and Histogram
Count
0
2
4
6
20
40
60
80
Value
Common sites gene−LOC105279461
1
2
3
4
5
6
7
8
9
RphaseB2
RphaseB1

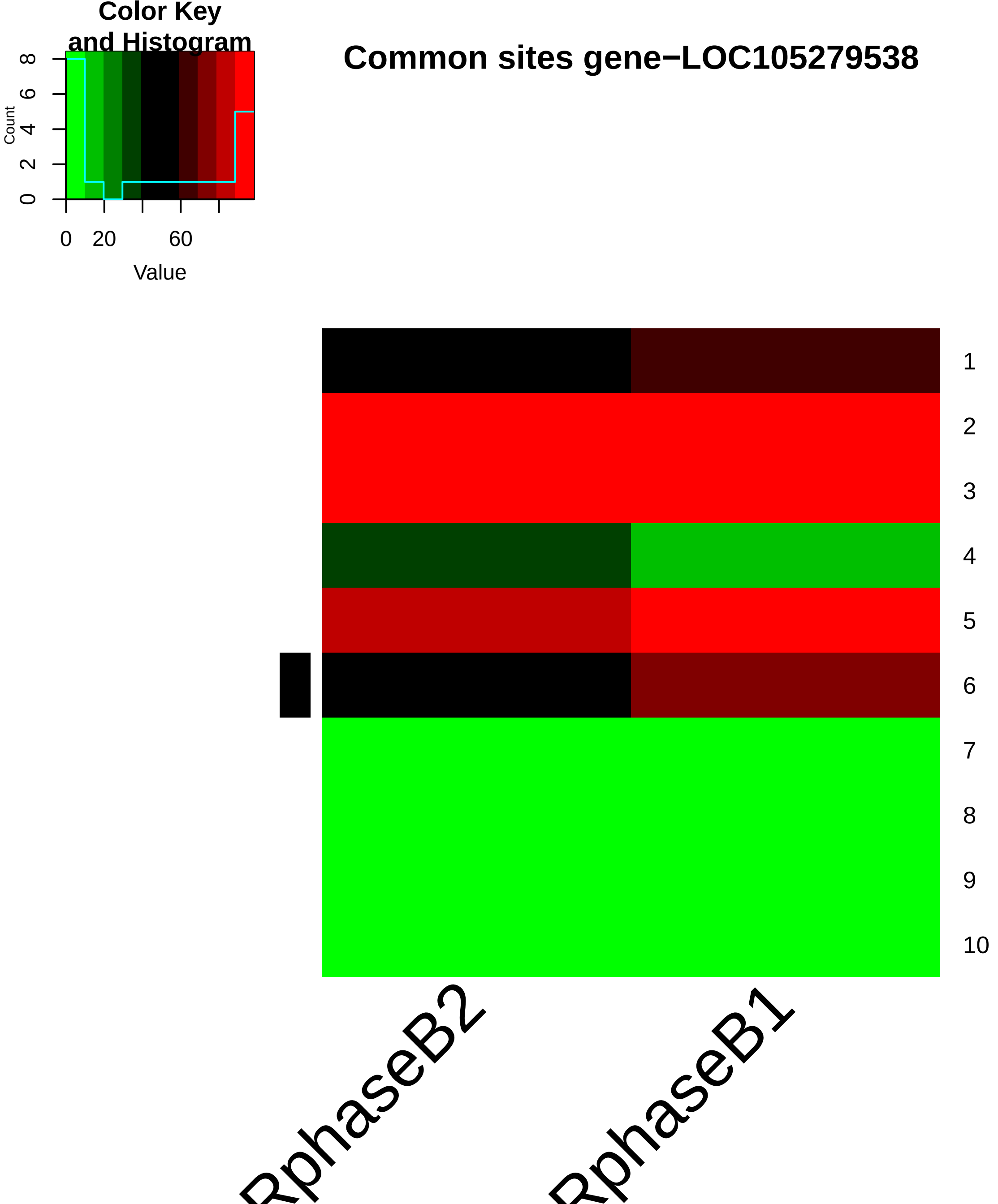

Color Key
and Histogram
Count
0
2
4
6
8
0
20
60
Value
Common sites gene−LOC105279538
1
2
3
4
5
6
7
8
9
10
RphaseB2
RphaseB1

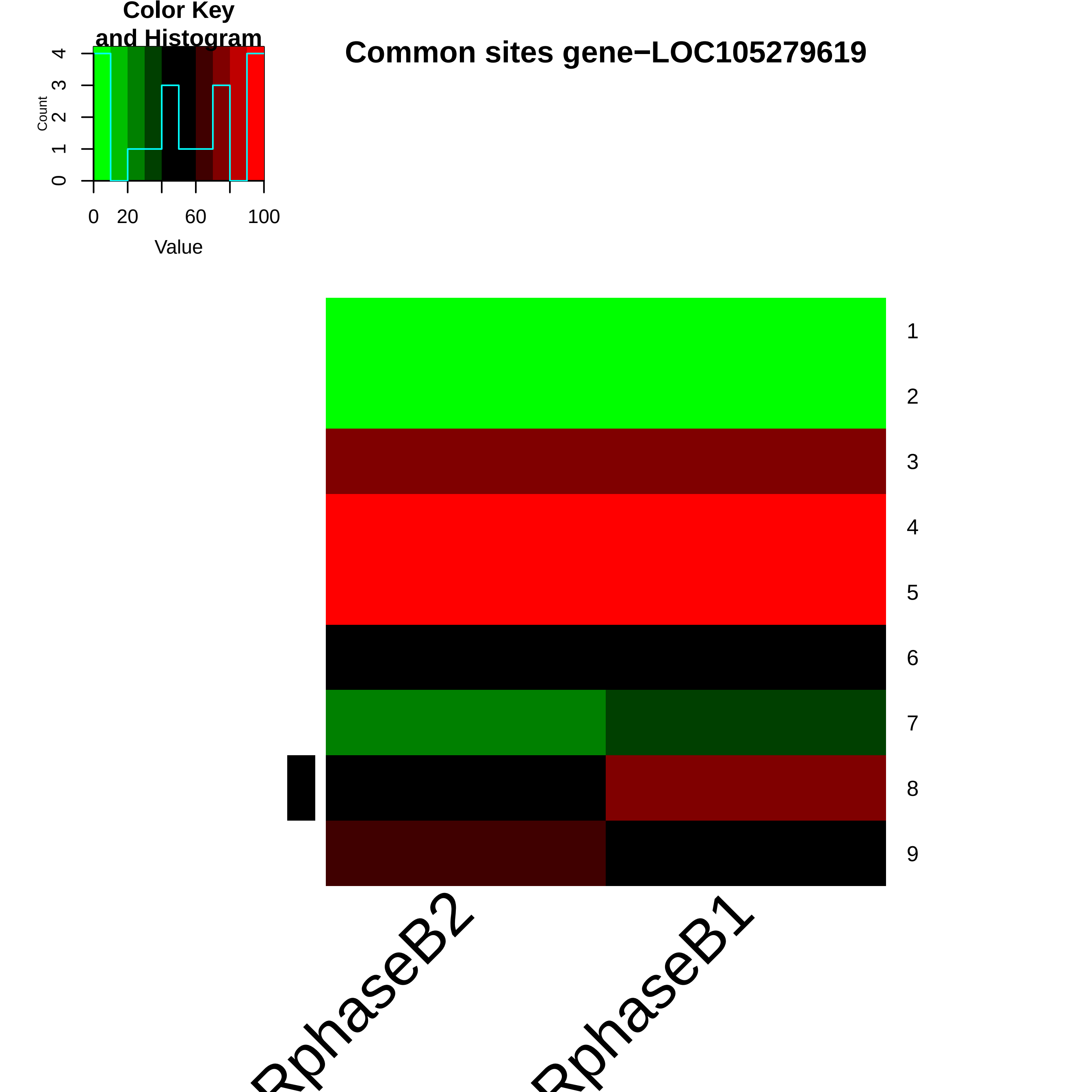
Color Key
and Histogram
Count
Value
0
20
60
100
0
1
2
3
4
Common sites gene−LOC105279619
1
2
3
4
5
6
7
8
9
RphaseB2
RphaseB1

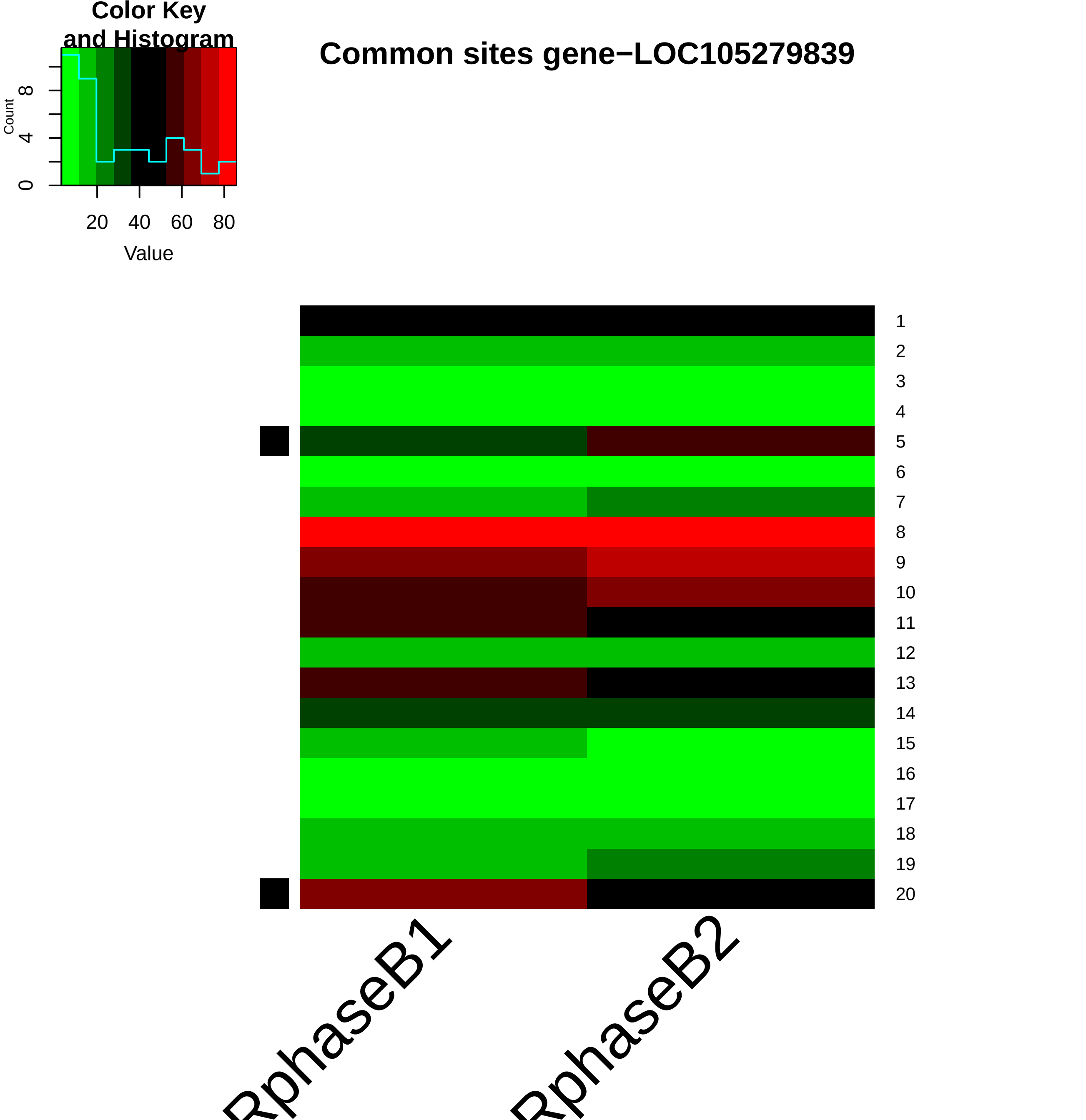

Color Key
and Histogram
Count
0
4
8
20
40
60
80
Value
Common sites gene−LOC105279839
RphaseB1
RphaseB2

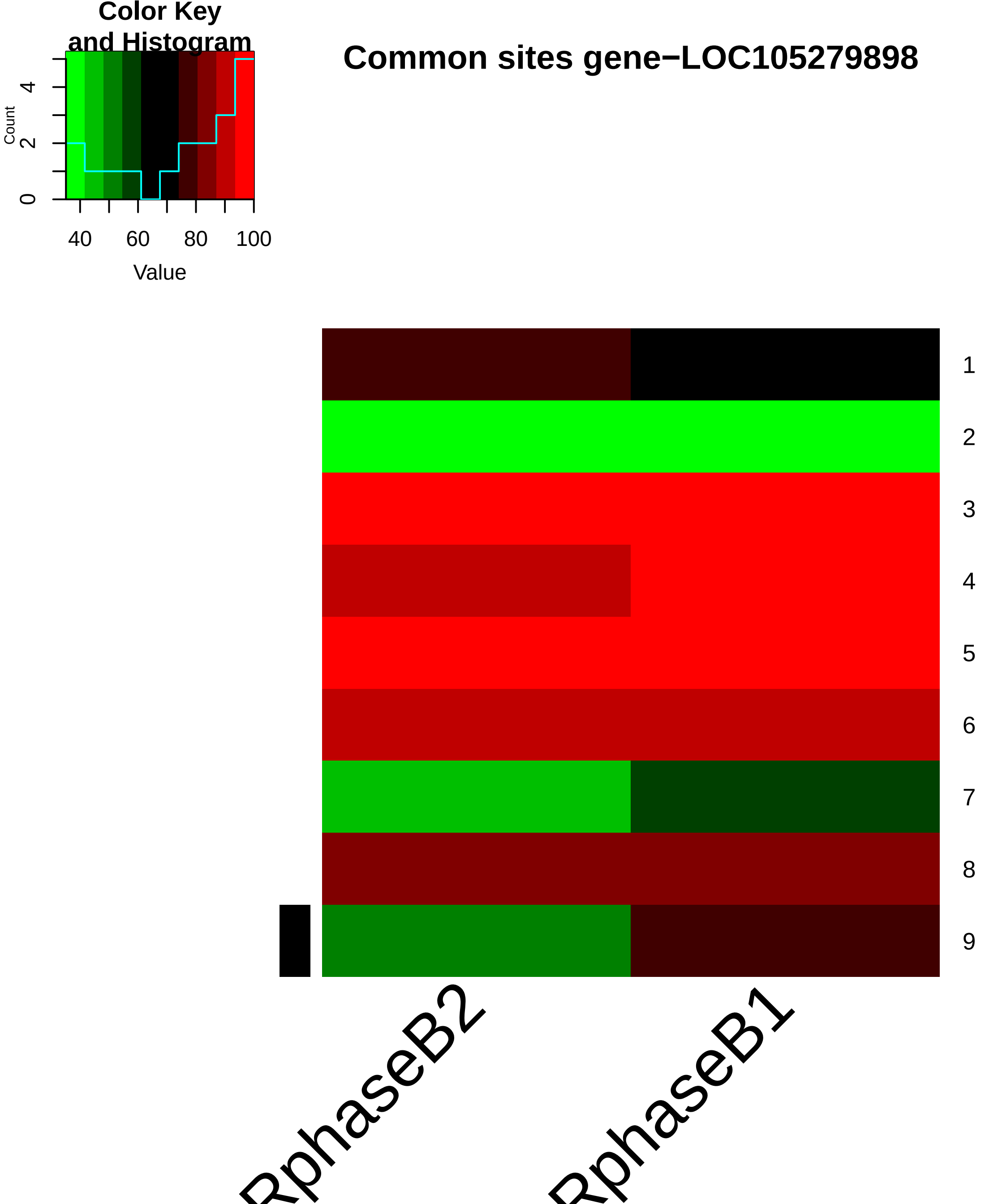

Color Key and Histogram
Count
0
2
4
40
60
80
100
Value
Common sites gene−LOC105279898
1
2
3
4
5
6
7
8
9
RphaseB2
RphaseB1

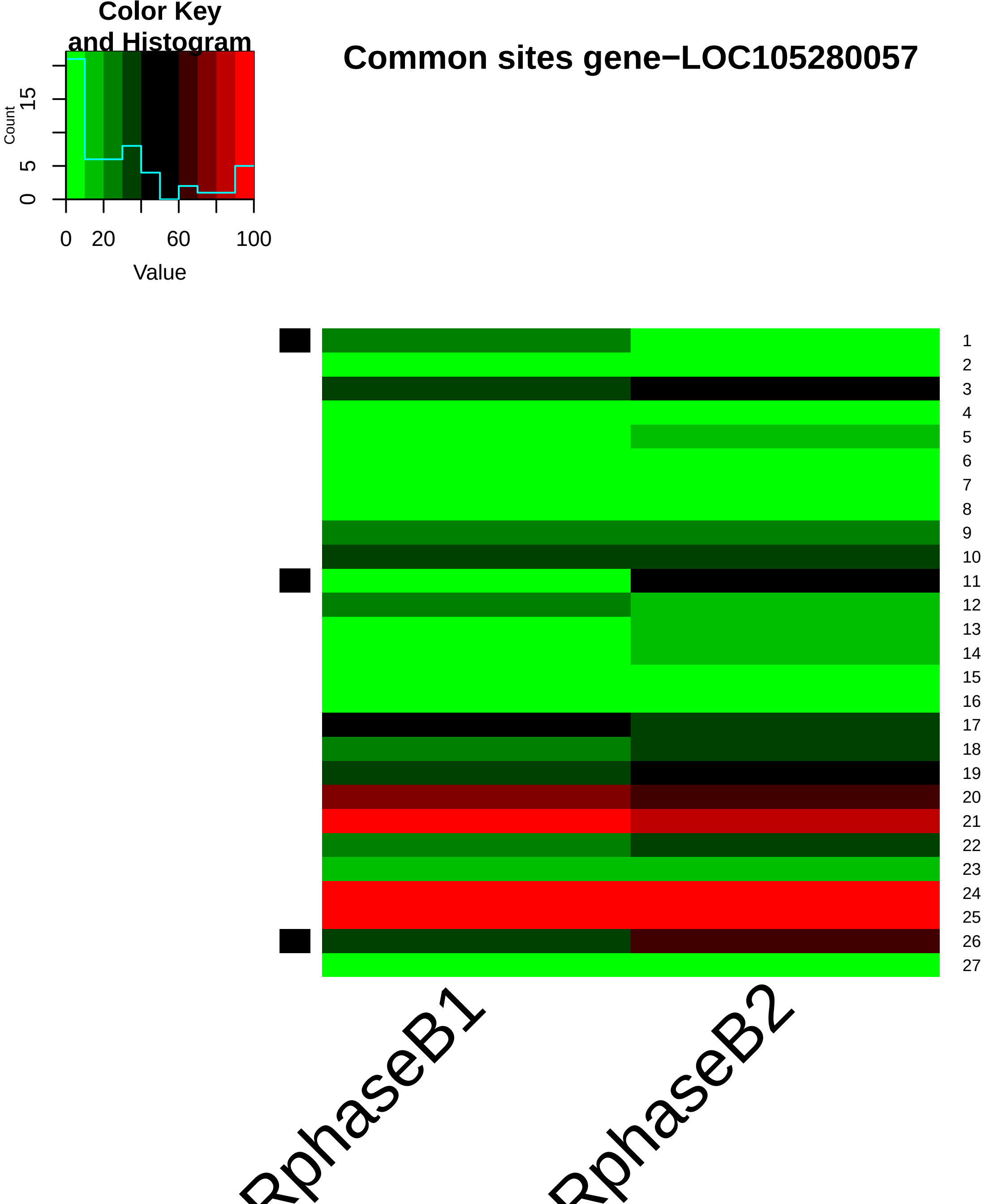

Color Key
and Histogram
Count
0
5
15
0
20
60
100
Value
Common sites gene−LOC105280057
RphaseB1
RphaseB2

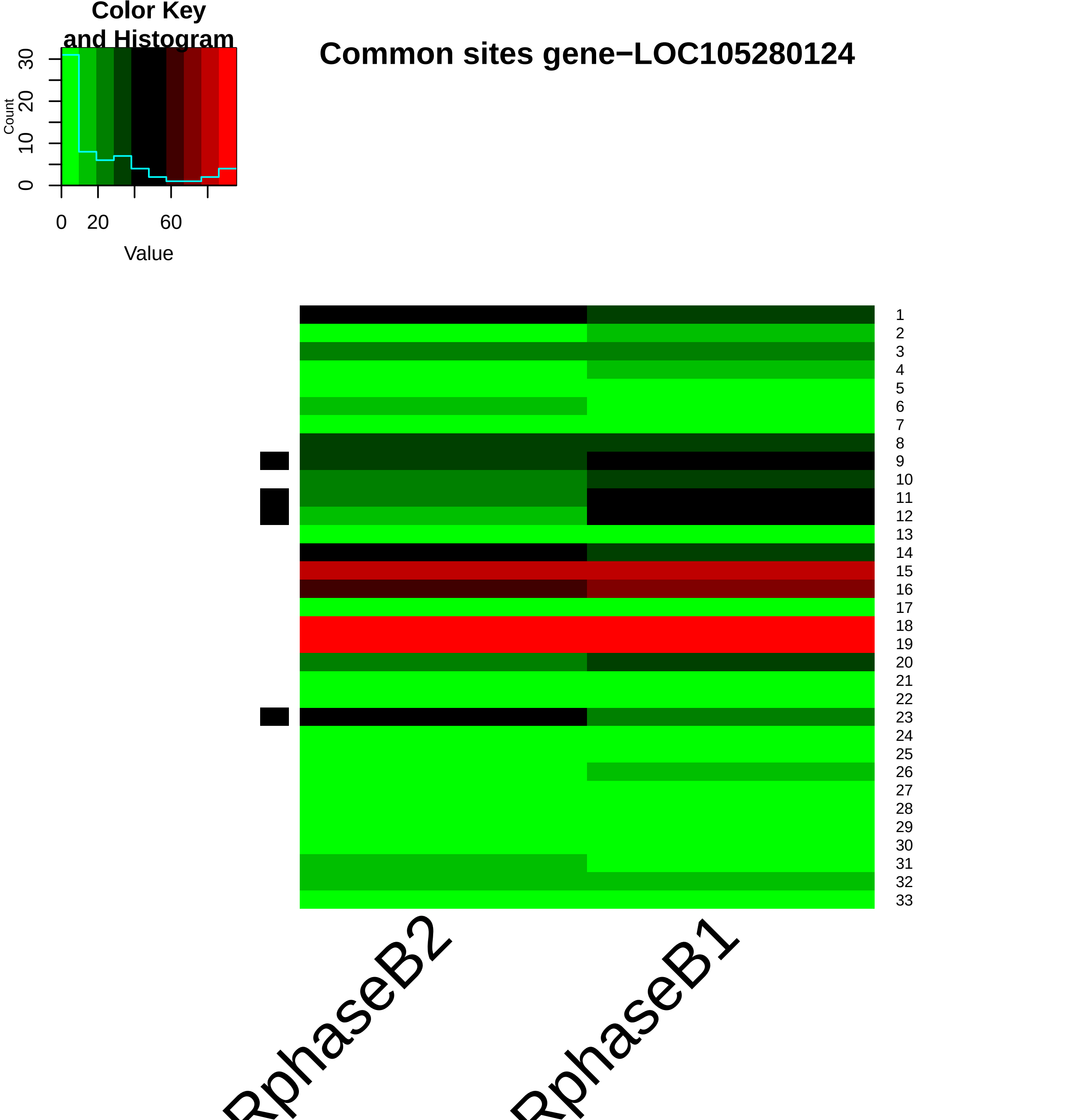

Color Key
and Histogram
Count
Value
0
20
60
0
10
20
30
Common sites gene−LOC105280124
RphaseB2
RphaseB1

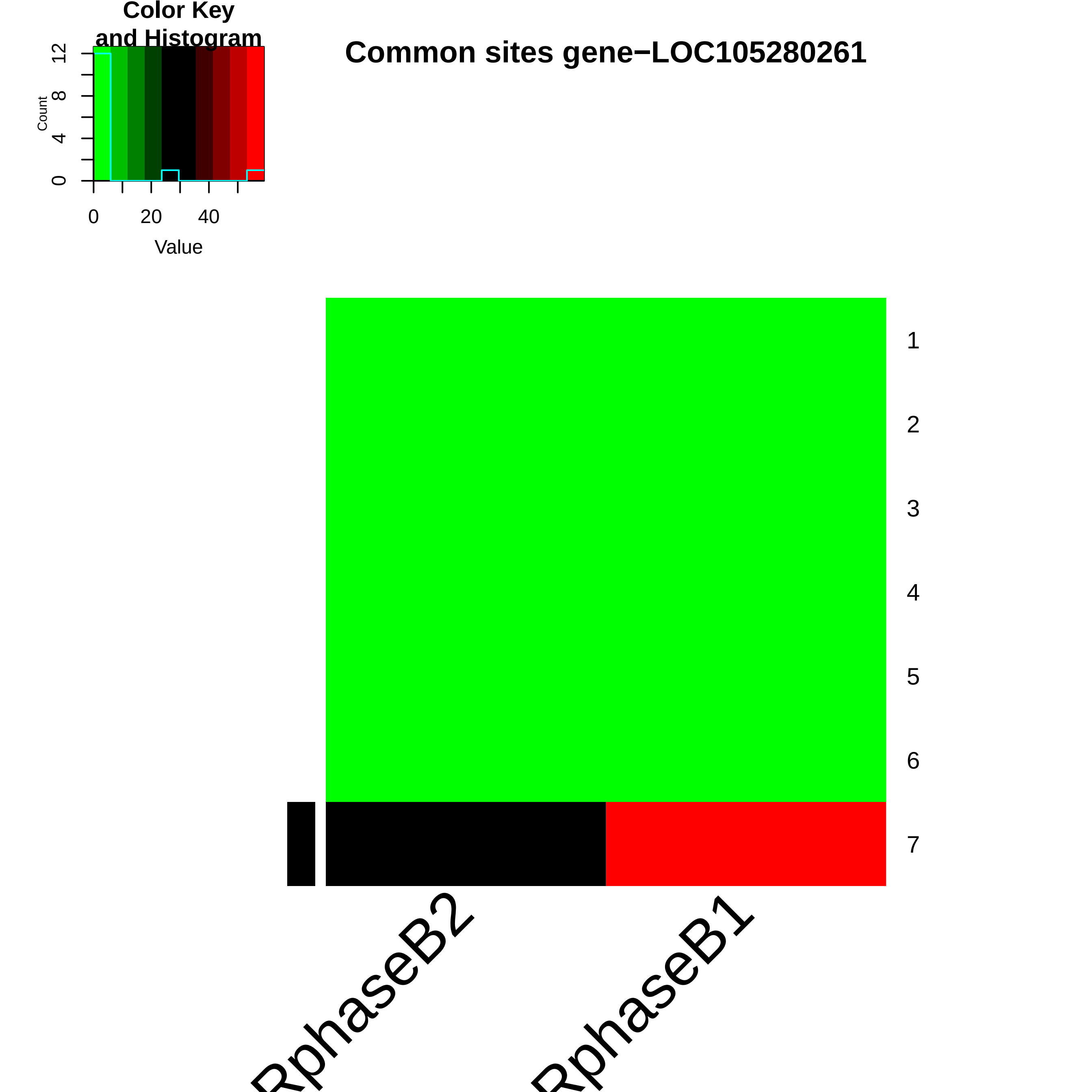

Color Key
and Histogram
Count
0
4
8
12
0
20
40
Value
Common sites gene−LOC105280261
1
2
3
4
5
6
7
RphaseB2
RphaseB1

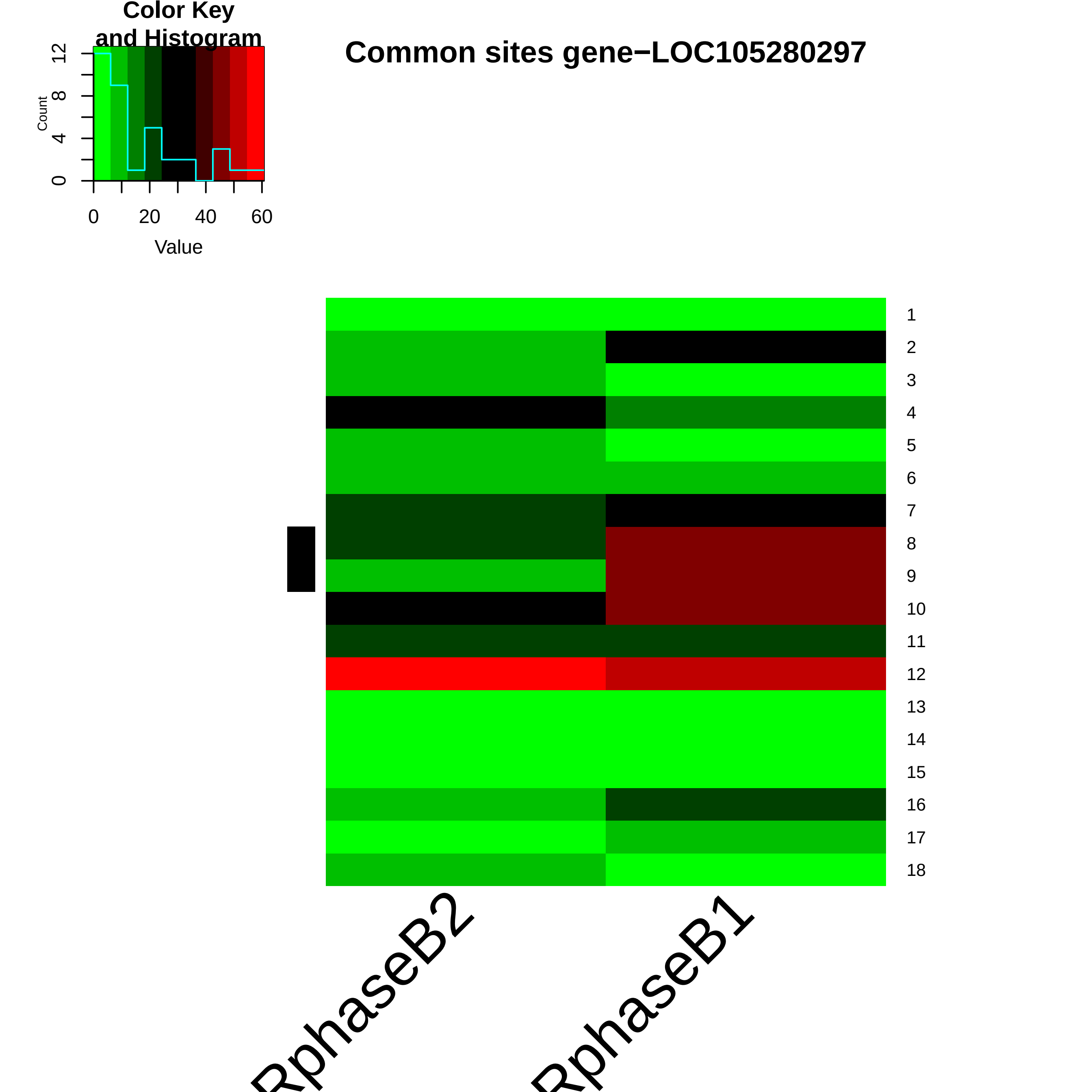

Color Key
and Histogram
Count
0
4
8
12
0
20
40
60
Value
Common sites gene−LOC105280297
RphaseB2
RphaseB1

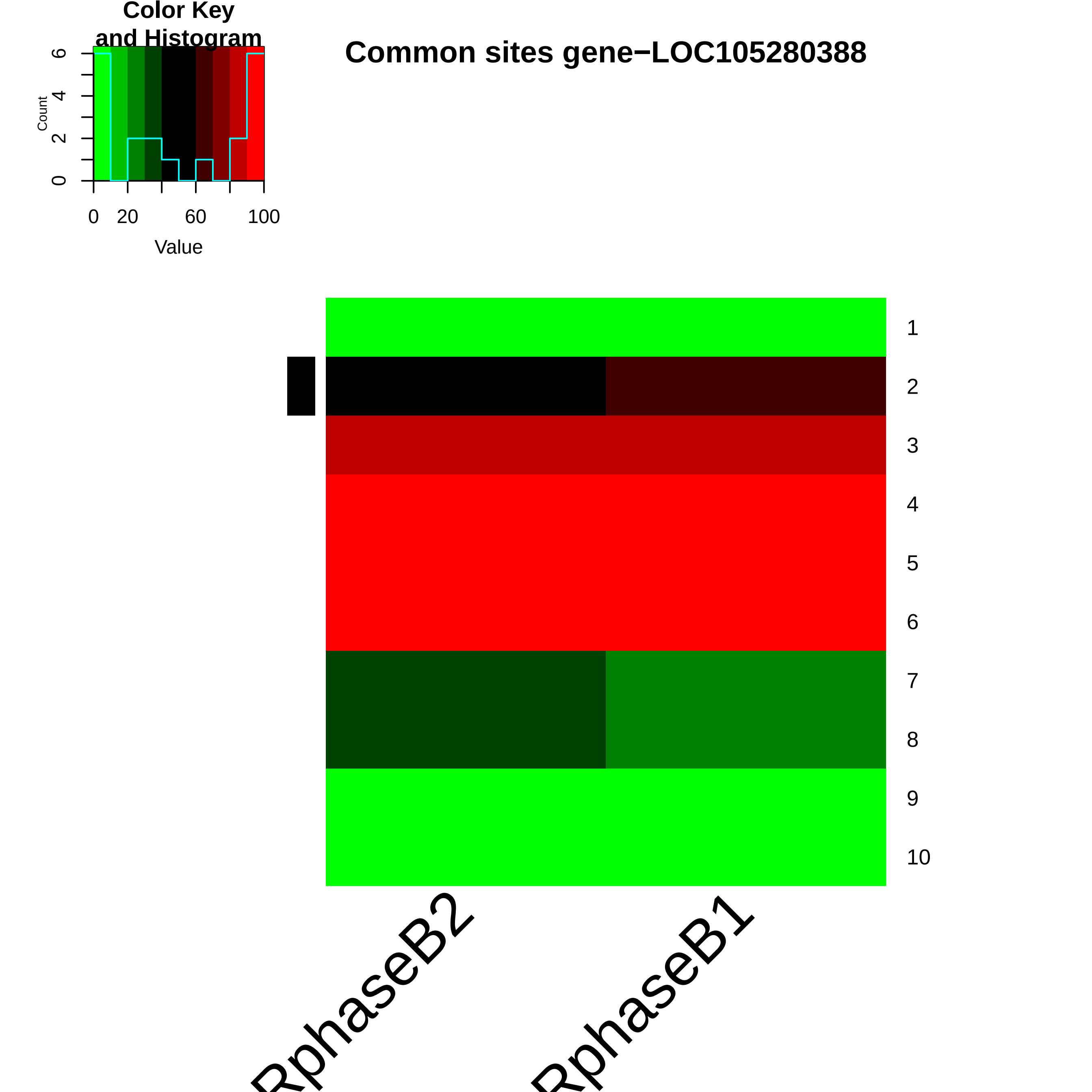

Color Key
and Histogram
Count
0
2
4
6
0
20
60
100
Value
Common sites gene−LOC105280388
1
2
3
4
5
6
7
8
9
10
RphaseB2
RphaseB1

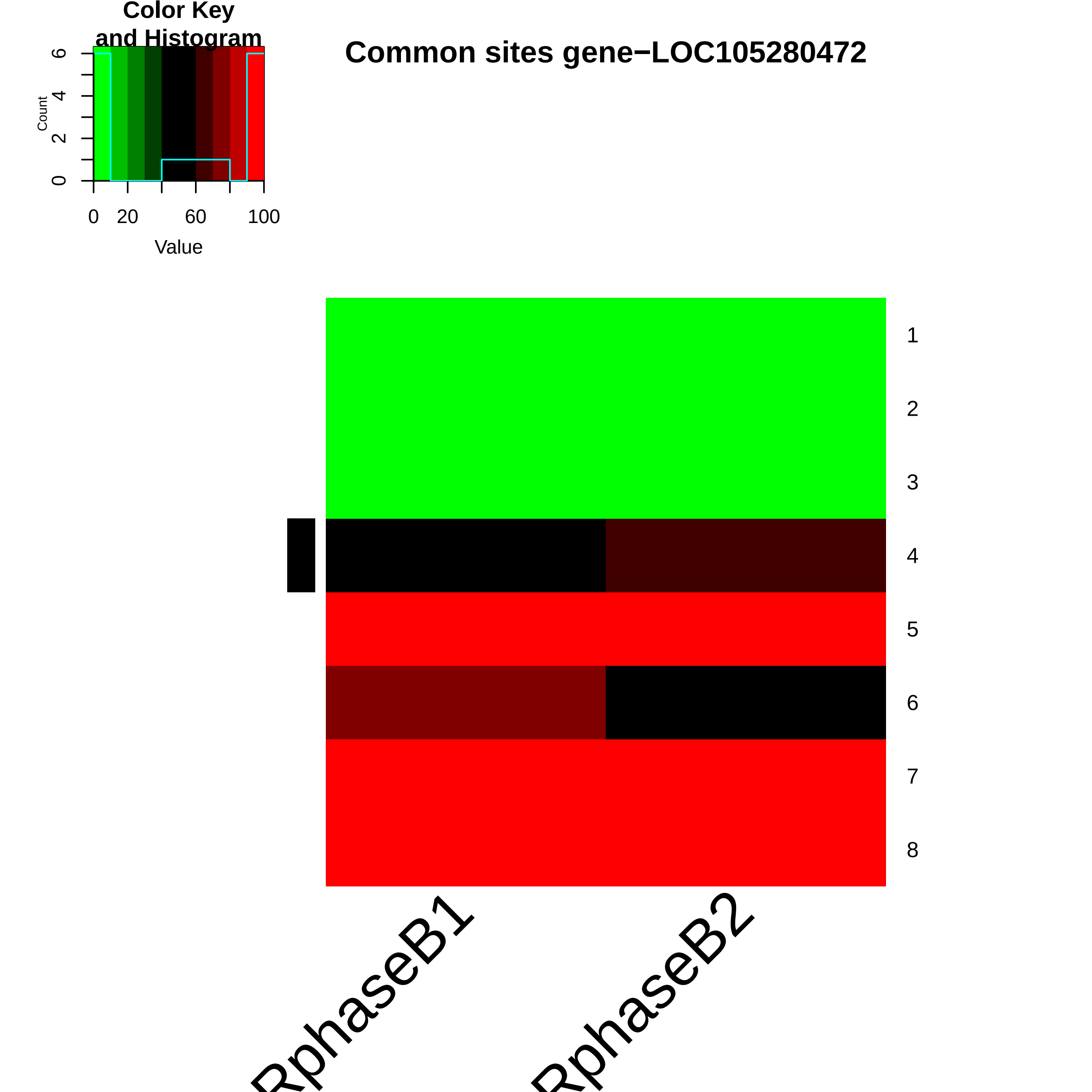

Color Key
and Histogram
Count
Value
0
20
60
100
0
2
4
6
Common sites gene−LOC105280472
1
2
3
4
5
6
7
8
RphaseB1
RphaseB2

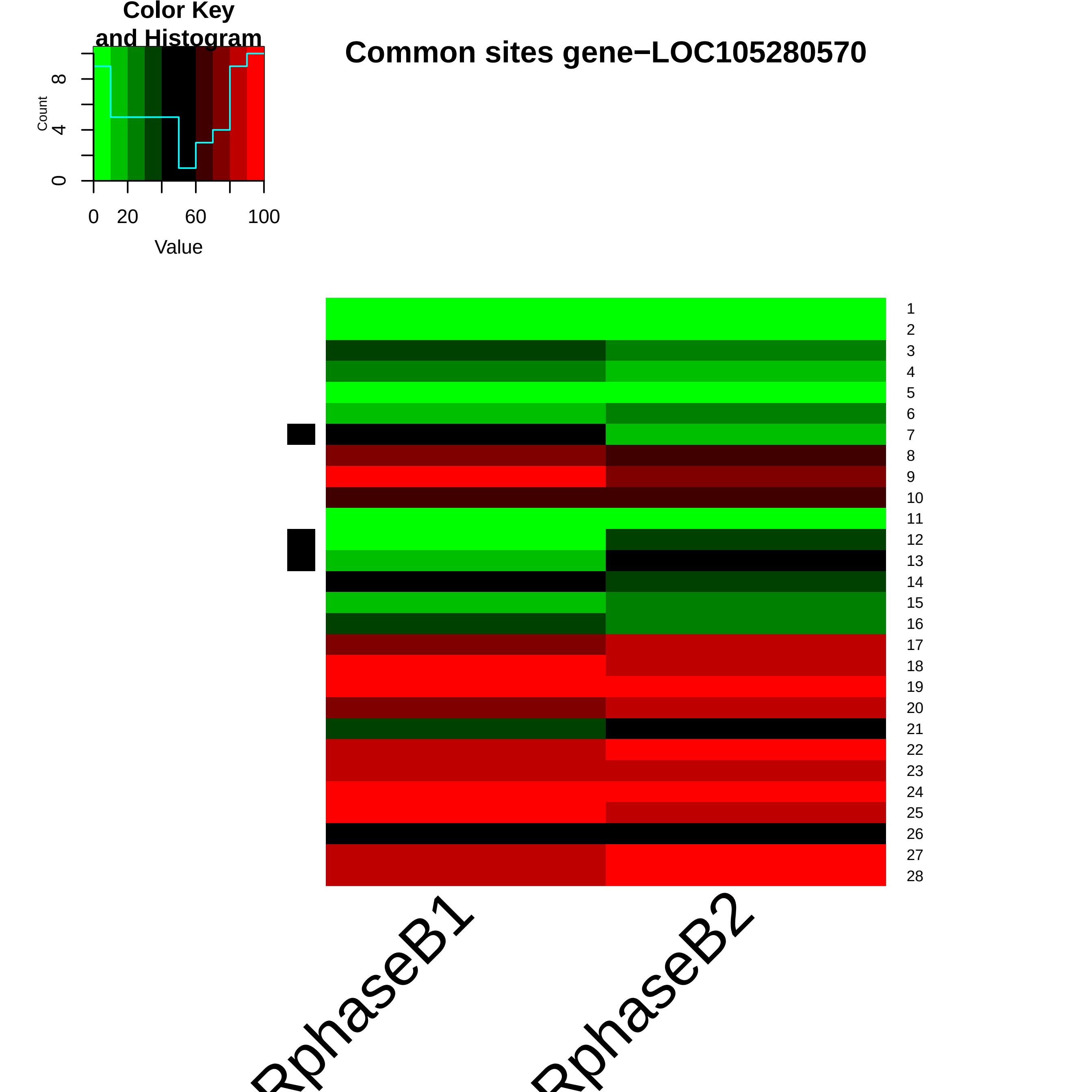

Color Key
and Histogram
Count
0
4
8
0
20
60
100
Value
Common sites gene−LOC105280570
RphaseB1
RphaseB2

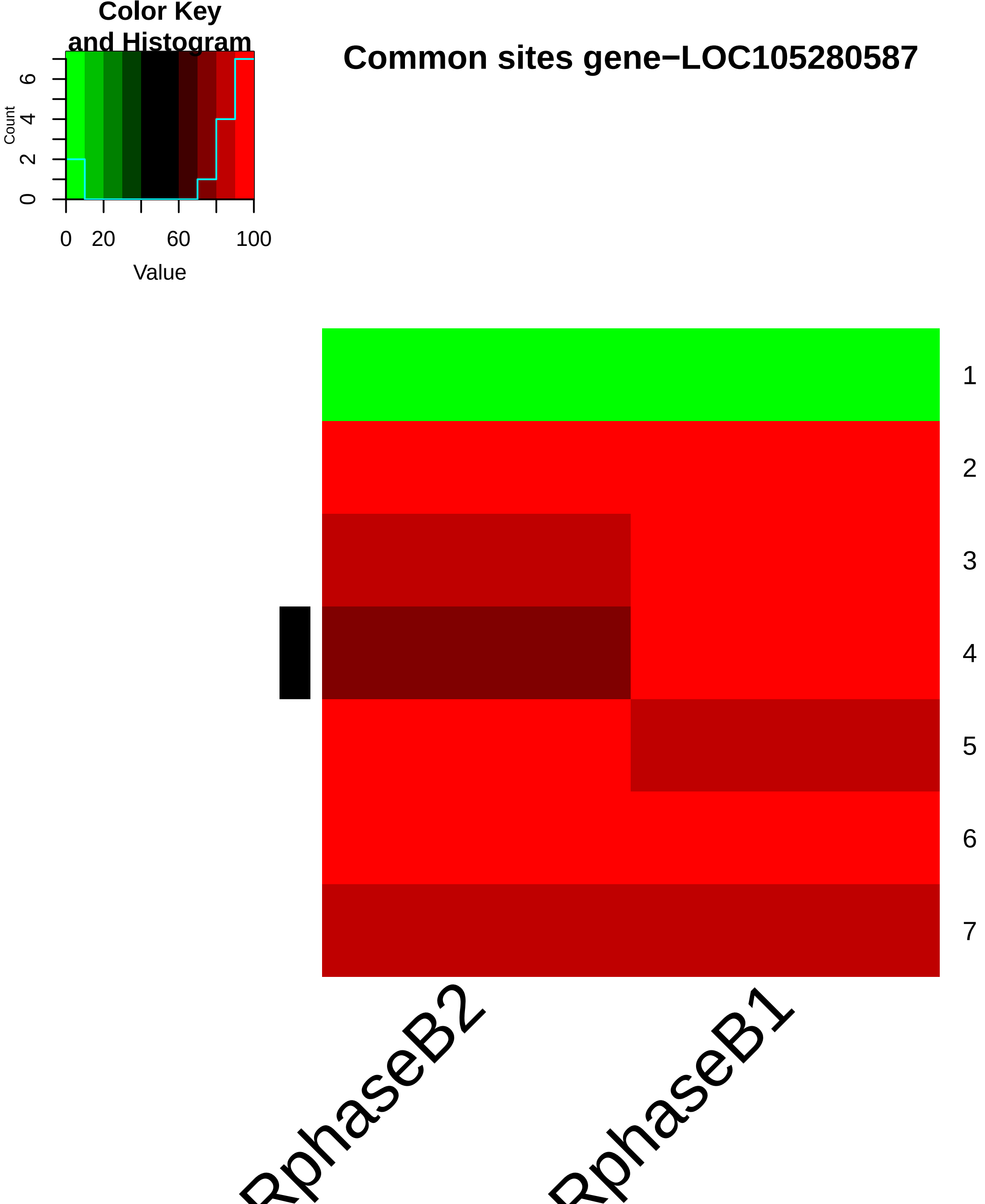
Color Key
and Histogram
Count
0
2
4
6
0
20
60
100
Value
Common sites gene−LOC105280587
1
2
3
4
5
6
7
RphaseB2
RphaseB1

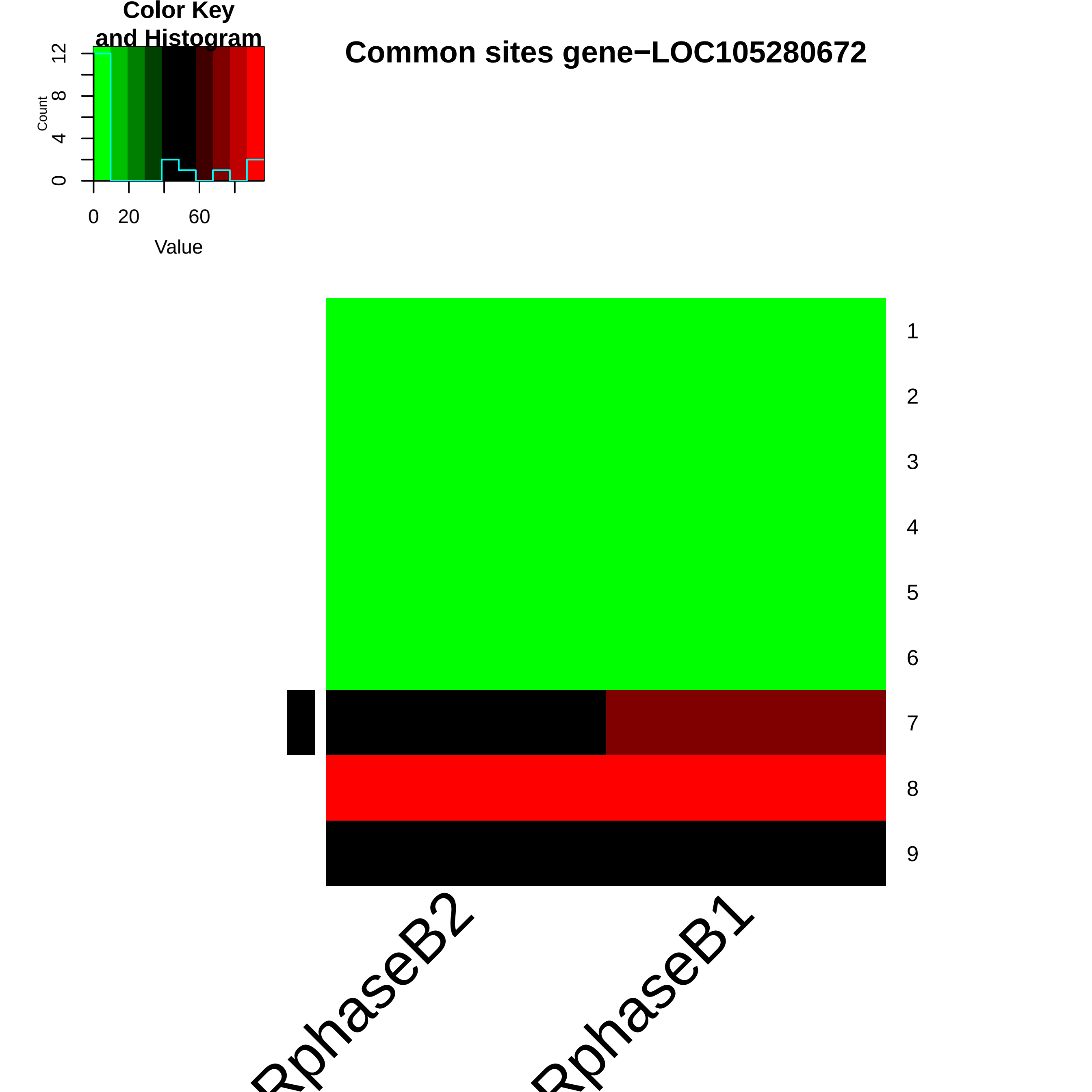

Color Key
and Histogram
Count
0
4
8
12
0
20
60
Value
Common sites gene−LOC105280672
1
2
3
4
5
6
7
8
9
RphaseB2
RphaseB1

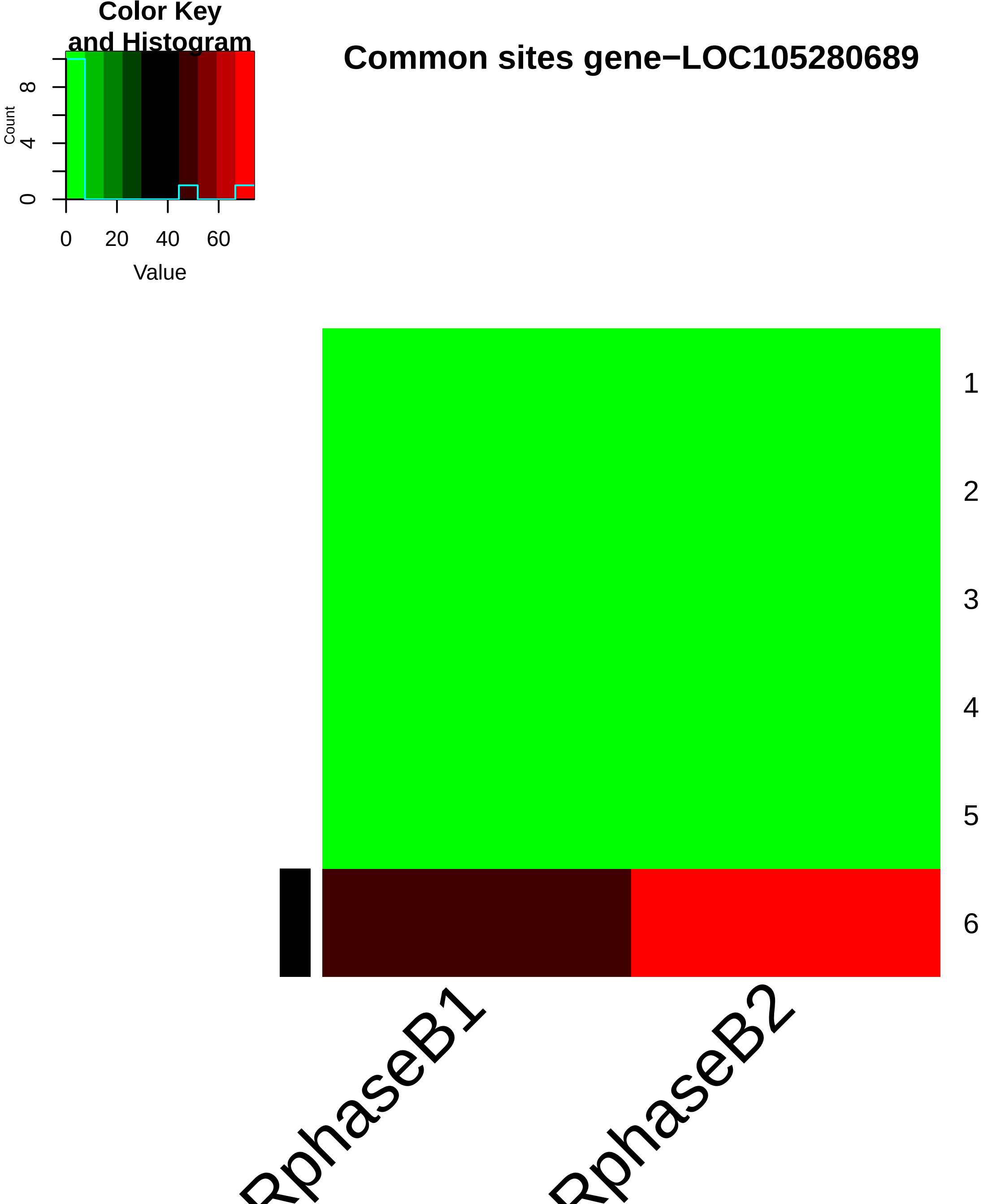

Color Key
and Histogram
Count
0
4
8
0
20
40
60
Value
Common sites gene−LOC105280689
1
2
3
4
5
6
RphaseB1
RphaseB2

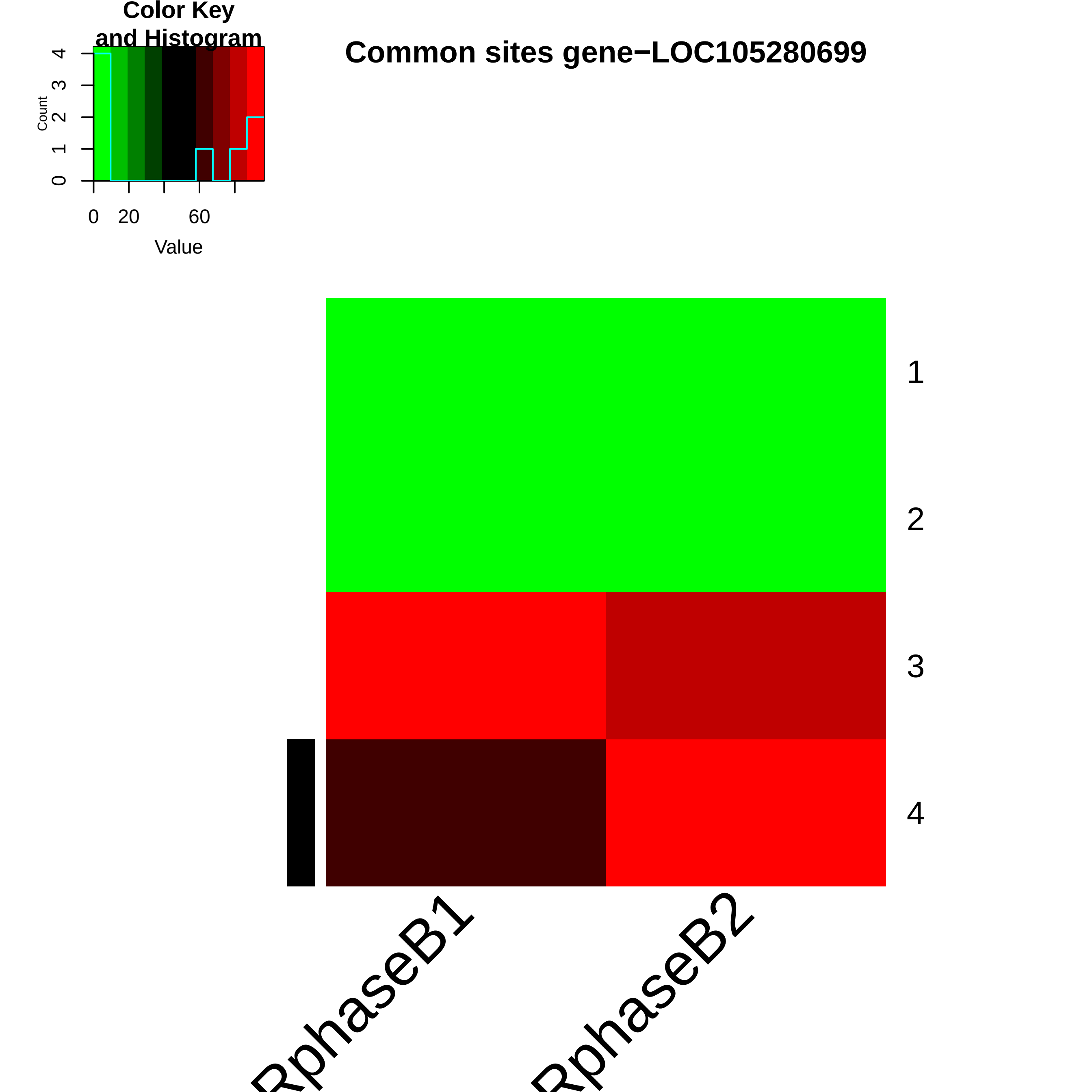

Color Key
and Histogram
Count
0
1
2
3
4
0
20
60
Value
Common sites gene−LOC105280699
1
2
3
4
RphaseB1
RphaseB2

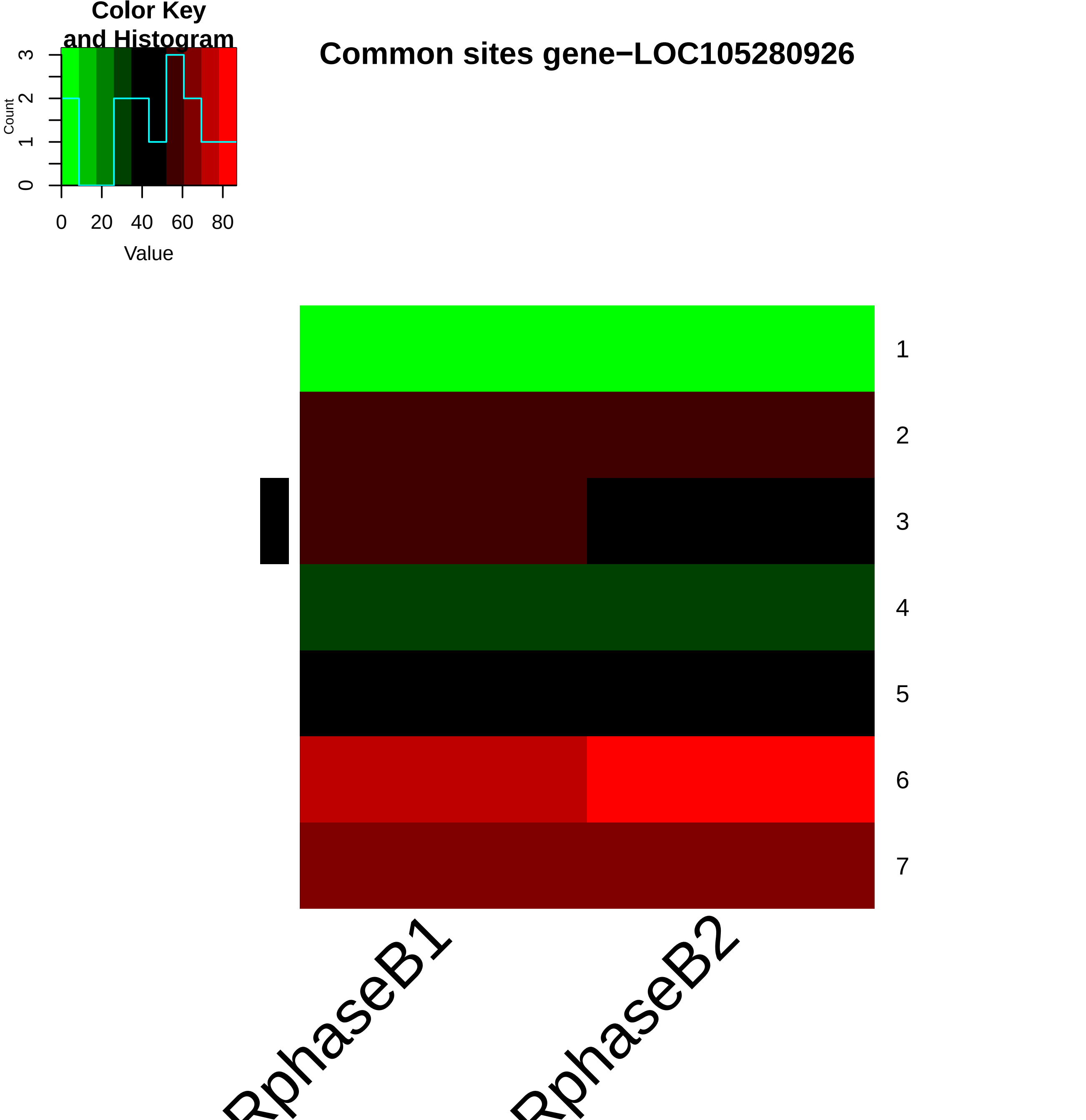
Color Key
and Histogram
Count
0
1
2
3
0
20
40
60
80
Value
Common sites gene−LOC105280926
1
2
3
4
5
6
7
RphaseB1
RphaseB2

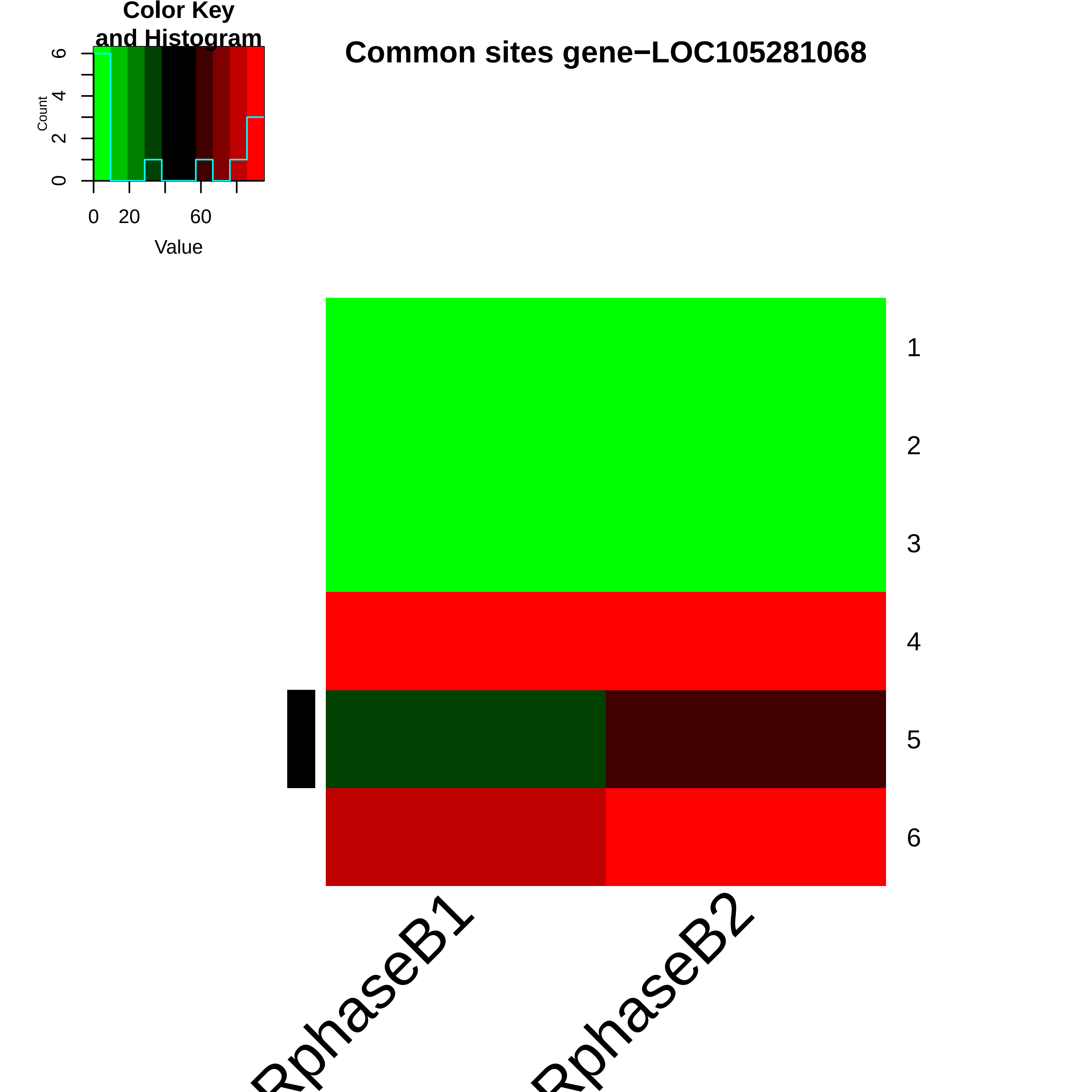

Color Key
and Histogram
Count
Value
0
20
60
0
2
4
6
Common sites gene−LOC105281068
1
2
3
4
5
6
RphaseB1
RphaseB2

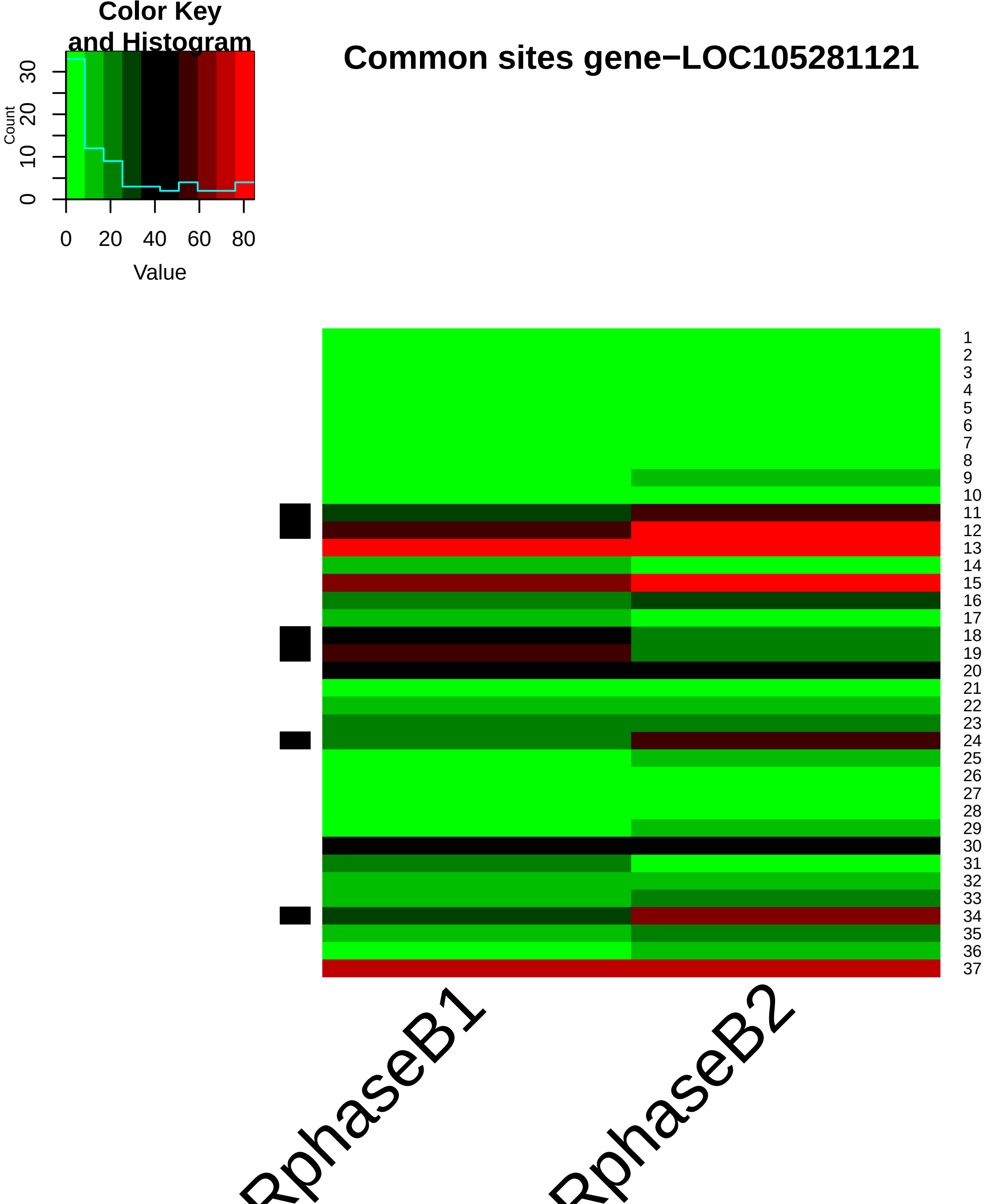

Color Key
and Histogram
Count
Value
0
10
20
30
40
60
80
Common sites gene−LOC105281121
RphaseB1
RphaseB2

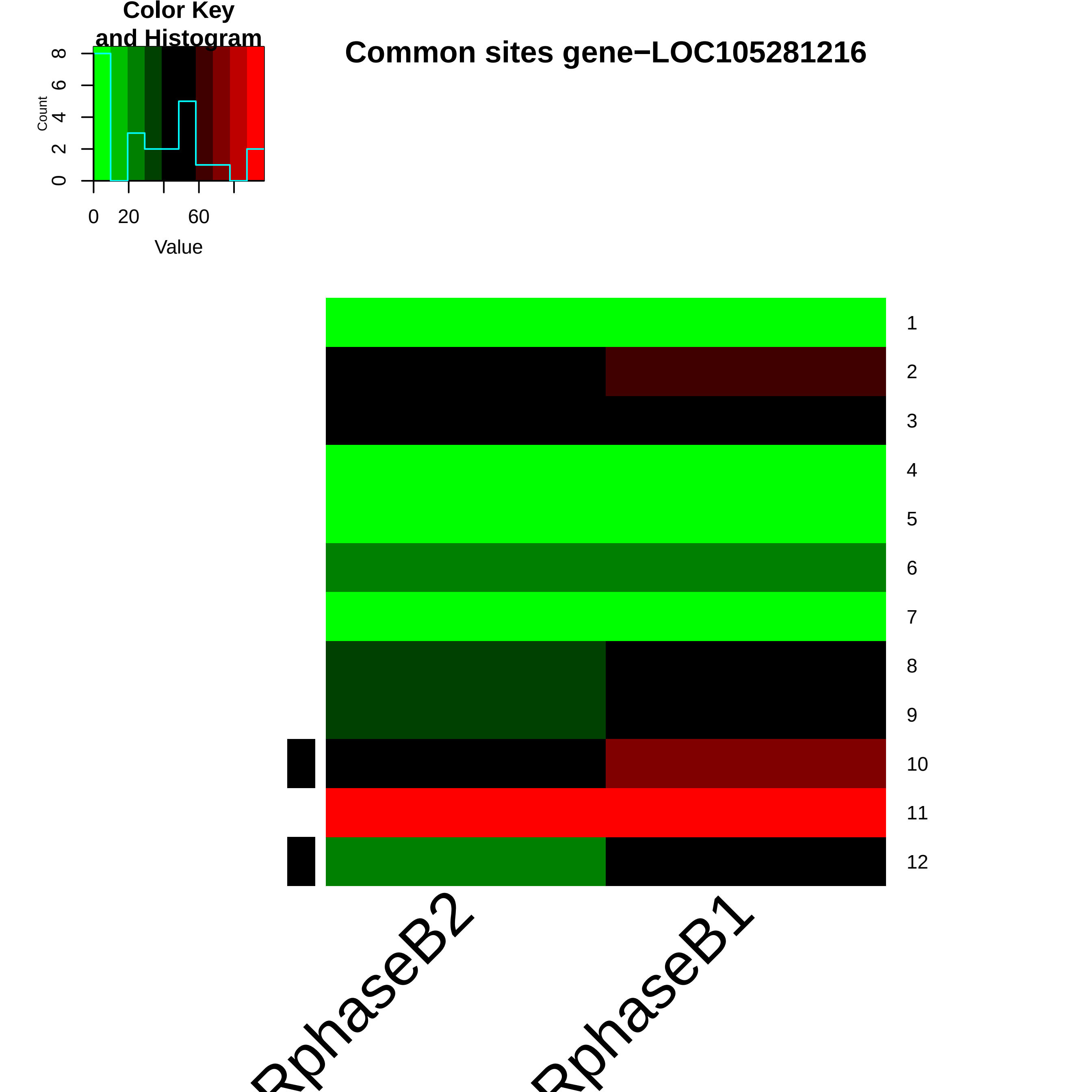

Color Key and Histogram
Count
Value
Common sites gene−LOC105281216
1
2
3
4
5
6
7
8
9
10
11
12
RphaseB2
RphaseB1

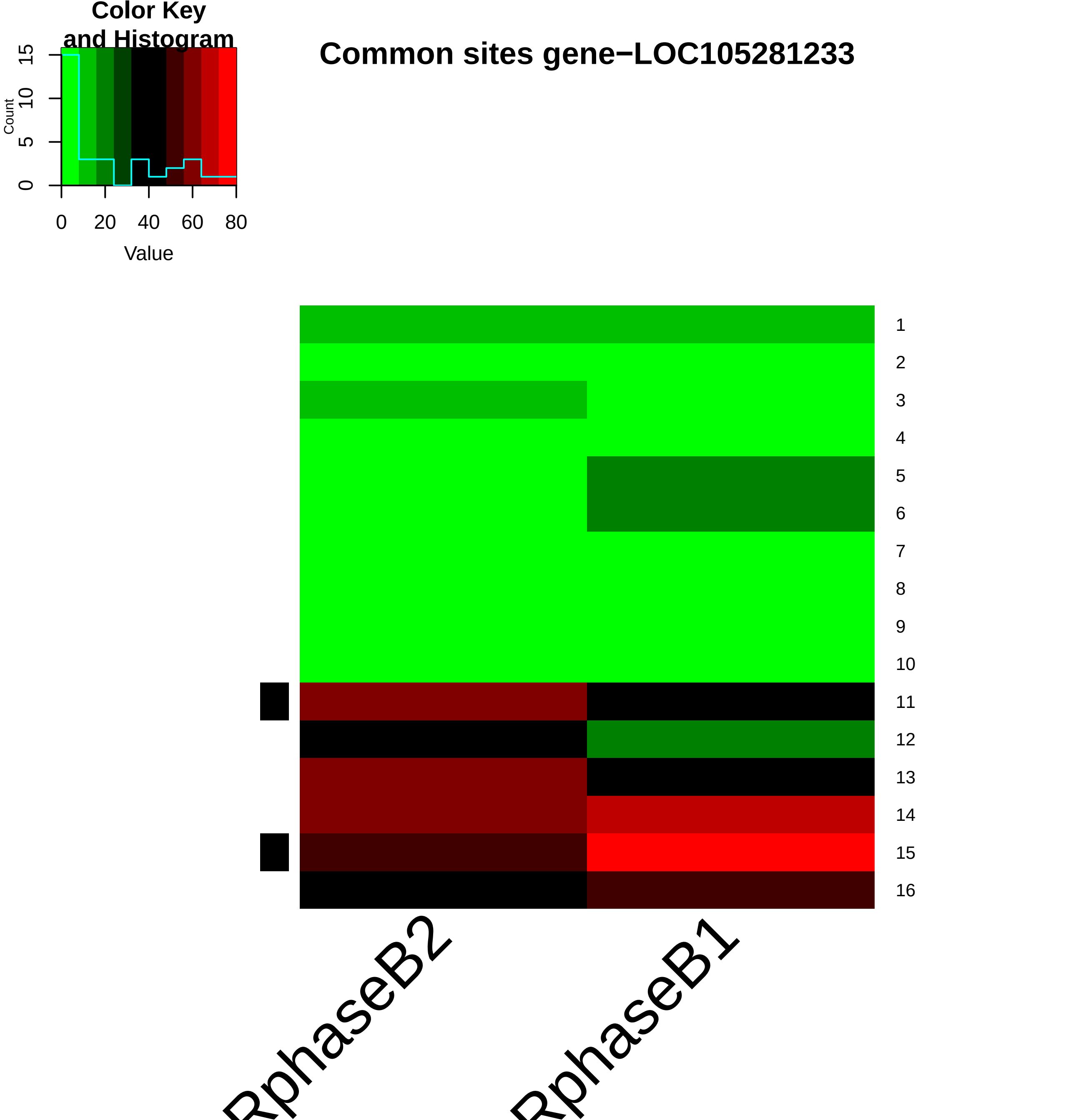

Color Key
and Histogram
Count
15
10
5
0
0
20
40
60
80
Value
Common sites gene−LOC105281233
RphaseB2
RphaseB1

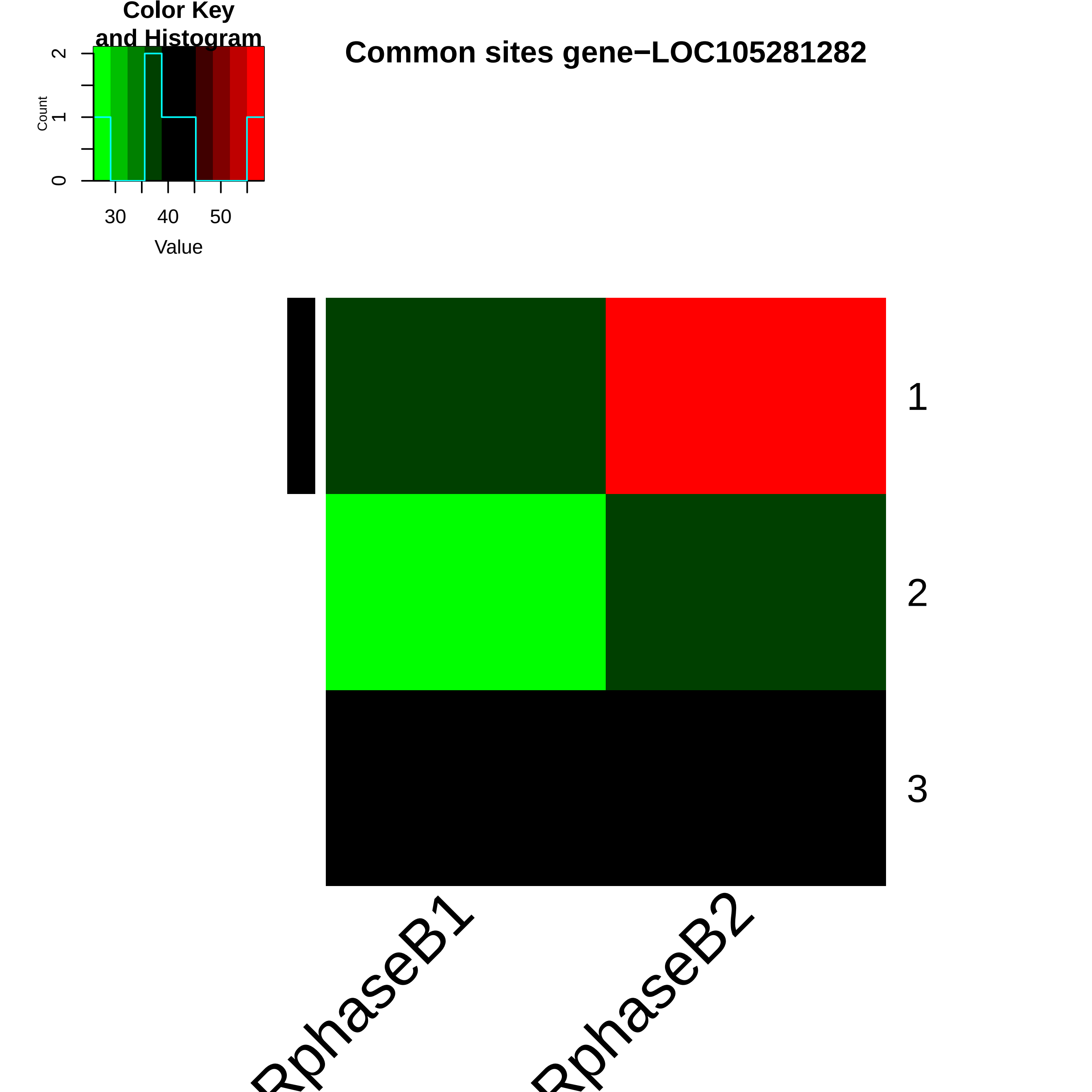

Color Key
and Histogram
Count
0
1
2
30
40
50
Value
Common sites gene−LOC105281282
1
2
3
RphaseB1
RphaseB2

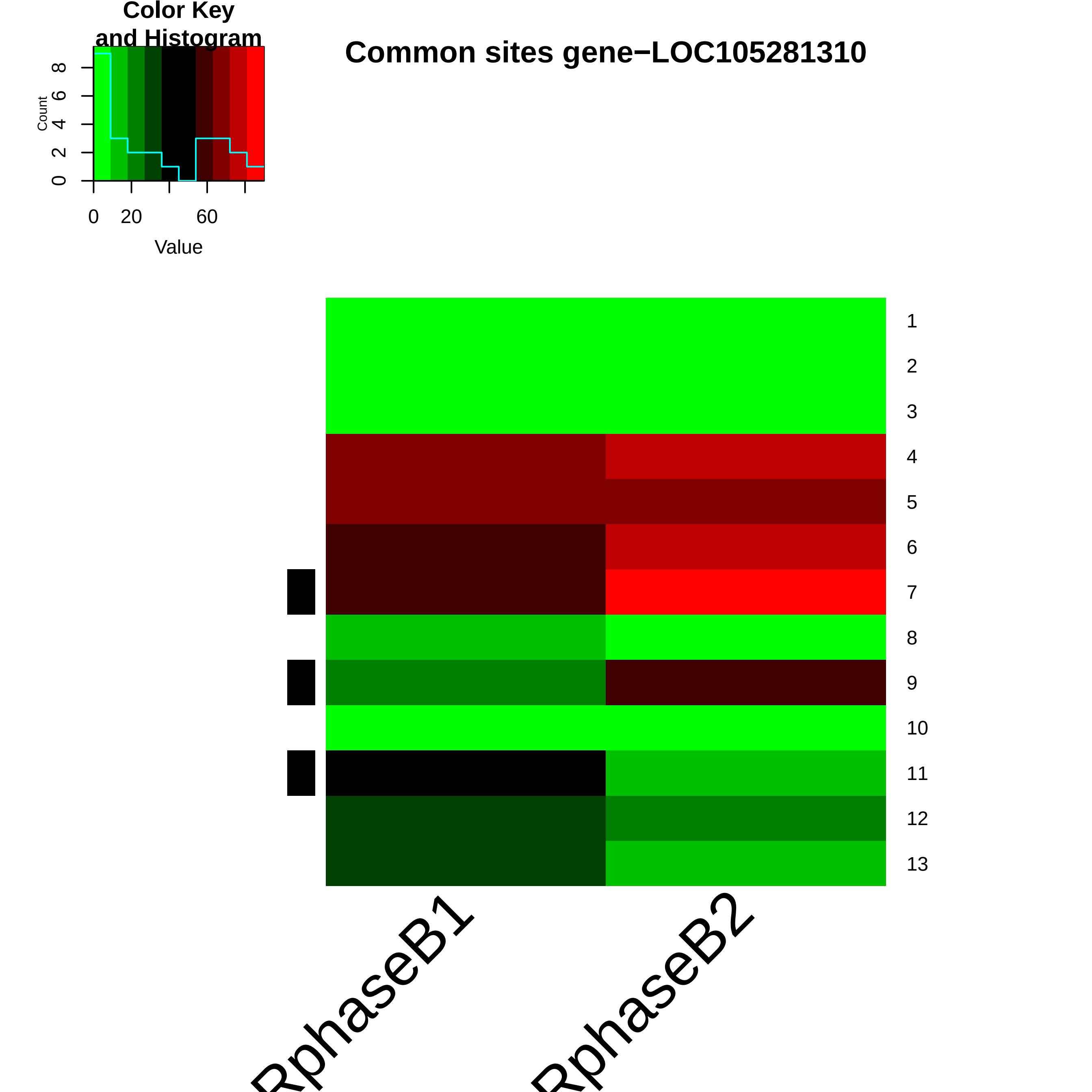

Color Key
and Histogram
Count
0
2
4
6
8
0
20
60
Value
Common sites gene−LOC105281310
1
2
3
4
5
6
7
8
9
10
11
12
13
RphaseB1
RphaseB2

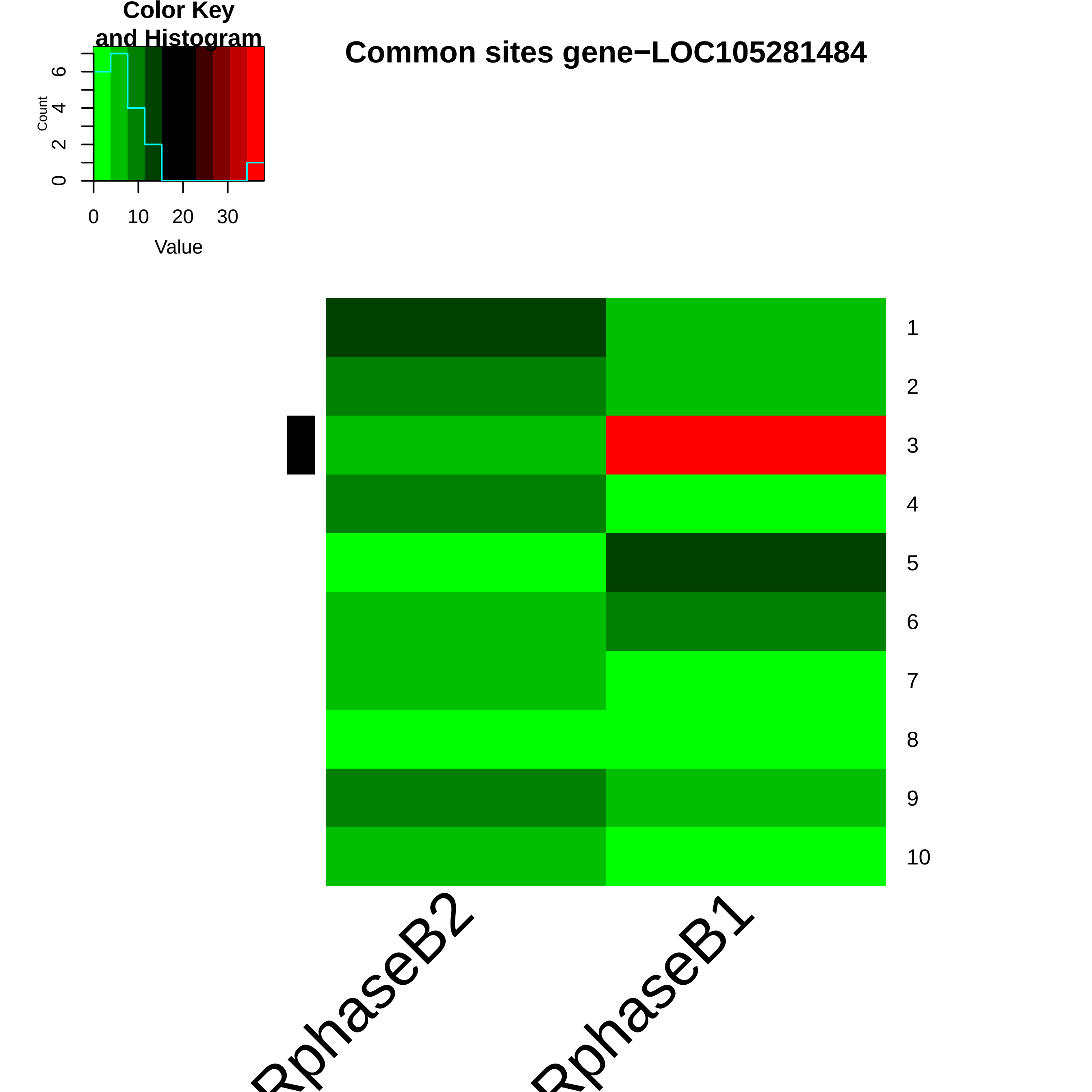
Color Key
and Histogram
Count
0
2
4
6
0
10
20
30
Value
Common sites gene−LOC105281484
1
2
3
4
5
6
7
8
9
10
RphaseB2
RphaseB1

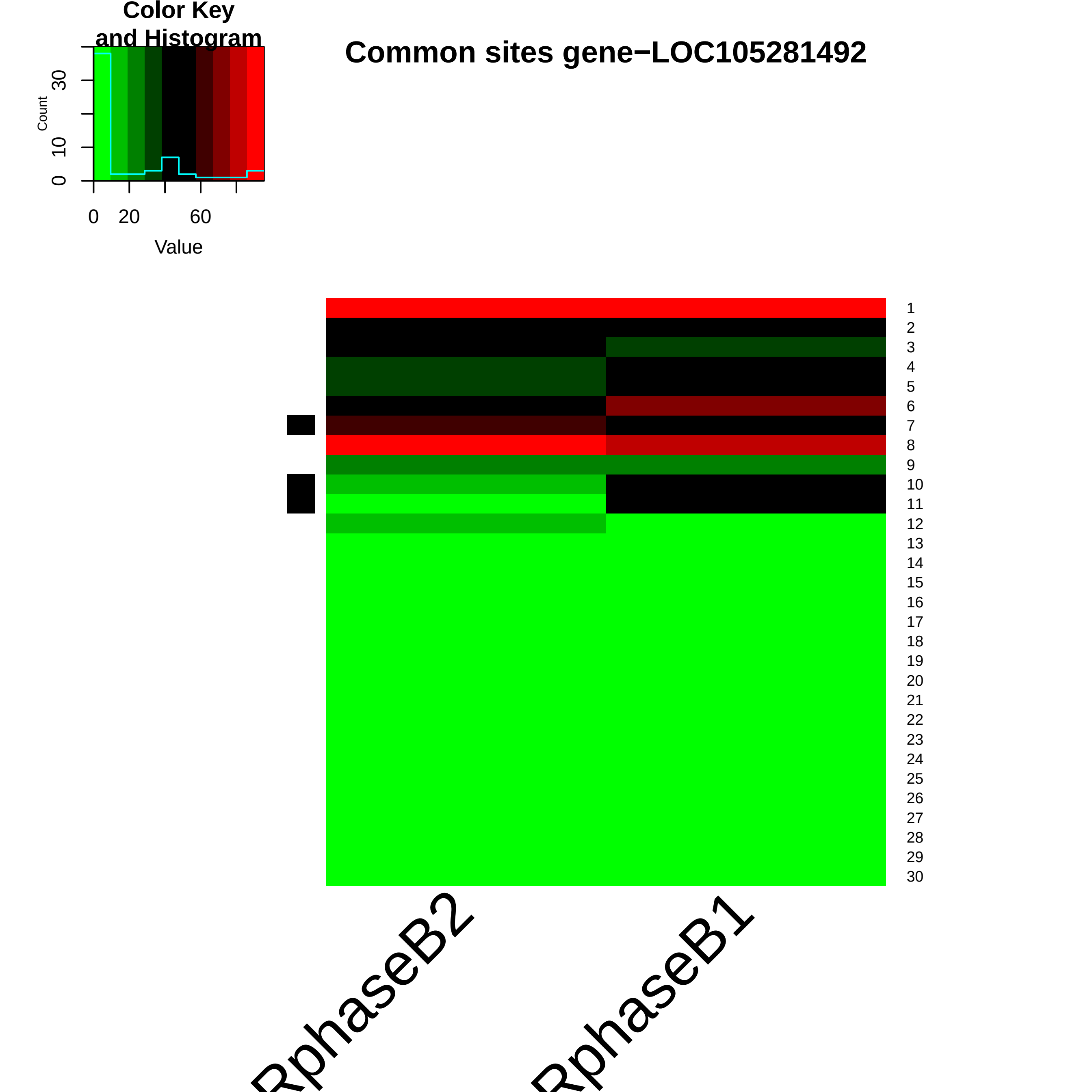
Color Key and Histogram
Count
0
10
30
0
20
60
Value
Common sites gene−LOC105281492
RphaseB2
RphaseB1

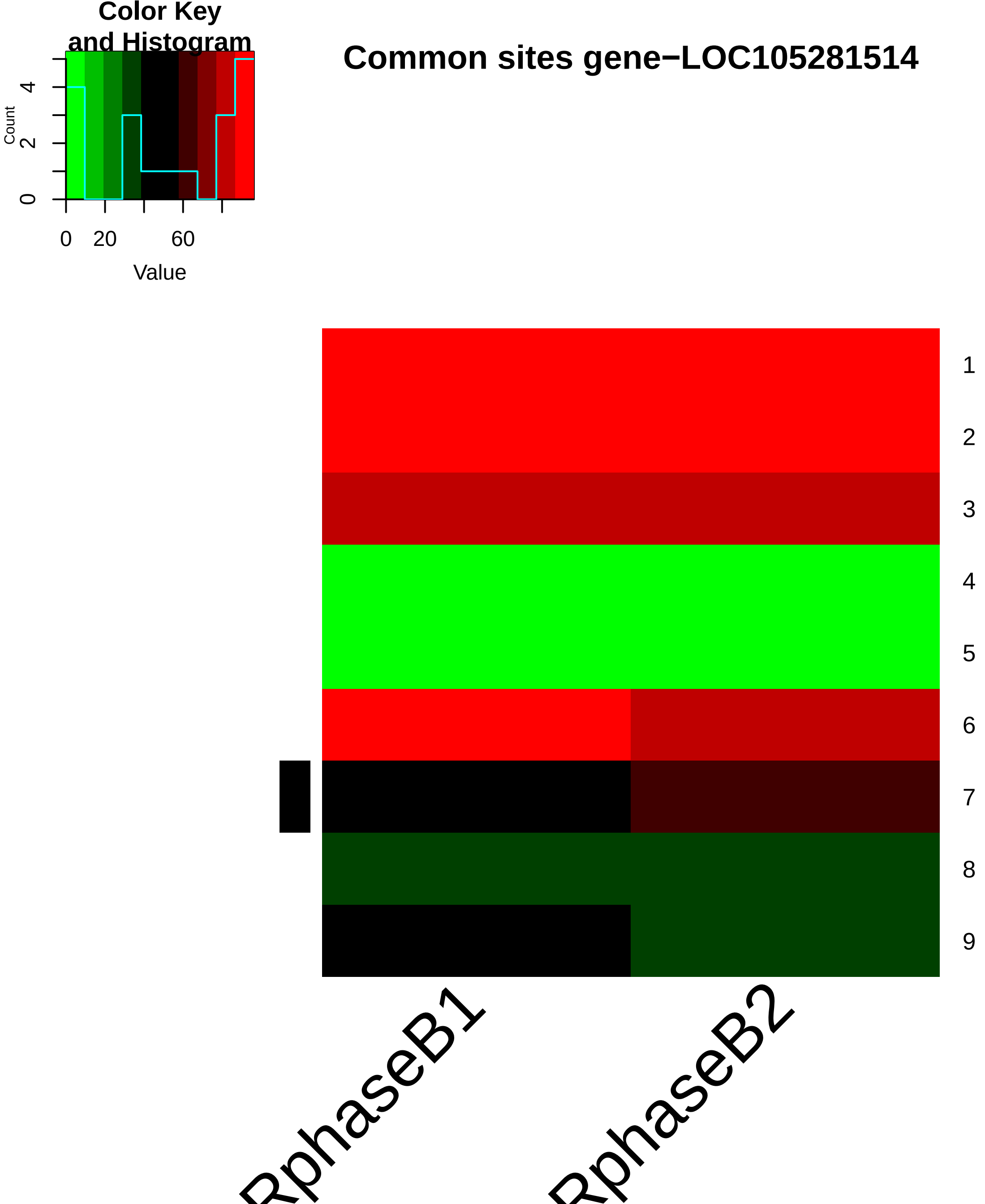
Color Key
and Histogram
Common sites gene−LOC105281514
Count
0
2
4
0
20
60
Value
1
2
3
4
5
6
7
8
9
RphaseB1
RphaseB2

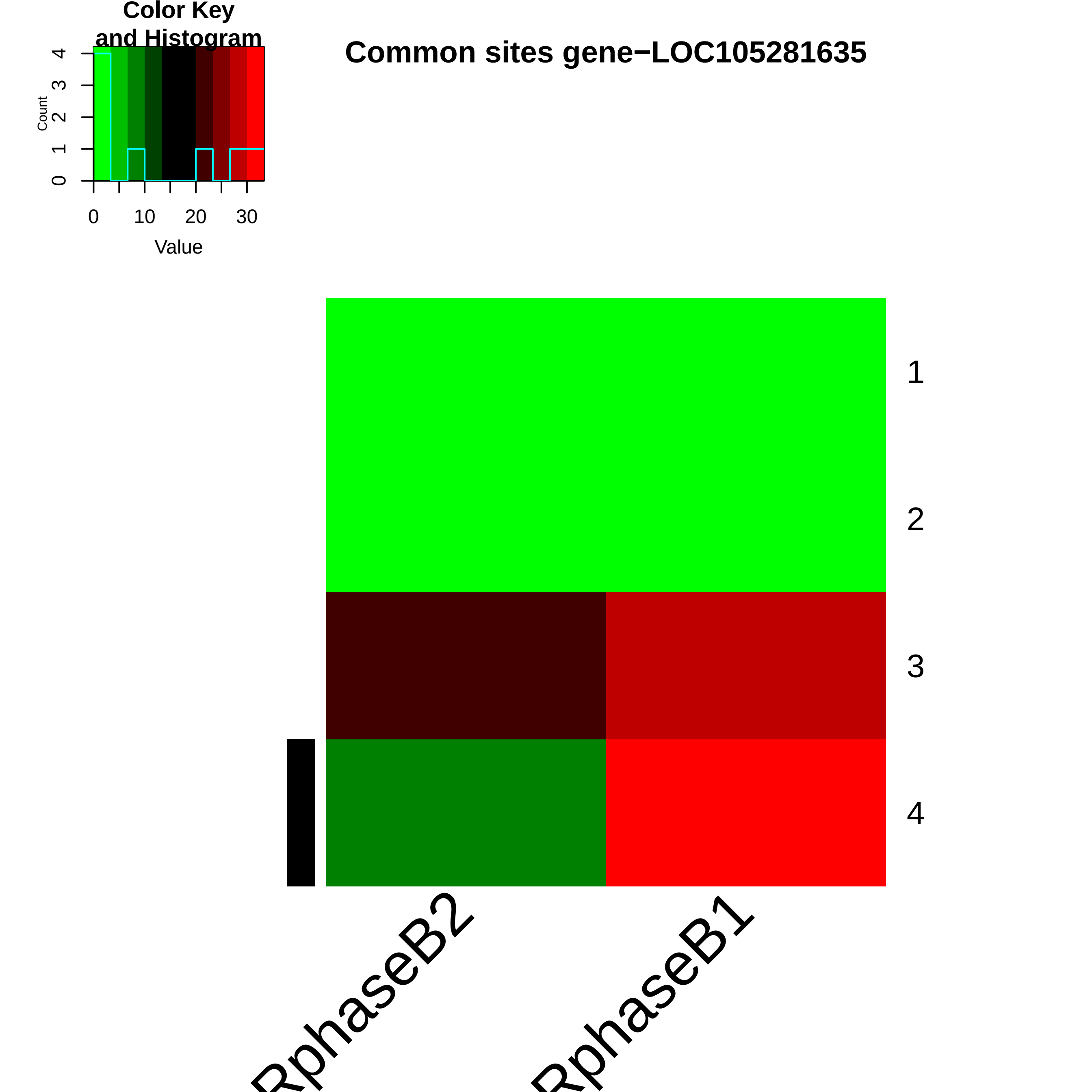

Color Key
and Histogram
Count
Value
0
10
20
30
0
1
2
3
4
Common sites gene−LOC105281635
1
2
3
4
RphaseB2
RphaseB1

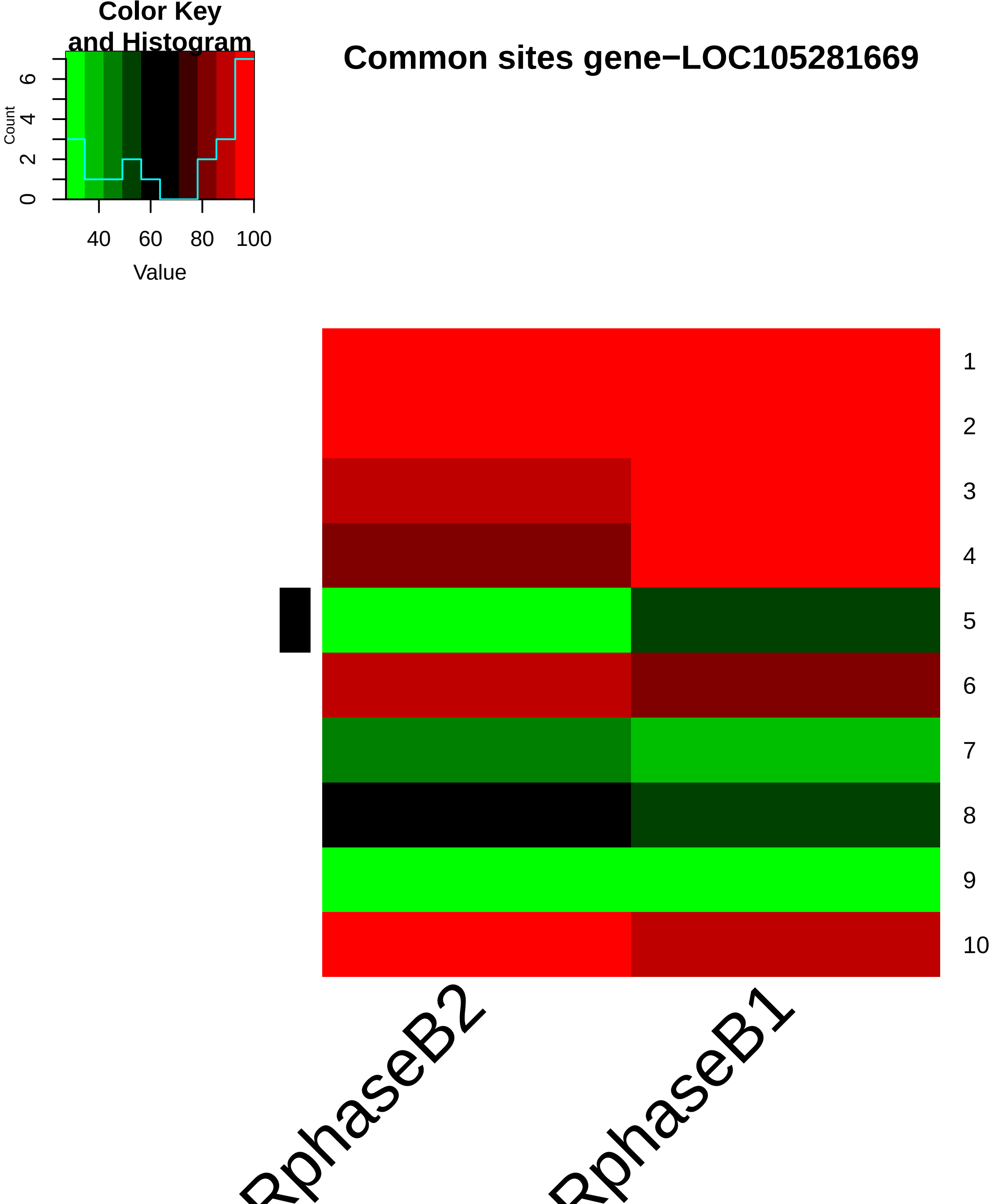
Color Key
and Histogram
Count
0
2
4
6
40
60
80
100
Value
Common sites gene−LOC105281669
1
2
3
4
5
6
7
8
9
10
RphaseB2
RphaseB1

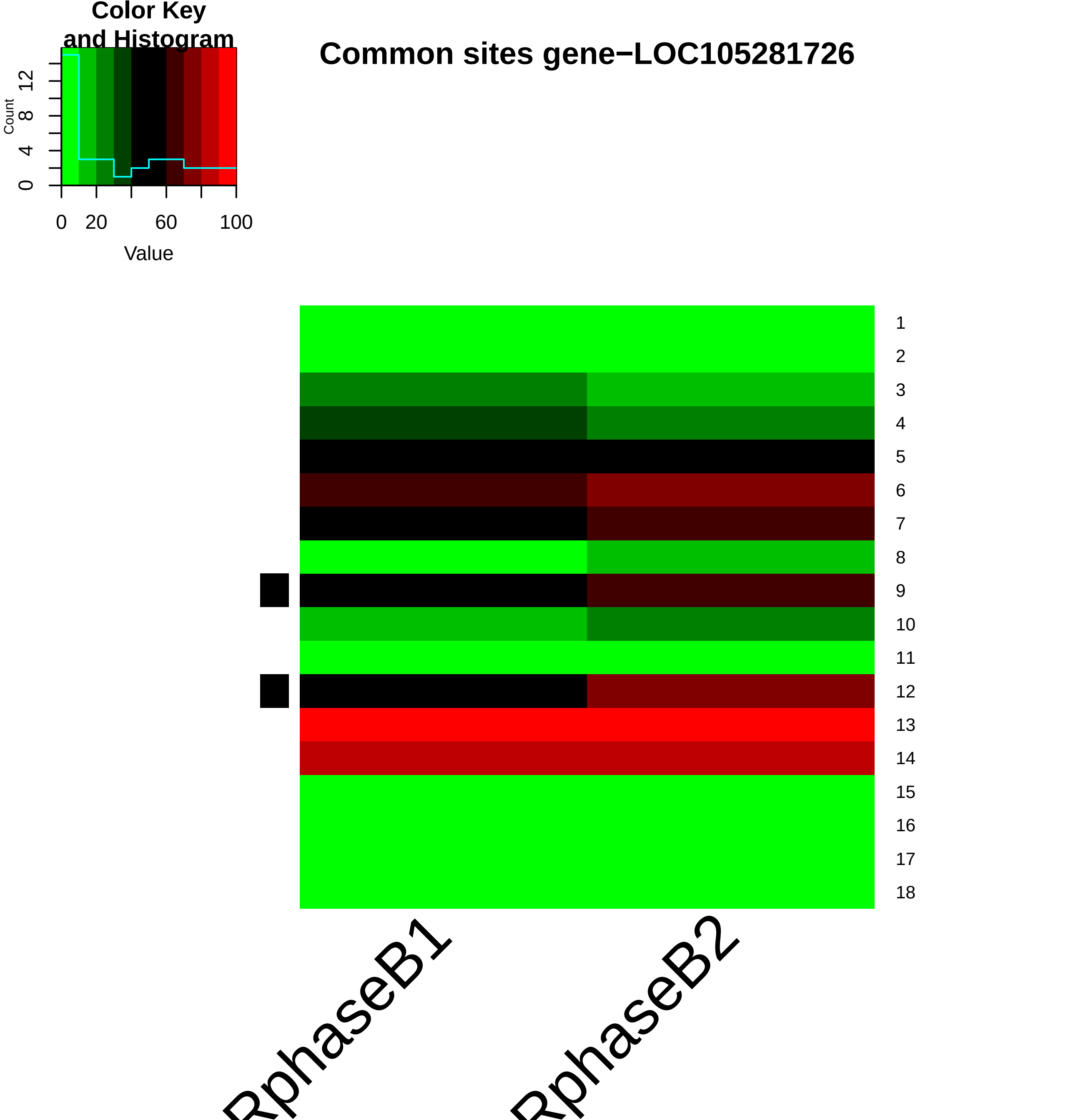
Color Key
and Histogram
Count
0
4
8
12
0
20
60
100
Value
Common sites gene−LOC105281726
RphaseB1
RphaseB2

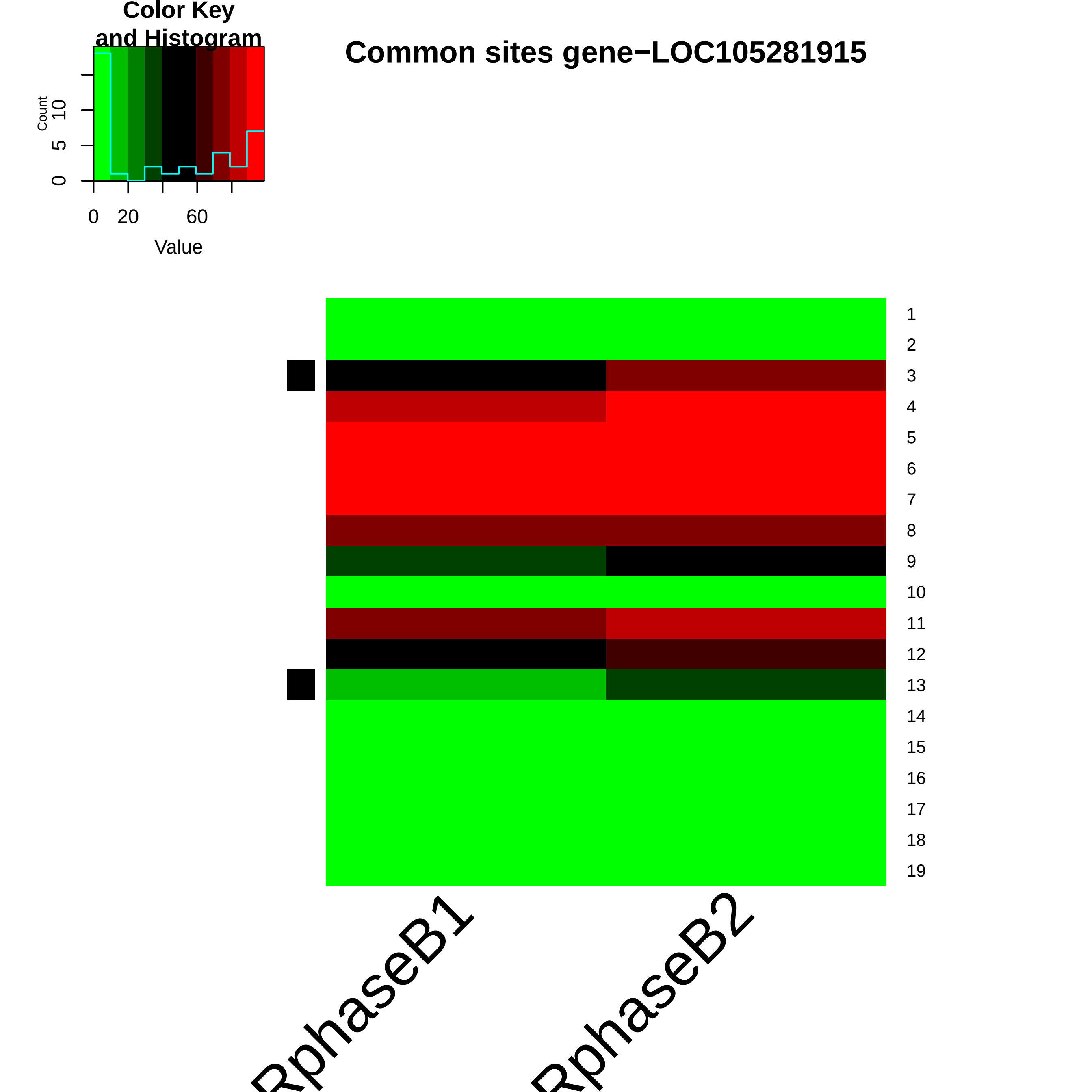

Color Key
and Histogram
Count
0
5
10
0
20
60
Value
Common sites gene−LOC105281915
RphaseB1
RphaseB2

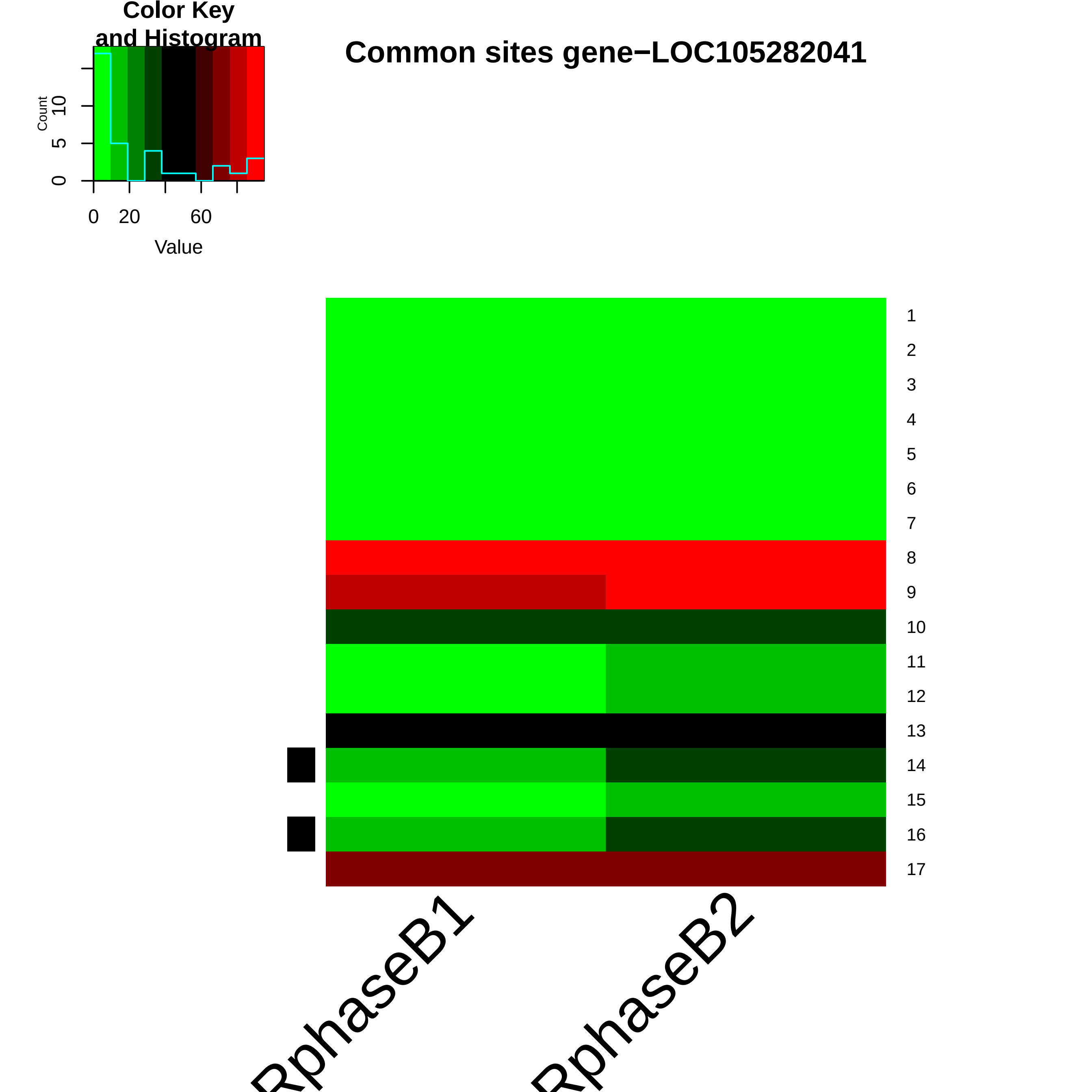

Color Key
and Histogram
Count
0
5
10
0
20
60
Value
Common sites gene−LOC105282041
RphaseB1
RphaseB2

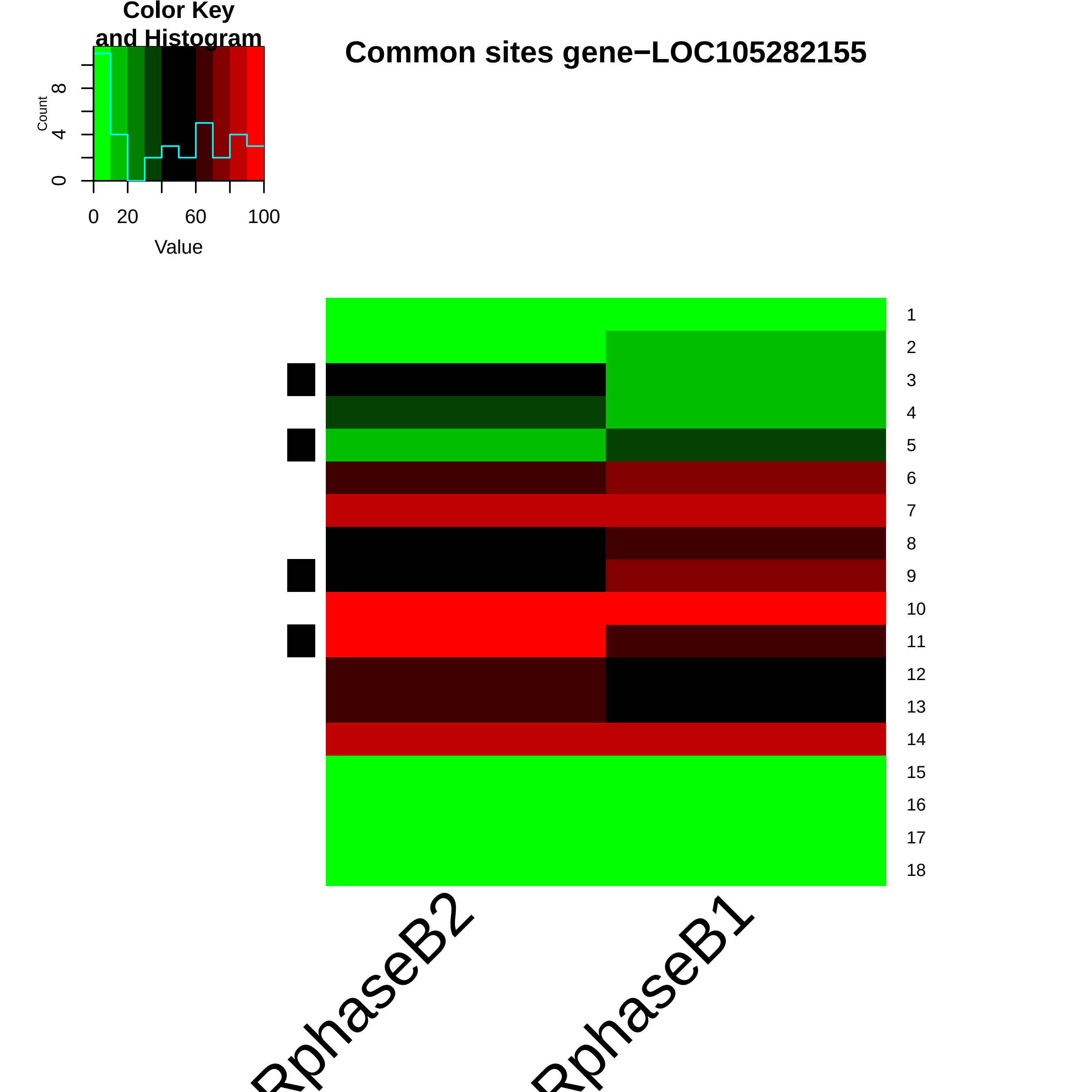

Color Key
and Histogram
Count
0
4
8
0
20
60
100
Value
Common sites gene−LOC105282155
RphaseB2
RphaseB1

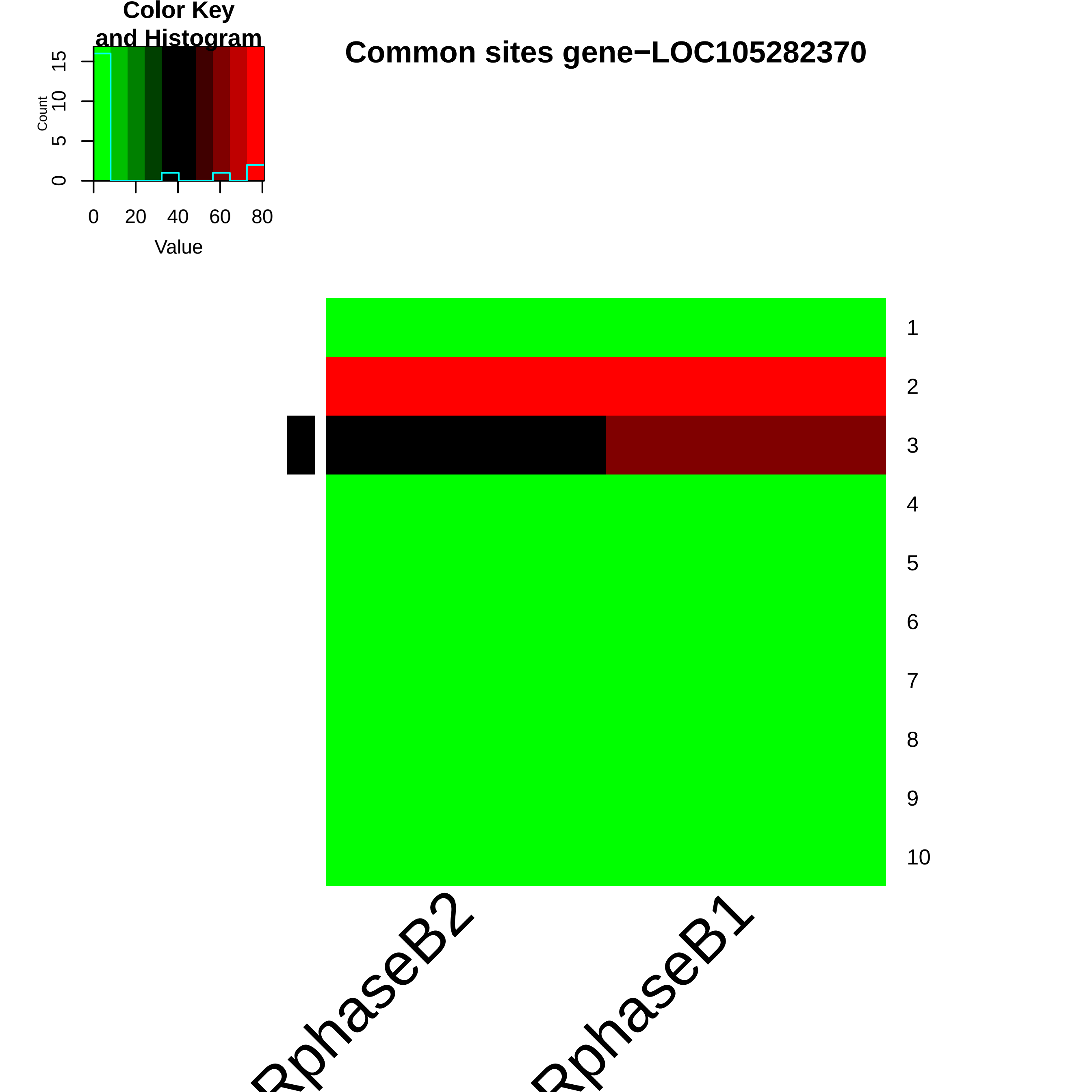

Color Key
and Histogram
Count
0
5
10
15
0
20
40
60
80
Value
Common sites gene−LOC105282370
1
2
3
4
5
6
7
8
9
10
RphaseB2
RphaseB1

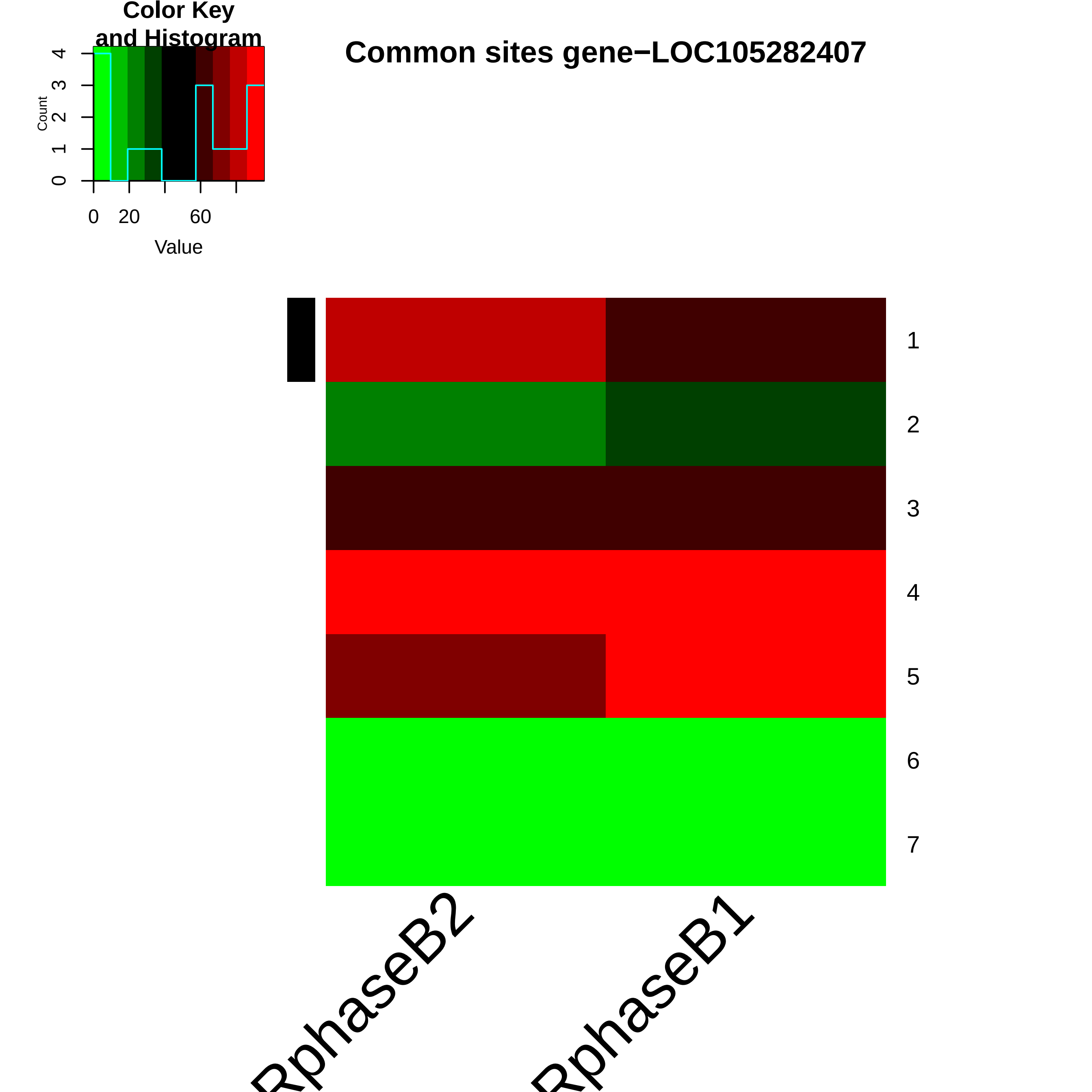

Color Key
and Histogram
Count
0
1
2
3
4
0
20
60
Value
Common sites gene−LOC105282407
1
2
3
4
5
6
7
RphaseB2
RphaseB1

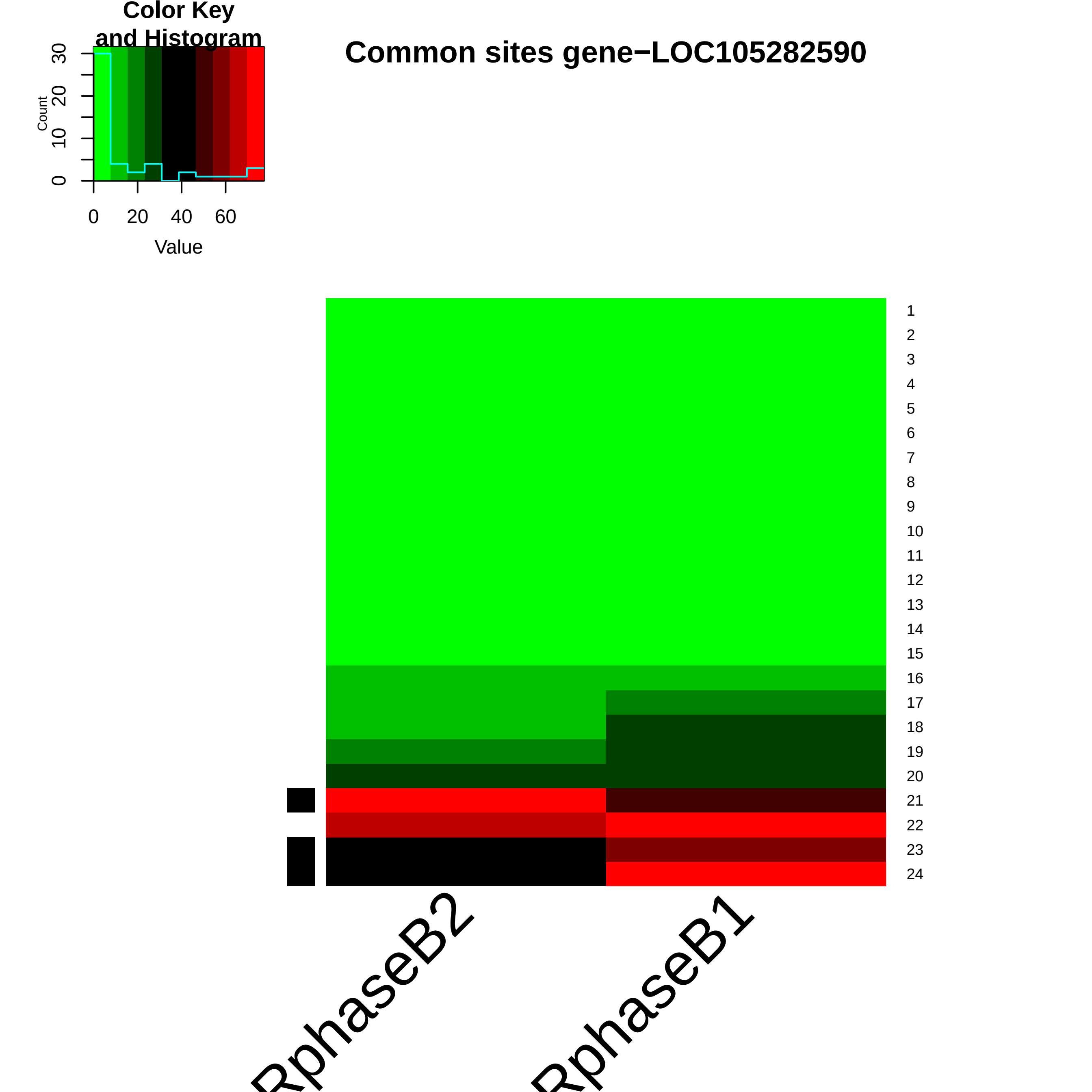

Color Key
and Histogram
Count
0
10
20
30
0
20
40
60
Value
Common sites gene−LOC105282590
RphaseB2
RphaseB1

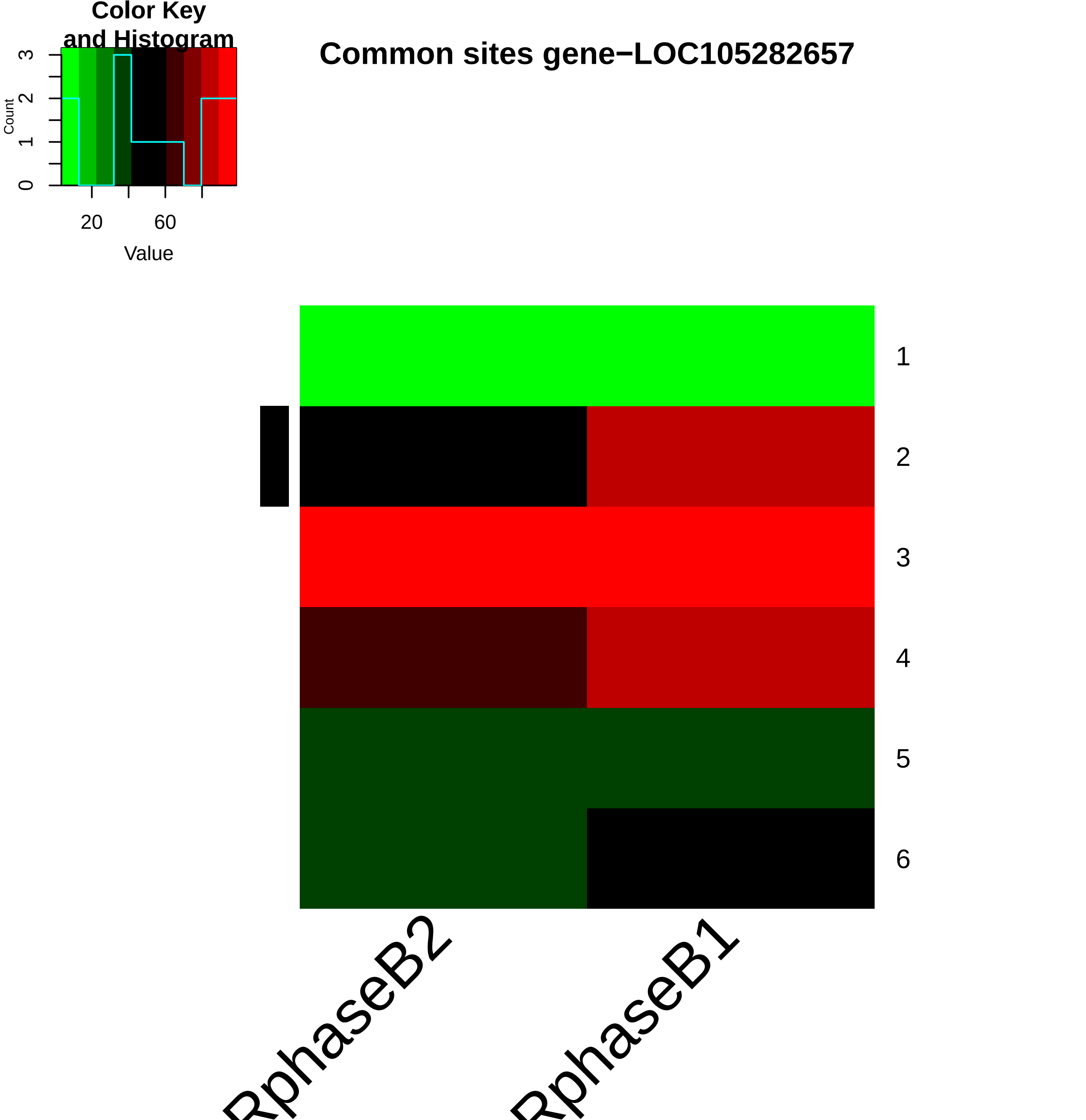
Color Key
and Histogram
Count
0
1
2
3
20
60
Value
Common sites gene−LOC105282657
1
2
3
4
5
6
RphaseB2
RphaseB1

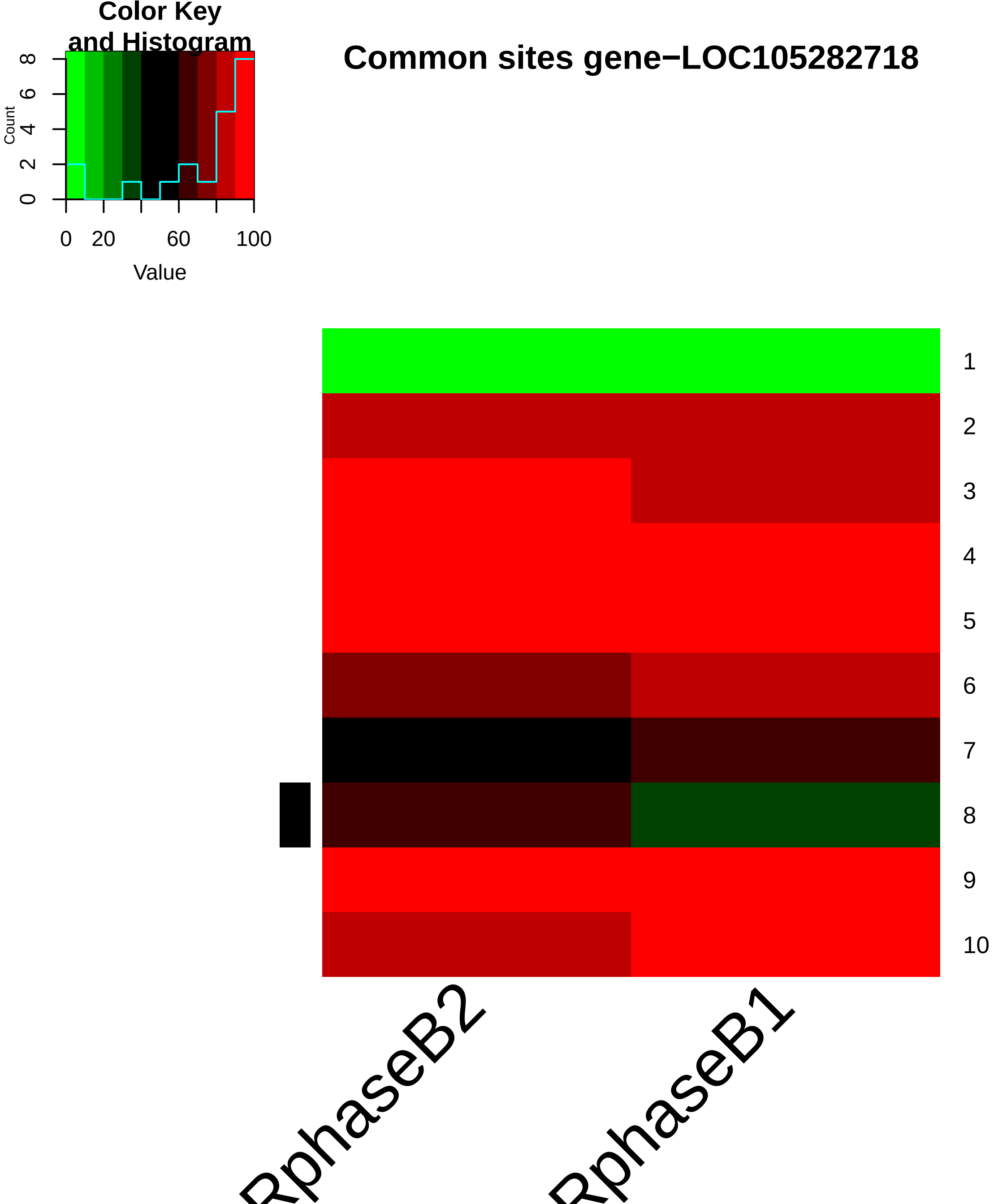
Color Key
and Histogram
Count
0
2
4
6
8
0
20
60
100
Value
Common sites gene−LOC105282718
1
2
3
4
5
6
7
8
9
10
RphaseB2
RphaseB1

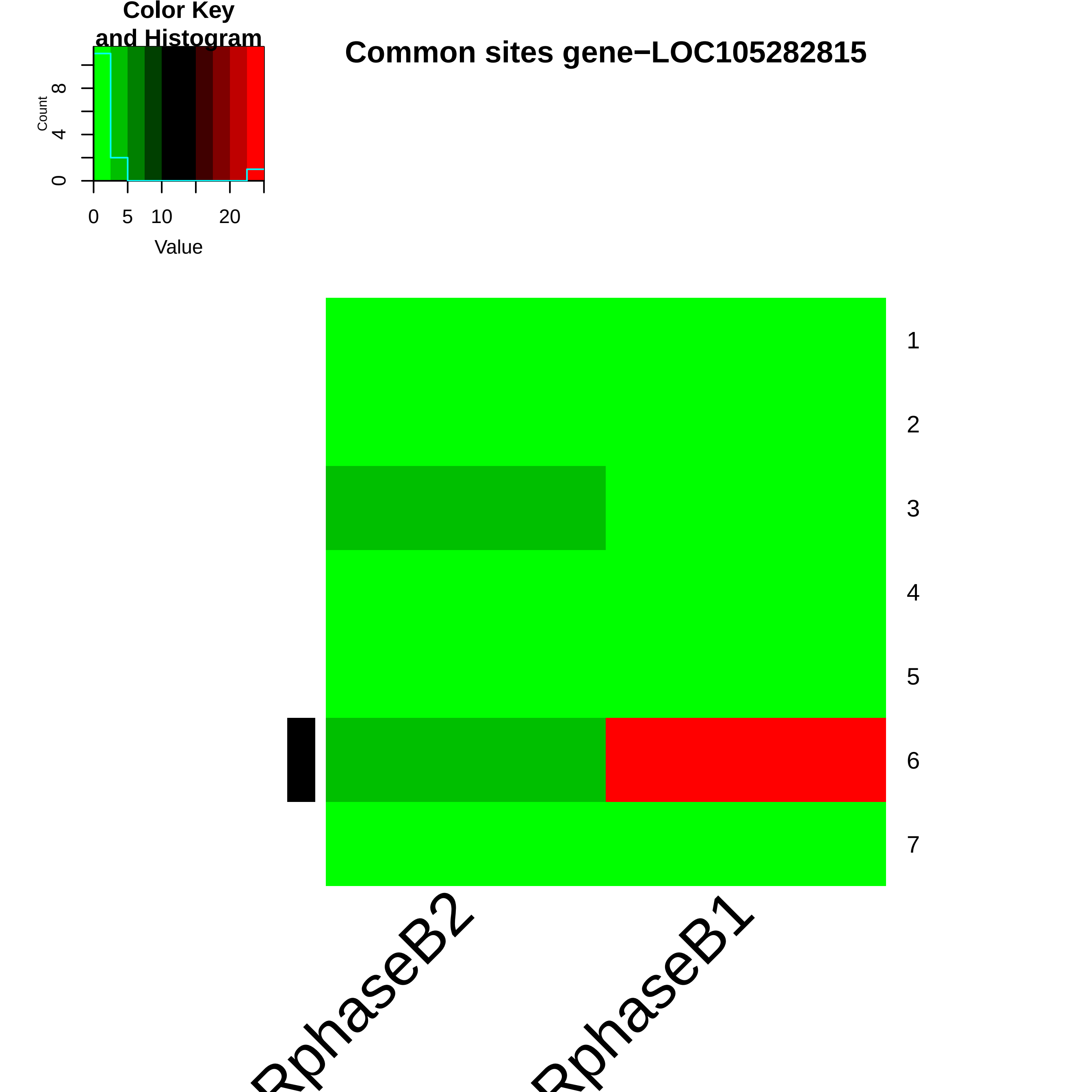
Color Key
and Histogram
Count
0
4
8
0
5
10
20
Value
Common sites gene−LOC105282815
1
2
3
4
5
6
7
RphaseB2
RphaseB1

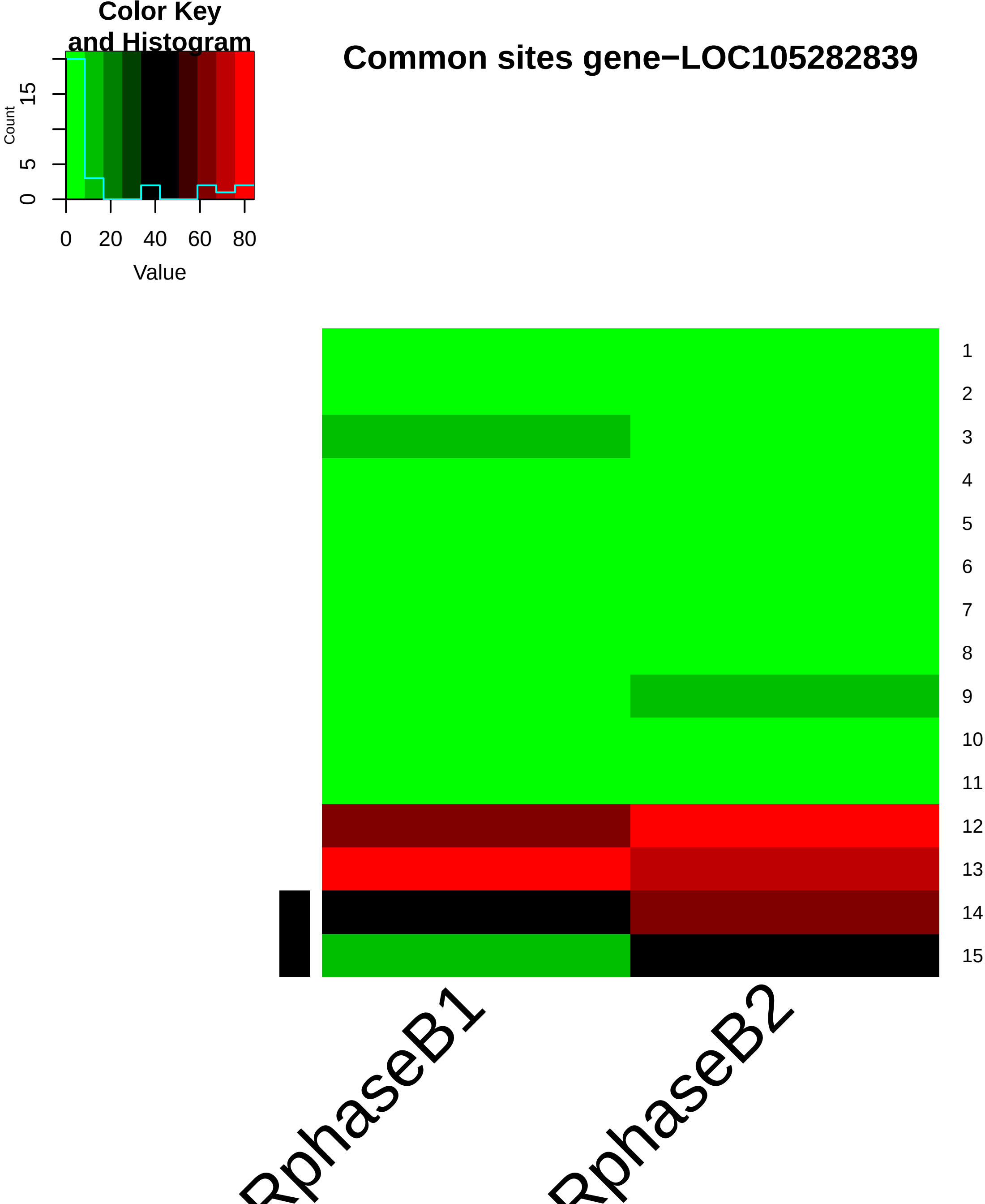
Color Key
and Histogram
Count
0
5
15
0
20
40
60
80
Value
Common sites gene−LOC105282839
RphaseB1
RphaseB2

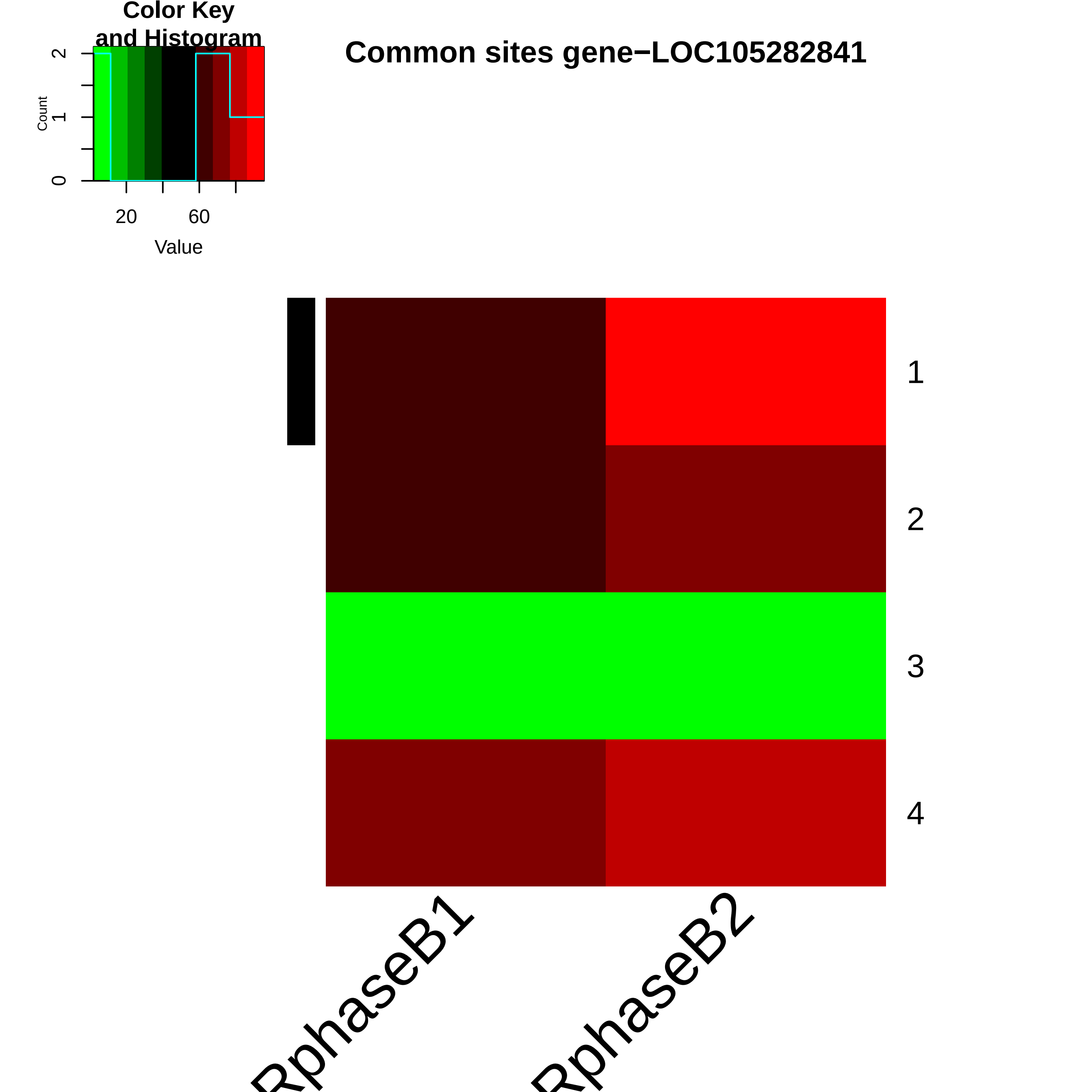

Color Key
and Histogram
Count
0
1
2
20
60
Value
Common sites gene−LOC105282841
1
2
3
4
RphaseB1
RphaseB2

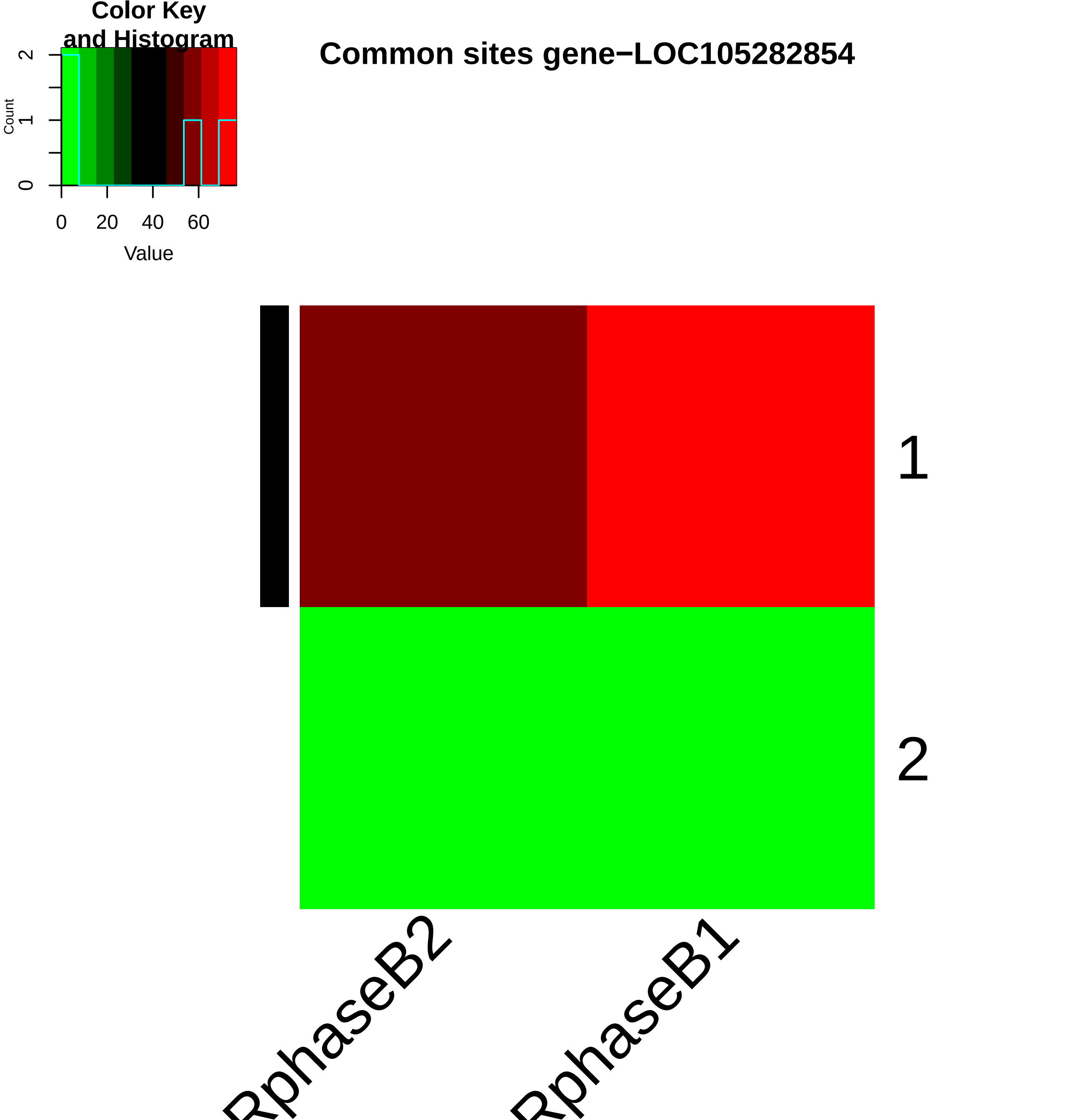
Color Key
and Histogram
Count
0
1
2
0
20
40
60
Value
Common sites gene−LOC105282854
1
2
RphaseB2
RphaseB1

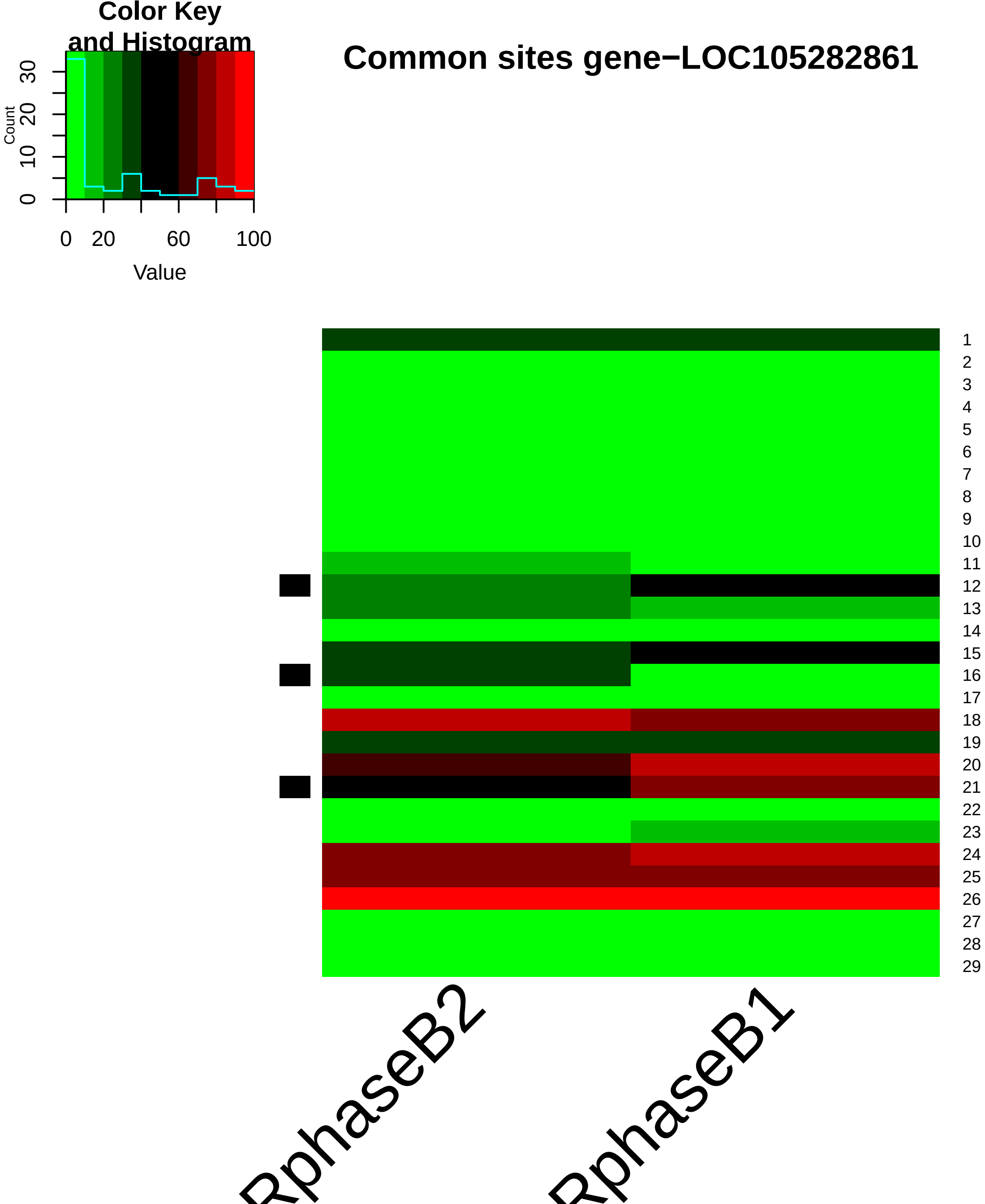

Color Key
and Histogram
Count
0
10
20
30
0
20
60
100
Value
Common sites gene−LOC105282861
RphaseB2
RphaseB1

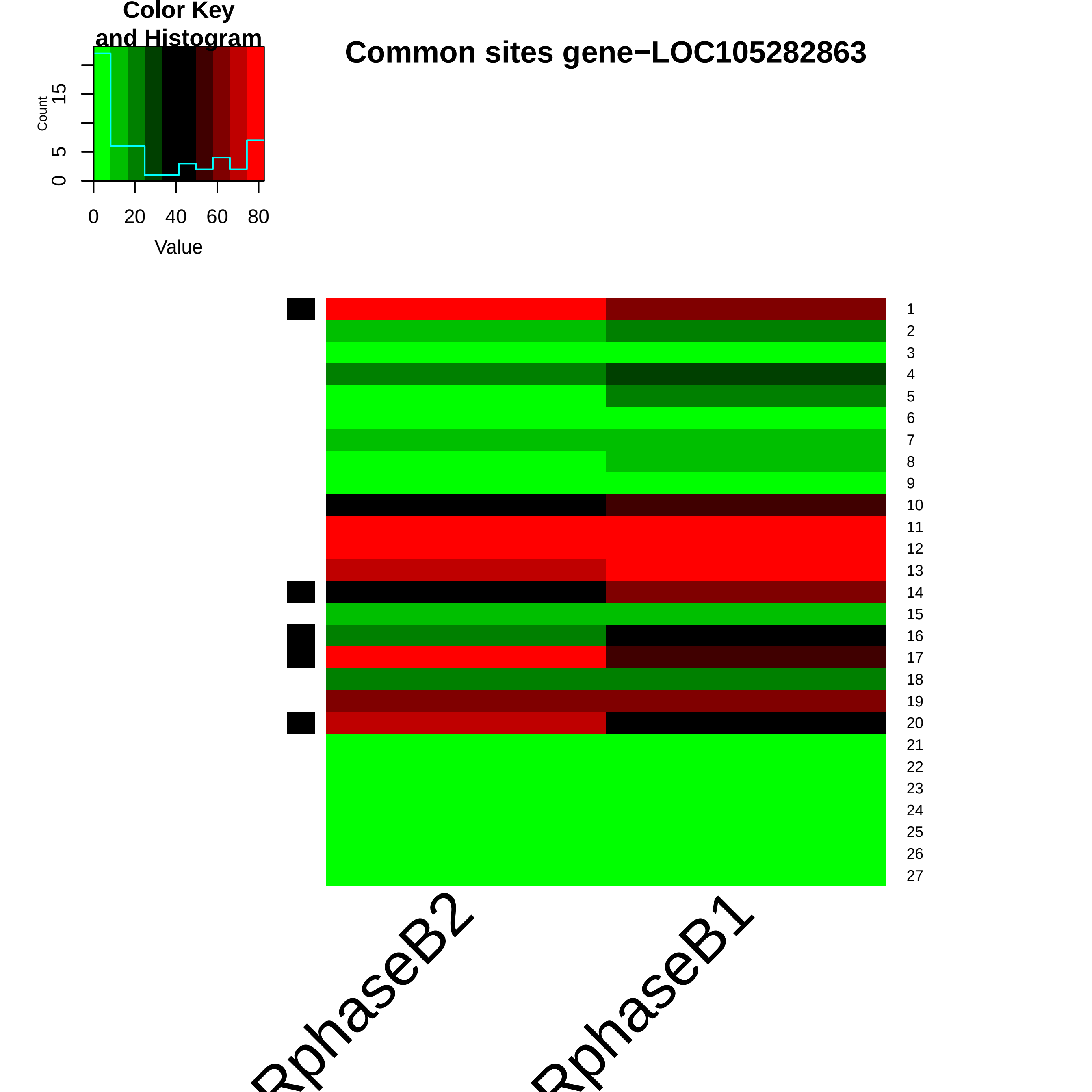

Color Key
and Histogram
Count
0
5
15
0
20
40
60
80
Value
Common sites gene−LOC105282863
RphaseB2
RphaseB1

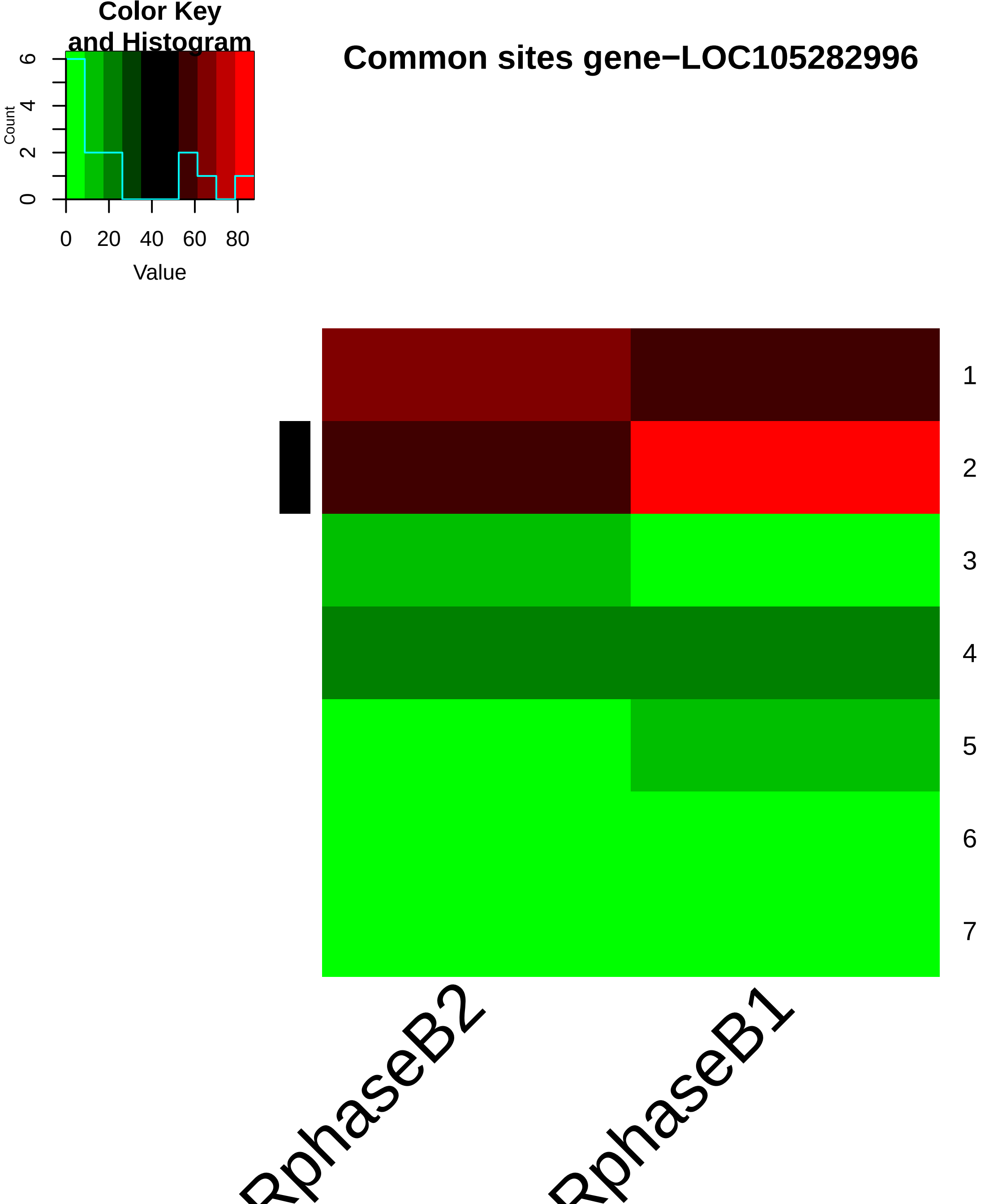

Color Key
and Histogram
Count
Value
0
20
40
60
80
0
2
4
6
Common sites gene−LOC105282996
1
2
3
4
5
6
7
RphaseB2
RphaseB1

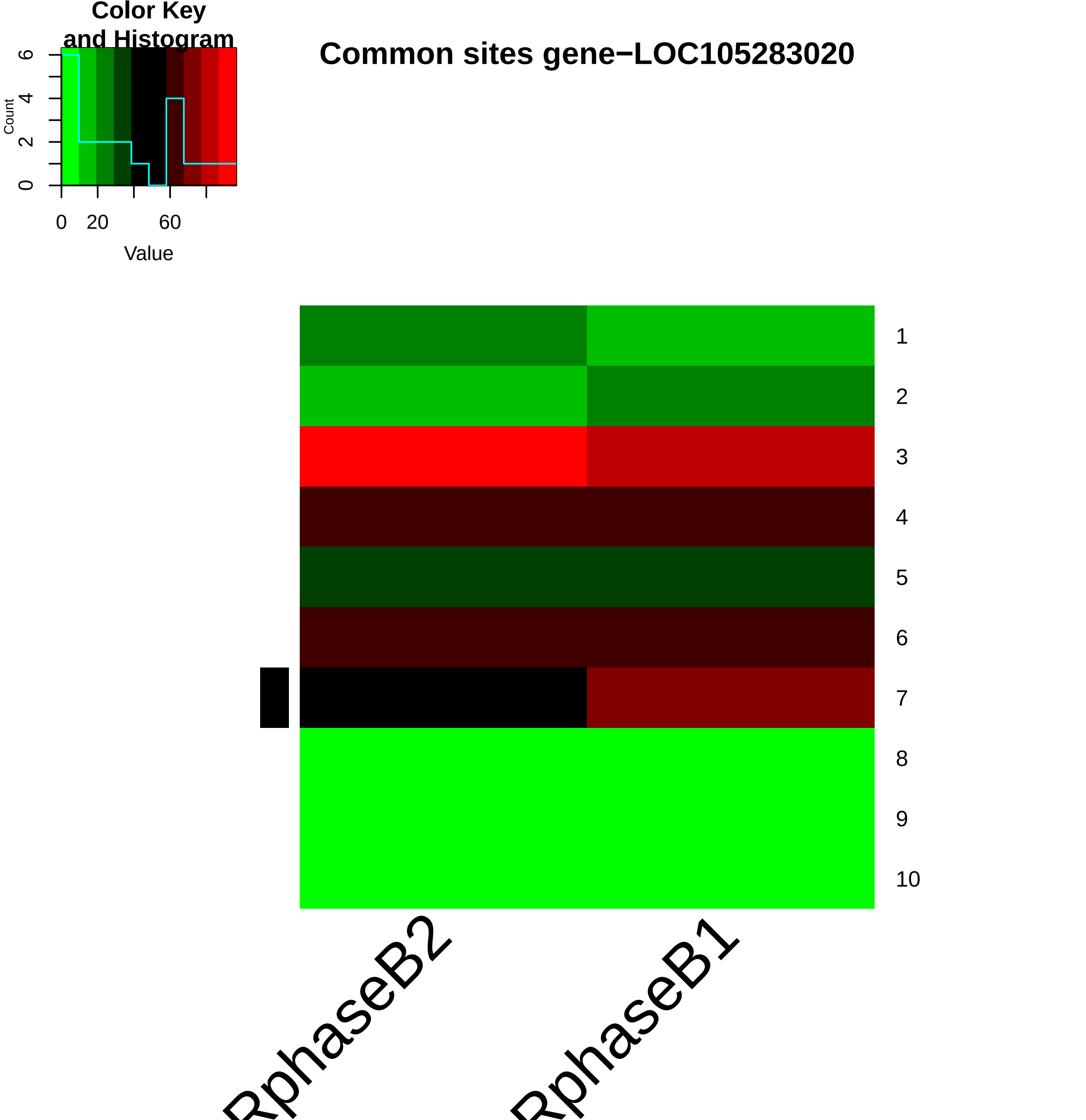
Color Key
and Histogram
Count
0
2
4
6
0
20
60
Value
Common sites gene−LOC105283020
1
2
3
4
5
6
7
8
9
10
RphaseB2
RphaseB1

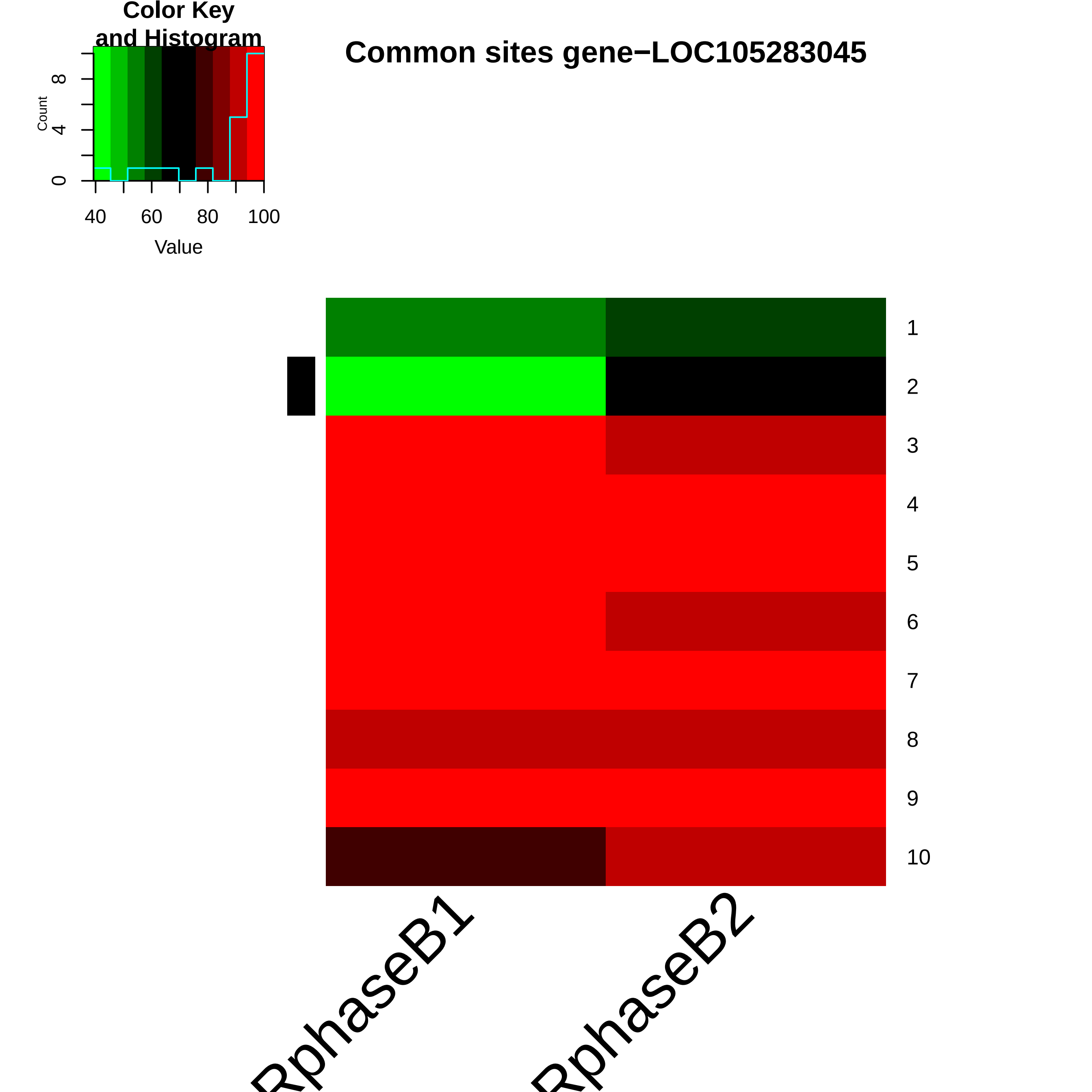
Color Key
and Histogram
Count
0
4
8
40
60
80
100
Value
Common sites gene−LOC105283045
1
2
3
4
5
6
7
8
9
10
RphaseB1
RphaseB2

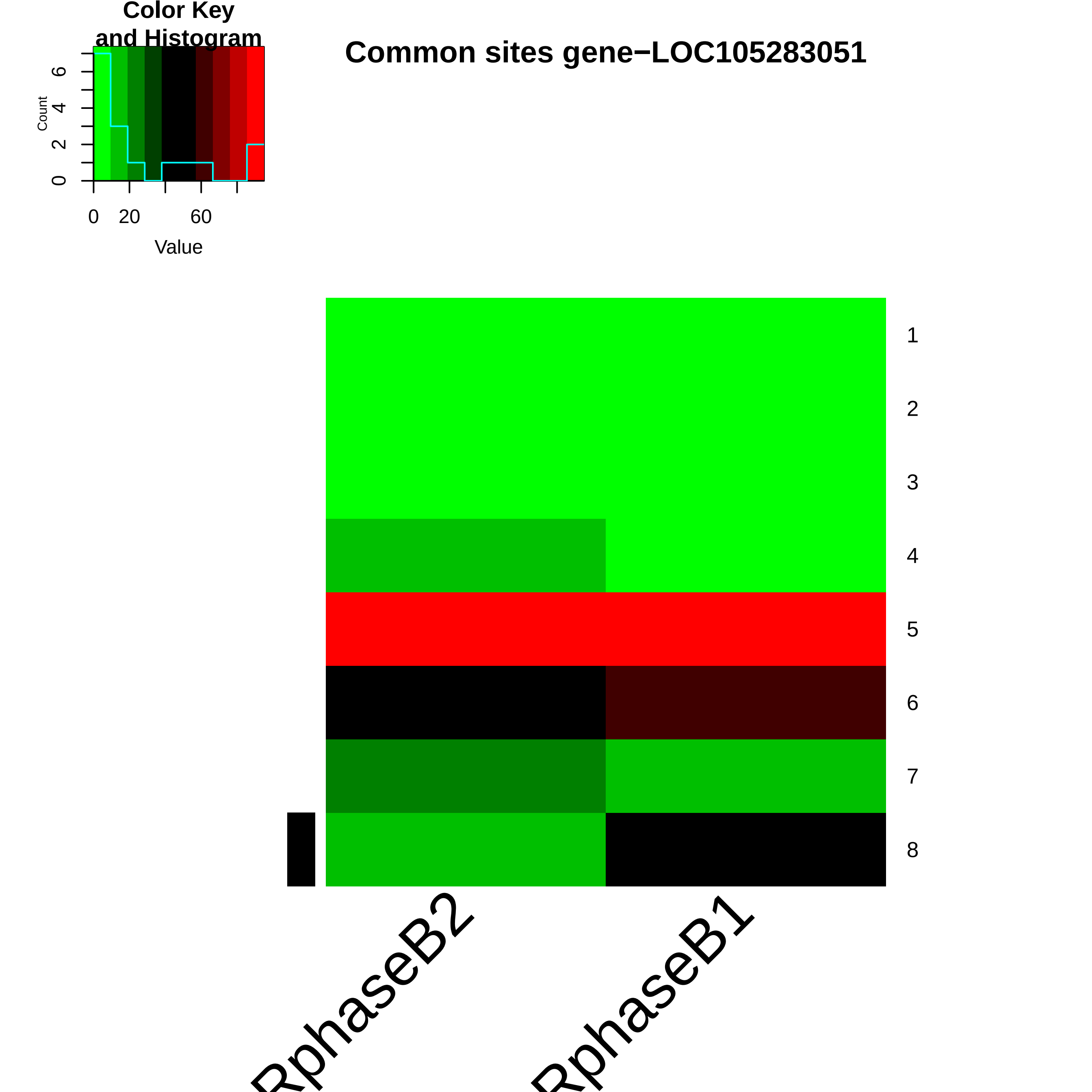
Color Key
and Histogram
Count
0
2
4
6
0
20
60
Value
Common sites gene−LOC105283051
1
2
3
4
5
6
7
8
RphaseB2
RphaseB1

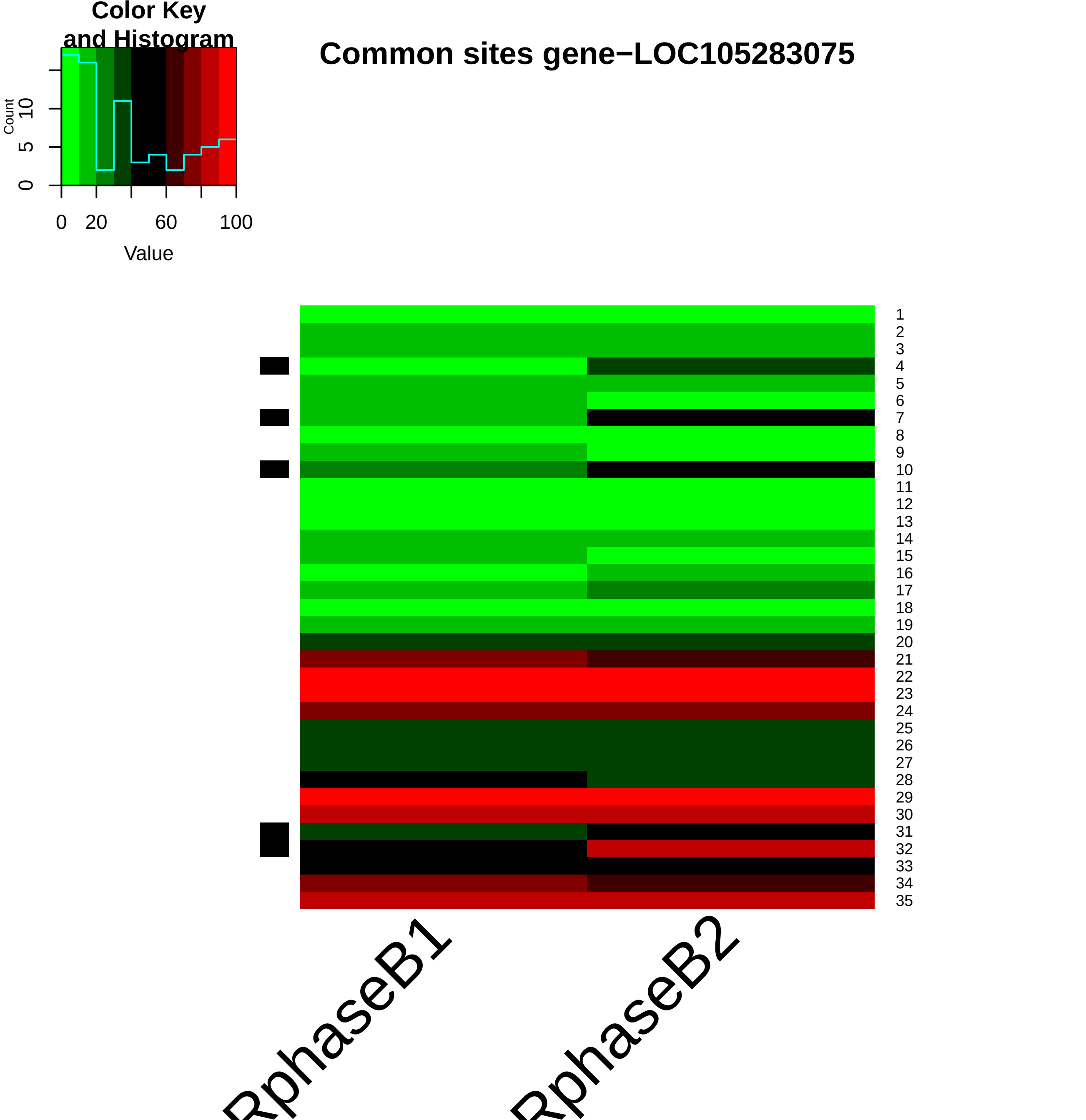

Color Key
and Histogram
Count
0
5
10
0
20
60
100
Value
Common sites gene−LOC105283075
RphaseB1
RphaseB2

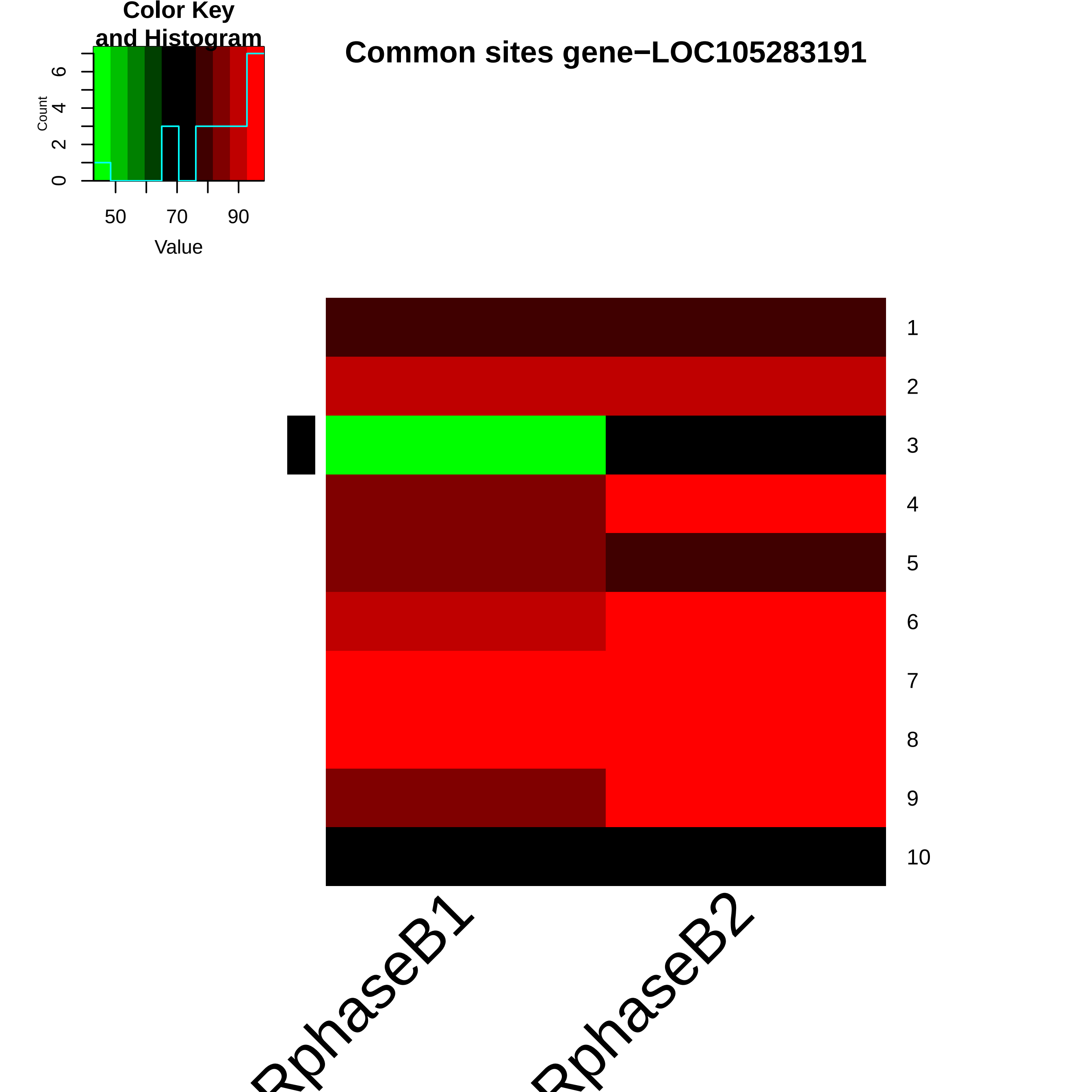
Color Key
and Histogram
Count
0
2
4
6
50
70
90
Value
Common sites gene−LOC105283191
1
2
3
4
5
6
7
8
9
10
RphaseB1
RphaseB2

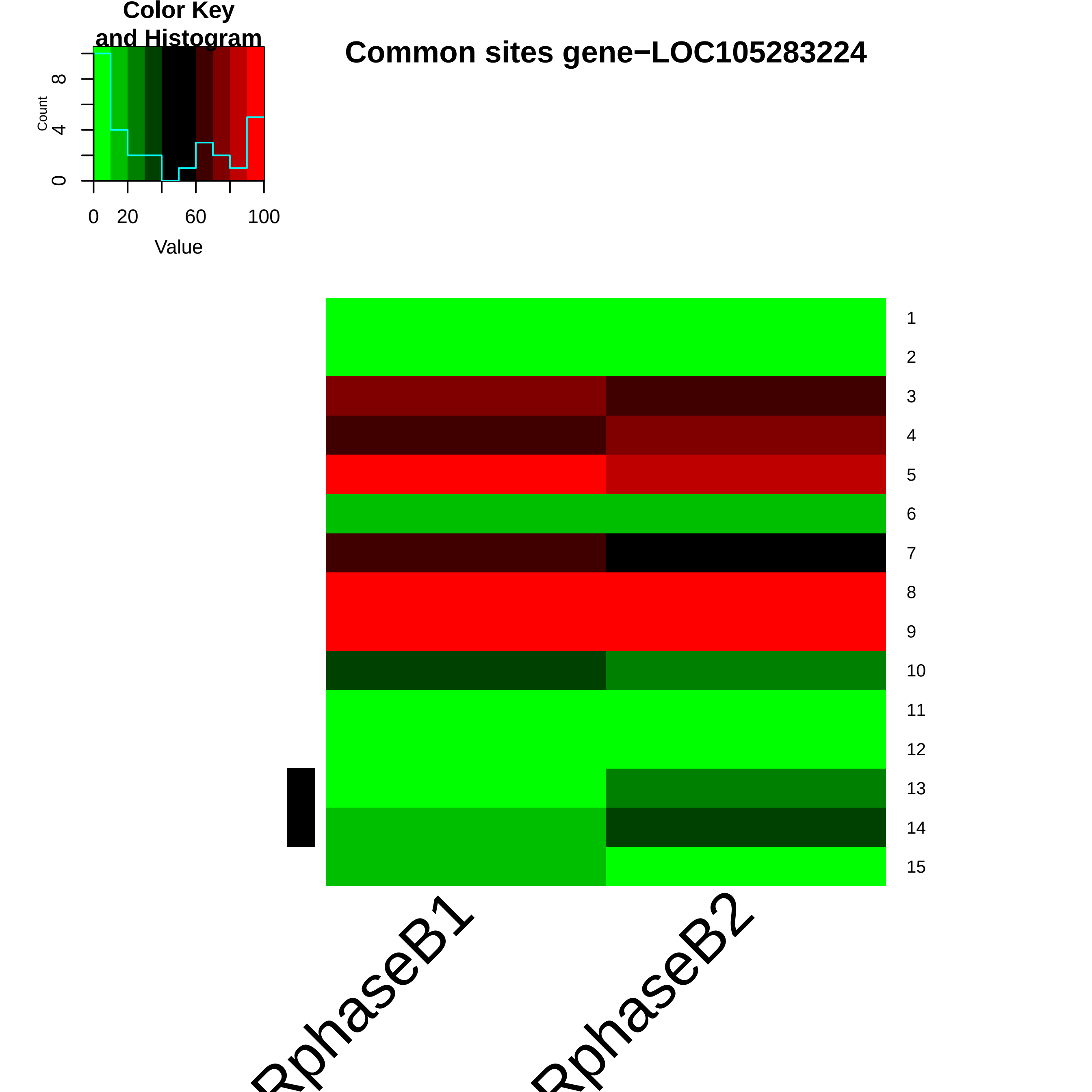
Color Key
and Histogram
Count
0
4
8
0
20
60
100
Value
Common sites gene−LOC105283224
RphaseB1
RphaseB2

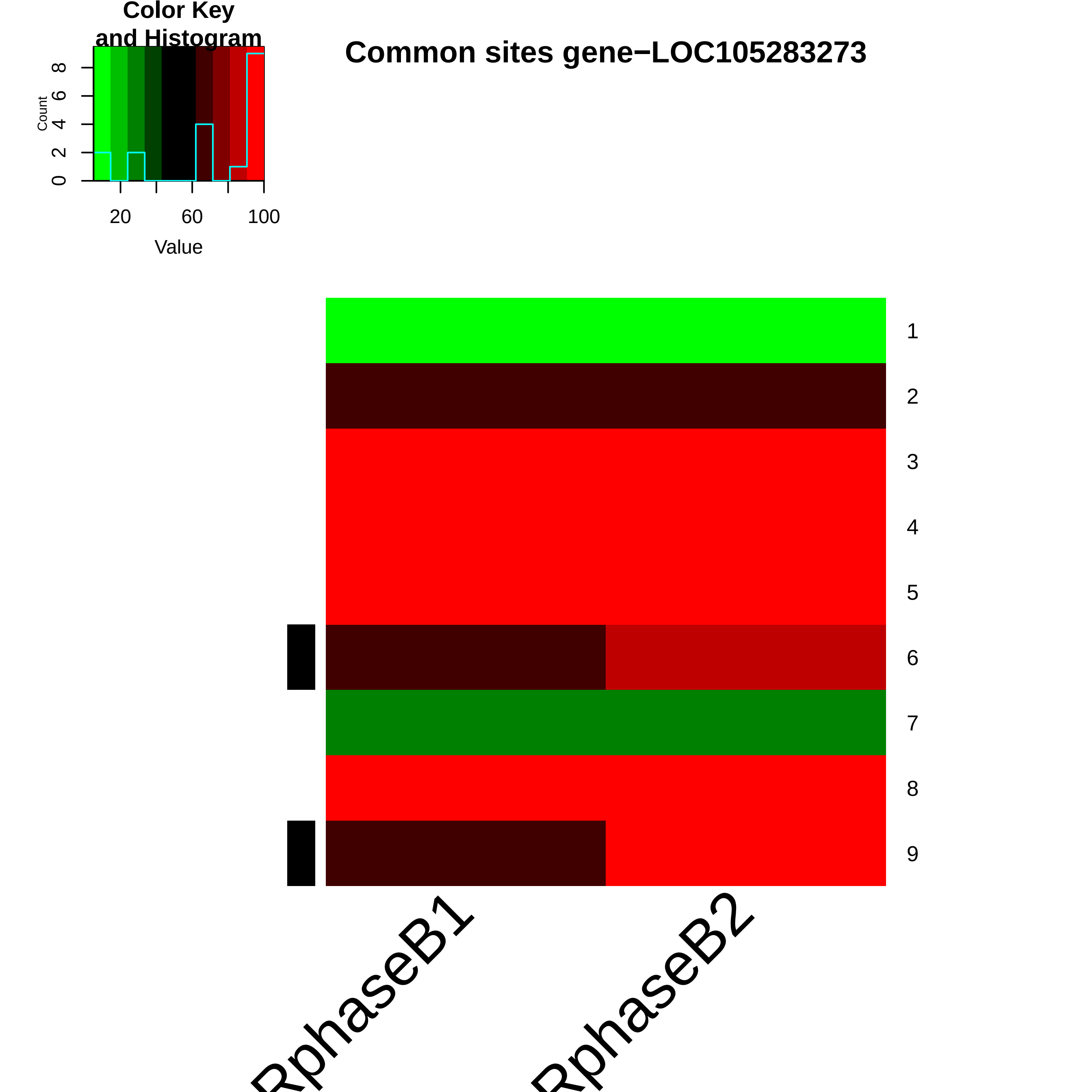
Color Key
and Histogram
Count
0
2
4
6
8
20
60
100
Value
Common sites gene−LOC105283273
1
2
3
4
5
6
7
8
9
RphaseB1
RphaseB2

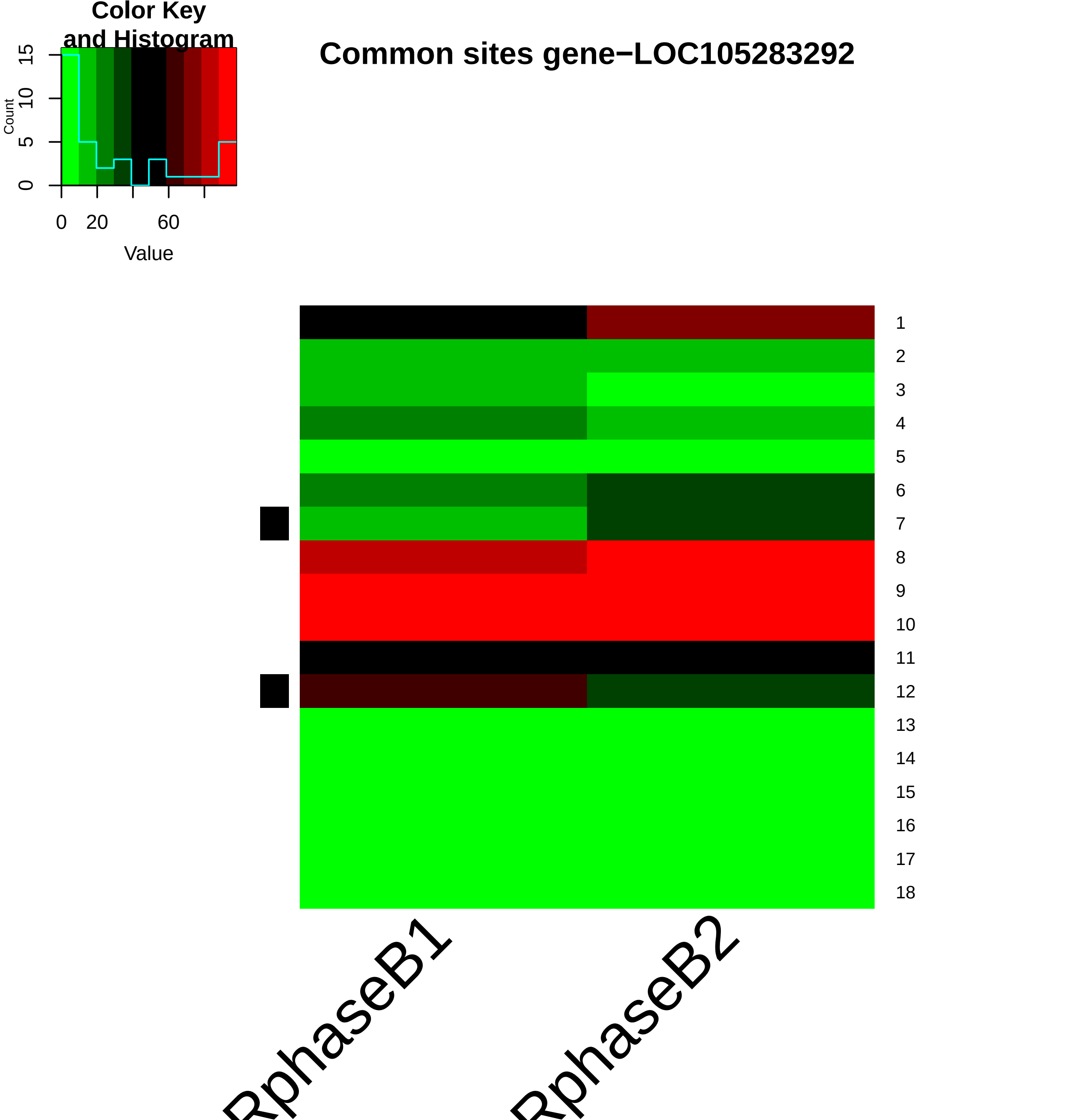
Color Key
and Histogram
Count
0
5
10
15
0
20
60
Value
Common sites gene−LOC105283292
RphaseB1
RphaseB2

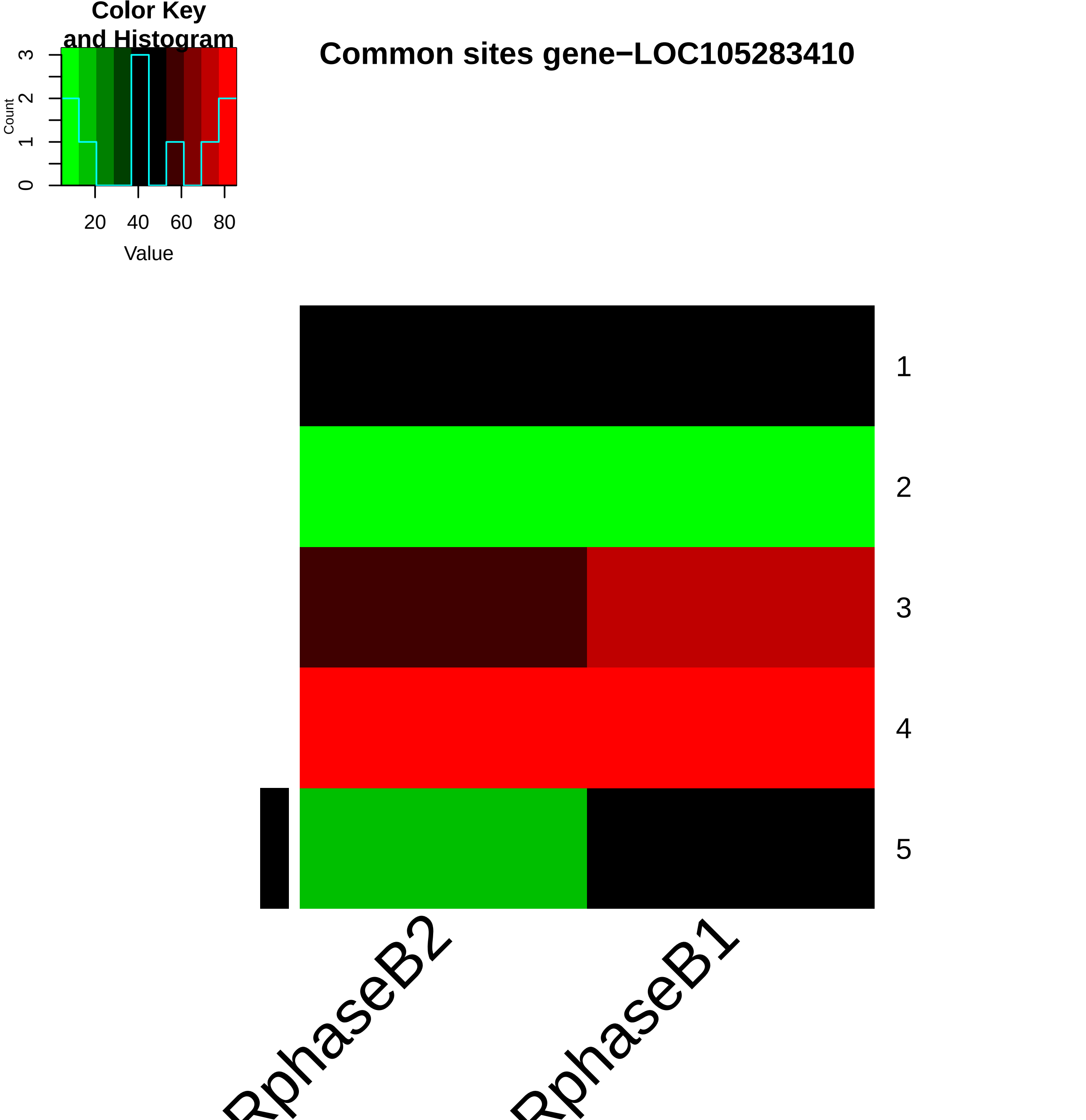

Color Key
and Histogram
Count
0
1
2
3
20
40
60
80
Value
Common sites gene−LOC105283410
1
2
3
4
5
RphaseB2
RphaseB1

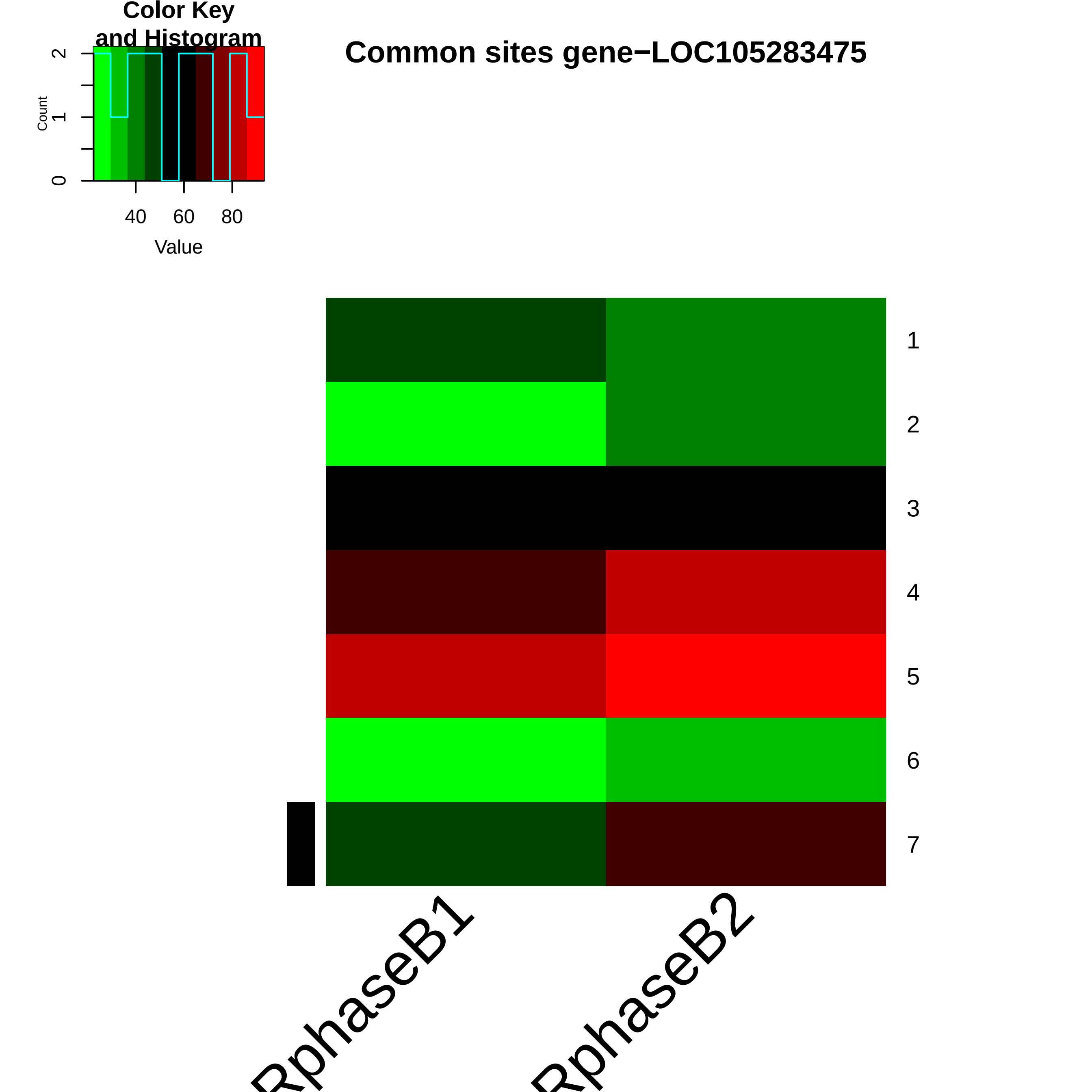

Color Key and Histogram
Count
Value
0
1
2
40
60
80
Common sites gene−LOC105283475
1
2
3
4
5
6
7
RphaseB1
RphaseB2

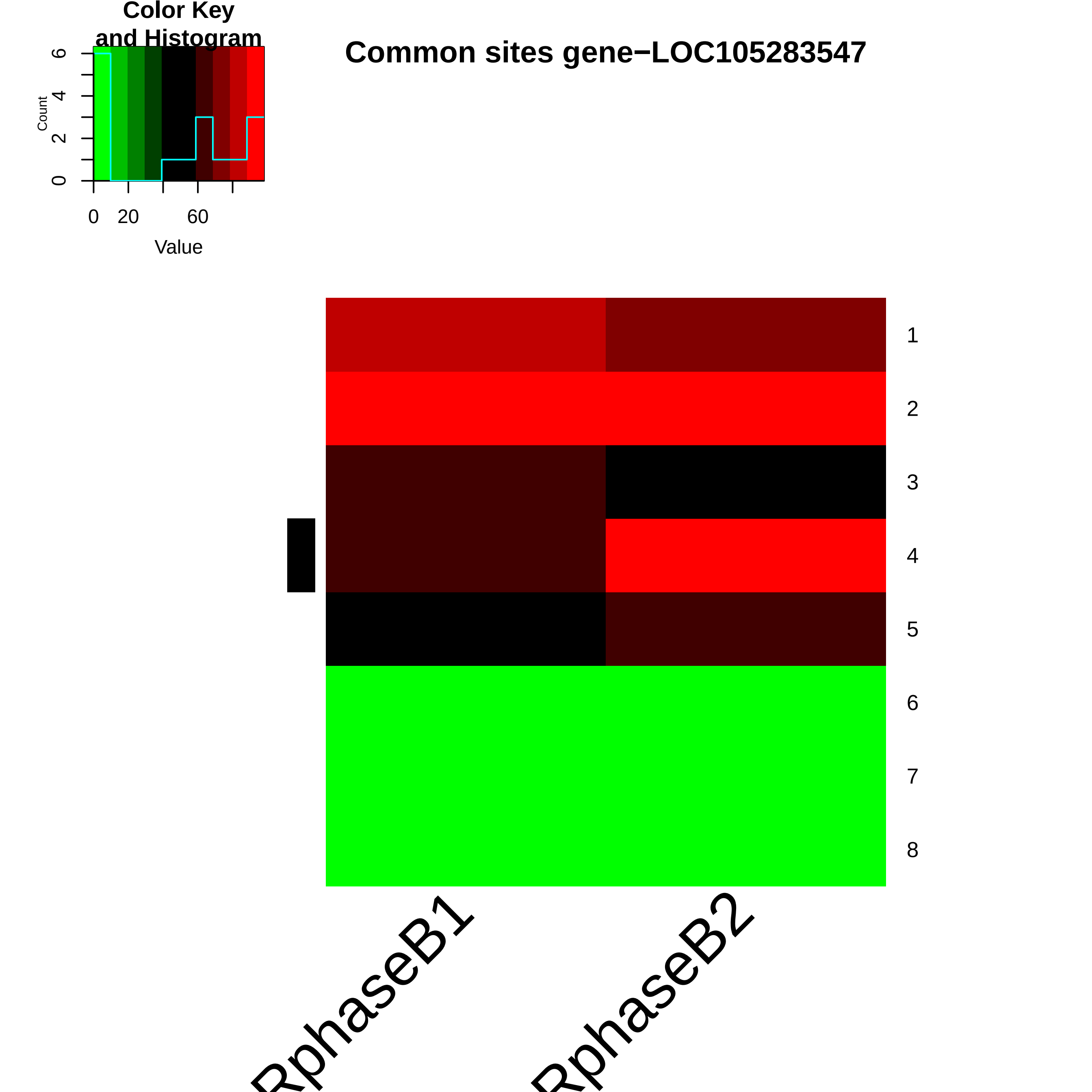

Color Key
and Histogram
Count
0
2
4
6
0
20
60
Value
Common sites gene−LOC105283547
1
2
3
4
5
6
7
8
RphaseB1
RphaseB2

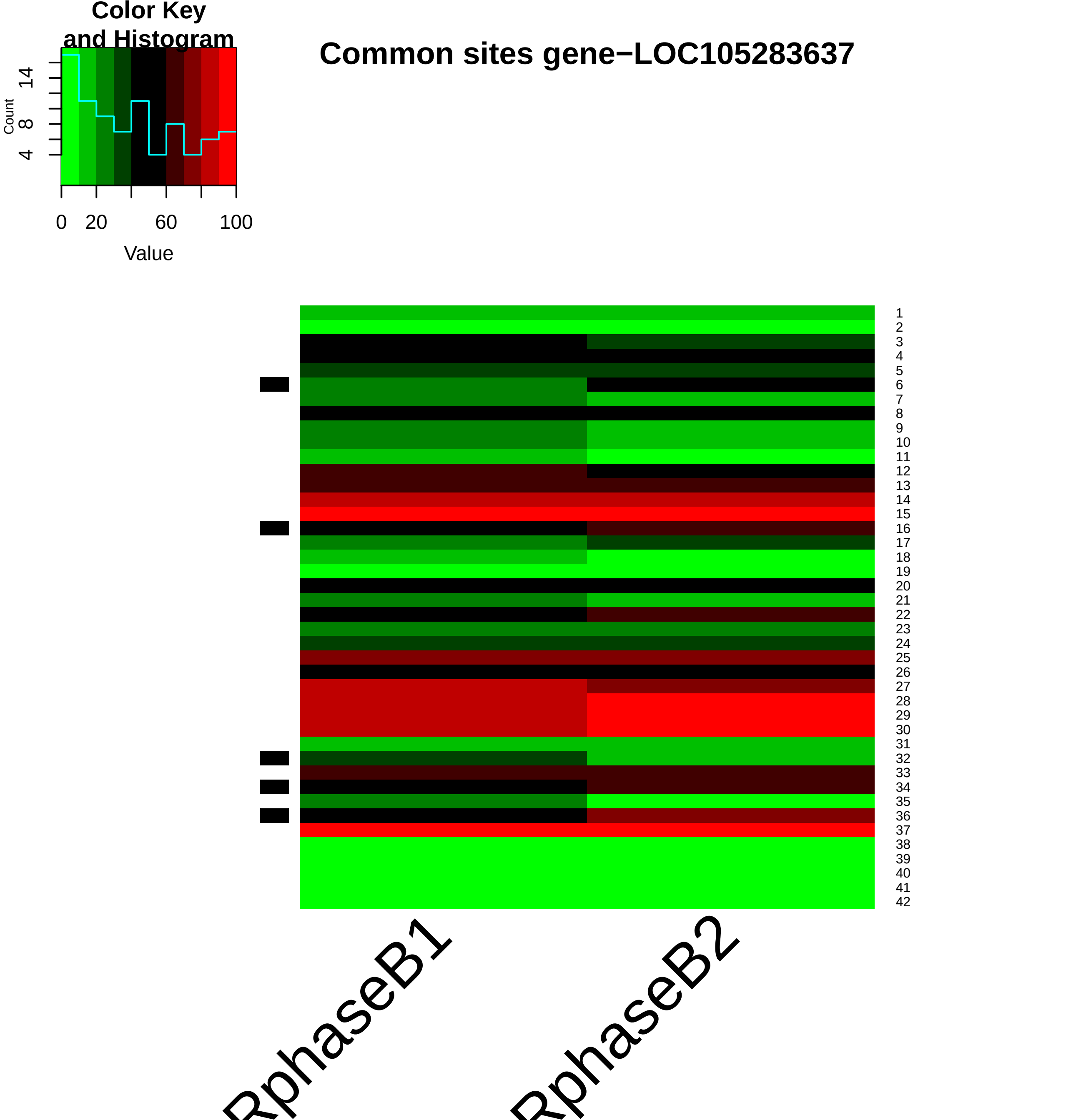
Color Key
and Histogram
Count
4
8
14
0
20
60
100
Value
Common sites gene−LOC105283637
RphaseB1
RphaseB2

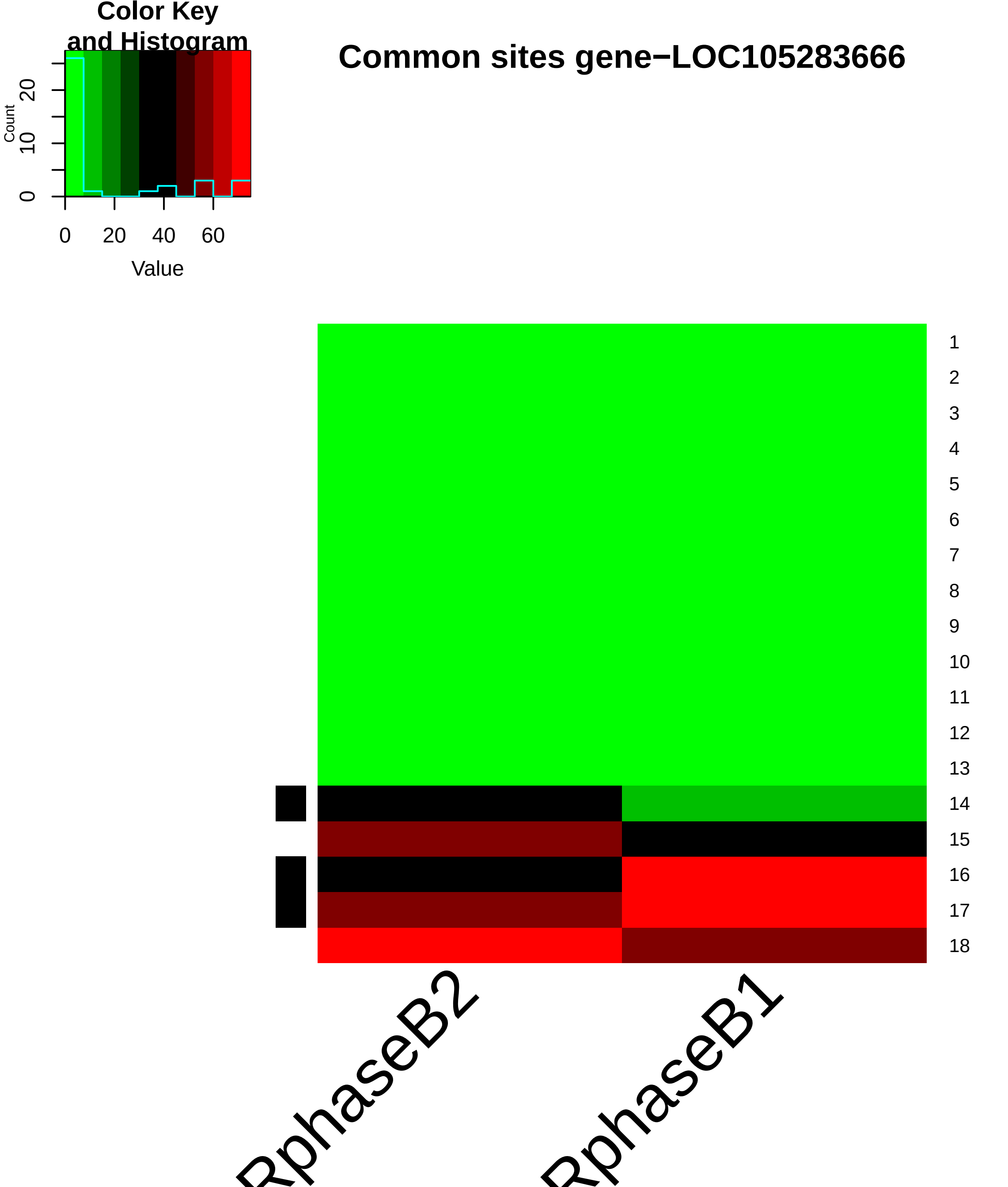
Color Key
and Histogram
Count
0
10
20
0
20
40
60
Value
Common sites gene−LOC105283666
RphaseB2
RphaseB1

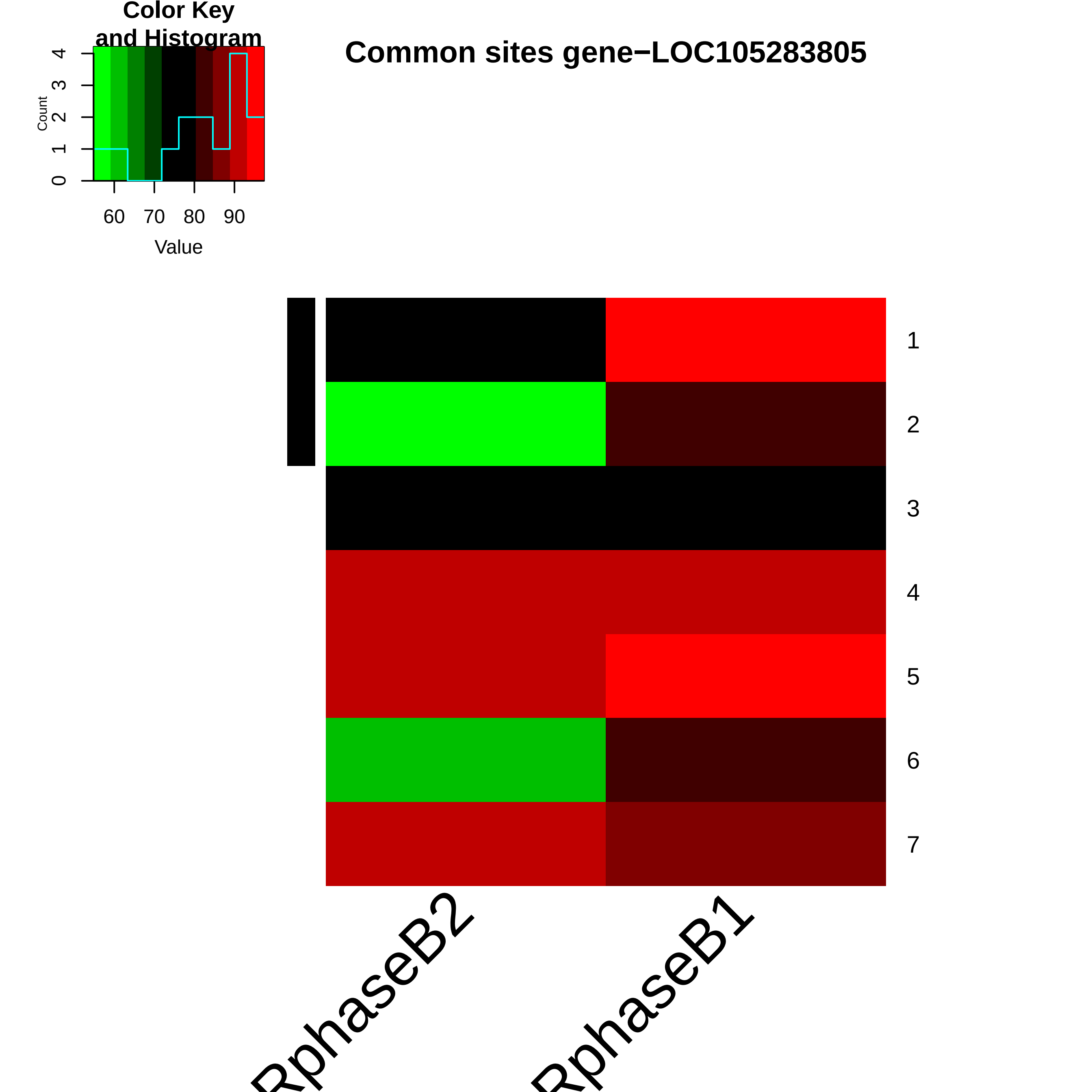
Color Key
and Histogram
Count
Value
60
70
80
90
0
1
2
3
4
Common sites gene−LOC105283805
1
2
3
4
5
6
7
RphaseB2
RphaseB1

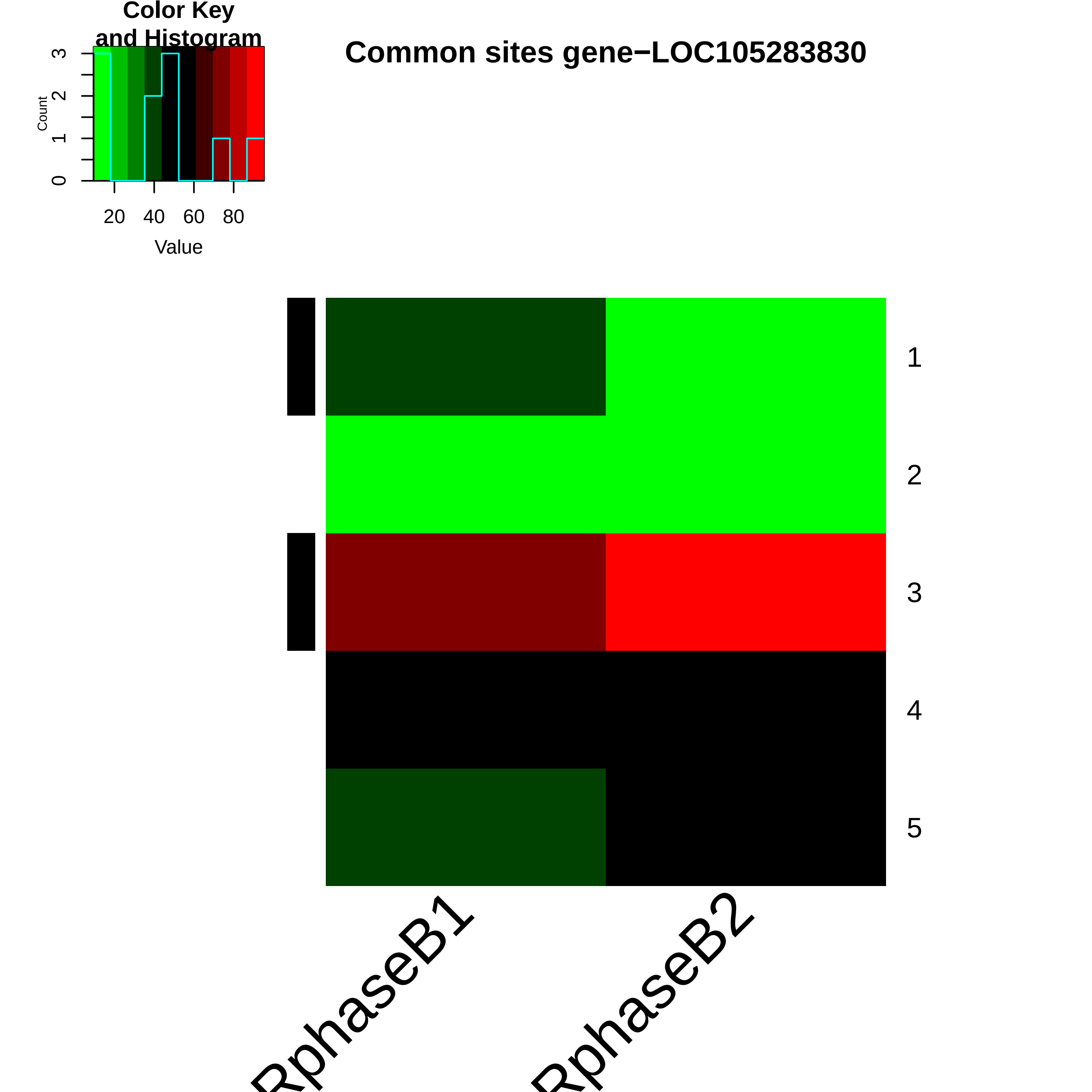

Color Key
and Histogram
Count
0
1
2
3
20
40
60
80
Value
Common sites gene−LOC105283830
1
2
3
4
5
RphaseB1
RphaseB2

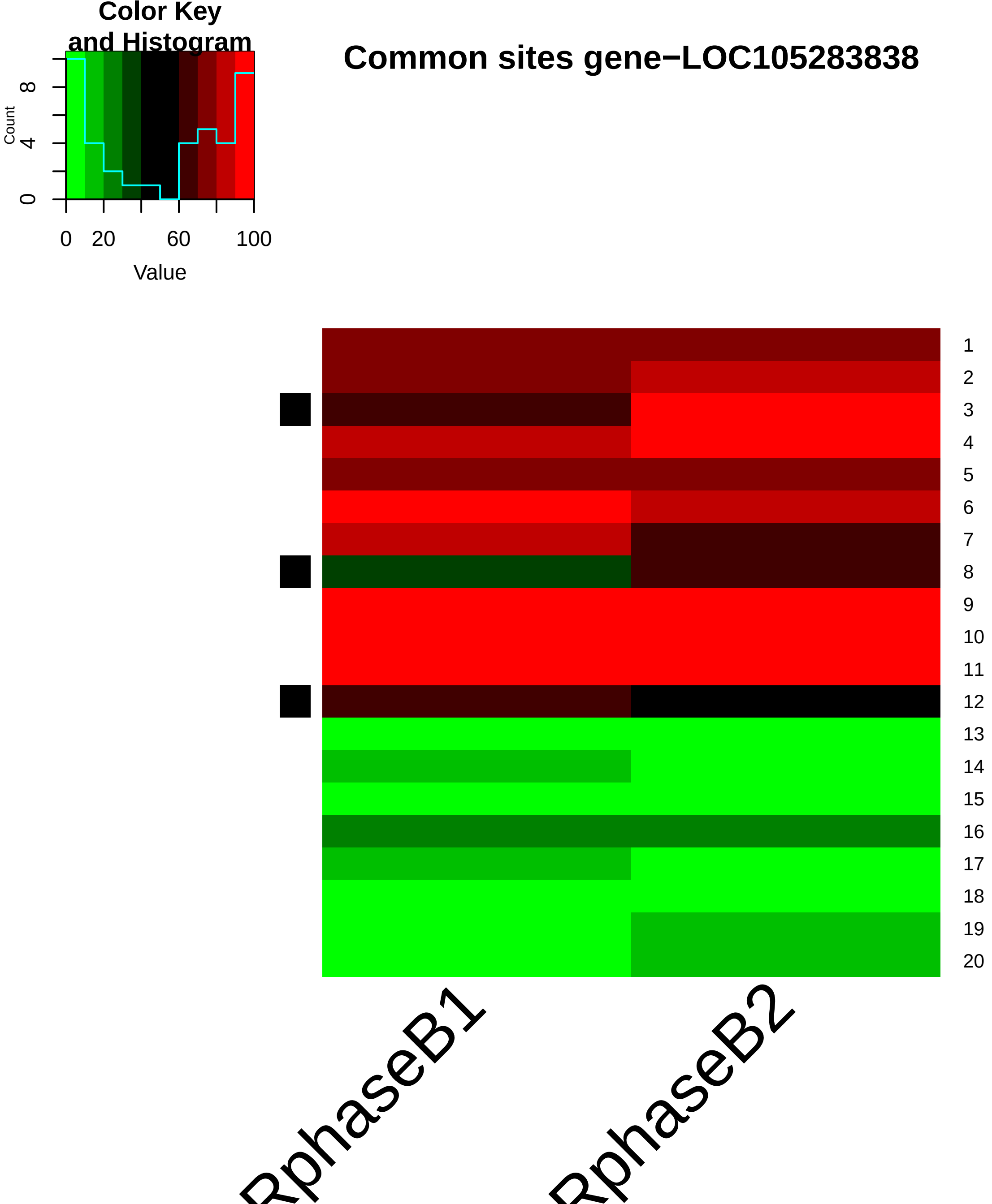

Color Key
and Histogram
Count
0
4
8
0
20
60
100
Value
Common sites gene−LOC105283838
RphaseB1
RphaseB2

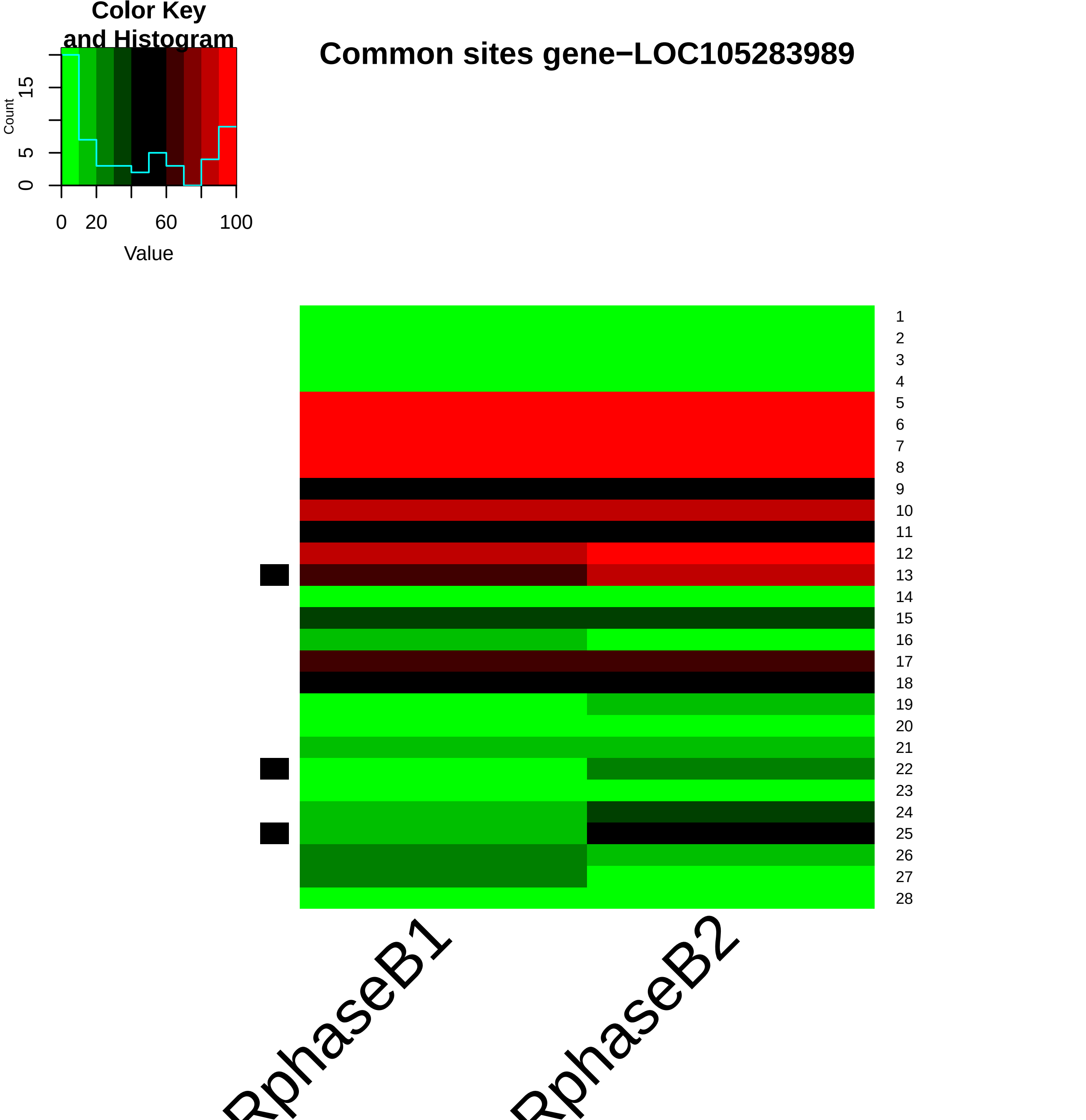
Color Key
and Histogram
Count
0
5
15
0
20
60
100
Value
Common sites gene−LOC105283989
RphaseB1
RphaseB2

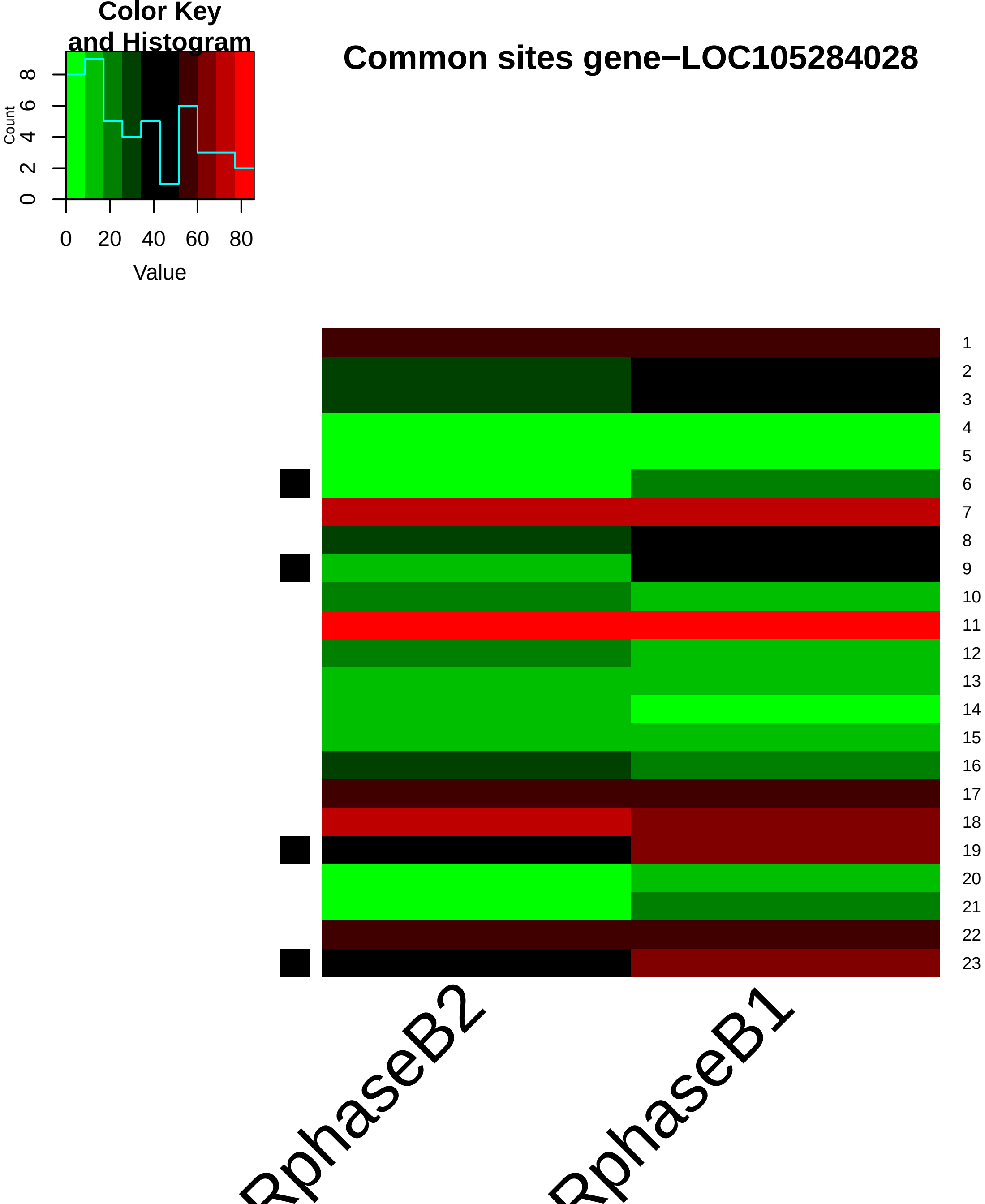

Color Key
and Histogram
Count
0
2
4
6
8
0
20
40
60
80
Value
Common sites gene−LOC105284028
RphaseB2
RphaseB1

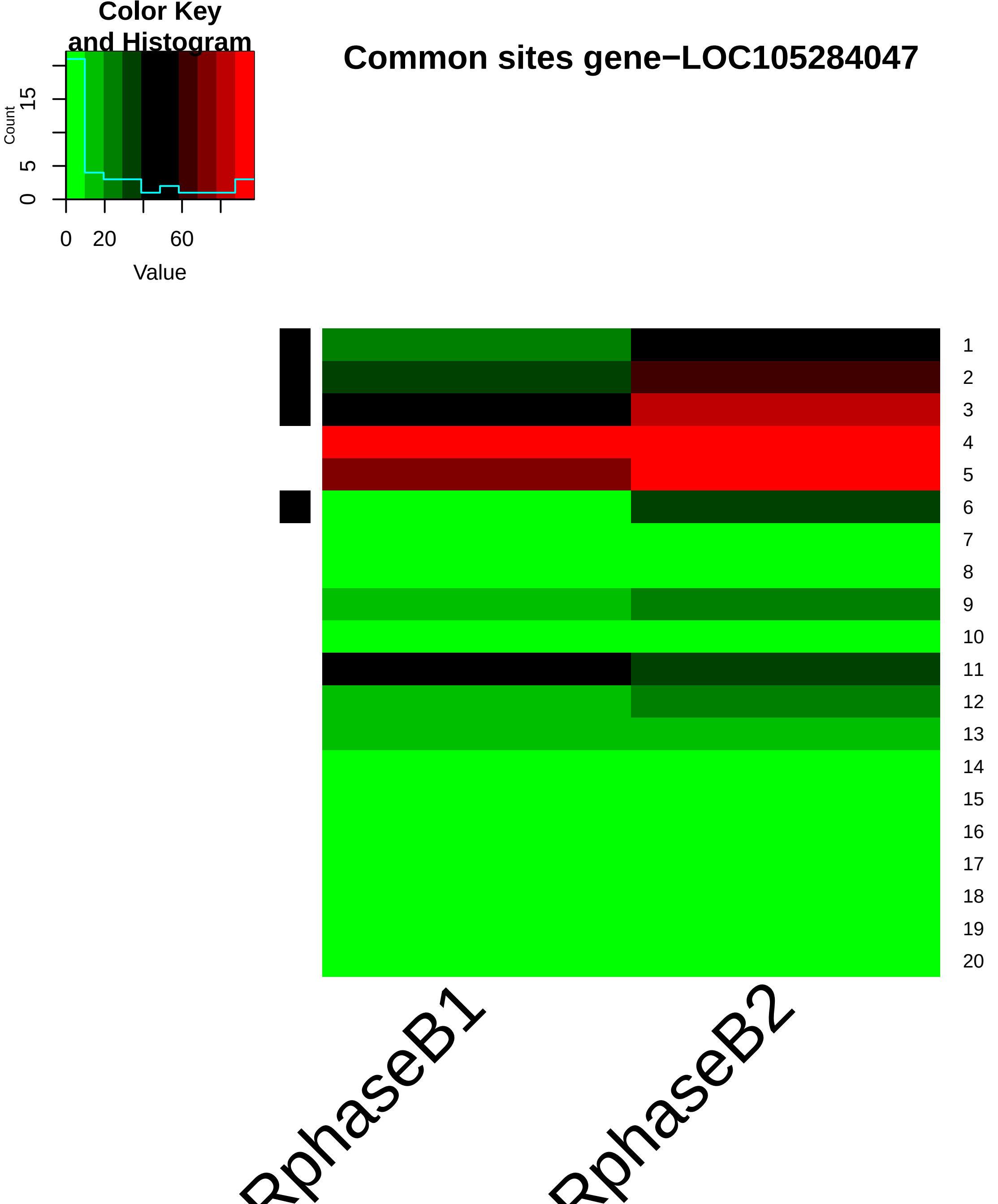
Color Key
and Histogram
Count
0
5
15
0
20
60
Value
Common sites gene−LOC105284047
RphaseB1
RphaseB2

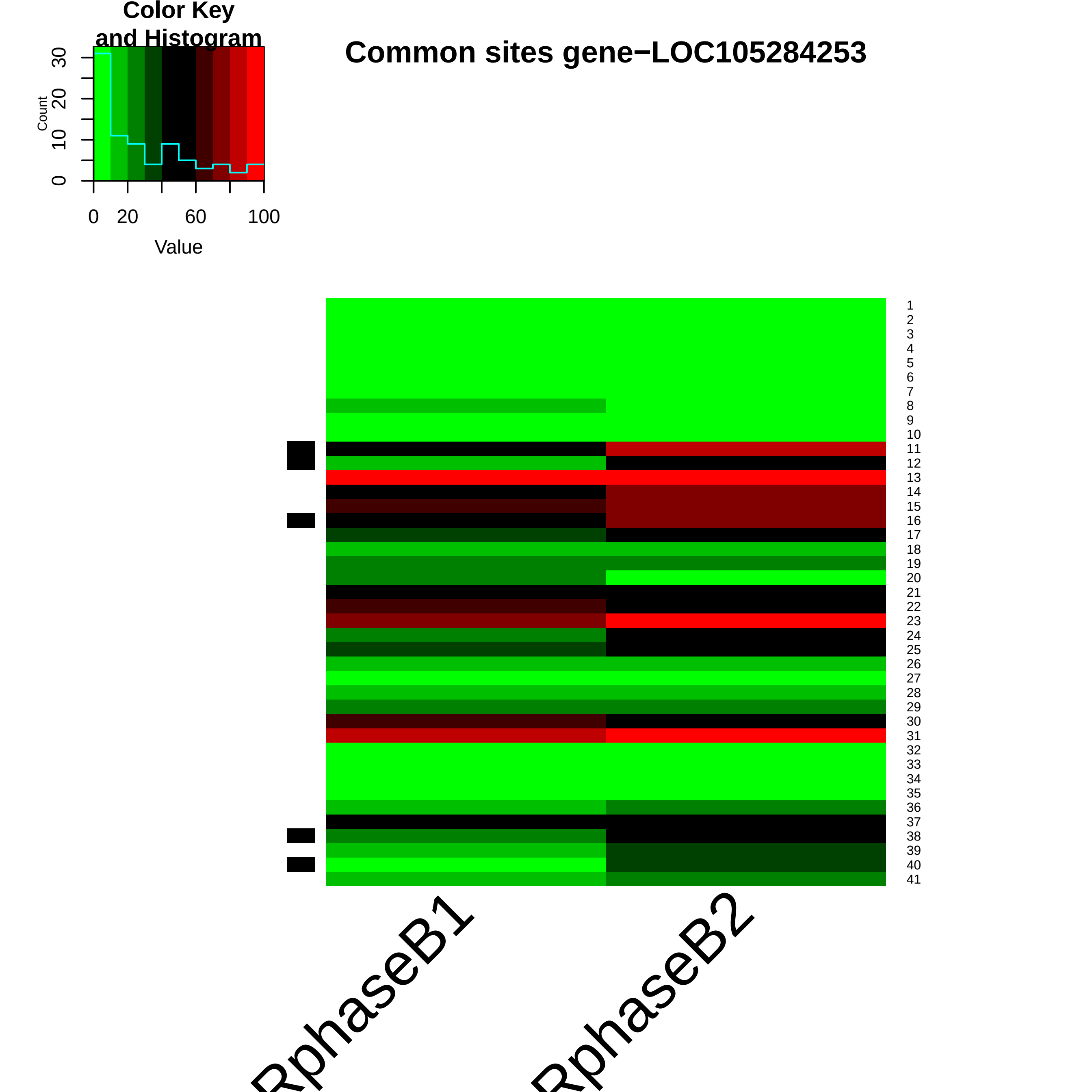

Color Key
and Histogram
Common sites gene−LOC105284253
Count
0
10
20
30
0
20
60
100
Value
RphaseB1
RphaseB2

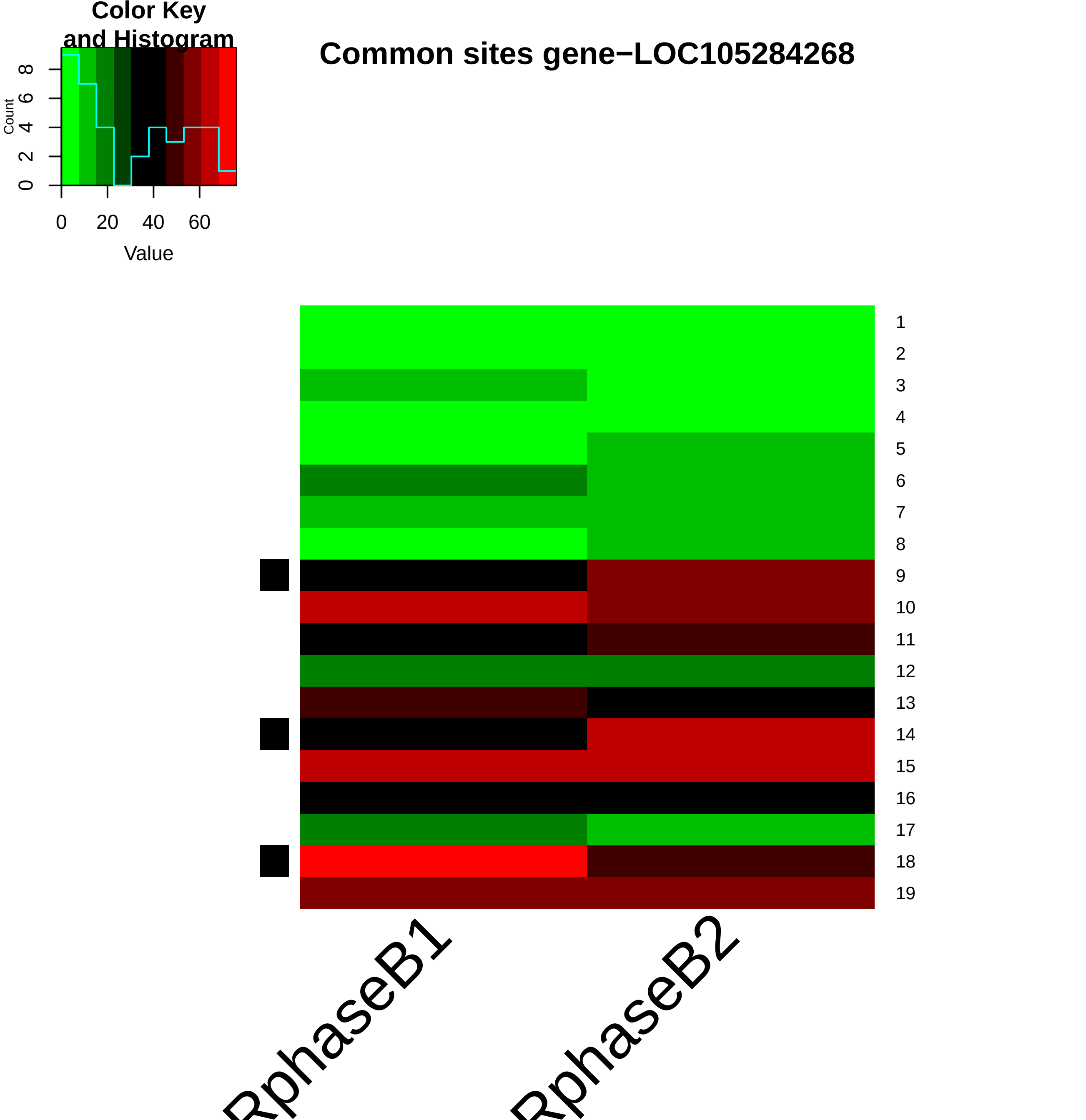
Color Key
and Histogram
Count
0
2
4
6
8
0
20
40
60
Value
Common sites gene−LOC105284268
RphaseB1
RphaseB2

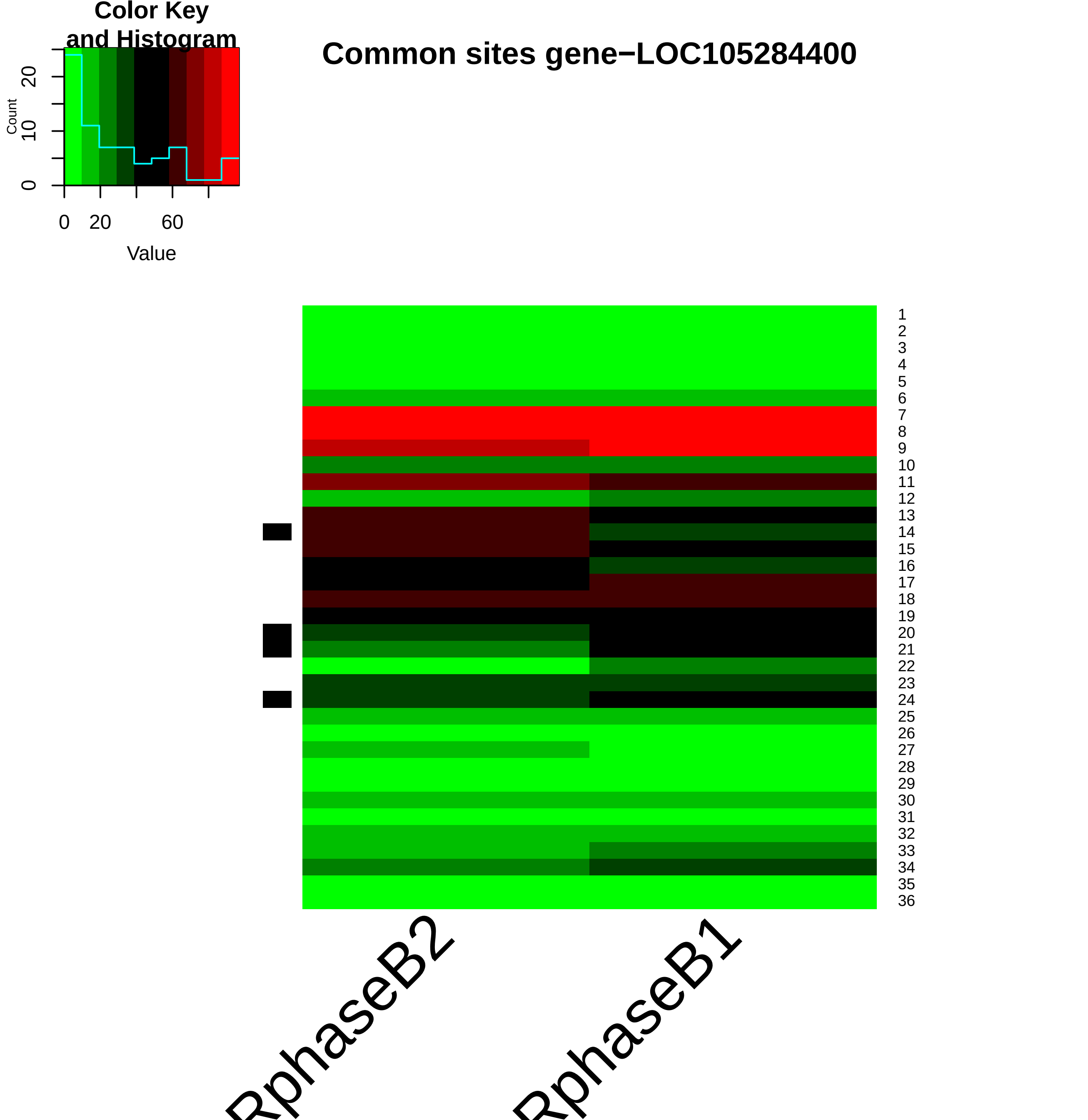
Color Key
and Histogram
Count
0
10
20
0
20
60
Value
Common sites gene−LOC105284400
RphaseB2
RphaseB1

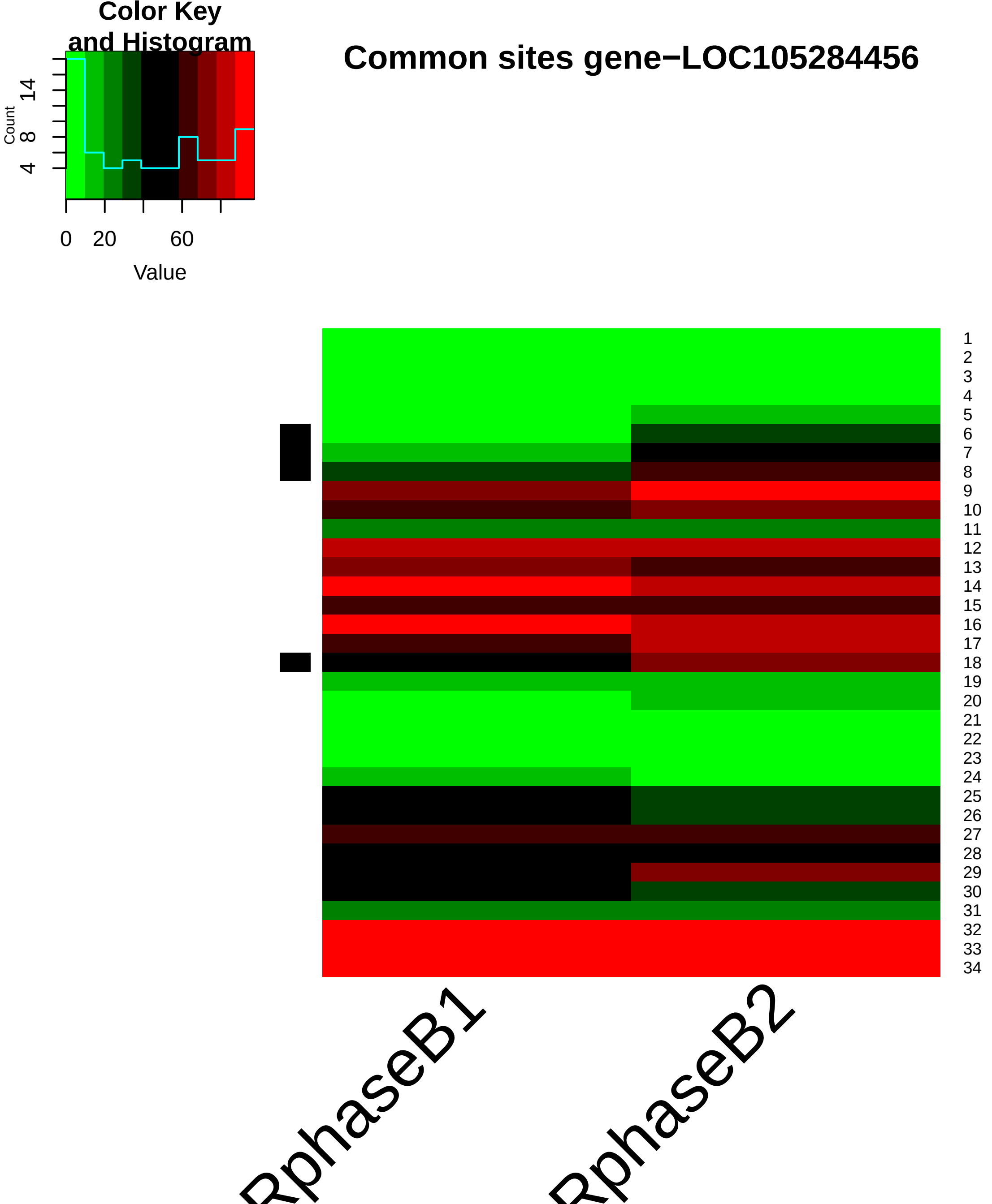
Color Key
and Histogram
Count
4
8
14
0
20
60
Value
Common sites gene−LOC105284456
RphaseB1
RphaseB2

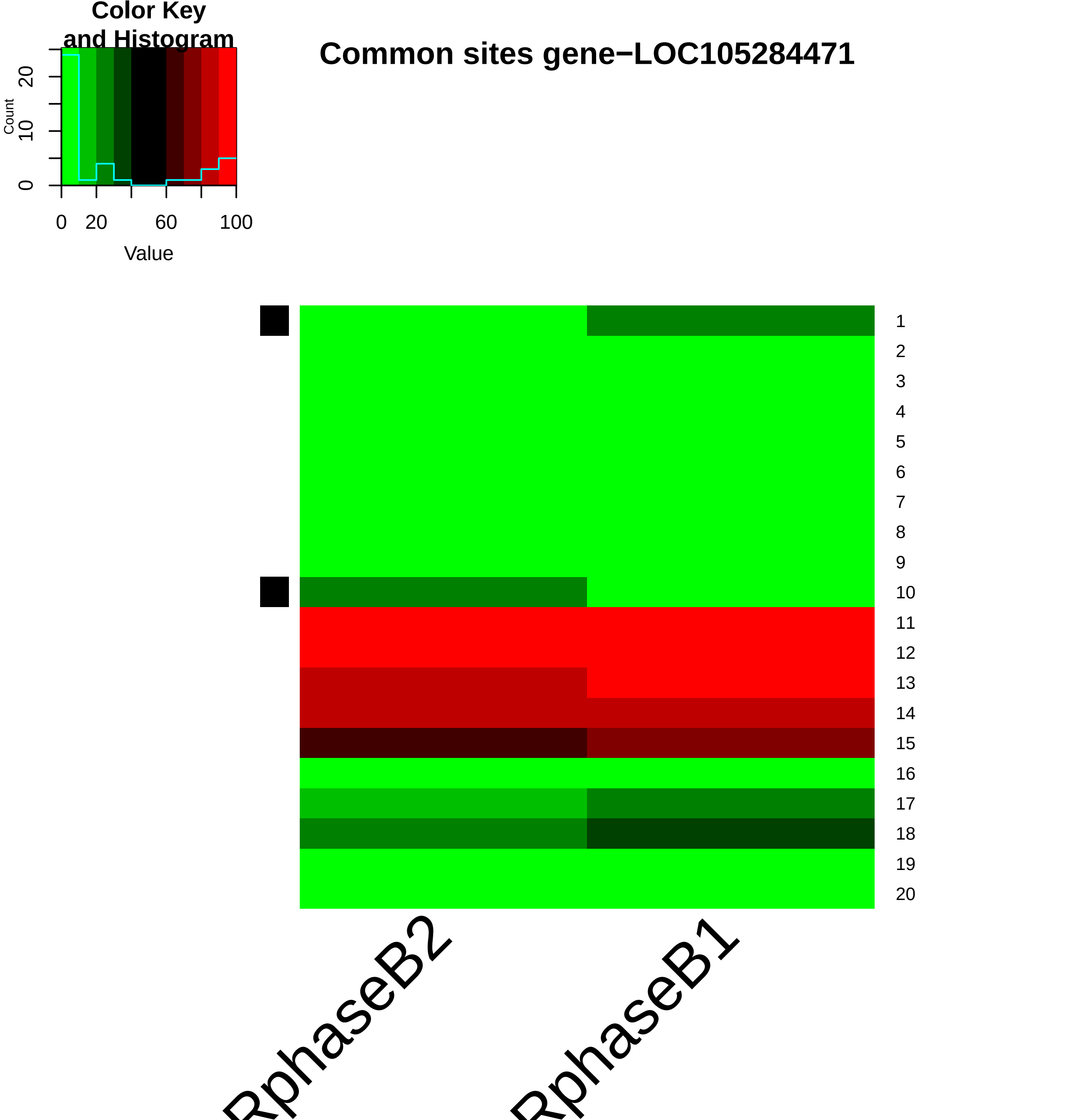
Color Key
and Histogram
Count
0
10
20
0
20
60
100
Value
Common sites gene−LOC105284471
RphaseB2
RphaseB1

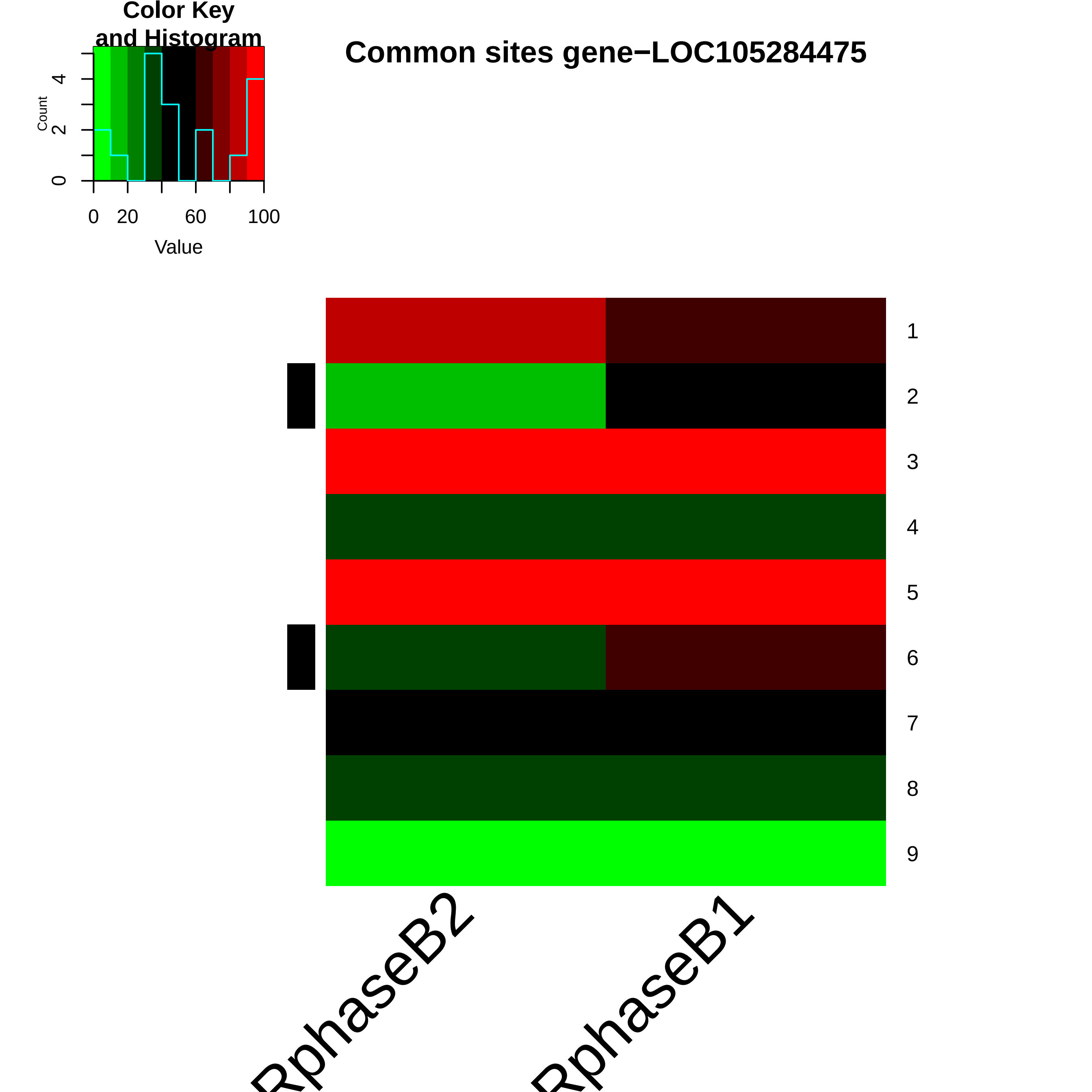
Color Key
and Histogram
Common sites gene−LOC105284475
Count
0
2
4
0
20
60
100
Value
1
2
3
4
5
6
7
8
9
RphaseB2
RphaseB1

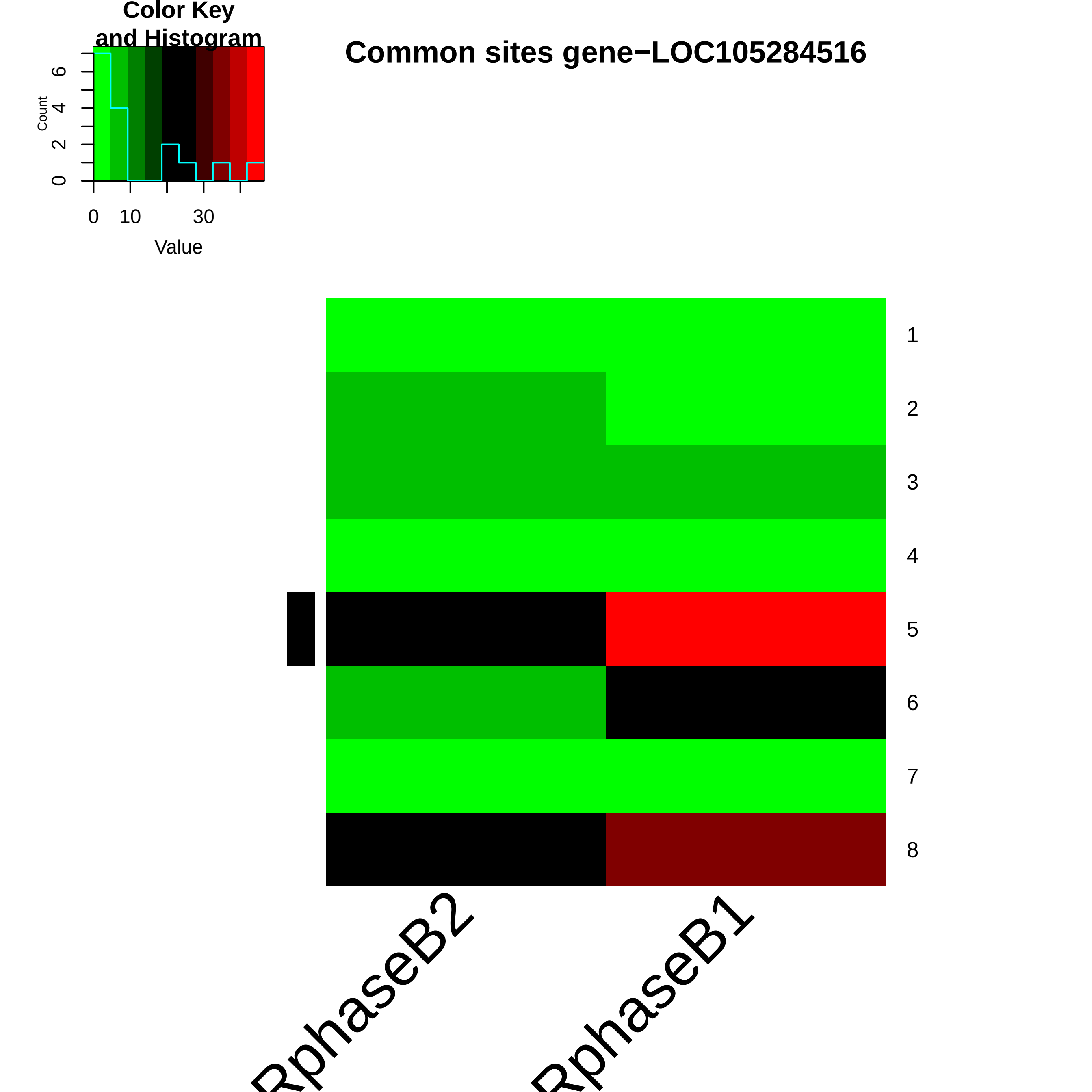
Color Key
and Histogram
Count
0
2
4
6
0
10
30
Value
Common sites gene−LOC105284516
1
2
3
4
5
6
7
8
RphaseB2
RphaseB1

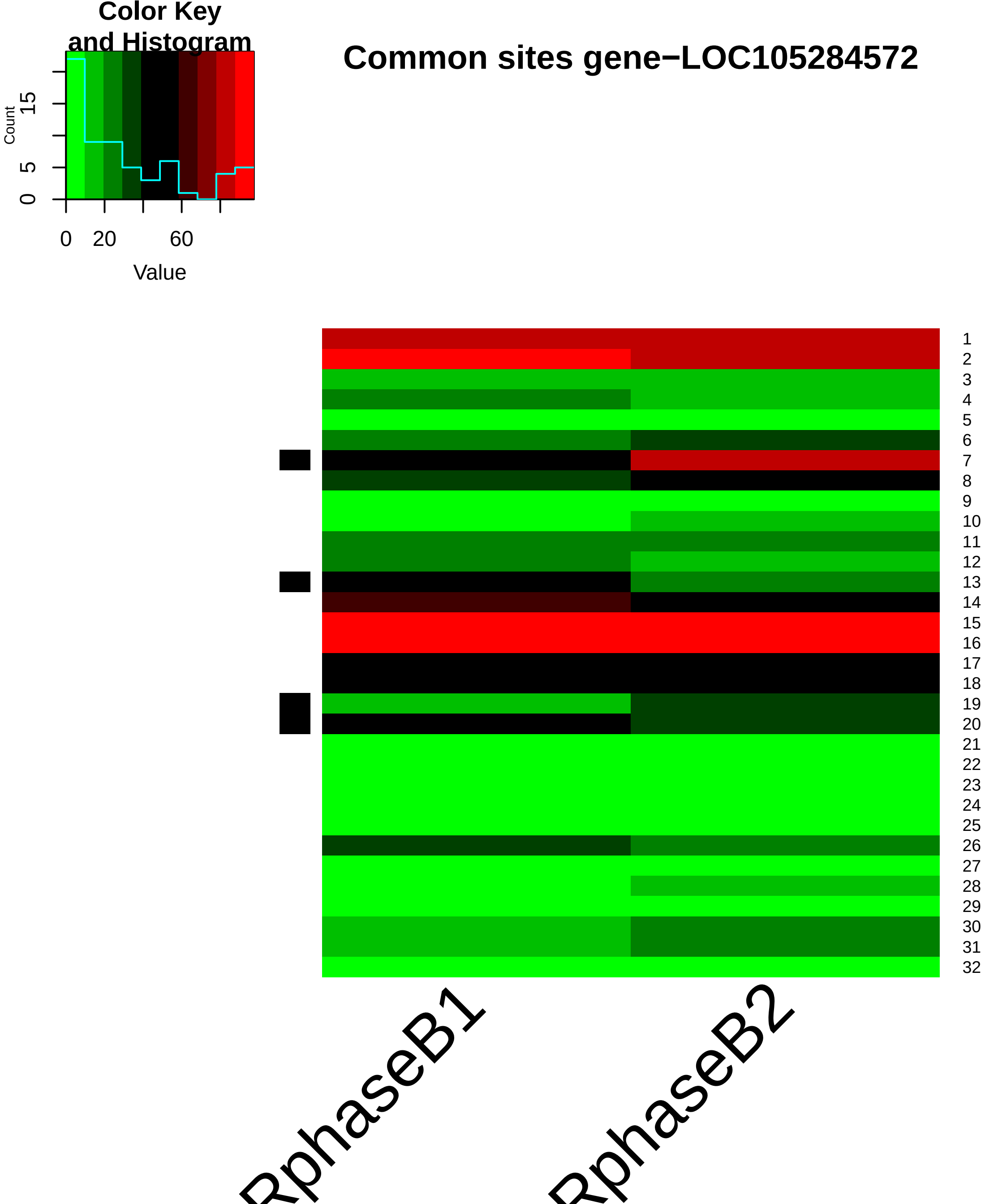
Color Key
and Histogram
Count
0
5
15
0
20
60
Value
Common sites gene−LOC105284572
RphaseB1
RphaseB2

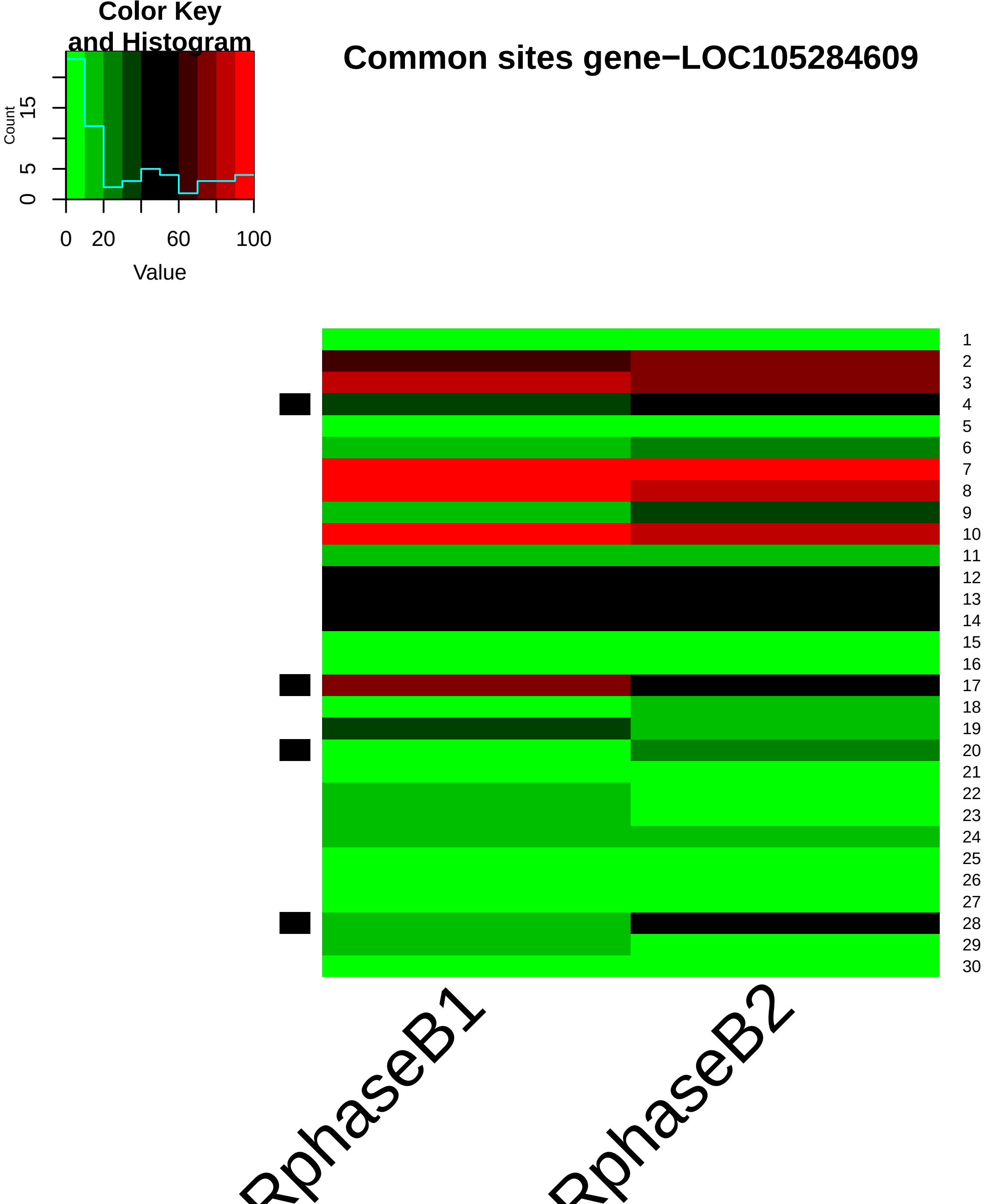
Color Key
and Histogram
Count
0
5
15
0
20
60
100
Value
Common sites gene−LOC105284609
RphaseB1
RphaseB2

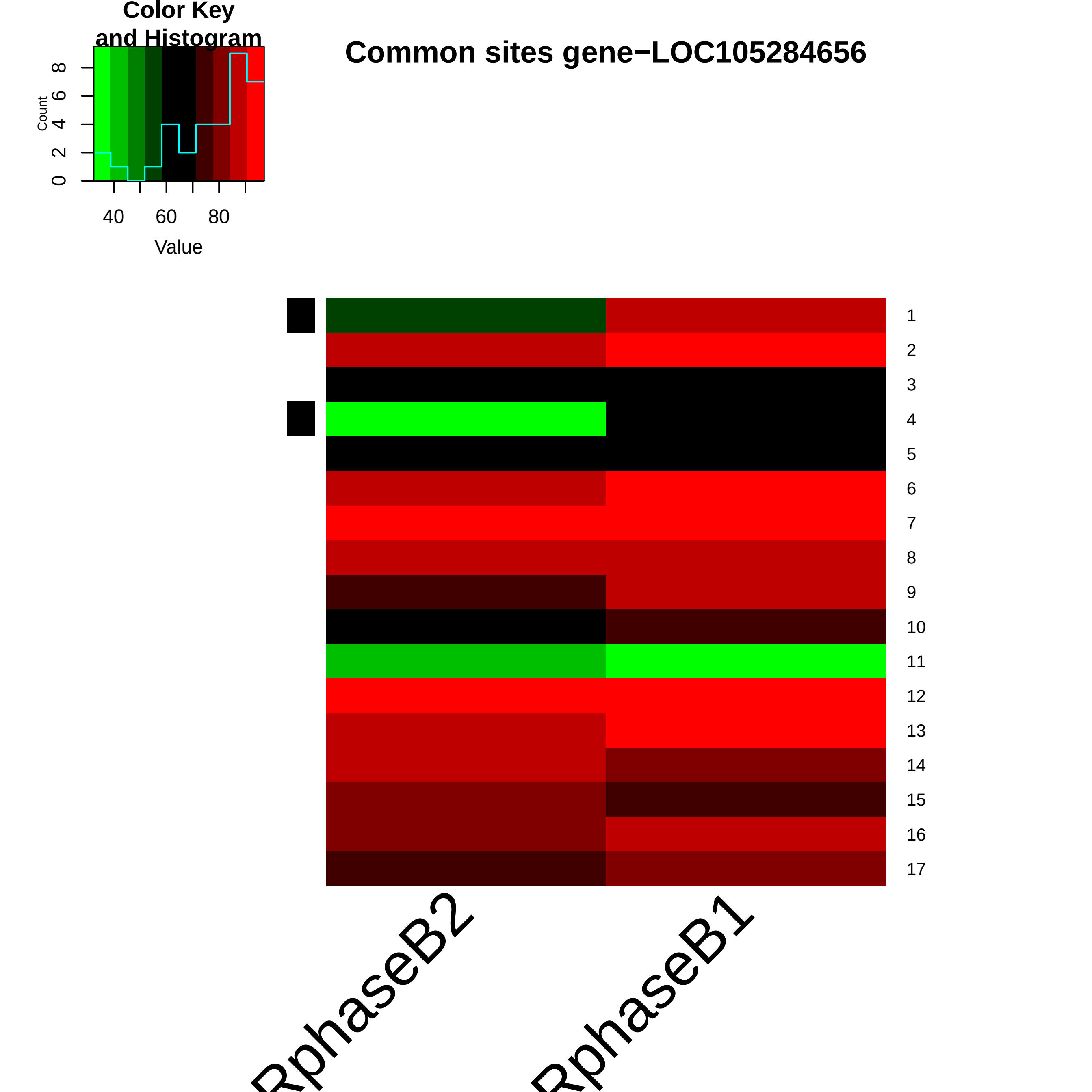

Color Key
and Histogram
Count
0
2
4
6
8
40
60
80
Value
Common sites gene−LOC105284656
RphaseB2
RphaseB1

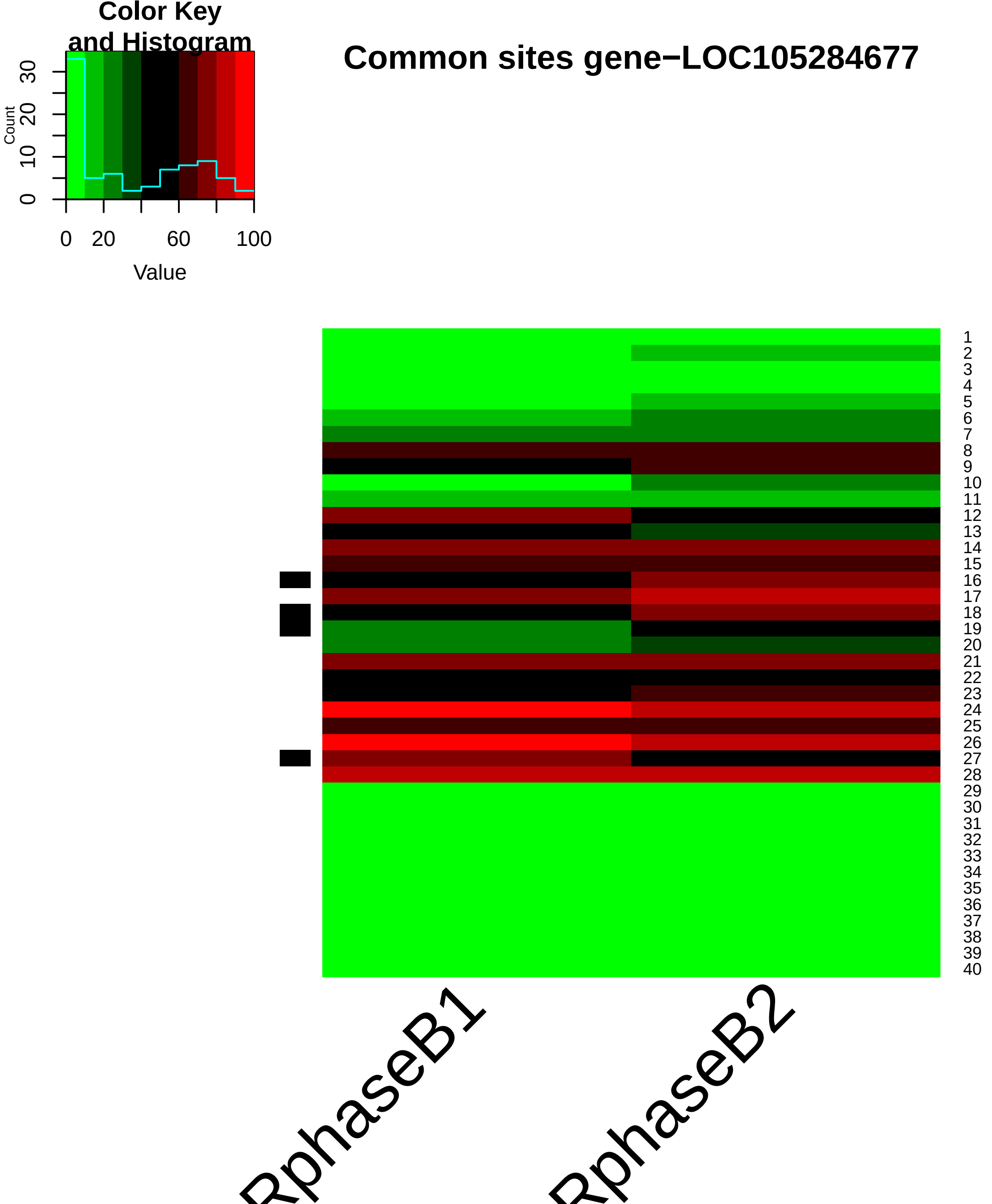
Color Key
and Histogram
Count
0
10
20
30
0
20
60
100
Value
Common sites gene−LOC105284677
RphaseB1
RphaseB2

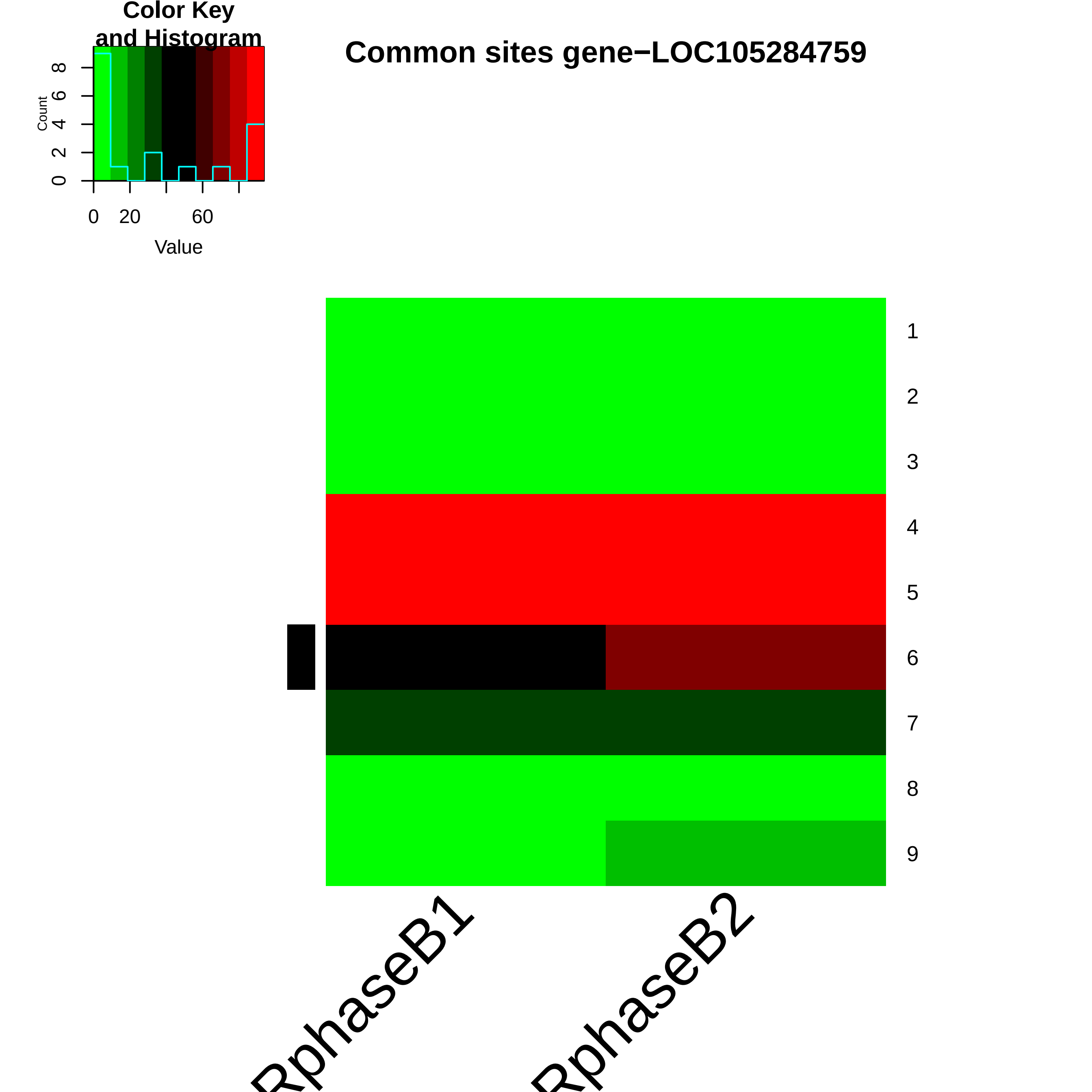

Color Key
and Histogram
Count
0
2
4
6
8
0
20
60
Value
Common sites gene−LOC105284759
1
2
3
4
5
6
7
8
9
RphaseB1
RphaseB2

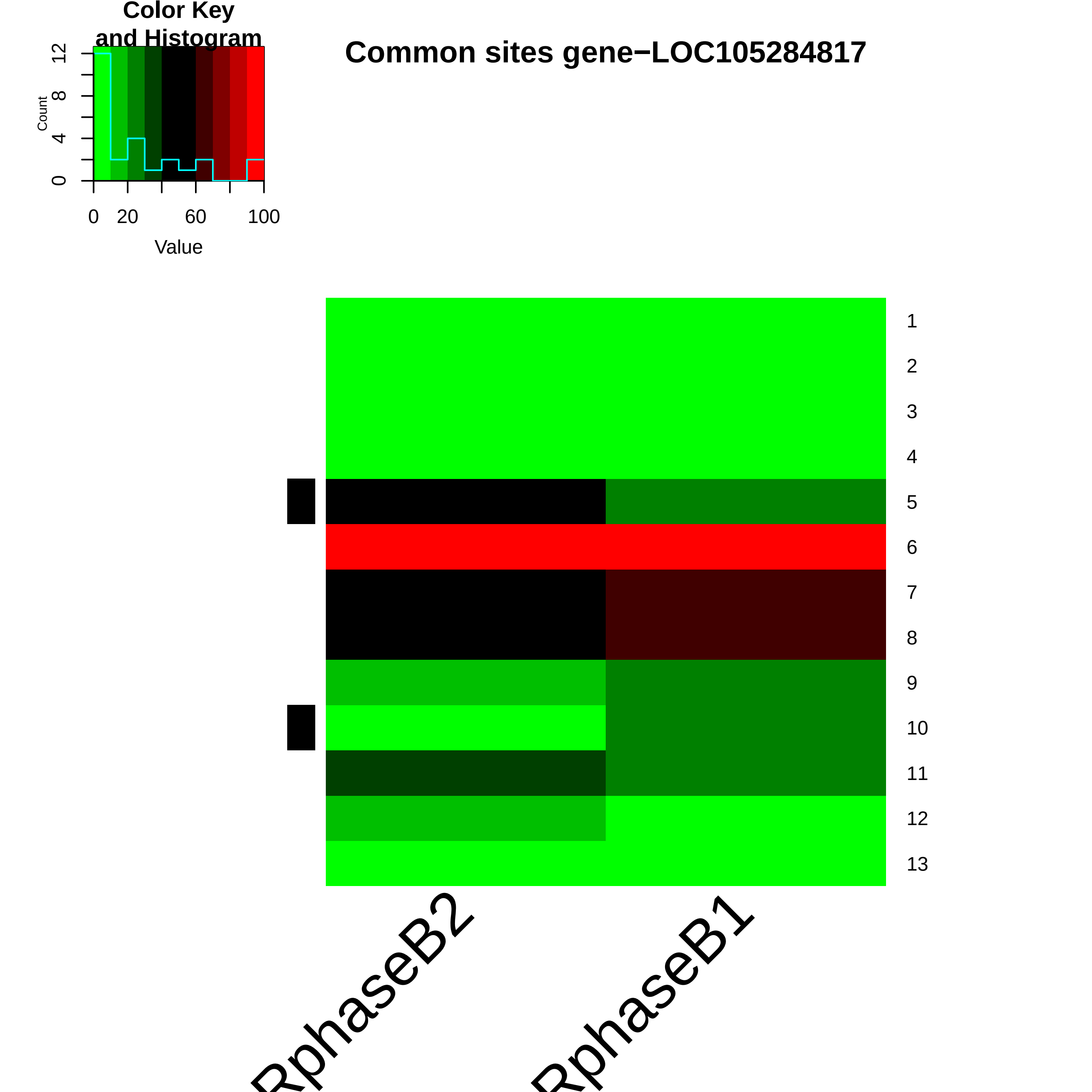

Color Key
and Histogram
Count
0
4
8
12
0
20
60
100
Value
Common sites gene−LOC105284817
1
2
3
4
5
6
7
8
9
10
11
12
13
RphaseB2
RphaseB1

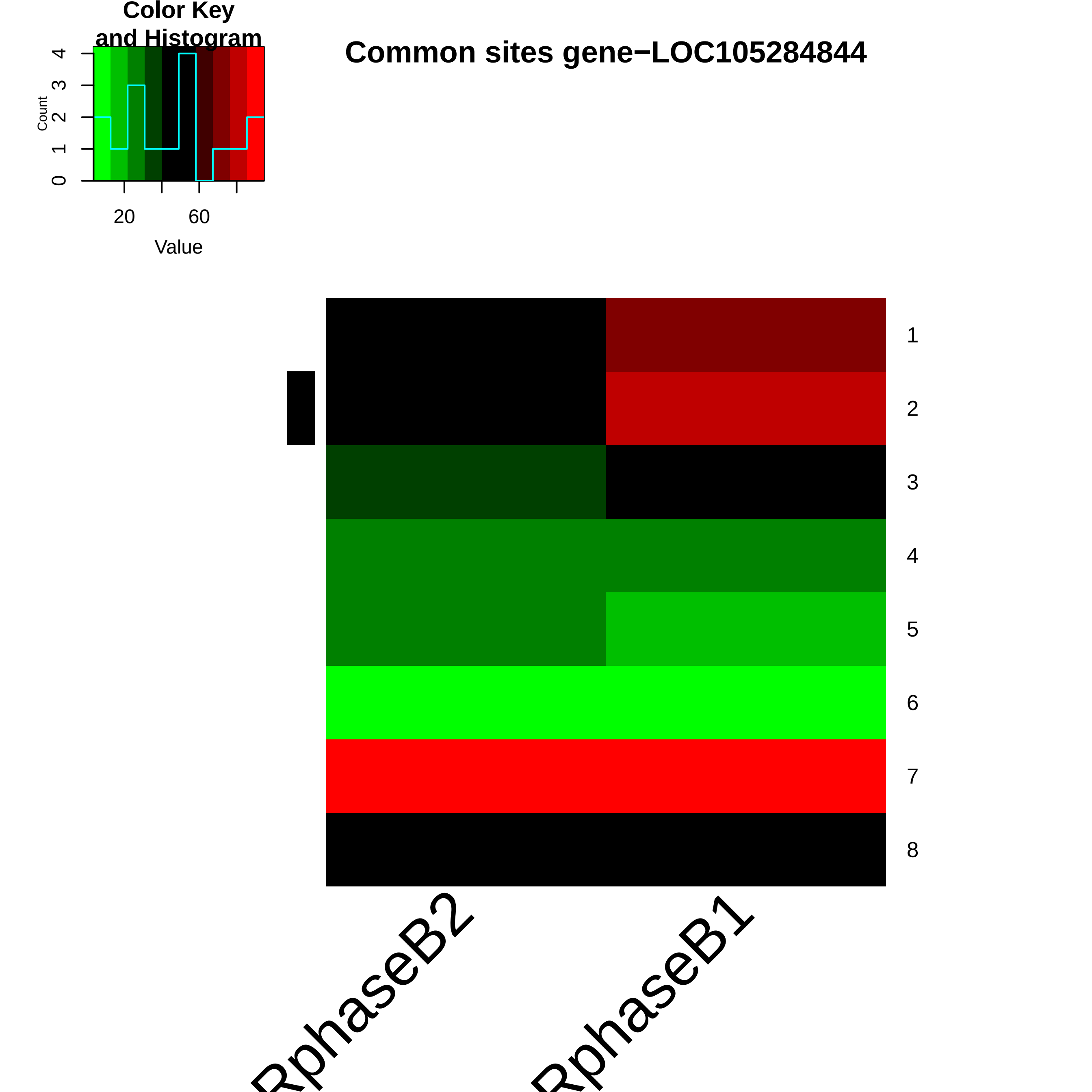
Color Key
and Histogram
Count
Value
0
1
2
3
4
20
60
Common sites gene−LOC105284844
1
2
3
4
5
6
7
8
RphaseB2
RphaseB1

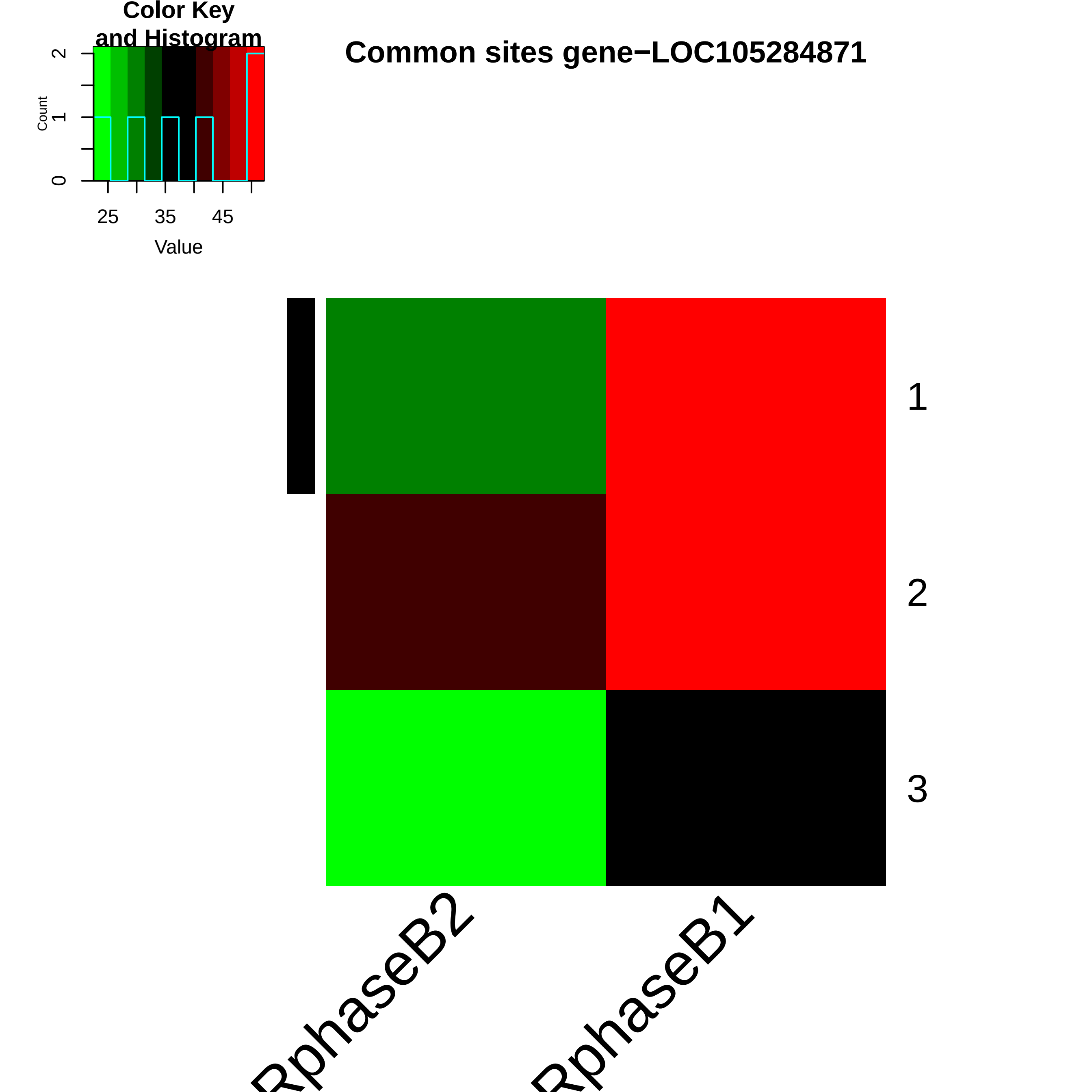

Color Key
and Histogram
Count
0
1
2
25
35
45
Value
Common sites gene−LOC105284871
1
2
3
RphaseB2
RphaseB1

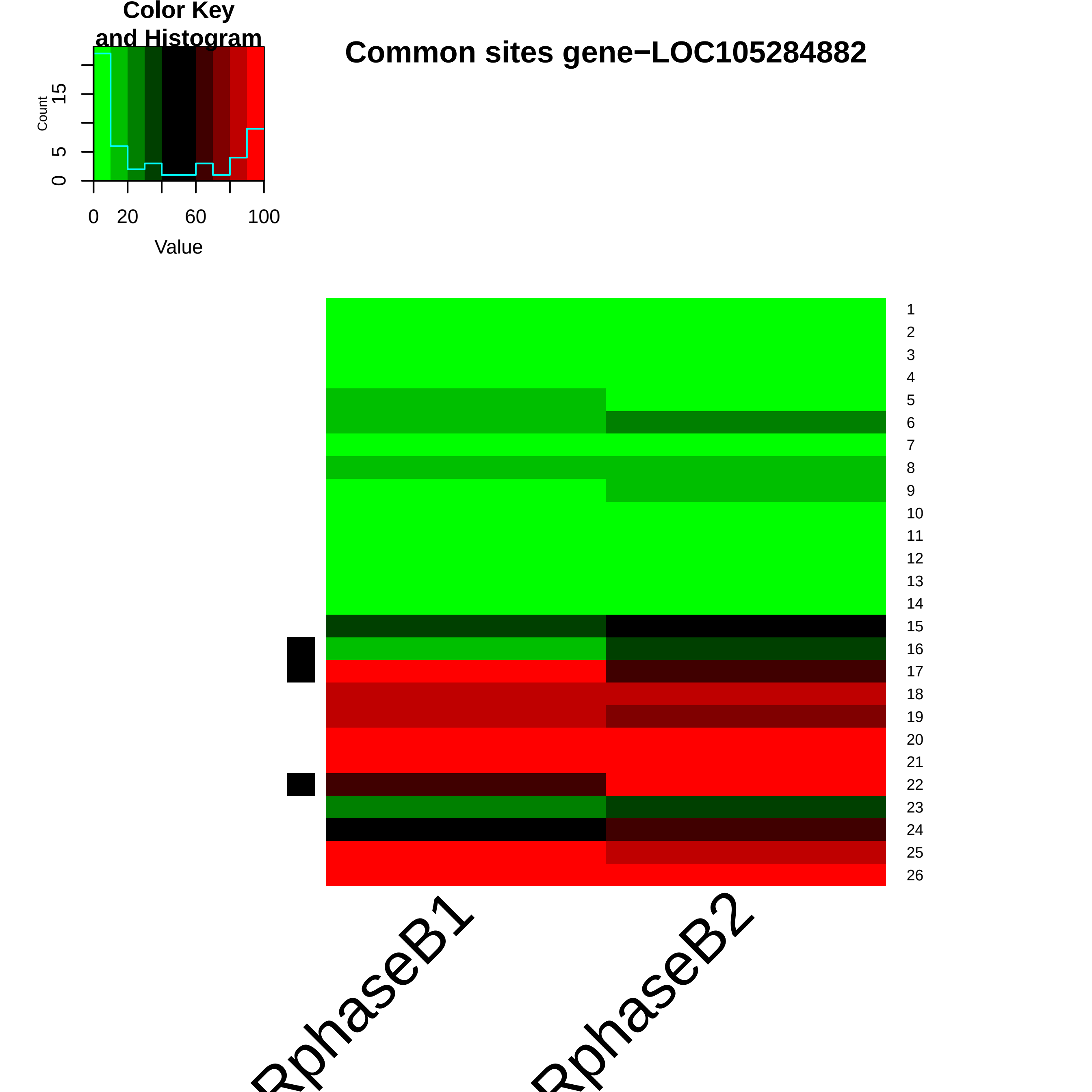

Color Key
and Histogram
Count
0
5
15
0
20
60
100
Value
Common sites gene−LOC105284882
RphaseB1
RphaseB2

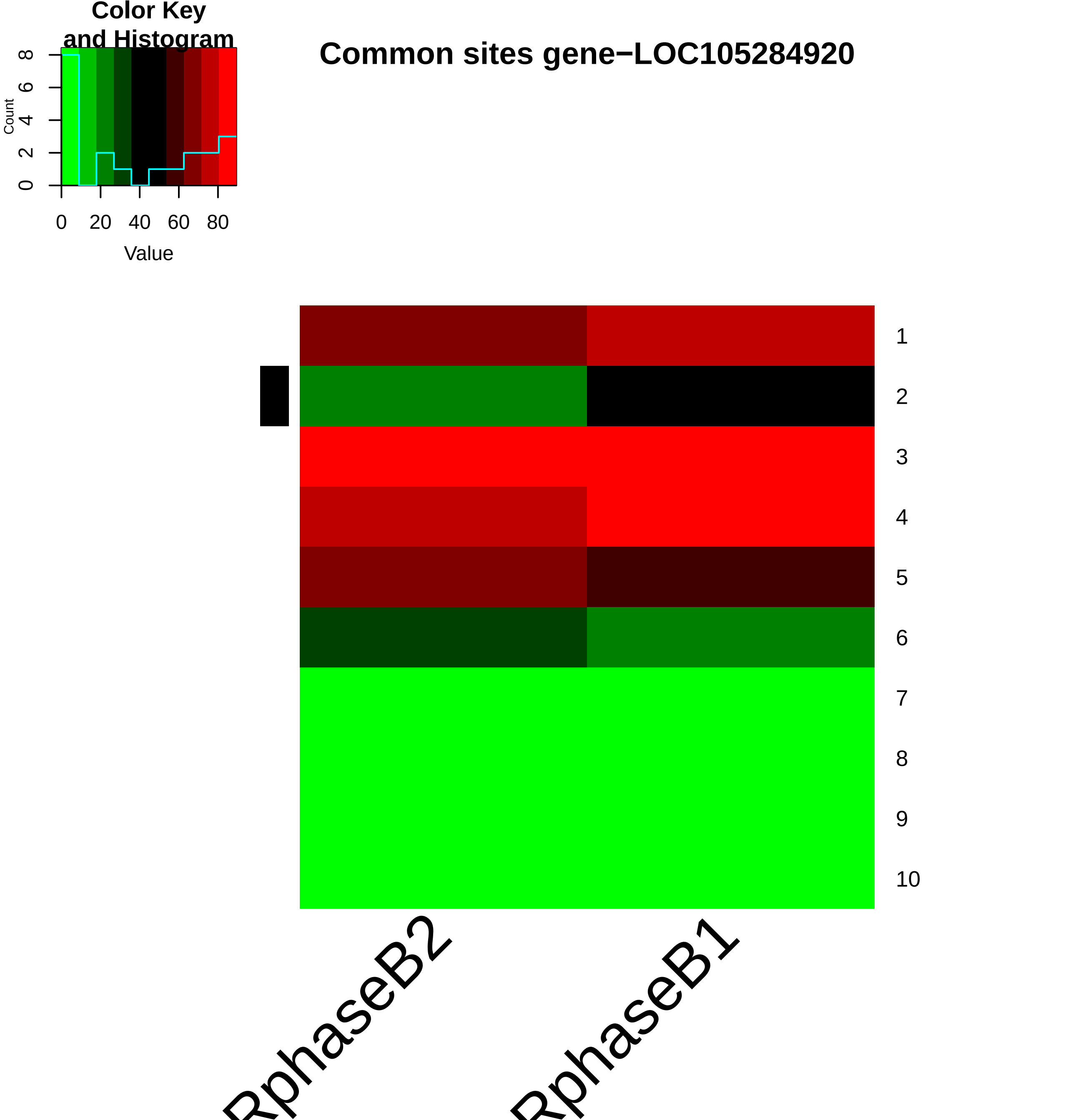
Color Key
and Histogram
Count
0
2
4
6
8
0
20
40
60
80
Value
Common sites gene−LOC105284920
1
2
3
4
5
6
7
8
9
10
RphaseB2
RphaseB1

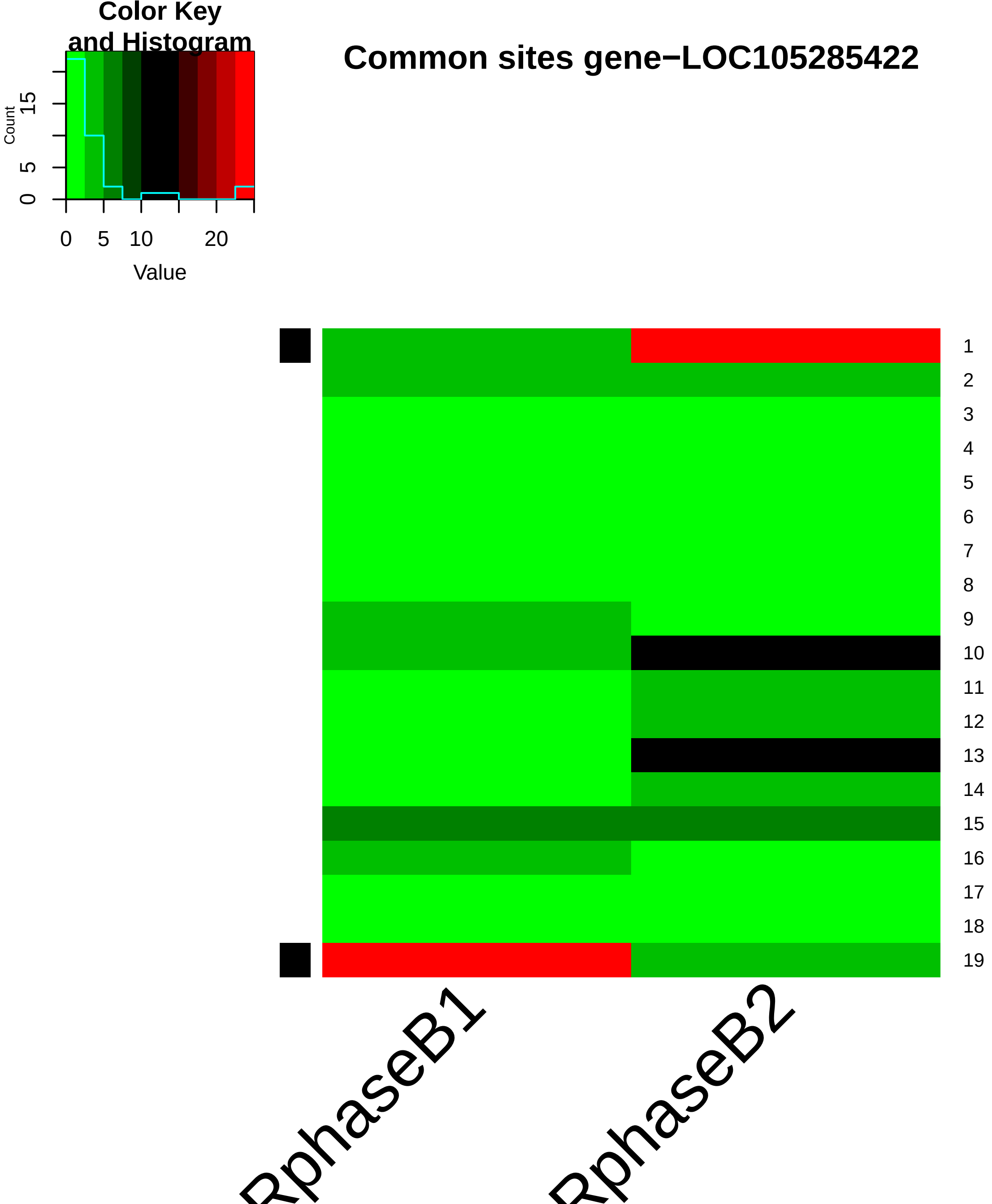

Color Key
and Histogram
Count
0
5
15
0
5
10
20
Value
Common sites gene−LOC105285422
RphaseB1
RphaseB2

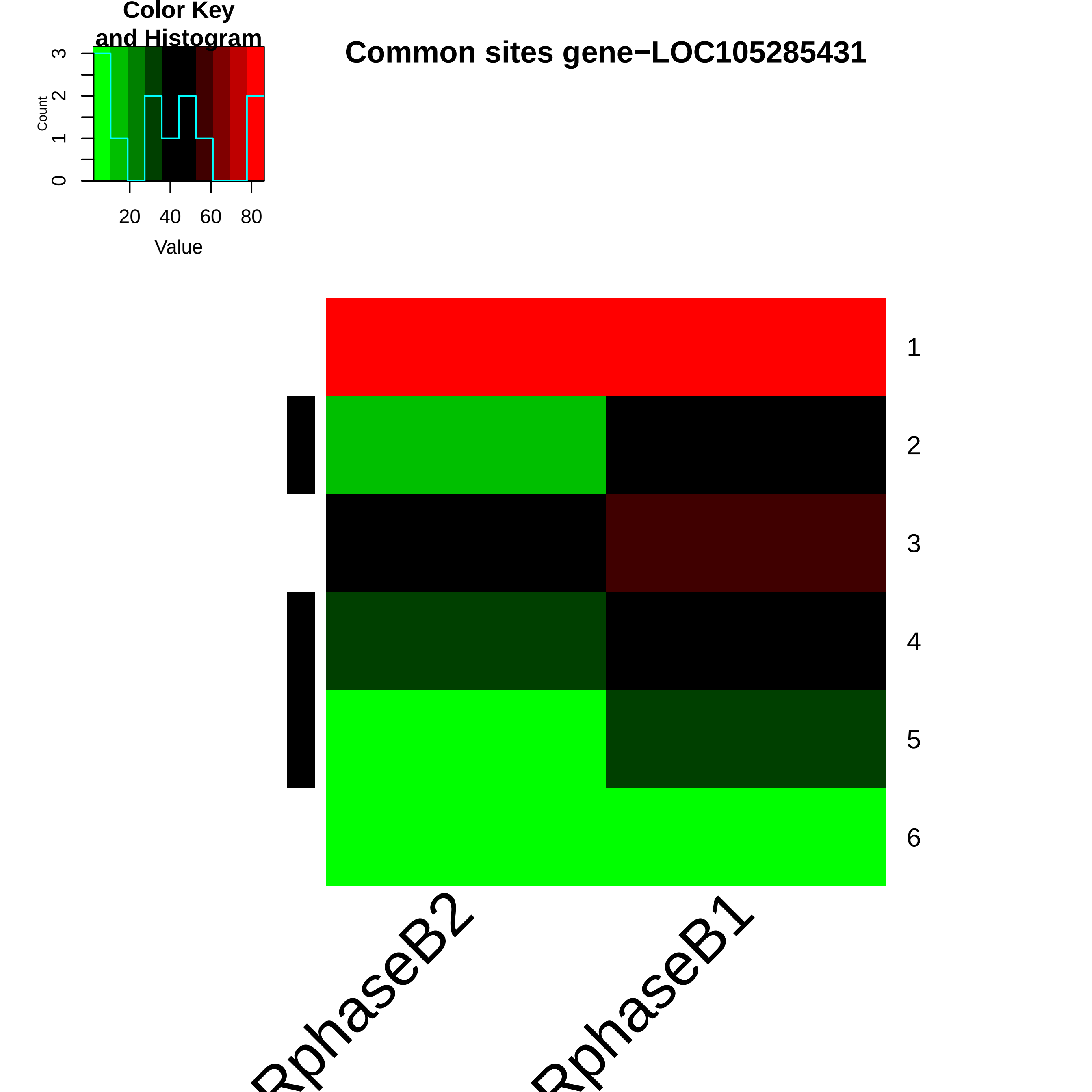
Color Key
and Histogram
Count
Value
20
40
60
80
0
1
2
3
Common sites gene−LOC105285431
1
2
3
4
5
6
RphaseB2
RphaseB1

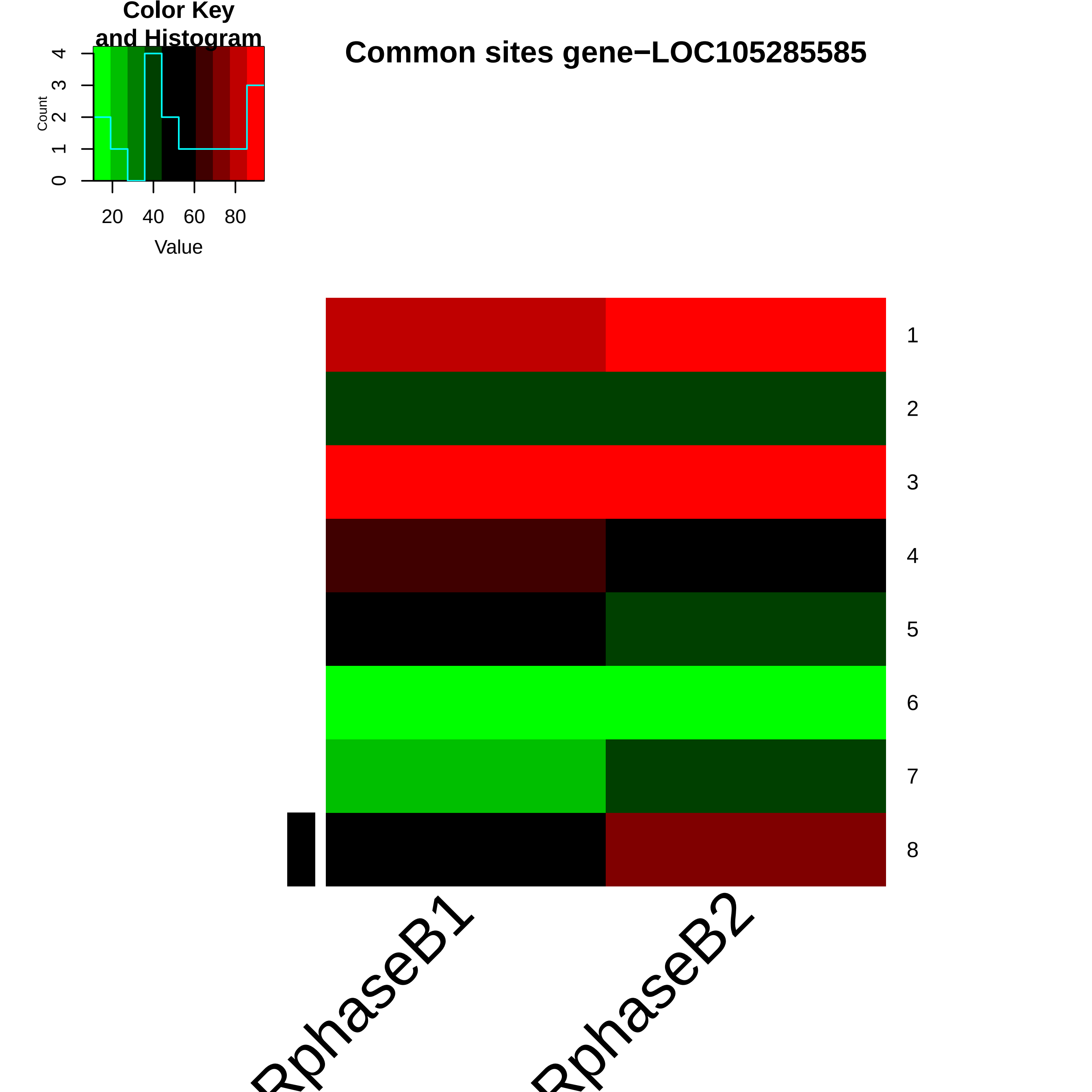

Color Key
and Histogram
Count
Value
0
1
2
3
4
20
40
60
80
Common sites gene−LOC105285585
1
2
3
4
5
6
7
8
RphaseB1
RphaseB2

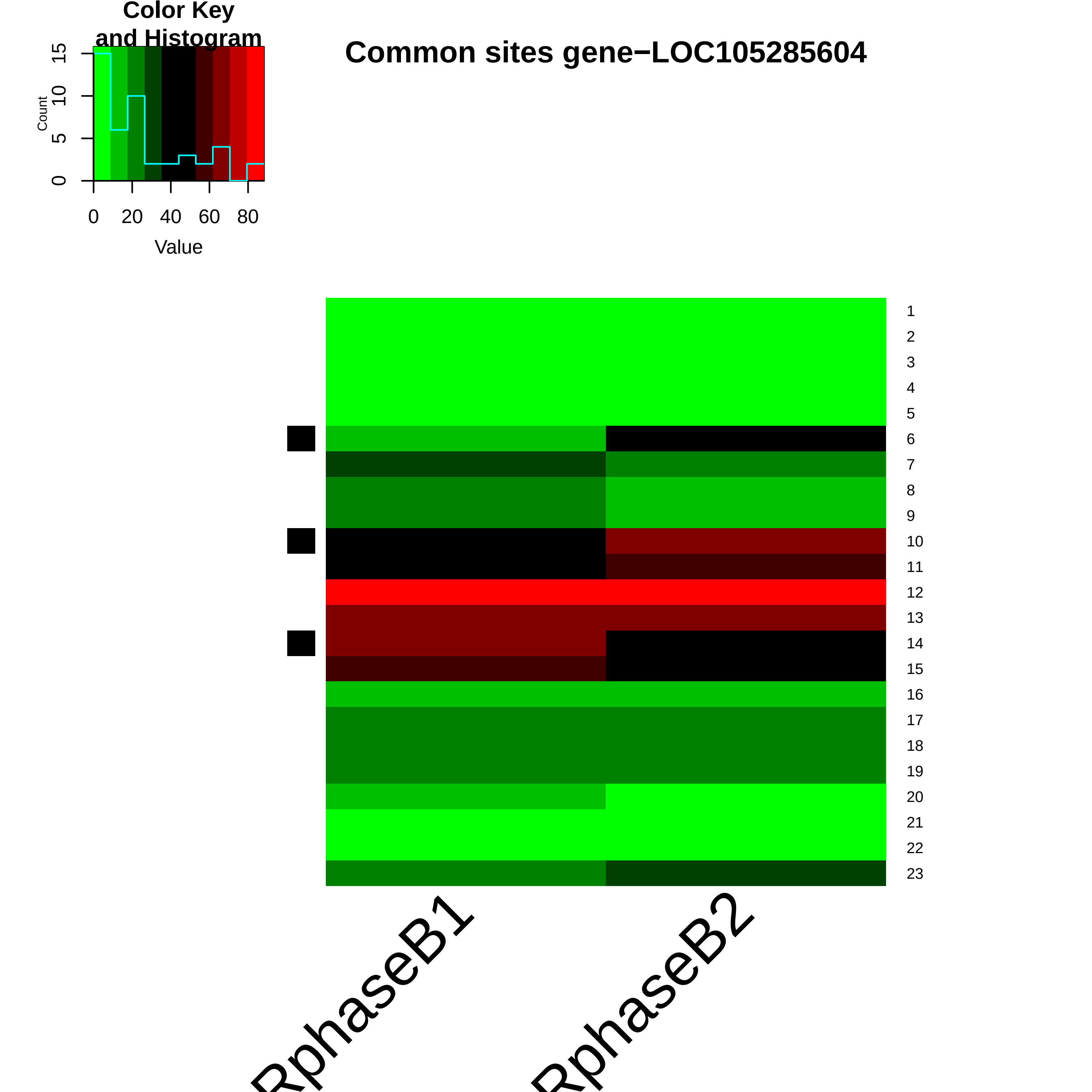
Color Key
and Histogram
Count
Value
0
5
10
15
0
20
40
60
80
Common sites gene−LOC105285604
RphaseB1
RphaseB2

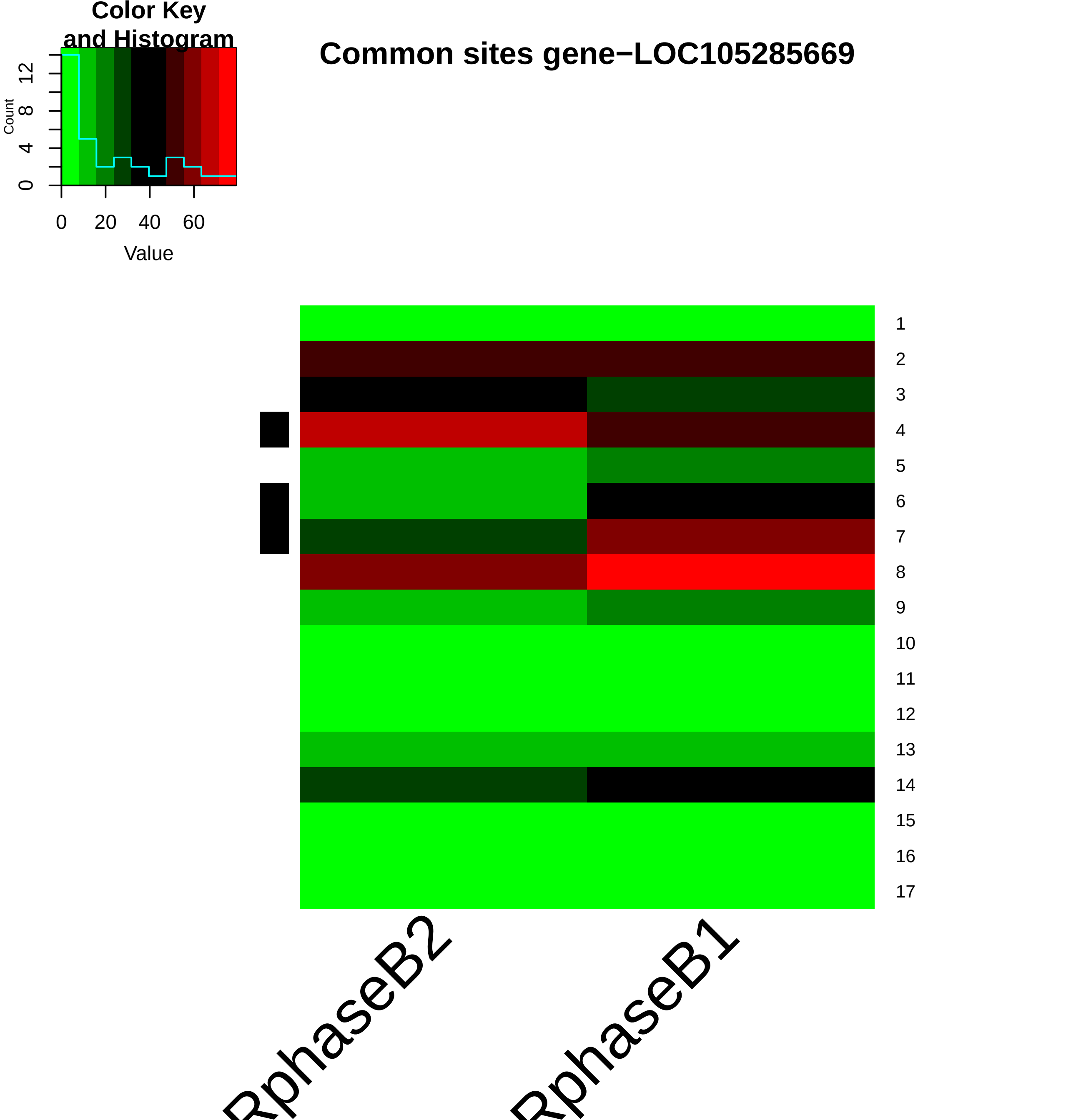

Color Key
and Histogram
Count
0
4
8
12
0
20
40
60
Value
Common sites gene−LOC105285669
RphaseB2
RphaseB1

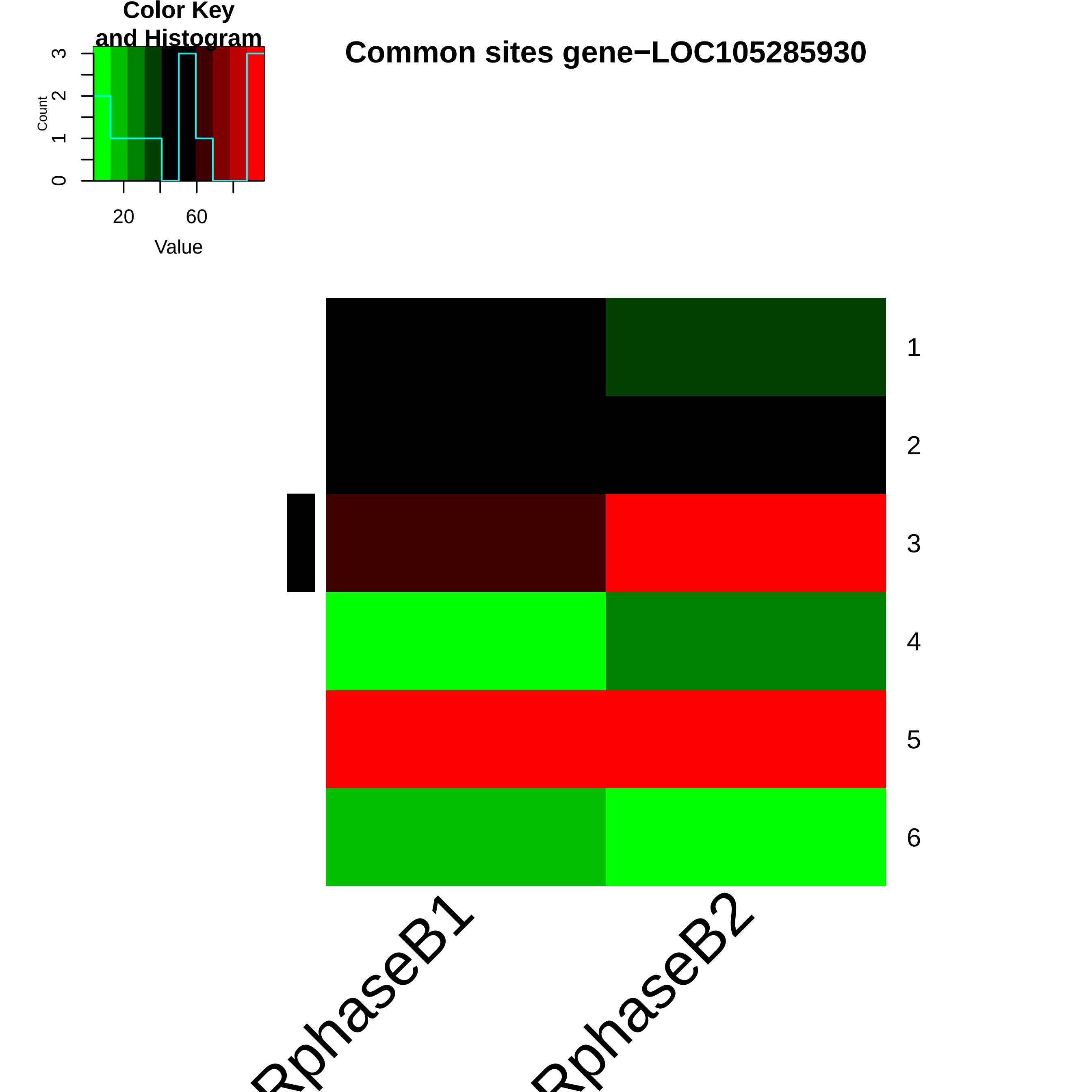

Color Key
and Histogram
Count
0
1
2
3
20
60
Value
Common sites gene−LOC105285930
1
2
3
4
5
6
RphaseB1
RphaseB2

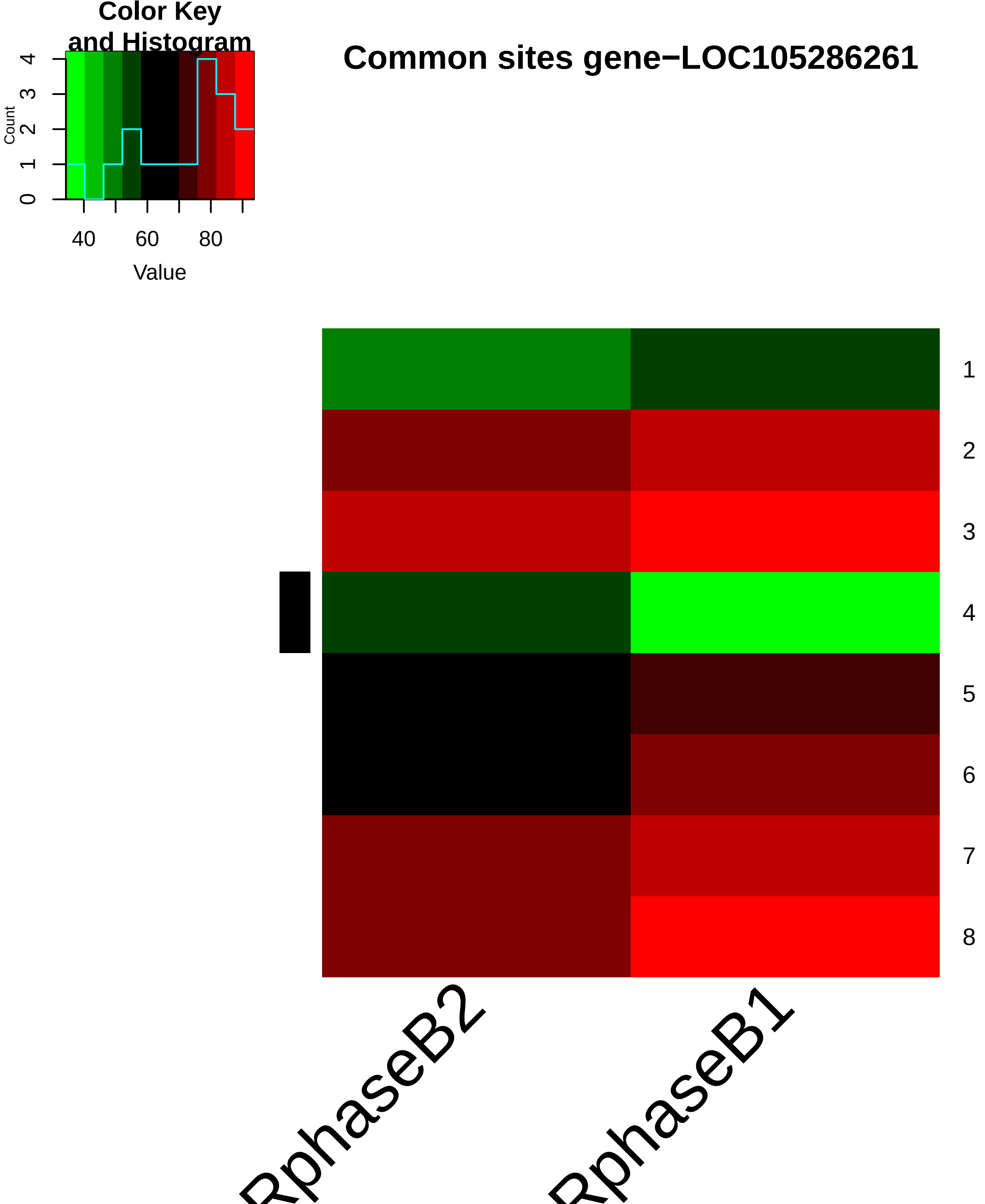

Color Key
and Histogram
Count
0
1
2
3
4
40
60
80
Value
Common sites gene−LOC105286261
1
2
3
4
5
6
7
8
RphaseB2
RphaseB1

### Common sites gene−LOC105286463

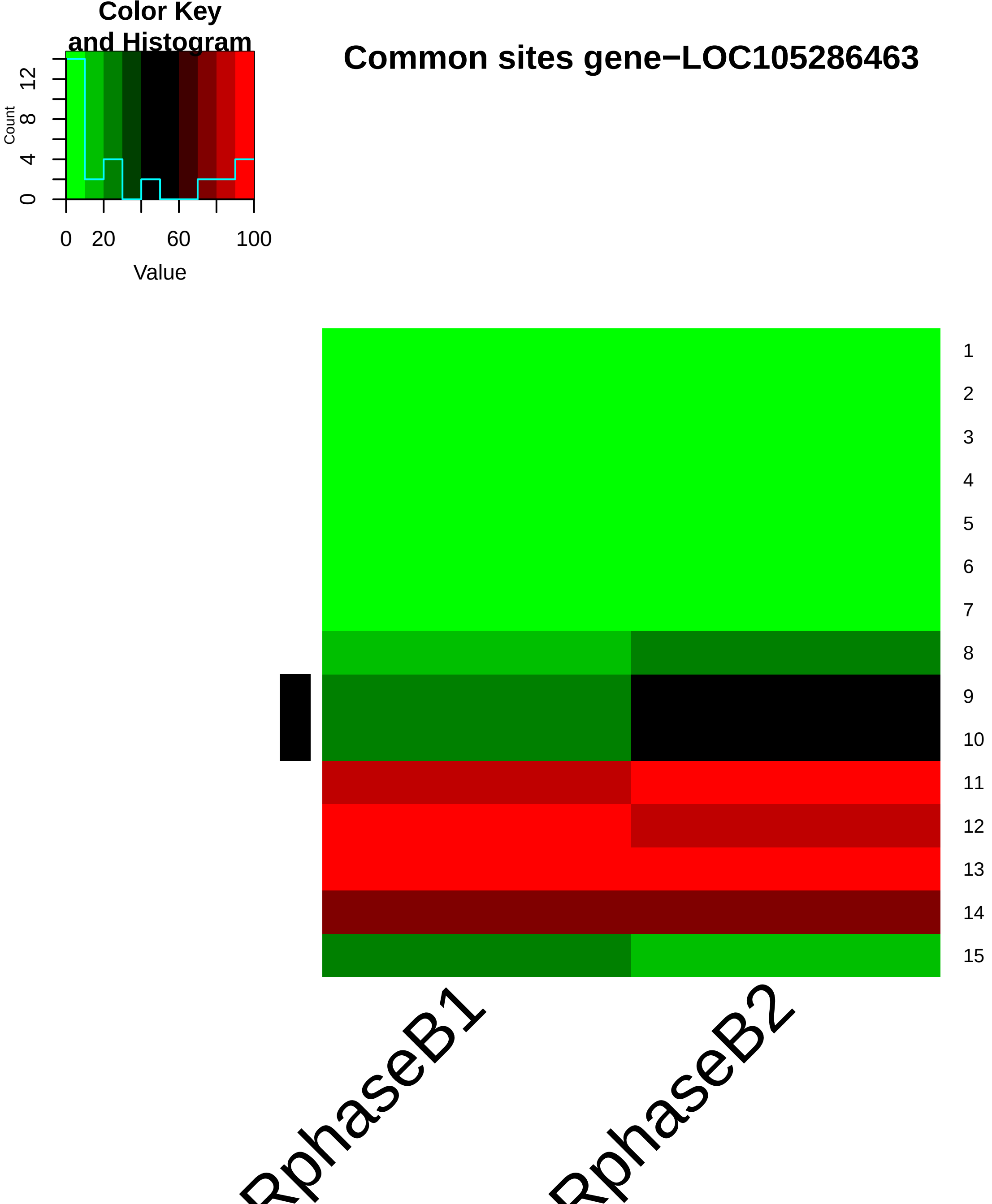

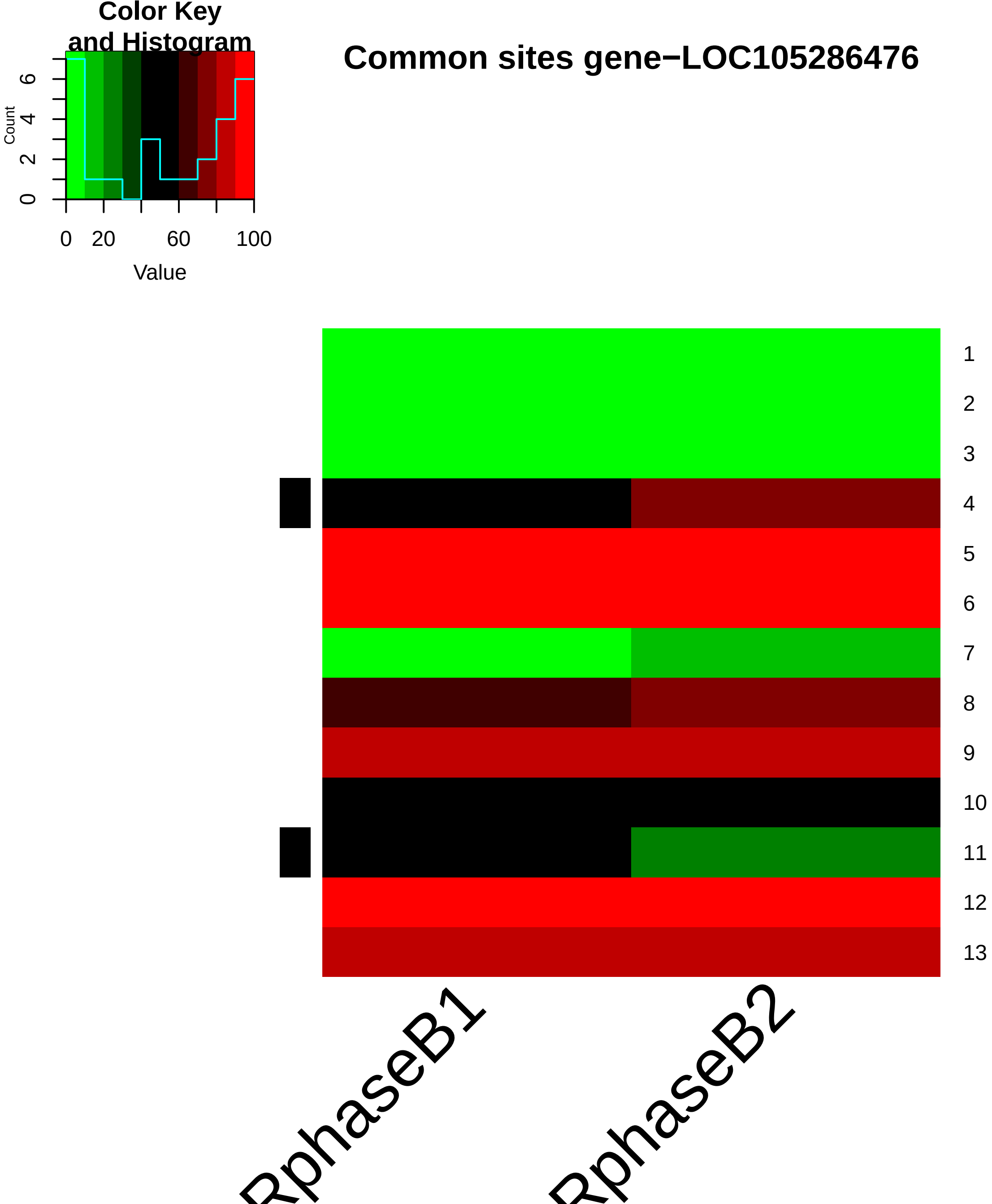
Color Key
and Histogram
Count
0
2
4
6
0
20
60
100
Value
Common sites gene−LOC105286476
1
2
3
4
5
6
7
8
9
10
11
12
13
RphaseB1
RphaseB2

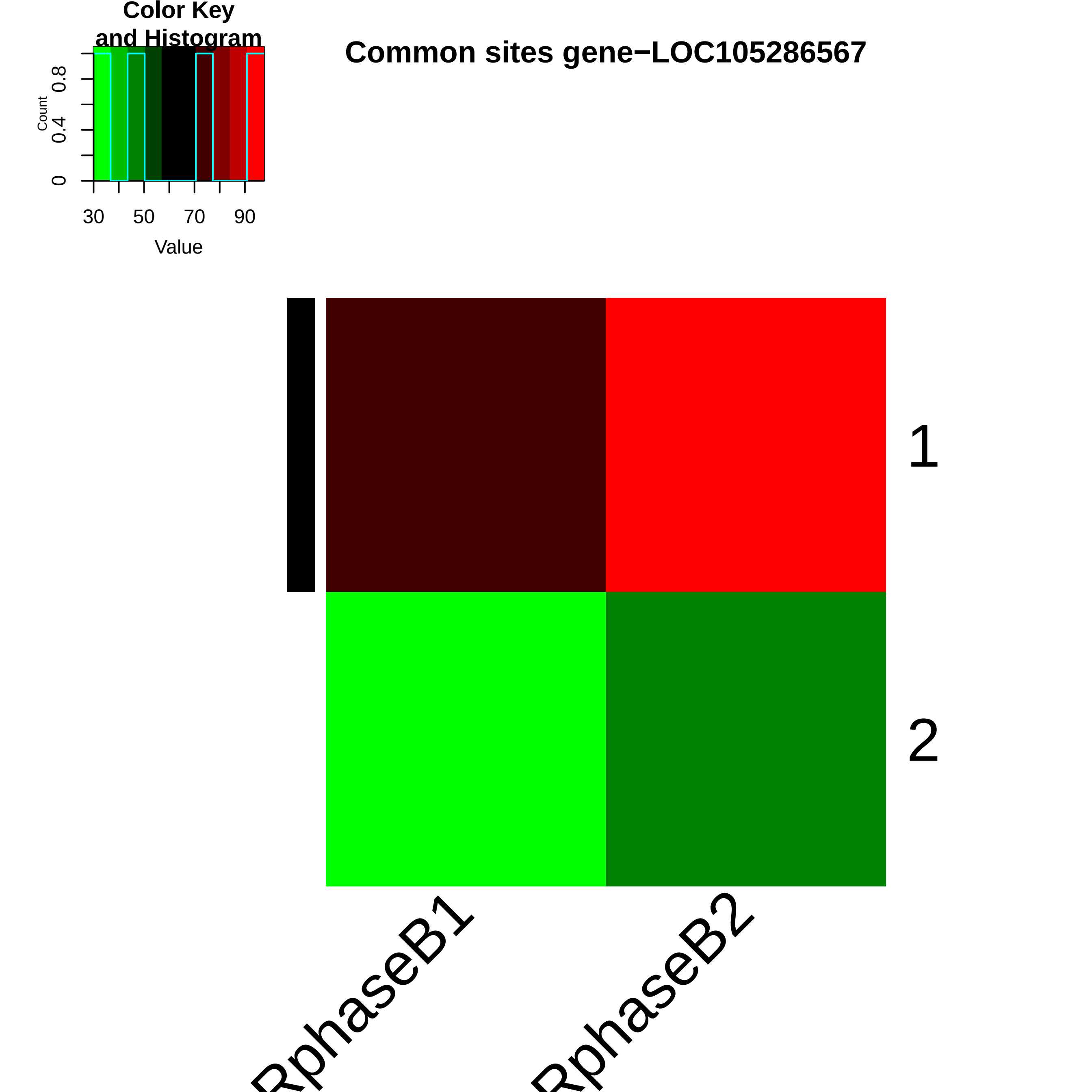

Color Key
and Histogram
Count
0
0.4
0.8
30
50
70
90
Value
Common sites gene−LOC105286567
1
2
RphaseB1
RphaseB2

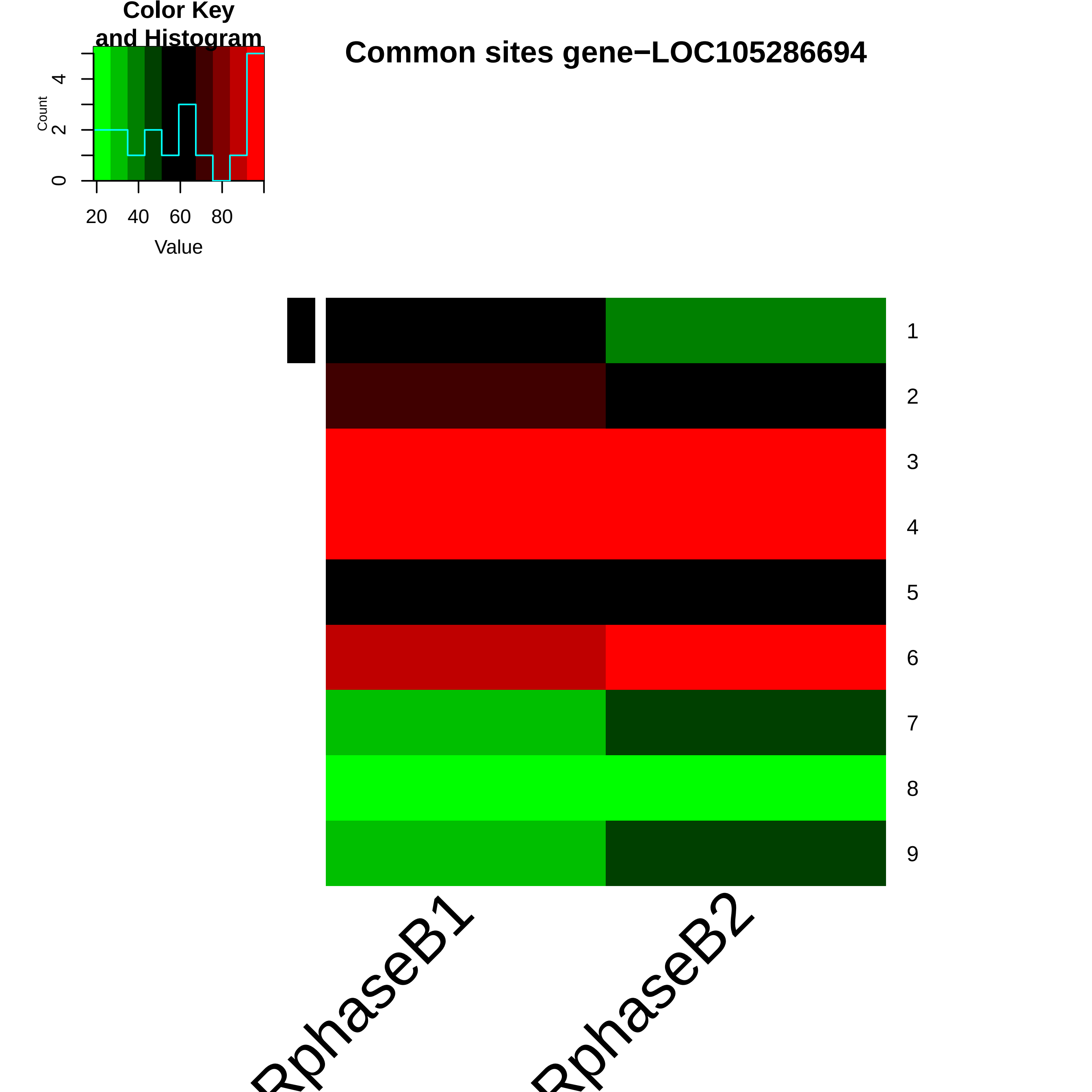

Color Key
and Histogram
Count
0
2
4
20
40
60
80
Value
Common sites gene−LOC105286694
1
2
3
4
5
6
7
8
9
RphaseB1
RphaseB2

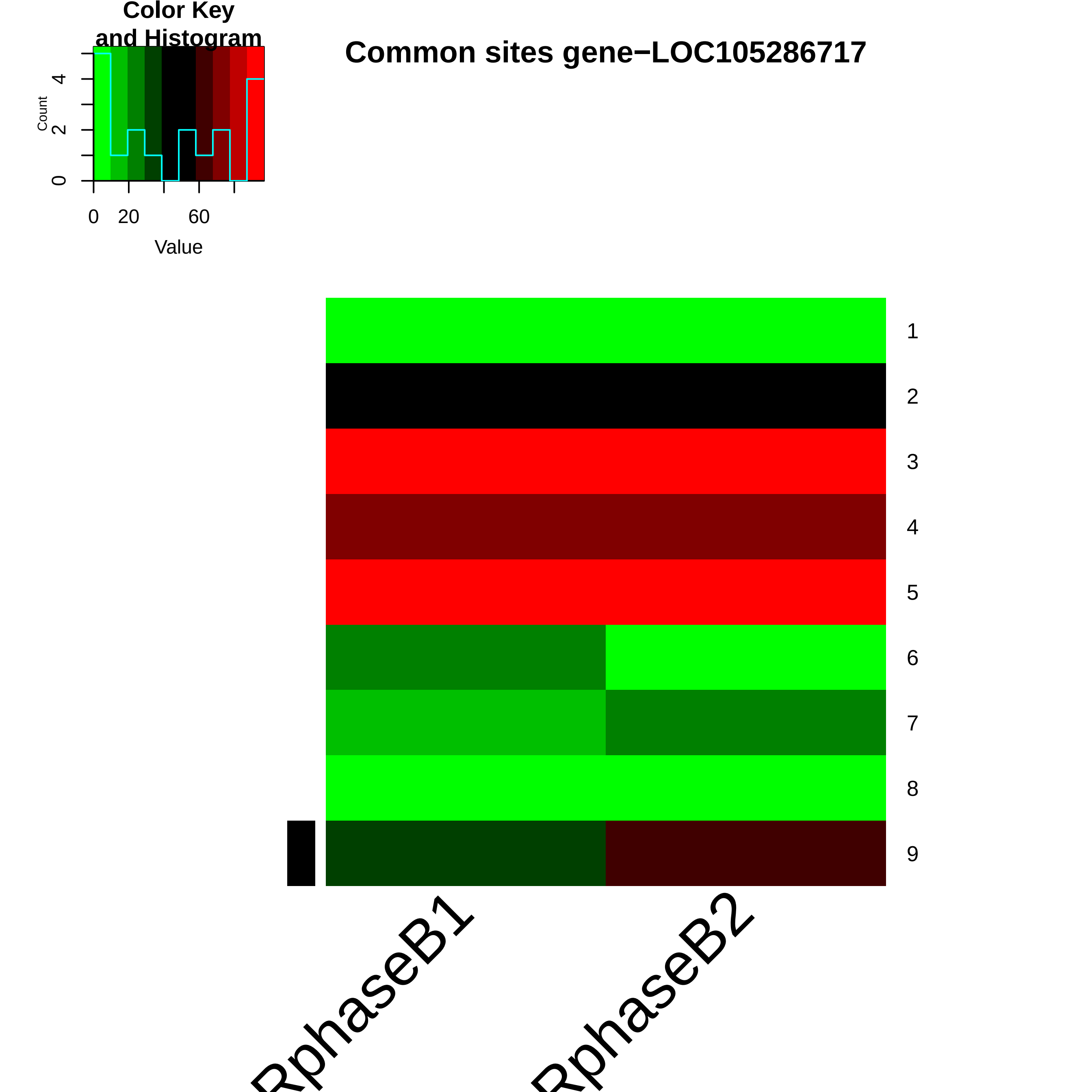
Color Key
and Histogram
Count
0
2
4
0
20
60
Value
Common sites gene−LOC105286717
1
2
3
4
5
6
7
8
9
RphaseB1
RphaseB2

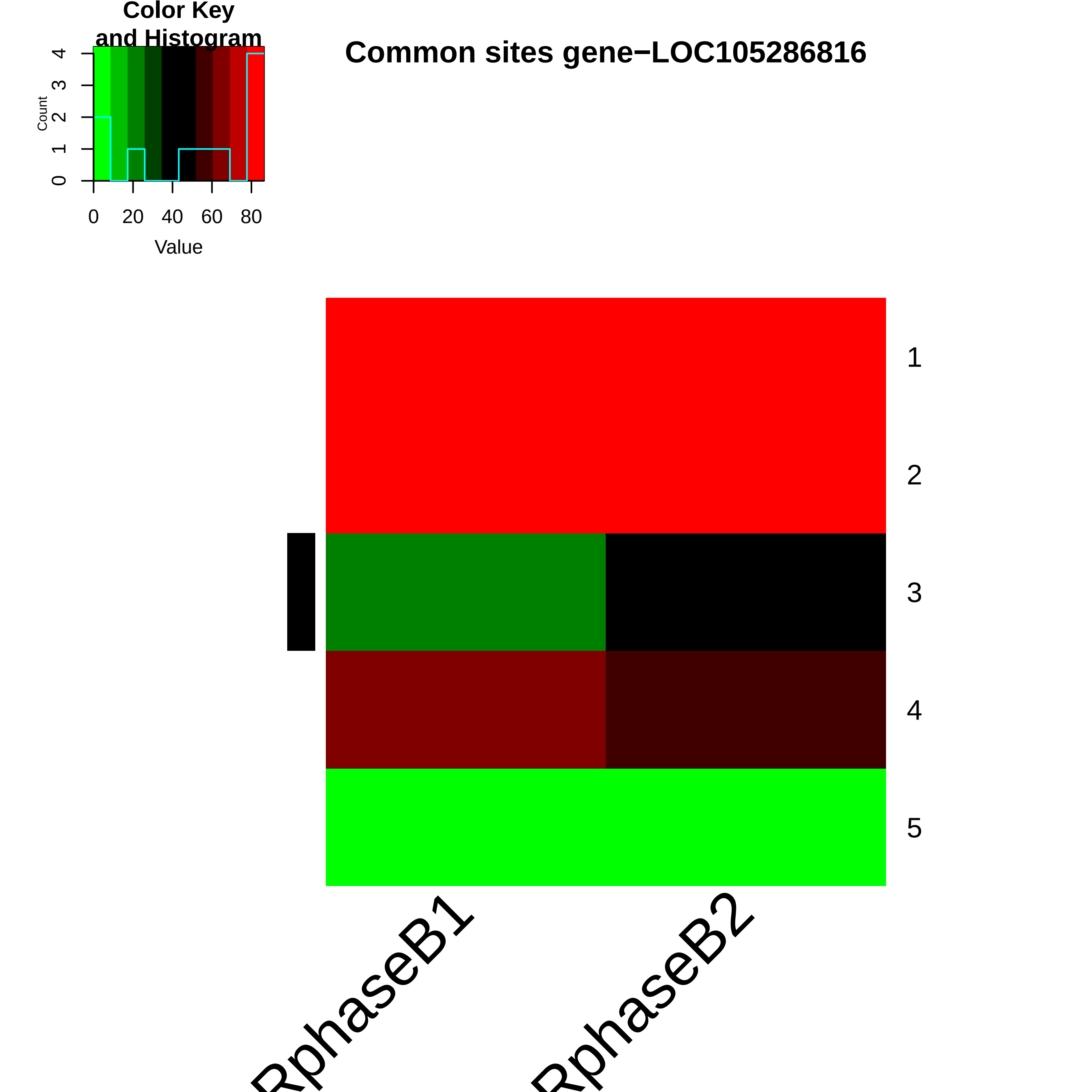

Color Key
and Histogram
Count
0
1
2
3
4
0
20
40
60
80
Value
Common sites gene−LOC105286816
1
2
3
4
5
RphaseB1
RphaseB2

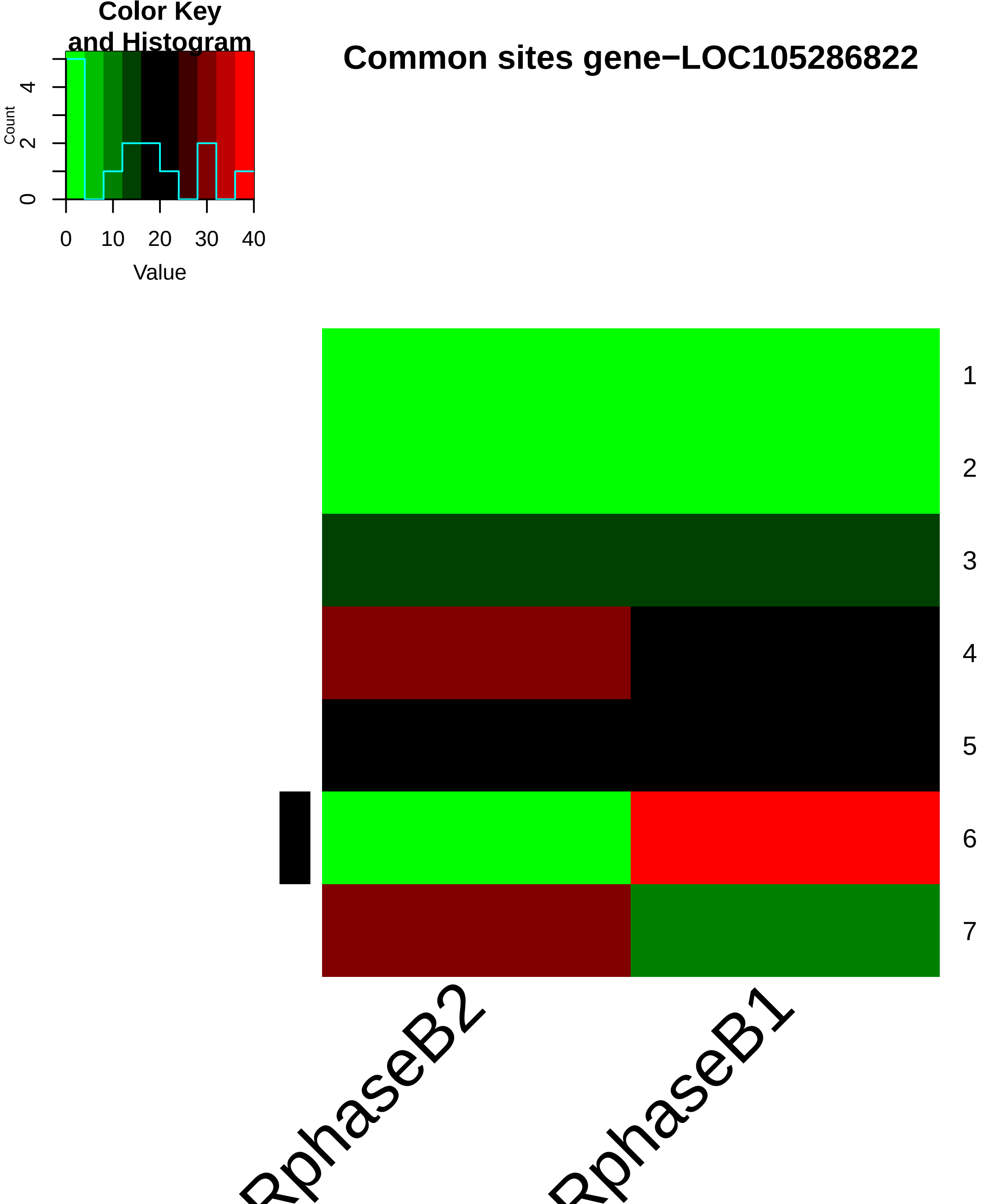
Color Key
and Histogram
Count
0
2
4
0
10
20
30
40
Value
Common sites gene−LOC105286822
1
2
3
4
5
6
7
RphaseB2
RphaseB1

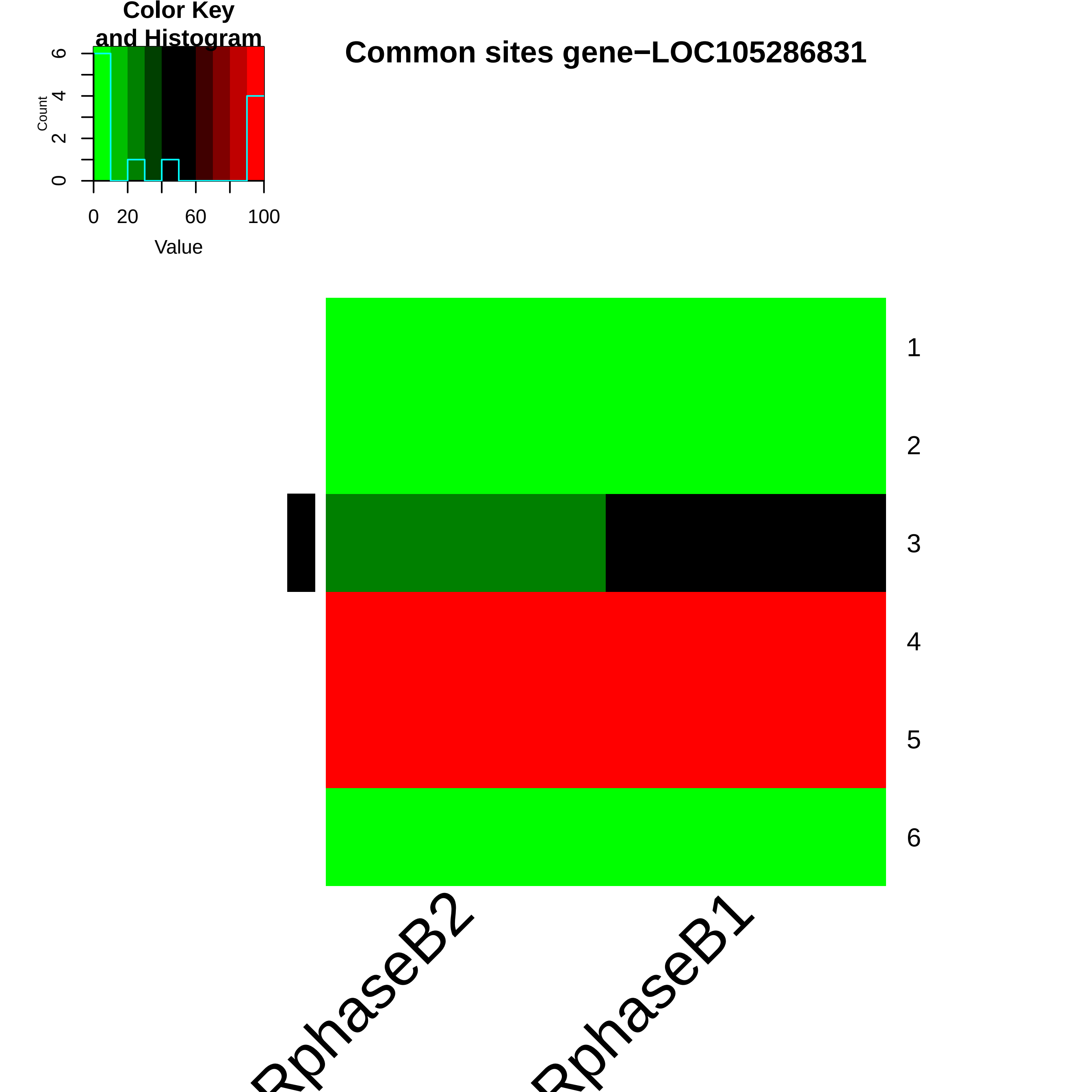

Color Key and Histogram
Count
Value
0
20
60
100
0
2
4
6
Common sites gene−LOC105286831
1
2
3
4
5
6
RphaseB2
RphaseB1

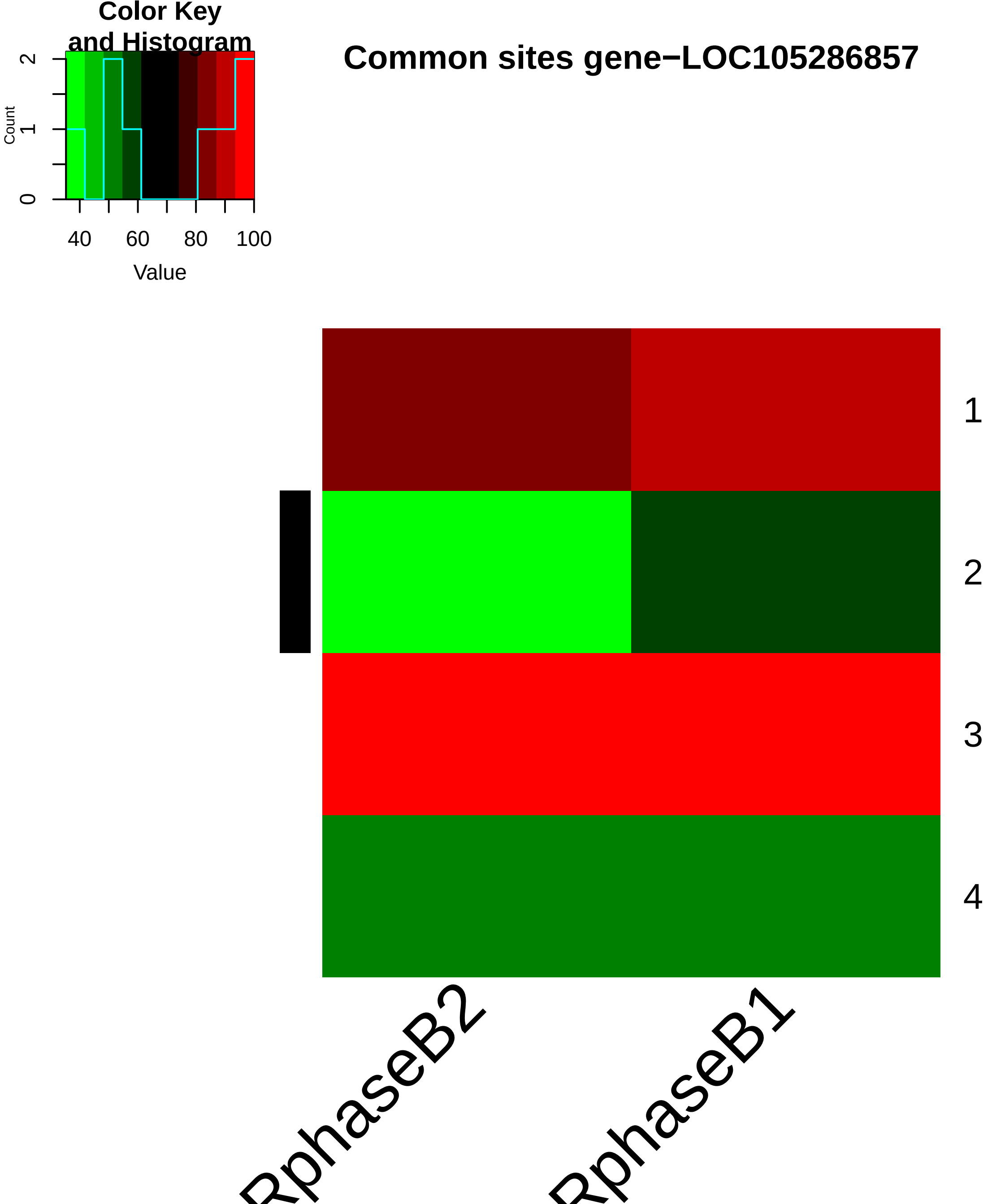
Color Key
and Histogram
Count
0
1
2
40
60
80
100
Value
Common sites gene−LOC105286857
1
2
3
4
RphaseB2
RphaseB1

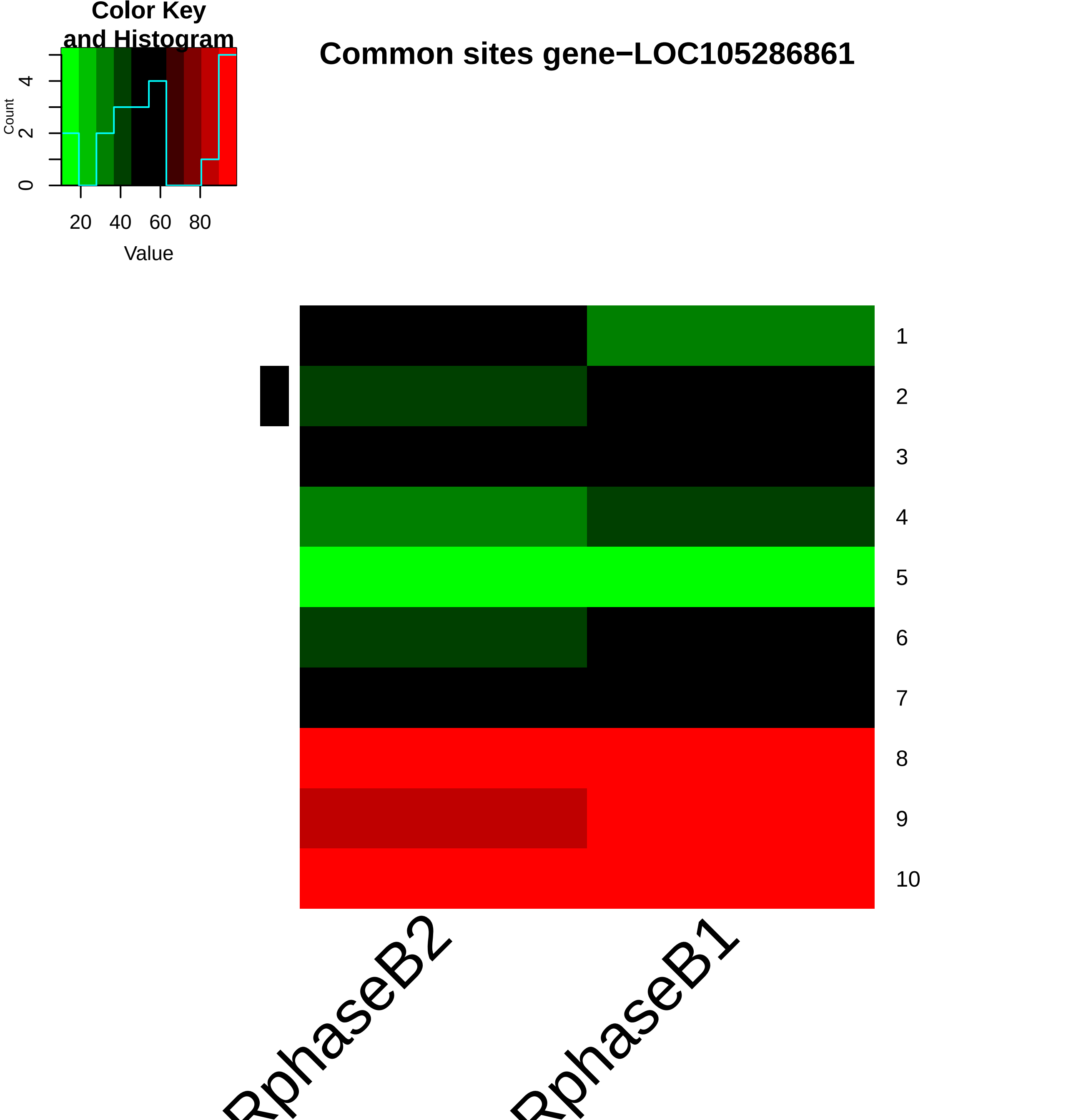

Color Key
and Histogram
Count
0
2
4
20
40
60
80
Value
Common sites gene−LOC105286861
1
2
3
4
5
6
7
8
9
10
RphaseB2
RphaseB1

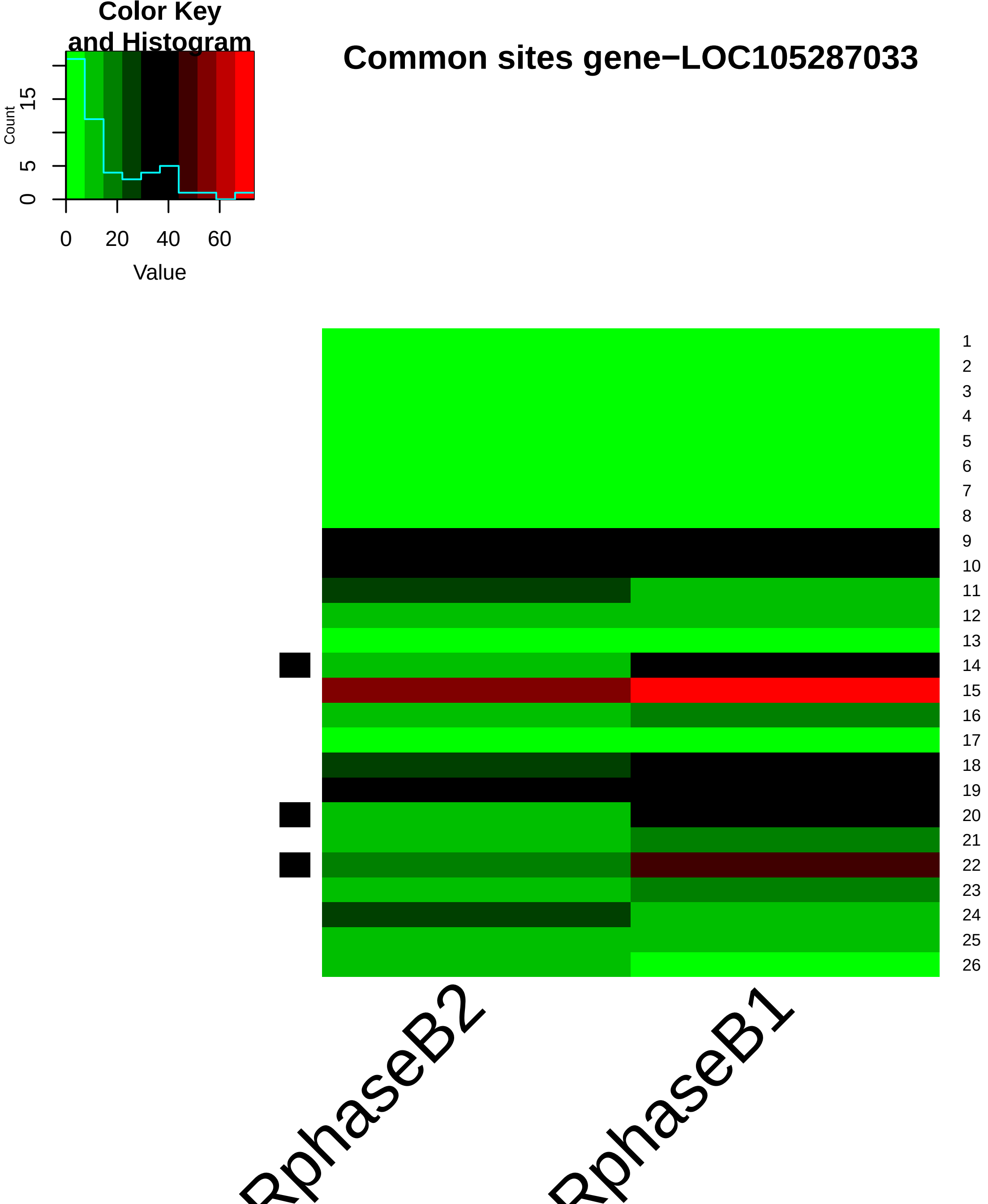
Color Key
and Histogram
Count
0
5
15
0
20
40
60
Value
Common sites gene−LOC105287033
RphaseB2
RphaseB1

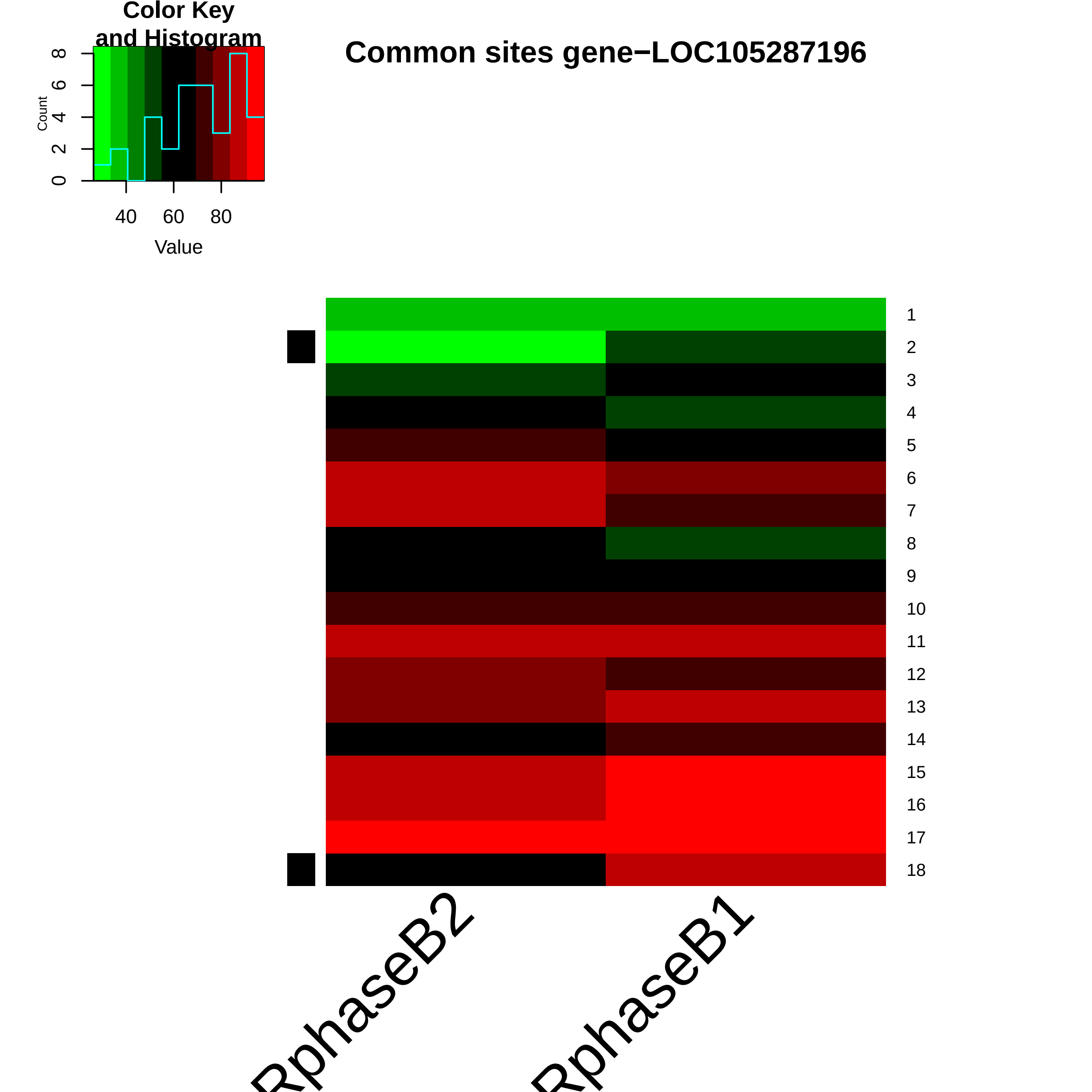

Color Key
and Histogram
Count
0
2
4
6
8
40
60
80
Value
Common sites gene−LOC105287196
RphaseB2
RphaseB1

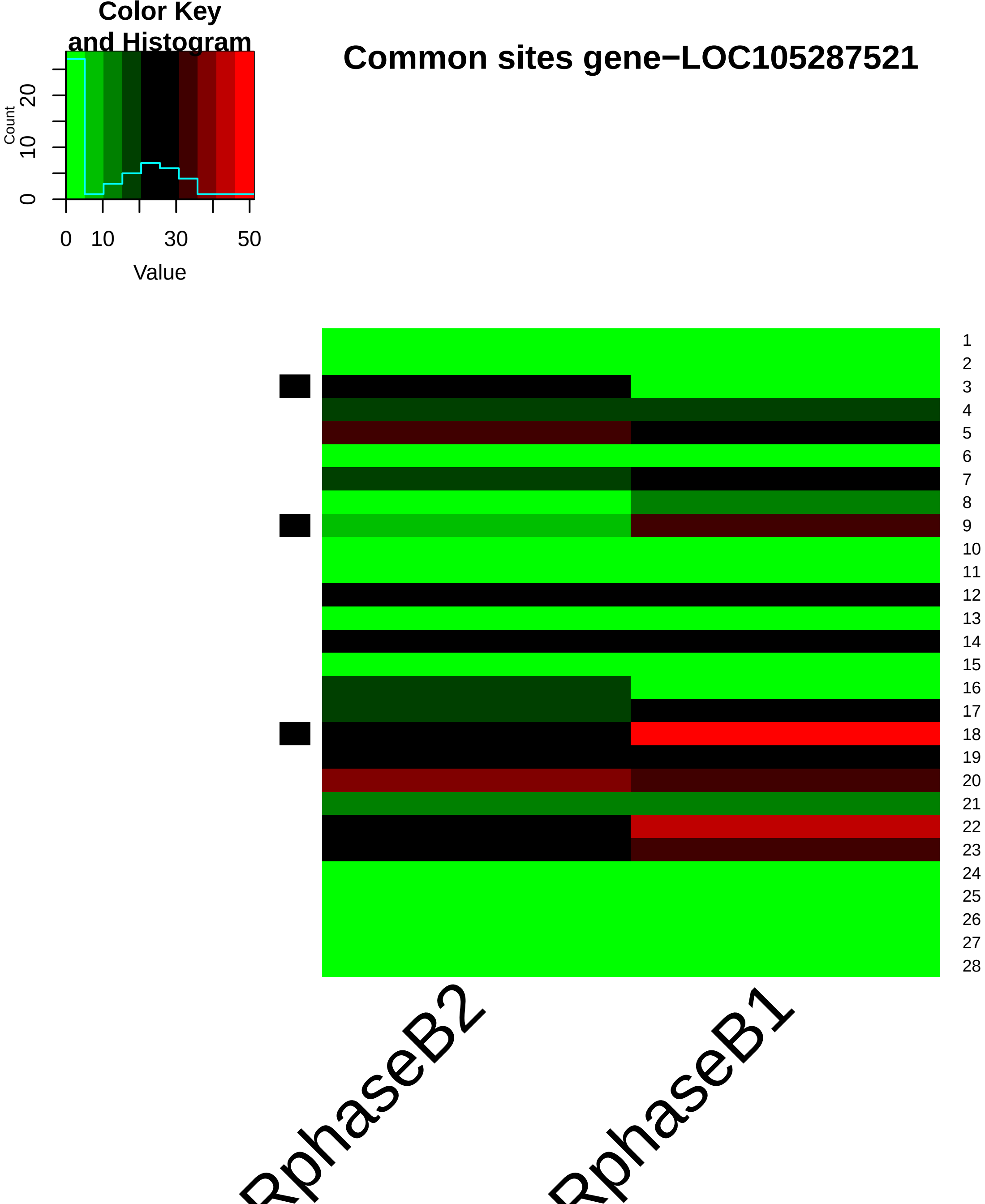
Color Key
and Histogram
Count
0
10
20
0
10
30
50
Value
Common sites gene−LOC105287521
RphaseB2
RphaseB1

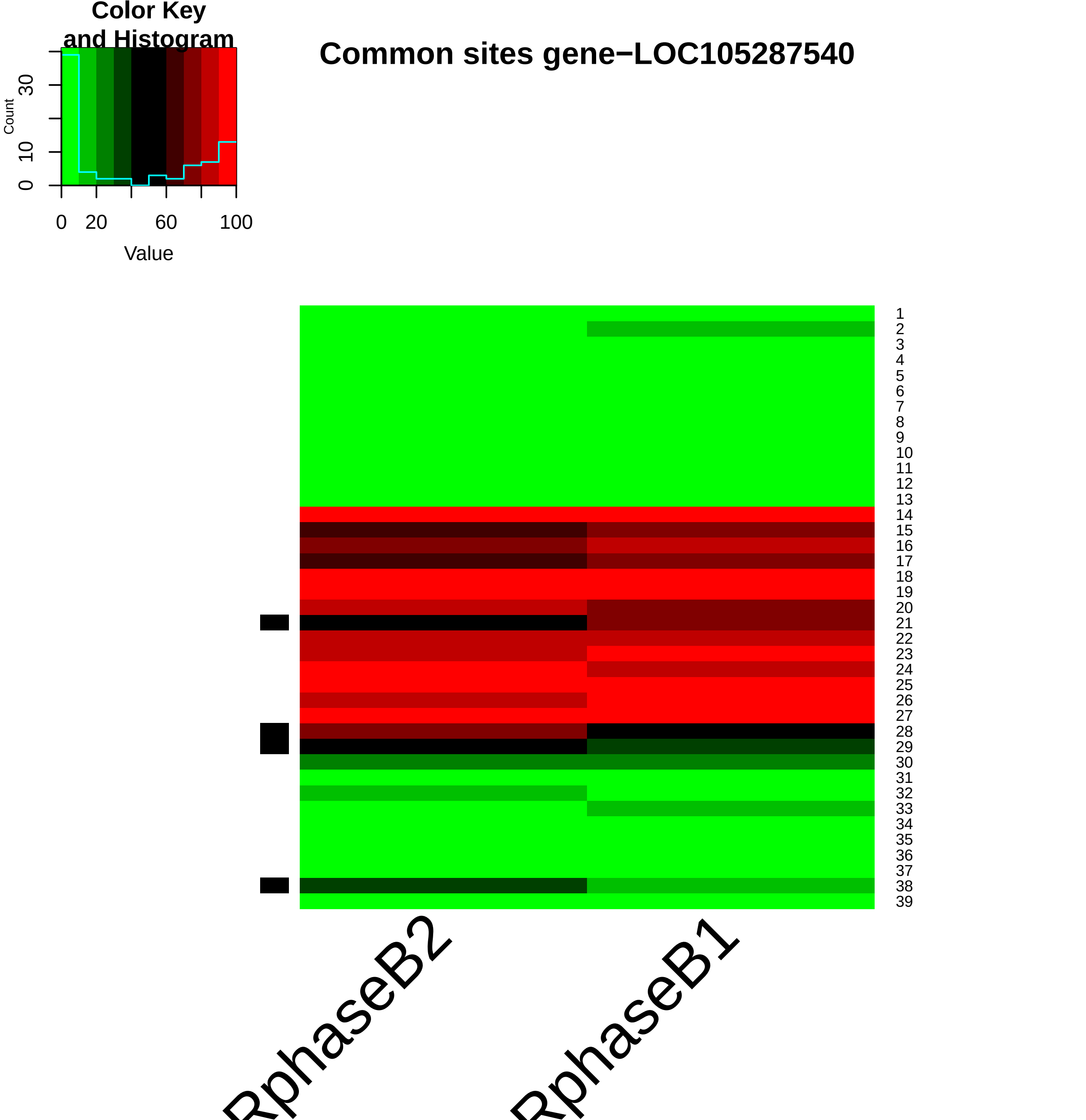
Color Key
and Histogram
Count
0
10
30
0
20
60
100
Value
Common sites gene−LOC105287540
RphaseB2
RphaseB1

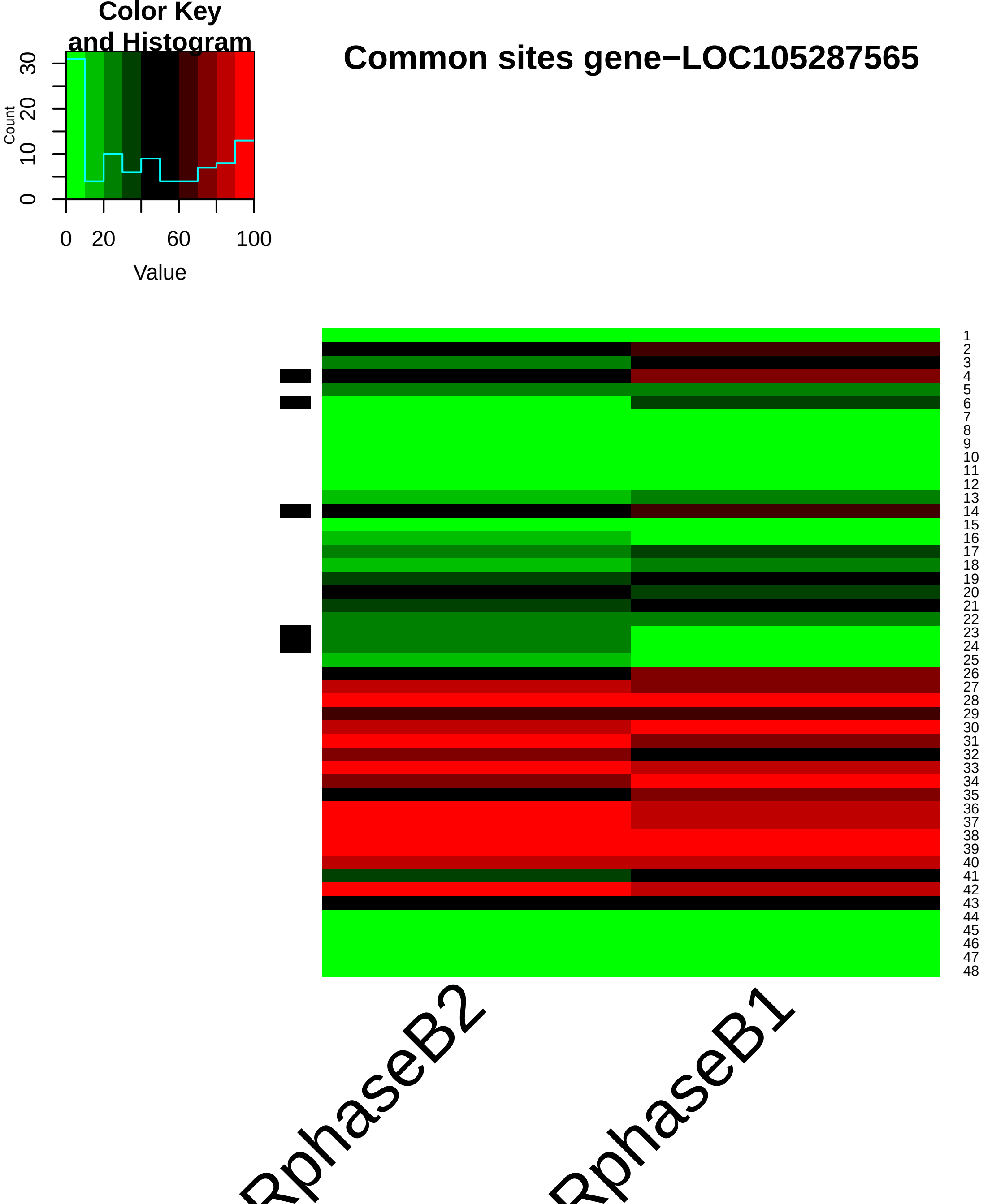
Color Key
and Histogram
Count
0
10
20
30
0
20
60
100
Value
Common sites gene−LOC105287565
RphaseB2
RphaseB1

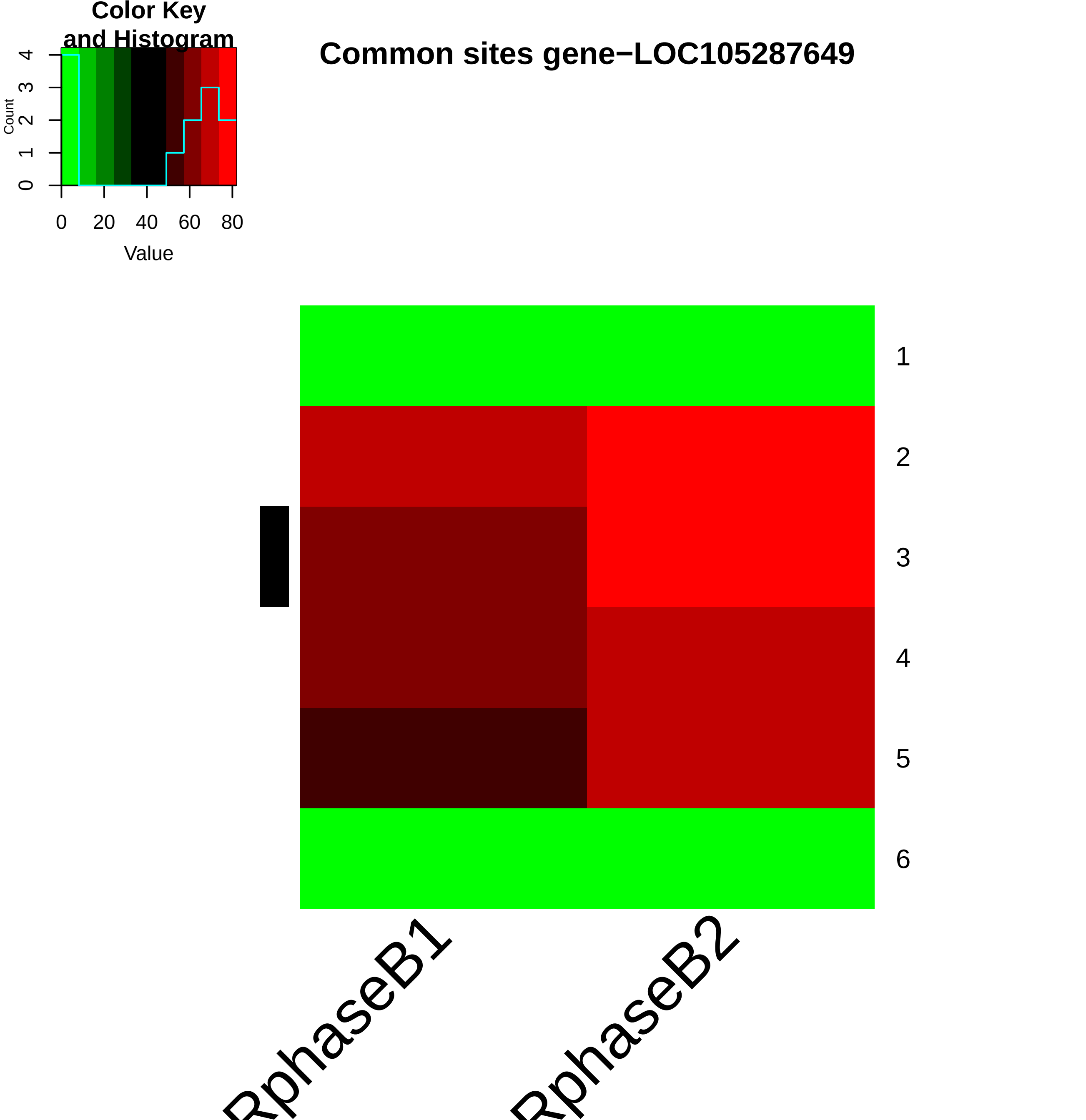
Color Key
and Histogram
Count
0
1
2
3
4
0
20
40
60
80
Value
Common sites gene−LOC105287649
1
2
3
4
5
6
RphaseB1
RphaseB2

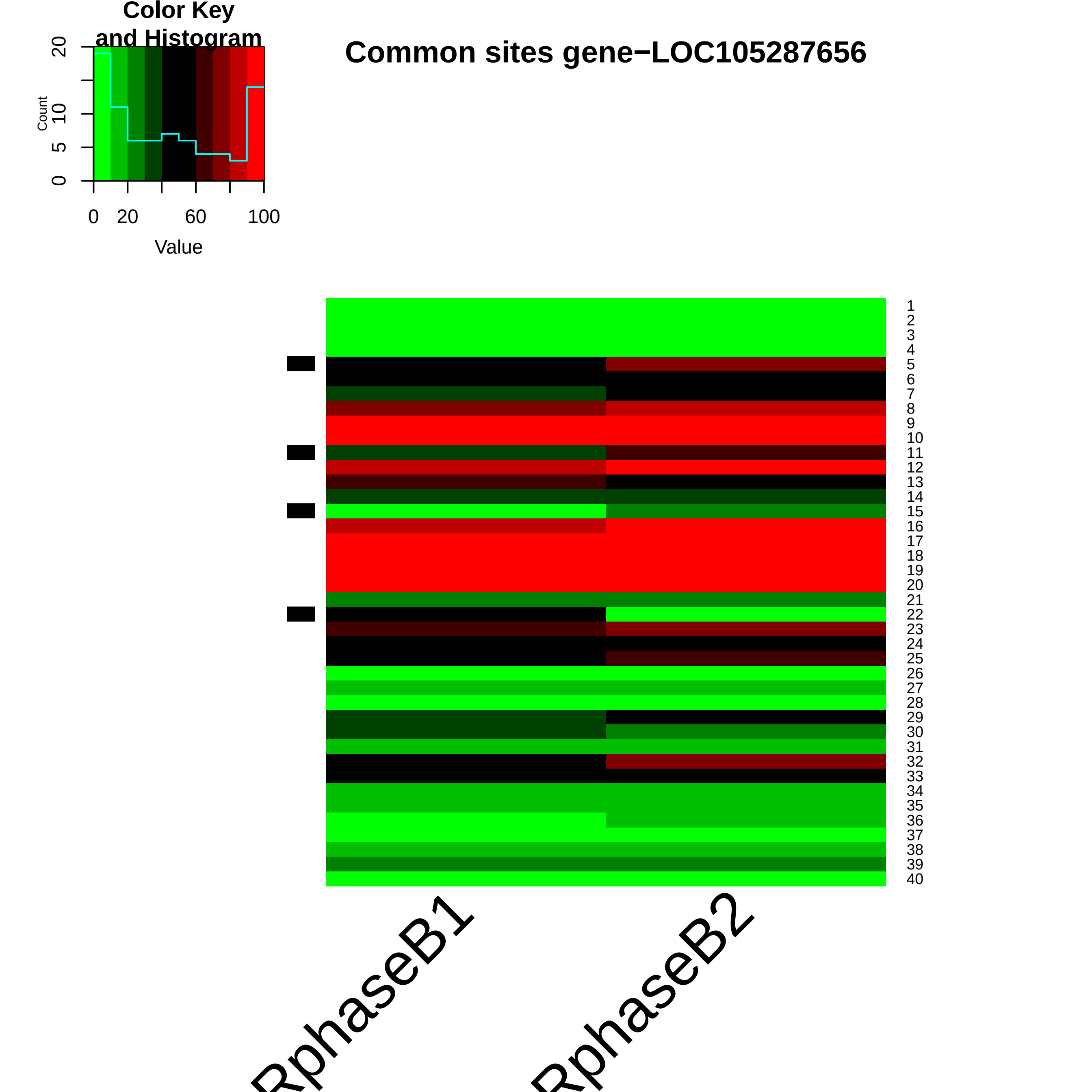
Color Key
and Histogram
Count
0
5
10
20
0
20
60
100
Value
Common sites gene−LOC105287656
RphaseB1
RphaseB2

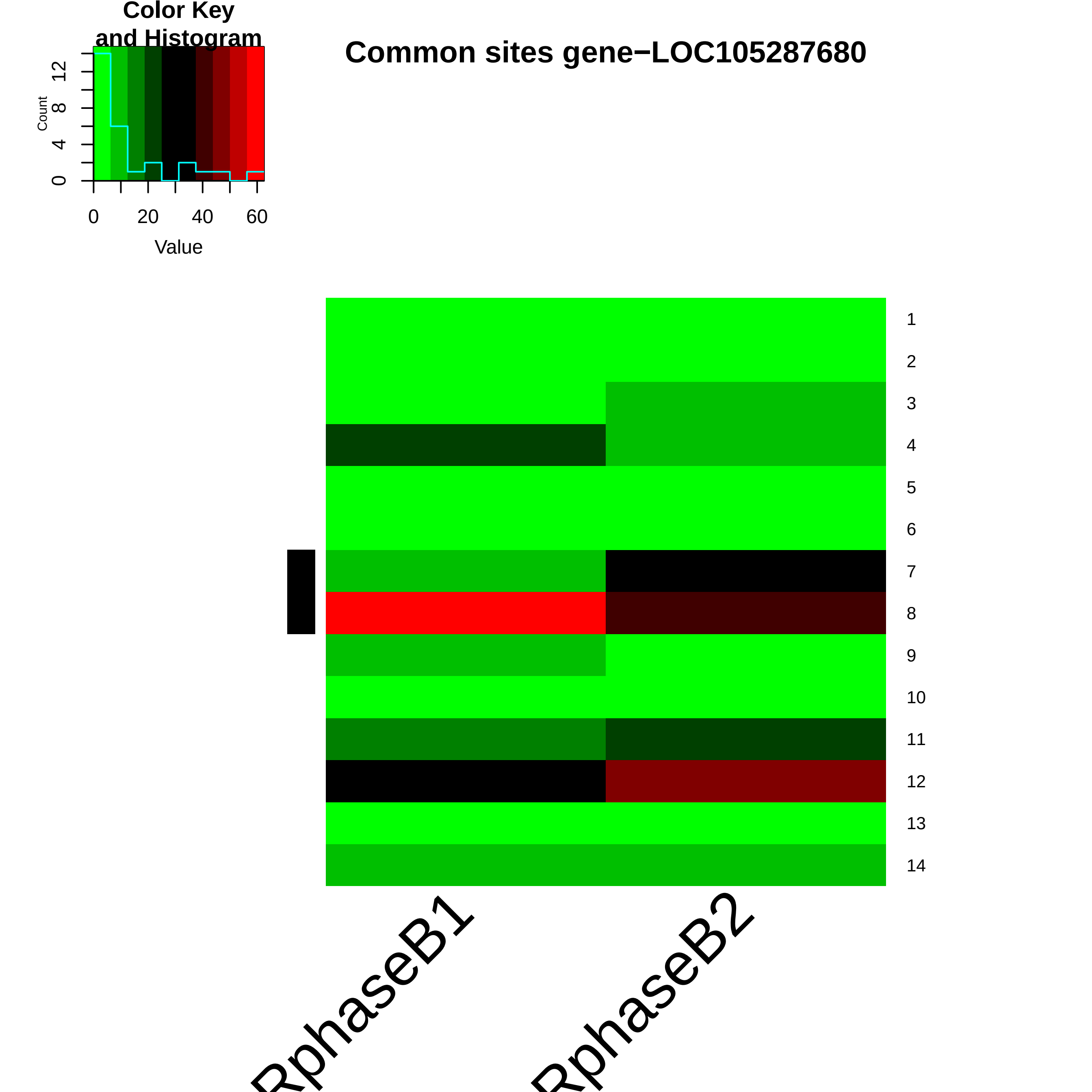

Color Key
and Histogram
Count
0
4
8
12
0
20
40
60
Value
Common sites gene−LOC105287680
1
2
3
4
5
6
7
8
9
10
11
12
13
14
RphaseB1
RphaseB2

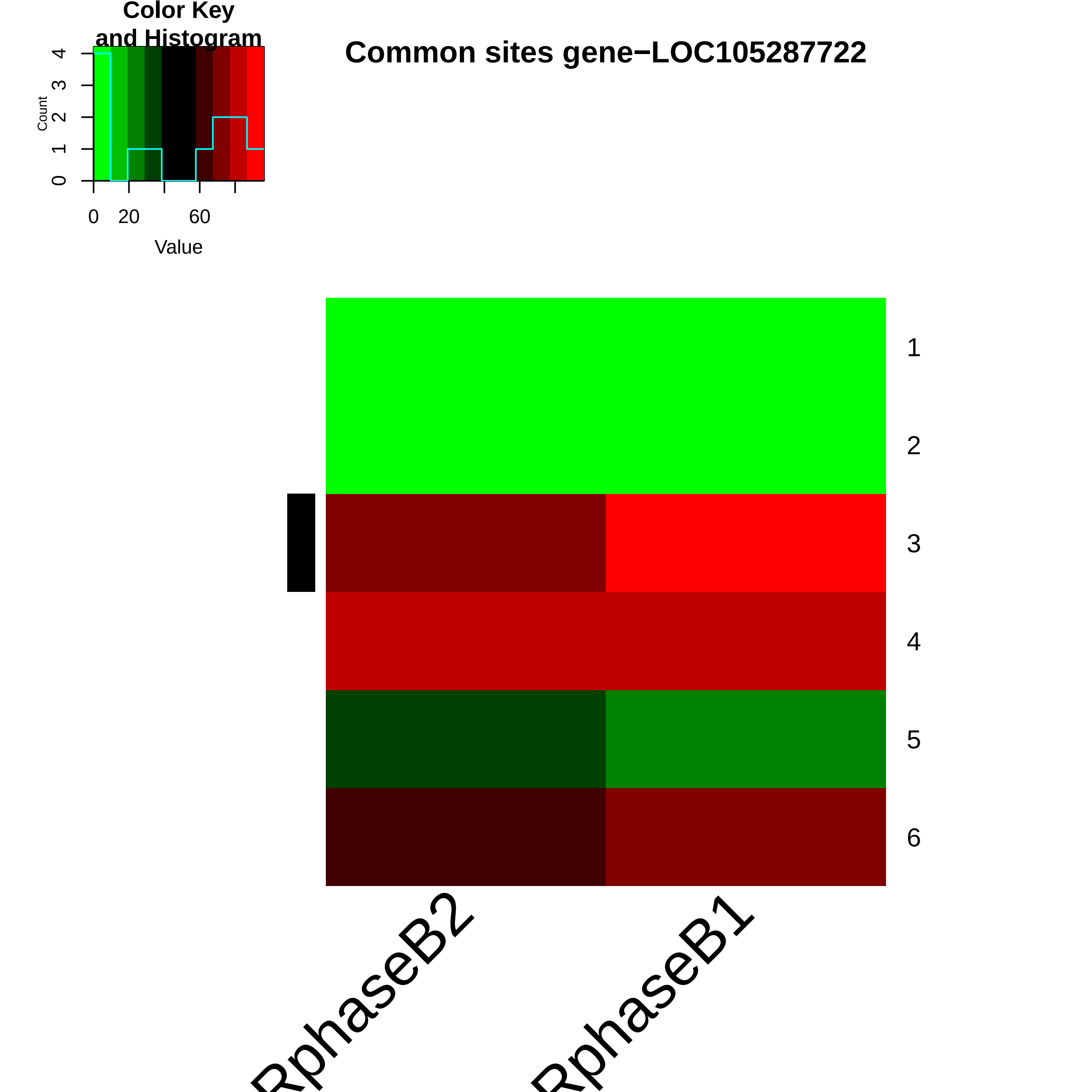

Color Key
and Histogram
Count
0
1
2
3
4
0
20
60
Value
Common sites gene−LOC105287722
1
2
3
4
5
6
RphaseB2
RphaseB1

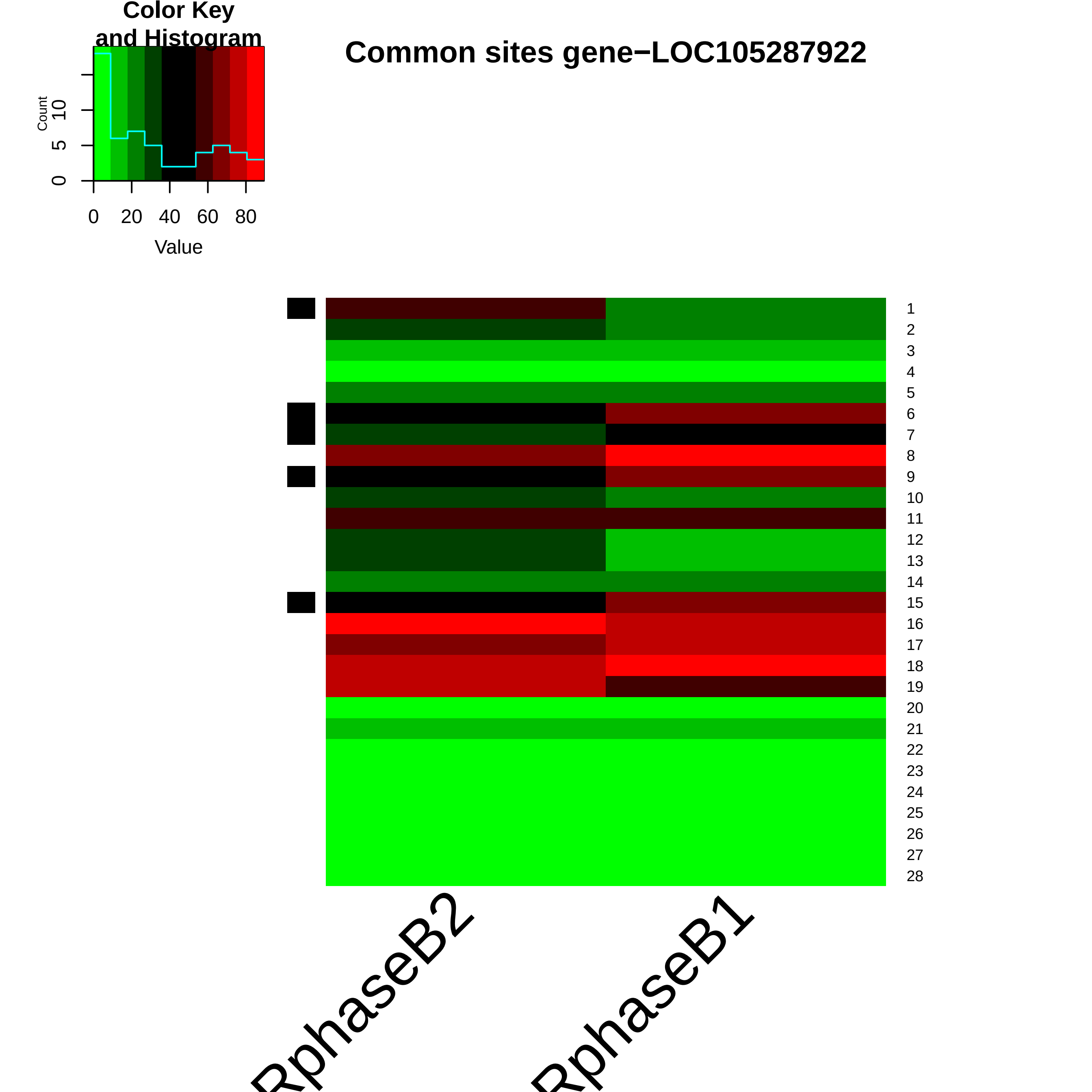
Color Key
and Histogram
Count
0
5
10
0
20
40
60
80
Value
Common sites gene−LOC105287922
RphaseB2
RphaseB1

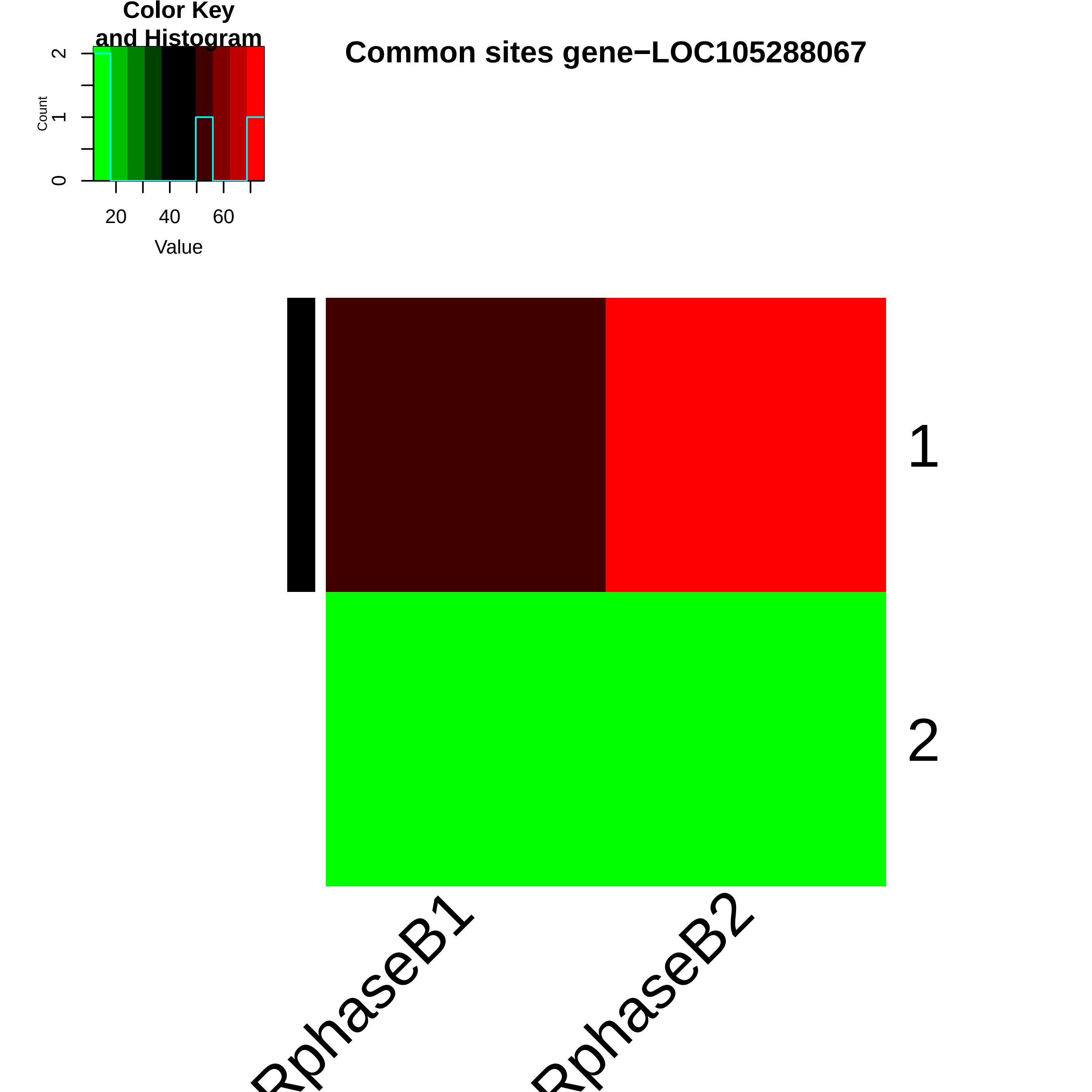

Color Key
and Histogram
Count
0
1
2
20
40
60
Value
Common sites gene−LOC105288067
1
2
RphaseB1
RphaseB2

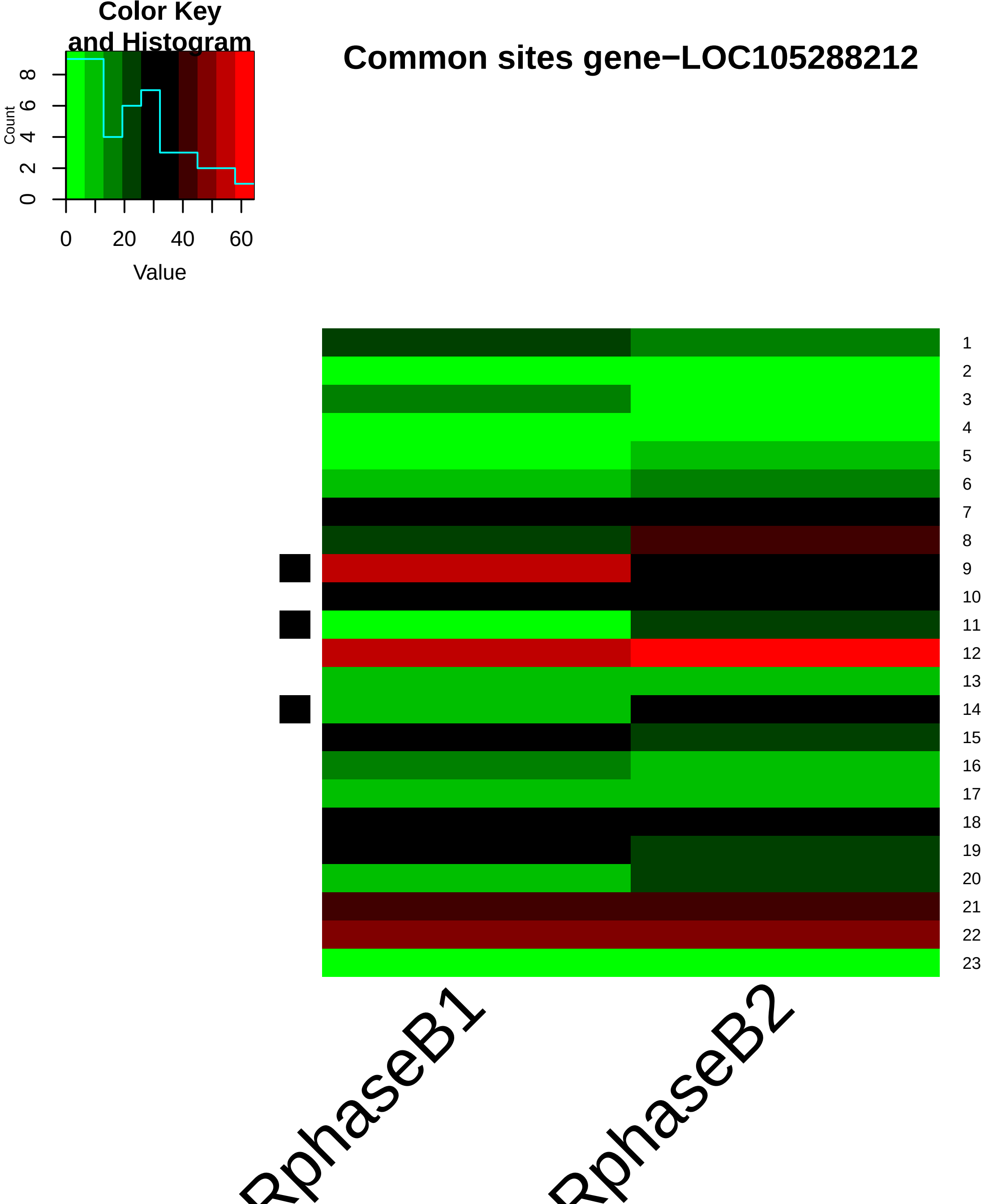
Color Key
and Histogram
Count
0
2
4
6
8
0
20
40
60
Value
Common sites gene−LOC105288212
RphaseB1
RphaseB2

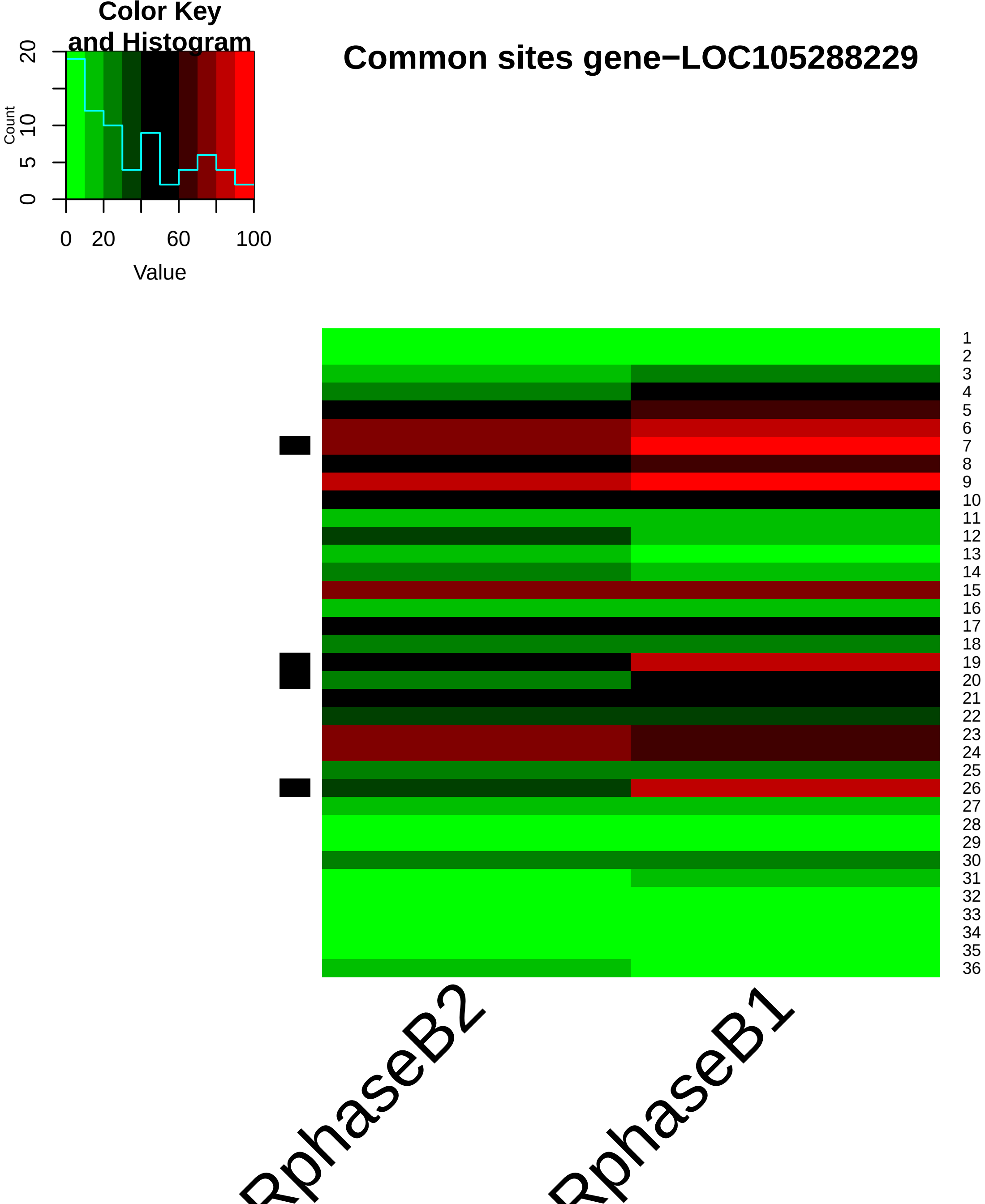

Color Key
and Histogram
Count
Value
Common sites gene−LOC105288229
RphaseB2
RphaseB1

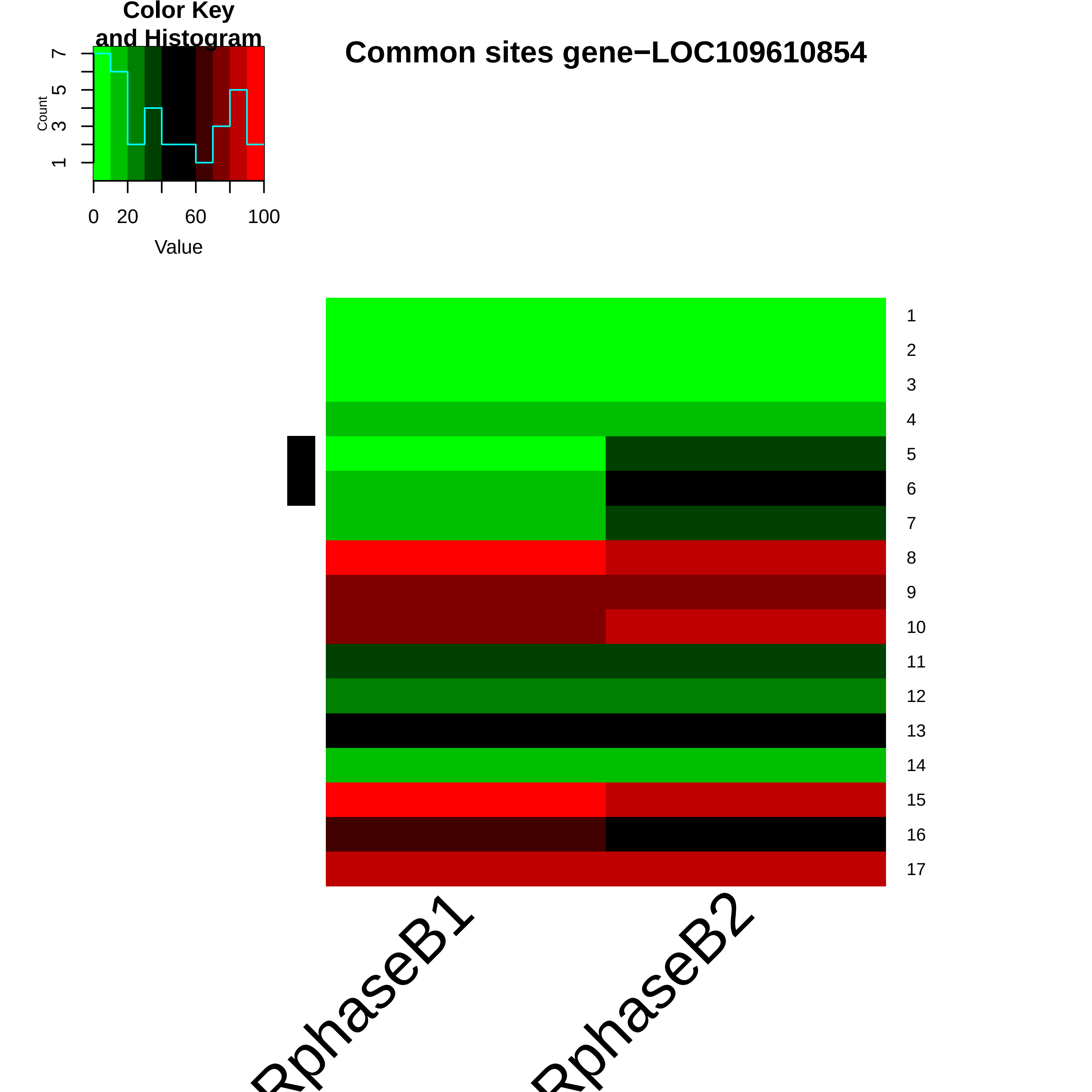
Color Key
and Histogram
Count
1
3
5
7
0
20
60
100
Value
Common sites gene−LOC109610854
RphaseB1
RphaseB2

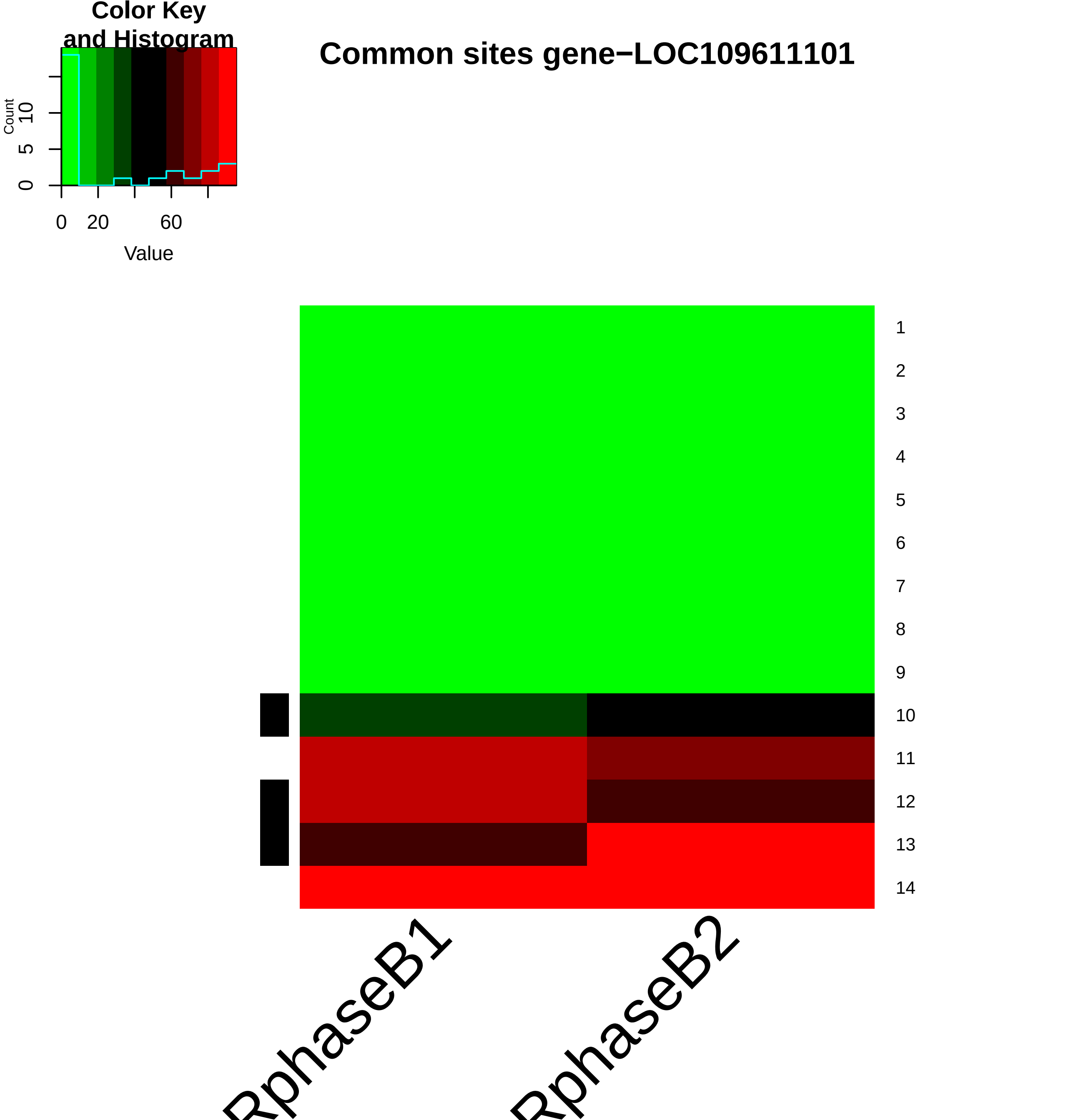
Color Key
and Histogram
Count
0
5
10
0
20
60
Value
Common sites gene−LOC109611101
1
2
3
4
5
6
7
8
9
10
11
12
13
14
RphaseB1
RphaseB2

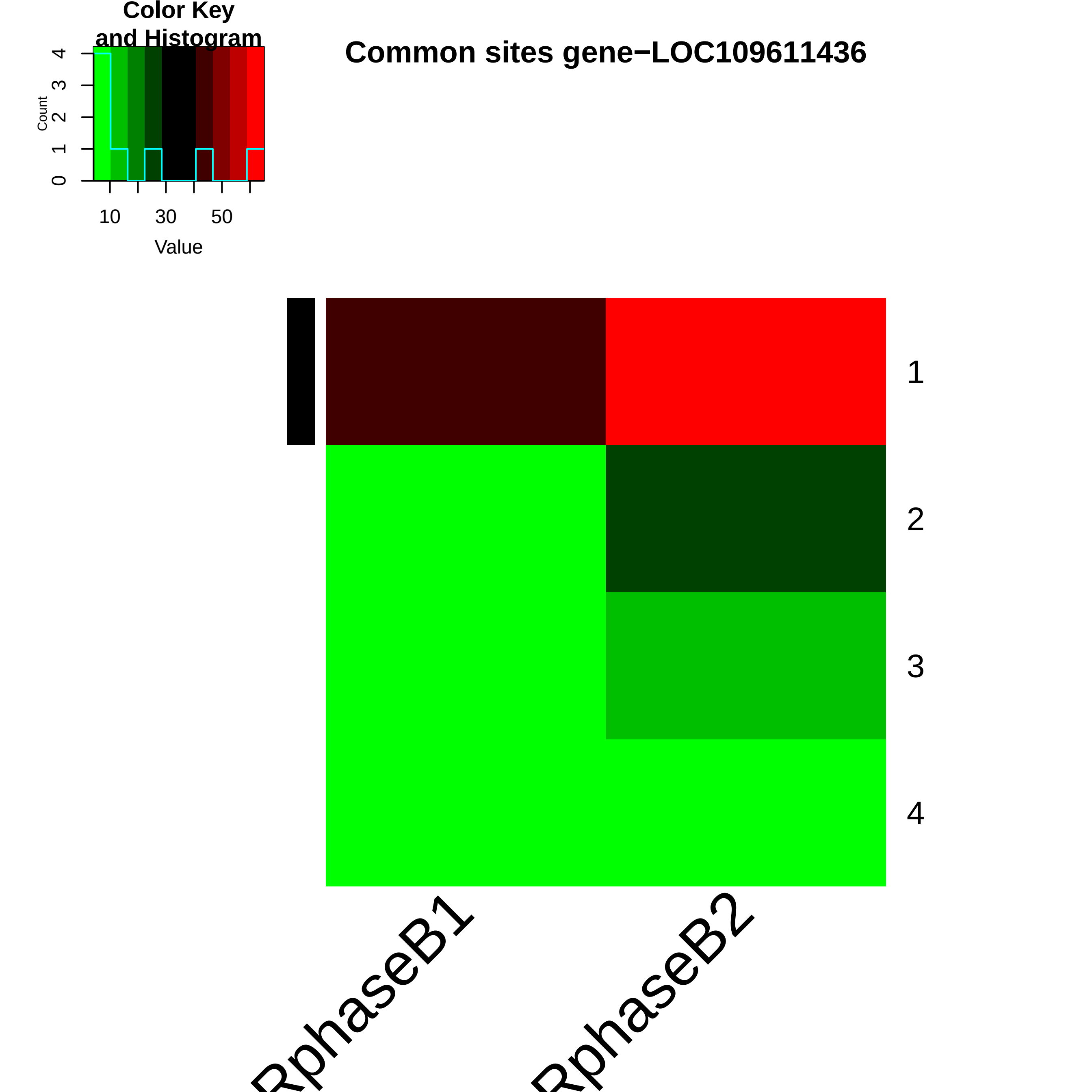

Color Key
and Histogram
Count
0
1
2
3
4
10
30
50
Value
Common sites gene−LOC109611436
1
2
3
4
RphaseB1
RphaseB2

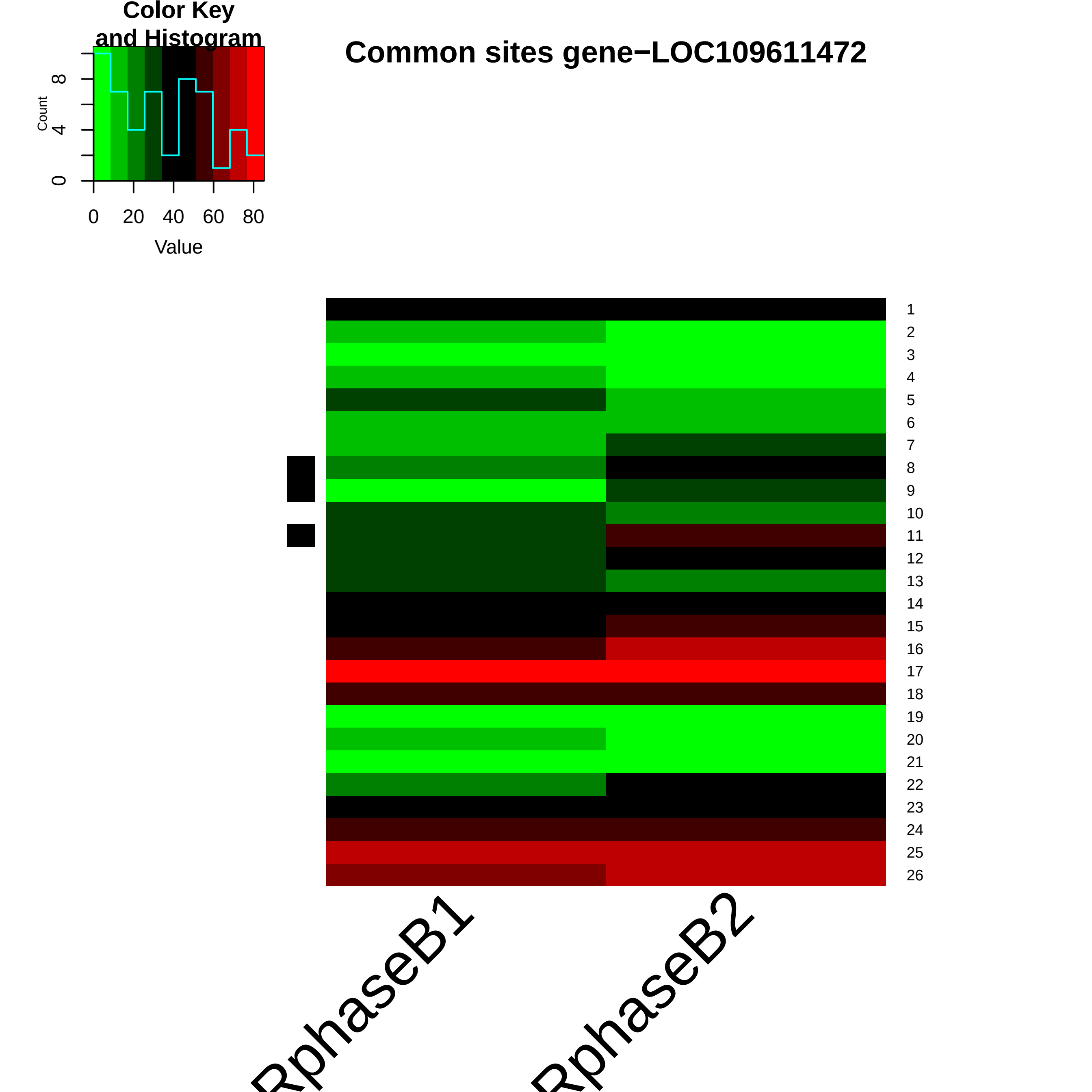

Color Key
and Histogram
Count
0
4
8
0
20
40
60
80
Value
Common sites gene−LOC109611472
RphaseB1
RphaseB2

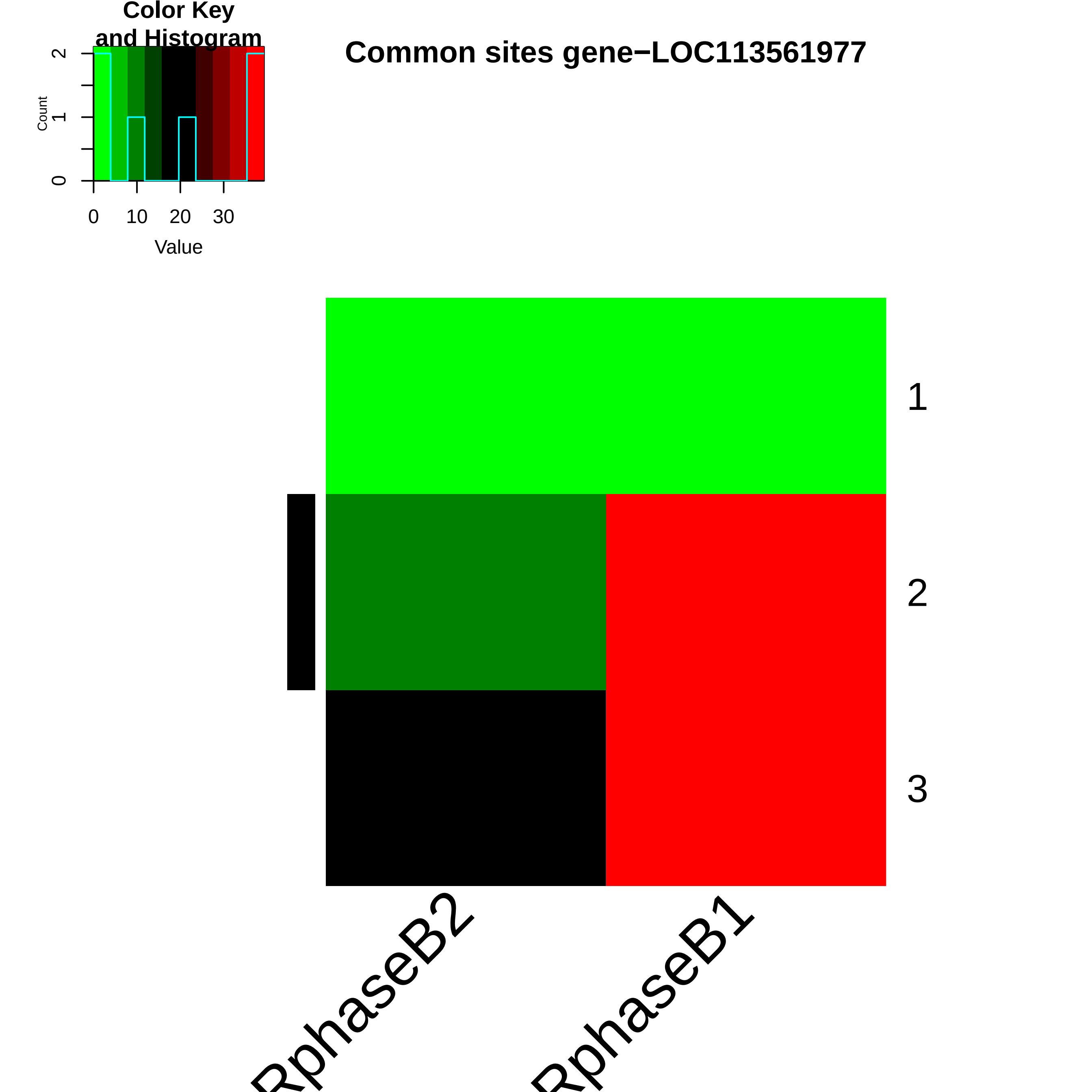

Color Key
and Histogram
Count
0
1
2
0
10
20
30
Value
Common sites gene−LOC113561977
1
2
3
RphaseB2
RphaseB1

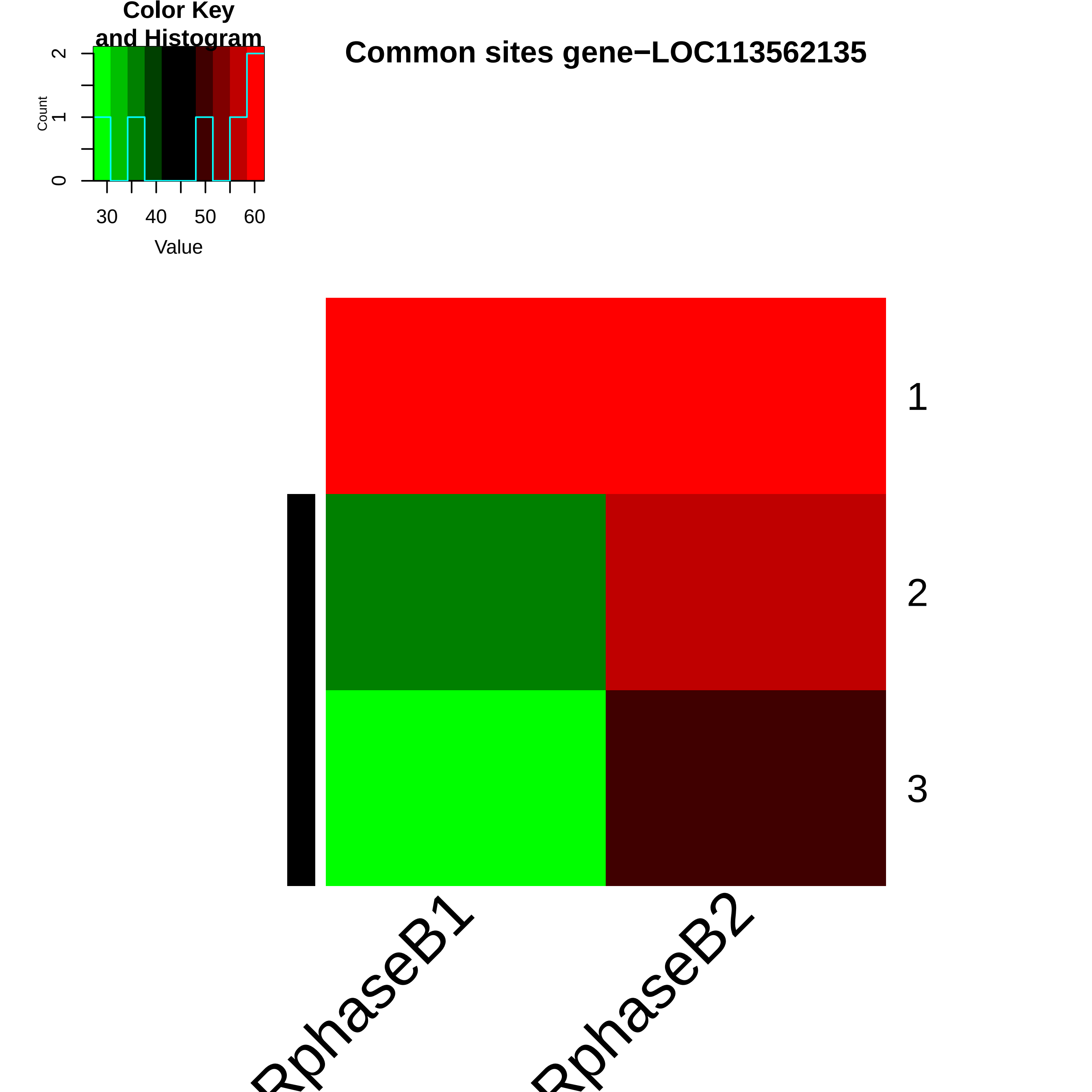

Color Key
and Histogram
Count
0
1
2
30
40
50
60
Value
Common sites gene−LOC113562135
1
2
3
RphaseB1
RphaseB2

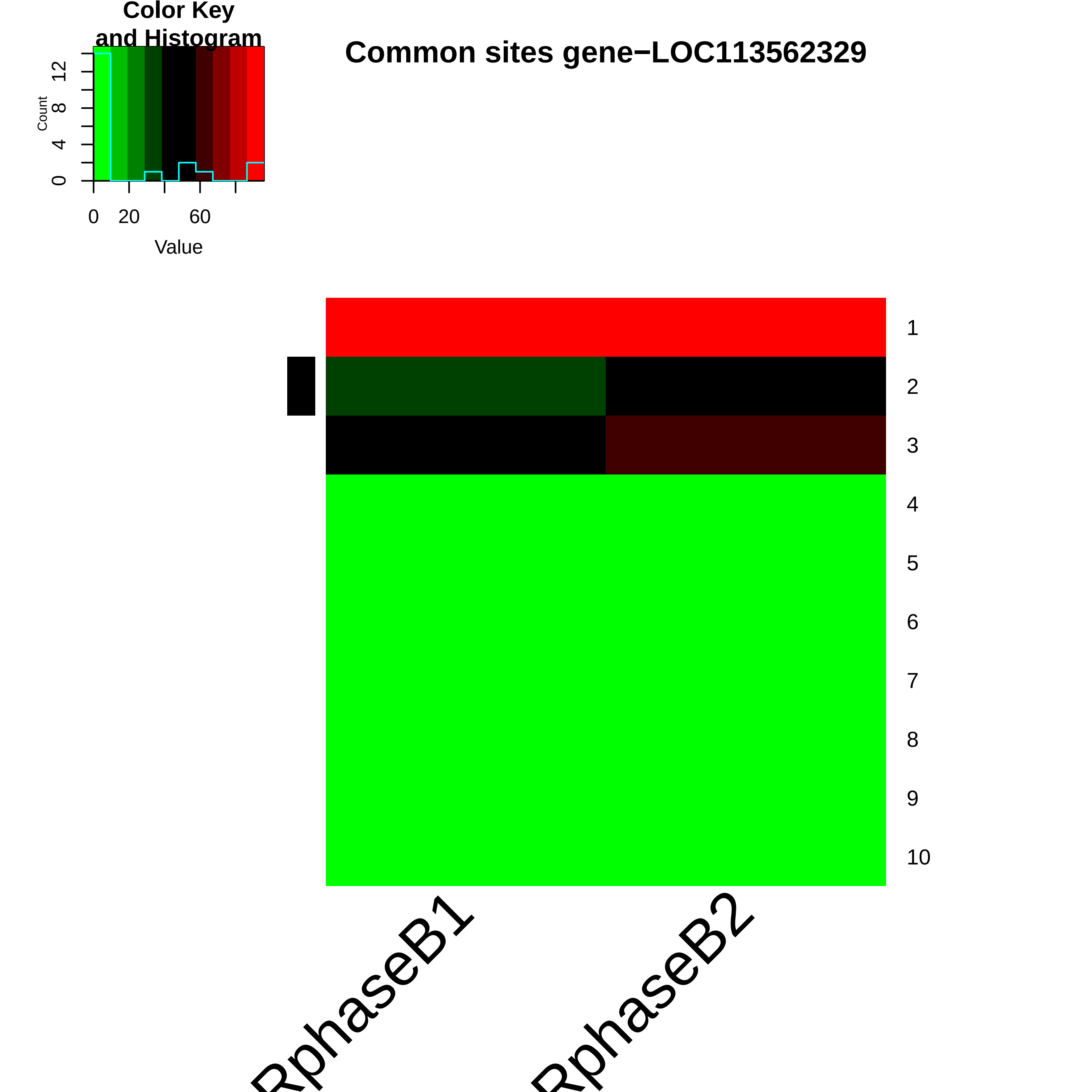
Color Key
and Histogram
Count
0
4
8
12
0
20
60
Value
Common sites gene−LOC113562329
1
2
3
4
5
6
7
8
9
10
RphaseB1
RphaseB2

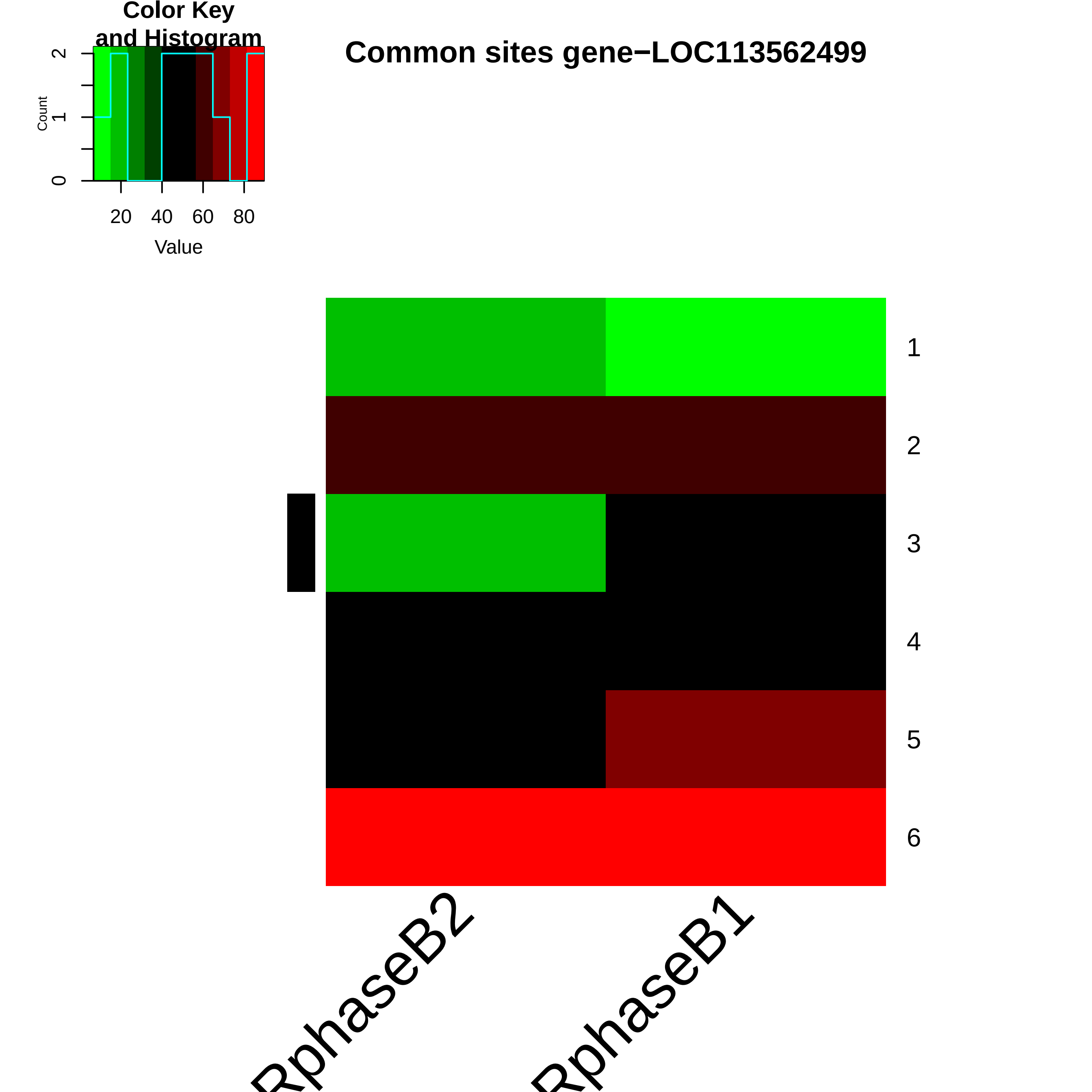

Color Key
and Histogram
Count
0
1
2
20
40
60
80
Value
Common sites gene−LOC113562499
1
2
3
4
5
6
RphaseB2
RphaseB1

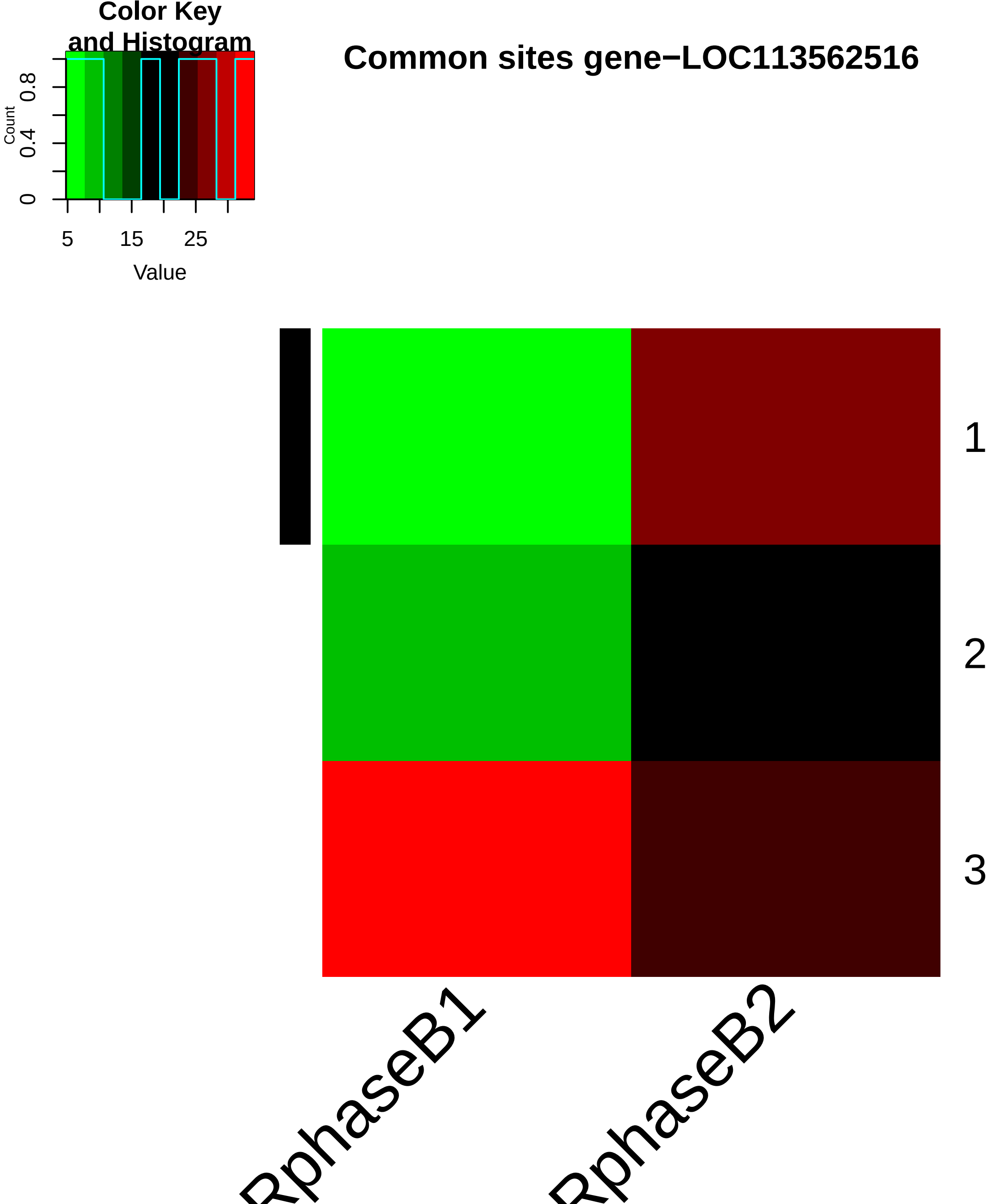

Color Key
and Histogram
Count
0
0.4
0.8
5
15
25
Value
Common sites gene−LOC113562516
1
2
3
RphaseB1
RphaseB2

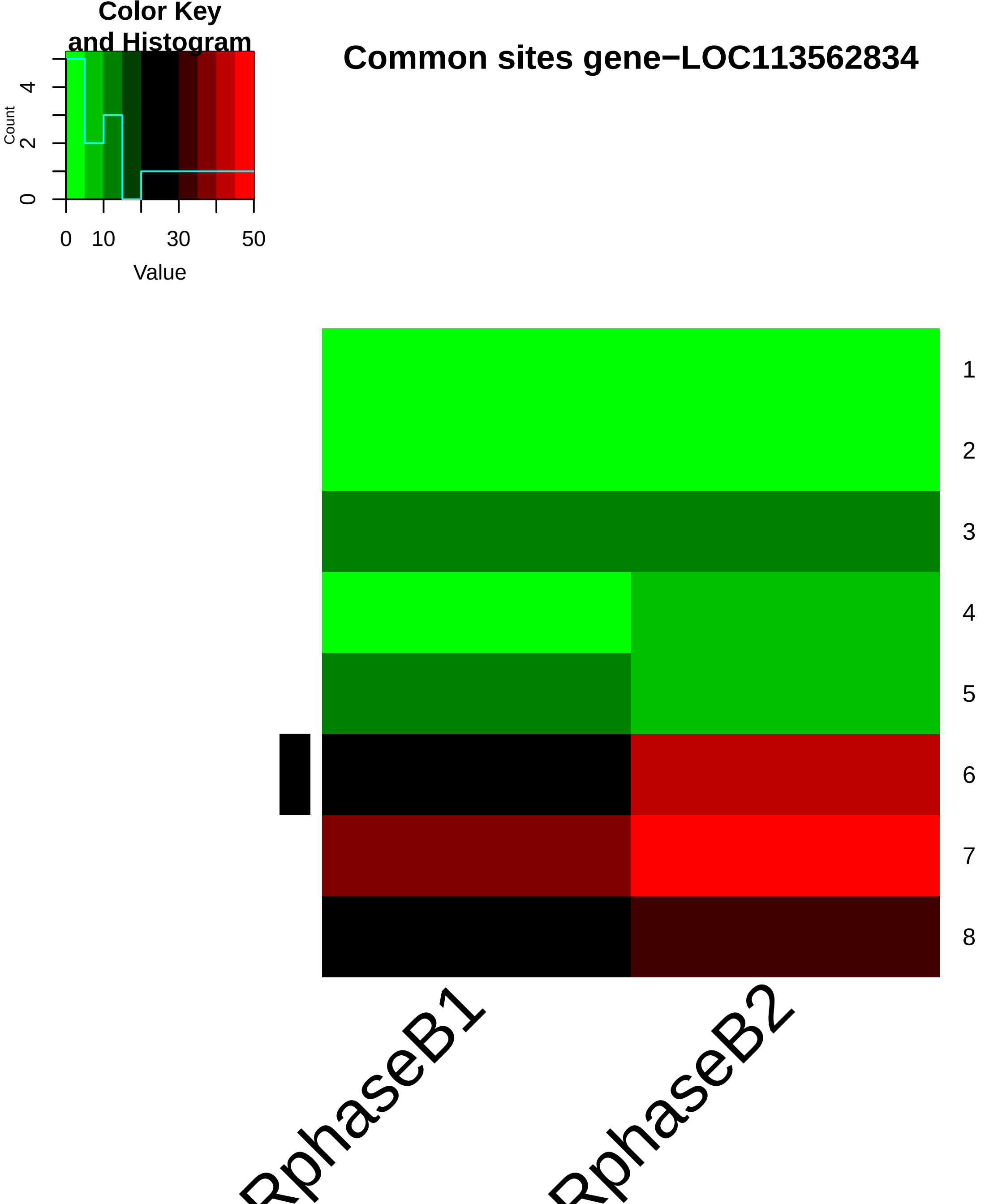
Color Key
and Histogram
Count
0
2
4
0
10
30
50
Value
Common sites gene−LOC113562834
1
2
3
4
5
6
7
8
RphaseB1
RphaseB2

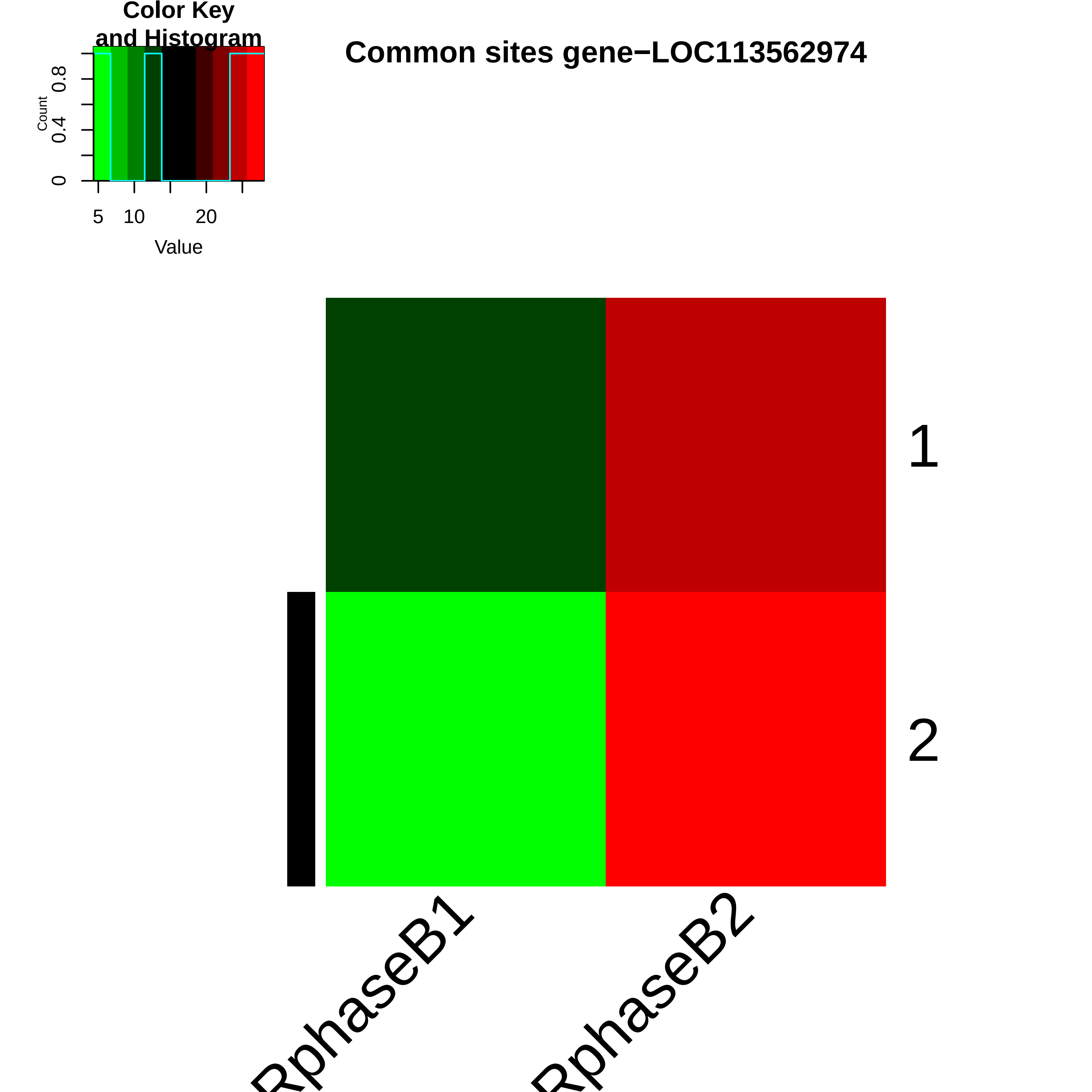

Color Key
and Histogram
Count
0
0.4
0.8
5
10
20
Value
Common sites gene−LOC113562974
1
2
RphaseB1
RphaseB2

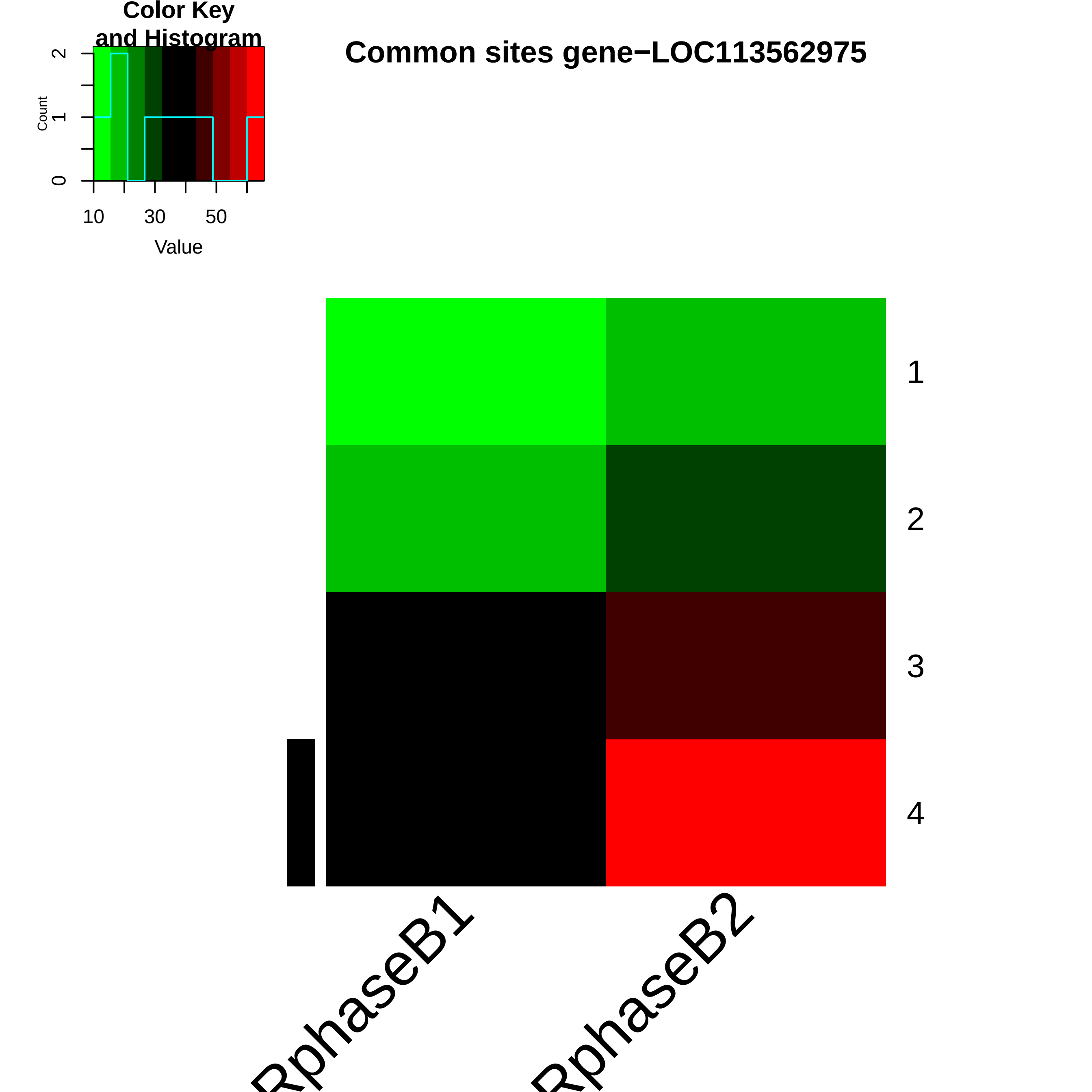

Color Key
and Histogram
Count
0
1
2
10
30
50
Value
Common sites gene−LOC113562975
1
2
3
4
RphaseB1
RphaseB2

```
Directory: OGL File: 0READMEogl

OGL - Ordered gene lists.

Input: studyhc annotation   maxgwidth minnbrdmsites
       dmgprp (= output of BWASPR:show_dmsg())

Output: files ogl-*.txt rnk-dmg-*.txt
         ogl-<study>_<sample>.txt ogl-<study>_<sample1>.vs.<sample2>.txt
         rnk-dmg--<study>_<sample1>.vs.<sample2>.txt rnk-dmg--<study>_<sample1>.vs.<sample2>.pdf
         wrt-<study>.txt

Notes: Output files ogl-<study>_<sample>.txt give tables for each sample with columns

        gene_ID gwidth #Sites #per10Kb %perSite %pNucl

       ordered by %pNucl.  If available, a link to an NCBI entry of gene_ID is inserted as second column.
       Abbreviations used: %perSite, percent methylation per site
                           %perNucl, percent methylation per nucleotide of the gene

       Output files ogl-<study>_<sample1>.vs.<sample2>.txt give tables for each comparison with columns

        gene_ID gwidth #Sites #per10Kb #dmSites #dmsp10kb %dmSites %pSite1 %pSite2 DMpSite ADMpSite DMpNucl ADMpNucl

       ordered by DMpSite.  If available, a link to an NCBI entry of gene_ID is inserted as second column.
       Abbreviations used: %dmSites, percent sites that are differentially methylated
                           %pSite1, average per site % methylation for sample1
                           %pSite2, average per site % methylation for sample2
                           DMpSite, average per site difference in % methylation between sample1 and sample2
                           ADMpSite, absolute value of DMpSite
                           DMpNucl, average per nucleotide difference in % methylation between sample1 and sample2
                           ADMpNucl, absolute value of DMpNucl

       Output files rnk-dmg-<study>_<sample1>.vs.<sample2>.txt are equivalent to files
        ogl-<study>_<sample1>.vs.<sample2>.txt but ordered by ADMpNucl.
       Output files rnk-dmg-<study>_<sample1>.vs.<sample2>.pdf provide visualization of the distribution
        of ADMpNucl values.

       Output file wrt-<study>.txt gives results of the Wilcoxon signed rank test comparing
        the %pSite1 and %pSite2 vectors

       In output directory OGL, tables are restricted to genes with gwidth <= maxgwidth and, for
        pairwise comparisons, #dmSites >= minnbrdmsites

       If either maxgwidth or minnbrdmsites is set, then output directory OGLall will show the full tables
        with all genes (for reference).
```

Directory: OGL File: ogl-Ob_LI_BCphaseA1.txt

| gene_ID | gene_link | gwidth | #Sites | #per10Kb | %perSite | %pNucl |
|---|---|---|---|---|---|---|
| gene-LOC105283830 | https://www.ncbi.nlm.nih.gov/gene/?term=LOC105283830 | 252 | 23 | 912.7 | 37.25 | 3.4 |
| gene-LOC105280739 | https://www.ncbi.nlm.nih.gov/gene/?term=LOC105280739 | 1991 | 123 | 617.78 | 54.86 | 3.39 |
| gene-LOC105286531 | https://www.ncbi.nlm.nih.gov/gene/?term=LOC105286531 | 372 | 35 | 940.86 | 34.73 | 3.27 |
| gene-LOC105274534 | https://www.ncbi.nlm.nih.gov/gene/?term=LOC105274534 | 1329 | 88 | 662.15 | 45.94 | 3.04 |
| gene-LOC105287811 | https://www.ncbi.nlm.nih.gov/gene/?term=LOC105287811 | 2155 | 123 | 570.77 | 52.68 | 3.01 |
| gene-LOC105285635 | https://www.ncbi.nlm.nih.gov/gene/?term=LOC105285635 | 1175 | 115 | 978.72 | 30.59 | 2.99 |
| gene-LOC105276360 | https://www.ncbi.nlm.nih.gov/gene/?term=LOC105276360 | 1466 | 131 | 893.59 | 33.23 | 2.97 |
| gene-LOC105278844 | https://www.ncbi.nlm.nih.gov/gene/?term=LOC105278844 | 1101 | 111 | 1008.17 | 29.43 | 2.97 |
| gene-LOC105286737 | https://www.ncbi.nlm.nih.gov/gene/?term=LOC105286737 | 2690 | 308 | 1144.98 | 25.86 | 2.96 |
| gene-LOC105278160 | https://www.ncbi.nlm.nih.gov/gene/?term=LOC105278160 | 1872 | 146 | 779.91 | 37.08 | 2.89 |
| gene-LOC105286801 | https://www.ncbi.nlm.nih.gov/gene/?term=LOC105286801 | 1102 | 84 | 762.25 | 37.74 | 2.88 |
| gene-LOC105278849 | https://www.ncbi.nlm.nih.gov/gene/?term=LOC105278849 | 776 | 94 | 1211.34 | 23.72 | 2.87 |
| gene-LOC105274711 | https://www.ncbi.nlm.nih.gov/gene/?term=LOC105274711 | 1655 | 100 | 604.23 | 46.56 | 2.81 |
| gene-LOC105276371 | https://www.ncbi.nlm.nih.gov/gene/?term=LOC105276371 | 871 | 55 | 631.46 | 44.03 | 2.78 |
| gene-LOC105286235 | https://www.ncbi.nlm.nih.gov/gene/?term=LOC105286235 | 1510 | 97 | 642.38 | 43.09 | 2.77 |
| gene-LOC113561532 | https://www.ncbi.nlm.nih.gov/gene/?term=LOC113561532 | 72 | 2 | 277.78 | 97.72 | 2.71 |
| gene-LOC105282149 | https://www.ncbi.nlm.nih.gov/gene/?term=LOC105282149 | 1787 | 116 | 649.13 | 41.07 | 2.67 |
| gene-LOC105287692 | https://www.ncbi.nlm.nih.gov/gene/?term=LOC105287692 | 1505 | 123 | 817.28 | 32.57 | 2.66 |
| gene-LOC105286269 | https://www.ncbi.nlm.nih.gov/gene/?term=LOC105286269 | 1401 | 68 | 485.37 | 54.59 | 2.65 |
| gene-LOC105275314 | https://www.ncbi.nlm.nih.gov/gene/?term=LOC105275314 | 1489 | 78 | 523.84 | 50.45 | 2.64 |
| gene-LOC105277254 | https://www.ncbi.nlm.nih.gov/gene/?term=LOC105277254 | 1377 | 108 | 784.31 | 33.57 | 2.63 |
| gene-LOC105274543 | https://www.ncbi.nlm.nih.gov/gene/?term=LOC105274543 | 1572 | 103 | 655.22 | 38.7 | 2.54 |
| gene-LOC105276439 | https://www.ncbi.nlm.nih.gov/gene/?term=LOC105276439 | 2382 | 141 | 591.94 | 42.03 | 2.49 |
| gene-LOC105282986 | https://www.ncbi.nlm.nih.gov/gene/?term=LOC105282986 | 1600 | 123 | 768.75 | 32.44 | 2.49 |

...
... only first 25 lines shown ...

Directory: OGL File: ogl-Ob_LI_BCphaseA1.vs.BCphaseA2.txt

| gene_ID | gene_link | gwidth | #Sites | #per10Kb | #dmSites | #dmsp10kb | %dmSites | %pSite1 | %pSite2 | DMpSite | ADMpSite | DMpNucl | ADMpNucl |
|---|---|---|---|---|---|---|---|---|---|---|---|---|---|
| gene-LOC105281173 | https://www.ncbi.nlm.nih.gov/gene/?term=LOC105281173 | 1595 | 25 | 156.74 | 1 | 6.27 | 4 | 64.76 | 61.8 | 2.96 | 2.96 | 0.05 | 0.05 |
| gene-LOC105278358 | https://www.ncbi.nlm.nih.gov/gene/?term=LOC105278358 | 2861 | 28 | 97.87 | 1 | 3.5 | 3.57 | 33.51 | 30.99 | 2.52 | 2.52 | 0.02 | 0.02 |
| gene-LOC109611283 | https://www.ncbi.nlm.nih.gov/gene/?term=LOC109611283 | 1451 | 36 | 248.1 | 1 | 6.89 | 2.78 | 37.85 | 35.62 | 2.23 | 2.23 | 0.06 | 0.06 |
| gene-LOC105286999 | https://www.ncbi.nlm.nih.gov/gene/?term=LOC105286999 | 2141 | 38 | 177.49 | 1 | 4.67 | 2.63 | 33.02 | 30.95 | 2.07 | 2.07 | 0.04 | 0.04 |
| gene-LOC105285202 | https://www.ncbi.nlm.nih.gov/gene/?term=LOC105285202 | 1838 | 33 | 179.54 | 1 | 5.44 | 3.03 | 20.53 | 19.01 | 1.52 | 1.52 | 0.03 | 0.03 |
| gene-LOC105282067 | https://www.ncbi.nlm.nih.gov/gene/?term=LOC105282067 | 5941 | 49 | 82.48 | 1 | 1.68 | 2.04 | 23.12 | 21.71 | 1.41 | 1.41 | 0.01 | 0.01 |
| gene-LOC105274449 | https://www.ncbi.nlm.nih.gov/gene/?term=LOC105274449 | 1549 | 42 | 271.14 | 1 | 6.46 | 2.38 | 55.88 | 54.57 | 1.32 | 1.32 | 0.04 | 0.04 |
| gene-LOC105278877 | https://www.ncbi.nlm.nih.gov/gene/?term=LOC105278877 | 1968 | 69 | 350.61 | 1 | 5.08 | 1.45 | 37.18 | 35.87 | 1.31 | 1.31 | 0.05 | 0.05 |
| gene-LOC105278228 | https://www.ncbi.nlm.nih.gov/gene/?term=LOC105278228 | 8610 | 146 | 169.57 | 1 | 1.16 | 0.68 | 23.97 | 22.69 | 1.27 | 1.27 | 0.02 | 0.02 |
| gene-LOC105285252 | https://www.ncbi.nlm.nih.gov/gene/?term=LOC105285252 | 7427 | 116 | 156.19 | 1 | 1.35 | 0.86 | 15.25 | 14.01 | 1.25 | 1.25 | 0.02 | 0.02 |
| gene-LOC105278282 | https://www.ncbi.nlm.nih.gov/gene/?term=LOC105278282 | 1775 | 43 | 242.25 | 1 | 5.63 | 2.33 | 46.59 | 45.35 | 1.24 | 1.24 | 0.03 | 0.03 |
| gene-LOC105284507 | https://www.ncbi.nlm.nih.gov/gene/?term=LOC105284507 | 5207 | 68 | 130.59 | 1 | 1.92 | 1.47 | 22.93 | 21.85 | 1.08 | 1.08 | 0.01 | 0.01 |
| gene-LOC105275653 | https://www.ncbi.nlm.nih.gov/gene/?term=LOC105275653 | 11251 | 236 | 209.76 | 1 | 0.89 | 0.42 | 9.52 | 8.44 | 1.07 | 1.07 | 0.02 | 0.02 |
| gene-LOC105275848 | https://www.ncbi.nlm.nih.gov/gene/?term=LOC105275848 | 3718 | 119 | 320.06 | 1 | 2.69 | 0.84 | 22.44 | 21.48 | 0.97 | 0.97 | 0.03 | 0.03 |
| gene-LOC105276272 | https://www.ncbi.nlm.nih.gov/gene/?term=LOC105276272 | 1965 | 25 | 127.23 | 1 | 5.09 | 4 | 43.93 | 43.09 | 0.84 | 0.84 | 0.01 | 0.01 |
| gene-LOC105275187 | https://www.ncbi.nlm.nih.gov/gene/?term=LOC105275187 | 5534 | 66 | 119.26 | 1 | 1.81 | 1.52 | 14.05 | 13.27 | 0.78 | 0.78 | 0.01 | 0.01 |
| gene-LOC105277435 | https://www.ncbi.nlm.nih.gov/gene/?term=LOC105277435 | 3643 | 150 | 411.75 | 1 | 2.74 | 0.67 | 12.77 | 12 | 0.77 | 0.77 | 0.03 | 0.03 |
| gene-LOC105282811 | https://www.ncbi.nlm.nih.gov/gene/?term=LOC105282811 | 5765 | 182 | 315.7 | 1 | 1.73 | 0.55 | 9.67 | 8.98 | 0.69 | 0.69 | 0.02 | 0.02 |
| gene-LOC105278632 | https://www.ncbi.nlm.nih.gov/gene/?term=LOC105278632 | 3624 | 119 | 328.37 | 1 | 2.76 | 0.84 | 15.91 | 15.28 | 0.63 | 0.63 | 0.02 | 0.02 |
| gene-LOC105276538 | https://www.ncbi.nlm.nih.gov/gene/?term=LOC105276538 | 1511 | 18 | 119.13 | 1 | 6.62 | 5.56 | 71.16 | 70.54 | 0.62 | 0.62 | 0.01 | 0.01 |
| gene-LOC105279664 | https://www.ncbi.nlm.nih.gov/gene/?term=LOC105279664 | 4023 | 56 | 139.2 | 1 | 2.49 | 1.79 | 21.21 | 20.62 | 0.59 | 0.59 | 0.01 | 0.01 |
| gene-LOC105280956 | https://www.ncbi.nlm.nih.gov/gene/?term=LOC105280956 | 2639 | 73 | 276.62 | 1 | 3.79 | 1.37 | 20.28 | 19.71 | 0.58 | 0.58 | 0.02 | 0.02 |
| gene-LOC105286515 | https://www.ncbi.nlm.nih.gov/gene/?term=LOC105286515 | 3751 | 107 | 285.26 | 1 | 2.67 | 0.93 | 10.47 | 9.92 | 0.55 | 0.55 | 0.02 | 0.02 |
| gene-LOC105276196 | https://www.ncbi.nlm.nih.gov/gene/?term=LOC105276196 | 2953 | 40 | 135.46 | 1 | 3.39 | 2.5 | 52.64 | 52.1 | 0.54 | 0.54 | 0.01 | 0.01 |
| gene-LOC105278454 | https://www.ncbi.nlm.nih.gov/gene/?term=LOC105278454 | 1972 | 36 | 182.56 | 1 | 5.07 | 2.78 | 47.84 | 47.3 | 0.54 | 0.54 | 0.01 | 0.01 |
| gene-LOC105275752 | https://www.ncbi.nlm.nih.gov/gene/?term=LOC105275752 | 3462 | 117 | 337.95 | 1 | 2.89 | 0.85 | 29.78 | 29.38 | 0.4 | 0.4 | 0.01 | 0.01 |
| gene-LOC105274706 | https://www.ncbi.nlm.nih.gov/gene/?term=LOC105274706 | 1446 | 51 | 352.7 | 1 | 6.92 | 1.96 | 37.41 | 37.03 | 0.38 | 0.38 | 0.01 | 0.01 |
| gene-LOC105280786 | https://www.ncbi.nlm.nih.gov/gene/?term=LOC105280786 | 2573 | 107 | 415.86 | 1 | 3.89 | 0.93 | 33.04 | 32.68 | 0.37 | 0.37 | 0.02 | 0.02 |
| gene-LOC105287417 | https://www.ncbi.nlm.nih.gov/gene/?term=LOC105287417 | 1025 | 28 | 273.17 | 1 | 9.76 | 3.57 | 42.43 | 42.07 | 0.36 | 0.36 | 0.01 | 0.01 |
| gene-LOC105276449 | https://www.ncbi.nlm.nih.gov/gene/?term=LOC105276449 | 1231 | 39 | 316.82 | 1 | 8.12 | 2.56 | 32.73 | 32.41 | 0.32 | 0.32 | 0.01 | 0.01 |
| gene-LOC105278534 | https://www.ncbi.nlm.nih.gov/gene/?term=LOC105278534 | 7364 | 166 | 225.42 | 1 | 1.36 | 0.6 | 18.8 | 18.53 | 0.26 | 0.26 | 0.01 | 0.01 |
| gene-LOC105286649 | https://www.ncbi.nlm.nih.gov/gene/?term=LOC105286649 | 14178 | 662 | 466.92 | 1 | 0.71 | 0.15 | 1.27 | 1.01 | 0.26 | 0.26 | 0.01 | 0.01 |
| gene-LOC105284774 | https://www.ncbi.nlm.nih.gov/gene/?term=LOC105284774 | 12089 | 74 | 61.21 | 1 | 0.83 | 1.35 | 19.79 | 19.53 | 0.25 | 0.25 | 0 | 0 |
| gene-LOC105283649 | https://www.ncbi.nlm.nih.gov/gene/?term=LOC105283649 | 1918 | 53 | 276.33 | 2 | 10.43 | 3.77 | 38.29 | 38.07 | 0.23 | 0.23 | 0.01 | 0.01 |
| gene-LOC105285807 | https://www.ncbi.nlm.nih.gov/gene/?term=LOC105285807 | 6063 | 174 | 286.99 | 1 | 1.65 | 0.57 | 7.5 | 7.29 | 0.22 | 0.22 | 0.01 | 0.01 |
| gene-LOC105282794 | https://www.ncbi.nlm.nih.gov/gene/?term=LOC105282794 | 4571 | 91 | 199.08 | 1 | 2.19 | 1.1 | 19.49 | 19.29 | 0.2 | 0.2 | 0 | 0 |
| gene-LOC105286274 | https://www.ncbi.nlm.nih.gov/gene/?term=LOC105286274 | 3817 | 181 | 474.19 | 1 | 2.62 | 0.55 | 15.32 | 15.14 | 0.18 | 0.18 | 0.01 | 0.01 |
| gene-LOC105288078 | https://www.ncbi.nlm.nih.gov/gene/?term=LOC105288078 | 19126 | 301 | 157.38 | 1 | 0.52 | 0.33 | 17.46 | 17.3 | 0.16 | 0.16 | 0 | 0 |
| gene-LOC105287095 | https://www.ncbi.nlm.nih.gov/gene/?term=LOC105287095 | 1435 | 52 | 362.37 | 1 | 6.97 | 1.92 | 30.47 | 30.36 | 0.12 | 0.12 | 0 | 0 |
| gene-LOC105284116 | https://www.ncbi.nlm.nih.gov/gene/?term=LOC105284116 | 3970 | 85 | 214.11 | 1 | 2.52 | 1.18 | 7.23 | 7.13 | 0.09 | 0.09 | 0 | 0 |
| gene-LOC105279038 | https://www.ncbi.nlm.nih.gov/gene/?term=LOC105279038 | 9453 | 298 | 315.24 | 1 | 1.06 | 0.34 | 9.03 | 8.95 | 0.08 | 0.08 | 0 | 0 |
| gene-LOC105281272 | https://www.ncbi.nlm.nih.gov/gene/?term=LOC105281272 | 5252 | 168 | 319.88 | 1 | 1.9 | 0.6 | 16.96 | 16.88 | 0.08 | 0.08 | 0 | 0 |
| gene-LOC105285742 | https://www.ncbi.nlm.nih.gov/gene/?term=LOC105285742 | 3446 | 91 | 264.07 | 1 | 2.9 | 1.1 | 26.72 | 26.67 | 0.05 | 0.05 | 0 | 0 |
| gene-LOC105276612 | https://www.ncbi.nlm.nih.gov/gene/?term=LOC105276612 | 4157 | 172 | 413.76 | 1 | 2.41 | 0.58 | 11.01 | 10.98 | 0.03 | 0.03 | 0 | 0 |
| gene-LOC105275327 | https://www.ncbi.nlm.nih.gov/gene/?term=LOC105275327 | 14245 | 413 | 289.93 | 1 | 0.7 | 0.24 | 7.72 | 7.75 | -0.03 | 0.03 | 0 | 0 |
| gene-LOC105280304 | https://www.ncbi.nlm.nih.gov/gene/?term=LOC105280304 | 8919 | 90 | 100.91 | 1 | 1.12 | 1.11 | 7.28 | 7.34 | -0.05 | 0.05 | 0 | 0 |
| gene-LOC105274534 | https://www.ncbi.nlm.nih.gov/gene/?term=LOC105274534 | 1329 | 53 | 398.8 | 1 | 7.52 | 1.89 | 47.05 | 47.11 | -0.06 | 0.06 | 0 | 0 |
| gene-LOC105278666 | https://www.ncbi.nlm.nih.gov/gene/?term=LOC105278666 | 10341 | 386 | 373.27 | 1 | 0.97 | 0.26 | 7.83 | 7.9 | -0.07 | 0.07 | 0 | 0 |
| gene-LOC105274531 | https://www.ncbi.nlm.nih.gov/gene/?term=LOC105274531 | 7774 | 252 | 324.16 | 1 | 1.29 | 0.4 | 11.91 | 11.99 | -0.08 | 0.08 | 0 | 0 |
| gene-LOC105276941 | https://www.ncbi.nlm.nih.gov/gene/?term=LOC105276941 | 15956 | 446 | 279.52 | 1 | 0.63 | 0.22 | 5.36 | 5.47 | -0.11 | 0.11 | 0 | 0 |
| gene-LOC105282825 | https://www.ncbi.nlm.nih.gov/gene/?term=LOC105282825 | 3919 | 80 | 204.13 | 1 | 2.55 | 1.25 | 13.05 | 13.16 | -0.12 | 0.12 | 0 | 0 |
| gene-LOC105286644 | https://www.ncbi.nlm.nih.gov/gene/?term=LOC105286644 | 1235 | 22 | 178.14 | 1 | 8.1 | 4.55 | 42.2 | 42.35 | -0.15 | 0.15 | 0 | 0 |
| gene-LOC105283989 | https://www.ncbi.nlm.nih.gov/gene/?term=LOC105283989 | 4948 | 25 | 50.53 | 1 | 2.02 | 4 | 45.66 | 45.85 | -0.19 | 0.19 | 0 | 0 |
| gene-LOC105278145 | https://www.ncbi.nlm.nih.gov/gene/?term=LOC105278145 | 4470 | 244 | 545.86 | 1 | 2.24 | 0.41 | 10.74 | 10.94 | -0.2 | 0.2 | -0.01 | 0.01 |
| gene-LOC105285534 | https://www.ncbi.nlm.nih.gov/gene/?term=LOC105285534 | 3944 | 211 | 534.99 | 1 | 2.54 | 0.47 | 11.95 | 12.16 | -0.21 | 0.21 | -0.01 | 0.01 |
| gene-LOC105276527 | https://www.ncbi.nlm.nih.gov/gene/?term=LOC105276527 | 7923 | 323 | 407.67 | 1 | 1.26 | 0.31 | 8.74 | 8.99 | -0.25 | 0.25 | -0.01 | 0.01 |
| gene-LOC105284117 | https://www.ncbi.nlm.nih.gov/gene/?term=LOC105284117 | 4340 | 64 | 147.47 | 1 | 2.3 | 1.56 | 10.83 | 11.09 | -0.25 | 0.25 | 0 | 0 |
| gene-LOC105274856 | https://www.ncbi.nlm.nih.gov/gene/?term=LOC105274856 | 5216 | 200 | 383.44 | 1 | 1.92 | 0.5 | 18.38 | 18.67 | -0.29 | 0.29 | -0.01 | 0.01 |
| gene-LOC105286223 | https://www.ncbi.nlm.nih.gov/gene/?term=LOC105286223 | 2225 | 79 | 355.06 | 1 | 4.49 | 1.27 | 30.58 | 30.96 | -0.38 | 0.38 | -0.01 | 0.01 |
| gene-LOC105285395 | https://www.ncbi.nlm.nih.gov/gene/?term=LOC105285395 | 8085 | 165 | 204.08 | 1 | 1.24 | 0.61 | 17.34 | 17.73 | -0.39 | 0.39 | -0.01 | 0.01 |

| | | | | | | | | | | | | | |
|---|---|---|---|---|---|---|---|---|---|---|---|---|---|
| gene-LOC105278538 | https://www.ncbi.nlm.nih.gov/gene/?term=LOC105278538 | 2150 | 76 | 353.49 | 1 | 4.65 | 1.32 | 18.23 | 18.64 | -0.41 | 0.41 | -0.01 | 0.01 |
| gene-LOC105283296 | https://www.ncbi.nlm.nih.gov/gene/?term=LOC105283296 | 2174 | 78 | 358.79 | 1 | 4.6 | 1.28 | 21.02 | 21.5 | -0.48 | 0.48 | -0.02 | 0.02 |
| gene-LOC105285406 | https://www.ncbi.nlm.nih.gov/gene/?term=LOC105285406 | 8015 | 39 | 48.66 | 1 | 1.25 | 2.56 | 33.07 | 33.55 | -0.48 | 0.48 | 0 | 0 |
| gene-LOC105275584 | https://www.ncbi.nlm.nih.gov/gene/?term=LOC105275584 | 3070 | 71 | 231.27 | 1 | 3.26 | 1.41 | 26.68 | 27.17 | -0.49 | 0.49 | -0.01 | 0.01 |
| gene-LOC105283943 | https://www.ncbi.nlm.nih.gov/gene/?term=LOC105283943 | 3559 | 48 | 134.87 | 1 | 2.81 | 2.08 | 17.72 | 18.23 | -0.51 | 0.51 | -0.01 | 0.01 |
| gene-LOC105281700 | https://www.ncbi.nlm.nih.gov/gene/?term=LOC105281700 | 2208 | 40 | 181.16 | 1 | 4.53 | 2.5 | 41.3 | 41.85 | -0.55 | 0.55 | -0.01 | 0.01 |
| gene-LOC105284355 | https://www.ncbi.nlm.nih.gov/gene/?term=LOC105284355 | 3904 | 135 | 345.8 | 1 | 2.56 | 0.74 | 22.88 | 23.48 | -0.59 | 0.59 | -0.02 | 0.02 |
| gene-LOC105275179 | https://www.ncbi.nlm.nih.gov/gene/?term=LOC105275179 | 3271 | 66 | 201.77 | 1 | 3.06 | 1.52 | 14.71 | 15.31 | -0.61 | 0.61 | -0.01 | 0.01 |
| gene-LOC105283728 | https://www.ncbi.nlm.nih.gov/gene/?term=LOC105283728 | 3256 | 103 | 316.34 | 2 | 6.14 | 1.94 | 21.65 | 22.28 | -0.62 | 0.62 | -0.02 | 0.02 |
| gene-LOC105286265 | https://www.ncbi.nlm.nih.gov/gene/?term=LOC105286265 | 2507 | 63 | 251.3 | 1 | 3.99 | 1.59 | 30.31 | 30.94 | -0.63 | 0.63 | -0.02 | 0.02 |
| gene-LOC105280410 | https://www.ncbi.nlm.nih.gov/gene/?term=LOC105280410 | 12580 | 235 | 186.8 | 1 | 0.79 | 0.43 | 20.46 | 21.13 | -0.67 | 0.67 | -0.01 | 0.01 |
| gene-LOC105275363 | https://www.ncbi.nlm.nih.gov/gene/?term=LOC105275363 | 3798 | 146 | 384.41 | 1 | 2.63 | 0.68 | 17.72 | 18.47 | -0.76 | 0.76 | -0.03 | 0.03 |
| gene-LOC105274851 | https://www.ncbi.nlm.nih.gov/gene/?term=LOC105274851 | 6616 | 122 | 184.4 | 1 | 1.51 | 0.82 | 10.53 | 11.32 | -0.79 | 0.79 | -0.01 | 0.01 |
| gene-LOC105287063 | https://www.ncbi.nlm.nih.gov/gene/?term=LOC105287063 | 1874 | 46 | 245.46 | 1 | 5.34 | 2.17 | 34.15 | 34.94 | -0.79 | 0.79 | -0.02 | 0.02 |
| gene-LOC105283987 | https://www.ncbi.nlm.nih.gov/gene/?term=LOC105283987 | 4841 | 64 | 132.2 | 1 | 2.07 | 1.56 | 32.11 | 32.92 | -0.81 | 0.81 | -0.01 | 0.01 |
| gene-LOC105279298 | https://www.ncbi.nlm.nih.gov/gene/?term=LOC105279298 | 2773 | 129 | 465.2 | 2 | 7.21 | 1.55 | 22.93 | 23.79 | -0.86 | 0.86 | -0.04 | 0.04 |
| gene-LOC105275460 | https://www.ncbi.nlm.nih.gov/gene/?term=LOC105275460 | 2473 | 107 | 432.67 | 1 | 4.04 | 0.93 | 13.53 | 14.43 | -0.91 | 0.91 | -0.04 | 0.04 |
| gene-LOC105275683 | https://www.ncbi.nlm.nih.gov/gene/?term=LOC105275683 | 4256 | 187 | 439.38 | 1 | 2.35 | 0.53 | 12 | 12.9 | -0.91 | 0.91 | -0.04 | 0.04 |
| gene-LOC105279657 | https://www.ncbi.nlm.nih.gov/gene/?term=LOC105279657 | 5878 | 79 | 134.4 | 1 | 1.7 | 1.27 | 19.77 | 20.7 | -0.93 | 0.93 | -0.01 | 0.01 |
| gene-LOC105275451 | https://www.ncbi.nlm.nih.gov/gene/?term=LOC105275451 | 6035 | 105 | 173.99 | 1 | 1.66 | 0.95 | 24.51 | 25.49 | -0.98 | 0.98 | -0.02 | 0.02 |
| gene-LOC105274590 | https://www.ncbi.nlm.nih.gov/gene/?term=LOC105274590 | 4516 | 61 | 135.08 | 1 | 2.21 | 1.64 | 27.27 | 28.28 | -1.01 | 1.01 | -0.01 | 0.01 |
| gene-LOC105285671 | https://www.ncbi.nlm.nih.gov/gene/?term=LOC105285671 | 2263 | 92 | 406.54 | 1 | 4.42 | 1.09 | 28.77 | 29.78 | -1.01 | 1.01 | -0.04 | 0.04 |
| gene-LOC105277216 | https://www.ncbi.nlm.nih.gov/gene/?term=LOC105277216 | 7627 | 76 | 99.65 | 1 | 1.31 | 1.32 | 7.73 | 8.77 | -1.04 | 1.04 | -0.01 | 0.01 |
| gene-LOC105275173 | https://www.ncbi.nlm.nih.gov/gene/?term=LOC105275173 | 4639 | 71 | 153.05 | 1 | 2.16 | 1.41 | 27.01 | 28.06 | -1.05 | 1.05 | -0.02 | 0.02 |
| gene-LOC105285179 | https://www.ncbi.nlm.nih.gov/gene/?term=LOC105285179 | 4191 | 148 | 353.14 | 1 | 2.39 | 0.68 | 17.81 | 18.87 | -1.05 | 1.05 | -0.04 | 0.04 |
| gene-LOC105281515 | https://www.ncbi.nlm.nih.gov/gene/?term=LOC105281515 | 8827 | 35 | 39.65 | 1 | 1.13 | 2.86 | 31.71 | 33.02 | -1.31 | 1.31 | -0.01 | 0.01 |
| gene-LOC105286452 | https://www.ncbi.nlm.nih.gov/gene/?term=LOC105286452 | 6888 | 55 | 79.85 | 1 | 1.45 | 1.82 | 32.52 | 33.96 | -1.44 | 1.44 | -0.01 | 0.01 |
| gene-LOC105276692 | https://www.ncbi.nlm.nih.gov/gene/?term=LOC105276692 | 1745 | 33 | 189.11 | 1 | 5.73 | 3.03 | 25.26 | 26.77 | -1.52 | 1.52 | -0.03 | 0.03 |
| gene-LOC105285055 | https://www.ncbi.nlm.nih.gov/gene/?term=LOC105285055 | 4108 | 197 | 479.55 | 1 | 2.43 | 0.51 | 13.57 | 15.12 | -1.55 | 1.55 | -0.07 | 0.07 |
| gene-LOC105275190 | https://www.ncbi.nlm.nih.gov/gene/?term=LOC105275190 | 3837 | 69 | 179.83 | 1 | 2.61 | 1.45 | 19.53 | 21.12 | -1.59 | 1.59 | -0.03 | 0.03 |
| gene-LOC105283592 | https://www.ncbi.nlm.nih.gov/gene/?term=LOC105283592 | 3479 | 66 | 189.71 | 1 | 2.87 | 1.52 | 25.49 | 27.08 | -1.59 | 1.59 | -0.03 | 0.03 |
| gene-LOC105279500 | https://www.ncbi.nlm.nih.gov/gene/?term=LOC105279500 | 3587 | 31 | 86.42 | 1 | 2.79 | 3.23 | 51.03 | 52.64 | -1.62 | 1.62 | -0.01 | 0.01 |
| gene-LOC105280774 | https://www.ncbi.nlm.nih.gov/gene/?term=LOC105280774 | 2793 | 97 | 347.3 | 1 | 3.58 | 1.03 | 31.81 | 33.47 | -1.66 | 1.66 | -0.06 | 0.06 |
| gene-LOC113561621 | https://www.ncbi.nlm.nih.gov/gene/?term=LOC113561621 | 2249 | 79 | 351.27 | 1 | 4.45 | 1.27 | 19.72 | 21.41 | -1.69 | 1.69 | -0.06 | 0.06 |
| gene-LOC105280641 | https://www.ncbi.nlm.nih.gov/gene/?term=LOC105280641 | 3682 | 53 | 143.94 | 1 | 2.72 | 1.89 | 17.07 | 18.78 | -1.71 | 1.71 | -0.02 | 0.02 |
| gene-LOC105275518 | https://www.ncbi.nlm.nih.gov/gene/?term=LOC105275518 | 2335 | 18 | 77.09 | 1 | 4.28 | 5.56 | 47.37 | 49.08 | -1.72 | 1.72 | -0.01 | 0.01 |
| gene-LOC105284184 | https://www.ncbi.nlm.nih.gov/gene/?term=LOC105284184 | 2535 | 67 | 264.3 | 1 | 3.94 | 1.49 | 41.64 | 43.45 | -1.81 | 1.81 | -0.05 | 0.05 |
| gene-LOC105278807 | https://www.ncbi.nlm.nih.gov/gene/?term=LOC105278807 | 2166 | 68 | 313.94 | 1 | 4.62 | 1.47 | 17.54 | 19.41 | -1.87 | 1.87 | -0.06 | 0.06 |
| gene-LOC105288066 | https://www.ncbi.nlm.nih.gov/gene/?term=LOC105288066 | 1964 | 16 | 81.47 | 1 | 5.09 | 6.25 | 41.07 | 43.1 | -2.03 | 2.03 | -0.02 | 0.02 |
| gene-LOC105282156 | https://www.ncbi.nlm.nih.gov/gene/?term=LOC105282156 | 2577 | 132 | 512.22 | 1 | 3.88 | 0.76 | 24.4 | 26.49 | -2.09 | 2.09 | -0.11 | 0.11 |
| gene-LOC105275656 | https://www.ncbi.nlm.nih.gov/gene/?term=LOC105275656 | 2803 | 127 | 453.09 | 1 | 3.57 | 0.79 | 26.43 | 28.64 | -2.2 | 2.2 | -0.1 | 0.1 |
| gene-LOC105283712 | https://www.ncbi.nlm.nih.gov/gene/?term=LOC105283712 | 1910 | 24 | 125.65 | 1 | 5.24 | 4.17 | 29.64 | 32.04 | -2.39 | 2.39 | -0.03 | 0.03 |
| gene-LOC105285695 | https://www.ncbi.nlm.nih.gov/gene/?term=LOC105285695 | 1371 | 62 | 452.22 | 1 | 7.29 | 1.61 | 23 | 25.77 | -2.77 | 2.77 | -0.13 | 0.13 |
| gene-LOC105278109 | https://www.ncbi.nlm.nih.gov/gene/?term=LOC105278109 | 4029 | 56 | 138.99 | 1 | 2.48 | 1.79 | 47.35 | 50.32 | -2.97 | 2.97 | -0.04 | 0.04 |
| gene-LOC105274900 | https://www.ncbi.nlm.nih.gov/gene/?term=LOC105274900 | 5561 | 47 | 84.52 | 2 | 3.6 | 4.26 | 31.62 | 34.67 | -3.05 | 3.05 | -0.03 | 0.03 |
| gene-LOC105285674 | https://www.ncbi.nlm.nih.gov/gene/?term=LOC105285674 | 3035 | 88 | 289.95 | 1 | 3.29 | 1.14 | 23.36 | 26.44 | -3.08 | 3.08 | -0.09 | 0.09 |
| gene-LOC105280348 | https://www.ncbi.nlm.nih.gov/gene/?term=LOC105280348 | 3797 | 50 | 131.68 | 1 | 2.63 | 2 | 25.73 | 28.9 | -3.17 | 3.17 | -0.04 | 0.04 |
| gene-LOC105279356 | https://www.ncbi.nlm.nih.gov/gene/?term=LOC105279356 | 1314 | 39 | 296.8 | 1 | 7.61 | 2.56 | 29.77 | 33.04 | -3.27 | 3.27 | -0.1 | 0.1 |
| gene-LOC105280972 | https://www.ncbi.nlm.nih.gov/gene/?term=LOC105280972 | 1532 | 26 | 169.71 | 1 | 6.53 | 3.85 | 20.04 | 23.88 | -3.84 | 3.84 | -0.07 | 0.07 |
| gene-LOC105281848 | https://www.ncbi.nlm.nih.gov/gene/?term=LOC105281848 | 2122 | 44 | 207.35 | 1 | 4.71 | 2.27 | 23.57 | 27.97 | -4.4 | 4.4 | -0.09 | 0.09 |
| gene-LOC105277215 | https://www.ncbi.nlm.nih.gov/gene/?term=LOC105277215 | 2619 | 20 | 76.37 | 1 | 3.82 | 5 | 28.06 | 32.55 | -4.49 | 4.49 | -0.03 | 0.03 |
| gene-LOC105279898 | https://www.ncbi.nlm.nih.gov/gene/?term=LOC105279898 | 1343 | 13 | 96.8 | 1 | 7.45 | 7.69 | 60.41 | 65.83 | -5.42 | 5.42 | -0.05 | 0.05 |
| gene-LOC105285514 | https://www.ncbi.nlm.nih.gov/gene/?term=LOC105285514 | 1277 | 8 | 62.65 | 1 | 7.83 | 12.5 | 82.36 | 88.52 | -6.16 | 6.16 | -0.04 | 0.04 |
| gene-LOC105276078 | https://www.ncbi.nlm.nih.gov/gene/?term=LOC105276078 | 1051 | 8 | 76.12 | 1 | 9.51 | 12.5 | 46.61 | 53.81 | -7.2 | 7.2 | -0.05 | 0.05 |
| gene-LOC105283552 | https://www.ncbi.nlm.nih.gov/gene/?term=LOC105283552 | 1562 | 27 | 172.86 | 1 | 6.4 | 3.7 | 56.95 | 65.52 | -8.57 | 8.57 | -0.15 | 0.15 |
| gene-LOC105285060 | https://www.ncbi.nlm.nih.gov/gene/?term=LOC105285060 | 1285 | 13 | 101.17 | 1 | 7.78 | 7.69 | 55.35 | 64 | -8.64 | 8.64 | -0.09 | 0.09 |
| gene-LOC105275708 | https://www.ncbi.nlm.nih.gov/gene/?term=LOC105275708 | 1772 | 10 | 56.43 | 1 | 5.64 | 10 | 41.45 | 50.31 | -8.85 | 8.85 | -0.05 | 0.05 |
| gene-LOC105275070 | https://www.ncbi.nlm.nih.gov/gene/?term=LOC105275070 | 5667 | 35 | 61.76 | 2 | 3.53 | 5.71 | 3.8 | 15.13 | -11.33 | 11.33 | -0.07 | 0.07 |
| gene-LOC113562345 | https://www.ncbi.nlm.nih.gov/gene/?term=LOC113562345 | 2290 | 62 | 270.74 | 15 | 65.5 | 24.19 | 17.91 | 32.18 | -14.27 | 14.27 | -0.39 | 0.39 |

Directory: OGL File: ogl-Ob_LI_BCphaseA1.vs.BCphaseB1.txt

| gene_ID | gene_link | gwidth | #Sites | #per10Kb | #dmSites | #dmsp10kb | %dmSites | %pSite1 | %pSite2 | DMpSite | ADMpSite | DMpNucl | ADMpNucl |
|---|---|---|---|---|---|---|---|---|---|---|---|---|---|
| gene-LOC105282885 | https://www.ncbi.nlm.nih.gov/gene/?term=LOC105282885 | 1870 | 41 | 219.25 | 1 | 5.35 | 2.44 | 37.52 | 34.93 | 2.59 | 2.59 | 0.06 | 0.06 |
| gene-LOC105281173 | https://www.ncbi.nlm.nih.gov/gene/?term=LOC105281173 | 1595 | 25 | 156.74 | 1 | 6.27 | 4 | 64.76 | 62.33 | 2.42 | 2.42 | 0.04 | 0.04 |
| gene-LOC105279528 | https://www.ncbi.nlm.nih.gov/gene/?term=LOC105279528 | 5802 | 46 | 79.28 | 1 | 1.72 | 2.17 | 17.18 | 14.84 | 2.33 | 2.33 | 0.02 | 0.02 |
| gene-LOC105285409 | https://www.ncbi.nlm.nih.gov/gene/?term=LOC105285409 | 5099 | 48 | 94.14 | 1 | 1.96 | 2.08 | 3.6 | 1.53 | 2.08 | 2.08 | 0.02 | 0.02 |
| gene-LOC105278454 | https://www.ncbi.nlm.nih.gov/gene/?term=LOC105278454 | 1972 | 37 | 187.63 | 1 | 5.07 | 2.7 | 46.8 | 45.07 | 1.73 | 1.73 | 0.03 | 0.03 |
| gene-LOC105285202 | https://www.ncbi.nlm.nih.gov/gene/?term=LOC105285202 | 1838 | 36 | 195.87 | 1 | 5.44 | 2.78 | 18.94 | 17.23 | 1.72 | 1.72 | 0.03 | 0.03 |
| gene-LOC105276196 | https://www.ncbi.nlm.nih.gov/gene/?term=LOC105276196 | 2953 | 54 | 182.86 | 1 | 3.39 | 1.85 | 40.59 | 39.19 | 1.4 | 1.4 | 0.03 | 0.03 |
| gene-LOC105283123 | https://www.ncbi.nlm.nih.gov/gene/?term=LOC105283123 | 1507 | 41 | 272.06 | 1 | 6.64 | 2.44 | 15.68 | 14.34 | 1.34 | 1.34 | 0.04 | 0.04 |
| gene-LOC105286265 | https://www.ncbi.nlm.nih.gov/gene/?term=LOC105286265 | 2507 | 64 | 255.29 | 1 | 3.99 | 1.56 | 30.65 | 29.46 | 1.19 | 1.19 | 0.03 | 0.03 |
| gene-LOC105288010 | https://www.ncbi.nlm.nih.gov/gene/?term=LOC105288010 | 3189 | 49 | 153.65 | 1 | 3.14 | 2.04 | 31.6 | 30.42 | 1.18 | 1.18 | 0.02 | 0.02 |
| gene-LOC105275653 | https://www.ncbi.nlm.nih.gov/gene/?term=LOC105275653 | 11251 | 262 | 232.87 | 1 | 0.89 | 0.38 | 9.15 | 8.02 | 1.13 | 1.13 | 0.03 | 0.03 |
| gene-LOC105274706 | https://www.ncbi.nlm.nih.gov/gene/?term=LOC105274706 | 1446 | 46 | 318.12 | 1 | 6.92 | 2.17 | 41.37 | 40.53 | 0.85 | 0.85 | 0.03 | 0.03 |
| gene-LOC105278282 | https://www.ncbi.nlm.nih.gov/gene/?term=LOC105278282 | 1775 | 47 | 264.79 | 1 | 5.63 | 2.13 | 43.73 | 42.89 | 0.84 | 0.84 | 0.02 | 0.02 |
| gene-LOC105280447 | https://www.ncbi.nlm.nih.gov/gene/?term=LOC105280447 | 2215 | 25 | 112.87 | 1 | 4.51 | 4 | 26.74 | 25.91 | 0.82 | 0.82 | 0.01 | 0.01 |
| gene-LOC105281047 | https://www.ncbi.nlm.nih.gov/gene/?term=LOC105281047 | 7313 | 117 | 159.99 | 1 | 1.37 | 0.85 | 10.66 | 9.84 | 0.82 | 0.82 | 0.01 | 0.01 |
| gene-LOC105278214 | https://www.ncbi.nlm.nih.gov/gene/?term=LOC105278214 | 3787 | 77 | 203.33 | 1 | 2.64 | 1.3 | 11.46 | 10.7 | 0.76 | 0.76 | 0.02 | 0.02 |
| gene-LOC105285583 | https://www.ncbi.nlm.nih.gov/gene/?term=LOC105285583 | 1947 | 49 | 251.67 | 1 | 5.14 | 2.04 | 29.41 | 28.65 | 0.76 | 0.76 | 0.02 | 0.02 |
| gene-LOC105278436 | https://www.ncbi.nlm.nih.gov/gene/?term=LOC105278436 | 2556 | 57 | 223 | 1 | 3.91 | 1.75 | 36.25 | 35.56 | 0.69 | 0.69 | 0.02 | 0.02 |
| gene-LOC105276429 | https://www.ncbi.nlm.nih.gov/gene/?term=LOC105276429 | 2901 | 112 | 386.07 | 1 | 3.45 | 0.89 | 26.41 | 25.73 | 0.68 | 0.68 | 0.03 | 0.03 |
| gene-LOC105285406 | https://www.ncbi.nlm.nih.gov/gene/?term=LOC105285406 | 8015 | 40 | 49.91 | 1 | 1.25 | 2.5 | 34.18 | 33.53 | 0.64 | 0.64 | 0 | 0 |
| gene-LOC105274449 | https://www.ncbi.nlm.nih.gov/gene/?term=LOC105274449 | 1549 | 43 | 277.6 | 1 | 6.46 | 2.33 | 54.59 | 53.98 | 0.6 | 0.6 | 0.02 | 0.02 |
| gene-LOC105281420 | https://www.ncbi.nlm.nih.gov/gene/?term=LOC105281420 | 3882 | 57 | 146.83 | 1 | 2.58 | 1.75 | 15.28 | 14.73 | 0.55 | 0.55 | 0.01 | 0.01 |
| gene-LOC105282811 | https://www.ncbi.nlm.nih.gov/gene/?term=LOC105282811 | 5765 | 203 | 352.12 | 1 | 1.73 | 0.49 | 9.79 | 9.26 | 0.53 | 0.53 | 0.02 | 0.02 |
| gene-LOC105277435 | https://www.ncbi.nlm.nih.gov/gene/?term=LOC105277435 | 3643 | 155 | 425.47 | 1 | 2.74 | 0.65 | 12.32 | 11.8 | 0.52 | 0.52 | 0.02 | 0.02 |
| gene-LOC105280786 | https://www.ncbi.nlm.nih.gov/gene/?term=LOC105280786 | 2573 | 120 | 466.38 | 1 | 3.89 | 0.83 | 29.63 | 29.12 | 0.52 | 0.52 | 0.02 | 0.02 |
| gene-LOC105284116 | https://www.ncbi.nlm.nih.gov/gene/?term=LOC105284116 | 3970 | 95 | 239.29 | 1 | 2.52 | 1.05 | 6.78 | 6.28 | 0.5 | 0.5 | 0.01 | 0.01 |
| gene-LOC105278877 | https://www.ncbi.nlm.nih.gov/gene/?term=LOC105278877 | 1968 | 77 | 391.26 | 1 | 5.08 | 1.3 | 32.7 | 32.23 | 0.46 | 0.46 | 0.02 | 0.02 |
| gene-LOC105284117 | https://www.ncbi.nlm.nih.gov/gene/?term=LOC105284117 | 4340 | 81 | 186.64 | 1 | 2.3 | 1.23 | 11.74 | 11.3 | 0.44 | 0.44 | 0.01 | 0.01 |
| gene-LOC105282600 | https://www.ncbi.nlm.nih.gov/gene/?term=LOC105282600 | 3715 | 57 | 153.43 | 1 | 2.69 | 1.75 | 29.16 | 28.81 | 0.35 | 0.35 | 0.01 | 0.01 |
| gene-LOC105285807 | https://www.ncbi.nlm.nih.gov/gene/?term=LOC105285807 | 6063 | 183 | 301.83 | 1 | 1.65 | 0.55 | 10.09 | 9.8 | 0.29 | 0.29 | 0.01 | 0.01 |
| gene-LOC105286515 | https://www.ncbi.nlm.nih.gov/gene/?term=LOC105286515 | 3751 | 103 | 274.59 | 1 | 2.67 | 0.97 | 10.73 | 10.44 | 0.29 | 0.29 | 0.01 | 0.01 |
| gene-LOC105279038 | https://www.ncbi.nlm.nih.gov/gene/?term=LOC105279038 | 9453 | 310 | 327.94 | 1 | 1.06 | 0.32 | 8.8 | 8.54 | 0.27 | 0.27 | 0.01 | 0.01 |
| gene-LOC105278666 | https://www.ncbi.nlm.nih.gov/gene/?term=LOC105278666 | 10341 | 405 | 391.64 | 1 | 0.97 | 0.25 | 7.09 | 6.88 | 0.22 | 0.22 | 0.01 | 0.01 |
| gene-LOC105284036 | https://www.ncbi.nlm.nih.gov/gene/?term=LOC105284036 | 3254 | 83 | 255.07 | 1 | 3.07 | 1.2 | 22.41 | 22.19 | 0.22 | 0.22 | 0.01 | 0.01 |
| gene-LOC105276901 | https://www.ncbi.nlm.nih.gov/gene/?term=LOC105276901 | 3254 | 95 | 291.95 | 1 | 3.07 | 1.05 | 23.05 | 22.87 | 0.18 | 0.18 | 0.01 | 0.01 |
| gene-LOC105278145 | https://www.ncbi.nlm.nih.gov/gene/?term=LOC105278145 | 4470 | 249 | 557.05 | 1 | 2.24 | 0.4 | 10.18 | 10.01 | 0.17 | 0.17 | 0.01 | 0.01 |
| gene-LOC105275752 | https://www.ncbi.nlm.nih.gov/gene/?term=LOC105275752 | 3462 | 124 | 358.17 | 1 | 2.89 | 0.81 | 28.83 | 28.67 | 0.16 | 0.16 | 0.01 | 0.01 |
| gene-LOC105279657 | https://www.ncbi.nlm.nih.gov/gene/?term=LOC105279657 | 5878 | 86 | 146.31 | 1 | 1.7 | 1.16 | 18.3 | 18.14 | 0.16 | 0.16 | 0 | 0 |
| gene-LOC105287095 | https://www.ncbi.nlm.nih.gov/gene/?term=LOC105287095 | 1435 | 61 | 425.09 | 1 | 6.97 | 1.64 | 26.05 | 25.92 | 0.14 | 0.14 | 0.01 | 0.01 |
| gene-LOC105275173 | https://www.ncbi.nlm.nih.gov/gene/?term=LOC105275173 | 4639 | 83 | 178.92 | 1 | 2.16 | 1.2 | 22.33 | 22.24 | 0.09 | 0.09 | 0 | 0 |
| gene-LOC105276059 | https://www.ncbi.nlm.nih.gov/gene/?term=LOC105276059 | 6256 | 134 | 214.19 | 1 | 1.6 | 0.75 | 10.93 | 10.85 | 0.08 | 0.08 | 0 | 0 |
| gene-LOC105279664 | https://www.ncbi.nlm.nih.gov/gene/?term=LOC105279664 | 4023 | 68 | 169.03 | 1 | 2.49 | 1.47 | 18.22 | 18.15 | 0.08 | 0.08 | 0 | 0 |
| gene-LOC105278228 | https://www.ncbi.nlm.nih.gov/gene/?term=LOC105278228 | 8610 | 165 | 191.64 | 1 | 1.16 | 0.61 | 22.79 | 22.73 | 0.06 | 0.06 | 0 | 0 |
| gene-LOC105282491 | https://www.ncbi.nlm.nih.gov/gene/?term=LOC105282491 | 4761 | 159 | 333.96 | 1 | 2.1 | 0.63 | 9.46 | 9.41 | 0.05 | 0.05 | 0 | 0 |
| gene-LOC105286274 | https://www.ncbi.nlm.nih.gov/gene/?term=LOC105286274 | 3817 | 177 | 463.71 | 1 | 2.62 | 0.56 | 15.66 | 15.62 | 0.04 | 0.04 | 0 | 0 |
| gene-LOC105275451 | https://www.ncbi.nlm.nih.gov/gene/?term=LOC105275451 | 6035 | 113 | 187.24 | 1 | 1.66 | 0.88 | 22.29 | 22.26 | 0.02 | 0.02 | 0 | 0 |
| gene-LOC105275327 | https://www.ncbi.nlm.nih.gov/gene/?term=LOC105275327 | 14245 | 477 | 334.85 | 1 | 0.7 | 0.21 | 7.15 | 7.14 | 0.01 | 0.01 | 0 | 0 |
| gene-LOC105282151 | https://www.ncbi.nlm.nih.gov/gene/?term=LOC105282151 | 1601 | 41 | 256.09 | 1 | 6.25 | 2.44 | 39.95 | 39.99 | -0.04 | 0.04 | 0 | 0 |
| gene-LOC105279778 | https://www.ncbi.nlm.nih.gov/gene/?term=LOC105279778 | 4347 | 139 | 319.76 | 1 | 2.3 | 0.72 | 9.36 | 9.42 | -0.06 | 0.06 | 0 | 0 |
| gene-LOC105274590 | https://www.ncbi.nlm.nih.gov/gene/?term=LOC105274590 | 4516 | 76 | 168.29 | 1 | 2.21 | 1.32 | 22.95 | 23.02 | -0.07 | 0.07 | 0 | 0 |
| gene-LOC105275170 | https://www.ncbi.nlm.nih.gov/gene/?term=LOC105275170 | 12918 | 337 | 260.88 | 1 | 0.77 | 0.3 | 2.06 | 2.13 | -0.07 | 0.07 | 0 | 0 |
| gene-LOC105280956 | https://www.ncbi.nlm.nih.gov/gene/?term=LOC105280956 | 2639 | 73 | 276.62 | 1 | 3.79 | 1.37 | 20.28 | 20.35 | -0.07 | 0.07 | 0 | 0 |
| gene-LOC105274531 | https://www.ncbi.nlm.nih.gov/gene/?term=LOC105274531 | 7774 | 254 | 326.73 | 1 | 1.29 | 0.39 | 11.9 | 12 | -0.1 | 0.1 | 0 | 0 |
| gene-LOC105274856 | https://www.ncbi.nlm.nih.gov/gene/?term=LOC105274856 | 5216 | 201 | 385.35 | 1 | 1.92 | 0.5 | 17.3 | 17.4 | -0.1 | 0.1 | 0 | 0 |
| gene-LOC105276538 | https://www.ncbi.nlm.nih.gov/gene/?term=LOC105276538 | 1511 | 18 | 119.13 | 1 | 6.62 | 5.56 | 71.72 | 71.83 | -0.11 | 0.11 | 0 | 0 |
| gene-LOC105284262 | https://www.ncbi.nlm.nih.gov/gene/?term=LOC105284262 | 6669 | 230 | 344.88 | 1 | 1.5 | 0.43 | 4.51 | 4.68 | -0.17 | 0.17 | -0.01 | 0.01 |
| gene-LOC105279500 | https://www.ncbi.nlm.nih.gov/gene/?term=LOC105279500 | 3587 | 32 | 89.21 | 1 | 2.79 | 3.12 | 49.74 | 49.96 | -0.21 | 0.21 | 0 | 0 |
| gene-LOC105275584 | https://www.ncbi.nlm.nih.gov/gene/?term=LOC105275584 | 3070 | 86 | 280.13 | 1 | 3.26 | 1.16 | 23.79 | 24 | -0.22 | 0.22 | -0.01 | 0.01 |
| gene-LOC105283649 | https://www.ncbi.nlm.nih.gov/gene/?term=LOC105283649 | 1918 | 53 | 276.33 | 1 | 5.21 | 1.89 | 38.29 | 38.55 | -0.25 | 0.25 | -0.01 | 0.01 |
| gene-LOC105280405 | https://www.ncbi.nlm.nih.gov/gene/?term=LOC105280405 | 5422 | 162 | 298.78 | 1 | 1.84 | 0.62 | 9.36 | 9.65 | -0.29 | 0.29 | -0.01 | 0.01 |

| | | | | | | | | | | | | |
|---|---|---|---|---|---|---|---|---|---|---|---|---|
| gene-LOC105285395 | https://www.ncbi.nlm.nih.gov/gene/?term=LOC105285395 | 8085 | 180 | 222.63 | 1 | 1.24 | 0.56 | 15.55 | 15.84 | -0.29 | 0.29 | -0.01 0.01 |
| gene-LOC105279298 | https://www.ncbi.nlm.nih.gov/gene/?term=LOC105279298 | 2773 | 134 | 483.23 | 1 | 3.61 | 0.75 | 22.72 | 23.03 | -0.3 | 0.3 | -0.01 0.01 |
| gene-LOC105282914 | https://www.ncbi.nlm.nih.gov/gene/?term=LOC105282914 | 5623 | 42 | 74.69 1 | 1.78 | 2.38 | 48.6 | 48.91 | -0.31 | 0.31 | 0 | 0 |
| gene-LOC105288078 | https://www.ncbi.nlm.nih.gov/gene/?term=LOC105288078 | 19126 | 313 | 163.65 | 1 | 0.52 | 0.32 | 17.33 | 17.64 | -0.31 | 0.31 | -0.01 0.01 |
| gene-LOC105284184 | https://www.ncbi.nlm.nih.gov/gene/?term=LOC105284184 | 2535 | 82 | 323.47 | 1 | 3.94 | 1.22 | 34.56 | 34.9 | -0.34 | 0.34 | -0.01 0.01 |
| gene-LOC105285534 | https://www.ncbi.nlm.nih.gov/gene/?term=LOC105285534 | 3944 | 221 | 560.34 | 1 | 2.54 | 0.45 | 11.41 | 11.78 | -0.37 | 0.37 | -0.02 0.02 |
| gene-LOC105278632 | https://www.ncbi.nlm.nih.gov/gene/?term=LOC105278632 | 3624 | 131 | 361.48 | 1 | 2.76 | 0.76 | 14.71 | 15.12 | -0.41 | 0.41 | -0.01 0.01 |
| gene-LOC105282947 | https://www.ncbi.nlm.nih.gov/gene/?term=LOC105282947 | 1162 | 45 | 387.26 | 1 | 8.61 | 2.22 | 32.13 | 32.56 | -0.42 | 0.42 | -0.02 0.02 |
| gene-LOC105276692 | https://www.ncbi.nlm.nih.gov/gene/?term=LOC105276692 | 1745 | 36 | 206.3 1 | 5.73 | 2.78 | 23.57 | 24.03 | -0.46 | 0.46 | -0.01 | 0.01 |
| gene-LOC105284355 | https://www.ncbi.nlm.nih.gov/gene/?term=LOC105284355 | 3904 | 154 | 394.47 | 1 | 2.56 | 0.65 | 20.37 | 20.87 | -0.5 | 0.5 | -0.02 0.02 |
| gene-LOC105278534 | https://www.ncbi.nlm.nih.gov/gene/?term=LOC105278534 | 7364 | 184 | 249.86 | 1 | 1.36 | 0.54 | 17.97 | 18.5 | -0.53 | 0.53 | -0.01 0.01 |
| gene-LOC105286770 | https://www.ncbi.nlm.nih.gov/gene/?term=LOC105286770 | 3055 | 42 | 137.48 | 1 | 3.27 | 2.38 | 8.53 | 9.07 | -0.55 | 0.55 | -0.01 0.01 |
| gene-LOC105276745 | https://www.ncbi.nlm.nih.gov/gene/?term=LOC105276745 | 3953 | 73 | 184.67 | 1 | 2.53 | 1.37 | 18.74 | 19.3 | -0.56 | 0.56 | -0.01 0.01 |
| gene-LOC105280002 | https://www.ncbi.nlm.nih.gov/gene/?term=LOC105280002 | 1940 | 28 | 144.33 | 1 | 5.15 | 3.57 | 33.09 | 33.65 | -0.57 | 0.57 | -0.01 0.01 |
| gene-LOC105278962 | https://www.ncbi.nlm.nih.gov/gene/?term=LOC105278962 | 4491 | 88 | 195.95 | 1 | 2.23 | 1.14 | 23.63 | 24.21 | -0.58 | 0.58 | -0.01 0.01 |
| gene-LOC105276527 | https://www.ncbi.nlm.nih.gov/gene/?term=LOC105276527 | 7923 | 345 | 435.44 | 1 | 1.26 | 0.29 | 8.28 | 8.94 | -0.66 | 0.66 | -0.03 0.03 |
| gene-LOC105277903 | https://www.ncbi.nlm.nih.gov/gene/?term=LOC105277903 | 1835 | 35 | 190.74 | 1 | 5.45 | 2.86 | 51.45 | 52.14 | -0.69 | 0.69 | -0.01 0.01 |
| gene-LOC105280641 | https://www.ncbi.nlm.nih.gov/gene/?term=LOC105280641 | 3682 | 62 | 168.39 | 1 | 2.72 | 1.61 | 16.36 | 17.05 | -0.69 | 0.69 | -0.01 0.01 |
| gene-LOC105285639 | https://www.ncbi.nlm.nih.gov/gene/?term=LOC105285639 | 10325 | 314 | 304.12 | 1 | 0.97 | 0.32 | 12 | 12.71 | -0.71 | 0.71 | -0.02 0.02 |
| gene-LOC105275190 | https://www.ncbi.nlm.nih.gov/gene/?term=LOC105275190 | 3837 | 72 | 187.65 | 1 | 2.61 | 1.39 | 18.78 | 19.54 | -0.76 | 0.76 | -0.01 0.01 |
| gene-LOC105285742 | https://www.ncbi.nlm.nih.gov/gene/?term=LOC105285742 | 3446 | 95 | 275.68 | 1 | 2.9 | 1.05 | 25.7 | 26.49 | -0.79 | 0.79 | -0.02 0.02 |
| gene-LOC105283728 | https://www.ncbi.nlm.nih.gov/gene/?term=LOC105283728 | 3256 | 111 | 340.91 | 1 | 3.07 | 0.9 | 20.22 | 21.04 | -0.81 | 0.81 | -0.03 0.03 |
| gene-LOC105275683 | https://www.ncbi.nlm.nih.gov/gene/?term=LOC105275683 | 4256 | 204 | 479.32 | 1 | 2.35 | 0.49 | 12.14 | 13.06 | -0.92 | 0.92 | -0.04 0.04 |
| gene-LOC105288066 | https://www.ncbi.nlm.nih.gov/gene/?term=LOC105288066 | 1964 | 25 | 127.29 | 1 | 5.09 | 4 | 27.34 | 28.29 | -0.95 | 0.95 | -0.01 0.01 |
| gene-LOC105283592 | https://www.ncbi.nlm.nih.gov/gene/?term=LOC105283592 | 3479 | 71 | 204.08 | 1 | 2.87 | 1.41 | 24.39 | 25.37 | -0.98 | 0.98 | -0.02 0.02 |
| gene-LOC105285671 | https://www.ncbi.nlm.nih.gov/gene/?term=LOC105285671 | 2263 | 95 | 419.8 1 | 4.42 | 1.05 | 27.86 | 28.85 | -1 | 1 | -0.04 | 0.04 |
| gene-LOC105277216 | https://www.ncbi.nlm.nih.gov/gene/?term=LOC105277216 | 7627 | 105 | 137.67 | 1 | 1.31 | 0.95 | 5.72 | 6.75 | -1.03 | 1.03 | -0.01 0.01 |
| gene-LOC105277631 | https://www.ncbi.nlm.nih.gov/gene/?term=LOC105277631 | 4387 | 285 | 649.65 | 1 | 2.28 | 0.35 | 9.88 | 10.92 | -1.04 | 1.04 | -0.07 0.07 |
| gene-LOC105281700 | https://www.ncbi.nlm.nih.gov/gene/?term=LOC105281700 | 2208 | 44 | 199.28 | 1 | 4.53 | 2.27 | 37.54 | 38.62 | -1.08 | 1.08 | -0.02 0.02 |
| gene-LOC105285060 | https://www.ncbi.nlm.nih.gov/gene/?term=LOC105285060 | 1285 | 16 | 124.51 | 1 | 7.78 | 6.25 | 50.64 | 51.74 | -1.1 | 1.1 | -0.01 0.01 |
| gene-LOC105275363 | https://www.ncbi.nlm.nih.gov/gene/?term=LOC105275363 | 3798 | 155 | 408.11 | 1 | 2.63 | 0.65 | 17.26 | 18.39 | -1.13 | 1.13 | -0.05 0.05 |
| gene-LOC105276704 | https://www.ncbi.nlm.nih.gov/gene/?term=LOC105276704 | 2063 | 84 | 407.17 | 2 | 9.69 | 2.38 | 12.84 | 14.02 | -1.18 | 1.18 | -0.05 0.05 |
| gene-LOC105275384 | https://www.ncbi.nlm.nih.gov/gene/?term=LOC105275384 | 1987 | 75 | 377.45 | 1 | 5.03 | 1.33 | 33.41 | 34.62 | -1.21 | 1.21 | -0.05 0.05 |
| gene-LOC105275460 | https://www.ncbi.nlm.nih.gov/gene/?term=LOC105275460 | 2473 | 116 | 469.07 | 1 | 4.04 | 0.86 | 12.57 | 13.81 | -1.24 | 1.24 | -0.06 0.06 |
| gene-LOC105285055 | https://www.ncbi.nlm.nih.gov/gene/?term=LOC105285055 | 4108 | 214 | 520.93 | 1 | 2.43 | 0.47 | 12.94 | 14.18 | -1.24 | 1.24 | -0.06 0.06 |
| gene-LOC105287417 | https://www.ncbi.nlm.nih.gov/gene/?term=LOC105287417 | 1025 | 33 | 321.95 | 1 | 9.76 | 3.03 | 36 | 37.27 | -1.27 | 1.27 | -0.04 0.04 |
| gene-LOC105282156 | https://www.ncbi.nlm.nih.gov/gene/?term=LOC105282156 | 2577 | 133 | 516.1 1 | 3.88 | 0.75 | 23.61 | 24.92 | -1.31 | 1.31 | -0.07 | 0.07 |
| gene-LOC105288036 | https://www.ncbi.nlm.nih.gov/gene/?term=LOC105288036 | 5110 | 198 | 387.48 | 1 | 1.96 | 0.51 | 16.95 | 18.26 | -1.31 | 1.31 | -0.05 0.05 |
| gene-LOC105283712 | https://www.ncbi.nlm.nih.gov/gene/?term=LOC105283712 | 1910 | 29 | 151.83 | 1 | 5.24 | 3.45 | 29.98 | 31.34 | -1.36 | 1.36 | -0.02 0.02 |
| gene-LOC105278260 | https://www.ncbi.nlm.nih.gov/gene/?term=LOC105278260 | 3240 | 42 | 129.63 | 1 | 3.09 | 2.38 | 12.8 | 14.28 | -1.47 | 1.47 | -0.02 0.02 |
| gene-LOC105285695 | https://www.ncbi.nlm.nih.gov/gene/?term=LOC105285695 | 1371 | 62 | 452.22 | 1 | 7.29 | 1.61 | 23 | 24.51 | -1.51 | 1.51 | -0.07 0.07 |
| gene-LOC105281515 | https://www.ncbi.nlm.nih.gov/gene/?term=LOC105281515 | 8827 | 39 | 44.18 1 | 1.13 | 2.56 | 34.44 | 35.97 | -1.53 | 1.53 | -0.01 | 0.01 |
| gene-LOC105278807 | https://www.ncbi.nlm.nih.gov/gene/?term=LOC105278807 | 2166 | 71 | 327.79 | 1 | 4.62 | 1.41 | 18.52 | 20.08 | -1.56 | 1.56 | -0.05 0.05 |
| gene-LOC105283943 | https://www.ncbi.nlm.nih.gov/gene/?term=LOC105283943 | 3559 | 59 | 165.78 | 1 | 2.81 | 1.69 | 20.25 | 21.81 | -1.56 | 1.56 | -0.03 0.03 |
| gene-LOC105286452 | https://www.ncbi.nlm.nih.gov/gene/?term=LOC105286452 | 6888 | 58 | 84.2 1 | 1.45 | 1.72 | 31.25 | 32.98 | -1.73 | 1.73 | -0.01 | 0.01 |
| gene-LOC105285674 | https://www.ncbi.nlm.nih.gov/gene/?term=LOC105285674 | 3035 | 91 | 299.84 | 2 | 6.59 | 2.2 | 23.1 | 24.91 | -1.81 | 1.81 | -0.05 0.05 |
| gene-LOC105282399 | https://www.ncbi.nlm.nih.gov/gene/?term=LOC105282399 | 2682 | 21 | 78.3 1 | 3.73 | 4.76 | 34.08 | 35.97 | -1.89 | 1.89 | -0.01 | 0.01 |
| gene-LOC105279356 | https://www.ncbi.nlm.nih.gov/gene/?term=LOC105279356 | 1314 | 42 | 319.63 | 1 | 7.61 | 2.38 | 28.83 | 30.95 | -2.12 | 2.12 | -0.07 0.07 |
| gene-LOC105280348 | https://www.ncbi.nlm.nih.gov/gene/?term=LOC105280348 | 3797 | 51 | 134.32 | 1 | 2.63 | 1.96 | 25.95 | 28.29 | -2.34 | 2.34 | -0.03 0.03 |
| gene-LOC105274900 | https://www.ncbi.nlm.nih.gov/gene/?term=LOC105274900 | 5561 | 56 | 100.7 2 | 3.6 | 3.57 | 26.85 | 29.47 | -2.62 | 2.62 | -0.03 | 0.03 |
| gene-LOC105283989 | https://www.ncbi.nlm.nih.gov/gene/?term=LOC105283989 | 4948 | 33 | 66.69 2 | 4.04 | 6.06 | 45.77 | 48.89 | -3.12 | 3.12 | -0.02 | 0.02 |
| gene-LOC105281848 | https://www.ncbi.nlm.nih.gov/gene/?term=LOC105281848 | 2122 | 47 | 221.49 | 1 | 4.71 | 2.13 | 22.74 | 25.94 | -3.2 | 3.2 | -0.07 0.07 |
| gene-LOC105280419 | https://www.ncbi.nlm.nih.gov/gene/?term=LOC105280419 | 1601 | 10 | 62.46 1 | 6.25 | 10 | 33.98 | 37.47 | -3.49 | 3.49 | -0.02 | 0.02 |
| gene-LOC105275518 | https://www.ncbi.nlm.nih.gov/gene/?term=LOC105275518 | 2335 | 20 | 85.65 1 | 4.28 | 5 | 47.29 | 50.84 | -3.55 | 3.55 | -0.03 | 0.03 |
| gene-LOC105277215 | https://www.ncbi.nlm.nih.gov/gene/?term=LOC105277215 | 2619 | 28 | 106.91 | 1 | 3.82 | 3.57 | 20.04 | 24.53 | -4.48 | 4.48 | -0.05 0.05 |
| gene-LOC105283552 | https://www.ncbi.nlm.nih.gov/gene/?term=LOC105283552 | 1562 | 33 | 211.27 | 1 | 6.4 | 3.03 | 54.5 | 59.97 | -5.47 | 5.47 | -0.12 0.12 |
| gene-LOC105279898 | https://www.ncbi.nlm.nih.gov/gene/?term=LOC105279898 | 1343 | 16 | 119.14 | 1 | 7.45 | 6.25 | 59.42 | 65.83 | -6.42 | 6.42 | -0.08 0.08 |
| gene-LOC105275708 | https://www.ncbi.nlm.nih.gov/gene/?term=LOC105275708 | 1772 | 13 | 73.36 1 | 5.64 | 7.69 | 46.34 | 53.52 | -7.18 | 7.18 | -0.05 | 0.05 |
| gene-LOC105285514 | https://www.ncbi.nlm.nih.gov/gene/?term=LOC105285514 | 1277 | 9 | 70.48 1 | 7.83 | 11.11 | 73.21 | 80.81 | -7.6 | 7.6 | -0.05 | 0.05 |
| gene-LOC105275070 | https://www.ncbi.nlm.nih.gov/gene/?term=LOC105275070 | 5667 | 36 | 63.53 2 | 3.53 | 5.56 | 3.69 | 13.97 | -10.28 | | 10.28 | -0.07 0.07 |
| gene-LOC113562345 | https://www.ncbi.nlm.nih.gov/gene/?term=LOC113562345 | 2290 | 62 | 270.74 | 14 | 61.14 | 22.58 | 17.91 | 32.35 | -14.44 | | 14.44 -0.39 0.39 |

Directory: OGL File: ogl-Ob_LI_BCphaseA1.vs.BCphaseB2.txt

| gene_ID | gene_link | gwidth | #Sites | #per10Kb | #dmSites | #dmsp10kb | %dmSites | %pSite1 | %pSite2 | DMpSite | ADMpSite | DMpNucl | ADMpNucl |
|---|---|---|---|---|---|---|---|---|---|---|---|---|---|
| gene-LOC105275343 | https://www.ncbi.nlm.nih.gov/gene/?term=LOC105275343 | 1746 | 4 | 22.91 | 2 | 11.45 | 50 | 47 | 32.01 | 14.99 | 14.99 | 0.03 | 0.03 |
| gene-LOC105280261 | https://www.ncbi.nlm.nih.gov/gene/?term=LOC105280261 | 1005 | 4 | 39.8 | 1 | 9.95 | 25 | 82.16 | 67.59 | 14.57 | 14.57 | 0.06 | 0.06 |
| gene-LOC113562975 | https://www.ncbi.nlm.nih.gov/gene/?term=LOC113562975 | 2751 | 3 | 10.91 | 1 | 3.64 | 33.33 | 46.98 | 32.86 | 14.12 | 14.12 | 0.02 | 0.02 |
| gene-LOC105278458 | https://www.ncbi.nlm.nih.gov/gene/?term=LOC105278458 | 216 | 3 | 138.89 | 1 | 46.3 | 33.33 | 20.87 | 10.63 | 10.24 | 10.24 | 0.14 | 0.14 |
| gene-LOC105281691 | https://www.ncbi.nlm.nih.gov/gene/?term=LOC105281691 | 2096 | 9 | 42.94 | 2 | 9.54 | 22.22 | 17 | 7.56 | 9.44 | 9.44 | 0.04 | 0.04 |
| gene-LOC105280926 | https://www.ncbi.nlm.nih.gov/gene/?term=LOC105280926 | 2139 | 6 | 28.05 | 2 | 9.35 | 33.33 | 47.28 | 37.87 | 9.41 | 9.41 | 0.03 | 0.03 |
| gene-LOC105287814 | https://www.ncbi.nlm.nih.gov/gene/?term=LOC105287814 | 892 | 7 | 78.48 | 1 | 11.21 | 14.29 | 37.51 | 29.17 | 8.34 | 8.34 | 0.07 | 0.07 |
| gene-LOC105283830 | https://www.ncbi.nlm.nih.gov/gene/?term=LOC105283830 | 252 | 13 | 515.87 | 1 | 39.68 | 7.69 | 42.02 | 33.83 | 8.19 | 8.19 | 0.42 | 0.42 |
| gene-LOC105285885 | https://www.ncbi.nlm.nih.gov/gene/?term=LOC105285885 | 1327 | 12 | 90.43 | 3 | 22.61 | 25 | 58.28 | 50.25 | 8.03 | 8.03 | 0.07 | 0.07 |
| gene-LOC105283963 | https://www.ncbi.nlm.nih.gov/gene/?term=LOC105283963 | 1330 | 8 | 60.15 | 1 | 7.52 | 12.5 | 51.82 | 43.92 | 7.9 | 7.9 | 0.05 | 0.05 |
| gene-LOC105275978 | https://www.ncbi.nlm.nih.gov/gene/?term=LOC105275978 | 3932 | 6 | 15.26 | 1 | 2.54 | 16.67 | 16.55 | 9.1 | 7.45 | 7.45 | 0.01 | 0.01 |
| gene-LOC105281507 | https://www.ncbi.nlm.nih.gov/gene/?term=LOC105281507 | 881 | 5 | 56.75 | 1 | 11.35 | 20 | 11.82 | 4.8 | 7.02 | 7.02 | 0.04 | 0.04 |
| gene-LOC105282405 | https://www.ncbi.nlm.nih.gov/gene/?term=LOC105282405 | 1310 | 31 | 236.64 | 3 | 22.9 | 9.68 | 51.18 | 44.58 | 6.6 | 6.6 | 0.16 | 0.16 |
| gene-LOC105280917 | https://www.ncbi.nlm.nih.gov/gene/?term=LOC105280917 | 2193 | 6 | 27.36 | 1 | 4.56 | 16.67 | 14.86 | 8.55 | 6.31 | 6.31 | 0.02 | 0.02 |
| gene-LOC105276706 | https://www.ncbi.nlm.nih.gov/gene/?term=LOC105276706 | 1440 | 16 | 111.11 | 1 | 6.94 | 6.25 | 82.64 | 76.36 | 6.28 | 6.28 | 0.07 | 0.07 |
| gene-LOC105281831 | https://www.ncbi.nlm.nih.gov/gene/?term=LOC105281831 | 1068 | 8 | 74.91 | 1 | 9.36 | 12.5 | 87.28 | 81.3 | 5.98 | 5.98 | 0.04 | 0.04 |
| gene-LOC113563560 | https://www.ncbi.nlm.nih.gov/gene/?term=LOC113563560 | 4945 | 5 | 10.11 | 1 | 2.02 | 20 | 86.62 | 80.66 | 5.96 | 5.96 | 0.01 | 0.01 |
| gene-LOC105283813 | https://www.ncbi.nlm.nih.gov/gene/?term=LOC105283813 | 1312 | 8 | 60.98 | 1 | 7.62 | 12.5 | 29.71 | 24.06 | 5.65 | 5.65 | 0.03 | 0.03 |
| gene-LOC105285051 | https://www.ncbi.nlm.nih.gov/gene/?term=LOC105285051 | 1357 | 11 | 81.06 | 3 | 22.11 | 27.27 | 46.19 | 40.59 | 5.6 | 5.6 | 0.05 | 0.05 |
| gene-LOC105278243 | https://www.ncbi.nlm.nih.gov/gene/?term=LOC105278243 | 1826 | 18 | 98.58 | 1 | 5.48 | 5.56 | 20.63 | 15.05 | 5.58 | 5.58 | 0.06 | 0.06 |
| gene-LOC105280891 | https://www.ncbi.nlm.nih.gov/gene/?term=LOC105280891 | 1844 | 26 | 141 | 4 | 21.69 | 15.38 | 25.44 | 20.09 | 5.35 | 5.35 | 0.08 | 0.08 |
| gene-LOC105284483 | https://www.ncbi.nlm.nih.gov/gene/?term=LOC105284483 | 2910 | 25 | 85.91 | 1 | 3.44 | 4 | 22.04 | 16.8 | 5.24 | 5.24 | 0.05 | 0.05 |
| gene-LOC105281226 | https://www.ncbi.nlm.nih.gov/gene/?term=LOC105281226 | 1308 | 6 | 45.87 | 1 | 7.65 | 16.67 | 62.79 | 57.58 | 5.21 | 5.21 | 0.02 | 0.02 |
| gene-LOC105280609 | https://www.ncbi.nlm.nih.gov/gene/?term=LOC105280609 | 1563 | 17 | 108.77 | 2 | 12.8 | 11.76 | 46.92 | 41.73 | 5.18 | 5.18 | 0.06 | 0.06 |

...
... only first 25 lines shown ...

Directory: OGL File: ogl-Ob_LI_BCphaseA1.vs.RphaseA1.txt

| gene_ID | gene_link | gwidth | #Sites | #per10Kb | #dmSites | #dmsp10kb | %dmSites | %pSite1 | %pSite2 | DMpSite | ADMpSite | DMpNucl | ADMpNucl |
|---|---|---|---|---|---|---|---|---|---|---|---|---|---|
| gene-LOC105286451 | https://www.ncbi.nlm.nih.gov/gene/?term=LOC105286451 | 1618 | 10 | 61.8 | 1 | 6.18 | 10 | 52.19 | 44.01 | 8.18 | 8.18 | 0.05 | 0.05 |
| gene-LOC105281173 | https://www.ncbi.nlm.nih.gov/gene/?term=LOC105281173 | 1595 | 25 | 156.74 | 1 | 6.27 | 4 | 64.76 | 61.03 | 3.73 | 3.73 | 0.06 | 0.06 |
| gene-LOC105276538 | https://www.ncbi.nlm.nih.gov/gene/?term=LOC105276538 | 1511 | 18 | 119.13 | 1 | 6.62 | 5.56 | 71.16 | 68.49 | 2.67 | 2.67 | 0.03 | 0.03 |
| gene-LOC105274449 | https://www.ncbi.nlm.nih.gov/gene/?term=LOC105274449 | 1549 | 43 | 277.6 | 1 | 6.46 | 2.33 | 54.59 | 52.01 | 2.58 | 2.58 | 0.07 | 0.07 |
| gene-LOC105284396 | https://www.ncbi.nlm.nih.gov/gene/?term=LOC105284396 | 1976 | 57 | 288.46 | 1 | 5.06 | 1.75 | 39.14 | 36.68 | 2.46 | 2.46 | 0.07 | 0.07 |
| gene-LOC105274706 | https://www.ncbi.nlm.nih.gov/gene/?term=LOC105274706 | 1446 | 48 | 331.95 | 1 | 6.92 | 2.08 | 39.02 | 37.1 | 1.92 | 1.92 | 0.06 | 0.06 |
| gene-LOC105285409 | https://www.ncbi.nlm.nih.gov/gene/?term=LOC105285409 | 5099 | 47 | 92.17 | 1 | 1.96 | 2.13 | 3.57 | 1.67 | 1.9 | 1.9 | 0.02 | 0.02 |
| gene-LOC105278877 | https://www.ncbi.nlm.nih.gov/gene/?term=LOC105278877 | 1968 | 78 | 396.34 | 1 | 5.08 | 1.28 | 32.36 | 30.87 | 1.48 | 1.48 | 0.06 | 0.06 |
| gene-LOC105275518 | https://www.ncbi.nlm.nih.gov/gene/?term=LOC105275518 | 2335 | 19 | 81.37 | 1 | 4.28 | 5.26 | 49.77 | 48.32 | 1.45 | 1.45 | 0.01 | 0.01 |
| gene-LOC105278538 | https://www.ncbi.nlm.nih.gov/gene/?term=LOC105278538 | 2150 | 75 | 348.84 | 1 | 4.65 | 1.33 | 18.53 | 17.09 | 1.44 | 1.44 | 0.05 | 0.05 |
| gene-LOC105278454 | https://www.ncbi.nlm.nih.gov/gene/?term=LOC105278454 | 1972 | 37 | 187.63 | 1 | 5.07 | 2.7 | 46.8 | 45.42 | 1.38 | 1.38 | 0.03 | 0.03 |
| gene-LOC105283649 | https://www.ncbi.nlm.nih.gov/gene/?term=LOC105283649 | 1918 | 53 | 276.33 | 2 | 10.43 | 3.77 | 38.29 | 37.03 | 1.27 | 1.27 | 0.04 | 0.04 |
| gene-LOC105279664 | https://www.ncbi.nlm.nih.gov/gene/?term=LOC105279664 | 4023 | 63 | 156.6 | 1 | 2.49 | 1.59 | 17.93 | 16.77 | 1.16 | 1.16 | 0.02 | 0.02 |
| gene-LOC105285202 | https://www.ncbi.nlm.nih.gov/gene/?term=LOC105285202 | 1838 | 35 | 190.42 | 1 | 5.44 | 2.86 | 19.49 | 18.4 | 1.08 | 1.08 | 0.02 | 0.02 |
| gene-LOC105276196 | https://www.ncbi.nlm.nih.gov/gene/?term=LOC105276196 | 2953 | 55 | 186.25 | 1 | 3.39 | 1.82 | 40.47 | 39.41 | 1.06 | 1.06 | 0.02 | 0.02 |
| gene-LOC105287095 | https://www.ncbi.nlm.nih.gov/gene/?term=LOC105287095 | 1435 | 60 | 418.12 | 1 | 6.97 | 1.67 | 26.49 | 25.42 | 1.06 | 1.06 | 0.04 | 0.04 |
| gene-LOC105282811 | https://www.ncbi.nlm.nih.gov/gene/?term=LOC105282811 | 5765 | 183 | 317.43 | 1 | 1.73 | 0.55 | 10.19 | 9.27 | 0.92 | 0.92 | 0.03 | 0.03 |
| gene-LOC105280786 | https://www.ncbi.nlm.nih.gov/gene/?term=LOC105280786 | 2573 | 115 | 446.95 | 1 | 3.89 | 0.87 | 29.1 | 28.31 | 0.79 | 0.79 | 0.04 | 0.04 |
| gene-LOC105281047 | https://www.ncbi.nlm.nih.gov/gene/?term=LOC105281047 | 7313 | 112 | 153.15 | 1 | 1.37 | 0.89 | 11.03 | 10.28 | 0.75 | 0.75 | 0.01 | 0.01 |
| gene-LOC105285807 | https://www.ncbi.nlm.nih.gov/gene/?term=LOC105285807 | 6063 | 180 | 296.88 | 1 | 1.65 | 0.56 | 10.12 | 9.41 | 0.72 | 0.72 | 0.02 | 0.02 |
| gene-LOC105282794 | https://www.ncbi.nlm.nih.gov/gene/?term=LOC105282794 | 4571 | 93 | 203.46 | 1 | 2.19 | 1.08 | 17.63 | 17.04 | 0.6 | 0.6 | 0.01 | 0.01 |
| gene-LOC105283123 | https://www.ncbi.nlm.nih.gov/gene/?term=LOC105283123 | 1507 | 35 | 232.25 | 1 | 6.64 | 2.86 | 18.15 | 17.56 | 0.59 | 0.59 | 0.01 | 0.01 |
| gene-LOC105275752 | https://www.ncbi.nlm.nih.gov/gene/?term=LOC105275752 | 3462 | 126 | 363.95 | 1 | 2.89 | 0.79 | 28.41 | 27.85 | 0.56 | 0.56 | 0.02 | 0.02 |
| gene-LOC105284116 | https://www.ncbi.nlm.nih.gov/gene/?term=LOC105284116 | 3970 | 90 | 226.7 | 1 | 2.52 | 1.11 | 7.07 | 6.57 | 0.5 | 0.5 | 0.01 | 0.01 |
| gene-LOC105279038 | https://www.ncbi.nlm.nih.gov/gene/?term=LOC105279038 | 9453 | 299 | 316.3 | 1 | 1.06 | 0.33 | 8.97 | 8.54 | 0.42 | 0.42 | 0.01 | 0.01 |
| gene-LOC105278666 | https://www.ncbi.nlm.nih.gov/gene/?term=LOC105278666 | 10341 | 393 | 380.04 | 1 | 0.97 | 0.25 | 7.76 | 7.36 | 0.4 | 0.4 | 0.02 | 0.02 |
| gene-LOC105280002 | https://www.ncbi.nlm.nih.gov/gene/?term=LOC105280002 | 1940 | 28 | 144.33 | 1 | 5.15 | 3.57 | 33.09 | 32.74 | 0.35 | 0.35 | 0.01 | 0.01 |
| gene-LOC105284117 | https://www.ncbi.nlm.nih.gov/gene/?term=LOC105284117 | 4340 | 82 | 188.94 | 1 | 2.3 | 1.22 | 10.79 | 10.47 | 0.33 | 0.33 | 0.01 | 0.01 |
| gene-LOC105274590 | https://www.ncbi.nlm.nih.gov/gene/?term=LOC105274590 | 4516 | 75 | 166.08 | 1 | 2.21 | 1.33 | 23.2 | 22.91 | 0.29 | 0.29 | 0 | 0 |
| gene-LOC105280819 | https://www.ncbi.nlm.nih.gov/gene/?term=LOC105280819 | 1639 | 41 | 250.15 | 1 | 6.1 | 2.44 | 40.58 | 40.31 | 0.27 | 0.27 | 0.01 | 0.01 |
| gene-LOC105288079 | https://www.ncbi.nlm.nih.gov/gene/?term=LOC105288079 | 3473 | 85 | 244.75 | 1 | 2.88 | 1.18 | 20.86 | 20.61 | 0.25 | 0.25 | 0.01 | 0.01 |
| gene-LOC105276901 | https://www.ncbi.nlm.nih.gov/gene/?term=LOC105276901 | 3254 | 96 | 295.02 | 1 | 3.07 | 1.04 | 22.83 | 22.72 | 0.1 | 0.1 | 0 | 0 |
| gene-LOC105278228 | https://www.ncbi.nlm.nih.gov/gene/?term=LOC105278228 | 8610 | 152 | 176.54 | 1 | 1.16 | 0.66 | 23.77 | 23.73 | 0.04 | 0.04 | 0 | 0 |
| gene-LOC105283728 | https://www.ncbi.nlm.nih.gov/gene/?term=LOC105283728 | 3256 | 113 | 347.05 | 2 | 6.14 | 1.77 | 19.87 | 19.82 | 0.04 | 0.04 | 0 | 0 |
| gene-LOC105278145 | https://www.ncbi.nlm.nih.gov/gene/?term=LOC105278145 | 4470 | 239 | 534.68 | 1 | 2.24 | 0.42 | 10.87 | 10.84 | 0.03 | 0.03 | 0 | 0 |
| gene-LOC105275327 | https://www.ncbi.nlm.nih.gov/gene/?term=LOC105275327 | 14245 | 461 | 323.62 | 1 | 0.7 | 0.22 | 7.38 | 7.42 | -0.03 | 0.03 | 0 | 0 |
| gene-LOC105286274 | https://www.ncbi.nlm.nih.gov/gene/?term=LOC105286274 | 3817 | 174 | 455.86 | 1 | 2.62 | 0.57 | 15.59 | 15.67 | -0.08 | 0.08 | 0 | 0 |
| gene-LOC105274856 | https://www.ncbi.nlm.nih.gov/gene/?term=LOC105274856 | 5216 | 201 | 385.35 | 1 | 1.92 | 0.5 | 17.77 | 17.88 | -0.11 | 0.11 | 0 | 0 |
| gene-LOC105278534 | https://www.ncbi.nlm.nih.gov/gene/?term=LOC105278534 | 7364 | 173 | 234.93 | 1 | 1.36 | 0.58 | 18.44 | 18.59 | -0.16 | 0.16 | 0 | 0 |
| gene-LOC105274851 | https://www.ncbi.nlm.nih.gov/gene/?term=LOC105274851 | 6616 | 133 | 201.03 | 1 | 1.51 | 0.75 | 10.15 | 10.35 | -0.2 | 0.2 | 0 | 0 |
| gene-LOC105286515 | https://www.ncbi.nlm.nih.gov/gene/?term=LOC105286515 | 3751 | 106 | 282.59 | 1 | 2.67 | 0.94 | 10.37 | 10.58 | -0.22 | 0.22 | -0.01 | 0.01 |
| gene-LOC105279500 | https://www.ncbi.nlm.nih.gov/gene/?term=LOC105279500 | 3587 | 29 | 80.85 | 1 | 2.79 | 3.45 | 48.07 | 48.34 | -0.27 | 0.27 | 0 | 0 |
| gene-LOC105285534 | https://www.ncbi.nlm.nih.gov/gene/?term=LOC105285534 | 3944 | 221 | 560.34 | 1 | 2.54 | 0.45 | 11.41 | 11.71 | -0.3 | 0.3 | -0.02 | 0.02 |
| gene-LOC105279298 | https://www.ncbi.nlm.nih.gov/gene/?term=LOC105279298 | 2773 | 135 | 486.84 | 1 | 3.61 | 0.74 | 22.57 | 22.9 | -0.33 | 0.33 | -0.02 | 0.02 |
| gene-LOC105288078 | https://www.ncbi.nlm.nih.gov/gene/?term=LOC105288078 | 19126 | 305 | 159.47 | 1 | 0.52 | 0.33 | 16.95 | 17.33 | -0.38 | 0.38 | -0.01 | 0.01 |
| gene-LOC105285406 | https://www.ncbi.nlm.nih.gov/gene/?term=LOC105285406 | 8015 | 39 | 48.66 | 1 | 1.25 | 2.56 | 33.07 | 33.49 | -0.42 | 0.42 | 0 | 0 |
| gene-LOC105274531 | https://www.ncbi.nlm.nih.gov/gene/?term=LOC105274531 | 7774 | 253 | 325.44 | 1 | 1.29 | 0.4 | 11.9 | 12.33 | -0.43 | 0.43 | -0.01 | 0.01 |
| gene-LOC105283712 | https://www.ncbi.nlm.nih.gov/gene/?term=LOC105283712 | 1910 | 29 | 151.83 | 1 | 5.24 | 3.45 | 29.98 | 30.42 | -0.44 | 0.44 | -0.01 | 0.01 |
| gene-LOC105275173 | https://www.ncbi.nlm.nih.gov/gene/?term=LOC105275173 | 4639 | 76 | 163.83 | 1 | 2.16 | 1.32 | 24.33 | 24.78 | -0.45 | 0.45 | -0.01 | 0.01 |
| gene-LOC105284184 | https://www.ncbi.nlm.nih.gov/gene/?term=LOC105284184 | 2535 | 80 | 315.58 | 1 | 3.94 | 1.25 | 34.85 | 35.33 | -0.47 | 0.47 | -0.01 | 0.01 |
| gene-LOC105286997 | https://www.ncbi.nlm.nih.gov/gene/?term=LOC105286997 | 2850 | 134 | 470.18 | 1 | 3.51 | 0.75 | 11.83 | 12.3 | -0.47 | 0.47 | -0.02 | 0.02 |
| gene-LOC105275898 | https://www.ncbi.nlm.nih.gov/gene/?term=LOC105275898 | 5358 | 132 | 246.36 | 1 | 1.87 | 0.76 | 12.84 | 13.34 | -0.5 | 0.5 | -0.01 | 0.01 |
| gene-LOC105280956 | https://www.ncbi.nlm.nih.gov/gene/?term=LOC105280956 | 2639 | 73 | 276.62 | 1 | 3.79 | 1.37 | 20.28 | 20.85 | -0.57 | 0.57 | -0.02 | 0.02 |
| gene-LOC105274534 | https://www.ncbi.nlm.nih.gov/gene/?term=LOC105274534 | 1329 | 57 | 428.89 | 1 | 7.52 | 1.75 | 43.75 | 44.38 | -0.63 | 0.63 | -0.03 | 0.03 |
| gene-LOC105275683 | https://www.ncbi.nlm.nih.gov/gene/?term=LOC105275683 | 4256 | 187 | 439.38 | 1 | 2.35 | 0.53 | 12.61 | 13.25 | -0.63 | 0.63 | -0.03 | 0.03 |
| gene-LOC105278497 | https://www.ncbi.nlm.nih.gov/gene/?term=LOC105278497 | 2072 | 97 | 468.15 | 1 | 4.83 | 1.03 | 18.56 | 19.21 | -0.64 | 0.64 | -0.03 | 0.03 |
| gene-LOC105276692 | https://www.ncbi.nlm.nih.gov/gene/?term=LOC105276692 | 1745 | 34 | 194.84 | 1 | 5.73 | 2.94 | 24.51 | 25.17 | -0.66 | 0.66 | -0.01 | 0.01 |
| gene-LOC105280641 | https://www.ncbi.nlm.nih.gov/gene/?term=LOC105280641 | 3682 | 55 | 149.38 | 1 | 2.72 | 1.82 | 18.35 | 19.01 | -0.66 | 0.66 | -0.01 | 0.01 |
| gene-LOC105285639 | https://www.ncbi.nlm.nih.gov/gene/?term=LOC105285639 | 10325 | 294 | 284.75 | 1 | 0.97 | 0.34 | 12.44 | 13.1 | -0.66 | 0.66 | -0.02 | 0.02 |
| gene-LOC105281700 | https://www.ncbi.nlm.nih.gov/gene/?term=LOC105281700 | 2208 | 45 | 203.8 | 1 | 4.53 | 2.22 | 38.83 | 39.5 | -0.67 | 0.67 | -0.01 | 0.01 |

| | | | | | | | | | | | | | |
|---|---|---|---|---|---|---|---|---|---|---|---|---|---|
| gene-LOC105288036 | https://www.ncbi.nlm.nih.gov/gene/?term=LOC105288036 | 5110 | 182 | 356.16 | 1 | 1.96 | 0.55 | 18.19 | 18.89 | -0.7 | 0.7 | -0.03 | 0.03 |
| gene-LOC105287063 | https://www.ncbi.nlm.nih.gov/gene/?term=LOC105287063 | 1874 | 46 | 245.46 | 1 | 5.34 | 2.17 | 34.15 | 34.88 | -0.73 | 0.73 | -0.02 | 0.02 |
| gene-LOC105279657 | https://www.ncbi.nlm.nih.gov/gene/?term=LOC105279657 | 5878 | 82 | 139.5 | 1 | 1.7 | 1.22 | 19.15 | 19.93 | -0.78 | 0.78 | -0.01 | 0.01 |
| gene-LOC105277856 | https://www.ncbi.nlm.nih.gov/gene/?term=LOC105277856 | 3442 | 28 | 81.35 | 1 | 2.91 | 3.57 | 36.24 | 37.06 | -0.82 | 0.82 | -0.01 | 0.01 |
| gene-LOC105275179 | https://www.ncbi.nlm.nih.gov/gene/?term=LOC105275179 | 3271 | 69 | 210.94 | 1 | 3.06 | 1.45 | 14.66 | 15.53 | -0.86 | 0.86 | -0.02 | 0.02 |
| gene-LOC105288237 | https://www.ncbi.nlm.nih.gov/gene/?term=LOC105288237 | 2226 | 45 | 202.16 | 2 | 8.98 | 4.44 | 36.5 | 37.4 | -0.9 | 0.9 | -0.02 | 0.02 |
| gene-LOC105275451 | https://www.ncbi.nlm.nih.gov/gene/?term=LOC105275451 | 6035 | 111 | 183.93 | 1 | 1.66 | 0.9 | 22.31 | 23.24 | -0.94 | 0.94 | -0.02 | 0.02 |
| gene-LOC105286265 | https://www.ncbi.nlm.nih.gov/gene/?term=LOC105286265 | 2507 | 65 | 259.27 | 1 | 3.99 | 1.54 | 31.58 | 32.53 | -0.95 | 0.95 | -0.02 | 0.02 |
| gene-LOC105275363 | https://www.ncbi.nlm.nih.gov/gene/?term=LOC105275363 | 3798 | 155 | 408.11 | 1 | 2.63 | 0.65 | 17.28 | 18.35 | -1.06 | 1.06 | -0.04 | 0.04 |
| gene-LOC105282156 | https://www.ncbi.nlm.nih.gov/gene/?term=LOC105282156 | 2577 | 128 | 496.7 | 1 | 3.88 | 0.78 | 23.38 | 24.45 | -1.07 | 1.07 | -0.05 | 0.05 |
| gene-LOC105282991 | https://www.ncbi.nlm.nih.gov/gene/?term=LOC105282991 | 5617 | 80 | 142.42 | 1 | 1.78 | 1.25 | 19.15 | 20.24 | -1.09 | 1.09 | -0.02 | 0.02 |
| gene-LOC105283943 | https://www.ncbi.nlm.nih.gov/gene/?term=LOC105283943 | 3559 | 49 | 137.68 | 1 | 2.81 | 2.04 | 22.13 | 23.23 | -1.1 | 1.1 | -0.02 | 0.02 |
| gene-LOC105285055 | https://www.ncbi.nlm.nih.gov/gene/?term=LOC105285055 | 4108 | 204 | 496.59 | 1 | 2.43 | 0.49 | 12.43 | 13.67 | -1.24 | 1.24 | -0.06 | 0.06 |
| gene-LOC105277216 | https://www.ncbi.nlm.nih.gov/gene/?term=LOC105277216 | 7627 | 91 | 119.31 | 1 | 1.31 | 1.1 | 6.55 | 7.88 | -1.33 | 1.33 | -0.02 | 0.02 |
| gene-LOC105286452 | https://www.ncbi.nlm.nih.gov/gene/?term=LOC105286452 | 6888 | 58 | 84.2 | 1 | 1.45 | 1.72 | 31.25 | 32.65 | -1.4 | 1.4 | -0.01 | 0.01 |
| gene-LOC105285695 | https://www.ncbi.nlm.nih.gov/gene/?term=LOC105285695 | 1371 | 61 | 444.93 | 1 | 7.29 | 1.64 | 22.52 | 23.97 | -1.45 | 1.45 | -0.06 | 0.06 |
| gene-LOC105283592 | https://www.ncbi.nlm.nih.gov/gene/?term=LOC105283592 | 3479 | 68 | 195.46 | 1 | 2.87 | 1.47 | 23.52 | 25 | -1.48 | 1.48 | -0.03 | 0.03 |
| gene-LOC105275190 | https://www.ncbi.nlm.nih.gov/gene/?term=LOC105275190 | 3837 | 65 | 169.4 | 1 | 2.61 | 1.54 | 21.02 | 22.59 | -1.58 | 1.58 | -0.03 | 0.03 |
| gene-LOC105280348 | https://www.ncbi.nlm.nih.gov/gene/?term=LOC105280348 | 3797 | 52 | 136.95 | 1 | 2.63 | 1.92 | 25.54 | 27.17 | -1.63 | 1.63 | -0.02 | 0.02 |
| gene-LOC105276704 | https://www.ncbi.nlm.nih.gov/gene/?term=LOC105276704 | 2063 | 80 | 387.78 | 1 | 4.85 | 1.25 | 13.36 | 15.07 | -1.71 | 1.71 | -0.07 | 0.07 |
| gene-LOC105275460 | https://www.ncbi.nlm.nih.gov/gene/?term=LOC105275460 | 2473 | 115 | 465.02 | 1 | 4.04 | 0.87 | 12.68 | 14.4 | -1.72 | 1.72 | -0.08 | 0.08 |
| gene-LOC105283989 | https://www.ncbi.nlm.nih.gov/gene/?term=LOC105283989 | 4948 | 30 | 60.63 | 1 | 2.02 | 3.33 | 45.76 | 47.5 | -1.74 | 1.74 | -0.01 | 0.01 |
| gene-LOC105285742 | https://www.ncbi.nlm.nih.gov/gene/?term=LOC105285742 | 3446 | 87 | 252.47 | 1 | 2.9 | 1.15 | 20.61 | 22.37 | -1.76 | 1.76 | -0.04 | 0.04 |
| gene-LOC105278436 | https://www.ncbi.nlm.nih.gov/gene/?term=LOC105278436 | 2556 | 56 | 219.09 | 1 | 3.91 | 1.79 | 36.89 | 38.91 | -2.02 | 2.02 | -0.04 | 0.04 |
| gene-LOC105278807 | https://www.ncbi.nlm.nih.gov/gene/?term=LOC105278807 | 2166 | 70 | 323.18 | 1 | 4.62 | 1.43 | 17.84 | 20.15 | -2.31 | 2.31 | -0.07 | 0.07 |
| gene-LOC105285671 | https://www.ncbi.nlm.nih.gov/gene/?term=LOC105285671 | 2263 | 90 | 397.7 | 1 | 4.42 | 1.11 | 29.29 | 31.61 | -2.32 | 2.32 | -0.09 | 0.09 |
| gene-LOC105279356 | https://www.ncbi.nlm.nih.gov/gene/?term=LOC105279356 | 1314 | 42 | 319.63 | 1 | 7.61 | 2.38 | 28.83 | 31.17 | -2.34 | 2.34 | -0.07 | 0.07 |
| gene-LOC105288066 | https://www.ncbi.nlm.nih.gov/gene/?term=LOC105288066 | 1964 | 24 | 122.2 | 1 | 5.09 | 4.17 | 27.7 | 30.29 | -2.59 | 2.59 | -0.03 | 0.03 |
| gene-LOC105285060 | https://www.ncbi.nlm.nih.gov/gene/?term=LOC105285060 | 1285 | 15 | 116.73 | 1 | 7.78 | 6.67 | 48.46 | 51.14 | -2.68 | 2.68 | -0.03 | 0.03 |
| gene-LOC105285674 | https://www.ncbi.nlm.nih.gov/gene/?term=LOC105285674 | 3035 | 88 | 289.95 | 2 | 6.59 | 2.27 | 23.51 | 26.27 | -2.76 | 2.76 | -0.08 | 0.08 |
| gene-LOC105281848 | https://www.ncbi.nlm.nih.gov/gene/?term=LOC105281848 | 2122 | 45 | 212.06 | 1 | 4.71 | 2.22 | 23.75 | 26.94 | -3.19 | 3.19 | -0.07 | 0.07 |
| gene-LOC105279898 | https://www.ncbi.nlm.nih.gov/gene/?term=LOC105279898 | 1343 | 13 | 96.8 | 1 | 7.45 | 7.69 | 65.18 | 68.44 | -3.25 | 3.25 | -0.03 | 0.03 |
| gene-LOC105283552 | https://www.ncbi.nlm.nih.gov/gene/?term=LOC105283552 | 1562 | 33 | 211.27 | 1 | 6.4 | 3.03 | 54.5 | 58 | -3.5 | 3.5 | -0.07 | 0.07 |
| gene-LOC105277903 | https://www.ncbi.nlm.nih.gov/gene/?term=LOC105277903 | 1835 | 35 | 190.74 | 1 | 5.45 | 2.86 | 51.45 | 55.03 | -3.59 | 3.59 | -0.07 | 0.07 |
| gene-LOC105274900 | https://www.ncbi.nlm.nih.gov/gene/?term=LOC105274900 | 5561 | 56 | 100.7 | 1 | 1.8 | 1.79 | 26.85 | 30.51 | -3.66 | 3.66 | -0.04 | 0.04 |
| gene-LOC105277215 | https://www.ncbi.nlm.nih.gov/gene/?term=LOC105277215 | 2619 | 27 | 103.09 | 1 | 3.82 | 3.7 | 20.79 | 25.43 | -4.64 | 4.64 | -0.05 | 0.05 |
| gene-LOC105275708 | https://www.ncbi.nlm.nih.gov/gene/?term=LOC105275708 | 1772 | 14 | 79.01 | 1 | 5.64 | 7.14 | 44.58 | 50.83 | -6.25 | 6.25 | -0.05 | 0.05 |
| gene-LOC105285514 | https://www.ncbi.nlm.nih.gov/gene/?term=LOC105285514 | 1277 | 8 | 62.65 | 1 | 7.83 | 12.5 | 70.46 | 79.23 | -8.77 | 8.77 | -0.05 | 0.05 |
| gene-LOC105275070 | https://www.ncbi.nlm.nih.gov/gene/?term=LOC105275070 | 5667 | 37 | 65.29 | 2 | 3.53 | 5.41 | 3.73 | 16.16 | -12.43 | 12.43 | -0.08 | 0.08 |
| gene-LOC113562345 | https://www.ncbi.nlm.nih.gov/gene/?term=LOC113562345 | 2290 | 61 | 266.38 | 10 | 43.67 | 16.39 | 17.52 | 31.67 | -14.16 | 14.16 | -0.38 | 0.38 |

Directory: OGL File: ogl-Ob_LI_BCphaseA1.vs.RphaseA2.txt

| gene_ID | gene_link | gwidth | #Sites | #per10Kb | #dmSites | #dmsp10kb | %dmSites | %pSite1 | %pSite2 | DMpSite | ADMpSite | DMpNucl | ADMpNucl |
|---|---|---|---|---|---|---|---|---|---|---|---|---|---|
| gene-LOC105277380 | https://www.ncbi.nlm.nih.gov/gene/?term=LOC105277380 | 1967 | 5 | 25.42 | 1 | 5.08 | 20 | 69.42 | 54.11 | 15.31 | 15.31 | 0.04 | 0.04 |
| gene-LOC105275859 | https://www.ncbi.nlm.nih.gov/gene/?term=LOC105275859 | 2331 | 25 | 107.25 | 1 | 4.29 | 4 | 53.95 | 51.3 | 2.65 | 2.65 | 0.03 | 0.03 |
| gene-LOC105285409 | https://www.ncbi.nlm.nih.gov/gene/?term=LOC105285409 | 5099 | 48 | 94.14 | 2 | 3.92 | 4.17 | 3.6 | 1.11 | 2.49 | 2.49 | 0.02 | 0.02 |
| gene-LOC105274449 | https://www.ncbi.nlm.nih.gov/gene/?term=LOC105274449 | 1549 | 42 | 271.14 | 1 | 6.46 | 2.38 | 55.88 | 53.68 | 2.2 | 2.2 | 0.06 | 0.06 |
| gene-LOC105281173 | https://www.ncbi.nlm.nih.gov/gene/?term=LOC105281173 | 1595 | 25 | 156.74 | 1 | 6.27 | 4 | 64.76 | 62.62 | 2.13 | 2.13 | 0.03 | 0.03 |
| gene-LOC105282885 | https://www.ncbi.nlm.nih.gov/gene/?term=LOC105282885 | 1870 | 42 | 224.6 | 1 | 5.35 | 2.38 | 36.63 | 34.78 | 1.84 | 1.84 | 0.04 | 0.04 |
| gene-LOC105276272 | https://www.ncbi.nlm.nih.gov/gene/?term=LOC105276272 | 1965 | 27 | 137.4 | 1 | 5.09 | 3.7 | 42.02 | 40.71 | 1.31 | 1.31 | 0.02 | 0.02 |
| gene-LOC105287095 | https://www.ncbi.nlm.nih.gov/gene/?term=LOC105287095 | 1435 | 61 | 425.09 | 1 | 6.97 | 1.64 | 26.05 | 24.77 | 1.29 | 1.29 | 0.05 | 0.05 |
| gene-LOC105285084 | https://www.ncbi.nlm.nih.gov/gene/?term=LOC105285084 | 3308 | 26 | 78.6 | 1 | 3.02 | 3.85 | 22.5 | 21.29 | 1.21 | 1.21 | 0.01 | 0.01 |
| gene-LOC105275848 | https://www.ncbi.nlm.nih.gov/gene/?term=LOC105275848 | 3718 | 130 | 349.65 | 1 | 2.69 | 0.77 | 21.27 | 20.2 | 1.07 | 1.07 | 0.04 | 0.04 |
| gene-LOC105281296 | https://www.ncbi.nlm.nih.gov/gene/?term=LOC105281296 | 2434 | 41 | 168.45 | 1 | 4.11 | 2.44 | 31.19 | 30.16 | 1.03 | 1.03 | 0.02 | 0.02 |
| gene-LOC105284507 | https://www.ncbi.nlm.nih.gov/gene/?term=LOC105284507 | 5207 | 82 | 157.48 | 1 | 1.92 | 1.22 | 21.61 | 20.58 | 1.03 | 1.03 | 0.02 | 0.02 |
| gene-LOC105285252 | https://www.ncbi.nlm.nih.gov/gene/?term=LOC105285252 | 7427 | 132 | 177.73 | 1 | 1.35 | 0.76 | 13.81 | 12.99 | 0.81 | 0.81 | 0.01 | 0.01 |
| gene-LOC105283123 | https://www.ncbi.nlm.nih.gov/gene/?term=LOC105283123 | 1507 | 43 | 285.34 | 1 | 6.64 | 2.33 | 17.05 | 16.29 | 0.76 | 0.76 | 0.02 | 0.02 |
| gene-LOC105284396 | https://www.ncbi.nlm.nih.gov/gene/?term=LOC105284396 | 1976 | 61 | 308.7 | 1 | 5.06 | 1.64 | 36.72 | 35.98 | 0.74 | 0.74 | 0.02 | 0.02 |
| gene-LOC105279664 | https://www.ncbi.nlm.nih.gov/gene/?term=LOC105279664 | 4023 | 72 | 178.97 | 1 | 2.49 | 1.39 | 17.48 | 16.74 | 0.73 | 0.73 | 0.01 | 0.01 |
| gene-LOC105278877 | https://www.ncbi.nlm.nih.gov/gene/?term=LOC105278877 | 1968 | 80 | 406.5 | 1 | 5.08 | 1.25 | 32.88 | 32.17 | 0.72 | 0.72 | 0.03 | 0.03 |
| gene-LOC105283264 | https://www.ncbi.nlm.nih.gov/gene/?term=LOC105283264 | 4005 | 88 | 219.73 | 1 | 2.5 | 1.14 | 24.81 | 24.09 | 0.72 | 0.72 | 0.02 | 0.02 |
| gene-LOC105282811 | https://www.ncbi.nlm.nih.gov/gene/?term=LOC105282811 | 5765 | 201 | 348.66 | 1 | 1.73 | 0.5 | 9.88 | 9.23 | 0.65 | 0.65 | 0.02 | 0.02 |
| gene-LOC105286515 | https://www.ncbi.nlm.nih.gov/gene/?term=LOC105286515 | 3751 | 116 | 309.25 | 1 | 2.67 | 0.86 | 11.85 | 11.25 | 0.6 | 0.6 | 0.02 | 0.02 |
| gene-LOC105280819 | https://www.ncbi.nlm.nih.gov/gene/?term=LOC105280819 | 1639 | 41 | 250.15 | 1 | 6.1 | 2.44 | 40.58 | 40.03 | 0.56 | 0.56 | 0.01 | 0.01 |
| gene-LOC105282794 | https://www.ncbi.nlm.nih.gov/gene/?term=LOC105282794 | 4571 | 98 | 214.4 | 1 | 2.19 | 1.02 | 18.44 | 17.88 | 0.56 | 0.56 | 0.01 | 0.01 |
| gene-LOC105281420 | https://www.ncbi.nlm.nih.gov/gene/?term=LOC105281420 | 3882 | 57 | 146.83 | 1 | 2.58 | 1.75 | 15.28 | 15.02 | 0.26 | 0.26 | 0 | 0 |
| gene-LOC105283649 | https://www.ncbi.nlm.nih.gov/gene/?term=LOC105283649 | 1918 | 53 | 276.33 | 2 | 10.43 | 3.77 | 38.29 | 38.03 | 0.26 | 0.26 | 0.01 | 0.01 |
| gene-LOC105286265 | https://www.ncbi.nlm.nih.gov/gene/?term=LOC105286265 | 2507 | 68 | 271.24 | 1 | 3.99 | 1.47 | 30.18 | 29.94 | 0.24 | 0.24 | 0.01 | 0.01 |
| gene-LOC105281834 | https://www.ncbi.nlm.nih.gov/gene/?term=LOC105281834 | 4711 | 204 | 433.03 | 1 | 2.12 | 0.49 | 10.43 | 10.29 | 0.14 | 0.14 | 0.01 | 0.01 |
| gene-LOC105278454 | https://www.ncbi.nlm.nih.gov/gene/?term=LOC105278454 | 1972 | 37 | 187.63 | 1 | 5.07 | 2.7 | 46.8 | 46.67 | 0.13 | 0.13 | 0 | 0 |
| gene-LOC105286649 | https://www.ncbi.nlm.nih.gov/gene/?term=LOC105286649 | 14178 | 701 | 494.43 | 1 | 0.71 | 0.14 | 1.28 | 1.15 | 0.13 | 0.13 | 0.01 | 0.01 |
| gene-LOC105276196 | https://www.ncbi.nlm.nih.gov/gene/?term=LOC105276196 | 2953 | 56 | 189.64 | 1 | 3.39 | 1.79 | 40.11 | 39.99 | 0.12 | 0.12 | 0 | 0 |
| gene-LOC105277273 | https://www.ncbi.nlm.nih.gov/gene/?term=LOC105277273 | 3711 | 151 | 406.9 | 1 | 2.69 | 0.66 | 12.76 | 12.67 | 0.09 | 0.09 | 0 | 0 |
| gene-LOC105278632 | https://www.ncbi.nlm.nih.gov/gene/?term=LOC105278632 | 3624 | 132 | 364.24 | 1 | 2.76 | 0.76 | 14.6 | 14.5 | 0.09 | 0.09 | 0 | 0 |
| gene-LOC105284116 | https://www.ncbi.nlm.nih.gov/gene/?term=LOC105284116 | 3970 | 96 | 241.81 | 1 | 2.52 | 1.04 | 6.8 | 6.72 | 0.08 | 0.08 | 0 | 0 |
| gene-LOC105278666 | https://www.ncbi.nlm.nih.gov/gene/?term=LOC105278666 | 10341 | 409 | 395.51 | 1 | 0.97 | 0.24 | 7.49 | 7.42 | 0.07 | 0.07 | 0 | 0 |
| gene-LOC105279528 | https://www.ncbi.nlm.nih.gov/gene/?term=LOC105279528 | 5802 | 46 | 79.28 | 1 | 1.72 | 2.17 | 15.76 | 15.72 | 0.04 | 0.04 | 0 | 0 |
| gene-LOC105280304 | https://www.ncbi.nlm.nih.gov/gene/?term=LOC105280304 | 8919 | 123 | 137.91 | 1 | 1.12 | 0.81 | 5.79 | 5.75 | 0.04 | 0.04 | 0 | 0 |
| gene-LOC105282914 | https://www.ncbi.nlm.nih.gov/gene/?term=LOC105282914 | 5623 | 42 | 74.69 | 1 | 1.78 | 2.38 | 48.6 | 48.57 | 0.03 | 0.03 | 0 | 0 |
| gene-LOC105274699 | https://www.ncbi.nlm.nih.gov/gene/?term=LOC105274699 | 2386 | 63 | 264.04 | 1 | 4.19 | 1.59 | 23.04 | 23.01 | 0.02 | 0.02 | 0 | 0 |
| gene-LOC105283296 | https://www.ncbi.nlm.nih.gov/gene/?term=LOC105283296 | 2174 | 89 | 409.38 | 1 | 4.6 | 1.12 | 19.56 | 19.55 | 0.01 | 0.01 | 0 | 0 |
| gene-LOC105276059 | https://www.ncbi.nlm.nih.gov/gene/?term=LOC105276059 | 6256 | 136 | 217.39 | 1 | 1.6 | 0.74 | 10.77 | 10.82 | -0.06 | 0.06 | 0 | 0 |
| gene-LOC105278538 | https://www.ncbi.nlm.nih.gov/gene/?term=LOC105278538 | 2150 | 83 | 386.05 | 1 | 4.65 | 1.2 | 18.97 | 19.05 | -0.08 | 0.08 | 0 | 0 |
| gene-LOC105284117 | https://www.ncbi.nlm.nih.gov/gene/?term=LOC105284117 | 4340 | 86 | 198.16 | 1 | 2.3 | 1.16 | 10.49 | 10.57 | -0.08 | 0.08 | 0 | 0 |
| gene-LOC105278145 | https://www.ncbi.nlm.nih.gov/gene/?term=LOC105278145 | 4470 | 256 | 572.71 | 1 | 2.24 | 0.39 | 10.38 | 10.47 | -0.09 | 0.09 | -0.01 | 0.01 |
| gene-LOC105284557 | https://www.ncbi.nlm.nih.gov/gene/?term=LOC105284557 | 1672 | 77 | 460.53 | 1 | 5.98 | 1.3 | 30.69 | 30.8 | -0.11 | 0.11 | -0.01 | 0.01 |
| gene-LOC105277761 | https://www.ncbi.nlm.nih.gov/gene/?term=LOC105277761 | 4648 | 150 | 322.72 | 1 | 2.15 | 0.67 | 14.73 | 14.87 | -0.13 | 0.13 | 0 | 0 |
| gene-LOC105284262 | https://www.ncbi.nlm.nih.gov/gene/?term=LOC105284262 | 6669 | 236 | 353.88 | 1 | 1.5 | 0.42 | 4.46 | 4.59 | -0.13 | 0.13 | 0 | 0 |
| gene-LOC105281272 | https://www.ncbi.nlm.nih.gov/gene/?term=LOC105281272 | 5252 | 179 | 340.82 | 1 | 1.9 | 0.56 | 16.36 | 16.51 | -0.15 | 0.15 | -0.01 | 0.01 |
| gene-LOC105285395 | https://www.ncbi.nlm.nih.gov/gene/?term=LOC105285395 | 8085 | 187 | 231.29 | 1 | 1.24 | 0.53 | 16.28 | 16.44 | -0.16 | 0.16 | 0 | 0 |
| gene-LOC105275187 | https://www.ncbi.nlm.nih.gov/gene/?term=LOC105275187 | 5534 | 75 | 135.53 | 1 | 1.81 | 1.33 | 14.15 | 14.34 | -0.18 | 0.18 | 0 | 0 |
| gene-LOC105276612 | https://www.ncbi.nlm.nih.gov/gene/?term=LOC105276612 | 4157 | 178 | 428.19 | 1 | 2.41 | 0.56 | 10.69 | 10.89 | -0.2 | 0.2 | -0.01 | 0.01 |
| gene-LOC105284036 | https://www.ncbi.nlm.nih.gov/gene/?term=LOC105284036 | 3254 | 83 | 255.07 | 1 | 3.07 | 1.2 | 22.52 | 22.73 | -0.2 | 0.2 | -0.01 | 0.01 |
| gene-LOC105285807 | https://www.ncbi.nlm.nih.gov/gene/?term=LOC105285807 | 6063 | 186 | 306.78 | 1 | 1.65 | 0.54 | 10.05 | 10.27 | -0.22 | 0.22 | -0.01 | 0.01 |
| gene-LOC105275327 | https://www.ncbi.nlm.nih.gov/gene/?term=LOC105275327 | 14245 | 479 | 336.26 | 1 | 0.7 | 0.21 | 7.11 | 7.34 | -0.23 | 0.23 | -0.01 | 0.01 |
| gene-LOC113562050 | https://www.ncbi.nlm.nih.gov/gene/?term=LOC113562050 | 17097 | 576 | 336.9 | 1 | 0.58 | 0.17 | 5.26 | 5.5 | -0.25 | 0.25 | -0.01 | 0.01 |
| gene-LOC105285202 | https://www.ncbi.nlm.nih.gov/gene/?term=LOC105285202 | 1838 | 36 | 195.87 | 1 | 5.44 | 2.78 | 18.94 | 19.21 | -0.27 | 0.27 | -0.01 | 0.01 |
| gene-LOC105274851 | https://www.ncbi.nlm.nih.gov/gene/?term=LOC105274851 | 6616 | 142 | 214.63 | 1 | 1.51 | 0.7 | 9.54 | 9.83 | -0.3 | 0.3 | -0.01 | 0.01 |
| gene-LOC105280104 | https://www.ncbi.nlm.nih.gov/gene/?term=LOC105280104 | 3960 | 120 | 303.03 | 1 | 2.53 | 0.83 | 14.59 | 14.9 | -0.31 | 0.31 | -0.01 | 0.01 |
| gene-LOC105288078 | https://www.ncbi.nlm.nih.gov/gene/?term=LOC105288078 | 19126 | 318 | 166.27 | 1 | 0.52 | 0.31 | 17.33 | 17.64 | -0.31 | 0.31 | -0.01 | 0.01 |
| gene-LOC105274531 | https://www.ncbi.nlm.nih.gov/gene/?term=LOC105274531 | 7774 | 257 | 330.59 | 1 | 1.29 | 0.39 | 11.81 | 12.14 | -0.32 | 0.32 | -0.01 | 0.01 |
| gene-LOC105278534 | https://www.ncbi.nlm.nih.gov/gene/?term=LOC105278534 | 7364 | 185 | 251.22 | 1 | 1.36 | 0.54 | 17.95 | 18.29 | -0.33 | 0.33 | -0.01 | 0.01 |
| gene-LOC105278384 | https://www.ncbi.nlm.nih.gov/gene/?term=LOC105278384 | 1390 | 33 | 237.41 | 1 | 7.19 | 3.03 | 45.97 | 46.32 | -0.35 | 0.35 | -0.01 | 0.01 |

| | | | | | | | | | | | | | |
|---|---|---|---|---|---|---|---|---|---|---|---|---|---|
| gene-LOC105279038 | https://www.ncbi.nlm.nih.gov/gene/?term=LOC105279038 | 9453 | 321 | 339.57 | 1 | 1.06 | 0.31 | 8.59 | 8.94 | -0.35 | 0.35 | -0.01 | 0.01 |
| gene-LOC105287704 | https://www.ncbi.nlm.nih.gov/gene/?term=LOC105287704 | 3019 | 125 | 414.04 | 1 | 3.31 | 0.8 | 21.19 | 21.56 | -0.37 | 0.37 | -0.02 | 0.02 |
| gene-LOC105278228 | https://www.ncbi.nlm.nih.gov/gene/?term=LOC105278228 | 8610 | 163 | 189.31 | 1 | 1.16 | 0.61 | 23.66 | 24.04 | -0.38 | 0.38 | -0.01 | 0.01 |
| gene-LOC105280956 | https://www.ncbi.nlm.nih.gov/gene/?term=LOC105280956 | 2639 | 73 | 276.62 | 1 | 3.79 | 1.37 | 20.28 | 20.69 | -0.4 | 0.4 | -0.01 | 0.01 |
| gene-LOC105284184 | https://www.ncbi.nlm.nih.gov/gene/?term=LOC105284184 | 2535 | 82 | 323.47 | 1 | 3.94 | 1.22 | 34.56 | 34.95 | -0.4 | 0.4 | -0.01 | 0.01 |
| gene-LOC105280786 | https://www.ncbi.nlm.nih.gov/gene/?term=LOC105280786 | 2573 | 122 | 474.15 | 1 | 3.89 | 0.82 | 29.15 | 29.56 | -0.41 | 0.41 | -0.02 | 0.02 |
| gene-LOC105285534 | https://www.ncbi.nlm.nih.gov/gene/?term=LOC105285534 | 3944 | 224 | 567.95 | 1 | 2.54 | 0.45 | 11.27 | 11.7 | -0.42 | 0.42 | -0.02 | 0.02 |
| gene-LOC105275451 | https://www.ncbi.nlm.nih.gov/gene/?term=LOC105275451 | 6035 | 118 | 195.53 | 1 | 1.66 | 0.85 | 22.19 | 22.63 | -0.44 | 0.44 | -0.01 | 0.01 |
| gene-LOC105288237 | https://www.ncbi.nlm.nih.gov/gene/?term=LOC105288237 | 2226 | 47 | 211.14 | 1 | 4.49 | 2.13 | 35.68 | 36.15 | -0.47 | 0.47 | -0.01 | 0.01 |
| gene-LOC105278283 | https://www.ncbi.nlm.nih.gov/gene/?term=LOC105278283 | 2317 | 130 | 561.07 | 1 | 4.32 | 0.77 | 24.9 | 25.39 | -0.48 | 0.48 | -0.03 | 0.03 |
| gene-LOC105284043 | https://www.ncbi.nlm.nih.gov/gene/?term=LOC105284043 | 5661 | 248 | 438.09 | 1 | 1.77 | 0.4 | 11.5 | 11.98 | -0.48 | 0.48 | -0.02 | 0.02 |
| gene-LOC105277435 | https://www.ncbi.nlm.nih.gov/gene/?term=LOC105277435 | 3643 | 158 | 433.71 | 1 | 2.74 | 0.63 | 12.84 | 13.36 | -0.52 | 0.52 | -0.02 | 0.02 |
| gene-LOC105282825 | https://www.ncbi.nlm.nih.gov/gene/?term=LOC105282825 | 3919 | 90 | 229.65 | 1 | 2.55 | 1.11 | 15.34 | 15.86 | -0.52 | 0.52 | -0.01 | 0.01 |
| gene-LOC105288079 | https://www.ncbi.nlm.nih.gov/gene/?term=LOC105288079 | 3473 | 89 | 256.26 | 1 | 2.88 | 1.12 | 19.96 | 20.49 | -0.52 | 0.52 | -0.01 | 0.01 |
| gene-LOC105276527 | https://www.ncbi.nlm.nih.gov/gene/?term=LOC105276527 | 7923 | 351 | 443.01 | 1 | 1.26 | 0.28 | 8.2 | 8.78 | -0.58 | 0.58 | -0.03 | 0.03 |
| gene-LOC105280410 | https://www.ncbi.nlm.nih.gov/gene/?term=LOC105280410 | 12580 | 260 | 206.68 | 1 | 0.79 | 0.38 | 19.03 | 19.62 | -0.59 | 0.59 | -0.01 | 0.01 |
| gene-LOC105275966 | https://www.ncbi.nlm.nih.gov/gene/?term=LOC105275966 | 3299 | 59 | 178.84 | 1 | 3.03 | 1.69 | 50.28 | 50.9 | -0.62 | 0.62 | -0.01 | 0.01 |
| gene-LOC105284355 | https://www.ncbi.nlm.nih.gov/gene/?term=LOC105284355 | 3904 | 147 | 376.54 | 1 | 2.56 | 0.68 | 21.32 | 21.98 | -0.67 | 0.67 | -0.03 | 0.03 |
| gene-LOC105286274 | https://www.ncbi.nlm.nih.gov/gene/?term=LOC105286274 | 3817 | 183 | 479.43 | 1 | 2.62 | 0.55 | 15.39 | 16.06 | -0.67 | 0.67 | -0.03 | 0.03 |
| gene-LOC105280641 | https://www.ncbi.nlm.nih.gov/gene/?term=LOC105280641 | 3682 | 66 | 179.25 | 1 | 2.72 | 1.52 | 15.43 | 16.12 | -0.68 | 0.68 | -0.01 | 0.01 |
| gene-LOC105286770 | https://www.ncbi.nlm.nih.gov/gene/?term=LOC105286770 | 3055 | 38 | 124.39 | 1 | 3.27 | 2.63 | 11.1 | 11.78 | -0.68 | 0.68 | -0.01 | 0.01 |
| gene-LOC105285179 | https://www.ncbi.nlm.nih.gov/gene/?term=LOC105285179 | 4191 | 156 | 372.23 | 1 | 2.39 | 0.64 | 17.53 | 18.23 | -0.69 | 0.69 | -0.03 | 0.03 |
| gene-LOC105283728 | https://www.ncbi.nlm.nih.gov/gene/?term=LOC105283728 | 3256 | 118 | 362.41 | 2 | 6.14 | 1.69 | 19.18 | 19.87 | -0.7 | 0.7 | -0.03 | 0.03 |
| gene-LOC105282600 | https://www.ncbi.nlm.nih.gov/gene/?term=LOC105282600 | 3715 | 65 | 174.97 | 1 | 2.69 | 1.54 | 26.29 | 27.03 | -0.73 | 0.73 | -0.01 | 0.01 |
| gene-LOC105276449 | https://www.ncbi.nlm.nih.gov/gene/?term=LOC105276449 | 1231 | 48 | 389.93 | 1 | 8.12 | 2.08 | 26.7 | 27.43 | -0.74 | 0.74 | -0.03 | 0.03 |
| gene-LOC105276917 | https://www.ncbi.nlm.nih.gov/gene/?term=LOC105276917 | 2336 | 26 | 111.3 1 | 4.28 | 3.85 | 36.57 | 37.3 | -0.74 | 0.74 | -0.01 | 0.01 | |
| gene-LOC105275752 | https://www.ncbi.nlm.nih.gov/gene/?term=LOC105275752 | 3462 | 123 | 355.29 | 1 | 2.89 | 0.81 | 28.75 | 29.5 | -0.75 | 0.75 | -0.03 | 0.03 |
| gene-LOC105283132 | https://www.ncbi.nlm.nih.gov/gene/?term=LOC105283132 | 1819 | 54 | 296.87 | 1 | 5.5 | 1.85 | 40.82 | 41.59 | -0.77 | 0.77 | -0.02 | 0.02 |
| gene-LOC105280774 | https://www.ncbi.nlm.nih.gov/gene/?term=LOC105280774 | 2793 | 117 | 418.9 1 | 3.58 | 0.85 | 27.44 | 28.24 | -0.79 | 0.79 | -0.03 | 0.03 | |
| gene-LOC105282067 | https://www.ncbi.nlm.nih.gov/gene/?term=LOC105282067 | 5941 | 54 | 90.89 1 | 1.68 | 1.85 | 23.11 | 23.91 | -0.8 | 0.8 | -0.01 | 0.01 | |
| gene-LOC105282563 | https://www.ncbi.nlm.nih.gov/gene/?term=LOC105282563 | 4353 | 146 | 335.4 1 | 2.3 | 0.68 | 15.5 | 16.32 | -0.83 | 0.83 | -0.03 | 0.03 | |
| gene-LOC105274856 | https://www.ncbi.nlm.nih.gov/gene/?term=LOC105274856 | 5216 | 208 | 398.77 | 1 | 1.92 | 0.48 | 18.49 | 19.33 | -0.84 | 0.84 | -0.03 | 0.03 |
| gene-LOC105274706 | https://www.ncbi.nlm.nih.gov/gene/?term=LOC105274706 | 1446 | 52 | 359.61 | 1 | 6.92 | 1.92 | 37.01 | 37.87 | -0.86 | 0.86 | -0.03 | 0.03 |
| gene-LOC105274773 | https://www.ncbi.nlm.nih.gov/gene/?term=LOC105274773 | 11034 | 167 | 151.35 | 1 | 0.91 | 0.6 | 16.37 | 17.25 | -0.88 | 0.88 | -0.01 | 0.01 |
| gene-LOC105275363 | https://www.ncbi.nlm.nih.gov/gene/?term=LOC105275363 | 3798 | 159 | 418.64 | 1 | 2.63 | 0.63 | 16.97 | 17.86 | -0.88 | 0.88 | -0.04 | 0.04 |
| gene-LOC105279657 | https://www.ncbi.nlm.nih.gov/gene/?term=LOC105279657 | 5878 | 84 | 142.91 | 1 | 1.7 | 1.19 | 18.69 | 19.56 | -0.88 | 0.88 | -0.01 | 0.01 |
| gene-LOC105274590 | https://www.ncbi.nlm.nih.gov/gene/?term=LOC105274590 | 4516 | 81 | 179.36 | 1 | 2.21 | 1.23 | 21.65 | 22.55 | -0.9 | 0.9 | -0.02 | 0.02 |
| gene-LOC105277216 | https://www.ncbi.nlm.nih.gov/gene/?term=LOC105277216 | 7627 | 89 | 116.69 | 1 | 1.31 | 1.12 | 6.69 | 7.62 | -0.93 | 0.93 | -0.01 | 0.01 |
| gene-LOC105280447 | https://www.ncbi.nlm.nih.gov/gene/?term=LOC105280447 | 2215 | 28 | 126.41 | 1 | 4.51 | 3.57 | 23.87 | 24.81 | -0.94 | 0.94 | -0.01 | 0.01 |
| gene-LOC105285406 | https://www.ncbi.nlm.nih.gov/gene/?term=LOC105285406 | 8015 | 42 | 52.4 1 | 1.25 | 2.38 | 35.2 | 36.16 | -0.96 | 0.96 | -0.01 | 0.01 | |
| gene-LOC105279298 | https://www.ncbi.nlm.nih.gov/gene/?term=LOC105279298 | 2773 | 138 | 497.66 | 2 | 7.21 | 1.45 | 22.22 | 23.22 | -1 | 1 | -0.05 | 0.05 |
| gene-LOC105282491 | https://www.ncbi.nlm.nih.gov/gene/?term=LOC105282491 | 4761 | 163 | 342.37 | 1 | 2.1 | 0.61 | 9.65 | 10.67 | -1.02 | 1.02 | -0.03 | 0.03 |
| gene-LOC105278497 | https://www.ncbi.nlm.nih.gov/gene/?term=LOC105278497 | 2072 | 100 | 482.63 | 1 | 4.83 | 1 | 18.86 | 19.92 | -1.06 | 1.06 | -0.05 | 0.05 |
| gene-LOC105275179 | https://www.ncbi.nlm.nih.gov/gene/?term=LOC105275179 | 3271 | 68 | 207.89 | 1 | 3.06 | 1.47 | 14.88 | 16 | -1.12 | 1.12 | -0.02 | 0.02 |
| gene-LOC105282947 | https://www.ncbi.nlm.nih.gov/gene/?term=LOC105282947 | 1162 | 49 | 421.69 | 1 | 8.61 | 2.04 | 29.51 | 30.69 | -1.17 | 1.17 | -0.05 | 0.05 |
| gene-LOC105278962 | https://www.ncbi.nlm.nih.gov/gene/?term=LOC105278962 | 4491 | 88 | 195.95 | 1 | 2.23 | 1.14 | 23.85 | 25.09 | -1.24 | 1.24 | -0.02 | 0.02 |
| gene-LOC105275683 | https://www.ncbi.nlm.nih.gov/gene/?term=LOC105275683 | 4256 | 203 | 476.97 | 1 | 2.35 | 0.49 | 12.81 | 14.05 | -1.25 | 1.25 | -0.06 | 0.06 |
| gene-LOC105283943 | https://www.ncbi.nlm.nih.gov/gene/?term=LOC105283943 | 3559 | 60 | 168.59 | 1 | 2.81 | 1.67 | 21.58 | 22.83 | -1.25 | 1.25 | -0.02 | 0.02 |
| gene-LOC105285742 | https://www.ncbi.nlm.nih.gov/gene/?term=LOC105285742 | 3446 | 100 | 290.19 | 1 | 2.9 | 1 | 24.69 | 25.96 | -1.27 | 1.27 | -0.04 | 0.04 |
| gene-LOC105281700 | https://www.ncbi.nlm.nih.gov/gene/?term=LOC105281700 | 2208 | 46 | 208.33 | 1 | 4.53 | 2.17 | 39.97 | 41.25 | -1.28 | 1.28 | -0.03 | 0.03 |
| gene-LOC105275460 | https://www.ncbi.nlm.nih.gov/gene/?term=LOC105275460 | 2473 | 112 | 452.89 | 1 | 4.04 | 0.89 | 12.89 | 14.18 | -1.3 | 1.3 | -0.06 | 0.06 |
| gene-LOC105278282 | https://www.ncbi.nlm.nih.gov/gene/?term=LOC105278282 | 1775 | 47 | 264.79 | 1 | 5.63 | 2.13 | 45.94 | 47.29 | -1.35 | 1.35 | -0.04 | 0.04 |
| gene-LOC105279500 | https://www.ncbi.nlm.nih.gov/gene/?term=LOC105279500 | 3587 | 33 | 92 1 | 2.79 | 3.03 | 49.6 | 51 | -1.4 | 1.4 | -0.01 | 0.01 | |
| gene-LOC105285055 | https://www.ncbi.nlm.nih.gov/gene/?term=LOC105285055 | 4108 | 217 | 528.24 | 1 | 2.43 | 0.46 | 12.76 | 14.18 | -1.42 | 1.42 | -0.07 | 0.07 |
| gene-LOC105277631 | https://www.ncbi.nlm.nih.gov/gene/?term=LOC105277631 | 4387 | 289 | 658.76 | 1 | 2.28 | 0.35 | 9.79 | 11.22 | -1.43 | 1.43 | -0.09 | 0.09 |
| gene-LOC105283592 | https://www.ncbi.nlm.nih.gov/gene/?term=LOC105283592 | 3479 | 71 | 204.08 | 1 | 2.87 | 1.41 | 24.39 | 25.83 | -1.45 | 1.45 | -0.03 | 0.03 |
| gene-LOC105275347 | https://www.ncbi.nlm.nih.gov/gene/?term=LOC105275347 | 2526 | 14 | 55.42 1 | 3.96 | 7.14 | 35.76 | 37.28 | -1.52 | 1.52 | -0.01 | 0.01 | |
| gene-LOC105276078 | https://www.ncbi.nlm.nih.gov/gene/?term=LOC105276078 | 1051 | 10 | 95.15 1 | 9.51 | 10 | 37.29 | 38.89 | -1.6 | 1.6 | -0.02 | 0.02 | |
| gene-LOC105287063 | https://www.ncbi.nlm.nih.gov/gene/?term=LOC105287063 | 1874 | 46 | 245.46 | 1 | 5.34 | 2.17 | 34.15 | 35.78 | -1.63 | 1.63 | -0.04 | 0.04 |
| gene-LOC105285695 | https://www.ncbi.nlm.nih.gov/gene/?term=LOC105285695 | 1371 | 62 | 452.22 | 1 | 7.29 | 1.61 | 23 | 24.72 | -1.72 | 1.72 | -0.08 | 0.08 |
| gene-LOC105279356 | https://www.ncbi.nlm.nih.gov/gene/?term=LOC105279356 | 1314 | 42 | 319.63 | 1 | 7.61 | 2.38 | 28.83 | 30.58 | -1.75 | 1.75 | -0.06 | 0.06 |
| gene-LOC105285671 | https://www.ncbi.nlm.nih.gov/gene/?term=LOC105285671 | 2263 | 95 | 419.8 1 | 4.42 | 1.05 | 27.86 | 29.64 | -1.78 | 1.78 | -0.07 | 0.07 | |
| gene-LOC105283989 | https://www.ncbi.nlm.nih.gov/gene/?term=LOC105283989 | 4948 | 34 | 68.71 2 | 4.04 | 5.88 | 44.42 | 46.21 | -1.79 | 1.79 | -0.01 | 0.01 | |
| gene-LOC105284704 | https://www.ncbi.nlm.nih.gov/gene/?term=LOC105284704 | 2400 | 65 | 270.83 | 1 | 4.17 | 1.54 | 25.06 | 26.85 | -1.79 | 1.79 | -0.05 | 0.05 |

| | | | | | | | | | | | | | |
|---|---|---|---|---|---|---|---|---|---|---|---|---|---|
| gene-LOC105276692 | https://www.ncbi.nlm.nih.gov/gene/?term=LOC105276692 | 1745 | 37 | 212.03 | 1 | 5.73 | 2.7 | 20.82 | 22.64 | -1.82 | 1.82 | -0.04 | 0.04 |
| gene-LOC105274900 | https://www.ncbi.nlm.nih.gov/gene/?term=LOC105274900 | 5561 | 56 | 100.7 | 2 | 3.6 | 3.57 | 26.85 | 28.69 | -1.84 | 1.84 | -0.02 | 0.02 |
| gene-LOC105278260 | https://www.ncbi.nlm.nih.gov/gene/?term=LOC105278260 | 3240 | 41 | 126.54 | 1 | 3.09 | 2.44 | 13.01 | 14.86 | -1.85 | 1.85 | -0.02 | 0.02 |
| gene-LOC105276745 | https://www.ncbi.nlm.nih.gov/gene/?term=LOC105276745 | 3953 | 73 | 184.67 | 1 | 2.53 | 1.37 | 21.09 | 23.02 | -1.93 | 1.93 | -0.04 | 0.04 |
| gene-LOC105278807 | https://www.ncbi.nlm.nih.gov/gene/?term=LOC105278807 | 2166 | 72 | 332.41 | 1 | 4.62 | 1.39 | 18.53 | 20.46 | -1.93 | 1.93 | -0.06 | 0.06 |
| gene-LOC105286223 | https://www.ncbi.nlm.nih.gov/gene/?term=LOC105286223 | 2225 | 83 | 373.03 | 1 | 4.49 | 1.2 | 30.2 | 32.18 | -1.98 | 1.98 | -0.07 | 0.07 |
| gene-LOC105286452 | https://www.ncbi.nlm.nih.gov/gene/?term=LOC105286452 | 6888 | 58 | 84.2 | 1 | 1.45 | 1.72 | 31.25 | 33.23 | -1.98 | 1.98 | -0.02 | 0.02 |
| gene-LOC105280348 | https://www.ncbi.nlm.nih.gov/gene/?term=LOC105280348 | 3797 | 52 | 136.95 | 1 | 2.63 | 1.92 | 25.54 | 27.53 | -1.99 | 1.99 | -0.03 | 0.03 |
| gene-LOC105282156 | https://www.ncbi.nlm.nih.gov/gene/?term=LOC105282156 | 2577 | 136 | 527.75 | 1 | 3.88 | 0.74 | 23.86 | 26.02 | -2.16 | 2.16 | -0.11 | 0.11 |
| gene-LOC105285583 | https://www.ncbi.nlm.nih.gov/gene/?term=LOC105285583 | 1947 | 51 | 261.94 | 1 | 5.14 | 1.96 | 28.26 | 30.43 | -2.17 | 2.17 | -0.06 | 0.06 |
| gene-LOC105282893 | https://www.ncbi.nlm.nih.gov/gene/?term=LOC105282893 | 2645 | 43 | 162.57 | 1 | 3.78 | 2.33 | 27.48 | 29.68 | -2.19 | 2.19 | -0.04 | 0.04 |
| gene-LOC105286644 | https://www.ncbi.nlm.nih.gov/gene/?term=LOC105286644 | 1235 | 22 | 178.14 | 1 | 8.1 | 4.55 | 42.2 | 44.4 | -2.2 | 2.2 | -0.04 | 0.04 |
| gene-LOC105280002 | https://www.ncbi.nlm.nih.gov/gene/?term=LOC105280002 | 1940 | 30 | 154.64 | 1 | 5.15 | 3.33 | 33.39 | 35.64 | -2.25 | 2.25 | -0.03 | 0.03 |
| gene-LOC105281515 | https://www.ncbi.nlm.nih.gov/gene/?term=LOC105281515 | 8827 | 41 | 46.45 | 1 | 1.13 | 2.44 | 32.87 | 35.23 | -2.37 | 2.37 | -0.01 | 0.01 |
| gene-LOC105274534 | https://www.ncbi.nlm.nih.gov/gene/?term=LOC105274534 | 1329 | 57 | 428.89 | 1 | 7.52 | 1.75 | 43.75 | 46.14 | -2.39 | 2.39 | -0.1 | 0.1 |
| gene-LOC105287417 | https://www.ncbi.nlm.nih.gov/gene/?term=LOC105287417 | 1025 | 33 | 321.95 | 1 | 9.76 | 3.03 | 36 | 38.42 | -2.42 | 2.42 | -0.08 | 0.08 |
| gene-LOC105275518 | https://www.ncbi.nlm.nih.gov/gene/?term=LOC105275518 | 2335 | 21 | 89.94 | 1 | 4.28 | 4.76 | 45.03 | 47.49 | -2.45 | 2.45 | -0.02 | 0.02 |
| gene-LOC105288066 | https://www.ncbi.nlm.nih.gov/gene/?term=LOC105288066 | 1964 | 26 | 132.38 | 1 | 5.09 | 3.85 | 26.28 | 28.74 | -2.45 | 2.45 | -0.03 | 0.03 |
| gene-LOC105287854 | https://www.ncbi.nlm.nih.gov/gene/?term=LOC105287854 | 3644 | 57 | 156.42 | 1 | 2.74 | 1.75 | 21.16 | 23.71 | -2.55 | 2.55 | -0.04 | 0.04 |
| gene-LOC105286502 | https://www.ncbi.nlm.nih.gov/gene/?term=LOC105286502 | 1286 | 44 | 342.15 | 1 | 7.78 | 2.27 | 23.97 | 26.78 | -2.81 | 2.81 | -0.1 | 0.1 |
| gene-LOC105276704 | https://www.ncbi.nlm.nih.gov/gene/?term=LOC105276704 | 2063 | 87 | 421.72 | 1 | 4.85 | 1.15 | 12.34 | 15.21 | -2.86 | 2.86 | -0.12 | 0.12 |
| gene-LOC105276538 | https://www.ncbi.nlm.nih.gov/gene/?term=LOC105276538 | 1511 | 19 | 125.74 | 1 | 6.62 | 5.26 | 72.68 | 75.55 | -2.87 | 2.87 | -0.04 | 0.04 |
| gene-LOC105277215 | https://www.ncbi.nlm.nih.gov/gene/?term=LOC105277215 | 2619 | 29 | 110.73 | 1 | 3.82 | 3.45 | 19.9 | 22.79 | -2.88 | 2.88 | -0.03 | 0.03 |
| gene-LOC105285674 | https://www.ncbi.nlm.nih.gov/gene/?term=LOC105285674 | 3035 | 91 | 299.84 | 2 | 6.59 | 2.2 | 23.1 | 26.01 | -2.91 | 2.91 | -0.09 | 0.09 |
| gene-LOC105283712 | https://www.ncbi.nlm.nih.gov/gene/?term=LOC105283712 | 1910 | 27 | 141.36 | 1 | 5.24 | 3.7 | 29.17 | 32.5 | -3.33 | 3.33 | -0.05 | 0.05 |
| gene-LOC105278109 | https://www.ncbi.nlm.nih.gov/gene/?term=LOC105278109 | 4029 | 62 | 153.88 | 1 | 2.48 | 1.61 | 43.07 | 46.49 | -3.42 | 3.42 | -0.05 | 0.05 |
| gene-LOC105282399 | https://www.ncbi.nlm.nih.gov/gene/?term=LOC105282399 | 2682 | 22 | 82.03 | 1 | 3.73 | 4.55 | 32.93 | 36.36 | -3.43 | 3.43 | -0.03 | 0.03 |
| gene-LOC113561805 | https://www.ncbi.nlm.nih.gov/gene/?term=LOC113561805 | 3337 | 64 | 191.79 | 1 | 3 | 1.56 | 37.95 | 41.4 | -3.45 | 3.45 | -0.07 | 0.07 |
| gene-LOC105275190 | https://www.ncbi.nlm.nih.gov/gene/?term=LOC105275190 | 3837 | 74 | 192.86 | 1 | 2.61 | 1.35 | 18.39 | 22.03 | -3.64 | 3.64 | -0.07 | 0.07 |
| gene-LOC105274778 | https://www.ncbi.nlm.nih.gov/gene/?term=LOC105274778 | 4662 | 38 | 81.51 | 1 | 2.15 | 2.63 | 44.96 | 48.85 | -3.88 | 3.88 | -0.03 | 0.03 |
| gene-LOC105281848 | https://www.ncbi.nlm.nih.gov/gene/?term=LOC105281848 | 2122 | 47 | 221.49 | 1 | 4.71 | 2.13 | 23.11 | 27.19 | -4.08 | 4.08 | -0.09 | 0.09 |
| gene-LOC105285060 | https://www.ncbi.nlm.nih.gov/gene/?term=LOC105285060 | 1285 | 15 | 116.73 | 1 | 7.78 | 6.67 | 48.46 | 52.7 | -4.25 | 4.25 | -0.05 | 0.05 |
| gene-LOC105277856 | https://www.ncbi.nlm.nih.gov/gene/?term=LOC105277856 | 3442 | 29 | 84.25 | 1 | 2.91 | 3.45 | 35.44 | 40.05 | -4.6 | 4.6 | -0.04 | 0.04 |
| gene-LOC105281039 | https://www.ncbi.nlm.nih.gov/gene/?term=LOC105281039 | 1471 | 13 | 88.38 | 1 | 6.8 | 7.69 | 36.12 | 40.81 | -4.7 | 4.7 | -0.04 | 0.04 |
| gene-LOC105286983 | https://www.ncbi.nlm.nih.gov/gene/?term=LOC105286983 | 16901 | 68 | 40.23 | 1 | 0.59 | 1.47 | 0.94 | 5.77 | -4.83 | 4.83 | -0.02 | 0.02 |
| gene-LOC105285514 | https://www.ncbi.nlm.nih.gov/gene/?term=LOC105285514 | 1277 | 9 | 70.48 | 1 | 7.83 | 11.11 | 73.21 | 78.44 | -5.23 | 5.23 | -0.04 | 0.04 |
| gene-LOC105284609 | https://www.ncbi.nlm.nih.gov/gene/?term=LOC105284609 | 5678 | 31 | 54.6 | 1 | 1.76 | 3.23 | 35.81 | 41.06 | -5.25 | 5.25 | -0.03 | 0.03 |
| gene-LOC105275708 | https://www.ncbi.nlm.nih.gov/gene/?term=LOC105275708 | 1772 | 14 | 79.01 | 1 | 5.64 | 7.14 | 44.58 | 50.56 | -5.98 | 5.98 | -0.05 | 0.05 |
| gene-LOC105279898 | https://www.ncbi.nlm.nih.gov/gene/?term=LOC105279898 | 1343 | 16 | 119.14 | 2 | 14.89 | 12.5 | 59.42 | 68.99 | -9.57 | 9.57 | -0.11 | 0.11 |
| gene-LOC105275070 | https://www.ncbi.nlm.nih.gov/gene/?term=LOC105275070 | 5667 | 38 | 67.05 | 3 | 5.29 | 7.89 | 3.63 | 13.45 | -9.82 | 9.82 | -0.07 | 0.07 |
| gene-LOC113562345 | https://www.ncbi.nlm.nih.gov/gene/?term=LOC113562345 | 2290 | 62 | 270.74 | 16 | 69.87 | 25.81 | 17.91 | 33 | -15.09 | 15.09 | -0.41 | 0.41 |

Directory: OGL File: ogl-Ob_LI_BCphaseA1.vs.RphaseB1.txt

| gene_ID | gene_link | gwidth | #Sites | #per10Kb | #dmSites | #dmsp10kb | %dmSites | %pSite1 | %pSite2 | DMpSite | ADMpSite | DMpNucl | ADMpNucl |
|---|---|---|---|---|---|---|---|---|---|---|---|---|---|
| gene-LOC113562975 | https://www.ncbi.nlm.nih.gov/gene/?term=LOC113562975 | 2751 | 3 | 10.91 | 1 | 3.64 | 33.33 | 46.98 | 30.96 | 16.02 | 16.02 | 0.02 | 0.02 |
| gene-LOC105284474 | https://www.ncbi.nlm.nih.gov/gene/?term=LOC105284474 | 479 | 2 | 41.75 | 1 | 20.88 | 50 | 33.34 | 19.05 | 14.29 | 14.29 | 0.06 | 0.06 |
| gene-LOC113562831 | https://www.ncbi.nlm.nih.gov/gene/?term=LOC113562831 | 574 | 2 | 34.84 | 1 | 17.42 | 50 | 18.75 | 6.94 | 11.8 | 11.8 | 0.04 | 0.04 |
| gene-LOC105288227 | https://www.ncbi.nlm.nih.gov/gene/?term=LOC105288227 | 1245 | 3 | 24.1 | 1 | 8.03 | 33.33 | 86.21 | 76.94 | 9.26 | 9.26 | 0.02 | 0.02 |
| gene-LOC113561934 | https://www.ncbi.nlm.nih.gov/gene/?term=LOC113561934 | 1252 | 16 | 127.8 | 4 | 31.95 | 25 | 36.29 | 28.08 | 8.21 | 8.21 | 0.1 | 0.1 |
| gene-LOC105276219 | https://www.ncbi.nlm.nih.gov/gene/?term=LOC105276219 | 916 | 6 | 65.5 | 1 | 10.92 | 16.67 | 47.07 | 39.74 | 7.33 | 7.33 | 0.05 | 0.05 |
| gene-LOC105275343 | https://www.ncbi.nlm.nih.gov/gene/?term=LOC105275343 | 1746 | 4 | 22.91 | 1 | 5.73 | 25 | 47 | 39.86 | 7.14 | 7.14 | 0.02 | 0.02 |
| gene-LOC105282405 | https://www.ncbi.nlm.nih.gov/gene/?term=LOC105282405 | 1310 | 31 | 236.64 | 2 | 15.27 | 6.45 | 51.18 | 44.54 | 6.64 | 6.64 | 0.16 | 0.16 |
| gene-LOC113562627 | https://www.ncbi.nlm.nih.gov/gene/?term=LOC113562627 | 397 | 10 | 251.89 | 1 | 25.19 | 10 | 77.44 | 70.89 | 6.56 | 6.56 | 0.17 | 0.17 |
| gene-LOC105285885 | https://www.ncbi.nlm.nih.gov/gene/?term=LOC105285885 | 1327 | 12 | 90.43 | 1 | 7.54 | 8.33 | 54.12 | 47.77 | 6.35 | 6.35 | 0.06 | 0.06 |
| gene-LOC105286816 | https://www.ncbi.nlm.nih.gov/gene/?term=LOC105286816 | 4880 | 9 | 18.44 | 1 | 2.05 | 11.11 | 37.19 | 31.02 | 6.17 | 6.17 | 0.01 | 0.01 |
| gene-LOC105281226 | https://www.ncbi.nlm.nih.gov/gene/?term=LOC105281226 | 1308 | 10 | 76.45 | 1 | 7.65 | 10 | 77.35 | 71.27 | 6.08 | 6.08 | 0.05 | 0.05 |
| gene-LOC113563111 | https://www.ncbi.nlm.nih.gov/gene/?term=LOC113563111 | 2455 | 35 | 142.57 | 2 | 8.15 | 5.71 | 26.92 | 21.34 | 5.57 | 5.57 | 0.08 | 0.08 |
| gene-LOC105283973 | https://www.ncbi.nlm.nih.gov/gene/?term=LOC105283973 | 1536 | 29 | 188.8 | 4 | 26.04 | 13.79 | 61.88 | 56.34 | 5.55 | 5.55 | 0.1 | 0.1 |
| gene-LOC105285241 | https://www.ncbi.nlm.nih.gov/gene/?term=LOC105285241 | 5118 | 25 | 48.85 | 2 | 3.91 | 8 | 30.97 | 25.5 | 5.47 | 5.47 | 0.03 | 0.03 |
| gene-LOC105284412 | https://www.ncbi.nlm.nih.gov/gene/?term=LOC105284412 | 829 | 11 | 132.69 | 1 | 12.06 | 9.09 | 56.74 | 51.39 | 5.35 | 5.35 | 0.07 | 0.07 |
| gene-LOC105278541 | https://www.ncbi.nlm.nih.gov/gene/?term=LOC105278541 | 1563 | 16 | 102.37 | 1 | 6.4 | 6.25 | 28.66 | 23.32 | 5.33 | 5.33 | 0.05 | 0.05 |
| gene-LOC105275539 | https://www.ncbi.nlm.nih.gov/gene/?term=LOC105275539 | 1272 | 23 | 180.82 | 2 | 15.72 | 8.7 | 20.8 | 15.48 | 5.32 | 5.32 | 0.1 | 0.1 |
| gene-LOC105281507 | https://www.ncbi.nlm.nih.gov/gene/?term=LOC105281507 | 881 | 5 | 56.75 | 1 | 11.35 | 20 | 11.82 | 6.67 | 5.15 | 5.15 | 0.03 | 0.03 |
| gene-LOC105282996 | https://www.ncbi.nlm.nih.gov/gene/?term=LOC105282996 | 3736 | 9 | 24.09 | 2 | 5.35 | 22.22 | 28.24 | 23.6 | 4.64 | 4.64 | 0.01 | 0.01 |
| gene-LOC105283988 | https://www.ncbi.nlm.nih.gov/gene/?term=LOC105283988 | 2510 | 14 | 55.78 | 1 | 3.98 | 7.14 | 40.87 | 36.23 | 4.64 | 4.64 | 0.03 | 0.03 |
| gene-LOC105285051 | https://www.ncbi.nlm.nih.gov/gene/?term=LOC105285051 | 1357 | 10 | 73.69 | 1 | 7.37 | 10 | 46.31 | 41.71 | 4.6 | 4.6 | 0.03 | 0.03 |
| gene-LOC105280926 | https://www.ncbi.nlm.nih.gov/gene/?term=LOC105280926 | 2139 | 7 | 32.73 | 1 | 4.68 | 14.29 | 49.74 | 45.26 | 4.48 | 4.48 | 0.01 | 0.01 |
| gene-LOC105284881 | https://www.ncbi.nlm.nih.gov/gene/?term=LOC105284881 | 2016 | 20 | 99.21 | 2 | 9.92 | 10 | 27.57 | 23.15 | 4.42 | 4.42 | 0.04 | 0.04 |

...

... only first 25 lines shown ...

Directory: OGL File: ogl-Ob_LI_BCphaseA1.vs.RphaseB2.txt

| gene_ID | gene_link | gwidth | #Sites | #per10Kb | #dmSites | #dmsp10kb | %dmSites | %pSite1 | %pSite2 | DMpSite | ADMpSite | DMpNucl | ADMpNucl |
|---|---|---|---|---|---|---|---|---|---|---|---|---|---|
| gene-LOC105288227 | https://www.ncbi.nlm.nih.gov/gene/?term=LOC105288227 | 1245 | 2 | 16.06 | 1 | 8.03 | 50 | 79.31 | 55.55 | 23.76 | 23.76 | 0.04 | 0.04 |
| gene-LOC105285431 | https://www.ncbi.nlm.nih.gov/gene/?term=LOC105285431 | 1550 | 5 | 32.26 | 2 | 12.9 | 40 | 49.86 | 35.35 | 14.51 | 14.51 | 0.05 | 0.05 |
| gene-LOC113562831 | https://www.ncbi.nlm.nih.gov/gene/?term=LOC113562831 | 574 | 2 | 34.84 | 1 | 17.42 | 50 | 18.75 | 5.88 | 12.87 | 12.87 | 0.04 | 0.04 |
| gene-LOC105287814 | https://www.ncbi.nlm.nih.gov/gene/?term=LOC105287814 | 892 | 5 | 56.05 | 2 | 22.42 | 40 | 36.02 | 23.39 | 12.63 | 12.63 | 0.07 | 0.07 |
| gene-LOC105283410 | https://www.ncbi.nlm.nih.gov/gene/?term=LOC105283410 | 1423 | 5 | 35.14 | 2 | 14.05 | 40 | 52.44 | 40.2 | 12.24 | 12.24 | 0.04 | 0.04 |
| gene-LOC105283241 | https://www.ncbi.nlm.nih.gov/gene/?term=LOC105283241 | 2011 | 4 | 19.89 | 1 | 4.97 | 25 | 39.29 | 29.03 | 10.26 | 10.26 | 0.02 | 0.02 |
| gene-LOC109611453 | https://www.ncbi.nlm.nih.gov/gene/?term=LOC109611453 | 3541 | 6 | 16.94 | 1 | 2.82 | 16.67 | 24.1 | 14.5 | 9.6 | 9.6 | 0.02 | 0.02 |
| gene-LOC113561934 | https://www.ncbi.nlm.nih.gov/gene/?term=LOC113561934 | 1252 | 15 | 119.81 | 2 | 15.97 | 13.33 | 36.71 | 27.17 | 9.54 | 9.54 | 0.11 | 0.11 |
| gene-LOC105276718 | https://www.ncbi.nlm.nih.gov/gene/?term=LOC105276718 | 1721 | 9 | 52.3 | 3 | 17.43 | 33.33 | 30.94 | 22.67 | 8.27 | 8.27 | 0.04 | 0.04 |
| gene-LOC105277368 | https://www.ncbi.nlm.nih.gov/gene/?term=LOC105277368 | 1735 | 7 | 40.35 | 1 | 5.76 | 14.29 | 19.45 | 12.67 | 6.78 | 6.78 | 0.03 | 0.03 |
| gene-LOC105286857 | https://www.ncbi.nlm.nih.gov/gene/?term=LOC105286857 | 1576 | 4 | 25.38 | 1 | 6.35 | 25 | 73.58 | 67.03 | 6.55 | 6.55 | 0.02 | 0.02 |
| gene-LOC105277862 | https://www.ncbi.nlm.nih.gov/gene/?term=LOC105277862 | 1490 | 7 | 46.98 | 1 | 6.71 | 14.29 | 61.97 | 55.64 | 6.33 | 6.33 | 0.03 | 0.03 |
| gene-LOC105282405 | https://www.ncbi.nlm.nih.gov/gene/?term=LOC105282405 | 1310 | 31 | 236.64 | 5 | 38.17 | 16.13 | 51.18 | 44.88 | 6.3 | 6.3 | 0.15 | 0.15 |
| gene-LOC105284899 | https://www.ncbi.nlm.nih.gov/gene/?term=LOC105284899 | 1779 | 14 | 78.7 | 1 | 5.62 | 7.14 | 40.04 | 33.95 | 6.09 | 6.09 | 0.05 | 0.05 |
| gene-LOC105275585 | https://www.ncbi.nlm.nih.gov/gene/?term=LOC105275585 | 2152 | 22 | 102.23 | 1 | 4.65 | 4.55 | 36.17 | 30.17 | 6 | 6 | 0.06 | 0.06 |
| gene-LOC105285429 | https://www.ncbi.nlm.nih.gov/gene/?term=LOC105285429 | 1324 | 8 | 60.42 | 1 | 7.55 | 12.5 | 53.59 | 47.67 | 5.92 | 5.92 | 0.04 | 0.04 |
| gene-LOC105277810 | https://www.ncbi.nlm.nih.gov/gene/?term=LOC105277810 | 1367 | 21 | 153.62 | 1 | 7.32 | 4.76 | 45.33 | 39.52 | 5.81 | 5.81 | 0.09 | 0.09 |
| gene-LOC113562901 | https://www.ncbi.nlm.nih.gov/gene/?term=LOC113562901 | 1531 | 20 | 130.63 | 4 | 26.13 | 20 | 50.7 | 45.01 | 5.69 | 5.69 | 0.07 | 0.07 |
| gene-LOC105279898 | https://www.ncbi.nlm.nih.gov/gene/?term=LOC105279898 | 1343 | 10 | 74.46 | 1 | 7.45 | 10 | 77.18 | 71.53 | 5.65 | 5.65 | 0.04 | 0.04 |
| gene-LOC105277745 | https://www.ncbi.nlm.nih.gov/gene/?term=LOC105277745 | 1365 | 8 | 58.61 | 1 | 7.33 | 12.5 | 67.79 | 62.35 | 5.44 | 5.44 | 0.03 | 0.03 |
| gene-LOC105286644 | https://www.ncbi.nlm.nih.gov/gene/?term=LOC105286644 | 1235 | 21 | 170.04 | 1 | 8.1 | 4.76 | 44.21 | 39.04 | 5.17 | 5.17 | 0.09 | 0.09 |
| gene-LOC105274625 | https://www.ncbi.nlm.nih.gov/gene/?term=LOC105274625 | 4523 | 33 | 72.96 | 1 | 2.21 | 3.03 | 24.56 | 19.42 | 5.14 | 5.14 | 0.04 | 0.04 |
| gene-LOC105280045 | https://www.ncbi.nlm.nih.gov/gene/?term=LOC105280045 | 1938 | 13 | 67.08 | 1 | 5.16 | 7.69 | 26.88 | 21.77 | 5.11 | 5.11 | 0.03 | 0.03 |
| gene-LOC105279461 | https://www.ncbi.nlm.nih.gov/gene/?term=LOC105279461 | 1298 | 7 | 53.93 | 2 | 15.41 | 28.57 | 67.83 | 63.08 | 4.75 | 4.75 | 0.03 | 0.03 |

...

... only first 25 lines shown ...

Directory: OGL File: ogl-Ob_LI_BCphaseA2.txt

| gene_ID | gene_link | gwidth | #Sites | #per10Kb | %perSite | %pNucl |
|---|---|---|---|---|---|---|
| gene-LOC113561532 | https://www.ncbi.nlm.nih.gov/gene/?term=LOC113561532 | 72 | 4 | 555.56 | 94.06 | 5.23 |
| gene-LOC105286531 | https://www.ncbi.nlm.nih.gov/gene/?term=LOC105286531 | 372 | 42 | 1129.03 | 38.35 | 4.33 |
| gene-LOC105280739 | https://www.ncbi.nlm.nih.gov/gene/?term=LOC105280739 | 1991 | 139 | 698.14 | 56.83 | 3.97 |
| gene-LOC105282149 | https://www.ncbi.nlm.nih.gov/gene/?term=LOC105282149 | 1787 | 156 | 872.97 | 45.44 | 3.97 |
| gene-LOC105286801 | https://www.ncbi.nlm.nih.gov/gene/?term=LOC105286801 | 1102 | 103 | 934.66 | 42.22 | 3.95 |
| gene-LOC105278849 | https://www.ncbi.nlm.nih.gov/gene/?term=LOC105278849 | 776 | 124 | 1597.94 | 24.65 | 3.94 |
| gene-LOC105285635 | https://www.ncbi.nlm.nih.gov/gene/?term=LOC105285635 | 1175 | 138 | 1174.47 | 31.91 | 3.75 |
| gene-LOC105274534 | https://www.ncbi.nlm.nih.gov/gene/?term=LOC105274534 | 1329 | 95 | 714.82 | 48.88 | 3.49 |
| gene-LOC105287811 | https://www.ncbi.nlm.nih.gov/gene/?term=LOC105287811 | 2155 | 151 | 700.7 | 49.56 | 3.47 |
| gene-LOC113562627 | https://www.ncbi.nlm.nih.gov/gene/?term=LOC113562627 | 397 | 20 | 503.78 | 68.12 | 3.43 |
| gene-LOC105286269 | https://www.ncbi.nlm.nih.gov/gene/?term=LOC105286269 | 1401 | 91 | 649.54 | 51.53 | 3.35 |
| gene-LOC105278844 | https://www.ncbi.nlm.nih.gov/gene/?term=LOC105278844 | 1101 | 109 | 990.01 | 33.65 | 3.33 |
| gene-LOC105278160 | https://www.ncbi.nlm.nih.gov/gene/?term=LOC105278160 | 1872 | 161 | 860.04 | 37.79 | 3.25 |
| gene-LOC105287677 | https://www.ncbi.nlm.nih.gov/gene/?term=LOC105287677 | 1588 | 123 | 774.56 | 41.72 | 3.23 |
| gene-LOC105286737 | https://www.ncbi.nlm.nih.gov/gene/?term=LOC105286737 | 2690 | 366 | 1360.59 | 23.68 | 3.22 |
| gene-LOC105286235 | https://www.ncbi.nlm.nih.gov/gene/?term=LOC105286235 | 1510 | 110 | 728.48 | 44.1 | 3.21 |
| gene-LOC105279052 | https://www.ncbi.nlm.nih.gov/gene/?term=LOC105279052 | 1130 | 139 | 1230.09 | 25.99 | 3.2 |
| gene-LOC105276371 | https://www.ncbi.nlm.nih.gov/gene/?term=LOC105276371 | 871 | 56 | 642.94 | 49.67 | 3.19 |
| gene-LOC105274711 | https://www.ncbi.nlm.nih.gov/gene/?term=LOC105274711 | 1655 | 109 | 658.61 | 47.63 | 3.14 |
| gene-LOC105278917 | https://www.ncbi.nlm.nih.gov/gene/?term=LOC105278917 | 2176 | 194 | 891.54 | 35.27 | 3.14 |
| gene-LOC105276360 | https://www.ncbi.nlm.nih.gov/gene/?term=LOC105276360 | 1466 | 153 | 1043.66 | 29.45 | 3.07 |
| gene-LOC105275231 | https://www.ncbi.nlm.nih.gov/gene/?term=LOC105275231 | 1270 | 83 | 653.54 | 45.54 | 2.98 |
| gene-LOC105275314 | https://www.ncbi.nlm.nih.gov/gene/?term=LOC105275314 | 1489 | 93 | 624.58 | 47.58 | 2.97 |
| gene-LOC105274543 | https://www.ncbi.nlm.nih.gov/gene/?term=LOC105274543 | 1572 | 127 | 807.89 | 35.56 | 2.87 |

...

... only first 25 lines shown ...

Directory: OGL File: ogl-Ob_LI_BCphaseA2.vs.BCphaseB1.txt

| gene_ID | gene_link | gwidth | #Sites | #per10Kb | #dmSites | #dmsp10kb | %dmSites | %pSite1 | %pSite2 | DMpSite | ADMpSite | DMpNucl | ADMpNucl |
|---|---|---|---|---|---|---|---|---|---|---|---|---|---|
| gene-LOC105275384 | https://www.ncbi.nlm.nih.gov/gene/?term=LOC105275384 | 1987 | 91 | 457.98 | 1 | 5.03 | 1.1 | 33.74 | 32.35 | 1.39 | 1.39 | 0.06 | 0.06 |
| gene-LOC105276429 | https://www.ncbi.nlm.nih.gov/gene/?term=LOC105276429 | 2901 | 120 | 413.65 | 1 | 3.45 | 0.83 | 23.61 | 22.3 | 1.32 | 1.32 | 0.05 | 0.05 |
| gene-LOC105277903 | https://www.ncbi.nlm.nih.gov/gene/?term=LOC105277903 | 1835 | 35 | 190.74 | 1 | 5.45 | 2.86 | 47.23 | 46.95 | 0.28 | 0.28 | 0.01 | 0.01 |
| gene-LOC105283296 | https://www.ncbi.nlm.nih.gov/gene/?term=LOC105283296 | 2174 | 92 | 423.18 | 1 | 4.6 | 1.09 | 19.15 | 18.97 | 0.18 | 0.18 | 0.01 | 0.01 |
| gene-LOC105280410 | https://www.ncbi.nlm.nih.gov/gene/?term=LOC105280410 | 12580 | 280 | 222.58 | 1 | 0.79 | 0.36 | 19.42 | 19.27 | 0.15 | 0.15 | 0 | 0 |
| gene-LOC105276452 | https://www.ncbi.nlm.nih.gov/gene/?term=LOC105276452 | 2917 | 262 | 898.18 | 1 | 3.43 | 0.38 | 10.28 | 10.37 | -0.08 | 0.08 | -0.01 | 0.01 |
| gene-LOC105288036 | https://www.ncbi.nlm.nih.gov/gene/?term=LOC105288036 | 5110 | 226 | 442.27 | 1 | 1.96 | 0.44 | 16.3 | 16.4 | -0.1 | 0.1 | 0 | 0 |
| gene-LOC105276385 | https://www.ncbi.nlm.nih.gov/gene/?term=LOC105276385 | 11575 | 494 | 426.78 | 1 | 0.86 | 0.2 | 5.31 | 5.56 | -0.25 | 0.25 | -0.01 | 0.01 |
| gene-LOC105281272 | https://www.ncbi.nlm.nih.gov/gene/?term=LOC105281272 | 5252 | 178 | 338.92 | 1 | 1.9 | 0.56 | 16.03 | 16.73 | -0.7 | 0.7 | -0.02 | 0.02 |
| gene-LOC105284507 | https://www.ncbi.nlm.nih.gov/gene/?term=LOC105284507 | 5207 | 94 | 180.53 | 1 | 1.92 | 1.06 | 20.48 | 21.18 | -0.7 | 0.7 | -0.01 | 0.01 |
| gene-LOC105288042 | https://www.ncbi.nlm.nih.gov/gene/?term=LOC105288042 | 3507 | 101 | 288 | 1 | 2.85 | 0.99 | 22.48 | 23.29 | -0.81 | 0.81 | -0.02 | 0.02 |
| gene-LOC105285252 | https://www.ncbi.nlm.nih.gov/gene/?term=LOC105285252 | 7427 | 129 | 173.69 | 1 | 1.35 | 0.78 | 12.88 | 13.9 | -1.02 | 1.02 | -0.02 | 0.02 |
| gene-LOC105283264 | https://www.ncbi.nlm.nih.gov/gene/?term=LOC105283264 | 4005 | 88 | 219.73 | 1 | 2.5 | 1.14 | 18.85 | 20.42 | -1.57 | 1.57 | -0.03 | 0.03 |

Directory: OGL File: ogl-Ob_LI_BCphaseA2.vs.BCphaseB2.txt

| gene_ID | gene_link | gwidth | #Sites | #per10Kb | #dmSites | #dmsp10kb | %dmSites | %pSite1 | %pSite2 | DMpSite | ADMpSite | DMpNucl | ADMpNucl |
|---|---|---|---|---|---|---|---|---|---|---|---|---|---|
| gene-LOC113562345 | https://www.ncbi.nlm.nih.gov/gene/?term=LOC113562345 | 2290 | 69 | 301.31 | 17 | 74.24 | 24.64 | 33.16 | 16.9 | 16.26 | 16.26 | 0.49 | 0.49 |
| gene-LOC105275708 | https://www.ncbi.nlm.nih.gov/gene/?term=LOC105275708 | 1772 | 10 | 56.43 | 1 | 5.64 | 10 | 50.31 | 39.56 | 10.75 | 10.75 | 0.06 | 0.06 |
| gene-LOC105275070 | https://www.ncbi.nlm.nih.gov/gene/?term=LOC105275070 | 5667 | 47 | 82.94 | 5 | 8.82 | 10.64 | 11.7 | 2 | 9.7 | 9.7 | 0.08 | 0.08 |
| gene-LOC105286694 | https://www.ncbi.nlm.nih.gov/gene/?term=LOC105286694 | 3409 | 11 | 32.27 | 1 | 2.93 | 9.09 | 63.21 | 54.71 | 8.5 | 8.5 | 0.03 | 0.03 |
| gene-LOC105276078 | https://www.ncbi.nlm.nih.gov/gene/?term=LOC105276078 | 1051 | 16 | 152.24 | 1 | 9.51 | 6.25 | 32.79 | 26.49 | 6.3 | 6.3 | 0.1 | 0.1 |
| gene-LOC105279898 | https://www.ncbi.nlm.nih.gov/gene/?term=LOC105279898 | 1343 | 14 | 104.24 | 1 | 7.45 | 7.14 | 61.13 | 54.89 | 6.24 | 6.24 | 0.07 | 0.07 |
| gene-LOC105282976 | https://www.ncbi.nlm.nih.gov/gene/?term=LOC105282976 | 1648 | 50 | 303.4 | 1 | 6.07 | 2 | 26.51 | 20.9 | 5.61 | 5.61 | 0.17 | 0.17 |
| gene-LOC105286302 | https://www.ncbi.nlm.nih.gov/gene/?term=LOC105286302 | 4536 | 18 | 39.68 | 1 | 2.2 | 5.56 | 22.71 | 17.52 | 5.2 | 5.2 | 0.02 | 0.02 |
| gene-LOC105275447 | https://www.ncbi.nlm.nih.gov/gene/?term=LOC105275447 | 1846 | 35 | 189.6 | 1 | 5.42 | 2.86 | 8.76 | 3.6 | 5.15 | 5.15 | 0.1 | 0.1 |
| gene-LOC105274900 | https://www.ncbi.nlm.nih.gov/gene/?term=LOC105274900 | 5561 | 52 | 93.51 | 2 | 3.6 | 3.85 | 36.59 | 32.11 | 4.48 | 4.48 | 0.04 | 0.04 |
| gene-LOC105286683 | https://www.ncbi.nlm.nih.gov/gene/?term=LOC105286683 | 1665 | 51 | 306.31 | 1 | 6.01 | 1.96 | 35.98 | 32.15 | 3.83 | 3.83 | 0.12 | 0.12 |
| gene-LOC105288063 | https://www.ncbi.nlm.nih.gov/gene/?term=LOC105288063 | 3400 | 53 | 155.88 | 1 | 2.94 | 1.89 | 16.83 | 13.02 | 3.81 | 3.81 | 0.06 | 0.06 |
| gene-LOC105283649 | https://www.ncbi.nlm.nih.gov/gene/?term=LOC105283649 | 1918 | 56 | 291.97 | 2 | 10.43 | 3.57 | 38.44 | 34.8 | 3.63 | 3.63 | 0.11 | 0.11 |
| gene-LOC113561805 | https://www.ncbi.nlm.nih.gov/gene/?term=LOC113561805 | 3337 | 77 | 230.75 | 1 | 3 | 1.3 | 37.31 | 34.13 | 3.18 | 3.18 | 0.07 | 0.07 |
| gene-LOC105285315 | https://www.ncbi.nlm.nih.gov/gene/?term=LOC105285315 | 5458 | 22 | 40.31 | 1 | 1.83 | 4.55 | 41.52 | 38.4 | 3.12 | 3.12 | 0.01 | 0.01 |
| gene-LOC105285674 | https://www.ncbi.nlm.nih.gov/gene/?term=LOC105285674 | 3035 | 93 | 306.43 | 2 | 6.59 | 2.15 | 27.41 | 24.3 | 3.11 | 3.11 | 0.1 | 0.1 |
| gene-LOC105281515 | https://www.ncbi.nlm.nih.gov/gene/?term=LOC105281515 | 8827 | 48 | 54.38 | 2 | 2.27 | 4.17 | 32.1 | 29.31 | 2.79 | 2.79 | 0.02 | 0.02 |
| gene-LOC105281420 | https://www.ncbi.nlm.nih.gov/gene/?term=LOC105281420 | 3882 | 55 | 141.68 | 1 | 2.58 | 1.82 | 15.69 | 12.96 | 2.73 | 2.73 | 0.04 | 0.04 |
| gene-LOC105280641 | https://www.ncbi.nlm.nih.gov/gene/?term=LOC105280641 | 3682 | 49 | 133.08 | 1 | 2.72 | 2.04 | 20.13 | 17.42 | 2.71 | 2.71 | 0.04 | 0.04 |
| gene-LOC105275347 | https://www.ncbi.nlm.nih.gov/gene/?term=LOC105275347 | 2526 | 15 | 59.38 | 1 | 3.96 | 6.67 | 31.33 | 28.89 | 2.44 | 2.44 | 0.01 | 0.01 |
| gene-LOC105275966 | https://www.ncbi.nlm.nih.gov/gene/?term=LOC105275966 | 3299 | 61 | 184.9 | 1 | 3.03 | 1.64 | 49.58 | 47.22 | 2.36 | 2.36 | 0.04 | 0.04 |
| gene-LOC105282156 | https://www.ncbi.nlm.nih.gov/gene/?term=LOC105282156 | 2577 | 136 | 527.75 | 1 | 3.88 | 0.74 | 26.99 | 24.63 | 2.36 | 2.36 | 0.12 | 0.12 |
| gene-LOC105276449 | https://www.ncbi.nlm.nih.gov/gene/?term=LOC105276449 | 1231 | 42 | 341.19 | 1 | 8.12 | 2.38 | 35.95 | 33.63 | 2.31 | 2.31 | 0.08 | 0.08 |
| gene-LOC105287648 | https://www.ncbi.nlm.nih.gov/gene/?term=LOC105287648 | 2205 | 27 | 122.45 | 1 | 4.54 | 3.7 | 41.48 | 39.19 | 2.29 | 2.29 | 0.03 | 0.03 |
| gene-LOC105278454 | https://www.ncbi.nlm.nih.gov/gene/?term=LOC105278454 | 1972 | 47 | 238.34 | 1 | 5.07 | 2.13 | 44.8 | 42.52 | 2.28 | 2.28 | 0.05 | 0.05 |
| gene-LOC105288066 | https://www.ncbi.nlm.nih.gov/gene/?term=LOC105288066 | 1964 | 21 | 106.92 | 1 | 5.09 | 4.76 | 46.21 | 43.96 | 2.25 | 2.25 | 0.02 | 0.02 |
| gene-LOC105275518 | https://www.ncbi.nlm.nih.gov/gene/?term=LOC105275518 | 2335 | 19 | 81.37 | 1 | 4.28 | 5.26 | 46.5 | 44.26 | 2.24 | 2.24 | 0.02 | 0.02 |
| gene-LOC105278109 | https://www.ncbi.nlm.nih.gov/gene/?term=LOC105278109 | 4029 | 65 | 161.33 | 1 | 2.48 | 1.54 | 48.15 | 45.97 | 2.19 | 2.19 | 0.04 | 0.04 |
| gene-LOC105282149 | https://www.ncbi.nlm.nih.gov/gene/?term=LOC105282149 | 1787 | 108 | 604.36 | 1 | 5.6 | 0.93 | 43.65 | 41.49 | 2.17 | 2.17 | 0.13 | 0.13 |
| gene-LOC105283592 | https://www.ncbi.nlm.nih.gov/gene/?term=LOC105283592 | 3479 | 71 | 204.08 | 1 | 2.87 | 1.41 | 26.49 | 24.49 | 2.01 | 2.01 | 0.04 | 0.04 |
| gene-LOC105286514 | https://www.ncbi.nlm.nih.gov/gene/?term=LOC105286514 | 2528 | 41 | 162.18 | 1 | 3.96 | 2.44 | 35.27 | 33.32 | 1.95 | 1.95 | 0.03 | 0.03 |
| gene-LOC105282970 | https://www.ncbi.nlm.nih.gov/gene/?term=LOC105282970 | 1529 | 39 | 255.07 | 1 | 6.54 | 2.56 | 54 | 52.14 | 1.86 | 1.86 | 0.05 | 0.05 |
| gene-LOC105282151 | https://www.ncbi.nlm.nih.gov/gene/?term=LOC105282151 | 1601 | 41 | 256.09 | 3 | 18.74 | 7.32 | 36.24 | 34.41 | 1.83 | 1.83 | 0.05 | 0.05 |
| gene-LOC105277856 | https://www.ncbi.nlm.nih.gov/gene/?term=LOC105277856 | 3442 | 34 | 98.78 | 1 | 2.91 | 2.94 | 37.69 | 35.88 | 1.81 | 1.81 | 0.02 | 0.02 |
| gene-LOC105286644 | https://www.ncbi.nlm.nih.gov/gene/?term=LOC105286644 | 1235 | 20 | 161.94 | 1 | 8.1 | 5 | 39.34 | 37.53 | 1.81 | 1.81 | 0.03 | 0.03 |
| gene-LOC105281848 | https://www.ncbi.nlm.nih.gov/gene/?term=LOC105281848 | 2122 | 56 | 263.9 | 1 | 4.71 | 1.79 | 24.08 | 22.28 | 1.8 | 1.8 | 0.05 | 0.05 |
| gene-LOC105280774 | https://www.ncbi.nlm.nih.gov/gene/?term=LOC105280774 | 2793 | 102 | 365.2 | 1 | 3.58 | 0.98 | 32.11 | 30.36 | 1.75 | 1.75 | 0.06 | 0.06 |
| gene-LOC105282658 | https://www.ncbi.nlm.nih.gov/gene/?term=LOC105282658 | 2421 | 65 | 268.48 | 1 | 4.13 | 1.54 | 21.38 | 19.92 | 1.46 | 1.46 | 0.04 | 0.04 |
| gene-LOC105280786 | https://www.ncbi.nlm.nih.gov/gene/?term=LOC105280786 | 2573 | 111 | 431.4 | 1 | 3.89 | 0.9 | 33.28 | 31.87 | 1.41 | 1.41 | 0.06 | 0.06 |
| gene-LOC105288010 | https://www.ncbi.nlm.nih.gov/gene/?term=LOC105288010 | 3189 | 39 | 122.3 | 1 | 3.14 | 2.56 | 37.68 | 36.3 | 1.38 | 1.38 | 0.02 | 0.02 |
| gene-LOC105280348 | https://www.ncbi.nlm.nih.gov/gene/?term=LOC105280348 | 3797 | 65 | 171.19 | 1 | 2.63 | 1.54 | 21.44 | 20.1 | 1.33 | 1.33 | 0.02 | 0.02 |
| gene-LOC105287095 | https://www.ncbi.nlm.nih.gov/gene/?term=LOC105287095 | 1435 | 57 | 397.21 | 1 | 6.97 | 1.75 | 29.87 | 28.54 | 1.33 | 1.33 | 0.05 | 0.05 |
| gene-LOC105277761 | https://www.ncbi.nlm.nih.gov/gene/?term=LOC105277761 | 4648 | 146 | 314.11 | 1 | 2.15 | 0.68 | 16.93 | 15.61 | 1.32 | 1.32 | 0.04 | 0.04 |
| gene-LOC105282600 | https://www.ncbi.nlm.nih.gov/gene/?term=LOC105282600 | 3715 | 53 | 142.66 | 1 | 2.69 | 1.89 | 32.07 | 30.77 | 1.3 | 1.3 | 0.02 | 0.02 |
| gene-LOC105276059 | https://www.ncbi.nlm.nih.gov/gene/?term=LOC105276059 | 6256 | 121 | 193.41 | 1 | 1.6 | 0.83 | 11.92 | 10.67 | 1.25 | 1.25 | 0.02 | 0.02 |
| gene-LOC105286190 | https://www.ncbi.nlm.nih.gov/gene/?term=LOC105286190 | 6447 | 298 | 462.23 | 1 | 1.55 | 0.34 | 8.47 | 7.3 | 1.16 | 1.16 | 0.05 | 0.05 |
| gene-LOC105279664 | https://www.ncbi.nlm.nih.gov/gene/?term=LOC105279664 | 4023 | 54 | 134.23 | 1 | 2.49 | 1.85 | 21.29 | 20.13 | 1.15 | 1.15 | 0.02 | 0.02 |
| gene-LOC105279298 | https://www.ncbi.nlm.nih.gov/gene/?term=LOC105279298 | 2773 | 128 | 461.59 | 2 | 7.21 | 1.56 | 25.02 | 23.92 | 1.1 | 1.1 | 0.05 | 0.05 |
| gene-LOC105285502 | https://www.ncbi.nlm.nih.gov/gene/?term=LOC105285502 | 9990 | 143 | 143.14 | 1 | 1 | 0.7 | 10.37 | 9.32 | 1.05 | 1.05 | 0.02 | 0.02 |
| gene-LOC105278497 | https://www.ncbi.nlm.nih.gov/gene/?term=LOC105278497 | 2072 | 94 | 453.67 | 1 | 4.83 | 1.06 | 21.1 | 20.09 | 1.01 | 1.01 | 0.05 | 0.05 |
| gene-LOC105274856 | https://www.ncbi.nlm.nih.gov/gene/?term=LOC105274856 | 5216 | 218 | 417.94 | 1 | 1.92 | 0.46 | 19.03 | 18.03 | 1 | 1 | 0.04 | 0.04 |
| gene-LOC105278807 | https://www.ncbi.nlm.nih.gov/gene/?term=LOC105278807 | 2166 | 73 | 337.03 | 1 | 4.62 | 1.37 | 18.28 | 17.29 | 0.99 | 0.99 | 0.03 | 0.03 |
| gene-LOC105284355 | https://www.ncbi.nlm.nih.gov/gene/?term=LOC105284355 | 3904 | 149 | 381.66 | 1 | 2.56 | 0.67 | 21.89 | 20.9 | 0.99 | 0.99 | 0.04 | 0.04 |
| gene-LOC105276704 | https://www.ncbi.nlm.nih.gov/gene/?term=LOC105276704 | 2063 | 98 | 475.04 | 1 | 4.85 | 1.02 | 13.8 | 12.82 | 0.98 | 0.98 | 0.05 | 0.05 |
| gene-LOC105278877 | https://www.ncbi.nlm.nih.gov/gene/?term=LOC105278877 | 1968 | 75 | 381.1 | 1 | 5.08 | 1.33 | 34.6 | 33.65 | 0.95 | 0.95 | 0.04 | 0.04 |
| gene-LOC105282794 | https://www.ncbi.nlm.nih.gov/gene/?term=LOC105282794 | 4571 | 93 | 203.46 | 1 | 2.19 | 1.08 | 18.93 | 18.03 | 0.9 | 0.9 | 0.02 | 0.02 |
| gene-LOC105288078 | https://www.ncbi.nlm.nih.gov/gene/?term=LOC105288078 | 19126 | 294 | 153.72 | 1 | 0.52 | 0.34 | 18.24 | 17.37 | 0.87 | 0.87 | 0.01 | 0.01 |
| gene-LOC105284184 | https://www.ncbi.nlm.nih.gov/gene/?term=LOC105284184 | 2535 | 85 | 335.31 | 1 | 3.94 | 1.18 | 34.37 | 33.53 | 0.83 | 0.83 | 0.03 | 0.03 |
| gene-LOC105285055 | https://www.ncbi.nlm.nih.gov/gene/?term=LOC105285055 | 4108 | 259 | 630.48 | 2 | 4.87 | 0.77 | 11.6 | 10.8 | 0.8 | 0.8 | 0.05 | 0.05 |
| gene-LOC105284329 | https://www.ncbi.nlm.nih.gov/gene/?term=LOC105284329 | 4051 | 50 | 123.43 | 1 | 2.47 | 2 | 19.81 | 19.02 | 0.79 | 0.79 | 0.01 | 0.01 |

| | | | | | | | | | | | | | |
|---|---|---|---|---|---|---|---|---|---|---|---|---|---|
| gene-LOC105276538 | https://www.ncbi.nlm.nih.gov/gene/?term=LOC105276538 | 1511 | 19 | 125.74 | 1 | 6.62 | 5.26 | 70.24 | 69.47 | 0.77 | 0.77 | 0.01 | 0.01 |
| gene-LOC105285202 | https://www.ncbi.nlm.nih.gov/gene/?term=LOC105285202 | 1838 | 66 | 359.09 | 1 | 5.44 | 1.52 | 25.35 | 24.58 | 0.77 | 0.77 | 0.03 | 0.03 |
| gene-LOC105275451 | https://www.ncbi.nlm.nih.gov/gene/?term=LOC105275451 | 6035 | 127 | 210.44 | 1 | 1.66 | 0.79 | 22.96 | 22.23 | 0.73 | 0.73 | 0.02 | 0.02 |
| gene-LOC105274452 | https://www.ncbi.nlm.nih.gov/gene/?term=LOC105274452 | 6595 | 261 | 395.75 | 1 | 1.52 | 0.38 | 13.63 | 12.94 | 0.69 | 0.69 | 0.03 | 0.03 |
| gene-LOC105283989 | https://www.ncbi.nlm.nih.gov/gene/?term=LOC105283989 | 4948 | 26 | 52.55 1 | 2.02 | 3.85 | 43.53 | 42.86 | 0.68 | 0.68 | 0 | 0 | |
| gene-LOC105280956 | https://www.ncbi.nlm.nih.gov/gene/?term=LOC105280956 | 2639 | 75 | 284.2 1 | 3.79 | 1.33 | 17.09 | 16.45 | 0.64 | 0.64 | 0.02 | 0.02 | |
| gene-LOC105279038 | https://www.ncbi.nlm.nih.gov/gene/?term=LOC105279038 | 9453 | 333 | 352.27 | 1 | 1.06 | 0.3 | 9.29 | 8.67 | 0.62 | 0.62 | 0.02 | 0.02 |
| gene-LOC105287063 | https://www.ncbi.nlm.nih.gov/gene/?term=LOC105287063 | 1874 | 48 | 256.14 | 1 | 5.34 | 2.08 | 30.52 | 29.9 | 0.62 | 0.62 | 0.02 | 0.02 |
| gene-LOC105274823 | https://www.ncbi.nlm.nih.gov/gene/?term=LOC105274823 | 4620 | 138 | 298.7 1 | 2.16 | 0.72 | 21.48 | 20.88 | 0.6 | 0.6 | 0.02 | 0.02 | |
| gene-LOC105280148 | https://www.ncbi.nlm.nih.gov/gene/?term=LOC105280148 | 3120 | 214 | 685.9 1 | 3.21 | 0.47 | 11.76 | 11.17 | 0.6 | 0.6 | 0.04 | 0.04 | |
| gene-LOC105280002 | https://www.ncbi.nlm.nih.gov/gene/?term=LOC105280002 | 1940 | 25 | 128.87 | 1 | 5.15 | 4 | 38.59 | 38.02 | 0.58 | 0.58 | 0.01 | 0.01 |
| gene-LOC105285807 | https://www.ncbi.nlm.nih.gov/gene/?term=LOC105285807 | 6063 | 218 | 359.56 | 1 | 1.65 | 0.46 | 7.08 | 6.5 | 0.58 | 0.58 | 0.02 | 0.02 |
| gene-LOC105286799 | https://www.ncbi.nlm.nih.gov/gene/?term=LOC105286799 | 3557 | 183 | 514.48 | 1 | 2.81 | 0.55 | 16.25 | 15.68 | 0.57 | 0.57 | 0.03 | 0.03 |
| gene-LOC105274851 | https://www.ncbi.nlm.nih.gov/gene/?term=LOC105274851 | 6616 | 128 | 193.47 | 1 | 1.51 | 0.78 | 11.71 | 11.16 | 0.54 | 0.54 | 0.01 | 0.01 |
| gene-LOC105275683 | https://www.ncbi.nlm.nih.gov/gene/?term=LOC105275683 | 4256 | 194 | 455.83 | 1 | 2.35 | 0.52 | 13.24 | 12.7 | 0.54 | 0.54 | 0.02 | 0.02 |
| gene-LOC105283943 | https://www.ncbi.nlm.nih.gov/gene/?term=LOC105283943 | 3559 | 58 | 162.97 | 1 | 2.81 | 1.72 | 14.89 | 14.35 | 0.54 | 0.54 | 0.01 | 0.01 |
| gene-LOC105278666 | https://www.ncbi.nlm.nih.gov/gene/?term=LOC105278666 | 10341 | 459 | 443.86 | 1 | 0.97 | 0.22 | 7.32 | 6.79 | 0.53 | 0.53 | 0.02 | 0.02 |
| gene-LOC105280723 | https://www.ncbi.nlm.nih.gov/gene/?term=LOC105280723 | 3572 | 170 | 475.92 | 1 | 2.8 | 0.59 | 16.88 | 16.35 | 0.53 | 0.53 | 0.03 | 0.03 |
| gene-LOC105285179 | https://www.ncbi.nlm.nih.gov/gene/?term=LOC105285179 | 4191 | 164 | 391.31 | 1 | 2.39 | 0.61 | 20.24 | 19.71 | 0.53 | 0.53 | 0.02 | 0.02 |
| gene-LOC105283123 | https://www.ncbi.nlm.nih.gov/gene/?term=LOC105283123 | 1507 | 52 | 345.06 | 1 | 6.64 | 1.92 | 17.86 | 17.35 | 0.52 | 0.52 | 0.02 | 0.02 |
| gene-LOC105285639 | https://www.ncbi.nlm.nih.gov/gene/?term=LOC105285639 | 10325 | 356 | 344.79 | 1 | 0.97 | 0.28 | 11.26 | 10.77 | 0.49 | 0.49 | 0.02 | 0.02 |
| gene-LOC105275848 | https://www.ncbi.nlm.nih.gov/gene/?term=LOC105275848 | 3718 | 139 | 373.86 | 1 | 2.69 | 0.72 | 22.47 | 21.99 | 0.48 | 0.48 | 0.02 | 0.02 |
| gene-LOC105285514 | https://www.ncbi.nlm.nih.gov/gene/?term=LOC105285514 | 1277 | 13 | 101.8 1 | 7.83 | 7.69 | 90.77 | 90.29 | 0.48 | 0.48 | 0 | 0 | |
| gene-LOC105280410 | https://www.ncbi.nlm.nih.gov/gene/?term=LOC105280410 | 12580 | 237 | 188.39 | 1 | 0.79 | 0.42 | 21.06 | 20.59 | 0.47 | 0.47 | 0.01 | 0.01 |
| gene-LOC105279657 | https://www.ncbi.nlm.nih.gov/gene/?term=LOC105279657 | 5878 | 86 | 146.31 | 1 | 1.7 | 1.16 | 19.59 | 19.13 | 0.46 | 0.46 | 0.01 | 0.01 |
| gene-LOC105283728 | https://www.ncbi.nlm.nih.gov/gene/?term=LOC105283728 | 3256 | 113 | 347.05 | 2 | 6.14 | 1.77 | 20.45 | 20.03 | 0.42 | 0.42 | 0.01 | 0.01 |
| gene-LOC113562050 | https://www.ncbi.nlm.nih.gov/gene/?term=LOC113562050 | 17097 | 583 | 341 1 | 0.58 | 0.17 | 5.52 | 5.1 | 0.42 | 0.42 | 0.01 | 0.01 | |
| gene-LOC105278145 | https://www.ncbi.nlm.nih.gov/gene/?term=LOC105278145 | 4470 | 241 | 539.15 | 1 | 2.24 | 0.41 | 10.99 | 10.59 | 0.4 | 0.4 | 0.02 | 0.02 |
| gene-LOC105286452 | https://www.ncbi.nlm.nih.gov/gene/?term=LOC105286452 | 6888 | 61 | 88.56 1 | 1.45 | 1.64 | 32.75 | 32.35 | 0.4 | 0.4 | 0 | 0 | |
| gene-LOC105276196 | https://www.ncbi.nlm.nih.gov/gene/?term=LOC105276196 | 2953 | 48 | 162.55 | 1 | 3.39 | 2.08 | 45.56 | 45.18 | 0.38 | 0.38 | 0.01 | 0.01 |
| gene-LOC105278632 | https://www.ncbi.nlm.nih.gov/gene/?term=LOC105278632 | 3624 | 129 | 355.96 | 1 | 2.76 | 0.78 | 14.81 | 14.44 | 0.37 | 0.37 | 0.01 | 0.01 |
| gene-LOC105285395 | https://www.ncbi.nlm.nih.gov/gene/?term=LOC105285395 | 8085 | 196 | 242.42 | 1 | 1.24 | 0.51 | 15.02 | 14.69 | 0.33 | 0.33 | 0.01 | 0.01 |
| gene-LOC105283712 | https://www.ncbi.nlm.nih.gov/gene/?term=LOC105283712 | 1910 | 31 | 162.3 1 | 5.24 | 3.23 | 30.95 | 30.64 | 0.31 | 0.31 | 0.01 | 0.01 | |
| gene-LOC105276941 | https://www.ncbi.nlm.nih.gov/gene/?term=LOC105276941 | 15956 | 551 | 345.32 | 1 | 0.63 | 0.18 | 5.1 | 4.82 | 0.28 | 0.28 | 0.01 | 0.01 |
| gene-LOC105275374 | https://www.ncbi.nlm.nih.gov/gene/?term=LOC105275374 | 11376 | 527 | 463.26 | 1 | 0.88 | 0.19 | 11.34 | 11.07 | 0.27 | 0.27 | 0.01 | 0.01 |
| gene-LOC105287417 | https://www.ncbi.nlm.nih.gov/gene/?term=LOC105287417 | 1025 | 28 | 273.17 | 1 | 9.76 | 3.57 | 42.07 | 41.82 | 0.26 | 0.26 | 0.01 | 0.01 |
| gene-LOC105276452 | https://www.ncbi.nlm.nih.gov/gene/?term=LOC105276452 | 2917 | 257 | 881.04 | 1 | 3.43 | 0.39 | 9.59 | 9.37 | 0.22 | 0.22 | 0.02 | 0.02 |
| gene-LOC105275327 | https://www.ncbi.nlm.nih.gov/gene/?term=LOC105275327 | 14245 | 493 | 346.09 | 1 | 0.7 | 0.2 | 6.9 | 6.7 | 0.19 | 0.19 | 0.01 | 0.01 |
| gene-LOC105287550 | https://www.ncbi.nlm.nih.gov/gene/?term=LOC105287550 | 15659 | 570 | 364.01 | 1 | 0.64 | 0.18 | 6.53 | 6.34 | 0.19 | 0.19 | 0.01 | 0.01 |
| gene-LOC105278282 | https://www.ncbi.nlm.nih.gov/gene/?term=LOC105278282 | 1775 | 56 | 315.49 | 1 | 5.63 | 1.79 | 35.74 | 35.56 | 0.18 | 0.18 | 0.01 | 0.01 |
| gene-LOC105281272 | https://www.ncbi.nlm.nih.gov/gene/?term=LOC105281272 | 5252 | 178 | 338.92 | 1 | 1.9 | 0.56 | 16.1 | 15.92 | 0.18 | 0.18 | 0.01 | 0.01 |
| gene-LOC105282491 | https://www.ncbi.nlm.nih.gov/gene/?term=LOC105282491 | 4761 | 165 | 346.57 | 1 | 2.1 | 0.61 | 8.62 | 8.44 | 0.18 | 0.18 | 0.01 | 0.01 |
| gene-LOC105282811 | https://www.ncbi.nlm.nih.gov/gene/?term=LOC105282811 | 5765 | 203 | 352.12 | 1 | 1.73 | 0.49 | 8.42 | 8.31 | 0.12 | 0.12 | 0 | 0 |
| gene-LOC105282825 | https://www.ncbi.nlm.nih.gov/gene/?term=LOC105282825 | 3919 | 80 | 204.13 | 1 | 2.55 | 1.25 | 12.57 | 12.46 | 0.11 | 0.11 | 0 | 0 |
| gene-LOC105274590 | https://www.ncbi.nlm.nih.gov/gene/?term=LOC105274590 | 4516 | 58 | 128.43 | 2 | 4.43 | 3.45 | 35.68 | 35.59 | 0.09 | 0.09 | 0 | 0 |
| gene-LOC105275363 | https://www.ncbi.nlm.nih.gov/gene/?term=LOC105275363 | 3798 | 156 | 410.74 | 1 | 2.63 | 0.64 | 17.23 | 17.17 | 0.07 | 0.07 | 0 | 0 |
| gene-LOC105283296 | https://www.ncbi.nlm.nih.gov/gene/?term=LOC105283296 | 2174 | 84 | 386.38 | 1 | 4.6 | 1.19 | 21.67 | 21.61 | 0.07 | 0.07 | 0 | 0 |
| gene-LOC105281700 | https://www.ncbi.nlm.nih.gov/gene/?term=LOC105281700 | 2208 | 37 | 167.57 | 1 | 4.53 | 2.7 | 39.7 | 39.67 | 0.04 | 0.04 | 0 | 0 |
| gene-LOC105274531 | https://www.ncbi.nlm.nih.gov/gene/?term=LOC105274531 | 7774 | 284 | 365.32 | 1 | 1.29 | 0.35 | 10.8 | 10.77 | 0.03 | 0.03 | 0 | 0 |
| gene-LOC105284043 | https://www.ncbi.nlm.nih.gov/gene/?term=LOC105284043 | 5661 | 279 | 492.85 | 1 | 1.77 | 0.36 | 10.73 | 10.7 | 0.03 | 0.03 | 0 | 0 |
| gene-LOC105275460 | https://www.ncbi.nlm.nih.gov/gene/?term=LOC105275460 | 2473 | 114 | 460.98 | 1 | 4.04 | 0.88 | 13.89 | 13.88 | 0.02 | 0.02 | 0 | 0 |
| gene-LOC105285406 | https://www.ncbi.nlm.nih.gov/gene/?term=LOC105285406 | 8015 | 59 | 73.61 1 | 1.25 | 1.69 | 26.4 | 26.39 | 0.01 | 0.01 | 0 | 0 | |
| gene-LOC105275179 | https://www.ncbi.nlm.nih.gov/gene/?term=LOC105275179 | 3271 | 73 | 223.17 | 1 | 3.06 | 1.37 | 20.99 | 20.99 | 0 | 0 | 0 | 0 |
| gene-LOC105278228 | https://www.ncbi.nlm.nih.gov/gene/?term=LOC105278228 | 8610 | 162 | 188.15 | 1 | 1.16 | 0.62 | 21.49 | 21.49 | 0 | 0 | 0 | 0 |
| gene-LOC105287764 | https://www.ncbi.nlm.nih.gov/gene/?term=LOC105287764 | 18359 | 396 | 215.7 1 | 0.54 | 0.25 | 2.87 | 2.88 | -0.01 | 0.01 | 0 | 0 | |
| gene-LOC105278283 | https://www.ncbi.nlm.nih.gov/gene/?term=LOC105278283 | 2317 | 125 | 539.49 | 1 | 4.32 | 0.8 | 28 | 28.03 | -0.02 | 0.02 | 0 | 0 |
| gene-LOC105282951 | https://www.ncbi.nlm.nih.gov/gene/?term=LOC105282951 | 3366 | 118 | 350.56 | 1 | 2.97 | 0.85 | 19.88 | 19.9 | -0.02 | 0.02 | 0 | 0 |
| gene-LOC105285533 | https://www.ncbi.nlm.nih.gov/gene/?term=LOC105285533 | 1640 | 60 | 365.85 | 1 | 6.1 | 1.67 | 26.06 | 26.09 | -0.03 | 0.03 | 0 | 0 |
| gene-LOC105276527 | https://www.ncbi.nlm.nih.gov/gene/?term=LOC105276527 | 7923 | 318 | 401.36 | 1 | 1.26 | 0.31 | 9.67 | 9.76 | -0.09 | 0.09 | 0 | 0 |
| gene-LOC105286649 | https://www.ncbi.nlm.nih.gov/gene/?term=LOC105286649 | 14178 | 826 | 582.59 | 1 | 0.71 | 0.12 | 0.92 | 1.04 | -0.12 | 0.12 | -0.01 | 0.01 |
| gene-LOC105282885 | https://www.ncbi.nlm.nih.gov/gene/?term=LOC105282885 | 1870 | 43 | 229.95 | 1 | 5.35 | 2.33 | 44.33 | 44.46 | -0.13 | 0.13 | 0 | 0 |
| gene-LOC105276462 | https://www.ncbi.nlm.nih.gov/gene/?term=LOC105276462 | 6561 | 278 | 423.72 | 1 | 1.52 | 0.36 | 7.35 | 7.5 | -0.15 | 0.15 | -0.01 | 0.01 |
| gene-LOC105282646 | https://www.ncbi.nlm.nih.gov/gene/?term=LOC105282646 | 3317 | 97 | 292.43 | 1 | 3.01 | 1.03 | 22.32 | 22.49 | -0.17 | 0.17 | -0.01 | 0.01 |
| gene-LOC105278691 | https://www.ncbi.nlm.nih.gov/gene/?term=LOC105278691 | 3871 | 147 | 379.75 | 1 | 2.58 | 0.68 | 14.28 | 14.5 | -0.22 | 0.22 | -0.01 | 0.01 |

| | | | | | | | | | | | | | |
|---|---|---|---|---|---|---|---|---|---|---|---|---|---|
| gene-LOC105280819 | https://www.ncbi.nlm.nih.gov/gene/?term=LOC105280819 | 1639 | 39 | 237.95 | 1 | 6.1 | 2.56 | 35.75 | 36.01 | -0.26 | 0.26 | -0.01 | 0.01 |
| gene-LOC105275187 | https://www.ncbi.nlm.nih.gov/gene/?term=LOC105275187 | 5534 | 70 | 126.49 | 1 | 1.81 | 1.43 | 14.08 | 14.45 | -0.36 | 0.36 | 0 | 0 |
| gene-LOC105275587 | https://www.ncbi.nlm.nih.gov/gene/?term=LOC105275587 | 2404 | 55 | 228.79 | 1 | 4.16 | 1.82 | 34.41 | 34.8 | -0.39 | 0.39 | -0.01 | 0.01 |
| gene-LOC105281312 | https://www.ncbi.nlm.nih.gov/gene/?term=LOC105281312 | 2551 | 72 | 282.24 | 1 | 3.92 | 1.39 | 29.66 | 30.06 | -0.4 | 0.4 | -0.01 | 0.01 |
| gene-LOC105286515 | https://www.ncbi.nlm.nih.gov/gene/?term=LOC105286515 | 3751 | 116 | 309.25 | 1 | 2.67 | 0.86 | 10.42 | 10.83 | -0.41 | 0.41 | -0.01 | 0.01 |
| gene-LOC105286999 | https://www.ncbi.nlm.nih.gov/gene/?term=LOC105286999 | 2141 | 41 | 191.5 | 1 | 4.67 | 2.44 | 37.42 | 37.83 | -0.41 | 0.41 | -0.01 | 0.01 |
| gene-LOC105286274 | https://www.ncbi.nlm.nih.gov/gene/?term=LOC105286274 | 3817 | 202 | 529.21 | 1 | 2.62 | 0.5 | 16.44 | 16.87 | -0.43 | 0.43 | -0.02 | 0.02 |
| gene-LOC105281754 | https://www.ncbi.nlm.nih.gov/gene/?term=LOC105281754 | 4300 | 117 | 272.09 | 1 | 2.33 | 0.85 | 16.59 | 17.09 | -0.5 | 0.5 | -0.01 | 0.01 |
| gene-LOC105283385 | https://www.ncbi.nlm.nih.gov/gene/?term=LOC105283385 | 9540 | 305 | 319.71 | 1 | 1.05 | 0.33 | 12.66 | 13.17 | -0.51 | 0.51 | -0.02 | 0.02 |
| gene-LOC113561621 | https://www.ncbi.nlm.nih.gov/gene/?term=LOC113561621 | 2249 | 78 | 346.82 | 1 | 4.45 | 1.28 | 20.85 | 21.38 | -0.53 | 0.53 | -0.02 | 0.02 |
| gene-LOC105278534 | https://www.ncbi.nlm.nih.gov/gene/?term=LOC105278534 | 7364 | 180 | 244.43 | 1 | 1.36 | 0.56 | 18.21 | 18.75 | -0.54 | 0.54 | -0.01 | 0.01 |
| gene-LOC105276272 | https://www.ncbi.nlm.nih.gov/gene/?term=LOC105276272 | 1965 | 36 | 183.21 | 1 | 5.09 | 2.78 | 32.08 | 32.69 | -0.61 | 0.61 | -0.01 | 0.01 |
| gene-LOC105281173 | https://www.ncbi.nlm.nih.gov/gene/?term=LOC105281173 | 1595 | 38 | 238.24 | 1 | 6.27 | 2.63 | 60.05 | 60.67 | -0.62 | 0.62 | -0.01 | 0.01 |
| gene-LOC105285742 | https://www.ncbi.nlm.nih.gov/gene/?term=LOC105285742 | 3446 | 107 | 310.5 | 1 | 2.9 | 0.93 | 21.42 | 22.06 | -0.64 | 0.64 | -0.02 | 0.02 |
| gene-LOC105287633 | https://www.ncbi.nlm.nih.gov/gene/?term=LOC105287633 | 1524 | 27 | 177.17 | 1 | 6.56 | 3.7 | 63.52 | 64.22 | -0.7 | 0.7 | -0.01 | 0.01 |
| gene-LOC105274534 | https://www.ncbi.nlm.nih.gov/gene/?term=LOC105274534 | 1329 | 54 | 406.32 | 1 | 7.52 | 1.85 | 45.08 | 45.82 | -0.75 | 0.75 | -0.03 | 0.03 |
| gene-LOC109611283 | https://www.ncbi.nlm.nih.gov/gene/?term=LOC109611283 | 1451 | 38 | 261.89 | 1 | 6.89 | 2.63 | 35.88 | 36.65 | -0.77 | 0.77 | -0.02 | 0.02 |
| gene-LOC105274706 | https://www.ncbi.nlm.nih.gov/gene/?term=LOC105274706 | 1446 | 55 | 380.36 | 1 | 6.92 | 1.82 | 37.25 | 38.23 | -0.98 | 0.98 | -0.04 | 0.04 |
| gene-LOC105274449 | https://www.ncbi.nlm.nih.gov/gene/?term=LOC105274449 | 1549 | 53 | 342.16 | 1 | 6.46 | 1.89 | 50.38 | 51.77 | -1.39 | 1.39 | -0.05 | 0.05 |
| gene-LOC105285252 | https://www.ncbi.nlm.nih.gov/gene/?term=LOC105285252 | 7427 | 117 | 157.53 | 1 | 1.35 | 0.85 | 13.64 | 15.21 | -1.57 | 1.57 | -0.02 | 0.02 |
| gene-LOC105284507 | https://www.ncbi.nlm.nih.gov/gene/?term=LOC105284507 | 5207 | 76 | 145.96 | 1 | 1.92 | 1.32 | 19.78 | 21.69 | -1.91 | 1.91 | -0.03 | 0.03 |
| gene-LOC105279500 | https://www.ncbi.nlm.nih.gov/gene/?term=LOC105279500 | 3587 | 33 | 92 | 1 | 2.79 | 3.03 | 44.93 | 47.01 | -2.08 | 2.08 | -0.02 | 0.02 |
| gene-LOC105282067 | https://www.ncbi.nlm.nih.gov/gene/?term=LOC105282067 | 5941 | 44 | 74.06 | 1 | 1.68 | 2.27 | 26.18 | 28.47 | -2.29 | 2.29 | -0.02 | 0.02 |
| gene-LOC105284502 | https://www.ncbi.nlm.nih.gov/gene/?term=LOC105284502 | 2157 | 19 | 88.09 | 1 | 4.64 | 5.26 | 36.83 | 43.93 | -7.1 | 7.1 | -0.06 | 0.06 |
| gene-LOC105284677 | https://www.ncbi.nlm.nih.gov/gene/?term=LOC105284677 | 1905 | 41 | 215.22 | 2 | 10.5 | 4.88 | 27.55 | 36.9 | -9.35 | 9.35 | -0.2 | 0.2 |

Directory: OGL File: ogl-Ob_LI_BCphaseA2.vs.RphaseA1.txt

| gene_ID | gene_link | gwidth | #Sites | #per10Kb | #dmSites | #dmsp10kb | %dmSites | %pSite1 | %pSite2 | DMpSite | ADMpSite | DMpNucl | ADMpNucl |
|---|---|---|---|---|---|---|---|---|---|---|---|---|---|
| gene-LOC105280604 | https://www.ncbi.nlm.nih.gov/gene/?term=LOC105280604 | 752 | 1 | 13.3 | 1 | 13.3 | 100 | 75 | 42.31 | 32.69 | 32.69 | 0.04 | 0.04 |
| gene-LOC105286178 | https://www.ncbi.nlm.nih.gov/gene/?term=LOC105286178 | 1590 | 1 | 6.29 | 1 | 6.29 | 100 | 78.38 | 56.76 | 21.62 | 21.62 | 0.01 | 0.01 |
| gene-LOC113561421 | https://www.ncbi.nlm.nih.gov/gene/?term=LOC113561421 | 907 | 1 | 11.03 | 1 | 11.03 | 100 | 53.57 | 32.65 | 20.92 | 20.92 | 0.02 | 0.02 |
| gene-LOC105286586 | https://www.ncbi.nlm.nih.gov/gene/?term=LOC105286586 | 1417 | 1 | 7.06 | 1 | 7.06 | 100 | 35.71 | 15.56 | 20.15 | 20.15 | 0.01 | 0.01 |
| gene-LOC105284346 | https://www.ncbi.nlm.nih.gov/gene/?term=LOC105284346 | 2002 | 2 | 9.99 | 1 | 5 | 50 | 28.11 | 14.29 | 13.83 | 13.83 | 0.01 | 0.01 |
| gene-LOC105283693 | https://www.ncbi.nlm.nih.gov/gene/?term=LOC105283693 | 1305 | 12 | 91.95 | 4 | 30.65 | 33.33 | 70.54 | 58.26 | 12.28 | 12.28 | 0.11 | 0.11 |
| gene-LOC113562901 | https://www.ncbi.nlm.nih.gov/gene/?term=LOC113562901 | 1531 | 22 | 143.7 | 8 | 52.25 | 36.36 | 60.1 | 48.53 | 11.57 | 11.57 | 0.17 | 0.17 |
| gene-LOC105281110 | https://www.ncbi.nlm.nih.gov/gene/?term=LOC105281110 | 960 | 5 | 52.08 | 1 | 10.42 | 20 | 48.85 | 39.79 | 9.06 | 9.06 | 0.05 | 0.05 |
| gene-LOC113562728 | https://www.ncbi.nlm.nih.gov/gene/?term=LOC113562728 | 1733 | 32 | 184.65 | 5 | 28.85 | 15.62 | 28.94 | 19.93 | 9 | 9 | 0.17 | 0.17 |
| gene-LOC105280683 | https://www.ncbi.nlm.nih.gov/gene/?term=LOC105280683 | 2171 | 21 | 96.73 | 4 | 18.42 | 19.05 | 54.85 | 46.7 | 8.15 | 8.15 | 0.08 | 0.08 |
| gene-LOC105284759 | https://www.ncbi.nlm.nih.gov/gene/?term=LOC105284759 | 1353 | 9 | 66.52 | 1 | 7.39 | 11.11 | 38.95 | 31.14 | 7.81 | 7.81 | 0.05 | 0.05 |
| gene-LOC105284521 | https://www.ncbi.nlm.nih.gov/gene/?term=LOC105284521 | 2105 | 14 | 66.51 | 1 | 4.75 | 7.14 | 62.65 | 55.05 | 7.59 | 7.59 | 0.05 | 0.05 |
| gene-LOC113562975 | https://www.ncbi.nlm.nih.gov/gene/?term=LOC113562975 | 2751 | 9 | 32.72 | 1 | 3.64 | 11.11 | 50.57 | 43.41 | 7.16 | 7.16 | 0.02 | 0.02 |
| gene-LOC105274834 | https://www.ncbi.nlm.nih.gov/gene/?term=LOC105274834 | 1248 | 9 | 72.12 | 3 | 24.04 | 33.33 | 53.69 | 46.65 | 7.03 | 7.03 | 0.05 | 0.05 |
| gene-LOC113561934 | https://www.ncbi.nlm.nih.gov/gene/?term=LOC113561934 | 1252 | 27 | 215.65 | 4 | 31.95 | 14.81 | 31.68 | 24.75 | 6.93 | 6.93 | 0.15 | 0.15 |
| gene-LOC105276591 | https://www.ncbi.nlm.nih.gov/gene/?term=LOC105276591 | 1863 | 7 | 37.57 | 1 | 5.37 | 14.29 | 44.34 | 37.79 | 6.54 | 6.54 | 0.02 | 0.02 |
| gene-LOC105279538 | https://www.ncbi.nlm.nih.gov/gene/?term=LOC105279538 | 889 | 17 | 191.23 | 2 | 22.5 | 11.76 | 36.14 | 29.61 | 6.53 | 6.53 | 0.12 | 0.12 |
| gene-LOC105275378 | https://www.ncbi.nlm.nih.gov/gene/?term=LOC105275378 | 1189 | 20 | 168.21 | 1 | 8.41 | 5 | 32.85 | 26.65 | 6.2 | 6.2 | 0.1 | 0.1 |
| gene-LOC105286263 | https://www.ncbi.nlm.nih.gov/gene/?term=LOC105286263 | 1867 | 23 | 123.19 | 4 | 21.42 | 17.39 | 38.1 | 31.96 | 6.14 | 6.14 | 0.08 | 0.08 |
| gene-LOC105280835 | https://www.ncbi.nlm.nih.gov/gene/?term=LOC105280835 | 1890 | 6 | 31.75 | 1 | 5.29 | 16.67 | 6.39 | 0.36 | 6.03 | 6.03 | 0.02 | 0.02 |
| gene-LOC105287955 | https://www.ncbi.nlm.nih.gov/gene/?term=LOC105287955 | 2486 | 12 | 48.27 | 1 | 4.02 | 8.33 | 91.37 | 85.34 | 6.03 | 6.03 | 0.03 | 0.03 |
| gene-LOC105286815 | https://www.ncbi.nlm.nih.gov/gene/?term=LOC105286815 | 1024 | 10 | 97.66 | 1 | 9.77 | 10 | 59.55 | 53.96 | 5.59 | 5.59 | 0.05 | 0.05 |
| gene-LOC105283475 | https://www.ncbi.nlm.nih.gov/gene/?term=LOC105283475 | 1721 | 7 | 40.67 | 1 | 5.81 | 14.29 | 61.36 | 55.92 | 5.44 | 5.44 | 0.02 | 0.02 |
| gene-LOC105284988 | https://www.ncbi.nlm.nih.gov/gene/?term=LOC105284988 | 3729 | 26 | 69.72 | 1 | 2.68 | 3.85 | 37.25 | 31.94 | 5.3 | 5.3 | 0.04 | 0.04 |

...
... only first 25 lines shown ...

Directory: OGL File: ogl-Ob_LI_BCphaseA2.vs.RphaseA2.txt

| gene_ID | gene_link | gwidth | #Sites | #per10Kb | #dmSites | #dmsp10kb | %dmSites | %pSite1 | %pSite2 | DMpSite | ADMpSite | DMpNucl | ADMpNucl |
|---|---|---|---|---|---|---|---|---|---|---|---|---|---|
| gene-LOC113563238 | https://www.ncbi.nlm.nih.gov/gene/?term=LOC113563238 | 2680 | 1 | 3.73 | 1 | 3.73 | 100 | 75 | 46.15 | 28.85 | 28.85 | 0.01 | 0.01 |
| gene-LOC105280604 | https://www.ncbi.nlm.nih.gov/gene/?term=LOC105280604 | 752 | 1 | 13.3 | 1 | 13.3 | 100 | 75 | 50 | 25 | 25 | 0.03 | 0.03 |
| gene-LOC105275861 | https://www.ncbi.nlm.nih.gov/gene/?term=LOC105275861 | 1979 | 7 | 35.37 | 1 | 5.05 | 14.29 | 39.97 | 30.1 | 9.87 | 9.87 | 0.03 | 0.03 |
| gene-LOC105281688 | https://www.ncbi.nlm.nih.gov/gene/?term=LOC105281688 | 1179 | 4 | 33.93 | 1 | 8.48 | 25 | 83.9 | 74.11 | 9.79 | 9.79 | 0.03 | 0.03 |
| gene-LOC105287647 | https://www.ncbi.nlm.nih.gov/gene/?term=LOC105287647 | 1949 | 10 | 51.31 | 1 | 5.13 | 10 | 41.7 | 35 | 6.7 | 6.7 | 0.03 | 0.03 |
| gene-LOC105287649 | https://www.ncbi.nlm.nih.gov/gene/?term=LOC105287649 | 1566 | 7 | 44.7 | 1 | 6.39 | 14.29 | 46.89 | 40.38 | 6.5 | 6.5 | 0.03 | 0.03 |
| gene-LOC105285051 | https://www.ncbi.nlm.nih.gov/gene/?term=LOC105285051 | 1357 | 16 | 117.91 | 2 | 14.74 | 12.5 | 34.98 | 29.29 | 5.69 | 5.69 | 0.07 | 0.07 |
| gene-LOC105281507 | https://www.ncbi.nlm.nih.gov/gene/?term=LOC105281507 | 881 | 7 | 79.46 | 1 | 11.35 | 14.29 | 24.99 | 19.48 | 5.51 | 5.51 | 0.04 | 0.04 |
| gene-LOC105280969 | https://www.ncbi.nlm.nih.gov/gene/?term=LOC105280969 | 1005 | 8 | 79.6 | 1 | 9.95 | 12.5 | 66.21 | 60.87 | 5.34 | 5.34 | 0.04 | 0.04 |
| gene-LOC105288141 | https://www.ncbi.nlm.nih.gov/gene/?term=LOC105288141 | 2845 | 27 | 94.9 | 1 | 3.51 | 3.7 | 26.78 | 22 | 4.78 | 4.78 | 0.05 | 0.05 |
| gene-LOC105280472 | https://www.ncbi.nlm.nih.gov/gene/?term=LOC105280472 | 1675 | 6 | 35.82 | 1 | 5.97 | 16.67 | 78.84 | 74.62 | 4.22 | 4.22 | 0.02 | 0.02 |
| gene-LOC105282975 | https://www.ncbi.nlm.nih.gov/gene/?term=LOC105282975 | 1500 | 22 | 146.67 | 2 | 13.33 | 9.09 | 49.9 | 46.07 | 3.84 | 3.84 | 0.06 | 0.06 |
| gene-LOC105283760 | https://www.ncbi.nlm.nih.gov/gene/?term=LOC105283760 | 1134 | 5 | 44.09 | 1 | 8.82 | 20 | 46 | 42.19 | 3.81 | 3.81 | 0.02 | 0.02 |
| gene-LOC113561391 | https://www.ncbi.nlm.nih.gov/gene/?term=LOC113561391 | 1463 | 12 | 82.02 | 1 | 6.84 | 8.33 | 17.64 | 13.97 | 3.68 | 3.68 | 0.03 | 0.03 |
| gene-LOC105275501 | https://www.ncbi.nlm.nih.gov/gene/?term=LOC105275501 | 1091 | 7 | 64.16 | 1 | 9.17 | 14.29 | 25.2 | 21.53 | 3.67 | 3.67 | 0.02 | 0.02 |
| gene-LOC105280589 | https://www.ncbi.nlm.nih.gov/gene/?term=LOC105280589 | 2884 | 47 | 162.97 | 2 | 6.93 | 4.26 | 32.58 | 28.94 | 3.64 | 3.64 | 0.06 | 0.06 |
| gene-LOC105280355 | https://www.ncbi.nlm.nih.gov/gene/?term=LOC105280355 | 1059 | 14 | 132.2 | 1 | 9.44 | 7.14 | 18.6 | 15.18 | 3.42 | 3.42 | 0.05 | 0.05 |
| gene-LOC105283851 | https://www.ncbi.nlm.nih.gov/gene/?term=LOC105283851 | 1886 | 9 | 47.72 | 1 | 5.3 | 11.11 | 44.96 | 41.54 | 3.42 | 3.42 | 0.02 | 0.02 |
| gene-LOC105279538 | https://www.ncbi.nlm.nih.gov/gene/?term=LOC105279538 | 889 | 20 | 224.97 | 2 | 22.5 | 10 | 30.72 | 27.36 | 3.36 | 3.36 | 0.08 | 0.08 |
| gene-LOC113562728 | https://www.ncbi.nlm.nih.gov/gene/?term=LOC113562728 | 1733 | 35 | 201.96 | 1 | 5.77 | 2.86 | 28.15 | 24.84 | 3.31 | 3.31 | 0.07 | 0.07 |
| gene-LOC113561448 | https://www.ncbi.nlm.nih.gov/gene/?term=LOC113561448 | 1279 | 13 | 101.64 | 1 | 7.82 | 7.69 | 66.97 | 63.82 | 3.15 | 3.15 | 0.03 | 0.03 |
| gene-LOC105276798 | https://www.ncbi.nlm.nih.gov/gene/?term=LOC105276798 | 3170 | 8 | 25.24 | 1 | 3.15 | 12.5 | 6.95 | 3.82 | 3.14 | 3.14 | 0.01 | 0.01 |
| gene-LOC105276840 | https://www.ncbi.nlm.nih.gov/gene/?term=LOC105276840 | 3299 | 11 | 33.34 | 1 | 3.03 | 9.09 | 6.96 | 3.82 | 3.14 | 3.14 | 0.01 | 0.01 |
| gene-LOC105286263 | https://www.ncbi.nlm.nih.gov/gene/?term=LOC105286263 | 1867 | 29 | 155.33 | 1 | 5.36 | 3.45 | 33.04 | 29.95 | 3.09 | 3.09 | 0.05 | 0.05 |

...

... only first 25 lines shown ...

Directory: OGL File: ogl-Ob_LI_BCphaseA2.vs.RphaseB1.txt

| gene_ID | gene_link | gwidth | #Sites | #per10Kb | #dmSites | #dmsp10kb | %dmSites | %pSite1 | %pSite2 | DMpSite | ADMpSite | DMpNucl | ADMpNucl |
|---|---|---|---|---|---|---|---|---|---|---|---|---|---|
| gene-LOC113562345 | https://www.ncbi.nlm.nih.gov/gene/?term=LOC113562345 | 2290 | 71 | 310.04 | 15 | 65.5 | 21.13 | 28.2 | 15.69 | 12.51 | 12.51 | 0.39 | 0.39 |
| gene-LOC105275070 | https://www.ncbi.nlm.nih.gov/gene/?term=LOC105275070 | 5667 | 53 | 93.52 | 4 | 7.06 | 7.55 | 16.59 | 5.92 | 10.67 | 10.67 | 0.1 | 0.1 |
| gene-LOC105275708 | https://www.ncbi.nlm.nih.gov/gene/?term=LOC105275708 | 1772 | 10 | 56.43 | 1 | 5.64 | 10 | 50.31 | 40.78 | 9.53 | 9.53 | 0.05 | 0.05 |
| gene-LOC105285060 | https://www.ncbi.nlm.nih.gov/gene/?term=LOC105285060 | 1285 | 13 | 101.17 | 1 | 7.78 | 7.69 | 56.31 | 50.43 | 5.87 | 5.87 | 0.06 | 0.06 |
| gene-LOC105285514 | https://www.ncbi.nlm.nih.gov/gene/?term=LOC105285514 | 1277 | 11 | 86.14 | 1 | 7.83 | 9.09 | 90.26 | 85.12 | 5.14 | 5.14 | 0.04 | 0.04 |
| gene-LOC105276078 | https://www.ncbi.nlm.nih.gov/gene/?term=LOC105276078 | 1051 | 14 | 133.21 | 1 | 9.51 | 7.14 | 31.96 | 27.06 | 4.9 | 4.9 | 0.07 | 0.07 |
| gene-LOC105275347 | https://www.ncbi.nlm.nih.gov/gene/?term=LOC105275347 | 2526 | 14 | 55.42 | 1 | 3.96 | 7.14 | 33.57 | 29.73 | 3.84 | 3.84 | 0.02 | 0.02 |
| gene-LOC105285695 | https://www.ncbi.nlm.nih.gov/gene/?term=LOC105285695 | 1371 | 74 | 539.75 | 1 | 7.29 | 1.35 | 27.97 | 24.61 | 3.36 | 3.36 | 0.18 | 0.18 |
| gene-LOC105283552 | https://www.ncbi.nlm.nih.gov/gene/?term=LOC105283552 | 1562 | 36 | 230.47 | 1 | 6.4 | 2.78 | 56.28 | 53.31 | 2.97 | 2.97 | 0.07 | 0.07 |
| gene-LOC105274900 | https://www.ncbi.nlm.nih.gov/gene/?term=LOC105274900 | 5561 | 51 | 91.71 | 2 | 3.6 | 3.92 | 39.2 | 36.36 | 2.84 | 2.84 | 0.03 | 0.03 |
| gene-LOC105285315 | https://www.ncbi.nlm.nih.gov/gene/?term=LOC105285315 | 5458 | 21 | 38.48 | 1 | 1.83 | 4.76 | 43.16 | 40.32 | 2.84 | 2.84 | 0.01 | 0.01 |
| gene-LOC105279898 | https://www.ncbi.nlm.nih.gov/gene/?term=LOC105279898 | 1343 | 13 | 96.8 | 1 | 7.45 | 7.69 | 58.71 | 56.01 | 2.69 | 2.69 | 0.03 | 0.03 |
| gene-LOC105287063 | https://www.ncbi.nlm.nih.gov/gene/?term=LOC105287063 | 1874 | 49 | 261.47 | 1 | 5.34 | 2.04 | 32.42 | 29.81 | 2.61 | 2.61 | 0.07 | 0.07 |
| gene-LOC105283712 | https://www.ncbi.nlm.nih.gov/gene/?term=LOC105283712 | 1910 | 29 | 151.83 | 1 | 5.24 | 3.45 | 33.08 | 30.49 | 2.59 | 2.59 | 0.04 | 0.04 |
| gene-LOC105284704 | https://www.ncbi.nlm.nih.gov/gene/?term=LOC105284704 | 2400 | 62 | 258.33 | 1 | 4.17 | 1.61 | 26.43 | 23.93 | 2.5 | 2.5 | 0.06 | 0.06 |
| gene-LOC105288066 | https://www.ncbi.nlm.nih.gov/gene/?term=LOC105288066 | 1964 | 23 | 117.11 | 1 | 5.09 | 4.35 | 46.39 | 44.06 | 2.34 | 2.34 | 0.03 | 0.03 |
| gene-LOC105276272 | https://www.ncbi.nlm.nih.gov/gene/?term=LOC105276272 | 1965 | 38 | 193.38 | 1 | 5.09 | 2.63 | 30.65 | 28.5 | 2.15 | 2.15 | 0.04 | 0.04 |
| gene-LOC105282156 | https://www.ncbi.nlm.nih.gov/gene/?term=LOC105282156 | 2577 | 128 | 496.7 | 1 | 3.88 | 0.78 | 28.45 | 26.37 | 2.08 | 2.08 | 0.1 | 0.1 |
| gene-LOC105280348 | https://www.ncbi.nlm.nih.gov/gene/?term=LOC105280348 | 3797 | 65 | 171.19 | 1 | 2.63 | 1.54 | 21.64 | 19.63 | 2.01 | 2.01 | 0.03 | 0.03 |
| gene-LOC105280774 | https://www.ncbi.nlm.nih.gov/gene/?term=LOC105280774 | 2793 | 101 | 361.62 | 1 | 3.58 | 0.99 | 32.38 | 30.39 | 1.99 | 1.99 | 0.07 | 0.07 |
| gene-LOC105281848 | https://www.ncbi.nlm.nih.gov/gene/?term=LOC105281848 | 2122 | 52 | 245.05 | 1 | 4.71 | 1.92 | 26.53 | 24.7 | 1.83 | 1.83 | 0.04 | 0.04 |
| gene-LOC105276704 | https://www.ncbi.nlm.nih.gov/gene/?term=LOC105276704 | 2063 | 89 | 431.41 | 1 | 4.85 | 1.12 | 14.86 | 13.07 | 1.79 | 1.79 | 0.08 | 0.08 |
| gene-LOC105280408 | https://www.ncbi.nlm.nih.gov/gene/?term=LOC105280408 | 5983 | 50 | 83.57 | 1 | 1.67 | 2 | 19.04 | 17.37 | 1.67 | 1.67 | 0.01 | 0.01 |
| gene-LOC105282149 | https://www.ncbi.nlm.nih.gov/gene/?term=LOC105282149 | 1787 | 105 | 587.58 | 1 | 5.6 | 0.95 | 43.64 | 41.97 | 1.67 | 1.67 | 0.1 | 0.1 |
| gene-LOC105280641 | https://www.ncbi.nlm.nih.gov/gene/?term=LOC105280641 | 3682 | 56 | 152.09 | 1 | 2.72 | 1.79 | 17.9 | 16.23 | 1.66 | 1.66 | 0.03 | 0.03 |
| gene-LOC105285674 | https://www.ncbi.nlm.nih.gov/gene/?term=LOC105285674 | 3035 | 93 | 306.43 | 1 | 3.29 | 1.08 | 26.91 | 25.27 | 1.64 | 1.64 | 0.05 | 0.05 |
| gene-LOC105284184 | https://www.ncbi.nlm.nih.gov/gene/?term=LOC105284184 | 2535 | 87 | 343.2 | 1 | 3.94 | 1.15 | 34.29 | 32.68 | 1.6 | 1.6 | 0.05 | 0.05 |
| gene-LOC105283123 | https://www.ncbi.nlm.nih.gov/gene/?term=LOC105283123 | 1507 | 51 | 338.42 | 1 | 6.64 | 1.96 | 16.3 | 14.72 | 1.58 | 1.58 | 0.05 | 0.05 |
| gene-LOC105283697 | https://www.ncbi.nlm.nih.gov/gene/?term=LOC105283697 | 5301 | 137 | 258.44 | 1 | 1.89 | 0.73 | 17.96 | 16.39 | 1.57 | 1.57 | 0.04 | 0.04 |
| gene-LOC105288010 | https://www.ncbi.nlm.nih.gov/gene/?term=LOC105288010 | 3189 | 51 | 159.92 | 1 | 3.14 | 1.96 | 34.39 | 32.85 | 1.54 | 1.54 | 0.02 | 0.02 |
| gene-LOC105277243 | https://www.ncbi.nlm.nih.gov/gene/?term=LOC105277243 | 2284 | 32 | 140.11 | 1 | 4.38 | 3.12 | 44.88 | 43.35 | 1.53 | 1.53 | 0.02 | 0.02 |
| gene-LOC105288079 | https://www.ncbi.nlm.nih.gov/gene/?term=LOC105288079 | 3473 | 92 | 264.9 | 1 | 2.88 | 1.09 | 19.48 | 18 | 1.49 | 1.49 | 0.04 | 0.04 |
| gene-LOC105276538 | https://www.ncbi.nlm.nih.gov/gene/?term=LOC105276538 | 1511 | 19 | 125.74 | 1 | 6.62 | 5.26 | 70.28 | 68.86 | 1.42 | 1.42 | 0.02 | 0.02 |
| gene-LOC105281515 | https://www.ncbi.nlm.nih.gov/gene/?term=LOC105281515 | 8827 | 48 | 54.38 | 2 | 2.27 | 4.17 | 32.1 | 30.7 | 1.4 | 1.4 | 0.01 | 0.01 |
| gene-LOC105280972 | https://www.ncbi.nlm.nih.gov/gene/?term=LOC105280972 | 1532 | 32 | 208.88 | 1 | 6.53 | 3.12 | 19.4 | 18.05 | 1.35 | 1.35 | 0.03 | 0.03 |
| gene-LOC105282885 | https://www.ncbi.nlm.nih.gov/gene/?term=LOC105282885 | 1870 | 38 | 203.21 | 1 | 5.35 | 2.63 | 45.14 | 43.81 | 1.33 | 1.33 | 0.03 | 0.03 |
| gene-LOC105279500 | https://www.ncbi.nlm.nih.gov/gene/?term=LOC105279500 | 3587 | 33 | 92 | 1 | 2.79 | 3.03 | 51.88 | 50.67 | 1.21 | 1.21 | 0.01 | 0.01 |
| gene-LOC105286265 | https://www.ncbi.nlm.nih.gov/gene/?term=LOC105286265 | 2507 | 64 | 255.29 | 1 | 3.99 | 1.56 | 30.69 | 29.49 | 1.21 | 1.21 | 0.03 | 0.03 |
| gene-LOC105274449 | https://www.ncbi.nlm.nih.gov/gene/?term=LOC105274449 | 1549 | 53 | 342.16 | 1 | 6.46 | 1.89 | 50.38 | 49.29 | 1.09 | 1.09 | 0.04 | 0.04 |
| gene-LOC105275518 | https://www.ncbi.nlm.nih.gov/gene/?term=LOC105275518 | 2335 | 19 | 81.37 | 1 | 4.28 | 5.26 | 51.76 | 50.67 | 1.09 | 1.09 | 0.01 | 0.01 |
| gene-LOC105280786 | https://www.ncbi.nlm.nih.gov/gene/?term=LOC105280786 | 2573 | 122 | 474.15 | 1 | 3.89 | 0.82 | 30.88 | 29.81 | 1.07 | 1.07 | 0.05 | 0.05 |
| gene-LOC105279664 | https://www.ncbi.nlm.nih.gov/gene/?term=LOC105279664 | 4023 | 57 | 141.69 | 1 | 2.49 | 1.75 | 22.82 | 21.77 | 1.05 | 1.05 | 0.01 | 0.01 |
| gene-LOC105287095 | https://www.ncbi.nlm.nih.gov/gene/?term=LOC105287095 | 1435 | 56 | 390.24 | 1 | 6.97 | 1.79 | 30.4 | 29.35 | 1.05 | 1.05 | 0.04 | 0.04 |
| gene-LOC105276449 | https://www.ncbi.nlm.nih.gov/gene/?term=LOC105276449 | 1231 | 43 | 349.31 | 1 | 8.12 | 2.33 | 30.8 | 29.78 | 1.02 | 1.02 | 0.04 | 0.04 |
| gene-LOC105278807 | https://www.ncbi.nlm.nih.gov/gene/?term=LOC105278807 | 2166 | 71 | 327.79 | 1 | 4.62 | 1.41 | 19.81 | 18.81 | 1 | 1 | 0.03 | 0.03 |
| gene-LOC105279298 | https://www.ncbi.nlm.nih.gov/gene/?term=LOC105279298 | 2773 | 123 | 443.56 | 1 | 3.61 | 0.81 | 24.88 | 23.88 | 1 | 1 | 0.04 | 0.04 |
| gene-LOC105274823 | https://www.ncbi.nlm.nih.gov/gene/?term=LOC105274823 | 4620 | 152 | 329 | 1 | 2.16 | 0.66 | 19.69 | 18.71 | 0.99 | 0.99 | 0.03 | 0.03 |
| gene-LOC105277215 | https://www.ncbi.nlm.nih.gov/gene/?term=LOC105277215 | 2619 | 23 | 87.82 | 1 | 3.82 | 4.35 | 35.64 | 34.71 | 0.93 | 0.93 | 0.01 | 0.01 |
| gene-LOC105278384 | https://www.ncbi.nlm.nih.gov/gene/?term=LOC105278384 | 1390 | 33 | 237.41 | 1 | 7.19 | 3.03 | 46.69 | 45.77 | 0.92 | 0.92 | 0.02 | 0.02 |
| gene-LOC105283838 | https://www.ncbi.nlm.nih.gov/gene/?term=LOC105283838 | 1748 | 21 | 120.14 | 1 | 5.72 | 4.76 | 46.37 | 45.45 | 0.92 | 0.92 | 0.01 | 0.01 |
| gene-LOC113561805 | https://www.ncbi.nlm.nih.gov/gene/?term=LOC113561805 | 3337 | 69 | 206.77 | 1 | 3 | 1.45 | 38.18 | 37.26 | 0.92 | 0.92 | 0.02 | 0.02 |
| gene-LOC105282794 | https://www.ncbi.nlm.nih.gov/gene/?term=LOC105282794 | 4571 | 99 | 216.58 | 1 | 2.19 | 1.01 | 18.66 | 17.76 | 0.9 | 0.9 | 0.02 | 0.02 |
| gene-LOC105286644 | https://www.ncbi.nlm.nih.gov/gene/?term=LOC105286644 | 1235 | 22 | 178.14 | 1 | 8.1 | 4.55 | 42.04 | 41.14 | 0.89 | 0.89 | 0.02 | 0.02 |
| gene-LOC105278109 | https://www.ncbi.nlm.nih.gov/gene/?term=LOC105278109 | 4029 | 65 | 161.33 | 1 | 2.48 | 1.54 | 46.32 | 45.45 | 0.88 | 0.88 | 0.01 | 0.01 |
| gene-LOC105278497 | https://www.ncbi.nlm.nih.gov/gene/?term=LOC105278497 | 2072 | 89 | 429.54 | 1 | 4.83 | 1.12 | 21.38 | 20.51 | 0.87 | 0.87 | 0.04 | 0.04 |
| gene-LOC105283592 | https://www.ncbi.nlm.nih.gov/gene/?term=LOC105283592 | 3479 | 75 | 215.58 | 1 | 2.87 | 1.33 | 25.19 | 24.33 | 0.86 | 0.86 | 0.02 | 0.02 |
| gene-LOC105283728 | https://www.ncbi.nlm.nih.gov/gene/?term=LOC105283728 | 3256 | 104 | 319.41 | 2 | 6.14 | 1.92 | 22.23 | 21.4 | 0.83 | 0.83 | 0.03 | 0.03 |
| gene-LOC105285055 | https://www.ncbi.nlm.nih.gov/gene/?term=LOC105285055 | 4108 | 241 | 586.66 | 2 | 4.87 | 0.83 | 12.15 | 11.34 | 0.81 | 0.81 | 0.05 | 0.05 |
| gene-LOC105278283 | https://www.ncbi.nlm.nih.gov/gene/?term=LOC105278283 | 2317 | 126 | 543.81 | 1 | 4.32 | 0.79 | 27.76 | 27.04 | 0.72 | 0.72 | 0.04 | 0.04 |
| gene-LOC105285179 | https://www.ncbi.nlm.nih.gov/gene/?term=LOC105285179 | 4191 | 178 | 424.72 | 1 | 2.39 | 0.56 | 17.79 | 17.08 | 0.71 | 0.71 | 0.03 | 0.03 |

| | | | | | | | | | | | | | |
|---|---|---|---|---|---|---|---|---|---|---|---|---|---|
| gene-LOC105276560 | https://www.ncbi.nlm.nih.gov/gene/?term=LOC105276560 | 18607 | 319 | 171.44 | 1 | 0.54 | 0.31 | 2.53 | 1.85 | 0.67 | 0.67 | 0.01 | 0.01 |
| gene-LOC105281312 | https://www.ncbi.nlm.nih.gov/gene/?term=LOC105281312 | 2551 | 68 | 266.56 | 1 | 3.92 | 1.47 | 31.46 | 30.8 | 0.65 | 0.65 | 0.02 | 0.02 |
| gene-LOC105286514 | https://www.ncbi.nlm.nih.gov/gene/?term=LOC105286514 | 2528 | 43 | 170.09 | 1 | 3.96 | 2.33 | 34.72 | 34.07 | 0.65 | 0.65 | 0.01 | 0.01 |
| gene-LOC105284355 | https://www.ncbi.nlm.nih.gov/gene/?term=LOC105284355 | 3904 | 143 | 366.29 | 1 | 2.56 | 0.7 | 22.99 | 22.35 | 0.64 | 0.64 | 0.02 | 0.02 |
| gene-LOC105278145 | https://www.ncbi.nlm.nih.gov/gene/?term=LOC105278145 | 4470 | 250 | 559.28 | 1 | 2.24 | 0.4 | 10.94 | 10.34 | 0.6 | 0.6 | 0.03 | 0.03 |
| gene-LOC105283989 | https://www.ncbi.nlm.nih.gov/gene/?term=LOC105283989 | 4948 | 25 | 50.53 1 | 2.02 | 4 | 46.26 | 45.67 | 0.59 | 0.59 | 0 | 0 | |
| gene-LOC105276059 | https://www.ncbi.nlm.nih.gov/gene/?term=LOC105276059 | 6256 | 123 | 196.61 | 1 | 1.6 | 0.81 | 12.44 | 11.86 | 0.58 | 0.58 | 0.01 | 0.01 |
| gene-LOC105280407 | https://www.ncbi.nlm.nih.gov/gene/?term=LOC105280407 | 10950 | 203 | 185.39 | 1 | 0.91 | 0.49 | 15.56 | 14.99 | 0.56 | 0.56 | 0.01 | 0.01 |
| gene-LOC105285671 | https://www.ncbi.nlm.nih.gov/gene/?term=LOC105285671 | 2263 | 99 | 437.47 | 1 | 4.42 | 1.01 | 30.99 | 30.44 | 0.55 | 0.55 | 0.02 | 0.02 |
| gene-LOC105285395 | https://www.ncbi.nlm.nih.gov/gene/?term=LOC105285395 | 8085 | 207 | 256.03 | 1 | 1.24 | 0.48 | 14.3 | 13.76 | 0.54 | 0.54 | 0.01 | 0.01 |
| gene-LOC105280819 | https://www.ncbi.nlm.nih.gov/gene/?term=LOC105280819 | 1639 | 39 | 237.95 | 1 | 6.1 | 2.56 | 35.99 | 35.46 | 0.53 | 0.53 | 0.01 | 0.01 |
| gene-LOC105275451 | https://www.ncbi.nlm.nih.gov/gene/?term=LOC105275451 | 6035 | 133 | 220.38 | 1 | 1.66 | 0.75 | 22.24 | 21.72 | 0.51 | 0.51 | 0.01 | 0.01 |
| gene-LOC105275460 | https://www.ncbi.nlm.nih.gov/gene/?term=LOC105275460 | 2473 | 118 | 477.15 | 1 | 4.04 | 0.85 | 13.46 | 12.95 | 0.51 | 0.51 | 0.02 | 0.02 |
| gene-LOC105278632 | https://www.ncbi.nlm.nih.gov/gene/?term=LOC105278632 | 3624 | 131 | 361.48 | 1 | 2.76 | 0.76 | 14.47 | 14.04 | 0.43 | 0.43 | 0.02 | 0.02 |
| gene-LOC105278534 | https://www.ncbi.nlm.nih.gov/gene/?term=LOC105278534 | 7364 | 180 | 244.43 | 1 | 1.36 | 0.56 | 18.01 | 17.6 | 0.41 | 0.41 | 0.01 | 0.01 |
| gene-LOC105280890 | https://www.ncbi.nlm.nih.gov/gene/?term=LOC105280890 | 1139 | 29 | 254.61 | 1 | 8.78 | 3.45 | 19.62 | 19.23 | 0.39 | 0.39 | 0.01 | 0.01 |
| gene-LOC105286999 | https://www.ncbi.nlm.nih.gov/gene/?term=LOC105286999 | 2141 | 44 | 205.51 | 1 | 4.67 | 2.27 | 35.31 | 34.92 | 0.39 | 0.39 | 0.01 | 0.01 |
| gene-LOC105276462 | https://www.ncbi.nlm.nih.gov/gene/?term=LOC105276462 | 6561 | 275 | 419.14 | 1 | 1.52 | 0.36 | 8.32 | 7.94 | 0.37 | 0.37 | 0.02 | 0.02 |
| gene-LOC105280410 | https://www.ncbi.nlm.nih.gov/gene/?term=LOC105280410 | 12580 | 265 | 210.65 | 1 | 0.79 | 0.38 | 19.92 | 19.57 | 0.35 | 0.35 | 0.01 | 0.01 |
| gene-LOC105283296 | https://www.ncbi.nlm.nih.gov/gene/?term=LOC105283296 | 2174 | 86 | 395.58 | 1 | 4.6 | 1.16 | 16.05 | 15.7 | 0.35 | 0.35 | 0.01 | 0.01 |
| gene-LOC105278666 | https://www.ncbi.nlm.nih.gov/gene/?term=LOC105278666 | 10341 | 456 | 440.96 | 1 | 0.97 | 0.22 | 7.52 | 7.19 | 0.34 | 0.34 | 0.01 | 0.01 |
| gene-LOC105279657 | https://www.ncbi.nlm.nih.gov/gene/?term=LOC105279657 | 5878 | 96 | 163.32 | 1 | 1.7 | 1.04 | 18.1 | 17.76 | 0.34 | 0.34 | 0.01 | 0.01 |
| gene-LOC105277435 | https://www.ncbi.nlm.nih.gov/gene/?term=LOC105277435 | 3643 | 175 | 480.37 | 1 | 2.74 | 0.57 | 11.3 | 10.97 | 0.33 | 0.33 | 0.02 | 0.02 |
| gene-LOC105284036 | https://www.ncbi.nlm.nih.gov/gene/?term=LOC105284036 | 3254 | 87 | 267.36 | 1 | 3.07 | 1.15 | 24.27 | 23.96 | 0.31 | 0.31 | 0.01 | 0.01 |
| gene-LOC105281272 | https://www.ncbi.nlm.nih.gov/gene/?term=LOC105281272 | 5252 | 175 | 333.21 | 1 | 1.9 | 0.57 | 16.26 | 15.96 | 0.3 | 0.3 | 0.01 | 0.01 |
| gene-LOC105285807 | https://www.ncbi.nlm.nih.gov/gene/?term=LOC105285807 | 6063 | 207 | 341.42 | 1 | 1.65 | 0.48 | 7.17 | 6.87 | 0.3 | 0.3 | 0.01 | 0.01 |
| gene-LOC105275187 | https://www.ncbi.nlm.nih.gov/gene/?term=LOC105275187 | 5534 | 69 | 124.68 | 1 | 1.81 | 1.45 | 14.14 | 13.85 | 0.29 | 0.29 | 0 | 0 |
| gene-LOC105282491 | https://www.ncbi.nlm.nih.gov/gene/?term=LOC105282491 | 4761 | 170 | 357.07 | 1 | 2.1 | 0.59 | 7.92 | 7.67 | 0.25 | 0.25 | 0.01 | 0.01 |
| gene-LOC105274706 | https://www.ncbi.nlm.nih.gov/gene/?term=LOC105274706 | 1446 | 50 | 345.78 | 1 | 6.92 | 2 | 39.23 | 39 | 0.23 | 0.23 | 0.01 | 0.01 |
| gene-LOC105283649 | https://www.ncbi.nlm.nih.gov/gene/?term=LOC105283649 | 1918 | 55 | 286.76 | 1 | 5.21 | 1.82 | 38.85 | 38.63 | 0.21 | 0.21 | 0.01 | 0.01 |
| gene-LOC105275374 | https://www.ncbi.nlm.nih.gov/gene/?term=LOC105275374 | 11376 | 525 | 461.5 1 | 0.88 | 0.19 | 10.23 | 10.06 | 0.17 | 0.17 | 0.01 | 0.01 | |
| gene-LOC105281700 | https://www.ncbi.nlm.nih.gov/gene/?term=LOC105281700 | 2208 | 38 | 172.1 1 | 4.53 | 2.63 | 39.22 | 39.09 | 0.14 | 0.14 | 0 | 0 | |
| gene-LOC105274531 | https://www.ncbi.nlm.nih.gov/gene/?term=LOC105274531 | 7774 | 283 | 364.03 | 1 | 1.29 | 0.35 | 10.82 | 10.73 | 0.09 | 0.09 | 0 | 0 |
| gene-LOC105282825 | https://www.ncbi.nlm.nih.gov/gene/?term=LOC105282825 | 3919 | 78 | 199.03 | 1 | 2.55 | 1.28 | 12.99 | 12.9 | 0.09 | 0.09 | 0 | 0 |
| gene-LOC105275179 | https://www.ncbi.nlm.nih.gov/gene/?term=LOC105275179 | 3271 | 71 | 217.06 | 1 | 3.06 | 1.41 | 16.78 | 16.7 | 0.08 | 0.08 | 0 | 0 |
| gene-LOC105279038 | https://www.ncbi.nlm.nih.gov/gene/?term=LOC105279038 | 9453 | 317 | 335.34 | 1 | 1.06 | 0.32 | 9.53 | 9.46 | 0.07 | 0.07 | 0 | 0 |
| gene-LOC105283987 | https://www.ncbi.nlm.nih.gov/gene/?term=LOC105283987 | 4841 | 71 | 146.66 | 1 | 2.07 | 1.41 | 32.31 | 32.24 | 0.07 | 0.07 | 0 | 0 |
| gene-LOC105284329 | https://www.ncbi.nlm.nih.gov/gene/?term=LOC105284329 | 4051 | 51 | 125.89 | 1 | 2.47 | 1.96 | 19.38 | 19.32 | 0.06 | 0.06 | 0 | 0 |
| gene-LOC105288078 | https://www.ncbi.nlm.nih.gov/gene/?term=LOC105288078 | 19126 | 307 | 160.51 | 1 | 0.52 | 0.33 | 16.81 | 16.78 | 0.03 | 0.03 | 0 | 0 |
| gene-LOC105276527 | https://www.ncbi.nlm.nih.gov/gene/?term=LOC105276527 | 7923 | 334 | 421.56 | 1 | 1.26 | 0.3 | 8.56 | 8.54 | 0.02 | 0.02 | 0 | 0 |
| gene-LOC105286452 | https://www.ncbi.nlm.nih.gov/gene/?term=LOC105286452 | 6888 | 68 | 98.72 1 | 1.45 | 1.47 | 28.88 | 28.89 | 0 | 0 | 0 | 0 | |
| gene-LOC105274856 | https://www.ncbi.nlm.nih.gov/gene/?term=LOC105274856 | 5216 | 212 | 406.44 | 1 | 1.92 | 0.47 | 18.3 | 18.31 | -0.01 | 0.01 | 0 | 0 |
| gene-LOC105275327 | https://www.ncbi.nlm.nih.gov/gene/?term=LOC105275327 | 14245 | 472 | 331.34 | 1 | 0.7 | 0.21 | 7.13 | 7.15 | -0.03 | 0.03 | 0 | 0 |
| gene-LOC105283943 | https://www.ncbi.nlm.nih.gov/gene/?term=LOC105283943 | 3559 | 56 | 157.35 | 1 | 2.81 | 1.79 | 16.7 | 16.73 | -0.03 | 0.03 | 0 | 0 |
| gene-LOC105284043 | https://www.ncbi.nlm.nih.gov/gene/?term=LOC105284043 | 5661 | 278 | 491.08 | 1 | 1.77 | 0.36 | 10.84 | 10.87 | -0.03 | 0.03 | 0 | 0 |
| gene-LOC105275683 | https://www.ncbi.nlm.nih.gov/gene/?term=LOC105275683 | 4256 | 191 | 448.78 | 1 | 2.35 | 0.52 | 12.17 | 12.21 | -0.04 | 0.04 | 0 | 0 |
| gene-LOC105277216 | https://www.ncbi.nlm.nih.gov/gene/?term=LOC105277216 | 7627 | 87 | 114.07 | 1 | 1.31 | 1.15 | 9.6 | 9.67 | -0.08 | 0.08 | 0 | 0 |
| gene-LOC105287417 | https://www.ncbi.nlm.nih.gov/gene/?term=LOC105287417 | 1025 | 28 | 273.17 | 1 | 9.76 | 3.57 | 42.07 | 42.23 | -0.16 | 0.16 | 0 | 0 |
| gene-LOC105282658 | https://www.ncbi.nlm.nih.gov/gene/?term=LOC105282658 | 2421 | 67 | 276.75 | 1 | 4.13 | 1.49 | 20.83 | 21.03 | -0.21 | 0.21 | -0.01 | 0.01 |
| gene-LOC105285981 | https://www.ncbi.nlm.nih.gov/gene/?term=LOC105285981 | 7666 | 183 | 238.72 | 1 | 1.3 | 0.55 | 18.71 | 18.92 | -0.21 | 0.21 | -0.01 | 0.01 |
| gene-LOC105278877 | https://www.ncbi.nlm.nih.gov/gene/?term=LOC105278877 | 1968 | 77 | 391.26 | 1 | 5.08 | 1.3 | 34.55 | 34.76 | -0.22 | 0.22 | -0.01 | 0.01 |
| gene-LOC105284774 | https://www.ncbi.nlm.nih.gov/gene/?term=LOC105284774 | 12089 | 81 | 67 1 | 0.83 | 1.23 | 21 | 21.27 | -0.27 | 0.27 | 0 | 0 | |
| gene-LOC105276196 | https://www.ncbi.nlm.nih.gov/gene/?term=LOC105276196 | 2953 | 45 | 152.39 | 1 | 3.39 | 2.22 | 48.59 | 48.91 | -0.31 | 0.31 | 0 | 0 |
| gene-LOC105274851 | https://www.ncbi.nlm.nih.gov/gene/?term=LOC105274851 | 6616 | 127 | 191.96 | 1 | 1.51 | 0.79 | 11.51 | 11.83 | -0.33 | 0.33 | -0.01 | 0.01 |
| gene-LOC105286649 | https://www.ncbi.nlm.nih.gov/gene/?term=LOC105286649 | 14178 | 785 | 553.67 | 1 | 0.71 | 0.13 | 0.94 | 1.3 | -0.36 | 0.36 | -0.02 | 0.02 |
| gene-LOC105285742 | https://www.ncbi.nlm.nih.gov/gene/?term=LOC105285742 | 3446 | 97 | 281.49 | 1 | 2.9 | 1.03 | 20.21 | 20.62 | -0.41 | 0.41 | -0.01 | 0.01 |
| gene-LOC105274590 | https://www.ncbi.nlm.nih.gov/gene/?term=LOC105274590 | 4516 | 64 | 141.72 | 2 | 4.43 | 3.12 | 32.42 | 32.94 | -0.51 | 0.51 | -0.01 | 0.01 |
| gene-LOC105284146 | https://www.ncbi.nlm.nih.gov/gene/?term=LOC105284146 | 2488 | 66 | 265.27 | 1 | 4.02 | 1.52 | 23.09 | 23.62 | -0.54 | 0.54 | -0.01 | 0.01 |
| gene-LOC105278228 | https://www.ncbi.nlm.nih.gov/gene/?term=LOC105278228 | 8610 | 169 | 196.28 | 1 | 1.16 | 0.59 | 21.96 | 22.51 | -0.55 | 0.55 | -0.01 | 0.01 |
| gene-LOC105280956 | https://www.ncbi.nlm.nih.gov/gene/?term=LOC105280956 | 2639 | 77 | 291.78 | 1 | 3.79 | 1.3 | 18.18 | 18.82 | -0.64 | 0.64 | -0.02 | 0.02 |
| gene-LOC105286515 | https://www.ncbi.nlm.nih.gov/gene/?term=LOC105286515 | 3751 | 113 | 301.25 | 1 | 2.67 | 0.88 | 10.68 | 11.31 | -0.64 | 0.64 | -0.02 | 0.02 |
| gene-LOC105278459 | https://www.ncbi.nlm.nih.gov/gene/?term=LOC105278459 | 2936 | 97 | 330.38 | 1 | 3.41 | 1.03 | 20.46 | 21.23 | -0.77 | 0.77 | -0.03 | 0.03 |
| gene-LOC105278454 | https://www.ncbi.nlm.nih.gov/gene/?term=LOC105278454 | 1972 | 41 | 207.91 | 1 | 5.07 | 2.44 | 43.16 | 44.07 | -0.91 | 0.91 | -0.02 | 0.02 |
| gene-LOC105287633 | https://www.ncbi.nlm.nih.gov/gene/?term=LOC105287633 | 1524 | 23 | 150.92 | 1 | 6.56 | 4.35 | 68.63 | 69.73 | -1.11 | 1.11 | -0.02 | 0.02 |

| | | | | | | | | | | | | | |
|---|---|---|---|---|---|---|---|---|---|---|---|---|---|
| gene-LOC105281173 | https://www.ncbi.nlm.nih.gov/gene/?term=LOC105281173 | 1595 | 39 | 244.51 | 1 | 6.27 | 2.56 | 60.96 | 62.08 | -1.12 | 1.12 | -0.03 | 0.03 |
| gene-LOC105278282 | https://www.ncbi.nlm.nih.gov/gene/?term=LOC105278282 | 1775 | 56 | 315.49 | 1 | 5.63 | 1.79 | 33.68 | 34.81 | -1.13 | 1.13 | -0.04 | 0.04 |
| gene-LOC105285252 | https://www.ncbi.nlm.nih.gov/gene/?term=LOC105285252 | 7427 | 124 | 166.96 | 2 | 2.69 | 1.61 | 12.83 | 14.05 | -1.22 | 1.22 | -0.02 | 0.02 |
| gene-LOC105274534 | https://www.ncbi.nlm.nih.gov/gene/?term=LOC105274534 | 1329 | 55 | 413.84 | 1 | 7.52 | 1.82 | 46.07 | 47.79 | -1.72 | 1.72 | -0.07 | 0.07 |
| gene-LOC105284507 | https://www.ncbi.nlm.nih.gov/gene/?term=LOC105284507 | 5207 | 86 | 165.16 | 1 | 1.92 | 1.16 | 21.7 | 23.52 | -1.83 | 1.83 | -0.03 | 0.03 |
| gene-LOC105284213 | https://www.ncbi.nlm.nih.gov/gene/?term=LOC105284213 | 868 | 15 | 172.81 | 1 | 11.52 | 6.67 | 15.38 | 18.37 | -2.99 | 2.99 | -0.05 | 0.05 |

Directory: OGL File: ogl-Ob_LI_BCphaseA2.vs.RphaseB2.txt

| gene_ID | gene_link | gwidth | #Sites | #per10Kb | #dmSites | #dmsp10kb | %dmSites | %pSite1 | %pSite2 | DMpSite | ADMpSite | DMpNucl | ADMpNucl |
|---|---|---|---|---|---|---|---|---|---|---|---|---|---|
| gene-LOC113562901 | https://www.ncbi.nlm.nih.gov/gene/?term=LOC113562901 | 1531 | 22 | 143.7 | 1 | 6.53 | 4.55 | 60.1 | 44.57 | 15.53 | 15.53 | 0.22 | 0.22 |
| gene-LOC105279898 | https://www.ncbi.nlm.nih.gov/gene/?term=LOC105279898 | 1343 | 10 | 74.46 | 1 | 7.45 | 10 | 84.43 | 71.53 | 12.89 | 12.89 | 0.1 | 0.1 |
| gene-LOC105275070 | https://www.ncbi.nlm.nih.gov/gene/?term=LOC105275070 | 5667 | 41 | 72.35 | 5 | 8.82 | 12.2 | 13.7 | 1.06 | 12.65 | 12.65 | 0.09 | 0.09 |
| gene-LOC113562345 | https://www.ncbi.nlm.nih.gov/gene/?term=LOC113562345 | 2290 | 67 | 292.58 | 11 | 48.03 | 16.42 | 29.88 | 17.32 | 12.57 | 12.57 | 0.37 | 0.37 |
| gene-LOC105286694 | https://www.ncbi.nlm.nih.gov/gene/?term=LOC105286694 | 3409 | 9 | 26.4 | 1 | 2.93 | 11.11 | 71.7 | 62.28 | 9.42 | 9.42 | 0.02 | 0.02 |
| gene-LOC105275708 | https://www.ncbi.nlm.nih.gov/gene/?term=LOC105275708 | 1772 | 10 | 56.43 | 1 | 5.64 | 10 | 50.31 | 44.22 | 6.09 | 6.09 | 0.03 | 0.03 |
| gene-LOC105282976 | https://www.ncbi.nlm.nih.gov/gene/?term=LOC105282976 | 1648 | 47 | 285.19 | 1 | 6.07 | 2.13 | 28.02 | 22.43 | 5.59 | 5.59 | 0.16 | 0.16 |
| gene-LOC105286644 | https://www.ncbi.nlm.nih.gov/gene/?term=LOC105286644 | 1235 | 22 | 178.14 | 1 | 8.1 | 4.55 | 42.35 | 37.26 | 5.09 | 5.09 | 0.09 | 0.09 |
| gene-LOC105285315 | https://www.ncbi.nlm.nih.gov/gene/?term=LOC105285315 | 5458 | 19 | 34.81 | 1 | 1.83 | 5.26 | 47.88 | 43.22 | 4.66 | 4.66 | 0.02 | 0.02 |
| gene-LOC105283552 | https://www.ncbi.nlm.nih.gov/gene/?term=LOC105283552 | 1562 | 36 | 230.47 | 1 | 6.4 | 2.78 | 56.28 | 51.7 | 4.58 | 4.58 | 0.11 | 0.11 |
| gene-LOC105276078 | https://www.ncbi.nlm.nih.gov/gene/?term=LOC105276078 | 1051 | 14 | 133.21 | 1 | 9.51 | 7.14 | 36.98 | 32.93 | 4.05 | 4.05 | 0.05 | 0.05 |
| gene-LOC105288063 | https://www.ncbi.nlm.nih.gov/gene/?term=LOC105288063 | 3400 | 43 | 126.47 | 1 | 2.94 | 2.33 | 18.5 | 14.57 | 3.92 | 3.92 | 0.05 | 0.05 |
| gene-LOC105278109 | https://www.ncbi.nlm.nih.gov/gene/?term=LOC105278109 | 4029 | 59 | 146.44 | 1 | 2.48 | 1.69 | 50.99 | 47.24 | 3.75 | 3.75 | 0.05 | 0.05 |
| gene-LOC105278215 | https://www.ncbi.nlm.nih.gov/gene/?term=LOC105278215 | 1700 | 29 | 170.59 | 1 | 5.88 | 3.45 | 57.89 | 54.66 | 3.23 | 3.23 | 0.06 | 0.06 |
| gene-LOC105275518 | https://www.ncbi.nlm.nih.gov/gene/?term=LOC105275518 | 2335 | 18 | 77.09 | 1 | 4.28 | 5.56 | 54.44 | 51.41 | 3.03 | 3.03 | 0.02 | 0.02 |
| gene-LOC105285674 | https://www.ncbi.nlm.nih.gov/gene/?term=LOC105285674 | 3035 | 93 | 306.43 | 2 | 6.59 | 2.15 | 26.74 | 23.71 | 3.03 | 3.03 | 0.09 | 0.09 |
| gene-LOC105285060 | https://www.ncbi.nlm.nih.gov/gene/?term=LOC105285060 | 1285 | 17 | 132.3 | 1 | 7.78 | 5.88 | 51.2 | 48.29 | 2.91 | 2.91 | 0.04 | 0.04 |
| gene-LOC105275347 | https://www.ncbi.nlm.nih.gov/gene/?term=LOC105275347 | 2526 | 14 | 55.42 | 1 | 3.96 | 7.14 | 33.57 | 30.77 | 2.8 | 2.8 | 0.02 | 0.02 |
| gene-LOC105274900 | https://www.ncbi.nlm.nih.gov/gene/?term=LOC105274900 | 5561 | 47 | 84.52 | 2 | 3.6 | 4.26 | 37.9 | 35.14 | 2.76 | 2.76 | 0.02 | 0.02 |
| gene-LOC105283649 | https://www.ncbi.nlm.nih.gov/gene/?term=LOC105283649 | 1918 | 54 | 281.54 | 1 | 5.21 | 1.85 | 37.63 | 35.03 | 2.6 | 2.6 | 0.07 | 0.07 |
| gene-LOC105279664 | https://www.ncbi.nlm.nih.gov/gene/?term=LOC105279664 | 4023 | 48 | 119.31 | 1 | 2.49 | 2.08 | 23.51 | 21.34 | 2.18 | 2.18 | 0.03 | 0.03 |
| gene-LOC105281515 | https://www.ncbi.nlm.nih.gov/gene/?term=LOC105281515 | 8827 | 38 | 43.05 | 1 | 1.13 | 2.63 | 31.29 | 29.13 | 2.16 | 2.16 | 0.01 | 0.01 |
| gene-LOC105276704 | https://www.ncbi.nlm.nih.gov/gene/?term=LOC105276704 | 2063 | 88 | 426.56 | 1 | 4.85 | 1.14 | 15.96 | 13.92 | 2.05 | 2.05 | 0.09 | 0.09 |
| gene-LOC105283123 | https://www.ncbi.nlm.nih.gov/gene/?term=LOC105283123 | 1507 | 44 | 291.97 | 1 | 6.64 | 2.27 | 19.91 | 17.87 | 2.05 | 2.05 | 0.06 | 0.06 |
| gene-LOC105281754 | https://www.ncbi.nlm.nih.gov/gene/?term=LOC105281754 | 4300 | 110 | 255.81 | 1 | 2.33 | 0.91 | 16.49 | 14.51 | 1.98 | 1.98 | 0.05 | 0.05 |
| gene-LOC113561805 | https://www.ncbi.nlm.nih.gov/gene/?term=LOC113561805 | 3337 | 68 | 203.78 | 1 | 3 | 1.47 | 35.02 | 33.08 | 1.94 | 1.94 | 0.04 | 0.04 |
| gene-LOC105285671 | https://www.ncbi.nlm.nih.gov/gene/?term=LOC105285671 | 2263 | 103 | 455.15 | 1 | 4.42 | 0.97 | 29.21 | 27.3 | 1.91 | 1.91 | 0.09 | 0.09 |
| gene-LOC105288066 | https://www.ncbi.nlm.nih.gov/gene/?term=LOC105288066 | 1964 | 20 | 101.83 | 1 | 5.09 | 5 | 44.28 | 42.38 | 1.9 | 1.9 | 0.02 | 0.02 |
| gene-LOC105282156 | https://www.ncbi.nlm.nih.gov/gene/?term=LOC105282156 | 2577 | 129 | 500.58 | 1 | 3.88 | 0.78 | 28.68 | 26.84 | 1.84 | 1.84 | 0.09 | 0.09 |
| gene-LOC105280408 | https://www.ncbi.nlm.nih.gov/gene/?term=LOC105280408 | 5983 | 47 | 78.56 | 1 | 1.67 | 2.13 | 19.12 | 17.33 | 1.78 | 1.78 | 0.01 | 0.01 |
| gene-LOC105282149 | https://www.ncbi.nlm.nih.gov/gene/?term=LOC105282149 | 1787 | 108 | 604.36 | 1 | 5.6 | 0.93 | 43.65 | 41.89 | 1.76 | 1.76 | 0.11 | 0.11 |
| gene-LOC105276190 | https://www.ncbi.nlm.nih.gov/gene/?term=LOC105276190 | 4839 | 82 | 169.46 | 1 | 2.07 | 1.22 | 11.63 | 9.92 | 1.71 | 1.71 | 0.03 | 0.03 |
| gene-LOC105282794 | https://www.ncbi.nlm.nih.gov/gene/?term=LOC105282794 | 4571 | 92 | 201.27 | 1 | 2.19 | 1.09 | 18.21 | 16.51 | 1.7 | 1.7 | 0.03 | 0.03 |
| gene-LOC105280786 | https://www.ncbi.nlm.nih.gov/gene/?term=LOC105280786 | 2573 | 113 | 439.18 | 1 | 3.89 | 0.88 | 32.35 | 30.66 | 1.69 | 1.69 | 0.07 | 0.07 |
| gene-LOC105282970 | https://www.ncbi.nlm.nih.gov/gene/?term=LOC105282970 | 1529 | 39 | 255.07 | 1 | 6.54 | 2.56 | 51.08 | 49.46 | 1.62 | 1.62 | 0.04 | 0.04 |
| gene-LOC105284184 | https://www.ncbi.nlm.nih.gov/gene/?term=LOC105284184 | 2535 | 79 | 311.64 | 1 | 3.94 | 1.27 | 37.02 | 35.48 | 1.54 | 1.54 | 0.05 | 0.05 |
| gene-LOC105277761 | https://www.ncbi.nlm.nih.gov/gene/?term=LOC105277761 | 4648 | 143 | 307.66 | 1 | 2.15 | 0.7 | 15.33 | 13.81 | 1.52 | 1.52 | 0.05 | 0.05 |
| gene-LOC105275966 | https://www.ncbi.nlm.nih.gov/gene/?term=LOC105275966 | 3299 | 53 | 160.65 | 1 | 3.03 | 1.89 | 54.5 | 52.99 | 1.51 | 1.51 | 0.02 | 0.02 |
| gene-LOC105278807 | https://www.ncbi.nlm.nih.gov/gene/?term=LOC105278807 | 2166 | 62 | 286.24 | 1 | 4.62 | 1.61 | 21.98 | 20.52 | 1.46 | 1.46 | 0.04 | 0.04 |
| gene-LOC105277565 | https://www.ncbi.nlm.nih.gov/gene/?term=LOC105277565 | 3216 | 169 | 525.5 | 1 | 3.11 | 0.59 | 20.65 | 19.2 | 1.45 | 1.45 | 0.08 | 0.08 |
| gene-LOC105283728 | https://www.ncbi.nlm.nih.gov/gene/?term=LOC105283728 | 3256 | 107 | 328.62 | 2 | 6.14 | 1.87 | 20.96 | 19.59 | 1.37 | 1.37 | 0.04 | 0.04 |
| gene-LOC105276059 | https://www.ncbi.nlm.nih.gov/gene/?term=LOC105276059 | 6256 | 121 | 193.41 | 1 | 1.6 | 0.83 | 10.95 | 9.59 | 1.36 | 1.36 | 0.03 | 0.03 |
| gene-LOC105283712 | https://www.ncbi.nlm.nih.gov/gene/?term=LOC105283712 | 1910 | 31 | 162.3 | 1 | 5.24 | 3.23 | 30.95 | 29.59 | 1.36 | 1.36 | 0.02 | 0.02 |
| gene-LOC105276875 | https://www.ncbi.nlm.nih.gov/gene/?term=LOC105276875 | 3151 | 109 | 345.92 | 1 | 3.17 | 0.92 | 25.45 | 24.19 | 1.26 | 1.26 | 0.04 | 0.04 |
| gene-LOC105278497 | https://www.ncbi.nlm.nih.gov/gene/?term=LOC105278497 | 2072 | 92 | 444.02 | 1 | 4.83 | 1.09 | 21.61 | 20.36 | 1.26 | 1.26 | 0.06 | 0.06 |
| gene-LOC105281173 | https://www.ncbi.nlm.nih.gov/gene/?term=LOC105281173 | 1595 | 38 | 238.24 | 1 | 6.27 | 2.63 | 60.05 | 58.85 | 1.2 | 1.2 | 0.03 | 0.03 |
| gene-LOC105279298 | https://www.ncbi.nlm.nih.gov/gene/?term=LOC105279298 | 2773 | 131 | 472.41 | 2 | 7.21 | 1.53 | 24.81 | 23.62 | 1.18 | 1.18 | 0.06 | 0.06 |
| gene-LOC105287063 | https://www.ncbi.nlm.nih.gov/gene/?term=LOC105287063 | 1874 | 48 | 256.14 | 1 | 5.34 | 2.08 | 31.5 | 30.33 | 1.17 | 1.17 | 0.03 | 0.03 |
| gene-LOC105281848 | https://www.ncbi.nlm.nih.gov/gene/?term=LOC105281848 | 2122 | 47 | 221.49 | 1 | 4.71 | 2.13 | 27.8 | 26.69 | 1.11 | 1.11 | 0.02 | 0.02 |
| gene-LOC105284355 | https://www.ncbi.nlm.nih.gov/gene/?term=LOC105284355 | 3904 | 143 | 366.29 | 1 | 2.56 | 0.7 | 22.85 | 21.77 | 1.08 | 1.08 | 0.04 | 0.04 |
| gene-LOC105274590 | https://www.ncbi.nlm.nih.gov/gene/?term=LOC105274590 | 4516 | 57 | 126.22 | 2 | 4.43 | 3.51 | 34.78 | 33.73 | 1.06 | 1.06 | 0.01 | 0.01 |
| gene-LOC105275451 | https://www.ncbi.nlm.nih.gov/gene/?term=LOC105275451 | 6035 | 118 | 195.53 | 1 | 1.66 | 0.85 | 23.45 | 22.41 | 1.04 | 1.04 | 0.02 | 0.02 |
| gene-LOC105285179 | https://www.ncbi.nlm.nih.gov/gene/?term=LOC105285179 | 4191 | 177 | 422.33 | 1 | 2.39 | 0.56 | 19.66 | 18.63 | 1.04 | 1.04 | 0.04 | 0.04 |
| gene-LOC105276272 | https://www.ncbi.nlm.nih.gov/gene/?term=LOC105276272 | 1965 | 34 | 173.03 | 1 | 5.09 | 2.94 | 33.29 | 32.32 | 0.98 | 0.98 | 0.02 | 0.02 |
| gene-LOC105279500 | https://www.ncbi.nlm.nih.gov/gene/?term=LOC105279500 | 3587 | 34 | 94.79 | 1 | 2.79 | 2.94 | 52.36 | 51.39 | 0.97 | 0.97 | 0.01 | 0.01 |
| gene-LOC105281272 | https://www.ncbi.nlm.nih.gov/gene/?term=LOC105281272 | 5252 | 156 | 297.03 | 1 | 1.9 | 0.64 | 18.91 | 17.94 | 0.97 | 0.97 | 0.03 | 0.03 |
| gene-LOC105280774 | https://www.ncbi.nlm.nih.gov/gene/?term=LOC105280774 | 2793 | 89 | 318.65 | 1 | 3.58 | 1.12 | 36.33 | 35.38 | 0.94 | 0.94 | 0.03 | 0.03 |
| gene-LOC105283987 | https://www.ncbi.nlm.nih.gov/gene/?term=LOC105283987 | 4841 | 64 | 132.2 | 1 | 2.07 | 1.56 | 30.28 | 29.39 | 0.89 | 0.89 | 0.01 | 0.01 |
| gene-LOC105287417 | https://www.ncbi.nlm.nih.gov/gene/?term=LOC105287417 | 1025 | 22 | 214.63 | 1 | 9.76 | 4.55 | 50.13 | 49.23 | 0.89 | 0.89 | 0.02 | 0.02 |
| gene-LOC105284036 | https://www.ncbi.nlm.nih.gov/gene/?term=LOC105284036 | 3254 | 96 | 295.02 | 1 | 3.07 | 1.04 | 22.78 | 21.95 | 0.83 | 0.83 | 0.02 | 0.02 |

| | | | | | | | | | | | | | |
|---|---|---|---|---|---|---|---|---|---|---|---|---|---|
| gene-LOC105276449 | https://www.ncbi.nlm.nih.gov/gene/?term=LOC105276449 | 1231 | 42 | 341.19 | 1 | 8.12 | 2.38 | 35.95 | 35.13 | 0.82 | 0.82 | 0.03 | 0.03 |
| gene-LOC105285055 | https://www.ncbi.nlm.nih.gov/gene/?term=LOC105285055 | 4108 | 255 | 620.74 | 2 | 4.87 | 0.78 | 11.66 | 10.91 | 0.76 | 0.76 | 0.05 | 0.05 |
| gene-LOC105285202 | https://www.ncbi.nlm.nih.gov/gene/?term=LOC105285202 | 1838 | 67 | 364.53 | 1 | 5.44 | 1.49 | 26.25 | 25.49 | 0.76 | 0.76 | 0.03 | 0.03 |
| gene-LOC105275363 | https://www.ncbi.nlm.nih.gov/gene/?term=LOC105275363 | 3798 | 150 | 394.94 | 1 | 2.63 | 0.67 | 18.12 | 17.37 | 0.75 | 0.75 | 0.03 | 0.03 |
| gene-LOC105283592 | https://www.ncbi.nlm.nih.gov/gene/?term=LOC105283592 | 3479 | 72 | 206.96 | 1 | 2.87 | 1.39 | 23.84 | 23.1 | 0.74 | 0.74 | 0.02 | 0.02 |
| gene-LOC105285395 | https://www.ncbi.nlm.nih.gov/gene/?term=LOC105285395 | 8085 | 185 | 228.82 | 1 | 1.24 | 0.54 | 16.99 | 16.27 | 0.72 | 0.72 | 0.02 | 0.02 |
| gene-LOC105287095 | https://www.ncbi.nlm.nih.gov/gene/?term=LOC105287095 | 1435 | 57 | 397.21 | 1 | 6.97 | 1.75 | 29.87 | 29.16 | 0.71 | 0.71 | 0.03 | 0.03 |
| gene-LOC113561621 | https://www.ncbi.nlm.nih.gov/gene/?term=LOC113561621 | 2249 | 82 | 364.61 | 1 | 4.45 | 1.22 | 20.63 | 19.95 | 0.68 | 0.68 | 0.02 | 0.02 |
| gene-LOC105285639 | https://www.ncbi.nlm.nih.gov/gene/?term=LOC105285639 | 10325 | 319 | 308.96 | 1 | 0.97 | 0.31 | 12.35 | 11.71 | 0.65 | 0.65 | 0.02 | 0.02 |
| gene-LOC105275374 | https://www.ncbi.nlm.nih.gov/gene/?term=LOC105275374 | 11376 | 489 | 429.85 | 1 | 0.88 | 0.2 | 11.7 | 11.06 | 0.64 | 0.64 | 0.03 | 0.03 |
| gene-LOC105274851 | https://www.ncbi.nlm.nih.gov/gene/?term=LOC105274851 | 6616 | 123 | 185.91 | 1 | 1.51 | 0.81 | 11.44 | 10.81 | 0.63 | 0.63 | 0.01 | 0.01 |
| gene-LOC105274452 | https://www.ncbi.nlm.nih.gov/gene/?term=LOC105274452 | 6595 | 252 | 382.11 | 1 | 1.52 | 0.4 | 13.74 | 13.13 | 0.61 | 0.61 | 0.02 | 0.02 |
| gene-LOC105282885 | https://www.ncbi.nlm.nih.gov/gene/?term=LOC105282885 | 1870 | 45 | 240.64 | 1 | 5.35 | 2.22 | 42.36 | 41.77 | 0.58 | 0.58 | 0.01 | 0.01 |
| gene-LOC105276692 | https://www.ncbi.nlm.nih.gov/gene/?term=LOC105276692 | 1745 | 29 | 166.19 | 1 | 5.73 | 3.45 | 25.51 | 24.96 | 0.55 | 0.55 | 0.01 | 0.01 |
| gene-LOC105278145 | https://www.ncbi.nlm.nih.gov/gene/?term=LOC105278145 | 4470 | 246 | 550.34 | 1 | 2.24 | 0.41 | 11.03 | 10.48 | 0.55 | 0.55 | 0.03 | 0.03 |
| gene-LOC105278283 | https://www.ncbi.nlm.nih.gov/gene/?term=LOC105278283 | 2317 | 125 | 539.49 | 1 | 4.32 | 0.8 | 28.1 | 27.55 | 0.55 | 0.55 | 0.03 | 0.03 |
| gene-LOC105284704 | https://www.ncbi.nlm.nih.gov/gene/?term=LOC105284704 | 2400 | 57 | 237.5 1 | 4.17 | 1.75 | 25.25 | 24.69 | 0.55 | 0.55 | 0.01 | 0.01 | |
| gene-LOC105282151 | https://www.ncbi.nlm.nih.gov/gene/?term=LOC105282151 | 1601 | 35 | 218.61 | 1 | 6.25 | 2.86 | 42.13 | 41.61 | 0.53 | 0.53 | 0.01 | 0.01 |
| gene-LOC105274706 | https://www.ncbi.nlm.nih.gov/gene/?term=LOC105274706 | 1446 | 55 | 380.36 | 1 | 6.92 | 1.82 | 37.05 | 36.54 | 0.5 | 0.5 | 0.02 | 0.02 |
| gene-LOC105280410 | https://www.ncbi.nlm.nih.gov/gene/?term=LOC105280410 | 12580 | 235 | 186.8 1 | 0.79 | 0.43 | 21.34 | 20.84 | 0.5 | 0.5 | 0.01 | 0.01 | |
| gene-LOC105274823 | https://www.ncbi.nlm.nih.gov/gene/?term=LOC105274823 | 4620 | 132 | 285.71 | 1 | 2.16 | 0.76 | 21.94 | 21.47 | 0.47 | 0.47 | 0.01 | 0.01 |
| gene-LOC105279038 | https://www.ncbi.nlm.nih.gov/gene/?term=LOC105279038 | 9453 | 306 | 323.71 | 1 | 1.06 | 0.33 | 10.13 | 9.67 | 0.46 | 0.46 | 0.01 | 0.01 |
| gene-LOC105274856 | https://www.ncbi.nlm.nih.gov/gene/?term=LOC105274856 | 5216 | 207 | 396.86 | 1 | 1.92 | 0.48 | 19.99 | 19.54 | 0.45 | 0.45 | 0.02 | 0.02 |
| gene-LOC105286514 | https://www.ncbi.nlm.nih.gov/gene/?term=LOC105286514 | 2528 | 44 | 174.05 | 1 | 3.96 | 2.27 | 33.92 | 33.47 | 0.45 | 0.45 | 0.01 | 0.01 |
| gene-LOC105278666 | https://www.ncbi.nlm.nih.gov/gene/?term=LOC105278666 | 10341 | 429 | 414.85 | 1 | 0.97 | 0.23 | 7.71 | 7.27 | 0.44 | 0.44 | 0.02 | 0.02 |
| gene-LOC105285406 | https://www.ncbi.nlm.nih.gov/gene/?term=LOC105285406 | 8015 | 60 | 74.86 1 | 1.25 | 1.67 | 25.96 | 25.52 | 0.44 | 0.44 | 0 | 0 | |
| gene-LOC105285807 | https://www.ncbi.nlm.nih.gov/gene/?term=LOC105285807 | 6063 | 214 | 352.96 | 1 | 1.65 | 0.47 | 7.55 | 7.11 | 0.44 | 0.44 | 0.02 | 0.02 |
| gene-LOC105278282 | https://www.ncbi.nlm.nih.gov/gene/?term=LOC105278282 | 1775 | 52 | 292.96 | 1 | 5.63 | 1.92 | 38.05 | 37.63 | 0.42 | 0.42 | 0.01 | 0.01 |
| gene-LOC105276612 | https://www.ncbi.nlm.nih.gov/gene/?term=LOC105276612 | 4157 | 181 | 435.41 | 1 | 2.41 | 0.55 | 10.16 | 9.76 | 0.4 | 0.4 | 0.02 | 0.02 |
| gene-LOC105276941 | https://www.ncbi.nlm.nih.gov/gene/?term=LOC105276941 | 15956 | 494 | 309.6 1 | 0.63 | 0.2 | 5.44 | 5.07 | 0.38 | 0.38 | 0.01 | 0.01 | |
| gene-LOC105277423 | https://www.ncbi.nlm.nih.gov/gene/?term=LOC105277423 | 2340 | 108 | 461.54 | 1 | 4.27 | 0.93 | 30.29 | 29.91 | 0.38 | 0.38 | 0.02 | 0.02 |
| gene-LOC105287689 | https://www.ncbi.nlm.nih.gov/gene/?term=LOC105287689 | 10867 | 331 | 304.59 | 1 | 0.92 | 0.3 | 9.53 | 9.17 | 0.36 | 0.36 | 0.01 | 0.01 |
| gene-LOC105276452 | https://www.ncbi.nlm.nih.gov/gene/?term=LOC105276452 | 2917 | 263 | 901.61 | 1 | 3.43 | 0.38 | 9.41 | 9.13 | 0.28 | 0.28 | 0.03 | 0.03 |
| gene-LOC105278454 | https://www.ncbi.nlm.nih.gov/gene/?term=LOC105278454 | 1972 | 44 | 223.12 | 1 | 5.07 | 2.27 | 46.65 | 46.38 | 0.26 | 0.26 | 0.01 | 0.01 |
| gene-LOC105278877 | https://www.ncbi.nlm.nih.gov/gene/?term=LOC105278877 | 1968 | 69 | 350.61 | 1 | 5.08 | 1.45 | 35.47 | 35.23 | 0.25 | 0.25 | 0.01 | 0.01 |
| gene-LOC105280407 | https://www.ncbi.nlm.nih.gov/gene/?term=LOC105280407 | 10950 | 189 | 172.6 1 | 0.91 | 0.53 | 16.21 | 15.97 | 0.24 | 0.24 | 0 | 0 | |
| gene-LOC105275460 | https://www.ncbi.nlm.nih.gov/gene/?term=LOC105275460 | 2473 | 116 | 469.07 | 1 | 4.04 | 0.86 | 13.65 | 13.42 | 0.23 | 0.23 | 0.01 | 0.01 |
| gene-LOC113562050 | https://www.ncbi.nlm.nih.gov/gene/?term=LOC113562050 | 17097 | 540 | 315.84 | 1 | 0.58 | 0.19 | 5.38 | 5.15 | 0.23 | 0.23 | 0.01 | 0.01 |
| gene-LOC105278534 | https://www.ncbi.nlm.nih.gov/gene/?term=LOC105278534 | 7364 | 175 | 237.64 | 1 | 1.36 | 0.57 | 18.73 | 18.51 | 0.22 | 0.22 | 0.01 | 0.01 |
| gene-LOC105286166 | https://www.ncbi.nlm.nih.gov/gene/?term=LOC105286166 | 3050 | 103 | 337.7 1 | 3.28 | 0.97 | 25.5 | 25.28 | 0.22 | 0.22 | 0.01 | 0.01 | |
| gene-LOC105274449 | https://www.ncbi.nlm.nih.gov/gene/?term=LOC105274449 | 1549 | 48 | 309.88 | 1 | 6.46 | 2.08 | 51.17 | 50.96 | 0.21 | 0.21 | 0.01 | 0.01 |
| gene-LOC105280956 | https://www.ncbi.nlm.nih.gov/gene/?term=LOC105280956 | 2639 | 69 | 261.46 | 1 | 3.79 | 1.45 | 18.51 | 18.31 | 0.21 | 0.21 | 0.01 | 0.01 |
| gene-LOC105275683 | https://www.ncbi.nlm.nih.gov/gene/?term=LOC105275683 | 4256 | 183 | 429.98 | 1 | 2.35 | 0.55 | 12.85 | 12.65 | 0.2 | 0.2 | 0.01 | 0.01 |
| gene-LOC105287764 | https://www.ncbi.nlm.nih.gov/gene/?term=LOC105287764 | 18359 | 375 | 204.26 | 1 | 0.54 | 0.27 | 3.25 | 3.09 | 0.17 | 0.17 | 0 | 0 |
| gene-LOC105282658 | https://www.ncbi.nlm.nih.gov/gene/?term=LOC105282658 | 2421 | 54 | 223.05 | 1 | 4.13 | 1.85 | 19.57 | 19.42 | 0.16 | 0.16 | 0 | 0 |
| gene-LOC105282825 | https://www.ncbi.nlm.nih.gov/gene/?term=LOC105282825 | 3919 | 69 | 176.07 | 1 | 2.55 | 1.45 | 13.52 | 13.36 | 0.16 | 0.16 | 0 | 0 |
| gene-LOC105276462 | https://www.ncbi.nlm.nih.gov/gene/?term=LOC105276462 | 6561 | 246 | 374.94 | 1 | 1.52 | 0.41 | 8.14 | 7.99 | 0.15 | 0.15 | 0.01 | 0.01 |
| gene-LOC105286999 | https://www.ncbi.nlm.nih.gov/gene/?term=LOC105286999 | 2141 | 42 | 196.17 | 1 | 4.67 | 2.38 | 36.17 | 36.05 | 0.12 | 0.12 | 0 | 0 |
| gene-LOC105282811 | https://www.ncbi.nlm.nih.gov/gene/?term=LOC105282811 | 5765 | 198 | 343.45 | 1 | 1.73 | 0.51 | 7.89 | 7.77 | 0.11 | 0.11 | 0 | 0 |
| gene-LOC105275327 | https://www.ncbi.nlm.nih.gov/gene/?term=LOC105275327 | 14245 | 460 | 322.92 | 1 | 0.7 | 0.22 | 7.25 | 7.2 | 0.05 | 0.05 | 0 | 0 |
| gene-LOC105282646 | https://www.ncbi.nlm.nih.gov/gene/?term=LOC105282646 | 3317 | 99 | 298.46 | 1 | 3.01 | 1.01 | 25.07 | 25.11 | -0.04 | 0.04 | 0 | 0 |
| gene-LOC105274531 | https://www.ncbi.nlm.nih.gov/gene/?term=LOC105274531 | 7774 | 285 | 366.61 | 1 | 1.29 | 0.35 | 10.62 | 10.74 | -0.11 | 0.11 | 0 | 0 |
| gene-LOC105280377 | https://www.ncbi.nlm.nih.gov/gene/?term=LOC105280377 | 2024 | 58 | 286.56 | 1 | 4.94 | 1.72 | 49.97 | 50.08 | -0.11 | 0.11 | 0 | 0 |
| gene-LOC105275848 | https://www.ncbi.nlm.nih.gov/gene/?term=LOC105275848 | 3718 | 133 | 357.72 | 1 | 2.69 | 0.75 | 21.71 | 21.82 | -0.12 | 0.12 | 0 | 0 |
| gene-LOC105276538 | https://www.ncbi.nlm.nih.gov/gene/?term=LOC105276538 | 1511 | 19 | 125.74 | 1 | 6.62 | 5.26 | 67.47 | 67.59 | -0.12 | 0.12 | 0 | 0 |
| gene-LOC105283296 | https://www.ncbi.nlm.nih.gov/gene/?term=LOC105283296 | 2174 | 84 | 386.38 | 1 | 4.6 | 1.19 | 19.37 | 19.49 | -0.12 | 0.12 | 0 | 0 |
| gene-LOC105278632 | https://www.ncbi.nlm.nih.gov/gene/?term=LOC105278632 | 3624 | 113 | 311.81 | 1 | 2.76 | 0.88 | 16.03 | 16.16 | -0.13 | 0.13 | 0 | 0 |
| gene-LOC105283943 | https://www.ncbi.nlm.nih.gov/gene/?term=LOC105283943 | 3559 | 42 | 118.01 | 1 | 2.81 | 2.38 | 14.31 | 14.48 | -0.17 | 0.17 | 0 | 0 |
| gene-LOC105275179 | https://www.ncbi.nlm.nih.gov/gene/?term=LOC105275179 | 3271 | 66 | 201.77 | 1 | 3.06 | 1.52 | 20.6 | 20.81 | -0.21 | 0.21 | 0 | 0 |
| gene-LOC105278649 | https://www.ncbi.nlm.nih.gov/gene/?term=LOC105278649 | 5616 | 161 | 286.68 | 1 | 1.78 | 0.62 | 11.64 | 11.9 | -0.26 | 0.26 | -0.01 | 0.01 |
| gene-LOC105276527 | https://www.ncbi.nlm.nih.gov/gene/?term=LOC105276527 | 7923 | 309 | 390 1 | 1.26 | 0.32 | 9.63 | 9.9 | -0.27 | 0.27 | -0.01 | 0.01 | |
| gene-LOC105286649 | https://www.ncbi.nlm.nih.gov/gene/?term=LOC105286649 | 14178 | 778 | 548.74 | 1 | 0.71 | 0.13 | 0.86 | 1.14 | -0.27 | 0.27 | -0.02 | 0.02 |
| gene-LOC105285514 | https://www.ncbi.nlm.nih.gov/gene/?term=LOC105285514 | 1277 | 14 | 109.63 | 1 | 7.83 | 7.14 | 85.59 | 85.88 | -0.28 | 0.28 | 0 | 0 |
| gene-LOC105286274 | https://www.ncbi.nlm.nih.gov/gene/?term=LOC105286274 | 3817 | 193 | 505.63 | 1 | 2.62 | 0.52 | 16.76 | 17.05 | -0.29 | 0.29 | -0.01 | 0.01 |

| | | | | | | | | | | | | | | | |
|---|---|---|---|---|---|---|---|---|---|---|---|---|---|---|---|
| gene-LOC105286515 | https://www.ncbi.nlm.nih.gov/gene/?term=LOC105286515 | 3751 | 99 | 263.93 | | 1 | 2.67 | 1.01 | 13.14 | 13.49 | -0.36 | 0.36 | -0.01 | 0.01 | |
| gene-LOC109611234 | https://www.ncbi.nlm.nih.gov/gene/?term=LOC109611234 | 3663 | 28 | 76.44 | 1 | 2.73 | 3.57 | 16.2 | 16.56 | -0.36 | 0.36 | 0 | 0 | | |
| gene-LOC105287633 | https://www.ncbi.nlm.nih.gov/gene/?term=LOC105287633 | 1524 | 27 | 177.17 | | 1 | 6.56 | 3.7 | 63.52 | 64 | -0.48 | 0.48 | -0.01 | 0.01 | |
| gene-LOC105286452 | https://www.ncbi.nlm.nih.gov/gene/?term=LOC105286452 | 6888 | 62 | 90.01 | 1 | 1.45 | 1.61 | 32.65 | 33.21 | -0.56 | 0.56 | -0.01 | 0.01 | | |
| gene-LOC105278108 | https://www.ncbi.nlm.nih.gov/gene/?term=LOC105278108 | 4392 | 44 | 100.18 | | 1 | 2.28 | 2.27 | 24.81 | 25.4 | -0.59 | 0.59 | -0.01 | 0.01 | |
| gene-LOC105275187 | https://www.ncbi.nlm.nih.gov/gene/?term=LOC105275187 | 5534 | 66 | 119.26 | | 1 | 1.81 | 1.52 | 11.92 | 12.68 | -0.77 | 0.77 | -0.01 | 0.01 | |
| gene-LOC105285252 | https://www.ncbi.nlm.nih.gov/gene/?term=LOC105285252 | 7427 | 112 | 150.8 | 2 | 2.69 | 1.79 | 14.17 | 15.17 | -1 | 1 | -0.02 | 0.02 | | |
| gene-LOC105281700 | https://www.ncbi.nlm.nih.gov/gene/?term=LOC105281700 | 2208 | 34 | 153.99 | | 1 | 4.53 | 2.94 | 35.91 | 36.92 | -1.02 | 1.02 | -0.02 | 0.02 | |
| gene-LOC105285860 | https://www.ncbi.nlm.nih.gov/gene/?term=LOC105285860 | 3110 | 52 | 167.2 | 1 | 3.22 | 1.92 | 20.92 | 22.04 | -1.12 | 1.12 | -0.02 | 0.02 | | |
| gene-LOC105284507 | https://www.ncbi.nlm.nih.gov/gene/?term=LOC105284507 | 5207 | 77 | 147.88 | | 1 | 1.92 | 1.3 | 22.93 | 24.13 | -1.2 | 1.2 | -0.02 | 0.02 | |
| gene-LOC105274534 | https://www.ncbi.nlm.nih.gov/gene/?term=LOC105274534 | 1329 | 56 | 421.37 | | 1 | 7.52 | 1.79 | 47.04 | 48.26 | -1.22 | 1.22 | -0.05 | 0.05 | |
| gene-LOC109611283 | https://www.ncbi.nlm.nih.gov/gene/?term=LOC109611283 | 1451 | 36 | 248.1 | 1 | 6.89 | 2.78 | 35.62 | 36.97 | -1.35 | 1.35 | -0.03 | 0.03 | | |
| gene-LOC105280819 | https://www.ncbi.nlm.nih.gov/gene/?term=LOC105280819 | 1639 | 37 | 225.75 | | 1 | 6.1 | 2.7 | 36.55 | 38.07 | -1.52 | 1.52 | -0.03 | 0.03 | |
| gene-LOC105282067 | https://www.ncbi.nlm.nih.gov/gene/?term=LOC105282067 | 5941 | 51 | 85.84 | 1 | 1.68 | 1.96 | 26.09 | 27.84 | -1.75 | 1.75 | -0.01 | 0.01 | | |
| gene-LOC105276196 | https://www.ncbi.nlm.nih.gov/gene/?term=LOC105276196 | 2953 | 48 | 162.55 | | 1 | 3.39 | 2.08 | 47.64 | 49.83 | -2.19 | 2.19 | -0.04 | 0.04 | |
| gene-LOC105283989 | https://www.ncbi.nlm.nih.gov/gene/?term=LOC105283989 | 4948 | 26 | 52.55 | 1 | 2.02 | 3.85 | 37.24 | 39.59 | -2.34 | 2.34 | -0.01 | 0.01 | | |
| gene-LOC105284677 | https://www.ncbi.nlm.nih.gov/gene/?term=LOC105284677 | 1905 | 39 | 204.72 | | 3 | 15.75 | 7.69 | 19.98 | 31.6 | -11.62 | | 11.62 | -0.24 | 0.24 |

Directory: OGL File: ogl-Ob_LI_BCphaseB1.txt

| gene_ID | gene_link | gwidth | #Sites | #per10Kb | %perSite | %pNucl |
|---|---|---|---|---|---|---|
| gene-LOC113561532 | https://www.ncbi.nlm.nih.gov/gene/?term=LOC113561532 | 72 | 5 | 694.44 | 94.44 | 6.56 |
| gene-LOC105286531 | https://www.ncbi.nlm.nih.gov/gene/?term=LOC105286531 | 372 | 43 | 1155.91 | 37.65 | 4.35 |
| gene-LOC105285635 | https://www.ncbi.nlm.nih.gov/gene/?term=LOC105285635 | 1175 | 148 | 1259.57 | 31.99 | 4.03 |
| gene-LOC105280739 | https://www.ncbi.nlm.nih.gov/gene/?term=LOC105280739 | 1991 | 143 | 718.23 | 54.78 | 3.93 |
| gene-LOC105286801 | https://www.ncbi.nlm.nih.gov/gene/?term=LOC105286801 | 1102 | 104 | 943.74 | 40.7 | 3.84 |
| gene-LOC113562627 | https://www.ncbi.nlm.nih.gov/gene/?term=LOC113562627 | 397 | 21 | 528.97 | 66.45 | 3.52 |
| gene-LOC105278160 | https://www.ncbi.nlm.nih.gov/gene/?term=LOC105278160 | 1872 | 163 | 870.73 | 39.22 | 3.42 |
| gene-LOC105274534 | https://www.ncbi.nlm.nih.gov/gene/?term=LOC105274534 | 1329 | 108 | 812.64 | 41.62 | 3.38 |
| gene-LOC105287811 | https://www.ncbi.nlm.nih.gov/gene/?term=LOC105287811 | 2155 | 152 | 705.34 | 47.67 | 3.36 |
| gene-LOC105278844 | https://www.ncbi.nlm.nih.gov/gene/?term=LOC105278844 | 1101 | 125 | 1135.33 | 29.16 | 3.31 |
| gene-LOC105278849 | https://www.ncbi.nlm.nih.gov/gene/?term=LOC105278849 | 776 | 101 | 1301.55 | 25.2 | 3.28 |
| gene-LOC105279052 | https://www.ncbi.nlm.nih.gov/gene/?term=LOC105279052 | 1130 | 139 | 1230.09 | 26.62 | 3.27 |
| gene-LOC105286235 | https://www.ncbi.nlm.nih.gov/gene/?term=LOC105286235 | 1510 | 111 | 735.1 | 44.43 | 3.27 |
| gene-LOC105286737 | https://www.ncbi.nlm.nih.gov/gene/?term=LOC105286737 | 2690 | 392 | 1457.25 | 22.38 | 3.26 |
| gene-LOC105287692 | https://www.ncbi.nlm.nih.gov/gene/?term=LOC105287692 | 1505 | 146 | 970.1 | 33.65 | 3.26 |
| gene-LOC105282149 | https://www.ncbi.nlm.nih.gov/gene/?term=LOC105282149 | 1787 | 146 | 817.01 | 39.37 | 3.22 |
| gene-LOC105274711 | https://www.ncbi.nlm.nih.gov/gene/?term=LOC105274711 | 1655 | 110 | 664.65 | 47.74 | 3.17 |
| gene-LOC105286269 | https://www.ncbi.nlm.nih.gov/gene/?term=LOC105286269 | 1401 | 85 | 606.71 | 51.17 | 3.1 |
| gene-LOC105277648 | https://www.ncbi.nlm.nih.gov/gene/?term=LOC105277648 | 722 | 40 | 554.02 | 55.5 | 3.07 |
| gene-LOC105282986 | https://www.ncbi.nlm.nih.gov/gene/?term=LOC105282986 | 1600 | 137 | 856.25 | 35.76 | 3.06 |
| gene-LOC105276360 | https://www.ncbi.nlm.nih.gov/gene/?term=LOC105276360 | 1466 | 162 | 1105.05 | 27 | 2.98 |
| gene-LOC105276371 | https://www.ncbi.nlm.nih.gov/gene/?term=LOC105276371 | 871 | 55 | 631.46 | 47.22 | 2.98 |
| gene-LOC105287677 | https://www.ncbi.nlm.nih.gov/gene/?term=LOC105287677 | 1588 | 118 | 743.07 | 39.84 | 2.96 |
| gene-LOC105274543 | https://www.ncbi.nlm.nih.gov/gene/?term=LOC105274543 | 1572 | 134 | 852.42 | 34.64 | 2.95 |

...
... only first 25 lines shown ...

Directory: OGL File: ogl-Ob_LI_BCphaseB1.vs.BCphaseB2.txt

| gene_ID | gene_link | gwidth | #Sites | #per10Kb | #dmSites | #dmsp10kb | %dmSites | %pSite1 | %pSite2 | DMpSite | ADMpSite | DMpNucl | ADMpNucl |
|---|---|---|---|---|---|---|---|---|---|---|---|---|---|
| gene-LOC113562345 | https://www.ncbi.nlm.nih.gov/gene/?term=LOC113562345 | 2290 | 64 | 279.48 | 15 | 65.5 | 23.44 | 32.83 | 17.93 | 14.9 | 14.9 | 0.42 | 0.42 |
| gene-LOC105275070 | https://www.ncbi.nlm.nih.gov/gene/?term=LOC105275070 | 5667 | 49 | 86.47 | 6 | 10.59 | 12.24 | 10.93 | 1.84 | 9.09 | 9.09 | 0.08 | 0.08 |
| gene-LOC105275708 | https://www.ncbi.nlm.nih.gov/gene/?term=LOC105275708 | 1772 | 15 | 84.65 | 1 | 5.64 | 6.67 | 46.38 | 38.5 | 7.88 | 7.88 | 0.07 | 0.07 |
| gene-LOC105281039 | https://www.ncbi.nlm.nih.gov/gene/?term=LOC105281039 | 1471 | 12 | 81.58 | 1 | 6.8 | 8.33 | 38.6 | 31.46 | 7.13 | 7.13 | 0.06 | 0.06 |
| gene-LOC105279898 | https://www.ncbi.nlm.nih.gov/gene/?term=LOC105279898 | 1343 | 13 | 96.8 | 1 | 7.45 | 7.69 | 64.63 | 59.11 | 5.52 | 5.52 | 0.05 | 0.05 |
| gene-LOC105275518 | https://www.ncbi.nlm.nih.gov/gene/?term=LOC105275518 | 2335 | 23 | 98.5 | 1 | 4.28 | 4.35 | 43.86 | 38.74 | 5.13 | 5.13 | 0.05 | 0.05 |
| gene-LOC105275447 | https://www.ncbi.nlm.nih.gov/gene/?term=LOC105275447 | 1846 | 32 | 173.35 | 1 | 5.42 | 3.12 | 8.01 | 3.17 | 4.84 | 4.84 | 0.08 | 0.08 |
| gene-LOC105274900 | https://www.ncbi.nlm.nih.gov/gene/?term=LOC105274900 | 5561 | 54 | 97.1 | 2 | 3.6 | 3.7 | 35.58 | 31 | 4.57 | 4.57 | 0.04 | 0.04 |
| gene-LOC105283649 | https://www.ncbi.nlm.nih.gov/gene/?term=LOC105283649 | 1918 | 56 | 291.97 | 1 | 5.21 | 1.79 | 39.64 | 36.17 | 3.47 | 3.47 | 0.1 | 0.1 |
| gene-LOC105281515 | https://www.ncbi.nlm.nih.gov/gene/?term=LOC105281515 | 8827 | 57 | 64.57 | 2 | 2.27 | 3.51 | 30.56 | 27.6 | 2.96 | 2.96 | 0.02 | 0.02 |
| gene-LOC105276538 | https://www.ncbi.nlm.nih.gov/gene/?term=LOC105276538 | 1511 | 17 | 112.51 | 1 | 6.62 | 5.88 | 72.11 | 69.24 | 2.87 | 2.87 | 0.03 | 0.03 |
| gene-LOC105285674 | https://www.ncbi.nlm.nih.gov/gene/?term=LOC105285674 | 3035 | 94 | 309.72 | 2 | 6.59 | 2.13 | 27.11 | 24.65 | 2.46 | 2.46 | 0.08 | 0.08 |
| gene-LOC105283989 | https://www.ncbi.nlm.nih.gov/gene/?term=LOC105283989 | 4948 | 31 | 62.65 | 1 | 2.02 | 3.23 | 46.67 | 44.28 | 2.39 | 2.39 | 0.01 | 0.01 |
| gene-LOC105285514 | https://www.ncbi.nlm.nih.gov/gene/?term=LOC105285514 | 1277 | 15 | 117.46 | 1 | 7.83 | 6.67 | 80.63 | 78.25 | 2.38 | 2.38 | 0.03 | 0.03 |
| gene-LOC105281420 | https://www.ncbi.nlm.nih.gov/gene/?term=LOC105281420 | 3882 | 67 | 172.59 | 1 | 2.58 | 1.49 | 15.32 | 12.95 | 2.37 | 2.37 | 0.04 | 0.04 |
| gene-LOC105279900 | https://www.ncbi.nlm.nih.gov/gene/?term=LOC105279900 | 2191 | 56 | 255.59 | 1 | 4.56 | 1.79 | 33.98 | 31.69 | 2.29 | 2.29 | 0.06 | 0.06 |
| gene-LOC105285315 | https://www.ncbi.nlm.nih.gov/gene/?term=LOC105285315 | 5458 | 26 | 47.64 | 1 | 1.83 | 3.85 | 38.49 | 36.2 | 2.29 | 2.29 | 0.01 | 0.01 |
| gene-LOC105276449 | https://www.ncbi.nlm.nih.gov/gene/?term=LOC105276449 | 1231 | 51 | 414.3 | 1 | 8.12 | 1.96 | 27.95 | 25.74 | 2.21 | 2.21 | 0.09 | 0.09 |
| gene-LOC105278877 | https://www.ncbi.nlm.nih.gov/gene/?term=LOC105278877 | 1968 | 82 | 416.67 | 1 | 5.08 | 1.22 | 32.75 | 30.76 | 1.99 | 1.99 | 0.08 | 0.08 |
| gene-LOC105283592 | https://www.ncbi.nlm.nih.gov/gene/?term=LOC105283592 | 3479 | 71 | 204.08 | 1 | 2.87 | 1.41 | 27.65 | 25.73 | 1.93 | 1.93 | 0.04 | 0.04 |
| gene-LOC105278454 | https://www.ncbi.nlm.nih.gov/gene/?term=LOC105278454 | 1972 | 48 | 243.41 | 1 | 5.07 | 2.08 | 42.9 | 41.15 | 1.75 | 1.75 | 0.04 | 0.04 |
| gene-LOC105279664 | https://www.ncbi.nlm.nih.gov/gene/?term=LOC105279664 | 4023 | 65 | 161.57 | 1 | 2.49 | 1.54 | 19.54 | 17.85 | 1.7 | 1.7 | 0.03 | 0.03 |
| gene-LOC105286514 | https://www.ncbi.nlm.nih.gov/gene/?term=LOC105286514 | 2528 | 46 | 181.96 | 1 | 3.96 | 2.17 | 37.97 | 36.35 | 1.61 | 1.61 | 0.03 | 0.03 |
| gene-LOC105283943 | https://www.ncbi.nlm.nih.gov/gene/?term=LOC105283943 | 3559 | 62 | 174.21 | 1 | 2.81 | 1.61 | 15.22 | 13.63 | 1.6 | 1.6 | 0.03 | 0.03 |
| gene-LOC105285583 | https://www.ncbi.nlm.nih.gov/gene/?term=LOC105285583 | 1947 | 45 | 231.12 | 1 | 5.14 | 2.22 | 29.78 | 28.25 | 1.53 | 1.53 | 0.04 | 0.04 |
| gene-LOC105282156 | https://www.ncbi.nlm.nih.gov/gene/?term=LOC105282156 | 2577 | 137 | 531.63 | 1 | 3.88 | 0.73 | 24.98 | 23.48 | 1.5 | 1.5 | 0.08 | 0.08 |
| gene-LOC105276635 | https://www.ncbi.nlm.nih.gov/gene/?term=LOC105276635 | 4962 | 33 | 66.51 | 1 | 2.02 | 3.03 | 19.08 | 17.59 | 1.49 | 1.49 | 0.01 | 0.01 |
| gene-LOC105287648 | https://www.ncbi.nlm.nih.gov/gene/?term=LOC105287648 | 2205 | 30 | 136.05 | 1 | 4.54 | 3.33 | 43.37 | 41.93 | 1.44 | 1.44 | 0.02 | 0.02 |
| gene-LOC105277903 | https://www.ncbi.nlm.nih.gov/gene/?term=LOC105277903 | 1835 | 37 | 201.63 | 1 | 5.45 | 2.7 | 54.03 | 52.65 | 1.39 | 1.39 | 0.03 | 0.03 |
| gene-LOC105287417 | https://www.ncbi.nlm.nih.gov/gene/?term=LOC105287417 | 1025 | 36 | 351.22 | 1 | 9.76 | 2.78 | 35.49 | 34.17 | 1.32 | 1.32 | 0.05 | 0.05 |
| gene-LOC105287854 | https://www.ncbi.nlm.nih.gov/gene/?term=LOC105287854 | 3644 | 67 | 183.86 | 1 | 2.74 | 1.49 | 16.53 | 15.24 | 1.29 | 1.29 | 0.02 | 0.02 |
| gene-LOC105282794 | https://www.ncbi.nlm.nih.gov/gene/?term=LOC105282794 | 4571 | 98 | 214.4 | 1 | 2.19 | 1.02 | 19.72 | 18.47 | 1.25 | 1.25 | 0.03 | 0.03 |
| gene-LOC105282991 | https://www.ncbi.nlm.nih.gov/gene/?term=LOC105282991 | 5617 | 105 | 186.93 | 1 | 1.78 | 0.95 | 19.71 | 18.51 | 1.19 | 1.19 | 0.02 | 0.02 |
| gene-LOC105280956 | https://www.ncbi.nlm.nih.gov/gene/?term=LOC105280956 | 2639 | 75 | 284.2 | 1 | 3.79 | 1.33 | 17.61 | 16.45 | 1.16 | 1.16 | 0.03 | 0.03 |
| gene-LOC105278632 | https://www.ncbi.nlm.nih.gov/gene/?term=LOC105278632 | 3624 | 126 | 347.68 | 1 | 2.76 | 0.79 | 16.08 | 14.93 | 1.15 | 1.15 | 0.04 | 0.04 |
| gene-LOC105288078 | https://www.ncbi.nlm.nih.gov/gene/?term=LOC105288078 | 19126 | 309 | 161.56 | 1 | 0.52 | 0.32 | 17.45 | 16.3 | 1.15 | 1.15 | 0.02 | 0.02 |
| gene-LOC105280786 | https://www.ncbi.nlm.nih.gov/gene/?term=LOC105280786 | 2573 | 119 | 462.5 | 1 | 3.89 | 0.84 | 30.85 | 29.73 | 1.12 | 1.12 | 0.05 | 0.05 |
| gene-LOC105287063 | https://www.ncbi.nlm.nih.gov/gene/?term=LOC105287063 | 1874 | 51 | 272.15 | 1 | 5.34 | 1.96 | 30.2 | 29.07 | 1.12 | 1.12 | 0.03 | 0.03 |
| gene-LOC105280641 | https://www.ncbi.nlm.nih.gov/gene/?term=LOC105280641 | 3682 | 55 | 149.38 | 1 | 2.72 | 1.82 | 19.12 | 18.02 | 1.1 | 1.1 | 0.02 | 0.02 |
| gene-LOC105280002 | https://www.ncbi.nlm.nih.gov/gene/?term=LOC105280002 | 1940 | 28 | 144.33 | 1 | 5.15 | 3.57 | 40.24 | 39.15 | 1.09 | 1.09 | 0.02 | 0.02 |
| gene-LOC105285202 | https://www.ncbi.nlm.nih.gov/gene/?term=LOC105285202 | 1838 | 63 | 342.76 | 1 | 5.44 | 1.59 | 23.99 | 22.95 | 1.05 | 1.05 | 0.04 | 0.04 |
| gene-LOC105284355 | https://www.ncbi.nlm.nih.gov/gene/?term=LOC105284355 | 3904 | 158 | 404.71 | 1 | 2.56 | 0.63 | 20.65 | 19.62 | 1.02 | 1.02 | 0.04 | 0.04 |
| gene-LOC105280348 | https://www.ncbi.nlm.nih.gov/gene/?term=LOC105280348 | 3797 | 76 | 200.16 | 1 | 2.63 | 1.32 | 20.16 | 19.19 | 0.97 | 0.97 | 0.02 | 0.02 |
| gene-LOC105288036 | https://www.ncbi.nlm.nih.gov/gene/?term=LOC105288036 | 5110 | 227 | 444.23 | 1 | 1.96 | 0.44 | 15.78 | 14.87 | 0.91 | 0.91 | 0.04 | 0.04 |
| gene-LOC105275384 | https://www.ncbi.nlm.nih.gov/gene/?term=LOC105275384 | 1987 | 92 | 463.01 | 1 | 5.03 | 1.09 | 32.77 | 31.88 | 0.89 | 0.89 | 0.04 | 0.04 |
| gene-LOC105288042 | https://www.ncbi.nlm.nih.gov/gene/?term=LOC105288042 | 3507 | 99 | 282.29 | 1 | 2.85 | 1.01 | 23.14 | 22.25 | 0.89 | 0.89 | 0.02 | 0.02 |
| gene-LOC105284117 | https://www.ncbi.nlm.nih.gov/gene/?term=LOC105284117 | 4340 | 94 | 216.59 | 1 | 2.3 | 1.06 | 11.08 | 10.26 | 0.83 | 0.83 | 0.02 | 0.02 |
| gene-LOC105275752 | https://www.ncbi.nlm.nih.gov/gene/?term=LOC105275752 | 3462 | 130 | 375.51 | 1 | 2.89 | 0.77 | 31.52 | 30.74 | 0.77 | 0.77 | 0.03 | 0.03 |
| gene-LOC105284116 | https://www.ncbi.nlm.nih.gov/gene/?term=LOC105284116 | 3970 | 107 | 269.52 | 1 | 2.52 | 0.93 | 7.59 | 6.83 | 0.75 | 0.75 | 0.02 | 0.02 |
| gene-LOC105276692 | https://www.ncbi.nlm.nih.gov/gene/?term=LOC105276692 | 1745 | 34 | 194.84 | 1 | 5.73 | 2.94 | 22.19 | 21.46 | 0.73 | 0.73 | 0.01 | 0.01 |
| gene-LOC105282600 | https://www.ncbi.nlm.nih.gov/gene/?term=LOC105282600 | 3715 | 50 | 134.59 | 1 | 2.69 | 2 | 33.68 | 32.96 | 0.73 | 0.73 | 0.01 | 0.01 |
| gene-LOC105285639 | https://www.ncbi.nlm.nih.gov/gene/?term=LOC105285639 | 10325 | 378 | 366.1 | 1 | 0.97 | 0.26 | 11.07 | 10.35 | 0.72 | 0.72 | 0.03 | 0.03 |
| gene-LOC105280405 | https://www.ncbi.nlm.nih.gov/gene/?term=LOC105280405 | 5422 | 161 | 296.94 | 1 | 1.84 | 0.62 | 10.76 | 10.05 | 0.71 | 0.71 | 0.02 | 0.02 |
| gene-LOC105288066 | https://www.ncbi.nlm.nih.gov/gene/?term=LOC105288066 | 1964 | 30 | 152.75 | 1 | 5.09 | 3.33 | 35.84 | 35.13 | 0.71 | 0.71 | 0.01 | 0.01 |
| gene-LOC105278807 | https://www.ncbi.nlm.nih.gov/gene/?term=LOC105278807 | 2166 | 80 | 369.34 | 1 | 4.62 | 1.25 | 18.57 | 17.87 | 0.69 | 0.69 | 0.03 | 0.03 |
| gene-LOC105281266 | https://www.ncbi.nlm.nih.gov/gene/?term=LOC105281266 | 4102 | 79 | 192.59 | 1 | 2.44 | 1.27 | 15.99 | 15.33 | 0.66 | 0.66 | 0.01 | 0.01 |
| gene-LOC105281848 | https://www.ncbi.nlm.nih.gov/gene/?term=LOC105281848 | 2122 | 60 | 282.75 | 1 | 4.71 | 1.67 | 20.32 | 19.71 | 0.62 | 0.62 | 0.02 | 0.02 |
| gene-LOC105278497 | https://www.ncbi.nlm.nih.gov/gene/?term=LOC105278497 | 2072 | 95 | 458.49 | 1 | 4.83 | 1.05 | 19.7 | 19.09 | 0.61 | 0.61 | 0.03 | 0.03 |
| gene-LOC105285055 | https://www.ncbi.nlm.nih.gov/gene/?term=LOC105285055 | 4108 | 253 | 615.87 | 1 | 2.43 | 0.4 | 12.35 | 11.74 | 0.61 | 0.61 | 0.04 | 0.04 |
| gene-LOC105282149 | https://www.ncbi.nlm.nih.gov/gene/?term=LOC105282149 | 1787 | 107 | 598.77 | 1 | 5.6 | 0.93 | 41.49 | 40.93 | 0.57 | 0.57 | 0.03 | 0.03 |

| | | | | | | | | | | | | | |
|---|---|---|---|---|---|---|---|---|---|---|---|---|---|
| gene-LOC105278962 | https://www.ncbi.nlm.nih.gov/gene/?term=LOC105278962 | 4491 | 106 | 236.03 | 1 | 2.23 | 0.94 | 20.64 | 20.08 | 0.56 | 0.56 | 0.01 | 0.01 |
| gene-LOC105288010 | https://www.ncbi.nlm.nih.gov/gene/?term=LOC105288010 | 3189 | 45 | 141.11 | 1 | 3.14 | 2.22 | 33.55 | 32.98 | 0.56 | 0.56 | 0.01 | 0.01 |
| gene-LOC105275187 | https://www.ncbi.nlm.nih.gov/gene/?term=LOC105275187 | 5534 | 74 | 133.72 | 1 | 1.81 | 1.35 | 16.17 | 15.62 | 0.54 | 0.54 | 0.01 | 0.01 |
| gene-LOC105283728 | https://www.ncbi.nlm.nih.gov/gene/?term=LOC105283728 | 3256 | 122 | 374.69 | 2 | 6.14 | 1.64 | 19.35 | 18.81 | 0.54 | 0.54 | 0.02 | 0.02 |
| gene-LOC105279298 | https://www.ncbi.nlm.nih.gov/gene/?term=LOC105279298 | 2773 | 132 | 476.02 | 2 | 7.21 | 1.52 | 23.66 | 23.13 | 0.53 | 0.53 | 0.03 | 0.03 |
| gene-LOC105275363 | https://www.ncbi.nlm.nih.gov/gene/?term=LOC105275363 | 3798 | 168 | 442.34 | 1 | 2.63 | 0.6 | 16.96 | 16.44 | 0.52 | 0.52 | 0.02 | 0.02 |
| gene-LOC105275683 | https://www.ncbi.nlm.nih.gov/gene/?term=LOC105275683 | 4256 | 205 | 481.67 | 1 | 2.35 | 0.49 | 13.27 | 12.75 | 0.52 | 0.52 | 0.02 | 0.02 |
| gene-LOC105278228 | https://www.ncbi.nlm.nih.gov/gene/?term=LOC105278228 | 8610 | 172 | 199.77 | 1 | 1.16 | 0.58 | 23.43 | 22.95 | 0.48 | 0.48 | 0.01 | 0.01 |
| gene-LOC105275170 | https://www.ncbi.nlm.nih.gov/gene/?term=LOC105275170 | 12918 | 298 | 230.69 | 1 | 0.77 | 0.34 | 2.35 | 1.88 | 0.47 | 0.47 | 0.01 | 0.01 |
| gene-LOC105286452 | https://www.ncbi.nlm.nih.gov/gene/?term=LOC105286452 | 6888 | 60 | 87.11 1 | 1.45 | 1.67 | 31.09 | 30.62 | 0.47 | 0.47 | 0 | 0 | |
| gene-LOC105282951 | https://www.ncbi.nlm.nih.gov/gene/?term=LOC105282951 | 3366 | 121 | 359.48 | 1 | 2.97 | 0.83 | 20.13 | 19.67 | 0.46 | 0.46 | 0.02 | 0.02 |
| gene-LOC105274856 | https://www.ncbi.nlm.nih.gov/gene/?term=LOC105274856 | 5216 | 222 | 425.61 | 1 | 1.92 | 0.45 | 17.35 | 16.9 | 0.45 | 0.45 | 0.02 | 0.02 |
| gene-LOC105276059 | https://www.ncbi.nlm.nih.gov/gene/?term=LOC105276059 | 6256 | 138 | 220.59 | 1 | 1.6 | 0.72 | 9.13 | 8.71 | 0.42 | 0.42 | 0.01 | 0.01 |
| gene-LOC105274531 | https://www.ncbi.nlm.nih.gov/gene/?term=LOC105274531 | 7774 | 291 | 374.32 | 1 | 1.29 | 0.34 | 10.83 | 10.42 | 0.41 | 0.41 | 0.02 | 0.02 |
| gene-LOC105282811 | https://www.ncbi.nlm.nih.gov/gene/?term=LOC105282811 | 5765 | 218 | 378.14 | 1 | 1.73 | 0.46 | 8.32 | 7.93 | 0.39 | 0.39 | 0.01 | 0.01 |
| gene-LOC105285395 | https://www.ncbi.nlm.nih.gov/gene/?term=LOC105285395 | 8085 | 208 | 257.27 | 1 | 1.24 | 0.48 | 14.72 | 14.34 | 0.39 | 0.39 | 0.01 | 0.01 |
| gene-LOC105274590 | https://www.ncbi.nlm.nih.gov/gene/?term=LOC105274590 | 4516 | 75 | 166.08 | 2 | 4.43 | 2.67 | 29.15 | 28.79 | 0.35 | 0.35 | 0.01 | 0.01 |
| gene-LOC105275451 | https://www.ncbi.nlm.nih.gov/gene/?term=LOC105275451 | 6035 | 136 | 225.35 | 1 | 1.66 | 0.74 | 21.08 | 20.75 | 0.33 | 0.33 | 0.01 | 0.01 |
| gene-LOC105280408 | https://www.ncbi.nlm.nih.gov/gene/?term=LOC105280408 | 5983 | 56 | 93.6 1 | 1.67 | 1.79 | 20.14 | 19.81 | 0.33 | 0.33 | 0 | 0 | |
| gene-LOC105277761 | https://www.ncbi.nlm.nih.gov/gene/?term=LOC105277761 | 4648 | 148 | 318.42 | 1 | 2.15 | 0.68 | 15.39 | 15.08 | 0.31 | 0.31 | 0.01 | 0.01 |
| gene-LOC105276683 | https://www.ncbi.nlm.nih.gov/gene/?term=LOC105276683 | 17000 | 137 | 80.59 1 | 0.59 | 0.73 | 6.59 | 6.28 | 0.3 | 0.3 | 0 | 0 | |
| gene-LOC105279038 | https://www.ncbi.nlm.nih.gov/gene/?term=LOC105279038 | 9453 | 363 | 384.01 | 1 | 1.06 | 0.28 | 8.55 | 8.25 | 0.3 | 0.3 | 0.01 | 0.01 |
| gene-LOC105285807 | https://www.ncbi.nlm.nih.gov/gene/?term=LOC105285807 | 6063 | 228 | 376.05 | 1 | 1.65 | 0.44 | 8.95 | 8.68 | 0.28 | 0.28 | 0.01 | 0.01 |
| gene-LOC105274452 | https://www.ncbi.nlm.nih.gov/gene/?term=LOC105274452 | 6595 | 260 | 394.24 | 1 | 1.52 | 0.38 | 13.07 | 12.82 | 0.25 | 0.25 | 0.01 | 0.01 |
| gene-LOC105276196 | https://www.ncbi.nlm.nih.gov/gene/?term=LOC105276196 | 2953 | 58 | 196.41 | 1 | 3.39 | 1.72 | 38.52 | 38.28 | 0.24 | 0.24 | 0 | 0 |
| gene-LOC105278666 | https://www.ncbi.nlm.nih.gov/gene/?term=LOC105278666 | 10341 | 477 | 461.27 | 1 | 0.97 | 0.21 | 6.78 | 6.53 | 0.24 | 0.24 | 0.01 | 0.01 |
| gene-LOC105276527 | https://www.ncbi.nlm.nih.gov/gene/?term=LOC105276527 | 7923 | 332 | 419.03 | 1 | 1.26 | 0.3 | 9.67 | 9.44 | 0.23 | 0.23 | 0.01 | 0.01 |
| gene-LOC105279657 | https://www.ncbi.nlm.nih.gov/gene/?term=LOC105279657 | 5878 | 88 | 149.71 | 1 | 1.7 | 1.14 | 19.35 | 19.14 | 0.21 | 0.21 | 0 | 0 |
| gene-LOC113562050 | https://www.ncbi.nlm.nih.gov/gene/?term=LOC113562050 | 17097 | 605 | 353.86 | 1 | 0.58 | 0.17 | 5.18 | 4.96 | 0.21 | 0.21 | 0.01 | 0.01 |
| gene-LOC105275327 | https://www.ncbi.nlm.nih.gov/gene/?term=LOC105275327 | 14245 | 543 | 381.19 | 1 | 0.7 | 0.18 | 6.62 | 6.43 | 0.19 | 0.19 | 0.01 | 0.01 |
| gene-LOC105282491 | https://www.ncbi.nlm.nih.gov/gene/?term=LOC105282491 | 4761 | 177 | 371.77 | 1 | 2.1 | 0.56 | 8.66 | 8.47 | 0.19 | 0.19 | 0.01 | 0.01 |
| gene-LOC105275460 | https://www.ncbi.nlm.nih.gov/gene/?term=LOC105275460 | 2473 | 123 | 497.37 | 1 | 4.04 | 0.81 | 13.15 | 12.97 | 0.18 | 0.18 | 0.01 | 0.01 |
| gene-LOC105282151 | https://www.ncbi.nlm.nih.gov/gene/?term=LOC105282151 | 1601 | 44 | 274.83 | 1 | 6.25 | 2.27 | 35.59 | 35.42 | 0.18 | 0.18 | 0 | 0 |
| gene-LOC105284262 | https://www.ncbi.nlm.nih.gov/gene/?term=LOC105284262 | 6669 | 235 | 352.38 | 1 | 1.5 | 0.43 | 4.64 | 4.46 | 0.18 | 0.18 | 0.01 | 0.01 |
| gene-LOC105287764 | https://www.ncbi.nlm.nih.gov/gene/?term=LOC105287764 | 18359 | 402 | 218.97 | 1 | 0.54 | 0.25 | 3.16 | 2.99 | 0.17 | 0.17 | 0 | 0 |
| gene-LOC105283712 | https://www.ncbi.nlm.nih.gov/gene/?term=LOC105283712 | 1910 | 36 | 188.48 | 1 | 5.24 | 2.78 | 29.2 | 29.04 | 0.16 | 0.16 | 0 | 0 |
| gene-LOC105284043 | https://www.ncbi.nlm.nih.gov/gene/?term=LOC105284043 | 5661 | 311 | 549.37 | 1 | 1.77 | 0.32 | 9.86 | 9.71 | 0.15 | 0.15 | 0.01 | 0.01 |
| gene-LOC105278145 | https://www.ncbi.nlm.nih.gov/gene/?term=LOC105278145 | 4470 | 235 | 525.73 | 1 | 2.24 | 0.43 | 10.67 | 10.54 | 0.14 | 0.14 | 0.01 | 0.01 |
| gene-LOC105284036 | https://www.ncbi.nlm.nih.gov/gene/?term=LOC105284036 | 3254 | 107 | 328.83 | 1 | 3.07 | 0.93 | 20.44 | 20.34 | 0.09 | 0.09 | 0 | 0 |
| gene-LOC105274823 | https://www.ncbi.nlm.nih.gov/gene/?term=LOC105274823 | 4620 | 146 | 316.02 | 1 | 2.16 | 0.68 | 20.43 | 20.37 | 0.06 | 0.06 | 0 | 0 |
| gene-LOC105281700 | https://www.ncbi.nlm.nih.gov/gene/?term=LOC105281700 | 2208 | 37 | 167.57 | 1 | 4.53 | 2.7 | 39.8 | 39.75 | 0.05 | 0.05 | 0 | 0 |
| gene-LOC105286515 | https://www.ncbi.nlm.nih.gov/gene/?term=LOC105286515 | 3751 | 106 | 282.59 | 1 | 2.67 | 0.94 | 10.25 | 10.22 | 0.04 | 0.04 | 0 | 0 |
| gene-LOC105280407 | https://www.ncbi.nlm.nih.gov/gene/?term=LOC105280407 | 10950 | 210 | 191.78 | 1 | 0.91 | 0.48 | 16.39 | 16.4 | -0.01 | 0.01 | 0 | 0 |
| gene-LOC105283123 | https://www.ncbi.nlm.nih.gov/gene/?term=LOC105283123 | 1507 | 55 | 364.96 | 1 | 6.64 | 1.82 | 18.11 | 18.13 | -0.02 | 0.02 | 0 | 0 |
| gene-LOC105274669 | https://www.ncbi.nlm.nih.gov/gene/?term=LOC105274669 | 8555 | 270 | 315.6 1 | 1.17 | 0.37 | 5.54 | 5.56 | -0.03 | 0.03 | 0 | 0 | |
| gene-LOC105278534 | https://www.ncbi.nlm.nih.gov/gene/?term=LOC105278534 | 7364 | 196 | 266.16 | 1 | 1.36 | 0.51 | 17.75 | 17.8 | -0.05 | 0.05 | 0 | 0 |
| gene-LOC105287550 | https://www.ncbi.nlm.nih.gov/gene/?term=LOC105287550 | 15659 | 616 | 393.38 | 1 | 0.64 | 0.16 | 5.82 | 5.86 | -0.05 | 0.05 | 0 | 0 |
| gene-LOC105275374 | https://www.ncbi.nlm.nih.gov/gene/?term=LOC105275374 | 11376 | 539 | 473.8 1 | 0.88 | 0.19 | 9.85 | 10 | -0.15 | 0.15 | -0.01 | 0.01 | |
| gene-LOC105278282 | https://www.ncbi.nlm.nih.gov/gene/?term=LOC105278282 | 1775 | 62 | 349.3 1 | 5.63 | 1.61 | 32.74 | 32.89 | -0.15 | 0.15 | -0.01 | 0.01 | |
| gene-LOC105285496 | https://www.ncbi.nlm.nih.gov/gene/?term=LOC105285496 | 4080 | 110 | 269.61 | 1 | 2.45 | 0.91 | 40.35 | 40.5 | -0.15 | 0.15 | 0 | 0 |
| gene-LOC105285742 | https://www.ncbi.nlm.nih.gov/gene/?term=LOC105285742 | 3446 | 123 | 356.94 | 1 | 2.9 | 0.81 | 19.52 | 19.69 | -0.16 | 0.16 | -0.01 | 0.01 |
| gene-LOC105280723 | https://www.ncbi.nlm.nih.gov/gene/?term=LOC105280723 | 3572 | 181 | 506.72 | 1 | 2.8 | 0.55 | 15.21 | 15.4 | -0.2 | 0.2 | -0.01 | 0.01 |
| gene-LOC105274851 | https://www.ncbi.nlm.nih.gov/gene/?term=LOC105274851 | 6616 | 147 | 222.19 | 1 | 1.51 | 0.68 | 9.12 | 9.33 | -0.21 | 0.21 | 0 | 0 |
| gene-LOC105284184 | https://www.ncbi.nlm.nih.gov/gene/?term=LOC105284184 | 2535 | 99 | 390.53 | 1 | 3.94 | 1.01 | 29.28 | 29.51 | -0.23 | 0.23 | -0.01 | 0.01 |
| gene-LOC105275587 | https://www.ncbi.nlm.nih.gov/gene/?term=LOC105275587 | 2404 | 55 | 228.79 | 1 | 4.16 | 1.82 | 36.62 | 36.89 | -0.27 | 0.27 | -0.01 | 0.01 |
| gene-LOC105286274 | https://www.ncbi.nlm.nih.gov/gene/?term=LOC105286274 | 3817 | 200 | 523.97 | 1 | 2.62 | 0.5 | 16.54 | 16.86 | -0.32 | 0.32 | -0.02 | 0.02 |
| gene-LOC105278691 | https://www.ncbi.nlm.nih.gov/gene/?term=LOC105278691 | 3871 | 168 | 434 1 | 2.58 | 0.6 | 14.64 | 14.98 | -0.34 | 0.34 | -0.01 | 0.01 | |
| gene-LOC105278283 | https://www.ncbi.nlm.nih.gov/gene/?term=LOC105278283 | 2317 | 139 | 599.91 | 1 | 4.32 | 0.72 | 25.04 | 25.46 | -0.41 | 0.41 | -0.02 | 0.02 |
| gene-LOC105285406 | https://www.ncbi.nlm.nih.gov/gene/?term=LOC105285406 | 8015 | 55 | 68.62 1 | 1.25 | 1.82 | 29.11 | 29.59 | -0.47 | 0.47 | 0 | 0 | |
| gene-LOC105276704 | https://www.ncbi.nlm.nih.gov/gene/?term=LOC105276704 | 2063 | 99 | 479.88 | 2 | 9.69 | 2.02 | 13.44 | 13.97 | -0.53 | 0.53 | -0.03 | 0.03 |
| gene-LOC105283967 | https://www.ncbi.nlm.nih.gov/gene/?term=LOC105283967 | 2393 | 53 | 221.48 | 1 | 4.18 | 1.89 | 45.17 | 45.77 | -0.59 | 0.59 | -0.01 | 0.01 |
| gene-LOC105285533 | https://www.ncbi.nlm.nih.gov/gene/?term=LOC105285533 | 1640 | 70 | 426.83 | 1 | 6.1 | 1.43 | 22.75 | 23.49 | -0.75 | 0.75 | -0.03 | 0.03 |
| gene-LOC105274534 | https://www.ncbi.nlm.nih.gov/gene/?term=LOC105274534 | 1329 | 60 | 451.47 | 1 | 7.52 | 1.67 | 40.57 | 41.38 | -0.82 | 0.82 | -0.04 | 0.04 |
| gene-LOC105274449 | https://www.ncbi.nlm.nih.gov/gene/?term=LOC105274449 | 1549 | 53 | 342.16 | 1 | 6.46 | 1.89 | 50.86 | 51.77 | -0.91 | 0.91 | -0.03 | 0.03 |

| | | | | | | | | | | | | | |
|---|---|---|---|---|---|---|---|---|---|---|---|---|---|
| gene-LOC105276429 | https://www.ncbi.nlm.nih.gov/gene/?term=LOC105276429 | 2901 | 108 | 372.29 | 1 | 3.45 | 0.93 | 24.11 | 25.23 | -1.12 | 1.12 | -0.04 | 0.04 |
| gene-LOC105281312 | https://www.ncbi.nlm.nih.gov/gene/?term=LOC105281312 | 2551 | 65 | 254.8 | 1 | 3.92 | 1.54 | 31.99 | 33.22 | -1.23 | 1.23 | -0.03 | 0.03 |
| gene-LOC105281173 | https://www.ncbi.nlm.nih.gov/gene/?term=LOC105281173 | 1595 | 38 | 238.24 | 1 | 6.27 | 2.63 | 59.28 | 60.67 | -1.38 | 1.38 | -0.03 | 0.03 |
| gene-LOC105274706 | https://www.ncbi.nlm.nih.gov/gene/?term=LOC105274706 | 1446 | 50 | 345.78 | 1 | 6.92 | 2 | 40.41 | 41.89 | -1.49 | 1.49 | -0.05 | 0.05 |
| gene-LOC105285409 | https://www.ncbi.nlm.nih.gov/gene/?term=LOC105285409 | 5099 | 58 | 113.75 | 1 | 1.96 | 1.72 | 1.58 | 3.25 | -1.67 | 1.67 | -0.02 | 0.02 |
| gene-LOC105287633 | https://www.ncbi.nlm.nih.gov/gene/?term=LOC105287633 | 1524 | 31 | 203.41 | 1 | 6.56 | 3.23 | 53.47 | 55.78 | -2.31 | 2.31 | -0.05 | 0.05 |
| gene-LOC105282914 | https://www.ncbi.nlm.nih.gov/gene/?term=LOC105282914 | 5623 | 41 | 72.91 | 1 | 1.78 | 2.44 | 49.33 | 52.21 | -2.88 | 2.88 | -0.02 | 0.02 |
| gene-LOC105282885 | https://www.ncbi.nlm.nih.gov/gene/?term=LOC105282885 | 1870 | 47 | 251.34 | 1 | 5.35 | 2.13 | 35.98 | 39.14 | -3.16 | 3.16 | -0.08 | 0.08 |
| gene-LOC105279500 | https://www.ncbi.nlm.nih.gov/gene/?term=LOC105279500 | 3587 | 33 | 92 | 1 | 2.79 | 3.03 | 43.74 | 47.01 | -3.27 | 3.27 | -0.03 | 0.03 |
| gene-LOC105284677 | https://www.ncbi.nlm.nih.gov/gene/?term=LOC105284677 | 1905 | 42 | 220.47 | 1 | 5.25 | 2.38 | 26.24 | 37.79 | -11.55 | 11.55 | -0.25 | 0.25 |

Directory: OGL File: ogl-Ob_LI_BCphaseB1.vs.RphaseA1.txt

| gene_ID | gene_link | gwidth | #Sites | #per10Kb | #dmSites | #dmsp10kb | %dmSites | %pSite1 | %pSite2 | DMpSite | ADMpSite | DMpNucl | ADMpNucl |
|---|---|---|---|---|---|---|---|---|---|---|---|---|---|
| gene-LOC105282854 | https://www.ncbi.nlm.nih.gov/gene/?term=LOC105282854 | 1825 | 6 | 32.88 | 1 | 5.48 | 16.67 | 52.41 | 41.89 | 10.52 | 10.52 | 0.03 | 0.03 |
| gene-LOC105286453 | https://www.ncbi.nlm.nih.gov/gene/?term=LOC105286453 | 1160 | 9 | 77.59 | 2 | 17.24 | 22.22 | 36.42 | 26.38 | 10.04 | 10.04 | 0.08 | 0.08 |
| gene-LOC105275378 | https://www.ncbi.nlm.nih.gov/gene/?term=LOC105275378 | 1189 | 21 | 176.62 | 3 | 25.23 | 14.29 | 38.9 | 30.15 | 8.75 | 8.75 | 0.15 | 0.15 |
| gene-LOC105286572 | https://www.ncbi.nlm.nih.gov/gene/?term=LOC105286572 | 2321 | 7 | 30.16 | 2 | 8.62 | 28.57 | 19.41 | 11.7 | 7.7 | 7.7 | 0.02 | 0.02 |
| gene-LOC105280699 | https://www.ncbi.nlm.nih.gov/gene/?term=LOC105280699 | 1563 | 5 | 31.99 | 1 | 6.4 | 20 | 31.61 | 25.04 | 6.57 | 6.57 | 0.02 | 0.02 |
| gene-LOC105275669 | https://www.ncbi.nlm.nih.gov/gene/?term=LOC105275669 | 1567 | 8 | 51.05 | 1 | 6.38 | 12.5 | 54.73 | 48.32 | 6.41 | 6.41 | 0.03 | 0.03 |
| gene-LOC105283227 | https://www.ncbi.nlm.nih.gov/gene/?term=LOC105283227 | 2373 | 12 | 50.57 | 2 | 8.43 | 16.67 | 43.46 | 37.45 | 6.01 | 6.01 | 0.03 | 0.03 |
| gene-LOC113563652 | https://www.ncbi.nlm.nih.gov/gene/?term=LOC113563652 | 1476 | 13 | 88.08 | 1 | 6.78 | 7.69 | 51.72 | 45.77 | 5.95 | 5.95 | 0.05 | 0.05 |
| gene-LOC113562135 | https://www.ncbi.nlm.nih.gov/gene/?term=LOC113562135 | 1413 | 9 | 63.69 | 1 | 7.08 | 11.11 | 40.16 | 34.54 | 5.62 | 5.62 | 0.04 | 0.04 |
| gene-LOC105284817 | https://www.ncbi.nlm.nih.gov/gene/?term=LOC105284817 | 2500 | 13 | 52 | 1 | 4 | 7.69 | 28.13 | 22.7 | 5.43 | 5.43 | 0.03 | 0.03 |
| gene-LOC105277491 | https://www.ncbi.nlm.nih.gov/gene/?term=LOC105277491 | 1647 | 11 | 66.79 | 1 | 6.07 | 9.09 | 19.05 | 13.86 | 5.19 | 5.19 | 0.03 | 0.03 |
| gene-LOC105280472 | https://www.ncbi.nlm.nih.gov/gene/?term=LOC105280472 | 1675 | 12 | 71.64 | 1 | 5.97 | 8.33 | 42.17 | 37.11 | 5.06 | 5.06 | 0.04 | 0.04 |
| gene-LOC105283785 | https://www.ncbi.nlm.nih.gov/gene/?term=LOC105283785 | 3447 | 33 | 95.74 | 1 | 2.9 | 3.03 | 19.56 | 14.53 | 5.03 | 5.03 | 0.05 | 0.05 |
| gene-LOC105286815 | https://www.ncbi.nlm.nih.gov/gene/?term=LOC105286815 | 1024 | 11 | 107.42 | 1 | 9.77 | 9.09 | 63.05 | 58.15 | 4.91 | 4.91 | 0.05 | 0.05 |
| gene-LOC105287814 | https://www.ncbi.nlm.nih.gov/gene/?term=LOC105287814 | 892 | 8 | 89.69 | 1 | 11.21 | 12.5 | 35.83 | 30.99 | 4.84 | 4.84 | 0.04 | 0.04 |
| gene-LOC105287978 | https://www.ncbi.nlm.nih.gov/gene/?term=LOC105287978 | 3017 | 7 | 23.2 | 1 | 3.31 | 14.29 | 17.4 | 12.85 | 4.56 | 4.56 | 0.01 | 0.01 |
| gene-LOC105276691 | https://www.ncbi.nlm.nih.gov/gene/?term=LOC105276691 | 2543 | 41 | 161.23 | 3 | 11.8 | 7.32 | 41.71 | 37.17 | 4.54 | 4.54 | 0.07 | 0.07 |
| gene-LOC105275518 | https://www.ncbi.nlm.nih.gov/gene/?term=LOC105275518 | 2335 | 26 | 111.35 | 2 | 8.57 | 7.69 | 47.97 | 43.47 | 4.5 | 4.5 | 0.05 | 0.05 |
| gene-LOC105282733 | https://www.ncbi.nlm.nih.gov/gene/?term=LOC105282733 | 3835 | 23 | 59.97 | 2 | 5.22 | 8.7 | 33.46 | 29 | 4.46 | 4.46 | 0.03 | 0.03 |
| gene-LOC105282407 | https://www.ncbi.nlm.nih.gov/gene/?term=LOC105282407 | 1210 | 12 | 99.17 | 1 | 8.26 | 8.33 | 50.3 | 45.91 | 4.39 | 4.39 | 0.04 | 0.04 |
| gene-LOC105276707 | https://www.ncbi.nlm.nih.gov/gene/?term=LOC105276707 | 1990 | 16 | 80.4 | 2 | 10.05 | 12.5 | 37.03 | 32.8 | 4.23 | 4.23 | 0.03 | 0.03 |
| gene-LOC105278080 | https://www.ncbi.nlm.nih.gov/gene/?term=LOC105278080 | 3258 | 21 | 64.46 | 1 | 3.07 | 4.76 | 41.67 | 37.47 | 4.21 | 4.21 | 0.03 | 0.03 |
| gene-LOC105285392 | https://www.ncbi.nlm.nih.gov/gene/?term=LOC105285392 | 1601 | 91 | 568.39 | 6 | 37.48 | 6.59 | 35.68 | 31.54 | 4.15 | 4.15 | 0.24 | 0.24 |
| gene-LOC105280372 | https://www.ncbi.nlm.nih.gov/gene/?term=LOC105280372 | 2159 | 19 | 88 | 1 | 4.63 | 5.26 | 55.74 | 51.64 | 4.1 | 4.1 | 0.04 | 0.04 |

...

... only first 25 lines shown ...

Directory: OGL File: ogl-Ob_LI_BCphaseB1.vs.RphaseA2.txt

| gene_ID | gene_link | gwidth | #Sites | #per10Kb | #dmSites | #dmsp10kb | %dmSites | %pSite1 | %pSite2 | DMpSite | ADMpSite | DMpNucl | ADMpNucl |
|---|---|---|---|---|---|---|---|---|---|---|---|---|---|
| gene-LOC105286572 | https://www.ncbi.nlm.nih.gov/gene/?term=LOC105286572 | 2321 | 8 | 34.47 | 1 | 4.31 | 12.5 | 20.96 | 13.96 | 7.01 | 7.01 | 0.02 | 0.02 |
| gene-LOC105283986 | https://www.ncbi.nlm.nih.gov/gene/?term=LOC105283986 | 1782 | 7 | 39.28 | 1 | 5.61 | 14.29 | 29.98 | 22.98 | 7 | 7 | 0.03 | 0.03 |
| gene-LOC113563238 | https://www.ncbi.nlm.nih.gov/gene/?term=LOC113563238 | 2680 | 4 | 14.93 | 1 | 3.73 | 25 | 22.36 | 15.49 | 6.86 | 6.86 | 0.01 | 0.01 |
| gene-LOC105280472 | https://www.ncbi.nlm.nih.gov/gene/?term=LOC105280472 | 1675 | 14 | 83.58 | 2 | 11.94 | 14.29 | 38.72 | 31.98 | 6.74 | 6.74 | 0.06 | 0.06 |
| gene-LOC105278607 | https://www.ncbi.nlm.nih.gov/gene/?term=LOC105278607 | 4407 | 29 | 65.8 | 2 | 4.54 | 6.9 | 11.68 | 5.33 | 6.35 | 6.35 | 0.04 | 0.04 |
| gene-LOC105283760 | https://www.ncbi.nlm.nih.gov/gene/?term=LOC105283760 | 1134 | 5 | 44.09 | 2 | 17.64 | 40 | 48.5 | 42.19 | 6.31 | 6.31 | 0.03 | 0.03 |
| gene-LOC105277405 | https://www.ncbi.nlm.nih.gov/gene/?term=LOC105277405 | 1255 | 4 | 31.87 | 1 | 7.97 | 25 | 25.11 | 19.1 | 6.02 | 6.02 | 0.02 | 0.02 |
| gene-LOC105287978 | https://www.ncbi.nlm.nih.gov/gene/?term=LOC105287978 | 3017 | 7 | 23.2 | 2 | 6.63 | 28.57 | 17.4 | 11.62 | 5.78 | 5.78 | 0.01 | 0.01 |
| gene-LOC105286453 | https://www.ncbi.nlm.nih.gov/gene/?term=LOC105286453 | 1160 | 14 | 120.69 | 2 | 17.24 | 14.29 | 32.33 | 26.68 | 5.65 | 5.65 | 0.07 | 0.07 |
| gene-LOC105286753 | https://www.ncbi.nlm.nih.gov/gene/?term=LOC105286753 | 3684 | 12 | 32.57 | 1 | 2.71 | 8.33 | 17.35 | 11.83 | 5.52 | 5.52 | 0.02 | 0.02 |
| gene-LOC105288020 | https://www.ncbi.nlm.nih.gov/gene/?term=LOC105288020 | 1568 | 11 | 70.15 | 1 | 6.38 | 9.09 | 35.44 | 30.49 | 4.95 | 4.95 | 0.03 | 0.03 |
| gene-LOC105274640 | https://www.ncbi.nlm.nih.gov/gene/?term=LOC105274640 | 2723 | 11 | 40.4 | 1 | 3.67 | 9.09 | 60.49 | 56.34 | 4.15 | 4.15 | 0.02 | 0.02 |
| gene-LOC105280969 | https://www.ncbi.nlm.nih.gov/gene/?term=LOC105280969 | 1005 | 14 | 139.3 | 1 | 9.95 | 7.14 | 39.18 | 35.04 | 4.14 | 4.14 | 0.06 | 0.06 |
| gene-LOC105285707 | https://www.ncbi.nlm.nih.gov/gene/?term=LOC105285707 | 2096 | 32 | 152.67 | 2 | 9.54 | 6.25 | 58.26 | 54.2 | 4.05 | 4.05 | 0.06 | 0.06 |
| gene-LOC105275378 | https://www.ncbi.nlm.nih.gov/gene/?term=LOC105275378 | 1189 | 26 | 218.67 | 2 | 16.82 | 7.69 | 46.76 | 43.04 | 3.73 | 3.73 | 0.08 | 0.08 |
| gene-LOC105275355 | https://www.ncbi.nlm.nih.gov/gene/?term=LOC105275355 | 3197 | 169 | 528.62 | 10 | 31.28 | 5.92 | 27.68 | 24.07 | 3.61 | 3.61 | 0.19 | 0.19 |
| gene-LOC105281081 | https://www.ncbi.nlm.nih.gov/gene/?term=LOC105281081 | 2358 | 15 | 63.61 | 1 | 4.24 | 6.67 | 19.8 | 16.63 | 3.16 | 3.16 | 0.02 | 0.02 |
| gene-LOC105280764 | https://www.ncbi.nlm.nih.gov/gene/?term=LOC105280764 | 1800 | 16 | 88.89 | 1 | 5.56 | 6.25 | 19.21 | 16.08 | 3.13 | 3.13 | 0.03 | 0.03 |
| gene-LOC105280001 | https://www.ncbi.nlm.nih.gov/gene/?term=LOC105280001 | 2566 | 23 | 89.63 | 1 | 3.9 | 4.35 | 24.54 | 21.43 | 3.11 | 3.11 | 0.03 | 0.03 |
| gene-LOC105284604 | https://www.ncbi.nlm.nih.gov/gene/?term=LOC105284604 | 1998 | 17 | 85.09 | 2 | 10.01 | 11.76 | 3.09 | 0 | 3.09 | 3.09 | 0.03 | 0.03 |
| gene-LOC105280448 | https://www.ncbi.nlm.nih.gov/gene/?term=LOC105280448 | 4000 | 29 | 72.5 | 2 | 5 | 6.9 | 18.2 | 15.16 | 3.04 | 3.04 | 0.02 | 0.02 |
| gene-LOC105277369 | https://www.ncbi.nlm.nih.gov/gene/?term=LOC105277369 | 2012 | 9 | 44.73 | 1 | 4.97 | 11.11 | 61.75 | 58.79 | 2.96 | 2.96 | 0.01 | 0.01 |
| gene-LOC105276691 | https://www.ncbi.nlm.nih.gov/gene/?term=LOC105276691 | 2543 | 44 | 173.02 | 3 | 11.8 | 6.82 | 39.83 | 36.94 | 2.89 | 2.89 | 0.05 | 0.05 |
| gene-LOC105280548 | https://www.ncbi.nlm.nih.gov/gene/?term=LOC105280548 | 1547 | 29 | 187.46 | 1 | 6.46 | 3.45 | 51.13 | 48.43 | 2.7 | 2.7 | 0.05 | 0.05 |

...
... only first 25 lines shown ...

Directory: OGL File: ogl-Ob_LI_BCphaseB1.vs.RphaseB1.txt

| gene_ID | gene_link | gwidth | #Sites | #per10Kb | #dmSites | #dmsp10kb | %dmSites | %pSite1 | %pSite2 | DMpSite | ADMpSite | DMpNucl | ADMpNucl |
|---|---|---|---|---|---|---|---|---|---|---|---|---|---|
| gene-LOC113562345 | https://www.ncbi.nlm.nih.gov/gene/?term=LOC113562345 | 2290 | 77 | 336.24 | 12 | 52.4 | 15.58 | 26.12 | 14.52 | 11.59 | 11.59 | 0.39 | 0.39 |
| gene-LOC105275070 | https://www.ncbi.nlm.nih.gov/gene/?term=LOC105275070 | 5667 | 61 | 107.64 | 2 | 3.53 | 3.28 | 14.18 | 5.29 | 8.9 | 8.9 | 0.1 | 0.1 |
| gene-LOC105275708 | https://www.ncbi.nlm.nih.gov/gene/?term=LOC105275708 | 1772 | 15 | 84.65 | 1 | 5.64 | 6.67 | 46.38 | 38.49 | 7.9 | 7.9 | 0.07 | 0.07 |
| gene-LOC105285514 | https://www.ncbi.nlm.nih.gov/gene/?term=LOC105285514 | 1277 | 11 | 86.14 | 1 | 7.83 | 9.09 | 92.13 | 85.12 | 7.01 | 7.01 | 0.06 | 0.06 |
| gene-LOC105275355 | https://www.ncbi.nlm.nih.gov/gene/?term=LOC105275355 | 3197 | 174 | 544.26 | 1 | 3.13 | 0.57 | 26.94 | 21.77 | 5.17 | 5.17 | 0.28 | 0.28 |
| gene-LOC105281039 | https://www.ncbi.nlm.nih.gov/gene/?term=LOC105281039 | 1471 | 14 | 95.17 | 1 | 6.8 | 7.14 | 44.63 | 40.12 | 4.51 | 4.51 | 0.04 | 0.04 |
| gene-LOC105275518 | https://www.ncbi.nlm.nih.gov/gene/?term=LOC105275518 | 2335 | 25 | 107.07 | 1 | 4.28 | 4 | 44.99 | 41.35 | 3.64 | 3.64 | 0.04 | 0.04 |
| gene-LOC105280419 | https://www.ncbi.nlm.nih.gov/gene/?term=LOC105280419 | 1601 | 11 | 68.71 | 1 | 6.25 | 9.09 | 46.06 | 42.48 | 3.57 | 3.57 | 0.02 | 0.02 |
| gene-LOC105283989 | https://www.ncbi.nlm.nih.gov/gene/?term=LOC105283989 | 4948 | 45 | 90.95 | 2 | 4.04 | 4.44 | 38.35 | 35.25 | 3.1 | 3.1 | 0.03 | 0.03 |
| gene-LOC105275384 | https://www.ncbi.nlm.nih.gov/gene/?term=LOC105275384 | 1987 | 92 | 463.01 | 1 | 5.03 | 1.09 | 34.18 | 31.19 | 2.98 | 2.98 | 0.14 | 0.14 |
| gene-LOC105276272 | https://www.ncbi.nlm.nih.gov/gene/?term=LOC105276272 | 1965 | 40 | 203.56 | 1 | 5.09 | 2.5 | 30.29 | 27.31 | 2.98 | 2.98 | 0.06 | 0.06 |
| gene-LOC105279900 | https://www.ncbi.nlm.nih.gov/gene/?term=LOC105279900 | 2191 | 56 | 255.59 | 1 | 4.56 | 1.79 | 31.89 | 28.91 | 2.98 | 2.98 | 0.08 | 0.08 |
| gene-LOC105285315 | https://www.ncbi.nlm.nih.gov/gene/?term=LOC105285315 | 5458 | 32 | 58.63 | 1 | 1.83 | 3.12 | 31.91 | 29.18 | 2.72 | 2.72 | 0.02 | 0.02 |
| gene-LOC105276611 | https://www.ncbi.nlm.nih.gov/gene/?term=LOC105276611 | 1731 | 36 | 207.97 | 1 | 5.78 | 2.78 | 30.25 | 27.6 | 2.65 | 2.65 | 0.06 | 0.06 |
| gene-LOC105283712 | https://www.ncbi.nlm.nih.gov/gene/?term=LOC105283712 | 1910 | 35 | 183.25 | 1 | 5.24 | 2.86 | 31.43 | 28.9 | 2.53 | 2.53 | 0.05 | 0.05 |
| gene-LOC105287063 | https://www.ncbi.nlm.nih.gov/gene/?term=LOC105287063 | 1874 | 50 | 266.81 | 1 | 5.34 | 2 | 31.85 | 29.62 | 2.23 | 2.23 | 0.06 | 0.06 |
| gene-LOC105274900 | https://www.ncbi.nlm.nih.gov/gene/?term=LOC105274900 | 5561 | 63 | 113.29 | 2 | 3.6 | 3.17 | 32.28 | 30.06 | 2.22 | 2.22 | 0.03 | 0.03 |
| gene-LOC105283552 | https://www.ncbi.nlm.nih.gov/gene/?term=LOC105283552 | 1562 | 41 | 262.48 | 1 | 6.4 | 2.44 | 53.31 | 51.1 | 2.21 | 2.21 | 0.06 | 0.06 |
| gene-LOC105279898 | https://www.ncbi.nlm.nih.gov/gene/?term=LOC105279898 | 1343 | 16 | 119.14 | 1 | 7.45 | 6.25 | 58.69 | 56.55 | 2.14 | 2.14 | 0.03 | 0.03 |
| gene-LOC105281515 | https://www.ncbi.nlm.nih.gov/gene/?term=LOC105281515 | 8827 | 55 | 62.31 | 2 | 2.27 | 3.64 | 34.93 | 32.95 | 1.98 | 1.98 | 0.01 | 0.01 |
| gene-LOC113561805 | https://www.ncbi.nlm.nih.gov/gene/?term=LOC113561805 | 3337 | 76 | 227.75 | 1 | 3 | 1.32 | 32.51 | 30.53 | 1.98 | 1.98 | 0.05 | 0.05 |
| gene-LOC105280704 | https://www.ncbi.nlm.nih.gov/gene/?term=LOC105280704 | 4266 | 147 | 344.59 | 1 | 2.34 | 0.68 | 18.45 | 16.63 | 1.81 | 1.81 | 0.06 | 0.06 |
| gene-LOC105280348 | https://www.ncbi.nlm.nih.gov/gene/?term=LOC105280348 | 3797 | 79 | 208.06 | 1 | 2.63 | 1.27 | 18.26 | 16.64 | 1.61 | 1.61 | 0.03 | 0.03 |
| gene-LOC105274449 | https://www.ncbi.nlm.nih.gov/gene/?term=LOC105274449 | 1549 | 53 | 342.16 | 1 | 6.46 | 1.89 | 50.86 | 49.29 | 1.57 | 1.57 | 0.05 | 0.05 |
| gene-LOC105287417 | https://www.ncbi.nlm.nih.gov/gene/?term=LOC105287417 | 1025 | 40 | 390.24 | 1 | 9.76 | 2.5 | 32.13 | 30.76 | 1.38 | 1.38 | 0.05 | 0.05 |
| gene-LOC105276692 | https://www.ncbi.nlm.nih.gov/gene/?term=LOC105276692 | 1745 | 35 | 200.57 | 1 | 5.73 | 2.86 | 24.46 | 23.1 | 1.36 | 1.36 | 0.03 | 0.03 |
| gene-LOC105276538 | https://www.ncbi.nlm.nih.gov/gene/?term=LOC105276538 | 1511 | 20 | 132.36 | 1 | 6.62 | 5 | 72.51 | 71.18 | 1.32 | 1.32 | 0.02 | 0.02 |
| gene-LOC105287095 | https://www.ncbi.nlm.nih.gov/gene/?term=LOC105287095 | 1435 | 70 | 487.8 | 1 | 6.97 | 1.43 | 25.15 | 23.89 | 1.26 | 1.26 | 0.06 | 0.06 |
| gene-LOC105288036 | https://www.ncbi.nlm.nih.gov/gene/?term=LOC105288036 | 5110 | 232 | 454.01 | 1 | 1.96 | 0.43 | 15.89 | 14.67 | 1.22 | 1.22 | 0.06 | 0.06 |
| gene-LOC105285060 | https://www.ncbi.nlm.nih.gov/gene/?term=LOC105285060 | 1285 | 16 | 124.51 | 1 | 7.78 | 6.25 | 42.41 | 41.27 | 1.13 | 1.13 | 0.01 | 0.01 |
| gene-LOC105278384 | https://www.ncbi.nlm.nih.gov/gene/?term=LOC105278384 | 1390 | 33 | 237.41 | 1 | 7.19 | 3.03 | 46.84 | 45.77 | 1.07 | 1.07 | 0.03 | 0.03 |
| gene-LOC105288079 | https://www.ncbi.nlm.nih.gov/gene/?term=LOC105288079 | 3473 | 110 | 316.73 | 1 | 2.88 | 0.91 | 18.5 | 17.43 | 1.07 | 1.07 | 0.03 | 0.03 |
| gene-LOC105280972 | https://www.ncbi.nlm.nih.gov/gene/?term=LOC105280972 | 1532 | 35 | 228.46 | 1 | 6.53 | 2.86 | 17.63 | 16.59 | 1.04 | 1.04 | 0.02 | 0.02 |
| gene-LOC105282794 | https://www.ncbi.nlm.nih.gov/gene/?term=LOC105282794 | 4571 | 110 | 240.65 | 1 | 2.19 | 0.91 | 18.14 | 17.16 | 0.98 | 0.98 | 0.02 | 0.02 |
| gene-LOC105281266 | https://www.ncbi.nlm.nih.gov/gene/?term=LOC105281266 | 4102 | 88 | 214.53 | 1 | 2.44 | 1.14 | 15.7 | 14.74 | 0.96 | 0.96 | 0.02 | 0.02 |
| gene-LOC105282156 | https://www.ncbi.nlm.nih.gov/gene/?term=LOC105282156 | 2577 | 132 | 512.22 | 1 | 3.88 | 0.76 | 26.63 | 25.67 | 0.96 | 0.96 | 0.05 | 0.05 |
| gene-LOC105285179 | https://www.ncbi.nlm.nih.gov/gene/?term=LOC105285179 | 4191 | 194 | 462.9 | 1 | 2.39 | 0.52 | 17.62 | 16.69 | 0.93 | 0.93 | 0.04 | 0.04 |
| gene-LOC105278632 | https://www.ncbi.nlm.nih.gov/gene/?term=LOC105278632 | 3624 | 139 | 383.55 | 1 | 2.76 | 0.72 | 14.57 | 13.67 | 0.9 | 0.9 | 0.03 | 0.03 |
| gene-LOC105283123 | https://www.ncbi.nlm.nih.gov/gene/?term=LOC105283123 | 1507 | 55 | 364.96 | 1 | 6.64 | 1.82 | 19.83 | 18.96 | 0.86 | 0.86 | 0.03 | 0.03 |
| gene-LOC105277215 | https://www.ncbi.nlm.nih.gov/gene/?term=LOC105277215 | 2619 | 29 | 110.73 | 1 | 3.82 | 3.45 | 25.46 | 24.64 | 0.82 | 0.82 | 0.01 | 0.01 |
| gene-LOC105281848 | https://www.ncbi.nlm.nih.gov/gene/?term=LOC105281848 | 2122 | 58 | 273.33 | 1 | 4.71 | 1.72 | 22.33 | 21.51 | 0.82 | 0.82 | 0.02 | 0.02 |
| gene-LOC105285055 | https://www.ncbi.nlm.nih.gov/gene/?term=LOC105285055 | 4108 | 248 | 603.7 | 1 | 2.43 | 0.4 | 12.58 | 11.79 | 0.79 | 0.79 | 0.05 | 0.05 |
| gene-LOC105285695 | https://www.ncbi.nlm.nih.gov/gene/?term=LOC105285695 | 1371 | 75 | 547.05 | 1 | 7.29 | 1.33 | 27.5 | 26.71 | 0.79 | 0.79 | 0.04 | 0.04 |
| gene-LOC105285674 | https://www.ncbi.nlm.nih.gov/gene/?term=LOC105285674 | 3035 | 109 | 359.14 | 1 | 3.29 | 0.92 | 24.18 | 23.43 | 0.75 | 0.75 | 0.03 | 0.03 |
| gene-LOC105275460 | https://www.ncbi.nlm.nih.gov/gene/?term=LOC105275460 | 2473 | 127 | 513.55 | 1 | 4.04 | 0.79 | 13.03 | 12.3 | 0.73 | 0.73 | 0.04 | 0.04 |
| gene-LOC105285639 | https://www.ncbi.nlm.nih.gov/gene/?term=LOC105285639 | 10325 | 372 | 360.29 | 2 | 1.94 | 0.54 | 11.5 | 10.78 | 0.72 | 0.72 | 0.03 | 0.03 |
| gene-LOC105283728 | https://www.ncbi.nlm.nih.gov/gene/?term=LOC105283728 | 3256 | 146 | 448.4 | 1 | 3.07 | 0.68 | 16.37 | 15.66 | 0.71 | 0.71 | 0.03 | 0.03 |
| gene-LOC105277435 | https://www.ncbi.nlm.nih.gov/gene/?term=LOC105277435 | 3643 | 179 | 491.35 | 1 | 2.74 | 0.56 | 10.99 | 10.3 | 0.7 | 0.7 | 0.03 | 0.03 |
| gene-LOC105285671 | https://www.ncbi.nlm.nih.gov/gene/?term=LOC105285671 | 2263 | 103 | 455.15 | 1 | 4.42 | 0.97 | 30.33 | 29.64 | 0.69 | 0.69 | 0.03 | 0.03 |
| gene-LOC105276449 | https://www.ncbi.nlm.nih.gov/gene/?term=LOC105276449 | 1231 | 55 | 446.79 | 1 | 8.12 | 1.82 | 22.22 | 21.54 | 0.67 | 0.67 | 0.03 | 0.03 |
| gene-LOC105284355 | https://www.ncbi.nlm.nih.gov/gene/?term=LOC105284355 | 3904 | 154 | 394.47 | 1 | 2.56 | 0.65 | 22.13 | 21.47 | 0.66 | 0.66 | 0.03 | 0.03 |
| gene-LOC105282480 | https://www.ncbi.nlm.nih.gov/gene/?term=LOC105282480 | 4177 | 166 | 397.41 | 1 | 2.39 | 0.6 | 12.65 | 12.01 | 0.65 | 0.65 | 0.03 | 0.03 |
| gene-LOC105276704 | https://www.ncbi.nlm.nih.gov/gene/?term=LOC105276704 | 2063 | 87 | 421.72 | 2 | 9.69 | 2.3 | 14.01 | 13.37 | 0.64 | 0.64 | 0.03 | 0.03 |
| gene-LOC105283943 | https://www.ncbi.nlm.nih.gov/gene/?term=LOC105283943 | 3559 | 69 | 193.87 | 1 | 2.81 | 1.45 | 20.61 | 20.02 | 0.59 | 0.59 | 0.01 | 0.01 |
| gene-LOC105276410 | https://www.ncbi.nlm.nih.gov/gene/?term=LOC105276410 | 11085 | 341 | 307.62 | 1 | 0.9 | 0.29 | 9.89 | 9.31 | 0.58 | 0.58 | 0.02 | 0.02 |
| gene-LOC105279664 | https://www.ncbi.nlm.nih.gov/gene/?term=LOC105279664 | 4023 | 67 | 166.54 | 1 | 2.49 | 1.49 | 18.98 | 18.43 | 0.55 | 0.55 | 0.01 | 0.01 |
| gene-LOC105274823 | https://www.ncbi.nlm.nih.gov/gene/?term=LOC105274823 | 4620 | 172 | 372.29 | 1 | 2.16 | 0.58 | 17.73 | 17.2 | 0.53 | 0.53 | 0.02 | 0.02 |
| gene-LOC105278497 | https://www.ncbi.nlm.nih.gov/gene/?term=LOC105278497 | 2072 | 92 | 444.02 | 1 | 4.83 | 1.09 | 20.29 | 19.78 | 0.51 | 0.51 | 0.02 | 0.02 |
| gene-LOC105280408 | https://www.ncbi.nlm.nih.gov/gene/?term=LOC105280408 | 5983 | 63 | 105.3 | 1 | 1.67 | 1.59 | 16.29 | 15.78 | 0.51 | 0.51 | 0.01 | 0.01 |
| gene-LOC105278534 | https://www.ncbi.nlm.nih.gov/gene/?term=LOC105278534 | 7364 | 204 | 277.02 | 1 | 1.36 | 0.49 | 17.28 | 16.79 | 0.5 | 0.5 | 0.01 | 0.01 |

| | | | | | | | | | | | | | |
|---|---|---|---|---|---|---|---|---|---|---|---|---|---|
| gene-LOC105279298 | https://www.ncbi.nlm.nih.gov/gene/?term=LOC105279298 | 2773 | 130 | 468.81 | 1 | 3.61 | 0.77 | 23.65 | 23.17 | 0.49 | 0.49 | 0.02 | 0.02 |
| gene-LOC105283649 | https://www.ncbi.nlm.nih.gov/gene/?term=LOC105283649 | 1918 | 57 | 297.18 | 1 | 5.21 | 1.75 | 40.29 | 39.8 | 0.48 | 0.48 | 0.01 | 0.01 |
| gene-LOC105286997 | https://www.ncbi.nlm.nih.gov/gene/?term=LOC105286997 | 2850 | 150 | 526.32 | 1 | 3.51 | 0.67 | 11.33 | 10.85 | 0.48 | 0.48 | 0.03 | 0.03 |
| gene-LOC105288078 | https://www.ncbi.nlm.nih.gov/gene/?term=LOC105288078 | 19126 | 350 | 183 | 1 | 0.52 | 0.29 | 15.9 | 15.43 | 0.47 | 0.47 | 0.01 | 0.01 |
| gene-LOC105277631 | https://www.ncbi.nlm.nih.gov/gene/?term=LOC105277631 | 4387 | 313 | 713.47 | 1 | 2.28 | 0.32 | 10.74 | 10.28 | 0.46 | 0.46 | 0.03 | 0.03 |
| gene-LOC105280641 | https://www.ncbi.nlm.nih.gov/gene/?term=LOC105280641 | 3682 | 84 | 228.14 | 1 | 2.72 | 1.19 | 13.99 | 13.53 | 0.46 | 0.46 | 0.01 | 0.01 |
| gene-LOC105275187 | https://www.ncbi.nlm.nih.gov/gene/?term=LOC105275187 | 5534 | 78 | 140.95 | 1 | 1.81 | 1.28 | 16.44 | 16.02 | 0.42 | 0.42 | 0.01 | 0.01 |
| gene-LOC105278807 | https://www.ncbi.nlm.nih.gov/gene/?term=LOC105278807 | 2166 | 75 | 346.26 | 1 | 4.62 | 1.33 | 20.48 | 20.06 | 0.41 | 0.41 | 0.01 | 0.01 |
| gene-LOC105286099 | https://www.ncbi.nlm.nih.gov/gene/?term=LOC105286099 | 5919 | 77 | 130.09 | 1 | 1.69 | 1.3 | 12.19 | 11.79 | 0.4 | 0.4 | 0.01 | 0.01 |
| gene-LOC105282491 | https://www.ncbi.nlm.nih.gov/gene/?term=LOC105282491 | 4761 | 193 | 405.38 | 1 | 2.1 | 0.52 | 8.12 | 7.73 | 0.38 | 0.38 | 0.02 | 0.02 |
| gene-LOC105276527 | https://www.ncbi.nlm.nih.gov/gene/?term=LOC105276527 | 7923 | 370 | 466.99 | 1 | 1.26 | 0.27 | 8.27 | 7.9 | 0.37 | 0.37 | 0.02 | 0.02 |
| gene-LOC105286514 | https://www.ncbi.nlm.nih.gov/gene/?term=LOC105286514 | 2528 | 51 | 201.74 | 1 | 3.96 | 1.96 | 35.82 | 35.46 | 0.36 | 0.36 | 0.01 | 0.01 |
| gene-LOC105275170 | https://www.ncbi.nlm.nih.gov/gene/?term=LOC105275170 | 12918 | 359 | 277.91 | 1 | 0.77 | 0.28 | 2.02 | 1.81 | 0.21 | 0.21 | 0.01 | 0.01 |
| gene-LOC105280407 | https://www.ncbi.nlm.nih.gov/gene/?term=LOC105280407 | 10950 | 227 | 207.31 | 1 | 0.91 | 0.44 | 15.19 | 14.98 | 0.21 | 0.21 | 0 | 0 |
| gene-LOC105279500 | https://www.ncbi.nlm.nih.gov/gene/?term=LOC105279500 | 3587 | 35 | 97.57 | 1 | 2.79 | 2.86 | 48.82 | 48.61 | 0.2 | 0.2 | 0 | 0 |
| gene-LOC105284262 | https://www.ncbi.nlm.nih.gov/gene/?term=LOC105284262 | 6669 | 241 | 361.37 | 1 | 1.5 | 0.41 | 5.35 | 5.16 | 0.19 | 0.19 | 0.01 | 0.01 |
| gene-LOC105288066 | https://www.ncbi.nlm.nih.gov/gene/?term=LOC105288066 | 1964 | 31 | 157.84 | 1 | 5.09 | 3.23 | 36.75 | 36.57 | 0.19 | 0.19 | 0 | 0 |
| gene-LOC105285742 | https://www.ncbi.nlm.nih.gov/gene/?term=LOC105285742 | 3446 | 118 | 342.43 | 1 | 2.9 | 0.85 | 17.32 | 17.14 | 0.18 | 0.18 | 0.01 | 0.01 |
| gene-LOC105285981 | https://www.ncbi.nlm.nih.gov/gene/?term=LOC105285981 | 7666 | 219 | 285.68 | 1 | 1.3 | 0.46 | 18.34 | 18.17 | 0.17 | 0.17 | 0 | 0 |
| gene-LOC105274531 | https://www.ncbi.nlm.nih.gov/gene/?term=LOC105274531 | 7774 | 295 | 379.47 | 1 | 1.29 | 0.34 | 10.65 | 10.48 | 0.16 | 0.16 | 0.01 | 0.01 |
| gene-LOC105278228 | https://www.ncbi.nlm.nih.gov/gene/?term=LOC105278228 | 8610 | 199 | 231.13 | 1 | 1.16 | 0.5 | 21.48 | 21.33 | 0.15 | 0.15 | 0 | 0 |
| gene-LOC105275451 | https://www.ncbi.nlm.nih.gov/gene/?term=LOC105275451 | 6035 | 142 | 235.29 | 1 | 1.66 | 0.7 | 20 | 19.9 | 0.1 | 0.1 | 0 | 0 |
| gene-LOC105283838 | https://www.ncbi.nlm.nih.gov/gene/?term=LOC105283838 | 1748 | 30 | 171.62 | 1 | 5.72 | 3.33 | 35.36 | 35.26 | 0.1 | 0.1 | 0 | 0 |
| gene-LOC105277216 | https://www.ncbi.nlm.nih.gov/gene/?term=LOC105277216 | 7627 | 117 | 153.4 | 1 | 1.31 | 0.85 | 6.74 | 6.65 | 0.09 | 0.09 | 0 | 0 |
| gene-LOC105278145 | https://www.ncbi.nlm.nih.gov/gene/?term=LOC105278145 | 4470 | 261 | 583.89 | 1 | 2.24 | 0.38 | 10.03 | 9.95 | 0.08 | 0.08 | 0 | 0 |
| gene-LOC105280002 | https://www.ncbi.nlm.nih.gov/gene/?term=LOC105280002 | 1940 | 32 | 164.95 | 1 | 5.15 | 3.12 | 34.33 | 34.27 | 0.06 | 0.06 | 0 | 0 |
| gene-LOC105284043 | https://www.ncbi.nlm.nih.gov/gene/?term=LOC105284043 | 5661 | 295 | 521.11 | 1 | 1.77 | 0.34 | 10.33 | 10.27 | 0.06 | 0.06 | 0 | 0 |
| gene-LOC105284184 | https://www.ncbi.nlm.nih.gov/gene/?term=LOC105284184 | 2535 | 102 | 402.37 | 1 | 3.94 | 0.98 | 30 | 29.94 | 0.06 | 0.06 | 0 | 0 |
| gene-LOC105282811 | https://www.ncbi.nlm.nih.gov/gene/?term=LOC105282811 | 5765 | 235 | 407.63 | 1 | 1.73 | 0.43 | 8.46 | 8.42 | 0.04 | 0.04 | 0 | 0 |
| gene-LOC105278283 | https://www.ncbi.nlm.nih.gov/gene/?term=LOC105278283 | 2317 | 138 | 595.6 | 1 | 4.32 | 0.72 | 25.04 | 25.03 | 0.02 | 0.02 | 0 | 0 |
| gene-LOC105278666 | https://www.ncbi.nlm.nih.gov/gene/?term=LOC105278666 | 10341 | 484 | 468.04 | 1 | 0.97 | 0.21 | 6.48 | 6.46 | 0.02 | 0.02 | 0 | 0 |
| gene-LOC105274706 | https://www.ncbi.nlm.nih.gov/gene/?term=LOC105274706 | 1446 | 46 | 318.12 | 1 | 6.92 | 2.17 | 42.4 | 42.39 | 0 | 0 | 0 | 0 |
| gene-LOC105276059 | https://www.ncbi.nlm.nih.gov/gene/?term=LOC105276059 | 6256 | 146 | 233.38 | 1 | 1.6 | 0.68 | 11.41 | 11.42 | 0 | 0 | 0 | 0 |
| gene-LOC105278877 | https://www.ncbi.nlm.nih.gov/gene/?term=LOC105278877 | 1968 | 88 | 447.15 | 1 | 5.08 | 1.14 | 30.96 | 31.01 | -0.05 | 0.05 | 0 | 0 |
| gene-LOC105287065 | https://www.ncbi.nlm.nih.gov/gene/?term=LOC105287065 | 4859 | 249 | 512.45 | 1 | 2.06 | 0.4 | 12.25 | 12.3 | -0.05 | 0.05 | 0 | 0 |
| gene-LOC105284774 | https://www.ncbi.nlm.nih.gov/gene/?term=LOC105284774 | 12089 | 100 | 82.72 | 1 | 0.83 | 1 | 19.53 | 19.59 | -0.06 | 0.06 | 0 | 0 |
| gene-LOC105275327 | https://www.ncbi.nlm.nih.gov/gene/?term=LOC105275327 | 14245 | 552 | 387.5 | 1 | 0.7 | 0.18 | 6.33 | 6.41 | -0.08 | 0.08 | 0 | 0 |
| gene-LOC105282600 | https://www.ncbi.nlm.nih.gov/gene/?term=LOC105282600 | 3715 | 60 | 161.51 | 1 | 2.69 | 1.67 | 28.86 | 28.93 | -0.08 | 0.08 | 0 | 0 |
| gene-LOC105286339 | https://www.ncbi.nlm.nih.gov/gene/?term=LOC105286339 | 5230 | 184 | 351.82 | 1 | 1.91 | 0.54 | 16.14 | 16.23 | -0.09 | 0.09 | 0 | 0 |
| gene-LOC105288010 | https://www.ncbi.nlm.nih.gov/gene/?term=LOC105288010 | 3189 | 52 | 163.06 | 1 | 3.14 | 1.92 | 33.66 | 33.76 | -0.1 | 0.1 | 0 | 0 |
| gene-LOC105285807 | https://www.ncbi.nlm.nih.gov/gene/?term=LOC105285807 | 6063 | 221 | 364.51 | 1 | 1.65 | 0.45 | 9.8 | 9.91 | -0.11 | 0.11 | 0 | 0 |
| gene-LOC105279038 | https://www.ncbi.nlm.nih.gov/gene/?term=LOC105279038 | 9453 | 346 | 366.02 | 1 | 1.06 | 0.29 | 8.85 | 9.03 | -0.18 | 0.18 | -0.01 | 0.01 |
| gene-LOC105288042 | https://www.ncbi.nlm.nih.gov/gene/?term=LOC105288042 | 3507 | 98 | 279.44 | 1 | 2.85 | 1.02 | 22.85 | 23.09 | -0.24 | 0.24 | -0.01 | 0.01 |
| gene-LOC105280956 | https://www.ncbi.nlm.nih.gov/gene/?term=LOC105280956 | 2639 | 81 | 306.93 | 1 | 3.79 | 1.23 | 18.56 | 18.86 | -0.29 | 0.29 | -0.01 | 0.01 |
| gene-LOC105281173 | https://www.ncbi.nlm.nih.gov/gene/?term=LOC105281173 | 1595 | 42 | 263.32 | 1 | 6.27 | 2.38 | 62.5 | 62.82 | -0.31 | 0.31 | -0.01 | 0.01 |
| gene-LOC105285004 | https://www.ncbi.nlm.nih.gov/gene/?term=LOC105285004 | 1334 | 35 | 262.37 | 1 | 7.5 | 2.86 | 19.44 | 19.75 | -0.31 | 0.31 | -0.01 | 0.01 |
| gene-LOC105278962 | https://www.ncbi.nlm.nih.gov/gene/?term=LOC105278962 | 4491 | 129 | 287.24 | 1 | 2.23 | 0.78 | 17.72 | 18.05 | -0.34 | 0.34 | -0.01 | 0.01 |
| gene-LOC105282110 | https://www.ncbi.nlm.nih.gov/gene/?term=LOC105282110 | 7748 | 148 | 191.02 | 1 | 1.29 | 0.68 | 11.12 | 11.53 | -0.42 | 0.42 | -0.01 | 0.01 |
| gene-LOC105276429 | https://www.ncbi.nlm.nih.gov/gene/?term=LOC105276429 | 2901 | 132 | 455.02 | 1 | 3.45 | 0.76 | 22.03 | 22.48 | -0.45 | 0.45 | -0.02 | 0.02 |
| gene-LOC105282825 | https://www.ncbi.nlm.nih.gov/gene/?term=LOC105282825 | 3919 | 98 | 250.06 | 1 | 2.55 | 1.02 | 13.43 | 13.91 | -0.48 | 0.48 | -0.01 | 0.01 |
| gene-LOC105274590 | https://www.ncbi.nlm.nih.gov/gene/?term=LOC105274590 | 4516 | 84 | 186.01 | 2 | 4.43 | 2.38 | 24.43 | 25 | -0.57 | 0.57 | -0.01 | 0.01 |
| gene-LOC105282151 | https://www.ncbi.nlm.nih.gov/gene/?term=LOC105282151 | 1601 | 43 | 268.58 | 1 | 6.25 | 2.33 | 38.6 | 39.18 | -0.58 | 0.58 | -0.02 | 0.02 |
| gene-LOC105285406 | https://www.ncbi.nlm.nih.gov/gene/?term=LOC105285406 | 8015 | 57 | 71.12 | 1 | 1.25 | 1.75 | 28.09 | 28.72 | -0.63 | 0.63 | 0 | 0 |
| gene-LOC105284036 | https://www.ncbi.nlm.nih.gov/gene/?term=LOC105284036 | 3254 | 103 | 316.53 | 1 | 3.07 | 0.97 | 23.72 | 24.37 | -0.65 | 0.65 | -0.02 | 0.02 |
| gene-LOC105286265 | https://www.ncbi.nlm.nih.gov/gene/?term=LOC105286265 | 2507 | 67 | 267.25 | 1 | 3.99 | 1.49 | 29.04 | 29.68 | -0.65 | 0.65 | -0.02 | 0.02 |
| gene-LOC105274856 | https://www.ncbi.nlm.nih.gov/gene/?term=LOC105274856 | 5216 | 222 | 425.61 | 1 | 1.92 | 0.45 | 16.84 | 17.52 | -0.68 | 0.68 | -0.03 | 0.03 |
| gene-LOC105276745 | https://www.ncbi.nlm.nih.gov/gene/?term=LOC105276745 | 3953 | 76 | 192.26 | 1 | 2.53 | 1.32 | 18.95 | 19.72 | -0.76 | 0.76 | -0.01 | 0.01 |
| gene-LOC105286515 | https://www.ncbi.nlm.nih.gov/gene/?term=LOC105286515 | 3751 | 111 | 295.92 | 1 | 2.67 | 0.9 | 10.2 | 10.99 | -0.8 | 0.8 | -0.02 | 0.02 |
| gene-LOC105279778 | https://www.ncbi.nlm.nih.gov/gene/?term=LOC105279778 | 4347 | 163 | 374.97 | 1 | 2.3 | 0.61 | 8.62 | 9.47 | -0.85 | 0.85 | -0.03 | 0.03 |
| gene-LOC105277903 | https://www.ncbi.nlm.nih.gov/gene/?term=LOC105277903 | 1835 | 38 | 207.08 | 1 | 5.45 | 2.63 | 52.47 | 53.46 | -0.99 | 0.99 | -0.02 | 0.02 |
| gene-LOC105283132 | https://www.ncbi.nlm.nih.gov/gene/?term=LOC105283132 | 1819 | 56 | 307.86 | 1 | 5.5 | 1.79 | 39.1 | 40.18 | -1.09 | 1.09 | -0.03 | 0.03 |
| gene-LOC105286770 | https://www.ncbi.nlm.nih.gov/gene/?term=LOC105286770 | 3055 | 49 | 160.39 | 1 | 3.27 | 2.04 | 11.46 | 12.59 | -1.13 | 1.13 | -0.02 | 0.02 |
| gene-LOC105276196 | https://www.ncbi.nlm.nih.gov/gene/?term=LOC105276196 | 2953 | 60 | 203.18 | 1 | 3.39 | 1.67 | 37.66 | 38.86 | -1.2 | 1.2 | -0.02 | 0.02 |
| gene-LOC105282914 | https://www.ncbi.nlm.nih.gov/gene/?term=LOC105282914 | 5623 | 43 | 76.47 | 1 | 1.78 | 2.33 | 48 | 49.79 | -1.79 | 1.79 | -0.01 | 0.01 |

| | | | | | | | | | | | | | |
|---|---|---|---|---|---|---|---|---|---|---|---|---|---|
| gene-LOC105278282 | https://www.ncbi.nlm.nih.gov/gene/?term=LOC105278282 | 1775 | 65 | 366.2 | 1 | 5.63 | 1.54 | 30.53 | 32.37 | -1.84 | 1.84 | -0.07 | 0.07 |
| gene-LOC105274534 | https://www.ncbi.nlm.nih.gov/gene/?term=LOC105274534 | 1329 | 59 | 443.94 | 1 | 7.52 | 1.69 | 41.22 | 43.19 | -1.98 | 1.98 | -0.09 | 0.09 |
| gene-LOC105282885 | https://www.ncbi.nlm.nih.gov/gene/?term=LOC105282885 | 1870 | 41 | 219.25 | 1 | 5.35 | 2.44 | 39.07 | 41.13 | -2.06 | 2.06 | -0.05 | 0.05 |
| gene-LOC105287633 | https://www.ncbi.nlm.nih.gov/gene/?term=LOC105287633 | 1524 | 25 | 164.04 | 1 | 6.56 | 4 | 61.97 | 64.15 | -2.18 | 2.18 | -0.04 | 0.04 |
| gene-LOC105279528 | https://www.ncbi.nlm.nih.gov/gene/?term=LOC105279528 | 5802 | 41 | 70.67 | 1 | 1.72 | 2.44 | 17.37 | 19.59 | -2.22 | 2.22 | -0.02 | 0.02 |
| gene-LOC105278454 | https://www.ncbi.nlm.nih.gov/gene/?term=LOC105278454 | 1972 | 41 | 207.91 | 1 | 5.07 | 2.44 | 41.77 | 44.02 | -2.26 | 2.26 | -0.05 | 0.05 |
| gene-LOC105280447 | https://www.ncbi.nlm.nih.gov/gene/?term=LOC105280447 | 2215 | 19 | 85.78 | 1 | 4.51 | 5.26 | 33.1 | 35.5 | -2.4 | 2.4 | -0.02 | 0.02 |
| gene-LOC105285409 | https://www.ncbi.nlm.nih.gov/gene/?term=LOC105285409 | 5099 | 66 | 129.44 | 1 | 1.96 | 1.52 | 2.12 | 4.53 | -2.41 | 2.41 | -0.03 | 0.03 |
| gene-LOC105284677 | https://www.ncbi.nlm.nih.gov/gene/?term=LOC105284677 | 1905 | 42 | 220.47 | 1 | 5.25 | 2.38 | 23.34 | 35.09 | -11.75 | 11.75 | -0.26 | 0.26 |

Directory: OGL File: ogl-Ob_LI_BCphaseB1.vs.RphaseB2.txt

| gene_ID | gene_link | gwidth | #Sites | #per10Kb | #dmSites | #dmsp10kb | %dmSites | %pSite1 | %pSite2 | DMpSite | ADMpSite | DMpNucl | ADMpNucl |
|---|---|---|---|---|---|---|---|---|---|---|---|---|---|
| gene-LOC113562345 | https://www.ncbi.nlm.nih.gov/gene/?term=LOC113562345 | 2290 | 66 | 288.21 | 12 | 52.4 | 18.18 | 30.39 | 17.58 | 12.82 | 12.82 | 0.37 | 0.37 |
| gene-LOC105275070 | https://www.ncbi.nlm.nih.gov/gene/?term=LOC105275070 | 5667 | 41 | 72.35 | 6 | 10.59 | 14.63 | 12.96 | 1.06 | 11.9 | 11.9 | 0.09 | 0.09 |
| gene-LOC105279898 | https://www.ncbi.nlm.nih.gov/gene/?term=LOC105279898 | 1343 | 10 | 74.46 | 1 | 7.45 | 10 | 83.43 | 71.53 | 11.89 | 11.89 | 0.09 | 0.09 |
| gene-LOC105275708 | https://www.ncbi.nlm.nih.gov/gene/?term=LOC105275708 | 1772 | 14 | 79.01 | 1 | 5.64 | 7.14 | 49.7 | 43.39 | 6.31 | 6.31 | 0.05 | 0.05 |
| gene-LOC105275518 | https://www.ncbi.nlm.nih.gov/gene/?term=LOC105275518 | 2335 | 20 | 85.65 | 1 | 4.28 | 5 | 58.2 | 52.79 | 5.42 | 5.42 | 0.05 | 0.05 |
| gene-LOC105283552 | https://www.ncbi.nlm.nih.gov/gene/?term=LOC105283552 | 1562 | 41 | 262.48 | 1 | 6.4 | 2.44 | 53.31 | 48.83 | 4.48 | 4.48 | 0.12 | 0.12 |
| gene-LOC105285315 | https://www.ncbi.nlm.nih.gov/gene/?term=LOC105285315 | 5458 | 19 | 34.81 | 1 | 1.83 | 5.26 | 47.14 | 43.22 | 3.93 | 3.93 | 0.01 | 0.01 |
| gene-LOC105287648 | https://www.ncbi.nlm.nih.gov/gene/?term=LOC105287648 | 2205 | 25 | 113.38 | 1 | 4.54 | 4 | 44.43 | 41.17 | 3.26 | 3.26 | 0.04 | 0.04 |
| gene-LOC105286683 | https://www.ncbi.nlm.nih.gov/gene/?term=LOC105286683 | 1665 | 55 | 330.33 | 1 | 6.01 | 1.82 | 34.95 | 31.78 | 3.17 | 3.17 | 0.1 | 0.1 |
| gene-LOC105274900 | https://www.ncbi.nlm.nih.gov/gene/?term=LOC105274900 | 5561 | 55 | 98.9 | 2 | 3.6 | 3.64 | 33.24 | 30.24 | 3 | 3 | 0.03 | 0.03 |
| gene-LOC105281515 | https://www.ncbi.nlm.nih.gov/gene/?term=LOC105281515 | 8827 | 40 | 45.32 | 2 | 2.27 | 5 | 31.53 | 28.62 | 2.91 | 2.91 | 0.01 | 0.01 |
| gene-LOC105285514 | https://www.ncbi.nlm.nih.gov/gene/?term=LOC105285514 | 1277 | 17 | 133.12 | 1 | 7.83 | 5.88 | 73.52 | 70.72 | 2.8 | 2.8 | 0.04 | 0.04 |
| gene-LOC105279664 | https://www.ncbi.nlm.nih.gov/gene/?term=LOC105279664 | 4023 | 54 | 134.23 | 1 | 2.49 | 1.85 | 21.88 | 19.13 | 2.74 | 2.74 | 0.04 | 0.04 |
| gene-LOC105282794 | https://www.ncbi.nlm.nih.gov/gene/?term=LOC105282794 | 4571 | 91 | 199.08 | 1 | 2.19 | 1.1 | 19.77 | 17.09 | 2.68 | 2.68 | 0.05 | 0.05 |
| gene-LOC105280104 | https://www.ncbi.nlm.nih.gov/gene/?term=LOC105280104 | 3960 | 135 | 340.91 | 1 | 2.53 | 0.74 | 18.4 | 16.2 | 2.2 | 2.2 | 0.07 | 0.07 |
| gene-LOC105278108 | https://www.ncbi.nlm.nih.gov/gene/?term=LOC105278108 | 4392 | 46 | 104.74 | 1 | 2.28 | 2.17 | 27.18 | 25.03 | 2.14 | 2.14 | 0.02 | 0.02 |
| gene-LOC105280641 | https://www.ncbi.nlm.nih.gov/gene/?term=LOC105280641 | 3682 | 50 | 135.8 | 1 | 2.72 | 2 | 17.97 | 16.08 | 1.89 | 1.89 | 0.03 | 0.03 |
| gene-LOC105285671 | https://www.ncbi.nlm.nih.gov/gene/?term=LOC105285671 | 2263 | 105 | 463.99 | 1 | 4.42 | 0.95 | 28.66 | 26.78 | 1.88 | 1.88 | 0.09 | 0.09 |
| gene-LOC105279356 | https://www.ncbi.nlm.nih.gov/gene/?term=LOC105279356 | 1314 | 43 | 327.25 | 1 | 7.61 | 2.33 | 32.17 | 30.3 | 1.87 | 1.87 | 0.06 | 0.06 |
| gene-LOC105287417 | https://www.ncbi.nlm.nih.gov/gene/?term=LOC105287417 | 1025 | 23 | 224.39 | 1 | 9.76 | 4.35 | 51.1 | 49.27 | 1.83 | 1.83 | 0.04 | 0.04 |
| gene-LOC105281419 | https://www.ncbi.nlm.nih.gov/gene/?term=LOC105281419 | 2366 | 80 | 338.12 | 1 | 4.23 | 1.25 | 22.59 | 20.79 | 1.81 | 1.81 | 0.06 | 0.06 |
| gene-LOC105285674 | https://www.ncbi.nlm.nih.gov/gene/?term=LOC105285674 | 3035 | 98 | 322.9 | 2 | 6.59 | 2.04 | 25.36 | 23.72 | 1.64 | 1.64 | 0.05 | 0.05 |
| gene-LOC105283728 | https://www.ncbi.nlm.nih.gov/gene/?term=LOC105283728 | 3256 | 115 | 353.19 | 1 | 3.07 | 0.87 | 19.83 | 18.35 | 1.48 | 1.48 | 0.05 | 0.05 |
| gene-LOC105288036 | https://www.ncbi.nlm.nih.gov/gene/?term=LOC105288036 | 5110 | 200 | 391.39 | 1 | 1.96 | 0.5 | 17.99 | 16.61 | 1.37 | 1.37 | 0.05 | 0.05 |
| gene-LOC105288042 | https://www.ncbi.nlm.nih.gov/gene/?term=LOC105288042 | 3507 | 89 | 253.78 | 1 | 2.85 | 1.12 | 22.75 | 21.38 | 1.37 | 1.37 | 0.03 | 0.03 |
| gene-LOC105280786 | https://www.ncbi.nlm.nih.gov/gene/?term=LOC105280786 | 2573 | 123 | 478.04 | 1 | 3.89 | 0.81 | 29.6 | 28.26 | 1.34 | 1.34 | 0.06 | 0.06 |
| gene-LOC113561621 | https://www.ncbi.nlm.nih.gov/gene/?term=LOC113561621 | 2249 | 83 | 369.05 | 1 | 4.45 | 1.2 | 21.13 | 19.82 | 1.31 | 1.31 | 0.05 | 0.05 |
| gene-LOC105285179 | https://www.ncbi.nlm.nih.gov/gene/?term=LOC105285179 | 4191 | 175 | 417.56 | 1 | 2.39 | 0.57 | 18.73 | 17.48 | 1.25 | 1.25 | 0.05 | 0.05 |
| gene-LOC113561805 | https://www.ncbi.nlm.nih.gov/gene/?term=LOC113561805 | 3337 | 65 | 194.79 | 1 | 3 | 1.54 | 35.44 | 34.26 | 1.18 | 1.18 | 0.02 | 0.02 |
| gene-LOC105276901 | https://www.ncbi.nlm.nih.gov/gene/?term=LOC105276901 | 3254 | 82 | 252 | 1 | 3.07 | 1.22 | 28 | 26.83 | 1.17 | 1.17 | 0.03 | 0.03 |
| gene-LOC105275363 | https://www.ncbi.nlm.nih.gov/gene/?term=LOC105275363 | 3798 | 160 | 421.27 | 1 | 2.63 | 0.62 | 17.46 | 16.32 | 1.14 | 1.14 | 0.05 | 0.05 |
| gene-LOC105282156 | https://www.ncbi.nlm.nih.gov/gene/?term=LOC105282156 | 2577 | 127 | 492.82 | 1 | 3.88 | 0.79 | 27.5 | 26.37 | 1.13 | 1.13 | 0.06 | 0.06 |
| gene-LOC105280408 | https://www.ncbi.nlm.nih.gov/gene/?term=LOC105280408 | 5983 | 56 | 93.6 | 1 | 1.67 | 1.79 | 17.1 | 16.06 | 1.03 | 1.03 | 0.01 | 0.01 |
| gene-LOC105284355 | https://www.ncbi.nlm.nih.gov/gene/?term=LOC105284355 | 3904 | 151 | 386.78 | 1 | 2.56 | 0.66 | 21.57 | 20.54 | 1.02 | 1.02 | 0.04 | 0.04 |
| gene-LOC105286099 | https://www.ncbi.nlm.nih.gov/gene/?term=LOC105286099 | 5919 | 72 | 121.64 | 1 | 1.69 | 1.39 | 10.67 | 9.65 | 1.01 | 1.01 | 0.01 | 0.01 |
| gene-LOC105274449 | https://www.ncbi.nlm.nih.gov/gene/?term=LOC105274449 | 1549 | 48 | 309.88 | 1 | 6.46 | 2.08 | 51.97 | 50.96 | 1 | 1 | 0.03 | 0.03 |
| gene-LOC105281848 | https://www.ncbi.nlm.nih.gov/gene/?term=LOC105281848 | 2122 | 47 | 221.49 | 1 | 4.71 | 2.13 | 27.57 | 26.57 | 1 | 1 | 0.02 | 0.02 |
| gene-LOC105285639 | https://www.ncbi.nlm.nih.gov/gene/?term=LOC105285639 | 10325 | 332 | 321.55 | 2 | 1.94 | 0.6 | 12.25 | 11.3 | 0.95 | 0.95 | 0.03 | 0.03 |
| gene-LOC105278877 | https://www.ncbi.nlm.nih.gov/gene/?term=LOC105278877 | 1968 | 69 | 350.61 | 1 | 5.08 | 1.45 | 36.22 | 35.3 | 0.91 | 0.91 | 0.03 | 0.03 |
| gene-LOC105276704 | https://www.ncbi.nlm.nih.gov/gene/?term=LOC105276704 | 2063 | 87 | 421.72 | 1 | 4.85 | 1.15 | 14.98 | 14.08 | 0.9 | 0.9 | 0.04 | 0.04 |
| gene-LOC105280447 | https://www.ncbi.nlm.nih.gov/gene/?term=LOC105280447 | 2215 | 21 | 94.81 | 1 | 4.51 | 4.76 | 30.62 | 29.74 | 0.88 | 0.88 | 0.01 | 0.01 |
| gene-LOC105278807 | https://www.ncbi.nlm.nih.gov/gene/?term=LOC105278807 | 2166 | 67 | 309.33 | 1 | 4.62 | 1.49 | 22.11 | 21.25 | 0.87 | 0.87 | 0.03 | 0.03 |
| gene-LOC105283592 | https://www.ncbi.nlm.nih.gov/gene/?term=LOC105283592 | 3479 | 79 | 227.08 | 1 | 2.87 | 1.27 | 22.25 | 21.39 | 0.85 | 0.85 | 0.02 | 0.02 |
| gene-LOC105276875 | https://www.ncbi.nlm.nih.gov/gene/?term=LOC105276875 | 3151 | 116 | 368.14 | 1 | 3.17 | 0.86 | 22.78 | 21.93 | 0.84 | 0.84 | 0.03 | 0.03 |
| gene-LOC105276059 | https://www.ncbi.nlm.nih.gov/gene/?term=LOC105276059 | 6256 | 132 | 211 | 1 | 1.6 | 0.76 | 10.3 | 9.47 | 0.83 | 0.83 | 0.02 | 0.02 |
| gene-LOC105283123 | https://www.ncbi.nlm.nih.gov/gene/?term=LOC105283123 | 1507 | 46 | 305.24 | 1 | 6.64 | 2.17 | 17.92 | 17.09 | 0.83 | 0.83 | 0.03 | 0.03 |
| gene-LOC105284043 | https://www.ncbi.nlm.nih.gov/gene/?term=LOC105284043 | 5661 | 294 | 519.34 | 1 | 1.77 | 0.34 | 10.31 | 9.51 | 0.8 | 0.8 | 0.04 | 0.04 |
| gene-LOC105278632 | https://www.ncbi.nlm.nih.gov/gene/?term=LOC105278632 | 3624 | 108 | 298.01 | 1 | 2.76 | 0.93 | 17.43 | 16.66 | 0.77 | 0.77 | 0.02 | 0.02 |
| gene-LOC105285395 | https://www.ncbi.nlm.nih.gov/gene/?term=LOC105285395 | 8085 | 196 | 242.42 | 1 | 1.24 | 0.51 | 16.42 | 15.66 | 0.76 | 0.76 | 0.02 | 0.02 |
| gene-LOC105280956 | https://www.ncbi.nlm.nih.gov/gene/?term=LOC105280956 | 2639 | 71 | 269.04 | 1 | 3.79 | 1.41 | 18.55 | 17.79 | 0.75 | 0.75 | 0.02 | 0.02 |
| gene-LOC105287095 | https://www.ncbi.nlm.nih.gov/gene/?term=LOC105287095 | 1435 | 58 | 404.18 | 1 | 6.97 | 1.72 | 29.38 | 28.65 | 0.73 | 0.73 | 0.03 | 0.03 |
| gene-LOC105278534 | https://www.ncbi.nlm.nih.gov/gene/?term=LOC105278534 | 7364 | 187 | 253.94 | 1 | 1.36 | 0.53 | 19.13 | 18.41 | 0.72 | 0.72 | 0.02 | 0.02 |
| gene-LOC105275584 | https://www.ncbi.nlm.nih.gov/gene/?term=LOC105275584 | 3070 | 89 | 289.9 | 1 | 3.26 | 1.12 | 19.48 | 18.76 | 0.71 | 0.71 | 0.02 | 0.02 |
| gene-LOC105278497 | https://www.ncbi.nlm.nih.gov/gene/?term=LOC105278497 | 2072 | 92 | 444.02 | 1 | 4.83 | 1.09 | 19.2 | 18.5 | 0.7 | 0.7 | 0.03 | 0.03 |
| gene-LOC105285055 | https://www.ncbi.nlm.nih.gov/gene/?term=LOC105285055 | 4108 | 256 | 623.17 | 1 | 2.43 | 0.39 | 12.1 | 11.41 | 0.69 | 0.69 | 0.04 | 0.04 |
| gene-LOC105284184 | https://www.ncbi.nlm.nih.gov/gene/?term=LOC105284184 | 2535 | 86 | 339.25 | 1 | 3.94 | 1.16 | 33.56 | 32.88 | 0.68 | 0.68 | 0.02 | 0.02 |
| gene-LOC105277423 | https://www.ncbi.nlm.nih.gov/gene/?term=LOC105277423 | 2340 | 106 | 452.99 | 1 | 4.27 | 0.94 | 29 | 28.34 | 0.66 | 0.66 | 0.03 | 0.03 |
| gene-LOC105275451 | https://www.ncbi.nlm.nih.gov/gene/?term=LOC105275451 | 6035 | 126 | 208.78 | 1 | 1.66 | 0.79 | 22.21 | 21.64 | 0.57 | 0.57 | 0.01 | 0.01 |
| gene-LOC105286452 | https://www.ncbi.nlm.nih.gov/gene/?term=LOC105286452 | 6888 | 59 | 85.66 | 1 | 1.45 | 1.69 | 32.42 | 31.88 | 0.55 | 0.55 | 0 | 0 |
| gene-LOC105281599 | https://www.ncbi.nlm.nih.gov/gene/?term=LOC105281599 | 3859 | 49 | 126.98 | 1 | 2.59 | 2.04 | 4.62 | 4.08 | 0.54 | 0.54 | 0.01 | 0.01 |

| | | | | | | | | | | | | | |
|---|---|---|---|---|---|---|---|---|---|---|---|---|---|
| gene-LOC105282811 | https://www.ncbi.nlm.nih.gov/gene/?term=LOC105282811 | 5765 | 215 | 372.94 | 1 | 1.73 | 0.47 | 8.61 | 8.16 | 0.45 | 0.45 | 0.02 | 0.02 |
| gene-LOC105275460 | https://www.ncbi.nlm.nih.gov/gene/?term=LOC105275460 | 2473 | 117 | 473.11 | 1 | 4.04 | 0.85 | 13.75 | 13.31 | 0.44 | 0.44 | 0.02 | 0.02 |
| gene-LOC105279298 | https://www.ncbi.nlm.nih.gov/gene/?term=LOC105279298 | 2773 | 133 | 479.62 | 2 | 7.21 | 1.5 | 23.5 | 23.06 | 0.44 | 0.44 | 0.02 | 0.02 |
| gene-LOC105285742 | https://www.ncbi.nlm.nih.gov/gene/?term=LOC105285742 | 3446 | 110 | 319.21 | 1 | 2.9 | 0.91 | 21.56 | 21.15 | 0.4 | 0.4 | 0.01 | 0.01 |
| gene-LOC105274706 | https://www.ncbi.nlm.nih.gov/gene/?term=LOC105274706 | 1446 | 50 | 345.78 | 1 | 6.92 | 2 | 40.48 | 40.09 | 0.39 | 0.39 | 0.01 | 0.01 |
| gene-LOC105280002 | https://www.ncbi.nlm.nih.gov/gene/?term=LOC105280002 | 1940 | 28 | 144.33 | 1 | 5.15 | 3.57 | 37.11 | 36.72 | 0.39 | 0.39 | 0.01 | 0.01 |
| gene-LOC105285406 | https://www.ncbi.nlm.nih.gov/gene/?term=LOC105285406 | 8015 | 53 | 66.13 1 | 1.25 | 1.89 | 28.7 | 28.32 | 0.39 | 0.39 | 0 | 0 | |
| gene-LOC105279038 | https://www.ncbi.nlm.nih.gov/gene/?term=LOC105279038 | 9453 | 314 | 332.17 | 1 | 1.06 | 0.32 | 9.5 | 9.12 | 0.38 | 0.38 | 0.01 | 0.01 |
| gene-LOC105280774 | https://www.ncbi.nlm.nih.gov/gene/?term=LOC105280774 | 2793 | 99 | 354.46 | 1 | 3.58 | 1.01 | 32.85 | 32.47 | 0.38 | 0.38 | 0.01 | 0.01 |
| gene-LOC105284036 | https://www.ncbi.nlm.nih.gov/gene/?term=LOC105284036 | 3254 | 102 | 313.46 | 1 | 3.07 | 0.98 | 22.68 | 22.3 | 0.38 | 0.38 | 0.01 | 0.01 |
| gene-LOC105274590 | https://www.ncbi.nlm.nih.gov/gene/?term=LOC105274590 | 4516 | 69 | 152.79 | 2 | 4.43 | 2.9 | 27.64 | 27.27 | 0.37 | 0.37 | 0.01 | 0.01 |
| gene-LOC105277079 | https://www.ncbi.nlm.nih.gov/gene/?term=LOC105277079 | 13009 | 273 | 209.85 | 1 | 0.77 | 0.37 | 9.7 | 9.33 | 0.37 | 0.37 | 0.01 | 0.01 |
| gene-LOC105283989 | https://www.ncbi.nlm.nih.gov/gene/?term=LOC105283989 | 4948 | 31 | 62.65 1 | 2.02 | 3.23 | 36.84 | 36.46 | 0.37 | 0.37 | 0 | 0 | |
| gene-LOC105278962 | https://www.ncbi.nlm.nih.gov/gene/?term=LOC105278962 | 4491 | 100 | 222.67 | 1 | 2.23 | 1 | 21.33 | 20.97 | 0.36 | 0.36 | 0.01 | 0.01 |
| gene-LOC105287764 | https://www.ncbi.nlm.nih.gov/gene/?term=LOC105287764 | 18359 | 394 | 214.61 | 1 | 0.54 | 0.25 | 3.28 | 2.97 | 0.31 | 0.31 | 0.01 | 0.01 |
| gene-LOC105281266 | https://www.ncbi.nlm.nih.gov/gene/?term=LOC105281266 | 4102 | 78 | 190.15 | 1 | 2.44 | 1.28 | 16.25 | 15.95 | 0.3 | 0.3 | 0.01 | 0.01 |
| gene-LOC105288066 | https://www.ncbi.nlm.nih.gov/gene/?term=LOC105288066 | 1964 | 28 | 142.57 | 1 | 5.09 | 3.57 | 32.79 | 32.53 | 0.26 | 0.26 | 0 | 0 |
| gene-LOC105283943 | https://www.ncbi.nlm.nih.gov/gene/?term=LOC105283943 | 3559 | 47 | 132.06 | 1 | 2.81 | 2.13 | 17.56 | 17.32 | 0.24 | 0.24 | 0 | 0 |
| gene-LOC105278145 | https://www.ncbi.nlm.nih.gov/gene/?term=LOC105278145 | 4470 | 235 | 525.73 | 1 | 2.24 | 0.43 | 10.78 | 10.55 | 0.23 | 0.23 | 0.01 | 0.01 |
| gene-LOC105284262 | https://www.ncbi.nlm.nih.gov/gene/?term=LOC105284262 | 6669 | 215 | 322.39 | 1 | 1.5 | 0.47 | 5.95 | 5.73 | 0.22 | 0.22 | 0.01 | 0.01 |
| gene-LOC105278666 | https://www.ncbi.nlm.nih.gov/gene/?term=LOC105278666 | 10341 | 444 | 429.36 | 1 | 0.97 | 0.23 | 6.98 | 6.79 | 0.19 | 0.19 | 0.01 | 0.01 |
| gene-LOC105282149 | https://www.ncbi.nlm.nih.gov/gene/?term=LOC105282149 | 1787 | 108 | 604.36 | 1 | 5.6 | 0.93 | 41.15 | 40.97 | 0.18 | 0.18 | 0.01 | 0.01 |
| gene-LOC105276527 | https://www.ncbi.nlm.nih.gov/gene/?term=LOC105276527 | 7923 | 325 | 410.2 1 | 1.26 | 0.31 | 9.54 | 9.36 | 0.17 | 0.17 | 0.01 | 0.01 | |
| gene-LOC105287550 | https://www.ncbi.nlm.nih.gov/gene/?term=LOC105287550 | 15659 | 590 | 376.78 | 1 | 0.64 | 0.17 | 6.31 | 6.15 | 0.16 | 0.16 | 0.01 | 0.01 |
| gene-LOC105276612 | https://www.ncbi.nlm.nih.gov/gene/?term=LOC105276612 | 4157 | 183 | 440.22 | 1 | 2.41 | 0.55 | 9.73 | 9.61 | 0.11 | 0.11 | 0.01 | 0.01 |
| gene-LOC105274531 | https://www.ncbi.nlm.nih.gov/gene/?term=LOC105274531 | 7774 | 291 | 374.32 | 1 | 1.29 | 0.34 | 10.67 | 10.57 | 0.1 | 0.1 | 0 | 0 |
| gene-LOC105275327 | https://www.ncbi.nlm.nih.gov/gene/?term=LOC105275327 | 14245 | 475 | 333.45 | 1 | 0.7 | 0.21 | 7.23 | 7.14 | 0.1 | 0.1 | 0 | 0 |
| gene-LOC105282110 | https://www.ncbi.nlm.nih.gov/gene/?term=LOC105282110 | 7748 | 119 | 153.59 | 1 | 1.29 | 0.84 | 13.03 | 12.95 | 0.08 | 0.08 | 0 | 0 |
| gene-LOC105285807 | https://www.ncbi.nlm.nih.gov/gene/?term=LOC105285807 | 6063 | 223 | 367.8 1 | 1.65 | 0.45 | 9.47 | 9.4 | 0.06 | 0.06 | 0 | 0 | |
| gene-LOC105278283 | https://www.ncbi.nlm.nih.gov/gene/?term=LOC105278283 | 2317 | 135 | 582.65 | 1 | 4.32 | 0.74 | 25.81 | 25.76 | 0.05 | 0.05 | 0 | 0 |
| gene-LOC105276538 | https://www.ncbi.nlm.nih.gov/gene/?term=LOC105276538 | 1511 | 17 | 112.51 | 1 | 6.62 | 5.88 | 69.95 | 69.91 | 0.04 | 0.04 | 0 | 0 |
| gene-LOC105287528 | https://www.ncbi.nlm.nih.gov/gene/?term=LOC105287528 | 14195 | 398 | 280.38 | 1 | 0.7 | 0.25 | 4.97 | 4.95 | 0.02 | 0.02 | 0 | 0 |
| gene-LOC105274823 | https://www.ncbi.nlm.nih.gov/gene/?term=LOC105274823 | 4620 | 138 | 298.7 1 | 2.16 | 0.72 | 21.27 | 21.26 | 0.01 | 0.01 | 0 | 0 | |
| gene-LOC105275127 | https://www.ncbi.nlm.nih.gov/gene/?term=LOC105275127 | 9377 | 295 | 314.6 1 | 1.07 | 0.34 | 7.39 | 7.38 | 0.01 | 0.01 | 0 | 0 | |
| gene-LOC105280407 | https://www.ncbi.nlm.nih.gov/gene/?term=LOC105280407 | 10950 | 207 | 189.04 | 1 | 0.91 | 0.48 | 16.49 | 16.48 | 0.01 | 0.01 | 0 | 0 |
| gene-LOC105275187 | https://www.ncbi.nlm.nih.gov/gene/?term=LOC105275187 | 5534 | 70 | 126.49 | 1 | 1.81 | 1.43 | 12.34 | 12.39 | -0.05 | 0.05 | 0 | 0 |
| gene-LOC105279500 | https://www.ncbi.nlm.nih.gov/gene/?term=LOC105279500 | 3587 | 36 | 100.36 | 1 | 2.79 | 2.78 | 49.66 | 49.71 | -0.05 | 0.05 | 0 | 0 |
| gene-LOC105276692 | https://www.ncbi.nlm.nih.gov/gene/?term=LOC105276692 | 1745 | 29 | 166.19 | 1 | 5.73 | 3.45 | 24.9 | 24.96 | -0.07 | 0.07 | 0 | 0 |
| gene-LOC105275374 | https://www.ncbi.nlm.nih.gov/gene/?term=LOC105275374 | 11376 | 502 | 441.28 | 1 | 0.88 | 0.2 | 10.16 | 10.25 | -0.09 | 0.09 | 0 | 0 |
| gene-LOC105281173 | https://www.ncbi.nlm.nih.gov/gene/?term=LOC105281173 | 1595 | 39 | 244.51 | 1 | 6.27 | 2.56 | 59.82 | 59.91 | -0.09 | 0.09 | 0 | 0 |
| gene-LOC105286274 | https://www.ncbi.nlm.nih.gov/gene/?term=LOC105286274 | 3817 | 191 | 500.39 | 1 | 2.62 | 0.52 | 17.09 | 17.23 | -0.14 | 0.14 | -0.01 | 0.01 |
| gene-LOC105278245 | https://www.ncbi.nlm.nih.gov/gene/?term=LOC105278245 | 6445 | 117 | 181.54 | 1 | 1.55 | 0.85 | 22.04 | 22.2 | -0.17 | 0.17 | 0 | 0 |
| gene-LOC105286514 | https://www.ncbi.nlm.nih.gov/gene/?term=LOC105286514 | 2528 | 48 | 189.87 | 1 | 3.96 | 2.08 | 36.18 | 36.38 | -0.2 | 0.2 | 0 | 0 |
| gene-LOC105275500 | https://www.ncbi.nlm.nih.gov/gene/?term=LOC105275500 | 10943 | 242 | 221.15 | 1 | 0.91 | 0.41 | 7.04 | 7.25 | -0.21 | 0.21 | 0 | 0 |
| gene-LOC105274856 | https://www.ncbi.nlm.nih.gov/gene/?term=LOC105274856 | 5216 | 210 | 402.61 | 1 | 1.92 | 0.48 | 18.25 | 18.52 | -0.26 | 0.26 | -0.01 | 0.01 |
| gene-LOC105277903 | https://www.ncbi.nlm.nih.gov/gene/?term=LOC105277903 | 1835 | 36 | 196.19 | 1 | 5.45 | 2.78 | 49.51 | 49.79 | -0.27 | 0.27 | -0.01 | 0.01 |
| gene-LOC105282491 | https://www.ncbi.nlm.nih.gov/gene/?term=LOC105282491 | 4761 | 165 | 346.57 | 1 | 2.1 | 0.61 | 8.1 | 8.48 | -0.38 | 0.38 | -0.01 | 0.01 |
| gene-LOC105274851 | https://www.ncbi.nlm.nih.gov/gene/?term=LOC105274851 | 6616 | 136 | 205.56 | 1 | 1.51 | 0.74 | 10.78 | 11.18 | -0.4 | 0.4 | -0.01 | 0.01 |
| gene-LOC105282067 | https://www.ncbi.nlm.nih.gov/gene/?term=LOC105282067 | 5941 | 49 | 82.48 1 | 1.68 | 2.04 | 26.22 | 26.65 | -0.43 | 0.43 | 0 | 0 | |
| gene-LOC105286515 | https://www.ncbi.nlm.nih.gov/gene/?term=LOC105286515 | 3751 | 102 | 271.93 | 1 | 2.67 | 0.98 | 12.76 | 13.19 | -0.43 | 0.43 | -0.01 | 0.01 |
| gene-LOC105282399 | https://www.ncbi.nlm.nih.gov/gene/?term=LOC105282399 | 2682 | 23 | 85.76 1 | 3.73 | 4.35 | 34.71 | 35.34 | -0.63 | 0.63 | -0.01 | 0.01 | |
| gene-LOC105288010 | https://www.ncbi.nlm.nih.gov/gene/?term=LOC105288010 | 3189 | 44 | 137.97 | 1 | 3.14 | 2.27 | 30.11 | 30.74 | -0.63 | 0.63 | -0.01 | 0.01 |
| gene-LOC105285060 | https://www.ncbi.nlm.nih.gov/gene/?term=LOC105285060 | 1285 | 18 | 140.08 | 1 | 7.78 | 5.56 | 44.89 | 45.6 | -0.72 | 0.72 | -0.01 | 0.01 |
| gene-LOC105278454 | https://www.ncbi.nlm.nih.gov/gene/?term=LOC105278454 | 1972 | 44 | 223.12 | 1 | 5.07 | 2.27 | 45.26 | 45.99 | -0.74 | 0.74 | -0.02 | 0.02 |
| gene-LOC105283712 | https://www.ncbi.nlm.nih.gov/gene/?term=LOC105283712 | 1910 | 35 | 183.25 | 1 | 5.24 | 2.86 | 26.72 | 27.47 | -0.75 | 0.75 | -0.01 | 0.01 |
| gene-LOC105286166 | https://www.ncbi.nlm.nih.gov/gene/?term=LOC105286166 | 3050 | 110 | 360.66 | 1 | 3.28 | 0.91 | 23.43 | 24.21 | -0.78 | 0.78 | -0.03 | 0.03 |
| gene-LOC105282151 | https://www.ncbi.nlm.nih.gov/gene/?term=LOC105282151 | 1601 | 39 | 243.6 1 | 6.25 | 2.56 | 40.09 | 41 | -0.91 | 0.91 | -0.02 | 0.02 | |
| gene-LOC105278282 | https://www.ncbi.nlm.nih.gov/gene/?term=LOC105278282 | 1775 | 56 | 315.49 | 1 | 5.63 | 1.79 | 37.29 | 38.34 | -1.05 | 1.05 | -0.03 | 0.03 |
| gene-LOC105280377 | https://www.ncbi.nlm.nih.gov/gene/?term=LOC105280377 | 2024 | 69 | 340.91 | 1 | 4.94 | 1.45 | 43.74 | 44.8 | -1.06 | 1.06 | -0.04 | 0.04 |
| gene-LOC105286999 | https://www.ncbi.nlm.nih.gov/gene/?term=LOC105286999 | 2141 | 46 | 214.85 | 1 | 4.67 | 2.17 | 31.8 | 32.92 | -1.12 | 1.12 | -0.02 | 0.02 |
| gene-LOC105276429 | https://www.ncbi.nlm.nih.gov/gene/?term=LOC105276429 | 2901 | 114 | 392.97 | 1 | 3.45 | 0.88 | 24.72 | 25.87 | -1.14 | 1.14 | -0.04 | 0.04 |
| gene-LOC105275966 | https://www.ncbi.nlm.nih.gov/gene/?term=LOC105275966 | 3299 | 59 | 178.84 | 1 | 3.03 | 1.69 | 49.71 | 51.14 | -1.42 | 1.42 | -0.03 | 0.03 |
| gene-LOC105274534 | https://www.ncbi.nlm.nih.gov/gene/?term=LOC105274534 | 1329 | 55 | 413.84 | 1 | 7.52 | 1.82 | 44.22 | 45.89 | -1.68 | 1.68 | -0.07 | 0.07 |
| gene-LOC105281700 | https://www.ncbi.nlm.nih.gov/gene/?term=LOC105281700 | 2208 | 32 | 144.93 | 1 | 4.53 | 3.12 | 37.39 | 39.09 | -1.7 | 1.7 | -0.02 | 0.02 |

| | | | | | | | | | | | | | | |
|---|---|---|---|---|---|---|---|---|---|---|---|---|---|---|
| gene-LOC105287633 | https://www.ncbi.nlm.nih.gov/gene/?term=LOC105287633 | 1524 | 31 | 203.41 | | 1 | 6.56 | 3.23 | 53.47 | 55.25 | -1.78 | 1.78 | -0.04 | 0.04 |
| gene-LOC105286770 | https://www.ncbi.nlm.nih.gov/gene/?term=LOC105286770 | 3055 | 30 | 98.2 | 1 | 3.27 | 3.33 | 17.03 | 18.87 | -1.84 | 1.84 | -0.02 | 0.02 | |
| gene-LOC105282885 | https://www.ncbi.nlm.nih.gov/gene/?term=LOC105282885 | 1870 | 47 | 251.34 | | 1 | 5.35 | 2.13 | 35.98 | 38.05 | -2.07 | 2.07 | -0.05 | 0.05 |
| gene-LOC105276196 | https://www.ncbi.nlm.nih.gov/gene/?term=LOC105276196 | 2953 | 60 | 203.18 | | 1 | 3.39 | 1.67 | 38.8 | 41.25 | -2.45 | 2.45 | -0.05 | 0.05 |
| gene-LOC105285409 | https://www.ncbi.nlm.nih.gov/gene/?term=LOC105285409 | 5099 | 49 | 96.1 | 2 | 3.92 | 4.08 | 2.16 | 4.82 | -2.66 | 2.66 | -0.03 | 0.03 | |
| gene-LOC105283492 | https://www.ncbi.nlm.nih.gov/gene/?term=LOC105283492 | 1535 | 10 | 65.15 | 1 | 6.51 | 10 | 34.7 | 42.8 | -8.1 | 8.1 | -0.05 | 0.05 | |

Directory: OGL File: ogl-Ob_LI_BCphaseB2.txt

| gene_ID | gene_link | gwidth | #Sites | #per10Kb | %perSite | %pNucl |
|---|---|---|---|---|---|---|
| gene-LOC105286531 | https://www.ncbi.nlm.nih.gov/gene/?term=LOC105286531 | 372 | 43 | 1155.91 | 35.45 | 4.1 |
| gene-LOC113561532 | https://www.ncbi.nlm.nih.gov/gene/?term=LOC113561532 | 72 | 3 | 416.67 | 98.48 | 4.1 |
| gene-LOC105282149 | https://www.ncbi.nlm.nih.gov/gene/?term=LOC105282149 | 1787 | 161 | 900.95 | 43.24 | 3.9 |
| gene-LOC113562627 | https://www.ncbi.nlm.nih.gov/gene/?term=LOC113562627 | 397 | 23 | 579.35 | 66.24 | 3.84 |
| gene-LOC105280739 | https://www.ncbi.nlm.nih.gov/gene/?term=LOC105280739 | 1991 | 131 | 657.96 | 55 | 3.62 |
| gene-LOC105274534 | https://www.ncbi.nlm.nih.gov/gene/?term=LOC105274534 | 1329 | 107 | 805.12 | 44.14 | 3.55 |
| gene-LOC105278849 | https://www.ncbi.nlm.nih.gov/gene/?term=LOC105278849 | 776 | 114 | 1469.07 | 23.06 | 3.39 |
| gene-LOC105285635 | https://www.ncbi.nlm.nih.gov/gene/?term=LOC105285635 | 1175 | 128 | 1089.36 | 30.64 | 3.34 |
| gene-LOC105286801 | https://www.ncbi.nlm.nih.gov/gene/?term=LOC105286801 | 1102 | 94 | 852.99 | 39.1 | 3.34 |
| gene-LOC105274711 | https://www.ncbi.nlm.nih.gov/gene/?term=LOC105274711 | 1655 | 118 | 712.99 | 45.69 | 3.26 |
| gene-LOC105285394 | https://www.ncbi.nlm.nih.gov/gene/?term=LOC105285394 | 1307 | 59 | 451.42 | 71.23 | 3.22 |
| gene-LOC105286269 | https://www.ncbi.nlm.nih.gov/gene/?term=LOC105286269 | 1401 | 89 | 635.26 | 49.97 | 3.17 |
| gene-LOC105279052 | https://www.ncbi.nlm.nih.gov/gene/?term=LOC105279052 | 1130 | 136 | 1203.54 | 25.81 | 3.11 |
| gene-LOC105287692 | https://www.ncbi.nlm.nih.gov/gene/?term=LOC105287692 | 1505 | 134 | 890.37 | 34.97 | 3.11 |
| gene-LOC105278844 | https://www.ncbi.nlm.nih.gov/gene/?term=LOC105278844 | 1101 | 115 | 1044.5 | 29.68 | 3.1 |
| gene-LOC105276360 | https://www.ncbi.nlm.nih.gov/gene/?term=LOC105276360 | 1466 | 159 | 1084.58 | 28.5 | 3.09 |
| gene-LOC105287811 | https://www.ncbi.nlm.nih.gov/gene/?term=LOC105287811 | 2155 | 136 | 631.09 | 47.8 | 3.02 |
| gene-LOC105286235 | https://www.ncbi.nlm.nih.gov/gene/?term=LOC105286235 | 1510 | 111 | 735.1 40.93 | 3.01 | |
| gene-LOC105286737 | https://www.ncbi.nlm.nih.gov/gene/?term=LOC105286737 | 2690 | 359 | 1334.57 | 22.22 | 2.97 |
| gene-LOC105288088 | https://www.ncbi.nlm.nih.gov/gene/?term=LOC105288088 | 1795 | 127 | 707.52 | 41.75 | 2.95 |
| gene-LOC105276371 | https://www.ncbi.nlm.nih.gov/gene/?term=LOC105276371 | 871 | 56 | 642.94 | 44.32 | 2.85 |
| gene-LOC105282986 | https://www.ncbi.nlm.nih.gov/gene/?term=LOC105282986 | 1600 | 142 | 887.5 32.15 | 2.85 | |
| gene-LOC105278160 | https://www.ncbi.nlm.nih.gov/gene/?term=LOC105278160 | 1872 | 155 | 827.99 | 34.29 | 2.84 |
| gene-LOC105275314 | https://www.ncbi.nlm.nih.gov/gene/?term=LOC105275314 | 1489 | 93 | 624.58 | 45.18 | 2.82 |

...
... only first 25 lines shown ...

Directory: OGL File: ogl-Ob_LI_BCphaseB2.vs.RphaseA1.txt

| gene_ID | gene_link | gwidth | #Sites | #per10Kb | #dmSites | #dmsp10kb | %dmSites | %pSite1 | %pSite2 | DMpSite | ADMpSite | DMpNucl | ADMpNucl |
|---|---|---|---|---|---|---|---|---|---|---|---|---|---|
| gene-LOC105282914 | https://www.ncbi.nlm.nih.gov/gene/?term=LOC105282914 | 5623 | 41 | 72.91 | 1 | 1.78 | 2.44 | 52.21 | 48.43 | 3.78 | 3.78 | 0.03 | 0.03 |
| gene-LOC105279500 | https://www.ncbi.nlm.nih.gov/gene/?term=LOC105279500 | 3587 | 30 | 83.64 | 1 | 2.79 | 3.33 | 48.41 | 46.03 | 2.38 | 2.38 | 0.02 | 0.02 |
| gene-LOC105284396 | https://www.ncbi.nlm.nih.gov/gene/?term=LOC105284396 | 1976 | 66 | 334.01 | 1 | 5.06 | 1.52 | 34.48 | 32.1 | 2.38 | 2.38 | 0.08 | 0.08 |
| gene-LOC105274706 | https://www.ncbi.nlm.nih.gov/gene/?term=LOC105274706 | 1446 | 50 | 345.78 | 1 | 6.92 | 2 | 39.84 | 37.9 | 1.94 | 1.94 | 0.07 | 0.07 |
| gene-LOC105274449 | https://www.ncbi.nlm.nih.gov/gene/?term=LOC105274449 | 1549 | 53 | 342.16 | 1 | 6.46 | 1.89 | 51.77 | 49.89 | 1.87 | 1.87 | 0.06 | 0.06 |
| gene-LOC105282885 | https://www.ncbi.nlm.nih.gov/gene/?term=LOC105282885 | 1870 | 46 | 245.99 | 1 | 5.35 | 2.17 | 37.06 | 35.54 | 1.52 | 1.52 | 0.04 | 0.04 |
| gene-LOC105276272 | https://www.ncbi.nlm.nih.gov/gene/?term=LOC105276272 | 1965 | 46 | 234.1 | 1 | 5.09 | 2.17 | 27.86 | 26.72 | 1.14 | 1.14 | 0.03 | 0.03 |
| gene-LOC105275587 | https://www.ncbi.nlm.nih.gov/gene/?term=LOC105275587 | 2404 | 57 | 237.1 | 1 | 4.16 | 1.75 | 37.35 | 36.26 | 1.09 | 1.09 | 0.03 | 0.03 |
| gene-LOC105283967 | https://www.ncbi.nlm.nih.gov/gene/?term=LOC105283967 | 2393 | 50 | 208.94 | 1 | 4.18 | 2 | 45.88 | 44.9 | 0.98 | 0.98 | 0.02 | 0.02 |
| gene-LOC105282970 | https://www.ncbi.nlm.nih.gov/gene/?term=LOC105282970 | 1529 | 40 | 261.61 | 1 | 6.54 | 2.5 | 50.96 | 50.14 | 0.82 | 0.82 | 0.02 | 0.02 |
| gene-LOC105281173 | https://www.ncbi.nlm.nih.gov/gene/?term=LOC105281173 | 1595 | 38 | 238.24 | 1 | 6.27 | 2.63 | 60.67 | 59.87 | 0.8 | 0.8 | 0.02 | 0.02 |
| gene-LOC105281312 | https://www.ncbi.nlm.nih.gov/gene/?term=LOC105281312 | 2551 | 65 | 254.8 | 1 | 3.92 | 1.54 | 33.11 | 32.31 | 0.8 | 0.8 | 0.02 | 0.02 |
| gene-LOC105278962 | https://www.ncbi.nlm.nih.gov/gene/?term=LOC105278962 | 4491 | 93 | 207.08 | 1 | 2.23 | 1.08 | 22.79 | 22.03 | 0.76 | 0.76 | 0.02 | 0.02 |
| gene-LOC105283712 | https://www.ncbi.nlm.nih.gov/gene/?term=LOC105283712 | 1910 | 36 | 188.48 | 1 | 5.24 | 2.78 | 26.26 | 25.56 | 0.7 | 0.7 | 0.01 | 0.01 |
| gene-LOC105280819 | https://www.ncbi.nlm.nih.gov/gene/?term=LOC105280819 | 1639 | 45 | 274.56 | 1 | 6.1 | 2.22 | 37.3 | 36.73 | 0.57 | 0.57 | 0.02 | 0.02 |
| gene-LOC105286274 | https://www.ncbi.nlm.nih.gov/gene/?term=LOC105286274 | 3817 | 187 | 489.91 | 1 | 2.62 | 0.53 | 16.7 | 16.2 | 0.51 | 0.51 | 0.02 | 0.02 |
| gene-LOC105284184 | https://www.ncbi.nlm.nih.gov/gene/?term=LOC105284184 | 2535 | 92 | 362.92 | 1 | 3.94 | 1.09 | 31.62 | 31.13 | 0.49 | 0.49 | 0.02 | 0.02 |
| gene-LOC105288042 | https://www.ncbi.nlm.nih.gov/gene/?term=LOC105288042 | 3507 | 98 | 279.44 | 1 | 2.85 | 1.02 | 22.45 | 21.99 | 0.46 | 0.46 | 0.01 | 0.01 |
| gene-LOC105282646 | https://www.ncbi.nlm.nih.gov/gene/?term=LOC105282646 | 3317 | 99 | 298.46 | 1 | 3.01 | 1.01 | 21.06 | 20.67 | 0.39 | 0.39 | 0.01 | 0.01 |
| gene-LOC105274590 | https://www.ncbi.nlm.nih.gov/gene/?term=LOC105274590 | 4516 | 74 | 163.86 | 1 | 2.21 | 1.35 | 30.44 | 30.06 | 0.38 | 0.38 | 0.01 | 0.01 |
| gene-LOC105276527 | https://www.ncbi.nlm.nih.gov/gene/?term=LOC105276527 | 7923 | 317 | 400.1 | 1 | 1.26 | 0.32 | 9.8 | 9.43 | 0.37 | 0.37 | 0.01 | 0.01 |
| gene-LOC105282951 | https://www.ncbi.nlm.nih.gov/gene/?term=LOC105282951 | 3366 | 120 | 356.51 | 1 | 2.97 | 0.83 | 19.9 | 19.57 | 0.33 | 0.33 | 0.01 | 0.01 |
| gene-LOC105276196 | https://www.ncbi.nlm.nih.gov/gene/?term=LOC105276196 | 2953 | 56 | 189.64 | 1 | 3.39 | 1.79 | 39.44 | 39.14 | 0.3 | 0.3 | 0.01 | 0.01 |
| gene-LOC105278534 | https://www.ncbi.nlm.nih.gov/gene/?term=LOC105278534 | 7364 | 185 | 251.22 | 2 | 2.72 | 1.08 | 18.38 | 18.12 | 0.25 | 0.25 | 0.01 | 0.01 |
| gene-LOC105280002 | https://www.ncbi.nlm.nih.gov/gene/?term=LOC105280002 | 1940 | 26 | 134.02 | 1 | 5.15 | 3.85 | 34.83 | 34.64 | 0.19 | 0.19 | 0 | 0 |
| gene-LOC105274534 | https://www.ncbi.nlm.nih.gov/gene/?term=LOC105274534 | 1329 | 59 | 443.94 | 1 | 7.52 | 1.69 | 42.17 | 42.03 | 0.14 | 0.14 | 0.01 | 0.01 |
| gene-LOC105278632 | https://www.ncbi.nlm.nih.gov/gene/?term=LOC105278632 | 3624 | 133 | 367 | 1 | 2.76 | 0.75 | 14.21 | 14.06 | 0.14 | 0.14 | 0.01 | 0.01 |
| gene-LOC105278691 | https://www.ncbi.nlm.nih.gov/gene/?term=LOC105278691 | 3871 | 151 | 390.08 | 1 | 2.58 | 0.66 | 14.98 | 14.84 | 0.14 | 0.14 | 0.01 | 0.01 |
| gene-LOC105282811 | https://www.ncbi.nlm.nih.gov/gene/?term=LOC105282811 | 5765 | 199 | 345.19 | 1 | 1.73 | 0.5 | 8.45 | 8.31 | 0.14 | 0.14 | 0 | 0 |
| gene-LOC105285807 | https://www.ncbi.nlm.nih.gov/gene/?term=LOC105285807 | 6063 | 217 | 357.91 | 1 | 1.65 | 0.46 | 8.91 | 8.79 | 0.12 | 0.12 | 0 | 0 |
| gene-LOC105281700 | https://www.ncbi.nlm.nih.gov/gene/?term=LOC105281700 | 2208 | 37 | 167.57 | 1 | 4.53 | 2.7 | 39.75 | 39.65 | 0.11 | 0.11 | 0 | 0 |
| gene-LOC105283728 | https://www.ncbi.nlm.nih.gov/gene/?term=LOC105283728 | 3256 | 122 | 374.69 | 2 | 6.14 | 1.64 | 18.7 | 18.6 | 0.1 | 0.1 | 0 | 0 |
| gene-LOC105276538 | https://www.ncbi.nlm.nih.gov/gene/?term=LOC105276538 | 1511 | 18 | 119.13 | 1 | 6.62 | 5.56 | 68.54 | 68.54 | 0 | 0 | 0 | 0 |
| gene-LOC105283123 | https://www.ncbi.nlm.nih.gov/gene/?term=LOC105283123 | 1507 | 47 | 311.88 | 1 | 6.64 | 2.13 | 17.1 | 17.12 | -0.02 | 0.02 | 0 | 0 |
| gene-LOC105286514 | https://www.ncbi.nlm.nih.gov/gene/?term=LOC105286514 | 2528 | 44 | 174.05 | 1 | 3.96 | 2.27 | 37.14 | 37.18 | -0.04 | 0.04 | 0 | 0 |
| gene-LOC105285179 | https://www.ncbi.nlm.nih.gov/gene/?term=LOC105285179 | 4191 | 176 | 419.95 | 1 | 2.39 | 0.57 | 19.46 | 19.52 | -0.06 | 0.06 | 0 | 0 |
| gene-LOC105285406 | https://www.ncbi.nlm.nih.gov/gene/?term=LOC105285406 | 8015 | 60 | 74.86 | 1 | 1.25 | 1.67 | 28.89 | 28.94 | -0.06 | 0.06 | 0 | 0 |
| gene-LOC105284043 | https://www.ncbi.nlm.nih.gov/gene/?term=LOC105284043 | 5661 | 295 | 521.11 | 1 | 1.77 | 0.34 | 9.91 | 9.97 | -0.07 | 0.07 | 0 | 0 |
| gene-LOC105287417 | https://www.ncbi.nlm.nih.gov/gene/?term=LOC105287417 | 1025 | 32 | 312.2 | 1 | 9.76 | 3.12 | 37.56 | 37.69 | -0.12 | 0.12 | 0 | 0 |
| gene-LOC113562050 | https://www.ncbi.nlm.nih.gov/gene/?term=LOC113562050 | 17097 | 587 | 343.34 | 1 | 0.58 | 0.17 | 4.91 | 5.05 | -0.13 | 0.13 | 0 | 0 |
| gene-LOC105275683 | https://www.ncbi.nlm.nih.gov/gene/?term=LOC105275683 | 4256 | 187 | 439.38 | 1 | 2.35 | 0.53 | 13.19 | 13.34 | -0.15 | 0.15 | -0.01 | 0.01 |
| gene-LOC105278228 | https://www.ncbi.nlm.nih.gov/gene/?term=LOC105278228 | 8610 | 154 | 178.86 | 1 | 1.16 | 0.65 | 23.95 | 24.14 | -0.2 | 0.2 | 0 | 0 |
| gene-LOC105274851 | https://www.ncbi.nlm.nih.gov/gene/?term=LOC105274851 | 6616 | 130 | 196.49 | 1 | 1.51 | 0.77 | 11.46 | 11.67 | -0.21 | 0.21 | 0 | 0 |
| gene-LOC105278666 | https://www.ncbi.nlm.nih.gov/gene/?term=LOC105278666 | 10341 | 471 | 455.47 | 1 | 0.97 | 0.21 | 6.62 | 6.83 | -0.21 | 0.21 | -0.01 | 0.01 |
| gene-LOC105275179 | https://www.ncbi.nlm.nih.gov/gene/?term=LOC105275179 | 3271 | 77 | 235.4 | 1 | 3.06 | 1.3 | 17.93 | 18.15 | -0.22 | 0.22 | -0.01 | 0.01 |
| gene-LOC105284355 | https://www.ncbi.nlm.nih.gov/gene/?term=LOC105284355 | 3904 | 162 | 414.96 | 1 | 2.56 | 0.62 | 19.15 | 19.36 | -0.22 | 0.22 | -0.01 | 0.01 |
| gene-LOC105286452 | https://www.ncbi.nlm.nih.gov/gene/?term=LOC105286452 | 6888 | 61 | 88.56 | 1 | 1.45 | 1.64 | 30.12 | 30.34 | -0.22 | 0.22 | 0 | 0 |
| gene-LOC105278283 | https://www.ncbi.nlm.nih.gov/gene/?term=LOC105278283 | 2317 | 139 | 599.91 | 1 | 4.32 | 0.72 | 25.21 | 25.46 | -0.25 | 0.25 | -0.02 | 0.02 |
| gene-LOC105275363 | https://www.ncbi.nlm.nih.gov/gene/?term=LOC105275363 | 3798 | 161 | 423.91 | 1 | 2.63 | 0.62 | 16.94 | 17.21 | -0.26 | 0.26 | -0.01 | 0.01 |
| gene-LOC105279038 | https://www.ncbi.nlm.nih.gov/gene/?term=LOC105279038 | 9453 | 340 | 359.67 | 1 | 1.06 | 0.29 | 8.66 | 8.92 | -0.26 | 0.26 | -0.01 | 0.01 |
| gene-LOC105288010 | https://www.ncbi.nlm.nih.gov/gene/?term=LOC105288010 | 3189 | 39 | 122.3 | 1 | 3.14 | 2.56 | 33.4 | 33.65 | -0.26 | 0.26 | 0 | 0 |
| gene-LOC105275374 | https://www.ncbi.nlm.nih.gov/gene/?term=LOC105275374 | 11376 | 488 | 428.97 | 1 | 0.88 | 0.2 | 10.37 | 10.64 | -0.27 | 0.27 | -0.01 | 0.01 |
| gene-LOC105275327 | https://www.ncbi.nlm.nih.gov/gene/?term=LOC105275327 | 14245 | 511 | 358.72 | 1 | 0.7 | 0.2 | 6.72 | 7.02 | -0.3 | 0.3 | -0.01 | 0.01 |
| gene-LOC105278497 | https://www.ncbi.nlm.nih.gov/gene/?term=LOC105278497 | 2072 | 96 | 463.32 | 1 | 4.83 | 1.04 | 18.89 | 19.19 | -0.31 | 0.31 | -0.01 | 0.01 |
| gene-LOC105275170 | https://www.ncbi.nlm.nih.gov/gene/?term=LOC105275170 | 12918 | 279 | 215.98 | 1 | 0.77 | 0.36 | 2.01 | 2.33 | -0.32 | 0.32 | -0.01 | 0.01 |
| gene-LOC105278282 | https://www.ncbi.nlm.nih.gov/gene/?term=LOC105278282 | 1775 | 51 | 287.32 | 1 | 5.63 | 1.96 | 40.43 | 40.76 | -0.33 | 0.33 | -0.01 | 0.01 |
| gene-LOC105282794 | https://www.ncbi.nlm.nih.gov/gene/?term=LOC105282794 | 4571 | 93 | 203.46 | 1 | 2.19 | 1.08 | 16.99 | 17.32 | -0.33 | 0.33 | -0.01 | 0.01 |
| gene-LOC105282149 | https://www.ncbi.nlm.nih.gov/gene/?term=LOC105282149 | 1787 | 103 | 576.39 | 1 | 5.6 | 0.97 | 40.98 | 41.32 | -0.34 | 0.34 | -0.02 | 0.02 |
| gene-LOC105274669 | https://www.ncbi.nlm.nih.gov/gene/?term=LOC105274669 | 8555 | 261 | 305.08 | 1 | 1.17 | 0.38 | 5.68 | 6.02 | -0.35 | 0.35 | -0.01 | 0.01 |
| gene-LOC105280774 | https://www.ncbi.nlm.nih.gov/gene/?term=LOC105280774 | 2793 | 117 | 418.9 | 1 | 3.58 | 0.85 | 27.14 | 27.51 | -0.36 | 0.36 | -0.02 | 0.02 |

| | | | | | | | | | | | | | |
|---|---|---|---|---|---|---|---|---|---|---|---|---|---|
| gene-LOC105277761 | https://www.ncbi.nlm.nih.gov/gene/?term=LOC105277761 | 4648 | 144 | 309.81 | 1 | 2.15 | 0.69 | 14.39 | 14.8 | -0.41 | 0.41 | -0.01 | 0.01 |
| gene-LOC105274531 | https://www.ncbi.nlm.nih.gov/gene/?term=LOC105274531 | 7774 | 275 | 353.74 | 1 | 1.29 | 0.36 | 10.93 | 11.37 | -0.43 | 0.43 | -0.02 | 0.02 |
| gene-LOC105274823 | https://www.ncbi.nlm.nih.gov/gene/?term=LOC105274823 | 4620 | 145 | 313.85 | 1 | 2.16 | 0.69 | 19.86 | 20.29 | -0.43 | 0.43 | -0.01 | 0.01 |
| gene-LOC105285639 | https://www.ncbi.nlm.nih.gov/gene/?term=LOC105285639 | 10325 | 344 | 333.17 | 1 | 0.97 | 0.29 | 11.23 | 11.66 | -0.43 | 0.43 | -0.01 | 0.01 |
| gene-LOC105287095 | https://www.ncbi.nlm.nih.gov/gene/?term=LOC105287095 | 1435 | 64 | 445.99 | 1 | 6.97 | 1.56 | 25.57 | 26.08 | -0.51 | 0.51 | -0.02 | 0.02 |
| gene-LOC105278145 | https://www.ncbi.nlm.nih.gov/gene/?term=LOC105278145 | 4470 | 235 | 525.73 | 2 | 4.47 | 0.85 | 10.77 | 11.29 | -0.52 | 0.52 | -0.03 | 0.03 |
| gene-LOC105275752 | https://www.ncbi.nlm.nih.gov/gene/?term=LOC105275752 | 3462 | 128 | 369.73 | 1 | 2.89 | 0.78 | 29.22 | 29.76 | -0.53 | 0.53 | -0.02 | 0.02 |
| gene-LOC105285742 | https://www.ncbi.nlm.nih.gov/gene/?term=LOC105285742 | 3446 | 105 | 304.7 | 1 | 2.9 | 0.95 | 17.29 | 17.85 | -0.55 | 0.55 | -0.02 | 0.02 |
| gene-LOC105288036 | https://www.ncbi.nlm.nih.gov/gene/?term=LOC105288036 | 5110 | 200 | 391.39 | 1 | 1.96 | 0.5 | 16.38 | 16.94 | -0.55 | 0.55 | -0.02 | 0.02 |
| gene-LOC105278649 | https://www.ncbi.nlm.nih.gov/gene/?term=LOC105278649 | 5616 | 137 | 243.95 | 1 | 1.78 | 0.73 | 13.17 | 13.73 | -0.56 | 0.56 | -0.01 | 0.01 |
| gene-LOC105275460 | https://www.ncbi.nlm.nih.gov/gene/?term=LOC105275460 | 2473 | 120 | 485.24 | 1 | 4.04 | 0.83 | 13.18 | 13.75 | -0.57 | 0.57 | -0.03 | 0.03 |
| gene-LOC105285055 | https://www.ncbi.nlm.nih.gov/gene/?term=LOC105285055 | 4108 | 256 | 623.17 | 1 | 2.43 | 0.39 | 10.36 | 10.97 | -0.61 | 0.61 | -0.04 | 0.04 |
| gene-LOC105282658 | https://www.ncbi.nlm.nih.gov/gene/?term=LOC105282658 | 2421 | 59 | 243.7 | 1 | 4.13 | 1.69 | 24.61 | 25.23 | -0.62 | 0.62 | -0.02 | 0.02 |
| gene-LOC105279298 | https://www.ncbi.nlm.nih.gov/gene/?term=LOC105279298 | 2773 | 130 | 468.81 | 2 | 7.21 | 1.54 | 23.75 | 24.38 | -0.63 | 0.63 | -0.03 | 0.03 |
| gene-LOC105282067 | https://www.ncbi.nlm.nih.gov/gene/?term=LOC105282067 | 5941 | 41 | 69.01 | 1 | 1.68 | 2.44 | 26.24 | 26.89 | -0.64 | 0.64 | 0 | 0 |
| gene-LOC105287063 | https://www.ncbi.nlm.nih.gov/gene/?term=LOC105287063 | 1874 | 49 | 261.47 | 1 | 5.34 | 2.04 | 29.55 | 30.23 | -0.68 | 0.68 | -0.02 | 0.02 |
| gene-LOC105277856 | https://www.ncbi.nlm.nih.gov/gene/?term=LOC105277856 | 3442 | 31 | 90.06 | 1 | 2.91 | 3.23 | 38.58 | 39.29 | -0.71 | 0.71 | -0.01 | 0.01 |
| gene-LOC105276059 | https://www.ncbi.nlm.nih.gov/gene/?term=LOC105276059 | 6256 | 129 | 206.2 | 1 | 1.6 | 0.78 | 9.24 | 9.96 | -0.72 | 0.72 | -0.01 | 0.01 |
| gene-LOC105279900 | https://www.ncbi.nlm.nih.gov/gene/?term=LOC105279900 | 2191 | 54 | 246.46 | 1 | 4.56 | 1.85 | 30.89 | 31.61 | -0.72 | 0.72 | -0.02 | 0.02 |
| gene-LOC105275451 | https://www.ncbi.nlm.nih.gov/gene/?term=LOC105275451 | 6035 | 123 | 203.81 | 1 | 1.66 | 0.81 | 22.94 | 23.69 | -0.75 | 0.75 | -0.02 | 0.02 |
| gene-LOC105274856 | https://www.ncbi.nlm.nih.gov/gene/?term=LOC105274856 | 5216 | 225 | 431.37 | 1 | 1.92 | 0.44 | 17.14 | 17.89 | -0.76 | 0.76 | -0.03 | 0.03 |
| gene-LOC105278384 | https://www.ncbi.nlm.nih.gov/gene/?term=LOC105278384 | 1390 | 35 | 251.8 | 1 | 7.19 | 2.86 | 45.32 | 46.11 | -0.79 | 0.79 | -0.02 | 0.02 |
| gene-LOC105280723 | https://www.ncbi.nlm.nih.gov/gene/?term=LOC105280723 | 3572 | 182 | 509.52 | 1 | 2.8 | 0.55 | 15.32 | 16.11 | -0.79 | 0.79 | -0.04 | 0.04 |
| gene-LOC105284116 | https://www.ncbi.nlm.nih.gov/gene/?term=LOC105284116 | 3970 | 107 | 269.52 | 1 | 2.52 | 0.93 | 7.18 | 8 | -0.82 | 0.82 | -0.02 | 0.02 |
| gene-LOC105284117 | https://www.ncbi.nlm.nih.gov/gene/?term=LOC105284117 | 4340 | 94 | 216.59 | 1 | 2.3 | 1.06 | 9.25 | 10.09 | -0.84 | 0.84 | -0.02 | 0.02 |
| gene-LOC105282491 | https://www.ncbi.nlm.nih.gov/gene/?term=LOC105282491 | 4761 | 174 | 365.47 | 1 | 2.1 | 0.57 | 8.54 | 9.39 | -0.85 | 0.85 | -0.03 | 0.03 |
| gene-LOC105275187 | https://www.ncbi.nlm.nih.gov/gene/?term=LOC105275187 | 5534 | 69 | 124.68 | 1 | 1.81 | 1.45 | 13.92 | 14.78 | -0.86 | 0.86 | -0.01 | 0.01 |
| gene-LOC105278877 | https://www.ncbi.nlm.nih.gov/gene/?term=LOC105278877 | 1968 | 82 | 416.67 | 1 | 5.08 | 1.22 | 30.66 | 31.6 | -0.94 | 0.94 | -0.04 | 0.04 |
| gene-LOC105278807 | https://www.ncbi.nlm.nih.gov/gene/?term=LOC105278807 | 2166 | 78 | 360.11 | 1 | 4.62 | 1.28 | 18.54 | 19.5 | -0.97 | 0.97 | -0.03 | 0.03 |
| gene-LOC105282991 | https://www.ncbi.nlm.nih.gov/gene/?term=LOC105282991 | 5617 | 101 | 179.81 | 1 | 1.78 | 0.99 | 19.27 | 20.29 | -1.02 | 1.02 | -0.02 | 0.02 |
| gene-LOC105279664 | https://www.ncbi.nlm.nih.gov/gene/?term=LOC105279664 | 4023 | 60 | 149.14 | 1 | 2.49 | 1.67 | 17.78 | 18.81 | -1.03 | 1.03 | -0.02 | 0.02 |
| gene-LOC105280348 | https://www.ncbi.nlm.nih.gov/gene/?term=LOC105280348 | 3797 | 71 | 186.99 | 1 | 2.63 | 1.41 | 18.59 | 19.68 | -1.09 | 1.09 | -0.02 | 0.02 |
| gene-LOC105282151 | https://www.ncbi.nlm.nih.gov/gene/?term=LOC105282151 | 1601 | 42 | 262.34 | 1 | 6.25 | 2.38 | 34.92 | 36.1 | -1.17 | 1.17 | -0.03 | 0.03 |
| gene-LOC105283989 | https://www.ncbi.nlm.nih.gov/gene/?term=LOC105283989 | 4948 | 27 | 54.57 | 1 | 2.02 | 3.7 | 44.98 | 46.16 | -1.18 | 1.18 | -0.01 | 0.01 |
| gene-LOC105287854 | https://www.ncbi.nlm.nih.gov/gene/?term=LOC105287854 | 3644 | 64 | 175.63 | 1 | 2.74 | 1.56 | 14.92 | 16.11 | -1.19 | 1.19 | -0.02 | 0.02 |
| gene-LOC105276692 | https://www.ncbi.nlm.nih.gov/gene/?term=LOC105276692 | 1745 | 33 | 189.11 | 1 | 5.73 | 3.03 | 21.96 | 23.19 | -1.23 | 1.23 | -0.02 | 0.02 |
| gene-LOC105288078 | https://www.ncbi.nlm.nih.gov/gene/?term=LOC105288078 | 19126 | 298 | 155.81 | 1 | 0.52 | 0.34 | 16.07 | 17.31 | -1.24 | 1.24 | -0.02 | 0.02 |
| gene-LOC105275347 | https://www.ncbi.nlm.nih.gov/gene/?term=LOC105275347 | 2526 | 16 | 63.34 | 1 | 3.96 | 6.25 | 40.9 | 42.25 | -1.34 | 1.34 | -0.01 | 0.01 |
| gene-LOC105280641 | https://www.ncbi.nlm.nih.gov/gene/?term=LOC105280641 | 3682 | 55 | 149.38 | 1 | 2.72 | 1.82 | 18.09 | 19.48 | -1.39 | 1.39 | -0.02 | 0.02 |
| gene-LOC105280956 | https://www.ncbi.nlm.nih.gov/gene/?term=LOC105280956 | 2639 | 72 | 272.83 | 1 | 3.79 | 1.39 | 17.08 | 18.51 | -1.43 | 1.43 | -0.04 | 0.04 |
| gene-LOC105281266 | https://www.ncbi.nlm.nih.gov/gene/?term=LOC105281266 | 4102 | 81 | 197.46 | 1 | 2.44 | 1.23 | 16.07 | 17.51 | -1.44 | 1.44 | -0.03 | 0.03 |
| gene-LOC105282156 | https://www.ncbi.nlm.nih.gov/gene/?term=LOC105282156 | 2577 | 133 | 516.1 | 1 | 3.88 | 0.75 | 22.7 | 24.15 | -1.45 | 1.45 | -0.07 | 0.07 |
| gene-LOC105287648 | https://www.ncbi.nlm.nih.gov/gene/?term=LOC105287648 | 2205 | 35 | 158.73 | 1 | 4.54 | 2.86 | 35.94 | 37.4 | -1.45 | 1.45 | -0.02 | 0.02 |
| gene-LOC105280786 | https://www.ncbi.nlm.nih.gov/gene/?term=LOC105280786 | 2573 | 109 | 423.63 | 1 | 3.89 | 0.92 | 28.63 | 30.24 | -1.61 | 1.61 | -0.07 | 0.07 |
| gene-LOC105275384 | https://www.ncbi.nlm.nih.gov/gene/?term=LOC105275384 | 1987 | 87 | 437.85 | 1 | 5.03 | 1.15 | 34.15 | 35.77 | -1.63 | 1.63 | -0.07 | 0.07 |
| gene-LOC105282600 | https://www.ncbi.nlm.nih.gov/gene/?term=LOC105282600 | 3715 | 52 | 139.97 | 1 | 2.69 | 1.92 | 31.36 | 33.08 | -1.72 | 1.72 | -0.02 | 0.02 |
| gene-LOC105283943 | https://www.ncbi.nlm.nih.gov/gene/?term=LOC105283943 | 3559 | 50 | 140.49 | 1 | 2.81 | 2 | 16.35 | 18.09 | -1.74 | 1.74 | -0.02 | 0.02 |
| gene-LOC105278436 | https://www.ncbi.nlm.nih.gov/gene/?term=LOC105278436 | 2556 | 54 | 211.27 | 1 | 3.91 | 1.85 | 41.69 | 43.45 | -1.76 | 1.76 | -0.04 | 0.04 |
| gene-LOC105288066 | https://www.ncbi.nlm.nih.gov/gene/?term=LOC105288066 | 1964 | 29 | 147.66 | 1 | 5.09 | 3.45 | 35.48 | 37.25 | -1.78 | 1.78 | -0.03 | 0.03 |
| gene-LOC105284329 | https://www.ncbi.nlm.nih.gov/gene/?term=LOC105284329 | 4051 | 50 | 123.43 | 1 | 2.47 | 2 | 17.13 | 19.12 | -1.99 | 1.99 | -0.02 | 0.02 |
| gene-LOC105285202 | https://www.ncbi.nlm.nih.gov/gene/?term=LOC105285202 | 1838 | 60 | 326.44 | 1 | 5.44 | 1.67 | 23.97 | 26.11 | -2.15 | 2.15 | -0.07 | 0.07 |
| gene-LOC105283592 | https://www.ncbi.nlm.nih.gov/gene/?term=LOC105283592 | 3479 | 64 | 183.96 | 1 | 2.87 | 1.56 | 24.19 | 26.35 | -2.16 | 2.16 | -0.04 | 0.04 |
| gene-LOC105278454 | https://www.ncbi.nlm.nih.gov/gene/?term=LOC105278454 | 1972 | 43 | 218.05 | 1 | 5.07 | 2.33 | 44.38 | 46.55 | -2.17 | 2.17 | -0.05 | 0.05 |
| gene-LOC105276449 | https://www.ncbi.nlm.nih.gov/gene/?term=LOC105276449 | 1231 | 45 | 365.56 | 1 | 8.12 | 2.22 | 27.05 | 29.32 | -2.27 | 2.27 | -0.08 | 0.08 |
| gene-LOC105283649 | https://www.ncbi.nlm.nih.gov/gene/?term=LOC105283649 | 1918 | 56 | 291.97 | 1 | 5.21 | 1.79 | 36.17 | 38.48 | -2.31 | 2.31 | -0.07 | 0.07 |
| gene-LOC105285674 | https://www.ncbi.nlm.nih.gov/gene/?term=LOC105285674 | 3035 | 84 | 276.77 | 2 | 6.59 | 2.38 | 24.69 | 27.77 | -3.08 | 3.08 | -0.09 | 0.09 |
| gene-LOC105277903 | https://www.ncbi.nlm.nih.gov/gene/?term=LOC105277903 | 1835 | 36 | 196.19 | 1 | 5.45 | 2.78 | 53.1 | 56.25 | -3.14 | 3.14 | -0.06 | 0.06 |
| gene-LOC105281515 | https://www.ncbi.nlm.nih.gov/gene/?term=LOC105281515 | 8827 | 60 | 67.97 | 2 | 2.27 | 3.33 | 26.32 | 29.67 | -3.35 | 3.35 | -0.02 | 0.02 |
| gene-LOC105281420 | https://www.ncbi.nlm.nih.gov/gene/?term=LOC105281420 | 3882 | 55 | 141.68 | 1 | 2.58 | 1.82 | 13.05 | 16.87 | -3.82 | 3.82 | -0.05 | 0.05 |
| gene-LOC105285514 | https://www.ncbi.nlm.nih.gov/gene/?term=LOC105285514 | 1277 | 11 | 86.14 | 1 | 7.83 | 9.09 | 70.96 | 74.98 | -4.02 | 4.02 | -0.03 | 0.03 |
| gene-LOC105274900 | https://www.ncbi.nlm.nih.gov/gene/?term=LOC105274900 | 5561 | 54 | 97.1 | 3 | 5.39 | 5.56 | 30 | 35.69 | -5.68 | 5.68 | -0.06 | 0.06 |
| gene-LOC105281039 | https://www.ncbi.nlm.nih.gov/gene/?term=LOC105281039 | 1471 | 11 | 74.78 | 1 | 6.8 | 9.09 | 29.26 | 35.1 | -5.84 | 5.84 | -0.04 | 0.04 |
| gene-LOC105279898 | https://www.ncbi.nlm.nih.gov/gene/?term=LOC105279898 | 1343 | 11 | 81.91 | 1 | 7.45 | 9.09 | 61.61 | 68.61 | -7 | 7 | -0.06 | 0.06 |
| gene-LOC105278567 | https://www.ncbi.nlm.nih.gov/gene/?term=LOC105278567 | 3627 | 29 | 79.96 | 1 | 2.76 | 3.45 | 11.32 | 18.71 | -7.39 | 7.39 | -0.06 | 0.06 |

| | | | | | | | | | | | | | |
|---|---|---|---|---|---|---|---|---|---|---|---|---|---|
| gene-LOC105275708 | https://www.ncbi.nlm.nih.gov/gene/?term=LOC105275708 | 1772 | 15 | 84.65 | 1 | 5.64 | 6.67 | 45.9 | 54.1 | -8.2 | 8.2 | -0.07 | 0.07 |
| gene-LOC105275070 | https://www.ncbi.nlm.nih.gov/gene/?term=LOC105275070 | 5667 | 40 | 70.58 | 6 | 10.59 | 15 | 2.13 | 14.38 | -12.25 | 12.25 | -0.09 | 0.09 |
| gene-LOC113562345 | https://www.ncbi.nlm.nih.gov/gene/?term=LOC113562345 | 2290 | 63 | 275.11 | 8 | 34.93 | 12.7 | 17.51 | 30.84 | -13.33 | 13.33 | -0.37 | 0.37 |

Directory: OGL File: ogl-Ob_LI_BCphaseB2.vs.RphaseA2.txt

| gene_ID | gene_link | gwidth | #Sites | #per10Kb | #dmSites | #dmsp10kb | %dmSites | %pSite1 | %pSite2 | DMpSite | ADMpSite | DMpNucl | ADMpNucl |
|---|---|---|---|---|---|---|---|---|---|---|---|---|---|
| gene-LOC105284677 | https://www.ncbi.nlm.nih.gov/gene/?term=LOC105284677 | 1905 | 43 | 225.72 | 2 | 10.5 | 4.65 | 38.99 | 29.38 | 9.62 | 9.62 | 0.22 | 0.22 |
| gene-LOC105282914 | https://www.ncbi.nlm.nih.gov/gene/?term=LOC105282914 | 5623 | 43 | 76.47 | 1 | 1.78 | 2.33 | 49.78 | 47.18 | 2.6 | 2.6 | 0.02 | 0.02 |
| gene-LOC105274449 | https://www.ncbi.nlm.nih.gov/gene/?term=LOC105274449 | 1549 | 53 | 342.16 | 1 | 6.46 | 1.89 | 51.77 | 49.51 | 2.26 | 2.26 | 0.08 | 0.08 |
| gene-LOC105285409 | https://www.ncbi.nlm.nih.gov/gene/?term=LOC105285409 | 5099 | 63 | 123.55 | 1 | 1.96 | 1.59 | 3.13 | 1.16 | 1.97 | 1.97 | 0.02 | 0.02 |
| gene-LOC105282885 | https://www.ncbi.nlm.nih.gov/gene/?term=LOC105282885 | 1870 | 49 | 262.03 | 1 | 5.35 | 2.04 | 39.47 | 37.7 | 1.77 | 1.77 | 0.05 | 0.05 |
| gene-LOC105279500 | https://www.ncbi.nlm.nih.gov/gene/?term=LOC105279500 | 3587 | 33 | 92 | 1 | 2.79 | 3.03 | 47.01 | 45.48 | 1.53 | 1.53 | 0.01 | 0.01 |
| gene-LOC105275743 | https://www.ncbi.nlm.nih.gov/gene/?term=LOC105275743 | 1782 | 76 | 426.49 | 1 | 5.61 | 1.32 | 23.56 | 22.24 | 1.32 | 1.32 | 0.06 | 0.06 |
| gene-LOC105281296 | https://www.ncbi.nlm.nih.gov/gene/?term=LOC105281296 | 2434 | 43 | 176.66 | 1 | 4.11 | 2.33 | 29.93 | 28.76 | 1.17 | 1.17 | 0.02 | 0.02 |
| gene-LOC105285252 | https://www.ncbi.nlm.nih.gov/gene/?term=LOC105285252 | 7427 | 125 | 168.3 | 1 | 1.35 | 0.8 | 14.66 | 13.59 | 1.08 | 1.08 | 0.02 | 0.02 |
| gene-LOC105284396 | https://www.ncbi.nlm.nih.gov/gene/?term=LOC105284396 | 1976 | 70 | 354.25 | 1 | 5.06 | 1.43 | 32.98 | 31.96 | 1.02 | 1.02 | 0.04 | 0.04 |
| gene-LOC105280819 | https://www.ncbi.nlm.nih.gov/gene/?term=LOC105280819 | 1639 | 43 | 262.36 | 1 | 6.1 | 2.33 | 39.04 | 38.17 | 0.87 | 0.87 | 0.02 | 0.02 |
| gene-LOC105288105 | https://www.ncbi.nlm.nih.gov/gene/?term=LOC105288105 | 1460 | 37 | 253.42 | 1 | 6.85 | 2.7 | 53 | 52.16 | 0.84 | 0.84 | 0.02 | 0.02 |
| gene-LOC105275587 | https://www.ncbi.nlm.nih.gov/gene/?term=LOC105275587 | 2404 | 57 | 237.1 | 1 | 4.16 | 1.75 | 36.63 | 35.9 | 0.73 | 0.73 | 0.02 | 0.02 |
| gene-LOC105276272 | https://www.ncbi.nlm.nih.gov/gene/?term=LOC105276272 | 1965 | 50 | 254.45 | 1 | 5.09 | 2 | 27.69 | 27.01 | 0.68 | 0.68 | 0.02 | 0.02 |
| gene-LOC105287633 | https://www.ncbi.nlm.nih.gov/gene/?term=LOC105287633 | 1524 | 35 | 229.66 | 1 | 6.56 | 2.86 | 49.8 | 49.12 | 0.68 | 0.68 | 0.02 | 0.02 |
| gene-LOC105281173 | https://www.ncbi.nlm.nih.gov/gene/?term=LOC105281173 | 1595 | 38 | 238.24 | 1 | 6.27 | 2.63 | 60.67 | 60.19 | 0.48 | 0.48 | 0.01 | 0.01 |
| gene-LOC105280304 | https://www.ncbi.nlm.nih.gov/gene/?term=LOC105280304 | 8919 | 115 | 128.94 | 1 | 1.12 | 0.87 | 7.4 | 6.97 | 0.43 | 0.43 | 0.01 | 0.01 |
| gene-LOC105284507 | https://www.ncbi.nlm.nih.gov/gene/?term=LOC105284507 | 5207 | 91 | 174.76 | 1 | 1.92 | 1.1 | 18.91 | 18.57 | 0.35 | 0.35 | 0.01 | 0.01 |
| gene-LOC105284329 | https://www.ncbi.nlm.nih.gov/gene/?term=LOC105284329 | 4051 | 53 | 130.83 | 1 | 2.47 | 1.89 | 19.36 | 19.04 | 0.32 | 0.32 | 0 | 0 |
| gene-LOC105283296 | https://www.ncbi.nlm.nih.gov/gene/?term=LOC105283296 | 2174 | 96 | 441.58 | 1 | 4.6 | 1.04 | 20.81 | 20.51 | 0.3 | 0.3 | 0.01 | 0.01 |
| gene-LOC105286770 | https://www.ncbi.nlm.nih.gov/gene/?term=LOC105286770 | 3055 | 38 | 124.39 | 1 | 3.27 | 2.63 | 15.65 | 15.48 | 0.17 | 0.17 | 0 | 0 |
| gene-LOC105287842 | https://www.ncbi.nlm.nih.gov/gene/?term=LOC105287842 | 2301 | 57 | 247.72 | 1 | 4.35 | 1.75 | 21.58 | 21.41 | 0.17 | 0.17 | 0 | 0 |
| gene-LOC105285496 | https://www.ncbi.nlm.nih.gov/gene/?term=LOC105285496 | 4080 | 112 | 274.51 | 1 | 2.45 | 0.89 | 40.26 | 40.12 | 0.14 | 0.14 | 0 | 0 |
| gene-LOC105278534 | https://www.ncbi.nlm.nih.gov/gene/?term=LOC105278534 | 7364 | 200 | 271.59 | 2 | 2.72 | 1 | 17.58 | 17.48 | 0.1 | 0.1 | 0 | 0 |
| gene-LOC105281312 | https://www.ncbi.nlm.nih.gov/gene/?term=LOC105281312 | 2551 | 74 | 290.08 | 1 | 3.92 | 1.35 | 29.31 | 29.21 | 0.1 | 0.1 | 0 | 0 |
| gene-LOC105282951 | https://www.ncbi.nlm.nih.gov/gene/?term=LOC105282951 | 3366 | 131 | 389.19 | 1 | 2.97 | 0.76 | 18.77 | 18.69 | 0.08 | 0.08 | 0 | 0 |
| gene-LOC105277273 | https://www.ncbi.nlm.nih.gov/gene/?term=LOC105277273 | 3711 | 169 | 455.4 | 1 | 2.69 | 0.59 | 12.24 | 12.17 | 0.06 | 0.06 | 0 | 0 |
| gene-LOC105281700 | https://www.ncbi.nlm.nih.gov/gene/?term=LOC105281700 | 2208 | 39 | 176.63 | 1 | 4.53 | 2.56 | 37.83 | 37.79 | 0.04 | 0.04 | 0 | 0 |
| gene-LOC105284184 | https://www.ncbi.nlm.nih.gov/gene/?term=LOC105284184 | 2535 | 102 | 402.37 | 1 | 3.94 | 0.98 | 28.97 | 28.93 | 0.03 | 0.03 | 0 | 0 |
| gene-LOC105285533 | https://www.ncbi.nlm.nih.gov/gene/?term=LOC105285533 | 1640 | 71 | 432.93 | 1 | 6.1 | 1.41 | 23.22 | 23.19 | 0.03 | 0.03 | 0 | 0 |
| gene-LOC105281754 | https://www.ncbi.nlm.nih.gov/gene/?term=LOC105281754 | 4300 | 122 | 283.72 | 1 | 2.33 | 0.82 | 16.75 | 16.74 | 0.01 | 0.01 | 0 | 0 |
| gene-LOC105274851 | https://www.ncbi.nlm.nih.gov/gene/?term=LOC105274851 | 6616 | 150 | 226.72 | 1 | 1.51 | 0.67 | 10.51 | 10.53 | -0.02 | 0.02 | 0 | 0 |
| gene-LOC105278691 | https://www.ncbi.nlm.nih.gov/gene/?term=LOC105278691 | 3871 | 172 | 444.33 | 1 | 2.58 | 0.58 | 14.5 | 14.55 | -0.05 | 0.05 | 0 | 0 |
| gene-LOC105286649 | https://www.ncbi.nlm.nih.gov/gene/?term=LOC105286649 | 14178 | 892 | 629.14 | 1 | 0.71 | 0.11 | 1.01 | 1.06 | -0.05 | 0.05 | 0 | 0 |
| gene-LOC105275848 | https://www.ncbi.nlm.nih.gov/gene/?term=LOC105275848 | 3718 | 149 | 400.75 | 1 | 2.69 | 0.67 | 21.41 | 21.47 | -0.06 | 0.06 | 0 | 0 |
| gene-LOC105282646 | https://www.ncbi.nlm.nih.gov/gene/?term=LOC105282646 | 3317 | 110 | 331.62 | 1 | 3.01 | 0.91 | 19.95 | 20.03 | -0.08 | 0.08 | 0 | 0 |
| gene-LOC105276462 | https://www.ncbi.nlm.nih.gov/gene/?term=LOC105276462 | 6561 | 296 | 451.15 | 1 | 1.52 | 0.34 | 7.73 | 7.83 | -0.1 | 0.1 | 0 | 0 |
| gene-LOC105276527 | https://www.ncbi.nlm.nih.gov/gene/?term=LOC105276527 | 7923 | 340 | 429.13 | 1 | 1.26 | 0.29 | 9.36 | 9.47 | -0.11 | 0.11 | 0 | 0 |
| gene-LOC105280408 | https://www.ncbi.nlm.nih.gov/gene/?term=LOC105280408 | 5983 | 60 | 100.28 | 1 | 1.67 | 1.67 | 21.93 | 22.03 | -0.11 | 0.11 | 0 | 0 |
| gene-LOC105280722 | https://www.ncbi.nlm.nih.gov/gene/?term=LOC105280722 | 3237 | 98 | 302.75 | 1 | 3.09 | 1.02 | 29.35 | 29.46 | -0.11 | 0.11 | 0 | 0 |
| gene-LOC105275179 | https://www.ncbi.nlm.nih.gov/gene/?term=LOC105275179 | 3271 | 85 | 259.86 | 1 | 3.06 | 1.18 | 19.41 | 19.53 | -0.12 | 0.12 | 0 | 0 |
| gene-LOC105284262 | https://www.ncbi.nlm.nih.gov/gene/?term=LOC105284262 | 6669 | 247 | 370.37 | 1 | 1.5 | 0.4 | 4.68 | 4.81 | -0.13 | 0.13 | 0 | 0 |
| gene-LOC105274823 | https://www.ncbi.nlm.nih.gov/gene/?term=LOC105274823 | 4620 | 148 | 320.35 | 1 | 2.16 | 0.68 | 20.3 | 20.45 | -0.14 | 0.14 | 0 | 0 |
| gene-LOC105278807 | https://www.ncbi.nlm.nih.gov/gene/?term=LOC105278807 | 2166 | 81 | 373.96 | 1 | 4.62 | 1.23 | 19.2 | 19.33 | -0.14 | 0.14 | -0.01 | 0.01 |
| gene-LOC105280410 | https://www.ncbi.nlm.nih.gov/gene/?term=LOC105280410 | 12580 | 263 | 209.06 | 1 | 0.79 | 0.38 | 19.71 | 19.86 | -0.15 | 0.15 | 0 | 0 |
| gene-LOC105286515 | https://www.ncbi.nlm.nih.gov/gene/?term=LOC105286515 | 3751 | 123 | 327.91 | 1 | 2.67 | 0.81 | 10.35 | 10.5 | -0.15 | 0.15 | 0 | 0 |
| gene-LOC105285179 | https://www.ncbi.nlm.nih.gov/gene/?term=LOC105285179 | 4191 | 183 | 436.65 | 1 | 2.39 | 0.55 | 19.23 | 19.39 | -0.16 | 0.16 | -0.01 | 0.01 |
| gene-LOC105282811 | https://www.ncbi.nlm.nih.gov/gene/?term=LOC105282811 | 5765 | 213 | 369.47 | 1 | 1.73 | 0.47 | 8.07 | 8.24 | -0.17 | 0.17 | -0.01 | 0.01 |
| gene-LOC105287550 | https://www.ncbi.nlm.nih.gov/gene/?term=LOC105287550 | 15659 | 656 | 418.93 | 1 | 0.64 | 0.15 | 6.08 | 6.26 | -0.18 | 0.18 | -0.01 | 0.01 |
| gene-LOC105287095 | https://www.ncbi.nlm.nih.gov/gene/?term=LOC105287095 | 1435 | 65 | 452.96 | 1 | 6.97 | 1.54 | 25.24 | 25.44 | -0.2 | 0.2 | -0.01 | 0.01 |
| gene-LOC105286274 | https://www.ncbi.nlm.nih.gov/gene/?term=LOC105286274 | 3817 | 206 | 539.69 | 1 | 2.62 | 0.49 | 16.61 | 16.82 | -0.21 | 0.21 | -0.01 | 0.01 |
| gene-LOC105274699 | https://www.ncbi.nlm.nih.gov/gene/?term=LOC105274699 | 2386 | 63 | 264.04 | 1 | 4.19 | 1.59 | 22.17 | 22.41 | -0.24 | 0.24 | -0.01 | 0.01 |
| gene-LOC105285406 | https://www.ncbi.nlm.nih.gov/gene/?term=LOC105285406 | 8015 | 59 | 73.61 | 1 | 1.25 | 1.69 | 29.55 | 29.82 | -0.27 | 0.27 | 0 | 0 |
| gene-LOC105276452 | https://www.ncbi.nlm.nih.gov/gene/?term=LOC105276452 | 2917 | 262 | 898.18 | 1 | 3.43 | 0.38 | 9.36 | 9.65 | -0.29 | 0.29 | -0.03 | 0.03 |
| gene-LOC105278145 | https://www.ncbi.nlm.nih.gov/gene/?term=LOC105278145 | 4470 | 252 | 563.76 | 1 | 2.24 | 0.4 | 10.24 | 10.56 | -0.32 | 0.32 | -0.02 | 0.02 |
| gene-LOC105274531 | https://www.ncbi.nlm.nih.gov/gene/?term=LOC105274531 | 7774 | 298 | 383.33 | 1 | 1.29 | 0.34 | 10.39 | 10.72 | -0.33 | 0.33 | -0.01 | 0.01 |
| gene-LOC105275327 | https://www.ncbi.nlm.nih.gov/gene/?term=LOC105275327 | 14245 | 568 | 398.74 | 1 | 0.7 | 0.18 | 6.19 | 6.55 | -0.36 | 0.36 | -0.01 | 0.01 |
| gene-LOC105282067 | https://www.ncbi.nlm.nih.gov/gene/?term=LOC105282067 | 5941 | 44 | 74.06 | 1 | 1.68 | 2.27 | 30.63 | 31 | -0.36 | 0.36 | 0 | 0 |
| gene-LOC105285163 | https://www.ncbi.nlm.nih.gov/gene/?term=LOC105285163 | 5061 | 85 | 167.95 | 1 | 1.98 | 1.18 | 19.84 | 20.21 | -0.37 | 0.37 | -0.01 | 0.01 |
| gene-LOC105278771 | https://www.ncbi.nlm.nih.gov/gene/?term=LOC105278771 | 10457 | 248 | 237.16 | 1 | 0.96 | 0.4 | 8.97 | 9.36 | -0.39 | 0.39 | -0.01 | 0.01 |

| | | | | | | | | | | | | | |
|---|---|---|---|---|---|---|---|---|---|---|---|---|---|
| gene-LOC105282794 | https://www.ncbi.nlm.nih.gov/gene/?term=LOC105282794 | 4571 | 100 | 218.77 | 1 | 2.19 | 1 | 16.99 | 17.37 | -0.39 | 0.39 | -0.01 | 0.01 |
| gene-LOC105283123 | https://www.ncbi.nlm.nih.gov/gene/?term=LOC105283123 | 1507 | 57 | 378.23 | 1 | 6.64 | 1.75 | 17.49 | 17.88 | -0.39 | 0.39 | -0.01 | 0.01 |
| gene-LOC105287309 | https://www.ncbi.nlm.nih.gov/gene/?term=LOC105287309 | 3553 | 96 | 270.19 | 1 | 2.81 | 1.04 | 7.77 | 8.16 | -0.39 | 0.39 | -0.01 | 0.01 |
| gene-LOC105288055 | https://www.ncbi.nlm.nih.gov/gene/?term=LOC105288055 | 9934 | 548 | 551.64 | 1 | 1.01 | 0.18 | 7.05 | 7.45 | -0.4 | 0.4 | -0.02 | 0.02 |
| gene-LOC105275374 | https://www.ncbi.nlm.nih.gov/gene/?term=LOC105275374 | 11376 | 567 | 498.42 | 1 | 0.88 | 0.18 | 10.6 | 11.02 | -0.41 | 0.41 | -0.02 | 0.02 |
| gene-LOC105285742 | https://www.ncbi.nlm.nih.gov/gene/?term=LOC105285742 | 3446 | 132 | 383.05 | 1 | 2.9 | 0.76 | 18.9 | 19.31 | -0.41 | 0.41 | -0.02 | 0.02 |
| gene-LOC105284043 | https://www.ncbi.nlm.nih.gov/gene/?term=LOC105284043 | 5661 | 325 | 574.1 | 1 | 1.77 | 0.31 | 9.91 | 10.34 | -0.43 | 0.43 | -0.02 | 0.02 |
| gene-LOC105278666 | https://www.ncbi.nlm.nih.gov/gene/?term=LOC105278666 | 10341 | 494 | 477.71 | 1 | 0.97 | 0.2 | 6.74 | 7.18 | -0.44 | 0.44 | -0.02 | 0.02 |
| gene-LOC105281272 | https://www.ncbi.nlm.nih.gov/gene/?term=LOC105281272 | 5252 | 203 | 386.52 | 1 | 1.9 | 0.49 | 14.55 | 14.99 | -0.44 | 0.44 | -0.02 | 0.02 |
| gene-LOC105278283 | https://www.ncbi.nlm.nih.gov/gene/?term=LOC105278283 | 2317 | 146 | 630.13 | 1 | 4.32 | 0.68 | 24.36 | 24.8 | -0.45 | 0.45 | -0.03 | 0.03 |
| gene-LOC105274669 | https://www.ncbi.nlm.nih.gov/gene/?term=LOC105274669 | 8555 | 293 | 342.49 | 1 | 1.17 | 0.34 | 5.34 | 5.82 | -0.48 | 0.48 | -0.02 | 0.02 |
| gene-LOC105275363 | https://www.ncbi.nlm.nih.gov/gene/?term=LOC105275363 | 3798 | 176 | 463.4 | 1 | 2.63 | 0.57 | 15.74 | 16.23 | -0.5 | 0.5 | -0.02 | 0.02 |
| gene-LOC105287648 | https://www.ncbi.nlm.nih.gov/gene/?term=LOC105287648 | 2205 | 35 | 158.73 | 1 | 4.54 | 2.86 | 35.94 | 36.45 | -0.5 | 0.5 | -0.01 | 0.01 |
| gene-LOC113562050 | https://www.ncbi.nlm.nih.gov/gene/?term=LOC113562050 | 17097 | 643 | 376.09 | 1 | 0.58 | 0.16 | 4.96 | 5.49 | -0.53 | 0.53 | -0.02 | 0.02 |
| gene-LOC105276059 | https://www.ncbi.nlm.nih.gov/gene/?term=LOC105276059 | 6256 | 145 | 231.78 | 1 | 1.6 | 0.69 | 8.95 | 9.49 | -0.54 | 0.54 | -0.01 | 0.01 |
| gene-LOC105275460 | https://www.ncbi.nlm.nih.gov/gene/?term=LOC105275460 | 2473 | 122 | 493.33 | 1 | 4.04 | 0.82 | 12.96 | 13.51 | -0.55 | 0.55 | -0.03 | 0.03 |
| gene-LOC105284036 | https://www.ncbi.nlm.nih.gov/gene/?term=LOC105284036 | 3254 | 112 | 344.19 | 1 | 3.07 | 0.89 | 21.95 | 22.5 | -0.55 | 0.55 | -0.02 | 0.02 |
| gene-LOC105283728 | https://www.ncbi.nlm.nih.gov/gene/?term=LOC105283728 | 3256 | 123 | 377.76 | 2 | 6.14 | 1.63 | 18.65 | 19.22 | -0.57 | 0.57 | -0.02 | 0.02 |
| gene-LOC105276875 | https://www.ncbi.nlm.nih.gov/gene/?term=LOC105276875 | 3151 | 124 | 393.53 | 1 | 3.17 | 0.81 | 21.3 | 21.9 | -0.6 | 0.6 | -0.02 | 0.02 |
| gene-LOC105274706 | https://www.ncbi.nlm.nih.gov/gene/?term=LOC105274706 | 1446 | 57 | 394.19 | 1 | 6.92 | 1.75 | 36.89 | 37.51 | -0.61 | 0.61 | -0.02 | 0.02 |
| gene-LOC105285502 | https://www.ncbi.nlm.nih.gov/gene/?term=LOC105285502 | 9990 | 164 | 164.16 | 1 | 1 | 0.61 | 8.55 | 9.16 | -0.62 | 0.62 | -0.01 | 0.01 |
| gene-LOC105285807 | https://www.ncbi.nlm.nih.gov/gene/?term=LOC105285807 | 6063 | 237 | 390.9 | 1 | 1.65 | 0.42 | 8.52 | 9.16 | -0.64 | 0.64 | -0.03 | 0.03 |
| gene-LOC105274452 | https://www.ncbi.nlm.nih.gov/gene/?term=LOC105274452 | 6595 | 279 | 423.05 | 1 | 1.52 | 0.36 | 12.34 | 12.98 | -0.65 | 0.65 | -0.03 | 0.03 |
| gene-LOC105276683 | https://www.ncbi.nlm.nih.gov/gene/?term=LOC105276683 | 17000 | 149 | 87.65 | 1 | 0.59 | 0.67 | 8.22 | 8.88 | -0.65 | 0.65 | -0.01 | 0.01 |
| gene-LOC105280723 | https://www.ncbi.nlm.nih.gov/gene/?term=LOC105280723 | 3572 | 185 | 517.92 | 1 | 2.8 | 0.54 | 15.07 | 15.74 | -0.67 | 0.67 | -0.03 | 0.03 |
| gene-LOC105286452 | https://www.ncbi.nlm.nih.gov/gene/?term=LOC105286452 | 6888 | 66 | 95.82 | 1 | 1.45 | 1.52 | 30.11 | 30.78 | -0.67 | 0.67 | -0.01 | 0.01 |
| gene-LOC105279657 | https://www.ncbi.nlm.nih.gov/gene/?term=LOC105279657 | 5878 | 91 | 154.81 | 1 | 1.7 | 1.1 | 18.9 | 19.62 | -0.72 | 0.72 | -0.01 | 0.01 |
| gene-LOC105275752 | https://www.ncbi.nlm.nih.gov/gene/?term=LOC105275752 | 3462 | 141 | 407.28 | 1 | 2.89 | 0.71 | 30.44 | 31.2 | -0.75 | 0.75 | -0.03 | 0.03 |
| gene-LOC105275451 | https://www.ncbi.nlm.nih.gov/gene/?term=LOC105275451 | 6035 | 142 | 235.29 | 1 | 1.66 | 0.7 | 20.66 | 21.45 | -0.78 | 0.78 | -0.02 | 0.02 |
| gene-LOC105280348 | https://www.ncbi.nlm.nih.gov/gene/?term=LOC105280348 | 3797 | 79 | 208.06 | 1 | 2.63 | 1.27 | 19.69 | 20.49 | -0.8 | 0.8 | -0.02 | 0.02 |
| gene-LOC105283967 | https://www.ncbi.nlm.nih.gov/gene/?term=LOC105283967 | 2393 | 60 | 250.73 | 1 | 4.18 | 1.67 | 41.56 | 42.36 | -0.8 | 0.8 | -0.02 | 0.02 |
| gene-LOC105285395 | https://www.ncbi.nlm.nih.gov/gene/?term=LOC105285395 | 8085 | 224 | 277.06 | 1 | 1.24 | 0.45 | 14.28 | 15.08 | -0.8 | 0.8 | -0.02 | 0.02 |
| gene-LOC105279038 | https://www.ncbi.nlm.nih.gov/gene/?term=LOC105279038 | 9453 | 369 | 390.35 | 1 | 1.06 | 0.27 | 8.12 | 8.94 | -0.82 | 0.82 | -0.03 | 0.03 |
| gene-LOC105275187 | https://www.ncbi.nlm.nih.gov/gene/?term=LOC105275187 | 5534 | 77 | 139.14 | 1 | 1.81 | 1.3 | 15.38 | 16.21 | -0.83 | 0.83 | -0.01 | 0.01 |
| gene-LOC105280407 | https://www.ncbi.nlm.nih.gov/gene/?term=LOC105280407 | 10950 | 218 | 199.09 | 1 | 0.91 | 0.46 | 16.66 | 17.49 | -0.83 | 0.83 | -0.02 | 0.02 |
| gene-LOC105278228 | https://www.ncbi.nlm.nih.gov/gene/?term=LOC105278228 | 8610 | 176 | 204.41 | 1 | 1.16 | 0.57 | 22.43 | 23.28 | -0.84 | 0.84 | -0.02 | 0.02 |
| gene-LOC105278632 | https://www.ncbi.nlm.nih.gov/gene/?term=LOC105278632 | 3624 | 141 | 389.07 | 1 | 2.76 | 0.71 | 13.56 | 14.44 | -0.87 | 0.87 | -0.03 | 0.03 |
| gene-LOC105285055 | https://www.ncbi.nlm.nih.gov/gene/?term=LOC105285055 | 4108 | 274 | 666.99 | 2 | 4.87 | 0.73 | 10.9 | 11.79 | -0.89 | 0.89 | -0.06 | 0.06 |
| gene-LOC105282491 | https://www.ncbi.nlm.nih.gov/gene/?term=LOC105282491 | 4761 | 184 | 386.47 | 1 | 2.1 | 0.54 | 8.25 | 9.16 | -0.91 | 0.91 | -0.04 | 0.04 |
| gene-LOC105282682 | https://www.ncbi.nlm.nih.gov/gene/?term=LOC105282682 | 4127 | 241 | 583.96 | 1 | 2.42 | 0.41 | 12.88 | 13.79 | -0.91 | 0.91 | -0.05 | 0.05 |
| gene-LOC105278877 | https://www.ncbi.nlm.nih.gov/gene/?term=LOC105278877 | 1968 | 86 | 436.99 | 1 | 5.08 | 1.16 | 31.6 | 32.52 | -0.92 | 0.92 | -0.04 | 0.04 |
| gene-LOC105285202 | https://www.ncbi.nlm.nih.gov/gene/?term=LOC105285202 | 1838 | 66 | 359.09 | 1 | 5.44 | 1.52 | 24.58 | 25.52 | -0.94 | 0.94 | -0.03 | 0.03 |
| gene-LOC105274590 | https://www.ncbi.nlm.nih.gov/gene/?term=LOC105274590 | 4516 | 80 | 177.15 | 2 | 4.43 | 2.5 | 29.75 | 30.69 | -0.95 | 0.95 | -0.02 | 0.02 |
| gene-LOC105276917 | https://www.ncbi.nlm.nih.gov/gene/?term=LOC105276917 | 2336 | 33 | 141.27 | 1 | 4.28 | 3.03 | 30.13 | 31.08 | -0.95 | 0.95 | -0.01 | 0.01 |
| gene-LOC105282825 | https://www.ncbi.nlm.nih.gov/gene/?term=LOC105282825 | 3919 | 98 | 250.06 | 1 | 2.55 | 1.02 | 14.18 | 15.14 | -0.96 | 0.96 | -0.02 | 0.02 |
| gene-LOC105284117 | https://www.ncbi.nlm.nih.gov/gene/?term=LOC105284117 | 4340 | 101 | 232.72 | 1 | 2.3 | 0.99 | 10.38 | 11.36 | -0.98 | 0.98 | -0.02 | 0.02 |
| gene-LOC105280774 | https://www.ncbi.nlm.nih.gov/gene/?term=LOC105280774 | 2793 | 122 | 436.81 | 1 | 3.58 | 0.82 | 26.14 | 27.14 | -0.99 | 0.99 | -0.04 | 0.04 |
| gene-LOC105281261 | https://www.ncbi.nlm.nih.gov/gene/?term=LOC105281261 | 5947 | 158 | 265.68 | 1 | 1.68 | 0.63 | 14.18 | 15.18 | -1 | 1 | -0.03 | 0.03 |
| gene-LOC105275683 | https://www.ncbi.nlm.nih.gov/gene/?term=LOC105275683 | 4256 | 210 | 493.42 | 1 | 2.35 | 0.48 | 13.47 | 14.47 | -1.01 | 1.01 | -0.05 | 0.05 |
| gene-LOC105278497 | https://www.ncbi.nlm.nih.gov/gene/?term=LOC105278497 | 2072 | 103 | 497.1 | 1 | 4.83 | 0.97 | 19.13 | 20.15 | -1.02 | 1.02 | -0.05 | 0.05 |
| gene-LOC105281848 | https://www.ncbi.nlm.nih.gov/gene/?term=LOC105281848 | 2122 | 64 | 301.6 | 1 | 4.71 | 1.56 | 21.13 | 22.15 | -1.02 | 1.02 | -0.03 | 0.03 |
| gene-LOC105285639 | https://www.ncbi.nlm.nih.gov/gene/?term=LOC105285639 | 10325 | 408 | 395.16 | 1 | 0.97 | 0.25 | 10.25 | 11.26 | -1.02 | 1.02 | -0.04 | 0.04 |
| gene-LOC105284116 | https://www.ncbi.nlm.nih.gov/gene/?term=LOC105284116 | 3970 | 113 | 284.63 | 1 | 2.52 | 0.88 | 7.21 | 8.25 | -1.03 | 1.03 | -0.03 | 0.03 |
| gene-LOC105276196 | https://www.ncbi.nlm.nih.gov/gene/?term=LOC105276196 | 2953 | 59 | 199.8 | 1 | 3.39 | 1.69 | 37.63 | 38.7 | -1.07 | 1.07 | -0.02 | 0.02 |
| gene-LOC105283943 | https://www.ncbi.nlm.nih.gov/gene/?term=LOC105283943 | 3559 | 64 | 179.83 | 1 | 2.81 | 1.56 | 14.69 | 15.79 | -1.1 | 1.1 | -0.02 | 0.02 |
| gene-LOC105276501 | https://www.ncbi.nlm.nih.gov/gene/?term=LOC105276501 | 6089 | 110 | 180.65 | 1 | 1.64 | 0.91 | 15.38 | 16.49 | -1.11 | 1.11 | -0.02 | 0.02 |
| gene-LOC105275347 | https://www.ncbi.nlm.nih.gov/gene/?term=LOC105275347 | 2526 | 20 | 79.18 | 1 | 3.96 | 5 | 37.1 | 38.21 | -1.12 | 1.12 | -0.01 | 0.01 |
| gene-LOC105285514 | https://www.ncbi.nlm.nih.gov/gene/?term=LOC105285514 | 1277 | 15 | 117.46 | 1 | 7.83 | 6.67 | 78.25 | 79.38 | -1.12 | 1.12 | -0.01 | 0.01 |
| gene-LOC105284355 | https://www.ncbi.nlm.nih.gov/gene/?term=LOC105284355 | 3904 | 165 | 422.64 | 1 | 2.56 | 0.61 | 19.1 | 20.26 | -1.16 | 1.16 | -0.05 | 0.05 |
| gene-LOC105277631 | https://www.ncbi.nlm.nih.gov/gene/?term=LOC105277631 | 4387 | 333 | 759.06 | 1 | 2.28 | 0.3 | 9.46 | 10.66 | -1.19 | 1.19 | -0.09 | 0.09 |
| gene-LOC105288078 | https://www.ncbi.nlm.nih.gov/gene/?term=LOC105288078 | 19126 | 332 | 173.59 | 1 | 0.52 | 0.3 | 16.16 | 17.36 | -1.2 | 1.2 | -0.02 | 0.02 |
| gene-LOC105277761 | https://www.ncbi.nlm.nih.gov/gene/?term=LOC105277761 | 4648 | 160 | 344.23 | 1 | 2.15 | 0.62 | 14.78 | 16.05 | -1.27 | 1.27 | -0.04 | 0.04 |
| gene-LOC105274856 | https://www.ncbi.nlm.nih.gov/gene/?term=LOC105274856 | 5216 | 232 | 444.79 | 1 | 1.92 | 0.43 | 17.14 | 18.43 | -1.29 | 1.29 | -0.06 | 0.06 |
| gene-LOC105283712 | https://www.ncbi.nlm.nih.gov/gene/?term=LOC105283712 | 1910 | 36 | 188.48 | 1 | 5.24 | 2.78 | 28.48 | 29.79 | -1.31 | 1.31 | -0.02 | 0.02 |

| | | | | | | | | | | | | | |
|---|---|---|---|---|---|---|---|---|---|---|---|---|---|
| gene-LOC105280641 | https://www.ncbi.nlm.nih.gov/gene/?term=LOC105280641 | 3682 | 61 | 165.67 | 1 | 2.72 | 1.64 | 16.75 | 18.07 | -1.32 | 1.32 | -0.02 | 0.02 |
| gene-LOC105282220 | https://www.ncbi.nlm.nih.gov/gene/?term=LOC105282220 | 2298 | 224 | 974.76 | 1 | 4.35 | 0.45 | 7.52 | 8.84 | -1.32 | 1.32 | -0.13 | 0.13 |
| gene-LOC105282991 | https://www.ncbi.nlm.nih.gov/gene/?term=LOC105282991 | 5617 | 112 | 199.39 | 1 | 1.78 | 0.89 | 19.26 | 20.58 | -1.32 | 1.32 | -0.03 | 0.03 |
| gene-LOC105274534 | https://www.ncbi.nlm.nih.gov/gene/?term=LOC105274534 | 1329 | 61 | 458.99 | 1 | 7.52 | 1.64 | 40.79 | 42.14 | -1.35 | 1.35 | -0.06 | 0.06 |
| gene-LOC105278962 | https://www.ncbi.nlm.nih.gov/gene/?term=LOC105278962 | 4491 | 107 | 238.25 | 1 | 2.23 | 0.93 | 20.7 | 22.05 | -1.35 | 1.35 | -0.03 | 0.03 |
| gene-LOC105280956 | https://www.ncbi.nlm.nih.gov/gene/?term=LOC105280956 | 2639 | 77 | 291.78 | 1 | 3.79 | 1.3 | 16.02 | 17.39 | -1.37 | 1.37 | -0.04 | 0.04 |
| gene-LOC105279664 | https://www.ncbi.nlm.nih.gov/gene/?term=LOC105279664 | 4023 | 74 | 183.94 | 1 | 2.49 | 1.35 | 16.49 | 17.94 | -1.45 | 1.45 | -0.03 | 0.03 |
| gene-LOC105279298 | https://www.ncbi.nlm.nih.gov/gene/?term=LOC105279298 | 2773 | 138 | 497.66 | 2 | 7.21 | 1.45 | 22.5 | 23.97 | -1.47 | 1.47 | -0.07 | 0.07 |
| gene-LOC105278109 | https://www.ncbi.nlm.nih.gov/gene/?term=LOC105278109 | 4029 | 72 | 178.7 | 1 | 2.48 | 1.39 | 42.29 | 43.78 | -1.48 | 1.48 | -0.03 | 0.03 |
| gene-LOC105286799 | https://www.ncbi.nlm.nih.gov/gene/?term=LOC105286799 | 3557 | 188 | 528.54 | 1 | 2.81 | 0.53 | 15.27 | 16.76 | -1.49 | 1.49 | -0.08 | 0.08 |
| gene-LOC105276704 | https://www.ncbi.nlm.nih.gov/gene/?term=LOC105276704 | 2063 | 101 | 489.58 | 2 | 9.69 | 1.98 | 13.1 | 14.6 | -1.5 | 1.5 | -0.07 | 0.07 |
| gene-LOC105282149 | https://www.ncbi.nlm.nih.gov/gene/?term=LOC105282149 | 1787 | 109 | 609.96 | 1 | 5.6 | 0.92 | 41.83 | 43.33 | -1.5 | 1.5 | -0.09 | 0.09 |
| gene-LOC105284557 | https://www.ncbi.nlm.nih.gov/gene/?term=LOC105284557 | 1672 | 84 | 502.39 | 1 | 5.98 | 1.19 | 28.16 | 29.69 | -1.53 | 1.53 | -0.08 | 0.08 |
| gene-LOC105282893 | https://www.ncbi.nlm.nih.gov/gene/?term=LOC105282893 | 2645 | 47 | 177.69 | 1 | 3.78 | 2.13 | 28.96 | 30.53 | -1.56 | 1.56 | -0.03 | 0.03 |
| gene-LOC105278282 | https://www.ncbi.nlm.nih.gov/gene/?term=LOC105278282 | 1775 | 64 | 360.56 | 1 | 5.63 | 1.56 | 33.91 | 35.49 | -1.58 | 1.58 | -0.06 | 0.06 |
| gene-LOC105283989 | https://www.ncbi.nlm.nih.gov/gene/?term=LOC105283989 | 4948 | 32 | 64.67 | 2 | 4.04 | 6.25 | 43.84 | 45.43 | -1.59 | 1.59 | -0.01 | 0.01 |
| gene-LOC105282970 | https://www.ncbi.nlm.nih.gov/gene/?term=LOC105282970 | 1529 | 40 | 261.61 | 1 | 6.54 | 2.5 | 50.96 | 52.56 | -1.6 | 1.6 | -0.04 | 0.04 |
| gene-LOC105278384 | https://www.ncbi.nlm.nih.gov/gene/?term=LOC105278384 | 1390 | 35 | 251.8 | 1 | 7.19 | 2.86 | 45.32 | 46.94 | -1.62 | 1.62 | -0.04 | 0.04 |
| gene-LOC105280002 | https://www.ncbi.nlm.nih.gov/gene/?term=LOC105280002 | 1940 | 28 | 144.33 | 1 | 5.15 | 3.57 | 39.15 | 40.87 | -1.73 | 1.73 | -0.02 | 0.02 |
| gene-LOC105275518 | https://www.ncbi.nlm.nih.gov/gene/?term=LOC105275518 | 2335 | 26 | 111.35 | 1 | 4.28 | 3.85 | 36.28 | 38.04 | -1.76 | 1.76 | -0.02 | 0.02 |
| gene-LOC105287854 | https://www.ncbi.nlm.nih.gov/gene/?term=LOC105287854 | 3644 | 66 | 181.12 | 1 | 2.74 | 1.52 | 15.87 | 17.63 | -1.76 | 1.76 | -0.03 | 0.03 |
| gene-LOC105282600 | https://www.ncbi.nlm.nih.gov/gene/?term=LOC105282600 | 3715 | 56 | 150.74 | 1 | 2.69 | 1.79 | 29.97 | 31.8 | -1.83 | 1.83 | -0.03 | 0.03 |
| gene-LOC105288010 | https://www.ncbi.nlm.nih.gov/gene/?term=LOC105288010 | 3189 | 45 | 141.11 | 1 | 3.14 | 2.22 | 34.76 | 36.59 | -1.83 | 1.83 | -0.03 | 0.03 |
| gene-LOC109611283 | https://www.ncbi.nlm.nih.gov/gene/?term=LOC109611283 | 1451 | 39 | 268.78 | 1 | 6.89 | 2.56 | 35.83 | 37.66 | -1.83 | 1.83 | -0.05 | 0.05 |
| gene-LOC105277856 | https://www.ncbi.nlm.nih.gov/gene/?term=LOC105277856 | 3442 | 37 | 107.5 | 1 | 2.91 | 2.7 | 37.9 | 39.79 | -1.89 | 1.89 | -0.02 | 0.02 |
| gene-LOC105282151 | https://www.ncbi.nlm.nih.gov/gene/?term=LOC105282151 | 1601 | 46 | 287.32 | 2 | 12.49 | 4.35 | 33.88 | 35.82 | -1.94 | 1.94 | -0.06 | 0.06 |
| gene-LOC105283592 | https://www.ncbi.nlm.nih.gov/gene/?term=LOC105283592 | 3479 | 72 | 206.96 | 1 | 2.87 | 1.39 | 25.41 | 27.37 | -1.95 | 1.95 | -0.04 | 0.04 |
| gene-LOC105280738 | https://www.ncbi.nlm.nih.gov/gene/?term=LOC105280738 | 4287 | 41 | 95.64 | 1 | 2.33 | 2.44 | 24.39 | 26.44 | -2.04 | 2.04 | -0.02 | 0.02 |
| gene-LOC105287063 | https://www.ncbi.nlm.nih.gov/gene/?term=LOC105287063 | 1874 | 54 | 288.15 | 1 | 5.34 | 1.85 | 27.46 | 29.52 | -2.06 | 2.06 | -0.06 | 0.06 |
| gene-LOC105275966 | https://www.ncbi.nlm.nih.gov/gene/?term=LOC105275966 | 3299 | 73 | 221.28 | 1 | 3.03 | 1.37 | 43.49 | 45.58 | -2.08 | 2.08 | -0.05 | 0.05 |
| gene-LOC105282156 | https://www.ncbi.nlm.nih.gov/gene/?term=LOC105282156 | 2577 | 145 | 562.67 | 1 | 3.88 | 0.69 | 23.3 | 25.57 | -2.27 | 2.27 | -0.13 | 0.13 |
| gene-LOC105286683 | https://www.ncbi.nlm.nih.gov/gene/?term=LOC105286683 | 1665 | 53 | 318.32 | 1 | 6.01 | 1.89 | 31.4 | 33.71 | -2.31 | 2.31 | -0.07 | 0.07 |
| gene-LOC105281420 | https://www.ncbi.nlm.nih.gov/gene/?term=LOC105281420 | 3882 | 67 | 172.59 | 1 | 2.58 | 1.49 | 12.82 | 15.24 | -2.42 | 2.42 | -0.04 | 0.04 |
| gene-LOC105285315 | https://www.ncbi.nlm.nih.gov/gene/?term=LOC105285315 | 5458 | 26 | 47.64 | 1 | 1.83 | 3.85 | 36.55 | 38.99 | -2.44 | 2.44 | -0.01 | 0.01 |
| gene-LOC105285583 | https://www.ncbi.nlm.nih.gov/gene/?term=LOC105285583 | 1947 | 47 | 241.4 | 1 | 5.14 | 2.13 | 27.05 | 29.68 | -2.63 | 2.63 | -0.06 | 0.06 |
| gene-LOC105276449 | https://www.ncbi.nlm.nih.gov/gene/?term=LOC105276449 | 1231 | 56 | 454.91 | 1 | 8.12 | 1.79 | 30.06 | 32.73 | -2.67 | 2.67 | -0.12 | 0.12 |
| gene-LOC105288063 | https://www.ncbi.nlm.nih.gov/gene/?term=LOC105288063 | 3400 | 57 | 167.65 | 1 | 2.94 | 1.75 | 12.44 | 15.19 | -2.75 | 2.75 | -0.05 | 0.05 |
| gene-LOC105280786 | https://www.ncbi.nlm.nih.gov/gene/?term=LOC105280786 | 2573 | 119 | 462.5 | 1 | 3.89 | 0.84 | 29.73 | 32.5 | -2.77 | 2.77 | -0.13 | 0.13 |
| gene-LOC105276680 | https://www.ncbi.nlm.nih.gov/gene/?term=LOC105276680 | 1977 | 35 | 177.04 | 1 | 5.06 | 2.86 | 58.22 | 61.02 | -2.8 | 2.8 | -0.05 | 0.05 |
| gene-LOC105278454 | https://www.ncbi.nlm.nih.gov/gene/?term=LOC105278454 | 1972 | 51 | 258.62 | 1 | 5.07 | 1.96 | 39.78 | 42.63 | -2.85 | 2.85 | -0.07 | 0.07 |
| gene-LOC105287417 | https://www.ncbi.nlm.nih.gov/gene/?term=LOC105287417 | 1025 | 36 | 351.22 | 1 | 9.76 | 2.78 | 34.17 | 37.2 | -3.03 | 3.03 | -0.11 | 0.11 |
| gene-LOC105283649 | https://www.ncbi.nlm.nih.gov/gene/?term=LOC105283649 | 1918 | 57 | 297.18 | 2 | 10.43 | 3.51 | 35.86 | 38.92 | -3.06 | 3.06 | -0.09 | 0.09 |
| gene-LOC105288066 | https://www.ncbi.nlm.nih.gov/gene/?term=LOC105288066 | 1964 | 32 | 162.93 | 1 | 5.09 | 3.12 | 32.93 | 36 | -3.07 | 3.07 | -0.05 | 0.05 |
| gene-LOC105284609 | https://www.ncbi.nlm.nih.gov/gene/?term=LOC105284609 | 5678 | 35 | 61.64 | 1 | 1.76 | 2.86 | 28.06 | 31.35 | -3.29 | 3.29 | -0.02 | 0.02 |
| gene-LOC105281515 | https://www.ncbi.nlm.nih.gov/gene/?term=LOC105281515 | 8827 | 54 | 61.18 | 2 | 2.27 | 3.7 | 27.93 | 31.25 | -3.31 | 3.31 | -0.02 | 0.02 |
| gene-LOC105276078 | https://www.ncbi.nlm.nih.gov/gene/?term=LOC105276078 | 1051 | 17 | 161.75 | 1 | 9.51 | 5.88 | 25.17 | 28.55 | -3.38 | 3.38 | -0.05 | 0.05 |
| gene-LOC105286514 | https://www.ncbi.nlm.nih.gov/gene/?term=LOC105286514 | 2528 | 47 | 185.92 | 1 | 3.96 | 2.13 | 35.58 | 39.1 | -3.52 | 3.52 | -0.07 | 0.07 |
| gene-LOC113561805 | https://www.ncbi.nlm.nih.gov/gene/?term=LOC113561805 | 3337 | 84 | 251.72 | 1 | 3 | 1.19 | 34.18 | 37.85 | -3.66 | 3.66 | -0.09 | 0.09 |
| gene-LOC105274900 | https://www.ncbi.nlm.nih.gov/gene/?term=LOC105274900 | 5561 | 55 | 98.9 | 2 | 3.6 | 3.64 | 30.44 | 34.13 | -3.69 | 3.69 | -0.04 | 0.04 |
| gene-LOC105280327 | https://www.ncbi.nlm.nih.gov/gene/?term=LOC105280327 | 2670 | 104 | 389.51 | 1 | 3.75 | 0.96 | 27.09 | 30.78 | -3.69 | 3.69 | -0.14 | 0.14 |
| gene-LOC105285674 | https://www.ncbi.nlm.nih.gov/gene/?term=LOC105285674 | 3035 | 97 | 319.6 | 2 | 6.59 | 2.06 | 24.1 | 27.79 | -3.69 | 3.69 | -0.12 | 0.12 |
| gene-LOC105279900 | https://www.ncbi.nlm.nih.gov/gene/?term=LOC105279900 | 2191 | 62 | 282.98 | 1 | 4.56 | 1.61 | 30.24 | 34.06 | -3.83 | 3.83 | -0.11 | 0.11 |
| gene-LOC105276635 | https://www.ncbi.nlm.nih.gov/gene/?term=LOC105276635 | 4962 | 40 | 80.61 | 1 | 2.02 | 2.5 | 23.53 | 27.39 | -3.86 | 3.86 | -0.03 | 0.03 |
| gene-LOC105286644 | https://www.ncbi.nlm.nih.gov/gene/?term=LOC105286644 | 1235 | 20 | 161.94 | 1 | 8.1 | 5 | 37.53 | 41.72 | -4.19 | 4.19 | -0.07 | 0.07 |
| gene-LOC105276538 | https://www.ncbi.nlm.nih.gov/gene/?term=LOC105276538 | 1511 | 19 | 125.74 | 1 | 6.62 | 5.26 | 69.47 | 74.28 | -4.82 | 4.82 | -0.06 | 0.06 |
| gene-LOC105279898 | https://www.ncbi.nlm.nih.gov/gene/?term=LOC105279898 | 1343 | 14 | 104.24 | 1 | 7.45 | 7.14 | 54.89 | 60.18 | -5.29 | 5.29 | -0.06 | 0.06 |
| gene-LOC105286983 | https://www.ncbi.nlm.nih.gov/gene/?term=LOC105286983 | 16901 | 64 | 37.87 | 2 | 1.18 | 3.12 | 0.67 | 5.99 | -5.32 | 5.32 | -0.02 | 0.02 |
| gene-LOC105275708 | https://www.ncbi.nlm.nih.gov/gene/?term=LOC105275708 | 1772 | 18 | 101.58 | 1 | 5.64 | 5.56 | 43.28 | 48.95 | -5.67 | 5.67 | -0.06 | 0.06 |
| gene-LOC105281039 | https://www.ncbi.nlm.nih.gov/gene/?term=LOC105281039 | 1471 | 12 | 81.58 | 1 | 6.8 | 8.33 | 31.46 | 37.55 | -6.08 | 6.08 | -0.05 | 0.05 |
| gene-LOC105275070 | https://www.ncbi.nlm.nih.gov/gene/?term=LOC105275070 | 5667 | 53 | 93.52 | 6 | 10.59 | 11.32 | 1.77 | 10.27 | -8.5 | 8.5 | -0.08 | 0.08 |
| gene-LOC113562901 | https://www.ncbi.nlm.nih.gov/gene/?term=LOC113562901 | 1531 | 22 | 143.7 | 3 | 19.6 | 13.64 | 45.03 | 59.16 | -14.13 | 14.13 | -0.2 | 0.2 |
| gene-LOC113562345 | https://www.ncbi.nlm.nih.gov/gene/?term=LOC113562345 | 2290 | 69 | 301.31 | 19 | 82.97 | 27.54 | 16.9 | 33.89 | -17 | 17 | -0.51 | 0.51 |
| gene-LOC105281282 | https://www.ncbi.nlm.nih.gov/gene/?term=LOC105281282 | 3250 | 3 | 9.23 | 1 | 3.08 | 33.33 | 23.25 | 65.16 | -41.92 | 41.92 | -0.04 | 0.04 |

Directory: OGL File: ogl-Ob_LI_BCphaseB2.vs.RphaseB1.txt

| gene_ID | gene_link | gwidth | #Sites | #per10Kb | #dmSites | #dmsp10kb | %dmSites | %pSite1 | %pSite2 | DMpSite | ADMpSite | DMpNucl | ADMpNucl |
|---|---|---|---|---|---|---|---|---|---|---|---|---|---|
| gene-LOC113562974 | https://www.ncbi.nlm.nih.gov/gene/?term=LOC113562974 | 1832 | 2 | 10.92 | 1 | 5.46 | 50 | 21.66 | 9 | 12.67 | 12.67 | 0.01 | 0.01 |
| gene-LOC105282841 | https://www.ncbi.nlm.nih.gov/gene/?term=LOC105282841 | 2047 | 5 | 24.43 | 1 | 4.89 | 20 | 67.42 | 55.09 | 12.33 | 12.33 | 0.03 | 0.03 |
| gene-LOC105278541 | https://www.ncbi.nlm.nih.gov/gene/?term=LOC105278541 | 1563 | 14 | 89.57 | 3 | 19.19 | 21.43 | 39.99 | 31.76 | 8.23 | 8.23 | 0.07 | 0.07 |
| gene-LOC105286822 | https://www.ncbi.nlm.nih.gov/gene/?term=LOC105286822 | 3538 | 6 | 16.96 | 1 | 2.83 | 16.67 | 18.11 | 10.53 | 7.58 | 7.58 | 0.01 | 0.01 |
| gene-LOC105287923 | https://www.ncbi.nlm.nih.gov/gene/?term=LOC105287923 | 1233 | 10 | 81.1 | 2 | 16.22 | 20 | 76.35 | 69.28 | 7.07 | 7.07 | 0.06 | 0.06 |
| gene-LOC105275454 | https://www.ncbi.nlm.nih.gov/gene/?term=LOC105275454 | 1145 | 13 | 113.54 | 3 | 26.2 | 23.08 | 55.74 | 49.23 | 6.51 | 6.51 | 0.07 | 0.07 |
| gene-LOC105284547 | https://www.ncbi.nlm.nih.gov/gene/?term=LOC105284547 | 2105 | 13 | 61.76 | 1 | 4.75 | 7.69 | 57.2 | 50.73 | 6.46 | 6.46 | 0.04 | 0.04 |
| gene-LOC105275176 | https://www.ncbi.nlm.nih.gov/gene/?term=LOC105275176 | 656 | 8 | 121.95 | 1 | 15.24 | 12.5 | 14.79 | 8.51 | 6.28 | 6.28 | 0.08 | 0.08 |
| gene-LOC105274672 | https://www.ncbi.nlm.nih.gov/gene/?term=LOC105274672 | 2878 | 15 | 52.12 | 2 | 6.95 | 13.33 | 40.13 | 34.21 | 5.92 | 5.92 | 0.03 | 0.03 |
| gene-LOC105284056 | https://www.ncbi.nlm.nih.gov/gene/?term=LOC105284056 | 1997 | 6 | 30.05 | 1 | 5.01 | 16.67 | 30.15 | 24.32 | 5.83 | 5.83 | 0.02 | 0.02 |
| gene-LOC105285334 | https://www.ncbi.nlm.nih.gov/gene/?term=LOC105285334 | 2363 | 33 | 139.65 | 4 | 16.93 | 12.12 | 24.89 | 19.32 | 5.57 | 5.57 | 0.08 | 0.08 |
| gene-LOC105285795 | https://www.ncbi.nlm.nih.gov/gene/?term=LOC105285795 | 1256 | 12 | 95.54 | 2 | 15.92 | 16.67 | 50.58 | 45.08 | 5.5 | 5.5 | 0.05 | 0.05 |
| gene-LOC105280473 | https://www.ncbi.nlm.nih.gov/gene/?term=LOC105280473 | 1816 | 6 | 33.04 | 1 | 5.51 | 16.67 | 88.83 | 83.46 | 5.36 | 5.36 | 0.02 | 0.02 |
| gene-LOC105276202 | https://www.ncbi.nlm.nih.gov/gene/?term=LOC105276202 | 1540 | 43 | 279.22 | 5 | 32.47 | 11.63 | 24.19 | 18.86 | 5.33 | 5.33 | 0.15 | 0.15 |
| gene-LOC105282371 | https://www.ncbi.nlm.nih.gov/gene/?term=LOC105282371 | 1722 | 8 | 46.46 | 1 | 5.81 | 12.5 | 31.57 | 26.4 | 5.17 | 5.17 | 0.02 | 0.02 |
| gene-LOC105275296 | https://www.ncbi.nlm.nih.gov/gene/?term=LOC105275296 | 2951 | 19 | 64.38 | 1 | 3.39 | 5.26 | 27.28 | 22.14 | 5.14 | 5.14 | 0.03 | 0.03 |
| gene-LOC105281716 | https://www.ncbi.nlm.nih.gov/gene/?term=LOC105281716 | 1802 | 8 | 44.4 | 1 | 5.55 | 12.5 | 36.4 | 31.28 | 5.11 | 5.11 | 0.02 | 0.02 |
| gene-LOC105284999 | https://www.ncbi.nlm.nih.gov/gene/?term=LOC105284999 | 1832 | 9 | 49.13 | 1 | 5.46 | 11.11 | 55.4 | 50.33 | 5.07 | 5.07 | 0.02 | 0.02 |
| gene-LOC105284475 | https://www.ncbi.nlm.nih.gov/gene/?term=LOC105284475 | 1364 | 11 | 80.65 | 1 | 7.33 | 9.09 | 56.71 | 51.84 | 4.87 | 4.87 | 0.04 | 0.04 |
| gene-LOC113563111 | https://www.ncbi.nlm.nih.gov/gene/?term=LOC113563111 | 2455 | 29 | 118.13 | 3 | 12.22 | 10.34 | 28.93 | 24.09 | 4.84 | 4.84 | 0.06 | 0.06 |
| gene-LOC105282041 | https://www.ncbi.nlm.nih.gov/gene/?term=LOC105282041 | 2041 | 14 | 68.59 | 2 | 9.8 | 14.29 | 31.5 | 26.67 | 4.83 | 4.83 | 0.03 | 0.03 |
| gene-LOC105278686 | https://www.ncbi.nlm.nih.gov/gene/?term=LOC105278686 | 2006 | 14 | 69.79 | 1 | 4.99 | 7.14 | 10.96 | 6.19 | 4.77 | 4.77 | 0.03 | 0.03 |
| gene-LOC105276171 | https://www.ncbi.nlm.nih.gov/gene/?term=LOC105276171 | 1370 | 14 | 102.19 | 2 | 14.6 | 14.29 | 17.56 | 12.89 | 4.67 | 4.67 | 0.05 | 0.05 |
| gene-LOC105277804 | https://www.ncbi.nlm.nih.gov/gene/?term=LOC105277804 | 1174 | 6 | 51.11 | 1 | 8.52 | 16.67 | 93.96 | 89.47 | 4.48 | 4.48 | 0.02 | 0.02 |

...
... only first 25 lines shown ...

Directory: OGL File: ogl-Ob_LI_BCphaseB2.vs.RphaseB2.txt

| gene_ID | gene_link | gwidth | #Sites | #per10Kb | #dmSites | #dmsp10kb | %dmSites | %pSite1 | %pSite2 | DMpSite | ADMpSite | DMpNucl | ADMpNucl |
|---|---|---|---|---|---|---|---|---|---|---|---|---|---|
| gene-LOC105286586 | https://www.ncbi.nlm.nih.gov/gene/?term=LOC105286586 | 1417 | 1 | 7.06 | 1 | 7.06 | 100 | 36.67 | 15.38 | 21.29 | 21.29 | 0.02 | 0.02 |
| gene-LOC113561765 | https://www.ncbi.nlm.nih.gov/gene/?term=LOC113561765 | 1822 | 4 | 21.95 | 1 | 5.49 | 25 | 29.77 | 18.68 | 11.1 | 11.1 | 0.02 | 0.02 |
| gene-LOC105275176 | https://www.ncbi.nlm.nih.gov/gene/?term=LOC105275176 | 656 | 8 | 121.95 | 2 | 30.49 | 25 | 14.79 | 6.32 | 8.47 | 8.47 | 0.1 | 0.1 |
| gene-LOC105286261 | https://www.ncbi.nlm.nih.gov/gene/?term=LOC105286261 | 1297 | 8 | 61.68 | 1 | 7.71 | 12.5 | 76.36 | 68.31 | 8.05 | 8.05 | 0.05 | 0.05 |
| gene-LOC105287216 | https://www.ncbi.nlm.nih.gov/gene/?term=LOC105287216 | 2142 | 6 | 28.01 | 1 | 4.67 | 16.67 | 27.53 | 19.82 | 7.72 | 7.72 | 0.02 | 0.02 |
| gene-LOC105283410 | https://www.ncbi.nlm.nih.gov/gene/?term=LOC105283410 | 1423 | 5 | 35.14 | 1 | 7.03 | 20 | 47.1 | 40.2 | 6.9 | 6.9 | 0.02 | 0.02 |
| gene-LOC105276075 | https://www.ncbi.nlm.nih.gov/gene/?term=LOC105276075 | 1440 | 10 | 69.44 | 1 | 6.94 | 10 | 39.46 | 33.04 | 6.42 | 6.42 | 0.04 | 0.04 |
| gene-LOC105282996 | https://www.ncbi.nlm.nih.gov/gene/?term=LOC105282996 | 3736 | 6 | 16.06 | 1 | 2.68 | 16.67 | 23.34 | 17.02 | 6.33 | 6.33 | 0.01 | 0.01 |
| gene-LOC105285229 | https://www.ncbi.nlm.nih.gov/gene/?term=LOC105285229 | 1396 | 5 | 35.82 | 1 | 7.16 | 20 | 23.52 | 17.61 | 5.91 | 5.91 | 0.02 | 0.02 |
| gene-LOC105277980 | https://www.ncbi.nlm.nih.gov/gene/?term=LOC105277980 | 2400 | 30 | 125 | 2 | 8.33 | 6.67 | 49.23 | 43.45 | 5.77 | 5.77 | 0.07 | 0.07 |
| gene-LOC105285203 | https://www.ncbi.nlm.nih.gov/gene/?term=LOC105285203 | 1020 | 27 | 264.71 | 2 | 19.61 | 7.41 | 30.05 | 24.28 | 5.76 | 5.76 | 0.15 | 0.15 |
| gene-LOC105281512 | https://www.ncbi.nlm.nih.gov/gene/?term=LOC105281512 | 2233 | 16 | 71.65 | 3 | 13.43 | 18.75 | 32.28 | 26.62 | 5.65 | 5.65 | 0.04 | 0.04 |
| gene-LOC105274857 | https://www.ncbi.nlm.nih.gov/gene/?term=LOC105274857 | 1569 | 4 | 25.49 | 1 | 6.37 | 25 | 58.2 | 52.63 | 5.57 | 5.57 | 0.01 | 0.01 |
| gene-LOC105285795 | https://www.ncbi.nlm.nih.gov/gene/?term=LOC105285795 | 1256 | 14 | 111.46 | 2 | 15.92 | 14.29 | 57.64 | 52.12 | 5.53 | 5.53 | 0.06 | 0.06 |
| gene-LOC105276569 | https://www.ncbi.nlm.nih.gov/gene/?term=LOC105276569 | 1963 | 14 | 71.32 | 1 | 5.09 | 7.14 | 23.68 | 18.21 | 5.47 | 5.47 | 0.04 | 0.04 |
| gene-LOC105281716 | https://www.ncbi.nlm.nih.gov/gene/?term=LOC105281716 | 1802 | 8 | 44.4 | 1 | 5.55 | 12.5 | 36.4 | 30.96 | 5.44 | 5.44 | 0.02 | 0.02 |
| gene-LOC105275503 | https://www.ncbi.nlm.nih.gov/gene/?term=LOC105275503 | 2000 | 4 | 20 | 1 | 5 | 25 | 38.28 | 33 | 5.28 | 5.28 | 0.01 | 0.01 |
| gene-LOC105276230 | https://www.ncbi.nlm.nih.gov/gene/?term=LOC105276230 | 3067 | 12 | 39.13 | 1 | 3.26 | 8.33 | 28.38 | 23.21 | 5.17 | 5.17 | 0.02 | 0.02 |
| gene-LOC105278606 | https://www.ncbi.nlm.nih.gov/gene/?term=LOC105278606 | 2650 | 5 | 18.87 | 1 | 3.77 | 20 | 6.01 | 0.87 | 5.14 | 5.14 | 0.01 | 0.01 |
| gene-LOC105283047 | https://www.ncbi.nlm.nih.gov/gene/?term=LOC105283047 | 1181 | 16 | 135.48 | 1 | 8.47 | 6.25 | 13.48 | 8.42 | 5.06 | 5.06 | 0.07 | 0.07 |
| gene-LOC105285669 | https://www.ncbi.nlm.nih.gov/gene/?term=LOC105285669 | 2193 | 16 | 72.96 | 1 | 4.56 | 6.25 | 26.19 | 21.17 | 5.02 | 5.02 | 0.04 | 0.04 |
| gene-LOC105285905 | https://www.ncbi.nlm.nih.gov/gene/?term=LOC105285905 | 1397 | 13 | 93.06 | 2 | 14.32 | 15.38 | 53.61 | 48.59 | 5.02 | 5.02 | 0.05 | 0.05 |
| gene-LOC105278108 | https://www.ncbi.nlm.nih.gov/gene/?term=LOC105278108 | 4392 | 46 | 104.74 | 1 | 2.28 | 2.17 | 29.79 | 24.81 | 4.98 | 4.98 | 0.05 | 0.05 |
| gene-LOC105286778 | https://www.ncbi.nlm.nih.gov/gene/?term=LOC105286778 | 6479 | 6 | 9.26 | 1 | 1.54 | 16.67 | 7.5 | 2.54 | 4.96 | 4.96 | 0 | 0 |

...
... only first 25 lines shown ...

Directory: OGL File: ogl-Ob_LI_RphaseA1.txt

| gene_ID | gene_link | gwidth | #Sites | #per10Kb | %perSite | %pNucl |
|---|---|---|---|---|---|---|
| gene-LOC113561532 | https://www.ncbi.nlm.nih.gov/gene/?term=LOC113561532 | 72 | 5 | 694.44 | 98.1 | 6.81 |
| gene-LOC105286531 | https://www.ncbi.nlm.nih.gov/gene/?term=LOC105286531 | 372 | 43 | 1155.91 | 36.48 | 4.22 |
| gene-LOC105280739 | https://www.ncbi.nlm.nih.gov/gene/?term=LOC105280739 | 1991 | 138 | 693.12 | 54.32 | 3.76 |
| gene-LOC105274534 | https://www.ncbi.nlm.nih.gov/gene/?term=LOC105274534 | 1329 | 104 | 782.54 | 47.62 | 3.73 |
| gene-LOC105285635 | https://www.ncbi.nlm.nih.gov/gene/?term=LOC105285635 | 1175 | 137 | 1165.96 | 30.78 | 3.59 |
| gene-LOC105287811 | https://www.ncbi.nlm.nih.gov/gene/?term=LOC105287811 | 2155 | 153 | 709.98 | 47.68 | 3.38 |
| gene-LOC105286235 | https://www.ncbi.nlm.nih.gov/gene/?term=LOC105286235 | 1510 | 118 | 781.46 | 42.69 | 3.34 |
| gene-LOC105286269 | https://www.ncbi.nlm.nih.gov/gene/?term=LOC105286269 | 1401 | 87 | 620.99 | 52.62 | 3.27 |
| gene-LOC105286737 | https://www.ncbi.nlm.nih.gov/gene/?term=LOC105286737 | 2690 | 365 | 1356.88 | 23.81 | 3.23 |
| gene-LOC105278844 | https://www.ncbi.nlm.nih.gov/gene/?term=LOC105278844 | 1101 | 109 | 990.01 | 32.11 | 3.18 |
| gene-LOC105282149 | https://www.ncbi.nlm.nih.gov/gene/?term=LOC105282149 | 1787 | 139 | 777.84 | 40.94 | 3.18 |
| gene-LOC105277648 | https://www.ncbi.nlm.nih.gov/gene/?term=LOC105277648 | 722 | 39 | 540.17 | 57.72 | 3.12 |
| gene-LOC105275314 | https://www.ncbi.nlm.nih.gov/gene/?term=LOC105275314 | 1489 | 97 | 651.44 | 47.77 | 3.11 |
| gene-LOC105279052 | https://www.ncbi.nlm.nih.gov/gene/?term=LOC105279052 | 1130 | 122 | 1079.65 | 28.5 | 3.08 |
| gene-LOC105276371 | https://www.ncbi.nlm.nih.gov/gene/?term=LOC105276371 | 871 | 55 | 631.46 | 48.54 | 3.07 |
| gene-LOC105278160 | https://www.ncbi.nlm.nih.gov/gene/?term=LOC105278160 | 1872 | 158 | 844.02 | 35.32 | 2.98 |
| gene-LOC105286801 | https://www.ncbi.nlm.nih.gov/gene/?term=LOC105286801 | 1102 | 86 | 780.4 | 38.13 | 2.98 |
| gene-LOC105276360 | https://www.ncbi.nlm.nih.gov/gene/?term=LOC105276360 | 1466 | 154 | 1050.48 | 27.76 | 2.92 |
| gene-LOC105287677 | https://www.ncbi.nlm.nih.gov/gene/?term=LOC105287677 | 1588 | 113 | 711.59 | 41.08 | 2.92 |
| gene-LOC105274711 | https://www.ncbi.nlm.nih.gov/gene/?term=LOC105274711 | 1655 | 102 | 616.31 | 46.87 | 2.89 |
| gene-LOC105278849 | https://www.ncbi.nlm.nih.gov/gene/?term=LOC105278849 | 776 | 101 | 1301.55 | 22.16 | 2.88 |
| gene-LOC105277254 | https://www.ncbi.nlm.nih.gov/gene/?term=LOC105277254 | 1377 | 112 | 813.36 | 35.34 | 2.87 |
| gene-LOC105285392 | https://www.ncbi.nlm.nih.gov/gene/?term=LOC105285392 | 1601 | 138 | 861.96 | 32.41 | 2.79 |
| gene-LOC105287692 | https://www.ncbi.nlm.nih.gov/gene/?term=LOC105287692 | 1505 | 120 | 797.34 | 34.96 | 2.79 |

...

... only first 25 lines shown ...

Directory: OGL File: ogl-Ob_LI_RphaseA1.vs.RphaseA2.txt

| gene_ID | gene_link | gwidth | #Sites | #per10Kb | #dmSites | #dmsp10kb | %dmSites | %pSite1 | %pSite2 | DMpSite | ADMpSite | DMpNucl | ADMpNucl |
|---|---|---|---|---|---|---|---|---|---|---|---|---|---|
| gene-LOC105283264 | https://www.ncbi.nlm.nih.gov/gene/?term=LOC105283264 | 4005 | 96 | 239.7 | 1 | 2.5 | 1.04 | 23.23 | 22.15 | 1.09 | 1.09 | 0.03 | 0.03 |
| gene-LOC105277903 | https://www.ncbi.nlm.nih.gov/gene/?term=LOC105277903 | 1835 | 39 | 212.53 | 1 | 5.45 | 2.56 | 54.3 | 53.51 | 0.79 | 0.79 | 0.02 | 0.02 |
| gene-LOC105283296 | https://www.ncbi.nlm.nih.gov/gene/?term=LOC105283296 | 2174 | 101 | 464.58 | 1 | 4.6 | 0.99 | 18.85 | 18.17 | 0.67 | 0.67 | 0.03 | 0.03 |
| gene-LOC105284557 | https://www.ncbi.nlm.nih.gov/gene/?term=LOC105284557 | 1672 | 84 | 502.39 | 1 | 5.98 | 1.19 | 28.63 | 28.17 | 0.46 | 0.46 | 0.02 | 0.02 |
| gene-LOC105280304 | https://www.ncbi.nlm.nih.gov/gene/?term=LOC105280304 | 8919 | 150 | 168.18 | 1 | 1.12 | 0.67 | 6.83 | 6.96 | -0.12 | 0.12 | 0 | 0 |
| gene-LOC105284507 | https://www.ncbi.nlm.nih.gov/gene/?term=LOC105284507 | 5207 | 112 | 215.1 | 1 | 1.92 | 0.89 | 18.79 | 19.06 | -0.28 | 0.28 | -0.01 | 0.01 |
| gene-LOC105275384 | https://www.ncbi.nlm.nih.gov/gene/?term=LOC105275384 | 1987 | 86 | 432.81 | 1 | 5.03 | 1.16 | 36.56 | 36.97 | -0.41 | 0.41 | -0.02 | 0.02 |
| gene-LOC105281272 | https://www.ncbi.nlm.nih.gov/gene/?term=LOC105281272 | 5252 | 199 | 378.9 | 1 | 1.9 | 0.5 | 14.62 | 15.07 | -0.45 | 0.45 | -0.02 | 0.02 |
| gene-LOC105288036 | https://www.ncbi.nlm.nih.gov/gene/?term=LOC105288036 | 5110 | 209 | 409 | 1 | 1.96 | 0.48 | 17.07 | 17.79 | -0.71 | 0.71 | -0.03 | 0.03 |
| gene-LOC105288042 | https://www.ncbi.nlm.nih.gov/gene/?term=LOC105288042 | 3507 | 111 | 316.51 | 1 | 2.85 | 0.9 | 22.13 | 23.4 | -1.28 | 1.28 | -0.04 | 0.04 |

Directory: OGL File: ogl-Ob_LI_RphaseA1.vs.RphaseB1.txt

| gene_ID | gene_link | gwidth | #Sites | #per10Kb | #dmSites | #dmsp10kb | %dmSites | %pSite1 | %pSite2 | DMpSite | ADMpSite | DMpNucl | ADMpNucl |
|---|---|---|---|---|---|---|---|---|---|---|---|---|---|
| gene-LOC105275070 | https://www.ncbi.nlm.nih.gov/gene/?term=LOC105275070 | 5667 | 56 | 98.82 | 3 | 5.29 | 5.36 | 17.83 | 5.85 | 11.98 | 11.98 | 0.12 | 0.12 |
| gene-LOC113562345 | https://www.ncbi.nlm.nih.gov/gene/?term=LOC113562345 | 2290 | 78 | 340.61 | 11 | 48.03 | 14.1 | 24.77 | 13.87 | 10.9 | 10.9 | 0.37 | 0.37 |
| gene-LOC105285514 | https://www.ncbi.nlm.nih.gov/gene/?term=LOC105285514 | 1277 | 7 | 54.82 | 1 | 7.83 | 14.29 | 90.55 | 80 | 10.54 | 10.54 | 0.06 | 0.06 |
| gene-LOC105275708 | https://www.ncbi.nlm.nih.gov/gene/?term=LOC105275708 | 1772 | 14 | 79.01 | 1 | 5.64 | 7.14 | 50.83 | 42.47 | 8.36 | 8.36 | 0.07 | 0.07 |
| gene-LOC105275355 | https://www.ncbi.nlm.nih.gov/gene/?term=LOC105275355 | 3197 | 174 | 544.26 | 1 | 3.13 | 0.57 | 25.86 | 21.8 | 4.06 | 4.06 | 0.22 | 0.22 |
| gene-LOC105281039 | https://www.ncbi.nlm.nih.gov/gene/?term=LOC105281039 | 1471 | 13 | 88.38 | 1 | 6.8 | 7.69 | 31.24 | 27.65 | 3.59 | 3.59 | 0.03 | 0.03 |
| gene-LOC105275384 | https://www.ncbi.nlm.nih.gov/gene/?term=LOC105275384 | 1987 | 88 | 442.88 | 1 | 5.03 | 1.14 | 36.35 | 32.79 | 3.56 | 3.56 | 0.16 | 0.16 |
| gene-LOC105275347 | https://www.ncbi.nlm.nih.gov/gene/?term=LOC105275347 | 2526 | 17 | 67.3 | 1 | 3.96 | 5.88 | 45.17 | 41.88 | 3.29 | 3.29 | 0.02 | 0.02 |
| gene-LOC105274900 | https://www.ncbi.nlm.nih.gov/gene/?term=LOC105274900 | 5561 | 61 | 109.69 | 3 | 5.39 | 4.92 | 33.5 | 30.47 | 3.02 | 3.02 | 0.03 | 0.03 |
| gene-LOC105287063 | https://www.ncbi.nlm.nih.gov/gene/?term=LOC105287063 | 1874 | 49 | 261.47 | 1 | 5.34 | 2.04 | 32.77 | 30.12 | 2.65 | 2.65 | 0.07 | 0.07 |
| gene-LOC105279898 | https://www.ncbi.nlm.nih.gov/gene/?term=LOC105279898 | 1343 | 15 | 111.69 | 1 | 7.45 | 6.67 | 61.47 | 58.93 | 2.54 | 2.54 | 0.03 | 0.03 |
| gene-LOC105281515 | https://www.ncbi.nlm.nih.gov/gene/?term=LOC105281515 | 8827 | 57 | 64.57 | 2 | 2.27 | 3.51 | 32.7 | 30.37 | 2.32 | 2.32 | 0.01 | 0.01 |
| gene-LOC113562102 | https://www.ncbi.nlm.nih.gov/gene/?term=LOC113562102 | 3433 | 63 | 183.51 | 1 | 2.91 | 1.59 | 17.89 | 15.59 | 2.3 | 2.3 | 0.04 | 0.04 |
| gene-LOC105276272 | https://www.ncbi.nlm.nih.gov/gene/?term=LOC105276272 | 1965 | 44 | 223.92 | 1 | 5.09 | 2.27 | 27.22 | 24.96 | 2.26 | 2.26 | 0.05 | 0.05 |
| gene-LOC105282991 | https://www.ncbi.nlm.nih.gov/gene/?term=LOC105282991 | 5617 | 103 | 183.37 | 1 | 1.78 | 0.97 | 18.36 | 16.13 | 2.23 | 2.23 | 0.04 | 0.04 |
| gene-LOC105285695 | https://www.ncbi.nlm.nih.gov/gene/?term=LOC105285695 | 1371 | 68 | 495.99 | 1 | 7.29 | 1.47 | 23.53 | 21.55 | 1.99 | 1.99 | 0.1 | 0.1 |
| gene-LOC105283712 | https://www.ncbi.nlm.nih.gov/gene/?term=LOC105283712 | 1910 | 35 | 183.25 | 1 | 5.24 | 2.86 | 28.45 | 26.47 | 1.98 | 1.98 | 0.04 | 0.04 |
| gene-LOC105285671 | https://www.ncbi.nlm.nih.gov/gene/?term=LOC105285671 | 2263 | 100 | 441.89 | 1 | 4.42 | 1 | 32.24 | 30.53 | 1.71 | 1.71 | 0.08 | 0.08 |
| gene-LOC105282652 | https://www.ncbi.nlm.nih.gov/gene/?term=LOC105282652 | 5065 | 85 | 167.82 | 1 | 1.97 | 1.18 | 16.28 | 14.6 | 1.68 | 1.68 | 0.03 | 0.03 |
| gene-LOC105276692 | https://www.ncbi.nlm.nih.gov/gene/?term=LOC105276692 | 1745 | 34 | 194.84 | 1 | 5.73 | 2.94 | 25.17 | 23.64 | 1.53 | 1.53 | 0.03 | 0.03 |
| gene-LOC105288066 | https://www.ncbi.nlm.nih.gov/gene/?term=LOC105288066 | 1964 | 30 | 152.75 | 1 | 5.09 | 3.33 | 38.49 | 37.01 | 1.48 | 1.48 | 0.02 | 0.02 |
| gene-LOC105286265 | https://www.ncbi.nlm.nih.gov/gene/?term=LOC105286265 | 2507 | 67 | 267.25 | 1 | 3.99 | 1.49 | 32.57 | 31.11 | 1.46 | 1.46 | 0.04 | 0.04 |
| gene-LOC113561805 | https://www.ncbi.nlm.nih.gov/gene/?term=LOC113561805 | 3337 | 75 | 224.75 | 1 | 3 | 1.33 | 34.35 | 32.91 | 1.44 | 1.44 | 0.03 | 0.03 |
| gene-LOC105280348 | https://www.ncbi.nlm.nih.gov/gene/?term=LOC105280348 | 3797 | 79 | 208.06 | 1 | 2.63 | 1.27 | 18.28 | 16.89 | 1.4 | 1.4 | 0.03 | 0.03 |
| gene-LOC105278807 | https://www.ncbi.nlm.nih.gov/gene/?term=LOC105278807 | 2166 | 74 | 341.64 | 1 | 4.62 | 1.35 | 21.42 | 20.03 | 1.39 | 1.39 | 0.05 | 0.05 |
| gene-LOC105282600 | https://www.ncbi.nlm.nih.gov/gene/?term=LOC105282600 | 3715 | 60 | 161.51 | 1 | 2.69 | 1.67 | 27.74 | 26.44 | 1.3 | 1.3 | 0.02 | 0.02 |
| gene-LOC105285060 | https://www.ncbi.nlm.nih.gov/gene/?term=LOC105285060 | 1285 | 16 | 124.51 | 1 | 7.78 | 6.25 | 42.55 | 41.27 | 1.27 | 1.27 | 0.02 | 0.02 |
| gene-LOC105285742 | https://www.ncbi.nlm.nih.gov/gene/?term=LOC105285742 | 3446 | 109 | 316.31 | 1 | 2.9 | 0.92 | 17.79 | 16.59 | 1.2 | 1.2 | 0.04 | 0.04 |
| gene-LOC105285674 | https://www.ncbi.nlm.nih.gov/gene/?term=LOC105285674 | 3035 | 100 | 329.49 | 2 | 6.59 | 2 | 25.96 | 24.78 | 1.18 | 1.18 | 0.04 | 0.04 |
| gene-LOC105279528 | https://www.ncbi.nlm.nih.gov/gene/?term=LOC105279528 | 5802 | 40 | 68.94 | 1 | 1.72 | 2.5 | 21.84 | 20.71 | 1.12 | 1.12 | 0.01 | 0.01 |
| gene-LOC105275187 | https://www.ncbi.nlm.nih.gov/gene/?term=LOC105275187 | 5534 | 73 | 131.91 | 1 | 1.81 | 1.37 | 14.7 | 13.62 | 1.08 | 1.08 | 0.01 | 0.01 |
| gene-LOC105283697 | https://www.ncbi.nlm.nih.gov/gene/?term=LOC105283697 | 5301 | 153 | 288.62 | 1 | 1.89 | 0.65 | 16.46 | 15.43 | 1.02 | 1.02 | 0.03 | 0.03 |
| gene-LOC105275460 | https://www.ncbi.nlm.nih.gov/gene/?term=LOC105275460 | 2473 | 126 | 509.5 | 1 | 4.04 | 0.79 | 13.41 | 12.4 | 1.01 | 1.01 | 0.05 | 0.05 |
| gene-LOC105276449 | https://www.ncbi.nlm.nih.gov/gene/?term=LOC105276449 | 1231 | 48 | 389.93 | 1 | 8.12 | 2.08 | 25.67 | 24.69 | 0.99 | 0.99 | 0.04 | 0.04 |
| gene-LOC105283552 | https://www.ncbi.nlm.nih.gov/gene/?term=LOC105283552 | 1562 | 40 | 256.08 | 1 | 6.4 | 2.5 | 50.43 | 49.44 | 0.99 | 0.99 | 0.03 | 0.03 |
| gene-LOC105274823 | https://www.ncbi.nlm.nih.gov/gene/?term=LOC105274823 | 4620 | 177 | 383.12 | 1 | 2.16 | 0.56 | 17.15 | 16.21 | 0.95 | 0.95 | 0.04 | 0.04 |
| gene-LOC105279900 | https://www.ncbi.nlm.nih.gov/gene/?term=LOC105279900 | 2191 | 55 | 251.03 | 1 | 4.56 | 1.82 | 30.23 | 29.28 | 0.95 | 0.95 | 0.02 | 0.02 |
| gene-LOC105288079 | https://www.ncbi.nlm.nih.gov/gene/?term=LOC105288079 | 3473 | 113 | 325.37 | 1 | 2.88 | 0.88 | 17.44 | 16.56 | 0.89 | 0.89 | 0.03 | 0.03 |
| gene-LOC105281848 | https://www.ncbi.nlm.nih.gov/gene/?term=LOC105281848 | 2122 | 57 | 268.61 | 1 | 4.71 | 1.75 | 23.82 | 22.98 | 0.84 | 0.84 | 0.02 | 0.02 |
| gene-LOC105282491 | https://www.ncbi.nlm.nih.gov/gene/?term=LOC105282491 | 4761 | 179 | 375.97 | 1 | 2.1 | 0.56 | 9.15 | 8.31 | 0.83 | 0.83 | 0.03 | 0.03 |
| gene-LOC105288036 | https://www.ncbi.nlm.nih.gov/gene/?term=LOC105288036 | 5110 | 200 | 391.39 | 1 | 1.96 | 0.5 | 17.43 | 16.6 | 0.83 | 0.83 | 0.03 | 0.03 |
| gene-LOC105282156 | https://www.ncbi.nlm.nih.gov/gene/?term=LOC105282156 | 2577 | 127 | 492.82 | 1 | 3.88 | 0.79 | 25.13 | 24.3 | 0.82 | 0.82 | 0.04 | 0.04 |
| gene-LOC105285981 | https://www.ncbi.nlm.nih.gov/gene/?term=LOC105285981 | 7666 | 179 | 233.5 | 1 | 1.3 | 0.56 | 19.94 | 19.14 | 0.81 | 0.81 | 0.02 | 0.02 |
| gene-LOC105283943 | https://www.ncbi.nlm.nih.gov/gene/?term=LOC105283943 | 3559 | 57 | 160.16 | 1 | 2.81 | 1.75 | 23.04 | 22.27 | 0.77 | 0.77 | 0.01 | 0.01 |
| gene-LOC105278283 | https://www.ncbi.nlm.nih.gov/gene/?term=LOC105278283 | 2317 | 142 | 612.86 | 1 | 4.32 | 0.7 | 24.95 | 24.22 | 0.73 | 0.73 | 0.04 | 0.04 |
| gene-LOC105282794 | https://www.ncbi.nlm.nih.gov/gene/?term=LOC105282794 | 4571 | 104 | 227.52 | 1 | 2.19 | 0.96 | 16.85 | 16.16 | 0.69 | 0.69 | 0.02 | 0.02 |
| gene-LOC105283989 | https://www.ncbi.nlm.nih.gov/gene/?term=LOC105283989 | 4948 | 32 | 64.67 | 1 | 2.02 | 3.12 | 46.57 | 45.88 | 0.69 | 0.69 | 0 | 0 |
| gene-LOC105288237 | https://www.ncbi.nlm.nih.gov/gene/?term=LOC105288237 | 2226 | 49 | 220.13 | 1 | 4.49 | 2.04 | 43.01 | 42.34 | 0.67 | 0.67 | 0.01 | 0.01 |
| gene-LOC105285055 | https://www.ncbi.nlm.nih.gov/gene/?term=LOC105285055 | 4108 | 253 | 615.87 | 1 | 2.43 | 0.4 | 10.93 | 10.3 | 0.63 | 0.63 | 0.04 | 0.04 |
| gene-LOC105275451 | https://www.ncbi.nlm.nih.gov/gene/?term=LOC105275451 | 6035 | 135 | 223.7 | 1 | 1.66 | 0.74 | 21.36 | 20.75 | 0.61 | 0.61 | 0.01 | 0.01 |
| gene-LOC105277215 | https://www.ncbi.nlm.nih.gov/gene/?term=LOC105277215 | 2619 | 29 | 110.73 | 1 | 3.82 | 3.45 | 25.25 | 24.64 | 0.61 | 0.61 | 0.01 | 0.01 |
| gene-LOC105274449 | https://www.ncbi.nlm.nih.gov/gene/?term=LOC105274449 | 1549 | 53 | 342.16 | 1 | 6.46 | 1.89 | 49.89 | 49.29 | 0.6 | 0.6 | 0.02 | 0.02 |
| gene-LOC105288078 | https://www.ncbi.nlm.nih.gov/gene/?term=LOC105288078 | 19126 | 337 | 176.2 | 1 | 0.52 | 0.3 | 15.64 | 15.05 | 0.59 | 0.59 | 0.01 | 0.01 |
| gene-LOC105280819 | https://www.ncbi.nlm.nih.gov/gene/?term=LOC105280819 | 1639 | 42 | 256.25 | 1 | 6.1 | 2.38 | 39.26 | 38.69 | 0.58 | 0.58 | 0.01 | 0.01 |
| gene-LOC105280641 | https://www.ncbi.nlm.nih.gov/gene/?term=LOC105280641 | 3682 | 75 | 203.69 | 1 | 2.72 | 1.33 | 13.21 | 12.66 | 0.55 | 0.55 | 0.01 | 0.01 |
| gene-LOC105285179 | https://www.ncbi.nlm.nih.gov/gene/?term=LOC105285179 | 4191 | 189 | 450.97 | 1 | 2.39 | 0.53 | 18.34 | 17.82 | 0.52 | 0.52 | 0.02 | 0.02 |
| gene-LOC105278145 | https://www.ncbi.nlm.nih.gov/gene/?term=LOC105278145 | 4470 | 252 | 563.76 | 2 | 4.47 | 0.79 | 10.98 | 10.51 | 0.48 | 0.48 | 0.03 | 0.03 |
| gene-LOC105274531 | https://www.ncbi.nlm.nih.gov/gene/?term=LOC105274531 | 7774 | 286 | 367.89 | 1 | 1.29 | 0.35 | 11.23 | 10.77 | 0.46 | 0.46 | 0.02 | 0.02 |
| gene-LOC105287417 | https://www.ncbi.nlm.nih.gov/gene/?term=LOC105287417 | 1025 | 32 | 312.2 | 1 | 9.76 | 3.12 | 37.69 | 37.23 | 0.46 | 0.46 | 0.01 | 0.01 |
| gene-LOC105277903 | https://www.ncbi.nlm.nih.gov/gene/?term=LOC105277903 | 1835 | 37 | 201.63 | 1 | 5.45 | 2.7 | 54.5 | 54.08 | 0.42 | 0.42 | 0.01 | 0.01 |

| | | | | | | | | | | | | | |
|---|---|---|---|---|---|---|---|---|---|---|---|---|---|
| gene-LOC105275363 | https://www.ncbi.nlm.nih.gov/gene/?term=LOC105275363 | 3798 | 179 | 471.3 | 1 | 2.63 | 0.56 | 15.94 | 15.54 | 0.4 | 0.4 | 0.02 | 0.02 |
| gene-LOC105286452 | https://www.ncbi.nlm.nih.gov/gene/?term=LOC105286452 | 6888 | 68 | 98.72 | 1 | 1.45 | 1.47 | 28.16 | 27.75 | 0.4 | 0.4 | 0 | 0 |
| gene-LOC105286355 | https://www.ncbi.nlm.nih.gov/gene/?term=LOC105286355 | 3245 | 149 | 459.17 | 1 | 3.08 | 0.67 | 21.45 | 21.07 | 0.38 | 0.38 | 0.02 | 0.02 |
| gene-LOC105283592 | https://www.ncbi.nlm.nih.gov/gene/?term=LOC105283592 | 3479 | 80 | 229.95 | 1 | 2.87 | 1.25 | 21.51 | 21.14 | 0.37 | 0.37 | 0.01 | 0.01 |
| gene-LOC105283838 | https://www.ncbi.nlm.nih.gov/gene/?term=LOC105283838 | 1748 | 30 | 171.62 | 1 | 5.72 | 3.33 | 33.04 | 32.68 | 0.36 | 0.36 | 0.01 | 0.01 |
| gene-LOC105278497 | https://www.ncbi.nlm.nih.gov/gene/?term=LOC105278497 | 2072 | 100 | 482.63 | 1 | 4.83 | 1 | 18.63 | 18.28 | 0.35 | 0.35 | 0.02 | 0.02 |
| gene-LOC105278384 | https://www.ncbi.nlm.nih.gov/gene/?term=LOC105278384 | 1390 | 33 | 237.41 | 1 | 7.19 | 3.03 | 46.11 | 45.77 | 0.34 | 0.34 | 0.01 | 0.01 |
| gene-LOC105285639 | https://www.ncbi.nlm.nih.gov/gene/?term=LOC105285639 | 10325 | 351 | 339.95 | 2 | 1.94 | 0.57 | 11.49 | 11.2 | 0.29 | 0.29 | 0.01 | 0.01 |
| gene-LOC105288010 | https://www.ncbi.nlm.nih.gov/gene/?term=LOC105288010 | 3189 | 52 | 163.06 | 1 | 3.14 | 1.92 | 29.32 | 29.04 | 0.28 | 0.28 | 0 | 0 |
| gene-LOC105279298 | https://www.ncbi.nlm.nih.gov/gene/?term=LOC105279298 | 2773 | 127 | 457.99 | 1 | 3.61 | 0.79 | 24.14 | 23.89 | 0.26 | 0.26 | 0.01 | 0.01 |
| gene-LOC105283123 | https://www.ncbi.nlm.nih.gov/gene/?term=LOC105283123 | 1507 | 47 | 311.88 | 1 | 6.64 | 2.13 | 20.21 | 19.96 | 0.25 | 0.25 | 0.01 | 0.01 |
| gene-LOC105284329 | https://www.ncbi.nlm.nih.gov/gene/?term=LOC105284329 | 4051 | 71 | 175.27 | 1 | 2.47 | 1.41 | 19.25 | 18.99 | 0.25 | 0.25 | 0 | 0 |
| gene-LOC105284184 | https://www.ncbi.nlm.nih.gov/gene/?term=LOC105284184 | 2535 | 92 | 362.92 | 1 | 3.94 | 1.09 | 31.61 | 31.41 | 0.2 | 0.2 | 0.01 | 0.01 |
| gene-LOC105286997 | https://www.ncbi.nlm.nih.gov/gene/?term=LOC105286997 | 2850 | 142 | 498.25 | 1 | 3.51 | 0.7 | 11.2 | 11.03 | 0.17 | 0.17 | 0.01 | 0.01 |
| gene-LOC105275170 | https://www.ncbi.nlm.nih.gov/gene/?term=LOC105275170 | 12918 | 350 | 270.94 | 1 | 0.77 | 0.29 | 1.94 | 1.8 | 0.15 | 0.15 | 0 | 0 |
| gene-LOC105275374 | https://www.ncbi.nlm.nih.gov/gene/?term=LOC105275374 | 11376 | 514 | 451.83 | 1 | 0.88 | 0.19 | 9.9 | 9.75 | 0.15 | 0.15 | 0.01 | 0.01 |
| gene-LOC105276059 | https://www.ncbi.nlm.nih.gov/gene/?term=LOC105276059 | 6256 | 134 | 214.19 | 1 | 1.6 | 0.75 | 11.07 | 10.91 | 0.15 | 0.15 | 0 | 0 |
| gene-LOC105283728 | https://www.ncbi.nlm.nih.gov/gene/?term=LOC105283728 | 3256 | 152 | 466.83 | 2 | 6.14 | 1.32 | 15.71 | 15.57 | 0.14 | 0.14 | 0.01 | 0.01 |
| gene-LOC105286099 | https://www.ncbi.nlm.nih.gov/gene/?term=LOC105286099 | 5919 | 70 | 118.26 | 1 | 1.69 | 1.43 | 12.4 | 12.26 | 0.14 | 0.14 | 0 | 0 |
| gene-LOC105278534 | https://www.ncbi.nlm.nih.gov/gene/?term=LOC105278534 | 7364 | 198 | 268.88 | 1 | 1.36 | 0.51 | 17.97 | 17.87 | 0.1 | 0.1 | 0 | 0 |
| gene-LOC105280774 | https://www.ncbi.nlm.nih.gov/gene/?term=LOC105280774 | 2793 | 132 | 472.61 | 1 | 3.58 | 0.76 | 25.2 | 25.1 | 0.1 | 0.1 | 0 | 0 |
| gene-LOC105282149 | https://www.ncbi.nlm.nih.gov/gene/?term=LOC105282149 | 1787 | 103 | 576.39 | 1 | 5.6 | 0.97 | 41.42 | 41.34 | 0.08 | 0.08 | 0 | 0 |
| gene-LOC105278666 | https://www.ncbi.nlm.nih.gov/gene/?term=LOC105278666 | 10341 | 474 | 458.37 | 1 | 0.97 | 0.21 | 6.99 | 6.95 | 0.05 | 0.05 | 0 | 0 |
| gene-LOC105280956 | https://www.ncbi.nlm.nih.gov/gene/?term=LOC105280956 | 2639 | 80 | 303.15 | 1 | 3.79 | 1.25 | 18.94 | 18.89 | 0.05 | 0.05 | 0 | 0 |
| gene-LOC105275327 | https://www.ncbi.nlm.nih.gov/gene/?term=LOC105275327 | 14245 | 529 | 371.36 | 1 | 0.7 | 0.19 | 6.66 | 6.63 | 0.03 | 0.03 | 0 | 0 |
| gene-LOC105284043 | https://www.ncbi.nlm.nih.gov/gene/?term=LOC105284043 | 5661 | 284 | 501.68 | 1 | 1.77 | 0.35 | 10.32 | 10.3 | 0.03 | 0.03 | 0 | 0 |
| gene-LOC105277216 | https://www.ncbi.nlm.nih.gov/gene/?term=LOC105277216 | 7627 | 106 | 138.98 | 1 | 1.31 | 0.94 | 7.36 | 7.34 | 0.02 | 0.02 | 0 | 0 |
| gene-LOC105279500 | https://www.ncbi.nlm.nih.gov/gene/?term=LOC105279500 | 3587 | 32 | 89.21 | 1 | 2.79 | 3.12 | 46.58 | 46.57 | 0.01 | 0.01 | 0 | 0 |
| gene-LOC105286515 | https://www.ncbi.nlm.nih.gov/gene/?term=LOC105286515 | 3751 | 117 | 311.92 | 1 | 2.67 | 0.85 | 10.68 | 10.68 | 0 | 0 | 0 | 0 |
| gene-LOC105278228 | https://www.ncbi.nlm.nih.gov/gene/?term=LOC105278228 | 8610 | 176 | 204.41 | 1 | 1.16 | 0.57 | 22.62 | 22.66 | -0.04 | 0.04 | 0 | 0 |
| gene-LOC105278877 | https://www.ncbi.nlm.nih.gov/gene/?term=LOC105278877 | 1968 | 87 | 442.07 | 1 | 5.08 | 1.15 | 29.15 | 29.21 | -0.06 | 0.06 | 0 | 0 |
| gene-LOC105287095 | https://www.ncbi.nlm.nih.gov/gene/?term=LOC105287095 | 1435 | 66 | 459.93 | 1 | 6.97 | 1.52 | 24.97 | 25.03 | -0.06 | 0.06 | 0 | 0 |
| gene-LOC105276527 | https://www.ncbi.nlm.nih.gov/gene/?term=LOC105276527 | 7923 | 338 | 426.61 | 1 | 1.26 | 0.3 | 8.29 | 8.43 | -0.14 | 0.14 | -0.01 | 0.01 |
| gene-LOC105279038 | https://www.ncbi.nlm.nih.gov/gene/?term=LOC105279038 | 9453 | 329 | 348.04 | 1 | 1.06 | 0.3 | 9.14 | 9.32 | -0.18 | 0.18 | -0.01 | 0.01 |
| gene-LOC105279664 | https://www.ncbi.nlm.nih.gov/gene/?term=LOC105279664 | 4023 | 64 | 159.09 | 1 | 2.49 | 1.56 | 17.75 | 17.94 | -0.19 | 0.19 | 0 | 0 |
| gene-LOC105282811 | https://www.ncbi.nlm.nih.gov/gene/?term=LOC105282811 | 5765 | 203 | 352.12 | 1 | 1.73 | 0.49 | 8.59 | 8.78 | -0.19 | 0.19 | -0.01 | 0.01 |
| gene-LOC105274856 | https://www.ncbi.nlm.nih.gov/gene/?term=LOC105274856 | 5216 | 228 | 437.12 | 1 | 1.92 | 0.44 | 17.7 | 17.9 | -0.2 | 0.2 | -0.01 | 0.01 |
| gene-LOC105277856 | https://www.ncbi.nlm.nih.gov/gene/?term=LOC105277856 | 3442 | 33 | 95.87 | 1 | 2.91 | 3.03 | 39.28 | 39.53 | -0.25 | 0.25 | 0 | 0 |
| gene-LOC105285807 | https://www.ncbi.nlm.nih.gov/gene/?term=LOC105285807 | 6063 | 215 | 354.61 | 1 | 1.65 | 0.47 | 8.65 | 8.94 | -0.28 | 0.28 | -0.01 | 0.01 |
| gene-LOC105274706 | https://www.ncbi.nlm.nih.gov/gene/?term=LOC105274706 | 1446 | 50 | 345.78 | 1 | 6.92 | 2 | 37.45 | 37.82 | -0.37 | 0.37 | -0.01 | 0.01 |
| gene-LOC105282825 | https://www.ncbi.nlm.nih.gov/gene/?term=LOC105282825 | 3919 | 96 | 244.96 | 1 | 2.55 | 1.04 | 14.53 | 14.91 | -0.38 | 0.38 | -0.01 | 0.01 |
| gene-LOC105278632 | https://www.ncbi.nlm.nih.gov/gene/?term=LOC105278632 | 3624 | 143 | 394.59 | 1 | 2.76 | 0.7 | 13.02 | 13.42 | -0.4 | 0.4 | -0.02 | 0.02 |
| gene-LOC105284355 | https://www.ncbi.nlm.nih.gov/gene/?term=LOC105284355 | 3904 | 155 | 397.03 | 1 | 2.56 | 0.65 | 20.33 | 20.86 | -0.53 | 0.53 | -0.02 | 0.02 |
| gene-LOC105274851 | https://www.ncbi.nlm.nih.gov/gene/?term=LOC105274851 | 6616 | 144 | 217.65 | 1 | 1.51 | 0.69 | 11.65 | 12.43 | -0.79 | 0.79 | -0.02 | 0.02 |
| gene-LOC105280002 | https://www.ncbi.nlm.nih.gov/gene/?term=LOC105280002 | 1940 | 31 | 159.79 | 1 | 5.15 | 3.23 | 31.38 | 32.27 | -0.88 | 0.88 | -0.01 | 0.01 |
| gene-LOC105282658 | https://www.ncbi.nlm.nih.gov/gene/?term=LOC105282658 | 2421 | 64 | 264.35 | 1 | 4.13 | 1.56 | 23.38 | 24.34 | -0.95 | 0.95 | -0.03 | 0.03 |
| gene-LOC105284036 | https://www.ncbi.nlm.nih.gov/gene/?term=LOC105284036 | 3254 | 98 | 301.17 | 1 | 3.07 | 1.02 | 22.14 | 23.09 | -0.95 | 0.95 | -0.03 | 0.03 |
| gene-LOC105279596 | https://www.ncbi.nlm.nih.gov/gene/?term=LOC105279596 | 7695 | 115 | 149.45 | 1 | 1.3 | 0.87 | 16.77 | 17.78 | -1 | 1 | -0.02 | 0.02 |
| gene-LOC105275518 | https://www.ncbi.nlm.nih.gov/gene/?term=LOC105275518 | 2335 | 23 | 98.5 | 1 | 4.28 | 4.35 | 45.78 | 46.9 | -1.12 | 1.12 | -0.01 | 0.01 |
| gene-LOC105274590 | https://www.ncbi.nlm.nih.gov/gene/?term=LOC105274590 | 4516 | 82 | 181.58 | 1 | 2.21 | 1.22 | 25.47 | 26.68 | -1.21 | 1.21 | -0.02 | 0.02 |
| gene-LOC105281173 | https://www.ncbi.nlm.nih.gov/gene/?term=LOC105281173 | 1595 | 38 | 238.24 | 1 | 6.27 | 2.63 | 59.87 | 61.09 | -1.22 | 1.22 | -0.03 | 0.03 |
| gene-LOC105283649 | https://www.ncbi.nlm.nih.gov/gene/?term=LOC105283649 | 1918 | 56 | 291.97 | 1 | 5.21 | 1.79 | 37.76 | 39.05 | -1.29 | 1.29 | -0.04 | 0.04 |
| gene-LOC105276196 | https://www.ncbi.nlm.nih.gov/gene/?term=LOC105276196 | 2953 | 57 | 193.02 | 1 | 3.39 | 1.75 | 39.57 | 40.89 | -1.32 | 1.32 | -0.03 | 0.03 |
| gene-LOC105274534 | https://www.ncbi.nlm.nih.gov/gene/?term=LOC105274534 | 1329 | 59 | 443.94 | 1 | 7.52 | 1.69 | 43.5 | 44.89 | -1.39 | 1.39 | -0.06 | 0.06 |
| gene-LOC105288042 | https://www.ncbi.nlm.nih.gov/gene/?term=LOC105288042 | 3507 | 100 | 285.14 | 1 | 2.85 | 1 | 20.77 | 22.23 | -1.46 | 1.46 | -0.04 | 0.04 |
| gene-LOC105278962 | https://www.ncbi.nlm.nih.gov/gene/?term=LOC105278962 | 4491 | 109 | 242.71 | 1 | 2.23 | 0.92 | 19.66 | 21.14 | -1.48 | 1.48 | -0.04 | 0.04 |
| gene-LOC105278282 | https://www.ncbi.nlm.nih.gov/gene/?term=LOC105278282 | 1775 | 51 | 287.32 | 1 | 5.63 | 1.96 | 39.13 | 40.69 | -1.57 | 1.57 | -0.05 | 0.05 |
| gene-LOC105286514 | https://www.ncbi.nlm.nih.gov/gene/?term=LOC105286514 | 2528 | 50 | 197.78 | 1 | 3.96 | 2 | 35.29 | 37.3 | -2.01 | 2.01 | -0.04 | 0.04 |
| gene-LOC105275571 | https://www.ncbi.nlm.nih.gov/gene/?term=LOC105275571 | 1926 | 30 | 155.76 | 1 | 5.19 | 3.33 | 42.71 | 44.75 | -2.04 | 2.04 | -0.03 | 0.03 |
| gene-LOC105276538 | https://www.ncbi.nlm.nih.gov/gene/?term=LOC105276538 | 1511 | 21 | 138.98 | 1 | 6.62 | 4.76 | 57.36 | 59.41 | -2.05 | 2.05 | -0.03 | 0.03 |
| gene-LOC105285004 | https://www.ncbi.nlm.nih.gov/gene/?term=LOC105285004 | 1334 | 37 | 277.36 | 1 | 7.5 | 2.7 | 16.38 | 18.79 | -2.41 | 2.41 | -0.07 | 0.07 |
| gene-LOC105282914 | https://www.ncbi.nlm.nih.gov/gene/?term=LOC105282914 | 5623 | 43 | 76.47 | 1 | 1.78 | 2.33 | 46.2 | 49.79 | -3.59 | 3.59 | -0.03 | 0.03 |

Directory: OGL File: ogl-Ob_LI_RphaseA1.vs.RphaseB2.txt

| gene_ID | gene_link | gwidth | #Sites | #per10Kb | #dmSites | #dmsp10kb | %dmSites | %pSite1 | %pSite2 | DMpSite | ADMpSite | DMpNucl | ADMpNucl |
|---|---|---|---|---|---|---|---|---|---|---|---|---|---|
| gene-LOC105275070 | https://www.ncbi.nlm.nih.gov/gene/?term=LOC105275070 | 5667 | 37 | 65.29 | 7 | 12.35 | 18.92 | 16.3 | 1.04 | 15.26 | 15.26 | 0.1 | 0.1 |
| gene-LOC113562345 | https://www.ncbi.nlm.nih.gov/gene/?term=LOC113562345 | 2290 | 67 | 292.58 | 6 | 26.2 | 8.96 | 28.84 | 16.72 | 12.12 | 12.12 | 0.35 | 0.35 |
| gene-LOC105279898 | https://www.ncbi.nlm.nih.gov/gene/?term=LOC105279898 | 1343 | 9 | 67.01 | 1 | 7.45 | 11.11 | 83.37 | 74.78 | 8.59 | 8.59 | 0.06 | 0.06 |
| gene-LOC105286694 | https://www.ncbi.nlm.nih.gov/gene/?term=LOC105286694 | 3409 | 8 | 23.47 | 1 | 2.93 | 12.5 | 66.22 | 57.86 | 8.36 | 8.36 | 0.02 | 0.02 |
| gene-LOC105275708 | https://www.ncbi.nlm.nih.gov/gene/?term=LOC105275708 | 1772 | 14 | 79.01 | 1 | 5.64 | 7.14 | 50.83 | 43.74 | 7.08 | 7.08 | 0.06 | 0.06 |
| gene-LOC105285514 | https://www.ncbi.nlm.nih.gov/gene/?term=LOC105285514 | 1277 | 14 | 109.63 | 1 | 7.83 | 7.14 | 69.69 | 65.1 | 4.59 | 4.59 | 0.05 | 0.05 |
| gene-LOC105283552 | https://www.ncbi.nlm.nih.gov/gene/?term=LOC105283552 | 1562 | 41 | 262.48 | 1 | 6.4 | 2.44 | 50.09 | 46.71 | 3.37 | 3.37 | 0.09 | 0.09 |
| gene-LOC105274900 | https://www.ncbi.nlm.nih.gov/gene/?term=LOC105274900 | 5561 | 55 | 98.9 | 3 | 5.39 | 5.45 | 32.79 | 29.47 | 3.31 | 3.31 | 0.03 | 0.03 |
| gene-LOC105281515 | https://www.ncbi.nlm.nih.gov/gene/?term=LOC105281515 | 8827 | 41 | 46.45 | 1 | 1.13 | 2.44 | 31.51 | 28.2 | 3.31 | 3.31 | 0.02 | 0.02 |
| gene-LOC105278436 | https://www.ncbi.nlm.nih.gov/gene/?term=LOC105278436 | 2556 | 54 | 211.27 | 1 | 3.91 | 1.85 | 40.35 | 37.22 | 3.13 | 3.13 | 0.07 | 0.07 |
| gene-LOC105285671 | https://www.ncbi.nlm.nih.gov/gene/?term=LOC105285671 | 2263 | 101 | 446.31 | 1 | 4.42 | 0.99 | 30.8 | 27.84 | 2.96 | 2.96 | 0.13 | 0.13 |
| gene-LOC105279356 | https://www.ncbi.nlm.nih.gov/gene/?term=LOC105279356 | 1314 | 43 | 327.25 | 1 | 7.61 | 2.33 | 32.62 | 30.3 | 2.31 | 2.31 | 0.08 | 0.08 |
| gene-LOC105285674 | https://www.ncbi.nlm.nih.gov/gene/?term=LOC105285674 | 3035 | 92 | 303.13 | 2 | 6.59 | 2.17 | 26.97 | 24.84 | 2.13 | 2.13 | 0.06 | 0.06 |
| gene-LOC105285202 | https://www.ncbi.nlm.nih.gov/gene/?term=LOC105285202 | 1838 | 61 | 331.88 | 1 | 5.44 | 1.64 | 26.9 | 24.8 | 2.1 | 2.1 | 0.07 | 0.07 |
| gene-LOC105275347 | https://www.ncbi.nlm.nih.gov/gene/?term=LOC105275347 | 2526 | 16 | 63.34 | 1 | 3.96 | 6.25 | 41.65 | 39.67 | 1.98 | 1.98 | 0.01 | 0.01 |
| gene-LOC105278807 | https://www.ncbi.nlm.nih.gov/gene/?term=LOC105278807 | 2166 | 65 | 300.09 | 1 | 4.62 | 1.54 | 22.17 | 20.43 | 1.74 | 1.74 | 0.05 | 0.05 |
| gene-LOC105280104 | https://www.ncbi.nlm.nih.gov/gene/?term=LOC105280104 | 3960 | 123 | 310.61 | 1 | 2.53 | 0.81 | 18.15 | 16.51 | 1.64 | 1.64 | 0.05 | 0.05 |
| gene-LOC105281266 | https://www.ncbi.nlm.nih.gov/gene/?term=LOC105281266 | 4102 | 78 | 190.15 | 1 | 2.44 | 1.28 | 18.58 | 17.01 | 1.57 | 1.57 | 0.03 | 0.03 |
| gene-LOC105286099 | https://www.ncbi.nlm.nih.gov/gene/?term=LOC105286099 | 5919 | 67 | 113.19 | 1 | 1.69 | 1.49 | 11.4 | 9.83 | 1.57 | 1.57 | 0.02 | 0.02 |
| gene-LOC105284329 | https://www.ncbi.nlm.nih.gov/gene/?term=LOC105284329 | 4051 | 52 | 128.36 | 1 | 2.47 | 1.92 | 19.88 | 18.38 | 1.5 | 1.5 | 0.02 | 0.02 |
| gene-LOC105282794 | https://www.ncbi.nlm.nih.gov/gene/?term=LOC105282794 | 4571 | 90 | 196.89 | 1 | 2.19 | 1.11 | 17.25 | 15.76 | 1.49 | 1.49 | 0.03 | 0.03 |
| gene-LOC105277903 | https://www.ncbi.nlm.nih.gov/gene/?term=LOC105277903 | 1835 | 36 | 196.19 | 1 | 5.45 | 2.78 | 51.23 | 49.79 | 1.45 | 1.45 | 0.03 | 0.03 |
| gene-LOC105280786 | https://www.ncbi.nlm.nih.gov/gene/?term=LOC105280786 | 2573 | 115 | 446.95 | 1 | 3.89 | 0.87 | 28.39 | 27.06 | 1.33 | 1.33 | 0.06 | 0.06 |
| gene-LOC105288066 | https://www.ncbi.nlm.nih.gov/gene/?term=LOC105288066 | 1964 | 27 | 137.47 | 1 | 5.09 | 3.7 | 34.07 | 32.8 | 1.27 | 1.27 | 0.02 | 0.02 |
| gene-LOC105275451 | https://www.ncbi.nlm.nih.gov/gene/?term=LOC105275451 | 6035 | 113 | 187.24 | 1 | 1.66 | 0.88 | 24.46 | 23.26 | 1.2 | 1.2 | 0.02 | 0.02 |
| gene-LOC105279664 | https://www.ncbi.nlm.nih.gov/gene/?term=LOC105279664 | 4023 | 53 | 131.74 | 1 | 2.49 | 1.89 | 19.07 | 17.88 | 1.19 | 1.19 | 0.02 | 0.02 |
| gene-LOC105283943 | https://www.ncbi.nlm.nih.gov/gene/?term=LOC105283943 | 3559 | 45 | 126.44 | 1 | 2.81 | 2.22 | 16.9 | 15.77 | 1.14 | 1.14 | 0.01 | 0.01 |
| gene-LOC113561805 | https://www.ncbi.nlm.nih.gov/gene/?term=LOC113561805 | 3337 | 67 | 200.78 | 1 | 3 | 1.49 | 36.13 | 35.02 | 1.11 | 1.11 | 0.02 | 0.02 |
| gene-LOC105275363 | https://www.ncbi.nlm.nih.gov/gene/?term=LOC105275363 | 3798 | 157 | 413.38 | 1 | 2.63 | 0.64 | 17.94 | 16.87 | 1.07 | 1.07 | 0.04 | 0.04 |
| gene-LOC105281173 | https://www.ncbi.nlm.nih.gov/gene/?term=LOC105281173 | 1595 | 38 | 238.24 | 1 | 6.27 | 2.63 | 59.87 | 58.85 | 1.02 | 1.02 | 0.02 | 0.02 |
| gene-LOC105276692 | https://www.ncbi.nlm.nih.gov/gene/?term=LOC105276692 | 1745 | 29 | 166.19 | 1 | 5.73 | 3.45 | 25.96 | 24.96 | 1 | 1 | 0.02 | 0.02 |
| gene-LOC105288078 | https://www.ncbi.nlm.nih.gov/gene/?term=LOC105288078 | 19126 | 286 | 149.53 | 1 | 0.52 | 0.35 | 17.66 | 16.67 | 0.99 | 0.99 | 0.01 | 0.01 |
| gene-LOC105280956 | https://www.ncbi.nlm.nih.gov/gene/?term=LOC105280956 | 2639 | 71 | 269.04 | 1 | 3.79 | 1.41 | 18.77 | 17.79 | 0.98 | 0.98 | 0.03 | 0.03 |
| gene-LOC105288036 | https://www.ncbi.nlm.nih.gov/gene/?term=LOC105288036 | 5110 | 181 | 354.21 | 1 | 1.96 | 0.55 | 18.99 | 18.01 | 0.98 | 0.98 | 0.03 | 0.03 |
| gene-LOC105283123 | https://www.ncbi.nlm.nih.gov/gene/?term=LOC105283123 | 1507 | 40 | 265.43 | 1 | 6.64 | 2.5 | 16.19 | 15.23 | 0.96 | 0.96 | 0.03 | 0.03 |
| gene-LOC113561621 | https://www.ncbi.nlm.nih.gov/gene/?term=LOC113561621 | 2249 | 76 | 337.93 | 1 | 4.45 | 1.32 | 22.44 | 21.49 | 0.95 | 0.95 | 0.03 | 0.03 |
| gene-LOC105276059 | https://www.ncbi.nlm.nih.gov/gene/?term=LOC105276059 | 6256 | 125 | 199.81 | 1 | 1.6 | 0.8 | 10.91 | 10 | 0.91 | 0.91 | 0.02 | 0.02 |
| gene-LOC105285060 | https://www.ncbi.nlm.nih.gov/gene/?term=LOC105285060 | 1285 | 16 | 124.51 | 1 | 7.78 | 6.25 | 47.94 | 47.04 | 0.9 | 0.9 | 0.01 | 0.01 |
| gene-LOC105283592 | https://www.ncbi.nlm.nih.gov/gene/?term=LOC105283592 | 3479 | 71 | 204.08 | 1 | 2.87 | 1.41 | 21.54 | 20.67 | 0.88 | 0.88 | 0.02 | 0.02 |
| gene-LOC105284184 | https://www.ncbi.nlm.nih.gov/gene/?term=LOC105284184 | 2535 | 80 | 315.58 | 1 | 3.94 | 1.25 | 35.73 | 34.87 | 0.86 | 0.86 | 0.03 | 0.03 |
| gene-LOC105276901 | https://www.ncbi.nlm.nih.gov/gene/?term=LOC105276901 | 3254 | 84 | 258.14 | 1 | 3.07 | 1.19 | 27.21 | 26.37 | 0.84 | 0.84 | 0.02 | 0.02 |
| gene-LOC105287854 | https://www.ncbi.nlm.nih.gov/gene/?term=LOC105287854 | 3644 | 49 | 134.47 | 1 | 2.74 | 2.04 | 24.37 | 23.53 | 0.84 | 0.84 | 0.01 | 0.01 |
| gene-LOC105274823 | https://www.ncbi.nlm.nih.gov/gene/?term=LOC105274823 | 4620 | 138 | 298.7 | 1 | 2.16 | 0.72 | 21.4 | 20.58 | 0.82 | 0.82 | 0.02 | 0.02 |
| gene-LOC105276612 | https://www.ncbi.nlm.nih.gov/gene/?term=LOC105276612 | 4157 | 178 | 428.19 | 1 | 2.41 | 0.56 | 10.68 | 9.88 | 0.8 | 0.8 | 0.03 | 0.03 |
| gene-LOC105284043 | https://www.ncbi.nlm.nih.gov/gene/?term=LOC105284043 | 5661 | 283 | 499.91 | 1 | 1.77 | 0.35 | 10.38 | 9.58 | 0.79 | 0.79 | 0.04 | 0.04 |
| gene-LOC105287417 | https://www.ncbi.nlm.nih.gov/gene/?term=LOC105287417 | 1025 | 19 | 185.37 | 1 | 9.76 | 5.26 | 59.44 | 58.65 | 0.79 | 0.79 | 0.01 | 0.01 |
| gene-LOC105282156 | https://www.ncbi.nlm.nih.gov/gene/?term=LOC105282156 | 2577 | 122 | 473.42 | 1 | 3.88 | 0.82 | 25.73 | 24.95 | 0.78 | 0.78 | 0.04 | 0.04 |
| gene-LOC105286452 | https://www.ncbi.nlm.nih.gov/gene/?term=LOC105286452 | 6888 | 64 | 92.92 | 1 | 1.45 | 1.56 | 30.17 | 29.46 | 0.72 | 0.72 | 0.01 | 0.01 |
| gene-LOC105283728 | https://www.ncbi.nlm.nih.gov/gene/?term=LOC105283728 | 3256 | 117 | 359.34 | 2 | 6.14 | 1.71 | 18.77 | 18.07 | 0.7 | 0.7 | 0.03 | 0.03 |
| gene-LOC105285534 | https://www.ncbi.nlm.nih.gov/gene/?term=LOC105285534 | 3944 | 232 | 588.24 | 1 | 2.54 | 0.43 | 11.59 | 10.9 | 0.69 | 0.69 | 0.04 | 0.04 |
| gene-LOC105275460 | https://www.ncbi.nlm.nih.gov/gene/?term=LOC105275460 | 2473 | 118 | 477.15 | 1 | 4.04 | 0.85 | 13.95 | 13.28 | 0.67 | 0.67 | 0.03 | 0.03 |
| gene-LOC105285639 | https://www.ncbi.nlm.nih.gov/gene/?term=LOC105285639 | 10325 | 312 | 302.18 | 2 | 1.94 | 0.64 | 12.89 | 12.24 | 0.65 | 0.65 | 0.02 | 0.02 |
| gene-LOC105285055 | https://www.ncbi.nlm.nih.gov/gene/?term=LOC105285055 | 4108 | 261 | 635.35 | 1 | 2.43 | 0.38 | 10.75 | 10.11 | 0.64 | 0.64 | 0.04 | 0.04 |
| gene-LOC105278534 | https://www.ncbi.nlm.nih.gov/gene/?term=LOC105278534 | 7364 | 176 | 239 | 2 | 2.72 | 1.14 | 19.64 | 19.01 | 0.63 | 0.63 | 0.02 | 0.02 |
| gene-LOC105281848 | https://www.ncbi.nlm.nih.gov/gene/?term=LOC105281848 | 2122 | 47 | 221.49 | 1 | 4.71 | 2.13 | 27.19 | 26.57 | 0.63 | 0.63 | 0.01 | 0.01 |
| gene-LOC105278283 | https://www.ncbi.nlm.nih.gov/gene/?term=LOC105278283 | 2317 | 138 | 595.6 | 1 | 4.32 | 0.72 | 25.64 | 25.05 | 0.59 | 0.59 | 0.04 | 0.04 |
| gene-LOC105279298 | https://www.ncbi.nlm.nih.gov/gene/?term=LOC105279298 | 2773 | 133 | 479.62 | 2 | 7.21 | 1.5 | 23.97 | 23.38 | 0.59 | 0.59 | 0.03 | 0.03 |
| gene-LOC105285179 | https://www.ncbi.nlm.nih.gov/gene/?term=LOC105285179 | 4191 | 171 | 408.02 | 1 | 2.39 | 0.58 | 19 | 18.45 | 0.55 | 0.55 | 0.02 | 0.02 |
| gene-LOC105278497 | https://www.ncbi.nlm.nih.gov/gene/?term=LOC105278497 | 2072 | 95 | 458.49 | 1 | 4.83 | 1.05 | 19.39 | 18.89 | 0.5 | 0.5 | 0.02 | 0.02 |
| gene-LOC105278108 | https://www.ncbi.nlm.nih.gov/gene/?term=LOC105278108 | 4392 | 43 | 97.91 | 1 | 2.28 | 2.33 | 26.73 | 26.25 | 0.49 | 0.49 | 0 | 0 |

| | | | | | | | | | | | | | |
|---|---|---|---|---|---|---|---|---|---|---|---|---|---|
| gene-LOC105278145 | https://www.ncbi.nlm.nih.gov/gene/?term=LOC105278145 | 4470 | 235 | 525.73 | 1 | 2.24 | 0.43 | 11.34 | 10.86 | 0.48 | 0.48 | 0.03 | 0.03 |
| gene-LOC105286997 | https://www.ncbi.nlm.nih.gov/gene/?term=LOC105286997 | 2850 | 106 | 371.93 | 1 | 3.51 | 0.94 | 12.07 | 11.62 | 0.46 | 0.46 | 0.02 | 0.02 |
| gene-LOC105275179 | https://www.ncbi.nlm.nih.gov/gene/?term=LOC105275179 | 3271 | 69 | 210.94 | 1 | 3.06 | 1.45 | 18.8 | 18.36 | 0.45 | 0.45 | 0.01 | 0.01 |
| gene-LOC105275374 | https://www.ncbi.nlm.nih.gov/gene/?term=LOC105275374 | 11376 | 454 | 399.09 | 1 | 0.88 | 0.22 | 10.82 | 10.43 | 0.38 | 0.38 | 0.02 | 0.02 |
| gene-LOC105287646 | https://www.ncbi.nlm.nih.gov/gene/?term=LOC105287646 | 3556 | 83 | 233.41 | 1 | 2.81 | 1.2 | 17.22 | 16.85 | 0.38 | 0.38 | 0.01 | 0.01 |
| gene-LOC105287550 | https://www.ncbi.nlm.nih.gov/gene/?term=LOC105287550 | 15659 | 549 | 350.6 | 1 | 0.64 | 0.18 | 6.33 | 5.97 | 0.37 | 0.37 | 0.01 | 0.01 |
| gene-LOC105285406 | https://www.ncbi.nlm.nih.gov/gene/?term=LOC105285406 | 8015 | 60 | 74.86 | 1 | 1.25 | 1.67 | 25.86 | 25.52 | 0.34 | 0.34 | 0 | 0 |
| gene-LOC105274856 | https://www.ncbi.nlm.nih.gov/gene/?term=LOC105274856 | 5216 | 209 | 400.69 | 1 | 1.92 | 0.48 | 18.76 | 18.44 | 0.31 | 0.31 | 0.01 | 0.01 |
| gene-LOC105274699 | https://www.ncbi.nlm.nih.gov/gene/?term=LOC105274699 | 2386 | 49 | 205.36 | 1 | 4.19 | 2.04 | 23.32 | 23.03 | 0.29 | 0.29 | 0.01 | 0.01 |
| gene-LOC105274531 | https://www.ncbi.nlm.nih.gov/gene/?term=LOC105274531 | 7774 | 273 | 351.17 | 1 | 1.29 | 0.37 | 11.43 | 11.16 | 0.27 | 0.27 | 0.01 | 0.01 |
| gene-LOC105283649 | https://www.ncbi.nlm.nih.gov/gene/?term=LOC105283649 | 1918 | 53 | 276.33 | 1 | 5.21 | 1.89 | 37.66 | 37.39 | 0.27 | 0.27 | 0.01 | 0.01 |
| gene-LOC105274851 | https://www.ncbi.nlm.nih.gov/gene/?term=LOC105274851 | 6616 | 129 | 194.98 | 1 | 1.51 | 0.78 | 11.16 | 10.94 | 0.22 | 0.22 | 0 | 0 |
| gene-LOC105279038 | https://www.ncbi.nlm.nih.gov/gene/?term=LOC105279038 | 9453 | 297 | 314.19 | 1 | 1.06 | 0.34 | 9.64 | 9.43 | 0.21 | 0.21 | 0.01 | 0.01 |
| gene-LOC105282149 | https://www.ncbi.nlm.nih.gov/gene/?term=LOC105282149 | 1787 | 103 | 576.39 | 1 | 5.6 | 0.97 | 41.32 | 41.12 | 0.2 | 0.2 | 0.01 | 0.01 |
| gene-LOC105275187 | https://www.ncbi.nlm.nih.gov/gene/?term=LOC105275187 | 5534 | 65 | 117.46 | 1 | 1.81 | 1.54 | 11.46 | 11.27 | 0.19 | 0.19 | 0 | 0 |
| gene-LOC105275327 | https://www.ncbi.nlm.nih.gov/gene/?term=LOC105275327 | 14245 | 458 | 321.52 | 1 | 0.7 | 0.22 | 7.5 | 7.33 | 0.17 | 0.17 | 0.01 | 0.01 |
| gene-LOC105278666 | https://www.ncbi.nlm.nih.gov/gene/?term=LOC105278666 | 10341 | 439 | 424.52 | 1 | 0.97 | 0.23 | 7.06 | 6.9 | 0.16 | 0.16 | 0.01 | 0.01 |
| gene-LOC105278649 | https://www.ncbi.nlm.nih.gov/gene/?term=LOC105278649 | 5616 | 138 | 245.73 | 1 | 1.78 | 0.72 | 13.54 | 13.42 | 0.12 | 0.12 | 0 | 0 |
| gene-LOC105288079 | https://www.ncbi.nlm.nih.gov/gene/?term=LOC105288079 | 3473 | 96 | 276.42 | 1 | 2.88 | 1.04 | 19.82 | 19.7 | 0.12 | 0.12 | 0 | 0 |
| gene-LOC105277856 | https://www.ncbi.nlm.nih.gov/gene/?term=LOC105277856 | 3442 | 29 | 84.25 | 1 | 2.91 | 3.45 | 32.81 | 32.75 | 0.06 | 0.06 | 0 | 0 |
| gene-LOC105284355 | https://www.ncbi.nlm.nih.gov/gene/?term=LOC105284355 | 3904 | 152 | 389.34 | 1 | 2.56 | 0.66 | 20.02 | 19.96 | 0.06 | 0.06 | 0 | 0 |
| gene-LOC105287528 | https://www.ncbi.nlm.nih.gov/gene/?term=LOC105287528 | 14195 | 375 | 264.18 | 1 | 0.7 | 0.27 | 5.12 | 5.07 | 0.05 | 0.05 | 0 | 0 |
| gene-LOC105282811 | https://www.ncbi.nlm.nih.gov/gene/?term=LOC105282811 | 5765 | 196 | 339.98 | 1 | 1.73 | 0.51 | 8.25 | 8.31 | -0.06 | 0.06 | 0 | 0 |
| gene-LOC105284704 | https://www.ncbi.nlm.nih.gov/gene/?term=LOC105284704 | 2400 | 59 | 245.83 | 1 | 4.17 | 1.69 | 22.63 | 22.7 | -0.07 | 0.07 | 0 | 0 |
| gene-LOC105274706 | https://www.ncbi.nlm.nih.gov/gene/?term=LOC105274706 | 1446 | 49 | 338.87 | 1 | 6.92 | 2.04 | 38.21 | 38.31 | -0.1 | 0.1 | 0 | 0 |
| gene-LOC105286274 | https://www.ncbi.nlm.nih.gov/gene/?term=LOC105286274 | 3817 | 179 | 468.95 | 1 | 2.62 | 0.56 | 16.95 | 17.06 | -0.11 | 0.11 | -0.01 | 0.01 |
| gene-LOC105278282 | https://www.ncbi.nlm.nih.gov/gene/?term=LOC105278282 | 1775 | 52 | 292.96 | 1 | 5.63 | 1.92 | 41.01 | 41.14 | -0.14 | 0.14 | 0 | 0 |
| gene-LOC105274590 | https://www.ncbi.nlm.nih.gov/gene/?term=LOC105274590 | 4516 | 67 | 148.36 | 1 | 2.21 | 1.49 | 28.5 | 28.65 | -0.15 | 0.15 | 0 | 0 |
| gene-LOC105282067 | https://www.ncbi.nlm.nih.gov/gene/?term=LOC105282067 | 5941 | 49 | 82.48 | 1 | 1.68 | 2.04 | 25.2 | 25.4 | -0.2 | 0.2 | 0 | 0 |
| gene-LOC105282658 | https://www.ncbi.nlm.nih.gov/gene/?term=LOC105282658 | 2421 | 52 | 214.79 | 1 | 4.13 | 1.92 | 21.81 | 22.01 | -0.2 | 0.2 | 0 | 0 |
| gene-LOC105276527 | https://www.ncbi.nlm.nih.gov/gene/?term=LOC105276527 | 7923 | 305 | 384.96 | 1 | 1.26 | 0.33 | 9.71 | 9.92 | -0.21 | 0.21 | -0.01 | 0.01 |
| gene-LOC105287095 | https://www.ncbi.nlm.nih.gov/gene/?term=LOC105287095 | 1435 | 58 | 404.18 | 1 | 6.97 | 1.72 | 28.41 | 28.65 | -0.24 | 0.24 | -0.01 | 0.01 |
| gene-LOC105288042 | https://www.ncbi.nlm.nih.gov/gene/?term=LOC105288042 | 3507 | 88 | 250.93 | 1 | 2.85 | 1.14 | 21.32 | 21.59 | -0.27 | 0.27 | -0.01 | 0.01 |
| gene-LOC105285807 | https://www.ncbi.nlm.nih.gov/gene/?term=LOC105285807 | 6063 | 213 | 351.31 | 1 | 1.65 | 0.47 | 9.29 | 9.56 | -0.28 | 0.28 | -0.01 | 0.01 |
| gene-LOC105286515 | https://www.ncbi.nlm.nih.gov/gene/?term=LOC105286515 | 3751 | 107 | 285.26 | 1 | 2.67 | 0.93 | 12.3 | 12.58 | -0.28 | 0.28 | -0.01 | 0.01 |
| gene-LOC105280774 | https://www.ncbi.nlm.nih.gov/gene/?term=LOC105280774 | 2793 | 100 | 358.04 | 1 | 3.58 | 1 | 31.9 | 32.19 | -0.29 | 0.29 | -0.01 | 0.01 |
| gene-LOC105278228 | https://www.ncbi.nlm.nih.gov/gene/?term=LOC105278228 | 8610 | 156 | 181.18 | 1 | 1.16 | 0.64 | 22.2 | 22.5 | -0.3 | 0.3 | -0.01 | 0.01 |
| gene-LOC105284036 | https://www.ncbi.nlm.nih.gov/gene/?term=LOC105284036 | 3254 | 96 | 295.02 | 1 | 3.07 | 1.04 | 23.11 | 23.46 | -0.35 | 0.35 | -0.01 | 0.01 |
| gene-LOC105280348 | https://www.ncbi.nlm.nih.gov/gene/?term=LOC105280348 | 3797 | 71 | 186.99 | 1 | 2.63 | 1.41 | 20.69 | 21.13 | -0.44 | 0.44 | -0.01 | 0.01 |
| gene-LOC105274449 | https://www.ncbi.nlm.nih.gov/gene/?term=LOC105274449 | 1549 | 48 | 309.88 | 1 | 6.46 | 2.08 | 50.45 | 50.96 | -0.51 | 0.51 | -0.02 | 0.02 |
| gene-LOC105278877 | https://www.ncbi.nlm.nih.gov/gene/?term=LOC105278877 | 1968 | 70 | 355.69 | 1 | 5.08 | 1.43 | 34.63 | 35.21 | -0.58 | 0.58 | -0.02 | 0.02 |
| gene-LOC105280377 | https://www.ncbi.nlm.nih.gov/gene/?term=LOC105280377 | 2024 | 65 | 321.15 | 1 | 4.94 | 1.54 | 47.38 | 47.97 | -0.59 | 0.59 | -0.02 | 0.02 |
| gene-LOC105278962 | https://www.ncbi.nlm.nih.gov/gene/?term=LOC105278962 | 4491 | 89 | 198.17 | 1 | 2.23 | 1.12 | 22.04 | 22.67 | -0.63 | 0.63 | -0.01 | 0.01 |
| gene-LOC105278454 | https://www.ncbi.nlm.nih.gov/gene/?term=LOC105278454 | 1972 | 40 | 202.84 | 1 | 5.07 | 2.5 | 48.64 | 49.28 | -0.64 | 0.64 | -0.01 | 0.01 |
| gene-LOC105280002 | https://www.ncbi.nlm.nih.gov/gene/?term=LOC105280002 | 1940 | 27 | 139.18 | 1 | 5.15 | 3.7 | 33.73 | 34.37 | -0.64 | 0.64 | -0.01 | 0.01 |
| gene-LOC105274534 | https://www.ncbi.nlm.nih.gov/gene/?term=LOC105274534 | 1329 | 57 | 428.89 | 1 | 7.52 | 1.75 | 46.71 | 47.65 | -0.94 | 0.94 | -0.04 | 0.04 |
| gene-LOC105280819 | https://www.ncbi.nlm.nih.gov/gene/?term=LOC105280819 | 1639 | 37 | 225.75 | 1 | 6.1 | 2.7 | 36.94 | 38.07 | -1.12 | 1.12 | -0.03 | 0.03 |
| gene-LOC105278384 | https://www.ncbi.nlm.nih.gov/gene/?term=LOC105278384 | 1390 | 35 | 251.8 | 1 | 7.19 | 2.86 | 48.96 | 50.26 | -1.3 | 1.3 | -0.03 | 0.03 |
| gene-LOC105283712 | https://www.ncbi.nlm.nih.gov/gene/?term=LOC105283712 | 1910 | 36 | 188.48 | 1 | 5.24 | 2.78 | 22.92 | 24.43 | -1.51 | 1.51 | -0.03 | 0.03 |
| gene-LOC105281700 | https://www.ncbi.nlm.nih.gov/gene/?term=LOC105281700 | 2208 | 32 | 144.93 | 1 | 4.53 | 3.12 | 37.54 | 39.09 | -1.55 | 1.55 | -0.02 | 0.02 |
| gene-LOC105276538 | https://www.ncbi.nlm.nih.gov/gene/?term=LOC105276538 | 1511 | 18 | 119.13 | 1 | 6.62 | 5.56 | 64.69 | 66.35 | -1.65 | 1.65 | -0.02 | 0.02 |
| gene-LOC105286514 | https://www.ncbi.nlm.nih.gov/gene/?term=LOC105286514 | 2528 | 46 | 181.96 | 1 | 3.96 | 2.17 | 35.96 | 37.79 | -1.83 | 1.83 | -0.03 | 0.03 |
| gene-LOC105280596 | https://www.ncbi.nlm.nih.gov/gene/?term=LOC105280596 | 3434 | 104 | 302.85 | 1 | 2.91 | 0.96 | 16.78 | 18.68 | -1.9 | 1.9 | -0.06 | 0.06 |
| gene-LOC105283989 | https://www.ncbi.nlm.nih.gov/gene/?term=LOC105283989 | 4948 | 25 | 50.53 | 1 | 2.02 | 4 | 35.77 | 37.79 | -2.02 | 2.02 | -0.01 | 0.01 |
| gene-LOC105286999 | https://www.ncbi.nlm.nih.gov/gene/?term=LOC105286999 | 2141 | 42 | 196.17 | 1 | 4.67 | 2.38 | 33.74 | 35.94 | -2.2 | 2.2 | -0.04 | 0.04 |
| gene-LOC105283967 | https://www.ncbi.nlm.nih.gov/gene/?term=LOC105283967 | 2393 | 40 | 167.15 | 1 | 4.18 | 2.5 | 52.37 | 54.65 | -2.28 | 2.28 | -0.04 | 0.04 |
| gene-LOC105276196 | https://www.ncbi.nlm.nih.gov/gene/?term=LOC105276196 | 2953 | 59 | 199.8 | 1 | 3.39 | 1.69 | 38.71 | 41.49 | -2.77 | 2.77 | -0.06 | 0.06 |
| gene-LOC105285409 | https://www.ncbi.nlm.nih.gov/gene/?term=LOC105285409 | 5099 | 49 | 96.1 | 1 | 1.96 | 2.04 | 1.72 | 4.82 | -3.09 | 3.09 | -0.03 | 0.03 |
| gene-LOC105283492 | https://www.ncbi.nlm.nih.gov/gene/?term=LOC105283492 | 1535 | 9 | 58.63 | 1 | 6.51 | 11.11 | 30.57 | 36.97 | -6.4 | 6.4 | -0.04 | 0.04 |

Directory: OGL File: ogl-Ob_LI_RphaseA2.txt

| gene_ID | gene_link | gwidth | #Sites | #per10Kb | %perSite | %pNucl |
|---|---|---|---|---|---|---|
| gene-LOC113561532 | https://www.ncbi.nlm.nih.gov/gene/?term=LOC113561532 | 72 | 6 | 833.33 | 95.64 | 7.97 |
| gene-LOC105282149 | https://www.ncbi.nlm.nih.gov/gene/?term=LOC105282149 | 1787 | 190 | 1063.23 | 44.1 | 4.69 |
| gene-LOC113562627 | https://www.ncbi.nlm.nih.gov/gene/?term=LOC113562627 | 397 | 26 | 654.91 | 67.37 | 4.41 |
| gene-LOC105285635 | https://www.ncbi.nlm.nih.gov/gene/?term=LOC105285635 | 1175 | 164 | 1395.74 | 29.93 | 4.18 |
| gene-LOC105283830 | https://www.ncbi.nlm.nih.gov/gene/?term=LOC105283830 | 252 | 24 | 952.38 | 43.77 | 4.17 |
| gene-LOC105286269 | https://www.ncbi.nlm.nih.gov/gene/?term=LOC105286269 | 1401 | 110 | 785.15 | 52.63 | 4.13 |
| gene-LOC105286531 | https://www.ncbi.nlm.nih.gov/gene/?term=LOC105286531 | 372 | 43 | 1155.91 | 35.68 | 4.12 |
| gene-LOC105280739 | https://www.ncbi.nlm.nih.gov/gene/?term=LOC105280739 | 1991 | 147 | 738.32 | 54.96 | 4.06 |
| gene-LOC105287811 | https://www.ncbi.nlm.nih.gov/gene/?term=LOC105287811 | 2155 | 191 | 886.31 | 45.74 | 4.05 |
| gene-LOC105278849 | https://www.ncbi.nlm.nih.gov/gene/?term=LOC105278849 | 776 | 127 | 1636.6 | 24.48 | 4.01 |
| gene-LOC105276360 | https://www.ncbi.nlm.nih.gov/gene/?term=LOC105276360 | 1466 | 203 | 1384.72 | 28.73 | 3.98 |
| gene-LOC105285394 | https://www.ncbi.nlm.nih.gov/gene/?term=LOC105285394 | 1307 | 74 | 566.18 | 69.6 | 3.94 |
| gene-LOC105286801 | https://www.ncbi.nlm.nih.gov/gene/?term=LOC105286801 | 1102 | 107 | 970.96 | 40.62 | 3.94 |
| gene-LOC105274534 | https://www.ncbi.nlm.nih.gov/gene/?term=LOC105274534 | 1329 | 119 | 895.41 | 42.8 | 3.83 |
| gene-LOC105275314 | https://www.ncbi.nlm.nih.gov/gene/?term=LOC105275314 | 1489 | 132 | 886.5 | 41.45 | 3.67 |
| gene-LOC105277648 | https://www.ncbi.nlm.nih.gov/gene/?term=LOC105277648 | 722 | 45 | 623.27 | 58.13 | 3.62 |
| gene-LOC105278160 | https://www.ncbi.nlm.nih.gov/gene/?term=LOC105278160 | 1872 | 184 | 982.91 | 36.84 | 3.62 |
| gene-LOC105279052 | https://www.ncbi.nlm.nih.gov/gene/?term=LOC105279052 | 1130 | 163 | 1442.48 | 24.87 | 3.59 |
| gene-LOC105274711 | https://www.ncbi.nlm.nih.gov/gene/?term=LOC105274711 | 1655 | 124 | 749.24 | 46.91 | 3.51 |
| gene-LOC105278844 | https://www.ncbi.nlm.nih.gov/gene/?term=LOC105278844 | 1101 | 139 | 1262.49 | 27.76 | 3.5 |
| gene-LOC105287677 | https://www.ncbi.nlm.nih.gov/gene/?term=LOC105287677 | 1588 | 143 | 900.5 | 38.14 | 3.43 |
| gene-LOC105286235 | https://www.ncbi.nlm.nih.gov/gene/?term=LOC105286235 | 1510 | 122 | 807.95 | 42.1 | 3.4 |
| gene-LOC105286737 | https://www.ncbi.nlm.nih.gov/gene/?term=LOC105286737 | 2690 | 404 | 1501.86 | 22.66 | 3.4 |
| gene-LOC105287692 | https://www.ncbi.nlm.nih.gov/gene/?term=LOC105287692 | 1505 | 156 | 1036.54 | 32.62 | 3.38 |

...

... only first 25 lines shown ...

Directory: OGL File: ogl-Ob_LI_RphaseA2.vs.RphaseB1.txt

| gene_ID | gene_link | gwidth | #Sites | #per10Kb | #dmSites | #dmsp10kb | %dmSites | %pSite1 | %pSite2 | DMpSite | ADMpSite | DMpNucl | ADMpNucl |
|---|---|---|---|---|---|---|---|---|---|---|---|---|---|
| gene-LOC113562974 | https://www.ncbi.nlm.nih.gov/gene/?term=LOC113562974 | 1832 | 2 | 10.92 | 1 | 5.46 | 50 | 57.94 | 9 | 48.94 | 48.94 | 0.05 | 0.05 |
| gene-LOC113562345 | https://www.ncbi.nlm.nih.gov/gene/?term=LOC113562345 | 2290 | 80 | 349.34 | 16 | 69.87 | 20 | 25.78 | 13.98 | 11.8 | 11.8 | 0.41 | 0.41 |
| gene-LOC105275070 | https://www.ncbi.nlm.nih.gov/gene/?term=LOC105275070 | 5667 | 64 | 112.93 | 6 | 10.59 | 9.38 | 14.32 | 5.2 | 9.13 | 9.13 | 0.1 | 0.1 |
| gene-LOC105275708 | https://www.ncbi.nlm.nih.gov/gene/?term=LOC105275708 | 1772 | 19 | 107.22 | 1 | 5.64 | 5.26 | 37.25 | 31.29 | 5.96 | 5.96 | 0.06 | 0.06 |
| gene-LOC105285514 | https://www.ncbi.nlm.nih.gov/gene/?term=LOC105285514 | 1277 | 11 | 86.14 | 1 | 7.83 | 9.09 | 90.74 | 85.12 | 5.62 | 5.62 | 0.05 | 0.05 |
| gene-LOC105286302 | https://www.ncbi.nlm.nih.gov/gene/?term=LOC105286302 | 4536 | 24 | 52.91 | 1 | 2.2 | 4.17 | 20.45 | 15.15 | 5.3 | 5.3 | 0.03 | 0.03 |
| gene-LOC105286983 | https://www.ncbi.nlm.nih.gov/gene/?term=LOC105286983 | 16901 | 70 | 41.42 | 1 | 0.59 | 1.43 | 6.21 | 1.14 | 5.06 | 5.06 | 0.02 | 0.02 |
| gene-LOC105281039 | https://www.ncbi.nlm.nih.gov/gene/?term=LOC105281039 | 1471 | 15 | 101.97 | 1 | 6.8 | 6.67 | 36.89 | 31.93 | 4.96 | 4.96 | 0.05 | 0.05 |
| gene-LOC105279900 | https://www.ncbi.nlm.nih.gov/gene/?term=LOC105279900 | 2191 | 56 | 255.59 | 1 | 4.56 | 1.79 | 33.63 | 28.91 | 4.72 | 4.72 | 0.12 | 0.12 |
| gene-LOC105284609 | https://www.ncbi.nlm.nih.gov/gene/?term=LOC105284609 | 5678 | 44 | 77.49 | 1 | 1.76 | 2.27 | 32.19 | 27.8 | 4.39 | 4.39 | 0.03 | 0.03 |
| gene-LOC105276538 | https://www.ncbi.nlm.nih.gov/gene/?term=LOC105276538 | 1511 | 23 | 152.22 | 1 | 6.62 | 4.35 | 69.63 | 65.38 | 4.25 | 4.25 | 0.06 | 0.06 |
| gene-LOC105285315 | https://www.ncbi.nlm.nih.gov/gene/?term=LOC105285315 | 5458 | 33 | 60.46 | 1 | 1.83 | 3.03 | 33.27 | 29.4 | 3.88 | 3.88 | 0.02 | 0.02 |
| gene-LOC105283712 | https://www.ncbi.nlm.nih.gov/gene/?term=LOC105283712 | 1910 | 33 | 172.77 | 1 | 5.24 | 3.03 | 32.87 | 29.11 | 3.76 | 3.76 | 0.06 | 0.06 |
| gene-LOC105282893 | https://www.ncbi.nlm.nih.gov/gene/?term=LOC105282893 | 2645 | 50 | 189.04 | 1 | 3.78 | 2 | 28.92 | 25.31 | 3.6 | 3.6 | 0.07 | 0.07 |
| gene-LOC105281515 | https://www.ncbi.nlm.nih.gov/gene/?term=LOC105281515 | 8827 | 56 | 63.44 | 2 | 2.27 | 3.57 | 36.05 | 32.61 | 3.44 | 3.44 | 0.02 | 0.02 |
| gene-LOC105287063 | https://www.ncbi.nlm.nih.gov/gene/?term=LOC105287063 | 1874 | 55 | 293.49 | 1 | 5.34 | 1.82 | 30.47 | 27.11 | 3.37 | 3.37 | 0.1 | 0.1 |
| gene-LOC105283457 | https://www.ncbi.nlm.nih.gov/gene/?term=LOC105283457 | 1882 | 49 | 260.36 | 1 | 5.31 | 2.04 | 35.19 | 32.03 | 3.16 | 3.16 | 0.08 | 0.08 |
| gene-LOC105287417 | https://www.ncbi.nlm.nih.gov/gene/?term=LOC105287417 | 1025 | 38 | 370.73 | 1 | 9.76 | 2.63 | 35.24 | 32.37 | 2.87 | 2.87 | 0.11 | 0.11 |
| gene-LOC105288066 | https://www.ncbi.nlm.nih.gov/gene/?term=LOC105288066 | 1964 | 31 | 157.84 | 1 | 5.09 | 3.23 | 39.38 | 36.57 | 2.81 | 2.81 | 0.04 | 0.04 |
| gene-LOC105284704 | https://www.ncbi.nlm.nih.gov/gene/?term=LOC105284704 | 2400 | 66 | 275 | 1 | 4.17 | 1.52 | 25.49 | 22.92 | 2.57 | 2.57 | 0.07 | 0.07 |
| gene-LOC105285583 | https://www.ncbi.nlm.nih.gov/gene/?term=LOC105285583 | 1947 | 62 | 318.44 | 1 | 5.14 | 1.61 | 27.07 | 24.61 | 2.46 | 2.46 | 0.08 | 0.08 |
| gene-LOC105275190 | https://www.ncbi.nlm.nih.gov/gene/?term=LOC105275190 | 3837 | 77 | 200.68 | 1 | 2.61 | 1.3 | 21.84 | 19.45 | 2.39 | 2.39 | 0.05 | 0.05 |
| gene-LOC113561805 | https://www.ncbi.nlm.nih.gov/gene/?term=LOC113561805 | 3337 | 73 | 218.76 | 1 | 3 | 1.37 | 38.02 | 35.67 | 2.34 | 2.34 | 0.05 | 0.05 |
| gene-LOC105279898 | https://www.ncbi.nlm.nih.gov/gene/?term=LOC105279898 | 1343 | 16 | 119.14 | 1 | 7.45 | 6.25 | 58.57 | 56.24 | 2.33 | 2.33 | 0.03 | 0.03 |
| gene-LOC105275518 | https://www.ncbi.nlm.nih.gov/gene/?term=LOC105275518 | 2335 | 28 | 119.91 | 1 | 4.28 | 3.57 | 41.74 | 39.46 | 2.28 | 2.28 | 0.03 | 0.03 |
| gene-LOC105285695 | https://www.ncbi.nlm.nih.gov/gene/?term=LOC105285695 | 1371 | 76 | 554.34 | 1 | 7.29 | 1.32 | 28.59 | 26.36 | 2.24 | 2.24 | 0.12 | 0.12 |
| gene-LOC105274706 | https://www.ncbi.nlm.nih.gov/gene/?term=LOC105274706 | 1446 | 56 | 387.28 | 1 | 6.92 | 1.79 | 39.8 | 37.63 | 2.17 | 2.17 | 0.08 | 0.08 |
| gene-LOC105288010 | https://www.ncbi.nlm.nih.gov/gene/?term=LOC105288010 | 3189 | 61 | 191.28 | 1 | 3.14 | 1.64 | 34.85 | 32.7 | 2.15 | 2.15 | 0.04 | 0.04 |
| gene-LOC105282156 | https://www.ncbi.nlm.nih.gov/gene/?term=LOC105282156 | 2577 | 134 | 519.98 | 1 | 3.88 | 0.75 | 27.52 | 25.46 | 2.06 | 2.06 | 0.11 | 0.11 |
| gene-LOC105285674 | https://www.ncbi.nlm.nih.gov/gene/?term=LOC105285674 | 3035 | 111 | 365.73 | 2 | 6.59 | 1.8 | 25.15 | 23.09 | 2.06 | 2.06 | 0.08 | 0.08 |
| gene-LOC105286644 | https://www.ncbi.nlm.nih.gov/gene/?term=LOC105286644 | 1235 | 23 | 186.23 | 1 | 8.1 | 4.35 | 43.78 | 41.75 | 2.04 | 2.04 | 0.04 | 0.04 |
| gene-LOC105285060 | https://www.ncbi.nlm.nih.gov/gene/?term=LOC105285060 | 1285 | 18 | 140.08 | 1 | 7.78 | 5.56 | 38.72 | 36.69 | 2.03 | 2.03 | 0.03 | 0.03 |
| gene-LOC105280786 | https://www.ncbi.nlm.nih.gov/gene/?term=LOC105280786 | 2573 | 143 | 555.77 | 1 | 3.89 | 0.7 | 27.63 | 25.63 | 2 | 2 | 0.11 | 0.11 |
| gene-LOC105276704 | https://www.ncbi.nlm.nih.gov/gene/?term=LOC105276704 | 2063 | 93 | 450.8 | 5 | 24.24 | 5.38 | 14.49 | 12.51 | 1.99 | 1.99 | 0.09 | 0.09 |
| gene-LOC105276272 | https://www.ncbi.nlm.nih.gov/gene/?term=LOC105276272 | 1965 | 47 | 239.19 | 1 | 5.09 | 2.13 | 25.43 | 23.44 | 1.98 | 1.98 | 0.05 | 0.05 |
| gene-LOC105286683 | https://www.ncbi.nlm.nih.gov/gene/?term=LOC105286683 | 1665 | 57 | 342.34 | 1 | 6.01 | 1.75 | 35.23 | 33.27 | 1.97 | 1.97 | 0.07 | 0.07 |
| gene-LOC105281848 | https://www.ncbi.nlm.nih.gov/gene/?term=LOC105281848 | 2122 | 60 | 282.75 | 1 | 4.71 | 1.67 | 24.48 | 22.53 | 1.95 | 1.95 | 0.06 | 0.06 |
| gene-LOC105274900 | https://www.ncbi.nlm.nih.gov/gene/?term=LOC105274900 | 5561 | 61 | 109.69 | 2 | 3.6 | 3.28 | 32.53 | 30.66 | 1.87 | 1.87 | 0.02 | 0.02 |
| gene-LOC105276078 | https://www.ncbi.nlm.nih.gov/gene/?term=LOC105276078 | 1051 | 18 | 171.27 | 1 | 9.51 | 5.56 | 28.41 | 26.59 | 1.83 | 1.83 | 0.03 | 0.03 |
| gene-LOC105280704 | https://www.ncbi.nlm.nih.gov/gene/?term=LOC105280704 | 4266 | 155 | 363.34 | 1 | 2.34 | 0.65 | 17.59 | 16.02 | 1.58 | 1.58 | 0.06 | 0.06 |
| gene-LOC105278108 | https://www.ncbi.nlm.nih.gov/gene/?term=LOC105278108 | 4392 | 57 | 129.78 | 1 | 2.28 | 1.75 | 25.91 | 24.39 | 1.52 | 1.52 | 0.02 | 0.02 |
| gene-LOC105278282 | https://www.ncbi.nlm.nih.gov/gene/?term=LOC105278282 | 1775 | 67 | 377.46 | 1 | 5.63 | 1.49 | 34.11 | 32.61 | 1.51 | 1.51 | 0.06 | 0.06 |
| gene-LOC105288079 | https://www.ncbi.nlm.nih.gov/gene/?term=LOC105288079 | 3473 | 112 | 322.49 | 1 | 2.88 | 0.89 | 18.94 | 17.43 | 1.51 | 1.51 | 0.05 | 0.05 |
| gene-LOC105276449 | https://www.ncbi.nlm.nih.gov/gene/?term=LOC105276449 | 1231 | 55 | 446.79 | 1 | 8.12 | 1.82 | 24.75 | 23.28 | 1.47 | 1.47 | 0.07 | 0.07 |
| gene-LOC105280348 | https://www.ncbi.nlm.nih.gov/gene/?term=LOC105280348 | 3797 | 78 | 205.43 | 1 | 2.63 | 1.28 | 18.55 | 17.12 | 1.44 | 1.44 | 0.03 | 0.03 |
| gene-LOC105277435 | https://www.ncbi.nlm.nih.gov/gene/?term=LOC105277435 | 3643 | 184 | 505.08 | 1 | 2.74 | 0.54 | 12.13 | 10.74 | 1.39 | 1.39 | 0.07 | 0.07 |
| gene-LOC105279298 | https://www.ncbi.nlm.nih.gov/gene/?term=LOC105279298 | 2773 | 134 | 483.23 | 2 | 7.21 | 1.49 | 24.2 | 22.81 | 1.39 | 1.39 | 0.07 | 0.07 |
| gene-LOC105282151 | https://www.ncbi.nlm.nih.gov/gene/?term=LOC105282151 | 1601 | 45 | 281.07 | 2 | 12.49 | 4.44 | 42.82 | 41.43 | 1.39 | 1.39 | 0.04 | 0.04 |
| gene-LOC105280002 | https://www.ncbi.nlm.nih.gov/gene/?term=LOC105280002 | 1940 | 34 | 175.26 | 1 | 5.15 | 2.94 | 33.64 | 32.26 | 1.38 | 1.38 | 0.02 | 0.02 |
| gene-LOC105282794 | https://www.ncbi.nlm.nih.gov/gene/?term=LOC105282794 | 4571 | 109 | 238.46 | 1 | 2.19 | 0.92 | 17.81 | 16.42 | 1.38 | 1.38 | 0.03 | 0.03 |
| gene-LOC105286514 | https://www.ncbi.nlm.nih.gov/gene/?term=LOC105286514 | 2528 | 55 | 217.56 | 1 | 3.96 | 1.82 | 37.15 | 35.79 | 1.36 | 1.36 | 0.03 | 0.03 |
| gene-LOC105285671 | https://www.ncbi.nlm.nih.gov/gene/?term=LOC105285671 | 2263 | 103 | 455.15 | 1 | 4.42 | 0.97 | 30.91 | 29.64 | 1.27 | 1.27 | 0.06 | 0.06 |
| gene-LOC105280722 | https://www.ncbi.nlm.nih.gov/gene/?term=LOC105280722 | 3237 | 93 | 287.3 | 1 | 3.09 | 1.08 | 27.81 | 26.63 | 1.18 | 1.18 | 0.03 | 0.03 |
| gene-LOC105279500 | https://www.ncbi.nlm.nih.gov/gene/?term=LOC105279500 | 3587 | 36 | 100.36 | 1 | 2.79 | 2.78 | 48.81 | 47.64 | 1.17 | 1.17 | 0.01 | 0.01 |
| gene-LOC105282682 | https://www.ncbi.nlm.nih.gov/gene/?term=LOC105282682 | 4127 | 240 | 581.54 | 1 | 2.42 | 0.42 | 13.64 | 12.51 | 1.14 | 1.14 | 0.07 | 0.07 |
| gene-LOC105288063 | https://www.ncbi.nlm.nih.gov/gene/?term=LOC105288063 | 3400 | 71 | 208.82 | 1 | 2.94 | 1.41 | 12.55 | 11.41 | 1.14 | 1.14 | 0.02 | 0.02 |
| gene-LOC105282600 | https://www.ncbi.nlm.nih.gov/gene/?term=LOC105282600 | 3715 | 67 | 180.35 | 1 | 2.69 | 1.49 | 27.6 | 26.48 | 1.12 | 1.12 | 0.02 | 0.02 |
| gene-LOC105283123 | https://www.ncbi.nlm.nih.gov/gene/?term=LOC105283123 | 1507 | 55 | 364.96 | 1 | 6.64 | 1.82 | 20.08 | 18.96 | 1.12 | 1.12 | 0.04 | 0.04 |
| gene-LOC105278071 | https://www.ncbi.nlm.nih.gov/gene/?term=LOC105278071 | 4307 | 61 | 141.63 | 1 | 2.32 | 1.64 | 11.12 | 10.02 | 1.1 | 1.1 | 0.02 | 0.02 |
| gene-LOC105285416 | https://www.ncbi.nlm.nih.gov/gene/?term=LOC105285416 | 3820 | 76 | 198.95 | 1 | 2.62 | 1.32 | 22.22 | 21.12 | 1.1 | 1.1 | 0.02 | 0.02 |

| | | | | | | | | | | | | | |
|---|---|---|---|---|---|---|---|---|---|---|---|---|---|
| gene-LOC105277856 | https://www.ncbi.nlm.nih.gov/gene/?term=LOC105277856 | 3442 | 36 | 104.59 | 1 | 2.91 | 2.78 | 37.81 | 36.75 | 1.06 | 1.06 | 0.01 | 0.01 |
| gene-LOC105280407 | https://www.ncbi.nlm.nih.gov/gene/?term=LOC105280407 | 10950 | 231 | 210.96 | 1 | 0.91 | 0.43 | 15.59 | 14.54 | 1.05 | 1.05 | 0.02 | 0.02 |
| gene-LOC105283838 | https://www.ncbi.nlm.nih.gov/gene/?term=LOC105283838 | 1748 | 33 | 188.79 | 1 | 5.72 | 3.03 | 33.49 | 32.48 | 1.01 | 1.01 | 0.02 | 0.02 |
| gene-LOC105276408 | https://www.ncbi.nlm.nih.gov/gene/?term=LOC105276408 | 5298 | 201 | 379.39 | 1 | 1.89 | 0.5 | 21.53 | 20.53 | 1 | 1 | 0.04 | 0.04 |
| gene-LOC105275460 | https://www.ncbi.nlm.nih.gov/gene/?term=LOC105275460 | 2473 | 123 | 497.37 | 1 | 4.04 | 0.81 | 13.29 | 12.32 | 0.97 | 0.97 | 0.05 | 0.05 |
| gene-LOC105280774 | https://www.ncbi.nlm.nih.gov/gene/?term=LOC105280774 | 2793 | 135 | 483.35 | 1 | 3.58 | 0.74 | 25.37 | 24.41 | 0.96 | 0.96 | 0.05 | 0.05 |
| gene-LOC105278283 | https://www.ncbi.nlm.nih.gov/gene/?term=LOC105278283 | 2317 | 145 | 625.81 | 1 | 4.32 | 0.69 | 24.77 | 23.82 | 0.95 | 0.95 | 0.06 | 0.06 |
| gene-LOC105284355 | https://www.ncbi.nlm.nih.gov/gene/?term=LOC105284355 | 3904 | 155 | 397.03 | 1 | 2.56 | 0.65 | 22.33 | 21.37 | 0.95 | 0.95 | 0.04 | 0.04 |
| gene-LOC105283592 | https://www.ncbi.nlm.nih.gov/gene/?term=LOC105283592 | 3479 | 87 | 250.07 | 1 | 2.87 | 1.15 | 22.28 | 21.36 | 0.92 | 0.92 | 0.02 | 0.02 |
| gene-LOC105285055 | https://www.ncbi.nlm.nih.gov/gene/?term=LOC105285055 | 4108 | 264 | 642.65 | 2 | 4.87 | 0.76 | 12 | 11.12 | 0.87 | 0.87 | 0.06 | 0.06 |
| gene-LOC105275347 | https://www.ncbi.nlm.nih.gov/gene/?term=LOC105275347 | 2526 | 21 | 83.14 1 | 3.96 | 4.76 | 35.82 | 34.96 | 0.86 | 0.86 | 0.01 | 0.01 | |
| gene-LOC105276969 | https://www.ncbi.nlm.nih.gov/gene/?term=LOC105276969 | 2997 | 104 | 347.01 | 1 | 3.34 | 0.96 | 11.67 | 10.81 | 0.86 | 0.86 | 0.03 | 0.03 |
| gene-LOC105277631 | https://www.ncbi.nlm.nih.gov/gene/?term=LOC105277631 | 4387 | 327 | 745.38 | 1 | 2.28 | 0.31 | 11.07 | 10.23 | 0.85 | 0.85 | 0.06 | 0.06 |
| gene-LOC105278807 | https://www.ncbi.nlm.nih.gov/gene/?term=LOC105278807 | 2166 | 77 | 355.49 | 1 | 4.62 | 1.3 | 21.89 | 21.08 | 0.81 | 0.81 | 0.03 | 0.03 |
| gene-LOC105274823 | https://www.ncbi.nlm.nih.gov/gene/?term=LOC105274823 | 4620 | 171 | 370.13 | 1 | 2.16 | 0.58 | 18.07 | 17.27 | 0.8 | 0.8 | 0.03 | 0.03 |
| gene-LOC105275179 | https://www.ncbi.nlm.nih.gov/gene/?term=LOC105275179 | 3271 | 85 | 259.86 | 1 | 3.06 | 1.18 | 16.68 | 15.92 | 0.76 | 0.76 | 0.02 | 0.02 |
| gene-LOC105280104 | https://www.ncbi.nlm.nih.gov/gene/?term=LOC105280104 | 3960 | 132 | 333.33 | 1 | 2.53 | 0.76 | 17.97 | 17.22 | 0.76 | 0.76 | 0.03 | 0.03 |
| gene-LOC105285742 | https://www.ncbi.nlm.nih.gov/gene/?term=LOC105285742 | 3446 | 122 | 354.03 | 1 | 2.9 | 0.82 | 17.63 | 16.88 | 0.76 | 0.76 | 0.03 | 0.03 |
| gene-LOC105278497 | https://www.ncbi.nlm.nih.gov/gene/?term=LOC105278497 | 2072 | 103 | 497.1 1 | 4.83 | 0.97 | 19.51 | 18.76 | 0.75 | 0.75 | 0.04 | 0.04 | |
| gene-LOC105281312 | https://www.ncbi.nlm.nih.gov/gene/?term=LOC105281312 | 2551 | 69 | 270.48 | 1 | 3.92 | 1.45 | 31.17 | 30.43 | 0.74 | 0.74 | 0.02 | 0.02 |
| gene-LOC105282491 | https://www.ncbi.nlm.nih.gov/gene/?term=LOC105282491 | 4761 | 191 | 401.18 | 1 | 2.1 | 0.52 | 9 | 8.31 | 0.69 | 0.69 | 0.03 | 0.03 |
| gene-LOC105283728 | https://www.ncbi.nlm.nih.gov/gene/?term=LOC105283728 | 3256 | 159 | 488.33 | 2 | 6.14 | 1.26 | 15.6 | 14.91 | 0.69 | 0.69 | 0.03 | 0.03 |
| gene-LOC105282825 | https://www.ncbi.nlm.nih.gov/gene/?term=LOC105282825 | 3919 | 102 | 260.27 | 1 | 2.55 | 0.98 | 14.91 | 14.23 | 0.68 | 0.68 | 0.02 | 0.02 |
| gene-LOC105283989 | https://www.ncbi.nlm.nih.gov/gene/?term=LOC105283989 | 4948 | 44 | 88.92 2 | 4.04 | 4.55 | 37.47 | 36.79 | 0.68 | 0.68 | 0.01 | 0.01 | |
| gene-LOC105283943 | https://www.ncbi.nlm.nih.gov/gene/?term=LOC105283943 | 3559 | 68 | 191.06 | 1 | 2.81 | 1.47 | 20.98 | 20.31 | 0.67 | 0.67 | 0.01 | 0.01 |
| gene-LOC105284557 | https://www.ncbi.nlm.nih.gov/gene/?term=LOC105284557 | 1672 | 91 | 544.26 | 1 | 5.98 | 1.1 | 27.82 | 27.15 | 0.67 | 0.67 | 0.04 | 0.04 |
| gene-LOC105275451 | https://www.ncbi.nlm.nih.gov/gene/?term=LOC105275451 | 6035 | 149 | 246.89 | 1 | 1.66 | 0.67 | 20.25 | 19.59 | 0.66 | 0.66 | 0.02 | 0.02 |
| gene-LOC105276560 | https://www.ncbi.nlm.nih.gov/gene/?term=LOC105276560 | 18607 | 384 | 206.37 | 3 | 1.61 | 0.78 | 2.53 | 1.87 | 0.66 | 0.66 | 0.01 | 0.01 |
| gene-LOC105288078 | https://www.ncbi.nlm.nih.gov/gene/?term=LOC105288078 | 19126 | 359 | 187.7 1 | 0.52 | 0.28 | 15.58 | 14.92 | 0.66 | 0.66 | 0.01 | 0.01 | |
| gene-LOC105287704 | https://www.ncbi.nlm.nih.gov/gene/?term=LOC105287704 | 3019 | 142 | 470.35 | 1 | 3.31 | 0.7 | 19.98 | 19.34 | 0.63 | 0.63 | 0.03 | 0.03 |
| gene-LOC105275683 | https://www.ncbi.nlm.nih.gov/gene/?term=LOC105275683 | 4256 | 204 | 479.32 | 1 | 2.35 | 0.49 | 13.38 | 12.77 | 0.61 | 0.61 | 0.03 | 0.03 |
| gene-LOC105277215 | https://www.ncbi.nlm.nih.gov/gene/?term=LOC105277215 | 2619 | 30 | 114.55 | 1 | 3.82 | 3.33 | 27.61 | 26.99 | 0.61 | 0.61 | 0.01 | 0.01 |
| gene-LOC105281296 | https://www.ncbi.nlm.nih.gov/gene/?term=LOC105281296 | 2434 | 48 | 197.21 | 1 | 4.11 | 2.08 | 28.14 | 27.55 | 0.59 | 0.59 | 0.01 | 0.01 |
| gene-LOC105278384 | https://www.ncbi.nlm.nih.gov/gene/?term=LOC105278384 | 1390 | 33 | 237.41 | 1 | 7.19 | 3.03 | 46.32 | 45.77 | 0.55 | 0.55 | 0.01 | 0.01 |
| gene-LOC105278632 | https://www.ncbi.nlm.nih.gov/gene/?term=LOC105278632 | 3624 | 150 | 413.91 | 1 | 2.76 | 0.67 | 13.57 | 13.05 | 0.53 | 0.53 | 0.02 | 0.02 |
| gene-LOC105274699 | https://www.ncbi.nlm.nih.gov/gene/?term=LOC105274699 | 2386 | 67 | 280.8 1 | 4.19 | 1.49 | 18.3 | 17.79 | 0.51 | 0.51 | 0.01 | 0.01 | |
| gene-LOC105285179 | https://www.ncbi.nlm.nih.gov/gene/?term=LOC105285179 | 4191 | 204 | 486.76 | 1 | 2.39 | 0.49 | 16.93 | 16.42 | 0.51 | 0.51 | 0.02 | 0.02 |
| gene-LOC105280641 | https://www.ncbi.nlm.nih.gov/gene/?term=LOC105280641 | 3682 | 78 | 211.84 | 1 | 2.72 | 1.28 | 13.91 | 13.42 | 0.49 | 0.49 | 0.01 | 0.01 |
| gene-LOC105275752 | https://www.ncbi.nlm.nih.gov/gene/?term=LOC105275752 | 3462 | 138 | 398.61 | 1 | 2.89 | 0.72 | 31.47 | 31.01 | 0.46 | 0.46 | 0.02 | 0.02 |
| gene-LOC105279038 | https://www.ncbi.nlm.nih.gov/gene/?term=LOC105279038 | 9453 | 345 | 364.96 | 1 | 1.06 | 0.29 | 9.49 | 9.03 | 0.45 | 0.45 | 0.02 | 0.02 |
| gene-LOC105280408 | https://www.ncbi.nlm.nih.gov/gene/?term=LOC105280408 | 5983 | 65 | 108.64 | 1 | 1.67 | 1.54 | 15.19 | 14.75 | 0.44 | 0.44 | 0 | 0 |
| gene-LOC105275363 | https://www.ncbi.nlm.nih.gov/gene/?term=LOC105275363 | 3798 | 212 | 558.19 | 1 | 2.63 | 0.47 | 14.03 | 13.61 | 0.43 | 0.43 | 0.02 | 0.02 |
| gene-LOC105285807 | https://www.ncbi.nlm.nih.gov/gene/?term=LOC105285807 | 6063 | 231 | 381 1 | 1.65 | 0.43 | 10.41 | 9.98 | 0.43 | 0.43 | 0.02 | 0.02 | |
| gene-LOC105274531 | https://www.ncbi.nlm.nih.gov/gene/?term=LOC105274531 | 7774 | 307 | 394.91 | 1 | 1.29 | 0.33 | 10.8 | 10.38 | 0.42 | 0.42 | 0.02 | 0.02 |
| gene-LOC105281272 | https://www.ncbi.nlm.nih.gov/gene/?term=LOC105281272 | 5252 | 201 | 382.71 | 1 | 1.9 | 0.5 | 15.32 | 14.89 | 0.42 | 0.42 | 0.02 | 0.02 |
| gene-LOC105275374 | https://www.ncbi.nlm.nih.gov/gene/?term=LOC105275374 | 11376 | 587 | 516 1 | 0.88 | 0.17 | 9.68 | 9.27 | 0.41 | 0.41 | 0.02 | 0.02 | |
| gene-LOC105285406 | https://www.ncbi.nlm.nih.gov/gene/?term=LOC105285406 | 8015 | 63 | 78.6 1 | 1.25 | 1.59 | 28.98 | 28.57 | 0.41 | 0.41 | 0 | 0 | |
| gene-LOC105274773 | https://www.ncbi.nlm.nih.gov/gene/?term=LOC105274773 | 11034 | 187 | 169.48 | 1 | 0.91 | 0.53 | 16.91 | 16.52 | 0.39 | 0.39 | 0.01 | 0.01 |
| gene-LOC105281700 | https://www.ncbi.nlm.nih.gov/gene/?term=LOC105281700 | 2208 | 43 | 194.75 | 1 | 4.53 | 2.33 | 36.88 | 36.49 | 0.39 | 0.39 | 0.01 | 0.01 |
| gene-LOC105284184 | https://www.ncbi.nlm.nih.gov/gene/?term=LOC105284184 | 2535 | 105 | 414.2 1 | 3.94 | 0.95 | 28.89 | 28.5 | 0.39 | 0.39 | 0.02 | 0.02 | |
| gene-LOC105276462 | https://www.ncbi.nlm.nih.gov/gene/?term=LOC105276462 | 6561 | 289 | 440.48 | 1 | 1.52 | 0.35 | 8.58 | 8.21 | 0.38 | 0.38 | 0.02 | 0.02 |
| gene-LOC105278145 | https://www.ncbi.nlm.nih.gov/gene/?term=LOC105278145 | 4470 | 274 | 612.98 | 1 | 2.24 | 0.36 | 10.24 | 9.87 | 0.37 | 0.37 | 0.02 | 0.02 |
| gene-LOC105284043 | https://www.ncbi.nlm.nih.gov/gene/?term=LOC105284043 | 5661 | 312 | 551.14 | 1 | 1.77 | 0.32 | 10.26 | 9.89 | 0.37 | 0.37 | 0.02 | 0.02 |
| gene-LOC105276527 | https://www.ncbi.nlm.nih.gov/gene/?term=LOC105276527 | 7923 | 381 | 480.88 | 1 | 1.26 | 0.26 | 8.09 | 7.73 | 0.36 | 0.36 | 0.02 | 0.02 |
| gene-LOC105278962 | https://www.ncbi.nlm.nih.gov/gene/?term=LOC105278962 | 4491 | 129 | 287.24 | 1 | 2.23 | 0.78 | 19.14 | 18.79 | 0.35 | 0.35 | 0.01 | 0.01 |
| gene-LOC105274534 | https://www.ncbi.nlm.nih.gov/gene/?term=LOC105274534 | 1329 | 60 | 451.47 | 1 | 7.52 | 1.67 | 44.47 | 44.14 | 0.32 | 0.32 | 0.01 | 0.01 |
| gene-LOC105285395 | https://www.ncbi.nlm.nih.gov/gene/?term=LOC105285395 | 8085 | 260 | 321.58 | 1 | 1.24 | 0.38 | 13.21 | 12.9 | 0.31 | 0.31 | 0.01 | 0.01 |
| gene-LOC105286265 | https://www.ncbi.nlm.nih.gov/gene/?term=LOC105286265 | 2507 | 68 | 271.24 | 1 | 3.99 | 1.47 | 31.01 | 30.72 | 0.29 | 0.29 | 0.01 | 0.01 |
| gene-LOC105278666 | https://www.ncbi.nlm.nih.gov/gene/?term=LOC105278666 | 10341 | 492 | 475.78 | 1 | 0.97 | 0.2 | 7.02 | 6.75 | 0.27 | 0.27 | 0.01 | 0.01 |
| gene-LOC105275187 | https://www.ncbi.nlm.nih.gov/gene/?term=LOC105275187 | 5534 | 87 | 157.21 | 1 | 1.81 | 1.15 | 15.13 | 14.88 | 0.25 | 0.25 | 0 | 0 |
| gene-LOC105274449 | https://www.ncbi.nlm.nih.gov/gene/?term=LOC105274449 | 1549 | 53 | 342.16 | 1 | 6.46 | 1.89 | 49.51 | 49.29 | 0.22 | 0.22 | 0.01 | 0.01 |
| gene-LOC105275327 | https://www.ncbi.nlm.nih.gov/gene/?term=LOC105275327 | 14245 | 550 | 386.1 1 | 0.7 | 0.18 | 6.58 | 6.38 | 0.2 | 0.2 | 0.01 | 0.01 | |
| gene-LOC105284262 | https://www.ncbi.nlm.nih.gov/gene/?term=LOC105284262 | 6669 | 248 | 371.87 | 1 | 1.5 | 0.4 | 5.25 | 5.05 | 0.2 | 0.2 | 0.01 | 0.01 |
| gene-LOC113562050 | https://www.ncbi.nlm.nih.gov/gene/?term=LOC113562050 | 17097 | 698 | 408.26 | 1 | 0.58 | 0.14 | 5.09 | 4.89 | 0.2 | 0.2 | 0.01 | 0.01 |

| | | | | | | | | | | | | | | |
|---|---|---|---|---|---|---|---|---|---|---|---|---|---|---|
| gene-LOC105278534 | https://www.ncbi.nlm.nih.gov/gene/?term=LOC105278534 | 7364 | 214 | 290.6 | 1 | 1.36 | 0.47 | 16.92 | 16.72 | 0.19 | 0.19 | 0.01 | 0.01 | |
| gene-LOC105274590 | https://www.ncbi.nlm.nih.gov/gene/?term=LOC105274590 | 4516 | 88 | 194.86 | | 2 | 4.43 | 2.27 | 24.95 | 24.81 | 0.14 | 0.14 | 0 | 0 |
| gene-LOC105274669 | https://www.ncbi.nlm.nih.gov/gene/?term=LOC105274669 | 8555 | 274 | 320.28 | | 2 | 2.34 | 0.73 | 5.92 | 5.78 | 0.14 | 0.14 | 0 | 0 |
| gene-LOC105274856 | https://www.ncbi.nlm.nih.gov/gene/?term=LOC105274856 | 5216 | 233 | 446.7 | 1 | 1.92 | 0.43 | 17.97 | 17.83 | 0.14 | 0.14 | 0.01 | 0.01 | |
| gene-LOC105286452 | https://www.ncbi.nlm.nih.gov/gene/?term=LOC105286452 | 6888 | 78 | 113.24 | | 1 | 1.45 | 1.28 | 26.06 | 25.92 | 0.14 | 0.14 | 0 | 0 |
| gene-LOC105286770 | https://www.ncbi.nlm.nih.gov/gene/?term=LOC105286770 | 3055 | 50 | 163.67 | | 1 | 3.27 | 2 | 14.75 | 14.61 | 0.14 | 0.14 | 0 | 0 |
| gene-LOC105280410 | https://www.ncbi.nlm.nih.gov/gene/?term=LOC105280410 | 12580 | 320 | 254.37 | | 1 | 0.79 | 0.31 | 17.81 | 17.7 | 0.11 | 0.11 | 0 | 0 |
| gene-LOC105287065 | https://www.ncbi.nlm.nih.gov/gene/?term=LOC105287065 | 4859 | 259 | 533.03 | | 1 | 2.06 | 0.39 | 12.11 | 12 | 0.11 | 0.11 | 0.01 | 0.01 |
| gene-LOC105278109 | https://www.ncbi.nlm.nih.gov/gene/?term=LOC105278109 | 4029 | 89 | 220.9 | 1 | 2.48 | 1.12 | 34.5 | 34.41 | 0.09 | 0.09 | 0 | 0 | |
| gene-LOC105282811 | https://www.ncbi.nlm.nih.gov/gene/?term=LOC105282811 | 5765 | 232 | 402.43 | | 1 | 1.73 | 0.43 | 8.6 | 8.51 | 0.09 | 0.09 | 0 | 0 |
| gene-LOC105287095 | https://www.ncbi.nlm.nih.gov/gene/?term=LOC105287095 | 1435 | 71 | 494.77 | | 1 | 6.97 | 1.41 | 23.64 | 23.55 | 0.09 | 0.09 | 0 | 0 |
| gene-LOC105276917 | https://www.ncbi.nlm.nih.gov/gene/?term=LOC105276917 | 2336 | 42 | 179.79 | | 1 | 4.28 | 2.38 | 28.76 | 28.68 | 0.08 | 0.08 | 0 | 0 |
| gene-LOC105276059 | https://www.ncbi.nlm.nih.gov/gene/?term=LOC105276059 | 6256 | 148 | 236.57 | | 1 | 1.6 | 0.68 | 11.97 | 11.9 | 0.07 | 0.07 | 0 | 0 |
| gene-LOC105283264 | https://www.ncbi.nlm.nih.gov/gene/?term=LOC105283264 | 4005 | 94 | 234.71 | | 1 | 2.5 | 1.06 | 22.97 | 22.91 | 0.07 | 0.07 | 0 | 0 |
| gene-LOC105283296 | https://www.ncbi.nlm.nih.gov/gene/?term=LOC105283296 | 2174 | 96 | 441.58 | | 1 | 4.6 | 1.04 | 14.24 | 14.2 | 0.04 | 0.04 | 0 | 0 |
| gene-LOC105280819 | https://www.ncbi.nlm.nih.gov/gene/?term=LOC105280819 | 1639 | 44 | 268.46 | | 1 | 6.1 | 2.27 | 37.71 | 37.7 | 0.02 | 0.02 | 0 | 0 |
| gene-LOC105277216 | https://www.ncbi.nlm.nih.gov/gene/?term=LOC105277216 | 7627 | 101 | 132.42 | | 1 | 1.31 | 0.99 | 8.48 | 8.48 | 0 | 0 | 0 | 0 |
| gene-LOC105279664 | https://www.ncbi.nlm.nih.gov/gene/?term=LOC105279664 | 4023 | 75 | 186.43 | | 1 | 2.49 | 1.33 | 19.05 | 19.05 | 0 | 0 | 0 | 0 |
| gene-LOC105279528 | https://www.ncbi.nlm.nih.gov/gene/?term=LOC105279528 | 5802 | 40 | 68.94 | 1 | 1.72 | 2.5 | 19.68 | 19.72 | -0.04 | 0.04 | 0 | 0 | |
| gene-LOC105277588 | https://www.ncbi.nlm.nih.gov/gene/?term=LOC105277588 | 3955 | 98 | 247.79 | | 1 | 2.53 | 1.02 | 8.55 | 8.61 | -0.06 | 0.06 | 0 | 0 |
| gene-LOC105277761 | https://www.ncbi.nlm.nih.gov/gene/?term=LOC105277761 | 4648 | 174 | 374.35 | | 1 | 2.15 | 0.57 | 14.23 | 14.3 | -0.07 | 0.07 | 0 | 0 |
| gene-LOC105279657 | https://www.ncbi.nlm.nih.gov/gene/?term=LOC105279657 | 5878 | 112 | 190.54 | | 1 | 1.7 | 0.89 | 16.48 | 16.55 | -0.07 | 0.07 | 0 | 0 |
| gene-LOC105284774 | https://www.ncbi.nlm.nih.gov/gene/?term=LOC105284774 | 12089 | 106 | 87.68 | 1 | 0.83 | 0.94 | 19.59 | 19.72 | -0.12 | 0.12 | 0 | 0 | |
| gene-LOC105276196 | https://www.ncbi.nlm.nih.gov/gene/?term=LOC105276196 | 2953 | 61 | 206.57 | | 1 | 3.39 | 1.64 | 38.98 | 39.11 | -0.13 | 0.13 | 0 | 0 |
| gene-LOC105284117 | https://www.ncbi.nlm.nih.gov/gene/?term=LOC105284117 | 4340 | 116 | 267.28 | | 1 | 2.3 | 0.86 | 11.19 | 11.33 | -0.14 | 0.14 | 0 | 0 |
| gene-LOC105286649 | https://www.ncbi.nlm.nih.gov/gene/?term=LOC105286649 | 14178 | 852 | 600.93 | | 1 | 0.71 | 0.12 | 1.32 | 1.49 | -0.16 | 0.16 | -0.01 | 0.01 |
| gene-LOC105284116 | https://www.ncbi.nlm.nih.gov/gene/?term=LOC105284116 | 3970 | 134 | 337.53 | | 1 | 2.52 | 0.75 | 7.83 | 8 | -0.17 | 0.17 | -0.01 | 0.01 |
| gene-LOC105287058 | https://www.ncbi.nlm.nih.gov/gene/?term=LOC105287058 | 4788 | 247 | 515.87 | | 1 | 2.09 | 0.4 | 8.85 | 9.06 | -0.2 | 0.2 | -0.01 | 0.01 |
| gene-LOC105278771 | https://www.ncbi.nlm.nih.gov/gene/?term=LOC105278771 | 10457 | 267 | 255.33 | | 1 | 0.96 | 0.37 | 9.06 | 9.27 | -0.21 | 0.21 | -0.01 | 0.01 |
| gene-LOC105286515 | https://www.ncbi.nlm.nih.gov/gene/?term=LOC105286515 | 3751 | 133 | 354.57 | | 1 | 2.67 | 0.75 | 10.29 | 10.56 | -0.27 | 0.27 | -0.01 | 0.01 |
| gene-LOC105285981 | https://www.ncbi.nlm.nih.gov/gene/?term=LOC105285981 | 7666 | 224 | 292.2 | 1 | 1.3 | 0.45 | 17.35 | 17.63 | -0.28 | 0.28 | -0.01 | 0.01 | |
| gene-LOC105280956 | https://www.ncbi.nlm.nih.gov/gene/?term=LOC105280956 | 2639 | 83 | 314.51 | | 1 | 3.79 | 1.2 | 18.28 | 18.58 | -0.3 | 0.3 | -0.01 | 0.01 |
| gene-LOC105283649 | https://www.ncbi.nlm.nih.gov/gene/?term=LOC105283649 | 1918 | 59 | 307.61 | | 2 | 10.43 | 3.39 | 41.12 | 41.42 | -0.31 | 0.31 | -0.01 | 0.01 |
| gene-LOC105285202 | https://www.ncbi.nlm.nih.gov/gene/?term=LOC105285202 | 1838 | 74 | 402.61 | | 1 | 5.44 | 1.35 | 23.68 | 24 | -0.32 | 0.32 | -0.01 | 0.01 |
| gene-LOC105283132 | https://www.ncbi.nlm.nih.gov/gene/?term=LOC105283132 | 1819 | 56 | 307.86 | | 1 | 5.5 | 1.79 | 39.81 | 40.18 | -0.38 | 0.38 | -0.01 | 0.01 |
| gene-LOC105280304 | https://www.ncbi.nlm.nih.gov/gene/?term=LOC105280304 | 8919 | 143 | 160.33 | | 1 | 1.12 | 0.7 | 6.95 | 7.37 | -0.42 | 0.42 | -0.01 | 0.01 |
| gene-LOC105278877 | https://www.ncbi.nlm.nih.gov/gene/?term=LOC105278877 | 1968 | 91 | 462.4 | 2 | 10.16 | 2.2 | 30.61 | 31.07 | -0.45 | 0.45 | -0.02 | 0.02 | |
| gene-LOC105284036 | https://www.ncbi.nlm.nih.gov/gene/?term=LOC105284036 | 3254 | 102 | 313.46 | | 1 | 3.07 | 0.98 | 24.26 | 24.78 | -0.52 | 0.52 | -0.02 | 0.02 |
| gene-LOC105286279 | https://www.ncbi.nlm.nih.gov/gene/?term=LOC105286279 | 5864 | 277 | 472.37 | | 1 | 1.71 | 0.36 | 2.15 | 2.72 | -0.57 | 0.57 | -0.03 | 0.03 |
| gene-LOC105274851 | https://www.ncbi.nlm.nih.gov/gene/?term=LOC105274851 | 6616 | 160 | 241.84 | | 1 | 1.51 | 0.62 | 10.97 | 11.63 | -0.66 | 0.66 | -0.02 | 0.02 |
| gene-LOC105278454 | https://www.ncbi.nlm.nih.gov/gene/?term=LOC105278454 | 1972 | 43 | 218.05 | | 1 | 5.07 | 2.33 | 42.19 | 42.86 | -0.67 | 0.67 | -0.01 | 0.01 |
| gene-LOC105275966 | https://www.ncbi.nlm.nih.gov/gene/?term=LOC105275966 | 3299 | 75 | 227.34 | | 1 | 3.03 | 1.33 | 43.86 | 44.6 | -0.74 | 0.74 | -0.02 | 0.02 |
| gene-LOC105284507 | https://www.ncbi.nlm.nih.gov/gene/?term=LOC105284507 | 5207 | 123 | 236.22 | | 1 | 1.92 | 0.81 | 18.34 | 19.09 | -0.75 | 0.75 | -0.02 | 0.02 |
| gene-LOC105280890 | https://www.ncbi.nlm.nih.gov/gene/?term=LOC105280890 | 1139 | 31 | 272.17 | | 1 | 8.78 | 3.23 | 17.12 | 17.99 | -0.87 | 0.87 | -0.02 | 0.02 |
| gene-LOC105281173 | https://www.ncbi.nlm.nih.gov/gene/?term=LOC105281173 | 1595 | 39 | 244.51 | | 1 | 6.27 | 2.56 | 61.21 | 62.08 | -0.87 | 0.87 | -0.02 | 0.02 |
| gene-LOC105282885 | https://www.ncbi.nlm.nih.gov/gene/?term=LOC105282885 | 1870 | 41 | 219.25 | | 1 | 5.35 | 2.44 | 40.22 | 41.13 | -0.91 | 0.91 | -0.02 | 0.02 |
| gene-LOC105285252 | https://www.ncbi.nlm.nih.gov/gene/?term=LOC105285252 | 7427 | 148 | 199.27 | | 1 | 1.35 | 0.68 | 11.91 | 12.93 | -1.01 | 1.01 | -0.02 | 0.02 |
| gene-LOC105287633 | https://www.ncbi.nlm.nih.gov/gene/?term=LOC105287633 | 1524 | 26 | 170.6 | 1 | 6.56 | 3.85 | 60.3 | 61.69 | -1.38 | 1.38 | -0.02 | 0.02 | |
| gene-LOC105284329 | https://www.ncbi.nlm.nih.gov/gene/?term=LOC105284329 | 4051 | 72 | 177.73 | | 1 | 2.47 | 1.39 | 18.86 | 20.42 | -1.56 | 1.56 | -0.03 | 0.03 |
| gene-LOC105275328 | https://www.ncbi.nlm.nih.gov/gene/?term=LOC105275328 | 2227 | 85 | 381.68 | | 1 | 4.49 | 1.18 | 17.9 | 19.55 | -1.65 | 1.65 | -0.06 | 0.06 |
| gene-LOC105282914 | https://www.ncbi.nlm.nih.gov/gene/?term=LOC105282914 | 5623 | 43 | 76.47 | 1 | 1.78 | 2.33 | 47.44 | 49.79 | -2.35 | 2.35 | -0.02 | 0.02 | |
| gene-LOC105284213 | https://www.ncbi.nlm.nih.gov/gene/?term=LOC105284213 | 868 | 23 | 264.98 | | 1 | 11.52 | 4.35 | 9.72 | 12.3 | -2.58 | 2.58 | -0.07 | 0.07 |
| gene-LOC105285409 | https://www.ncbi.nlm.nih.gov/gene/?term=LOC105285409 | 5099 | 67 | 131.4 | 2 | 3.92 | 2.99 | 1.72 | 4.47 | -2.75 | 2.75 | -0.04 | 0.04 | |

Directory: OGL File: ogl-Ob_LI_RphaseA2.vs.RphaseB2.txt

| gene_ID | gene_link | gwidth | #Sites | #per10Kb | #dmSites | #dmsp10kb | %dmSites | %pSite1 | %pSite2 | DMpSite | ADMpSite | DMpNucl | ADMpNucl |
|---|---|---|---|---|---|---|---|---|---|---|---|---|---|
| gene-LOC113562901 | https://www.ncbi.nlm.nih.gov/gene/?term=LOC113562901 | 1531 | 24 | 156.76 | 3 | 19.6 | 12.5 | 59.3 | 45.31 | 14 | 14 | 0.22 | 0.22 |
| gene-LOC113562345 | https://www.ncbi.nlm.nih.gov/gene/?term=LOC113562345 | 2290 | 69 | 301.31 | 14 | 61.14 | 20.29 | 29.74 | 16.81 | 12.93 | 12.93 | 0.39 | 0.39 |
| gene-LOC105275070 | https://www.ncbi.nlm.nih.gov/gene/?term=LOC105275070 | 5667 | 42 | 74.11 | 5 | 8.82 | 11.9 | 13.23 | 1.03 | 12.2 | 12.2 | 0.09 | 0.09 |
| gene-LOC105279898 | https://www.ncbi.nlm.nih.gov/gene/?term=LOC105279898 | 1343 | 10 | 74.46 | 1 | 7.45 | 10 | 83.16 | 71.53 | 11.63 | 11.63 | 0.09 | 0.09 |
| gene-LOC105286644 | https://www.ncbi.nlm.nih.gov/gene/?term=LOC105286644 | 1235 | 22 | 178.14 | 1 | 8.1 | 4.55 | 44.4 | 37.26 | 7.14 | 7.14 | 0.13 | 0.13 |
| gene-LOC105275708 | https://www.ncbi.nlm.nih.gov/gene/?term=LOC105275708 | 1772 | 15 | 84.65 | 1 | 5.64 | 6.67 | 47.19 | 40.83 | 6.36 | 6.36 | 0.05 | 0.05 |
| gene-LOC105275447 | https://www.ncbi.nlm.nih.gov/gene/?term=LOC105275447 | 1846 | 31 | 167.93 | 1 | 5.42 | 3.23 | 9.13 | 2.78 | 6.34 | 6.34 | 0.11 | 0.11 |
| gene-LOC105286694 | https://www.ncbi.nlm.nih.gov/gene/?term=LOC105286694 | 3409 | 12 | 35.2 | 1 | 2.93 | 8.33 | 52.59 | 46.71 | 5.88 | 5.88 | 0.02 | 0.02 |
| gene-LOC105282388 | https://www.ncbi.nlm.nih.gov/gene/?term=LOC105282388 | 1661 | 34 | 204.7 | 1 | 6.02 | 2.94 | 48.95 | 44.6 | 4.35 | 4.35 | 0.09 | 0.09 |
| gene-LOC105287417 | https://www.ncbi.nlm.nih.gov/gene/?term=LOC105287417 | 1025 | 23 | 224.39 | 1 | 9.76 | 4.35 | 53.58 | 49.27 | 4.31 | 4.31 | 0.1 | 0.1 |
| gene-LOC105282893 | https://www.ncbi.nlm.nih.gov/gene/?term=LOC105282893 | 2645 | 47 | 177.69 | 1 | 3.78 | 2.13 | 30.08 | 25.9 | 4.18 | 4.18 | 0.07 | 0.07 |
| gene-LOC105276538 | https://www.ncbi.nlm.nih.gov/gene/?term=LOC105276538 | 1511 | 20 | 132.36 | 1 | 6.62 | 5 | 73.31 | 69.21 | 4.1 | 4.1 | 0.05 | 0.05 |
| gene-LOC105285583 | https://www.ncbi.nlm.nih.gov/gene/?term=LOC105285583 | 1947 | 49 | 251.67 | 1 | 5.14 | 2.04 | 30.89 | 26.83 | 4.06 | 4.06 | 0.1 | 0.1 |
| gene-LOC105280555 | https://www.ncbi.nlm.nih.gov/gene/?term=LOC105280555 | 1496 | 31 | 207.22 | 1 | 6.68 | 3.23 | 61.05 | 57.2 | 3.85 | 3.85 | 0.08 | 0.08 |
| gene-LOC105274778 | https://www.ncbi.nlm.nih.gov/gene/?term=LOC105274778 | 4662 | 40 | 85.8 | 1 | 2.15 | 2.5 | 47.99 | 44.37 | 3.62 | 3.62 | 0.03 | 0.03 |
| gene-LOC105284609 | https://www.ncbi.nlm.nih.gov/gene/?term=LOC105284609 | 5678 | 33 | 58.12 | 1 | 1.76 | 3.03 | 34 | 30.46 | 3.54 | 3.54 | 0.02 | 0.02 |
| gene-LOC105285315 | https://www.ncbi.nlm.nih.gov/gene/?term=LOC105285315 | 5458 | 19 | 34.81 | 1 | 1.83 | 5.26 | 46.71 | 43.22 | 3.49 | 3.49 | 0.01 | 0.01 |
| gene-LOC105275518 | https://www.ncbi.nlm.nih.gov/gene/?term=LOC105275518 | 2335 | 24 | 102.78 | 1 | 4.28 | 4.17 | 49.73 | 46.6 | 3.13 | 3.13 | 0.03 | 0.03 |
| gene-LOC105286683 | https://www.ncbi.nlm.nih.gov/gene/?term=LOC105286683 | 1665 | 57 | 342.34 | 2 | 12.01 | 3.51 | 34.89 | 31.76 | 3.13 | 3.13 | 0.11 | 0.11 |
| gene-LOC105274706 | https://www.ncbi.nlm.nih.gov/gene/?term=LOC105274706 | 1446 | 56 | 387.28 | 1 | 6.92 | 1.79 | 39.74 | 36.7 | 3.05 | 3.05 | 0.12 | 0.12 |
| gene-LOC105285674 | https://www.ncbi.nlm.nih.gov/gene/?term=LOC105285674 | 3035 | 102 | 336.08 | 2 | 6.59 | 1.96 | 26.23 | 23.21 | 3.03 | 3.03 | 0.1 | 0.1 |
| gene-LOC105278215 | https://www.ncbi.nlm.nih.gov/gene/?term=LOC105278215 | 1700 | 29 | 170.59 | 1 | 5.88 | 3.45 | 57.59 | 54.66 | 2.93 | 2.93 | 0.05 | 0.05 |
| gene-LOC105278109 | https://www.ncbi.nlm.nih.gov/gene/?term=LOC105278109 | 4029 | 67 | 166.29 | 1 | 2.48 | 1.49 | 44.53 | 41.68 | 2.86 | 2.86 | 0.05 | 0.05 |
| gene-LOC105281515 | https://www.ncbi.nlm.nih.gov/gene/?term=LOC105281515 | 8827 | 42 | 47.58 | 1 | 1.13 | 2.38 | 30.33 | 27.53 | 2.81 | 2.81 | 0.01 | 0.01 |
| gene-LOC105280786 | https://www.ncbi.nlm.nih.gov/gene/?term=LOC105280786 | 2573 | 127 | 493.59 | 1 | 3.89 | 0.79 | 30.11 | 27.41 | 2.7 | 2.7 | 0.13 | 0.13 |
| gene-LOC105287063 | https://www.ncbi.nlm.nih.gov/gene/?term=LOC105287063 | 1874 | 48 | 256.14 | 1 | 5.34 | 2.08 | 32.76 | 30.33 | 2.44 | 2.44 | 0.06 | 0.06 |
| gene-LOC105287854 | https://www.ncbi.nlm.nih.gov/gene/?term=LOC105287854 | 3644 | 54 | 148.19 | 1 | 2.74 | 1.85 | 25.58 | 23.15 | 2.43 | 2.43 | 0.04 | 0.04 |
| gene-LOC105282794 | https://www.ncbi.nlm.nih.gov/gene/?term=LOC105282794 | 4571 | 95 | 207.83 | 1 | 2.19 | 1.05 | 18.64 | 16.23 | 2.41 | 2.41 | 0.05 | 0.05 |
| gene-LOC105285671 | https://www.ncbi.nlm.nih.gov/gene/?term=LOC105285671 | 2263 | 106 | 468.4 | 1 | 4.42 | 0.94 | 28.93 | 26.53 | 2.4 | 2.4 | 0.11 | 0.11 |
| gene-LOC105276917 | https://www.ncbi.nlm.nih.gov/gene/?term=LOC105276917 | 2336 | 38 | 162.67 | 1 | 4.28 | 2.63 | 28.71 | 26.35 | 2.36 | 2.36 | 0.04 | 0.04 |
| gene-LOC105274900 | https://www.ncbi.nlm.nih.gov/gene/?term=LOC105274900 | 5561 | 56 | 100.7 | 2 | 3.6 | 3.57 | 32.09 | 29.76 | 2.33 | 2.33 | 0.02 | 0.02 |
| gene-LOC105276704 | https://www.ncbi.nlm.nih.gov/gene/?term=LOC105276704 | 2063 | 88 | 426.56 | 1 | 4.85 | 1.14 | 16.14 | 13.92 | 2.23 | 2.23 | 0.1 | 0.1 |
| gene-LOC105279664 | https://www.ncbi.nlm.nih.gov/gene/?term=LOC105279664 | 4023 | 56 | 139.2 | 1 | 2.49 | 1.79 | 21.11 | 18.99 | 2.12 | 2.12 | 0.03 | 0.03 |
| gene-LOC105280104 | https://www.ncbi.nlm.nih.gov/gene/?term=LOC105280104 | 3960 | 139 | 351.01 | 1 | 2.53 | 0.72 | 17.44 | 15.39 | 2.05 | 2.05 | 0.07 | 0.07 |
| gene-LOC105283132 | https://www.ncbi.nlm.nih.gov/gene/?term=LOC105283132 | 1819 | 57 | 313.36 | 1 | 5.5 | 1.75 | 39.92 | 37.88 | 2.04 | 2.04 | 0.06 | 0.06 |
| gene-LOC105277856 | https://www.ncbi.nlm.nih.gov/gene/?term=LOC105277856 | 3442 | 35 | 101.69 | 1 | 2.91 | 2.86 | 35 | 32.98 | 2.02 | 2.02 | 0.02 | 0.02 |
| gene-LOC105278108 | https://www.ncbi.nlm.nih.gov/gene/?term=LOC105278108 | 4392 | 51 | 116.12 | 1 | 2.28 | 1.96 | 25.15 | 23.14 | 2.01 | 2.01 | 0.02 | 0.02 |
| gene-LOC105280641 | https://www.ncbi.nlm.nih.gov/gene/?term=LOC105280641 | 3682 | 53 | 143.94 | 1 | 2.72 | 1.89 | 17.27 | 15.26 | 2.01 | 2.01 | 0.03 | 0.03 |
| gene-LOC105280377 | https://www.ncbi.nlm.nih.gov/gene/?term=LOC105280377 | 2024 | 70 | 345.85 | 1 | 4.94 | 1.43 | 48.1 | 46.12 | 1.98 | 1.98 | 0.07 | 0.07 |
| gene-LOC105282156 | https://www.ncbi.nlm.nih.gov/gene/?term=LOC105282156 | 2577 | 130 | 504.46 | 1 | 3.88 | 0.77 | 28.69 | 26.74 | 1.95 | 1.95 | 0.1 | 0.1 |
| gene-LOC113561805 | https://www.ncbi.nlm.nih.gov/gene/?term=LOC113561805 | 3337 | 71 | 212.77 | 1 | 3 | 1.41 | 36.21 | 34.33 | 1.88 | 1.88 | 0.04 | 0.04 |
| gene-LOC105282600 | https://www.ncbi.nlm.nih.gov/gene/?term=LOC105282600 | 3715 | 56 | 150.74 | 1 | 2.69 | 1.79 | 27.25 | 25.37 | 1.87 | 1.87 | 0.03 | 0.03 |
| gene-LOC105282947 | https://www.ncbi.nlm.nih.gov/gene/?term=LOC105282947 | 1162 | 41 | 352.84 | 1 | 8.61 | 2.44 | 36.57 | 34.72 | 1.85 | 1.85 | 0.07 | 0.07 |
| gene-LOC105287646 | https://www.ncbi.nlm.nih.gov/gene/?term=LOC105287646 | 3556 | 90 | 253.09 | 1 | 2.81 | 1.11 | 18.4 | 16.59 | 1.81 | 1.81 | 0.05 | 0.05 |
| gene-LOC105276190 | https://www.ncbi.nlm.nih.gov/gene/?term=LOC105276190 | 4839 | 93 | 192.19 | 1 | 2.07 | 1.08 | 11.07 | 9.37 | 1.7 | 1.7 | 0.03 | 0.03 |
| gene-LOC105281420 | https://www.ncbi.nlm.nih.gov/gene/?term=LOC105281420 | 3882 | 62 | 159.71 | 1 | 2.58 | 1.61 | 15.69 | 14 | 1.7 | 1.7 | 0.03 | 0.03 |
| gene-LOC105284117 | https://www.ncbi.nlm.nih.gov/gene/?term=LOC105284117 | 4340 | 65 | 149.77 | 1 | 2.3 | 1.54 | 12.82 | 11.26 | 1.56 | 1.56 | 0.02 | 0.02 |
| gene-LOC105287648 | https://www.ncbi.nlm.nih.gov/gene/?term=LOC105287648 | 2205 | 26 | 117.91 | 1 | 4.54 | 3.85 | 41.13 | 39.59 | 1.54 | 1.54 | 0.02 | 0.02 |
| gene-LOC105288063 | https://www.ncbi.nlm.nih.gov/gene/?term=LOC105288063 | 3400 | 47 | 138.24 | 1 | 2.94 | 2.13 | 15.4 | 13.86 | 1.54 | 1.54 | 0.02 | 0.02 |
| gene-LOC105283182 | https://www.ncbi.nlm.nih.gov/gene/?term=LOC105283182 | 3585 | 131 | 365.41 | 1 | 2.79 | 0.76 | 14.16 | 12.63 | 1.52 | 1.52 | 0.06 | 0.06 |
| gene-LOC105283649 | https://www.ncbi.nlm.nih.gov/gene/?term=LOC105283649 | 1918 | 56 | 291.97 | 2 | 10.43 | 3.57 | 37.98 | 36.48 | 1.5 | 1.5 | 0.04 | 0.04 |
| gene-LOC105275347 | https://www.ncbi.nlm.nih.gov/gene/?term=LOC105275347 | 2526 | 19 | 75.22 | 1 | 3.96 | 5.26 | 38.97 | 37.49 | 1.48 | 1.48 | 0.01 | 0.01 |
| gene-LOC105276692 | https://www.ncbi.nlm.nih.gov/gene/?term=LOC105276692 | 1745 | 35 | 200.57 | 1 | 5.73 | 2.86 | 22.12 | 20.68 | 1.44 | 1.44 | 0.03 | 0.03 |
| gene-LOC105278282 | https://www.ncbi.nlm.nih.gov/gene/?term=LOC105278282 | 1775 | 56 | 315.49 | 1 | 5.63 | 1.79 | 39.77 | 38.34 | 1.43 | 1.43 | 0.05 | 0.05 |
| gene-LOC105282970 | https://www.ncbi.nlm.nih.gov/gene/?term=LOC105282970 | 1529 | 40 | 261.61 | 1 | 6.54 | 2.5 | 49.63 | 48.22 | 1.41 | 1.41 | 0.04 | 0.04 |
| gene-LOC105281272 | https://www.ncbi.nlm.nih.gov/gene/?term=LOC105281272 | 5252 | 164 | 312.26 | 1 | 1.9 | 0.61 | 19.36 | 17.96 | 1.4 | 1.4 | 0.04 | 0.04 |
| gene-LOC105281532 | https://www.ncbi.nlm.nih.gov/gene/?term=LOC105281532 | 7793 | 81 | 103.94 | 1 | 1.28 | 1.23 | 12.92 | 11.51 | 1.4 | 1.4 | 0.01 | 0.01 |
| gene-LOC105276875 | https://www.ncbi.nlm.nih.gov/gene/?term=LOC105276875 | 3151 | 118 | 374.48 | 1 | 3.17 | 0.85 | 23.91 | 22.53 | 1.38 | 1.38 | 0.05 | 0.05 |
| gene-LOC105284704 | https://www.ncbi.nlm.nih.gov/gene/?term=LOC105284704 | 2400 | 62 | 258.33 | 1 | 4.17 | 1.61 | 24.3 | 22.92 | 1.38 | 1.38 | 0.04 | 0.04 |
| gene-LOC105276449 | https://www.ncbi.nlm.nih.gov/gene/?term=LOC105276449 | 1231 | 50 | 406.17 | 1 | 8.12 | 2 | 32.79 | 31.42 | 1.37 | 1.37 | 0.06 | 0.06 |

| | | | | | | | | | | | | | |
|---|---|---|---|---|---|---|---|---|---|---|---|---|---|
| gene-LOC105284355 | https://www.ncbi.nlm.nih.gov/gene/?term=LOC105284355 | 3904 | 153 | 391.91 | 1 | 2.56 | 0.65 | 21.82 | 20.47 | 1.35 | 1.35 | 0.05 | 0.05 |
| gene-LOC105274590 | https://www.ncbi.nlm.nih.gov/gene/?term=LOC105274590 | 4516 | 71 | 157.22 | 2 | 4.43 | 2.82 | 29.16 | 27.84 | 1.32 | 1.32 | 0.02 | 0.02 |
| gene-LOC105283728 | https://www.ncbi.nlm.nih.gov/gene/?term=LOC105283728 | 3256 | 120 | 368.55 | 2 | 6.14 | 1.67 | 19.01 | 17.69 | 1.32 | 1.32 | 0.05 | 0.05 |
| gene-LOC105283123 | https://www.ncbi.nlm.nih.gov/gene/?term=LOC105283123 | 1507 | 48 | 318.51 | 1 | 6.64 | 2.08 | 17.79 | 16.48 | 1.31 | 1.31 | 0.04 | 0.04 |
| gene-LOC105281419 | https://www.ncbi.nlm.nih.gov/gene/?term=LOC105281419 | 2366 | 83 | 350.8 | 1 | 4.23 | 1.2 | 23.42 | 22.13 | 1.3 | 1.3 | 0.05 | 0.05 |
| gene-LOC105282151 | https://www.ncbi.nlm.nih.gov/gene/?term=LOC105282151 | 1601 | 39 | 243.6 | 2 | 12.49 | 5.13 | 42.25 | 41 | 1.25 | 1.25 | 0.03 | 0.03 |
| gene-LOC105283712 | https://www.ncbi.nlm.nih.gov/gene/?term=LOC105283712 | 1910 | 36 | 188.48 | 1 | 5.24 | 2.78 | 27.71 | 26.46 | 1.25 | 1.25 | 0.02 | 0.02 |
| gene-LOC105281848 | https://www.ncbi.nlm.nih.gov/gene/?term=LOC105281848 | 2122 | 48 | 226.2 | 1 | 4.71 | 2.08 | 27.66 | 26.43 | 1.23 | 1.23 | 0.03 | 0.03 |
| gene-LOC105288079 | https://www.ncbi.nlm.nih.gov/gene/?term=LOC105288079 | 3473 | 101 | 290.81 | 1 | 2.88 | 0.99 | 20.37 | 19.15 | 1.21 | 1.21 | 0.04 | 0.04 |
| gene-LOC105277238 | https://www.ncbi.nlm.nih.gov/gene/?term=LOC105277238 | 2708 | 88 | 324.96 | 1 | 3.69 | 1.14 | 27.16 | 25.96 | 1.2 | 1.2 | 0.04 | 0.04 |
| gene-LOC105278497 | https://www.ncbi.nlm.nih.gov/gene/?term=LOC105278497 | 2072 | 100 | 482.63 | 1 | 4.83 | 1 | 19.97 | 18.78 | 1.2 | 1.2 | 0.06 | 0.06 |
| gene-LOC105288066 | https://www.ncbi.nlm.nih.gov/gene/?term=LOC105288066 | 1964 | 30 | 152.75 | 1 | 5.09 | 3.33 | 32.32 | 31.12 | 1.2 | 1.2 | 0.02 | 0.02 |
| gene-LOC105277761 | https://www.ncbi.nlm.nih.gov/gene/?term=LOC105277761 | 4648 | 150 | 322.72 | 1 | 2.15 | 0.67 | 14.38 | 13.19 | 1.19 | 1.19 | 0.04 | 0.04 |
| gene-LOC105287720 | https://www.ncbi.nlm.nih.gov/gene/?term=LOC105287720 | 4843 | 99 | 204.42 | 1 | 2.06 | 1.01 | 16.39 | 15.24 | 1.16 | 1.16 | 0.02 | 0.02 |
| gene-LOC105277631 | https://www.ncbi.nlm.nih.gov/gene/?term=LOC105277631 | 4387 | 304 | 692.96 | 1 | 2.28 | 0.33 | 10.76 | 9.61 | 1.15 | 1.15 | 0.08 | 0.08 |
| gene-LOC105281173 | https://www.ncbi.nlm.nih.gov/gene/?term=LOC105281173 | 1595 | 39 | 244.51 | 1 | 6.27 | 2.56 | 61.03 | 59.91 | 1.13 | 1.13 | 0.03 | 0.03 |
| gene-LOC105284184 | https://www.ncbi.nlm.nih.gov/gene/?term=LOC105284184 | 2535 | 86 | 339.25 | 1 | 3.94 | 1.16 | 34.02 | 32.88 | 1.13 | 1.13 | 0.04 | 0.04 |
| gene-LOC105275363 | https://www.ncbi.nlm.nih.gov/gene/?term=LOC105275363 | 3798 | 165 | 434.44 | 1 | 2.63 | 0.61 | 17.38 | 16.26 | 1.12 | 1.12 | 0.05 | 0.05 |
| gene-LOC105279298 | https://www.ncbi.nlm.nih.gov/gene/?term=LOC105279298 | 2773 | 136 | 490.44 | 2 | 7.21 | 1.47 | 23.98 | 22.86 | 1.11 | 1.11 | 0.05 | 0.05 |
| gene-LOC113561621 | https://www.ncbi.nlm.nih.gov/gene/?term=LOC113561621 | 2249 | 86 | 382.39 | 1 | 4.45 | 1.16 | 20.44 | 19.32 | 1.11 | 1.11 | 0.04 | 0.04 |
| gene-LOC105279500 | https://www.ncbi.nlm.nih.gov/gene/?term=LOC105279500 | 3587 | 37 | 103.15 | 1 | 2.79 | 2.7 | 50.42 | 49.31 | 1.1 | 1.1 | 0.01 | 0.01 |
| gene-LOC105285639 | https://www.ncbi.nlm.nih.gov/gene/?term=LOC105285639 | 10325 | 349 | 338.01 | 1 | 0.97 | 0.29 | 13.03 | 11.95 | 1.08 | 1.08 | 0.04 | 0.04 |
| gene-LOC105275451 | https://www.ncbi.nlm.nih.gov/gene/?term=LOC105275451 | 6035 | 130 | 215.41 | 1 | 1.66 | 0.77 | 22.16 | 21.09 | 1.07 | 1.07 | 0.02 | 0.02 |
| gene-LOC105288010 | https://www.ncbi.nlm.nih.gov/gene/?term=LOC105288010 | 3189 | 51 | 159.92 | 1 | 3.14 | 1.96 | 29.8 | 28.73 | 1.07 | 1.07 | 0.02 | 0.02 |
| gene-LOC105284043 | https://www.ncbi.nlm.nih.gov/gene/?term=LOC105284043 | 5661 | 308 | 544.07 | 1 | 1.77 | 0.32 | 10.53 | 9.46 | 1.06 | 1.06 | 0.06 | 0.06 |
| gene-LOC105278283 | https://www.ncbi.nlm.nih.gov/gene/?term=LOC105278283 | 2317 | 142 | 612.86 | 1 | 4.32 | 0.7 | 25.61 | 24.56 | 1.05 | 1.05 | 0.06 | 0.06 |
| gene-LOC105282149 | https://www.ncbi.nlm.nih.gov/gene/?term=LOC105282149 | 1787 | 110 | 615.56 | 1 | 5.6 | 0.91 | 43.04 | 42 | 1.04 | 1.04 | 0.06 | 0.06 |
| gene-LOC105285742 | https://www.ncbi.nlm.nih.gov/gene/?term=LOC105285742 | 3446 | 123 | 356.94 | 1 | 2.9 | 0.81 | 20.56 | 19.51 | 1.04 | 1.04 | 0.04 | 0.04 |
| gene-LOC105286514 | https://www.ncbi.nlm.nih.gov/gene/?term=LOC105286514 | 2528 | 49 | 193.83 | 1 | 3.96 | 2.04 | 36.75 | 35.71 | 1.04 | 1.04 | 0.02 | 0.02 |
| gene-LOC105283592 | https://www.ncbi.nlm.nih.gov/gene/?term=LOC105283592 | 3479 | 80 | 229.95 | 1 | 2.87 | 1.25 | 23.33 | 22.32 | 1.01 | 1.01 | 0.02 | 0.02 |
| gene-LOC109611283 | https://www.ncbi.nlm.nih.gov/gene/?term=LOC109611283 | 1451 | 37 | 255 | 1 | 6.89 | 2.7 | 37.11 | 36.1 | 1 | 1 | 0.03 | 0.03 |
| gene-LOC105285395 | https://www.ncbi.nlm.nih.gov/gene/?term=LOC105285395 | 8085 | 218 | 269.64 | 1 | 1.24 | 0.46 | 17.01 | 16.05 | 0.96 | 0.96 | 0.03 | 0.03 |
| gene-LOC105278807 | https://www.ncbi.nlm.nih.gov/gene/?term=LOC105278807 | 2166 | 67 | 309.33 | 1 | 4.62 | 1.49 | 22.84 | 21.92 | 0.93 | 0.93 | 0.03 | 0.03 |
| gene-LOC105282682 | https://www.ncbi.nlm.nih.gov/gene/?term=LOC105282682 | 4127 | 223 | 540.34 | 1 | 2.42 | 0.45 | 13.69 | 12.76 | 0.93 | 0.93 | 0.05 | 0.05 |
| gene-LOC105278962 | https://www.ncbi.nlm.nih.gov/gene/?term=LOC105278962 | 4491 | 104 | 231.57 | 1 | 2.23 | 0.96 | 22.01 | 21.09 | 0.92 | 0.92 | 0.02 | 0.02 |
| gene-LOC105274856 | https://www.ncbi.nlm.nih.gov/gene/?term=LOC105274856 | 5216 | 216 | 414.11 | 1 | 1.92 | 0.46 | 19.8 | 18.89 | 0.91 | 0.91 | 0.04 | 0.04 |
| gene-LOC105283987 | https://www.ncbi.nlm.nih.gov/gene/?term=LOC105283987 | 4841 | 68 | 140.47 | 1 | 2.07 | 1.47 | 29.57 | 28.67 | 0.91 | 0.91 | 0.01 | 0.01 |
| gene-LOC105279038 | https://www.ncbi.nlm.nih.gov/gene/?term=LOC105279038 | 9453 | 313 | 331.11 | 1 | 1.06 | 0.32 | 10.35 | 9.46 | 0.9 | 0.9 | 0.03 | 0.03 |
| gene-LOC105285406 | https://www.ncbi.nlm.nih.gov/gene/?term=LOC105285406 | 8015 | 60 | 74.86 | 1 | 1.25 | 1.67 | 26.39 | 25.52 | 0.87 | 0.87 | 0.01 | 0.01 |
| gene-LOC105276059 | https://www.ncbi.nlm.nih.gov/gene/?term=LOC105276059 | 6256 | 133 | 212.6 | 1 | 1.6 | 0.75 | 10.26 | 9.4 | 0.86 | 0.86 | 0.02 | 0.02 |
| gene-LOC105280408 | https://www.ncbi.nlm.nih.gov/gene/?term=LOC105280408 | 5983 | 58 | 96.94 | 1 | 1.67 | 1.72 | 16.75 | 15.89 | 0.86 | 0.86 | 0.01 | 0.01 |
| gene-LOC105280956 | https://www.ncbi.nlm.nih.gov/gene/?term=LOC105280956 | 2639 | 71 | 269.04 | 1 | 3.79 | 1.41 | 18.65 | 17.79 | 0.86 | 0.86 | 0.02 | 0.02 |
| gene-LOC105280407 | https://www.ncbi.nlm.nih.gov/gene/?term=LOC105280407 | 10950 | 211 | 192.69 | 1 | 0.91 | 0.47 | 16.65 | 15.81 | 0.84 | 0.84 | 0.02 | 0.02 |
| gene-LOC105285055 | https://www.ncbi.nlm.nih.gov/gene/?term=LOC105285055 | 4108 | 276 | 671.86 | 2 | 4.87 | 0.72 | 11.41 | 10.6 | 0.8 | 0.8 | 0.05 | 0.05 |
| gene-LOC105280002 | https://www.ncbi.nlm.nih.gov/gene/?term=LOC105280002 | 1940 | 28 | 144.33 | 1 | 5.15 | 3.57 | 37.5 | 36.72 | 0.79 | 0.79 | 0.01 | 0.01 |
| gene-LOC105284036 | https://www.ncbi.nlm.nih.gov/gene/?term=LOC105284036 | 3254 | 108 | 331.9 | 1 | 3.07 | 0.93 | 22.88 | 22.09 | 0.79 | 0.79 | 0.03 | 0.03 |
| gene-LOC105275460 | https://www.ncbi.nlm.nih.gov/gene/?term=LOC105275460 | 2473 | 117 | 473.11 | 1 | 4.04 | 0.85 | 14.05 | 13.26 | 0.78 | 0.78 | 0.04 | 0.04 |
| gene-LOC105280774 | https://www.ncbi.nlm.nih.gov/gene/?term=LOC105280774 | 2793 | 100 | 358.04 | 1 | 3.58 | 1 | 32.96 | 32.18 | 0.78 | 0.78 | 0.03 | 0.03 |
| gene-LOC105285534 | https://www.ncbi.nlm.nih.gov/gene/?term=LOC105285534 | 3944 | 237 | 600.91 | 1 | 2.54 | 0.42 | 11.45 | 10.67 | 0.78 | 0.78 | 0.05 | 0.05 |
| gene-LOC105285060 | https://www.ncbi.nlm.nih.gov/gene/?term=LOC105285060 | 1285 | 21 | 163.42 | 1 | 7.78 | 4.76 | 39.84 | 39.09 | 0.75 | 0.75 | 0.01 | 0.01 |
| gene-LOC105284116 | https://www.ncbi.nlm.nih.gov/gene/?term=LOC105284116 | 3970 | 78 | 196.47 | 1 | 2.52 | 1.28 | 7.09 | 6.36 | 0.73 | 0.73 | 0.01 | 0.01 |
| gene-LOC105274534 | https://www.ncbi.nlm.nih.gov/gene/?term=LOC105274534 | 1329 | 57 | 428.89 | 1 | 7.52 | 1.75 | 48.36 | 47.65 | 0.71 | 0.71 | 0.03 | 0.03 |
| gene-LOC105282067 | https://www.ncbi.nlm.nih.gov/gene/?term=LOC105282067 | 5941 | 54 | 90.89 | 1 | 1.68 | 1.85 | 28.74 | 28.06 | 0.67 | 0.67 | 0.01 | 0.01 |
| gene-LOC105285514 | https://www.ncbi.nlm.nih.gov/gene/?term=LOC105285514 | 1277 | 17 | 133.12 | 1 | 7.83 | 5.88 | 71.39 | 70.72 | 0.67 | 0.67 | 0.01 | 0.01 |
| gene-LOC105274699 | https://www.ncbi.nlm.nih.gov/gene/?term=LOC105274699 | 2386 | 52 | 217.94 | 1 | 4.19 | 1.92 | 24.82 | 24.15 | 0.66 | 0.66 | 0.01 | 0.01 |
| gene-LOC105284557 | https://www.ncbi.nlm.nih.gov/gene/?term=LOC105284557 | 1672 | 84 | 502.39 | 1 | 5.98 | 1.19 | 29.36 | 28.7 | 0.66 | 0.66 | 0.03 | 0.03 |
| gene-LOC105275966 | https://www.ncbi.nlm.nih.gov/gene/?term=LOC105275966 | 3299 | 64 | 194 | 1 | 3.03 | 1.56 | 48.15 | 47.49 | 0.65 | 0.65 | 0.01 | 0.01 |
| gene-LOC105282399 | https://www.ncbi.nlm.nih.gov/gene/?term=LOC105282399 | 2682 | 24 | 89.49 | 1 | 3.73 | 4.17 | 35.91 | 35.25 | 0.65 | 0.65 | 0.01 | 0.01 |
| gene-LOC105275683 | https://www.ncbi.nlm.nih.gov/gene/?term=LOC105275683 | 4256 | 193 | 453.48 | 1 | 2.35 | 0.52 | 13.76 | 13.13 | 0.63 | 0.63 | 0.03 | 0.03 |
| gene-LOC105285179 | https://www.ncbi.nlm.nih.gov/gene/?term=LOC105285179 | 4191 | 185 | 441.42 | 1 | 2.39 | 0.54 | 18.94 | 18.34 | 0.6 | 0.6 | 0.03 | 0.03 |
| gene-LOC105283943 | https://www.ncbi.nlm.nih.gov/gene/?term=LOC105283943 | 3559 | 48 | 134.87 | 1 | 2.81 | 2.08 | 17.55 | 16.96 | 0.59 | 0.59 | 0.01 | 0.01 |
| gene-LOC105275374 | https://www.ncbi.nlm.nih.gov/gene/?term=LOC105275374 | 11376 | 523 | 459.74 | 1 | 0.88 | 0.19 | 11.54 | 10.97 | 0.57 | 0.57 | 0.03 | 0.03 |
| gene-LOC105276612 | https://www.ncbi.nlm.nih.gov/gene/?term=LOC105276612 | 4157 | 187 | 449.84 | 1 | 2.41 | 0.53 | 10.04 | 9.47 | 0.57 | 0.57 | 0.03 | 0.03 |
| gene-LOC105282825 | https://www.ncbi.nlm.nih.gov/gene/?term=LOC105282825 | 3919 | 85 | 216.89 | 1 | 2.55 | 1.18 | 14.44 | 13.87 | 0.57 | 0.57 | 0.01 | 0.01 |

| | | | | | | | | | | | | | | |
|---|---|---|---|---|---|---|---|---|---|---|---|---|---|---|
| gene-LOC105274823 | https://www.ncbi.nlm.nih.gov/gene/?term=LOC105274823 | 4620 | 141 | 305.19 | | 1 | 2.16 | 0.71 | 21.37 | 20.81 | 0.56 | 0.56 | 0.02 | 0.02 |
| gene-LOC105278534 | https://www.ncbi.nlm.nih.gov/gene/?term=LOC105278534 | 7364 | 197 | 267.52 | | 2 | 2.72 | 1.02 | 18.17 | 17.62 | 0.55 | 0.55 | 0.01 | 0.01 |
| gene-LOC105281599 | https://www.ncbi.nlm.nih.gov/gene/?term=LOC105281599 | 3859 | 49 | 126.98 | | 1 | 2.59 | 2.04 | 4.6 | 4.08 | 0.53 | 0.53 | 0.01 | 0.01 |
| gene-LOC105275127 | https://www.ncbi.nlm.nih.gov/gene/?term=LOC105275127 | 9377 | 301 | 321 | 1 | 1.07 | 0.33 | 7.86 | 7.34 | 0.52 | 0.52 | 0.02 | 0.02 | |
| gene-LOC105280410 | https://www.ncbi.nlm.nih.gov/gene/?term=LOC105280410 | 12580 | 264 | 209.86 | | 1 | 0.79 | 0.38 | 19.99 | 19.51 | 0.48 | 0.48 | 0.01 | 0.01 |
| gene-LOC105285202 | https://www.ncbi.nlm.nih.gov/gene/?term=LOC105285202 | 1838 | 70 | 380.85 | | 1 | 5.44 | 1.43 | 25.94 | 25.46 | 0.48 | 0.48 | 0.02 | 0.02 |
| gene-LOC105278454 | https://www.ncbi.nlm.nih.gov/gene/?term=LOC105278454 | 1972 | 46 | 233.27 | | 1 | 5.07 | 2.17 | 44.92 | 44.47 | 0.46 | 0.46 | 0.01 | 0.01 |
| gene-LOC105275179 | https://www.ncbi.nlm.nih.gov/gene/?term=LOC105275179 | 3271 | 72 | 220.12 | | 1 | 3.06 | 1.39 | 21.5 | 21.05 | 0.45 | 0.45 | 0.01 | 0.01 |
| gene-LOC105282491 | https://www.ncbi.nlm.nih.gov/gene/?term=LOC105282491 | 4761 | 170 | 357.07 | | 1 | 2.1 | 0.59 | 10.04 | 9.59 | 0.45 | 0.45 | 0.02 | 0.02 |
| gene-LOC105276078 | https://www.ncbi.nlm.nih.gov/gene/?term=LOC105276078 | 1051 | 14 | 133.21 | | 1 | 9.51 | 7.14 | 33.32 | 32.93 | 0.39 | 0.39 | 0.01 | 0.01 |
| gene-LOC105285860 | https://www.ncbi.nlm.nih.gov/gene/?term=LOC105285860 | 3110 | 55 | 176.85 | | 1 | 3.22 | 1.82 | 24.17 | 23.77 | 0.39 | 0.39 | 0.01 | 0.01 |
| gene-LOC105275327 | https://www.ncbi.nlm.nih.gov/gene/?term=LOC105275327 | 14245 | 492 | 345.38 | | 1 | 0.7 | 0.2 | 7.28 | 6.9 | 0.38 | 0.38 | 0.01 | 0.01 |
| gene-LOC105278666 | https://www.ncbi.nlm.nih.gov/gene/?term=LOC105278666 | 10341 | 451 | 436.13 | | 1 | 0.97 | 0.22 | 7.32 | 6.95 | 0.38 | 0.38 | 0.02 | 0.02 |
| gene-LOC105274531 | https://www.ncbi.nlm.nih.gov/gene/?term=LOC105274531 | 7774 | 301 | 387.19 | | 1 | 1.29 | 0.33 | 10.99 | 10.62 | 0.37 | 0.37 | 0.01 | 0.01 |
| gene-LOC105282811 | https://www.ncbi.nlm.nih.gov/gene/?term=LOC105282811 | 5765 | 213 | 369.47 | | 1 | 1.73 | 0.47 | 8.63 | 8.26 | 0.37 | 0.37 | 0.01 | 0.01 |
| gene-LOC105276452 | https://www.ncbi.nlm.nih.gov/gene/?term=LOC105276452 | 2917 | 269 | 922.18 | | 1 | 3.43 | 0.37 | 9.31 | 8.95 | 0.36 | 0.36 | 0.03 | 0.03 |
| gene-LOC105278145 | https://www.ncbi.nlm.nih.gov/gene/?term=LOC105278145 | 4470 | 251 | 561.52 | | 1 | 2.24 | 0.4 | 10.77 | 10.43 | 0.34 | 0.34 | 0.02 | 0.02 |
| gene-LOC105287371 | https://www.ncbi.nlm.nih.gov/gene/?term=LOC105287371 | 8224 | 351 | 426.8 | 1 | 1.22 | 0.28 | 7.67 | 7.34 | 0.33 | 0.33 | 0.01 | 0.01 | |
| gene-LOC105287528 | https://www.ncbi.nlm.nih.gov/gene/?term=LOC105287528 | 14195 | 419 | 295.17 | | 1 | 0.7 | 0.24 | 5.19 | 4.86 | 0.33 | 0.33 | 0.01 | 0.01 |
| gene-LOC105286274 | https://www.ncbi.nlm.nih.gov/gene/?term=LOC105286274 | 3817 | 193 | 505.63 | | 1 | 2.62 | 0.52 | 17.37 | 17.05 | 0.32 | 0.32 | 0.02 | 0.02 |
| gene-LOC105286452 | https://www.ncbi.nlm.nih.gov/gene/?term=LOC105286452 | 6888 | 67 | 97.27 | 1 | 1.45 | 1.49 | 31.04 | 30.73 | 0.31 | 0.31 | 0 | 0 | |
| gene-LOC105287550 | https://www.ncbi.nlm.nih.gov/gene/?term=LOC105287550 | 15659 | 613 | 391.47 | | 1 | 0.64 | 0.16 | 6.43 | 6.12 | 0.31 | 0.31 | 0.01 | 0.01 |
| gene-LOC105286166 | https://www.ncbi.nlm.nih.gov/gene/?term=LOC105286166 | 3050 | 110 | 360.66 | | 1 | 3.28 | 0.91 | 24.5 | 24.21 | 0.29 | 0.29 | 0.01 | 0.01 |
| gene-LOC113562050 | https://www.ncbi.nlm.nih.gov/gene/?term=LOC113562050 | 17097 | 576 | 336.9 | 1 | 0.58 | 0.17 | 5.24 | 4.95 | 0.29 | 0.29 | 0.01 | 0.01 | |
| gene-LOC105278877 | https://www.ncbi.nlm.nih.gov/gene/?term=LOC105278877 | 1968 | 72 | 365.85 | | 1 | 5.08 | 1.39 | 36.04 | 35.76 | 0.28 | 0.28 | 0.01 | 0.01 |
| gene-LOC105277079 | https://www.ncbi.nlm.nih.gov/gene/?term=LOC105277079 | 13009 | 288 | 221.39 | | 1 | 0.77 | 0.35 | 9.86 | 9.59 | 0.27 | 0.27 | 0.01 | 0.01 |
| gene-LOC105278245 | https://www.ncbi.nlm.nih.gov/gene/?term=LOC105278245 | 6445 | 117 | 181.54 | | 1 | 1.55 | 0.85 | 22.48 | 22.2 | 0.27 | 0.27 | 0 | 0 |
| gene-LOC105284262 | https://www.ncbi.nlm.nih.gov/gene/?term=LOC105284262 | 6669 | 225 | 337.38 | | 1 | 1.5 | 0.44 | 6.17 | 5.94 | 0.23 | 0.23 | 0.01 | 0.01 |
| gene-LOC105278632 | https://www.ncbi.nlm.nih.gov/gene/?term=LOC105278632 | 3624 | 120 | 331.13 | | 1 | 2.76 | 0.83 | 15.73 | 15.53 | 0.2 | 0.2 | 0.01 | 0.01 |
| gene-LOC105285807 | https://www.ncbi.nlm.nih.gov/gene/?term=LOC105285807 | 6063 | 227 | 374.4 | 1 | 1.65 | 0.44 | 9.73 | 9.55 | 0.18 | 0.18 | 0.01 | 0.01 | |
| gene-LOC105276272 | https://www.ncbi.nlm.nih.gov/gene/?term=LOC105276272 | 1965 | 48 | 244.27 | | 1 | 5.09 | 2.08 | 27.74 | 27.58 | 0.17 | 0.17 | 0 | 0 |
| gene-LOC105276527 | https://www.ncbi.nlm.nih.gov/gene/?term=LOC105276527 | 7923 | 329 | 415.25 | | 1 | 1.26 | 0.3 | 9.51 | 9.36 | 0.15 | 0.15 | 0.01 | 0.01 |
| gene-LOC105274669 | https://www.ncbi.nlm.nih.gov/gene/?term=LOC105274669 | 8555 | 256 | 299.24 | | 1 | 1.17 | 0.39 | 6.43 | 6.35 | 0.08 | 0.08 | 0 | 0 |
| gene-LOC105275752 | https://www.ncbi.nlm.nih.gov/gene/?term=LOC105275752 | 3462 | 132 | 381.28 | | 1 | 2.89 | 0.76 | 32.74 | 32.68 | 0.06 | 0.06 | 0 | 0 |
| gene-LOC105286515 | https://www.ncbi.nlm.nih.gov/gene/?term=LOC105286515 | 3751 | 114 | 303.92 | | 1 | 2.67 | 0.88 | 11.87 | 11.89 | -0.02 | 0.02 | 0 | 0 |
| gene-LOC105283296 | https://www.ncbi.nlm.nih.gov/gene/?term=LOC105283296 | 2174 | 94 | 432.38 | | 1 | 4.6 | 1.06 | 18.21 | 18.24 | -0.03 | 0.03 | 0 | 0 |
| gene-LOC105287695 | https://www.ncbi.nlm.nih.gov/gene/?term=LOC105287695 | 6106 | 201 | 329.18 | | 1 | 1.64 | 0.5 | 6.09 | 6.16 | -0.06 | 0.06 | 0 | 0 |
| gene-LOC105274851 | https://www.ncbi.nlm.nih.gov/gene/?term=LOC105274851 | 6616 | 139 | 210.1 | 1 | 1.51 | 0.72 | 11.55 | 11.65 | -0.1 | 0.1 | 0 | 0 | |
| gene-LOC105275187 | https://www.ncbi.nlm.nih.gov/gene/?term=LOC105275187 | 5534 | 71 | 128.3 | 1 | 1.81 | 1.41 | 12.33 | 12.43 | -0.1 | 0.1 | 0 | 0 | |
| gene-LOC105286649 | https://www.ncbi.nlm.nih.gov/gene/?term=LOC105286649 | 14178 | 817 | 576.24 | | 1 | 0.71 | 0.12 | 1 | 1.1 | -0.1 | 0.1 | -0.01 | 0.01 |
| gene-LOC105282646 | https://www.ncbi.nlm.nih.gov/gene/?term=LOC105282646 | 3317 | 116 | 349.71 | | 1 | 3.01 | 0.86 | 24.81 | 24.92 | -0.11 | 0.11 | 0 | 0 |
| gene-LOC105283806 | https://www.ncbi.nlm.nih.gov/gene/?term=LOC105283806 | 10501 | 74 | 70.47 | 1 | 0.95 | 1.35 | 16.11 | 16.23 | -0.12 | 0.12 | 0 | 0 | |
| gene-LOC105284507 | https://www.ncbi.nlm.nih.gov/gene/?term=LOC105284507 | 5207 | 87 | 167.08 | | 1 | 1.92 | 1.15 | 21.48 | 21.62 | -0.13 | 0.13 | 0 | 0 |
| gene-LOC105275848 | https://www.ncbi.nlm.nih.gov/gene/?term=LOC105275848 | 3718 | 144 | 387.31 | | 1 | 2.69 | 0.69 | 20.8 | 20.95 | -0.15 | 0.15 | -0.01 | 0.01 |
| gene-LOC105280704 | https://www.ncbi.nlm.nih.gov/gene/?term=LOC105280704 | 4266 | 134 | 314.11 | | 1 | 2.34 | 0.75 | 19.89 | 20.04 | -0.15 | 0.15 | 0 | 0 |
| gene-LOC105280348 | https://www.ncbi.nlm.nih.gov/gene/?term=LOC105280348 | 3797 | 70 | 184.36 | | 1 | 2.63 | 1.43 | 20.64 | 20.81 | -0.17 | 0.17 | 0 | 0 |
| gene-LOC105283967 | https://www.ncbi.nlm.nih.gov/gene/?term=LOC105283967 | 2393 | 47 | 196.41 | | 1 | 4.18 | 2.13 | 48.9 | 49.11 | -0.21 | 0.21 | 0 | 0 |
| gene-LOC105285163 | https://www.ncbi.nlm.nih.gov/gene/?term=LOC105285163 | 5061 | 87 | 171.9 | 1 | 1.98 | 1.15 | 21.7 | 21.95 | -0.26 | 0.26 | 0 | 0 | |
| gene-LOC105287633 | https://www.ncbi.nlm.nih.gov/gene/?term=LOC105287633 | 1524 | 34 | 223.1 | 1 | 6.56 | 2.94 | 50.71 | 50.96 | -0.26 | 0.26 | -0.01 | 0.01 | |
| gene-LOC105281312 | https://www.ncbi.nlm.nih.gov/gene/?term=LOC105281312 | 2551 | 63 | 246.96 | | 1 | 3.92 | 1.59 | 34.27 | 34.56 | -0.29 | 0.29 | -0.01 | 0.01 |
| gene-LOC105286999 | https://www.ncbi.nlm.nih.gov/gene/?term=LOC105286999 | 2141 | 44 | 205.51 | | 1 | 4.67 | 2.27 | 34.01 | 34.41 | -0.41 | 0.41 | -0.01 | 0.01 |
| gene-LOC105287095 | https://www.ncbi.nlm.nih.gov/gene/?term=LOC105287095 | 1435 | 58 | 404.18 | | 1 | 6.97 | 1.72 | 28.05 | 28.65 | -0.6 | 0.6 | -0.02 | 0.02 |
| gene-LOC105274449 | https://www.ncbi.nlm.nih.gov/gene/?term=LOC105274449 | 1549 | 48 | 309.88 | | 1 | 6.46 | 2.08 | 50.3 | 50.96 | -0.66 | 0.66 | -0.02 | 0.02 |
| gene-LOC105285252 | https://www.ncbi.nlm.nih.gov/gene/?term=LOC105285252 | 7427 | 117 | 157.53 | | 1 | 1.35 | 0.85 | 14.46 | 15.16 | -0.7 | 0.7 | -0.01 | 0.01 |
| gene-LOC105281296 | https://www.ncbi.nlm.nih.gov/gene/?term=LOC105281296 | 2434 | 37 | 152.01 | | 1 | 4.11 | 2.7 | 30.9 | 31.72 | -0.83 | 0.83 | -0.01 | 0.01 |
| gene-LOC105280596 | https://www.ncbi.nlm.nih.gov/gene/?term=LOC105280596 | 3434 | 113 | 329.06 | | 1 | 2.91 | 0.88 | 17.62 | 18.56 | -0.94 | 0.94 | -0.03 | 0.03 |
| gene-LOC105282885 | https://www.ncbi.nlm.nih.gov/gene/?term=LOC105282885 | 1870 | 48 | 256.68 | | 1 | 5.35 | 2.08 | 38.27 | 39.26 | -0.99 | 0.99 | -0.03 | 0.03 |
| gene-LOC105278384 | https://www.ncbi.nlm.nih.gov/gene/?term=LOC105278384 | 1390 | 36 | 258.99 | | 1 | 7.19 | 2.78 | 50.65 | 51.65 | -1 | 1 | -0.03 | 0.03 |
| gene-LOC105286770 | https://www.ncbi.nlm.nih.gov/gene/?term=LOC105286770 | 3055 | 33 | 108.02 | | 1 | 3.27 | 3.03 | 17.27 | 18.34 | -1.07 | 1.07 | -0.01 | 0.01 |
| gene-LOC105280819 | https://www.ncbi.nlm.nih.gov/gene/?term=LOC105280819 | 1639 | 38 | 231.85 | | 1 | 6.1 | 2.63 | 35.58 | 37.06 | -1.49 | 1.49 | -0.03 | 0.03 |
| gene-LOC105283989 | https://www.ncbi.nlm.nih.gov/gene/?term=LOC105283989 | 4948 | 31 | 62.65 | 2 | 4.04 | 6.45 | 34.97 | 36.46 | -1.49 | 1.49 | -0.01 | 0.01 | |
| gene-LOC105276196 | https://www.ncbi.nlm.nih.gov/gene/?term=LOC105276196 | 2953 | 62 | 209.96 | | 1 | 3.39 | 1.61 | 36.95 | 38.53 | -1.58 | 1.58 | -0.03 | 0.03 |
| gene-LOC105281700 | https://www.ncbi.nlm.nih.gov/gene/?term=LOC105281700 | 2208 | 34 | 153.99 | | 1 | 4.53 | 2.94 | 35.23 | 36.92 | -1.69 | 1.69 | -0.03 | 0.03 |
| gene-LOC105284047 | https://www.ncbi.nlm.nih.gov/gene/?term=LOC105284047 | 1212 | 24 | 198.02 | | 1 | 8.25 | 4.17 | 25.81 | 27.68 | -1.87 | 1.87 | -0.04 | 0.04 |

| | | | | | | | | | | | | | | | |
|---|---|---|---|---|---|---|---|---|---|---|---|---|---|---|---|
| gene-LOC105285409 | https://www.ncbi.nlm.nih.gov/gene/?term=LOC105285409 | 5099 | 49 | 96.1 | 2 | 3.92 | 4.08 | 1.55 | 4.82 | -3.27 | 3.27 | -0.03 | 0.03 | | |
| gene-LOC105283492 | https://www.ncbi.nlm.nih.gov/gene/?term=LOC105283492 | 1535 | 10 | 65.15 | 1 | 6.51 | 10 | 37.74 | 42.8 | -5.06 | 5.06 | -0.03 | 0.03 | | |
| gene-LOC105284677 | https://www.ncbi.nlm.nih.gov/gene/?term=LOC105284677 | 1905 | 42 | 220.47 | | 1 | 5.25 | 2.38 | 23.81 | 34.99 | -11.18 | | 11.18 | -0.25 | 0.25 |

Directory: OGL File: ogl-Ob_LI_RphaseB1.txt

| gene_ID | gene_link | gwidth | #Sites | #per10Kb | %perSite | %pNucl |
|---|---|---|---|---|---|---|
| gene-LOC113561532 | https://www.ncbi.nlm.nih.gov/gene/?term=LOC113561532 | 72 | 5 | 694.44 | 95.54 | 6.64 |
| gene-LOC105286531 | https://www.ncbi.nlm.nih.gov/gene/?term=LOC105286531 | 372 | 40 | 1075.27 | 35.13 | 3.78 |
| gene-LOC105280739 | https://www.ncbi.nlm.nih.gov/gene/?term=LOC105280739 | 1991 | 138 | 693.12 | 53.97 | 3.74 |
| gene-LOC113562627 | https://www.ncbi.nlm.nih.gov/gene/?term=LOC113562627 | 397 | 22 | 554.16 | 65.67 | 3.64 |
| gene-LOC105274534 | https://www.ncbi.nlm.nih.gov/gene/?term=LOC105274534 | 1329 | 102 | 767.49 | 45.61 | 3.5 |
| gene-LOC105286801 | https://www.ncbi.nlm.nih.gov/gene/?term=LOC105286801 | 1102 | 101 | 916.52 | 37.9 | 3.47 |
| gene-LOC105285635 | https://www.ncbi.nlm.nih.gov/gene/?term=LOC105285635 | 1175 | 127 | 1080.85 | 31.75 | 3.43 |
| gene-LOC105286269 | https://www.ncbi.nlm.nih.gov/gene/?term=LOC105286269 | 1401 | 95 | 678.09 | 50.19 | 3.4 |
| gene-LOC105276360 | https://www.ncbi.nlm.nih.gov/gene/?term=LOC105276360 | 1466 | 169 | 1152.8 | 29.3 | 3.38 |
| gene-LOC105287811 | https://www.ncbi.nlm.nih.gov/gene/?term=LOC105287811 | 2155 | 135 | 626.45 | 53.84 | 3.37 |
| gene-LOC105278844 | https://www.ncbi.nlm.nih.gov/gene/?term=LOC105278844 | 1101 | 115 | 1044.5 | 30.86 | 3.22 |
| gene-LOC105278849 | https://www.ncbi.nlm.nih.gov/gene/?term=LOC105278849 | 776 | 114 | 1469.07 | 21.62 | 3.18 |
| gene-LOC105279052 | https://www.ncbi.nlm.nih.gov/gene/?term=LOC105279052 | 1130 | 127 | 1123.89 | 27.56 | 3.1 |
| gene-LOC105278160 | https://www.ncbi.nlm.nih.gov/gene/?term=LOC105278160 | 1872 | 168 | 897.44 | 34.28 | 3.08 |
| gene-LOC105285392 | https://www.ncbi.nlm.nih.gov/gene/?term=LOC105285392 | 1601 | 134 | 836.98 | 36.82 | 3.08 |
| gene-LOC105274711 | https://www.ncbi.nlm.nih.gov/gene/?term=LOC105274711 | 1655 | 113 | 682.78 | 44.97 | 3.07 |
| gene-LOC105286235 | https://www.ncbi.nlm.nih.gov/gene/?term=LOC105286235 | 1510 | 105 | 695.36 | 44.05 | 3.06 |
| gene-LOC105286737 | https://www.ncbi.nlm.nih.gov/gene/?term=LOC105286737 | 2690 | 368 | 1368.03 | 22.4 | 3.06 |
| gene-LOC105278549 | https://www.ncbi.nlm.nih.gov/gene/?term=LOC105278549 | 1771 | 117 | 660.64 | 46.07 | 3.04 |
| gene-LOC105282149 | https://www.ncbi.nlm.nih.gov/gene/?term=LOC105282149 | 1787 | 134 | 749.86 | 40.5 | 3.04 |
| gene-LOC105287692 | https://www.ncbi.nlm.nih.gov/gene/?term=LOC105287692 | 1505 | 137 | 910.3 | 32.53 | 2.96 |
| gene-LOC105275314 | https://www.ncbi.nlm.nih.gov/gene/?term=LOC105275314 | 1489 | 104 | 698.46 | 41.67 | 2.91 |
| gene-LOC105277648 | https://www.ncbi.nlm.nih.gov/gene/?term=LOC105277648 | 722 | 39 | 540.17 | 53.41 | 2.89 |
| gene-LOC105288088 | https://www.ncbi.nlm.nih.gov/gene/?term=LOC105288088 | 1795 | 123 | 685.24 | 42.1 | 2.89 |

...

... only first 25 lines shown ...

Directory: OGL File: ogl-Ob_LI_RphaseB1.vs.RphaseB2.txt

| gene_ID | gene_link | gwidth | #Sites | #per10Kb | #dmSites | #dmsp10kb | %dmSites | %pSite1 | %pSite2 | DMpSite | ADMpSite | DMpNucl | ADMpNucl |
|---|---|---|---|---|---|---|---|---|---|---|---|---|---|
| gene-LOC105285431 | https://www.ncbi.nlm.nih.gov/gene/?term=LOC105285431 | 1550 | 6 | 38.71 | 3 | 19.35 | 50 | 46.72 | 30.29 | 16.43 | 16.43 | 0.06 | 0.06 |
| gene-LOC105284871 | https://www.ncbi.nlm.nih.gov/gene/?term=LOC105284871 | 1878 | 3 | 15.97 | 1 | 5.32 | 33.33 | 46.42 | 30.96 | 15.47 | 15.47 | 0.02 | 0.02 |
| gene-LOC113561977 | https://www.ncbi.nlm.nih.gov/gene/?term=LOC113561977 | 1058 | 3 | 28.36 | 1 | 9.45 | 33.33 | 25 | 10.74 | 14.26 | 14.26 | 0.04 | 0.04 |
| gene-LOC105277491 | https://www.ncbi.nlm.nih.gov/gene/?term=LOC105277491 | 1647 | 12 | 72.86 | 2 | 12.14 | 16.67 | 21.62 | 11.32 | 10.3 | 10.3 | 0.08 | 0.08 |
| gene-LOC105279429 | https://www.ncbi.nlm.nih.gov/gene/?term=LOC105279429 | 1741 | 10 | 57.44 | 1 | 5.74 | 10 | 23.71 | 13.86 | 9.85 | 9.85 | 0.06 | 0.06 |
| gene-LOC105283805 | https://www.ncbi.nlm.nih.gov/gene/?term=LOC105283805 | 2008 | 7 | 34.86 | 2 | 9.96 | 28.57 | 87.5 | 77.76 | 9.74 | 9.74 | 0.03 | 0.03 |
| gene-LOC105282854 | https://www.ncbi.nlm.nih.gov/gene/?term=LOC105282854 | 1825 | 2 | 10.96 | 1 | 5.48 | 50 | 38.24 | 28.94 | 9.3 | 9.3 | 0.01 | 0.01 |
| gene-LOC105281216 | https://www.ncbi.nlm.nih.gov/gene/?term=LOC105281216 | 3765 | 12 | 31.87 | 2 | 5.31 | 16.67 | 39.32 | 30.3 | 9.02 | 9.02 | 0.03 | 0.03 |
| gene-LOC105283410 | https://www.ncbi.nlm.nih.gov/gene/?term=LOC105283410 | 1423 | 5 | 35.14 | 1 | 7.03 | 20 | 49.1 | 40.2 | 8.9 | 8.9 | 0.03 | 0.03 |
| gene-LOC113561900 | https://www.ncbi.nlm.nih.gov/gene/?term=LOC113561900 | 629 | 11 | 174.88 | 1 | 15.9 | 9.09 | 33.39 | 25.51 | 7.88 | 7.88 | 0.14 | 0.14 |
| gene-LOC105281635 | https://www.ncbi.nlm.nih.gov/gene/?term=LOC105281635 | 1508 | 4 | 26.53 | 1 | 6.63 | 25 | 15.47 | 7.61 | 7.86 | 7.86 | 0.02 | 0.02 |
| gene-LOC105286857 | https://www.ncbi.nlm.nih.gov/gene/?term=LOC105286857 | 1576 | 4 | 25.38 | 1 | 6.35 | 25 | 74.57 | 67.03 | 7.54 | 7.54 | 0.02 | 0.02 |
| gene-LOC105282657 | https://www.ncbi.nlm.nih.gov/gene/?term=LOC105282657 | 1409 | 6 | 42.58 | 1 | 7.1 | 16.67 | 58.16 | 50.81 | 7.34 | 7.34 | 0.03 | 0.03 |
| gene-LOC105277862 | https://www.ncbi.nlm.nih.gov/gene/?term=LOC105277862 | 1490 | 7 | 46.98 | 1 | 6.71 | 14.29 | 62.25 | 55.64 | 6.6 | 6.6 | 0.03 | 0.03 |
| gene-LOC105286261 | https://www.ncbi.nlm.nih.gov/gene/?term=LOC105286261 | 1297 | 8 | 61.68 | 1 | 7.71 | 12.5 | 74.9 | 68.31 | 6.59 | 6.59 | 0.04 | 0.04 |
| gene-LOC113562499 | https://www.ncbi.nlm.nih.gov/gene/?term=LOC113562499 | 491 | 6 | 122.2 | 1 | 20.37 | 16.67 | 53.38 | 46.94 | 6.45 | 6.45 | 0.08 | 0.08 |
| gene-LOC105276718 | https://www.ncbi.nlm.nih.gov/gene/?term=LOC105276718 | 1721 | 10 | 58.11 | 1 | 5.81 | 10 | 28.17 | 21.77 | 6.4 | 6.4 | 0.04 | 0.04 |
| gene-LOC105274857 | https://www.ncbi.nlm.nih.gov/gene/?term=LOC105274857 | 1569 | 5 | 31.87 | 1 | 6.37 | 20 | 57.81 | 51.63 | 6.18 | 6.18 | 0.02 | 0.02 |
| gene-LOC105280261 | https://www.ncbi.nlm.nih.gov/gene/?term=LOC105280261 | 1005 | 7 | 69.65 | 1 | 9.95 | 14.29 | 8.97 | 3.43 | 5.54 | 5.54 | 0.04 | 0.04 |
| gene-LOC105284028 | https://www.ncbi.nlm.nih.gov/gene/?term=LOC105284028 | 2536 | 23 | 90.69 | 4 | 15.77 | 17.39 | 36.1 | 30.67 | 5.43 | 5.43 | 0.05 | 0.05 |
| gene-LOC105287722 | https://www.ncbi.nlm.nih.gov/gene/?term=LOC105287722 | 3481 | 6 | 17.24 | 1 | 2.87 | 16.67 | 47.51 | 42.11 | 5.39 | 5.39 | 0.01 | 0.01 |
| gene-LOC105279461 | https://www.ncbi.nlm.nih.gov/gene/?term=LOC105279461 | 1298 | 9 | 69.34 | 1 | 7.7 | 11.11 | 73.75 | 68.65 | 5.1 | 5.1 | 0.04 | 0.04 |
| gene-LOC105284656 | https://www.ncbi.nlm.nih.gov/gene/?term=LOC105284656 | 1312 | 17 | 129.57 | 2 | 15.24 | 11.76 | 79.02 | 73.94 | 5.08 | 5.08 | 0.07 | 0.07 |
| gene-LOC105285670 | https://www.ncbi.nlm.nih.gov/gene/?term=LOC105285670 | 1477 | 21 | 142.18 | 1 | 6.77 | 4.76 | 49.06 | 44.02 | 5.03 | 5.03 | 0.07 | 0.07 |

...

... only first 25 lines shown ...

Directory: OGL File: ogl-Ob_LI_RphaseB2.txt

| gene_ID | gene_link | gwidth | #Sites | #per10Kb | %perSite | %pNucl |
|---|---|---|---|---|---|---|
| gene-LOC113561532 | https://www.ncbi.nlm.nih.gov/gene/?term=LOC113561532 | 72 | 4 | 555.56 | 94.52 | 5.25 |
| gene-LOC105286531 | https://www.ncbi.nlm.nih.gov/gene/?term=LOC105286531 | 372 | 41 | 1102.15 | 36.36 | 4.01 |
| gene-LOC105286801 | https://www.ncbi.nlm.nih.gov/gene/?term=LOC105286801 | 1102 | 95 | 862.07 | 43.07 | 3.71 |
| gene-LOC105282149 | https://www.ncbi.nlm.nih.gov/gene/?term=LOC105282149 | 1787 | 157 | 878.57 | 41.23 | 3.62 |
| gene-LOC105280739 | https://www.ncbi.nlm.nih.gov/gene/?term=LOC105280739 | 1991 | 127 | 637.87 | 54.82 | 3.5 |
| gene-LOC105278849 | https://www.ncbi.nlm.nih.gov/gene/?term=LOC105278849 | 776 | 113 | 1456.19 | 23.54 | 3.43 |
| gene-LOC105285635 | https://www.ncbi.nlm.nih.gov/gene/?term=LOC105285635 | 1175 | 123 | 1046.81 | 32.53 | 3.4 |
| gene-LOC105276360 | https://www.ncbi.nlm.nih.gov/gene/?term=LOC105276360 | 1466 | 141 | 961.8 | 35 | 3.37 |
| gene-LOC105274534 | https://www.ncbi.nlm.nih.gov/gene/?term=LOC105274534 | 1329 | 89 | 669.68 | 48.75 | 3.26 |
| gene-LOC113562627 | https://www.ncbi.nlm.nih.gov/gene/?term=LOC113562627 | 397 | 19 | 478.59 | 66.86 | 3.2 |
| gene-LOC105287811 | https://www.ncbi.nlm.nih.gov/gene/?term=LOC105287811 | 2155 | 147 | 682.13 | 46.69 | 3.19 |
| gene-LOC105274711 | https://www.ncbi.nlm.nih.gov/gene/?term=LOC105274711 | 1655 | 111 | 670.69 | 47.04 | 3.15 |
| gene-LOC105278160 | https://www.ncbi.nlm.nih.gov/gene/?term=LOC105278160 | 1872 | 159 | 849.36 | 36.67 | 3.11 |
| gene-LOC105278844 | https://www.ncbi.nlm.nih.gov/gene/?term=LOC105278844 | 1101 | 111 | 1008.17 | 30.45 | 3.07 |
| gene-LOC105276371 | https://www.ncbi.nlm.nih.gov/gene/?term=LOC105276371 | 871 | 57 | 654.42 | 46.14 | 3.02 |
| gene-LOC105287692 | https://www.ncbi.nlm.nih.gov/gene/?term=LOC105287692 | 1505 | 129 | 857.14 | 34.93 | 2.99 |
| gene-LOC105279052 | https://www.ncbi.nlm.nih.gov/gene/?term=LOC105279052 | 1130 | 112 | 991.15 | 29.29 | 2.9 |
| gene-LOC105286235 | https://www.ncbi.nlm.nih.gov/gene/?term=LOC105286235 | 1510 | 96 | 635.76 | 45.62 | 2.9 |
| gene-LOC105286737 | https://www.ncbi.nlm.nih.gov/gene/?term=LOC105286737 | 2690 | 337 | 1252.79 | 22.93 | 2.87 |
| gene-LOC105275314 | https://www.ncbi.nlm.nih.gov/gene/?term=LOC105275314 | 1489 | 89 | 597.72 | 47.67 | 2.85 |
| gene-LOC105283717 | https://www.ncbi.nlm.nih.gov/gene/?term=LOC105283717 | 1373 | 112 | 815.73 | 34.69 | 2.83 |
| gene-LOC105287677 | https://www.ncbi.nlm.nih.gov/gene/?term=LOC105287677 | 1588 | 115 | 724.18 | 38.65 | 2.8 |
| gene-LOC105286269 | https://www.ncbi.nlm.nih.gov/gene/?term=LOC105286269 | 1401 | 68 | 485.37 | 55.58 | 2.7 |
| gene-LOC105275897 | https://www.ncbi.nlm.nih.gov/gene/?term=LOC105275897 | 1684 | 116 | 688.84 | 38.05 | 2.62 |

...

... only first 25 lines shown ...

Directory: OGL File: rnk-dmg-Ob_LI_BCphaseA1.vs.BCphaseA2.txt

| gene_ID | gene_link | gwidth | #Sites | #per10Kb | #dmSites | #dmsp10kb | %dmSites | %pSite1 | %pSite2 | DMpSite | ADMpSite | DMpNucl | ADMpNucl |
|---|---|---|---|---|---|---|---|---|---|---|---|---|---|
| gene-LOC113562345 | https://www.ncbi.nlm.nih.gov/gene/?term=LOC113562345 | 2290 | 62 | 270.74 | 15 | 65.5 | 24.19 | 17.91 | 32.18 | -14.27 | 14.27 | -0.39 | 0.39 |
| gene-LOC105283552 | https://www.ncbi.nlm.nih.gov/gene/?term=LOC105283552 | 1562 | 27 | 172.86 | 1 | 6.4 | 3.7 | 56.95 | 65.52 | -8.57 | 8.57 | -0.15 | 0.15 |
| gene-LOC105285695 | https://www.ncbi.nlm.nih.gov/gene/?term=LOC105285695 | 1371 | 62 | 452.22 | 1 | 7.29 | 1.61 | 23 | 25.77 | -2.77 | 2.77 | -0.13 | 0.13 |
| gene-LOC105282156 | https://www.ncbi.nlm.nih.gov/gene/?term=LOC105282156 | 2577 | 132 | 512.22 | 1 | 3.88 | 0.76 | 24.4 | 26.49 | -2.09 | 2.09 | -0.11 | 0.11 |
| gene-LOC105275656 | https://www.ncbi.nlm.nih.gov/gene/?term=LOC105275656 | 2803 | 127 | 453.09 | 1 | 3.57 | 0.79 | 26.43 | 28.64 | -2.2 | 2.2 | -0.1 | 0.1 |
| gene-LOC105279356 | https://www.ncbi.nlm.nih.gov/gene/?term=LOC105279356 | 1314 | 39 | 296.8 | 1 | 7.61 | 2.56 | 29.77 | 33.04 | -3.27 | 3.27 | -0.1 | 0.1 |
| gene-LOC105285674 | https://www.ncbi.nlm.nih.gov/gene/?term=LOC105285674 | 3035 | 88 | 289.95 | 1 | 3.29 | 1.14 | 23.36 | 26.44 | -3.08 | 3.08 | -0.09 | 0.09 |
| gene-LOC105281848 | https://www.ncbi.nlm.nih.gov/gene/?term=LOC105281848 | 2122 | 44 | 207.35 | 1 | 4.71 | 2.27 | 23.57 | 27.97 | -4.4 | 4.4 | -0.09 | 0.09 |
| gene-LOC105285060 | https://www.ncbi.nlm.nih.gov/gene/?term=LOC105285060 | 1285 | 13 | 101.17 | 1 | 7.78 | 7.69 | 55.35 | 64 | -8.64 | 8.64 | -0.09 | 0.09 |
| gene-LOC105285055 | https://www.ncbi.nlm.nih.gov/gene/?term=LOC105285055 | 4108 | 197 | 479.55 | 1 | 2.43 | 0.51 | 13.57 | 15.12 | -1.55 | 1.55 | -0.07 | 0.07 |
| gene-LOC105280972 | https://www.ncbi.nlm.nih.gov/gene/?term=LOC105280972 | 1532 | 26 | 169.71 | 1 | 6.53 | 3.85 | 20.04 | 23.88 | -3.84 | 3.84 | -0.07 | 0.07 |
| gene-LOC105275070 | https://www.ncbi.nlm.nih.gov/gene/?term=LOC105275070 | 5667 | 35 | 61.76 | 2 | 3.53 | 5.71 | 3.8 | 15.13 | -11.33 | 11.33 | -0.07 | 0.07 |
| gene-LOC109611283 | https://www.ncbi.nlm.nih.gov/gene/?term=LOC109611283 | 1451 | 36 | 248.1 | 1 | 6.89 | 2.78 | 37.85 | 35.62 | 2.23 | 2.23 | 0.06 | 0.06 |
| gene-LOC105280774 | https://www.ncbi.nlm.nih.gov/gene/?term=LOC105280774 | 2793 | 97 | 347.3 | 1 | 3.58 | 1.03 | 31.81 | 33.47 | -1.66 | 1.66 | -0.06 | 0.06 |
| gene-LOC113561621 | https://www.ncbi.nlm.nih.gov/gene/?term=LOC113561621 | 2249 | 79 | 351.27 | 1 | 4.45 | 1.27 | 19.72 | 21.41 | -1.69 | 1.69 | -0.06 | 0.06 |
| gene-LOC105278807 | https://www.ncbi.nlm.nih.gov/gene/?term=LOC105278807 | 2166 | 68 | 313.94 | 1 | 4.62 | 1.47 | 17.54 | 19.41 | -1.87 | 1.87 | -0.06 | 0.06 |
| gene-LOC105281173 | https://www.ncbi.nlm.nih.gov/gene/?term=LOC105281173 | 1595 | 25 | 156.74 | 1 | 6.27 | 4 | 64.76 | 61.8 | 2.96 | 2.96 | 0.05 | 0.05 |
| gene-LOC105278877 | https://www.ncbi.nlm.nih.gov/gene/?term=LOC105278877 | 1968 | 69 | 350.61 | 1 | 5.08 | 1.45 | 37.18 | 35.87 | 1.31 | 1.31 | 0.05 | 0.05 |
| gene-LOC105284184 | https://www.ncbi.nlm.nih.gov/gene/?term=LOC105284184 | 2535 | 67 | 264.3 | 1 | 3.94 | 1.49 | 41.64 | 43.45 | -1.81 | 1.81 | -0.05 | 0.05 |
| gene-LOC105279898 | https://www.ncbi.nlm.nih.gov/gene/?term=LOC105279898 | 1343 | 13 | 96.8 | 1 | 7.45 | 7.69 | 60.41 | 65.83 | -5.42 | 5.42 | -0.05 | 0.05 |
| gene-LOC105276078 | https://www.ncbi.nlm.nih.gov/gene/?term=LOC105276078 | 1051 | 8 | 76.12 | 1 | 9.51 | 12.5 | 46.61 | 53.81 | -7.2 | 7.2 | -0.05 | 0.05 |
| gene-LOC105275708 | https://www.ncbi.nlm.nih.gov/gene/?term=LOC105275708 | 1772 | 10 | 56.43 | 1 | 5.64 | 10 | 41.45 | 50.31 | -8.85 | 8.85 | -0.05 | 0.05 |
| gene-LOC105286999 | https://www.ncbi.nlm.nih.gov/gene/?term=LOC105286999 | 2141 | 38 | 177.49 | 1 | 4.67 | 2.63 | 33.02 | 30.95 | 2.07 | 2.07 | 0.04 | 0.04 |
| gene-LOC105274449 | https://www.ncbi.nlm.nih.gov/gene/?term=LOC105274449 | 1549 | 42 | 271.14 | 1 | 6.46 | 2.38 | 55.88 | 54.57 | 1.32 | 1.32 | 0.04 | 0.04 |
| gene-LOC105279298 | https://www.ncbi.nlm.nih.gov/gene/?term=LOC105279298 | 2773 | 129 | 465.2 | 2 | 7.21 | 1.55 | 22.93 | 23.79 | -0.86 | 0.86 | -0.04 | 0.04 |
| gene-LOC105275460 | https://www.ncbi.nlm.nih.gov/gene/?term=LOC105275460 | 2473 | 107 | 432.67 | 1 | 4.04 | 0.93 | 13.53 | 14.43 | -0.91 | 0.91 | -0.04 | 0.04 |
| gene-LOC105275683 | https://www.ncbi.nlm.nih.gov/gene/?term=LOC105275683 | 4256 | 187 | 439.38 | 1 | 2.35 | 0.53 | 12 | 12.9 | -0.91 | 0.91 | -0.04 | 0.04 |
| gene-LOC105285671 | https://www.ncbi.nlm.nih.gov/gene/?term=LOC105285671 | 2263 | 92 | 406.54 | 1 | 4.42 | 1.09 | 28.77 | 29.78 | -1.01 | 1.01 | -0.04 | 0.04 |
| gene-LOC105285179 | https://www.ncbi.nlm.nih.gov/gene/?term=LOC105285179 | 4191 | 148 | 353.14 | 1 | 2.39 | 0.68 | 17.81 | 18.87 | -1.05 | 1.05 | -0.04 | 0.04 |
| gene-LOC105278109 | https://www.ncbi.nlm.nih.gov/gene/?term=LOC105278109 | 4029 | 56 | 138.99 | 1 | 2.48 | 1.79 | 47.35 | 50.32 | -2.97 | 2.97 | -0.04 | 0.04 |
| gene-LOC105280348 | https://www.ncbi.nlm.nih.gov/gene/?term=LOC105280348 | 3797 | 50 | 131.68 | 1 | 2.63 | 2 | 25.73 | 28.9 | -3.17 | 3.17 | -0.04 | 0.04 |
| gene-LOC105285514 | https://www.ncbi.nlm.nih.gov/gene/?term=LOC105285514 | 1277 | 8 | 62.65 | 1 | 7.83 | 12.5 | 82.36 | 88.52 | -6.16 | 6.16 | -0.04 | 0.04 |
| gene-LOC105285202 | https://www.ncbi.nlm.nih.gov/gene/?term=LOC105285202 | 1838 | 33 | 179.54 | 1 | 5.44 | 3.03 | 20.53 | 19.01 | 1.52 | 1.52 | 0.03 | 0.03 |
| gene-LOC105278282 | https://www.ncbi.nlm.nih.gov/gene/?term=LOC105278282 | 1775 | 43 | 242.25 | 1 | 5.63 | 2.33 | 46.59 | 45.35 | 1.24 | 1.24 | 0.03 | 0.03 |
| gene-LOC105275848 | https://www.ncbi.nlm.nih.gov/gene/?term=LOC105275848 | 3718 | 119 | 320.06 | 1 | 2.69 | 0.84 | 22.44 | 21.48 | 0.97 | 0.97 | 0.03 | 0.03 |
| gene-LOC105277435 | https://www.ncbi.nlm.nih.gov/gene/?term=LOC105277435 | 3643 | 150 | 411.75 | 1 | 2.74 | 0.67 | 12.77 | 12 | 0.77 | 0.77 | 0.03 | 0.03 |
| gene-LOC105275363 | https://www.ncbi.nlm.nih.gov/gene/?term=LOC105275363 | 3798 | 146 | 384.41 | 1 | 2.63 | 0.68 | 17.72 | 18.47 | -0.76 | 0.76 | -0.03 | 0.03 |
| gene-LOC105276692 | https://www.ncbi.nlm.nih.gov/gene/?term=LOC105276692 | 1745 | 33 | 189.11 | 1 | 5.73 | 3.03 | 25.26 | 26.77 | -1.52 | 1.52 | -0.03 | 0.03 |
| gene-LOC105275190 | https://www.ncbi.nlm.nih.gov/gene/?term=LOC105275190 | 3837 | 69 | 179.83 | 1 | 2.61 | 1.45 | 19.53 | 21.12 | -1.59 | 1.59 | -0.03 | 0.03 |
| gene-LOC105283592 | https://www.ncbi.nlm.nih.gov/gene/?term=LOC105283592 | 3479 | 66 | 189.71 | 1 | 2.87 | 1.52 | 25.49 | 27.08 | -1.59 | 1.59 | -0.03 | 0.03 |
| gene-LOC105283712 | https://www.ncbi.nlm.nih.gov/gene/?term=LOC105283712 | 1910 | 24 | 125.65 | 1 | 5.24 | 4.17 | 29.64 | 32.04 | -2.39 | 2.39 | -0.03 | 0.03 |
| gene-LOC105274900 | https://www.ncbi.nlm.nih.gov/gene/?term=LOC105274900 | 5561 | 47 | 84.52 | 2 | 3.6 | 4.26 | 31.62 | 34.67 | -3.05 | 3.05 | -0.03 | 0.03 |
| gene-LOC105277215 | https://www.ncbi.nlm.nih.gov/gene/?term=LOC105277215 | 2619 | 20 | 76.37 | 1 | 3.82 | 5 | 28.06 | 32.55 | -4.49 | 4.49 | -0.03 | 0.03 |
| gene-LOC105278358 | https://www.ncbi.nlm.nih.gov/gene/?term=LOC105278358 | 2861 | 28 | 97.87 | 1 | 3.5 | 3.57 | 33.51 | 30.99 | 2.52 | 2.52 | 0.02 | 0.02 |
| gene-LOC105278228 | https://www.ncbi.nlm.nih.gov/gene/?term=LOC105278228 | 8610 | 146 | 169.57 | 1 | 1.16 | 0.68 | 23.97 | 22.69 | 1.27 | 1.27 | 0.02 | 0.02 |
| gene-LOC105285252 | https://www.ncbi.nlm.nih.gov/gene/?term=LOC105285252 | 7427 | 116 | 156.19 | 1 | 1.35 | 0.86 | 15.25 | 14.01 | 1.25 | 1.25 | 0.02 | 0.02 |
| gene-LOC105275653 | https://www.ncbi.nlm.nih.gov/gene/?term=LOC105275653 | 11251 | 236 | 209.76 | 1 | 0.89 | 0.42 | 9.52 | 8.44 | 1.07 | 1.07 | 0.02 | 0.02 |
| gene-LOC105282811 | https://www.ncbi.nlm.nih.gov/gene/?term=LOC105282811 | 5765 | 182 | 315.7 | 1 | 1.73 | 0.55 | 9.67 | 8.98 | 0.69 | 0.69 | 0.02 | 0.02 |
| gene-LOC105278632 | https://www.ncbi.nlm.nih.gov/gene/?term=LOC105278632 | 3624 | 119 | 328.37 | 1 | 2.76 | 0.84 | 15.91 | 15.28 | 0.63 | 0.63 | 0.02 | 0.02 |
| gene-LOC105280956 | https://www.ncbi.nlm.nih.gov/gene/?term=LOC105280956 | 2639 | 73 | 276.62 | 1 | 3.79 | 1.37 | 20.28 | 19.71 | 0.58 | 0.58 | 0.02 | 0.02 |
| gene-LOC105286515 | https://www.ncbi.nlm.nih.gov/gene/?term=LOC105286515 | 3751 | 107 | 285.26 | 1 | 2.67 | 0.93 | 10.47 | 9.92 | 0.55 | 0.55 | 0.02 | 0.02 |
| gene-LOC105280786 | https://www.ncbi.nlm.nih.gov/gene/?term=LOC105280786 | 2573 | 107 | 415.86 | 1 | 3.89 | 0.93 | 33.04 | 32.68 | 0.37 | 0.37 | 0.02 | 0.02 |
| gene-LOC105283296 | https://www.ncbi.nlm.nih.gov/gene/?term=LOC105283296 | 2174 | 78 | 358.79 | 1 | 4.6 | 1.28 | 21.02 | 21.5 | -0.48 | 0.48 | -0.02 | 0.02 |
| gene-LOC105284355 | https://www.ncbi.nlm.nih.gov/gene/?term=LOC105284355 | 3904 | 135 | 345.8 | 1 | 2.56 | 0.74 | 22.88 | 23.48 | -0.59 | 0.59 | -0.02 | 0.02 |
| gene-LOC105283728 | https://www.ncbi.nlm.nih.gov/gene/?term=LOC105283728 | 3256 | 103 | 316.34 | 2 | 6.14 | 1.94 | 21.65 | 22.28 | -0.62 | 0.62 | -0.02 | 0.02 |
| gene-LOC105286265 | https://www.ncbi.nlm.nih.gov/gene/?term=LOC105286265 | 2507 | 63 | 251.3 | 1 | 3.99 | 1.59 | 30.31 | 30.94 | -0.63 | 0.63 | -0.02 | 0.02 |
| gene-LOC105287063 | https://www.ncbi.nlm.nih.gov/gene/?term=LOC105287063 | 1874 | 46 | 245.46 | 1 | 5.34 | 2.17 | 34.15 | 34.94 | -0.79 | 0.79 | -0.02 | 0.02 |
| gene-LOC105275451 | https://www.ncbi.nlm.nih.gov/gene/?term=LOC105275451 | 6035 | 105 | 173.99 | 1 | 1.66 | 0.95 | 24.51 | 25.49 | -0.98 | 0.98 | -0.02 | 0.02 |
| gene-LOC105275173 | https://www.ncbi.nlm.nih.gov/gene/?term=LOC105275173 | 4639 | 71 | 153.05 | 1 | 2.16 | 1.41 | 27.01 | 28.06 | -1.05 | 1.05 | -0.02 | 0.02 |
| gene-LOC105280641 | https://www.ncbi.nlm.nih.gov/gene/?term=LOC105280641 | 3682 | 53 | 143.94 | 1 | 2.72 | 1.89 | 17.07 | 18.78 | -1.71 | 1.71 | -0.02 | 0.02 |

| | | | | | | | | | | | | | |
|---|---|---|---|---|---|---|---|---|---|---|---|---|---|
| gene-LOC105288066 | https://www.ncbi.nlm.nih.gov/gene/?term=LOC105288066 | 1964 | 16 | 81.47 | 1 | 5.09 | 6.25 | 41.07 | 43.1 | -2.03 | 2.03 | -0.02 | 0.02 |
| gene-LOC105282067 | https://www.ncbi.nlm.nih.gov/gene/?term=LOC105282067 | 5941 | 49 | 82.48 | 1 | 1.68 | 2.04 | 23.12 | 21.71 | 1.41 | 1.41 | 0.01 | 0.01 |
| gene-LOC105284507 | https://www.ncbi.nlm.nih.gov/gene/?term=LOC105284507 | 5207 | 68 | 130.59 | 1 | 1.92 | 1.47 | 22.93 | 21.85 | 1.08 | 1.08 | 0.01 | 0.01 |
| gene-LOC105276272 | https://www.ncbi.nlm.nih.gov/gene/?term=LOC105276272 | 1965 | 25 | 127.23 | 1 | 5.09 | 4 | 43.93 | 43.09 | 0.84 | 0.84 | 0.01 | 0.01 |
| gene-LOC105275187 | https://www.ncbi.nlm.nih.gov/gene/?term=LOC105275187 | 5534 | 66 | 119.26 | 1 | 1.81 | 1.52 | 14.05 | 13.27 | 0.78 | 0.78 | 0.01 | 0.01 |
| gene-LOC105276538 | https://www.ncbi.nlm.nih.gov/gene/?term=LOC105276538 | 1511 | 18 | 119.13 | 1 | 6.62 | 5.56 | 71.16 | 70.54 | 0.62 | 0.62 | 0.01 | 0.01 |
| gene-LOC105279664 | https://www.ncbi.nlm.nih.gov/gene/?term=LOC105279664 | 4023 | 56 | 139.2 | 1 | 2.49 | 1.79 | 21.21 | 20.62 | 0.59 | 0.59 | 0.01 | 0.01 |
| gene-LOC105276196 | https://www.ncbi.nlm.nih.gov/gene/?term=LOC105276196 | 2953 | 40 | 135.46 | 1 | 3.39 | 2.5 | 52.64 | 52.1 | 0.54 | 0.54 | 0.01 | 0.01 |
| gene-LOC105278454 | https://www.ncbi.nlm.nih.gov/gene/?term=LOC105278454 | 1972 | 36 | 182.56 | 1 | 5.07 | 2.78 | 47.84 | 47.3 | 0.54 | 0.54 | 0.01 | 0.01 |
| gene-LOC105275752 | https://www.ncbi.nlm.nih.gov/gene/?term=LOC105275752 | 3462 | 117 | 337.95 | 1 | 2.89 | 0.85 | 29.78 | 29.38 | 0.4 | 0.4 | 0.01 | 0.01 |
| gene-LOC105274706 | https://www.ncbi.nlm.nih.gov/gene/?term=LOC105274706 | 1446 | 51 | 352.7 | 1 | 6.92 | 1.96 | 37.41 | 37.03 | 0.38 | 0.38 | 0.01 | 0.01 |
| gene-LOC105287417 | https://www.ncbi.nlm.nih.gov/gene/?term=LOC105287417 | 1025 | 28 | 273.17 | 1 | 9.76 | 3.57 | 42.43 | 42.07 | 0.36 | 0.36 | 0.01 | 0.01 |
| gene-LOC105276449 | https://www.ncbi.nlm.nih.gov/gene/?term=LOC105276449 | 1231 | 39 | 316.82 | 1 | 8.12 | 2.56 | 32.73 | 32.41 | 0.32 | 0.32 | 0.01 | 0.01 |
| gene-LOC105278534 | https://www.ncbi.nlm.nih.gov/gene/?term=LOC105278534 | 7364 | 166 | 225.42 | 1 | 1.36 | 0.6 | 18.8 | 18.53 | 0.26 | 0.26 | 0.01 | 0.01 |
| gene-LOC105286649 | https://www.ncbi.nlm.nih.gov/gene/?term=LOC105286649 | 14178 | 662 | 466.92 | 1 | 0.71 | 0.15 | 1.27 | 1.01 | 0.26 | 0.26 | 0.01 | 0.01 |
| gene-LOC105283649 | https://www.ncbi.nlm.nih.gov/gene/?term=LOC105283649 | 1918 | 53 | 276.33 | 2 | 10.43 | 3.77 | 38.29 | 38.07 | 0.23 | 0.23 | 0.01 | 0.01 |
| gene-LOC105285807 | https://www.ncbi.nlm.nih.gov/gene/?term=LOC105285807 | 6063 | 174 | 286.99 | 1 | 1.65 | 0.57 | 7.5 | 7.29 | 0.22 | 0.22 | 0.01 | 0.01 |
| gene-LOC105286274 | https://www.ncbi.nlm.nih.gov/gene/?term=LOC105286274 | 3817 | 181 | 474.19 | 1 | 2.62 | 0.55 | 15.32 | 15.14 | 0.18 | 0.18 | 0.01 | 0.01 |
| gene-LOC105278145 | https://www.ncbi.nlm.nih.gov/gene/?term=LOC105278145 | 4470 | 244 | 545.86 | 1 | 2.24 | 0.41 | 10.74 | 10.94 | -0.2 | 0.2 | -0.01 | 0.01 |
| gene-LOC105285534 | https://www.ncbi.nlm.nih.gov/gene/?term=LOC105285534 | 3944 | 211 | 534.99 | 1 | 2.54 | 0.47 | 11.95 | 12.16 | -0.21 | 0.21 | -0.01 | 0.01 |
| gene-LOC105276527 | https://www.ncbi.nlm.nih.gov/gene/?term=LOC105276527 | 7923 | 323 | 407.67 | 1 | 1.26 | 0.31 | 8.74 | 8.99 | -0.25 | 0.25 | -0.01 | 0.01 |
| gene-LOC105274856 | https://www.ncbi.nlm.nih.gov/gene/?term=LOC105274856 | 5216 | 200 | 383.44 | 1 | 1.92 | 0.5 | 18.38 | 18.67 | -0.29 | 0.29 | -0.01 | 0.01 |
| gene-LOC105286223 | https://www.ncbi.nlm.nih.gov/gene/?term=LOC105286223 | 2225 | 79 | 355.06 | 1 | 4.49 | 1.27 | 30.58 | 30.96 | -0.38 | 0.38 | -0.01 | 0.01 |
| gene-LOC105285395 | https://www.ncbi.nlm.nih.gov/gene/?term=LOC105285395 | 8085 | 165 | 204.08 | 1 | 1.24 | 0.61 | 17.34 | 17.73 | -0.39 | 0.39 | -0.01 | 0.01 |
| gene-LOC105278538 | https://www.ncbi.nlm.nih.gov/gene/?term=LOC105278538 | 2150 | 76 | 353.49 | 1 | 4.65 | 1.32 | 18.23 | 18.64 | -0.41 | 0.41 | -0.01 | 0.01 |
| gene-LOC105275584 | https://www.ncbi.nlm.nih.gov/gene/?term=LOC105275584 | 3070 | 71 | 231.27 | 1 | 3.26 | 1.41 | 26.68 | 27.17 | -0.49 | 0.49 | -0.01 | 0.01 |
| gene-LOC105283943 | https://www.ncbi.nlm.nih.gov/gene/?term=LOC105283943 | 3559 | 48 | 134.87 | 1 | 2.81 | 2.08 | 17.72 | 18.23 | -0.51 | 0.51 | -0.01 | 0.01 |
| gene-LOC105281700 | https://www.ncbi.nlm.nih.gov/gene/?term=LOC105281700 | 2208 | 40 | 181.16 | 1 | 4.53 | 2.5 | 41.3 | 41.85 | -0.55 | 0.55 | -0.01 | 0.01 |
| gene-LOC105275179 | https://www.ncbi.nlm.nih.gov/gene/?term=LOC105275179 | 3271 | 66 | 201.77 | 1 | 3.06 | 1.52 | 14.71 | 15.31 | -0.61 | 0.61 | -0.01 | 0.01 |
| gene-LOC105280410 | https://www.ncbi.nlm.nih.gov/gene/?term=LOC105280410 | 12580 | 235 | 186.8 | 1 | 0.79 | 0.43 | 20.46 | 21.13 | -0.67 | 0.67 | -0.01 | 0.01 |
| gene-LOC105274851 | https://www.ncbi.nlm.nih.gov/gene/?term=LOC105274851 | 6616 | 122 | 184.4 | 1 | 1.51 | 0.82 | 10.53 | 11.32 | -0.79 | 0.79 | -0.01 | 0.01 |
| gene-LOC105283987 | https://www.ncbi.nlm.nih.gov/gene/?term=LOC105283987 | 4841 | 64 | 132.2 | 1 | 2.07 | 1.56 | 32.11 | 32.92 | -0.81 | 0.81 | -0.01 | 0.01 |
| gene-LOC105279657 | https://www.ncbi.nlm.nih.gov/gene/?term=LOC105279657 | 5878 | 79 | 134.4 | 1 | 1.7 | 1.27 | 19.77 | 20.7 | -0.93 | 0.93 | -0.01 | 0.01 |
| gene-LOC105274590 | https://www.ncbi.nlm.nih.gov/gene/?term=LOC105274590 | 4516 | 61 | 135.08 | 1 | 2.21 | 1.64 | 27.27 | 28.28 | -1.01 | 1.01 | -0.01 | 0.01 |
| gene-LOC105277216 | https://www.ncbi.nlm.nih.gov/gene/?term=LOC105277216 | 7627 | 76 | 99.65 | 1 | 1.31 | 1.32 | 7.73 | 8.77 | -1.04 | 1.04 | -0.01 | 0.01 |
| gene-LOC105281515 | https://www.ncbi.nlm.nih.gov/gene/?term=LOC105281515 | 8827 | 35 | 39.65 | 1 | 1.13 | 2.86 | 31.71 | 33.02 | -1.31 | 1.31 | -0.01 | 0.01 |
| gene-LOC105286452 | https://www.ncbi.nlm.nih.gov/gene/?term=LOC105286452 | 6888 | 55 | 79.85 | 1 | 1.45 | 1.82 | 32.52 | 33.96 | -1.44 | 1.44 | -0.01 | 0.01 |
| gene-LOC105279500 | https://www.ncbi.nlm.nih.gov/gene/?term=LOC105279500 | 3587 | 31 | 86.42 | 1 | 2.79 | 3.23 | 51.03 | 52.64 | -1.62 | 1.62 | -0.01 | 0.01 |
| gene-LOC105275518 | https://www.ncbi.nlm.nih.gov/gene/?term=LOC105275518 | 2335 | 18 | 77.09 | 1 | 4.28 | 5.56 | 47.37 | 49.08 | -1.72 | 1.72 | -0.01 | 0.01 |
| gene-LOC105284774 | https://www.ncbi.nlm.nih.gov/gene/?term=LOC105284774 | 12089 | 74 | 61.21 | 1 | 0.83 | 1.35 | 19.79 | 19.53 | 0.25 | 0.25 | 0 | 0 |
| gene-LOC105282794 | https://www.ncbi.nlm.nih.gov/gene/?term=LOC105282794 | 4571 | 91 | 199.08 | 1 | 2.19 | 1.1 | 19.49 | 19.29 | 0.2 | 0.2 | 0 | 0 |
| gene-LOC105288078 | https://www.ncbi.nlm.nih.gov/gene/?term=LOC105288078 | 19126 | 301 | 157.38 | 1 | 0.52 | 0.33 | 17.46 | 17.3 | 0.16 | 0.16 | 0 | 0 |
| gene-LOC105287095 | https://www.ncbi.nlm.nih.gov/gene/?term=LOC105287095 | 1435 | 52 | 362.37 | 1 | 6.97 | 1.92 | 30.47 | 30.36 | 0.12 | 0.12 | 0 | 0 |
| gene-LOC105284116 | https://www.ncbi.nlm.nih.gov/gene/?term=LOC105284116 | 3970 | 85 | 214.11 | 1 | 2.52 | 1.18 | 7.23 | 7.13 | 0.09 | 0.09 | 0 | 0 |
| gene-LOC105279038 | https://www.ncbi.nlm.nih.gov/gene/?term=LOC105279038 | 9453 | 298 | 315.24 | 1 | 1.06 | 0.34 | 9.03 | 8.95 | 0.08 | 0.08 | 0 | 0 |
| gene-LOC105281272 | https://www.ncbi.nlm.nih.gov/gene/?term=LOC105281272 | 5252 | 168 | 319.88 | 1 | 1.9 | 0.6 | 16.96 | 16.88 | 0.08 | 0.08 | 0 | 0 |
| gene-LOC105285742 | https://www.ncbi.nlm.nih.gov/gene/?term=LOC105285742 | 3446 | 91 | 264.07 | 1 | 2.9 | 1.1 | 26.72 | 26.67 | 0.05 | 0.05 | 0 | 0 |
| gene-LOC105276612 | https://www.ncbi.nlm.nih.gov/gene/?term=LOC105276612 | 4157 | 172 | 413.76 | 1 | 2.41 | 0.58 | 11.01 | 10.98 | 0.03 | 0.03 | 0 | 0 |
| gene-LOC105275327 | https://www.ncbi.nlm.nih.gov/gene/?term=LOC105275327 | 14245 | 413 | 289.93 | 1 | 0.7 | 0.24 | 7.72 | 7.75 | -0.03 | 0.03 | 0 | 0 |
| gene-LOC105280304 | https://www.ncbi.nlm.nih.gov/gene/?term=LOC105280304 | 8919 | 90 | 100.91 | 1 | 1.12 | 1.11 | 7.28 | 7.34 | -0.05 | 0.05 | 0 | 0 |
| gene-LOC105274534 | https://www.ncbi.nlm.nih.gov/gene/?term=LOC105274534 | 1329 | 53 | 398.8 | 1 | 7.52 | 1.89 | 47.05 | 47.11 | -0.06 | 0.06 | 0 | 0 |
| gene-LOC105278666 | https://www.ncbi.nlm.nih.gov/gene/?term=LOC105278666 | 10341 | 386 | 373.27 | 1 | 0.97 | 0.26 | 7.83 | 7.9 | -0.07 | 0.07 | 0 | 0 |
| gene-LOC105274531 | https://www.ncbi.nlm.nih.gov/gene/?term=LOC105274531 | 7774 | 252 | 324.16 | 1 | 1.29 | 0.4 | 11.91 | 11.99 | -0.08 | 0.08 | 0 | 0 |
| gene-LOC105276941 | https://www.ncbi.nlm.nih.gov/gene/?term=LOC105276941 | 15956 | 446 | 279.52 | 1 | 0.63 | 0.22 | 5.36 | 5.47 | -0.11 | 0.11 | 0 | 0 |
| gene-LOC105282825 | https://www.ncbi.nlm.nih.gov/gene/?term=LOC105282825 | 3919 | 80 | 204.13 | 1 | 2.55 | 1.25 | 13.05 | 13.16 | -0.12 | 0.12 | 0 | 0 |
| gene-LOC105286644 | https://www.ncbi.nlm.nih.gov/gene/?term=LOC105286644 | 1235 | 22 | 178.14 | 1 | 8.1 | 4.55 | 42.2 | 42.35 | -0.15 | 0.15 | 0 | 0 |
| gene-LOC105283989 | https://www.ncbi.nlm.nih.gov/gene/?term=LOC105283989 | 4948 | 25 | 50.53 | 1 | 2.02 | 4 | 45.66 | 45.85 | -0.19 | 0.19 | 0 | 0 |
| gene-LOC105284117 | https://www.ncbi.nlm.nih.gov/gene/?term=LOC105284117 | 4340 | 64 | 147.47 | 1 | 2.3 | 1.56 | 10.83 | 11.09 | -0.25 | 0.25 | 0 | 0 |
| gene-LOC105285406 | https://www.ncbi.nlm.nih.gov/gene/?term=LOC105285406 | 8015 | 39 | 48.66 | 1 | 1.25 | 2.56 | 33.07 | 33.55 | -0.48 | 0.48 | 0 | 0 |

Directory: OGL File: rnk-dmg-Ob_LI_BCphaseA1.vs.BCphaseB1.txt

| gene_ID | gene_link | gwidth | #Sites | #per10Kb | #dmSites | #dmsp10kb | %dmSites | %pSite1 | %pSite2 | DMpSite | ADMpSite | DMpNucl | ADMpNucl |
|---|---|---|---|---|---|---|---|---|---|---|---|---|---|
| gene-LOC113562345 | https://www.ncbi.nlm.nih.gov/gene/?term=LOC113562345 | 2290 | 62 | 270.74 | 14 | 61.14 | 22.58 | 17.91 | 32.35 | -14.44 | 14.44 | -0.39 | 0.39 |
| gene-LOC105283552 | https://www.ncbi.nlm.nih.gov/gene/?term=LOC105283552 | 1562 | 33 | 211.27 | 1 | 6.4 | 3.03 | 54.5 | 59.97 | -5.47 | 5.47 | -0.12 | 0.12 |
| gene-LOC105279898 | https://www.ncbi.nlm.nih.gov/gene/?term=LOC105279898 | 1343 | 16 | 119.14 | 1 | 7.45 | 6.25 | 59.42 | 65.83 | -6.42 | 6.42 | -0.08 | 0.08 |
| gene-LOC105277631 | https://www.ncbi.nlm.nih.gov/gene/?term=LOC105277631 | 4387 | 285 | 649.65 | 1 | 2.28 | 0.35 | 9.88 | 10.92 | -1.04 | 1.04 | -0.07 | 0.07 |
| gene-LOC105282156 | https://www.ncbi.nlm.nih.gov/gene/?term=LOC105282156 | 2577 | 133 | 516.1 | 1 | 3.88 | 0.75 | 23.61 | 24.92 | -1.31 | 1.31 | -0.07 | 0.07 |
| gene-LOC105285695 | https://www.ncbi.nlm.nih.gov/gene/?term=LOC105285695 | 1371 | 62 | 452.22 | 1 | 7.29 | 1.61 | 23 | 24.51 | -1.51 | 1.51 | -0.07 | 0.07 |
| gene-LOC105279356 | https://www.ncbi.nlm.nih.gov/gene/?term=LOC105279356 | 1314 | 42 | 319.63 | 1 | 7.61 | 2.38 | 28.83 | 30.95 | -2.12 | 2.12 | -0.07 | 0.07 |
| gene-LOC105281848 | https://www.ncbi.nlm.nih.gov/gene/?term=LOC105281848 | 2122 | 47 | 221.49 | 1 | 4.71 | 2.13 | 22.74 | 25.94 | -3.2 | 3.2 | -0.07 | 0.07 |
| gene-LOC105275070 | https://www.ncbi.nlm.nih.gov/gene/?term=LOC105275070 | 5667 | 36 | 63.53 | 2 | 3.53 | 5.56 | 3.69 | 13.97 | -10.28 | 10.28 | -0.07 | 0.07 |
| gene-LOC105282885 | https://www.ncbi.nlm.nih.gov/gene/?term=LOC105282885 | 1870 | 41 | 219.25 | 1 | 5.35 | 2.44 | 37.52 | 34.93 | 2.59 | 2.59 | 0.06 | 0.06 |
| gene-LOC105275460 | https://www.ncbi.nlm.nih.gov/gene/?term=LOC105275460 | 2473 | 116 | 469.07 | 1 | 4.04 | 0.86 | 12.57 | 13.81 | -1.24 | 1.24 | -0.06 | 0.06 |
| gene-LOC105285055 | https://www.ncbi.nlm.nih.gov/gene/?term=LOC105285055 | 4108 | 214 | 520.93 | 1 | 2.43 | 0.47 | 12.94 | 14.18 | -1.24 | 1.24 | -0.06 | 0.06 |
| gene-LOC105275363 | https://www.ncbi.nlm.nih.gov/gene/?term=LOC105275363 | 3798 | 155 | 408.11 | 1 | 2.63 | 0.65 | 17.26 | 18.39 | -1.13 | 1.13 | -0.05 | 0.05 |
| gene-LOC105276704 | https://www.ncbi.nlm.nih.gov/gene/?term=LOC105276704 | 2063 | 84 | 407.17 | 2 | 9.69 | 2.38 | 12.84 | 14.02 | -1.18 | 1.18 | -0.05 | 0.05 |
| gene-LOC105275384 | https://www.ncbi.nlm.nih.gov/gene/?term=LOC105275384 | 1987 | 75 | 377.45 | 1 | 5.03 | 1.33 | 33.41 | 34.62 | -1.21 | 1.21 | -0.05 | 0.05 |
| gene-LOC105288036 | https://www.ncbi.nlm.nih.gov/gene/?term=LOC105288036 | 5110 | 198 | 387.48 | 1 | 1.96 | 0.51 | 16.95 | 18.26 | -1.31 | 1.31 | -0.05 | 0.05 |
| gene-LOC105278807 | https://www.ncbi.nlm.nih.gov/gene/?term=LOC105278807 | 2166 | 71 | 327.79 | 1 | 4.62 | 1.41 | 18.52 | 20.08 | -1.56 | 1.56 | -0.05 | 0.05 |
| gene-LOC105285674 | https://www.ncbi.nlm.nih.gov/gene/?term=LOC105285674 | 3035 | 91 | 299.84 | 2 | 6.59 | 2.2 | 23.1 | 24.91 | -1.81 | 1.81 | -0.05 | 0.05 |
| gene-LOC105277215 | https://www.ncbi.nlm.nih.gov/gene/?term=LOC105277215 | 2619 | 28 | 106.91 | 1 | 3.82 | 3.57 | 20.04 | 24.53 | -4.48 | 4.48 | -0.05 | 0.05 |
| gene-LOC105275708 | https://www.ncbi.nlm.nih.gov/gene/?term=LOC105275708 | 1772 | 13 | 73.36 | 1 | 5.64 | 7.69 | 46.34 | 53.52 | -7.18 | 7.18 | -0.05 | 0.05 |
| gene-LOC105285514 | https://www.ncbi.nlm.nih.gov/gene/?term=LOC105285514 | 1277 | 9 | 70.48 | 1 | 7.83 | 11.11 | 73.21 | 80.81 | -7.6 | 7.6 | -0.05 | 0.05 |
| gene-LOC105281173 | https://www.ncbi.nlm.nih.gov/gene/?term=LOC105281173 | 1595 | 25 | 156.74 | 1 | 6.27 | 4 | 64.76 | 62.33 | 2.42 | 2.42 | 0.04 | 0.04 |
| gene-LOC105283123 | https://www.ncbi.nlm.nih.gov/gene/?term=LOC105283123 | 1507 | 41 | 272.06 | 1 | 6.64 | 2.44 | 15.68 | 14.34 | 1.34 | 1.34 | 0.04 | 0.04 |
| gene-LOC105275683 | https://www.ncbi.nlm.nih.gov/gene/?term=LOC105275683 | 4256 | 204 | 479.32 | 1 | 2.35 | 0.49 | 12.14 | 13.06 | -0.92 | 0.92 | -0.04 | 0.04 |
| gene-LOC105285671 | https://www.ncbi.nlm.nih.gov/gene/?term=LOC105285671 | 2263 | 95 | 419.8 | 1 | 4.42 | 1.05 | 27.86 | 28.85 | -1 | 1 | -0.04 | 0.04 |
| gene-LOC105287417 | https://www.ncbi.nlm.nih.gov/gene/?term=LOC105287417 | 1025 | 33 | 321.95 | 1 | 9.76 | 3.03 | 36 | 37.27 | -1.27 | 1.27 | -0.04 | 0.04 |
| gene-LOC105278454 | https://www.ncbi.nlm.nih.gov/gene/?term=LOC105278454 | 1972 | 37 | 187.63 | 1 | 5.07 | 2.7 | 46.8 | 45.07 | 1.73 | 1.73 | 0.03 | 0.03 |
| gene-LOC105285202 | https://www.ncbi.nlm.nih.gov/gene/?term=LOC105285202 | 1838 | 36 | 195.87 | 1 | 5.44 | 2.78 | 18.94 | 17.23 | 1.72 | 1.72 | 0.03 | 0.03 |
| gene-LOC105276196 | https://www.ncbi.nlm.nih.gov/gene/?term=LOC105276196 | 2953 | 54 | 182.86 | 1 | 3.39 | 1.85 | 40.59 | 39.19 | 1.4 | 1.4 | 0.03 | 0.03 |
| gene-LOC105286265 | https://www.ncbi.nlm.nih.gov/gene/?term=LOC105286265 | 2507 | 64 | 255.29 | 1 | 3.99 | 1.56 | 30.65 | 29.46 | 1.19 | 1.19 | 0.03 | 0.03 |
| gene-LOC105275653 | https://www.ncbi.nlm.nih.gov/gene/?term=LOC105275653 | 11251 | 262 | 232.87 | 1 | 0.89 | 0.38 | 9.15 | 8.02 | 1.13 | 1.13 | 0.03 | 0.03 |
| gene-LOC105274706 | https://www.ncbi.nlm.nih.gov/gene/?term=LOC105274706 | 1446 | 46 | 318.12 | 1 | 6.92 | 2.17 | 41.37 | 40.53 | 0.85 | 0.85 | 0.03 | 0.03 |
| gene-LOC105276429 | https://www.ncbi.nlm.nih.gov/gene/?term=LOC105276429 | 2901 | 112 | 386.07 | 1 | 3.45 | 0.89 | 26.41 | 25.73 | 0.68 | 0.68 | 0.03 | 0.03 |
| gene-LOC105276527 | https://www.ncbi.nlm.nih.gov/gene/?term=LOC105276527 | 7923 | 345 | 435.44 | 1 | 1.26 | 0.29 | 8.28 | 8.94 | -0.66 | 0.66 | -0.03 | 0.03 |
| gene-LOC105283728 | https://www.ncbi.nlm.nih.gov/gene/?term=LOC105283728 | 3256 | 111 | 340.91 | 1 | 3.07 | 0.9 | 20.22 | 21.04 | -0.81 | 0.81 | -0.03 | 0.03 |
| gene-LOC105283943 | https://www.ncbi.nlm.nih.gov/gene/?term=LOC105283943 | 3559 | 59 | 165.78 | 1 | 2.81 | 1.69 | 20.25 | 21.81 | -1.56 | 1.56 | -0.03 | 0.03 |
| gene-LOC105280348 | https://www.ncbi.nlm.nih.gov/gene/?term=LOC105280348 | 3797 | 51 | 134.32 | 1 | 2.63 | 1.96 | 25.95 | 28.29 | -2.34 | 2.34 | -0.03 | 0.03 |
| gene-LOC105274900 | https://www.ncbi.nlm.nih.gov/gene/?term=LOC105274900 | 5561 | 56 | 100.7 | 2 | 3.6 | 3.57 | 26.85 | 29.47 | -2.62 | 2.62 | -0.03 | 0.03 |
| gene-LOC105275518 | https://www.ncbi.nlm.nih.gov/gene/?term=LOC105275518 | 2335 | 20 | 85.65 | 1 | 4.28 | 5 | 47.29 | 50.84 | -3.55 | 3.55 | -0.03 | 0.03 |
| gene-LOC105279528 | https://www.ncbi.nlm.nih.gov/gene/?term=LOC105279528 | 5802 | 46 | 79.28 | 1 | 1.72 | 2.17 | 17.18 | 14.84 | 2.33 | 2.33 | 0.02 | 0.02 |
| gene-LOC105285409 | https://www.ncbi.nlm.nih.gov/gene/?term=LOC105285409 | 5099 | 48 | 94.14 | 1 | 1.96 | 2.08 | 3.6 | 1.53 | 2.08 | 2.08 | 0.02 | 0.02 |
| gene-LOC105288010 | https://www.ncbi.nlm.nih.gov/gene/?term=LOC105288010 | 3189 | 49 | 153.65 | 1 | 3.14 | 2.04 | 31.6 | 30.42 | 1.18 | 1.18 | 0.02 | 0.02 |
| gene-LOC105278282 | https://www.ncbi.nlm.nih.gov/gene/?term=LOC105278282 | 1775 | 47 | 264.79 | 1 | 5.63 | 2.13 | 43.73 | 42.89 | 0.84 | 0.84 | 0.02 | 0.02 |
| gene-LOC105278214 | https://www.ncbi.nlm.nih.gov/gene/?term=LOC105278214 | 3787 | 77 | 203.33 | 1 | 2.64 | 1.3 | 11.46 | 10.7 | 0.76 | 0.76 | 0.02 | 0.02 |
| gene-LOC105285583 | https://www.ncbi.nlm.nih.gov/gene/?term=LOC105285583 | 1947 | 49 | 251.67 | 1 | 5.14 | 2.04 | 29.41 | 28.65 | 0.76 | 0.76 | 0.02 | 0.02 |
| gene-LOC105278436 | https://www.ncbi.nlm.nih.gov/gene/?term=LOC105278436 | 2556 | 57 | 223 | 1 | 3.91 | 1.75 | 36.25 | 35.56 | 0.69 | 0.69 | 0.02 | 0.02 |
| gene-LOC105274449 | https://www.ncbi.nlm.nih.gov/gene/?term=LOC105274449 | 1549 | 43 | 277.6 | 1 | 6.46 | 2.33 | 54.59 | 53.98 | 0.6 | 0.6 | 0.02 | 0.02 |
| gene-LOC105282811 | https://www.ncbi.nlm.nih.gov/gene/?term=LOC105282811 | 5765 | 203 | 352.12 | 1 | 1.73 | 0.49 | 9.79 | 9.26 | 0.53 | 0.53 | 0.02 | 0.02 |
| gene-LOC105277435 | https://www.ncbi.nlm.nih.gov/gene/?term=LOC105277435 | 3643 | 155 | 425.47 | 1 | 2.74 | 0.65 | 12.32 | 11.8 | 0.52 | 0.52 | 0.02 | 0.02 |
| gene-LOC105280786 | https://www.ncbi.nlm.nih.gov/gene/?term=LOC105280786 | 2573 | 120 | 466.38 | 1 | 3.89 | 0.83 | 29.63 | 29.12 | 0.52 | 0.52 | 0.02 | 0.02 |
| gene-LOC105278877 | https://www.ncbi.nlm.nih.gov/gene/?term=LOC105278877 | 1968 | 77 | 391.26 | 1 | 5.08 | 1.3 | 32.7 | 32.23 | 0.46 | 0.46 | 0.02 | 0.02 |
| gene-LOC105285534 | https://www.ncbi.nlm.nih.gov/gene/?term=LOC105285534 | 3944 | 221 | 560.34 | 1 | 2.54 | 0.45 | 11.41 | 11.78 | -0.37 | 0.37 | -0.02 | 0.02 |
| gene-LOC105282947 | https://www.ncbi.nlm.nih.gov/gene/?term=LOC105282947 | 1162 | 45 | 387.26 | 1 | 8.61 | 2.22 | 32.13 | 32.56 | -0.42 | 0.42 | -0.02 | 0.02 |
| gene-LOC105284355 | https://www.ncbi.nlm.nih.gov/gene/?term=LOC105284355 | 3904 | 154 | 394.47 | 1 | 2.56 | 0.65 | 20.37 | 20.87 | -0.5 | 0.5 | -0.02 | 0.02 |
| gene-LOC105285639 | https://www.ncbi.nlm.nih.gov/gene/?term=LOC105285639 | 10325 | 314 | 304.12 | 1 | 0.97 | 0.32 | 12 | 12.71 | -0.71 | 0.71 | -0.02 | 0.02 |
| gene-LOC105285742 | https://www.ncbi.nlm.nih.gov/gene/?term=LOC105285742 | 3446 | 95 | 275.68 | 1 | 2.9 | 1.05 | 25.7 | 26.49 | -0.79 | 0.79 | -0.02 | 0.02 |
| gene-LOC105283592 | https://www.ncbi.nlm.nih.gov/gene/?term=LOC105283592 | 3479 | 71 | 204.08 | 1 | 2.87 | 1.41 | 24.39 | 25.37 | -0.98 | 0.98 | -0.02 | 0.02 |
| gene-LOC105281700 | https://www.ncbi.nlm.nih.gov/gene/?term=LOC105281700 | 2208 | 44 | 199.28 | 1 | 4.53 | 2.27 | 37.54 | 38.62 | -1.08 | 1.08 | -0.02 | 0.02 |
| gene-LOC105283712 | https://www.ncbi.nlm.nih.gov/gene/?term=LOC105283712 | 1910 | 29 | 151.83 | 1 | 5.24 | 3.45 | 29.98 | 31.34 | -1.36 | 1.36 | -0.02 | 0.02 |
| gene-LOC105278260 | https://www.ncbi.nlm.nih.gov/gene/?term=LOC105278260 | 3240 | 42 | 129.63 | 1 | 3.09 | 2.38 | 12.8 | 14.28 | -1.47 | 1.47 | -0.02 | 0.02 |

| | | | | | | | | | | | | | |
|---|---|---|---|---|---|---|---|---|---|---|---|---|---|
| gene-LOC105283989 | https://www.ncbi.nlm.nih.gov/gene/?term=LOC105283989 | 4948 | 33 | 66.69 | 2 | 4.04 | 6.06 | 45.77 | 48.89 | -3.12 | 3.12 | -0.02 | 0.02 |
| gene-LOC105280419 | https://www.ncbi.nlm.nih.gov/gene/?term=LOC105280419 | 1601 | 10 | 62.46 | 1 | 6.25 | 10 | 33.98 | 37.47 | -3.49 | 3.49 | -0.02 | 0.02 |
| gene-LOC105280447 | https://www.ncbi.nlm.nih.gov/gene/?term=LOC105280447 | 2215 | 25 | 112.87 | 1 | 4.51 | 4 | 26.74 | 25.91 | 0.82 | 0.82 | 0.01 | 0.01 |
| gene-LOC105281047 | https://www.ncbi.nlm.nih.gov/gene/?term=LOC105281047 | 7313 | 117 | 159.99 | 1 | 1.37 | 0.85 | 10.66 | 9.84 | 0.82 | 0.82 | 0.01 | 0.01 |
| gene-LOC105281420 | https://www.ncbi.nlm.nih.gov/gene/?term=LOC105281420 | 3882 | 57 | 146.83 | 1 | 2.58 | 1.75 | 15.28 | 14.73 | 0.55 | 0.55 | 0.01 | 0.01 |
| gene-LOC105284116 | https://www.ncbi.nlm.nih.gov/gene/?term=LOC105284116 | 3970 | 95 | 239.29 | 1 | 2.52 | 1.05 | 6.78 | 6.28 | 0.5 | 0.5 | 0.01 | 0.01 |
| gene-LOC105284117 | https://www.ncbi.nlm.nih.gov/gene/?term=LOC105284117 | 4340 | 81 | 186.64 | 1 | 2.3 | 1.23 | 11.74 | 11.3 | 0.44 | 0.44 | 0.01 | 0.01 |
| gene-LOC105282600 | https://www.ncbi.nlm.nih.gov/gene/?term=LOC105282600 | 3715 | 57 | 153.43 | 1 | 2.69 | 1.75 | 29.16 | 28.81 | 0.35 | 0.35 | 0.01 | 0.01 |
| gene-LOC105285807 | https://www.ncbi.nlm.nih.gov/gene/?term=LOC105285807 | 6063 | 183 | 301.83 | 1 | 1.65 | 0.55 | 10.09 | 9.8 | 0.29 | 0.29 | 0.01 | 0.01 |
| gene-LOC105286515 | https://www.ncbi.nlm.nih.gov/gene/?term=LOC105286515 | 3751 | 103 | 274.59 | 1 | 2.67 | 0.97 | 10.73 | 10.44 | 0.29 | 0.29 | 0.01 | 0.01 |
| gene-LOC105279038 | https://www.ncbi.nlm.nih.gov/gene/?term=LOC105279038 | 9453 | 310 | 327.94 | 1 | 1.06 | 0.32 | 8.8 | 8.54 | 0.27 | 0.27 | 0.01 | 0.01 |
| gene-LOC105278666 | https://www.ncbi.nlm.nih.gov/gene/?term=LOC105278666 | 10341 | 405 | 391.64 | 1 | 0.97 | 0.25 | 7.09 | 6.88 | 0.22 | 0.22 | 0.01 | 0.01 |
| gene-LOC105284036 | https://www.ncbi.nlm.nih.gov/gene/?term=LOC105284036 | 3254 | 83 | 255.07 | 1 | 3.07 | 1.2 | 22.41 | 22.19 | 0.22 | 0.22 | 0.01 | 0.01 |
| gene-LOC105276901 | https://www.ncbi.nlm.nih.gov/gene/?term=LOC105276901 | 3254 | 95 | 291.95 | 1 | 3.07 | 1.05 | 23.05 | 22.87 | 0.18 | 0.18 | 0.01 | 0.01 |
| gene-LOC105278145 | https://www.ncbi.nlm.nih.gov/gene/?term=LOC105278145 | 4470 | 249 | 557.05 | 1 | 2.24 | 0.4 | 10.18 | 10.01 | 0.17 | 0.17 | 0.01 | 0.01 |
| gene-LOC105275752 | https://www.ncbi.nlm.nih.gov/gene/?term=LOC105275752 | 3462 | 124 | 358.17 | 1 | 2.89 | 0.81 | 28.83 | 28.67 | 0.16 | 0.16 | 0.01 | 0.01 |
| gene-LOC105287095 | https://www.ncbi.nlm.nih.gov/gene/?term=LOC105287095 | 1435 | 61 | 425.09 | 1 | 6.97 | 1.64 | 26.05 | 25.92 | 0.14 | 0.14 | 0.01 | 0.01 |
| gene-LOC105284262 | https://www.ncbi.nlm.nih.gov/gene/?term=LOC105284262 | 6669 | 230 | 344.88 | 1 | 1.5 | 0.43 | 4.51 | 4.68 | -0.17 | 0.17 | -0.01 | 0.01 |
| gene-LOC105275584 | https://www.ncbi.nlm.nih.gov/gene/?term=LOC105275584 | 3070 | 86 | 280.13 | 1 | 3.26 | 1.16 | 23.79 | 24 | -0.22 | 0.22 | -0.01 | 0.01 |
| gene-LOC105283649 | https://www.ncbi.nlm.nih.gov/gene/?term=LOC105283649 | 1918 | 53 | 276.33 | 1 | 5.21 | 1.89 | 38.29 | 38.55 | -0.25 | 0.25 | -0.01 | 0.01 |
| gene-LOC105280405 | https://www.ncbi.nlm.nih.gov/gene/?term=LOC105280405 | 5422 | 162 | 298.78 | 1 | 1.84 | 0.62 | 9.36 | 9.65 | -0.29 | 0.29 | -0.01 | 0.01 |
| gene-LOC105285395 | https://www.ncbi.nlm.nih.gov/gene/?term=LOC105285395 | 8085 | 180 | 222.63 | 1 | 1.24 | 0.56 | 15.55 | 15.84 | -0.29 | 0.29 | -0.01 | 0.01 |
| gene-LOC105279298 | https://www.ncbi.nlm.nih.gov/gene/?term=LOC105279298 | 2773 | 134 | 483.23 | 1 | 3.61 | 0.75 | 22.72 | 23.03 | -0.3 | 0.3 | -0.01 | 0.01 |
| gene-LOC105288078 | https://www.ncbi.nlm.nih.gov/gene/?term=LOC105288078 | 19126 | 313 | 163.65 | 1 | 0.52 | 0.32 | 17.33 | 17.64 | -0.31 | 0.31 | -0.01 | 0.01 |
| gene-LOC105284184 | https://www.ncbi.nlm.nih.gov/gene/?term=LOC105284184 | 2535 | 82 | 323.47 | 1 | 3.94 | 1.22 | 34.56 | 34.9 | -0.34 | 0.34 | -0.01 | 0.01 |
| gene-LOC105278632 | https://www.ncbi.nlm.nih.gov/gene/?term=LOC105278632 | 3624 | 131 | 361.48 | 1 | 2.76 | 0.76 | 14.71 | 15.12 | -0.41 | 0.41 | -0.01 | 0.01 |
| gene-LOC105276692 | https://www.ncbi.nlm.nih.gov/gene/?term=LOC105276692 | 1745 | 36 | 206.3 | 1 | 5.73 | 2.78 | 23.57 | 24.03 | -0.46 | 0.46 | -0.01 | 0.01 |
| gene-LOC105278534 | https://www.ncbi.nlm.nih.gov/gene/?term=LOC105278534 | 7364 | 184 | 249.86 | 1 | 1.36 | 0.54 | 17.97 | 18.5 | -0.53 | 0.53 | -0.01 | 0.01 |
| gene-LOC105286770 | https://www.ncbi.nlm.nih.gov/gene/?term=LOC105286770 | 3055 | 42 | 137.48 | 1 | 3.27 | 2.38 | 8.53 | 9.07 | -0.55 | 0.55 | -0.01 | 0.01 |
| gene-LOC105276745 | https://www.ncbi.nlm.nih.gov/gene/?term=LOC105276745 | 3953 | 73 | 184.67 | 1 | 2.53 | 1.37 | 18.74 | 19.3 | -0.56 | 0.56 | -0.01 | 0.01 |
| gene-LOC105280002 | https://www.ncbi.nlm.nih.gov/gene/?term=LOC105280002 | 1940 | 28 | 144.33 | 1 | 5.15 | 3.57 | 33.09 | 33.65 | -0.57 | 0.57 | -0.01 | 0.01 |
| gene-LOC105278962 | https://www.ncbi.nlm.nih.gov/gene/?term=LOC105278962 | 4491 | 88 | 195.95 | 1 | 2.23 | 1.14 | 23.63 | 24.21 | -0.58 | 0.58 | -0.01 | 0.01 |
| gene-LOC105277903 | https://www.ncbi.nlm.nih.gov/gene/?term=LOC105277903 | 1835 | 35 | 190.74 | 1 | 5.45 | 2.86 | 51.45 | 52.14 | -0.69 | 0.69 | -0.01 | 0.01 |
| gene-LOC105280641 | https://www.ncbi.nlm.nih.gov/gene/?term=LOC105280641 | 3682 | 62 | 168.39 | 1 | 2.72 | 1.61 | 16.36 | 17.05 | -0.69 | 0.69 | -0.01 | 0.01 |
| gene-LOC105275190 | https://www.ncbi.nlm.nih.gov/gene/?term=LOC105275190 | 3837 | 72 | 187.65 | 1 | 2.61 | 1.39 | 18.78 | 19.54 | -0.76 | 0.76 | -0.01 | 0.01 |
| gene-LOC105288066 | https://www.ncbi.nlm.nih.gov/gene/?term=LOC105288066 | 1964 | 25 | 127.29 | 1 | 5.09 | 4 | 27.34 | 28.29 | -0.95 | 0.95 | -0.01 | 0.01 |
| gene-LOC105277216 | https://www.ncbi.nlm.nih.gov/gene/?term=LOC105277216 | 7627 | 105 | 137.67 | 1 | 1.31 | 0.95 | 5.72 | 6.75 | -1.03 | 1.03 | -0.01 | 0.01 |
| gene-LOC105285060 | https://www.ncbi.nlm.nih.gov/gene/?term=LOC105285060 | 1285 | 16 | 124.51 | 1 | 7.78 | 6.25 | 50.64 | 51.74 | -1.1 | 1.1 | -0.01 | 0.01 |
| gene-LOC105281515 | https://www.ncbi.nlm.nih.gov/gene/?term=LOC105281515 | 8827 | 39 | 44.18 | 1 | 1.13 | 2.56 | 34.44 | 35.97 | -1.53 | 1.53 | -0.01 | 0.01 |
| gene-LOC105286452 | https://www.ncbi.nlm.nih.gov/gene/?term=LOC105286452 | 6888 | 58 | 84.2 | 1 | 1.45 | 1.72 | 31.25 | 32.98 | -1.73 | 1.73 | -0.01 | 0.01 |
| gene-LOC105282399 | https://www.ncbi.nlm.nih.gov/gene/?term=LOC105282399 | 2682 | 21 | 78.3 | 1 | 3.73 | 4.76 | 34.08 | 35.97 | -1.89 | 1.89 | -0.01 | 0.01 |
| gene-LOC105285406 | https://www.ncbi.nlm.nih.gov/gene/?term=LOC105285406 | 8015 | 40 | 49.91 | 1 | 1.25 | 2.5 | 34.18 | 33.53 | 0.64 | 0.64 | 0 | 0 |
| gene-LOC105279657 | https://www.ncbi.nlm.nih.gov/gene/?term=LOC105279657 | 5878 | 86 | 146.31 | 1 | 1.7 | 1.16 | 18.3 | 18.14 | 0.16 | 0.16 | 0 | 0 |
| gene-LOC105275173 | https://www.ncbi.nlm.nih.gov/gene/?term=LOC105275173 | 4639 | 83 | 178.92 | 1 | 2.16 | 1.2 | 22.33 | 22.24 | 0.09 | 0.09 | 0 | 0 |
| gene-LOC105276059 | https://www.ncbi.nlm.nih.gov/gene/?term=LOC105276059 | 6256 | 134 | 214.19 | 1 | 1.6 | 0.75 | 10.93 | 10.85 | 0.08 | 0.08 | 0 | 0 |
| gene-LOC105279664 | https://www.ncbi.nlm.nih.gov/gene/?term=LOC105279664 | 4023 | 68 | 169.03 | 1 | 2.49 | 1.47 | 18.22 | 18.15 | 0.08 | 0.08 | 0 | 0 |
| gene-LOC105278228 | https://www.ncbi.nlm.nih.gov/gene/?term=LOC105278228 | 8610 | 165 | 191.64 | 1 | 1.16 | 0.61 | 22.79 | 22.73 | 0.06 | 0.06 | 0 | 0 |
| gene-LOC105282491 | https://www.ncbi.nlm.nih.gov/gene/?term=LOC105282491 | 4761 | 159 | 333.96 | 1 | 2.1 | 0.63 | 9.46 | 9.41 | 0.05 | 0.05 | 0 | 0 |
| gene-LOC105286274 | https://www.ncbi.nlm.nih.gov/gene/?term=LOC105286274 | 3817 | 177 | 463.71 | 1 | 2.62 | 0.56 | 15.66 | 15.62 | 0.04 | 0.04 | 0 | 0 |
| gene-LOC105275451 | https://www.ncbi.nlm.nih.gov/gene/?term=LOC105275451 | 6035 | 113 | 187.24 | 1 | 1.66 | 0.88 | 22.29 | 22.26 | 0.02 | 0.02 | 0 | 0 |
| gene-LOC105275327 | https://www.ncbi.nlm.nih.gov/gene/?term=LOC105275327 | 14245 | 477 | 334.85 | 1 | 0.7 | 0.21 | 7.15 | 7.14 | 0.01 | 0.01 | 0 | 0 |
| gene-LOC105282151 | https://www.ncbi.nlm.nih.gov/gene/?term=LOC105282151 | 1601 | 41 | 256.09 | 1 | 6.25 | 2.44 | 39.95 | 39.99 | -0.04 | 0.04 | 0 | 0 |
| gene-LOC105279778 | https://www.ncbi.nlm.nih.gov/gene/?term=LOC105279778 | 4347 | 139 | 319.76 | 1 | 2.3 | 0.72 | 9.36 | 9.42 | -0.06 | 0.06 | 0 | 0 |
| gene-LOC105274590 | https://www.ncbi.nlm.nih.gov/gene/?term=LOC105274590 | 4516 | 76 | 168.29 | 1 | 2.21 | 1.32 | 22.95 | 23.02 | -0.07 | 0.07 | 0 | 0 |
| gene-LOC105275170 | https://www.ncbi.nlm.nih.gov/gene/?term=LOC105275170 | 12918 | 337 | 260.88 | 1 | 0.77 | 0.3 | 2.06 | 2.13 | -0.07 | 0.07 | 0 | 0 |
| gene-LOC105280956 | https://www.ncbi.nlm.nih.gov/gene/?term=LOC105280956 | 2639 | 73 | 276.62 | 1 | 3.79 | 1.37 | 20.28 | 20.35 | -0.07 | 0.07 | 0 | 0 |
| gene-LOC105274531 | https://www.ncbi.nlm.nih.gov/gene/?term=LOC105274531 | 7774 | 254 | 326.73 | 1 | 1.29 | 0.39 | 11.9 | 12 | -0.1 | 0.1 | 0 | 0 |
| gene-LOC105274856 | https://www.ncbi.nlm.nih.gov/gene/?term=LOC105274856 | 5216 | 201 | 385.35 | 1 | 1.92 | 0.5 | 17.3 | 17.4 | -0.1 | 0.1 | 0 | 0 |
| gene-LOC105276538 | https://www.ncbi.nlm.nih.gov/gene/?term=LOC105276538 | 1511 | 18 | 119.13 | 1 | 6.62 | 5.56 | 71.72 | 71.83 | -0.11 | 0.11 | 0 | 0 |
| gene-LOC105279500 | https://www.ncbi.nlm.nih.gov/gene/?term=LOC105279500 | 3587 | 32 | 89.21 | 1 | 2.79 | 3.12 | 49.74 | 49.96 | -0.21 | 0.21 | 0 | 0 |
| gene-LOC105282914 | https://www.ncbi.nlm.nih.gov/gene/?term=LOC105282914 | 5623 | 42 | 74.69 | 1 | 1.78 | 2.38 | 48.6 | 48.91 | -0.31 | 0.31 | 0 | 0 |

Directory: OGL File: rnk-dmg-Ob_LI_BCphaseA1.vs.BCphaseB2.txt

| gene_ID | gene_link | gwidth | #Sites | #per10Kb | #dmSites | #dmsp10kb | %dmSites | %pSite1 | %pSite2 | DMpSite | ADMpSite | DMpNucl | ADMpNucl |
|---|---|---|---|---|---|---|---|---|---|---|---|---|---|
| gene-LOC105283830 | https://www.ncbi.nlm.nih.gov/gene/?term=LOC105283830 | 252 | 13 | 515.87 | 1 | 39.68 | 7.69 | 42.02 | 33.83 | 8.19 | 8.19 | 0.42 | 0.42 |
| gene-LOC105278849 | https://www.ncbi.nlm.nih.gov/gene/?term=LOC105278849 | 776 | 75 | 966.49 | 2 | 25.77 | 2.67 | 24.26 | 22.31 | 1.95 | 1.95 | 0.19 | 0.19 |
| gene-LOC105282405 | https://www.ncbi.nlm.nih.gov/gene/?term=LOC105282405 | 1310 | 31 | 236.64 | 3 | 22.9 | 9.68 | 51.18 | 44.58 | 6.6 | 6.6 | 0.16 | 0.16 |
| gene-LOC105278458 | https://www.ncbi.nlm.nih.gov/gene/?term=LOC105278458 | 216 | 3 | 138.89 | 1 | 46.3 | 33.33 | 20.87 | 10.63 | 10.24 | 10.24 | 0.14 | 0.14 |
| gene-LOC105287856 | https://www.ncbi.nlm.nih.gov/gene/?term=LOC105287856 | 2090 | 116 | 555.02 | 3 | 14.35 | 2.59 | 12.68 | 14.95 | -2.27 | 2.27 | -0.13 | 0.13 |
| gene-LOC105284168 | https://www.ncbi.nlm.nih.gov/gene/?term=LOC105284168 | 1400 | 72 | 514.29 | 3 | 21.43 | 4.17 | 21.98 | 24.55 | -2.57 | 2.57 | -0.13 | 0.13 |
| gene-LOC105287662 | https://www.ncbi.nlm.nih.gov/gene/?term=LOC105287662 | 1704 | 39 | 228.87 | 3 | 17.61 | 7.69 | 30.88 | 36.7 | -5.82 | 5.82 | -0.13 | 0.13 |
| gene-LOC113562627 | https://www.ncbi.nlm.nih.gov/gene/?term=LOC113562627 | 397 | 10 | 251.89 | 2 | 50.38 | 20 | 77.44 | 72.85 | 4.6 | 4.6 | 0.12 | 0.12 |
| gene-LOC105276679 | https://www.ncbi.nlm.nih.gov/gene/?term=LOC105276679 | 2005 | 65 | 324.19 | 4 | 19.95 | 6.15 | 39.83 | 36.18 | 3.65 | 3.65 | 0.12 | 0.12 |
| gene-LOC105283016 | https://www.ncbi.nlm.nih.gov/gene/?term=LOC105283016 | 2324 | 111 | 477.62 | 3 | 12.91 | 2.7 | 23.61 | 21.2 | 2.41 | 2.41 | 0.12 | 0.12 |
| gene-LOC105282986 | https://www.ncbi.nlm.nih.gov/gene/?term=LOC105282986 | 1600 | 81 | 506.25 | 2 | 12.5 | 2.47 | 30.91 | 28.82 | 2.08 | 2.08 | 0.11 | 0.11 |
| gene-LOC105274711 | https://www.ncbi.nlm.nih.gov/gene/?term=LOC105274711 | 1655 | 87 | 525.68 | 3 | 18.13 | 3.45 | 45.85 | 43.79 | 2.06 | 2.06 | 0.11 | 0.11 |
| gene-LOC105283717 | https://www.ncbi.nlm.nih.gov/gene/?term=LOC105283717 | 1373 | 83 | 604.52 | 2 | 14.57 | 2.41 | 24.03 | 25.8 | -1.76 | 1.76 | -0.11 | 0.11 |
| gene-LOC105286845 | https://www.ncbi.nlm.nih.gov/gene/?term=LOC105286845 | 1159 | 59 | 509.06 | 3 | 25.88 | 5.08 | 17.66 | 19.8 | -2.14 | 2.14 | -0.11 | 0.11 |
| gene-LOC105287895 | https://www.ncbi.nlm.nih.gov/gene/?term=LOC105287895 | 2424 | 50 | 206.27 | 2 | 8.25 | 4 | 28.82 | 34.37 | -5.56 | 5.56 | -0.11 | 0.11 |
| gene-LOC105277648 | https://www.ncbi.nlm.nih.gov/gene/?term=LOC105277648 | 722 | 26 | 360.11 | 1 | 13.85 | 3.85 | 56.47 | 53.69 | 2.78 | 2.78 | 0.1 | 0.1 |
| gene-LOC105280786 | https://www.ncbi.nlm.nih.gov/gene/?term=LOC105280786 | 2573 | 109 | 423.63 | 3 | 11.66 | 2.75 | 32.44 | 30.15 | 2.29 | 2.29 | 0.1 | 0.1 |
| gene-LOC105275484 | https://www.ncbi.nlm.nih.gov/gene/?term=LOC105275484 | 931 | 33 | 354.46 | 1 | 10.74 | 3.03 | 34.17 | 37.06 | -2.89 | 2.89 | -0.1 | 0.1 |
| gene-LOC105287565 | https://www.ncbi.nlm.nih.gov/gene/?term=LOC105287565 | 1735 | 49 | 282.42 | 5 | 28.82 | 10.2 | 38.14 | 41.53 | -3.39 | 3.39 | -0.1 | 0.1 |
| gene-LOC105283649 | https://www.ncbi.nlm.nih.gov/gene/?term=LOC105283649 | 1918 | 53 | 276.33 | 4 | 20.86 | 7.55 | 38.29 | 35.16 | 3.13 | 3.13 | 0.09 | 0.09 |
| gene-LOC105275761 | https://www.ncbi.nlm.nih.gov/gene/?term=LOC105275761 | 2635 | 98 | 371.92 | 4 | 15.18 | 4.08 | 30.05 | 27.74 | 2.31 | 2.31 | 0.09 | 0.09 |
| gene-LOC105279335 | https://www.ncbi.nlm.nih.gov/gene/?term=LOC105279335 | 737 | 33 | 447.76 | 1 | 13.57 | 3.03 | 19.48 | 17.37 | 2.11 | 2.11 | 0.09 | 0.09 |
| gene-LOC105279035 | https://www.ncbi.nlm.nih.gov/gene/?term=LOC105279035 | 2108 | 145 | 687.86 | 5 | 23.72 | 3.45 | 18.41 | 17.12 | 1.29 | 1.29 | 0.09 | 0.09 |
| gene-LOC105278737 | https://www.ncbi.nlm.nih.gov/gene/?term=LOC105278737 | 1346 | 89 | 661.22 | 1 | 7.43 | 1.12 | 18.69 | 20.11 | -1.41 | 1.41 | -0.09 | 0.09 |

...
... only first 25 lines shown ...

Directory: OGL File: rnk-dmg-Ob_LI_BCphaseA1.vs.RphaseA1.txt

| gene_ID | gene_link | gwidth | #Sites | #per10Kb | #dmSites | #dmsp10kb | %dmSites | %pSite1 | %pSite2 | DMpSite | ADMpSite | DMpNucl | ADMpNucl |
|---|---|---|---|---|---|---|---|---|---|---|---|---|---|
| gene-LOC113562345 | https://www.ncbi.nlm.nih.gov/gene/?term=LOC113562345 | 2290 | 61 | 266.38 | 10 | 43.67 | 16.39 | 17.52 | 31.67 | -14.16 | 14.16 | -0.38 | 0.38 |
| gene-LOC105285671 | https://www.ncbi.nlm.nih.gov/gene/?term=LOC105285671 | 2263 | 90 | 397.7 | 1 | 4.42 | 1.11 | 29.29 | 31.61 | -2.32 | 2.32 | -0.09 | 0.09 |
| gene-LOC105275460 | https://www.ncbi.nlm.nih.gov/gene/?term=LOC105275460 | 2473 | 115 | 465.02 | 1 | 4.04 | 0.87 | 12.68 | 14.4 | -1.72 | 1.72 | -0.08 | 0.08 |
| gene-LOC105285674 | https://www.ncbi.nlm.nih.gov/gene/?term=LOC105285674 | 3035 | 88 | 289.95 | 2 | 6.59 | 2.27 | 23.51 | 26.27 | -2.76 | 2.76 | -0.08 | 0.08 |
| gene-LOC105275070 | https://www.ncbi.nlm.nih.gov/gene/?term=LOC105275070 | 5667 | 37 | 65.29 | 2 | 3.53 | 5.41 | 3.73 | 16.16 | -12.43 | 12.43 | -0.08 | 0.08 |
| gene-LOC105274449 | https://www.ncbi.nlm.nih.gov/gene/?term=LOC105274449 | 1549 | 43 | 277.6 | 1 | 6.46 | 2.33 | 54.59 | 52.01 | 2.58 | 2.58 | 0.07 | 0.07 |
| gene-LOC105284396 | https://www.ncbi.nlm.nih.gov/gene/?term=LOC105284396 | 1976 | 57 | 288.46 | 1 | 5.06 | 1.75 | 39.14 | 36.68 | 2.46 | 2.46 | 0.07 | 0.07 |
| gene-LOC105276704 | https://www.ncbi.nlm.nih.gov/gene/?term=LOC105276704 | 2063 | 80 | 387.78 | 1 | 4.85 | 1.25 | 13.36 | 15.07 | -1.71 | 1.71 | -0.07 | 0.07 |
| gene-LOC105278807 | https://www.ncbi.nlm.nih.gov/gene/?term=LOC105278807 | 2166 | 70 | 323.18 | 1 | 4.62 | 1.43 | 17.84 | 20.15 | -2.31 | 2.31 | -0.07 | 0.07 |
| gene-LOC105279356 | https://www.ncbi.nlm.nih.gov/gene/?term=LOC105279356 | 1314 | 42 | 319.63 | 1 | 7.61 | 2.38 | 28.83 | 31.17 | -2.34 | 2.34 | -0.07 | 0.07 |
| gene-LOC105281848 | https://www.ncbi.nlm.nih.gov/gene/?term=LOC105281848 | 2122 | 45 | 212.06 | 1 | 4.71 | 2.22 | 23.75 | 26.94 | -3.19 | 3.19 | -0.07 | 0.07 |
| gene-LOC105283552 | https://www.ncbi.nlm.nih.gov/gene/?term=LOC105283552 | 1562 | 33 | 211.27 | 1 | 6.4 | 3.03 | 54.5 | 58 | -3.5 | 3.5 | -0.07 | 0.07 |
| gene-LOC105277903 | https://www.ncbi.nlm.nih.gov/gene/?term=LOC105277903 | 1835 | 35 | 190.74 | 1 | 5.45 | 2.86 | 51.45 | 55.03 | -3.59 | 3.59 | -0.07 | 0.07 |
| gene-LOC105281173 | https://www.ncbi.nlm.nih.gov/gene/?term=LOC105281173 | 1595 | 25 | 156.74 | 1 | 6.27 | 4 | 64.76 | 61.03 | 3.73 | 3.73 | 0.06 | 0.06 |
| gene-LOC105274706 | https://www.ncbi.nlm.nih.gov/gene/?term=LOC105274706 | 1446 | 48 | 331.95 | 1 | 6.92 | 2.08 | 39.02 | 37.1 | 1.92 | 1.92 | 0.06 | 0.06 |
| gene-LOC105278877 | https://www.ncbi.nlm.nih.gov/gene/?term=LOC105278877 | 1968 | 78 | 396.34 | 1 | 5.08 | 1.28 | 32.36 | 30.87 | 1.48 | 1.48 | 0.06 | 0.06 |
| gene-LOC105285055 | https://www.ncbi.nlm.nih.gov/gene/?term=LOC105285055 | 4108 | 204 | 496.59 | 1 | 2.43 | 0.49 | 12.43 | 13.67 | -1.24 | 1.24 | -0.06 | 0.06 |
| gene-LOC105285695 | https://www.ncbi.nlm.nih.gov/gene/?term=LOC105285695 | 1371 | 61 | 444.93 | 1 | 7.29 | 1.64 | 22.52 | 23.97 | -1.45 | 1.45 | -0.06 | 0.06 |
| gene-LOC105286451 | https://www.ncbi.nlm.nih.gov/gene/?term=LOC105286451 | 1618 | 10 | 61.8 | 1 | 6.18 | 10 | 52.19 | 44.01 | 8.18 | 8.18 | 0.05 | 0.05 |
| gene-LOC105278538 | https://www.ncbi.nlm.nih.gov/gene/?term=LOC105278538 | 2150 | 75 | 348.84 | 1 | 4.65 | 1.33 | 18.53 | 17.09 | 1.44 | 1.44 | 0.05 | 0.05 |
| gene-LOC105282156 | https://www.ncbi.nlm.nih.gov/gene/?term=LOC105282156 | 2577 | 128 | 496.7 | 1 | 3.88 | 0.78 | 23.38 | 24.45 | -1.07 | 1.07 | -0.05 | 0.05 |
| gene-LOC105277215 | https://www.ncbi.nlm.nih.gov/gene/?term=LOC105277215 | 2619 | 27 | 103.09 | 1 | 3.82 | 3.7 | 20.79 | 25.43 | -4.64 | 4.64 | -0.05 | 0.05 |
| gene-LOC105275708 | https://www.ncbi.nlm.nih.gov/gene/?term=LOC105275708 | 1772 | 14 | 79.01 | 1 | 5.64 | 7.14 | 44.58 | 50.83 | -6.25 | 6.25 | -0.05 | 0.05 |
| gene-LOC105285514 | https://www.ncbi.nlm.nih.gov/gene/?term=LOC105285514 | 1277 | 8 | 62.65 | 1 | 7.83 | 12.5 | 70.46 | 79.23 | -8.77 | 8.77 | -0.05 | 0.05 |
| gene-LOC105283649 | https://www.ncbi.nlm.nih.gov/gene/?term=LOC105283649 | 1918 | 53 | 276.33 | 2 | 10.43 | 3.77 | 38.29 | 37.03 | 1.27 | 1.27 | 0.04 | 0.04 |
| gene-LOC105287095 | https://www.ncbi.nlm.nih.gov/gene/?term=LOC105287095 | 1435 | 60 | 418.12 | 1 | 6.97 | 1.67 | 26.49 | 25.42 | 1.06 | 1.06 | 0.04 | 0.04 |
| gene-LOC105280786 | https://www.ncbi.nlm.nih.gov/gene/?term=LOC105280786 | 2573 | 115 | 446.95 | 1 | 3.89 | 0.87 | 29.1 | 28.31 | 0.79 | 0.79 | 0.04 | 0.04 |
| gene-LOC105275363 | https://www.ncbi.nlm.nih.gov/gene/?term=LOC105275363 | 3798 | 155 | 408.11 | 1 | 2.63 | 0.65 | 17.28 | 18.35 | -1.06 | 1.06 | -0.04 | 0.04 |
| gene-LOC105285742 | https://www.ncbi.nlm.nih.gov/gene/?term=LOC105285742 | 3446 | 87 | 252.47 | 1 | 2.9 | 1.15 | 20.61 | 22.37 | -1.76 | 1.76 | -0.04 | 0.04 |
| gene-LOC105278436 | https://www.ncbi.nlm.nih.gov/gene/?term=LOC105278436 | 2556 | 56 | 219.09 | 1 | 3.91 | 1.79 | 36.89 | 38.91 | -2.02 | 2.02 | -0.04 | 0.04 |
| gene-LOC105274900 | https://www.ncbi.nlm.nih.gov/gene/?term=LOC105274900 | 5561 | 56 | 100.7 | 1 | 1.8 | 1.79 | 26.85 | 30.51 | -3.66 | 3.66 | -0.04 | 0.04 |
| gene-LOC105276538 | https://www.ncbi.nlm.nih.gov/gene/?term=LOC105276538 | 1511 | 18 | 119.13 | 1 | 6.62 | 5.56 | 71.16 | 68.49 | 2.67 | 2.67 | 0.03 | 0.03 |
| gene-LOC105278454 | https://www.ncbi.nlm.nih.gov/gene/?term=LOC105278454 | 1972 | 37 | 187.63 | 1 | 5.07 | 2.7 | 46.8 | 45.42 | 1.38 | 1.38 | 0.03 | 0.03 |
| gene-LOC105282811 | https://www.ncbi.nlm.nih.gov/gene/?term=LOC105282811 | 5765 | 183 | 317.43 | 1 | 1.73 | 0.55 | 10.19 | 9.27 | 0.92 | 0.92 | 0.03 | 0.03 |
| gene-LOC105274534 | https://www.ncbi.nlm.nih.gov/gene/?term=LOC105274534 | 1329 | 57 | 428.89 | 1 | 7.52 | 1.75 | 43.75 | 44.38 | -0.63 | 0.63 | -0.03 | 0.03 |
| gene-LOC105275683 | https://www.ncbi.nlm.nih.gov/gene/?term=LOC105275683 | 4256 | 187 | 439.38 | 1 | 2.35 | 0.53 | 12.61 | 13.25 | -0.63 | 0.63 | -0.03 | 0.03 |
| gene-LOC105278497 | https://www.ncbi.nlm.nih.gov/gene/?term=LOC105278497 | 2072 | 97 | 468.15 | 1 | 4.83 | 1.03 | 18.56 | 19.21 | -0.64 | 0.64 | -0.03 | 0.03 |
| gene-LOC105288036 | https://www.ncbi.nlm.nih.gov/gene/?term=LOC105288036 | 5110 | 182 | 356.16 | 1 | 1.96 | 0.55 | 18.19 | 18.89 | -0.7 | 0.7 | -0.03 | 0.03 |
| gene-LOC105283592 | https://www.ncbi.nlm.nih.gov/gene/?term=LOC105283592 | 3479 | 68 | 195.46 | 1 | 2.87 | 1.47 | 23.52 | 25 | -1.48 | 1.48 | -0.03 | 0.03 |
| gene-LOC105275190 | https://www.ncbi.nlm.nih.gov/gene/?term=LOC105275190 | 3837 | 65 | 169.4 | 1 | 2.61 | 1.54 | 21.02 | 22.59 | -1.58 | 1.58 | -0.03 | 0.03 |
| gene-LOC105288066 | https://www.ncbi.nlm.nih.gov/gene/?term=LOC105288066 | 1964 | 24 | 122.2 | 1 | 5.09 | 4.17 | 27.7 | 30.29 | -2.59 | 2.59 | -0.03 | 0.03 |
| gene-LOC105285060 | https://www.ncbi.nlm.nih.gov/gene/?term=LOC105285060 | 1285 | 15 | 116.73 | 1 | 7.78 | 6.67 | 48.46 | 51.14 | -2.68 | 2.68 | -0.03 | 0.03 |
| gene-LOC105279898 | https://www.ncbi.nlm.nih.gov/gene/?term=LOC105279898 | 1343 | 13 | 96.8 | 1 | 7.45 | 7.69 | 65.18 | 68.44 | -3.25 | 3.25 | -0.03 | 0.03 |
| gene-LOC105285409 | https://www.ncbi.nlm.nih.gov/gene/?term=LOC105285409 | 5099 | 47 | 92.17 | 1 | 1.96 | 2.13 | 3.57 | 1.67 | 1.9 | 1.9 | 0.02 | 0.02 |
| gene-LOC105279664 | https://www.ncbi.nlm.nih.gov/gene/?term=LOC105279664 | 4023 | 63 | 156.6 | 1 | 2.49 | 1.59 | 17.93 | 16.77 | 1.16 | 1.16 | 0.02 | 0.02 |
| gene-LOC105285202 | https://www.ncbi.nlm.nih.gov/gene/?term=LOC105285202 | 1838 | 35 | 190.42 | 1 | 5.44 | 2.86 | 19.49 | 18.4 | 1.08 | 1.08 | 0.02 | 0.02 |
| gene-LOC105276196 | https://www.ncbi.nlm.nih.gov/gene/?term=LOC105276196 | 2953 | 55 | 186.25 | 1 | 3.39 | 1.82 | 40.47 | 39.41 | 1.06 | 1.06 | 0.02 | 0.02 |
| gene-LOC105285807 | https://www.ncbi.nlm.nih.gov/gene/?term=LOC105285807 | 6063 | 180 | 296.88 | 1 | 1.65 | 0.56 | 10.12 | 9.41 | 0.72 | 0.72 | 0.02 | 0.02 |
| gene-LOC105275752 | https://www.ncbi.nlm.nih.gov/gene/?term=LOC105275752 | 3462 | 126 | 363.95 | 1 | 2.89 | 0.79 | 28.41 | 27.85 | 0.56 | 0.56 | 0.02 | 0.02 |
| gene-LOC105278666 | https://www.ncbi.nlm.nih.gov/gene/?term=LOC105278666 | 10341 | 393 | 380.04 | 1 | 0.97 | 0.25 | 7.76 | 7.36 | 0.4 | 0.4 | 0.02 | 0.02 |
| gene-LOC105285534 | https://www.ncbi.nlm.nih.gov/gene/?term=LOC105285534 | 3944 | 221 | 560.34 | 1 | 2.54 | 0.45 | 11.41 | 11.71 | -0.3 | 0.3 | -0.02 | 0.02 |
| gene-LOC105279298 | https://www.ncbi.nlm.nih.gov/gene/?term=LOC105279298 | 2773 | 135 | 486.84 | 1 | 3.61 | 0.74 | 22.57 | 22.9 | -0.33 | 0.33 | -0.02 | 0.02 |
| gene-LOC105286997 | https://www.ncbi.nlm.nih.gov/gene/?term=LOC105286997 | 2850 | 134 | 470.18 | 1 | 3.51 | 0.75 | 11.83 | 12.3 | -0.47 | 0.47 | -0.02 | 0.02 |
| gene-LOC105280956 | https://www.ncbi.nlm.nih.gov/gene/?term=LOC105280956 | 2639 | 73 | 276.62 | 1 | 3.79 | 1.37 | 20.28 | 20.85 | -0.57 | 0.57 | -0.02 | 0.02 |
| gene-LOC105285639 | https://www.ncbi.nlm.nih.gov/gene/?term=LOC105285639 | 10325 | 294 | 284.75 | 1 | 0.97 | 0.34 | 12.44 | 13.1 | -0.66 | 0.66 | -0.02 | 0.02 |
| gene-LOC105287063 | https://www.ncbi.nlm.nih.gov/gene/?term=LOC105287063 | 1874 | 46 | 245.46 | 1 | 5.34 | 2.17 | 34.15 | 34.88 | -0.73 | 0.73 | -0.02 | 0.02 |
| gene-LOC105275179 | https://www.ncbi.nlm.nih.gov/gene/?term=LOC105275179 | 3271 | 69 | 210.94 | 1 | 3.06 | 1.45 | 14.66 | 15.53 | -0.86 | 0.86 | -0.02 | 0.02 |
| gene-LOC105288237 | https://www.ncbi.nlm.nih.gov/gene/?term=LOC105288237 | 2226 | 45 | 202.16 | 2 | 8.98 | 4.44 | 36.5 | 37.4 | -0.9 | 0.9 | -0.02 | 0.02 |
| gene-LOC105275451 | https://www.ncbi.nlm.nih.gov/gene/?term=LOC105275451 | 6035 | 111 | 183.93 | 1 | 1.66 | 0.9 | 22.31 | 23.24 | -0.94 | 0.94 | -0.02 | 0.02 |
| gene-LOC105286265 | https://www.ncbi.nlm.nih.gov/gene/?term=LOC105286265 | 2507 | 65 | 259.27 | 1 | 3.99 | 1.54 | 31.58 | 32.53 | -0.95 | 0.95 | -0.02 | 0.02 |

| | | | | | | | | | | | | | | |
|---|---|---|---|---|---|---|---|---|---|---|---|---|---|---|
| gene-LOC105282991 | https://www.ncbi.nlm.nih.gov/gene/?term=LOC105282991 | 5617 | 80 | 142.42 | | 1 | 1.78 | 1.25 | 19.15 | 20.24 | -1.09 | 1.09 | -0.02 | 0.02 |
| gene-LOC105283943 | https://www.ncbi.nlm.nih.gov/gene/?term=LOC105283943 | 3559 | 49 | 137.68 | | 1 | 2.81 | 2.04 | 22.13 | 23.23 | -1.1 | 1.1 | -0.02 | 0.02 |
| gene-LOC105277216 | https://www.ncbi.nlm.nih.gov/gene/?term=LOC105277216 | 7627 | 91 | 119.31 | | 1 | 1.31 | 1.1 | 6.55 | 7.88 | -1.33 | 1.33 | -0.02 | 0.02 |
| gene-LOC105280348 | https://www.ncbi.nlm.nih.gov/gene/?term=LOC105280348 | 3797 | 52 | 136.95 | | 1 | 2.63 | 1.92 | 25.54 | 27.17 | -1.63 | 1.63 | -0.02 | 0.02 |
| gene-LOC105275518 | https://www.ncbi.nlm.nih.gov/gene/?term=LOC105275518 | 2335 | 19 | 81.37 | 1 | 4.28 | 5.26 | 49.77 | 48.32 | 1.45 | 1.45 | 0.01 | 0.01 | |
| gene-LOC105281047 | https://www.ncbi.nlm.nih.gov/gene/?term=LOC105281047 | 7313 | 112 | 153.15 | | 1 | 1.37 | 0.89 | 11.03 | 10.28 | 0.75 | 0.75 | 0.01 | 0.01 |
| gene-LOC105282794 | https://www.ncbi.nlm.nih.gov/gene/?term=LOC105282794 | 4571 | 93 | 203.46 | | 1 | 2.19 | 1.08 | 17.63 | 17.04 | 0.6 | 0.6 | 0.01 | 0.01 |
| gene-LOC105283123 | https://www.ncbi.nlm.nih.gov/gene/?term=LOC105283123 | 1507 | 35 | 232.25 | | 1 | 6.64 | 2.86 | 18.15 | 17.56 | 0.59 | 0.59 | 0.01 | 0.01 |
| gene-LOC105284116 | https://www.ncbi.nlm.nih.gov/gene/?term=LOC105284116 | 3970 | 90 | 226.7 | 1 | 2.52 | 1.11 | 7.07 | 6.57 | 0.5 | 0.5 | 0.01 | 0.01 | |
| gene-LOC105279038 | https://www.ncbi.nlm.nih.gov/gene/?term=LOC105279038 | 9453 | 299 | 316.3 | 1 | 1.06 | 0.33 | 8.97 | 8.54 | 0.42 | 0.42 | 0.01 | 0.01 | |
| gene-LOC105280002 | https://www.ncbi.nlm.nih.gov/gene/?term=LOC105280002 | 1940 | 28 | 144.33 | | 1 | 5.15 | 3.57 | 33.09 | 32.74 | 0.35 | 0.35 | 0.01 | 0.01 |
| gene-LOC105284117 | https://www.ncbi.nlm.nih.gov/gene/?term=LOC105284117 | 4340 | 82 | 188.94 | | 1 | 2.3 | 1.22 | 10.79 | 10.47 | 0.33 | 0.33 | 0.01 | 0.01 |
| gene-LOC105280819 | https://www.ncbi.nlm.nih.gov/gene/?term=LOC105280819 | 1639 | 41 | 250.15 | | 1 | 6.1 | 2.44 | 40.58 | 40.31 | 0.27 | 0.27 | 0.01 | 0.01 |
| gene-LOC105288079 | https://www.ncbi.nlm.nih.gov/gene/?term=LOC105288079 | 3473 | 85 | 244.75 | | 1 | 2.88 | 1.18 | 20.86 | 20.61 | 0.25 | 0.25 | 0.01 | 0.01 |
| gene-LOC105286515 | https://www.ncbi.nlm.nih.gov/gene/?term=LOC105286515 | 3751 | 106 | 282.59 | | 1 | 2.67 | 0.94 | 10.37 | 10.58 | -0.22 | 0.22 | -0.01 | 0.01 |
| gene-LOC105288078 | https://www.ncbi.nlm.nih.gov/gene/?term=LOC105288078 | 19126 | 305 | 159.47 | | 1 | 0.52 | 0.33 | 16.95 | 17.33 | -0.38 | 0.38 | -0.01 | 0.01 |
| gene-LOC105274531 | https://www.ncbi.nlm.nih.gov/gene/?term=LOC105274531 | 7774 | 253 | 325.44 | | 1 | 1.29 | 0.4 | 11.9 | 12.33 | -0.43 | 0.43 | -0.01 | 0.01 |
| gene-LOC105283712 | https://www.ncbi.nlm.nih.gov/gene/?term=LOC105283712 | 1910 | 29 | 151.83 | | 1 | 5.24 | 3.45 | 29.98 | 30.42 | -0.44 | 0.44 | -0.01 | 0.01 |
| gene-LOC105275173 | https://www.ncbi.nlm.nih.gov/gene/?term=LOC105275173 | 4639 | 76 | 163.83 | | 1 | 2.16 | 1.32 | 24.33 | 24.78 | -0.45 | 0.45 | -0.01 | 0.01 |
| gene-LOC105284184 | https://www.ncbi.nlm.nih.gov/gene/?term=LOC105284184 | 2535 | 80 | 315.58 | | 1 | 3.94 | 1.25 | 34.85 | 35.33 | -0.47 | 0.47 | -0.01 | 0.01 |
| gene-LOC105275898 | https://www.ncbi.nlm.nih.gov/gene/?term=LOC105275898 | 5358 | 132 | 246.36 | | 1 | 1.87 | 0.76 | 12.84 | 13.34 | -0.5 | 0.5 | -0.01 | 0.01 |
| gene-LOC105276692 | https://www.ncbi.nlm.nih.gov/gene/?term=LOC105276692 | 1745 | 34 | 194.84 | | 1 | 5.73 | 2.94 | 24.51 | 25.17 | -0.66 | 0.66 | -0.01 | 0.01 |
| gene-LOC105280641 | https://www.ncbi.nlm.nih.gov/gene/?term=LOC105280641 | 3682 | 55 | 149.38 | | 1 | 2.72 | 1.82 | 18.35 | 19.01 | -0.66 | 0.66 | -0.01 | 0.01 |
| gene-LOC105281700 | https://www.ncbi.nlm.nih.gov/gene/?term=LOC105281700 | 2208 | 45 | 203.8 | 1 | 4.53 | 2.22 | 38.83 | 39.5 | -0.67 | 0.67 | -0.01 | 0.01 | |
| gene-LOC105279657 | https://www.ncbi.nlm.nih.gov/gene/?term=LOC105279657 | 5878 | 82 | 139.5 | 1 | 1.7 | 1.22 | 19.15 | 19.93 | -0.78 | 0.78 | -0.01 | 0.01 | |
| gene-LOC105277856 | https://www.ncbi.nlm.nih.gov/gene/?term=LOC105277856 | 3442 | 28 | 81.35 | 1 | 2.91 | 3.57 | 36.24 | 37.06 | -0.82 | 0.82 | -0.01 | 0.01 | |
| gene-LOC105286452 | https://www.ncbi.nlm.nih.gov/gene/?term=LOC105286452 | 6888 | 58 | 84.2 | 1 | 1.45 | 1.72 | 31.25 | 32.65 | -1.4 | 1.4 | -0.01 | 0.01 | |
| gene-LOC105283989 | https://www.ncbi.nlm.nih.gov/gene/?term=LOC105283989 | 4948 | 30 | 60.63 | 1 | 2.02 | 3.33 | 45.76 | 47.5 | -1.74 | 1.74 | -0.01 | 0.01 | |
| gene-LOC105274590 | https://www.ncbi.nlm.nih.gov/gene/?term=LOC105274590 | 4516 | 75 | 166.08 | | 1 | 2.21 | 1.33 | 23.2 | 22.91 | 0.29 | 0.29 | 0 | 0 |
| gene-LOC105276901 | https://www.ncbi.nlm.nih.gov/gene/?term=LOC105276901 | 3254 | 96 | 295.02 | | 1 | 3.07 | 1.04 | 22.83 | 22.72 | 0.1 | 0.1 | 0 | 0 |
| gene-LOC105278228 | https://www.ncbi.nlm.nih.gov/gene/?term=LOC105278228 | 8610 | 152 | 176.54 | | 1 | 1.16 | 0.66 | 23.77 | 23.73 | 0.04 | 0.04 | 0 | 0 |
| gene-LOC105283728 | https://www.ncbi.nlm.nih.gov/gene/?term=LOC105283728 | 3256 | 113 | 347.05 | | 2 | 6.14 | 1.77 | 19.87 | 19.82 | 0.04 | 0.04 | 0 | 0 |
| gene-LOC105278145 | https://www.ncbi.nlm.nih.gov/gene/?term=LOC105278145 | 4470 | 239 | 534.68 | | 1 | 2.24 | 0.42 | 10.87 | 10.84 | 0.03 | 0.03 | 0 | 0 |
| gene-LOC105275327 | https://www.ncbi.nlm.nih.gov/gene/?term=LOC105275327 | 14245 | 461 | 323.62 | | 1 | 0.7 | 0.22 | 7.38 | 7.42 | -0.03 | 0.03 | 0 | 0 |
| gene-LOC105286274 | https://www.ncbi.nlm.nih.gov/gene/?term=LOC105286274 | 3817 | 174 | 455.86 | | 1 | 2.62 | 0.57 | 15.59 | 15.67 | -0.08 | 0.08 | 0 | 0 |
| gene-LOC105274856 | https://www.ncbi.nlm.nih.gov/gene/?term=LOC105274856 | 5216 | 201 | 385.35 | | 1 | 1.92 | 0.5 | 17.77 | 17.88 | -0.11 | 0.11 | 0 | 0 |
| gene-LOC105278534 | https://www.ncbi.nlm.nih.gov/gene/?term=LOC105278534 | 7364 | 173 | 234.93 | | 1 | 1.36 | 0.58 | 18.44 | 18.59 | -0.16 | 0.16 | 0 | 0 |
| gene-LOC105274851 | https://www.ncbi.nlm.nih.gov/gene/?term=LOC105274851 | 6616 | 133 | 201.03 | | 1 | 1.51 | 0.75 | 10.15 | 10.35 | -0.2 | 0.2 | 0 | 0 |
| gene-LOC105279500 | https://www.ncbi.nlm.nih.gov/gene/?term=LOC105279500 | 3587 | 29 | 80.85 | 1 | 2.79 | 3.45 | 48.07 | 48.34 | -0.27 | 0.27 | 0 | 0 | |
| gene-LOC105285406 | https://www.ncbi.nlm.nih.gov/gene/?term=LOC105285406 | 8015 | 39 | 48.66 | 1 | 1.25 | 2.56 | 33.07 | 33.49 | -0.42 | 0.42 | 0 | 0 | |

Directory: OGL File: rnk-dmg-Ob_LI_BCphaseA1.vs.RphaseA2.txt

| gene_ID | gene_link | gwidth | #Sites | #per10Kb | #dmSites | #dmsp10kb | %dmSites | %pSite1 | %pSite2 | DMpSite | ADMpSite | DMpNucl | ADMpNucl |
|---|---|---|---|---|---|---|---|---|---|---|---|---|---|
| gene-LOC113562345 | https://www.ncbi.nlm.nih.gov/gene/?term=LOC113562345 | 2290 | 62 | 270.74 | 16 | 69.87 | 25.81 | 17.91 | 33 | -15.09 | 15.09 | -0.41 | 0.41 |
| gene-LOC105276704 | https://www.ncbi.nlm.nih.gov/gene/?term=LOC105276704 | 2063 | 87 | 421.72 | 1 | 4.85 | 1.15 | 12.34 | 15.21 | -2.86 | 2.86 | -0.12 | 0.12 |
| gene-LOC105282156 | https://www.ncbi.nlm.nih.gov/gene/?term=LOC105282156 | 2577 | 136 | 527.75 | 1 | 3.88 | 0.74 | 23.86 | 26.02 | -2.16 | 2.16 | -0.11 | 0.11 |
| gene-LOC105279898 | https://www.ncbi.nlm.nih.gov/gene/?term=LOC105279898 | 1343 | 16 | 119.14 | 2 | 14.89 | 12.5 | 59.42 | 68.99 | -9.57 | 9.57 | -0.11 | 0.11 |
| gene-LOC105274534 | https://www.ncbi.nlm.nih.gov/gene/?term=LOC105274534 | 1329 | 57 | 428.89 | 1 | 7.52 | 1.75 | 43.75 | 46.14 | -2.39 | 2.39 | -0.1 | 0.1 |
| gene-LOC105286502 | https://www.ncbi.nlm.nih.gov/gene/?term=LOC105286502 | 1286 | 44 | 342.15 | 1 | 7.78 | 2.27 | 23.97 | 26.78 | -2.81 | 2.81 | -0.1 | 0.1 |
| gene-LOC105277631 | https://www.ncbi.nlm.nih.gov/gene/?term=LOC105277631 | 4387 | 289 | 658.76 | 1 | 2.28 | 0.35 | 9.79 | 11.22 | -1.43 | 1.43 | -0.09 | 0.09 |
| gene-LOC105285674 | https://www.ncbi.nlm.nih.gov/gene/?term=LOC105285674 | 3035 | 91 | 299.84 | 2 | 6.59 | 2.2 | 23.1 | 26.01 | -2.91 | 2.91 | -0.09 | 0.09 |
| gene-LOC105281848 | https://www.ncbi.nlm.nih.gov/gene/?term=LOC105281848 | 2122 | 47 | 221.49 | 1 | 4.71 | 2.13 | 23.11 | 27.19 | -4.08 | 4.08 | -0.09 | 0.09 |
| gene-LOC105285695 | https://www.ncbi.nlm.nih.gov/gene/?term=LOC105285695 | 1371 | 62 | 452.22 | 1 | 7.29 | 1.61 | 23 | 24.72 | -1.72 | 1.72 | -0.08 | 0.08 |
| gene-LOC105287417 | https://www.ncbi.nlm.nih.gov/gene/?term=LOC105287417 | 1025 | 33 | 321.95 | 1 | 9.76 | 3.03 | 36 | 38.42 | -2.42 | 2.42 | -0.08 | 0.08 |
| gene-LOC105285055 | https://www.ncbi.nlm.nih.gov/gene/?term=LOC105285055 | 4108 | 217 | 528.24 | 1 | 2.43 | 0.46 | 12.76 | 14.18 | -1.42 | 1.42 | -0.07 | 0.07 |
| gene-LOC105285671 | https://www.ncbi.nlm.nih.gov/gene/?term=LOC105285671 | 2263 | 95 | 419.8 | 1 | 4.42 | 1.05 | 27.86 | 29.64 | -1.78 | 1.78 | -0.07 | 0.07 |
| gene-LOC105286223 | https://www.ncbi.nlm.nih.gov/gene/?term=LOC105286223 | 2225 | 83 | 373.03 | 1 | 4.49 | 1.2 | 30.2 | 32.18 | -1.98 | 1.98 | -0.07 | 0.07 |
| gene-LOC113561805 | https://www.ncbi.nlm.nih.gov/gene/?term=LOC113561805 | 3337 | 64 | 191.79 | 1 | 3 | 1.56 | 37.95 | 41.4 | -3.45 | 3.45 | -0.07 | 0.07 |
| gene-LOC105275190 | https://www.ncbi.nlm.nih.gov/gene/?term=LOC105275190 | 3837 | 74 | 192.86 | 1 | 2.61 | 1.35 | 18.39 | 22.03 | -3.64 | 3.64 | -0.07 | 0.07 |
| gene-LOC105275070 | https://www.ncbi.nlm.nih.gov/gene/?term=LOC105275070 | 5667 | 38 | 67.05 | 3 | 5.29 | 7.89 | 3.63 | 13.45 | -9.82 | 9.82 | -0.07 | 0.07 |
| gene-LOC105274449 | https://www.ncbi.nlm.nih.gov/gene/?term=LOC105274449 | 1549 | 42 | 271.14 | 1 | 6.46 | 2.38 | 55.88 | 53.68 | 2.2 | 2.2 | 0.06 | 0.06 |
| gene-LOC105275683 | https://www.ncbi.nlm.nih.gov/gene/?term=LOC105275683 | 4256 | 203 | 476.97 | 1 | 2.35 | 0.49 | 12.81 | 14.05 | -1.25 | 1.25 | -0.06 | 0.06 |
| gene-LOC105275460 | https://www.ncbi.nlm.nih.gov/gene/?term=LOC105275460 | 2473 | 112 | 452.89 | 1 | 4.04 | 0.89 | 12.89 | 14.18 | -1.3 | 1.3 | -0.06 | 0.06 |
| gene-LOC105279356 | https://www.ncbi.nlm.nih.gov/gene/?term=LOC105279356 | 1314 | 42 | 319.63 | 1 | 7.61 | 2.38 | 28.83 | 30.58 | -1.75 | 1.75 | -0.06 | 0.06 |
| gene-LOC105278807 | https://www.ncbi.nlm.nih.gov/gene/?term=LOC105278807 | 2166 | 72 | 332.41 | 1 | 4.62 | 1.39 | 18.53 | 20.46 | -1.93 | 1.93 | -0.06 | 0.06 |
| gene-LOC105285583 | https://www.ncbi.nlm.nih.gov/gene/?term=LOC105285583 | 1947 | 51 | 261.94 | 1 | 5.14 | 1.96 | 28.26 | 30.43 | -2.17 | 2.17 | -0.06 | 0.06 |
| gene-LOC105287095 | https://www.ncbi.nlm.nih.gov/gene/?term=LOC105287095 | 1435 | 61 | 425.09 | 1 | 6.97 | 1.64 | 26.05 | 24.77 | 1.29 | 1.29 | 0.05 | 0.05 |
| gene-LOC105279298 | https://www.ncbi.nlm.nih.gov/gene/?term=LOC105279298 | 2773 | 138 | 497.66 | 2 | 7.21 | 1.45 | 22.22 | 23.22 | -1 | 1 | -0.05 | 0.05 |
| gene-LOC105278497 | https://www.ncbi.nlm.nih.gov/gene/?term=LOC105278497 | 2072 | 100 | 482.63 | 1 | 4.83 | 1 | 18.86 | 19.92 | -1.06 | 1.06 | -0.05 | 0.05 |
| gene-LOC105282947 | https://www.ncbi.nlm.nih.gov/gene/?term=LOC105282947 | 1162 | 49 | 421.69 | 1 | 8.61 | 2.04 | 29.51 | 30.69 | -1.17 | 1.17 | -0.05 | 0.05 |
| gene-LOC105284704 | https://www.ncbi.nlm.nih.gov/gene/?term=LOC105284704 | 2400 | 65 | 270.83 | 1 | 4.17 | 1.54 | 25.06 | 26.85 | -1.79 | 1.79 | -0.05 | 0.05 |
| gene-LOC105283712 | https://www.ncbi.nlm.nih.gov/gene/?term=LOC105283712 | 1910 | 27 | 141.36 | 1 | 5.24 | 3.7 | 29.17 | 32.5 | -3.33 | 3.33 | -0.05 | 0.05 |
| gene-LOC105278109 | https://www.ncbi.nlm.nih.gov/gene/?term=LOC105278109 | 4029 | 62 | 153.88 | 1 | 2.48 | 1.61 | 43.07 | 46.49 | -3.42 | 3.42 | -0.05 | 0.05 |
| gene-LOC105285060 | https://www.ncbi.nlm.nih.gov/gene/?term=LOC105285060 | 1285 | 15 | 116.73 | 1 | 7.78 | 6.67 | 48.46 | 52.7 | -4.25 | 4.25 | -0.05 | 0.05 |
| gene-LOC105275708 | https://www.ncbi.nlm.nih.gov/gene/?term=LOC105275708 | 1772 | 14 | 79.01 | 1 | 5.64 | 7.14 | 44.58 | 50.56 | -5.98 | 5.98 | -0.05 | 0.05 |
| gene-LOC105277380 | https://www.ncbi.nlm.nih.gov/gene/?term=LOC105277380 | 1967 | 5 | 25.42 | 1 | 5.08 | 20 | 69.42 | 54.11 | 15.31 | 15.31 | 0.04 | 0.04 |
| gene-LOC105282885 | https://www.ncbi.nlm.nih.gov/gene/?term=LOC105282885 | 1870 | 42 | 224.6 | 1 | 5.35 | 2.38 | 36.63 | 34.78 | 1.84 | 1.84 | 0.04 | 0.04 |
| gene-LOC105275848 | https://www.ncbi.nlm.nih.gov/gene/?term=LOC105275848 | 3718 | 130 | 349.65 | 1 | 2.69 | 0.77 | 21.27 | 20.2 | 1.07 | 1.07 | 0.04 | 0.04 |
| gene-LOC105275363 | https://www.ncbi.nlm.nih.gov/gene/?term=LOC105275363 | 3798 | 159 | 418.64 | 1 | 2.63 | 0.63 | 16.97 | 17.86 | -0.88 | 0.88 | -0.04 | 0.04 |
| gene-LOC105285742 | https://www.ncbi.nlm.nih.gov/gene/?term=LOC105285742 | 3446 | 100 | 290.19 | 1 | 2.9 | 1 | 24.69 | 25.96 | -1.27 | 1.27 | -0.04 | 0.04 |
| gene-LOC105278282 | https://www.ncbi.nlm.nih.gov/gene/?term=LOC105278282 | 1775 | 47 | 264.79 | 1 | 5.63 | 2.13 | 45.94 | 47.29 | -1.35 | 1.35 | -0.04 | 0.04 |
| gene-LOC105287063 | https://www.ncbi.nlm.nih.gov/gene/?term=LOC105287063 | 1874 | 46 | 245.46 | 1 | 5.34 | 2.17 | 34.15 | 35.78 | -1.63 | 1.63 | -0.04 | 0.04 |
| gene-LOC105276692 | https://www.ncbi.nlm.nih.gov/gene/?term=LOC105276692 | 1745 | 37 | 212.03 | 1 | 5.73 | 2.7 | 20.82 | 22.64 | -1.82 | 1.82 | -0.04 | 0.04 |
| gene-LOC105276745 | https://www.ncbi.nlm.nih.gov/gene/?term=LOC105276745 | 3953 | 73 | 184.67 | 1 | 2.53 | 1.37 | 21.09 | 23.02 | -1.93 | 1.93 | -0.04 | 0.04 |
| gene-LOC105282893 | https://www.ncbi.nlm.nih.gov/gene/?term=LOC105282893 | 2645 | 43 | 162.57 | 1 | 3.78 | 2.33 | 27.48 | 29.68 | -2.19 | 2.19 | -0.04 | 0.04 |
| gene-LOC105286644 | https://www.ncbi.nlm.nih.gov/gene/?term=LOC105286644 | 1235 | 22 | 178.14 | 1 | 8.1 | 4.55 | 42.2 | 44.4 | -2.2 | 2.2 | -0.04 | 0.04 |
| gene-LOC105287854 | https://www.ncbi.nlm.nih.gov/gene/?term=LOC105287854 | 3644 | 57 | 156.42 | 1 | 2.74 | 1.75 | 21.16 | 23.71 | -2.55 | 2.55 | -0.04 | 0.04 |
| gene-LOC105276538 | https://www.ncbi.nlm.nih.gov/gene/?term=LOC105276538 | 1511 | 19 | 125.74 | 1 | 6.62 | 5.26 | 72.68 | 75.55 | -2.87 | 2.87 | -0.04 | 0.04 |
| gene-LOC105277856 | https://www.ncbi.nlm.nih.gov/gene/?term=LOC105277856 | 3442 | 29 | 84.25 | 1 | 2.91 | 3.45 | 35.44 | 40.05 | -4.6 | 4.6 | -0.04 | 0.04 |
| gene-LOC105281039 | https://www.ncbi.nlm.nih.gov/gene/?term=LOC105281039 | 1471 | 13 | 88.38 | 1 | 6.8 | 7.69 | 36.12 | 40.81 | -4.7 | 4.7 | -0.04 | 0.04 |
| gene-LOC105285514 | https://www.ncbi.nlm.nih.gov/gene/?term=LOC105285514 | 1277 | 9 | 70.48 | 1 | 7.83 | 11.11 | 73.21 | 78.44 | -5.23 | 5.23 | -0.04 | 0.04 |
| gene-LOC105275859 | https://www.ncbi.nlm.nih.gov/gene/?term=LOC105275859 | 2331 | 25 | 107.25 | 1 | 4.29 | 4 | 53.95 | 51.3 | 2.65 | 2.65 | 0.03 | 0.03 |
| gene-LOC105281173 | https://www.ncbi.nlm.nih.gov/gene/?term=LOC105281173 | 1595 | 25 | 156.74 | 1 | 6.27 | 4 | 64.76 | 62.62 | 2.13 | 2.13 | 0.03 | 0.03 |
| gene-LOC105278877 | https://www.ncbi.nlm.nih.gov/gene/?term=LOC105278877 | 1968 | 80 | 406.5 | 1 | 5.08 | 1.25 | 32.88 | 32.17 | 0.72 | 0.72 | 0.03 | 0.03 |
| gene-LOC105278283 | https://www.ncbi.nlm.nih.gov/gene/?term=LOC105278283 | 2317 | 130 | 561.07 | 1 | 4.32 | 0.77 | 24.9 | 25.39 | -0.48 | 0.48 | -0.03 | 0.03 |
| gene-LOC105276527 | https://www.ncbi.nlm.nih.gov/gene/?term=LOC105276527 | 7923 | 351 | 443.01 | 1 | 1.26 | 0.28 | 8.2 | 8.78 | -0.58 | 0.58 | -0.03 | 0.03 |
| gene-LOC105284355 | https://www.ncbi.nlm.nih.gov/gene/?term=LOC105284355 | 3904 | 147 | 376.54 | 1 | 2.56 | 0.68 | 21.32 | 21.98 | -0.67 | 0.67 | -0.03 | 0.03 |
| gene-LOC105286274 | https://www.ncbi.nlm.nih.gov/gene/?term=LOC105286274 | 3817 | 183 | 479.43 | 1 | 2.62 | 0.55 | 15.39 | 16.06 | -0.67 | 0.67 | -0.03 | 0.03 |
| gene-LOC105285179 | https://www.ncbi.nlm.nih.gov/gene/?term=LOC105285179 | 4191 | 156 | 372.23 | 1 | 2.39 | 0.64 | 17.53 | 18.23 | -0.69 | 0.69 | -0.03 | 0.03 |
| gene-LOC105283728 | https://www.ncbi.nlm.nih.gov/gene/?term=LOC105283728 | 3256 | 118 | 362.41 | 2 | 6.14 | 1.69 | 19.18 | 19.87 | -0.7 | 0.7 | -0.03 | 0.03 |
| gene-LOC105276449 | https://www.ncbi.nlm.nih.gov/gene/?term=LOC105276449 | 1231 | 48 | 389.93 | 1 | 8.12 | 2.08 | 26.7 | 27.43 | -0.74 | 0.74 | -0.03 | 0.03 |
| gene-LOC105275752 | https://www.ncbi.nlm.nih.gov/gene/?term=LOC105275752 | 3462 | 123 | 355.29 | 1 | 2.89 | 0.81 | 28.75 | 29.5 | -0.75 | 0.75 | -0.03 | 0.03 |
| gene-LOC105280774 | https://www.ncbi.nlm.nih.gov/gene/?term=LOC105280774 | 2793 | 117 | 418.9 | 1 | 3.58 | 0.85 | 27.44 | 28.24 | -0.79 | 0.79 | -0.03 | 0.03 |

| | | | | | | | | | | | | | |
|---|---|---|---|---|---|---|---|---|---|---|---|---|---|
| gene-LOC105282563 | https://www.ncbi.nlm.nih.gov/gene/?term=LOC105282563 | 4353 | 146 | 335.4 | 1 | 2.3 | 0.68 | 15.5 | 16.32 | -0.83 | 0.83 | -0.03 | 0.03 |
| gene-LOC105274856 | https://www.ncbi.nlm.nih.gov/gene/?term=LOC105274856 | 5216 | 208 | 398.77 | 1 | 1.92 | 0.48 | 18.49 | 19.33 | -0.84 | 0.84 | -0.03 | 0.03 |
| gene-LOC105274706 | https://www.ncbi.nlm.nih.gov/gene/?term=LOC105274706 | 1446 | 52 | 359.61 | 1 | 6.92 | 1.92 | 37.01 | 37.87 | -0.86 | 0.86 | -0.03 | 0.03 |
| gene-LOC105282491 | https://www.ncbi.nlm.nih.gov/gene/?term=LOC105282491 | 4761 | 163 | 342.37 | 1 | 2.1 | 0.61 | 9.65 | 10.67 | -1.02 | 1.02 | -0.03 | 0.03 |
| gene-LOC105281700 | https://www.ncbi.nlm.nih.gov/gene/?term=LOC105281700 | 2208 | 46 | 208.33 | 1 | 4.53 | 2.17 | 39.97 | 41.25 | -1.28 | 1.28 | -0.03 | 0.03 |
| gene-LOC105283592 | https://www.ncbi.nlm.nih.gov/gene/?term=LOC105283592 | 3479 | 71 | 204.08 | 1 | 2.87 | 1.41 | 24.39 | 25.83 | -1.45 | 1.45 | -0.03 | 0.03 |
| gene-LOC105280348 | https://www.ncbi.nlm.nih.gov/gene/?term=LOC105280348 | 3797 | 52 | 136.95 | 1 | 2.63 | 1.92 | 25.54 | 27.53 | -1.99 | 1.99 | -0.03 | 0.03 |
| gene-LOC105280002 | https://www.ncbi.nlm.nih.gov/gene/?term=LOC105280002 | 1940 | 30 | 154.64 | 1 | 5.15 | 3.33 | 33.39 | 35.64 | -2.25 | 2.25 | -0.03 | 0.03 |
| gene-LOC105288066 | https://www.ncbi.nlm.nih.gov/gene/?term=LOC105288066 | 1964 | 26 | 132.38 | 1 | 5.09 | 3.85 | 26.28 | 28.74 | -2.45 | 2.45 | -0.03 | 0.03 |
| gene-LOC105277215 | https://www.ncbi.nlm.nih.gov/gene/?term=LOC105277215 | 2619 | 29 | 110.73 | 1 | 3.82 | 3.45 | 19.9 | 22.79 | -2.88 | 2.88 | -0.03 | 0.03 |
| gene-LOC105282399 | https://www.ncbi.nlm.nih.gov/gene/?term=LOC105282399 | 2682 | 22 | 82.03 | 1 | 3.73 | 4.55 | 32.93 | 36.36 | -3.43 | 3.43 | -0.03 | 0.03 |
| gene-LOC105274778 | https://www.ncbi.nlm.nih.gov/gene/?term=LOC105274778 | 4662 | 38 | 81.51 | 1 | 2.15 | 2.63 | 44.96 | 48.85 | -3.88 | 3.88 | -0.03 | 0.03 |
| gene-LOC105284609 | https://www.ncbi.nlm.nih.gov/gene/?term=LOC105284609 | 5678 | 31 | 54.6 | 1 | 1.76 | 3.23 | 35.81 | 41.06 | -5.25 | 5.25 | -0.03 | 0.03 |
| gene-LOC105285409 | https://www.ncbi.nlm.nih.gov/gene/?term=LOC105285409 | 5099 | 48 | 94.14 | 2 | 3.92 | 4.17 | 3.6 | 1.11 | 2.49 | 2.49 | 0.02 | 0.02 |
| gene-LOC105276272 | https://www.ncbi.nlm.nih.gov/gene/?term=LOC105276272 | 1965 | 27 | 137.4 | 1 | 5.09 | 3.7 | 42.02 | 40.71 | 1.31 | 1.31 | 0.02 | 0.02 |
| gene-LOC105281296 | https://www.ncbi.nlm.nih.gov/gene/?term=LOC105281296 | 2434 | 41 | 168.45 | 1 | 4.11 | 2.44 | 31.19 | 30.16 | 1.03 | 1.03 | 0.02 | 0.02 |
| gene-LOC105284507 | https://www.ncbi.nlm.nih.gov/gene/?term=LOC105284507 | 5207 | 82 | 157.48 | 1 | 1.92 | 1.22 | 21.61 | 20.58 | 1.03 | 1.03 | 0.02 | 0.02 |
| gene-LOC105283123 | https://www.ncbi.nlm.nih.gov/gene/?term=LOC105283123 | 1507 | 43 | 285.34 | 1 | 6.64 | 2.33 | 17.05 | 16.29 | 0.76 | 0.76 | 0.02 | 0.02 |
| gene-LOC105284396 | https://www.ncbi.nlm.nih.gov/gene/?term=LOC105284396 | 1976 | 61 | 308.7 | 1 | 5.06 | 1.64 | 36.72 | 35.98 | 0.74 | 0.74 | 0.02 | 0.02 |
| gene-LOC105283264 | https://www.ncbi.nlm.nih.gov/gene/?term=LOC105283264 | 4005 | 88 | 219.73 | 1 | 2.5 | 1.14 | 24.81 | 24.09 | 0.72 | 0.72 | 0.02 | 0.02 |
| gene-LOC105282811 | https://www.ncbi.nlm.nih.gov/gene/?term=LOC105282811 | 5765 | 201 | 348.66 | 1 | 1.73 | 0.5 | 9.88 | 9.23 | 0.65 | 0.65 | 0.02 | 0.02 |
| gene-LOC105286515 | https://www.ncbi.nlm.nih.gov/gene/?term=LOC105286515 | 3751 | 116 | 309.25 | 1 | 2.67 | 0.86 | 11.85 | 11.25 | 0.6 | 0.6 | 0.02 | 0.02 |
| gene-LOC105287704 | https://www.ncbi.nlm.nih.gov/gene/?term=LOC105287704 | 3019 | 125 | 414.04 | 1 | 3.31 | 0.8 | 21.19 | 21.56 | -0.37 | 0.37 | -0.02 | 0.02 |
| gene-LOC105280786 | https://www.ncbi.nlm.nih.gov/gene/?term=LOC105280786 | 2573 | 122 | 474.15 | 1 | 3.89 | 0.82 | 29.15 | 29.56 | -0.41 | 0.41 | -0.02 | 0.02 |
| gene-LOC105285534 | https://www.ncbi.nlm.nih.gov/gene/?term=LOC105285534 | 3944 | 224 | 567.95 | 1 | 2.54 | 0.45 | 11.27 | 11.7 | -0.42 | 0.42 | -0.02 | 0.02 |
| gene-LOC105284043 | https://www.ncbi.nlm.nih.gov/gene/?term=LOC105284043 | 5661 | 248 | 438.09 | 1 | 1.77 | 0.4 | 11.5 | 11.98 | -0.48 | 0.48 | -0.02 | 0.02 |
| gene-LOC105277435 | https://www.ncbi.nlm.nih.gov/gene/?term=LOC105277435 | 3643 | 158 | 433.71 | 1 | 2.74 | 0.63 | 12.84 | 13.36 | -0.52 | 0.52 | -0.02 | 0.02 |
| gene-LOC105283132 | https://www.ncbi.nlm.nih.gov/gene/?term=LOC105283132 | 1819 | 54 | 296.87 | 1 | 5.5 | 1.85 | 40.82 | 41.59 | -0.77 | 0.77 | -0.02 | 0.02 |
| gene-LOC105274590 | https://www.ncbi.nlm.nih.gov/gene/?term=LOC105274590 | 4516 | 81 | 179.36 | 1 | 2.21 | 1.23 | 21.65 | 22.55 | -0.9 | 0.9 | -0.02 | 0.02 |
| gene-LOC105275179 | https://www.ncbi.nlm.nih.gov/gene/?term=LOC105275179 | 3271 | 68 | 207.89 | 1 | 3.06 | 1.47 | 14.88 | 16 | -1.12 | 1.12 | -0.02 | 0.02 |
| gene-LOC105278962 | https://www.ncbi.nlm.nih.gov/gene/?term=LOC105278962 | 4491 | 88 | 195.95 | 1 | 2.23 | 1.14 | 23.85 | 25.09 | -1.24 | 1.24 | -0.02 | 0.02 |
| gene-LOC105283943 | https://www.ncbi.nlm.nih.gov/gene/?term=LOC105283943 | 3559 | 60 | 168.59 | 1 | 2.81 | 1.67 | 21.58 | 22.83 | -1.25 | 1.25 | -0.02 | 0.02 |
| gene-LOC105276078 | https://www.ncbi.nlm.nih.gov/gene/?term=LOC105276078 | 1051 | 10 | 95.15 | 1 | 9.51 | 10 | 37.29 | 38.89 | -1.6 | 1.6 | -0.02 | 0.02 |
| gene-LOC105274900 | https://www.ncbi.nlm.nih.gov/gene/?term=LOC105274900 | 5561 | 56 | 100.7 | 2 | 3.6 | 3.57 | 26.85 | 28.69 | -1.84 | 1.84 | -0.02 | 0.02 |
| gene-LOC105278260 | https://www.ncbi.nlm.nih.gov/gene/?term=LOC105278260 | 3240 | 41 | 126.54 | 1 | 3.09 | 2.44 | 13.01 | 14.86 | -1.85 | 1.85 | -0.02 | 0.02 |
| gene-LOC105286452 | https://www.ncbi.nlm.nih.gov/gene/?term=LOC105286452 | 6888 | 58 | 84.2 | 1 | 1.45 | 1.72 | 31.25 | 33.23 | -1.98 | 1.98 | -0.02 | 0.02 |
| gene-LOC105275518 | https://www.ncbi.nlm.nih.gov/gene/?term=LOC105275518 | 2335 | 21 | 89.94 | 1 | 4.28 | 4.76 | 45.03 | 47.49 | -2.45 | 2.45 | -0.02 | 0.02 |
| gene-LOC105286983 | https://www.ncbi.nlm.nih.gov/gene/?term=LOC105286983 | 16901 | 68 | 40.23 | 1 | 0.59 | 1.47 | 0.94 | 5.77 | -4.83 | 4.83 | -0.02 | 0.02 |
| gene-LOC105285084 | https://www.ncbi.nlm.nih.gov/gene/?term=LOC105285084 | 3308 | 26 | 78.6 | 1 | 3.02 | 3.85 | 22.5 | 21.29 | 1.21 | 1.21 | 0.01 | 0.01 |
| gene-LOC105285252 | https://www.ncbi.nlm.nih.gov/gene/?term=LOC105285252 | 7427 | 132 | 177.73 | 1 | 1.35 | 0.76 | 13.81 | 12.99 | 0.81 | 0.81 | 0.01 | 0.01 |
| gene-LOC105279664 | https://www.ncbi.nlm.nih.gov/gene/?term=LOC105279664 | 4023 | 72 | 178.97 | 1 | 2.49 | 1.39 | 17.48 | 16.74 | 0.73 | 0.73 | 0.01 | 0.01 |
| gene-LOC105280819 | https://www.ncbi.nlm.nih.gov/gene/?term=LOC105280819 | 1639 | 41 | 250.15 | 1 | 6.1 | 2.44 | 40.58 | 40.03 | 0.56 | 0.56 | 0.01 | 0.01 |
| gene-LOC105282794 | https://www.ncbi.nlm.nih.gov/gene/?term=LOC105282794 | 4571 | 98 | 214.4 | 1 | 2.19 | 1.02 | 18.44 | 17.88 | 0.56 | 0.56 | 0.01 | 0.01 |
| gene-LOC105283649 | https://www.ncbi.nlm.nih.gov/gene/?term=LOC105283649 | 1918 | 53 | 276.33 | 2 | 10.43 | 3.77 | 38.29 | 38.03 | 0.26 | 0.26 | 0.01 | 0.01 |
| gene-LOC105286265 | https://www.ncbi.nlm.nih.gov/gene/?term=LOC105286265 | 2507 | 68 | 271.24 | 1 | 3.99 | 1.47 | 30.18 | 29.94 | 0.24 | 0.24 | 0.01 | 0.01 |
| gene-LOC105281834 | https://www.ncbi.nlm.nih.gov/gene/?term=LOC105281834 | 4711 | 204 | 433.03 | 1 | 2.12 | 0.49 | 10.43 | 10.29 | 0.14 | 0.14 | 0.01 | 0.01 |
| gene-LOC105286649 | https://www.ncbi.nlm.nih.gov/gene/?term=LOC105286649 | 14178 | 701 | 494.43 | 1 | 0.71 | 0.14 | 1.28 | 1.15 | 0.13 | 0.13 | 0.01 | 0.01 |
| gene-LOC105278145 | https://www.ncbi.nlm.nih.gov/gene/?term=LOC105278145 | 4470 | 256 | 572.71 | 1 | 2.24 | 0.39 | 10.38 | 10.47 | -0.09 | 0.09 | -0.01 | 0.01 |
| gene-LOC105284557 | https://www.ncbi.nlm.nih.gov/gene/?term=LOC105284557 | 1672 | 77 | 460.53 | 1 | 5.98 | 1.3 | 30.69 | 30.8 | -0.11 | 0.11 | -0.01 | 0.01 |
| gene-LOC105281272 | https://www.ncbi.nlm.nih.gov/gene/?term=LOC105281272 | 5252 | 179 | 340.82 | 1 | 1.9 | 0.56 | 16.36 | 16.51 | -0.15 | 0.15 | -0.01 | 0.01 |
| gene-LOC105276612 | https://www.ncbi.nlm.nih.gov/gene/?term=LOC105276612 | 4157 | 178 | 428.19 | 1 | 2.41 | 0.56 | 10.69 | 10.89 | -0.2 | 0.2 | -0.01 | 0.01 |
| gene-LOC105284036 | https://www.ncbi.nlm.nih.gov/gene/?term=LOC105284036 | 3254 | 83 | 255.07 | 1 | 3.07 | 1.2 | 22.52 | 22.73 | -0.2 | 0.2 | -0.01 | 0.01 |
| gene-LOC105285807 | https://www.ncbi.nlm.nih.gov/gene/?term=LOC105285807 | 6063 | 186 | 306.78 | 1 | 1.65 | 0.54 | 10.05 | 10.27 | -0.22 | 0.22 | -0.01 | 0.01 |
| gene-LOC105275327 | https://www.ncbi.nlm.nih.gov/gene/?term=LOC105275327 | 14245 | 479 | 336.26 | 1 | 0.7 | 0.21 | 7.11 | 7.34 | -0.23 | 0.23 | -0.01 | 0.01 |
| gene-LOC113562050 | https://www.ncbi.nlm.nih.gov/gene/?term=LOC113562050 | 17097 | 576 | 336.9 | 1 | 0.58 | 0.17 | 5.26 | 5.5 | -0.25 | 0.25 | -0.01 | 0.01 |
| gene-LOC105285202 | https://www.ncbi.nlm.nih.gov/gene/?term=LOC105285202 | 1838 | 36 | 195.87 | 1 | 5.44 | 2.78 | 18.94 | 19.21 | -0.27 | 0.27 | -0.01 | 0.01 |
| gene-LOC105274851 | https://www.ncbi.nlm.nih.gov/gene/?term=LOC105274851 | 6616 | 142 | 214.63 | 1 | 1.51 | 0.7 | 9.54 | 9.83 | -0.3 | 0.3 | -0.01 | 0.01 |
| gene-LOC105280104 | https://www.ncbi.nlm.nih.gov/gene/?term=LOC105280104 | 3960 | 120 | 303.03 | 1 | 2.53 | 0.83 | 14.59 | 14.9 | -0.31 | 0.31 | -0.01 | 0.01 |
| gene-LOC105288078 | https://www.ncbi.nlm.nih.gov/gene/?term=LOC105288078 | 19126 | 318 | 166.27 | 1 | 0.52 | 0.31 | 17.33 | 17.64 | -0.31 | 0.31 | -0.01 | 0.01 |
| gene-LOC105274531 | https://www.ncbi.nlm.nih.gov/gene/?term=LOC105274531 | 7774 | 257 | 330.59 | 1 | 1.29 | 0.39 | 11.81 | 12.14 | -0.32 | 0.32 | -0.01 | 0.01 |
| gene-LOC105278534 | https://www.ncbi.nlm.nih.gov/gene/?term=LOC105278534 | 7364 | 185 | 251.22 | 1 | 1.36 | 0.54 | 17.95 | 18.29 | -0.33 | 0.33 | -0.01 | 0.01 |
| gene-LOC105278384 | https://www.ncbi.nlm.nih.gov/gene/?term=LOC105278384 | 1390 | 33 | 237.41 | 1 | 7.19 | 3.03 | 45.97 | 46.32 | -0.35 | 0.35 | -0.01 | 0.01 |
| gene-LOC105279038 | https://www.ncbi.nlm.nih.gov/gene/?term=LOC105279038 | 9453 | 321 | 339.57 | 1 | 1.06 | 0.31 | 8.59 | 8.94 | -0.35 | 0.35 | -0.01 | 0.01 |
| gene-LOC105278228 | https://www.ncbi.nlm.nih.gov/gene/?term=LOC105278228 | 8610 | 163 | 189.31 | 1 | 1.16 | 0.61 | 23.66 | 24.04 | -0.38 | 0.38 | -0.01 | 0.01 |

| | | | | | | | | | | | | | |
|---|---|---|---|---|---|---|---|---|---|---|---|---|---|
| gene-LOC105280956 | https://www.ncbi.nlm.nih.gov/gene/?term=LOC105280956 | 2639 | 73 | 276.62 | 1 | 3.79 | 1.37 | 20.28 | 20.69 | -0.4 | 0.4 | -0.01 | 0.01 |
| gene-LOC105284184 | https://www.ncbi.nlm.nih.gov/gene/?term=LOC105284184 | 2535 | 82 | 323.47 | 1 | 3.94 | 1.22 | 34.56 | 34.95 | -0.4 | 0.4 | -0.01 | 0.01 |
| gene-LOC105275451 | https://www.ncbi.nlm.nih.gov/gene/?term=LOC105275451 | 6035 | 118 | 195.53 | 1 | 1.66 | 0.85 | 22.19 | 22.63 | -0.44 | 0.44 | -0.01 | 0.01 |
| gene-LOC105288237 | https://www.ncbi.nlm.nih.gov/gene/?term=LOC105288237 | 2226 | 47 | 211.14 | 1 | 4.49 | 2.13 | 35.68 | 36.15 | -0.47 | 0.47 | -0.01 | 0.01 |
| gene-LOC105282825 | https://www.ncbi.nlm.nih.gov/gene/?term=LOC105282825 | 3919 | 90 | 229.65 | 1 | 2.55 | 1.11 | 15.34 | 15.86 | -0.52 | 0.52 | -0.01 | 0.01 |
| gene-LOC105288079 | https://www.ncbi.nlm.nih.gov/gene/?term=LOC105288079 | 3473 | 89 | 256.26 | 1 | 2.88 | 1.12 | 19.96 | 20.49 | -0.52 | 0.52 | -0.01 | 0.01 |
| gene-LOC105280410 | https://www.ncbi.nlm.nih.gov/gene/?term=LOC105280410 | 12580 | 260 | 206.68 | 1 | 0.79 | 0.38 | 19.03 | 19.62 | -0.59 | 0.59 | -0.01 | 0.01 |
| gene-LOC105275966 | https://www.ncbi.nlm.nih.gov/gene/?term=LOC105275966 | 3299 | 59 | 178.84 | 1 | 3.03 | 1.69 | 50.28 | 50.9 | -0.62 | 0.62 | -0.01 | 0.01 |
| gene-LOC105280641 | https://www.ncbi.nlm.nih.gov/gene/?term=LOC105280641 | 3682 | 66 | 179.25 | 1 | 2.72 | 1.52 | 15.43 | 16.12 | -0.68 | 0.68 | -0.01 | 0.01 |
| gene-LOC105286770 | https://www.ncbi.nlm.nih.gov/gene/?term=LOC105286770 | 3055 | 38 | 124.39 | 1 | 3.27 | 2.63 | 11.1 | 11.78 | -0.68 | 0.68 | -0.01 | 0.01 |
| gene-LOC105282600 | https://www.ncbi.nlm.nih.gov/gene/?term=LOC105282600 | 3715 | 65 | 174.97 | 1 | 2.69 | 1.54 | 26.29 | 27.03 | -0.73 | 0.73 | -0.01 | 0.01 |
| gene-LOC105276917 | https://www.ncbi.nlm.nih.gov/gene/?term=LOC105276917 | 2336 | 26 | 111.3 | 1 | 4.28 | 3.85 | 36.57 | 37.3 | -0.74 | 0.74 | -0.01 | 0.01 |
| gene-LOC105282067 | https://www.ncbi.nlm.nih.gov/gene/?term=LOC105282067 | 5941 | 54 | 90.89 | 1 | 1.68 | 1.85 | 23.11 | 23.91 | -0.8 | 0.8 | -0.01 | 0.01 |
| gene-LOC105274773 | https://www.ncbi.nlm.nih.gov/gene/?term=LOC105274773 | 11034 | 167 | 151.35 | 1 | 0.91 | 0.6 | 16.37 | 17.25 | -0.88 | 0.88 | -0.01 | 0.01 |
| gene-LOC105279657 | https://www.ncbi.nlm.nih.gov/gene/?term=LOC105279657 | 5878 | 84 | 142.91 | 1 | 1.7 | 1.19 | 18.69 | 19.56 | -0.88 | 0.88 | -0.01 | 0.01 |
| gene-LOC105277216 | https://www.ncbi.nlm.nih.gov/gene/?term=LOC105277216 | 7627 | 89 | 116.69 | 1 | 1.31 | 1.12 | 6.69 | 7.62 | -0.93 | 0.93 | -0.01 | 0.01 |
| gene-LOC105280447 | https://www.ncbi.nlm.nih.gov/gene/?term=LOC105280447 | 2215 | 28 | 126.41 | 1 | 4.51 | 3.57 | 23.87 | 24.81 | -0.94 | 0.94 | -0.01 | 0.01 |
| gene-LOC105285406 | https://www.ncbi.nlm.nih.gov/gene/?term=LOC105285406 | 8015 | 42 | 52.4 | 1 | 1.25 | 2.38 | 35.2 | 36.16 | -0.96 | 0.96 | -0.01 | 0.01 |
| gene-LOC105279500 | https://www.ncbi.nlm.nih.gov/gene/?term=LOC105279500 | 3587 | 33 | 92 | 1 | 2.79 | 3.03 | 49.6 | 51 | -1.4 | 1.4 | -0.01 | 0.01 |
| gene-LOC105275347 | https://www.ncbi.nlm.nih.gov/gene/?term=LOC105275347 | 2526 | 14 | 55.42 | 1 | 3.96 | 7.14 | 35.76 | 37.28 | -1.52 | 1.52 | -0.01 | 0.01 |
| gene-LOC105283989 | https://www.ncbi.nlm.nih.gov/gene/?term=LOC105283989 | 4948 | 34 | 68.71 | 2 | 4.04 | 5.88 | 44.42 | 46.21 | -1.79 | 1.79 | -0.01 | 0.01 |
| gene-LOC105281515 | https://www.ncbi.nlm.nih.gov/gene/?term=LOC105281515 | 8827 | 41 | 46.45 | 1 | 1.13 | 2.44 | 32.87 | 35.23 | -2.37 | 2.37 | -0.01 | 0.01 |
| gene-LOC105281420 | https://www.ncbi.nlm.nih.gov/gene/?term=LOC105281420 | 3882 | 57 | 146.83 | 1 | 2.58 | 1.75 | 15.28 | 15.02 | 0.26 | 0.26 | 0 | 0 |
| gene-LOC105278454 | https://www.ncbi.nlm.nih.gov/gene/?term=LOC105278454 | 1972 | 37 | 187.63 | 1 | 5.07 | 2.7 | 46.8 | 46.67 | 0.13 | 0.13 | 0 | 0 |
| gene-LOC105276196 | https://www.ncbi.nlm.nih.gov/gene/?term=LOC105276196 | 2953 | 56 | 189.64 | 1 | 3.39 | 1.79 | 40.11 | 39.99 | 0.12 | 0.12 | 0 | 0 |
| gene-LOC105277273 | https://www.ncbi.nlm.nih.gov/gene/?term=LOC105277273 | 3711 | 151 | 406.9 | 1 | 2.69 | 0.66 | 12.76 | 12.67 | 0.09 | 0.09 | 0 | 0 |
| gene-LOC105278632 | https://www.ncbi.nlm.nih.gov/gene/?term=LOC105278632 | 3624 | 132 | 364.24 | 1 | 2.76 | 0.76 | 14.6 | 14.5 | 0.09 | 0.09 | 0 | 0 |
| gene-LOC105284116 | https://www.ncbi.nlm.nih.gov/gene/?term=LOC105284116 | 3970 | 96 | 241.81 | 1 | 2.52 | 1.04 | 6.8 | 6.72 | 0.08 | 0.08 | 0 | 0 |
| gene-LOC105278666 | https://www.ncbi.nlm.nih.gov/gene/?term=LOC105278666 | 10341 | 409 | 395.51 | 1 | 0.97 | 0.24 | 7.49 | 7.42 | 0.07 | 0.07 | 0 | 0 |
| gene-LOC105279528 | https://www.ncbi.nlm.nih.gov/gene/?term=LOC105279528 | 5802 | 46 | 79.28 | 1 | 1.72 | 2.17 | 15.76 | 15.72 | 0.04 | 0.04 | 0 | 0 |
| gene-LOC105280304 | https://www.ncbi.nlm.nih.gov/gene/?term=LOC105280304 | 8919 | 123 | 137.91 | 1 | 1.12 | 0.81 | 5.79 | 5.75 | 0.04 | 0.04 | 0 | 0 |
| gene-LOC105282914 | https://www.ncbi.nlm.nih.gov/gene/?term=LOC105282914 | 5623 | 42 | 74.69 | 1 | 1.78 | 2.38 | 48.6 | 48.57 | 0.03 | 0.03 | 0 | 0 |
| gene-LOC105274699 | https://www.ncbi.nlm.nih.gov/gene/?term=LOC105274699 | 2386 | 63 | 264.04 | 1 | 4.19 | 1.59 | 23.04 | 23.01 | 0.02 | 0.02 | 0 | 0 |
| gene-LOC105283296 | https://www.ncbi.nlm.nih.gov/gene/?term=LOC105283296 | 2174 | 89 | 409.38 | 1 | 4.6 | 1.12 | 19.56 | 19.55 | 0.01 | 0.01 | 0 | 0 |
| gene-LOC105276059 | https://www.ncbi.nlm.nih.gov/gene/?term=LOC105276059 | 6256 | 136 | 217.39 | 1 | 1.6 | 0.74 | 10.77 | 10.82 | -0.06 | 0.06 | 0 | 0 |
| gene-LOC105278538 | https://www.ncbi.nlm.nih.gov/gene/?term=LOC105278538 | 2150 | 83 | 386.05 | 1 | 4.65 | 1.2 | 18.97 | 19.05 | -0.08 | 0.08 | 0 | 0 |
| gene-LOC105284117 | https://www.ncbi.nlm.nih.gov/gene/?term=LOC105284117 | 4340 | 86 | 198.16 | 1 | 2.3 | 1.16 | 10.49 | 10.57 | -0.08 | 0.08 | 0 | 0 |
| gene-LOC105277761 | https://www.ncbi.nlm.nih.gov/gene/?term=LOC105277761 | 4648 | 150 | 322.72 | 1 | 2.15 | 0.67 | 14.73 | 14.87 | -0.13 | 0.13 | 0 | 0 |
| gene-LOC105284262 | https://www.ncbi.nlm.nih.gov/gene/?term=LOC105284262 | 6669 | 236 | 353.88 | 1 | 1.5 | 0.42 | 4.46 | 4.59 | -0.13 | 0.13 | 0 | 0 |
| gene-LOC105285395 | https://www.ncbi.nlm.nih.gov/gene/?term=LOC105285395 | 8085 | 187 | 231.29 | 1 | 1.24 | 0.53 | 16.28 | 16.44 | -0.16 | 0.16 | 0 | 0 |
| gene-LOC105275187 | https://www.ncbi.nlm.nih.gov/gene/?term=LOC105275187 | 5534 | 75 | 135.53 | 1 | 1.81 | 1.33 | 14.15 | 14.34 | -0.18 | 0.18 | 0 | 0 |

Directory: OGL File: rnk-dmg-Ob_LI_BCphaseA1.vs.RphaseB1.txt

| gene_ID | gene_link | gwidth | #Sites | #per10Kb | #dmSites | #dmsp10kb | %dmSites | %pSite1 | %pSite2 | DMpSite | ADMpSite | DMpNucl | ADMpNucl |
|---|---|---|---|---|---|---|---|---|---|---|---|---|---|
| gene-LOC113562627 | https://www.ncbi.nlm.nih.gov/gene/?term=LOC113562627 | 397 | 10 | 251.89 | 1 | 25.19 | 10 | 77.44 | 70.89 | 6.56 | 6.56 | 0.17 | 0.17 |
| gene-LOC105282405 | https://www.ncbi.nlm.nih.gov/gene/?term=LOC105282405 | 1310 | 31 | 236.64 | 2 | 15.27 | 6.45 | 51.18 | 44.54 | 6.64 | 6.64 | 0.16 | 0.16 |
| gene-LOC113561900 | https://www.ncbi.nlm.nih.gov/gene/?term=LOC113561900 | 629 | 10 | 158.98 | 2 | 31.8 | 20 | 21.61 | 31.23 | -9.62 | 9.62 | -0.15 | 0.15 |
| gene-LOC105288193 | https://www.ncbi.nlm.nih.gov/gene/?term=LOC105288193 | 1495 | 63 | 421.4 | 4 | 26.76 | 6.35 | 33.59 | 30.52 | 3.07 | 3.07 | 0.13 | 0.13 |
| gene-LOC105279335 | https://www.ncbi.nlm.nih.gov/gene/?term=LOC105279335 | 737 | 34 | 461.33 | 2 | 27.14 | 5.88 | 18.91 | 16.04 | 2.87 | 2.87 | 0.13 | 0.13 |
| gene-LOC113562894 | https://www.ncbi.nlm.nih.gov/gene/?term=LOC113562894 | 781 | 34 | 435.34 | 1 | 12.8 | 2.94 | 12.48 | 15.26 | -2.78 | 2.78 | -0.12 | 0.12 |
| gene-LOC105283861 | https://www.ncbi.nlm.nih.gov/gene/?term=LOC105283861 | 1298 | 19 | 146.38 | 3 | 23.11 | 15.79 | 35.76 | 43.84 | -8.08 | 8.08 | -0.12 | 0.12 |
| gene-LOC105278160 | https://www.ncbi.nlm.nih.gov/gene/?term=LOC105278160 | 1872 | 85 | 454.06 | 2 | 10.68 | 2.35 | 38.22 | 35.85 | 2.37 | 2.37 | 0.11 | 0.11 |
| gene-LOC105285213 | https://www.ncbi.nlm.nih.gov/gene/?term=LOC105285213 | 1491 | 37 | 248.16 | 4 | 26.83 | 10.81 | 13.74 | 18.03 | -4.29 | 4.29 | -0.11 | 0.11 |
| gene-LOC113561934 | https://www.ncbi.nlm.nih.gov/gene/?term=LOC113561934 | 1252 | 16 | 127.8 | 4 | 31.95 | 25 | 36.29 | 28.08 | 8.21 | 8.21 | 0.1 | 0.1 |
| gene-LOC105283973 | https://www.ncbi.nlm.nih.gov/gene/?term=LOC105283973 | 1536 | 29 | 188.8 | 4 | 26.04 | 13.79 | 61.88 | 56.34 | 5.55 | 5.55 | 0.1 | 0.1 |
| gene-LOC105275539 | https://www.ncbi.nlm.nih.gov/gene/?term=LOC105275539 | 1272 | 23 | 180.82 | 2 | 15.72 | 8.7 | 20.8 | 15.48 | 5.32 | 5.32 | 0.1 | 0.1 |
| gene-LOC105275761 | https://www.ncbi.nlm.nih.gov/gene/?term=LOC105275761 | 2635 | 101 | 383.3 | 6 | 22.77 | 5.94 | 29.65 | 26.96 | 2.69 | 2.69 | 0.1 | 0.1 |
| gene-LOC105274555 | https://www.ncbi.nlm.nih.gov/gene/?term=LOC105274555 | 1395 | 56 | 401.43 | 2 | 14.34 | 3.57 | 36.48 | 34.09 | 2.39 | 2.39 | 0.1 | 0.1 |
| gene-LOC105287692 | https://www.ncbi.nlm.nih.gov/gene/?term=LOC105287692 | 1505 | 99 | 657.81 | 3 | 19.93 | 3.03 | 29.69 | 28.16 | 1.52 | 1.52 | 0.1 | 0.1 |
| gene-LOC109610897 | https://www.ncbi.nlm.nih.gov/gene/?term=LOC109610897 | 1561 | 110 | 704.68 | 3 | 19.22 | 2.73 | 15.43 | 14.07 | 1.37 | 1.37 | 0.1 | 0.1 |
| gene-LOC105280710 | https://www.ncbi.nlm.nih.gov/gene/?term=LOC105280710 | 1215 | 69 | 567.9 | 5 | 41.15 | 7.25 | 20.73 | 22.43 | -1.7 | 1.7 | -0.1 | 0.1 |
| gene-LOC105286235 | https://www.ncbi.nlm.nih.gov/gene/?term=LOC105286235 | 1510 | 76 | 503.31 | 3 | 19.87 | 3.95 | 37.9 | 39.89 | -1.99 | 1.99 | -0.1 | 0.1 |
| gene-LOC105287465 | https://www.ncbi.nlm.nih.gov/gene/?term=LOC105287465 | 2477 | 92 | 371.42 | 7 | 28.26 | 7.61 | 40.81 | 43.43 | -2.62 | 2.62 | -0.1 | 0.1 |
| gene-LOC105278537 | https://www.ncbi.nlm.nih.gov/gene/?term=LOC105278537 | 1660 | 60 | 361.45 | 5 | 30.12 | 8.33 | 29.09 | 31.82 | -2.73 | 2.73 | -0.1 | 0.1 |
| gene-LOC105279087 | https://www.ncbi.nlm.nih.gov/gene/?term=LOC105279087 | 1077 | 28 | 259.98 | 1 | 9.29 | 3.57 | 53 | 56.81 | -3.81 | 3.81 | -0.1 | 0.1 |
| gene-LOC105276115 | https://www.ncbi.nlm.nih.gov/gene/?term=LOC105276115 | 1520 | 38 | 250 | 3 | 19.74 | 7.89 | 23.97 | 28.04 | -4.07 | 4.07 | -0.1 | 0.1 |
| gene-LOC105286358 | https://www.ncbi.nlm.nih.gov/gene/?term=LOC105286358 | 1489 | 54 | 362.66 | 3 | 20.15 | 5.56 | 34.15 | 31.79 | 2.36 | 2.36 | 0.09 | 0.09 |
| gene-LOC105280310 | https://www.ncbi.nlm.nih.gov/gene/?term=LOC105280310 | 1916 | 75 | 391.44 | 1 | 5.22 | 1.33 | 34.06 | 31.75 | 2.31 | 2.31 | 0.09 | 0.09 |

...

... only first 25 lines shown ...

Directory: OGL File: rnk-dmg-Ob_LI_BCphaseA1.vs.RphaseB2.txt

| gene_ID | gene_link | gwidth | #Sites | #per10Kb | #dmSites | #dmsp10kb | %dmSites | %pSite1 | %pSite2 | DMpSite | ADMpSite | DMpNucl | ADMpNucl |
|---|---|---|---|---|---|---|---|---|---|---|---|---|---|
| gene-LOC105282405 | https://www.ncbi.nlm.nih.gov/gene/?term=LOC105282405 | 1310 | 31 | 236.64 | 5 | 38.17 | 16.13 | 51.18 | 44.88 | 6.3 | 6.3 | 0.15 | 0.15 |
| gene-LOC105276585 | https://www.ncbi.nlm.nih.gov/gene/?term=LOC105276585 | 1534 | 113 | 736.64 | 1 | 6.52 | 0.88 | 7.26 | 5.41 | 1.85 | 1.85 | 0.14 | 0.14 |
| gene-LOC105285203 | https://www.ncbi.nlm.nih.gov/gene/?term=LOC105285203 | 1020 | 27 | 264.71 | 1 | 9.8 | 3.7 | 30.19 | 25.46 | 4.73 | 4.73 | 0.13 | 0.13 |
| gene-LOC105278849 | https://www.ncbi.nlm.nih.gov/gene/?term=LOC105278849 | 776 | 74 | 953.61 | 2 | 25.77 | 2.7 | 24.36 | 22.99 | 1.37 | 1.37 | 0.13 | 0.13 |
| gene-LOC105278266 | https://www.ncbi.nlm.nih.gov/gene/?term=LOC105278266 | 2442 | 108 | 442.26 | 4 | 16.38 | 3.7 | 25.68 | 28.72 | -3.05 | 3.05 | -0.13 | 0.13 |
| gene-LOC113562894 | https://www.ncbi.nlm.nih.gov/gene/?term=LOC113562894 | 781 | 34 | 435.34 | 2 | 25.61 | 5.88 | 11.76 | 14.52 | -2.76 | 2.76 | -0.12 | 0.12 |
| gene-LOC105278458 | https://www.ncbi.nlm.nih.gov/gene/?term=LOC105278458 | 216 | 3 | 138.89 | 1 | 46.3 | 33.33 | 20.87 | 29.54 | -8.67 | 8.67 | -0.12 | 0.12 |
| gene-LOC113561934 | https://www.ncbi.nlm.nih.gov/gene/?term=LOC113561934 | 1252 | 15 | 119.81 | 2 | 15.97 | 13.33 | 36.71 | 27.17 | 9.54 | 9.54 | 0.11 | 0.11 |
| gene-LOC105278234 | https://www.ncbi.nlm.nih.gov/gene/?term=LOC105278234 | 2068 | 106 | 512.57 | 1 | 4.84 | 0.94 | 10.74 | 8.52 | 2.21 | 2.21 | 0.11 | 0.11 |
| gene-LOC105278873 | https://www.ncbi.nlm.nih.gov/gene/?term=LOC105278873 | 1897 | 143 | 753.82 | 4 | 21.09 | 2.8 | 19.98 | 21.44 | -1.47 | 1.47 | -0.11 | 0.11 |
| gene-LOC105280710 | https://www.ncbi.nlm.nih.gov/gene/?term=LOC105280710 | 1215 | 64 | 526.75 | 1 | 8.23 | 1.56 | 22.3 | 24.34 | -2.04 | 2.04 | -0.11 | 0.11 |
| gene-LOC105287856 | https://www.ncbi.nlm.nih.gov/gene/?term=LOC105287856 | 2090 | 113 | 540.67 | 6 | 28.71 | 5.31 | 12.28 | 14.37 | -2.09 | 2.09 | -0.11 | 0.11 |
| gene-LOC105278537 | https://www.ncbi.nlm.nih.gov/gene/?term=LOC105278537 | 1660 | 62 | 373.49 | 1 | 6.02 | 1.61 | 29.42 | 32.25 | -2.83 | 2.83 | -0.11 | 0.11 |
| gene-LOC105279863 | https://www.ncbi.nlm.nih.gov/gene/?term=LOC105279863 | 1204 | 18 | 149.5 | 4 | 33.22 | 22.22 | 11.07 | 18.73 | -7.66 | 7.66 | -0.11 | 0.11 |
| gene-LOC105278824 | https://www.ncbi.nlm.nih.gov/gene/?term=LOC105278824 | 1230 | 61 | 495.93 | 1 | 8.13 | 1.64 | 24.95 | 22.9 | 2.05 | 2.05 | 0.1 | 0.1 |
| gene-LOC105279346 | https://www.ncbi.nlm.nih.gov/gene/?term=LOC105279346 | 4649 | 229 | 492.58 | 9 | 19.36 | 3.93 | 14.52 | 16.49 | -1.97 | 1.97 | -0.1 | 0.1 |
| gene-LOC105277398 | https://www.ncbi.nlm.nih.gov/gene/?term=LOC105277398 | 1963 | 59 | 300.56 | 5 | 25.47 | 8.47 | 36.03 | 39.34 | -3.31 | 3.31 | -0.1 | 0.1 |
| gene-LOC105285207 | https://www.ncbi.nlm.nih.gov/gene/?term=LOC105285207 | 3030 | 64 | 211.22 | 5 | 16.5 | 7.81 | 20.3 | 25.24 | -4.94 | 4.94 | -0.1 | 0.1 |
| gene-LOC105277810 | https://www.ncbi.nlm.nih.gov/gene/?term=LOC105277810 | 1367 | 21 | 153.62 | 1 | 7.32 | 4.76 | 45.33 | 39.52 | 5.81 | 5.81 | 0.09 | 0.09 |
| gene-LOC105286644 | https://www.ncbi.nlm.nih.gov/gene/?term=LOC105286644 | 1235 | 21 | 170.04 | 1 | 8.1 | 4.76 | 44.21 | 39.04 | 5.17 | 5.17 | 0.09 | 0.09 |
| gene-LOC105276679 | https://www.ncbi.nlm.nih.gov/gene/?term=LOC105276679 | 2005 | 67 | 334.16 | 1 | 4.99 | 1.49 | 37.86 | 35.03 | 2.83 | 2.83 | 0.09 | 0.09 |
| gene-LOC105279487 | https://www.ncbi.nlm.nih.gov/gene/?term=LOC105279487 | 2117 | 82 | 387.34 | 2 | 9.45 | 2.44 | 17.82 | 15.61 | 2.21 | 2.21 | 0.09 | 0.09 |
| gene-LOC105288104 | https://www.ncbi.nlm.nih.gov/gene/?term=LOC105288104 | 2069 | 92 | 444.66 | 2 | 9.67 | 2.17 | 27.34 | 25.28 | 2.06 | 2.06 | 0.09 | 0.09 |
| gene-LOC105280786 | https://www.ncbi.nlm.nih.gov/gene/?term=LOC105280786 | 2573 | 115 | 446.95 | 1 | 3.89 | 0.87 | 30.29 | 28.35 | 1.94 | 1.94 | 0.09 | 0.09 |

...

... only first 25 lines shown ...

Directory: OGL File: rnk-dmg-Ob_LI_BCphaseA2.vs.BCphaseB1.txt

| gene_ID | gene_link | gwidth | #Sites | #per10Kb | #dmSites | #dmsp10kb | %dmSites | %pSite1 | %pSite2 | DMpSite | ADMpSite | DMpNucl | ADMpNucl |
|---|---|---|---|---|---|---|---|---|---|---|---|---|---|
| gene-LOC105275384 | https://www.ncbi.nlm.nih.gov/gene/?term=LOC105275384 | 1987 | 91 | 457.98 | 1 | 5.03 | 1.1 | 33.74 | 32.35 | 1.39 | 1.39 | 0.06 | 0.06 |
| gene-LOC105276429 | https://www.ncbi.nlm.nih.gov/gene/?term=LOC105276429 | 2901 | 120 | 413.65 | 1 | 3.45 | 0.83 | 23.61 | 22.3 | 1.32 | 1.32 | 0.05 | 0.05 |
| gene-LOC105283264 | https://www.ncbi.nlm.nih.gov/gene/?term=LOC105283264 | 4005 | 88 | 219.73 | 1 | 2.5 | 1.14 | 18.85 | 20.42 | -1.57 | 1.57 | -0.03 | 0.03 |
| gene-LOC105281272 | https://www.ncbi.nlm.nih.gov/gene/?term=LOC105281272 | 5252 | 178 | 338.92 | 1 | 1.9 | 0.56 | 16.03 | 16.73 | -0.7 | 0.7 | -0.02 | 0.02 |
| gene-LOC105288042 | https://www.ncbi.nlm.nih.gov/gene/?term=LOC105288042 | 3507 | 101 | 288 | 1 | 2.85 | 0.99 | 22.48 | 23.29 | -0.81 | 0.81 | -0.02 | 0.02 |
| gene-LOC105285252 | https://www.ncbi.nlm.nih.gov/gene/?term=LOC105285252 | 7427 | 129 | 173.69 | 1 | 1.35 | 0.78 | 12.88 | 13.9 | -1.02 | 1.02 | -0.02 | 0.02 |
| gene-LOC105277903 | https://www.ncbi.nlm.nih.gov/gene/?term=LOC105277903 | 1835 | 35 | 190.74 | 1 | 5.45 | 2.86 | 47.23 | 46.95 | 0.28 | 0.28 | 0.01 | 0.01 |
| gene-LOC105283296 | https://www.ncbi.nlm.nih.gov/gene/?term=LOC105283296 | 2174 | 92 | 423.18 | 1 | 4.6 | 1.09 | 19.15 | 18.97 | 0.18 | 0.18 | 0.01 | 0.01 |
| gene-LOC105276452 | https://www.ncbi.nlm.nih.gov/gene/?term=LOC105276452 | 2917 | 262 | 898.18 | 1 | 3.43 | 0.38 | 10.28 | 10.37 | -0.08 | 0.08 | -0.01 | 0.01 |
| gene-LOC105276385 | https://www.ncbi.nlm.nih.gov/gene/?term=LOC105276385 | 11575 | 494 | 426.78 | 1 | 0.86 | 0.2 | 5.31 | 5.56 | -0.25 | 0.25 | -0.01 | 0.01 |
| gene-LOC105284507 | https://www.ncbi.nlm.nih.gov/gene/?term=LOC105284507 | 5207 | 94 | 180.53 | 1 | 1.92 | 1.06 | 20.48 | 21.18 | -0.7 | 0.7 | -0.01 | 0.01 |
| gene-LOC105280410 | https://www.ncbi.nlm.nih.gov/gene/?term=LOC105280410 | 12580 | 280 | 222.58 | 1 | 0.79 | 0.36 | 19.42 | 19.27 | 0.15 | 0.15 | 0 | 0 |
| gene-LOC105288036 | https://www.ncbi.nlm.nih.gov/gene/?term=LOC105288036 | 5110 | 226 | 442.27 | 1 | 1.96 | 0.44 | 16.3 | 16.4 | -0.1 | 0.1 | 0 | 0 |

Directory: OGL File: rnk-dmg-Ob_LI_BCphaseA2.vs.BCphaseB2.txt

| gene_ID | gene_link | gwidth | #Sites | #per10Kb | #dmSites | #dmsp10kb | %dmSites | %pSite1 | %pSite2 | DMpSite | ADMpSite | DMpNucl | ADMpNucl |
|---|---|---|---|---|---|---|---|---|---|---|---|---|---|
| gene-LOC113562345 | https://www.ncbi.nlm.nih.gov/gene/?term=LOC113562345 | 2290 | 69 | 301.31 | 17 | 74.24 | 24.64 | 33.16 | 16.9 | 16.26 | 16.26 | 0.49 | 0.49 |
| gene-LOC105284677 | https://www.ncbi.nlm.nih.gov/gene/?term=LOC105284677 | 1905 | 41 | 215.22 | 2 | 10.5 | 4.88 | 27.55 | 36.9 | -9.35 | 9.35 | -0.2 | 0.2 |
| gene-LOC105282976 | https://www.ncbi.nlm.nih.gov/gene/?term=LOC105282976 | 1648 | 50 | 303.4 | 1 | 6.07 | 2 | 26.51 | 20.9 | 5.61 | 5.61 | 0.17 | 0.17 |
| gene-LOC105282149 | https://www.ncbi.nlm.nih.gov/gene/?term=LOC105282149 | 1787 | 108 | 604.36 | 1 | 5.6 | 0.93 | 43.65 | 41.49 | 2.17 | 2.17 | 0.13 | 0.13 |
| gene-LOC105286683 | https://www.ncbi.nlm.nih.gov/gene/?term=LOC105286683 | 1665 | 51 | 306.31 | 1 | 6.01 | 1.96 | 35.98 | 32.15 | 3.83 | 3.83 | 0.12 | 0.12 |
| gene-LOC105282156 | https://www.ncbi.nlm.nih.gov/gene/?term=LOC105282156 | 2577 | 136 | 527.75 | 1 | 3.88 | 0.74 | 26.99 | 24.63 | 2.36 | 2.36 | 0.12 | 0.12 |
| gene-LOC105283649 | https://www.ncbi.nlm.nih.gov/gene/?term=LOC105283649 | 1918 | 56 | 291.97 | 2 | 10.43 | 3.57 | 38.44 | 34.8 | 3.63 | 3.63 | 0.11 | 0.11 |
| gene-LOC105276078 | https://www.ncbi.nlm.nih.gov/gene/?term=LOC105276078 | 1051 | 16 | 152.24 | 1 | 9.51 | 6.25 | 32.79 | 26.49 | 6.3 | 6.3 | 0.1 | 0.1 |
| gene-LOC105275447 | https://www.ncbi.nlm.nih.gov/gene/?term=LOC105275447 | 1846 | 35 | 189.6 | 1 | 5.42 | 2.86 | 8.76 | 3.6 | 5.15 | 5.15 | 0.1 | 0.1 |
| gene-LOC105285674 | https://www.ncbi.nlm.nih.gov/gene/?term=LOC105285674 | 3035 | 93 | 306.43 | 2 | 6.59 | 2.15 | 27.41 | 24.3 | 3.11 | 3.11 | 0.1 | 0.1 |
| gene-LOC105275070 | https://www.ncbi.nlm.nih.gov/gene/?term=LOC105275070 | 5667 | 47 | 82.94 | 5 | 8.82 | 10.64 | 11.7 | 2 | 9.7 | 9.7 | 0.08 | 0.08 |
| gene-LOC105276449 | https://www.ncbi.nlm.nih.gov/gene/?term=LOC105276449 | 1231 | 42 | 341.19 | 1 | 8.12 | 2.38 | 35.95 | 33.63 | 2.31 | 2.31 | 0.08 | 0.08 |
| gene-LOC105279898 | https://www.ncbi.nlm.nih.gov/gene/?term=LOC105279898 | 1343 | 14 | 104.24 | 1 | 7.45 | 7.14 | 61.13 | 54.89 | 6.24 | 6.24 | 0.07 | 0.07 |
| gene-LOC113561805 | https://www.ncbi.nlm.nih.gov/gene/?term=LOC113561805 | 3337 | 77 | 230.75 | 1 | 3 | 1.3 | 37.31 | 34.13 | 3.18 | 3.18 | 0.07 | 0.07 |
| gene-LOC105275708 | https://www.ncbi.nlm.nih.gov/gene/?term=LOC105275708 | 1772 | 10 | 56.43 | 1 | 5.64 | 10 | 50.31 | 39.56 | 10.75 | 10.75 | 0.06 | 0.06 |
| gene-LOC105288063 | https://www.ncbi.nlm.nih.gov/gene/?term=LOC105288063 | 3400 | 53 | 155.88 | 1 | 2.94 | 1.89 | 16.83 | 13.02 | 3.81 | 3.81 | 0.06 | 0.06 |
| gene-LOC105280774 | https://www.ncbi.nlm.nih.gov/gene/?term=LOC105280774 | 2793 | 102 | 365.2 | 1 | 3.58 | 0.98 | 32.11 | 30.36 | 1.75 | 1.75 | 0.06 | 0.06 |
| gene-LOC105280786 | https://www.ncbi.nlm.nih.gov/gene/?term=LOC105280786 | 2573 | 111 | 431.4 | 1 | 3.89 | 0.9 | 33.28 | 31.87 | 1.41 | 1.41 | 0.06 | 0.06 |
| gene-LOC105284502 | https://www.ncbi.nlm.nih.gov/gene/?term=LOC105284502 | 2157 | 19 | 88.09 | 1 | 4.64 | 5.26 | 36.83 | 43.93 | -7.1 | 7.1 | -0.06 | 0.06 |
| gene-LOC105278454 | https://www.ncbi.nlm.nih.gov/gene/?term=LOC105278454 | 1972 | 47 | 238.34 | 1 | 5.07 | 2.13 | 44.8 | 42.52 | 2.28 | 2.28 | 0.05 | 0.05 |
| gene-LOC105282970 | https://www.ncbi.nlm.nih.gov/gene/?term=LOC105282970 | 1529 | 39 | 255.07 | 1 | 6.54 | 2.56 | 54 | 52.14 | 1.86 | 1.86 | 0.05 | 0.05 |
| gene-LOC105282151 | https://www.ncbi.nlm.nih.gov/gene/?term=LOC105282151 | 1601 | 41 | 256.09 | 3 | 18.74 | 7.32 | 36.24 | 34.41 | 1.83 | 1.83 | 0.05 | 0.05 |
| gene-LOC105281848 | https://www.ncbi.nlm.nih.gov/gene/?term=LOC105281848 | 2122 | 56 | 263.9 | 1 | 4.71 | 1.79 | 24.08 | 22.28 | 1.8 | 1.8 | 0.05 | 0.05 |
| gene-LOC105287095 | https://www.ncbi.nlm.nih.gov/gene/?term=LOC105287095 | 1435 | 57 | 397.21 | 1 | 6.97 | 1.75 | 29.87 | 28.54 | 1.33 | 1.33 | 0.05 | 0.05 |
| gene-LOC105286190 | https://www.ncbi.nlm.nih.gov/gene/?term=LOC105286190 | 6447 | 298 | 462.23 | 1 | 1.55 | 0.34 | 8.47 | 7.3 | 1.16 | 1.16 | 0.05 | 0.05 |
| gene-LOC105279298 | https://www.ncbi.nlm.nih.gov/gene/?term=LOC105279298 | 2773 | 128 | 461.59 | 2 | 7.21 | 1.56 | 25.02 | 23.92 | 1.1 | 1.1 | 0.05 | 0.05 |
| gene-LOC105278497 | https://www.ncbi.nlm.nih.gov/gene/?term=LOC105278497 | 2072 | 94 | 453.67 | 1 | 4.83 | 1.06 | 21.1 | 20.09 | 1.01 | 1.01 | 0.05 | 0.05 |
| gene-LOC105276704 | https://www.ncbi.nlm.nih.gov/gene/?term=LOC105276704 | 2063 | 98 | 475.04 | 1 | 4.85 | 1.02 | 13.8 | 12.82 | 0.98 | 0.98 | 0.05 | 0.05 |
| gene-LOC105285055 | https://www.ncbi.nlm.nih.gov/gene/?term=LOC105285055 | 4108 | 259 | 630.48 | 2 | 4.87 | 0.77 | 11.6 | 10.8 | 0.8 | 0.8 | 0.05 | 0.05 |
| gene-LOC105274449 | https://www.ncbi.nlm.nih.gov/gene/?term=LOC105274449 | 1549 | 53 | 342.16 | 1 | 6.46 | 1.89 | 50.38 | 51.77 | -1.39 | 1.39 | -0.05 | 0.05 |
| gene-LOC105274900 | https://www.ncbi.nlm.nih.gov/gene/?term=LOC105274900 | 5561 | 52 | 93.51 | 2 | 3.6 | 3.85 | 36.59 | 32.11 | 4.48 | 4.48 | 0.04 | 0.04 |
| gene-LOC105281420 | https://www.ncbi.nlm.nih.gov/gene/?term=LOC105281420 | 3882 | 55 | 141.68 | 1 | 2.58 | 1.82 | 15.69 | 12.96 | 2.73 | 2.73 | 0.04 | 0.04 |
| gene-LOC105280641 | https://www.ncbi.nlm.nih.gov/gene/?term=LOC105280641 | 3682 | 49 | 133.08 | 1 | 2.72 | 2.04 | 20.13 | 17.42 | 2.71 | 2.71 | 0.04 | 0.04 |
| gene-LOC105275966 | https://www.ncbi.nlm.nih.gov/gene/?term=LOC105275966 | 3299 | 61 | 184.9 | 1 | 3.03 | 1.64 | 49.58 | 47.22 | 2.36 | 2.36 | 0.04 | 0.04 |
| gene-LOC105278109 | https://www.ncbi.nlm.nih.gov/gene/?term=LOC105278109 | 4029 | 65 | 161.33 | 1 | 2.48 | 1.54 | 48.15 | 45.97 | 2.19 | 2.19 | 0.04 | 0.04 |
| gene-LOC105283592 | https://www.ncbi.nlm.nih.gov/gene/?term=LOC105283592 | 3479 | 71 | 204.08 | 1 | 2.87 | 1.41 | 26.49 | 24.49 | 2.01 | 2.01 | 0.04 | 0.04 |
| gene-LOC105282658 | https://www.ncbi.nlm.nih.gov/gene/?term=LOC105282658 | 2421 | 65 | 268.48 | 1 | 4.13 | 1.54 | 21.38 | 19.92 | 1.46 | 1.46 | 0.04 | 0.04 |
| gene-LOC105277761 | https://www.ncbi.nlm.nih.gov/gene/?term=LOC105277761 | 4648 | 146 | 314.11 | 1 | 2.15 | 0.68 | 16.93 | 15.61 | 1.32 | 1.32 | 0.04 | 0.04 |
| gene-LOC105274856 | https://www.ncbi.nlm.nih.gov/gene/?term=LOC105274856 | 5216 | 218 | 417.94 | 1 | 1.92 | 0.46 | 19.03 | 18.03 | 1 | 1 | 0.04 | 0.04 |
| gene-LOC105284355 | https://www.ncbi.nlm.nih.gov/gene/?term=LOC105284355 | 3904 | 149 | 381.66 | 1 | 2.56 | 0.67 | 21.89 | 20.9 | 0.99 | 0.99 | 0.04 | 0.04 |
| gene-LOC105278877 | https://www.ncbi.nlm.nih.gov/gene/?term=LOC105278877 | 1968 | 75 | 381.1 | 1 | 5.08 | 1.33 | 34.6 | 33.65 | 0.95 | 0.95 | 0.04 | 0.04 |
| gene-LOC105280148 | https://www.ncbi.nlm.nih.gov/gene/?term=LOC105280148 | 3120 | 214 | 685.9 | 1 | 3.21 | 0.47 | 11.76 | 11.17 | 0.6 | 0.6 | 0.04 | 0.04 |
| gene-LOC105274706 | https://www.ncbi.nlm.nih.gov/gene/?term=LOC105274706 | 1446 | 55 | 380.36 | 1 | 6.92 | 1.82 | 37.25 | 38.23 | -0.98 | 0.98 | -0.04 | 0.04 |
| gene-LOC105286694 | https://www.ncbi.nlm.nih.gov/gene/?term=LOC105286694 | 3409 | 11 | 32.27 | 1 | 2.93 | 9.09 | 63.21 | 54.71 | 8.5 | 8.5 | 0.03 | 0.03 |
| gene-LOC105287648 | https://www.ncbi.nlm.nih.gov/gene/?term=LOC105287648 | 2205 | 27 | 122.45 | 1 | 4.54 | 3.7 | 41.48 | 39.19 | 2.29 | 2.29 | 0.03 | 0.03 |
| gene-LOC105286514 | https://www.ncbi.nlm.nih.gov/gene/?term=LOC105286514 | 2528 | 41 | 162.18 | 1 | 3.96 | 2.44 | 35.27 | 33.32 | 1.95 | 1.95 | 0.03 | 0.03 |
| gene-LOC105286644 | https://www.ncbi.nlm.nih.gov/gene/?term=LOC105286644 | 1235 | 20 | 161.94 | 1 | 8.1 | 5 | 39.34 | 37.53 | 1.81 | 1.81 | 0.03 | 0.03 |
| gene-LOC105278807 | https://www.ncbi.nlm.nih.gov/gene/?term=LOC105278807 | 2166 | 73 | 337.03 | 1 | 4.62 | 1.37 | 18.28 | 17.29 | 0.99 | 0.99 | 0.03 | 0.03 |
| gene-LOC105284184 | https://www.ncbi.nlm.nih.gov/gene/?term=LOC105284184 | 2535 | 85 | 335.31 | 1 | 3.94 | 1.18 | 34.37 | 33.53 | 0.83 | 0.83 | 0.03 | 0.03 |
| gene-LOC105285202 | https://www.ncbi.nlm.nih.gov/gene/?term=LOC105285202 | 1838 | 66 | 359.09 | 1 | 5.44 | 1.52 | 25.35 | 24.58 | 0.77 | 0.77 | 0.03 | 0.03 |
| gene-LOC105274452 | https://www.ncbi.nlm.nih.gov/gene/?term=LOC105274452 | 6595 | 261 | 395.75 | 1 | 1.52 | 0.38 | 13.63 | 12.94 | 0.69 | 0.69 | 0.03 | 0.03 |
| gene-LOC105286799 | https://www.ncbi.nlm.nih.gov/gene/?term=LOC105286799 | 3557 | 183 | 514.48 | 1 | 2.81 | 0.55 | 16.25 | 15.68 | 0.57 | 0.57 | 0.03 | 0.03 |
| gene-LOC105280723 | https://www.ncbi.nlm.nih.gov/gene/?term=LOC105280723 | 3572 | 170 | 475.92 | 1 | 2.8 | 0.59 | 16.88 | 16.35 | 0.53 | 0.53 | 0.03 | 0.03 |
| gene-LOC105274534 | https://www.ncbi.nlm.nih.gov/gene/?term=LOC105274534 | 1329 | 54 | 406.32 | 1 | 7.52 | 1.85 | 45.08 | 45.82 | -0.75 | 0.75 | -0.03 | 0.03 |
| gene-LOC105284507 | https://www.ncbi.nlm.nih.gov/gene/?term=LOC105284507 | 5207 | 76 | 145.96 | 1 | 1.92 | 1.32 | 19.78 | 21.69 | -1.91 | 1.91 | -0.03 | 0.03 |
| gene-LOC105286302 | https://www.ncbi.nlm.nih.gov/gene/?term=LOC105286302 | 4536 | 18 | 39.68 | 1 | 2.2 | 5.56 | 22.71 | 17.52 | 5.2 | 5.2 | 0.02 | 0.02 |
| gene-LOC105281515 | https://www.ncbi.nlm.nih.gov/gene/?term=LOC105281515 | 8827 | 48 | 54.38 | 2 | 2.27 | 4.17 | 32.1 | 29.31 | 2.79 | 2.79 | 0.02 | 0.02 |
| gene-LOC105288066 | https://www.ncbi.nlm.nih.gov/gene/?term=LOC105288066 | 1964 | 21 | 106.92 | 1 | 5.09 | 4.76 | 46.21 | 43.96 | 2.25 | 2.25 | 0.02 | 0.02 |
| gene-LOC105275518 | https://www.ncbi.nlm.nih.gov/gene/?term=LOC105275518 | 2335 | 19 | 81.37 | 1 | 4.28 | 5.26 | 46.5 | 44.26 | 2.24 | 2.24 | 0.02 | 0.02 |
| gene-LOC105277856 | https://www.ncbi.nlm.nih.gov/gene/?term=LOC105277856 | 3442 | 34 | 98.78 | 1 | 2.91 | 2.94 | 37.69 | 35.88 | 1.81 | 1.81 | 0.02 | 0.02 |

| | | | | | | | | | | | | | | |
|---|---|---|---|---|---|---|---|---|---|---|---|---|---|---|
| gene-LOC105288010 | https://www.ncbi.nlm.nih.gov/gene/?term=LOC105288010 | 3189 | 39 | 122.3 | 1 | 3.14 | 2.56 | 37.68 | 36.3 | 1.38 | 1.38 | 0.02 | 0.02 | |
| gene-LOC105280348 | https://www.ncbi.nlm.nih.gov/gene/?term=LOC105280348 | 3797 | 65 | 171.19 | | 1 | 2.63 | 1.54 | 21.44 | 20.1 | 1.33 | 1.33 | 0.02 | 0.02 |
| gene-LOC105282600 | https://www.ncbi.nlm.nih.gov/gene/?term=LOC105282600 | 3715 | 53 | 142.66 | | 1 | 2.69 | 1.89 | 32.07 | 30.77 | 1.3 | 1.3 | 0.02 | 0.02 |
| gene-LOC105276059 | https://www.ncbi.nlm.nih.gov/gene/?term=LOC105276059 | 6256 | 121 | 193.41 | | 1 | 1.6 | 0.83 | 11.92 | 10.67 | 1.25 | 1.25 | 0.02 | 0.02 |
| gene-LOC105279664 | https://www.ncbi.nlm.nih.gov/gene/?term=LOC105279664 | 4023 | 54 | 134.23 | | 1 | 2.49 | 1.85 | 21.29 | 20.13 | 1.15 | 1.15 | 0.02 | 0.02 |
| gene-LOC105285502 | https://www.ncbi.nlm.nih.gov/gene/?term=LOC105285502 | 9990 | 143 | 143.14 | | 1 | 1 | 0.7 | 10.37 | 9.32 | 1.05 | 1.05 | 0.02 | 0.02 |
| gene-LOC105282794 | https://www.ncbi.nlm.nih.gov/gene/?term=LOC105282794 | 4571 | 93 | 203.46 | | 1 | 2.19 | 1.08 | 18.93 | 18.03 | 0.9 | 0.9 | 0.02 | 0.02 |
| gene-LOC105275451 | https://www.ncbi.nlm.nih.gov/gene/?term=LOC105275451 | 6035 | 127 | 210.44 | | 1 | 1.66 | 0.79 | 22.96 | 22.23 | 0.73 | 0.73 | 0.02 | 0.02 |
| gene-LOC105280956 | https://www.ncbi.nlm.nih.gov/gene/?term=LOC105280956 | 2639 | 75 | 284.2 | 1 | 3.79 | 1.33 | 17.09 | 16.45 | 0.64 | 0.64 | 0.02 | 0.02 | |
| gene-LOC105279038 | https://www.ncbi.nlm.nih.gov/gene/?term=LOC105279038 | 9453 | 333 | 352.27 | | 1 | 1.06 | 0.3 | 9.29 | 8.67 | 0.62 | 0.62 | 0.02 | 0.02 |
| gene-LOC105287063 | https://www.ncbi.nlm.nih.gov/gene/?term=LOC105287063 | 1874 | 48 | 256.14 | | 1 | 5.34 | 2.08 | 30.52 | 29.9 | 0.62 | 0.62 | 0.02 | 0.02 |
| gene-LOC105274823 | https://www.ncbi.nlm.nih.gov/gene/?term=LOC105274823 | 4620 | 138 | 298.7 | 1 | 2.16 | 0.72 | 21.48 | 20.88 | 0.6 | 0.6 | 0.02 | 0.02 | |
| gene-LOC105285807 | https://www.ncbi.nlm.nih.gov/gene/?term=LOC105285807 | 6063 | 218 | 359.56 | | 1 | 1.65 | 0.46 | 7.08 | 6.5 | 0.58 | 0.58 | 0.02 | 0.02 |
| gene-LOC105275683 | https://www.ncbi.nlm.nih.gov/gene/?term=LOC105275683 | 4256 | 194 | 455.83 | | 1 | 2.35 | 0.52 | 13.24 | 12.7 | 0.54 | 0.54 | 0.02 | 0.02 |
| gene-LOC105278666 | https://www.ncbi.nlm.nih.gov/gene/?term=LOC105278666 | 10341 | 459 | 443.86 | | 1 | 0.97 | 0.22 | 7.32 | 6.79 | 0.53 | 0.53 | 0.02 | 0.02 |
| gene-LOC105285179 | https://www.ncbi.nlm.nih.gov/gene/?term=LOC105285179 | 4191 | 164 | 391.31 | | 1 | 2.39 | 0.61 | 20.24 | 19.71 | 0.53 | 0.53 | 0.02 | 0.02 |
| gene-LOC105283123 | https://www.ncbi.nlm.nih.gov/gene/?term=LOC105283123 | 1507 | 52 | 345.06 | | 1 | 6.64 | 1.92 | 17.86 | 17.35 | 0.52 | 0.52 | 0.02 | 0.02 |
| gene-LOC105285639 | https://www.ncbi.nlm.nih.gov/gene/?term=LOC105285639 | 10325 | 356 | 344.79 | | 1 | 0.97 | 0.28 | 11.26 | 10.77 | 0.49 | 0.49 | 0.02 | 0.02 |
| gene-LOC105275848 | https://www.ncbi.nlm.nih.gov/gene/?term=LOC105275848 | 3718 | 139 | 373.86 | | 1 | 2.69 | 0.72 | 22.47 | 21.99 | 0.48 | 0.48 | 0.02 | 0.02 |
| gene-LOC105278145 | https://www.ncbi.nlm.nih.gov/gene/?term=LOC105278145 | 4470 | 241 | 539.15 | | 1 | 2.24 | 0.41 | 10.99 | 10.59 | 0.4 | 0.4 | 0.02 | 0.02 |
| gene-LOC105276452 | https://www.ncbi.nlm.nih.gov/gene/?term=LOC105276452 | 2917 | 257 | 881.04 | | 1 | 3.43 | 0.39 | 9.59 | 9.37 | 0.22 | 0.22 | 0.02 | 0.02 |
| gene-LOC105286274 | https://www.ncbi.nlm.nih.gov/gene/?term=LOC105286274 | 3817 | 202 | 529.21 | | 1 | 2.62 | 0.5 | 16.44 | 16.87 | -0.43 | 0.43 | -0.02 | 0.02 |
| gene-LOC105283385 | https://www.ncbi.nlm.nih.gov/gene/?term=LOC105283385 | 9540 | 305 | 319.71 | | 1 | 1.05 | 0.33 | 12.66 | 13.17 | -0.51 | 0.51 | -0.02 | 0.02 |
| gene-LOC113561621 | https://www.ncbi.nlm.nih.gov/gene/?term=LOC113561621 | 2249 | 78 | 346.82 | | 1 | 4.45 | 1.28 | 20.85 | 21.38 | -0.53 | 0.53 | -0.02 | 0.02 |
| gene-LOC105285742 | https://www.ncbi.nlm.nih.gov/gene/?term=LOC105285742 | 3446 | 107 | 310.5 | 1 | 2.9 | 0.93 | 21.42 | 22.06 | -0.64 | 0.64 | -0.02 | 0.02 | |
| gene-LOC109611283 | https://www.ncbi.nlm.nih.gov/gene/?term=LOC109611283 | 1451 | 38 | 261.89 | | 1 | 6.89 | 2.63 | 35.88 | 36.65 | -0.77 | 0.77 | -0.02 | 0.02 |
| gene-LOC105285252 | https://www.ncbi.nlm.nih.gov/gene/?term=LOC105285252 | 7427 | 117 | 157.53 | | 1 | 1.35 | 0.85 | 13.64 | 15.21 | -1.57 | 1.57 | -0.02 | 0.02 |
| gene-LOC105279500 | https://www.ncbi.nlm.nih.gov/gene/?term=LOC105279500 | 3587 | 33 | 92 | 1 | 2.79 | 3.03 | 44.93 | 47.01 | -2.08 | 2.08 | -0.02 | 0.02 | |
| gene-LOC105282067 | https://www.ncbi.nlm.nih.gov/gene/?term=LOC105282067 | 5941 | 44 | 74.06 | 1 | 1.68 | 2.27 | 26.18 | 28.47 | -2.29 | 2.29 | -0.02 | 0.02 | |
| gene-LOC105285315 | https://www.ncbi.nlm.nih.gov/gene/?term=LOC105285315 | 5458 | 22 | 40.31 | 1 | 1.83 | 4.55 | 41.52 | 38.4 | 3.12 | 3.12 | 0.01 | 0.01 | |
| gene-LOC105275347 | https://www.ncbi.nlm.nih.gov/gene/?term=LOC105275347 | 2526 | 15 | 59.38 | 1 | 3.96 | 6.67 | 31.33 | 28.89 | 2.44 | 2.44 | 0.01 | 0.01 | |
| gene-LOC105288078 | https://www.ncbi.nlm.nih.gov/gene/?term=LOC105288078 | 19126 | 294 | 153.72 | | 1 | 0.52 | 0.34 | 18.24 | 17.37 | 0.87 | 0.87 | 0.01 | 0.01 |
| gene-LOC105284329 | https://www.ncbi.nlm.nih.gov/gene/?term=LOC105284329 | 4051 | 50 | 123.43 | | 1 | 2.47 | 2 | 19.81 | 19.02 | 0.79 | 0.79 | 0.01 | 0.01 |
| gene-LOC105276538 | https://www.ncbi.nlm.nih.gov/gene/?term=LOC105276538 | 1511 | 19 | 125.74 | | 1 | 6.62 | 5.26 | 70.24 | 69.47 | 0.77 | 0.77 | 0.01 | 0.01 |
| gene-LOC105280002 | https://www.ncbi.nlm.nih.gov/gene/?term=LOC105280002 | 1940 | 25 | 128.87 | | 1 | 5.15 | 4 | 38.59 | 38.02 | 0.58 | 0.58 | 0.01 | 0.01 |
| gene-LOC105274851 | https://www.ncbi.nlm.nih.gov/gene/?term=LOC105274851 | 6616 | 128 | 193.47 | | 1 | 1.51 | 0.78 | 11.71 | 11.16 | 0.54 | 0.54 | 0.01 | 0.01 |
| gene-LOC105283943 | https://www.ncbi.nlm.nih.gov/gene/?term=LOC105283943 | 3559 | 58 | 162.97 | | 1 | 2.81 | 1.72 | 14.89 | 14.35 | 0.54 | 0.54 | 0.01 | 0.01 |
| gene-LOC105280410 | https://www.ncbi.nlm.nih.gov/gene/?term=LOC105280410 | 12580 | 237 | 188.39 | | 1 | 0.79 | 0.42 | 21.06 | 20.59 | 0.47 | 0.47 | 0.01 | 0.01 |
| gene-LOC105279657 | https://www.ncbi.nlm.nih.gov/gene/?term=LOC105279657 | 5878 | 86 | 146.31 | | 1 | 1.7 | 1.16 | 19.59 | 19.13 | 0.46 | 0.46 | 0.01 | 0.01 |
| gene-LOC105283728 | https://www.ncbi.nlm.nih.gov/gene/?term=LOC105283728 | 3256 | 113 | 347.05 | | 2 | 6.14 | 1.77 | 20.45 | 20.03 | 0.42 | 0.42 | 0.01 | 0.01 |
| gene-LOC113562050 | https://www.ncbi.nlm.nih.gov/gene/?term=LOC113562050 | 17097 | 583 | 341 | 1 | 0.58 | 0.17 | 5.52 | 5.1 | 0.42 | 0.42 | 0.01 | 0.01 | |
| gene-LOC105276196 | https://www.ncbi.nlm.nih.gov/gene/?term=LOC105276196 | 2953 | 48 | 162.55 | | 1 | 3.39 | 2.08 | 45.56 | 45.18 | 0.38 | 0.38 | 0.01 | 0.01 |
| gene-LOC105278632 | https://www.ncbi.nlm.nih.gov/gene/?term=LOC105278632 | 3624 | 129 | 355.96 | | 1 | 2.76 | 0.78 | 14.81 | 14.44 | 0.37 | 0.37 | 0.01 | 0.01 |
| gene-LOC105285395 | https://www.ncbi.nlm.nih.gov/gene/?term=LOC105285395 | 8085 | 196 | 242.42 | | 1 | 1.24 | 0.51 | 15.02 | 14.69 | 0.33 | 0.33 | 0.01 | 0.01 |
| gene-LOC105283712 | https://www.ncbi.nlm.nih.gov/gene/?term=LOC105283712 | 1910 | 31 | 162.3 | 1 | 5.24 | 3.23 | 30.95 | 30.64 | 0.31 | 0.31 | 0.01 | 0.01 | |
| gene-LOC105276941 | https://www.ncbi.nlm.nih.gov/gene/?term=LOC105276941 | 15956 | 551 | 345.32 | | 1 | 0.63 | 0.18 | 5.1 | 4.82 | 0.28 | 0.28 | 0.01 | 0.01 |
| gene-LOC105275374 | https://www.ncbi.nlm.nih.gov/gene/?term=LOC105275374 | 11376 | 527 | 463.26 | | 1 | 0.88 | 0.19 | 11.34 | 11.07 | 0.27 | 0.27 | 0.01 | 0.01 |
| gene-LOC105287417 | https://www.ncbi.nlm.nih.gov/gene/?term=LOC105287417 | 1025 | 28 | 273.17 | | 1 | 9.76 | 3.57 | 42.07 | 41.82 | 0.26 | 0.26 | 0.01 | 0.01 |
| gene-LOC105275327 | https://www.ncbi.nlm.nih.gov/gene/?term=LOC105275327 | 14245 | 493 | 346.09 | | 1 | 0.7 | 0.2 | 6.9 | 6.7 | 0.19 | 0.19 | 0.01 | 0.01 |
| gene-LOC105287550 | https://www.ncbi.nlm.nih.gov/gene/?term=LOC105287550 | 15659 | 570 | 364.01 | | 1 | 0.64 | 0.18 | 6.53 | 6.34 | 0.19 | 0.19 | 0.01 | 0.01 |
| gene-LOC105278282 | https://www.ncbi.nlm.nih.gov/gene/?term=LOC105278282 | 1775 | 56 | 315.49 | | 1 | 5.63 | 1.79 | 35.74 | 35.56 | 0.18 | 0.18 | 0.01 | 0.01 |
| gene-LOC105281272 | https://www.ncbi.nlm.nih.gov/gene/?term=LOC105281272 | 5252 | 178 | 338.92 | | 1 | 1.9 | 0.56 | 16.1 | 15.92 | 0.18 | 0.18 | 0.01 | 0.01 |
| gene-LOC105282491 | https://www.ncbi.nlm.nih.gov/gene/?term=LOC105282491 | 4761 | 165 | 346.57 | | 1 | 2.1 | 0.61 | 8.62 | 8.44 | 0.18 | 0.18 | 0.01 | 0.01 |
| gene-LOC105286649 | https://www.ncbi.nlm.nih.gov/gene/?term=LOC105286649 | 14178 | 826 | 582.59 | | 1 | 0.71 | 0.12 | 0.92 | 1.04 | -0.12 | 0.12 | -0.01 | 0.01 |
| gene-LOC105276462 | https://www.ncbi.nlm.nih.gov/gene/?term=LOC105276462 | 6561 | 278 | 423.72 | | 1 | 1.52 | 0.36 | 7.35 | 7.5 | -0.15 | 0.15 | -0.01 | 0.01 |
| gene-LOC105282646 | https://www.ncbi.nlm.nih.gov/gene/?term=LOC105282646 | 3317 | 97 | 292.43 | | 1 | 3.01 | 1.03 | 22.32 | 22.49 | -0.17 | 0.17 | -0.01 | 0.01 |
| gene-LOC105278691 | https://www.ncbi.nlm.nih.gov/gene/?term=LOC105278691 | 3871 | 147 | 379.75 | | 1 | 2.58 | 0.68 | 14.28 | 14.5 | -0.22 | 0.22 | -0.01 | 0.01 |
| gene-LOC105280819 | https://www.ncbi.nlm.nih.gov/gene/?term=LOC105280819 | 1639 | 39 | 237.95 | | 1 | 6.1 | 2.56 | 35.75 | 36.01 | -0.26 | 0.26 | -0.01 | 0.01 |
| gene-LOC105275587 | https://www.ncbi.nlm.nih.gov/gene/?term=LOC105275587 | 2404 | 55 | 228.79 | | 1 | 4.16 | 1.82 | 34.41 | 34.8 | -0.39 | 0.39 | -0.01 | 0.01 |
| gene-LOC105281312 | https://www.ncbi.nlm.nih.gov/gene/?term=LOC105281312 | 2551 | 72 | 282.24 | | 1 | 3.92 | 1.39 | 29.66 | 30.06 | -0.4 | 0.4 | -0.01 | 0.01 |
| gene-LOC105286515 | https://www.ncbi.nlm.nih.gov/gene/?term=LOC105286515 | 3751 | 116 | 309.25 | | 1 | 2.67 | 0.86 | 10.42 | 10.83 | -0.41 | 0.41 | -0.01 | 0.01 |
| gene-LOC105286999 | https://www.ncbi.nlm.nih.gov/gene/?term=LOC105286999 | 2141 | 41 | 191.5 | 1 | 4.67 | 2.44 | 37.42 | 37.83 | -0.41 | 0.41 | -0.01 | 0.01 | |
| gene-LOC105281754 | https://www.ncbi.nlm.nih.gov/gene/?term=LOC105281754 | 4300 | 117 | 272.09 | | 1 | 2.33 | 0.85 | 16.59 | 17.09 | -0.5 | 0.5 | -0.01 | 0.01 |
| gene-LOC105278534 | https://www.ncbi.nlm.nih.gov/gene/?term=LOC105278534 | 7364 | 180 | 244.43 | | 1 | 1.36 | 0.56 | 18.21 | 18.75 | -0.54 | 0.54 | -0.01 | 0.01 |

| | | | | | | | | | | | | | | |
|---|---|---|---|---|---|---|---|---|---|---|---|---|---|---|
| gene-LOC105276272 | https://www.ncbi.nlm.nih.gov/gene/?term=LOC105276272 | 1965 | 36 | 183.21 | | 1 | 5.09 | 2.78 | 32.08 | 32.69 | -0.61 | 0.61 | -0.01 | 0.01 |
| gene-LOC105281173 | https://www.ncbi.nlm.nih.gov/gene/?term=LOC105281173 | 1595 | 38 | 238.24 | | 1 | 6.27 | 2.63 | 60.05 | 60.67 | -0.62 | 0.62 | -0.01 | 0.01 |
| gene-LOC105287633 | https://www.ncbi.nlm.nih.gov/gene/?term=LOC105287633 | 1524 | 27 | 177.17 | | 1 | 6.56 | 3.7 | 63.52 | 64.22 | -0.7 | 0.7 | -0.01 | 0.01 |
| gene-LOC105283989 | https://www.ncbi.nlm.nih.gov/gene/?term=LOC105283989 | 4948 | 26 | 52.55 | 1 | 2.02 | 3.85 | 43.53 | 42.86 | 0.68 | 0.68 | 0 | 0 | |
| gene-LOC105285514 | https://www.ncbi.nlm.nih.gov/gene/?term=LOC105285514 | 1277 | 13 | 101.8 | 1 | 7.83 | 7.69 | 90.77 | 90.29 | 0.48 | 0.48 | 0 | 0 | |
| gene-LOC105286452 | https://www.ncbi.nlm.nih.gov/gene/?term=LOC105286452 | 6888 | 61 | 88.56 | 1 | 1.45 | 1.64 | 32.75 | 32.35 | 0.4 | 0.4 | 0 | 0 | |
| gene-LOC105282811 | https://www.ncbi.nlm.nih.gov/gene/?term=LOC105282811 | 5765 | 203 | 352.12 | | 1 | 1.73 | 0.49 | 8.42 | 8.31 | 0.12 | 0.12 | 0 | 0 |
| gene-LOC105282825 | https://www.ncbi.nlm.nih.gov/gene/?term=LOC105282825 | 3919 | 80 | 204.13 | | 1 | 2.55 | 1.25 | 12.57 | 12.46 | 0.11 | 0.11 | 0 | 0 |
| gene-LOC105274590 | https://www.ncbi.nlm.nih.gov/gene/?term=LOC105274590 | 4516 | 58 | 128.43 | | 2 | 4.43 | 3.45 | 35.68 | 35.59 | 0.09 | 0.09 | 0 | 0 |
| gene-LOC105275363 | https://www.ncbi.nlm.nih.gov/gene/?term=LOC105275363 | 3798 | 156 | 410.74 | | 1 | 2.63 | 0.64 | 17.23 | 17.17 | 0.07 | 0.07 | 0 | 0 |
| gene-LOC105283296 | https://www.ncbi.nlm.nih.gov/gene/?term=LOC105283296 | 2174 | 84 | 386.38 | | 1 | 4.6 | 1.19 | 21.67 | 21.61 | 0.07 | 0.07 | 0 | 0 |
| gene-LOC105281700 | https://www.ncbi.nlm.nih.gov/gene/?term=LOC105281700 | 2208 | 37 | 167.57 | | 1 | 4.53 | 2.7 | 39.7 | 39.67 | 0.04 | 0.04 | 0 | 0 |
| gene-LOC105274531 | https://www.ncbi.nlm.nih.gov/gene/?term=LOC105274531 | 7774 | 284 | 365.32 | | 1 | 1.29 | 0.35 | 10.8 | 10.77 | 0.03 | 0.03 | 0 | 0 |
| gene-LOC105284043 | https://www.ncbi.nlm.nih.gov/gene/?term=LOC105284043 | 5661 | 279 | 492.85 | | 1 | 1.77 | 0.36 | 10.73 | 10.7 | 0.03 | 0.03 | 0 | 0 |
| gene-LOC105275460 | https://www.ncbi.nlm.nih.gov/gene/?term=LOC105275460 | 2473 | 114 | 460.98 | | 1 | 4.04 | 0.88 | 13.89 | 13.88 | 0.02 | 0.02 | 0 | 0 |
| gene-LOC105285406 | https://www.ncbi.nlm.nih.gov/gene/?term=LOC105285406 | 8015 | 59 | 73.61 | 1 | 1.25 | 1.69 | 26.4 | 26.39 | 0.01 | 0.01 | 0 | 0 | |
| gene-LOC105275179 | https://www.ncbi.nlm.nih.gov/gene/?term=LOC105275179 | 3271 | 73 | 223.17 | | 1 | 3.06 | 1.37 | 20.99 | 20.99 | 0 | 0 | 0 | 0 |
| gene-LOC105278228 | https://www.ncbi.nlm.nih.gov/gene/?term=LOC105278228 | 8610 | 162 | 188.15 | | 1 | 1.16 | 0.62 | 21.49 | 21.49 | 0 | 0 | 0 | 0 |
| gene-LOC105287764 | https://www.ncbi.nlm.nih.gov/gene/?term=LOC105287764 | 18359 | 396 | 215.7 | 1 | 0.54 | 0.25 | 2.87 | 2.88 | -0.01 | 0.01 | 0 | 0 | |
| gene-LOC105278283 | https://www.ncbi.nlm.nih.gov/gene/?term=LOC105278283 | 2317 | 125 | 539.49 | | 1 | 4.32 | 0.8 | 28 | 28.03 | -0.02 | 0.02 | 0 | 0 |
| gene-LOC105282951 | https://www.ncbi.nlm.nih.gov/gene/?term=LOC105282951 | 3366 | 118 | 350.56 | | 1 | 2.97 | 0.85 | 19.88 | 19.9 | -0.02 | 0.02 | 0 | 0 |
| gene-LOC105285533 | https://www.ncbi.nlm.nih.gov/gene/?term=LOC105285533 | 1640 | 60 | 365.85 | | 1 | 6.1 | 1.67 | 26.06 | 26.09 | -0.03 | 0.03 | 0 | 0 |
| gene-LOC105276527 | https://www.ncbi.nlm.nih.gov/gene/?term=LOC105276527 | 7923 | 318 | 401.36 | | 1 | 1.26 | 0.31 | 9.67 | 9.76 | -0.09 | 0.09 | 0 | 0 |
| gene-LOC105282885 | https://www.ncbi.nlm.nih.gov/gene/?term=LOC105282885 | 1870 | 43 | 229.95 | | 1 | 5.35 | 2.33 | 44.33 | 44.46 | -0.13 | 0.13 | 0 | 0 |
| gene-LOC105275187 | https://www.ncbi.nlm.nih.gov/gene/?term=LOC105275187 | 5534 | 70 | 126.49 | | 1 | 1.81 | 1.43 | 14.08 | 14.45 | -0.36 | 0.36 | 0 | 0 |

Directory: OGL File: rnk-dmg-Ob_LI_BCphaseA2.vs.RphaseA1.txt

| gene_ID | gene_link | gwidth | #Sites | #per10Kb | #dmSites | #dmsp10kb | %dmSites | %pSite1 | %pSite2 | DMpSite | ADMpSite | DMpNucl | ADMpNucl |
|---|---|---|---|---|---|---|---|---|---|---|---|---|---|
| gene-LOC105278458 | https://www.ncbi.nlm.nih.gov/gene/?term=LOC105278458 | 216 | 3 | 138.89 | 1 | 46.3 | 33.33 | 10.57 | 24.69 | -14.12 | 14.12 | -0.2 | 0.2 |
| gene-LOC105286531 | https://www.ncbi.nlm.nih.gov/gene/?term=LOC105286531 | 372 | 23 | 618.28 | 1 | 26.88 | 4.35 | 39.65 | 36.5 | 3.15 | 3.15 | 0.19 | 0.19 |
| gene-LOC113562901 | https://www.ncbi.nlm.nih.gov/gene/?term=LOC113562901 | 1531 | 22 | 143.7 | 8 | 52.25 | 36.36 | 60.1 | 48.53 | 11.57 | 11.57 | 0.17 | 0.17 |
| gene-LOC113562728 | https://www.ncbi.nlm.nih.gov/gene/?term=LOC113562728 | 1733 | 32 | 184.65 | 5 | 28.85 | 15.62 | 28.94 | 19.93 | 9 | 9 | 0.17 | 0.17 |
| gene-LOC109611487 | https://www.ncbi.nlm.nih.gov/gene/?term=LOC109611487 | 437 | 32 | 732.27 | 1 | 22.88 | 3.12 | 16.38 | 14.11 | 2.27 | 2.27 | 0.17 | 0.17 |
| gene-LOC105285392 | https://www.ncbi.nlm.nih.gov/gene/?term=LOC105285392 | 1601 | 82 | 512.18 | 3 | 18.74 | 3.66 | 38.11 | 34.95 | 3.16 | 3.16 | 0.16 | 0.16 |
| gene-LOC113561934 | https://www.ncbi.nlm.nih.gov/gene/?term=LOC113561934 | 1252 | 27 | 215.65 | 4 | 31.95 | 14.81 | 31.68 | 24.75 | 6.93 | 6.93 | 0.15 | 0.15 |
| gene-LOC105275833 | https://www.ncbi.nlm.nih.gov/gene/?term=LOC105275833 | 981 | 39 | 397.55 | 2 | 20.39 | 5.13 | 40.15 | 36.28 | 3.87 | 3.87 | 0.15 | 0.15 |
| gene-LOC105284411 | https://www.ncbi.nlm.nih.gov/gene/?term=LOC105284411 | 1795 | 63 | 350.97 | 4 | 22.28 | 6.35 | 36.82 | 33 | 3.82 | 3.82 | 0.13 | 0.13 |
| gene-LOC105286320 | https://www.ncbi.nlm.nih.gov/gene/?term=LOC105286320 | 1659 | 60 | 361.66 | 4 | 24.11 | 6.67 | 41.91 | 38.27 | 3.64 | 3.64 | 0.13 | 0.13 |
| gene-LOC105275355 | https://www.ncbi.nlm.nih.gov/gene/?term=LOC105275355 | 3197 | 163 | 509.85 | 7 | 21.9 | 4.29 | 24.89 | 27.47 | -2.59 | 2.59 | -0.13 | 0.13 |
| gene-LOC113561930 | https://www.ncbi.nlm.nih.gov/gene/?term=LOC113561930 | 422 | 20 | 473.93 | 1 | 23.7 | 5 | 10.9 | 13.66 | -2.76 | 2.76 | -0.13 | 0.13 |
| gene-LOC105279538 | https://www.ncbi.nlm.nih.gov/gene/?term=LOC105279538 | 889 | 17 | 191.23 | 2 | 22.5 | 11.76 | 36.14 | 29.61 | 6.53 | 6.53 | 0.12 | 0.12 |
| gene-LOC105281325 | https://www.ncbi.nlm.nih.gov/gene/?term=LOC105281325 | 942 | 29 | 307.86 | 1 | 10.62 | 3.45 | 25.29 | 21.51 | 3.78 | 3.78 | 0.12 | 0.12 |
| gene-LOC105275231 | https://www.ncbi.nlm.nih.gov/gene/?term=LOC105275231 | 1270 | 43 | 338.58 | 1 | 7.87 | 2.33 | 43.15 | 39.57 | 3.59 | 3.59 | 0.12 | 0.12 |
| gene-LOC105286946 | https://www.ncbi.nlm.nih.gov/gene/?term=LOC105286946 | 1228 | 49 | 399.02 | 3 | 24.43 | 6.12 | 34.75 | 31.69 | 3.06 | 3.06 | 0.12 | 0.12 |
| gene-LOC105276563 | https://www.ncbi.nlm.nih.gov/gene/?term=LOC105276563 | 1958 | 115 | 587.33 | 6 | 30.64 | 5.22 | 18.73 | 16.62 | 2.11 | 2.11 | 0.12 | 0.12 |
| gene-LOC105278873 | https://www.ncbi.nlm.nih.gov/gene/?term=LOC105278873 | 1897 | 146 | 769.64 | 6 | 31.63 | 4.11 | 21.31 | 19.69 | 1.62 | 1.62 | 0.12 | 0.12 |
| gene-LOC105283693 | https://www.ncbi.nlm.nih.gov/gene/?term=LOC105283693 | 1305 | 12 | 91.95 | 4 | 30.65 | 33.33 | 70.54 | 58.26 | 12.28 | 12.28 | 0.11 | 0.11 |
| gene-LOC105277390 | https://www.ncbi.nlm.nih.gov/gene/?term=LOC105277390 | 2567 | 94 | 366.19 | 3 | 11.69 | 3.19 | 39.27 | 36.36 | 2.9 | 2.9 | 0.11 | 0.11 |
| gene-LOC105275378 | https://www.ncbi.nlm.nih.gov/gene/?term=LOC105275378 | 1189 | 20 | 168.21 | 1 | 8.41 | 5 | 32.85 | 26.65 | 6.2 | 6.2 | 0.1 | 0.1 |
| gene-LOC105276679 | https://www.ncbi.nlm.nih.gov/gene/?term=LOC105276679 | 2005 | 64 | 319.2 | 2 | 9.98 | 3.12 | 39.86 | 36.61 | 3.25 | 3.25 | 0.1 | 0.1 |
| gene-LOC105285812 | https://www.ncbi.nlm.nih.gov/gene/?term=LOC105285812 | 1413 | 41 | 290.16 | 1 | 7.08 | 2.44 | 25.66 | 22.39 | 3.27 | 3.27 | 0.09 | 0.09 |
| gene-LOC105279437 | https://www.ncbi.nlm.nih.gov/gene/?term=LOC105279437 | 1069 | 41 | 383.54 | 1 | 9.35 | 2.44 | 22.95 | 20.6 | 2.35 | 2.35 | 0.09 | 0.09 |

...

... only first 25 lines shown ...

Directory: OGL File: rnk-dmg-Ob_LI_BCphaseA2.vs.RphaseA2.txt

| gene_ID | gene_link | gwidth | #Sites | #per10Kb | #dmSites | #dmsp10kb | %dmSites | %pSite1 | %pSite2 | DMpSite | ADMpSite | DMpNucl | ADMpNucl |
|---|---|---|---|---|---|---|---|---|---|---|---|---|---|
| gene-Trnam-cau-2 | https://www.ncbi.nlm.nih.gov/gene/?term=Trnam-cau | 72 | 1 | 138.89 | 1 | 138.89 | 100 | 0 | 24 | -24 | 24 | -0.33 | 0.33 |
| gene-LOC113561930 | https://www.ncbi.nlm.nih.gov/gene/?term=LOC113561930 | 422 | 20 | 473.93 | 2 | 47.39 | 10 | 13.1 | 16.23 | -3.12 | 3.12 | -0.15 | 0.15 |
| gene-LOC109611472 | https://www.ncbi.nlm.nih.gov/gene/?term=LOC109611472 | 1351 | 27 | 199.85 | 2 | 14.8 | 7.41 | 32.9 | 39.85 | -6.95 | 6.95 | -0.14 | 0.14 |
| gene-LOC105286212 | https://www.ncbi.nlm.nih.gov/gene/?term=LOC105286212 | 1022 | 36 | 352.25 | 1 | 9.78 | 2.78 | 37.91 | 41.67 | -3.76 | 3.76 | -0.13 | 0.13 |
| gene-LOC105282572 | https://www.ncbi.nlm.nih.gov/gene/?term=LOC105282572 | 847 | 37 | 436.84 | 1 | 11.81 | 2.7 | 30.33 | 33.17 | -2.84 | 2.84 | -0.12 | 0.12 |
| gene-LOC105287692 | https://www.ncbi.nlm.nih.gov/gene/?term=LOC105287692 | 1505 | 100 | 664.45 | 3 | 19.93 | 3 | 27.86 | 29.51 | -1.66 | 1.66 | -0.11 | 0.11 |
| gene-LOC105278847 | https://www.ncbi.nlm.nih.gov/gene/?term=LOC105278847 | 1851 | 112 | 605.08 | 1 | 5.4 | 0.89 | 10.12 | 11.9 | -1.78 | 1.78 | -0.11 | 0.11 |
| gene-LOC105276920 | https://www.ncbi.nlm.nih.gov/gene/?term=LOC105276920 | 2623 | 95 | 362.18 | 1 | 3.81 | 1.05 | 19.9 | 22.94 | -3.04 | 3.04 | -0.11 | 0.11 |
| gene-LOC105274524 | https://www.ncbi.nlm.nih.gov/gene/?term=LOC105274524 | 1126 | 38 | 337.48 | 1 | 8.88 | 2.63 | 28.9 | 32.2 | -3.3 | 3.3 | -0.11 | 0.11 |
| gene-LOC105284411 | https://www.ncbi.nlm.nih.gov/gene/?term=LOC105284411 | 1795 | 63 | 350.97 | 1 | 5.57 | 1.59 | 35.7 | 32.95 | 2.74 | 2.74 | 0.1 | 0.1 |
| gene-LOC105276360 | https://www.ncbi.nlm.nih.gov/gene/?term=LOC105276360 | 1466 | 108 | 736.7 | 1 | 6.82 | 0.93 | 29.44 | 30.83 | -1.39 | 1.39 | -0.1 | 0.1 |
| gene-LOC105276423 | https://www.ncbi.nlm.nih.gov/gene/?term=LOC105276423 | 1689 | 77 | 455.89 | 1 | 5.92 | 1.3 | 16.63 | 18.76 | -2.13 | 2.13 | -0.1 | 0.1 |
| gene-LOC105287677 | https://www.ncbi.nlm.nih.gov/gene/?term=LOC105287677 | 1588 | 75 | 472.29 | 1 | 6.3 | 1.33 | 40.32 | 38.45 | 1.87 | 1.87 | 0.09 | 0.09 |
| gene-LOC105278824 | https://www.ncbi.nlm.nih.gov/gene/?term=LOC105278824 | 1230 | 61 | 495.93 | 1 | 8.13 | 1.64 | 27.78 | 29.67 | -1.89 | 1.89 | -0.09 | 0.09 |
| gene-LOC105275307 | https://www.ncbi.nlm.nih.gov/gene/?term=LOC105275307 | 1420 | 53 | 373.24 | 2 | 14.08 | 3.77 | 40.01 | 42.31 | -2.31 | 2.31 | -0.09 | 0.09 |
| gene-LOC105280327 | https://www.ncbi.nlm.nih.gov/gene/?term=LOC105280327 | 2670 | 100 | 374.53 | 1 | 3.75 | 1 | 29.14 | 31.51 | -2.37 | 2.37 | -0.09 | 0.09 |
| gene-LOC105281038 | https://www.ncbi.nlm.nih.gov/gene/?term=LOC105281038 | 2221 | 78 | 351.19 | 1 | 4.5 | 1.28 | 24.56 | 27.03 | -2.46 | 2.46 | -0.09 | 0.09 |
| gene-LOC105277012 | https://www.ncbi.nlm.nih.gov/gene/?term=LOC105277012 | 1494 | 52 | 348.06 | 3 | 20.08 | 5.77 | 24.71 | 27.23 | -2.52 | 2.52 | -0.09 | 0.09 |
| gene-LOC105281009 | https://www.ncbi.nlm.nih.gov/gene/?term=LOC105281009 | 1562 | 42 | 268.89 | 3 | 19.21 | 7.14 | 50.63 | 54.06 | -3.42 | 3.42 | -0.09 | 0.09 |
| gene-LOC105284400 | https://www.ncbi.nlm.nih.gov/gene/?term=LOC105284400 | 1906 | 44 | 230.85 | 3 | 15.74 | 6.82 | 27.71 | 31.42 | -3.71 | 3.71 | -0.09 | 0.09 |
| gene-LOC105278074 | https://www.ncbi.nlm.nih.gov/gene/?term=LOC105278074 | 623 | 11 | 176.57 | 2 | 32.1 | 18.18 | 11.57 | 16.88 | -5.31 | 5.31 | -0.09 | 0.09 |
| gene-LOC105281691 | https://www.ncbi.nlm.nih.gov/gene/?term=LOC105281691 | 2096 | 13 | 62.02 | 4 | 19.08 | 30.77 | 7.23 | 22.12 | -14.89 | 14.89 | -0.09 | 0.09 |
| gene-LOC105279538 | https://www.ncbi.nlm.nih.gov/gene/?term=LOC105279538 | 889 | 20 | 224.97 | 2 | 22.5 | 10 | 30.72 | 27.36 | 3.36 | 3.36 | 0.08 | 0.08 |
| gene-LOC113561393 | https://www.ncbi.nlm.nih.gov/gene/?term=LOC113561393 | 1872 | 63 | 336.54 | 2 | 10.68 | 3.17 | 21.1 | 18.81 | 2.28 | 2.28 | 0.08 | 0.08 |

...
... only first 25 lines shown ...

Directory: OGL File: rnk-dmg-Ob_LI_BCphaseA2.vs.RphaseB1.txt

| gene_ID | gene_link | gwidth | #Sites | #per10Kb | #dmSites | #dmsp10kb | %dmSites | %pSite1 | %pSite2 | DMpSite | ADMpSite | DMpNucl | ADMpNucl |
|---|---|---|---|---|---|---|---|---|---|---|---|---|---|
| gene-LOC113562345 | https://www.ncbi.nlm.nih.gov/gene/?term=LOC113562345 | 2290 | 71 | 310.04 | 15 | 65.5 | 21.13 | 28.2 | 15.69 | 12.51 | 12.51 | 0.39 | 0.39 |
| gene-LOC105285695 | https://www.ncbi.nlm.nih.gov/gene/?term=LOC105285695 | 1371 | 74 | 539.75 | 1 | 7.29 | 1.35 | 27.97 | 24.61 | 3.36 | 3.36 | 0.18 | 0.18 |
| gene-LOC105275070 | https://www.ncbi.nlm.nih.gov/gene/?term=LOC105275070 | 5667 | 53 | 93.52 | 4 | 7.06 | 7.55 | 16.59 | 5.92 | 10.67 | 10.67 | 0.1 | 0.1 |
| gene-LOC105282156 | https://www.ncbi.nlm.nih.gov/gene/?term=LOC105282156 | 2577 | 128 | 496.7 | 1 | 3.88 | 0.78 | 28.45 | 26.37 | 2.08 | 2.08 | 0.1 | 0.1 |
| gene-LOC105282149 | https://www.ncbi.nlm.nih.gov/gene/?term=LOC105282149 | 1787 | 105 | 587.58 | 1 | 5.6 | 0.95 | 43.64 | 41.97 | 1.67 | 1.67 | 0.1 | 0.1 |
| gene-LOC105276704 | https://www.ncbi.nlm.nih.gov/gene/?term=LOC105276704 | 2063 | 89 | 431.41 | 1 | 4.85 | 1.12 | 14.86 | 13.07 | 1.79 | 1.79 | 0.08 | 0.08 |
| gene-LOC105276078 | https://www.ncbi.nlm.nih.gov/gene/?term=LOC105276078 | 1051 | 14 | 133.21 | 1 | 9.51 | 7.14 | 31.96 | 27.06 | 4.9 | 4.9 | 0.07 | 0.07 |
| gene-LOC105283552 | https://www.ncbi.nlm.nih.gov/gene/?term=LOC105283552 | 1562 | 36 | 230.47 | 1 | 6.4 | 2.78 | 56.28 | 53.31 | 2.97 | 2.97 | 0.07 | 0.07 |
| gene-LOC105287063 | https://www.ncbi.nlm.nih.gov/gene/?term=LOC105287063 | 1874 | 49 | 261.47 | 1 | 5.34 | 2.04 | 32.42 | 29.81 | 2.61 | 2.61 | 0.07 | 0.07 |
| gene-LOC105280774 | https://www.ncbi.nlm.nih.gov/gene/?term=LOC105280774 | 2793 | 101 | 361.62 | 1 | 3.58 | 0.99 | 32.38 | 30.39 | 1.99 | 1.99 | 0.07 | 0.07 |
| gene-LOC105274534 | https://www.ncbi.nlm.nih.gov/gene/?term=LOC105274534 | 1329 | 55 | 413.84 | 1 | 7.52 | 1.82 | 46.07 | 47.79 | -1.72 | 1.72 | -0.07 | 0.07 |
| gene-LOC105285060 | https://www.ncbi.nlm.nih.gov/gene/?term=LOC105285060 | 1285 | 13 | 101.17 | 1 | 7.78 | 7.69 | 56.31 | 50.43 | 5.87 | 5.87 | 0.06 | 0.06 |
| gene-LOC105284704 | https://www.ncbi.nlm.nih.gov/gene/?term=LOC105284704 | 2400 | 62 | 258.33 | 1 | 4.17 | 1.61 | 26.43 | 23.93 | 2.5 | 2.5 | 0.06 | 0.06 |
| gene-LOC105275708 | https://www.ncbi.nlm.nih.gov/gene/?term=LOC105275708 | 1772 | 10 | 56.43 | 1 | 5.64 | 10 | 50.31 | 40.78 | 9.53 | 9.53 | 0.05 | 0.05 |
| gene-LOC105285674 | https://www.ncbi.nlm.nih.gov/gene/?term=LOC105285674 | 3035 | 93 | 306.43 | 1 | 3.29 | 1.08 | 26.91 | 25.27 | 1.64 | 1.64 | 0.05 | 0.05 |
| gene-LOC105284184 | https://www.ncbi.nlm.nih.gov/gene/?term=LOC105284184 | 2535 | 87 | 343.2 | 1 | 3.94 | 1.15 | 34.29 | 32.68 | 1.6 | 1.6 | 0.05 | 0.05 |
| gene-LOC105283123 | https://www.ncbi.nlm.nih.gov/gene/?term=LOC105283123 | 1507 | 51 | 338.42 | 1 | 6.64 | 1.96 | 16.3 | 14.72 | 1.58 | 1.58 | 0.05 | 0.05 |
| gene-LOC105280786 | https://www.ncbi.nlm.nih.gov/gene/?term=LOC105280786 | 2573 | 122 | 474.15 | 1 | 3.89 | 0.82 | 30.88 | 29.81 | 1.07 | 1.07 | 0.05 | 0.05 |
| gene-LOC105285055 | https://www.ncbi.nlm.nih.gov/gene/?term=LOC105285055 | 4108 | 241 | 586.66 | 2 | 4.87 | 0.83 | 12.15 | 11.34 | 0.81 | 0.81 | 0.05 | 0.05 |
| gene-LOC105284213 | https://www.ncbi.nlm.nih.gov/gene/?term=LOC105284213 | 868 | 15 | 172.81 | 1 | 11.52 | 6.67 | 15.38 | 18.37 | -2.99 | 2.99 | -0.05 | 0.05 |
| gene-LOC105285514 | https://www.ncbi.nlm.nih.gov/gene/?term=LOC105285514 | 1277 | 11 | 86.14 | 1 | 7.83 | 9.09 | 90.26 | 85.12 | 5.14 | 5.14 | 0.04 | 0.04 |
| gene-LOC105283712 | https://www.ncbi.nlm.nih.gov/gene/?term=LOC105283712 | 1910 | 29 | 151.83 | 1 | 5.24 | 3.45 | 33.08 | 30.49 | 2.59 | 2.59 | 0.04 | 0.04 |
| gene-LOC105276272 | https://www.ncbi.nlm.nih.gov/gene/?term=LOC105276272 | 1965 | 38 | 193.38 | 1 | 5.09 | 2.63 | 30.65 | 28.5 | 2.15 | 2.15 | 0.04 | 0.04 |
| gene-LOC105281848 | https://www.ncbi.nlm.nih.gov/gene/?term=LOC105281848 | 2122 | 52 | 245.05 | 1 | 4.71 | 1.92 | 26.53 | 24.7 | 1.83 | 1.83 | 0.04 | 0.04 |
| gene-LOC105283697 | https://www.ncbi.nlm.nih.gov/gene/?term=LOC105283697 | 5301 | 137 | 258.44 | 1 | 1.89 | 0.73 | 17.96 | 16.39 | 1.57 | 1.57 | 0.04 | 0.04 |
| gene-LOC105288079 | https://www.ncbi.nlm.nih.gov/gene/?term=LOC105288079 | 3473 | 92 | 264.9 | 1 | 2.88 | 1.09 | 19.48 | 18 | 1.49 | 1.49 | 0.04 | 0.04 |
| gene-LOC105274449 | https://www.ncbi.nlm.nih.gov/gene/?term=LOC105274449 | 1549 | 53 | 342.16 | 1 | 6.46 | 1.89 | 50.38 | 49.29 | 1.09 | 1.09 | 0.04 | 0.04 |
| gene-LOC105287095 | https://www.ncbi.nlm.nih.gov/gene/?term=LOC105287095 | 1435 | 56 | 390.24 | 1 | 6.97 | 1.79 | 30.4 | 29.35 | 1.05 | 1.05 | 0.04 | 0.04 |
| gene-LOC105276449 | https://www.ncbi.nlm.nih.gov/gene/?term=LOC105276449 | 1231 | 43 | 349.31 | 1 | 8.12 | 2.33 | 30.8 | 29.78 | 1.02 | 1.02 | 0.04 | 0.04 |
| gene-LOC105279298 | https://www.ncbi.nlm.nih.gov/gene/?term=LOC105279298 | 2773 | 123 | 443.56 | 1 | 3.61 | 0.81 | 24.88 | 23.88 | 1 | 1 | 0.04 | 0.04 |
| gene-LOC105278497 | https://www.ncbi.nlm.nih.gov/gene/?term=LOC105278497 | 2072 | 89 | 429.54 | 1 | 4.83 | 1.12 | 21.38 | 20.51 | 0.87 | 0.87 | 0.04 | 0.04 |
| gene-LOC105278283 | https://www.ncbi.nlm.nih.gov/gene/?term=LOC105278283 | 2317 | 126 | 543.81 | 1 | 4.32 | 0.79 | 27.76 | 27.04 | 0.72 | 0.72 | 0.04 | 0.04 |
| gene-LOC105278282 | https://www.ncbi.nlm.nih.gov/gene/?term=LOC105278282 | 1775 | 56 | 315.49 | 1 | 5.63 | 1.79 | 33.68 | 34.81 | -1.13 | 1.13 | -0.04 | 0.04 |
| gene-LOC105274900 | https://www.ncbi.nlm.nih.gov/gene/?term=LOC105274900 | 5561 | 51 | 91.71 | 2 | 3.6 | 3.92 | 39.2 | 36.36 | 2.84 | 2.84 | 0.03 | 0.03 |
| gene-LOC105279898 | https://www.ncbi.nlm.nih.gov/gene/?term=LOC105279898 | 1343 | 13 | 96.8 | 1 | 7.45 | 7.69 | 58.71 | 56.01 | 2.69 | 2.69 | 0.03 | 0.03 |
| gene-LOC105288066 | https://www.ncbi.nlm.nih.gov/gene/?term=LOC105288066 | 1964 | 23 | 117.11 | 1 | 5.09 | 4.35 | 46.39 | 44.06 | 2.34 | 2.34 | 0.03 | 0.03 |
| gene-LOC105280348 | https://www.ncbi.nlm.nih.gov/gene/?term=LOC105280348 | 3797 | 65 | 171.19 | 1 | 2.63 | 1.54 | 21.64 | 19.63 | 2.01 | 2.01 | 0.03 | 0.03 |
| gene-LOC105280641 | https://www.ncbi.nlm.nih.gov/gene/?term=LOC105280641 | 3682 | 56 | 152.09 | 1 | 2.72 | 1.79 | 17.9 | 16.23 | 1.66 | 1.66 | 0.03 | 0.03 |
| gene-LOC105280972 | https://www.ncbi.nlm.nih.gov/gene/?term=LOC105280972 | 1532 | 32 | 208.88 | 1 | 6.53 | 3.12 | 19.4 | 18.05 | 1.35 | 1.35 | 0.03 | 0.03 |
| gene-LOC105282885 | https://www.ncbi.nlm.nih.gov/gene/?term=LOC105282885 | 1870 | 38 | 203.21 | 1 | 5.35 | 2.63 | 45.14 | 43.81 | 1.33 | 1.33 | 0.03 | 0.03 |
| gene-LOC105286265 | https://www.ncbi.nlm.nih.gov/gene/?term=LOC105286265 | 2507 | 64 | 255.29 | 1 | 3.99 | 1.56 | 30.69 | 29.49 | 1.21 | 1.21 | 0.03 | 0.03 |
| gene-LOC105278807 | https://www.ncbi.nlm.nih.gov/gene/?term=LOC105278807 | 2166 | 71 | 327.79 | 1 | 4.62 | 1.41 | 19.81 | 18.81 | 1 | 1 | 0.03 | 0.03 |
| gene-LOC105274823 | https://www.ncbi.nlm.nih.gov/gene/?term=LOC105274823 | 4620 | 152 | 329 | 1 | 2.16 | 0.66 | 19.69 | 18.71 | 0.99 | 0.99 | 0.03 | 0.03 |
| gene-LOC105283728 | https://www.ncbi.nlm.nih.gov/gene/?term=LOC105283728 | 3256 | 104 | 319.41 | 2 | 6.14 | 1.92 | 22.23 | 21.4 | 0.83 | 0.83 | 0.03 | 0.03 |
| gene-LOC105285179 | https://www.ncbi.nlm.nih.gov/gene/?term=LOC105285179 | 4191 | 178 | 424.72 | 1 | 2.39 | 0.56 | 17.79 | 17.08 | 0.71 | 0.71 | 0.03 | 0.03 |
| gene-LOC105278145 | https://www.ncbi.nlm.nih.gov/gene/?term=LOC105278145 | 4470 | 250 | 559.28 | 1 | 2.24 | 0.4 | 10.94 | 10.34 | 0.6 | 0.6 | 0.03 | 0.03 |
| gene-LOC105278459 | https://www.ncbi.nlm.nih.gov/gene/?term=LOC105278459 | 2936 | 97 | 330.38 | 1 | 3.41 | 1.03 | 20.46 | 21.23 | -0.77 | 0.77 | -0.03 | 0.03 |
| gene-LOC105281173 | https://www.ncbi.nlm.nih.gov/gene/?term=LOC105281173 | 1595 | 39 | 244.51 | 1 | 6.27 | 2.56 | 60.96 | 62.08 | -1.12 | 1.12 | -0.03 | 0.03 |
| gene-LOC105284507 | https://www.ncbi.nlm.nih.gov/gene/?term=LOC105284507 | 5207 | 86 | 165.16 | 1 | 1.92 | 1.16 | 21.7 | 23.52 | -1.83 | 1.83 | -0.03 | 0.03 |
| gene-LOC105275347 | https://www.ncbi.nlm.nih.gov/gene/?term=LOC105275347 | 2526 | 14 | 55.42 | 1 | 3.96 | 7.14 | 33.57 | 29.73 | 3.84 | 3.84 | 0.02 | 0.02 |
| gene-LOC105288010 | https://www.ncbi.nlm.nih.gov/gene/?term=LOC105288010 | 3189 | 51 | 159.92 | 1 | 3.14 | 1.96 | 34.39 | 32.85 | 1.54 | 1.54 | 0.02 | 0.02 |
| gene-LOC105277243 | https://www.ncbi.nlm.nih.gov/gene/?term=LOC105277243 | 2284 | 32 | 140.11 | 1 | 4.38 | 3.12 | 44.88 | 43.35 | 1.53 | 1.53 | 0.02 | 0.02 |
| gene-LOC105276538 | https://www.ncbi.nlm.nih.gov/gene/?term=LOC105276538 | 1511 | 19 | 125.74 | 1 | 6.62 | 5.26 | 70.28 | 68.86 | 1.42 | 1.42 | 0.02 | 0.02 |
| gene-LOC105278384 | https://www.ncbi.nlm.nih.gov/gene/?term=LOC105278384 | 1390 | 33 | 237.41 | 1 | 7.19 | 3.03 | 46.69 | 45.77 | 0.92 | 0.92 | 0.02 | 0.02 |
| gene-LOC113561805 | https://www.ncbi.nlm.nih.gov/gene/?term=LOC113561805 | 3337 | 69 | 206.77 | 1 | 3 | 1.45 | 38.18 | 37.26 | 0.92 | 0.92 | 0.02 | 0.02 |
| gene-LOC105282794 | https://www.ncbi.nlm.nih.gov/gene/?term=LOC105282794 | 4571 | 99 | 216.58 | 1 | 2.19 | 1.01 | 18.66 | 17.76 | 0.9 | 0.9 | 0.02 | 0.02 |
| gene-LOC105286644 | https://www.ncbi.nlm.nih.gov/gene/?term=LOC105286644 | 1235 | 22 | 178.14 | 1 | 8.1 | 4.55 | 42.04 | 41.14 | 0.89 | 0.89 | 0.02 | 0.02 |
| gene-LOC105283592 | https://www.ncbi.nlm.nih.gov/gene/?term=LOC105283592 | 3479 | 75 | 215.58 | 1 | 2.87 | 1.33 | 25.19 | 24.33 | 0.86 | 0.86 | 0.02 | 0.02 |
| gene-LOC105281312 | https://www.ncbi.nlm.nih.gov/gene/?term=LOC105281312 | 2551 | 68 | 266.56 | 1 | 3.92 | 1.47 | 31.46 | 30.8 | 0.65 | 0.65 | 0.02 | 0.02 |
| gene-LOC105284355 | https://www.ncbi.nlm.nih.gov/gene/?term=LOC105284355 | 3904 | 143 | 366.29 | 1 | 2.56 | 0.7 | 22.99 | 22.35 | 0.64 | 0.64 | 0.02 | 0.02 |

| | | | | | | | | | | | | | |
|---|---|---|---|---|---|---|---|---|---|---|---|---|---|
| gene-LOC105285671 | https://www.ncbi.nlm.nih.gov/gene/?term=LOC105285671 | 2263 | 99 | 437.47 | 1 | 4.42 | 1.01 | 30.99 | 30.44 | 0.55 | 0.55 | 0.02 | 0.02 |
| gene-LOC105275460 | https://www.ncbi.nlm.nih.gov/gene/?term=LOC105275460 | 2473 | 118 | 477.15 | 1 | 4.04 | 0.85 | 13.46 | 12.95 | 0.51 | 0.51 | 0.02 | 0.02 |
| gene-LOC105278632 | https://www.ncbi.nlm.nih.gov/gene/?term=LOC105278632 | 3624 | 131 | 361.48 | 1 | 2.76 | 0.76 | 14.47 | 14.04 | 0.43 | 0.43 | 0.02 | 0.02 |
| gene-LOC105276462 | https://www.ncbi.nlm.nih.gov/gene/?term=LOC105276462 | 6561 | 275 | 419.14 | 1 | 1.52 | 0.36 | 8.32 | 7.94 | 0.37 | 0.37 | 0.02 | 0.02 |
| gene-LOC105277435 | https://www.ncbi.nlm.nih.gov/gene/?term=LOC105277435 | 3643 | 175 | 480.37 | 1 | 2.74 | 0.57 | 11.3 | 10.97 | 0.33 | 0.33 | 0.02 | 0.02 |
| gene-LOC105286649 | https://www.ncbi.nlm.nih.gov/gene/?term=LOC105286649 | 14178 | 785 | 553.67 | 1 | 0.71 | 0.13 | 0.94 | 1.3 | -0.36 | 0.36 | -0.02 | 0.02 |
| gene-LOC105280956 | https://www.ncbi.nlm.nih.gov/gene/?term=LOC105280956 | 2639 | 77 | 291.78 | 1 | 3.79 | 1.3 | 18.18 | 18.82 | -0.64 | 0.64 | -0.02 | 0.02 |
| gene-LOC105286515 | https://www.ncbi.nlm.nih.gov/gene/?term=LOC105286515 | 3751 | 113 | 301.25 | 1 | 2.67 | 0.88 | 10.68 | 11.31 | -0.64 | 0.64 | -0.02 | 0.02 |
| gene-LOC105278454 | https://www.ncbi.nlm.nih.gov/gene/?term=LOC105278454 | 1972 | 41 | 207.91 | 1 | 5.07 | 2.44 | 43.16 | 44.07 | -0.91 | 0.91 | -0.02 | 0.02 |
| gene-LOC105287633 | https://www.ncbi.nlm.nih.gov/gene/?term=LOC105287633 | 1524 | 23 | 150.92 | 1 | 6.56 | 4.35 | 68.63 | 69.73 | -1.11 | 1.11 | -0.02 | 0.02 |
| gene-LOC105285252 | https://www.ncbi.nlm.nih.gov/gene/?term=LOC105285252 | 7427 | 124 | 166.96 | 2 | 2.69 | 1.61 | 12.83 | 14.05 | -1.22 | 1.22 | -0.02 | 0.02 |
| gene-LOC105285315 | https://www.ncbi.nlm.nih.gov/gene/?term=LOC105285315 | 5458 | 21 | 38.48 1 | 1.83 | 4.76 | 43.16 | 40.32 | 2.84 | 2.84 | 0.01 | 0.01 | |
| gene-LOC105280408 | https://www.ncbi.nlm.nih.gov/gene/?term=LOC105280408 | 5983 | 50 | 83.57 1 | 1.67 | 2 | 19.04 | 17.37 | 1.67 | 1.67 | 0.01 | 0.01 | |
| gene-LOC105281515 | https://www.ncbi.nlm.nih.gov/gene/?term=LOC105281515 | 8827 | 48 | 54.38 2 | 2.27 | 4.17 | 32.1 | 30.7 | 1.4 | 1.4 | 0.01 | 0.01 | |
| gene-LOC105279500 | https://www.ncbi.nlm.nih.gov/gene/?term=LOC105279500 | 3587 | 33 | 92 1 | 2.79 | 3.03 | 51.88 | 50.67 | 1.21 | 1.21 | 0.01 | 0.01 | |
| gene-LOC105275518 | https://www.ncbi.nlm.nih.gov/gene/?term=LOC105275518 | 2335 | 19 | 81.37 1 | 4.28 | 5.26 | 51.76 | 50.67 | 1.09 | 1.09 | 0.01 | 0.01 | |
| gene-LOC105279664 | https://www.ncbi.nlm.nih.gov/gene/?term=LOC105279664 | 4023 | 57 | 141.69 | 1 | 2.49 | 1.75 | 22.82 | 21.77 | 1.05 | 1.05 | 0.01 | 0.01 |
| gene-LOC105277215 | https://www.ncbi.nlm.nih.gov/gene/?term=LOC105277215 | 2619 | 23 | 87.82 1 | 3.82 | 4.35 | 35.64 | 34.71 | 0.93 | 0.93 | 0.01 | 0.01 | |
| gene-LOC105283838 | https://www.ncbi.nlm.nih.gov/gene/?term=LOC105283838 | 1748 | 21 | 120.14 | 1 | 5.72 | 4.76 | 46.37 | 45.45 | 0.92 | 0.92 | 0.01 | 0.01 |
| gene-LOC105278109 | https://www.ncbi.nlm.nih.gov/gene/?term=LOC105278109 | 4029 | 65 | 161.33 | 1 | 2.48 | 1.54 | 46.32 | 45.45 | 0.88 | 0.88 | 0.01 | 0.01 |
| gene-LOC105276560 | https://www.ncbi.nlm.nih.gov/gene/?term=LOC105276560 | 18607 | 319 | 171.44 | 1 | 0.54 | 0.31 | 2.53 | 1.85 | 0.67 | 0.67 | 0.01 | 0.01 |
| gene-LOC105286514 | https://www.ncbi.nlm.nih.gov/gene/?term=LOC105286514 | 2528 | 43 | 170.09 | 1 | 3.96 | 2.33 | 34.72 | 34.07 | 0.65 | 0.65 | 0.01 | 0.01 |
| gene-LOC105276059 | https://www.ncbi.nlm.nih.gov/gene/?term=LOC105276059 | 6256 | 123 | 196.61 | 1 | 1.6 | 0.81 | 12.44 | 11.86 | 0.58 | 0.58 | 0.01 | 0.01 |
| gene-LOC105280407 | https://www.ncbi.nlm.nih.gov/gene/?term=LOC105280407 | 10950 | 203 | 185.39 | 1 | 0.91 | 0.49 | 15.56 | 14.99 | 0.56 | 0.56 | 0.01 | 0.01 |
| gene-LOC105285395 | https://www.ncbi.nlm.nih.gov/gene/?term=LOC105285395 | 8085 | 207 | 256.03 | 1 | 1.24 | 0.48 | 14.3 | 13.76 | 0.54 | 0.54 | 0.01 | 0.01 |
| gene-LOC105280819 | https://www.ncbi.nlm.nih.gov/gene/?term=LOC105280819 | 1639 | 39 | 237.95 | 1 | 6.1 | 2.56 | 35.99 | 35.46 | 0.53 | 0.53 | 0.01 | 0.01 |
| gene-LOC105275451 | https://www.ncbi.nlm.nih.gov/gene/?term=LOC105275451 | 6035 | 133 | 220.38 | 1 | 1.66 | 0.75 | 22.24 | 21.72 | 0.51 | 0.51 | 0.01 | 0.01 |
| gene-LOC105278534 | https://www.ncbi.nlm.nih.gov/gene/?term=LOC105278534 | 7364 | 180 | 244.43 | 1 | 1.36 | 0.56 | 18.01 | 17.6 | 0.41 | 0.41 | 0.01 | 0.01 |
| gene-LOC105280890 | https://www.ncbi.nlm.nih.gov/gene/?term=LOC105280890 | 1139 | 29 | 254.61 | 1 | 8.78 | 3.45 | 19.62 | 19.23 | 0.39 | 0.39 | 0.01 | 0.01 |
| gene-LOC105286999 | https://www.ncbi.nlm.nih.gov/gene/?term=LOC105286999 | 2141 | 44 | 205.51 | 1 | 4.67 | 2.27 | 35.31 | 34.92 | 0.39 | 0.39 | 0.01 | 0.01 |
| gene-LOC105280410 | https://www.ncbi.nlm.nih.gov/gene/?term=LOC105280410 | 12580 | 265 | 210.65 | 1 | 0.79 | 0.38 | 19.92 | 19.57 | 0.35 | 0.35 | 0.01 | 0.01 |
| gene-LOC105283296 | https://www.ncbi.nlm.nih.gov/gene/?term=LOC105283296 | 2174 | 86 | 395.58 | 1 | 4.6 | 1.16 | 16.05 | 15.7 | 0.35 | 0.35 | 0.01 | 0.01 |
| gene-LOC105278666 | https://www.ncbi.nlm.nih.gov/gene/?term=LOC105278666 | 10341 | 456 | 440.96 | 1 | 0.97 | 0.22 | 7.52 | 7.19 | 0.34 | 0.34 | 0.01 | 0.01 |
| gene-LOC105279657 | https://www.ncbi.nlm.nih.gov/gene/?term=LOC105279657 | 5878 | 96 | 163.32 | 1 | 1.7 | 1.04 | 18.1 | 17.76 | 0.34 | 0.34 | 0.01 | 0.01 |
| gene-LOC105284036 | https://www.ncbi.nlm.nih.gov/gene/?term=LOC105284036 | 3254 | 87 | 267.36 | 1 | 3.07 | 1.15 | 24.27 | 23.96 | 0.31 | 0.31 | 0.01 | 0.01 |
| gene-LOC105281272 | https://www.ncbi.nlm.nih.gov/gene/?term=LOC105281272 | 5252 | 175 | 333.21 | 1 | 1.9 | 0.57 | 16.26 | 15.96 | 0.3 | 0.3 | 0.01 | 0.01 |
| gene-LOC105285807 | https://www.ncbi.nlm.nih.gov/gene/?term=LOC105285807 | 6063 | 207 | 341.42 | 1 | 1.65 | 0.48 | 7.17 | 6.87 | 0.3 | 0.3 | 0.01 | 0.01 |
| gene-LOC105282491 | https://www.ncbi.nlm.nih.gov/gene/?term=LOC105282491 | 4761 | 170 | 357.07 | 1 | 2.1 | 0.59 | 7.92 | 7.67 | 0.25 | 0.25 | 0.01 | 0.01 |
| gene-LOC105274706 | https://www.ncbi.nlm.nih.gov/gene/?term=LOC105274706 | 1446 | 50 | 345.78 | 1 | 6.92 | 2 | 39.23 | 39 | 0.23 | 0.23 | 0.01 | 0.01 |
| gene-LOC105283649 | https://www.ncbi.nlm.nih.gov/gene/?term=LOC105283649 | 1918 | 55 | 286.76 | 1 | 5.21 | 1.82 | 38.85 | 38.63 | 0.21 | 0.21 | 0.01 | 0.01 |
| gene-LOC105275374 | https://www.ncbi.nlm.nih.gov/gene/?term=LOC105275374 | 11376 | 525 | 461.5 1 | 0.88 | 0.19 | 10.23 | 10.06 | 0.17 | 0.17 | 0.01 | 0.01 | |
| gene-LOC105282658 | https://www.ncbi.nlm.nih.gov/gene/?term=LOC105282658 | 2421 | 67 | 276.75 | 1 | 4.13 | 1.49 | 20.83 | 21.03 | -0.21 | 0.21 | -0.01 | 0.01 |
| gene-LOC105285981 | https://www.ncbi.nlm.nih.gov/gene/?term=LOC105285981 | 7666 | 183 | 238.72 | 1 | 1.3 | 0.55 | 18.71 | 18.92 | -0.21 | 0.21 | -0.01 | 0.01 |
| gene-LOC105278877 | https://www.ncbi.nlm.nih.gov/gene/?term=LOC105278877 | 1968 | 77 | 391.26 | 1 | 5.08 | 1.3 | 34.55 | 34.76 | -0.22 | 0.22 | -0.01 | 0.01 |
| gene-LOC105274851 | https://www.ncbi.nlm.nih.gov/gene/?term=LOC105274851 | 6616 | 127 | 191.96 | 1 | 1.51 | 0.79 | 11.51 | 11.83 | -0.33 | 0.33 | -0.01 | 0.01 |
| gene-LOC105285742 | https://www.ncbi.nlm.nih.gov/gene/?term=LOC105285742 | 3446 | 97 | 281.49 | 1 | 2.9 | 1.03 | 20.21 | 20.62 | -0.41 | 0.41 | -0.01 | 0.01 |
| gene-LOC105274590 | https://www.ncbi.nlm.nih.gov/gene/?term=LOC105274590 | 4516 | 64 | 141.72 | 2 | 4.43 | 3.12 | 32.42 | 32.94 | -0.51 | 0.51 | -0.01 | 0.01 |
| gene-LOC105284146 | https://www.ncbi.nlm.nih.gov/gene/?term=LOC105284146 | 2488 | 66 | 265.27 | 1 | 4.02 | 1.52 | 23.09 | 23.62 | -0.54 | 0.54 | -0.01 | 0.01 |
| gene-LOC105278228 | https://www.ncbi.nlm.nih.gov/gene/?term=LOC105278228 | 8610 | 169 | 196.28 | 1 | 1.16 | 0.59 | 21.96 | 22.51 | -0.55 | 0.55 | -0.01 | 0.01 |
| gene-LOC105283989 | https://www.ncbi.nlm.nih.gov/gene/?term=LOC105283989 | 4948 | 25 | 50.53 1 | 2.02 | 4 | 46.26 | 45.67 | 0.59 | 0.59 | 0 | 0 | |
| gene-LOC105275187 | https://www.ncbi.nlm.nih.gov/gene/?term=LOC105275187 | 5534 | 69 | 124.68 | 1 | 1.81 | 1.45 | 14.14 | 13.85 | 0.29 | 0.29 | 0 | 0 |
| gene-LOC105281700 | https://www.ncbi.nlm.nih.gov/gene/?term=LOC105281700 | 2208 | 38 | 172.1 1 | 4.53 | 2.63 | 39.22 | 39.09 | 0.14 | 0.14 | 0 | 0 | |
| gene-LOC105274531 | https://www.ncbi.nlm.nih.gov/gene/?term=LOC105274531 | 7774 | 283 | 364.03 | 1 | 1.29 | 0.35 | 10.82 | 10.73 | 0.09 | 0.09 | 0 | 0 |
| gene-LOC105282825 | https://www.ncbi.nlm.nih.gov/gene/?term=LOC105282825 | 3919 | 78 | 199.03 | 1 | 2.55 | 1.28 | 12.99 | 12.9 | 0.09 | 0.09 | 0 | 0 |
| gene-LOC105275179 | https://www.ncbi.nlm.nih.gov/gene/?term=LOC105275179 | 3271 | 71 | 217.06 | 1 | 3.06 | 1.41 | 16.78 | 16.7 | 0.08 | 0.08 | 0 | 0 |
| gene-LOC105279038 | https://www.ncbi.nlm.nih.gov/gene/?term=LOC105279038 | 9453 | 317 | 335.34 | 1 | 1.06 | 0.32 | 9.53 | 9.46 | 0.07 | 0.07 | 0 | 0 |
| gene-LOC105283987 | https://www.ncbi.nlm.nih.gov/gene/?term=LOC105283987 | 4841 | 71 | 146.66 | 1 | 2.07 | 1.41 | 32.31 | 32.24 | 0.07 | 0.07 | 0 | 0 |
| gene-LOC105284329 | https://www.ncbi.nlm.nih.gov/gene/?term=LOC105284329 | 4051 | 51 | 125.89 | 1 | 2.47 | 1.96 | 19.38 | 19.32 | 0.06 | 0.06 | 0 | 0 |
| gene-LOC105288078 | https://www.ncbi.nlm.nih.gov/gene/?term=LOC105288078 | 19126 | 307 | 160.51 | 1 | 0.52 | 0.33 | 16.81 | 16.78 | 0.03 | 0.03 | 0 | 0 |
| gene-LOC105276527 | https://www.ncbi.nlm.nih.gov/gene/?term=LOC105276527 | 7923 | 334 | 421.56 | 1 | 1.26 | 0.3 | 8.56 | 8.54 | 0.02 | 0.02 | 0 | 0 |
| gene-LOC105286452 | https://www.ncbi.nlm.nih.gov/gene/?term=LOC105286452 | 6888 | 68 | 98.72 1 | 1.45 | 1.47 | 28.88 | 28.89 | 0 | 0 | 0 | 0 | |
| gene-LOC105274856 | https://www.ncbi.nlm.nih.gov/gene/?term=LOC105274856 | 5216 | 212 | 406.44 | 1 | 1.92 | 0.47 | 18.3 | 18.31 | -0.01 | 0.01 | 0 | 0 |
| gene-LOC105275327 | https://www.ncbi.nlm.nih.gov/gene/?term=LOC105275327 | 14245 | 472 | 331.34 | 1 | 0.7 | 0.21 | 7.13 | 7.15 | -0.03 | 0.03 | 0 | 0 |
| gene-LOC105283943 | https://www.ncbi.nlm.nih.gov/gene/?term=LOC105283943 | 3559 | 56 | 157.35 | 1 | 2.81 | 1.79 | 16.7 | 16.73 | -0.03 | 0.03 | 0 | 0 |

| | | | | | | | | | | | | | | |
|---|---|---|---|---|---|---|---|---|---|---|---|---|---|---|
| gene-LOC105284043 | https://www.ncbi.nlm.nih.gov/gene/?term=LOC105284043 | 5661 | 278 | 491.08 | | 1 | 1.77 | 0.36 | 10.84 | 10.87 | -0.03 | 0.03 | 0 | 0 |
| gene-LOC105275683 | https://www.ncbi.nlm.nih.gov/gene/?term=LOC105275683 | 4256 | 191 | 448.78 | | 1 | 2.35 | 0.52 | 12.17 | 12.21 | -0.04 | 0.04 | 0 | 0 |
| gene-LOC105277216 | https://www.ncbi.nlm.nih.gov/gene/?term=LOC105277216 | 7627 | 87 | 114.07 | | 1 | 1.31 | 1.15 | 9.6 | 9.67 | -0.08 | 0.08 | 0 | 0 |
| gene-LOC105287417 | https://www.ncbi.nlm.nih.gov/gene/?term=LOC105287417 | 1025 | 28 | 273.17 | | 1 | 9.76 | 3.57 | 42.07 | 42.23 | -0.16 | 0.16 | 0 | 0 |
| gene-LOC105284774 | https://www.ncbi.nlm.nih.gov/gene/?term=LOC105284774 | 12089 | 81 | 67 | 1 | 0.83 | 1.23 | 21 | 21.27 | -0.27 | 0.27 | 0 | 0 | |
| gene-LOC105276196 | https://www.ncbi.nlm.nih.gov/gene/?term=LOC105276196 | 2953 | 45 | 152.39 | | 1 | 3.39 | 2.22 | 48.59 | 48.91 | -0.31 | 0.31 | 0 | 0 |

Directory: OGL File: rnk-dmg-Ob_LI_BCphaseA2.vs.RphaseB2.txt

| gene_ID | gene_link | gwidth | #Sites | #per10Kb | #dmSites | #dmsp10kb | %dmSites | %pSite1 | %pSite2 | DMpSite | ADMpSite | DMpNucl | ADMpNucl |
|---|---|---|---|---|---|---|---|---|---|---|---|---|---|
| gene-LOC113562345 | https://www.ncbi.nlm.nih.gov/gene/?term=LOC113562345 | 2290 | 67 | 292.58 | 11 | 48.03 | 16.42 | 29.88 | 17.32 | 12.57 | 12.57 | 0.37 | 0.37 |
| gene-LOC105284677 | https://www.ncbi.nlm.nih.gov/gene/?term=LOC105284677 | 1905 | 39 | 204.72 | 3 | 15.75 | 7.69 | 19.98 | 31.6 | -11.62 | 11.62 | -0.24 | 0.24 |
| gene-LOC113562901 | https://www.ncbi.nlm.nih.gov/gene/?term=LOC113562901 | 1531 | 22 | 143.7 | 1 | 6.53 | 4.55 | 60.1 | 44.57 | 15.53 | 15.53 | 0.22 | 0.22 |
| gene-LOC105282976 | https://www.ncbi.nlm.nih.gov/gene/?term=LOC105282976 | 1648 | 47 | 285.19 | 1 | 6.07 | 2.13 | 28.02 | 22.43 | 5.59 | 5.59 | 0.16 | 0.16 |
| gene-LOC105283552 | https://www.ncbi.nlm.nih.gov/gene/?term=LOC105283552 | 1562 | 36 | 230.47 | 1 | 6.4 | 2.78 | 56.28 | 51.7 | 4.58 | 4.58 | 0.11 | 0.11 |
| gene-LOC105282149 | https://www.ncbi.nlm.nih.gov/gene/?term=LOC105282149 | 1787 | 108 | 604.36 | 1 | 5.6 | 0.93 | 43.65 | 41.89 | 1.76 | 1.76 | 0.11 | 0.11 |
| gene-LOC105279898 | https://www.ncbi.nlm.nih.gov/gene/?term=LOC105279898 | 1343 | 10 | 74.46 | 1 | 7.45 | 10 | 84.43 | 71.53 | 12.89 | 12.89 | 0.1 | 0.1 |
| gene-LOC105275070 | https://www.ncbi.nlm.nih.gov/gene/?term=LOC105275070 | 5667 | 41 | 72.35 | 5 | 8.82 | 12.2 | 13.7 | 1.06 | 12.65 | 12.65 | 0.09 | 0.09 |
| gene-LOC105286644 | https://www.ncbi.nlm.nih.gov/gene/?term=LOC105286644 | 1235 | 22 | 178.14 | 1 | 8.1 | 4.55 | 42.35 | 37.26 | 5.09 | 5.09 | 0.09 | 0.09 |
| gene-LOC105285674 | https://www.ncbi.nlm.nih.gov/gene/?term=LOC105285674 | 3035 | 93 | 306.43 | 2 | 6.59 | 2.15 | 26.74 | 23.71 | 3.03 | 3.03 | 0.09 | 0.09 |
| gene-LOC105276704 | https://www.ncbi.nlm.nih.gov/gene/?term=LOC105276704 | 2063 | 88 | 426.56 | 1 | 4.85 | 1.14 | 15.96 | 13.92 | 2.05 | 2.05 | 0.09 | 0.09 |
| gene-LOC105285671 | https://www.ncbi.nlm.nih.gov/gene/?term=LOC105285671 | 2263 | 103 | 455.15 | 1 | 4.42 | 0.97 | 29.21 | 27.3 | 1.91 | 1.91 | 0.09 | 0.09 |
| gene-LOC105282156 | https://www.ncbi.nlm.nih.gov/gene/?term=LOC105282156 | 2577 | 129 | 500.58 | 1 | 3.88 | 0.78 | 28.68 | 26.84 | 1.84 | 1.84 | 0.09 | 0.09 |
| gene-LOC105277565 | https://www.ncbi.nlm.nih.gov/gene/?term=LOC105277565 | 3216 | 169 | 525.5 | 1 | 3.11 | 0.59 | 20.65 | 19.2 | 1.45 | 1.45 | 0.08 | 0.08 |
| gene-LOC105283649 | https://www.ncbi.nlm.nih.gov/gene/?term=LOC105283649 | 1918 | 54 | 281.54 | 1 | 5.21 | 1.85 | 37.63 | 35.03 | 2.6 | 2.6 | 0.07 | 0.07 |
| gene-LOC105280786 | https://www.ncbi.nlm.nih.gov/gene/?term=LOC105280786 | 2573 | 113 | 439.18 | 1 | 3.89 | 0.88 | 32.35 | 30.66 | 1.69 | 1.69 | 0.07 | 0.07 |
| gene-LOC105278215 | https://www.ncbi.nlm.nih.gov/gene/?term=LOC105278215 | 1700 | 29 | 170.59 | 1 | 5.88 | 3.45 | 57.89 | 54.66 | 3.23 | 3.23 | 0.06 | 0.06 |
| gene-LOC105283123 | https://www.ncbi.nlm.nih.gov/gene/?term=LOC105283123 | 1507 | 44 | 291.97 | 1 | 6.64 | 2.27 | 19.91 | 17.87 | 2.05 | 2.05 | 0.06 | 0.06 |
| gene-LOC105278497 | https://www.ncbi.nlm.nih.gov/gene/?term=LOC105278497 | 2072 | 92 | 444.02 | 1 | 4.83 | 1.09 | 21.61 | 20.36 | 1.26 | 1.26 | 0.06 | 0.06 |
| gene-LOC105279298 | https://www.ncbi.nlm.nih.gov/gene/?term=LOC105279298 | 2773 | 131 | 472.41 | 2 | 7.21 | 1.53 | 24.81 | 23.62 | 1.18 | 1.18 | 0.06 | 0.06 |
| gene-LOC105276078 | https://www.ncbi.nlm.nih.gov/gene/?term=LOC105276078 | 1051 | 14 | 133.21 | 1 | 9.51 | 7.14 | 36.98 | 32.93 | 4.05 | 4.05 | 0.05 | 0.05 |
| gene-LOC105288063 | https://www.ncbi.nlm.nih.gov/gene/?term=LOC105288063 | 3400 | 43 | 126.47 | 1 | 2.94 | 2.33 | 18.5 | 14.57 | 3.92 | 3.92 | 0.05 | 0.05 |
| gene-LOC105278109 | https://www.ncbi.nlm.nih.gov/gene/?term=LOC105278109 | 4029 | 59 | 146.44 | 1 | 2.48 | 1.69 | 50.99 | 47.24 | 3.75 | 3.75 | 0.05 | 0.05 |
| gene-LOC105281754 | https://www.ncbi.nlm.nih.gov/gene/?term=LOC105281754 | 4300 | 110 | 255.81 | 1 | 2.33 | 0.91 | 16.49 | 14.51 | 1.98 | 1.98 | 0.05 | 0.05 |
| gene-LOC105284184 | https://www.ncbi.nlm.nih.gov/gene/?term=LOC105284184 | 2535 | 79 | 311.64 | 1 | 3.94 | 1.27 | 37.02 | 35.48 | 1.54 | 1.54 | 0.05 | 0.05 |
| gene-LOC105277761 | https://www.ncbi.nlm.nih.gov/gene/?term=LOC105277761 | 4648 | 143 | 307.66 | 1 | 2.15 | 0.7 | 15.33 | 13.81 | 1.52 | 1.52 | 0.05 | 0.05 |
| gene-LOC105285055 | https://www.ncbi.nlm.nih.gov/gene/?term=LOC105285055 | 4108 | 255 | 620.74 | 2 | 4.87 | 0.78 | 11.66 | 10.91 | 0.76 | 0.76 | 0.05 | 0.05 |
| gene-LOC105274534 | https://www.ncbi.nlm.nih.gov/gene/?term=LOC105274534 | 1329 | 56 | 421.37 | 1 | 7.52 | 1.79 | 47.04 | 48.26 | -1.22 | 1.22 | -0.05 | 0.05 |
| gene-LOC105285060 | https://www.ncbi.nlm.nih.gov/gene/?term=LOC105285060 | 1285 | 17 | 132.3 | 1 | 7.78 | 5.88 | 51.2 | 48.29 | 2.91 | 2.91 | 0.04 | 0.04 |
| gene-LOC113561805 | https://www.ncbi.nlm.nih.gov/gene/?term=LOC113561805 | 3337 | 68 | 203.78 | 1 | 3 | 1.47 | 35.02 | 33.08 | 1.94 | 1.94 | 0.04 | 0.04 |
| gene-LOC105282970 | https://www.ncbi.nlm.nih.gov/gene/?term=LOC105282970 | 1529 | 39 | 255.07 | 1 | 6.54 | 2.56 | 51.08 | 49.46 | 1.62 | 1.62 | 0.04 | 0.04 |
| gene-LOC105278807 | https://www.ncbi.nlm.nih.gov/gene/?term=LOC105278807 | 2166 | 62 | 286.24 | 1 | 4.62 | 1.61 | 21.98 | 20.52 | 1.46 | 1.46 | 0.04 | 0.04 |
| gene-LOC105283728 | https://www.ncbi.nlm.nih.gov/gene/?term=LOC105283728 | 3256 | 107 | 328.62 | 2 | 6.14 | 1.87 | 20.96 | 19.59 | 1.37 | 1.37 | 0.04 | 0.04 |
| gene-LOC105276875 | https://www.ncbi.nlm.nih.gov/gene/?term=LOC105276875 | 3151 | 109 | 345.92 | 1 | 3.17 | 0.92 | 25.45 | 24.19 | 1.26 | 1.26 | 0.04 | 0.04 |
| gene-LOC105284355 | https://www.ncbi.nlm.nih.gov/gene/?term=LOC105284355 | 3904 | 143 | 366.29 | 1 | 2.56 | 0.7 | 22.85 | 21.77 | 1.08 | 1.08 | 0.04 | 0.04 |
| gene-LOC105285179 | https://www.ncbi.nlm.nih.gov/gene/?term=LOC105285179 | 4191 | 177 | 422.33 | 1 | 2.39 | 0.56 | 19.66 | 18.63 | 1.04 | 1.04 | 0.04 | 0.04 |
| gene-LOC105276196 | https://www.ncbi.nlm.nih.gov/gene/?term=LOC105276196 | 2953 | 48 | 162.55 | 1 | 3.39 | 2.08 | 47.64 | 49.83 | -2.19 | 2.19 | -0.04 | 0.04 |
| gene-LOC105275708 | https://www.ncbi.nlm.nih.gov/gene/?term=LOC105275708 | 1772 | 10 | 56.43 | 1 | 5.64 | 10 | 50.31 | 44.22 | 6.09 | 6.09 | 0.03 | 0.03 |
| gene-LOC105279664 | https://www.ncbi.nlm.nih.gov/gene/?term=LOC105279664 | 4023 | 48 | 119.31 | 1 | 2.49 | 2.08 | 23.51 | 21.34 | 2.18 | 2.18 | 0.03 | 0.03 |
| gene-LOC105276190 | https://www.ncbi.nlm.nih.gov/gene/?term=LOC105276190 | 4839 | 82 | 169.46 | 1 | 2.07 | 1.22 | 11.63 | 9.92 | 1.71 | 1.71 | 0.03 | 0.03 |
| gene-LOC105282794 | https://www.ncbi.nlm.nih.gov/gene/?term=LOC105282794 | 4571 | 92 | 201.27 | 1 | 2.19 | 1.09 | 18.21 | 16.51 | 1.7 | 1.7 | 0.03 | 0.03 |
| gene-LOC105276059 | https://www.ncbi.nlm.nih.gov/gene/?term=LOC105276059 | 6256 | 121 | 193.41 | 1 | 1.6 | 0.83 | 10.95 | 9.59 | 1.36 | 1.36 | 0.03 | 0.03 |
| gene-LOC105281173 | https://www.ncbi.nlm.nih.gov/gene/?term=LOC105281173 | 1595 | 38 | 238.24 | 1 | 6.27 | 2.63 | 60.05 | 58.85 | 1.2 | 1.2 | 0.03 | 0.03 |
| gene-LOC105287063 | https://www.ncbi.nlm.nih.gov/gene/?term=LOC105287063 | 1874 | 48 | 256.14 | 1 | 5.34 | 2.08 | 31.5 | 30.33 | 1.17 | 1.17 | 0.03 | 0.03 |
| gene-LOC105281272 | https://www.ncbi.nlm.nih.gov/gene/?term=LOC105281272 | 5252 | 156 | 297.03 | 1 | 1.9 | 0.64 | 18.91 | 17.94 | 0.97 | 0.97 | 0.03 | 0.03 |
| gene-LOC105280774 | https://www.ncbi.nlm.nih.gov/gene/?term=LOC105280774 | 2793 | 89 | 318.65 | 1 | 3.58 | 1.12 | 36.33 | 35.38 | 0.94 | 0.94 | 0.03 | 0.03 |
| gene-LOC105276449 | https://www.ncbi.nlm.nih.gov/gene/?term=LOC105276449 | 1231 | 42 | 341.19 | 1 | 8.12 | 2.38 | 35.95 | 35.13 | 0.82 | 0.82 | 0.03 | 0.03 |
| gene-LOC105285202 | https://www.ncbi.nlm.nih.gov/gene/?term=LOC105285202 | 1838 | 67 | 364.53 | 1 | 5.44 | 1.49 | 26.25 | 25.49 | 0.76 | 0.76 | 0.03 | 0.03 |
| gene-LOC105275363 | https://www.ncbi.nlm.nih.gov/gene/?term=LOC105275363 | 3798 | 150 | 394.94 | 1 | 2.63 | 0.67 | 18.12 | 17.37 | 0.75 | 0.75 | 0.03 | 0.03 |
| gene-LOC105287095 | https://www.ncbi.nlm.nih.gov/gene/?term=LOC105287095 | 1435 | 57 | 397.21 | 1 | 6.97 | 1.75 | 29.87 | 29.16 | 0.71 | 0.71 | 0.03 | 0.03 |
| gene-LOC105275374 | https://www.ncbi.nlm.nih.gov/gene/?term=LOC105275374 | 11376 | 489 | 429.85 | 1 | 0.88 | 0.2 | 11.7 | 11.06 | 0.64 | 0.64 | 0.03 | 0.03 |
| gene-LOC105278145 | https://www.ncbi.nlm.nih.gov/gene/?term=LOC105278145 | 4470 | 246 | 550.34 | 1 | 2.24 | 0.41 | 11.03 | 10.48 | 0.55 | 0.55 | 0.03 | 0.03 |
| gene-LOC105278283 | https://www.ncbi.nlm.nih.gov/gene/?term=LOC105278283 | 2317 | 125 | 539.49 | 1 | 4.32 | 0.8 | 28.1 | 27.55 | 0.55 | 0.55 | 0.03 | 0.03 |
| gene-LOC105276452 | https://www.ncbi.nlm.nih.gov/gene/?term=LOC105276452 | 2917 | 263 | 901.61 | 1 | 3.43 | 0.38 | 9.41 | 9.13 | 0.28 | 0.28 | 0.03 | 0.03 |
| gene-LOC109611283 | https://www.ncbi.nlm.nih.gov/gene/?term=LOC109611283 | 1451 | 36 | 248.1 | 1 | 6.89 | 2.78 | 35.62 | 36.97 | -1.35 | 1.35 | -0.03 | 0.03 |
| gene-LOC105280819 | https://www.ncbi.nlm.nih.gov/gene/?term=LOC105280819 | 1639 | 37 | 225.75 | 1 | 6.1 | 2.7 | 36.55 | 38.07 | -1.52 | 1.52 | -0.03 | 0.03 |
| gene-LOC105286694 | https://www.ncbi.nlm.nih.gov/gene/?term=LOC105286694 | 3409 | 9 | 26.4 | 1 | 2.93 | 11.11 | 71.7 | 62.28 | 9.42 | 9.42 | 0.02 | 0.02 |
| gene-LOC105285315 | https://www.ncbi.nlm.nih.gov/gene/?term=LOC105285315 | 5458 | 19 | 34.81 | 1 | 1.83 | 5.26 | 47.88 | 43.22 | 4.66 | 4.66 | 0.02 | 0.02 |
| gene-LOC105275518 | https://www.ncbi.nlm.nih.gov/gene/?term=LOC105275518 | 2335 | 18 | 77.09 | 1 | 4.28 | 5.56 | 54.44 | 51.41 | 3.03 | 3.03 | 0.02 | 0.02 |
| gene-LOC105275347 | https://www.ncbi.nlm.nih.gov/gene/?term=LOC105275347 | 2526 | 14 | 55.42 | 1 | 3.96 | 7.14 | 33.57 | 30.77 | 2.8 | 2.8 | 0.02 | 0.02 |

| | | | | | | | | | | | | | | |
|---|---|---|---|---|---|---|---|---|---|---|---|---|---|---|
| gene-LOC105274900 | https://www.ncbi.nlm.nih.gov/gene/?term=LOC105274900 | 5561 | 47 | 84.52 | 2 | 3.6 | 4.26 | 37.9 | 35.14 | 2.76 | 2.76 | 0.02 | 0.02 | |
| gene-LOC105288066 | https://www.ncbi.nlm.nih.gov/gene/?term=LOC105288066 | 1964 | 20 | 101.83 | | 1 | 5.09 | 5 | 44.28 | 42.38 | 1.9 | 1.9 | 0.02 | 0.02 |
| gene-LOC105275966 | https://www.ncbi.nlm.nih.gov/gene/?term=LOC105275966 | 3299 | 53 | 160.65 | | 1 | 3.03 | 1.89 | 54.5 | 52.99 | 1.51 | 1.51 | 0.02 | 0.02 |
| gene-LOC105283712 | https://www.ncbi.nlm.nih.gov/gene/?term=LOC105283712 | 1910 | 31 | 162.3 | 1 | 5.24 | 3.23 | 30.95 | 29.59 | 1.36 | 1.36 | 0.02 | 0.02 | |
| gene-LOC105281848 | https://www.ncbi.nlm.nih.gov/gene/?term=LOC105281848 | 2122 | 47 | 221.49 | | 1 | 4.71 | 2.13 | 27.8 | 26.69 | 1.11 | 1.11 | 0.02 | 0.02 |
| gene-LOC105275451 | https://www.ncbi.nlm.nih.gov/gene/?term=LOC105275451 | 6035 | 118 | 195.53 | | 1 | 1.66 | 0.85 | 23.45 | 22.41 | 1.04 | 1.04 | 0.02 | 0.02 |
| gene-LOC105276272 | https://www.ncbi.nlm.nih.gov/gene/?term=LOC105276272 | 1965 | 34 | 173.03 | | 1 | 5.09 | 2.94 | 33.29 | 32.32 | 0.98 | 0.98 | 0.02 | 0.02 |
| gene-LOC105287417 | https://www.ncbi.nlm.nih.gov/gene/?term=LOC105287417 | 1025 | 22 | 214.63 | | 1 | 9.76 | 4.55 | 50.13 | 49.23 | 0.89 | 0.89 | 0.02 | 0.02 |
| gene-LOC105284036 | https://www.ncbi.nlm.nih.gov/gene/?term=LOC105284036 | 3254 | 96 | 295.02 | | 1 | 3.07 | 1.04 | 22.78 | 21.95 | 0.83 | 0.83 | 0.02 | 0.02 |
| gene-LOC105283592 | https://www.ncbi.nlm.nih.gov/gene/?term=LOC105283592 | 3479 | 72 | 206.96 | | 1 | 2.87 | 1.39 | 23.84 | 23.1 | 0.74 | 0.74 | 0.02 | 0.02 |
| gene-LOC105285395 | https://www.ncbi.nlm.nih.gov/gene/?term=LOC105285395 | 8085 | 185 | 228.82 | | 1 | 1.24 | 0.54 | 16.99 | 16.27 | 0.72 | 0.72 | 0.02 | 0.02 |
| gene-LOC113561621 | https://www.ncbi.nlm.nih.gov/gene/?term=LOC113561621 | 2249 | 82 | 364.61 | | 1 | 4.45 | 1.22 | 20.63 | 19.95 | 0.68 | 0.68 | 0.02 | 0.02 |
| gene-LOC105285639 | https://www.ncbi.nlm.nih.gov/gene/?term=LOC105285639 | 10325 | 319 | 308.96 | | 1 | 0.97 | 0.31 | 12.35 | 11.71 | 0.65 | 0.65 | 0.02 | 0.02 |
| gene-LOC105274452 | https://www.ncbi.nlm.nih.gov/gene/?term=LOC105274452 | 6595 | 252 | 382.11 | | 1 | 1.52 | 0.4 | 13.74 | 13.13 | 0.61 | 0.61 | 0.02 | 0.02 |
| gene-LOC105274706 | https://www.ncbi.nlm.nih.gov/gene/?term=LOC105274706 | 1446 | 55 | 380.36 | | 1 | 6.92 | 1.82 | 37.05 | 36.54 | 0.5 | 0.5 | 0.02 | 0.02 |
| gene-LOC105274856 | https://www.ncbi.nlm.nih.gov/gene/?term=LOC105274856 | 5216 | 207 | 396.86 | | 1 | 1.92 | 0.48 | 19.99 | 19.54 | 0.45 | 0.45 | 0.02 | 0.02 |
| gene-LOC105278666 | https://www.ncbi.nlm.nih.gov/gene/?term=LOC105278666 | 10341 | 429 | 414.85 | | 1 | 0.97 | 0.23 | 7.71 | 7.27 | 0.44 | 0.44 | 0.02 | 0.02 |
| gene-LOC105285807 | https://www.ncbi.nlm.nih.gov/gene/?term=LOC105285807 | 6063 | 214 | 352.96 | | 1 | 1.65 | 0.47 | 7.55 | 7.11 | 0.44 | 0.44 | 0.02 | 0.02 |
| gene-LOC105276612 | https://www.ncbi.nlm.nih.gov/gene/?term=LOC105276612 | 4157 | 181 | 435.41 | | 1 | 2.41 | 0.55 | 10.16 | 9.76 | 0.4 | 0.4 | 0.02 | 0.02 |
| gene-LOC105277423 | https://www.ncbi.nlm.nih.gov/gene/?term=LOC105277423 | 2340 | 108 | 461.54 | | 1 | 4.27 | 0.93 | 30.29 | 29.91 | 0.38 | 0.38 | 0.02 | 0.02 |
| gene-LOC105286649 | https://www.ncbi.nlm.nih.gov/gene/?term=LOC105286649 | 14178 | 778 | 548.74 | | 1 | 0.71 | 0.13 | 0.86 | 1.14 | -0.27 | 0.27 | -0.02 | 0.02 |
| gene-LOC105285252 | https://www.ncbi.nlm.nih.gov/gene/?term=LOC105285252 | 7427 | 112 | 150.8 | 2 | 2.69 | 1.79 | 14.17 | 15.17 | -1 | 1 | -0.02 | 0.02 | |
| gene-LOC105281700 | https://www.ncbi.nlm.nih.gov/gene/?term=LOC105281700 | 2208 | 34 | 153.99 | | 1 | 4.53 | 2.94 | 35.91 | 36.92 | -1.02 | 1.02 | -0.02 | 0.02 |
| gene-LOC105285860 | https://www.ncbi.nlm.nih.gov/gene/?term=LOC105285860 | 3110 | 52 | 167.2 | 1 | 3.22 | 1.92 | 20.92 | 22.04 | -1.12 | 1.12 | -0.02 | 0.02 | |
| gene-LOC105284507 | https://www.ncbi.nlm.nih.gov/gene/?term=LOC105284507 | 5207 | 77 | 147.88 | | 1 | 1.92 | 1.3 | 22.93 | 24.13 | -1.2 | 1.2 | -0.02 | 0.02 |
| gene-LOC105281515 | https://www.ncbi.nlm.nih.gov/gene/?term=LOC105281515 | 8827 | 38 | 43.05 | 1 | 1.13 | 2.63 | 31.29 | 29.13 | 2.16 | 2.16 | 0.01 | 0.01 | |
| gene-LOC105280408 | https://www.ncbi.nlm.nih.gov/gene/?term=LOC105280408 | 5983 | 47 | 78.56 | 1 | 1.67 | 2.13 | 19.12 | 17.33 | 1.78 | 1.78 | 0.01 | 0.01 | |
| gene-LOC105274590 | https://www.ncbi.nlm.nih.gov/gene/?term=LOC105274590 | 4516 | 57 | 126.22 | | 2 | 4.43 | 3.51 | 34.78 | 33.73 | 1.06 | 1.06 | 0.01 | 0.01 |
| gene-LOC105279500 | https://www.ncbi.nlm.nih.gov/gene/?term=LOC105279500 | 3587 | 34 | 94.79 | 1 | 2.79 | 2.94 | 52.36 | 51.39 | 0.97 | 0.97 | 0.01 | 0.01 | |
| gene-LOC105283987 | https://www.ncbi.nlm.nih.gov/gene/?term=LOC105283987 | 4841 | 64 | 132.2 | 1 | 2.07 | 1.56 | 30.28 | 29.39 | 0.89 | 0.89 | 0.01 | 0.01 | |
| gene-LOC105274851 | https://www.ncbi.nlm.nih.gov/gene/?term=LOC105274851 | 6616 | 123 | 185.91 | | 1 | 1.51 | 0.81 | 11.44 | 10.81 | 0.63 | 0.63 | 0.01 | 0.01 |
| gene-LOC105282885 | https://www.ncbi.nlm.nih.gov/gene/?term=LOC105282885 | 1870 | 45 | 240.64 | | 1 | 5.35 | 2.22 | 42.36 | 41.77 | 0.58 | 0.58 | 0.01 | 0.01 |
| gene-LOC105276692 | https://www.ncbi.nlm.nih.gov/gene/?term=LOC105276692 | 1745 | 29 | 166.19 | | 1 | 5.73 | 3.45 | 25.51 | 24.96 | 0.55 | 0.55 | 0.01 | 0.01 |
| gene-LOC105284704 | https://www.ncbi.nlm.nih.gov/gene/?term=LOC105284704 | 2400 | 57 | 237.5 | 1 | 4.17 | 1.75 | 25.25 | 24.69 | 0.55 | 0.55 | 0.01 | 0.01 | |
| gene-LOC105282151 | https://www.ncbi.nlm.nih.gov/gene/?term=LOC105282151 | 1601 | 35 | 218.61 | | 1 | 6.25 | 2.86 | 42.13 | 41.61 | 0.53 | 0.53 | 0.01 | 0.01 |
| gene-LOC105280410 | https://www.ncbi.nlm.nih.gov/gene/?term=LOC105280410 | 12580 | 235 | 186.8 | 1 | 0.79 | 0.43 | 21.34 | 20.84 | 0.5 | 0.5 | 0.01 | 0.01 | |
| gene-LOC105274823 | https://www.ncbi.nlm.nih.gov/gene/?term=LOC105274823 | 4620 | 132 | 285.71 | | 1 | 2.16 | 0.76 | 21.94 | 21.47 | 0.47 | 0.47 | 0.01 | 0.01 |
| gene-LOC105279038 | https://www.ncbi.nlm.nih.gov/gene/?term=LOC105279038 | 9453 | 306 | 323.71 | | 1 | 1.06 | 0.33 | 10.13 | 9.67 | 0.46 | 0.46 | 0.01 | 0.01 |
| gene-LOC105286514 | https://www.ncbi.nlm.nih.gov/gene/?term=LOC105286514 | 2528 | 44 | 174.05 | | 1 | 3.96 | 2.27 | 33.92 | 33.47 | 0.45 | 0.45 | 0.01 | 0.01 |
| gene-LOC105278282 | https://www.ncbi.nlm.nih.gov/gene/?term=LOC105278282 | 1775 | 52 | 292.96 | | 1 | 5.63 | 1.92 | 38.05 | 37.63 | 0.42 | 0.42 | 0.01 | 0.01 |
| gene-LOC105276941 | https://www.ncbi.nlm.nih.gov/gene/?term=LOC105276941 | 15956 | 494 | 309.6 | 1 | 0.63 | 0.2 | 5.44 | 5.07 | 0.38 | 0.38 | 0.01 | 0.01 | |
| gene-LOC105287689 | https://www.ncbi.nlm.nih.gov/gene/?term=LOC105287689 | 10867 | 331 | 304.59 | | 1 | 0.92 | 0.3 | 9.53 | 9.17 | 0.36 | 0.36 | 0.01 | 0.01 |
| gene-LOC105278454 | https://www.ncbi.nlm.nih.gov/gene/?term=LOC105278454 | 1972 | 44 | 223.12 | | 1 | 5.07 | 2.27 | 46.65 | 46.38 | 0.26 | 0.26 | 0.01 | 0.01 |
| gene-LOC105278877 | https://www.ncbi.nlm.nih.gov/gene/?term=LOC105278877 | 1968 | 69 | 350.61 | | 1 | 5.08 | 1.45 | 35.47 | 35.23 | 0.25 | 0.25 | 0.01 | 0.01 |
| gene-LOC105275460 | https://www.ncbi.nlm.nih.gov/gene/?term=LOC105275460 | 2473 | 116 | 469.07 | | 1 | 4.04 | 0.86 | 13.65 | 13.42 | 0.23 | 0.23 | 0.01 | 0.01 |
| gene-LOC113562050 | https://www.ncbi.nlm.nih.gov/gene/?term=LOC113562050 | 17097 | 540 | 315.84 | | 1 | 0.58 | 0.19 | 5.38 | 5.15 | 0.23 | 0.23 | 0.01 | 0.01 |
| gene-LOC105278534 | https://www.ncbi.nlm.nih.gov/gene/?term=LOC105278534 | 7364 | 175 | 237.64 | | 1 | 1.36 | 0.57 | 18.73 | 18.51 | 0.22 | 0.22 | 0.01 | 0.01 |
| gene-LOC105286166 | https://www.ncbi.nlm.nih.gov/gene/?term=LOC105286166 | 3050 | 103 | 337.7 | 1 | 3.28 | 0.97 | 25.5 | 25.28 | 0.22 | 0.22 | 0.01 | 0.01 | |
| gene-LOC105274449 | https://www.ncbi.nlm.nih.gov/gene/?term=LOC105274449 | 1549 | 48 | 309.88 | | 1 | 6.46 | 2.08 | 51.17 | 50.96 | 0.21 | 0.21 | 0.01 | 0.01 |
| gene-LOC105280956 | https://www.ncbi.nlm.nih.gov/gene/?term=LOC105280956 | 2639 | 69 | 261.46 | | 1 | 3.79 | 1.45 | 18.51 | 18.31 | 0.21 | 0.21 | 0.01 | 0.01 |
| gene-LOC105275683 | https://www.ncbi.nlm.nih.gov/gene/?term=LOC105275683 | 4256 | 183 | 429.98 | | 1 | 2.35 | 0.55 | 12.85 | 12.65 | 0.2 | 0.2 | 0.01 | 0.01 |
| gene-LOC105276462 | https://www.ncbi.nlm.nih.gov/gene/?term=LOC105276462 | 6561 | 246 | 374.94 | | 1 | 1.52 | 0.41 | 8.14 | 7.99 | 0.15 | 0.15 | 0.01 | 0.01 |
| gene-LOC105278649 | https://www.ncbi.nlm.nih.gov/gene/?term=LOC105278649 | 5616 | 161 | 286.68 | | 1 | 1.78 | 0.62 | 11.64 | 11.9 | -0.26 | 0.26 | -0.01 | 0.01 |
| gene-LOC105276527 | https://www.ncbi.nlm.nih.gov/gene/?term=LOC105276527 | 7923 | 309 | 390 | 1 | 1.26 | 0.32 | 9.63 | 9.9 | -0.27 | 0.27 | -0.01 | 0.01 | |
| gene-LOC105286274 | https://www.ncbi.nlm.nih.gov/gene/?term=LOC105286274 | 3817 | 193 | 505.63 | | 1 | 2.62 | 0.52 | 16.76 | 17.05 | -0.29 | 0.29 | -0.01 | 0.01 |
| gene-LOC105286515 | https://www.ncbi.nlm.nih.gov/gene/?term=LOC105286515 | 3751 | 99 | 263.93 | | 1 | 2.67 | 1.01 | 13.14 | 13.49 | -0.36 | 0.36 | -0.01 | 0.01 |
| gene-LOC105287633 | https://www.ncbi.nlm.nih.gov/gene/?term=LOC105287633 | 1524 | 27 | 177.17 | | 1 | 6.56 | 3.7 | 63.52 | 64 | -0.48 | 0.48 | -0.01 | 0.01 |
| gene-LOC105286452 | https://www.ncbi.nlm.nih.gov/gene/?term=LOC105286452 | 6888 | 62 | 90.01 | 1 | 1.45 | 1.61 | 32.65 | 33.21 | -0.56 | 0.56 | -0.01 | 0.01 | |
| gene-LOC105278108 | https://www.ncbi.nlm.nih.gov/gene/?term=LOC105278108 | 4392 | 44 | 100.18 | | 1 | 2.28 | 2.27 | 24.81 | 25.4 | -0.59 | 0.59 | -0.01 | 0.01 |
| gene-LOC105275187 | https://www.ncbi.nlm.nih.gov/gene/?term=LOC105275187 | 5534 | 66 | 119.26 | | 1 | 1.81 | 1.52 | 11.92 | 12.68 | -0.77 | 0.77 | -0.01 | 0.01 |
| gene-LOC105282067 | https://www.ncbi.nlm.nih.gov/gene/?term=LOC105282067 | 5941 | 51 | 85.84 | 1 | 1.68 | 1.96 | 26.09 | 27.84 | -1.75 | 1.75 | -0.01 | 0.01 | |
| gene-LOC105283989 | https://www.ncbi.nlm.nih.gov/gene/?term=LOC105283989 | 4948 | 26 | 52.55 | 1 | 2.02 | 3.85 | 37.24 | 39.59 | -2.34 | 2.34 | -0.01 | 0.01 | |
| gene-LOC105285406 | https://www.ncbi.nlm.nih.gov/gene/?term=LOC105285406 | 8015 | 60 | 74.86 | 1 | 1.25 | 1.67 | 25.96 | 25.52 | 0.44 | 0.44 | 0 | 0 | |
| gene-LOC105280407 | https://www.ncbi.nlm.nih.gov/gene/?term=LOC105280407 | 10950 | 189 | 172.6 | 1 | 0.91 | 0.53 | 16.21 | 15.97 | 0.24 | 0.24 | 0 | 0 | |

| | | | | | | | | | | | | | |
|---|---|---|---|---|---|---|---|---|---|---|---|---|---|
| gene-LOC105287764 | https://www.ncbi.nlm.nih.gov/gene/?term=LOC105287764 | 18359 | 375 | 204.26 | 1 | 0.54 | 0.27 | 3.25 | 3.09 | 0.17 | 0.17 | 0 | 0 |
| gene-LOC105282658 | https://www.ncbi.nlm.nih.gov/gene/?term=LOC105282658 | 2421 | 54 | 223.05 | 1 | 4.13 | 1.85 | 19.57 | 19.42 | 0.16 | 0.16 | 0 | 0 |
| gene-LOC105282825 | https://www.ncbi.nlm.nih.gov/gene/?term=LOC105282825 | 3919 | 69 | 176.07 | 1 | 2.55 | 1.45 | 13.52 | 13.36 | 0.16 | 0.16 | 0 | 0 |
| gene-LOC105286999 | https://www.ncbi.nlm.nih.gov/gene/?term=LOC105286999 | 2141 | 42 | 196.17 | 1 | 4.67 | 2.38 | 36.17 | 36.05 | 0.12 | 0.12 | 0 | 0 |
| gene-LOC105282811 | https://www.ncbi.nlm.nih.gov/gene/?term=LOC105282811 | 5765 | 198 | 343.45 | 1 | 1.73 | 0.51 | 7.89 | 7.77 | 0.11 | 0.11 | 0 | 0 |
| gene-LOC105275327 | https://www.ncbi.nlm.nih.gov/gene/?term=LOC105275327 | 14245 | 460 | 322.92 | 1 | 0.7 | 0.22 | 7.25 | 7.2 | 0.05 | 0.05 | 0 | 0 |
| gene-LOC105282646 | https://www.ncbi.nlm.nih.gov/gene/?term=LOC105282646 | 3317 | 99 | 298.46 | 1 | 3.01 | 1.01 | 25.07 | 25.11 | -0.04 | 0.04 | 0 | 0 |
| gene-LOC105274531 | https://www.ncbi.nlm.nih.gov/gene/?term=LOC105274531 | 7774 | 285 | 366.61 | 1 | 1.29 | 0.35 | 10.62 | 10.74 | -0.11 | 0.11 | 0 | 0 |
| gene-LOC105280377 | https://www.ncbi.nlm.nih.gov/gene/?term=LOC105280377 | 2024 | 58 | 286.56 | 1 | 4.94 | 1.72 | 49.97 | 50.08 | -0.11 | 0.11 | 0 | 0 |
| gene-LOC105275848 | https://www.ncbi.nlm.nih.gov/gene/?term=LOC105275848 | 3718 | 133 | 357.72 | 1 | 2.69 | 0.75 | 21.71 | 21.82 | -0.12 | 0.12 | 0 | 0 |
| gene-LOC105276538 | https://www.ncbi.nlm.nih.gov/gene/?term=LOC105276538 | 1511 | 19 | 125.74 | 1 | 6.62 | 5.26 | 67.47 | 67.59 | -0.12 | 0.12 | 0 | 0 |
| gene-LOC105283296 | https://www.ncbi.nlm.nih.gov/gene/?term=LOC105283296 | 2174 | 84 | 386.38 | 1 | 4.6 | 1.19 | 19.37 | 19.49 | -0.12 | 0.12 | 0 | 0 |
| gene-LOC105278632 | https://www.ncbi.nlm.nih.gov/gene/?term=LOC105278632 | 3624 | 113 | 311.81 | 1 | 2.76 | 0.88 | 16.03 | 16.16 | -0.13 | 0.13 | 0 | 0 |
| gene-LOC105283943 | https://www.ncbi.nlm.nih.gov/gene/?term=LOC105283943 | 3559 | 42 | 118.01 | 1 | 2.81 | 2.38 | 14.31 | 14.48 | -0.17 | 0.17 | 0 | 0 |
| gene-LOC105275179 | https://www.ncbi.nlm.nih.gov/gene/?term=LOC105275179 | 3271 | 66 | 201.77 | 1 | 3.06 | 1.52 | 20.6 | 20.81 | -0.21 | 0.21 | 0 | 0 |
| gene-LOC105285514 | https://www.ncbi.nlm.nih.gov/gene/?term=LOC105285514 | 1277 | 14 | 109.63 | 1 | 7.83 | 7.14 | 85.59 | 85.88 | -0.28 | 0.28 | 0 | 0 |
| gene-LOC109611234 | https://www.ncbi.nlm.nih.gov/gene/?term=LOC109611234 | 3663 | 28 | 76.44 1 | 2.73 | 3.57 | 16.2 | 16.56 | -0.36 | 0.36 | 0 | 0 | |

Directory: OGL File: rnk-dmg-Ob_LI_BCphaseB1.vs.BCphaseB2.txt

| gene_ID | gene_link | gwidth | #Sites | #per10Kb | #dmSites | #dmsp10kb | %dmSites | %pSite1 | %pSite2 | DMpSite | ADMpSite | DMpNucl | ADMpNucl |
|---|---|---|---|---|---|---|---|---|---|---|---|---|---|
| gene-LOC113562345 | https://www.ncbi.nlm.nih.gov/gene/?term=LOC113562345 | 2290 | 64 | 279.48 | 15 | 65.5 | 23.44 | 32.83 | 17.93 | 14.9 | 14.9 | 0.42 | 0.42 |
| gene-LOC105284677 | https://www.ncbi.nlm.nih.gov/gene/?term=LOC105284677 | 1905 | 42 | 220.47 | 1 | 5.25 | 2.38 | 26.24 | 37.79 | -11.55 | 11.55 | -0.25 | 0.25 |
| gene-LOC105283649 | https://www.ncbi.nlm.nih.gov/gene/?term=LOC105283649 | 1918 | 56 | 291.97 | 1 | 5.21 | 1.79 | 39.64 | 36.17 | 3.47 | 3.47 | 0.1 | 0.1 |
| gene-LOC105276449 | https://www.ncbi.nlm.nih.gov/gene/?term=LOC105276449 | 1231 | 51 | 414.3 | 1 | 8.12 | 1.96 | 27.95 | 25.74 | 2.21 | 2.21 | 0.09 | 0.09 |
| gene-LOC105275070 | https://www.ncbi.nlm.nih.gov/gene/?term=LOC105275070 | 5667 | 49 | 86.47 | 6 | 10.59 | 12.24 | 10.93 | 1.84 | 9.09 | 9.09 | 0.08 | 0.08 |
| gene-LOC105275447 | https://www.ncbi.nlm.nih.gov/gene/?term=LOC105275447 | 1846 | 32 | 173.35 | 1 | 5.42 | 3.12 | 8.01 | 3.17 | 4.84 | 4.84 | 0.08 | 0.08 |
| gene-LOC105285674 | https://www.ncbi.nlm.nih.gov/gene/?term=LOC105285674 | 3035 | 94 | 309.72 | 2 | 6.59 | 2.13 | 27.11 | 24.65 | 2.46 | 2.46 | 0.08 | 0.08 |
| gene-LOC105278877 | https://www.ncbi.nlm.nih.gov/gene/?term=LOC105278877 | 1968 | 82 | 416.67 | 1 | 5.08 | 1.22 | 32.75 | 30.76 | 1.99 | 1.99 | 0.08 | 0.08 |
| gene-LOC105282156 | https://www.ncbi.nlm.nih.gov/gene/?term=LOC105282156 | 2577 | 137 | 531.63 | 1 | 3.88 | 0.73 | 24.98 | 23.48 | 1.5 | 1.5 | 0.08 | 0.08 |
| gene-LOC105282885 | https://www.ncbi.nlm.nih.gov/gene/?term=LOC105282885 | 1870 | 47 | 251.34 | 1 | 5.35 | 2.13 | 35.98 | 39.14 | -3.16 | 3.16 | -0.08 | 0.08 |
| gene-LOC105275708 | https://www.ncbi.nlm.nih.gov/gene/?term=LOC105275708 | 1772 | 15 | 84.65 | 1 | 5.64 | 6.67 | 46.38 | 38.5 | 7.88 | 7.88 | 0.07 | 0.07 |
| gene-LOC105281039 | https://www.ncbi.nlm.nih.gov/gene/?term=LOC105281039 | 1471 | 12 | 81.58 | 1 | 6.8 | 8.33 | 38.6 | 31.46 | 7.13 | 7.13 | 0.06 | 0.06 |
| gene-LOC105279900 | https://www.ncbi.nlm.nih.gov/gene/?term=LOC105279900 | 2191 | 56 | 255.59 | 1 | 4.56 | 1.79 | 33.98 | 31.69 | 2.29 | 2.29 | 0.06 | 0.06 |
| gene-LOC105279898 | https://www.ncbi.nlm.nih.gov/gene/?term=LOC105279898 | 1343 | 13 | 96.8 | 1 | 7.45 | 7.69 | 64.63 | 59.11 | 5.52 | 5.52 | 0.05 | 0.05 |
| gene-LOC105275518 | https://www.ncbi.nlm.nih.gov/gene/?term=LOC105275518 | 2335 | 23 | 98.5 | 1 | 4.28 | 4.35 | 43.86 | 38.74 | 5.13 | 5.13 | 0.05 | 0.05 |
| gene-LOC105287417 | https://www.ncbi.nlm.nih.gov/gene/?term=LOC105287417 | 1025 | 36 | 351.22 | 1 | 9.76 | 2.78 | 35.49 | 34.17 | 1.32 | 1.32 | 0.05 | 0.05 |
| gene-LOC105280786 | https://www.ncbi.nlm.nih.gov/gene/?term=LOC105280786 | 2573 | 119 | 462.5 | 1 | 3.89 | 0.84 | 30.85 | 29.73 | 1.12 | 1.12 | 0.05 | 0.05 |
| gene-LOC105274706 | https://www.ncbi.nlm.nih.gov/gene/?term=LOC105274706 | 1446 | 50 | 345.78 | 1 | 6.92 | 2 | 40.41 | 41.89 | -1.49 | 1.49 | -0.05 | 0.05 |
| gene-LOC105287633 | https://www.ncbi.nlm.nih.gov/gene/?term=LOC105287633 | 1524 | 31 | 203.41 | 1 | 6.56 | 3.23 | 53.47 | 55.78 | -2.31 | 2.31 | -0.05 | 0.05 |
| gene-LOC105274900 | https://www.ncbi.nlm.nih.gov/gene/?term=LOC105274900 | 5561 | 54 | 97.1 | 2 | 3.6 | 3.7 | 35.58 | 31 | 4.57 | 4.57 | 0.04 | 0.04 |
| gene-LOC105281420 | https://www.ncbi.nlm.nih.gov/gene/?term=LOC105281420 | 3882 | 67 | 172.59 | 1 | 2.58 | 1.49 | 15.32 | 12.95 | 2.37 | 2.37 | 0.04 | 0.04 |
| gene-LOC105283592 | https://www.ncbi.nlm.nih.gov/gene/?term=LOC105283592 | 3479 | 71 | 204.08 | 1 | 2.87 | 1.41 | 27.65 | 25.73 | 1.93 | 1.93 | 0.04 | 0.04 |
| gene-LOC105278454 | https://www.ncbi.nlm.nih.gov/gene/?term=LOC105278454 | 1972 | 48 | 243.41 | 1 | 5.07 | 2.08 | 42.9 | 41.15 | 1.75 | 1.75 | 0.04 | 0.04 |
| gene-LOC105285583 | https://www.ncbi.nlm.nih.gov/gene/?term=LOC105285583 | 1947 | 45 | 231.12 | 1 | 5.14 | 2.22 | 29.78 | 28.25 | 1.53 | 1.53 | 0.04 | 0.04 |
| gene-LOC105278632 | https://www.ncbi.nlm.nih.gov/gene/?term=LOC105278632 | 3624 | 126 | 347.68 | 1 | 2.76 | 0.79 | 16.08 | 14.93 | 1.15 | 1.15 | 0.04 | 0.04 |
| gene-LOC105285202 | https://www.ncbi.nlm.nih.gov/gene/?term=LOC105285202 | 1838 | 63 | 342.76 | 1 | 5.44 | 1.59 | 23.99 | 22.95 | 1.05 | 1.05 | 0.04 | 0.04 |
| gene-LOC105284355 | https://www.ncbi.nlm.nih.gov/gene/?term=LOC105284355 | 3904 | 158 | 404.71 | 1 | 2.56 | 0.63 | 20.65 | 19.62 | 1.02 | 1.02 | 0.04 | 0.04 |
| gene-LOC105288036 | https://www.ncbi.nlm.nih.gov/gene/?term=LOC105288036 | 5110 | 227 | 444.23 | 1 | 1.96 | 0.44 | 15.78 | 14.87 | 0.91 | 0.91 | 0.04 | 0.04 |
| gene-LOC105275384 | https://www.ncbi.nlm.nih.gov/gene/?term=LOC105275384 | 1987 | 92 | 463.01 | 1 | 5.03 | 1.09 | 32.77 | 31.88 | 0.89 | 0.89 | 0.04 | 0.04 |
| gene-LOC105285055 | https://www.ncbi.nlm.nih.gov/gene/?term=LOC105285055 | 4108 | 253 | 615.87 | 1 | 2.43 | 0.4 | 12.35 | 11.74 | 0.61 | 0.61 | 0.04 | 0.04 |
| gene-LOC105274534 | https://www.ncbi.nlm.nih.gov/gene/?term=LOC105274534 | 1329 | 60 | 451.47 | 1 | 7.52 | 1.67 | 40.57 | 41.38 | -0.82 | 0.82 | -0.04 | 0.04 |
| gene-LOC105276429 | https://www.ncbi.nlm.nih.gov/gene/?term=LOC105276429 | 2901 | 108 | 372.29 | 1 | 3.45 | 0.93 | 24.11 | 25.23 | -1.12 | 1.12 | -0.04 | 0.04 |
| gene-LOC105276538 | https://www.ncbi.nlm.nih.gov/gene/?term=LOC105276538 | 1511 | 17 | 112.51 | 1 | 6.62 | 5.88 | 72.11 | 69.24 | 2.87 | 2.87 | 0.03 | 0.03 |
| gene-LOC105285514 | https://www.ncbi.nlm.nih.gov/gene/?term=LOC105285514 | 1277 | 15 | 117.46 | 1 | 7.83 | 6.67 | 80.63 | 78.25 | 2.38 | 2.38 | 0.03 | 0.03 |
| gene-LOC105279664 | https://www.ncbi.nlm.nih.gov/gene/?term=LOC105279664 | 4023 | 65 | 161.57 | 1 | 2.49 | 1.54 | 19.54 | 17.85 | 1.7 | 1.7 | 0.03 | 0.03 |
| gene-LOC105286514 | https://www.ncbi.nlm.nih.gov/gene/?term=LOC105286514 | 2528 | 46 | 181.96 | 1 | 3.96 | 2.17 | 37.97 | 36.35 | 1.61 | 1.61 | 0.03 | 0.03 |
| gene-LOC105283943 | https://www.ncbi.nlm.nih.gov/gene/?term=LOC105283943 | 3559 | 62 | 174.21 | 1 | 2.81 | 1.61 | 15.22 | 13.63 | 1.6 | 1.6 | 0.03 | 0.03 |
| gene-LOC105277903 | https://www.ncbi.nlm.nih.gov/gene/?term=LOC105277903 | 1835 | 37 | 201.63 | 1 | 5.45 | 2.7 | 54.03 | 52.65 | 1.39 | 1.39 | 0.03 | 0.03 |
| gene-LOC105282794 | https://www.ncbi.nlm.nih.gov/gene/?term=LOC105282794 | 4571 | 98 | 214.4 | 1 | 2.19 | 1.02 | 19.72 | 18.47 | 1.25 | 1.25 | 0.03 | 0.03 |
| gene-LOC105280956 | https://www.ncbi.nlm.nih.gov/gene/?term=LOC105280956 | 2639 | 75 | 284.2 | 1 | 3.79 | 1.33 | 17.61 | 16.45 | 1.16 | 1.16 | 0.03 | 0.03 |
| gene-LOC105287063 | https://www.ncbi.nlm.nih.gov/gene/?term=LOC105287063 | 1874 | 51 | 272.15 | 1 | 5.34 | 1.96 | 30.2 | 29.07 | 1.12 | 1.12 | 0.03 | 0.03 |
| gene-LOC105275752 | https://www.ncbi.nlm.nih.gov/gene/?term=LOC105275752 | 3462 | 130 | 375.51 | 1 | 2.89 | 0.77 | 31.52 | 30.74 | 0.77 | 0.77 | 0.03 | 0.03 |
| gene-LOC105285639 | https://www.ncbi.nlm.nih.gov/gene/?term=LOC105285639 | 10325 | 378 | 366.1 | 1 | 0.97 | 0.26 | 11.07 | 10.35 | 0.72 | 0.72 | 0.03 | 0.03 |
| gene-LOC105278807 | https://www.ncbi.nlm.nih.gov/gene/?term=LOC105278807 | 2166 | 80 | 369.34 | 1 | 4.62 | 1.25 | 18.57 | 17.87 | 0.69 | 0.69 | 0.03 | 0.03 |
| gene-LOC105278497 | https://www.ncbi.nlm.nih.gov/gene/?term=LOC105278497 | 2072 | 95 | 458.49 | 1 | 4.83 | 1.05 | 19.7 | 19.09 | 0.61 | 0.61 | 0.03 | 0.03 |
| gene-LOC105282149 | https://www.ncbi.nlm.nih.gov/gene/?term=LOC105282149 | 1787 | 107 | 598.77 | 1 | 5.6 | 0.93 | 41.49 | 40.93 | 0.57 | 0.57 | 0.03 | 0.03 |
| gene-LOC105279298 | https://www.ncbi.nlm.nih.gov/gene/?term=LOC105279298 | 2773 | 132 | 476.02 | 2 | 7.21 | 1.52 | 23.66 | 23.13 | 0.53 | 0.53 | 0.03 | 0.03 |
| gene-LOC105276704 | https://www.ncbi.nlm.nih.gov/gene/?term=LOC105276704 | 2063 | 99 | 479.88 | 2 | 9.69 | 2.02 | 13.44 | 13.97 | -0.53 | 0.53 | -0.03 | 0.03 |
| gene-LOC105285533 | https://www.ncbi.nlm.nih.gov/gene/?term=LOC105285533 | 1640 | 70 | 426.83 | 1 | 6.1 | 1.43 | 22.75 | 23.49 | -0.75 | 0.75 | -0.03 | 0.03 |
| gene-LOC105274449 | https://www.ncbi.nlm.nih.gov/gene/?term=LOC105274449 | 1549 | 53 | 342.16 | 1 | 6.46 | 1.89 | 50.86 | 51.77 | -0.91 | 0.91 | -0.03 | 0.03 |
| gene-LOC105281312 | https://www.ncbi.nlm.nih.gov/gene/?term=LOC105281312 | 2551 | 65 | 254.8 | 1 | 3.92 | 1.54 | 31.99 | 33.22 | -1.23 | 1.23 | -0.03 | 0.03 |
| gene-LOC105281173 | https://www.ncbi.nlm.nih.gov/gene/?term=LOC105281173 | 1595 | 38 | 238.24 | 1 | 6.27 | 2.63 | 59.28 | 60.67 | -1.38 | 1.38 | -0.03 | 0.03 |
| gene-LOC105279500 | https://www.ncbi.nlm.nih.gov/gene/?term=LOC105279500 | 3587 | 33 | 92 | 1 | 2.79 | 3.03 | 43.74 | 47.01 | -3.27 | 3.27 | -0.03 | 0.03 |
| gene-LOC105281515 | https://www.ncbi.nlm.nih.gov/gene/?term=LOC105281515 | 8827 | 57 | 64.57 | 2 | 2.27 | 3.51 | 30.56 | 27.6 | 2.96 | 2.96 | 0.02 | 0.02 |
| gene-LOC105287648 | https://www.ncbi.nlm.nih.gov/gene/?term=LOC105287648 | 2205 | 30 | 136.05 | 1 | 4.54 | 3.33 | 43.37 | 41.93 | 1.44 | 1.44 | 0.02 | 0.02 |
| gene-LOC105287854 | https://www.ncbi.nlm.nih.gov/gene/?term=LOC105287854 | 3644 | 67 | 183.86 | 1 | 2.74 | 1.49 | 16.53 | 15.24 | 1.29 | 1.29 | 0.02 | 0.02 |
| gene-LOC105282991 | https://www.ncbi.nlm.nih.gov/gene/?term=LOC105282991 | 5617 | 105 | 186.93 | 1 | 1.78 | 0.95 | 19.71 | 18.51 | 1.19 | 1.19 | 0.02 | 0.02 |
| gene-LOC105288078 | https://www.ncbi.nlm.nih.gov/gene/?term=LOC105288078 | 19126 | 309 | 161.56 | 1 | 0.52 | 0.32 | 17.45 | 16.3 | 1.15 | 1.15 | 0.02 | 0.02 |
| gene-LOC105280641 | https://www.ncbi.nlm.nih.gov/gene/?term=LOC105280641 | 3682 | 55 | 149.38 | 1 | 2.72 | 1.82 | 19.12 | 18.02 | 1.1 | 1.1 | 0.02 | 0.02 |
| gene-LOC105280002 | https://www.ncbi.nlm.nih.gov/gene/?term=LOC105280002 | 1940 | 28 | 144.33 | 1 | 5.15 | 3.57 | 40.24 | 39.15 | 1.09 | 1.09 | 0.02 | 0.02 |

| | | | | | | | | | | | | | |
|---|---|---|---|---|---|---|---|---|---|---|---|---|---|
| gene-LOC105280348 | https://www.ncbi.nlm.nih.gov/gene/?term=LOC105280348 | 3797 | 76 | 200.16 | 1 | 2.63 | 1.32 | 20.16 | 19.19 | 0.97 | 0.97 | 0.02 | 0.02 |
| gene-LOC105288042 | https://www.ncbi.nlm.nih.gov/gene/?term=LOC105288042 | 3507 | 99 | 282.29 | 1 | 2.85 | 1.01 | 23.14 | 22.25 | 0.89 | 0.89 | 0.02 | 0.02 |
| gene-LOC105284117 | https://www.ncbi.nlm.nih.gov/gene/?term=LOC105284117 | 4340 | 94 | 216.59 | 1 | 2.3 | 1.06 | 11.08 | 10.26 | 0.83 | 0.83 | 0.02 | 0.02 |
| gene-LOC105284116 | https://www.ncbi.nlm.nih.gov/gene/?term=LOC105284116 | 3970 | 107 | 269.52 | 1 | 2.52 | 0.93 | 7.59 | 6.83 | 0.75 | 0.75 | 0.02 | 0.02 |
| gene-LOC105280405 | https://www.ncbi.nlm.nih.gov/gene/?term=LOC105280405 | 5422 | 161 | 296.94 | 1 | 1.84 | 0.62 | 10.76 | 10.05 | 0.71 | 0.71 | 0.02 | 0.02 |
| gene-LOC105281848 | https://www.ncbi.nlm.nih.gov/gene/?term=LOC105281848 | 2122 | 60 | 282.75 | 1 | 4.71 | 1.67 | 20.32 | 19.71 | 0.62 | 0.62 | 0.02 | 0.02 |
| gene-LOC105283728 | https://www.ncbi.nlm.nih.gov/gene/?term=LOC105283728 | 3256 | 122 | 374.69 | 2 | 6.14 | 1.64 | 19.35 | 18.81 | 0.54 | 0.54 | 0.02 | 0.02 |
| gene-LOC105275363 | https://www.ncbi.nlm.nih.gov/gene/?term=LOC105275363 | 3798 | 168 | 442.34 | 1 | 2.63 | 0.6 | 16.96 | 16.44 | 0.52 | 0.52 | 0.02 | 0.02 |
| gene-LOC105275683 | https://www.ncbi.nlm.nih.gov/gene/?term=LOC105275683 | 4256 | 205 | 481.67 | 1 | 2.35 | 0.49 | 13.27 | 12.75 | 0.52 | 0.52 | 0.02 | 0.02 |
| gene-LOC105282951 | https://www.ncbi.nlm.nih.gov/gene/?term=LOC105282951 | 3366 | 121 | 359.48 | 1 | 2.97 | 0.83 | 20.13 | 19.67 | 0.46 | 0.46 | 0.02 | 0.02 |
| gene-LOC105274856 | https://www.ncbi.nlm.nih.gov/gene/?term=LOC105274856 | 5216 | 222 | 425.61 | 1 | 1.92 | 0.45 | 17.35 | 16.9 | 0.45 | 0.45 | 0.02 | 0.02 |
| gene-LOC105274531 | https://www.ncbi.nlm.nih.gov/gene/?term=LOC105274531 | 7774 | 291 | 374.32 | 1 | 1.29 | 0.34 | 10.83 | 10.42 | 0.41 | 0.41 | 0.02 | 0.02 |
| gene-LOC105286274 | https://www.ncbi.nlm.nih.gov/gene/?term=LOC105286274 | 3817 | 200 | 523.97 | 1 | 2.62 | 0.5 | 16.54 | 16.86 | -0.32 | 0.32 | -0.02 | 0.02 |
| gene-LOC105278283 | https://www.ncbi.nlm.nih.gov/gene/?term=LOC105278283 | 2317 | 139 | 599.91 | 1 | 4.32 | 0.72 | 25.04 | 25.46 | -0.41 | 0.41 | -0.02 | 0.02 |
| gene-LOC105285409 | https://www.ncbi.nlm.nih.gov/gene/?term=LOC105285409 | 5099 | 58 | 113.75 | 1 | 1.96 | 1.72 | 1.58 | 3.25 | -1.67 | 1.67 | -0.02 | 0.02 |
| gene-LOC105282914 | https://www.ncbi.nlm.nih.gov/gene/?term=LOC105282914 | 5623 | 41 | 72.91 1 | 1.78 | 2.44 | 49.33 | 52.21 | -2.88 | 2.88 | -0.02 | 0.02 | |
| gene-LOC105283989 | https://www.ncbi.nlm.nih.gov/gene/?term=LOC105283989 | 4948 | 31 | 62.65 1 | 2.02 | 3.23 | 46.67 | 44.28 | 2.39 | 2.39 | 0.01 | 0.01 | |
| gene-LOC105285315 | https://www.ncbi.nlm.nih.gov/gene/?term=LOC105285315 | 5458 | 26 | 47.64 1 | 1.83 | 3.85 | 38.49 | 36.2 | 2.29 | 2.29 | 0.01 | 0.01 | |
| gene-LOC105276635 | https://www.ncbi.nlm.nih.gov/gene/?term=LOC105276635 | 4962 | 33 | 66.51 1 | 2.02 | 3.03 | 19.08 | 17.59 | 1.49 | 1.49 | 0.01 | 0.01 | |
| gene-LOC105276692 | https://www.ncbi.nlm.nih.gov/gene/?term=LOC105276692 | 1745 | 34 | 194.84 | 1 | 5.73 | 2.94 | 22.19 | 21.46 | 0.73 | 0.73 | 0.01 | 0.01 |
| gene-LOC105282600 | https://www.ncbi.nlm.nih.gov/gene/?term=LOC105282600 | 3715 | 50 | 134.59 | 1 | 2.69 | 2 | 33.68 | 32.96 | 0.73 | 0.73 | 0.01 | 0.01 |
| gene-LOC105288066 | https://www.ncbi.nlm.nih.gov/gene/?term=LOC105288066 | 1964 | 30 | 152.75 | 1 | 5.09 | 3.33 | 35.84 | 35.13 | 0.71 | 0.71 | 0.01 | 0.01 |
| gene-LOC105281266 | https://www.ncbi.nlm.nih.gov/gene/?term=LOC105281266 | 4102 | 79 | 192.59 | 1 | 2.44 | 1.27 | 15.99 | 15.33 | 0.66 | 0.66 | 0.01 | 0.01 |
| gene-LOC105278962 | https://www.ncbi.nlm.nih.gov/gene/?term=LOC105278962 | 4491 | 106 | 236.03 | 1 | 2.23 | 0.94 | 20.64 | 20.08 | 0.56 | 0.56 | 0.01 | 0.01 |
| gene-LOC105288010 | https://www.ncbi.nlm.nih.gov/gene/?term=LOC105288010 | 3189 | 45 | 141.11 | 1 | 3.14 | 2.22 | 33.55 | 32.98 | 0.56 | 0.56 | 0.01 | 0.01 |
| gene-LOC105275187 | https://www.ncbi.nlm.nih.gov/gene/?term=LOC105275187 | 5534 | 74 | 133.72 | 1 | 1.81 | 1.35 | 16.17 | 15.62 | 0.54 | 0.54 | 0.01 | 0.01 |
| gene-LOC105278228 | https://www.ncbi.nlm.nih.gov/gene/?term=LOC105278228 | 8610 | 172 | 199.77 | 1 | 1.16 | 0.58 | 23.43 | 22.95 | 0.48 | 0.48 | 0.01 | 0.01 |
| gene-LOC105275170 | https://www.ncbi.nlm.nih.gov/gene/?term=LOC105275170 | 12918 | 298 | 230.69 | 1 | 0.77 | 0.34 | 2.35 | 1.88 | 0.47 | 0.47 | 0.01 | 0.01 |
| gene-LOC105276059 | https://www.ncbi.nlm.nih.gov/gene/?term=LOC105276059 | 6256 | 138 | 220.59 | 1 | 1.6 | 0.72 | 9.13 | 8.71 | 0.42 | 0.42 | 0.01 | 0.01 |
| gene-LOC105282811 | https://www.ncbi.nlm.nih.gov/gene/?term=LOC105282811 | 5765 | 218 | 378.14 | 1 | 1.73 | 0.46 | 8.32 | 7.93 | 0.39 | 0.39 | 0.01 | 0.01 |
| gene-LOC105285395 | https://www.ncbi.nlm.nih.gov/gene/?term=LOC105285395 | 8085 | 208 | 257.27 | 1 | 1.24 | 0.48 | 14.72 | 14.34 | 0.39 | 0.39 | 0.01 | 0.01 |
| gene-LOC105274590 | https://www.ncbi.nlm.nih.gov/gene/?term=LOC105274590 | 4516 | 75 | 166.08 | 2 | 4.43 | 2.67 | 29.15 | 28.79 | 0.35 | 0.35 | 0.01 | 0.01 |
| gene-LOC105275451 | https://www.ncbi.nlm.nih.gov/gene/?term=LOC105275451 | 6035 | 136 | 225.35 | 1 | 1.66 | 0.74 | 21.08 | 20.75 | 0.33 | 0.33 | 0.01 | 0.01 |
| gene-LOC105277761 | https://www.ncbi.nlm.nih.gov/gene/?term=LOC105277761 | 4648 | 148 | 318.42 | 1 | 2.15 | 0.68 | 15.39 | 15.08 | 0.31 | 0.31 | 0.01 | 0.01 |
| gene-LOC105279038 | https://www.ncbi.nlm.nih.gov/gene/?term=LOC105279038 | 9453 | 363 | 384.01 | 1 | 1.06 | 0.28 | 8.55 | 8.25 | 0.3 | 0.3 | 0.01 | 0.01 |
| gene-LOC105285807 | https://www.ncbi.nlm.nih.gov/gene/?term=LOC105285807 | 6063 | 228 | 376.05 | 1 | 1.65 | 0.44 | 8.95 | 8.68 | 0.28 | 0.28 | 0.01 | 0.01 |
| gene-LOC105274452 | https://www.ncbi.nlm.nih.gov/gene/?term=LOC105274452 | 6595 | 260 | 394.24 | 1 | 1.52 | 0.38 | 13.07 | 12.82 | 0.25 | 0.25 | 0.01 | 0.01 |
| gene-LOC105278666 | https://www.ncbi.nlm.nih.gov/gene/?term=LOC105278666 | 10341 | 477 | 461.27 | 1 | 0.97 | 0.21 | 6.78 | 6.53 | 0.24 | 0.24 | 0.01 | 0.01 |
| gene-LOC105276527 | https://www.ncbi.nlm.nih.gov/gene/?term=LOC105276527 | 7923 | 332 | 419.03 | 1 | 1.26 | 0.3 | 9.67 | 9.44 | 0.23 | 0.23 | 0.01 | 0.01 |
| gene-LOC113562050 | https://www.ncbi.nlm.nih.gov/gene/?term=LOC113562050 | 17097 | 605 | 353.86 | 1 | 0.58 | 0.17 | 5.18 | 4.96 | 0.21 | 0.21 | 0.01 | 0.01 |
| gene-LOC105275327 | https://www.ncbi.nlm.nih.gov/gene/?term=LOC105275327 | 14245 | 543 | 381.19 | 1 | 0.7 | 0.18 | 6.62 | 6.43 | 0.19 | 0.19 | 0.01 | 0.01 |
| gene-LOC105282491 | https://www.ncbi.nlm.nih.gov/gene/?term=LOC105282491 | 4761 | 177 | 371.77 | 1 | 2.1 | 0.56 | 8.66 | 8.47 | 0.19 | 0.19 | 0.01 | 0.01 |
| gene-LOC105275460 | https://www.ncbi.nlm.nih.gov/gene/?term=LOC105275460 | 2473 | 123 | 497.37 | 1 | 4.04 | 0.81 | 13.15 | 12.97 | 0.18 | 0.18 | 0.01 | 0.01 |
| gene-LOC105284262 | https://www.ncbi.nlm.nih.gov/gene/?term=LOC105284262 | 6669 | 235 | 352.38 | 1 | 1.5 | 0.43 | 4.64 | 4.46 | 0.18 | 0.18 | 0.01 | 0.01 |
| gene-LOC105284043 | https://www.ncbi.nlm.nih.gov/gene/?term=LOC105284043 | 5661 | 311 | 549.37 | 1 | 1.77 | 0.32 | 9.86 | 9.71 | 0.15 | 0.15 | 0.01 | 0.01 |
| gene-LOC105278145 | https://www.ncbi.nlm.nih.gov/gene/?term=LOC105278145 | 4470 | 235 | 525.73 | 1 | 2.24 | 0.43 | 10.67 | 10.54 | 0.14 | 0.14 | 0.01 | 0.01 |
| gene-LOC105275374 | https://www.ncbi.nlm.nih.gov/gene/?term=LOC105275374 | 11376 | 539 | 473.8 1 | 0.88 | 0.19 | 9.85 | 10 | -0.15 | 0.15 | -0.01 | 0.01 | |
| gene-LOC105278282 | https://www.ncbi.nlm.nih.gov/gene/?term=LOC105278282 | 1775 | 62 | 349.3 1 | 5.63 | 1.61 | 32.74 | 32.89 | -0.15 | 0.15 | -0.01 | 0.01 | |
| gene-LOC105285742 | https://www.ncbi.nlm.nih.gov/gene/?term=LOC105285742 | 3446 | 123 | 356.94 | 1 | 2.9 | 0.81 | 19.52 | 19.69 | -0.16 | 0.16 | -0.01 | 0.01 |
| gene-LOC105280723 | https://www.ncbi.nlm.nih.gov/gene/?term=LOC105280723 | 3572 | 181 | 506.72 | 1 | 2.8 | 0.55 | 15.21 | 15.4 | -0.2 | 0.2 | -0.01 | 0.01 |
| gene-LOC105284184 | https://www.ncbi.nlm.nih.gov/gene/?term=LOC105284184 | 2535 | 99 | 390.53 | 1 | 3.94 | 1.01 | 29.28 | 29.51 | -0.23 | 0.23 | -0.01 | 0.01 |
| gene-LOC105275587 | https://www.ncbi.nlm.nih.gov/gene/?term=LOC105275587 | 2404 | 55 | 228.79 | 1 | 4.16 | 1.82 | 36.62 | 36.89 | -0.27 | 0.27 | -0.01 | 0.01 |
| gene-LOC105278691 | https://www.ncbi.nlm.nih.gov/gene/?term=LOC105278691 | 3871 | 168 | 434 1 | 2.58 | 0.6 | 14.64 | 14.98 | -0.34 | 0.34 | -0.01 | 0.01 | |
| gene-LOC105283967 | https://www.ncbi.nlm.nih.gov/gene/?term=LOC105283967 | 2393 | 53 | 221.48 | 1 | 4.18 | 1.89 | 45.17 | 45.77 | -0.59 | 0.59 | -0.01 | 0.01 |
| gene-LOC105286452 | https://www.ncbi.nlm.nih.gov/gene/?term=LOC105286452 | 6888 | 60 | 87.11 1 | 1.45 | 1.67 | 31.09 | 30.62 | 0.47 | 0.47 | 0 | 0 | |
| gene-LOC105280408 | https://www.ncbi.nlm.nih.gov/gene/?term=LOC105280408 | 5983 | 56 | 93.6 1 | 1.67 | 1.79 | 20.14 | 19.81 | 0.33 | 0.33 | 0 | 0 | |
| gene-LOC105276683 | https://www.ncbi.nlm.nih.gov/gene/?term=LOC105276683 | 17000 | 137 | 80.59 1 | 0.59 | 0.73 | 6.59 | 6.28 | 0.3 | 0.3 | 0 | 0 | |
| gene-LOC105276196 | https://www.ncbi.nlm.nih.gov/gene/?term=LOC105276196 | 2953 | 58 | 196.41 | 1 | 3.39 | 1.72 | 38.52 | 38.28 | 0.24 | 0.24 | 0 | 0 |
| gene-LOC105279657 | https://www.ncbi.nlm.nih.gov/gene/?term=LOC105279657 | 5878 | 88 | 149.71 | 1 | 1.7 | 1.14 | 19.35 | 19.14 | 0.21 | 0.21 | 0 | 0 |
| gene-LOC105282151 | https://www.ncbi.nlm.nih.gov/gene/?term=LOC105282151 | 1601 | 44 | 274.83 | 1 | 6.25 | 2.27 | 35.59 | 35.42 | 0.18 | 0.18 | 0 | 0 |
| gene-LOC105287764 | https://www.ncbi.nlm.nih.gov/gene/?term=LOC105287764 | 18359 | 402 | 218.97 | 1 | 0.54 | 0.25 | 3.16 | 2.99 | 0.17 | 0.17 | 0 | 0 |
| gene-LOC105283712 | https://www.ncbi.nlm.nih.gov/gene/?term=LOC105283712 | 1910 | 36 | 188.48 | 1 | 5.24 | 2.78 | 29.2 | 29.04 | 0.16 | 0.16 | 0 | 0 |
| gene-LOC105284036 | https://www.ncbi.nlm.nih.gov/gene/?term=LOC105284036 | 3254 | 107 | 328.83 | 1 | 3.07 | 0.93 | 20.44 | 20.34 | 0.09 | 0.09 | 0 | 0 |
| gene-LOC105274823 | https://www.ncbi.nlm.nih.gov/gene/?term=LOC105274823 | 4620 | 146 | 316.02 | 1 | 2.16 | 0.68 | 20.43 | 20.37 | 0.06 | 0.06 | 0 | 0 |

| | | | | | | | | | | | | | | |
|---|---|---|---|---|---|---|---|---|---|---|---|---|---|---|
| gene-LOC105281700 | https://www.ncbi.nlm.nih.gov/gene/?term=LOC105281700 | 2208 | 37 | 167.57 | | 1 | 4.53 | 2.7 | 39.8 | 39.75 | 0.05 | 0.05 | 0 | 0 |
| gene-LOC105286515 | https://www.ncbi.nlm.nih.gov/gene/?term=LOC105286515 | 3751 | 106 | 282.59 | | 1 | 2.67 | 0.94 | 10.25 | 10.22 | 0.04 | 0.04 | 0 | 0 |
| gene-LOC105280407 | https://www.ncbi.nlm.nih.gov/gene/?term=LOC105280407 | 10950 | 210 | 191.78 | | 1 | 0.91 | 0.48 | 16.39 | 16.4 | -0.01 | 0.01 | 0 | 0 |
| gene-LOC105283123 | https://www.ncbi.nlm.nih.gov/gene/?term=LOC105283123 | 1507 | 55 | 364.96 | | 1 | 6.64 | 1.82 | 18.11 | 18.13 | -0.02 | 0.02 | 0 | 0 |
| gene-LOC105274669 | https://www.ncbi.nlm.nih.gov/gene/?term=LOC105274669 | 8555 | 270 | 315.6 | 1 | 1.17 | 0.37 | 5.54 | 5.56 | -0.03 | 0.03 | 0 | 0 | |
| gene-LOC105278534 | https://www.ncbi.nlm.nih.gov/gene/?term=LOC105278534 | 7364 | 196 | 266.16 | | 1 | 1.36 | 0.51 | 17.75 | 17.8 | -0.05 | 0.05 | 0 | 0 |
| gene-LOC105287550 | https://www.ncbi.nlm.nih.gov/gene/?term=LOC105287550 | 15659 | 616 | 393.38 | | 1 | 0.64 | 0.16 | 5.82 | 5.86 | -0.05 | 0.05 | 0 | 0 |
| gene-LOC105285496 | https://www.ncbi.nlm.nih.gov/gene/?term=LOC105285496 | 4080 | 110 | 269.61 | | 1 | 2.45 | 0.91 | 40.35 | 40.5 | -0.15 | 0.15 | 0 | 0 |
| gene-LOC105274851 | https://www.ncbi.nlm.nih.gov/gene/?term=LOC105274851 | 6616 | 147 | 222.19 | | 1 | 1.51 | 0.68 | 9.12 | 9.33 | -0.21 | 0.21 | 0 | 0 |
| gene-LOC105285406 | https://www.ncbi.nlm.nih.gov/gene/?term=LOC105285406 | 8015 | 55 | 68.62 | 1 | 1.25 | 1.82 | 29.11 | 29.59 | -0.47 | 0.47 | 0 | 0 | |

Directory: OGL File: rnk-dmg-Ob_LI_BCphaseB1.vs.RphaseA1.txt

| gene_ID | gene_link | gwidth | #Sites | #per10Kb | #dmSites | #dmsp10kb | %dmSites | %pSite1 | %pSite2 | DMpSite | ADMpSite | DMpNucl | ADMpNucl |
|---|---|---|---|---|---|---|---|---|---|---|---|---|---|
| gene-LOC105285392 | https://www.ncbi.nlm.nih.gov/gene/?term=LOC105285392 | 1601 | 91 | 568.39 | 6 | 37.48 | 6.59 | 35.68 | 31.54 | 4.15 | 4.15 | 0.24 | 0.24 |
| gene-LOC113561930 | https://www.ncbi.nlm.nih.gov/gene/?term=LOC113561930 | 422 | 22 | 521.33 | 1 | 23.7 | 4.55 | 10.08 | 13.72 | -3.64 | 3.64 | -0.19 | 0.19 |
| gene-LOC105275378 | https://www.ncbi.nlm.nih.gov/gene/?term=LOC105275378 | 1189 | 21 | 176.62 | 3 | 25.23 | 14.29 | 38.9 | 30.15 | 8.75 | 8.75 | 0.15 | 0.15 |
| gene-LOC105278153 | https://www.ncbi.nlm.nih.gov/gene/?term=LOC105278153 | 550 | 23 | 418.18 | 3 | 54.55 | 13.04 | 16.52 | 13.29 | 3.23 | 3.23 | 0.14 | 0.14 |
| gene-LOC109610897 | https://www.ncbi.nlm.nih.gov/gene/?term=LOC109610897 | 1561 | 124 | 794.36 | 2 | 12.81 | 1.61 | 14.34 | 16.15 | -1.8 | 1.8 | -0.14 | 0.14 |
| gene-LOC105286645 | https://www.ncbi.nlm.nih.gov/gene/?term=LOC105286645 | 1499 | 55 | 366.91 | 1 | 6.67 | 1.82 | 37.78 | 41.38 | -3.6 | 3.6 | -0.13 | 0.13 |
| gene-LOC105274528 | https://www.ncbi.nlm.nih.gov/gene/?term=LOC105274528 | 2564 | 155 | 604.52 | 3 | 11.7 | 1.94 | 18.68 | 16.63 | 2.05 | 2.05 | 0.12 | 0.12 |
| gene-LOC105278849 | https://www.ncbi.nlm.nih.gov/gene/?term=LOC105278849 | 776 | 75 | 966.49 | 1 | 12.89 | 1.33 | 24.51 | 23.23 | 1.28 | 1.28 | 0.12 | 0.12 |
| gene-LOC105284174 | https://www.ncbi.nlm.nih.gov/gene/?term=LOC105284174 | 1627 | 83 | 510.14 | 1 | 6.15 | 1.2 | 19.97 | 22.34 | -2.36 | 2.36 | -0.12 | 0.12 |
| gene-LOC105286269 | https://www.ncbi.nlm.nih.gov/gene/?term=LOC105286269 | 1401 | 51 | 364.03 | 4 | 28.55 | 7.84 | 50.68 | 53.95 | -3.27 | 3.27 | -0.12 | 0.12 |
| gene-LOC105283717 | https://www.ncbi.nlm.nih.gov/gene/?term=LOC105283717 | 1373 | 87 | 633.65 | 2 | 14.57 | 2.3 | 29.33 | 27.67 | 1.66 | 1.66 | 0.11 | 0.11 |
| gene-LOC105285635 | https://www.ncbi.nlm.nih.gov/gene/?term=LOC105285635 | 1175 | 92 | 782.98 | 1 | 8.51 | 1.09 | 28.39 | 26.97 | 1.42 | 1.42 | 0.11 | 0.11 |
| gene-LOC105288242 | https://www.ncbi.nlm.nih.gov/gene/?term=LOC105288242 | 1260 | 36 | 285.71 | 1 | 7.94 | 2.78 | 42.4 | 46.4 | -4 | 4 | -0.11 | 0.11 |
| gene-LOC113562457 | https://www.ncbi.nlm.nih.gov/gene/?term=LOC113562457 | 739 | 11 | 148.85 | 2 | 27.06 | 18.18 | 47.39 | 54.48 | -7.08 | 7.08 | -0.11 | 0.11 |
| gene-LOC105275214 | https://www.ncbi.nlm.nih.gov/gene/?term=LOC105275214 | 1299 | 44 | 338.72 | 1 | 7.7 | 2.27 | 33.78 | 30.91 | 2.87 | 2.87 | 0.1 | 0.1 |
| gene-LOC105276563 | https://www.ncbi.nlm.nih.gov/gene/?term=LOC105276563 | 1958 | 116 | 592.44 | 5 | 25.54 | 4.31 | 17.76 | 16.02 | 1.74 | 1.74 | 0.1 | 0.1 |
| gene-LOC105285716 | https://www.ncbi.nlm.nih.gov/gene/?term=LOC105285716 | 1138 | 36 | 316.34 | 1 | 8.79 | 2.78 | 36.22 | 39.41 | -3.19 | 3.19 | -0.1 | 0.1 |
| gene-LOC105280327 | https://www.ncbi.nlm.nih.gov/gene/?term=LOC105280327 | 2670 | 109 | 408.24 | 6 | 22.47 | 5.5 | 29.28 | 27.03 | 2.25 | 2.25 | 0.09 | 0.09 |
| gene-LOC105274711 | https://www.ncbi.nlm.nih.gov/gene/?term=LOC105274711 | 1655 | 86 | 519.64 | 2 | 12.08 | 2.33 | 46.42 | 44.73 | 1.69 | 1.69 | 0.09 | 0.09 |
| gene-LOC105278873 | https://www.ncbi.nlm.nih.gov/gene/?term=LOC105278873 | 1897 | 149 | 785.45 | 6 | 31.63 | 4.03 | 20.63 | 19.44 | 1.19 | 1.19 | 0.09 | 0.09 |
| gene-LOC105287690 | https://www.ncbi.nlm.nih.gov/gene/?term=LOC105287690 | 1378 | 108 | 783.74 | 3 | 21.77 | 2.78 | 18.67 | 17.5 | 1.16 | 1.16 | 0.09 | 0.09 |
| gene-LOC105287942 | https://www.ncbi.nlm.nih.gov/gene/?term=LOC105287942 | 2147 | 78 | 363.3 | 5 | 23.29 | 6.41 | 31.04 | 33.39 | -2.35 | 2.35 | -0.09 | 0.09 |
| gene-LOC105286212 | https://www.ncbi.nlm.nih.gov/gene/?term=LOC105286212 | 1022 | 32 | 313.11 | 1 | 9.78 | 3.12 | 35.65 | 38.39 | -2.74 | 2.74 | -0.09 | 0.09 |
| gene-LOC105286107 | https://www.ncbi.nlm.nih.gov/gene/?term=LOC105286107 | 1902 | 47 | 247.11 | 2 | 10.52 | 4.26 | 14.16 | 17.81 | -3.65 | 3.65 | -0.09 | 0.09 |

...
... only first 25 lines shown ...

Directory: OGL File: rnk-dmg-Ob_LI_BCphaseB1.vs.RphaseA2.txt

| gene_ID | gene_link | gwidth | #Sites | #per10Kb | #dmSites | #dmsp10kb | %dmSites | %pSite1 | %pSite2 | DMpSite | ADMpSite | DMpNucl | ADMpNucl |
|---|---|---|---|---|---|---|---|---|---|---|---|---|---|
| gene-LOC109611487 | https://www.ncbi.nlm.nih.gov/gene/?term=LOC109611487 | 437 | 32 | 732.27 | 1 | 22.88 | 3.12 | 14.09 | 17.07 | -2.98 | 2.98 | -0.22 | 0.22 |
| gene-LOC105275355 | https://www.ncbi.nlm.nih.gov/gene/?term=LOC105275355 | 3197 | 169 | 528.62 | 10 | 31.28 | 5.92 | 27.68 | 24.07 | 3.61 | 3.61 | 0.19 | 0.19 |
| gene-LOC105286212 | https://www.ncbi.nlm.nih.gov/gene/?term=LOC105286212 | 1022 | 39 | 381.6 | 3 | 29.35 | 7.69 | 33.25 | 37.99 | -4.73 | 4.73 | -0.18 | 0.18 |
| gene-LOC105276451 | https://www.ncbi.nlm.nih.gov/gene/?term=LOC105276451 | 1360 | 40 | 294.12 | 3 | 22.06 | 7.5 | 35.85 | 41.18 | -5.34 | 5.34 | -0.16 | 0.16 |
| gene-LOC105279049 | https://www.ncbi.nlm.nih.gov/gene/?term=LOC105279049 | 1285 | 100 | 778.21 | 1 | 7.78 | 1 | 16.79 | 18.78 | -1.99 | 1.99 | -0.15 | 0.15 |
| gene-LOC105282220 | https://www.ncbi.nlm.nih.gov/gene/?term=LOC105282220 | 2298 | 236 | 1026.98 | 3 | 13.05 | 1.27 | 7.12 | 8.51 | -1.39 | 1.39 | -0.14 | 0.14 |
| gene-LOC105275307 | https://www.ncbi.nlm.nih.gov/gene/?term=LOC105275307 | 1420 | 54 | 380.28 | 2 | 14.08 | 3.7 | 38.11 | 41.77 | -3.67 | 3.67 | -0.14 | 0.14 |
| gene-LOC109610897 | https://www.ncbi.nlm.nih.gov/gene/?term=LOC109610897 | 1561 | 147 | 941.7 | 2 | 12.81 | 1.36 | 13.43 | 14.76 | -1.34 | 1.34 | -0.13 | 0.13 |
| gene-LOC105281172 | https://www.ncbi.nlm.nih.gov/gene/?term=LOC105281172 | 1310 | 30 | 229.01 | 3 | 22.9 | 10 | 42.61 | 48.18 | -5.57 | 5.57 | -0.13 | 0.13 |
| gene-LOC105285628 | https://www.ncbi.nlm.nih.gov/gene/?term=LOC105285628 | 1033 | 22 | 212.97 | 3 | 29.04 | 13.64 | 32.6 | 38.87 | -6.27 | 6.27 | -0.13 | 0.13 |
| gene-LOC105278849 | https://www.ncbi.nlm.nih.gov/gene/?term=LOC105278849 | 776 | 76 | 979.38 | 1 | 12.89 | 1.32 | 24.69 | 23.43 | 1.26 | 1.26 | 0.12 | 0.12 |
| gene-LOC105279035 | https://www.ncbi.nlm.nih.gov/gene/?term=LOC105279035 | 2108 | 166 | 787.48 | 4 | 18.98 | 2.41 | 18.4 | 19.91 | -1.5 | 1.5 | -0.12 | 0.12 |
| gene-LOC105278396 | https://www.ncbi.nlm.nih.gov/gene/?term=LOC105278396 | 3012 | 190 | 630.81 | 5 | 16.6 | 2.63 | 24.21 | 26.15 | -1.94 | 1.94 | -0.12 | 0.12 |
| gene-LOC105285716 | https://www.ncbi.nlm.nih.gov/gene/?term=LOC105285716 | 1138 | 42 | 369.07 | 3 | 26.36 | 7.14 | 36.53 | 39.66 | -3.13 | 3.13 | -0.12 | 0.12 |
| gene-LOC113562901 | https://www.ncbi.nlm.nih.gov/gene/?term=LOC113562901 | 1531 | 25 | 163.29 | 3 | 19.6 | 12 | 50.23 | 57.46 | -7.23 | 7.23 | -0.12 | 0.12 |
| gene-LOC105284174 | https://www.ncbi.nlm.nih.gov/gene/?term=LOC105284174 | 1627 | 92 | 565.46 | 1 | 6.15 | 1.09 | 19.2 | 21.11 | -1.92 | 1.92 | -0.11 | 0.11 |
| gene-LOC113561930 | https://www.ncbi.nlm.nih.gov/gene/?term=LOC113561930 | 422 | 23 | 545.02 | 1 | 23.7 | 4.35 | 12.86 | 14.92 | -2.06 | 2.06 | -0.11 | 0.11 |
| gene-LOC105274542 | https://www.ncbi.nlm.nih.gov/gene/?term=LOC105274542 | 1266 | 61 | 481.83 | 2 | 15.8 | 3.28 | 29.9 | 32.16 | -2.26 | 2.26 | -0.11 | 0.11 |
| gene-LOC105285056 | https://www.ncbi.nlm.nih.gov/gene/?term=LOC105285056 | 1849 | 88 | 475.93 | 1 | 5.41 | 1.14 | 26.08 | 28.38 | -2.3 | 2.3 | -0.11 | 0.11 |
| gene-LOC105283847 | https://www.ncbi.nlm.nih.gov/gene/?term=LOC105283847 | 1590 | 74 | 465.41 | 2 | 12.58 | 2.7 | 33.62 | 36.08 | -2.47 | 2.47 | -0.11 | 0.11 |
| gene-LOC105274524 | https://www.ncbi.nlm.nih.gov/gene/?term=LOC105274524 | 1126 | 43 | 381.88 | 2 | 17.76 | 4.65 | 27.35 | 30.21 | -2.85 | 2.85 | -0.11 | 0.11 |
| gene-LOC105278282 | https://www.ncbi.nlm.nih.gov/gene/?term=LOC105278282 | 1775 | 68 | 383.1 | 3 | 16.9 | 4.41 | 31.98 | 34.93 | -2.95 | 2.95 | -0.11 | 0.11 |
| gene-LOC105286269 | https://www.ncbi.nlm.nih.gov/gene/?term=LOC105286269 | 1401 | 51 | 364.03 | 1 | 7.14 | 1.96 | 50.68 | 53.75 | -3.07 | 3.07 | -0.11 | 0.11 |
| gene-LOC105280117 | https://www.ncbi.nlm.nih.gov/gene/?term=LOC105280117 | 1817 | 63 | 346.73 | 2 | 11.01 | 3.17 | 46.36 | 49.66 | -3.3 | 3.3 | -0.11 | 0.11 |

...
... only first 25 lines shown ...

Directory: OGL File: rnk-dmg-Ob_LI_BCphaseB1.vs.RphaseB1.txt

| gene_ID | gene_link | gwidth | #Sites | #per10Kb | #dmSites | #dmsp10kb | %dmSites | %pSite1 | %pSite2 | DMpSite | ADMpSite | DMpNucl | ADMpNucl |
|---|---|---|---|---|---|---|---|---|---|---|---|---|---|
| gene-LOC113562345 | https://www.ncbi.nlm.nih.gov/gene/?term=LOC113562345 | 2290 | 77 | 336.24 | 12 | 52.4 | 15.58 | 26.12 | 14.52 | 11.59 | 11.59 | 0.39 | 0.39 |
| gene-LOC105275355 | https://www.ncbi.nlm.nih.gov/gene/?term=LOC105275355 | 3197 | 174 | 544.26 | 1 | 3.13 | 0.57 | 26.94 | 21.77 | 5.17 | 5.17 | 0.28 | 0.28 |
| gene-LOC105284677 | https://www.ncbi.nlm.nih.gov/gene/?term=LOC105284677 | 1905 | 42 | 220.47 | 1 | 5.25 | 2.38 | 23.34 | 35.09 | -11.75 | 11.75 | -0.26 | 0.26 |
| gene-LOC105275384 | https://www.ncbi.nlm.nih.gov/gene/?term=LOC105275384 | 1987 | 92 | 463.01 | 1 | 5.03 | 1.09 | 34.18 | 31.19 | 2.98 | 2.98 | 0.14 | 0.14 |
| gene-LOC105275070 | https://www.ncbi.nlm.nih.gov/gene/?term=LOC105275070 | 5667 | 61 | 107.64 | 2 | 3.53 | 3.28 | 14.18 | 5.29 | 8.9 | 8.9 | 0.1 | 0.1 |
| gene-LOC105274534 | https://www.ncbi.nlm.nih.gov/gene/?term=LOC105274534 | 1329 | 59 | 443.94 | 1 | 7.52 | 1.69 | 41.22 | 43.19 | -1.98 | 1.98 | -0.09 | 0.09 |
| gene-LOC105279900 | https://www.ncbi.nlm.nih.gov/gene/?term=LOC105279900 | 2191 | 56 | 255.59 | 1 | 4.56 | 1.79 | 31.89 | 28.91 | 2.98 | 2.98 | 0.08 | 0.08 |
| gene-LOC105275708 | https://www.ncbi.nlm.nih.gov/gene/?term=LOC105275708 | 1772 | 15 | 84.65 | 1 | 5.64 | 6.67 | 46.38 | 38.49 | 7.9 | 7.9 | 0.07 | 0.07 |
| gene-LOC105278282 | https://www.ncbi.nlm.nih.gov/gene/?term=LOC105278282 | 1775 | 65 | 366.2 | 1 | 5.63 | 1.54 | 30.53 | 32.37 | -1.84 | 1.84 | -0.07 | 0.07 |
| gene-LOC105285514 | https://www.ncbi.nlm.nih.gov/gene/?term=LOC105285514 | 1277 | 11 | 86.14 | 1 | 7.83 | 9.09 | 92.13 | 85.12 | 7.01 | 7.01 | 0.06 | 0.06 |
| gene-LOC105276272 | https://www.ncbi.nlm.nih.gov/gene/?term=LOC105276272 | 1965 | 40 | 203.56 | 1 | 5.09 | 2.5 | 30.29 | 27.31 | 2.98 | 2.98 | 0.06 | 0.06 |
| gene-LOC105276611 | https://www.ncbi.nlm.nih.gov/gene/?term=LOC105276611 | 1731 | 36 | 207.97 | 1 | 5.78 | 2.78 | 30.25 | 27.6 | 2.65 | 2.65 | 0.06 | 0.06 |
| gene-LOC105287063 | https://www.ncbi.nlm.nih.gov/gene/?term=LOC105287063 | 1874 | 50 | 266.81 | 1 | 5.34 | 2 | 31.85 | 29.62 | 2.23 | 2.23 | 0.06 | 0.06 |
| gene-LOC105283552 | https://www.ncbi.nlm.nih.gov/gene/?term=LOC105283552 | 1562 | 41 | 262.48 | 1 | 6.4 | 2.44 | 53.31 | 51.1 | 2.21 | 2.21 | 0.06 | 0.06 |
| gene-LOC105280704 | https://www.ncbi.nlm.nih.gov/gene/?term=LOC105280704 | 4266 | 147 | 344.59 | 1 | 2.34 | 0.68 | 18.45 | 16.63 | 1.81 | 1.81 | 0.06 | 0.06 |
| gene-LOC105287095 | https://www.ncbi.nlm.nih.gov/gene/?term=LOC105287095 | 1435 | 70 | 487.8 | 1 | 6.97 | 1.43 | 25.15 | 23.89 | 1.26 | 1.26 | 0.06 | 0.06 |
| gene-LOC105288036 | https://www.ncbi.nlm.nih.gov/gene/?term=LOC105288036 | 5110 | 232 | 454.01 | 1 | 1.96 | 0.43 | 15.89 | 14.67 | 1.22 | 1.22 | 0.06 | 0.06 |
| gene-LOC105283712 | https://www.ncbi.nlm.nih.gov/gene/?term=LOC105283712 | 1910 | 35 | 183.25 | 1 | 5.24 | 2.86 | 31.43 | 28.9 | 2.53 | 2.53 | 0.05 | 0.05 |
| gene-LOC113561805 | https://www.ncbi.nlm.nih.gov/gene/?term=LOC113561805 | 3337 | 76 | 227.75 | 1 | 3 | 1.32 | 32.51 | 30.53 | 1.98 | 1.98 | 0.05 | 0.05 |
| gene-LOC105274449 | https://www.ncbi.nlm.nih.gov/gene/?term=LOC105274449 | 1549 | 53 | 342.16 | 1 | 6.46 | 1.89 | 50.86 | 49.29 | 1.57 | 1.57 | 0.05 | 0.05 |
| gene-LOC105287417 | https://www.ncbi.nlm.nih.gov/gene/?term=LOC105287417 | 1025 | 40 | 390.24 | 1 | 9.76 | 2.5 | 32.13 | 30.76 | 1.38 | 1.38 | 0.05 | 0.05 |
| gene-LOC105282156 | https://www.ncbi.nlm.nih.gov/gene/?term=LOC105282156 | 2577 | 132 | 512.22 | 1 | 3.88 | 0.76 | 26.63 | 25.67 | 0.96 | 0.96 | 0.05 | 0.05 |
| gene-LOC105285055 | https://www.ncbi.nlm.nih.gov/gene/?term=LOC105285055 | 4108 | 248 | 603.7 | 1 | 2.43 | 0.4 | 12.58 | 11.79 | 0.79 | 0.79 | 0.05 | 0.05 |
| gene-LOC105282885 | https://www.ncbi.nlm.nih.gov/gene/?term=LOC105282885 | 1870 | 41 | 219.25 | 1 | 5.35 | 2.44 | 39.07 | 41.13 | -2.06 | 2.06 | -0.05 | 0.05 |
| gene-LOC105278454 | https://www.ncbi.nlm.nih.gov/gene/?term=LOC105278454 | 1972 | 41 | 207.91 | 1 | 5.07 | 2.44 | 41.77 | 44.02 | -2.26 | 2.26 | -0.05 | 0.05 |
| gene-LOC105281039 | https://www.ncbi.nlm.nih.gov/gene/?term=LOC105281039 | 1471 | 14 | 95.17 | 1 | 6.8 | 7.14 | 44.63 | 40.12 | 4.51 | 4.51 | 0.04 | 0.04 |
| gene-LOC105275518 | https://www.ncbi.nlm.nih.gov/gene/?term=LOC105275518 | 2335 | 25 | 107.07 | 1 | 4.28 | 4 | 44.99 | 41.35 | 3.64 | 3.64 | 0.04 | 0.04 |
| gene-LOC105285179 | https://www.ncbi.nlm.nih.gov/gene/?term=LOC105285179 | 4191 | 194 | 462.9 | 1 | 2.39 | 0.52 | 17.62 | 16.69 | 0.93 | 0.93 | 0.04 | 0.04 |
| gene-LOC105285695 | https://www.ncbi.nlm.nih.gov/gene/?term=LOC105285695 | 1371 | 75 | 547.05 | 1 | 7.29 | 1.33 | 27.5 | 26.71 | 0.79 | 0.79 | 0.04 | 0.04 |
| gene-LOC105275460 | https://www.ncbi.nlm.nih.gov/gene/?term=LOC105275460 | 2473 | 127 | 513.55 | 1 | 4.04 | 0.79 | 13.03 | 12.3 | 0.73 | 0.73 | 0.04 | 0.04 |
| gene-LOC105287633 | https://www.ncbi.nlm.nih.gov/gene/?term=LOC105287633 | 1524 | 25 | 164.04 | 1 | 6.56 | 4 | 61.97 | 64.15 | -2.18 | 2.18 | -0.04 | 0.04 |
| gene-LOC105283989 | https://www.ncbi.nlm.nih.gov/gene/?term=LOC105283989 | 4948 | 45 | 90.95 | 2 | 4.04 | 4.44 | 38.35 | 35.25 | 3.1 | 3.1 | 0.03 | 0.03 |
| gene-LOC105274900 | https://www.ncbi.nlm.nih.gov/gene/?term=LOC105274900 | 5561 | 63 | 113.29 | 2 | 3.6 | 3.17 | 32.28 | 30.06 | 2.22 | 2.22 | 0.03 | 0.03 |
| gene-LOC105279898 | https://www.ncbi.nlm.nih.gov/gene/?term=LOC105279898 | 1343 | 16 | 119.14 | 1 | 7.45 | 6.25 | 58.69 | 56.55 | 2.14 | 2.14 | 0.03 | 0.03 |
| gene-LOC105280348 | https://www.ncbi.nlm.nih.gov/gene/?term=LOC105280348 | 3797 | 79 | 208.06 | 1 | 2.63 | 1.27 | 18.26 | 16.64 | 1.61 | 1.61 | 0.03 | 0.03 |
| gene-LOC105276692 | https://www.ncbi.nlm.nih.gov/gene/?term=LOC105276692 | 1745 | 35 | 200.57 | 1 | 5.73 | 2.86 | 24.46 | 23.1 | 1.36 | 1.36 | 0.03 | 0.03 |
| gene-LOC105278384 | https://www.ncbi.nlm.nih.gov/gene/?term=LOC105278384 | 1390 | 33 | 237.41 | 1 | 7.19 | 3.03 | 46.84 | 45.77 | 1.07 | 1.07 | 0.03 | 0.03 |
| gene-LOC105288079 | https://www.ncbi.nlm.nih.gov/gene/?term=LOC105288079 | 3473 | 110 | 316.73 | 1 | 2.88 | 0.91 | 18.5 | 17.43 | 1.07 | 1.07 | 0.03 | 0.03 |
| gene-LOC105278632 | https://www.ncbi.nlm.nih.gov/gene/?term=LOC105278632 | 3624 | 139 | 383.55 | 1 | 2.76 | 0.72 | 14.57 | 13.67 | 0.9 | 0.9 | 0.03 | 0.03 |
| gene-LOC105283123 | https://www.ncbi.nlm.nih.gov/gene/?term=LOC105283123 | 1507 | 55 | 364.96 | 1 | 6.64 | 1.82 | 19.83 | 18.96 | 0.86 | 0.86 | 0.03 | 0.03 |
| gene-LOC105285674 | https://www.ncbi.nlm.nih.gov/gene/?term=LOC105285674 | 3035 | 109 | 359.14 | 1 | 3.29 | 0.92 | 24.18 | 23.43 | 0.75 | 0.75 | 0.03 | 0.03 |
| gene-LOC105285639 | https://www.ncbi.nlm.nih.gov/gene/?term=LOC105285639 | 10325 | 372 | 360.29 | 2 | 1.94 | 0.54 | 11.5 | 10.78 | 0.72 | 0.72 | 0.03 | 0.03 |
| gene-LOC105283728 | https://www.ncbi.nlm.nih.gov/gene/?term=LOC105283728 | 3256 | 146 | 448.4 | 1 | 3.07 | 0.68 | 16.37 | 15.66 | 0.71 | 0.71 | 0.03 | 0.03 |
| gene-LOC105277435 | https://www.ncbi.nlm.nih.gov/gene/?term=LOC105277435 | 3643 | 179 | 491.35 | 1 | 2.74 | 0.56 | 10.99 | 10.3 | 0.7 | 0.7 | 0.03 | 0.03 |
| gene-LOC105285671 | https://www.ncbi.nlm.nih.gov/gene/?term=LOC105285671 | 2263 | 103 | 455.15 | 1 | 4.42 | 0.97 | 30.33 | 29.64 | 0.69 | 0.69 | 0.03 | 0.03 |
| gene-LOC105276449 | https://www.ncbi.nlm.nih.gov/gene/?term=LOC105276449 | 1231 | 55 | 446.79 | 1 | 8.12 | 1.82 | 22.22 | 21.54 | 0.67 | 0.67 | 0.03 | 0.03 |
| gene-LOC105284355 | https://www.ncbi.nlm.nih.gov/gene/?term=LOC105284355 | 3904 | 154 | 394.47 | 1 | 2.56 | 0.65 | 22.13 | 21.47 | 0.66 | 0.66 | 0.03 | 0.03 |
| gene-LOC105282480 | https://www.ncbi.nlm.nih.gov/gene/?term=LOC105282480 | 4177 | 166 | 397.41 | 1 | 2.39 | 0.6 | 12.65 | 12.01 | 0.65 | 0.65 | 0.03 | 0.03 |
| gene-LOC105276704 | https://www.ncbi.nlm.nih.gov/gene/?term=LOC105276704 | 2063 | 87 | 421.72 | 2 | 9.69 | 2.3 | 14.01 | 13.37 | 0.64 | 0.64 | 0.03 | 0.03 |
| gene-LOC105286997 | https://www.ncbi.nlm.nih.gov/gene/?term=LOC105286997 | 2850 | 150 | 526.32 | 1 | 3.51 | 0.67 | 11.33 | 10.85 | 0.48 | 0.48 | 0.03 | 0.03 |
| gene-LOC105277631 | https://www.ncbi.nlm.nih.gov/gene/?term=LOC105277631 | 4387 | 313 | 713.47 | 1 | 2.28 | 0.32 | 10.74 | 10.28 | 0.46 | 0.46 | 0.03 | 0.03 |
| gene-LOC105274856 | https://www.ncbi.nlm.nih.gov/gene/?term=LOC105274856 | 5216 | 222 | 425.61 | 1 | 1.92 | 0.45 | 16.84 | 17.52 | -0.68 | 0.68 | -0.03 | 0.03 |
| gene-LOC105279778 | https://www.ncbi.nlm.nih.gov/gene/?term=LOC105279778 | 4347 | 163 | 374.97 | 1 | 2.3 | 0.61 | 8.62 | 9.47 | -0.85 | 0.85 | -0.03 | 0.03 |
| gene-LOC105283132 | https://www.ncbi.nlm.nih.gov/gene/?term=LOC105283132 | 1819 | 56 | 307.86 | 1 | 5.5 | 1.79 | 39.1 | 40.18 | -1.09 | 1.09 | -0.03 | 0.03 |
| gene-LOC105285409 | https://www.ncbi.nlm.nih.gov/gene/?term=LOC105285409 | 5099 | 66 | 129.44 | 1 | 1.96 | 1.52 | 2.12 | 4.53 | -2.41 | 2.41 | -0.03 | 0.03 |
| gene-LOC105280419 | https://www.ncbi.nlm.nih.gov/gene/?term=LOC105280419 | 1601 | 11 | 68.71 | 1 | 6.25 | 9.09 | 46.06 | 42.48 | 3.57 | 3.57 | 0.02 | 0.02 |
| gene-LOC105285315 | https://www.ncbi.nlm.nih.gov/gene/?term=LOC105285315 | 5458 | 32 | 58.63 | 1 | 1.83 | 3.12 | 31.91 | 29.18 | 2.72 | 2.72 | 0.02 | 0.02 |
| gene-LOC105276538 | https://www.ncbi.nlm.nih.gov/gene/?term=LOC105276538 | 1511 | 20 | 132.36 | 1 | 6.62 | 5 | 72.51 | 71.18 | 1.32 | 1.32 | 0.02 | 0.02 |
| gene-LOC105280972 | https://www.ncbi.nlm.nih.gov/gene/?term=LOC105280972 | 1532 | 35 | 228.46 | 1 | 6.53 | 2.86 | 17.63 | 16.59 | 1.04 | 1.04 | 0.02 | 0.02 |
| gene-LOC105282794 | https://www.ncbi.nlm.nih.gov/gene/?term=LOC105282794 | 4571 | 110 | 240.65 | 1 | 2.19 | 0.91 | 18.14 | 17.16 | 0.98 | 0.98 | 0.02 | 0.02 |

| | | | | | | | | | | | | | |
|---|---|---|---|---|---|---|---|---|---|---|---|---|---|
| gene-LOC105281266 | https://www.ncbi.nlm.nih.gov/gene/?term=LOC105281266 | 4102 | 88 | 214.53 | 1 | 2.44 | 1.14 | 15.7 | 14.74 | 0.96 | 0.96 | 0.02 | 0.02 |
| gene-LOC105281848 | https://www.ncbi.nlm.nih.gov/gene/?term=LOC105281848 | 2122 | 58 | 273.33 | 1 | 4.71 | 1.72 | 22.33 | 21.51 | 0.82 | 0.82 | 0.02 | 0.02 |
| gene-LOC105276410 | https://www.ncbi.nlm.nih.gov/gene/?term=LOC105276410 | 11085 | 341 | 307.62 | 1 | 0.9 | 0.29 | 9.89 | 9.31 | 0.58 | 0.58 | 0.02 | 0.02 |
| gene-LOC105274823 | https://www.ncbi.nlm.nih.gov/gene/?term=LOC105274823 | 4620 | 172 | 372.29 | 1 | 2.16 | 0.58 | 17.73 | 17.2 | 0.53 | 0.53 | 0.02 | 0.02 |
| gene-LOC105278497 | https://www.ncbi.nlm.nih.gov/gene/?term=LOC105278497 | 2072 | 92 | 444.02 | 1 | 4.83 | 1.09 | 20.29 | 19.78 | 0.51 | 0.51 | 0.02 | 0.02 |
| gene-LOC105279298 | https://www.ncbi.nlm.nih.gov/gene/?term=LOC105279298 | 2773 | 130 | 468.81 | 1 | 3.61 | 0.77 | 23.65 | 23.17 | 0.49 | 0.49 | 0.02 | 0.02 |
| gene-LOC105282491 | https://www.ncbi.nlm.nih.gov/gene/?term=LOC105282491 | 4761 | 193 | 405.38 | 1 | 2.1 | 0.52 | 8.12 | 7.73 | 0.38 | 0.38 | 0.02 | 0.02 |
| gene-LOC105276527 | https://www.ncbi.nlm.nih.gov/gene/?term=LOC105276527 | 7923 | 370 | 466.99 | 1 | 1.26 | 0.27 | 8.27 | 7.9 | 0.37 | 0.37 | 0.02 | 0.02 |
| gene-LOC105276429 | https://www.ncbi.nlm.nih.gov/gene/?term=LOC105276429 | 2901 | 132 | 455.02 | 1 | 3.45 | 0.76 | 22.03 | 22.48 | -0.45 | 0.45 | -0.02 | 0.02 |
| gene-LOC105282151 | https://www.ncbi.nlm.nih.gov/gene/?term=LOC105282151 | 1601 | 43 | 268.58 | 1 | 6.25 | 2.33 | 38.6 | 39.18 | -0.58 | 0.58 | -0.02 | 0.02 |
| gene-LOC105284036 | https://www.ncbi.nlm.nih.gov/gene/?term=LOC105284036 | 3254 | 103 | 316.53 | 1 | 3.07 | 0.97 | 23.72 | 24.37 | -0.65 | 0.65 | -0.02 | 0.02 |
| gene-LOC105286265 | https://www.ncbi.nlm.nih.gov/gene/?term=LOC105286265 | 2507 | 67 | 267.25 | 1 | 3.99 | 1.49 | 29.04 | 29.68 | -0.65 | 0.65 | -0.02 | 0.02 |
| gene-LOC105286515 | https://www.ncbi.nlm.nih.gov/gene/?term=LOC105286515 | 3751 | 111 | 295.92 | 1 | 2.67 | 0.9 | 10.2 | 10.99 | -0.8 | 0.8 | -0.02 | 0.02 |
| gene-LOC105277903 | https://www.ncbi.nlm.nih.gov/gene/?term=LOC105277903 | 1835 | 38 | 207.08 | 1 | 5.45 | 2.63 | 52.47 | 53.46 | -0.99 | 0.99 | -0.02 | 0.02 |
| gene-LOC105286770 | https://www.ncbi.nlm.nih.gov/gene/?term=LOC105286770 | 3055 | 49 | 160.39 | 1 | 3.27 | 2.04 | 11.46 | 12.59 | -1.13 | 1.13 | -0.02 | 0.02 |
| gene-LOC105276196 | https://www.ncbi.nlm.nih.gov/gene/?term=LOC105276196 | 2953 | 60 | 203.18 | 1 | 3.39 | 1.67 | 37.66 | 38.86 | -1.2 | 1.2 | -0.02 | 0.02 |
| gene-LOC105279528 | https://www.ncbi.nlm.nih.gov/gene/?term=LOC105279528 | 5802 | 41 | 70.67 1 | 1.72 | 2.44 | 17.37 | 19.59 | -2.22 | 2.22 | -0.02 | 0.02 | |
| gene-LOC105280447 | https://www.ncbi.nlm.nih.gov/gene/?term=LOC105280447 | 2215 | 19 | 85.78 1 | 4.51 | 5.26 | 33.1 | 35.5 | -2.4 | 2.4 | -0.02 | 0.02 | |
| gene-LOC105281515 | https://www.ncbi.nlm.nih.gov/gene/?term=LOC105281515 | 8827 | 55 | 62.31 2 | 2.27 | 3.64 | 34.93 | 32.95 | 1.98 | 1.98 | 0.01 | 0.01 | |
| gene-LOC105285060 | https://www.ncbi.nlm.nih.gov/gene/?term=LOC105285060 | 1285 | 16 | 124.51 | 1 | 7.78 | 6.25 | 42.41 | 41.27 | 1.13 | 1.13 | 0.01 | 0.01 |
| gene-LOC105277215 | https://www.ncbi.nlm.nih.gov/gene/?term=LOC105277215 | 2619 | 29 | 110.73 | 1 | 3.82 | 3.45 | 25.46 | 24.64 | 0.82 | 0.82 | 0.01 | 0.01 |
| gene-LOC105283943 | https://www.ncbi.nlm.nih.gov/gene/?term=LOC105283943 | 3559 | 69 | 193.87 | 1 | 2.81 | 1.45 | 20.61 | 20.02 | 0.59 | 0.59 | 0.01 | 0.01 |
| gene-LOC105279664 | https://www.ncbi.nlm.nih.gov/gene/?term=LOC105279664 | 4023 | 67 | 166.54 | 1 | 2.49 | 1.49 | 18.98 | 18.43 | 0.55 | 0.55 | 0.01 | 0.01 |
| gene-LOC105280408 | https://www.ncbi.nlm.nih.gov/gene/?term=LOC105280408 | 5983 | 63 | 105.3 1 | 1.67 | 1.59 | 16.29 | 15.78 | 0.51 | 0.51 | 0.01 | 0.01 | |
| gene-LOC105278534 | https://www.ncbi.nlm.nih.gov/gene/?term=LOC105278534 | 7364 | 204 | 277.02 | 1 | 1.36 | 0.49 | 17.28 | 16.79 | 0.5 | 0.5 | 0.01 | 0.01 |
| gene-LOC105283649 | https://www.ncbi.nlm.nih.gov/gene/?term=LOC105283649 | 1918 | 57 | 297.18 | 1 | 5.21 | 1.75 | 40.29 | 39.8 | 0.48 | 0.48 | 0.01 | 0.01 |
| gene-LOC105288078 | https://www.ncbi.nlm.nih.gov/gene/?term=LOC105288078 | 19126 | 350 | 183 1 | 0.52 | 0.29 | 15.9 | 15.43 | 0.47 | 0.47 | 0.01 | 0.01 | |
| gene-LOC105280641 | https://www.ncbi.nlm.nih.gov/gene/?term=LOC105280641 | 3682 | 84 | 228.14 | 1 | 2.72 | 1.19 | 13.99 | 13.53 | 0.46 | 0.46 | 0.01 | 0.01 |
| gene-LOC105275187 | https://www.ncbi.nlm.nih.gov/gene/?term=LOC105275187 | 5534 | 78 | 140.95 | 1 | 1.81 | 1.28 | 16.44 | 16.02 | 0.42 | 0.42 | 0.01 | 0.01 |
| gene-LOC105278807 | https://www.ncbi.nlm.nih.gov/gene/?term=LOC105278807 | 2166 | 75 | 346.26 | 1 | 4.62 | 1.33 | 20.48 | 20.06 | 0.41 | 0.41 | 0.01 | 0.01 |
| gene-LOC105286099 | https://www.ncbi.nlm.nih.gov/gene/?term=LOC105286099 | 5919 | 77 | 130.09 | 1 | 1.69 | 1.3 | 12.19 | 11.79 | 0.4 | 0.4 | 0.01 | 0.01 |
| gene-LOC105286514 | https://www.ncbi.nlm.nih.gov/gene/?term=LOC105286514 | 2528 | 51 | 201.74 | 1 | 3.96 | 1.96 | 35.82 | 35.46 | 0.36 | 0.36 | 0.01 | 0.01 |
| gene-LOC105275170 | https://www.ncbi.nlm.nih.gov/gene/?term=LOC105275170 | 12918 | 359 | 277.91 | 1 | 0.77 | 0.28 | 2.02 | 1.81 | 0.21 | 0.21 | 0.01 | 0.01 |
| gene-LOC105284262 | https://www.ncbi.nlm.nih.gov/gene/?term=LOC105284262 | 6669 | 241 | 361.37 | 1 | 1.5 | 0.41 | 5.35 | 5.16 | 0.19 | 0.19 | 0.01 | 0.01 |
| gene-LOC105285742 | https://www.ncbi.nlm.nih.gov/gene/?term=LOC105285742 | 3446 | 118 | 342.43 | 1 | 2.9 | 0.85 | 17.32 | 17.14 | 0.18 | 0.18 | 0.01 | 0.01 |
| gene-LOC105274531 | https://www.ncbi.nlm.nih.gov/gene/?term=LOC105274531 | 7774 | 295 | 379.47 | 1 | 1.29 | 0.34 | 10.65 | 10.48 | 0.16 | 0.16 | 0.01 | 0.01 |
| gene-LOC105279038 | https://www.ncbi.nlm.nih.gov/gene/?term=LOC105279038 | 9453 | 346 | 366.02 | 1 | 1.06 | 0.29 | 8.85 | 9.03 | -0.18 | 0.18 | -0.01 | 0.01 |
| gene-LOC105288042 | https://www.ncbi.nlm.nih.gov/gene/?term=LOC105288042 | 3507 | 98 | 279.44 | 1 | 2.85 | 1.02 | 22.85 | 23.09 | -0.24 | 0.24 | -0.01 | 0.01 |
| gene-LOC105280956 | https://www.ncbi.nlm.nih.gov/gene/?term=LOC105280956 | 2639 | 81 | 306.93 | 1 | 3.79 | 1.23 | 18.56 | 18.86 | -0.29 | 0.29 | -0.01 | 0.01 |
| gene-LOC105281173 | https://www.ncbi.nlm.nih.gov/gene/?term=LOC105281173 | 1595 | 42 | 263.32 | 1 | 6.27 | 2.38 | 62.5 | 62.82 | -0.31 | 0.31 | -0.01 | 0.01 |
| gene-LOC105285004 | https://www.ncbi.nlm.nih.gov/gene/?term=LOC105285004 | 1334 | 35 | 262.37 | 1 | 7.5 | 2.86 | 19.44 | 19.75 | -0.31 | 0.31 | -0.01 | 0.01 |
| gene-LOC105278962 | https://www.ncbi.nlm.nih.gov/gene/?term=LOC105278962 | 4491 | 129 | 287.24 | 1 | 2.23 | 0.78 | 17.72 | 18.05 | -0.34 | 0.34 | -0.01 | 0.01 |
| gene-LOC105282110 | https://www.ncbi.nlm.nih.gov/gene/?term=LOC105282110 | 7748 | 148 | 191.02 | 1 | 1.29 | 0.68 | 11.12 | 11.53 | -0.42 | 0.42 | -0.01 | 0.01 |
| gene-LOC105282825 | https://www.ncbi.nlm.nih.gov/gene/?term=LOC105282825 | 3919 | 98 | 250.06 | 1 | 2.55 | 1.02 | 13.43 | 13.91 | -0.48 | 0.48 | -0.01 | 0.01 |
| gene-LOC105274590 | https://www.ncbi.nlm.nih.gov/gene/?term=LOC105274590 | 4516 | 84 | 186.01 | 2 | 4.43 | 2.38 | 24.43 | 25 | -0.57 | 0.57 | -0.01 | 0.01 |
| gene-LOC105276745 | https://www.ncbi.nlm.nih.gov/gene/?term=LOC105276745 | 3953 | 76 | 192.26 | 1 | 2.53 | 1.32 | 18.95 | 19.72 | -0.76 | 0.76 | -0.01 | 0.01 |
| gene-LOC105282914 | https://www.ncbi.nlm.nih.gov/gene/?term=LOC105282914 | 5623 | 43 | 76.47 1 | 1.78 | 2.33 | 48 | 49.79 | -1.79 | 1.79 | -0.01 | 0.01 | |
| gene-LOC105280407 | https://www.ncbi.nlm.nih.gov/gene/?term=LOC105280407 | 10950 | 227 | 207.31 | 1 | 0.91 | 0.44 | 15.19 | 14.98 | 0.21 | 0.21 | 0 | 0 |
| gene-LOC105279500 | https://www.ncbi.nlm.nih.gov/gene/?term=LOC105279500 | 3587 | 35 | 97.57 1 | 2.79 | 2.86 | 48.82 | 48.61 | 0.2 | 0.2 | 0 | 0 | |
| gene-LOC105288066 | https://www.ncbi.nlm.nih.gov/gene/?term=LOC105288066 | 1964 | 31 | 157.84 | 1 | 5.09 | 3.23 | 36.75 | 36.57 | 0.19 | 0.19 | 0 | 0 |
| gene-LOC105285981 | https://www.ncbi.nlm.nih.gov/gene/?term=LOC105285981 | 7666 | 219 | 285.68 | 1 | 1.3 | 0.46 | 18.34 | 18.17 | 0.17 | 0.17 | 0 | 0 |
| gene-LOC105278228 | https://www.ncbi.nlm.nih.gov/gene/?term=LOC105278228 | 8610 | 199 | 231.13 | 1 | 1.16 | 0.5 | 21.48 | 21.33 | 0.15 | 0.15 | 0 | 0 |
| gene-LOC105275451 | https://www.ncbi.nlm.nih.gov/gene/?term=LOC105275451 | 6035 | 142 | 235.29 | 1 | 1.66 | 0.7 | 20 | 19.9 | 0.1 | 0.1 | 0 | 0 |
| gene-LOC105283838 | https://www.ncbi.nlm.nih.gov/gene/?term=LOC105283838 | 1748 | 30 | 171.62 | 1 | 5.72 | 3.33 | 35.36 | 35.26 | 0.1 | 0.1 | 0 | 0 |
| gene-LOC105277216 | https://www.ncbi.nlm.nih.gov/gene/?term=LOC105277216 | 7627 | 117 | 153.4 1 | 1.31 | 0.85 | 6.74 | 6.65 | 0.09 | 0.09 | 0 | 0 | |
| gene-LOC105278145 | https://www.ncbi.nlm.nih.gov/gene/?term=LOC105278145 | 4470 | 261 | 583.89 | 1 | 2.24 | 0.38 | 10.03 | 9.95 | 0.08 | 0.08 | 0 | 0 |
| gene-LOC105280002 | https://www.ncbi.nlm.nih.gov/gene/?term=LOC105280002 | 1940 | 32 | 164.95 | 1 | 5.15 | 3.12 | 34.33 | 34.27 | 0.06 | 0.06 | 0 | 0 |
| gene-LOC105284043 | https://www.ncbi.nlm.nih.gov/gene/?term=LOC105284043 | 5661 | 295 | 521.11 | 1 | 1.77 | 0.34 | 10.33 | 10.27 | 0.06 | 0.06 | 0 | 0 |
| gene-LOC105284184 | https://www.ncbi.nlm.nih.gov/gene/?term=LOC105284184 | 2535 | 102 | 402.37 | 1 | 3.94 | 0.98 | 30 | 29.94 | 0.06 | 0.06 | 0 | 0 |
| gene-LOC105282811 | https://www.ncbi.nlm.nih.gov/gene/?term=LOC105282811 | 5765 | 235 | 407.63 | 1 | 1.73 | 0.43 | 8.46 | 8.42 | 0.04 | 0.04 | 0 | 0 |
| gene-LOC105278283 | https://www.ncbi.nlm.nih.gov/gene/?term=LOC105278283 | 2317 | 138 | 595.6 1 | 4.32 | 0.72 | 25.04 | 25.03 | 0.02 | 0.02 | 0 | 0 | |
| gene-LOC105278666 | https://www.ncbi.nlm.nih.gov/gene/?term=LOC105278666 | 10341 | 484 | 468.04 | 1 | 0.97 | 0.21 | 6.48 | 6.46 | 0.02 | 0.02 | 0 | 0 |
| gene-LOC105274706 | https://www.ncbi.nlm.nih.gov/gene/?term=LOC105274706 | 1446 | 46 | 318.12 | 1 | 6.92 | 2.17 | 42.4 | 42.39 | 0 | 0 | 0 | 0 |
| gene-LOC105276059 | https://www.ncbi.nlm.nih.gov/gene/?term=LOC105276059 | 6256 | 146 | 233.38 | 1 | 1.6 | 0.68 | 11.41 | 11.42 | 0 | 0 | 0 | 0 |

| | | | | | | | | | | | | | | |
|---|---|---|---|---|---|---|---|---|---|---|---|---|---|---|
| gene-LOC105278877 | https://www.ncbi.nlm.nih.gov/gene/?term=LOC105278877 | 1968 | 88 | 447.15 | | 1 | 5.08 | 1.14 | 30.96 | 31.01 | -0.05 | 0.05 | 0 | 0 |
| gene-LOC105287065 | https://www.ncbi.nlm.nih.gov/gene/?term=LOC105287065 | 4859 | 249 | 512.45 | | 1 | 2.06 | 0.4 | 12.25 | 12.3 | -0.05 | 0.05 | 0 | 0 |
| gene-LOC105284774 | https://www.ncbi.nlm.nih.gov/gene/?term=LOC105284774 | 12089 | 100 | 82.72 | 1 | 0.83 | 1 | 19.53 | 19.59 | -0.06 | 0.06 | 0 | 0 | |
| gene-LOC105275327 | https://www.ncbi.nlm.nih.gov/gene/?term=LOC105275327 | 14245 | 552 | 387.5 | 1 | 0.7 | 0.18 | 6.33 | 6.41 | -0.08 | 0.08 | 0 | 0 | |
| gene-LOC105282600 | https://www.ncbi.nlm.nih.gov/gene/?term=LOC105282600 | 3715 | 60 | 161.51 | | 1 | 2.69 | 1.67 | 28.86 | 28.93 | -0.08 | 0.08 | 0 | 0 |
| gene-LOC105286339 | https://www.ncbi.nlm.nih.gov/gene/?term=LOC105286339 | 5230 | 184 | 351.82 | | 1 | 1.91 | 0.54 | 16.14 | 16.23 | -0.09 | 0.09 | 0 | 0 |
| gene-LOC105288010 | https://www.ncbi.nlm.nih.gov/gene/?term=LOC105288010 | 3189 | 52 | 163.06 | | 1 | 3.14 | 1.92 | 33.66 | 33.76 | -0.1 | 0.1 | 0 | 0 |
| gene-LOC105285807 | https://www.ncbi.nlm.nih.gov/gene/?term=LOC105285807 | 6063 | 221 | 364.51 | | 1 | 1.65 | 0.45 | 9.8 | 9.91 | -0.11 | 0.11 | 0 | 0 |
| gene-LOC105285406 | https://www.ncbi.nlm.nih.gov/gene/?term=LOC105285406 | 8015 | 57 | 71.12 | 1 | 1.25 | 1.75 | 28.09 | 28.72 | -0.63 | 0.63 | 0 | 0 | |

Directory: OGL File: rnk-dmg-Ob_LI_BCphaseB1.vs.RphaseB2.txt

| gene_ID | gene_link | gwidth | #Sites | #per10Kb | #dmSites | #dmsp10kb | %dmSites | %pSite1 | %pSite2 | DMpSite | ADMpSite | DMpNucl | ADMpNucl |
|---|---|---|---|---|---|---|---|---|---|---|---|---|---|
| gene-LOC113562345 | https://www.ncbi.nlm.nih.gov/gene/?term=LOC113562345 | 2290 | 66 | 288.21 | 12 | 52.4 | 18.18 | 30.39 | 17.58 | 12.82 | 12.82 | 0.37 | 0.37 |
| gene-LOC105283552 | https://www.ncbi.nlm.nih.gov/gene/?term=LOC105283552 | 1562 | 41 | 262.48 | 1 | 6.4 | 2.44 | 53.31 | 48.83 | 4.48 | 4.48 | 0.12 | 0.12 |
| gene-LOC105286683 | https://www.ncbi.nlm.nih.gov/gene/?term=LOC105286683 | 1665 | 55 | 330.33 | 1 | 6.01 | 1.82 | 34.95 | 31.78 | 3.17 | 3.17 | 0.1 | 0.1 |
| gene-LOC105275070 | https://www.ncbi.nlm.nih.gov/gene/?term=LOC105275070 | 5667 | 41 | 72.35 | 6 | 10.59 | 14.63 | 12.96 | 1.06 | 11.9 | 11.9 | 0.09 | 0.09 |
| gene-LOC105279898 | https://www.ncbi.nlm.nih.gov/gene/?term=LOC105279898 | 1343 | 10 | 74.46 | 1 | 7.45 | 10 | 83.43 | 71.53 | 11.89 | 11.89 | 0.09 | 0.09 |
| gene-LOC105285671 | https://www.ncbi.nlm.nih.gov/gene/?term=LOC105285671 | 2263 | 105 | 463.99 | 1 | 4.42 | 0.95 | 28.66 | 26.78 | 1.88 | 1.88 | 0.09 | 0.09 |
| gene-LOC105280104 | https://www.ncbi.nlm.nih.gov/gene/?term=LOC105280104 | 3960 | 135 | 340.91 | 1 | 2.53 | 0.74 | 18.4 | 16.2 | 2.2 | 2.2 | 0.07 | 0.07 |
| gene-LOC105274534 | https://www.ncbi.nlm.nih.gov/gene/?term=LOC105274534 | 1329 | 55 | 413.84 | 1 | 7.52 | 1.82 | 44.22 | 45.89 | -1.68 | 1.68 | -0.07 | 0.07 |
| gene-LOC105279356 | https://www.ncbi.nlm.nih.gov/gene/?term=LOC105279356 | 1314 | 43 | 327.25 | 1 | 7.61 | 2.33 | 32.17 | 30.3 | 1.87 | 1.87 | 0.06 | 0.06 |
| gene-LOC105281419 | https://www.ncbi.nlm.nih.gov/gene/?term=LOC105281419 | 2366 | 80 | 338.12 | 1 | 4.23 | 1.25 | 22.59 | 20.79 | 1.81 | 1.81 | 0.06 | 0.06 |
| gene-LOC105280786 | https://www.ncbi.nlm.nih.gov/gene/?term=LOC105280786 | 2573 | 123 | 478.04 | 1 | 3.89 | 0.81 | 29.6 | 28.26 | 1.34 | 1.34 | 0.06 | 0.06 |
| gene-LOC105282156 | https://www.ncbi.nlm.nih.gov/gene/?term=LOC105282156 | 2577 | 127 | 492.82 | 1 | 3.88 | 0.79 | 27.5 | 26.37 | 1.13 | 1.13 | 0.06 | 0.06 |
| gene-LOC105275708 | https://www.ncbi.nlm.nih.gov/gene/?term=LOC105275708 | 1772 | 14 | 79.01 | 1 | 5.64 | 7.14 | 49.7 | 43.39 | 6.31 | 6.31 | 0.05 | 0.05 |
| gene-LOC105275518 | https://www.ncbi.nlm.nih.gov/gene/?term=LOC105275518 | 2335 | 20 | 85.65 | 1 | 4.28 | 5 | 58.2 | 52.79 | 5.42 | 5.42 | 0.05 | 0.05 |
| gene-LOC105282794 | https://www.ncbi.nlm.nih.gov/gene/?term=LOC105282794 | 4571 | 91 | 199.08 | 1 | 2.19 | 1.1 | 19.77 | 17.09 | 2.68 | 2.68 | 0.05 | 0.05 |
| gene-LOC105285674 | https://www.ncbi.nlm.nih.gov/gene/?term=LOC105285674 | 3035 | 98 | 322.9 | 2 | 6.59 | 2.04 | 25.36 | 23.72 | 1.64 | 1.64 | 0.05 | 0.05 |
| gene-LOC105283728 | https://www.ncbi.nlm.nih.gov/gene/?term=LOC105283728 | 3256 | 115 | 353.19 | 1 | 3.07 | 0.87 | 19.83 | 18.35 | 1.48 | 1.48 | 0.05 | 0.05 |
| gene-LOC105288036 | https://www.ncbi.nlm.nih.gov/gene/?term=LOC105288036 | 5110 | 200 | 391.39 | 1 | 1.96 | 0.5 | 17.99 | 16.61 | 1.37 | 1.37 | 0.05 | 0.05 |
| gene-LOC113561621 | https://www.ncbi.nlm.nih.gov/gene/?term=LOC113561621 | 2249 | 83 | 369.05 | 1 | 4.45 | 1.2 | 21.13 | 19.82 | 1.31 | 1.31 | 0.05 | 0.05 |
| gene-LOC105285179 | https://www.ncbi.nlm.nih.gov/gene/?term=LOC105285179 | 4191 | 175 | 417.56 | 1 | 2.39 | 0.57 | 18.73 | 17.48 | 1.25 | 1.25 | 0.05 | 0.05 |
| gene-LOC105275363 | https://www.ncbi.nlm.nih.gov/gene/?term=LOC105275363 | 3798 | 160 | 421.27 | 1 | 2.63 | 0.62 | 17.46 | 16.32 | 1.14 | 1.14 | 0.05 | 0.05 |
| gene-LOC105282885 | https://www.ncbi.nlm.nih.gov/gene/?term=LOC105282885 | 1870 | 47 | 251.34 | 1 | 5.35 | 2.13 | 35.98 | 38.05 | -2.07 | 2.07 | -0.05 | 0.05 |
| gene-LOC105276196 | https://www.ncbi.nlm.nih.gov/gene/?term=LOC105276196 | 2953 | 60 | 203.18 | 1 | 3.39 | 1.67 | 38.8 | 41.25 | -2.45 | 2.45 | -0.05 | 0.05 |
| gene-LOC105283492 | https://www.ncbi.nlm.nih.gov/gene/?term=LOC105283492 | 1535 | 10 | 65.15 | 1 | 6.51 | 10 | 34.7 | 42.8 | -8.1 | 8.1 | -0.05 | 0.05 |
| gene-LOC105287648 | https://www.ncbi.nlm.nih.gov/gene/?term=LOC105287648 | 2205 | 25 | 113.38 | 1 | 4.54 | 4 | 44.43 | 41.17 | 3.26 | 3.26 | 0.04 | 0.04 |
| gene-LOC105285514 | https://www.ncbi.nlm.nih.gov/gene/?term=LOC105285514 | 1277 | 17 | 133.12 | 1 | 7.83 | 5.88 | 73.52 | 70.72 | 2.8 | 2.8 | 0.04 | 0.04 |
| gene-LOC105279664 | https://www.ncbi.nlm.nih.gov/gene/?term=LOC105279664 | 4023 | 54 | 134.23 | 1 | 2.49 | 1.85 | 21.88 | 19.13 | 2.74 | 2.74 | 0.04 | 0.04 |
| gene-LOC105287417 | https://www.ncbi.nlm.nih.gov/gene/?term=LOC105287417 | 1025 | 23 | 224.39 | 1 | 9.76 | 4.35 | 51.1 | 49.27 | 1.83 | 1.83 | 0.04 | 0.04 |
| gene-LOC105284355 | https://www.ncbi.nlm.nih.gov/gene/?term=LOC105284355 | 3904 | 151 | 386.78 | 1 | 2.56 | 0.66 | 21.57 | 20.54 | 1.02 | 1.02 | 0.04 | 0.04 |
| gene-LOC105276704 | https://www.ncbi.nlm.nih.gov/gene/?term=LOC105276704 | 2063 | 87 | 421.72 | 1 | 4.85 | 1.15 | 14.98 | 14.08 | 0.9 | 0.9 | 0.04 | 0.04 |
| gene-LOC105284043 | https://www.ncbi.nlm.nih.gov/gene/?term=LOC105284043 | 5661 | 294 | 519.34 | 1 | 1.77 | 0.34 | 10.31 | 9.51 | 0.8 | 0.8 | 0.04 | 0.04 |
| gene-LOC105285055 | https://www.ncbi.nlm.nih.gov/gene/?term=LOC105285055 | 4108 | 256 | 623.17 | 1 | 2.43 | 0.39 | 12.1 | 11.41 | 0.69 | 0.69 | 0.04 | 0.04 |
| gene-LOC105280377 | https://www.ncbi.nlm.nih.gov/gene/?term=LOC105280377 | 2024 | 69 | 340.91 | 1 | 4.94 | 1.45 | 43.74 | 44.8 | -1.06 | 1.06 | -0.04 | 0.04 |
| gene-LOC105276429 | https://www.ncbi.nlm.nih.gov/gene/?term=LOC105276429 | 2901 | 114 | 392.97 | 1 | 3.45 | 0.88 | 24.72 | 25.87 | -1.14 | 1.14 | -0.04 | 0.04 |
| gene-LOC105287633 | https://www.ncbi.nlm.nih.gov/gene/?term=LOC105287633 | 1524 | 31 | 203.41 | 1 | 6.56 | 3.23 | 53.47 | 55.25 | -1.78 | 1.78 | -0.04 | 0.04 |
| gene-LOC105274900 | https://www.ncbi.nlm.nih.gov/gene/?term=LOC105274900 | 5561 | 55 | 98.9 | 2 | 3.6 | 3.64 | 33.24 | 30.24 | 3 | 3 | 0.03 | 0.03 |
| gene-LOC105280641 | https://www.ncbi.nlm.nih.gov/gene/?term=LOC105280641 | 3682 | 50 | 135.8 | 1 | 2.72 | 2 | 17.97 | 16.08 | 1.89 | 1.89 | 0.03 | 0.03 |
| gene-LOC105288042 | https://www.ncbi.nlm.nih.gov/gene/?term=LOC105288042 | 3507 | 89 | 253.78 | 1 | 2.85 | 1.12 | 22.75 | 21.38 | 1.37 | 1.37 | 0.03 | 0.03 |
| gene-LOC105276901 | https://www.ncbi.nlm.nih.gov/gene/?term=LOC105276901 | 3254 | 82 | 252 | 1 | 3.07 | 1.22 | 28 | 26.83 | 1.17 | 1.17 | 0.03 | 0.03 |
| gene-LOC105274449 | https://www.ncbi.nlm.nih.gov/gene/?term=LOC105274449 | 1549 | 48 | 309.88 | 1 | 6.46 | 2.08 | 51.97 | 50.96 | 1 | 1 | 0.03 | 0.03 |
| gene-LOC105285639 | https://www.ncbi.nlm.nih.gov/gene/?term=LOC105285639 | 10325 | 332 | 321.55 | 2 | 1.94 | 0.6 | 12.25 | 11.3 | 0.95 | 0.95 | 0.03 | 0.03 |
| gene-LOC105278877 | https://www.ncbi.nlm.nih.gov/gene/?term=LOC105278877 | 1968 | 69 | 350.61 | 1 | 5.08 | 1.45 | 36.22 | 35.3 | 0.91 | 0.91 | 0.03 | 0.03 |
| gene-LOC105278807 | https://www.ncbi.nlm.nih.gov/gene/?term=LOC105278807 | 2166 | 67 | 309.33 | 1 | 4.62 | 1.49 | 22.11 | 21.25 | 0.87 | 0.87 | 0.03 | 0.03 |
| gene-LOC105276875 | https://www.ncbi.nlm.nih.gov/gene/?term=LOC105276875 | 3151 | 116 | 368.14 | 1 | 3.17 | 0.86 | 22.78 | 21.93 | 0.84 | 0.84 | 0.03 | 0.03 |
| gene-LOC105283123 | https://www.ncbi.nlm.nih.gov/gene/?term=LOC105283123 | 1507 | 46 | 305.24 | 1 | 6.64 | 2.17 | 17.92 | 17.09 | 0.83 | 0.83 | 0.03 | 0.03 |
| gene-LOC105287095 | https://www.ncbi.nlm.nih.gov/gene/?term=LOC105287095 | 1435 | 58 | 404.18 | 1 | 6.97 | 1.72 | 29.38 | 28.65 | 0.73 | 0.73 | 0.03 | 0.03 |
| gene-LOC105278497 | https://www.ncbi.nlm.nih.gov/gene/?term=LOC105278497 | 2072 | 92 | 444.02 | 1 | 4.83 | 1.09 | 19.2 | 18.5 | 0.7 | 0.7 | 0.03 | 0.03 |
| gene-LOC105277423 | https://www.ncbi.nlm.nih.gov/gene/?term=LOC105277423 | 2340 | 106 | 452.99 | 1 | 4.27 | 0.94 | 29 | 28.34 | 0.66 | 0.66 | 0.03 | 0.03 |
| gene-LOC105286166 | https://www.ncbi.nlm.nih.gov/gene/?term=LOC105286166 | 3050 | 110 | 360.66 | 1 | 3.28 | 0.91 | 23.43 | 24.21 | -0.78 | 0.78 | -0.03 | 0.03 |
| gene-LOC105278282 | https://www.ncbi.nlm.nih.gov/gene/?term=LOC105278282 | 1775 | 56 | 315.49 | 1 | 5.63 | 1.79 | 37.29 | 38.34 | -1.05 | 1.05 | -0.03 | 0.03 |
| gene-LOC105275966 | https://www.ncbi.nlm.nih.gov/gene/?term=LOC105275966 | 3299 | 59 | 178.84 | 1 | 3.03 | 1.69 | 49.71 | 51.14 | -1.42 | 1.42 | -0.03 | 0.03 |
| gene-LOC105285409 | https://www.ncbi.nlm.nih.gov/gene/?term=LOC105285409 | 5099 | 49 | 96.1 | 2 | 3.92 | 4.08 | 2.16 | 4.82 | -2.66 | 2.66 | -0.03 | 0.03 |
| gene-LOC105278108 | https://www.ncbi.nlm.nih.gov/gene/?term=LOC105278108 | 4392 | 46 | 104.74 | 1 | 2.28 | 2.17 | 27.18 | 25.03 | 2.14 | 2.14 | 0.02 | 0.02 |
| gene-LOC113561805 | https://www.ncbi.nlm.nih.gov/gene/?term=LOC113561805 | 3337 | 65 | 194.79 | 1 | 3 | 1.54 | 35.44 | 34.26 | 1.18 | 1.18 | 0.02 | 0.02 |
| gene-LOC105281848 | https://www.ncbi.nlm.nih.gov/gene/?term=LOC105281848 | 2122 | 47 | 221.49 | 1 | 4.71 | 2.13 | 27.57 | 26.57 | 1 | 1 | 0.02 | 0.02 |
| gene-LOC105283592 | https://www.ncbi.nlm.nih.gov/gene/?term=LOC105283592 | 3479 | 79 | 227.08 | 1 | 2.87 | 1.27 | 22.25 | 21.39 | 0.85 | 0.85 | 0.02 | 0.02 |
| gene-LOC105276059 | https://www.ncbi.nlm.nih.gov/gene/?term=LOC105276059 | 6256 | 132 | 211 | 1 | 1.6 | 0.76 | 10.3 | 9.47 | 0.83 | 0.83 | 0.02 | 0.02 |
| gene-LOC105278632 | https://www.ncbi.nlm.nih.gov/gene/?term=LOC105278632 | 3624 | 108 | 298.01 | 1 | 2.76 | 0.93 | 17.43 | 16.66 | 0.77 | 0.77 | 0.02 | 0.02 |
| gene-LOC105285395 | https://www.ncbi.nlm.nih.gov/gene/?term=LOC105285395 | 8085 | 196 | 242.42 | 1 | 1.24 | 0.51 | 16.42 | 15.66 | 0.76 | 0.76 | 0.02 | 0.02 |
| gene-LOC105280956 | https://www.ncbi.nlm.nih.gov/gene/?term=LOC105280956 | 2639 | 71 | 269.04 | 1 | 3.79 | 1.41 | 18.55 | 17.79 | 0.75 | 0.75 | 0.02 | 0.02 |

| | | | | | | | | | | | | | |
|---|---|---|---|---|---|---|---|---|---|---|---|---|---|
| gene-LOC105278534 | https://www.ncbi.nlm.nih.gov/gene/?term=LOC105278534 | 7364 | 187 | 253.94 | 1 | 1.36 | 0.53 | 19.13 | 18.41 | 0.72 | 0.72 | 0.02 | 0.02 |
| gene-LOC105275584 | https://www.ncbi.nlm.nih.gov/gene/?term=LOC105275584 | 3070 | 89 | 289.9 | 1 | 3.26 | 1.12 | 19.48 | 18.76 | 0.71 | 0.71 | 0.02 | 0.02 |
| gene-LOC105284184 | https://www.ncbi.nlm.nih.gov/gene/?term=LOC105284184 | 2535 | 86 | 339.25 | 1 | 3.94 | 1.16 | 33.56 | 32.88 | 0.68 | 0.68 | 0.02 | 0.02 |
| gene-LOC105282811 | https://www.ncbi.nlm.nih.gov/gene/?term=LOC105282811 | 5765 | 215 | 372.94 | 1 | 1.73 | 0.47 | 8.61 | 8.16 | 0.45 | 0.45 | 0.02 | 0.02 |
| gene-LOC105275460 | https://www.ncbi.nlm.nih.gov/gene/?term=LOC105275460 | 2473 | 117 | 473.11 | 1 | 4.04 | 0.85 | 13.75 | 13.31 | 0.44 | 0.44 | 0.02 | 0.02 |
| gene-LOC105279298 | https://www.ncbi.nlm.nih.gov/gene/?term=LOC105279298 | 2773 | 133 | 479.62 | 2 | 7.21 | 1.5 | 23.5 | 23.06 | 0.44 | 0.44 | 0.02 | 0.02 |
| gene-LOC105278454 | https://www.ncbi.nlm.nih.gov/gene/?term=LOC105278454 | 1972 | 44 | 223.12 | 1 | 5.07 | 2.27 | 45.26 | 45.99 | -0.74 | 0.74 | -0.02 | 0.02 |
| gene-LOC105282151 | https://www.ncbi.nlm.nih.gov/gene/?term=LOC105282151 | 1601 | 39 | 243.6 | 1 | 6.25 | 2.56 | 40.09 | 41 | -0.91 | 0.91 | -0.02 | 0.02 |
| gene-LOC105286999 | https://www.ncbi.nlm.nih.gov/gene/?term=LOC105286999 | 2141 | 46 | 214.85 | 1 | 4.67 | 2.17 | 31.8 | 32.92 | -1.12 | 1.12 | -0.02 | 0.02 |
| gene-LOC105281700 | https://www.ncbi.nlm.nih.gov/gene/?term=LOC105281700 | 2208 | 32 | 144.93 | 1 | 4.53 | 3.12 | 37.39 | 39.09 | -1.7 | 1.7 | -0.02 | 0.02 |
| gene-LOC105286770 | https://www.ncbi.nlm.nih.gov/gene/?term=LOC105286770 | 3055 | 30 | 98.2 | 1 | 3.27 | 3.33 | 17.03 | 18.87 | -1.84 | 1.84 | -0.02 | 0.02 |
| gene-LOC105285315 | https://www.ncbi.nlm.nih.gov/gene/?term=LOC105285315 | 5458 | 19 | 34.81 | 1 | 1.83 | 5.26 | 47.14 | 43.22 | 3.93 | 3.93 | 0.01 | 0.01 |
| gene-LOC105281515 | https://www.ncbi.nlm.nih.gov/gene/?term=LOC105281515 | 8827 | 40 | 45.32 | 2 | 2.27 | 5 | 31.53 | 28.62 | 2.91 | 2.91 | 0.01 | 0.01 |
| gene-LOC105280408 | https://www.ncbi.nlm.nih.gov/gene/?term=LOC105280408 | 5983 | 56 | 93.6 | 1 | 1.67 | 1.79 | 17.1 | 16.06 | 1.03 | 1.03 | 0.01 | 0.01 |
| gene-LOC105286099 | https://www.ncbi.nlm.nih.gov/gene/?term=LOC105286099 | 5919 | 72 | 121.64 | 1 | 1.69 | 1.39 | 10.67 | 9.65 | 1.01 | 1.01 | 0.01 | 0.01 |
| gene-LOC105280447 | https://www.ncbi.nlm.nih.gov/gene/?term=LOC105280447 | 2215 | 21 | 94.81 | 1 | 4.51 | 4.76 | 30.62 | 29.74 | 0.88 | 0.88 | 0.01 | 0.01 |
| gene-LOC105275451 | https://www.ncbi.nlm.nih.gov/gene/?term=LOC105275451 | 6035 | 126 | 208.78 | 1 | 1.66 | 0.79 | 22.21 | 21.64 | 0.57 | 0.57 | 0.01 | 0.01 |
| gene-LOC105281599 | https://www.ncbi.nlm.nih.gov/gene/?term=LOC105281599 | 3859 | 49 | 126.98 | 1 | 2.59 | 2.04 | 4.62 | 4.08 | 0.54 | 0.54 | 0.01 | 0.01 |
| gene-LOC105285742 | https://www.ncbi.nlm.nih.gov/gene/?term=LOC105285742 | 3446 | 110 | 319.21 | 1 | 2.9 | 0.91 | 21.56 | 21.15 | 0.4 | 0.4 | 0.01 | 0.01 |
| gene-LOC105274706 | https://www.ncbi.nlm.nih.gov/gene/?term=LOC105274706 | 1446 | 50 | 345.78 | 1 | 6.92 | 2 | 40.48 | 40.09 | 0.39 | 0.39 | 0.01 | 0.01 |
| gene-LOC105280002 | https://www.ncbi.nlm.nih.gov/gene/?term=LOC105280002 | 1940 | 28 | 144.33 | 1 | 5.15 | 3.57 | 37.11 | 36.72 | 0.39 | 0.39 | 0.01 | 0.01 |
| gene-LOC105279038 | https://www.ncbi.nlm.nih.gov/gene/?term=LOC105279038 | 9453 | 314 | 332.17 | 1 | 1.06 | 0.32 | 9.5 | 9.12 | 0.38 | 0.38 | 0.01 | 0.01 |
| gene-LOC105280774 | https://www.ncbi.nlm.nih.gov/gene/?term=LOC105280774 | 2793 | 99 | 354.46 | 1 | 3.58 | 1.01 | 32.85 | 32.47 | 0.38 | 0.38 | 0.01 | 0.01 |
| gene-LOC105284036 | https://www.ncbi.nlm.nih.gov/gene/?term=LOC105284036 | 3254 | 102 | 313.46 | 1 | 3.07 | 0.98 | 22.68 | 22.3 | 0.38 | 0.38 | 0.01 | 0.01 |
| gene-LOC105274590 | https://www.ncbi.nlm.nih.gov/gene/?term=LOC105274590 | 4516 | 69 | 152.79 | 2 | 4.43 | 2.9 | 27.64 | 27.27 | 0.37 | 0.37 | 0.01 | 0.01 |
| gene-LOC105277079 | https://www.ncbi.nlm.nih.gov/gene/?term=LOC105277079 | 13009 | 273 | 209.85 | 1 | 0.77 | 0.37 | 9.7 | 9.33 | 0.37 | 0.37 | 0.01 | 0.01 |
| gene-LOC105278962 | https://www.ncbi.nlm.nih.gov/gene/?term=LOC105278962 | 4491 | 100 | 222.67 | 1 | 2.23 | 1 | 21.33 | 20.97 | 0.36 | 0.36 | 0.01 | 0.01 |
| gene-LOC105287764 | https://www.ncbi.nlm.nih.gov/gene/?term=LOC105287764 | 18359 | 394 | 214.61 | 1 | 0.54 | 0.25 | 3.28 | 2.97 | 0.31 | 0.31 | 0.01 | 0.01 |
| gene-LOC105281266 | https://www.ncbi.nlm.nih.gov/gene/?term=LOC105281266 | 4102 | 78 | 190.15 | 1 | 2.44 | 1.28 | 16.25 | 15.95 | 0.3 | 0.3 | 0.01 | 0.01 |
| gene-LOC105278145 | https://www.ncbi.nlm.nih.gov/gene/?term=LOC105278145 | 4470 | 235 | 525.73 | 1 | 2.24 | 0.43 | 10.78 | 10.55 | 0.23 | 0.23 | 0.01 | 0.01 |
| gene-LOC105284262 | https://www.ncbi.nlm.nih.gov/gene/?term=LOC105284262 | 6669 | 215 | 322.39 | 1 | 1.5 | 0.47 | 5.95 | 5.73 | 0.22 | 0.22 | 0.01 | 0.01 |
| gene-LOC105278666 | https://www.ncbi.nlm.nih.gov/gene/?term=LOC105278666 | 10341 | 444 | 429.36 | 1 | 0.97 | 0.23 | 6.98 | 6.79 | 0.19 | 0.19 | 0.01 | 0.01 |
| gene-LOC105282149 | https://www.ncbi.nlm.nih.gov/gene/?term=LOC105282149 | 1787 | 108 | 604.36 | 1 | 5.6 | 0.93 | 41.15 | 40.97 | 0.18 | 0.18 | 0.01 | 0.01 |
| gene-LOC105276527 | https://www.ncbi.nlm.nih.gov/gene/?term=LOC105276527 | 7923 | 325 | 410.2 | 1 | 1.26 | 0.31 | 9.54 | 9.36 | 0.17 | 0.17 | 0.01 | 0.01 |
| gene-LOC105287550 | https://www.ncbi.nlm.nih.gov/gene/?term=LOC105287550 | 15659 | 590 | 376.78 | 1 | 0.64 | 0.17 | 6.31 | 6.15 | 0.16 | 0.16 | 0.01 | 0.01 |
| gene-LOC105276612 | https://www.ncbi.nlm.nih.gov/gene/?term=LOC105276612 | 4157 | 183 | 440.22 | 1 | 2.41 | 0.55 | 9.73 | 9.61 | 0.11 | 0.11 | 0.01 | 0.01 |
| gene-LOC105286274 | https://www.ncbi.nlm.nih.gov/gene/?term=LOC105286274 | 3817 | 191 | 500.39 | 1 | 2.62 | 0.52 | 17.09 | 17.23 | -0.14 | 0.14 | -0.01 | 0.01 |
| gene-LOC105274856 | https://www.ncbi.nlm.nih.gov/gene/?term=LOC105274856 | 5216 | 210 | 402.61 | 1 | 1.92 | 0.48 | 18.25 | 18.52 | -0.26 | 0.26 | -0.01 | 0.01 |
| gene-LOC105277903 | https://www.ncbi.nlm.nih.gov/gene/?term=LOC105277903 | 1835 | 36 | 196.19 | 1 | 5.45 | 2.78 | 49.51 | 49.79 | -0.27 | 0.27 | -0.01 | 0.01 |
| gene-LOC105282491 | https://www.ncbi.nlm.nih.gov/gene/?term=LOC105282491 | 4761 | 165 | 346.57 | 1 | 2.1 | 0.61 | 8.1 | 8.48 | -0.38 | 0.38 | -0.01 | 0.01 |
| gene-LOC105274851 | https://www.ncbi.nlm.nih.gov/gene/?term=LOC105274851 | 6616 | 136 | 205.56 | 1 | 1.51 | 0.74 | 10.78 | 11.18 | -0.4 | 0.4 | -0.01 | 0.01 |
| gene-LOC105286515 | https://www.ncbi.nlm.nih.gov/gene/?term=LOC105286515 | 3751 | 102 | 271.93 | 1 | 2.67 | 0.98 | 12.76 | 13.19 | -0.43 | 0.43 | -0.01 | 0.01 |
| gene-LOC105282399 | https://www.ncbi.nlm.nih.gov/gene/?term=LOC105282399 | 2682 | 23 | 85.76 | 1 | 3.73 | 4.35 | 34.71 | 35.34 | -0.63 | 0.63 | -0.01 | 0.01 |
| gene-LOC105288010 | https://www.ncbi.nlm.nih.gov/gene/?term=LOC105288010 | 3189 | 44 | 137.97 | 1 | 3.14 | 2.27 | 30.11 | 30.74 | -0.63 | 0.63 | -0.01 | 0.01 |
| gene-LOC105285060 | https://www.ncbi.nlm.nih.gov/gene/?term=LOC105285060 | 1285 | 18 | 140.08 | 1 | 7.78 | 5.56 | 44.89 | 45.6 | -0.72 | 0.72 | -0.01 | 0.01 |
| gene-LOC105283712 | https://www.ncbi.nlm.nih.gov/gene/?term=LOC105283712 | 1910 | 35 | 183.25 | 1 | 5.24 | 2.86 | 26.72 | 27.47 | -0.75 | 0.75 | -0.01 | 0.01 |
| gene-LOC105286452 | https://www.ncbi.nlm.nih.gov/gene/?term=LOC105286452 | 6888 | 59 | 85.66 | 1 | 1.45 | 1.69 | 32.42 | 31.88 | 0.55 | 0.55 | 0 | 0 |
| gene-LOC105285406 | https://www.ncbi.nlm.nih.gov/gene/?term=LOC105285406 | 8015 | 53 | 66.13 | 1 | 1.25 | 1.89 | 28.7 | 28.32 | 0.39 | 0.39 | 0 | 0 |
| gene-LOC105283989 | https://www.ncbi.nlm.nih.gov/gene/?term=LOC105283989 | 4948 | 31 | 62.65 | 1 | 2.02 | 3.23 | 36.84 | 36.46 | 0.37 | 0.37 | 0 | 0 |
| gene-LOC105288066 | https://www.ncbi.nlm.nih.gov/gene/?term=LOC105288066 | 1964 | 28 | 142.57 | 1 | 5.09 | 3.57 | 32.79 | 32.53 | 0.26 | 0.26 | 0 | 0 |
| gene-LOC105283943 | https://www.ncbi.nlm.nih.gov/gene/?term=LOC105283943 | 3559 | 47 | 132.06 | 1 | 2.81 | 2.13 | 17.56 | 17.32 | 0.24 | 0.24 | 0 | 0 |
| gene-LOC105274531 | https://www.ncbi.nlm.nih.gov/gene/?term=LOC105274531 | 7774 | 291 | 374.32 | 1 | 1.29 | 0.34 | 10.67 | 10.57 | 0.1 | 0.1 | 0 | 0 |
| gene-LOC105275327 | https://www.ncbi.nlm.nih.gov/gene/?term=LOC105275327 | 14245 | 475 | 333.45 | 1 | 0.7 | 0.21 | 7.23 | 7.14 | 0.1 | 0.1 | 0 | 0 |
| gene-LOC105282110 | https://www.ncbi.nlm.nih.gov/gene/?term=LOC105282110 | 7748 | 119 | 153.59 | 1 | 1.29 | 0.84 | 13.03 | 12.95 | 0.08 | 0.08 | 0 | 0 |
| gene-LOC105285807 | https://www.ncbi.nlm.nih.gov/gene/?term=LOC105285807 | 6063 | 223 | 367.8 | 1 | 1.65 | 0.45 | 9.47 | 9.4 | 0.06 | 0.06 | 0 | 0 |
| gene-LOC105278283 | https://www.ncbi.nlm.nih.gov/gene/?term=LOC105278283 | 2317 | 135 | 582.65 | 1 | 4.32 | 0.74 | 25.81 | 25.76 | 0.05 | 0.05 | 0 | 0 |
| gene-LOC105276538 | https://www.ncbi.nlm.nih.gov/gene/?term=LOC105276538 | 1511 | 17 | 112.51 | 1 | 6.62 | 5.88 | 69.95 | 69.91 | 0.04 | 0.04 | 0 | 0 |
| gene-LOC105287528 | https://www.ncbi.nlm.nih.gov/gene/?term=LOC105287528 | 14195 | 398 | 280.38 | 1 | 0.7 | 0.25 | 4.97 | 4.95 | 0.02 | 0.02 | 0 | 0 |
| gene-LOC105274823 | https://www.ncbi.nlm.nih.gov/gene/?term=LOC105274823 | 4620 | 138 | 298.7 | 1 | 2.16 | 0.72 | 21.27 | 21.26 | 0.01 | 0.01 | 0 | 0 |
| gene-LOC105275127 | https://www.ncbi.nlm.nih.gov/gene/?term=LOC105275127 | 9377 | 295 | 314.6 | 1 | 1.07 | 0.34 | 7.39 | 7.38 | 0.01 | 0.01 | 0 | 0 |
| gene-LOC105280407 | https://www.ncbi.nlm.nih.gov/gene/?term=LOC105280407 | 10950 | 207 | 189.04 | 1 | 0.91 | 0.48 | 16.49 | 16.48 | 0.01 | 0.01 | 0 | 0 |
| gene-LOC105275187 | https://www.ncbi.nlm.nih.gov/gene/?term=LOC105275187 | 5534 | 70 | 126.49 | 1 | 1.81 | 1.43 | 12.34 | 12.39 | -0.05 | 0.05 | 0 | 0 |
| gene-LOC105279500 | https://www.ncbi.nlm.nih.gov/gene/?term=LOC105279500 | 3587 | 36 | 100.36 | 1 | 2.79 | 2.78 | 49.66 | 49.71 | -0.05 | 0.05 | 0 | 0 |
| gene-LOC105276692 | https://www.ncbi.nlm.nih.gov/gene/?term=LOC105276692 | 1745 | 29 | 166.19 | 1 | 5.73 | 3.45 | 24.9 | 24.96 | -0.07 | 0.07 | 0 | 0 |

| | | | | | | | | | | | | | | |
|---|---|---|---|---|---|---|---|---|---|---|---|---|---|---|
| gene-LOC105275374 | https://www.ncbi.nlm.nih.gov/gene/?term=LOC105275374 | 11376 | 502 | 441.28 | | 1 | 0.88 | 0.2 | 10.16 | 10.25 | -0.09 | 0.09 | 0 | 0 |
| gene-LOC105281173 | https://www.ncbi.nlm.nih.gov/gene/?term=LOC105281173 | 1595 | 39 | 244.51 | | 1 | 6.27 | 2.56 | 59.82 | 59.91 | -0.09 | 0.09 | 0 | 0 |
| gene-LOC105278245 | https://www.ncbi.nlm.nih.gov/gene/?term=LOC105278245 | 6445 | 117 | 181.54 | | 1 | 1.55 | 0.85 | 22.04 | 22.2 | -0.17 | 0.17 | 0 | 0 |
| gene-LOC105286514 | https://www.ncbi.nlm.nih.gov/gene/?term=LOC105286514 | 2528 | 48 | 189.87 | | 1 | 3.96 | 2.08 | 36.18 | 36.38 | -0.2 | 0.2 | 0 | 0 |
| gene-LOC105275500 | https://www.ncbi.nlm.nih.gov/gene/?term=LOC105275500 | 10943 | 242 | 221.15 | | 1 | 0.91 | 0.41 | 7.04 | 7.25 | -0.21 | 0.21 | 0 | 0 |
| gene-LOC105282067 | https://www.ncbi.nlm.nih.gov/gene/?term=LOC105282067 | 5941 | 49 | 82.48 | 1 | 1.68 | 2.04 | 26.22 | 26.65 | -0.43 | 0.43 | 0 | 0 | |

Directory: OGL File: rnk-dmg-Ob_LI_BCphaseB2.vs.RphaseA1.txt

| gene_ID | gene_link | gwidth | #Sites | #per10Kb | #dmSites | #dmsp10kb | %dmSites | %pSite1 | %pSite2 | DMpSite | ADMpSite | DMpNucl | ADMpNucl |
|---|---|---|---|---|---|---|---|---|---|---|---|---|---|
| gene-LOC113562345 | https://www.ncbi.nlm.nih.gov/gene/?term=LOC113562345 | 2290 | 63 | 275.11 | 8 | 34.93 | 12.7 | 17.51 | 30.84 | -13.33 | 13.33 | -0.37 | 0.37 |
| gene-LOC105285674 | https://www.ncbi.nlm.nih.gov/gene/?term=LOC105285674 | 3035 | 84 | 276.77 | 2 | 6.59 | 2.38 | 24.69 | 27.77 | -3.08 | 3.08 | -0.09 | 0.09 |
| gene-LOC105275070 | https://www.ncbi.nlm.nih.gov/gene/?term=LOC105275070 | 5667 | 40 | 70.58 | 6 | 10.59 | 15 | 2.13 | 14.38 | -12.25 | 12.25 | -0.09 | 0.09 |
| gene-LOC105284396 | https://www.ncbi.nlm.nih.gov/gene/?term=LOC105284396 | 1976 | 66 | 334.01 | 1 | 5.06 | 1.52 | 34.48 | 32.1 | 2.38 | 2.38 | 0.08 | 0.08 |
| gene-LOC105276449 | https://www.ncbi.nlm.nih.gov/gene/?term=LOC105276449 | 1231 | 45 | 365.56 | 1 | 8.12 | 2.22 | 27.05 | 29.32 | -2.27 | 2.27 | -0.08 | 0.08 |
| gene-LOC105274706 | https://www.ncbi.nlm.nih.gov/gene/?term=LOC105274706 | 1446 | 50 | 345.78 | 1 | 6.92 | 2 | 39.84 | 37.9 | 1.94 | 1.94 | 0.07 | 0.07 |
| gene-LOC105282156 | https://www.ncbi.nlm.nih.gov/gene/?term=LOC105282156 | 2577 | 133 | 516.1 | 1 | 3.88 | 0.75 | 22.7 | 24.15 | -1.45 | 1.45 | -0.07 | 0.07 |
| gene-LOC105280786 | https://www.ncbi.nlm.nih.gov/gene/?term=LOC105280786 | 2573 | 109 | 423.63 | 1 | 3.89 | 0.92 | 28.63 | 30.24 | -1.61 | 1.61 | -0.07 | 0.07 |
| gene-LOC105275384 | https://www.ncbi.nlm.nih.gov/gene/?term=LOC105275384 | 1987 | 87 | 437.85 | 1 | 5.03 | 1.15 | 34.15 | 35.77 | -1.63 | 1.63 | -0.07 | 0.07 |
| gene-LOC105285202 | https://www.ncbi.nlm.nih.gov/gene/?term=LOC105285202 | 1838 | 60 | 326.44 | 1 | 5.44 | 1.67 | 23.97 | 26.11 | -2.15 | 2.15 | -0.07 | 0.07 |
| gene-LOC105283649 | https://www.ncbi.nlm.nih.gov/gene/?term=LOC105283649 | 1918 | 56 | 291.97 | 1 | 5.21 | 1.79 | 36.17 | 38.48 | -2.31 | 2.31 | -0.07 | 0.07 |
| gene-LOC105275708 | https://www.ncbi.nlm.nih.gov/gene/?term=LOC105275708 | 1772 | 15 | 84.65 | 1 | 5.64 | 6.67 | 45.9 | 54.1 | -8.2 | 8.2 | -0.07 | 0.07 |
| gene-LOC105274449 | https://www.ncbi.nlm.nih.gov/gene/?term=LOC105274449 | 1549 | 53 | 342.16 | 1 | 6.46 | 1.89 | 51.77 | 49.89 | 1.87 | 1.87 | 0.06 | 0.06 |
| gene-LOC105277903 | https://www.ncbi.nlm.nih.gov/gene/?term=LOC105277903 | 1835 | 36 | 196.19 | 1 | 5.45 | 2.78 | 53.1 | 56.25 | -3.14 | 3.14 | -0.06 | 0.06 |
| gene-LOC105274900 | https://www.ncbi.nlm.nih.gov/gene/?term=LOC105274900 | 5561 | 54 | 97.1 | 3 | 5.39 | 5.56 | 30 | 35.69 | -5.68 | 5.68 | -0.06 | 0.06 |
| gene-LOC105279898 | https://www.ncbi.nlm.nih.gov/gene/?term=LOC105279898 | 1343 | 11 | 81.91 | 1 | 7.45 | 9.09 | 61.61 | 68.61 | -7 | 7 | -0.06 | 0.06 |
| gene-LOC105278567 | https://www.ncbi.nlm.nih.gov/gene/?term=LOC105278567 | 3627 | 29 | 79.96 | 1 | 2.76 | 3.45 | 11.32 | 18.71 | -7.39 | 7.39 | -0.06 | 0.06 |
| gene-LOC105278454 | https://www.ncbi.nlm.nih.gov/gene/?term=LOC105278454 | 1972 | 43 | 218.05 | 1 | 5.07 | 2.33 | 44.38 | 46.55 | -2.17 | 2.17 | -0.05 | 0.05 |
| gene-LOC105281420 | https://www.ncbi.nlm.nih.gov/gene/?term=LOC105281420 | 3882 | 55 | 141.68 | 1 | 2.58 | 1.82 | 13.05 | 16.87 | -3.82 | 3.82 | -0.05 | 0.05 |
| gene-LOC105282885 | https://www.ncbi.nlm.nih.gov/gene/?term=LOC105282885 | 1870 | 46 | 245.99 | 1 | 5.35 | 2.17 | 37.06 | 35.54 | 1.52 | 1.52 | 0.04 | 0.04 |
| gene-LOC105285055 | https://www.ncbi.nlm.nih.gov/gene/?term=LOC105285055 | 4108 | 256 | 623.17 | 1 | 2.43 | 0.39 | 10.36 | 10.97 | -0.61 | 0.61 | -0.04 | 0.04 |
| gene-LOC105280723 | https://www.ncbi.nlm.nih.gov/gene/?term=LOC105280723 | 3572 | 182 | 509.52 | 1 | 2.8 | 0.55 | 15.32 | 16.11 | -0.79 | 0.79 | -0.04 | 0.04 |
| gene-LOC105278877 | https://www.ncbi.nlm.nih.gov/gene/?term=LOC105278877 | 1968 | 82 | 416.67 | 1 | 5.08 | 1.22 | 30.66 | 31.6 | -0.94 | 0.94 | -0.04 | 0.04 |
| gene-LOC105280956 | https://www.ncbi.nlm.nih.gov/gene/?term=LOC105280956 | 2639 | 72 | 272.83 | 1 | 3.79 | 1.39 | 17.08 | 18.51 | -1.43 | 1.43 | -0.04 | 0.04 |
| gene-LOC105278436 | https://www.ncbi.nlm.nih.gov/gene/?term=LOC105278436 | 2556 | 54 | 211.27 | 1 | 3.91 | 1.85 | 41.69 | 43.45 | -1.76 | 1.76 | -0.04 | 0.04 |
| gene-LOC105283592 | https://www.ncbi.nlm.nih.gov/gene/?term=LOC105283592 | 3479 | 64 | 183.96 | 1 | 2.87 | 1.56 | 24.19 | 26.35 | -2.16 | 2.16 | -0.04 | 0.04 |
| gene-LOC105281039 | https://www.ncbi.nlm.nih.gov/gene/?term=LOC105281039 | 1471 | 11 | 74.78 | 1 | 6.8 | 9.09 | 29.26 | 35.1 | -5.84 | 5.84 | -0.04 | 0.04 |
| gene-LOC105282914 | https://www.ncbi.nlm.nih.gov/gene/?term=LOC105282914 | 5623 | 41 | 72.91 | 1 | 1.78 | 2.44 | 52.21 | 48.43 | 3.78 | 3.78 | 0.03 | 0.03 |
| gene-LOC105276272 | https://www.ncbi.nlm.nih.gov/gene/?term=LOC105276272 | 1965 | 46 | 234.1 | 1 | 5.09 | 2.17 | 27.86 | 26.72 | 1.14 | 1.14 | 0.03 | 0.03 |
| gene-LOC105275587 | https://www.ncbi.nlm.nih.gov/gene/?term=LOC105275587 | 2404 | 57 | 237.1 | 1 | 4.16 | 1.75 | 37.35 | 36.26 | 1.09 | 1.09 | 0.03 | 0.03 |
| gene-LOC105278145 | https://www.ncbi.nlm.nih.gov/gene/?term=LOC105278145 | 4470 | 235 | 525.73 | 2 | 4.47 | 0.85 | 10.77 | 11.29 | -0.52 | 0.52 | -0.03 | 0.03 |
| gene-LOC105275460 | https://www.ncbi.nlm.nih.gov/gene/?term=LOC105275460 | 2473 | 120 | 485.24 | 1 | 4.04 | 0.83 | 13.18 | 13.75 | -0.57 | 0.57 | -0.03 | 0.03 |
| gene-LOC105279298 | https://www.ncbi.nlm.nih.gov/gene/?term=LOC105279298 | 2773 | 130 | 468.81 | 2 | 7.21 | 1.54 | 23.75 | 24.38 | -0.63 | 0.63 | -0.03 | 0.03 |
| gene-LOC105274856 | https://www.ncbi.nlm.nih.gov/gene/?term=LOC105274856 | 5216 | 225 | 431.37 | 1 | 1.92 | 0.44 | 17.14 | 17.89 | -0.76 | 0.76 | -0.03 | 0.03 |
| gene-LOC105282491 | https://www.ncbi.nlm.nih.gov/gene/?term=LOC105282491 | 4761 | 174 | 365.47 | 1 | 2.1 | 0.57 | 8.54 | 9.39 | -0.85 | 0.85 | -0.03 | 0.03 |
| gene-LOC105278807 | https://www.ncbi.nlm.nih.gov/gene/?term=LOC105278807 | 2166 | 78 | 360.11 | 1 | 4.62 | 1.28 | 18.54 | 19.5 | -0.97 | 0.97 | -0.03 | 0.03 |
| gene-LOC105282151 | https://www.ncbi.nlm.nih.gov/gene/?term=LOC105282151 | 1601 | 42 | 262.34 | 1 | 6.25 | 2.38 | 34.92 | 36.1 | -1.17 | 1.17 | -0.03 | 0.03 |
| gene-LOC105281266 | https://www.ncbi.nlm.nih.gov/gene/?term=LOC105281266 | 4102 | 81 | 197.46 | 1 | 2.44 | 1.23 | 16.07 | 17.51 | -1.44 | 1.44 | -0.03 | 0.03 |
| gene-LOC105288066 | https://www.ncbi.nlm.nih.gov/gene/?term=LOC105288066 | 1964 | 29 | 147.66 | 1 | 5.09 | 3.45 | 35.48 | 37.25 | -1.78 | 1.78 | -0.03 | 0.03 |
| gene-LOC105285514 | https://www.ncbi.nlm.nih.gov/gene/?term=LOC105285514 | 1277 | 11 | 86.14 | 1 | 7.83 | 9.09 | 70.96 | 74.98 | -4.02 | 4.02 | -0.03 | 0.03 |
| gene-LOC105279500 | https://www.ncbi.nlm.nih.gov/gene/?term=LOC105279500 | 3587 | 30 | 83.64 | 1 | 2.79 | 3.33 | 48.41 | 46.03 | 2.38 | 2.38 | 0.02 | 0.02 |
| gene-LOC105283967 | https://www.ncbi.nlm.nih.gov/gene/?term=LOC105283967 | 2393 | 50 | 208.94 | 1 | 4.18 | 2 | 45.88 | 44.9 | 0.98 | 0.98 | 0.02 | 0.02 |
| gene-LOC105282970 | https://www.ncbi.nlm.nih.gov/gene/?term=LOC105282970 | 1529 | 40 | 261.61 | 1 | 6.54 | 2.5 | 50.96 | 50.14 | 0.82 | 0.82 | 0.02 | 0.02 |
| gene-LOC105281173 | https://www.ncbi.nlm.nih.gov/gene/?term=LOC105281173 | 1595 | 38 | 238.24 | 1 | 6.27 | 2.63 | 60.67 | 59.87 | 0.8 | 0.8 | 0.02 | 0.02 |
| gene-LOC105281312 | https://www.ncbi.nlm.nih.gov/gene/?term=LOC105281312 | 2551 | 65 | 254.8 | 1 | 3.92 | 1.54 | 33.11 | 32.31 | 0.8 | 0.8 | 0.02 | 0.02 |
| gene-LOC105278962 | https://www.ncbi.nlm.nih.gov/gene/?term=LOC105278962 | 4491 | 93 | 207.08 | 1 | 2.23 | 1.08 | 22.79 | 22.03 | 0.76 | 0.76 | 0.02 | 0.02 |
| gene-LOC105280819 | https://www.ncbi.nlm.nih.gov/gene/?term=LOC105280819 | 1639 | 45 | 274.56 | 1 | 6.1 | 2.22 | 37.3 | 36.73 | 0.57 | 0.57 | 0.02 | 0.02 |
| gene-LOC105286274 | https://www.ncbi.nlm.nih.gov/gene/?term=LOC105286274 | 3817 | 187 | 489.91 | 1 | 2.62 | 0.53 | 16.7 | 16.2 | 0.51 | 0.51 | 0.02 | 0.02 |
| gene-LOC105284184 | https://www.ncbi.nlm.nih.gov/gene/?term=LOC105284184 | 2535 | 92 | 362.92 | 1 | 3.94 | 1.09 | 31.62 | 31.13 | 0.49 | 0.49 | 0.02 | 0.02 |
| gene-LOC105278283 | https://www.ncbi.nlm.nih.gov/gene/?term=LOC105278283 | 2317 | 139 | 599.91 | 1 | 4.32 | 0.72 | 25.21 | 25.46 | -0.25 | 0.25 | -0.02 | 0.02 |
| gene-LOC105282149 | https://www.ncbi.nlm.nih.gov/gene/?term=LOC105282149 | 1787 | 103 | 576.39 | 1 | 5.6 | 0.97 | 40.98 | 41.32 | -0.34 | 0.34 | -0.02 | 0.02 |
| gene-LOC105280774 | https://www.ncbi.nlm.nih.gov/gene/?term=LOC105280774 | 2793 | 117 | 418.9 | 1 | 3.58 | 0.85 | 27.14 | 27.51 | -0.36 | 0.36 | -0.02 | 0.02 |
| gene-LOC105274531 | https://www.ncbi.nlm.nih.gov/gene/?term=LOC105274531 | 7774 | 275 | 353.74 | 1 | 1.29 | 0.36 | 10.93 | 11.37 | -0.43 | 0.43 | -0.02 | 0.02 |
| gene-LOC105287095 | https://www.ncbi.nlm.nih.gov/gene/?term=LOC105287095 | 1435 | 64 | 445.99 | 1 | 6.97 | 1.56 | 25.57 | 26.08 | -0.51 | 0.51 | -0.02 | 0.02 |
| gene-LOC105275752 | https://www.ncbi.nlm.nih.gov/gene/?term=LOC105275752 | 3462 | 128 | 369.73 | 1 | 2.89 | 0.78 | 29.22 | 29.76 | -0.53 | 0.53 | -0.02 | 0.02 |
| gene-LOC105285742 | https://www.ncbi.nlm.nih.gov/gene/?term=LOC105285742 | 3446 | 105 | 304.7 | 1 | 2.9 | 0.95 | 17.29 | 17.85 | -0.55 | 0.55 | -0.02 | 0.02 |
| gene-LOC105288036 | https://www.ncbi.nlm.nih.gov/gene/?term=LOC105288036 | 5110 | 200 | 391.39 | 1 | 1.96 | 0.5 | 16.38 | 16.94 | -0.55 | 0.55 | -0.02 | 0.02 |
| gene-LOC105282658 | https://www.ncbi.nlm.nih.gov/gene/?term=LOC105282658 | 2421 | 59 | 243.7 | 1 | 4.13 | 1.69 | 24.61 | 25.23 | -0.62 | 0.62 | -0.02 | 0.02 |
| gene-LOC105287063 | https://www.ncbi.nlm.nih.gov/gene/?term=LOC105287063 | 1874 | 49 | 261.47 | 1 | 5.34 | 2.04 | 29.55 | 30.23 | -0.68 | 0.68 | -0.02 | 0.02 |
| gene-LOC105279900 | https://www.ncbi.nlm.nih.gov/gene/?term=LOC105279900 | 2191 | 54 | 246.46 | 1 | 4.56 | 1.85 | 30.89 | 31.61 | -0.72 | 0.72 | -0.02 | 0.02 |

| | | | | | | | | | | | | | |
|---|---|---|---|---|---|---|---|---|---|---|---|---|---|
| gene-LOC105275451 | https://www.ncbi.nlm.nih.gov/gene/?term=LOC105275451 | 6035 | 123 | 203.81 | 1 | 1.66 | 0.81 | 22.94 | 23.69 | -0.75 | 0.75 | -0.02 | 0.02 |
| gene-LOC105278384 | https://www.ncbi.nlm.nih.gov/gene/?term=LOC105278384 | 1390 | 35 | 251.8 1 | 7.19 | 2.86 | 45.32 | 46.11 | -0.79 | 0.79 | -0.02 | 0.02 | |
| gene-LOC105284116 | https://www.ncbi.nlm.nih.gov/gene/?term=LOC105284116 | 3970 | 107 | 269.52 | 1 | 2.52 | 0.93 | 7.18 | 8 | -0.82 | 0.82 | -0.02 | 0.02 |
| gene-LOC105284117 | https://www.ncbi.nlm.nih.gov/gene/?term=LOC105284117 | 4340 | 94 | 216.59 | 1 | 2.3 | 1.06 | 9.25 | 10.09 | -0.84 | 0.84 | -0.02 | 0.02 |
| gene-LOC105282991 | https://www.ncbi.nlm.nih.gov/gene/?term=LOC105282991 | 5617 | 101 | 179.81 | 1 | 1.78 | 0.99 | 19.27 | 20.29 | -1.02 | 1.02 | -0.02 | 0.02 |
| gene-LOC105279664 | https://www.ncbi.nlm.nih.gov/gene/?term=LOC105279664 | 4023 | 60 | 149.14 | 1 | 2.49 | 1.67 | 17.78 | 18.81 | -1.03 | 1.03 | -0.02 | 0.02 |
| gene-LOC105280348 | https://www.ncbi.nlm.nih.gov/gene/?term=LOC105280348 | 3797 | 71 | 186.99 | 1 | 2.63 | 1.41 | 18.59 | 19.68 | -1.09 | 1.09 | -0.02 | 0.02 |
| gene-LOC105287854 | https://www.ncbi.nlm.nih.gov/gene/?term=LOC105287854 | 3644 | 64 | 175.63 | 1 | 2.74 | 1.56 | 14.92 | 16.11 | -1.19 | 1.19 | -0.02 | 0.02 |
| gene-LOC105276692 | https://www.ncbi.nlm.nih.gov/gene/?term=LOC105276692 | 1745 | 33 | 189.11 | 1 | 5.73 | 3.03 | 21.96 | 23.19 | -1.23 | 1.23 | -0.02 | 0.02 |
| gene-LOC105288078 | https://www.ncbi.nlm.nih.gov/gene/?term=LOC105288078 | 19126 | 298 | 155.81 | 1 | 0.52 | 0.34 | 16.07 | 17.31 | -1.24 | 1.24 | -0.02 | 0.02 |
| gene-LOC105280641 | https://www.ncbi.nlm.nih.gov/gene/?term=LOC105280641 | 3682 | 55 | 149.38 | 1 | 2.72 | 1.82 | 18.09 | 19.48 | -1.39 | 1.39 | -0.02 | 0.02 |
| gene-LOC105287648 | https://www.ncbi.nlm.nih.gov/gene/?term=LOC105287648 | 2205 | 35 | 158.73 | 1 | 4.54 | 2.86 | 35.94 | 37.4 | -1.45 | 1.45 | -0.02 | 0.02 |
| gene-LOC105282600 | https://www.ncbi.nlm.nih.gov/gene/?term=LOC105282600 | 3715 | 52 | 139.97 | 1 | 2.69 | 1.92 | 31.36 | 33.08 | -1.72 | 1.72 | -0.02 | 0.02 |
| gene-LOC105283943 | https://www.ncbi.nlm.nih.gov/gene/?term=LOC105283943 | 3559 | 50 | 140.49 | 1 | 2.81 | 2 | 16.35 | 18.09 | -1.74 | 1.74 | -0.02 | 0.02 |
| gene-LOC105284329 | https://www.ncbi.nlm.nih.gov/gene/?term=LOC105284329 | 4051 | 50 | 123.43 | 1 | 2.47 | 2 | 17.13 | 19.12 | -1.99 | 1.99 | -0.02 | 0.02 |
| gene-LOC105281515 | https://www.ncbi.nlm.nih.gov/gene/?term=LOC105281515 | 8827 | 60 | 67.97 2 | 2.27 | 3.33 | 26.32 | 29.67 | -3.35 | 3.35 | -0.02 | 0.02 | |
| gene-LOC105283712 | https://www.ncbi.nlm.nih.gov/gene/?term=LOC105283712 | 1910 | 36 | 188.48 | 1 | 5.24 | 2.78 | 26.26 | 25.56 | 0.7 | 0.7 | 0.01 | 0.01 |
| gene-LOC105288042 | https://www.ncbi.nlm.nih.gov/gene/?term=LOC105288042 | 3507 | 98 | 279.44 | 1 | 2.85 | 1.02 | 22.45 | 21.99 | 0.46 | 0.46 | 0.01 | 0.01 |
| gene-LOC105282646 | https://www.ncbi.nlm.nih.gov/gene/?term=LOC105282646 | 3317 | 99 | 298.46 | 1 | 3.01 | 1.01 | 21.06 | 20.67 | 0.39 | 0.39 | 0.01 | 0.01 |
| gene-LOC105274590 | https://www.ncbi.nlm.nih.gov/gene/?term=LOC105274590 | 4516 | 74 | 163.86 | 1 | 2.21 | 1.35 | 30.44 | 30.06 | 0.38 | 0.38 | 0.01 | 0.01 |
| gene-LOC105276527 | https://www.ncbi.nlm.nih.gov/gene/?term=LOC105276527 | 7923 | 317 | 400.1 1 | 1.26 | 0.32 | 9.8 | 9.43 | 0.37 | 0.37 | 0.01 | 0.01 | |
| gene-LOC105282951 | https://www.ncbi.nlm.nih.gov/gene/?term=LOC105282951 | 3366 | 120 | 356.51 | 1 | 2.97 | 0.83 | 19.9 | 19.57 | 0.33 | 0.33 | 0.01 | 0.01 |
| gene-LOC105276196 | https://www.ncbi.nlm.nih.gov/gene/?term=LOC105276196 | 2953 | 56 | 189.64 | 1 | 3.39 | 1.79 | 39.44 | 39.14 | 0.3 | 0.3 | 0.01 | 0.01 |
| gene-LOC105278534 | https://www.ncbi.nlm.nih.gov/gene/?term=LOC105278534 | 7364 | 185 | 251.22 | 2 | 2.72 | 1.08 | 18.38 | 18.12 | 0.25 | 0.25 | 0.01 | 0.01 |
| gene-LOC105274534 | https://www.ncbi.nlm.nih.gov/gene/?term=LOC105274534 | 1329 | 59 | 443.94 | 1 | 7.52 | 1.69 | 42.17 | 42.03 | 0.14 | 0.14 | 0.01 | 0.01 |
| gene-LOC105278632 | https://www.ncbi.nlm.nih.gov/gene/?term=LOC105278632 | 3624 | 133 | 367 1 | 2.76 | 0.75 | 14.21 | 14.06 | 0.14 | 0.14 | 0.01 | 0.01 | |
| gene-LOC105278691 | https://www.ncbi.nlm.nih.gov/gene/?term=LOC105278691 | 3871 | 151 | 390.08 | 1 | 2.58 | 0.66 | 14.98 | 14.84 | 0.14 | 0.14 | 0.01 | 0.01 |
| gene-LOC105275683 | https://www.ncbi.nlm.nih.gov/gene/?term=LOC105275683 | 4256 | 187 | 439.38 | 1 | 2.35 | 0.53 | 13.19 | 13.34 | -0.15 | 0.15 | -0.01 | 0.01 |
| gene-LOC105278666 | https://www.ncbi.nlm.nih.gov/gene/?term=LOC105278666 | 10341 | 471 | 455.47 | 1 | 0.97 | 0.21 | 6.62 | 6.83 | -0.21 | 0.21 | -0.01 | 0.01 |
| gene-LOC105275179 | https://www.ncbi.nlm.nih.gov/gene/?term=LOC105275179 | 3271 | 77 | 235.4 1 | 3.06 | 1.3 | 17.93 | 18.15 | -0.22 | 0.22 | -0.01 | 0.01 | |
| gene-LOC105284355 | https://www.ncbi.nlm.nih.gov/gene/?term=LOC105284355 | 3904 | 162 | 414.96 | 1 | 2.56 | 0.62 | 19.15 | 19.36 | -0.22 | 0.22 | -0.01 | 0.01 |
| gene-LOC105275363 | https://www.ncbi.nlm.nih.gov/gene/?term=LOC105275363 | 3798 | 161 | 423.91 | 1 | 2.63 | 0.62 | 16.94 | 17.21 | -0.26 | 0.26 | -0.01 | 0.01 |
| gene-LOC105279038 | https://www.ncbi.nlm.nih.gov/gene/?term=LOC105279038 | 9453 | 340 | 359.67 | 1 | 1.06 | 0.29 | 8.66 | 8.92 | -0.26 | 0.26 | -0.01 | 0.01 |
| gene-LOC105275374 | https://www.ncbi.nlm.nih.gov/gene/?term=LOC105275374 | 11376 | 488 | 428.97 | 1 | 0.88 | 0.2 | 10.37 | 10.64 | -0.27 | 0.27 | -0.01 | 0.01 |
| gene-LOC105275327 | https://www.ncbi.nlm.nih.gov/gene/?term=LOC105275327 | 14245 | 511 | 358.72 | 1 | 0.7 | 0.2 | 6.72 | 7.02 | -0.3 | 0.3 | -0.01 | 0.01 |
| gene-LOC105278497 | https://www.ncbi.nlm.nih.gov/gene/?term=LOC105278497 | 2072 | 96 | 463.32 | 1 | 4.83 | 1.04 | 18.89 | 19.19 | -0.31 | 0.31 | -0.01 | 0.01 |
| gene-LOC105275170 | https://www.ncbi.nlm.nih.gov/gene/?term=LOC105275170 | 12918 | 279 | 215.98 | 1 | 0.77 | 0.36 | 2.01 | 2.33 | -0.32 | 0.32 | -0.01 | 0.01 |
| gene-LOC105278282 | https://www.ncbi.nlm.nih.gov/gene/?term=LOC105278282 | 1775 | 51 | 287.32 | 1 | 5.63 | 1.96 | 40.43 | 40.76 | -0.33 | 0.33 | -0.01 | 0.01 |
| gene-LOC105282794 | https://www.ncbi.nlm.nih.gov/gene/?term=LOC105282794 | 4571 | 93 | 203.46 | 1 | 2.19 | 1.08 | 16.99 | 17.32 | -0.33 | 0.33 | -0.01 | 0.01 |
| gene-LOC105274669 | https://www.ncbi.nlm.nih.gov/gene/?term=LOC105274669 | 8555 | 261 | 305.08 | 1 | 1.17 | 0.38 | 5.68 | 6.02 | -0.35 | 0.35 | -0.01 | 0.01 |
| gene-LOC105277761 | https://www.ncbi.nlm.nih.gov/gene/?term=LOC105277761 | 4648 | 144 | 309.81 | 1 | 2.15 | 0.69 | 14.39 | 14.8 | -0.41 | 0.41 | -0.01 | 0.01 |
| gene-LOC105274823 | https://www.ncbi.nlm.nih.gov/gene/?term=LOC105274823 | 4620 | 145 | 313.85 | 1 | 2.16 | 0.69 | 19.86 | 20.29 | -0.43 | 0.43 | -0.01 | 0.01 |
| gene-LOC105285639 | https://www.ncbi.nlm.nih.gov/gene/?term=LOC105285639 | 10325 | 344 | 333.17 | 1 | 0.97 | 0.29 | 11.23 | 11.66 | -0.43 | 0.43 | -0.01 | 0.01 |
| gene-LOC105278649 | https://www.ncbi.nlm.nih.gov/gene/?term=LOC105278649 | 5616 | 137 | 243.95 | 1 | 1.78 | 0.73 | 13.17 | 13.73 | -0.56 | 0.56 | -0.01 | 0.01 |
| gene-LOC105277856 | https://www.ncbi.nlm.nih.gov/gene/?term=LOC105277856 | 3442 | 31 | 90.06 1 | 2.91 | 3.23 | 38.58 | 39.29 | -0.71 | 0.71 | -0.01 | 0.01 | |
| gene-LOC105276059 | https://www.ncbi.nlm.nih.gov/gene/?term=LOC105276059 | 6256 | 129 | 206.2 1 | 1.6 | 0.78 | 9.24 | 9.96 | -0.72 | 0.72 | -0.01 | 0.01 | |
| gene-LOC105275187 | https://www.ncbi.nlm.nih.gov/gene/?term=LOC105275187 | 5534 | 69 | 124.68 | 1 | 1.81 | 1.45 | 13.92 | 14.78 | -0.86 | 0.86 | -0.01 | 0.01 |
| gene-LOC105283989 | https://www.ncbi.nlm.nih.gov/gene/?term=LOC105283989 | 4948 | 27 | 54.57 1 | 2.02 | 3.7 | 44.98 | 46.16 | -1.18 | 1.18 | -0.01 | 0.01 | |
| gene-LOC105275347 | https://www.ncbi.nlm.nih.gov/gene/?term=LOC105275347 | 2526 | 16 | 63.34 1 | 3.96 | 6.25 | 40.9 | 42.25 | -1.34 | 1.34 | -0.01 | 0.01 | |
| gene-LOC105280002 | https://www.ncbi.nlm.nih.gov/gene/?term=LOC105280002 | 1940 | 26 | 134.02 | 1 | 5.15 | 3.85 | 34.83 | 34.64 | 0.19 | 0.19 | 0 | 0 |
| gene-LOC105282811 | https://www.ncbi.nlm.nih.gov/gene/?term=LOC105282811 | 5765 | 199 | 345.19 | 1 | 1.73 | 0.5 | 8.45 | 8.31 | 0.14 | 0.14 | 0 | 0 |
| gene-LOC105285807 | https://www.ncbi.nlm.nih.gov/gene/?term=LOC105285807 | 6063 | 217 | 357.91 | 1 | 1.65 | 0.46 | 8.91 | 8.79 | 0.12 | 0.12 | 0 | 0 |
| gene-LOC105281700 | https://www.ncbi.nlm.nih.gov/gene/?term=LOC105281700 | 2208 | 37 | 167.57 | 1 | 4.53 | 2.7 | 39.75 | 39.65 | 0.11 | 0.11 | 0 | 0 |
| gene-LOC105283728 | https://www.ncbi.nlm.nih.gov/gene/?term=LOC105283728 | 3256 | 122 | 374.69 | 2 | 6.14 | 1.64 | 18.7 | 18.6 | 0.1 | 0.1 | 0 | 0 |
| gene-LOC105276538 | https://www.ncbi.nlm.nih.gov/gene/?term=LOC105276538 | 1511 | 18 | 119.13 | 1 | 6.62 | 5.56 | 68.54 | 68.54 | 0 | 0 | 0 | 0 |
| gene-LOC105283123 | https://www.ncbi.nlm.nih.gov/gene/?term=LOC105283123 | 1507 | 47 | 311.88 | 1 | 6.64 | 2.13 | 17.1 | 17.12 | -0.02 | 0.02 | 0 | 0 |
| gene-LOC105286514 | https://www.ncbi.nlm.nih.gov/gene/?term=LOC105286514 | 2528 | 44 | 174.05 | 1 | 3.96 | 2.27 | 37.14 | 37.18 | -0.04 | 0.04 | 0 | 0 |
| gene-LOC105285179 | https://www.ncbi.nlm.nih.gov/gene/?term=LOC105285179 | 4191 | 176 | 419.95 | 1 | 2.39 | 0.57 | 19.46 | 19.52 | -0.06 | 0.06 | 0 | 0 |
| gene-LOC105285406 | https://www.ncbi.nlm.nih.gov/gene/?term=LOC105285406 | 8015 | 60 | 74.86 1 | 1.25 | 1.67 | 28.89 | 28.94 | -0.06 | 0.06 | 0 | 0 | |
| gene-LOC105284043 | https://www.ncbi.nlm.nih.gov/gene/?term=LOC105284043 | 5661 | 295 | 521.11 | 1 | 1.77 | 0.34 | 9.91 | 9.97 | -0.07 | 0.07 | 0 | 0 |
| gene-LOC105287417 | https://www.ncbi.nlm.nih.gov/gene/?term=LOC105287417 | 1025 | 32 | 312.2 1 | 9.76 | 3.12 | 37.56 | 37.69 | -0.12 | 0.12 | 0 | 0 | |
| gene-LOC113562050 | https://www.ncbi.nlm.nih.gov/gene/?term=LOC113562050 | 17097 | 587 | 343.34 | 1 | 0.58 | 0.17 | 4.91 | 5.05 | -0.13 | 0.13 | 0 | 0 |
| gene-LOC105278228 | https://www.ncbi.nlm.nih.gov/gene/?term=LOC105278228 | 8610 | 154 | 178.86 | 1 | 1.16 | 0.65 | 23.95 | 24.14 | -0.2 | 0.2 | 0 | 0 |
| gene-LOC105274851 | https://www.ncbi.nlm.nih.gov/gene/?term=LOC105274851 | 6616 | 130 | 196.49 | 1 | 1.51 | 0.77 | 11.46 | 11.67 | -0.21 | 0.21 | 0 | 0 |

| | | | | | | | | | | | | | |
|---|---|---|---|---|---|---|---|---|---|---|---|---|---|
| gene-LOC105286452 | https://www.ncbi.nlm.nih.gov/gene/?term=LOC105286452 | 6888 | 61 | 88.56 | 1 | 1.45 | 1.64 | 30.12 | 30.34 | -0.22 | 0.22 | 0 | 0 |
| gene-LOC105288010 | https://www.ncbi.nlm.nih.gov/gene/?term=LOC105288010 | 3189 | 39 | 122.3 | 1 | 3.14 | 2.56 | 33.4 | 33.65 | -0.26 | 0.26 | 0 | 0 |
| gene-LOC105282067 | https://www.ncbi.nlm.nih.gov/gene/?term=LOC105282067 | 5941 | 41 | 69.01 | 1 | 1.68 | 2.44 | 26.24 | 26.89 | -0.64 | 0.64 | 0 | 0 |

Directory: OGL File: rnk-dmg-Ob_LI_BCphaseB2.vs.RphaseA2.txt

| gene_ID | gene_link | gwidth | #Sites | #per10Kb | #dmSites | #dmsp10kb | %dmSites | %pSite1 | %pSite2 | DMpSite | ADMpSite | DMpNucl | ADMpNucl |
|---|---|---|---|---|---|---|---|---|---|---|---|---|---|
| gene-LOC113562345 | https://www.ncbi.nlm.nih.gov/gene/?term=LOC113562345 | 2290 | 69 | 301.31 | 19 | 82.97 | 27.54 | 16.9 | 33.89 | -17 | 17 | -0.51 | 0.51 |
| gene-LOC105284677 | https://www.ncbi.nlm.nih.gov/gene/?term=LOC105284677 | 1905 | 43 | 225.72 | 2 | 10.5 | 4.65 | 38.99 | 29.38 | 9.62 | 9.62 | 0.22 | 0.22 |
| gene-LOC113562901 | https://www.ncbi.nlm.nih.gov/gene/?term=LOC113562901 | 1531 | 22 | 143.7 | 3 | 19.6 | 13.64 | 45.03 | 59.16 | -14.13 | 14.13 | -0.2 | 0.2 |
| gene-LOC105280327 | https://www.ncbi.nlm.nih.gov/gene/?term=LOC105280327 | 2670 | 104 | 389.51 | 1 | 3.75 | 0.96 | 27.09 | 30.78 | -3.69 | 3.69 | -0.14 | 0.14 |
| gene-LOC105282220 | https://www.ncbi.nlm.nih.gov/gene/?term=LOC105282220 | 2298 | 224 | 974.76 | 1 | 4.35 | 0.45 | 7.52 | 8.84 | -1.32 | 1.32 | -0.13 | 0.13 |
| gene-LOC105282156 | https://www.ncbi.nlm.nih.gov/gene/?term=LOC105282156 | 2577 | 145 | 562.67 | 1 | 3.88 | 0.69 | 23.3 | 25.57 | -2.27 | 2.27 | -0.13 | 0.13 |
| gene-LOC105280786 | https://www.ncbi.nlm.nih.gov/gene/?term=LOC105280786 | 2573 | 119 | 462.5 | 1 | 3.89 | 0.84 | 29.73 | 32.5 | -2.77 | 2.77 | -0.13 | 0.13 |
| gene-LOC105276449 | https://www.ncbi.nlm.nih.gov/gene/?term=LOC105276449 | 1231 | 56 | 454.91 | 1 | 8.12 | 1.79 | 30.06 | 32.73 | -2.67 | 2.67 | -0.12 | 0.12 |
| gene-LOC105285674 | https://www.ncbi.nlm.nih.gov/gene/?term=LOC105285674 | 3035 | 97 | 319.6 | 2 | 6.59 | 2.06 | 24.1 | 27.79 | -3.69 | 3.69 | -0.12 | 0.12 |
| gene-LOC105287417 | https://www.ncbi.nlm.nih.gov/gene/?term=LOC105287417 | 1025 | 36 | 351.22 | 1 | 9.76 | 2.78 | 34.17 | 37.2 | -3.03 | 3.03 | -0.11 | 0.11 |
| gene-LOC105279900 | https://www.ncbi.nlm.nih.gov/gene/?term=LOC105279900 | 2191 | 62 | 282.98 | 1 | 4.56 | 1.61 | 30.24 | 34.06 | -3.83 | 3.83 | -0.11 | 0.11 |
| gene-LOC105277631 | https://www.ncbi.nlm.nih.gov/gene/?term=LOC105277631 | 4387 | 333 | 759.06 | 1 | 2.28 | 0.3 | 9.46 | 10.66 | -1.19 | 1.19 | -0.09 | 0.09 |
| gene-LOC105282149 | https://www.ncbi.nlm.nih.gov/gene/?term=LOC105282149 | 1787 | 109 | 609.96 | 1 | 5.6 | 0.92 | 41.83 | 43.33 | -1.5 | 1.5 | -0.09 | 0.09 |
| gene-LOC105283649 | https://www.ncbi.nlm.nih.gov/gene/?term=LOC105283649 | 1918 | 57 | 297.18 | 2 | 10.43 | 3.51 | 35.86 | 38.92 | -3.06 | 3.06 | -0.09 | 0.09 |
| gene-LOC113561805 | https://www.ncbi.nlm.nih.gov/gene/?term=LOC113561805 | 3337 | 84 | 251.72 | 1 | 3 | 1.19 | 34.18 | 37.85 | -3.66 | 3.66 | -0.09 | 0.09 |
| gene-LOC105274449 | https://www.ncbi.nlm.nih.gov/gene/?term=LOC105274449 | 1549 | 53 | 342.16 | 1 | 6.46 | 1.89 | 51.77 | 49.51 | 2.26 | 2.26 | 0.08 | 0.08 |
| gene-LOC105286799 | https://www.ncbi.nlm.nih.gov/gene/?term=LOC105286799 | 3557 | 188 | 528.54 | 1 | 2.81 | 0.53 | 15.27 | 16.76 | -1.49 | 1.49 | -0.08 | 0.08 |
| gene-LOC105284557 | https://www.ncbi.nlm.nih.gov/gene/?term=LOC105284557 | 1672 | 84 | 502.39 | 1 | 5.98 | 1.19 | 28.16 | 29.69 | -1.53 | 1.53 | -0.08 | 0.08 |
| gene-LOC105275070 | https://www.ncbi.nlm.nih.gov/gene/?term=LOC105275070 | 5667 | 53 | 93.52 | 6 | 10.59 | 11.32 | 1.77 | 10.27 | -8.5 | 8.5 | -0.08 | 0.08 |
| gene-LOC105279298 | https://www.ncbi.nlm.nih.gov/gene/?term=LOC105279298 | 2773 | 138 | 497.66 | 2 | 7.21 | 1.45 | 22.5 | 23.97 | -1.47 | 1.47 | -0.07 | 0.07 |
| gene-LOC105276704 | https://www.ncbi.nlm.nih.gov/gene/?term=LOC105276704 | 2063 | 101 | 489.58 | 2 | 9.69 | 1.98 | 13.1 | 14.6 | -1.5 | 1.5 | -0.07 | 0.07 |
| gene-LOC105286683 | https://www.ncbi.nlm.nih.gov/gene/?term=LOC105286683 | 1665 | 53 | 318.32 | 1 | 6.01 | 1.89 | 31.4 | 33.71 | -2.31 | 2.31 | -0.07 | 0.07 |
| gene-LOC105278454 | https://www.ncbi.nlm.nih.gov/gene/?term=LOC105278454 | 1972 | 51 | 258.62 | 1 | 5.07 | 1.96 | 39.78 | 42.63 | -2.85 | 2.85 | -0.07 | 0.07 |
| gene-LOC105286514 | https://www.ncbi.nlm.nih.gov/gene/?term=LOC105286514 | 2528 | 47 | 185.92 | 1 | 3.96 | 2.13 | 35.58 | 39.1 | -3.52 | 3.52 | -0.07 | 0.07 |
| gene-LOC105286644 | https://www.ncbi.nlm.nih.gov/gene/?term=LOC105286644 | 1235 | 20 | 161.94 | 1 | 8.1 | 5 | 37.53 | 41.72 | -4.19 | 4.19 | -0.07 | 0.07 |
| gene-LOC105275743 | https://www.ncbi.nlm.nih.gov/gene/?term=LOC105275743 | 1782 | 76 | 426.49 | 1 | 5.61 | 1.32 | 23.56 | 22.24 | 1.32 | 1.32 | 0.06 | 0.06 |
| gene-LOC105285055 | https://www.ncbi.nlm.nih.gov/gene/?term=LOC105285055 | 4108 | 274 | 666.99 | 2 | 4.87 | 0.73 | 10.9 | 11.79 | -0.89 | 0.89 | -0.06 | 0.06 |
| gene-LOC105274856 | https://www.ncbi.nlm.nih.gov/gene/?term=LOC105274856 | 5216 | 232 | 444.79 | 1 | 1.92 | 0.43 | 17.14 | 18.43 | -1.29 | 1.29 | -0.06 | 0.06 |
| gene-LOC105274534 | https://www.ncbi.nlm.nih.gov/gene/?term=LOC105274534 | 1329 | 61 | 458.99 | 1 | 7.52 | 1.64 | 40.79 | 42.14 | -1.35 | 1.35 | -0.06 | 0.06 |
| gene-LOC105278282 | https://www.ncbi.nlm.nih.gov/gene/?term=LOC105278282 | 1775 | 64 | 360.56 | 1 | 5.63 | 1.56 | 33.91 | 35.49 | -1.58 | 1.58 | -0.06 | 0.06 |
| gene-LOC105282151 | https://www.ncbi.nlm.nih.gov/gene/?term=LOC105282151 | 1601 | 46 | 287.32 | 2 | 12.49 | 4.35 | 33.88 | 35.82 | -1.94 | 1.94 | -0.06 | 0.06 |
| gene-LOC105287063 | https://www.ncbi.nlm.nih.gov/gene/?term=LOC105287063 | 1874 | 54 | 288.15 | 1 | 5.34 | 1.85 | 27.46 | 29.52 | -2.06 | 2.06 | -0.06 | 0.06 |
| gene-LOC105285583 | https://www.ncbi.nlm.nih.gov/gene/?term=LOC105285583 | 1947 | 47 | 241.4 | 1 | 5.14 | 2.13 | 27.05 | 29.68 | -2.63 | 2.63 | -0.06 | 0.06 |
| gene-LOC105276538 | https://www.ncbi.nlm.nih.gov/gene/?term=LOC105276538 | 1511 | 19 | 125.74 | 1 | 6.62 | 5.26 | 69.47 | 74.28 | -4.82 | 4.82 | -0.06 | 0.06 |
| gene-LOC105279898 | https://www.ncbi.nlm.nih.gov/gene/?term=LOC105279898 | 1343 | 14 | 104.24 | 1 | 7.45 | 7.14 | 54.89 | 60.18 | -5.29 | 5.29 | -0.06 | 0.06 |
| gene-LOC105275708 | https://www.ncbi.nlm.nih.gov/gene/?term=LOC105275708 | 1772 | 18 | 101.58 | 1 | 5.64 | 5.56 | 43.28 | 48.95 | -5.67 | 5.67 | -0.06 | 0.06 |
| gene-LOC105282885 | https://www.ncbi.nlm.nih.gov/gene/?term=LOC105282885 | 1870 | 49 | 262.03 | 1 | 5.35 | 2.04 | 39.47 | 37.7 | 1.77 | 1.77 | 0.05 | 0.05 |
| gene-LOC105282682 | https://www.ncbi.nlm.nih.gov/gene/?term=LOC105282682 | 4127 | 241 | 583.96 | 1 | 2.42 | 0.41 | 12.88 | 13.79 | -0.91 | 0.91 | -0.05 | 0.05 |
| gene-LOC105275683 | https://www.ncbi.nlm.nih.gov/gene/?term=LOC105275683 | 4256 | 210 | 493.42 | 1 | 2.35 | 0.48 | 13.47 | 14.47 | -1.01 | 1.01 | -0.05 | 0.05 |
| gene-LOC105278497 | https://www.ncbi.nlm.nih.gov/gene/?term=LOC105278497 | 2072 | 103 | 497.1 | 1 | 4.83 | 0.97 | 19.13 | 20.15 | -1.02 | 1.02 | -0.05 | 0.05 |
| gene-LOC105284355 | https://www.ncbi.nlm.nih.gov/gene/?term=LOC105284355 | 3904 | 165 | 422.64 | 1 | 2.56 | 0.61 | 19.1 | 20.26 | -1.16 | 1.16 | -0.05 | 0.05 |
| gene-LOC109611283 | https://www.ncbi.nlm.nih.gov/gene/?term=LOC109611283 | 1451 | 39 | 268.78 | 1 | 6.89 | 2.56 | 35.83 | 37.66 | -1.83 | 1.83 | -0.05 | 0.05 |
| gene-LOC105275966 | https://www.ncbi.nlm.nih.gov/gene/?term=LOC105275966 | 3299 | 73 | 221.28 | 1 | 3.03 | 1.37 | 43.49 | 45.58 | -2.08 | 2.08 | -0.05 | 0.05 |
| gene-LOC105288063 | https://www.ncbi.nlm.nih.gov/gene/?term=LOC105288063 | 3400 | 57 | 167.65 | 1 | 2.94 | 1.75 | 12.44 | 15.19 | -2.75 | 2.75 | -0.05 | 0.05 |
| gene-LOC105276680 | https://www.ncbi.nlm.nih.gov/gene/?term=LOC105276680 | 1977 | 35 | 177.04 | 1 | 5.06 | 2.86 | 58.22 | 61.02 | -2.8 | 2.8 | -0.05 | 0.05 |
| gene-LOC105288066 | https://www.ncbi.nlm.nih.gov/gene/?term=LOC105288066 | 1964 | 32 | 162.93 | 1 | 5.09 | 3.12 | 32.93 | 36 | -3.07 | 3.07 | -0.05 | 0.05 |
| gene-LOC105276078 | https://www.ncbi.nlm.nih.gov/gene/?term=LOC105276078 | 1051 | 17 | 161.75 | 1 | 9.51 | 5.88 | 25.17 | 28.55 | -3.38 | 3.38 | -0.05 | 0.05 |
| gene-LOC105281039 | https://www.ncbi.nlm.nih.gov/gene/?term=LOC105281039 | 1471 | 12 | 81.58 | 1 | 6.8 | 8.33 | 31.46 | 37.55 | -6.08 | 6.08 | -0.05 | 0.05 |
| gene-LOC105284396 | https://www.ncbi.nlm.nih.gov/gene/?term=LOC105284396 | 1976 | 70 | 354.25 | 1 | 5.06 | 1.43 | 32.98 | 31.96 | 1.02 | 1.02 | 0.04 | 0.04 |
| gene-LOC105282491 | https://www.ncbi.nlm.nih.gov/gene/?term=LOC105282491 | 4761 | 184 | 386.47 | 1 | 2.1 | 0.54 | 8.25 | 9.16 | -0.91 | 0.91 | -0.04 | 0.04 |
| gene-LOC105278877 | https://www.ncbi.nlm.nih.gov/gene/?term=LOC105278877 | 1968 | 86 | 436.99 | 1 | 5.08 | 1.16 | 31.6 | 32.52 | -0.92 | 0.92 | -0.04 | 0.04 |
| gene-LOC105280774 | https://www.ncbi.nlm.nih.gov/gene/?term=LOC105280774 | 2793 | 122 | 436.81 | 1 | 3.58 | 0.82 | 26.14 | 27.14 | -0.99 | 0.99 | -0.04 | 0.04 |
| gene-LOC105285639 | https://www.ncbi.nlm.nih.gov/gene/?term=LOC105285639 | 10325 | 408 | 395.16 | 1 | 0.97 | 0.25 | 10.25 | 11.26 | -1.02 | 1.02 | -0.04 | 0.04 |
| gene-LOC105277761 | https://www.ncbi.nlm.nih.gov/gene/?term=LOC105277761 | 4648 | 160 | 344.23 | 1 | 2.15 | 0.62 | 14.78 | 16.05 | -1.27 | 1.27 | -0.04 | 0.04 |
| gene-LOC105280956 | https://www.ncbi.nlm.nih.gov/gene/?term=LOC105280956 | 2639 | 77 | 291.78 | 1 | 3.79 | 1.3 | 16.02 | 17.39 | -1.37 | 1.37 | -0.04 | 0.04 |
| gene-LOC105282970 | https://www.ncbi.nlm.nih.gov/gene/?term=LOC105282970 | 1529 | 40 | 261.61 | 1 | 6.54 | 2.5 | 50.96 | 52.56 | -1.6 | 1.6 | -0.04 | 0.04 |
| gene-LOC105278384 | https://www.ncbi.nlm.nih.gov/gene/?term=LOC105278384 | 1390 | 35 | 251.8 | 1 | 7.19 | 2.86 | 45.32 | 46.94 | -1.62 | 1.62 | -0.04 | 0.04 |
| gene-LOC105283592 | https://www.ncbi.nlm.nih.gov/gene/?term=LOC105283592 | 3479 | 72 | 206.96 | 1 | 2.87 | 1.39 | 25.41 | 27.37 | -1.95 | 1.95 | -0.04 | 0.04 |
| gene-LOC105281420 | https://www.ncbi.nlm.nih.gov/gene/?term=LOC105281420 | 3882 | 67 | 172.59 | 1 | 2.58 | 1.49 | 12.82 | 15.24 | -2.42 | 2.42 | -0.04 | 0.04 |
| gene-LOC105274900 | https://www.ncbi.nlm.nih.gov/gene/?term=LOC105274900 | 5561 | 55 | 98.9 | 2 | 3.6 | 3.64 | 30.44 | 34.13 | -3.69 | 3.69 | -0.04 | 0.04 |

| | | | | | | | | | | | | | | |
|---|---|---|---|---|---|---|---|---|---|---|---|---|---|---|
| gene-LOC105281282 | https://www.ncbi.nlm.nih.gov/gene/?term=LOC105281282 | 3250 | 3 | 9.23 | 1 | 3.08 | 33.33 | 23.25 | 65.16 | -41.92 | | 41.92 | -0.04 | 0.04 |
| gene-LOC105276452 | https://www.ncbi.nlm.nih.gov/gene/?term=LOC105276452 | 2917 | 262 | 898.18 | | 1 | 3.43 | 0.38 | 9.36 | 9.65 | -0.29 | 0.29 | -0.03 | 0.03 |
| gene-LOC105278283 | https://www.ncbi.nlm.nih.gov/gene/?term=LOC105278283 | 2317 | 146 | 630.13 | | 1 | 4.32 | 0.68 | 24.36 | 24.8 | -0.45 | 0.45 | -0.03 | 0.03 |
| gene-LOC105275460 | https://www.ncbi.nlm.nih.gov/gene/?term=LOC105275460 | 2473 | 122 | 493.33 | | 1 | 4.04 | 0.82 | 12.96 | 13.51 | -0.55 | 0.55 | -0.03 | 0.03 |
| gene-LOC105285807 | https://www.ncbi.nlm.nih.gov/gene/?term=LOC105285807 | 6063 | 237 | 390.9 | 1 | 1.65 | 0.42 | 8.52 | 9.16 | -0.64 | 0.64 | -0.03 | 0.03 | |
| gene-LOC105274452 | https://www.ncbi.nlm.nih.gov/gene/?term=LOC105274452 | 6595 | 279 | 423.05 | | 1 | 1.52 | 0.36 | 12.34 | 12.98 | -0.65 | 0.65 | -0.03 | 0.03 |
| gene-LOC105280723 | https://www.ncbi.nlm.nih.gov/gene/?term=LOC105280723 | 3572 | 185 | 517.92 | | 1 | 2.8 | 0.54 | 15.07 | 15.74 | -0.67 | 0.67 | -0.03 | 0.03 |
| gene-LOC105275752 | https://www.ncbi.nlm.nih.gov/gene/?term=LOC105275752 | 3462 | 141 | 407.28 | | 1 | 2.89 | 0.71 | 30.44 | 31.2 | -0.75 | 0.75 | -0.03 | 0.03 |
| gene-LOC105279038 | https://www.ncbi.nlm.nih.gov/gene/?term=LOC105279038 | 9453 | 369 | 390.35 | | 1 | 1.06 | 0.27 | 8.12 | 8.94 | -0.82 | 0.82 | -0.03 | 0.03 |
| gene-LOC105278632 | https://www.ncbi.nlm.nih.gov/gene/?term=LOC105278632 | 3624 | 141 | 389.07 | | 1 | 2.76 | 0.71 | 13.56 | 14.44 | -0.87 | 0.87 | -0.03 | 0.03 |
| gene-LOC105285202 | https://www.ncbi.nlm.nih.gov/gene/?term=LOC105285202 | 1838 | 66 | 359.09 | | 1 | 5.44 | 1.52 | 24.58 | 25.52 | -0.94 | 0.94 | -0.03 | 0.03 |
| gene-LOC105281261 | https://www.ncbi.nlm.nih.gov/gene/?term=LOC105281261 | 5947 | 158 | 265.68 | | 1 | 1.68 | 0.63 | 14.18 | 15.18 | -1 | 1 | -0.03 | 0.03 |
| gene-LOC105281848 | https://www.ncbi.nlm.nih.gov/gene/?term=LOC105281848 | 2122 | 64 | 301.6 | 1 | 4.71 | 1.56 | 21.13 | 22.15 | -1.02 | 1.02 | -0.03 | 0.03 | |
| gene-LOC105284116 | https://www.ncbi.nlm.nih.gov/gene/?term=LOC105284116 | 3970 | 113 | 284.63 | | 1 | 2.52 | 0.88 | 7.21 | 8.25 | -1.03 | 1.03 | -0.03 | 0.03 |
| gene-LOC105282991 | https://www.ncbi.nlm.nih.gov/gene/?term=LOC105282991 | 5617 | 112 | 199.39 | | 1 | 1.78 | 0.89 | 19.26 | 20.58 | -1.32 | 1.32 | -0.03 | 0.03 |
| gene-LOC105278962 | https://www.ncbi.nlm.nih.gov/gene/?term=LOC105278962 | 4491 | 107 | 238.25 | | 1 | 2.23 | 0.93 | 20.7 | 22.05 | -1.35 | 1.35 | -0.03 | 0.03 |
| gene-LOC105279664 | https://www.ncbi.nlm.nih.gov/gene/?term=LOC105279664 | 4023 | 74 | 183.94 | | 1 | 2.49 | 1.35 | 16.49 | 17.94 | -1.45 | 1.45 | -0.03 | 0.03 |
| gene-LOC105278109 | https://www.ncbi.nlm.nih.gov/gene/?term=LOC105278109 | 4029 | 72 | 178.7 | 1 | 2.48 | 1.39 | 42.29 | 43.78 | -1.48 | 1.48 | -0.03 | 0.03 | |
| gene-LOC105282893 | https://www.ncbi.nlm.nih.gov/gene/?term=LOC105282893 | 2645 | 47 | 177.69 | | 1 | 3.78 | 2.13 | 28.96 | 30.53 | -1.56 | 1.56 | -0.03 | 0.03 |
| gene-LOC105287854 | https://www.ncbi.nlm.nih.gov/gene/?term=LOC105287854 | 3644 | 66 | 181.12 | | 1 | 2.74 | 1.52 | 15.87 | 17.63 | -1.76 | 1.76 | -0.03 | 0.03 |
| gene-LOC105282600 | https://www.ncbi.nlm.nih.gov/gene/?term=LOC105282600 | 3715 | 56 | 150.74 | | 1 | 2.69 | 1.79 | 29.97 | 31.8 | -1.83 | 1.83 | -0.03 | 0.03 |
| gene-LOC105288010 | https://www.ncbi.nlm.nih.gov/gene/?term=LOC105288010 | 3189 | 45 | 141.11 | | 1 | 3.14 | 2.22 | 34.76 | 36.59 | -1.83 | 1.83 | -0.03 | 0.03 |
| gene-LOC105276635 | https://www.ncbi.nlm.nih.gov/gene/?term=LOC105276635 | 4962 | 40 | 80.61 | 1 | 2.02 | 2.5 | 23.53 | 27.39 | -3.86 | 3.86 | -0.03 | 0.03 | |
| gene-LOC105282914 | https://www.ncbi.nlm.nih.gov/gene/?term=LOC105282914 | 5623 | 43 | 76.47 | 1 | 1.78 | 2.33 | 49.78 | 47.18 | 2.6 | 2.6 | 0.02 | 0.02 | |
| gene-LOC105285409 | https://www.ncbi.nlm.nih.gov/gene/?term=LOC105285409 | 5099 | 63 | 123.55 | | 1 | 1.96 | 1.59 | 3.13 | 1.16 | 1.97 | 1.97 | 0.02 | 0.02 |
| gene-LOC105281296 | https://www.ncbi.nlm.nih.gov/gene/?term=LOC105281296 | 2434 | 43 | 176.66 | | 1 | 4.11 | 2.33 | 29.93 | 28.76 | 1.17 | 1.17 | 0.02 | 0.02 |
| gene-LOC105285252 | https://www.ncbi.nlm.nih.gov/gene/?term=LOC105285252 | 7427 | 125 | 168.3 | 1 | 1.35 | 0.8 | 14.66 | 13.59 | 1.08 | 1.08 | 0.02 | 0.02 | |
| gene-LOC105280819 | https://www.ncbi.nlm.nih.gov/gene/?term=LOC105280819 | 1639 | 43 | 262.36 | | 1 | 6.1 | 2.33 | 39.04 | 38.17 | 0.87 | 0.87 | 0.02 | 0.02 |
| gene-LOC105288105 | https://www.ncbi.nlm.nih.gov/gene/?term=LOC105288105 | 1460 | 37 | 253.42 | | 1 | 6.85 | 2.7 | 53 | 52.16 | 0.84 | 0.84 | 0.02 | 0.02 |
| gene-LOC105275587 | https://www.ncbi.nlm.nih.gov/gene/?term=LOC105275587 | 2404 | 57 | 237.1 | 1 | 4.16 | 1.75 | 36.63 | 35.9 | 0.73 | 0.73 | 0.02 | 0.02 | |
| gene-LOC105276272 | https://www.ncbi.nlm.nih.gov/gene/?term=LOC105276272 | 1965 | 50 | 254.45 | | 1 | 5.09 | 2 | 27.69 | 27.01 | 0.68 | 0.68 | 0.02 | 0.02 |
| gene-LOC105287633 | https://www.ncbi.nlm.nih.gov/gene/?term=LOC105287633 | 1524 | 35 | 229.66 | | 1 | 6.56 | 2.86 | 49.8 | 49.12 | 0.68 | 0.68 | 0.02 | 0.02 |
| gene-LOC105278145 | https://www.ncbi.nlm.nih.gov/gene/?term=LOC105278145 | 4470 | 252 | 563.76 | | 1 | 2.24 | 0.4 | 10.24 | 10.56 | -0.32 | 0.32 | -0.02 | 0.02 |
| gene-LOC105288055 | https://www.ncbi.nlm.nih.gov/gene/?term=LOC105288055 | 9934 | 548 | 551.64 | | 1 | 1.01 | 0.18 | 7.05 | 7.45 | -0.4 | 0.4 | -0.02 | 0.02 |
| gene-LOC105275374 | https://www.ncbi.nlm.nih.gov/gene/?term=LOC105275374 | 11376 | 567 | 498.42 | | 1 | 0.88 | 0.18 | 10.6 | 11.02 | -0.41 | 0.41 | -0.02 | 0.02 |
| gene-LOC105285742 | https://www.ncbi.nlm.nih.gov/gene/?term=LOC105285742 | 3446 | 132 | 383.05 | | 1 | 2.9 | 0.76 | 18.9 | 19.31 | -0.41 | 0.41 | -0.02 | 0.02 |
| gene-LOC105284043 | https://www.ncbi.nlm.nih.gov/gene/?term=LOC105284043 | 5661 | 325 | 574.1 | 1 | 1.77 | 0.31 | 9.91 | 10.34 | -0.43 | 0.43 | -0.02 | 0.02 | |
| gene-LOC105278666 | https://www.ncbi.nlm.nih.gov/gene/?term=LOC105278666 | 10341 | 494 | 477.71 | | 1 | 0.97 | 0.2 | 6.74 | 7.18 | -0.44 | 0.44 | -0.02 | 0.02 |
| gene-LOC105281272 | https://www.ncbi.nlm.nih.gov/gene/?term=LOC105281272 | 5252 | 203 | 386.52 | | 1 | 1.9 | 0.49 | 14.55 | 14.99 | -0.44 | 0.44 | -0.02 | 0.02 |
| gene-LOC105274669 | https://www.ncbi.nlm.nih.gov/gene/?term=LOC105274669 | 8555 | 293 | 342.49 | | 1 | 1.17 | 0.34 | 5.34 | 5.82 | -0.48 | 0.48 | -0.02 | 0.02 |
| gene-LOC105275363 | https://www.ncbi.nlm.nih.gov/gene/?term=LOC105275363 | 3798 | 176 | 463.4 | 1 | 2.63 | 0.57 | 15.74 | 16.23 | -0.5 | 0.5 | -0.02 | 0.02 | |
| gene-LOC113562050 | https://www.ncbi.nlm.nih.gov/gene/?term=LOC113562050 | 17097 | 643 | 376.09 | | 1 | 0.58 | 0.16 | 4.96 | 5.49 | -0.53 | 0.53 | -0.02 | 0.02 |
| gene-LOC105284036 | https://www.ncbi.nlm.nih.gov/gene/?term=LOC105284036 | 3254 | 112 | 344.19 | | 1 | 3.07 | 0.89 | 21.95 | 22.5 | -0.55 | 0.55 | -0.02 | 0.02 |
| gene-LOC105283728 | https://www.ncbi.nlm.nih.gov/gene/?term=LOC105283728 | 3256 | 123 | 377.76 | | 2 | 6.14 | 1.63 | 18.65 | 19.22 | -0.57 | 0.57 | -0.02 | 0.02 |
| gene-LOC105276875 | https://www.ncbi.nlm.nih.gov/gene/?term=LOC105276875 | 3151 | 124 | 393.53 | | 1 | 3.17 | 0.81 | 21.3 | 21.9 | -0.6 | 0.6 | -0.02 | 0.02 |
| gene-LOC105274706 | https://www.ncbi.nlm.nih.gov/gene/?term=LOC105274706 | 1446 | 57 | 394.19 | | 1 | 6.92 | 1.75 | 36.89 | 37.51 | -0.61 | 0.61 | -0.02 | 0.02 |
| gene-LOC105275451 | https://www.ncbi.nlm.nih.gov/gene/?term=LOC105275451 | 6035 | 142 | 235.29 | | 1 | 1.66 | 0.7 | 20.66 | 21.45 | -0.78 | 0.78 | -0.02 | 0.02 |
| gene-LOC105280348 | https://www.ncbi.nlm.nih.gov/gene/?term=LOC105280348 | 3797 | 79 | 208.06 | | 1 | 2.63 | 1.27 | 19.69 | 20.49 | -0.8 | 0.8 | -0.02 | 0.02 |
| gene-LOC105283967 | https://www.ncbi.nlm.nih.gov/gene/?term=LOC105283967 | 2393 | 60 | 250.73 | | 1 | 4.18 | 1.67 | 41.56 | 42.36 | -0.8 | 0.8 | -0.02 | 0.02 |
| gene-LOC105285395 | https://www.ncbi.nlm.nih.gov/gene/?term=LOC105285395 | 8085 | 224 | 277.06 | | 1 | 1.24 | 0.45 | 14.28 | 15.08 | -0.8 | 0.8 | -0.02 | 0.02 |
| gene-LOC105280407 | https://www.ncbi.nlm.nih.gov/gene/?term=LOC105280407 | 10950 | 218 | 199.09 | | 1 | 0.91 | 0.46 | 16.66 | 17.49 | -0.83 | 0.83 | -0.02 | 0.02 |
| gene-LOC105278228 | https://www.ncbi.nlm.nih.gov/gene/?term=LOC105278228 | 8610 | 176 | 204.41 | | 1 | 1.16 | 0.57 | 22.43 | 23.28 | -0.84 | 0.84 | -0.02 | 0.02 |
| gene-LOC105274590 | https://www.ncbi.nlm.nih.gov/gene/?term=LOC105274590 | 4516 | 80 | 177.15 | | 2 | 4.43 | 2.5 | 29.75 | 30.69 | -0.95 | 0.95 | -0.02 | 0.02 |
| gene-LOC105282825 | https://www.ncbi.nlm.nih.gov/gene/?term=LOC105282825 | 3919 | 98 | 250.06 | | 1 | 2.55 | 1.02 | 14.18 | 15.14 | -0.96 | 0.96 | -0.02 | 0.02 |
| gene-LOC105284117 | https://www.ncbi.nlm.nih.gov/gene/?term=LOC105284117 | 4340 | 101 | 232.72 | | 1 | 2.3 | 0.99 | 10.38 | 11.36 | -0.98 | 0.98 | -0.02 | 0.02 |
| gene-LOC105276196 | https://www.ncbi.nlm.nih.gov/gene/?term=LOC105276196 | 2953 | 59 | 199.8 | 1 | 3.39 | 1.69 | 37.63 | 38.7 | -1.07 | 1.07 | -0.02 | 0.02 | |
| gene-LOC105283943 | https://www.ncbi.nlm.nih.gov/gene/?term=LOC105283943 | 3559 | 64 | 179.83 | | 1 | 2.81 | 1.56 | 14.69 | 15.79 | -1.1 | 1.1 | -0.02 | 0.02 |
| gene-LOC105276501 | https://www.ncbi.nlm.nih.gov/gene/?term=LOC105276501 | 6089 | 110 | 180.65 | | 1 | 1.64 | 0.91 | 15.38 | 16.49 | -1.11 | 1.11 | -0.02 | 0.02 |
| gene-LOC105288078 | https://www.ncbi.nlm.nih.gov/gene/?term=LOC105288078 | 19126 | 332 | 173.59 | | 1 | 0.52 | 0.3 | 16.16 | 17.36 | -1.2 | 1.2 | -0.02 | 0.02 |
| gene-LOC105283712 | https://www.ncbi.nlm.nih.gov/gene/?term=LOC105283712 | 1910 | 36 | 188.48 | | 1 | 5.24 | 2.78 | 28.48 | 29.79 | -1.31 | 1.31 | -0.02 | 0.02 |
| gene-LOC105280641 | https://www.ncbi.nlm.nih.gov/gene/?term=LOC105280641 | 3682 | 61 | 165.67 | | 1 | 2.72 | 1.64 | 16.75 | 18.07 | -1.32 | 1.32 | -0.02 | 0.02 |
| gene-LOC105280002 | https://www.ncbi.nlm.nih.gov/gene/?term=LOC105280002 | 1940 | 28 | 144.33 | | 1 | 5.15 | 3.57 | 39.15 | 40.87 | -1.73 | 1.73 | -0.02 | 0.02 |
| gene-LOC105275518 | https://www.ncbi.nlm.nih.gov/gene/?term=LOC105275518 | 2335 | 26 | 111.35 | | 1 | 4.28 | 3.85 | 36.28 | 38.04 | -1.76 | 1.76 | -0.02 | 0.02 |
| gene-LOC105277856 | https://www.ncbi.nlm.nih.gov/gene/?term=LOC105277856 | 3442 | 37 | 107.5 | 1 | 2.91 | 2.7 | 37.9 | 39.79 | -1.89 | 1.89 | -0.02 | 0.02 | |

| | | | | | | | | | | | | | | |
|---|---|---|---|---|---|---|---|---|---|---|---|---|---|---|
| gene-LOC105280738 | https://www.ncbi.nlm.nih.gov/gene/?term=LOC105280738 | 4287 | 41 | 95.64 | 1 | 2.33 | 2.44 | 24.39 | 26.44 | -2.04 | 2.04 | -0.02 | 0.02 | |
| gene-LOC105284609 | https://www.ncbi.nlm.nih.gov/gene/?term=LOC105284609 | 5678 | 35 | 61.64 | 1 | 1.76 | 2.86 | 28.06 | 31.35 | -3.29 | 3.29 | -0.02 | 0.02 | |
| gene-LOC105281515 | https://www.ncbi.nlm.nih.gov/gene/?term=LOC105281515 | 8827 | 54 | 61.18 | 2 | 2.27 | 3.7 | 27.93 | 31.25 | -3.31 | 3.31 | -0.02 | 0.02 | |
| gene-LOC105286983 | https://www.ncbi.nlm.nih.gov/gene/?term=LOC105286983 | 16901 | 64 | 37.87 | 2 | 1.18 | 3.12 | 0.67 | 5.99 | -5.32 | 5.32 | -0.02 | 0.02 | |
| gene-LOC105279500 | https://www.ncbi.nlm.nih.gov/gene/?term=LOC105279500 | 3587 | 33 | 92 | 1 | 2.79 | 3.03 | 47.01 | 45.48 | 1.53 | 1.53 | 0.01 | 0.01 | |
| gene-LOC105281173 | https://www.ncbi.nlm.nih.gov/gene/?term=LOC105281173 | 1595 | 38 | 238.24 | | 1 | 6.27 | 2.63 | 60.67 | 60.19 | 0.48 | 0.48 | 0.01 | 0.01 |
| gene-LOC105280304 | https://www.ncbi.nlm.nih.gov/gene/?term=LOC105280304 | 8919 | 115 | 128.94 | | 1 | 1.12 | 0.87 | 7.4 | 6.97 | 0.43 | 0.43 | 0.01 | 0.01 |
| gene-LOC105284507 | https://www.ncbi.nlm.nih.gov/gene/?term=LOC105284507 | 5207 | 91 | 174.76 | | 1 | 1.92 | 1.1 | 18.91 | 18.57 | 0.35 | 0.35 | 0.01 | 0.01 |
| gene-LOC105283296 | https://www.ncbi.nlm.nih.gov/gene/?term=LOC105283296 | 2174 | 96 | 441.58 | | 1 | 4.6 | 1.04 | 20.81 | 20.51 | 0.3 | 0.3 | 0.01 | 0.01 |
| gene-LOC105278807 | https://www.ncbi.nlm.nih.gov/gene/?term=LOC105278807 | 2166 | 81 | 373.96 | | 1 | 4.62 | 1.23 | 19.2 | 19.33 | -0.14 | 0.14 | -0.01 | 0.01 |
| gene-LOC105285179 | https://www.ncbi.nlm.nih.gov/gene/?term=LOC105285179 | 4191 | 183 | 436.65 | | 1 | 2.39 | 0.55 | 19.23 | 19.39 | -0.16 | 0.16 | -0.01 | 0.01 |
| gene-LOC105282811 | https://www.ncbi.nlm.nih.gov/gene/?term=LOC105282811 | 5765 | 213 | 369.47 | | 1 | 1.73 | 0.47 | 8.07 | 8.24 | -0.17 | 0.17 | -0.01 | 0.01 |
| gene-LOC105287550 | https://www.ncbi.nlm.nih.gov/gene/?term=LOC105287550 | 15659 | 656 | 418.93 | | 1 | 0.64 | 0.15 | 6.08 | 6.26 | -0.18 | 0.18 | -0.01 | 0.01 |
| gene-LOC105287095 | https://www.ncbi.nlm.nih.gov/gene/?term=LOC105287095 | 1435 | 65 | 452.96 | | 1 | 6.97 | 1.54 | 25.24 | 25.44 | -0.2 | 0.2 | -0.01 | 0.01 |
| gene-LOC105286274 | https://www.ncbi.nlm.nih.gov/gene/?term=LOC105286274 | 3817 | 206 | 539.69 | | 1 | 2.62 | 0.49 | 16.61 | 16.82 | -0.21 | 0.21 | -0.01 | 0.01 |
| gene-LOC105274699 | https://www.ncbi.nlm.nih.gov/gene/?term=LOC105274699 | 2386 | 63 | 264.04 | | 1 | 4.19 | 1.59 | 22.17 | 22.41 | -0.24 | 0.24 | -0.01 | 0.01 |
| gene-LOC105274531 | https://www.ncbi.nlm.nih.gov/gene/?term=LOC105274531 | 7774 | 298 | 383.33 | | 1 | 1.29 | 0.34 | 10.39 | 10.72 | -0.33 | 0.33 | -0.01 | 0.01 |
| gene-LOC105275327 | https://www.ncbi.nlm.nih.gov/gene/?term=LOC105275327 | 14245 | 568 | 398.74 | | 1 | 0.7 | 0.18 | 6.19 | 6.55 | -0.36 | 0.36 | -0.01 | 0.01 |
| gene-LOC105285163 | https://www.ncbi.nlm.nih.gov/gene/?term=LOC105285163 | 5061 | 85 | 167.95 | | 1 | 1.98 | 1.18 | 19.84 | 20.21 | -0.37 | 0.37 | -0.01 | 0.01 |
| gene-LOC105278771 | https://www.ncbi.nlm.nih.gov/gene/?term=LOC105278771 | 10457 | 248 | 237.16 | | 1 | 0.96 | 0.4 | 8.97 | 9.36 | -0.39 | 0.39 | -0.01 | 0.01 |
| gene-LOC105282794 | https://www.ncbi.nlm.nih.gov/gene/?term=LOC105282794 | 4571 | 100 | 218.77 | | 1 | 2.19 | 1 | 16.99 | 17.37 | -0.39 | 0.39 | -0.01 | 0.01 |
| gene-LOC105283123 | https://www.ncbi.nlm.nih.gov/gene/?term=LOC105283123 | 1507 | 57 | 378.23 | | 1 | 6.64 | 1.75 | 17.49 | 17.88 | -0.39 | 0.39 | -0.01 | 0.01 |
| gene-LOC105287309 | https://www.ncbi.nlm.nih.gov/gene/?term=LOC105287309 | 3553 | 96 | 270.19 | | 1 | 2.81 | 1.04 | 7.77 | 8.16 | -0.39 | 0.39 | -0.01 | 0.01 |
| gene-LOC105287648 | https://www.ncbi.nlm.nih.gov/gene/?term=LOC105287648 | 2205 | 35 | 158.73 | | 1 | 4.54 | 2.86 | 35.94 | 36.45 | -0.5 | 0.5 | -0.01 | 0.01 |
| gene-LOC105276059 | https://www.ncbi.nlm.nih.gov/gene/?term=LOC105276059 | 6256 | 145 | 231.78 | | 1 | 1.6 | 0.69 | 8.95 | 9.49 | -0.54 | 0.54 | -0.01 | 0.01 |
| gene-LOC105285502 | https://www.ncbi.nlm.nih.gov/gene/?term=LOC105285502 | 9990 | 164 | 164.16 | | 1 | 1 | 0.61 | 8.55 | 9.16 | -0.62 | 0.62 | -0.01 | 0.01 |
| gene-LOC105276683 | https://www.ncbi.nlm.nih.gov/gene/?term=LOC105276683 | 17000 | 149 | 87.65 | 1 | 0.59 | 0.67 | 8.22 | 8.88 | -0.65 | 0.65 | -0.01 | 0.01 | |
| gene-LOC105286452 | https://www.ncbi.nlm.nih.gov/gene/?term=LOC105286452 | 6888 | 66 | 95.82 | 1 | 1.45 | 1.52 | 30.11 | 30.78 | -0.67 | 0.67 | -0.01 | 0.01 | |
| gene-LOC105279657 | https://www.ncbi.nlm.nih.gov/gene/?term=LOC105279657 | 5878 | 91 | 154.81 | | 1 | 1.7 | 1.1 | 18.9 | 19.62 | -0.72 | 0.72 | -0.01 | 0.01 |
| gene-LOC105275187 | https://www.ncbi.nlm.nih.gov/gene/?term=LOC105275187 | 5534 | 77 | 139.14 | | 1 | 1.81 | 1.3 | 15.38 | 16.21 | -0.83 | 0.83 | -0.01 | 0.01 |
| gene-LOC105276917 | https://www.ncbi.nlm.nih.gov/gene/?term=LOC105276917 | 2336 | 33 | 141.27 | | 1 | 4.28 | 3.03 | 30.13 | 31.08 | -0.95 | 0.95 | -0.01 | 0.01 |
| gene-LOC105275347 | https://www.ncbi.nlm.nih.gov/gene/?term=LOC105275347 | 2526 | 20 | 79.18 | 1 | 3.96 | 5 | 37.1 | 38.21 | -1.12 | 1.12 | -0.01 | 0.01 | |
| gene-LOC105285514 | https://www.ncbi.nlm.nih.gov/gene/?term=LOC105285514 | 1277 | 15 | 117.46 | | 1 | 7.83 | 6.67 | 78.25 | 79.38 | -1.12 | 1.12 | -0.01 | 0.01 |
| gene-LOC105283989 | https://www.ncbi.nlm.nih.gov/gene/?term=LOC105283989 | 4948 | 32 | 64.67 | 2 | 4.04 | 6.25 | 43.84 | 45.43 | -1.59 | 1.59 | -0.01 | 0.01 | |
| gene-LOC105285315 | https://www.ncbi.nlm.nih.gov/gene/?term=LOC105285315 | 5458 | 26 | 47.64 | 1 | 1.83 | 3.85 | 36.55 | 38.99 | -2.44 | 2.44 | -0.01 | 0.01 | |
| gene-LOC105284329 | https://www.ncbi.nlm.nih.gov/gene/?term=LOC105284329 | 4051 | 53 | 130.83 | | 1 | 2.47 | 1.89 | 19.36 | 19.04 | 0.32 | 0.32 | 0 | 0 |
| gene-LOC105286770 | https://www.ncbi.nlm.nih.gov/gene/?term=LOC105286770 | 3055 | 38 | 124.39 | | 1 | 3.27 | 2.63 | 15.65 | 15.48 | 0.17 | 0.17 | 0 | 0 |
| gene-LOC105287842 | https://www.ncbi.nlm.nih.gov/gene/?term=LOC105287842 | 2301 | 57 | 247.72 | | 1 | 4.35 | 1.75 | 21.58 | 21.41 | 0.17 | 0.17 | 0 | 0 |
| gene-LOC105285496 | https://www.ncbi.nlm.nih.gov/gene/?term=LOC105285496 | 4080 | 112 | 274.51 | | 1 | 2.45 | 0.89 | 40.26 | 40.12 | 0.14 | 0.14 | 0 | 0 |
| gene-LOC105278534 | https://www.ncbi.nlm.nih.gov/gene/?term=LOC105278534 | 7364 | 200 | 271.59 | | 2 | 2.72 | 1 | 17.58 | 17.48 | 0.1 | 0.1 | 0 | 0 |
| gene-LOC105281312 | https://www.ncbi.nlm.nih.gov/gene/?term=LOC105281312 | 2551 | 74 | 290.08 | | 1 | 3.92 | 1.35 | 29.31 | 29.21 | 0.1 | 0.1 | 0 | 0 |
| gene-LOC105282951 | https://www.ncbi.nlm.nih.gov/gene/?term=LOC105282951 | 3366 | 131 | 389.19 | | 1 | 2.97 | 0.76 | 18.77 | 18.69 | 0.08 | 0.08 | 0 | 0 |
| gene-LOC105277273 | https://www.ncbi.nlm.nih.gov/gene/?term=LOC105277273 | 3711 | 169 | 455.4 | 1 | 2.69 | 0.59 | 12.24 | 12.17 | 0.06 | 0.06 | 0 | 0 | |
| gene-LOC105281700 | https://www.ncbi.nlm.nih.gov/gene/?term=LOC105281700 | 2208 | 39 | 176.63 | | 1 | 4.53 | 2.56 | 37.83 | 37.79 | 0.04 | 0.04 | 0 | 0 |
| gene-LOC105284184 | https://www.ncbi.nlm.nih.gov/gene/?term=LOC105284184 | 2535 | 102 | 402.37 | | 1 | 3.94 | 0.98 | 28.97 | 28.93 | 0.03 | 0.03 | 0 | 0 |
| gene-LOC105285533 | https://www.ncbi.nlm.nih.gov/gene/?term=LOC105285533 | 1640 | 71 | 432.93 | | 1 | 6.1 | 1.41 | 23.22 | 23.19 | 0.03 | 0.03 | 0 | 0 |
| gene-LOC105281754 | https://www.ncbi.nlm.nih.gov/gene/?term=LOC105281754 | 4300 | 122 | 283.72 | | 1 | 2.33 | 0.82 | 16.75 | 16.74 | 0.01 | 0.01 | 0 | 0 |
| gene-LOC105274851 | https://www.ncbi.nlm.nih.gov/gene/?term=LOC105274851 | 6616 | 150 | 226.72 | | 1 | 1.51 | 0.67 | 10.51 | 10.53 | -0.02 | 0.02 | 0 | 0 |
| gene-LOC105278691 | https://www.ncbi.nlm.nih.gov/gene/?term=LOC105278691 | 3871 | 172 | 444.33 | | 1 | 2.58 | 0.58 | 14.5 | 14.55 | -0.05 | 0.05 | 0 | 0 |
| gene-LOC105286649 | https://www.ncbi.nlm.nih.gov/gene/?term=LOC105286649 | 14178 | 892 | 629.14 | | 1 | 0.71 | 0.11 | 1.01 | 1.06 | -0.05 | 0.05 | 0 | 0 |
| gene-LOC105275848 | https://www.ncbi.nlm.nih.gov/gene/?term=LOC105275848 | 3718 | 149 | 400.75 | | 1 | 2.69 | 0.67 | 21.41 | 21.47 | -0.06 | 0.06 | 0 | 0 |
| gene-LOC105282646 | https://www.ncbi.nlm.nih.gov/gene/?term=LOC105282646 | 3317 | 110 | 331.62 | | 1 | 3.01 | 0.91 | 19.95 | 20.03 | -0.08 | 0.08 | 0 | 0 |
| gene-LOC105276462 | https://www.ncbi.nlm.nih.gov/gene/?term=LOC105276462 | 6561 | 296 | 451.15 | | 1 | 1.52 | 0.34 | 7.73 | 7.83 | -0.1 | 0.1 | 0 | 0 |
| gene-LOC105276527 | https://www.ncbi.nlm.nih.gov/gene/?term=LOC105276527 | 7923 | 340 | 429.13 | | 1 | 1.26 | 0.29 | 9.36 | 9.47 | -0.11 | 0.11 | 0 | 0 |
| gene-LOC105280408 | https://www.ncbi.nlm.nih.gov/gene/?term=LOC105280408 | 5983 | 60 | 100.28 | | 1 | 1.67 | 1.67 | 21.93 | 22.03 | -0.11 | 0.11 | 0 | 0 |
| gene-LOC105280722 | https://www.ncbi.nlm.nih.gov/gene/?term=LOC105280722 | 3237 | 98 | 302.75 | | 1 | 3.09 | 1.02 | 29.35 | 29.46 | -0.11 | 0.11 | 0 | 0 |
| gene-LOC105275179 | https://www.ncbi.nlm.nih.gov/gene/?term=LOC105275179 | 3271 | 85 | 259.86 | | 1 | 3.06 | 1.18 | 19.41 | 19.53 | -0.12 | 0.12 | 0 | 0 |
| gene-LOC105284262 | https://www.ncbi.nlm.nih.gov/gene/?term=LOC105284262 | 6669 | 247 | 370.37 | | 1 | 1.5 | 0.4 | 4.68 | 4.81 | -0.13 | 0.13 | 0 | 0 |
| gene-LOC105274823 | https://www.ncbi.nlm.nih.gov/gene/?term=LOC105274823 | 4620 | 148 | 320.35 | | 1 | 2.16 | 0.68 | 20.3 | 20.45 | -0.14 | 0.14 | 0 | 0 |
| gene-LOC105280410 | https://www.ncbi.nlm.nih.gov/gene/?term=LOC105280410 | 12580 | 263 | 209.06 | | 1 | 0.79 | 0.38 | 19.71 | 19.86 | -0.15 | 0.15 | 0 | 0 |
| gene-LOC105286515 | https://www.ncbi.nlm.nih.gov/gene/?term=LOC105286515 | 3751 | 123 | 327.91 | | 1 | 2.67 | 0.81 | 10.35 | 10.5 | -0.15 | 0.15 | 0 | 0 |
| gene-LOC105285406 | https://www.ncbi.nlm.nih.gov/gene/?term=LOC105285406 | 8015 | 59 | 73.61 | 1 | 1.25 | 1.69 | 29.55 | 29.82 | -0.27 | 0.27 | 0 | 0 | |
| gene-LOC105282067 | https://www.ncbi.nlm.nih.gov/gene/?term=LOC105282067 | 5941 | 44 | 74.06 | 1 | 1.68 | 2.27 | 30.63 | 31 | -0.36 | 0.36 | 0 | 0 | |

Directory: OGL File: rnk-dmg-Ob_LI_BCphaseB2.vs.RphaseB1.txt

| gene_ID | gene_link | gwidth | #Sites | #per10Kb | #dmSites | #dmsp10kb | %dmSites | %pSite1 | %pSite2 | DMpSite | ADMpSite | DMpNucl | ADMpNucl |
|---|---|---|---|---|---|---|---|---|---|---|---|---|---|
| gene-LOC105283830 | https://www.ncbi.nlm.nih.gov/gene/?term=LOC105283830 | 252 | 14 | 555.56 | 1 | 39.68 | 7.14 | 34.01 | 40.39 | -6.38 | 6.38 | -0.35 | 0.35 |
| gene-LOC105288193 | https://www.ncbi.nlm.nih.gov/gene/?term=LOC105288193 | 1495 | 85 | 568.56 | 5 | 33.44 | 5.88 | 31.62 | 28.31 | 3.31 | 3.31 | 0.19 | 0.19 |
| gene-LOC105275833 | https://www.ncbi.nlm.nih.gov/gene/?term=LOC105275833 | 981 | 41 | 417.94 | 1 | 10.19 | 2.44 | 35.54 | 39.29 | -3.75 | 3.75 | -0.16 | 0.16 |
| gene-LOC105276202 | https://www.ncbi.nlm.nih.gov/gene/?term=LOC105276202 | 1540 | 43 | 279.22 | 5 | 32.47 | 11.63 | 24.19 | 18.86 | 5.33 | 5.33 | 0.15 | 0.15 |
| gene-LOC113562453 | https://www.ncbi.nlm.nih.gov/gene/?term=LOC113562453 | 763 | 32 | 419.4 | 2 | 26.21 | 6.25 | 26.93 | 30.48 | -3.55 | 3.55 | -0.15 | 0.15 |
| gene-LOC105287692 | https://www.ncbi.nlm.nih.gov/gene/?term=LOC105287692 | 1505 | 97 | 644.52 | 5 | 33.22 | 5.15 | 29.7 | 27.6 | 2.1 | 2.1 | 0.14 | 0.14 |
| gene-LOC105278849 | https://www.ncbi.nlm.nih.gov/gene/?term=LOC105278849 | 776 | 75 | 966.49 | 3 | 38.66 | 4 | 22.31 | 21.08 | 1.24 | 1.24 | 0.12 | 0.12 |
| gene-LOC105283267 | https://www.ncbi.nlm.nih.gov/gene/?term=LOC105283267 | 2571 | 73 | 283.94 | 5 | 19.45 | 6.85 | 39.09 | 43.39 | -4.29 | 4.29 | -0.12 | 0.12 |
| gene-LOC105276115 | https://www.ncbi.nlm.nih.gov/gene/?term=LOC105276115 | 1520 | 36 | 236.84 | 3 | 19.74 | 8.33 | 20.99 | 26.1 | -5.11 | 5.11 | -0.12 | 0.12 |
| gene-LOC105280891 | https://www.ncbi.nlm.nih.gov/gene/?term=LOC105280891 | 1844 | 36 | 195.23 | 2 | 10.85 | 5.56 | 18.81 | 24.77 | -5.96 | 5.96 | -0.12 | 0.12 |
| gene-LOC105285004 | https://www.ncbi.nlm.nih.gov/gene/?term=LOC105285004 | 1334 | 33 | 247.38 | 3 | 22.49 | 9.09 | 22.51 | 18.04 | 4.47 | 4.47 | 0.11 | 0.11 |
| gene-LOC105277371 | https://www.ncbi.nlm.nih.gov/gene/?term=LOC105277371 | 1816 | 74 | 407.49 | 2 | 11.01 | 2.7 | 20.36 | 17.6 | 2.76 | 2.76 | 0.11 | 0.11 |
| gene-LOC105286845 | https://www.ncbi.nlm.nih.gov/gene/?term=LOC105286845 | 1159 | 81 | 698.88 | 1 | 8.63 | 1.23 | 14.6 | 12.97 | 1.63 | 1.63 | 0.11 | 0.11 |
| gene-LOC105284093 | https://www.ncbi.nlm.nih.gov/gene/?term=LOC105284093 | 809 | 59 | 729.3 | 1 | 12.36 | 1.69 | 8.36 | 9.93 | -1.57 | 1.57 | -0.11 | 0.11 |
| gene-LOC105280710 | https://www.ncbi.nlm.nih.gov/gene/?term=LOC105280710 | 1215 | 68 | 559.67 | 2 | 16.46 | 2.94 | 20.85 | 22.76 | -1.91 | 1.91 | -0.11 | 0.11 |
| gene-LOC105285203 | https://www.ncbi.nlm.nih.gov/gene/?term=LOC105285203 | 1020 | 27 | 264.71 | 1 | 9.8 | 3.7 | 30.05 | 26.28 | 3.77 | 3.77 | 0.1 | 0.1 |
| gene-LOC105284168 | https://www.ncbi.nlm.nih.gov/gene/?term=LOC105284168 | 1400 | 76 | 542.86 | 2 | 14.29 | 2.63 | 24.73 | 22.83 | 1.9 | 1.9 | 0.1 | 0.1 |
| gene-LOC105283909 | https://www.ncbi.nlm.nih.gov/gene/?term=LOC105283909 | 2234 | 112 | 501.34 | 2 | 8.95 | 1.79 | 10.52 | 12.48 | -1.96 | 1.96 | -0.1 | 0.1 |
| gene-LOC105284189 | https://www.ncbi.nlm.nih.gov/gene/?term=LOC105284189 | 2601 | 130 | 499.81 | 3 | 11.53 | 2.31 | 14.87 | 16.94 | -2.07 | 2.07 | -0.1 | 0.1 |
| gene-LOC105275897 | https://www.ncbi.nlm.nih.gov/gene/?term=LOC105275897 | 1684 | 72 | 427.55 | 1 | 5.94 | 1.39 | 36.2 | 38.44 | -2.24 | 2.24 | -0.1 | 0.1 |
| gene-LOC105277345 | https://www.ncbi.nlm.nih.gov/gene/?term=LOC105277345 | 2151 | 82 | 381.22 | 1 | 4.65 | 1.22 | 16.59 | 19.1 | -2.51 | 2.51 | -0.1 | 0.1 |
| gene-LOC105278537 | https://www.ncbi.nlm.nih.gov/gene/?term=LOC105278537 | 1660 | 62 | 373.49 | 4 | 24.1 | 6.45 | 28.42 | 31.22 | -2.8 | 2.8 | -0.1 | 0.1 |
| gene-LOC105284046 | https://www.ncbi.nlm.nih.gov/gene/?term=LOC105284046 | 1027 | 28 | 272.64 | 4 | 38.95 | 14.29 | 19.04 | 15.74 | 3.3 | 3.3 | 0.09 | 0.09 |
| gene-LOC105282209 | https://www.ncbi.nlm.nih.gov/gene/?term=LOC105282209 | 1041 | 35 | 336.22 | 1 | 9.61 | 2.86 | 31.01 | 28.36 | 2.65 | 2.65 | 0.09 | 0.09 |

...

... only first 25 lines shown ...

Directory: OGL File: rnk-dmg-Ob_LI_BCphaseB2.vs.RphaseB2.txt

| gene_ID | gene_link | gwidth | #Sites | #per10Kb | #dmSites | #dmsp10kb | %dmSites | %pSite1 | %pSite2 | DMpSite | ADMpSite | DMpNucl | ADMpNucl |
|---|---|---|---|---|---|---|---|---|---|---|---|---|---|
| gene-LOC105278458 | https://www.ncbi.nlm.nih.gov/gene/?term=LOC105278458 | 216 | 3 | 138.89 | 1 | 46.3 | 33.33 | 10.63 | 29.54 | -18.91 | 18.91 | -0.26 | 0.26 |
| gene-LOC105283830 | https://www.ncbi.nlm.nih.gov/gene/?term=LOC105283830 | 252 | 5 | 198.41 | 1 | 39.68 | 20 | 31.26 | 44.09 | -12.83 | 12.83 | -0.25 | 0.25 |
| gene-LOC105285203 | https://www.ncbi.nlm.nih.gov/gene/?term=LOC105285203 | 1020 | 27 | 264.71 | 2 | 19.61 | 7.41 | 30.05 | 24.28 | 5.76 | 5.76 | 0.15 | 0.15 |
| gene-LOC105275897 | https://www.ncbi.nlm.nih.gov/gene/?term=LOC105275897 | 1684 | 72 | 427.55 | 1 | 5.94 | 1.39 | 36.2 | 39.56 | -3.37 | 3.37 | -0.14 | 0.14 |
| gene-LOC105278938 | https://www.ncbi.nlm.nih.gov/gene/?term=LOC105278938 | 1818 | 110 | 605.06 | 1 | 5.5 | 0.91 | 20.68 | 22.88 | -2.2 | 2.2 | -0.13 | 0.13 |
| gene-LOC105276563 | https://www.ncbi.nlm.nih.gov/gene/?term=LOC105276563 | 1958 | 113 | 577.12 | 6 | 30.64 | 5.31 | 19.41 | 17.48 | 1.93 | 1.93 | 0.11 | 0.11 |
| gene-LOC105287692 | https://www.ncbi.nlm.nih.gov/gene/?term=LOC105287692 | 1505 | 89 | 591.36 | 5 | 33.22 | 5.62 | 31.34 | 29.41 | 1.93 | 1.93 | 0.11 | 0.11 |
| gene-LOC105278737 | https://www.ncbi.nlm.nih.gov/gene/?term=LOC105278737 | 1346 | 98 | 728.08 | 2 | 14.86 | 2.04 | 19.42 | 17.93 | 1.49 | 1.49 | 0.11 | 0.11 |
| gene-LOC105278873 | https://www.ncbi.nlm.nih.gov/gene/?term=LOC105278873 | 1897 | 147 | 774.91 | 4 | 21.09 | 2.72 | 19.42 | 20.81 | -1.39 | 1.39 | -0.11 | 0.11 |
| gene-LOC105279049 | https://www.ncbi.nlm.nih.gov/gene/?term=LOC105279049 | 1285 | 90 | 700.39 | 1 | 7.78 | 1.11 | 17.36 | 18.97 | -1.61 | 1.61 | -0.11 | 0.11 |
| gene-LOC105280710 | https://www.ncbi.nlm.nih.gov/gene/?term=LOC105280710 | 1215 | 64 | 526.75 | 2 | 16.46 | 3.12 | 22.16 | 24.28 | -2.12 | 2.12 | -0.11 | 0.11 |
| gene-LOC105274670 | https://www.ncbi.nlm.nih.gov/gene/?term=LOC105274670 | 1306 | 39 | 298.62 | 2 | 15.31 | 5.13 | 33.71 | 37.28 | -3.56 | 3.56 | -0.11 | 0.11 |
| gene-LOC105284320 | https://www.ncbi.nlm.nih.gov/gene/?term=LOC105284320 | 390 | 10 | 256.41 | 1 | 25.64 | 10 | 21.93 | 26.26 | -4.33 | 4.33 | -0.11 | 0.11 |
| gene-LOC105275176 | https://www.ncbi.nlm.nih.gov/gene/?term=LOC105275176 | 656 | 8 | 121.95 | 2 | 30.49 | 25 | 14.79 | 6.32 | 8.47 | 8.47 | 0.1 | 0.1 |
| gene-LOC105283552 | https://www.ncbi.nlm.nih.gov/gene/?term=LOC105283552 | 1562 | 37 | 236.88 | 1 | 6.4 | 2.7 | 55 | 50.64 | 4.36 | 4.36 | 0.1 | 0.1 |
| gene-LOC105285914 | https://www.ncbi.nlm.nih.gov/gene/?term=LOC105285914 | 867 | 28 | 322.95 | 1 | 11.53 | 3.57 | 16.16 | 12.99 | 3.17 | 3.17 | 0.1 | 0.1 |
| gene-LOC105284253 | https://www.ncbi.nlm.nih.gov/gene/?term=LOC105284253 | 1255 | 42 | 334.66 | 1 | 7.97 | 2.38 | 28.76 | 31.67 | -2.91 | 2.91 | -0.1 | 0.1 |
| gene-LOC105282481 | https://www.ncbi.nlm.nih.gov/gene/?term=LOC105282481 | 1900 | 59 | 310.53 | 2 | 10.53 | 3.39 | 33.33 | 36.46 | -3.13 | 3.13 | -0.1 | 0.1 |
| gene-LOC105274938 | https://www.ncbi.nlm.nih.gov/gene/?term=LOC105274938 | 1964 | 54 | 274.95 | 4 | 20.37 | 7.41 | 27.19 | 30.79 | -3.6 | 3.6 | -0.1 | 0.1 |
| gene-LOC105275219 | https://www.ncbi.nlm.nih.gov/gene/?term=LOC105275219 | 997 | 18 | 180.54 | 1 | 10.03 | 5.56 | 37.08 | 42.66 | -5.58 | 5.58 | -0.1 | 0.1 |
| gene-LOC105278083 | https://www.ncbi.nlm.nih.gov/gene/?term=LOC105278083 | 1605 | 38 | 236.76 | 1 | 6.23 | 2.63 | 32.5 | 28.84 | 3.66 | 3.66 | 0.09 | 0.09 |
| gene-LOC105281704 | https://www.ncbi.nlm.nih.gov/gene/?term=LOC105281704 | 2195 | 57 | 259.68 | 3 | 13.67 | 5.26 | 54.26 | 50.77 | 3.49 | 3.49 | 0.09 | 0.09 |
| gene-LOC105277371 | https://www.ncbi.nlm.nih.gov/gene/?term=LOC105277371 | 1816 | 71 | 390.97 | 1 | 5.51 | 1.41 | 20.94 | 18.75 | 2.2 | 2.2 | 0.09 | 0.09 |
| gene-LOC105277398 | https://www.ncbi.nlm.nih.gov/gene/?term=LOC105277398 | 1963 | 70 | 356.6 | 3 | 15.28 | 4.29 | 35.51 | 37.95 | -2.44 | 2.44 | -0.09 | 0.09 |

...

... only first 25 lines shown ...

Directory: OGL File: rnk-dmg-Ob_LI_RphaseA1.vs.RphaseA2.txt

| gene_ID | gene_link | gwidth | #Sites | #per10Kb | #dmSites | #dmsp10kb | %dmSites | %pSite1 | %pSite2 | DMpSite | ADMpSite | DMpNucl | ADMpNucl |
|---|---|---|---|---|---|---|---|---|---|---|---|---|---|
| gene-LOC105288042 | https://www.ncbi.nlm.nih.gov/gene/?term=LOC105288042 | 3507 | 111 | 316.51 | 1 | 2.85 | 0.9 | 22.13 | 23.4 | -1.28 | 1.28 | -0.04 | 0.04 |
| gene-LOC105283264 | https://www.ncbi.nlm.nih.gov/gene/?term=LOC105283264 | 4005 | 96 | 239.7 | 1 | 2.5 | 1.04 | 23.23 | 22.15 | 1.09 | 1.09 | 0.03 | 0.03 |
| gene-LOC105283296 | https://www.ncbi.nlm.nih.gov/gene/?term=LOC105283296 | 2174 | 101 | 464.58 | 1 | 4.6 | 0.99 | 18.85 | 18.17 | 0.67 | 0.67 | 0.03 | 0.03 |
| gene-LOC105288036 | https://www.ncbi.nlm.nih.gov/gene/?term=LOC105288036 | 5110 | 209 | 409 | 1 | 1.96 | 0.48 | 17.07 | 17.79 | -0.71 | 0.71 | -0.03 | 0.03 |
| gene-LOC105277903 | https://www.ncbi.nlm.nih.gov/gene/?term=LOC105277903 | 1835 | 39 | 212.53 | 1 | 5.45 | 2.56 | 54.3 | 53.51 | 0.79 | 0.79 | 0.02 | 0.02 |
| gene-LOC105284557 | https://www.ncbi.nlm.nih.gov/gene/?term=LOC105284557 | 1672 | 84 | 502.39 | 1 | 5.98 | 1.19 | 28.63 | 28.17 | 0.46 | 0.46 | 0.02 | 0.02 |
| gene-LOC105275384 | https://www.ncbi.nlm.nih.gov/gene/?term=LOC105275384 | 1987 | 86 | 432.81 | 1 | 5.03 | 1.16 | 36.56 | 36.97 | -0.41 | 0.41 | -0.02 | 0.02 |
| gene-LOC105281272 | https://www.ncbi.nlm.nih.gov/gene/?term=LOC105281272 | 5252 | 199 | 378.9 | 1 | 1.9 | 0.5 | 14.62 | 15.07 | -0.45 | 0.45 | -0.02 | 0.02 |
| gene-LOC105284507 | https://www.ncbi.nlm.nih.gov/gene/?term=LOC105284507 | 5207 | 112 | 215.1 | 1 | 1.92 | 0.89 | 18.79 | 19.06 | -0.28 | 0.28 | -0.01 | 0.01 |
| gene-LOC105280304 | https://www.ncbi.nlm.nih.gov/gene/?term=LOC105280304 | 8919 | 150 | 168.18 | 1 | 1.12 | 0.67 | 6.83 | 6.96 | -0.12 | 0.12 | 0 | 0 |

Directory: OGL File: rnk-dmg-Ob_LI_RphaseA1.vs.RphaseB1.txt

| gene_ID | gene_link | gwidth | #Sites | #per10Kb | #dmSites | #dmsp10kb | %dmSites | %pSite1 | %pSite2 | DMpSite | ADMpSite | DMpNucl | ADMpNucl |
|---|---|---|---|---|---|---|---|---|---|---|---|---|---|
| gene-LOC113562345 | https://www.ncbi.nlm.nih.gov/gene/?term=LOC113562345 | 2290 | 78 | 340.61 | 11 | 48.03 | 14.1 | 24.77 | 13.87 | 10.9 | 10.9 | 0.37 | 0.37 |
| gene-LOC105275355 | https://www.ncbi.nlm.nih.gov/gene/?term=LOC105275355 | 3197 | 174 | 544.26 | 1 | 3.13 | 0.57 | 25.86 | 21.8 | 4.06 | 4.06 | 0.22 | 0.22 |
| gene-LOC105275384 | https://www.ncbi.nlm.nih.gov/gene/?term=LOC105275384 | 1987 | 88 | 442.88 | 1 | 5.03 | 1.14 | 36.35 | 32.79 | 3.56 | 3.56 | 0.16 | 0.16 |
| gene-LOC105275070 | https://www.ncbi.nlm.nih.gov/gene/?term=LOC105275070 | 5667 | 56 | 98.82 | 3 | 5.29 | 5.36 | 17.83 | 5.85 | 11.98 | 11.98 | 0.12 | 0.12 |
| gene-LOC105285695 | https://www.ncbi.nlm.nih.gov/gene/?term=LOC105285695 | 1371 | 68 | 495.99 | 1 | 7.29 | 1.47 | 23.53 | 21.55 | 1.99 | 1.99 | 0.1 | 0.1 |
| gene-LOC105285671 | https://www.ncbi.nlm.nih.gov/gene/?term=LOC105285671 | 2263 | 100 | 441.89 | 1 | 4.42 | 1 | 32.24 | 30.53 | 1.71 | 1.71 | 0.08 | 0.08 |
| gene-LOC105275708 | https://www.ncbi.nlm.nih.gov/gene/?term=LOC105275708 | 1772 | 14 | 79.01 | 1 | 5.64 | 7.14 | 50.83 | 42.47 | 8.36 | 8.36 | 0.07 | 0.07 |
| gene-LOC105287063 | https://www.ncbi.nlm.nih.gov/gene/?term=LOC105287063 | 1874 | 49 | 261.47 | 1 | 5.34 | 2.04 | 32.77 | 30.12 | 2.65 | 2.65 | 0.07 | 0.07 |
| gene-LOC105285004 | https://www.ncbi.nlm.nih.gov/gene/?term=LOC105285004 | 1334 | 37 | 277.36 | 1 | 7.5 | 2.7 | 16.38 | 18.79 | -2.41 | 2.41 | -0.07 | 0.07 |
| gene-LOC105285514 | https://www.ncbi.nlm.nih.gov/gene/?term=LOC105285514 | 1277 | 7 | 54.82 | 1 | 7.83 | 14.29 | 90.55 | 80 | 10.54 | 10.54 | 0.06 | 0.06 |
| gene-LOC105274534 | https://www.ncbi.nlm.nih.gov/gene/?term=LOC105274534 | 1329 | 59 | 443.94 | 1 | 7.52 | 1.69 | 43.5 | 44.89 | -1.39 | 1.39 | -0.06 | 0.06 |
| gene-LOC105276272 | https://www.ncbi.nlm.nih.gov/gene/?term=LOC105276272 | 1965 | 44 | 223.92 | 1 | 5.09 | 2.27 | 27.22 | 24.96 | 2.26 | 2.26 | 0.05 | 0.05 |
| gene-LOC105278807 | https://www.ncbi.nlm.nih.gov/gene/?term=LOC105278807 | 2166 | 74 | 341.64 | 1 | 4.62 | 1.35 | 21.42 | 20.03 | 1.39 | 1.39 | 0.05 | 0.05 |
| gene-LOC105275460 | https://www.ncbi.nlm.nih.gov/gene/?term=LOC105275460 | 2473 | 126 | 509.5 | 1 | 4.04 | 0.79 | 13.41 | 12.4 | 1.01 | 1.01 | 0.05 | 0.05 |
| gene-LOC105278282 | https://www.ncbi.nlm.nih.gov/gene/?term=LOC105278282 | 1775 | 51 | 287.32 | 1 | 5.63 | 1.96 | 39.13 | 40.69 | -1.57 | 1.57 | -0.05 | 0.05 |
| gene-LOC113562102 | https://www.ncbi.nlm.nih.gov/gene/?term=LOC113562102 | 3433 | 63 | 183.51 | 1 | 2.91 | 1.59 | 17.89 | 15.59 | 2.3 | 2.3 | 0.04 | 0.04 |
| gene-LOC105282991 | https://www.ncbi.nlm.nih.gov/gene/?term=LOC105282991 | 5617 | 103 | 183.37 | 1 | 1.78 | 0.97 | 18.36 | 16.13 | 2.23 | 2.23 | 0.04 | 0.04 |
| gene-LOC105283712 | https://www.ncbi.nlm.nih.gov/gene/?term=LOC105283712 | 1910 | 35 | 183.25 | 1 | 5.24 | 2.86 | 28.45 | 26.47 | 1.98 | 1.98 | 0.04 | 0.04 |
| gene-LOC105286265 | https://www.ncbi.nlm.nih.gov/gene/?term=LOC105286265 | 2507 | 67 | 267.25 | 1 | 3.99 | 1.49 | 32.57 | 31.11 | 1.46 | 1.46 | 0.04 | 0.04 |
| gene-LOC105285742 | https://www.ncbi.nlm.nih.gov/gene/?term=LOC105285742 | 3446 | 109 | 316.31 | 1 | 2.9 | 0.92 | 17.79 | 16.59 | 1.2 | 1.2 | 0.04 | 0.04 |
| gene-LOC105285674 | https://www.ncbi.nlm.nih.gov/gene/?term=LOC105285674 | 3035 | 100 | 329.49 | 2 | 6.59 | 2 | 25.96 | 24.78 | 1.18 | 1.18 | 0.04 | 0.04 |
| gene-LOC105276449 | https://www.ncbi.nlm.nih.gov/gene/?term=LOC105276449 | 1231 | 48 | 389.93 | 1 | 8.12 | 2.08 | 25.67 | 24.69 | 0.99 | 0.99 | 0.04 | 0.04 |
| gene-LOC105274823 | https://www.ncbi.nlm.nih.gov/gene/?term=LOC105274823 | 4620 | 177 | 383.12 | 1 | 2.16 | 0.56 | 17.15 | 16.21 | 0.95 | 0.95 | 0.04 | 0.04 |
| gene-LOC105282156 | https://www.ncbi.nlm.nih.gov/gene/?term=LOC105282156 | 2577 | 127 | 492.82 | 1 | 3.88 | 0.79 | 25.13 | 24.3 | 0.82 | 0.82 | 0.04 | 0.04 |
| gene-LOC105278283 | https://www.ncbi.nlm.nih.gov/gene/?term=LOC105278283 | 2317 | 142 | 612.86 | 1 | 4.32 | 0.7 | 24.95 | 24.22 | 0.73 | 0.73 | 0.04 | 0.04 |
| gene-LOC105285055 | https://www.ncbi.nlm.nih.gov/gene/?term=LOC105285055 | 4108 | 253 | 615.87 | 1 | 2.43 | 0.4 | 10.93 | 10.3 | 0.63 | 0.63 | 0.04 | 0.04 |
| gene-LOC105283649 | https://www.ncbi.nlm.nih.gov/gene/?term=LOC105283649 | 1918 | 56 | 291.97 | 1 | 5.21 | 1.79 | 37.76 | 39.05 | -1.29 | 1.29 | -0.04 | 0.04 |
| gene-LOC105288042 | https://www.ncbi.nlm.nih.gov/gene/?term=LOC105288042 | 3507 | 100 | 285.14 | 1 | 2.85 | 1 | 20.77 | 22.23 | -1.46 | 1.46 | -0.04 | 0.04 |
| gene-LOC105278962 | https://www.ncbi.nlm.nih.gov/gene/?term=LOC105278962 | 4491 | 109 | 242.71 | 1 | 2.23 | 0.92 | 19.66 | 21.14 | -1.48 | 1.48 | -0.04 | 0.04 |
| gene-LOC105286514 | https://www.ncbi.nlm.nih.gov/gene/?term=LOC105286514 | 2528 | 50 | 197.78 | 1 | 3.96 | 2 | 35.29 | 37.3 | -2.01 | 2.01 | -0.04 | 0.04 |
| gene-LOC105281039 | https://www.ncbi.nlm.nih.gov/gene/?term=LOC105281039 | 1471 | 13 | 88.38 | 1 | 6.8 | 7.69 | 31.24 | 27.65 | 3.59 | 3.59 | 0.03 | 0.03 |
| gene-LOC105274900 | https://www.ncbi.nlm.nih.gov/gene/?term=LOC105274900 | 5561 | 61 | 109.69 | 3 | 5.39 | 4.92 | 33.5 | 30.47 | 3.02 | 3.02 | 0.03 | 0.03 |
| gene-LOC105279898 | https://www.ncbi.nlm.nih.gov/gene/?term=LOC105279898 | 1343 | 15 | 111.69 | 1 | 7.45 | 6.67 | 61.47 | 58.93 | 2.54 | 2.54 | 0.03 | 0.03 |
| gene-LOC105282652 | https://www.ncbi.nlm.nih.gov/gene/?term=LOC105282652 | 5065 | 85 | 167.82 | 1 | 1.97 | 1.18 | 16.28 | 14.6 | 1.68 | 1.68 | 0.03 | 0.03 |
| gene-LOC105276692 | https://www.ncbi.nlm.nih.gov/gene/?term=LOC105276692 | 1745 | 34 | 194.84 | 1 | 5.73 | 2.94 | 25.17 | 23.64 | 1.53 | 1.53 | 0.03 | 0.03 |
| gene-LOC113561805 | https://www.ncbi.nlm.nih.gov/gene/?term=LOC113561805 | 3337 | 75 | 224.75 | 1 | 3 | 1.33 | 34.35 | 32.91 | 1.44 | 1.44 | 0.03 | 0.03 |
| gene-LOC105280348 | https://www.ncbi.nlm.nih.gov/gene/?term=LOC105280348 | 3797 | 79 | 208.06 | 1 | 2.63 | 1.27 | 18.28 | 16.89 | 1.4 | 1.4 | 0.03 | 0.03 |
| gene-LOC105283697 | https://www.ncbi.nlm.nih.gov/gene/?term=LOC105283697 | 5301 | 153 | 288.62 | 1 | 1.89 | 0.65 | 16.46 | 15.43 | 1.02 | 1.02 | 0.03 | 0.03 |
| gene-LOC105283552 | https://www.ncbi.nlm.nih.gov/gene/?term=LOC105283552 | 1562 | 40 | 256.08 | 1 | 6.4 | 2.5 | 50.43 | 49.44 | 0.99 | 0.99 | 0.03 | 0.03 |
| gene-LOC105288079 | https://www.ncbi.nlm.nih.gov/gene/?term=LOC105288079 | 3473 | 113 | 325.37 | 1 | 2.88 | 0.88 | 17.44 | 16.56 | 0.89 | 0.89 | 0.03 | 0.03 |
| gene-LOC105282491 | https://www.ncbi.nlm.nih.gov/gene/?term=LOC105282491 | 4761 | 179 | 375.97 | 1 | 2.1 | 0.56 | 9.15 | 8.31 | 0.83 | 0.83 | 0.03 | 0.03 |
| gene-LOC105288036 | https://www.ncbi.nlm.nih.gov/gene/?term=LOC105288036 | 5110 | 200 | 391.39 | 1 | 1.96 | 0.5 | 17.43 | 16.6 | 0.83 | 0.83 | 0.03 | 0.03 |
| gene-LOC105278145 | https://www.ncbi.nlm.nih.gov/gene/?term=LOC105278145 | 4470 | 252 | 563.76 | 2 | 4.47 | 0.79 | 10.98 | 10.51 | 0.48 | 0.48 | 0.03 | 0.03 |
| gene-LOC105282658 | https://www.ncbi.nlm.nih.gov/gene/?term=LOC105282658 | 2421 | 64 | 264.35 | 1 | 4.13 | 1.56 | 23.38 | 24.34 | -0.95 | 0.95 | -0.03 | 0.03 |
| gene-LOC105284036 | https://www.ncbi.nlm.nih.gov/gene/?term=LOC105284036 | 3254 | 98 | 301.17 | 1 | 3.07 | 1.02 | 22.14 | 23.09 | -0.95 | 0.95 | -0.03 | 0.03 |
| gene-LOC105281173 | https://www.ncbi.nlm.nih.gov/gene/?term=LOC105281173 | 1595 | 38 | 238.24 | 1 | 6.27 | 2.63 | 59.87 | 61.09 | -1.22 | 1.22 | -0.03 | 0.03 |
| gene-LOC105276196 | https://www.ncbi.nlm.nih.gov/gene/?term=LOC105276196 | 2953 | 57 | 193.02 | 1 | 3.39 | 1.75 | 39.57 | 40.89 | -1.32 | 1.32 | -0.03 | 0.03 |
| gene-LOC105275571 | https://www.ncbi.nlm.nih.gov/gene/?term=LOC105275571 | 1926 | 30 | 155.76 | 1 | 5.19 | 3.33 | 42.71 | 44.75 | -2.04 | 2.04 | -0.03 | 0.03 |
| gene-LOC105276538 | https://www.ncbi.nlm.nih.gov/gene/?term=LOC105276538 | 1511 | 21 | 138.98 | 1 | 6.62 | 4.76 | 57.36 | 59.41 | -2.05 | 2.05 | -0.03 | 0.03 |
| gene-LOC105282914 | https://www.ncbi.nlm.nih.gov/gene/?term=LOC105282914 | 5623 | 43 | 76.47 | 1 | 1.78 | 2.33 | 46.2 | 49.79 | -3.59 | 3.59 | -0.03 | 0.03 |
| gene-LOC105275347 | https://www.ncbi.nlm.nih.gov/gene/?term=LOC105275347 | 2526 | 17 | 67.3 | 1 | 3.96 | 5.88 | 45.17 | 41.88 | 3.29 | 3.29 | 0.02 | 0.02 |
| gene-LOC105288066 | https://www.ncbi.nlm.nih.gov/gene/?term=LOC105288066 | 1964 | 30 | 152.75 | 1 | 5.09 | 3.33 | 38.49 | 37.01 | 1.48 | 1.48 | 0.02 | 0.02 |
| gene-LOC105282600 | https://www.ncbi.nlm.nih.gov/gene/?term=LOC105282600 | 3715 | 60 | 161.51 | 1 | 2.69 | 1.67 | 27.74 | 26.44 | 1.3 | 1.3 | 0.02 | 0.02 |
| gene-LOC105285060 | https://www.ncbi.nlm.nih.gov/gene/?term=LOC105285060 | 1285 | 16 | 124.51 | 1 | 7.78 | 6.25 | 42.55 | 41.27 | 1.27 | 1.27 | 0.02 | 0.02 |
| gene-LOC105279900 | https://www.ncbi.nlm.nih.gov/gene/?term=LOC105279900 | 2191 | 55 | 251.03 | 1 | 4.56 | 1.82 | 30.23 | 29.28 | 0.95 | 0.95 | 0.02 | 0.02 |
| gene-LOC105281848 | https://www.ncbi.nlm.nih.gov/gene/?term=LOC105281848 | 2122 | 57 | 268.61 | 1 | 4.71 | 1.75 | 23.82 | 22.98 | 0.84 | 0.84 | 0.02 | 0.02 |
| gene-LOC105285981 | https://www.ncbi.nlm.nih.gov/gene/?term=LOC105285981 | 7666 | 179 | 233.5 | 1 | 1.3 | 0.56 | 19.94 | 19.14 | 0.81 | 0.81 | 0.02 | 0.02 |
| gene-LOC105282794 | https://www.ncbi.nlm.nih.gov/gene/?term=LOC105282794 | 4571 | 104 | 227.52 | 1 | 2.19 | 0.96 | 16.85 | 16.16 | 0.69 | 0.69 | 0.02 | 0.02 |
| gene-LOC105274449 | https://www.ncbi.nlm.nih.gov/gene/?term=LOC105274449 | 1549 | 53 | 342.16 | 1 | 6.46 | 1.89 | 49.89 | 49.29 | 0.6 | 0.6 | 0.02 | 0.02 |
| gene-LOC105285179 | https://www.ncbi.nlm.nih.gov/gene/?term=LOC105285179 | 4191 | 189 | 450.97 | 1 | 2.39 | 0.53 | 18.34 | 17.82 | 0.52 | 0.52 | 0.02 | 0.02 |

| | | | | | | | | | | | | | |
|---|---|---|---|---|---|---|---|---|---|---|---|---|---|
| gene-LOC105274531 | https://www.ncbi.nlm.nih.gov/gene/?term=LOC105274531 | 7774 | 286 | 367.89 | 1 | 1.29 | 0.35 | 11.23 | 10.77 | 0.46 | 0.46 | 0.02 | 0.02 |
| gene-LOC105275363 | https://www.ncbi.nlm.nih.gov/gene/?term=LOC105275363 | 3798 | 179 | 471.3 1 | 2.63 | 0.56 | 15.94 | 15.54 | 0.4 | 0.4 | 0.02 | 0.02 | |
| gene-LOC105286355 | https://www.ncbi.nlm.nih.gov/gene/?term=LOC105286355 | 3245 | 149 | 459.17 | 1 | 3.08 | 0.67 | 21.45 | 21.07 | 0.38 | 0.38 | 0.02 | 0.02 |
| gene-LOC105278497 | https://www.ncbi.nlm.nih.gov/gene/?term=LOC105278497 | 2072 | 100 | 482.63 | 1 | 4.83 | 1 | 18.63 | 18.28 | 0.35 | 0.35 | 0.02 | 0.02 |
| gene-LOC105278632 | https://www.ncbi.nlm.nih.gov/gene/?term=LOC105278632 | 3624 | 143 | 394.59 | 1 | 2.76 | 0.7 | 13.02 | 13.42 | -0.4 | 0.4 | -0.02 | 0.02 |
| gene-LOC105284355 | https://www.ncbi.nlm.nih.gov/gene/?term=LOC105284355 | 3904 | 155 | 397.03 | 1 | 2.56 | 0.65 | 20.33 | 20.86 | -0.53 | 0.53 | -0.02 | 0.02 |
| gene-LOC105274851 | https://www.ncbi.nlm.nih.gov/gene/?term=LOC105274851 | 6616 | 144 | 217.65 | 1 | 1.51 | 0.69 | 11.65 | 12.43 | -0.79 | 0.79 | -0.02 | 0.02 |
| gene-LOC105279596 | https://www.ncbi.nlm.nih.gov/gene/?term=LOC105279596 | 7695 | 115 | 149.45 | 1 | 1.3 | 0.87 | 16.77 | 17.78 | -1 | 1 | -0.02 | 0.02 |
| gene-LOC105274590 | https://www.ncbi.nlm.nih.gov/gene/?term=LOC105274590 | 4516 | 82 | 181.58 | 1 | 2.21 | 1.22 | 25.47 | 26.68 | -1.21 | 1.21 | -0.02 | 0.02 |
| gene-LOC105281515 | https://www.ncbi.nlm.nih.gov/gene/?term=LOC105281515 | 8827 | 57 | 64.57 2 | 2.27 | 3.51 | 32.7 | 30.37 | 2.32 | 2.32 | 0.01 | 0.01 | |
| gene-LOC105279528 | https://www.ncbi.nlm.nih.gov/gene/?term=LOC105279528 | 5802 | 40 | 68.94 1 | 1.72 | 2.5 | 21.84 | 20.71 | 1.12 | 1.12 | 0.01 | 0.01 | |
| gene-LOC105275187 | https://www.ncbi.nlm.nih.gov/gene/?term=LOC105275187 | 5534 | 73 | 131.91 | 1 | 1.81 | 1.37 | 14.7 | 13.62 | 1.08 | 1.08 | 0.01 | 0.01 |
| gene-LOC105283943 | https://www.ncbi.nlm.nih.gov/gene/?term=LOC105283943 | 3559 | 57 | 160.16 | 1 | 2.81 | 1.75 | 23.04 | 22.27 | 0.77 | 0.77 | 0.01 | 0.01 |
| gene-LOC105288237 | https://www.ncbi.nlm.nih.gov/gene/?term=LOC105288237 | 2226 | 49 | 220.13 | 1 | 4.49 | 2.04 | 43.01 | 42.34 | 0.67 | 0.67 | 0.01 | 0.01 |
| gene-LOC105275451 | https://www.ncbi.nlm.nih.gov/gene/?term=LOC105275451 | 6035 | 135 | 223.7 1 | 1.66 | 0.74 | 21.36 | 20.75 | 0.61 | 0.61 | 0.01 | 0.01 | |
| gene-LOC105277215 | https://www.ncbi.nlm.nih.gov/gene/?term=LOC105277215 | 2619 | 29 | 110.73 | 1 | 3.82 | 3.45 | 25.25 | 24.64 | 0.61 | 0.61 | 0.01 | 0.01 |
| gene-LOC105288078 | https://www.ncbi.nlm.nih.gov/gene/?term=LOC105288078 | 19126 | 337 | 176.2 1 | 0.52 | 0.3 | 15.64 | 15.05 | 0.59 | 0.59 | 0.01 | 0.01 | |
| gene-LOC105280819 | https://www.ncbi.nlm.nih.gov/gene/?term=LOC105280819 | 1639 | 42 | 256.25 | 1 | 6.1 | 2.38 | 39.26 | 38.69 | 0.58 | 0.58 | 0.01 | 0.01 |
| gene-LOC105280641 | https://www.ncbi.nlm.nih.gov/gene/?term=LOC105280641 | 3682 | 75 | 203.69 | 1 | 2.72 | 1.33 | 13.21 | 12.66 | 0.55 | 0.55 | 0.01 | 0.01 |
| gene-LOC105287417 | https://www.ncbi.nlm.nih.gov/gene/?term=LOC105287417 | 1025 | 32 | 312.2 1 | 9.76 | 3.12 | 37.69 | 37.23 | 0.46 | 0.46 | 0.01 | 0.01 | |
| gene-LOC105277903 | https://www.ncbi.nlm.nih.gov/gene/?term=LOC105277903 | 1835 | 37 | 201.63 | 1 | 5.45 | 2.7 | 54.5 | 54.08 | 0.42 | 0.42 | 0.01 | 0.01 |
| gene-LOC105283592 | https://www.ncbi.nlm.nih.gov/gene/?term=LOC105283592 | 3479 | 80 | 229.95 | 1 | 2.87 | 1.25 | 21.51 | 21.14 | 0.37 | 0.37 | 0.01 | 0.01 |
| gene-LOC105283838 | https://www.ncbi.nlm.nih.gov/gene/?term=LOC105283838 | 1748 | 30 | 171.62 | 1 | 5.72 | 3.33 | 33.04 | 32.68 | 0.36 | 0.36 | 0.01 | 0.01 |
| gene-LOC105278384 | https://www.ncbi.nlm.nih.gov/gene/?term=LOC105278384 | 1390 | 33 | 237.41 | 1 | 7.19 | 3.03 | 46.11 | 45.77 | 0.34 | 0.34 | 0.01 | 0.01 |
| gene-LOC105285639 | https://www.ncbi.nlm.nih.gov/gene/?term=LOC105285639 | 10325 | 351 | 339.95 | 2 | 1.94 | 0.57 | 11.49 | 11.2 | 0.29 | 0.29 | 0.01 | 0.01 |
| gene-LOC105279298 | https://www.ncbi.nlm.nih.gov/gene/?term=LOC105279298 | 2773 | 127 | 457.99 | 1 | 3.61 | 0.79 | 24.14 | 23.89 | 0.26 | 0.26 | 0.01 | 0.01 |
| gene-LOC105283123 | https://www.ncbi.nlm.nih.gov/gene/?term=LOC105283123 | 1507 | 47 | 311.88 | 1 | 6.64 | 2.13 | 20.21 | 19.96 | 0.25 | 0.25 | 0.01 | 0.01 |
| gene-LOC105284184 | https://www.ncbi.nlm.nih.gov/gene/?term=LOC105284184 | 2535 | 92 | 362.92 | 1 | 3.94 | 1.09 | 31.61 | 31.41 | 0.2 | 0.2 | 0.01 | 0.01 |
| gene-LOC105286997 | https://www.ncbi.nlm.nih.gov/gene/?term=LOC105286997 | 2850 | 142 | 498.25 | 1 | 3.51 | 0.7 | 11.2 | 11.03 | 0.17 | 0.17 | 0.01 | 0.01 |
| gene-LOC105275374 | https://www.ncbi.nlm.nih.gov/gene/?term=LOC105275374 | 11376 | 514 | 451.83 | 1 | 0.88 | 0.19 | 9.9 | 9.75 | 0.15 | 0.15 | 0.01 | 0.01 |
| gene-LOC105283728 | https://www.ncbi.nlm.nih.gov/gene/?term=LOC105283728 | 3256 | 152 | 466.83 | 2 | 6.14 | 1.32 | 15.71 | 15.57 | 0.14 | 0.14 | 0.01 | 0.01 |
| gene-LOC105276527 | https://www.ncbi.nlm.nih.gov/gene/?term=LOC105276527 | 7923 | 338 | 426.61 | 1 | 1.26 | 0.3 | 8.29 | 8.43 | -0.14 | 0.14 | -0.01 | 0.01 |
| gene-LOC105279038 | https://www.ncbi.nlm.nih.gov/gene/?term=LOC105279038 | 9453 | 329 | 348.04 | 1 | 1.06 | 0.3 | 9.14 | 9.32 | -0.18 | 0.18 | -0.01 | 0.01 |
| gene-LOC105282811 | https://www.ncbi.nlm.nih.gov/gene/?term=LOC105282811 | 5765 | 203 | 352.12 | 1 | 1.73 | 0.49 | 8.59 | 8.78 | -0.19 | 0.19 | -0.01 | 0.01 |
| gene-LOC105274856 | https://www.ncbi.nlm.nih.gov/gene/?term=LOC105274856 | 5216 | 228 | 437.12 | 1 | 1.92 | 0.44 | 17.7 | 17.9 | -0.2 | 0.2 | -0.01 | 0.01 |
| gene-LOC105285807 | https://www.ncbi.nlm.nih.gov/gene/?term=LOC105285807 | 6063 | 215 | 354.61 | 1 | 1.65 | 0.47 | 8.65 | 8.94 | -0.28 | 0.28 | -0.01 | 0.01 |
| gene-LOC105274706 | https://www.ncbi.nlm.nih.gov/gene/?term=LOC105274706 | 1446 | 50 | 345.78 | 1 | 6.92 | 2 | 37.45 | 37.82 | -0.37 | 0.37 | -0.01 | 0.01 |
| gene-LOC105282825 | https://www.ncbi.nlm.nih.gov/gene/?term=LOC105282825 | 3919 | 96 | 244.96 | 1 | 2.55 | 1.04 | 14.53 | 14.91 | -0.38 | 0.38 | -0.01 | 0.01 |
| gene-LOC105280002 | https://www.ncbi.nlm.nih.gov/gene/?term=LOC105280002 | 1940 | 31 | 159.79 | 1 | 5.15 | 3.23 | 31.38 | 32.27 | -0.88 | 0.88 | -0.01 | 0.01 |
| gene-LOC105275518 | https://www.ncbi.nlm.nih.gov/gene/?term=LOC105275518 | 2335 | 23 | 98.5 1 | 4.28 | 4.35 | 45.78 | 46.9 | -1.12 | 1.12 | -0.01 | 0.01 | |
| gene-LOC105283989 | https://www.ncbi.nlm.nih.gov/gene/?term=LOC105283989 | 4948 | 32 | 64.67 1 | 2.02 | 3.12 | 46.57 | 45.88 | 0.69 | 0.69 | 0 | 0 | |
| gene-LOC105286452 | https://www.ncbi.nlm.nih.gov/gene/?term=LOC105286452 | 6888 | 68 | 98.72 1 | 1.45 | 1.47 | 28.16 | 27.75 | 0.4 | 0.4 | 0 | 0 | |
| gene-LOC105288010 | https://www.ncbi.nlm.nih.gov/gene/?term=LOC105288010 | 3189 | 52 | 163.06 | 1 | 3.14 | 1.92 | 29.32 | 29.04 | 0.28 | 0.28 | 0 | 0 |
| gene-LOC105284329 | https://www.ncbi.nlm.nih.gov/gene/?term=LOC105284329 | 4051 | 71 | 175.27 | 1 | 2.47 | 1.41 | 19.25 | 18.99 | 0.25 | 0.25 | 0 | 0 |
| gene-LOC105275170 | https://www.ncbi.nlm.nih.gov/gene/?term=LOC105275170 | 12918 | 350 | 270.94 | 1 | 0.77 | 0.29 | 1.94 | 1.8 | 0.15 | 0.15 | 0 | 0 |
| gene-LOC105276059 | https://www.ncbi.nlm.nih.gov/gene/?term=LOC105276059 | 6256 | 134 | 214.19 | 1 | 1.6 | 0.75 | 11.07 | 10.91 | 0.15 | 0.15 | 0 | 0 |
| gene-LOC105286099 | https://www.ncbi.nlm.nih.gov/gene/?term=LOC105286099 | 5919 | 70 | 118.26 | 1 | 1.69 | 1.43 | 12.4 | 12.26 | 0.14 | 0.14 | 0 | 0 |
| gene-LOC105278534 | https://www.ncbi.nlm.nih.gov/gene/?term=LOC105278534 | 7364 | 198 | 268.88 | 1 | 1.36 | 0.51 | 17.97 | 17.87 | 0.1 | 0.1 | 0 | 0 |
| gene-LOC105280774 | https://www.ncbi.nlm.nih.gov/gene/?term=LOC105280774 | 2793 | 132 | 472.61 | 1 | 3.58 | 0.76 | 25.2 | 25.1 | 0.1 | 0.1 | 0 | 0 |
| gene-LOC105282149 | https://www.ncbi.nlm.nih.gov/gene/?term=LOC105282149 | 1787 | 103 | 576.39 | 1 | 5.6 | 0.97 | 41.42 | 41.34 | 0.08 | 0.08 | 0 | 0 |
| gene-LOC105278666 | https://www.ncbi.nlm.nih.gov/gene/?term=LOC105278666 | 10341 | 474 | 458.37 | 1 | 0.97 | 0.21 | 6.99 | 6.95 | 0.05 | 0.05 | 0 | 0 |
| gene-LOC105280956 | https://www.ncbi.nlm.nih.gov/gene/?term=LOC105280956 | 2639 | 80 | 303.15 | 1 | 3.79 | 1.25 | 18.94 | 18.89 | 0.05 | 0.05 | 0 | 0 |
| gene-LOC105275327 | https://www.ncbi.nlm.nih.gov/gene/?term=LOC105275327 | 14245 | 529 | 371.36 | 1 | 0.7 | 0.19 | 6.66 | 6.63 | 0.03 | 0.03 | 0 | 0 |
| gene-LOC105284043 | https://www.ncbi.nlm.nih.gov/gene/?term=LOC105284043 | 5661 | 284 | 501.68 | 1 | 1.77 | 0.35 | 10.32 | 10.3 | 0.03 | 0.03 | 0 | 0 |
| gene-LOC105277216 | https://www.ncbi.nlm.nih.gov/gene/?term=LOC105277216 | 7627 | 106 | 138.98 | 1 | 1.31 | 0.94 | 7.36 | 7.34 | 0.02 | 0.02 | 0 | 0 |
| gene-LOC105279500 | https://www.ncbi.nlm.nih.gov/gene/?term=LOC105279500 | 3587 | 32 | 89.21 1 | 2.79 | 3.12 | 46.58 | 46.57 | 0.01 | 0.01 | 0 | 0 | |
| gene-LOC105286515 | https://www.ncbi.nlm.nih.gov/gene/?term=LOC105286515 | 3751 | 117 | 311.92 | 1 | 2.67 | 0.85 | 10.68 | 10.68 | 0 | 0 | 0 | 0 |
| gene-LOC105278228 | https://www.ncbi.nlm.nih.gov/gene/?term=LOC105278228 | 8610 | 176 | 204.41 | 1 | 1.16 | 0.57 | 22.62 | 22.66 | -0.04 | 0.04 | 0 | 0 |
| gene-LOC105278877 | https://www.ncbi.nlm.nih.gov/gene/?term=LOC105278877 | 1968 | 87 | 442.07 | 1 | 5.08 | 1.15 | 29.15 | 29.21 | -0.06 | 0.06 | 0 | 0 |
| gene-LOC105287095 | https://www.ncbi.nlm.nih.gov/gene/?term=LOC105287095 | 1435 | 66 | 459.93 | 1 | 6.97 | 1.52 | 24.97 | 25.03 | -0.06 | 0.06 | 0 | 0 |
| gene-LOC105279664 | https://www.ncbi.nlm.nih.gov/gene/?term=LOC105279664 | 4023 | 64 | 159.09 | 1 | 2.49 | 1.56 | 17.75 | 17.94 | -0.19 | 0.19 | 0 | 0 |
| gene-LOC105277856 | https://www.ncbi.nlm.nih.gov/gene/?term=LOC105277856 | 3442 | 33 | 95.87 1 | 2.91 | 3.03 | 39.28 | 39.53 | -0.25 | 0.25 | 0 | 0 | |

Directory: OGL File: rnk-dmg-Ob_LI_RphaseA1.vs.RphaseB2.txt

| gene_ID | gene_link | gwidth | #Sites | #per10Kb | #dmSites | #dmsp10kb | %dmSites | %pSite1 | %pSite2 | DMpSite | ADMpSite | DMpNucl | ADMpNucl |
|---|---|---|---|---|---|---|---|---|---|---|---|---|---|
| gene-LOC113562345 | https://www.ncbi.nlm.nih.gov/gene/?term=LOC113562345 | 2290 | 67 | 292.58 | 6 | 26.2 | 8.96 | 28.84 | 16.72 | 12.12 | 12.12 | 0.35 | 0.35 |
| gene-LOC105285671 | https://www.ncbi.nlm.nih.gov/gene/?term=LOC105285671 | 2263 | 101 | 446.31 | 1 | 4.42 | 0.99 | 30.8 | 27.84 | 2.96 | 2.96 | 0.13 | 0.13 |
| gene-LOC105275070 | https://www.ncbi.nlm.nih.gov/gene/?term=LOC105275070 | 5667 | 37 | 65.29 | 7 | 12.35 | 18.92 | 16.3 | 1.04 | 15.26 | 15.26 | 0.1 | 0.1 |
| gene-LOC105283552 | https://www.ncbi.nlm.nih.gov/gene/?term=LOC105283552 | 1562 | 41 | 262.48 | 1 | 6.4 | 2.44 | 50.09 | 46.71 | 3.37 | 3.37 | 0.09 | 0.09 |
| gene-LOC105279356 | https://www.ncbi.nlm.nih.gov/gene/?term=LOC105279356 | 1314 | 43 | 327.25 | 1 | 7.61 | 2.33 | 32.62 | 30.3 | 2.31 | 2.31 | 0.08 | 0.08 |
| gene-LOC105278436 | https://www.ncbi.nlm.nih.gov/gene/?term=LOC105278436 | 2556 | 54 | 211.27 | 1 | 3.91 | 1.85 | 40.35 | 37.22 | 3.13 | 3.13 | 0.07 | 0.07 |
| gene-LOC105285202 | https://www.ncbi.nlm.nih.gov/gene/?term=LOC105285202 | 1838 | 61 | 331.88 | 1 | 5.44 | 1.64 | 26.9 | 24.8 | 2.1 | 2.1 | 0.07 | 0.07 |
| gene-LOC105279898 | https://www.ncbi.nlm.nih.gov/gene/?term=LOC105279898 | 1343 | 9 | 67.01 | 1 | 7.45 | 11.11 | 83.37 | 74.78 | 8.59 | 8.59 | 0.06 | 0.06 |
| gene-LOC105275708 | https://www.ncbi.nlm.nih.gov/gene/?term=LOC105275708 | 1772 | 14 | 79.01 | 1 | 5.64 | 7.14 | 50.83 | 43.74 | 7.08 | 7.08 | 0.06 | 0.06 |
| gene-LOC105285674 | https://www.ncbi.nlm.nih.gov/gene/?term=LOC105285674 | 3035 | 92 | 303.13 | 2 | 6.59 | 2.17 | 26.97 | 24.84 | 2.13 | 2.13 | 0.06 | 0.06 |
| gene-LOC105280786 | https://www.ncbi.nlm.nih.gov/gene/?term=LOC105280786 | 2573 | 115 | 446.95 | 1 | 3.89 | 0.87 | 28.39 | 27.06 | 1.33 | 1.33 | 0.06 | 0.06 |
| gene-LOC105280596 | https://www.ncbi.nlm.nih.gov/gene/?term=LOC105280596 | 3434 | 104 | 302.85 | 1 | 2.91 | 0.96 | 16.78 | 18.68 | -1.9 | 1.9 | -0.06 | 0.06 |
| gene-LOC105276196 | https://www.ncbi.nlm.nih.gov/gene/?term=LOC105276196 | 2953 | 59 | 199.8 | 1 | 3.39 | 1.69 | 38.71 | 41.49 | -2.77 | 2.77 | -0.06 | 0.06 |
| gene-LOC105285514 | https://www.ncbi.nlm.nih.gov/gene/?term=LOC105285514 | 1277 | 14 | 109.63 | 1 | 7.83 | 7.14 | 69.69 | 65.1 | 4.59 | 4.59 | 0.05 | 0.05 |
| gene-LOC105278807 | https://www.ncbi.nlm.nih.gov/gene/?term=LOC105278807 | 2166 | 65 | 300.09 | 1 | 4.62 | 1.54 | 22.17 | 20.43 | 1.74 | 1.74 | 0.05 | 0.05 |
| gene-LOC105280104 | https://www.ncbi.nlm.nih.gov/gene/?term=LOC105280104 | 3960 | 123 | 310.61 | 1 | 2.53 | 0.81 | 18.15 | 16.51 | 1.64 | 1.64 | 0.05 | 0.05 |
| gene-LOC105275363 | https://www.ncbi.nlm.nih.gov/gene/?term=LOC105275363 | 3798 | 157 | 413.38 | 1 | 2.63 | 0.64 | 17.94 | 16.87 | 1.07 | 1.07 | 0.04 | 0.04 |
| gene-LOC105284043 | https://www.ncbi.nlm.nih.gov/gene/?term=LOC105284043 | 5661 | 283 | 499.91 | 1 | 1.77 | 0.35 | 10.38 | 9.58 | 0.79 | 0.79 | 0.04 | 0.04 |
| gene-LOC105282156 | https://www.ncbi.nlm.nih.gov/gene/?term=LOC105282156 | 2577 | 122 | 473.42 | 1 | 3.88 | 0.82 | 25.73 | 24.95 | 0.78 | 0.78 | 0.04 | 0.04 |
| gene-LOC105285534 | https://www.ncbi.nlm.nih.gov/gene/?term=LOC105285534 | 3944 | 232 | 588.24 | 1 | 2.54 | 0.43 | 11.59 | 10.9 | 0.69 | 0.69 | 0.04 | 0.04 |
| gene-LOC105285055 | https://www.ncbi.nlm.nih.gov/gene/?term=LOC105285055 | 4108 | 261 | 635.35 | 1 | 2.43 | 0.38 | 10.75 | 10.11 | 0.64 | 0.64 | 0.04 | 0.04 |
| gene-LOC105278283 | https://www.ncbi.nlm.nih.gov/gene/?term=LOC105278283 | 2317 | 138 | 595.6 | 1 | 4.32 | 0.72 | 25.64 | 25.05 | 0.59 | 0.59 | 0.04 | 0.04 |
| gene-LOC105274534 | https://www.ncbi.nlm.nih.gov/gene/?term=LOC105274534 | 1329 | 57 | 428.89 | 1 | 7.52 | 1.75 | 46.71 | 47.65 | -0.94 | 0.94 | -0.04 | 0.04 |
| gene-LOC105286999 | https://www.ncbi.nlm.nih.gov/gene/?term=LOC105286999 | 2141 | 42 | 196.17 | 1 | 4.67 | 2.38 | 33.74 | 35.94 | -2.2 | 2.2 | -0.04 | 0.04 |
| gene-LOC105283967 | https://www.ncbi.nlm.nih.gov/gene/?term=LOC105283967 | 2393 | 40 | 167.15 | 1 | 4.18 | 2.5 | 52.37 | 54.65 | -2.28 | 2.28 | -0.04 | 0.04 |
| gene-LOC105283492 | https://www.ncbi.nlm.nih.gov/gene/?term=LOC105283492 | 1535 | 9 | 58.63 | 1 | 6.51 | 11.11 | 30.57 | 36.97 | -6.4 | 6.4 | -0.04 | 0.04 |
| gene-LOC105274900 | https://www.ncbi.nlm.nih.gov/gene/?term=LOC105274900 | 5561 | 55 | 98.9 | 3 | 5.39 | 5.45 | 32.79 | 29.47 | 3.31 | 3.31 | 0.03 | 0.03 |
| gene-LOC105281266 | https://www.ncbi.nlm.nih.gov/gene/?term=LOC105281266 | 4102 | 78 | 190.15 | 1 | 2.44 | 1.28 | 18.58 | 17.01 | 1.57 | 1.57 | 0.03 | 0.03 |
| gene-LOC105282794 | https://www.ncbi.nlm.nih.gov/gene/?term=LOC105282794 | 4571 | 90 | 196.89 | 1 | 2.19 | 1.11 | 17.25 | 15.76 | 1.49 | 1.49 | 0.03 | 0.03 |
| gene-LOC105277903 | https://www.ncbi.nlm.nih.gov/gene/?term=LOC105277903 | 1835 | 36 | 196.19 | 1 | 5.45 | 2.78 | 51.23 | 49.79 | 1.45 | 1.45 | 0.03 | 0.03 |
| gene-LOC105280956 | https://www.ncbi.nlm.nih.gov/gene/?term=LOC105280956 | 2639 | 71 | 269.04 | 1 | 3.79 | 1.41 | 18.77 | 17.79 | 0.98 | 0.98 | 0.03 | 0.03 |
| gene-LOC105288036 | https://www.ncbi.nlm.nih.gov/gene/?term=LOC105288036 | 5110 | 181 | 354.21 | 1 | 1.96 | 0.55 | 18.99 | 18.01 | 0.98 | 0.98 | 0.03 | 0.03 |
| gene-LOC105283123 | https://www.ncbi.nlm.nih.gov/gene/?term=LOC105283123 | 1507 | 40 | 265.43 | 1 | 6.64 | 2.5 | 16.19 | 15.23 | 0.96 | 0.96 | 0.03 | 0.03 |
| gene-LOC113561621 | https://www.ncbi.nlm.nih.gov/gene/?term=LOC113561621 | 2249 | 76 | 337.93 | 1 | 4.45 | 1.32 | 22.44 | 21.49 | 0.95 | 0.95 | 0.03 | 0.03 |
| gene-LOC105284184 | https://www.ncbi.nlm.nih.gov/gene/?term=LOC105284184 | 2535 | 80 | 315.58 | 1 | 3.94 | 1.25 | 35.73 | 34.87 | 0.86 | 0.86 | 0.03 | 0.03 |
| gene-LOC105276612 | https://www.ncbi.nlm.nih.gov/gene/?term=LOC105276612 | 4157 | 178 | 428.19 | 1 | 2.41 | 0.56 | 10.68 | 9.88 | 0.8 | 0.8 | 0.03 | 0.03 |
| gene-LOC105283728 | https://www.ncbi.nlm.nih.gov/gene/?term=LOC105283728 | 3256 | 117 | 359.34 | 2 | 6.14 | 1.71 | 18.77 | 18.07 | 0.7 | 0.7 | 0.03 | 0.03 |
| gene-LOC105275460 | https://www.ncbi.nlm.nih.gov/gene/?term=LOC105275460 | 2473 | 118 | 477.15 | 1 | 4.04 | 0.85 | 13.95 | 13.28 | 0.67 | 0.67 | 0.03 | 0.03 |
| gene-LOC105279298 | https://www.ncbi.nlm.nih.gov/gene/?term=LOC105279298 | 2773 | 133 | 479.62 | 2 | 7.21 | 1.5 | 23.97 | 23.38 | 0.59 | 0.59 | 0.03 | 0.03 |
| gene-LOC105278145 | https://www.ncbi.nlm.nih.gov/gene/?term=LOC105278145 | 4470 | 235 | 525.73 | 1 | 2.24 | 0.43 | 11.34 | 10.86 | 0.48 | 0.48 | 0.03 | 0.03 |
| gene-LOC105280819 | https://www.ncbi.nlm.nih.gov/gene/?term=LOC105280819 | 1639 | 37 | 225.75 | 1 | 6.1 | 2.7 | 36.94 | 38.07 | -1.12 | 1.12 | -0.03 | 0.03 |
| gene-LOC105278384 | https://www.ncbi.nlm.nih.gov/gene/?term=LOC105278384 | 1390 | 35 | 251.8 | 1 | 7.19 | 2.86 | 48.96 | 50.26 | -1.3 | 1.3 | -0.03 | 0.03 |
| gene-LOC105283712 | https://www.ncbi.nlm.nih.gov/gene/?term=LOC105283712 | 1910 | 36 | 188.48 | 1 | 5.24 | 2.78 | 22.92 | 24.43 | -1.51 | 1.51 | -0.03 | 0.03 |
| gene-LOC105286514 | https://www.ncbi.nlm.nih.gov/gene/?term=LOC105286514 | 2528 | 46 | 181.96 | 1 | 3.96 | 2.17 | 35.96 | 37.79 | -1.83 | 1.83 | -0.03 | 0.03 |
| gene-LOC105285409 | https://www.ncbi.nlm.nih.gov/gene/?term=LOC105285409 | 5099 | 49 | 96.1 | 1 | 1.96 | 2.04 | 1.72 | 4.82 | -3.09 | 3.09 | -0.03 | 0.03 |
| gene-LOC105286694 | https://www.ncbi.nlm.nih.gov/gene/?term=LOC105286694 | 3409 | 8 | 23.47 | 1 | 2.93 | 12.5 | 66.22 | 57.86 | 8.36 | 8.36 | 0.02 | 0.02 |
| gene-LOC105281515 | https://www.ncbi.nlm.nih.gov/gene/?term=LOC105281515 | 8827 | 41 | 46.45 | 1 | 1.13 | 2.44 | 31.51 | 28.2 | 3.31 | 3.31 | 0.02 | 0.02 |
| gene-LOC105286099 | https://www.ncbi.nlm.nih.gov/gene/?term=LOC105286099 | 5919 | 67 | 113.19 | 1 | 1.69 | 1.49 | 11.4 | 9.83 | 1.57 | 1.57 | 0.02 | 0.02 |
| gene-LOC105284329 | https://www.ncbi.nlm.nih.gov/gene/?term=LOC105284329 | 4051 | 52 | 128.36 | 1 | 2.47 | 1.92 | 19.88 | 18.38 | 1.5 | 1.5 | 0.02 | 0.02 |
| gene-LOC105288066 | https://www.ncbi.nlm.nih.gov/gene/?term=LOC105288066 | 1964 | 27 | 137.47 | 1 | 5.09 | 3.7 | 34.07 | 32.8 | 1.27 | 1.27 | 0.02 | 0.02 |
| gene-LOC105275451 | https://www.ncbi.nlm.nih.gov/gene/?term=LOC105275451 | 6035 | 113 | 187.24 | 1 | 1.66 | 0.88 | 24.46 | 23.26 | 1.2 | 1.2 | 0.02 | 0.02 |
| gene-LOC105279664 | https://www.ncbi.nlm.nih.gov/gene/?term=LOC105279664 | 4023 | 53 | 131.74 | 1 | 2.49 | 1.89 | 19.07 | 17.88 | 1.19 | 1.19 | 0.02 | 0.02 |
| gene-LOC113561805 | https://www.ncbi.nlm.nih.gov/gene/?term=LOC113561805 | 3337 | 67 | 200.78 | 1 | 3 | 1.49 | 36.13 | 35.02 | 1.11 | 1.11 | 0.02 | 0.02 |
| gene-LOC105281173 | https://www.ncbi.nlm.nih.gov/gene/?term=LOC105281173 | 1595 | 38 | 238.24 | 1 | 6.27 | 2.63 | 59.87 | 58.85 | 1.02 | 1.02 | 0.02 | 0.02 |
| gene-LOC105276692 | https://www.ncbi.nlm.nih.gov/gene/?term=LOC105276692 | 1745 | 29 | 166.19 | 1 | 5.73 | 3.45 | 25.96 | 24.96 | 1 | 1 | 0.02 | 0.02 |
| gene-LOC105276059 | https://www.ncbi.nlm.nih.gov/gene/?term=LOC105276059 | 6256 | 125 | 199.81 | 1 | 1.6 | 0.8 | 10.91 | 10 | 0.91 | 0.91 | 0.02 | 0.02 |
| gene-LOC105283592 | https://www.ncbi.nlm.nih.gov/gene/?term=LOC105283592 | 3479 | 71 | 204.08 | 1 | 2.87 | 1.41 | 21.54 | 20.67 | 0.88 | 0.88 | 0.02 | 0.02 |
| gene-LOC105276901 | https://www.ncbi.nlm.nih.gov/gene/?term=LOC105276901 | 3254 | 84 | 258.14 | 1 | 3.07 | 1.19 | 27.21 | 26.37 | 0.84 | 0.84 | 0.02 | 0.02 |
| gene-LOC105274823 | https://www.ncbi.nlm.nih.gov/gene/?term=LOC105274823 | 4620 | 138 | 298.7 | 1 | 2.16 | 0.72 | 21.4 | 20.58 | 0.82 | 0.82 | 0.02 | 0.02 |
| gene-LOC105285639 | https://www.ncbi.nlm.nih.gov/gene/?term=LOC105285639 | 10325 | 312 | 302.18 | 2 | 1.94 | 0.64 | 12.89 | 12.24 | 0.65 | 0.65 | 0.02 | 0.02 |

| | | | | | | | | | | | | | |
|---|---|---|---|---|---|---|---|---|---|---|---|---|---|
| gene-LOC105278534 | https://www.ncbi.nlm.nih.gov/gene/?term=LOC105278534 | 7364 | 176 | 239 | 2 | 2.72 | 1.14 | 19.64 | 19.01 | 0.63 | 0.63 | 0.02 | 0.02 |
| gene-LOC105285179 | https://www.ncbi.nlm.nih.gov/gene/?term=LOC105285179 | 4191 | 171 | 408.02 | 1 | 2.39 | 0.58 | 19 | 18.45 | 0.55 | 0.55 | 0.02 | 0.02 |
| gene-LOC105278497 | https://www.ncbi.nlm.nih.gov/gene/?term=LOC105278497 | 2072 | 95 | 458.49 | 1 | 4.83 | 1.05 | 19.39 | 18.89 | 0.5 | 0.5 | 0.02 | 0.02 |
| gene-LOC105286997 | https://www.ncbi.nlm.nih.gov/gene/?term=LOC105286997 | 2850 | 106 | 371.93 | 1 | 3.51 | 0.94 | 12.07 | 11.62 | 0.46 | 0.46 | 0.02 | 0.02 |
| gene-LOC105275374 | https://www.ncbi.nlm.nih.gov/gene/?term=LOC105275374 | 11376 | 454 | 399.09 | 1 | 0.88 | 0.22 | 10.82 | 10.43 | 0.38 | 0.38 | 0.02 | 0.02 |
| gene-LOC105274449 | https://www.ncbi.nlm.nih.gov/gene/?term=LOC105274449 | 1549 | 48 | 309.88 | 1 | 6.46 | 2.08 | 50.45 | 50.96 | -0.51 | 0.51 | -0.02 | 0.02 |
| gene-LOC105278877 | https://www.ncbi.nlm.nih.gov/gene/?term=LOC105278877 | 1968 | 70 | 355.69 | 1 | 5.08 | 1.43 | 34.63 | 35.21 | -0.58 | 0.58 | -0.02 | 0.02 |
| gene-LOC105280377 | https://www.ncbi.nlm.nih.gov/gene/?term=LOC105280377 | 2024 | 65 | 321.15 | 1 | 4.94 | 1.54 | 47.38 | 47.97 | -0.59 | 0.59 | -0.02 | 0.02 |
| gene-LOC105281700 | https://www.ncbi.nlm.nih.gov/gene/?term=LOC105281700 | 2208 | 32 | 144.93 | 1 | 4.53 | 3.12 | 37.54 | 39.09 | -1.55 | 1.55 | -0.02 | 0.02 |
| gene-LOC105276538 | https://www.ncbi.nlm.nih.gov/gene/?term=LOC105276538 | 1511 | 18 | 119.13 | 1 | 6.62 | 5.56 | 64.69 | 66.35 | -1.65 | 1.65 | -0.02 | 0.02 |
| gene-LOC105275347 | https://www.ncbi.nlm.nih.gov/gene/?term=LOC105275347 | 2526 | 16 | 63.34 | 1 | 3.96 | 6.25 | 41.65 | 39.67 | 1.98 | 1.98 | 0.01 | 0.01 |
| gene-LOC105283943 | https://www.ncbi.nlm.nih.gov/gene/?term=LOC105283943 | 3559 | 45 | 126.44 | 1 | 2.81 | 2.22 | 16.9 | 15.77 | 1.14 | 1.14 | 0.01 | 0.01 |
| gene-LOC105288078 | https://www.ncbi.nlm.nih.gov/gene/?term=LOC105288078 | 19126 | 286 | 149.53 | 1 | 0.52 | 0.35 | 17.66 | 16.67 | 0.99 | 0.99 | 0.01 | 0.01 |
| gene-LOC105285060 | https://www.ncbi.nlm.nih.gov/gene/?term=LOC105285060 | 1285 | 16 | 124.51 | 1 | 7.78 | 6.25 | 47.94 | 47.04 | 0.9 | 0.9 | 0.01 | 0.01 |
| gene-LOC105287854 | https://www.ncbi.nlm.nih.gov/gene/?term=LOC105287854 | 3644 | 49 | 134.47 | 1 | 2.74 | 2.04 | 24.37 | 23.53 | 0.84 | 0.84 | 0.01 | 0.01 |
| gene-LOC105287417 | https://www.ncbi.nlm.nih.gov/gene/?term=LOC105287417 | 1025 | 19 | 185.37 | 1 | 9.76 | 5.26 | 59.44 | 58.65 | 0.79 | 0.79 | 0.01 | 0.01 |
| gene-LOC105286452 | https://www.ncbi.nlm.nih.gov/gene/?term=LOC105286452 | 6888 | 64 | 92.92 | 1 | 1.45 | 1.56 | 30.17 | 29.46 | 0.72 | 0.72 | 0.01 | 0.01 |
| gene-LOC105281848 | https://www.ncbi.nlm.nih.gov/gene/?term=LOC105281848 | 2122 | 47 | 221.49 | 1 | 4.71 | 2.13 | 27.19 | 26.57 | 0.63 | 0.63 | 0.01 | 0.01 |
| gene-LOC105275179 | https://www.ncbi.nlm.nih.gov/gene/?term=LOC105275179 | 3271 | 69 | 210.94 | 1 | 3.06 | 1.45 | 18.8 | 18.36 | 0.45 | 0.45 | 0.01 | 0.01 |
| gene-LOC105287646 | https://www.ncbi.nlm.nih.gov/gene/?term=LOC105287646 | 3556 | 83 | 233.41 | 1 | 2.81 | 1.2 | 17.22 | 16.85 | 0.38 | 0.38 | 0.01 | 0.01 |
| gene-LOC105287550 | https://www.ncbi.nlm.nih.gov/gene/?term=LOC105287550 | 15659 | 549 | 350.6 | 1 | 0.64 | 0.18 | 6.33 | 5.97 | 0.37 | 0.37 | 0.01 | 0.01 |
| gene-LOC105274856 | https://www.ncbi.nlm.nih.gov/gene/?term=LOC105274856 | 5216 | 209 | 400.69 | 1 | 1.92 | 0.48 | 18.76 | 18.44 | 0.31 | 0.31 | 0.01 | 0.01 |
| gene-LOC105274699 | https://www.ncbi.nlm.nih.gov/gene/?term=LOC105274699 | 2386 | 49 | 205.36 | 1 | 4.19 | 2.04 | 23.32 | 23.03 | 0.29 | 0.29 | 0.01 | 0.01 |
| gene-LOC105274531 | https://www.ncbi.nlm.nih.gov/gene/?term=LOC105274531 | 7774 | 273 | 351.17 | 1 | 1.29 | 0.37 | 11.43 | 11.16 | 0.27 | 0.27 | 0.01 | 0.01 |
| gene-LOC105283649 | https://www.ncbi.nlm.nih.gov/gene/?term=LOC105283649 | 1918 | 53 | 276.33 | 1 | 5.21 | 1.89 | 37.66 | 37.39 | 0.27 | 0.27 | 0.01 | 0.01 |
| gene-LOC105279038 | https://www.ncbi.nlm.nih.gov/gene/?term=LOC105279038 | 9453 | 297 | 314.19 | 1 | 1.06 | 0.34 | 9.64 | 9.43 | 0.21 | 0.21 | 0.01 | 0.01 |
| gene-LOC105282149 | https://www.ncbi.nlm.nih.gov/gene/?term=LOC105282149 | 1787 | 103 | 576.39 | 1 | 5.6 | 0.97 | 41.32 | 41.12 | 0.2 | 0.2 | 0.01 | 0.01 |
| gene-LOC105275327 | https://www.ncbi.nlm.nih.gov/gene/?term=LOC105275327 | 14245 | 458 | 321.52 | 1 | 0.7 | 0.22 | 7.5 | 7.33 | 0.17 | 0.17 | 0.01 | 0.01 |
| gene-LOC105278666 | https://www.ncbi.nlm.nih.gov/gene/?term=LOC105278666 | 10341 | 439 | 424.52 | 1 | 0.97 | 0.23 | 7.06 | 6.9 | 0.16 | 0.16 | 0.01 | 0.01 |
| gene-LOC105286274 | https://www.ncbi.nlm.nih.gov/gene/?term=LOC105286274 | 3817 | 179 | 468.95 | 1 | 2.62 | 0.56 | 16.95 | 17.06 | -0.11 | 0.11 | -0.01 | 0.01 |
| gene-LOC105276527 | https://www.ncbi.nlm.nih.gov/gene/?term=LOC105276527 | 7923 | 305 | 384.96 | 1 | 1.26 | 0.33 | 9.71 | 9.92 | -0.21 | 0.21 | -0.01 | 0.01 |
| gene-LOC105287095 | https://www.ncbi.nlm.nih.gov/gene/?term=LOC105287095 | 1435 | 58 | 404.18 | 1 | 6.97 | 1.72 | 28.41 | 28.65 | -0.24 | 0.24 | -0.01 | 0.01 |
| gene-LOC105288042 | https://www.ncbi.nlm.nih.gov/gene/?term=LOC105288042 | 3507 | 88 | 250.93 | 1 | 2.85 | 1.14 | 21.32 | 21.59 | -0.27 | 0.27 | -0.01 | 0.01 |
| gene-LOC105285807 | https://www.ncbi.nlm.nih.gov/gene/?term=LOC105285807 | 6063 | 213 | 351.31 | 1 | 1.65 | 0.47 | 9.29 | 9.56 | -0.28 | 0.28 | -0.01 | 0.01 |
| gene-LOC105286515 | https://www.ncbi.nlm.nih.gov/gene/?term=LOC105286515 | 3751 | 107 | 285.26 | 1 | 2.67 | 0.93 | 12.3 | 12.58 | -0.28 | 0.28 | -0.01 | 0.01 |
| gene-LOC105280774 | https://www.ncbi.nlm.nih.gov/gene/?term=LOC105280774 | 2793 | 100 | 358.04 | 1 | 3.58 | 1 | 31.9 | 32.19 | -0.29 | 0.29 | -0.01 | 0.01 |
| gene-LOC105278228 | https://www.ncbi.nlm.nih.gov/gene/?term=LOC105278228 | 8610 | 156 | 181.18 | 1 | 1.16 | 0.64 | 22.2 | 22.5 | -0.3 | 0.3 | -0.01 | 0.01 |
| gene-LOC105284036 | https://www.ncbi.nlm.nih.gov/gene/?term=LOC105284036 | 3254 | 96 | 295.02 | 1 | 3.07 | 1.04 | 23.11 | 23.46 | -0.35 | 0.35 | -0.01 | 0.01 |
| gene-LOC105280348 | https://www.ncbi.nlm.nih.gov/gene/?term=LOC105280348 | 3797 | 71 | 186.99 | 1 | 2.63 | 1.41 | 20.69 | 21.13 | -0.44 | 0.44 | -0.01 | 0.01 |
| gene-LOC105278962 | https://www.ncbi.nlm.nih.gov/gene/?term=LOC105278962 | 4491 | 89 | 198.17 | 1 | 2.23 | 1.12 | 22.04 | 22.67 | -0.63 | 0.63 | -0.01 | 0.01 |
| gene-LOC105278454 | https://www.ncbi.nlm.nih.gov/gene/?term=LOC105278454 | 1972 | 40 | 202.84 | 1 | 5.07 | 2.5 | 48.64 | 49.28 | -0.64 | 0.64 | -0.01 | 0.01 |
| gene-LOC105280002 | https://www.ncbi.nlm.nih.gov/gene/?term=LOC105280002 | 1940 | 27 | 139.18 | 1 | 5.15 | 3.7 | 33.73 | 34.37 | -0.64 | 0.64 | -0.01 | 0.01 |
| gene-LOC105283989 | https://www.ncbi.nlm.nih.gov/gene/?term=LOC105283989 | 4948 | 25 | 50.53 | 1 | 2.02 | 4 | 35.77 | 37.79 | -2.02 | 2.02 | -0.01 | 0.01 |
| gene-LOC105278108 | https://www.ncbi.nlm.nih.gov/gene/?term=LOC105278108 | 4392 | 43 | 97.91 | 1 | 2.28 | 2.33 | 26.73 | 26.25 | 0.49 | 0.49 | 0 | 0 |
| gene-LOC105285406 | https://www.ncbi.nlm.nih.gov/gene/?term=LOC105285406 | 8015 | 60 | 74.86 | 1 | 1.25 | 1.67 | 25.86 | 25.52 | 0.34 | 0.34 | 0 | 0 |
| gene-LOC105274851 | https://www.ncbi.nlm.nih.gov/gene/?term=LOC105274851 | 6616 | 129 | 194.98 | 1 | 1.51 | 0.78 | 11.16 | 10.94 | 0.22 | 0.22 | 0 | 0 |
| gene-LOC105275187 | https://www.ncbi.nlm.nih.gov/gene/?term=LOC105275187 | 5534 | 65 | 117.46 | 1 | 1.81 | 1.54 | 11.46 | 11.27 | 0.19 | 0.19 | 0 | 0 |
| gene-LOC105278649 | https://www.ncbi.nlm.nih.gov/gene/?term=LOC105278649 | 5616 | 138 | 245.73 | 1 | 1.78 | 0.72 | 13.54 | 13.42 | 0.12 | 0.12 | 0 | 0 |
| gene-LOC105288079 | https://www.ncbi.nlm.nih.gov/gene/?term=LOC105288079 | 3473 | 96 | 276.42 | 1 | 2.88 | 1.04 | 19.82 | 19.7 | 0.12 | 0.12 | 0 | 0 |
| gene-LOC105277856 | https://www.ncbi.nlm.nih.gov/gene/?term=LOC105277856 | 3442 | 29 | 84.25 | 1 | 2.91 | 3.45 | 32.81 | 32.75 | 0.06 | 0.06 | 0 | 0 |
| gene-LOC105284355 | https://www.ncbi.nlm.nih.gov/gene/?term=LOC105284355 | 3904 | 152 | 389.34 | 1 | 2.56 | 0.66 | 20.02 | 19.96 | 0.06 | 0.06 | 0 | 0 |
| gene-LOC105287528 | https://www.ncbi.nlm.nih.gov/gene/?term=LOC105287528 | 14195 | 375 | 264.18 | 1 | 0.7 | 0.27 | 5.12 | 5.07 | 0.05 | 0.05 | 0 | 0 |
| gene-LOC105282811 | https://www.ncbi.nlm.nih.gov/gene/?term=LOC105282811 | 5765 | 196 | 339.98 | 1 | 1.73 | 0.51 | 8.25 | 8.31 | -0.06 | 0.06 | 0 | 0 |
| gene-LOC105284704 | https://www.ncbi.nlm.nih.gov/gene/?term=LOC105284704 | 2400 | 59 | 245.83 | 1 | 4.17 | 1.69 | 22.63 | 22.7 | -0.07 | 0.07 | 0 | 0 |
| gene-LOC105274706 | https://www.ncbi.nlm.nih.gov/gene/?term=LOC105274706 | 1446 | 49 | 338.87 | 1 | 6.92 | 2.04 | 38.21 | 38.31 | -0.1 | 0.1 | 0 | 0 |
| gene-LOC105278282 | https://www.ncbi.nlm.nih.gov/gene/?term=LOC105278282 | 1775 | 52 | 292.96 | 1 | 5.63 | 1.92 | 41.01 | 41.14 | -0.14 | 0.14 | 0 | 0 |
| gene-LOC105274590 | https://www.ncbi.nlm.nih.gov/gene/?term=LOC105274590 | 4516 | 67 | 148.36 | 1 | 2.21 | 1.49 | 28.5 | 28.65 | -0.15 | 0.15 | 0 | 0 |
| gene-LOC105282067 | https://www.ncbi.nlm.nih.gov/gene/?term=LOC105282067 | 5941 | 49 | 82.48 | 1 | 1.68 | 2.04 | 25.2 | 25.4 | -0.2 | 0.2 | 0 | 0 |
| gene-LOC105282658 | https://www.ncbi.nlm.nih.gov/gene/?term=LOC105282658 | 2421 | 52 | 214.79 | 1 | 4.13 | 1.92 | 21.81 | 22.01 | -0.2 | 0.2 | 0 | 0 |

Directory: OGL File: rnk-dmg-Ob_LI_RphaseA2.vs.RphaseB1.txt

| gene_ID | gene_link | gwidth | #Sites | #per10Kb | #dmSites | #dmsp10kb | %dmSites | %pSite1 | %pSite2 | DMpSite | ADMpSite | DMpNucl | ADMpNucl |
|---|---|---|---|---|---|---|---|---|---|---|---|---|---|
| gene-LOC113562345 | https://www.ncbi.nlm.nih.gov/gene/?term=LOC113562345 | 2290 | 80 | 349.34 | 16 | 69.87 | 20 | 25.78 | 13.98 | 11.8 | 11.8 | 0.41 | 0.41 |
| gene-LOC105279900 | https://www.ncbi.nlm.nih.gov/gene/?term=LOC105279900 | 2191 | 56 | 255.59 | 1 | 4.56 | 1.79 | 33.63 | 28.91 | 4.72 | 4.72 | 0.12 | 0.12 |
| gene-LOC105285695 | https://www.ncbi.nlm.nih.gov/gene/?term=LOC105285695 | 1371 | 76 | 554.34 | 1 | 7.29 | 1.32 | 28.59 | 26.36 | 2.24 | 2.24 | 0.12 | 0.12 |
| gene-LOC105287417 | https://www.ncbi.nlm.nih.gov/gene/?term=LOC105287417 | 1025 | 38 | 370.73 | 1 | 9.76 | 2.63 | 35.24 | 32.37 | 2.87 | 2.87 | 0.11 | 0.11 |
| gene-LOC105282156 | https://www.ncbi.nlm.nih.gov/gene/?term=LOC105282156 | 2577 | 134 | 519.98 | 1 | 3.88 | 0.75 | 27.52 | 25.46 | 2.06 | 2.06 | 0.11 | 0.11 |
| gene-LOC105280786 | https://www.ncbi.nlm.nih.gov/gene/?term=LOC105280786 | 2573 | 143 | 555.77 | 1 | 3.89 | 0.7 | 27.63 | 25.63 | 2 | 2 | 0.11 | 0.11 |
| gene-LOC105275070 | https://www.ncbi.nlm.nih.gov/gene/?term=LOC105275070 | 5667 | 64 | 112.93 | 6 | 10.59 | 9.38 | 14.32 | 5.2 | 9.13 | 9.13 | 0.1 | 0.1 |
| gene-LOC105287063 | https://www.ncbi.nlm.nih.gov/gene/?term=LOC105287063 | 1874 | 55 | 293.49 | 1 | 5.34 | 1.82 | 30.47 | 27.11 | 3.37 | 3.37 | 0.1 | 0.1 |
| gene-LOC105276704 | https://www.ncbi.nlm.nih.gov/gene/?term=LOC105276704 | 2063 | 93 | 450.8 | 5 | 24.24 | 5.38 | 14.49 | 12.51 | 1.99 | 1.99 | 0.09 | 0.09 |
| gene-LOC105283457 | https://www.ncbi.nlm.nih.gov/gene/?term=LOC105283457 | 1882 | 49 | 260.36 | 1 | 5.31 | 2.04 | 35.19 | 32.03 | 3.16 | 3.16 | 0.08 | 0.08 |
| gene-LOC105285583 | https://www.ncbi.nlm.nih.gov/gene/?term=LOC105285583 | 1947 | 62 | 318.44 | 1 | 5.14 | 1.61 | 27.07 | 24.61 | 2.46 | 2.46 | 0.08 | 0.08 |
| gene-LOC105274706 | https://www.ncbi.nlm.nih.gov/gene/?term=LOC105274706 | 1446 | 56 | 387.28 | 1 | 6.92 | 1.79 | 39.8 | 37.63 | 2.17 | 2.17 | 0.08 | 0.08 |
| gene-LOC105285674 | https://www.ncbi.nlm.nih.gov/gene/?term=LOC105285674 | 3035 | 111 | 365.73 | 2 | 6.59 | 1.8 | 25.15 | 23.09 | 2.06 | 2.06 | 0.08 | 0.08 |
| gene-LOC105282893 | https://www.ncbi.nlm.nih.gov/gene/?term=LOC105282893 | 2645 | 50 | 189.04 | 1 | 3.78 | 2 | 28.92 | 25.31 | 3.6 | 3.6 | 0.07 | 0.07 |
| gene-LOC105284704 | https://www.ncbi.nlm.nih.gov/gene/?term=LOC105284704 | 2400 | 66 | 275 | 1 | 4.17 | 1.52 | 25.49 | 22.92 | 2.57 | 2.57 | 0.07 | 0.07 |
| gene-LOC105286683 | https://www.ncbi.nlm.nih.gov/gene/?term=LOC105286683 | 1665 | 57 | 342.34 | 1 | 6.01 | 1.75 | 35.23 | 33.27 | 1.97 | 1.97 | 0.07 | 0.07 |
| gene-LOC105276449 | https://www.ncbi.nlm.nih.gov/gene/?term=LOC105276449 | 1231 | 55 | 446.79 | 1 | 8.12 | 1.82 | 24.75 | 23.28 | 1.47 | 1.47 | 0.07 | 0.07 |
| gene-LOC105277435 | https://www.ncbi.nlm.nih.gov/gene/?term=LOC105277435 | 3643 | 184 | 505.08 | 1 | 2.74 | 0.54 | 12.13 | 10.74 | 1.39 | 1.39 | 0.07 | 0.07 |
| gene-LOC105279298 | https://www.ncbi.nlm.nih.gov/gene/?term=LOC105279298 | 2773 | 134 | 483.23 | 2 | 7.21 | 1.49 | 24.2 | 22.81 | 1.39 | 1.39 | 0.07 | 0.07 |
| gene-LOC105282682 | https://www.ncbi.nlm.nih.gov/gene/?term=LOC105282682 | 4127 | 240 | 581.54 | 1 | 2.42 | 0.42 | 13.64 | 12.51 | 1.14 | 1.14 | 0.07 | 0.07 |
| gene-LOC105284213 | https://www.ncbi.nlm.nih.gov/gene/?term=LOC105284213 | 868 | 23 | 264.98 | 1 | 11.52 | 4.35 | 9.72 | 12.3 | -2.58 | 2.58 | -0.07 | 0.07 |
| gene-LOC105275708 | https://www.ncbi.nlm.nih.gov/gene/?term=LOC105275708 | 1772 | 19 | 107.22 | 1 | 5.64 | 5.26 | 37.25 | 31.29 | 5.96 | 5.96 | 0.06 | 0.06 |
| gene-LOC105276538 | https://www.ncbi.nlm.nih.gov/gene/?term=LOC105276538 | 1511 | 23 | 152.22 | 1 | 6.62 | 4.35 | 69.63 | 65.38 | 4.25 | 4.25 | 0.06 | 0.06 |
| gene-LOC105283712 | https://www.ncbi.nlm.nih.gov/gene/?term=LOC105283712 | 1910 | 33 | 172.77 | 1 | 5.24 | 3.03 | 32.87 | 29.11 | 3.76 | 3.76 | 0.06 | 0.06 |
| gene-LOC105281848 | https://www.ncbi.nlm.nih.gov/gene/?term=LOC105281848 | 2122 | 60 | 282.75 | 1 | 4.71 | 1.67 | 24.48 | 22.53 | 1.95 | 1.95 | 0.06 | 0.06 |
| gene-LOC105280704 | https://www.ncbi.nlm.nih.gov/gene/?term=LOC105280704 | 4266 | 155 | 363.34 | 1 | 2.34 | 0.65 | 17.59 | 16.02 | 1.58 | 1.58 | 0.06 | 0.06 |
| gene-LOC105278282 | https://www.ncbi.nlm.nih.gov/gene/?term=LOC105278282 | 1775 | 67 | 377.46 | 1 | 5.63 | 1.49 | 34.11 | 32.61 | 1.51 | 1.51 | 0.06 | 0.06 |
| gene-LOC105285671 | https://www.ncbi.nlm.nih.gov/gene/?term=LOC105285671 | 2263 | 103 | 455.15 | 1 | 4.42 | 0.97 | 30.91 | 29.64 | 1.27 | 1.27 | 0.06 | 0.06 |
| gene-LOC105278283 | https://www.ncbi.nlm.nih.gov/gene/?term=LOC105278283 | 2317 | 145 | 625.81 | 1 | 4.32 | 0.69 | 24.77 | 23.82 | 0.95 | 0.95 | 0.06 | 0.06 |
| gene-LOC105285055 | https://www.ncbi.nlm.nih.gov/gene/?term=LOC105285055 | 4108 | 264 | 642.65 | 2 | 4.87 | 0.76 | 12 | 11.12 | 0.87 | 0.87 | 0.06 | 0.06 |
| gene-LOC105277631 | https://www.ncbi.nlm.nih.gov/gene/?term=LOC105277631 | 4387 | 327 | 745.38 | 1 | 2.28 | 0.31 | 11.07 | 10.23 | 0.85 | 0.85 | 0.06 | 0.06 |
| gene-LOC105275328 | https://www.ncbi.nlm.nih.gov/gene/?term=LOC105275328 | 2227 | 85 | 381.68 | 1 | 4.49 | 1.18 | 17.9 | 19.55 | -1.65 | 1.65 | -0.06 | 0.06 |
| gene-LOC113562974 | https://www.ncbi.nlm.nih.gov/gene/?term=LOC113562974 | 1832 | 2 | 10.92 | 1 | 5.46 | 50 | 57.94 | 9 | 48.94 | 48.94 | 0.05 | 0.05 |
| gene-LOC105285514 | https://www.ncbi.nlm.nih.gov/gene/?term=LOC105285514 | 1277 | 11 | 86.14 | 1 | 7.83 | 9.09 | 90.74 | 85.12 | 5.62 | 5.62 | 0.05 | 0.05 |
| gene-LOC105281039 | https://www.ncbi.nlm.nih.gov/gene/?term=LOC105281039 | 1471 | 15 | 101.97 | 1 | 6.8 | 6.67 | 36.89 | 31.93 | 4.96 | 4.96 | 0.05 | 0.05 |
| gene-LOC105275190 | https://www.ncbi.nlm.nih.gov/gene/?term=LOC105275190 | 3837 | 77 | 200.68 | 1 | 2.61 | 1.3 | 21.84 | 19.45 | 2.39 | 2.39 | 0.05 | 0.05 |
| gene-LOC113561805 | https://www.ncbi.nlm.nih.gov/gene/?term=LOC113561805 | 3337 | 73 | 218.76 | 1 | 3 | 1.37 | 38.02 | 35.67 | 2.34 | 2.34 | 0.05 | 0.05 |
| gene-LOC105276272 | https://www.ncbi.nlm.nih.gov/gene/?term=LOC105276272 | 1965 | 47 | 239.19 | 1 | 5.09 | 2.13 | 25.43 | 23.44 | 1.98 | 1.98 | 0.05 | 0.05 |
| gene-LOC105288079 | https://www.ncbi.nlm.nih.gov/gene/?term=LOC105288079 | 3473 | 112 | 322.49 | 1 | 2.88 | 0.89 | 18.94 | 17.43 | 1.51 | 1.51 | 0.05 | 0.05 |
| gene-LOC105275460 | https://www.ncbi.nlm.nih.gov/gene/?term=LOC105275460 | 2473 | 123 | 497.37 | 1 | 4.04 | 0.81 | 13.29 | 12.32 | 0.97 | 0.97 | 0.05 | 0.05 |
| gene-LOC105280774 | https://www.ncbi.nlm.nih.gov/gene/?term=LOC105280774 | 2793 | 135 | 483.35 | 1 | 3.58 | 0.74 | 25.37 | 24.41 | 0.96 | 0.96 | 0.05 | 0.05 |
| gene-LOC105288066 | https://www.ncbi.nlm.nih.gov/gene/?term=LOC105288066 | 1964 | 31 | 157.84 | 1 | 5.09 | 3.23 | 39.38 | 36.57 | 2.81 | 2.81 | 0.04 | 0.04 |
| gene-LOC105288010 | https://www.ncbi.nlm.nih.gov/gene/?term=LOC105288010 | 3189 | 61 | 191.28 | 1 | 3.14 | 1.64 | 34.85 | 32.7 | 2.15 | 2.15 | 0.04 | 0.04 |
| gene-LOC105286644 | https://www.ncbi.nlm.nih.gov/gene/?term=LOC105286644 | 1235 | 23 | 186.23 | 1 | 8.1 | 4.35 | 43.78 | 41.75 | 2.04 | 2.04 | 0.04 | 0.04 |
| gene-LOC105282151 | https://www.ncbi.nlm.nih.gov/gene/?term=LOC105282151 | 1601 | 45 | 281.07 | 2 | 12.49 | 4.44 | 42.82 | 41.43 | 1.39 | 1.39 | 0.04 | 0.04 |
| gene-LOC105283123 | https://www.ncbi.nlm.nih.gov/gene/?term=LOC105283123 | 1507 | 55 | 364.96 | 1 | 6.64 | 1.82 | 20.08 | 18.96 | 1.12 | 1.12 | 0.04 | 0.04 |
| gene-LOC105276408 | https://www.ncbi.nlm.nih.gov/gene/?term=LOC105276408 | 5298 | 201 | 379.39 | 1 | 1.89 | 0.5 | 21.53 | 20.53 | 1 | 1 | 0.04 | 0.04 |
| gene-LOC105284355 | https://www.ncbi.nlm.nih.gov/gene/?term=LOC105284355 | 3904 | 155 | 397.03 | 1 | 2.56 | 0.65 | 22.33 | 21.37 | 0.95 | 0.95 | 0.04 | 0.04 |
| gene-LOC105278497 | https://www.ncbi.nlm.nih.gov/gene/?term=LOC105278497 | 2072 | 103 | 497.1 | 1 | 4.83 | 0.97 | 19.51 | 18.76 | 0.75 | 0.75 | 0.04 | 0.04 |
| gene-LOC105284557 | https://www.ncbi.nlm.nih.gov/gene/?term=LOC105284557 | 1672 | 91 | 544.26 | 1 | 5.98 | 1.1 | 27.82 | 27.15 | 0.67 | 0.67 | 0.04 | 0.04 |
| gene-LOC105285409 | https://www.ncbi.nlm.nih.gov/gene/?term=LOC105285409 | 5099 | 67 | 131.4 | 2 | 3.92 | 2.99 | 1.72 | 4.47 | -2.75 | 2.75 | -0.04 | 0.04 |
| gene-LOC105286302 | https://www.ncbi.nlm.nih.gov/gene/?term=LOC105286302 | 4536 | 24 | 52.91 | 1 | 2.2 | 4.17 | 20.45 | 15.15 | 5.3 | 5.3 | 0.03 | 0.03 |
| gene-LOC105284609 | https://www.ncbi.nlm.nih.gov/gene/?term=LOC105284609 | 5678 | 44 | 77.49 | 1 | 1.76 | 2.27 | 32.19 | 27.8 | 4.39 | 4.39 | 0.03 | 0.03 |
| gene-LOC105279898 | https://www.ncbi.nlm.nih.gov/gene/?term=LOC105279898 | 1343 | 16 | 119.14 | 1 | 7.45 | 6.25 | 58.57 | 56.24 | 2.33 | 2.33 | 0.03 | 0.03 |
| gene-LOC105275518 | https://www.ncbi.nlm.nih.gov/gene/?term=LOC105275518 | 2335 | 28 | 119.91 | 1 | 4.28 | 3.57 | 41.74 | 39.46 | 2.28 | 2.28 | 0.03 | 0.03 |
| gene-LOC105285060 | https://www.ncbi.nlm.nih.gov/gene/?term=LOC105285060 | 1285 | 18 | 140.08 | 1 | 7.78 | 5.56 | 38.72 | 36.69 | 2.03 | 2.03 | 0.03 | 0.03 |
| gene-LOC105276078 | https://www.ncbi.nlm.nih.gov/gene/?term=LOC105276078 | 1051 | 18 | 171.27 | 1 | 9.51 | 5.56 | 28.41 | 26.59 | 1.83 | 1.83 | 0.03 | 0.03 |
| gene-LOC105280348 | https://www.ncbi.nlm.nih.gov/gene/?term=LOC105280348 | 3797 | 78 | 205.43 | 1 | 2.63 | 1.28 | 18.55 | 17.12 | 1.44 | 1.44 | 0.03 | 0.03 |
| gene-LOC105282794 | https://www.ncbi.nlm.nih.gov/gene/?term=LOC105282794 | 4571 | 109 | 238.46 | 1 | 2.19 | 0.92 | 17.81 | 16.42 | 1.38 | 1.38 | 0.03 | 0.03 |
| gene-LOC105286514 | https://www.ncbi.nlm.nih.gov/gene/?term=LOC105286514 | 2528 | 55 | 217.56 | 1 | 3.96 | 1.82 | 37.15 | 35.79 | 1.36 | 1.36 | 0.03 | 0.03 |

| | | | | | | | | | | | | | |
|---|---|---|---|---|---|---|---|---|---|---|---|---|---|
| gene-LOC105280722 | https://www.ncbi.nlm.nih.gov/gene/?term=LOC105280722 | 3237 | 93 | 287.3 1 | 3.09 | 1.08 | 27.81 | 26.63 | 1.18 | 1.18 | 0.03 | 0.03 | |
| gene-LOC105276969 | https://www.ncbi.nlm.nih.gov/gene/?term=LOC105276969 | 2997 | 104 | 347.01 | 1 | 3.34 | 0.96 | 11.67 | 10.81 | 0.86 | 0.86 | 0.03 | 0.03 |
| gene-LOC105278807 | https://www.ncbi.nlm.nih.gov/gene/?term=LOC105278807 | 2166 | 77 | 355.49 | 1 | 4.62 | 1.3 | 21.89 | 21.08 | 0.81 | 0.81 | 0.03 | 0.03 |
| gene-LOC105274823 | https://www.ncbi.nlm.nih.gov/gene/?term=LOC105274823 | 4620 | 171 | 370.13 | 1 | 2.16 | 0.58 | 18.07 | 17.27 | 0.8 | 0.8 | 0.03 | 0.03 |
| gene-LOC105280104 | https://www.ncbi.nlm.nih.gov/gene/?term=LOC105280104 | 3960 | 132 | 333.33 | 1 | 2.53 | 0.76 | 17.97 | 17.22 | 0.76 | 0.76 | 0.03 | 0.03 |
| gene-LOC105285742 | https://www.ncbi.nlm.nih.gov/gene/?term=LOC105285742 | 3446 | 122 | 354.03 | 1 | 2.9 | 0.82 | 17.63 | 16.88 | 0.76 | 0.76 | 0.03 | 0.03 |
| gene-LOC105282491 | https://www.ncbi.nlm.nih.gov/gene/?term=LOC105282491 | 4761 | 191 | 401.18 | 1 | 2.1 | 0.52 | 9 | 8.31 | 0.69 | 0.69 | 0.03 | 0.03 |
| gene-LOC105283728 | https://www.ncbi.nlm.nih.gov/gene/?term=LOC105283728 | 3256 | 159 | 488.33 | 2 | 6.14 | 1.26 | 15.6 | 14.91 | 0.69 | 0.69 | 0.03 | 0.03 |
| gene-LOC105287704 | https://www.ncbi.nlm.nih.gov/gene/?term=LOC105287704 | 3019 | 142 | 470.35 | 1 | 3.31 | 0.7 | 19.98 | 19.34 | 0.63 | 0.63 | 0.03 | 0.03 |
| gene-LOC105275683 | https://www.ncbi.nlm.nih.gov/gene/?term=LOC105275683 | 4256 | 204 | 479.32 | 1 | 2.35 | 0.49 | 13.38 | 12.77 | 0.61 | 0.61 | 0.03 | 0.03 |
| gene-LOC105286279 | https://www.ncbi.nlm.nih.gov/gene/?term=LOC105286279 | 5864 | 277 | 472.37 | 1 | 1.71 | 0.36 | 2.15 | 2.72 | -0.57 | 0.57 | -0.03 | 0.03 |
| gene-LOC105284329 | https://www.ncbi.nlm.nih.gov/gene/?term=LOC105284329 | 4051 | 72 | 177.73 | 1 | 2.47 | 1.39 | 18.86 | 20.42 | -1.56 | 1.56 | -0.03 | 0.03 |
| gene-LOC105286983 | https://www.ncbi.nlm.nih.gov/gene/?term=LOC105286983 | 16901 | 70 | 41.42 1 | 0.59 | 1.43 | 6.21 | 1.14 | 5.06 | 5.06 | 0.02 | 0.02 | |
| gene-LOC105285315 | https://www.ncbi.nlm.nih.gov/gene/?term=LOC105285315 | 5458 | 33 | 60.46 1 | 1.83 | 3.03 | 33.27 | 29.4 | 3.88 | 3.88 | 0.02 | 0.02 | |
| gene-LOC105281515 | https://www.ncbi.nlm.nih.gov/gene/?term=LOC105281515 | 8827 | 56 | 63.44 2 | 2.27 | 3.57 | 36.05 | 32.61 | 3.44 | 3.44 | 0.02 | 0.02 | |
| gene-LOC105274900 | https://www.ncbi.nlm.nih.gov/gene/?term=LOC105274900 | 5561 | 61 | 109.69 | 2 | 3.6 | 3.28 | 32.53 | 30.66 | 1.87 | 1.87 | 0.02 | 0.02 |
| gene-LOC105278108 | https://www.ncbi.nlm.nih.gov/gene/?term=LOC105278108 | 4392 | 57 | 129.78 | 1 | 2.28 | 1.75 | 25.91 | 24.39 | 1.52 | 1.52 | 0.02 | 0.02 |
| gene-LOC105280002 | https://www.ncbi.nlm.nih.gov/gene/?term=LOC105280002 | 1940 | 34 | 175.26 | 1 | 5.15 | 2.94 | 33.64 | 32.26 | 1.38 | 1.38 | 0.02 | 0.02 |
| gene-LOC105288063 | https://www.ncbi.nlm.nih.gov/gene/?term=LOC105288063 | 3400 | 71 | 208.82 | 1 | 2.94 | 1.41 | 12.55 | 11.41 | 1.14 | 1.14 | 0.02 | 0.02 |
| gene-LOC105282600 | https://www.ncbi.nlm.nih.gov/gene/?term=LOC105282600 | 3715 | 67 | 180.35 | 1 | 2.69 | 1.49 | 27.6 | 26.48 | 1.12 | 1.12 | 0.02 | 0.02 |
| gene-LOC105278071 | https://www.ncbi.nlm.nih.gov/gene/?term=LOC105278071 | 4307 | 61 | 141.63 | 1 | 2.32 | 1.64 | 11.12 | 10.02 | 1.1 | 1.1 | 0.02 | 0.02 |
| gene-LOC105285416 | https://www.ncbi.nlm.nih.gov/gene/?term=LOC105285416 | 3820 | 76 | 198.95 | 1 | 2.62 | 1.32 | 22.22 | 21.12 | 1.1 | 1.1 | 0.02 | 0.02 |
| gene-LOC105280407 | https://www.ncbi.nlm.nih.gov/gene/?term=LOC105280407 | 10950 | 231 | 210.96 | 1 | 0.91 | 0.43 | 15.59 | 14.54 | 1.05 | 1.05 | 0.02 | 0.02 |
| gene-LOC105283838 | https://www.ncbi.nlm.nih.gov/gene/?term=LOC105283838 | 1748 | 33 | 188.79 | 1 | 5.72 | 3.03 | 33.49 | 32.48 | 1.01 | 1.01 | 0.02 | 0.02 |
| gene-LOC105283592 | https://www.ncbi.nlm.nih.gov/gene/?term=LOC105283592 | 3479 | 87 | 250.07 | 1 | 2.87 | 1.15 | 22.28 | 21.36 | 0.92 | 0.92 | 0.02 | 0.02 |
| gene-LOC105275179 | https://www.ncbi.nlm.nih.gov/gene/?term=LOC105275179 | 3271 | 85 | 259.86 | 1 | 3.06 | 1.18 | 16.68 | 15.92 | 0.76 | 0.76 | 0.02 | 0.02 |
| gene-LOC105281312 | https://www.ncbi.nlm.nih.gov/gene/?term=LOC105281312 | 2551 | 69 | 270.48 | 1 | 3.92 | 1.45 | 31.17 | 30.43 | 0.74 | 0.74 | 0.02 | 0.02 |
| gene-LOC105282825 | https://www.ncbi.nlm.nih.gov/gene/?term=LOC105282825 | 3919 | 102 | 260.27 | 1 | 2.55 | 0.98 | 14.91 | 14.23 | 0.68 | 0.68 | 0.02 | 0.02 |
| gene-LOC105275451 | https://www.ncbi.nlm.nih.gov/gene/?term=LOC105275451 | 6035 | 149 | 246.89 | 1 | 1.66 | 0.67 | 20.25 | 19.59 | 0.66 | 0.66 | 0.02 | 0.02 |
| gene-LOC105278632 | https://www.ncbi.nlm.nih.gov/gene/?term=LOC105278632 | 3624 | 150 | 413.91 | 1 | 2.76 | 0.67 | 13.57 | 13.05 | 0.53 | 0.53 | 0.02 | 0.02 |
| gene-LOC105285179 | https://www.ncbi.nlm.nih.gov/gene/?term=LOC105285179 | 4191 | 204 | 486.76 | 1 | 2.39 | 0.49 | 16.93 | 16.42 | 0.51 | 0.51 | 0.02 | 0.02 |
| gene-LOC105275752 | https://www.ncbi.nlm.nih.gov/gene/?term=LOC105275752 | 3462 | 138 | 398.61 | 1 | 2.89 | 0.72 | 31.47 | 31.01 | 0.46 | 0.46 | 0.02 | 0.02 |
| gene-LOC105279038 | https://www.ncbi.nlm.nih.gov/gene/?term=LOC105279038 | 9453 | 345 | 364.96 | 1 | 1.06 | 0.29 | 9.49 | 9.03 | 0.45 | 0.45 | 0.02 | 0.02 |
| gene-LOC105275363 | https://www.ncbi.nlm.nih.gov/gene/?term=LOC105275363 | 3798 | 212 | 558.19 | 1 | 2.63 | 0.47 | 14.03 | 13.61 | 0.43 | 0.43 | 0.02 | 0.02 |
| gene-LOC105285807 | https://www.ncbi.nlm.nih.gov/gene/?term=LOC105285807 | 6063 | 231 | 381 1 | 1.65 | 0.43 | 10.41 | 9.98 | 0.43 | 0.43 | 0.02 | 0.02 | |
| gene-LOC105274531 | https://www.ncbi.nlm.nih.gov/gene/?term=LOC105274531 | 7774 | 307 | 394.91 | 1 | 1.29 | 0.33 | 10.8 | 10.38 | 0.42 | 0.42 | 0.02 | 0.02 |
| gene-LOC105281272 | https://www.ncbi.nlm.nih.gov/gene/?term=LOC105281272 | 5252 | 201 | 382.71 | 1 | 1.9 | 0.5 | 15.32 | 14.89 | 0.42 | 0.42 | 0.02 | 0.02 |
| gene-LOC105275374 | https://www.ncbi.nlm.nih.gov/gene/?term=LOC105275374 | 11376 | 587 | 516 1 | 0.88 | 0.17 | 9.68 | 9.27 | 0.41 | 0.41 | 0.02 | 0.02 | |
| gene-LOC105284184 | https://www.ncbi.nlm.nih.gov/gene/?term=LOC105284184 | 2535 | 105 | 414.2 1 | 3.94 | 0.95 | 28.89 | 28.5 | 0.39 | 0.39 | 0.02 | 0.02 | |
| gene-LOC105276462 | https://www.ncbi.nlm.nih.gov/gene/?term=LOC105276462 | 6561 | 289 | 440.48 | 1 | 1.52 | 0.35 | 8.58 | 8.21 | 0.38 | 0.38 | 0.02 | 0.02 |
| gene-LOC105278145 | https://www.ncbi.nlm.nih.gov/gene/?term=LOC105278145 | 4470 | 274 | 612.98 | 1 | 2.24 | 0.36 | 10.24 | 9.87 | 0.37 | 0.37 | 0.02 | 0.02 |
| gene-LOC105284043 | https://www.ncbi.nlm.nih.gov/gene/?term=LOC105284043 | 5661 | 312 | 551.14 | 1 | 1.77 | 0.32 | 10.26 | 9.89 | 0.37 | 0.37 | 0.02 | 0.02 |
| gene-LOC105276527 | https://www.ncbi.nlm.nih.gov/gene/?term=LOC105276527 | 7923 | 381 | 480.88 | 1 | 1.26 | 0.26 | 8.09 | 7.73 | 0.36 | 0.36 | 0.02 | 0.02 |
| gene-LOC105278877 | https://www.ncbi.nlm.nih.gov/gene/?term=LOC105278877 | 1968 | 91 | 462.4 2 | 10.16 | 2.2 | 30.61 | 31.07 | -0.45 | 0.45 | -0.02 | 0.02 | |
| gene-LOC105284036 | https://www.ncbi.nlm.nih.gov/gene/?term=LOC105284036 | 3254 | 102 | 313.46 | 1 | 3.07 | 0.98 | 24.26 | 24.78 | -0.52 | 0.52 | -0.02 | 0.02 |
| gene-LOC105274851 | https://www.ncbi.nlm.nih.gov/gene/?term=LOC105274851 | 6616 | 160 | 241.84 | 1 | 1.51 | 0.62 | 10.97 | 11.63 | -0.66 | 0.66 | -0.02 | 0.02 |
| gene-LOC105275966 | https://www.ncbi.nlm.nih.gov/gene/?term=LOC105275966 | 3299 | 75 | 227.34 | 1 | 3.03 | 1.33 | 43.86 | 44.6 | -0.74 | 0.74 | -0.02 | 0.02 |
| gene-LOC105284507 | https://www.ncbi.nlm.nih.gov/gene/?term=LOC105284507 | 5207 | 123 | 236.22 | 1 | 1.92 | 0.81 | 18.34 | 19.09 | -0.75 | 0.75 | -0.02 | 0.02 |
| gene-LOC105280890 | https://www.ncbi.nlm.nih.gov/gene/?term=LOC105280890 | 1139 | 31 | 272.17 | 1 | 8.78 | 3.23 | 17.12 | 17.99 | -0.87 | 0.87 | -0.02 | 0.02 |
| gene-LOC105281173 | https://www.ncbi.nlm.nih.gov/gene/?term=LOC105281173 | 1595 | 39 | 244.51 | 1 | 6.27 | 2.56 | 61.21 | 62.08 | -0.87 | 0.87 | -0.02 | 0.02 |
| gene-LOC105282885 | https://www.ncbi.nlm.nih.gov/gene/?term=LOC105282885 | 1870 | 41 | 219.25 | 1 | 5.35 | 2.44 | 40.22 | 41.13 | -0.91 | 0.91 | -0.02 | 0.02 |
| gene-LOC105285252 | https://www.ncbi.nlm.nih.gov/gene/?term=LOC105285252 | 7427 | 148 | 199.27 | 1 | 1.35 | 0.68 | 11.91 | 12.93 | -1.01 | 1.01 | -0.02 | 0.02 |
| gene-LOC105287633 | https://www.ncbi.nlm.nih.gov/gene/?term=LOC105287633 | 1524 | 26 | 170.6 1 | 6.56 | 3.85 | 60.3 | 61.69 | -1.38 | 1.38 | -0.02 | 0.02 | |
| gene-LOC105282914 | https://www.ncbi.nlm.nih.gov/gene/?term=LOC105282914 | 5623 | 43 | 76.47 1 | 1.78 | 2.33 | 47.44 | 49.79 | -2.35 | 2.35 | -0.02 | 0.02 | |
| gene-LOC105279500 | https://www.ncbi.nlm.nih.gov/gene/?term=LOC105279500 | 3587 | 36 | 100.36 | 1 | 2.79 | 2.78 | 48.81 | 47.64 | 1.17 | 1.17 | 0.01 | 0.01 |
| gene-LOC105277856 | https://www.ncbi.nlm.nih.gov/gene/?term=LOC105277856 | 3442 | 36 | 104.59 | 1 | 2.91 | 2.78 | 37.81 | 36.75 | 1.06 | 1.06 | 0.01 | 0.01 |
| gene-LOC105275347 | https://www.ncbi.nlm.nih.gov/gene/?term=LOC105275347 | 2526 | 21 | 83.14 1 | 3.96 | 4.76 | 35.82 | 34.96 | 0.86 | 0.86 | 0.01 | 0.01 | |
| gene-LOC105283989 | https://www.ncbi.nlm.nih.gov/gene/?term=LOC105283989 | 4948 | 44 | 88.92 2 | 4.04 | 4.55 | 37.47 | 36.79 | 0.68 | 0.68 | 0.01 | 0.01 | |
| gene-LOC105283943 | https://www.ncbi.nlm.nih.gov/gene/?term=LOC105283943 | 3559 | 68 | 191.06 | 1 | 2.81 | 1.47 | 20.98 | 20.31 | 0.67 | 0.67 | 0.01 | 0.01 |
| gene-LOC105276560 | https://www.ncbi.nlm.nih.gov/gene/?term=LOC105276560 | 18607 | 384 | 206.37 | 3 | 1.61 | 0.78 | 2.53 | 1.87 | 0.66 | 0.66 | 0.01 | 0.01 |
| gene-LOC105288078 | https://www.ncbi.nlm.nih.gov/gene/?term=LOC105288078 | 19126 | 359 | 187.7 1 | 0.52 | 0.28 | 15.58 | 14.92 | 0.66 | 0.66 | 0.01 | 0.01 | |
| gene-LOC105277215 | https://www.ncbi.nlm.nih.gov/gene/?term=LOC105277215 | 2619 | 30 | 114.55 | 1 | 3.82 | 3.33 | 27.61 | 26.99 | 0.61 | 0.61 | 0.01 | 0.01 |
| gene-LOC105281296 | https://www.ncbi.nlm.nih.gov/gene/?term=LOC105281296 | 2434 | 48 | 197.21 | 1 | 4.11 | 2.08 | 28.14 | 27.55 | 0.59 | 0.59 | 0.01 | 0.01 |
| gene-LOC105278384 | https://www.ncbi.nlm.nih.gov/gene/?term=LOC105278384 | 1390 | 33 | 237.41 | 1 | 7.19 | 3.03 | 46.32 | 45.77 | 0.55 | 0.55 | 0.01 | 0.01 |

| | | | | | | | | | | | | | |
|---|---|---|---|---|---|---|---|---|---|---|---|---|---|
| gene-LOC105274699 | https://www.ncbi.nlm.nih.gov/gene/?term=LOC105274699 | 2386 | 67 | 280.8 | 1 | 4.19 | 1.49 | 18.3 | 17.79 | 0.51 | 0.51 | 0.01 | 0.01 |
| gene-LOC105280641 | https://www.ncbi.nlm.nih.gov/gene/?term=LOC105280641 | 3682 | 78 | 211.84 | 1 | 2.72 | 1.28 | 13.91 | 13.42 | 0.49 | 0.49 | 0.01 | 0.01 |
| gene-LOC105274773 | https://www.ncbi.nlm.nih.gov/gene/?term=LOC105274773 | 11034 | 187 | 169.48 | 1 | 0.91 | 0.53 | 16.91 | 16.52 | 0.39 | 0.39 | 0.01 | 0.01 |
| gene-LOC105281700 | https://www.ncbi.nlm.nih.gov/gene/?term=LOC105281700 | 2208 | 43 | 194.75 | 1 | 4.53 | 2.33 | 36.88 | 36.49 | 0.39 | 0.39 | 0.01 | 0.01 |
| gene-LOC105278962 | https://www.ncbi.nlm.nih.gov/gene/?term=LOC105278962 | 4491 | 129 | 287.24 | 1 | 2.23 | 0.78 | 19.14 | 18.79 | 0.35 | 0.35 | 0.01 | 0.01 |
| gene-LOC105274534 | https://www.ncbi.nlm.nih.gov/gene/?term=LOC105274534 | 1329 | 60 | 451.47 | 1 | 7.52 | 1.67 | 44.47 | 44.14 | 0.32 | 0.32 | 0.01 | 0.01 |
| gene-LOC105285395 | https://www.ncbi.nlm.nih.gov/gene/?term=LOC105285395 | 8085 | 260 | 321.58 | 1 | 1.24 | 0.38 | 13.21 | 12.9 | 0.31 | 0.31 | 0.01 | 0.01 |
| gene-LOC105286265 | https://www.ncbi.nlm.nih.gov/gene/?term=LOC105286265 | 2507 | 68 | 271.24 | 1 | 3.99 | 1.47 | 31.01 | 30.72 | 0.29 | 0.29 | 0.01 | 0.01 |
| gene-LOC105278666 | https://www.ncbi.nlm.nih.gov/gene/?term=LOC105278666 | 10341 | 492 | 475.78 | 1 | 0.97 | 0.2 | 7.02 | 6.75 | 0.27 | 0.27 | 0.01 | 0.01 |
| gene-LOC105274449 | https://www.ncbi.nlm.nih.gov/gene/?term=LOC105274449 | 1549 | 53 | 342.16 | 1 | 6.46 | 1.89 | 49.51 | 49.29 | 0.22 | 0.22 | 0.01 | 0.01 |
| gene-LOC105275327 | https://www.ncbi.nlm.nih.gov/gene/?term=LOC105275327 | 14245 | 550 | 386.1 | 1 | 0.7 | 0.18 | 6.58 | 6.38 | 0.2 | 0.2 | 0.01 | 0.01 |
| gene-LOC105284262 | https://www.ncbi.nlm.nih.gov/gene/?term=LOC105284262 | 6669 | 248 | 371.87 | 1 | 1.5 | 0.4 | 5.25 | 5.05 | 0.2 | 0.2 | 0.01 | 0.01 |
| gene-LOC113562050 | https://www.ncbi.nlm.nih.gov/gene/?term=LOC113562050 | 17097 | 698 | 408.26 | 1 | 0.58 | 0.14 | 5.09 | 4.89 | 0.2 | 0.2 | 0.01 | 0.01 |
| gene-LOC105278534 | https://www.ncbi.nlm.nih.gov/gene/?term=LOC105278534 | 7364 | 214 | 290.6 | 1 | 1.36 | 0.47 | 16.92 | 16.72 | 0.19 | 0.19 | 0.01 | 0.01 |
| gene-LOC105274856 | https://www.ncbi.nlm.nih.gov/gene/?term=LOC105274856 | 5216 | 233 | 446.7 | 1 | 1.92 | 0.43 | 17.97 | 17.83 | 0.14 | 0.14 | 0.01 | 0.01 |
| gene-LOC105287065 | https://www.ncbi.nlm.nih.gov/gene/?term=LOC105287065 | 4859 | 259 | 533.03 | 1 | 2.06 | 0.39 | 12.11 | 12 | 0.11 | 0.11 | 0.01 | 0.01 |
| gene-LOC105286649 | https://www.ncbi.nlm.nih.gov/gene/?term=LOC105286649 | 14178 | 852 | 600.93 | 1 | 0.71 | 0.12 | 1.32 | 1.49 | -0.16 | 0.16 | -0.01 | 0.01 |
| gene-LOC105284116 | https://www.ncbi.nlm.nih.gov/gene/?term=LOC105284116 | 3970 | 134 | 337.53 | 1 | 2.52 | 0.75 | 7.83 | 8 | -0.17 | 0.17 | -0.01 | 0.01 |
| gene-LOC105287058 | https://www.ncbi.nlm.nih.gov/gene/?term=LOC105287058 | 4788 | 247 | 515.87 | 1 | 2.09 | 0.4 | 8.85 | 9.06 | -0.2 | 0.2 | -0.01 | 0.01 |
| gene-LOC105278771 | https://www.ncbi.nlm.nih.gov/gene/?term=LOC105278771 | 10457 | 267 | 255.33 | 1 | 0.96 | 0.37 | 9.06 | 9.27 | -0.21 | 0.21 | -0.01 | 0.01 |
| gene-LOC105286515 | https://www.ncbi.nlm.nih.gov/gene/?term=LOC105286515 | 3751 | 133 | 354.57 | 1 | 2.67 | 0.75 | 10.29 | 10.56 | -0.27 | 0.27 | -0.01 | 0.01 |
| gene-LOC105285981 | https://www.ncbi.nlm.nih.gov/gene/?term=LOC105285981 | 7666 | 224 | 292.2 | 1 | 1.3 | 0.45 | 17.35 | 17.63 | -0.28 | 0.28 | -0.01 | 0.01 |
| gene-LOC105280956 | https://www.ncbi.nlm.nih.gov/gene/?term=LOC105280956 | 2639 | 83 | 314.51 | 1 | 3.79 | 1.2 | 18.28 | 18.58 | -0.3 | 0.3 | -0.01 | 0.01 |
| gene-LOC105283649 | https://www.ncbi.nlm.nih.gov/gene/?term=LOC105283649 | 1918 | 59 | 307.61 | 2 | 10.43 | 3.39 | 41.12 | 41.42 | -0.31 | 0.31 | -0.01 | 0.01 |
| gene-LOC105285202 | https://www.ncbi.nlm.nih.gov/gene/?term=LOC105285202 | 1838 | 74 | 402.61 | 1 | 5.44 | 1.35 | 23.68 | 24 | -0.32 | 0.32 | -0.01 | 0.01 |
| gene-LOC105283132 | https://www.ncbi.nlm.nih.gov/gene/?term=LOC105283132 | 1819 | 56 | 307.86 | 1 | 5.5 | 1.79 | 39.81 | 40.18 | -0.38 | 0.38 | -0.01 | 0.01 |
| gene-LOC105280304 | https://www.ncbi.nlm.nih.gov/gene/?term=LOC105280304 | 8919 | 143 | 160.33 | 1 | 1.12 | 0.7 | 6.95 | 7.37 | -0.42 | 0.42 | -0.01 | 0.01 |
| gene-LOC105278454 | https://www.ncbi.nlm.nih.gov/gene/?term=LOC105278454 | 1972 | 43 | 218.05 | 1 | 5.07 | 2.33 | 42.19 | 42.86 | -0.67 | 0.67 | -0.01 | 0.01 |
| gene-LOC105280408 | https://www.ncbi.nlm.nih.gov/gene/?term=LOC105280408 | 5983 | 65 | 108.64 | 1 | 1.67 | 1.54 | 15.19 | 14.75 | 0.44 | 0.44 | 0 | 0 |
| gene-LOC105285406 | https://www.ncbi.nlm.nih.gov/gene/?term=LOC105285406 | 8015 | 63 | 78.6 | 1 | 1.25 | 1.59 | 28.98 | 28.57 | 0.41 | 0.41 | 0 | 0 |
| gene-LOC105275187 | https://www.ncbi.nlm.nih.gov/gene/?term=LOC105275187 | 5534 | 87 | 157.21 | 1 | 1.81 | 1.15 | 15.13 | 14.88 | 0.25 | 0.25 | 0 | 0 |
| gene-LOC105274590 | https://www.ncbi.nlm.nih.gov/gene/?term=LOC105274590 | 4516 | 88 | 194.86 | 2 | 4.43 | 2.27 | 24.95 | 24.81 | 0.14 | 0.14 | 0 | 0 |
| gene-LOC105274669 | https://www.ncbi.nlm.nih.gov/gene/?term=LOC105274669 | 8555 | 274 | 320.28 | 2 | 2.34 | 0.73 | 5.92 | 5.78 | 0.14 | 0.14 | 0 | 0 |
| gene-LOC105286452 | https://www.ncbi.nlm.nih.gov/gene/?term=LOC105286452 | 6888 | 78 | 113.24 | 1 | 1.45 | 1.28 | 26.06 | 25.92 | 0.14 | 0.14 | 0 | 0 |
| gene-LOC105286770 | https://www.ncbi.nlm.nih.gov/gene/?term=LOC105286770 | 3055 | 50 | 163.67 | 1 | 3.27 | 2 | 14.75 | 14.61 | 0.14 | 0.14 | 0 | 0 |
| gene-LOC105280410 | https://www.ncbi.nlm.nih.gov/gene/?term=LOC105280410 | 12580 | 320 | 254.37 | 1 | 0.79 | 0.31 | 17.81 | 17.7 | 0.11 | 0.11 | 0 | 0 |
| gene-LOC105278109 | https://www.ncbi.nlm.nih.gov/gene/?term=LOC105278109 | 4029 | 89 | 220.9 | 1 | 2.48 | 1.12 | 34.5 | 34.41 | 0.09 | 0.09 | 0 | 0 |
| gene-LOC105282811 | https://www.ncbi.nlm.nih.gov/gene/?term=LOC105282811 | 5765 | 232 | 402.43 | 1 | 1.73 | 0.43 | 8.6 | 8.51 | 0.09 | 0.09 | 0 | 0 |
| gene-LOC105287095 | https://www.ncbi.nlm.nih.gov/gene/?term=LOC105287095 | 1435 | 71 | 494.77 | 1 | 6.97 | 1.41 | 23.64 | 23.55 | 0.09 | 0.09 | 0 | 0 |
| gene-LOC105276917 | https://www.ncbi.nlm.nih.gov/gene/?term=LOC105276917 | 2336 | 42 | 179.79 | 1 | 4.28 | 2.38 | 28.76 | 28.68 | 0.08 | 0.08 | 0 | 0 |
| gene-LOC105276059 | https://www.ncbi.nlm.nih.gov/gene/?term=LOC105276059 | 6256 | 148 | 236.57 | 1 | 1.6 | 0.68 | 11.97 | 11.9 | 0.07 | 0.07 | 0 | 0 |
| gene-LOC105283264 | https://www.ncbi.nlm.nih.gov/gene/?term=LOC105283264 | 4005 | 94 | 234.71 | 1 | 2.5 | 1.06 | 22.97 | 22.91 | 0.07 | 0.07 | 0 | 0 |
| gene-LOC105283296 | https://www.ncbi.nlm.nih.gov/gene/?term=LOC105283296 | 2174 | 96 | 441.58 | 1 | 4.6 | 1.04 | 14.24 | 14.2 | 0.04 | 0.04 | 0 | 0 |
| gene-LOC105280819 | https://www.ncbi.nlm.nih.gov/gene/?term=LOC105280819 | 1639 | 44 | 268.46 | 1 | 6.1 | 2.27 | 37.71 | 37.7 | 0.02 | 0.02 | 0 | 0 |
| gene-LOC105277216 | https://www.ncbi.nlm.nih.gov/gene/?term=LOC105277216 | 7627 | 101 | 132.42 | 1 | 1.31 | 0.99 | 8.48 | 8.48 | 0 | 0 | 0 | 0 |
| gene-LOC105279664 | https://www.ncbi.nlm.nih.gov/gene/?term=LOC105279664 | 4023 | 75 | 186.43 | 1 | 2.49 | 1.33 | 19.05 | 19.05 | 0 | 0 | 0 | 0 |
| gene-LOC105279528 | https://www.ncbi.nlm.nih.gov/gene/?term=LOC105279528 | 5802 | 40 | 68.94 | 1 | 1.72 | 2.5 | 19.68 | 19.72 | -0.04 | 0.04 | 0 | 0 |
| gene-LOC105277588 | https://www.ncbi.nlm.nih.gov/gene/?term=LOC105277588 | 3955 | 98 | 247.79 | 1 | 2.53 | 1.02 | 8.55 | 8.61 | -0.06 | 0.06 | 0 | 0 |
| gene-LOC105277761 | https://www.ncbi.nlm.nih.gov/gene/?term=LOC105277761 | 4648 | 174 | 374.35 | 1 | 2.15 | 0.57 | 14.23 | 14.3 | -0.07 | 0.07 | 0 | 0 |
| gene-LOC105279657 | https://www.ncbi.nlm.nih.gov/gene/?term=LOC105279657 | 5878 | 112 | 190.54 | 1 | 1.7 | 0.89 | 16.48 | 16.55 | -0.07 | 0.07 | 0 | 0 |
| gene-LOC105284774 | https://www.ncbi.nlm.nih.gov/gene/?term=LOC105284774 | 12089 | 106 | 87.68 | 1 | 0.83 | 0.94 | 19.59 | 19.72 | -0.12 | 0.12 | 0 | 0 |
| gene-LOC105276196 | https://www.ncbi.nlm.nih.gov/gene/?term=LOC105276196 | 2953 | 61 | 206.57 | 1 | 3.39 | 1.64 | 38.98 | 39.11 | -0.13 | 0.13 | 0 | 0 |
| gene-LOC105284117 | https://www.ncbi.nlm.nih.gov/gene/?term=LOC105284117 | 4340 | 116 | 267.28 | 1 | 2.3 | 0.86 | 11.19 | 11.33 | -0.14 | 0.14 | 0 | 0 |

Directory: OGL File: rnk-dmg-Ob_LI_RphaseA2.vs.RphaseB2.txt

| gene_ID | gene_link | gwidth | #Sites | #per10Kb | #dmSites | #dmsp10kb | %dmSites | %pSite1 | %pSite2 | DMpSite | ADMpSite | DMpNucl | ADMpNucl |
|---|---|---|---|---|---|---|---|---|---|---|---|---|---|
| gene-LOC113562345 | https://www.ncbi.nlm.nih.gov/gene/?term=LOC113562345 | 2290 | 69 | 301.31 | 14 | 61.14 | 20.29 | 29.74 | 16.81 | 12.93 | 12.93 | 0.39 | 0.39 |
| gene-LOC105284677 | https://www.ncbi.nlm.nih.gov/gene/?term=LOC105284677 | 1905 | 42 | 220.47 | 1 | 5.25 | 2.38 | 23.81 | 34.99 | -11.18 | 11.18 | -0.25 | 0.25 |
| gene-LOC113562901 | https://www.ncbi.nlm.nih.gov/gene/?term=LOC113562901 | 1531 | 24 | 156.76 | 3 | 19.6 | 12.5 | 59.3 | 45.31 | 14 | 14 | 0.22 | 0.22 |
| gene-LOC105286644 | https://www.ncbi.nlm.nih.gov/gene/?term=LOC105286644 | 1235 | 22 | 178.14 | 1 | 8.1 | 4.55 | 44.4 | 37.26 | 7.14 | 7.14 | 0.13 | 0.13 |
| gene-LOC105280786 | https://www.ncbi.nlm.nih.gov/gene/?term=LOC105280786 | 2573 | 127 | 493.59 | 1 | 3.89 | 0.79 | 30.11 | 27.41 | 2.7 | 2.7 | 0.13 | 0.13 |
| gene-LOC105274706 | https://www.ncbi.nlm.nih.gov/gene/?term=LOC105274706 | 1446 | 56 | 387.28 | 1 | 6.92 | 1.79 | 39.74 | 36.7 | 3.05 | 3.05 | 0.12 | 0.12 |
| gene-LOC105275447 | https://www.ncbi.nlm.nih.gov/gene/?term=LOC105275447 | 1846 | 31 | 167.93 | 1 | 5.42 | 3.23 | 9.13 | 2.78 | 6.34 | 6.34 | 0.11 | 0.11 |
| gene-LOC105286683 | https://www.ncbi.nlm.nih.gov/gene/?term=LOC105286683 | 1665 | 57 | 342.34 | 2 | 12.01 | 3.51 | 34.89 | 31.76 | 3.13 | 3.13 | 0.11 | 0.11 |
| gene-LOC105285671 | https://www.ncbi.nlm.nih.gov/gene/?term=LOC105285671 | 2263 | 106 | 468.4 | 1 | 4.42 | 0.94 | 28.93 | 26.53 | 2.4 | 2.4 | 0.11 | 0.11 |
| gene-LOC105287417 | https://www.ncbi.nlm.nih.gov/gene/?term=LOC105287417 | 1025 | 23 | 224.39 | 1 | 9.76 | 4.35 | 53.58 | 49.27 | 4.31 | 4.31 | 0.1 | 0.1 |
| gene-LOC105285583 | https://www.ncbi.nlm.nih.gov/gene/?term=LOC105285583 | 1947 | 49 | 251.67 | 1 | 5.14 | 2.04 | 30.89 | 26.83 | 4.06 | 4.06 | 0.1 | 0.1 |
| gene-LOC105285674 | https://www.ncbi.nlm.nih.gov/gene/?term=LOC105285674 | 3035 | 102 | 336.08 | 2 | 6.59 | 1.96 | 26.23 | 23.21 | 3.03 | 3.03 | 0.1 | 0.1 |
| gene-LOC105276704 | https://www.ncbi.nlm.nih.gov/gene/?term=LOC105276704 | 2063 | 88 | 426.56 | 1 | 4.85 | 1.14 | 16.14 | 13.92 | 2.23 | 2.23 | 0.1 | 0.1 |
| gene-LOC105282156 | https://www.ncbi.nlm.nih.gov/gene/?term=LOC105282156 | 2577 | 130 | 504.46 | 1 | 3.88 | 0.77 | 28.69 | 26.74 | 1.95 | 1.95 | 0.1 | 0.1 |
| gene-LOC105275070 | https://www.ncbi.nlm.nih.gov/gene/?term=LOC105275070 | 5667 | 42 | 74.11 | 5 | 8.82 | 11.9 | 13.23 | 1.03 | 12.2 | 12.2 | 0.09 | 0.09 |
| gene-LOC105279898 | https://www.ncbi.nlm.nih.gov/gene/?term=LOC105279898 | 1343 | 10 | 74.46 | 1 | 7.45 | 10 | 83.16 | 71.53 | 11.63 | 11.63 | 0.09 | 0.09 |
| gene-LOC105282388 | https://www.ncbi.nlm.nih.gov/gene/?term=LOC105282388 | 1661 | 34 | 204.7 | 1 | 6.02 | 2.94 | 48.95 | 44.6 | 4.35 | 4.35 | 0.09 | 0.09 |
| gene-LOC105280555 | https://www.ncbi.nlm.nih.gov/gene/?term=LOC105280555 | 1496 | 31 | 207.22 | 1 | 6.68 | 3.23 | 61.05 | 57.2 | 3.85 | 3.85 | 0.08 | 0.08 |
| gene-LOC105277631 | https://www.ncbi.nlm.nih.gov/gene/?term=LOC105277631 | 4387 | 304 | 692.96 | 1 | 2.28 | 0.33 | 10.76 | 9.61 | 1.15 | 1.15 | 0.08 | 0.08 |
| gene-LOC105282893 | https://www.ncbi.nlm.nih.gov/gene/?term=LOC105282893 | 2645 | 47 | 177.69 | 1 | 3.78 | 2.13 | 30.08 | 25.9 | 4.18 | 4.18 | 0.07 | 0.07 |
| gene-LOC105280104 | https://www.ncbi.nlm.nih.gov/gene/?term=LOC105280104 | 3960 | 139 | 351.01 | 1 | 2.53 | 0.72 | 17.44 | 15.39 | 2.05 | 2.05 | 0.07 | 0.07 |
| gene-LOC105280377 | https://www.ncbi.nlm.nih.gov/gene/?term=LOC105280377 | 2024 | 70 | 345.85 | 1 | 4.94 | 1.43 | 48.1 | 46.12 | 1.98 | 1.98 | 0.07 | 0.07 |
| gene-LOC105282947 | https://www.ncbi.nlm.nih.gov/gene/?term=LOC105282947 | 1162 | 41 | 352.84 | 1 | 8.61 | 2.44 | 36.57 | 34.72 | 1.85 | 1.85 | 0.07 | 0.07 |
| gene-LOC105287063 | https://www.ncbi.nlm.nih.gov/gene/?term=LOC105287063 | 1874 | 48 | 256.14 | 1 | 5.34 | 2.08 | 32.76 | 30.33 | 2.44 | 2.44 | 0.06 | 0.06 |
| gene-LOC105283132 | https://www.ncbi.nlm.nih.gov/gene/?term=LOC105283132 | 1819 | 57 | 313.36 | 1 | 5.5 | 1.75 | 39.92 | 37.88 | 2.04 | 2.04 | 0.06 | 0.06 |
| gene-LOC105283182 | https://www.ncbi.nlm.nih.gov/gene/?term=LOC105283182 | 3585 | 131 | 365.41 | 1 | 2.79 | 0.76 | 14.16 | 12.63 | 1.52 | 1.52 | 0.06 | 0.06 |
| gene-LOC105276449 | https://www.ncbi.nlm.nih.gov/gene/?term=LOC105276449 | 1231 | 50 | 406.17 | 1 | 8.12 | 2 | 32.79 | 31.42 | 1.37 | 1.37 | 0.06 | 0.06 |
| gene-LOC105278497 | https://www.ncbi.nlm.nih.gov/gene/?term=LOC105278497 | 2072 | 100 | 482.63 | 1 | 4.83 | 1 | 19.97 | 18.78 | 1.2 | 1.2 | 0.06 | 0.06 |
| gene-LOC105284043 | https://www.ncbi.nlm.nih.gov/gene/?term=LOC105284043 | 5661 | 308 | 544.07 | 1 | 1.77 | 0.32 | 10.53 | 9.46 | 1.06 | 1.06 | 0.06 | 0.06 |
| gene-LOC105278283 | https://www.ncbi.nlm.nih.gov/gene/?term=LOC105278283 | 2317 | 142 | 612.86 | 1 | 4.32 | 0.7 | 25.61 | 24.56 | 1.05 | 1.05 | 0.06 | 0.06 |
| gene-LOC105282149 | https://www.ncbi.nlm.nih.gov/gene/?term=LOC105282149 | 1787 | 110 | 615.56 | 1 | 5.6 | 0.91 | 43.04 | 42 | 1.04 | 1.04 | 0.06 | 0.06 |
| gene-LOC105275708 | https://www.ncbi.nlm.nih.gov/gene/?term=LOC105275708 | 1772 | 15 | 84.65 | 1 | 5.64 | 6.67 | 47.19 | 40.83 | 6.36 | 6.36 | 0.05 | 0.05 |
| gene-LOC105276538 | https://www.ncbi.nlm.nih.gov/gene/?term=LOC105276538 | 1511 | 20 | 132.36 | 1 | 6.62 | 5 | 73.31 | 69.21 | 4.1 | 4.1 | 0.05 | 0.05 |
| gene-LOC105278215 | https://www.ncbi.nlm.nih.gov/gene/?term=LOC105278215 | 1700 | 29 | 170.59 | 1 | 5.88 | 3.45 | 57.59 | 54.66 | 2.93 | 2.93 | 0.05 | 0.05 |
| gene-LOC105278109 | https://www.ncbi.nlm.nih.gov/gene/?term=LOC105278109 | 4029 | 67 | 166.29 | 1 | 2.48 | 1.49 | 44.53 | 41.68 | 2.86 | 2.86 | 0.05 | 0.05 |
| gene-LOC105282794 | https://www.ncbi.nlm.nih.gov/gene/?term=LOC105282794 | 4571 | 95 | 207.83 | 1 | 2.19 | 1.05 | 18.64 | 16.23 | 2.41 | 2.41 | 0.05 | 0.05 |
| gene-LOC105287646 | https://www.ncbi.nlm.nih.gov/gene/?term=LOC105287646 | 3556 | 90 | 253.09 | 1 | 2.81 | 1.11 | 18.4 | 16.59 | 1.81 | 1.81 | 0.05 | 0.05 |
| gene-LOC105278282 | https://www.ncbi.nlm.nih.gov/gene/?term=LOC105278282 | 1775 | 56 | 315.49 | 1 | 5.63 | 1.79 | 39.77 | 38.34 | 1.43 | 1.43 | 0.05 | 0.05 |
| gene-LOC105276875 | https://www.ncbi.nlm.nih.gov/gene/?term=LOC105276875 | 3151 | 118 | 374.48 | 1 | 3.17 | 0.85 | 23.91 | 22.53 | 1.38 | 1.38 | 0.05 | 0.05 |
| gene-LOC105284355 | https://www.ncbi.nlm.nih.gov/gene/?term=LOC105284355 | 3904 | 153 | 391.91 | 1 | 2.56 | 0.65 | 21.82 | 20.47 | 1.35 | 1.35 | 0.05 | 0.05 |
| gene-LOC105283728 | https://www.ncbi.nlm.nih.gov/gene/?term=LOC105283728 | 3256 | 120 | 368.55 | 2 | 6.14 | 1.67 | 19.01 | 17.69 | 1.32 | 1.32 | 0.05 | 0.05 |
| gene-LOC105281419 | https://www.ncbi.nlm.nih.gov/gene/?term=LOC105281419 | 2366 | 83 | 350.8 | 1 | 4.23 | 1.2 | 23.42 | 22.13 | 1.3 | 1.3 | 0.05 | 0.05 |
| gene-LOC105275363 | https://www.ncbi.nlm.nih.gov/gene/?term=LOC105275363 | 3798 | 165 | 434.44 | 1 | 2.63 | 0.61 | 17.38 | 16.26 | 1.12 | 1.12 | 0.05 | 0.05 |
| gene-LOC105279298 | https://www.ncbi.nlm.nih.gov/gene/?term=LOC105279298 | 2773 | 136 | 490.44 | 2 | 7.21 | 1.47 | 23.98 | 22.86 | 1.11 | 1.11 | 0.05 | 0.05 |
| gene-LOC105282682 | https://www.ncbi.nlm.nih.gov/gene/?term=LOC105282682 | 4127 | 223 | 540.34 | 1 | 2.42 | 0.45 | 13.69 | 12.76 | 0.93 | 0.93 | 0.05 | 0.05 |
| gene-LOC105285055 | https://www.ncbi.nlm.nih.gov/gene/?term=LOC105285055 | 4108 | 276 | 671.86 | 2 | 4.87 | 0.72 | 11.41 | 10.6 | 0.8 | 0.8 | 0.05 | 0.05 |
| gene-LOC105285534 | https://www.ncbi.nlm.nih.gov/gene/?term=LOC105285534 | 3944 | 237 | 600.91 | 1 | 2.54 | 0.42 | 11.45 | 10.67 | 0.78 | 0.78 | 0.05 | 0.05 |
| gene-LOC105287854 | https://www.ncbi.nlm.nih.gov/gene/?term=LOC105287854 | 3644 | 54 | 148.19 | 1 | 2.74 | 1.85 | 25.58 | 23.15 | 2.43 | 2.43 | 0.04 | 0.04 |
| gene-LOC105276917 | https://www.ncbi.nlm.nih.gov/gene/?term=LOC105276917 | 2336 | 38 | 162.67 | 1 | 4.28 | 2.63 | 28.71 | 26.35 | 2.36 | 2.36 | 0.04 | 0.04 |
| gene-LOC113561805 | https://www.ncbi.nlm.nih.gov/gene/?term=LOC113561805 | 3337 | 71 | 212.77 | 1 | 3 | 1.41 | 36.21 | 34.33 | 1.88 | 1.88 | 0.04 | 0.04 |
| gene-LOC105283649 | https://www.ncbi.nlm.nih.gov/gene/?term=LOC105283649 | 1918 | 56 | 291.97 | 2 | 10.43 | 3.57 | 37.98 | 36.48 | 1.5 | 1.5 | 0.04 | 0.04 |
| gene-LOC105282970 | https://www.ncbi.nlm.nih.gov/gene/?term=LOC105282970 | 1529 | 40 | 261.61 | 1 | 6.54 | 2.5 | 49.63 | 48.22 | 1.41 | 1.41 | 0.04 | 0.04 |
| gene-LOC105281272 | https://www.ncbi.nlm.nih.gov/gene/?term=LOC105281272 | 5252 | 164 | 312.26 | 1 | 1.9 | 0.61 | 19.36 | 17.96 | 1.4 | 1.4 | 0.04 | 0.04 |
| gene-LOC105284704 | https://www.ncbi.nlm.nih.gov/gene/?term=LOC105284704 | 2400 | 62 | 258.33 | 1 | 4.17 | 1.61 | 24.3 | 22.92 | 1.38 | 1.38 | 0.04 | 0.04 |
| gene-LOC105283123 | https://www.ncbi.nlm.nih.gov/gene/?term=LOC105283123 | 1507 | 48 | 318.51 | 1 | 6.64 | 2.08 | 17.79 | 16.48 | 1.31 | 1.31 | 0.04 | 0.04 |
| gene-LOC105288079 | https://www.ncbi.nlm.nih.gov/gene/?term=LOC105288079 | 3473 | 101 | 290.81 | 1 | 2.88 | 0.99 | 20.37 | 19.15 | 1.21 | 1.21 | 0.04 | 0.04 |
| gene-LOC105277238 | https://www.ncbi.nlm.nih.gov/gene/?term=LOC105277238 | 2708 | 88 | 324.96 | 1 | 3.69 | 1.14 | 27.16 | 25.96 | 1.2 | 1.2 | 0.04 | 0.04 |
| gene-LOC105277761 | https://www.ncbi.nlm.nih.gov/gene/?term=LOC105277761 | 4648 | 150 | 322.72 | 1 | 2.15 | 0.67 | 14.38 | 13.19 | 1.19 | 1.19 | 0.04 | 0.04 |
| gene-LOC105284184 | https://www.ncbi.nlm.nih.gov/gene/?term=LOC105284184 | 2535 | 86 | 339.25 | 1 | 3.94 | 1.16 | 34.02 | 32.88 | 1.13 | 1.13 | 0.04 | 0.04 |
| gene-LOC113561621 | https://www.ncbi.nlm.nih.gov/gene/?term=LOC113561621 | 2249 | 86 | 382.39 | 1 | 4.45 | 1.16 | 20.44 | 19.32 | 1.11 | 1.11 | 0.04 | 0.04 |

| | | | | | | | | | | | | | |
|---|---|---|---|---|---|---|---|---|---|---|---|---|---|
| gene-LOC105285639 | https://www.ncbi.nlm.nih.gov/gene/?term=LOC105285639 | 10325 | 349 | 338.01 | 1 | 0.97 | 0.29 | 13.03 | 11.95 | 1.08 | 1.08 | 0.04 | 0.04 |
| gene-LOC105285742 | https://www.ncbi.nlm.nih.gov/gene/?term=LOC105285742 | 3446 | 123 | 356.94 | 1 | 2.9 | 0.81 | 20.56 | 19.51 | 1.04 | 1.04 | 0.04 | 0.04 |
| gene-LOC105274856 | https://www.ncbi.nlm.nih.gov/gene/?term=LOC105274856 | 5216 | 216 | 414.11 | 1 | 1.92 | 0.46 | 19.8 | 18.89 | 0.91 | 0.91 | 0.04 | 0.04 |
| gene-LOC105275460 | https://www.ncbi.nlm.nih.gov/gene/?term=LOC105275460 | 2473 | 117 | 473.11 | 1 | 4.04 | 0.85 | 14.05 | 13.26 | 0.78 | 0.78 | 0.04 | 0.04 |
| gene-LOC105284047 | https://www.ncbi.nlm.nih.gov/gene/?term=LOC105284047 | 1212 | 24 | 198.02 | 1 | 8.25 | 4.17 | 25.81 | 27.68 | -1.87 | 1.87 | -0.04 | 0.04 |
| gene-LOC105274778 | https://www.ncbi.nlm.nih.gov/gene/?term=LOC105274778 | 4662 | 40 | 85.8 1 | 2.15 | 2.5 | 47.99 | 44.37 | 3.62 | 3.62 | 0.03 | 0.03 | |
| gene-LOC105275518 | https://www.ncbi.nlm.nih.gov/gene/?term=LOC105275518 | 2335 | 24 | 102.78 | 1 | 4.28 | 4.17 | 49.73 | 46.6 | 3.13 | 3.13 | 0.03 | 0.03 |
| gene-LOC105279664 | https://www.ncbi.nlm.nih.gov/gene/?term=LOC105279664 | 4023 | 56 | 139.2 1 | 2.49 | 1.79 | 21.11 | 18.99 | 2.12 | 2.12 | 0.03 | 0.03 | |
| gene-LOC105280641 | https://www.ncbi.nlm.nih.gov/gene/?term=LOC105280641 | 3682 | 53 | 143.94 | 1 | 2.72 | 1.89 | 17.27 | 15.26 | 2.01 | 2.01 | 0.03 | 0.03 |
| gene-LOC105282600 | https://www.ncbi.nlm.nih.gov/gene/?term=LOC105282600 | 3715 | 56 | 150.74 | 1 | 2.69 | 1.79 | 27.25 | 25.37 | 1.87 | 1.87 | 0.03 | 0.03 |
| gene-LOC105276190 | https://www.ncbi.nlm.nih.gov/gene/?term=LOC105276190 | 4839 | 93 | 192.19 | 1 | 2.07 | 1.08 | 11.07 | 9.37 | 1.7 | 1.7 | 0.03 | 0.03 |
| gene-LOC105281420 | https://www.ncbi.nlm.nih.gov/gene/?term=LOC105281420 | 3882 | 62 | 159.71 | 1 | 2.58 | 1.61 | 15.69 | 14 | 1.7 | 1.7 | 0.03 | 0.03 |
| gene-LOC105276692 | https://www.ncbi.nlm.nih.gov/gene/?term=LOC105276692 | 1745 | 35 | 200.57 | 1 | 5.73 | 2.86 | 22.12 | 20.68 | 1.44 | 1.44 | 0.03 | 0.03 |
| gene-LOC105282151 | https://www.ncbi.nlm.nih.gov/gene/?term=LOC105282151 | 1601 | 39 | 243.6 2 | 12.49 | 5.13 | 42.25 | 41 | 1.25 | 1.25 | 0.03 | 0.03 | |
| gene-LOC105281848 | https://www.ncbi.nlm.nih.gov/gene/?term=LOC105281848 | 2122 | 48 | 226.2 1 | 4.71 | 2.08 | 27.66 | 26.43 | 1.23 | 1.23 | 0.03 | 0.03 | |
| gene-LOC105281173 | https://www.ncbi.nlm.nih.gov/gene/?term=LOC105281173 | 1595 | 39 | 244.51 | 1 | 6.27 | 2.56 | 61.03 | 59.91 | 1.13 | 1.13 | 0.03 | 0.03 |
| gene-LOC109611283 | https://www.ncbi.nlm.nih.gov/gene/?term=LOC109611283 | 1451 | 37 | 255 1 | 6.89 | 2.7 | 37.11 | 36.1 | 1 | 1 | 0.03 | 0.03 | |
| gene-LOC105285395 | https://www.ncbi.nlm.nih.gov/gene/?term=LOC105285395 | 8085 | 218 | 269.64 | 1 | 1.24 | 0.46 | 17.01 | 16.05 | 0.96 | 0.96 | 0.03 | 0.03 |
| gene-LOC105278807 | https://www.ncbi.nlm.nih.gov/gene/?term=LOC105278807 | 2166 | 67 | 309.33 | 1 | 4.62 | 1.49 | 22.84 | 21.92 | 0.93 | 0.93 | 0.03 | 0.03 |
| gene-LOC105279038 | https://www.ncbi.nlm.nih.gov/gene/?term=LOC105279038 | 9453 | 313 | 331.11 | 1 | 1.06 | 0.32 | 10.35 | 9.46 | 0.9 | 0.9 | 0.03 | 0.03 |
| gene-LOC105284036 | https://www.ncbi.nlm.nih.gov/gene/?term=LOC105284036 | 3254 | 108 | 331.9 1 | 3.07 | 0.93 | 22.88 | 22.09 | 0.79 | 0.79 | 0.03 | 0.03 | |
| gene-LOC105280774 | https://www.ncbi.nlm.nih.gov/gene/?term=LOC105280774 | 2793 | 100 | 358.04 | 1 | 3.58 | 1 | 32.96 | 32.18 | 0.78 | 0.78 | 0.03 | 0.03 |
| gene-LOC105274534 | https://www.ncbi.nlm.nih.gov/gene/?term=LOC105274534 | 1329 | 57 | 428.89 | 1 | 7.52 | 1.75 | 48.36 | 47.65 | 0.71 | 0.71 | 0.03 | 0.03 |
| gene-LOC105284557 | https://www.ncbi.nlm.nih.gov/gene/?term=LOC105284557 | 1672 | 84 | 502.39 | 1 | 5.98 | 1.19 | 29.36 | 28.7 | 0.66 | 0.66 | 0.03 | 0.03 |
| gene-LOC105275683 | https://www.ncbi.nlm.nih.gov/gene/?term=LOC105275683 | 4256 | 193 | 453.48 | 1 | 2.35 | 0.52 | 13.76 | 13.13 | 0.63 | 0.63 | 0.03 | 0.03 |
| gene-LOC105285179 | https://www.ncbi.nlm.nih.gov/gene/?term=LOC105285179 | 4191 | 185 | 441.42 | 1 | 2.39 | 0.54 | 18.94 | 18.34 | 0.6 | 0.6 | 0.03 | 0.03 |
| gene-LOC105275374 | https://www.ncbi.nlm.nih.gov/gene/?term=LOC105275374 | 11376 | 523 | 459.74 | 1 | 0.88 | 0.19 | 11.54 | 10.97 | 0.57 | 0.57 | 0.03 | 0.03 |
| gene-LOC105276612 | https://www.ncbi.nlm.nih.gov/gene/?term=LOC105276612 | 4157 | 187 | 449.84 | 1 | 2.41 | 0.53 | 10.04 | 9.47 | 0.57 | 0.57 | 0.03 | 0.03 |
| gene-LOC105276452 | https://www.ncbi.nlm.nih.gov/gene/?term=LOC105276452 | 2917 | 269 | 922.18 | 1 | 3.43 | 0.37 | 9.31 | 8.95 | 0.36 | 0.36 | 0.03 | 0.03 |
| gene-LOC105280596 | https://www.ncbi.nlm.nih.gov/gene/?term=LOC105280596 | 3434 | 113 | 329.06 | 1 | 2.91 | 0.88 | 17.62 | 18.56 | -0.94 | 0.94 | -0.03 | 0.03 |
| gene-LOC105282885 | https://www.ncbi.nlm.nih.gov/gene/?term=LOC105282885 | 1870 | 48 | 256.68 | 1 | 5.35 | 2.08 | 38.27 | 39.26 | -0.99 | 0.99 | -0.03 | 0.03 |
| gene-LOC105278384 | https://www.ncbi.nlm.nih.gov/gene/?term=LOC105278384 | 1390 | 36 | 258.99 | 1 | 7.19 | 2.78 | 50.65 | 51.65 | -1 | 1 | -0.03 | 0.03 |
| gene-LOC105280819 | https://www.ncbi.nlm.nih.gov/gene/?term=LOC105280819 | 1639 | 38 | 231.85 | 1 | 6.1 | 2.63 | 35.58 | 37.06 | -1.49 | 1.49 | -0.03 | 0.03 |
| gene-LOC105276196 | https://www.ncbi.nlm.nih.gov/gene/?term=LOC105276196 | 2953 | 62 | 209.96 | 1 | 3.39 | 1.61 | 36.95 | 38.53 | -1.58 | 1.58 | -0.03 | 0.03 |
| gene-LOC105281700 | https://www.ncbi.nlm.nih.gov/gene/?term=LOC105281700 | 2208 | 34 | 153.99 | 1 | 4.53 | 2.94 | 35.23 | 36.92 | -1.69 | 1.69 | -0.03 | 0.03 |
| gene-LOC105285409 | https://www.ncbi.nlm.nih.gov/gene/?term=LOC105285409 | 5099 | 49 | 96.1 2 | 3.92 | 4.08 | 1.55 | 4.82 | -3.27 | 3.27 | -0.03 | 0.03 | |
| gene-LOC105283492 | https://www.ncbi.nlm.nih.gov/gene/?term=LOC105283492 | 1535 | 10 | 65.15 1 | 6.51 | 10 | 37.74 | 42.8 | -5.06 | 5.06 | -0.03 | 0.03 | |
| gene-LOC105286694 | https://www.ncbi.nlm.nih.gov/gene/?term=LOC105286694 | 3409 | 12 | 35.2 1 | 2.93 | 8.33 | 52.59 | 46.71 | 5.88 | 5.88 | 0.02 | 0.02 | |
| gene-LOC105284609 | https://www.ncbi.nlm.nih.gov/gene/?term=LOC105284609 | 5678 | 33 | 58.12 1 | 1.76 | 3.03 | 34 | 30.46 | 3.54 | 3.54 | 0.02 | 0.02 | |
| gene-LOC105274900 | https://www.ncbi.nlm.nih.gov/gene/?term=LOC105274900 | 5561 | 56 | 100.7 2 | 3.6 | 3.57 | 32.09 | 29.76 | 2.33 | 2.33 | 0.02 | 0.02 | |
| gene-LOC105277856 | https://www.ncbi.nlm.nih.gov/gene/?term=LOC105277856 | 3442 | 35 | 101.69 | 1 | 2.91 | 2.86 | 35 | 32.98 | 2.02 | 2.02 | 0.02 | 0.02 |
| gene-LOC105278108 | https://www.ncbi.nlm.nih.gov/gene/?term=LOC105278108 | 4392 | 51 | 116.12 | 1 | 2.28 | 1.96 | 25.15 | 23.14 | 2.01 | 2.01 | 0.02 | 0.02 |
| gene-LOC105284117 | https://www.ncbi.nlm.nih.gov/gene/?term=LOC105284117 | 4340 | 65 | 149.77 | 1 | 2.3 | 1.54 | 12.82 | 11.26 | 1.56 | 1.56 | 0.02 | 0.02 |
| gene-LOC105287648 | https://www.ncbi.nlm.nih.gov/gene/?term=LOC105287648 | 2205 | 26 | 117.91 | 1 | 4.54 | 3.85 | 41.13 | 39.59 | 1.54 | 1.54 | 0.02 | 0.02 |
| gene-LOC105288063 | https://www.ncbi.nlm.nih.gov/gene/?term=LOC105288063 | 3400 | 47 | 138.24 | 1 | 2.94 | 2.13 | 15.4 | 13.86 | 1.54 | 1.54 | 0.02 | 0.02 |
| gene-LOC105274590 | https://www.ncbi.nlm.nih.gov/gene/?term=LOC105274590 | 4516 | 71 | 157.22 | 2 | 4.43 | 2.82 | 29.16 | 27.84 | 1.32 | 1.32 | 0.02 | 0.02 |
| gene-LOC105283712 | https://www.ncbi.nlm.nih.gov/gene/?term=LOC105283712 | 1910 | 36 | 188.48 | 1 | 5.24 | 2.78 | 27.71 | 26.46 | 1.25 | 1.25 | 0.02 | 0.02 |
| gene-LOC105288066 | https://www.ncbi.nlm.nih.gov/gene/?term=LOC105288066 | 1964 | 30 | 152.75 | 1 | 5.09 | 3.33 | 32.32 | 31.12 | 1.2 | 1.2 | 0.02 | 0.02 |
| gene-LOC105287720 | https://www.ncbi.nlm.nih.gov/gene/?term=LOC105287720 | 4843 | 99 | 204.42 | 1 | 2.06 | 1.01 | 16.39 | 15.24 | 1.16 | 1.16 | 0.02 | 0.02 |
| gene-LOC105275451 | https://www.ncbi.nlm.nih.gov/gene/?term=LOC105275451 | 6035 | 130 | 215.41 | 1 | 1.66 | 0.77 | 22.16 | 21.09 | 1.07 | 1.07 | 0.02 | 0.02 |
| gene-LOC105288010 | https://www.ncbi.nlm.nih.gov/gene/?term=LOC105288010 | 3189 | 51 | 159.92 | 1 | 3.14 | 1.96 | 29.8 | 28.73 | 1.07 | 1.07 | 0.02 | 0.02 |
| gene-LOC105286514 | https://www.ncbi.nlm.nih.gov/gene/?term=LOC105286514 | 2528 | 49 | 193.83 | 1 | 3.96 | 2.04 | 36.75 | 35.71 | 1.04 | 1.04 | 0.02 | 0.02 |
| gene-LOC105283592 | https://www.ncbi.nlm.nih.gov/gene/?term=LOC105283592 | 3479 | 80 | 229.95 | 1 | 2.87 | 1.25 | 23.33 | 22.32 | 1.01 | 1.01 | 0.02 | 0.02 |
| gene-LOC105278962 | https://www.ncbi.nlm.nih.gov/gene/?term=LOC105278962 | 4491 | 104 | 231.57 | 1 | 2.23 | 0.96 | 22.01 | 21.09 | 0.92 | 0.92 | 0.02 | 0.02 |
| gene-LOC105276059 | https://www.ncbi.nlm.nih.gov/gene/?term=LOC105276059 | 6256 | 133 | 212.6 1 | 1.6 | 0.75 | 10.26 | 9.4 | 0.86 | 0.86 | 0.02 | 0.02 | |
| gene-LOC105280956 | https://www.ncbi.nlm.nih.gov/gene/?term=LOC105280956 | 2639 | 71 | 269.04 | 1 | 3.79 | 1.41 | 18.65 | 17.79 | 0.86 | 0.86 | 0.02 | 0.02 |
| gene-LOC105280407 | https://www.ncbi.nlm.nih.gov/gene/?term=LOC105280407 | 10950 | 211 | 192.69 | 1 | 0.91 | 0.47 | 16.65 | 15.81 | 0.84 | 0.84 | 0.02 | 0.02 |
| gene-LOC105274823 | https://www.ncbi.nlm.nih.gov/gene/?term=LOC105274823 | 4620 | 141 | 305.19 | 1 | 2.16 | 0.71 | 21.37 | 20.81 | 0.56 | 0.56 | 0.02 | 0.02 |
| gene-LOC105275127 | https://www.ncbi.nlm.nih.gov/gene/?term=LOC105275127 | 9377 | 301 | 321 1 | 1.07 | 0.33 | 7.86 | 7.34 | 0.52 | 0.52 | 0.02 | 0.02 | |
| gene-LOC105285202 | https://www.ncbi.nlm.nih.gov/gene/?term=LOC105285202 | 1838 | 70 | 380.85 | 1 | 5.44 | 1.43 | 25.94 | 25.46 | 0.48 | 0.48 | 0.02 | 0.02 |
| gene-LOC105282491 | https://www.ncbi.nlm.nih.gov/gene/?term=LOC105282491 | 4761 | 170 | 357.07 | 1 | 2.1 | 0.59 | 10.04 | 9.59 | 0.45 | 0.45 | 0.02 | 0.02 |
| gene-LOC105278666 | https://www.ncbi.nlm.nih.gov/gene/?term=LOC105278666 | 10341 | 451 | 436.13 | 1 | 0.97 | 0.22 | 7.32 | 6.95 | 0.38 | 0.38 | 0.02 | 0.02 |
| gene-LOC105278145 | https://www.ncbi.nlm.nih.gov/gene/?term=LOC105278145 | 4470 | 251 | 561.52 | 1 | 2.24 | 0.4 | 10.77 | 10.43 | 0.34 | 0.34 | 0.02 | 0.02 |
| gene-LOC105286274 | https://www.ncbi.nlm.nih.gov/gene/?term=LOC105286274 | 3817 | 193 | 505.63 | 1 | 2.62 | 0.52 | 17.37 | 17.05 | 0.32 | 0.32 | 0.02 | 0.02 |

| | | | | | | | | | | | | | |
|---|---|---|---|---|---|---|---|---|---|---|---|---|---|
| gene-LOC105287095 | https://www.ncbi.nlm.nih.gov/gene/?term=LOC105287095 | 1435 | 58 | 404.18 | 1 | 6.97 | 1.72 | 28.05 | 28.65 | -0.6 | 0.6 | -0.02 | 0.02 |
| gene-LOC105274449 | https://www.ncbi.nlm.nih.gov/gene/?term=LOC105274449 | 1549 | 48 | 309.88 | 1 | 6.46 | 2.08 | 50.3 | 50.96 | -0.66 | 0.66 | -0.02 | 0.02 |
| gene-LOC105285315 | https://www.ncbi.nlm.nih.gov/gene/?term=LOC105285315 | 5458 | 19 | 34.81 | 1 | 1.83 | 5.26 | 46.71 | 43.22 | 3.49 | 3.49 | 0.01 | 0.01 |
| gene-LOC105281515 | https://www.ncbi.nlm.nih.gov/gene/?term=LOC105281515 | 8827 | 42 | 47.58 | 1 | 1.13 | 2.38 | 30.33 | 27.53 | 2.81 | 2.81 | 0.01 | 0.01 |
| gene-LOC105275347 | https://www.ncbi.nlm.nih.gov/gene/?term=LOC105275347 | 2526 | 19 | 75.22 | 1 | 3.96 | 5.26 | 38.97 | 37.49 | 1.48 | 1.48 | 0.01 | 0.01 |
| gene-LOC105281532 | https://www.ncbi.nlm.nih.gov/gene/?term=LOC105281532 | 7793 | 81 | 103.94 | 1 | 1.28 | 1.23 | 12.92 | 11.51 | 1.4 | 1.4 | 0.01 | 0.01 |
| gene-LOC105279500 | https://www.ncbi.nlm.nih.gov/gene/?term=LOC105279500 | 3587 | 37 | 103.15 | 1 | 2.79 | 2.7 | 50.42 | 49.31 | 1.1 | 1.1 | 0.01 | 0.01 |
| gene-LOC105283987 | https://www.ncbi.nlm.nih.gov/gene/?term=LOC105283987 | 4841 | 68 | 140.47 | 1 | 2.07 | 1.47 | 29.57 | 28.67 | 0.91 | 0.91 | 0.01 | 0.01 |
| gene-LOC105285406 | https://www.ncbi.nlm.nih.gov/gene/?term=LOC105285406 | 8015 | 60 | 74.86 | 1 | 1.25 | 1.67 | 26.39 | 25.52 | 0.87 | 0.87 | 0.01 | 0.01 |
| gene-LOC105280408 | https://www.ncbi.nlm.nih.gov/gene/?term=LOC105280408 | 5983 | 58 | 96.94 | 1 | 1.67 | 1.72 | 16.75 | 15.89 | 0.86 | 0.86 | 0.01 | 0.01 |
| gene-LOC105280002 | https://www.ncbi.nlm.nih.gov/gene/?term=LOC105280002 | 1940 | 28 | 144.33 | 1 | 5.15 | 3.57 | 37.5 | 36.72 | 0.79 | 0.79 | 0.01 | 0.01 |
| gene-LOC105285060 | https://www.ncbi.nlm.nih.gov/gene/?term=LOC105285060 | 1285 | 21 | 163.42 | 1 | 7.78 | 4.76 | 39.84 | 39.09 | 0.75 | 0.75 | 0.01 | 0.01 |
| gene-LOC105284116 | https://www.ncbi.nlm.nih.gov/gene/?term=LOC105284116 | 3970 | 78 | 196.47 | 1 | 2.52 | 1.28 | 7.09 | 6.36 | 0.73 | 0.73 | 0.01 | 0.01 |
| gene-LOC105282067 | https://www.ncbi.nlm.nih.gov/gene/?term=LOC105282067 | 5941 | 54 | 90.89 | 1 | 1.68 | 1.85 | 28.74 | 28.06 | 0.67 | 0.67 | 0.01 | 0.01 |
| gene-LOC105285514 | https://www.ncbi.nlm.nih.gov/gene/?term=LOC105285514 | 1277 | 17 | 133.12 | 1 | 7.83 | 5.88 | 71.39 | 70.72 | 0.67 | 0.67 | 0.01 | 0.01 |
| gene-LOC105274699 | https://www.ncbi.nlm.nih.gov/gene/?term=LOC105274699 | 2386 | 52 | 217.94 | 1 | 4.19 | 1.92 | 24.82 | 24.15 | 0.66 | 0.66 | 0.01 | 0.01 |
| gene-LOC105275966 | https://www.ncbi.nlm.nih.gov/gene/?term=LOC105275966 | 3299 | 64 | 194 | 1 | 3.03 | 1.56 | 48.15 | 47.49 | 0.65 | 0.65 | 0.01 | 0.01 |
| gene-LOC105282399 | https://www.ncbi.nlm.nih.gov/gene/?term=LOC105282399 | 2682 | 24 | 89.49 | 1 | 3.73 | 4.17 | 35.91 | 35.25 | 0.65 | 0.65 | 0.01 | 0.01 |
| gene-LOC105283943 | https://www.ncbi.nlm.nih.gov/gene/?term=LOC105283943 | 3559 | 48 | 134.87 | 1 | 2.81 | 2.08 | 17.55 | 16.96 | 0.59 | 0.59 | 0.01 | 0.01 |
| gene-LOC105282825 | https://www.ncbi.nlm.nih.gov/gene/?term=LOC105282825 | 3919 | 85 | 216.89 | 1 | 2.55 | 1.18 | 14.44 | 13.87 | 0.57 | 0.57 | 0.01 | 0.01 |
| gene-LOC105278534 | https://www.ncbi.nlm.nih.gov/gene/?term=LOC105278534 | 7364 | 197 | 267.52 | 2 | 2.72 | 1.02 | 18.17 | 17.62 | 0.55 | 0.55 | 0.01 | 0.01 |
| gene-LOC105281599 | https://www.ncbi.nlm.nih.gov/gene/?term=LOC105281599 | 3859 | 49 | 126.98 | 1 | 2.59 | 2.04 | 4.6 | 4.08 | 0.53 | 0.53 | 0.01 | 0.01 |
| gene-LOC105280410 | https://www.ncbi.nlm.nih.gov/gene/?term=LOC105280410 | 12580 | 264 | 209.86 | 1 | 0.79 | 0.38 | 19.99 | 19.51 | 0.48 | 0.48 | 0.01 | 0.01 |
| gene-LOC105278454 | https://www.ncbi.nlm.nih.gov/gene/?term=LOC105278454 | 1972 | 46 | 233.27 | 1 | 5.07 | 2.17 | 44.92 | 44.47 | 0.46 | 0.46 | 0.01 | 0.01 |
| gene-LOC105275179 | https://www.ncbi.nlm.nih.gov/gene/?term=LOC105275179 | 3271 | 72 | 220.12 | 1 | 3.06 | 1.39 | 21.5 | 21.05 | 0.45 | 0.45 | 0.01 | 0.01 |
| gene-LOC105276078 | https://www.ncbi.nlm.nih.gov/gene/?term=LOC105276078 | 1051 | 14 | 133.21 | 1 | 9.51 | 7.14 | 33.32 | 32.93 | 0.39 | 0.39 | 0.01 | 0.01 |
| gene-LOC105285860 | https://www.ncbi.nlm.nih.gov/gene/?term=LOC105285860 | 3110 | 55 | 176.85 | 1 | 3.22 | 1.82 | 24.17 | 23.77 | 0.39 | 0.39 | 0.01 | 0.01 |
| gene-LOC105275327 | https://www.ncbi.nlm.nih.gov/gene/?term=LOC105275327 | 14245 | 492 | 345.38 | 1 | 0.7 | 0.2 | 7.28 | 6.9 | 0.38 | 0.38 | 0.01 | 0.01 |
| gene-LOC105274531 | https://www.ncbi.nlm.nih.gov/gene/?term=LOC105274531 | 7774 | 301 | 387.19 | 1 | 1.29 | 0.33 | 10.99 | 10.62 | 0.37 | 0.37 | 0.01 | 0.01 |
| gene-LOC105282811 | https://www.ncbi.nlm.nih.gov/gene/?term=LOC105282811 | 5765 | 213 | 369.47 | 1 | 1.73 | 0.47 | 8.63 | 8.26 | 0.37 | 0.37 | 0.01 | 0.01 |
| gene-LOC105287371 | https://www.ncbi.nlm.nih.gov/gene/?term=LOC105287371 | 8224 | 351 | 426.8 | 1 | 1.22 | 0.28 | 7.67 | 7.34 | 0.33 | 0.33 | 0.01 | 0.01 |
| gene-LOC105287528 | https://www.ncbi.nlm.nih.gov/gene/?term=LOC105287528 | 14195 | 419 | 295.17 | 1 | 0.7 | 0.24 | 5.19 | 4.86 | 0.33 | 0.33 | 0.01 | 0.01 |
| gene-LOC105287550 | https://www.ncbi.nlm.nih.gov/gene/?term=LOC105287550 | 15659 | 613 | 391.47 | 1 | 0.64 | 0.16 | 6.43 | 6.12 | 0.31 | 0.31 | 0.01 | 0.01 |
| gene-LOC105286166 | https://www.ncbi.nlm.nih.gov/gene/?term=LOC105286166 | 3050 | 110 | 360.66 | 1 | 3.28 | 0.91 | 24.5 | 24.21 | 0.29 | 0.29 | 0.01 | 0.01 |
| gene-LOC113562050 | https://www.ncbi.nlm.nih.gov/gene/?term=LOC113562050 | 17097 | 576 | 336.9 | 1 | 0.58 | 0.17 | 5.24 | 4.95 | 0.29 | 0.29 | 0.01 | 0.01 |
| gene-LOC105278877 | https://www.ncbi.nlm.nih.gov/gene/?term=LOC105278877 | 1968 | 72 | 365.85 | 1 | 5.08 | 1.39 | 36.04 | 35.76 | 0.28 | 0.28 | 0.01 | 0.01 |
| gene-LOC105277079 | https://www.ncbi.nlm.nih.gov/gene/?term=LOC105277079 | 13009 | 288 | 221.39 | 1 | 0.77 | 0.35 | 9.86 | 9.59 | 0.27 | 0.27 | 0.01 | 0.01 |
| gene-LOC105284262 | https://www.ncbi.nlm.nih.gov/gene/?term=LOC105284262 | 6669 | 225 | 337.38 | 1 | 1.5 | 0.44 | 6.17 | 5.94 | 0.23 | 0.23 | 0.01 | 0.01 |
| gene-LOC105278632 | https://www.ncbi.nlm.nih.gov/gene/?term=LOC105278632 | 3624 | 120 | 331.13 | 1 | 2.76 | 0.83 | 15.73 | 15.53 | 0.2 | 0.2 | 0.01 | 0.01 |
| gene-LOC105285807 | https://www.ncbi.nlm.nih.gov/gene/?term=LOC105285807 | 6063 | 227 | 374.4 | 1 | 1.65 | 0.44 | 9.73 | 9.55 | 0.18 | 0.18 | 0.01 | 0.01 |
| gene-LOC105276527 | https://www.ncbi.nlm.nih.gov/gene/?term=LOC105276527 | 7923 | 329 | 415.25 | 1 | 1.26 | 0.3 | 9.51 | 9.36 | 0.15 | 0.15 | 0.01 | 0.01 |
| gene-LOC105286649 | https://www.ncbi.nlm.nih.gov/gene/?term=LOC105286649 | 14178 | 817 | 576.24 | 1 | 0.71 | 0.12 | 1 | 1.1 | -0.1 | 0.1 | -0.01 | 0.01 |
| gene-LOC105275848 | https://www.ncbi.nlm.nih.gov/gene/?term=LOC105275848 | 3718 | 144 | 387.31 | 1 | 2.69 | 0.69 | 20.8 | 20.95 | -0.15 | 0.15 | -0.01 | 0.01 |
| gene-LOC105287633 | https://www.ncbi.nlm.nih.gov/gene/?term=LOC105287633 | 1524 | 34 | 223.1 | 1 | 6.56 | 2.94 | 50.71 | 50.96 | -0.26 | 0.26 | -0.01 | 0.01 |
| gene-LOC105281312 | https://www.ncbi.nlm.nih.gov/gene/?term=LOC105281312 | 2551 | 63 | 246.96 | 1 | 3.92 | 1.59 | 34.27 | 34.56 | -0.29 | 0.29 | -0.01 | 0.01 |
| gene-LOC105286999 | https://www.ncbi.nlm.nih.gov/gene/?term=LOC105286999 | 2141 | 44 | 205.51 | 1 | 4.67 | 2.27 | 34.01 | 34.41 | -0.41 | 0.41 | -0.01 | 0.01 |
| gene-LOC105285252 | https://www.ncbi.nlm.nih.gov/gene/?term=LOC105285252 | 7427 | 117 | 157.53 | 1 | 1.35 | 0.85 | 14.46 | 15.16 | -0.7 | 0.7 | -0.01 | 0.01 |
| gene-LOC105281296 | https://www.ncbi.nlm.nih.gov/gene/?term=LOC105281296 | 2434 | 37 | 152.01 | 1 | 4.11 | 2.7 | 30.9 | 31.72 | -0.83 | 0.83 | -0.01 | 0.01 |
| gene-LOC105286770 | https://www.ncbi.nlm.nih.gov/gene/?term=LOC105286770 | 3055 | 33 | 108.02 | 1 | 3.27 | 3.03 | 17.27 | 18.34 | -1.07 | 1.07 | -0.01 | 0.01 |
| gene-LOC105283989 | https://www.ncbi.nlm.nih.gov/gene/?term=LOC105283989 | 4948 | 31 | 62.65 | 2 | 4.04 | 6.45 | 34.97 | 36.46 | -1.49 | 1.49 | -0.01 | 0.01 |
| gene-LOC105286452 | https://www.ncbi.nlm.nih.gov/gene/?term=LOC105286452 | 6888 | 67 | 97.27 | 1 | 1.45 | 1.49 | 31.04 | 30.73 | 0.31 | 0.31 | 0 | 0 |
| gene-LOC105278245 | https://www.ncbi.nlm.nih.gov/gene/?term=LOC105278245 | 6445 | 117 | 181.54 | 1 | 1.55 | 0.85 | 22.48 | 22.2 | 0.27 | 0.27 | 0 | 0 |
| gene-LOC105276272 | https://www.ncbi.nlm.nih.gov/gene/?term=LOC105276272 | 1965 | 48 | 244.27 | 1 | 5.09 | 2.08 | 27.74 | 27.58 | 0.17 | 0.17 | 0 | 0 |
| gene-LOC105274669 | https://www.ncbi.nlm.nih.gov/gene/?term=LOC105274669 | 8555 | 256 | 299.24 | 1 | 1.17 | 0.39 | 6.43 | 6.35 | 0.08 | 0.08 | 0 | 0 |
| gene-LOC105275752 | https://www.ncbi.nlm.nih.gov/gene/?term=LOC105275752 | 3462 | 132 | 381.28 | 1 | 2.89 | 0.76 | 32.74 | 32.68 | 0.06 | 0.06 | 0 | 0 |
| gene-LOC105286515 | https://www.ncbi.nlm.nih.gov/gene/?term=LOC105286515 | 3751 | 114 | 303.92 | 1 | 2.67 | 0.88 | 11.87 | 11.89 | -0.02 | 0.02 | 0 | 0 |
| gene-LOC105283296 | https://www.ncbi.nlm.nih.gov/gene/?term=LOC105283296 | 2174 | 94 | 432.38 | 1 | 4.6 | 1.06 | 18.21 | 18.24 | -0.03 | 0.03 | 0 | 0 |
| gene-LOC105287695 | https://www.ncbi.nlm.nih.gov/gene/?term=LOC105287695 | 6106 | 201 | 329.18 | 1 | 1.64 | 0.5 | 6.09 | 6.16 | -0.06 | 0.06 | 0 | 0 |
| gene-LOC105274851 | https://www.ncbi.nlm.nih.gov/gene/?term=LOC105274851 | 6616 | 139 | 210.1 | 1 | 1.51 | 0.72 | 11.55 | 11.65 | -0.1 | 0.1 | 0 | 0 |
| gene-LOC105275187 | https://www.ncbi.nlm.nih.gov/gene/?term=LOC105275187 | 5534 | 71 | 128.3 | 1 | 1.81 | 1.41 | 12.33 | 12.43 | -0.1 | 0.1 | 0 | 0 |
| gene-LOC105282646 | https://www.ncbi.nlm.nih.gov/gene/?term=LOC105282646 | 3317 | 116 | 349.71 | 1 | 3.01 | 0.86 | 24.81 | 24.92 | -0.11 | 0.11 | 0 | 0 |
| gene-LOC105283806 | https://www.ncbi.nlm.nih.gov/gene/?term=LOC105283806 | 10501 | 74 | 70.47 | 1 | 0.95 | 1.35 | 16.11 | 16.23 | -0.12 | 0.12 | 0 | 0 |
| gene-LOC105284507 | https://www.ncbi.nlm.nih.gov/gene/?term=LOC105284507 | 5207 | 87 | 167.08 | 1 | 1.92 | 1.15 | 21.48 | 21.62 | -0.13 | 0.13 | 0 | 0 |
| gene-LOC105280704 | https://www.ncbi.nlm.nih.gov/gene/?term=LOC105280704 | 4266 | 134 | 314.11 | 1 | 2.34 | 0.75 | 19.89 | 20.04 | -0.15 | 0.15 | 0 | 0 |

| | | | | | | | | | | | | | |
|---|---|---|---|---|---|---|---|---|---|---|---|---|---|
| gene-LOC105280348 | https://www.ncbi.nlm.nih.gov/gene/?term=LOC105280348 | 3797 | 70 | 184.36 | 1 | 2.63 | 1.43 | 20.64 | 20.81 | -0.17 | 0.17 | 0 | 0 |
| gene-LOC105283967 | https://www.ncbi.nlm.nih.gov/gene/?term=LOC105283967 | 2393 | 47 | 196.41 | 1 | 4.18 | 2.13 | 48.9 | 49.11 | -0.21 | 0.21 | 0 | 0 |
| gene-LOC105285163 | https://www.ncbi.nlm.nih.gov/gene/?term=LOC105285163 | 5061 | 87 | 171.9 | 1 | 1.98 | 1.15 | 21.7 | 21.95 | -0.26 | 0.26 | 0 | 0 |

Directory: OGL File: rnk-dmg-Ob_LI_RphaseB1.vs.RphaseB2.txt

| gene_ID | gene_link | gwidth | #Sites | #per10Kb | #dmSites | #dmsp10kb | %dmSites | %pSite1 | %pSite2 | DMpSite | ADMpSite | DMpNucl | ADMpNucl |
|---|---|---|---|---|---|---|---|---|---|---|---|---|---|
| gene-LOC105278849 | https://www.ncbi.nlm.nih.gov/gene/?term=LOC105278849 | 776 | 74 | 953.61 | 2 | 25.77 | 2.7 | 20.95 | 22.99 | -2.05 | 2.05 | -0.2 | 0.2 |
| gene-LOC113562453 | https://www.ncbi.nlm.nih.gov/gene/?term=LOC113562453 | 763 | 32 | 419.4 | 2 | 26.21 | 6.25 | 30.48 | 26.18 | 4.3 | 4.3 | 0.18 | 0.18 |
| gene-LOC105284093 | https://www.ncbi.nlm.nih.gov/gene/?term=LOC105284093 | 809 | 60 | 741.66 | 2 | 24.72 | 3.33 | 10.26 | 7.81 | 2.45 | 2.45 | 0.18 | 0.18 |
| gene-LOC105284253 | https://www.ncbi.nlm.nih.gov/gene/?term=LOC105284253 | 1255 | 41 | 326.69 | 5 | 39.84 | 12.2 | 26.29 | 31.86 | -5.58 | 5.58 | -0.18 | 0.18 |
| gene-LOC105284320 | https://www.ncbi.nlm.nih.gov/gene/?term=LOC105284320 | 390 | 11 | 282.05 | 1 | 25.64 | 9.09 | 22.16 | 27.96 | -5.81 | 5.81 | -0.16 | 0.16 |
| gene-LOC105288193 | https://www.ncbi.nlm.nih.gov/gene/?term=LOC105288193 | 1495 | 76 | 508.36 | 5 | 33.44 | 6.58 | 30.07 | 33 | -2.93 | 2.93 | -0.15 | 0.15 |
| gene-LOC113561900 | https://www.ncbi.nlm.nih.gov/gene/?term=LOC113561900 | 629 | 11 | 174.88 | 1 | 15.9 | 9.09 | 33.39 | 25.51 | 7.88 | 7.88 | 0.14 | 0.14 |
| gene-LOC105282481 | https://www.ncbi.nlm.nih.gov/gene/?term=LOC105282481 | 1900 | 60 | 315.79 | 4 | 21.05 | 6.67 | 31.49 | 36.02 | -4.52 | 4.52 | -0.14 | 0.14 |
| gene-LOC105284189 | https://www.ncbi.nlm.nih.gov/gene/?term=LOC105284189 | 2601 | 133 | 511.34 | 3 | 11.53 | 2.26 | 17.44 | 14.95 | 2.49 | 2.49 | 0.13 | 0.13 |
| gene-LOC105278160 | https://www.ncbi.nlm.nih.gov/gene/?term=LOC105278160 | 1872 | 91 | 486.11 | 1 | 5.34 | 1.1 | 34.58 | 37.24 | -2.67 | 2.67 | -0.13 | 0.13 |
| gene-LOC105277345 | https://www.ncbi.nlm.nih.gov/gene/?term=LOC105277345 | 2151 | 77 | 357.97 | 3 | 13.95 | 3.9 | 18.53 | 15.19 | 3.34 | 3.34 | 0.12 | 0.12 |
| gene-LOC105276563 | https://www.ncbi.nlm.nih.gov/gene/?term=LOC105276563 | 1958 | 114 | 582.23 | 5 | 25.54 | 4.39 | 18.98 | 16.96 | 2.02 | 2.02 | 0.12 | 0.12 |
| gene-LOC105279346 | https://www.ncbi.nlm.nih.gov/gene/?term=LOC105279346 | 4649 | 252 | 542.05 | 6 | 12.91 | 2.38 | 14.38 | 16.52 | -2.15 | 2.15 | -0.12 | 0.12 |
| gene-LOC105278458 | https://www.ncbi.nlm.nih.gov/gene/?term=LOC105278458 | 216 | 3 | 138.89 | 1 | 46.3 | 33.33 | 20.6 | 29.54 | -8.94 | 8.94 | -0.12 | 0.12 |
| gene-LOC105285203 | https://www.ncbi.nlm.nih.gov/gene/?term=LOC105285203 | 1020 | 32 | 313.73 | 2 | 19.61 | 6.25 | 29.55 | 25.89 | 3.66 | 3.66 | 0.11 | 0.11 |
| gene-LOC105278025 | https://www.ncbi.nlm.nih.gov/gene/?term=LOC105278025 | 1625 | 61 | 375.38 | 2 | 12.31 | 3.28 | 30.61 | 33.53 | -2.91 | 2.91 | -0.11 | 0.11 |
| gene-LOC105286845 | https://www.ncbi.nlm.nih.gov/gene/?term=LOC105286845 | 1159 | 78 | 672.99 | 2 | 17.26 | 2.56 | 11.01 | 12.57 | -1.56 | 1.56 | -0.1 | 0.1 |
| gene-LOC105278938 | https://www.ncbi.nlm.nih.gov/gene/?term=LOC105278938 | 1818 | 110 | 605.06 | 4 | 22 | 3.64 | 19.53 | 21.23 | -1.71 | 1.71 | -0.1 | 0.1 |
| gene-LOC105284047 | https://www.ncbi.nlm.nih.gov/gene/?term=LOC105284047 | 1212 | 20 | 165.02 | 4 | 33 | 20 | 20.52 | 26.47 | -5.95 | 5.95 | -0.1 | 0.1 |
| gene-LOC105285713 | https://www.ncbi.nlm.nih.gov/gene/?term=LOC105285713 | 1048 | 39 | 372.14 | 2 | 19.08 | 5.13 | 46.18 | 43.72 | 2.46 | 2.46 | 0.09 | 0.09 |
| gene-LOC105274542 | https://www.ncbi.nlm.nih.gov/gene/?term=LOC105274542 | 1266 | 52 | 410.74 | 2 | 15.8 | 3.85 | 30.12 | 27.91 | 2.21 | 2.21 | 0.09 | 0.09 |
| gene-LOC105279487 | https://www.ncbi.nlm.nih.gov/gene/?term=LOC105279487 | 2117 | 101 | 477.09 | 2 | 9.45 | 1.98 | 14.93 | 12.99 | 1.94 | 1.94 | 0.09 | 0.09 |
| gene-LOC105277091 | https://www.ncbi.nlm.nih.gov/gene/?term=LOC105277091 | 902 | 43 | 476.72 | 2 | 22.17 | 4.65 | 23.8 | 21.99 | 1.82 | 1.82 | 0.09 | 0.09 |
| gene-LOC105285392 | https://www.ncbi.nlm.nih.gov/gene/?term=LOC105285392 | 1601 | 82 | 512.18 | 9 | 56.21 | 10.98 | 37.61 | 35.85 | 1.76 | 1.76 | 0.09 | 0.09 |

...
... only first 25 lines shown ...

```
Directory: OGL File: wrt-Ob_LI.txt

... comparing  BCphaseA1.vs.BCphaseA2 ...

Number of genes <=  20000  and with #dmsites >=  1 :  119
sample1     sum of percent methylation:  3154.84     average:    26.51
sample2     sum of percent methylation:  3274.53     average:    27.52

sum difference:  -119.65     average difference:  -1.0055

      Wilcoxon signed rank test with continuity correction

data:  pw_summary$`%pSite1` and pw_summary$`%pSite2`
V = 2075, p-value = 7e-05
alternative hypothesis: true location shift is not equal to 0

... comparing  BCphaseA1.vs.BCphaseB1 ...

Number of genes <=  20000  and with #dmsites >=  1 :  121
sample1     sum of percent methylation:  2984.97     average:    24.67
sample2     sum of percent methylation:  3074.12     average:    25.41

sum difference:   -89.12     average difference:  -0.7365

      Wilcoxon signed rank test with continuity correction
...
... only first 25 lines shown ...
```

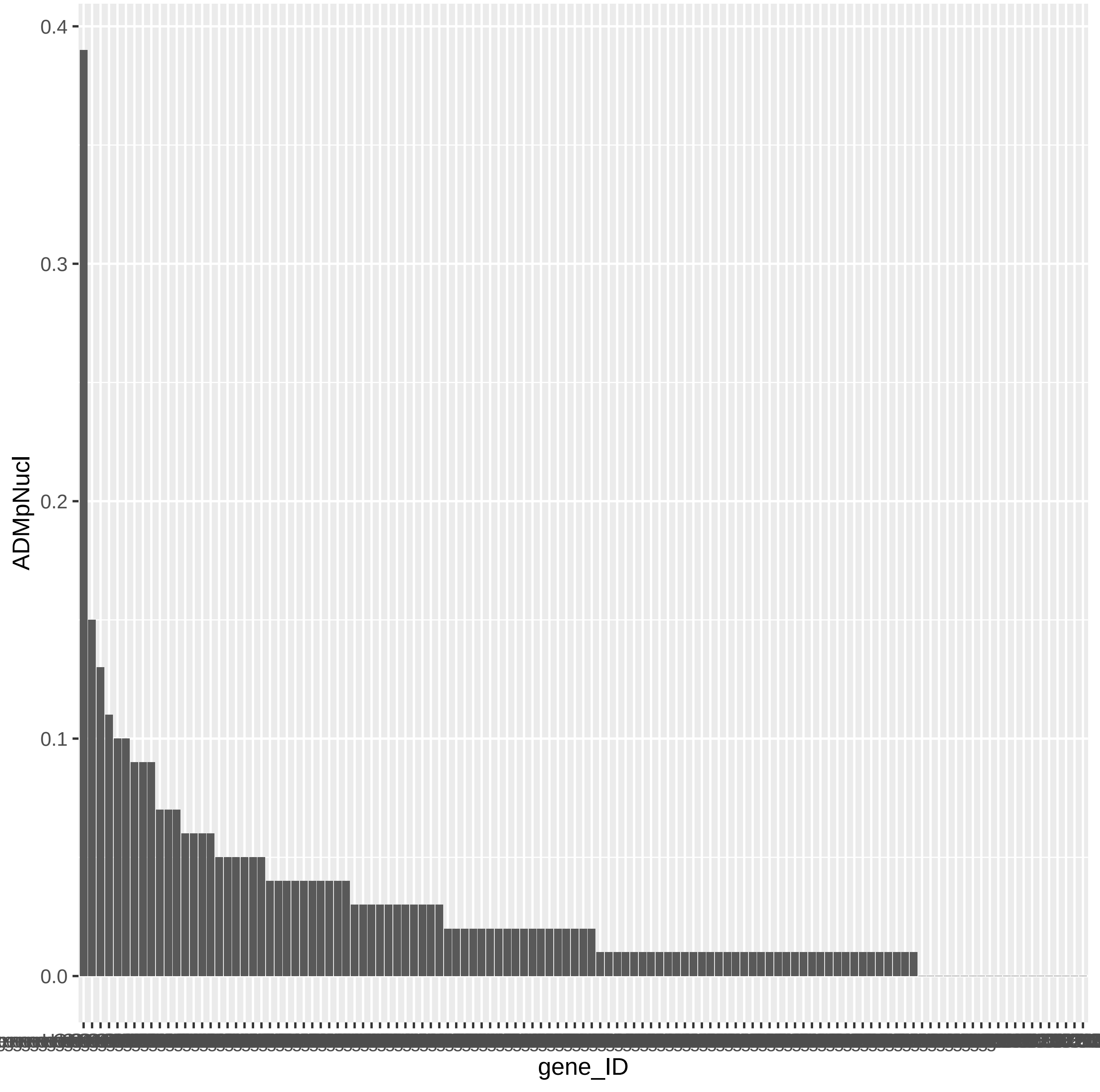

0.4
0.3
0.2
0.1
0.0
ADMpNucl
gene_ID

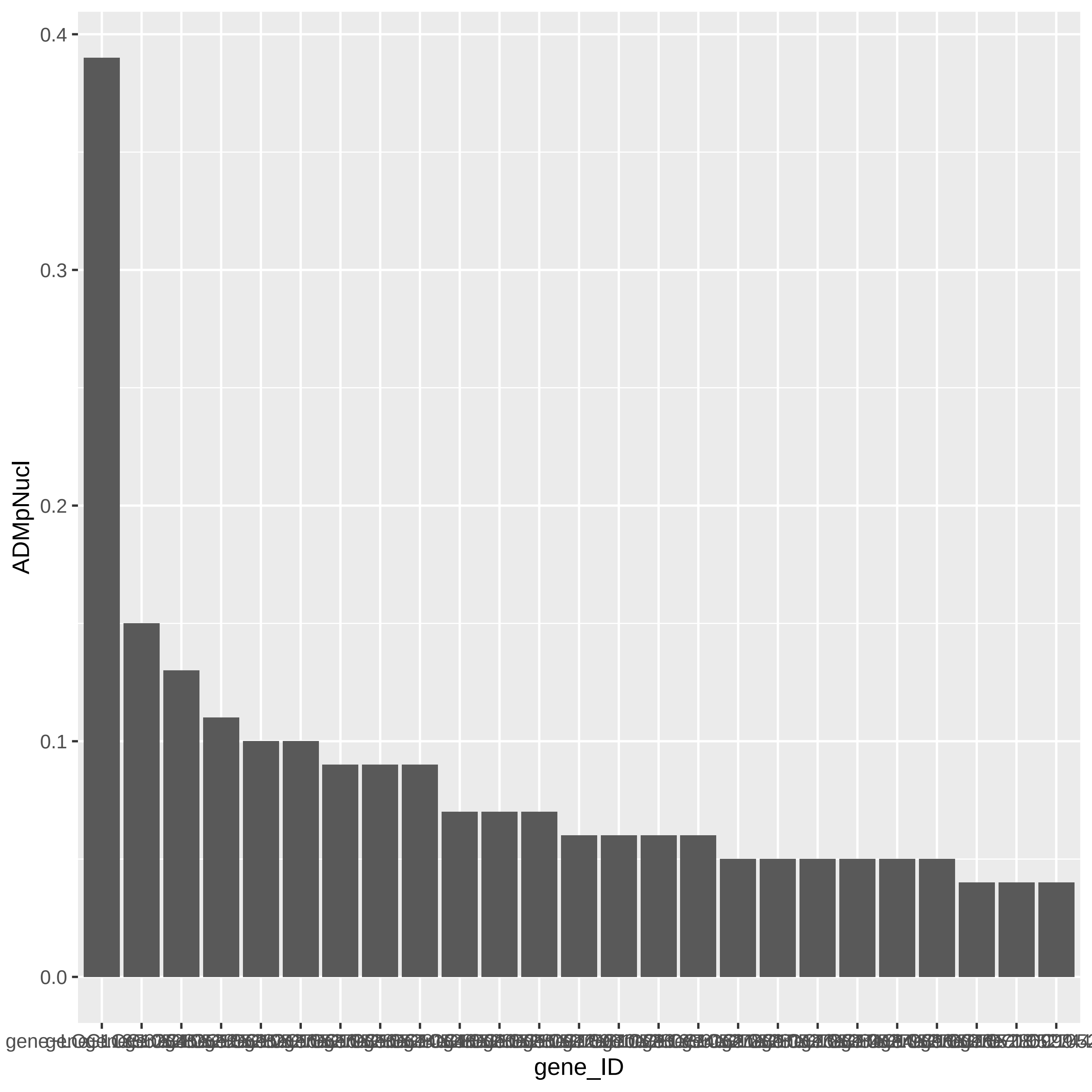
0.4
0.3
0.2
0.1
0.0
ADMpNucl
gene_ID

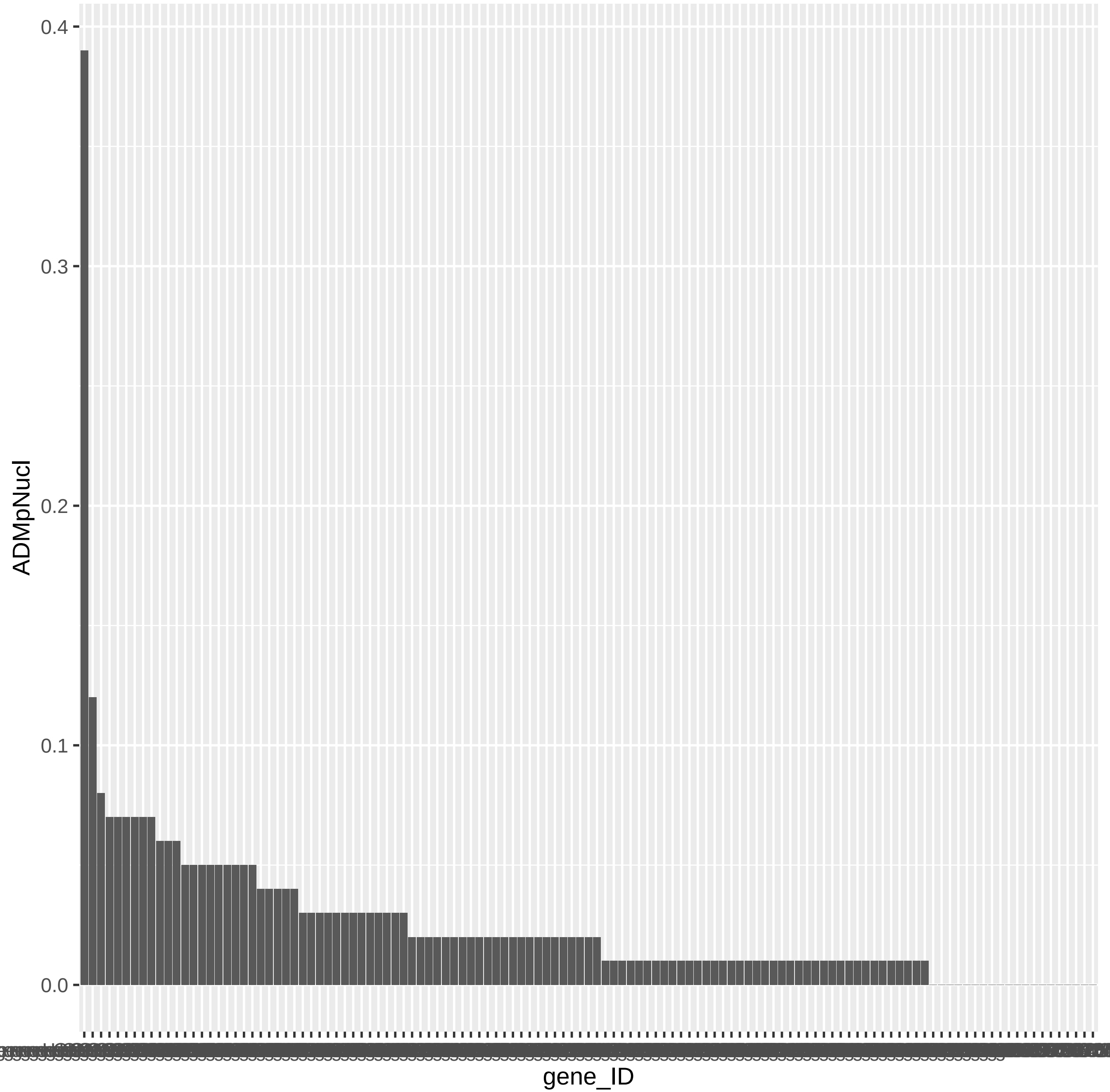

0.4
0.3
0.2
0.1
0.0
ADMpNucl
gene_ID

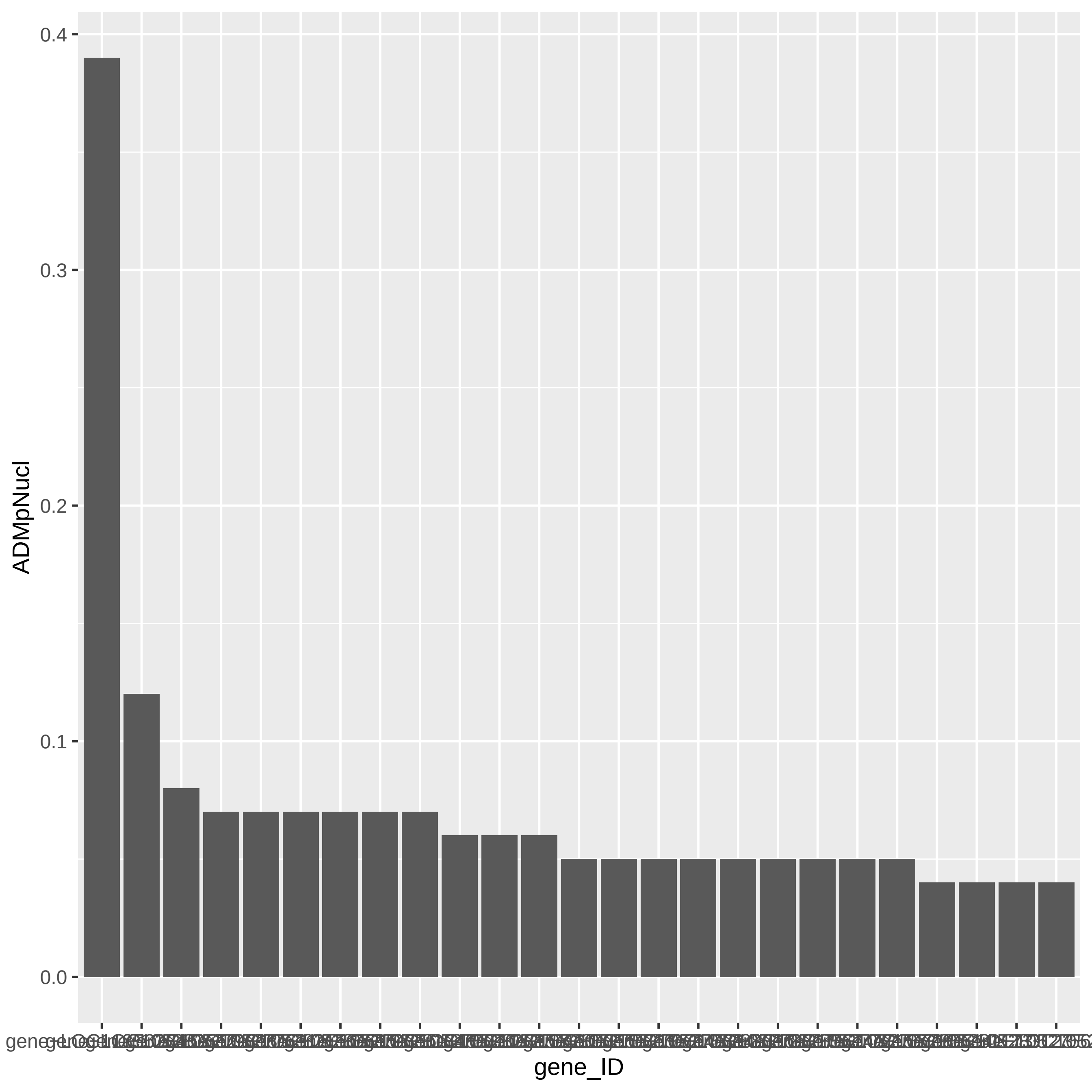

0.4
0.3
0.2
0.1
0.0
ADMpNucl
gene_ID

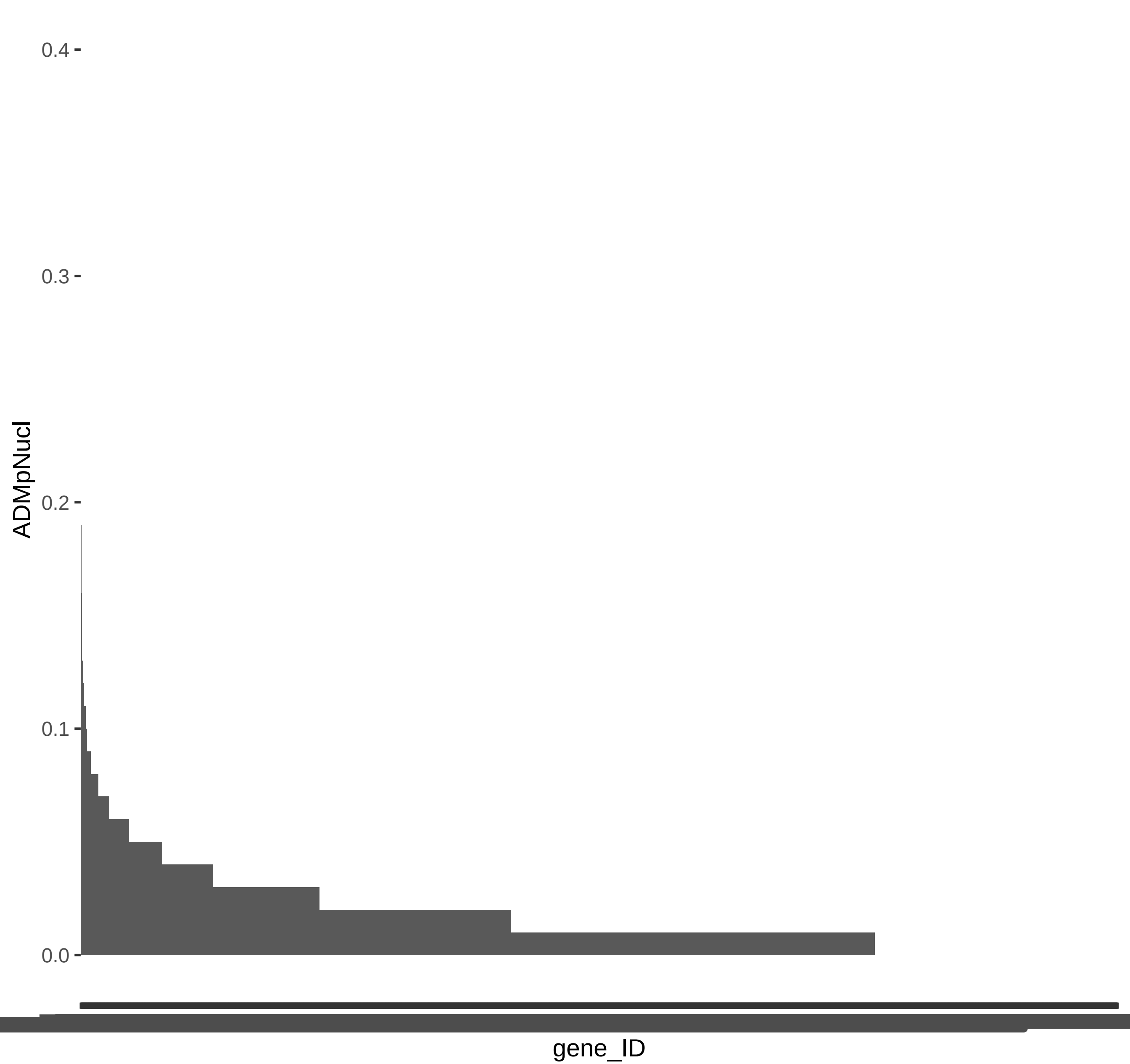

0.4
0.3
0.2
0.1
0.0
ADMpNucl
gene_ID

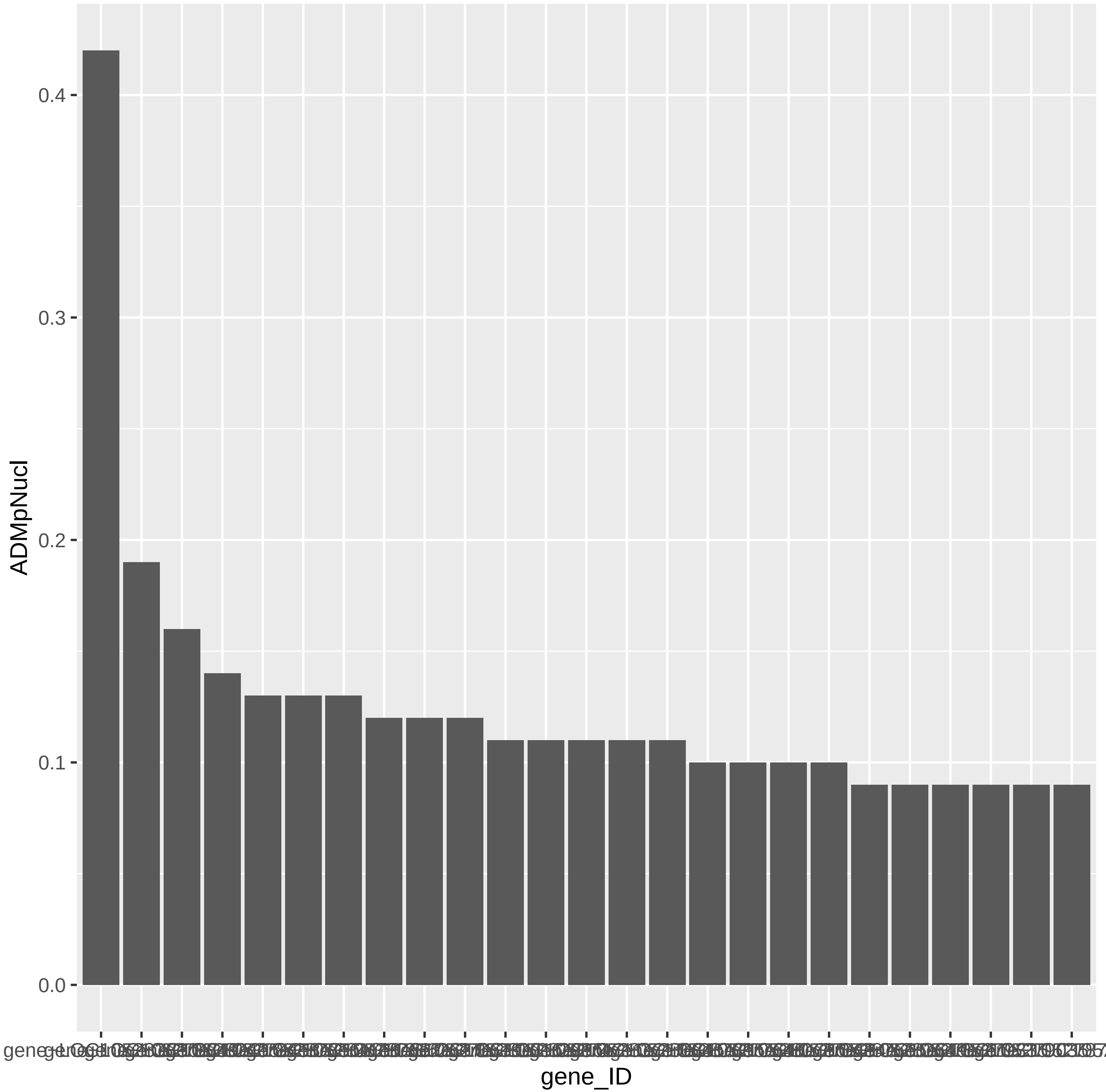
ADMpNucl
0.4
0.3
0.2
0.1
0.0
gene_ID

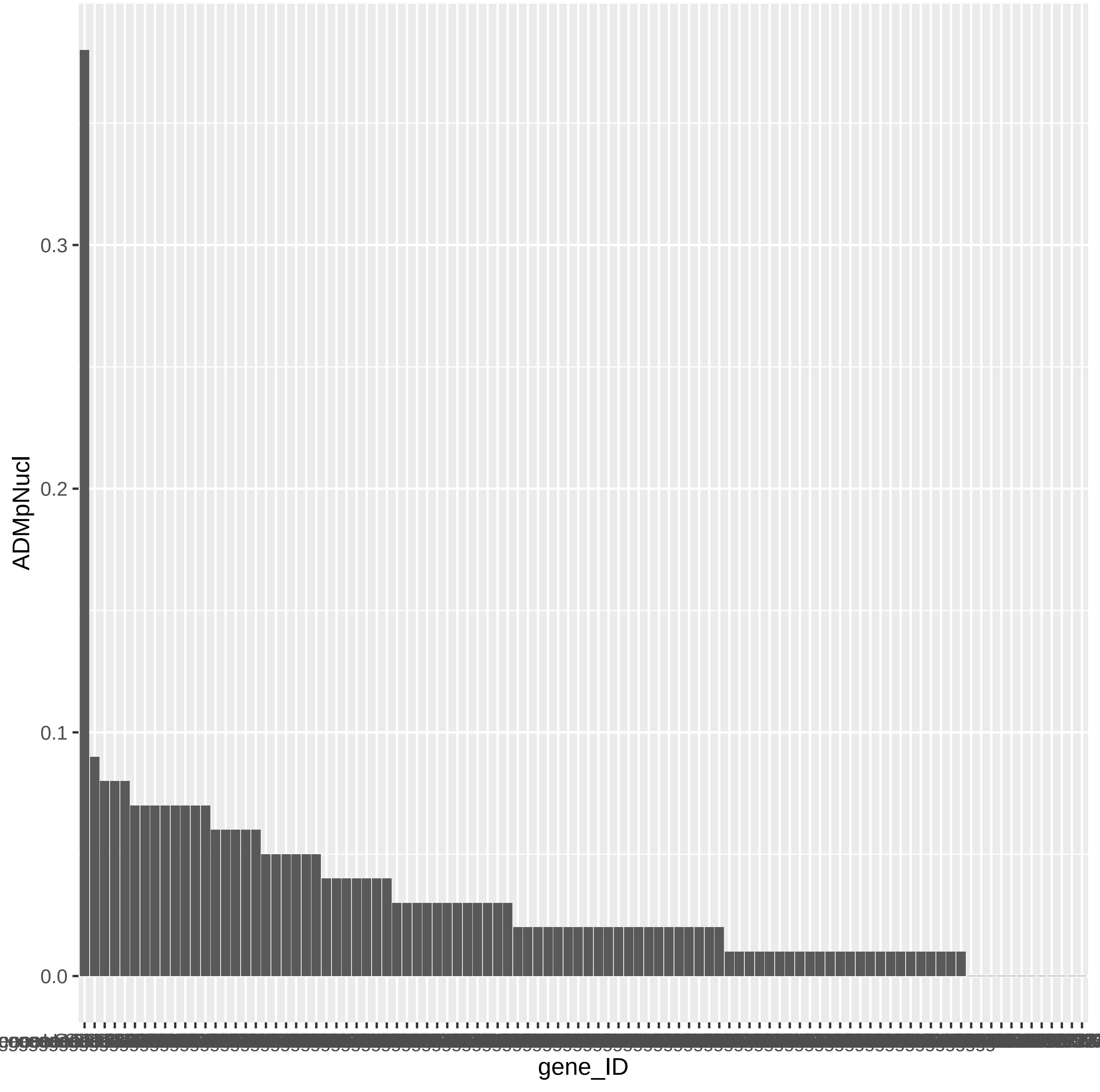

ADMpNucl
0.3
0.2
0.1
0.0
gene_ID

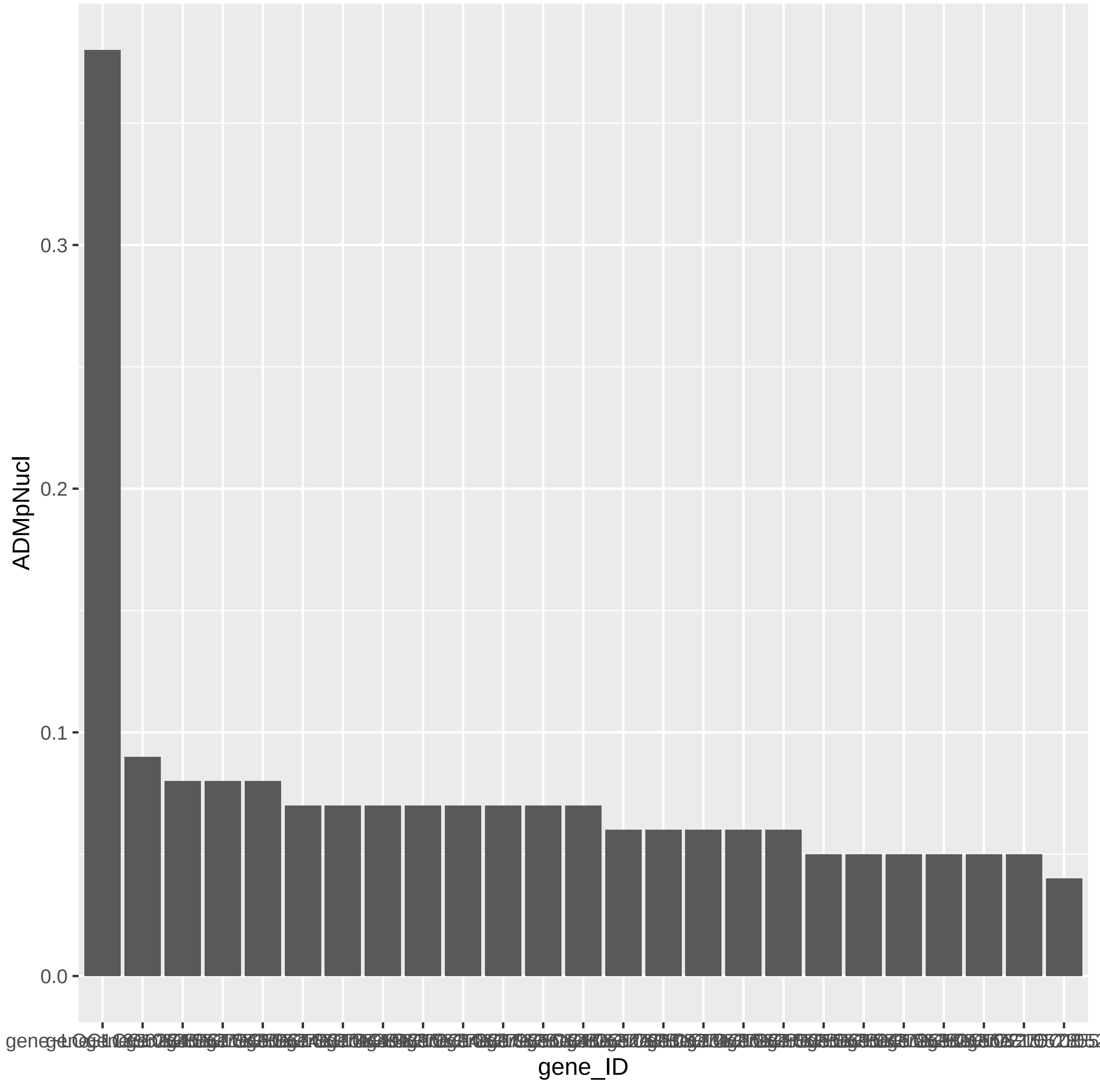

ADMpNucl
0.0
0.1
0.2
0.3
gene_ID

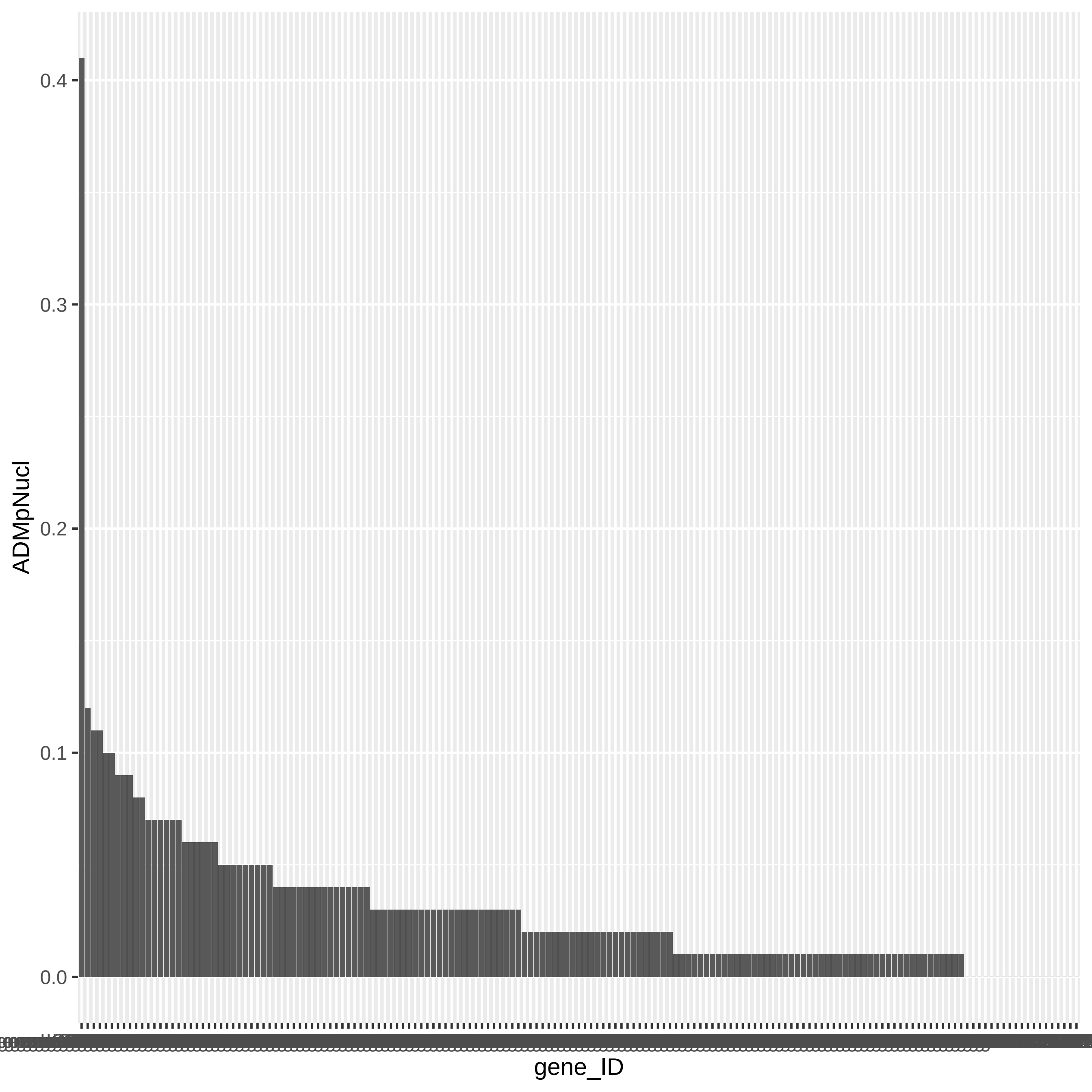
ADMpNucl
0.4
0.3
0.2
0.1
0.0
gene_ID

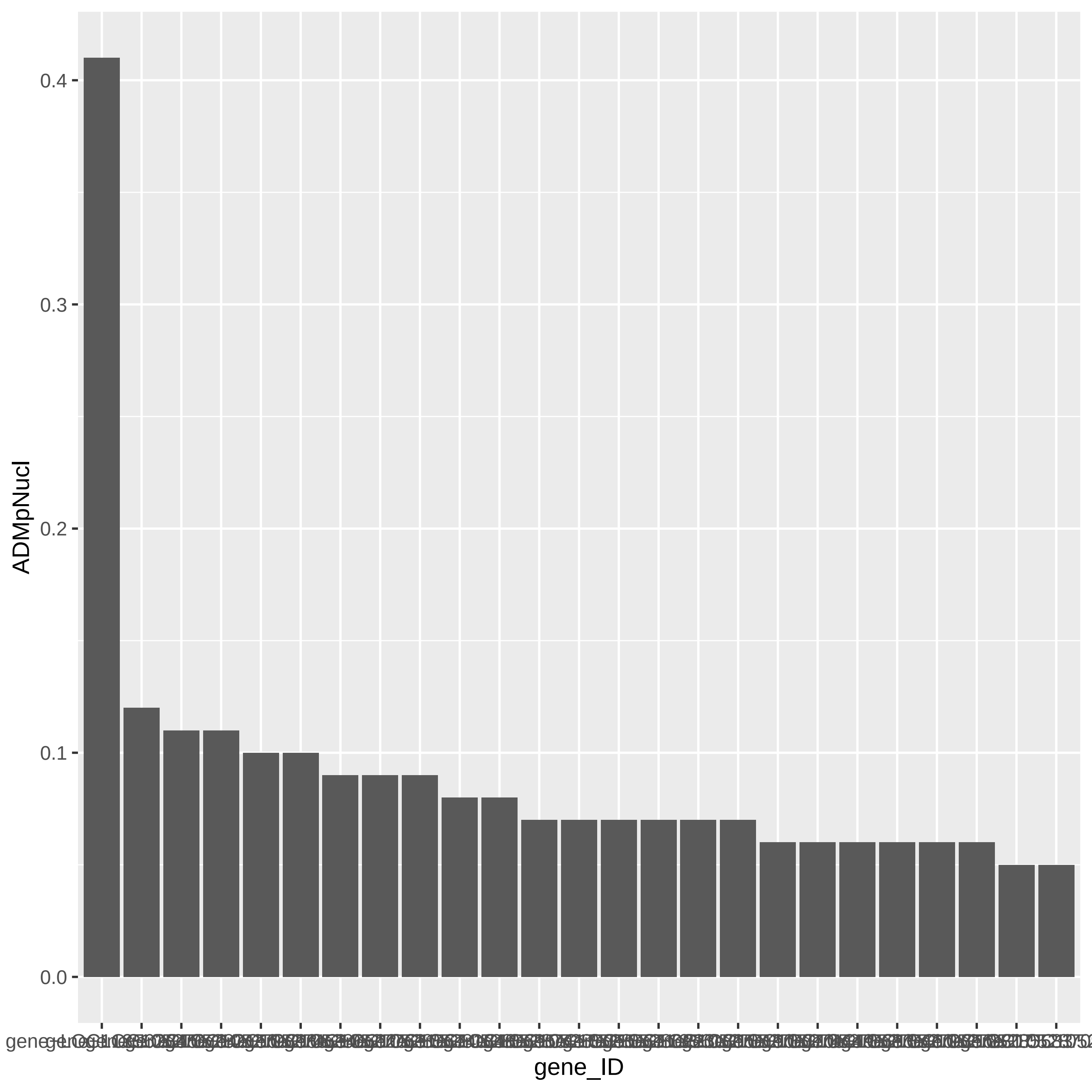

0.4
0.3
0.2
0.1
0.0
ADMpNucl
gene_ID

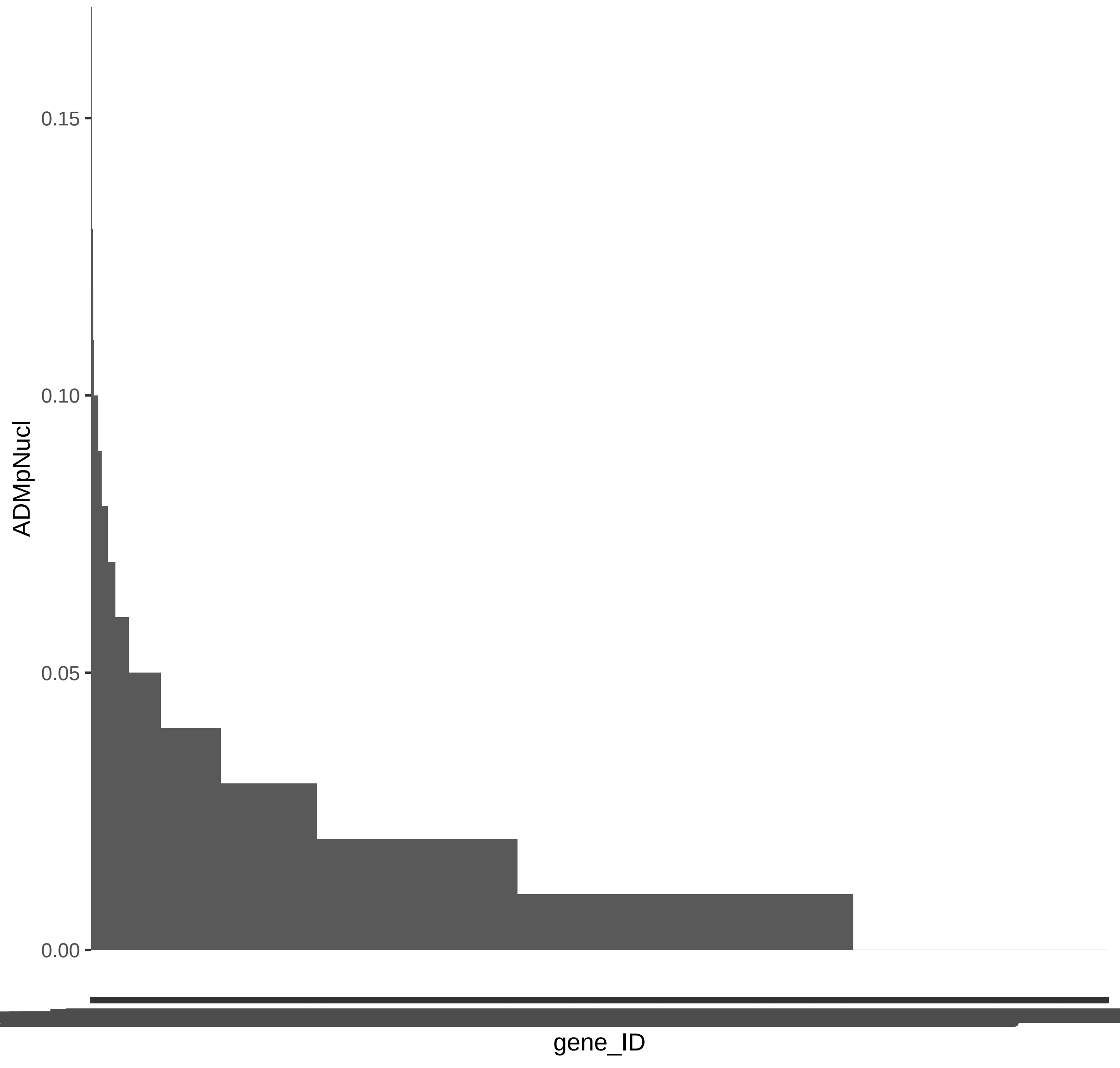
0.15
0.10
0.05
0.00
ADMpNucl
gene_ID

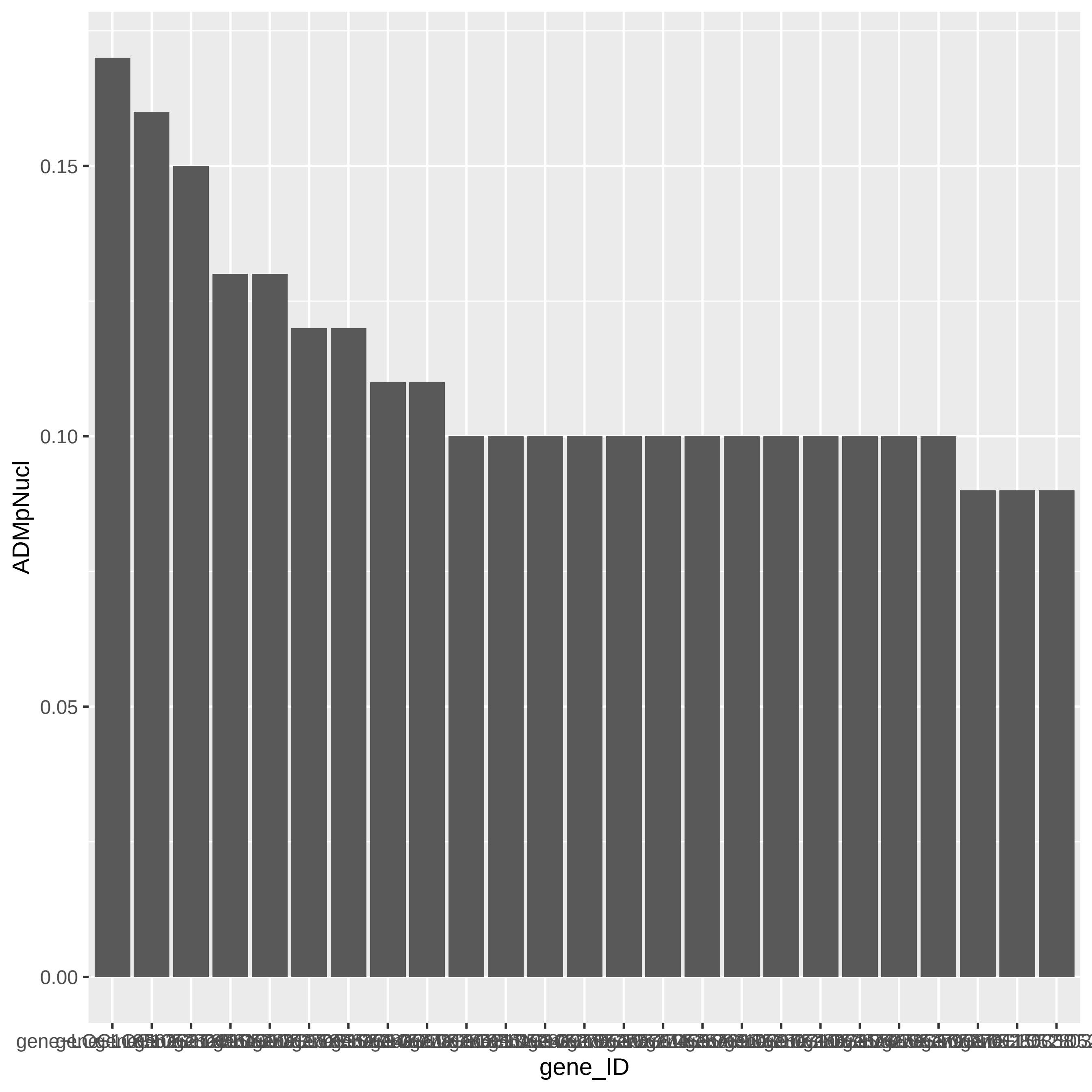

0.15
0.10
0.05
0.00
ADMpNucl
gene_ID

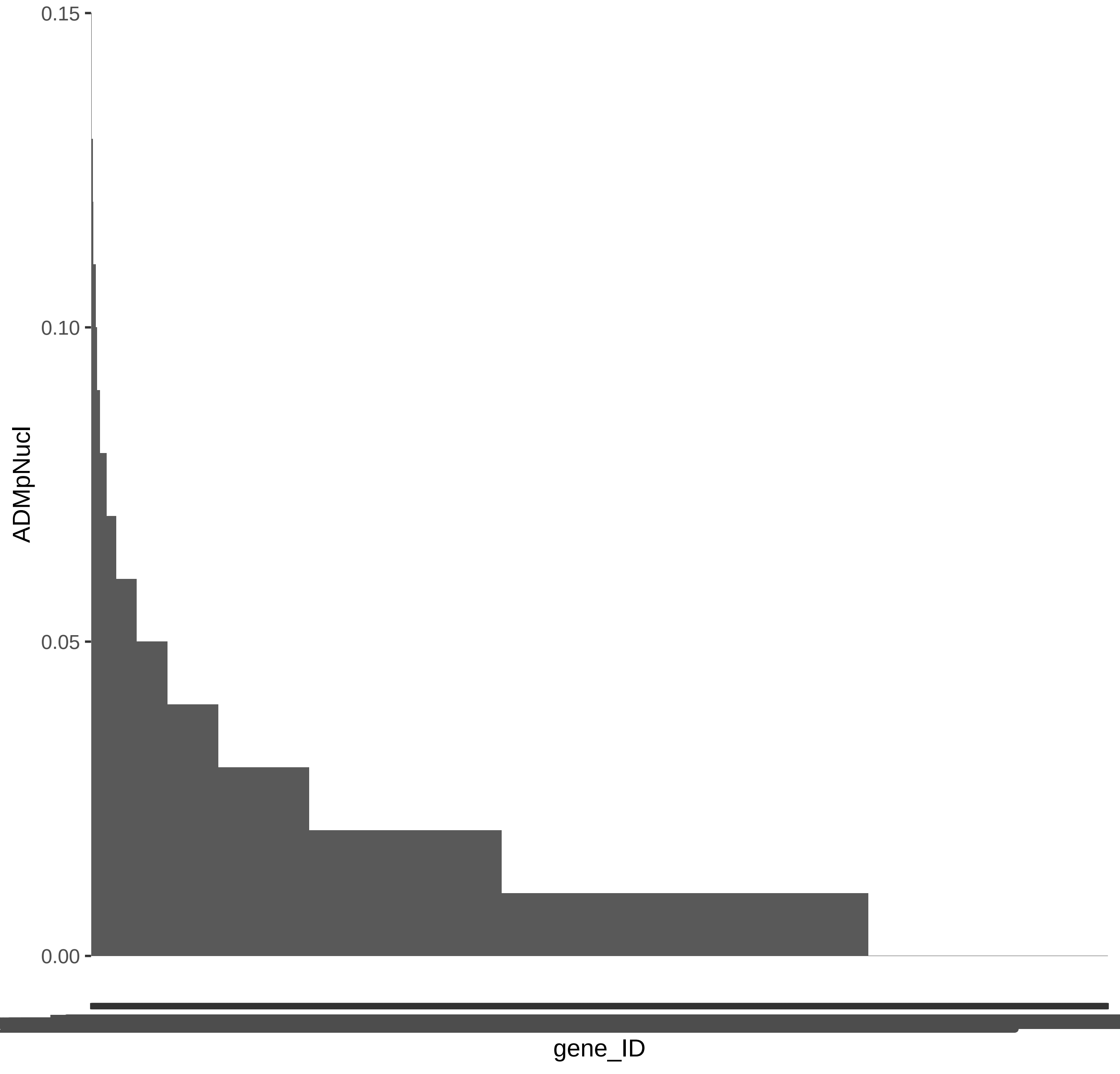
0.15
0.10
0.05
0.00
ADMpNucl
gene_ID

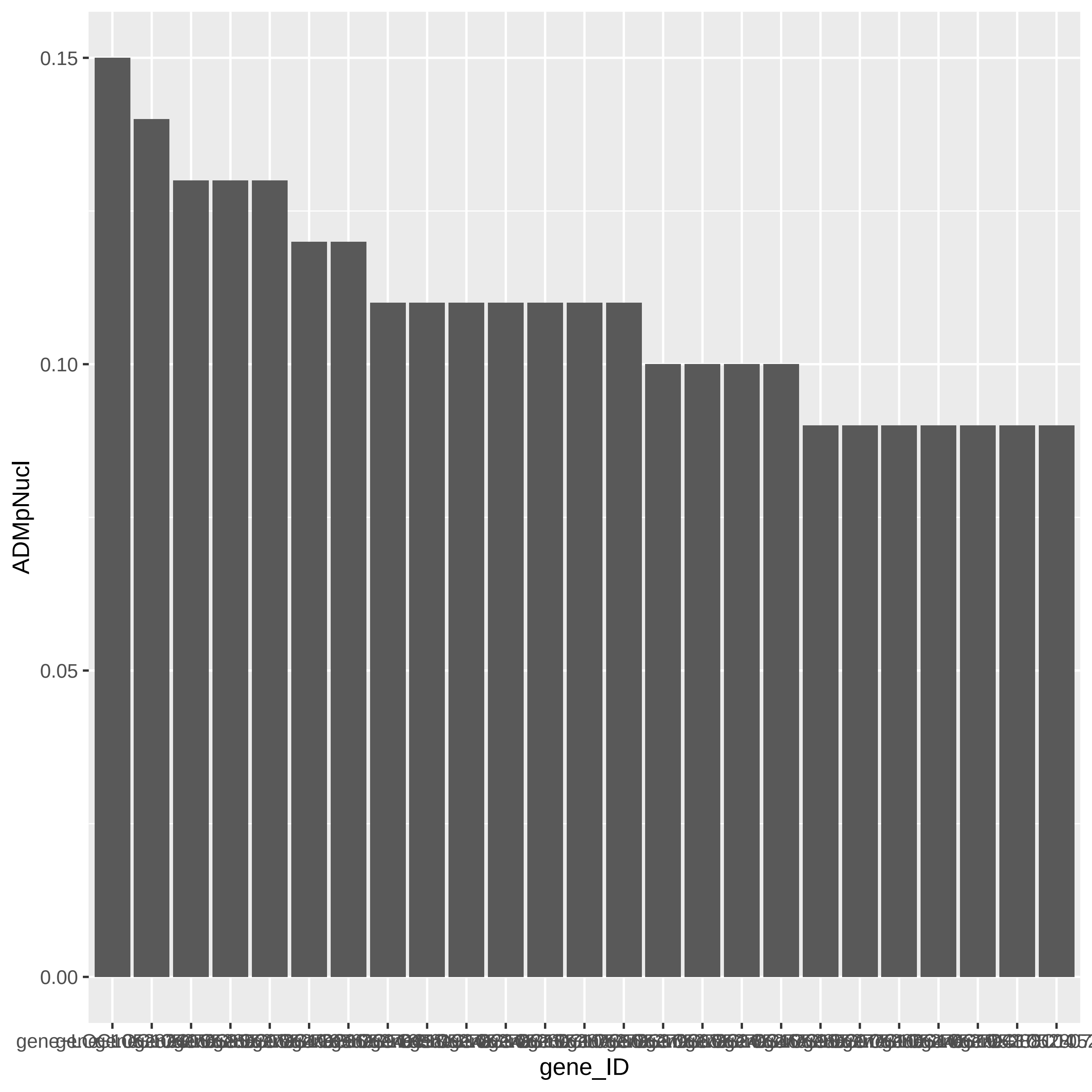

0.15
0.10
0.05
0.00
ADMpNucl
gene_ID

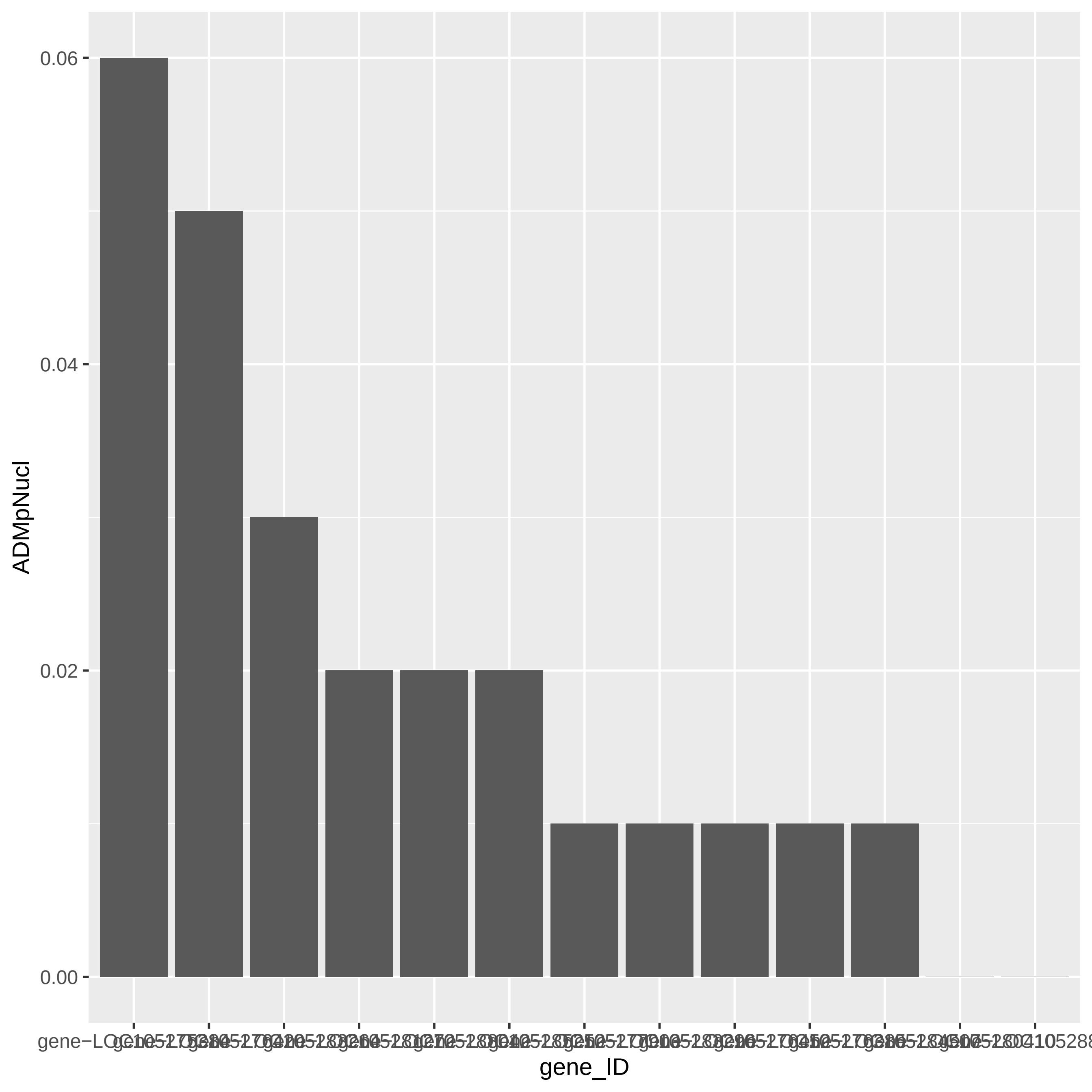
0.06
0.04
0.02
0.00
ADMpNucl
gene_ID

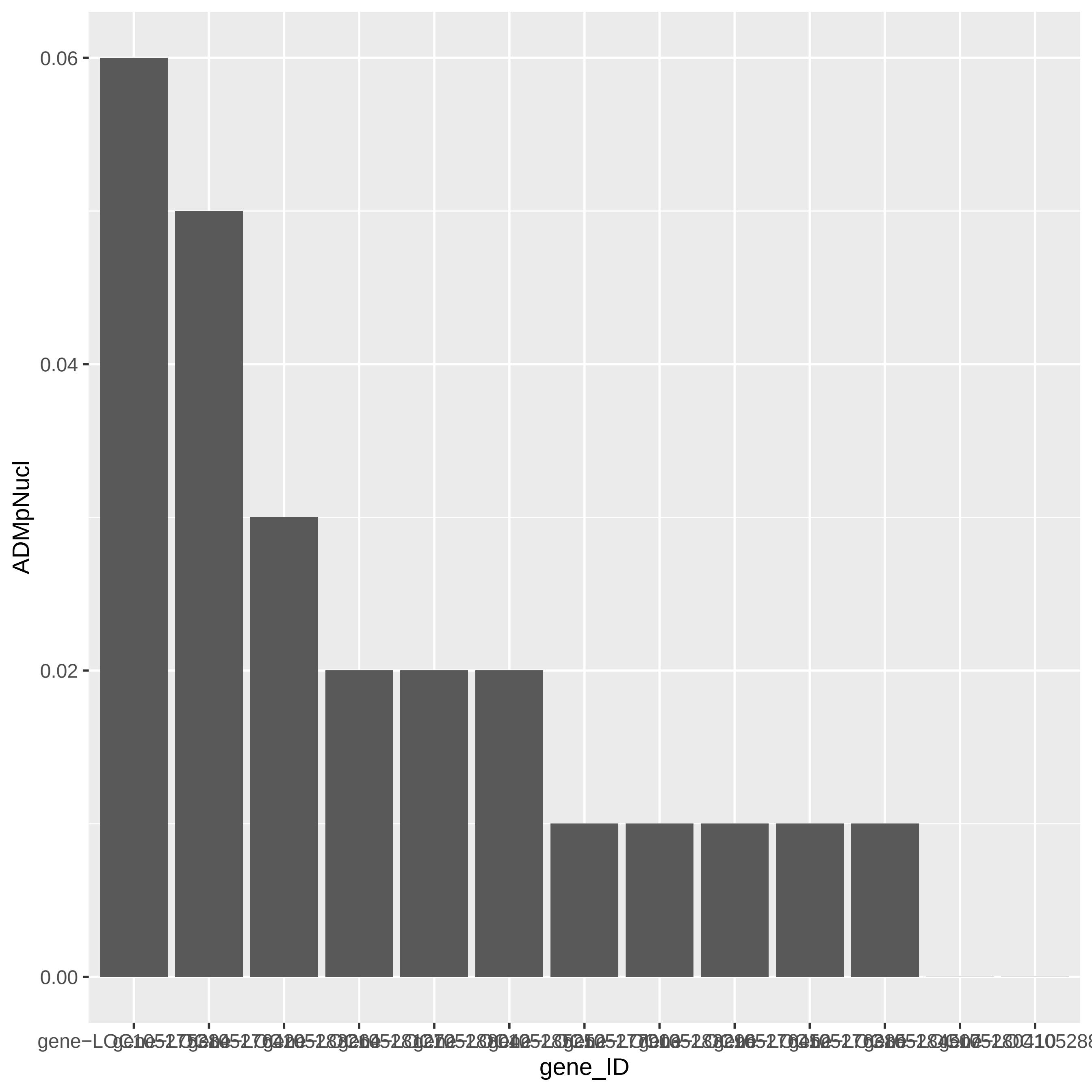

0.06
0.04
0.02
0.00
ADMpNucl
gene_ID

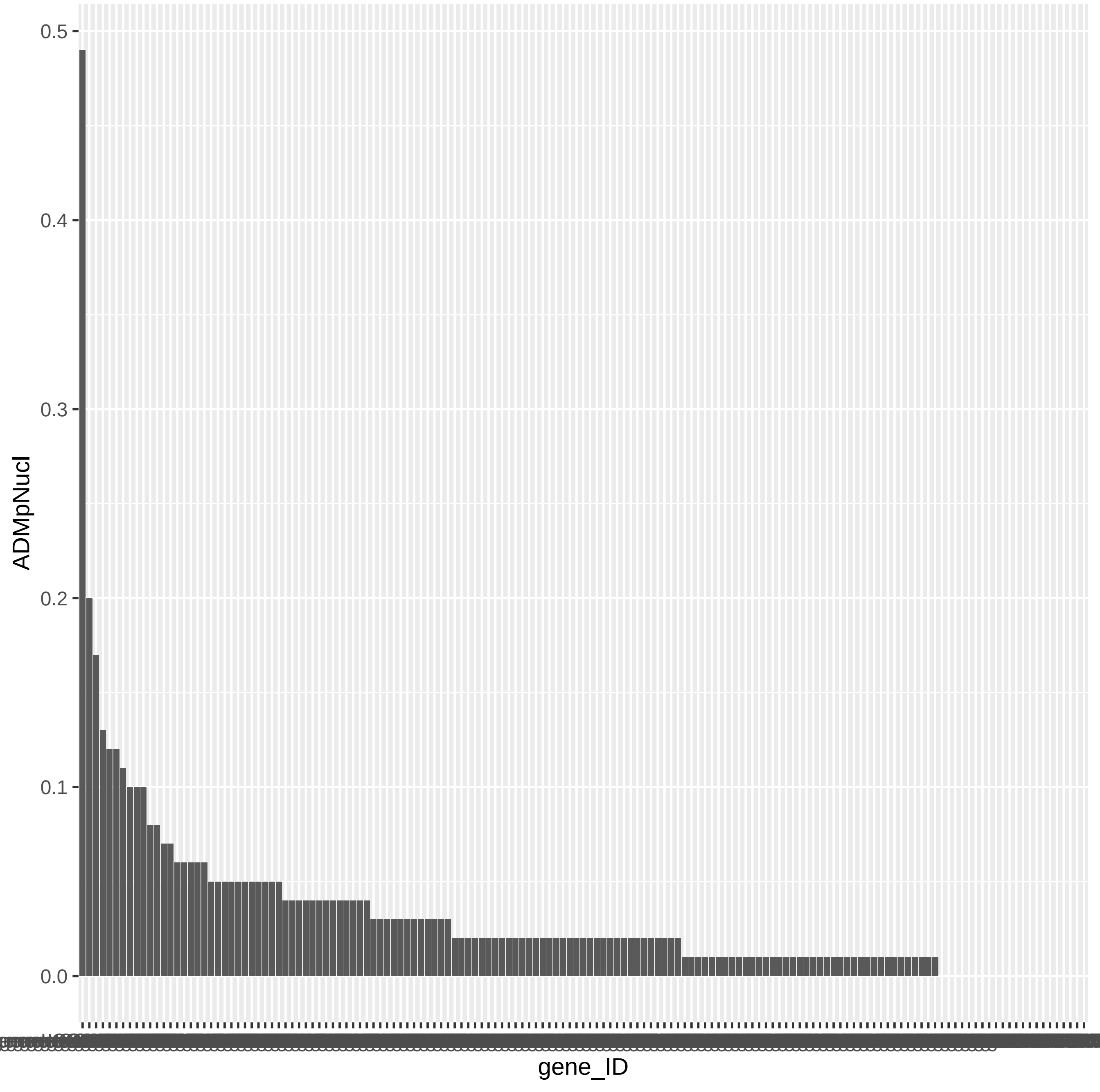
0.5
0.4
0.3
0.2
0.1
0.0
ADMpNucl
gene_ID

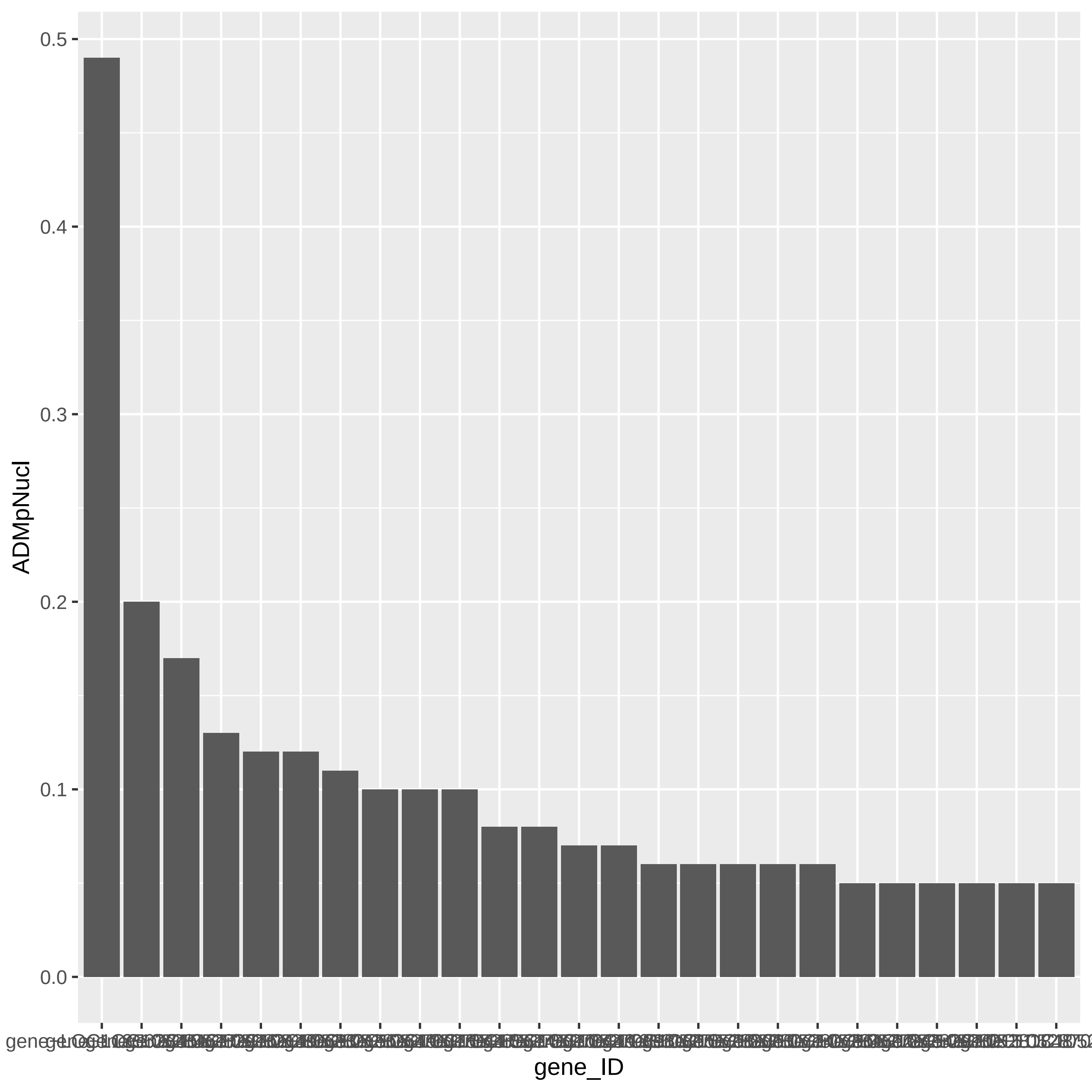

ADMpNucl
gene_ID
0.0
0.1
0.2
0.3
0.4
0.5

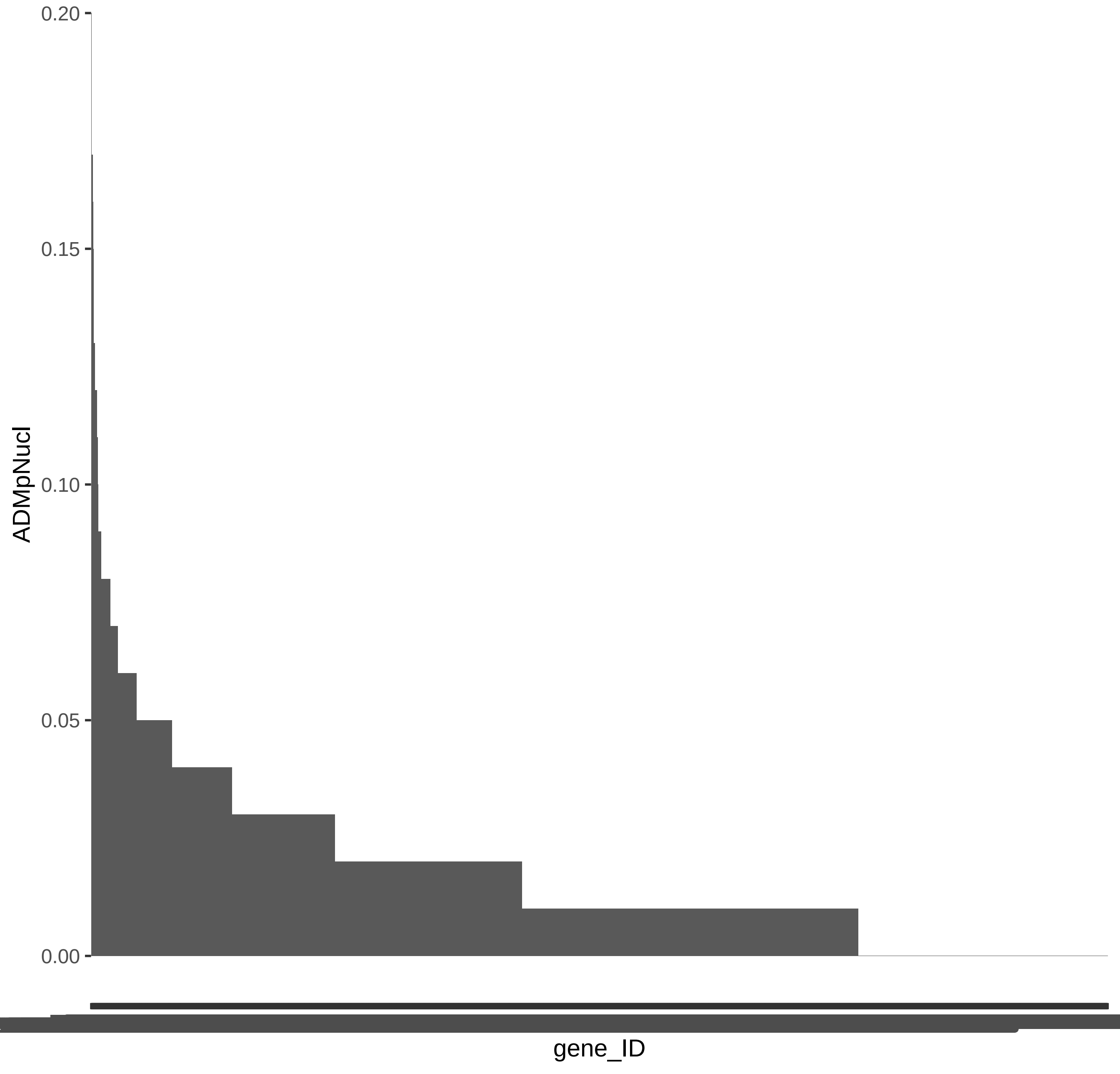
0.20
0.15
0.10
0.05
0.00
ADMpNucl
gene_ID

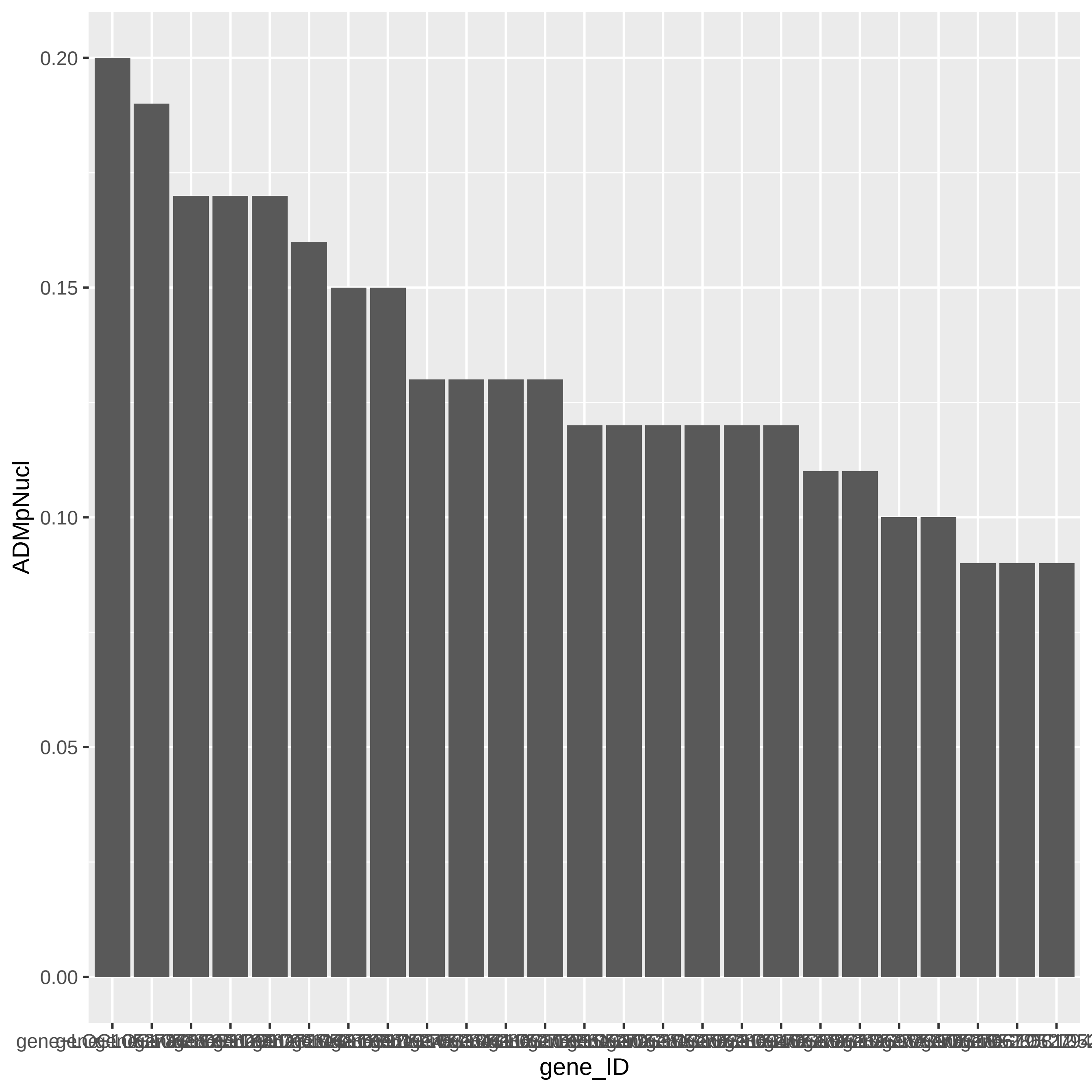
0.20
0.15
0.10
0.05
0.00
ADMpNucl
gene_ID

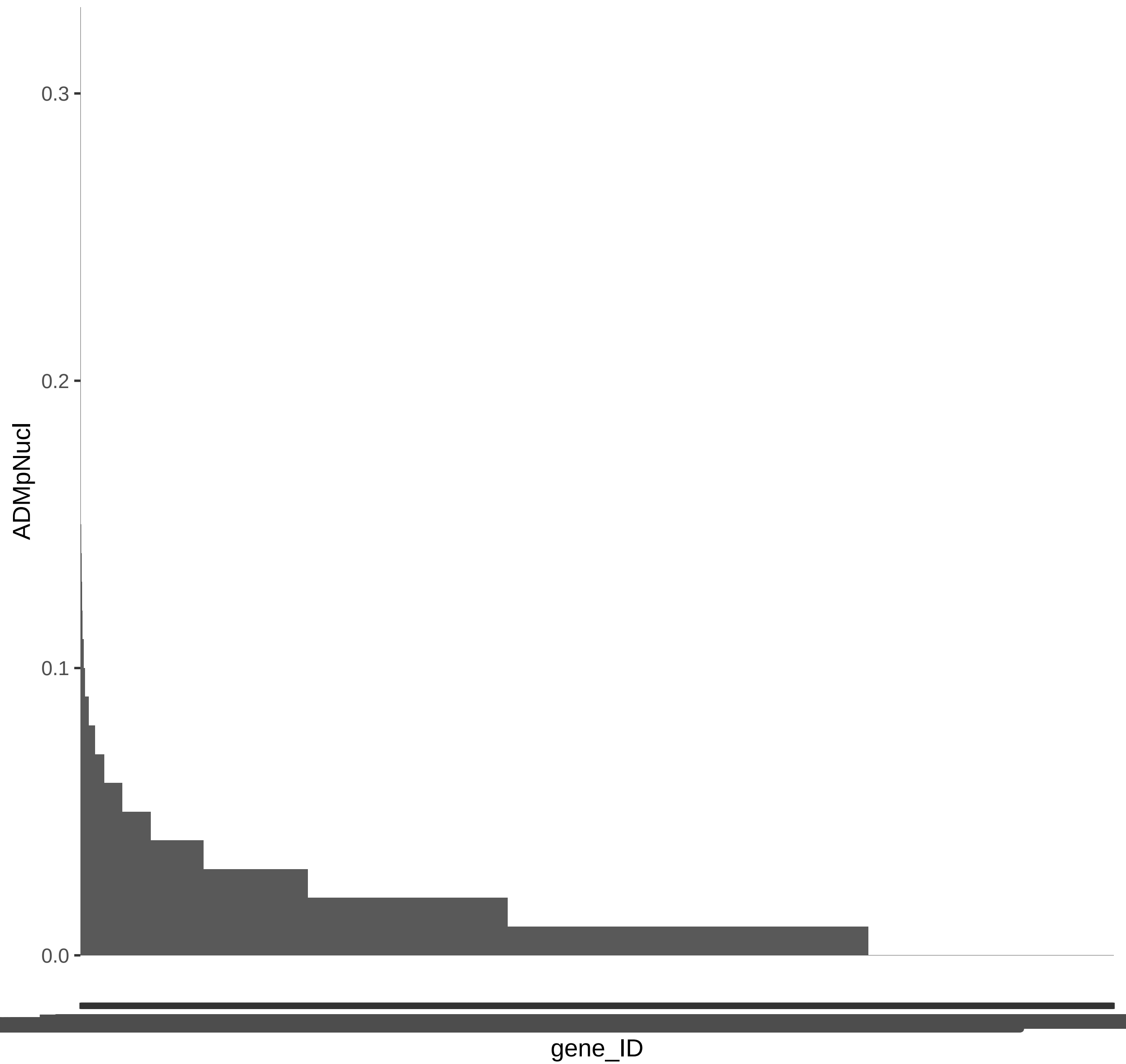
0.3
0.2
0.1
0.0
ADMpNucl
gene_ID

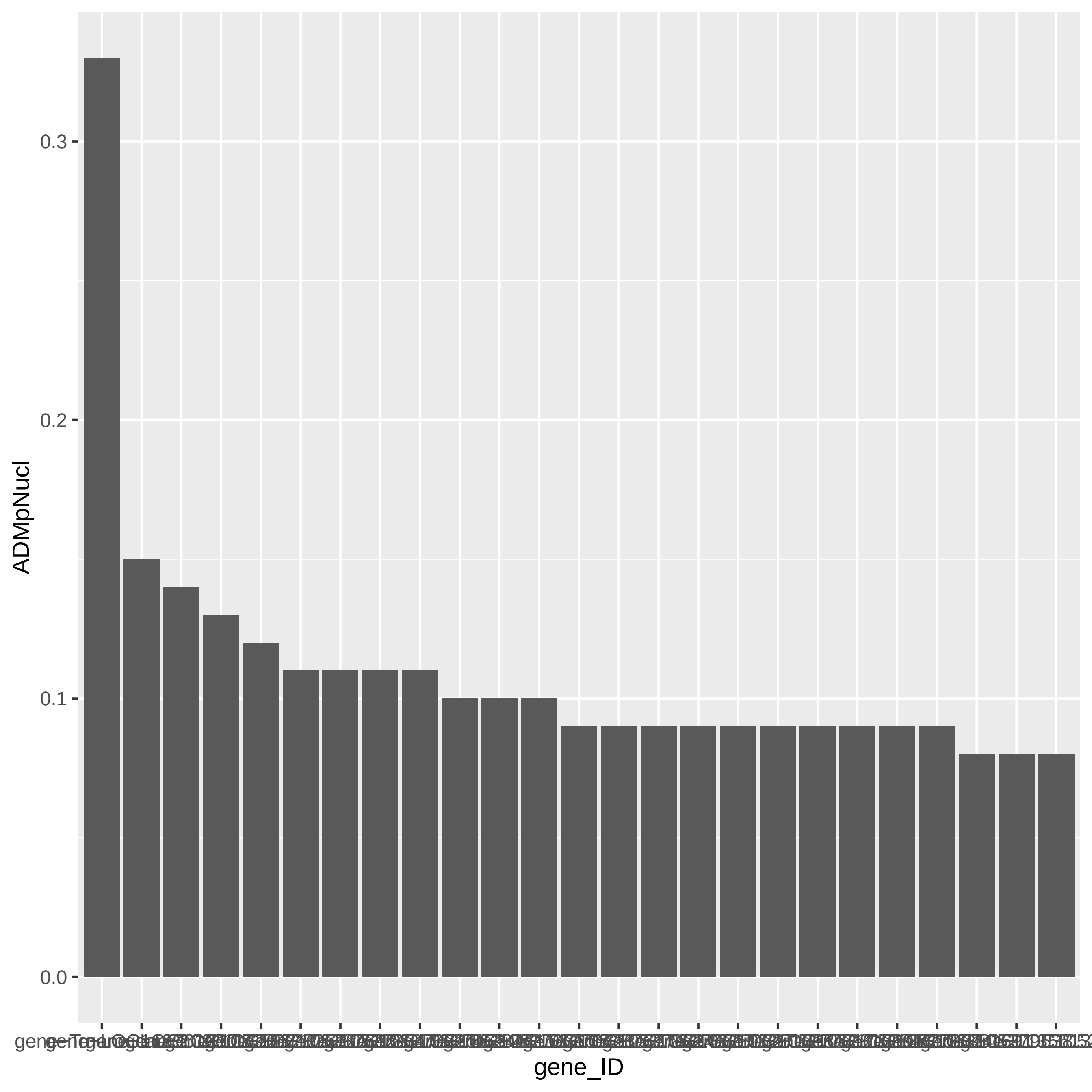

ADMpNucl
0.3
0.2
0.1
0.0
gene_ID

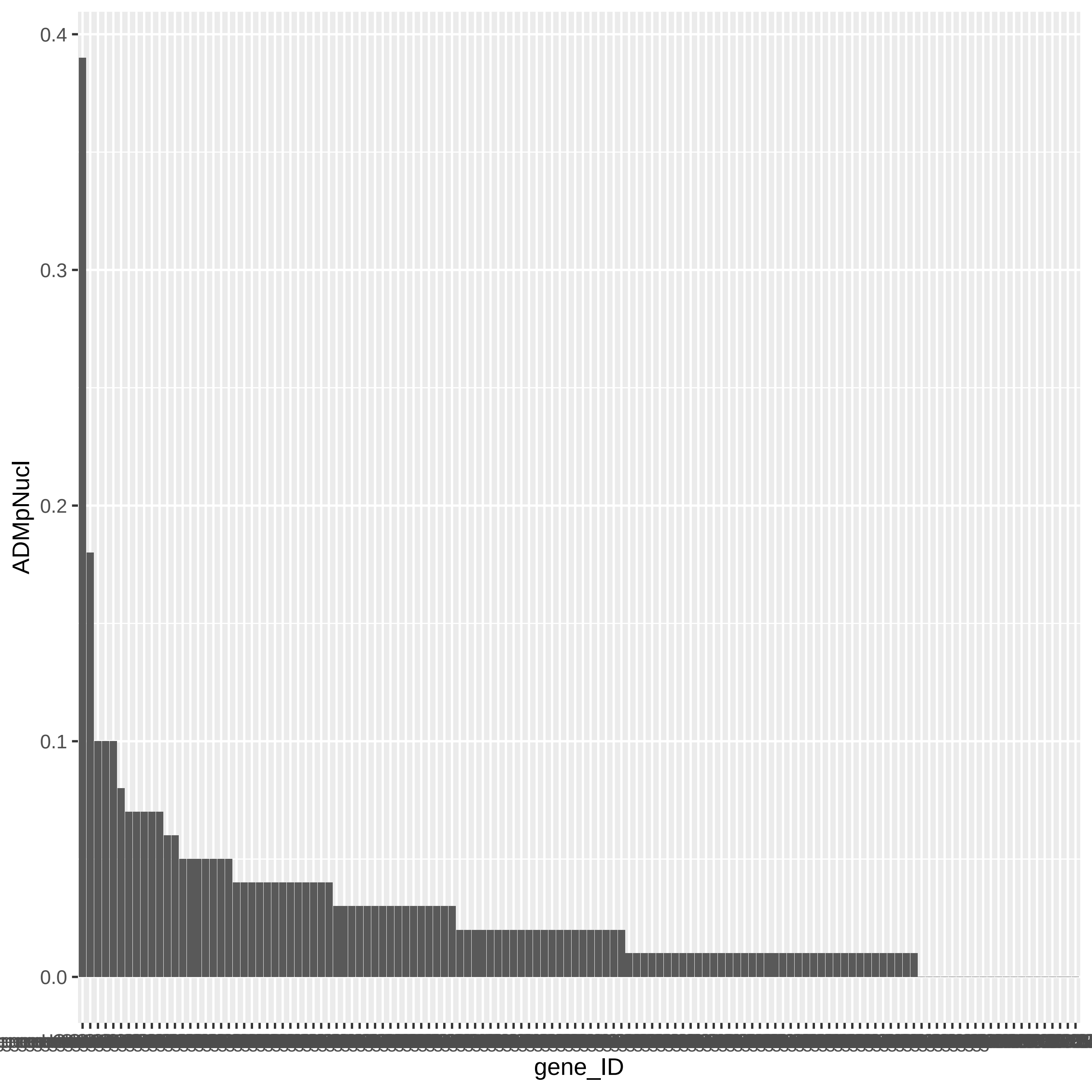

ADMpNucl
0.4
0.3
0.2
0.1
0.0
gene_ID

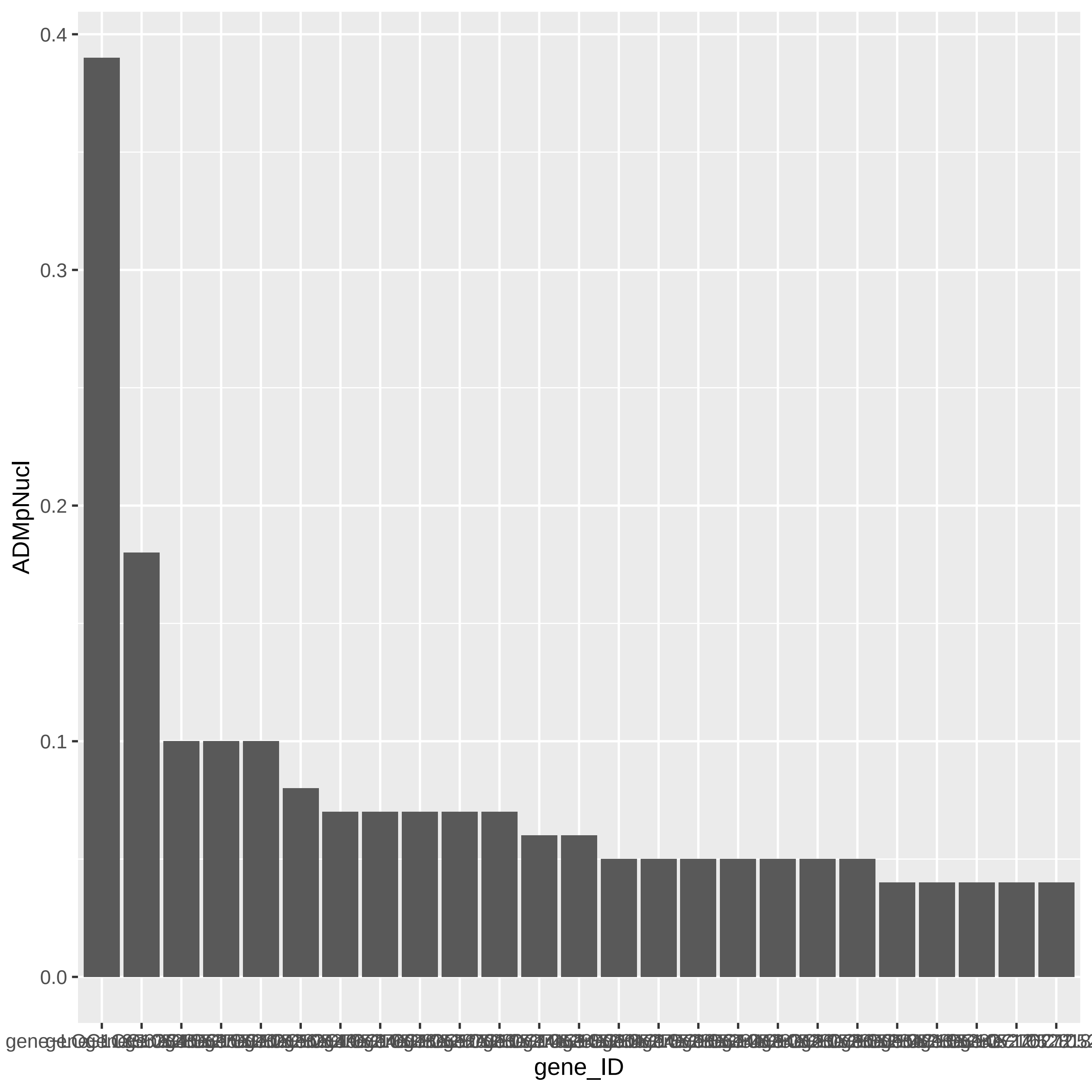

ADMpNucl
0.4
0.3
0.2
0.1
0.0
gene_ID

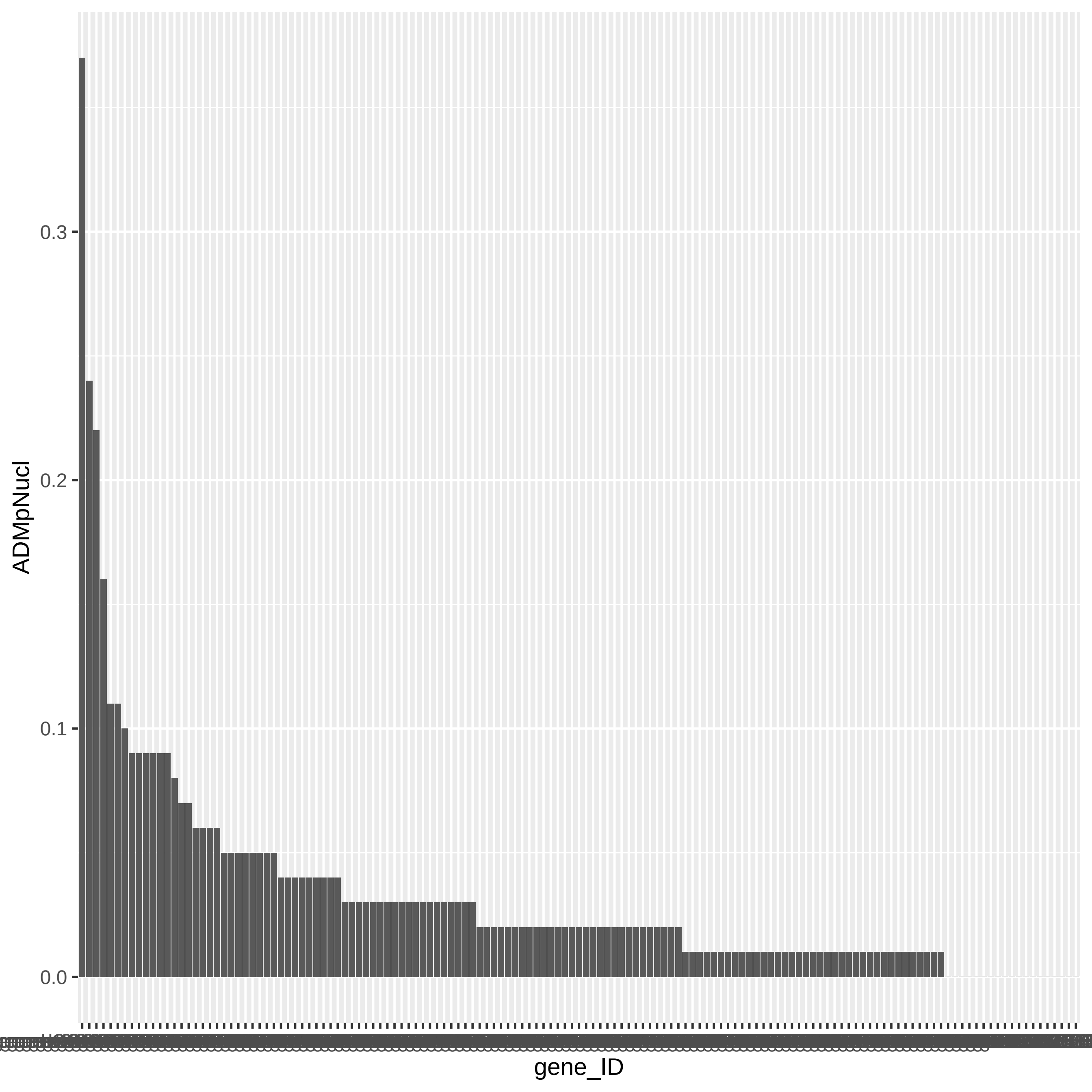
ADMpNucl
0.3
0.2
0.1
0.0
gene_ID

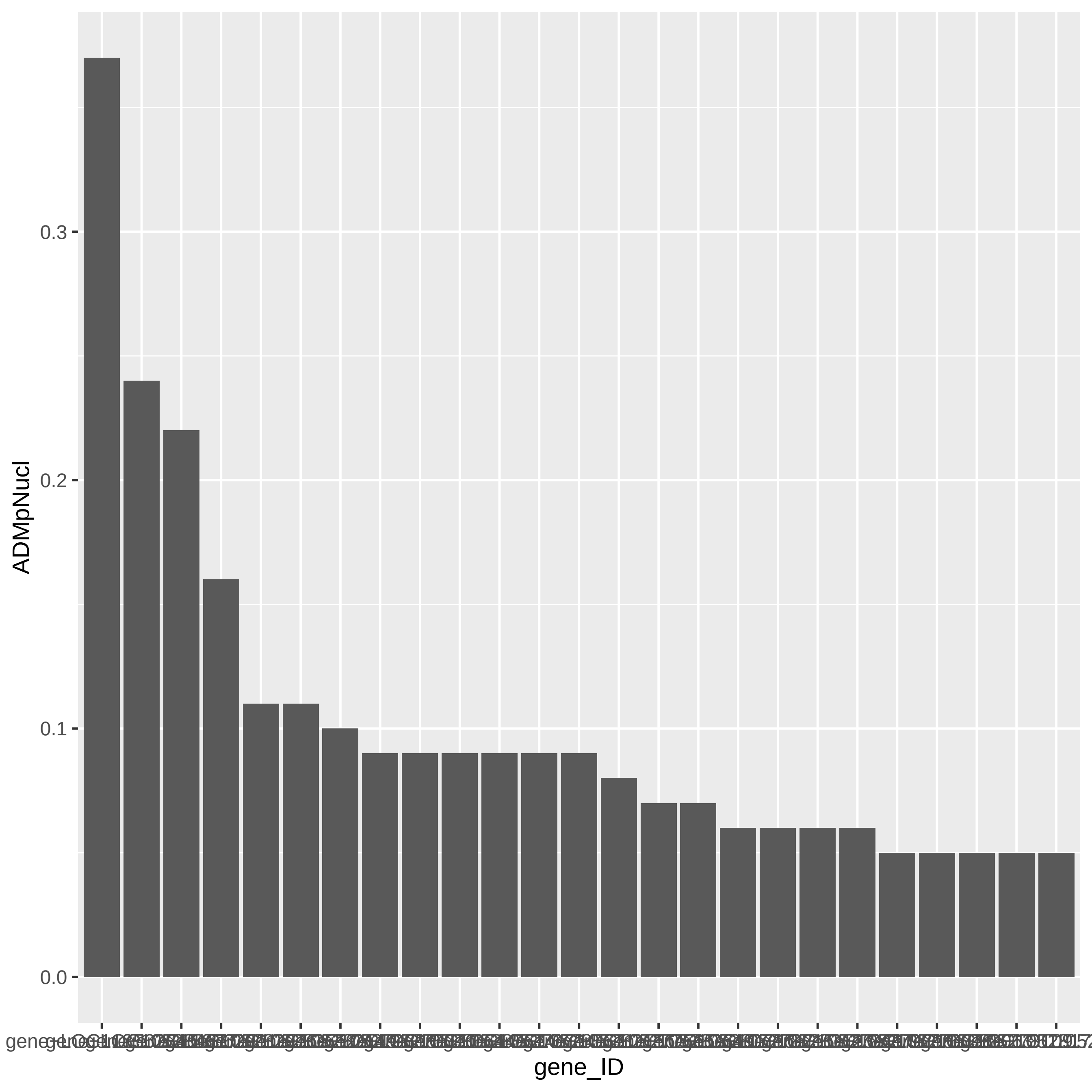

ADMpNucl
0.0
0.1
0.2
0.3
gene_ID

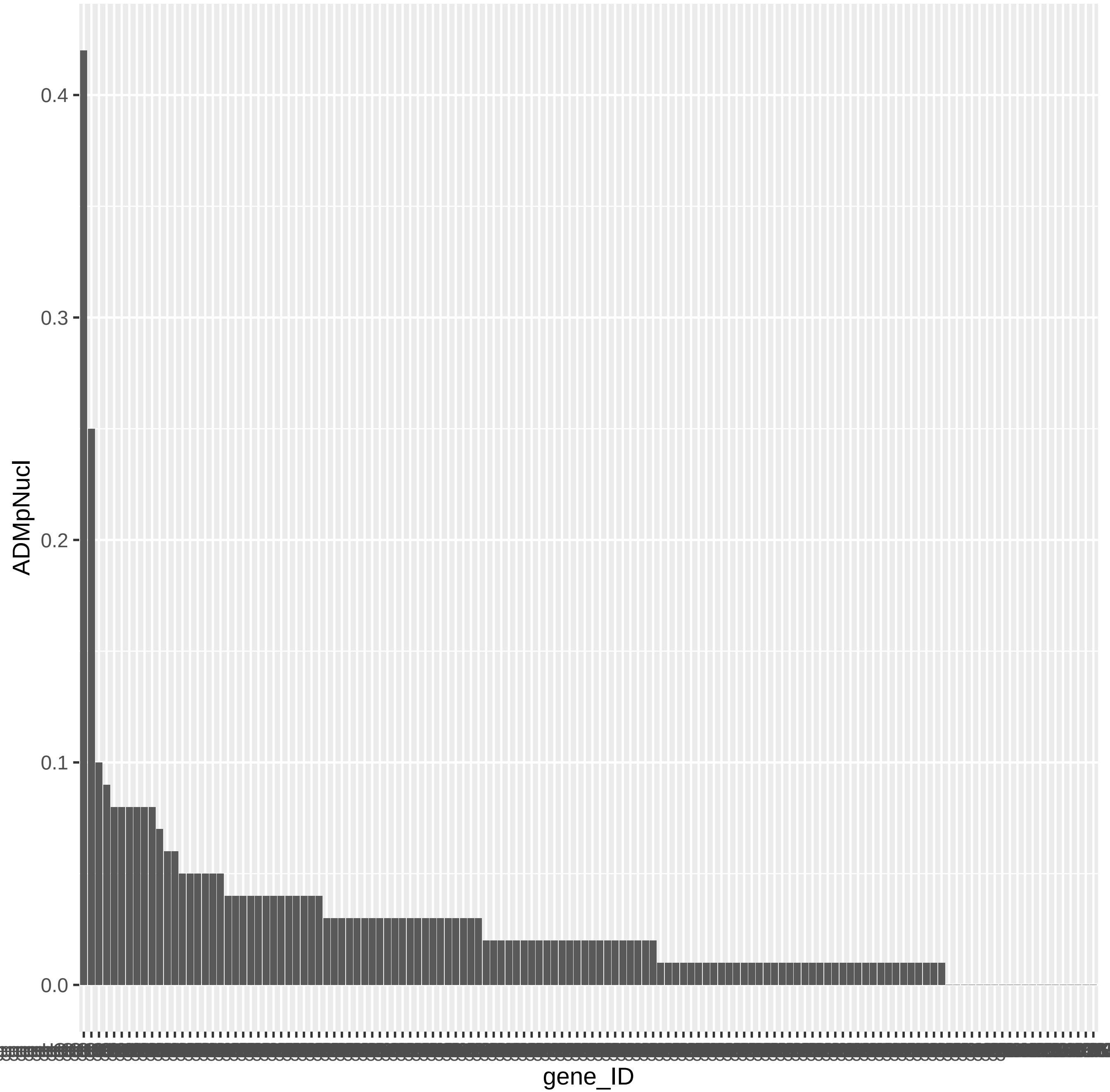

0.4
0.3
0.2
0.1
0.0
ADMpNucl
gene_ID

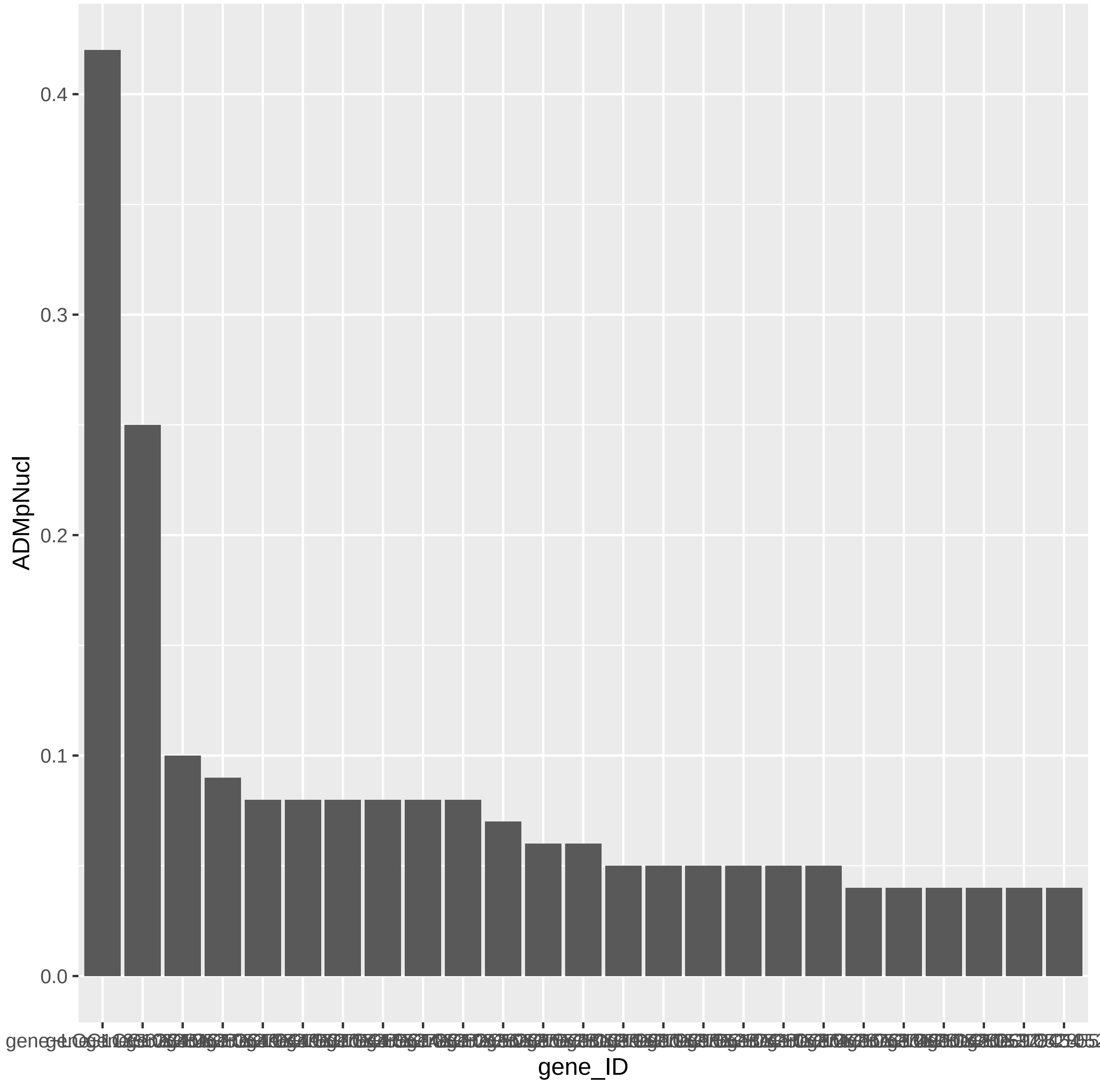

ADMpNucl
gene_ID
0.0
0.1
0.2
0.3
0.4

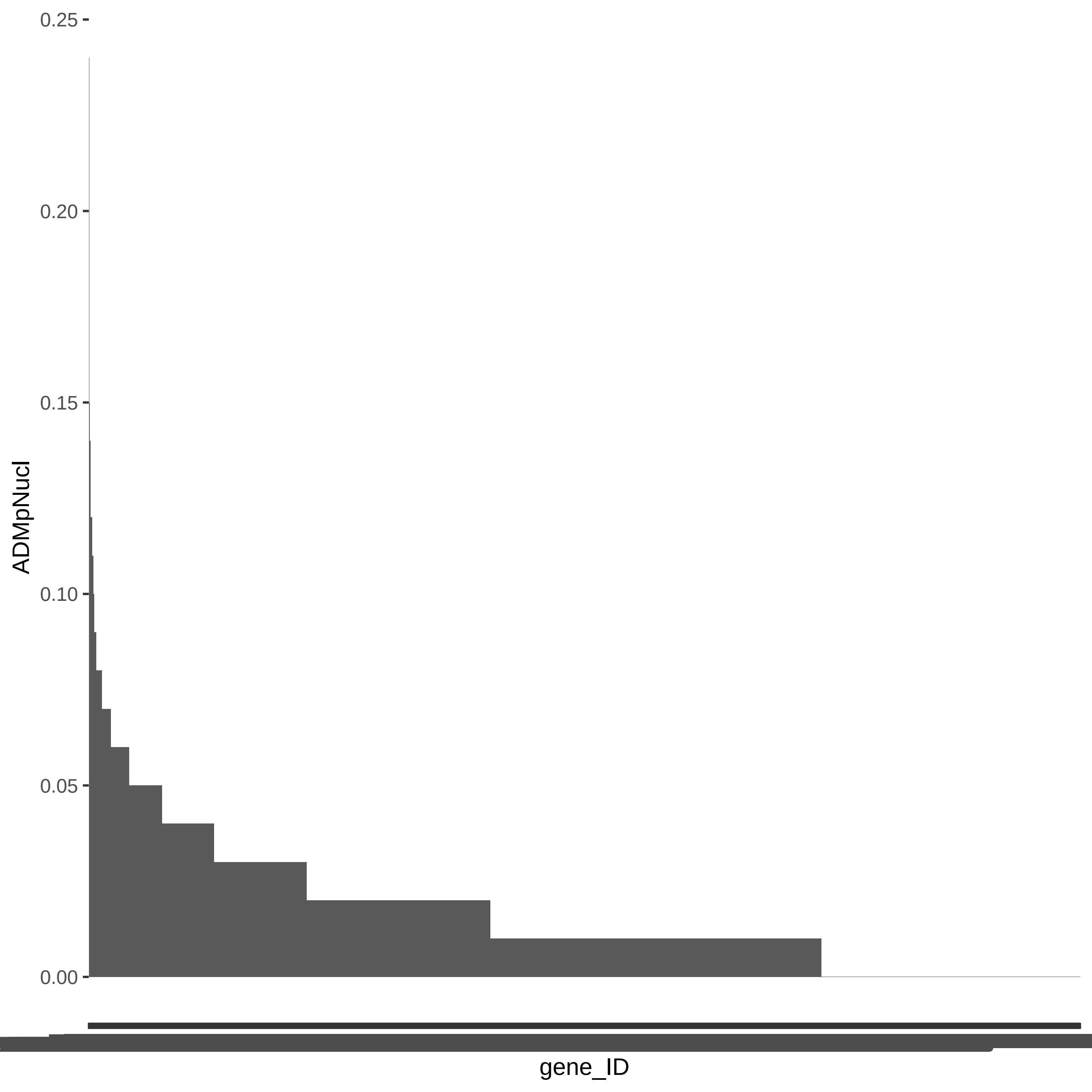

0.25
0.20
0.15
0.10
0.05
0.00
ADMpNucl
gene_ID

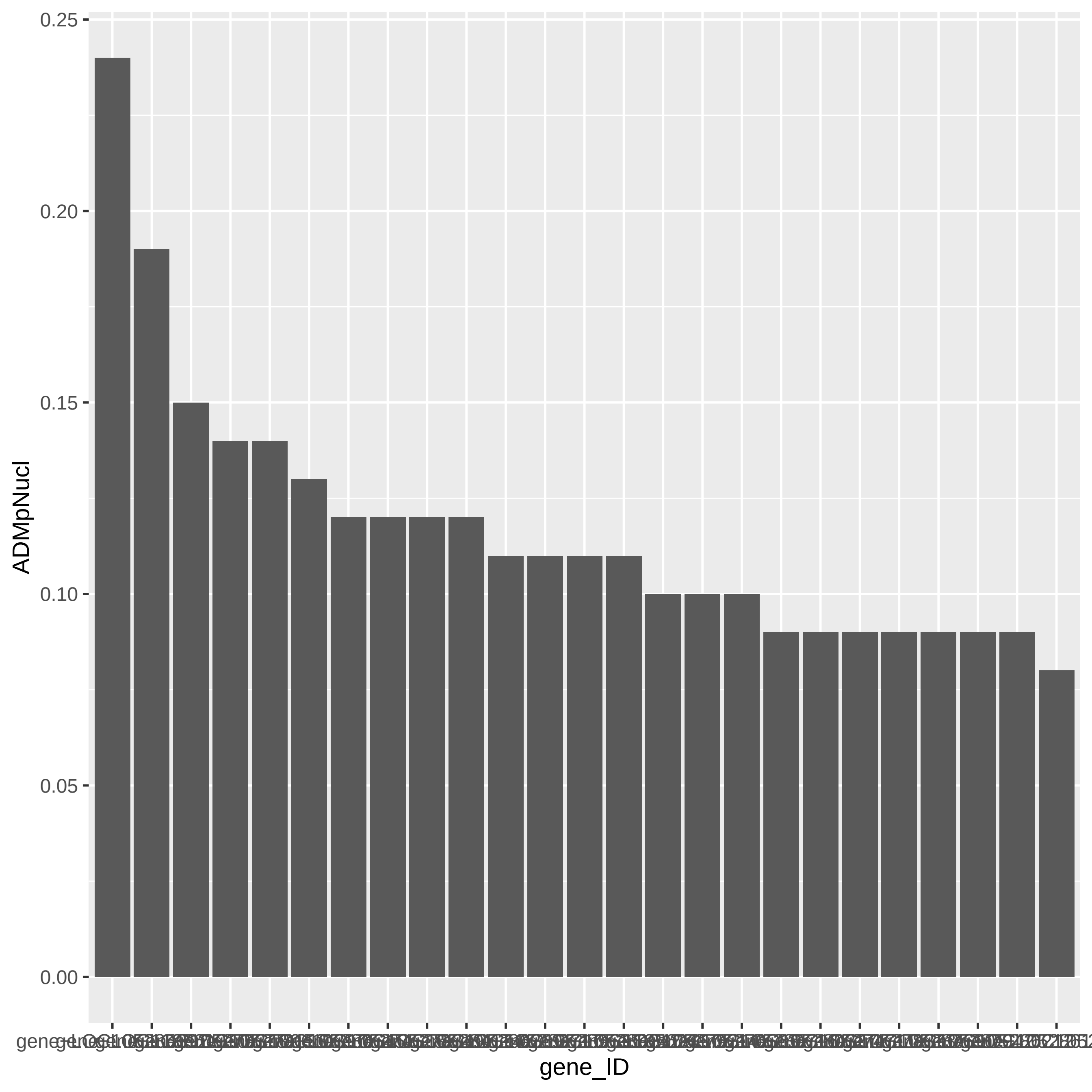

0.25
0.20
0.15
0.10
0.05
0.00
ADMpNucl
gene_ID

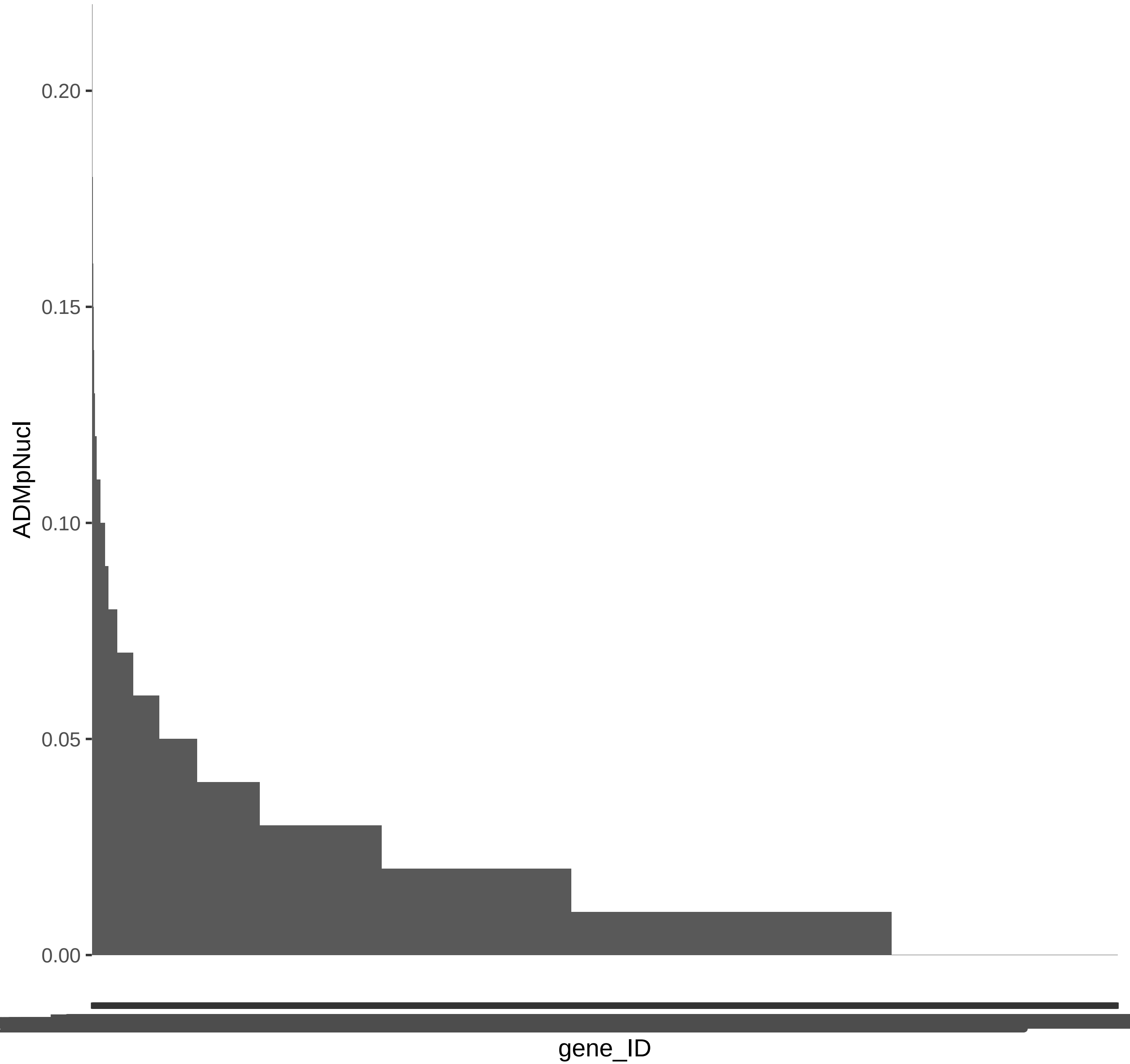

0.20
0.15
0.10
0.05
0.00
ADMpNucl
gene_ID

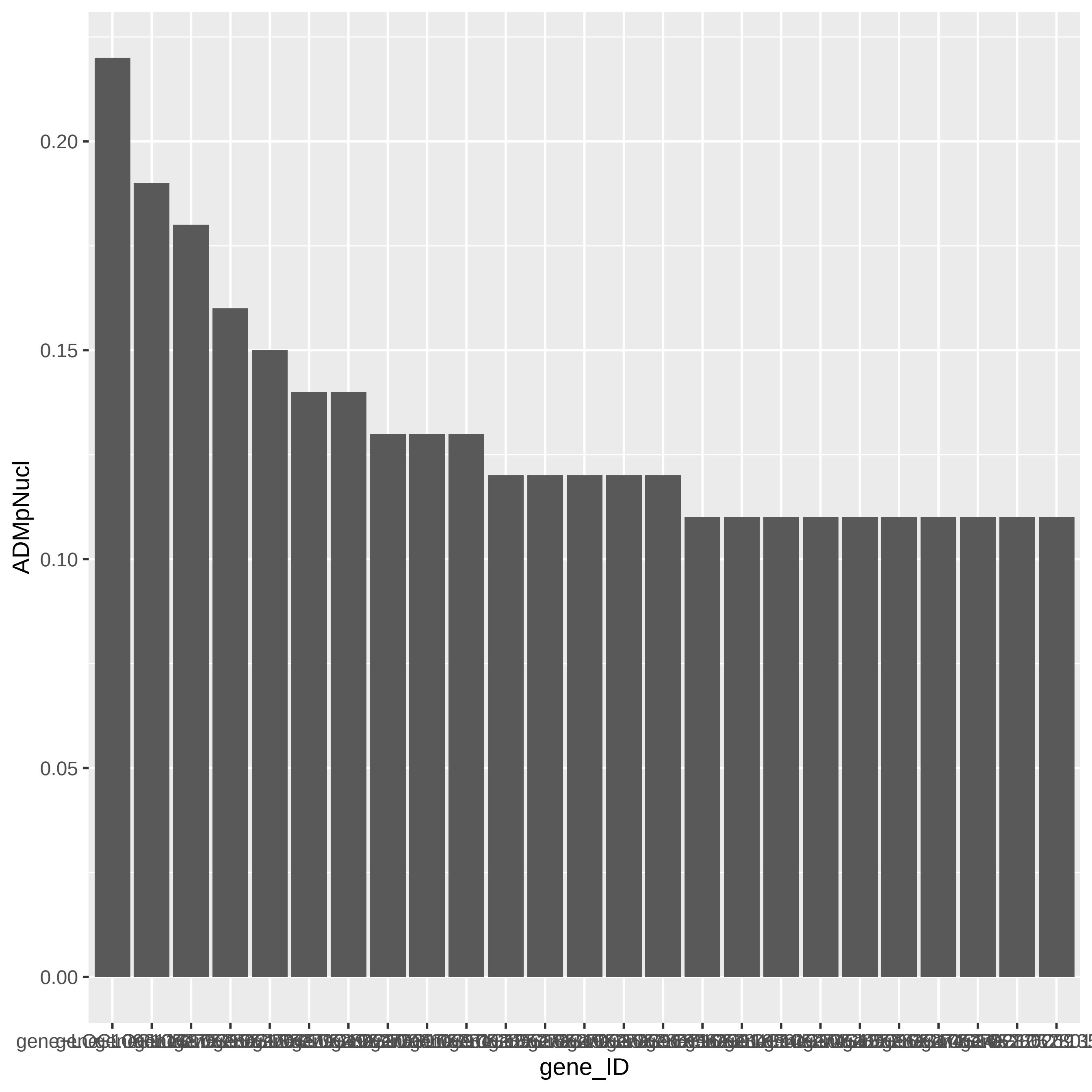

ADMpNucl
0.00
0.05
0.10
0.15
0.20
gene_ID

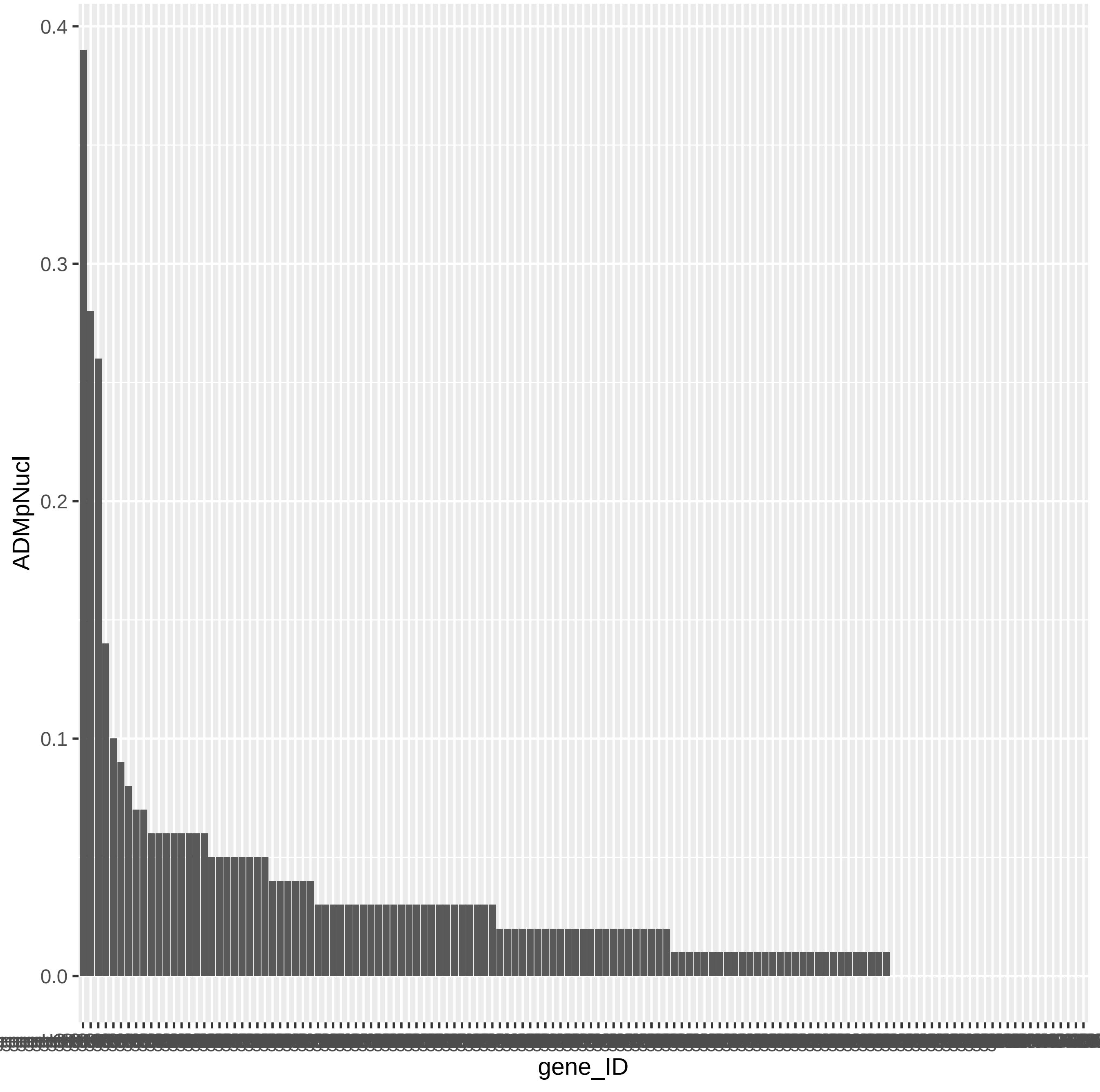

0.4
0.3
0.2
0.1
0.0
ADMpNucl
gene_ID

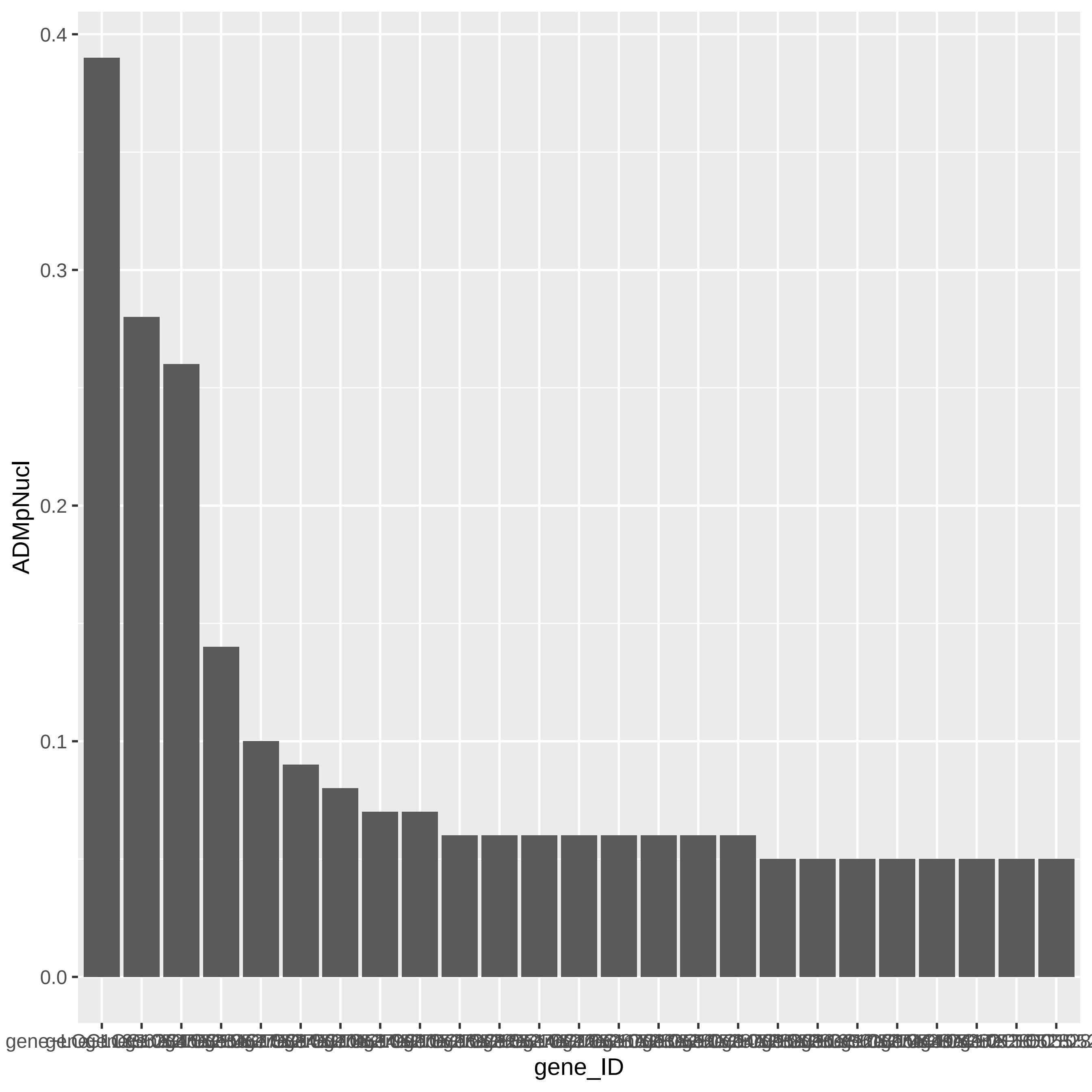

0.4
0.3
0.2
0.1
0.0
ADMpNucl
gene_ID

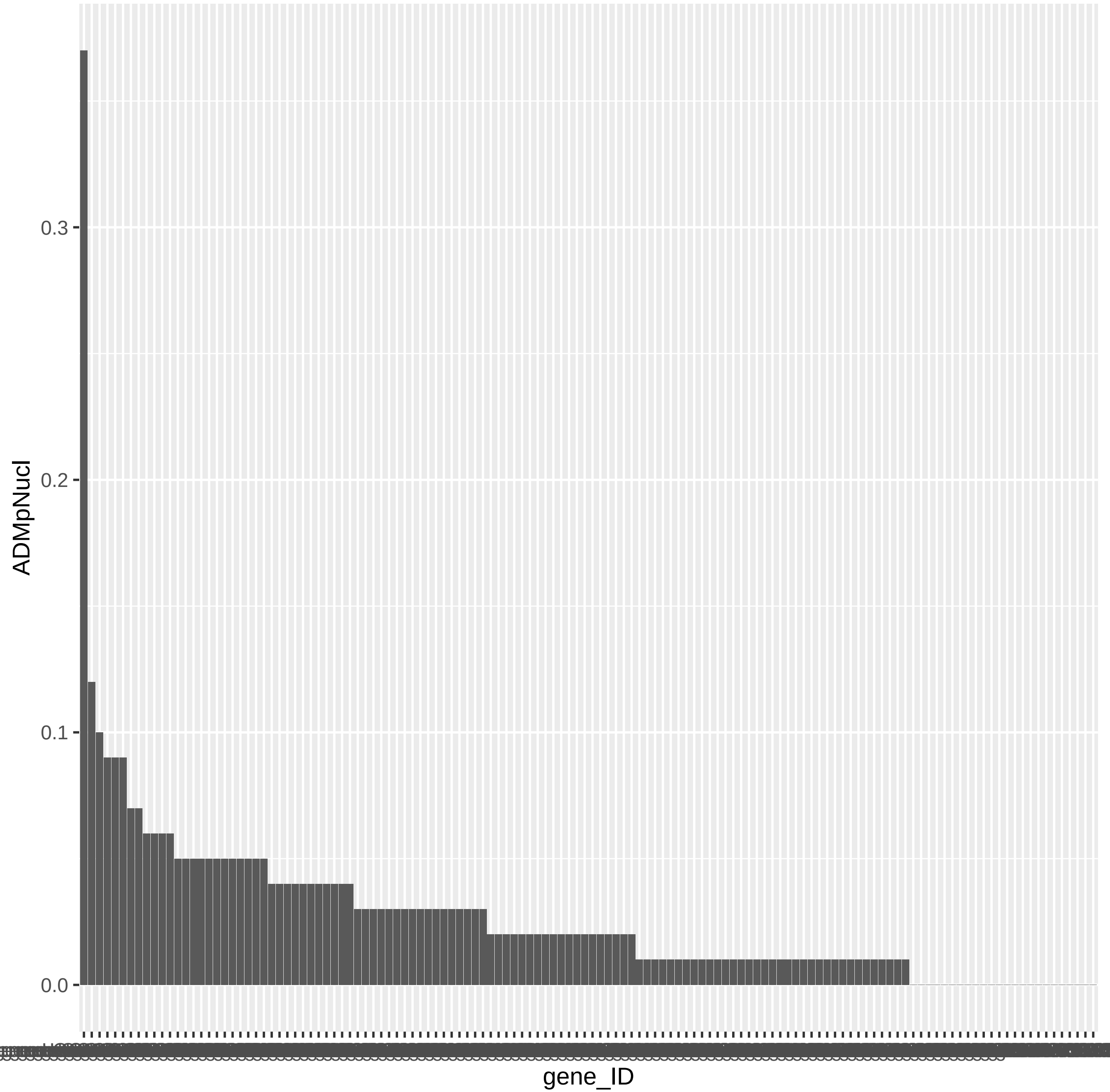
ADMpNucl
0.3
0.2
0.1
0.0
gene_ID

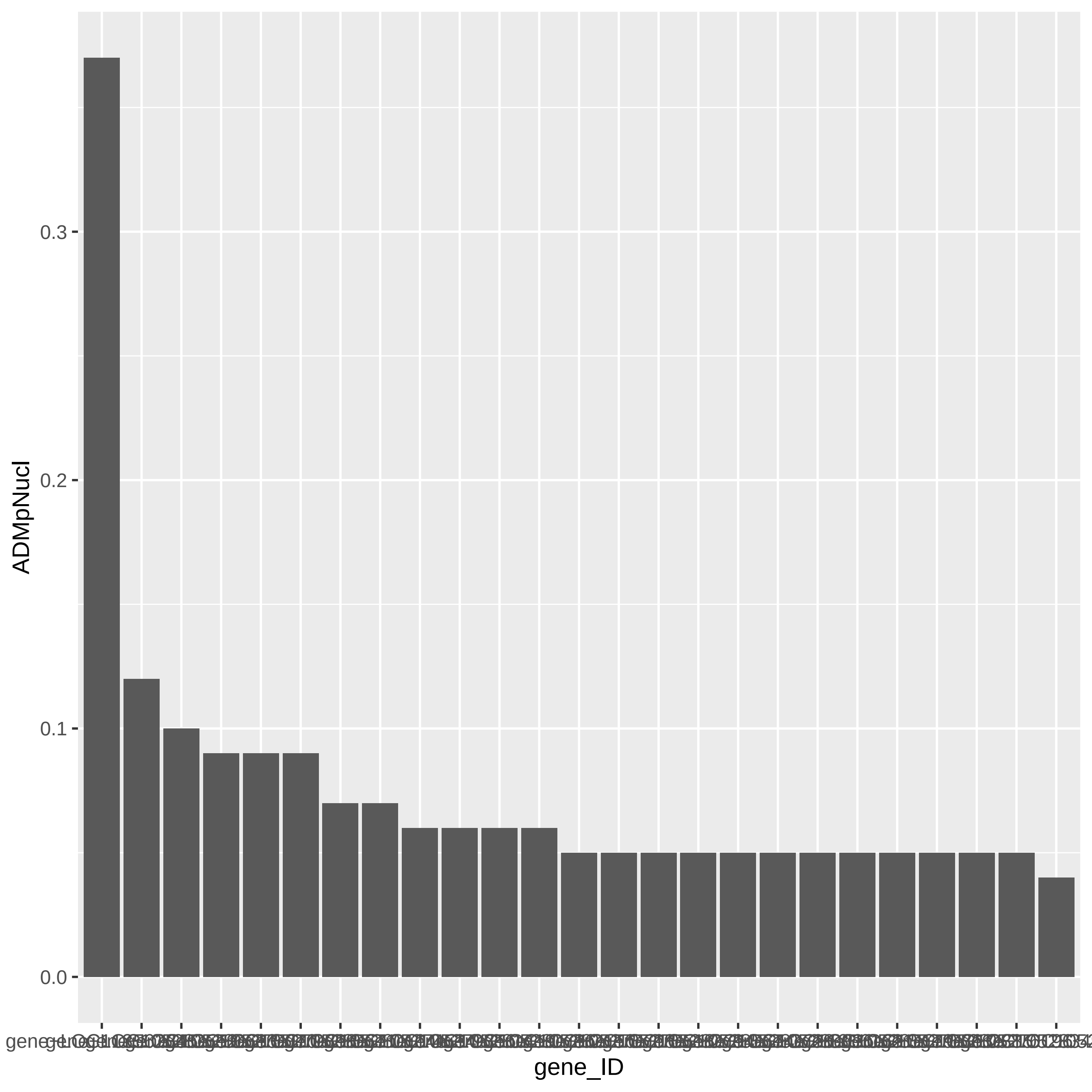

0.3
0.2
0.1
0.0
ADMpNucl
gene_ID

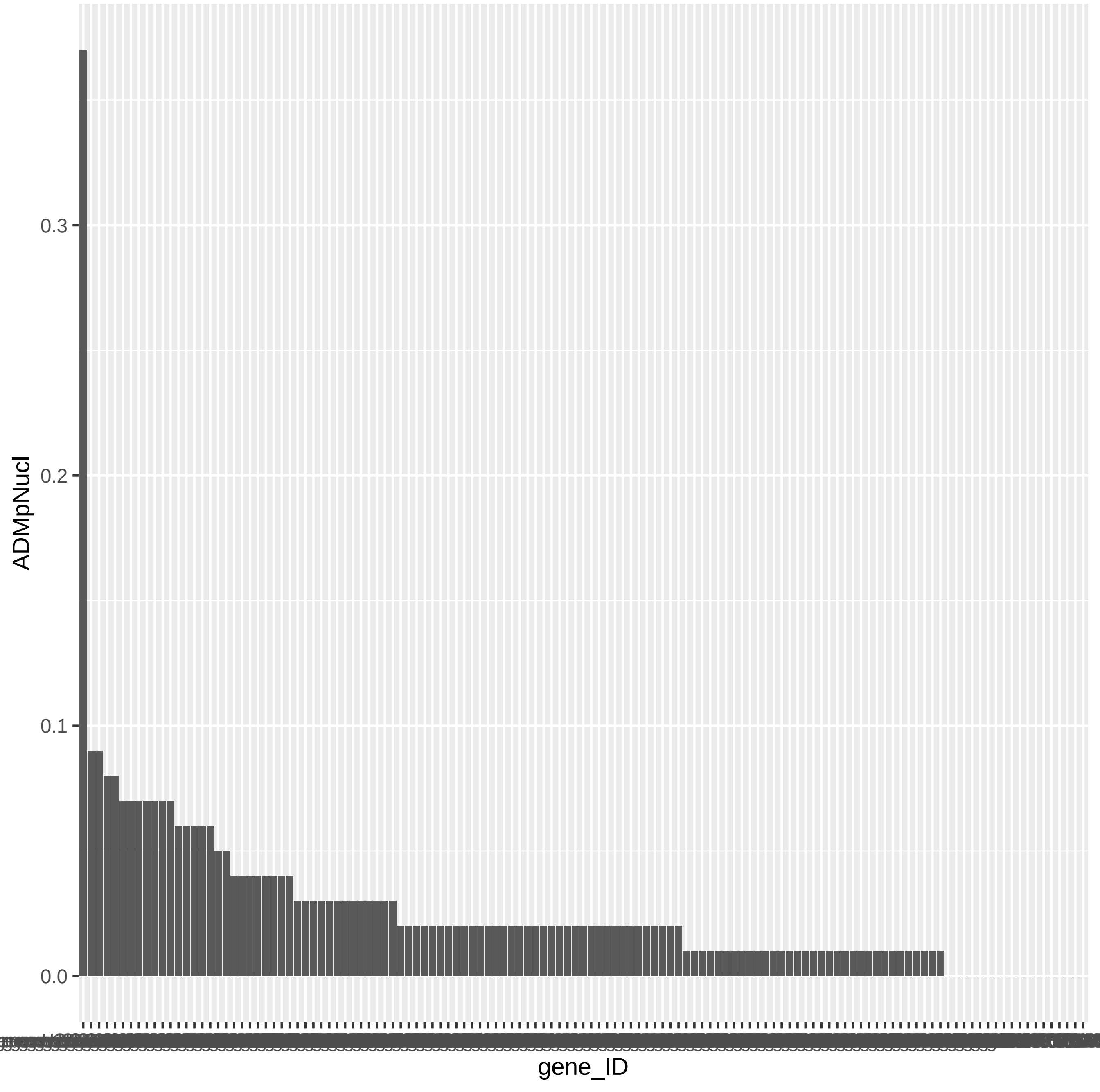

ADMpNucl
0.3
0.2
0.1
0.0
gene_ID

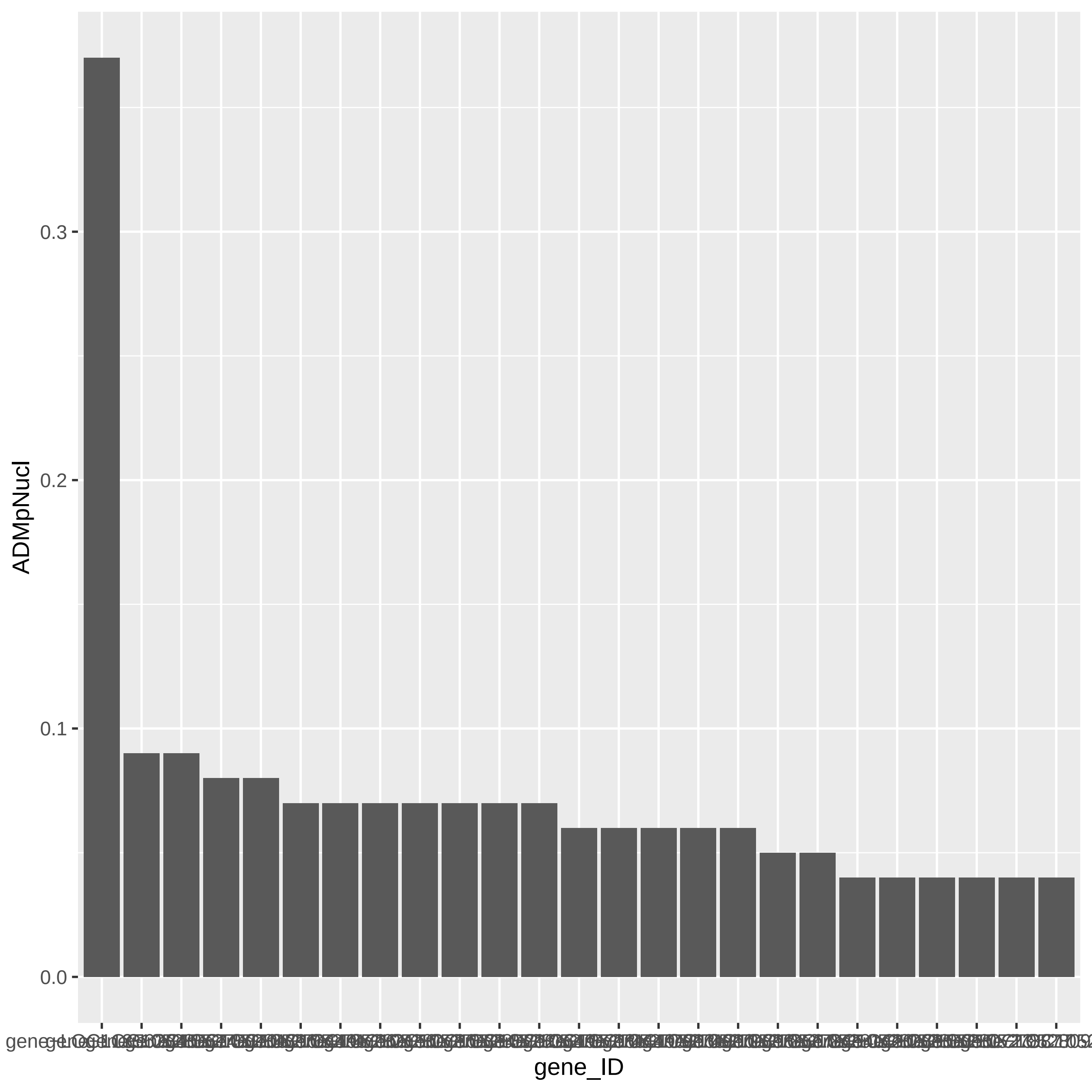
ADMpNucl
0.3
0.2
0.1
0.0
gene_ID

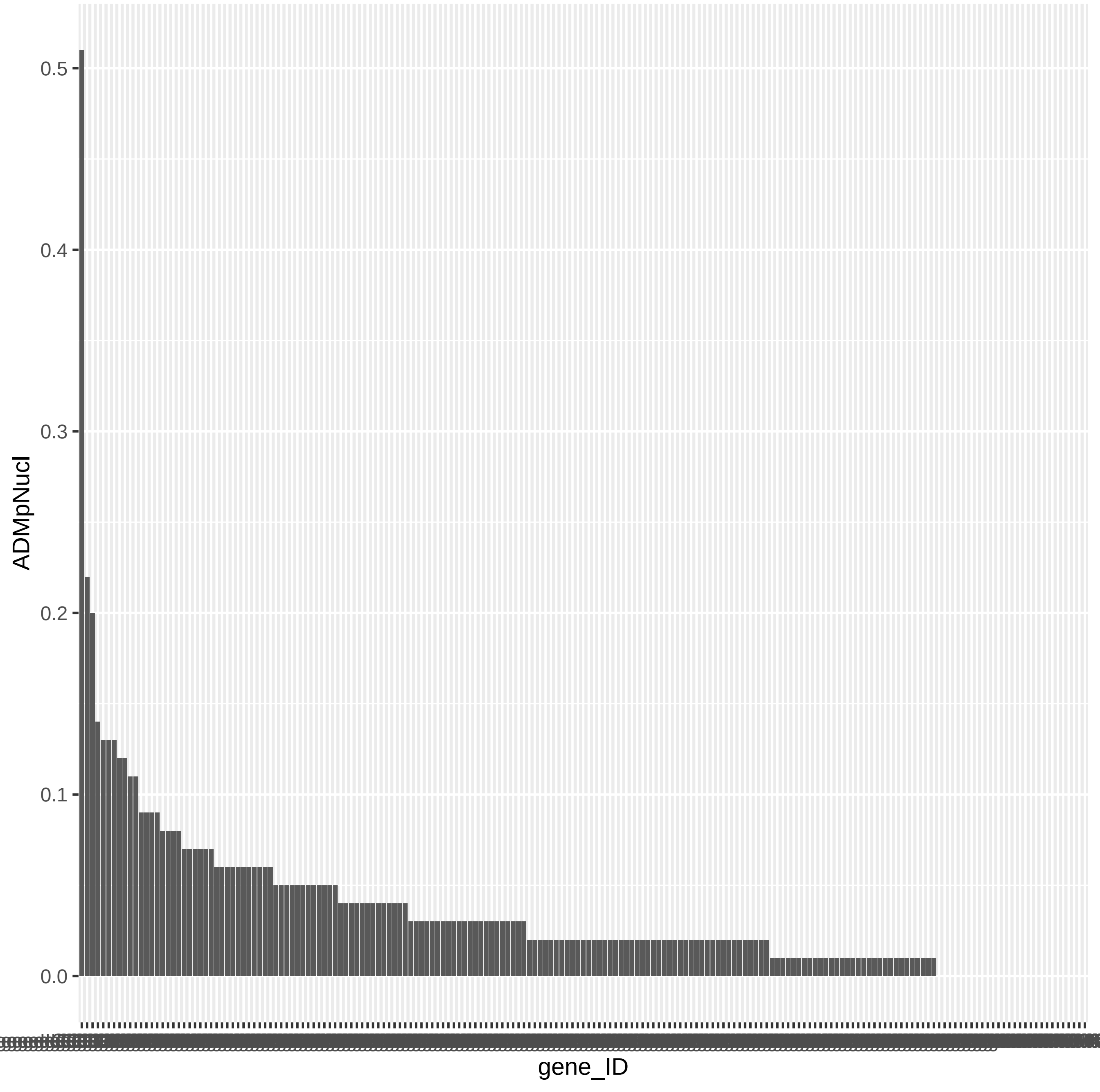

0.5
0.4
0.3
0.2
0.1
0.0
ADMpNucl
gene_ID

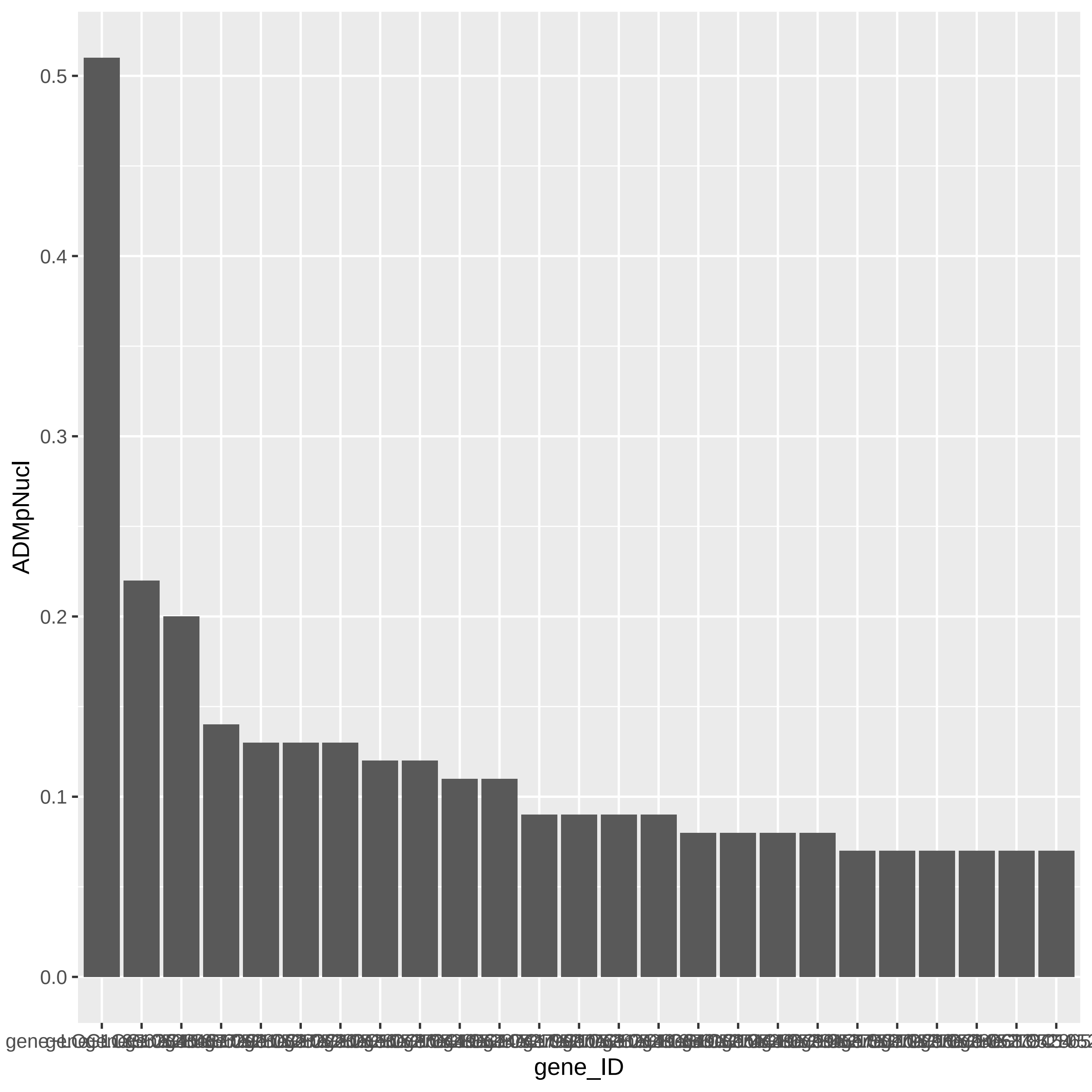

ADMpNucl
0.5
0.4
0.3
0.2
0.1
0.0
gene_ID

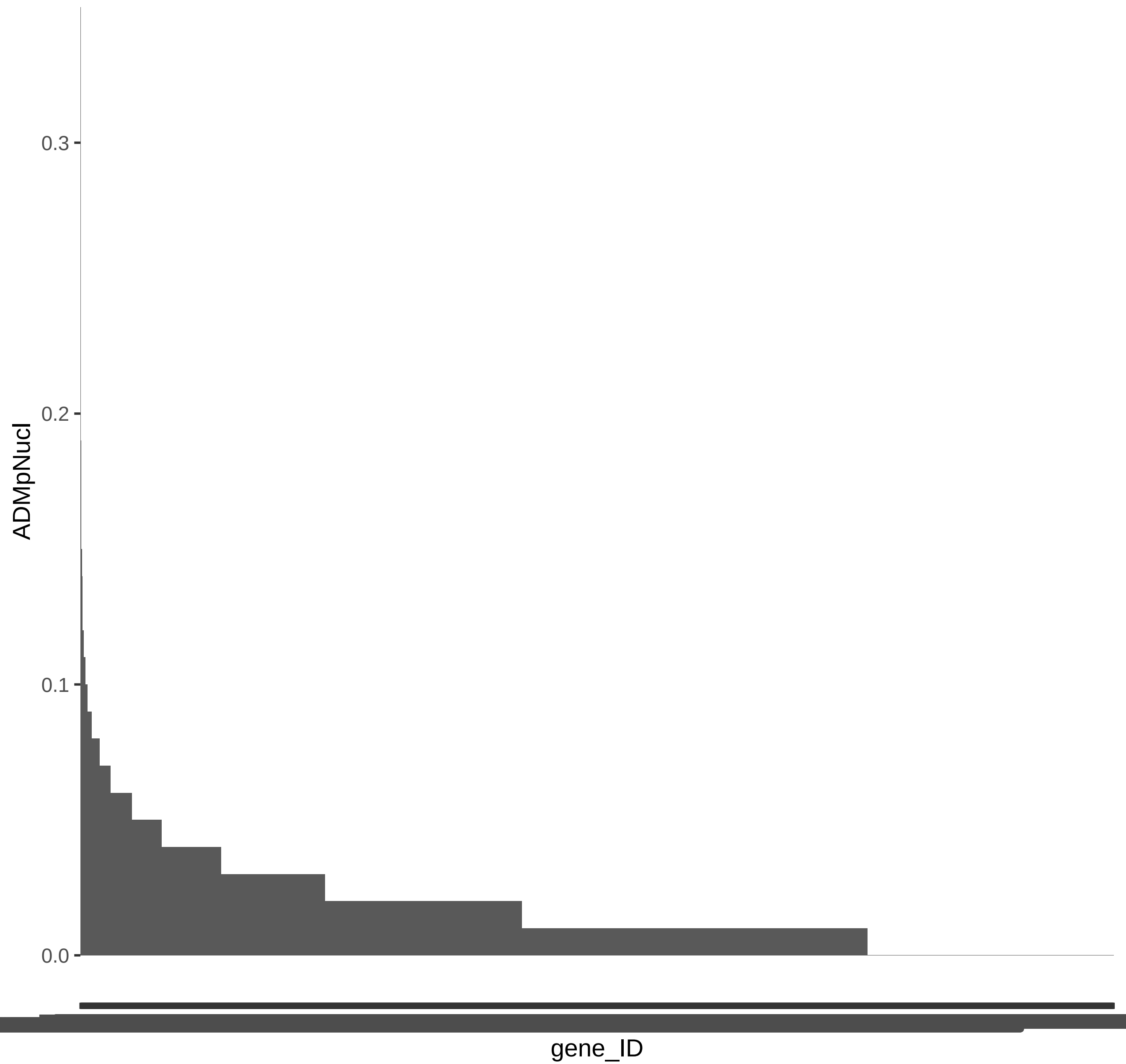
0.3
0.2
0.1
0.0
ADMpNucl
gene_ID

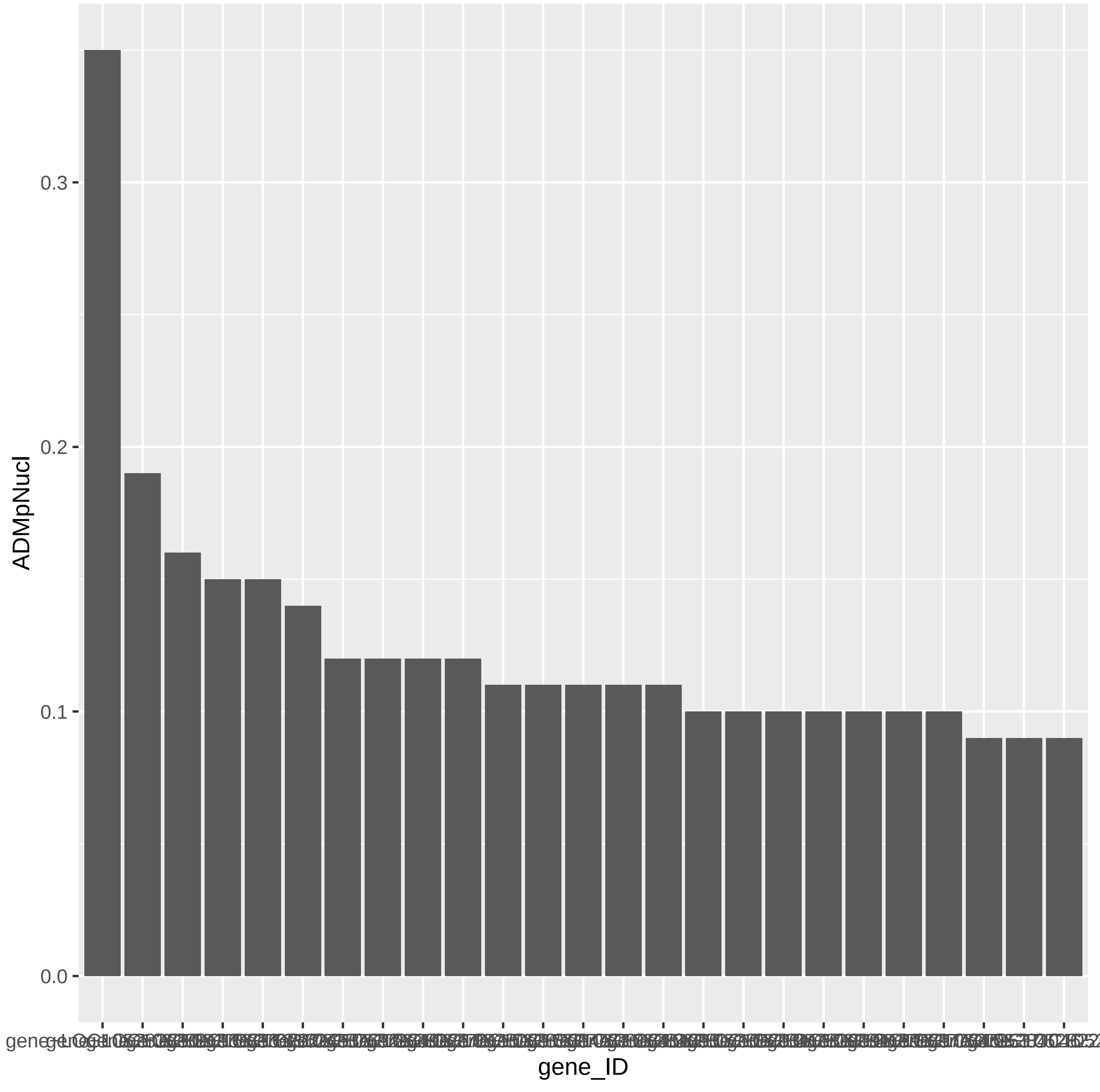

ADMpNucl
0.3
0.2
0.1
0.0
gene_ID

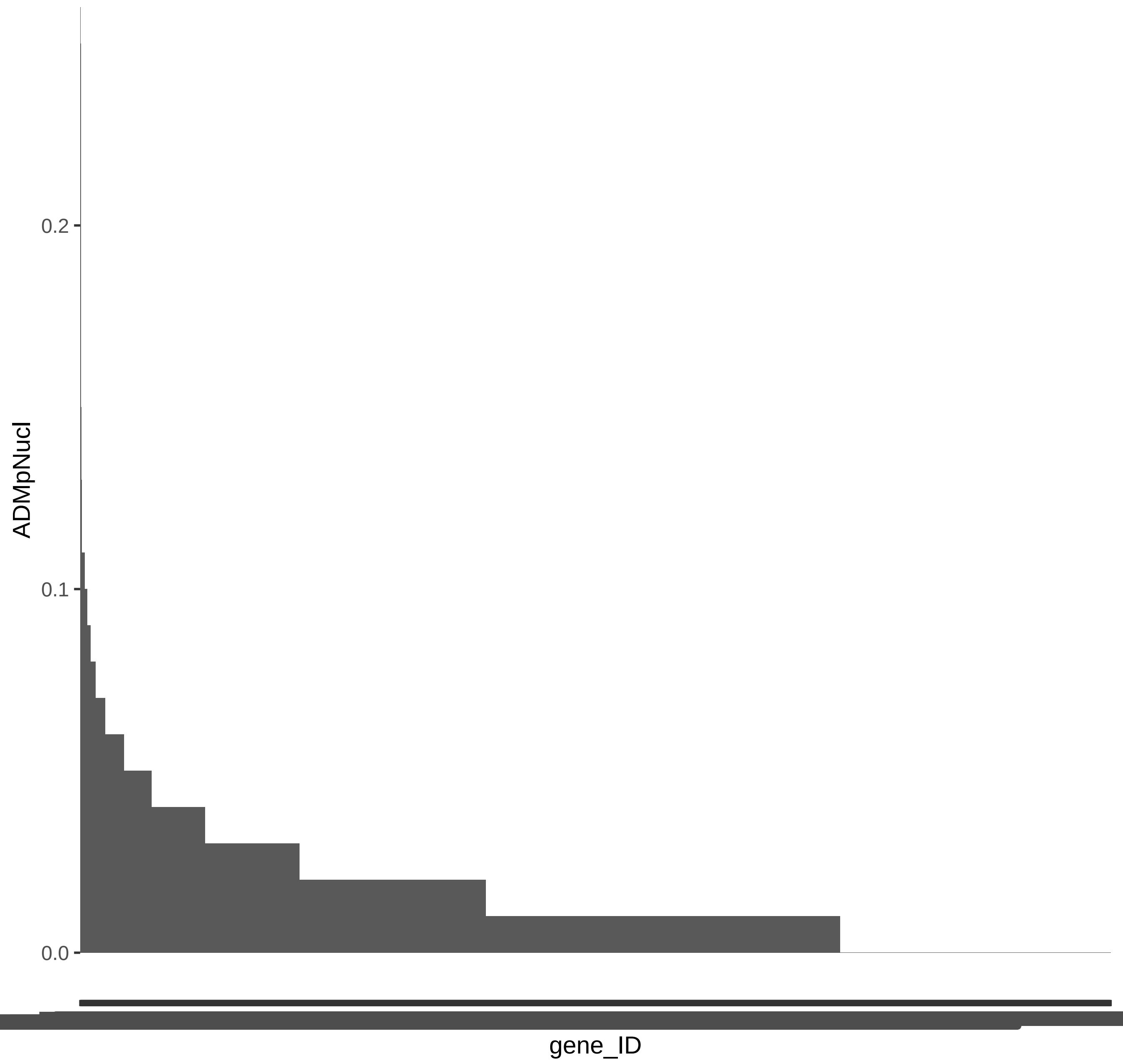

0.2
0.1
0.0
ADMpNucl
gene_ID

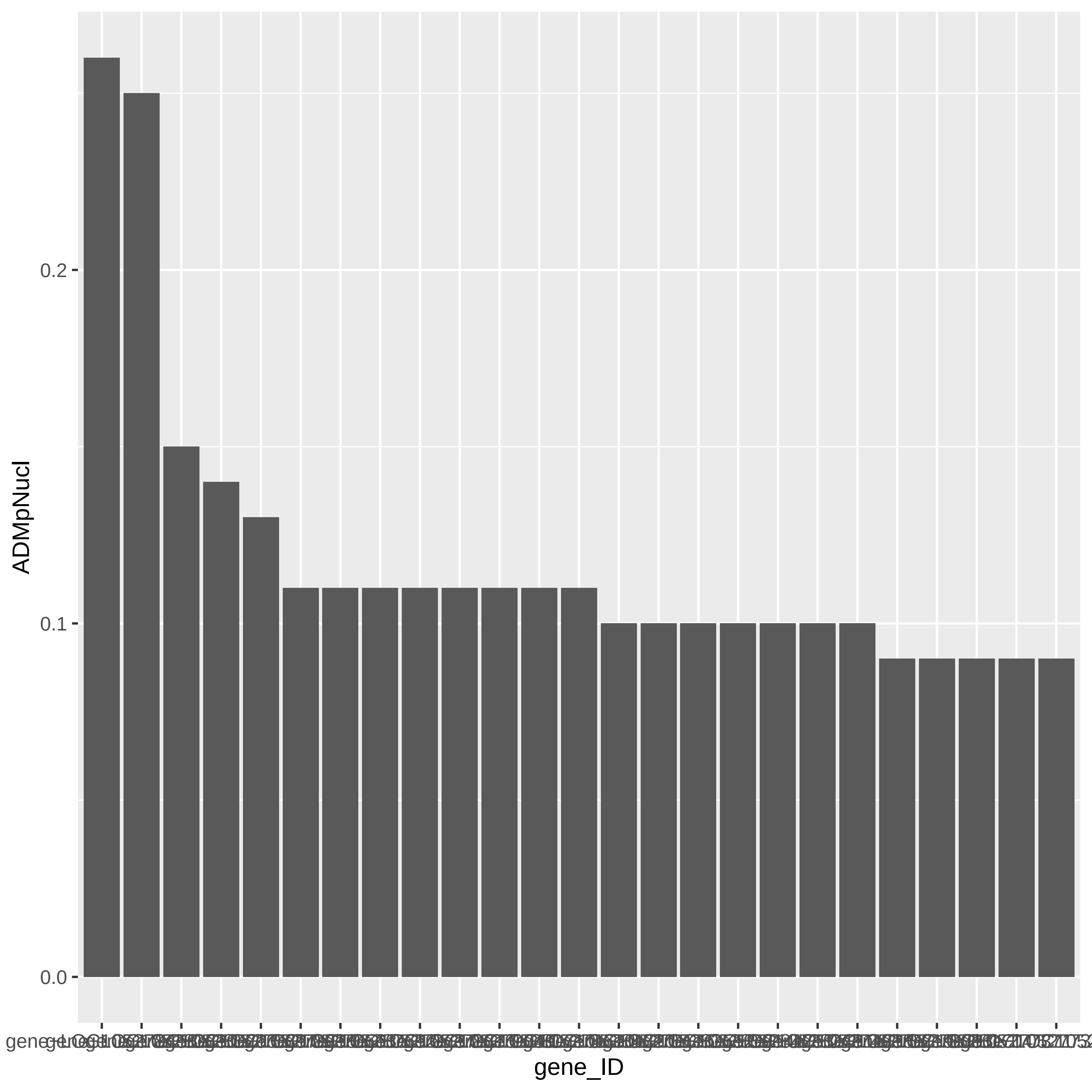
ADMpNucl
0.0
0.1
0.2
gene_ID

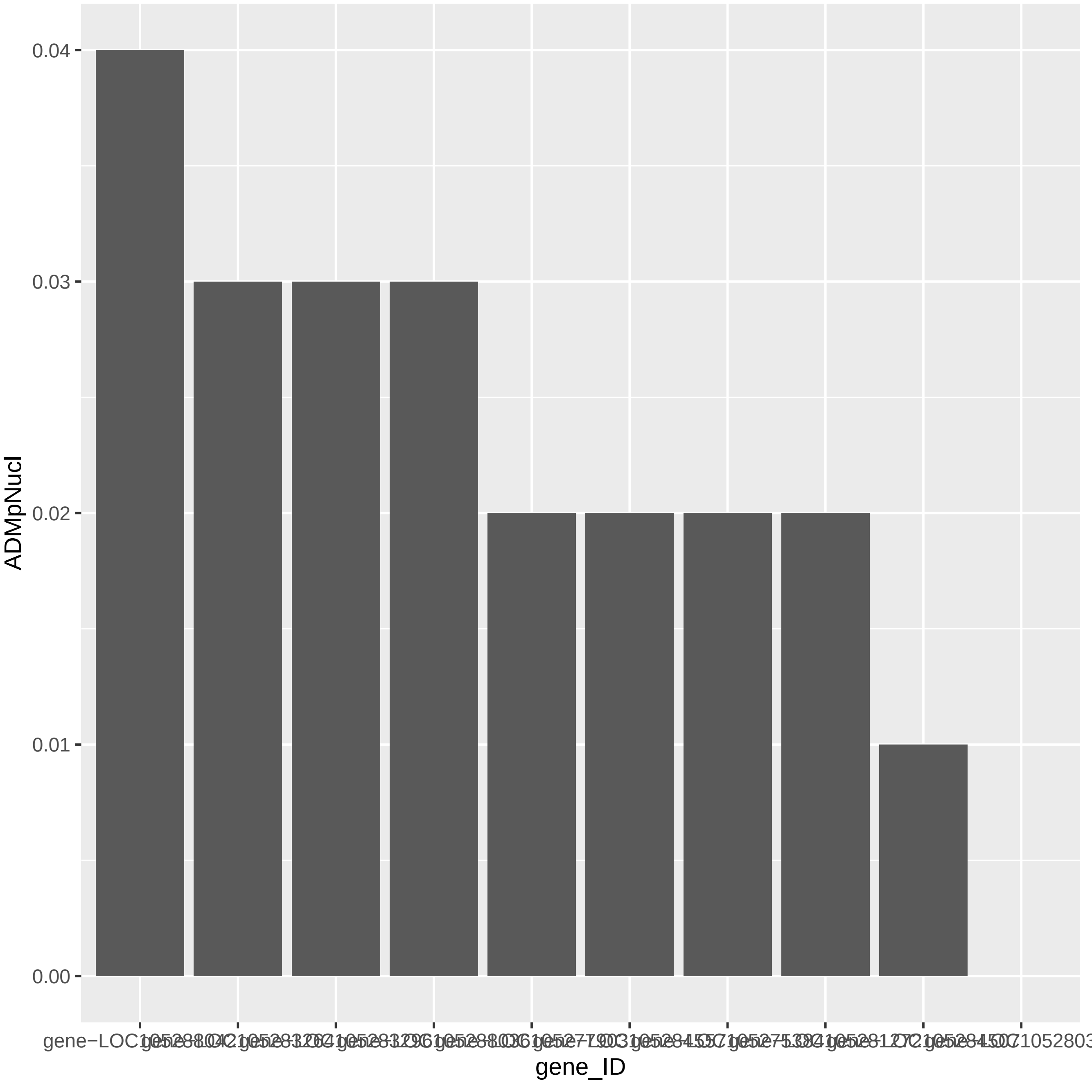
0.04
0.03
0.02
0.01
0.00
ADMpNucl
gene_ID

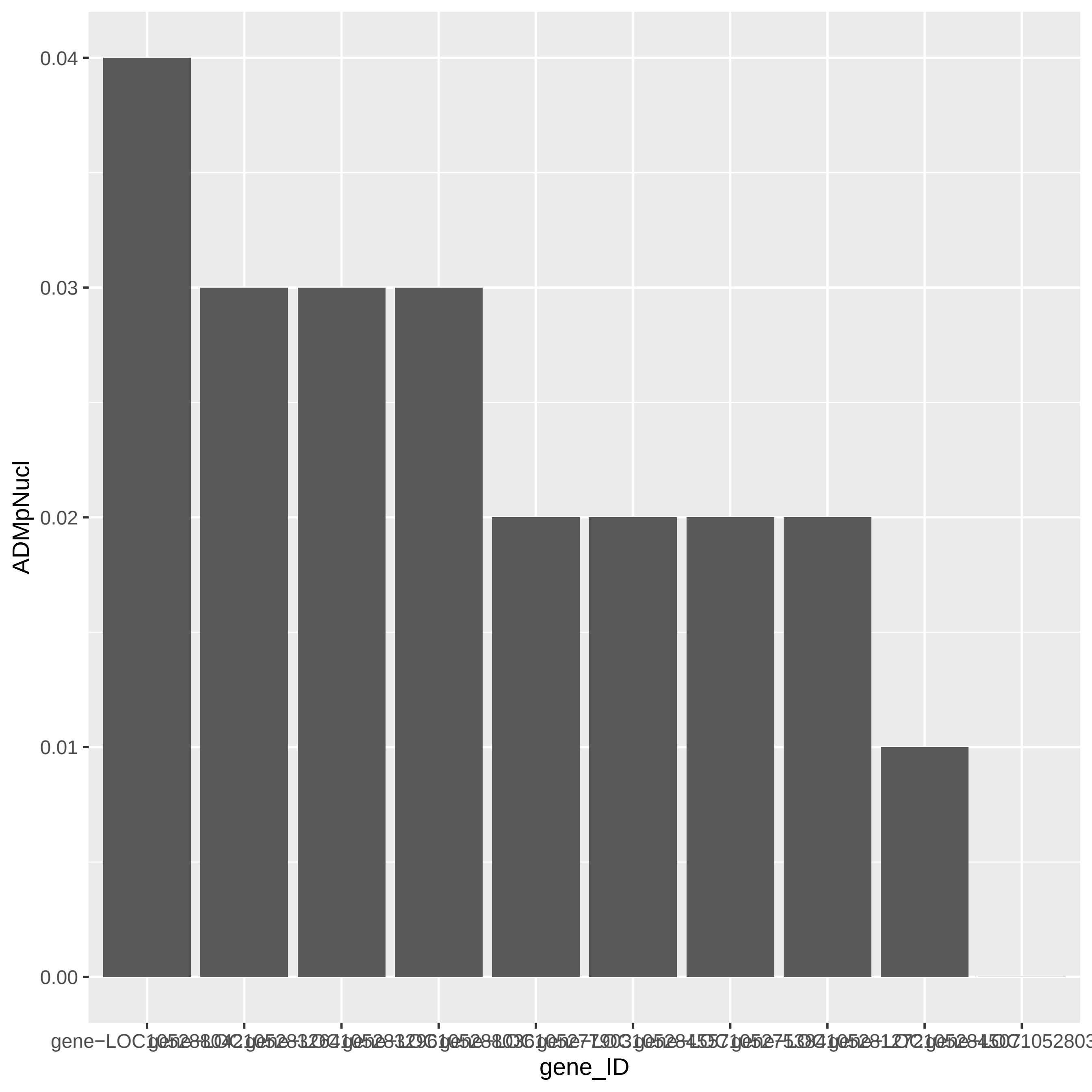

0.04
0.03
0.02
0.01
0.00
ADMpNucl
gene_ID

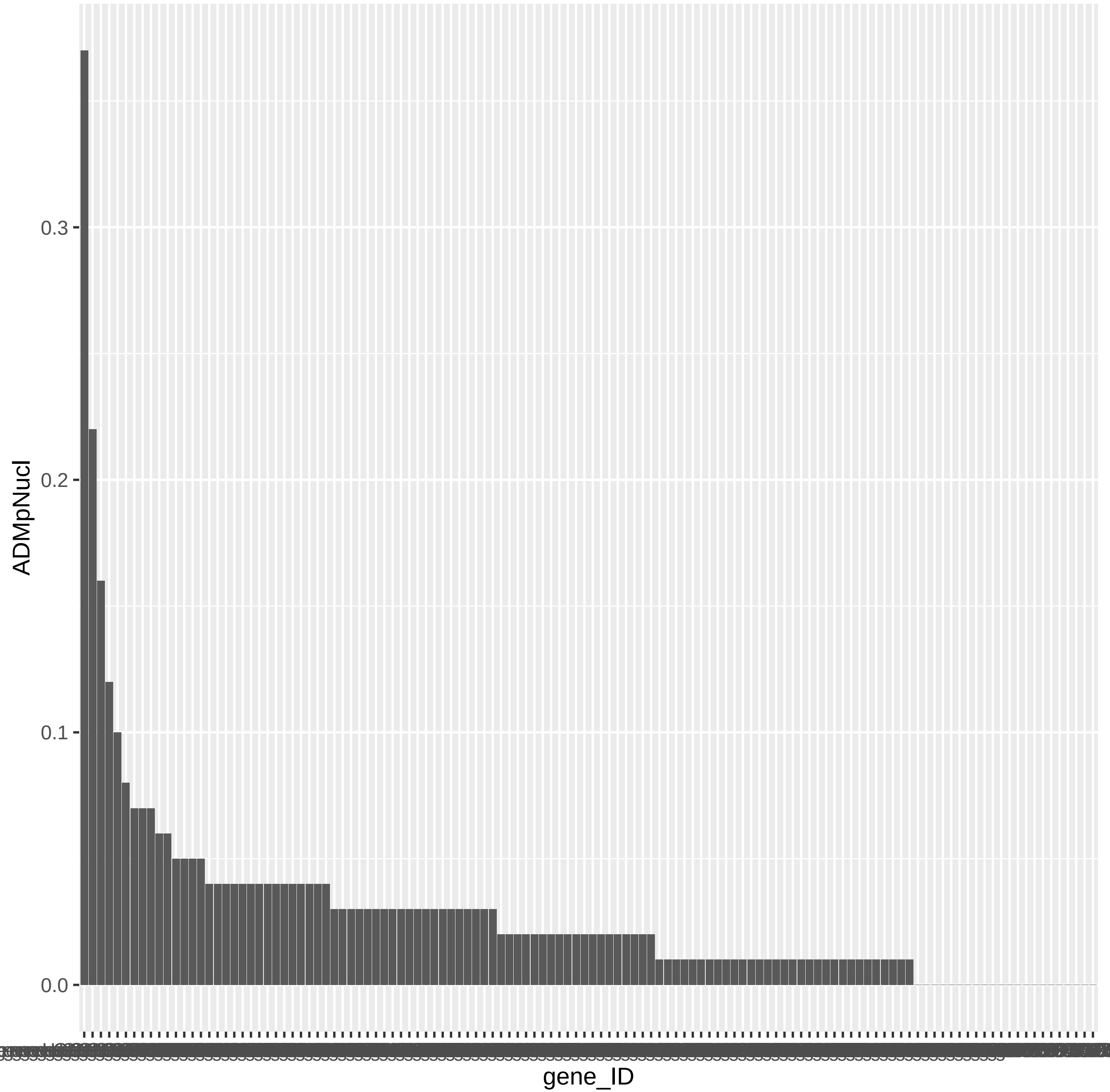

0.3
0.2
0.1
0.0
ADMpNucl
gene_ID

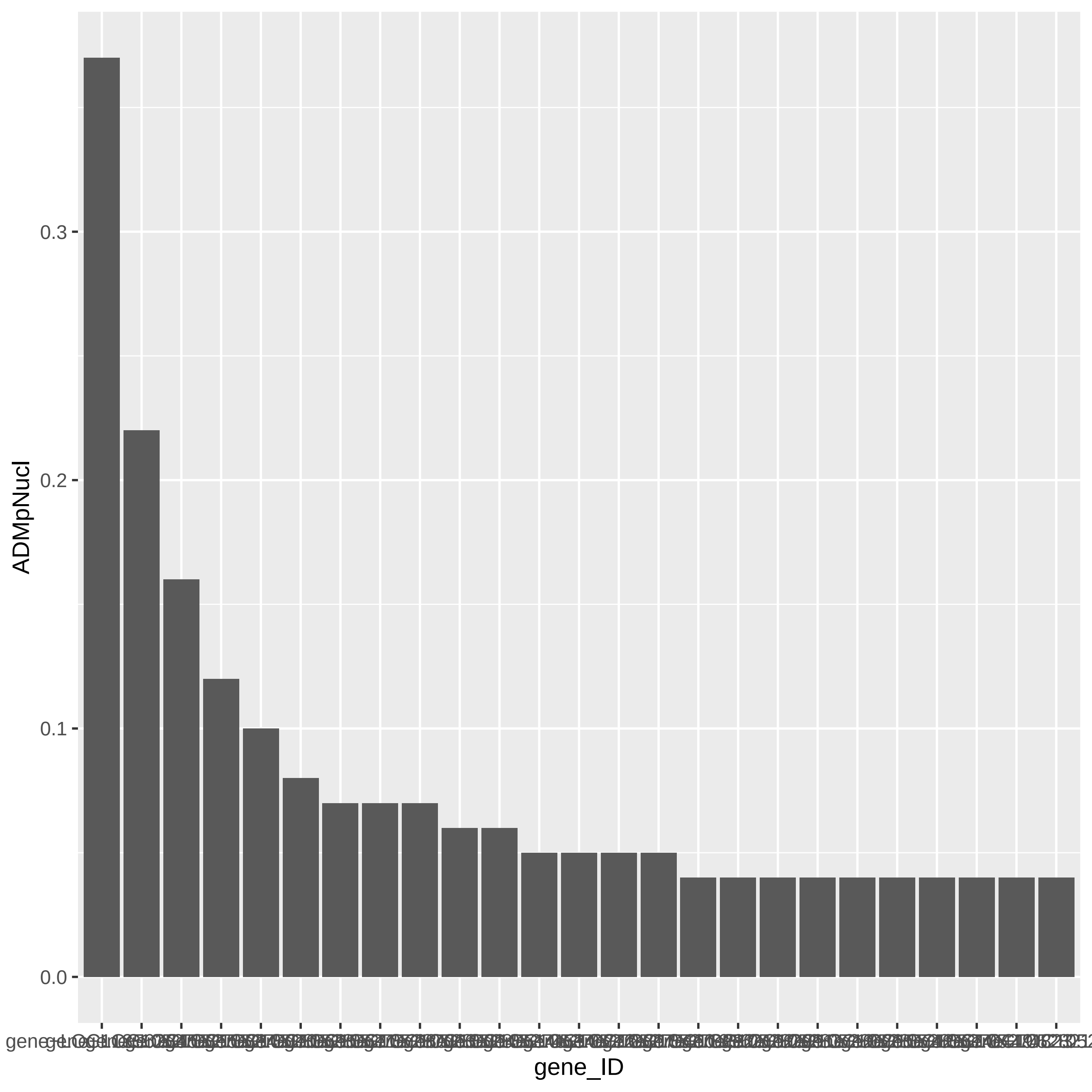

0.3
0.2
0.1
0.0
ADMpNucl
gene_ID

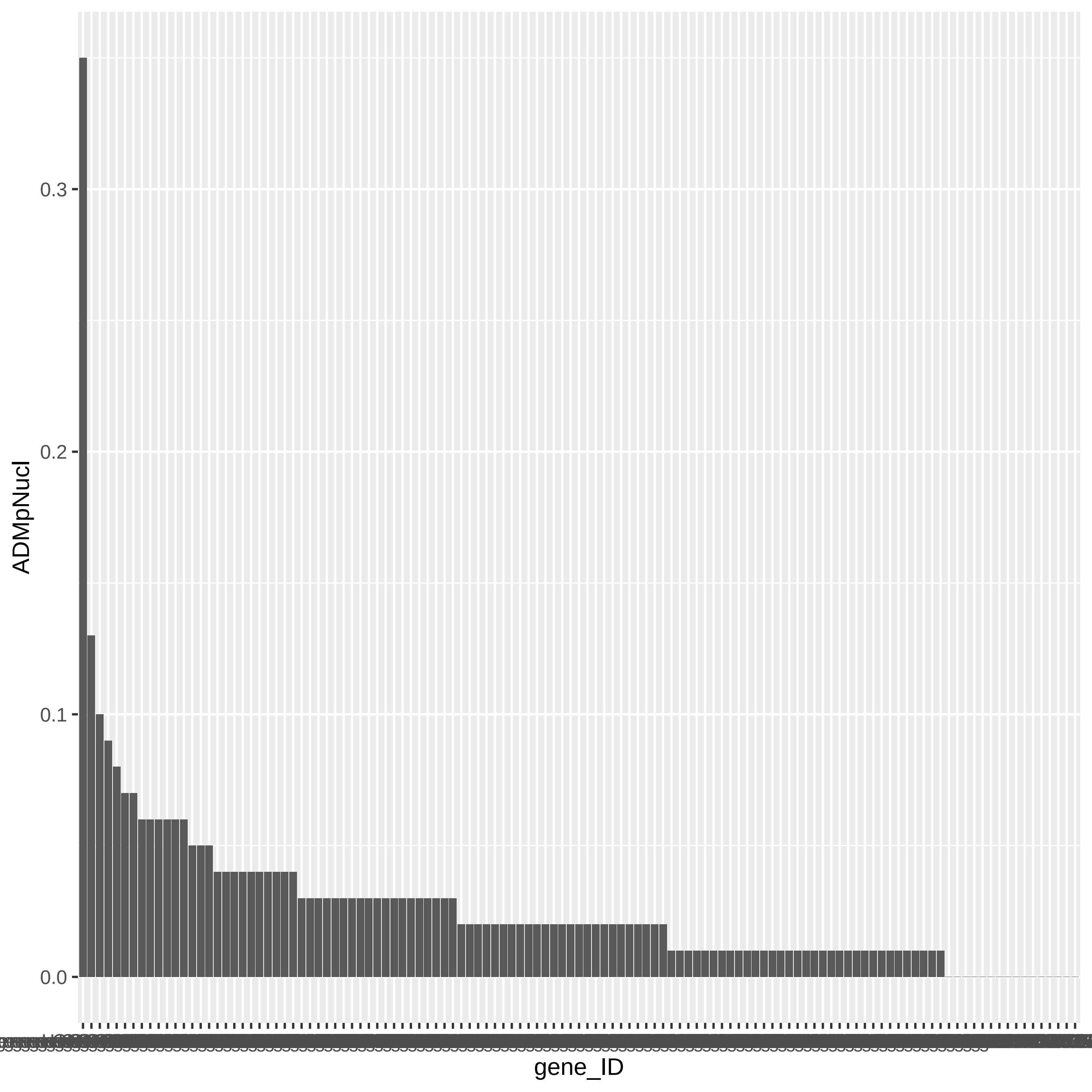

0.3
0.2
0.1
0.0
ADMpNucl
gene_ID

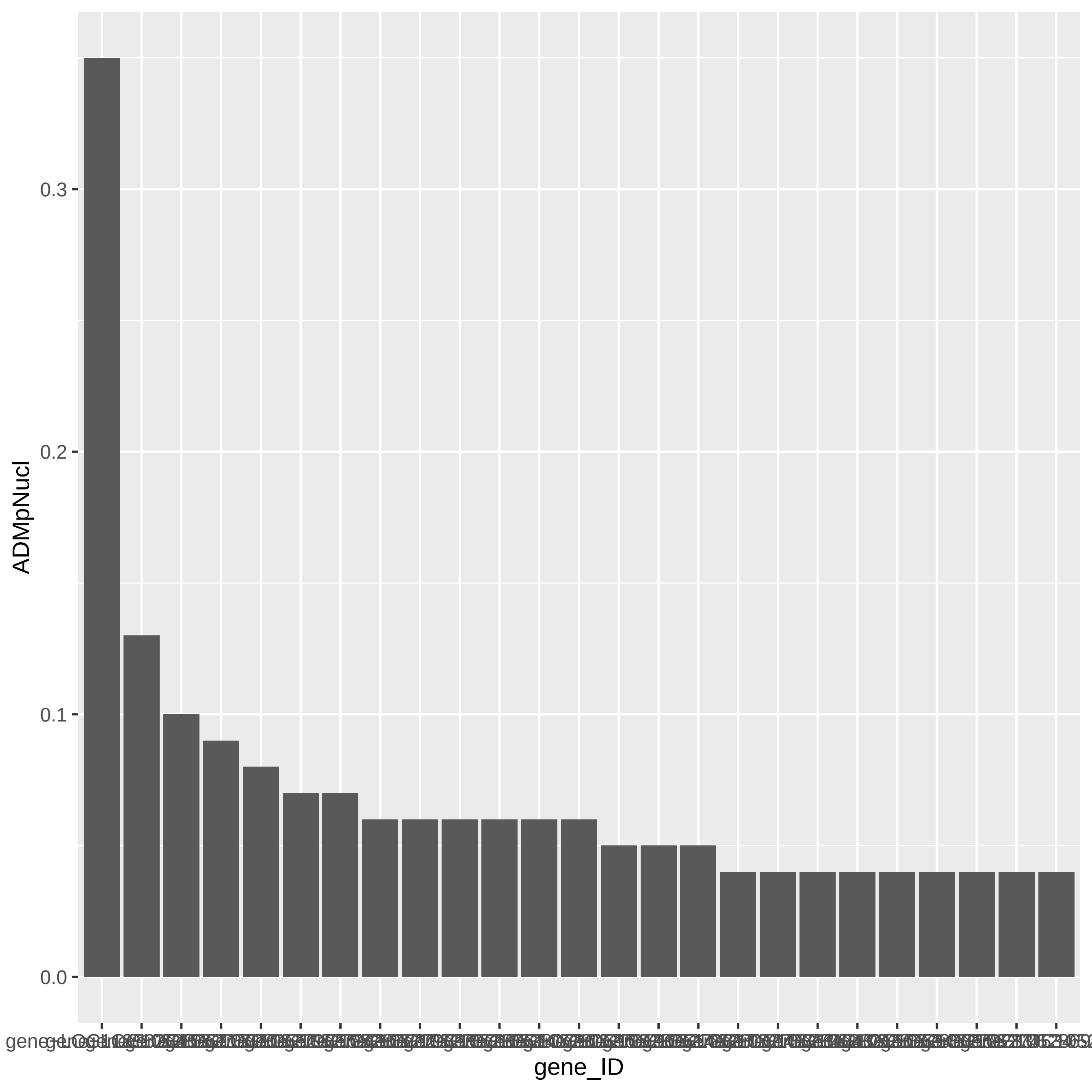

ADMpNucl
0.3
0.2
0.1
0.0
gene_ID

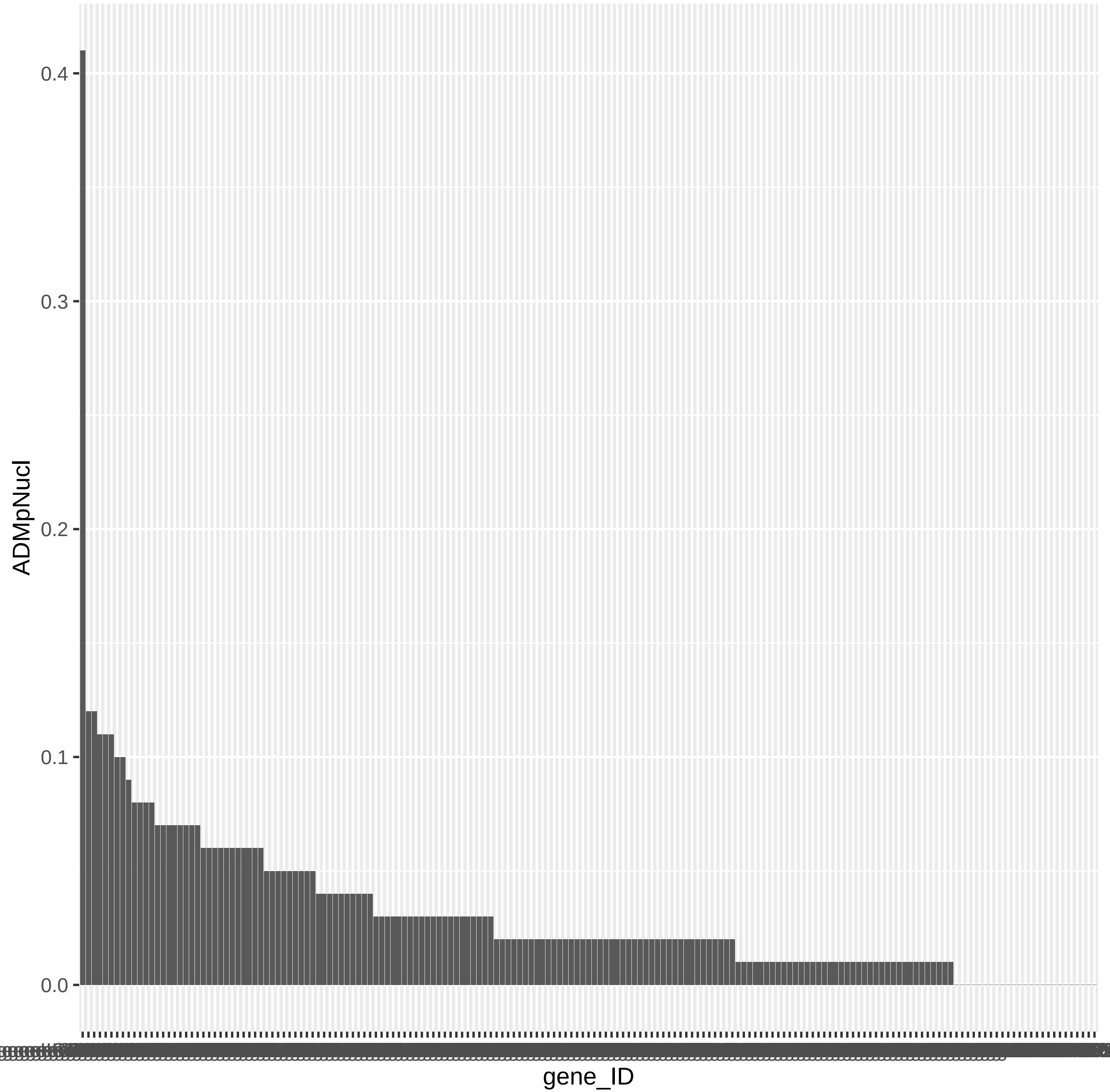

0.4
0.3
0.2
0.1
0.0
ADMpNucl
gene_ID

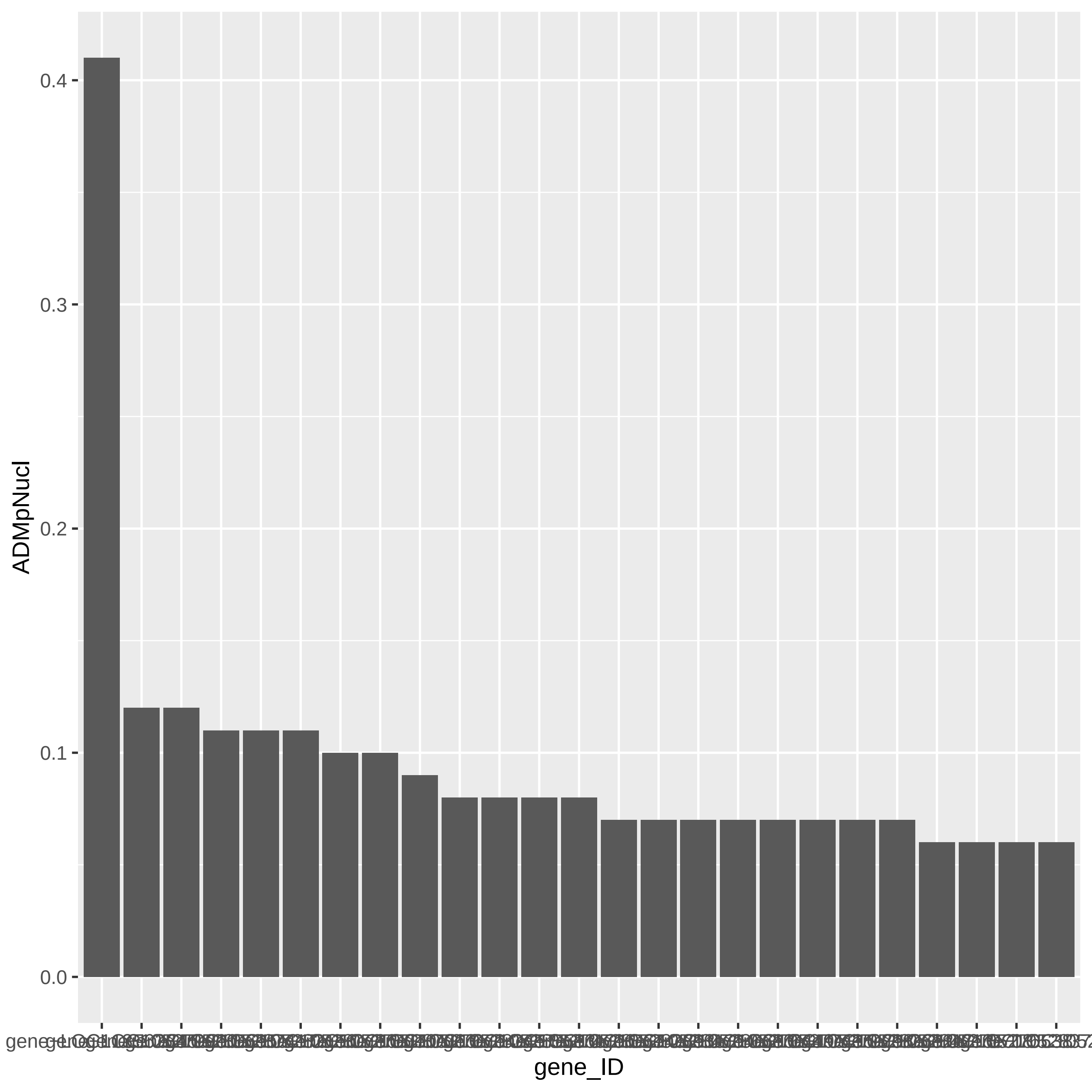
ADMpNucl
0.0
0.1
0.2
0.3
0.4
gene_ID

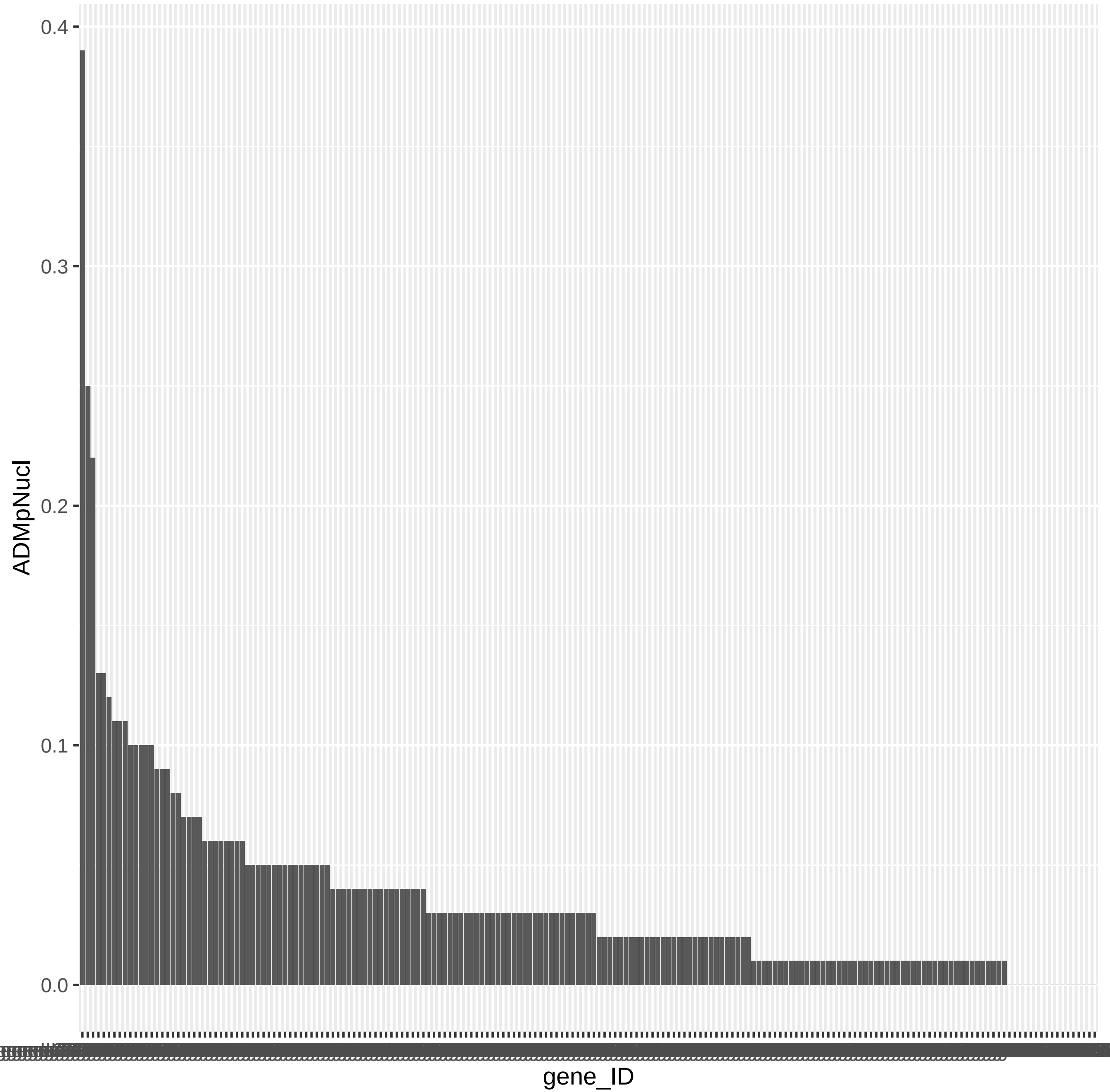

ADMpNucl
0.4
0.3
0.2
0.1
0.0
gene_ID

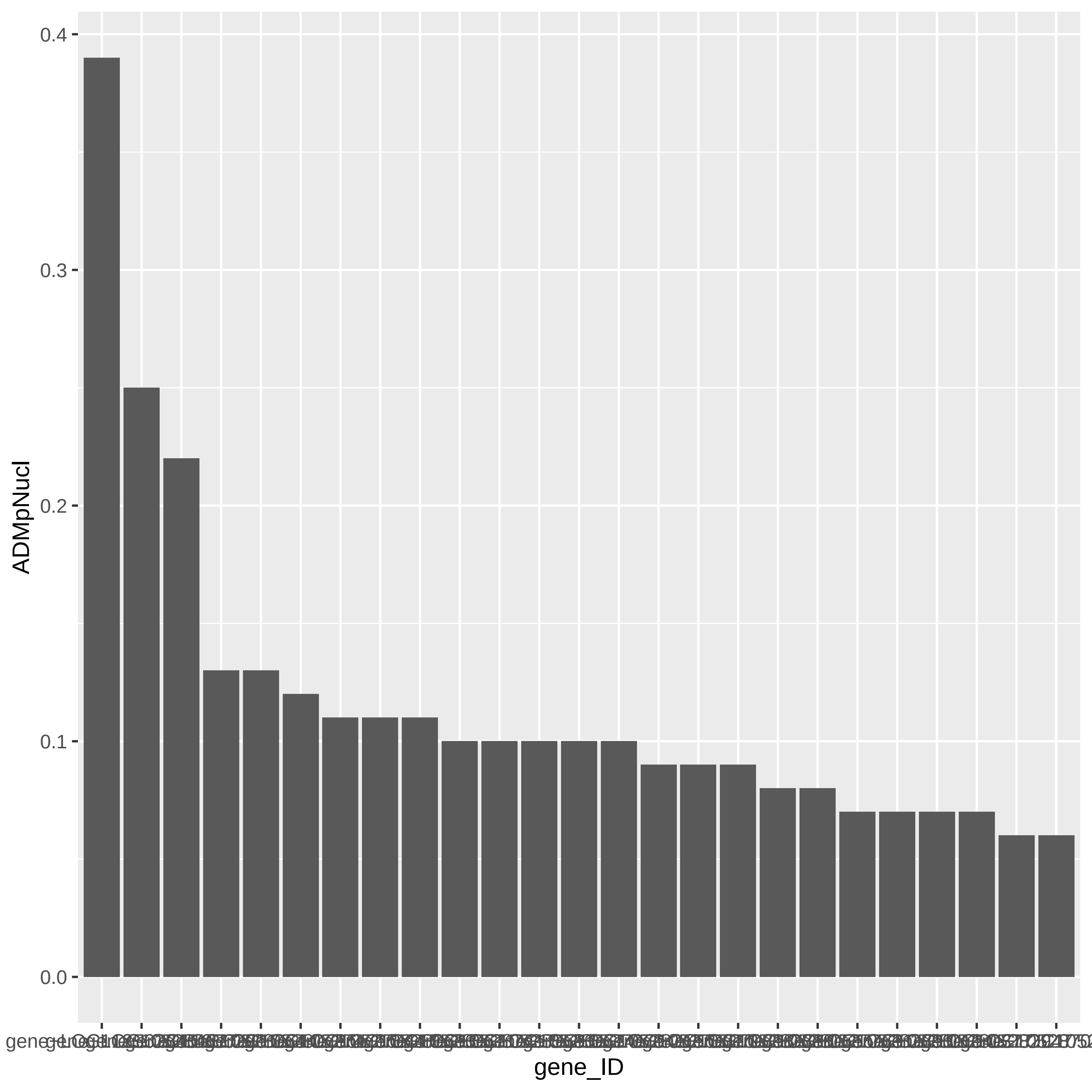

0.4
0.3
0.2
0.1
0.0
ADMpNucl
gene_ID

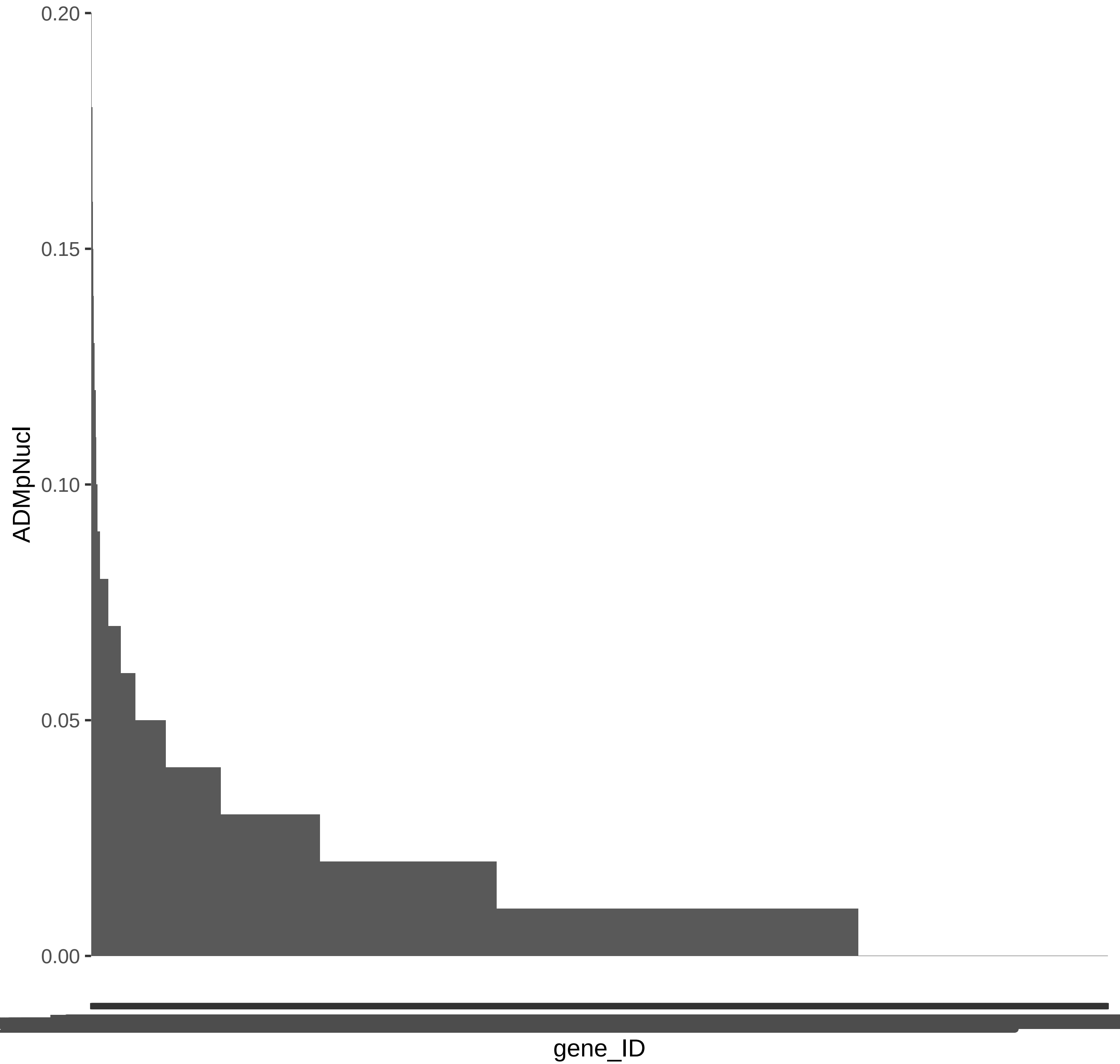

0.20
0.15
0.10
0.05
0.00
ADMpNucl
gene_ID

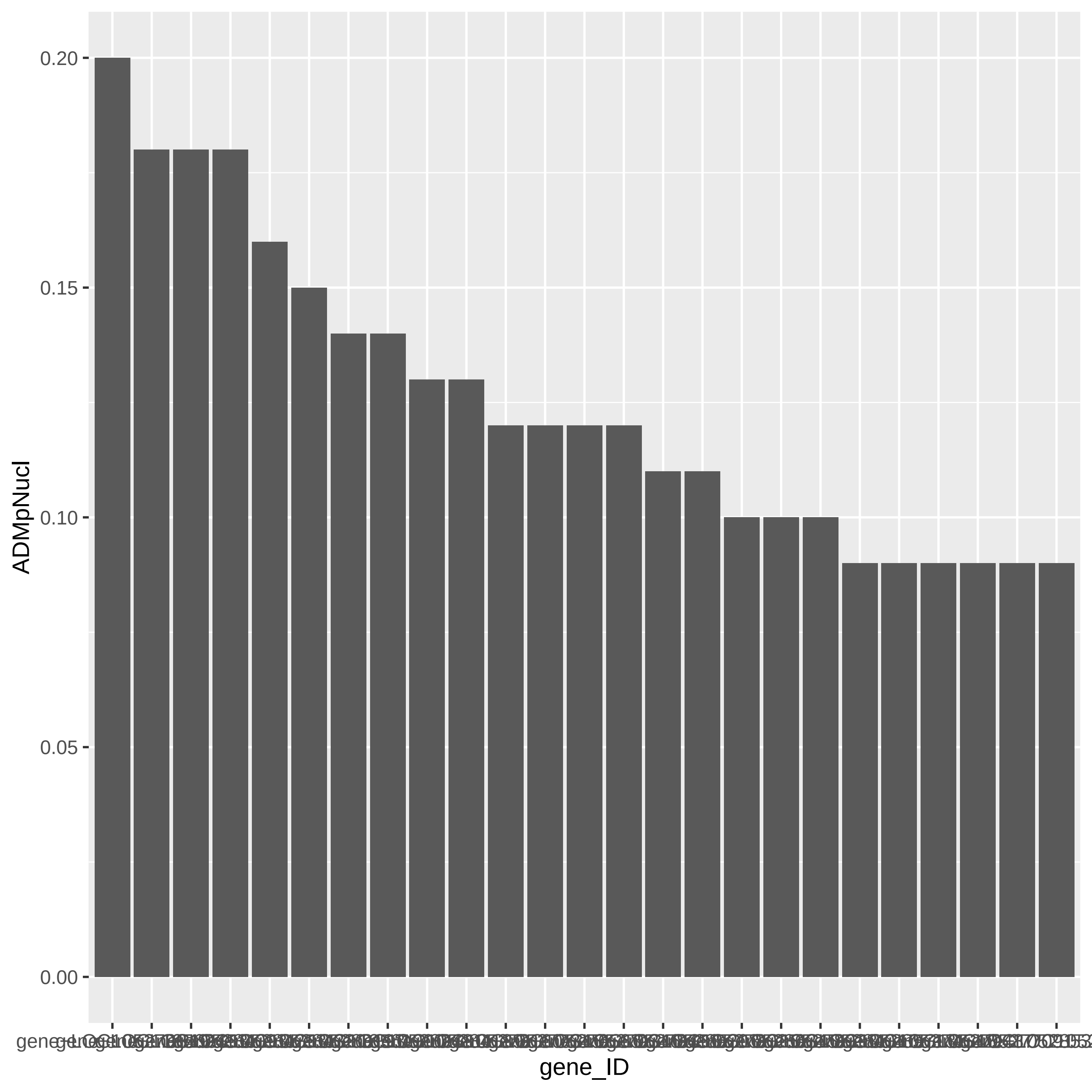

0.20
0.15
0.10
0.05
0.00
ADMpNucl
gene_ID

```
Directory: OGLall File: 0READMEogl

OGL - Ordered gene lists.

Input: studyhc annotation   maxgwidth minnbrdmsites
       dmgprp (= output of BWASPR:show_dmsg())

Output: files ogl-*.txt rnk-dmg-*.txt
         ogl-<study>_<sample>.txt ogl-<study>_<sample1>.vs.<sample2>.txt
         rnk-dmg--<study>_<sample1>.vs.<sample2>.txt rnk-dmg--<study>_<sample1>.vs.<sample2>.pdf
         wrt-<study>.txt

Notes: Output files ogl-<study>_<sample>.txt give tables for each sample with columns

        gene_ID gwidth #Sites #per10Kb %perSite %pNucl

       ordered by %pNucl.  If available, a link to an NCBI entry of gene_ID is inserted as second column.
       Abbreviations used: %perSite, percent methylation per site
                           %perNucl, percent methylation per nucleotide of the gene

       Output files ogl-<study>_<sample1>.vs.<sample2>.txt give tables for each comparison with columns

        gene_ID gwidth #Sites #per10Kb #dmSites #dmsp10kb %dmSites %pSite1 %pSite2 DMpSite ADMpSite DMpNucl ADMpNucl

       ordered by DMpSite.  If available, a link to an NCBI entry of gene_ID is inserted as second column.
       Abbreviations used: %dmSites, percent sites that are differentially methylated
                           %pSite1, average per site % methylation for sample1
                           %pSite2, average per site % methylation for sample2
                           DMpSite, average per site difference in % methylation between sample1 and sample2
                           ADMpSite, absolute value of DMpSite
                           DMpNucl, average per nucleotide difference in % methylation between sample1 and sample2
                           ADMpNucl, absolute value of DMpNucl

       Output files rnk-dmg-<study>_<sample1>.vs.<sample2>.txt are equivalent to files
        ogl-<study>_<sample1>.vs.<sample2>.txt but ordered by ADMpNucl.
       Output files rnk-dmg-<study>_<sample1>.vs.<sample2>.pdf provide visualization of the distribution
        of ADMpNucl values.

       Output file wrt-<study>.txt gives results of the Wilcoxon signed rank test comparing
        the %pSite1 and %pSite2 vectors

       In output directory OGL, tables are restricted to genes with gwidth <= maxgwidth and, for
        pairwise comparisons, #dmSites >= minnbrdmsites

       If either maxgwidth or minnbrdmsites is set, then output directory OGLall will show the full tables
        with all genes (for reference).
```

Directory: OGLall File: ogl-Ob_LI_BCphaseA1.txt

| gene_ID | gene_link | gwidth | #Sites | #per10Kb | %perSite | %pNucl |
|---|---|---|---|---|---|---|
| gene-LOC105283830 | https://www.ncbi.nlm.nih.gov/gene/?term=LOC105283830 | 252 | 23 | 912.7 | 37.25 | 3.4 |
| gene-LOC105280739 | https://www.ncbi.nlm.nih.gov/gene/?term=LOC105280739 | 1991 | 123 | 617.78 | 54.86 | 3.39 |
| gene-LOC105286531 | https://www.ncbi.nlm.nih.gov/gene/?term=LOC105286531 | 372 | 35 | 940.86 | 34.73 | 3.27 |
| gene-LOC105274534 | https://www.ncbi.nlm.nih.gov/gene/?term=LOC105274534 | 1329 | 88 | 662.15 | 45.94 | 3.04 |
| gene-LOC105287811 | https://www.ncbi.nlm.nih.gov/gene/?term=LOC105287811 | 2155 | 123 | 570.77 | 52.68 | 3.01 |
| gene-LOC105285635 | https://www.ncbi.nlm.nih.gov/gene/?term=LOC105285635 | 1175 | 115 | 978.72 | 30.59 | 2.99 |
| gene-LOC105276360 | https://www.ncbi.nlm.nih.gov/gene/?term=LOC105276360 | 1466 | 131 | 893.59 | 33.23 | 2.97 |
| gene-LOC105278844 | https://www.ncbi.nlm.nih.gov/gene/?term=LOC105278844 | 1101 | 111 | 1008.17 | 29.43 | 2.97 |
| gene-LOC105286737 | https://www.ncbi.nlm.nih.gov/gene/?term=LOC105286737 | 2690 | 308 | 1144.98 | 25.86 | 2.96 |
| gene-LOC105278160 | https://www.ncbi.nlm.nih.gov/gene/?term=LOC105278160 | 1872 | 146 | 779.91 | 37.08 | 2.89 |
| gene-LOC105286801 | https://www.ncbi.nlm.nih.gov/gene/?term=LOC105286801 | 1102 | 84 | 762.25 | 37.74 | 2.88 |
| gene-LOC105278849 | https://www.ncbi.nlm.nih.gov/gene/?term=LOC105278849 | 776 | 94 | 1211.34 | 23.72 | 2.87 |
| gene-LOC105274711 | https://www.ncbi.nlm.nih.gov/gene/?term=LOC105274711 | 1655 | 100 | 604.23 | 46.56 | 2.81 |
| gene-LOC105276371 | https://www.ncbi.nlm.nih.gov/gene/?term=LOC105276371 | 871 | 55 | 631.46 | 44.03 | 2.78 |
| gene-LOC105286235 | https://www.ncbi.nlm.nih.gov/gene/?term=LOC105286235 | 1510 | 97 | 642.38 | 43.09 | 2.77 |
| gene-LOC113561532 | https://www.ncbi.nlm.nih.gov/gene/?term=LOC113561532 | 72 | 2 | 277.78 | 97.72 | 2.71 |
| gene-LOC105282149 | https://www.ncbi.nlm.nih.gov/gene/?term=LOC105282149 | 1787 | 116 | 649.13 | 41.07 | 2.67 |
| gene-LOC105287692 | https://www.ncbi.nlm.nih.gov/gene/?term=LOC105287692 | 1505 | 123 | 817.28 | 32.57 | 2.66 |
| gene-LOC105286269 | https://www.ncbi.nlm.nih.gov/gene/?term=LOC105286269 | 1401 | 68 | 485.37 | 54.59 | 2.65 |
| gene-LOC105275314 | https://www.ncbi.nlm.nih.gov/gene/?term=LOC105275314 | 1489 | 78 | 523.84 | 50.45 | 2.64 |
| gene-LOC105277254 | https://www.ncbi.nlm.nih.gov/gene/?term=LOC105277254 | 1377 | 108 | 784.31 | 33.57 | 2.63 |
| gene-LOC105274543 | https://www.ncbi.nlm.nih.gov/gene/?term=LOC105274543 | 1572 | 103 | 655.22 | 38.7 | 2.54 |
| gene-LOC105276439 | https://www.ncbi.nlm.nih.gov/gene/?term=LOC105276439 | 2382 | 141 | 591.94 | 42.03 | 2.49 |
| gene-LOC105282986 | https://www.ncbi.nlm.nih.gov/gene/?term=LOC105282986 | 1600 | 123 | 768.75 | 32.44 | 2.49 |

...

... only first 25 lines shown ...

Directory: OGLall File: ogl-Ob_LI_BCphaseA1.vs.BCphaseA2.txt

| gene_ID | gene_link | gwidth | #Sites | #per10Kb | #dmSites | #dmsp10kb | %dmSites | %pSite1 | %pSite2 | DMpSite | ADMpSite | DMpNucl | ADMpNucl |
|---|---|---|---|---|---|---|---|---|---|---|---|---|---|
| gene-LOC105281173 | https://www.ncbi.nlm.nih.gov/gene/?term=LOC105281173 | 1595 | 25 | 156.74 | 1 | 6.27 | 4 | 64.76 | 61.8 | 2.96 | 2.96 | 0.05 | 0.05 |
| gene-LOC105278358 | https://www.ncbi.nlm.nih.gov/gene/?term=LOC105278358 | 2861 | 28 | 97.87 | 1 | 3.5 | 3.57 | 33.51 | 30.99 | 2.52 | 2.52 | 0.02 | 0.02 |
| gene-LOC109611283 | https://www.ncbi.nlm.nih.gov/gene/?term=LOC109611283 | 1451 | 36 | 248.1 | 1 | 6.89 | 2.78 | 37.85 | 35.62 | 2.23 | 2.23 | 0.06 | 0.06 |
| gene-LOC105286999 | https://www.ncbi.nlm.nih.gov/gene/?term=LOC105286999 | 2141 | 38 | 177.49 | 1 | 4.67 | 2.63 | 33.02 | 30.95 | 2.07 | 2.07 | 0.04 | 0.04 |
| gene-LOC105285202 | https://www.ncbi.nlm.nih.gov/gene/?term=LOC105285202 | 1838 | 33 | 179.54 | 1 | 5.44 | 3.03 | 20.53 | 19.01 | 1.52 | 1.52 | 0.03 | 0.03 |
| gene-LOC105282067 | https://www.ncbi.nlm.nih.gov/gene/?term=LOC105282067 | 5941 | 49 | 82.48 | 1 | 1.68 | 2.04 | 23.12 | 21.71 | 1.41 | 1.41 | 0.01 | 0.01 |
| gene-LOC105274449 | https://www.ncbi.nlm.nih.gov/gene/?term=LOC105274449 | 1549 | 42 | 271.14 | 1 | 6.46 | 2.38 | 55.88 | 54.57 | 1.32 | 1.32 | 0.04 | 0.04 |
| gene-LOC105278877 | https://www.ncbi.nlm.nih.gov/gene/?term=LOC105278877 | 1968 | 69 | 350.61 | 1 | 5.08 | 1.45 | 37.18 | 35.87 | 1.31 | 1.31 | 0.05 | 0.05 |
| gene-LOC105278228 | https://www.ncbi.nlm.nih.gov/gene/?term=LOC105278228 | 8610 | 146 | 169.57 | 1 | 1.16 | 0.68 | 23.97 | 22.69 | 1.27 | 1.27 | 0.02 | 0.02 |
| gene-LOC105285252 | https://www.ncbi.nlm.nih.gov/gene/?term=LOC105285252 | 7427 | 116 | 156.19 | 1 | 1.35 | 0.86 | 15.25 | 14.01 | 1.25 | 1.25 | 0.02 | 0.02 |
| gene-LOC105278282 | https://www.ncbi.nlm.nih.gov/gene/?term=LOC105278282 | 1775 | 43 | 242.25 | 1 | 5.63 | 2.33 | 46.59 | 45.35 | 1.24 | 1.24 | 0.03 | 0.03 |
| gene-LOC105284507 | https://www.ncbi.nlm.nih.gov/gene/?term=LOC105284507 | 5207 | 68 | 130.59 | 1 | 1.92 | 1.47 | 22.93 | 21.85 | 1.08 | 1.08 | 0.01 | 0.01 |
| gene-LOC105275653 | https://www.ncbi.nlm.nih.gov/gene/?term=LOC105275653 | 11251 | 236 | 209.76 | 1 | 0.89 | 0.42 | 9.52 | 8.44 | 1.07 | 1.07 | 0.02 | 0.02 |
| gene-LOC105275848 | https://www.ncbi.nlm.nih.gov/gene/?term=LOC105275848 | 3718 | 119 | 320.06 | 1 | 2.69 | 0.84 | 22.44 | 21.48 | 0.97 | 0.97 | 0.03 | 0.03 |
| gene-LOC105276272 | https://www.ncbi.nlm.nih.gov/gene/?term=LOC105276272 | 1965 | 25 | 127.23 | 1 | 5.09 | 4 | 43.93 | 43.09 | 0.84 | 0.84 | 0.01 | 0.01 |
| gene-LOC105275187 | https://www.ncbi.nlm.nih.gov/gene/?term=LOC105275187 | 5534 | 66 | 119.26 | 1 | 1.81 | 1.52 | 14.05 | 13.27 | 0.78 | 0.78 | 0.01 | 0.01 |
| gene-LOC105277435 | https://www.ncbi.nlm.nih.gov/gene/?term=LOC105277435 | 3643 | 150 | 411.75 | 1 | 2.74 | 0.67 | 12.77 | 12 | 0.77 | 0.77 | 0.03 | 0.03 |
| gene-LOC105282811 | https://www.ncbi.nlm.nih.gov/gene/?term=LOC105282811 | 5765 | 182 | 315.7 | 1 | 1.73 | 0.55 | 9.67 | 8.98 | 0.69 | 0.69 | 0.02 | 0.02 |
| gene-LOC105278632 | https://www.ncbi.nlm.nih.gov/gene/?term=LOC105278632 | 3624 | 119 | 328.37 | 1 | 2.76 | 0.84 | 15.91 | 15.28 | 0.63 | 0.63 | 0.02 | 0.02 |
| gene-LOC105276538 | https://www.ncbi.nlm.nih.gov/gene/?term=LOC105276538 | 1511 | 18 | 119.13 | 1 | 6.62 | 5.56 | 71.16 | 70.54 | 0.62 | 0.62 | 0.01 | 0.01 |
| gene-LOC105279664 | https://www.ncbi.nlm.nih.gov/gene/?term=LOC105279664 | 4023 | 56 | 139.2 | 1 | 2.49 | 1.79 | 21.21 | 20.62 | 0.59 | 0.59 | 0.01 | 0.01 |
| gene-LOC105280956 | https://www.ncbi.nlm.nih.gov/gene/?term=LOC105280956 | 2639 | 73 | 276.62 | 1 | 3.79 | 1.37 | 20.28 | 19.71 | 0.58 | 0.58 | 0.02 | 0.02 |
| gene-LOC105286515 | https://www.ncbi.nlm.nih.gov/gene/?term=LOC105286515 | 3751 | 107 | 285.26 | 1 | 2.67 | 0.93 | 10.47 | 9.92 | 0.55 | 0.55 | 0.02 | 0.02 |
| gene-LOC105276196 | https://www.ncbi.nlm.nih.gov/gene/?term=LOC105276196 | 2953 | 40 | 135.46 | 1 | 3.39 | 2.5 | 52.64 | 52.1 | 0.54 | 0.54 | 0.01 | 0.01 |
| gene-LOC105278454 | https://www.ncbi.nlm.nih.gov/gene/?term=LOC105278454 | 1972 | 36 | 182.56 | 1 | 5.07 | 2.78 | 47.84 | 47.3 | 0.54 | 0.54 | 0.01 | 0.01 |
| gene-LOC105275752 | https://www.ncbi.nlm.nih.gov/gene/?term=LOC105275752 | 3462 | 117 | 337.95 | 1 | 2.89 | 0.85 | 29.78 | 29.38 | 0.4 | 0.4 | 0.01 | 0.01 |
| gene-LOC105274706 | https://www.ncbi.nlm.nih.gov/gene/?term=LOC105274706 | 1446 | 51 | 352.7 | 1 | 6.92 | 1.96 | 37.41 | 37.03 | 0.38 | 0.38 | 0.01 | 0.01 |
| gene-LOC105280786 | https://www.ncbi.nlm.nih.gov/gene/?term=LOC105280786 | 2573 | 107 | 415.86 | 1 | 3.89 | 0.93 | 33.04 | 32.68 | 0.37 | 0.37 | 0.02 | 0.02 |
| gene-LOC105287417 | https://www.ncbi.nlm.nih.gov/gene/?term=LOC105287417 | 1025 | 28 | 273.17 | 1 | 9.76 | 3.57 | 42.43 | 42.07 | 0.36 | 0.36 | 0.01 | 0.01 |
| gene-LOC105276449 | https://www.ncbi.nlm.nih.gov/gene/?term=LOC105276449 | 1231 | 39 | 316.82 | 1 | 8.12 | 2.56 | 32.73 | 32.41 | 0.32 | 0.32 | 0.01 | 0.01 |
| gene-LOC105278534 | https://www.ncbi.nlm.nih.gov/gene/?term=LOC105278534 | 7364 | 166 | 225.42 | 1 | 1.36 | 0.6 | 18.8 | 18.53 | 0.26 | 0.26 | 0.01 | 0.01 |
| gene-LOC105286649 | https://www.ncbi.nlm.nih.gov/gene/?term=LOC105286649 | 14178 | 662 | 466.92 | 1 | 0.71 | 0.15 | 1.27 | 1.01 | 0.26 | 0.26 | 0.01 | 0.01 |
| gene-LOC105284774 | https://www.ncbi.nlm.nih.gov/gene/?term=LOC105284774 | 12089 | 74 | 61.21 | 1 | 0.83 | 1.35 | 19.79 | 19.53 | 0.25 | 0.25 | 0 | 0 |
| gene-LOC105283649 | https://www.ncbi.nlm.nih.gov/gene/?term=LOC105283649 | 1918 | 53 | 276.33 | 2 | 10.43 | 3.77 | 38.29 | 38.07 | 0.23 | 0.23 | 0.01 | 0.01 |
| gene-LOC105285807 | https://www.ncbi.nlm.nih.gov/gene/?term=LOC105285807 | 6063 | 174 | 286.99 | 1 | 1.65 | 0.57 | 7.5 | 7.29 | 0.22 | 0.22 | 0.01 | 0.01 |
| gene-LOC105282794 | https://www.ncbi.nlm.nih.gov/gene/?term=LOC105282794 | 4571 | 91 | 199.08 | 1 | 2.19 | 1.1 | 19.49 | 19.29 | 0.2 | 0.2 | 0 | 0 |
| gene-LOC105286274 | https://www.ncbi.nlm.nih.gov/gene/?term=LOC105286274 | 3817 | 181 | 474.19 | 1 | 2.62 | 0.55 | 15.32 | 15.14 | 0.18 | 0.18 | 0.01 | 0.01 |
| gene-LOC105288078 | https://www.ncbi.nlm.nih.gov/gene/?term=LOC105288078 | 19126 | 301 | 157.38 | 1 | 0.52 | 0.33 | 17.46 | 17.3 | 0.16 | 0.16 | 0 | 0 |
| gene-LOC105277866 | https://www.ncbi.nlm.nih.gov/gene/?term=LOC105277866 | 33045 | 370 | 111.97 | 1 | 0.3 | 0.27 | 3.21 | 3.09 | 0.13 | 0.13 | 0 | 0 |
| gene-LOC105287095 | https://www.ncbi.nlm.nih.gov/gene/?term=LOC105287095 | 1435 | 52 | 362.37 | 1 | 6.97 | 1.92 | 30.47 | 30.36 | 0.12 | 0.12 | 0 | 0 |
| gene-LOC105284116 | https://www.ncbi.nlm.nih.gov/gene/?term=LOC105284116 | 3970 | 85 | 214.11 | 1 | 2.52 | 1.18 | 7.23 | 7.13 | 0.09 | 0.09 | 0 | 0 |
| gene-LOC105279038 | https://www.ncbi.nlm.nih.gov/gene/?term=LOC105279038 | 9453 | 298 | 315.24 | 1 | 1.06 | 0.34 | 9.03 | 8.95 | 0.08 | 0.08 | 0 | 0 |
| gene-LOC105280265 | https://www.ncbi.nlm.nih.gov/gene/?term=LOC105280265 | 85984 | 2570 | 298.89 | 1 | 0.12 | 0.04 | 1.12 | 1.04 | 0.08 | 0.08 | 0 | 0 |
| gene-LOC105281272 | https://www.ncbi.nlm.nih.gov/gene/?term=LOC105281272 | 5252 | 168 | 319.88 | 1 | 1.9 | 0.6 | 16.96 | 16.88 | 0.08 | 0.08 | 0 | 0 |
| gene-LOC105278455 | https://www.ncbi.nlm.nih.gov/gene/?term=LOC105278455 | 27181 | 800 | 294.32 | 1 | 0.37 | 0.12 | 7.96 | 7.89 | 0.07 | 0.07 | 0 | 0 |
| gene-LOC105285742 | https://www.ncbi.nlm.nih.gov/gene/?term=LOC105285742 | 3446 | 91 | 264.07 | 1 | 2.9 | 1.1 | 26.72 | 26.67 | 0.05 | 0.05 | 0 | 0 |
| gene-LOC105276612 | https://www.ncbi.nlm.nih.gov/gene/?term=LOC105276612 | 4157 | 172 | 413.76 | 1 | 2.41 | 0.58 | 11.01 | 10.98 | 0.03 | 0.03 | 0 | 0 |
| gene-LOC105277125 | https://www.ncbi.nlm.nih.gov/gene/?term=LOC105277125 | 51079 | 1589 | 311.09 | 1 | 0.2 | 0.06 | 1.64 | 1.63 | 0 | 0 | 0 | 0 |
| gene-LOC105286194 | https://www.ncbi.nlm.nih.gov/gene/?term=LOC105286194 | 152619 | 1870 | 122.53 | 1 | 0.07 | 0.05 | 6.39 | 6.41 | -0.02 | 0.02 | 0 | 0 |
| gene-LOC105275327 | https://www.ncbi.nlm.nih.gov/gene/?term=LOC105275327 | 14245 | 413 | 289.93 | 1 | 0.7 | 0.24 | 7.72 | 7.75 | -0.03 | 0.03 | 0 | 0 |
| gene-LOC105280304 | https://www.ncbi.nlm.nih.gov/gene/?term=LOC105280304 | 8919 | 90 | 100.91 | 1 | 1.12 | 1.11 | 7.28 | 7.34 | -0.05 | 0.05 | 0 | 0 |
| gene-LOC105274534 | https://www.ncbi.nlm.nih.gov/gene/?term=LOC105274534 | 1329 | 53 | 398.8 | 1 | 7.52 | 1.89 | 47.05 | 47.11 | -0.06 | 0.06 | 0 | 0 |
| gene-LOC105278666 | https://www.ncbi.nlm.nih.gov/gene/?term=LOC105278666 | 10341 | 386 | 373.27 | 1 | 0.97 | 0.26 | 7.83 | 7.9 | -0.07 | 0.07 | 0 | 0 |
| gene-LOC105274531 | https://www.ncbi.nlm.nih.gov/gene/?term=LOC105274531 | 7774 | 252 | 324.16 | 1 | 1.29 | 0.4 | 11.91 | 11.99 | -0.08 | 0.08 | 0 | 0 |
| gene-LOC105276941 | https://www.ncbi.nlm.nih.gov/gene/?term=LOC105276941 | 15956 | 446 | 279.52 | 1 | 0.63 | 0.22 | 5.36 | 5.47 | -0.11 | 0.11 | 0 | 0 |
| gene-LOC105282825 | https://www.ncbi.nlm.nih.gov/gene/?term=LOC105282825 | 3919 | 80 | 204.13 | 1 | 2.55 | 1.25 | 13.05 | 13.16 | -0.12 | 0.12 | 0 | 0 |
| gene-LOC105286644 | https://www.ncbi.nlm.nih.gov/gene/?term=LOC105286644 | 1235 | 22 | 178.14 | 1 | 8.1 | 4.55 | 42.2 | 42.35 | -0.15 | 0.15 | 0 | 0 |
| gene-LOC105283989 | https://www.ncbi.nlm.nih.gov/gene/?term=LOC105283989 | 4948 | 25 | 50.53 | 1 | 2.02 | 4 | 45.66 | 45.85 | -0.19 | 0.19 | 0 | 0 |
| gene-LOC105278145 | https://www.ncbi.nlm.nih.gov/gene/?term=LOC105278145 | 4470 | 244 | 545.86 | 1 | 2.24 | 0.41 | 10.74 | 10.94 | -0.2 | 0.2 | -0.01 | 0.01 |
| gene-LOC105285534 | https://www.ncbi.nlm.nih.gov/gene/?term=LOC105285534 | 3944 | 211 | 534.99 | 1 | 2.54 | 0.47 | 11.95 | 12.16 | -0.21 | 0.21 | -0.01 | 0.01 |

| | | | | | | | | | | | | | |
|---|---|---|---|---|---|---|---|---|---|---|---|---|---|
| gene-LOC105276527 | https://www.ncbi.nlm.nih.gov/gene/?term=LOC105276527 | 7923 | 323 | 407.67 | 1 | 1.26 | 0.31 | 8.74 | 8.99 | -0.25 | 0.25 | -0.01 | 0.01 |
| gene-LOC105284117 | https://www.ncbi.nlm.nih.gov/gene/?term=LOC105284117 | 4340 | 64 | 147.47 | 1 | 2.3 | 1.56 | 10.83 | 11.09 | -0.25 | 0.25 | 0 | 0 |
| gene-LOC105274856 | https://www.ncbi.nlm.nih.gov/gene/?term=LOC105274856 | 5216 | 200 | 383.44 | 1 | 1.92 | 0.5 | 18.38 | 18.67 | -0.29 | 0.29 | -0.01 | 0.01 |
| gene-LOC105286223 | https://www.ncbi.nlm.nih.gov/gene/?term=LOC105286223 | 2225 | 79 | 355.06 | 1 | 4.49 | 1.27 | 30.58 | 30.96 | -0.38 | 0.38 | -0.01 | 0.01 |
| gene-LOC105285395 | https://www.ncbi.nlm.nih.gov/gene/?term=LOC105285395 | 8085 | 165 | 204.08 | 1 | 1.24 | 0.61 | 17.34 | 17.73 | -0.39 | 0.39 | -0.01 | 0.01 |
| gene-LOC105278538 | https://www.ncbi.nlm.nih.gov/gene/?term=LOC105278538 | 2150 | 76 | 353.49 | 1 | 4.65 | 1.32 | 18.23 | 18.64 | -0.41 | 0.41 | -0.01 | 0.01 |
| gene-LOC105283296 | https://www.ncbi.nlm.nih.gov/gene/?term=LOC105283296 | 2174 | 78 | 358.79 | 1 | 4.6 | 1.28 | 21.02 | 21.5 | -0.48 | 0.48 | -0.02 | 0.02 |
| gene-LOC105285406 | https://www.ncbi.nlm.nih.gov/gene/?term=LOC105285406 | 8015 | 39 | 48.66 | 1 | 1.25 | 2.56 | 33.07 | 33.55 | -0.48 | 0.48 | 0 | 0 |
| gene-LOC105275584 | https://www.ncbi.nlm.nih.gov/gene/?term=LOC105275584 | 3070 | 71 | 231.27 | 1 | 3.26 | 1.41 | 26.68 | 27.17 | -0.49 | 0.49 | -0.01 | 0.01 |
| gene-LOC105283943 | https://www.ncbi.nlm.nih.gov/gene/?term=LOC105283943 | 3559 | 48 | 134.87 | 1 | 2.81 | 2.08 | 17.72 | 18.23 | -0.51 | 0.51 | -0.01 | 0.01 |
| gene-LOC105281700 | https://www.ncbi.nlm.nih.gov/gene/?term=LOC105281700 | 2208 | 40 | 181.16 | 1 | 4.53 | 2.5 | 41.3 | 41.85 | -0.55 | 0.55 | -0.01 | 0.01 |
| gene-LOC105284355 | https://www.ncbi.nlm.nih.gov/gene/?term=LOC105284355 | 3904 | 135 | 345.8 | 1 | 2.56 | 0.74 | 22.88 | 23.48 | -0.59 | 0.59 | -0.02 | 0.02 |
| gene-LOC105275179 | https://www.ncbi.nlm.nih.gov/gene/?term=LOC105275179 | 3271 | 66 | 201.77 | 1 | 3.06 | 1.52 | 14.71 | 15.31 | -0.61 | 0.61 | -0.01 | 0.01 |
| gene-LOC105283728 | https://www.ncbi.nlm.nih.gov/gene/?term=LOC105283728 | 3256 | 103 | 316.34 | 2 | 6.14 | 1.94 | 21.65 | 22.28 | -0.62 | 0.62 | -0.02 | 0.02 |
| gene-LOC105286265 | https://www.ncbi.nlm.nih.gov/gene/?term=LOC105286265 | 2507 | 63 | 251.3 | 1 | 3.99 | 1.59 | 30.31 | 30.94 | -0.63 | 0.63 | -0.02 | 0.02 |
| gene-LOC105280410 | https://www.ncbi.nlm.nih.gov/gene/?term=LOC105280410 | 12580 | 235 | 186.8 | 1 | 0.79 | 0.43 | 20.46 | 21.13 | -0.67 | 0.67 | -0.01 | 0.01 |
| gene-LOC105275363 | https://www.ncbi.nlm.nih.gov/gene/?term=LOC105275363 | 3798 | 146 | 384.41 | 1 | 2.63 | 0.68 | 17.72 | 18.47 | -0.76 | 0.76 | -0.03 | 0.03 |
| gene-LOC105274851 | https://www.ncbi.nlm.nih.gov/gene/?term=LOC105274851 | 6616 | 122 | 184.4 | 1 | 1.51 | 0.82 | 10.53 | 11.32 | -0.79 | 0.79 | -0.01 | 0.01 |
| gene-LOC105287063 | https://www.ncbi.nlm.nih.gov/gene/?term=LOC105287063 | 1874 | 46 | 245.46 | 1 | 5.34 | 2.17 | 34.15 | 34.94 | -0.79 | 0.79 | -0.02 | 0.02 |
| gene-LOC105283987 | https://www.ncbi.nlm.nih.gov/gene/?term=LOC105283987 | 4841 | 64 | 132.2 | 1 | 2.07 | 1.56 | 32.11 | 32.92 | -0.81 | 0.81 | -0.01 | 0.01 |
| gene-LOC105279298 | https://www.ncbi.nlm.nih.gov/gene/?term=LOC105279298 | 2773 | 129 | 465.2 | 2 | 7.21 | 1.55 | 22.93 | 23.79 | -0.86 | 0.86 | -0.04 | 0.04 |
| gene-LOC105275460 | https://www.ncbi.nlm.nih.gov/gene/?term=LOC105275460 | 2473 | 107 | 432.67 | 1 | 4.04 | 0.93 | 13.53 | 14.43 | -0.91 | 0.91 | -0.04 | 0.04 |
| gene-LOC105275683 | https://www.ncbi.nlm.nih.gov/gene/?term=LOC105275683 | 4256 | 187 | 439.38 | 1 | 2.35 | 0.53 | 12 | 12.9 | -0.91 | 0.91 | -0.04 | 0.04 |
| gene-LOC105279657 | https://www.ncbi.nlm.nih.gov/gene/?term=LOC105279657 | 5878 | 79 | 134.4 | 1 | 1.7 | 1.27 | 19.77 | 20.7 | -0.93 | 0.93 | -0.01 | 0.01 |
| gene-LOC105275451 | https://www.ncbi.nlm.nih.gov/gene/?term=LOC105275451 | 6035 | 105 | 173.99 | 1 | 1.66 | 0.95 | 24.51 | 25.49 | -0.98 | 0.98 | -0.02 | 0.02 |
| gene-LOC105274590 | https://www.ncbi.nlm.nih.gov/gene/?term=LOC105274590 | 4516 | 61 | 135.08 | 1 | 2.21 | 1.64 | 27.27 | 28.28 | -1.01 | 1.01 | -0.01 | 0.01 |
| gene-LOC105285671 | https://www.ncbi.nlm.nih.gov/gene/?term=LOC105285671 | 2263 | 92 | 406.54 | 1 | 4.42 | 1.09 | 28.77 | 29.78 | -1.01 | 1.01 | -0.04 | 0.04 |
| gene-LOC105277216 | https://www.ncbi.nlm.nih.gov/gene/?term=LOC105277216 | 7627 | 76 | 99.65 | 1 | 1.31 | 1.32 | 7.73 | 8.77 | -1.04 | 1.04 | -0.01 | 0.01 |
| gene-LOC105275173 | https://www.ncbi.nlm.nih.gov/gene/?term=LOC105275173 | 4639 | 71 | 153.05 | 1 | 2.16 | 1.41 | 27.01 | 28.06 | -1.05 | 1.05 | -0.02 | 0.02 |
| gene-LOC105285179 | https://www.ncbi.nlm.nih.gov/gene/?term=LOC105285179 | 4191 | 148 | 353.14 | 1 | 2.39 | 0.68 | 17.81 | 18.87 | -1.05 | 1.05 | -0.04 | 0.04 |
| gene-LOC105281515 | https://www.ncbi.nlm.nih.gov/gene/?term=LOC105281515 | 8827 | 35 | 39.65 | 1 | 1.13 | 2.86 | 31.71 | 33.02 | -1.31 | 1.31 | -0.01 | 0.01 |
| gene-LOC105286452 | https://www.ncbi.nlm.nih.gov/gene/?term=LOC105286452 | 6888 | 55 | 79.85 | 1 | 1.45 | 1.82 | 32.52 | 33.96 | -1.44 | 1.44 | -0.01 | 0.01 |
| gene-LOC105276692 | https://www.ncbi.nlm.nih.gov/gene/?term=LOC105276692 | 1745 | 33 | 189.11 | 1 | 5.73 | 3.03 | 25.26 | 26.77 | -1.52 | 1.52 | -0.03 | 0.03 |
| gene-LOC105285055 | https://www.ncbi.nlm.nih.gov/gene/?term=LOC105285055 | 4108 | 197 | 479.55 | 1 | 2.43 | 0.51 | 13.57 | 15.12 | -1.55 | 1.55 | -0.07 | 0.07 |
| gene-LOC105275190 | https://www.ncbi.nlm.nih.gov/gene/?term=LOC105275190 | 3837 | 69 | 179.83 | 1 | 2.61 | 1.45 | 19.53 | 21.12 | -1.59 | 1.59 | -0.03 | 0.03 |
| gene-LOC105283592 | https://www.ncbi.nlm.nih.gov/gene/?term=LOC105283592 | 3479 | 66 | 189.71 | 1 | 2.87 | 1.52 | 25.49 | 27.08 | -1.59 | 1.59 | -0.03 | 0.03 |
| gene-LOC105279500 | https://www.ncbi.nlm.nih.gov/gene/?term=LOC105279500 | 3587 | 31 | 86.42 | 1 | 2.79 | 3.23 | 51.03 | 52.64 | -1.62 | 1.62 | -0.01 | 0.01 |
| gene-LOC105280774 | https://www.ncbi.nlm.nih.gov/gene/?term=LOC105280774 | 2793 | 97 | 347.3 | 1 | 3.58 | 1.03 | 31.81 | 33.47 | -1.66 | 1.66 | -0.06 | 0.06 |
| gene-LOC113561621 | https://www.ncbi.nlm.nih.gov/gene/?term=LOC113561621 | 2249 | 79 | 351.27 | 1 | 4.45 | 1.27 | 19.72 | 21.41 | -1.69 | 1.69 | -0.06 | 0.06 |
| gene-LOC105280641 | https://www.ncbi.nlm.nih.gov/gene/?term=LOC105280641 | 3682 | 53 | 143.94 | 1 | 2.72 | 1.89 | 17.07 | 18.78 | -1.71 | 1.71 | -0.02 | 0.02 |
| gene-LOC105275518 | https://www.ncbi.nlm.nih.gov/gene/?term=LOC105275518 | 2335 | 18 | 77.09 | 1 | 4.28 | 5.56 | 47.37 | 49.08 | -1.72 | 1.72 | -0.01 | 0.01 |
| gene-LOC105284184 | https://www.ncbi.nlm.nih.gov/gene/?term=LOC105284184 | 2535 | 67 | 264.3 | 1 | 3.94 | 1.49 | 41.64 | 43.45 | -1.81 | 1.81 | -0.05 | 0.05 |
| gene-LOC105278807 | https://www.ncbi.nlm.nih.gov/gene/?term=LOC105278807 | 2166 | 68 | 313.94 | 1 | 4.62 | 1.47 | 17.54 | 19.41 | -1.87 | 1.87 | -0.06 | 0.06 |
| gene-LOC105288066 | https://www.ncbi.nlm.nih.gov/gene/?term=LOC105288066 | 1964 | 16 | 81.47 | 1 | 5.09 | 6.25 | 41.07 | 43.1 | -2.03 | 2.03 | -0.02 | 0.02 |
| gene-LOC105282156 | https://www.ncbi.nlm.nih.gov/gene/?term=LOC105282156 | 2577 | 132 | 512.22 | 1 | 3.88 | 0.76 | 24.4 | 26.49 | -2.09 | 2.09 | -0.11 | 0.11 |
| gene-LOC105275656 | https://www.ncbi.nlm.nih.gov/gene/?term=LOC105275656 | 2803 | 127 | 453.09 | 1 | 3.57 | 0.79 | 26.43 | 28.64 | -2.2 | 2.2 | -0.1 | 0.1 |
| gene-LOC105283712 | https://www.ncbi.nlm.nih.gov/gene/?term=LOC105283712 | 1910 | 24 | 125.65 | 1 | 5.24 | 4.17 | 29.64 | 32.04 | -2.39 | 2.39 | -0.03 | 0.03 |
| gene-LOC105285695 | https://www.ncbi.nlm.nih.gov/gene/?term=LOC105285695 | 1371 | 62 | 452.22 | 1 | 7.29 | 1.61 | 23 | 25.77 | -2.77 | 2.77 | -0.13 | 0.13 |
| gene-LOC105278109 | https://www.ncbi.nlm.nih.gov/gene/?term=LOC105278109 | 4029 | 56 | 138.99 | 1 | 2.48 | 1.79 | 47.35 | 50.32 | -2.97 | 2.97 | -0.04 | 0.04 |
| gene-LOC105274900 | https://www.ncbi.nlm.nih.gov/gene/?term=LOC105274900 | 5561 | 47 | 84.52 | 2 | 3.6 | 4.26 | 31.62 | 34.67 | -3.05 | 3.05 | -0.03 | 0.03 |
| gene-LOC105285674 | https://www.ncbi.nlm.nih.gov/gene/?term=LOC105285674 | 3035 | 88 | 289.95 | 1 | 3.29 | 1.14 | 23.36 | 26.44 | -3.08 | 3.08 | -0.09 | 0.09 |
| gene-LOC105280348 | https://www.ncbi.nlm.nih.gov/gene/?term=LOC105280348 | 3797 | 50 | 131.68 | 1 | 2.63 | 2 | 25.73 | 28.9 | -3.17 | 3.17 | -0.04 | 0.04 |
| gene-LOC105279356 | https://www.ncbi.nlm.nih.gov/gene/?term=LOC105279356 | 1314 | 39 | 296.8 | 1 | 7.61 | 2.56 | 29.77 | 33.04 | -3.27 | 3.27 | -0.1 | 0.1 |
| gene-LOC105280972 | https://www.ncbi.nlm.nih.gov/gene/?term=LOC105280972 | 1532 | 26 | 169.71 | 1 | 6.53 | 3.85 | 20.04 | 23.88 | -3.84 | 3.84 | -0.07 | 0.07 |
| gene-LOC105281848 | https://www.ncbi.nlm.nih.gov/gene/?term=LOC105281848 | 2122 | 44 | 207.35 | 1 | 4.71 | 2.27 | 23.57 | 27.97 | -4.4 | 4.4 | -0.09 | 0.09 |
| gene-LOC105277215 | https://www.ncbi.nlm.nih.gov/gene/?term=LOC105277215 | 2619 | 20 | 76.37 | 1 | 3.82 | 5 | 28.06 | 32.55 | -4.49 | 4.49 | -0.03 | 0.03 |
| gene-LOC105279898 | https://www.ncbi.nlm.nih.gov/gene/?term=LOC105279898 | 1343 | 13 | 96.8 | 1 | 7.45 | 7.69 | 60.41 | 65.83 | -5.42 | 5.42 | -0.05 | 0.05 |
| gene-LOC105285514 | https://www.ncbi.nlm.nih.gov/gene/?term=LOC105285514 | 1277 | 8 | 62.65 | 1 | 7.83 | 12.5 | 82.36 | 88.52 | -6.16 | 6.16 | -0.04 | 0.04 |
| gene-LOC105276078 | https://www.ncbi.nlm.nih.gov/gene/?term=LOC105276078 | 1051 | 8 | 76.12 | 1 | 9.51 | 12.5 | 46.61 | 53.81 | -7.2 | 7.2 | -0.05 | 0.05 |
| gene-LOC105283552 | https://www.ncbi.nlm.nih.gov/gene/?term=LOC105283552 | 1562 | 27 | 172.86 | 1 | 6.4 | 3.7 | 56.95 | 65.52 | -8.57 | 8.57 | -0.15 | 0.15 |
| gene-LOC105285060 | https://www.ncbi.nlm.nih.gov/gene/?term=LOC105285060 | 1285 | 13 | 101.17 | 1 | 7.78 | 7.69 | 55.35 | 64 | -8.64 | 8.64 | -0.09 | 0.09 |
| gene-LOC105275708 | https://www.ncbi.nlm.nih.gov/gene/?term=LOC105275708 | 1772 | 10 | 56.43 | 1 | 5.64 | 10 | 41.45 | 50.31 | -8.85 | 8.85 | -0.05 | 0.05 |
| gene-LOC105275070 | https://www.ncbi.nlm.nih.gov/gene/?term=LOC105275070 | 5667 | 35 | 61.76 | 2 | 3.53 | 5.71 | 3.8 | 15.13 | -11.33 | 11.33 | -0.07 | 0.07 |
| gene-LOC113562345 | https://www.ncbi.nlm.nih.gov/gene/?term=LOC113562345 | 2290 | 62 | 270.74 | 15 | 65.5 | 24.19 | 17.91 | 32.18 | -14.27 | 14.27 | -0.39 | 0.39 |

Directory: OGLall File: ogl-Ob_LI_BCphaseA1.vs.BCphaseB1.txt

| gene_ID | gene_link | gwidth | #Sites | #per10Kb | #dmSites | #dmsp10kb | %dmSites | %pSite1 | %pSite2 | DMpSite | ADMpSite | DMpNucl | ADMpNucl |
|---|---|---|---|---|---|---|---|---|---|---|---|---|---|
| gene-LOC105282885 | https://www.ncbi.nlm.nih.gov/gene/?term=LOC105282885 | 1870 | 41 | 219.25 | 1 | 5.35 | 2.44 | 37.52 | 34.93 | 2.59 | 2.59 | 0.06 | 0.06 |
| gene-LOC105281173 | https://www.ncbi.nlm.nih.gov/gene/?term=LOC105281173 | 1595 | 25 | 156.74 | 1 | 6.27 | 4 | 64.76 | 62.33 | 2.42 | 2.42 | 0.04 | 0.04 |
| gene-LOC105279528 | https://www.ncbi.nlm.nih.gov/gene/?term=LOC105279528 | 5802 | 46 | 79.28 | 1 | 1.72 | 2.17 | 17.18 | 14.84 | 2.33 | 2.33 | 0.02 | 0.02 |
| gene-LOC105285409 | https://www.ncbi.nlm.nih.gov/gene/?term=LOC105285409 | 5099 | 48 | 94.14 | 1 | 1.96 | 2.08 | 3.6 | 1.53 | 2.08 | 2.08 | 0.02 | 0.02 |
| gene-LOC105278454 | https://www.ncbi.nlm.nih.gov/gene/?term=LOC105278454 | 1972 | 37 | 187.63 | 1 | 5.07 | 2.7 | 46.8 | 45.07 | 1.73 | 1.73 | 0.03 | 0.03 |
| gene-LOC105285202 | https://www.ncbi.nlm.nih.gov/gene/?term=LOC105285202 | 1838 | 36 | 195.87 | 1 | 5.44 | 2.78 | 18.94 | 17.23 | 1.72 | 1.72 | 0.03 | 0.03 |
| gene-LOC105276196 | https://www.ncbi.nlm.nih.gov/gene/?term=LOC105276196 | 2953 | 54 | 182.86 | 1 | 3.39 | 1.85 | 40.59 | 39.19 | 1.4 | 1.4 | 0.03 | 0.03 |
| gene-LOC105283123 | https://www.ncbi.nlm.nih.gov/gene/?term=LOC105283123 | 1507 | 41 | 272.06 | 1 | 6.64 | 2.44 | 15.68 | 14.34 | 1.34 | 1.34 | 0.04 | 0.04 |
| gene-LOC105286265 | https://www.ncbi.nlm.nih.gov/gene/?term=LOC105286265 | 2507 | 64 | 255.29 | 1 | 3.99 | 1.56 | 30.65 | 29.46 | 1.19 | 1.19 | 0.03 | 0.03 |
| gene-LOC105288010 | https://www.ncbi.nlm.nih.gov/gene/?term=LOC105288010 | 3189 | 49 | 153.65 | 1 | 3.14 | 2.04 | 31.6 | 30.42 | 1.18 | 1.18 | 0.02 | 0.02 |
| gene-LOC105275653 | https://www.ncbi.nlm.nih.gov/gene/?term=LOC105275653 | 11251 | 262 | 232.87 | 1 | 0.89 | 0.38 | 9.15 | 8.02 | 1.13 | 1.13 | 0.03 | 0.03 |
| gene-LOC105274706 | https://www.ncbi.nlm.nih.gov/gene/?term=LOC105274706 | 1446 | 46 | 318.12 | 1 | 6.92 | 2.17 | 41.37 | 40.53 | 0.85 | 0.85 | 0.03 | 0.03 |
| gene-LOC105278282 | https://www.ncbi.nlm.nih.gov/gene/?term=LOC105278282 | 1775 | 47 | 264.79 | 1 | 5.63 | 2.13 | 43.73 | 42.89 | 0.84 | 0.84 | 0.02 | 0.02 |
| gene-LOC105280447 | https://www.ncbi.nlm.nih.gov/gene/?term=LOC105280447 | 2215 | 25 | 112.87 | 1 | 4.51 | 4 | 26.74 | 25.91 | 0.82 | 0.82 | 0.01 | 0.01 |
| gene-LOC105281047 | https://www.ncbi.nlm.nih.gov/gene/?term=LOC105281047 | 7313 | 117 | 159.99 | 1 | 1.37 | 0.85 | 10.66 | 9.84 | 0.82 | 0.82 | 0.01 | 0.01 |
| gene-LOC105278214 | https://www.ncbi.nlm.nih.gov/gene/?term=LOC105278214 | 3787 | 77 | 203.33 | 1 | 2.64 | 1.3 | 11.46 | 10.7 | 0.76 | 0.76 | 0.02 | 0.02 |
| gene-LOC105285583 | https://www.ncbi.nlm.nih.gov/gene/?term=LOC105285583 | 1947 | 49 | 251.67 | 1 | 5.14 | 2.04 | 29.41 | 28.65 | 0.76 | 0.76 | 0.02 | 0.02 |
| gene-LOC105278436 | https://www.ncbi.nlm.nih.gov/gene/?term=LOC105278436 | 2556 | 57 | 223 | 1 | 3.91 | 1.75 | 36.25 | 35.56 | 0.69 | 0.69 | 0.02 | 0.02 |
| gene-LOC105276429 | https://www.ncbi.nlm.nih.gov/gene/?term=LOC105276429 | 2901 | 112 | 386.07 | 1 | 3.45 | 0.89 | 26.41 | 25.73 | 0.68 | 0.68 | 0.03 | 0.03 |
| gene-LOC105285406 | https://www.ncbi.nlm.nih.gov/gene/?term=LOC105285406 | 8015 | 40 | 49.91 | 1 | 1.25 | 2.5 | 34.18 | 33.53 | 0.64 | 0.64 | 0 | 0 |
| gene-LOC105274449 | https://www.ncbi.nlm.nih.gov/gene/?term=LOC105274449 | 1549 | 43 | 277.6 | 1 | 6.46 | 2.33 | 54.59 | 53.98 | 0.6 | 0.6 | 0.02 | 0.02 |
| gene-LOC105281420 | https://www.ncbi.nlm.nih.gov/gene/?term=LOC105281420 | 3882 | 57 | 146.83 | 1 | 2.58 | 1.75 | 15.28 | 14.73 | 0.55 | 0.55 | 0.01 | 0.01 |
| gene-LOC105282811 | https://www.ncbi.nlm.nih.gov/gene/?term=LOC105282811 | 5765 | 203 | 352.12 | 1 | 1.73 | 0.49 | 9.79 | 9.26 | 0.53 | 0.53 | 0.02 | 0.02 |
| gene-LOC105277435 | https://www.ncbi.nlm.nih.gov/gene/?term=LOC105277435 | 3643 | 155 | 425.47 | 1 | 2.74 | 0.65 | 12.32 | 11.8 | 0.52 | 0.52 | 0.02 | 0.02 |
| gene-LOC105280786 | https://www.ncbi.nlm.nih.gov/gene/?term=LOC105280786 | 2573 | 120 | 466.38 | 1 | 3.89 | 0.83 | 29.63 | 29.12 | 0.52 | 0.52 | 0.02 | 0.02 |
| gene-LOC105284116 | https://www.ncbi.nlm.nih.gov/gene/?term=LOC105284116 | 3970 | 95 | 239.29 | 1 | 2.52 | 1.05 | 6.78 | 6.28 | 0.5 | 0.5 | 0.01 | 0.01 |
| gene-LOC105278877 | https://www.ncbi.nlm.nih.gov/gene/?term=LOC105278877 | 1968 | 77 | 391.26 | 1 | 5.08 | 1.3 | 32.7 | 32.23 | 0.46 | 0.46 | 0.02 | 0.02 |
| gene-LOC105284117 | https://www.ncbi.nlm.nih.gov/gene/?term=LOC105284117 | 4340 | 81 | 186.64 | 1 | 2.3 | 1.23 | 11.74 | 11.3 | 0.44 | 0.44 | 0.01 | 0.01 |
| gene-LOC105282600 | https://www.ncbi.nlm.nih.gov/gene/?term=LOC105282600 | 3715 | 57 | 153.43 | 1 | 2.69 | 1.75 | 29.16 | 28.81 | 0.35 | 0.35 | 0.01 | 0.01 |
| gene-LOC105278455 | https://www.ncbi.nlm.nih.gov/gene/?term=LOC105278455 | 27181 | 854 | 314.19 | 1 | 0.37 | 0.12 | 7.49 | 7.16 | 0.33 | 0.33 | 0.01 | 0.01 |
| gene-LOC105285807 | https://www.ncbi.nlm.nih.gov/gene/?term=LOC105285807 | 6063 | 183 | 301.83 | 1 | 1.65 | 0.55 | 10.09 | 9.8 | 0.29 | 0.29 | 0.01 | 0.01 |
| gene-LOC105286515 | https://www.ncbi.nlm.nih.gov/gene/?term=LOC105286515 | 3751 | 103 | 274.59 | 1 | 2.67 | 0.97 | 10.73 | 10.44 | 0.29 | 0.29 | 0.01 | 0.01 |
| gene-LOC105277866 | https://www.ncbi.nlm.nih.gov/gene/?term=LOC105277866 | 33045 | 442 | 133.76 | 1 | 0.3 | 0.23 | 3.08 | 2.82 | 0.27 | 0.27 | 0 | 0 |
| gene-LOC105279038 | https://www.ncbi.nlm.nih.gov/gene/?term=LOC105279038 | 9453 | 310 | 327.94 | 1 | 1.06 | 0.32 | 8.8 | 8.54 | 0.27 | 0.27 | 0.01 | 0.01 |
| gene-LOC105278666 | https://www.ncbi.nlm.nih.gov/gene/?term=LOC105278666 | 10341 | 405 | 391.64 | 1 | 0.97 | 0.25 | 7.09 | 6.88 | 0.22 | 0.22 | 0.01 | 0.01 |
| gene-LOC105284036 | https://www.ncbi.nlm.nih.gov/gene/?term=LOC105284036 | 3254 | 83 | 255.07 | 1 | 3.07 | 1.2 | 22.41 | 22.19 | 0.22 | 0.22 | 0.01 | 0.01 |
| gene-LOC105276901 | https://www.ncbi.nlm.nih.gov/gene/?term=LOC105276901 | 3254 | 95 | 291.95 | 1 | 3.07 | 1.05 | 23.05 | 22.87 | 0.18 | 0.18 | 0.01 | 0.01 |
| gene-LOC105278145 | https://www.ncbi.nlm.nih.gov/gene/?term=LOC105278145 | 4470 | 249 | 557.05 | 1 | 2.24 | 0.4 | 10.18 | 10.01 | 0.17 | 0.17 | 0.01 | 0.01 |
| gene-LOC105275752 | https://www.ncbi.nlm.nih.gov/gene/?term=LOC105275752 | 3462 | 124 | 358.17 | 1 | 2.89 | 0.81 | 28.83 | 28.67 | 0.16 | 0.16 | 0.01 | 0.01 |
| gene-LOC105279657 | https://www.ncbi.nlm.nih.gov/gene/?term=LOC105279657 | 5878 | 86 | 146.31 | 1 | 1.7 | 1.16 | 18.3 | 18.14 | 0.16 | 0.16 | 0 | 0 |
| gene-LOC105286194 | https://www.ncbi.nlm.nih.gov/gene/?term=LOC105286194 | 152619 | 2028 | 132.88 | 1 | 0.07 | 0.05 | 6.64 | 6.49 | 0.14 | 0.14 | 0 | 0 |
| gene-LOC105287095 | https://www.ncbi.nlm.nih.gov/gene/?term=LOC105287095 | 1435 | 61 | 425.09 | 1 | 6.97 | 1.64 | 26.05 | 25.92 | 0.14 | 0.14 | 0.01 | 0.01 |
| gene-LOC105275173 | https://www.ncbi.nlm.nih.gov/gene/?term=LOC105275173 | 4639 | 83 | 178.92 | 1 | 2.16 | 1.2 | 22.33 | 22.24 | 0.09 | 0.09 | 0 | 0 |
| gene-LOC105276059 | https://www.ncbi.nlm.nih.gov/gene/?term=LOC105276059 | 6256 | 134 | 214.19 | 1 | 1.6 | 0.75 | 10.93 | 10.85 | 0.08 | 0.08 | 0 | 0 |
| gene-LOC105279664 | https://www.ncbi.nlm.nih.gov/gene/?term=LOC105279664 | 4023 | 68 | 169.03 | 1 | 2.49 | 1.47 | 18.22 | 18.15 | 0.08 | 0.08 | 0 | 0 |
| gene-LOC105288161 | https://www.ncbi.nlm.nih.gov/gene/?term=LOC105288161 | 26065 | 287 | 110.11 | 1 | 0.38 | 0.35 | 12.1 | 12.03 | 0.07 | 0.07 | 0 | 0 |
| gene-LOC105278228 | https://www.ncbi.nlm.nih.gov/gene/?term=LOC105278228 | 8610 | 165 | 191.64 | 1 | 1.16 | 0.61 | 22.79 | 22.73 | 0.06 | 0.06 | 0 | 0 |
| gene-LOC105282491 | https://www.ncbi.nlm.nih.gov/gene/?term=LOC105282491 | 4761 | 159 | 333.96 | 1 | 2.1 | 0.63 | 9.46 | 9.41 | 0.05 | 0.05 | 0 | 0 |
| gene-LOC105286274 | https://www.ncbi.nlm.nih.gov/gene/?term=LOC105286274 | 3817 | 177 | 463.71 | 1 | 2.62 | 0.56 | 15.66 | 15.62 | 0.04 | 0.04 | 0 | 0 |
| gene-LOC105275451 | https://www.ncbi.nlm.nih.gov/gene/?term=LOC105275451 | 6035 | 113 | 187.24 | 1 | 1.66 | 0.88 | 22.29 | 22.26 | 0.02 | 0.02 | 0 | 0 |
| gene-LOC105275327 | https://www.ncbi.nlm.nih.gov/gene/?term=LOC105275327 | 14245 | 477 | 334.85 | 1 | 0.7 | 0.21 | 7.15 | 7.14 | 0.01 | 0.01 | 0 | 0 |
| gene-LOC105280265 | https://www.ncbi.nlm.nih.gov/gene/?term=LOC105280265 | 85984 | 2825 | 328.55 | 1 | 0.12 | 0.04 | 1.07 | 1.08 | -0.01 | 0.01 | 0 | 0 |
| gene-LOC105282151 | https://www.ncbi.nlm.nih.gov/gene/?term=LOC105282151 | 1601 | 41 | 256.09 | 1 | 6.25 | 2.44 | 39.95 | 39.99 | -0.04 | 0.04 | 0 | 0 |
| gene-LOC105279778 | https://www.ncbi.nlm.nih.gov/gene/?term=LOC105279778 | 4347 | 139 | 319.76 | 1 | 2.3 | 0.72 | 9.36 | 9.42 | -0.06 | 0.06 | 0 | 0 |
| gene-LOC105274590 | https://www.ncbi.nlm.nih.gov/gene/?term=LOC105274590 | 4516 | 76 | 168.29 | 1 | 2.21 | 1.32 | 22.95 | 23.02 | -0.07 | 0.07 | 0 | 0 |
| gene-LOC105275170 | https://www.ncbi.nlm.nih.gov/gene/?term=LOC105275170 | 12918 | 337 | 260.88 | 1 | 0.77 | 0.3 | 2.06 | 2.13 | -0.07 | 0.07 | 0 | 0 |
| gene-LOC105280956 | https://www.ncbi.nlm.nih.gov/gene/?term=LOC105280956 | 2639 | 73 | 276.62 | 1 | 3.79 | 1.37 | 20.28 | 20.35 | -0.07 | 0.07 | 0 | 0 |
| gene-LOC105274531 | https://www.ncbi.nlm.nih.gov/gene/?term=LOC105274531 | 7774 | 254 | 326.73 | 1 | 1.29 | 0.39 | 11.9 | 12 | -0.1 | 0.1 | 0 | 0 |
| gene-LOC105274856 | https://www.ncbi.nlm.nih.gov/gene/?term=LOC105274856 | 5216 | 201 | 385.35 | 1 | 1.92 | 0.5 | 17.3 | 17.4 | -0.1 | 0.1 | 0 | 0 |
| gene-LOC105279865 | https://www.ncbi.nlm.nih.gov/gene/?term=LOC105279865 | 62373 | 1507 | 241.61 | 1 | 0.16 | 0.07 | 1.62 | 1.72 | -0.1 | 0.1 | 0 | 0 |

| | | | | | | | | | | | | | |
|---|---|---|---|---|---|---|---|---|---|---|---|---|---|
| gene-LOC105276538 | https://www.ncbi.nlm.nih.gov/gene/?term=LOC105276538 | 1511 | 18 | 119.13 | 1 | 6.62 | 5.56 | 71.72 | 71.83 | -0.11 | 0.11 | 0 | 0 |
| gene-LOC105284262 | https://www.ncbi.nlm.nih.gov/gene/?term=LOC105284262 | 6669 | 230 | 344.88 | 1 | 1.5 | 0.43 | 4.51 | 4.68 | -0.17 | 0.17 | -0.01 | 0.01 |
| gene-LOC105279500 | https://www.ncbi.nlm.nih.gov/gene/?term=LOC105279500 | 3587 | 32 | 89.21 | 1 | 2.79 | 3.12 | 49.74 | 49.96 | -0.21 | 0.21 | 0 | 0 |
| gene-LOC105275584 | https://www.ncbi.nlm.nih.gov/gene/?term=LOC105275584 | 3070 | 86 | 280.13 | 1 | 3.26 | 1.16 | 23.79 | 24 | -0.22 | 0.22 | -0.01 | 0.01 |
| gene-LOC105283649 | https://www.ncbi.nlm.nih.gov/gene/?term=LOC105283649 | 1918 | 53 | 276.33 | 1 | 5.21 | 1.89 | 38.29 | 38.55 | -0.25 | 0.25 | -0.01 | 0.01 |
| gene-LOC105280405 | https://www.ncbi.nlm.nih.gov/gene/?term=LOC105280405 | 5422 | 162 | 298.78 | 1 | 1.84 | 0.62 | 9.36 | 9.65 | -0.29 | 0.29 | -0.01 | 0.01 |
| gene-LOC105285395 | https://www.ncbi.nlm.nih.gov/gene/?term=LOC105285395 | 8085 | 180 | 222.63 | 1 | 1.24 | 0.56 | 15.55 | 15.84 | -0.29 | 0.29 | -0.01 | 0.01 |
| gene-LOC105279298 | https://www.ncbi.nlm.nih.gov/gene/?term=LOC105279298 | 2773 | 134 | 483.23 | 1 | 3.61 | 0.75 | 22.72 | 23.03 | -0.3 | 0.3 | -0.01 | 0.01 |
| gene-LOC105282914 | https://www.ncbi.nlm.nih.gov/gene/?term=LOC105282914 | 5623 | 42 | 74.69 | 1 | 1.78 | 2.38 | 48.6 | 48.91 | -0.31 | 0.31 | 0 | 0 |
| gene-LOC105288078 | https://www.ncbi.nlm.nih.gov/gene/?term=LOC105288078 | 19126 | 313 | 163.65 | 1 | 0.52 | 0.32 | 17.33 | 17.64 | -0.31 | 0.31 | -0.01 | 0.01 |
| gene-LOC105284184 | https://www.ncbi.nlm.nih.gov/gene/?term=LOC105284184 | 2535 | 82 | 323.47 | 1 | 3.94 | 1.22 | 34.56 | 34.9 | -0.34 | 0.34 | -0.01 | 0.01 |
| gene-LOC105285534 | https://www.ncbi.nlm.nih.gov/gene/?term=LOC105285534 | 3944 | 221 | 560.34 | 1 | 2.54 | 0.45 | 11.41 | 11.78 | -0.37 | 0.37 | -0.02 | 0.02 |
| gene-LOC105278632 | https://www.ncbi.nlm.nih.gov/gene/?term=LOC105278632 | 3624 | 131 | 361.48 | 1 | 2.76 | 0.76 | 14.71 | 15.12 | -0.41 | 0.41 | -0.01 | 0.01 |
| gene-LOC105282947 | https://www.ncbi.nlm.nih.gov/gene/?term=LOC105282947 | 1162 | 45 | 387.26 | 1 | 8.61 | 2.22 | 32.13 | 32.56 | -0.42 | 0.42 | -0.02 | 0.02 |
| gene-LOC105276692 | https://www.ncbi.nlm.nih.gov/gene/?term=LOC105276692 | 1745 | 36 | 206.3 | 1 | 5.73 | 2.78 | 23.57 | 24.03 | -0.46 | 0.46 | -0.01 | 0.01 |
| gene-LOC105284355 | https://www.ncbi.nlm.nih.gov/gene/?term=LOC105284355 | 3904 | 154 | 394.47 | 1 | 2.56 | 0.65 | 20.37 | 20.87 | -0.5 | 0.5 | -0.02 | 0.02 |
| gene-LOC105278534 | https://www.ncbi.nlm.nih.gov/gene/?term=LOC105278534 | 7364 | 184 | 249.86 | 1 | 1.36 | 0.54 | 17.97 | 18.5 | -0.53 | 0.53 | -0.01 | 0.01 |
| gene-LOC105286770 | https://www.ncbi.nlm.nih.gov/gene/?term=LOC105286770 | 3055 | 42 | 137.48 | 1 | 3.27 | 2.38 | 8.53 | 9.07 | -0.55 | 0.55 | -0.01 | 0.01 |
| gene-LOC105276745 | https://www.ncbi.nlm.nih.gov/gene/?term=LOC105276745 | 3953 | 73 | 184.67 | 1 | 2.53 | 1.37 | 18.74 | 19.3 | -0.56 | 0.56 | -0.01 | 0.01 |
| gene-LOC105280002 | https://www.ncbi.nlm.nih.gov/gene/?term=LOC105280002 | 1940 | 28 | 144.33 | 1 | 5.15 | 3.57 | 33.09 | 33.65 | -0.57 | 0.57 | -0.01 | 0.01 |
| gene-LOC105278962 | https://www.ncbi.nlm.nih.gov/gene/?term=LOC105278962 | 4491 | 88 | 195.95 | 1 | 2.23 | 1.14 | 23.63 | 24.21 | -0.58 | 0.58 | -0.01 | 0.01 |
| gene-LOC105276527 | https://www.ncbi.nlm.nih.gov/gene/?term=LOC105276527 | 7923 | 345 | 435.44 | 1 | 1.26 | 0.29 | 8.28 | 8.94 | -0.66 | 0.66 | -0.03 | 0.03 |
| gene-LOC105277903 | https://www.ncbi.nlm.nih.gov/gene/?term=LOC105277903 | 1835 | 35 | 190.74 | 1 | 5.45 | 2.86 | 51.45 | 52.14 | -0.69 | 0.69 | -0.01 | 0.01 |
| gene-LOC105280641 | https://www.ncbi.nlm.nih.gov/gene/?term=LOC105280641 | 3682 | 62 | 168.39 | 1 | 2.72 | 1.61 | 16.36 | 17.05 | -0.69 | 0.69 | -0.01 | 0.01 |
| gene-LOC105285639 | https://www.ncbi.nlm.nih.gov/gene/?term=LOC105285639 | 10325 | 314 | 304.12 | 1 | 0.97 | 0.32 | 12 | 12.71 | -0.71 | 0.71 | -0.02 | 0.02 |
| gene-LOC105275190 | https://www.ncbi.nlm.nih.gov/gene/?term=LOC105275190 | 3837 | 72 | 187.65 | 1 | 2.61 | 1.39 | 18.78 | 19.54 | -0.76 | 0.76 | -0.01 | 0.01 |
| gene-LOC105285742 | https://www.ncbi.nlm.nih.gov/gene/?term=LOC105285742 | 3446 | 95 | 275.68 | 1 | 2.9 | 1.05 | 25.7 | 26.49 | -0.79 | 0.79 | -0.02 | 0.02 |
| gene-LOC105283728 | https://www.ncbi.nlm.nih.gov/gene/?term=LOC105283728 | 3256 | 111 | 340.91 | 1 | 3.07 | 0.9 | 20.22 | 21.04 | -0.81 | 0.81 | -0.03 | 0.03 |
| gene-LOC105275683 | https://www.ncbi.nlm.nih.gov/gene/?term=LOC105275683 | 4256 | 204 | 479.32 | 1 | 2.35 | 0.49 | 12.14 | 13.06 | -0.92 | 0.92 | -0.04 | 0.04 |
| gene-LOC105288066 | https://www.ncbi.nlm.nih.gov/gene/?term=LOC105288066 | 1964 | 25 | 127.29 | 1 | 5.09 | 4 | 27.34 | 28.29 | -0.95 | 0.95 | -0.01 | 0.01 |
| gene-LOC105283592 | https://www.ncbi.nlm.nih.gov/gene/?term=LOC105283592 | 3479 | 71 | 204.08 | 1 | 2.87 | 1.41 | 24.39 | 25.37 | -0.98 | 0.98 | -0.02 | 0.02 |
| gene-LOC105285671 | https://www.ncbi.nlm.nih.gov/gene/?term=LOC105285671 | 2263 | 95 | 419.8 | 1 | 4.42 | 1.05 | 27.86 | 28.85 | -1 | 1 | -0.04 | 0.04 |
| gene-LOC105277216 | https://www.ncbi.nlm.nih.gov/gene/?term=LOC105277216 | 7627 | 105 | 137.67 | 1 | 1.31 | 0.95 | 5.72 | 6.75 | -1.03 | 1.03 | -0.01 | 0.01 |
| gene-LOC105277631 | https://www.ncbi.nlm.nih.gov/gene/?term=LOC105277631 | 4387 | 285 | 649.65 | 1 | 2.28 | 0.35 | 9.88 | 10.92 | -1.04 | 1.04 | -0.07 | 0.07 |
| gene-LOC105281700 | https://www.ncbi.nlm.nih.gov/gene/?term=LOC105281700 | 2208 | 44 | 199.28 | 1 | 4.53 | 2.27 | 37.54 | 38.62 | -1.08 | 1.08 | -0.02 | 0.02 |
| gene-LOC105285060 | https://www.ncbi.nlm.nih.gov/gene/?term=LOC105285060 | 1285 | 16 | 124.51 | 1 | 7.78 | 6.25 | 50.64 | 51.74 | -1.1 | 1.1 | -0.01 | 0.01 |
| gene-LOC105275363 | https://www.ncbi.nlm.nih.gov/gene/?term=LOC105275363 | 3798 | 155 | 408.11 | 1 | 2.63 | 0.65 | 17.26 | 18.39 | -1.13 | 1.13 | -0.05 | 0.05 |
| gene-LOC105276704 | https://www.ncbi.nlm.nih.gov/gene/?term=LOC105276704 | 2063 | 84 | 407.17 | 2 | 9.69 | 2.38 | 12.84 | 14.02 | -1.18 | 1.18 | -0.05 | 0.05 |
| gene-LOC105275384 | https://www.ncbi.nlm.nih.gov/gene/?term=LOC105275384 | 1987 | 75 | 377.45 | 1 | 5.03 | 1.33 | 33.41 | 34.62 | -1.21 | 1.21 | -0.05 | 0.05 |
| gene-LOC105275460 | https://www.ncbi.nlm.nih.gov/gene/?term=LOC105275460 | 2473 | 116 | 469.07 | 1 | 4.04 | 0.86 | 12.57 | 13.81 | -1.24 | 1.24 | -0.06 | 0.06 |
| gene-LOC105285055 | https://www.ncbi.nlm.nih.gov/gene/?term=LOC105285055 | 4108 | 214 | 520.93 | 1 | 2.43 | 0.47 | 12.94 | 14.18 | -1.24 | 1.24 | -0.06 | 0.06 |
| gene-LOC105287417 | https://www.ncbi.nlm.nih.gov/gene/?term=LOC105287417 | 1025 | 33 | 321.95 | 1 | 9.76 | 3.03 | 36 | 37.27 | -1.27 | 1.27 | -0.04 | 0.04 |
| gene-LOC105282156 | https://www.ncbi.nlm.nih.gov/gene/?term=LOC105282156 | 2577 | 133 | 516.1 | 1 | 3.88 | 0.75 | 23.61 | 24.92 | -1.31 | 1.31 | -0.07 | 0.07 |
| gene-LOC105288036 | https://www.ncbi.nlm.nih.gov/gene/?term=LOC105288036 | 5110 | 198 | 387.48 | 1 | 1.96 | 0.51 | 16.95 | 18.26 | -1.31 | 1.31 | -0.05 | 0.05 |
| gene-LOC105283712 | https://www.ncbi.nlm.nih.gov/gene/?term=LOC105283712 | 1910 | 29 | 151.83 | 1 | 5.24 | 3.45 | 29.98 | 31.34 | -1.36 | 1.36 | -0.02 | 0.02 |
| gene-LOC105278260 | https://www.ncbi.nlm.nih.gov/gene/?term=LOC105278260 | 3240 | 42 | 129.63 | 1 | 3.09 | 2.38 | 12.8 | 14.28 | -1.47 | 1.47 | -0.02 | 0.02 |
| gene-LOC105285695 | https://www.ncbi.nlm.nih.gov/gene/?term=LOC105285695 | 1371 | 62 | 452.22 | 1 | 7.29 | 1.61 | 23 | 24.51 | -1.51 | 1.51 | -0.07 | 0.07 |
| gene-LOC105281515 | https://www.ncbi.nlm.nih.gov/gene/?term=LOC105281515 | 8827 | 39 | 44.18 | 1 | 1.13 | 2.56 | 34.44 | 35.97 | -1.53 | 1.53 | -0.01 | 0.01 |
| gene-LOC105278807 | https://www.ncbi.nlm.nih.gov/gene/?term=LOC105278807 | 2166 | 71 | 327.79 | 1 | 4.62 | 1.41 | 18.52 | 20.08 | -1.56 | 1.56 | -0.05 | 0.05 |
| gene-LOC105283943 | https://www.ncbi.nlm.nih.gov/gene/?term=LOC105283943 | 3559 | 59 | 165.78 | 1 | 2.81 | 1.69 | 20.25 | 21.81 | -1.56 | 1.56 | -0.03 | 0.03 |
| gene-LOC105286452 | https://www.ncbi.nlm.nih.gov/gene/?term=LOC105286452 | 6888 | 58 | 84.2 | 1 | 1.45 | 1.72 | 31.25 | 32.98 | -1.73 | 1.73 | -0.01 | 0.01 |
| gene-LOC105285674 | https://www.ncbi.nlm.nih.gov/gene/?term=LOC105285674 | 3035 | 91 | 299.84 | 2 | 6.59 | 2.2 | 23.1 | 24.91 | -1.81 | 1.81 | -0.05 | 0.05 |
| gene-LOC105282399 | https://www.ncbi.nlm.nih.gov/gene/?term=LOC105282399 | 2682 | 21 | 78.3 | 1 | 3.73 | 4.76 | 34.08 | 35.97 | -1.89 | 1.89 | -0.01 | 0.01 |
| gene-LOC105279356 | https://www.ncbi.nlm.nih.gov/gene/?term=LOC105279356 | 1314 | 42 | 319.63 | 1 | 7.61 | 2.38 | 28.83 | 30.95 | -2.12 | 2.12 | -0.07 | 0.07 |
| gene-LOC105280348 | https://www.ncbi.nlm.nih.gov/gene/?term=LOC105280348 | 3797 | 51 | 134.32 | 1 | 2.63 | 1.96 | 25.95 | 28.29 | -2.34 | 2.34 | -0.03 | 0.03 |
| gene-LOC105274900 | https://www.ncbi.nlm.nih.gov/gene/?term=LOC105274900 | 5561 | 56 | 100.7 | 2 | 3.6 | 3.57 | 26.85 | 29.47 | -2.62 | 2.62 | -0.03 | 0.03 |
| gene-LOC105283989 | https://www.ncbi.nlm.nih.gov/gene/?term=LOC105283989 | 4948 | 33 | 66.69 | 2 | 4.04 | 6.06 | 45.77 | 48.89 | -3.12 | 3.12 | -0.02 | 0.02 |
| gene-LOC105281848 | https://www.ncbi.nlm.nih.gov/gene/?term=LOC105281848 | 2122 | 47 | 221.49 | 1 | 4.71 | 2.13 | 22.74 | 25.94 | -3.2 | 3.2 | -0.07 | 0.07 |
| gene-LOC105280419 | https://www.ncbi.nlm.nih.gov/gene/?term=LOC105280419 | 1601 | 10 | 62.46 | 1 | 6.25 | 10 | 33.98 | 37.47 | -3.49 | 3.49 | -0.02 | 0.02 |
| gene-LOC105275518 | https://www.ncbi.nlm.nih.gov/gene/?term=LOC105275518 | 2335 | 20 | 85.65 | 1 | 4.28 | 5 | 47.29 | 50.84 | -3.55 | 3.55 | -0.03 | 0.03 |
| gene-LOC105277215 | https://www.ncbi.nlm.nih.gov/gene/?term=LOC105277215 | 2619 | 28 | 106.91 | 1 | 3.82 | 3.57 | 20.04 | 24.53 | -4.48 | 4.48 | -0.05 | 0.05 |
| gene-LOC105283552 | https://www.ncbi.nlm.nih.gov/gene/?term=LOC105283552 | 1562 | 33 | 211.27 | 1 | 6.4 | 3.03 | 54.5 | 59.97 | -5.47 | 5.47 | -0.12 | 0.12 |
| gene-LOC105279898 | https://www.ncbi.nlm.nih.gov/gene/?term=LOC105279898 | 1343 | 16 | 119.14 | 1 | 7.45 | 6.25 | 59.42 | 65.83 | -6.42 | 6.42 | -0.08 | 0.08 |
| gene-LOC105275708 | https://www.ncbi.nlm.nih.gov/gene/?term=LOC105275708 | 1772 | 13 | 73.36 | 1 | 5.64 | 7.69 | 46.34 | 53.52 | -7.18 | 7.18 | -0.05 | 0.05 |

| | | | | | | | | | | | | | | | |
|---|---|---|---|---|---|---|---|---|---|---|---|---|---|---|---|
| gene-LOC105285514 | https://www.ncbi.nlm.nih.gov/gene/?term=LOC105285514 | 1277 | 9 | 70.48 | 1 | 7.83 | 11.11 | 73.21 | 80.81 | -7.6 | 7.6 | -0.05 | 0.05 | | |
| gene-LOC105275070 | https://www.ncbi.nlm.nih.gov/gene/?term=LOC105275070 | 5667 | 36 | 63.53 | 2 | 3.53 | 5.56 | 3.69 | 13.97 | -10.28 | | 10.28 | -0.07 | 0.07 | |
| gene-LOC113562345 | https://www.ncbi.nlm.nih.gov/gene/?term=LOC113562345 | 2290 | 62 | 270.74 | | 14 | 61.14 | 22.58 | 17.91 | 32.35 | -14.44 | | 14.44 | -0.39 | 0.39 |

Directory: OGLall File: ogl-Ob_LI_BCphaseA1.vs.BCphaseB2.txt

| gene_ID | gene_link | gwidth | #Sites | #per10Kb | #dmSites | #dmsp10kb | %dmSites | %pSite1 | %pSite2 | DMpSite | ADMpSite | DMpNucl | ADMpNucl |
|---|---|---|---|---|---|---|---|---|---|---|---|---|---|
| gene-LOC105275343 | https://www.ncbi.nlm.nih.gov/gene/?term=LOC105275343 | 1746 | 4 | 22.91 | 2 | 11.45 | 50 | 47 | 32.01 | 14.99 | 14.99 | 0.03 | 0.03 |
| gene-LOC105280261 | https://www.ncbi.nlm.nih.gov/gene/?term=LOC105280261 | 1005 | 4 | 39.8 | 1 | 9.95 | 25 | 82.16 | 67.59 | 14.57 | 14.57 | 0.06 | 0.06 |
| gene-LOC113562975 | https://www.ncbi.nlm.nih.gov/gene/?term=LOC113562975 | 2751 | 3 | 10.91 | 1 | 3.64 | 33.33 | 46.98 | 32.86 | 14.12 | 14.12 | 0.02 | 0.02 |
| gene-LOC105278458 | https://www.ncbi.nlm.nih.gov/gene/?term=LOC105278458 | 216 | 3 | 138.89 | 1 | 46.3 | 33.33 | 20.87 | 10.63 | 10.24 | 10.24 | 0.14 | 0.14 |
| gene-LOC105281691 | https://www.ncbi.nlm.nih.gov/gene/?term=LOC105281691 | 2096 | 9 | 42.94 | 2 | 9.54 | 22.22 | 17 | 7.56 | 9.44 | 9.44 | 0.04 | 0.04 |
| gene-LOC105280926 | https://www.ncbi.nlm.nih.gov/gene/?term=LOC105280926 | 2139 | 6 | 28.05 | 2 | 9.35 | 33.33 | 47.28 | 37.87 | 9.41 | 9.41 | 0.03 | 0.03 |
| gene-LOC105287814 | https://www.ncbi.nlm.nih.gov/gene/?term=LOC105287814 | 892 | 7 | 78.48 | 1 | 11.21 | 14.29 | 37.51 | 29.17 | 8.34 | 8.34 | 0.07 | 0.07 |
| gene-LOC105283830 | https://www.ncbi.nlm.nih.gov/gene/?term=LOC105283830 | 252 | 13 | 515.87 | 1 | 39.68 | 7.69 | 42.02 | 33.83 | 8.19 | 8.19 | 0.42 | 0.42 |
| gene-LOC105285885 | https://www.ncbi.nlm.nih.gov/gene/?term=LOC105285885 | 1327 | 12 | 90.43 | 3 | 22.61 | 25 | 58.28 | 50.25 | 8.03 | 8.03 | 0.07 | 0.07 |
| gene-LOC105283963 | https://www.ncbi.nlm.nih.gov/gene/?term=LOC105283963 | 1330 | 8 | 60.15 | 1 | 7.52 | 12.5 | 51.82 | 43.92 | 7.9 | 7.9 | 0.05 | 0.05 |
| gene-LOC105275978 | https://www.ncbi.nlm.nih.gov/gene/?term=LOC105275978 | 3932 | 6 | 15.26 | 1 | 2.54 | 16.67 | 16.55 | 9.1 | 7.45 | 7.45 | 0.01 | 0.01 |
| gene-LOC105281507 | https://www.ncbi.nlm.nih.gov/gene/?term=LOC105281507 | 881 | 5 | 56.75 | 1 | 11.35 | 20 | 11.82 | 4.8 | 7.02 | 7.02 | 0.04 | 0.04 |
| gene-LOC105282405 | https://www.ncbi.nlm.nih.gov/gene/?term=LOC105282405 | 1310 | 31 | 236.64 | 3 | 22.9 | 9.68 | 51.18 | 44.58 | 6.6 | 6.6 | 0.16 | 0.16 |
| gene-LOC105280917 | https://www.ncbi.nlm.nih.gov/gene/?term=LOC105280917 | 2193 | 6 | 27.36 | 1 | 4.56 | 16.67 | 14.86 | 8.55 | 6.31 | 6.31 | 0.02 | 0.02 |
| gene-LOC105276706 | https://www.ncbi.nlm.nih.gov/gene/?term=LOC105276706 | 1440 | 16 | 111.11 | 1 | 6.94 | 6.25 | 82.64 | 76.36 | 6.28 | 6.28 | 0.07 | 0.07 |
| gene-LOC105281831 | https://www.ncbi.nlm.nih.gov/gene/?term=LOC105281831 | 1068 | 8 | 74.91 | 1 | 9.36 | 12.5 | 87.28 | 81.3 | 5.98 | 5.98 | 0.04 | 0.04 |
| gene-LOC113563560 | https://www.ncbi.nlm.nih.gov/gene/?term=LOC113563560 | 4945 | 5 | 10.11 | 1 | 2.02 | 20 | 86.62 | 80.66 | 5.96 | 5.96 | 0.01 | 0.01 |
| gene-LOC105283813 | https://www.ncbi.nlm.nih.gov/gene/?term=LOC105283813 | 1312 | 8 | 60.98 | 1 | 7.62 | 12.5 | 29.71 | 24.06 | 5.65 | 5.65 | 0.03 | 0.03 |
| gene-LOC105285051 | https://www.ncbi.nlm.nih.gov/gene/?term=LOC105285051 | 1357 | 11 | 81.06 | 3 | 22.11 | 27.27 | 46.19 | 40.59 | 5.6 | 5.6 | 0.05 | 0.05 |
| gene-LOC105278243 | https://www.ncbi.nlm.nih.gov/gene/?term=LOC105278243 | 1826 | 18 | 98.58 | 1 | 5.48 | 5.56 | 20.63 | 15.05 | 5.58 | 5.58 | 0.06 | 0.06 |
| gene-LOC105280891 | https://www.ncbi.nlm.nih.gov/gene/?term=LOC105280891 | 1844 | 26 | 141 | 4 | 21.69 | 15.38 | 25.44 | 20.09 | 5.35 | 5.35 | 0.08 | 0.08 |
| gene-LOC105284483 | https://www.ncbi.nlm.nih.gov/gene/?term=LOC105284483 | 2910 | 25 | 85.91 | 1 | 3.44 | 4 | 22.04 | 16.8 | 5.24 | 5.24 | 0.05 | 0.05 |
| gene-LOC105281226 | https://www.ncbi.nlm.nih.gov/gene/?term=LOC105281226 | 1308 | 6 | 45.87 | 1 | 7.65 | 16.67 | 62.79 | 57.58 | 5.21 | 5.21 | 0.02 | 0.02 |
| gene-LOC105280609 | https://www.ncbi.nlm.nih.gov/gene/?term=LOC105280609 | 1563 | 17 | 108.77 | 2 | 12.8 | 11.76 | 46.92 | 41.73 | 5.18 | 5.18 | 0.06 | 0.06 |

...
... only first 25 lines shown ...

Directory: OGLall File: ogl-Ob_LI_BCphaseA1.vs.RphaseA1.txt

| gene_ID | gene_link | gwidth | #Sites | #per10Kb | #dmSites | #dmsp10kb | %dmSites | %pSite1 | %pSite2 | DMpSite | ADMpSite | DMpNucl | ADMpNucl |
|---|---|---|---|---|---|---|---|---|---|---|---|---|---|
| gene-LOC105286451 | https://www.ncbi.nlm.nih.gov/gene/?term=LOC105286451 | 1618 | 10 | 61.8 | 1 | 6.18 | 10 | 52.19 | 44.01 | 8.18 | 8.18 | 0.05 | 0.05 |
| gene-LOC105281173 | https://www.ncbi.nlm.nih.gov/gene/?term=LOC105281173 | 1595 | 25 | 156.74 | 1 | 6.27 | 4 | 64.76 | 61.03 | 3.73 | 3.73 | 0.06 | 0.06 |
| gene-LOC105276538 | https://www.ncbi.nlm.nih.gov/gene/?term=LOC105276538 | 1511 | 18 | 119.13 | 1 | 6.62 | 5.56 | 71.16 | 68.49 | 2.67 | 2.67 | 0.03 | 0.03 |
| gene-LOC105274449 | https://www.ncbi.nlm.nih.gov/gene/?term=LOC105274449 | 1549 | 43 | 277.6 | 1 | 6.46 | 2.33 | 54.59 | 52.01 | 2.58 | 2.58 | 0.07 | 0.07 |
| gene-LOC105284396 | https://www.ncbi.nlm.nih.gov/gene/?term=LOC105284396 | 1976 | 57 | 288.46 | 1 | 5.06 | 1.75 | 39.14 | 36.68 | 2.46 | 2.46 | 0.07 | 0.07 |
| gene-LOC105274706 | https://www.ncbi.nlm.nih.gov/gene/?term=LOC105274706 | 1446 | 48 | 331.95 | 1 | 6.92 | 2.08 | 39.02 | 37.1 | 1.92 | 1.92 | 0.06 | 0.06 |
| gene-LOC105285409 | https://www.ncbi.nlm.nih.gov/gene/?term=LOC105285409 | 5099 | 47 | 92.17 | 1 | 1.96 | 2.13 | 3.57 | 1.67 | 1.9 | 1.9 | 0.02 | 0.02 |
| gene-LOC105278877 | https://www.ncbi.nlm.nih.gov/gene/?term=LOC105278877 | 1968 | 78 | 396.34 | 1 | 5.08 | 1.28 | 32.36 | 30.87 | 1.48 | 1.48 | 0.06 | 0.06 |
| gene-LOC105275518 | https://www.ncbi.nlm.nih.gov/gene/?term=LOC105275518 | 2335 | 19 | 81.37 | 1 | 4.28 | 5.26 | 49.77 | 48.32 | 1.45 | 1.45 | 0.01 | 0.01 |
| gene-LOC105278538 | https://www.ncbi.nlm.nih.gov/gene/?term=LOC105278538 | 2150 | 75 | 348.84 | 1 | 4.65 | 1.33 | 18.53 | 17.09 | 1.44 | 1.44 | 0.05 | 0.05 |
| gene-LOC105278454 | https://www.ncbi.nlm.nih.gov/gene/?term=LOC105278454 | 1972 | 37 | 187.63 | 1 | 5.07 | 2.7 | 46.8 | 45.42 | 1.38 | 1.38 | 0.03 | 0.03 |
| gene-LOC105283649 | https://www.ncbi.nlm.nih.gov/gene/?term=LOC105283649 | 1918 | 53 | 276.33 | 2 | 10.43 | 3.77 | 38.29 | 37.03 | 1.27 | 1.27 | 0.04 | 0.04 |
| gene-LOC105279664 | https://www.ncbi.nlm.nih.gov/gene/?term=LOC105279664 | 4023 | 63 | 156.6 | 1 | 2.49 | 1.59 | 17.93 | 16.77 | 1.16 | 1.16 | 0.02 | 0.02 |
| gene-LOC105285202 | https://www.ncbi.nlm.nih.gov/gene/?term=LOC105285202 | 1838 | 35 | 190.42 | 1 | 5.44 | 2.86 | 19.49 | 18.4 | 1.08 | 1.08 | 0.02 | 0.02 |
| gene-LOC105276196 | https://www.ncbi.nlm.nih.gov/gene/?term=LOC105276196 | 2953 | 55 | 186.25 | 1 | 3.39 | 1.82 | 40.47 | 39.41 | 1.06 | 1.06 | 0.02 | 0.02 |
| gene-LOC105287095 | https://www.ncbi.nlm.nih.gov/gene/?term=LOC105287095 | 1435 | 60 | 418.12 | 1 | 6.97 | 1.67 | 26.49 | 25.42 | 1.06 | 1.06 | 0.04 | 0.04 |
| gene-LOC105282811 | https://www.ncbi.nlm.nih.gov/gene/?term=LOC105282811 | 5765 | 183 | 317.43 | 1 | 1.73 | 0.55 | 10.19 | 9.27 | 0.92 | 0.92 | 0.03 | 0.03 |
| gene-LOC105280786 | https://www.ncbi.nlm.nih.gov/gene/?term=LOC105280786 | 2573 | 115 | 446.95 | 1 | 3.89 | 0.87 | 29.1 | 28.31 | 0.79 | 0.79 | 0.04 | 0.04 |
| gene-LOC105281047 | https://www.ncbi.nlm.nih.gov/gene/?term=LOC105281047 | 7313 | 112 | 153.15 | 1 | 1.37 | 0.89 | 11.03 | 10.28 | 0.75 | 0.75 | 0.01 | 0.01 |
| gene-LOC105285807 | https://www.ncbi.nlm.nih.gov/gene/?term=LOC105285807 | 6063 | 180 | 296.88 | 1 | 1.65 | 0.56 | 10.12 | 9.41 | 0.72 | 0.72 | 0.02 | 0.02 |
| gene-LOC105282794 | https://www.ncbi.nlm.nih.gov/gene/?term=LOC105282794 | 4571 | 93 | 203.46 | 1 | 2.19 | 1.08 | 17.63 | 17.04 | 0.6 | 0.6 | 0.01 | 0.01 |
| gene-LOC105283123 | https://www.ncbi.nlm.nih.gov/gene/?term=LOC105283123 | 1507 | 35 | 232.25 | 1 | 6.64 | 2.86 | 18.15 | 17.56 | 0.59 | 0.59 | 0.01 | 0.01 |
| gene-LOC105275752 | https://www.ncbi.nlm.nih.gov/gene/?term=LOC105275752 | 3462 | 126 | 363.95 | 1 | 2.89 | 0.79 | 28.41 | 27.85 | 0.56 | 0.56 | 0.02 | 0.02 |
| gene-LOC105284116 | https://www.ncbi.nlm.nih.gov/gene/?term=LOC105284116 | 3970 | 90 | 226.7 | 1 | 2.52 | 1.11 | 7.07 | 6.57 | 0.5 | 0.5 | 0.01 | 0.01 |
| gene-LOC105279038 | https://www.ncbi.nlm.nih.gov/gene/?term=LOC105279038 | 9453 | 299 | 316.3 | 1 | 1.06 | 0.33 | 8.97 | 8.54 | 0.42 | 0.42 | 0.01 | 0.01 |
| gene-LOC105278666 | https://www.ncbi.nlm.nih.gov/gene/?term=LOC105278666 | 10341 | 393 | 380.04 | 1 | 0.97 | 0.25 | 7.76 | 7.36 | 0.4 | 0.4 | 0.02 | 0.02 |
| gene-LOC105280002 | https://www.ncbi.nlm.nih.gov/gene/?term=LOC105280002 | 1940 | 28 | 144.33 | 1 | 5.15 | 3.57 | 33.09 | 32.74 | 0.35 | 0.35 | 0.01 | 0.01 |
| gene-LOC105284117 | https://www.ncbi.nlm.nih.gov/gene/?term=LOC105284117 | 4340 | 82 | 188.94 | 1 | 2.3 | 1.22 | 10.79 | 10.47 | 0.33 | 0.33 | 0.01 | 0.01 |
| gene-LOC105274590 | https://www.ncbi.nlm.nih.gov/gene/?term=LOC105274590 | 4516 | 75 | 166.08 | 1 | 2.21 | 1.33 | 23.2 | 22.91 | 0.29 | 0.29 | 0 | 0 |
| gene-LOC105280819 | https://www.ncbi.nlm.nih.gov/gene/?term=LOC105280819 | 1639 | 41 | 250.15 | 1 | 6.1 | 2.44 | 40.58 | 40.31 | 0.27 | 0.27 | 0.01 | 0.01 |
| gene-LOC105288079 | https://www.ncbi.nlm.nih.gov/gene/?term=LOC105288079 | 3473 | 85 | 244.75 | 1 | 2.88 | 1.18 | 20.86 | 20.61 | 0.25 | 0.25 | 0.01 | 0.01 |
| gene-LOC105286194 | https://www.ncbi.nlm.nih.gov/gene/?term=LOC105286194 | 152619 | 1939 | 127.05 | 1 | 0.07 | 0.05 | 6.35 | 6.21 | 0.14 | 0.14 | 0 | 0 |
| gene-LOC105276901 | https://www.ncbi.nlm.nih.gov/gene/?term=LOC105276901 | 3254 | 96 | 295.02 | 1 | 3.07 | 1.04 | 22.83 | 22.72 | 0.1 | 0.1 | 0 | 0 |
| gene-LOC105277866 | https://www.ncbi.nlm.nih.gov/gene/?term=LOC105277866 | 33045 | 403 | 121.95 | 1 | 0.3 | 0.25 | 3.32 | 3.22 | 0.1 | 0.1 | 0 | 0 |
| gene-LOC105280265 | https://www.ncbi.nlm.nih.gov/gene/?term=LOC105280265 | 85984 | 2645 | 307.62 | 1 | 0.12 | 0.04 | 1.12 | 1.06 | 0.07 | 0.07 | 0 | 0 |
| gene-LOC105278455 | https://www.ncbi.nlm.nih.gov/gene/?term=LOC105278455 | 27181 | 836 | 307.57 | 1 | 0.37 | 0.12 | 7.62 | 7.57 | 0.05 | 0.05 | 0 | 0 |
| gene-LOC105278228 | https://www.ncbi.nlm.nih.gov/gene/?term=LOC105278228 | 8610 | 152 | 176.54 | 1 | 1.16 | 0.66 | 23.77 | 23.73 | 0.04 | 0.04 | 0 | 0 |
| gene-LOC105283728 | https://www.ncbi.nlm.nih.gov/gene/?term=LOC105283728 | 3256 | 113 | 347.05 | 2 | 6.14 | 1.77 | 19.87 | 19.82 | 0.04 | 0.04 | 0 | 0 |
| gene-LOC105278145 | https://www.ncbi.nlm.nih.gov/gene/?term=LOC105278145 | 4470 | 239 | 534.68 | 1 | 2.24 | 0.42 | 10.87 | 10.84 | 0.03 | 0.03 | 0 | 0 |
| gene-LOC105275327 | https://www.ncbi.nlm.nih.gov/gene/?term=LOC105275327 | 14245 | 461 | 323.62 | 1 | 0.7 | 0.22 | 7.38 | 7.42 | -0.03 | 0.03 | 0 | 0 |
| gene-LOC105286274 | https://www.ncbi.nlm.nih.gov/gene/?term=LOC105286274 | 3817 | 174 | 455.86 | 1 | 2.62 | 0.57 | 15.59 | 15.67 | -0.08 | 0.08 | 0 | 0 |
| gene-LOC105274856 | https://www.ncbi.nlm.nih.gov/gene/?term=LOC105274856 | 5216 | 201 | 385.35 | 1 | 1.92 | 0.5 | 17.77 | 17.88 | -0.11 | 0.11 | 0 | 0 |
| gene-LOC105278534 | https://www.ncbi.nlm.nih.gov/gene/?term=LOC105278534 | 7364 | 173 | 234.93 | 1 | 1.36 | 0.58 | 18.44 | 18.59 | -0.16 | 0.16 | 0 | 0 |
| gene-LOC105274851 | https://www.ncbi.nlm.nih.gov/gene/?term=LOC105274851 | 6616 | 133 | 201.03 | 1 | 1.51 | 0.75 | 10.15 | 10.35 | -0.2 | 0.2 | 0 | 0 |
| gene-LOC105286515 | https://www.ncbi.nlm.nih.gov/gene/?term=LOC105286515 | 3751 | 106 | 282.59 | 1 | 2.67 | 0.94 | 10.37 | 10.58 | -0.22 | 0.22 | -0.01 | 0.01 |
| gene-LOC105279500 | https://www.ncbi.nlm.nih.gov/gene/?term=LOC105279500 | 3587 | 29 | 80.85 | 1 | 2.79 | 3.45 | 48.07 | 48.34 | -0.27 | 0.27 | 0 | 0 |
| gene-LOC105285534 | https://www.ncbi.nlm.nih.gov/gene/?term=LOC105285534 | 3944 | 221 | 560.34 | 1 | 2.54 | 0.45 | 11.41 | 11.71 | -0.3 | 0.3 | -0.02 | 0.02 |
| gene-LOC105279298 | https://www.ncbi.nlm.nih.gov/gene/?term=LOC105279298 | 2773 | 135 | 486.84 | 1 | 3.61 | 0.74 | 22.57 | 22.9 | -0.33 | 0.33 | -0.02 | 0.02 |
| gene-LOC105288078 | https://www.ncbi.nlm.nih.gov/gene/?term=LOC105288078 | 19126 | 305 | 159.47 | 1 | 0.52 | 0.33 | 16.95 | 17.33 | -0.38 | 0.38 | -0.01 | 0.01 |
| gene-LOC105285406 | https://www.ncbi.nlm.nih.gov/gene/?term=LOC105285406 | 8015 | 39 | 48.66 | 1 | 1.25 | 2.56 | 33.07 | 33.49 | -0.42 | 0.42 | 0 | 0 |
| gene-LOC105274531 | https://www.ncbi.nlm.nih.gov/gene/?term=LOC105274531 | 7774 | 253 | 325.44 | 1 | 1.29 | 0.4 | 11.9 | 12.33 | -0.43 | 0.43 | -0.01 | 0.01 |
| gene-LOC105283712 | https://www.ncbi.nlm.nih.gov/gene/?term=LOC105283712 | 1910 | 29 | 151.83 | 1 | 5.24 | 3.45 | 29.98 | 30.42 | -0.44 | 0.44 | -0.01 | 0.01 |
| gene-LOC105275173 | https://www.ncbi.nlm.nih.gov/gene/?term=LOC105275173 | 4639 | 76 | 163.83 | 1 | 2.16 | 1.32 | 24.33 | 24.78 | -0.45 | 0.45 | -0.01 | 0.01 |
| gene-LOC105284184 | https://www.ncbi.nlm.nih.gov/gene/?term=LOC105284184 | 2535 | 80 | 315.58 | 1 | 3.94 | 1.25 | 34.85 | 35.33 | -0.47 | 0.47 | -0.01 | 0.01 |
| gene-LOC105286997 | https://www.ncbi.nlm.nih.gov/gene/?term=LOC105286997 | 2850 | 134 | 470.18 | 1 | 3.51 | 0.75 | 11.83 | 12.3 | -0.47 | 0.47 | -0.02 | 0.02 |
| gene-LOC105275898 | https://www.ncbi.nlm.nih.gov/gene/?term=LOC105275898 | 5358 | 132 | 246.36 | 1 | 1.87 | 0.76 | 12.84 | 13.34 | -0.5 | 0.5 | -0.01 | 0.01 |
| gene-LOC105280956 | https://www.ncbi.nlm.nih.gov/gene/?term=LOC105280956 | 2639 | 73 | 276.62 | 1 | 3.79 | 1.37 | 20.28 | 20.85 | -0.57 | 0.57 | -0.02 | 0.02 |
| gene-LOC105274534 | https://www.ncbi.nlm.nih.gov/gene/?term=LOC105274534 | 1329 | 57 | 428.89 | 1 | 7.52 | 1.75 | 43.75 | 44.38 | -0.63 | 0.63 | -0.03 | 0.03 |
| gene-LOC105275683 | https://www.ncbi.nlm.nih.gov/gene/?term=LOC105275683 | 4256 | 187 | 439.38 | 1 | 2.35 | 0.53 | 12.61 | 13.25 | -0.63 | 0.63 | -0.03 | 0.03 |
| gene-LOC105278497 | https://www.ncbi.nlm.nih.gov/gene/?term=LOC105278497 | 2072 | 97 | 468.15 | 1 | 4.83 | 1.03 | 18.56 | 19.21 | -0.64 | 0.64 | -0.03 | 0.03 |

| | | | | | | | | | | | | | | |
|---|---|---|---|---|---|---|---|---|---|---|---|---|---|---|
| gene-LOC105276692 | https://www.ncbi.nlm.nih.gov/gene/?term=LOC105276692 | 1745 | 34 | 194.84 | 1 | 5.73 | 2.94 | 24.51 | 25.17 | -0.66 | 0.66 | -0.01 | 0.01 | |
| gene-LOC105280641 | https://www.ncbi.nlm.nih.gov/gene/?term=LOC105280641 | 3682 | 55 | 149.38 | 1 | 2.72 | 1.82 | 18.35 | 19.01 | -0.66 | 0.66 | -0.01 | 0.01 | |
| gene-LOC105285639 | https://www.ncbi.nlm.nih.gov/gene/?term=LOC105285639 | 10325 | 294 | 284.75 | 1 | 0.97 | 0.34 | 12.44 | 13.1 | -0.66 | 0.66 | -0.02 | 0.02 | |
| gene-LOC105281700 | https://www.ncbi.nlm.nih.gov/gene/?term=LOC105281700 | 2208 | 45 | 203.8 1 | 4.53 | 2.22 | 38.83 | 39.5 | -0.67 | 0.67 | -0.01 | 0.01 | | |
| gene-LOC105288036 | https://www.ncbi.nlm.nih.gov/gene/?term=LOC105288036 | 5110 | 182 | 356.16 | 1 | 1.96 | 0.55 | 18.19 | 18.89 | -0.7 | 0.7 | -0.03 | 0.03 | |
| gene-LOC105287063 | https://www.ncbi.nlm.nih.gov/gene/?term=LOC105287063 | 1874 | 46 | 245.46 | 1 | 5.34 | 2.17 | 34.15 | 34.88 | -0.73 | 0.73 | -0.02 | 0.02 | |
| gene-LOC105279657 | https://www.ncbi.nlm.nih.gov/gene/?term=LOC105279657 | 5878 | 82 | 139.5 1 | 1.7 | 1.22 | 19.15 | 19.93 | -0.78 | 0.78 | -0.01 | 0.01 | | |
| gene-LOC105277856 | https://www.ncbi.nlm.nih.gov/gene/?term=LOC105277856 | 3442 | 28 | 81.35 1 | 2.91 | 3.57 | 36.24 | 37.06 | -0.82 | 0.82 | -0.01 | 0.01 | | |
| gene-LOC105275179 | https://www.ncbi.nlm.nih.gov/gene/?term=LOC105275179 | 3271 | 69 | 210.94 | 1 | 3.06 | 1.45 | 14.66 | 15.53 | -0.86 | 0.86 | -0.02 | 0.02 | |
| gene-LOC105288237 | https://www.ncbi.nlm.nih.gov/gene/?term=LOC105288237 | 2226 | 45 | 202.16 | 2 | 8.98 | 4.44 | 36.5 | 37.4 | -0.9 | 0.9 | -0.02 | 0.02 | |
| gene-LOC105275451 | https://www.ncbi.nlm.nih.gov/gene/?term=LOC105275451 | 6035 | 111 | 183.93 | 1 | 1.66 | 0.9 | 22.31 | 23.24 | -0.94 | 0.94 | -0.02 | 0.02 | |
| gene-LOC105286265 | https://www.ncbi.nlm.nih.gov/gene/?term=LOC105286265 | 2507 | 65 | 259.27 | 1 | 3.99 | 1.54 | 31.58 | 32.53 | -0.95 | 0.95 | -0.02 | 0.02 | |
| gene-LOC105275363 | https://www.ncbi.nlm.nih.gov/gene/?term=LOC105275363 | 3798 | 155 | 408.11 | 1 | 2.63 | 0.65 | 17.28 | 18.35 | -1.06 | 1.06 | -0.04 | 0.04 | |
| gene-LOC105282156 | https://www.ncbi.nlm.nih.gov/gene/?term=LOC105282156 | 2577 | 128 | 496.7 1 | 3.88 | 0.78 | 23.38 | 24.45 | -1.07 | 1.07 | -0.05 | 0.05 | | |
| gene-LOC105282991 | https://www.ncbi.nlm.nih.gov/gene/?term=LOC105282991 | 5617 | 80 | 142.42 | 1 | 1.78 | 1.25 | 19.15 | 20.24 | -1.09 | 1.09 | -0.02 | 0.02 | |
| gene-LOC105283943 | https://www.ncbi.nlm.nih.gov/gene/?term=LOC105283943 | 3559 | 49 | 137.68 | 1 | 2.81 | 2.04 | 22.13 | 23.23 | -1.1 | 1.1 | -0.02 | 0.02 | |
| gene-LOC105285055 | https://www.ncbi.nlm.nih.gov/gene/?term=LOC105285055 | 4108 | 204 | 496.59 | 1 | 2.43 | 0.49 | 12.43 | 13.67 | -1.24 | 1.24 | -0.06 | 0.06 | |
| gene-LOC105277216 | https://www.ncbi.nlm.nih.gov/gene/?term=LOC105277216 | 7627 | 91 | 119.31 | 1 | 1.31 | 1.1 | 6.55 | 7.88 | -1.33 | 1.33 | -0.02 | 0.02 | |
| gene-LOC105286452 | https://www.ncbi.nlm.nih.gov/gene/?term=LOC105286452 | 6888 | 58 | 84.2 1 | 1.45 | 1.72 | 31.25 | 32.65 | -1.4 | 1.4 | -0.01 | 0.01 | | |
| gene-LOC105285695 | https://www.ncbi.nlm.nih.gov/gene/?term=LOC105285695 | 1371 | 61 | 444.93 | 1 | 7.29 | 1.64 | 22.52 | 23.97 | -1.45 | 1.45 | -0.06 | 0.06 | |
| gene-LOC105283592 | https://www.ncbi.nlm.nih.gov/gene/?term=LOC105283592 | 3479 | 68 | 195.46 | 1 | 2.87 | 1.47 | 23.52 | 25 | -1.48 | 1.48 | -0.03 | 0.03 | |
| gene-LOC105275190 | https://www.ncbi.nlm.nih.gov/gene/?term=LOC105275190 | 3837 | 65 | 169.4 1 | 2.61 | 1.54 | 21.02 | 22.59 | -1.58 | 1.58 | -0.03 | 0.03 | | |
| gene-LOC105280348 | https://www.ncbi.nlm.nih.gov/gene/?term=LOC105280348 | 3797 | 52 | 136.95 | 1 | 2.63 | 1.92 | 25.54 | 27.17 | -1.63 | 1.63 | -0.02 | 0.02 | |
| gene-LOC105276704 | https://www.ncbi.nlm.nih.gov/gene/?term=LOC105276704 | 2063 | 80 | 387.78 | 1 | 4.85 | 1.25 | 13.36 | 15.07 | -1.71 | 1.71 | -0.07 | 0.07 | |
| gene-LOC105275460 | https://www.ncbi.nlm.nih.gov/gene/?term=LOC105275460 | 2473 | 115 | 465.02 | 1 | 4.04 | 0.87 | 12.68 | 14.4 | -1.72 | 1.72 | -0.08 | 0.08 | |
| gene-LOC105283989 | https://www.ncbi.nlm.nih.gov/gene/?term=LOC105283989 | 4948 | 30 | 60.63 1 | 2.02 | 3.33 | 45.76 | 47.5 | -1.74 | 1.74 | -0.01 | 0.01 | | |
| gene-LOC105285742 | https://www.ncbi.nlm.nih.gov/gene/?term=LOC105285742 | 3446 | 87 | 252.47 | 1 | 2.9 | 1.15 | 20.61 | 22.37 | -1.76 | 1.76 | -0.04 | 0.04 | |
| gene-LOC105278436 | https://www.ncbi.nlm.nih.gov/gene/?term=LOC105278436 | 2556 | 56 | 219.09 | 1 | 3.91 | 1.79 | 36.89 | 38.91 | -2.02 | 2.02 | -0.04 | 0.04 | |
| gene-LOC105278807 | https://www.ncbi.nlm.nih.gov/gene/?term=LOC105278807 | 2166 | 70 | 323.18 | 1 | 4.62 | 1.43 | 17.84 | 20.15 | -2.31 | 2.31 | -0.07 | 0.07 | |
| gene-LOC105285671 | https://www.ncbi.nlm.nih.gov/gene/?term=LOC105285671 | 2263 | 90 | 397.7 1 | 4.42 | 1.11 | 29.29 | 31.61 | -2.32 | 2.32 | -0.09 | 0.09 | | |
| gene-LOC105279356 | https://www.ncbi.nlm.nih.gov/gene/?term=LOC105279356 | 1314 | 42 | 319.63 | 1 | 7.61 | 2.38 | 28.83 | 31.17 | -2.34 | 2.34 | -0.07 | 0.07 | |
| gene-LOC105288066 | https://www.ncbi.nlm.nih.gov/gene/?term=LOC105288066 | 1964 | 24 | 122.2 1 | 5.09 | 4.17 | 27.7 | 30.29 | -2.59 | 2.59 | -0.03 | 0.03 | | |
| gene-LOC105285060 | https://www.ncbi.nlm.nih.gov/gene/?term=LOC105285060 | 1285 | 15 | 116.73 | 1 | 7.78 | 6.67 | 48.46 | 51.14 | -2.68 | 2.68 | -0.03 | 0.03 | |
| gene-LOC105285674 | https://www.ncbi.nlm.nih.gov/gene/?term=LOC105285674 | 3035 | 88 | 289.95 | 2 | 6.59 | 2.27 | 23.51 | 26.27 | -2.76 | 2.76 | -0.08 | 0.08 | |
| gene-LOC105281848 | https://www.ncbi.nlm.nih.gov/gene/?term=LOC105281848 | 2122 | 45 | 212.06 | 1 | 4.71 | 2.22 | 23.75 | 26.94 | -3.19 | 3.19 | -0.07 | 0.07 | |
| gene-LOC105279898 | https://www.ncbi.nlm.nih.gov/gene/?term=LOC105279898 | 1343 | 13 | 96.8 1 | 7.45 | 7.69 | 65.18 | 68.44 | -3.25 | 3.25 | -0.03 | 0.03 | | |
| gene-LOC105283552 | https://www.ncbi.nlm.nih.gov/gene/?term=LOC105283552 | 1562 | 33 | 211.27 | 1 | 6.4 | 3.03 | 54.5 | 58 | -3.5 | 3.5 | -0.07 | 0.07 | |
| gene-LOC105277903 | https://www.ncbi.nlm.nih.gov/gene/?term=LOC105277903 | 1835 | 35 | 190.74 | 1 | 5.45 | 2.86 | 51.45 | 55.03 | -3.59 | 3.59 | -0.07 | 0.07 | |
| gene-LOC105274900 | https://www.ncbi.nlm.nih.gov/gene/?term=LOC105274900 | 5561 | 56 | 100.7 1 | 1.8 | 1.79 | 26.85 | 30.51 | -3.66 | 3.66 | -0.04 | 0.04 | | |
| gene-LOC105277215 | https://www.ncbi.nlm.nih.gov/gene/?term=LOC105277215 | 2619 | 27 | 103.09 | 1 | 3.82 | 3.7 | 20.79 | 25.43 | -4.64 | 4.64 | -0.05 | 0.05 | |
| gene-LOC105275708 | https://www.ncbi.nlm.nih.gov/gene/?term=LOC105275708 | 1772 | 14 | 79.01 1 | 5.64 | 7.14 | 44.58 | 50.83 | -6.25 | 6.25 | -0.05 | 0.05 | | |
| gene-LOC105285514 | https://www.ncbi.nlm.nih.gov/gene/?term=LOC105285514 | 1277 | 8 | 62.65 1 | 7.83 | 12.5 | 70.46 | 79.23 | -8.77 | 8.77 | -0.05 | 0.05 | | |
| gene-LOC105275070 | https://www.ncbi.nlm.nih.gov/gene/?term=LOC105275070 | 5667 | 37 | 65.29 2 | 3.53 | 5.41 | 3.73 | 16.16 | -12.43 | | 12.43 | -0.08 | 0.08 | |
| gene-LOC113562345 | https://www.ncbi.nlm.nih.gov/gene/?term=LOC113562345 | 2290 | 61 | 266.38 | 10 | 43.67 | 16.39 | 17.52 | 31.67 | -14.16 | | 14.16 | -0.38 | 0.38 |

Directory: OGLall File: ogl-Ob_LI_BCphaseA1.vs.RphaseA2.txt

| gene_ID | gene_link | gwidth | #Sites | #per10Kb | #dmSites | #dmsp10kb | %dmSites | %pSite1 | %pSite2 | DMpSite | ADMpSite | DMpNucl | ADMpNucl |
|---|---|---|---|---|---|---|---|---|---|---|---|---|---|
| gene-LOC105277380 | https://www.ncbi.nlm.nih.gov/gene/?term=LOC105277380 | 1967 | 5 | 25.42 | 1 | 5.08 | 20 | 69.42 | 54.11 | 15.31 | 15.31 | 0.04 | 0.04 |
| gene-LOC105275859 | https://www.ncbi.nlm.nih.gov/gene/?term=LOC105275859 | 2331 | 25 | 107.25 | 1 | 4.29 | 4 | 53.95 | 51.3 | 2.65 | 2.65 | 0.03 | 0.03 |
| gene-LOC105285409 | https://www.ncbi.nlm.nih.gov/gene/?term=LOC105285409 | 5099 | 48 | 94.14 | 2 | 3.92 | 4.17 | 3.6 | 1.11 | 2.49 | 2.49 | 0.02 | 0.02 |
| gene-LOC105274449 | https://www.ncbi.nlm.nih.gov/gene/?term=LOC105274449 | 1549 | 42 | 271.14 | 1 | 6.46 | 2.38 | 55.88 | 53.68 | 2.2 | 2.2 | 0.06 | 0.06 |
| gene-LOC105281173 | https://www.ncbi.nlm.nih.gov/gene/?term=LOC105281173 | 1595 | 25 | 156.74 | 1 | 6.27 | 4 | 64.76 | 62.62 | 2.13 | 2.13 | 0.03 | 0.03 |
| gene-LOC105282885 | https://www.ncbi.nlm.nih.gov/gene/?term=LOC105282885 | 1870 | 42 | 224.6 | 1 | 5.35 | 2.38 | 36.63 | 34.78 | 1.84 | 1.84 | 0.04 | 0.04 |
| gene-LOC105276272 | https://www.ncbi.nlm.nih.gov/gene/?term=LOC105276272 | 1965 | 27 | 137.4 | 1 | 5.09 | 3.7 | 42.02 | 40.71 | 1.31 | 1.31 | 0.02 | 0.02 |
| gene-LOC105287095 | https://www.ncbi.nlm.nih.gov/gene/?term=LOC105287095 | 1435 | 61 | 425.09 | 1 | 6.97 | 1.64 | 26.05 | 24.77 | 1.29 | 1.29 | 0.05 | 0.05 |
| gene-LOC105285084 | https://www.ncbi.nlm.nih.gov/gene/?term=LOC105285084 | 3308 | 26 | 78.6 | 1 | 3.02 | 3.85 | 22.5 | 21.29 | 1.21 | 1.21 | 0.01 | 0.01 |
| gene-LOC105275848 | https://www.ncbi.nlm.nih.gov/gene/?term=LOC105275848 | 3718 | 130 | 349.65 | 1 | 2.69 | 0.77 | 21.27 | 20.2 | 1.07 | 1.07 | 0.04 | 0.04 |
| gene-LOC105281296 | https://www.ncbi.nlm.nih.gov/gene/?term=LOC105281296 | 2434 | 41 | 168.45 | 1 | 4.11 | 2.44 | 31.19 | 30.16 | 1.03 | 1.03 | 0.02 | 0.02 |
| gene-LOC105284507 | https://www.ncbi.nlm.nih.gov/gene/?term=LOC105284507 | 5207 | 82 | 157.48 | 1 | 1.92 | 1.22 | 21.61 | 20.58 | 1.03 | 1.03 | 0.02 | 0.02 |
| gene-LOC105285252 | https://www.ncbi.nlm.nih.gov/gene/?term=LOC105285252 | 7427 | 132 | 177.73 | 1 | 1.35 | 0.76 | 13.81 | 12.99 | 0.81 | 0.81 | 0.01 | 0.01 |
| gene-LOC105283123 | https://www.ncbi.nlm.nih.gov/gene/?term=LOC105283123 | 1507 | 43 | 285.34 | 1 | 6.64 | 2.33 | 17.05 | 16.29 | 0.76 | 0.76 | 0.02 | 0.02 |
| gene-LOC105284396 | https://www.ncbi.nlm.nih.gov/gene/?term=LOC105284396 | 1976 | 61 | 308.7 | 1 | 5.06 | 1.64 | 36.72 | 35.98 | 0.74 | 0.74 | 0.02 | 0.02 |
| gene-LOC105279664 | https://www.ncbi.nlm.nih.gov/gene/?term=LOC105279664 | 4023 | 72 | 178.97 | 1 | 2.49 | 1.39 | 17.48 | 16.74 | 0.73 | 0.73 | 0.01 | 0.01 |
| gene-LOC105278877 | https://www.ncbi.nlm.nih.gov/gene/?term=LOC105278877 | 1968 | 80 | 406.5 | 1 | 5.08 | 1.25 | 32.88 | 32.17 | 0.72 | 0.72 | 0.03 | 0.03 |
| gene-LOC105283264 | https://www.ncbi.nlm.nih.gov/gene/?term=LOC105283264 | 4005 | 88 | 219.73 | 1 | 2.5 | 1.14 | 24.81 | 24.09 | 0.72 | 0.72 | 0.02 | 0.02 |
| gene-LOC105282811 | https://www.ncbi.nlm.nih.gov/gene/?term=LOC105282811 | 5765 | 201 | 348.66 | 1 | 1.73 | 0.5 | 9.88 | 9.23 | 0.65 | 0.65 | 0.02 | 0.02 |
| gene-LOC105286515 | https://www.ncbi.nlm.nih.gov/gene/?term=LOC105286515 | 3751 | 116 | 309.25 | 1 | 2.67 | 0.86 | 11.85 | 11.25 | 0.6 | 0.6 | 0.02 | 0.02 |
| gene-LOC105280819 | https://www.ncbi.nlm.nih.gov/gene/?term=LOC105280819 | 1639 | 41 | 250.15 | 1 | 6.1 | 2.44 | 40.58 | 40.03 | 0.56 | 0.56 | 0.01 | 0.01 |
| gene-LOC105282794 | https://www.ncbi.nlm.nih.gov/gene/?term=LOC105282794 | 4571 | 98 | 214.4 | 1 | 2.19 | 1.02 | 18.44 | 17.88 | 0.56 | 0.56 | 0.01 | 0.01 |
| gene-LOC105281420 | https://www.ncbi.nlm.nih.gov/gene/?term=LOC105281420 | 3882 | 57 | 146.83 | 1 | 2.58 | 1.75 | 15.28 | 15.02 | 0.26 | 0.26 | 0 | 0 |
| gene-LOC105283649 | https://www.ncbi.nlm.nih.gov/gene/?term=LOC105283649 | 1918 | 53 | 276.33 | 2 | 10.43 | 3.77 | 38.29 | 38.03 | 0.26 | 0.26 | 0.01 | 0.01 |
| gene-LOC105286265 | https://www.ncbi.nlm.nih.gov/gene/?term=LOC105286265 | 2507 | 68 | 271.24 | 1 | 3.99 | 1.47 | 30.18 | 29.94 | 0.24 | 0.24 | 0.01 | 0.01 |
| gene-LOC105281834 | https://www.ncbi.nlm.nih.gov/gene/?term=LOC105281834 | 4711 | 204 | 433.03 | 1 | 2.12 | 0.49 | 10.43 | 10.29 | 0.14 | 0.14 | 0.01 | 0.01 |
| gene-LOC105278454 | https://www.ncbi.nlm.nih.gov/gene/?term=LOC105278454 | 1972 | 37 | 187.63 | 1 | 5.07 | 2.7 | 46.8 | 46.67 | 0.13 | 0.13 | 0 | 0 |
| gene-LOC105278455 | https://www.ncbi.nlm.nih.gov/gene/?term=LOC105278455 | 27181 | 875 | 321.92 | 1 | 0.37 | 0.11 | 7.33 | 7.2 | 0.13 | 0.13 | 0 | 0 |
| gene-LOC105286649 | https://www.ncbi.nlm.nih.gov/gene/?term=LOC105286649 | 14178 | 701 | 494.43 | 1 | 0.71 | 0.14 | 1.28 | 1.15 | 0.13 | 0.13 | 0.01 | 0.01 |
| gene-LOC105276196 | https://www.ncbi.nlm.nih.gov/gene/?term=LOC105276196 | 2953 | 56 | 189.64 | 1 | 3.39 | 1.79 | 40.11 | 39.99 | 0.12 | 0.12 | 0 | 0 |
| gene-LOC105277273 | https://www.ncbi.nlm.nih.gov/gene/?term=LOC105277273 | 3711 | 151 | 406.9 | 1 | 2.69 | 0.66 | 12.76 | 12.67 | 0.09 | 0.09 | 0 | 0 |
| gene-LOC105278632 | https://www.ncbi.nlm.nih.gov/gene/?term=LOC105278632 | 3624 | 132 | 364.24 | 1 | 2.76 | 0.76 | 14.6 | 14.5 | 0.09 | 0.09 | 0 | 0 |
| gene-LOC105284116 | https://www.ncbi.nlm.nih.gov/gene/?term=LOC105284116 | 3970 | 96 | 241.81 | 1 | 2.52 | 1.04 | 6.8 | 6.72 | 0.08 | 0.08 | 0 | 0 |
| gene-LOC105278666 | https://www.ncbi.nlm.nih.gov/gene/?term=LOC105278666 | 10341 | 409 | 395.51 | 1 | 0.97 | 0.24 | 7.49 | 7.42 | 0.07 | 0.07 | 0 | 0 |
| gene-LOC105279528 | https://www.ncbi.nlm.nih.gov/gene/?term=LOC105279528 | 5802 | 46 | 79.28 | 1 | 1.72 | 2.17 | 15.76 | 15.72 | 0.04 | 0.04 | 0 | 0 |
| gene-LOC105280304 | https://www.ncbi.nlm.nih.gov/gene/?term=LOC105280304 | 8919 | 123 | 137.91 | 1 | 1.12 | 0.81 | 5.79 | 5.75 | 0.04 | 0.04 | 0 | 0 |
| gene-LOC105282914 | https://www.ncbi.nlm.nih.gov/gene/?term=LOC105282914 | 5623 | 42 | 74.69 | 1 | 1.78 | 2.38 | 48.6 | 48.57 | 0.03 | 0.03 | 0 | 0 |
| gene-LOC105274699 | https://www.ncbi.nlm.nih.gov/gene/?term=LOC105274699 | 2386 | 63 | 264.04 | 1 | 4.19 | 1.59 | 23.04 | 23.01 | 0.02 | 0.02 | 0 | 0 |
| gene-LOC105283296 | https://www.ncbi.nlm.nih.gov/gene/?term=LOC105283296 | 2174 | 89 | 409.38 | 1 | 4.6 | 1.12 | 19.56 | 19.55 | 0.01 | 0.01 | 0 | 0 |
| gene-LOC105277866 | https://www.ncbi.nlm.nih.gov/gene/?term=LOC105277866 | 33045 | 447 | 135.27 | 1 | 0.3 | 0.22 | 3.08 | 3.1 | -0.02 | 0.02 | 0 | 0 |
| gene-LOC105280265 | https://www.ncbi.nlm.nih.gov/gene/?term=LOC105280265 | 85984 | 2872 | 334.02 | 1 | 0.12 | 0.03 | 1.07 | 1.1 | -0.02 | 0.02 | 0 | 0 |
| gene-LOC105277125 | https://www.ncbi.nlm.nih.gov/gene/?term=LOC105277125 | 51079 | 1743 | 341.24 | 1 | 0.2 | 0.06 | 1.56 | 1.61 | -0.05 | 0.05 | 0 | 0 |
| gene-LOC105276059 | https://www.ncbi.nlm.nih.gov/gene/?term=LOC105276059 | 6256 | 136 | 217.39 | 1 | 1.6 | 0.74 | 10.77 | 10.82 | -0.06 | 0.06 | 0 | 0 |
| gene-LOC105286332 | https://www.ncbi.nlm.nih.gov/gene/?term=LOC105286332 | 27577 | 507 | 183.85 | 1 | 0.36 | 0.2 | 7.74 | 7.81 | -0.07 | 0.07 | 0 | 0 |
| gene-LOC105278538 | https://www.ncbi.nlm.nih.gov/gene/?term=LOC105278538 | 2150 | 83 | 386.05 | 1 | 4.65 | 1.2 | 18.97 | 19.05 | -0.08 | 0.08 | 0 | 0 |
| gene-LOC105284117 | https://www.ncbi.nlm.nih.gov/gene/?term=LOC105284117 | 4340 | 86 | 198.16 | 1 | 2.3 | 1.16 | 10.49 | 10.57 | -0.08 | 0.08 | 0 | 0 |
| gene-LOC105278145 | https://www.ncbi.nlm.nih.gov/gene/?term=LOC105278145 | 4470 | 256 | 572.71 | 1 | 2.24 | 0.39 | 10.38 | 10.47 | -0.09 | 0.09 | -0.01 | 0.01 |
| gene-LOC105284557 | https://www.ncbi.nlm.nih.gov/gene/?term=LOC105284557 | 1672 | 77 | 460.53 | 1 | 5.98 | 1.3 | 30.69 | 30.8 | -0.11 | 0.11 | -0.01 | 0.01 |
| gene-LOC105277761 | https://www.ncbi.nlm.nih.gov/gene/?term=LOC105277761 | 4648 | 150 | 322.72 | 1 | 2.15 | 0.67 | 14.73 | 14.87 | -0.13 | 0.13 | 0 | 0 |
| gene-LOC105284262 | https://www.ncbi.nlm.nih.gov/gene/?term=LOC105284262 | 6669 | 236 | 353.88 | 1 | 1.5 | 0.42 | 4.46 | 4.59 | -0.13 | 0.13 | 0 | 0 |
| gene-LOC105281272 | https://www.ncbi.nlm.nih.gov/gene/?term=LOC105281272 | 5252 | 179 | 340.82 | 1 | 1.9 | 0.56 | 16.36 | 16.51 | -0.15 | 0.15 | -0.01 | 0.01 |
| gene-LOC105285395 | https://www.ncbi.nlm.nih.gov/gene/?term=LOC105285395 | 8085 | 187 | 231.29 | 1 | 1.24 | 0.53 | 16.28 | 16.44 | -0.16 | 0.16 | 0 | 0 |
| gene-LOC105275187 | https://www.ncbi.nlm.nih.gov/gene/?term=LOC105275187 | 5534 | 75 | 135.53 | 1 | 1.81 | 1.33 | 14.15 | 14.34 | -0.18 | 0.18 | 0 | 0 |
| gene-LOC105276612 | https://www.ncbi.nlm.nih.gov/gene/?term=LOC105276612 | 4157 | 178 | 428.19 | 1 | 2.41 | 0.56 | 10.69 | 10.89 | -0.2 | 0.2 | -0.01 | 0.01 |
| gene-LOC105284036 | https://www.ncbi.nlm.nih.gov/gene/?term=LOC105284036 | 3254 | 83 | 255.07 | 1 | 3.07 | 1.2 | 22.52 | 22.73 | -0.2 | 0.2 | -0.01 | 0.01 |
| gene-LOC105286194 | https://www.ncbi.nlm.nih.gov/gene/?term=LOC105286194 | 152619 | 2093 | 137.14 | 1 | 0.07 | 0.05 | 6.53 | 6.74 | -0.21 | 0.21 | 0 | 0 |
| gene-LOC105285807 | https://www.ncbi.nlm.nih.gov/gene/?term=LOC105285807 | 6063 | 186 | 306.78 | 1 | 1.65 | 0.54 | 10.05 | 10.27 | -0.22 | 0.22 | -0.01 | 0.01 |
| gene-LOC105275327 | https://www.ncbi.nlm.nih.gov/gene/?term=LOC105275327 | 14245 | 479 | 336.26 | 1 | 0.7 | 0.21 | 7.11 | 7.34 | -0.23 | 0.23 | -0.01 | 0.01 |
| gene-LOC105288161 | https://www.ncbi.nlm.nih.gov/gene/?term=LOC105288161 | 26065 | 302 | 115.86 | 1 | 0.38 | 0.33 | 11.58 | 11.82 | -0.24 | 0.24 | 0 | 0 |
| gene-LOC113562050 | https://www.ncbi.nlm.nih.gov/gene/?term=LOC113562050 | 17097 | 576 | 336.9 | 1 | 0.58 | 0.17 | 5.26 | 5.5 | -0.25 | 0.25 | -0.01 | 0.01 |

| | | | | | | | | | | | | | |
|---|---|---|---|---|---|---|---|---|---|---|---|---|---|
| gene-LOC105285202 | https://www.ncbi.nlm.nih.gov/gene/?term=LOC105285202 | 1838 | 36 | 195.87 | 1 | 5.44 | 2.78 | 18.94 | 19.21 | -0.27 | 0.27 | -0.01 | 0.01 |
| gene-LOC105274851 | https://www.ncbi.nlm.nih.gov/gene/?term=LOC105274851 | 6616 | 142 | 214.63 | 1 | 1.51 | 0.7 | 9.54 | 9.83 | -0.3 | 0.3 | -0.01 | 0.01 |
| gene-LOC105280104 | https://www.ncbi.nlm.nih.gov/gene/?term=LOC105280104 | 3960 | 120 | 303.03 | 1 | 2.53 | 0.83 | 14.59 | 14.9 | -0.31 | 0.31 | -0.01 | 0.01 |
| gene-LOC105288078 | https://www.ncbi.nlm.nih.gov/gene/?term=LOC105288078 | 19126 | 318 | 166.27 | 1 | 0.52 | 0.31 | 17.33 | 17.64 | -0.31 | 0.31 | -0.01 | 0.01 |
| gene-LOC105274531 | https://www.ncbi.nlm.nih.gov/gene/?term=LOC105274531 | 7774 | 257 | 330.59 | 1 | 1.29 | 0.39 | 11.81 | 12.14 | -0.32 | 0.32 | -0.01 | 0.01 |
| gene-LOC105278534 | https://www.ncbi.nlm.nih.gov/gene/?term=LOC105278534 | 7364 | 185 | 251.22 | 1 | 1.36 | 0.54 | 17.95 | 18.29 | -0.33 | 0.33 | -0.01 | 0.01 |
| gene-LOC105278384 | https://www.ncbi.nlm.nih.gov/gene/?term=LOC105278384 | 1390 | 33 | 237.41 | 1 | 7.19 | 3.03 | 45.97 | 46.32 | -0.35 | 0.35 | -0.01 | 0.01 |
| gene-LOC105279038 | https://www.ncbi.nlm.nih.gov/gene/?term=LOC105279038 | 9453 | 321 | 339.57 | 1 | 1.06 | 0.31 | 8.59 | 8.94 | -0.35 | 0.35 | -0.01 | 0.01 |
| gene-LOC105287704 | https://www.ncbi.nlm.nih.gov/gene/?term=LOC105287704 | 3019 | 125 | 414.04 | 1 | 3.31 | 0.8 | 21.19 | 21.56 | -0.37 | 0.37 | -0.02 | 0.02 |
| gene-LOC105278228 | https://www.ncbi.nlm.nih.gov/gene/?term=LOC105278228 | 8610 | 163 | 189.31 | 1 | 1.16 | 0.61 | 23.66 | 24.04 | -0.38 | 0.38 | -0.01 | 0.01 |
| gene-LOC105280956 | https://www.ncbi.nlm.nih.gov/gene/?term=LOC105280956 | 2639 | 73 | 276.62 | 1 | 3.79 | 1.37 | 20.28 | 20.69 | -0.4 | 0.4 | -0.01 | 0.01 |
| gene-LOC105284184 | https://www.ncbi.nlm.nih.gov/gene/?term=LOC105284184 | 2535 | 82 | 323.47 | 1 | 3.94 | 1.22 | 34.56 | 34.95 | -0.4 | 0.4 | -0.01 | 0.01 |
| gene-LOC105280786 | https://www.ncbi.nlm.nih.gov/gene/?term=LOC105280786 | 2573 | 122 | 474.15 | 1 | 3.89 | 0.82 | 29.15 | 29.56 | -0.41 | 0.41 | -0.02 | 0.02 |
| gene-LOC105285534 | https://www.ncbi.nlm.nih.gov/gene/?term=LOC105285534 | 3944 | 224 | 567.95 | 1 | 2.54 | 0.45 | 11.27 | 11.7 | -0.42 | 0.42 | -0.02 | 0.02 |
| gene-LOC105275451 | https://www.ncbi.nlm.nih.gov/gene/?term=LOC105275451 | 6035 | 118 | 195.53 | 1 | 1.66 | 0.85 | 22.19 | 22.63 | -0.44 | 0.44 | -0.01 | 0.01 |
| gene-LOC105288237 | https://www.ncbi.nlm.nih.gov/gene/?term=LOC105288237 | 2226 | 47 | 211.14 | 1 | 4.49 | 2.13 | 35.68 | 36.15 | -0.47 | 0.47 | -0.01 | 0.01 |
| gene-LOC105278283 | https://www.ncbi.nlm.nih.gov/gene/?term=LOC105278283 | 2317 | 130 | 561.07 | 1 | 4.32 | 0.77 | 24.9 | 25.39 | -0.48 | 0.48 | -0.03 | 0.03 |
| gene-LOC105284043 | https://www.ncbi.nlm.nih.gov/gene/?term=LOC105284043 | 5661 | 248 | 438.09 | 1 | 1.77 | 0.4 | 11.5 | 11.98 | -0.48 | 0.48 | -0.02 | 0.02 |
| gene-LOC105277435 | https://www.ncbi.nlm.nih.gov/gene/?term=LOC105277435 | 3643 | 158 | 433.71 | 1 | 2.74 | 0.63 | 12.84 | 13.36 | -0.52 | 0.52 | -0.02 | 0.02 |
| gene-LOC105282825 | https://www.ncbi.nlm.nih.gov/gene/?term=LOC105282825 | 3919 | 90 | 229.65 | 1 | 2.55 | 1.11 | 15.34 | 15.86 | -0.52 | 0.52 | -0.01 | 0.01 |
| gene-LOC105288079 | https://www.ncbi.nlm.nih.gov/gene/?term=LOC105288079 | 3473 | 89 | 256.26 | 1 | 2.88 | 1.12 | 19.96 | 20.49 | -0.52 | 0.52 | -0.01 | 0.01 |
| gene-LOC105276527 | https://www.ncbi.nlm.nih.gov/gene/?term=LOC105276527 | 7923 | 351 | 443.01 | 1 | 1.26 | 0.28 | 8.2 | 8.78 | -0.58 | 0.58 | -0.03 | 0.03 |
| gene-LOC105280410 | https://www.ncbi.nlm.nih.gov/gene/?term=LOC105280410 | 12580 | 260 | 206.68 | 1 | 0.79 | 0.38 | 19.03 | 19.62 | -0.59 | 0.59 | -0.01 | 0.01 |
| gene-LOC105275966 | https://www.ncbi.nlm.nih.gov/gene/?term=LOC105275966 | 3299 | 59 | 178.84 | 1 | 3.03 | 1.69 | 50.28 | 50.9 | -0.62 | 0.62 | -0.01 | 0.01 |
| gene-LOC105284355 | https://www.ncbi.nlm.nih.gov/gene/?term=LOC105284355 | 3904 | 147 | 376.54 | 1 | 2.56 | 0.68 | 21.32 | 21.98 | -0.67 | 0.67 | -0.03 | 0.03 |
| gene-LOC105286274 | https://www.ncbi.nlm.nih.gov/gene/?term=LOC105286274 | 3817 | 183 | 479.43 | 1 | 2.62 | 0.55 | 15.39 | 16.06 | -0.67 | 0.67 | -0.03 | 0.03 |
| gene-LOC105280641 | https://www.ncbi.nlm.nih.gov/gene/?term=LOC105280641 | 3682 | 66 | 179.25 | 1 | 2.72 | 1.52 | 15.43 | 16.12 | -0.68 | 0.68 | -0.01 | 0.01 |
| gene-LOC105286770 | https://www.ncbi.nlm.nih.gov/gene/?term=LOC105286770 | 3055 | 38 | 124.39 | 1 | 3.27 | 2.63 | 11.1 | 11.78 | -0.68 | 0.68 | -0.01 | 0.01 |
| gene-LOC105285179 | https://www.ncbi.nlm.nih.gov/gene/?term=LOC105285179 | 4191 | 156 | 372.23 | 1 | 2.39 | 0.64 | 17.53 | 18.23 | -0.69 | 0.69 | -0.03 | 0.03 |
| gene-LOC105283728 | https://www.ncbi.nlm.nih.gov/gene/?term=LOC105283728 | 3256 | 118 | 362.41 | 2 | 6.14 | 1.69 | 19.18 | 19.87 | -0.7 | 0.7 | -0.03 | 0.03 |
| gene-LOC105282600 | https://www.ncbi.nlm.nih.gov/gene/?term=LOC105282600 | 3715 | 65 | 174.97 | 1 | 2.69 | 1.54 | 26.29 | 27.03 | -0.73 | 0.73 | -0.01 | 0.01 |
| gene-LOC105276449 | https://www.ncbi.nlm.nih.gov/gene/?term=LOC105276449 | 1231 | 48 | 389.93 | 1 | 8.12 | 2.08 | 26.7 | 27.43 | -0.74 | 0.74 | -0.03 | 0.03 |
| gene-LOC105276917 | https://www.ncbi.nlm.nih.gov/gene/?term=LOC105276917 | 2336 | 26 | 111.3 1 | 4.28 | 3.85 | 36.57 | 37.3 | -0.74 | 0.74 | -0.01 | 0.01 | |
| gene-LOC105275752 | https://www.ncbi.nlm.nih.gov/gene/?term=LOC105275752 | 3462 | 123 | 355.29 | 1 | 2.89 | 0.81 | 28.75 | 29.5 | -0.75 | 0.75 | -0.03 | 0.03 |
| gene-LOC105283132 | https://www.ncbi.nlm.nih.gov/gene/?term=LOC105283132 | 1819 | 54 | 296.87 | 1 | 5.5 | 1.85 | 40.82 | 41.59 | -0.77 | 0.77 | -0.02 | 0.02 |
| gene-LOC105280774 | https://www.ncbi.nlm.nih.gov/gene/?term=LOC105280774 | 2793 | 117 | 418.9 1 | 3.58 | 0.85 | 27.44 | 28.24 | -0.79 | 0.79 | -0.03 | 0.03 | |
| gene-LOC105282067 | https://www.ncbi.nlm.nih.gov/gene/?term=LOC105282067 | 5941 | 54 | 90.89 1 | 1.68 | 1.85 | 23.11 | 23.91 | -0.8 | 0.8 | -0.01 | 0.01 | |
| gene-LOC105282563 | https://www.ncbi.nlm.nih.gov/gene/?term=LOC105282563 | 4353 | 146 | 335.4 1 | 2.3 | 0.68 | 15.5 | 16.32 | -0.83 | 0.83 | -0.03 | 0.03 | |
| gene-LOC105274856 | https://www.ncbi.nlm.nih.gov/gene/?term=LOC105274856 | 5216 | 208 | 398.77 | 1 | 1.92 | 0.48 | 18.49 | 19.33 | -0.84 | 0.84 | -0.03 | 0.03 |
| gene-LOC105274706 | https://www.ncbi.nlm.nih.gov/gene/?term=LOC105274706 | 1446 | 52 | 359.61 | 1 | 6.92 | 1.92 | 37.01 | 37.87 | -0.86 | 0.86 | -0.03 | 0.03 |
| gene-LOC105274773 | https://www.ncbi.nlm.nih.gov/gene/?term=LOC105274773 | 11034 | 167 | 151.35 | 1 | 0.91 | 0.6 | 16.37 | 17.25 | -0.88 | 0.88 | -0.01 | 0.01 |
| gene-LOC105275363 | https://www.ncbi.nlm.nih.gov/gene/?term=LOC105275363 | 3798 | 159 | 418.64 | 1 | 2.63 | 0.63 | 16.97 | 17.86 | -0.88 | 0.88 | -0.04 | 0.04 |
| gene-LOC105279657 | https://www.ncbi.nlm.nih.gov/gene/?term=LOC105279657 | 5878 | 84 | 142.91 | 1 | 1.7 | 1.19 | 18.69 | 19.56 | -0.88 | 0.88 | -0.01 | 0.01 |
| gene-LOC105274590 | https://www.ncbi.nlm.nih.gov/gene/?term=LOC105274590 | 4516 | 81 | 179.36 | 1 | 2.21 | 1.23 | 21.65 | 22.55 | -0.9 | 0.9 | -0.02 | 0.02 |
| gene-LOC105277216 | https://www.ncbi.nlm.nih.gov/gene/?term=LOC105277216 | 7627 | 89 | 116.69 | 1 | 1.31 | 1.12 | 6.69 | 7.62 | -0.93 | 0.93 | -0.01 | 0.01 |
| gene-LOC105280447 | https://www.ncbi.nlm.nih.gov/gene/?term=LOC105280447 | 2215 | 28 | 126.41 | 1 | 4.51 | 3.57 | 23.87 | 24.81 | -0.94 | 0.94 | -0.01 | 0.01 |
| gene-LOC105285406 | https://www.ncbi.nlm.nih.gov/gene/?term=LOC105285406 | 8015 | 42 | 52.4 1 | 1.25 | 2.38 | 35.2 | 36.16 | -0.96 | 0.96 | -0.01 | 0.01 | |
| gene-LOC105279298 | https://www.ncbi.nlm.nih.gov/gene/?term=LOC105279298 | 2773 | 138 | 497.66 | 2 | 7.21 | 1.45 | 22.22 | 23.22 | -1 | 1 | -0.05 | 0.05 |
| gene-LOC105282491 | https://www.ncbi.nlm.nih.gov/gene/?term=LOC105282491 | 4761 | 163 | 342.37 | 1 | 2.1 | 0.61 | 9.65 | 10.67 | -1.02 | 1.02 | -0.03 | 0.03 |
| gene-LOC105278497 | https://www.ncbi.nlm.nih.gov/gene/?term=LOC105278497 | 2072 | 100 | 482.63 | 1 | 4.83 | 1 | 18.86 | 19.92 | -1.06 | 1.06 | -0.05 | 0.05 |
| gene-LOC105275179 | https://www.ncbi.nlm.nih.gov/gene/?term=LOC105275179 | 3271 | 68 | 207.89 | 1 | 3.06 | 1.47 | 14.88 | 16 | -1.12 | 1.12 | -0.02 | 0.02 |
| gene-LOC105282947 | https://www.ncbi.nlm.nih.gov/gene/?term=LOC105282947 | 1162 | 49 | 421.69 | 1 | 8.61 | 2.04 | 29.51 | 30.69 | -1.17 | 1.17 | -0.05 | 0.05 |
| gene-LOC105278962 | https://www.ncbi.nlm.nih.gov/gene/?term=LOC105278962 | 4491 | 88 | 195.95 | 1 | 2.23 | 1.14 | 23.85 | 25.09 | -1.24 | 1.24 | -0.02 | 0.02 |
| gene-LOC105275683 | https://www.ncbi.nlm.nih.gov/gene/?term=LOC105275683 | 4256 | 203 | 476.97 | 1 | 2.35 | 0.49 | 12.81 | 14.05 | -1.25 | 1.25 | -0.06 | 0.06 |
| gene-LOC105283943 | https://www.ncbi.nlm.nih.gov/gene/?term=LOC105283943 | 3559 | 60 | 168.59 | 1 | 2.81 | 1.67 | 21.58 | 22.83 | -1.25 | 1.25 | -0.02 | 0.02 |
| gene-LOC105285742 | https://www.ncbi.nlm.nih.gov/gene/?term=LOC105285742 | 3446 | 100 | 290.19 | 1 | 2.9 | 1 | 24.69 | 25.96 | -1.27 | 1.27 | -0.04 | 0.04 |
| gene-LOC105281700 | https://www.ncbi.nlm.nih.gov/gene/?term=LOC105281700 | 2208 | 46 | 208.33 | 1 | 4.53 | 2.17 | 39.97 | 41.25 | -1.28 | 1.28 | -0.03 | 0.03 |
| gene-LOC105275460 | https://www.ncbi.nlm.nih.gov/gene/?term=LOC105275460 | 2473 | 112 | 452.89 | 1 | 4.04 | 0.89 | 12.89 | 14.18 | -1.3 | 1.3 | -0.06 | 0.06 |
| gene-LOC105278282 | https://www.ncbi.nlm.nih.gov/gene/?term=LOC105278282 | 1775 | 47 | 264.79 | 1 | 5.63 | 2.13 | 45.94 | 47.29 | -1.35 | 1.35 | -0.04 | 0.04 |
| gene-LOC105279500 | https://www.ncbi.nlm.nih.gov/gene/?term=LOC105279500 | 3587 | 33 | 92 1 | 2.79 | 3.03 | 49.6 | 51 | -1.4 | 1.4 | -0.01 | 0.01 | |
| gene-LOC105285055 | https://www.ncbi.nlm.nih.gov/gene/?term=LOC105285055 | 4108 | 217 | 528.24 | 1 | 2.43 | 0.46 | 12.76 | 14.18 | -1.42 | 1.42 | -0.07 | 0.07 |
| gene-LOC105277631 | https://www.ncbi.nlm.nih.gov/gene/?term=LOC105277631 | 4387 | 289 | 658.76 | 1 | 2.28 | 0.35 | 9.79 | 11.22 | -1.43 | 1.43 | -0.09 | 0.09 |
| gene-LOC105283592 | https://www.ncbi.nlm.nih.gov/gene/?term=LOC105283592 | 3479 | 71 | 204.08 | 1 | 2.87 | 1.41 | 24.39 | 25.83 | -1.45 | 1.45 | -0.03 | 0.03 |
| gene-LOC105275347 | https://www.ncbi.nlm.nih.gov/gene/?term=LOC105275347 | 2526 | 14 | 55.42 1 | 3.96 | 7.14 | 35.76 | 37.28 | -1.52 | 1.52 | -0.01 | 0.01 | |

| | | | | | | | | | | | | | |
|---|---|---|---|---|---|---|---|---|---|---|---|---|---|
| gene-LOC105276078 | https://www.ncbi.nlm.nih.gov/gene/?term=LOC105276078 | 1051 | 10 | 95.15 | 1 | 9.51 | 10 | 37.29 | 38.89 | -1.6 | 1.6 | -0.02 | 0.02 |
| gene-LOC105287063 | https://www.ncbi.nlm.nih.gov/gene/?term=LOC105287063 | 1874 | 46 | 245.46 | 1 | 5.34 | 2.17 | 34.15 | 35.78 | -1.63 | 1.63 | -0.04 | 0.04 |
| gene-LOC105285695 | https://www.ncbi.nlm.nih.gov/gene/?term=LOC105285695 | 1371 | 62 | 452.22 | 1 | 7.29 | 1.61 | 23 | 24.72 | -1.72 | 1.72 | -0.08 | 0.08 |
| gene-LOC105279356 | https://www.ncbi.nlm.nih.gov/gene/?term=LOC105279356 | 1314 | 42 | 319.63 | 1 | 7.61 | 2.38 | 28.83 | 30.58 | -1.75 | 1.75 | -0.06 | 0.06 |
| gene-LOC105285671 | https://www.ncbi.nlm.nih.gov/gene/?term=LOC105285671 | 2263 | 95 | 419.8 | 1 | 4.42 | 1.05 | 27.86 | 29.64 | -1.78 | 1.78 | -0.07 | 0.07 |
| gene-LOC105283989 | https://www.ncbi.nlm.nih.gov/gene/?term=LOC105283989 | 4948 | 34 | 68.71 | 2 | 4.04 | 5.88 | 44.42 | 46.21 | -1.79 | 1.79 | -0.01 | 0.01 |
| gene-LOC105284704 | https://www.ncbi.nlm.nih.gov/gene/?term=LOC105284704 | 2400 | 65 | 270.83 | 1 | 4.17 | 1.54 | 25.06 | 26.85 | -1.79 | 1.79 | -0.05 | 0.05 |
| gene-LOC105276692 | https://www.ncbi.nlm.nih.gov/gene/?term=LOC105276692 | 1745 | 37 | 212.03 | 1 | 5.73 | 2.7 | 20.82 | 22.64 | -1.82 | 1.82 | -0.04 | 0.04 |
| gene-LOC105274900 | https://www.ncbi.nlm.nih.gov/gene/?term=LOC105274900 | 5561 | 56 | 100.7 | 2 | 3.6 | 3.57 | 26.85 | 28.69 | -1.84 | 1.84 | -0.02 | 0.02 |
| gene-LOC105278260 | https://www.ncbi.nlm.nih.gov/gene/?term=LOC105278260 | 3240 | 41 | 126.54 | 1 | 3.09 | 2.44 | 13.01 | 14.86 | -1.85 | 1.85 | -0.02 | 0.02 |
| gene-LOC105276745 | https://www.ncbi.nlm.nih.gov/gene/?term=LOC105276745 | 3953 | 73 | 184.67 | 1 | 2.53 | 1.37 | 21.09 | 23.02 | -1.93 | 1.93 | -0.04 | 0.04 |
| gene-LOC105278807 | https://www.ncbi.nlm.nih.gov/gene/?term=LOC105278807 | 2166 | 72 | 332.41 | 1 | 4.62 | 1.39 | 18.53 | 20.46 | -1.93 | 1.93 | -0.06 | 0.06 |
| gene-LOC105286223 | https://www.ncbi.nlm.nih.gov/gene/?term=LOC105286223 | 2225 | 83 | 373.03 | 1 | 4.49 | 1.2 | 30.2 | 32.18 | -1.98 | 1.98 | -0.07 | 0.07 |
| gene-LOC105286452 | https://www.ncbi.nlm.nih.gov/gene/?term=LOC105286452 | 6888 | 58 | 84.2 | 1 | 1.45 | 1.72 | 31.25 | 33.23 | -1.98 | 1.98 | -0.02 | 0.02 |
| gene-LOC105280348 | https://www.ncbi.nlm.nih.gov/gene/?term=LOC105280348 | 3797 | 52 | 136.95 | 1 | 2.63 | 1.92 | 25.54 | 27.53 | -1.99 | 1.99 | -0.03 | 0.03 |
| gene-LOC105282156 | https://www.ncbi.nlm.nih.gov/gene/?term=LOC105282156 | 2577 | 136 | 527.75 | 1 | 3.88 | 0.74 | 23.86 | 26.02 | -2.16 | 2.16 | -0.11 | 0.11 |
| gene-LOC105285583 | https://www.ncbi.nlm.nih.gov/gene/?term=LOC105285583 | 1947 | 51 | 261.94 | 1 | 5.14 | 1.96 | 28.26 | 30.43 | -2.17 | 2.17 | -0.06 | 0.06 |
| gene-LOC105282893 | https://www.ncbi.nlm.nih.gov/gene/?term=LOC105282893 | 2645 | 43 | 162.57 | 1 | 3.78 | 2.33 | 27.48 | 29.68 | -2.19 | 2.19 | -0.04 | 0.04 |
| gene-LOC105286644 | https://www.ncbi.nlm.nih.gov/gene/?term=LOC105286644 | 1235 | 22 | 178.14 | 1 | 8.1 | 4.55 | 42.2 | 44.4 | -2.2 | 2.2 | -0.04 | 0.04 |
| gene-LOC105280002 | https://www.ncbi.nlm.nih.gov/gene/?term=LOC105280002 | 1940 | 30 | 154.64 | 1 | 5.15 | 3.33 | 33.39 | 35.64 | -2.25 | 2.25 | -0.03 | 0.03 |
| gene-LOC105281515 | https://www.ncbi.nlm.nih.gov/gene/?term=LOC105281515 | 8827 | 41 | 46.45 | 1 | 1.13 | 2.44 | 32.87 | 35.23 | -2.37 | 2.37 | -0.01 | 0.01 |
| gene-LOC105274534 | https://www.ncbi.nlm.nih.gov/gene/?term=LOC105274534 | 1329 | 57 | 428.89 | 1 | 7.52 | 1.75 | 43.75 | 46.14 | -2.39 | 2.39 | -0.1 | 0.1 |
| gene-LOC105287417 | https://www.ncbi.nlm.nih.gov/gene/?term=LOC105287417 | 1025 | 33 | 321.95 | 1 | 9.76 | 3.03 | 36 | 38.42 | -2.42 | 2.42 | -0.08 | 0.08 |
| gene-LOC105275518 | https://www.ncbi.nlm.nih.gov/gene/?term=LOC105275518 | 2335 | 21 | 89.94 | 1 | 4.28 | 4.76 | 45.03 | 47.49 | -2.45 | 2.45 | -0.02 | 0.02 |
| gene-LOC105288066 | https://www.ncbi.nlm.nih.gov/gene/?term=LOC105288066 | 1964 | 26 | 132.38 | 1 | 5.09 | 3.85 | 26.28 | 28.74 | -2.45 | 2.45 | -0.03 | 0.03 |
| gene-LOC105287854 | https://www.ncbi.nlm.nih.gov/gene/?term=LOC105287854 | 3644 | 57 | 156.42 | 1 | 2.74 | 1.75 | 21.16 | 23.71 | -2.55 | 2.55 | -0.04 | 0.04 |
| gene-LOC105286502 | https://www.ncbi.nlm.nih.gov/gene/?term=LOC105286502 | 1286 | 44 | 342.15 | 1 | 7.78 | 2.27 | 23.97 | 26.78 | -2.81 | 2.81 | -0.1 | 0.1 |
| gene-LOC105276704 | https://www.ncbi.nlm.nih.gov/gene/?term=LOC105276704 | 2063 | 87 | 421.72 | 1 | 4.85 | 1.15 | 12.34 | 15.21 | -2.86 | 2.86 | -0.12 | 0.12 |
| gene-LOC105276538 | https://www.ncbi.nlm.nih.gov/gene/?term=LOC105276538 | 1511 | 19 | 125.74 | 1 | 6.62 | 5.26 | 72.68 | 75.55 | -2.87 | 2.87 | -0.04 | 0.04 |
| gene-LOC105277215 | https://www.ncbi.nlm.nih.gov/gene/?term=LOC105277215 | 2619 | 29 | 110.73 | 1 | 3.82 | 3.45 | 19.9 | 22.79 | -2.88 | 2.88 | -0.03 | 0.03 |
| gene-LOC105285674 | https://www.ncbi.nlm.nih.gov/gene/?term=LOC105285674 | 3035 | 91 | 299.84 | 2 | 6.59 | 2.2 | 23.1 | 26.01 | -2.91 | 2.91 | -0.09 | 0.09 |
| gene-LOC105283712 | https://www.ncbi.nlm.nih.gov/gene/?term=LOC105283712 | 1910 | 27 | 141.36 | 1 | 5.24 | 3.7 | 29.17 | 32.5 | -3.33 | 3.33 | -0.05 | 0.05 |
| gene-LOC105278109 | https://www.ncbi.nlm.nih.gov/gene/?term=LOC105278109 | 4029 | 62 | 153.88 | 1 | 2.48 | 1.61 | 43.07 | 46.49 | -3.42 | 3.42 | -0.05 | 0.05 |
| gene-LOC105282399 | https://www.ncbi.nlm.nih.gov/gene/?term=LOC105282399 | 2682 | 22 | 82.03 | 1 | 3.73 | 4.55 | 32.93 | 36.36 | -3.43 | 3.43 | -0.03 | 0.03 |
| gene-LOC113561805 | https://www.ncbi.nlm.nih.gov/gene/?term=LOC113561805 | 3337 | 64 | 191.79 | 1 | 3 | 1.56 | 37.95 | 41.4 | -3.45 | 3.45 | -0.07 | 0.07 |
| gene-LOC105275190 | https://www.ncbi.nlm.nih.gov/gene/?term=LOC105275190 | 3837 | 74 | 192.86 | 1 | 2.61 | 1.35 | 18.39 | 22.03 | -3.64 | 3.64 | -0.07 | 0.07 |
| gene-LOC105274778 | https://www.ncbi.nlm.nih.gov/gene/?term=LOC105274778 | 4662 | 38 | 81.51 | 1 | 2.15 | 2.63 | 44.96 | 48.85 | -3.88 | 3.88 | -0.03 | 0.03 |
| gene-LOC105281848 | https://www.ncbi.nlm.nih.gov/gene/?term=LOC105281848 | 2122 | 47 | 221.49 | 1 | 4.71 | 2.13 | 23.11 | 27.19 | -4.08 | 4.08 | -0.09 | 0.09 |
| gene-LOC105285060 | https://www.ncbi.nlm.nih.gov/gene/?term=LOC105285060 | 1285 | 15 | 116.73 | 1 | 7.78 | 6.67 | 48.46 | 52.7 | -4.25 | 4.25 | -0.05 | 0.05 |
| gene-LOC105277856 | https://www.ncbi.nlm.nih.gov/gene/?term=LOC105277856 | 3442 | 29 | 84.25 | 1 | 2.91 | 3.45 | 35.44 | 40.05 | -4.6 | 4.6 | -0.04 | 0.04 |
| gene-LOC105281039 | https://www.ncbi.nlm.nih.gov/gene/?term=LOC105281039 | 1471 | 13 | 88.38 | 1 | 6.8 | 7.69 | 36.12 | 40.81 | -4.7 | 4.7 | -0.04 | 0.04 |
| gene-LOC105286983 | https://www.ncbi.nlm.nih.gov/gene/?term=LOC105286983 | 16901 | 68 | 40.23 | 1 | 0.59 | 1.47 | 0.94 | 5.77 | -4.83 | 4.83 | -0.02 | 0.02 |
| gene-LOC105285514 | https://www.ncbi.nlm.nih.gov/gene/?term=LOC105285514 | 1277 | 9 | 70.48 | 1 | 7.83 | 11.11 | 73.21 | 78.44 | -5.23 | 5.23 | -0.04 | 0.04 |
| gene-LOC105284609 | https://www.ncbi.nlm.nih.gov/gene/?term=LOC105284609 | 5678 | 31 | 54.6 | 1 | 1.76 | 3.23 | 35.81 | 41.06 | -5.25 | 5.25 | -0.03 | 0.03 |
| gene-LOC105275708 | https://www.ncbi.nlm.nih.gov/gene/?term=LOC105275708 | 1772 | 14 | 79.01 | 1 | 5.64 | 7.14 | 44.58 | 50.56 | -5.98 | 5.98 | -0.05 | 0.05 |
| gene-LOC105279898 | https://www.ncbi.nlm.nih.gov/gene/?term=LOC105279898 | 1343 | 16 | 119.14 | 2 | 14.89 | 12.5 | 59.42 | 68.99 | -9.57 | 9.57 | -0.11 | 0.11 |
| gene-LOC105275070 | https://www.ncbi.nlm.nih.gov/gene/?term=LOC105275070 | 5667 | 38 | 67.05 | 3 | 5.29 | 7.89 | 3.63 | 13.45 | -9.82 | 9.82 | -0.07 | 0.07 |
| gene-LOC113562345 | https://www.ncbi.nlm.nih.gov/gene/?term=LOC113562345 | 2290 | 62 | 270.74 | 16 | 69.87 | 25.81 | 17.91 | 33 | -15.09 | 15.09 | -0.41 | 0.41 |

Directory: OGLall File: ogl-Ob_LI_BCphaseA1.vs.RphaseB1.txt

| gene_ID | gene_link | gwidth | #Sites | #per10Kb | #dmSites | #dmsp10kb | %dmSites | %pSite1 | %pSite2 | DMpSite | ADMpSite | DMpNucl | ADMpNucl |
|---|---|---|---|---|---|---|---|---|---|---|---|---|---|
| gene-LOC113562975 | https://www.ncbi.nlm.nih.gov/gene/?term=LOC113562975 | 2751 | 3 | 10.91 | 1 | 3.64 | 33.33 | 46.98 | 30.96 | 16.02 | 16.02 | 0.02 | 0.02 |
| gene-LOC105284474 | https://www.ncbi.nlm.nih.gov/gene/?term=LOC105284474 | 479 | 2 | 41.75 | 1 | 20.88 | 50 | 33.34 | 19.05 | 14.29 | 14.29 | 0.06 | 0.06 |
| gene-LOC113562831 | https://www.ncbi.nlm.nih.gov/gene/?term=LOC113562831 | 574 | 2 | 34.84 | 1 | 17.42 | 50 | 18.75 | 6.94 | 11.8 | 11.8 | 0.04 | 0.04 |
| gene-LOC105288227 | https://www.ncbi.nlm.nih.gov/gene/?term=LOC105288227 | 1245 | 3 | 24.1 | 1 | 8.03 | 33.33 | 86.21 | 76.94 | 9.26 | 9.26 | 0.02 | 0.02 |
| gene-LOC113561934 | https://www.ncbi.nlm.nih.gov/gene/?term=LOC113561934 | 1252 | 16 | 127.8 | 4 | 31.95 | 25 | 36.29 | 28.08 | 8.21 | 8.21 | 0.1 | 0.1 |
| gene-LOC105276219 | https://www.ncbi.nlm.nih.gov/gene/?term=LOC105276219 | 916 | 6 | 65.5 | 1 | 10.92 | 16.67 | 47.07 | 39.74 | 7.33 | 7.33 | 0.05 | 0.05 |
| gene-LOC105275343 | https://www.ncbi.nlm.nih.gov/gene/?term=LOC105275343 | 1746 | 4 | 22.91 | 1 | 5.73 | 25 | 47 | 39.86 | 7.14 | 7.14 | 0.02 | 0.02 |
| gene-LOC105282405 | https://www.ncbi.nlm.nih.gov/gene/?term=LOC105282405 | 1310 | 31 | 236.64 | 2 | 15.27 | 6.45 | 51.18 | 44.54 | 6.64 | 6.64 | 0.16 | 0.16 |
| gene-LOC113562627 | https://www.ncbi.nlm.nih.gov/gene/?term=LOC113562627 | 397 | 10 | 251.89 | 1 | 25.19 | 10 | 77.44 | 70.89 | 6.56 | 6.56 | 0.17 | 0.17 |
| gene-LOC105285885 | https://www.ncbi.nlm.nih.gov/gene/?term=LOC105285885 | 1327 | 12 | 90.43 | 1 | 7.54 | 8.33 | 54.12 | 47.77 | 6.35 | 6.35 | 0.06 | 0.06 |
| gene-LOC105286816 | https://www.ncbi.nlm.nih.gov/gene/?term=LOC105286816 | 4880 | 9 | 18.44 | 1 | 2.05 | 11.11 | 37.19 | 31.02 | 6.17 | 6.17 | 0.01 | 0.01 |
| gene-LOC105281226 | https://www.ncbi.nlm.nih.gov/gene/?term=LOC105281226 | 1308 | 10 | 76.45 | 1 | 7.65 | 10 | 77.35 | 71.27 | 6.08 | 6.08 | 0.05 | 0.05 |
| gene-LOC113563111 | https://www.ncbi.nlm.nih.gov/gene/?term=LOC113563111 | 2455 | 35 | 142.57 | 2 | 8.15 | 5.71 | 26.92 | 21.34 | 5.57 | 5.57 | 0.08 | 0.08 |
| gene-LOC105283973 | https://www.ncbi.nlm.nih.gov/gene/?term=LOC105283973 | 1536 | 29 | 188.8 | 4 | 26.04 | 13.79 | 61.88 | 56.34 | 5.55 | 5.55 | 0.1 | 0.1 |
| gene-LOC105285241 | https://www.ncbi.nlm.nih.gov/gene/?term=LOC105285241 | 5118 | 25 | 48.85 | 2 | 3.91 | 8 | 30.97 | 25.5 | 5.47 | 5.47 | 0.03 | 0.03 |
| gene-LOC105284412 | https://www.ncbi.nlm.nih.gov/gene/?term=LOC105284412 | 829 | 11 | 132.69 | 1 | 12.06 | 9.09 | 56.74 | 51.39 | 5.35 | 5.35 | 0.07 | 0.07 |
| gene-LOC105278541 | https://www.ncbi.nlm.nih.gov/gene/?term=LOC105278541 | 1563 | 16 | 102.37 | 1 | 6.4 | 6.25 | 28.66 | 23.32 | 5.33 | 5.33 | 0.05 | 0.05 |
| gene-LOC105275539 | https://www.ncbi.nlm.nih.gov/gene/?term=LOC105275539 | 1272 | 23 | 180.82 | 2 | 15.72 | 8.7 | 20.8 | 15.48 | 5.32 | 5.32 | 0.1 | 0.1 |
| gene-LOC105281507 | https://www.ncbi.nlm.nih.gov/gene/?term=LOC105281507 | 881 | 5 | 56.75 | 1 | 11.35 | 20 | 11.82 | 6.67 | 5.15 | 5.15 | 0.03 | 0.03 |
| gene-LOC105282996 | https://www.ncbi.nlm.nih.gov/gene/?term=LOC105282996 | 3736 | 9 | 24.09 | 2 | 5.35 | 22.22 | 28.24 | 23.6 | 4.64 | 4.64 | 0.01 | 0.01 |
| gene-LOC105283988 | https://www.ncbi.nlm.nih.gov/gene/?term=LOC105283988 | 2510 | 14 | 55.78 | 1 | 3.98 | 7.14 | 40.87 | 36.23 | 4.64 | 4.64 | 0.03 | 0.03 |
| gene-LOC105285051 | https://www.ncbi.nlm.nih.gov/gene/?term=LOC105285051 | 1357 | 10 | 73.69 | 1 | 7.37 | 10 | 46.31 | 41.71 | 4.6 | 4.6 | 0.03 | 0.03 |
| gene-LOC105280926 | https://www.ncbi.nlm.nih.gov/gene/?term=LOC105280926 | 2139 | 7 | 32.73 | 1 | 4.68 | 14.29 | 49.74 | 45.26 | 4.48 | 4.48 | 0.01 | 0.01 |
| gene-LOC105284881 | https://www.ncbi.nlm.nih.gov/gene/?term=LOC105284881 | 2016 | 20 | 99.21 | 2 | 9.92 | 10 | 27.57 | 23.15 | 4.42 | 4.42 | 0.04 | 0.04 |

...
... only first 25 lines shown ...

Directory: OGLall File: ogl-Ob_LI_BCphaseA1.vs.RphaseB2.txt

| gene_ID | gene_link | gwidth | #Sites | #per10Kb | #dmSites | #dmsp10kb | %dmSites | %pSite1 | %pSite2 | DMpSite | ADMpSite | DMpNucl | ADMpNucl |
|---|---|---|---|---|---|---|---|---|---|---|---|---|---|
| gene-LOC105288227 | https://www.ncbi.nlm.nih.gov/gene/?term=LOC105288227 | 1245 | 2 | 16.06 | 1 | 8.03 | 50 | 79.31 | 55.55 | 23.76 | 23.76 | 0.04 | 0.04 |
| gene-LOC105285431 | https://www.ncbi.nlm.nih.gov/gene/?term=LOC105285431 | 1550 | 5 | 32.26 | 2 | 12.9 | 40 | 49.86 | 35.35 | 14.51 | 14.51 | 0.05 | 0.05 |
| gene-LOC113562831 | https://www.ncbi.nlm.nih.gov/gene/?term=LOC113562831 | 574 | 2 | 34.84 | 1 | 17.42 | 50 | 18.75 | 5.88 | 12.87 | 12.87 | 0.04 | 0.04 |
| gene-LOC105287814 | https://www.ncbi.nlm.nih.gov/gene/?term=LOC105287814 | 892 | 5 | 56.05 | 2 | 22.42 | 40 | 36.02 | 23.39 | 12.63 | 12.63 | 0.07 | 0.07 |
| gene-LOC105283410 | https://www.ncbi.nlm.nih.gov/gene/?term=LOC105283410 | 1423 | 5 | 35.14 | 2 | 14.05 | 40 | 52.44 | 40.2 | 12.24 | 12.24 | 0.04 | 0.04 |
| gene-LOC105283241 | https://www.ncbi.nlm.nih.gov/gene/?term=LOC105283241 | 2011 | 4 | 19.89 | 1 | 4.97 | 25 | 39.29 | 29.03 | 10.26 | 10.26 | 0.02 | 0.02 |
| gene-LOC109611453 | https://www.ncbi.nlm.nih.gov/gene/?term=LOC109611453 | 3541 | 6 | 16.94 | 1 | 2.82 | 16.67 | 24.1 | 14.5 | 9.6 | 9.6 | 0.02 | 0.02 |
| gene-LOC113561934 | https://www.ncbi.nlm.nih.gov/gene/?term=LOC113561934 | 1252 | 15 | 119.81 | 2 | 15.97 | 13.33 | 36.71 | 27.17 | 9.54 | 9.54 | 0.11 | 0.11 |
| gene-LOC105276718 | https://www.ncbi.nlm.nih.gov/gene/?term=LOC105276718 | 1721 | 9 | 52.3 | 3 | 17.43 | 33.33 | 30.94 | 22.67 | 8.27 | 8.27 | 0.04 | 0.04 |
| gene-LOC105277368 | https://www.ncbi.nlm.nih.gov/gene/?term=LOC105277368 | 1735 | 7 | 40.35 | 1 | 5.76 | 14.29 | 19.45 | 12.67 | 6.78 | 6.78 | 0.03 | 0.03 |
| gene-LOC105286857 | https://www.ncbi.nlm.nih.gov/gene/?term=LOC105286857 | 1576 | 4 | 25.38 | 1 | 6.35 | 25 | 73.58 | 67.03 | 6.55 | 6.55 | 0.02 | 0.02 |
| gene-LOC105277862 | https://www.ncbi.nlm.nih.gov/gene/?term=LOC105277862 | 1490 | 7 | 46.98 | 1 | 6.71 | 14.29 | 61.97 | 55.64 | 6.33 | 6.33 | 0.03 | 0.03 |
| gene-LOC105282405 | https://www.ncbi.nlm.nih.gov/gene/?term=LOC105282405 | 1310 | 31 | 236.64 | 5 | 38.17 | 16.13 | 51.18 | 44.88 | 6.3 | 6.3 | 0.15 | 0.15 |
| gene-LOC105284899 | https://www.ncbi.nlm.nih.gov/gene/?term=LOC105284899 | 1779 | 14 | 78.7 | 1 | 5.62 | 7.14 | 40.04 | 33.95 | 6.09 | 6.09 | 0.05 | 0.05 |
| gene-LOC105275585 | https://www.ncbi.nlm.nih.gov/gene/?term=LOC105275585 | 2152 | 22 | 102.23 | 1 | 4.65 | 4.55 | 36.17 | 30.17 | 6 | 6 | 0.06 | 0.06 |
| gene-LOC105285429 | https://www.ncbi.nlm.nih.gov/gene/?term=LOC105285429 | 1324 | 8 | 60.42 | 1 | 7.55 | 12.5 | 53.59 | 47.67 | 5.92 | 5.92 | 0.04 | 0.04 |
| gene-LOC105277810 | https://www.ncbi.nlm.nih.gov/gene/?term=LOC105277810 | 1367 | 21 | 153.62 | 1 | 7.32 | 4.76 | 45.33 | 39.52 | 5.81 | 5.81 | 0.09 | 0.09 |
| gene-LOC113562901 | https://www.ncbi.nlm.nih.gov/gene/?term=LOC113562901 | 1531 | 20 | 130.63 | 4 | 26.13 | 20 | 50.7 | 45.01 | 5.69 | 5.69 | 0.07 | 0.07 |
| gene-LOC105279898 | https://www.ncbi.nlm.nih.gov/gene/?term=LOC105279898 | 1343 | 10 | 74.46 | 1 | 7.45 | 10 | 77.18 | 71.53 | 5.65 | 5.65 | 0.04 | 0.04 |
| gene-LOC105277745 | https://www.ncbi.nlm.nih.gov/gene/?term=LOC105277745 | 1365 | 8 | 58.61 | 1 | 7.33 | 12.5 | 67.79 | 62.35 | 5.44 | 5.44 | 0.03 | 0.03 |
| gene-LOC105286644 | https://www.ncbi.nlm.nih.gov/gene/?term=LOC105286644 | 1235 | 21 | 170.04 | 1 | 8.1 | 4.76 | 44.21 | 39.04 | 5.17 | 5.17 | 0.09 | 0.09 |
| gene-LOC105274625 | https://www.ncbi.nlm.nih.gov/gene/?term=LOC105274625 | 4523 | 33 | 72.96 | 1 | 2.21 | 3.03 | 24.56 | 19.42 | 5.14 | 5.14 | 0.04 | 0.04 |
| gene-LOC105280045 | https://www.ncbi.nlm.nih.gov/gene/?term=LOC105280045 | 1938 | 13 | 67.08 | 1 | 5.16 | 7.69 | 26.88 | 21.77 | 5.11 | 5.11 | 0.03 | 0.03 |
| gene-LOC105279461 | https://www.ncbi.nlm.nih.gov/gene/?term=LOC105279461 | 1298 | 7 | 53.93 | 2 | 15.41 | 28.57 | 67.83 | 63.08 | 4.75 | 4.75 | 0.03 | 0.03 |

...
... only first 25 lines shown ...

Directory: OGLall File: ogl-Ob_LI_BCphaseA2.txt

| gene_ID | gene_link | gwidth | #Sites | #per10Kb | %perSite | %pNucl |
|---|---|---|---|---|---|---|
| gene-LOC113561532 | https://www.ncbi.nlm.nih.gov/gene/?term=LOC113561532 | 72 | 4 | 555.56 | 94.06 | 5.23 |
| gene-LOC105286531 | https://www.ncbi.nlm.nih.gov/gene/?term=LOC105286531 | 372 | 42 | 1129.03 | 38.35 | 4.33 |
| gene-LOC105280739 | https://www.ncbi.nlm.nih.gov/gene/?term=LOC105280739 | 1991 | 139 | 698.14 | 56.83 | 3.97 |
| gene-LOC105282149 | https://www.ncbi.nlm.nih.gov/gene/?term=LOC105282149 | 1787 | 156 | 872.97 | 45.44 | 3.97 |
| gene-LOC105286801 | https://www.ncbi.nlm.nih.gov/gene/?term=LOC105286801 | 1102 | 103 | 934.66 | 42.22 | 3.95 |
| gene-LOC105278849 | https://www.ncbi.nlm.nih.gov/gene/?term=LOC105278849 | 776 | 124 | 1597.94 | 24.65 | 3.94 |
| gene-LOC105285635 | https://www.ncbi.nlm.nih.gov/gene/?term=LOC105285635 | 1175 | 138 | 1174.47 | 31.91 | 3.75 |
| gene-LOC105274534 | https://www.ncbi.nlm.nih.gov/gene/?term=LOC105274534 | 1329 | 95 | 714.82 | 48.88 | 3.49 |
| gene-LOC105287811 | https://www.ncbi.nlm.nih.gov/gene/?term=LOC105287811 | 2155 | 151 | 700.7 | 49.56 | 3.47 |
| gene-LOC113562627 | https://www.ncbi.nlm.nih.gov/gene/?term=LOC113562627 | 397 | 20 | 503.78 | 68.12 | 3.43 |
| gene-LOC105286269 | https://www.ncbi.nlm.nih.gov/gene/?term=LOC105286269 | 1401 | 91 | 649.54 | 51.53 | 3.35 |
| gene-LOC105278844 | https://www.ncbi.nlm.nih.gov/gene/?term=LOC105278844 | 1101 | 109 | 990.01 | 33.65 | 3.33 |
| gene-LOC105278160 | https://www.ncbi.nlm.nih.gov/gene/?term=LOC105278160 | 1872 | 161 | 860.04 | 37.79 | 3.25 |
| gene-LOC105287677 | https://www.ncbi.nlm.nih.gov/gene/?term=LOC105287677 | 1588 | 123 | 774.56 | 41.72 | 3.23 |
| gene-LOC105286737 | https://www.ncbi.nlm.nih.gov/gene/?term=LOC105286737 | 2690 | 366 | 1360.59 | 23.68 | 3.22 |
| gene-LOC105286235 | https://www.ncbi.nlm.nih.gov/gene/?term=LOC105286235 | 1510 | 110 | 728.48 | 44.1 | 3.21 |
| gene-LOC105279052 | https://www.ncbi.nlm.nih.gov/gene/?term=LOC105279052 | 1130 | 139 | 1230.09 | 25.99 | 3.2 |
| gene-LOC105276371 | https://www.ncbi.nlm.nih.gov/gene/?term=LOC105276371 | 871 | 56 | 642.94 | 49.67 | 3.19 |
| gene-LOC105274711 | https://www.ncbi.nlm.nih.gov/gene/?term=LOC105274711 | 1655 | 109 | 658.61 | 47.63 | 3.14 |
| gene-LOC105278917 | https://www.ncbi.nlm.nih.gov/gene/?term=LOC105278917 | 2176 | 194 | 891.54 | 35.27 | 3.14 |
| gene-LOC105276360 | https://www.ncbi.nlm.nih.gov/gene/?term=LOC105276360 | 1466 | 153 | 1043.66 | 29.45 | 3.07 |
| gene-LOC105275231 | https://www.ncbi.nlm.nih.gov/gene/?term=LOC105275231 | 1270 | 83 | 653.54 | 45.54 | 2.98 |
| gene-LOC105275314 | https://www.ncbi.nlm.nih.gov/gene/?term=LOC105275314 | 1489 | 93 | 624.58 | 47.58 | 2.97 |
| gene-LOC105274543 | https://www.ncbi.nlm.nih.gov/gene/?term=LOC105274543 | 1572 | 127 | 807.89 | 35.56 | 2.87 |

...
... only first 25 lines shown ...

Directory: OGLall File: ogl-Ob_LI_BCphaseA2.vs.BCphaseB1.txt

| gene_ID | gene_link | gwidth | #Sites | #per10Kb | #dmSites | #dmsp10kb | %dmSites | %pSite1 | %pSite2 | DMpSite | ADMpSite | DMpNucl | ADMpNucl |
|---|---|---|---|---|---|---|---|---|---|---|---|---|---|
| gene-LOC105275384 | https://www.ncbi.nlm.nih.gov/gene/?term=LOC105275384 | 1987 | 91 | 457.98 | 1 | 5.03 | 1.1 | 33.74 | 32.35 | 1.39 | 1.39 | 0.06 | 0.06 |
| gene-LOC105276429 | https://www.ncbi.nlm.nih.gov/gene/?term=LOC105276429 | 2901 | 120 | 413.65 | 1 | 3.45 | 0.83 | 23.61 | 22.3 | 1.32 | 1.32 | 0.05 | 0.05 |
| gene-LOC105277903 | https://www.ncbi.nlm.nih.gov/gene/?term=LOC105277903 | 1835 | 35 | 190.74 | 1 | 5.45 | 2.86 | 47.23 | 46.95 | 0.28 | 0.28 | 0.01 | 0.01 |
| gene-LOC105283296 | https://www.ncbi.nlm.nih.gov/gene/?term=LOC105283296 | 2174 | 92 | 423.18 | 1 | 4.6 | 1.09 | 19.15 | 18.97 | 0.18 | 0.18 | 0.01 | 0.01 |
| gene-LOC105280410 | https://www.ncbi.nlm.nih.gov/gene/?term=LOC105280410 | 12580 | 280 | 222.58 | 1 | 0.79 | 0.36 | 19.42 | 19.27 | 0.15 | 0.15 | 0 | 0 |
| gene-LOC105277125 | https://www.ncbi.nlm.nih.gov/gene/?term=LOC105277125 | 51079 | 1879 | 367.86 | 1 | 0.2 | 0.05 | 1.58 | 1.64 | -0.06 | 0.06 | 0 | 0 |
| gene-LOC105276452 | https://www.ncbi.nlm.nih.gov/gene/?term=LOC105276452 | 2917 | 262 | 898.18 | 1 | 3.43 | 0.38 | 10.28 | 10.37 | -0.08 | 0.08 | -0.01 | 0.01 |
| gene-LOC105288036 | https://www.ncbi.nlm.nih.gov/gene/?term=LOC105288036 | 5110 | 226 | 442.27 | 1 | 1.96 | 0.44 | 16.3 | 16.4 | -0.1 | 0.1 | 0 | 0 |
| gene-LOC105276385 | https://www.ncbi.nlm.nih.gov/gene/?term=LOC105276385 | 11575 | 494 | 426.78 | 1 | 0.86 | 0.2 | 5.31 | 5.56 | -0.25 | 0.25 | -0.01 | 0.01 |
| gene-LOC105281272 | https://www.ncbi.nlm.nih.gov/gene/?term=LOC105281272 | 5252 | 178 | 338.92 | 1 | 1.9 | 0.56 | 16.03 | 16.73 | -0.7 | 0.7 | -0.02 | 0.02 |
| gene-LOC105284507 | https://www.ncbi.nlm.nih.gov/gene/?term=LOC105284507 | 5207 | 94 | 180.53 | 1 | 1.92 | 1.06 | 20.48 | 21.18 | -0.7 | 0.7 | -0.01 | 0.01 |
| gene-LOC105288042 | https://www.ncbi.nlm.nih.gov/gene/?term=LOC105288042 | 3507 | 101 | 288 | 1 | 2.85 | 0.99 | 22.48 | 23.29 | -0.81 | 0.81 | -0.02 | 0.02 |
| gene-LOC105285252 | https://www.ncbi.nlm.nih.gov/gene/?term=LOC105285252 | 7427 | 129 | 173.69 | 1 | 1.35 | 0.78 | 12.88 | 13.9 | -1.02 | 1.02 | -0.02 | 0.02 |
| gene-LOC105283264 | https://www.ncbi.nlm.nih.gov/gene/?term=LOC105283264 | 4005 | 88 | 219.73 | 1 | 2.5 | 1.14 | 18.85 | 20.42 | -1.57 | 1.57 | -0.03 | 0.03 |

Directory: OGLall File: ogl-Ob_LI_BCphaseA2.vs.BCphaseB2.txt

| gene_ID | gene_link | gwidth | #Sites | #per10Kb | #dmSites | #dmsp10kb | %dmSites | %pSite1 | %pSite2 | DMpSite | ADMpSite | DMpNucl | ADMpNucl |
|---|---|---|---|---|---|---|---|---|---|---|---|---|---|
| gene-LOC113562345 | https://www.ncbi.nlm.nih.gov/gene/?term=LOC113562345 | 2290 | 69 | 301.31 | 17 | 74.24 | 24.64 | 33.16 | 16.9 | 16.26 | 16.26 | 0.49 | 0.49 |
| gene-LOC105275708 | https://www.ncbi.nlm.nih.gov/gene/?term=LOC105275708 | 1772 | 10 | 56.43 | 1 | 5.64 | 10 | 50.31 | 39.56 | 10.75 | 10.75 | 0.06 | 0.06 |
| gene-LOC105275070 | https://www.ncbi.nlm.nih.gov/gene/?term=LOC105275070 | 5667 | 47 | 82.94 | 5 | 8.82 | 10.64 | 11.7 | 2 | 9.7 | 9.7 | 0.08 | 0.08 |
| gene-LOC105286694 | https://www.ncbi.nlm.nih.gov/gene/?term=LOC105286694 | 3409 | 11 | 32.27 | 1 | 2.93 | 9.09 | 63.21 | 54.71 | 8.5 | 8.5 | 0.03 | 0.03 |
| gene-LOC105276078 | https://www.ncbi.nlm.nih.gov/gene/?term=LOC105276078 | 1051 | 16 | 152.24 | 1 | 9.51 | 6.25 | 32.79 | 26.49 | 6.3 | 6.3 | 0.1 | 0.1 |
| gene-LOC105279898 | https://www.ncbi.nlm.nih.gov/gene/?term=LOC105279898 | 1343 | 14 | 104.24 | 1 | 7.45 | 7.14 | 61.13 | 54.89 | 6.24 | 6.24 | 0.07 | 0.07 |
| gene-LOC105282976 | https://www.ncbi.nlm.nih.gov/gene/?term=LOC105282976 | 1648 | 50 | 303.4 | 1 | 6.07 | 2 | 26.51 | 20.9 | 5.61 | 5.61 | 0.17 | 0.17 |
| gene-LOC105286302 | https://www.ncbi.nlm.nih.gov/gene/?term=LOC105286302 | 4536 | 18 | 39.68 | 1 | 2.2 | 5.56 | 22.71 | 17.52 | 5.2 | 5.2 | 0.02 | 0.02 |
| gene-LOC105275447 | https://www.ncbi.nlm.nih.gov/gene/?term=LOC105275447 | 1846 | 35 | 189.6 | 1 | 5.42 | 2.86 | 8.76 | 3.6 | 5.15 | 5.15 | 0.1 | 0.1 |
| gene-LOC105274900 | https://www.ncbi.nlm.nih.gov/gene/?term=LOC105274900 | 5561 | 52 | 93.51 | 2 | 3.6 | 3.85 | 36.59 | 32.11 | 4.48 | 4.48 | 0.04 | 0.04 |
| gene-LOC105286683 | https://www.ncbi.nlm.nih.gov/gene/?term=LOC105286683 | 1665 | 51 | 306.31 | 1 | 6.01 | 1.96 | 35.98 | 32.15 | 3.83 | 3.83 | 0.12 | 0.12 |
| gene-LOC105288063 | https://www.ncbi.nlm.nih.gov/gene/?term=LOC105288063 | 3400 | 53 | 155.88 | 1 | 2.94 | 1.89 | 16.83 | 13.02 | 3.81 | 3.81 | 0.06 | 0.06 |
| gene-LOC105283649 | https://www.ncbi.nlm.nih.gov/gene/?term=LOC105283649 | 1918 | 56 | 291.97 | 2 | 10.43 | 3.57 | 38.44 | 34.8 | 3.63 | 3.63 | 0.11 | 0.11 |
| gene-LOC113561805 | https://www.ncbi.nlm.nih.gov/gene/?term=LOC113561805 | 3337 | 77 | 230.75 | 1 | 3 | 1.3 | 37.31 | 34.13 | 3.18 | 3.18 | 0.07 | 0.07 |
| gene-LOC105285315 | https://www.ncbi.nlm.nih.gov/gene/?term=LOC105285315 | 5458 | 22 | 40.31 | 1 | 1.83 | 4.55 | 41.52 | 38.4 | 3.12 | 3.12 | 0.01 | 0.01 |
| gene-LOC105285674 | https://www.ncbi.nlm.nih.gov/gene/?term=LOC105285674 | 3035 | 93 | 306.43 | 2 | 6.59 | 2.15 | 27.41 | 24.3 | 3.11 | 3.11 | 0.1 | 0.1 |
| gene-LOC105281515 | https://www.ncbi.nlm.nih.gov/gene/?term=LOC105281515 | 8827 | 48 | 54.38 | 2 | 2.27 | 4.17 | 32.1 | 29.31 | 2.79 | 2.79 | 0.02 | 0.02 |
| gene-LOC105281420 | https://www.ncbi.nlm.nih.gov/gene/?term=LOC105281420 | 3882 | 55 | 141.68 | 1 | 2.58 | 1.82 | 15.69 | 12.96 | 2.73 | 2.73 | 0.04 | 0.04 |
| gene-LOC105280641 | https://www.ncbi.nlm.nih.gov/gene/?term=LOC105280641 | 3682 | 49 | 133.08 | 1 | 2.72 | 2.04 | 20.13 | 17.42 | 2.71 | 2.71 | 0.04 | 0.04 |
| gene-LOC105275347 | https://www.ncbi.nlm.nih.gov/gene/?term=LOC105275347 | 2526 | 15 | 59.38 | 1 | 3.96 | 6.67 | 31.33 | 28.89 | 2.44 | 2.44 | 0.01 | 0.01 |
| gene-LOC105275966 | https://www.ncbi.nlm.nih.gov/gene/?term=LOC105275966 | 3299 | 61 | 184.9 | 1 | 3.03 | 1.64 | 49.58 | 47.22 | 2.36 | 2.36 | 0.04 | 0.04 |
| gene-LOC105282156 | https://www.ncbi.nlm.nih.gov/gene/?term=LOC105282156 | 2577 | 136 | 527.75 | 1 | 3.88 | 0.74 | 26.99 | 24.63 | 2.36 | 2.36 | 0.12 | 0.12 |
| gene-LOC105276449 | https://www.ncbi.nlm.nih.gov/gene/?term=LOC105276449 | 1231 | 42 | 341.19 | 1 | 8.12 | 2.38 | 35.95 | 33.63 | 2.31 | 2.31 | 0.08 | 0.08 |
| gene-LOC105287648 | https://www.ncbi.nlm.nih.gov/gene/?term=LOC105287648 | 2205 | 27 | 122.45 | 1 | 4.54 | 3.7 | 41.48 | 39.19 | 2.29 | 2.29 | 0.03 | 0.03 |
| gene-LOC105278454 | https://www.ncbi.nlm.nih.gov/gene/?term=LOC105278454 | 1972 | 47 | 238.34 | 1 | 5.07 | 2.13 | 44.8 | 42.52 | 2.28 | 2.28 | 0.05 | 0.05 |
| gene-LOC105288066 | https://www.ncbi.nlm.nih.gov/gene/?term=LOC105288066 | 1964 | 21 | 106.92 | 1 | 5.09 | 4.76 | 46.21 | 43.96 | 2.25 | 2.25 | 0.02 | 0.02 |
| gene-LOC105275518 | https://www.ncbi.nlm.nih.gov/gene/?term=LOC105275518 | 2335 | 19 | 81.37 | 1 | 4.28 | 5.26 | 46.5 | 44.26 | 2.24 | 2.24 | 0.02 | 0.02 |
| gene-LOC105278109 | https://www.ncbi.nlm.nih.gov/gene/?term=LOC105278109 | 4029 | 65 | 161.33 | 1 | 2.48 | 1.54 | 48.15 | 45.97 | 2.19 | 2.19 | 0.04 | 0.04 |
| gene-LOC105282149 | https://www.ncbi.nlm.nih.gov/gene/?term=LOC105282149 | 1787 | 108 | 604.36 | 1 | 5.6 | 0.93 | 43.65 | 41.49 | 2.17 | 2.17 | 0.13 | 0.13 |
| gene-LOC105283592 | https://www.ncbi.nlm.nih.gov/gene/?term=LOC105283592 | 3479 | 71 | 204.08 | 1 | 2.87 | 1.41 | 26.49 | 24.49 | 2.01 | 2.01 | 0.04 | 0.04 |
| gene-LOC105286514 | https://www.ncbi.nlm.nih.gov/gene/?term=LOC105286514 | 2528 | 41 | 162.18 | 1 | 3.96 | 2.44 | 35.27 | 33.32 | 1.95 | 1.95 | 0.03 | 0.03 |
| gene-LOC105282970 | https://www.ncbi.nlm.nih.gov/gene/?term=LOC105282970 | 1529 | 39 | 255.07 | 1 | 6.54 | 2.56 | 54 | 52.14 | 1.86 | 1.86 | 0.05 | 0.05 |
| gene-LOC105282151 | https://www.ncbi.nlm.nih.gov/gene/?term=LOC105282151 | 1601 | 41 | 256.09 | 3 | 18.74 | 7.32 | 36.24 | 34.41 | 1.83 | 1.83 | 0.05 | 0.05 |
| gene-LOC105277856 | https://www.ncbi.nlm.nih.gov/gene/?term=LOC105277856 | 3442 | 34 | 98.78 | 1 | 2.91 | 2.94 | 37.69 | 35.88 | 1.81 | 1.81 | 0.02 | 0.02 |
| gene-LOC105286644 | https://www.ncbi.nlm.nih.gov/gene/?term=LOC105286644 | 1235 | 20 | 161.94 | 1 | 8.1 | 5 | 39.34 | 37.53 | 1.81 | 1.81 | 0.03 | 0.03 |
| gene-LOC105281848 | https://www.ncbi.nlm.nih.gov/gene/?term=LOC105281848 | 2122 | 56 | 263.9 | 1 | 4.71 | 1.79 | 24.08 | 22.28 | 1.8 | 1.8 | 0.05 | 0.05 |
| gene-LOC105280774 | https://www.ncbi.nlm.nih.gov/gene/?term=LOC105280774 | 2793 | 102 | 365.2 | 1 | 3.58 | 0.98 | 32.11 | 30.36 | 1.75 | 1.75 | 0.06 | 0.06 |
| gene-LOC105282658 | https://www.ncbi.nlm.nih.gov/gene/?term=LOC105282658 | 2421 | 65 | 268.48 | 1 | 4.13 | 1.54 | 21.38 | 19.92 | 1.46 | 1.46 | 0.04 | 0.04 |
| gene-LOC105280786 | https://www.ncbi.nlm.nih.gov/gene/?term=LOC105280786 | 2573 | 111 | 431.4 | 1 | 3.89 | 0.9 | 33.28 | 31.87 | 1.41 | 1.41 | 0.06 | 0.06 |
| gene-LOC105288010 | https://www.ncbi.nlm.nih.gov/gene/?term=LOC105288010 | 3189 | 39 | 122.3 | 1 | 3.14 | 2.56 | 37.68 | 36.3 | 1.38 | 1.38 | 0.02 | 0.02 |
| gene-LOC105280348 | https://www.ncbi.nlm.nih.gov/gene/?term=LOC105280348 | 3797 | 65 | 171.19 | 1 | 2.63 | 1.54 | 21.44 | 20.1 | 1.33 | 1.33 | 0.02 | 0.02 |
| gene-LOC105287095 | https://www.ncbi.nlm.nih.gov/gene/?term=LOC105287095 | 1435 | 57 | 397.21 | 1 | 6.97 | 1.75 | 29.87 | 28.54 | 1.33 | 1.33 | 0.05 | 0.05 |
| gene-LOC105277761 | https://www.ncbi.nlm.nih.gov/gene/?term=LOC105277761 | 4648 | 146 | 314.11 | 1 | 2.15 | 0.68 | 16.93 | 15.61 | 1.32 | 1.32 | 0.04 | 0.04 |
| gene-LOC105282600 | https://www.ncbi.nlm.nih.gov/gene/?term=LOC105282600 | 3715 | 53 | 142.66 | 1 | 2.69 | 1.89 | 32.07 | 30.77 | 1.3 | 1.3 | 0.02 | 0.02 |
| gene-LOC105276059 | https://www.ncbi.nlm.nih.gov/gene/?term=LOC105276059 | 6256 | 121 | 193.41 | 1 | 1.6 | 0.83 | 11.92 | 10.67 | 1.25 | 1.25 | 0.02 | 0.02 |
| gene-LOC105286190 | https://www.ncbi.nlm.nih.gov/gene/?term=LOC105286190 | 6447 | 298 | 462.23 | 1 | 1.55 | 0.34 | 8.47 | 7.3 | 1.16 | 1.16 | 0.05 | 0.05 |
| gene-LOC105279664 | https://www.ncbi.nlm.nih.gov/gene/?term=LOC105279664 | 4023 | 54 | 134.23 | 1 | 2.49 | 1.85 | 21.29 | 20.13 | 1.15 | 1.15 | 0.02 | 0.02 |
| gene-LOC105279298 | https://www.ncbi.nlm.nih.gov/gene/?term=LOC105279298 | 2773 | 128 | 461.59 | 2 | 7.21 | 1.56 | 25.02 | 23.92 | 1.1 | 1.1 | 0.05 | 0.05 |
| gene-LOC105285502 | https://www.ncbi.nlm.nih.gov/gene/?term=LOC105285502 | 9990 | 143 | 143.14 | 1 | 1 | 0.7 | 10.37 | 9.32 | 1.05 | 1.05 | 0.02 | 0.02 |
| gene-LOC105278497 | https://www.ncbi.nlm.nih.gov/gene/?term=LOC105278497 | 2072 | 94 | 453.67 | 1 | 4.83 | 1.06 | 21.1 | 20.09 | 1.01 | 1.01 | 0.05 | 0.05 |
| gene-LOC105274856 | https://www.ncbi.nlm.nih.gov/gene/?term=LOC105274856 | 5216 | 218 | 417.94 | 1 | 1.92 | 0.46 | 19.03 | 18.03 | 1 | 1 | 0.04 | 0.04 |
| gene-LOC105278807 | https://www.ncbi.nlm.nih.gov/gene/?term=LOC105278807 | 2166 | 73 | 337.03 | 1 | 4.62 | 1.37 | 18.28 | 17.29 | 0.99 | 0.99 | 0.03 | 0.03 |
| gene-LOC105284355 | https://www.ncbi.nlm.nih.gov/gene/?term=LOC105284355 | 3904 | 149 | 381.66 | 1 | 2.56 | 0.67 | 21.89 | 20.9 | 0.99 | 0.99 | 0.04 | 0.04 |
| gene-LOC105276704 | https://www.ncbi.nlm.nih.gov/gene/?term=LOC105276704 | 2063 | 98 | 475.04 | 1 | 4.85 | 1.02 | 13.8 | 12.82 | 0.98 | 0.98 | 0.05 | 0.05 |
| gene-LOC105278877 | https://www.ncbi.nlm.nih.gov/gene/?term=LOC105278877 | 1968 | 75 | 381.1 | 1 | 5.08 | 1.33 | 34.6 | 33.65 | 0.95 | 0.95 | 0.04 | 0.04 |
| gene-LOC105282794 | https://www.ncbi.nlm.nih.gov/gene/?term=LOC105282794 | 4571 | 93 | 203.46 | 1 | 2.19 | 1.08 | 18.93 | 18.03 | 0.9 | 0.9 | 0.02 | 0.02 |
| gene-LOC105288078 | https://www.ncbi.nlm.nih.gov/gene/?term=LOC105288078 | 19126 | 294 | 153.72 | 1 | 0.52 | 0.34 | 18.24 | 17.37 | 0.87 | 0.87 | 0.01 | 0.01 |
| gene-LOC105284184 | https://www.ncbi.nlm.nih.gov/gene/?term=LOC105284184 | 2535 | 85 | 335.31 | 1 | 3.94 | 1.18 | 34.37 | 33.53 | 0.83 | 0.83 | 0.03 | 0.03 |
| gene-LOC105285055 | https://www.ncbi.nlm.nih.gov/gene/?term=LOC105285055 | 4108 | 259 | 630.48 | 2 | 4.87 | 0.77 | 11.6 | 10.8 | 0.8 | 0.8 | 0.05 | 0.05 |
| gene-LOC105284329 | https://www.ncbi.nlm.nih.gov/gene/?term=LOC105284329 | 4051 | 50 | 123.43 | 1 | 2.47 | 2 | 19.81 | 19.02 | 0.79 | 0.79 | 0.01 | 0.01 |

| | | | | | | | | | | | | | | | |
|---|---|---|---|---|---|---|---|---|---|---|---|---|---|---|---|
| gene-LOC105276538 | https://www.ncbi.nlm.nih.gov/gene/?term=LOC105276538 | 1511 | 19 | 125.74 | | 1 | 6.62 | 5.26 | 70.24 | 69.47 | 0.77 | 0.77 | 0.01 | 0.01 | |
| gene-LOC105285202 | https://www.ncbi.nlm.nih.gov/gene/?term=LOC105285202 | 1838 | 66 | 359.09 | | 1 | 5.44 | 1.52 | 25.35 | 24.58 | 0.77 | 0.77 | 0.03 | 0.03 | |
| gene-LOC105275451 | https://www.ncbi.nlm.nih.gov/gene/?term=LOC105275451 | 6035 | 127 | 210.44 | | 1 | 1.66 | 0.79 | 22.96 | 22.23 | 0.73 | 0.73 | 0.02 | 0.02 | |
| gene-LOC105274452 | https://www.ncbi.nlm.nih.gov/gene/?term=LOC105274452 | 6595 | 261 | 395.75 | | 1 | 1.52 | 0.38 | 13.63 | 12.94 | 0.69 | 0.69 | 0.03 | 0.03 | |
| gene-LOC105283989 | https://www.ncbi.nlm.nih.gov/gene/?term=LOC105283989 | 4948 | 26 | 52.55 | 1 | 2.02 | 3.85 | 43.53 | 42.86 | 0.68 | 0.68 | 0 | 0 | | |
| gene-LOC105280956 | https://www.ncbi.nlm.nih.gov/gene/?term=LOC105280956 | 2639 | 75 | 284.2 | 1 | 3.79 | 1.33 | 17.09 | 16.45 | 0.64 | 0.64 | 0.02 | 0.02 | | |
| gene-LOC105279038 | https://www.ncbi.nlm.nih.gov/gene/?term=LOC105279038 | 9453 | 333 | 352.27 | | 1 | 1.06 | 0.3 | 9.29 | 8.67 | 0.62 | 0.62 | 0.02 | 0.02 | |
| gene-LOC105287063 | https://www.ncbi.nlm.nih.gov/gene/?term=LOC105287063 | 1874 | 48 | 256.14 | | 1 | 5.34 | 2.08 | 30.52 | 29.9 | 0.62 | 0.62 | 0.02 | 0.02 | |
| gene-LOC105274823 | https://www.ncbi.nlm.nih.gov/gene/?term=LOC105274823 | 4620 | 138 | 298.7 | 1 | 2.16 | 0.72 | 21.48 | 20.88 | 0.6 | 0.6 | 0.02 | 0.02 | | |
| gene-LOC105280148 | https://www.ncbi.nlm.nih.gov/gene/?term=LOC105280148 | 3120 | 214 | 685.9 | 1 | 3.21 | 0.47 | 11.76 | 11.17 | 0.6 | 0.6 | 0.04 | 0.04 | | |
| gene-LOC105280002 | https://www.ncbi.nlm.nih.gov/gene/?term=LOC105280002 | 1940 | 25 | 128.87 | | 1 | 5.15 | 4 | 38.59 | 38.02 | 0.58 | 0.58 | 0.01 | 0.01 | |
| gene-LOC105285807 | https://www.ncbi.nlm.nih.gov/gene/?term=LOC105285807 | 6063 | 218 | 359.56 | | 1 | 1.65 | 0.46 | 7.08 | 6.5 | 0.58 | 0.58 | 0.02 | 0.02 | |
| gene-LOC105286799 | https://www.ncbi.nlm.nih.gov/gene/?term=LOC105286799 | 3557 | 183 | 514.48 | | 1 | 2.81 | 0.55 | 16.25 | 15.68 | 0.57 | 0.57 | 0.03 | 0.03 | |
| gene-LOC105274851 | https://www.ncbi.nlm.nih.gov/gene/?term=LOC105274851 | 6616 | 128 | 193.47 | | 1 | 1.51 | 0.78 | 11.71 | 11.16 | 0.54 | 0.54 | 0.01 | 0.01 | |
| gene-LOC105275683 | https://www.ncbi.nlm.nih.gov/gene/?term=LOC105275683 | 4256 | 194 | 455.83 | | 1 | 2.35 | 0.52 | 13.24 | 12.7 | 0.54 | 0.54 | 0.02 | 0.02 | |
| gene-LOC105283943 | https://www.ncbi.nlm.nih.gov/gene/?term=LOC105283943 | 3559 | 58 | 162.97 | | 1 | 2.81 | 1.72 | 14.89 | 14.35 | 0.54 | 0.54 | 0.01 | 0.01 | |
| gene-LOC105278666 | https://www.ncbi.nlm.nih.gov/gene/?term=LOC105278666 | 10341 | 459 | 443.86 | | 1 | 0.97 | 0.22 | 7.32 | 6.79 | 0.53 | 0.53 | 0.02 | 0.02 | |
| gene-LOC105280723 | https://www.ncbi.nlm.nih.gov/gene/?term=LOC105280723 | 3572 | 170 | 475.92 | | 1 | 2.8 | 0.59 | 16.88 | 16.35 | 0.53 | 0.53 | 0.03 | 0.03 | |
| gene-LOC105285179 | https://www.ncbi.nlm.nih.gov/gene/?term=LOC105285179 | 4191 | 164 | 391.31 | | 1 | 2.39 | 0.61 | 20.24 | 19.71 | 0.53 | 0.53 | 0.02 | 0.02 | |
| gene-LOC105283123 | https://www.ncbi.nlm.nih.gov/gene/?term=LOC105283123 | 1507 | 52 | 345.06 | | 1 | 6.64 | 1.92 | 17.86 | 17.35 | 0.52 | 0.52 | 0.02 | 0.02 | |
| gene-LOC105285639 | https://www.ncbi.nlm.nih.gov/gene/?term=LOC105285639 | 10325 | 356 | 344.79 | | 1 | 0.97 | 0.28 | 11.26 | 10.77 | 0.49 | 0.49 | 0.02 | 0.02 | |
| gene-LOC105275848 | https://www.ncbi.nlm.nih.gov/gene/?term=LOC105275848 | 3718 | 139 | 373.86 | | 1 | 2.69 | 0.72 | 22.47 | 21.99 | 0.48 | 0.48 | 0.02 | 0.02 | |
| gene-LOC105285514 | https://www.ncbi.nlm.nih.gov/gene/?term=LOC105285514 | 1277 | 13 | 101.8 | 1 | 7.83 | 7.69 | 90.77 | 90.29 | 0.48 | 0.48 | 0 | 0 | | |
| gene-LOC105280410 | https://www.ncbi.nlm.nih.gov/gene/?term=LOC105280410 | 12580 | 237 | 188.39 | | 1 | 0.79 | 0.42 | 21.06 | 20.59 | 0.47 | 0.47 | 0.01 | 0.01 | |
| gene-LOC105279657 | https://www.ncbi.nlm.nih.gov/gene/?term=LOC105279657 | 5878 | 86 | 146.31 | | 1 | 1.7 | 1.16 | 19.59 | 19.13 | 0.46 | 0.46 | 0.01 | 0.01 | |
| gene-LOC105283728 | https://www.ncbi.nlm.nih.gov/gene/?term=LOC105283728 | 3256 | 113 | 347.05 | | 2 | 6.14 | 1.77 | 20.45 | 20.03 | 0.42 | 0.42 | 0.01 | 0.01 | |
| gene-LOC113562050 | https://www.ncbi.nlm.nih.gov/gene/?term=LOC113562050 | 17097 | 583 | 341 | 1 | 0.58 | 0.17 | 5.52 | 5.1 | 0.42 | 0.42 | 0.01 | 0.01 | | |
| gene-LOC105278145 | https://www.ncbi.nlm.nih.gov/gene/?term=LOC105278145 | 4470 | 241 | 539.15 | | 1 | 2.24 | 0.41 | 10.99 | 10.59 | 0.4 | 0.4 | 0.02 | 0.02 | |
| gene-LOC105286452 | https://www.ncbi.nlm.nih.gov/gene/?term=LOC105286452 | 6888 | 61 | 88.56 | 1 | 1.45 | 1.64 | 32.75 | 32.35 | 0.4 | 0.4 | 0 | 0 | | |
| gene-LOC105276196 | https://www.ncbi.nlm.nih.gov/gene/?term=LOC105276196 | 2953 | 48 | 162.55 | | 1 | 3.39 | 2.08 | 45.56 | 45.18 | 0.38 | 0.38 | 0.01 | 0.01 | |
| gene-LOC105278632 | https://www.ncbi.nlm.nih.gov/gene/?term=LOC105278632 | 3624 | 129 | 355.96 | | 1 | 2.76 | 0.78 | 14.81 | 14.44 | 0.37 | 0.37 | 0.01 | 0.01 | |
| gene-LOC105285395 | https://www.ncbi.nlm.nih.gov/gene/?term=LOC105285395 | 8085 | 196 | 242.42 | | 1 | 1.24 | 0.51 | 15.02 | 14.69 | 0.33 | 0.33 | 0.01 | 0.01 | |
| gene-LOC105278455 | https://www.ncbi.nlm.nih.gov/gene/?term=LOC105278455 | 27181 | 925 | 340.31 | | 1 | 0.37 | 0.11 | 7.49 | 7.17 | 0.32 | 0.32 | 0.01 | 0.01 | |
| gene-LOC105283712 | https://www.ncbi.nlm.nih.gov/gene/?term=LOC105283712 | 1910 | 31 | 162.3 | 1 | 5.24 | 3.23 | 30.95 | 30.64 | 0.31 | 0.31 | 0.01 | 0.01 | | |
| gene-LOC105276941 | https://www.ncbi.nlm.nih.gov/gene/?term=LOC105276941 | 15956 | 551 | 345.32 | | 1 | 0.63 | 0.18 | 5.1 | 4.82 | 0.28 | 0.28 | 0.01 | 0.01 | |
| gene-LOC105275374 | https://www.ncbi.nlm.nih.gov/gene/?term=LOC105275374 | 11376 | 527 | 463.26 | | 1 | 0.88 | 0.19 | 11.34 | 11.07 | 0.27 | 0.27 | 0.01 | 0.01 | |
| gene-LOC105287417 | https://www.ncbi.nlm.nih.gov/gene/?term=LOC105287417 | 1025 | 28 | 273.17 | | 1 | 9.76 | 3.57 | 42.07 | 41.82 | 0.26 | 0.26 | 0.01 | 0.01 | |
| gene-LOC105286194 | https://www.ncbi.nlm.nih.gov/gene/?term=LOC105286194 | 152619 | | 2166 | 141.92 | | 1 | 0.07 | 0.05 | 6.19 | 5.95 | 0.23 | 0.23 | 0 | 0 |
| gene-LOC105276452 | https://www.ncbi.nlm.nih.gov/gene/?term=LOC105276452 | 2917 | 257 | 881.04 | | 1 | 3.43 | 0.39 | 9.59 | 9.37 | 0.22 | 0.22 | 0.02 | 0.02 | |
| gene-LOC105275327 | https://www.ncbi.nlm.nih.gov/gene/?term=LOC105275327 | 14245 | 493 | 346.09 | | 1 | 0.7 | 0.2 | 6.9 | 6.7 | 0.19 | 0.19 | 0.01 | 0.01 | |
| gene-LOC105287550 | https://www.ncbi.nlm.nih.gov/gene/?term=LOC105287550 | 15659 | 570 | 364.01 | | 1 | 0.64 | 0.18 | 6.53 | 6.34 | 0.19 | 0.19 | 0.01 | 0.01 | |
| gene-LOC105278282 | https://www.ncbi.nlm.nih.gov/gene/?term=LOC105278282 | 1775 | 56 | 315.49 | | 1 | 5.63 | 1.79 | 35.74 | 35.56 | 0.18 | 0.18 | 0.01 | 0.01 | |
| gene-LOC105281272 | https://www.ncbi.nlm.nih.gov/gene/?term=LOC105281272 | 5252 | 178 | 338.92 | | 1 | 1.9 | 0.56 | 16.1 | 15.92 | 0.18 | 0.18 | 0.01 | 0.01 | |
| gene-LOC105282491 | https://www.ncbi.nlm.nih.gov/gene/?term=LOC105282491 | 4761 | 165 | 346.57 | | 1 | 2.1 | 0.61 | 8.62 | 8.44 | 0.18 | 0.18 | 0.01 | 0.01 | |
| gene-LOC105282811 | https://www.ncbi.nlm.nih.gov/gene/?term=LOC105282811 | 5765 | 203 | 352.12 | | 1 | 1.73 | 0.49 | 8.42 | 8.31 | 0.12 | 0.12 | 0 | 0 | |
| gene-LOC105282825 | https://www.ncbi.nlm.nih.gov/gene/?term=LOC105282825 | 3919 | 80 | 204.13 | | 1 | 2.55 | 1.25 | 12.57 | 12.46 | 0.11 | 0.11 | 0 | 0 | |
| gene-LOC105274590 | https://www.ncbi.nlm.nih.gov/gene/?term=LOC105274590 | 4516 | 58 | 128.43 | | 2 | 4.43 | 3.45 | 35.68 | 35.59 | 0.09 | 0.09 | 0 | 0 | |
| gene-LOC105275363 | https://www.ncbi.nlm.nih.gov/gene/?term=LOC105275363 | 3798 | 156 | 410.74 | | 1 | 2.63 | 0.64 | 17.23 | 17.17 | 0.07 | 0.07 | 0 | 0 | |
| gene-LOC105283296 | https://www.ncbi.nlm.nih.gov/gene/?term=LOC105283296 | 2174 | 84 | 386.38 | | 1 | 4.6 | 1.19 | 21.67 | 21.61 | 0.07 | 0.07 | 0 | 0 | |
| gene-LOC105281700 | https://www.ncbi.nlm.nih.gov/gene/?term=LOC105281700 | 2208 | 37 | 167.57 | | 1 | 4.53 | 2.7 | 39.7 | 39.67 | 0.04 | 0.04 | 0 | 0 | |
| gene-LOC105274531 | https://www.ncbi.nlm.nih.gov/gene/?term=LOC105274531 | 7774 | 284 | 365.32 | | 1 | 1.29 | 0.35 | 10.8 | 10.77 | 0.03 | 0.03 | 0 | 0 | |
| gene-LOC105284043 | https://www.ncbi.nlm.nih.gov/gene/?term=LOC105284043 | 5661 | 279 | 492.85 | | 1 | 1.77 | 0.36 | 10.73 | 10.7 | 0.03 | 0.03 | 0 | 0 | |
| gene-LOC105275460 | https://www.ncbi.nlm.nih.gov/gene/?term=LOC105275460 | 2473 | 114 | 460.98 | | 1 | 4.04 | 0.88 | 13.89 | 13.88 | 0.02 | 0.02 | 0 | 0 | |
| gene-LOC105277866 | https://www.ncbi.nlm.nih.gov/gene/?term=LOC105277866 | 33045 | 378 | 114.39 | | 1 | 0.3 | 0.26 | 3.22 | 3.21 | 0.01 | 0.01 | 0 | 0 | |
| gene-LOC105285406 | https://www.ncbi.nlm.nih.gov/gene/?term=LOC105285406 | 8015 | 59 | 73.61 | 1 | 1.25 | 1.69 | 26.4 | 26.39 | 0.01 | 0.01 | 0 | 0 | | |
| gene-LOC105275179 | https://www.ncbi.nlm.nih.gov/gene/?term=LOC105275179 | 3271 | 73 | 223.17 | | 1 | 3.06 | 1.37 | 20.99 | 20.99 | 0 | 0 | 0 | 0 | |
| gene-LOC105278228 | https://www.ncbi.nlm.nih.gov/gene/?term=LOC105278228 | 8610 | 162 | 188.15 | | 1 | 1.16 | 0.62 | 21.49 | 21.49 | 0 | 0 | 0 | 0 | |
| gene-LOC105287764 | https://www.ncbi.nlm.nih.gov/gene/?term=LOC105287764 | 18359 | 396 | 215.7 | 1 | 0.54 | 0.25 | 2.87 | 2.88 | -0.01 | 0.01 | 0 | 0 | | |
| gene-LOC105278283 | https://www.ncbi.nlm.nih.gov/gene/?term=LOC105278283 | 2317 | 125 | 539.49 | | 1 | 4.32 | 0.8 | 28 | 28.03 | -0.02 | 0.02 | 0 | 0 | |
| gene-LOC105280265 | https://www.ncbi.nlm.nih.gov/gene/?term=LOC105280265 | 85984 | 3074 | 357.51 | | 1 | 0.12 | 0.03 | 1.06 | 1.08 | -0.02 | 0.02 | 0 | 0 | |
| gene-LOC105282951 | https://www.ncbi.nlm.nih.gov/gene/?term=LOC105282951 | 3366 | 118 | 350.56 | | 1 | 2.97 | 0.85 | 19.88 | 19.9 | -0.02 | 0.02 | 0 | 0 | |
| gene-LOC105285533 | https://www.ncbi.nlm.nih.gov/gene/?term=LOC105285533 | 1640 | 60 | 365.85 | | 1 | 6.1 | 1.67 | 26.06 | 26.09 | -0.03 | 0.03 | 0 | 0 | |
| gene-LOC105280513 | https://www.ncbi.nlm.nih.gov/gene/?term=LOC105280513 | 69542 | 1246 | 179.17 | | 1 | 0.14 | 0.08 | 0.53 | 0.59 | -0.06 | 0.06 | 0 | 0 | |
| gene-LOC105277125 | https://www.ncbi.nlm.nih.gov/gene/?term=LOC105277125 | 51079 | 1856 | 363.36 | | 1 | 0.2 | 0.05 | 1.57 | 1.64 | -0.07 | 0.07 | 0 | 0 | |

| | | | | | | | | | | | | | |
|---|---|---|---|---|---|---|---|---|---|---|---|---|---|
| gene-LOC105276527 | https://www.ncbi.nlm.nih.gov/gene/?term=LOC105276527 | 7923 | 318 | 401.36 | 1 | 1.26 | 0.31 | 9.67 | 9.76 | -0.09 | 0.09 | 0 | 0 |
| gene-LOC105286649 | https://www.ncbi.nlm.nih.gov/gene/?term=LOC105286649 | 14178 | 826 | 582.59 | 1 | 0.71 | 0.12 | 0.92 | 1.04 | -0.12 | 0.12 | -0.01 | 0.01 |
| gene-LOC105282885 | https://www.ncbi.nlm.nih.gov/gene/?term=LOC105282885 | 1870 | 43 | 229.95 | 1 | 5.35 | 2.33 | 44.33 | 44.46 | -0.13 | 0.13 | 0 | 0 |
| gene-LOC105276462 | https://www.ncbi.nlm.nih.gov/gene/?term=LOC105276462 | 6561 | 278 | 423.72 | 1 | 1.52 | 0.36 | 7.35 | 7.5 | -0.15 | 0.15 | -0.01 | 0.01 |
| gene-LOC105282646 | https://www.ncbi.nlm.nih.gov/gene/?term=LOC105282646 | 3317 | 97 | 292.43 | 1 | 3.01 | 1.03 | 22.32 | 22.49 | -0.17 | 0.17 | -0.01 | 0.01 |
| gene-LOC105278691 | https://www.ncbi.nlm.nih.gov/gene/?term=LOC105278691 | 3871 | 147 | 379.75 | 1 | 2.58 | 0.68 | 14.28 | 14.5 | -0.22 | 0.22 | -0.01 | 0.01 |
| gene-LOC105280819 | https://www.ncbi.nlm.nih.gov/gene/?term=LOC105280819 | 1639 | 39 | 237.95 | 1 | 6.1 | 2.56 | 35.75 | 36.01 | -0.26 | 0.26 | -0.01 | 0.01 |
| gene-LOC105275187 | https://www.ncbi.nlm.nih.gov/gene/?term=LOC105275187 | 5534 | 70 | 126.49 | 1 | 1.81 | 1.43 | 14.08 | 14.45 | -0.36 | 0.36 | 0 | 0 |
| gene-LOC105275587 | https://www.ncbi.nlm.nih.gov/gene/?term=LOC105275587 | 2404 | 55 | 228.79 | 1 | 4.16 | 1.82 | 34.41 | 34.8 | -0.39 | 0.39 | -0.01 | 0.01 |
| gene-LOC105281312 | https://www.ncbi.nlm.nih.gov/gene/?term=LOC105281312 | 2551 | 72 | 282.24 | 1 | 3.92 | 1.39 | 29.66 | 30.06 | -0.4 | 0.4 | -0.01 | 0.01 |
| gene-LOC105286515 | https://www.ncbi.nlm.nih.gov/gene/?term=LOC105286515 | 3751 | 116 | 309.25 | 1 | 2.67 | 0.86 | 10.42 | 10.83 | -0.41 | 0.41 | -0.01 | 0.01 |
| gene-LOC105286999 | https://www.ncbi.nlm.nih.gov/gene/?term=LOC105286999 | 2141 | 41 | 191.5 | 1 | 4.67 | 2.44 | 37.42 | 37.83 | -0.41 | 0.41 | -0.01 | 0.01 |
| gene-LOC105286274 | https://www.ncbi.nlm.nih.gov/gene/?term=LOC105286274 | 3817 | 202 | 529.21 | 1 | 2.62 | 0.5 | 16.44 | 16.87 | -0.43 | 0.43 | -0.02 | 0.02 |
| gene-LOC105281754 | https://www.ncbi.nlm.nih.gov/gene/?term=LOC105281754 | 4300 | 117 | 272.09 | 1 | 2.33 | 0.85 | 16.59 | 17.09 | -0.5 | 0.5 | -0.01 | 0.01 |
| gene-LOC105283385 | https://www.ncbi.nlm.nih.gov/gene/?term=LOC105283385 | 9540 | 305 | 319.71 | 1 | 1.05 | 0.33 | 12.66 | 13.17 | -0.51 | 0.51 | -0.02 | 0.02 |
| gene-LOC113561621 | https://www.ncbi.nlm.nih.gov/gene/?term=LOC113561621 | 2249 | 78 | 346.82 | 1 | 4.45 | 1.28 | 20.85 | 21.38 | -0.53 | 0.53 | -0.02 | 0.02 |
| gene-LOC105278534 | https://www.ncbi.nlm.nih.gov/gene/?term=LOC105278534 | 7364 | 180 | 244.43 | 1 | 1.36 | 0.56 | 18.21 | 18.75 | -0.54 | 0.54 | -0.01 | 0.01 |
| gene-LOC105276272 | https://www.ncbi.nlm.nih.gov/gene/?term=LOC105276272 | 1965 | 36 | 183.21 | 1 | 5.09 | 2.78 | 32.08 | 32.69 | -0.61 | 0.61 | -0.01 | 0.01 |
| gene-LOC105281173 | https://www.ncbi.nlm.nih.gov/gene/?term=LOC105281173 | 1595 | 38 | 238.24 | 1 | 6.27 | 2.63 | 60.05 | 60.67 | -0.62 | 0.62 | -0.01 | 0.01 |
| gene-LOC105285742 | https://www.ncbi.nlm.nih.gov/gene/?term=LOC105285742 | 3446 | 107 | 310.5 | 1 | 2.9 | 0.93 | 21.42 | 22.06 | -0.64 | 0.64 | -0.02 | 0.02 |
| gene-LOC105287633 | https://www.ncbi.nlm.nih.gov/gene/?term=LOC105287633 | 1524 | 27 | 177.17 | 1 | 6.56 | 3.7 | 63.52 | 64.22 | -0.7 | 0.7 | -0.01 | 0.01 |
| gene-LOC105274534 | https://www.ncbi.nlm.nih.gov/gene/?term=LOC105274534 | 1329 | 54 | 406.32 | 1 | 7.52 | 1.85 | 45.08 | 45.82 | -0.75 | 0.75 | -0.03 | 0.03 |
| gene-LOC109611283 | https://www.ncbi.nlm.nih.gov/gene/?term=LOC109611283 | 1451 | 38 | 261.89 | 1 | 6.89 | 2.63 | 35.88 | 36.65 | -0.77 | 0.77 | -0.02 | 0.02 |
| gene-LOC105274706 | https://www.ncbi.nlm.nih.gov/gene/?term=LOC105274706 | 1446 | 55 | 380.36 | 1 | 6.92 | 1.82 | 37.25 | 38.23 | -0.98 | 0.98 | -0.04 | 0.04 |
| gene-LOC105274449 | https://www.ncbi.nlm.nih.gov/gene/?term=LOC105274449 | 1549 | 53 | 342.16 | 1 | 6.46 | 1.89 | 50.38 | 51.77 | -1.39 | 1.39 | -0.05 | 0.05 |
| gene-LOC105285252 | https://www.ncbi.nlm.nih.gov/gene/?term=LOC105285252 | 7427 | 117 | 157.53 | 1 | 1.35 | 0.85 | 13.64 | 15.21 | -1.57 | 1.57 | -0.02 | 0.02 |
| gene-LOC105284507 | https://www.ncbi.nlm.nih.gov/gene/?term=LOC105284507 | 5207 | 76 | 145.96 | 1 | 1.92 | 1.32 | 19.78 | 21.69 | -1.91 | 1.91 | -0.03 | 0.03 |
| gene-LOC105279500 | https://www.ncbi.nlm.nih.gov/gene/?term=LOC105279500 | 3587 | 33 | 92 | 1 | 2.79 | 3.03 | 44.93 | 47.01 | -2.08 | 2.08 | -0.02 | 0.02 |
| gene-LOC105282067 | https://www.ncbi.nlm.nih.gov/gene/?term=LOC105282067 | 5941 | 44 | 74.06 | 1 | 1.68 | 2.27 | 26.18 | 28.47 | -2.29 | 2.29 | -0.02 | 0.02 |
| gene-LOC105284502 | https://www.ncbi.nlm.nih.gov/gene/?term=LOC105284502 | 2157 | 19 | 88.09 | 1 | 4.64 | 5.26 | 36.83 | 43.93 | -7.1 | 7.1 | -0.06 | 0.06 |
| gene-LOC105284677 | https://www.ncbi.nlm.nih.gov/gene/?term=LOC105284677 | 1905 | 41 | 215.22 | 2 | 10.5 | 4.88 | 27.55 | 36.9 | -9.35 | 9.35 | -0.2 | 0.2 |

Directory: OGLall File: ogl-Ob_LI_BCphaseA2.vs.RphaseA1.txt

| gene_ID | gene_link | gwidth | #Sites | #per10Kb | #dmSites | #dmsp10kb | %dmSites | %pSite1 | %pSite2 | DMpSite | ADMpSite | DMpNucl | ADMpNucl |
|---|---|---|---|---|---|---|---|---|---|---|---|---|---|
| gene-LOC105280604 | https://www.ncbi.nlm.nih.gov/gene/?term=LOC105280604 | 752 | 1 | 13.3 | 1 | 13.3 | 100 | 75 | 42.31 | 32.69 | 32.69 | 0.04 | 0.04 |
| gene-LOC105286178 | https://www.ncbi.nlm.nih.gov/gene/?term=LOC105286178 | 1590 | 1 | 6.29 | 1 | 6.29 | 100 | 78.38 | 56.76 | 21.62 | 21.62 | 0.01 | 0.01 |
| gene-LOC113561421 | https://www.ncbi.nlm.nih.gov/gene/?term=LOC113561421 | 907 | 1 | 11.03 | 1 | 11.03 | 100 | 53.57 | 32.65 | 20.92 | 20.92 | 0.02 | 0.02 |
| gene-LOC105286586 | https://www.ncbi.nlm.nih.gov/gene/?term=LOC105286586 | 1417 | 1 | 7.06 | 1 | 7.06 | 100 | 35.71 | 15.56 | 20.15 | 20.15 | 0.01 | 0.01 |
| gene-LOC105284346 | https://www.ncbi.nlm.nih.gov/gene/?term=LOC105284346 | 2002 | 2 | 9.99 | 1 | 5 | 50 | 28.11 | 14.29 | 13.83 | 13.83 | 0.01 | 0.01 |
| gene-LOC105283693 | https://www.ncbi.nlm.nih.gov/gene/?term=LOC105283693 | 1305 | 12 | 91.95 | 4 | 30.65 | 33.33 | 70.54 | 58.26 | 12.28 | 12.28 | 0.11 | 0.11 |
| gene-LOC113562901 | https://www.ncbi.nlm.nih.gov/gene/?term=LOC113562901 | 1531 | 22 | 143.7 | 8 | 52.25 | 36.36 | 60.1 | 48.53 | 11.57 | 11.57 | 0.17 | 0.17 |
| gene-LOC105281110 | https://www.ncbi.nlm.nih.gov/gene/?term=LOC105281110 | 960 | 5 | 52.08 | 1 | 10.42 | 20 | 48.85 | 39.79 | 9.06 | 9.06 | 0.05 | 0.05 |
| gene-LOC113562728 | https://www.ncbi.nlm.nih.gov/gene/?term=LOC113562728 | 1733 | 32 | 184.65 | 5 | 28.85 | 15.62 | 28.94 | 19.93 | 9 | 9 | 0.17 | 0.17 |
| gene-LOC105280683 | https://www.ncbi.nlm.nih.gov/gene/?term=LOC105280683 | 2171 | 21 | 96.73 | 4 | 18.42 | 19.05 | 54.85 | 46.7 | 8.15 | 8.15 | 0.08 | 0.08 |
| gene-LOC105284759 | https://www.ncbi.nlm.nih.gov/gene/?term=LOC105284759 | 1353 | 9 | 66.52 | 1 | 7.39 | 11.11 | 38.95 | 31.14 | 7.81 | 7.81 | 0.05 | 0.05 |
| gene-LOC105284521 | https://www.ncbi.nlm.nih.gov/gene/?term=LOC105284521 | 2105 | 14 | 66.51 | 1 | 4.75 | 7.14 | 62.65 | 55.05 | 7.59 | 7.59 | 0.05 | 0.05 |
| gene-LOC113562975 | https://www.ncbi.nlm.nih.gov/gene/?term=LOC113562975 | 2751 | 9 | 32.72 | 1 | 3.64 | 11.11 | 50.57 | 43.41 | 7.16 | 7.16 | 0.02 | 0.02 |
| gene-LOC105274834 | https://www.ncbi.nlm.nih.gov/gene/?term=LOC105274834 | 1248 | 9 | 72.12 | 3 | 24.04 | 33.33 | 53.69 | 46.65 | 7.03 | 7.03 | 0.05 | 0.05 |
| gene-LOC113561934 | https://www.ncbi.nlm.nih.gov/gene/?term=LOC113561934 | 1252 | 27 | 215.65 | 4 | 31.95 | 14.81 | 31.68 | 24.75 | 6.93 | 6.93 | 0.15 | 0.15 |
| gene-LOC105276591 | https://www.ncbi.nlm.nih.gov/gene/?term=LOC105276591 | 1863 | 7 | 37.57 | 1 | 5.37 | 14.29 | 44.34 | 37.79 | 6.54 | 6.54 | 0.02 | 0.02 |
| gene-LOC105279538 | https://www.ncbi.nlm.nih.gov/gene/?term=LOC105279538 | 889 | 17 | 191.23 | 2 | 22.5 | 11.76 | 36.14 | 29.61 | 6.53 | 6.53 | 0.12 | 0.12 |
| gene-LOC105275378 | https://www.ncbi.nlm.nih.gov/gene/?term=LOC105275378 | 1189 | 20 | 168.21 | 1 | 8.41 | 5 | 32.85 | 26.65 | 6.2 | 6.2 | 0.1 | 0.1 |
| gene-LOC105286263 | https://www.ncbi.nlm.nih.gov/gene/?term=LOC105286263 | 1867 | 23 | 123.19 | 4 | 21.42 | 17.39 | 38.1 | 31.96 | 6.14 | 6.14 | 0.08 | 0.08 |
| gene-LOC105280835 | https://www.ncbi.nlm.nih.gov/gene/?term=LOC105280835 | 1890 | 6 | 31.75 | 1 | 5.29 | 16.67 | 6.39 | 0.36 | 6.03 | 6.03 | 0.02 | 0.02 |
| gene-LOC105287955 | https://www.ncbi.nlm.nih.gov/gene/?term=LOC105287955 | 2486 | 12 | 48.27 | 1 | 4.02 | 8.33 | 91.37 | 85.34 | 6.03 | 6.03 | 0.03 | 0.03 |
| gene-LOC105286815 | https://www.ncbi.nlm.nih.gov/gene/?term=LOC105286815 | 1024 | 10 | 97.66 | 1 | 9.77 | 10 | 59.55 | 53.96 | 5.59 | 5.59 | 0.05 | 0.05 |
| gene-LOC105283475 | https://www.ncbi.nlm.nih.gov/gene/?term=LOC105283475 | 1721 | 7 | 40.67 | 1 | 5.81 | 14.29 | 61.36 | 55.92 | 5.44 | 5.44 | 0.02 | 0.02 |
| gene-LOC105284988 | https://www.ncbi.nlm.nih.gov/gene/?term=LOC105284988 | 3729 | 26 | 69.72 | 1 | 2.68 | 3.85 | 37.25 | 31.94 | 5.3 | 5.3 | 0.04 | 0.04 |

...

... only first 25 lines shown ...

Directory: OGLall File: ogl-Ob_LI_BCphaseA2.vs.RphaseA2.txt

| gene_ID | gene_link | gwidth | #Sites | #per10Kb | #dmSites | #dmsp10kb | %dmSites | %pSite1 | %pSite2 | DMpSite | ADMpSite | DMpNucl | ADMpNucl |
|---|---|---|---|---|---|---|---|---|---|---|---|---|---|
| gene-LOC113563238 | https://www.ncbi.nlm.nih.gov/gene/?term=LOC113563238 | 2680 | 1 | 3.73 | 1 | 3.73 | 100 | 75 | 46.15 | 28.85 | 28.85 | 0.01 | 0.01 |
| gene-LOC105280604 | https://www.ncbi.nlm.nih.gov/gene/?term=LOC105280604 | 752 | 1 | 13.3 | 1 | 13.3 | 100 | 75 | 50 | 25 | 25 | 0.03 | 0.03 |
| gene-LOC105275861 | https://www.ncbi.nlm.nih.gov/gene/?term=LOC105275861 | 1979 | 7 | 35.37 | 1 | 5.05 | 14.29 | 39.97 | 30.1 | 9.87 | 9.87 | 0.03 | 0.03 |
| gene-LOC105281688 | https://www.ncbi.nlm.nih.gov/gene/?term=LOC105281688 | 1179 | 4 | 33.93 | 1 | 8.48 | 25 | 83.9 | 74.11 | 9.79 | 9.79 | 0.03 | 0.03 |
| gene-LOC105287647 | https://www.ncbi.nlm.nih.gov/gene/?term=LOC105287647 | 1949 | 10 | 51.31 | 1 | 5.13 | 10 | 41.7 | 35 | 6.7 | 6.7 | 0.03 | 0.03 |
| gene-LOC105287649 | https://www.ncbi.nlm.nih.gov/gene/?term=LOC105287649 | 1566 | 7 | 44.7 | 1 | 6.39 | 14.29 | 46.89 | 40.38 | 6.5 | 6.5 | 0.03 | 0.03 |
| gene-LOC105285051 | https://www.ncbi.nlm.nih.gov/gene/?term=LOC105285051 | 1357 | 16 | 117.91 | 2 | 14.74 | 12.5 | 34.98 | 29.29 | 5.69 | 5.69 | 0.07 | 0.07 |
| gene-LOC105281507 | https://www.ncbi.nlm.nih.gov/gene/?term=LOC105281507 | 881 | 7 | 79.46 | 1 | 11.35 | 14.29 | 24.99 | 19.48 | 5.51 | 5.51 | 0.04 | 0.04 |
| gene-LOC105280969 | https://www.ncbi.nlm.nih.gov/gene/?term=LOC105280969 | 1005 | 8 | 79.6 | 1 | 9.95 | 12.5 | 66.21 | 60.87 | 5.34 | 5.34 | 0.04 | 0.04 |
| gene-LOC105288141 | https://www.ncbi.nlm.nih.gov/gene/?term=LOC105288141 | 2845 | 27 | 94.9 | 1 | 3.51 | 3.7 | 26.78 | 22 | 4.78 | 4.78 | 0.05 | 0.05 |
| gene-LOC105280472 | https://www.ncbi.nlm.nih.gov/gene/?term=LOC105280472 | 1675 | 6 | 35.82 | 1 | 5.97 | 16.67 | 78.84 | 74.62 | 4.22 | 4.22 | 0.02 | 0.02 |
| gene-LOC105282975 | https://www.ncbi.nlm.nih.gov/gene/?term=LOC105282975 | 1500 | 22 | 146.67 | 2 | 13.33 | 9.09 | 49.9 | 46.07 | 3.84 | 3.84 | 0.06 | 0.06 |
| gene-LOC105283760 | https://www.ncbi.nlm.nih.gov/gene/?term=LOC105283760 | 1134 | 5 | 44.09 | 1 | 8.82 | 20 | 46 | 42.19 | 3.81 | 3.81 | 0.02 | 0.02 |
| gene-LOC113561391 | https://www.ncbi.nlm.nih.gov/gene/?term=LOC113561391 | 1463 | 12 | 82.02 | 1 | 6.84 | 8.33 | 17.64 | 13.97 | 3.68 | 3.68 | 0.03 | 0.03 |
| gene-LOC105275501 | https://www.ncbi.nlm.nih.gov/gene/?term=LOC105275501 | 1091 | 7 | 64.16 | 1 | 9.17 | 14.29 | 25.2 | 21.53 | 3.67 | 3.67 | 0.02 | 0.02 |
| gene-LOC105280589 | https://www.ncbi.nlm.nih.gov/gene/?term=LOC105280589 | 2884 | 47 | 162.97 | 2 | 6.93 | 4.26 | 32.58 | 28.94 | 3.64 | 3.64 | 0.06 | 0.06 |
| gene-LOC105280355 | https://www.ncbi.nlm.nih.gov/gene/?term=LOC105280355 | 1059 | 14 | 132.2 | 1 | 9.44 | 7.14 | 18.6 | 15.18 | 3.42 | 3.42 | 0.05 | 0.05 |
| gene-LOC105283851 | https://www.ncbi.nlm.nih.gov/gene/?term=LOC105283851 | 1886 | 9 | 47.72 | 1 | 5.3 | 11.11 | 44.96 | 41.54 | 3.42 | 3.42 | 0.02 | 0.02 |
| gene-LOC105279538 | https://www.ncbi.nlm.nih.gov/gene/?term=LOC105279538 | 889 | 20 | 224.97 | 2 | 22.5 | 10 | 30.72 | 27.36 | 3.36 | 3.36 | 0.08 | 0.08 |
| gene-LOC113562728 | https://www.ncbi.nlm.nih.gov/gene/?term=LOC113562728 | 1733 | 35 | 201.96 | 1 | 5.77 | 2.86 | 28.15 | 24.84 | 3.31 | 3.31 | 0.07 | 0.07 |
| gene-LOC113561448 | https://www.ncbi.nlm.nih.gov/gene/?term=LOC113561448 | 1279 | 13 | 101.64 | 1 | 7.82 | 7.69 | 66.97 | 63.82 | 3.15 | 3.15 | 0.03 | 0.03 |
| gene-LOC105276798 | https://www.ncbi.nlm.nih.gov/gene/?term=LOC105276798 | 3170 | 8 | 25.24 | 1 | 3.15 | 12.5 | 6.95 | 3.82 | 3.14 | 3.14 | 0.01 | 0.01 |
| gene-LOC105276840 | https://www.ncbi.nlm.nih.gov/gene/?term=LOC105276840 | 3299 | 11 | 33.34 | 1 | 3.03 | 9.09 | 6.96 | 3.82 | 3.14 | 3.14 | 0.01 | 0.01 |
| gene-LOC105286263 | https://www.ncbi.nlm.nih.gov/gene/?term=LOC105286263 | 1867 | 29 | 155.33 | 1 | 5.36 | 3.45 | 33.04 | 29.95 | 3.09 | 3.09 | 0.05 | 0.05 |

...
... only first 25 lines shown ...

Directory: OGLall File: ogl-Ob_LI_BCphaseA2.vs.RphaseB1.txt

| gene_ID | gene_link | gwidth | #Sites | #per10Kb | #dmSites | #dmsp10kb | %dmSites | %pSite1 | %pSite2 | DMpSite | ADMpSite | DMpNucl | ADMpNucl |
|---|---|---|---|---|---|---|---|---|---|---|---|---|---|
| gene-LOC113562345 | https://www.ncbi.nlm.nih.gov/gene/?term=LOC113562345 | 2290 | 71 | 310.04 | 15 | 65.5 | 21.13 | 28.2 | 15.69 | 12.51 | 12.51 | 0.39 | 0.39 |
| gene-LOC105275070 | https://www.ncbi.nlm.nih.gov/gene/?term=LOC105275070 | 5667 | 53 | 93.52 | 4 | 7.06 | 7.55 | 16.59 | 5.92 | 10.67 | 10.67 | 0.1 | 0.1 |
| gene-LOC105275708 | https://www.ncbi.nlm.nih.gov/gene/?term=LOC105275708 | 1772 | 10 | 56.43 | 1 | 5.64 | 10 | 50.31 | 40.78 | 9.53 | 9.53 | 0.05 | 0.05 |
| gene-LOC105285060 | https://www.ncbi.nlm.nih.gov/gene/?term=LOC105285060 | 1285 | 13 | 101.17 | 1 | 7.78 | 7.69 | 56.31 | 50.43 | 5.87 | 5.87 | 0.06 | 0.06 |
| gene-LOC105285514 | https://www.ncbi.nlm.nih.gov/gene/?term=LOC105285514 | 1277 | 11 | 86.14 | 1 | 7.83 | 9.09 | 90.26 | 85.12 | 5.14 | 5.14 | 0.04 | 0.04 |
| gene-LOC105276078 | https://www.ncbi.nlm.nih.gov/gene/?term=LOC105276078 | 1051 | 14 | 133.21 | 1 | 9.51 | 7.14 | 31.96 | 27.06 | 4.9 | 4.9 | 0.07 | 0.07 |
| gene-LOC105275347 | https://www.ncbi.nlm.nih.gov/gene/?term=LOC105275347 | 2526 | 14 | 55.42 | 1 | 3.96 | 7.14 | 33.57 | 29.73 | 3.84 | 3.84 | 0.02 | 0.02 |
| gene-LOC105285695 | https://www.ncbi.nlm.nih.gov/gene/?term=LOC105285695 | 1371 | 74 | 539.75 | 1 | 7.29 | 1.35 | 27.97 | 24.61 | 3.36 | 3.36 | 0.18 | 0.18 |
| gene-LOC105283552 | https://www.ncbi.nlm.nih.gov/gene/?term=LOC105283552 | 1562 | 36 | 230.47 | 1 | 6.4 | 2.78 | 56.28 | 53.31 | 2.97 | 2.97 | 0.07 | 0.07 |
| gene-LOC105274900 | https://www.ncbi.nlm.nih.gov/gene/?term=LOC105274900 | 5561 | 51 | 91.71 | 2 | 3.6 | 3.92 | 39.2 | 36.36 | 2.84 | 2.84 | 0.03 | 0.03 |
| gene-LOC105285315 | https://www.ncbi.nlm.nih.gov/gene/?term=LOC105285315 | 5458 | 21 | 38.48 | 1 | 1.83 | 4.76 | 43.16 | 40.32 | 2.84 | 2.84 | 0.01 | 0.01 |
| gene-LOC105279898 | https://www.ncbi.nlm.nih.gov/gene/?term=LOC105279898 | 1343 | 13 | 96.8 | 1 | 7.45 | 7.69 | 58.71 | 56.01 | 2.69 | 2.69 | 0.03 | 0.03 |
| gene-LOC105287063 | https://www.ncbi.nlm.nih.gov/gene/?term=LOC105287063 | 1874 | 49 | 261.47 | 1 | 5.34 | 2.04 | 32.42 | 29.81 | 2.61 | 2.61 | 0.07 | 0.07 |
| gene-LOC105283712 | https://www.ncbi.nlm.nih.gov/gene/?term=LOC105283712 | 1910 | 29 | 151.83 | 1 | 5.24 | 3.45 | 33.08 | 30.49 | 2.59 | 2.59 | 0.04 | 0.04 |
| gene-LOC105284704 | https://www.ncbi.nlm.nih.gov/gene/?term=LOC105284704 | 2400 | 62 | 258.33 | 1 | 4.17 | 1.61 | 26.43 | 23.93 | 2.5 | 2.5 | 0.06 | 0.06 |
| gene-LOC105288066 | https://www.ncbi.nlm.nih.gov/gene/?term=LOC105288066 | 1964 | 23 | 117.11 | 1 | 5.09 | 4.35 | 46.39 | 44.06 | 2.34 | 2.34 | 0.03 | 0.03 |
| gene-LOC105276272 | https://www.ncbi.nlm.nih.gov/gene/?term=LOC105276272 | 1965 | 38 | 193.38 | 1 | 5.09 | 2.63 | 30.65 | 28.5 | 2.15 | 2.15 | 0.04 | 0.04 |
| gene-LOC105282156 | https://www.ncbi.nlm.nih.gov/gene/?term=LOC105282156 | 2577 | 128 | 496.7 | 1 | 3.88 | 0.78 | 28.45 | 26.37 | 2.08 | 2.08 | 0.1 | 0.1 |
| gene-LOC105280348 | https://www.ncbi.nlm.nih.gov/gene/?term=LOC105280348 | 3797 | 65 | 171.19 | 1 | 2.63 | 1.54 | 21.64 | 19.63 | 2.01 | 2.01 | 0.03 | 0.03 |
| gene-LOC105280774 | https://www.ncbi.nlm.nih.gov/gene/?term=LOC105280774 | 2793 | 101 | 361.62 | 1 | 3.58 | 0.99 | 32.38 | 30.39 | 1.99 | 1.99 | 0.07 | 0.07 |
| gene-LOC105281848 | https://www.ncbi.nlm.nih.gov/gene/?term=LOC105281848 | 2122 | 52 | 245.05 | 1 | 4.71 | 1.92 | 26.53 | 24.7 | 1.83 | 1.83 | 0.04 | 0.04 |
| gene-LOC105276704 | https://www.ncbi.nlm.nih.gov/gene/?term=LOC105276704 | 2063 | 89 | 431.41 | 1 | 4.85 | 1.12 | 14.86 | 13.07 | 1.79 | 1.79 | 0.08 | 0.08 |
| gene-LOC105280408 | https://www.ncbi.nlm.nih.gov/gene/?term=LOC105280408 | 5983 | 50 | 83.57 | 1 | 1.67 | 2 | 19.04 | 17.37 | 1.67 | 1.67 | 0.01 | 0.01 |
| gene-LOC105282149 | https://www.ncbi.nlm.nih.gov/gene/?term=LOC105282149 | 1787 | 105 | 587.58 | 1 | 5.6 | 0.95 | 43.64 | 41.97 | 1.67 | 1.67 | 0.1 | 0.1 |
| gene-LOC105280641 | https://www.ncbi.nlm.nih.gov/gene/?term=LOC105280641 | 3682 | 56 | 152.09 | 1 | 2.72 | 1.79 | 17.9 | 16.23 | 1.66 | 1.66 | 0.03 | 0.03 |
| gene-LOC105285674 | https://www.ncbi.nlm.nih.gov/gene/?term=LOC105285674 | 3035 | 93 | 306.43 | 1 | 3.29 | 1.08 | 26.91 | 25.27 | 1.64 | 1.64 | 0.05 | 0.05 |
| gene-LOC105284184 | https://www.ncbi.nlm.nih.gov/gene/?term=LOC105284184 | 2535 | 87 | 343.2 | 1 | 3.94 | 1.15 | 34.29 | 32.68 | 1.6 | 1.6 | 0.05 | 0.05 |
| gene-LOC105283123 | https://www.ncbi.nlm.nih.gov/gene/?term=LOC105283123 | 1507 | 51 | 338.42 | 1 | 6.64 | 1.96 | 16.3 | 14.72 | 1.58 | 1.58 | 0.05 | 0.05 |
| gene-LOC105283697 | https://www.ncbi.nlm.nih.gov/gene/?term=LOC105283697 | 5301 | 137 | 258.44 | 1 | 1.89 | 0.73 | 17.96 | 16.39 | 1.57 | 1.57 | 0.04 | 0.04 |
| gene-LOC105288010 | https://www.ncbi.nlm.nih.gov/gene/?term=LOC105288010 | 3189 | 51 | 159.92 | 1 | 3.14 | 1.96 | 34.39 | 32.85 | 1.54 | 1.54 | 0.02 | 0.02 |
| gene-LOC105277243 | https://www.ncbi.nlm.nih.gov/gene/?term=LOC105277243 | 2284 | 32 | 140.11 | 1 | 4.38 | 3.12 | 44.88 | 43.35 | 1.53 | 1.53 | 0.02 | 0.02 |
| gene-LOC105288079 | https://www.ncbi.nlm.nih.gov/gene/?term=LOC105288079 | 3473 | 92 | 264.9 | 1 | 2.88 | 1.09 | 19.48 | 18 | 1.49 | 1.49 | 0.04 | 0.04 |
| gene-LOC105276538 | https://www.ncbi.nlm.nih.gov/gene/?term=LOC105276538 | 1511 | 19 | 125.74 | 1 | 6.62 | 5.26 | 70.28 | 68.86 | 1.42 | 1.42 | 0.02 | 0.02 |
| gene-LOC105281515 | https://www.ncbi.nlm.nih.gov/gene/?term=LOC105281515 | 8827 | 48 | 54.38 | 2 | 2.27 | 4.17 | 32.1 | 30.7 | 1.4 | 1.4 | 0.01 | 0.01 |
| gene-LOC105280972 | https://www.ncbi.nlm.nih.gov/gene/?term=LOC105280972 | 1532 | 32 | 208.88 | 1 | 6.53 | 3.12 | 19.4 | 18.05 | 1.35 | 1.35 | 0.03 | 0.03 |
| gene-LOC105282885 | https://www.ncbi.nlm.nih.gov/gene/?term=LOC105282885 | 1870 | 38 | 203.21 | 1 | 5.35 | 2.63 | 45.14 | 43.81 | 1.33 | 1.33 | 0.03 | 0.03 |
| gene-LOC105279500 | https://www.ncbi.nlm.nih.gov/gene/?term=LOC105279500 | 3587 | 33 | 92 | 1 | 2.79 | 3.03 | 51.88 | 50.67 | 1.21 | 1.21 | 0.01 | 0.01 |
| gene-LOC105286265 | https://www.ncbi.nlm.nih.gov/gene/?term=LOC105286265 | 2507 | 64 | 255.29 | 1 | 3.99 | 1.56 | 30.69 | 29.49 | 1.21 | 1.21 | 0.03 | 0.03 |
| gene-LOC105274449 | https://www.ncbi.nlm.nih.gov/gene/?term=LOC105274449 | 1549 | 53 | 342.16 | 1 | 6.46 | 1.89 | 50.38 | 49.29 | 1.09 | 1.09 | 0.04 | 0.04 |
| gene-LOC105275518 | https://www.ncbi.nlm.nih.gov/gene/?term=LOC105275518 | 2335 | 19 | 81.37 | 1 | 4.28 | 5.26 | 51.76 | 50.67 | 1.09 | 1.09 | 0.01 | 0.01 |
| gene-LOC105280786 | https://www.ncbi.nlm.nih.gov/gene/?term=LOC105280786 | 2573 | 122 | 474.15 | 1 | 3.89 | 0.82 | 30.88 | 29.81 | 1.07 | 1.07 | 0.05 | 0.05 |
| gene-LOC105279664 | https://www.ncbi.nlm.nih.gov/gene/?term=LOC105279664 | 4023 | 57 | 141.69 | 1 | 2.49 | 1.75 | 22.82 | 21.77 | 1.05 | 1.05 | 0.01 | 0.01 |
| gene-LOC105287095 | https://www.ncbi.nlm.nih.gov/gene/?term=LOC105287095 | 1435 | 56 | 390.24 | 1 | 6.97 | 1.79 | 30.4 | 29.35 | 1.05 | 1.05 | 0.04 | 0.04 |
| gene-LOC105276449 | https://www.ncbi.nlm.nih.gov/gene/?term=LOC105276449 | 1231 | 43 | 349.31 | 1 | 8.12 | 2.33 | 30.8 | 29.78 | 1.02 | 1.02 | 0.04 | 0.04 |
| gene-LOC105278807 | https://www.ncbi.nlm.nih.gov/gene/?term=LOC105278807 | 2166 | 71 | 327.79 | 1 | 4.62 | 1.41 | 19.81 | 18.81 | 1 | 1 | 0.03 | 0.03 |
| gene-LOC105279298 | https://www.ncbi.nlm.nih.gov/gene/?term=LOC105279298 | 2773 | 123 | 443.56 | 1 | 3.61 | 0.81 | 24.88 | 23.88 | 1 | 1 | 0.04 | 0.04 |
| gene-LOC105274823 | https://www.ncbi.nlm.nih.gov/gene/?term=LOC105274823 | 4620 | 152 | 329 | 1 | 2.16 | 0.66 | 19.69 | 18.71 | 0.99 | 0.99 | 0.03 | 0.03 |
| gene-LOC105277215 | https://www.ncbi.nlm.nih.gov/gene/?term=LOC105277215 | 2619 | 23 | 87.82 | 1 | 3.82 | 4.35 | 35.64 | 34.71 | 0.93 | 0.93 | 0.01 | 0.01 |
| gene-LOC105278384 | https://www.ncbi.nlm.nih.gov/gene/?term=LOC105278384 | 1390 | 33 | 237.41 | 1 | 7.19 | 3.03 | 46.69 | 45.77 | 0.92 | 0.92 | 0.02 | 0.02 |
| gene-LOC105283838 | https://www.ncbi.nlm.nih.gov/gene/?term=LOC105283838 | 1748 | 21 | 120.14 | 1 | 5.72 | 4.76 | 46.37 | 45.45 | 0.92 | 0.92 | 0.01 | 0.01 |
| gene-LOC113561805 | https://www.ncbi.nlm.nih.gov/gene/?term=LOC113561805 | 3337 | 69 | 206.77 | 1 | 3 | 1.45 | 38.18 | 37.26 | 0.92 | 0.92 | 0.02 | 0.02 |
| gene-LOC105282794 | https://www.ncbi.nlm.nih.gov/gene/?term=LOC105282794 | 4571 | 99 | 216.58 | 1 | 2.19 | 1.01 | 18.66 | 17.76 | 0.9 | 0.9 | 0.02 | 0.02 |
| gene-LOC105286644 | https://www.ncbi.nlm.nih.gov/gene/?term=LOC105286644 | 1235 | 22 | 178.14 | 1 | 8.1 | 4.55 | 42.04 | 41.14 | 0.89 | 0.89 | 0.02 | 0.02 |
| gene-LOC105278109 | https://www.ncbi.nlm.nih.gov/gene/?term=LOC105278109 | 4029 | 65 | 161.33 | 1 | 2.48 | 1.54 | 46.32 | 45.45 | 0.88 | 0.88 | 0.01 | 0.01 |
| gene-LOC105278497 | https://www.ncbi.nlm.nih.gov/gene/?term=LOC105278497 | 2072 | 89 | 429.54 | 1 | 4.83 | 1.12 | 21.38 | 20.51 | 0.87 | 0.87 | 0.04 | 0.04 |
| gene-LOC105283592 | https://www.ncbi.nlm.nih.gov/gene/?term=LOC105283592 | 3479 | 75 | 215.58 | 1 | 2.87 | 1.33 | 25.19 | 24.33 | 0.86 | 0.86 | 0.02 | 0.02 |
| gene-LOC105283728 | https://www.ncbi.nlm.nih.gov/gene/?term=LOC105283728 | 3256 | 104 | 319.41 | 2 | 6.14 | 1.92 | 22.23 | 21.4 | 0.83 | 0.83 | 0.03 | 0.03 |
| gene-LOC105285055 | https://www.ncbi.nlm.nih.gov/gene/?term=LOC105285055 | 4108 | 241 | 586.66 | 2 | 4.87 | 0.83 | 12.15 | 11.34 | 0.81 | 0.81 | 0.05 | 0.05 |
| gene-LOC105278283 | https://www.ncbi.nlm.nih.gov/gene/?term=LOC105278283 | 2317 | 126 | 543.81 | 1 | 4.32 | 0.79 | 27.76 | 27.04 | 0.72 | 0.72 | 0.04 | 0.04 |
| gene-LOC105285179 | https://www.ncbi.nlm.nih.gov/gene/?term=LOC105285179 | 4191 | 178 | 424.72 | 1 | 2.39 | 0.56 | 17.79 | 17.08 | 0.71 | 0.71 | 0.03 | 0.03 |

| | | | | | | | | | | | | | | |
|---|---|---|---|---|---|---|---|---|---|---|---|---|---|---|
| gene-LOC105276560 | https://www.ncbi.nlm.nih.gov/gene/?term=LOC105276560 | 18607 | 319 | 171.44 | 1 | 0.54 | 0.31 | 2.53 | 1.85 | 0.67 | 0.67 | 0.01 | 0.01 | |
| gene-LOC105281312 | https://www.ncbi.nlm.nih.gov/gene/?term=LOC105281312 | 2551 | 68 | 266.56 | 1 | 3.92 | 1.47 | 31.46 | 30.8 | 0.65 | 0.65 | 0.02 | 0.02 | |
| gene-LOC105286514 | https://www.ncbi.nlm.nih.gov/gene/?term=LOC105286514 | 2528 | 43 | 170.09 | 1 | 3.96 | 2.33 | 34.72 | 34.07 | 0.65 | 0.65 | 0.01 | 0.01 | |
| gene-LOC105284355 | https://www.ncbi.nlm.nih.gov/gene/?term=LOC105284355 | 3904 | 143 | 366.29 | 1 | 2.56 | 0.7 | 22.99 | 22.35 | 0.64 | 0.64 | 0.02 | 0.02 | |
| gene-LOC105278145 | https://www.ncbi.nlm.nih.gov/gene/?term=LOC105278145 | 4470 | 250 | 559.28 | 1 | 2.24 | 0.4 | 10.94 | 10.34 | 0.6 | 0.6 | 0.03 | 0.03 | |
| gene-LOC105283989 | https://www.ncbi.nlm.nih.gov/gene/?term=LOC105283989 | 4948 | 25 | 50.53 1 | 2.02 | 4 | 46.26 | 45.67 | 0.59 | 0.59 | 0 | 0 | | |
| gene-LOC105276059 | https://www.ncbi.nlm.nih.gov/gene/?term=LOC105276059 | 6256 | 123 | 196.61 | 1 | 1.6 | 0.81 | 12.44 | 11.86 | 0.58 | 0.58 | 0.01 | 0.01 | |
| gene-LOC105280407 | https://www.ncbi.nlm.nih.gov/gene/?term=LOC105280407 | 10950 | 203 | 185.39 | 1 | 0.91 | 0.49 | 15.56 | 14.99 | 0.56 | 0.56 | 0.01 | 0.01 | |
| gene-LOC105285671 | https://www.ncbi.nlm.nih.gov/gene/?term=LOC105285671 | 2263 | 99 | 437.47 | 1 | 4.42 | 1.01 | 30.99 | 30.44 | 0.55 | 0.55 | 0.02 | 0.02 | |
| gene-LOC105285395 | https://www.ncbi.nlm.nih.gov/gene/?term=LOC105285395 | 8085 | 207 | 256.03 | 1 | 1.24 | 0.48 | 14.3 | 13.76 | 0.54 | 0.54 | 0.01 | 0.01 | |
| gene-LOC105280819 | https://www.ncbi.nlm.nih.gov/gene/?term=LOC105280819 | 1639 | 39 | 237.95 | 1 | 6.1 | 2.56 | 35.99 | 35.46 | 0.53 | 0.53 | 0.01 | 0.01 | |
| gene-LOC105275451 | https://www.ncbi.nlm.nih.gov/gene/?term=LOC105275451 | 6035 | 133 | 220.38 | 1 | 1.66 | 0.75 | 22.24 | 21.72 | 0.51 | 0.51 | 0.01 | 0.01 | |
| gene-LOC105275460 | https://www.ncbi.nlm.nih.gov/gene/?term=LOC105275460 | 2473 | 118 | 477.15 | 1 | 4.04 | 0.85 | 13.46 | 12.95 | 0.51 | 0.51 | 0.02 | 0.02 | |
| gene-LOC105278632 | https://www.ncbi.nlm.nih.gov/gene/?term=LOC105278632 | 3624 | 131 | 361.48 | 1 | 2.76 | 0.76 | 14.47 | 14.04 | 0.43 | 0.43 | 0.02 | 0.02 | |
| gene-LOC105278534 | https://www.ncbi.nlm.nih.gov/gene/?term=LOC105278534 | 7364 | 180 | 244.43 | 1 | 1.36 | 0.56 | 18.01 | 17.6 | 0.41 | 0.41 | 0.01 | 0.01 | |
| gene-LOC105280890 | https://www.ncbi.nlm.nih.gov/gene/?term=LOC105280890 | 1139 | 29 | 254.61 | 1 | 8.78 | 3.45 | 19.62 | 19.23 | 0.39 | 0.39 | 0.01 | 0.01 | |
| gene-LOC105286999 | https://www.ncbi.nlm.nih.gov/gene/?term=LOC105286999 | 2141 | 44 | 205.51 | 1 | 4.67 | 2.27 | 35.31 | 34.92 | 0.39 | 0.39 | 0.01 | 0.01 | |
| gene-LOC105276462 | https://www.ncbi.nlm.nih.gov/gene/?term=LOC105276462 | 6561 | 275 | 419.14 | 1 | 1.52 | 0.36 | 8.32 | 7.94 | 0.37 | 0.37 | 0.02 | 0.02 | |
| gene-LOC105280410 | https://www.ncbi.nlm.nih.gov/gene/?term=LOC105280410 | 12580 | 265 | 210.65 | 1 | 0.79 | 0.38 | 19.92 | 19.57 | 0.35 | 0.35 | 0.01 | 0.01 | |
| gene-LOC105283296 | https://www.ncbi.nlm.nih.gov/gene/?term=LOC105283296 | 2174 | 86 | 395.58 | 1 | 4.6 | 1.16 | 16.05 | 15.7 | 0.35 | 0.35 | 0.01 | 0.01 | |
| gene-LOC105278666 | https://www.ncbi.nlm.nih.gov/gene/?term=LOC105278666 | 10341 | 456 | 440.96 | 1 | 0.97 | 0.22 | 7.52 | 7.19 | 0.34 | 0.34 | 0.01 | 0.01 | |
| gene-LOC105279657 | https://www.ncbi.nlm.nih.gov/gene/?term=LOC105279657 | 5878 | 96 | 163.32 | 1 | 1.7 | 1.04 | 18.1 | 17.76 | 0.34 | 0.34 | 0.01 | 0.01 | |
| gene-LOC105277435 | https://www.ncbi.nlm.nih.gov/gene/?term=LOC105277435 | 3643 | 175 | 480.37 | 1 | 2.74 | 0.57 | 11.3 | 10.97 | 0.33 | 0.33 | 0.02 | 0.02 | |
| gene-LOC105284036 | https://www.ncbi.nlm.nih.gov/gene/?term=LOC105284036 | 3254 | 87 | 267.36 | 1 | 3.07 | 1.15 | 24.27 | 23.96 | 0.31 | 0.31 | 0.01 | 0.01 | |
| gene-LOC105281272 | https://www.ncbi.nlm.nih.gov/gene/?term=LOC105281272 | 5252 | 175 | 333.21 | 1 | 1.9 | 0.57 | 16.26 | 15.96 | 0.3 | 0.3 | 0.01 | 0.01 | |
| gene-LOC105285807 | https://www.ncbi.nlm.nih.gov/gene/?term=LOC105285807 | 6063 | 207 | 341.42 | 1 | 1.65 | 0.48 | 7.17 | 6.87 | 0.3 | 0.3 | 0.01 | 0.01 | |
| gene-LOC105275187 | https://www.ncbi.nlm.nih.gov/gene/?term=LOC105275187 | 5534 | 69 | 124.68 | 1 | 1.81 | 1.45 | 14.14 | 13.85 | 0.29 | 0.29 | 0 | 0 | |
| gene-LOC105282491 | https://www.ncbi.nlm.nih.gov/gene/?term=LOC105282491 | 4761 | 170 | 357.07 | 1 | 2.1 | 0.59 | 7.92 | 7.67 | 0.25 | 0.25 | 0.01 | 0.01 | |
| gene-LOC105274706 | https://www.ncbi.nlm.nih.gov/gene/?term=LOC105274706 | 1446 | 50 | 345.78 | 1 | 6.92 | 2 | 39.23 | 39 | 0.23 | 0.23 | 0.01 | 0.01 | |
| gene-LOC105283649 | https://www.ncbi.nlm.nih.gov/gene/?term=LOC105283649 | 1918 | 55 | 286.76 | 1 | 5.21 | 1.82 | 38.85 | 38.63 | 0.21 | 0.21 | 0.01 | 0.01 | |
| gene-LOC105275374 | https://www.ncbi.nlm.nih.gov/gene/?term=LOC105275374 | 11376 | 525 | 461.5 1 | 0.88 | 0.19 | 10.23 | 10.06 | 0.17 | 0.17 | 0.01 | 0.01 | | |
| gene-LOC105281700 | https://www.ncbi.nlm.nih.gov/gene/?term=LOC105281700 | 2208 | 38 | 172.1 1 | 4.53 | 2.63 | 39.22 | 39.09 | 0.14 | 0.14 | 0 | 0 | | |
| gene-LOC105274531 | https://www.ncbi.nlm.nih.gov/gene/?term=LOC105274531 | 7774 | 283 | 364.03 | 1 | 1.29 | 0.35 | 10.82 | 10.73 | 0.09 | 0.09 | 0 | 0 | |
| gene-LOC105282825 | https://www.ncbi.nlm.nih.gov/gene/?term=LOC105282825 | 3919 | 78 | 199.03 | 1 | 2.55 | 1.28 | 12.99 | 12.9 | 0.09 | 0.09 | 0 | 0 | |
| gene-LOC105275179 | https://www.ncbi.nlm.nih.gov/gene/?term=LOC105275179 | 3271 | 71 | 217.06 | 1 | 3.06 | 1.41 | 16.78 | 16.7 | 0.08 | 0.08 | 0 | 0 | |
| gene-LOC105279038 | https://www.ncbi.nlm.nih.gov/gene/?term=LOC105279038 | 9453 | 317 | 335.34 | 1 | 1.06 | 0.32 | 9.53 | 9.46 | 0.07 | 0.07 | 0 | 0 | |
| gene-LOC105283987 | https://www.ncbi.nlm.nih.gov/gene/?term=LOC105283987 | 4841 | 71 | 146.66 | 1 | 2.07 | 1.41 | 32.31 | 32.24 | 0.07 | 0.07 | 0 | 0 | |
| gene-LOC105284329 | https://www.ncbi.nlm.nih.gov/gene/?term=LOC105284329 | 4051 | 51 | 125.89 | 1 | 2.47 | 1.96 | 19.38 | 19.32 | 0.06 | 0.06 | 0 | 0 | |
| gene-LOC105286194 | https://www.ncbi.nlm.nih.gov/gene/?term=LOC105286194 | 152619 | | 2202 144.28 | | 1 | 0.07 | 0.05 | 6.22 | 6.17 | 0.04 | 0.04 | 0 | 0 |
| gene-LOC105288078 | https://www.ncbi.nlm.nih.gov/gene/?term=LOC105288078 | 19126 | 307 | 160.51 | 1 | 0.52 | 0.33 | 16.81 | 16.78 | 0.03 | 0.03 | 0 | 0 | |
| gene-LOC105276527 | https://www.ncbi.nlm.nih.gov/gene/?term=LOC105276527 | 7923 | 334 | 421.56 | 1 | 1.26 | 0.3 | 8.56 | 8.54 | 0.02 | 0.02 | 0 | 0 | |
| gene-LOC105286452 | https://www.ncbi.nlm.nih.gov/gene/?term=LOC105286452 | 6888 | 68 | 98.72 1 | 1.45 | 1.47 | 28.88 | 28.89 | 0 | 0 | 0 | 0 | | |
| gene-LOC105274856 | https://www.ncbi.nlm.nih.gov/gene/?term=LOC105274856 | 5216 | 212 | 406.44 | 1 | 1.92 | 0.47 | 18.3 | 18.31 | -0.01 | 0.01 | 0 | 0 | |
| gene-LOC105275327 | https://www.ncbi.nlm.nih.gov/gene/?term=LOC105275327 | 14245 | 472 | 331.34 | 1 | 0.7 | 0.21 | 7.13 | 7.15 | -0.03 | 0.03 | 0 | 0 | |
| gene-LOC105283943 | https://www.ncbi.nlm.nih.gov/gene/?term=LOC105283943 | 3559 | 56 | 157.35 | 1 | 2.81 | 1.79 | 16.7 | 16.73 | -0.03 | 0.03 | 0 | 0 | |
| gene-LOC105284043 | https://www.ncbi.nlm.nih.gov/gene/?term=LOC105284043 | 5661 | 278 | 491.08 | 1 | 1.77 | 0.36 | 10.84 | 10.87 | -0.03 | 0.03 | 0 | 0 | |
| gene-LOC105275683 | https://www.ncbi.nlm.nih.gov/gene/?term=LOC105275683 | 4256 | 191 | 448.78 | 1 | 2.35 | 0.52 | 12.17 | 12.21 | -0.04 | 0.04 | 0 | 0 | |
| gene-LOC105277866 | https://www.ncbi.nlm.nih.gov/gene/?term=LOC105277866 | 33045 | 409 | 123.77 | 1 | 0.3 | 0.24 | 3.12 | 3.19 | -0.07 | 0.07 | 0 | 0 | |
| gene-LOC105277216 | https://www.ncbi.nlm.nih.gov/gene/?term=LOC105277216 | 7627 | 87 | 114.07 | 1 | 1.31 | 1.15 | 9.6 | 9.67 | -0.08 | 0.08 | 0 | 0 | |
| gene-LOC105277125 | https://www.ncbi.nlm.nih.gov/gene/?term=LOC105277125 | 51079 | 1802 | 352.79 | 1 | 0.2 | 0.06 | 1.61 | 1.7 | -0.09 | 0.09 | 0 | 0 | |
| gene-LOC105278455 | https://www.ncbi.nlm.nih.gov/gene/?term=LOC105278455 | 27181 | 915 | 336.63 | 1 | 0.37 | 0.11 | 7.11 | 7.23 | -0.12 | 0.12 | 0 | 0 | |
| gene-LOC105280265 | https://www.ncbi.nlm.nih.gov/gene/?term=LOC105280265 | 85984 | 3009 | 349.95 | 1 | 0.12 | 0.03 | 1.05 | 1.18 | -0.12 | 0.12 | 0 | 0 | |
| gene-LOC105287417 | https://www.ncbi.nlm.nih.gov/gene/?term=LOC105287417 | 1025 | 28 | 273.17 | 1 | 9.76 | 3.57 | 42.07 | 42.23 | -0.16 | 0.16 | 0 | 0 | |
| gene-LOC105282658 | https://www.ncbi.nlm.nih.gov/gene/?term=LOC105282658 | 2421 | 67 | 276.75 | 1 | 4.13 | 1.49 | 20.83 | 21.03 | -0.21 | 0.21 | -0.01 | 0.01 | |
| gene-LOC105285981 | https://www.ncbi.nlm.nih.gov/gene/?term=LOC105285981 | 7666 | 183 | 238.72 | 1 | 1.3 | 0.55 | 18.71 | 18.92 | -0.21 | 0.21 | -0.01 | 0.01 | |
| gene-LOC105278877 | https://www.ncbi.nlm.nih.gov/gene/?term=LOC105278877 | 1968 | 77 | 391.26 | 1 | 5.08 | 1.3 | 34.55 | 34.76 | -0.22 | 0.22 | -0.01 | 0.01 | |
| gene-LOC105284147 | https://www.ncbi.nlm.nih.gov/gene/?term=LOC105284147 | 77820 | 534 | 68.62 1 | 0.13 | 0.19 | 9 | 9.26 | -0.26 | 0.26 | 0 | 0 | | |
| gene-LOC105284774 | https://www.ncbi.nlm.nih.gov/gene/?term=LOC105284774 | 12089 | 81 | 67 1 | 0.83 | 1.23 | 21 | 21.27 | -0.27 | 0.27 | 0 | 0 | | |
| gene-LOC105276196 | https://www.ncbi.nlm.nih.gov/gene/?term=LOC105276196 | 2953 | 45 | 152.39 | 1 | 3.39 | 2.22 | 48.59 | 48.91 | -0.31 | 0.31 | 0 | 0 | |
| gene-LOC105274851 | https://www.ncbi.nlm.nih.gov/gene/?term=LOC105274851 | 6616 | 127 | 191.96 | 1 | 1.51 | 0.79 | 11.51 | 11.83 | -0.33 | 0.33 | -0.01 | 0.01 | |
| gene-LOC105286649 | https://www.ncbi.nlm.nih.gov/gene/?term=LOC105286649 | 14178 | 785 | 553.67 | 1 | 0.71 | 0.13 | 0.94 | 1.3 | -0.36 | 0.36 | -0.02 | 0.02 | |
| gene-LOC105285742 | https://www.ncbi.nlm.nih.gov/gene/?term=LOC105285742 | 3446 | 97 | 281.49 | 1 | 2.9 | 1.03 | 20.21 | 20.62 | -0.41 | 0.41 | -0.01 | 0.01 | |
| gene-LOC105274590 | https://www.ncbi.nlm.nih.gov/gene/?term=LOC105274590 | 4516 | 64 | 141.72 | 2 | 4.43 | 3.12 | 32.42 | 32.94 | -0.51 | 0.51 | -0.01 | 0.01 | |
| gene-LOC105284146 | https://www.ncbi.nlm.nih.gov/gene/?term=LOC105284146 | 2488 | 66 | 265.27 | 1 | 4.02 | 1.52 | 23.09 | 23.62 | -0.54 | 0.54 | -0.01 | 0.01 | |

| | | | | | | | | | | | | | |
|---|---|---|---|---|---|---|---|---|---|---|---|---|---|
| gene-LOC105278228 | https://www.ncbi.nlm.nih.gov/gene/?term=LOC105278228 | 8610 | 169 | 196.28 | 1 | 1.16 | 0.59 | 21.96 | 22.51 | -0.55 | 0.55 | -0.01 | 0.01 |
| gene-LOC105280956 | https://www.ncbi.nlm.nih.gov/gene/?term=LOC105280956 | 2639 | 77 | 291.78 | 1 | 3.79 | 1.3 | 18.18 | 18.82 | -0.64 | 0.64 | -0.02 | 0.02 |
| gene-LOC105286515 | https://www.ncbi.nlm.nih.gov/gene/?term=LOC105286515 | 3751 | 113 | 301.25 | 1 | 2.67 | 0.88 | 10.68 | 11.31 | -0.64 | 0.64 | -0.02 | 0.02 |
| gene-LOC105278459 | https://www.ncbi.nlm.nih.gov/gene/?term=LOC105278459 | 2936 | 97 | 330.38 | 1 | 3.41 | 1.03 | 20.46 | 21.23 | -0.77 | 0.77 | -0.03 | 0.03 |
| gene-LOC105278454 | https://www.ncbi.nlm.nih.gov/gene/?term=LOC105278454 | 1972 | 41 | 207.91 | 1 | 5.07 | 2.44 | 43.16 | 44.07 | -0.91 | 0.91 | -0.02 | 0.02 |
| gene-LOC105287633 | https://www.ncbi.nlm.nih.gov/gene/?term=LOC105287633 | 1524 | 23 | 150.92 | 1 | 6.56 | 4.35 | 68.63 | 69.73 | -1.11 | 1.11 | -0.02 | 0.02 |
| gene-LOC105281173 | https://www.ncbi.nlm.nih.gov/gene/?term=LOC105281173 | 1595 | 39 | 244.51 | 1 | 6.27 | 2.56 | 60.96 | 62.08 | -1.12 | 1.12 | -0.03 | 0.03 |
| gene-LOC105278282 | https://www.ncbi.nlm.nih.gov/gene/?term=LOC105278282 | 1775 | 56 | 315.49 | 1 | 5.63 | 1.79 | 33.68 | 34.81 | -1.13 | 1.13 | -0.04 | 0.04 |
| gene-LOC105285252 | https://www.ncbi.nlm.nih.gov/gene/?term=LOC105285252 | 7427 | 124 | 166.96 | 2 | 2.69 | 1.61 | 12.83 | 14.05 | -1.22 | 1.22 | -0.02 | 0.02 |
| gene-LOC105274534 | https://www.ncbi.nlm.nih.gov/gene/?term=LOC105274534 | 1329 | 55 | 413.84 | 1 | 7.52 | 1.82 | 46.07 | 47.79 | -1.72 | 1.72 | -0.07 | 0.07 |
| gene-LOC105284507 | https://www.ncbi.nlm.nih.gov/gene/?term=LOC105284507 | 5207 | 86 | 165.16 | 1 | 1.92 | 1.16 | 21.7 | 23.52 | -1.83 | 1.83 | -0.03 | 0.03 |
| gene-LOC105284213 | https://www.ncbi.nlm.nih.gov/gene/?term=LOC105284213 | 868 | 15 | 172.81 | 1 | 11.52 | 6.67 | 15.38 | 18.37 | -2.99 | 2.99 | -0.05 | 0.05 |

Directory: OGLall File: ogl-Ob_LI_BCphaseA2.vs.RphaseB2.txt

| gene_ID | gene_link | gwidth | #Sites | #per10Kb | #dmSites | #dmsp10kb | %dmSites | %pSite1 | %pSite2 | DMpSite | ADMpSite | DMpNucl | ADMpNucl |
|---|---|---|---|---|---|---|---|---|---|---|---|---|---|
| gene-LOC113562901 | https://www.ncbi.nlm.nih.gov/gene/?term=LOC113562901 | 1531 | 22 | 143.7 | 1 | 6.53 | 4.55 | 60.1 | 44.57 | 15.53 | 15.53 | 0.22 | 0.22 |
| gene-LOC105279898 | https://www.ncbi.nlm.nih.gov/gene/?term=LOC105279898 | 1343 | 10 | 74.46 | 1 | 7.45 | 10 | 84.43 | 71.53 | 12.89 | 12.89 | 0.1 | 0.1 |
| gene-LOC105275070 | https://www.ncbi.nlm.nih.gov/gene/?term=LOC105275070 | 5667 | 41 | 72.35 | 5 | 8.82 | 12.2 | 13.7 | 1.06 | 12.65 | 12.65 | 0.09 | 0.09 |
| gene-LOC113562345 | https://www.ncbi.nlm.nih.gov/gene/?term=LOC113562345 | 2290 | 67 | 292.58 | 11 | 48.03 | 16.42 | 29.88 | 17.32 | 12.57 | 12.57 | 0.37 | 0.37 |
| gene-LOC105286694 | https://www.ncbi.nlm.nih.gov/gene/?term=LOC105286694 | 3409 | 9 | 26.4 | 1 | 2.93 | 11.11 | 71.7 | 62.28 | 9.42 | 9.42 | 0.02 | 0.02 |
| gene-LOC105275708 | https://www.ncbi.nlm.nih.gov/gene/?term=LOC105275708 | 1772 | 10 | 56.43 | 1 | 5.64 | 10 | 50.31 | 44.22 | 6.09 | 6.09 | 0.03 | 0.03 |
| gene-LOC105282976 | https://www.ncbi.nlm.nih.gov/gene/?term=LOC105282976 | 1648 | 47 | 285.19 | 1 | 6.07 | 2.13 | 28.02 | 22.43 | 5.59 | 5.59 | 0.16 | 0.16 |
| gene-LOC105286644 | https://www.ncbi.nlm.nih.gov/gene/?term=LOC105286644 | 1235 | 22 | 178.14 | 1 | 8.1 | 4.55 | 42.35 | 37.26 | 5.09 | 5.09 | 0.09 | 0.09 |
| gene-LOC105285315 | https://www.ncbi.nlm.nih.gov/gene/?term=LOC105285315 | 5458 | 19 | 34.81 | 1 | 1.83 | 5.26 | 47.88 | 43.22 | 4.66 | 4.66 | 0.02 | 0.02 |
| gene-LOC105283552 | https://www.ncbi.nlm.nih.gov/gene/?term=LOC105283552 | 1562 | 36 | 230.47 | 1 | 6.4 | 2.78 | 56.28 | 51.7 | 4.58 | 4.58 | 0.11 | 0.11 |
| gene-LOC105276078 | https://www.ncbi.nlm.nih.gov/gene/?term=LOC105276078 | 1051 | 14 | 133.21 | 1 | 9.51 | 7.14 | 36.98 | 32.93 | 4.05 | 4.05 | 0.05 | 0.05 |
| gene-LOC105288063 | https://www.ncbi.nlm.nih.gov/gene/?term=LOC105288063 | 3400 | 43 | 126.47 | 1 | 2.94 | 2.33 | 18.5 | 14.57 | 3.92 | 3.92 | 0.05 | 0.05 |
| gene-LOC105278109 | https://www.ncbi.nlm.nih.gov/gene/?term=LOC105278109 | 4029 | 59 | 146.44 | 1 | 2.48 | 1.69 | 50.99 | 47.24 | 3.75 | 3.75 | 0.05 | 0.05 |
| gene-LOC105278215 | https://www.ncbi.nlm.nih.gov/gene/?term=LOC105278215 | 1700 | 29 | 170.59 | 1 | 5.88 | 3.45 | 57.89 | 54.66 | 3.23 | 3.23 | 0.06 | 0.06 |
| gene-LOC105275518 | https://www.ncbi.nlm.nih.gov/gene/?term=LOC105275518 | 2335 | 18 | 77.09 | 1 | 4.28 | 5.56 | 54.44 | 51.41 | 3.03 | 3.03 | 0.02 | 0.02 |
| gene-LOC105285674 | https://www.ncbi.nlm.nih.gov/gene/?term=LOC105285674 | 3035 | 93 | 306.43 | 2 | 6.59 | 2.15 | 26.74 | 23.71 | 3.03 | 3.03 | 0.09 | 0.09 |
| gene-LOC105285060 | https://www.ncbi.nlm.nih.gov/gene/?term=LOC105285060 | 1285 | 17 | 132.3 | 1 | 7.78 | 5.88 | 51.2 | 48.29 | 2.91 | 2.91 | 0.04 | 0.04 |
| gene-LOC105275347 | https://www.ncbi.nlm.nih.gov/gene/?term=LOC105275347 | 2526 | 14 | 55.42 | 1 | 3.96 | 7.14 | 33.57 | 30.77 | 2.8 | 2.8 | 0.02 | 0.02 |
| gene-LOC105274900 | https://www.ncbi.nlm.nih.gov/gene/?term=LOC105274900 | 5561 | 47 | 84.52 | 2 | 3.6 | 4.26 | 37.9 | 35.14 | 2.76 | 2.76 | 0.02 | 0.02 |
| gene-LOC105283649 | https://www.ncbi.nlm.nih.gov/gene/?term=LOC105283649 | 1918 | 54 | 281.54 | 1 | 5.21 | 1.85 | 37.63 | 35.03 | 2.6 | 2.6 | 0.07 | 0.07 |
| gene-LOC105279664 | https://www.ncbi.nlm.nih.gov/gene/?term=LOC105279664 | 4023 | 48 | 119.31 | 1 | 2.49 | 2.08 | 23.51 | 21.34 | 2.18 | 2.18 | 0.03 | 0.03 |
| gene-LOC105281515 | https://www.ncbi.nlm.nih.gov/gene/?term=LOC105281515 | 8827 | 38 | 43.05 | 1 | 1.13 | 2.63 | 31.29 | 29.13 | 2.16 | 2.16 | 0.01 | 0.01 |
| gene-LOC105276704 | https://www.ncbi.nlm.nih.gov/gene/?term=LOC105276704 | 2063 | 88 | 426.56 | 1 | 4.85 | 1.14 | 15.96 | 13.92 | 2.05 | 2.05 | 0.09 | 0.09 |
| gene-LOC105283123 | https://www.ncbi.nlm.nih.gov/gene/?term=LOC105283123 | 1507 | 44 | 291.97 | 1 | 6.64 | 2.27 | 19.91 | 17.87 | 2.05 | 2.05 | 0.06 | 0.06 |
| gene-LOC105281754 | https://www.ncbi.nlm.nih.gov/gene/?term=LOC105281754 | 4300 | 110 | 255.81 | 1 | 2.33 | 0.91 | 16.49 | 14.51 | 1.98 | 1.98 | 0.05 | 0.05 |
| gene-LOC113561805 | https://www.ncbi.nlm.nih.gov/gene/?term=LOC113561805 | 3337 | 68 | 203.78 | 1 | 3 | 1.47 | 35.02 | 33.08 | 1.94 | 1.94 | 0.04 | 0.04 |
| gene-LOC105285671 | https://www.ncbi.nlm.nih.gov/gene/?term=LOC105285671 | 2263 | 103 | 455.15 | 1 | 4.42 | 0.97 | 29.21 | 27.3 | 1.91 | 1.91 | 0.09 | 0.09 |
| gene-LOC105288066 | https://www.ncbi.nlm.nih.gov/gene/?term=LOC105288066 | 1964 | 20 | 101.83 | 1 | 5.09 | 5 | 44.28 | 42.38 | 1.9 | 1.9 | 0.02 | 0.02 |
| gene-LOC105282156 | https://www.ncbi.nlm.nih.gov/gene/?term=LOC105282156 | 2577 | 129 | 500.58 | 1 | 3.88 | 0.78 | 28.68 | 26.84 | 1.84 | 1.84 | 0.09 | 0.09 |
| gene-LOC105280408 | https://www.ncbi.nlm.nih.gov/gene/?term=LOC105280408 | 5983 | 47 | 78.56 | 1 | 1.67 | 2.13 | 19.12 | 17.33 | 1.78 | 1.78 | 0.01 | 0.01 |
| gene-LOC105282149 | https://www.ncbi.nlm.nih.gov/gene/?term=LOC105282149 | 1787 | 108 | 604.36 | 1 | 5.6 | 0.93 | 43.65 | 41.89 | 1.76 | 1.76 | 0.11 | 0.11 |
| gene-LOC105276190 | https://www.ncbi.nlm.nih.gov/gene/?term=LOC105276190 | 4839 | 82 | 169.46 | 1 | 2.07 | 1.22 | 11.63 | 9.92 | 1.71 | 1.71 | 0.03 | 0.03 |
| gene-LOC105282794 | https://www.ncbi.nlm.nih.gov/gene/?term=LOC105282794 | 4571 | 92 | 201.27 | 1 | 2.19 | 1.09 | 18.21 | 16.51 | 1.7 | 1.7 | 0.03 | 0.03 |
| gene-LOC105280786 | https://www.ncbi.nlm.nih.gov/gene/?term=LOC105280786 | 2573 | 113 | 439.18 | 1 | 3.89 | 0.88 | 32.35 | 30.66 | 1.69 | 1.69 | 0.07 | 0.07 |
| gene-LOC105282970 | https://www.ncbi.nlm.nih.gov/gene/?term=LOC105282970 | 1529 | 39 | 255.07 | 1 | 6.54 | 2.56 | 51.08 | 49.46 | 1.62 | 1.62 | 0.04 | 0.04 |
| gene-LOC105284184 | https://www.ncbi.nlm.nih.gov/gene/?term=LOC105284184 | 2535 | 79 | 311.64 | 1 | 3.94 | 1.27 | 37.02 | 35.48 | 1.54 | 1.54 | 0.05 | 0.05 |
| gene-LOC105277761 | https://www.ncbi.nlm.nih.gov/gene/?term=LOC105277761 | 4648 | 143 | 307.66 | 1 | 2.15 | 0.7 | 15.33 | 13.81 | 1.52 | 1.52 | 0.05 | 0.05 |
| gene-LOC105275966 | https://www.ncbi.nlm.nih.gov/gene/?term=LOC105275966 | 3299 | 53 | 160.65 | 1 | 3.03 | 1.89 | 54.5 | 52.99 | 1.51 | 1.51 | 0.02 | 0.02 |
| gene-LOC105278807 | https://www.ncbi.nlm.nih.gov/gene/?term=LOC105278807 | 2166 | 62 | 286.24 | 1 | 4.62 | 1.61 | 21.98 | 20.52 | 1.46 | 1.46 | 0.04 | 0.04 |
| gene-LOC105277565 | https://www.ncbi.nlm.nih.gov/gene/?term=LOC105277565 | 3216 | 169 | 525.5 | 1 | 3.11 | 0.59 | 20.65 | 19.2 | 1.45 | 1.45 | 0.08 | 0.08 |
| gene-LOC105283728 | https://www.ncbi.nlm.nih.gov/gene/?term=LOC105283728 | 3256 | 107 | 328.62 | 2 | 6.14 | 1.87 | 20.96 | 19.59 | 1.37 | 1.37 | 0.04 | 0.04 |
| gene-LOC105276059 | https://www.ncbi.nlm.nih.gov/gene/?term=LOC105276059 | 6256 | 121 | 193.41 | 1 | 1.6 | 0.83 | 10.95 | 9.59 | 1.36 | 1.36 | 0.03 | 0.03 |
| gene-LOC105283712 | https://www.ncbi.nlm.nih.gov/gene/?term=LOC105283712 | 1910 | 31 | 162.3 | 1 | 5.24 | 3.23 | 30.95 | 29.59 | 1.36 | 1.36 | 0.02 | 0.02 |
| gene-LOC105276875 | https://www.ncbi.nlm.nih.gov/gene/?term=LOC105276875 | 3151 | 109 | 345.92 | 1 | 3.17 | 0.92 | 25.45 | 24.19 | 1.26 | 1.26 | 0.04 | 0.04 |
| gene-LOC105278497 | https://www.ncbi.nlm.nih.gov/gene/?term=LOC105278497 | 2072 | 92 | 444.02 | 1 | 4.83 | 1.09 | 21.61 | 20.36 | 1.26 | 1.26 | 0.06 | 0.06 |
| gene-LOC105281173 | https://www.ncbi.nlm.nih.gov/gene/?term=LOC105281173 | 1595 | 38 | 238.24 | 1 | 6.27 | 2.63 | 60.05 | 58.85 | 1.2 | 1.2 | 0.03 | 0.03 |
| gene-LOC105279298 | https://www.ncbi.nlm.nih.gov/gene/?term=LOC105279298 | 2773 | 131 | 472.41 | 2 | 7.21 | 1.53 | 24.81 | 23.62 | 1.18 | 1.18 | 0.06 | 0.06 |
| gene-LOC105287063 | https://www.ncbi.nlm.nih.gov/gene/?term=LOC105287063 | 1874 | 48 | 256.14 | 1 | 5.34 | 2.08 | 31.5 | 30.33 | 1.17 | 1.17 | 0.03 | 0.03 |
| gene-LOC105281848 | https://www.ncbi.nlm.nih.gov/gene/?term=LOC105281848 | 2122 | 47 | 221.49 | 1 | 4.71 | 2.13 | 27.8 | 26.69 | 1.11 | 1.11 | 0.02 | 0.02 |
| gene-LOC105284355 | https://www.ncbi.nlm.nih.gov/gene/?term=LOC105284355 | 3904 | 143 | 366.29 | 1 | 2.56 | 0.7 | 22.85 | 21.77 | 1.08 | 1.08 | 0.04 | 0.04 |
| gene-LOC105274590 | https://www.ncbi.nlm.nih.gov/gene/?term=LOC105274590 | 4516 | 57 | 126.22 | 2 | 4.43 | 3.51 | 34.78 | 33.73 | 1.06 | 1.06 | 0.01 | 0.01 |
| gene-LOC105275451 | https://www.ncbi.nlm.nih.gov/gene/?term=LOC105275451 | 6035 | 118 | 195.53 | 1 | 1.66 | 0.85 | 23.45 | 22.41 | 1.04 | 1.04 | 0.02 | 0.02 |
| gene-LOC105285179 | https://www.ncbi.nlm.nih.gov/gene/?term=LOC105285179 | 4191 | 177 | 422.33 | 1 | 2.39 | 0.56 | 19.66 | 18.63 | 1.04 | 1.04 | 0.04 | 0.04 |
| gene-LOC105276272 | https://www.ncbi.nlm.nih.gov/gene/?term=LOC105276272 | 1965 | 34 | 173.03 | 1 | 5.09 | 2.94 | 33.29 | 32.32 | 0.98 | 0.98 | 0.02 | 0.02 |
| gene-LOC105279500 | https://www.ncbi.nlm.nih.gov/gene/?term=LOC105279500 | 3587 | 34 | 94.79 | 1 | 2.79 | 2.94 | 52.36 | 51.39 | 0.97 | 0.97 | 0.01 | 0.01 |
| gene-LOC105281272 | https://www.ncbi.nlm.nih.gov/gene/?term=LOC105281272 | 5252 | 156 | 297.03 | 1 | 1.9 | 0.64 | 18.91 | 17.94 | 0.97 | 0.97 | 0.03 | 0.03 |
| gene-LOC105280774 | https://www.ncbi.nlm.nih.gov/gene/?term=LOC105280774 | 2793 | 89 | 318.65 | 1 | 3.58 | 1.12 | 36.33 | 35.38 | 0.94 | 0.94 | 0.03 | 0.03 |
| gene-LOC105283987 | https://www.ncbi.nlm.nih.gov/gene/?term=LOC105283987 | 4841 | 64 | 132.2 | 1 | 2.07 | 1.56 | 30.28 | 29.39 | 0.89 | 0.89 | 0.01 | 0.01 |
| gene-LOC105287417 | https://www.ncbi.nlm.nih.gov/gene/?term=LOC105287417 | 1025 | 22 | 214.63 | 1 | 9.76 | 4.55 | 50.13 | 49.23 | 0.89 | 0.89 | 0.02 | 0.02 |
| gene-LOC105284036 | https://www.ncbi.nlm.nih.gov/gene/?term=LOC105284036 | 3254 | 96 | 295.02 | 1 | 3.07 | 1.04 | 22.78 | 21.95 | 0.83 | 0.83 | 0.02 | 0.02 |

| | | | | | | | | | | | | | | |
|---|---|---|---|---|---|---|---|---|---|---|---|---|---|---|
| gene-LOC105276449 | https://www.ncbi.nlm.nih.gov/gene/?term=LOC105276449 | 1231 | 42 | 341.19 | 1 | 8.12 | 2.38 | 35.95 | 35.13 | 0.82 | 0.82 | 0.03 | 0.03 | |
| gene-LOC105285055 | https://www.ncbi.nlm.nih.gov/gene/?term=LOC105285055 | 4108 | 255 | 620.74 | 2 | 4.87 | 0.78 | 11.66 | 10.91 | 0.76 | 0.76 | 0.05 | 0.05 | |
| gene-LOC105285202 | https://www.ncbi.nlm.nih.gov/gene/?term=LOC105285202 | 1838 | 67 | 364.53 | 1 | 5.44 | 1.49 | 26.25 | 25.49 | 0.76 | 0.76 | 0.03 | 0.03 | |
| gene-LOC105275363 | https://www.ncbi.nlm.nih.gov/gene/?term=LOC105275363 | 3798 | 150 | 394.94 | 1 | 2.63 | 0.67 | 18.12 | 17.37 | 0.75 | 0.75 | 0.03 | 0.03 | |
| gene-LOC105283592 | https://www.ncbi.nlm.nih.gov/gene/?term=LOC105283592 | 3479 | 72 | 206.96 | 1 | 2.87 | 1.39 | 23.84 | 23.1 | 0.74 | 0.74 | 0.02 | 0.02 | |
| gene-LOC105285395 | https://www.ncbi.nlm.nih.gov/gene/?term=LOC105285395 | 8085 | 185 | 228.82 | 1 | 1.24 | 0.54 | 16.99 | 16.27 | 0.72 | 0.72 | 0.02 | 0.02 | |
| gene-LOC105287095 | https://www.ncbi.nlm.nih.gov/gene/?term=LOC105287095 | 1435 | 57 | 397.21 | 1 | 6.97 | 1.75 | 29.87 | 29.16 | 0.71 | 0.71 | 0.03 | 0.03 | |
| gene-LOC113561621 | https://www.ncbi.nlm.nih.gov/gene/?term=LOC113561621 | 2249 | 82 | 364.61 | 1 | 4.45 | 1.22 | 20.63 | 19.95 | 0.68 | 0.68 | 0.02 | 0.02 | |
| gene-LOC105285639 | https://www.ncbi.nlm.nih.gov/gene/?term=LOC105285639 | 10325 | 319 | 308.96 | 1 | 0.97 | 0.31 | 12.35 | 11.71 | 0.65 | 0.65 | 0.02 | 0.02 | |
| gene-LOC105275374 | https://www.ncbi.nlm.nih.gov/gene/?term=LOC105275374 | 11376 | 489 | 429.85 | 1 | 0.88 | 0.2 | 11.7 | 11.06 | 0.64 | 0.64 | 0.03 | 0.03 | |
| gene-LOC105274851 | https://www.ncbi.nlm.nih.gov/gene/?term=LOC105274851 | 6616 | 123 | 185.91 | 1 | 1.51 | 0.81 | 11.44 | 10.81 | 0.63 | 0.63 | 0.01 | 0.01 | |
| gene-LOC105274452 | https://www.ncbi.nlm.nih.gov/gene/?term=LOC105274452 | 6595 | 252 | 382.11 | 1 | 1.52 | 0.4 | 13.74 | 13.13 | 0.61 | 0.61 | 0.02 | 0.02 | |
| gene-LOC105282885 | https://www.ncbi.nlm.nih.gov/gene/?term=LOC105282885 | 1870 | 45 | 240.64 | 1 | 5.35 | 2.22 | 42.36 | 41.77 | 0.58 | 0.58 | 0.01 | 0.01 | |
| gene-LOC105276692 | https://www.ncbi.nlm.nih.gov/gene/?term=LOC105276692 | 1745 | 29 | 166.19 | 1 | 5.73 | 3.45 | 25.51 | 24.96 | 0.55 | 0.55 | 0.01 | 0.01 | |
| gene-LOC105278145 | https://www.ncbi.nlm.nih.gov/gene/?term=LOC105278145 | 4470 | 246 | 550.34 | 1 | 2.24 | 0.41 | 11.03 | 10.48 | 0.55 | 0.55 | 0.03 | 0.03 | |
| gene-LOC105278283 | https://www.ncbi.nlm.nih.gov/gene/?term=LOC105278283 | 2317 | 125 | 539.49 | 1 | 4.32 | 0.8 | 28.1 | 27.55 | 0.55 | 0.55 | 0.03 | 0.03 | |
| gene-LOC105284704 | https://www.ncbi.nlm.nih.gov/gene/?term=LOC105284704 | 2400 | 57 | 237.5 1 | 4.17 | 1.75 | 25.25 | 24.69 | 0.55 | 0.55 | 0.01 | 0.01 | | |
| gene-LOC105282151 | https://www.ncbi.nlm.nih.gov/gene/?term=LOC105282151 | 1601 | 35 | 218.61 | 1 | 6.25 | 2.86 | 42.13 | 41.61 | 0.53 | 0.53 | 0.01 | 0.01 | |
| gene-LOC105274706 | https://www.ncbi.nlm.nih.gov/gene/?term=LOC105274706 | 1446 | 55 | 380.36 | 1 | 6.92 | 1.82 | 37.05 | 36.54 | 0.5 | 0.5 | 0.02 | 0.02 | |
| gene-LOC105280410 | https://www.ncbi.nlm.nih.gov/gene/?term=LOC105280410 | 12580 | 235 | 186.8 1 | 0.79 | 0.43 | 21.34 | 20.84 | 0.5 | 0.5 | 0.01 | 0.01 | | |
| gene-LOC105274823 | https://www.ncbi.nlm.nih.gov/gene/?term=LOC105274823 | 4620 | 132 | 285.71 | 1 | 2.16 | 0.76 | 21.94 | 21.47 | 0.47 | 0.47 | 0.01 | 0.01 | |
| gene-LOC105279038 | https://www.ncbi.nlm.nih.gov/gene/?term=LOC105279038 | 9453 | 306 | 323.71 | 1 | 1.06 | 0.33 | 10.13 | 9.67 | 0.46 | 0.46 | 0.01 | 0.01 | |
| gene-LOC105274856 | https://www.ncbi.nlm.nih.gov/gene/?term=LOC105274856 | 5216 | 207 | 396.86 | 1 | 1.92 | 0.48 | 19.99 | 19.54 | 0.45 | 0.45 | 0.02 | 0.02 | |
| gene-LOC105286514 | https://www.ncbi.nlm.nih.gov/gene/?term=LOC105286514 | 2528 | 44 | 174.05 | 1 | 3.96 | 2.27 | 33.92 | 33.47 | 0.45 | 0.45 | 0.01 | 0.01 | |
| gene-LOC105278666 | https://www.ncbi.nlm.nih.gov/gene/?term=LOC105278666 | 10341 | 429 | 414.85 | 1 | 0.97 | 0.23 | 7.71 | 7.27 | 0.44 | 0.44 | 0.02 | 0.02 | |
| gene-LOC105285406 | https://www.ncbi.nlm.nih.gov/gene/?term=LOC105285406 | 8015 | 60 | 74.86 1 | 1.25 | 1.67 | 25.96 | 25.52 | 0.44 | 0.44 | 0 | 0 | | |
| gene-LOC105285807 | https://www.ncbi.nlm.nih.gov/gene/?term=LOC105285807 | 6063 | 214 | 352.96 | 1 | 1.65 | 0.47 | 7.55 | 7.11 | 0.44 | 0.44 | 0.02 | 0.02 | |
| gene-LOC105278282 | https://www.ncbi.nlm.nih.gov/gene/?term=LOC105278282 | 1775 | 52 | 292.96 | 1 | 5.63 | 1.92 | 38.05 | 37.63 | 0.42 | 0.42 | 0.01 | 0.01 | |
| gene-LOC105276612 | https://www.ncbi.nlm.nih.gov/gene/?term=LOC105276612 | 4157 | 181 | 435.41 | 1 | 2.41 | 0.55 | 10.16 | 9.76 | 0.4 | 0.4 | 0.02 | 0.02 | |
| gene-LOC105276941 | https://www.ncbi.nlm.nih.gov/gene/?term=LOC105276941 | 15956 | 494 | 309.6 1 | 0.63 | 0.2 | 5.44 | 5.07 | 0.38 | 0.38 | 0.01 | 0.01 | | |
| gene-LOC105277423 | https://www.ncbi.nlm.nih.gov/gene/?term=LOC105277423 | 2340 | 108 | 461.54 | 1 | 4.27 | 0.93 | 30.29 | 29.91 | 0.38 | 0.38 | 0.02 | 0.02 | |
| gene-LOC105287689 | https://www.ncbi.nlm.nih.gov/gene/?term=LOC105287689 | 10867 | 331 | 304.59 | 1 | 0.92 | 0.3 | 9.53 | 9.17 | 0.36 | 0.36 | 0.01 | 0.01 | |
| gene-LOC105286194 | https://www.ncbi.nlm.nih.gov/gene/?term=LOC105286194 | 152619 | | 2040 | 133.67 | 1 | 0.07 | 0.05 | 6.36 | 6.07 | 0.3 | 0.3 | 0 | 0 |
| gene-LOC105276452 | https://www.ncbi.nlm.nih.gov/gene/?term=LOC105276452 | 2917 | 263 | 901.61 | 1 | 3.43 | 0.38 | 9.41 | 9.13 | 0.28 | 0.28 | 0.03 | 0.03 | |
| gene-LOC105278454 | https://www.ncbi.nlm.nih.gov/gene/?term=LOC105278454 | 1972 | 44 | 223.12 | 1 | 5.07 | 2.27 | 46.65 | 46.38 | 0.26 | 0.26 | 0.01 | 0.01 | |
| gene-LOC105278877 | https://www.ncbi.nlm.nih.gov/gene/?term=LOC105278877 | 1968 | 69 | 350.61 | 1 | 5.08 | 1.45 | 35.47 | 35.23 | 0.25 | 0.25 | 0.01 | 0.01 | |
| gene-LOC105280407 | https://www.ncbi.nlm.nih.gov/gene/?term=LOC105280407 | 10950 | 189 | 172.6 1 | 0.91 | 0.53 | 16.21 | 15.97 | 0.24 | 0.24 | 0 | 0 | | |
| gene-LOC105275460 | https://www.ncbi.nlm.nih.gov/gene/?term=LOC105275460 | 2473 | 116 | 469.07 | 1 | 4.04 | 0.86 | 13.65 | 13.42 | 0.23 | 0.23 | 0.01 | 0.01 | |
| gene-LOC113562050 | https://www.ncbi.nlm.nih.gov/gene/?term=LOC113562050 | 17097 | 540 | 315.84 | 1 | 0.58 | 0.19 | 5.38 | 5.15 | 0.23 | 0.23 | 0.01 | 0.01 | |
| gene-LOC105278534 | https://www.ncbi.nlm.nih.gov/gene/?term=LOC105278534 | 7364 | 175 | 237.64 | 1 | 1.36 | 0.57 | 18.73 | 18.51 | 0.22 | 0.22 | 0.01 | 0.01 | |
| gene-LOC105286166 | https://www.ncbi.nlm.nih.gov/gene/?term=LOC105286166 | 3050 | 103 | 337.7 1 | 3.28 | 0.97 | 25.5 | 25.28 | 0.22 | 0.22 | 0.01 | 0.01 | | |
| gene-LOC105274449 | https://www.ncbi.nlm.nih.gov/gene/?term=LOC105274449 | 1549 | 48 | 309.88 | 1 | 6.46 | 2.08 | 51.17 | 50.96 | 0.21 | 0.21 | 0.01 | 0.01 | |
| gene-LOC105280956 | https://www.ncbi.nlm.nih.gov/gene/?term=LOC105280956 | 2639 | 69 | 261.46 | 1 | 3.79 | 1.45 | 18.51 | 18.31 | 0.21 | 0.21 | 0.01 | 0.01 | |
| gene-LOC105275683 | https://www.ncbi.nlm.nih.gov/gene/?term=LOC105275683 | 4256 | 183 | 429.98 | 1 | 2.35 | 0.55 | 12.85 | 12.65 | 0.2 | 0.2 | 0.01 | 0.01 | |
| gene-LOC105287764 | https://www.ncbi.nlm.nih.gov/gene/?term=LOC105287764 | 18359 | 375 | 204.26 | 1 | 0.54 | 0.27 | 3.25 | 3.09 | 0.17 | 0.17 | 0 | 0 | |
| gene-LOC105282658 | https://www.ncbi.nlm.nih.gov/gene/?term=LOC105282658 | 2421 | 54 | 223.05 | 1 | 4.13 | 1.85 | 19.57 | 19.42 | 0.16 | 0.16 | 0 | 0 | |
| gene-LOC105282825 | https://www.ncbi.nlm.nih.gov/gene/?term=LOC105282825 | 3919 | 69 | 176.07 | 1 | 2.55 | 1.45 | 13.52 | 13.36 | 0.16 | 0.16 | 0 | 0 | |
| gene-LOC105276462 | https://www.ncbi.nlm.nih.gov/gene/?term=LOC105276462 | 6561 | 246 | 374.94 | 1 | 1.52 | 0.41 | 8.14 | 7.99 | 0.15 | 0.15 | 0.01 | 0.01 | |
| gene-LOC105278455 | https://www.ncbi.nlm.nih.gov/gene/?term=LOC105278455 | 27181 | 882 | 324.49 | 1 | 0.37 | 0.11 | 7.78 | 7.66 | 0.12 | 0.12 | 0 | 0 | |
| gene-LOC105286999 | https://www.ncbi.nlm.nih.gov/gene/?term=LOC105286999 | 2141 | 42 | 196.17 | 1 | 4.67 | 2.38 | 36.17 | 36.05 | 0.12 | 0.12 | 0 | 0 | |
| gene-LOC105282811 | https://www.ncbi.nlm.nih.gov/gene/?term=LOC105282811 | 5765 | 198 | 343.45 | 1 | 1.73 | 0.51 | 7.89 | 7.77 | 0.11 | 0.11 | 0 | 0 | |
| gene-LOC105275327 | https://www.ncbi.nlm.nih.gov/gene/?term=LOC105275327 | 14245 | 460 | 322.92 | 1 | 0.7 | 0.22 | 7.25 | 7.2 | 0.05 | 0.05 | 0 | 0 | |
| gene-LOC105280265 | https://www.ncbi.nlm.nih.gov/gene/?term=LOC105280265 | 85984 | 2858 | 332.39 | 1 | 0.12 | 0.03 | 1.09 | 1.08 | 0.01 | 0.01 | 0 | 0 | |
| gene-LOC105277125 | https://www.ncbi.nlm.nih.gov/gene/?term=LOC105277125 | 51079 | 1693 | 331.45 | 1 | 0.2 | 0.06 | 1.64 | 1.64 | 0 | 0 | 0 | 0 | |
| gene-LOC105277866 | https://www.ncbi.nlm.nih.gov/gene/?term=LOC105277866 | 33045 | 352 | 106.52 | 1 | 0.3 | 0.28 | 3.39 | 3.44 | -0.04 | 0.04 | 0 | 0 | |
| gene-LOC105282646 | https://www.ncbi.nlm.nih.gov/gene/?term=LOC105282646 | 3317 | 99 | 298.46 | 1 | 3.01 | 1.01 | 25.07 | 25.11 | -0.04 | 0.04 | 0 | 0 | |
| gene-LOC105274531 | https://www.ncbi.nlm.nih.gov/gene/?term=LOC105274531 | 7774 | 285 | 366.61 | 1 | 1.29 | 0.35 | 10.62 | 10.74 | -0.11 | 0.11 | 0 | 0 | |
| gene-LOC105280377 | https://www.ncbi.nlm.nih.gov/gene/?term=LOC105280377 | 2024 | 58 | 286.56 | 1 | 4.94 | 1.72 | 49.97 | 50.08 | -0.11 | 0.11 | 0 | 0 | |
| gene-LOC105275848 | https://www.ncbi.nlm.nih.gov/gene/?term=LOC105275848 | 3718 | 133 | 357.72 | 1 | 2.69 | 0.75 | 21.71 | 21.82 | -0.12 | 0.12 | 0 | 0 | |
| gene-LOC105276538 | https://www.ncbi.nlm.nih.gov/gene/?term=LOC105276538 | 1511 | 19 | 125.74 | 1 | 6.62 | 5.26 | 67.47 | 67.59 | -0.12 | 0.12 | 0 | 0 | |
| gene-LOC105283296 | https://www.ncbi.nlm.nih.gov/gene/?term=LOC105283296 | 2174 | 84 | 386.38 | 1 | 4.6 | 1.19 | 19.37 | 19.49 | -0.12 | 0.12 | 0 | 0 | |
| gene-LOC105278632 | https://www.ncbi.nlm.nih.gov/gene/?term=LOC105278632 | 3624 | 113 | 311.81 | 1 | 2.76 | 0.88 | 16.03 | 16.16 | -0.13 | 0.13 | 0 | 0 | |
| gene-LOC105283943 | https://www.ncbi.nlm.nih.gov/gene/?term=LOC105283943 | 3559 | 42 | 118.01 | 1 | 2.81 | 2.38 | 14.31 | 14.48 | -0.17 | 0.17 | 0 | 0 | |
| gene-LOC105275179 | https://www.ncbi.nlm.nih.gov/gene/?term=LOC105275179 | 3271 | 66 | 201.77 | 1 | 3.06 | 1.52 | 20.6 | 20.81 | -0.21 | 0.21 | 0 | 0 | |

| | | | | | | | | | | | | | | | |
|---|---|---|---|---|---|---|---|---|---|---|---|---|---|---|---|
| gene-LOC105278649 | https://www.ncbi.nlm.nih.gov/gene/?term=LOC105278649 | 5616 | 161 | 286.68 | | 1 | 1.78 | 0.62 | 11.64 | 11.9 | -0.26 | 0.26 | -0.01 | 0.01 | |
| gene-LOC105276527 | https://www.ncbi.nlm.nih.gov/gene/?term=LOC105276527 | 7923 | 309 | 390 | 1 | 1.26 | 0.32 | 9.63 | 9.9 | -0.27 | 0.27 | -0.01 | 0.01 | | |
| gene-LOC105286649 | https://www.ncbi.nlm.nih.gov/gene/?term=LOC105286649 | 14178 | 778 | 548.74 | | 1 | 0.71 | 0.13 | 0.86 | 1.14 | -0.27 | 0.27 | -0.02 | 0.02 | |
| gene-LOC105285514 | https://www.ncbi.nlm.nih.gov/gene/?term=LOC105285514 | 1277 | 14 | 109.63 | | 1 | 7.83 | 7.14 | 85.59 | 85.88 | -0.28 | 0.28 | 0 | 0 | |
| gene-LOC105286274 | https://www.ncbi.nlm.nih.gov/gene/?term=LOC105286274 | 3817 | 193 | 505.63 | | 1 | 2.62 | 0.52 | 16.76 | 17.05 | -0.29 | 0.29 | -0.01 | 0.01 | |
| gene-LOC105286515 | https://www.ncbi.nlm.nih.gov/gene/?term=LOC105286515 | 3751 | 99 | 263.93 | | 1 | 2.67 | 1.01 | 13.14 | 13.49 | -0.36 | 0.36 | -0.01 | 0.01 | |
| gene-LOC109611234 | https://www.ncbi.nlm.nih.gov/gene/?term=LOC109611234 | 3663 | 28 | 76.44 | 1 | 2.73 | 3.57 | 16.2 | 16.56 | -0.36 | 0.36 | 0 | 0 | | |
| gene-LOC105287633 | https://www.ncbi.nlm.nih.gov/gene/?term=LOC105287633 | 1524 | 27 | 177.17 | | 1 | 6.56 | 3.7 | 63.52 | 64 | -0.48 | 0.48 | -0.01 | 0.01 | |
| gene-LOC105286452 | https://www.ncbi.nlm.nih.gov/gene/?term=LOC105286452 | 6888 | 62 | 90.01 | 1 | 1.45 | 1.61 | 32.65 | 33.21 | -0.56 | 0.56 | -0.01 | 0.01 | | |
| gene-LOC105278108 | https://www.ncbi.nlm.nih.gov/gene/?term=LOC105278108 | 4392 | 44 | 100.18 | | 1 | 2.28 | 2.27 | 24.81 | 25.4 | -0.59 | 0.59 | -0.01 | 0.01 | |
| gene-LOC105275187 | https://www.ncbi.nlm.nih.gov/gene/?term=LOC105275187 | 5534 | 66 | 119.26 | | 1 | 1.81 | 1.52 | 11.92 | 12.68 | -0.77 | 0.77 | -0.01 | 0.01 | |
| gene-LOC105285252 | https://www.ncbi.nlm.nih.gov/gene/?term=LOC105285252 | 7427 | 112 | 150.8 | 2 | 2.69 | 1.79 | 14.17 | 15.17 | -1 | 1 | -0.02 | 0.02 | | |
| gene-LOC105281700 | https://www.ncbi.nlm.nih.gov/gene/?term=LOC105281700 | 2208 | 34 | 153.99 | | 1 | 4.53 | 2.94 | 35.91 | 36.92 | -1.02 | 1.02 | -0.02 | 0.02 | |
| gene-LOC105285860 | https://www.ncbi.nlm.nih.gov/gene/?term=LOC105285860 | 3110 | 52 | 167.2 | 1 | 3.22 | 1.92 | 20.92 | 22.04 | -1.12 | 1.12 | -0.02 | 0.02 | | |
| gene-LOC105284507 | https://www.ncbi.nlm.nih.gov/gene/?term=LOC105284507 | 5207 | 77 | 147.88 | | 1 | 1.92 | 1.3 | 22.93 | 24.13 | -1.2 | 1.2 | -0.02 | 0.02 | |
| gene-LOC105274534 | https://www.ncbi.nlm.nih.gov/gene/?term=LOC105274534 | 1329 | 56 | 421.37 | | 1 | 7.52 | 1.79 | 47.04 | 48.26 | -1.22 | 1.22 | -0.05 | 0.05 | |
| gene-LOC109611283 | https://www.ncbi.nlm.nih.gov/gene/?term=LOC109611283 | 1451 | 36 | 248.1 | 1 | 6.89 | 2.78 | 35.62 | 36.97 | -1.35 | 1.35 | -0.03 | 0.03 | | |
| gene-LOC105280819 | https://www.ncbi.nlm.nih.gov/gene/?term=LOC105280819 | 1639 | 37 | 225.75 | | 1 | 6.1 | 2.7 | 36.55 | 38.07 | -1.52 | 1.52 | -0.03 | 0.03 | |
| gene-LOC105282067 | https://www.ncbi.nlm.nih.gov/gene/?term=LOC105282067 | 5941 | 51 | 85.84 | 1 | 1.68 | 1.96 | 26.09 | 27.84 | -1.75 | 1.75 | -0.01 | 0.01 | | |
| gene-LOC105276196 | https://www.ncbi.nlm.nih.gov/gene/?term=LOC105276196 | 2953 | 48 | 162.55 | | 1 | 3.39 | 2.08 | 47.64 | 49.83 | -2.19 | 2.19 | -0.04 | 0.04 | |
| gene-LOC105283989 | https://www.ncbi.nlm.nih.gov/gene/?term=LOC105283989 | 4948 | 26 | 52.55 | 1 | 2.02 | 3.85 | 37.24 | 39.59 | -2.34 | 2.34 | -0.01 | 0.01 | | |
| gene-LOC105284677 | https://www.ncbi.nlm.nih.gov/gene/?term=LOC105284677 | 1905 | 39 | 204.72 | | 3 | 15.75 | 7.69 | 19.98 | 31.6 | -11.62 | | 11.62 | -0.24 | 0.24 |

Directory: OGLall File: ogl-Ob_LI_BCphaseB1.txt

| gene_ID | gene_link | gwidth | #Sites | #per10Kb | %perSite | %pNucl |
|---|---|---|---|---|---|---|
| gene-LOC113561532 | https://www.ncbi.nlm.nih.gov/gene/?term=LOC113561532 | 72 | 5 | 694.44 | 94.44 | 6.56 |
| gene-LOC105286531 | https://www.ncbi.nlm.nih.gov/gene/?term=LOC105286531 | 372 | 43 | 1155.91 | 37.65 | 4.35 |
| gene-LOC105285635 | https://www.ncbi.nlm.nih.gov/gene/?term=LOC105285635 | 1175 | 148 | 1259.57 | 31.99 | 4.03 |
| gene-LOC105280739 | https://www.ncbi.nlm.nih.gov/gene/?term=LOC105280739 | 1991 | 143 | 718.23 | 54.78 | 3.93 |
| gene-LOC105286801 | https://www.ncbi.nlm.nih.gov/gene/?term=LOC105286801 | 1102 | 104 | 943.74 | 40.7 | 3.84 |
| gene-LOC113562627 | https://www.ncbi.nlm.nih.gov/gene/?term=LOC113562627 | 397 | 21 | 528.97 | 66.45 | 3.52 |
| gene-LOC105278160 | https://www.ncbi.nlm.nih.gov/gene/?term=LOC105278160 | 1872 | 163 | 870.73 | 39.22 | 3.42 |
| gene-LOC105274534 | https://www.ncbi.nlm.nih.gov/gene/?term=LOC105274534 | 1329 | 108 | 812.64 | 41.62 | 3.38 |
| gene-LOC105287811 | https://www.ncbi.nlm.nih.gov/gene/?term=LOC105287811 | 2155 | 152 | 705.34 | 47.67 | 3.36 |
| gene-LOC105278844 | https://www.ncbi.nlm.nih.gov/gene/?term=LOC105278844 | 1101 | 125 | 1135.33 | 29.16 | 3.31 |
| gene-LOC105278849 | https://www.ncbi.nlm.nih.gov/gene/?term=LOC105278849 | 776 | 101 | 1301.55 | 25.2 | 3.28 |
| gene-LOC105279052 | https://www.ncbi.nlm.nih.gov/gene/?term=LOC105279052 | 1130 | 139 | 1230.09 | 26.62 | 3.27 |
| gene-LOC105286235 | https://www.ncbi.nlm.nih.gov/gene/?term=LOC105286235 | 1510 | 111 | 735.1 | 44.43 | 3.27 |
| gene-LOC105286737 | https://www.ncbi.nlm.nih.gov/gene/?term=LOC105286737 | 2690 | 392 | 1457.25 | 22.38 | 3.26 |
| gene-LOC105287692 | https://www.ncbi.nlm.nih.gov/gene/?term=LOC105287692 | 1505 | 146 | 970.1 | 33.65 | 3.26 |
| gene-LOC105282149 | https://www.ncbi.nlm.nih.gov/gene/?term=LOC105282149 | 1787 | 146 | 817.01 | 39.37 | 3.22 |
| gene-LOC105274711 | https://www.ncbi.nlm.nih.gov/gene/?term=LOC105274711 | 1655 | 110 | 664.65 | 47.74 | 3.17 |
| gene-LOC105286269 | https://www.ncbi.nlm.nih.gov/gene/?term=LOC105286269 | 1401 | 85 | 606.71 | 51.17 | 3.1 |
| gene-LOC105277648 | https://www.ncbi.nlm.nih.gov/gene/?term=LOC105277648 | 722 | 40 | 554.02 | 55.5 | 3.07 |
| gene-LOC105282986 | https://www.ncbi.nlm.nih.gov/gene/?term=LOC105282986 | 1600 | 137 | 856.25 | 35.76 | 3.06 |
| gene-LOC105276360 | https://www.ncbi.nlm.nih.gov/gene/?term=LOC105276360 | 1466 | 162 | 1105.05 | 27 | 2.98 |
| gene-LOC105276371 | https://www.ncbi.nlm.nih.gov/gene/?term=LOC105276371 | 871 | 55 | 631.46 | 47.22 | 2.98 |
| gene-LOC105287677 | https://www.ncbi.nlm.nih.gov/gene/?term=LOC105287677 | 1588 | 118 | 743.07 | 39.84 | 2.96 |
| gene-LOC105274543 | https://www.ncbi.nlm.nih.gov/gene/?term=LOC105274543 | 1572 | 134 | 852.42 | 34.64 | 2.95 |

...

... only first 25 lines shown ...

Directory: OGLall File: ogl-Ob_LI_BCphaseB1.vs.BCphaseB2.txt

| gene_ID | gene_link | gwidth | #Sites | #per10Kb | #dmSites | #dmsp10kb | %dmSites | %pSite1 | %pSite2 | DMpSite | ADMpSite | DMpNucl | ADMpNucl |
|---|---|---|---|---|---|---|---|---|---|---|---|---|---|
| gene-LOC113562345 | https://www.ncbi.nlm.nih.gov/gene/?term=LOC113562345 | 2290 | 64 | 279.48 | 15 | 65.5 | 23.44 | 32.83 | 17.93 | 14.9 | 14.9 | 0.42 | 0.42 |
| gene-LOC105275070 | https://www.ncbi.nlm.nih.gov/gene/?term=LOC105275070 | 5667 | 49 | 86.47 | 6 | 10.59 | 12.24 | 10.93 | 1.84 | 9.09 | 9.09 | 0.08 | 0.08 |
| gene-LOC105275708 | https://www.ncbi.nlm.nih.gov/gene/?term=LOC105275708 | 1772 | 15 | 84.65 | 1 | 5.64 | 6.67 | 46.38 | 38.5 | 7.88 | 7.88 | 0.07 | 0.07 |
| gene-LOC105281039 | https://www.ncbi.nlm.nih.gov/gene/?term=LOC105281039 | 1471 | 12 | 81.58 | 1 | 6.8 | 8.33 | 38.6 | 31.46 | 7.13 | 7.13 | 0.06 | 0.06 |
| gene-LOC105279898 | https://www.ncbi.nlm.nih.gov/gene/?term=LOC105279898 | 1343 | 13 | 96.8 | 1 | 7.45 | 7.69 | 64.63 | 59.11 | 5.52 | 5.52 | 0.05 | 0.05 |
| gene-LOC105275518 | https://www.ncbi.nlm.nih.gov/gene/?term=LOC105275518 | 2335 | 23 | 98.5 | 1 | 4.28 | 4.35 | 43.86 | 38.74 | 5.13 | 5.13 | 0.05 | 0.05 |
| gene-LOC105275447 | https://www.ncbi.nlm.nih.gov/gene/?term=LOC105275447 | 1846 | 32 | 173.35 | 1 | 5.42 | 3.12 | 8.01 | 3.17 | 4.84 | 4.84 | 0.08 | 0.08 |
| gene-LOC105274900 | https://www.ncbi.nlm.nih.gov/gene/?term=LOC105274900 | 5561 | 54 | 97.1 | 2 | 3.6 | 3.7 | 35.58 | 31 | 4.57 | 4.57 | 0.04 | 0.04 |
| gene-LOC105283649 | https://www.ncbi.nlm.nih.gov/gene/?term=LOC105283649 | 1918 | 56 | 291.97 | 1 | 5.21 | 1.79 | 39.64 | 36.17 | 3.47 | 3.47 | 0.1 | 0.1 |
| gene-LOC105281515 | https://www.ncbi.nlm.nih.gov/gene/?term=LOC105281515 | 8827 | 57 | 64.57 | 2 | 2.27 | 3.51 | 30.56 | 27.6 | 2.96 | 2.96 | 0.02 | 0.02 |
| gene-LOC105276538 | https://www.ncbi.nlm.nih.gov/gene/?term=LOC105276538 | 1511 | 17 | 112.51 | 1 | 6.62 | 5.88 | 72.11 | 69.24 | 2.87 | 2.87 | 0.03 | 0.03 |
| gene-LOC105285674 | https://www.ncbi.nlm.nih.gov/gene/?term=LOC105285674 | 3035 | 94 | 309.72 | 2 | 6.59 | 2.13 | 27.11 | 24.65 | 2.46 | 2.46 | 0.08 | 0.08 |
| gene-LOC105283989 | https://www.ncbi.nlm.nih.gov/gene/?term=LOC105283989 | 4948 | 31 | 62.65 | 1 | 2.02 | 3.23 | 46.67 | 44.28 | 2.39 | 2.39 | 0.01 | 0.01 |
| gene-LOC105285514 | https://www.ncbi.nlm.nih.gov/gene/?term=LOC105285514 | 1277 | 15 | 117.46 | 1 | 7.83 | 6.67 | 80.63 | 78.25 | 2.38 | 2.38 | 0.03 | 0.03 |
| gene-LOC105281420 | https://www.ncbi.nlm.nih.gov/gene/?term=LOC105281420 | 3882 | 67 | 172.59 | 1 | 2.58 | 1.49 | 15.32 | 12.95 | 2.37 | 2.37 | 0.04 | 0.04 |
| gene-LOC105279900 | https://www.ncbi.nlm.nih.gov/gene/?term=LOC105279900 | 2191 | 56 | 255.59 | 1 | 4.56 | 1.79 | 33.98 | 31.69 | 2.29 | 2.29 | 0.06 | 0.06 |
| gene-LOC105285315 | https://www.ncbi.nlm.nih.gov/gene/?term=LOC105285315 | 5458 | 26 | 47.64 | 1 | 1.83 | 3.85 | 38.49 | 36.2 | 2.29 | 2.29 | 0.01 | 0.01 |
| gene-LOC105276449 | https://www.ncbi.nlm.nih.gov/gene/?term=LOC105276449 | 1231 | 51 | 414.3 | 1 | 8.12 | 1.96 | 27.95 | 25.74 | 2.21 | 2.21 | 0.09 | 0.09 |
| gene-LOC105278877 | https://www.ncbi.nlm.nih.gov/gene/?term=LOC105278877 | 1968 | 82 | 416.67 | 1 | 5.08 | 1.22 | 32.75 | 30.76 | 1.99 | 1.99 | 0.08 | 0.08 |
| gene-LOC105283592 | https://www.ncbi.nlm.nih.gov/gene/?term=LOC105283592 | 3479 | 71 | 204.08 | 1 | 2.87 | 1.41 | 27.65 | 25.73 | 1.93 | 1.93 | 0.04 | 0.04 |
| gene-LOC105278454 | https://www.ncbi.nlm.nih.gov/gene/?term=LOC105278454 | 1972 | 48 | 243.41 | 1 | 5.07 | 2.08 | 42.9 | 41.15 | 1.75 | 1.75 | 0.04 | 0.04 |
| gene-LOC105279664 | https://www.ncbi.nlm.nih.gov/gene/?term=LOC105279664 | 4023 | 65 | 161.57 | 1 | 2.49 | 1.54 | 19.54 | 17.85 | 1.7 | 1.7 | 0.03 | 0.03 |
| gene-LOC105286514 | https://www.ncbi.nlm.nih.gov/gene/?term=LOC105286514 | 2528 | 46 | 181.96 | 1 | 3.96 | 2.17 | 37.97 | 36.35 | 1.61 | 1.61 | 0.03 | 0.03 |
| gene-LOC105283943 | https://www.ncbi.nlm.nih.gov/gene/?term=LOC105283943 | 3559 | 62 | 174.21 | 1 | 2.81 | 1.61 | 15.22 | 13.63 | 1.6 | 1.6 | 0.03 | 0.03 |
| gene-LOC105285583 | https://www.ncbi.nlm.nih.gov/gene/?term=LOC105285583 | 1947 | 45 | 231.12 | 1 | 5.14 | 2.22 | 29.78 | 28.25 | 1.53 | 1.53 | 0.04 | 0.04 |
| gene-LOC105282156 | https://www.ncbi.nlm.nih.gov/gene/?term=LOC105282156 | 2577 | 137 | 531.63 | 1 | 3.88 | 0.73 | 24.98 | 23.48 | 1.5 | 1.5 | 0.08 | 0.08 |
| gene-LOC105276635 | https://www.ncbi.nlm.nih.gov/gene/?term=LOC105276635 | 4962 | 33 | 66.51 | 1 | 2.02 | 3.03 | 19.08 | 17.59 | 1.49 | 1.49 | 0.01 | 0.01 |
| gene-LOC105287648 | https://www.ncbi.nlm.nih.gov/gene/?term=LOC105287648 | 2205 | 30 | 136.05 | 1 | 4.54 | 3.33 | 43.37 | 41.93 | 1.44 | 1.44 | 0.02 | 0.02 |
| gene-LOC105277903 | https://www.ncbi.nlm.nih.gov/gene/?term=LOC105277903 | 1835 | 37 | 201.63 | 1 | 5.45 | 2.7 | 54.03 | 52.65 | 1.39 | 1.39 | 0.03 | 0.03 |
| gene-LOC105287417 | https://www.ncbi.nlm.nih.gov/gene/?term=LOC105287417 | 1025 | 36 | 351.22 | 1 | 9.76 | 2.78 | 35.49 | 34.17 | 1.32 | 1.32 | 0.05 | 0.05 |
| gene-LOC105287854 | https://www.ncbi.nlm.nih.gov/gene/?term=LOC105287854 | 3644 | 67 | 183.86 | 1 | 2.74 | 1.49 | 16.53 | 15.24 | 1.29 | 1.29 | 0.02 | 0.02 |
| gene-LOC105282794 | https://www.ncbi.nlm.nih.gov/gene/?term=LOC105282794 | 4571 | 98 | 214.4 | 1 | 2.19 | 1.02 | 19.72 | 18.47 | 1.25 | 1.25 | 0.03 | 0.03 |
| gene-LOC105282991 | https://www.ncbi.nlm.nih.gov/gene/?term=LOC105282991 | 5617 | 105 | 186.93 | 1 | 1.78 | 0.95 | 19.71 | 18.51 | 1.19 | 1.19 | 0.02 | 0.02 |
| gene-LOC105280956 | https://www.ncbi.nlm.nih.gov/gene/?term=LOC105280956 | 2639 | 75 | 284.2 | 1 | 3.79 | 1.33 | 17.61 | 16.45 | 1.16 | 1.16 | 0.03 | 0.03 |
| gene-LOC105278632 | https://www.ncbi.nlm.nih.gov/gene/?term=LOC105278632 | 3624 | 126 | 347.68 | 1 | 2.76 | 0.79 | 16.08 | 14.93 | 1.15 | 1.15 | 0.04 | 0.04 |
| gene-LOC105288078 | https://www.ncbi.nlm.nih.gov/gene/?term=LOC105288078 | 19126 | 309 | 161.56 | 1 | 0.52 | 0.32 | 17.45 | 16.3 | 1.15 | 1.15 | 0.02 | 0.02 |
| gene-LOC105280786 | https://www.ncbi.nlm.nih.gov/gene/?term=LOC105280786 | 2573 | 119 | 462.5 | 1 | 3.89 | 0.84 | 30.85 | 29.73 | 1.12 | 1.12 | 0.05 | 0.05 |
| gene-LOC105287063 | https://www.ncbi.nlm.nih.gov/gene/?term=LOC105287063 | 1874 | 51 | 272.15 | 1 | 5.34 | 1.96 | 30.2 | 29.07 | 1.12 | 1.12 | 0.03 | 0.03 |
| gene-LOC105280641 | https://www.ncbi.nlm.nih.gov/gene/?term=LOC105280641 | 3682 | 55 | 149.38 | 1 | 2.72 | 1.82 | 19.12 | 18.02 | 1.1 | 1.1 | 0.02 | 0.02 |
| gene-LOC105280002 | https://www.ncbi.nlm.nih.gov/gene/?term=LOC105280002 | 1940 | 28 | 144.33 | 1 | 5.15 | 3.57 | 40.24 | 39.15 | 1.09 | 1.09 | 0.02 | 0.02 |
| gene-LOC105285202 | https://www.ncbi.nlm.nih.gov/gene/?term=LOC105285202 | 1838 | 63 | 342.76 | 1 | 5.44 | 1.59 | 23.99 | 22.95 | 1.05 | 1.05 | 0.04 | 0.04 |
| gene-LOC105284355 | https://www.ncbi.nlm.nih.gov/gene/?term=LOC105284355 | 3904 | 158 | 404.71 | 1 | 2.56 | 0.63 | 20.65 | 19.62 | 1.02 | 1.02 | 0.04 | 0.04 |
| gene-LOC105280348 | https://www.ncbi.nlm.nih.gov/gene/?term=LOC105280348 | 3797 | 76 | 200.16 | 1 | 2.63 | 1.32 | 20.16 | 19.19 | 0.97 | 0.97 | 0.02 | 0.02 |
| gene-LOC105288036 | https://www.ncbi.nlm.nih.gov/gene/?term=LOC105288036 | 5110 | 227 | 444.23 | 1 | 1.96 | 0.44 | 15.78 | 14.87 | 0.91 | 0.91 | 0.04 | 0.04 |
| gene-LOC105275384 | https://www.ncbi.nlm.nih.gov/gene/?term=LOC105275384 | 1987 | 92 | 463.01 | 1 | 5.03 | 1.09 | 32.77 | 31.88 | 0.89 | 0.89 | 0.04 | 0.04 |
| gene-LOC105288042 | https://www.ncbi.nlm.nih.gov/gene/?term=LOC105288042 | 3507 | 99 | 282.29 | 1 | 2.85 | 1.01 | 23.14 | 22.25 | 0.89 | 0.89 | 0.02 | 0.02 |
| gene-LOC105284117 | https://www.ncbi.nlm.nih.gov/gene/?term=LOC105284117 | 4340 | 94 | 216.59 | 1 | 2.3 | 1.06 | 11.08 | 10.26 | 0.83 | 0.83 | 0.02 | 0.02 |
| gene-LOC105275752 | https://www.ncbi.nlm.nih.gov/gene/?term=LOC105275752 | 3462 | 130 | 375.51 | 1 | 2.89 | 0.77 | 31.52 | 30.74 | 0.77 | 0.77 | 0.03 | 0.03 |
| gene-LOC105284116 | https://www.ncbi.nlm.nih.gov/gene/?term=LOC105284116 | 3970 | 107 | 269.52 | 1 | 2.52 | 0.93 | 7.59 | 6.83 | 0.75 | 0.75 | 0.02 | 0.02 |
| gene-LOC105276692 | https://www.ncbi.nlm.nih.gov/gene/?term=LOC105276692 | 1745 | 34 | 194.84 | 1 | 5.73 | 2.94 | 22.19 | 21.46 | 0.73 | 0.73 | 0.01 | 0.01 |
| gene-LOC105282600 | https://www.ncbi.nlm.nih.gov/gene/?term=LOC105282600 | 3715 | 50 | 134.59 | 1 | 2.69 | 2 | 33.68 | 32.96 | 0.73 | 0.73 | 0.01 | 0.01 |
| gene-LOC105285639 | https://www.ncbi.nlm.nih.gov/gene/?term=LOC105285639 | 10325 | 378 | 366.1 | 1 | 0.97 | 0.26 | 11.07 | 10.35 | 0.72 | 0.72 | 0.03 | 0.03 |
| gene-LOC105280405 | https://www.ncbi.nlm.nih.gov/gene/?term=LOC105280405 | 5422 | 161 | 296.94 | 1 | 1.84 | 0.62 | 10.76 | 10.05 | 0.71 | 0.71 | 0.02 | 0.02 |
| gene-LOC105288066 | https://www.ncbi.nlm.nih.gov/gene/?term=LOC105288066 | 1964 | 30 | 152.75 | 1 | 5.09 | 3.33 | 35.84 | 35.13 | 0.71 | 0.71 | 0.01 | 0.01 |
| gene-LOC105278807 | https://www.ncbi.nlm.nih.gov/gene/?term=LOC105278807 | 2166 | 80 | 369.34 | 1 | 4.62 | 1.25 | 18.57 | 17.87 | 0.69 | 0.69 | 0.03 | 0.03 |
| gene-LOC105281266 | https://www.ncbi.nlm.nih.gov/gene/?term=LOC105281266 | 4102 | 79 | 192.59 | 1 | 2.44 | 1.27 | 15.99 | 15.33 | 0.66 | 0.66 | 0.01 | 0.01 |
| gene-LOC105281848 | https://www.ncbi.nlm.nih.gov/gene/?term=LOC105281848 | 2122 | 60 | 282.75 | 1 | 4.71 | 1.67 | 20.32 | 19.71 | 0.62 | 0.62 | 0.02 | 0.02 |
| gene-LOC105278497 | https://www.ncbi.nlm.nih.gov/gene/?term=LOC105278497 | 2072 | 95 | 458.49 | 1 | 4.83 | 1.05 | 19.7 | 19.09 | 0.61 | 0.61 | 0.03 | 0.03 |
| gene-LOC105285055 | https://www.ncbi.nlm.nih.gov/gene/?term=LOC105285055 | 4108 | 253 | 615.87 | 1 | 2.43 | 0.4 | 12.35 | 11.74 | 0.61 | 0.61 | 0.04 | 0.04 |
| gene-LOC105282149 | https://www.ncbi.nlm.nih.gov/gene/?term=LOC105282149 | 1787 | 107 | 598.77 | 1 | 5.6 | 0.93 | 41.49 | 40.93 | 0.57 | 0.57 | 0.03 | 0.03 |

| | | | | | | | | | | | | | | |
|---|---|---|---|---|---|---|---|---|---|---|---|---|---|---|
| gene-LOC105278962 | https://www.ncbi.nlm.nih.gov/gene/?term=LOC105278962 | 4491 | 106 | 236.03 | 1 | 2.23 | 0.94 | 20.64 | 20.08 | 0.56 | 0.56 | 0.01 | 0.01 | |
| gene-LOC105288010 | https://www.ncbi.nlm.nih.gov/gene/?term=LOC105288010 | 3189 | 45 | 141.11 | 1 | 3.14 | 2.22 | 33.55 | 32.98 | 0.56 | 0.56 | 0.01 | 0.01 | |
| gene-LOC105275187 | https://www.ncbi.nlm.nih.gov/gene/?term=LOC105275187 | 5534 | 74 | 133.72 | 1 | 1.81 | 1.35 | 16.17 | 15.62 | 0.54 | 0.54 | 0.01 | 0.01 | |
| gene-LOC105283728 | https://www.ncbi.nlm.nih.gov/gene/?term=LOC105283728 | 3256 | 122 | 374.69 | 2 | 6.14 | 1.64 | 19.35 | 18.81 | 0.54 | 0.54 | 0.02 | 0.02 | |
| gene-LOC105279298 | https://www.ncbi.nlm.nih.gov/gene/?term=LOC105279298 | 2773 | 132 | 476.02 | 2 | 7.21 | 1.52 | 23.66 | 23.13 | 0.53 | 0.53 | 0.03 | 0.03 | |
| gene-LOC105275363 | https://www.ncbi.nlm.nih.gov/gene/?term=LOC105275363 | 3798 | 168 | 442.34 | 1 | 2.63 | 0.6 | 16.96 | 16.44 | 0.52 | 0.52 | 0.02 | 0.02 | |
| gene-LOC105275683 | https://www.ncbi.nlm.nih.gov/gene/?term=LOC105275683 | 4256 | 205 | 481.67 | 1 | 2.35 | 0.49 | 13.27 | 12.75 | 0.52 | 0.52 | 0.02 | 0.02 | |
| gene-LOC105278228 | https://www.ncbi.nlm.nih.gov/gene/?term=LOC105278228 | 8610 | 172 | 199.77 | 1 | 1.16 | 0.58 | 23.43 | 22.95 | 0.48 | 0.48 | 0.01 | 0.01 | |
| gene-LOC105275170 | https://www.ncbi.nlm.nih.gov/gene/?term=LOC105275170 | 12918 | 298 | 230.69 | 1 | 0.77 | 0.34 | 2.35 | 1.88 | 0.47 | 0.47 | 0.01 | 0.01 | |
| gene-LOC105286452 | https://www.ncbi.nlm.nih.gov/gene/?term=LOC105286452 | 6888 | 60 | 87.11 1 | 1.45 | 1.67 | 31.09 | 30.62 | 0.47 | 0.47 | 0 | 0 | | |
| gene-LOC105282951 | https://www.ncbi.nlm.nih.gov/gene/?term=LOC105282951 | 3366 | 121 | 359.48 | 1 | 2.97 | 0.83 | 20.13 | 19.67 | 0.46 | 0.46 | 0.02 | 0.02 | |
| gene-LOC105274856 | https://www.ncbi.nlm.nih.gov/gene/?term=LOC105274856 | 5216 | 222 | 425.61 | 1 | 1.92 | 0.45 | 17.35 | 16.9 | 0.45 | 0.45 | 0.02 | 0.02 | |
| gene-LOC105276059 | https://www.ncbi.nlm.nih.gov/gene/?term=LOC105276059 | 6256 | 138 | 220.59 | 1 | 1.6 | 0.72 | 9.13 | 8.71 | 0.42 | 0.42 | 0.01 | 0.01 | |
| gene-LOC105274531 | https://www.ncbi.nlm.nih.gov/gene/?term=LOC105274531 | 7774 | 291 | 374.32 | 1 | 1.29 | 0.34 | 10.83 | 10.42 | 0.41 | 0.41 | 0.02 | 0.02 | |
| gene-LOC105282811 | https://www.ncbi.nlm.nih.gov/gene/?term=LOC105282811 | 5765 | 218 | 378.14 | 1 | 1.73 | 0.46 | 8.32 | 7.93 | 0.39 | 0.39 | 0.01 | 0.01 | |
| gene-LOC105285395 | https://www.ncbi.nlm.nih.gov/gene/?term=LOC105285395 | 8085 | 208 | 257.27 | 1 | 1.24 | 0.48 | 14.72 | 14.34 | 0.39 | 0.39 | 0.01 | 0.01 | |
| gene-LOC105274590 | https://www.ncbi.nlm.nih.gov/gene/?term=LOC105274590 | 4516 | 75 | 166.08 | 2 | 4.43 | 2.67 | 29.15 | 28.79 | 0.35 | 0.35 | 0.01 | 0.01 | |
| gene-LOC105275451 | https://www.ncbi.nlm.nih.gov/gene/?term=LOC105275451 | 6035 | 136 | 225.35 | 1 | 1.66 | 0.74 | 21.08 | 20.75 | 0.33 | 0.33 | 0.01 | 0.01 | |
| gene-LOC105280408 | https://www.ncbi.nlm.nih.gov/gene/?term=LOC105280408 | 5983 | 56 | 93.6 1 | 1.67 | 1.79 | 20.14 | 19.81 | 0.33 | 0.33 | 0 | 0 | | |
| gene-LOC105277761 | https://www.ncbi.nlm.nih.gov/gene/?term=LOC105277761 | 4648 | 148 | 318.42 | 1 | 2.15 | 0.68 | 15.39 | 15.08 | 0.31 | 0.31 | 0.01 | 0.01 | |
| gene-LOC105276683 | https://www.ncbi.nlm.nih.gov/gene/?term=LOC105276683 | 17000 | 137 | 80.59 1 | 0.59 | 0.73 | 6.59 | 6.28 | 0.3 | 0.3 | 0 | 0 | | |
| gene-LOC105279038 | https://www.ncbi.nlm.nih.gov/gene/?term=LOC105279038 | 9453 | 363 | 384.01 | 1 | 1.06 | 0.28 | 8.55 | 8.25 | 0.3 | 0.3 | 0.01 | 0.01 | |
| gene-LOC105285807 | https://www.ncbi.nlm.nih.gov/gene/?term=LOC105285807 | 6063 | 228 | 376.05 | 1 | 1.65 | 0.44 | 8.95 | 8.68 | 0.28 | 0.28 | 0.01 | 0.01 | |
| gene-LOC105274452 | https://www.ncbi.nlm.nih.gov/gene/?term=LOC105274452 | 6595 | 260 | 394.24 | 1 | 1.52 | 0.38 | 13.07 | 12.82 | 0.25 | 0.25 | 0.01 | 0.01 | |
| gene-LOC105276196 | https://www.ncbi.nlm.nih.gov/gene/?term=LOC105276196 | 2953 | 58 | 196.41 | 1 | 3.39 | 1.72 | 38.52 | 38.28 | 0.24 | 0.24 | 0 | 0 | |
| gene-LOC105278666 | https://www.ncbi.nlm.nih.gov/gene/?term=LOC105278666 | 10341 | 477 | 461.27 | 1 | 0.97 | 0.21 | 6.78 | 6.53 | 0.24 | 0.24 | 0.01 | 0.01 | |
| gene-LOC105276527 | https://www.ncbi.nlm.nih.gov/gene/?term=LOC105276527 | 7923 | 332 | 419.03 | 1 | 1.26 | 0.3 | 9.67 | 9.44 | 0.23 | 0.23 | 0.01 | 0.01 | |
| gene-LOC105288161 | https://www.ncbi.nlm.nih.gov/gene/?term=LOC105288161 | 26065 | 321 | 123.15 | 1 | 0.38 | 0.31 | 11.9 | 11.67 | 0.23 | 0.23 | 0 | 0 | |
| gene-LOC105279657 | https://www.ncbi.nlm.nih.gov/gene/?term=LOC105279657 | 5878 | 88 | 149.71 | 1 | 1.7 | 1.14 | 19.35 | 19.14 | 0.21 | 0.21 | 0 | 0 | |
| gene-LOC113562050 | https://www.ncbi.nlm.nih.gov/gene/?term=LOC113562050 | 17097 | 605 | 353.86 | 1 | 0.58 | 0.17 | 5.18 | 4.96 | 0.21 | 0.21 | 0.01 | 0.01 | |
| gene-LOC105275327 | https://www.ncbi.nlm.nih.gov/gene/?term=LOC105275327 | 14245 | 543 | 381.19 | 1 | 0.7 | 0.18 | 6.62 | 6.43 | 0.19 | 0.19 | 0.01 | 0.01 | |
| gene-LOC105282491 | https://www.ncbi.nlm.nih.gov/gene/?term=LOC105282491 | 4761 | 177 | 371.77 | 1 | 2.1 | 0.56 | 8.66 | 8.47 | 0.19 | 0.19 | 0.01 | 0.01 | |
| gene-LOC105275460 | https://www.ncbi.nlm.nih.gov/gene/?term=LOC105275460 | 2473 | 123 | 497.37 | 1 | 4.04 | 0.81 | 13.15 | 12.97 | 0.18 | 0.18 | 0.01 | 0.01 | |
| gene-LOC105282151 | https://www.ncbi.nlm.nih.gov/gene/?term=LOC105282151 | 1601 | 44 | 274.83 | 1 | 6.25 | 2.27 | 35.59 | 35.42 | 0.18 | 0.18 | 0 | 0 | |
| gene-LOC105284262 | https://www.ncbi.nlm.nih.gov/gene/?term=LOC105284262 | 6669 | 235 | 352.38 | 1 | 1.5 | 0.43 | 4.64 | 4.46 | 0.18 | 0.18 | 0.01 | 0.01 | |
| gene-LOC105287764 | https://www.ncbi.nlm.nih.gov/gene/?term=LOC105287764 | 18359 | 402 | 218.97 | 1 | 0.54 | 0.25 | 3.16 | 2.99 | 0.17 | 0.17 | 0 | 0 | |
| gene-LOC105283712 | https://www.ncbi.nlm.nih.gov/gene/?term=LOC105283712 | 1910 | 36 | 188.48 | 1 | 5.24 | 2.78 | 29.2 | 29.04 | 0.16 | 0.16 | 0 | 0 | |
| gene-LOC105284043 | https://www.ncbi.nlm.nih.gov/gene/?term=LOC105284043 | 5661 | 311 | 549.37 | 1 | 1.77 | 0.32 | 9.86 | 9.71 | 0.15 | 0.15 | 0.01 | 0.01 | |
| gene-LOC105278145 | https://www.ncbi.nlm.nih.gov/gene/?term=LOC105278145 | 4470 | 235 | 525.73 | 1 | 2.24 | 0.43 | 10.67 | 10.54 | 0.14 | 0.14 | 0.01 | 0.01 | |
| gene-LOC105284036 | https://www.ncbi.nlm.nih.gov/gene/?term=LOC105284036 | 3254 | 107 | 328.83 | 1 | 3.07 | 0.93 | 20.44 | 20.34 | 0.09 | 0.09 | 0 | 0 | |
| gene-LOC105274823 | https://www.ncbi.nlm.nih.gov/gene/?term=LOC105274823 | 4620 | 146 | 316.02 | 1 | 2.16 | 0.68 | 20.43 | 20.37 | 0.06 | 0.06 | 0 | 0 | |
| gene-LOC105280265 | https://www.ncbi.nlm.nih.gov/gene/?term=LOC105280265 | 85984 | 3274 | 380.77 | 1 | 0.12 | 0.03 | 1.1 | 1.05 | 0.06 | 0.06 | 0 | 0 | |
| gene-LOC105280513 | https://www.ncbi.nlm.nih.gov/gene/?term=LOC105280513 | 69542 | 1377 | 198.01 | 1 | 0.14 | 0.07 | 0.72 | 0.66 | 0.06 | 0.06 | 0 | 0 | |
| gene-LOC105286194 | https://www.ncbi.nlm.nih.gov/gene/?term=LOC105286194 | 152619 | | 2206 144.54 | | 1 | 0.07 | 0.05 | 6.36 | 6.3 | 0.06 | 0.06 | 0 | 0 |
| gene-LOC105281700 | https://www.ncbi.nlm.nih.gov/gene/?term=LOC105281700 | 2208 | 37 | 167.57 | 1 | 4.53 | 2.7 | 39.8 | 39.75 | 0.05 | 0.05 | 0 | 0 | |
| gene-LOC105286515 | https://www.ncbi.nlm.nih.gov/gene/?term=LOC105286515 | 3751 | 106 | 282.59 | 1 | 2.67 | 0.94 | 10.25 | 10.22 | 0.04 | 0.04 | 0 | 0 | |
| gene-LOC105278455 | https://www.ncbi.nlm.nih.gov/gene/?term=LOC105278455 | 27181 | 985 | 362.39 | 1 | 0.37 | 0.1 | 6.75 | 6.75 | 0 | 0 | 0 | 0 | |
| gene-LOC105280407 | https://www.ncbi.nlm.nih.gov/gene/?term=LOC105280407 | 10950 | 210 | 191.78 | 1 | 0.91 | 0.48 | 16.39 | 16.4 | -0.01 | 0.01 | 0 | 0 | |
| gene-LOC105283123 | https://www.ncbi.nlm.nih.gov/gene/?term=LOC105283123 | 1507 | 55 | 364.96 | 1 | 6.64 | 1.82 | 18.11 | 18.13 | -0.02 | 0.02 | 0 | 0 | |
| gene-LOC105274669 | https://www.ncbi.nlm.nih.gov/gene/?term=LOC105274669 | 8555 | 270 | 315.6 1 | 1.17 | 0.37 | 5.54 | 5.56 | -0.03 | 0.03 | 0 | 0 | | |
| gene-LOC105278534 | https://www.ncbi.nlm.nih.gov/gene/?term=LOC105278534 | 7364 | 196 | 266.16 | 1 | 1.36 | 0.51 | 17.75 | 17.8 | -0.05 | 0.05 | 0 | 0 | |
| gene-LOC105287550 | https://www.ncbi.nlm.nih.gov/gene/?term=LOC105287550 | 15659 | 616 | 393.38 | 1 | 0.64 | 0.16 | 5.82 | 5.86 | -0.05 | 0.05 | 0 | 0 | |
| gene-LOC105275374 | https://www.ncbi.nlm.nih.gov/gene/?term=LOC105275374 | 11376 | 539 | 473.8 1 | 0.88 | 0.19 | 9.85 | 10 | -0.15 | 0.15 | -0.01 | 0.01 | | |
| gene-LOC105278282 | https://www.ncbi.nlm.nih.gov/gene/?term=LOC105278282 | 1775 | 62 | 349.3 1 | 5.63 | 1.61 | 32.74 | 32.89 | -0.15 | 0.15 | -0.01 | 0.01 | | |
| gene-LOC105285496 | https://www.ncbi.nlm.nih.gov/gene/?term=LOC105285496 | 4080 | 110 | 269.61 | 1 | 2.45 | 0.91 | 40.35 | 40.5 | -0.15 | 0.15 | 0 | 0 | |
| gene-LOC105285742 | https://www.ncbi.nlm.nih.gov/gene/?term=LOC105285742 | 3446 | 123 | 356.94 | 1 | 2.9 | 0.81 | 19.52 | 19.69 | -0.16 | 0.16 | -0.01 | 0.01 | |
| gene-LOC105280723 | https://www.ncbi.nlm.nih.gov/gene/?term=LOC105280723 | 3572 | 181 | 506.72 | 1 | 2.8 | 0.55 | 15.21 | 15.4 | -0.2 | 0.2 | -0.01 | 0.01 | |
| gene-LOC105274851 | https://www.ncbi.nlm.nih.gov/gene/?term=LOC105274851 | 6616 | 147 | 222.19 | 1 | 1.51 | 0.68 | 9.12 | 9.33 | -0.21 | 0.21 | 0 | 0 | |
| gene-LOC105284184 | https://www.ncbi.nlm.nih.gov/gene/?term=LOC105284184 | 2535 | 99 | 390.53 | 1 | 3.94 | 1.01 | 29.28 | 29.51 | -0.23 | 0.23 | -0.01 | 0.01 | |
| gene-LOC105275587 | https://www.ncbi.nlm.nih.gov/gene/?term=LOC105275587 | 2404 | 55 | 228.79 | 1 | 4.16 | 1.82 | 36.62 | 36.89 | -0.27 | 0.27 | -0.01 | 0.01 | |
| gene-LOC105277866 | https://www.ncbi.nlm.nih.gov/gene/?term=LOC105277866 | 33045 | 423 | 128.01 | 1 | 0.3 | 0.24 | 3 | 3.27 | -0.28 | 0.28 | 0 | 0 | |
| gene-LOC105286274 | https://www.ncbi.nlm.nih.gov/gene/?term=LOC105286274 | 3817 | 200 | 523.97 | 1 | 2.62 | 0.5 | 16.54 | 16.86 | -0.32 | 0.32 | -0.02 | 0.02 | |
| gene-LOC105278691 | https://www.ncbi.nlm.nih.gov/gene/?term=LOC105278691 | 3871 | 168 | 434 1 | 2.58 | 0.6 | 14.64 | 14.98 | -0.34 | 0.34 | -0.01 | 0.01 | | |
| gene-LOC105278283 | https://www.ncbi.nlm.nih.gov/gene/?term=LOC105278283 | 2317 | 139 | 599.91 | 1 | 4.32 | 0.72 | 25.04 | 25.46 | -0.41 | 0.41 | -0.02 | 0.02 | |

| | | | | | | | | | | | | | |
|---|---|---|---|---|---|---|---|---|---|---|---|---|---|
| gene-LOC105285406 | https://www.ncbi.nlm.nih.gov/gene/?term=LOC105285406 | 8015 | 55 | 68.62 | 1 | 1.25 | 1.82 | 29.11 | 29.59 | -0.47 | 0.47 | 0 | 0 |
| gene-LOC105276704 | https://www.ncbi.nlm.nih.gov/gene/?term=LOC105276704 | 2063 | 99 | 479.88 | 2 | 9.69 | 2.02 | 13.44 | 13.97 | -0.53 | 0.53 | -0.03 | 0.03 |
| gene-LOC105283967 | https://www.ncbi.nlm.nih.gov/gene/?term=LOC105283967 | 2393 | 53 | 221.48 | 1 | 4.18 | 1.89 | 45.17 | 45.77 | -0.59 | 0.59 | -0.01 | 0.01 |
| gene-LOC105285533 | https://www.ncbi.nlm.nih.gov/gene/?term=LOC105285533 | 1640 | 70 | 426.83 | 1 | 6.1 | 1.43 | 22.75 | 23.49 | -0.75 | 0.75 | -0.03 | 0.03 |
| gene-LOC105274534 | https://www.ncbi.nlm.nih.gov/gene/?term=LOC105274534 | 1329 | 60 | 451.47 | 1 | 7.52 | 1.67 | 40.57 | 41.38 | -0.82 | 0.82 | -0.04 | 0.04 |
| gene-LOC105274449 | https://www.ncbi.nlm.nih.gov/gene/?term=LOC105274449 | 1549 | 53 | 342.16 | 1 | 6.46 | 1.89 | 50.86 | 51.77 | -0.91 | 0.91 | -0.03 | 0.03 |
| gene-LOC105276429 | https://www.ncbi.nlm.nih.gov/gene/?term=LOC105276429 | 2901 | 108 | 372.29 | 1 | 3.45 | 0.93 | 24.11 | 25.23 | -1.12 | 1.12 | -0.04 | 0.04 |
| gene-LOC105281312 | https://www.ncbi.nlm.nih.gov/gene/?term=LOC105281312 | 2551 | 65 | 254.8 | 1 | 3.92 | 1.54 | 31.99 | 33.22 | -1.23 | 1.23 | -0.03 | 0.03 |
| gene-LOC105281173 | https://www.ncbi.nlm.nih.gov/gene/?term=LOC105281173 | 1595 | 38 | 238.24 | 1 | 6.27 | 2.63 | 59.28 | 60.67 | -1.38 | 1.38 | -0.03 | 0.03 |
| gene-LOC105274706 | https://www.ncbi.nlm.nih.gov/gene/?term=LOC105274706 | 1446 | 50 | 345.78 | 1 | 6.92 | 2 | 40.41 | 41.89 | -1.49 | 1.49 | -0.05 | 0.05 |
| gene-LOC105285409 | https://www.ncbi.nlm.nih.gov/gene/?term=LOC105285409 | 5099 | 58 | 113.75 | 1 | 1.96 | 1.72 | 1.58 | 3.25 | -1.67 | 1.67 | -0.02 | 0.02 |
| gene-LOC105287633 | https://www.ncbi.nlm.nih.gov/gene/?term=LOC105287633 | 1524 | 31 | 203.41 | 1 | 6.56 | 3.23 | 53.47 | 55.78 | -2.31 | 2.31 | -0.05 | 0.05 |
| gene-LOC105282914 | https://www.ncbi.nlm.nih.gov/gene/?term=LOC105282914 | 5623 | 41 | 72.91 | 1 | 1.78 | 2.44 | 49.33 | 52.21 | -2.88 | 2.88 | -0.02 | 0.02 |
| gene-LOC105282885 | https://www.ncbi.nlm.nih.gov/gene/?term=LOC105282885 | 1870 | 47 | 251.34 | 1 | 5.35 | 2.13 | 35.98 | 39.14 | -3.16 | 3.16 | -0.08 | 0.08 |
| gene-LOC105279500 | https://www.ncbi.nlm.nih.gov/gene/?term=LOC105279500 | 3587 | 33 | 92 | 1 | 2.79 | 3.03 | 43.74 | 47.01 | -3.27 | 3.27 | -0.03 | 0.03 |
| gene-LOC105284677 | https://www.ncbi.nlm.nih.gov/gene/?term=LOC105284677 | 1905 | 42 | 220.47 | 1 | 5.25 | 2.38 | 26.24 | 37.79 | -11.55 | 11.55 | -0.25 | 0.25 |

Directory: OGLall File: ogl-Ob_LI_BCphaseB1.vs.RphaseA1.txt

| gene_ID | gene_link | gwidth | #Sites | #per10Kb | #dmSites | #dmsp10kb | %dmSites | %pSite1 | %pSite2 | DMpSite | ADMpSite | DMpNucl | ADMpNucl |
|---|---|---|---|---|---|---|---|---|---|---|---|---|---|
| gene-LOC105282854 | https://www.ncbi.nlm.nih.gov/gene/?term=LOC105282854 | 1825 | 6 | 32.88 | 1 | 5.48 | 16.67 | 52.41 | 41.89 | 10.52 | 10.52 | 0.03 | 0.03 |
| gene-LOC105286453 | https://www.ncbi.nlm.nih.gov/gene/?term=LOC105286453 | 1160 | 9 | 77.59 | 2 | 17.24 | 22.22 | 36.42 | 26.38 | 10.04 | 10.04 | 0.08 | 0.08 |
| gene-LOC105275378 | https://www.ncbi.nlm.nih.gov/gene/?term=LOC105275378 | 1189 | 21 | 176.62 | 3 | 25.23 | 14.29 | 38.9 | 30.15 | 8.75 | 8.75 | 0.15 | 0.15 |
| gene-LOC105286572 | https://www.ncbi.nlm.nih.gov/gene/?term=LOC105286572 | 2321 | 7 | 30.16 | 2 | 8.62 | 28.57 | 19.41 | 11.7 | 7.7 | 7.7 | 0.02 | 0.02 |
| gene-LOC105280699 | https://www.ncbi.nlm.nih.gov/gene/?term=LOC105280699 | 1563 | 5 | 31.99 | 1 | 6.4 | 20 | 31.61 | 25.04 | 6.57 | 6.57 | 0.02 | 0.02 |
| gene-LOC105275669 | https://www.ncbi.nlm.nih.gov/gene/?term=LOC105275669 | 1567 | 8 | 51.05 | 1 | 6.38 | 12.5 | 54.73 | 48.32 | 6.41 | 6.41 | 0.03 | 0.03 |
| gene-LOC105283227 | https://www.ncbi.nlm.nih.gov/gene/?term=LOC105283227 | 2373 | 12 | 50.57 | 2 | 8.43 | 16.67 | 43.46 | 37.45 | 6.01 | 6.01 | 0.03 | 0.03 |
| gene-LOC113563652 | https://www.ncbi.nlm.nih.gov/gene/?term=LOC113563652 | 1476 | 13 | 88.08 | 1 | 6.78 | 7.69 | 51.72 | 45.77 | 5.95 | 5.95 | 0.05 | 0.05 |
| gene-LOC113562135 | https://www.ncbi.nlm.nih.gov/gene/?term=LOC113562135 | 1413 | 9 | 63.69 | 1 | 7.08 | 11.11 | 40.16 | 34.54 | 5.62 | 5.62 | 0.04 | 0.04 |
| gene-LOC105284817 | https://www.ncbi.nlm.nih.gov/gene/?term=LOC105284817 | 2500 | 13 | 52 | 1 | 4 | 7.69 | 28.13 | 22.7 | 5.43 | 5.43 | 0.03 | 0.03 |
| gene-LOC105277491 | https://www.ncbi.nlm.nih.gov/gene/?term=LOC105277491 | 1647 | 11 | 66.79 | 1 | 6.07 | 9.09 | 19.05 | 13.86 | 5.19 | 5.19 | 0.03 | 0.03 |
| gene-LOC105280472 | https://www.ncbi.nlm.nih.gov/gene/?term=LOC105280472 | 1675 | 12 | 71.64 | 1 | 5.97 | 8.33 | 42.17 | 37.11 | 5.06 | 5.06 | 0.04 | 0.04 |
| gene-LOC105283785 | https://www.ncbi.nlm.nih.gov/gene/?term=LOC105283785 | 3447 | 33 | 95.74 | 1 | 2.9 | 3.03 | 19.56 | 14.53 | 5.03 | 5.03 | 0.05 | 0.05 |
| gene-LOC105286815 | https://www.ncbi.nlm.nih.gov/gene/?term=LOC105286815 | 1024 | 11 | 107.42 | 1 | 9.77 | 9.09 | 63.05 | 58.15 | 4.91 | 4.91 | 0.05 | 0.05 |
| gene-LOC105287814 | https://www.ncbi.nlm.nih.gov/gene/?term=LOC105287814 | 892 | 8 | 89.69 | 1 | 11.21 | 12.5 | 35.83 | 30.99 | 4.84 | 4.84 | 0.04 | 0.04 |
| gene-LOC105287978 | https://www.ncbi.nlm.nih.gov/gene/?term=LOC105287978 | 3017 | 7 | 23.2 | 1 | 3.31 | 14.29 | 17.4 | 12.85 | 4.56 | 4.56 | 0.01 | 0.01 |
| gene-LOC105276691 | https://www.ncbi.nlm.nih.gov/gene/?term=LOC105276691 | 2543 | 41 | 161.23 | 3 | 11.8 | 7.32 | 41.71 | 37.17 | 4.54 | 4.54 | 0.07 | 0.07 |
| gene-LOC105275518 | https://www.ncbi.nlm.nih.gov/gene/?term=LOC105275518 | 2335 | 26 | 111.35 | 2 | 8.57 | 7.69 | 47.97 | 43.47 | 4.5 | 4.5 | 0.05 | 0.05 |
| gene-LOC105282733 | https://www.ncbi.nlm.nih.gov/gene/?term=LOC105282733 | 3835 | 23 | 59.97 | 2 | 5.22 | 8.7 | 33.46 | 29 | 4.46 | 4.46 | 0.03 | 0.03 |
| gene-LOC105282407 | https://www.ncbi.nlm.nih.gov/gene/?term=LOC105282407 | 1210 | 12 | 99.17 | 1 | 8.26 | 8.33 | 50.3 | 45.91 | 4.39 | 4.39 | 0.04 | 0.04 |
| gene-LOC105276707 | https://www.ncbi.nlm.nih.gov/gene/?term=LOC105276707 | 1990 | 16 | 80.4 | 2 | 10.05 | 12.5 | 37.03 | 32.8 | 4.23 | 4.23 | 0.03 | 0.03 |
| gene-LOC105278080 | https://www.ncbi.nlm.nih.gov/gene/?term=LOC105278080 | 3258 | 21 | 64.46 | 1 | 3.07 | 4.76 | 41.67 | 37.47 | 4.21 | 4.21 | 0.03 | 0.03 |
| gene-LOC105285392 | https://www.ncbi.nlm.nih.gov/gene/?term=LOC105285392 | 1601 | 91 | 568.39 | 6 | 37.48 | 6.59 | 35.68 | 31.54 | 4.15 | 4.15 | 0.24 | 0.24 |
| gene-LOC105280372 | https://www.ncbi.nlm.nih.gov/gene/?term=LOC105280372 | 2159 | 19 | 88 | 1 | 4.63 | 5.26 | 55.74 | 51.64 | 4.1 | 4.1 | 0.04 | 0.04 |

...
... only first 25 lines shown ...

Directory: OGLall File: ogl-Ob_LI_BCphaseB1.vs.RphaseA2.txt

| gene_ID | gene_link | gwidth | #Sites | #per10Kb | #dmSites | #dmsp10kb | %dmSites | %pSite1 | %pSite2 | DMpSite | ADMpSite | DMpNucl | ADMpNucl |
|---|---|---|---|---|---|---|---|---|---|---|---|---|---|
| gene-LOC105286572 | https://www.ncbi.nlm.nih.gov/gene/?term=LOC105286572 | 2321 | 8 | 34.47 | 1 | 4.31 | 12.5 | 20.96 | 13.96 | 7.01 | 7.01 | 0.02 | 0.02 |
| gene-LOC105283986 | https://www.ncbi.nlm.nih.gov/gene/?term=LOC105283986 | 1782 | 7 | 39.28 | 1 | 5.61 | 14.29 | 29.98 | 22.98 | 7 | 7 | 0.03 | 0.03 |
| gene-LOC113563238 | https://www.ncbi.nlm.nih.gov/gene/?term=LOC113563238 | 2680 | 4 | 14.93 | 1 | 3.73 | 25 | 22.36 | 15.49 | 6.86 | 6.86 | 0.01 | 0.01 |
| gene-LOC105280472 | https://www.ncbi.nlm.nih.gov/gene/?term=LOC105280472 | 1675 | 14 | 83.58 | 2 | 11.94 | 14.29 | 38.72 | 31.98 | 6.74 | 6.74 | 0.06 | 0.06 |
| gene-LOC105278607 | https://www.ncbi.nlm.nih.gov/gene/?term=LOC105278607 | 4407 | 29 | 65.8 | 2 | 4.54 | 6.9 | 11.68 | 5.33 | 6.35 | 6.35 | 0.04 | 0.04 |
| gene-LOC105283760 | https://www.ncbi.nlm.nih.gov/gene/?term=LOC105283760 | 1134 | 5 | 44.09 | 2 | 17.64 | 40 | 48.5 | 42.19 | 6.31 | 6.31 | 0.03 | 0.03 |
| gene-LOC105277405 | https://www.ncbi.nlm.nih.gov/gene/?term=LOC105277405 | 1255 | 4 | 31.87 | 1 | 7.97 | 25 | 25.11 | 19.1 | 6.02 | 6.02 | 0.02 | 0.02 |
| gene-LOC105287978 | https://www.ncbi.nlm.nih.gov/gene/?term=LOC105287978 | 3017 | 7 | 23.2 | 2 | 6.63 | 28.57 | 17.4 | 11.62 | 5.78 | 5.78 | 0.01 | 0.01 |
| gene-LOC105286453 | https://www.ncbi.nlm.nih.gov/gene/?term=LOC105286453 | 1160 | 14 | 120.69 | 2 | 17.24 | 14.29 | 32.33 | 26.68 | 5.65 | 5.65 | 0.07 | 0.07 |
| gene-LOC105286753 | https://www.ncbi.nlm.nih.gov/gene/?term=LOC105286753 | 3684 | 12 | 32.57 | 1 | 2.71 | 8.33 | 17.35 | 11.83 | 5.52 | 5.52 | 0.02 | 0.02 |
| gene-LOC105288020 | https://www.ncbi.nlm.nih.gov/gene/?term=LOC105288020 | 1568 | 11 | 70.15 | 1 | 6.38 | 9.09 | 35.44 | 30.49 | 4.95 | 4.95 | 0.03 | 0.03 |
| gene-LOC105274640 | https://www.ncbi.nlm.nih.gov/gene/?term=LOC105274640 | 2723 | 11 | 40.4 | 1 | 3.67 | 9.09 | 60.49 | 56.34 | 4.15 | 4.15 | 0.02 | 0.02 |
| gene-LOC105280969 | https://www.ncbi.nlm.nih.gov/gene/?term=LOC105280969 | 1005 | 14 | 139.3 | 1 | 9.95 | 7.14 | 39.18 | 35.04 | 4.14 | 4.14 | 0.06 | 0.06 |
| gene-LOC105285707 | https://www.ncbi.nlm.nih.gov/gene/?term=LOC105285707 | 2096 | 32 | 152.67 | 2 | 9.54 | 6.25 | 58.26 | 54.2 | 4.05 | 4.05 | 0.06 | 0.06 |
| gene-LOC105275378 | https://www.ncbi.nlm.nih.gov/gene/?term=LOC105275378 | 1189 | 26 | 218.67 | 2 | 16.82 | 7.69 | 46.76 | 43.04 | 3.73 | 3.73 | 0.08 | 0.08 |
| gene-LOC105275355 | https://www.ncbi.nlm.nih.gov/gene/?term=LOC105275355 | 3197 | 169 | 528.62 | 10 | 31.28 | 5.92 | 27.68 | 24.07 | 3.61 | 3.61 | 0.19 | 0.19 |
| gene-LOC105281081 | https://www.ncbi.nlm.nih.gov/gene/?term=LOC105281081 | 2358 | 15 | 63.61 | 1 | 4.24 | 6.67 | 19.8 | 16.63 | 3.16 | 3.16 | 0.02 | 0.02 |
| gene-LOC105280764 | https://www.ncbi.nlm.nih.gov/gene/?term=LOC105280764 | 1800 | 16 | 88.89 | 1 | 5.56 | 6.25 | 19.21 | 16.08 | 3.13 | 3.13 | 0.03 | 0.03 |
| gene-LOC105280001 | https://www.ncbi.nlm.nih.gov/gene/?term=LOC105280001 | 2566 | 23 | 89.63 | 1 | 3.9 | 4.35 | 24.54 | 21.43 | 3.11 | 3.11 | 0.03 | 0.03 |
| gene-LOC105284604 | https://www.ncbi.nlm.nih.gov/gene/?term=LOC105284604 | 1998 | 17 | 85.09 | 2 | 10.01 | 11.76 | 3.09 | 0 | 3.09 | 3.09 | 0.03 | 0.03 |
| gene-LOC105280448 | https://www.ncbi.nlm.nih.gov/gene/?term=LOC105280448 | 4000 | 29 | 72.5 | 2 | 5 | 6.9 | 18.2 | 15.16 | 3.04 | 3.04 | 0.02 | 0.02 |
| gene-LOC105277369 | https://www.ncbi.nlm.nih.gov/gene/?term=LOC105277369 | 2012 | 9 | 44.73 | 1 | 4.97 | 11.11 | 61.75 | 58.79 | 2.96 | 2.96 | 0.01 | 0.01 |
| gene-LOC105276691 | https://www.ncbi.nlm.nih.gov/gene/?term=LOC105276691 | 2543 | 44 | 173.02 | 3 | 11.8 | 6.82 | 39.83 | 36.94 | 2.89 | 2.89 | 0.05 | 0.05 |
| gene-LOC105280548 | https://www.ncbi.nlm.nih.gov/gene/?term=LOC105280548 | 1547 | 29 | 187.46 | 1 | 6.46 | 3.45 | 51.13 | 48.43 | 2.7 | 2.7 | 0.05 | 0.05 |

...
... only first 25 lines shown ...

Directory: OGLall File: ogl-Ob_LI_BCphaseB1.vs.RphaseB1.txt

| gene_ID | gene_link | gwidth | #Sites | #per10Kb | #dmSites | #dmsp10kb | %dmSites | %pSite1 | %pSite2 | DMpSite | ADMpSite | DMpNucl | ADMpNucl |
|---|---|---|---|---|---|---|---|---|---|---|---|---|---|
| gene-LOC113562345 | https://www.ncbi.nlm.nih.gov/gene/?term=LOC113562345 | 2290 | 77 | 336.24 | 12 | 52.4 | 15.58 | 26.12 | 14.52 | 11.59 | 11.59 | 0.39 | 0.39 |
| gene-LOC105275070 | https://www.ncbi.nlm.nih.gov/gene/?term=LOC105275070 | 5667 | 61 | 107.64 | 2 | 3.53 | 3.28 | 14.18 | 5.29 | 8.9 | 8.9 | 0.1 | 0.1 |
| gene-LOC105275708 | https://www.ncbi.nlm.nih.gov/gene/?term=LOC105275708 | 1772 | 15 | 84.65 | 1 | 5.64 | 6.67 | 46.38 | 38.49 | 7.9 | 7.9 | 0.07 | 0.07 |
| gene-LOC105285514 | https://www.ncbi.nlm.nih.gov/gene/?term=LOC105285514 | 1277 | 11 | 86.14 | 1 | 7.83 | 9.09 | 92.13 | 85.12 | 7.01 | 7.01 | 0.06 | 0.06 |
| gene-LOC105275355 | https://www.ncbi.nlm.nih.gov/gene/?term=LOC105275355 | 3197 | 174 | 544.26 | 1 | 3.13 | 0.57 | 26.94 | 21.77 | 5.17 | 5.17 | 0.28 | 0.28 |
| gene-LOC105281039 | https://www.ncbi.nlm.nih.gov/gene/?term=LOC105281039 | 1471 | 14 | 95.17 | 1 | 6.8 | 7.14 | 44.63 | 40.12 | 4.51 | 4.51 | 0.04 | 0.04 |
| gene-LOC105275518 | https://www.ncbi.nlm.nih.gov/gene/?term=LOC105275518 | 2335 | 25 | 107.07 | 1 | 4.28 | 4 | 44.99 | 41.35 | 3.64 | 3.64 | 0.04 | 0.04 |
| gene-LOC105280419 | https://www.ncbi.nlm.nih.gov/gene/?term=LOC105280419 | 1601 | 11 | 68.71 | 1 | 6.25 | 9.09 | 46.06 | 42.48 | 3.57 | 3.57 | 0.02 | 0.02 |
| gene-LOC105283989 | https://www.ncbi.nlm.nih.gov/gene/?term=LOC105283989 | 4948 | 45 | 90.95 | 2 | 4.04 | 4.44 | 38.35 | 35.25 | 3.1 | 3.1 | 0.03 | 0.03 |
| gene-LOC105275384 | https://www.ncbi.nlm.nih.gov/gene/?term=LOC105275384 | 1987 | 92 | 463.01 | 1 | 5.03 | 1.09 | 34.18 | 31.19 | 2.98 | 2.98 | 0.14 | 0.14 |
| gene-LOC105276272 | https://www.ncbi.nlm.nih.gov/gene/?term=LOC105276272 | 1965 | 40 | 203.56 | 1 | 5.09 | 2.5 | 30.29 | 27.31 | 2.98 | 2.98 | 0.06 | 0.06 |
| gene-LOC105279900 | https://www.ncbi.nlm.nih.gov/gene/?term=LOC105279900 | 2191 | 56 | 255.59 | 1 | 4.56 | 1.79 | 31.89 | 28.91 | 2.98 | 2.98 | 0.08 | 0.08 |
| gene-LOC105285315 | https://www.ncbi.nlm.nih.gov/gene/?term=LOC105285315 | 5458 | 32 | 58.63 | 1 | 1.83 | 3.12 | 31.91 | 29.18 | 2.72 | 2.72 | 0.02 | 0.02 |
| gene-LOC105276611 | https://www.ncbi.nlm.nih.gov/gene/?term=LOC105276611 | 1731 | 36 | 207.97 | 1 | 5.78 | 2.78 | 30.25 | 27.6 | 2.65 | 2.65 | 0.06 | 0.06 |
| gene-LOC105283712 | https://www.ncbi.nlm.nih.gov/gene/?term=LOC105283712 | 1910 | 35 | 183.25 | 1 | 5.24 | 2.86 | 31.43 | 28.9 | 2.53 | 2.53 | 0.05 | 0.05 |
| gene-LOC105287063 | https://www.ncbi.nlm.nih.gov/gene/?term=LOC105287063 | 1874 | 50 | 266.81 | 1 | 5.34 | 2 | 31.85 | 29.62 | 2.23 | 2.23 | 0.06 | 0.06 |
| gene-LOC105274900 | https://www.ncbi.nlm.nih.gov/gene/?term=LOC105274900 | 5561 | 63 | 113.29 | 2 | 3.6 | 3.17 | 32.28 | 30.06 | 2.22 | 2.22 | 0.03 | 0.03 |
| gene-LOC105283552 | https://www.ncbi.nlm.nih.gov/gene/?term=LOC105283552 | 1562 | 41 | 262.48 | 1 | 6.4 | 2.44 | 53.31 | 51.1 | 2.21 | 2.21 | 0.06 | 0.06 |
| gene-LOC105279898 | https://www.ncbi.nlm.nih.gov/gene/?term=LOC105279898 | 1343 | 16 | 119.14 | 1 | 7.45 | 6.25 | 58.69 | 56.55 | 2.14 | 2.14 | 0.03 | 0.03 |
| gene-LOC105281515 | https://www.ncbi.nlm.nih.gov/gene/?term=LOC105281515 | 8827 | 55 | 62.31 | 2 | 2.27 | 3.64 | 34.93 | 32.95 | 1.98 | 1.98 | 0.01 | 0.01 |
| gene-LOC113561805 | https://www.ncbi.nlm.nih.gov/gene/?term=LOC113561805 | 3337 | 76 | 227.75 | 1 | 3 | 1.32 | 32.51 | 30.53 | 1.98 | 1.98 | 0.05 | 0.05 |
| gene-LOC105280704 | https://www.ncbi.nlm.nih.gov/gene/?term=LOC105280704 | 4266 | 147 | 344.59 | 1 | 2.34 | 0.68 | 18.45 | 16.63 | 1.81 | 1.81 | 0.06 | 0.06 |
| gene-LOC105280348 | https://www.ncbi.nlm.nih.gov/gene/?term=LOC105280348 | 3797 | 79 | 208.06 | 1 | 2.63 | 1.27 | 18.26 | 16.64 | 1.61 | 1.61 | 0.03 | 0.03 |
| gene-LOC105274449 | https://www.ncbi.nlm.nih.gov/gene/?term=LOC105274449 | 1549 | 53 | 342.16 | 1 | 6.46 | 1.89 | 50.86 | 49.29 | 1.57 | 1.57 | 0.05 | 0.05 |
| gene-LOC105287417 | https://www.ncbi.nlm.nih.gov/gene/?term=LOC105287417 | 1025 | 40 | 390.24 | 1 | 9.76 | 2.5 | 32.13 | 30.76 | 1.38 | 1.38 | 0.05 | 0.05 |
| gene-LOC105276692 | https://www.ncbi.nlm.nih.gov/gene/?term=LOC105276692 | 1745 | 35 | 200.57 | 1 | 5.73 | 2.86 | 24.46 | 23.1 | 1.36 | 1.36 | 0.03 | 0.03 |
| gene-LOC105276538 | https://www.ncbi.nlm.nih.gov/gene/?term=LOC105276538 | 1511 | 20 | 132.36 | 1 | 6.62 | 5 | 72.51 | 71.18 | 1.32 | 1.32 | 0.02 | 0.02 |
| gene-LOC105287095 | https://www.ncbi.nlm.nih.gov/gene/?term=LOC105287095 | 1435 | 70 | 487.8 | 1 | 6.97 | 1.43 | 25.15 | 23.89 | 1.26 | 1.26 | 0.06 | 0.06 |
| gene-LOC105288036 | https://www.ncbi.nlm.nih.gov/gene/?term=LOC105288036 | 5110 | 232 | 454.01 | 1 | 1.96 | 0.43 | 15.89 | 14.67 | 1.22 | 1.22 | 0.06 | 0.06 |
| gene-LOC105285060 | https://www.ncbi.nlm.nih.gov/gene/?term=LOC105285060 | 1285 | 16 | 124.51 | 1 | 7.78 | 6.25 | 42.41 | 41.27 | 1.13 | 1.13 | 0.01 | 0.01 |
| gene-LOC105278384 | https://www.ncbi.nlm.nih.gov/gene/?term=LOC105278384 | 1390 | 33 | 237.41 | 1 | 7.19 | 3.03 | 46.84 | 45.77 | 1.07 | 1.07 | 0.03 | 0.03 |
| gene-LOC105288079 | https://www.ncbi.nlm.nih.gov/gene/?term=LOC105288079 | 3473 | 110 | 316.73 | 1 | 2.88 | 0.91 | 18.5 | 17.43 | 1.07 | 1.07 | 0.03 | 0.03 |
| gene-LOC105280972 | https://www.ncbi.nlm.nih.gov/gene/?term=LOC105280972 | 1532 | 35 | 228.46 | 1 | 6.53 | 2.86 | 17.63 | 16.59 | 1.04 | 1.04 | 0.02 | 0.02 |
| gene-LOC105282794 | https://www.ncbi.nlm.nih.gov/gene/?term=LOC105282794 | 4571 | 110 | 240.65 | 1 | 2.19 | 0.91 | 18.14 | 17.16 | 0.98 | 0.98 | 0.02 | 0.02 |
| gene-LOC105281266 | https://www.ncbi.nlm.nih.gov/gene/?term=LOC105281266 | 4102 | 88 | 214.53 | 1 | 2.44 | 1.14 | 15.7 | 14.74 | 0.96 | 0.96 | 0.02 | 0.02 |
| gene-LOC105282156 | https://www.ncbi.nlm.nih.gov/gene/?term=LOC105282156 | 2577 | 132 | 512.22 | 1 | 3.88 | 0.76 | 26.63 | 25.67 | 0.96 | 0.96 | 0.05 | 0.05 |
| gene-LOC105285179 | https://www.ncbi.nlm.nih.gov/gene/?term=LOC105285179 | 4191 | 194 | 462.9 | 1 | 2.39 | 0.52 | 17.62 | 16.69 | 0.93 | 0.93 | 0.04 | 0.04 |
| gene-LOC105278632 | https://www.ncbi.nlm.nih.gov/gene/?term=LOC105278632 | 3624 | 139 | 383.55 | 1 | 2.76 | 0.72 | 14.57 | 13.67 | 0.9 | 0.9 | 0.03 | 0.03 |
| gene-LOC105283123 | https://www.ncbi.nlm.nih.gov/gene/?term=LOC105283123 | 1507 | 55 | 364.96 | 1 | 6.64 | 1.82 | 19.83 | 18.96 | 0.86 | 0.86 | 0.03 | 0.03 |
| gene-LOC105277215 | https://www.ncbi.nlm.nih.gov/gene/?term=LOC105277215 | 2619 | 29 | 110.73 | 1 | 3.82 | 3.45 | 25.46 | 24.64 | 0.82 | 0.82 | 0.01 | 0.01 |
| gene-LOC105281848 | https://www.ncbi.nlm.nih.gov/gene/?term=LOC105281848 | 2122 | 58 | 273.33 | 1 | 4.71 | 1.72 | 22.33 | 21.51 | 0.82 | 0.82 | 0.02 | 0.02 |
| gene-LOC105285055 | https://www.ncbi.nlm.nih.gov/gene/?term=LOC105285055 | 4108 | 248 | 603.7 | 1 | 2.43 | 0.4 | 12.58 | 11.79 | 0.79 | 0.79 | 0.05 | 0.05 |
| gene-LOC105285695 | https://www.ncbi.nlm.nih.gov/gene/?term=LOC105285695 | 1371 | 75 | 547.05 | 1 | 7.29 | 1.33 | 27.5 | 26.71 | 0.79 | 0.79 | 0.04 | 0.04 |
| gene-LOC105285674 | https://www.ncbi.nlm.nih.gov/gene/?term=LOC105285674 | 3035 | 109 | 359.14 | 1 | 3.29 | 0.92 | 24.18 | 23.43 | 0.75 | 0.75 | 0.03 | 0.03 |
| gene-LOC105275460 | https://www.ncbi.nlm.nih.gov/gene/?term=LOC105275460 | 2473 | 127 | 513.55 | 1 | 4.04 | 0.79 | 13.03 | 12.3 | 0.73 | 0.73 | 0.04 | 0.04 |
| gene-LOC105285639 | https://www.ncbi.nlm.nih.gov/gene/?term=LOC105285639 | 10325 | 372 | 360.29 | 2 | 1.94 | 0.54 | 11.5 | 10.78 | 0.72 | 0.72 | 0.03 | 0.03 |
| gene-LOC105283728 | https://www.ncbi.nlm.nih.gov/gene/?term=LOC105283728 | 3256 | 146 | 448.4 | 1 | 3.07 | 0.68 | 16.37 | 15.66 | 0.71 | 0.71 | 0.03 | 0.03 |
| gene-LOC105277435 | https://www.ncbi.nlm.nih.gov/gene/?term=LOC105277435 | 3643 | 179 | 491.35 | 1 | 2.74 | 0.56 | 10.99 | 10.3 | 0.7 | 0.7 | 0.03 | 0.03 |
| gene-LOC105285671 | https://www.ncbi.nlm.nih.gov/gene/?term=LOC105285671 | 2263 | 103 | 455.15 | 1 | 4.42 | 0.97 | 30.33 | 29.64 | 0.69 | 0.69 | 0.03 | 0.03 |
| gene-LOC105276449 | https://www.ncbi.nlm.nih.gov/gene/?term=LOC105276449 | 1231 | 55 | 446.79 | 1 | 8.12 | 1.82 | 22.22 | 21.54 | 0.67 | 0.67 | 0.03 | 0.03 |
| gene-LOC105284355 | https://www.ncbi.nlm.nih.gov/gene/?term=LOC105284355 | 3904 | 154 | 394.47 | 1 | 2.56 | 0.65 | 22.13 | 21.47 | 0.66 | 0.66 | 0.03 | 0.03 |
| gene-LOC105282480 | https://www.ncbi.nlm.nih.gov/gene/?term=LOC105282480 | 4177 | 166 | 397.41 | 1 | 2.39 | 0.6 | 12.65 | 12.01 | 0.65 | 0.65 | 0.03 | 0.03 |
| gene-LOC105276704 | https://www.ncbi.nlm.nih.gov/gene/?term=LOC105276704 | 2063 | 87 | 421.72 | 2 | 9.69 | 2.3 | 14.01 | 13.37 | 0.64 | 0.64 | 0.03 | 0.03 |
| gene-LOC105283943 | https://www.ncbi.nlm.nih.gov/gene/?term=LOC105283943 | 3559 | 69 | 193.87 | 1 | 2.81 | 1.45 | 20.61 | 20.02 | 0.59 | 0.59 | 0.01 | 0.01 |
| gene-LOC105276410 | https://www.ncbi.nlm.nih.gov/gene/?term=LOC105276410 | 11085 | 341 | 307.62 | 1 | 0.9 | 0.29 | 9.89 | 9.31 | 0.58 | 0.58 | 0.02 | 0.02 |
| gene-LOC105279664 | https://www.ncbi.nlm.nih.gov/gene/?term=LOC105279664 | 4023 | 67 | 166.54 | 1 | 2.49 | 1.49 | 18.98 | 18.43 | 0.55 | 0.55 | 0.01 | 0.01 |
| gene-LOC105274823 | https://www.ncbi.nlm.nih.gov/gene/?term=LOC105274823 | 4620 | 172 | 372.29 | 1 | 2.16 | 0.58 | 17.73 | 17.2 | 0.53 | 0.53 | 0.02 | 0.02 |
| gene-LOC105278497 | https://www.ncbi.nlm.nih.gov/gene/?term=LOC105278497 | 2072 | 92 | 444.02 | 1 | 4.83 | 1.09 | 20.29 | 19.78 | 0.51 | 0.51 | 0.02 | 0.02 |
| gene-LOC105280408 | https://www.ncbi.nlm.nih.gov/gene/?term=LOC105280408 | 5983 | 63 | 105.3 | 1 | 1.67 | 1.59 | 16.29 | 15.78 | 0.51 | 0.51 | 0.01 | 0.01 |
| gene-LOC105278534 | https://www.ncbi.nlm.nih.gov/gene/?term=LOC105278534 | 7364 | 204 | 277.02 | 1 | 1.36 | 0.49 | 17.28 | 16.79 | 0.5 | 0.5 | 0.01 | 0.01 |

| | | | | | | | | | | | | | |
|---|---|---|---|---|---|---|---|---|---|---|---|---|---|
| gene-LOC105279298 | https://www.ncbi.nlm.nih.gov/gene/?term=LOC105279298 | 2773 | 130 | 468.81 | 1 | 3.61 | 0.77 | 23.65 | 23.17 | 0.49 | 0.49 | 0.02 | 0.02 |
| gene-LOC105283649 | https://www.ncbi.nlm.nih.gov/gene/?term=LOC105283649 | 1918 | 57 | 297.18 | 1 | 5.21 | 1.75 | 40.29 | 39.8 | 0.48 | 0.48 | 0.01 | 0.01 |
| gene-LOC105286997 | https://www.ncbi.nlm.nih.gov/gene/?term=LOC105286997 | 2850 | 150 | 526.32 | 1 | 3.51 | 0.67 | 11.33 | 10.85 | 0.48 | 0.48 | 0.03 | 0.03 |
| gene-LOC105288078 | https://www.ncbi.nlm.nih.gov/gene/?term=LOC105288078 | 19126 | 350 | 183 | 1 | 0.52 | 0.29 | 15.9 | 15.43 | 0.47 | 0.47 | 0.01 | 0.01 |
| gene-LOC105277631 | https://www.ncbi.nlm.nih.gov/gene/?term=LOC105277631 | 4387 | 313 | 713.47 | 1 | 2.28 | 0.32 | 10.74 | 10.28 | 0.46 | 0.46 | 0.03 | 0.03 |
| gene-LOC105280641 | https://www.ncbi.nlm.nih.gov/gene/?term=LOC105280641 | 3682 | 84 | 228.14 | 1 | 2.72 | 1.19 | 13.99 | 13.53 | 0.46 | 0.46 | 0.01 | 0.01 |
| gene-LOC105275187 | https://www.ncbi.nlm.nih.gov/gene/?term=LOC105275187 | 5534 | 78 | 140.95 | 1 | 1.81 | 1.28 | 16.44 | 16.02 | 0.42 | 0.42 | 0.01 | 0.01 |
| gene-LOC105278807 | https://www.ncbi.nlm.nih.gov/gene/?term=LOC105278807 | 2166 | 75 | 346.26 | 1 | 4.62 | 1.33 | 20.48 | 20.06 | 0.41 | 0.41 | 0.01 | 0.01 |
| gene-LOC105286099 | https://www.ncbi.nlm.nih.gov/gene/?term=LOC105286099 | 5919 | 77 | 130.09 | 1 | 1.69 | 1.3 | 12.19 | 11.79 | 0.4 | 0.4 | 0.01 | 0.01 |
| gene-LOC105282491 | https://www.ncbi.nlm.nih.gov/gene/?term=LOC105282491 | 4761 | 193 | 405.38 | 1 | 2.1 | 0.52 | 8.12 | 7.73 | 0.38 | 0.38 | 0.02 | 0.02 |
| gene-LOC105276527 | https://www.ncbi.nlm.nih.gov/gene/?term=LOC105276527 | 7923 | 370 | 466.99 | 1 | 1.26 | 0.27 | 8.27 | 7.9 | 0.37 | 0.37 | 0.02 | 0.02 |
| gene-LOC105286514 | https://www.ncbi.nlm.nih.gov/gene/?term=LOC105286514 | 2528 | 51 | 201.74 | 1 | 3.96 | 1.96 | 35.82 | 35.46 | 0.36 | 0.36 | 0.01 | 0.01 |
| gene-LOC105275170 | https://www.ncbi.nlm.nih.gov/gene/?term=LOC105275170 | 12918 | 359 | 277.91 | 1 | 0.77 | 0.28 | 2.02 | 1.81 | 0.21 | 0.21 | 0.01 | 0.01 |
| gene-LOC105280407 | https://www.ncbi.nlm.nih.gov/gene/?term=LOC105280407 | 10950 | 227 | 207.31 | 1 | 0.91 | 0.44 | 15.19 | 14.98 | 0.21 | 0.21 | 0 | 0 |
| gene-LOC105279500 | https://www.ncbi.nlm.nih.gov/gene/?term=LOC105279500 | 3587 | 35 | 97.57 | 1 | 2.79 | 2.86 | 48.82 | 48.61 | 0.2 | 0.2 | 0 | 0 |
| gene-LOC105284262 | https://www.ncbi.nlm.nih.gov/gene/?term=LOC105284262 | 6669 | 241 | 361.37 | 1 | 1.5 | 0.41 | 5.35 | 5.16 | 0.19 | 0.19 | 0.01 | 0.01 |
| gene-LOC105288066 | https://www.ncbi.nlm.nih.gov/gene/?term=LOC105288066 | 1964 | 31 | 157.84 | 1 | 5.09 | 3.23 | 36.75 | 36.57 | 0.19 | 0.19 | 0 | 0 |
| gene-LOC105285742 | https://www.ncbi.nlm.nih.gov/gene/?term=LOC105285742 | 3446 | 118 | 342.43 | 1 | 2.9 | 0.85 | 17.32 | 17.14 | 0.18 | 0.18 | 0.01 | 0.01 |
| gene-LOC105285981 | https://www.ncbi.nlm.nih.gov/gene/?term=LOC105285981 | 7666 | 219 | 285.68 | 1 | 1.3 | 0.46 | 18.34 | 18.17 | 0.17 | 0.17 | 0 | 0 |
| gene-LOC105274531 | https://www.ncbi.nlm.nih.gov/gene/?term=LOC105274531 | 7774 | 295 | 379.47 | 1 | 1.29 | 0.34 | 10.65 | 10.48 | 0.16 | 0.16 | 0.01 | 0.01 |
| gene-LOC105278228 | https://www.ncbi.nlm.nih.gov/gene/?term=LOC105278228 | 8610 | 199 | 231.13 | 1 | 1.16 | 0.5 | 21.48 | 21.33 | 0.15 | 0.15 | 0 | 0 |
| gene-LOC105275451 | https://www.ncbi.nlm.nih.gov/gene/?term=LOC105275451 | 6035 | 142 | 235.29 | 1 | 1.66 | 0.7 | 20 | 19.9 | 0.1 | 0.1 | 0 | 0 |
| gene-LOC105283838 | https://www.ncbi.nlm.nih.gov/gene/?term=LOC105283838 | 1748 | 30 | 171.62 | 1 | 5.72 | 3.33 | 35.36 | 35.26 | 0.1 | 0.1 | 0 | 0 |
| gene-LOC105277216 | https://www.ncbi.nlm.nih.gov/gene/?term=LOC105277216 | 7627 | 117 | 153.4 | 1 | 1.31 | 0.85 | 6.74 | 6.65 | 0.09 | 0 | 0 | |
| gene-LOC105278145 | https://www.ncbi.nlm.nih.gov/gene/?term=LOC105278145 | 4470 | 261 | 583.89 | 1 | 2.24 | 0.38 | 10.03 | 9.95 | 0.08 | 0.08 | 0 | 0 |
| gene-LOC105280002 | https://www.ncbi.nlm.nih.gov/gene/?term=LOC105280002 | 1940 | 32 | 164.95 | 1 | 5.15 | 3.12 | 34.33 | 34.27 | 0.06 | 0.06 | 0 | 0 |
| gene-LOC105284043 | https://www.ncbi.nlm.nih.gov/gene/?term=LOC105284043 | 5661 | 295 | 521.11 | 1 | 1.77 | 0.34 | 10.33 | 10.27 | 0.06 | 0.06 | 0 | 0 |
| gene-LOC105284184 | https://www.ncbi.nlm.nih.gov/gene/?term=LOC105284184 | 2535 | 102 | 402.37 | 1 | 3.94 | 0.98 | 30 | 29.94 | 0.06 | 0.06 | 0 | 0 |
| gene-LOC105282811 | https://www.ncbi.nlm.nih.gov/gene/?term=LOC105282811 | 5765 | 235 | 407.63 | 1 | 1.73 | 0.43 | 8.46 | 8.42 | 0.04 | 0.04 | 0 | 0 |
| gene-LOC105278283 | https://www.ncbi.nlm.nih.gov/gene/?term=LOC105278283 | 2317 | 138 | 595.6 | 1 | 4.32 | 0.72 | 25.04 | 25.03 | 0.02 | 0.02 | 0 | 0 |
| gene-LOC105278666 | https://www.ncbi.nlm.nih.gov/gene/?term=LOC105278666 | 10341 | 484 | 468.04 | 1 | 0.97 | 0.21 | 6.48 | 6.46 | 0.02 | 0.02 | 0 | 0 |
| gene-LOC105274706 | https://www.ncbi.nlm.nih.gov/gene/?term=LOC105274706 | 1446 | 46 | 318.12 | 1 | 6.92 | 2.17 | 42.4 | 42.39 | 0 | 0 | 0 | 0 |
| gene-LOC105276059 | https://www.ncbi.nlm.nih.gov/gene/?term=LOC105276059 | 6256 | 146 | 233.38 | 1 | 1.6 | 0.68 | 11.41 | 11.42 | 0 | 0 | 0 | 0 |
| gene-LOC105280265 | https://www.ncbi.nlm.nih.gov/gene/?term=LOC105280265 | 85984 | 3391 | 394.38 | 1 | 0.12 | 0.03 | 1.11 | 1.13 | -0.03 | 0.03 | 0 | 0 |
| gene-LOC105278877 | https://www.ncbi.nlm.nih.gov/gene/?term=LOC105278877 | 1968 | 88 | 447.15 | 1 | 5.08 | 1.14 | 30.96 | 31.01 | -0.05 | 0.05 | 0 | 0 |
| gene-LOC105287065 | https://www.ncbi.nlm.nih.gov/gene/?term=LOC105287065 | 4859 | 249 | 512.45 | 1 | 2.06 | 0.4 | 12.25 | 12.3 | -0.05 | 0.05 | 0 | 0 |
| gene-LOC105284774 | https://www.ncbi.nlm.nih.gov/gene/?term=LOC105284774 | 12089 | 100 | 82.72 | 1 | 0.83 | 1 | 19.53 | 19.59 | -0.06 | 0.06 | 0 | 0 |
| gene-LOC105275327 | https://www.ncbi.nlm.nih.gov/gene/?term=LOC105275327 | 14245 | 552 | 387.5 | 1 | 0.7 | 0.18 | 6.33 | 6.41 | -0.08 | 0.08 | 0 | 0 |
| gene-LOC105282600 | https://www.ncbi.nlm.nih.gov/gene/?term=LOC105282600 | 3715 | 60 | 161.51 | 1 | 2.69 | 1.67 | 28.86 | 28.93 | -0.08 | 0.08 | 0 | 0 |
| gene-LOC105286339 | https://www.ncbi.nlm.nih.gov/gene/?term=LOC105286339 | 5230 | 184 | 351.82 | 1 | 1.91 | 0.54 | 16.14 | 16.23 | -0.09 | 0.09 | 0 | 0 |
| gene-LOC105288010 | https://www.ncbi.nlm.nih.gov/gene/?term=LOC105288010 | 3189 | 52 | 163.06 | 1 | 3.14 | 1.92 | 33.66 | 33.76 | -0.1 | 0.1 | 0 | 0 |
| gene-LOC105285807 | https://www.ncbi.nlm.nih.gov/gene/?term=LOC105285807 | 6063 | 221 | 364.51 | 1 | 1.65 | 0.45 | 9.8 | 9.91 | -0.11 | 0.11 | 0 | 0 |
| gene-LOC105286194 | https://www.ncbi.nlm.nih.gov/gene/?term=LOC105286194 | 152619 | 2446 | 160.27 | 1 | 0.07 | 0.04 | 6.09 | 6.23 | -0.14 | 0.14 | 0 | 0 |
| gene-LOC105279038 | https://www.ncbi.nlm.nih.gov/gene/?term=LOC105279038 | 9453 | 346 | 366.02 | 1 | 1.06 | 0.29 | 8.85 | 9.03 | -0.18 | 0.18 | -0.01 | 0.01 |
| gene-LOC105277866 | https://www.ncbi.nlm.nih.gov/gene/?term=LOC105277866 | 33045 | 544 | 164.62 | 1 | 0.3 | 0.18 | 2.82 | 3.03 | -0.21 | 0.21 | 0 | 0 |
| gene-LOC105288042 | https://www.ncbi.nlm.nih.gov/gene/?term=LOC105288042 | 3507 | 98 | 279.44 | 1 | 2.85 | 1.02 | 22.85 | 23.09 | -0.24 | 0.24 | -0.01 | 0.01 |
| gene-LOC105280956 | https://www.ncbi.nlm.nih.gov/gene/?term=LOC105280956 | 2639 | 81 | 306.93 | 1 | 3.79 | 1.23 | 18.56 | 18.86 | -0.29 | 0.29 | -0.01 | 0.01 |
| gene-LOC105281173 | https://www.ncbi.nlm.nih.gov/gene/?term=LOC105281173 | 1595 | 42 | 263.32 | 1 | 6.27 | 2.38 | 62.5 | 62.82 | -0.31 | 0.31 | -0.01 | 0.01 |
| gene-LOC105285004 | https://www.ncbi.nlm.nih.gov/gene/?term=LOC105285004 | 1334 | 35 | 262.37 | 1 | 7.5 | 2.86 | 19.44 | 19.75 | -0.31 | 0.31 | -0.01 | 0.01 |
| gene-LOC105278962 | https://www.ncbi.nlm.nih.gov/gene/?term=LOC105278962 | 4491 | 129 | 287.24 | 1 | 2.23 | 0.78 | 17.72 | 18.05 | -0.34 | 0.34 | -0.01 | 0.01 |
| gene-LOC105278455 | https://www.ncbi.nlm.nih.gov/gene/?term=LOC105278455 | 27181 | 996 | 366.43 | 1 | 0.37 | 0.1 | 6.32 | 6.68 | -0.36 | 0.36 | -0.01 | 0.01 |
| gene-LOC105282110 | https://www.ncbi.nlm.nih.gov/gene/?term=LOC105282110 | 7748 | 148 | 191.02 | 1 | 1.29 | 0.68 | 11.12 | 11.53 | -0.42 | 0.42 | -0.01 | 0.01 |
| gene-LOC105276429 | https://www.ncbi.nlm.nih.gov/gene/?term=LOC105276429 | 2901 | 132 | 455.02 | 1 | 3.45 | 0.76 | 22.03 | 22.48 | -0.45 | 0.45 | -0.02 | 0.02 |
| gene-LOC105282825 | https://www.ncbi.nlm.nih.gov/gene/?term=LOC105282825 | 3919 | 98 | 250.06 | 1 | 2.55 | 1.02 | 13.43 | 13.91 | -0.48 | 0.48 | -0.01 | 0.01 |
| gene-LOC105274590 | https://www.ncbi.nlm.nih.gov/gene/?term=LOC105274590 | 4516 | 84 | 186.01 | 2 | 4.43 | 2.38 | 24.43 | 25 | -0.57 | 0.57 | -0.01 | 0.01 |
| gene-LOC105282151 | https://www.ncbi.nlm.nih.gov/gene/?term=LOC105282151 | 1601 | 43 | 268.58 | 1 | 6.25 | 2.33 | 38.6 | 39.18 | -0.58 | 0.58 | -0.02 | 0.02 |
| gene-LOC105285406 | https://www.ncbi.nlm.nih.gov/gene/?term=LOC105285406 | 8015 | 57 | 71.12 | 1 | 1.25 | 1.75 | 28.09 | 28.72 | -0.63 | 0.63 | 0 | 0 |
| gene-LOC105284036 | https://www.ncbi.nlm.nih.gov/gene/?term=LOC105284036 | 3254 | 103 | 316.53 | 1 | 3.07 | 0.97 | 23.72 | 24.37 | -0.65 | 0.65 | -0.02 | 0.02 |
| gene-LOC105286265 | https://www.ncbi.nlm.nih.gov/gene/?term=LOC105286265 | 2507 | 67 | 267.25 | 1 | 3.99 | 1.49 | 29.04 | 29.68 | -0.65 | 0.65 | -0.02 | 0.02 |
| gene-LOC105274856 | https://www.ncbi.nlm.nih.gov/gene/?term=LOC105274856 | 5216 | 222 | 425.61 | 1 | 1.92 | 0.45 | 16.84 | 17.52 | -0.68 | 0.68 | -0.03 | 0.03 |
| gene-LOC105276745 | https://www.ncbi.nlm.nih.gov/gene/?term=LOC105276745 | 3953 | 76 | 192.26 | 1 | 2.53 | 1.32 | 18.95 | 19.72 | -0.76 | 0.76 | -0.01 | 0.01 |
| gene-LOC105286515 | https://www.ncbi.nlm.nih.gov/gene/?term=LOC105286515 | 3751 | 111 | 295.92 | 1 | 2.67 | 0.9 | 10.2 | 10.99 | -0.8 | 0.8 | -0.02 | 0.02 |
| gene-LOC105279778 | https://www.ncbi.nlm.nih.gov/gene/?term=LOC105279778 | 4347 | 163 | 374.97 | 1 | 2.3 | 0.61 | 8.62 | 9.47 | -0.85 | 0.85 | -0.03 | 0.03 |
| gene-LOC105277903 | https://www.ncbi.nlm.nih.gov/gene/?term=LOC105277903 | 1835 | 38 | 207.08 | 1 | 5.45 | 2.63 | 52.47 | 53.46 | -0.99 | 0.99 | -0.02 | 0.02 |

| | | | | | | | | | | | | | |
|---|---|---|---|---|---|---|---|---|---|---|---|---|---|
| gene-LOC105283132 | https://www.ncbi.nlm.nih.gov/gene/?term=LOC105283132 | 1819 | 56 | 307.86 | 1 | 5.5 | 1.79 | 39.1 | 40.18 | -1.09 | 1.09 | -0.03 | 0.03 |
| gene-LOC105286770 | https://www.ncbi.nlm.nih.gov/gene/?term=LOC105286770 | 3055 | 49 | 160.39 | 1 | 3.27 | 2.04 | 11.46 | 12.59 | -1.13 | 1.13 | -0.02 | 0.02 |
| gene-LOC105276196 | https://www.ncbi.nlm.nih.gov/gene/?term=LOC105276196 | 2953 | 60 | 203.18 | 1 | 3.39 | 1.67 | 37.66 | 38.86 | -1.2 | 1.2 | -0.02 | 0.02 |
| gene-LOC105282914 | https://www.ncbi.nlm.nih.gov/gene/?term=LOC105282914 | 5623 | 43 | 76.47 | 1 | 1.78 | 2.33 | 48 | 49.79 | -1.79 | 1.79 | -0.01 | 0.01 |
| gene-LOC105278282 | https://www.ncbi.nlm.nih.gov/gene/?term=LOC105278282 | 1775 | 65 | 366.2 | 1 | 5.63 | 1.54 | 30.53 | 32.37 | -1.84 | 1.84 | -0.07 | 0.07 |
| gene-LOC105274534 | https://www.ncbi.nlm.nih.gov/gene/?term=LOC105274534 | 1329 | 59 | 443.94 | 1 | 7.52 | 1.69 | 41.22 | 43.19 | -1.98 | 1.98 | -0.09 | 0.09 |
| gene-LOC105282885 | https://www.ncbi.nlm.nih.gov/gene/?term=LOC105282885 | 1870 | 41 | 219.25 | 1 | 5.35 | 2.44 | 39.07 | 41.13 | -2.06 | 2.06 | -0.05 | 0.05 |
| gene-LOC105287633 | https://www.ncbi.nlm.nih.gov/gene/?term=LOC105287633 | 1524 | 25 | 164.04 | 1 | 6.56 | 4 | 61.97 | 64.15 | -2.18 | 2.18 | -0.04 | 0.04 |
| gene-LOC105279528 | https://www.ncbi.nlm.nih.gov/gene/?term=LOC105279528 | 5802 | 41 | 70.67 | 1 | 1.72 | 2.44 | 17.37 | 19.59 | -2.22 | 2.22 | -0.02 | 0.02 |
| gene-LOC105278454 | https://www.ncbi.nlm.nih.gov/gene/?term=LOC105278454 | 1972 | 41 | 207.91 | 1 | 5.07 | 2.44 | 41.77 | 44.02 | -2.26 | 2.26 | -0.05 | 0.05 |
| gene-LOC105280447 | https://www.ncbi.nlm.nih.gov/gene/?term=LOC105280447 | 2215 | 19 | 85.78 | 1 | 4.51 | 5.26 | 33.1 | 35.5 | -2.4 | 2.4 | -0.02 | 0.02 |
| gene-LOC105285409 | https://www.ncbi.nlm.nih.gov/gene/?term=LOC105285409 | 5099 | 66 | 129.44 | 1 | 1.96 | 1.52 | 2.12 | 4.53 | -2.41 | 2.41 | -0.03 | 0.03 |
| gene-LOC105284677 | https://www.ncbi.nlm.nih.gov/gene/?term=LOC105284677 | 1905 | 42 | 220.47 | 1 | 5.25 | 2.38 | 23.34 | 35.09 | -11.75 | 11.75 | -0.26 | 0.26 |

Directory: OGLall File: ogl-Ob_LI_BCphaseB1.vs.RphaseB2.txt

| gene_ID | gene_link | gwidth | #Sites | #per10Kb | #dmSites | #dmsp10kb | %dmSites | %pSite1 | %pSite2 | DMpSite | ADMpSite | DMpNucl | ADMpNucl |
|---|---|---|---|---|---|---|---|---|---|---|---|---|---|
| gene-LOC113562345 | https://www.ncbi.nlm.nih.gov/gene/?term=LOC113562345 | 2290 | 66 | 288.21 | 12 | 52.4 | 18.18 | 30.39 | 17.58 | 12.82 | 12.82 | 0.37 | 0.37 |
| gene-LOC105275070 | https://www.ncbi.nlm.nih.gov/gene/?term=LOC105275070 | 5667 | 41 | 72.35 | 6 | 10.59 | 14.63 | 12.96 | 1.06 | 11.9 | 11.9 | 0.09 | 0.09 |
| gene-LOC105279898 | https://www.ncbi.nlm.nih.gov/gene/?term=LOC105279898 | 1343 | 10 | 74.46 | 1 | 7.45 | 10 | 83.43 | 71.53 | 11.89 | 11.89 | 0.09 | 0.09 |
| gene-LOC105275708 | https://www.ncbi.nlm.nih.gov/gene/?term=LOC105275708 | 1772 | 14 | 79.01 | 1 | 5.64 | 7.14 | 49.7 | 43.39 | 6.31 | 6.31 | 0.05 | 0.05 |
| gene-LOC105275518 | https://www.ncbi.nlm.nih.gov/gene/?term=LOC105275518 | 2335 | 20 | 85.65 | 1 | 4.28 | 5 | 58.2 | 52.79 | 5.42 | 5.42 | 0.05 | 0.05 |
| gene-LOC105283552 | https://www.ncbi.nlm.nih.gov/gene/?term=LOC105283552 | 1562 | 41 | 262.48 | 1 | 6.4 | 2.44 | 53.31 | 48.83 | 4.48 | 4.48 | 0.12 | 0.12 |
| gene-LOC105285315 | https://www.ncbi.nlm.nih.gov/gene/?term=LOC105285315 | 5458 | 19 | 34.81 | 1 | 1.83 | 5.26 | 47.14 | 43.22 | 3.93 | 3.93 | 0.01 | 0.01 |
| gene-LOC105287648 | https://www.ncbi.nlm.nih.gov/gene/?term=LOC105287648 | 2205 | 25 | 113.38 | 1 | 4.54 | 4 | 44.43 | 41.17 | 3.26 | 3.26 | 0.04 | 0.04 |
| gene-LOC105286683 | https://www.ncbi.nlm.nih.gov/gene/?term=LOC105286683 | 1665 | 55 | 330.33 | 1 | 6.01 | 1.82 | 34.95 | 31.78 | 3.17 | 3.17 | 0.1 | 0.1 |
| gene-LOC105274900 | https://www.ncbi.nlm.nih.gov/gene/?term=LOC105274900 | 5561 | 55 | 98.9 | 2 | 3.6 | 3.64 | 33.24 | 30.24 | 3 | 3 | 0.03 | 0.03 |
| gene-LOC105281515 | https://www.ncbi.nlm.nih.gov/gene/?term=LOC105281515 | 8827 | 40 | 45.32 | 2 | 2.27 | 5 | 31.53 | 28.62 | 2.91 | 2.91 | 0.01 | 0.01 |
| gene-LOC105285514 | https://www.ncbi.nlm.nih.gov/gene/?term=LOC105285514 | 1277 | 17 | 133.12 | 1 | 7.83 | 5.88 | 73.52 | 70.72 | 2.8 | 2.8 | 0.04 | 0.04 |
| gene-LOC105279664 | https://www.ncbi.nlm.nih.gov/gene/?term=LOC105279664 | 4023 | 54 | 134.23 | 1 | 2.49 | 1.85 | 21.88 | 19.13 | 2.74 | 2.74 | 0.04 | 0.04 |
| gene-LOC105282794 | https://www.ncbi.nlm.nih.gov/gene/?term=LOC105282794 | 4571 | 91 | 199.08 | 1 | 2.19 | 1.1 | 19.77 | 17.09 | 2.68 | 2.68 | 0.05 | 0.05 |
| gene-LOC105280104 | https://www.ncbi.nlm.nih.gov/gene/?term=LOC105280104 | 3960 | 135 | 340.91 | 1 | 2.53 | 0.74 | 18.4 | 16.2 | 2.2 | 2.2 | 0.07 | 0.07 |
| gene-LOC105278108 | https://www.ncbi.nlm.nih.gov/gene/?term=LOC105278108 | 4392 | 46 | 104.74 | 1 | 2.28 | 2.17 | 27.18 | 25.03 | 2.14 | 2.14 | 0.02 | 0.02 |
| gene-LOC105280641 | https://www.ncbi.nlm.nih.gov/gene/?term=LOC105280641 | 3682 | 50 | 135.8 | 1 | 2.72 | 2 | 17.97 | 16.08 | 1.89 | 1.89 | 0.03 | 0.03 |
| gene-LOC105285671 | https://www.ncbi.nlm.nih.gov/gene/?term=LOC105285671 | 2263 | 105 | 463.99 | 1 | 4.42 | 0.95 | 28.66 | 26.78 | 1.88 | 1.88 | 0.09 | 0.09 |
| gene-LOC105279356 | https://www.ncbi.nlm.nih.gov/gene/?term=LOC105279356 | 1314 | 43 | 327.25 | 1 | 7.61 | 2.33 | 32.17 | 30.3 | 1.87 | 1.87 | 0.06 | 0.06 |
| gene-LOC105287417 | https://www.ncbi.nlm.nih.gov/gene/?term=LOC105287417 | 1025 | 23 | 224.39 | 1 | 9.76 | 4.35 | 51.1 | 49.27 | 1.83 | 1.83 | 0.04 | 0.04 |
| gene-LOC105281419 | https://www.ncbi.nlm.nih.gov/gene/?term=LOC105281419 | 2366 | 80 | 338.12 | 1 | 4.23 | 1.25 | 22.59 | 20.79 | 1.81 | 1.81 | 0.06 | 0.06 |
| gene-LOC105285674 | https://www.ncbi.nlm.nih.gov/gene/?term=LOC105285674 | 3035 | 98 | 322.9 | 2 | 6.59 | 2.04 | 25.36 | 23.72 | 1.64 | 1.64 | 0.05 | 0.05 |
| gene-LOC105283728 | https://www.ncbi.nlm.nih.gov/gene/?term=LOC105283728 | 3256 | 115 | 353.19 | 1 | 3.07 | 0.87 | 19.83 | 18.35 | 1.48 | 1.48 | 0.05 | 0.05 |
| gene-LOC105288036 | https://www.ncbi.nlm.nih.gov/gene/?term=LOC105288036 | 5110 | 200 | 391.39 | 1 | 1.96 | 0.5 | 17.99 | 16.61 | 1.37 | 1.37 | 0.05 | 0.05 |
| gene-LOC105288042 | https://www.ncbi.nlm.nih.gov/gene/?term=LOC105288042 | 3507 | 89 | 253.78 | 1 | 2.85 | 1.12 | 22.75 | 21.38 | 1.37 | 1.37 | 0.03 | 0.03 |
| gene-LOC105280786 | https://www.ncbi.nlm.nih.gov/gene/?term=LOC105280786 | 2573 | 123 | 478.04 | 1 | 3.89 | 0.81 | 29.6 | 28.26 | 1.34 | 1.34 | 0.06 | 0.06 |
| gene-LOC113561621 | https://www.ncbi.nlm.nih.gov/gene/?term=LOC113561621 | 2249 | 83 | 369.05 | 1 | 4.45 | 1.2 | 21.13 | 19.82 | 1.31 | 1.31 | 0.05 | 0.05 |
| gene-LOC105285179 | https://www.ncbi.nlm.nih.gov/gene/?term=LOC105285179 | 4191 | 175 | 417.56 | 1 | 2.39 | 0.57 | 18.73 | 17.48 | 1.25 | 1.25 | 0.05 | 0.05 |
| gene-LOC113561805 | https://www.ncbi.nlm.nih.gov/gene/?term=LOC113561805 | 3337 | 65 | 194.79 | 1 | 3 | 1.54 | 35.44 | 34.26 | 1.18 | 1.18 | 0.02 | 0.02 |
| gene-LOC105276901 | https://www.ncbi.nlm.nih.gov/gene/?term=LOC105276901 | 3254 | 82 | 252 | 1 | 3.07 | 1.22 | 28 | 26.83 | 1.17 | 1.17 | 0.03 | 0.03 |
| gene-LOC105275363 | https://www.ncbi.nlm.nih.gov/gene/?term=LOC105275363 | 3798 | 160 | 421.27 | 1 | 2.63 | 0.62 | 17.46 | 16.32 | 1.14 | 1.14 | 0.05 | 0.05 |
| gene-LOC105282156 | https://www.ncbi.nlm.nih.gov/gene/?term=LOC105282156 | 2577 | 127 | 492.82 | 1 | 3.88 | 0.79 | 27.5 | 26.37 | 1.13 | 1.13 | 0.06 | 0.06 |
| gene-LOC105280408 | https://www.ncbi.nlm.nih.gov/gene/?term=LOC105280408 | 5983 | 56 | 93.6 | 1 | 1.67 | 1.79 | 17.1 | 16.06 | 1.03 | 1.03 | 0.01 | 0.01 |
| gene-LOC105284355 | https://www.ncbi.nlm.nih.gov/gene/?term=LOC105284355 | 3904 | 151 | 386.78 | 1 | 2.56 | 0.66 | 21.57 | 20.54 | 1.02 | 1.02 | 0.04 | 0.04 |
| gene-LOC105286099 | https://www.ncbi.nlm.nih.gov/gene/?term=LOC105286099 | 5919 | 72 | 121.64 | 1 | 1.69 | 1.39 | 10.67 | 9.65 | 1.01 | 1.01 | 0.01 | 0.01 |
| gene-LOC105274449 | https://www.ncbi.nlm.nih.gov/gene/?term=LOC105274449 | 1549 | 48 | 309.88 | 1 | 6.46 | 2.08 | 51.97 | 50.96 | 1 | 1 | 0.03 | 0.03 |
| gene-LOC105281848 | https://www.ncbi.nlm.nih.gov/gene/?term=LOC105281848 | 2122 | 47 | 221.49 | 1 | 4.71 | 2.13 | 27.57 | 26.57 | 1 | 1 | 0.02 | 0.02 |
| gene-LOC105285639 | https://www.ncbi.nlm.nih.gov/gene/?term=LOC105285639 | 10325 | 332 | 321.55 | 2 | 1.94 | 0.6 | 12.25 | 11.3 | 0.95 | 0.95 | 0.03 | 0.03 |
| gene-LOC105278877 | https://www.ncbi.nlm.nih.gov/gene/?term=LOC105278877 | 1968 | 69 | 350.61 | 1 | 5.08 | 1.45 | 36.22 | 35.3 | 0.91 | 0.91 | 0.03 | 0.03 |
| gene-LOC105276704 | https://www.ncbi.nlm.nih.gov/gene/?term=LOC105276704 | 2063 | 87 | 421.72 | 1 | 4.85 | 1.15 | 14.98 | 14.08 | 0.9 | 0.9 | 0.04 | 0.04 |
| gene-LOC105280447 | https://www.ncbi.nlm.nih.gov/gene/?term=LOC105280447 | 2215 | 21 | 94.81 | 1 | 4.51 | 4.76 | 30.62 | 29.74 | 0.88 | 0.88 | 0.01 | 0.01 |
| gene-LOC105278807 | https://www.ncbi.nlm.nih.gov/gene/?term=LOC105278807 | 2166 | 67 | 309.33 | 1 | 4.62 | 1.49 | 22.11 | 21.25 | 0.87 | 0.87 | 0.03 | 0.03 |
| gene-LOC105283592 | https://www.ncbi.nlm.nih.gov/gene/?term=LOC105283592 | 3479 | 79 | 227.08 | 1 | 2.87 | 1.27 | 22.25 | 21.39 | 0.85 | 0.85 | 0.02 | 0.02 |
| gene-LOC105276875 | https://www.ncbi.nlm.nih.gov/gene/?term=LOC105276875 | 3151 | 116 | 368.14 | 1 | 3.17 | 0.86 | 22.78 | 21.93 | 0.84 | 0.84 | 0.03 | 0.03 |
| gene-LOC105276059 | https://www.ncbi.nlm.nih.gov/gene/?term=LOC105276059 | 6256 | 132 | 211 | 1 | 1.6 | 0.76 | 10.3 | 9.47 | 0.83 | 0.83 | 0.02 | 0.02 |
| gene-LOC105283123 | https://www.ncbi.nlm.nih.gov/gene/?term=LOC105283123 | 1507 | 46 | 305.24 | 1 | 6.64 | 2.17 | 17.92 | 17.09 | 0.83 | 0.83 | 0.03 | 0.03 |
| gene-LOC105284043 | https://www.ncbi.nlm.nih.gov/gene/?term=LOC105284043 | 5661 | 294 | 519.34 | 1 | 1.77 | 0.34 | 10.31 | 9.51 | 0.8 | 0.8 | 0.04 | 0.04 |
| gene-LOC105278632 | https://www.ncbi.nlm.nih.gov/gene/?term=LOC105278632 | 3624 | 108 | 298.01 | 1 | 2.76 | 0.93 | 17.43 | 16.66 | 0.77 | 0.77 | 0.02 | 0.02 |
| gene-LOC105285395 | https://www.ncbi.nlm.nih.gov/gene/?term=LOC105285395 | 8085 | 196 | 242.42 | 1 | 1.24 | 0.51 | 16.42 | 15.66 | 0.76 | 0.76 | 0.02 | 0.02 |
| gene-LOC105280956 | https://www.ncbi.nlm.nih.gov/gene/?term=LOC105280956 | 2639 | 71 | 269.04 | 1 | 3.79 | 1.41 | 18.55 | 17.79 | 0.75 | 0.75 | 0.02 | 0.02 |
| gene-LOC105287095 | https://www.ncbi.nlm.nih.gov/gene/?term=LOC105287095 | 1435 | 58 | 404.18 | 1 | 6.97 | 1.72 | 29.38 | 28.65 | 0.73 | 0.73 | 0.03 | 0.03 |
| gene-LOC105278534 | https://www.ncbi.nlm.nih.gov/gene/?term=LOC105278534 | 7364 | 187 | 253.94 | 1 | 1.36 | 0.53 | 19.13 | 18.41 | 0.72 | 0.72 | 0.02 | 0.02 |
| gene-LOC105275584 | https://www.ncbi.nlm.nih.gov/gene/?term=LOC105275584 | 3070 | 89 | 289.9 | 1 | 3.26 | 1.12 | 19.48 | 18.76 | 0.71 | 0.71 | 0.02 | 0.02 |
| gene-LOC105278497 | https://www.ncbi.nlm.nih.gov/gene/?term=LOC105278497 | 2072 | 92 | 444.02 | 1 | 4.83 | 1.09 | 19.2 | 18.5 | 0.7 | 0.7 | 0.03 | 0.03 |
| gene-LOC105285055 | https://www.ncbi.nlm.nih.gov/gene/?term=LOC105285055 | 4108 | 256 | 623.17 | 1 | 2.43 | 0.39 | 12.1 | 11.41 | 0.69 | 0.69 | 0.04 | 0.04 |
| gene-LOC105284184 | https://www.ncbi.nlm.nih.gov/gene/?term=LOC105284184 | 2535 | 86 | 339.25 | 1 | 3.94 | 1.16 | 33.56 | 32.88 | 0.68 | 0.68 | 0.02 | 0.02 |
| gene-LOC105277423 | https://www.ncbi.nlm.nih.gov/gene/?term=LOC105277423 | 2340 | 106 | 452.99 | 1 | 4.27 | 0.94 | 29 | 28.34 | 0.66 | 0.66 | 0.03 | 0.03 |
| gene-LOC105275451 | https://www.ncbi.nlm.nih.gov/gene/?term=LOC105275451 | 6035 | 126 | 208.78 | 1 | 1.66 | 0.79 | 22.21 | 21.64 | 0.57 | 0.57 | 0.01 | 0.01 |
| gene-LOC105286452 | https://www.ncbi.nlm.nih.gov/gene/?term=LOC105286452 | 6888 | 59 | 85.66 | 1 | 1.45 | 1.69 | 32.42 | 31.88 | 0.55 | 0.55 | 0 | 0 |
| gene-LOC105281599 | https://www.ncbi.nlm.nih.gov/gene/?term=LOC105281599 | 3859 | 49 | 126.98 | 1 | 2.59 | 2.04 | 4.62 | 4.08 | 0.54 | 0.54 | 0.01 | 0.01 |

| | | | | | | | | | | | | | | |
|---|---|---|---|---|---|---|---|---|---|---|---|---|---|---|
| gene-LOC105282811 | https://www.ncbi.nlm.nih.gov/gene/?term=LOC105282811 | 5765 | 215 | 372.94 | 1 | 1.73 | 0.47 | 8.61 | 8.16 | 0.45 | 0.45 | 0.02 | 0.02 | |
| gene-LOC105275460 | https://www.ncbi.nlm.nih.gov/gene/?term=LOC105275460 | 2473 | 117 | 473.11 | 1 | 4.04 | 0.85 | 13.75 | 13.31 | 0.44 | 0.44 | 0.02 | 0.02 | |
| gene-LOC105279298 | https://www.ncbi.nlm.nih.gov/gene/?term=LOC105279298 | 2773 | 133 | 479.62 | 2 | 7.21 | 1.5 | 23.5 | 23.06 | 0.44 | 0.44 | 0.02 | 0.02 | |
| gene-LOC105285742 | https://www.ncbi.nlm.nih.gov/gene/?term=LOC105285742 | 3446 | 110 | 319.21 | 1 | 2.9 | 0.91 | 21.56 | 21.15 | 0.4 | 0.4 | 0.01 | 0.01 | |
| gene-LOC105274706 | https://www.ncbi.nlm.nih.gov/gene/?term=LOC105274706 | 1446 | 50 | 345.78 | 1 | 6.92 | 2 | 40.48 | 40.09 | 0.39 | 0.39 | 0.01 | 0.01 | |
| gene-LOC105280002 | https://www.ncbi.nlm.nih.gov/gene/?term=LOC105280002 | 1940 | 28 | 144.33 | 1 | 5.15 | 3.57 | 37.11 | 36.72 | 0.39 | 0.39 | 0.01 | 0.01 | |
| gene-LOC105285406 | https://www.ncbi.nlm.nih.gov/gene/?term=LOC105285406 | 8015 | 53 | 66.13 1 | 1.25 | 1.89 | 28.7 | 28.32 | 0.39 | 0.39 | 0 | 0 | | |
| gene-LOC105279038 | https://www.ncbi.nlm.nih.gov/gene/?term=LOC105279038 | 9453 | 314 | 332.17 | 1 | 1.06 | 0.32 | 9.5 | 9.12 | 0.38 | 0.38 | 0.01 | 0.01 | |
| gene-LOC105280774 | https://www.ncbi.nlm.nih.gov/gene/?term=LOC105280774 | 2793 | 99 | 354.46 | 1 | 3.58 | 1.01 | 32.85 | 32.47 | 0.38 | 0.38 | 0.01 | 0.01 | |
| gene-LOC105284036 | https://www.ncbi.nlm.nih.gov/gene/?term=LOC105284036 | 3254 | 102 | 313.46 | 1 | 3.07 | 0.98 | 22.68 | 22.3 | 0.38 | 0.38 | 0.01 | 0.01 | |
| gene-LOC105274590 | https://www.ncbi.nlm.nih.gov/gene/?term=LOC105274590 | 4516 | 69 | 152.79 | 2 | 4.43 | 2.9 | 27.64 | 27.27 | 0.37 | 0.37 | 0.01 | 0.01 | |
| gene-LOC105277079 | https://www.ncbi.nlm.nih.gov/gene/?term=LOC105277079 | 13009 | 273 | 209.85 | 1 | 0.77 | 0.37 | 9.7 | 9.33 | 0.37 | 0.37 | 0.01 | 0.01 | |
| gene-LOC105283989 | https://www.ncbi.nlm.nih.gov/gene/?term=LOC105283989 | 4948 | 31 | 62.65 1 | 2.02 | 3.23 | 36.84 | 36.46 | 0.37 | 0.37 | 0 | 0 | | |
| gene-LOC105278962 | https://www.ncbi.nlm.nih.gov/gene/?term=LOC105278962 | 4491 | 100 | 222.67 | 1 | 2.23 | 1 | 21.33 | 20.97 | 0.36 | 0.36 | 0.01 | 0.01 | |
| gene-LOC105287764 | https://www.ncbi.nlm.nih.gov/gene/?term=LOC105287764 | 18359 | 394 | 214.61 | 1 | 0.54 | 0.25 | 3.28 | 2.97 | 0.31 | 0.31 | 0.01 | 0.01 | |
| gene-LOC105281266 | https://www.ncbi.nlm.nih.gov/gene/?term=LOC105281266 | 4102 | 78 | 190.15 | 1 | 2.44 | 1.28 | 16.25 | 15.95 | 0.3 | 0.3 | 0.01 | 0.01 | |
| gene-LOC105288066 | https://www.ncbi.nlm.nih.gov/gene/?term=LOC105288066 | 1964 | 28 | 142.57 | 1 | 5.09 | 3.57 | 32.79 | 32.53 | 0.26 | 0.26 | 0 | 0 | |
| gene-LOC105283943 | https://www.ncbi.nlm.nih.gov/gene/?term=LOC105283943 | 3559 | 47 | 132.06 | 1 | 2.81 | 2.13 | 17.56 | 17.32 | 0.24 | 0.24 | 0 | 0 | |
| gene-LOC105278145 | https://www.ncbi.nlm.nih.gov/gene/?term=LOC105278145 | 4470 | 235 | 525.73 | 1 | 2.24 | 0.43 | 10.78 | 10.55 | 0.23 | 0.23 | 0.01 | 0.01 | |
| gene-LOC105284262 | https://www.ncbi.nlm.nih.gov/gene/?term=LOC105284262 | 6669 | 215 | 322.39 | 1 | 1.5 | 0.47 | 5.95 | 5.73 | 0.22 | 0.22 | 0.01 | 0.01 | |
| gene-LOC105278666 | https://www.ncbi.nlm.nih.gov/gene/?term=LOC105278666 | 10341 | 444 | 429.36 | 1 | 0.97 | 0.23 | 6.98 | 6.79 | 0.19 | 0.19 | 0.01 | 0.01 | |
| gene-LOC105282149 | https://www.ncbi.nlm.nih.gov/gene/?term=LOC105282149 | 1787 | 108 | 604.36 | 1 | 5.6 | 0.93 | 41.15 | 40.97 | 0.18 | 0.18 | 0.01 | 0.01 | |
| gene-LOC105276527 | https://www.ncbi.nlm.nih.gov/gene/?term=LOC105276527 | 7923 | 325 | 410.2 1 | 1.26 | 0.31 | 9.54 | 9.36 | 0.17 | 0.17 | 0.01 | 0.01 | | |
| gene-LOC105287550 | https://www.ncbi.nlm.nih.gov/gene/?term=LOC105287550 | 15659 | 590 | 376.78 | 1 | 0.64 | 0.17 | 6.31 | 6.15 | 0.16 | 0.16 | 0.01 | 0.01 | |
| gene-LOC105286194 | https://www.ncbi.nlm.nih.gov/gene/?term=LOC105286194 | 152619 | | 2124 | 139.17 | 1 | 0.07 | 0.05 | 6.36 | 6.23 | 0.14 | 0.14 | 0 | 0 |
| gene-LOC105276612 | https://www.ncbi.nlm.nih.gov/gene/?term=LOC105276612 | 4157 | 183 | 440.22 | 1 | 2.41 | 0.55 | 9.73 | 9.61 | 0.11 | 0.11 | 0.01 | 0.01 | |
| gene-LOC105274531 | https://www.ncbi.nlm.nih.gov/gene/?term=LOC105274531 | 7774 | 291 | 374.32 | 1 | 1.29 | 0.34 | 10.67 | 10.57 | 0.1 | 0.1 | 0 | 0 | |
| gene-LOC105275327 | https://www.ncbi.nlm.nih.gov/gene/?term=LOC105275327 | 14245 | 475 | 333.45 | 1 | 0.7 | 0.21 | 7.23 | 7.14 | 0.1 | 0.1 | 0 | 0 | |
| gene-LOC105280265 | https://www.ncbi.nlm.nih.gov/gene/?term=LOC105280265 | 85984 | 3003 | 349.25 | 1 | 0.12 | 0.03 | 1.15 | 1.07 | 0.08 | 0.08 | 0 | 0 | |
| gene-LOC105282110 | https://www.ncbi.nlm.nih.gov/gene/?term=LOC105282110 | 7748 | 119 | 153.59 | 1 | 1.29 | 0.84 | 13.03 | 12.95 | 0.08 | 0.08 | 0 | 0 | |
| gene-LOC105285807 | https://www.ncbi.nlm.nih.gov/gene/?term=LOC105285807 | 6063 | 223 | 367.8 1 | 1.65 | 0.45 | 9.47 | 9.4 | 0.06 | 0.06 | 0 | 0 | | |
| gene-LOC105278283 | https://www.ncbi.nlm.nih.gov/gene/?term=LOC105278283 | 2317 | 135 | 582.65 | 1 | 4.32 | 0.74 | 25.81 | 25.76 | 0.05 | 0.05 | 0 | 0 | |
| gene-LOC105276538 | https://www.ncbi.nlm.nih.gov/gene/?term=LOC105276538 | 1511 | 17 | 112.51 | 1 | 6.62 | 5.88 | 69.95 | 69.91 | 0.04 | 0.04 | 0 | 0 | |
| gene-LOC105287528 | https://www.ncbi.nlm.nih.gov/gene/?term=LOC105287528 | 14195 | 398 | 280.38 | 1 | 0.7 | 0.25 | 4.97 | 4.95 | 0.02 | 0.02 | 0 | 0 | |
| gene-LOC105274823 | https://www.ncbi.nlm.nih.gov/gene/?term=LOC105274823 | 4620 | 138 | 298.7 1 | 2.16 | 0.72 | 21.27 | 21.26 | 0.01 | 0.01 | 0 | 0 | | |
| gene-LOC105275127 | https://www.ncbi.nlm.nih.gov/gene/?term=LOC105275127 | 9377 | 295 | 314.6 1 | 1.07 | 0.34 | 7.39 | 7.38 | 0.01 | 0.01 | 0 | 0 | | |
| gene-LOC105280407 | https://www.ncbi.nlm.nih.gov/gene/?term=LOC105280407 | 10950 | 207 | 189.04 | 1 | 0.91 | 0.48 | 16.49 | 16.48 | 0.01 | 0.01 | 0 | 0 | |
| gene-LOC105275187 | https://www.ncbi.nlm.nih.gov/gene/?term=LOC105275187 | 5534 | 70 | 126.49 | 1 | 1.81 | 1.43 | 12.34 | 12.39 | -0.05 | 0.05 | 0 | 0 | |
| gene-LOC105279500 | https://www.ncbi.nlm.nih.gov/gene/?term=LOC105279500 | 3587 | 36 | 100.36 | 1 | 2.79 | 2.78 | 49.66 | 49.71 | -0.05 | 0.05 | 0 | 0 | |
| gene-LOC105276692 | https://www.ncbi.nlm.nih.gov/gene/?term=LOC105276692 | 1745 | 29 | 166.19 | 1 | 5.73 | 3.45 | 24.9 | 24.96 | -0.07 | 0.07 | 0 | 0 | |
| gene-LOC105275374 | https://www.ncbi.nlm.nih.gov/gene/?term=LOC105275374 | 11376 | 502 | 441.28 | 1 | 0.88 | 0.2 | 10.16 | 10.25 | -0.09 | 0.09 | 0 | 0 | |
| gene-LOC105281173 | https://www.ncbi.nlm.nih.gov/gene/?term=LOC105281173 | 1595 | 39 | 244.51 | 1 | 6.27 | 2.56 | 59.82 | 59.91 | -0.09 | 0.09 | 0 | 0 | |
| gene-LOC105286274 | https://www.ncbi.nlm.nih.gov/gene/?term=LOC105286274 | 3817 | 191 | 500.39 | 1 | 2.62 | 0.52 | 17.09 | 17.23 | -0.14 | 0.14 | -0.01 | 0.01 | |
| gene-LOC105278245 | https://www.ncbi.nlm.nih.gov/gene/?term=LOC105278245 | 6445 | 117 | 181.54 | 1 | 1.55 | 0.85 | 22.04 | 22.2 | -0.17 | 0.17 | 0 | 0 | |
| gene-LOC105278455 | https://www.ncbi.nlm.nih.gov/gene/?term=LOC105278455 | 27181 | 921 | 338.84 | 1 | 0.37 | 0.11 | 7.15 | 7.32 | -0.18 | 0.18 | -0.01 | 0.01 | |
| gene-LOC105277866 | https://www.ncbi.nlm.nih.gov/gene/?term=LOC105277866 | 33045 | 390 | 118.02 | 1 | 0.3 | 0.26 | 3.22 | 3.41 | -0.19 | 0.19 | 0 | 0 | |
| gene-LOC105286514 | https://www.ncbi.nlm.nih.gov/gene/?term=LOC105286514 | 2528 | 48 | 189.87 | 1 | 3.96 | 2.08 | 36.18 | 36.38 | -0.2 | 0.2 | 0 | 0 | |
| gene-LOC105275500 | https://www.ncbi.nlm.nih.gov/gene/?term=LOC105275500 | 10943 | 242 | 221.15 | 1 | 0.91 | 0.41 | 7.04 | 7.25 | -0.21 | 0.21 | 0 | 0 | |
| gene-LOC105274856 | https://www.ncbi.nlm.nih.gov/gene/?term=LOC105274856 | 5216 | 210 | 402.61 | 1 | 1.92 | 0.48 | 18.25 | 18.52 | -0.26 | 0.26 | -0.01 | 0.01 | |
| gene-LOC105277903 | https://www.ncbi.nlm.nih.gov/gene/?term=LOC105277903 | 1835 | 36 | 196.19 | 1 | 5.45 | 2.78 | 49.51 | 49.79 | -0.27 | 0.27 | -0.01 | 0.01 | |
| gene-LOC105282491 | https://www.ncbi.nlm.nih.gov/gene/?term=LOC105282491 | 4761 | 165 | 346.57 | 1 | 2.1 | 0.61 | 8.1 | 8.48 | -0.38 | 0.38 | -0.01 | 0.01 | |
| gene-LOC105274851 | https://www.ncbi.nlm.nih.gov/gene/?term=LOC105274851 | 6616 | 136 | 205.56 | 1 | 1.51 | 0.74 | 10.78 | 11.18 | -0.4 | 0.4 | -0.01 | 0.01 | |
| gene-LOC105282067 | https://www.ncbi.nlm.nih.gov/gene/?term=LOC105282067 | 5941 | 49 | 82.48 1 | 1.68 | 2.04 | 26.22 | 26.65 | -0.43 | 0.43 | 0 | 0 | | |
| gene-LOC105286515 | https://www.ncbi.nlm.nih.gov/gene/?term=LOC105286515 | 3751 | 102 | 271.93 | 1 | 2.67 | 0.98 | 12.76 | 13.19 | -0.43 | 0.43 | -0.01 | 0.01 | |
| gene-LOC105282399 | https://www.ncbi.nlm.nih.gov/gene/?term=LOC105282399 | 2682 | 23 | 85.76 1 | 3.73 | 4.35 | 34.71 | 35.34 | -0.63 | 0.63 | -0.01 | 0.01 | | |
| gene-LOC105288010 | https://www.ncbi.nlm.nih.gov/gene/?term=LOC105288010 | 3189 | 44 | 137.97 | 1 | 3.14 | 2.27 | 30.11 | 30.74 | -0.63 | 0.63 | -0.01 | 0.01 | |
| gene-LOC105285060 | https://www.ncbi.nlm.nih.gov/gene/?term=LOC105285060 | 1285 | 18 | 140.08 | 1 | 7.78 | 5.56 | 44.89 | 45.6 | -0.72 | 0.72 | -0.01 | 0.01 | |
| gene-LOC105278454 | https://www.ncbi.nlm.nih.gov/gene/?term=LOC105278454 | 1972 | 44 | 223.12 | 1 | 5.07 | 2.27 | 45.26 | 45.99 | -0.74 | 0.74 | -0.02 | 0.02 | |
| gene-LOC105283712 | https://www.ncbi.nlm.nih.gov/gene/?term=LOC105283712 | 1910 | 35 | 183.25 | 1 | 5.24 | 2.86 | 26.72 | 27.47 | -0.75 | 0.75 | -0.01 | 0.01 | |
| gene-LOC105286166 | https://www.ncbi.nlm.nih.gov/gene/?term=LOC105286166 | 3050 | 110 | 360.66 | 1 | 3.28 | 0.91 | 23.43 | 24.21 | -0.78 | 0.78 | -0.03 | 0.03 | |
| gene-LOC105282151 | https://www.ncbi.nlm.nih.gov/gene/?term=LOC105282151 | 1601 | 39 | 243.6 1 | 6.25 | 2.56 | 40.09 | 41 | -0.91 | 0.91 | -0.02 | 0.02 | | |
| gene-LOC105278282 | https://www.ncbi.nlm.nih.gov/gene/?term=LOC105278282 | 1775 | 56 | 315.49 | 1 | 5.63 | 1.79 | 37.29 | 38.34 | -1.05 | 1.05 | -0.03 | 0.03 | |
| gene-LOC105280377 | https://www.ncbi.nlm.nih.gov/gene/?term=LOC105280377 | 2024 | 69 | 340.91 | 1 | 4.94 | 1.45 | 43.74 | 44.8 | -1.06 | 1.06 | -0.04 | 0.04 | |
| gene-LOC105286999 | https://www.ncbi.nlm.nih.gov/gene/?term=LOC105286999 | 2141 | 46 | 214.85 | 1 | 4.67 | 2.17 | 31.8 | 32.92 | -1.12 | 1.12 | -0.02 | 0.02 | |

| | | | | | | | | | | | | | |
|---|---|---|---|---|---|---|---|---|---|---|---|---|---|
| gene-LOC105276429 | https://www.ncbi.nlm.nih.gov/gene/?term=LOC105276429 | 2901 | 114 | 392.97 | 1 | 3.45 | 0.88 | 24.72 | 25.87 | -1.14 | 1.14 | -0.04 | 0.04 |
| gene-LOC105275966 | https://www.ncbi.nlm.nih.gov/gene/?term=LOC105275966 | 3299 | 59 | 178.84 | 1 | 3.03 | 1.69 | 49.71 | 51.14 | -1.42 | 1.42 | -0.03 | 0.03 |
| gene-LOC105274534 | https://www.ncbi.nlm.nih.gov/gene/?term=LOC105274534 | 1329 | 55 | 413.84 | 1 | 7.52 | 1.82 | 44.22 | 45.89 | -1.68 | 1.68 | -0.07 | 0.07 |
| gene-LOC105281700 | https://www.ncbi.nlm.nih.gov/gene/?term=LOC105281700 | 2208 | 32 | 144.93 | 1 | 4.53 | 3.12 | 37.39 | 39.09 | -1.7 | 1.7 | -0.02 | 0.02 |
| gene-LOC105287633 | https://www.ncbi.nlm.nih.gov/gene/?term=LOC105287633 | 1524 | 31 | 203.41 | 1 | 6.56 | 3.23 | 53.47 | 55.25 | -1.78 | 1.78 | -0.04 | 0.04 |
| gene-LOC105286770 | https://www.ncbi.nlm.nih.gov/gene/?term=LOC105286770 | 3055 | 30 | 98.2 | 1 | 3.27 | 3.33 | 17.03 | 18.87 | -1.84 | 1.84 | -0.02 | 0.02 |
| gene-LOC105282885 | https://www.ncbi.nlm.nih.gov/gene/?term=LOC105282885 | 1870 | 47 | 251.34 | 1 | 5.35 | 2.13 | 35.98 | 38.05 | -2.07 | 2.07 | -0.05 | 0.05 |
| gene-LOC105276196 | https://www.ncbi.nlm.nih.gov/gene/?term=LOC105276196 | 2953 | 60 | 203.18 | 1 | 3.39 | 1.67 | 38.8 | 41.25 | -2.45 | 2.45 | -0.05 | 0.05 |
| gene-LOC105285409 | https://www.ncbi.nlm.nih.gov/gene/?term=LOC105285409 | 5099 | 49 | 96.1 | 2 | 3.92 | 4.08 | 2.16 | 4.82 | -2.66 | 2.66 | -0.03 | 0.03 |
| gene-LOC105283492 | https://www.ncbi.nlm.nih.gov/gene/?term=LOC105283492 | 1535 | 10 | 65.15 | 1 | 6.51 | 10 | 34.7 | 42.8 | -8.1 | 8.1 | -0.05 | 0.05 |

Directory: OGLall File: ogl-Ob_LI_BCphaseB2.txt

| gene_ID | gene_link | gwidth | #Sites | #per10Kb | %perSite | %pNucl |
|---|---|---|---|---|---|---|
| gene-LOC105286531 | https://www.ncbi.nlm.nih.gov/gene/?term=LOC105286531 | 372 | 43 | 1155.91 | 35.45 | 4.1 |
| gene-LOC113561532 | https://www.ncbi.nlm.nih.gov/gene/?term=LOC113561532 | 72 | 3 | 416.67 | 98.48 | 4.1 |
| gene-LOC105282149 | https://www.ncbi.nlm.nih.gov/gene/?term=LOC105282149 | 1787 | 161 | 900.95 | 43.24 | 3.9 |
| gene-LOC113562627 | https://www.ncbi.nlm.nih.gov/gene/?term=LOC113562627 | 397 | 23 | 579.35 | 66.24 | 3.84 |
| gene-LOC105280739 | https://www.ncbi.nlm.nih.gov/gene/?term=LOC105280739 | 1991 | 131 | 657.96 | 55 | 3.62 |
| gene-LOC105274534 | https://www.ncbi.nlm.nih.gov/gene/?term=LOC105274534 | 1329 | 107 | 805.12 | 44.14 | 3.55 |
| gene-LOC105278849 | https://www.ncbi.nlm.nih.gov/gene/?term=LOC105278849 | 776 | 114 | 1469.07 | 23.06 | 3.39 |
| gene-LOC105285635 | https://www.ncbi.nlm.nih.gov/gene/?term=LOC105285635 | 1175 | 128 | 1089.36 | 30.64 | 3.34 |
| gene-LOC105286801 | https://www.ncbi.nlm.nih.gov/gene/?term=LOC105286801 | 1102 | 94 | 852.99 | 39.1 | 3.34 |
| gene-LOC105274711 | https://www.ncbi.nlm.nih.gov/gene/?term=LOC105274711 | 1655 | 118 | 712.99 | 45.69 | 3.26 |
| gene-LOC105285394 | https://www.ncbi.nlm.nih.gov/gene/?term=LOC105285394 | 1307 | 59 | 451.42 | 71.23 | 3.22 |
| gene-LOC105286269 | https://www.ncbi.nlm.nih.gov/gene/?term=LOC105286269 | 1401 | 89 | 635.26 | 49.97 | 3.17 |
| gene-LOC105279052 | https://www.ncbi.nlm.nih.gov/gene/?term=LOC105279052 | 1130 | 136 | 1203.54 | 25.81 | 3.11 |
| gene-LOC105287692 | https://www.ncbi.nlm.nih.gov/gene/?term=LOC105287692 | 1505 | 134 | 890.37 | 34.97 | 3.11 |
| gene-LOC105278844 | https://www.ncbi.nlm.nih.gov/gene/?term=LOC105278844 | 1101 | 115 | 1044.5 | 29.68 | 3.1 |
| gene-LOC105276360 | https://www.ncbi.nlm.nih.gov/gene/?term=LOC105276360 | 1466 | 159 | 1084.58 | 28.5 | 3.09 |
| gene-LOC105287811 | https://www.ncbi.nlm.nih.gov/gene/?term=LOC105287811 | 2155 | 136 | 631.09 | 47.8 | 3.02 |
| gene-LOC105286235 | https://www.ncbi.nlm.nih.gov/gene/?term=LOC105286235 | 1510 | 111 | 735.1 | 40.93 | 3.01 |
| gene-LOC105286737 | https://www.ncbi.nlm.nih.gov/gene/?term=LOC105286737 | 2690 | 359 | 1334.57 | 22.22 | 2.97 |
| gene-LOC105288088 | https://www.ncbi.nlm.nih.gov/gene/?term=LOC105288088 | 1795 | 127 | 707.52 | 41.75 | 2.95 |
| gene-LOC105276371 | https://www.ncbi.nlm.nih.gov/gene/?term=LOC105276371 | 871 | 56 | 642.94 | 44.32 | 2.85 |
| gene-LOC105282986 | https://www.ncbi.nlm.nih.gov/gene/?term=LOC105282986 | 1600 | 142 | 887.5 | 32.15 | 2.85 |
| gene-LOC105278160 | https://www.ncbi.nlm.nih.gov/gene/?term=LOC105278160 | 1872 | 155 | 827.99 | 34.29 | 2.84 |
| gene-LOC105275314 | https://www.ncbi.nlm.nih.gov/gene/?term=LOC105275314 | 1489 | 93 | 624.58 | 45.18 | 2.82 |

...

... only first 25 lines shown ...

Directory: OGLall File: ogl-Ob_LI_BCphaseB2.vs.RphaseA1.txt

| gene_ID | gene_link | gwidth | #Sites | #per10Kb | #dmSites | #dmsp10kb | %dmSites | %pSite1 | %pSite2 | DMpSite | ADMpSite | DMpNucl | ADMpNucl |
|---|---|---|---|---|---|---|---|---|---|---|---|---|---|
| gene-LOC105282914 | https://www.ncbi.nlm.nih.gov/gene/?term=LOC105282914 | 5623 | 41 | 72.91 | 1 | 1.78 | 2.44 | 52.21 | 48.43 | 3.78 | 3.78 | 0.03 | 0.03 |
| gene-LOC105279500 | https://www.ncbi.nlm.nih.gov/gene/?term=LOC105279500 | 3587 | 30 | 83.64 | 1 | 2.79 | 3.33 | 48.41 | 46.03 | 2.38 | 2.38 | 0.02 | 0.02 |
| gene-LOC105284396 | https://www.ncbi.nlm.nih.gov/gene/?term=LOC105284396 | 1976 | 66 | 334.01 | 1 | 5.06 | 1.52 | 34.48 | 32.1 | 2.38 | 2.38 | 0.08 | 0.08 |
| gene-LOC105274706 | https://www.ncbi.nlm.nih.gov/gene/?term=LOC105274706 | 1446 | 50 | 345.78 | 1 | 6.92 | 2 | 39.84 | 37.9 | 1.94 | 1.94 | 0.07 | 0.07 |
| gene-LOC105274449 | https://www.ncbi.nlm.nih.gov/gene/?term=LOC105274449 | 1549 | 53 | 342.16 | 1 | 6.46 | 1.89 | 51.77 | 49.89 | 1.87 | 1.87 | 0.06 | 0.06 |
| gene-LOC105282885 | https://www.ncbi.nlm.nih.gov/gene/?term=LOC105282885 | 1870 | 46 | 245.99 | 1 | 5.35 | 2.17 | 37.06 | 35.54 | 1.52 | 1.52 | 0.04 | 0.04 |
| gene-LOC105276272 | https://www.ncbi.nlm.nih.gov/gene/?term=LOC105276272 | 1965 | 46 | 234.1 | 1 | 5.09 | 2.17 | 27.86 | 26.72 | 1.14 | 1.14 | 0.03 | 0.03 |
| gene-LOC105275587 | https://www.ncbi.nlm.nih.gov/gene/?term=LOC105275587 | 2404 | 57 | 237.1 | 1 | 4.16 | 1.75 | 37.35 | 36.26 | 1.09 | 1.09 | 0.03 | 0.03 |
| gene-LOC105283967 | https://www.ncbi.nlm.nih.gov/gene/?term=LOC105283967 | 2393 | 50 | 208.94 | 1 | 4.18 | 2 | 45.88 | 44.9 | 0.98 | 0.98 | 0.02 | 0.02 |
| gene-LOC105282970 | https://www.ncbi.nlm.nih.gov/gene/?term=LOC105282970 | 1529 | 40 | 261.61 | 1 | 6.54 | 2.5 | 50.96 | 50.14 | 0.82 | 0.82 | 0.02 | 0.02 |
| gene-LOC105281173 | https://www.ncbi.nlm.nih.gov/gene/?term=LOC105281173 | 1595 | 38 | 238.24 | 1 | 6.27 | 2.63 | 60.67 | 59.87 | 0.8 | 0.8 | 0.02 | 0.02 |
| gene-LOC105281312 | https://www.ncbi.nlm.nih.gov/gene/?term=LOC105281312 | 2551 | 65 | 254.8 | 1 | 3.92 | 1.54 | 33.11 | 32.31 | 0.8 | 0.8 | 0.02 | 0.02 |
| gene-LOC105278962 | https://www.ncbi.nlm.nih.gov/gene/?term=LOC105278962 | 4491 | 93 | 207.08 | 1 | 2.23 | 1.08 | 22.79 | 22.03 | 0.76 | 0.76 | 0.02 | 0.02 |
| gene-LOC105283712 | https://www.ncbi.nlm.nih.gov/gene/?term=LOC105283712 | 1910 | 36 | 188.48 | 1 | 5.24 | 2.78 | 26.26 | 25.56 | 0.7 | 0.7 | 0.01 | 0.01 |
| gene-LOC105280819 | https://www.ncbi.nlm.nih.gov/gene/?term=LOC105280819 | 1639 | 45 | 274.56 | 1 | 6.1 | 2.22 | 37.3 | 36.73 | 0.57 | 0.57 | 0.02 | 0.02 |
| gene-LOC105286274 | https://www.ncbi.nlm.nih.gov/gene/?term=LOC105286274 | 3817 | 187 | 489.91 | 1 | 2.62 | 0.53 | 16.7 | 16.2 | 0.51 | 0.51 | 0.02 | 0.02 |
| gene-LOC105284184 | https://www.ncbi.nlm.nih.gov/gene/?term=LOC105284184 | 2535 | 92 | 362.92 | 1 | 3.94 | 1.09 | 31.62 | 31.13 | 0.49 | 0.49 | 0.02 | 0.02 |
| gene-LOC105288042 | https://www.ncbi.nlm.nih.gov/gene/?term=LOC105288042 | 3507 | 98 | 279.44 | 1 | 2.85 | 1.02 | 22.45 | 21.99 | 0.46 | 0.46 | 0.01 | 0.01 |
| gene-LOC105282646 | https://www.ncbi.nlm.nih.gov/gene/?term=LOC105282646 | 3317 | 99 | 298.46 | 1 | 3.01 | 1.01 | 21.06 | 20.67 | 0.39 | 0.39 | 0.01 | 0.01 |
| gene-LOC105274590 | https://www.ncbi.nlm.nih.gov/gene/?term=LOC105274590 | 4516 | 74 | 163.86 | 1 | 2.21 | 1.35 | 30.44 | 30.06 | 0.38 | 0.38 | 0.01 | 0.01 |
| gene-LOC105276527 | https://www.ncbi.nlm.nih.gov/gene/?term=LOC105276527 | 7923 | 317 | 400.1 | 1 | 1.26 | 0.32 | 9.8 | 9.43 | 0.37 | 0.37 | 0.01 | 0.01 |
| gene-LOC105282951 | https://www.ncbi.nlm.nih.gov/gene/?term=LOC105282951 | 3366 | 120 | 356.51 | 1 | 2.97 | 0.83 | 19.9 | 19.57 | 0.33 | 0.33 | 0.01 | 0.01 |
| gene-LOC105276196 | https://www.ncbi.nlm.nih.gov/gene/?term=LOC105276196 | 2953 | 56 | 189.64 | 1 | 3.39 | 1.79 | 39.44 | 39.14 | 0.3 | 0.3 | 0.01 | 0.01 |
| gene-LOC105278534 | https://www.ncbi.nlm.nih.gov/gene/?term=LOC105278534 | 7364 | 185 | 251.22 | 2 | 2.72 | 1.08 | 18.38 | 18.12 | 0.25 | 0.25 | 0.01 | 0.01 |
| gene-LOC105280002 | https://www.ncbi.nlm.nih.gov/gene/?term=LOC105280002 | 1940 | 26 | 134.02 | 1 | 5.15 | 3.85 | 34.83 | 34.64 | 0.19 | 0.19 | 0 | 0 |
| gene-LOC105274534 | https://www.ncbi.nlm.nih.gov/gene/?term=LOC105274534 | 1329 | 59 | 443.94 | 1 | 7.52 | 1.69 | 42.17 | 42.03 | 0.14 | 0.14 | 0.01 | 0.01 |
| gene-LOC105278632 | https://www.ncbi.nlm.nih.gov/gene/?term=LOC105278632 | 3624 | 133 | 367 | 1 | 2.76 | 0.75 | 14.21 | 14.06 | 0.14 | 0.14 | 0.01 | 0.01 |
| gene-LOC105278691 | https://www.ncbi.nlm.nih.gov/gene/?term=LOC105278691 | 3871 | 151 | 390.08 | 1 | 2.58 | 0.66 | 14.98 | 14.84 | 0.14 | 0.14 | 0.01 | 0.01 |
| gene-LOC105282811 | https://www.ncbi.nlm.nih.gov/gene/?term=LOC105282811 | 5765 | 199 | 345.19 | 1 | 1.73 | 0.5 | 8.45 | 8.31 | 0.14 | 0.14 | 0 | 0 |
| gene-LOC105285807 | https://www.ncbi.nlm.nih.gov/gene/?term=LOC105285807 | 6063 | 217 | 357.91 | 1 | 1.65 | 0.46 | 8.91 | 8.79 | 0.12 | 0.12 | 0 | 0 |
| gene-LOC105281700 | https://www.ncbi.nlm.nih.gov/gene/?term=LOC105281700 | 2208 | 37 | 167.57 | 1 | 4.53 | 2.7 | 39.75 | 39.65 | 0.11 | 0.11 | 0 | 0 |
| gene-LOC105283728 | https://www.ncbi.nlm.nih.gov/gene/?term=LOC105283728 | 3256 | 122 | 374.69 | 2 | 6.14 | 1.64 | 18.7 | 18.6 | 0.1 | 0.1 | 0 | 0 |
| gene-LOC105276538 | https://www.ncbi.nlm.nih.gov/gene/?term=LOC105276538 | 1511 | 18 | 119.13 | 1 | 6.62 | 5.56 | 68.54 | 68.54 | 0 | 0 | 0 | 0 |
| gene-LOC105280265 | https://www.ncbi.nlm.nih.gov/gene/?term=LOC105280265 | 85984 | 2983 | 346.93 | 1 | 0.12 | 0.03 | 1.1 | 1.12 | -0.02 | 0.02 | 0 | 0 |
| gene-LOC105283123 | https://www.ncbi.nlm.nih.gov/gene/?term=LOC105283123 | 1507 | 47 | 311.88 | 1 | 6.64 | 2.13 | 17.1 | 17.12 | -0.02 | 0.02 | 0 | 0 |
| gene-LOC105286514 | https://www.ncbi.nlm.nih.gov/gene/?term=LOC105286514 | 2528 | 44 | 174.05 | 1 | 3.96 | 2.27 | 37.14 | 37.18 | -0.04 | 0.04 | 0 | 0 |
| gene-LOC105280513 | https://www.ncbi.nlm.nih.gov/gene/?term=LOC105280513 | 69542 | 1333 | 191.68 | 1 | 0.14 | 0.08 | 0.66 | 0.72 | -0.05 | 0.05 | 0 | 0 |
| gene-LOC105285179 | https://www.ncbi.nlm.nih.gov/gene/?term=LOC105285179 | 4191 | 176 | 419.95 | 1 | 2.39 | 0.57 | 19.46 | 19.52 | -0.06 | 0.06 | 0 | 0 |
| gene-LOC105285406 | https://www.ncbi.nlm.nih.gov/gene/?term=LOC105285406 | 8015 | 60 | 74.86 | 1 | 1.25 | 1.67 | 28.89 | 28.94 | -0.06 | 0.06 | 0 | 0 |
| gene-LOC105284043 | https://www.ncbi.nlm.nih.gov/gene/?term=LOC105284043 | 5661 | 295 | 521.11 | 1 | 1.77 | 0.34 | 9.91 | 9.97 | -0.07 | 0.07 | 0 | 0 |
| gene-LOC105280421 | https://www.ncbi.nlm.nih.gov/gene/?term=LOC105280421 | 54824 | 1624 | 296.22 | 1 | 0.18 | 0.06 | 2.3 | 2.37 | -0.08 | 0.08 | 0 | 0 |
| gene-LOC105277866 | https://www.ncbi.nlm.nih.gov/gene/?term=LOC105277866 | 33045 | 394 | 119.23 | 1 | 0.3 | 0.25 | 3.34 | 3.43 | -0.09 | 0.09 | 0 | 0 |
| gene-LOC105286194 | https://www.ncbi.nlm.nih.gov/gene/?term=LOC105286194 | 152619 | 2103 | 137.79 | 1 | 0.07 | 0.05 | 6.13 | 6.22 | -0.09 | 0.09 | 0 | 0 |
| gene-LOC105287417 | https://www.ncbi.nlm.nih.gov/gene/?term=LOC105287417 | 1025 | 32 | 312.2 | 1 | 9.76 | 3.12 | 37.56 | 37.69 | -0.12 | 0.12 | 0 | 0 |
| gene-LOC113562050 | https://www.ncbi.nlm.nih.gov/gene/?term=LOC113562050 | 17097 | 587 | 343.34 | 1 | 0.58 | 0.17 | 4.91 | 5.05 | -0.13 | 0.13 | 0 | 0 |
| gene-LOC105275683 | https://www.ncbi.nlm.nih.gov/gene/?term=LOC105275683 | 4256 | 187 | 439.38 | 1 | 2.35 | 0.53 | 13.19 | 13.34 | -0.15 | 0.15 | -0.01 | 0.01 |
| gene-LOC105278228 | https://www.ncbi.nlm.nih.gov/gene/?term=LOC105278228 | 8610 | 154 | 178.86 | 1 | 1.16 | 0.65 | 23.95 | 24.14 | -0.2 | 0.2 | 0 | 0 |
| gene-LOC105274851 | https://www.ncbi.nlm.nih.gov/gene/?term=LOC105274851 | 6616 | 130 | 196.49 | 1 | 1.51 | 0.77 | 11.46 | 11.67 | -0.21 | 0.21 | 0 | 0 |
| gene-LOC105278666 | https://www.ncbi.nlm.nih.gov/gene/?term=LOC105278666 | 10341 | 471 | 455.47 | 1 | 0.97 | 0.21 | 6.62 | 6.83 | -0.21 | 0.21 | -0.01 | 0.01 |
| gene-LOC105275179 | https://www.ncbi.nlm.nih.gov/gene/?term=LOC105275179 | 3271 | 77 | 235.4 | 1 | 3.06 | 1.3 | 17.93 | 18.15 | -0.22 | 0.22 | -0.01 | 0.01 |
| gene-LOC105284355 | https://www.ncbi.nlm.nih.gov/gene/?term=LOC105284355 | 3904 | 162 | 414.96 | 1 | 2.56 | 0.62 | 19.15 | 19.36 | -0.22 | 0.22 | -0.01 | 0.01 |
| gene-LOC105286452 | https://www.ncbi.nlm.nih.gov/gene/?term=LOC105286452 | 6888 | 61 | 88.56 | 1 | 1.45 | 1.64 | 30.12 | 30.34 | -0.22 | 0.22 | 0 | 0 |
| gene-LOC105278283 | https://www.ncbi.nlm.nih.gov/gene/?term=LOC105278283 | 2317 | 139 | 599.91 | 1 | 4.32 | 0.72 | 25.21 | 25.46 | -0.25 | 0.25 | -0.02 | 0.02 |
| gene-LOC105275363 | https://www.ncbi.nlm.nih.gov/gene/?term=LOC105275363 | 3798 | 161 | 423.91 | 1 | 2.63 | 0.62 | 16.94 | 17.21 | -0.26 | 0.26 | -0.01 | 0.01 |
| gene-LOC105279038 | https://www.ncbi.nlm.nih.gov/gene/?term=LOC105279038 | 9453 | 340 | 359.67 | 1 | 1.06 | 0.29 | 8.66 | 8.92 | -0.26 | 0.26 | -0.01 | 0.01 |
| gene-LOC105288010 | https://www.ncbi.nlm.nih.gov/gene/?term=LOC105288010 | 3189 | 39 | 122.3 | 1 | 3.14 | 2.56 | 33.4 | 33.65 | -0.26 | 0.26 | 0 | 0 |
| gene-LOC105275374 | https://www.ncbi.nlm.nih.gov/gene/?term=LOC105275374 | 11376 | 488 | 428.97 | 1 | 0.88 | 0.2 | 10.37 | 10.64 | -0.27 | 0.27 | -0.01 | 0.01 |
| gene-LOC105275327 | https://www.ncbi.nlm.nih.gov/gene/?term=LOC105275327 | 14245 | 511 | 358.72 | 1 | 0.7 | 0.2 | 6.72 | 7.02 | -0.3 | 0.3 | -0.01 | 0.01 |
| gene-LOC105278497 | https://www.ncbi.nlm.nih.gov/gene/?term=LOC105278497 | 2072 | 96 | 463.32 | 1 | 4.83 | 1.04 | 18.89 | 19.19 | -0.31 | 0.31 | -0.01 | 0.01 |
| gene-LOC105275170 | https://www.ncbi.nlm.nih.gov/gene/?term=LOC105275170 | 12918 | 279 | 215.98 | 1 | 0.77 | 0.36 | 2.01 | 2.33 | -0.32 | 0.32 | -0.01 | 0.01 |

| | | | | | | | | | | | | | |
|---|---|---|---|---|---|---|---|---|---|---|---|---|---|
| gene-LOC105278282 | https://www.ncbi.nlm.nih.gov/gene/?term=LOC105278282 | 1775 | 51 | 287.32 | 1 | 5.63 | 1.96 | 40.43 | 40.76 | -0.33 | 0.33 | -0.01 | 0.01 |
| gene-LOC105282794 | https://www.ncbi.nlm.nih.gov/gene/?term=LOC105282794 | 4571 | 93 | 203.46 | 1 | 2.19 | 1.08 | 16.99 | 17.32 | -0.33 | 0.33 | -0.01 | 0.01 |
| gene-LOC105282149 | https://www.ncbi.nlm.nih.gov/gene/?term=LOC105282149 | 1787 | 103 | 576.39 | 1 | 5.6 | 0.97 | 40.98 | 41.32 | -0.34 | 0.34 | -0.02 | 0.02 |
| gene-LOC105274669 | https://www.ncbi.nlm.nih.gov/gene/?term=LOC105274669 | 8555 | 261 | 305.08 | 1 | 1.17 | 0.38 | 5.68 | 6.02 | -0.35 | 0.35 | -0.01 | 0.01 |
| gene-LOC105278455 | https://www.ncbi.nlm.nih.gov/gene/?term=LOC105278455 | 27181 | 922 | 339.21 | 1 | 0.37 | 0.11 | 7.06 | 7.41 | -0.35 | 0.35 | -0.01 | 0.01 |
| gene-LOC105280774 | https://www.ncbi.nlm.nih.gov/gene/?term=LOC105280774 | 2793 | 117 | 418.9 | 1 | 3.58 | 0.85 | 27.14 | 27.51 | -0.36 | 0.36 | -0.02 | 0.02 |
| gene-LOC105277761 | https://www.ncbi.nlm.nih.gov/gene/?term=LOC105277761 | 4648 | 144 | 309.81 | 1 | 2.15 | 0.69 | 14.39 | 14.8 | -0.41 | 0.41 | -0.01 | 0.01 |
| gene-LOC105274531 | https://www.ncbi.nlm.nih.gov/gene/?term=LOC105274531 | 7774 | 275 | 353.74 | 1 | 1.29 | 0.36 | 10.93 | 11.37 | -0.43 | 0.43 | -0.02 | 0.02 |
| gene-LOC105274823 | https://www.ncbi.nlm.nih.gov/gene/?term=LOC105274823 | 4620 | 145 | 313.85 | 1 | 2.16 | 0.69 | 19.86 | 20.29 | -0.43 | 0.43 | -0.01 | 0.01 |
| gene-LOC105285639 | https://www.ncbi.nlm.nih.gov/gene/?term=LOC105285639 | 10325 | 344 | 333.17 | 1 | 0.97 | 0.29 | 11.23 | 11.66 | -0.43 | 0.43 | -0.01 | 0.01 |
| gene-LOC105287095 | https://www.ncbi.nlm.nih.gov/gene/?term=LOC105287095 | 1435 | 64 | 445.99 | 1 | 6.97 | 1.56 | 25.57 | 26.08 | -0.51 | 0.51 | -0.02 | 0.02 |
| gene-LOC105278145 | https://www.ncbi.nlm.nih.gov/gene/?term=LOC105278145 | 4470 | 235 | 525.73 | 2 | 4.47 | 0.85 | 10.77 | 11.29 | -0.52 | 0.52 | -0.03 | 0.03 |
| gene-LOC105275752 | https://www.ncbi.nlm.nih.gov/gene/?term=LOC105275752 | 3462 | 128 | 369.73 | 1 | 2.89 | 0.78 | 29.22 | 29.76 | -0.53 | 0.53 | -0.02 | 0.02 |
| gene-LOC105285742 | https://www.ncbi.nlm.nih.gov/gene/?term=LOC105285742 | 3446 | 105 | 304.7 | 1 | 2.9 | 0.95 | 17.29 | 17.85 | -0.55 | 0.55 | -0.02 | 0.02 |
| gene-LOC105288036 | https://www.ncbi.nlm.nih.gov/gene/?term=LOC105288036 | 5110 | 200 | 391.39 | 1 | 1.96 | 0.5 | 16.38 | 16.94 | -0.55 | 0.55 | -0.02 | 0.02 |
| gene-LOC105278649 | https://www.ncbi.nlm.nih.gov/gene/?term=LOC105278649 | 5616 | 137 | 243.95 | 1 | 1.78 | 0.73 | 13.17 | 13.73 | -0.56 | 0.56 | -0.01 | 0.01 |
| gene-LOC105275460 | https://www.ncbi.nlm.nih.gov/gene/?term=LOC105275460 | 2473 | 120 | 485.24 | 1 | 4.04 | 0.83 | 13.18 | 13.75 | -0.57 | 0.57 | -0.03 | 0.03 |
| gene-LOC105285055 | https://www.ncbi.nlm.nih.gov/gene/?term=LOC105285055 | 4108 | 256 | 623.17 | 1 | 2.43 | 0.39 | 10.36 | 10.97 | -0.61 | 0.61 | -0.04 | 0.04 |
| gene-LOC105282658 | https://www.ncbi.nlm.nih.gov/gene/?term=LOC105282658 | 2421 | 59 | 243.7 | 1 | 4.13 | 1.69 | 24.61 | 25.23 | -0.62 | 0.62 | -0.02 | 0.02 |
| gene-LOC105279298 | https://www.ncbi.nlm.nih.gov/gene/?term=LOC105279298 | 2773 | 130 | 468.81 | 2 | 7.21 | 1.54 | 23.75 | 24.38 | -0.63 | 0.63 | -0.03 | 0.03 |
| gene-LOC105282067 | https://www.ncbi.nlm.nih.gov/gene/?term=LOC105282067 | 5941 | 41 | 69.01 | 1 | 1.68 | 2.44 | 26.24 | 26.89 | -0.64 | 0.64 | 0 | 0 |
| gene-LOC105287063 | https://www.ncbi.nlm.nih.gov/gene/?term=LOC105287063 | 1874 | 49 | 261.47 | 1 | 5.34 | 2.04 | 29.55 | 30.23 | -0.68 | 0.68 | -0.02 | 0.02 |
| gene-LOC105277856 | https://www.ncbi.nlm.nih.gov/gene/?term=LOC105277856 | 3442 | 31 | 90.06 | 1 | 2.91 | 3.23 | 38.58 | 39.29 | -0.71 | 0.71 | -0.01 | 0.01 |
| gene-LOC105276059 | https://www.ncbi.nlm.nih.gov/gene/?term=LOC105276059 | 6256 | 129 | 206.2 | 1 | 1.6 | 0.78 | 9.24 | 9.96 | -0.72 | 0.72 | -0.01 | 0.01 |
| gene-LOC105279900 | https://www.ncbi.nlm.nih.gov/gene/?term=LOC105279900 | 2191 | 54 | 246.46 | 1 | 4.56 | 1.85 | 30.89 | 31.61 | -0.72 | 0.72 | -0.02 | 0.02 |
| gene-LOC105275451 | https://www.ncbi.nlm.nih.gov/gene/?term=LOC105275451 | 6035 | 123 | 203.81 | 1 | 1.66 | 0.81 | 22.94 | 23.69 | -0.75 | 0.75 | -0.02 | 0.02 |
| gene-LOC105274856 | https://www.ncbi.nlm.nih.gov/gene/?term=LOC105274856 | 5216 | 225 | 431.37 | 1 | 1.92 | 0.44 | 17.14 | 17.89 | -0.76 | 0.76 | -0.03 | 0.03 |
| gene-LOC105278384 | https://www.ncbi.nlm.nih.gov/gene/?term=LOC105278384 | 1390 | 35 | 251.8 | 1 | 7.19 | 2.86 | 45.32 | 46.11 | -0.79 | 0.79 | -0.02 | 0.02 |
| gene-LOC105280723 | https://www.ncbi.nlm.nih.gov/gene/?term=LOC105280723 | 3572 | 182 | 509.52 | 1 | 2.8 | 0.55 | 15.32 | 16.11 | -0.79 | 0.79 | -0.04 | 0.04 |
| gene-LOC105284116 | https://www.ncbi.nlm.nih.gov/gene/?term=LOC105284116 | 3970 | 107 | 269.52 | 1 | 2.52 | 0.93 | 7.18 | 8 | -0.82 | 0.82 | -0.02 | 0.02 |
| gene-LOC105284117 | https://www.ncbi.nlm.nih.gov/gene/?term=LOC105284117 | 4340 | 94 | 216.59 | 1 | 2.3 | 1.06 | 9.25 | 10.09 | -0.84 | 0.84 | -0.02 | 0.02 |
| gene-LOC105282491 | https://www.ncbi.nlm.nih.gov/gene/?term=LOC105282491 | 4761 | 174 | 365.47 | 1 | 2.1 | 0.57 | 8.54 | 9.39 | -0.85 | 0.85 | -0.03 | 0.03 |
| gene-LOC105275187 | https://www.ncbi.nlm.nih.gov/gene/?term=LOC105275187 | 5534 | 69 | 124.68 | 1 | 1.81 | 1.45 | 13.92 | 14.78 | -0.86 | 0.86 | -0.01 | 0.01 |
| gene-LOC105278877 | https://www.ncbi.nlm.nih.gov/gene/?term=LOC105278877 | 1968 | 82 | 416.67 | 1 | 5.08 | 1.22 | 30.66 | 31.6 | -0.94 | 0.94 | -0.04 | 0.04 |
| gene-LOC105278807 | https://www.ncbi.nlm.nih.gov/gene/?term=LOC105278807 | 2166 | 78 | 360.11 | 1 | 4.62 | 1.28 | 18.54 | 19.5 | -0.97 | 0.97 | -0.03 | 0.03 |
| gene-LOC105282991 | https://www.ncbi.nlm.nih.gov/gene/?term=LOC105282991 | 5617 | 101 | 179.81 | 1 | 1.78 | 0.99 | 19.27 | 20.29 | -1.02 | 1.02 | -0.02 | 0.02 |
| gene-LOC105279664 | https://www.ncbi.nlm.nih.gov/gene/?term=LOC105279664 | 4023 | 60 | 149.14 | 1 | 2.49 | 1.67 | 17.78 | 18.81 | -1.03 | 1.03 | -0.02 | 0.02 |
| gene-LOC105280348 | https://www.ncbi.nlm.nih.gov/gene/?term=LOC105280348 | 3797 | 71 | 186.99 | 1 | 2.63 | 1.41 | 18.59 | 19.68 | -1.09 | 1.09 | -0.02 | 0.02 |
| gene-LOC105282151 | https://www.ncbi.nlm.nih.gov/gene/?term=LOC105282151 | 1601 | 42 | 262.34 | 1 | 6.25 | 2.38 | 34.92 | 36.1 | -1.17 | 1.17 | -0.03 | 0.03 |
| gene-LOC105283989 | https://www.ncbi.nlm.nih.gov/gene/?term=LOC105283989 | 4948 | 27 | 54.57 | 1 | 2.02 | 3.7 | 44.98 | 46.16 | -1.18 | 1.18 | -0.01 | 0.01 |
| gene-LOC105287854 | https://www.ncbi.nlm.nih.gov/gene/?term=LOC105287854 | 3644 | 64 | 175.63 | 1 | 2.74 | 1.56 | 14.92 | 16.11 | -1.19 | 1.19 | -0.02 | 0.02 |
| gene-LOC105276692 | https://www.ncbi.nlm.nih.gov/gene/?term=LOC105276692 | 1745 | 33 | 189.11 | 1 | 5.73 | 3.03 | 21.96 | 23.19 | -1.23 | 1.23 | -0.02 | 0.02 |
| gene-LOC105288078 | https://www.ncbi.nlm.nih.gov/gene/?term=LOC105288078 | 19126 | 298 | 155.81 | 1 | 0.52 | 0.34 | 16.07 | 17.31 | -1.24 | 1.24 | -0.02 | 0.02 |
| gene-LOC105275347 | https://www.ncbi.nlm.nih.gov/gene/?term=LOC105275347 | 2526 | 16 | 63.34 | 1 | 3.96 | 6.25 | 40.9 | 42.25 | -1.34 | 1.34 | -0.01 | 0.01 |
| gene-LOC105280641 | https://www.ncbi.nlm.nih.gov/gene/?term=LOC105280641 | 3682 | 55 | 149.38 | 1 | 2.72 | 1.82 | 18.09 | 19.48 | -1.39 | 1.39 | -0.02 | 0.02 |
| gene-LOC105280956 | https://www.ncbi.nlm.nih.gov/gene/?term=LOC105280956 | 2639 | 72 | 272.83 | 1 | 3.79 | 1.39 | 17.08 | 18.51 | -1.43 | 1.43 | -0.04 | 0.04 |
| gene-LOC105281266 | https://www.ncbi.nlm.nih.gov/gene/?term=LOC105281266 | 4102 | 81 | 197.46 | 1 | 2.44 | 1.23 | 16.07 | 17.51 | -1.44 | 1.44 | -0.03 | 0.03 |
| gene-LOC105282156 | https://www.ncbi.nlm.nih.gov/gene/?term=LOC105282156 | 2577 | 133 | 516.1 | 1 | 3.88 | 0.75 | 22.7 | 24.15 | -1.45 | 1.45 | -0.07 | 0.07 |
| gene-LOC105287648 | https://www.ncbi.nlm.nih.gov/gene/?term=LOC105287648 | 2205 | 35 | 158.73 | 1 | 4.54 | 2.86 | 35.94 | 37.4 | -1.45 | 1.45 | -0.02 | 0.02 |
| gene-LOC105280786 | https://www.ncbi.nlm.nih.gov/gene/?term=LOC105280786 | 2573 | 109 | 423.63 | 1 | 3.89 | 0.92 | 28.63 | 30.24 | -1.61 | 1.61 | -0.07 | 0.07 |
| gene-LOC105275384 | https://www.ncbi.nlm.nih.gov/gene/?term=LOC105275384 | 1987 | 87 | 437.85 | 1 | 5.03 | 1.15 | 34.15 | 35.77 | -1.63 | 1.63 | -0.07 | 0.07 |
| gene-LOC105282600 | https://www.ncbi.nlm.nih.gov/gene/?term=LOC105282600 | 3715 | 52 | 139.97 | 1 | 2.69 | 1.92 | 31.36 | 33.08 | -1.72 | 1.72 | -0.02 | 0.02 |
| gene-LOC105283943 | https://www.ncbi.nlm.nih.gov/gene/?term=LOC105283943 | 3559 | 50 | 140.49 | 1 | 2.81 | 2 | 16.35 | 18.09 | -1.74 | 1.74 | -0.02 | 0.02 |
| gene-LOC105278436 | https://www.ncbi.nlm.nih.gov/gene/?term=LOC105278436 | 2556 | 54 | 211.27 | 1 | 3.91 | 1.85 | 41.69 | 43.45 | -1.76 | 1.76 | -0.04 | 0.04 |
| gene-LOC105288066 | https://www.ncbi.nlm.nih.gov/gene/?term=LOC105288066 | 1964 | 29 | 147.66 | 1 | 5.09 | 3.45 | 35.48 | 37.25 | -1.78 | 1.78 | -0.03 | 0.03 |
| gene-LOC105284329 | https://www.ncbi.nlm.nih.gov/gene/?term=LOC105284329 | 4051 | 50 | 123.43 | 1 | 2.47 | 2 | 17.13 | 19.12 | -1.99 | 1.99 | -0.02 | 0.02 |
| gene-LOC105285202 | https://www.ncbi.nlm.nih.gov/gene/?term=LOC105285202 | 1838 | 60 | 326.44 | 1 | 5.44 | 1.67 | 23.97 | 26.11 | -2.15 | 2.15 | -0.07 | 0.07 |
| gene-LOC105283592 | https://www.ncbi.nlm.nih.gov/gene/?term=LOC105283592 | 3479 | 64 | 183.96 | 1 | 2.87 | 1.56 | 24.19 | 26.35 | -2.16 | 2.16 | -0.04 | 0.04 |
| gene-LOC105278454 | https://www.ncbi.nlm.nih.gov/gene/?term=LOC105278454 | 1972 | 43 | 218.05 | 1 | 5.07 | 2.33 | 44.38 | 46.55 | -2.17 | 2.17 | -0.05 | 0.05 |
| gene-LOC105276449 | https://www.ncbi.nlm.nih.gov/gene/?term=LOC105276449 | 1231 | 45 | 365.56 | 1 | 8.12 | 2.22 | 27.05 | 29.32 | -2.27 | 2.27 | -0.08 | 0.08 |
| gene-LOC105283649 | https://www.ncbi.nlm.nih.gov/gene/?term=LOC105283649 | 1918 | 56 | 291.97 | 1 | 5.21 | 1.79 | 36.17 | 38.48 | -2.31 | 2.31 | -0.07 | 0.07 |
| gene-LOC105285674 | https://www.ncbi.nlm.nih.gov/gene/?term=LOC105285674 | 3035 | 84 | 276.77 | 2 | 6.59 | 2.38 | 24.69 | 27.77 | -3.08 | 3.08 | -0.09 | 0.09 |
| gene-LOC105277903 | https://www.ncbi.nlm.nih.gov/gene/?term=LOC105277903 | 1835 | 36 | 196.19 | 1 | 5.45 | 2.78 | 53.1 | 56.25 | -3.14 | 3.14 | -0.06 | 0.06 |
| gene-LOC105281515 | https://www.ncbi.nlm.nih.gov/gene/?term=LOC105281515 | 8827 | 60 | 67.97 | 2 | 2.27 | 3.33 | 26.32 | 29.67 | -3.35 | 3.35 | -0.02 | 0.02 |

| | | | | | | | | | | | | | | | |
|---|---|---|---|---|---|---|---|---|---|---|---|---|---|---|---|
| gene-LOC105281420 | https://www.ncbi.nlm.nih.gov/gene/?term=LOC105281420 | 3882 | 55 | 141.68 | | 1 | 2.58 | 1.82 | 13.05 | 16.87 | -3.82 | 3.82 | -0.05 | 0.05 | |
| gene-LOC105285514 | https://www.ncbi.nlm.nih.gov/gene/?term=LOC105285514 | 1277 | 11 | 86.14 | 1 | 7.83 | 9.09 | 70.96 | 74.98 | -4.02 | 4.02 | -0.03 | 0.03 | | |
| gene-LOC105274900 | https://www.ncbi.nlm.nih.gov/gene/?term=LOC105274900 | 5561 | 54 | 97.1 | 3 | 5.39 | 5.56 | 30 | 35.69 | -5.68 | 5.68 | -0.06 | 0.06 | | |
| gene-LOC105281039 | https://www.ncbi.nlm.nih.gov/gene/?term=LOC105281039 | 1471 | 11 | 74.78 | 1 | 6.8 | 9.09 | 29.26 | 35.1 | -5.84 | 5.84 | -0.04 | 0.04 | | |
| gene-LOC105279898 | https://www.ncbi.nlm.nih.gov/gene/?term=LOC105279898 | 1343 | 11 | 81.91 | 1 | 7.45 | 9.09 | 61.61 | 68.61 | -7 | 7 | -0.06 | 0.06 | | |
| gene-LOC105278567 | https://www.ncbi.nlm.nih.gov/gene/?term=LOC105278567 | 3627 | 29 | 79.96 | 1 | 2.76 | 3.45 | 11.32 | 18.71 | -7.39 | 7.39 | -0.06 | 0.06 | | |
| gene-LOC105275708 | https://www.ncbi.nlm.nih.gov/gene/?term=LOC105275708 | 1772 | 15 | 84.65 | 1 | 5.64 | 6.67 | 45.9 | 54.1 | -8.2 | 8.2 | -0.07 | 0.07 | | |
| gene-LOC105275070 | https://www.ncbi.nlm.nih.gov/gene/?term=LOC105275070 | 5667 | 40 | 70.58 | 6 | 10.59 | 15 | 2.13 | 14.38 | -12.25 | | 12.25 | -0.09 | 0.09 | |
| gene-LOC113562345 | https://www.ncbi.nlm.nih.gov/gene/?term=LOC113562345 | 2290 | 63 | 275.11 | | 8 | 34.93 | 12.7 | 17.51 | 30.84 | -13.33 | | 13.33 | -0.37 | 0.37 |

Directory: OGLall File: ogl-Ob_LI_BCphaseB2.vs.RphaseA2.txt

| gene_ID | gene_link | gwidth | #Sites | #per10Kb | #dmSites | #dmsp10kb | %dmSites | %pSite1 | %pSite2 | DMpSite | ADMpSite | DMpNucl | ADMpNucl |
|---|---|---|---|---|---|---|---|---|---|---|---|---|---|
| gene-LOC105284677 | https://www.ncbi.nlm.nih.gov/gene/?term=LOC105284677 | 1905 | 43 | 225.72 | 2 | 10.5 | 4.65 | 38.99 | 29.38 | 9.62 | 9.62 | 0.22 | 0.22 |
| gene-LOC105282914 | https://www.ncbi.nlm.nih.gov/gene/?term=LOC105282914 | 5623 | 43 | 76.47 | 1 | 1.78 | 2.33 | 49.78 | 47.18 | 2.6 | 2.6 | 0.02 | 0.02 |
| gene-LOC105274449 | https://www.ncbi.nlm.nih.gov/gene/?term=LOC105274449 | 1549 | 53 | 342.16 | 1 | 6.46 | 1.89 | 51.77 | 49.51 | 2.26 | 2.26 | 0.08 | 0.08 |
| gene-LOC105285409 | https://www.ncbi.nlm.nih.gov/gene/?term=LOC105285409 | 5099 | 63 | 123.55 | 1 | 1.96 | 1.59 | 3.13 | 1.16 | 1.97 | 1.97 | 0.02 | 0.02 |
| gene-LOC105282885 | https://www.ncbi.nlm.nih.gov/gene/?term=LOC105282885 | 1870 | 49 | 262.03 | 1 | 5.35 | 2.04 | 39.47 | 37.7 | 1.77 | 1.77 | 0.05 | 0.05 |
| gene-LOC105279500 | https://www.ncbi.nlm.nih.gov/gene/?term=LOC105279500 | 3587 | 33 | 92 | 1 | 2.79 | 3.03 | 47.01 | 45.48 | 1.53 | 1.53 | 0.01 | 0.01 |
| gene-LOC105275743 | https://www.ncbi.nlm.nih.gov/gene/?term=LOC105275743 | 1782 | 76 | 426.49 | 1 | 5.61 | 1.32 | 23.56 | 22.24 | 1.32 | 1.32 | 0.06 | 0.06 |
| gene-LOC105281296 | https://www.ncbi.nlm.nih.gov/gene/?term=LOC105281296 | 2434 | 43 | 176.66 | 1 | 4.11 | 2.33 | 29.93 | 28.76 | 1.17 | 1.17 | 0.02 | 0.02 |
| gene-LOC105285252 | https://www.ncbi.nlm.nih.gov/gene/?term=LOC105285252 | 7427 | 125 | 168.3 | 1 | 1.35 | 0.8 | 14.66 | 13.59 | 1.08 | 1.08 | 0.02 | 0.02 |
| gene-LOC105284396 | https://www.ncbi.nlm.nih.gov/gene/?term=LOC105284396 | 1976 | 70 | 354.25 | 1 | 5.06 | 1.43 | 32.98 | 31.96 | 1.02 | 1.02 | 0.04 | 0.04 |
| gene-LOC105280819 | https://www.ncbi.nlm.nih.gov/gene/?term=LOC105280819 | 1639 | 43 | 262.36 | 1 | 6.1 | 2.33 | 39.04 | 38.17 | 0.87 | 0.87 | 0.02 | 0.02 |
| gene-LOC105288105 | https://www.ncbi.nlm.nih.gov/gene/?term=LOC105288105 | 1460 | 37 | 253.42 | 1 | 6.85 | 2.7 | 53 | 52.16 | 0.84 | 0.84 | 0.02 | 0.02 |
| gene-LOC105275587 | https://www.ncbi.nlm.nih.gov/gene/?term=LOC105275587 | 2404 | 57 | 237.1 | 1 | 4.16 | 1.75 | 36.63 | 35.9 | 0.73 | 0.73 | 0.02 | 0.02 |
| gene-LOC105276272 | https://www.ncbi.nlm.nih.gov/gene/?term=LOC105276272 | 1965 | 50 | 254.45 | 1 | 5.09 | 2 | 27.69 | 27.01 | 0.68 | 0.68 | 0.02 | 0.02 |
| gene-LOC105287633 | https://www.ncbi.nlm.nih.gov/gene/?term=LOC105287633 | 1524 | 35 | 229.66 | 1 | 6.56 | 2.86 | 49.8 | 49.12 | 0.68 | 0.68 | 0.02 | 0.02 |
| gene-LOC105281173 | https://www.ncbi.nlm.nih.gov/gene/?term=LOC105281173 | 1595 | 38 | 238.24 | 1 | 6.27 | 2.63 | 60.67 | 60.19 | 0.48 | 0.48 | 0.01 | 0.01 |
| gene-LOC105280304 | https://www.ncbi.nlm.nih.gov/gene/?term=LOC105280304 | 8919 | 115 | 128.94 | 1 | 1.12 | 0.87 | 7.4 | 6.97 | 0.43 | 0.43 | 0.01 | 0.01 |
| gene-LOC105284507 | https://www.ncbi.nlm.nih.gov/gene/?term=LOC105284507 | 5207 | 91 | 174.76 | 1 | 1.92 | 1.1 | 18.91 | 18.57 | 0.35 | 0.35 | 0.01 | 0.01 |
| gene-LOC105284329 | https://www.ncbi.nlm.nih.gov/gene/?term=LOC105284329 | 4051 | 53 | 130.83 | 1 | 2.47 | 1.89 | 19.36 | 19.04 | 0.32 | 0.32 | 0 | 0 |
| gene-LOC105283296 | https://www.ncbi.nlm.nih.gov/gene/?term=LOC105283296 | 2174 | 96 | 441.58 | 1 | 4.6 | 1.04 | 20.81 | 20.51 | 0.3 | 0.3 | 0.01 | 0.01 |
| gene-LOC105286770 | https://www.ncbi.nlm.nih.gov/gene/?term=LOC105286770 | 3055 | 38 | 124.39 | 1 | 3.27 | 2.63 | 15.65 | 15.48 | 0.17 | 0.17 | 0 | 0 |
| gene-LOC105287842 | https://www.ncbi.nlm.nih.gov/gene/?term=LOC105287842 | 2301 | 57 | 247.72 | 1 | 4.35 | 1.75 | 21.58 | 21.41 | 0.17 | 0.17 | 0 | 0 |
| gene-LOC105285496 | https://www.ncbi.nlm.nih.gov/gene/?term=LOC105285496 | 4080 | 112 | 274.51 | 1 | 2.45 | 0.89 | 40.26 | 40.12 | 0.14 | 0.14 | 0 | 0 |
| gene-LOC105278534 | https://www.ncbi.nlm.nih.gov/gene/?term=LOC105278534 | 7364 | 200 | 271.59 | 2 | 2.72 | 1 | 17.58 | 17.48 | 0.1 | 0.1 | 0 | 0 |
| gene-LOC105281312 | https://www.ncbi.nlm.nih.gov/gene/?term=LOC105281312 | 2551 | 74 | 290.08 | 1 | 3.92 | 1.35 | 29.31 | 29.21 | 0.1 | 0.1 | 0 | 0 |
| gene-LOC105282951 | https://www.ncbi.nlm.nih.gov/gene/?term=LOC105282951 | 3366 | 131 | 389.19 | 1 | 2.97 | 0.76 | 18.77 | 18.69 | 0.08 | 0.08 | 0 | 0 |
| gene-LOC105277273 | https://www.ncbi.nlm.nih.gov/gene/?term=LOC105277273 | 3711 | 169 | 455.4 | 1 | 2.69 | 0.59 | 12.24 | 12.17 | 0.06 | 0.06 | 0 | 0 |
| gene-LOC105281700 | https://www.ncbi.nlm.nih.gov/gene/?term=LOC105281700 | 2208 | 39 | 176.63 | 1 | 4.53 | 2.56 | 37.83 | 37.79 | 0.04 | 0.04 | 0 | 0 |
| gene-LOC105284184 | https://www.ncbi.nlm.nih.gov/gene/?term=LOC105284184 | 2535 | 102 | 402.37 | 1 | 3.94 | 0.98 | 28.97 | 28.93 | 0.03 | 0.03 | 0 | 0 |
| gene-LOC105285533 | https://www.ncbi.nlm.nih.gov/gene/?term=LOC105285533 | 1640 | 71 | 432.93 | 1 | 6.1 | 1.41 | 23.22 | 23.19 | 0.03 | 0.03 | 0 | 0 |
| gene-LOC105281754 | https://www.ncbi.nlm.nih.gov/gene/?term=LOC105281754 | 4300 | 122 | 283.72 | 1 | 2.33 | 0.82 | 16.75 | 16.74 | 0.01 | 0.01 | 0 | 0 |
| gene-LOC105285838 | https://www.ncbi.nlm.nih.gov/gene/?term=LOC105285838 | 21225 | 249 | 117.31 | 1 | 0.47 | 0.4 | 4.78 | 4.78 | 0 | 0 | 0 | 0 |
| gene-LOC105274851 | https://www.ncbi.nlm.nih.gov/gene/?term=LOC105274851 | 6616 | 150 | 226.72 | 1 | 1.51 | 0.67 | 10.51 | 10.53 | -0.02 | 0.02 | 0 | 0 |
| gene-LOC105277866 | https://www.ncbi.nlm.nih.gov/gene/?term=LOC105277866 | 33045 | 432 | 130.73 | 1 | 0.3 | 0.23 | 3.25 | 3.28 | -0.03 | 0.03 | 0 | 0 |
| gene-LOC105280806 | https://www.ncbi.nlm.nih.gov/gene/?term=LOC105280806 | 357459 | 15148 | 423.77 | 1 | 0.03 | 0.01 | 0.61 | 0.64 | -0.03 | 0.03 | 0 | 0 |
| gene-LOC105278691 | https://www.ncbi.nlm.nih.gov/gene/?term=LOC105278691 | 3871 | 172 | 444.33 | 1 | 2.58 | 0.58 | 14.5 | 14.55 | -0.05 | 0.05 | 0 | 0 |
| gene-LOC105280265 | https://www.ncbi.nlm.nih.gov/gene/?term=LOC105280265 | 85984 | 3467 | 403.21 | 1 | 0.12 | 0.03 | 1.03 | 1.08 | -0.05 | 0.05 | 0 | 0 |
| gene-LOC105286649 | https://www.ncbi.nlm.nih.gov/gene/?term=LOC105286649 | 14178 | 892 | 629.14 | 1 | 0.71 | 0.11 | 1.01 | 1.06 | -0.05 | 0.05 | 0 | 0 |
| gene-LOC105275848 | https://www.ncbi.nlm.nih.gov/gene/?term=LOC105275848 | 3718 | 149 | 400.75 | 1 | 2.69 | 0.67 | 21.41 | 21.47 | -0.06 | 0.06 | 0 | 0 |
| gene-LOC105277125 | https://www.ncbi.nlm.nih.gov/gene/?term=LOC105277125 | 51079 | 2082 | 407.6 | 1 | 0.2 | 0.05 | 1.65 | 1.71 | -0.06 | 0.06 | 0 | 0 |
| gene-LOC105280513 | https://www.ncbi.nlm.nih.gov/gene/?term=LOC105280513 | 69542 | 1415 | 203.47 | 1 | 0.14 | 0.07 | 0.66 | 0.73 | -0.06 | 0.06 | 0 | 0 |
| gene-LOC105282646 | https://www.ncbi.nlm.nih.gov/gene/?term=LOC105282646 | 3317 | 110 | 331.62 | 1 | 3.01 | 0.91 | 19.95 | 20.03 | -0.08 | 0.08 | 0 | 0 |
| gene-LOC105276462 | https://www.ncbi.nlm.nih.gov/gene/?term=LOC105276462 | 6561 | 296 | 451.15 | 1 | 1.52 | 0.34 | 7.73 | 7.83 | -0.1 | 0.1 | 0 | 0 |
| gene-LOC105276527 | https://www.ncbi.nlm.nih.gov/gene/?term=LOC105276527 | 7923 | 340 | 429.13 | 1 | 1.26 | 0.29 | 9.36 | 9.47 | -0.11 | 0.11 | 0 | 0 |
| gene-LOC105280408 | https://www.ncbi.nlm.nih.gov/gene/?term=LOC105280408 | 5983 | 60 | 100.28 | 1 | 1.67 | 1.67 | 21.93 | 22.03 | -0.11 | 0.11 | 0 | 0 |
| gene-LOC105280421 | https://www.ncbi.nlm.nih.gov/gene/?term=LOC105280421 | 54824 | 1900 | 346.56 | 1 | 0.18 | 0.05 | 2.13 | 2.24 | -0.11 | 0.11 | 0 | 0 |
| gene-LOC105280722 | https://www.ncbi.nlm.nih.gov/gene/?term=LOC105280722 | 3237 | 98 | 302.75 | 1 | 3.09 | 1.02 | 29.35 | 29.46 | -0.11 | 0.11 | 0 | 0 |
| gene-LOC105275179 | https://www.ncbi.nlm.nih.gov/gene/?term=LOC105275179 | 3271 | 85 | 259.86 | 1 | 3.06 | 1.18 | 19.41 | 19.53 | -0.12 | 0.12 | 0 | 0 |
| gene-LOC105284262 | https://www.ncbi.nlm.nih.gov/gene/?term=LOC105284262 | 6669 | 247 | 370.37 | 1 | 1.5 | 0.4 | 4.68 | 4.81 | -0.13 | 0.13 | 0 | 0 |
| gene-LOC105274823 | https://www.ncbi.nlm.nih.gov/gene/?term=LOC105274823 | 4620 | 148 | 320.35 | 1 | 2.16 | 0.68 | 20.3 | 20.45 | -0.14 | 0.14 | 0 | 0 |
| gene-LOC105278807 | https://www.ncbi.nlm.nih.gov/gene/?term=LOC105278807 | 2166 | 81 | 373.96 | 1 | 4.62 | 1.23 | 19.2 | 19.33 | -0.14 | 0.14 | -0.01 | 0.01 |
| gene-LOC105280410 | https://www.ncbi.nlm.nih.gov/gene/?term=LOC105280410 | 12580 | 263 | 209.06 | 1 | 0.79 | 0.38 | 19.71 | 19.86 | -0.15 | 0.15 | 0 | 0 |
| gene-LOC105286515 | https://www.ncbi.nlm.nih.gov/gene/?term=LOC105286515 | 3751 | 123 | 327.91 | 1 | 2.67 | 0.81 | 10.35 | 10.5 | -0.15 | 0.15 | 0 | 0 |
| gene-LOC105285179 | https://www.ncbi.nlm.nih.gov/gene/?term=LOC105285179 | 4191 | 183 | 436.65 | 1 | 2.39 | 0.55 | 19.23 | 19.39 | -0.16 | 0.16 | -0.01 | 0.01 |
| gene-LOC105282811 | https://www.ncbi.nlm.nih.gov/gene/?term=LOC105282811 | 5765 | 213 | 369.47 | 1 | 1.73 | 0.47 | 8.07 | 8.24 | -0.17 | 0.17 | -0.01 | 0.01 |
| gene-LOC105287550 | https://www.ncbi.nlm.nih.gov/gene/?term=LOC105287550 | 15659 | 656 | 418.93 | 1 | 0.64 | 0.15 | 6.08 | 6.26 | -0.18 | 0.18 | -0.01 | 0.01 |
| gene-LOC105287095 | https://www.ncbi.nlm.nih.gov/gene/?term=LOC105287095 | 1435 | 65 | 452.96 | 1 | 6.97 | 1.54 | 25.24 | 25.44 | -0.2 | 0.2 | -0.01 | 0.01 |
| gene-LOC105286274 | https://www.ncbi.nlm.nih.gov/gene/?term=LOC105286274 | 3817 | 206 | 539.69 | 1 | 2.62 | 0.49 | 16.61 | 16.82 | -0.21 | 0.21 | -0.01 | 0.01 |
| gene-LOC105278455 | https://www.ncbi.nlm.nih.gov/gene/?term=LOC105278455 | 27181 | 1027 | 377.84 | 2 | 0.74 | 0.19 | 6.56 | 6.78 | -0.22 | 0.22 | -0.01 | 0.01 |
| gene-LOC105274699 | https://www.ncbi.nlm.nih.gov/gene/?term=LOC105274699 | 2386 | 63 | 264.04 | 1 | 4.19 | 1.59 | 22.17 | 22.41 | -0.24 | 0.24 | -0.01 | 0.01 |

| | | | | | | | | | | | | | | |
|---|---|---|---|---|---|---|---|---|---|---|---|---|---|---|
| gene-LOC105285406 | https://www.ncbi.nlm.nih.gov/gene/?term=LOC105285406 | 8015 | 59 | 73.61 1 | 1.25 | 1.69 | 29.55 | 29.82 | -0.27 | 0.27 | 0 | 0 | | |
| gene-LOC105276452 | https://www.ncbi.nlm.nih.gov/gene/?term=LOC105276452 | 2917 | 262 | 898.18 | 1 | 3.43 | 0.38 | 9.36 | 9.65 | -0.29 | 0.29 | -0.03 | 0.03 | |
| gene-LOC105278145 | https://www.ncbi.nlm.nih.gov/gene/?term=LOC105278145 | 4470 | 252 | 563.76 | 1 | 2.24 | 0.4 | 10.24 | 10.56 | -0.32 | 0.32 | -0.02 | 0.02 | |
| gene-LOC105286194 | https://www.ncbi.nlm.nih.gov/gene/?term=LOC105286194 | 152619 | | 2371 155.35 | | 1 | 0.07 | 0.04 | 6.01 | 6.33 | -0.32 | 0.32 | 0 | 0 |
| gene-LOC105274531 | https://www.ncbi.nlm.nih.gov/gene/?term=LOC105274531 | 7774 | 298 | 383.33 | 1 | 1.29 | 0.34 | 10.39 | 10.72 | -0.33 | 0.33 | -0.01 | 0.01 | |
| gene-LOC105275327 | https://www.ncbi.nlm.nih.gov/gene/?term=LOC105275327 | 14245 | 568 | 398.74 | 1 | 0.7 | 0.18 | 6.19 | 6.55 | -0.36 | 0.36 | -0.01 | 0.01 | |
| gene-LOC105282067 | https://www.ncbi.nlm.nih.gov/gene/?term=LOC105282067 | 5941 | 44 | 74.06 1 | 1.68 | 2.27 | 30.63 | 31 | -0.36 | 0.36 | 0 | 0 | | |
| gene-LOC105285163 | https://www.ncbi.nlm.nih.gov/gene/?term=LOC105285163 | 5061 | 85 | 167.95 | 1 | 1.98 | 1.18 | 19.84 | 20.21 | -0.37 | 0.37 | -0.01 | 0.01 | |
| gene-LOC105278771 | https://www.ncbi.nlm.nih.gov/gene/?term=LOC105278771 | 10457 | 248 | 237.16 | 1 | 0.96 | 0.4 | 8.97 | 9.36 | -0.39 | 0.39 | -0.01 | 0.01 | |
| gene-LOC105282794 | https://www.ncbi.nlm.nih.gov/gene/?term=LOC105282794 | 4571 | 100 | 218.77 | 1 | 2.19 | 1 | 16.99 | 17.37 | -0.39 | 0.39 | -0.01 | 0.01 | |
| gene-LOC105283123 | https://www.ncbi.nlm.nih.gov/gene/?term=LOC105283123 | 1507 | 57 | 378.23 | 1 | 6.64 | 1.75 | 17.49 | 17.88 | -0.39 | 0.39 | -0.01 | 0.01 | |
| gene-LOC105287309 | https://www.ncbi.nlm.nih.gov/gene/?term=LOC105287309 | 3553 | 96 | 270.19 | 1 | 2.81 | 1.04 | 7.77 | 8.16 | -0.39 | 0.39 | -0.01 | 0.01 | |
| gene-LOC105288055 | https://www.ncbi.nlm.nih.gov/gene/?term=LOC105288055 | 9934 | 548 | 551.64 | 1 | 1.01 | 0.18 | 7.05 | 7.45 | -0.4 | 0.4 | -0.02 | 0.02 | |
| gene-LOC105275374 | https://www.ncbi.nlm.nih.gov/gene/?term=LOC105275374 | 11376 | 567 | 498.42 | 1 | 0.88 | 0.18 | 10.6 | 11.02 | -0.41 | 0.41 | -0.02 | 0.02 | |
| gene-LOC105285742 | https://www.ncbi.nlm.nih.gov/gene/?term=LOC105285742 | 3446 | 132 | 383.05 | 1 | 2.9 | 0.76 | 18.9 | 19.31 | -0.41 | 0.41 | -0.02 | 0.02 | |
| gene-LOC105284043 | https://www.ncbi.nlm.nih.gov/gene/?term=LOC105284043 | 5661 | 325 | 574.1 1 | 1.77 | 0.31 | 9.91 | 10.34 | -0.43 | 0.43 | -0.02 | 0.02 | | |
| gene-LOC105278666 | https://www.ncbi.nlm.nih.gov/gene/?term=LOC105278666 | 10341 | 494 | 477.71 | 1 | 0.97 | 0.2 | 6.74 | 7.18 | -0.44 | 0.44 | -0.02 | 0.02 | |
| gene-LOC105281272 | https://www.ncbi.nlm.nih.gov/gene/?term=LOC105281272 | 5252 | 203 | 386.52 | 1 | 1.9 | 0.49 | 14.55 | 14.99 | -0.44 | 0.44 | -0.02 | 0.02 | |
| gene-LOC105278283 | https://www.ncbi.nlm.nih.gov/gene/?term=LOC105278283 | 2317 | 146 | 630.13 | 1 | 4.32 | 0.68 | 24.36 | 24.8 | -0.45 | 0.45 | -0.03 | 0.03 | |
| gene-LOC105274669 | https://www.ncbi.nlm.nih.gov/gene/?term=LOC105274669 | 8555 | 293 | 342.49 | 1 | 1.17 | 0.34 | 5.34 | 5.82 | -0.48 | 0.48 | -0.02 | 0.02 | |
| gene-LOC105275363 | https://www.ncbi.nlm.nih.gov/gene/?term=LOC105275363 | 3798 | 176 | 463.4 1 | 2.63 | 0.57 | 15.74 | 16.23 | -0.5 | 0.5 | -0.02 | 0.02 | | |
| gene-LOC105287648 | https://www.ncbi.nlm.nih.gov/gene/?term=LOC105287648 | 2205 | 35 | 158.73 | 1 | 4.54 | 2.86 | 35.94 | 36.45 | -0.5 | 0.5 | -0.01 | 0.01 | |
| gene-LOC113562050 | https://www.ncbi.nlm.nih.gov/gene/?term=LOC113562050 | 17097 | 643 | 376.09 | 1 | 0.58 | 0.16 | 4.96 | 5.49 | -0.53 | 0.53 | -0.02 | 0.02 | |
| gene-LOC105276059 | https://www.ncbi.nlm.nih.gov/gene/?term=LOC105276059 | 6256 | 145 | 231.78 | 1 | 1.6 | 0.69 | 8.95 | 9.49 | -0.54 | 0.54 | -0.01 | 0.01 | |
| gene-LOC105275460 | https://www.ncbi.nlm.nih.gov/gene/?term=LOC105275460 | 2473 | 122 | 493.33 | 1 | 4.04 | 0.82 | 12.96 | 13.51 | -0.55 | 0.55 | -0.03 | 0.03 | |
| gene-LOC105284036 | https://www.ncbi.nlm.nih.gov/gene/?term=LOC105284036 | 3254 | 112 | 344.19 | 1 | 3.07 | 0.89 | 21.95 | 22.5 | -0.55 | 0.55 | -0.02 | 0.02 | |
| gene-LOC105283728 | https://www.ncbi.nlm.nih.gov/gene/?term=LOC105283728 | 3256 | 123 | 377.76 | 2 | 6.14 | 1.63 | 18.65 | 19.22 | -0.57 | 0.57 | -0.02 | 0.02 | |
| gene-LOC105276875 | https://www.ncbi.nlm.nih.gov/gene/?term=LOC105276875 | 3151 | 124 | 393.53 | 1 | 3.17 | 0.81 | 21.3 | 21.9 | -0.6 | 0.6 | -0.02 | 0.02 | |
| gene-LOC105274706 | https://www.ncbi.nlm.nih.gov/gene/?term=LOC105274706 | 1446 | 57 | 394.19 | 1 | 6.92 | 1.75 | 36.89 | 37.51 | -0.61 | 0.61 | -0.02 | 0.02 | |
| gene-LOC105285502 | https://www.ncbi.nlm.nih.gov/gene/?term=LOC105285502 | 9990 | 164 | 164.16 | 1 | 1 | 0.61 | 8.55 | 9.16 | -0.62 | 0.62 | -0.01 | 0.01 | |
| gene-LOC105285807 | https://www.ncbi.nlm.nih.gov/gene/?term=LOC105285807 | 6063 | 237 | 390.9 1 | 1.65 | 0.42 | 8.52 | 9.16 | -0.64 | 0.64 | -0.03 | 0.03 | | |
| gene-LOC105274452 | https://www.ncbi.nlm.nih.gov/gene/?term=LOC105274452 | 6595 | 279 | 423.05 | 1 | 1.52 | 0.36 | 12.34 | 12.98 | -0.65 | 0.65 | -0.03 | 0.03 | |
| gene-LOC105276683 | https://www.ncbi.nlm.nih.gov/gene/?term=LOC105276683 | 17000 | 149 | 87.65 1 | 0.59 | 0.67 | 8.22 | 8.88 | -0.65 | 0.65 | -0.01 | 0.01 | | |
| gene-LOC105280723 | https://www.ncbi.nlm.nih.gov/gene/?term=LOC105280723 | 3572 | 185 | 517.92 | 1 | 2.8 | 0.54 | 15.07 | 15.74 | -0.67 | 0.67 | -0.03 | 0.03 | |
| gene-LOC105286452 | https://www.ncbi.nlm.nih.gov/gene/?term=LOC105286452 | 6888 | 66 | 95.82 1 | 1.45 | 1.52 | 30.11 | 30.78 | -0.67 | 0.67 | -0.01 | 0.01 | | |
| gene-LOC105279657 | https://www.ncbi.nlm.nih.gov/gene/?term=LOC105279657 | 5878 | 91 | 154.81 | 1 | 1.7 | 1.1 | 18.9 | 19.62 | -0.72 | 0.72 | -0.01 | 0.01 | |
| gene-LOC105288161 | https://www.ncbi.nlm.nih.gov/gene/?term=LOC105288161 | 26065 | 356 | 136.58 | 1 | 0.38 | 0.28 | 10.97 | 11.71 | -0.74 | 0.74 | -0.01 | 0.01 | |
| gene-LOC105275752 | https://www.ncbi.nlm.nih.gov/gene/?term=LOC105275752 | 3462 | 141 | 407.28 | 1 | 2.89 | 0.71 | 30.44 | 31.2 | -0.75 | 0.75 | -0.03 | 0.03 | |
| gene-LOC105275451 | https://www.ncbi.nlm.nih.gov/gene/?term=LOC105275451 | 6035 | 142 | 235.29 | 1 | 1.66 | 0.7 | 20.66 | 21.45 | -0.78 | 0.78 | -0.02 | 0.02 | |
| gene-LOC105280348 | https://www.ncbi.nlm.nih.gov/gene/?term=LOC105280348 | 3797 | 79 | 208.06 | 1 | 2.63 | 1.27 | 19.69 | 20.49 | -0.8 | 0.8 | -0.02 | 0.02 | |
| gene-LOC105283967 | https://www.ncbi.nlm.nih.gov/gene/?term=LOC105283967 | 2393 | 60 | 250.73 | 1 | 4.18 | 1.67 | 41.56 | 42.36 | -0.8 | 0.8 | -0.02 | 0.02 | |
| gene-LOC105285395 | https://www.ncbi.nlm.nih.gov/gene/?term=LOC105285395 | 8085 | 224 | 277.06 | 1 | 1.24 | 0.45 | 14.28 | 15.08 | -0.8 | 0.8 | -0.02 | 0.02 | |
| gene-LOC105279038 | https://www.ncbi.nlm.nih.gov/gene/?term=LOC105279038 | 9453 | 369 | 390.35 | 1 | 1.06 | 0.27 | 8.12 | 8.94 | -0.82 | 0.82 | -0.03 | 0.03 | |
| gene-LOC105286332 | https://www.ncbi.nlm.nih.gov/gene/?term=LOC105286332 | 27577 | 532 | 192.91 | 1 | 0.36 | 0.19 | 7.3 | 8.12 | -0.82 | 0.82 | -0.02 | 0.02 | |
| gene-LOC105275187 | https://www.ncbi.nlm.nih.gov/gene/?term=LOC105275187 | 5534 | 77 | 139.14 | 1 | 1.81 | 1.3 | 15.38 | 16.21 | -0.83 | 0.83 | -0.01 | 0.01 | |
| gene-LOC105280407 | https://www.ncbi.nlm.nih.gov/gene/?term=LOC105280407 | 10950 | 218 | 199.09 | 1 | 0.91 | 0.46 | 16.66 | 17.49 | -0.83 | 0.83 | -0.02 | 0.02 | |
| gene-LOC105278228 | https://www.ncbi.nlm.nih.gov/gene/?term=LOC105278228 | 8610 | 176 | 204.41 | 1 | 1.16 | 0.57 | 22.43 | 23.28 | -0.84 | 0.84 | -0.02 | 0.02 | |
| gene-LOC105278632 | https://www.ncbi.nlm.nih.gov/gene/?term=LOC105278632 | 3624 | 141 | 389.07 | 1 | 2.76 | 0.71 | 13.56 | 14.44 | -0.87 | 0.87 | -0.03 | 0.03 | |
| gene-LOC105285055 | https://www.ncbi.nlm.nih.gov/gene/?term=LOC105285055 | 4108 | 274 | 666.99 | 2 | 4.87 | 0.73 | 10.9 | 11.79 | -0.89 | 0.89 | -0.06 | 0.06 | |
| gene-LOC105282491 | https://www.ncbi.nlm.nih.gov/gene/?term=LOC105282491 | 4761 | 184 | 386.47 | 1 | 2.1 | 0.54 | 8.25 | 9.16 | -0.91 | 0.91 | -0.04 | 0.04 | |
| gene-LOC105282682 | https://www.ncbi.nlm.nih.gov/gene/?term=LOC105282682 | 4127 | 241 | 583.96 | 1 | 2.42 | 0.41 | 12.88 | 13.79 | -0.91 | 0.91 | -0.05 | 0.05 | |
| gene-LOC105278877 | https://www.ncbi.nlm.nih.gov/gene/?term=LOC105278877 | 1968 | 86 | 436.99 | 1 | 5.08 | 1.16 | 31.6 | 32.52 | -0.92 | 0.92 | -0.04 | 0.04 | |
| gene-LOC105285202 | https://www.ncbi.nlm.nih.gov/gene/?term=LOC105285202 | 1838 | 66 | 359.09 | 1 | 5.44 | 1.52 | 24.58 | 25.52 | -0.94 | 0.94 | -0.03 | 0.03 | |
| gene-LOC105274590 | https://www.ncbi.nlm.nih.gov/gene/?term=LOC105274590 | 4516 | 80 | 177.15 | 2 | 4.43 | 2.5 | 29.75 | 30.69 | -0.95 | 0.95 | -0.02 | 0.02 | |
| gene-LOC105276917 | https://www.ncbi.nlm.nih.gov/gene/?term=LOC105276917 | 2336 | 33 | 141.27 | 1 | 4.28 | 3.03 | 30.13 | 31.08 | -0.95 | 0.95 | -0.01 | 0.01 | |
| gene-LOC105282825 | https://www.ncbi.nlm.nih.gov/gene/?term=LOC105282825 | 3919 | 98 | 250.06 | 1 | 2.55 | 1.02 | 14.18 | 15.14 | -0.96 | 0.96 | -0.02 | 0.02 | |
| gene-LOC105284117 | https://www.ncbi.nlm.nih.gov/gene/?term=LOC105284117 | 4340 | 101 | 232.72 | 1 | 2.3 | 0.99 | 10.38 | 11.36 | -0.98 | 0.98 | -0.02 | 0.02 | |
| gene-LOC105280774 | https://www.ncbi.nlm.nih.gov/gene/?term=LOC105280774 | 2793 | 122 | 436.81 | 1 | 3.58 | 0.82 | 26.14 | 27.14 | -0.99 | 0.99 | -0.04 | 0.04 | |
| gene-LOC105281261 | https://www.ncbi.nlm.nih.gov/gene/?term=LOC105281261 | 5947 | 158 | 265.68 | 1 | 1.68 | 0.63 | 14.18 | 15.18 | -1 | 1 | -0.03 | 0.03 | |
| gene-LOC105275683 | https://www.ncbi.nlm.nih.gov/gene/?term=LOC105275683 | 4256 | 210 | 493.42 | 1 | 2.35 | 0.48 | 13.47 | 14.47 | -1.01 | 1.01 | -0.05 | 0.05 | |
| gene-LOC105278497 | https://www.ncbi.nlm.nih.gov/gene/?term=LOC105278497 | 2072 | 103 | 497.1 1 | 4.83 | 0.97 | 19.13 | 20.15 | -1.02 | 1.02 | -0.05 | 0.05 | | |
| gene-LOC105281848 | https://www.ncbi.nlm.nih.gov/gene/?term=LOC105281848 | 2122 | 64 | 301.6 1 | 4.71 | 1.56 | 21.13 | 22.15 | -1.02 | 1.02 | -0.03 | 0.03 | | |
| gene-LOC105285639 | https://www.ncbi.nlm.nih.gov/gene/?term=LOC105285639 | 10325 | 408 | 395.16 | 1 | 0.97 | 0.25 | 10.25 | 11.26 | -1.02 | 1.02 | -0.04 | 0.04 | |
| gene-LOC105284116 | https://www.ncbi.nlm.nih.gov/gene/?term=LOC105284116 | 3970 | 113 | 284.63 | 1 | 2.52 | 0.88 | 7.21 | 8.25 | -1.03 | 1.03 | -0.03 | 0.03 | |

| | | | | | | | | | | | | | |
|---|---|---|---|---|---|---|---|---|---|---|---|---|---|
| gene-LOC105276196 | https://www.ncbi.nlm.nih.gov/gene/?term=LOC105276196 | 2953 | 59 | 199.8 | 1 | 3.39 | 1.69 | 37.63 | 38.7 | -1.07 | 1.07 | -0.02 | 0.02 |
| gene-LOC105283943 | https://www.ncbi.nlm.nih.gov/gene/?term=LOC105283943 | 3559 | 64 | 179.83 | 1 | 2.81 | 1.56 | 14.69 | 15.79 | -1.1 | 1.1 | -0.02 | 0.02 |
| gene-LOC105276501 | https://www.ncbi.nlm.nih.gov/gene/?term=LOC105276501 | 6089 | 110 | 180.65 | 1 | 1.64 | 0.91 | 15.38 | 16.49 | -1.11 | 1.11 | -0.02 | 0.02 |
| gene-LOC105275347 | https://www.ncbi.nlm.nih.gov/gene/?term=LOC105275347 | 2526 | 20 | 79.18 | 1 | 3.96 | 5 | 37.1 | 38.21 | -1.12 | 1.12 | -0.01 | 0.01 |
| gene-LOC105285514 | https://www.ncbi.nlm.nih.gov/gene/?term=LOC105285514 | 1277 | 15 | 117.46 | 1 | 7.83 | 6.67 | 78.25 | 79.38 | -1.12 | 1.12 | -0.01 | 0.01 |
| gene-LOC105284355 | https://www.ncbi.nlm.nih.gov/gene/?term=LOC105284355 | 3904 | 165 | 422.64 | 1 | 2.56 | 0.61 | 19.1 | 20.26 | -1.16 | 1.16 | -0.05 | 0.05 |
| gene-LOC105277631 | https://www.ncbi.nlm.nih.gov/gene/?term=LOC105277631 | 4387 | 333 | 759.06 | 1 | 2.28 | 0.3 | 9.46 | 10.66 | -1.19 | 1.19 | -0.09 | 0.09 |
| gene-LOC105288078 | https://www.ncbi.nlm.nih.gov/gene/?term=LOC105288078 | 19126 | 332 | 173.59 | 1 | 0.52 | 0.3 | 16.16 | 17.36 | -1.2 | 1.2 | -0.02 | 0.02 |
| gene-LOC105277761 | https://www.ncbi.nlm.nih.gov/gene/?term=LOC105277761 | 4648 | 160 | 344.23 | 1 | 2.15 | 0.62 | 14.78 | 16.05 | -1.27 | 1.27 | -0.04 | 0.04 |
| gene-LOC105274856 | https://www.ncbi.nlm.nih.gov/gene/?term=LOC105274856 | 5216 | 232 | 444.79 | 1 | 1.92 | 0.43 | 17.14 | 18.43 | -1.29 | 1.29 | -0.06 | 0.06 |
| gene-LOC105283712 | https://www.ncbi.nlm.nih.gov/gene/?term=LOC105283712 | 1910 | 36 | 188.48 | 1 | 5.24 | 2.78 | 28.48 | 29.79 | -1.31 | 1.31 | -0.02 | 0.02 |
| gene-LOC105280641 | https://www.ncbi.nlm.nih.gov/gene/?term=LOC105280641 | 3682 | 61 | 165.67 | 1 | 2.72 | 1.64 | 16.75 | 18.07 | -1.32 | 1.32 | -0.02 | 0.02 |
| gene-LOC105282220 | https://www.ncbi.nlm.nih.gov/gene/?term=LOC105282220 | 2298 | 224 | 974.76 | 1 | 4.35 | 0.45 | 7.52 | 8.84 | -1.32 | 1.32 | -0.13 | 0.13 |
| gene-LOC105282991 | https://www.ncbi.nlm.nih.gov/gene/?term=LOC105282991 | 5617 | 112 | 199.39 | 1 | 1.78 | 0.89 | 19.26 | 20.58 | -1.32 | 1.32 | -0.03 | 0.03 |
| gene-LOC105274534 | https://www.ncbi.nlm.nih.gov/gene/?term=LOC105274534 | 1329 | 61 | 458.99 | 1 | 7.52 | 1.64 | 40.79 | 42.14 | -1.35 | 1.35 | -0.06 | 0.06 |
| gene-LOC105278962 | https://www.ncbi.nlm.nih.gov/gene/?term=LOC105278962 | 4491 | 107 | 238.25 | 1 | 2.23 | 0.93 | 20.7 | 22.05 | -1.35 | 1.35 | -0.03 | 0.03 |
| gene-LOC105280956 | https://www.ncbi.nlm.nih.gov/gene/?term=LOC105280956 | 2639 | 77 | 291.78 | 1 | 3.79 | 1.3 | 16.02 | 17.39 | -1.37 | 1.37 | -0.04 | 0.04 |
| gene-LOC105279664 | https://www.ncbi.nlm.nih.gov/gene/?term=LOC105279664 | 4023 | 74 | 183.94 | 1 | 2.49 | 1.35 | 16.49 | 17.94 | -1.45 | 1.45 | -0.03 | 0.03 |
| gene-LOC105279298 | https://www.ncbi.nlm.nih.gov/gene/?term=LOC105279298 | 2773 | 138 | 497.66 | 2 | 7.21 | 1.45 | 22.5 | 23.97 | -1.47 | 1.47 | -0.07 | 0.07 |
| gene-LOC105278109 | https://www.ncbi.nlm.nih.gov/gene/?term=LOC105278109 | 4029 | 72 | 178.7 | 1 | 2.48 | 1.39 | 42.29 | 43.78 | -1.48 | 1.48 | -0.03 | 0.03 |
| gene-LOC105286799 | https://www.ncbi.nlm.nih.gov/gene/?term=LOC105286799 | 3557 | 188 | 528.54 | 1 | 2.81 | 0.53 | 15.27 | 16.76 | -1.49 | 1.49 | -0.08 | 0.08 |
| gene-LOC105276704 | https://www.ncbi.nlm.nih.gov/gene/?term=LOC105276704 | 2063 | 101 | 489.58 | 2 | 9.69 | 1.98 | 13.1 | 14.6 | -1.5 | 1.5 | -0.07 | 0.07 |
| gene-LOC105282149 | https://www.ncbi.nlm.nih.gov/gene/?term=LOC105282149 | 1787 | 109 | 609.96 | 1 | 5.6 | 0.92 | 41.83 | 43.33 | -1.5 | 1.5 | -0.09 | 0.09 |
| gene-LOC105284557 | https://www.ncbi.nlm.nih.gov/gene/?term=LOC105284557 | 1672 | 84 | 502.39 | 1 | 5.98 | 1.19 | 28.16 | 29.69 | -1.53 | 1.53 | -0.08 | 0.08 |
| gene-LOC105282893 | https://www.ncbi.nlm.nih.gov/gene/?term=LOC105282893 | 2645 | 47 | 177.69 | 1 | 3.78 | 2.13 | 28.96 | 30.53 | -1.56 | 1.56 | -0.03 | 0.03 |
| gene-LOC105278282 | https://www.ncbi.nlm.nih.gov/gene/?term=LOC105278282 | 1775 | 64 | 360.56 | 1 | 5.63 | 1.56 | 33.91 | 35.49 | -1.58 | 1.58 | -0.06 | 0.06 |
| gene-LOC105283989 | https://www.ncbi.nlm.nih.gov/gene/?term=LOC105283989 | 4948 | 32 | 64.67 | 2 | 4.04 | 6.25 | 43.84 | 45.43 | -1.59 | 1.59 | -0.01 | 0.01 |
| gene-LOC105282970 | https://www.ncbi.nlm.nih.gov/gene/?term=LOC105282970 | 1529 | 40 | 261.61 | 1 | 6.54 | 2.5 | 50.96 | 52.56 | -1.6 | 1.6 | -0.04 | 0.04 |
| gene-LOC105278384 | https://www.ncbi.nlm.nih.gov/gene/?term=LOC105278384 | 1390 | 35 | 251.8 | 1 | 7.19 | 2.86 | 45.32 | 46.94 | -1.62 | 1.62 | -0.04 | 0.04 |
| gene-LOC105280002 | https://www.ncbi.nlm.nih.gov/gene/?term=LOC105280002 | 1940 | 28 | 144.33 | 1 | 5.15 | 3.57 | 39.15 | 40.87 | -1.73 | 1.73 | -0.02 | 0.02 |
| gene-LOC105275518 | https://www.ncbi.nlm.nih.gov/gene/?term=LOC105275518 | 2335 | 26 | 111.35 | 1 | 4.28 | 3.85 | 36.28 | 38.04 | -1.76 | 1.76 | -0.02 | 0.02 |
| gene-LOC105287854 | https://www.ncbi.nlm.nih.gov/gene/?term=LOC105287854 | 3644 | 66 | 181.12 | 1 | 2.74 | 1.52 | 15.87 | 17.63 | -1.76 | 1.76 | -0.03 | 0.03 |
| gene-LOC105282600 | https://www.ncbi.nlm.nih.gov/gene/?term=LOC105282600 | 3715 | 56 | 150.74 | 1 | 2.69 | 1.79 | 29.97 | 31.8 | -1.83 | 1.83 | -0.03 | 0.03 |
| gene-LOC105288010 | https://www.ncbi.nlm.nih.gov/gene/?term=LOC105288010 | 3189 | 45 | 141.11 | 1 | 3.14 | 2.22 | 34.76 | 36.59 | -1.83 | 1.83 | -0.03 | 0.03 |
| gene-LOC109611283 | https://www.ncbi.nlm.nih.gov/gene/?term=LOC109611283 | 1451 | 39 | 268.78 | 1 | 6.89 | 2.56 | 35.83 | 37.66 | -1.83 | 1.83 | -0.05 | 0.05 |
| gene-LOC105277856 | https://www.ncbi.nlm.nih.gov/gene/?term=LOC105277856 | 3442 | 37 | 107.5 | 1 | 2.91 | 2.7 | 37.9 | 39.79 | -1.89 | 1.89 | -0.02 | 0.02 |
| gene-LOC105282151 | https://www.ncbi.nlm.nih.gov/gene/?term=LOC105282151 | 1601 | 46 | 287.32 | 2 | 12.49 | 4.35 | 33.88 | 35.82 | -1.94 | 1.94 | -0.06 | 0.06 |
| gene-LOC105283592 | https://www.ncbi.nlm.nih.gov/gene/?term=LOC105283592 | 3479 | 72 | 206.96 | 1 | 2.87 | 1.39 | 25.41 | 27.37 | -1.95 | 1.95 | -0.04 | 0.04 |
| gene-LOC105280738 | https://www.ncbi.nlm.nih.gov/gene/?term=LOC105280738 | 4287 | 41 | 95.64 | 1 | 2.33 | 2.44 | 24.39 | 26.44 | -2.04 | 2.04 | -0.02 | 0.02 |
| gene-LOC105287063 | https://www.ncbi.nlm.nih.gov/gene/?term=LOC105287063 | 1874 | 54 | 288.15 | 1 | 5.34 | 1.85 | 27.46 | 29.52 | -2.06 | 2.06 | -0.06 | 0.06 |
| gene-LOC105275966 | https://www.ncbi.nlm.nih.gov/gene/?term=LOC105275966 | 3299 | 73 | 221.28 | 1 | 3.03 | 1.37 | 43.49 | 45.58 | -2.08 | 2.08 | -0.05 | 0.05 |
| gene-LOC105282156 | https://www.ncbi.nlm.nih.gov/gene/?term=LOC105282156 | 2577 | 145 | 562.67 | 1 | 3.88 | 0.69 | 23.3 | 25.57 | -2.27 | 2.27 | -0.13 | 0.13 |
| gene-LOC105286683 | https://www.ncbi.nlm.nih.gov/gene/?term=LOC105286683 | 1665 | 53 | 318.32 | 1 | 6.01 | 1.89 | 31.4 | 33.71 | -2.31 | 2.31 | -0.07 | 0.07 |
| gene-LOC105281420 | https://www.ncbi.nlm.nih.gov/gene/?term=LOC105281420 | 3882 | 67 | 172.59 | 1 | 2.58 | 1.49 | 12.82 | 15.24 | -2.42 | 2.42 | -0.04 | 0.04 |
| gene-LOC105285315 | https://www.ncbi.nlm.nih.gov/gene/?term=LOC105285315 | 5458 | 26 | 47.64 | 1 | 1.83 | 3.85 | 36.55 | 38.99 | -2.44 | 2.44 | -0.01 | 0.01 |
| gene-LOC105285583 | https://www.ncbi.nlm.nih.gov/gene/?term=LOC105285583 | 1947 | 47 | 241.4 | 1 | 5.14 | 2.13 | 27.05 | 29.68 | -2.63 | 2.63 | -0.06 | 0.06 |
| gene-LOC105276449 | https://www.ncbi.nlm.nih.gov/gene/?term=LOC105276449 | 1231 | 56 | 454.91 | 1 | 8.12 | 1.79 | 30.06 | 32.73 | -2.67 | 2.67 | -0.12 | 0.12 |
| gene-LOC105288063 | https://www.ncbi.nlm.nih.gov/gene/?term=LOC105288063 | 3400 | 57 | 167.65 | 1 | 2.94 | 1.75 | 12.44 | 15.19 | -2.75 | 2.75 | -0.05 | 0.05 |
| gene-LOC105280786 | https://www.ncbi.nlm.nih.gov/gene/?term=LOC105280786 | 2573 | 119 | 462.5 | 1 | 3.89 | 0.84 | 29.73 | 32.5 | -2.77 | 2.77 | -0.13 | 0.13 |
| gene-LOC105276680 | https://www.ncbi.nlm.nih.gov/gene/?term=LOC105276680 | 1977 | 35 | 177.04 | 1 | 5.06 | 2.86 | 58.22 | 61.02 | -2.8 | 2.8 | -0.05 | 0.05 |
| gene-LOC105278454 | https://www.ncbi.nlm.nih.gov/gene/?term=LOC105278454 | 1972 | 51 | 258.62 | 1 | 5.07 | 1.96 | 39.78 | 42.63 | -2.85 | 2.85 | -0.07 | 0.07 |
| gene-LOC105287417 | https://www.ncbi.nlm.nih.gov/gene/?term=LOC105287417 | 1025 | 36 | 351.22 | 1 | 9.76 | 2.78 | 34.17 | 37.2 | -3.03 | 3.03 | -0.11 | 0.11 |
| gene-LOC105283649 | https://www.ncbi.nlm.nih.gov/gene/?term=LOC105283649 | 1918 | 57 | 297.18 | 2 | 10.43 | 3.51 | 35.86 | 38.92 | -3.06 | 3.06 | -0.09 | 0.09 |
| gene-LOC105288066 | https://www.ncbi.nlm.nih.gov/gene/?term=LOC105288066 | 1964 | 32 | 162.93 | 1 | 5.09 | 3.12 | 32.93 | 36 | -3.07 | 3.07 | -0.05 | 0.05 |
| gene-LOC105284609 | https://www.ncbi.nlm.nih.gov/gene/?term=LOC105284609 | 5678 | 35 | 61.64 | 1 | 1.76 | 2.86 | 28.06 | 31.35 | -3.29 | 3.29 | -0.02 | 0.02 |
| gene-LOC105281515 | https://www.ncbi.nlm.nih.gov/gene/?term=LOC105281515 | 8827 | 54 | 61.18 | 2 | 2.27 | 3.7 | 27.93 | 31.25 | -3.31 | 3.31 | -0.02 | 0.02 |
| gene-LOC105276078 | https://www.ncbi.nlm.nih.gov/gene/?term=LOC105276078 | 1051 | 17 | 161.75 | 1 | 9.51 | 5.88 | 25.17 | 28.55 | -3.38 | 3.38 | -0.05 | 0.05 |
| gene-LOC105286514 | https://www.ncbi.nlm.nih.gov/gene/?term=LOC105286514 | 2528 | 47 | 185.92 | 1 | 3.96 | 2.13 | 35.58 | 39.1 | -3.52 | 3.52 | -0.07 | 0.07 |
| gene-LOC113561805 | https://www.ncbi.nlm.nih.gov/gene/?term=LOC113561805 | 3337 | 84 | 251.72 | 1 | 3 | 1.19 | 34.18 | 37.85 | -3.66 | 3.66 | -0.09 | 0.09 |
| gene-LOC105274900 | https://www.ncbi.nlm.nih.gov/gene/?term=LOC105274900 | 5561 | 55 | 98.9 | 2 | 3.6 | 3.64 | 30.44 | 34.13 | -3.69 | 3.69 | -0.04 | 0.04 |
| gene-LOC105280327 | https://www.ncbi.nlm.nih.gov/gene/?term=LOC105280327 | 2670 | 104 | 389.51 | 1 | 3.75 | 0.96 | 27.09 | 30.78 | -3.69 | 3.69 | -0.14 | 0.14 |
| gene-LOC105285674 | https://www.ncbi.nlm.nih.gov/gene/?term=LOC105285674 | 3035 | 97 | 319.6 | 2 | 6.59 | 2.06 | 24.1 | 27.79 | -3.69 | 3.69 | -0.12 | 0.12 |
| gene-LOC105279900 | https://www.ncbi.nlm.nih.gov/gene/?term=LOC105279900 | 2191 | 62 | 282.98 | 1 | 4.56 | 1.61 | 30.24 | 34.06 | -3.83 | 3.83 | -0.11 | 0.11 |
| gene-LOC105276635 | https://www.ncbi.nlm.nih.gov/gene/?term=LOC105276635 | 4962 | 40 | 80.61 | 1 | 2.02 | 2.5 | 23.53 | 27.39 | -3.86 | 3.86 | -0.03 | 0.03 |

| | | | | | | | | | | | | | |
|---|---|---|---|---|---|---|---|---|---|---|---|---|---|
| gene-LOC105286644 | https://www.ncbi.nlm.nih.gov/gene/?term=LOC105286644 | 1235 | 20 | 161.94 | 1 | 8.1 | 5 | 37.53 | 41.72 | -4.19 | 4.19 | -0.07 | 0.07 |
| gene-LOC105276538 | https://www.ncbi.nlm.nih.gov/gene/?term=LOC105276538 | 1511 | 19 | 125.74 | 1 | 6.62 | 5.26 | 69.47 | 74.28 | -4.82 | 4.82 | -0.06 | 0.06 |
| gene-LOC105279898 | https://www.ncbi.nlm.nih.gov/gene/?term=LOC105279898 | 1343 | 14 | 104.24 | 1 | 7.45 | 7.14 | 54.89 | 60.18 | -5.29 | 5.29 | -0.06 | 0.06 |
| gene-LOC105286983 | https://www.ncbi.nlm.nih.gov/gene/?term=LOC105286983 | 16901 | 64 | 37.87 | 2 | 1.18 | 3.12 | 0.67 | 5.99 | -5.32 | 5.32 | -0.02 | 0.02 |
| gene-LOC105275708 | https://www.ncbi.nlm.nih.gov/gene/?term=LOC105275708 | 1772 | 18 | 101.58 | 1 | 5.64 | 5.56 | 43.28 | 48.95 | -5.67 | 5.67 | -0.06 | 0.06 |
| gene-LOC105281039 | https://www.ncbi.nlm.nih.gov/gene/?term=LOC105281039 | 1471 | 12 | 81.58 | 1 | 6.8 | 8.33 | 31.46 | 37.55 | -6.08 | 6.08 | -0.05 | 0.05 |
| gene-LOC105275070 | https://www.ncbi.nlm.nih.gov/gene/?term=LOC105275070 | 5667 | 53 | 93.52 | 6 | 10.59 | 11.32 | 1.77 | 10.27 | -8.5 | 8.5 | -0.08 | 0.08 |
| gene-LOC113562901 | https://www.ncbi.nlm.nih.gov/gene/?term=LOC113562901 | 1531 | 22 | 143.7 | 3 | 19.6 | 13.64 | 45.03 | 59.16 | -14.13 | 14.13 | -0.2 | 0.2 |
| gene-LOC113562345 | https://www.ncbi.nlm.nih.gov/gene/?term=LOC113562345 | 2290 | 69 | 301.31 | 19 | 82.97 | 27.54 | 16.9 | 33.89 | -17 | 17 | -0.51 | 0.51 |
| gene-LOC105281282 | https://www.ncbi.nlm.nih.gov/gene/?term=LOC105281282 | 3250 | 3 | 9.23 | 1 | 3.08 | 33.33 | 23.25 | 65.16 | -41.92 | 41.92 | -0.04 | 0.04 |

Directory: OGLall File: ogl-Ob_LI_BCphaseB2.vs.RphaseB1.txt

| gene_ID | gene_link | gwidth | #Sites | #per10Kb | #dmSites | #dmsp10kb | %dmSites | %pSite1 | %pSite2 | DMpSite | ADMpSite | DMpNucl | ADMpNucl |
|---|---|---|---|---|---|---|---|---|---|---|---|---|---|
| gene-LOC113562974 | https://www.ncbi.nlm.nih.gov/gene/?term=LOC113562974 | 1832 | 2 | 10.92 | 1 | 5.46 | 50 | 21.66 | 9 | 12.67 | 12.67 | 0.01 | 0.01 |
| gene-LOC105282841 | https://www.ncbi.nlm.nih.gov/gene/?term=LOC105282841 | 2047 | 5 | 24.43 | 1 | 4.89 | 20 | 67.42 | 55.09 | 12.33 | 12.33 | 0.03 | 0.03 |
| gene-LOC105278541 | https://www.ncbi.nlm.nih.gov/gene/?term=LOC105278541 | 1563 | 14 | 89.57 | 3 | 19.19 | 21.43 | 39.99 | 31.76 | 8.23 | 8.23 | 0.07 | 0.07 |
| gene-LOC105286822 | https://www.ncbi.nlm.nih.gov/gene/?term=LOC105286822 | 3538 | 6 | 16.96 | 1 | 2.83 | 16.67 | 18.11 | 10.53 | 7.58 | 7.58 | 0.01 | 0.01 |
| gene-LOC105287923 | https://www.ncbi.nlm.nih.gov/gene/?term=LOC105287923 | 1233 | 10 | 81.1 | 2 | 16.22 | 20 | 76.35 | 69.28 | 7.07 | 7.07 | 0.06 | 0.06 |
| gene-LOC105275454 | https://www.ncbi.nlm.nih.gov/gene/?term=LOC105275454 | 1145 | 13 | 113.54 | 3 | 26.2 | 23.08 | 55.74 | 49.23 | 6.51 | 6.51 | 0.07 | 0.07 |
| gene-LOC105284547 | https://www.ncbi.nlm.nih.gov/gene/?term=LOC105284547 | 2105 | 13 | 61.76 | 1 | 4.75 | 7.69 | 57.2 | 50.73 | 6.46 | 6.46 | 0.04 | 0.04 |
| gene-LOC105275176 | https://www.ncbi.nlm.nih.gov/gene/?term=LOC105275176 | 656 | 8 | 121.95 | 1 | 15.24 | 12.5 | 14.79 | 8.51 | 6.28 | 6.28 | 0.08 | 0.08 |
| gene-LOC105274672 | https://www.ncbi.nlm.nih.gov/gene/?term=LOC105274672 | 2878 | 15 | 52.12 | 2 | 6.95 | 13.33 | 40.13 | 34.21 | 5.92 | 5.92 | 0.03 | 0.03 |
| gene-LOC105284056 | https://www.ncbi.nlm.nih.gov/gene/?term=LOC105284056 | 1997 | 6 | 30.05 | 1 | 5.01 | 16.67 | 30.15 | 24.32 | 5.83 | 5.83 | 0.02 | 0.02 |
| gene-LOC105285334 | https://www.ncbi.nlm.nih.gov/gene/?term=LOC105285334 | 2363 | 33 | 139.65 | 4 | 16.93 | 12.12 | 24.89 | 19.32 | 5.57 | 5.57 | 0.08 | 0.08 |
| gene-LOC105285795 | https://www.ncbi.nlm.nih.gov/gene/?term=LOC105285795 | 1256 | 12 | 95.54 | 2 | 15.92 | 16.67 | 50.58 | 45.08 | 5.5 | 5.5 | 0.05 | 0.05 |
| gene-LOC105280473 | https://www.ncbi.nlm.nih.gov/gene/?term=LOC105280473 | 1816 | 6 | 33.04 | 1 | 5.51 | 16.67 | 88.83 | 83.46 | 5.36 | 5.36 | 0.02 | 0.02 |
| gene-LOC105276202 | https://www.ncbi.nlm.nih.gov/gene/?term=LOC105276202 | 1540 | 43 | 279.22 | 5 | 32.47 | 11.63 | 24.19 | 18.86 | 5.33 | 5.33 | 0.15 | 0.15 |
| gene-LOC105282371 | https://www.ncbi.nlm.nih.gov/gene/?term=LOC105282371 | 1722 | 8 | 46.46 | 1 | 5.81 | 12.5 | 31.57 | 26.4 | 5.17 | 5.17 | 0.02 | 0.02 |
| gene-LOC105275296 | https://www.ncbi.nlm.nih.gov/gene/?term=LOC105275296 | 2951 | 19 | 64.38 | 1 | 3.39 | 5.26 | 27.28 | 22.14 | 5.14 | 5.14 | 0.03 | 0.03 |
| gene-LOC105281716 | https://www.ncbi.nlm.nih.gov/gene/?term=LOC105281716 | 1802 | 8 | 44.4 | 1 | 5.55 | 12.5 | 36.4 | 31.28 | 5.11 | 5.11 | 0.02 | 0.02 |
| gene-LOC105284999 | https://www.ncbi.nlm.nih.gov/gene/?term=LOC105284999 | 1832 | 9 | 49.13 | 1 | 5.46 | 11.11 | 55.4 | 50.33 | 5.07 | 5.07 | 0.02 | 0.02 |
| gene-LOC105284475 | https://www.ncbi.nlm.nih.gov/gene/?term=LOC105284475 | 1364 | 11 | 80.65 | 1 | 7.33 | 9.09 | 56.71 | 51.84 | 4.87 | 4.87 | 0.04 | 0.04 |
| gene-LOC113563111 | https://www.ncbi.nlm.nih.gov/gene/?term=LOC113563111 | 2455 | 29 | 118.13 | 3 | 12.22 | 10.34 | 28.93 | 24.09 | 4.84 | 4.84 | 0.06 | 0.06 |
| gene-LOC105282041 | https://www.ncbi.nlm.nih.gov/gene/?term=LOC105282041 | 2041 | 14 | 68.59 | 2 | 9.8 | 14.29 | 31.5 | 26.67 | 4.83 | 4.83 | 0.03 | 0.03 |
| gene-LOC105278686 | https://www.ncbi.nlm.nih.gov/gene/?term=LOC105278686 | 2006 | 14 | 69.79 | 1 | 4.99 | 7.14 | 10.96 | 6.19 | 4.77 | 4.77 | 0.03 | 0.03 |
| gene-LOC105276171 | https://www.ncbi.nlm.nih.gov/gene/?term=LOC105276171 | 1370 | 14 | 102.19 | 2 | 14.6 | 14.29 | 17.56 | 12.89 | 4.67 | 4.67 | 0.05 | 0.05 |
| gene-LOC105277804 | https://www.ncbi.nlm.nih.gov/gene/?term=LOC105277804 | 1174 | 6 | 51.11 | 1 | 8.52 | 16.67 | 93.96 | 89.47 | 4.48 | 4.48 | 0.02 | 0.02 |

...

... only first 25 lines shown ...

Directory: OGLall File: ogl-Ob_LI_BCphaseB2.vs.RphaseB2.txt

| gene_ID | gene_link | gwidth | #Sites | #per10Kb | #dmSites | #dmsp10kb | %dmSites | %pSite1 | %pSite2 | DMpSite | ADMpSite | DMpNucl | ADMpNucl |
|---|---|---|---|---|---|---|---|---|---|---|---|---|---|
| gene-LOC105286586 | https://www.ncbi.nlm.nih.gov/gene/?term=LOC105286586 | 1417 | 1 | 7.06 | 1 | 7.06 | 100 | 36.67 | 15.38 | 21.29 | 21.29 | 0.02 | 0.02 |
| gene-LOC113561765 | https://www.ncbi.nlm.nih.gov/gene/?term=LOC113561765 | 1822 | 4 | 21.95 | 1 | 5.49 | 25 | 29.77 | 18.68 | 11.1 | 11.1 | 0.02 | 0.02 |
| gene-LOC105275176 | https://www.ncbi.nlm.nih.gov/gene/?term=LOC105275176 | 656 | 8 | 121.95 | 2 | 30.49 | 25 | 14.79 | 6.32 | 8.47 | 8.47 | 0.1 | 0.1 |
| gene-LOC105286261 | https://www.ncbi.nlm.nih.gov/gene/?term=LOC105286261 | 1297 | 8 | 61.68 | 1 | 7.71 | 12.5 | 76.36 | 68.31 | 8.05 | 8.05 | 0.05 | 0.05 |
| gene-LOC105287216 | https://www.ncbi.nlm.nih.gov/gene/?term=LOC105287216 | 2142 | 6 | 28.01 | 1 | 4.67 | 16.67 | 27.53 | 19.82 | 7.72 | 7.72 | 0.02 | 0.02 |
| gene-LOC105283410 | https://www.ncbi.nlm.nih.gov/gene/?term=LOC105283410 | 1423 | 5 | 35.14 | 1 | 7.03 | 20 | 47.1 | 40.2 | 6.9 | 6.9 | 0.02 | 0.02 |
| gene-LOC105276075 | https://www.ncbi.nlm.nih.gov/gene/?term=LOC105276075 | 1440 | 10 | 69.44 | 1 | 6.94 | 10 | 39.46 | 33.04 | 6.42 | 6.42 | 0.04 | 0.04 |
| gene-LOC105282996 | https://www.ncbi.nlm.nih.gov/gene/?term=LOC105282996 | 3736 | 6 | 16.06 | 1 | 2.68 | 16.67 | 23.34 | 17.02 | 6.33 | 6.33 | 0.01 | 0.01 |
| gene-LOC105285229 | https://www.ncbi.nlm.nih.gov/gene/?term=LOC105285229 | 1396 | 5 | 35.82 | 1 | 7.16 | 20 | 23.52 | 17.61 | 5.91 | 5.91 | 0.02 | 0.02 |
| gene-LOC105277980 | https://www.ncbi.nlm.nih.gov/gene/?term=LOC105277980 | 2400 | 30 | 125 | 2 | 8.33 | 6.67 | 49.23 | 43.45 | 5.77 | 5.77 | 0.07 | 0.07 |
| gene-LOC105285203 | https://www.ncbi.nlm.nih.gov/gene/?term=LOC105285203 | 1020 | 27 | 264.71 | 2 | 19.61 | 7.41 | 30.05 | 24.28 | 5.76 | 5.76 | 0.15 | 0.15 |
| gene-LOC105281512 | https://www.ncbi.nlm.nih.gov/gene/?term=LOC105281512 | 2233 | 16 | 71.65 | 3 | 13.43 | 18.75 | 32.28 | 26.62 | 5.65 | 5.65 | 0.04 | 0.04 |
| gene-LOC105274857 | https://www.ncbi.nlm.nih.gov/gene/?term=LOC105274857 | 1569 | 4 | 25.49 | 1 | 6.37 | 25 | 58.2 | 52.63 | 5.57 | 5.57 | 0.01 | 0.01 |
| gene-LOC105285795 | https://www.ncbi.nlm.nih.gov/gene/?term=LOC105285795 | 1256 | 14 | 111.46 | 2 | 15.92 | 14.29 | 57.64 | 52.12 | 5.53 | 5.53 | 0.06 | 0.06 |
| gene-LOC105276569 | https://www.ncbi.nlm.nih.gov/gene/?term=LOC105276569 | 1963 | 14 | 71.32 | 1 | 5.09 | 7.14 | 23.68 | 18.21 | 5.47 | 5.47 | 0.04 | 0.04 |
| gene-LOC105281716 | https://www.ncbi.nlm.nih.gov/gene/?term=LOC105281716 | 1802 | 8 | 44.4 | 1 | 5.55 | 12.5 | 36.4 | 30.96 | 5.44 | 5.44 | 0.02 | 0.02 |
| gene-LOC105275503 | https://www.ncbi.nlm.nih.gov/gene/?term=LOC105275503 | 2000 | 4 | 20 | 1 | 5 | 25 | 38.28 | 33 | 5.28 | 5.28 | 0.01 | 0.01 |
| gene-LOC105276230 | https://www.ncbi.nlm.nih.gov/gene/?term=LOC105276230 | 3067 | 12 | 39.13 | 1 | 3.26 | 8.33 | 28.38 | 23.21 | 5.17 | 5.17 | 0.02 | 0.02 |
| gene-LOC105278606 | https://www.ncbi.nlm.nih.gov/gene/?term=LOC105278606 | 2650 | 5 | 18.87 | 1 | 3.77 | 20 | 6.01 | 0.87 | 5.14 | 5.14 | 0.01 | 0.01 |
| gene-LOC105283047 | https://www.ncbi.nlm.nih.gov/gene/?term=LOC105283047 | 1181 | 16 | 135.48 | 1 | 8.47 | 6.25 | 13.48 | 8.42 | 5.06 | 5.06 | 0.07 | 0.07 |
| gene-LOC105285669 | https://www.ncbi.nlm.nih.gov/gene/?term=LOC105285669 | 2193 | 16 | 72.96 | 1 | 4.56 | 6.25 | 26.19 | 21.17 | 5.02 | 5.02 | 0.04 | 0.04 |
| gene-LOC105285905 | https://www.ncbi.nlm.nih.gov/gene/?term=LOC105285905 | 1397 | 13 | 93.06 | 2 | 14.32 | 15.38 | 53.61 | 48.59 | 5.02 | 5.02 | 0.05 | 0.05 |
| gene-LOC105278108 | https://www.ncbi.nlm.nih.gov/gene/?term=LOC105278108 | 4392 | 46 | 104.74 | 1 | 2.28 | 2.17 | 29.79 | 24.81 | 4.98 | 4.98 | 0.05 | 0.05 |
| gene-LOC105286778 | https://www.ncbi.nlm.nih.gov/gene/?term=LOC105286778 | 6479 | 6 | 9.26 | 1 | 1.54 | 16.67 | 7.5 | 2.54 | 4.96 | 4.96 | 0 | 0 |

...
... only first 25 lines shown ...

Directory: OGLall File: ogl-Ob_LI_RphaseA1.txt

| gene_ID | gene_link | gwidth | #Sites | #per10Kb | %perSite | %pNucl |
|---|---|---|---|---|---|---|
| gene-LOC113561532 | https://www.ncbi.nlm.nih.gov/gene/?term=LOC113561532 | 72 | 5 | 694.44 | 98.1 | 6.81 |
| gene-LOC105286531 | https://www.ncbi.nlm.nih.gov/gene/?term=LOC105286531 | 372 | 43 | 1155.91 | 36.48 | 4.22 |
| gene-LOC105280739 | https://www.ncbi.nlm.nih.gov/gene/?term=LOC105280739 | 1991 | 138 | 693.12 | 54.32 | 3.76 |
| gene-LOC105274534 | https://www.ncbi.nlm.nih.gov/gene/?term=LOC105274534 | 1329 | 104 | 782.54 | 47.62 | 3.73 |
| gene-LOC105285635 | https://www.ncbi.nlm.nih.gov/gene/?term=LOC105285635 | 1175 | 137 | 1165.96 | 30.78 | 3.59 |
| gene-LOC105287811 | https://www.ncbi.nlm.nih.gov/gene/?term=LOC105287811 | 2155 | 153 | 709.98 | 47.68 | 3.38 |
| gene-LOC105286235 | https://www.ncbi.nlm.nih.gov/gene/?term=LOC105286235 | 1510 | 118 | 781.46 | 42.69 | 3.34 |
| gene-LOC105286269 | https://www.ncbi.nlm.nih.gov/gene/?term=LOC105286269 | 1401 | 87 | 620.99 | 52.62 | 3.27 |
| gene-LOC105286737 | https://www.ncbi.nlm.nih.gov/gene/?term=LOC105286737 | 2690 | 365 | 1356.88 | 23.81 | 3.23 |
| gene-LOC105278844 | https://www.ncbi.nlm.nih.gov/gene/?term=LOC105278844 | 1101 | 109 | 990.01 | 32.11 | 3.18 |
| gene-LOC105282149 | https://www.ncbi.nlm.nih.gov/gene/?term=LOC105282149 | 1787 | 139 | 777.84 | 40.94 | 3.18 |
| gene-LOC105277648 | https://www.ncbi.nlm.nih.gov/gene/?term=LOC105277648 | 722 | 39 | 540.17 | 57.72 | 3.12 |
| gene-LOC105275314 | https://www.ncbi.nlm.nih.gov/gene/?term=LOC105275314 | 1489 | 97 | 651.44 | 47.77 | 3.11 |
| gene-LOC105279052 | https://www.ncbi.nlm.nih.gov/gene/?term=LOC105279052 | 1130 | 122 | 1079.65 | 28.5 | 3.08 |
| gene-LOC105276371 | https://www.ncbi.nlm.nih.gov/gene/?term=LOC105276371 | 871 | 55 | 631.46 | 48.54 | 3.07 |
| gene-LOC105278160 | https://www.ncbi.nlm.nih.gov/gene/?term=LOC105278160 | 1872 | 158 | 844.02 | 35.32 | 2.98 |
| gene-LOC105286801 | https://www.ncbi.nlm.nih.gov/gene/?term=LOC105286801 | 1102 | 86 | 780.4 | 38.13 | 2.98 |
| gene-LOC105276360 | https://www.ncbi.nlm.nih.gov/gene/?term=LOC105276360 | 1466 | 154 | 1050.48 | 27.76 | 2.92 |
| gene-LOC105287677 | https://www.ncbi.nlm.nih.gov/gene/?term=LOC105287677 | 1588 | 113 | 711.59 | 41.08 | 2.92 |
| gene-LOC105274711 | https://www.ncbi.nlm.nih.gov/gene/?term=LOC105274711 | 1655 | 102 | 616.31 | 46.87 | 2.89 |
| gene-LOC105278849 | https://www.ncbi.nlm.nih.gov/gene/?term=LOC105278849 | 776 | 101 | 1301.55 | 22.16 | 2.88 |
| gene-LOC105277254 | https://www.ncbi.nlm.nih.gov/gene/?term=LOC105277254 | 1377 | 112 | 813.36 | 35.34 | 2.87 |
| gene-LOC105285392 | https://www.ncbi.nlm.nih.gov/gene/?term=LOC105285392 | 1601 | 138 | 861.96 | 32.41 | 2.79 |
| gene-LOC105287692 | https://www.ncbi.nlm.nih.gov/gene/?term=LOC105287692 | 1505 | 120 | 797.34 | 34.96 | 2.79 |

...

... only first 25 lines shown ...

Directory: OGLall File: ogl-Ob_LI_RphaseA1.vs.RphaseA2.txt

| gene_ID | gene_link | gwidth | #Sites | #per10Kb | #dmSites | #dmsp10kb | %dmSites | %pSite1 | %pSite2 | DMpSite | ADMpSite | DMpNucl | ADMpNucl |
|---|---|---|---|---|---|---|---|---|---|---|---|---|---|
| gene-LOC105283264 | https://www.ncbi.nlm.nih.gov/gene/?term=LOC105283264 | 4005 | 96 | 239.7 | 1 | 2.5 | 1.04 | 23.23 | 22.15 | 1.09 | 1.09 | 0.03 | 0.03 |
| gene-LOC105277903 | https://www.ncbi.nlm.nih.gov/gene/?term=LOC105277903 | 1835 | 39 | 212.53 | 1 | 5.45 | 2.56 | 54.3 | 53.51 | 0.79 | 0.79 | 0.02 | 0.02 |
| gene-LOC105283296 | https://www.ncbi.nlm.nih.gov/gene/?term=LOC105283296 | 2174 | 101 | 464.58 | 1 | 4.6 | 0.99 | 18.85 | 18.17 | 0.67 | 0.67 | 0.03 | 0.03 |
| gene-LOC105284557 | https://www.ncbi.nlm.nih.gov/gene/?term=LOC105284557 | 1672 | 84 | 502.39 | 1 | 5.98 | 1.19 | 28.63 | 28.17 | 0.46 | 0.46 | 0.02 | 0.02 |
| gene-LOC105277125 | https://www.ncbi.nlm.nih.gov/gene/?term=LOC105277125 | 51079 | 2004 | 392.33 | 1 | 0.2 | 0.05 | 1.65 | 1.59 | 0.06 | 0.06 | 0 | 0 |
| gene-LOC105280304 | https://www.ncbi.nlm.nih.gov/gene/?term=LOC105280304 | 8919 | 150 | 168.18 | 1 | 1.12 | 0.67 | 6.83 | 6.96 | -0.12 | 0.12 | 0 | 0 |
| gene-LOC105284507 | https://www.ncbi.nlm.nih.gov/gene/?term=LOC105284507 | 5207 | 112 | 215.1 | 1 | 1.92 | 0.89 | 18.79 | 19.06 | -0.28 | 0.28 | -0.01 | 0.01 |
| gene-LOC105275384 | https://www.ncbi.nlm.nih.gov/gene/?term=LOC105275384 | 1987 | 86 | 432.81 | 1 | 5.03 | 1.16 | 36.56 | 36.97 | -0.41 | 0.41 | -0.02 | 0.02 |
| gene-LOC105281272 | https://www.ncbi.nlm.nih.gov/gene/?term=LOC105281272 | 5252 | 199 | 378.9 | 1 | 1.9 | 0.5 | 14.62 | 15.07 | -0.45 | 0.45 | -0.02 | 0.02 |
| gene-LOC105288036 | https://www.ncbi.nlm.nih.gov/gene/?term=LOC105288036 | 5110 | 209 | 409 | 1 | 1.96 | 0.48 | 17.07 | 17.79 | -0.71 | 0.71 | -0.03 | 0.03 |
| gene-LOC105288042 | https://www.ncbi.nlm.nih.gov/gene/?term=LOC105288042 | 3507 | 111 | 316.51 | 1 | 2.85 | 0.9 | 22.13 | 23.4 | -1.28 | 1.28 | -0.04 | 0.04 |

Directory: OGLall File: ogl-Ob_LI_RphaseA1.vs.RphaseB1.txt

| gene_ID | gene_link | gwidth | #Sites | #per10Kb | #dmSites | #dmsp10kb | %dmSites | %pSite1 | %pSite2 | DMpSite | ADMpSite | DMpNucl | ADMpNucl |
|---|---|---|---|---|---|---|---|---|---|---|---|---|---|
| gene-LOC105275070 | https://www.ncbi.nlm.nih.gov/gene/?term=LOC105275070 | 5667 | 56 | 98.82 | 3 | 5.29 | 5.36 | 17.83 | 5.85 | 11.98 | 11.98 | 0.12 | 0.12 |
| gene-LOC113562345 | https://www.ncbi.nlm.nih.gov/gene/?term=LOC113562345 | 2290 | 78 | 340.61 | 11 | 48.03 | 14.1 | 24.77 | 13.87 | 10.9 | 10.9 | 0.37 | 0.37 |
| gene-LOC105285514 | https://www.ncbi.nlm.nih.gov/gene/?term=LOC105285514 | 1277 | 7 | 54.82 | 1 | 7.83 | 14.29 | 90.55 | 80 | 10.54 | 10.54 | 0.06 | 0.06 |
| gene-LOC105275708 | https://www.ncbi.nlm.nih.gov/gene/?term=LOC105275708 | 1772 | 14 | 79.01 | 1 | 5.64 | 7.14 | 50.83 | 42.47 | 8.36 | 8.36 | 0.07 | 0.07 |
| gene-LOC105275355 | https://www.ncbi.nlm.nih.gov/gene/?term=LOC105275355 | 3197 | 174 | 544.26 | 1 | 3.13 | 0.57 | 25.86 | 21.8 | 4.06 | 4.06 | 0.22 | 0.22 |
| gene-LOC105281039 | https://www.ncbi.nlm.nih.gov/gene/?term=LOC105281039 | 1471 | 13 | 88.38 | 1 | 6.8 | 7.69 | 31.24 | 27.65 | 3.59 | 3.59 | 0.03 | 0.03 |
| gene-LOC105275384 | https://www.ncbi.nlm.nih.gov/gene/?term=LOC105275384 | 1987 | 88 | 442.88 | 1 | 5.03 | 1.14 | 36.35 | 32.79 | 3.56 | 3.56 | 0.16 | 0.16 |
| gene-LOC105275347 | https://www.ncbi.nlm.nih.gov/gene/?term=LOC105275347 | 2526 | 17 | 67.3 | 1 | 3.96 | 5.88 | 45.17 | 41.88 | 3.29 | 3.29 | 0.02 | 0.02 |
| gene-LOC105274900 | https://www.ncbi.nlm.nih.gov/gene/?term=LOC105274900 | 5561 | 61 | 109.69 | 3 | 5.39 | 4.92 | 33.5 | 30.47 | 3.02 | 3.02 | 0.03 | 0.03 |
| gene-LOC105287063 | https://www.ncbi.nlm.nih.gov/gene/?term=LOC105287063 | 1874 | 49 | 261.47 | 1 | 5.34 | 2.04 | 32.77 | 30.12 | 2.65 | 2.65 | 0.07 | 0.07 |
| gene-LOC105279898 | https://www.ncbi.nlm.nih.gov/gene/?term=LOC105279898 | 1343 | 15 | 111.69 | 1 | 7.45 | 6.67 | 61.47 | 58.93 | 2.54 | 2.54 | 0.03 | 0.03 |
| gene-LOC105281515 | https://www.ncbi.nlm.nih.gov/gene/?term=LOC105281515 | 8827 | 57 | 64.57 | 2 | 2.27 | 3.51 | 32.7 | 30.37 | 2.32 | 2.32 | 0.01 | 0.01 |
| gene-LOC113562102 | https://www.ncbi.nlm.nih.gov/gene/?term=LOC113562102 | 3433 | 63 | 183.51 | 1 | 2.91 | 1.59 | 17.89 | 15.59 | 2.3 | 2.3 | 0.04 | 0.04 |
| gene-LOC105276272 | https://www.ncbi.nlm.nih.gov/gene/?term=LOC105276272 | 1965 | 44 | 223.92 | 1 | 5.09 | 2.27 | 27.22 | 24.96 | 2.26 | 2.26 | 0.05 | 0.05 |
| gene-LOC105282991 | https://www.ncbi.nlm.nih.gov/gene/?term=LOC105282991 | 5617 | 103 | 183.37 | 1 | 1.78 | 0.97 | 18.36 | 16.13 | 2.23 | 2.23 | 0.04 | 0.04 |
| gene-LOC105285695 | https://www.ncbi.nlm.nih.gov/gene/?term=LOC105285695 | 1371 | 68 | 495.99 | 1 | 7.29 | 1.47 | 23.53 | 21.55 | 1.99 | 1.99 | 0.1 | 0.1 |
| gene-LOC105283712 | https://www.ncbi.nlm.nih.gov/gene/?term=LOC105283712 | 1910 | 35 | 183.25 | 1 | 5.24 | 2.86 | 28.45 | 26.47 | 1.98 | 1.98 | 0.04 | 0.04 |
| gene-LOC105285671 | https://www.ncbi.nlm.nih.gov/gene/?term=LOC105285671 | 2263 | 100 | 441.89 | 1 | 4.42 | 1 | 32.24 | 30.53 | 1.71 | 1.71 | 0.08 | 0.08 |
| gene-LOC105282652 | https://www.ncbi.nlm.nih.gov/gene/?term=LOC105282652 | 5065 | 85 | 167.82 | 1 | 1.97 | 1.18 | 16.28 | 14.6 | 1.68 | 1.68 | 0.03 | 0.03 |
| gene-LOC105276692 | https://www.ncbi.nlm.nih.gov/gene/?term=LOC105276692 | 1745 | 34 | 194.84 | 1 | 5.73 | 2.94 | 25.17 | 23.64 | 1.53 | 1.53 | 0.03 | 0.03 |
| gene-LOC105288066 | https://www.ncbi.nlm.nih.gov/gene/?term=LOC105288066 | 1964 | 30 | 152.75 | 1 | 5.09 | 3.33 | 38.49 | 37.01 | 1.48 | 1.48 | 0.02 | 0.02 |
| gene-LOC105286265 | https://www.ncbi.nlm.nih.gov/gene/?term=LOC105286265 | 2507 | 67 | 267.25 | 1 | 3.99 | 1.49 | 32.57 | 31.11 | 1.46 | 1.46 | 0.04 | 0.04 |
| gene-LOC113561805 | https://www.ncbi.nlm.nih.gov/gene/?term=LOC113561805 | 3337 | 75 | 224.75 | 1 | 3 | 1.33 | 34.35 | 32.91 | 1.44 | 1.44 | 0.03 | 0.03 |
| gene-LOC105280348 | https://www.ncbi.nlm.nih.gov/gene/?term=LOC105280348 | 3797 | 79 | 208.06 | 1 | 2.63 | 1.27 | 18.28 | 16.89 | 1.4 | 1.4 | 0.03 | 0.03 |
| gene-LOC105278807 | https://www.ncbi.nlm.nih.gov/gene/?term=LOC105278807 | 2166 | 74 | 341.64 | 1 | 4.62 | 1.35 | 21.42 | 20.03 | 1.39 | 1.39 | 0.05 | 0.05 |
| gene-LOC105282600 | https://www.ncbi.nlm.nih.gov/gene/?term=LOC105282600 | 3715 | 60 | 161.51 | 1 | 2.69 | 1.67 | 27.74 | 26.44 | 1.3 | 1.3 | 0.02 | 0.02 |
| gene-LOC105285060 | https://www.ncbi.nlm.nih.gov/gene/?term=LOC105285060 | 1285 | 16 | 124.51 | 1 | 7.78 | 6.25 | 42.55 | 41.27 | 1.27 | 1.27 | 0.02 | 0.02 |
| gene-LOC105285742 | https://www.ncbi.nlm.nih.gov/gene/?term=LOC105285742 | 3446 | 109 | 316.31 | 1 | 2.9 | 0.92 | 17.79 | 16.59 | 1.2 | 1.2 | 0.04 | 0.04 |
| gene-LOC105285674 | https://www.ncbi.nlm.nih.gov/gene/?term=LOC105285674 | 3035 | 100 | 329.49 | 2 | 6.59 | 2 | 25.96 | 24.78 | 1.18 | 1.18 | 0.04 | 0.04 |
| gene-LOC105279528 | https://www.ncbi.nlm.nih.gov/gene/?term=LOC105279528 | 5802 | 40 | 68.94 | 1 | 1.72 | 2.5 | 21.84 | 20.71 | 1.12 | 1.12 | 0.01 | 0.01 |
| gene-LOC105275187 | https://www.ncbi.nlm.nih.gov/gene/?term=LOC105275187 | 5534 | 73 | 131.91 | 1 | 1.81 | 1.37 | 14.7 | 13.62 | 1.08 | 1.08 | 0.01 | 0.01 |
| gene-LOC105283697 | https://www.ncbi.nlm.nih.gov/gene/?term=LOC105283697 | 5301 | 153 | 288.62 | 1 | 1.89 | 0.65 | 16.46 | 15.43 | 1.02 | 1.02 | 0.03 | 0.03 |
| gene-LOC105275460 | https://www.ncbi.nlm.nih.gov/gene/?term=LOC105275460 | 2473 | 126 | 509.5 | 1 | 4.04 | 0.79 | 13.41 | 12.4 | 1.01 | 1.01 | 0.05 | 0.05 |
| gene-LOC105276449 | https://www.ncbi.nlm.nih.gov/gene/?term=LOC105276449 | 1231 | 48 | 389.93 | 1 | 8.12 | 2.08 | 25.67 | 24.69 | 0.99 | 0.99 | 0.04 | 0.04 |
| gene-LOC105283552 | https://www.ncbi.nlm.nih.gov/gene/?term=LOC105283552 | 1562 | 40 | 256.08 | 1 | 6.4 | 2.5 | 50.43 | 49.44 | 0.99 | 0.99 | 0.03 | 0.03 |
| gene-LOC105274823 | https://www.ncbi.nlm.nih.gov/gene/?term=LOC105274823 | 4620 | 177 | 383.12 | 1 | 2.16 | 0.56 | 17.15 | 16.21 | 0.95 | 0.95 | 0.04 | 0.04 |
| gene-LOC105279900 | https://www.ncbi.nlm.nih.gov/gene/?term=LOC105279900 | 2191 | 55 | 251.03 | 1 | 4.56 | 1.82 | 30.23 | 29.28 | 0.95 | 0.95 | 0.02 | 0.02 |
| gene-LOC105288079 | https://www.ncbi.nlm.nih.gov/gene/?term=LOC105288079 | 3473 | 113 | 325.37 | 1 | 2.88 | 0.88 | 17.44 | 16.56 | 0.89 | 0.89 | 0.03 | 0.03 |
| gene-LOC105281848 | https://www.ncbi.nlm.nih.gov/gene/?term=LOC105281848 | 2122 | 57 | 268.61 | 1 | 4.71 | 1.75 | 23.82 | 22.98 | 0.84 | 0.84 | 0.02 | 0.02 |
| gene-LOC105282491 | https://www.ncbi.nlm.nih.gov/gene/?term=LOC105282491 | 4761 | 179 | 375.97 | 1 | 2.1 | 0.56 | 9.15 | 8.31 | 0.83 | 0.83 | 0.03 | 0.03 |
| gene-LOC105288036 | https://www.ncbi.nlm.nih.gov/gene/?term=LOC105288036 | 5110 | 200 | 391.39 | 1 | 1.96 | 0.5 | 17.43 | 16.6 | 0.83 | 0.83 | 0.03 | 0.03 |
| gene-LOC105282156 | https://www.ncbi.nlm.nih.gov/gene/?term=LOC105282156 | 2577 | 127 | 492.82 | 1 | 3.88 | 0.79 | 25.13 | 24.3 | 0.82 | 0.82 | 0.04 | 0.04 |
| gene-LOC105285981 | https://www.ncbi.nlm.nih.gov/gene/?term=LOC105285981 | 7666 | 179 | 233.5 | 1 | 1.3 | 0.56 | 19.94 | 19.14 | 0.81 | 0.81 | 0.02 | 0.02 |
| gene-LOC105283943 | https://www.ncbi.nlm.nih.gov/gene/?term=LOC105283943 | 3559 | 57 | 160.16 | 1 | 2.81 | 1.75 | 23.04 | 22.27 | 0.77 | 0.77 | 0.01 | 0.01 |
| gene-LOC105278283 | https://www.ncbi.nlm.nih.gov/gene/?term=LOC105278283 | 2317 | 142 | 612.86 | 1 | 4.32 | 0.7 | 24.95 | 24.22 | 0.73 | 0.73 | 0.04 | 0.04 |
| gene-LOC105282794 | https://www.ncbi.nlm.nih.gov/gene/?term=LOC105282794 | 4571 | 104 | 227.52 | 1 | 2.19 | 0.96 | 16.85 | 16.16 | 0.69 | 0.69 | 0.02 | 0.02 |
| gene-LOC105283989 | https://www.ncbi.nlm.nih.gov/gene/?term=LOC105283989 | 4948 | 32 | 64.67 | 1 | 2.02 | 3.12 | 46.57 | 45.88 | 0.69 | 0.69 | 0 | 0 |
| gene-LOC105288237 | https://www.ncbi.nlm.nih.gov/gene/?term=LOC105288237 | 2226 | 49 | 220.13 | 1 | 4.49 | 2.04 | 43.01 | 42.34 | 0.67 | 0.67 | 0.01 | 0.01 |
| gene-LOC105285055 | https://www.ncbi.nlm.nih.gov/gene/?term=LOC105285055 | 4108 | 253 | 615.87 | 1 | 2.43 | 0.4 | 10.93 | 10.3 | 0.63 | 0.63 | 0.04 | 0.04 |
| gene-LOC105275451 | https://www.ncbi.nlm.nih.gov/gene/?term=LOC105275451 | 6035 | 135 | 223.7 | 1 | 1.66 | 0.74 | 21.36 | 20.75 | 0.61 | 0.61 | 0.01 | 0.01 |
| gene-LOC105277215 | https://www.ncbi.nlm.nih.gov/gene/?term=LOC105277215 | 2619 | 29 | 110.73 | 1 | 3.82 | 3.45 | 25.25 | 24.64 | 0.61 | 0.61 | 0.01 | 0.01 |
| gene-LOC105274449 | https://www.ncbi.nlm.nih.gov/gene/?term=LOC105274449 | 1549 | 53 | 342.16 | 1 | 6.46 | 1.89 | 49.89 | 49.29 | 0.6 | 0.6 | 0.02 | 0.02 |
| gene-LOC105288078 | https://www.ncbi.nlm.nih.gov/gene/?term=LOC105288078 | 19126 | 337 | 176.2 | 1 | 0.52 | 0.3 | 15.64 | 15.05 | 0.59 | 0.59 | 0.01 | 0.01 |
| gene-LOC105280819 | https://www.ncbi.nlm.nih.gov/gene/?term=LOC105280819 | 1639 | 42 | 256.25 | 1 | 6.1 | 2.38 | 39.26 | 38.69 | 0.58 | 0.58 | 0.01 | 0.01 |
| gene-LOC105280641 | https://www.ncbi.nlm.nih.gov/gene/?term=LOC105280641 | 3682 | 75 | 203.69 | 1 | 2.72 | 1.33 | 13.21 | 12.66 | 0.55 | 0.55 | 0.01 | 0.01 |
| gene-LOC105285179 | https://www.ncbi.nlm.nih.gov/gene/?term=LOC105285179 | 4191 | 189 | 450.97 | 1 | 2.39 | 0.53 | 18.34 | 17.82 | 0.52 | 0.52 | 0.02 | 0.02 |
| gene-LOC105278145 | https://www.ncbi.nlm.nih.gov/gene/?term=LOC105278145 | 4470 | 252 | 563.76 | 2 | 4.47 | 0.79 | 10.98 | 10.51 | 0.48 | 0.48 | 0.03 | 0.03 |
| gene-LOC105274531 | https://www.ncbi.nlm.nih.gov/gene/?term=LOC105274531 | 7774 | 286 | 367.89 | 1 | 1.29 | 0.35 | 11.23 | 10.77 | 0.46 | 0.46 | 0.02 | 0.02 |
| gene-LOC105287417 | https://www.ncbi.nlm.nih.gov/gene/?term=LOC105287417 | 1025 | 32 | 312.2 | 1 | 9.76 | 3.12 | 37.69 | 37.23 | 0.46 | 0.46 | 0.01 | 0.01 |
| gene-LOC105277903 | https://www.ncbi.nlm.nih.gov/gene/?term=LOC105277903 | 1835 | 37 | 201.63 | 1 | 5.45 | 2.7 | 54.5 | 54.08 | 0.42 | 0.42 | 0.01 | 0.01 |

| | | | | | | | | | | | | | |
|---|---|---|---|---|---|---|---|---|---|---|---|---|---|
| gene-LOC105275363 | https://www.ncbi.nlm.nih.gov/gene/?term=LOC105275363 | 3798 | 179 | 471.3 | 1 | 2.63 | 0.56 | 15.94 | 15.54 | 0.4 | 0.4 | 0.02 | 0.02 |
| gene-LOC105286452 | https://www.ncbi.nlm.nih.gov/gene/?term=LOC105286452 | 6888 | 68 | 98.72 | 1 | 1.45 | 1.47 | 28.16 | 27.75 | 0.4 | 0.4 | 0 | 0 |
| gene-LOC105286355 | https://www.ncbi.nlm.nih.gov/gene/?term=LOC105286355 | 3245 | 149 | 459.17 | 1 | 3.08 | 0.67 | 21.45 | 21.07 | 0.38 | 0.38 | 0.02 | 0.02 |
| gene-LOC105283592 | https://www.ncbi.nlm.nih.gov/gene/?term=LOC105283592 | 3479 | 80 | 229.95 | 1 | 2.87 | 1.25 | 21.51 | 21.14 | 0.37 | 0.37 | 0.01 | 0.01 |
| gene-LOC105283838 | https://www.ncbi.nlm.nih.gov/gene/?term=LOC105283838 | 1748 | 30 | 171.62 | 1 | 5.72 | 3.33 | 33.04 | 32.68 | 0.36 | 0.36 | 0.01 | 0.01 |
| gene-LOC105278497 | https://www.ncbi.nlm.nih.gov/gene/?term=LOC105278497 | 2072 | 100 | 482.63 | 1 | 4.83 | 1 | 18.63 | 18.28 | 0.35 | 0.35 | 0.02 | 0.02 |
| gene-LOC105278384 | https://www.ncbi.nlm.nih.gov/gene/?term=LOC105278384 | 1390 | 33 | 237.41 | 1 | 7.19 | 3.03 | 46.11 | 45.77 | 0.34 | 0.34 | 0.01 | 0.01 |
| gene-LOC105285639 | https://www.ncbi.nlm.nih.gov/gene/?term=LOC105285639 | 10325 | 351 | 339.95 | 2 | 1.94 | 0.57 | 11.49 | 11.2 | 0.29 | 0.29 | 0.01 | 0.01 |
| gene-LOC105288010 | https://www.ncbi.nlm.nih.gov/gene/?term=LOC105288010 | 3189 | 52 | 163.06 | 1 | 3.14 | 1.92 | 29.32 | 29.04 | 0.28 | 0.28 | 0 | 0 |
| gene-LOC105279298 | https://www.ncbi.nlm.nih.gov/gene/?term=LOC105279298 | 2773 | 127 | 457.99 | 1 | 3.61 | 0.79 | 24.14 | 23.89 | 0.26 | 0.26 | 0.01 | 0.01 |
| gene-LOC105283123 | https://www.ncbi.nlm.nih.gov/gene/?term=LOC105283123 | 1507 | 47 | 311.88 | 1 | 6.64 | 2.13 | 20.21 | 19.96 | 0.25 | 0.25 | 0.01 | 0.01 |
| gene-LOC105284329 | https://www.ncbi.nlm.nih.gov/gene/?term=LOC105284329 | 4051 | 71 | 175.27 | 1 | 2.47 | 1.41 | 19.25 | 18.99 | 0.25 | 0.25 | 0 | 0 |
| gene-LOC105284184 | https://www.ncbi.nlm.nih.gov/gene/?term=LOC105284184 | 2535 | 92 | 362.92 | 1 | 3.94 | 1.09 | 31.61 | 31.41 | 0.2 | 0.2 | 0.01 | 0.01 |
| gene-LOC105286997 | https://www.ncbi.nlm.nih.gov/gene/?term=LOC105286997 | 2850 | 142 | 498.25 | 1 | 3.51 | 0.7 | 11.2 | 11.03 | 0.17 | 0.17 | 0.01 | 0.01 |
| gene-LOC105275170 | https://www.ncbi.nlm.nih.gov/gene/?term=LOC105275170 | 12918 | 350 | 270.94 | 1 | 0.77 | 0.29 | 1.94 | 1.8 | 0.15 | 0.15 | 0 | 0 |
| gene-LOC105275374 | https://www.ncbi.nlm.nih.gov/gene/?term=LOC105275374 | 11376 | 514 | 451.83 | 1 | 0.88 | 0.19 | 9.9 | 9.75 | 0.15 | 0.15 | 0.01 | 0.01 |
| gene-LOC105276059 | https://www.ncbi.nlm.nih.gov/gene/?term=LOC105276059 | 6256 | 134 | 214.19 | 1 | 1.6 | 0.75 | 11.07 | 10.91 | 0.15 | 0.15 | 0 | 0 |
| gene-LOC105283728 | https://www.ncbi.nlm.nih.gov/gene/?term=LOC105283728 | 3256 | 152 | 466.83 | 2 | 6.14 | 1.32 | 15.71 | 15.57 | 0.14 | 0.14 | 0.01 | 0.01 |
| gene-LOC105286099 | https://www.ncbi.nlm.nih.gov/gene/?term=LOC105286099 | 5919 | 70 | 118.26 | 1 | 1.69 | 1.43 | 12.4 | 12.26 | 0.14 | 0.14 | 0 | 0 |
| gene-LOC105278534 | https://www.ncbi.nlm.nih.gov/gene/?term=LOC105278534 | 7364 | 198 | 268.88 | 1 | 1.36 | 0.51 | 17.97 | 17.87 | 0.1 | 0.1 | 0 | 0 |
| gene-LOC105280774 | https://www.ncbi.nlm.nih.gov/gene/?term=LOC105280774 | 2793 | 132 | 472.61 | 1 | 3.58 | 0.76 | 25.2 | 25.1 | 0.1 | 0.1 | 0 | 0 |
| gene-LOC105282149 | https://www.ncbi.nlm.nih.gov/gene/?term=LOC105282149 | 1787 | 103 | 576.39 | 1 | 5.6 | 0.97 | 41.42 | 41.34 | 0.08 | 0.08 | 0 | 0 |
| gene-LOC105278666 | https://www.ncbi.nlm.nih.gov/gene/?term=LOC105278666 | 10341 | 474 | 458.37 | 1 | 0.97 | 0.21 | 6.99 | 6.95 | 0.05 | 0.05 | 0 | 0 |
| gene-LOC105280956 | https://www.ncbi.nlm.nih.gov/gene/?term=LOC105280956 | 2639 | 80 | 303.15 | 1 | 3.79 | 1.25 | 18.94 | 18.89 | 0.05 | 0.05 | 0 | 0 |
| gene-LOC105275327 | https://www.ncbi.nlm.nih.gov/gene/?term=LOC105275327 | 14245 | 529 | 371.36 | 1 | 0.7 | 0.19 | 6.66 | 6.63 | 0.03 | 0.03 | 0 | 0 |
| gene-LOC105284043 | https://www.ncbi.nlm.nih.gov/gene/?term=LOC105284043 | 5661 | 284 | 501.68 | 1 | 1.77 | 0.35 | 10.32 | 10.3 | 0.03 | 0.03 | 0 | 0 |
| gene-LOC105277216 | https://www.ncbi.nlm.nih.gov/gene/?term=LOC105277216 | 7627 | 106 | 138.98 | 1 | 1.31 | 0.94 | 7.36 | 7.34 | 0.02 | 0.02 | 0 | 0 |
| gene-LOC105277866 | https://www.ncbi.nlm.nih.gov/gene/?term=LOC105277866 | 33045 | 482 | 145.86 | 1 | 0.3 | 0.21 | 3.05 | 3.04 | 0.02 | 0.02 | 0 | 0 |
| gene-LOC105279500 | https://www.ncbi.nlm.nih.gov/gene/?term=LOC105279500 | 3587 | 32 | 89.21 | 1 | 2.79 | 3.12 | 46.58 | 46.57 | 0.01 | 0.01 | 0 | 0 |
| gene-LOC105286515 | https://www.ncbi.nlm.nih.gov/gene/?term=LOC105286515 | 3751 | 117 | 311.92 | 1 | 2.67 | 0.85 | 10.68 | 10.68 | 0 | 0 | 0 | 0 |
| gene-LOC105278228 | https://www.ncbi.nlm.nih.gov/gene/?term=LOC105278228 | 8610 | 176 | 204.41 | 1 | 1.16 | 0.57 | 22.62 | 22.66 | -0.04 | 0.04 | 0 | 0 |
| gene-LOC105278877 | https://www.ncbi.nlm.nih.gov/gene/?term=LOC105278877 | 1968 | 87 | 442.07 | 1 | 5.08 | 1.15 | 29.15 | 29.21 | -0.06 | 0.06 | 0 | 0 |
| gene-LOC105287095 | https://www.ncbi.nlm.nih.gov/gene/?term=LOC105287095 | 1435 | 66 | 459.93 | 1 | 6.97 | 1.52 | 24.97 | 25.03 | -0.06 | 0.06 | 0 | 0 |
| gene-LOC105286194 | https://www.ncbi.nlm.nih.gov/gene/?term=LOC105286194 | 152619 | 2310 | 151.36 | 1 | 0.07 | 0.04 | 5.89 | 6 | -0.11 | 0.11 | 0 | 0 |
| gene-LOC105276527 | https://www.ncbi.nlm.nih.gov/gene/?term=LOC105276527 | 7923 | 338 | 426.61 | 1 | 1.26 | 0.3 | 8.29 | 8.43 | -0.14 | 0.14 | -0.01 | 0.01 |
| gene-LOC105279038 | https://www.ncbi.nlm.nih.gov/gene/?term=LOC105279038 | 9453 | 329 | 348.04 | 1 | 1.06 | 0.3 | 9.14 | 9.32 | -0.18 | 0.18 | -0.01 | 0.01 |
| gene-LOC105279664 | https://www.ncbi.nlm.nih.gov/gene/?term=LOC105279664 | 4023 | 64 | 159.09 | 1 | 2.49 | 1.56 | 17.75 | 17.94 | -0.19 | 0.19 | 0 | 0 |
| gene-LOC105282811 | https://www.ncbi.nlm.nih.gov/gene/?term=LOC105282811 | 5765 | 203 | 352.12 | 1 | 1.73 | 0.49 | 8.59 | 8.78 | -0.19 | 0.19 | -0.01 | 0.01 |
| gene-LOC105274856 | https://www.ncbi.nlm.nih.gov/gene/?term=LOC105274856 | 5216 | 228 | 437.12 | 1 | 1.92 | 0.44 | 17.7 | 17.9 | -0.2 | 0.2 | -0.01 | 0.01 |
| gene-LOC105277856 | https://www.ncbi.nlm.nih.gov/gene/?term=LOC105277856 | 3442 | 33 | 95.87 | 1 | 2.91 | 3.03 | 39.28 | 39.53 | -0.25 | 0.25 | 0 | 0 |
| gene-LOC105285807 | https://www.ncbi.nlm.nih.gov/gene/?term=LOC105285807 | 6063 | 215 | 354.61 | 1 | 1.65 | 0.47 | 8.65 | 8.94 | -0.28 | 0.28 | -0.01 | 0.01 |
| gene-LOC105274706 | https://www.ncbi.nlm.nih.gov/gene/?term=LOC105274706 | 1446 | 50 | 345.78 | 1 | 6.92 | 2 | 37.45 | 37.82 | -0.37 | 0.37 | -0.01 | 0.01 |
| gene-LOC105282825 | https://www.ncbi.nlm.nih.gov/gene/?term=LOC105282825 | 3919 | 96 | 244.96 | 1 | 2.55 | 1.04 | 14.53 | 14.91 | -0.38 | 0.38 | -0.01 | 0.01 |
| gene-LOC105278632 | https://www.ncbi.nlm.nih.gov/gene/?term=LOC105278632 | 3624 | 143 | 394.59 | 1 | 2.76 | 0.7 | 13.02 | 13.42 | -0.4 | 0.4 | -0.02 | 0.02 |
| gene-LOC105284355 | https://www.ncbi.nlm.nih.gov/gene/?term=LOC105284355 | 3904 | 155 | 397.03 | 1 | 2.56 | 0.65 | 20.33 | 20.86 | -0.53 | 0.53 | -0.02 | 0.02 |
| gene-LOC105274851 | https://www.ncbi.nlm.nih.gov/gene/?term=LOC105274851 | 6616 | 144 | 217.65 | 1 | 1.51 | 0.69 | 11.65 | 12.43 | -0.79 | 0.79 | -0.02 | 0.02 |
| gene-LOC105280002 | https://www.ncbi.nlm.nih.gov/gene/?term=LOC105280002 | 1940 | 31 | 159.79 | 1 | 5.15 | 3.23 | 31.38 | 32.27 | -0.88 | 0.88 | -0.01 | 0.01 |
| gene-LOC105282658 | https://www.ncbi.nlm.nih.gov/gene/?term=LOC105282658 | 2421 | 64 | 264.35 | 1 | 4.13 | 1.56 | 23.38 | 24.34 | -0.95 | 0.95 | -0.03 | 0.03 |
| gene-LOC105284036 | https://www.ncbi.nlm.nih.gov/gene/?term=LOC105284036 | 3254 | 98 | 301.17 | 1 | 3.07 | 1.02 | 22.14 | 23.09 | -0.95 | 0.95 | -0.03 | 0.03 |
| gene-LOC105279596 | https://www.ncbi.nlm.nih.gov/gene/?term=LOC105279596 | 7695 | 115 | 149.45 | 1 | 1.3 | 0.87 | 16.77 | 17.78 | -1 | 1 | -0.02 | 0.02 |
| gene-LOC105275518 | https://www.ncbi.nlm.nih.gov/gene/?term=LOC105275518 | 2335 | 23 | 98.5 | 1 | 4.28 | 4.35 | 45.78 | 46.9 | -1.12 | 1.12 | -0.01 | 0.01 |
| gene-LOC105274590 | https://www.ncbi.nlm.nih.gov/gene/?term=LOC105274590 | 4516 | 82 | 181.58 | 1 | 2.21 | 1.22 | 25.47 | 26.68 | -1.21 | 1.21 | -0.02 | 0.02 |
| gene-LOC105281173 | https://www.ncbi.nlm.nih.gov/gene/?term=LOC105281173 | 1595 | 38 | 238.24 | 1 | 6.27 | 2.63 | 59.87 | 61.09 | -1.22 | 1.22 | -0.03 | 0.03 |
| gene-LOC105283649 | https://www.ncbi.nlm.nih.gov/gene/?term=LOC105283649 | 1918 | 56 | 291.97 | 1 | 5.21 | 1.79 | 37.76 | 39.05 | -1.29 | 1.29 | -0.04 | 0.04 |
| gene-LOC105276196 | https://www.ncbi.nlm.nih.gov/gene/?term=LOC105276196 | 2953 | 57 | 193.02 | 1 | 3.39 | 1.75 | 39.57 | 40.89 | -1.32 | 1.32 | -0.03 | 0.03 |
| gene-LOC105274534 | https://www.ncbi.nlm.nih.gov/gene/?term=LOC105274534 | 1329 | 59 | 443.94 | 1 | 7.52 | 1.69 | 43.5 | 44.89 | -1.39 | 1.39 | -0.06 | 0.06 |
| gene-LOC105288042 | https://www.ncbi.nlm.nih.gov/gene/?term=LOC105288042 | 3507 | 100 | 285.14 | 1 | 2.85 | 1 | 20.77 | 22.23 | -1.46 | 1.46 | -0.04 | 0.04 |
| gene-LOC105278962 | https://www.ncbi.nlm.nih.gov/gene/?term=LOC105278962 | 4491 | 109 | 242.71 | 1 | 2.23 | 0.92 | 19.66 | 21.14 | -1.48 | 1.48 | -0.04 | 0.04 |
| gene-LOC105278282 | https://www.ncbi.nlm.nih.gov/gene/?term=LOC105278282 | 1775 | 51 | 287.32 | 1 | 5.63 | 1.96 | 39.13 | 40.69 | -1.57 | 1.57 | -0.05 | 0.05 |
| gene-LOC105286514 | https://www.ncbi.nlm.nih.gov/gene/?term=LOC105286514 | 2528 | 50 | 197.78 | 1 | 3.96 | 2 | 35.29 | 37.3 | -2.01 | 2.01 | -0.04 | 0.04 |
| gene-LOC105275571 | https://www.ncbi.nlm.nih.gov/gene/?term=LOC105275571 | 1926 | 30 | 155.76 | 1 | 5.19 | 3.33 | 42.71 | 44.75 | -2.04 | 2.04 | -0.03 | 0.03 |
| gene-LOC105276538 | https://www.ncbi.nlm.nih.gov/gene/?term=LOC105276538 | 1511 | 21 | 138.98 | 1 | 6.62 | 4.76 | 57.36 | 59.41 | -2.05 | 2.05 | -0.03 | 0.03 |
| gene-LOC105285004 | https://www.ncbi.nlm.nih.gov/gene/?term=LOC105285004 | 1334 | 37 | 277.36 | 1 | 7.5 | 2.7 | 16.38 | 18.79 | -2.41 | 2.41 | -0.07 | 0.07 |
| gene-LOC105282914 | https://www.ncbi.nlm.nih.gov/gene/?term=LOC105282914 | 5623 | 43 | 76.47 | 1 | 1.78 | 2.33 | 46.2 | 49.79 | -3.59 | 3.59 | -0.03 | 0.03 |

Directory: OGLall File: ogl-Ob_LI_RphaseA1.vs.RphaseB2.txt

| gene_ID | gene_link | gwidth | #Sites | #per10Kb | #dmSites | #dmsp10kb | %dmSites | %pSite1 | %pSite2 | DMpSite | ADMpSite | DMpNucl | ADMpNucl |
|---|---|---|---|---|---|---|---|---|---|---|---|---|---|
| gene-LOC105275070 | https://www.ncbi.nlm.nih.gov/gene/?term=LOC105275070 | 5667 | 37 | 65.29 | 7 | 12.35 | 18.92 | 16.3 | 1.04 | 15.26 | 15.26 | 0.1 | 0.1 |
| gene-LOC113562345 | https://www.ncbi.nlm.nih.gov/gene/?term=LOC113562345 | 2290 | 67 | 292.58 | 6 | 26.2 | 8.96 | 28.84 | 16.72 | 12.12 | 12.12 | 0.35 | 0.35 |
| gene-LOC105279898 | https://www.ncbi.nlm.nih.gov/gene/?term=LOC105279898 | 1343 | 9 | 67.01 | 1 | 7.45 | 11.11 | 83.37 | 74.78 | 8.59 | 8.59 | 0.06 | 0.06 |
| gene-LOC105286694 | https://www.ncbi.nlm.nih.gov/gene/?term=LOC105286694 | 3409 | 8 | 23.47 | 1 | 2.93 | 12.5 | 66.22 | 57.86 | 8.36 | 8.36 | 0.02 | 0.02 |
| gene-LOC105275708 | https://www.ncbi.nlm.nih.gov/gene/?term=LOC105275708 | 1772 | 14 | 79.01 | 1 | 5.64 | 7.14 | 50.83 | 43.74 | 7.08 | 7.08 | 0.06 | 0.06 |
| gene-LOC105285514 | https://www.ncbi.nlm.nih.gov/gene/?term=LOC105285514 | 1277 | 14 | 109.63 | 1 | 7.83 | 7.14 | 69.69 | 65.1 | 4.59 | 4.59 | 0.05 | 0.05 |
| gene-LOC105283552 | https://www.ncbi.nlm.nih.gov/gene/?term=LOC105283552 | 1562 | 41 | 262.48 | 1 | 6.4 | 2.44 | 50.09 | 46.71 | 3.37 | 3.37 | 0.09 | 0.09 |
| gene-LOC105274900 | https://www.ncbi.nlm.nih.gov/gene/?term=LOC105274900 | 5561 | 55 | 98.9 | 3 | 5.39 | 5.45 | 32.79 | 29.47 | 3.31 | 3.31 | 0.03 | 0.03 |
| gene-LOC105281515 | https://www.ncbi.nlm.nih.gov/gene/?term=LOC105281515 | 8827 | 41 | 46.45 | 1 | 1.13 | 2.44 | 31.51 | 28.2 | 3.31 | 3.31 | 0.02 | 0.02 |
| gene-LOC105278436 | https://www.ncbi.nlm.nih.gov/gene/?term=LOC105278436 | 2556 | 54 | 211.27 | 1 | 3.91 | 1.85 | 40.35 | 37.22 | 3.13 | 3.13 | 0.07 | 0.07 |
| gene-LOC105285671 | https://www.ncbi.nlm.nih.gov/gene/?term=LOC105285671 | 2263 | 101 | 446.31 | 1 | 4.42 | 0.99 | 30.8 | 27.84 | 2.96 | 2.96 | 0.13 | 0.13 |
| gene-LOC105279356 | https://www.ncbi.nlm.nih.gov/gene/?term=LOC105279356 | 1314 | 43 | 327.25 | 1 | 7.61 | 2.33 | 32.62 | 30.3 | 2.31 | 2.31 | 0.08 | 0.08 |
| gene-LOC105285674 | https://www.ncbi.nlm.nih.gov/gene/?term=LOC105285674 | 3035 | 92 | 303.13 | 2 | 6.59 | 2.17 | 26.97 | 24.84 | 2.13 | 2.13 | 0.06 | 0.06 |
| gene-LOC105285202 | https://www.ncbi.nlm.nih.gov/gene/?term=LOC105285202 | 1838 | 61 | 331.88 | 1 | 5.44 | 1.64 | 26.9 | 24.8 | 2.1 | 2.1 | 0.07 | 0.07 |
| gene-LOC105275347 | https://www.ncbi.nlm.nih.gov/gene/?term=LOC105275347 | 2526 | 16 | 63.34 | 1 | 3.96 | 6.25 | 41.65 | 39.67 | 1.98 | 1.98 | 0.01 | 0.01 |
| gene-LOC105278807 | https://www.ncbi.nlm.nih.gov/gene/?term=LOC105278807 | 2166 | 65 | 300.09 | 1 | 4.62 | 1.54 | 22.17 | 20.43 | 1.74 | 1.74 | 0.05 | 0.05 |
| gene-LOC105280104 | https://www.ncbi.nlm.nih.gov/gene/?term=LOC105280104 | 3960 | 123 | 310.61 | 1 | 2.53 | 0.81 | 18.15 | 16.51 | 1.64 | 1.64 | 0.05 | 0.05 |
| gene-LOC105281266 | https://www.ncbi.nlm.nih.gov/gene/?term=LOC105281266 | 4102 | 78 | 190.15 | 1 | 2.44 | 1.28 | 18.58 | 17.01 | 1.57 | 1.57 | 0.03 | 0.03 |
| gene-LOC105286099 | https://www.ncbi.nlm.nih.gov/gene/?term=LOC105286099 | 5919 | 67 | 113.19 | 1 | 1.69 | 1.49 | 11.4 | 9.83 | 1.57 | 1.57 | 0.02 | 0.02 |
| gene-LOC105284329 | https://www.ncbi.nlm.nih.gov/gene/?term=LOC105284329 | 4051 | 52 | 128.36 | 1 | 2.47 | 1.92 | 19.88 | 18.38 | 1.5 | 1.5 | 0.02 | 0.02 |
| gene-LOC105282794 | https://www.ncbi.nlm.nih.gov/gene/?term=LOC105282794 | 4571 | 90 | 196.89 | 1 | 2.19 | 1.11 | 17.25 | 15.76 | 1.49 | 1.49 | 0.03 | 0.03 |
| gene-LOC105277903 | https://www.ncbi.nlm.nih.gov/gene/?term=LOC105277903 | 1835 | 36 | 196.19 | 1 | 5.45 | 2.78 | 51.23 | 49.79 | 1.45 | 1.45 | 0.03 | 0.03 |
| gene-LOC105280786 | https://www.ncbi.nlm.nih.gov/gene/?term=LOC105280786 | 2573 | 115 | 446.95 | 1 | 3.89 | 0.87 | 28.39 | 27.06 | 1.33 | 1.33 | 0.06 | 0.06 |
| gene-LOC105288066 | https://www.ncbi.nlm.nih.gov/gene/?term=LOC105288066 | 1964 | 27 | 137.47 | 1 | 5.09 | 3.7 | 34.07 | 32.8 | 1.27 | 1.27 | 0.02 | 0.02 |
| gene-LOC105275451 | https://www.ncbi.nlm.nih.gov/gene/?term=LOC105275451 | 6035 | 113 | 187.24 | 1 | 1.66 | 0.88 | 24.46 | 23.26 | 1.2 | 1.2 | 0.02 | 0.02 |
| gene-LOC105279664 | https://www.ncbi.nlm.nih.gov/gene/?term=LOC105279664 | 4023 | 53 | 131.74 | 1 | 2.49 | 1.89 | 19.07 | 17.88 | 1.19 | 1.19 | 0.02 | 0.02 |
| gene-LOC105283943 | https://www.ncbi.nlm.nih.gov/gene/?term=LOC105283943 | 3559 | 45 | 126.44 | 1 | 2.81 | 2.22 | 16.9 | 15.77 | 1.14 | 1.14 | 0.01 | 0.01 |
| gene-LOC113561805 | https://www.ncbi.nlm.nih.gov/gene/?term=LOC113561805 | 3337 | 67 | 200.78 | 1 | 3 | 1.49 | 36.13 | 35.02 | 1.11 | 1.11 | 0.02 | 0.02 |
| gene-LOC105275363 | https://www.ncbi.nlm.nih.gov/gene/?term=LOC105275363 | 3798 | 157 | 413.38 | 1 | 2.63 | 0.64 | 17.94 | 16.87 | 1.07 | 1.07 | 0.04 | 0.04 |
| gene-LOC105281173 | https://www.ncbi.nlm.nih.gov/gene/?term=LOC105281173 | 1595 | 38 | 238.24 | 1 | 6.27 | 2.63 | 59.87 | 58.85 | 1.02 | 1.02 | 0.02 | 0.02 |
| gene-LOC105276692 | https://www.ncbi.nlm.nih.gov/gene/?term=LOC105276692 | 1745 | 29 | 166.19 | 1 | 5.73 | 3.45 | 25.96 | 24.96 | 1 | 1 | 0.02 | 0.02 |
| gene-LOC105288078 | https://www.ncbi.nlm.nih.gov/gene/?term=LOC105288078 | 19126 | 286 | 149.53 | 1 | 0.52 | 0.35 | 17.66 | 16.67 | 0.99 | 0.99 | 0.01 | 0.01 |
| gene-LOC105280956 | https://www.ncbi.nlm.nih.gov/gene/?term=LOC105280956 | 2639 | 71 | 269.04 | 1 | 3.79 | 1.41 | 18.77 | 17.79 | 0.98 | 0.98 | 0.03 | 0.03 |
| gene-LOC105288036 | https://www.ncbi.nlm.nih.gov/gene/?term=LOC105288036 | 5110 | 181 | 354.21 | 1 | 1.96 | 0.55 | 18.99 | 18.01 | 0.98 | 0.98 | 0.03 | 0.03 |
| gene-LOC105283123 | https://www.ncbi.nlm.nih.gov/gene/?term=LOC105283123 | 1507 | 40 | 265.43 | 1 | 6.64 | 2.5 | 16.19 | 15.23 | 0.96 | 0.96 | 0.03 | 0.03 |
| gene-LOC113561621 | https://www.ncbi.nlm.nih.gov/gene/?term=LOC113561621 | 2249 | 76 | 337.93 | 1 | 4.45 | 1.32 | 22.44 | 21.49 | 0.95 | 0.95 | 0.03 | 0.03 |
| gene-LOC105276059 | https://www.ncbi.nlm.nih.gov/gene/?term=LOC105276059 | 6256 | 125 | 199.81 | 1 | 1.6 | 0.8 | 10.91 | 10 | 0.91 | 0.91 | 0.02 | 0.02 |
| gene-LOC105285060 | https://www.ncbi.nlm.nih.gov/gene/?term=LOC105285060 | 1285 | 16 | 124.51 | 1 | 7.78 | 6.25 | 47.94 | 47.04 | 0.9 | 0.9 | 0.01 | 0.01 |
| gene-LOC105283592 | https://www.ncbi.nlm.nih.gov/gene/?term=LOC105283592 | 3479 | 71 | 204.08 | 1 | 2.87 | 1.41 | 21.54 | 20.67 | 0.88 | 0.88 | 0.02 | 0.02 |
| gene-LOC105284184 | https://www.ncbi.nlm.nih.gov/gene/?term=LOC105284184 | 2535 | 80 | 315.58 | 1 | 3.94 | 1.25 | 35.73 | 34.87 | 0.86 | 0.86 | 0.03 | 0.03 |
| gene-LOC105276901 | https://www.ncbi.nlm.nih.gov/gene/?term=LOC105276901 | 3254 | 84 | 258.14 | 1 | 3.07 | 1.19 | 27.21 | 26.37 | 0.84 | 0.84 | 0.02 | 0.02 |
| gene-LOC105287854 | https://www.ncbi.nlm.nih.gov/gene/?term=LOC105287854 | 3644 | 49 | 134.47 | 1 | 2.74 | 2.04 | 24.37 | 23.53 | 0.84 | 0.84 | 0.01 | 0.01 |
| gene-LOC105274823 | https://www.ncbi.nlm.nih.gov/gene/?term=LOC105274823 | 4620 | 138 | 298.7 | 1 | 2.16 | 0.72 | 21.4 | 20.58 | 0.82 | 0.82 | 0.02 | 0.02 |
| gene-LOC105276612 | https://www.ncbi.nlm.nih.gov/gene/?term=LOC105276612 | 4157 | 178 | 428.19 | 1 | 2.41 | 0.56 | 10.68 | 9.88 | 0.8 | 0.8 | 0.03 | 0.03 |
| gene-LOC105284043 | https://www.ncbi.nlm.nih.gov/gene/?term=LOC105284043 | 5661 | 283 | 499.91 | 1 | 1.77 | 0.35 | 10.38 | 9.58 | 0.79 | 0.79 | 0.04 | 0.04 |
| gene-LOC105287417 | https://www.ncbi.nlm.nih.gov/gene/?term=LOC105287417 | 1025 | 19 | 185.37 | 1 | 9.76 | 5.26 | 59.44 | 58.65 | 0.79 | 0.79 | 0.01 | 0.01 |
| gene-LOC105282156 | https://www.ncbi.nlm.nih.gov/gene/?term=LOC105282156 | 2577 | 122 | 473.42 | 1 | 3.88 | 0.82 | 25.73 | 24.95 | 0.78 | 0.78 | 0.04 | 0.04 |
| gene-LOC105286452 | https://www.ncbi.nlm.nih.gov/gene/?term=LOC105286452 | 6888 | 64 | 92.92 | 1 | 1.45 | 1.56 | 30.17 | 29.46 | 0.72 | 0.72 | 0.01 | 0.01 |
| gene-LOC105283728 | https://www.ncbi.nlm.nih.gov/gene/?term=LOC105283728 | 3256 | 117 | 359.34 | 2 | 6.14 | 1.71 | 18.77 | 18.07 | 0.7 | 0.7 | 0.03 | 0.03 |
| gene-LOC105285534 | https://www.ncbi.nlm.nih.gov/gene/?term=LOC105285534 | 3944 | 232 | 588.24 | 1 | 2.54 | 0.43 | 11.59 | 10.9 | 0.69 | 0.69 | 0.04 | 0.04 |
| gene-LOC105275460 | https://www.ncbi.nlm.nih.gov/gene/?term=LOC105275460 | 2473 | 118 | 477.15 | 1 | 4.04 | 0.85 | 13.95 | 13.28 | 0.67 | 0.67 | 0.03 | 0.03 |
| gene-LOC105285639 | https://www.ncbi.nlm.nih.gov/gene/?term=LOC105285639 | 10325 | 312 | 302.18 | 2 | 1.94 | 0.64 | 12.89 | 12.24 | 0.65 | 0.65 | 0.02 | 0.02 |
| gene-LOC105285055 | https://www.ncbi.nlm.nih.gov/gene/?term=LOC105285055 | 4108 | 261 | 635.35 | 1 | 2.43 | 0.38 | 10.75 | 10.11 | 0.64 | 0.64 | 0.04 | 0.04 |
| gene-LOC105278534 | https://www.ncbi.nlm.nih.gov/gene/?term=LOC105278534 | 7364 | 176 | 239 | 2 | 2.72 | 1.14 | 19.64 | 19.01 | 0.63 | 0.63 | 0.02 | 0.02 |
| gene-LOC105281848 | https://www.ncbi.nlm.nih.gov/gene/?term=LOC105281848 | 2122 | 47 | 221.49 | 1 | 4.71 | 2.13 | 27.19 | 26.57 | 0.63 | 0.63 | 0.01 | 0.01 |
| gene-LOC105278283 | https://www.ncbi.nlm.nih.gov/gene/?term=LOC105278283 | 2317 | 138 | 595.6 | 1 | 4.32 | 0.72 | 25.64 | 25.05 | 0.59 | 0.59 | 0.04 | 0.04 |
| gene-LOC105279298 | https://www.ncbi.nlm.nih.gov/gene/?term=LOC105279298 | 2773 | 133 | 479.62 | 2 | 7.21 | 1.5 | 23.97 | 23.38 | 0.59 | 0.59 | 0.03 | 0.03 |
| gene-LOC105285179 | https://www.ncbi.nlm.nih.gov/gene/?term=LOC105285179 | 4191 | 171 | 408.02 | 1 | 2.39 | 0.58 | 19 | 18.45 | 0.55 | 0.55 | 0.02 | 0.02 |
| gene-LOC105278497 | https://www.ncbi.nlm.nih.gov/gene/?term=LOC105278497 | 2072 | 95 | 458.49 | 1 | 4.83 | 1.05 | 19.39 | 18.89 | 0.5 | 0.5 | 0.02 | 0.02 |
| gene-LOC105278108 | https://www.ncbi.nlm.nih.gov/gene/?term=LOC105278108 | 4392 | 43 | 97.91 | 1 | 2.28 | 2.33 | 26.73 | 26.25 | 0.49 | 0.49 | 0 | 0 |

| | | | | | | | | | | | | | |
|---|---|---|---|---|---|---|---|---|---|---|---|---|---|
| gene-LOC105278145 | https://www.ncbi.nlm.nih.gov/gene/?term=LOC105278145 | 4470 | 235 | 525.73 | 1 | 2.24 | 0.43 | 11.34 | 10.86 | 0.48 | 0.48 | 0.03 | 0.03 |
| gene-LOC105286997 | https://www.ncbi.nlm.nih.gov/gene/?term=LOC105286997 | 2850 | 106 | 371.93 | 1 | 3.51 | 0.94 | 12.07 | 11.62 | 0.46 | 0.46 | 0.02 | 0.02 |
| gene-LOC105275179 | https://www.ncbi.nlm.nih.gov/gene/?term=LOC105275179 | 3271 | 69 | 210.94 | 1 | 3.06 | 1.45 | 18.8 | 18.36 | 0.45 | 0.45 | 0.01 | 0.01 |
| gene-LOC105275374 | https://www.ncbi.nlm.nih.gov/gene/?term=LOC105275374 | 11376 | 454 | 399.09 | 1 | 0.88 | 0.22 | 10.82 | 10.43 | 0.38 | 0.38 | 0.02 | 0.02 |
| gene-LOC105287646 | https://www.ncbi.nlm.nih.gov/gene/?term=LOC105287646 | 3556 | 83 | 233.41 | 1 | 2.81 | 1.2 | 17.22 | 16.85 | 0.38 | 0.38 | 0.01 | 0.01 |
| gene-LOC105287550 | https://www.ncbi.nlm.nih.gov/gene/?term=LOC105287550 | 15659 | 549 | 350.6 | 1 | 0.64 | 0.18 | 6.33 | 5.97 | 0.37 | 0.37 | 0.01 | 0.01 |
| gene-LOC105285406 | https://www.ncbi.nlm.nih.gov/gene/?term=LOC105285406 | 8015 | 60 | 74.86 | 1 | 1.25 | 1.67 | 25.86 | 25.52 | 0.34 | 0.34 | 0 | 0 |
| gene-LOC105274856 | https://www.ncbi.nlm.nih.gov/gene/?term=LOC105274856 | 5216 | 209 | 400.69 | 1 | 1.92 | 0.48 | 18.76 | 18.44 | 0.31 | 0.31 | 0.01 | 0.01 |
| gene-LOC105274699 | https://www.ncbi.nlm.nih.gov/gene/?term=LOC105274699 | 2386 | 49 | 205.36 | 1 | 4.19 | 2.04 | 23.32 | 23.03 | 0.29 | 0.29 | 0.01 | 0.01 |
| gene-LOC105274531 | https://www.ncbi.nlm.nih.gov/gene/?term=LOC105274531 | 7774 | 273 | 351.17 | 1 | 1.29 | 0.37 | 11.43 | 11.16 | 0.27 | 0.27 | 0.01 | 0.01 |
| gene-LOC105283649 | https://www.ncbi.nlm.nih.gov/gene/?term=LOC105283649 | 1918 | 53 | 276.33 | 1 | 5.21 | 1.89 | 37.66 | 37.39 | 0.27 | 0.27 | 0.01 | 0.01 |
| gene-LOC105274851 | https://www.ncbi.nlm.nih.gov/gene/?term=LOC105274851 | 6616 | 129 | 194.98 | 1 | 1.51 | 0.78 | 11.16 | 10.94 | 0.22 | 0.22 | 0 | 0 |
| gene-LOC105279038 | https://www.ncbi.nlm.nih.gov/gene/?term=LOC105279038 | 9453 | 297 | 314.19 | 1 | 1.06 | 0.34 | 9.64 | 9.43 | 0.21 | 0.21 | 0.01 | 0.01 |
| gene-LOC105282149 | https://www.ncbi.nlm.nih.gov/gene/?term=LOC105282149 | 1787 | 103 | 576.39 | 1 | 5.6 | 0.97 | 41.32 | 41.12 | 0.2 | 0.2 | 0.01 | 0.01 |
| gene-LOC105275187 | https://www.ncbi.nlm.nih.gov/gene/?term=LOC105275187 | 5534 | 65 | 117.46 | 1 | 1.81 | 1.54 | 11.46 | 11.27 | 0.19 | 0.19 | 0 | 0 |
| gene-LOC105275327 | https://www.ncbi.nlm.nih.gov/gene/?term=LOC105275327 | 14245 | 458 | 321.52 | 1 | 0.7 | 0.22 | 7.5 | 7.33 | 0.17 | 0.17 | 0.01 | 0.01 |
| gene-LOC105278666 | https://www.ncbi.nlm.nih.gov/gene/?term=LOC105278666 | 10341 | 439 | 424.52 | 1 | 0.97 | 0.23 | 7.06 | 6.9 | 0.16 | 0.16 | 0.01 | 0.01 |
| gene-LOC105286194 | https://www.ncbi.nlm.nih.gov/gene/?term=LOC105286194 | 152619 | 1987 | 130.19 | 1 | 0.07 | 0.05 | 6.27 | 6.13 | 0.14 | 0.14 | 0 | 0 |
| gene-LOC105278649 | https://www.ncbi.nlm.nih.gov/gene/?term=LOC105278649 | 5616 | 138 | 245.73 | 1 | 1.78 | 0.72 | 13.54 | 13.42 | 0.12 | 0.12 | 0 | 0 |
| gene-LOC105288079 | https://www.ncbi.nlm.nih.gov/gene/?term=LOC105288079 | 3473 | 96 | 276.42 | 1 | 2.88 | 1.04 | 19.82 | 19.7 | 0.12 | 0.12 | 0 | 0 |
| gene-LOC105278455 | https://www.ncbi.nlm.nih.gov/gene/?term=LOC105278455 | 27181 | 875 | 321.92 | 1 | 0.37 | 0.11 | 7.69 | 7.59 | 0.1 | 0.1 | 0 | 0 |
| gene-LOC105277856 | https://www.ncbi.nlm.nih.gov/gene/?term=LOC105277856 | 3442 | 29 | 84.25 | 1 | 2.91 | 3.45 | 32.81 | 32.75 | 0.06 | 0.06 | 0 | 0 |
| gene-LOC105284355 | https://www.ncbi.nlm.nih.gov/gene/?term=LOC105284355 | 3904 | 152 | 389.34 | 1 | 2.56 | 0.66 | 20.02 | 19.96 | 0.06 | 0.06 | 0 | 0 |
| gene-LOC105287528 | https://www.ncbi.nlm.nih.gov/gene/?term=LOC105287528 | 14195 | 375 | 264.18 | 1 | 0.7 | 0.27 | 5.12 | 5.07 | 0.05 | 0.05 | 0 | 0 |
| gene-LOC105277866 | https://www.ncbi.nlm.nih.gov/gene/?term=LOC105277866 | 33045 | 368 | 111.36 | 1 | 0.3 | 0.27 | 3.48 | 3.54 | -0.06 | 0.06 | 0 | 0 |
| gene-LOC105282811 | https://www.ncbi.nlm.nih.gov/gene/?term=LOC105282811 | 5765 | 196 | 339.98 | 1 | 1.73 | 0.51 | 8.25 | 8.31 | -0.06 | 0.06 | 0 | 0 |
| gene-LOC105284704 | https://www.ncbi.nlm.nih.gov/gene/?term=LOC105284704 | 2400 | 59 | 245.83 | 1 | 4.17 | 1.69 | 22.63 | 22.7 | -0.07 | 0.07 | 0 | 0 |
| gene-LOC105274706 | https://www.ncbi.nlm.nih.gov/gene/?term=LOC105274706 | 1446 | 49 | 338.87 | 1 | 6.92 | 2.04 | 38.21 | 38.31 | -0.1 | 0.1 | 0 | 0 |
| gene-LOC105286274 | https://www.ncbi.nlm.nih.gov/gene/?term=LOC105286274 | 3817 | 179 | 468.95 | 1 | 2.62 | 0.56 | 16.95 | 17.06 | -0.11 | 0.11 | -0.01 | 0.01 |
| gene-LOC105278282 | https://www.ncbi.nlm.nih.gov/gene/?term=LOC105278282 | 1775 | 52 | 292.96 | 1 | 5.63 | 1.92 | 41.01 | 41.14 | -0.14 | 0.14 | 0 | 0 |
| gene-LOC105274590 | https://www.ncbi.nlm.nih.gov/gene/?term=LOC105274590 | 4516 | 67 | 148.36 | 1 | 2.21 | 1.49 | 28.5 | 28.65 | -0.15 | 0.15 | 0 | 0 |
| gene-LOC105282067 | https://www.ncbi.nlm.nih.gov/gene/?term=LOC105282067 | 5941 | 49 | 82.48 | 1 | 1.68 | 2.04 | 25.2 | 25.4 | -0.2 | 0.2 | 0 | 0 |
| gene-LOC105282658 | https://www.ncbi.nlm.nih.gov/gene/?term=LOC105282658 | 2421 | 52 | 214.79 | 1 | 4.13 | 1.92 | 21.81 | 22.01 | -0.2 | 0.2 | 0 | 0 |
| gene-LOC105276527 | https://www.ncbi.nlm.nih.gov/gene/?term=LOC105276527 | 7923 | 305 | 384.96 | 1 | 1.26 | 0.33 | 9.71 | 9.92 | -0.21 | 0.21 | -0.01 | 0.01 |
| gene-LOC105287095 | https://www.ncbi.nlm.nih.gov/gene/?term=LOC105287095 | 1435 | 58 | 404.18 | 1 | 6.97 | 1.72 | 28.41 | 28.65 | -0.24 | 0.24 | -0.01 | 0.01 |
| gene-LOC105288042 | https://www.ncbi.nlm.nih.gov/gene/?term=LOC105288042 | 3507 | 88 | 250.93 | 1 | 2.85 | 1.14 | 21.32 | 21.59 | -0.27 | 0.27 | -0.01 | 0.01 |
| gene-LOC105285807 | https://www.ncbi.nlm.nih.gov/gene/?term=LOC105285807 | 6063 | 213 | 351.31 | 1 | 1.65 | 0.47 | 9.29 | 9.56 | -0.28 | 0.28 | -0.01 | 0.01 |
| gene-LOC105286515 | https://www.ncbi.nlm.nih.gov/gene/?term=LOC105286515 | 3751 | 107 | 285.26 | 1 | 2.67 | 0.93 | 12.3 | 12.58 | -0.28 | 0.28 | -0.01 | 0.01 |
| gene-LOC105280774 | https://www.ncbi.nlm.nih.gov/gene/?term=LOC105280774 | 2793 | 100 | 358.04 | 1 | 3.58 | 1 | 31.9 | 32.19 | -0.29 | 0.29 | -0.01 | 0.01 |
| gene-LOC105278228 | https://www.ncbi.nlm.nih.gov/gene/?term=LOC105278228 | 8610 | 156 | 181.18 | 1 | 1.16 | 0.64 | 22.2 | 22.5 | -0.3 | 0.3 | -0.01 | 0.01 |
| gene-LOC105284036 | https://www.ncbi.nlm.nih.gov/gene/?term=LOC105284036 | 3254 | 96 | 295.02 | 1 | 3.07 | 1.04 | 23.11 | 23.46 | -0.35 | 0.35 | -0.01 | 0.01 |
| gene-LOC105280348 | https://www.ncbi.nlm.nih.gov/gene/?term=LOC105280348 | 3797 | 71 | 186.99 | 1 | 2.63 | 1.41 | 20.69 | 21.13 | -0.44 | 0.44 | -0.01 | 0.01 |
| gene-LOC105274449 | https://www.ncbi.nlm.nih.gov/gene/?term=LOC105274449 | 1549 | 48 | 309.88 | 1 | 6.46 | 2.08 | 50.45 | 50.96 | -0.51 | 0.51 | -0.02 | 0.02 |
| gene-LOC105278877 | https://www.ncbi.nlm.nih.gov/gene/?term=LOC105278877 | 1968 | 70 | 355.69 | 1 | 5.08 | 1.43 | 34.63 | 35.21 | -0.58 | 0.58 | -0.02 | 0.02 |
| gene-LOC105280377 | https://www.ncbi.nlm.nih.gov/gene/?term=LOC105280377 | 2024 | 65 | 321.15 | 1 | 4.94 | 1.54 | 47.38 | 47.97 | -0.59 | 0.59 | -0.02 | 0.02 |
| gene-LOC105278962 | https://www.ncbi.nlm.nih.gov/gene/?term=LOC105278962 | 4491 | 89 | 198.17 | 1 | 2.23 | 1.12 | 22.04 | 22.67 | -0.63 | 0.63 | -0.01 | 0.01 |
| gene-LOC105278454 | https://www.ncbi.nlm.nih.gov/gene/?term=LOC105278454 | 1972 | 40 | 202.84 | 1 | 5.07 | 2.5 | 48.64 | 49.28 | -0.64 | 0.64 | -0.01 | 0.01 |
| gene-LOC105280002 | https://www.ncbi.nlm.nih.gov/gene/?term=LOC105280002 | 1940 | 27 | 139.18 | 1 | 5.15 | 3.7 | 33.73 | 34.37 | -0.64 | 0.64 | -0.01 | 0.01 |
| gene-LOC105274534 | https://www.ncbi.nlm.nih.gov/gene/?term=LOC105274534 | 1329 | 57 | 428.89 | 1 | 7.52 | 1.75 | 46.71 | 47.65 | -0.94 | 0.94 | -0.04 | 0.04 |
| gene-LOC105280819 | https://www.ncbi.nlm.nih.gov/gene/?term=LOC105280819 | 1639 | 37 | 225.75 | 1 | 6.1 | 2.7 | 36.94 | 38.07 | -1.12 | 1.12 | -0.03 | 0.03 |
| gene-LOC105278384 | https://www.ncbi.nlm.nih.gov/gene/?term=LOC105278384 | 1390 | 35 | 251.8 | 1 | 7.19 | 2.86 | 48.96 | 50.26 | -1.3 | 1.3 | -0.03 | 0.03 |
| gene-LOC105283712 | https://www.ncbi.nlm.nih.gov/gene/?term=LOC105283712 | 1910 | 36 | 188.48 | 1 | 5.24 | 2.78 | 22.92 | 24.43 | -1.51 | 1.51 | -0.03 | 0.03 |
| gene-LOC105281700 | https://www.ncbi.nlm.nih.gov/gene/?term=LOC105281700 | 2208 | 32 | 144.93 | 1 | 4.53 | 3.12 | 37.54 | 39.09 | -1.55 | 1.55 | -0.02 | 0.02 |
| gene-LOC105276538 | https://www.ncbi.nlm.nih.gov/gene/?term=LOC105276538 | 1511 | 18 | 119.13 | 1 | 6.62 | 5.56 | 64.69 | 66.35 | -1.65 | 1.65 | -0.02 | 0.02 |
| gene-LOC105286514 | https://www.ncbi.nlm.nih.gov/gene/?term=LOC105286514 | 2528 | 46 | 181.96 | 1 | 3.96 | 2.17 | 35.96 | 37.79 | -1.83 | 1.83 | -0.03 | 0.03 |
| gene-LOC105280596 | https://www.ncbi.nlm.nih.gov/gene/?term=LOC105280596 | 3434 | 104 | 302.85 | 1 | 2.91 | 0.96 | 16.78 | 18.68 | -1.9 | 1.9 | -0.06 | 0.06 |
| gene-LOC105283989 | https://www.ncbi.nlm.nih.gov/gene/?term=LOC105283989 | 4948 | 25 | 50.53 | 1 | 2.02 | 4 | 35.77 | 37.79 | -2.02 | 2.02 | -0.01 | 0.01 |
| gene-LOC105286999 | https://www.ncbi.nlm.nih.gov/gene/?term=LOC105286999 | 2141 | 42 | 196.17 | 1 | 4.67 | 2.38 | 33.74 | 35.94 | -2.2 | 2.2 | -0.04 | 0.04 |
| gene-LOC105283967 | https://www.ncbi.nlm.nih.gov/gene/?term=LOC105283967 | 2393 | 40 | 167.15 | 1 | 4.18 | 2.5 | 52.37 | 54.65 | -2.28 | 2.28 | -0.04 | 0.04 |
| gene-LOC105276196 | https://www.ncbi.nlm.nih.gov/gene/?term=LOC105276196 | 2953 | 59 | 199.8 | 1 | 3.39 | 1.69 | 38.71 | 41.49 | -2.77 | 2.77 | -0.06 | 0.06 |
| gene-LOC105285409 | https://www.ncbi.nlm.nih.gov/gene/?term=LOC105285409 | 5099 | 49 | 96.1 | 1 | 1.96 | 2.04 | 1.72 | 4.82 | -3.09 | 3.09 | -0.03 | 0.03 |
| gene-LOC105283492 | https://www.ncbi.nlm.nih.gov/gene/?term=LOC105283492 | 1535 | 9 | 58.63 | 1 | 6.51 | 11.11 | 30.57 | 36.97 | -6.4 | 6.4 | -0.04 | 0.04 |

Directory: OGLall File: ogl-Ob_LI_RphaseA2.txt

| gene_ID | gene_link | gwidth | #Sites | #per10Kb | %perSite | %pNucl |
|---|---|---|---|---|---|---|
| gene-LOC113561532 | https://www.ncbi.nlm.nih.gov/gene/?term=LOC113561532 | 72 | 6 | 833.33 | 95.64 | 7.97 |
| gene-LOC105282149 | https://www.ncbi.nlm.nih.gov/gene/?term=LOC105282149 | 1787 | 190 | 1063.23 | 44.1 | 4.69 |
| gene-LOC113562627 | https://www.ncbi.nlm.nih.gov/gene/?term=LOC113562627 | 397 | 26 | 654.91 | 67.37 | 4.41 |
| gene-LOC105285635 | https://www.ncbi.nlm.nih.gov/gene/?term=LOC105285635 | 1175 | 164 | 1395.74 | 29.93 | 4.18 |
| gene-LOC105283830 | https://www.ncbi.nlm.nih.gov/gene/?term=LOC105283830 | 252 | 24 | 952.38 | 43.77 | 4.17 |
| gene-LOC105286269 | https://www.ncbi.nlm.nih.gov/gene/?term=LOC105286269 | 1401 | 110 | 785.15 | 52.63 | 4.13 |
| gene-LOC105286531 | https://www.ncbi.nlm.nih.gov/gene/?term=LOC105286531 | 372 | 43 | 1155.91 | 35.68 | 4.12 |
| gene-LOC105280739 | https://www.ncbi.nlm.nih.gov/gene/?term=LOC105280739 | 1991 | 147 | 738.32 | 54.96 | 4.06 |
| gene-LOC105287811 | https://www.ncbi.nlm.nih.gov/gene/?term=LOC105287811 | 2155 | 191 | 886.31 | 45.74 | 4.05 |
| gene-LOC105278849 | https://www.ncbi.nlm.nih.gov/gene/?term=LOC105278849 | 776 | 127 | 1636.6 | 24.48 | 4.01 |
| gene-LOC105276360 | https://www.ncbi.nlm.nih.gov/gene/?term=LOC105276360 | 1466 | 203 | 1384.72 | 28.73 | 3.98 |
| gene-LOC105285394 | https://www.ncbi.nlm.nih.gov/gene/?term=LOC105285394 | 1307 | 74 | 566.18 | 69.6 | 3.94 |
| gene-LOC105286801 | https://www.ncbi.nlm.nih.gov/gene/?term=LOC105286801 | 1102 | 107 | 970.96 | 40.62 | 3.94 |
| gene-LOC105274534 | https://www.ncbi.nlm.nih.gov/gene/?term=LOC105274534 | 1329 | 119 | 895.41 | 42.8 | 3.83 |
| gene-LOC105275314 | https://www.ncbi.nlm.nih.gov/gene/?term=LOC105275314 | 1489 | 132 | 886.5 | 41.45 | 3.67 |
| gene-LOC105277648 | https://www.ncbi.nlm.nih.gov/gene/?term=LOC105277648 | 722 | 45 | 623.27 | 58.13 | 3.62 |
| gene-LOC105278160 | https://www.ncbi.nlm.nih.gov/gene/?term=LOC105278160 | 1872 | 184 | 982.91 | 36.84 | 3.62 |
| gene-LOC105279052 | https://www.ncbi.nlm.nih.gov/gene/?term=LOC105279052 | 1130 | 163 | 1442.48 | 24.87 | 3.59 |
| gene-LOC105274711 | https://www.ncbi.nlm.nih.gov/gene/?term=LOC105274711 | 1655 | 124 | 749.24 | 46.91 | 3.51 |
| gene-LOC105278844 | https://www.ncbi.nlm.nih.gov/gene/?term=LOC105278844 | 1101 | 139 | 1262.49 | 27.76 | 3.5 |
| gene-LOC105287677 | https://www.ncbi.nlm.nih.gov/gene/?term=LOC105287677 | 1588 | 143 | 900.5 | 38.14 | 3.43 |
| gene-LOC105286235 | https://www.ncbi.nlm.nih.gov/gene/?term=LOC105286235 | 1510 | 122 | 807.95 | 42.1 | 3.4 |
| gene-LOC105286737 | https://www.ncbi.nlm.nih.gov/gene/?term=LOC105286737 | 2690 | 404 | 1501.86 | 22.66 | 3.4 |
| gene-LOC105287692 | https://www.ncbi.nlm.nih.gov/gene/?term=LOC105287692 | 1505 | 156 | 1036.54 | 32.62 | 3.38 |

...
... only first 25 lines shown ...

Directory: OGLall File: ogl-Ob_LI_RphaseA2.vs.RphaseB1.txt

| gene_ID | gene_link | gwidth | #Sites | #per10Kb | #dmSites | #dmsp10kb | %dmSites | %pSite1 | %pSite2 | DMpSite | ADMpSite | DMpNucl | ADMpNucl |
|---|---|---|---|---|---|---|---|---|---|---|---|---|---|
| gene-LOC113562974 | https://www.ncbi.nlm.nih.gov/gene/?term=LOC113562974 | 1832 | 2 | 10.92 | 1 | 5.46 | 50 | 57.94 | 9 | 48.94 | 48.94 | 0.05 | 0.05 |
| gene-LOC113562345 | https://www.ncbi.nlm.nih.gov/gene/?term=LOC113562345 | 2290 | 80 | 349.34 | 16 | 69.87 | 20 | 25.78 | 13.98 | 11.8 | 11.8 | 0.41 | 0.41 |
| gene-LOC105275070 | https://www.ncbi.nlm.nih.gov/gene/?term=LOC105275070 | 5667 | 64 | 112.93 | 6 | 10.59 | 9.38 | 14.32 | 5.2 | 9.13 | 9.13 | 0.1 | 0.1 |
| gene-LOC105275708 | https://www.ncbi.nlm.nih.gov/gene/?term=LOC105275708 | 1772 | 19 | 107.22 | 1 | 5.64 | 5.26 | 37.25 | 31.29 | 5.96 | 5.96 | 0.06 | 0.06 |
| gene-LOC105285514 | https://www.ncbi.nlm.nih.gov/gene/?term=LOC105285514 | 1277 | 11 | 86.14 | 1 | 7.83 | 9.09 | 90.74 | 85.12 | 5.62 | 5.62 | 0.05 | 0.05 |
| gene-LOC105286302 | https://www.ncbi.nlm.nih.gov/gene/?term=LOC105286302 | 4536 | 24 | 52.91 | 1 | 2.2 | 4.17 | 20.45 | 15.15 | 5.3 | 5.3 | 0.03 | 0.03 |
| gene-LOC105286983 | https://www.ncbi.nlm.nih.gov/gene/?term=LOC105286983 | 16901 | 70 | 41.42 | 1 | 0.59 | 1.43 | 6.21 | 1.14 | 5.06 | 5.06 | 0.02 | 0.02 |
| gene-LOC105281039 | https://www.ncbi.nlm.nih.gov/gene/?term=LOC105281039 | 1471 | 15 | 101.97 | 1 | 6.8 | 6.67 | 36.89 | 31.93 | 4.96 | 4.96 | 0.05 | 0.05 |
| gene-LOC105279900 | https://www.ncbi.nlm.nih.gov/gene/?term=LOC105279900 | 2191 | 56 | 255.59 | 1 | 4.56 | 1.79 | 33.63 | 28.91 | 4.72 | 4.72 | 0.12 | 0.12 |
| gene-LOC105284609 | https://www.ncbi.nlm.nih.gov/gene/?term=LOC105284609 | 5678 | 44 | 77.49 | 1 | 1.76 | 2.27 | 32.19 | 27.8 | 4.39 | 4.39 | 0.03 | 0.03 |
| gene-LOC105276538 | https://www.ncbi.nlm.nih.gov/gene/?term=LOC105276538 | 1511 | 23 | 152.22 | 1 | 6.62 | 4.35 | 69.63 | 65.38 | 4.25 | 4.25 | 0.06 | 0.06 |
| gene-LOC105285315 | https://www.ncbi.nlm.nih.gov/gene/?term=LOC105285315 | 5458 | 33 | 60.46 | 1 | 1.83 | 3.03 | 33.27 | 29.4 | 3.88 | 3.88 | 0.02 | 0.02 |
| gene-LOC105283712 | https://www.ncbi.nlm.nih.gov/gene/?term=LOC105283712 | 1910 | 33 | 172.77 | 1 | 5.24 | 3.03 | 32.87 | 29.11 | 3.76 | 3.76 | 0.06 | 0.06 |
| gene-LOC105282893 | https://www.ncbi.nlm.nih.gov/gene/?term=LOC105282893 | 2645 | 50 | 189.04 | 1 | 3.78 | 2 | 28.92 | 25.31 | 3.6 | 3.6 | 0.07 | 0.07 |
| gene-LOC105281515 | https://www.ncbi.nlm.nih.gov/gene/?term=LOC105281515 | 8827 | 56 | 63.44 | 2 | 2.27 | 3.57 | 36.05 | 32.61 | 3.44 | 3.44 | 0.02 | 0.02 |
| gene-LOC105287063 | https://www.ncbi.nlm.nih.gov/gene/?term=LOC105287063 | 1874 | 55 | 293.49 | 1 | 5.34 | 1.82 | 30.47 | 27.11 | 3.37 | 3.37 | 0.1 | 0.1 |
| gene-LOC105283457 | https://www.ncbi.nlm.nih.gov/gene/?term=LOC105283457 | 1882 | 49 | 260.36 | 1 | 5.31 | 2.04 | 35.19 | 32.03 | 3.16 | 3.16 | 0.08 | 0.08 |
| gene-LOC105287417 | https://www.ncbi.nlm.nih.gov/gene/?term=LOC105287417 | 1025 | 38 | 370.73 | 1 | 9.76 | 2.63 | 35.24 | 32.37 | 2.87 | 2.87 | 0.11 | 0.11 |
| gene-LOC105288066 | https://www.ncbi.nlm.nih.gov/gene/?term=LOC105288066 | 1964 | 31 | 157.84 | 1 | 5.09 | 3.23 | 39.38 | 36.57 | 2.81 | 2.81 | 0.04 | 0.04 |
| gene-LOC105284704 | https://www.ncbi.nlm.nih.gov/gene/?term=LOC105284704 | 2400 | 66 | 275 | 1 | 4.17 | 1.52 | 25.49 | 22.92 | 2.57 | 2.57 | 0.07 | 0.07 |
| gene-LOC105285583 | https://www.ncbi.nlm.nih.gov/gene/?term=LOC105285583 | 1947 | 62 | 318.44 | 1 | 5.14 | 1.61 | 27.07 | 24.61 | 2.46 | 2.46 | 0.08 | 0.08 |
| gene-LOC105275190 | https://www.ncbi.nlm.nih.gov/gene/?term=LOC105275190 | 3837 | 77 | 200.68 | 1 | 2.61 | 1.3 | 21.84 | 19.45 | 2.39 | 2.39 | 0.05 | 0.05 |
| gene-LOC113561805 | https://www.ncbi.nlm.nih.gov/gene/?term=LOC113561805 | 3337 | 73 | 218.76 | 1 | 3 | 1.37 | 38.02 | 35.67 | 2.34 | 2.34 | 0.05 | 0.05 |
| gene-LOC105279898 | https://www.ncbi.nlm.nih.gov/gene/?term=LOC105279898 | 1343 | 16 | 119.14 | 1 | 7.45 | 6.25 | 58.57 | 56.24 | 2.33 | 2.33 | 0.03 | 0.03 |
| gene-LOC105275518 | https://www.ncbi.nlm.nih.gov/gene/?term=LOC105275518 | 2335 | 28 | 119.91 | 1 | 4.28 | 3.57 | 41.74 | 39.46 | 2.28 | 2.28 | 0.03 | 0.03 |
| gene-LOC105285695 | https://www.ncbi.nlm.nih.gov/gene/?term=LOC105285695 | 1371 | 76 | 554.34 | 1 | 7.29 | 1.32 | 28.59 | 26.36 | 2.24 | 2.24 | 0.12 | 0.12 |
| gene-LOC105274706 | https://www.ncbi.nlm.nih.gov/gene/?term=LOC105274706 | 1446 | 56 | 387.28 | 1 | 6.92 | 1.79 | 39.8 | 37.63 | 2.17 | 2.17 | 0.08 | 0.08 |
| gene-LOC105288010 | https://www.ncbi.nlm.nih.gov/gene/?term=LOC105288010 | 3189 | 61 | 191.28 | 1 | 3.14 | 1.64 | 34.85 | 32.7 | 2.15 | 2.15 | 0.04 | 0.04 |
| gene-LOC105282156 | https://www.ncbi.nlm.nih.gov/gene/?term=LOC105282156 | 2577 | 134 | 519.98 | 1 | 3.88 | 0.75 | 27.52 | 25.46 | 2.06 | 2.06 | 0.11 | 0.11 |
| gene-LOC105285674 | https://www.ncbi.nlm.nih.gov/gene/?term=LOC105285674 | 3035 | 111 | 365.73 | 2 | 6.59 | 1.8 | 25.15 | 23.09 | 2.06 | 2.06 | 0.08 | 0.08 |
| gene-LOC105286644 | https://www.ncbi.nlm.nih.gov/gene/?term=LOC105286644 | 1235 | 23 | 186.23 | 1 | 8.1 | 4.35 | 43.78 | 41.75 | 2.04 | 2.04 | 0.04 | 0.04 |
| gene-LOC105285060 | https://www.ncbi.nlm.nih.gov/gene/?term=LOC105285060 | 1285 | 18 | 140.08 | 1 | 7.78 | 5.56 | 38.72 | 36.69 | 2.03 | 2.03 | 0.03 | 0.03 |
| gene-LOC105280786 | https://www.ncbi.nlm.nih.gov/gene/?term=LOC105280786 | 2573 | 143 | 555.77 | 1 | 3.89 | 0.7 | 27.63 | 25.63 | 2 | 2 | 0.11 | 0.11 |
| gene-LOC105276704 | https://www.ncbi.nlm.nih.gov/gene/?term=LOC105276704 | 2063 | 93 | 450.8 | 5 | 24.24 | 5.38 | 14.49 | 12.51 | 1.99 | 1.99 | 0.09 | 0.09 |
| gene-LOC105276272 | https://www.ncbi.nlm.nih.gov/gene/?term=LOC105276272 | 1965 | 47 | 239.19 | 1 | 5.09 | 2.13 | 25.43 | 23.44 | 1.98 | 1.98 | 0.05 | 0.05 |
| gene-LOC105286683 | https://www.ncbi.nlm.nih.gov/gene/?term=LOC105286683 | 1665 | 57 | 342.34 | 1 | 6.01 | 1.75 | 35.23 | 33.27 | 1.97 | 1.97 | 0.07 | 0.07 |
| gene-LOC105281848 | https://www.ncbi.nlm.nih.gov/gene/?term=LOC105281848 | 2122 | 60 | 282.75 | 1 | 4.71 | 1.67 | 24.48 | 22.53 | 1.95 | 1.95 | 0.06 | 0.06 |
| gene-LOC105274900 | https://www.ncbi.nlm.nih.gov/gene/?term=LOC105274900 | 5561 | 61 | 109.69 | 2 | 3.6 | 3.28 | 32.53 | 30.66 | 1.87 | 1.87 | 0.02 | 0.02 |
| gene-LOC105276078 | https://www.ncbi.nlm.nih.gov/gene/?term=LOC105276078 | 1051 | 18 | 171.27 | 1 | 9.51 | 5.56 | 28.41 | 26.59 | 1.83 | 1.83 | 0.03 | 0.03 |
| gene-LOC105280704 | https://www.ncbi.nlm.nih.gov/gene/?term=LOC105280704 | 4266 | 155 | 363.34 | 1 | 2.34 | 0.65 | 17.59 | 16.02 | 1.58 | 1.58 | 0.06 | 0.06 |
| gene-LOC105278108 | https://www.ncbi.nlm.nih.gov/gene/?term=LOC105278108 | 4392 | 57 | 129.78 | 1 | 2.28 | 1.75 | 25.91 | 24.39 | 1.52 | 1.52 | 0.02 | 0.02 |
| gene-LOC105278282 | https://www.ncbi.nlm.nih.gov/gene/?term=LOC105278282 | 1775 | 67 | 377.46 | 1 | 5.63 | 1.49 | 34.11 | 32.61 | 1.51 | 1.51 | 0.06 | 0.06 |
| gene-LOC105288079 | https://www.ncbi.nlm.nih.gov/gene/?term=LOC105288079 | 3473 | 112 | 322.49 | 1 | 2.88 | 0.89 | 18.94 | 17.43 | 1.51 | 1.51 | 0.05 | 0.05 |
| gene-LOC105276449 | https://www.ncbi.nlm.nih.gov/gene/?term=LOC105276449 | 1231 | 55 | 446.79 | 1 | 8.12 | 1.82 | 24.75 | 23.28 | 1.47 | 1.47 | 0.07 | 0.07 |
| gene-LOC105280348 | https://www.ncbi.nlm.nih.gov/gene/?term=LOC105280348 | 3797 | 78 | 205.43 | 1 | 2.63 | 1.28 | 18.55 | 17.12 | 1.44 | 1.44 | 0.03 | 0.03 |
| gene-LOC105277435 | https://www.ncbi.nlm.nih.gov/gene/?term=LOC105277435 | 3643 | 184 | 505.08 | 1 | 2.74 | 0.54 | 12.13 | 10.74 | 1.39 | 1.39 | 0.07 | 0.07 |
| gene-LOC105279298 | https://www.ncbi.nlm.nih.gov/gene/?term=LOC105279298 | 2773 | 134 | 483.23 | 2 | 7.21 | 1.49 | 24.2 | 22.81 | 1.39 | 1.39 | 0.07 | 0.07 |
| gene-LOC105282151 | https://www.ncbi.nlm.nih.gov/gene/?term=LOC105282151 | 1601 | 45 | 281.07 | 2 | 12.49 | 4.44 | 42.82 | 41.43 | 1.39 | 1.39 | 0.04 | 0.04 |
| gene-LOC105280002 | https://www.ncbi.nlm.nih.gov/gene/?term=LOC105280002 | 1940 | 34 | 175.26 | 1 | 5.15 | 2.94 | 33.64 | 32.26 | 1.38 | 1.38 | 0.02 | 0.02 |
| gene-LOC105282794 | https://www.ncbi.nlm.nih.gov/gene/?term=LOC105282794 | 4571 | 109 | 238.46 | 1 | 2.19 | 0.92 | 17.81 | 16.42 | 1.38 | 1.38 | 0.03 | 0.03 |
| gene-LOC105286514 | https://www.ncbi.nlm.nih.gov/gene/?term=LOC105286514 | 2528 | 55 | 217.56 | 1 | 3.96 | 1.82 | 37.15 | 35.79 | 1.36 | 1.36 | 0.03 | 0.03 |
| gene-LOC105285671 | https://www.ncbi.nlm.nih.gov/gene/?term=LOC105285671 | 2263 | 103 | 455.15 | 1 | 4.42 | 0.97 | 30.91 | 29.64 | 1.27 | 1.27 | 0.06 | 0.06 |
| gene-LOC105280722 | https://www.ncbi.nlm.nih.gov/gene/?term=LOC105280722 | 3237 | 93 | 287.3 | 1 | 3.09 | 1.08 | 27.81 | 26.63 | 1.18 | 1.18 | 0.03 | 0.03 |
| gene-LOC105279500 | https://www.ncbi.nlm.nih.gov/gene/?term=LOC105279500 | 3587 | 36 | 100.36 | 1 | 2.79 | 2.78 | 48.81 | 47.64 | 1.17 | 1.17 | 0.01 | 0.01 |
| gene-LOC105282682 | https://www.ncbi.nlm.nih.gov/gene/?term=LOC105282682 | 4127 | 240 | 581.54 | 1 | 2.42 | 0.42 | 13.64 | 12.51 | 1.14 | 1.14 | 0.07 | 0.07 |
| gene-LOC105288063 | https://www.ncbi.nlm.nih.gov/gene/?term=LOC105288063 | 3400 | 71 | 208.82 | 1 | 2.94 | 1.41 | 12.55 | 11.41 | 1.14 | 1.14 | 0.02 | 0.02 |
| gene-LOC105282600 | https://www.ncbi.nlm.nih.gov/gene/?term=LOC105282600 | 3715 | 67 | 180.35 | 1 | 2.69 | 1.49 | 27.6 | 26.48 | 1.12 | 1.12 | 0.02 | 0.02 |
| gene-LOC105283123 | https://www.ncbi.nlm.nih.gov/gene/?term=LOC105283123 | 1507 | 55 | 364.96 | 1 | 6.64 | 1.82 | 20.08 | 18.96 | 1.12 | 1.12 | 0.04 | 0.04 |
| gene-LOC105278071 | https://www.ncbi.nlm.nih.gov/gene/?term=LOC105278071 | 4307 | 61 | 141.63 | 1 | 2.32 | 1.64 | 11.12 | 10.02 | 1.1 | 1.1 | 0.02 | 0.02 |
| gene-LOC105285416 | https://www.ncbi.nlm.nih.gov/gene/?term=LOC105285416 | 3820 | 76 | 198.95 | 1 | 2.62 | 1.32 | 22.22 | 21.12 | 1.1 | 1.1 | 0.02 | 0.02 |

| | | | | | | | | | | | | | |
|---|---|---|---|---|---|---|---|---|---|---|---|---|---|
| gene-LOC105277856 | https://www.ncbi.nlm.nih.gov/gene/?term=LOC105277856 | 3442 | 36 | 104.59 | 1 | 2.91 | 2.78 | 37.81 | 36.75 | 1.06 | 1.06 | 0.01 | 0.01 |
| gene-LOC105280407 | https://www.ncbi.nlm.nih.gov/gene/?term=LOC105280407 | 10950 | 231 | 210.96 | 1 | 0.91 | 0.43 | 15.59 | 14.54 | 1.05 | 1.05 | 0.02 | 0.02 |
| gene-LOC105283838 | https://www.ncbi.nlm.nih.gov/gene/?term=LOC105283838 | 1748 | 33 | 188.79 | 1 | 5.72 | 3.03 | 33.49 | 32.48 | 1.01 | 1.01 | 0.02 | 0.02 |
| gene-LOC105276408 | https://www.ncbi.nlm.nih.gov/gene/?term=LOC105276408 | 5298 | 201 | 379.39 | 1 | 1.89 | 0.5 | 21.53 | 20.53 | 1 | 1 | 0.04 | 0.04 |
| gene-LOC105275460 | https://www.ncbi.nlm.nih.gov/gene/?term=LOC105275460 | 2473 | 123 | 497.37 | 1 | 4.04 | 0.81 | 13.29 | 12.32 | 0.97 | 0.97 | 0.05 | 0.05 |
| gene-LOC105280774 | https://www.ncbi.nlm.nih.gov/gene/?term=LOC105280774 | 2793 | 135 | 483.35 | 1 | 3.58 | 0.74 | 25.37 | 24.41 | 0.96 | 0.96 | 0.05 | 0.05 |
| gene-LOC105278283 | https://www.ncbi.nlm.nih.gov/gene/?term=LOC105278283 | 2317 | 145 | 625.81 | 1 | 4.32 | 0.69 | 24.77 | 23.82 | 0.95 | 0.95 | 0.06 | 0.06 |
| gene-LOC105284355 | https://www.ncbi.nlm.nih.gov/gene/?term=LOC105284355 | 3904 | 155 | 397.03 | 1 | 2.56 | 0.65 | 22.33 | 21.37 | 0.95 | 0.95 | 0.04 | 0.04 |
| gene-LOC105283592 | https://www.ncbi.nlm.nih.gov/gene/?term=LOC105283592 | 3479 | 87 | 250.07 | 1 | 2.87 | 1.15 | 22.28 | 21.36 | 0.92 | 0.92 | 0.02 | 0.02 |
| gene-LOC105285055 | https://www.ncbi.nlm.nih.gov/gene/?term=LOC105285055 | 4108 | 264 | 642.65 | 2 | 4.87 | 0.76 | 12 | 11.12 | 0.87 | 0.87 | 0.06 | 0.06 |
| gene-LOC105275347 | https://www.ncbi.nlm.nih.gov/gene/?term=LOC105275347 | 2526 | 21 | 83.14 1 | 3.96 | 4.76 | 35.82 | 34.96 | 0.86 | 0.86 | 0.01 | 0.01 | |
| gene-LOC105276969 | https://www.ncbi.nlm.nih.gov/gene/?term=LOC105276969 | 2997 | 104 | 347.01 | 1 | 3.34 | 0.96 | 11.67 | 10.81 | 0.86 | 0.86 | 0.03 | 0.03 |
| gene-LOC105277631 | https://www.ncbi.nlm.nih.gov/gene/?term=LOC105277631 | 4387 | 327 | 745.38 | 1 | 2.28 | 0.31 | 11.07 | 10.23 | 0.85 | 0.85 | 0.06 | 0.06 |
| gene-LOC105278807 | https://www.ncbi.nlm.nih.gov/gene/?term=LOC105278807 | 2166 | 77 | 355.49 | 1 | 4.62 | 1.3 | 21.89 | 21.08 | 0.81 | 0.81 | 0.03 | 0.03 |
| gene-LOC105274823 | https://www.ncbi.nlm.nih.gov/gene/?term=LOC105274823 | 4620 | 171 | 370.13 | 1 | 2.16 | 0.58 | 18.07 | 17.27 | 0.8 | 0.8 | 0.03 | 0.03 |
| gene-LOC105275179 | https://www.ncbi.nlm.nih.gov/gene/?term=LOC105275179 | 3271 | 85 | 259.86 | 1 | 3.06 | 1.18 | 16.68 | 15.92 | 0.76 | 0.76 | 0.02 | 0.02 |
| gene-LOC105280104 | https://www.ncbi.nlm.nih.gov/gene/?term=LOC105280104 | 3960 | 132 | 333.33 | 1 | 2.53 | 0.76 | 17.97 | 17.22 | 0.76 | 0.76 | 0.03 | 0.03 |
| gene-LOC105285742 | https://www.ncbi.nlm.nih.gov/gene/?term=LOC105285742 | 3446 | 122 | 354.03 | 1 | 2.9 | 0.82 | 17.63 | 16.88 | 0.76 | 0.76 | 0.03 | 0.03 |
| gene-LOC105278497 | https://www.ncbi.nlm.nih.gov/gene/?term=LOC105278497 | 2072 | 103 | 497.1 1 | 4.83 | 0.97 | 19.51 | 18.76 | 0.75 | 0.75 | 0.04 | 0.04 | |
| gene-LOC105281312 | https://www.ncbi.nlm.nih.gov/gene/?term=LOC105281312 | 2551 | 69 | 270.48 | 1 | 3.92 | 1.45 | 31.17 | 30.43 | 0.74 | 0.74 | 0.02 | 0.02 |
| gene-LOC105282491 | https://www.ncbi.nlm.nih.gov/gene/?term=LOC105282491 | 4761 | 191 | 401.18 | 1 | 2.1 | 0.52 | 9 | 8.31 | 0.69 | 0.69 | 0.03 | 0.03 |
| gene-LOC105283728 | https://www.ncbi.nlm.nih.gov/gene/?term=LOC105283728 | 3256 | 159 | 488.33 | 2 | 6.14 | 1.26 | 15.6 | 14.91 | 0.69 | 0.69 | 0.03 | 0.03 |
| gene-LOC105282825 | https://www.ncbi.nlm.nih.gov/gene/?term=LOC105282825 | 3919 | 102 | 260.27 | 1 | 2.55 | 0.98 | 14.91 | 14.23 | 0.68 | 0.68 | 0.02 | 0.02 |
| gene-LOC105283989 | https://www.ncbi.nlm.nih.gov/gene/?term=LOC105283989 | 4948 | 44 | 88.92 2 | 4.04 | 4.55 | 37.47 | 36.79 | 0.68 | 0.68 | 0.01 | 0.01 | |
| gene-LOC105283943 | https://www.ncbi.nlm.nih.gov/gene/?term=LOC105283943 | 3559 | 68 | 191.06 | 1 | 2.81 | 1.47 | 20.98 | 20.31 | 0.67 | 0.67 | 0.01 | 0.01 |
| gene-LOC105284557 | https://www.ncbi.nlm.nih.gov/gene/?term=LOC105284557 | 1672 | 91 | 544.26 | 1 | 5.98 | 1.1 | 27.82 | 27.15 | 0.67 | 0.67 | 0.04 | 0.04 |
| gene-LOC105275451 | https://www.ncbi.nlm.nih.gov/gene/?term=LOC105275451 | 6035 | 149 | 246.89 | 1 | 1.66 | 0.67 | 20.25 | 19.59 | 0.66 | 0.66 | 0.02 | 0.02 |
| gene-LOC105276560 | https://www.ncbi.nlm.nih.gov/gene/?term=LOC105276560 | 18607 | 384 | 206.37 | 3 | 1.61 | 0.78 | 2.53 | 1.87 | 0.66 | 0.66 | 0.01 | 0.01 |
| gene-LOC105288078 | https://www.ncbi.nlm.nih.gov/gene/?term=LOC105288078 | 19126 | 359 | 187.7 1 | 0.52 | 0.28 | 15.58 | 14.92 | 0.66 | 0.66 | 0.01 | 0.01 | |
| gene-LOC105287704 | https://www.ncbi.nlm.nih.gov/gene/?term=LOC105287704 | 3019 | 142 | 470.35 | 1 | 3.31 | 0.7 | 19.98 | 19.34 | 0.63 | 0.63 | 0.03 | 0.03 |
| gene-LOC105275683 | https://www.ncbi.nlm.nih.gov/gene/?term=LOC105275683 | 4256 | 204 | 479.32 | 1 | 2.35 | 0.49 | 13.38 | 12.77 | 0.61 | 0.61 | 0.03 | 0.03 |
| gene-LOC105277215 | https://www.ncbi.nlm.nih.gov/gene/?term=LOC105277215 | 2619 | 30 | 114.55 | 1 | 3.82 | 3.33 | 27.61 | 26.99 | 0.61 | 0.61 | 0.01 | 0.01 |
| gene-LOC105281296 | https://www.ncbi.nlm.nih.gov/gene/?term=LOC105281296 | 2434 | 48 | 197.21 | 1 | 4.11 | 2.08 | 28.14 | 27.55 | 0.59 | 0.59 | 0.01 | 0.01 |
| gene-LOC105278384 | https://www.ncbi.nlm.nih.gov/gene/?term=LOC105278384 | 1390 | 33 | 237.41 | 1 | 7.19 | 3.03 | 46.32 | 45.77 | 0.55 | 0.55 | 0.01 | 0.01 |
| gene-LOC105278632 | https://www.ncbi.nlm.nih.gov/gene/?term=LOC105278632 | 3624 | 150 | 413.91 | 1 | 2.76 | 0.67 | 13.57 | 13.05 | 0.53 | 0.53 | 0.02 | 0.02 |
| gene-LOC105274699 | https://www.ncbi.nlm.nih.gov/gene/?term=LOC105274699 | 2386 | 67 | 280.8 1 | 4.19 | 1.49 | 18.3 | 17.79 | 0.51 | 0.51 | 0.01 | 0.01 | |
| gene-LOC105285179 | https://www.ncbi.nlm.nih.gov/gene/?term=LOC105285179 | 4191 | 204 | 486.76 | 1 | 2.39 | 0.49 | 16.93 | 16.42 | 0.51 | 0.51 | 0.02 | 0.02 |
| gene-LOC105280641 | https://www.ncbi.nlm.nih.gov/gene/?term=LOC105280641 | 3682 | 78 | 211.84 | 1 | 2.72 | 1.28 | 13.91 | 13.42 | 0.49 | 0.49 | 0.01 | 0.01 |
| gene-LOC105275752 | https://www.ncbi.nlm.nih.gov/gene/?term=LOC105275752 | 3462 | 138 | 398.61 | 1 | 2.89 | 0.72 | 31.47 | 31.01 | 0.46 | 0.46 | 0.02 | 0.02 |
| gene-LOC105279038 | https://www.ncbi.nlm.nih.gov/gene/?term=LOC105279038 | 9453 | 345 | 364.96 | 1 | 1.06 | 0.29 | 9.49 | 9.03 | 0.45 | 0.45 | 0.02 | 0.02 |
| gene-LOC105280408 | https://www.ncbi.nlm.nih.gov/gene/?term=LOC105280408 | 5983 | 65 | 108.64 | 1 | 1.67 | 1.54 | 15.19 | 14.75 | 0.44 | 0.44 | 0 | 0 |
| gene-LOC105275363 | https://www.ncbi.nlm.nih.gov/gene/?term=LOC105275363 | 3798 | 212 | 558.19 | 1 | 2.63 | 0.47 | 14.03 | 13.61 | 0.43 | 0.43 | 0.02 | 0.02 |
| gene-LOC105285807 | https://www.ncbi.nlm.nih.gov/gene/?term=LOC105285807 | 6063 | 231 | 381 1 | 1.65 | 0.43 | 10.41 | 9.98 | 0.43 | 0.43 | 0.02 | 0.02 | |
| gene-LOC105274531 | https://www.ncbi.nlm.nih.gov/gene/?term=LOC105274531 | 7774 | 307 | 394.91 | 1 | 1.29 | 0.33 | 10.8 | 10.38 | 0.42 | 0.42 | 0.02 | 0.02 |
| gene-LOC105281272 | https://www.ncbi.nlm.nih.gov/gene/?term=LOC105281272 | 5252 | 201 | 382.71 | 1 | 1.9 | 0.5 | 15.32 | 14.89 | 0.42 | 0.42 | 0.02 | 0.02 |
| gene-LOC105275374 | https://www.ncbi.nlm.nih.gov/gene/?term=LOC105275374 | 11376 | 587 | 516 1 | 0.88 | 0.17 | 9.68 | 9.27 | 0.41 | 0.41 | 0.02 | 0.02 | |
| gene-LOC105285406 | https://www.ncbi.nlm.nih.gov/gene/?term=LOC105285406 | 8015 | 63 | 78.6 1 | 1.25 | 1.59 | 28.98 | 28.57 | 0.41 | 0.41 | 0 | 0 | |
| gene-LOC105274773 | https://www.ncbi.nlm.nih.gov/gene/?term=LOC105274773 | 11034 | 187 | 169.48 | 1 | 0.91 | 0.53 | 16.91 | 16.52 | 0.39 | 0.39 | 0.01 | 0.01 |
| gene-LOC105281700 | https://www.ncbi.nlm.nih.gov/gene/?term=LOC105281700 | 2208 | 43 | 194.75 | 1 | 4.53 | 2.33 | 36.88 | 36.49 | 0.39 | 0.39 | 0.01 | 0.01 |
| gene-LOC105284184 | https://www.ncbi.nlm.nih.gov/gene/?term=LOC105284184 | 2535 | 105 | 414.2 1 | 3.94 | 0.95 | 28.89 | 28.5 | 0.39 | 0.39 | 0.02 | 0.02 | |
| gene-LOC105276462 | https://www.ncbi.nlm.nih.gov/gene/?term=LOC105276462 | 6561 | 289 | 440.48 | 1 | 1.52 | 0.35 | 8.58 | 8.21 | 0.38 | 0.38 | 0.02 | 0.02 |
| gene-LOC105278145 | https://www.ncbi.nlm.nih.gov/gene/?term=LOC105278145 | 4470 | 274 | 612.98 | 1 | 2.24 | 0.36 | 10.24 | 9.87 | 0.37 | 0.37 | 0.02 | 0.02 |
| gene-LOC105284043 | https://www.ncbi.nlm.nih.gov/gene/?term=LOC105284043 | 5661 | 312 | 551.14 | 1 | 1.77 | 0.32 | 10.26 | 9.89 | 0.37 | 0.37 | 0.02 | 0.02 |
| gene-LOC105276527 | https://www.ncbi.nlm.nih.gov/gene/?term=LOC105276527 | 7923 | 381 | 480.88 | 1 | 1.26 | 0.26 | 8.09 | 7.73 | 0.36 | 0.36 | 0.02 | 0.02 |
| gene-LOC105278962 | https://www.ncbi.nlm.nih.gov/gene/?term=LOC105278962 | 4491 | 129 | 287.24 | 1 | 2.23 | 0.78 | 19.14 | 18.79 | 0.35 | 0.35 | 0.01 | 0.01 |
| gene-LOC105274534 | https://www.ncbi.nlm.nih.gov/gene/?term=LOC105274534 | 1329 | 60 | 451.47 | 1 | 7.52 | 1.67 | 44.47 | 44.14 | 0.32 | 0.32 | 0.01 | 0.01 |
| gene-LOC105286332 | https://www.ncbi.nlm.nih.gov/gene/?term=LOC105286332 | 27577 | 573 | 207.78 | 1 | 0.36 | 0.17 | 7.89 | 7.57 | 0.32 | 0.32 | 0.01 | 0.01 |
| gene-LOC105285395 | https://www.ncbi.nlm.nih.gov/gene/?term=LOC105285395 | 8085 | 260 | 321.58 | 1 | 1.24 | 0.38 | 13.21 | 12.9 | 0.31 | 0.31 | 0.01 | 0.01 |
| gene-LOC105286265 | https://www.ncbi.nlm.nih.gov/gene/?term=LOC105286265 | 2507 | 68 | 271.24 | 1 | 3.99 | 1.47 | 31.01 | 30.72 | 0.29 | 0.29 | 0.01 | 0.01 |
| gene-LOC105278666 | https://www.ncbi.nlm.nih.gov/gene/?term=LOC105278666 | 10341 | 492 | 475.78 | 1 | 0.97 | 0.2 | 7.02 | 6.75 | 0.27 | 0.27 | 0.01 | 0.01 |
| gene-LOC105275187 | https://www.ncbi.nlm.nih.gov/gene/?term=LOC105275187 | 5534 | 87 | 157.21 | 1 | 1.81 | 1.15 | 15.13 | 14.88 | 0.25 | 0.25 | 0 | 0 |
| gene-LOC105274449 | https://www.ncbi.nlm.nih.gov/gene/?term=LOC105274449 | 1549 | 53 | 342.16 | 1 | 6.46 | 1.89 | 49.51 | 49.29 | 0.22 | 0.22 | 0.01 | 0.01 |
| gene-LOC105275327 | https://www.ncbi.nlm.nih.gov/gene/?term=LOC105275327 | 14245 | 550 | 386.1 1 | 0.7 | 0.18 | 6.58 | 6.38 | 0.2 | 0.2 | 0.01 | 0.01 | |
| gene-LOC105284262 | https://www.ncbi.nlm.nih.gov/gene/?term=LOC105284262 | 6669 | 248 | 371.87 | 1 | 1.5 | 0.4 | 5.25 | 5.05 | 0.2 | 0.2 | 0.01 | 0.01 |

| | | | | | | | | | | | | | |
|---|---|---|---|---|---|---|---|---|---|---|---|---|---|
| gene-LOC113562050 | https://www.ncbi.nlm.nih.gov/gene/?term=LOC113562050 | 17097 | 698 | 408.26 | 1 | 0.58 | 0.14 | 5.09 | 4.89 | 0.2 | 0.2 | 0.01 | 0.01 |
| gene-LOC105278534 | https://www.ncbi.nlm.nih.gov/gene/?term=LOC105278534 | 7364 | 214 | 290.6 | 1 | 1.36 | 0.47 | 16.92 | 16.72 | 0.19 | 0.19 | 0.01 | 0.01 |
| gene-LOC105286194 | https://www.ncbi.nlm.nih.gov/gene/?term=LOC105286194 | 152619 | 2577 | 168.85 | 1 | 0.07 | 0.04 | 6.51 | 6.31 | 0.19 | 0.19 | 0 | 0 |
| gene-LOC105274590 | https://www.ncbi.nlm.nih.gov/gene/?term=LOC105274590 | 4516 | 88 | 194.86 | 2 | 4.43 | 2.27 | 24.95 | 24.81 | 0.14 | 0.14 | 0 | 0 |
| gene-LOC105274669 | https://www.ncbi.nlm.nih.gov/gene/?term=LOC105274669 | 8555 | 274 | 320.28 | 2 | 2.34 | 0.73 | 5.92 | 5.78 | 0.14 | 0.14 | 0 | 0 |
| gene-LOC105274856 | https://www.ncbi.nlm.nih.gov/gene/?term=LOC105274856 | 5216 | 233 | 446.7 | 1 | 1.92 | 0.43 | 17.97 | 17.83 | 0.14 | 0.14 | 0.01 | 0.01 |
| gene-LOC105277866 | https://www.ncbi.nlm.nih.gov/gene/?term=LOC105277866 | 33045 | 539 | 163.11 | 1 | 0.3 | 0.19 | 3.2 | 3.07 | 0.14 | 0.14 | 0 | 0 |
| gene-LOC105286452 | https://www.ncbi.nlm.nih.gov/gene/?term=LOC105286452 | 6888 | 78 | 113.24 | 1 | 1.45 | 1.28 | 26.06 | 25.92 | 0.14 | 0.14 | 0 | 0 |
| gene-LOC105286770 | https://www.ncbi.nlm.nih.gov/gene/?term=LOC105286770 | 3055 | 50 | 163.67 | 1 | 3.27 | 2 | 14.75 | 14.61 | 0.14 | 0.14 | 0 | 0 |
| gene-LOC105280410 | https://www.ncbi.nlm.nih.gov/gene/?term=LOC105280410 | 12580 | 320 | 254.37 | 1 | 0.79 | 0.31 | 17.81 | 17.7 | 0.11 | 0.11 | 0 | 0 |
| gene-LOC105287065 | https://www.ncbi.nlm.nih.gov/gene/?term=LOC105287065 | 4859 | 259 | 533.03 | 1 | 2.06 | 0.39 | 12.11 | 12 | 0.11 | 0.11 | 0.01 | 0.01 |
| gene-LOC105278109 | https://www.ncbi.nlm.nih.gov/gene/?term=LOC105278109 | 4029 | 89 | 220.9 | 1 | 2.48 | 1.12 | 34.5 | 34.41 | 0.09 | 0.09 | 0 | 0 |
| gene-LOC105282811 | https://www.ncbi.nlm.nih.gov/gene/?term=LOC105282811 | 5765 | 232 | 402.43 | 1 | 1.73 | 0.43 | 8.6 | 8.51 | 0.09 | 0.09 | 0 | 0 |
| gene-LOC105287095 | https://www.ncbi.nlm.nih.gov/gene/?term=LOC105287095 | 1435 | 71 | 494.77 | 1 | 6.97 | 1.41 | 23.64 | 23.55 | 0.09 | 0.09 | 0 | 0 |
| gene-LOC105276917 | https://www.ncbi.nlm.nih.gov/gene/?term=LOC105276917 | 2336 | 42 | 179.79 | 1 | 4.28 | 2.38 | 28.76 | 28.68 | 0.08 | 0.08 | 0 | 0 |
| gene-LOC105276059 | https://www.ncbi.nlm.nih.gov/gene/?term=LOC105276059 | 6256 | 148 | 236.57 | 1 | 1.6 | 0.68 | 11.97 | 11.9 | 0.07 | 0.07 | 0 | 0 |
| gene-LOC105283264 | https://www.ncbi.nlm.nih.gov/gene/?term=LOC105283264 | 4005 | 94 | 234.71 | 1 | 2.5 | 1.06 | 22.97 | 22.91 | 0.07 | 0.07 | 0 | 0 |
| gene-LOC105283296 | https://www.ncbi.nlm.nih.gov/gene/?term=LOC105283296 | 2174 | 96 | 441.58 | 1 | 4.6 | 1.04 | 14.24 | 14.2 | 0.04 | 0.04 | 0 | 0 |
| gene-LOC105280819 | https://www.ncbi.nlm.nih.gov/gene/?term=LOC105280819 | 1639 | 44 | 268.46 | 1 | 6.1 | 2.27 | 37.71 | 37.7 | 0.02 | 0.02 | 0 | 0 |
| gene-LOC105280513 | https://www.ncbi.nlm.nih.gov/gene/?term=LOC105280513 | 69542 | 1608 | 231.23 | 1 | 0.14 | 0.06 | 0.74 | 0.72 | 0.01 | 0.01 | 0 | 0 |
| gene-LOC105277216 | https://www.ncbi.nlm.nih.gov/gene/?term=LOC105277216 | 7627 | 101 | 132.42 | 1 | 1.31 | 0.99 | 8.48 | 8.48 | 0 | 0 | 0 | 0 |
| gene-LOC105279664 | https://www.ncbi.nlm.nih.gov/gene/?term=LOC105279664 | 4023 | 75 | 186.43 | 1 | 2.49 | 1.33 | 19.05 | 19.05 | 0 | 0 | 0 | 0 |
| gene-LOC105277125 | https://www.ncbi.nlm.nih.gov/gene/?term=LOC105277125 | 51079 | 2092 | 409.56 | 1 | 0.2 | 0.05 | 1.63 | 1.65 | -0.02 | 0.02 | 0 | 0 |
| gene-LOC105279528 | https://www.ncbi.nlm.nih.gov/gene/?term=LOC105279528 | 5802 | 40 | 68.94 | 1 | 1.72 | 2.5 | 19.68 | 19.72 | -0.04 | 0.04 | 0 | 0 |
| gene-LOC105280265 | https://www.ncbi.nlm.nih.gov/gene/?term=LOC105280265 | 85984 | 3486 | 405.42 | 1 | 0.12 | 0.03 | 1.07 | 1.12 | -0.04 | 0.04 | 0 | 0 |
| gene-LOC105277588 | https://www.ncbi.nlm.nih.gov/gene/?term=LOC105277588 | 3955 | 98 | 247.79 | 1 | 2.53 | 1.02 | 8.55 | 8.61 | -0.06 | 0.06 | 0 | 0 |
| gene-LOC105277761 | https://www.ncbi.nlm.nih.gov/gene/?term=LOC105277761 | 4648 | 174 | 374.35 | 1 | 2.15 | 0.57 | 14.23 | 14.3 | -0.07 | 0.07 | 0 | 0 |
| gene-LOC105279657 | https://www.ncbi.nlm.nih.gov/gene/?term=LOC105279657 | 5878 | 112 | 190.54 | 1 | 1.7 | 0.89 | 16.48 | 16.55 | -0.07 | 0.07 | 0 | 0 |
| gene-LOC105284774 | https://www.ncbi.nlm.nih.gov/gene/?term=LOC105284774 | 12089 | 106 | 87.68 | 1 | 0.83 | 0.94 | 19.59 | 19.72 | -0.12 | 0.12 | 0 | 0 |
| gene-LOC105276196 | https://www.ncbi.nlm.nih.gov/gene/?term=LOC105276196 | 2953 | 61 | 206.57 | 1 | 3.39 | 1.64 | 38.98 | 39.11 | -0.13 | 0.13 | 0 | 0 |
| gene-LOC105284117 | https://www.ncbi.nlm.nih.gov/gene/?term=LOC105284117 | 4340 | 116 | 267.28 | 1 | 2.3 | 0.86 | 11.19 | 11.33 | -0.14 | 0.14 | 0 | 0 |
| gene-LOC105286649 | https://www.ncbi.nlm.nih.gov/gene/?term=LOC105286649 | 14178 | 852 | 600.93 | 1 | 0.71 | 0.12 | 1.32 | 1.49 | -0.16 | 0.16 | -0.01 | 0.01 |
| gene-LOC105284116 | https://www.ncbi.nlm.nih.gov/gene/?term=LOC105284116 | 3970 | 134 | 337.53 | 1 | 2.52 | 0.75 | 7.83 | 8 | -0.17 | 0.17 | -0.01 | 0.01 |
| gene-LOC105278455 | https://www.ncbi.nlm.nih.gov/gene/?term=LOC105278455 | 27181 | 1036 | 381.15 | 1 | 0.37 | 0.1 | 6.32 | 6.5 | -0.18 | 0.18 | -0.01 | 0.01 |
| gene-LOC105287058 | https://www.ncbi.nlm.nih.gov/gene/?term=LOC105287058 | 4788 | 247 | 515.87 | 1 | 2.09 | 0.4 | 8.85 | 9.06 | -0.2 | 0.2 | -0.01 | 0.01 |
| gene-LOC105278771 | https://www.ncbi.nlm.nih.gov/gene/?term=LOC105278771 | 10457 | 267 | 255.33 | 1 | 0.96 | 0.37 | 9.06 | 9.27 | -0.21 | 0.21 | -0.01 | 0.01 |
| gene-LOC105286515 | https://www.ncbi.nlm.nih.gov/gene/?term=LOC105286515 | 3751 | 133 | 354.57 | 1 | 2.67 | 0.75 | 10.29 | 10.56 | -0.27 | 0.27 | -0.01 | 0.01 |
| gene-LOC105285981 | https://www.ncbi.nlm.nih.gov/gene/?term=LOC105285981 | 7666 | 224 | 292.2 | 1 | 1.3 | 0.45 | 17.35 | 17.63 | -0.28 | 0.28 | -0.01 | 0.01 |
| gene-LOC105280956 | https://www.ncbi.nlm.nih.gov/gene/?term=LOC105280956 | 2639 | 83 | 314.51 | 1 | 3.79 | 1.2 | 18.28 | 18.58 | -0.3 | 0.3 | -0.01 | 0.01 |
| gene-LOC105283649 | https://www.ncbi.nlm.nih.gov/gene/?term=LOC105283649 | 1918 | 59 | 307.61 | 2 | 10.43 | 3.39 | 41.12 | 41.42 | -0.31 | 0.31 | -0.01 | 0.01 |
| gene-LOC105285202 | https://www.ncbi.nlm.nih.gov/gene/?term=LOC105285202 | 1838 | 74 | 402.61 | 1 | 5.44 | 1.35 | 23.68 | 24 | -0.32 | 0.32 | -0.01 | 0.01 |
| gene-LOC105283132 | https://www.ncbi.nlm.nih.gov/gene/?term=LOC105283132 | 1819 | 56 | 307.86 | 1 | 5.5 | 1.79 | 39.81 | 40.18 | -0.38 | 0.38 | -0.01 | 0.01 |
| gene-LOC105280304 | https://www.ncbi.nlm.nih.gov/gene/?term=LOC105280304 | 8919 | 143 | 160.33 | 1 | 1.12 | 0.7 | 6.95 | 7.37 | -0.42 | 0.42 | -0.01 | 0.01 |
| gene-LOC105278877 | https://www.ncbi.nlm.nih.gov/gene/?term=LOC105278877 | 1968 | 91 | 462.4 | 2 | 10.16 | 2.2 | 30.61 | 31.07 | -0.45 | 0.45 | -0.02 | 0.02 |
| gene-LOC105284036 | https://www.ncbi.nlm.nih.gov/gene/?term=LOC105284036 | 3254 | 102 | 313.46 | 1 | 3.07 | 0.98 | 24.26 | 24.78 | -0.52 | 0.52 | -0.02 | 0.02 |
| gene-LOC105286279 | https://www.ncbi.nlm.nih.gov/gene/?term=LOC105286279 | 5864 | 277 | 472.37 | 1 | 1.71 | 0.36 | 2.15 | 2.72 | -0.57 | 0.57 | -0.03 | 0.03 |
| gene-LOC105274851 | https://www.ncbi.nlm.nih.gov/gene/?term=LOC105274851 | 6616 | 160 | 241.84 | 1 | 1.51 | 0.62 | 10.97 | 11.63 | -0.66 | 0.66 | -0.02 | 0.02 |
| gene-LOC105278454 | https://www.ncbi.nlm.nih.gov/gene/?term=LOC105278454 | 1972 | 43 | 218.05 | 1 | 5.07 | 2.33 | 42.19 | 42.86 | -0.67 | 0.67 | -0.01 | 0.01 |
| gene-LOC105275966 | https://www.ncbi.nlm.nih.gov/gene/?term=LOC105275966 | 3299 | 75 | 227.34 | 1 | 3.03 | 1.33 | 43.86 | 44.6 | -0.74 | 0.74 | -0.02 | 0.02 |
| gene-LOC105284507 | https://www.ncbi.nlm.nih.gov/gene/?term=LOC105284507 | 5207 | 123 | 236.22 | 1 | 1.92 | 0.81 | 18.34 | 19.09 | -0.75 | 0.75 | -0.02 | 0.02 |
| gene-LOC105280890 | https://www.ncbi.nlm.nih.gov/gene/?term=LOC105280890 | 1139 | 31 | 272.17 | 1 | 8.78 | 3.23 | 17.12 | 17.99 | -0.87 | 0.87 | -0.02 | 0.02 |
| gene-LOC105281173 | https://www.ncbi.nlm.nih.gov/gene/?term=LOC105281173 | 1595 | 39 | 244.51 | 1 | 6.27 | 2.56 | 61.21 | 62.08 | -0.87 | 0.87 | -0.02 | 0.02 |
| gene-LOC105282885 | https://www.ncbi.nlm.nih.gov/gene/?term=LOC105282885 | 1870 | 41 | 219.25 | 1 | 5.35 | 2.44 | 40.22 | 41.13 | -0.91 | 0.91 | -0.02 | 0.02 |
| gene-LOC105285252 | https://www.ncbi.nlm.nih.gov/gene/?term=LOC105285252 | 7427 | 148 | 199.27 | 1 | 1.35 | 0.68 | 11.91 | 12.93 | -1.01 | 1.01 | -0.02 | 0.02 |
| gene-LOC105287633 | https://www.ncbi.nlm.nih.gov/gene/?term=LOC105287633 | 1524 | 26 | 170.6 | 1 | 6.56 | 3.85 | 60.3 | 61.69 | -1.38 | 1.38 | -0.02 | 0.02 |
| gene-LOC105284329 | https://www.ncbi.nlm.nih.gov/gene/?term=LOC105284329 | 4051 | 72 | 177.73 | 1 | 2.47 | 1.39 | 18.86 | 20.42 | -1.56 | 1.56 | -0.03 | 0.03 |
| gene-LOC105275328 | https://www.ncbi.nlm.nih.gov/gene/?term=LOC105275328 | 2227 | 85 | 381.68 | 1 | 4.49 | 1.18 | 17.9 | 19.55 | -1.65 | 1.65 | -0.06 | 0.06 |
| gene-LOC105282914 | https://www.ncbi.nlm.nih.gov/gene/?term=LOC105282914 | 5623 | 43 | 76.47 | 1 | 1.78 | 2.33 | 47.44 | 49.79 | -2.35 | 2.35 | -0.02 | 0.02 |
| gene-LOC105284213 | https://www.ncbi.nlm.nih.gov/gene/?term=LOC105284213 | 868 | 23 | 264.98 | 1 | 11.52 | 4.35 | 9.72 | 12.3 | -2.58 | 2.58 | -0.07 | 0.07 |
| gene-LOC105285409 | https://www.ncbi.nlm.nih.gov/gene/?term=LOC105285409 | 5099 | 67 | 131.4 | 2 | 3.92 | 2.99 | 1.72 | 4.47 | -2.75 | 2.75 | -0.04 | 0.04 |

Directory: OGLall File: ogl-Ob_LI_RphaseA2.vs.RphaseB2.txt

| gene_ID | gene_link | gwidth | #Sites | #per10Kb | #dmSites | #dmsp10kb | %dmSites | %pSite1 | %pSite2 | DMpSite | ADMpSite | DMpNucl | ADMpNucl |
|---|---|---|---|---|---|---|---|---|---|---|---|---|---|
| gene-LOC113562901 | https://www.ncbi.nlm.nih.gov/gene/?term=LOC113562901 | 1531 | 24 | 156.76 | 3 | 19.6 | 12.5 | 59.3 | 45.31 | 14 | 14 | 0.22 | 0.22 |
| gene-LOC113562345 | https://www.ncbi.nlm.nih.gov/gene/?term=LOC113562345 | 2290 | 69 | 301.31 | 14 | 61.14 | 20.29 | 29.74 | 16.81 | 12.93 | 12.93 | 0.39 | 0.39 |
| gene-LOC105275070 | https://www.ncbi.nlm.nih.gov/gene/?term=LOC105275070 | 5667 | 42 | 74.11 | 5 | 8.82 | 11.9 | 13.23 | 1.03 | 12.2 | 12.2 | 0.09 | 0.09 |
| gene-LOC105279898 | https://www.ncbi.nlm.nih.gov/gene/?term=LOC105279898 | 1343 | 10 | 74.46 | 1 | 7.45 | 10 | 83.16 | 71.53 | 11.63 | 11.63 | 0.09 | 0.09 |
| gene-LOC105286644 | https://www.ncbi.nlm.nih.gov/gene/?term=LOC105286644 | 1235 | 22 | 178.14 | 1 | 8.1 | 4.55 | 44.4 | 37.26 | 7.14 | 7.14 | 0.13 | 0.13 |
| gene-LOC105275708 | https://www.ncbi.nlm.nih.gov/gene/?term=LOC105275708 | 1772 | 15 | 84.65 | 1 | 5.64 | 6.67 | 47.19 | 40.83 | 6.36 | 6.36 | 0.05 | 0.05 |
| gene-LOC105275447 | https://www.ncbi.nlm.nih.gov/gene/?term=LOC105275447 | 1846 | 31 | 167.93 | 1 | 5.42 | 3.23 | 9.13 | 2.78 | 6.34 | 6.34 | 0.11 | 0.11 |
| gene-LOC105286694 | https://www.ncbi.nlm.nih.gov/gene/?term=LOC105286694 | 3409 | 12 | 35.2 | 1 | 2.93 | 8.33 | 52.59 | 46.71 | 5.88 | 5.88 | 0.02 | 0.02 |
| gene-LOC105282388 | https://www.ncbi.nlm.nih.gov/gene/?term=LOC105282388 | 1661 | 34 | 204.7 | 1 | 6.02 | 2.94 | 48.95 | 44.6 | 4.35 | 4.35 | 0.09 | 0.09 |
| gene-LOC105287417 | https://www.ncbi.nlm.nih.gov/gene/?term=LOC105287417 | 1025 | 23 | 224.39 | 1 | 9.76 | 4.35 | 53.58 | 49.27 | 4.31 | 4.31 | 0.1 | 0.1 |
| gene-LOC105282893 | https://www.ncbi.nlm.nih.gov/gene/?term=LOC105282893 | 2645 | 47 | 177.69 | 1 | 3.78 | 2.13 | 30.08 | 25.9 | 4.18 | 4.18 | 0.07 | 0.07 |
| gene-LOC105276538 | https://www.ncbi.nlm.nih.gov/gene/?term=LOC105276538 | 1511 | 20 | 132.36 | 1 | 6.62 | 5 | 73.31 | 69.21 | 4.1 | 4.1 | 0.05 | 0.05 |
| gene-LOC105285583 | https://www.ncbi.nlm.nih.gov/gene/?term=LOC105285583 | 1947 | 49 | 251.67 | 1 | 5.14 | 2.04 | 30.89 | 26.83 | 4.06 | 4.06 | 0.1 | 0.1 |
| gene-LOC105280555 | https://www.ncbi.nlm.nih.gov/gene/?term=LOC105280555 | 1496 | 31 | 207.22 | 1 | 6.68 | 3.23 | 61.05 | 57.2 | 3.85 | 3.85 | 0.08 | 0.08 |
| gene-LOC105274778 | https://www.ncbi.nlm.nih.gov/gene/?term=LOC105274778 | 4662 | 40 | 85.8 | 1 | 2.15 | 2.5 | 47.99 | 44.37 | 3.62 | 3.62 | 0.03 | 0.03 |
| gene-LOC105284609 | https://www.ncbi.nlm.nih.gov/gene/?term=LOC105284609 | 5678 | 33 | 58.12 | 1 | 1.76 | 3.03 | 34 | 30.46 | 3.54 | 3.54 | 0.02 | 0.02 |
| gene-LOC105285315 | https://www.ncbi.nlm.nih.gov/gene/?term=LOC105285315 | 5458 | 19 | 34.81 | 1 | 1.83 | 5.26 | 46.71 | 43.22 | 3.49 | 3.49 | 0.01 | 0.01 |
| gene-LOC105275518 | https://www.ncbi.nlm.nih.gov/gene/?term=LOC105275518 | 2335 | 24 | 102.78 | 1 | 4.28 | 4.17 | 49.73 | 46.6 | 3.13 | 3.13 | 0.03 | 0.03 |
| gene-LOC105286683 | https://www.ncbi.nlm.nih.gov/gene/?term=LOC105286683 | 1665 | 57 | 342.34 | 2 | 12.01 | 3.51 | 34.89 | 31.76 | 3.13 | 3.13 | 0.11 | 0.11 |
| gene-LOC105274706 | https://www.ncbi.nlm.nih.gov/gene/?term=LOC105274706 | 1446 | 56 | 387.28 | 1 | 6.92 | 1.79 | 39.74 | 36.7 | 3.05 | 3.05 | 0.12 | 0.12 |
| gene-LOC105285674 | https://www.ncbi.nlm.nih.gov/gene/?term=LOC105285674 | 3035 | 102 | 336.08 | 2 | 6.59 | 1.96 | 26.23 | 23.21 | 3.03 | 3.03 | 0.1 | 0.1 |
| gene-LOC105278215 | https://www.ncbi.nlm.nih.gov/gene/?term=LOC105278215 | 1700 | 29 | 170.59 | 1 | 5.88 | 3.45 | 57.59 | 54.66 | 2.93 | 2.93 | 0.05 | 0.05 |
| gene-LOC105278109 | https://www.ncbi.nlm.nih.gov/gene/?term=LOC105278109 | 4029 | 67 | 166.29 | 1 | 2.48 | 1.49 | 44.53 | 41.68 | 2.86 | 2.86 | 0.05 | 0.05 |
| gene-LOC105281515 | https://www.ncbi.nlm.nih.gov/gene/?term=LOC105281515 | 8827 | 42 | 47.58 | 1 | 1.13 | 2.38 | 30.33 | 27.53 | 2.81 | 2.81 | 0.01 | 0.01 |
| gene-LOC105280786 | https://www.ncbi.nlm.nih.gov/gene/?term=LOC105280786 | 2573 | 127 | 493.59 | 1 | 3.89 | 0.79 | 30.11 | 27.41 | 2.7 | 2.7 | 0.13 | 0.13 |
| gene-LOC105287063 | https://www.ncbi.nlm.nih.gov/gene/?term=LOC105287063 | 1874 | 48 | 256.14 | 1 | 5.34 | 2.08 | 32.76 | 30.33 | 2.44 | 2.44 | 0.06 | 0.06 |
| gene-LOC105287854 | https://www.ncbi.nlm.nih.gov/gene/?term=LOC105287854 | 3644 | 54 | 148.19 | 1 | 2.74 | 1.85 | 25.58 | 23.15 | 2.43 | 2.43 | 0.04 | 0.04 |
| gene-LOC105282794 | https://www.ncbi.nlm.nih.gov/gene/?term=LOC105282794 | 4571 | 95 | 207.83 | 1 | 2.19 | 1.05 | 18.64 | 16.23 | 2.41 | 2.41 | 0.05 | 0.05 |
| gene-LOC105285671 | https://www.ncbi.nlm.nih.gov/gene/?term=LOC105285671 | 2263 | 106 | 468.4 | 1 | 4.42 | 0.94 | 28.93 | 26.53 | 2.4 | 2.4 | 0.11 | 0.11 |
| gene-LOC105276917 | https://www.ncbi.nlm.nih.gov/gene/?term=LOC105276917 | 2336 | 38 | 162.67 | 1 | 4.28 | 2.63 | 28.71 | 26.35 | 2.36 | 2.36 | 0.04 | 0.04 |
| gene-LOC105274900 | https://www.ncbi.nlm.nih.gov/gene/?term=LOC105274900 | 5561 | 56 | 100.7 | 2 | 3.6 | 3.57 | 32.09 | 29.76 | 2.33 | 2.33 | 0.02 | 0.02 |
| gene-LOC105276704 | https://www.ncbi.nlm.nih.gov/gene/?term=LOC105276704 | 2063 | 88 | 426.56 | 1 | 4.85 | 1.14 | 16.14 | 13.92 | 2.23 | 2.23 | 0.1 | 0.1 |
| gene-LOC105279664 | https://www.ncbi.nlm.nih.gov/gene/?term=LOC105279664 | 4023 | 56 | 139.2 | 1 | 2.49 | 1.79 | 21.11 | 18.99 | 2.12 | 2.12 | 0.03 | 0.03 |
| gene-LOC105280104 | https://www.ncbi.nlm.nih.gov/gene/?term=LOC105280104 | 3960 | 139 | 351.01 | 1 | 2.53 | 0.72 | 17.44 | 15.39 | 2.05 | 2.05 | 0.07 | 0.07 |
| gene-LOC105283132 | https://www.ncbi.nlm.nih.gov/gene/?term=LOC105283132 | 1819 | 57 | 313.36 | 1 | 5.5 | 1.75 | 39.92 | 37.88 | 2.04 | 2.04 | 0.06 | 0.06 |
| gene-LOC105277856 | https://www.ncbi.nlm.nih.gov/gene/?term=LOC105277856 | 3442 | 35 | 101.69 | 1 | 2.91 | 2.86 | 35 | 32.98 | 2.02 | 2.02 | 0.02 | 0.02 |
| gene-LOC105278108 | https://www.ncbi.nlm.nih.gov/gene/?term=LOC105278108 | 4392 | 51 | 116.12 | 1 | 2.28 | 1.96 | 25.15 | 23.14 | 2.01 | 2.01 | 0.02 | 0.02 |
| gene-LOC105280641 | https://www.ncbi.nlm.nih.gov/gene/?term=LOC105280641 | 3682 | 53 | 143.94 | 1 | 2.72 | 1.89 | 17.27 | 15.26 | 2.01 | 2.01 | 0.03 | 0.03 |
| gene-LOC105280377 | https://www.ncbi.nlm.nih.gov/gene/?term=LOC105280377 | 2024 | 70 | 345.85 | 1 | 4.94 | 1.43 | 48.1 | 46.12 | 1.98 | 1.98 | 0.07 | 0.07 |
| gene-LOC105282156 | https://www.ncbi.nlm.nih.gov/gene/?term=LOC105282156 | 2577 | 130 | 504.46 | 1 | 3.88 | 0.77 | 28.69 | 26.74 | 1.95 | 1.95 | 0.1 | 0.1 |
| gene-LOC113561805 | https://www.ncbi.nlm.nih.gov/gene/?term=LOC113561805 | 3337 | 71 | 212.77 | 1 | 3 | 1.41 | 36.21 | 34.33 | 1.88 | 1.88 | 0.04 | 0.04 |
| gene-LOC105282600 | https://www.ncbi.nlm.nih.gov/gene/?term=LOC105282600 | 3715 | 56 | 150.74 | 1 | 2.69 | 1.79 | 27.25 | 25.37 | 1.87 | 1.87 | 0.03 | 0.03 |
| gene-LOC105282947 | https://www.ncbi.nlm.nih.gov/gene/?term=LOC105282947 | 1162 | 41 | 352.84 | 1 | 8.61 | 2.44 | 36.57 | 34.72 | 1.85 | 1.85 | 0.07 | 0.07 |
| gene-LOC105287646 | https://www.ncbi.nlm.nih.gov/gene/?term=LOC105287646 | 3556 | 90 | 253.09 | 1 | 2.81 | 1.11 | 18.4 | 16.59 | 1.81 | 1.81 | 0.05 | 0.05 |
| gene-LOC105276190 | https://www.ncbi.nlm.nih.gov/gene/?term=LOC105276190 | 4839 | 93 | 192.19 | 1 | 2.07 | 1.08 | 11.07 | 9.37 | 1.7 | 1.7 | 0.03 | 0.03 |
| gene-LOC105281420 | https://www.ncbi.nlm.nih.gov/gene/?term=LOC105281420 | 3882 | 62 | 159.71 | 1 | 2.58 | 1.61 | 15.69 | 14 | 1.7 | 1.7 | 0.03 | 0.03 |
| gene-LOC105284117 | https://www.ncbi.nlm.nih.gov/gene/?term=LOC105284117 | 4340 | 65 | 149.77 | 1 | 2.3 | 1.54 | 12.82 | 11.26 | 1.56 | 1.56 | 0.02 | 0.02 |
| gene-LOC105287648 | https://www.ncbi.nlm.nih.gov/gene/?term=LOC105287648 | 2205 | 26 | 117.91 | 1 | 4.54 | 3.85 | 41.13 | 39.59 | 1.54 | 1.54 | 0.02 | 0.02 |
| gene-LOC105288063 | https://www.ncbi.nlm.nih.gov/gene/?term=LOC105288063 | 3400 | 47 | 138.24 | 1 | 2.94 | 2.13 | 15.4 | 13.86 | 1.54 | 1.54 | 0.02 | 0.02 |
| gene-LOC105283182 | https://www.ncbi.nlm.nih.gov/gene/?term=LOC105283182 | 3585 | 131 | 365.41 | 1 | 2.79 | 0.76 | 14.16 | 12.63 | 1.52 | 1.52 | 0.06 | 0.06 |
| gene-LOC105283649 | https://www.ncbi.nlm.nih.gov/gene/?term=LOC105283649 | 1918 | 56 | 291.97 | 2 | 10.43 | 3.57 | 37.98 | 36.48 | 1.5 | 1.5 | 0.04 | 0.04 |
| gene-LOC105275347 | https://www.ncbi.nlm.nih.gov/gene/?term=LOC105275347 | 2526 | 19 | 75.22 | 1 | 3.96 | 5.26 | 38.97 | 37.49 | 1.48 | 1.48 | 0.01 | 0.01 |
| gene-LOC105276692 | https://www.ncbi.nlm.nih.gov/gene/?term=LOC105276692 | 1745 | 35 | 200.57 | 1 | 5.73 | 2.86 | 22.12 | 20.68 | 1.44 | 1.44 | 0.03 | 0.03 |
| gene-LOC105278282 | https://www.ncbi.nlm.nih.gov/gene/?term=LOC105278282 | 1775 | 56 | 315.49 | 1 | 5.63 | 1.79 | 39.77 | 38.34 | 1.43 | 1.43 | 0.05 | 0.05 |
| gene-LOC105282970 | https://www.ncbi.nlm.nih.gov/gene/?term=LOC105282970 | 1529 | 40 | 261.61 | 1 | 6.54 | 2.5 | 49.63 | 48.22 | 1.41 | 1.41 | 0.04 | 0.04 |
| gene-LOC105281272 | https://www.ncbi.nlm.nih.gov/gene/?term=LOC105281272 | 5252 | 164 | 312.26 | 1 | 1.9 | 0.61 | 19.36 | 17.96 | 1.4 | 1.4 | 0.04 | 0.04 |
| gene-LOC105281532 | https://www.ncbi.nlm.nih.gov/gene/?term=LOC105281532 | 7793 | 81 | 103.94 | 1 | 1.28 | 1.23 | 12.92 | 11.51 | 1.4 | 1.4 | 0.01 | 0.01 |
| gene-LOC105276875 | https://www.ncbi.nlm.nih.gov/gene/?term=LOC105276875 | 3151 | 118 | 374.48 | 1 | 3.17 | 0.85 | 23.91 | 22.53 | 1.38 | 1.38 | 0.05 | 0.05 |
| gene-LOC105284704 | https://www.ncbi.nlm.nih.gov/gene/?term=LOC105284704 | 2400 | 62 | 258.33 | 1 | 4.17 | 1.61 | 24.3 | 22.92 | 1.38 | 1.38 | 0.04 | 0.04 |
| gene-LOC105276449 | https://www.ncbi.nlm.nih.gov/gene/?term=LOC105276449 | 1231 | 50 | 406.17 | 1 | 8.12 | 2 | 32.79 | 31.42 | 1.37 | 1.37 | 0.06 | 0.06 |

| | | | | | | | | | | | | | |
|---|---|---|---|---|---|---|---|---|---|---|---|---|---|
| gene-LOC105284355 | https://www.ncbi.nlm.nih.gov/gene/?term=LOC105284355 | 3904 | 153 | 391.91 | 1 | 2.56 | 0.65 | 21.82 | 20.47 | 1.35 | 1.35 | 0.05 | 0.05 |
| gene-LOC105274590 | https://www.ncbi.nlm.nih.gov/gene/?term=LOC105274590 | 4516 | 71 | 157.22 | 2 | 4.43 | 2.82 | 29.16 | 27.84 | 1.32 | 1.32 | 0.02 | 0.02 |
| gene-LOC105283728 | https://www.ncbi.nlm.nih.gov/gene/?term=LOC105283728 | 3256 | 120 | 368.55 | 2 | 6.14 | 1.67 | 19.01 | 17.69 | 1.32 | 1.32 | 0.05 | 0.05 |
| gene-LOC105283123 | https://www.ncbi.nlm.nih.gov/gene/?term=LOC105283123 | 1507 | 48 | 318.51 | 1 | 6.64 | 2.08 | 17.79 | 16.48 | 1.31 | 1.31 | 0.04 | 0.04 |
| gene-LOC105281419 | https://www.ncbi.nlm.nih.gov/gene/?term=LOC105281419 | 2366 | 83 | 350.8 1 | 4.23 | 1.2 | 23.42 | 22.13 | 1.3 | 1.3 | 0.05 | 0.05 | |
| gene-LOC105282151 | https://www.ncbi.nlm.nih.gov/gene/?term=LOC105282151 | 1601 | 39 | 243.6 2 | 12.49 | 5.13 | 42.25 | 41 | 1.25 | 1.25 | 0.03 | 0.03 | |
| gene-LOC105283712 | https://www.ncbi.nlm.nih.gov/gene/?term=LOC105283712 | 1910 | 36 | 188.48 | 1 | 5.24 | 2.78 | 27.71 | 26.46 | 1.25 | 1.25 | 0.02 | 0.02 |
| gene-LOC105281848 | https://www.ncbi.nlm.nih.gov/gene/?term=LOC105281848 | 2122 | 48 | 226.2 1 | 4.71 | 2.08 | 27.66 | 26.43 | 1.23 | 1.23 | 0.03 | 0.03 | |
| gene-LOC105288079 | https://www.ncbi.nlm.nih.gov/gene/?term=LOC105288079 | 3473 | 101 | 290.81 | 1 | 2.88 | 0.99 | 20.37 | 19.15 | 1.21 | 1.21 | 0.04 | 0.04 |
| gene-LOC105277238 | https://www.ncbi.nlm.nih.gov/gene/?term=LOC105277238 | 2708 | 88 | 324.96 | 1 | 3.69 | 1.14 | 27.16 | 25.96 | 1.2 | 1.2 | 0.04 | 0.04 |
| gene-LOC105278497 | https://www.ncbi.nlm.nih.gov/gene/?term=LOC105278497 | 2072 | 100 | 482.63 | 1 | 4.83 | 1 | 19.97 | 18.78 | 1.2 | 1.2 | 0.06 | 0.06 |
| gene-LOC105288066 | https://www.ncbi.nlm.nih.gov/gene/?term=LOC105288066 | 1964 | 30 | 152.75 | 1 | 5.09 | 3.33 | 32.32 | 31.12 | 1.2 | 1.2 | 0.02 | 0.02 |
| gene-LOC105277761 | https://www.ncbi.nlm.nih.gov/gene/?term=LOC105277761 | 4648 | 150 | 322.72 | 1 | 2.15 | 0.67 | 14.38 | 13.19 | 1.19 | 1.19 | 0.04 | 0.04 |
| gene-LOC105287720 | https://www.ncbi.nlm.nih.gov/gene/?term=LOC105287720 | 4843 | 99 | 204.42 | 1 | 2.06 | 1.01 | 16.39 | 15.24 | 1.16 | 1.16 | 0.02 | 0.02 |
| gene-LOC105277631 | https://www.ncbi.nlm.nih.gov/gene/?term=LOC105277631 | 4387 | 304 | 692.96 | 1 | 2.28 | 0.33 | 10.76 | 9.61 | 1.15 | 1.15 | 0.08 | 0.08 |
| gene-LOC105281173 | https://www.ncbi.nlm.nih.gov/gene/?term=LOC105281173 | 1595 | 39 | 244.51 | 1 | 6.27 | 2.56 | 61.03 | 59.91 | 1.13 | 1.13 | 0.03 | 0.03 |
| gene-LOC105284184 | https://www.ncbi.nlm.nih.gov/gene/?term=LOC105284184 | 2535 | 86 | 339.25 | 1 | 3.94 | 1.16 | 34.02 | 32.88 | 1.13 | 1.13 | 0.04 | 0.04 |
| gene-LOC105275363 | https://www.ncbi.nlm.nih.gov/gene/?term=LOC105275363 | 3798 | 165 | 434.44 | 1 | 2.63 | 0.61 | 17.38 | 16.26 | 1.12 | 1.12 | 0.05 | 0.05 |
| gene-LOC105279298 | https://www.ncbi.nlm.nih.gov/gene/?term=LOC105279298 | 2773 | 136 | 490.44 | 2 | 7.21 | 1.47 | 23.98 | 22.86 | 1.11 | 1.11 | 0.05 | 0.05 |
| gene-LOC113561621 | https://www.ncbi.nlm.nih.gov/gene/?term=LOC113561621 | 2249 | 86 | 382.39 | 1 | 4.45 | 1.16 | 20.44 | 19.32 | 1.11 | 1.11 | 0.04 | 0.04 |
| gene-LOC105279500 | https://www.ncbi.nlm.nih.gov/gene/?term=LOC105279500 | 3587 | 37 | 103.15 | 1 | 2.79 | 2.7 | 50.42 | 49.31 | 1.1 | 1.1 | 0.01 | 0.01 |
| gene-LOC105285639 | https://www.ncbi.nlm.nih.gov/gene/?term=LOC105285639 | 10325 | 349 | 338.01 | 1 | 0.97 | 0.29 | 13.03 | 11.95 | 1.08 | 1.08 | 0.04 | 0.04 |
| gene-LOC105275451 | https://www.ncbi.nlm.nih.gov/gene/?term=LOC105275451 | 6035 | 130 | 215.41 | 1 | 1.66 | 0.77 | 22.16 | 21.09 | 1.07 | 1.07 | 0.02 | 0.02 |
| gene-LOC105288010 | https://www.ncbi.nlm.nih.gov/gene/?term=LOC105288010 | 3189 | 51 | 159.92 | 1 | 3.14 | 1.96 | 29.8 | 28.73 | 1.07 | 1.07 | 0.02 | 0.02 |
| gene-LOC105284043 | https://www.ncbi.nlm.nih.gov/gene/?term=LOC105284043 | 5661 | 308 | 544.07 | 1 | 1.77 | 0.32 | 10.53 | 9.46 | 1.06 | 1.06 | 0.06 | 0.06 |
| gene-LOC105278283 | https://www.ncbi.nlm.nih.gov/gene/?term=LOC105278283 | 2317 | 142 | 612.86 | 1 | 4.32 | 0.7 | 25.61 | 24.56 | 1.05 | 1.05 | 0.06 | 0.06 |
| gene-LOC105282149 | https://www.ncbi.nlm.nih.gov/gene/?term=LOC105282149 | 1787 | 110 | 615.56 | 1 | 5.6 | 0.91 | 43.04 | 42 | 1.04 | 1.04 | 0.06 | 0.06 |
| gene-LOC105285742 | https://www.ncbi.nlm.nih.gov/gene/?term=LOC105285742 | 3446 | 123 | 356.94 | 1 | 2.9 | 0.81 | 20.56 | 19.51 | 1.04 | 1.04 | 0.04 | 0.04 |
| gene-LOC105286514 | https://www.ncbi.nlm.nih.gov/gene/?term=LOC105286514 | 2528 | 49 | 193.83 | 1 | 3.96 | 2.04 | 36.75 | 35.71 | 1.04 | 1.04 | 0.02 | 0.02 |
| gene-LOC105283592 | https://www.ncbi.nlm.nih.gov/gene/?term=LOC105283592 | 3479 | 80 | 229.95 | 1 | 2.87 | 1.25 | 23.33 | 22.32 | 1.01 | 1.01 | 0.02 | 0.02 |
| gene-LOC109611283 | https://www.ncbi.nlm.nih.gov/gene/?term=LOC109611283 | 1451 | 37 | 255 1 | 6.89 | 2.7 | 37.11 | 36.1 | 1 | 1 | 0.03 | 0.03 | |
| gene-LOC105285395 | https://www.ncbi.nlm.nih.gov/gene/?term=LOC105285395 | 8085 | 218 | 269.64 | 1 | 1.24 | 0.46 | 17.01 | 16.05 | 0.96 | 0.96 | 0.03 | 0.03 |
| gene-LOC105278807 | https://www.ncbi.nlm.nih.gov/gene/?term=LOC105278807 | 2166 | 67 | 309.33 | 1 | 4.62 | 1.49 | 22.84 | 21.92 | 0.93 | 0.93 | 0.03 | 0.03 |
| gene-LOC105282682 | https://www.ncbi.nlm.nih.gov/gene/?term=LOC105282682 | 4127 | 223 | 540.34 | 1 | 2.42 | 0.45 | 13.69 | 12.76 | 0.93 | 0.93 | 0.05 | 0.05 |
| gene-LOC105278962 | https://www.ncbi.nlm.nih.gov/gene/?term=LOC105278962 | 4491 | 104 | 231.57 | 1 | 2.23 | 0.96 | 22.01 | 21.09 | 0.92 | 0.92 | 0.02 | 0.02 |
| gene-LOC105274856 | https://www.ncbi.nlm.nih.gov/gene/?term=LOC105274856 | 5216 | 216 | 414.11 | 1 | 1.92 | 0.46 | 19.8 | 18.89 | 0.91 | 0.91 | 0.04 | 0.04 |
| gene-LOC105283987 | https://www.ncbi.nlm.nih.gov/gene/?term=LOC105283987 | 4841 | 68 | 140.47 | 1 | 2.07 | 1.47 | 29.57 | 28.67 | 0.91 | 0.91 | 0.01 | 0.01 |
| gene-LOC105279038 | https://www.ncbi.nlm.nih.gov/gene/?term=LOC105279038 | 9453 | 313 | 331.11 | 1 | 1.06 | 0.32 | 10.35 | 9.46 | 0.9 | 0.9 | 0.03 | 0.03 |
| gene-LOC105285406 | https://www.ncbi.nlm.nih.gov/gene/?term=LOC105285406 | 8015 | 60 | 74.86 1 | 1.25 | 1.67 | 26.39 | 25.52 | 0.87 | 0.87 | 0.01 | 0.01 | |
| gene-LOC105276059 | https://www.ncbi.nlm.nih.gov/gene/?term=LOC105276059 | 6256 | 133 | 212.6 1 | 1.6 | 0.75 | 10.26 | 9.4 | 0.86 | 0.86 | 0.02 | 0.02 | |
| gene-LOC105280408 | https://www.ncbi.nlm.nih.gov/gene/?term=LOC105280408 | 5983 | 58 | 96.94 1 | 1.67 | 1.72 | 16.75 | 15.89 | 0.86 | 0.86 | 0.01 | 0.01 | |
| gene-LOC105280956 | https://www.ncbi.nlm.nih.gov/gene/?term=LOC105280956 | 2639 | 71 | 269.04 | 1 | 3.79 | 1.41 | 18.65 | 17.79 | 0.86 | 0.86 | 0.02 | 0.02 |
| gene-LOC105280407 | https://www.ncbi.nlm.nih.gov/gene/?term=LOC105280407 | 10950 | 211 | 192.69 | 1 | 0.91 | 0.47 | 16.65 | 15.81 | 0.84 | 0.84 | 0.02 | 0.02 |
| gene-LOC105285055 | https://www.ncbi.nlm.nih.gov/gene/?term=LOC105285055 | 4108 | 276 | 671.86 | 2 | 4.87 | 0.72 | 11.41 | 10.6 | 0.8 | 0.8 | 0.05 | 0.05 |
| gene-LOC105280002 | https://www.ncbi.nlm.nih.gov/gene/?term=LOC105280002 | 1940 | 28 | 144.33 | 1 | 5.15 | 3.57 | 37.5 | 36.72 | 0.79 | 0.79 | 0.01 | 0.01 |
| gene-LOC105284036 | https://www.ncbi.nlm.nih.gov/gene/?term=LOC105284036 | 3254 | 108 | 331.9 1 | 3.07 | 0.93 | 22.88 | 22.09 | 0.79 | 0.79 | 0.03 | 0.03 | |
| gene-LOC105275460 | https://www.ncbi.nlm.nih.gov/gene/?term=LOC105275460 | 2473 | 117 | 473.11 | 1 | 4.04 | 0.85 | 14.05 | 13.26 | 0.78 | 0.78 | 0.04 | 0.04 |
| gene-LOC105280774 | https://www.ncbi.nlm.nih.gov/gene/?term=LOC105280774 | 2793 | 100 | 358.04 | 1 | 3.58 | 1 | 32.96 | 32.18 | 0.78 | 0.78 | 0.03 | 0.03 |
| gene-LOC105285534 | https://www.ncbi.nlm.nih.gov/gene/?term=LOC105285534 | 3944 | 237 | 600.91 | 1 | 2.54 | 0.42 | 11.45 | 10.67 | 0.78 | 0.78 | 0.05 | 0.05 |
| gene-LOC105285060 | https://www.ncbi.nlm.nih.gov/gene/?term=LOC105285060 | 1285 | 21 | 163.42 | 1 | 7.78 | 4.76 | 39.84 | 39.09 | 0.75 | 0.75 | 0.01 | 0.01 |
| gene-LOC105284116 | https://www.ncbi.nlm.nih.gov/gene/?term=LOC105284116 | 3970 | 78 | 196.47 | 1 | 2.52 | 1.28 | 7.09 | 6.36 | 0.73 | 0.73 | 0.01 | 0.01 |
| gene-LOC105274534 | https://www.ncbi.nlm.nih.gov/gene/?term=LOC105274534 | 1329 | 57 | 428.89 | 1 | 7.52 | 1.75 | 48.36 | 47.65 | 0.71 | 0.71 | 0.03 | 0.03 |
| gene-LOC105282067 | https://www.ncbi.nlm.nih.gov/gene/?term=LOC105282067 | 5941 | 54 | 90.89 1 | 1.68 | 1.85 | 28.74 | 28.06 | 0.67 | 0.67 | 0.01 | 0.01 | |
| gene-LOC105285514 | https://www.ncbi.nlm.nih.gov/gene/?term=LOC105285514 | 1277 | 17 | 133.12 | 1 | 7.83 | 5.88 | 71.39 | 70.72 | 0.67 | 0.67 | 0.01 | 0.01 |
| gene-LOC105274699 | https://www.ncbi.nlm.nih.gov/gene/?term=LOC105274699 | 2386 | 52 | 217.94 | 1 | 4.19 | 1.92 | 24.82 | 24.15 | 0.66 | 0.66 | 0.01 | 0.01 |
| gene-LOC105284557 | https://www.ncbi.nlm.nih.gov/gene/?term=LOC105284557 | 1672 | 84 | 502.39 | 1 | 5.98 | 1.19 | 29.36 | 28.7 | 0.66 | 0.66 | 0.03 | 0.03 |
| gene-LOC105275966 | https://www.ncbi.nlm.nih.gov/gene/?term=LOC105275966 | 3299 | 64 | 194 1 | 3.03 | 1.56 | 48.15 | 47.49 | 0.65 | 0.65 | 0.01 | 0.01 | |
| gene-LOC105282399 | https://www.ncbi.nlm.nih.gov/gene/?term=LOC105282399 | 2682 | 24 | 89.49 1 | 3.73 | 4.17 | 35.91 | 35.25 | 0.65 | 0.65 | 0.01 | 0.01 | |
| gene-LOC105275683 | https://www.ncbi.nlm.nih.gov/gene/?term=LOC105275683 | 4256 | 193 | 453.48 | 1 | 2.35 | 0.52 | 13.76 | 13.13 | 0.63 | 0.63 | 0.03 | 0.03 |
| gene-LOC105285179 | https://www.ncbi.nlm.nih.gov/gene/?term=LOC105285179 | 4191 | 185 | 441.42 | 1 | 2.39 | 0.54 | 18.94 | 18.34 | 0.6 | 0.6 | 0.03 | 0.03 |
| gene-LOC105283943 | https://www.ncbi.nlm.nih.gov/gene/?term=LOC105283943 | 3559 | 48 | 134.87 | 1 | 2.81 | 2.08 | 17.55 | 16.96 | 0.59 | 0.59 | 0.01 | 0.01 |
| gene-LOC105275374 | https://www.ncbi.nlm.nih.gov/gene/?term=LOC105275374 | 11376 | 523 | 459.74 | 1 | 0.88 | 0.19 | 11.54 | 10.97 | 0.57 | 0.57 | 0.03 | 0.03 |
| gene-LOC105276612 | https://www.ncbi.nlm.nih.gov/gene/?term=LOC105276612 | 4157 | 187 | 449.84 | 1 | 2.41 | 0.53 | 10.04 | 9.47 | 0.57 | 0.57 | 0.03 | 0.03 |
| gene-LOC105282825 | https://www.ncbi.nlm.nih.gov/gene/?term=LOC105282825 | 3919 | 85 | 216.89 | 1 | 2.55 | 1.18 | 14.44 | 13.87 | 0.57 | 0.57 | 0.01 | 0.01 |

| | | | | | | | | | | | | | | |
|---|---|---|---|---|---|---|---|---|---|---|---|---|---|---|
| gene-LOC105274823 | https://www.ncbi.nlm.nih.gov/gene/?term=LOC105274823 | 4620 | 141 | 305.19 | | 1 | 2.16 | 0.71 | 21.37 | 20.81 | 0.56 | 0.56 | 0.02 | 0.02 |
| gene-LOC105278534 | https://www.ncbi.nlm.nih.gov/gene/?term=LOC105278534 | 7364 | 197 | 267.52 | | 2 | 2.72 | 1.02 | 18.17 | 17.62 | 0.55 | 0.55 | 0.01 | 0.01 |
| gene-LOC105281599 | https://www.ncbi.nlm.nih.gov/gene/?term=LOC105281599 | 3859 | 49 | 126.98 | | 1 | 2.59 | 2.04 | 4.6 | 4.08 | 0.53 | 0.53 | 0.01 | 0.01 |
| gene-LOC105275127 | https://www.ncbi.nlm.nih.gov/gene/?term=LOC105275127 | 9377 | 301 | 321 | 1 | 1.07 | 0.33 | 7.86 | 7.34 | 0.52 | 0.52 | 0.02 | 0.02 | |
| gene-LOC105280410 | https://www.ncbi.nlm.nih.gov/gene/?term=LOC105280410 | 12580 | 264 | 209.86 | | 1 | 0.79 | 0.38 | 19.99 | 19.51 | 0.48 | 0.48 | 0.01 | 0.01 |
| gene-LOC105285202 | https://www.ncbi.nlm.nih.gov/gene/?term=LOC105285202 | 1838 | 70 | 380.85 | | 1 | 5.44 | 1.43 | 25.94 | 25.46 | 0.48 | 0.48 | 0.02 | 0.02 |
| gene-LOC105278454 | https://www.ncbi.nlm.nih.gov/gene/?term=LOC105278454 | 1972 | 46 | 233.27 | | 1 | 5.07 | 2.17 | 44.92 | 44.47 | 0.46 | 0.46 | 0.01 | 0.01 |
| gene-LOC105275179 | https://www.ncbi.nlm.nih.gov/gene/?term=LOC105275179 | 3271 | 72 | 220.12 | | 1 | 3.06 | 1.39 | 21.5 | 21.05 | 0.45 | 0.45 | 0.01 | 0.01 |
| gene-LOC105282491 | https://www.ncbi.nlm.nih.gov/gene/?term=LOC105282491 | 4761 | 170 | 357.07 | | 1 | 2.1 | 0.59 | 10.04 | 9.59 | 0.45 | 0.45 | 0.02 | 0.02 |
| gene-LOC105276078 | https://www.ncbi.nlm.nih.gov/gene/?term=LOC105276078 | 1051 | 14 | 133.21 | | 1 | 9.51 | 7.14 | 33.32 | 32.93 | 0.39 | 0.39 | 0.01 | 0.01 |
| gene-LOC105285860 | https://www.ncbi.nlm.nih.gov/gene/?term=LOC105285860 | 3110 | 55 | 176.85 | | 1 | 3.22 | 1.82 | 24.17 | 23.77 | 0.39 | 0.39 | 0.01 | 0.01 |
| gene-LOC105275327 | https://www.ncbi.nlm.nih.gov/gene/?term=LOC105275327 | 14245 | 492 | 345.38 | | 1 | 0.7 | 0.2 | 7.28 | 6.9 | 0.38 | 0.38 | 0.01 | 0.01 |
| gene-LOC105278666 | https://www.ncbi.nlm.nih.gov/gene/?term=LOC105278666 | 10341 | 451 | 436.13 | | 1 | 0.97 | 0.22 | 7.32 | 6.95 | 0.38 | 0.38 | 0.02 | 0.02 |
| gene-LOC105286194 | https://www.ncbi.nlm.nih.gov/gene/?term=LOC105286194 | 152619 | | 2242 | 146.9 | 1 | 0.07 | 0.04 | 6.47 | 6.09 | 0.38 | 0.38 | 0.01 | 0.01 |
| gene-LOC105274531 | https://www.ncbi.nlm.nih.gov/gene/?term=LOC105274531 | 7774 | 301 | 387.19 | | 1 | 1.29 | 0.33 | 10.99 | 10.62 | 0.37 | 0.37 | 0.01 | 0.01 |
| gene-LOC105282811 | https://www.ncbi.nlm.nih.gov/gene/?term=LOC105282811 | 5765 | 213 | 369.47 | | 1 | 1.73 | 0.47 | 8.63 | 8.26 | 0.37 | 0.37 | 0.01 | 0.01 |
| gene-LOC105276452 | https://www.ncbi.nlm.nih.gov/gene/?term=LOC105276452 | 2917 | 269 | 922.18 | | 1 | 3.43 | 0.37 | 9.31 | 8.95 | 0.36 | 0.36 | 0.03 | 0.03 |
| gene-LOC105278145 | https://www.ncbi.nlm.nih.gov/gene/?term=LOC105278145 | 4470 | 251 | 561.52 | | 1 | 2.24 | 0.4 | 10.77 | 10.43 | 0.34 | 0.34 | 0.02 | 0.02 |
| gene-LOC105287371 | https://www.ncbi.nlm.nih.gov/gene/?term=LOC105287371 | 8224 | 351 | 426.8 | 1 | 1.22 | 0.28 | 7.67 | 7.34 | 0.33 | 0.33 | 0.01 | 0.01 | |
| gene-LOC105287528 | https://www.ncbi.nlm.nih.gov/gene/?term=LOC105287528 | 14195 | 419 | 295.17 | | 1 | 0.7 | 0.24 | 5.19 | 4.86 | 0.33 | 0.33 | 0.01 | 0.01 |
| gene-LOC105286274 | https://www.ncbi.nlm.nih.gov/gene/?term=LOC105286274 | 3817 | 193 | 505.63 | | 1 | 2.62 | 0.52 | 17.37 | 17.05 | 0.32 | 0.32 | 0.02 | 0.02 |
| gene-LOC105286452 | https://www.ncbi.nlm.nih.gov/gene/?term=LOC105286452 | 6888 | 67 | 97.27 | 1 | 1.45 | 1.49 | 31.04 | 30.73 | 0.31 | 0.31 | 0 | 0 | |
| gene-LOC105287550 | https://www.ncbi.nlm.nih.gov/gene/?term=LOC105287550 | 15659 | 613 | 391.47 | | 1 | 0.64 | 0.16 | 6.43 | 6.12 | 0.31 | 0.31 | 0.01 | 0.01 |
| gene-LOC105286166 | https://www.ncbi.nlm.nih.gov/gene/?term=LOC105286166 | 3050 | 110 | 360.66 | | 1 | 3.28 | 0.91 | 24.5 | 24.21 | 0.29 | 0.29 | 0.01 | 0.01 |
| gene-LOC113562050 | https://www.ncbi.nlm.nih.gov/gene/?term=LOC113562050 | 17097 | 576 | 336.9 | 1 | 0.58 | 0.17 | 5.24 | 4.95 | 0.29 | 0.29 | 0.01 | 0.01 | |
| gene-LOC105278877 | https://www.ncbi.nlm.nih.gov/gene/?term=LOC105278877 | 1968 | 72 | 365.85 | | 1 | 5.08 | 1.39 | 36.04 | 35.76 | 0.28 | 0.28 | 0.01 | 0.01 |
| gene-LOC105277079 | https://www.ncbi.nlm.nih.gov/gene/?term=LOC105277079 | 13009 | 288 | 221.39 | | 1 | 0.77 | 0.35 | 9.86 | 9.59 | 0.27 | 0.27 | 0.01 | 0.01 |
| gene-LOC105278245 | https://www.ncbi.nlm.nih.gov/gene/?term=LOC105278245 | 6445 | 117 | 181.54 | | 1 | 1.55 | 0.85 | 22.48 | 22.2 | 0.27 | 0.27 | 0 | 0 |
| gene-LOC105284262 | https://www.ncbi.nlm.nih.gov/gene/?term=LOC105284262 | 6669 | 225 | 337.38 | | 1 | 1.5 | 0.44 | 6.17 | 5.94 | 0.23 | 0.23 | 0.01 | 0.01 |
| gene-LOC105278632 | https://www.ncbi.nlm.nih.gov/gene/?term=LOC105278632 | 3624 | 120 | 331.13 | | 1 | 2.76 | 0.83 | 15.73 | 15.53 | 0.2 | 0.2 | 0.01 | 0.01 |
| gene-LOC105285807 | https://www.ncbi.nlm.nih.gov/gene/?term=LOC105285807 | 6063 | 227 | 374.4 | 1 | 1.65 | 0.44 | 9.73 | 9.55 | 0.18 | 0.18 | 0.01 | 0.01 | |
| gene-LOC105276272 | https://www.ncbi.nlm.nih.gov/gene/?term=LOC105276272 | 1965 | 48 | 244.27 | | 1 | 5.09 | 2.08 | 27.74 | 27.58 | 0.17 | 0.17 | 0 | 0 |
| gene-LOC105276527 | https://www.ncbi.nlm.nih.gov/gene/?term=LOC105276527 | 7923 | 329 | 415.25 | | 1 | 1.26 | 0.3 | 9.51 | 9.36 | 0.15 | 0.15 | 0.01 | 0.01 |
| gene-LOC105277125 | https://www.ncbi.nlm.nih.gov/gene/?term=LOC105277125 | 51079 | 1858 | 363.75 | | 1 | 0.2 | 0.05 | 1.73 | 1.6 | 0.13 | 0.13 | 0 | 0 |
| gene-LOC105277866 | https://www.ncbi.nlm.nih.gov/gene/?term=LOC105277866 | 33045 | 400 | 121.05 | | 1 | 0.3 | 0.25 | 3.44 | 3.35 | 0.1 | 0.1 | 0 | 0 |
| gene-LOC105280265 | https://www.ncbi.nlm.nih.gov/gene/?term=LOC105280265 | 85984 | 3130 | 364.02 | | 1 | 0.12 | 0.03 | 1.14 | 1.05 | 0.09 | 0.09 | 0 | 0 |
| gene-LOC105274669 | https://www.ncbi.nlm.nih.gov/gene/?term=LOC105274669 | 8555 | 256 | 299.24 | | 1 | 1.17 | 0.39 | 6.43 | 6.35 | 0.08 | 0.08 | 0 | 0 |
| gene-LOC105275752 | https://www.ncbi.nlm.nih.gov/gene/?term=LOC105275752 | 3462 | 132 | 381.28 | | 1 | 2.89 | 0.76 | 32.74 | 32.68 | 0.06 | 0.06 | 0 | 0 |
| gene-LOC105278455 | https://www.ncbi.nlm.nih.gov/gene/?term=LOC105278455 | 27181 | 961 | 353.56 | | 1 | 0.37 | 0.1 | 7.16 | 7.14 | 0.02 | 0.02 | 0 | 0 |
| gene-LOC105286515 | https://www.ncbi.nlm.nih.gov/gene/?term=LOC105286515 | 3751 | 114 | 303.92 | | 1 | 2.67 | 0.88 | 11.87 | 11.89 | -0.02 | 0.02 | 0 | 0 |
| gene-LOC105283296 | https://www.ncbi.nlm.nih.gov/gene/?term=LOC105283296 | 2174 | 94 | 432.38 | | 1 | 4.6 | 1.06 | 18.21 | 18.24 | -0.03 | 0.03 | 0 | 0 |
| gene-LOC105287695 | https://www.ncbi.nlm.nih.gov/gene/?term=LOC105287695 | 6106 | 201 | 329.18 | | 1 | 1.64 | 0.5 | 6.09 | 6.16 | -0.06 | 0.06 | 0 | 0 |
| gene-LOC105274851 | https://www.ncbi.nlm.nih.gov/gene/?term=LOC105274851 | 6616 | 139 | 210.1 | 1 | 1.51 | 0.72 | 11.55 | 11.65 | -0.1 | 0.1 | 0 | 0 | |
| gene-LOC105275187 | https://www.ncbi.nlm.nih.gov/gene/?term=LOC105275187 | 5534 | 71 | 128.3 | 1 | 1.81 | 1.41 | 12.33 | 12.43 | -0.1 | 0.1 | 0 | 0 | |
| gene-LOC105286649 | https://www.ncbi.nlm.nih.gov/gene/?term=LOC105286649 | 14178 | 817 | 576.24 | | 1 | 0.71 | 0.12 | 1 | 1.1 | -0.1 | 0.1 | -0.01 | 0.01 |
| gene-LOC105282646 | https://www.ncbi.nlm.nih.gov/gene/?term=LOC105282646 | 3317 | 116 | 349.71 | | 1 | 3.01 | 0.86 | 24.81 | 24.92 | -0.11 | 0.11 | 0 | 0 |
| gene-LOC105283806 | https://www.ncbi.nlm.nih.gov/gene/?term=LOC105283806 | 10501 | 74 | 70.47 | 1 | 0.95 | 1.35 | 16.11 | 16.23 | -0.12 | 0.12 | 0 | 0 | |
| gene-LOC105284507 | https://www.ncbi.nlm.nih.gov/gene/?term=LOC105284507 | 5207 | 87 | 167.08 | | 1 | 1.92 | 1.15 | 21.48 | 21.62 | -0.13 | 0.13 | 0 | 0 |
| gene-LOC105275848 | https://www.ncbi.nlm.nih.gov/gene/?term=LOC105275848 | 3718 | 144 | 387.31 | | 1 | 2.69 | 0.69 | 20.8 | 20.95 | -0.15 | 0.15 | -0.01 | 0.01 |
| gene-LOC105280704 | https://www.ncbi.nlm.nih.gov/gene/?term=LOC105280704 | 4266 | 134 | 314.11 | | 1 | 2.34 | 0.75 | 19.89 | 20.04 | -0.15 | 0.15 | 0 | 0 |
| gene-LOC105280348 | https://www.ncbi.nlm.nih.gov/gene/?term=LOC105280348 | 3797 | 70 | 184.36 | | 1 | 2.63 | 1.43 | 20.64 | 20.81 | -0.17 | 0.17 | 0 | 0 |
| gene-LOC105283967 | https://www.ncbi.nlm.nih.gov/gene/?term=LOC105283967 | 2393 | 47 | 196.41 | | 1 | 4.18 | 2.13 | 48.9 | 49.11 | -0.21 | 0.21 | 0 | 0 |
| gene-LOC105285163 | https://www.ncbi.nlm.nih.gov/gene/?term=LOC105285163 | 5061 | 87 | 171.9 | 1 | 1.98 | 1.15 | 21.7 | 21.95 | -0.26 | 0.26 | 0 | 0 | |
| gene-LOC105287633 | https://www.ncbi.nlm.nih.gov/gene/?term=LOC105287633 | 1524 | 34 | 223.1 | 1 | 6.56 | 2.94 | 50.71 | 50.96 | -0.26 | 0.26 | -0.01 | 0.01 | |
| gene-LOC105281312 | https://www.ncbi.nlm.nih.gov/gene/?term=LOC105281312 | 2551 | 63 | 246.96 | | 1 | 3.92 | 1.59 | 34.27 | 34.56 | -0.29 | 0.29 | -0.01 | 0.01 |
| gene-LOC105286999 | https://www.ncbi.nlm.nih.gov/gene/?term=LOC105286999 | 2141 | 44 | 205.51 | | 1 | 4.67 | 2.27 | 34.01 | 34.41 | -0.41 | 0.41 | -0.01 | 0.01 |
| gene-LOC105287095 | https://www.ncbi.nlm.nih.gov/gene/?term=LOC105287095 | 1435 | 58 | 404.18 | | 1 | 6.97 | 1.72 | 28.05 | 28.65 | -0.6 | 0.6 | -0.02 | 0.02 |
| gene-LOC105274449 | https://www.ncbi.nlm.nih.gov/gene/?term=LOC105274449 | 1549 | 48 | 309.88 | | 1 | 6.46 | 2.08 | 50.3 | 50.96 | -0.66 | 0.66 | -0.02 | 0.02 |
| gene-LOC105285252 | https://www.ncbi.nlm.nih.gov/gene/?term=LOC105285252 | 7427 | 117 | 157.53 | | 1 | 1.35 | 0.85 | 14.46 | 15.16 | -0.7 | 0.7 | -0.01 | 0.01 |
| gene-LOC105281296 | https://www.ncbi.nlm.nih.gov/gene/?term=LOC105281296 | 2434 | 37 | 152.01 | | 1 | 4.11 | 2.7 | 30.9 | 31.72 | -0.83 | 0.83 | -0.01 | 0.01 |
| gene-LOC105280596 | https://www.ncbi.nlm.nih.gov/gene/?term=LOC105280596 | 3434 | 113 | 329.06 | | 1 | 2.91 | 0.88 | 17.62 | 18.56 | -0.94 | 0.94 | -0.03 | 0.03 |
| gene-LOC105282885 | https://www.ncbi.nlm.nih.gov/gene/?term=LOC105282885 | 1870 | 48 | 256.68 | | 1 | 5.35 | 2.08 | 38.27 | 39.26 | -0.99 | 0.99 | -0.03 | 0.03 |
| gene-LOC105278384 | https://www.ncbi.nlm.nih.gov/gene/?term=LOC105278384 | 1390 | 36 | 258.99 | | 1 | 7.19 | 2.78 | 50.65 | 51.65 | -1 | 1 | -0.03 | 0.03 |
| gene-LOC105286770 | https://www.ncbi.nlm.nih.gov/gene/?term=LOC105286770 | 3055 | 33 | 108.02 | | 1 | 3.27 | 3.03 | 17.27 | 18.34 | -1.07 | 1.07 | -0.01 | 0.01 |

| | | | | | | | | | | | | | | | |
|---|---|---|---|---|---|---|---|---|---|---|---|---|---|---|---|
| gene-LOC105280819 | https://www.ncbi.nlm.nih.gov/gene/?term=LOC105280819 | 1639 | 38 | 231.85 | | 1 | 6.1 | 2.63 | 35.58 | 37.06 | -1.49 | 1.49 | -0.03 | 0.03 | |
| gene-LOC105283989 | https://www.ncbi.nlm.nih.gov/gene/?term=LOC105283989 | 4948 | 31 | 62.65 | 2 | 4.04 | 6.45 | 34.97 | 36.46 | -1.49 | 1.49 | -0.01 | 0.01 | | |
| gene-LOC105276196 | https://www.ncbi.nlm.nih.gov/gene/?term=LOC105276196 | 2953 | 62 | 209.96 | | 1 | 3.39 | 1.61 | 36.95 | 38.53 | -1.58 | 1.58 | -0.03 | 0.03 | |
| gene-LOC105281700 | https://www.ncbi.nlm.nih.gov/gene/?term=LOC105281700 | 2208 | 34 | 153.99 | | 1 | 4.53 | 2.94 | 35.23 | 36.92 | -1.69 | 1.69 | -0.03 | 0.03 | |
| gene-LOC105284047 | https://www.ncbi.nlm.nih.gov/gene/?term=LOC105284047 | 1212 | 24 | 198.02 | | 1 | 8.25 | 4.17 | 25.81 | 27.68 | -1.87 | 1.87 | -0.04 | 0.04 | |
| gene-LOC105285409 | https://www.ncbi.nlm.nih.gov/gene/?term=LOC105285409 | 5099 | 49 | 96.1 | 2 | 3.92 | 4.08 | 1.55 | 4.82 | -3.27 | 3.27 | -0.03 | 0.03 | | |
| gene-LOC105283492 | https://www.ncbi.nlm.nih.gov/gene/?term=LOC105283492 | 1535 | 10 | 65.15 | 1 | 6.51 | 10 | 37.74 | 42.8 | -5.06 | 5.06 | -0.03 | 0.03 | | |
| gene-LOC105284677 | https://www.ncbi.nlm.nih.gov/gene/?term=LOC105284677 | 1905 | 42 | 220.47 | | 1 | 5.25 | 2.38 | 23.81 | 34.99 | -11.18 | | 11.18 | -0.25 | 0.25 |

Directory: OGLall File: ogl-Ob_LI_RphaseB1.txt

| gene_ID | gene_link | gwidth | #Sites | #per10Kb | %perSite | %pNucl |
|---|---|---|---|---|---|---|
| gene-LOC113561532 | https://www.ncbi.nlm.nih.gov/gene/?term=LOC113561532 | 72 | 5 | 694.44 | 95.54 | 6.64 |
| gene-LOC105286531 | https://www.ncbi.nlm.nih.gov/gene/?term=LOC105286531 | 372 | 40 | 1075.27 | 35.13 | 3.78 |
| gene-LOC105280739 | https://www.ncbi.nlm.nih.gov/gene/?term=LOC105280739 | 1991 | 138 | 693.12 | 53.97 | 3.74 |
| gene-LOC113562627 | https://www.ncbi.nlm.nih.gov/gene/?term=LOC113562627 | 397 | 22 | 554.16 | 65.67 | 3.64 |
| gene-LOC105274534 | https://www.ncbi.nlm.nih.gov/gene/?term=LOC105274534 | 1329 | 102 | 767.49 | 45.61 | 3.5 |
| gene-LOC105286801 | https://www.ncbi.nlm.nih.gov/gene/?term=LOC105286801 | 1102 | 101 | 916.52 | 37.9 | 3.47 |
| gene-LOC105285635 | https://www.ncbi.nlm.nih.gov/gene/?term=LOC105285635 | 1175 | 127 | 1080.85 | 31.75 | 3.43 |
| gene-LOC105286269 | https://www.ncbi.nlm.nih.gov/gene/?term=LOC105286269 | 1401 | 95 | 678.09 | 50.19 | 3.4 |
| gene-LOC105276360 | https://www.ncbi.nlm.nih.gov/gene/?term=LOC105276360 | 1466 | 169 | 1152.8 | 29.3 | 3.38 |
| gene-LOC105287811 | https://www.ncbi.nlm.nih.gov/gene/?term=LOC105287811 | 2155 | 135 | 626.45 | 53.84 | 3.37 |
| gene-LOC105278844 | https://www.ncbi.nlm.nih.gov/gene/?term=LOC105278844 | 1101 | 115 | 1044.5 | 30.86 | 3.22 |
| gene-LOC105278849 | https://www.ncbi.nlm.nih.gov/gene/?term=LOC105278849 | 776 | 114 | 1469.07 | 21.62 | 3.18 |
| gene-LOC105279052 | https://www.ncbi.nlm.nih.gov/gene/?term=LOC105279052 | 1130 | 127 | 1123.89 | 27.56 | 3.1 |
| gene-LOC105278160 | https://www.ncbi.nlm.nih.gov/gene/?term=LOC105278160 | 1872 | 168 | 897.44 | 34.28 | 3.08 |
| gene-LOC105285392 | https://www.ncbi.nlm.nih.gov/gene/?term=LOC105285392 | 1601 | 134 | 836.98 | 36.82 | 3.08 |
| gene-LOC105274711 | https://www.ncbi.nlm.nih.gov/gene/?term=LOC105274711 | 1655 | 113 | 682.78 | 44.97 | 3.07 |
| gene-LOC105286235 | https://www.ncbi.nlm.nih.gov/gene/?term=LOC105286235 | 1510 | 105 | 695.36 | 44.05 | 3.06 |
| gene-LOC105286737 | https://www.ncbi.nlm.nih.gov/gene/?term=LOC105286737 | 2690 | 368 | 1368.03 | 22.4 | 3.06 |
| gene-LOC105278549 | https://www.ncbi.nlm.nih.gov/gene/?term=LOC105278549 | 1771 | 117 | 660.64 | 46.07 | 3.04 |
| gene-LOC105282149 | https://www.ncbi.nlm.nih.gov/gene/?term=LOC105282149 | 1787 | 134 | 749.86 | 40.5 | 3.04 |
| gene-LOC105287692 | https://www.ncbi.nlm.nih.gov/gene/?term=LOC105287692 | 1505 | 137 | 910.3 | 32.53 | 2.96 |
| gene-LOC105275314 | https://www.ncbi.nlm.nih.gov/gene/?term=LOC105275314 | 1489 | 104 | 698.46 | 41.67 | 2.91 |
| gene-LOC105277648 | https://www.ncbi.nlm.nih.gov/gene/?term=LOC105277648 | 722 | 39 | 540.17 | 53.41 | 2.89 |
| gene-LOC105288088 | https://www.ncbi.nlm.nih.gov/gene/?term=LOC105288088 | 1795 | 123 | 685.24 | 42.1 | 2.89 |

...

... only first 25 lines shown ...

Directory: OGLall File: ogl-Ob_LI_RphaseB1.vs.RphaseB2.txt

| gene_ID | gene_link | gwidth | #Sites | #per10Kb | #dmSites | #dmsp10kb | %dmSites | %pSite1 | %pSite2 | DMpSite | ADMpSite | DMpNucl | ADMpNucl |
|---|---|---|---|---|---|---|---|---|---|---|---|---|---|
| gene-LOC105285431 | https://www.ncbi.nlm.nih.gov/gene/?term=LOC105285431 | 1550 | 6 | 38.71 | 3 | 19.35 | 50 | 46.72 | 30.29 | 16.43 | 16.43 | 0.06 | 0.06 |
| gene-LOC105284871 | https://www.ncbi.nlm.nih.gov/gene/?term=LOC105284871 | 1878 | 3 | 15.97 | 1 | 5.32 | 33.33 | 46.42 | 30.96 | 15.47 | 15.47 | 0.02 | 0.02 |
| gene-LOC113561977 | https://www.ncbi.nlm.nih.gov/gene/?term=LOC113561977 | 1058 | 3 | 28.36 | 1 | 9.45 | 33.33 | 25 | 10.74 | 14.26 | 14.26 | 0.04 | 0.04 |
| gene-LOC105277491 | https://www.ncbi.nlm.nih.gov/gene/?term=LOC105277491 | 1647 | 12 | 72.86 | 2 | 12.14 | 16.67 | 21.62 | 11.32 | 10.3 | 10.3 | 0.08 | 0.08 |
| gene-LOC105279429 | https://www.ncbi.nlm.nih.gov/gene/?term=LOC105279429 | 1741 | 10 | 57.44 | 1 | 5.74 | 10 | 23.71 | 13.86 | 9.85 | 9.85 | 0.06 | 0.06 |
| gene-LOC105283805 | https://www.ncbi.nlm.nih.gov/gene/?term=LOC105283805 | 2008 | 7 | 34.86 | 2 | 9.96 | 28.57 | 87.5 | 77.76 | 9.74 | 9.74 | 0.03 | 0.03 |
| gene-LOC105282854 | https://www.ncbi.nlm.nih.gov/gene/?term=LOC105282854 | 1825 | 2 | 10.96 | 1 | 5.48 | 50 | 38.24 | 28.94 | 9.3 | 9.3 | 0.01 | 0.01 |
| gene-LOC105281216 | https://www.ncbi.nlm.nih.gov/gene/?term=LOC105281216 | 3765 | 12 | 31.87 | 2 | 5.31 | 16.67 | 39.32 | 30.3 | 9.02 | 9.02 | 0.03 | 0.03 |
| gene-LOC105283410 | https://www.ncbi.nlm.nih.gov/gene/?term=LOC105283410 | 1423 | 5 | 35.14 | 1 | 7.03 | 20 | 49.1 | 40.2 | 8.9 | 8.9 | 0.03 | 0.03 |
| gene-LOC113561900 | https://www.ncbi.nlm.nih.gov/gene/?term=LOC113561900 | 629 | 11 | 174.88 | 1 | 15.9 | 9.09 | 33.39 | 25.51 | 7.88 | 7.88 | 0.14 | 0.14 |
| gene-LOC105281635 | https://www.ncbi.nlm.nih.gov/gene/?term=LOC105281635 | 1508 | 4 | 26.53 | 1 | 6.63 | 25 | 15.47 | 7.61 | 7.86 | 7.86 | 0.02 | 0.02 |
| gene-LOC105286857 | https://www.ncbi.nlm.nih.gov/gene/?term=LOC105286857 | 1576 | 4 | 25.38 | 1 | 6.35 | 25 | 74.57 | 67.03 | 7.54 | 7.54 | 0.02 | 0.02 |
| gene-LOC105282657 | https://www.ncbi.nlm.nih.gov/gene/?term=LOC105282657 | 1409 | 6 | 42.58 | 1 | 7.1 | 16.67 | 58.16 | 50.81 | 7.34 | 7.34 | 0.03 | 0.03 |
| gene-LOC105277862 | https://www.ncbi.nlm.nih.gov/gene/?term=LOC105277862 | 1490 | 7 | 46.98 | 1 | 6.71 | 14.29 | 62.25 | 55.64 | 6.6 | 6.6 | 0.03 | 0.03 |
| gene-LOC105286261 | https://www.ncbi.nlm.nih.gov/gene/?term=LOC105286261 | 1297 | 8 | 61.68 | 1 | 7.71 | 12.5 | 74.9 | 68.31 | 6.59 | 6.59 | 0.04 | 0.04 |
| gene-LOC113562499 | https://www.ncbi.nlm.nih.gov/gene/?term=LOC113562499 | 491 | 6 | 122.2 | 1 | 20.37 | 16.67 | 53.38 | 46.94 | 6.45 | 6.45 | 0.08 | 0.08 |
| gene-LOC105276718 | https://www.ncbi.nlm.nih.gov/gene/?term=LOC105276718 | 1721 | 10 | 58.11 | 1 | 5.81 | 10 | 28.17 | 21.77 | 6.4 | 6.4 | 0.04 | 0.04 |
| gene-LOC105274857 | https://www.ncbi.nlm.nih.gov/gene/?term=LOC105274857 | 1569 | 5 | 31.87 | 1 | 6.37 | 20 | 57.81 | 51.63 | 6.18 | 6.18 | 0.02 | 0.02 |
| gene-LOC105280261 | https://www.ncbi.nlm.nih.gov/gene/?term=LOC105280261 | 1005 | 7 | 69.65 | 1 | 9.95 | 14.29 | 8.97 | 3.43 | 5.54 | 5.54 | 0.04 | 0.04 |
| gene-LOC105284028 | https://www.ncbi.nlm.nih.gov/gene/?term=LOC105284028 | 2536 | 23 | 90.69 | 4 | 15.77 | 17.39 | 36.1 | 30.67 | 5.43 | 5.43 | 0.05 | 0.05 |
| gene-LOC105287722 | https://www.ncbi.nlm.nih.gov/gene/?term=LOC105287722 | 3481 | 6 | 17.24 | 1 | 2.87 | 16.67 | 47.51 | 42.11 | 5.39 | 5.39 | 0.01 | 0.01 |
| gene-LOC105279461 | https://www.ncbi.nlm.nih.gov/gene/?term=LOC105279461 | 1298 | 9 | 69.34 | 1 | 7.7 | 11.11 | 73.75 | 68.65 | 5.1 | 5.1 | 0.04 | 0.04 |
| gene-LOC105284656 | https://www.ncbi.nlm.nih.gov/gene/?term=LOC105284656 | 1312 | 17 | 129.57 | 2 | 15.24 | 11.76 | 79.02 | 73.94 | 5.08 | 5.08 | 0.07 | 0.07 |
| gene-LOC105285670 | https://www.ncbi.nlm.nih.gov/gene/?term=LOC105285670 | 1477 | 21 | 142.18 | 1 | 6.77 | 4.76 | 49.06 | 44.02 | 5.03 | 5.03 | 0.07 | 0.07 |

...

... only first 25 lines shown ...

Directory: OGLall File: ogl-Ob_LI_RphaseB2.txt

| gene_ID | gene_link | gwidth | #Sites | #per10Kb | %perSite | %pNucl |
|---|---|---|---|---|---|---|
| gene-LOC113561532 | https://www.ncbi.nlm.nih.gov/gene/?term=LOC113561532 | 72 | 4 | 555.56 | 94.52 | 5.25 |
| gene-LOC105286531 | https://www.ncbi.nlm.nih.gov/gene/?term=LOC105286531 | 372 | 41 | 1102.15 | 36.36 | 4.01 |
| gene-LOC105286801 | https://www.ncbi.nlm.nih.gov/gene/?term=LOC105286801 | 1102 | 95 | 862.07 | 43.07 | 3.71 |
| gene-LOC105282149 | https://www.ncbi.nlm.nih.gov/gene/?term=LOC105282149 | 1787 | 157 | 878.57 | 41.23 | 3.62 |
| gene-LOC105280739 | https://www.ncbi.nlm.nih.gov/gene/?term=LOC105280739 | 1991 | 127 | 637.87 | 54.82 | 3.5 |
| gene-LOC105278849 | https://www.ncbi.nlm.nih.gov/gene/?term=LOC105278849 | 776 | 113 | 1456.19 | 23.54 | 3.43 |
| gene-LOC105285635 | https://www.ncbi.nlm.nih.gov/gene/?term=LOC105285635 | 1175 | 123 | 1046.81 | 32.53 | 3.4 |
| gene-LOC105276360 | https://www.ncbi.nlm.nih.gov/gene/?term=LOC105276360 | 1466 | 141 | 961.8 | 35 | 3.37 |
| gene-LOC105274534 | https://www.ncbi.nlm.nih.gov/gene/?term=LOC105274534 | 1329 | 89 | 669.68 | 48.75 | 3.26 |
| gene-LOC113562627 | https://www.ncbi.nlm.nih.gov/gene/?term=LOC113562627 | 397 | 19 | 478.59 | 66.86 | 3.2 |
| gene-LOC105287811 | https://www.ncbi.nlm.nih.gov/gene/?term=LOC105287811 | 2155 | 147 | 682.13 | 46.69 | 3.19 |
| gene-LOC105274711 | https://www.ncbi.nlm.nih.gov/gene/?term=LOC105274711 | 1655 | 111 | 670.69 | 47.04 | 3.15 |
| gene-LOC105278160 | https://www.ncbi.nlm.nih.gov/gene/?term=LOC105278160 | 1872 | 159 | 849.36 | 36.67 | 3.11 |
| gene-LOC105278844 | https://www.ncbi.nlm.nih.gov/gene/?term=LOC105278844 | 1101 | 111 | 1008.17 | 30.45 | 3.07 |
| gene-LOC105276371 | https://www.ncbi.nlm.nih.gov/gene/?term=LOC105276371 | 871 | 57 | 654.42 | 46.14 | 3.02 |
| gene-LOC105287692 | https://www.ncbi.nlm.nih.gov/gene/?term=LOC105287692 | 1505 | 129 | 857.14 | 34.93 | 2.99 |
| gene-LOC105279052 | https://www.ncbi.nlm.nih.gov/gene/?term=LOC105279052 | 1130 | 112 | 991.15 | 29.29 | 2.9 |
| gene-LOC105286235 | https://www.ncbi.nlm.nih.gov/gene/?term=LOC105286235 | 1510 | 96 | 635.76 | 45.62 | 2.9 |
| gene-LOC105286737 | https://www.ncbi.nlm.nih.gov/gene/?term=LOC105286737 | 2690 | 337 | 1252.79 | 22.93 | 2.87 |
| gene-LOC105275314 | https://www.ncbi.nlm.nih.gov/gene/?term=LOC105275314 | 1489 | 89 | 597.72 | 47.67 | 2.85 |
| gene-LOC105283717 | https://www.ncbi.nlm.nih.gov/gene/?term=LOC105283717 | 1373 | 112 | 815.73 | 34.69 | 2.83 |
| gene-LOC105287677 | https://www.ncbi.nlm.nih.gov/gene/?term=LOC105287677 | 1588 | 115 | 724.18 | 38.65 | 2.8 |
| gene-LOC105286269 | https://www.ncbi.nlm.nih.gov/gene/?term=LOC105286269 | 1401 | 68 | 485.37 | 55.58 | 2.7 |
| gene-LOC105275897 | https://www.ncbi.nlm.nih.gov/gene/?term=LOC105275897 | 1684 | 116 | 688.84 | 38.05 | 2.62 |

...

... only first 25 lines shown ...

Directory: OGLall File: rnk-dmg-Ob_LI_BCphaseA1.vs.BCphaseA2.txt

| gene_ID | gene_link | gwidth | #Sites | #per10Kb | #dmSites | #dmsp10kb | %dmSites | %pSite1 | %pSite2 | DMpSite | ADMpSite | DMpNucl | ADMpNucl |
|---|---|---|---|---|---|---|---|---|---|---|---|---|---|
| gene-LOC113562345 | https://www.ncbi.nlm.nih.gov/gene/?term=LOC113562345 | 2290 | 62 | 270.74 | 15 | 65.5 | 24.19 | 17.91 | 32.18 | -14.27 | 14.27 | -0.39 | 0.39 |
| gene-LOC105283552 | https://www.ncbi.nlm.nih.gov/gene/?term=LOC105283552 | 1562 | 27 | 172.86 | 1 | 6.4 | 3.7 | 56.95 | 65.52 | -8.57 | 8.57 | -0.15 | 0.15 |
| gene-LOC105285695 | https://www.ncbi.nlm.nih.gov/gene/?term=LOC105285695 | 1371 | 62 | 452.22 | 1 | 7.29 | 1.61 | 23 | 25.77 | -2.77 | 2.77 | -0.13 | 0.13 |
| gene-LOC105282156 | https://www.ncbi.nlm.nih.gov/gene/?term=LOC105282156 | 2577 | 132 | 512.22 | 1 | 3.88 | 0.76 | 24.4 | 26.49 | -2.09 | 2.09 | -0.11 | 0.11 |
| gene-LOC105275656 | https://www.ncbi.nlm.nih.gov/gene/?term=LOC105275656 | 2803 | 127 | 453.09 | 1 | 3.57 | 0.79 | 26.43 | 28.64 | -2.2 | 2.2 | -0.1 | 0.1 |
| gene-LOC105279356 | https://www.ncbi.nlm.nih.gov/gene/?term=LOC105279356 | 1314 | 39 | 296.8 | 1 | 7.61 | 2.56 | 29.77 | 33.04 | -3.27 | 3.27 | -0.1 | 0.1 |
| gene-LOC105285674 | https://www.ncbi.nlm.nih.gov/gene/?term=LOC105285674 | 3035 | 88 | 289.95 | 1 | 3.29 | 1.14 | 23.36 | 26.44 | -3.08 | 3.08 | -0.09 | 0.09 |
| gene-LOC105281848 | https://www.ncbi.nlm.nih.gov/gene/?term=LOC105281848 | 2122 | 44 | 207.35 | 1 | 4.71 | 2.27 | 23.57 | 27.97 | -4.4 | 4.4 | -0.09 | 0.09 |
| gene-LOC105285060 | https://www.ncbi.nlm.nih.gov/gene/?term=LOC105285060 | 1285 | 13 | 101.17 | 1 | 7.78 | 7.69 | 55.35 | 64 | -8.64 | 8.64 | -0.09 | 0.09 |
| gene-LOC105285055 | https://www.ncbi.nlm.nih.gov/gene/?term=LOC105285055 | 4108 | 197 | 479.55 | 1 | 2.43 | 0.51 | 13.57 | 15.12 | -1.55 | 1.55 | -0.07 | 0.07 |
| gene-LOC105280972 | https://www.ncbi.nlm.nih.gov/gene/?term=LOC105280972 | 1532 | 26 | 169.71 | 1 | 6.53 | 3.85 | 20.04 | 23.88 | -3.84 | 3.84 | -0.07 | 0.07 |
| gene-LOC105275070 | https://www.ncbi.nlm.nih.gov/gene/?term=LOC105275070 | 5667 | 35 | 61.76 | 2 | 3.53 | 5.71 | 3.8 | 15.13 | -11.33 | 11.33 | -0.07 | 0.07 |
| gene-LOC109611283 | https://www.ncbi.nlm.nih.gov/gene/?term=LOC109611283 | 1451 | 36 | 248.1 | 1 | 6.89 | 2.78 | 37.85 | 35.62 | 2.23 | 2.23 | 0.06 | 0.06 |
| gene-LOC105280774 | https://www.ncbi.nlm.nih.gov/gene/?term=LOC105280774 | 2793 | 97 | 347.3 | 1 | 3.58 | 1.03 | 31.81 | 33.47 | -1.66 | 1.66 | -0.06 | 0.06 |
| gene-LOC113561621 | https://www.ncbi.nlm.nih.gov/gene/?term=LOC113561621 | 2249 | 79 | 351.27 | 1 | 4.45 | 1.27 | 19.72 | 21.41 | -1.69 | 1.69 | -0.06 | 0.06 |
| gene-LOC105278807 | https://www.ncbi.nlm.nih.gov/gene/?term=LOC105278807 | 2166 | 68 | 313.94 | 1 | 4.62 | 1.47 | 17.54 | 19.41 | -1.87 | 1.87 | -0.06 | 0.06 |
| gene-LOC105281173 | https://www.ncbi.nlm.nih.gov/gene/?term=LOC105281173 | 1595 | 25 | 156.74 | 1 | 6.27 | 4 | 64.76 | 61.8 | 2.96 | 2.96 | 0.05 | 0.05 |
| gene-LOC105278877 | https://www.ncbi.nlm.nih.gov/gene/?term=LOC105278877 | 1968 | 69 | 350.61 | 1 | 5.08 | 1.45 | 37.18 | 35.87 | 1.31 | 1.31 | 0.05 | 0.05 |
| gene-LOC105284184 | https://www.ncbi.nlm.nih.gov/gene/?term=LOC105284184 | 2535 | 67 | 264.3 | 1 | 3.94 | 1.49 | 41.64 | 43.45 | -1.81 | 1.81 | -0.05 | 0.05 |
| gene-LOC105279898 | https://www.ncbi.nlm.nih.gov/gene/?term=LOC105279898 | 1343 | 13 | 96.8 | 1 | 7.45 | 7.69 | 60.41 | 65.83 | -5.42 | 5.42 | -0.05 | 0.05 |
| gene-LOC105276078 | https://www.ncbi.nlm.nih.gov/gene/?term=LOC105276078 | 1051 | 8 | 76.12 | 1 | 9.51 | 12.5 | 46.61 | 53.81 | -7.2 | 7.2 | -0.05 | 0.05 |
| gene-LOC105275708 | https://www.ncbi.nlm.nih.gov/gene/?term=LOC105275708 | 1772 | 10 | 56.43 | 1 | 5.64 | 10 | 41.45 | 50.31 | -8.85 | 8.85 | -0.05 | 0.05 |
| gene-LOC105286999 | https://www.ncbi.nlm.nih.gov/gene/?term=LOC105286999 | 2141 | 38 | 177.49 | 1 | 4.67 | 2.63 | 33.02 | 30.95 | 2.07 | 2.07 | 0.04 | 0.04 |
| gene-LOC105274449 | https://www.ncbi.nlm.nih.gov/gene/?term=LOC105274449 | 1549 | 42 | 271.14 | 1 | 6.46 | 2.38 | 55.88 | 54.57 | 1.32 | 1.32 | 0.04 | 0.04 |
| gene-LOC105279298 | https://www.ncbi.nlm.nih.gov/gene/?term=LOC105279298 | 2773 | 129 | 465.2 | 2 | 7.21 | 1.55 | 22.93 | 23.79 | -0.86 | 0.86 | -0.04 | 0.04 |
| gene-LOC105275460 | https://www.ncbi.nlm.nih.gov/gene/?term=LOC105275460 | 2473 | 107 | 432.67 | 1 | 4.04 | 0.93 | 13.53 | 14.43 | -0.91 | 0.91 | -0.04 | 0.04 |
| gene-LOC105275683 | https://www.ncbi.nlm.nih.gov/gene/?term=LOC105275683 | 4256 | 187 | 439.38 | 1 | 2.35 | 0.53 | 12 | 12.9 | -0.91 | 0.91 | -0.04 | 0.04 |
| gene-LOC105285671 | https://www.ncbi.nlm.nih.gov/gene/?term=LOC105285671 | 2263 | 92 | 406.54 | 1 | 4.42 | 1.09 | 28.77 | 29.78 | -1.01 | 1.01 | -0.04 | 0.04 |
| gene-LOC105285179 | https://www.ncbi.nlm.nih.gov/gene/?term=LOC105285179 | 4191 | 148 | 353.14 | 1 | 2.39 | 0.68 | 17.81 | 18.87 | -1.05 | 1.05 | -0.04 | 0.04 |
| gene-LOC105278109 | https://www.ncbi.nlm.nih.gov/gene/?term=LOC105278109 | 4029 | 56 | 138.99 | 1 | 2.48 | 1.79 | 47.35 | 50.32 | -2.97 | 2.97 | -0.04 | 0.04 |
| gene-LOC105280348 | https://www.ncbi.nlm.nih.gov/gene/?term=LOC105280348 | 3797 | 50 | 131.68 | 1 | 2.63 | 2 | 25.73 | 28.9 | -3.17 | 3.17 | -0.04 | 0.04 |
| gene-LOC105285514 | https://www.ncbi.nlm.nih.gov/gene/?term=LOC105285514 | 1277 | 8 | 62.65 | 1 | 7.83 | 12.5 | 82.36 | 88.52 | -6.16 | 6.16 | -0.04 | 0.04 |
| gene-LOC105285202 | https://www.ncbi.nlm.nih.gov/gene/?term=LOC105285202 | 1838 | 33 | 179.54 | 1 | 5.44 | 3.03 | 20.53 | 19.01 | 1.52 | 1.52 | 0.03 | 0.03 |
| gene-LOC105278282 | https://www.ncbi.nlm.nih.gov/gene/?term=LOC105278282 | 1775 | 43 | 242.25 | 1 | 5.63 | 2.33 | 46.59 | 45.35 | 1.24 | 1.24 | 0.03 | 0.03 |
| gene-LOC105275848 | https://www.ncbi.nlm.nih.gov/gene/?term=LOC105275848 | 3718 | 119 | 320.06 | 1 | 2.69 | 0.84 | 22.44 | 21.48 | 0.97 | 0.97 | 0.03 | 0.03 |
| gene-LOC105277435 | https://www.ncbi.nlm.nih.gov/gene/?term=LOC105277435 | 3643 | 150 | 411.75 | 1 | 2.74 | 0.67 | 12.77 | 12 | 0.77 | 0.77 | 0.03 | 0.03 |
| gene-LOC105275363 | https://www.ncbi.nlm.nih.gov/gene/?term=LOC105275363 | 3798 | 146 | 384.41 | 1 | 2.63 | 0.68 | 17.72 | 18.47 | -0.76 | 0.76 | -0.03 | 0.03 |
| gene-LOC105276692 | https://www.ncbi.nlm.nih.gov/gene/?term=LOC105276692 | 1745 | 33 | 189.11 | 1 | 5.73 | 3.03 | 25.26 | 26.77 | -1.52 | 1.52 | -0.03 | 0.03 |
| gene-LOC105275190 | https://www.ncbi.nlm.nih.gov/gene/?term=LOC105275190 | 3837 | 69 | 179.83 | 1 | 2.61 | 1.45 | 19.53 | 21.12 | -1.59 | 1.59 | -0.03 | 0.03 |
| gene-LOC105283592 | https://www.ncbi.nlm.nih.gov/gene/?term=LOC105283592 | 3479 | 66 | 189.71 | 1 | 2.87 | 1.52 | 25.49 | 27.08 | -1.59 | 1.59 | -0.03 | 0.03 |
| gene-LOC105283712 | https://www.ncbi.nlm.nih.gov/gene/?term=LOC105283712 | 1910 | 24 | 125.65 | 1 | 5.24 | 4.17 | 29.64 | 32.04 | -2.39 | 2.39 | -0.03 | 0.03 |
| gene-LOC105274900 | https://www.ncbi.nlm.nih.gov/gene/?term=LOC105274900 | 5561 | 47 | 84.52 | 2 | 3.6 | 4.26 | 31.62 | 34.67 | -3.05 | 3.05 | -0.03 | 0.03 |
| gene-LOC105277215 | https://www.ncbi.nlm.nih.gov/gene/?term=LOC105277215 | 2619 | 20 | 76.37 | 1 | 3.82 | 5 | 28.06 | 32.55 | -4.49 | 4.49 | -0.03 | 0.03 |
| gene-LOC105278358 | https://www.ncbi.nlm.nih.gov/gene/?term=LOC105278358 | 2861 | 28 | 97.87 | 1 | 3.5 | 3.57 | 33.51 | 30.99 | 2.52 | 2.52 | 0.02 | 0.02 |
| gene-LOC105278228 | https://www.ncbi.nlm.nih.gov/gene/?term=LOC105278228 | 8610 | 146 | 169.57 | 1 | 1.16 | 0.68 | 23.97 | 22.69 | 1.27 | 1.27 | 0.02 | 0.02 |
| gene-LOC105285252 | https://www.ncbi.nlm.nih.gov/gene/?term=LOC105285252 | 7427 | 116 | 156.19 | 1 | 1.35 | 0.86 | 15.25 | 14.01 | 1.25 | 1.25 | 0.02 | 0.02 |
| gene-LOC105275653 | https://www.ncbi.nlm.nih.gov/gene/?term=LOC105275653 | 11251 | 236 | 209.76 | 1 | 0.89 | 0.42 | 9.52 | 8.44 | 1.07 | 1.07 | 0.02 | 0.02 |
| gene-LOC105282811 | https://www.ncbi.nlm.nih.gov/gene/?term=LOC105282811 | 5765 | 182 | 315.7 | 1 | 1.73 | 0.55 | 9.67 | 8.98 | 0.69 | 0.69 | 0.02 | 0.02 |
| gene-LOC105278632 | https://www.ncbi.nlm.nih.gov/gene/?term=LOC105278632 | 3624 | 119 | 328.37 | 1 | 2.76 | 0.84 | 15.91 | 15.28 | 0.63 | 0.63 | 0.02 | 0.02 |
| gene-LOC105280956 | https://www.ncbi.nlm.nih.gov/gene/?term=LOC105280956 | 2639 | 73 | 276.62 | 1 | 3.79 | 1.37 | 20.28 | 19.71 | 0.58 | 0.58 | 0.02 | 0.02 |
| gene-LOC105286515 | https://www.ncbi.nlm.nih.gov/gene/?term=LOC105286515 | 3751 | 107 | 285.26 | 1 | 2.67 | 0.93 | 10.47 | 9.92 | 0.55 | 0.55 | 0.02 | 0.02 |
| gene-LOC105280786 | https://www.ncbi.nlm.nih.gov/gene/?term=LOC105280786 | 2573 | 107 | 415.86 | 1 | 3.89 | 0.93 | 33.04 | 32.68 | 0.37 | 0.37 | 0.02 | 0.02 |
| gene-LOC105283296 | https://www.ncbi.nlm.nih.gov/gene/?term=LOC105283296 | 2174 | 78 | 358.79 | 1 | 4.6 | 1.28 | 21.02 | 21.5 | -0.48 | 0.48 | -0.02 | 0.02 |
| gene-LOC105284355 | https://www.ncbi.nlm.nih.gov/gene/?term=LOC105284355 | 3904 | 135 | 345.8 | 1 | 2.56 | 0.74 | 22.88 | 23.48 | -0.59 | 0.59 | -0.02 | 0.02 |
| gene-LOC105283728 | https://www.ncbi.nlm.nih.gov/gene/?term=LOC105283728 | 3256 | 103 | 316.34 | 2 | 6.14 | 1.94 | 21.65 | 22.28 | -0.62 | 0.62 | -0.02 | 0.02 |
| gene-LOC105286265 | https://www.ncbi.nlm.nih.gov/gene/?term=LOC105286265 | 2507 | 63 | 251.3 | 1 | 3.99 | 1.59 | 30.31 | 30.94 | -0.63 | 0.63 | -0.02 | 0.02 |
| gene-LOC105287063 | https://www.ncbi.nlm.nih.gov/gene/?term=LOC105287063 | 1874 | 46 | 245.46 | 1 | 5.34 | 2.17 | 34.15 | 34.94 | -0.79 | 0.79 | -0.02 | 0.02 |
| gene-LOC105275451 | https://www.ncbi.nlm.nih.gov/gene/?term=LOC105275451 | 6035 | 105 | 173.99 | 1 | 1.66 | 0.95 | 24.51 | 25.49 | -0.98 | 0.98 | -0.02 | 0.02 |
| gene-LOC105275173 | https://www.ncbi.nlm.nih.gov/gene/?term=LOC105275173 | 4639 | 71 | 153.05 | 1 | 2.16 | 1.41 | 27.01 | 28.06 | -1.05 | 1.05 | -0.02 | 0.02 |
| gene-LOC105280641 | https://www.ncbi.nlm.nih.gov/gene/?term=LOC105280641 | 3682 | 53 | 143.94 | 1 | 2.72 | 1.89 | 17.07 | 18.78 | -1.71 | 1.71 | -0.02 | 0.02 |

| | | | | | | | | | | | | | |
|---|---|---|---|---|---|---|---|---|---|---|---|---|---|
| gene-LOC105288066 | https://www.ncbi.nlm.nih.gov/gene/?term=LOC105288066 | 1964 | 16 | 81.47 | 1 | 5.09 | 6.25 | 41.07 | 43.1 | -2.03 | 2.03 | -0.02 | 0.02 |
| gene-LOC105282067 | https://www.ncbi.nlm.nih.gov/gene/?term=LOC105282067 | 5941 | 49 | 82.48 | 1 | 1.68 | 2.04 | 23.12 | 21.71 | 1.41 | 1.41 | 0.01 | 0.01 |
| gene-LOC105284507 | https://www.ncbi.nlm.nih.gov/gene/?term=LOC105284507 | 5207 | 68 | 130.59 | 1 | 1.92 | 1.47 | 22.93 | 21.85 | 1.08 | 1.08 | 0.01 | 0.01 |
| gene-LOC105276272 | https://www.ncbi.nlm.nih.gov/gene/?term=LOC105276272 | 1965 | 25 | 127.23 | 1 | 5.09 | 4 | 43.93 | 43.09 | 0.84 | 0.84 | 0.01 | 0.01 |
| gene-LOC105275187 | https://www.ncbi.nlm.nih.gov/gene/?term=LOC105275187 | 5534 | 66 | 119.26 | 1 | 1.81 | 1.52 | 14.05 | 13.27 | 0.78 | 0.78 | 0.01 | 0.01 |
| gene-LOC105276538 | https://www.ncbi.nlm.nih.gov/gene/?term=LOC105276538 | 1511 | 18 | 119.13 | 1 | 6.62 | 5.56 | 71.16 | 70.54 | 0.62 | 0.62 | 0.01 | 0.01 |
| gene-LOC105279664 | https://www.ncbi.nlm.nih.gov/gene/?term=LOC105279664 | 4023 | 56 | 139.2 | 1 | 2.49 | 1.79 | 21.21 | 20.62 | 0.59 | 0.59 | 0.01 | 0.01 |
| gene-LOC105276196 | https://www.ncbi.nlm.nih.gov/gene/?term=LOC105276196 | 2953 | 40 | 135.46 | 1 | 3.39 | 2.5 | 52.64 | 52.1 | 0.54 | 0.54 | 0.01 | 0.01 |
| gene-LOC105278454 | https://www.ncbi.nlm.nih.gov/gene/?term=LOC105278454 | 1972 | 36 | 182.56 | 1 | 5.07 | 2.78 | 47.84 | 47.3 | 0.54 | 0.54 | 0.01 | 0.01 |
| gene-LOC105275752 | https://www.ncbi.nlm.nih.gov/gene/?term=LOC105275752 | 3462 | 117 | 337.95 | 1 | 2.89 | 0.85 | 29.78 | 29.38 | 0.4 | 0.4 | 0.01 | 0.01 |
| gene-LOC105274706 | https://www.ncbi.nlm.nih.gov/gene/?term=LOC105274706 | 1446 | 51 | 352.7 | 1 | 6.92 | 1.96 | 37.41 | 37.03 | 0.38 | 0.38 | 0.01 | 0.01 |
| gene-LOC105287417 | https://www.ncbi.nlm.nih.gov/gene/?term=LOC105287417 | 1025 | 28 | 273.17 | 1 | 9.76 | 3.57 | 42.43 | 42.07 | 0.36 | 0.36 | 0.01 | 0.01 |
| gene-LOC105276449 | https://www.ncbi.nlm.nih.gov/gene/?term=LOC105276449 | 1231 | 39 | 316.82 | 1 | 8.12 | 2.56 | 32.73 | 32.41 | 0.32 | 0.32 | 0.01 | 0.01 |
| gene-LOC105278534 | https://www.ncbi.nlm.nih.gov/gene/?term=LOC105278534 | 7364 | 166 | 225.42 | 1 | 1.36 | 0.6 | 18.8 | 18.53 | 0.26 | 0.26 | 0.01 | 0.01 |
| gene-LOC105286649 | https://www.ncbi.nlm.nih.gov/gene/?term=LOC105286649 | 14178 | 662 | 466.92 | 1 | 0.71 | 0.15 | 1.27 | 1.01 | 0.26 | 0.26 | 0.01 | 0.01 |
| gene-LOC105283649 | https://www.ncbi.nlm.nih.gov/gene/?term=LOC105283649 | 1918 | 53 | 276.33 | 2 | 10.43 | 3.77 | 38.29 | 38.07 | 0.23 | 0.23 | 0.01 | 0.01 |
| gene-LOC105285807 | https://www.ncbi.nlm.nih.gov/gene/?term=LOC105285807 | 6063 | 174 | 286.99 | 1 | 1.65 | 0.57 | 7.5 | 7.29 | 0.22 | 0.22 | 0.01 | 0.01 |
| gene-LOC105286274 | https://www.ncbi.nlm.nih.gov/gene/?term=LOC105286274 | 3817 | 181 | 474.19 | 1 | 2.62 | 0.55 | 15.32 | 15.14 | 0.18 | 0.18 | 0.01 | 0.01 |
| gene-LOC105278145 | https://www.ncbi.nlm.nih.gov/gene/?term=LOC105278145 | 4470 | 244 | 545.86 | 1 | 2.24 | 0.41 | 10.74 | 10.94 | -0.2 | 0.2 | -0.01 | 0.01 |
| gene-LOC105285534 | https://www.ncbi.nlm.nih.gov/gene/?term=LOC105285534 | 3944 | 211 | 534.99 | 1 | 2.54 | 0.47 | 11.95 | 12.16 | -0.21 | 0.21 | -0.01 | 0.01 |
| gene-LOC105276527 | https://www.ncbi.nlm.nih.gov/gene/?term=LOC105276527 | 7923 | 323 | 407.67 | 1 | 1.26 | 0.31 | 8.74 | 8.99 | -0.25 | 0.25 | -0.01 | 0.01 |
| gene-LOC105274856 | https://www.ncbi.nlm.nih.gov/gene/?term=LOC105274856 | 5216 | 200 | 383.44 | 1 | 1.92 | 0.5 | 18.38 | 18.67 | -0.29 | 0.29 | -0.01 | 0.01 |
| gene-LOC105286223 | https://www.ncbi.nlm.nih.gov/gene/?term=LOC105286223 | 2225 | 79 | 355.06 | 1 | 4.49 | 1.27 | 30.58 | 30.96 | -0.38 | 0.38 | -0.01 | 0.01 |
| gene-LOC105285395 | https://www.ncbi.nlm.nih.gov/gene/?term=LOC105285395 | 8085 | 165 | 204.08 | 1 | 1.24 | 0.61 | 17.34 | 17.73 | -0.39 | 0.39 | -0.01 | 0.01 |
| gene-LOC105278538 | https://www.ncbi.nlm.nih.gov/gene/?term=LOC105278538 | 2150 | 76 | 353.49 | 1 | 4.65 | 1.32 | 18.23 | 18.64 | -0.41 | 0.41 | -0.01 | 0.01 |
| gene-LOC105275584 | https://www.ncbi.nlm.nih.gov/gene/?term=LOC105275584 | 3070 | 71 | 231.27 | 1 | 3.26 | 1.41 | 26.68 | 27.17 | -0.49 | 0.49 | -0.01 | 0.01 |
| gene-LOC105283943 | https://www.ncbi.nlm.nih.gov/gene/?term=LOC105283943 | 3559 | 48 | 134.87 | 1 | 2.81 | 2.08 | 17.72 | 18.23 | -0.51 | 0.51 | -0.01 | 0.01 |
| gene-LOC105281700 | https://www.ncbi.nlm.nih.gov/gene/?term=LOC105281700 | 2208 | 40 | 181.16 | 1 | 4.53 | 2.5 | 41.3 | 41.85 | -0.55 | 0.55 | -0.01 | 0.01 |
| gene-LOC105275179 | https://www.ncbi.nlm.nih.gov/gene/?term=LOC105275179 | 3271 | 66 | 201.77 | 1 | 3.06 | 1.52 | 14.71 | 15.31 | -0.61 | 0.61 | -0.01 | 0.01 |
| gene-LOC105280410 | https://www.ncbi.nlm.nih.gov/gene/?term=LOC105280410 | 12580 | 235 | 186.8 | 1 | 0.79 | 0.43 | 20.46 | 21.13 | -0.67 | 0.67 | -0.01 | 0.01 |
| gene-LOC105274851 | https://www.ncbi.nlm.nih.gov/gene/?term=LOC105274851 | 6616 | 122 | 184.4 | 1 | 1.51 | 0.82 | 10.53 | 11.32 | -0.79 | 0.79 | -0.01 | 0.01 |
| gene-LOC105283987 | https://www.ncbi.nlm.nih.gov/gene/?term=LOC105283987 | 4841 | 64 | 132.2 | 1 | 2.07 | 1.56 | 32.11 | 32.92 | -0.81 | 0.81 | -0.01 | 0.01 |
| gene-LOC105279657 | https://www.ncbi.nlm.nih.gov/gene/?term=LOC105279657 | 5878 | 79 | 134.4 | 1 | 1.7 | 1.27 | 19.77 | 20.7 | -0.93 | 0.93 | -0.01 | 0.01 |
| gene-LOC105274590 | https://www.ncbi.nlm.nih.gov/gene/?term=LOC105274590 | 4516 | 61 | 135.08 | 1 | 2.21 | 1.64 | 27.27 | 28.28 | -1.01 | 1.01 | -0.01 | 0.01 |
| gene-LOC105277216 | https://www.ncbi.nlm.nih.gov/gene/?term=LOC105277216 | 7627 | 76 | 99.65 | 1 | 1.31 | 1.32 | 7.73 | 8.77 | -1.04 | 1.04 | -0.01 | 0.01 |
| gene-LOC105281515 | https://www.ncbi.nlm.nih.gov/gene/?term=LOC105281515 | 8827 | 35 | 39.65 | 1 | 1.13 | 2.86 | 31.71 | 33.02 | -1.31 | 1.31 | -0.01 | 0.01 |
| gene-LOC105286452 | https://www.ncbi.nlm.nih.gov/gene/?term=LOC105286452 | 6888 | 55 | 79.85 | 1 | 1.45 | 1.82 | 32.52 | 33.96 | -1.44 | 1.44 | -0.01 | 0.01 |
| gene-LOC105279500 | https://www.ncbi.nlm.nih.gov/gene/?term=LOC105279500 | 3587 | 31 | 86.42 | 1 | 2.79 | 3.23 | 51.03 | 52.64 | -1.62 | 1.62 | -0.01 | 0.01 |
| gene-LOC105275518 | https://www.ncbi.nlm.nih.gov/gene/?term=LOC105275518 | 2335 | 18 | 77.09 | 1 | 4.28 | 5.56 | 47.37 | 49.08 | -1.72 | 1.72 | -0.01 | 0.01 |
| gene-LOC105284774 | https://www.ncbi.nlm.nih.gov/gene/?term=LOC105284774 | 12089 | 74 | 61.21 | 1 | 0.83 | 1.35 | 19.79 | 19.53 | 0.25 | 0.25 | 0 | 0 |
| gene-LOC105282794 | https://www.ncbi.nlm.nih.gov/gene/?term=LOC105282794 | 4571 | 91 | 199.08 | 1 | 2.19 | 1.1 | 19.49 | 19.29 | 0.2 | 0.2 | 0 | 0 |
| gene-LOC105288078 | https://www.ncbi.nlm.nih.gov/gene/?term=LOC105288078 | 19126 | 301 | 157.38 | 1 | 0.52 | 0.33 | 17.46 | 17.3 | 0.16 | 0.16 | 0 | 0 |
| gene-LOC105277866 | https://www.ncbi.nlm.nih.gov/gene/?term=LOC105277866 | 33045 | 370 | 111.97 | 1 | 0.3 | 0.27 | 3.21 | 3.09 | 0.13 | 0.13 | 0 | 0 |
| gene-LOC105287095 | https://www.ncbi.nlm.nih.gov/gene/?term=LOC105287095 | 1435 | 52 | 362.37 | 1 | 6.97 | 1.92 | 30.47 | 30.36 | 0.12 | 0.12 | 0 | 0 |
| gene-LOC105284116 | https://www.ncbi.nlm.nih.gov/gene/?term=LOC105284116 | 3970 | 85 | 214.11 | 1 | 2.52 | 1.18 | 7.23 | 7.13 | 0.09 | 0.09 | 0 | 0 |
| gene-LOC105279038 | https://www.ncbi.nlm.nih.gov/gene/?term=LOC105279038 | 9453 | 298 | 315.24 | 1 | 1.06 | 0.34 | 9.03 | 8.95 | 0.08 | 0.08 | 0 | 0 |
| gene-LOC105280265 | https://www.ncbi.nlm.nih.gov/gene/?term=LOC105280265 | 85984 | 2570 | 298.89 | 1 | 0.12 | 0.04 | 1.12 | 1.04 | 0.08 | 0.08 | 0 | 0 |
| gene-LOC105281272 | https://www.ncbi.nlm.nih.gov/gene/?term=LOC105281272 | 5252 | 168 | 319.88 | 1 | 1.9 | 0.6 | 16.96 | 16.88 | 0.08 | 0.08 | 0 | 0 |
| gene-LOC105278455 | https://www.ncbi.nlm.nih.gov/gene/?term=LOC105278455 | 27181 | 800 | 294.32 | 1 | 0.37 | 0.12 | 7.96 | 7.89 | 0.07 | 0.07 | 0 | 0 |
| gene-LOC105285742 | https://www.ncbi.nlm.nih.gov/gene/?term=LOC105285742 | 3446 | 91 | 264.07 | 1 | 2.9 | 1.1 | 26.72 | 26.67 | 0.05 | 0.05 | 0 | 0 |
| gene-LOC105276612 | https://www.ncbi.nlm.nih.gov/gene/?term=LOC105276612 | 4157 | 172 | 413.76 | 1 | 2.41 | 0.58 | 11.01 | 10.98 | 0.03 | 0.03 | 0 | 0 |
| gene-LOC105277125 | https://www.ncbi.nlm.nih.gov/gene/?term=LOC105277125 | 51079 | 1589 | 311.09 | 1 | 0.2 | 0.06 | 1.64 | 1.63 | 0 | 0 | 0 | 0 |
| gene-LOC105286194 | https://www.ncbi.nlm.nih.gov/gene/?term=LOC105286194 | 152619 | 1870 | 122.53 | 1 | 0.07 | 0.05 | 6.39 | 6.41 | -0.02 | 0.02 | 0 | 0 |
| gene-LOC105275327 | https://www.ncbi.nlm.nih.gov/gene/?term=LOC105275327 | 14245 | 413 | 289.93 | 1 | 0.7 | 0.24 | 7.72 | 7.75 | -0.03 | 0.03 | 0 | 0 |
| gene-LOC105280304 | https://www.ncbi.nlm.nih.gov/gene/?term=LOC105280304 | 8919 | 90 | 100.91 | 1 | 1.12 | 1.11 | 7.28 | 7.34 | -0.05 | 0.05 | 0 | 0 |
| gene-LOC105274534 | https://www.ncbi.nlm.nih.gov/gene/?term=LOC105274534 | 1329 | 53 | 398.8 | 1 | 7.52 | 1.89 | 47.05 | 47.11 | -0.06 | 0.06 | 0 | 0 |
| gene-LOC105278666 | https://www.ncbi.nlm.nih.gov/gene/?term=LOC105278666 | 10341 | 386 | 373.27 | 1 | 0.97 | 0.26 | 7.83 | 7.9 | -0.07 | 0.07 | 0 | 0 |
| gene-LOC105274531 | https://www.ncbi.nlm.nih.gov/gene/?term=LOC105274531 | 7774 | 252 | 324.16 | 1 | 1.29 | 0.4 | 11.91 | 11.99 | -0.08 | 0.08 | 0 | 0 |
| gene-LOC105276941 | https://www.ncbi.nlm.nih.gov/gene/?term=LOC105276941 | 15956 | 446 | 279.52 | 1 | 0.63 | 0.22 | 5.36 | 5.47 | -0.11 | 0.11 | 0 | 0 |
| gene-LOC105282825 | https://www.ncbi.nlm.nih.gov/gene/?term=LOC105282825 | 3919 | 80 | 204.13 | 1 | 2.55 | 1.25 | 13.05 | 13.16 | -0.12 | 0.12 | 0 | 0 |
| gene-LOC105286644 | https://www.ncbi.nlm.nih.gov/gene/?term=LOC105286644 | 1235 | 22 | 178.14 | 1 | 8.1 | 4.55 | 42.2 | 42.35 | -0.15 | 0.15 | 0 | 0 |
| gene-LOC105283989 | https://www.ncbi.nlm.nih.gov/gene/?term=LOC105283989 | 4948 | 25 | 50.53 | 1 | 2.02 | 4 | 45.66 | 45.85 | -0.19 | 0.19 | 0 | 0 |
| gene-LOC105284117 | https://www.ncbi.nlm.nih.gov/gene/?term=LOC105284117 | 4340 | 64 | 147.47 | 1 | 2.3 | 1.56 | 10.83 | 11.09 | -0.25 | 0.25 | 0 | 0 |
| gene-LOC105285406 | https://www.ncbi.nlm.nih.gov/gene/?term=LOC105285406 | 8015 | 39 | 48.66 | 1 | 1.25 | 2.56 | 33.07 | 33.55 | -0.48 | 0.48 | 0 | 0 |

Directory: OGLall File: rnk-dmg-Ob_LI_BCphaseA1.vs.BCphaseB1.txt

| gene_ID | gene_link | gwidth | #Sites | #per10Kb | #dmSites | #dmsp10kb | %dmSites | %pSite1 | %pSite2 | DMpSite | ADMpSite | DMpNucl | ADMpNucl |
|---|---|---|---|---|---|---|---|---|---|---|---|---|---|
| gene-LOC113562345 | https://www.ncbi.nlm.nih.gov/gene/?term=LOC113562345 | 2290 | 62 | 270.74 | 14 | 61.14 | 22.58 | 17.91 | 32.35 | -14.44 | 14.44 | -0.39 | 0.39 |
| gene-LOC105283552 | https://www.ncbi.nlm.nih.gov/gene/?term=LOC105283552 | 1562 | 33 | 211.27 | 1 | 6.4 | 3.03 | 54.5 | 59.97 | -5.47 | 5.47 | -0.12 | 0.12 |
| gene-LOC105279898 | https://www.ncbi.nlm.nih.gov/gene/?term=LOC105279898 | 1343 | 16 | 119.14 | 1 | 7.45 | 6.25 | 59.42 | 65.83 | -6.42 | 6.42 | -0.08 | 0.08 |
| gene-LOC105277631 | https://www.ncbi.nlm.nih.gov/gene/?term=LOC105277631 | 4387 | 285 | 649.65 | 1 | 2.28 | 0.35 | 9.88 | 10.92 | -1.04 | 1.04 | -0.07 | 0.07 |
| gene-LOC105282156 | https://www.ncbi.nlm.nih.gov/gene/?term=LOC105282156 | 2577 | 133 | 516.1 | 1 | 3.88 | 0.75 | 23.61 | 24.92 | -1.31 | 1.31 | -0.07 | 0.07 |
| gene-LOC105285695 | https://www.ncbi.nlm.nih.gov/gene/?term=LOC105285695 | 1371 | 62 | 452.22 | 1 | 7.29 | 1.61 | 23 | 24.51 | -1.51 | 1.51 | -0.07 | 0.07 |
| gene-LOC105279356 | https://www.ncbi.nlm.nih.gov/gene/?term=LOC105279356 | 1314 | 42 | 319.63 | 1 | 7.61 | 2.38 | 28.83 | 30.95 | -2.12 | 2.12 | -0.07 | 0.07 |
| gene-LOC105281848 | https://www.ncbi.nlm.nih.gov/gene/?term=LOC105281848 | 2122 | 47 | 221.49 | 1 | 4.71 | 2.13 | 22.74 | 25.94 | -3.2 | 3.2 | -0.07 | 0.07 |
| gene-LOC105275070 | https://www.ncbi.nlm.nih.gov/gene/?term=LOC105275070 | 5667 | 36 | 63.53 | 2 | 3.53 | 5.56 | 3.69 | 13.97 | -10.28 | 10.28 | -0.07 | 0.07 |
| gene-LOC105282885 | https://www.ncbi.nlm.nih.gov/gene/?term=LOC105282885 | 1870 | 41 | 219.25 | 1 | 5.35 | 2.44 | 37.52 | 34.93 | 2.59 | 2.59 | 0.06 | 0.06 |
| gene-LOC105275460 | https://www.ncbi.nlm.nih.gov/gene/?term=LOC105275460 | 2473 | 116 | 469.07 | 1 | 4.04 | 0.86 | 12.57 | 13.81 | -1.24 | 1.24 | -0.06 | 0.06 |
| gene-LOC105285055 | https://www.ncbi.nlm.nih.gov/gene/?term=LOC105285055 | 4108 | 214 | 520.93 | 1 | 2.43 | 0.47 | 12.94 | 14.18 | -1.24 | 1.24 | -0.06 | 0.06 |
| gene-LOC105275363 | https://www.ncbi.nlm.nih.gov/gene/?term=LOC105275363 | 3798 | 155 | 408.11 | 1 | 2.63 | 0.65 | 17.26 | 18.39 | -1.13 | 1.13 | -0.05 | 0.05 |
| gene-LOC105276704 | https://www.ncbi.nlm.nih.gov/gene/?term=LOC105276704 | 2063 | 84 | 407.17 | 2 | 9.69 | 2.38 | 12.84 | 14.02 | -1.18 | 1.18 | -0.05 | 0.05 |
| gene-LOC105275384 | https://www.ncbi.nlm.nih.gov/gene/?term=LOC105275384 | 1987 | 75 | 377.45 | 1 | 5.03 | 1.33 | 33.41 | 34.62 | -1.21 | 1.21 | -0.05 | 0.05 |
| gene-LOC105288036 | https://www.ncbi.nlm.nih.gov/gene/?term=LOC105288036 | 5110 | 198 | 387.48 | 1 | 1.96 | 0.51 | 16.95 | 18.26 | -1.31 | 1.31 | -0.05 | 0.05 |
| gene-LOC105278807 | https://www.ncbi.nlm.nih.gov/gene/?term=LOC105278807 | 2166 | 71 | 327.79 | 1 | 4.62 | 1.41 | 18.52 | 20.08 | -1.56 | 1.56 | -0.05 | 0.05 |
| gene-LOC105285674 | https://www.ncbi.nlm.nih.gov/gene/?term=LOC105285674 | 3035 | 91 | 299.84 | 2 | 6.59 | 2.2 | 23.1 | 24.91 | -1.81 | 1.81 | -0.05 | 0.05 |
| gene-LOC105277215 | https://www.ncbi.nlm.nih.gov/gene/?term=LOC105277215 | 2619 | 28 | 106.91 | 1 | 3.82 | 3.57 | 20.04 | 24.53 | -4.48 | 4.48 | -0.05 | 0.05 |
| gene-LOC105275708 | https://www.ncbi.nlm.nih.gov/gene/?term=LOC105275708 | 1772 | 13 | 73.36 | 1 | 5.64 | 7.69 | 46.34 | 53.52 | -7.18 | 7.18 | -0.05 | 0.05 |
| gene-LOC105285514 | https://www.ncbi.nlm.nih.gov/gene/?term=LOC105285514 | 1277 | 9 | 70.48 | 1 | 7.83 | 11.11 | 73.21 | 80.81 | -7.6 | 7.6 | -0.05 | 0.05 |
| gene-LOC105281173 | https://www.ncbi.nlm.nih.gov/gene/?term=LOC105281173 | 1595 | 25 | 156.74 | 1 | 6.27 | 4 | 64.76 | 62.33 | 2.42 | 2.42 | 0.04 | 0.04 |
| gene-LOC105283123 | https://www.ncbi.nlm.nih.gov/gene/?term=LOC105283123 | 1507 | 41 | 272.06 | 1 | 6.64 | 2.44 | 15.68 | 14.34 | 1.34 | 1.34 | 0.04 | 0.04 |
| gene-LOC105275683 | https://www.ncbi.nlm.nih.gov/gene/?term=LOC105275683 | 4256 | 204 | 479.32 | 1 | 2.35 | 0.49 | 12.14 | 13.06 | -0.92 | 0.92 | -0.04 | 0.04 |
| gene-LOC105285671 | https://www.ncbi.nlm.nih.gov/gene/?term=LOC105285671 | 2263 | 95 | 419.8 | 1 | 4.42 | 1.05 | 27.86 | 28.85 | -1 | 1 | -0.04 | 0.04 |
| gene-LOC105287417 | https://www.ncbi.nlm.nih.gov/gene/?term=LOC105287417 | 1025 | 33 | 321.95 | 1 | 9.76 | 3.03 | 36 | 37.27 | -1.27 | 1.27 | -0.04 | 0.04 |
| gene-LOC105278454 | https://www.ncbi.nlm.nih.gov/gene/?term=LOC105278454 | 1972 | 37 | 187.63 | 1 | 5.07 | 2.7 | 46.8 | 45.07 | 1.73 | 1.73 | 0.03 | 0.03 |
| gene-LOC105285202 | https://www.ncbi.nlm.nih.gov/gene/?term=LOC105285202 | 1838 | 36 | 195.87 | 1 | 5.44 | 2.78 | 18.94 | 17.23 | 1.72 | 1.72 | 0.03 | 0.03 |
| gene-LOC105276196 | https://www.ncbi.nlm.nih.gov/gene/?term=LOC105276196 | 2953 | 54 | 182.86 | 1 | 3.39 | 1.85 | 40.59 | 39.19 | 1.4 | 1.4 | 0.03 | 0.03 |
| gene-LOC105286265 | https://www.ncbi.nlm.nih.gov/gene/?term=LOC105286265 | 2507 | 64 | 255.29 | 1 | 3.99 | 1.56 | 30.65 | 29.46 | 1.19 | 1.19 | 0.03 | 0.03 |
| gene-LOC105275653 | https://www.ncbi.nlm.nih.gov/gene/?term=LOC105275653 | 11251 | 262 | 232.87 | 1 | 0.89 | 0.38 | 9.15 | 8.02 | 1.13 | 1.13 | 0.03 | 0.03 |
| gene-LOC105274706 | https://www.ncbi.nlm.nih.gov/gene/?term=LOC105274706 | 1446 | 46 | 318.12 | 1 | 6.92 | 2.17 | 41.37 | 40.53 | 0.85 | 0.85 | 0.03 | 0.03 |
| gene-LOC105276429 | https://www.ncbi.nlm.nih.gov/gene/?term=LOC105276429 | 2901 | 112 | 386.07 | 1 | 3.45 | 0.89 | 26.41 | 25.73 | 0.68 | 0.68 | 0.03 | 0.03 |
| gene-LOC105276527 | https://www.ncbi.nlm.nih.gov/gene/?term=LOC105276527 | 7923 | 345 | 435.44 | 1 | 1.26 | 0.29 | 8.28 | 8.94 | -0.66 | 0.66 | -0.03 | 0.03 |
| gene-LOC105283728 | https://www.ncbi.nlm.nih.gov/gene/?term=LOC105283728 | 3256 | 111 | 340.91 | 1 | 3.07 | 0.9 | 20.22 | 21.04 | -0.81 | 0.81 | -0.03 | 0.03 |
| gene-LOC105283943 | https://www.ncbi.nlm.nih.gov/gene/?term=LOC105283943 | 3559 | 59 | 165.78 | 1 | 2.81 | 1.69 | 20.25 | 21.81 | -1.56 | 1.56 | -0.03 | 0.03 |
| gene-LOC105280348 | https://www.ncbi.nlm.nih.gov/gene/?term=LOC105280348 | 3797 | 51 | 134.32 | 1 | 2.63 | 1.96 | 25.95 | 28.29 | -2.34 | 2.34 | -0.03 | 0.03 |
| gene-LOC105274900 | https://www.ncbi.nlm.nih.gov/gene/?term=LOC105274900 | 5561 | 56 | 100.7 | 2 | 3.6 | 3.57 | 26.85 | 29.47 | -2.62 | 2.62 | -0.03 | 0.03 |
| gene-LOC105275518 | https://www.ncbi.nlm.nih.gov/gene/?term=LOC105275518 | 2335 | 20 | 85.65 | 1 | 4.28 | 5 | 47.29 | 50.84 | -3.55 | 3.55 | -0.03 | 0.03 |
| gene-LOC105279528 | https://www.ncbi.nlm.nih.gov/gene/?term=LOC105279528 | 5802 | 46 | 79.28 | 1 | 1.72 | 2.17 | 17.18 | 14.84 | 2.33 | 2.33 | 0.02 | 0.02 |
| gene-LOC105285409 | https://www.ncbi.nlm.nih.gov/gene/?term=LOC105285409 | 5099 | 48 | 94.14 | 1 | 1.96 | 2.08 | 3.6 | 1.53 | 2.08 | 2.08 | 0.02 | 0.02 |
| gene-LOC105288010 | https://www.ncbi.nlm.nih.gov/gene/?term=LOC105288010 | 3189 | 49 | 153.65 | 1 | 3.14 | 2.04 | 31.6 | 30.42 | 1.18 | 1.18 | 0.02 | 0.02 |
| gene-LOC105278282 | https://www.ncbi.nlm.nih.gov/gene/?term=LOC105278282 | 1775 | 47 | 264.79 | 1 | 5.63 | 2.13 | 43.73 | 42.89 | 0.84 | 0.84 | 0.02 | 0.02 |
| gene-LOC105278214 | https://www.ncbi.nlm.nih.gov/gene/?term=LOC105278214 | 3787 | 77 | 203.33 | 1 | 2.64 | 1.3 | 11.46 | 10.7 | 0.76 | 0.76 | 0.02 | 0.02 |
| gene-LOC105285583 | https://www.ncbi.nlm.nih.gov/gene/?term=LOC105285583 | 1947 | 49 | 251.67 | 1 | 5.14 | 2.04 | 29.41 | 28.65 | 0.76 | 0.76 | 0.02 | 0.02 |
| gene-LOC105278436 | https://www.ncbi.nlm.nih.gov/gene/?term=LOC105278436 | 2556 | 57 | 223 | 1 | 3.91 | 1.75 | 36.25 | 35.56 | 0.69 | 0.69 | 0.02 | 0.02 |
| gene-LOC105274449 | https://www.ncbi.nlm.nih.gov/gene/?term=LOC105274449 | 1549 | 43 | 277.6 | 1 | 6.46 | 2.33 | 54.59 | 53.98 | 0.6 | 0.6 | 0.02 | 0.02 |
| gene-LOC105282811 | https://www.ncbi.nlm.nih.gov/gene/?term=LOC105282811 | 5765 | 203 | 352.12 | 1 | 1.73 | 0.49 | 9.79 | 9.26 | 0.53 | 0.53 | 0.02 | 0.02 |
| gene-LOC105277435 | https://www.ncbi.nlm.nih.gov/gene/?term=LOC105277435 | 3643 | 155 | 425.47 | 1 | 2.74 | 0.65 | 12.32 | 11.8 | 0.52 | 0.52 | 0.02 | 0.02 |
| gene-LOC105280786 | https://www.ncbi.nlm.nih.gov/gene/?term=LOC105280786 | 2573 | 120 | 466.38 | 1 | 3.89 | 0.83 | 29.63 | 29.12 | 0.52 | 0.52 | 0.02 | 0.02 |
| gene-LOC105278877 | https://www.ncbi.nlm.nih.gov/gene/?term=LOC105278877 | 1968 | 77 | 391.26 | 1 | 5.08 | 1.3 | 32.7 | 32.23 | 0.46 | 0.46 | 0.02 | 0.02 |
| gene-LOC105285534 | https://www.ncbi.nlm.nih.gov/gene/?term=LOC105285534 | 3944 | 221 | 560.34 | 1 | 2.54 | 0.45 | 11.41 | 11.78 | -0.37 | 0.37 | -0.02 | 0.02 |
| gene-LOC105282947 | https://www.ncbi.nlm.nih.gov/gene/?term=LOC105282947 | 1162 | 45 | 387.26 | 1 | 8.61 | 2.22 | 32.13 | 32.56 | -0.42 | 0.42 | -0.02 | 0.02 |
| gene-LOC105284355 | https://www.ncbi.nlm.nih.gov/gene/?term=LOC105284355 | 3904 | 154 | 394.47 | 1 | 2.56 | 0.65 | 20.37 | 20.87 | -0.5 | 0.5 | -0.02 | 0.02 |
| gene-LOC105285639 | https://www.ncbi.nlm.nih.gov/gene/?term=LOC105285639 | 10325 | 314 | 304.12 | 1 | 0.97 | 0.32 | 12 | 12.71 | -0.71 | 0.71 | -0.02 | 0.02 |
| gene-LOC105285742 | https://www.ncbi.nlm.nih.gov/gene/?term=LOC105285742 | 3446 | 95 | 275.68 | 1 | 2.9 | 1.05 | 25.7 | 26.49 | -0.79 | 0.79 | -0.02 | 0.02 |
| gene-LOC105283592 | https://www.ncbi.nlm.nih.gov/gene/?term=LOC105283592 | 3479 | 71 | 204.08 | 1 | 2.87 | 1.41 | 24.39 | 25.37 | -0.98 | 0.98 | -0.02 | 0.02 |
| gene-LOC105281700 | https://www.ncbi.nlm.nih.gov/gene/?term=LOC105281700 | 2208 | 44 | 199.28 | 1 | 4.53 | 2.27 | 37.54 | 38.62 | -1.08 | 1.08 | -0.02 | 0.02 |
| gene-LOC105283712 | https://www.ncbi.nlm.nih.gov/gene/?term=LOC105283712 | 1910 | 29 | 151.83 | 1 | 5.24 | 3.45 | 29.98 | 31.34 | -1.36 | 1.36 | -0.02 | 0.02 |
| gene-LOC105278260 | https://www.ncbi.nlm.nih.gov/gene/?term=LOC105278260 | 3240 | 42 | 129.63 | 1 | 3.09 | 2.38 | 12.8 | 14.28 | -1.47 | 1.47 | -0.02 | 0.02 |

| | | | | | | | | | | | | | |
|---|---|---|---|---|---|---|---|---|---|---|---|---|---|
| gene-LOC105283989 | https://www.ncbi.nlm.nih.gov/gene/?term=LOC105283989 | 4948 | 33 | 66.69 | 2 | 4.04 | 6.06 | 45.77 | 48.89 | -3.12 | 3.12 | -0.02 | 0.02 |
| gene-LOC105280419 | https://www.ncbi.nlm.nih.gov/gene/?term=LOC105280419 | 1601 | 10 | 62.46 | 1 | 6.25 | 10 | 33.98 | 37.47 | -3.49 | 3.49 | -0.02 | 0.02 |
| gene-LOC105280447 | https://www.ncbi.nlm.nih.gov/gene/?term=LOC105280447 | 2215 | 25 | 112.87 | 1 | 4.51 | 4 | 26.74 | 25.91 | 0.82 | 0.82 | 0.01 | 0.01 |
| gene-LOC105281047 | https://www.ncbi.nlm.nih.gov/gene/?term=LOC105281047 | 7313 | 117 | 159.99 | 1 | 1.37 | 0.85 | 10.66 | 9.84 | 0.82 | 0.82 | 0.01 | 0.01 |
| gene-LOC105281420 | https://www.ncbi.nlm.nih.gov/gene/?term=LOC105281420 | 3882 | 57 | 146.83 | 1 | 2.58 | 1.75 | 15.28 | 14.73 | 0.55 | 0.55 | 0.01 | 0.01 |
| gene-LOC105284116 | https://www.ncbi.nlm.nih.gov/gene/?term=LOC105284116 | 3970 | 95 | 239.29 | 1 | 2.52 | 1.05 | 6.78 | 6.28 | 0.5 | 0.5 | 0.01 | 0.01 |
| gene-LOC105284117 | https://www.ncbi.nlm.nih.gov/gene/?term=LOC105284117 | 4340 | 81 | 186.64 | 1 | 2.3 | 1.23 | 11.74 | 11.3 | 0.44 | 0.44 | 0.01 | 0.01 |
| gene-LOC105282600 | https://www.ncbi.nlm.nih.gov/gene/?term=LOC105282600 | 3715 | 57 | 153.43 | 1 | 2.69 | 1.75 | 29.16 | 28.81 | 0.35 | 0.35 | 0.01 | 0.01 |
| gene-LOC105278455 | https://www.ncbi.nlm.nih.gov/gene/?term=LOC105278455 | 27181 | 854 | 314.19 | 1 | 0.37 | 0.12 | 7.49 | 7.16 | 0.33 | 0.33 | 0.01 | 0.01 |
| gene-LOC105285807 | https://www.ncbi.nlm.nih.gov/gene/?term=LOC105285807 | 6063 | 183 | 301.83 | 1 | 1.65 | 0.55 | 10.09 | 9.8 | 0.29 | 0.29 | 0.01 | 0.01 |
| gene-LOC105286515 | https://www.ncbi.nlm.nih.gov/gene/?term=LOC105286515 | 3751 | 103 | 274.59 | 1 | 2.67 | 0.97 | 10.73 | 10.44 | 0.29 | 0.29 | 0.01 | 0.01 |
| gene-LOC105279038 | https://www.ncbi.nlm.nih.gov/gene/?term=LOC105279038 | 9453 | 310 | 327.94 | 1 | 1.06 | 0.32 | 8.8 | 8.54 | 0.27 | 0.27 | 0.01 | 0.01 |
| gene-LOC105278666 | https://www.ncbi.nlm.nih.gov/gene/?term=LOC105278666 | 10341 | 405 | 391.64 | 1 | 0.97 | 0.25 | 7.09 | 6.88 | 0.22 | 0.22 | 0.01 | 0.01 |
| gene-LOC105284036 | https://www.ncbi.nlm.nih.gov/gene/?term=LOC105284036 | 3254 | 83 | 255.07 | 1 | 3.07 | 1.2 | 22.41 | 22.19 | 0.22 | 0.22 | 0.01 | 0.01 |
| gene-LOC105276901 | https://www.ncbi.nlm.nih.gov/gene/?term=LOC105276901 | 3254 | 95 | 291.95 | 1 | 3.07 | 1.05 | 23.05 | 22.87 | 0.18 | 0.18 | 0.01 | 0.01 |
| gene-LOC105278145 | https://www.ncbi.nlm.nih.gov/gene/?term=LOC105278145 | 4470 | 249 | 557.05 | 1 | 2.24 | 0.4 | 10.18 | 10.01 | 0.17 | 0.17 | 0.01 | 0.01 |
| gene-LOC105275752 | https://www.ncbi.nlm.nih.gov/gene/?term=LOC105275752 | 3462 | 124 | 358.17 | 1 | 2.89 | 0.81 | 28.83 | 28.67 | 0.16 | 0.16 | 0.01 | 0.01 |
| gene-LOC105287095 | https://www.ncbi.nlm.nih.gov/gene/?term=LOC105287095 | 1435 | 61 | 425.09 | 1 | 6.97 | 1.64 | 26.05 | 25.92 | 0.14 | 0.14 | 0.01 | 0.01 |
| gene-LOC105284262 | https://www.ncbi.nlm.nih.gov/gene/?term=LOC105284262 | 6669 | 230 | 344.88 | 1 | 1.5 | 0.43 | 4.51 | 4.68 | -0.17 | 0.17 | -0.01 | 0.01 |
| gene-LOC105275584 | https://www.ncbi.nlm.nih.gov/gene/?term=LOC105275584 | 3070 | 86 | 280.13 | 1 | 3.26 | 1.16 | 23.79 | 24 | -0.22 | 0.22 | -0.01 | 0.01 |
| gene-LOC105283649 | https://www.ncbi.nlm.nih.gov/gene/?term=LOC105283649 | 1918 | 53 | 276.33 | 1 | 5.21 | 1.89 | 38.29 | 38.55 | -0.25 | 0.25 | -0.01 | 0.01 |
| gene-LOC105280405 | https://www.ncbi.nlm.nih.gov/gene/?term=LOC105280405 | 5422 | 162 | 298.78 | 1 | 1.84 | 0.62 | 9.36 | 9.65 | -0.29 | 0.29 | -0.01 | 0.01 |
| gene-LOC105285395 | https://www.ncbi.nlm.nih.gov/gene/?term=LOC105285395 | 8085 | 180 | 222.63 | 1 | 1.24 | 0.56 | 15.55 | 15.84 | -0.29 | 0.29 | -0.01 | 0.01 |
| gene-LOC105279298 | https://www.ncbi.nlm.nih.gov/gene/?term=LOC105279298 | 2773 | 134 | 483.23 | 1 | 3.61 | 0.75 | 22.72 | 23.03 | -0.3 | 0.3 | -0.01 | 0.01 |
| gene-LOC105288078 | https://www.ncbi.nlm.nih.gov/gene/?term=LOC105288078 | 19126 | 313 | 163.65 | 1 | 0.52 | 0.32 | 17.33 | 17.64 | -0.31 | 0.31 | -0.01 | 0.01 |
| gene-LOC105284184 | https://www.ncbi.nlm.nih.gov/gene/?term=LOC105284184 | 2535 | 82 | 323.47 | 1 | 3.94 | 1.22 | 34.56 | 34.9 | -0.34 | 0.34 | -0.01 | 0.01 |
| gene-LOC105278632 | https://www.ncbi.nlm.nih.gov/gene/?term=LOC105278632 | 3624 | 131 | 361.48 | 1 | 2.76 | 0.76 | 14.71 | 15.12 | -0.41 | 0.41 | -0.01 | 0.01 |
| gene-LOC105276692 | https://www.ncbi.nlm.nih.gov/gene/?term=LOC105276692 | 1745 | 36 | 206.3 | 1 | 5.73 | 2.78 | 23.57 | 24.03 | -0.46 | 0.46 | -0.01 | 0.01 |
| gene-LOC105278534 | https://www.ncbi.nlm.nih.gov/gene/?term=LOC105278534 | 7364 | 184 | 249.86 | 1 | 1.36 | 0.54 | 17.97 | 18.5 | -0.53 | 0.53 | -0.01 | 0.01 |
| gene-LOC105286770 | https://www.ncbi.nlm.nih.gov/gene/?term=LOC105286770 | 3055 | 42 | 137.48 | 1 | 3.27 | 2.38 | 8.53 | 9.07 | -0.55 | 0.55 | -0.01 | 0.01 |
| gene-LOC105276745 | https://www.ncbi.nlm.nih.gov/gene/?term=LOC105276745 | 3953 | 73 | 184.67 | 1 | 2.53 | 1.37 | 18.74 | 19.3 | -0.56 | 0.56 | -0.01 | 0.01 |
| gene-LOC105280002 | https://www.ncbi.nlm.nih.gov/gene/?term=LOC105280002 | 1940 | 28 | 144.33 | 1 | 5.15 | 3.57 | 33.09 | 33.65 | -0.57 | 0.57 | -0.01 | 0.01 |
| gene-LOC105278962 | https://www.ncbi.nlm.nih.gov/gene/?term=LOC105278962 | 4491 | 88 | 195.95 | 1 | 2.23 | 1.14 | 23.63 | 24.21 | -0.58 | 0.58 | -0.01 | 0.01 |
| gene-LOC105277903 | https://www.ncbi.nlm.nih.gov/gene/?term=LOC105277903 | 1835 | 35 | 190.74 | 1 | 5.45 | 2.86 | 51.45 | 52.14 | -0.69 | 0.69 | -0.01 | 0.01 |
| gene-LOC105280641 | https://www.ncbi.nlm.nih.gov/gene/?term=LOC105280641 | 3682 | 62 | 168.39 | 1 | 2.72 | 1.61 | 16.36 | 17.05 | -0.69 | 0.69 | -0.01 | 0.01 |
| gene-LOC105275190 | https://www.ncbi.nlm.nih.gov/gene/?term=LOC105275190 | 3837 | 72 | 187.65 | 1 | 2.61 | 1.39 | 18.78 | 19.54 | -0.76 | 0.76 | -0.01 | 0.01 |
| gene-LOC105288066 | https://www.ncbi.nlm.nih.gov/gene/?term=LOC105288066 | 1964 | 25 | 127.29 | 1 | 5.09 | 4 | 27.34 | 28.29 | -0.95 | 0.95 | -0.01 | 0.01 |
| gene-LOC105277216 | https://www.ncbi.nlm.nih.gov/gene/?term=LOC105277216 | 7627 | 105 | 137.67 | 1 | 1.31 | 0.95 | 5.72 | 6.75 | -1.03 | 1.03 | -0.01 | 0.01 |
| gene-LOC105285060 | https://www.ncbi.nlm.nih.gov/gene/?term=LOC105285060 | 1285 | 16 | 124.51 | 1 | 7.78 | 6.25 | 50.64 | 51.74 | -1.1 | 1.1 | -0.01 | 0.01 |
| gene-LOC105281515 | https://www.ncbi.nlm.nih.gov/gene/?term=LOC105281515 | 8827 | 39 | 44.18 | 1 | 1.13 | 2.56 | 34.44 | 35.97 | -1.53 | 1.53 | -0.01 | 0.01 |
| gene-LOC105286452 | https://www.ncbi.nlm.nih.gov/gene/?term=LOC105286452 | 6888 | 58 | 84.2 | 1 | 1.45 | 1.72 | 31.25 | 32.98 | -1.73 | 1.73 | -0.01 | 0.01 |
| gene-LOC105282399 | https://www.ncbi.nlm.nih.gov/gene/?term=LOC105282399 | 2682 | 21 | 78.3 | 1 | 3.73 | 4.76 | 34.08 | 35.97 | -1.89 | 1.89 | -0.01 | 0.01 |
| gene-LOC105285406 | https://www.ncbi.nlm.nih.gov/gene/?term=LOC105285406 | 8015 | 40 | 49.91 | 1 | 1.25 | 2.5 | 34.18 | 33.53 | 0.64 | 0.64 | 0 | 0 |
| gene-LOC105277866 | https://www.ncbi.nlm.nih.gov/gene/?term=LOC105277866 | 33045 | 442 | 133.76 | 1 | 0.3 | 0.23 | 3.08 | 2.82 | 0.27 | 0.27 | 0 | 0 |
| gene-LOC105279657 | https://www.ncbi.nlm.nih.gov/gene/?term=LOC105279657 | 5878 | 86 | 146.31 | 1 | 1.7 | 1.16 | 18.3 | 18.14 | 0.16 | 0.16 | 0 | 0 |
| gene-LOC105286194 | https://www.ncbi.nlm.nih.gov/gene/?term=LOC105286194 | 152619 | 2028 | 132.88 | 1 | 0.07 | 0.05 | 6.64 | 6.49 | 0.14 | 0.14 | 0 | 0 |
| gene-LOC105275173 | https://www.ncbi.nlm.nih.gov/gene/?term=LOC105275173 | 4639 | 83 | 178.92 | 1 | 2.16 | 1.2 | 22.33 | 22.24 | 0.09 | 0.09 | 0 | 0 |
| gene-LOC105276059 | https://www.ncbi.nlm.nih.gov/gene/?term=LOC105276059 | 6256 | 134 | 214.19 | 1 | 1.6 | 0.75 | 10.93 | 10.85 | 0.08 | 0.08 | 0 | 0 |
| gene-LOC105279664 | https://www.ncbi.nlm.nih.gov/gene/?term=LOC105279664 | 4023 | 68 | 169.03 | 1 | 2.49 | 1.47 | 18.22 | 18.15 | 0.08 | 0.08 | 0 | 0 |
| gene-LOC105288161 | https://www.ncbi.nlm.nih.gov/gene/?term=LOC105288161 | 26065 | 287 | 110.11 | 1 | 0.38 | 0.35 | 12.1 | 12.03 | 0.07 | 0.07 | 0 | 0 |
| gene-LOC105278228 | https://www.ncbi.nlm.nih.gov/gene/?term=LOC105278228 | 8610 | 165 | 191.64 | 1 | 1.16 | 0.61 | 22.79 | 22.73 | 0.06 | 0.06 | 0 | 0 |
| gene-LOC105282491 | https://www.ncbi.nlm.nih.gov/gene/?term=LOC105282491 | 4761 | 159 | 333.96 | 1 | 2.1 | 0.63 | 9.46 | 9.41 | 0.05 | 0.05 | 0 | 0 |
| gene-LOC105286274 | https://www.ncbi.nlm.nih.gov/gene/?term=LOC105286274 | 3817 | 177 | 463.71 | 1 | 2.62 | 0.56 | 15.66 | 15.62 | 0.04 | 0.04 | 0 | 0 |
| gene-LOC105275451 | https://www.ncbi.nlm.nih.gov/gene/?term=LOC105275451 | 6035 | 113 | 187.24 | 1 | 1.66 | 0.88 | 22.29 | 22.26 | 0.02 | 0.02 | 0 | 0 |
| gene-LOC105275327 | https://www.ncbi.nlm.nih.gov/gene/?term=LOC105275327 | 14245 | 477 | 334.85 | 1 | 0.7 | 0.21 | 7.15 | 7.14 | 0.01 | 0.01 | 0 | 0 |
| gene-LOC105280265 | https://www.ncbi.nlm.nih.gov/gene/?term=LOC105280265 | 85984 | 2825 | 328.55 | 1 | 0.12 | 0.04 | 1.07 | 1.08 | -0.01 | 0.01 | 0 | 0 |
| gene-LOC105282151 | https://www.ncbi.nlm.nih.gov/gene/?term=LOC105282151 | 1601 | 41 | 256.09 | 1 | 6.25 | 2.44 | 39.95 | 39.99 | -0.04 | 0.04 | 0 | 0 |
| gene-LOC105279778 | https://www.ncbi.nlm.nih.gov/gene/?term=LOC105279778 | 4347 | 139 | 319.76 | 1 | 2.3 | 0.72 | 9.36 | 9.42 | -0.06 | 0.06 | 0 | 0 |
| gene-LOC105274590 | https://www.ncbi.nlm.nih.gov/gene/?term=LOC105274590 | 4516 | 76 | 168.29 | 1 | 2.21 | 1.32 | 22.95 | 23.02 | -0.07 | 0.07 | 0 | 0 |
| gene-LOC105275170 | https://www.ncbi.nlm.nih.gov/gene/?term=LOC105275170 | 12918 | 337 | 260.88 | 1 | 0.77 | 0.3 | 2.06 | 2.13 | -0.07 | 0.07 | 0 | 0 |
| gene-LOC105280956 | https://www.ncbi.nlm.nih.gov/gene/?term=LOC105280956 | 2639 | 73 | 276.62 | 1 | 3.79 | 1.37 | 20.28 | 20.35 | -0.07 | 0.07 | 0 | 0 |
| gene-LOC105274531 | https://www.ncbi.nlm.nih.gov/gene/?term=LOC105274531 | 7774 | 254 | 326.73 | 1 | 1.29 | 0.39 | 11.9 | 12 | -0.1 | 0.1 | 0 | 0 |
| gene-LOC105274856 | https://www.ncbi.nlm.nih.gov/gene/?term=LOC105274856 | 5216 | 201 | 385.35 | 1 | 1.92 | 0.5 | 17.3 | 17.4 | -0.1 | 0.1 | 0 | 0 |
| gene-LOC105279865 | https://www.ncbi.nlm.nih.gov/gene/?term=LOC105279865 | 62373 | 1507 | 241.61 | 1 | 0.16 | 0.07 | 1.62 | 1.72 | -0.1 | 0.1 | 0 | 0 |

| | | | | | | | | | | | | | |
|---|---|---|---|---|---|---|---|---|---|---|---|---|---|
| gene-LOC105276538 | https://www.ncbi.nlm.nih.gov/gene/?term=LOC105276538 | 1511 | 18 | 119.13 | 1 | 6.62 | 5.56 | 71.72 | 71.83 | -0.11 | 0.11 | 0 | 0 |
| gene-LOC105279500 | https://www.ncbi.nlm.nih.gov/gene/?term=LOC105279500 | 3587 | 32 | 89.21 | 1 | 2.79 | 3.12 | 49.74 | 49.96 | -0.21 | 0.21 | 0 | 0 |
| gene-LOC105282914 | https://www.ncbi.nlm.nih.gov/gene/?term=LOC105282914 | 5623 | 42 | 74.69 | 1 | 1.78 | 2.38 | 48.6 | 48.91 | -0.31 | 0.31 | 0 | 0 |

Directory: OGLall File: rnk-dmg-Ob_LI_BCphaseA1.vs.BCphaseB2.txt

| gene_ID | gene_link | gwidth | #Sites | #per10Kb | #dmSites | #dmsp10kb | %dmSites | %pSite1 | %pSite2 | DMpSite | ADMpSite | DMpNucl | ADMpNucl |
|---|---|---|---|---|---|---|---|---|---|---|---|---|---|
| gene-LOC105283830 | https://www.ncbi.nlm.nih.gov/gene/?term=LOC105283830 | 252 | 13 | 515.87 | 1 | 39.68 | 7.69 | 42.02 | 33.83 | 8.19 | 8.19 | 0.42 | 0.42 |
| gene-LOC105278849 | https://www.ncbi.nlm.nih.gov/gene/?term=LOC105278849 | 776 | 75 | 966.49 | 2 | 25.77 | 2.67 | 24.26 | 22.31 | 1.95 | 1.95 | 0.19 | 0.19 |
| gene-LOC105282405 | https://www.ncbi.nlm.nih.gov/gene/?term=LOC105282405 | 1310 | 31 | 236.64 | 3 | 22.9 | 9.68 | 51.18 | 44.58 | 6.6 | 6.6 | 0.16 | 0.16 |
| gene-LOC105278458 | https://www.ncbi.nlm.nih.gov/gene/?term=LOC105278458 | 216 | 3 | 138.89 | 1 | 46.3 | 33.33 | 20.87 | 10.63 | 10.24 | 10.24 | 0.14 | 0.14 |
| gene-LOC105287856 | https://www.ncbi.nlm.nih.gov/gene/?term=LOC105287856 | 2090 | 116 | 555.02 | 3 | 14.35 | 2.59 | 12.68 | 14.95 | -2.27 | 2.27 | -0.13 | 0.13 |
| gene-LOC105284168 | https://www.ncbi.nlm.nih.gov/gene/?term=LOC105284168 | 1400 | 72 | 514.29 | 3 | 21.43 | 4.17 | 21.98 | 24.55 | -2.57 | 2.57 | -0.13 | 0.13 |
| gene-LOC105287662 | https://www.ncbi.nlm.nih.gov/gene/?term=LOC105287662 | 1704 | 39 | 228.87 | 3 | 17.61 | 7.69 | 30.88 | 36.7 | -5.82 | 5.82 | -0.13 | 0.13 |
| gene-LOC113562627 | https://www.ncbi.nlm.nih.gov/gene/?term=LOC113562627 | 397 | 10 | 251.89 | 2 | 50.38 | 20 | 77.44 | 72.85 | 4.6 | 4.6 | 0.12 | 0.12 |
| gene-LOC105276679 | https://www.ncbi.nlm.nih.gov/gene/?term=LOC105276679 | 2005 | 65 | 324.19 | 4 | 19.95 | 6.15 | 39.83 | 36.18 | 3.65 | 3.65 | 0.12 | 0.12 |
| gene-LOC105283016 | https://www.ncbi.nlm.nih.gov/gene/?term=LOC105283016 | 2324 | 111 | 477.62 | 3 | 12.91 | 2.7 | 23.61 | 21.2 | 2.41 | 2.41 | 0.12 | 0.12 |
| gene-LOC105282986 | https://www.ncbi.nlm.nih.gov/gene/?term=LOC105282986 | 1600 | 81 | 506.25 | 2 | 12.5 | 2.47 | 30.91 | 28.82 | 2.08 | 2.08 | 0.11 | 0.11 |
| gene-LOC105274711 | https://www.ncbi.nlm.nih.gov/gene/?term=LOC105274711 | 1655 | 87 | 525.68 | 3 | 18.13 | 3.45 | 45.85 | 43.79 | 2.06 | 2.06 | 0.11 | 0.11 |
| gene-LOC105283717 | https://www.ncbi.nlm.nih.gov/gene/?term=LOC105283717 | 1373 | 83 | 604.52 | 2 | 14.57 | 2.41 | 24.03 | 25.8 | -1.76 | 1.76 | -0.11 | 0.11 |
| gene-LOC105286845 | https://www.ncbi.nlm.nih.gov/gene/?term=LOC105286845 | 1159 | 59 | 509.06 | 3 | 25.88 | 5.08 | 17.66 | 19.8 | -2.14 | 2.14 | -0.11 | 0.11 |
| gene-LOC105287895 | https://www.ncbi.nlm.nih.gov/gene/?term=LOC105287895 | 2424 | 50 | 206.27 | 2 | 8.25 | 4 | 28.82 | 34.37 | -5.56 | 5.56 | -0.11 | 0.11 |
| gene-LOC105277648 | https://www.ncbi.nlm.nih.gov/gene/?term=LOC105277648 | 722 | 26 | 360.11 | 1 | 13.85 | 3.85 | 56.47 | 53.69 | 2.78 | 2.78 | 0.1 | 0.1 |
| gene-LOC105280786 | https://www.ncbi.nlm.nih.gov/gene/?term=LOC105280786 | 2573 | 109 | 423.63 | 3 | 11.66 | 2.75 | 32.44 | 30.15 | 2.29 | 2.29 | 0.1 | 0.1 |
| gene-LOC105275484 | https://www.ncbi.nlm.nih.gov/gene/?term=LOC105275484 | 931 | 33 | 354.46 | 1 | 10.74 | 3.03 | 34.17 | 37.06 | -2.89 | 2.89 | -0.1 | 0.1 |
| gene-LOC105287565 | https://www.ncbi.nlm.nih.gov/gene/?term=LOC105287565 | 1735 | 49 | 282.42 | 5 | 28.82 | 10.2 | 38.14 | 41.53 | -3.39 | 3.39 | -0.1 | 0.1 |
| gene-LOC105283649 | https://www.ncbi.nlm.nih.gov/gene/?term=LOC105283649 | 1918 | 53 | 276.33 | 4 | 20.86 | 7.55 | 38.29 | 35.16 | 3.13 | 3.13 | 0.09 | 0.09 |
| gene-LOC105275761 | https://www.ncbi.nlm.nih.gov/gene/?term=LOC105275761 | 2635 | 98 | 371.92 | 4 | 15.18 | 4.08 | 30.05 | 27.74 | 2.31 | 2.31 | 0.09 | 0.09 |
| gene-LOC105279335 | https://www.ncbi.nlm.nih.gov/gene/?term=LOC105279335 | 737 | 33 | 447.76 | 1 | 13.57 | 3.03 | 19.48 | 17.37 | 2.11 | 2.11 | 0.09 | 0.09 |
| gene-LOC105279035 | https://www.ncbi.nlm.nih.gov/gene/?term=LOC105279035 | 2108 | 145 | 687.86 | 5 | 23.72 | 3.45 | 18.41 | 17.12 | 1.29 | 1.29 | 0.09 | 0.09 |
| gene-LOC105278737 | https://www.ncbi.nlm.nih.gov/gene/?term=LOC105278737 | 1346 | 89 | 661.22 | 1 | 7.43 | 1.12 | 18.69 | 20.11 | -1.41 | 1.41 | -0.09 | 0.09 |

...
... only first 25 lines shown ...

Directory: OGLall File: rnk-dmg-Ob_LI_BCphaseA1.vs.RphaseA1.txt

| gene_ID | gene_link | gwidth | #Sites | #per10Kb | #dmSites | #dmsp10kb | %dmSites | %pSite1 | %pSite2 | DMpSite | ADMpSite | DMpNucl | ADMpNucl |
|---|---|---|---|---|---|---|---|---|---|---|---|---|---|
| gene-LOC113562345 | https://www.ncbi.nlm.nih.gov/gene/?term=LOC113562345 | 2290 | 61 | 266.38 | 10 | 43.67 | 16.39 | 17.52 | 31.67 | -14.16 | 14.16 | -0.38 | 0.38 |
| gene-LOC105285671 | https://www.ncbi.nlm.nih.gov/gene/?term=LOC105285671 | 2263 | 90 | 397.7 | 1 | 4.42 | 1.11 | 29.29 | 31.61 | -2.32 | 2.32 | -0.09 | 0.09 |
| gene-LOC105275460 | https://www.ncbi.nlm.nih.gov/gene/?term=LOC105275460 | 2473 | 115 | 465.02 | 1 | 4.04 | 0.87 | 12.68 | 14.4 | -1.72 | 1.72 | -0.08 | 0.08 |
| gene-LOC105285674 | https://www.ncbi.nlm.nih.gov/gene/?term=LOC105285674 | 3035 | 88 | 289.95 | 2 | 6.59 | 2.27 | 23.51 | 26.27 | -2.76 | 2.76 | -0.08 | 0.08 |
| gene-LOC105275070 | https://www.ncbi.nlm.nih.gov/gene/?term=LOC105275070 | 5667 | 37 | 65.29 | 2 | 3.53 | 5.41 | 3.73 | 16.16 | -12.43 | 12.43 | -0.08 | 0.08 |
| gene-LOC105274449 | https://www.ncbi.nlm.nih.gov/gene/?term=LOC105274449 | 1549 | 43 | 277.6 | 1 | 6.46 | 2.33 | 54.59 | 52.01 | 2.58 | 2.58 | 0.07 | 0.07 |
| gene-LOC105284396 | https://www.ncbi.nlm.nih.gov/gene/?term=LOC105284396 | 1976 | 57 | 288.46 | 1 | 5.06 | 1.75 | 39.14 | 36.68 | 2.46 | 2.46 | 0.07 | 0.07 |
| gene-LOC105276704 | https://www.ncbi.nlm.nih.gov/gene/?term=LOC105276704 | 2063 | 80 | 387.78 | 1 | 4.85 | 1.25 | 13.36 | 15.07 | -1.71 | 1.71 | -0.07 | 0.07 |
| gene-LOC105278807 | https://www.ncbi.nlm.nih.gov/gene/?term=LOC105278807 | 2166 | 70 | 323.18 | 1 | 4.62 | 1.43 | 17.84 | 20.15 | -2.31 | 2.31 | -0.07 | 0.07 |
| gene-LOC105279356 | https://www.ncbi.nlm.nih.gov/gene/?term=LOC105279356 | 1314 | 42 | 319.63 | 1 | 7.61 | 2.38 | 28.83 | 31.17 | -2.34 | 2.34 | -0.07 | 0.07 |
| gene-LOC105281848 | https://www.ncbi.nlm.nih.gov/gene/?term=LOC105281848 | 2122 | 45 | 212.06 | 1 | 4.71 | 2.22 | 23.75 | 26.94 | -3.19 | 3.19 | -0.07 | 0.07 |
| gene-LOC105283552 | https://www.ncbi.nlm.nih.gov/gene/?term=LOC105283552 | 1562 | 33 | 211.27 | 1 | 6.4 | 3.03 | 54.5 | 58 | -3.5 | 3.5 | -0.07 | 0.07 |
| gene-LOC105277903 | https://www.ncbi.nlm.nih.gov/gene/?term=LOC105277903 | 1835 | 35 | 190.74 | 1 | 5.45 | 2.86 | 51.45 | 55.03 | -3.59 | 3.59 | -0.07 | 0.07 |
| gene-LOC105281173 | https://www.ncbi.nlm.nih.gov/gene/?term=LOC105281173 | 1595 | 25 | 156.74 | 1 | 6.27 | 4 | 64.76 | 61.03 | 3.73 | 3.73 | 0.06 | 0.06 |
| gene-LOC105274706 | https://www.ncbi.nlm.nih.gov/gene/?term=LOC105274706 | 1446 | 48 | 331.95 | 1 | 6.92 | 2.08 | 39.02 | 37.1 | 1.92 | 1.92 | 0.06 | 0.06 |
| gene-LOC105278877 | https://www.ncbi.nlm.nih.gov/gene/?term=LOC105278877 | 1968 | 78 | 396.34 | 1 | 5.08 | 1.28 | 32.36 | 30.87 | 1.48 | 1.48 | 0.06 | 0.06 |
| gene-LOC105285055 | https://www.ncbi.nlm.nih.gov/gene/?term=LOC105285055 | 4108 | 204 | 496.59 | 1 | 2.43 | 0.49 | 12.43 | 13.67 | -1.24 | 1.24 | -0.06 | 0.06 |
| gene-LOC105285695 | https://www.ncbi.nlm.nih.gov/gene/?term=LOC105285695 | 1371 | 61 | 444.93 | 1 | 7.29 | 1.64 | 22.52 | 23.97 | -1.45 | 1.45 | -0.06 | 0.06 |
| gene-LOC105286451 | https://www.ncbi.nlm.nih.gov/gene/?term=LOC105286451 | 1618 | 10 | 61.8 | 1 | 6.18 | 10 | 52.19 | 44.01 | 8.18 | 8.18 | 0.05 | 0.05 |
| gene-LOC105278538 | https://www.ncbi.nlm.nih.gov/gene/?term=LOC105278538 | 2150 | 75 | 348.84 | 1 | 4.65 | 1.33 | 18.53 | 17.09 | 1.44 | 1.44 | 0.05 | 0.05 |
| gene-LOC105282156 | https://www.ncbi.nlm.nih.gov/gene/?term=LOC105282156 | 2577 | 128 | 496.7 | 1 | 3.88 | 0.78 | 23.38 | 24.45 | -1.07 | 1.07 | -0.05 | 0.05 |
| gene-LOC105277215 | https://www.ncbi.nlm.nih.gov/gene/?term=LOC105277215 | 2619 | 27 | 103.09 | 1 | 3.82 | 3.7 | 20.79 | 25.43 | -4.64 | 4.64 | -0.05 | 0.05 |
| gene-LOC105275708 | https://www.ncbi.nlm.nih.gov/gene/?term=LOC105275708 | 1772 | 14 | 79.01 | 1 | 5.64 | 7.14 | 44.58 | 50.83 | -6.25 | 6.25 | -0.05 | 0.05 |
| gene-LOC105285514 | https://www.ncbi.nlm.nih.gov/gene/?term=LOC105285514 | 1277 | 8 | 62.65 | 1 | 7.83 | 12.5 | 70.46 | 79.23 | -8.77 | 8.77 | -0.05 | 0.05 |
| gene-LOC105283649 | https://www.ncbi.nlm.nih.gov/gene/?term=LOC105283649 | 1918 | 53 | 276.33 | 2 | 10.43 | 3.77 | 38.29 | 37.03 | 1.27 | 1.27 | 0.04 | 0.04 |
| gene-LOC105287095 | https://www.ncbi.nlm.nih.gov/gene/?term=LOC105287095 | 1435 | 60 | 418.12 | 1 | 6.97 | 1.67 | 26.49 | 25.42 | 1.06 | 1.06 | 0.04 | 0.04 |
| gene-LOC105280786 | https://www.ncbi.nlm.nih.gov/gene/?term=LOC105280786 | 2573 | 115 | 446.95 | 1 | 3.89 | 0.87 | 29.1 | 28.31 | 0.79 | 0.79 | 0.04 | 0.04 |
| gene-LOC105275363 | https://www.ncbi.nlm.nih.gov/gene/?term=LOC105275363 | 3798 | 155 | 408.11 | 1 | 2.63 | 0.65 | 17.28 | 18.35 | -1.06 | 1.06 | -0.04 | 0.04 |
| gene-LOC105285742 | https://www.ncbi.nlm.nih.gov/gene/?term=LOC105285742 | 3446 | 87 | 252.47 | 1 | 2.9 | 1.15 | 20.61 | 22.37 | -1.76 | 1.76 | -0.04 | 0.04 |
| gene-LOC105278436 | https://www.ncbi.nlm.nih.gov/gene/?term=LOC105278436 | 2556 | 56 | 219.09 | 1 | 3.91 | 1.79 | 36.89 | 38.91 | -2.02 | 2.02 | -0.04 | 0.04 |
| gene-LOC105274900 | https://www.ncbi.nlm.nih.gov/gene/?term=LOC105274900 | 5561 | 56 | 100.7 | 1 | 1.8 | 1.79 | 26.85 | 30.51 | -3.66 | 3.66 | -0.04 | 0.04 |
| gene-LOC105276538 | https://www.ncbi.nlm.nih.gov/gene/?term=LOC105276538 | 1511 | 18 | 119.13 | 1 | 6.62 | 5.56 | 71.16 | 68.49 | 2.67 | 2.67 | 0.03 | 0.03 |
| gene-LOC105278454 | https://www.ncbi.nlm.nih.gov/gene/?term=LOC105278454 | 1972 | 37 | 187.63 | 1 | 5.07 | 2.7 | 46.8 | 45.42 | 1.38 | 1.38 | 0.03 | 0.03 |
| gene-LOC105282811 | https://www.ncbi.nlm.nih.gov/gene/?term=LOC105282811 | 5765 | 183 | 317.43 | 1 | 1.73 | 0.55 | 10.19 | 9.27 | 0.92 | 0.92 | 0.03 | 0.03 |
| gene-LOC105274534 | https://www.ncbi.nlm.nih.gov/gene/?term=LOC105274534 | 1329 | 57 | 428.89 | 1 | 7.52 | 1.75 | 43.75 | 44.38 | -0.63 | 0.63 | -0.03 | 0.03 |
| gene-LOC105275683 | https://www.ncbi.nlm.nih.gov/gene/?term=LOC105275683 | 4256 | 187 | 439.38 | 1 | 2.35 | 0.53 | 12.61 | 13.25 | -0.63 | 0.63 | -0.03 | 0.03 |
| gene-LOC105278497 | https://www.ncbi.nlm.nih.gov/gene/?term=LOC105278497 | 2072 | 97 | 468.15 | 1 | 4.83 | 1.03 | 18.56 | 19.21 | -0.64 | 0.64 | -0.03 | 0.03 |
| gene-LOC105288036 | https://www.ncbi.nlm.nih.gov/gene/?term=LOC105288036 | 5110 | 182 | 356.16 | 1 | 1.96 | 0.55 | 18.19 | 18.89 | -0.7 | 0.7 | -0.03 | 0.03 |
| gene-LOC105283592 | https://www.ncbi.nlm.nih.gov/gene/?term=LOC105283592 | 3479 | 68 | 195.46 | 1 | 2.87 | 1.47 | 23.52 | 25 | -1.48 | 1.48 | -0.03 | 0.03 |
| gene-LOC105275190 | https://www.ncbi.nlm.nih.gov/gene/?term=LOC105275190 | 3837 | 65 | 169.4 | 1 | 2.61 | 1.54 | 21.02 | 22.59 | -1.58 | 1.58 | -0.03 | 0.03 |
| gene-LOC105288066 | https://www.ncbi.nlm.nih.gov/gene/?term=LOC105288066 | 1964 | 24 | 122.2 | 1 | 5.09 | 4.17 | 27.7 | 30.29 | -2.59 | 2.59 | -0.03 | 0.03 |
| gene-LOC105285060 | https://www.ncbi.nlm.nih.gov/gene/?term=LOC105285060 | 1285 | 15 | 116.73 | 1 | 7.78 | 6.67 | 48.46 | 51.14 | -2.68 | 2.68 | -0.03 | 0.03 |
| gene-LOC105279898 | https://www.ncbi.nlm.nih.gov/gene/?term=LOC105279898 | 1343 | 13 | 96.8 | 1 | 7.45 | 7.69 | 65.18 | 68.44 | -3.25 | 3.25 | -0.03 | 0.03 |
| gene-LOC105285409 | https://www.ncbi.nlm.nih.gov/gene/?term=LOC105285409 | 5099 | 47 | 92.17 | 1 | 1.96 | 2.13 | 3.57 | 1.67 | 1.9 | 1.9 | 0.02 | 0.02 |
| gene-LOC105279664 | https://www.ncbi.nlm.nih.gov/gene/?term=LOC105279664 | 4023 | 63 | 156.6 | 1 | 2.49 | 1.59 | 17.93 | 16.77 | 1.16 | 1.16 | 0.02 | 0.02 |
| gene-LOC105285202 | https://www.ncbi.nlm.nih.gov/gene/?term=LOC105285202 | 1838 | 35 | 190.42 | 1 | 5.44 | 2.86 | 19.49 | 18.4 | 1.08 | 1.08 | 0.02 | 0.02 |
| gene-LOC105276196 | https://www.ncbi.nlm.nih.gov/gene/?term=LOC105276196 | 2953 | 55 | 186.25 | 1 | 3.39 | 1.82 | 40.47 | 39.41 | 1.06 | 1.06 | 0.02 | 0.02 |
| gene-LOC105285807 | https://www.ncbi.nlm.nih.gov/gene/?term=LOC105285807 | 6063 | 180 | 296.88 | 1 | 1.65 | 0.56 | 10.12 | 9.41 | 0.72 | 0.72 | 0.02 | 0.02 |
| gene-LOC105275752 | https://www.ncbi.nlm.nih.gov/gene/?term=LOC105275752 | 3462 | 126 | 363.95 | 1 | 2.89 | 0.79 | 28.41 | 27.85 | 0.56 | 0.56 | 0.02 | 0.02 |
| gene-LOC105278666 | https://www.ncbi.nlm.nih.gov/gene/?term=LOC105278666 | 10341 | 393 | 380.04 | 1 | 0.97 | 0.25 | 7.76 | 7.36 | 0.4 | 0.4 | 0.02 | 0.02 |
| gene-LOC105285534 | https://www.ncbi.nlm.nih.gov/gene/?term=LOC105285534 | 3944 | 221 | 560.34 | 1 | 2.54 | 0.45 | 11.41 | 11.71 | -0.3 | 0.3 | -0.02 | 0.02 |
| gene-LOC105279298 | https://www.ncbi.nlm.nih.gov/gene/?term=LOC105279298 | 2773 | 135 | 486.84 | 1 | 3.61 | 0.74 | 22.57 | 22.9 | -0.33 | 0.33 | -0.02 | 0.02 |
| gene-LOC105286997 | https://www.ncbi.nlm.nih.gov/gene/?term=LOC105286997 | 2850 | 134 | 470.18 | 1 | 3.51 | 0.75 | 11.83 | 12.3 | -0.47 | 0.47 | -0.02 | 0.02 |
| gene-LOC105280956 | https://www.ncbi.nlm.nih.gov/gene/?term=LOC105280956 | 2639 | 73 | 276.62 | 1 | 3.79 | 1.37 | 20.28 | 20.85 | -0.57 | 0.57 | -0.02 | 0.02 |
| gene-LOC105285639 | https://www.ncbi.nlm.nih.gov/gene/?term=LOC105285639 | 10325 | 294 | 284.75 | 1 | 0.97 | 0.34 | 12.44 | 13.1 | -0.66 | 0.66 | -0.02 | 0.02 |
| gene-LOC105287063 | https://www.ncbi.nlm.nih.gov/gene/?term=LOC105287063 | 1874 | 46 | 245.46 | 1 | 5.34 | 2.17 | 34.15 | 34.88 | -0.73 | 0.73 | -0.02 | 0.02 |
| gene-LOC105275179 | https://www.ncbi.nlm.nih.gov/gene/?term=LOC105275179 | 3271 | 69 | 210.94 | 1 | 3.06 | 1.45 | 14.66 | 15.53 | -0.86 | 0.86 | -0.02 | 0.02 |
| gene-LOC105288237 | https://www.ncbi.nlm.nih.gov/gene/?term=LOC105288237 | 2226 | 45 | 202.16 | 2 | 8.98 | 4.44 | 36.5 | 37.4 | -0.9 | 0.9 | -0.02 | 0.02 |
| gene-LOC105275451 | https://www.ncbi.nlm.nih.gov/gene/?term=LOC105275451 | 6035 | 111 | 183.93 | 1 | 1.66 | 0.9 | 22.31 | 23.24 | -0.94 | 0.94 | -0.02 | 0.02 |
| gene-LOC105286265 | https://www.ncbi.nlm.nih.gov/gene/?term=LOC105286265 | 2507 | 65 | 259.27 | 1 | 3.99 | 1.54 | 31.58 | 32.53 | -0.95 | 0.95 | -0.02 | 0.02 |

| | | | | | | | | | | | | | | | |
|---|---|---|---|---|---|---|---|---|---|---|---|---|---|---|---|
| gene-LOC105282991 | https://www.ncbi.nlm.nih.gov/gene/?term=LOC105282991 | 5617 | 80 | 142.42 | | 1 | 1.78 | 1.25 | 19.15 | 20.24 | -1.09 | 1.09 | -0.02 | 0.02 | |
| gene-LOC105283943 | https://www.ncbi.nlm.nih.gov/gene/?term=LOC105283943 | 3559 | 49 | 137.68 | | 1 | 2.81 | 2.04 | 22.13 | 23.23 | -1.1 | 1.1 | -0.02 | 0.02 | |
| gene-LOC105277216 | https://www.ncbi.nlm.nih.gov/gene/?term=LOC105277216 | 7627 | 91 | 119.31 | | 1 | 1.31 | 1.1 | 6.55 | 7.88 | -1.33 | 1.33 | -0.02 | 0.02 | |
| gene-LOC105280348 | https://www.ncbi.nlm.nih.gov/gene/?term=LOC105280348 | 3797 | 52 | 136.95 | | 1 | 2.63 | 1.92 | 25.54 | 27.17 | -1.63 | 1.63 | -0.02 | 0.02 | |
| gene-LOC105275518 | https://www.ncbi.nlm.nih.gov/gene/?term=LOC105275518 | 2335 | 19 | 81.37 | 1 | 4.28 | 5.26 | 49.77 | 48.32 | 1.45 | 1.45 | 0.01 | 0.01 | | |
| gene-LOC105281047 | https://www.ncbi.nlm.nih.gov/gene/?term=LOC105281047 | 7313 | 112 | 153.15 | | 1 | 1.37 | 0.89 | 11.03 | 10.28 | 0.75 | 0.75 | 0.01 | 0.01 | |
| gene-LOC105282794 | https://www.ncbi.nlm.nih.gov/gene/?term=LOC105282794 | 4571 | 93 | 203.46 | | 1 | 2.19 | 1.08 | 17.63 | 17.04 | 0.6 | 0.6 | 0.01 | 0.01 | |
| gene-LOC105283123 | https://www.ncbi.nlm.nih.gov/gene/?term=LOC105283123 | 1507 | 35 | 232.25 | | 1 | 6.64 | 2.86 | 18.15 | 17.56 | 0.59 | 0.59 | 0.01 | 0.01 | |
| gene-LOC105284116 | https://www.ncbi.nlm.nih.gov/gene/?term=LOC105284116 | 3970 | 90 | 226.7 | 1 | 2.52 | 1.11 | 7.07 | 6.57 | 0.5 | 0.5 | 0.01 | 0.01 | | |
| gene-LOC105279038 | https://www.ncbi.nlm.nih.gov/gene/?term=LOC105279038 | 9453 | 299 | 316.3 | 1 | 1.06 | 0.33 | 8.97 | 8.54 | 0.42 | 0.42 | 0.01 | 0.01 | | |
| gene-LOC105280002 | https://www.ncbi.nlm.nih.gov/gene/?term=LOC105280002 | 1940 | 28 | 144.33 | | 1 | 5.15 | 3.57 | 33.09 | 32.74 | 0.35 | 0.35 | 0.01 | 0.01 | |
| gene-LOC105284117 | https://www.ncbi.nlm.nih.gov/gene/?term=LOC105284117 | 4340 | 82 | 188.94 | | 1 | 2.3 | 1.22 | 10.79 | 10.47 | 0.33 | 0.33 | 0.01 | 0.01 | |
| gene-LOC105280819 | https://www.ncbi.nlm.nih.gov/gene/?term=LOC105280819 | 1639 | 41 | 250.15 | | 1 | 6.1 | 2.44 | 40.58 | 40.31 | 0.27 | 0.27 | 0.01 | 0.01 | |
| gene-LOC105288079 | https://www.ncbi.nlm.nih.gov/gene/?term=LOC105288079 | 3473 | 85 | 244.75 | | 1 | 2.88 | 1.18 | 20.86 | 20.61 | 0.25 | 0.25 | 0.01 | 0.01 | |
| gene-LOC105286515 | https://www.ncbi.nlm.nih.gov/gene/?term=LOC105286515 | 3751 | 106 | 282.59 | | 1 | 2.67 | 0.94 | 10.37 | 10.58 | -0.22 | 0.22 | -0.01 | 0.01 | |
| gene-LOC105288078 | https://www.ncbi.nlm.nih.gov/gene/?term=LOC105288078 | 19126 | 305 | 159.47 | | 1 | 0.52 | 0.33 | 16.95 | 17.33 | -0.38 | 0.38 | -0.01 | 0.01 | |
| gene-LOC105274531 | https://www.ncbi.nlm.nih.gov/gene/?term=LOC105274531 | 7774 | 253 | 325.44 | | 1 | 1.29 | 0.4 | 11.9 | 12.33 | -0.43 | 0.43 | -0.01 | 0.01 | |
| gene-LOC105283712 | https://www.ncbi.nlm.nih.gov/gene/?term=LOC105283712 | 1910 | 29 | 151.83 | | 1 | 5.24 | 3.45 | 29.98 | 30.42 | -0.44 | 0.44 | -0.01 | 0.01 | |
| gene-LOC105275173 | https://www.ncbi.nlm.nih.gov/gene/?term=LOC105275173 | 4639 | 76 | 163.83 | | 1 | 2.16 | 1.32 | 24.33 | 24.78 | -0.45 | 0.45 | -0.01 | 0.01 | |
| gene-LOC105284184 | https://www.ncbi.nlm.nih.gov/gene/?term=LOC105284184 | 2535 | 80 | 315.58 | | 1 | 3.94 | 1.25 | 34.85 | 35.33 | -0.47 | 0.47 | -0.01 | 0.01 | |
| gene-LOC105275898 | https://www.ncbi.nlm.nih.gov/gene/?term=LOC105275898 | 5358 | 132 | 246.36 | | 1 | 1.87 | 0.76 | 12.84 | 13.34 | -0.5 | 0.5 | -0.01 | 0.01 | |
| gene-LOC105276692 | https://www.ncbi.nlm.nih.gov/gene/?term=LOC105276692 | 1745 | 34 | 194.84 | | 1 | 5.73 | 2.94 | 24.51 | 25.17 | -0.66 | 0.66 | -0.01 | 0.01 | |
| gene-LOC105280641 | https://www.ncbi.nlm.nih.gov/gene/?term=LOC105280641 | 3682 | 55 | 149.38 | | 1 | 2.72 | 1.82 | 18.35 | 19.01 | -0.66 | 0.66 | -0.01 | 0.01 | |
| gene-LOC105281700 | https://www.ncbi.nlm.nih.gov/gene/?term=LOC105281700 | 2208 | 45 | 203.8 | 1 | 4.53 | 2.22 | 38.83 | 39.5 | -0.67 | 0.67 | -0.01 | 0.01 | | |
| gene-LOC105279657 | https://www.ncbi.nlm.nih.gov/gene/?term=LOC105279657 | 5878 | 82 | 139.5 | 1 | 1.7 | 1.22 | 19.15 | 19.93 | -0.78 | 0.78 | -0.01 | 0.01 | | |
| gene-LOC105277856 | https://www.ncbi.nlm.nih.gov/gene/?term=LOC105277856 | 3442 | 28 | 81.35 | 1 | 2.91 | 3.57 | 36.24 | 37.06 | -0.82 | 0.82 | -0.01 | 0.01 | | |
| gene-LOC105286452 | https://www.ncbi.nlm.nih.gov/gene/?term=LOC105286452 | 6888 | 58 | 84.2 | 1 | 1.45 | 1.72 | 31.25 | 32.65 | -1.4 | 1.4 | -0.01 | 0.01 | | |
| gene-LOC105283989 | https://www.ncbi.nlm.nih.gov/gene/?term=LOC105283989 | 4948 | 30 | 60.63 | 1 | 2.02 | 3.33 | 45.76 | 47.5 | -1.74 | 1.74 | -0.01 | 0.01 | | |
| gene-LOC105274590 | https://www.ncbi.nlm.nih.gov/gene/?term=LOC105274590 | 4516 | 75 | 166.08 | | 1 | 2.21 | 1.33 | 23.2 | 22.91 | 0.29 | 0.29 | 0 | 0 | |
| gene-LOC105286194 | https://www.ncbi.nlm.nih.gov/gene/?term=LOC105286194 | 152619 | | 1939 | 127.05 | | 1 | 0.07 | 0.05 | 6.35 | 6.21 | 0.14 | 0.14 | 0 | 0 |
| gene-LOC105276901 | https://www.ncbi.nlm.nih.gov/gene/?term=LOC105276901 | 3254 | 96 | 295.02 | | 1 | 3.07 | 1.04 | 22.83 | 22.72 | 0.1 | 0.1 | 0 | 0 | |
| gene-LOC105277866 | https://www.ncbi.nlm.nih.gov/gene/?term=LOC105277866 | 33045 | 403 | 121.95 | | 1 | 0.3 | 0.25 | 3.32 | 3.22 | 0.1 | 0.1 | 0 | 0 | |
| gene-LOC105280265 | https://www.ncbi.nlm.nih.gov/gene/?term=LOC105280265 | 85984 | 2645 | 307.62 | | 1 | 0.12 | 0.04 | 1.12 | 1.06 | 0.07 | 0.07 | 0 | 0 | |
| gene-LOC105278455 | https://www.ncbi.nlm.nih.gov/gene/?term=LOC105278455 | 27181 | 836 | 307.57 | | 1 | 0.37 | 0.12 | 7.62 | 7.57 | 0.05 | 0.05 | 0 | 0 | |
| gene-LOC105278228 | https://www.ncbi.nlm.nih.gov/gene/?term=LOC105278228 | 8610 | 152 | 176.54 | | 1 | 1.16 | 0.66 | 23.77 | 23.73 | 0.04 | 0.04 | 0 | 0 | |
| gene-LOC105283728 | https://www.ncbi.nlm.nih.gov/gene/?term=LOC105283728 | 3256 | 113 | 347.05 | | 2 | 6.14 | 1.77 | 19.87 | 19.82 | 0.04 | 0.04 | 0 | 0 | |
| gene-LOC105278145 | https://www.ncbi.nlm.nih.gov/gene/?term=LOC105278145 | 4470 | 239 | 534.68 | | 1 | 2.24 | 0.42 | 10.87 | 10.84 | 0.03 | 0.03 | 0 | 0 | |
| gene-LOC105275327 | https://www.ncbi.nlm.nih.gov/gene/?term=LOC105275327 | 14245 | 461 | 323.62 | | 1 | 0.7 | 0.22 | 7.38 | 7.42 | -0.03 | 0.03 | 0 | 0 | |
| gene-LOC105286274 | https://www.ncbi.nlm.nih.gov/gene/?term=LOC105286274 | 3817 | 174 | 455.86 | | 1 | 2.62 | 0.57 | 15.59 | 15.67 | -0.08 | 0.08 | 0 | 0 | |
| gene-LOC105274856 | https://www.ncbi.nlm.nih.gov/gene/?term=LOC105274856 | 5216 | 201 | 385.35 | | 1 | 1.92 | 0.5 | 17.77 | 17.88 | -0.11 | 0.11 | 0 | 0 | |
| gene-LOC105278534 | https://www.ncbi.nlm.nih.gov/gene/?term=LOC105278534 | 7364 | 173 | 234.93 | | 1 | 1.36 | 0.58 | 18.44 | 18.59 | -0.16 | 0.16 | 0 | 0 | |
| gene-LOC105274851 | https://www.ncbi.nlm.nih.gov/gene/?term=LOC105274851 | 6616 | 133 | 201.03 | | 1 | 1.51 | 0.75 | 10.15 | 10.35 | -0.2 | 0.2 | 0 | 0 | |
| gene-LOC105279500 | https://www.ncbi.nlm.nih.gov/gene/?term=LOC105279500 | 3587 | 29 | 80.85 | 1 | 2.79 | 3.45 | 48.07 | 48.34 | -0.27 | 0.27 | 0 | 0 | | |
| gene-LOC105285406 | https://www.ncbi.nlm.nih.gov/gene/?term=LOC105285406 | 8015 | 39 | 48.66 | 1 | 1.25 | 2.56 | 33.07 | 33.49 | -0.42 | 0.42 | 0 | 0 | | |

Directory: OGLall File: rnk-dmg-Ob_LI_BCphaseA1.vs.RphaseA2.txt

| gene_ID | gene_link | gwidth | #Sites | #per10Kb | #dmSites | #dmsp10kb | %dmSites | %pSite1 | %pSite2 | DMpSite | ADMpSite | DMpNucl | ADMpNucl |
|---|---|---|---|---|---|---|---|---|---|---|---|---|---|
| gene-LOC113562345 | https://www.ncbi.nlm.nih.gov/gene/?term=LOC113562345 | 2290 | 62 | 270.74 | 16 | 69.87 | 25.81 | 17.91 | 33 | -15.09 | 15.09 | -0.41 | 0.41 |
| gene-LOC105276704 | https://www.ncbi.nlm.nih.gov/gene/?term=LOC105276704 | 2063 | 87 | 421.72 | 1 | 4.85 | 1.15 | 12.34 | 15.21 | -2.86 | 2.86 | -0.12 | 0.12 |
| gene-LOC105282156 | https://www.ncbi.nlm.nih.gov/gene/?term=LOC105282156 | 2577 | 136 | 527.75 | 1 | 3.88 | 0.74 | 23.86 | 26.02 | -2.16 | 2.16 | -0.11 | 0.11 |
| gene-LOC105279898 | https://www.ncbi.nlm.nih.gov/gene/?term=LOC105279898 | 1343 | 16 | 119.14 | 2 | 14.89 | 12.5 | 59.42 | 68.99 | -9.57 | 9.57 | -0.11 | 0.11 |
| gene-LOC105274534 | https://www.ncbi.nlm.nih.gov/gene/?term=LOC105274534 | 1329 | 57 | 428.89 | 1 | 7.52 | 1.75 | 43.75 | 46.14 | -2.39 | 2.39 | -0.1 | 0.1 |
| gene-LOC105286502 | https://www.ncbi.nlm.nih.gov/gene/?term=LOC105286502 | 1286 | 44 | 342.15 | 1 | 7.78 | 2.27 | 23.97 | 26.78 | -2.81 | 2.81 | -0.1 | 0.1 |
| gene-LOC105277631 | https://www.ncbi.nlm.nih.gov/gene/?term=LOC105277631 | 4387 | 289 | 658.76 | 1 | 2.28 | 0.35 | 9.79 | 11.22 | -1.43 | 1.43 | -0.09 | 0.09 |
| gene-LOC105285674 | https://www.ncbi.nlm.nih.gov/gene/?term=LOC105285674 | 3035 | 91 | 299.84 | 2 | 6.59 | 2.2 | 23.1 | 26.01 | -2.91 | 2.91 | -0.09 | 0.09 |
| gene-LOC105281848 | https://www.ncbi.nlm.nih.gov/gene/?term=LOC105281848 | 2122 | 47 | 221.49 | 1 | 4.71 | 2.13 | 23.11 | 27.19 | -4.08 | 4.08 | -0.09 | 0.09 |
| gene-LOC105285695 | https://www.ncbi.nlm.nih.gov/gene/?term=LOC105285695 | 1371 | 62 | 452.22 | 1 | 7.29 | 1.61 | 23 | 24.72 | -1.72 | 1.72 | -0.08 | 0.08 |
| gene-LOC105287417 | https://www.ncbi.nlm.nih.gov/gene/?term=LOC105287417 | 1025 | 33 | 321.95 | 1 | 9.76 | 3.03 | 36 | 38.42 | -2.42 | 2.42 | -0.08 | 0.08 |
| gene-LOC105285055 | https://www.ncbi.nlm.nih.gov/gene/?term=LOC105285055 | 4108 | 217 | 528.24 | 1 | 2.43 | 0.46 | 12.76 | 14.18 | -1.42 | 1.42 | -0.07 | 0.07 |
| gene-LOC105285671 | https://www.ncbi.nlm.nih.gov/gene/?term=LOC105285671 | 2263 | 95 | 419.8 | 1 | 4.42 | 1.05 | 27.86 | 29.64 | -1.78 | 1.78 | -0.07 | 0.07 |
| gene-LOC105286223 | https://www.ncbi.nlm.nih.gov/gene/?term=LOC105286223 | 2225 | 83 | 373.03 | 1 | 4.49 | 1.2 | 30.2 | 32.18 | -1.98 | 1.98 | -0.07 | 0.07 |
| gene-LOC113561805 | https://www.ncbi.nlm.nih.gov/gene/?term=LOC113561805 | 3337 | 64 | 191.79 | 1 | 3 | 1.56 | 37.95 | 41.4 | -3.45 | 3.45 | -0.07 | 0.07 |
| gene-LOC105275190 | https://www.ncbi.nlm.nih.gov/gene/?term=LOC105275190 | 3837 | 74 | 192.86 | 1 | 2.61 | 1.35 | 18.39 | 22.03 | -3.64 | 3.64 | -0.07 | 0.07 |
| gene-LOC105275070 | https://www.ncbi.nlm.nih.gov/gene/?term=LOC105275070 | 5667 | 38 | 67.05 | 3 | 5.29 | 7.89 | 3.63 | 13.45 | -9.82 | 9.82 | -0.07 | 0.07 |
| gene-LOC105274449 | https://www.ncbi.nlm.nih.gov/gene/?term=LOC105274449 | 1549 | 42 | 271.14 | 1 | 6.46 | 2.38 | 55.88 | 53.68 | 2.2 | 2.2 | 0.06 | 0.06 |
| gene-LOC105275683 | https://www.ncbi.nlm.nih.gov/gene/?term=LOC105275683 | 4256 | 203 | 476.97 | 1 | 2.35 | 0.49 | 12.81 | 14.05 | -1.25 | 1.25 | -0.06 | 0.06 |
| gene-LOC105275460 | https://www.ncbi.nlm.nih.gov/gene/?term=LOC105275460 | 2473 | 112 | 452.89 | 1 | 4.04 | 0.89 | 12.89 | 14.18 | -1.3 | 1.3 | -0.06 | 0.06 |
| gene-LOC105279356 | https://www.ncbi.nlm.nih.gov/gene/?term=LOC105279356 | 1314 | 42 | 319.63 | 1 | 7.61 | 2.38 | 28.83 | 30.58 | -1.75 | 1.75 | -0.06 | 0.06 |
| gene-LOC105278807 | https://www.ncbi.nlm.nih.gov/gene/?term=LOC105278807 | 2166 | 72 | 332.41 | 1 | 4.62 | 1.39 | 18.53 | 20.46 | -1.93 | 1.93 | -0.06 | 0.06 |
| gene-LOC105285583 | https://www.ncbi.nlm.nih.gov/gene/?term=LOC105285583 | 1947 | 51 | 261.94 | 1 | 5.14 | 1.96 | 28.26 | 30.43 | -2.17 | 2.17 | -0.06 | 0.06 |
| gene-LOC105287095 | https://www.ncbi.nlm.nih.gov/gene/?term=LOC105287095 | 1435 | 61 | 425.09 | 1 | 6.97 | 1.64 | 26.05 | 24.77 | 1.29 | 1.29 | 0.05 | 0.05 |
| gene-LOC105279298 | https://www.ncbi.nlm.nih.gov/gene/?term=LOC105279298 | 2773 | 138 | 497.66 | 2 | 7.21 | 1.45 | 22.22 | 23.22 | -1 | 1 | -0.05 | 0.05 |
| gene-LOC105278497 | https://www.ncbi.nlm.nih.gov/gene/?term=LOC105278497 | 2072 | 100 | 482.63 | 1 | 4.83 | 1 | 18.86 | 19.92 | -1.06 | 1.06 | -0.05 | 0.05 |
| gene-LOC105282947 | https://www.ncbi.nlm.nih.gov/gene/?term=LOC105282947 | 1162 | 49 | 421.69 | 1 | 8.61 | 2.04 | 29.51 | 30.69 | -1.17 | 1.17 | -0.05 | 0.05 |
| gene-LOC105284704 | https://www.ncbi.nlm.nih.gov/gene/?term=LOC105284704 | 2400 | 65 | 270.83 | 1 | 4.17 | 1.54 | 25.06 | 26.85 | -1.79 | 1.79 | -0.05 | 0.05 |
| gene-LOC105283712 | https://www.ncbi.nlm.nih.gov/gene/?term=LOC105283712 | 1910 | 27 | 141.36 | 1 | 5.24 | 3.7 | 29.17 | 32.5 | -3.33 | 3.33 | -0.05 | 0.05 |
| gene-LOC105278109 | https://www.ncbi.nlm.nih.gov/gene/?term=LOC105278109 | 4029 | 62 | 153.88 | 1 | 2.48 | 1.61 | 43.07 | 46.49 | -3.42 | 3.42 | -0.05 | 0.05 |
| gene-LOC105285060 | https://www.ncbi.nlm.nih.gov/gene/?term=LOC105285060 | 1285 | 15 | 116.73 | 1 | 7.78 | 6.67 | 48.46 | 52.7 | -4.25 | 4.25 | -0.05 | 0.05 |
| gene-LOC105275708 | https://www.ncbi.nlm.nih.gov/gene/?term=LOC105275708 | 1772 | 14 | 79.01 | 1 | 5.64 | 7.14 | 44.58 | 50.56 | -5.98 | 5.98 | -0.05 | 0.05 |
| gene-LOC105277380 | https://www.ncbi.nlm.nih.gov/gene/?term=LOC105277380 | 1967 | 5 | 25.42 | 1 | 5.08 | 20 | 69.42 | 54.11 | 15.31 | 15.31 | 0.04 | 0.04 |
| gene-LOC105282885 | https://www.ncbi.nlm.nih.gov/gene/?term=LOC105282885 | 1870 | 42 | 224.6 | 1 | 5.35 | 2.38 | 36.63 | 34.78 | 1.84 | 1.84 | 0.04 | 0.04 |
| gene-LOC105275848 | https://www.ncbi.nlm.nih.gov/gene/?term=LOC105275848 | 3718 | 130 | 349.65 | 1 | 2.69 | 0.77 | 21.27 | 20.2 | 1.07 | 1.07 | 0.04 | 0.04 |
| gene-LOC105275363 | https://www.ncbi.nlm.nih.gov/gene/?term=LOC105275363 | 3798 | 159 | 418.64 | 1 | 2.63 | 0.63 | 16.97 | 17.86 | -0.88 | 0.88 | -0.04 | 0.04 |
| gene-LOC105285742 | https://www.ncbi.nlm.nih.gov/gene/?term=LOC105285742 | 3446 | 100 | 290.19 | 1 | 2.9 | 1 | 24.69 | 25.96 | -1.27 | 1.27 | -0.04 | 0.04 |
| gene-LOC105278282 | https://www.ncbi.nlm.nih.gov/gene/?term=LOC105278282 | 1775 | 47 | 264.79 | 1 | 5.63 | 2.13 | 45.94 | 47.29 | -1.35 | 1.35 | -0.04 | 0.04 |
| gene-LOC105287063 | https://www.ncbi.nlm.nih.gov/gene/?term=LOC105287063 | 1874 | 46 | 245.46 | 1 | 5.34 | 2.17 | 34.15 | 35.78 | -1.63 | 1.63 | -0.04 | 0.04 |
| gene-LOC105276692 | https://www.ncbi.nlm.nih.gov/gene/?term=LOC105276692 | 1745 | 37 | 212.03 | 1 | 5.73 | 2.7 | 20.82 | 22.64 | -1.82 | 1.82 | -0.04 | 0.04 |
| gene-LOC105276745 | https://www.ncbi.nlm.nih.gov/gene/?term=LOC105276745 | 3953 | 73 | 184.67 | 1 | 2.53 | 1.37 | 21.09 | 23.02 | -1.93 | 1.93 | -0.04 | 0.04 |
| gene-LOC105282893 | https://www.ncbi.nlm.nih.gov/gene/?term=LOC105282893 | 2645 | 43 | 162.57 | 1 | 3.78 | 2.33 | 27.48 | 29.68 | -2.19 | 2.19 | -0.04 | 0.04 |
| gene-LOC105286644 | https://www.ncbi.nlm.nih.gov/gene/?term=LOC105286644 | 1235 | 22 | 178.14 | 1 | 8.1 | 4.55 | 42.2 | 44.4 | -2.2 | 2.2 | -0.04 | 0.04 |
| gene-LOC105287854 | https://www.ncbi.nlm.nih.gov/gene/?term=LOC105287854 | 3644 | 57 | 156.42 | 1 | 2.74 | 1.75 | 21.16 | 23.71 | -2.55 | 2.55 | -0.04 | 0.04 |
| gene-LOC105276538 | https://www.ncbi.nlm.nih.gov/gene/?term=LOC105276538 | 1511 | 19 | 125.74 | 1 | 6.62 | 5.26 | 72.68 | 75.55 | -2.87 | 2.87 | -0.04 | 0.04 |
| gene-LOC105277856 | https://www.ncbi.nlm.nih.gov/gene/?term=LOC105277856 | 3442 | 29 | 84.25 | 1 | 2.91 | 3.45 | 35.44 | 40.05 | -4.6 | 4.6 | -0.04 | 0.04 |
| gene-LOC105281039 | https://www.ncbi.nlm.nih.gov/gene/?term=LOC105281039 | 1471 | 13 | 88.38 | 1 | 6.8 | 7.69 | 36.12 | 40.81 | -4.7 | 4.7 | -0.04 | 0.04 |
| gene-LOC105285514 | https://www.ncbi.nlm.nih.gov/gene/?term=LOC105285514 | 1277 | 9 | 70.48 | 1 | 7.83 | 11.11 | 73.21 | 78.44 | -5.23 | 5.23 | -0.04 | 0.04 |
| gene-LOC105275859 | https://www.ncbi.nlm.nih.gov/gene/?term=LOC105275859 | 2331 | 25 | 107.25 | 1 | 4.29 | 4 | 53.95 | 51.3 | 2.65 | 2.65 | 0.03 | 0.03 |
| gene-LOC105281173 | https://www.ncbi.nlm.nih.gov/gene/?term=LOC105281173 | 1595 | 25 | 156.74 | 1 | 6.27 | 4 | 64.76 | 62.62 | 2.13 | 2.13 | 0.03 | 0.03 |
| gene-LOC105278877 | https://www.ncbi.nlm.nih.gov/gene/?term=LOC105278877 | 1968 | 80 | 406.5 | 1 | 5.08 | 1.25 | 32.88 | 32.17 | 0.72 | 0.72 | 0.03 | 0.03 |
| gene-LOC105278283 | https://www.ncbi.nlm.nih.gov/gene/?term=LOC105278283 | 2317 | 130 | 561.07 | 1 | 4.32 | 0.77 | 24.9 | 25.39 | -0.48 | 0.48 | -0.03 | 0.03 |
| gene-LOC105276527 | https://www.ncbi.nlm.nih.gov/gene/?term=LOC105276527 | 7923 | 351 | 443.01 | 1 | 1.26 | 0.28 | 8.2 | 8.78 | -0.58 | 0.58 | -0.03 | 0.03 |
| gene-LOC105284355 | https://www.ncbi.nlm.nih.gov/gene/?term=LOC105284355 | 3904 | 147 | 376.54 | 1 | 2.56 | 0.68 | 21.32 | 21.98 | -0.67 | 0.67 | -0.03 | 0.03 |
| gene-LOC105286274 | https://www.ncbi.nlm.nih.gov/gene/?term=LOC105286274 | 3817 | 183 | 479.43 | 1 | 2.62 | 0.55 | 15.39 | 16.06 | -0.67 | 0.67 | -0.03 | 0.03 |
| gene-LOC105285179 | https://www.ncbi.nlm.nih.gov/gene/?term=LOC105285179 | 4191 | 156 | 372.23 | 1 | 2.39 | 0.64 | 17.53 | 18.23 | -0.69 | 0.69 | -0.03 | 0.03 |
| gene-LOC105283728 | https://www.ncbi.nlm.nih.gov/gene/?term=LOC105283728 | 3256 | 118 | 362.41 | 2 | 6.14 | 1.69 | 19.18 | 19.87 | -0.7 | 0.7 | -0.03 | 0.03 |
| gene-LOC105276449 | https://www.ncbi.nlm.nih.gov/gene/?term=LOC105276449 | 1231 | 48 | 389.93 | 1 | 8.12 | 2.08 | 26.7 | 27.43 | -0.74 | 0.74 | -0.03 | 0.03 |
| gene-LOC105275752 | https://www.ncbi.nlm.nih.gov/gene/?term=LOC105275752 | 3462 | 123 | 355.29 | 1 | 2.89 | 0.81 | 28.75 | 29.5 | -0.75 | 0.75 | -0.03 | 0.03 |
| gene-LOC105280774 | https://www.ncbi.nlm.nih.gov/gene/?term=LOC105280774 | 2793 | 117 | 418.9 | 1 | 3.58 | 0.85 | 27.44 | 28.24 | -0.79 | 0.79 | -0.03 | 0.03 |

| | | | | | | | | | | | | | |
|---|---|---|---|---|---|---|---|---|---|---|---|---|---|
| gene-LOC105282563 | https://www.ncbi.nlm.nih.gov/gene/?term=LOC105282563 | 4353 | 146 | 335.4 | 1 | 2.3 | 0.68 | 15.5 | 16.32 | -0.83 | 0.83 | -0.03 | 0.03 |
| gene-LOC105274856 | https://www.ncbi.nlm.nih.gov/gene/?term=LOC105274856 | 5216 | 208 | 398.77 | 1 | 1.92 | 0.48 | 18.49 | 19.33 | -0.84 | 0.84 | -0.03 | 0.03 |
| gene-LOC105274706 | https://www.ncbi.nlm.nih.gov/gene/?term=LOC105274706 | 1446 | 52 | 359.61 | 1 | 6.92 | 1.92 | 37.01 | 37.87 | -0.86 | 0.86 | -0.03 | 0.03 |
| gene-LOC105282491 | https://www.ncbi.nlm.nih.gov/gene/?term=LOC105282491 | 4761 | 163 | 342.37 | 1 | 2.1 | 0.61 | 9.65 | 10.67 | -1.02 | 1.02 | -0.03 | 0.03 |
| gene-LOC105281700 | https://www.ncbi.nlm.nih.gov/gene/?term=LOC105281700 | 2208 | 46 | 208.33 | 1 | 4.53 | 2.17 | 39.97 | 41.25 | -1.28 | 1.28 | -0.03 | 0.03 |
| gene-LOC105283592 | https://www.ncbi.nlm.nih.gov/gene/?term=LOC105283592 | 3479 | 71 | 204.08 | 1 | 2.87 | 1.41 | 24.39 | 25.83 | -1.45 | 1.45 | -0.03 | 0.03 |
| gene-LOC105280348 | https://www.ncbi.nlm.nih.gov/gene/?term=LOC105280348 | 3797 | 52 | 136.95 | 1 | 2.63 | 1.92 | 25.54 | 27.53 | -1.99 | 1.99 | -0.03 | 0.03 |
| gene-LOC105280002 | https://www.ncbi.nlm.nih.gov/gene/?term=LOC105280002 | 1940 | 30 | 154.64 | 1 | 5.15 | 3.33 | 33.39 | 35.64 | -2.25 | 2.25 | -0.03 | 0.03 |
| gene-LOC105288066 | https://www.ncbi.nlm.nih.gov/gene/?term=LOC105288066 | 1964 | 26 | 132.38 | 1 | 5.09 | 3.85 | 26.28 | 28.74 | -2.45 | 2.45 | -0.03 | 0.03 |
| gene-LOC105277215 | https://www.ncbi.nlm.nih.gov/gene/?term=LOC105277215 | 2619 | 29 | 110.73 | 1 | 3.82 | 3.45 | 19.9 | 22.79 | -2.88 | 2.88 | -0.03 | 0.03 |
| gene-LOC105282399 | https://www.ncbi.nlm.nih.gov/gene/?term=LOC105282399 | 2682 | 22 | 82.03 | 1 | 3.73 | 4.55 | 32.93 | 36.36 | -3.43 | 3.43 | -0.03 | 0.03 |
| gene-LOC105274778 | https://www.ncbi.nlm.nih.gov/gene/?term=LOC105274778 | 4662 | 38 | 81.51 | 1 | 2.15 | 2.63 | 44.96 | 48.85 | -3.88 | 3.88 | -0.03 | 0.03 |
| gene-LOC105284609 | https://www.ncbi.nlm.nih.gov/gene/?term=LOC105284609 | 5678 | 31 | 54.6 | 1 | 1.76 | 3.23 | 35.81 | 41.06 | -5.25 | 5.25 | -0.03 | 0.03 |
| gene-LOC105285409 | https://www.ncbi.nlm.nih.gov/gene/?term=LOC105285409 | 5099 | 48 | 94.14 | 2 | 3.92 | 4.17 | 3.6 | 1.11 | 2.49 | 2.49 | 0.02 | 0.02 |
| gene-LOC105276272 | https://www.ncbi.nlm.nih.gov/gene/?term=LOC105276272 | 1965 | 27 | 137.4 | 1 | 5.09 | 3.7 | 42.02 | 40.71 | 1.31 | 1.31 | 0.02 | 0.02 |
| gene-LOC105281296 | https://www.ncbi.nlm.nih.gov/gene/?term=LOC105281296 | 2434 | 41 | 168.45 | 1 | 4.11 | 2.44 | 31.19 | 30.16 | 1.03 | 1.03 | 0.02 | 0.02 |
| gene-LOC105284507 | https://www.ncbi.nlm.nih.gov/gene/?term=LOC105284507 | 5207 | 82 | 157.48 | 1 | 1.92 | 1.22 | 21.61 | 20.58 | 1.03 | 1.03 | 0.02 | 0.02 |
| gene-LOC105283123 | https://www.ncbi.nlm.nih.gov/gene/?term=LOC105283123 | 1507 | 43 | 285.34 | 1 | 6.64 | 2.33 | 17.05 | 16.29 | 0.76 | 0.76 | 0.02 | 0.02 |
| gene-LOC105284396 | https://www.ncbi.nlm.nih.gov/gene/?term=LOC105284396 | 1976 | 61 | 308.7 | 1 | 5.06 | 1.64 | 36.72 | 35.98 | 0.74 | 0.74 | 0.02 | 0.02 |
| gene-LOC105283264 | https://www.ncbi.nlm.nih.gov/gene/?term=LOC105283264 | 4005 | 88 | 219.73 | 1 | 2.5 | 1.14 | 24.81 | 24.09 | 0.72 | 0.72 | 0.02 | 0.02 |
| gene-LOC105282811 | https://www.ncbi.nlm.nih.gov/gene/?term=LOC105282811 | 5765 | 201 | 348.66 | 1 | 1.73 | 0.5 | 9.88 | 9.23 | 0.65 | 0.65 | 0.02 | 0.02 |
| gene-LOC105286515 | https://www.ncbi.nlm.nih.gov/gene/?term=LOC105286515 | 3751 | 116 | 309.25 | 1 | 2.67 | 0.86 | 11.85 | 11.25 | 0.6 | 0.6 | 0.02 | 0.02 |
| gene-LOC105287704 | https://www.ncbi.nlm.nih.gov/gene/?term=LOC105287704 | 3019 | 125 | 414.04 | 1 | 3.31 | 0.8 | 21.19 | 21.56 | -0.37 | 0.37 | -0.02 | 0.02 |
| gene-LOC105280786 | https://www.ncbi.nlm.nih.gov/gene/?term=LOC105280786 | 2573 | 122 | 474.15 | 1 | 3.89 | 0.82 | 29.15 | 29.56 | -0.41 | 0.41 | -0.02 | 0.02 |
| gene-LOC105285534 | https://www.ncbi.nlm.nih.gov/gene/?term=LOC105285534 | 3944 | 224 | 567.95 | 1 | 2.54 | 0.45 | 11.27 | 11.7 | -0.42 | 0.42 | -0.02 | 0.02 |
| gene-LOC105284043 | https://www.ncbi.nlm.nih.gov/gene/?term=LOC105284043 | 5661 | 248 | 438.09 | 1 | 1.77 | 0.4 | 11.5 | 11.98 | -0.48 | 0.48 | -0.02 | 0.02 |
| gene-LOC105277435 | https://www.ncbi.nlm.nih.gov/gene/?term=LOC105277435 | 3643 | 158 | 433.71 | 1 | 2.74 | 0.63 | 12.84 | 13.36 | -0.52 | 0.52 | -0.02 | 0.02 |
| gene-LOC105283132 | https://www.ncbi.nlm.nih.gov/gene/?term=LOC105283132 | 1819 | 54 | 296.87 | 1 | 5.5 | 1.85 | 40.82 | 41.59 | -0.77 | 0.77 | -0.02 | 0.02 |
| gene-LOC105274590 | https://www.ncbi.nlm.nih.gov/gene/?term=LOC105274590 | 4516 | 81 | 179.36 | 1 | 2.21 | 1.23 | 21.65 | 22.55 | -0.9 | 0.9 | -0.02 | 0.02 |
| gene-LOC105275179 | https://www.ncbi.nlm.nih.gov/gene/?term=LOC105275179 | 3271 | 68 | 207.89 | 1 | 3.06 | 1.47 | 14.88 | 16 | -1.12 | 1.12 | -0.02 | 0.02 |
| gene-LOC105278962 | https://www.ncbi.nlm.nih.gov/gene/?term=LOC105278962 | 4491 | 88 | 195.95 | 1 | 2.23 | 1.14 | 23.85 | 25.09 | -1.24 | 1.24 | -0.02 | 0.02 |
| gene-LOC105283943 | https://www.ncbi.nlm.nih.gov/gene/?term=LOC105283943 | 3559 | 60 | 168.59 | 1 | 2.81 | 1.67 | 21.58 | 22.83 | -1.25 | 1.25 | -0.02 | 0.02 |
| gene-LOC105276078 | https://www.ncbi.nlm.nih.gov/gene/?term=LOC105276078 | 1051 | 10 | 95.15 | 1 | 9.51 | 10 | 37.29 | 38.89 | -1.6 | 1.6 | -0.02 | 0.02 |
| gene-LOC105274900 | https://www.ncbi.nlm.nih.gov/gene/?term=LOC105274900 | 5561 | 56 | 100.7 | 2 | 3.6 | 3.57 | 26.85 | 28.69 | -1.84 | 1.84 | -0.02 | 0.02 |
| gene-LOC105278260 | https://www.ncbi.nlm.nih.gov/gene/?term=LOC105278260 | 3240 | 41 | 126.54 | 1 | 3.09 | 2.44 | 13.01 | 14.86 | -1.85 | 1.85 | -0.02 | 0.02 |
| gene-LOC105286452 | https://www.ncbi.nlm.nih.gov/gene/?term=LOC105286452 | 6888 | 58 | 84.2 | 1 | 1.45 | 1.72 | 31.25 | 33.23 | -1.98 | 1.98 | -0.02 | 0.02 |
| gene-LOC105275518 | https://www.ncbi.nlm.nih.gov/gene/?term=LOC105275518 | 2335 | 21 | 89.94 | 1 | 4.28 | 4.76 | 45.03 | 47.49 | -2.45 | 2.45 | -0.02 | 0.02 |
| gene-LOC105286983 | https://www.ncbi.nlm.nih.gov/gene/?term=LOC105286983 | 16901 | 68 | 40.23 | 1 | 0.59 | 1.47 | 0.94 | 5.77 | -4.83 | 4.83 | -0.02 | 0.02 |
| gene-LOC105285084 | https://www.ncbi.nlm.nih.gov/gene/?term=LOC105285084 | 3308 | 26 | 78.6 | 1 | 3.02 | 3.85 | 22.5 | 21.29 | 1.21 | 1.21 | 0.01 | 0.01 |
| gene-LOC105285252 | https://www.ncbi.nlm.nih.gov/gene/?term=LOC105285252 | 7427 | 132 | 177.73 | 1 | 1.35 | 0.76 | 13.81 | 12.99 | 0.81 | 0.81 | 0.01 | 0.01 |
| gene-LOC105279664 | https://www.ncbi.nlm.nih.gov/gene/?term=LOC105279664 | 4023 | 72 | 178.97 | 1 | 2.49 | 1.39 | 17.48 | 16.74 | 0.73 | 0.73 | 0.01 | 0.01 |
| gene-LOC105280819 | https://www.ncbi.nlm.nih.gov/gene/?term=LOC105280819 | 1639 | 41 | 250.15 | 1 | 6.1 | 2.44 | 40.58 | 40.03 | 0.56 | 0.56 | 0.01 | 0.01 |
| gene-LOC105282794 | https://www.ncbi.nlm.nih.gov/gene/?term=LOC105282794 | 4571 | 98 | 214.4 | 1 | 2.19 | 1.02 | 18.44 | 17.88 | 0.56 | 0.56 | 0.01 | 0.01 |
| gene-LOC105283649 | https://www.ncbi.nlm.nih.gov/gene/?term=LOC105283649 | 1918 | 53 | 276.33 | 2 | 10.43 | 3.77 | 38.29 | 38.03 | 0.26 | 0.26 | 0.01 | 0.01 |
| gene-LOC105286265 | https://www.ncbi.nlm.nih.gov/gene/?term=LOC105286265 | 2507 | 68 | 271.24 | 1 | 3.99 | 1.47 | 30.18 | 29.94 | 0.24 | 0.24 | 0.01 | 0.01 |
| gene-LOC105281834 | https://www.ncbi.nlm.nih.gov/gene/?term=LOC105281834 | 4711 | 204 | 433.03 | 1 | 2.12 | 0.49 | 10.43 | 10.29 | 0.14 | 0.14 | 0.01 | 0.01 |
| gene-LOC105286649 | https://www.ncbi.nlm.nih.gov/gene/?term=LOC105286649 | 14178 | 701 | 494.43 | 1 | 0.71 | 0.14 | 1.28 | 1.15 | 0.13 | 0.13 | 0.01 | 0.01 |
| gene-LOC105278145 | https://www.ncbi.nlm.nih.gov/gene/?term=LOC105278145 | 4470 | 256 | 572.71 | 1 | 2.24 | 0.39 | 10.38 | 10.47 | -0.09 | 0.09 | -0.01 | 0.01 |
| gene-LOC105284557 | https://www.ncbi.nlm.nih.gov/gene/?term=LOC105284557 | 1672 | 77 | 460.53 | 1 | 5.98 | 1.3 | 30.69 | 30.8 | -0.11 | 0.11 | -0.01 | 0.01 |
| gene-LOC105281272 | https://www.ncbi.nlm.nih.gov/gene/?term=LOC105281272 | 5252 | 179 | 340.82 | 1 | 1.9 | 0.56 | 16.36 | 16.51 | -0.15 | 0.15 | -0.01 | 0.01 |
| gene-LOC105276612 | https://www.ncbi.nlm.nih.gov/gene/?term=LOC105276612 | 4157 | 178 | 428.19 | 1 | 2.41 | 0.56 | 10.69 | 10.89 | -0.2 | 0.2 | -0.01 | 0.01 |
| gene-LOC105284036 | https://www.ncbi.nlm.nih.gov/gene/?term=LOC105284036 | 3254 | 83 | 255.07 | 1 | 3.07 | 1.2 | 22.52 | 22.73 | -0.2 | 0.2 | -0.01 | 0.01 |
| gene-LOC105285807 | https://www.ncbi.nlm.nih.gov/gene/?term=LOC105285807 | 6063 | 186 | 306.78 | 1 | 1.65 | 0.54 | 10.05 | 10.27 | -0.22 | 0.22 | -0.01 | 0.01 |
| gene-LOC105275327 | https://www.ncbi.nlm.nih.gov/gene/?term=LOC105275327 | 14245 | 479 | 336.26 | 1 | 0.7 | 0.21 | 7.11 | 7.34 | -0.23 | 0.23 | -0.01 | 0.01 |
| gene-LOC113562050 | https://www.ncbi.nlm.nih.gov/gene/?term=LOC113562050 | 17097 | 576 | 336.9 | 1 | 0.58 | 0.17 | 5.26 | 5.5 | -0.25 | 0.25 | -0.01 | 0.01 |
| gene-LOC105285202 | https://www.ncbi.nlm.nih.gov/gene/?term=LOC105285202 | 1838 | 36 | 195.87 | 1 | 5.44 | 2.78 | 18.94 | 19.21 | -0.27 | 0.27 | -0.01 | 0.01 |
| gene-LOC105274851 | https://www.ncbi.nlm.nih.gov/gene/?term=LOC105274851 | 6616 | 142 | 214.63 | 1 | 1.51 | 0.7 | 9.54 | 9.83 | -0.3 | 0.3 | -0.01 | 0.01 |
| gene-LOC105280104 | https://www.ncbi.nlm.nih.gov/gene/?term=LOC105280104 | 3960 | 120 | 303.03 | 1 | 2.53 | 0.83 | 14.59 | 14.9 | -0.31 | 0.31 | -0.01 | 0.01 |
| gene-LOC105288078 | https://www.ncbi.nlm.nih.gov/gene/?term=LOC105288078 | 19126 | 318 | 166.27 | 1 | 0.52 | 0.31 | 17.33 | 17.64 | -0.31 | 0.31 | -0.01 | 0.01 |
| gene-LOC105274531 | https://www.ncbi.nlm.nih.gov/gene/?term=LOC105274531 | 7774 | 257 | 330.59 | 1 | 1.29 | 0.39 | 11.81 | 12.14 | -0.32 | 0.32 | -0.01 | 0.01 |
| gene-LOC105278534 | https://www.ncbi.nlm.nih.gov/gene/?term=LOC105278534 | 7364 | 185 | 251.22 | 1 | 1.36 | 0.54 | 17.95 | 18.29 | -0.33 | 0.33 | -0.01 | 0.01 |
| gene-LOC105278384 | https://www.ncbi.nlm.nih.gov/gene/?term=LOC105278384 | 1390 | 33 | 237.41 | 1 | 7.19 | 3.03 | 45.97 | 46.32 | -0.35 | 0.35 | -0.01 | 0.01 |
| gene-LOC105279038 | https://www.ncbi.nlm.nih.gov/gene/?term=LOC105279038 | 9453 | 321 | 339.57 | 1 | 1.06 | 0.31 | 8.59 | 8.94 | -0.35 | 0.35 | -0.01 | 0.01 |
| gene-LOC105278228 | https://www.ncbi.nlm.nih.gov/gene/?term=LOC105278228 | 8610 | 163 | 189.31 | 1 | 1.16 | 0.61 | 23.66 | 24.04 | -0.38 | 0.38 | -0.01 | 0.01 |

| | | | | | | | | | | | | | |
|---|---|---|---|---|---|---|---|---|---|---|---|---|---|
| gene-LOC105280956 | https://www.ncbi.nlm.nih.gov/gene/?term=LOC105280956 | 2639 | 73 | 276.62 | 1 | 3.79 | 1.37 | 20.28 | 20.69 | -0.4 | 0.4 | -0.01 | 0.01 |
| gene-LOC105284184 | https://www.ncbi.nlm.nih.gov/gene/?term=LOC105284184 | 2535 | 82 | 323.47 | 1 | 3.94 | 1.22 | 34.56 | 34.95 | -0.4 | 0.4 | -0.01 | 0.01 |
| gene-LOC105275451 | https://www.ncbi.nlm.nih.gov/gene/?term=LOC105275451 | 6035 | 118 | 195.53 | 1 | 1.66 | 0.85 | 22.19 | 22.63 | -0.44 | 0.44 | -0.01 | 0.01 |
| gene-LOC105288237 | https://www.ncbi.nlm.nih.gov/gene/?term=LOC105288237 | 2226 | 47 | 211.14 | 1 | 4.49 | 2.13 | 35.68 | 36.15 | -0.47 | 0.47 | -0.01 | 0.01 |
| gene-LOC105282825 | https://www.ncbi.nlm.nih.gov/gene/?term=LOC105282825 | 3919 | 90 | 229.65 | 1 | 2.55 | 1.11 | 15.34 | 15.86 | -0.52 | 0.52 | -0.01 | 0.01 |
| gene-LOC105288079 | https://www.ncbi.nlm.nih.gov/gene/?term=LOC105288079 | 3473 | 89 | 256.26 | 1 | 2.88 | 1.12 | 19.96 | 20.49 | -0.52 | 0.52 | -0.01 | 0.01 |
| gene-LOC105280410 | https://www.ncbi.nlm.nih.gov/gene/?term=LOC105280410 | 12580 | 260 | 206.68 | 1 | 0.79 | 0.38 | 19.03 | 19.62 | -0.59 | 0.59 | -0.01 | 0.01 |
| gene-LOC105275966 | https://www.ncbi.nlm.nih.gov/gene/?term=LOC105275966 | 3299 | 59 | 178.84 | 1 | 3.03 | 1.69 | 50.28 | 50.9 | -0.62 | 0.62 | -0.01 | 0.01 |
| gene-LOC105280641 | https://www.ncbi.nlm.nih.gov/gene/?term=LOC105280641 | 3682 | 66 | 179.25 | 1 | 2.72 | 1.52 | 15.43 | 16.12 | -0.68 | 0.68 | -0.01 | 0.01 |
| gene-LOC105286770 | https://www.ncbi.nlm.nih.gov/gene/?term=LOC105286770 | 3055 | 38 | 124.39 | 1 | 3.27 | 2.63 | 11.1 | 11.78 | -0.68 | 0.68 | -0.01 | 0.01 |
| gene-LOC105282600 | https://www.ncbi.nlm.nih.gov/gene/?term=LOC105282600 | 3715 | 65 | 174.97 | 1 | 2.69 | 1.54 | 26.29 | 27.03 | -0.73 | 0.73 | -0.01 | 0.01 |
| gene-LOC105276917 | https://www.ncbi.nlm.nih.gov/gene/?term=LOC105276917 | 2336 | 26 | 111.3 | 1 | 4.28 | 3.85 | 36.57 | 37.3 | -0.74 | 0.74 | -0.01 | 0.01 |
| gene-LOC105282067 | https://www.ncbi.nlm.nih.gov/gene/?term=LOC105282067 | 5941 | 54 | 90.89 | 1 | 1.68 | 1.85 | 23.11 | 23.91 | -0.8 | 0.8 | -0.01 | 0.01 |
| gene-LOC105274773 | https://www.ncbi.nlm.nih.gov/gene/?term=LOC105274773 | 11034 | 167 | 151.35 | 1 | 0.91 | 0.6 | 16.37 | 17.25 | -0.88 | 0.88 | -0.01 | 0.01 |
| gene-LOC105279657 | https://www.ncbi.nlm.nih.gov/gene/?term=LOC105279657 | 5878 | 84 | 142.91 | 1 | 1.7 | 1.19 | 18.69 | 19.56 | -0.88 | 0.88 | -0.01 | 0.01 |
| gene-LOC105277216 | https://www.ncbi.nlm.nih.gov/gene/?term=LOC105277216 | 7627 | 89 | 116.69 | 1 | 1.31 | 1.12 | 6.69 | 7.62 | -0.93 | 0.93 | -0.01 | 0.01 |
| gene-LOC105280447 | https://www.ncbi.nlm.nih.gov/gene/?term=LOC105280447 | 2215 | 28 | 126.41 | 1 | 4.51 | 3.57 | 23.87 | 24.81 | -0.94 | 0.94 | -0.01 | 0.01 |
| gene-LOC105285406 | https://www.ncbi.nlm.nih.gov/gene/?term=LOC105285406 | 8015 | 42 | 52.4 | 1 | 1.25 | 2.38 | 35.2 | 36.16 | -0.96 | 0.96 | -0.01 | 0.01 |
| gene-LOC105279500 | https://www.ncbi.nlm.nih.gov/gene/?term=LOC105279500 | 3587 | 33 | 92 | 1 | 2.79 | 3.03 | 49.6 | 51 | -1.4 | 1.4 | -0.01 | 0.01 |
| gene-LOC105275347 | https://www.ncbi.nlm.nih.gov/gene/?term=LOC105275347 | 2526 | 14 | 55.42 | 1 | 3.96 | 7.14 | 35.76 | 37.28 | -1.52 | 1.52 | -0.01 | 0.01 |
| gene-LOC105283989 | https://www.ncbi.nlm.nih.gov/gene/?term=LOC105283989 | 4948 | 34 | 68.71 | 2 | 4.04 | 5.88 | 44.42 | 46.21 | -1.79 | 1.79 | -0.01 | 0.01 |
| gene-LOC105281515 | https://www.ncbi.nlm.nih.gov/gene/?term=LOC105281515 | 8827 | 41 | 46.45 | 1 | 1.13 | 2.44 | 32.87 | 35.23 | -2.37 | 2.37 | -0.01 | 0.01 |
| gene-LOC105281420 | https://www.ncbi.nlm.nih.gov/gene/?term=LOC105281420 | 3882 | 57 | 146.83 | 1 | 2.58 | 1.75 | 15.28 | 15.02 | 0.26 | 0.26 | 0 | 0 |
| gene-LOC105278454 | https://www.ncbi.nlm.nih.gov/gene/?term=LOC105278454 | 1972 | 37 | 187.63 | 1 | 5.07 | 2.7 | 46.8 | 46.67 | 0.13 | 0.13 | 0 | 0 |
| gene-LOC105278455 | https://www.ncbi.nlm.nih.gov/gene/?term=LOC105278455 | 27181 | 875 | 321.92 | 1 | 0.37 | 0.11 | 7.33 | 7.2 | 0.13 | 0.13 | 0 | 0 |
| gene-LOC105276196 | https://www.ncbi.nlm.nih.gov/gene/?term=LOC105276196 | 2953 | 56 | 189.64 | 1 | 3.39 | 1.79 | 40.11 | 39.99 | 0.12 | 0.12 | 0 | 0 |
| gene-LOC105277273 | https://www.ncbi.nlm.nih.gov/gene/?term=LOC105277273 | 3711 | 151 | 406.9 | 1 | 2.69 | 0.66 | 12.76 | 12.67 | 0.09 | 0.09 | 0 | 0 |
| gene-LOC105278632 | https://www.ncbi.nlm.nih.gov/gene/?term=LOC105278632 | 3624 | 132 | 364.24 | 1 | 2.76 | 0.76 | 14.6 | 14.5 | 0.09 | 0.09 | 0 | 0 |
| gene-LOC105284116 | https://www.ncbi.nlm.nih.gov/gene/?term=LOC105284116 | 3970 | 96 | 241.81 | 1 | 2.52 | 1.04 | 6.8 | 6.72 | 0.08 | 0.08 | 0 | 0 |
| gene-LOC105278666 | https://www.ncbi.nlm.nih.gov/gene/?term=LOC105278666 | 10341 | 409 | 395.51 | 1 | 0.97 | 0.24 | 7.49 | 7.42 | 0.07 | 0.07 | 0 | 0 |
| gene-LOC105279528 | https://www.ncbi.nlm.nih.gov/gene/?term=LOC105279528 | 5802 | 46 | 79.28 | 1 | 1.72 | 2.17 | 15.76 | 15.72 | 0.04 | 0.04 | 0 | 0 |
| gene-LOC105280304 | https://www.ncbi.nlm.nih.gov/gene/?term=LOC105280304 | 8919 | 123 | 137.91 | 1 | 1.12 | 0.81 | 5.79 | 5.75 | 0.04 | 0.04 | 0 | 0 |
| gene-LOC105282914 | https://www.ncbi.nlm.nih.gov/gene/?term=LOC105282914 | 5623 | 42 | 74.69 | 1 | 1.78 | 2.38 | 48.6 | 48.57 | 0.03 | 0.03 | 0 | 0 |
| gene-LOC105274699 | https://www.ncbi.nlm.nih.gov/gene/?term=LOC105274699 | 2386 | 63 | 264.04 | 1 | 4.19 | 1.59 | 23.04 | 23.01 | 0.02 | 0.02 | 0 | 0 |
| gene-LOC105283296 | https://www.ncbi.nlm.nih.gov/gene/?term=LOC105283296 | 2174 | 89 | 409.38 | 1 | 4.6 | 1.12 | 19.56 | 19.55 | 0.01 | 0.01 | 0 | 0 |
| gene-LOC105277866 | https://www.ncbi.nlm.nih.gov/gene/?term=LOC105277866 | 33045 | 447 | 135.27 | 1 | 0.3 | 0.22 | 3.08 | 3.1 | -0.02 | 0.02 | 0 | 0 |
| gene-LOC105280265 | https://www.ncbi.nlm.nih.gov/gene/?term=LOC105280265 | 85984 | 2872 | 334.02 | 1 | 0.12 | 0.03 | 1.07 | 1.1 | -0.02 | 0.02 | 0 | 0 |
| gene-LOC105277125 | https://www.ncbi.nlm.nih.gov/gene/?term=LOC105277125 | 51079 | 1743 | 341.24 | 1 | 0.2 | 0.06 | 1.56 | 1.61 | -0.05 | 0.05 | 0 | 0 |
| gene-LOC105276059 | https://www.ncbi.nlm.nih.gov/gene/?term=LOC105276059 | 6256 | 136 | 217.39 | 1 | 1.6 | 0.74 | 10.77 | 10.82 | -0.06 | 0.06 | 0 | 0 |
| gene-LOC105286332 | https://www.ncbi.nlm.nih.gov/gene/?term=LOC105286332 | 27577 | 507 | 183.85 | 1 | 0.36 | 0.2 | 7.74 | 7.81 | -0.07 | 0.07 | 0 | 0 |
| gene-LOC105278538 | https://www.ncbi.nlm.nih.gov/gene/?term=LOC105278538 | 2150 | 83 | 386.05 | 1 | 4.65 | 1.2 | 18.97 | 19.05 | -0.08 | 0.08 | 0 | 0 |
| gene-LOC105284117 | https://www.ncbi.nlm.nih.gov/gene/?term=LOC105284117 | 4340 | 86 | 198.16 | 1 | 2.3 | 1.16 | 10.49 | 10.57 | -0.08 | 0.08 | 0 | 0 |
| gene-LOC105277761 | https://www.ncbi.nlm.nih.gov/gene/?term=LOC105277761 | 4648 | 150 | 322.72 | 1 | 2.15 | 0.67 | 14.73 | 14.87 | -0.13 | 0.13 | 0 | 0 |
| gene-LOC105284262 | https://www.ncbi.nlm.nih.gov/gene/?term=LOC105284262 | 6669 | 236 | 353.88 | 1 | 1.5 | 0.42 | 4.46 | 4.59 | -0.13 | 0.13 | 0 | 0 |
| gene-LOC105285395 | https://www.ncbi.nlm.nih.gov/gene/?term=LOC105285395 | 8085 | 187 | 231.29 | 1 | 1.24 | 0.53 | 16.28 | 16.44 | -0.16 | 0.16 | 0 | 0 |
| gene-LOC105275187 | https://www.ncbi.nlm.nih.gov/gene/?term=LOC105275187 | 5534 | 75 | 135.53 | 1 | 1.81 | 1.33 | 14.15 | 14.34 | -0.18 | 0.18 | 0 | 0 |
| gene-LOC105286194 | https://www.ncbi.nlm.nih.gov/gene/?term=LOC105286194 | 152619 | 2093 | 137.14 | 1 | 0.07 | 0.05 | 6.53 | 6.74 | -0.21 | 0.21 | 0 | 0 |
| gene-LOC105288161 | https://www.ncbi.nlm.nih.gov/gene/?term=LOC105288161 | 26065 | 302 | 115.86 | 1 | 0.38 | 0.33 | 11.58 | 11.82 | -0.24 | 0.24 | 0 | 0 |

Directory: OGLall File: rnk-dmg-Ob_LI_BCphaseA1.vs.RphaseB1.txt

| gene_ID | gene_link | gwidth | #Sites | #per10Kb | #dmSites | #dmsp10kb | %dmSites | %pSite1 | %pSite2 | DMpSite | ADMpSite | DMpNucl | ADMpNucl |
|---|---|---|---|---|---|---|---|---|---|---|---|---|---|
| gene-LOC113562627 | https://www.ncbi.nlm.nih.gov/gene/?term=LOC113562627 | 397 | 10 | 251.89 | 1 | 25.19 | 10 | 77.44 | 70.89 | 6.56 | 6.56 | 0.17 | 0.17 |
| gene-LOC105282405 | https://www.ncbi.nlm.nih.gov/gene/?term=LOC105282405 | 1310 | 31 | 236.64 | 2 | 15.27 | 6.45 | 51.18 | 44.54 | 6.64 | 6.64 | 0.16 | 0.16 |
| gene-LOC113561900 | https://www.ncbi.nlm.nih.gov/gene/?term=LOC113561900 | 629 | 10 | 158.98 | 2 | 31.8 | 20 | 21.61 | 31.23 | -9.62 | 9.62 | -0.15 | 0.15 |
| gene-LOC105288193 | https://www.ncbi.nlm.nih.gov/gene/?term=LOC105288193 | 1495 | 63 | 421.4 | 4 | 26.76 | 6.35 | 33.59 | 30.52 | 3.07 | 3.07 | 0.13 | 0.13 |
| gene-LOC105279335 | https://www.ncbi.nlm.nih.gov/gene/?term=LOC105279335 | 737 | 34 | 461.33 | 2 | 27.14 | 5.88 | 18.91 | 16.04 | 2.87 | 2.87 | 0.13 | 0.13 |
| gene-LOC113562894 | https://www.ncbi.nlm.nih.gov/gene/?term=LOC113562894 | 781 | 34 | 435.34 | 1 | 12.8 | 2.94 | 12.48 | 15.26 | -2.78 | 2.78 | -0.12 | 0.12 |
| gene-LOC105283861 | https://www.ncbi.nlm.nih.gov/gene/?term=LOC105283861 | 1298 | 19 | 146.38 | 3 | 23.11 | 15.79 | 35.76 | 43.84 | -8.08 | 8.08 | -0.12 | 0.12 |
| gene-LOC105278160 | https://www.ncbi.nlm.nih.gov/gene/?term=LOC105278160 | 1872 | 85 | 454.06 | 2 | 10.68 | 2.35 | 38.22 | 35.85 | 2.37 | 2.37 | 0.11 | 0.11 |
| gene-LOC105285213 | https://www.ncbi.nlm.nih.gov/gene/?term=LOC105285213 | 1491 | 37 | 248.16 | 4 | 26.83 | 10.81 | 13.74 | 18.03 | -4.29 | 4.29 | -0.11 | 0.11 |
| gene-LOC113561934 | https://www.ncbi.nlm.nih.gov/gene/?term=LOC113561934 | 1252 | 16 | 127.8 | 4 | 31.95 | 25 | 36.29 | 28.08 | 8.21 | 8.21 | 0.1 | 0.1 |
| gene-LOC105283973 | https://www.ncbi.nlm.nih.gov/gene/?term=LOC105283973 | 1536 | 29 | 188.8 | 4 | 26.04 | 13.79 | 61.88 | 56.34 | 5.55 | 5.55 | 0.1 | 0.1 |
| gene-LOC105275539 | https://www.ncbi.nlm.nih.gov/gene/?term=LOC105275539 | 1272 | 23 | 180.82 | 2 | 15.72 | 8.7 | 20.8 | 15.48 | 5.32 | 5.32 | 0.1 | 0.1 |
| gene-LOC105275761 | https://www.ncbi.nlm.nih.gov/gene/?term=LOC105275761 | 2635 | 101 | 383.3 | 6 | 22.77 | 5.94 | 29.65 | 26.96 | 2.69 | 2.69 | 0.1 | 0.1 |
| gene-LOC105274555 | https://www.ncbi.nlm.nih.gov/gene/?term=LOC105274555 | 1395 | 56 | 401.43 | 2 | 14.34 | 3.57 | 36.48 | 34.09 | 2.39 | 2.39 | 0.1 | 0.1 |
| gene-LOC105287692 | https://www.ncbi.nlm.nih.gov/gene/?term=LOC105287692 | 1505 | 99 | 657.81 | 3 | 19.93 | 3.03 | 29.69 | 28.16 | 1.52 | 1.52 | 0.1 | 0.1 |
| gene-LOC109610897 | https://www.ncbi.nlm.nih.gov/gene/?term=LOC109610897 | 1561 | 110 | 704.68 | 3 | 19.22 | 2.73 | 15.43 | 14.07 | 1.37 | 1.37 | 0.1 | 0.1 |
| gene-LOC105280710 | https://www.ncbi.nlm.nih.gov/gene/?term=LOC105280710 | 1215 | 69 | 567.9 | 5 | 41.15 | 7.25 | 20.73 | 22.43 | -1.7 | 1.7 | -0.1 | 0.1 |
| gene-LOC105286235 | https://www.ncbi.nlm.nih.gov/gene/?term=LOC105286235 | 1510 | 76 | 503.31 | 3 | 19.87 | 3.95 | 37.9 | 39.89 | -1.99 | 1.99 | -0.1 | 0.1 |
| gene-LOC105287465 | https://www.ncbi.nlm.nih.gov/gene/?term=LOC105287465 | 2477 | 92 | 371.42 | 7 | 28.26 | 7.61 | 40.81 | 43.43 | -2.62 | 2.62 | -0.1 | 0.1 |
| gene-LOC105278537 | https://www.ncbi.nlm.nih.gov/gene/?term=LOC105278537 | 1660 | 60 | 361.45 | 5 | 30.12 | 8.33 | 29.09 | 31.82 | -2.73 | 2.73 | -0.1 | 0.1 |
| gene-LOC105279087 | https://www.ncbi.nlm.nih.gov/gene/?term=LOC105279087 | 1077 | 28 | 259.98 | 1 | 9.29 | 3.57 | 53 | 56.81 | -3.81 | 3.81 | -0.1 | 0.1 |
| gene-LOC105276115 | https://www.ncbi.nlm.nih.gov/gene/?term=LOC105276115 | 1520 | 38 | 250 | 3 | 19.74 | 7.89 | 23.97 | 28.04 | -4.07 | 4.07 | -0.1 | 0.1 |
| gene-LOC105286358 | https://www.ncbi.nlm.nih.gov/gene/?term=LOC105286358 | 1489 | 54 | 362.66 | 3 | 20.15 | 5.56 | 34.15 | 31.79 | 2.36 | 2.36 | 0.09 | 0.09 |
| gene-LOC105280310 | https://www.ncbi.nlm.nih.gov/gene/?term=LOC105280310 | 1916 | 75 | 391.44 | 1 | 5.22 | 1.33 | 34.06 | 31.75 | 2.31 | 2.31 | 0.09 | 0.09 |

...

... only first 25 lines shown ...

Directory: OGLall File: rnk-dmg-Ob_LI_BCphaseA1.vs.RphaseB2.txt

| gene_ID | gene_link | gwidth | #Sites | #per10Kb | #dmSites | #dmsp10kb | %dmSites | %pSite1 | %pSite2 | DMpSite | ADMpSite | DMpNucl | ADMpNucl |
|---|---|---|---|---|---|---|---|---|---|---|---|---|---|
| gene-LOC105282405 | https://www.ncbi.nlm.nih.gov/gene/?term=LOC105282405 | 1310 | 31 | 236.64 | 5 | 38.17 | 16.13 | 51.18 | 44.88 | 6.3 | 6.3 | 0.15 | 0.15 |
| gene-LOC105276585 | https://www.ncbi.nlm.nih.gov/gene/?term=LOC105276585 | 1534 | 113 | 736.64 | 1 | 6.52 | 0.88 | 7.26 | 5.41 | 1.85 | 1.85 | 0.14 | 0.14 |
| gene-LOC105285203 | https://www.ncbi.nlm.nih.gov/gene/?term=LOC105285203 | 1020 | 27 | 264.71 | 1 | 9.8 | 3.7 | 30.19 | 25.46 | 4.73 | 4.73 | 0.13 | 0.13 |
| gene-LOC105278849 | https://www.ncbi.nlm.nih.gov/gene/?term=LOC105278849 | 776 | 74 | 953.61 | 2 | 25.77 | 2.7 | 24.36 | 22.99 | 1.37 | 1.37 | 0.13 | 0.13 |
| gene-LOC105278266 | https://www.ncbi.nlm.nih.gov/gene/?term=LOC105278266 | 2442 | 108 | 442.26 | 4 | 16.38 | 3.7 | 25.68 | 28.72 | -3.05 | 3.05 | -0.13 | 0.13 |
| gene-LOC113562894 | https://www.ncbi.nlm.nih.gov/gene/?term=LOC113562894 | 781 | 34 | 435.34 | 2 | 25.61 | 5.88 | 11.76 | 14.52 | -2.76 | 2.76 | -0.12 | 0.12 |
| gene-LOC105278458 | https://www.ncbi.nlm.nih.gov/gene/?term=LOC105278458 | 216 | 3 | 138.89 | 1 | 46.3 | 33.33 | 20.87 | 29.54 | -8.67 | 8.67 | -0.12 | 0.12 |
| gene-LOC113561934 | https://www.ncbi.nlm.nih.gov/gene/?term=LOC113561934 | 1252 | 15 | 119.81 | 2 | 15.97 | 13.33 | 36.71 | 27.17 | 9.54 | 9.54 | 0.11 | 0.11 |
| gene-LOC105278234 | https://www.ncbi.nlm.nih.gov/gene/?term=LOC105278234 | 2068 | 106 | 512.57 | 1 | 4.84 | 0.94 | 10.74 | 8.52 | 2.21 | 2.21 | 0.11 | 0.11 |
| gene-LOC105278873 | https://www.ncbi.nlm.nih.gov/gene/?term=LOC105278873 | 1897 | 143 | 753.82 | 4 | 21.09 | 2.8 | 19.98 | 21.44 | -1.47 | 1.47 | -0.11 | 0.11 |
| gene-LOC105280710 | https://www.ncbi.nlm.nih.gov/gene/?term=LOC105280710 | 1215 | 64 | 526.75 | 1 | 8.23 | 1.56 | 22.3 | 24.34 | -2.04 | 2.04 | -0.11 | 0.11 |
| gene-LOC105287856 | https://www.ncbi.nlm.nih.gov/gene/?term=LOC105287856 | 2090 | 113 | 540.67 | 6 | 28.71 | 5.31 | 12.28 | 14.37 | -2.09 | 2.09 | -0.11 | 0.11 |
| gene-LOC105278537 | https://www.ncbi.nlm.nih.gov/gene/?term=LOC105278537 | 1660 | 62 | 373.49 | 1 | 6.02 | 1.61 | 29.42 | 32.25 | -2.83 | 2.83 | -0.11 | 0.11 |
| gene-LOC105279863 | https://www.ncbi.nlm.nih.gov/gene/?term=LOC105279863 | 1204 | 18 | 149.5 | 4 | 33.22 | 22.22 | 11.07 | 18.73 | -7.66 | 7.66 | -0.11 | 0.11 |
| gene-LOC105278824 | https://www.ncbi.nlm.nih.gov/gene/?term=LOC105278824 | 1230 | 61 | 495.93 | 1 | 8.13 | 1.64 | 24.95 | 22.9 | 2.05 | 2.05 | 0.1 | 0.1 |
| gene-LOC105279346 | https://www.ncbi.nlm.nih.gov/gene/?term=LOC105279346 | 4649 | 229 | 492.58 | 9 | 19.36 | 3.93 | 14.52 | 16.49 | -1.97 | 1.97 | -0.1 | 0.1 |
| gene-LOC105277398 | https://www.ncbi.nlm.nih.gov/gene/?term=LOC105277398 | 1963 | 59 | 300.56 | 5 | 25.47 | 8.47 | 36.03 | 39.34 | -3.31 | 3.31 | -0.1 | 0.1 |
| gene-LOC105285207 | https://www.ncbi.nlm.nih.gov/gene/?term=LOC105285207 | 3030 | 64 | 211.22 | 5 | 16.5 | 7.81 | 20.3 | 25.24 | -4.94 | 4.94 | -0.1 | 0.1 |
| gene-LOC105277810 | https://www.ncbi.nlm.nih.gov/gene/?term=LOC105277810 | 1367 | 21 | 153.62 | 1 | 7.32 | 4.76 | 45.33 | 39.52 | 5.81 | 5.81 | 0.09 | 0.09 |
| gene-LOC105286644 | https://www.ncbi.nlm.nih.gov/gene/?term=LOC105286644 | 1235 | 21 | 170.04 | 1 | 8.1 | 4.76 | 44.21 | 39.04 | 5.17 | 5.17 | 0.09 | 0.09 |
| gene-LOC105276679 | https://www.ncbi.nlm.nih.gov/gene/?term=LOC105276679 | 2005 | 67 | 334.16 | 1 | 4.99 | 1.49 | 37.86 | 35.03 | 2.83 | 2.83 | 0.09 | 0.09 |
| gene-LOC105279487 | https://www.ncbi.nlm.nih.gov/gene/?term=LOC105279487 | 2117 | 82 | 387.34 | 2 | 9.45 | 2.44 | 17.82 | 15.61 | 2.21 | 2.21 | 0.09 | 0.09 |
| gene-LOC105288104 | https://www.ncbi.nlm.nih.gov/gene/?term=LOC105288104 | 2069 | 92 | 444.66 | 2 | 9.67 | 2.17 | 27.34 | 25.28 | 2.06 | 2.06 | 0.09 | 0.09 |
| gene-LOC105280786 | https://www.ncbi.nlm.nih.gov/gene/?term=LOC105280786 | 2573 | 115 | 446.95 | 1 | 3.89 | 0.87 | 30.29 | 28.35 | 1.94 | 1.94 | 0.09 | 0.09 |

...

... only first 25 lines shown ...

Directory: OGLall File: rnk-dmg-Ob_LI_BCphaseA2.vs.BCphaseB1.txt

| gene_ID | gene_link | gwidth | #Sites | #per10Kb | #dmSites | #dmsp10kb | %dmSites | %pSite1 | %pSite2 | DMpSite | ADMpSite | DMpNucl | ADMpNucl |
|---|---|---|---|---|---|---|---|---|---|---|---|---|---|
| gene-LOC105275384 | https://www.ncbi.nlm.nih.gov/gene/?term=LOC105275384 | 1987 | 91 | 457.98 | 1 | 5.03 | 1.1 | 33.74 | 32.35 | 1.39 | 1.39 | 0.06 | 0.06 |
| gene-LOC105276429 | https://www.ncbi.nlm.nih.gov/gene/?term=LOC105276429 | 2901 | 120 | 413.65 | 1 | 3.45 | 0.83 | 23.61 | 22.3 | 1.32 | 1.32 | 0.05 | 0.05 |
| gene-LOC105283264 | https://www.ncbi.nlm.nih.gov/gene/?term=LOC105283264 | 4005 | 88 | 219.73 | 1 | 2.5 | 1.14 | 18.85 | 20.42 | -1.57 | 1.57 | -0.03 | 0.03 |
| gene-LOC105281272 | https://www.ncbi.nlm.nih.gov/gene/?term=LOC105281272 | 5252 | 178 | 338.92 | 1 | 1.9 | 0.56 | 16.03 | 16.73 | -0.7 | 0.7 | -0.02 | 0.02 |
| gene-LOC105288042 | https://www.ncbi.nlm.nih.gov/gene/?term=LOC105288042 | 3507 | 101 | 288 | 1 | 2.85 | 0.99 | 22.48 | 23.29 | -0.81 | 0.81 | -0.02 | 0.02 |
| gene-LOC105285252 | https://www.ncbi.nlm.nih.gov/gene/?term=LOC105285252 | 7427 | 129 | 173.69 | 1 | 1.35 | 0.78 | 12.88 | 13.9 | -1.02 | 1.02 | -0.02 | 0.02 |
| gene-LOC105277903 | https://www.ncbi.nlm.nih.gov/gene/?term=LOC105277903 | 1835 | 35 | 190.74 | 1 | 5.45 | 2.86 | 47.23 | 46.95 | 0.28 | 0.28 | 0.01 | 0.01 |
| gene-LOC105283296 | https://www.ncbi.nlm.nih.gov/gene/?term=LOC105283296 | 2174 | 92 | 423.18 | 1 | 4.6 | 1.09 | 19.15 | 18.97 | 0.18 | 0.18 | 0.01 | 0.01 |
| gene-LOC105276452 | https://www.ncbi.nlm.nih.gov/gene/?term=LOC105276452 | 2917 | 262 | 898.18 | 1 | 3.43 | 0.38 | 10.28 | 10.37 | -0.08 | 0.08 | -0.01 | 0.01 |
| gene-LOC105276385 | https://www.ncbi.nlm.nih.gov/gene/?term=LOC105276385 | 11575 | 494 | 426.78 | 1 | 0.86 | 0.2 | 5.31 | 5.56 | -0.25 | 0.25 | -0.01 | 0.01 |
| gene-LOC105284507 | https://www.ncbi.nlm.nih.gov/gene/?term=LOC105284507 | 5207 | 94 | 180.53 | 1 | 1.92 | 1.06 | 20.48 | 21.18 | -0.7 | 0.7 | -0.01 | 0.01 |
| gene-LOC105280410 | https://www.ncbi.nlm.nih.gov/gene/?term=LOC105280410 | 12580 | 280 | 222.58 | 1 | 0.79 | 0.36 | 19.42 | 19.27 | 0.15 | 0.15 | 0 | 0 |
| gene-LOC105277125 | https://www.ncbi.nlm.nih.gov/gene/?term=LOC105277125 | 51079 | 1879 | 367.86 | 1 | 0.2 | 0.05 | 1.58 | 1.64 | -0.06 | 0.06 | 0 | 0 |
| gene-LOC105288036 | https://www.ncbi.nlm.nih.gov/gene/?term=LOC105288036 | 5110 | 226 | 442.27 | 1 | 1.96 | 0.44 | 16.3 | 16.4 | -0.1 | 0.1 | 0 | 0 |

Directory: OGLall File: rnk-dmg-Ob_LI_BCphaseA2.vs.BCphaseB2.txt

| gene_ID | gene_link | gwidth | #Sites | #per10Kb | #dmSites | #dmsp10kb | %dmSites | %pSite1 | %pSite2 | DMpSite | ADMpSite | DMpNucl | ADMpNucl |
|---|---|---|---|---|---|---|---|---|---|---|---|---|---|
| gene-LOC113562345 | https://www.ncbi.nlm.nih.gov/gene/?term=LOC113562345 | 2290 | 69 | 301.31 | 17 | 74.24 | 24.64 | 33.16 | 16.9 | 16.26 | 16.26 | 0.49 | 0.49 |
| gene-LOC105284677 | https://www.ncbi.nlm.nih.gov/gene/?term=LOC105284677 | 1905 | 41 | 215.22 | 2 | 10.5 | 4.88 | 27.55 | 36.9 | -9.35 | 9.35 | -0.2 | 0.2 |
| gene-LOC105282976 | https://www.ncbi.nlm.nih.gov/gene/?term=LOC105282976 | 1648 | 50 | 303.4 | 1 | 6.07 | 2 | 26.51 | 20.9 | 5.61 | 5.61 | 0.17 | 0.17 |
| gene-LOC105282149 | https://www.ncbi.nlm.nih.gov/gene/?term=LOC105282149 | 1787 | 108 | 604.36 | 1 | 5.6 | 0.93 | 43.65 | 41.49 | 2.17 | 2.17 | 0.13 | 0.13 |
| gene-LOC105286683 | https://www.ncbi.nlm.nih.gov/gene/?term=LOC105286683 | 1665 | 51 | 306.31 | 1 | 6.01 | 1.96 | 35.98 | 32.15 | 3.83 | 3.83 | 0.12 | 0.12 |
| gene-LOC105282156 | https://www.ncbi.nlm.nih.gov/gene/?term=LOC105282156 | 2577 | 136 | 527.75 | 1 | 3.88 | 0.74 | 26.99 | 24.63 | 2.36 | 2.36 | 0.12 | 0.12 |
| gene-LOC105283649 | https://www.ncbi.nlm.nih.gov/gene/?term=LOC105283649 | 1918 | 56 | 291.97 | 2 | 10.43 | 3.57 | 38.44 | 34.8 | 3.63 | 3.63 | 0.11 | 0.11 |
| gene-LOC105276078 | https://www.ncbi.nlm.nih.gov/gene/?term=LOC105276078 | 1051 | 16 | 152.24 | 1 | 9.51 | 6.25 | 32.79 | 26.49 | 6.3 | 6.3 | 0.1 | 0.1 |
| gene-LOC105275447 | https://www.ncbi.nlm.nih.gov/gene/?term=LOC105275447 | 1846 | 35 | 189.6 | 1 | 5.42 | 2.86 | 8.76 | 3.6 | 5.15 | 5.15 | 0.1 | 0.1 |
| gene-LOC105285674 | https://www.ncbi.nlm.nih.gov/gene/?term=LOC105285674 | 3035 | 93 | 306.43 | 2 | 6.59 | 2.15 | 27.41 | 24.3 | 3.11 | 3.11 | 0.1 | 0.1 |
| gene-LOC105275070 | https://www.ncbi.nlm.nih.gov/gene/?term=LOC105275070 | 5667 | 47 | 82.94 | 5 | 8.82 | 10.64 | 11.7 | 2 | 9.7 | 9.7 | 0.08 | 0.08 |
| gene-LOC105276449 | https://www.ncbi.nlm.nih.gov/gene/?term=LOC105276449 | 1231 | 42 | 341.19 | 1 | 8.12 | 2.38 | 35.95 | 33.63 | 2.31 | 2.31 | 0.08 | 0.08 |
| gene-LOC105279898 | https://www.ncbi.nlm.nih.gov/gene/?term=LOC105279898 | 1343 | 14 | 104.24 | 1 | 7.45 | 7.14 | 61.13 | 54.89 | 6.24 | 6.24 | 0.07 | 0.07 |
| gene-LOC113561805 | https://www.ncbi.nlm.nih.gov/gene/?term=LOC113561805 | 3337 | 77 | 230.75 | 1 | 3 | 1.3 | 37.31 | 34.13 | 3.18 | 3.18 | 0.07 | 0.07 |
| gene-LOC105275708 | https://www.ncbi.nlm.nih.gov/gene/?term=LOC105275708 | 1772 | 10 | 56.43 | 1 | 5.64 | 10 | 50.31 | 39.56 | 10.75 | 10.75 | 0.06 | 0.06 |
| gene-LOC105288063 | https://www.ncbi.nlm.nih.gov/gene/?term=LOC105288063 | 3400 | 53 | 155.88 | 1 | 2.94 | 1.89 | 16.83 | 13.02 | 3.81 | 3.81 | 0.06 | 0.06 |
| gene-LOC105280774 | https://www.ncbi.nlm.nih.gov/gene/?term=LOC105280774 | 2793 | 102 | 365.2 | 1 | 3.58 | 0.98 | 32.11 | 30.36 | 1.75 | 1.75 | 0.06 | 0.06 |
| gene-LOC105280786 | https://www.ncbi.nlm.nih.gov/gene/?term=LOC105280786 | 2573 | 111 | 431.4 | 1 | 3.89 | 0.9 | 33.28 | 31.87 | 1.41 | 1.41 | 0.06 | 0.06 |
| gene-LOC105284502 | https://www.ncbi.nlm.nih.gov/gene/?term=LOC105284502 | 2157 | 19 | 88.09 | 1 | 4.64 | 5.26 | 36.83 | 43.93 | -7.1 | 7.1 | -0.06 | 0.06 |
| gene-LOC105278454 | https://www.ncbi.nlm.nih.gov/gene/?term=LOC105278454 | 1972 | 47 | 238.34 | 1 | 5.07 | 2.13 | 44.8 | 42.52 | 2.28 | 2.28 | 0.05 | 0.05 |
| gene-LOC105282970 | https://www.ncbi.nlm.nih.gov/gene/?term=LOC105282970 | 1529 | 39 | 255.07 | 1 | 6.54 | 2.56 | 54 | 52.14 | 1.86 | 1.86 | 0.05 | 0.05 |
| gene-LOC105282151 | https://www.ncbi.nlm.nih.gov/gene/?term=LOC105282151 | 1601 | 41 | 256.09 | 3 | 18.74 | 7.32 | 36.24 | 34.41 | 1.83 | 1.83 | 0.05 | 0.05 |
| gene-LOC105281848 | https://www.ncbi.nlm.nih.gov/gene/?term=LOC105281848 | 2122 | 56 | 263.9 | 1 | 4.71 | 1.79 | 24.08 | 22.28 | 1.8 | 1.8 | 0.05 | 0.05 |
| gene-LOC105287095 | https://www.ncbi.nlm.nih.gov/gene/?term=LOC105287095 | 1435 | 57 | 397.21 | 1 | 6.97 | 1.75 | 29.87 | 28.54 | 1.33 | 1.33 | 0.05 | 0.05 |
| gene-LOC105286190 | https://www.ncbi.nlm.nih.gov/gene/?term=LOC105286190 | 6447 | 298 | 462.23 | 1 | 1.55 | 0.34 | 8.47 | 7.3 | 1.16 | 1.16 | 0.05 | 0.05 |
| gene-LOC105279298 | https://www.ncbi.nlm.nih.gov/gene/?term=LOC105279298 | 2773 | 128 | 461.59 | 2 | 7.21 | 1.56 | 25.02 | 23.92 | 1.1 | 1.1 | 0.05 | 0.05 |
| gene-LOC105278497 | https://www.ncbi.nlm.nih.gov/gene/?term=LOC105278497 | 2072 | 94 | 453.67 | 1 | 4.83 | 1.06 | 21.1 | 20.09 | 1.01 | 1.01 | 0.05 | 0.05 |
| gene-LOC105276704 | https://www.ncbi.nlm.nih.gov/gene/?term=LOC105276704 | 2063 | 98 | 475.04 | 1 | 4.85 | 1.02 | 13.8 | 12.82 | 0.98 | 0.98 | 0.05 | 0.05 |
| gene-LOC105285055 | https://www.ncbi.nlm.nih.gov/gene/?term=LOC105285055 | 4108 | 259 | 630.48 | 2 | 4.87 | 0.77 | 11.6 | 10.8 | 0.8 | 0.8 | 0.05 | 0.05 |
| gene-LOC105274449 | https://www.ncbi.nlm.nih.gov/gene/?term=LOC105274449 | 1549 | 53 | 342.16 | 1 | 6.46 | 1.89 | 50.38 | 51.77 | -1.39 | 1.39 | -0.05 | 0.05 |
| gene-LOC105274900 | https://www.ncbi.nlm.nih.gov/gene/?term=LOC105274900 | 5561 | 52 | 93.51 | 2 | 3.6 | 3.85 | 36.59 | 32.11 | 4.48 | 4.48 | 0.04 | 0.04 |
| gene-LOC105281420 | https://www.ncbi.nlm.nih.gov/gene/?term=LOC105281420 | 3882 | 55 | 141.68 | 1 | 2.58 | 1.82 | 15.69 | 12.96 | 2.73 | 2.73 | 0.04 | 0.04 |
| gene-LOC105280641 | https://www.ncbi.nlm.nih.gov/gene/?term=LOC105280641 | 3682 | 49 | 133.08 | 1 | 2.72 | 2.04 | 20.13 | 17.42 | 2.71 | 2.71 | 0.04 | 0.04 |
| gene-LOC105275966 | https://www.ncbi.nlm.nih.gov/gene/?term=LOC105275966 | 3299 | 61 | 184.9 | 1 | 3.03 | 1.64 | 49.58 | 47.22 | 2.36 | 2.36 | 0.04 | 0.04 |
| gene-LOC105278109 | https://www.ncbi.nlm.nih.gov/gene/?term=LOC105278109 | 4029 | 65 | 161.33 | 1 | 2.48 | 1.54 | 48.15 | 45.97 | 2.19 | 2.19 | 0.04 | 0.04 |
| gene-LOC105283592 | https://www.ncbi.nlm.nih.gov/gene/?term=LOC105283592 | 3479 | 71 | 204.08 | 1 | 2.87 | 1.41 | 26.49 | 24.49 | 2.01 | 2.01 | 0.04 | 0.04 |
| gene-LOC105282658 | https://www.ncbi.nlm.nih.gov/gene/?term=LOC105282658 | 2421 | 65 | 268.48 | 1 | 4.13 | 1.54 | 21.38 | 19.92 | 1.46 | 1.46 | 0.04 | 0.04 |
| gene-LOC105277761 | https://www.ncbi.nlm.nih.gov/gene/?term=LOC105277761 | 4648 | 146 | 314.11 | 1 | 2.15 | 0.68 | 16.93 | 15.61 | 1.32 | 1.32 | 0.04 | 0.04 |
| gene-LOC105274856 | https://www.ncbi.nlm.nih.gov/gene/?term=LOC105274856 | 5216 | 218 | 417.94 | 1 | 1.92 | 0.46 | 19.03 | 18.03 | 1 | 1 | 0.04 | 0.04 |
| gene-LOC105284355 | https://www.ncbi.nlm.nih.gov/gene/?term=LOC105284355 | 3904 | 149 | 381.66 | 1 | 2.56 | 0.67 | 21.89 | 20.9 | 0.99 | 0.99 | 0.04 | 0.04 |
| gene-LOC105278877 | https://www.ncbi.nlm.nih.gov/gene/?term=LOC105278877 | 1968 | 75 | 381.1 | 1 | 5.08 | 1.33 | 34.6 | 33.65 | 0.95 | 0.95 | 0.04 | 0.04 |
| gene-LOC105280148 | https://www.ncbi.nlm.nih.gov/gene/?term=LOC105280148 | 3120 | 214 | 685.9 | 1 | 3.21 | 0.47 | 11.76 | 11.17 | 0.6 | 0.6 | 0.04 | 0.04 |
| gene-LOC105274706 | https://www.ncbi.nlm.nih.gov/gene/?term=LOC105274706 | 1446 | 55 | 380.36 | 1 | 6.92 | 1.82 | 37.25 | 38.23 | -0.98 | 0.98 | -0.04 | 0.04 |
| gene-LOC105286694 | https://www.ncbi.nlm.nih.gov/gene/?term=LOC105286694 | 3409 | 11 | 32.27 | 1 | 2.93 | 9.09 | 63.21 | 54.71 | 8.5 | 8.5 | 0.03 | 0.03 |
| gene-LOC105287648 | https://www.ncbi.nlm.nih.gov/gene/?term=LOC105287648 | 2205 | 27 | 122.45 | 1 | 4.54 | 3.7 | 41.48 | 39.19 | 2.29 | 2.29 | 0.03 | 0.03 |
| gene-LOC105286514 | https://www.ncbi.nlm.nih.gov/gene/?term=LOC105286514 | 2528 | 41 | 162.18 | 1 | 3.96 | 2.44 | 35.27 | 33.32 | 1.95 | 1.95 | 0.03 | 0.03 |
| gene-LOC105286644 | https://www.ncbi.nlm.nih.gov/gene/?term=LOC105286644 | 1235 | 20 | 161.94 | 1 | 8.1 | 5 | 39.34 | 37.53 | 1.81 | 1.81 | 0.03 | 0.03 |
| gene-LOC105278807 | https://www.ncbi.nlm.nih.gov/gene/?term=LOC105278807 | 2166 | 73 | 337.03 | 1 | 4.62 | 1.37 | 18.28 | 17.29 | 0.99 | 0.99 | 0.03 | 0.03 |
| gene-LOC105284184 | https://www.ncbi.nlm.nih.gov/gene/?term=LOC105284184 | 2535 | 85 | 335.31 | 1 | 3.94 | 1.18 | 34.37 | 33.53 | 0.83 | 0.83 | 0.03 | 0.03 |
| gene-LOC105285202 | https://www.ncbi.nlm.nih.gov/gene/?term=LOC105285202 | 1838 | 66 | 359.09 | 1 | 5.44 | 1.52 | 25.35 | 24.58 | 0.77 | 0.77 | 0.03 | 0.03 |
| gene-LOC105274452 | https://www.ncbi.nlm.nih.gov/gene/?term=LOC105274452 | 6595 | 261 | 395.75 | 1 | 1.52 | 0.38 | 13.63 | 12.94 | 0.69 | 0.69 | 0.03 | 0.03 |
| gene-LOC105286799 | https://www.ncbi.nlm.nih.gov/gene/?term=LOC105286799 | 3557 | 183 | 514.48 | 1 | 2.81 | 0.55 | 16.25 | 15.68 | 0.57 | 0.57 | 0.03 | 0.03 |
| gene-LOC105280723 | https://www.ncbi.nlm.nih.gov/gene/?term=LOC105280723 | 3572 | 170 | 475.92 | 1 | 2.8 | 0.59 | 16.88 | 16.35 | 0.53 | 0.53 | 0.03 | 0.03 |
| gene-LOC105274534 | https://www.ncbi.nlm.nih.gov/gene/?term=LOC105274534 | 1329 | 54 | 406.32 | 1 | 7.52 | 1.85 | 45.08 | 45.82 | -0.75 | 0.75 | -0.03 | 0.03 |
| gene-LOC105284507 | https://www.ncbi.nlm.nih.gov/gene/?term=LOC105284507 | 5207 | 76 | 145.96 | 1 | 1.92 | 1.32 | 19.78 | 21.69 | -1.91 | 1.91 | -0.03 | 0.03 |
| gene-LOC105286302 | https://www.ncbi.nlm.nih.gov/gene/?term=LOC105286302 | 4536 | 18 | 39.68 | 1 | 2.2 | 5.56 | 22.71 | 17.52 | 5.2 | 5.2 | 0.02 | 0.02 |
| gene-LOC105281515 | https://www.ncbi.nlm.nih.gov/gene/?term=LOC105281515 | 8827 | 48 | 54.38 | 2 | 2.27 | 4.17 | 32.1 | 29.31 | 2.79 | 2.79 | 0.02 | 0.02 |
| gene-LOC105288066 | https://www.ncbi.nlm.nih.gov/gene/?term=LOC105288066 | 1964 | 21 | 106.92 | 1 | 5.09 | 4.76 | 46.21 | 43.96 | 2.25 | 2.25 | 0.02 | 0.02 |
| gene-LOC105275518 | https://www.ncbi.nlm.nih.gov/gene/?term=LOC105275518 | 2335 | 19 | 81.37 | 1 | 4.28 | 5.26 | 46.5 | 44.26 | 2.24 | 2.24 | 0.02 | 0.02 |
| gene-LOC105277856 | https://www.ncbi.nlm.nih.gov/gene/?term=LOC105277856 | 3442 | 34 | 98.78 | 1 | 2.91 | 2.94 | 37.69 | 35.88 | 1.81 | 1.81 | 0.02 | 0.02 |

| | | | | | | | | | | | | | |
|---|---|---|---|---|---|---|---|---|---|---|---|---|---|
| gene-LOC105288010 | https://www.ncbi.nlm.nih.gov/gene/?term=LOC105288010 | 3189 | 39 | 122.3 | 1 | 3.14 | 2.56 | 37.68 | 36.3 | 1.38 | 1.38 | 0.02 | 0.02 |
| gene-LOC105280348 | https://www.ncbi.nlm.nih.gov/gene/?term=LOC105280348 | 3797 | 65 | 171.19 | 1 | 2.63 | 1.54 | 21.44 | 20.1 | 1.33 | 1.33 | 0.02 | 0.02 |
| gene-LOC105282600 | https://www.ncbi.nlm.nih.gov/gene/?term=LOC105282600 | 3715 | 53 | 142.66 | 1 | 2.69 | 1.89 | 32.07 | 30.77 | 1.3 | 1.3 | 0.02 | 0.02 |
| gene-LOC105276059 | https://www.ncbi.nlm.nih.gov/gene/?term=LOC105276059 | 6256 | 121 | 193.41 | 1 | 1.6 | 0.83 | 11.92 | 10.67 | 1.25 | 1.25 | 0.02 | 0.02 |
| gene-LOC105279664 | https://www.ncbi.nlm.nih.gov/gene/?term=LOC105279664 | 4023 | 54 | 134.23 | 1 | 2.49 | 1.85 | 21.29 | 20.13 | 1.15 | 1.15 | 0.02 | 0.02 |
| gene-LOC105285502 | https://www.ncbi.nlm.nih.gov/gene/?term=LOC105285502 | 9990 | 143 | 143.14 | 1 | 1 | 0.7 | 10.37 | 9.32 | 1.05 | 1.05 | 0.02 | 0.02 |
| gene-LOC105282794 | https://www.ncbi.nlm.nih.gov/gene/?term=LOC105282794 | 4571 | 93 | 203.46 | 1 | 2.19 | 1.08 | 18.93 | 18.03 | 0.9 | 0.9 | 0.02 | 0.02 |
| gene-LOC105275451 | https://www.ncbi.nlm.nih.gov/gene/?term=LOC105275451 | 6035 | 127 | 210.44 | 1 | 1.66 | 0.79 | 22.96 | 22.23 | 0.73 | 0.73 | 0.02 | 0.02 |
| gene-LOC105280956 | https://www.ncbi.nlm.nih.gov/gene/?term=LOC105280956 | 2639 | 75 | 284.2 | 1 | 3.79 | 1.33 | 17.09 | 16.45 | 0.64 | 0.64 | 0.02 | 0.02 |
| gene-LOC105279038 | https://www.ncbi.nlm.nih.gov/gene/?term=LOC105279038 | 9453 | 333 | 352.27 | 1 | 1.06 | 0.3 | 9.29 | 8.67 | 0.62 | 0.62 | 0.02 | 0.02 |
| gene-LOC105287063 | https://www.ncbi.nlm.nih.gov/gene/?term=LOC105287063 | 1874 | 48 | 256.14 | 1 | 5.34 | 2.08 | 30.52 | 29.9 | 0.62 | 0.62 | 0.02 | 0.02 |
| gene-LOC105274823 | https://www.ncbi.nlm.nih.gov/gene/?term=LOC105274823 | 4620 | 138 | 298.7 | 1 | 2.16 | 0.72 | 21.48 | 20.88 | 0.6 | 0.6 | 0.02 | 0.02 |
| gene-LOC105285807 | https://www.ncbi.nlm.nih.gov/gene/?term=LOC105285807 | 6063 | 218 | 359.56 | 1 | 1.65 | 0.46 | 7.08 | 6.5 | 0.58 | 0.58 | 0.02 | 0.02 |
| gene-LOC105275683 | https://www.ncbi.nlm.nih.gov/gene/?term=LOC105275683 | 4256 | 194 | 455.83 | 1 | 2.35 | 0.52 | 13.24 | 12.7 | 0.54 | 0.54 | 0.02 | 0.02 |
| gene-LOC105278666 | https://www.ncbi.nlm.nih.gov/gene/?term=LOC105278666 | 10341 | 459 | 443.86 | 1 | 0.97 | 0.22 | 7.32 | 6.79 | 0.53 | 0.53 | 0.02 | 0.02 |
| gene-LOC105285179 | https://www.ncbi.nlm.nih.gov/gene/?term=LOC105285179 | 4191 | 164 | 391.31 | 1 | 2.39 | 0.61 | 20.24 | 19.71 | 0.53 | 0.53 | 0.02 | 0.02 |
| gene-LOC105283123 | https://www.ncbi.nlm.nih.gov/gene/?term=LOC105283123 | 1507 | 52 | 345.06 | 1 | 6.64 | 1.92 | 17.86 | 17.35 | 0.52 | 0.52 | 0.02 | 0.02 |
| gene-LOC105285639 | https://www.ncbi.nlm.nih.gov/gene/?term=LOC105285639 | 10325 | 356 | 344.79 | 1 | 0.97 | 0.28 | 11.26 | 10.77 | 0.49 | 0.49 | 0.02 | 0.02 |
| gene-LOC105275848 | https://www.ncbi.nlm.nih.gov/gene/?term=LOC105275848 | 3718 | 139 | 373.86 | 1 | 2.69 | 0.72 | 22.47 | 21.99 | 0.48 | 0.48 | 0.02 | 0.02 |
| gene-LOC105278145 | https://www.ncbi.nlm.nih.gov/gene/?term=LOC105278145 | 4470 | 241 | 539.15 | 1 | 2.24 | 0.41 | 10.99 | 10.59 | 0.4 | 0.4 | 0.02 | 0.02 |
| gene-LOC105276452 | https://www.ncbi.nlm.nih.gov/gene/?term=LOC105276452 | 2917 | 257 | 881.04 | 1 | 3.43 | 0.39 | 9.59 | 9.37 | 0.22 | 0.22 | 0.02 | 0.02 |
| gene-LOC105286274 | https://www.ncbi.nlm.nih.gov/gene/?term=LOC105286274 | 3817 | 202 | 529.21 | 1 | 2.62 | 0.5 | 16.44 | 16.87 | -0.43 | 0.43 | -0.02 | 0.02 |
| gene-LOC105283385 | https://www.ncbi.nlm.nih.gov/gene/?term=LOC105283385 | 9540 | 305 | 319.71 | 1 | 1.05 | 0.33 | 12.66 | 13.17 | -0.51 | 0.51 | -0.02 | 0.02 |
| gene-LOC113561621 | https://www.ncbi.nlm.nih.gov/gene/?term=LOC113561621 | 2249 | 78 | 346.82 | 1 | 4.45 | 1.28 | 20.85 | 21.38 | -0.53 | 0.53 | -0.02 | 0.02 |
| gene-LOC105285742 | https://www.ncbi.nlm.nih.gov/gene/?term=LOC105285742 | 3446 | 107 | 310.5 | 1 | 2.9 | 0.93 | 21.42 | 22.06 | -0.64 | 0.64 | -0.02 | 0.02 |
| gene-LOC109611283 | https://www.ncbi.nlm.nih.gov/gene/?term=LOC109611283 | 1451 | 38 | 261.89 | 1 | 6.89 | 2.63 | 35.88 | 36.65 | -0.77 | 0.77 | -0.02 | 0.02 |
| gene-LOC105285252 | https://www.ncbi.nlm.nih.gov/gene/?term=LOC105285252 | 7427 | 117 | 157.53 | 1 | 1.35 | 0.85 | 13.64 | 15.21 | -1.57 | 1.57 | -0.02 | 0.02 |
| gene-LOC105279500 | https://www.ncbi.nlm.nih.gov/gene/?term=LOC105279500 | 3587 | 33 | 92 | 1 | 2.79 | 3.03 | 44.93 | 47.01 | -2.08 | 2.08 | -0.02 | 0.02 |
| gene-LOC105282067 | https://www.ncbi.nlm.nih.gov/gene/?term=LOC105282067 | 5941 | 44 | 74.06 | 1 | 1.68 | 2.27 | 26.18 | 28.47 | -2.29 | 2.29 | -0.02 | 0.02 |
| gene-LOC105285315 | https://www.ncbi.nlm.nih.gov/gene/?term=LOC105285315 | 5458 | 22 | 40.31 | 1 | 1.83 | 4.55 | 41.52 | 38.4 | 3.12 | 3.12 | 0.01 | 0.01 |
| gene-LOC105275347 | https://www.ncbi.nlm.nih.gov/gene/?term=LOC105275347 | 2526 | 15 | 59.38 | 1 | 3.96 | 6.67 | 31.33 | 28.89 | 2.44 | 2.44 | 0.01 | 0.01 |
| gene-LOC105288078 | https://www.ncbi.nlm.nih.gov/gene/?term=LOC105288078 | 19126 | 294 | 153.72 | 1 | 0.52 | 0.34 | 18.24 | 17.37 | 0.87 | 0.87 | 0.01 | 0.01 |
| gene-LOC105284329 | https://www.ncbi.nlm.nih.gov/gene/?term=LOC105284329 | 4051 | 50 | 123.43 | 1 | 2.47 | 2 | 19.81 | 19.02 | 0.79 | 0.79 | 0.01 | 0.01 |
| gene-LOC105276538 | https://www.ncbi.nlm.nih.gov/gene/?term=LOC105276538 | 1511 | 19 | 125.74 | 1 | 6.62 | 5.26 | 70.24 | 69.47 | 0.77 | 0.77 | 0.01 | 0.01 |
| gene-LOC105280002 | https://www.ncbi.nlm.nih.gov/gene/?term=LOC105280002 | 1940 | 25 | 128.87 | 1 | 5.15 | 4 | 38.59 | 38.02 | 0.58 | 0.58 | 0.01 | 0.01 |
| gene-LOC105274851 | https://www.ncbi.nlm.nih.gov/gene/?term=LOC105274851 | 6616 | 128 | 193.47 | 1 | 1.51 | 0.78 | 11.71 | 11.16 | 0.54 | 0.54 | 0.01 | 0.01 |
| gene-LOC105283943 | https://www.ncbi.nlm.nih.gov/gene/?term=LOC105283943 | 3559 | 58 | 162.97 | 1 | 2.81 | 1.72 | 14.89 | 14.35 | 0.54 | 0.54 | 0.01 | 0.01 |
| gene-LOC105280410 | https://www.ncbi.nlm.nih.gov/gene/?term=LOC105280410 | 12580 | 237 | 188.39 | 1 | 0.79 | 0.42 | 21.06 | 20.59 | 0.47 | 0.47 | 0.01 | 0.01 |
| gene-LOC105279657 | https://www.ncbi.nlm.nih.gov/gene/?term=LOC105279657 | 5878 | 86 | 146.31 | 1 | 1.7 | 1.16 | 19.59 | 19.13 | 0.46 | 0.46 | 0.01 | 0.01 |
| gene-LOC105283728 | https://www.ncbi.nlm.nih.gov/gene/?term=LOC105283728 | 3256 | 113 | 347.05 | 2 | 6.14 | 1.77 | 20.45 | 20.03 | 0.42 | 0.42 | 0.01 | 0.01 |
| gene-LOC113562050 | https://www.ncbi.nlm.nih.gov/gene/?term=LOC113562050 | 17097 | 583 | 341 | 1 | 0.58 | 0.17 | 5.52 | 5.1 | 0.42 | 0.42 | 0.01 | 0.01 |
| gene-LOC105276196 | https://www.ncbi.nlm.nih.gov/gene/?term=LOC105276196 | 2953 | 48 | 162.55 | 1 | 3.39 | 2.08 | 45.56 | 45.18 | 0.38 | 0.38 | 0.01 | 0.01 |
| gene-LOC105278632 | https://www.ncbi.nlm.nih.gov/gene/?term=LOC105278632 | 3624 | 129 | 355.96 | 1 | 2.76 | 0.78 | 14.81 | 14.44 | 0.37 | 0.37 | 0.01 | 0.01 |
| gene-LOC105285395 | https://www.ncbi.nlm.nih.gov/gene/?term=LOC105285395 | 8085 | 196 | 242.42 | 1 | 1.24 | 0.51 | 15.02 | 14.69 | 0.33 | 0.33 | 0.01 | 0.01 |
| gene-LOC105278455 | https://www.ncbi.nlm.nih.gov/gene/?term=LOC105278455 | 27181 | 925 | 340.31 | 1 | 0.37 | 0.11 | 7.49 | 7.17 | 0.32 | 0.32 | 0.01 | 0.01 |
| gene-LOC105283712 | https://www.ncbi.nlm.nih.gov/gene/?term=LOC105283712 | 1910 | 31 | 162.3 | 1 | 5.24 | 3.23 | 30.95 | 30.64 | 0.31 | 0.31 | 0.01 | 0.01 |
| gene-LOC105276941 | https://www.ncbi.nlm.nih.gov/gene/?term=LOC105276941 | 15956 | 551 | 345.32 | 1 | 0.63 | 0.18 | 5.1 | 4.82 | 0.28 | 0.28 | 0.01 | 0.01 |
| gene-LOC105275374 | https://www.ncbi.nlm.nih.gov/gene/?term=LOC105275374 | 11376 | 527 | 463.26 | 1 | 0.88 | 0.19 | 11.34 | 11.07 | 0.27 | 0.27 | 0.01 | 0.01 |
| gene-LOC105287417 | https://www.ncbi.nlm.nih.gov/gene/?term=LOC105287417 | 1025 | 28 | 273.17 | 1 | 9.76 | 3.57 | 42.07 | 41.82 | 0.26 | 0.26 | 0.01 | 0.01 |
| gene-LOC105275327 | https://www.ncbi.nlm.nih.gov/gene/?term=LOC105275327 | 14245 | 493 | 346.09 | 1 | 0.7 | 0.2 | 6.9 | 6.7 | 0.19 | 0.19 | 0.01 | 0.01 |
| gene-LOC105287550 | https://www.ncbi.nlm.nih.gov/gene/?term=LOC105287550 | 15659 | 570 | 364.01 | 1 | 0.64 | 0.18 | 6.53 | 6.34 | 0.19 | 0.19 | 0.01 | 0.01 |
| gene-LOC105278282 | https://www.ncbi.nlm.nih.gov/gene/?term=LOC105278282 | 1775 | 56 | 315.49 | 1 | 5.63 | 1.79 | 35.74 | 35.56 | 0.18 | 0.18 | 0.01 | 0.01 |
| gene-LOC105281272 | https://www.ncbi.nlm.nih.gov/gene/?term=LOC105281272 | 5252 | 178 | 338.92 | 1 | 1.9 | 0.56 | 16.1 | 15.92 | 0.18 | 0.18 | 0.01 | 0.01 |
| gene-LOC105282491 | https://www.ncbi.nlm.nih.gov/gene/?term=LOC105282491 | 4761 | 165 | 346.57 | 1 | 2.1 | 0.61 | 8.62 | 8.44 | 0.18 | 0.18 | 0.01 | 0.01 |
| gene-LOC105286649 | https://www.ncbi.nlm.nih.gov/gene/?term=LOC105286649 | 14178 | 826 | 582.59 | 1 | 0.71 | 0.12 | 0.92 | 1.04 | -0.12 | 0.12 | -0.01 | 0.01 |
| gene-LOC105276462 | https://www.ncbi.nlm.nih.gov/gene/?term=LOC105276462 | 6561 | 278 | 423.72 | 1 | 1.52 | 0.36 | 7.35 | 7.5 | -0.15 | 0.15 | -0.01 | 0.01 |
| gene-LOC105282646 | https://www.ncbi.nlm.nih.gov/gene/?term=LOC105282646 | 3317 | 97 | 292.43 | 1 | 3.01 | 1.03 | 22.32 | 22.49 | -0.17 | 0.17 | -0.01 | 0.01 |
| gene-LOC105278691 | https://www.ncbi.nlm.nih.gov/gene/?term=LOC105278691 | 3871 | 147 | 379.75 | 1 | 2.58 | 0.68 | 14.28 | 14.5 | -0.22 | 0.22 | -0.01 | 0.01 |
| gene-LOC105280819 | https://www.ncbi.nlm.nih.gov/gene/?term=LOC105280819 | 1639 | 39 | 237.95 | 1 | 6.1 | 2.56 | 35.75 | 36.01 | -0.26 | 0.26 | -0.01 | 0.01 |
| gene-LOC105275587 | https://www.ncbi.nlm.nih.gov/gene/?term=LOC105275587 | 2404 | 55 | 228.79 | 1 | 4.16 | 1.82 | 34.41 | 34.8 | -0.39 | 0.39 | -0.01 | 0.01 |
| gene-LOC105281312 | https://www.ncbi.nlm.nih.gov/gene/?term=LOC105281312 | 2551 | 72 | 282.24 | 1 | 3.92 | 1.39 | 29.66 | 30.06 | -0.4 | 0.4 | -0.01 | 0.01 |
| gene-LOC105286515 | https://www.ncbi.nlm.nih.gov/gene/?term=LOC105286515 | 3751 | 116 | 309.25 | 1 | 2.67 | 0.86 | 10.42 | 10.83 | -0.41 | 0.41 | -0.01 | 0.01 |
| gene-LOC105286999 | https://www.ncbi.nlm.nih.gov/gene/?term=LOC105286999 | 2141 | 41 | 191.5 | 1 | 4.67 | 2.44 | 37.42 | 37.83 | -0.41 | 0.41 | -0.01 | 0.01 |
| gene-LOC105281754 | https://www.ncbi.nlm.nih.gov/gene/?term=LOC105281754 | 4300 | 117 | 272.09 | 1 | 2.33 | 0.85 | 16.59 | 17.09 | -0.5 | 0.5 | -0.01 | 0.01 |

| | | | | | | | | | | | | | |
|---|---|---|---|---|---|---|---|---|---|---|---|---|---|
| gene-LOC105278534 | https://www.ncbi.nlm.nih.gov/gene/?term=LOC105278534 | 7364 | 180 | 244.43 | 1 | 1.36 | 0.56 | 18.21 | 18.75 | -0.54 | 0.54 | -0.01 | 0.01 |
| gene-LOC105276272 | https://www.ncbi.nlm.nih.gov/gene/?term=LOC105276272 | 1965 | 36 | 183.21 | 1 | 5.09 | 2.78 | 32.08 | 32.69 | -0.61 | 0.61 | -0.01 | 0.01 |
| gene-LOC105281173 | https://www.ncbi.nlm.nih.gov/gene/?term=LOC105281173 | 1595 | 38 | 238.24 | 1 | 6.27 | 2.63 | 60.05 | 60.67 | -0.62 | 0.62 | -0.01 | 0.01 |
| gene-LOC105287633 | https://www.ncbi.nlm.nih.gov/gene/?term=LOC105287633 | 1524 | 27 | 177.17 | 1 | 6.56 | 3.7 | 63.52 | 64.22 | -0.7 | 0.7 | -0.01 | 0.01 |
| gene-LOC105283989 | https://www.ncbi.nlm.nih.gov/gene/?term=LOC105283989 | 4948 | 26 | 52.55 | 1 | 2.02 | 3.85 | 43.53 | 42.86 | 0.68 | 0.68 | 0 | 0 |
| gene-LOC105285514 | https://www.ncbi.nlm.nih.gov/gene/?term=LOC105285514 | 1277 | 13 | 101.8 | 1 | 7.83 | 7.69 | 90.77 | 90.29 | 0.48 | 0.48 | 0 | 0 |
| gene-LOC105286452 | https://www.ncbi.nlm.nih.gov/gene/?term=LOC105286452 | 6888 | 61 | 88.56 | 1 | 1.45 | 1.64 | 32.75 | 32.35 | 0.4 | 0.4 | 0 | 0 |
| gene-LOC105286194 | https://www.ncbi.nlm.nih.gov/gene/?term=LOC105286194 | 152619 | 2166 | 141.92 | 1 | 0.07 | 0.05 | 6.19 | 5.95 | 0.23 | 0.23 | 0 | 0 |
| gene-LOC105282811 | https://www.ncbi.nlm.nih.gov/gene/?term=LOC105282811 | 5765 | 203 | 352.12 | 1 | 1.73 | 0.49 | 8.42 | 8.31 | 0.12 | 0.12 | 0 | 0 |
| gene-LOC105282825 | https://www.ncbi.nlm.nih.gov/gene/?term=LOC105282825 | 3919 | 80 | 204.13 | 1 | 2.55 | 1.25 | 12.57 | 12.46 | 0.11 | 0.11 | 0 | 0 |
| gene-LOC105274590 | https://www.ncbi.nlm.nih.gov/gene/?term=LOC105274590 | 4516 | 58 | 128.43 | 2 | 4.43 | 3.45 | 35.68 | 35.59 | 0.09 | 0.09 | 0 | 0 |
| gene-LOC105275363 | https://www.ncbi.nlm.nih.gov/gene/?term=LOC105275363 | 3798 | 156 | 410.74 | 1 | 2.63 | 0.64 | 17.23 | 17.17 | 0.07 | 0.07 | 0 | 0 |
| gene-LOC105283296 | https://www.ncbi.nlm.nih.gov/gene/?term=LOC105283296 | 2174 | 84 | 386.38 | 1 | 4.6 | 1.19 | 21.67 | 21.61 | 0.07 | 0.07 | 0 | 0 |
| gene-LOC105281700 | https://www.ncbi.nlm.nih.gov/gene/?term=LOC105281700 | 2208 | 37 | 167.57 | 1 | 4.53 | 2.7 | 39.7 | 39.67 | 0.04 | 0.04 | 0 | 0 |
| gene-LOC105274531 | https://www.ncbi.nlm.nih.gov/gene/?term=LOC105274531 | 7774 | 284 | 365.32 | 1 | 1.29 | 0.35 | 10.8 | 10.77 | 0.03 | 0.03 | 0 | 0 |
| gene-LOC105284043 | https://www.ncbi.nlm.nih.gov/gene/?term=LOC105284043 | 5661 | 279 | 492.85 | 1 | 1.77 | 0.36 | 10.73 | 10.7 | 0.03 | 0.03 | 0 | 0 |
| gene-LOC105275460 | https://www.ncbi.nlm.nih.gov/gene/?term=LOC105275460 | 2473 | 114 | 460.98 | 1 | 4.04 | 0.88 | 13.89 | 13.88 | 0.02 | 0.02 | 0 | 0 |
| gene-LOC105277866 | https://www.ncbi.nlm.nih.gov/gene/?term=LOC105277866 | 33045 | 378 | 114.39 | 1 | 0.3 | 0.26 | 3.22 | 3.21 | 0.01 | 0.01 | 0 | 0 |
| gene-LOC105285406 | https://www.ncbi.nlm.nih.gov/gene/?term=LOC105285406 | 8015 | 59 | 73.61 | 1 | 1.25 | 1.69 | 26.4 | 26.39 | 0.01 | 0.01 | 0 | 0 |
| gene-LOC105275179 | https://www.ncbi.nlm.nih.gov/gene/?term=LOC105275179 | 3271 | 73 | 223.17 | 1 | 3.06 | 1.37 | 20.99 | 20.99 | 0 | 0 | 0 | 0 |
| gene-LOC105278228 | https://www.ncbi.nlm.nih.gov/gene/?term=LOC105278228 | 8610 | 162 | 188.15 | 1 | 1.16 | 0.62 | 21.49 | 21.49 | 0 | 0 | 0 | 0 |
| gene-LOC105287764 | https://www.ncbi.nlm.nih.gov/gene/?term=LOC105287764 | 18359 | 396 | 215.7 | 1 | 0.54 | 0.25 | 2.87 | 2.88 | -0.01 | 0.01 | 0 | 0 |
| gene-LOC105278283 | https://www.ncbi.nlm.nih.gov/gene/?term=LOC105278283 | 2317 | 125 | 539.49 | 1 | 4.32 | 0.8 | 28 | 28.03 | -0.02 | 0.02 | 0 | 0 |
| gene-LOC105280265 | https://www.ncbi.nlm.nih.gov/gene/?term=LOC105280265 | 85984 | 3074 | 357.51 | 1 | 0.12 | 0.03 | 1.06 | 1.08 | -0.02 | 0.02 | 0 | 0 |
| gene-LOC105282951 | https://www.ncbi.nlm.nih.gov/gene/?term=LOC105282951 | 3366 | 118 | 350.56 | 1 | 2.97 | 0.85 | 19.88 | 19.9 | -0.02 | 0.02 | 0 | 0 |
| gene-LOC105285533 | https://www.ncbi.nlm.nih.gov/gene/?term=LOC105285533 | 1640 | 60 | 365.85 | 1 | 6.1 | 1.67 | 26.06 | 26.09 | -0.03 | 0.03 | 0 | 0 |
| gene-LOC105280513 | https://www.ncbi.nlm.nih.gov/gene/?term=LOC105280513 | 69542 | 1246 | 179.17 | 1 | 0.14 | 0.08 | 0.53 | 0.59 | -0.06 | 0.06 | 0 | 0 |
| gene-LOC105277125 | https://www.ncbi.nlm.nih.gov/gene/?term=LOC105277125 | 51079 | 1856 | 363.36 | 1 | 0.2 | 0.05 | 1.57 | 1.64 | -0.07 | 0.07 | 0 | 0 |
| gene-LOC105276527 | https://www.ncbi.nlm.nih.gov/gene/?term=LOC105276527 | 7923 | 318 | 401.36 | 1 | 1.26 | 0.31 | 9.67 | 9.76 | -0.09 | 0.09 | 0 | 0 |
| gene-LOC105282885 | https://www.ncbi.nlm.nih.gov/gene/?term=LOC105282885 | 1870 | 43 | 229.95 | 1 | 5.35 | 2.33 | 44.33 | 44.46 | -0.13 | 0.13 | 0 | 0 |
| gene-LOC105275187 | https://www.ncbi.nlm.nih.gov/gene/?term=LOC105275187 | 5534 | 70 | 126.49 | 1 | 1.81 | 1.43 | 14.08 | 14.45 | -0.36 | 0.36 | 0 | 0 |

Directory: OGLall File: rnk-dmg-Ob_LI_BCphaseA2.vs.RphaseA1.txt

| gene_ID | gene_link | gwidth | #Sites | #per10Kb | #dmSites | #dmsp10kb | %dmSites | %pSite1 | %pSite2 | DMpSite | ADMpSite | DMpNucl | ADMpNucl |
|---|---|---|---|---|---|---|---|---|---|---|---|---|---|
| gene-LOC105278458 | https://www.ncbi.nlm.nih.gov/gene/?term=LOC105278458 | 216 | 3 | 138.89 | 1 | 46.3 | 33.33 | 10.57 | 24.69 | -14.12 | 14.12 | -0.2 | 0.2 |
| gene-LOC105286531 | https://www.ncbi.nlm.nih.gov/gene/?term=LOC105286531 | 372 | 23 | 618.28 | 1 | 26.88 | 4.35 | 39.65 | 36.5 | 3.15 | 3.15 | 0.19 | 0.19 |
| gene-LOC113562901 | https://www.ncbi.nlm.nih.gov/gene/?term=LOC113562901 | 1531 | 22 | 143.7 | 8 | 52.25 | 36.36 | 60.1 | 48.53 | 11.57 | 11.57 | 0.17 | 0.17 |
| gene-LOC113562728 | https://www.ncbi.nlm.nih.gov/gene/?term=LOC113562728 | 1733 | 32 | 184.65 | 5 | 28.85 | 15.62 | 28.94 | 19.93 | 9 | 9 | 0.17 | 0.17 |
| gene-LOC109611487 | https://www.ncbi.nlm.nih.gov/gene/?term=LOC109611487 | 437 | 32 | 732.27 | 1 | 22.88 | 3.12 | 16.38 | 14.11 | 2.27 | 2.27 | 0.17 | 0.17 |
| gene-LOC105285392 | https://www.ncbi.nlm.nih.gov/gene/?term=LOC105285392 | 1601 | 82 | 512.18 | 3 | 18.74 | 3.66 | 38.11 | 34.95 | 3.16 | 3.16 | 0.16 | 0.16 |
| gene-LOC113561934 | https://www.ncbi.nlm.nih.gov/gene/?term=LOC113561934 | 1252 | 27 | 215.65 | 4 | 31.95 | 14.81 | 31.68 | 24.75 | 6.93 | 6.93 | 0.15 | 0.15 |
| gene-LOC105275833 | https://www.ncbi.nlm.nih.gov/gene/?term=LOC105275833 | 981 | 39 | 397.55 | 2 | 20.39 | 5.13 | 40.15 | 36.28 | 3.87 | 3.87 | 0.15 | 0.15 |
| gene-LOC105284411 | https://www.ncbi.nlm.nih.gov/gene/?term=LOC105284411 | 1795 | 63 | 350.97 | 4 | 22.28 | 6.35 | 36.82 | 33 | 3.82 | 3.82 | 0.13 | 0.13 |
| gene-LOC105286320 | https://www.ncbi.nlm.nih.gov/gene/?term=LOC105286320 | 1659 | 60 | 361.66 | 4 | 24.11 | 6.67 | 41.91 | 38.27 | 3.64 | 3.64 | 0.13 | 0.13 |
| gene-LOC105275355 | https://www.ncbi.nlm.nih.gov/gene/?term=LOC105275355 | 3197 | 163 | 509.85 | 7 | 21.9 | 4.29 | 24.89 | 27.47 | -2.59 | 2.59 | -0.13 | 0.13 |
| gene-LOC113561930 | https://www.ncbi.nlm.nih.gov/gene/?term=LOC113561930 | 422 | 20 | 473.93 | 1 | 23.7 | 5 | 10.9 | 13.66 | -2.76 | 2.76 | -0.13 | 0.13 |
| gene-LOC105279538 | https://www.ncbi.nlm.nih.gov/gene/?term=LOC105279538 | 889 | 17 | 191.23 | 2 | 22.5 | 11.76 | 36.14 | 29.61 | 6.53 | 6.53 | 0.12 | 0.12 |
| gene-LOC105281325 | https://www.ncbi.nlm.nih.gov/gene/?term=LOC105281325 | 942 | 29 | 307.86 | 1 | 10.62 | 3.45 | 25.29 | 21.51 | 3.78 | 3.78 | 0.12 | 0.12 |
| gene-LOC105275231 | https://www.ncbi.nlm.nih.gov/gene/?term=LOC105275231 | 1270 | 43 | 338.58 | 1 | 7.87 | 2.33 | 43.15 | 39.57 | 3.59 | 3.59 | 0.12 | 0.12 |
| gene-LOC105286946 | https://www.ncbi.nlm.nih.gov/gene/?term=LOC105286946 | 1228 | 49 | 399.02 | 3 | 24.43 | 6.12 | 34.75 | 31.69 | 3.06 | 3.06 | 0.12 | 0.12 |
| gene-LOC105276563 | https://www.ncbi.nlm.nih.gov/gene/?term=LOC105276563 | 1958 | 115 | 587.33 | 6 | 30.64 | 5.22 | 18.73 | 16.62 | 2.11 | 2.11 | 0.12 | 0.12 |
| gene-LOC105278873 | https://www.ncbi.nlm.nih.gov/gene/?term=LOC105278873 | 1897 | 146 | 769.64 | 6 | 31.63 | 4.11 | 21.31 | 19.69 | 1.62 | 1.62 | 0.12 | 0.12 |
| gene-LOC105283693 | https://www.ncbi.nlm.nih.gov/gene/?term=LOC105283693 | 1305 | 12 | 91.95 | 4 | 30.65 | 33.33 | 70.54 | 58.26 | 12.28 | 12.28 | 0.11 | 0.11 |
| gene-LOC105277390 | https://www.ncbi.nlm.nih.gov/gene/?term=LOC105277390 | 2567 | 94 | 366.19 | 3 | 11.69 | 3.19 | 39.27 | 36.36 | 2.9 | 2.9 | 0.11 | 0.11 |
| gene-LOC105275378 | https://www.ncbi.nlm.nih.gov/gene/?term=LOC105275378 | 1189 | 20 | 168.21 | 1 | 8.41 | 5 | 32.85 | 26.65 | 6.2 | 6.2 | 0.1 | 0.1 |
| gene-LOC105276679 | https://www.ncbi.nlm.nih.gov/gene/?term=LOC105276679 | 2005 | 64 | 319.2 | 2 | 9.98 | 3.12 | 39.86 | 36.61 | 3.25 | 3.25 | 0.1 | 0.1 |
| gene-LOC105285812 | https://www.ncbi.nlm.nih.gov/gene/?term=LOC105285812 | 1413 | 41 | 290.16 | 1 | 7.08 | 2.44 | 25.66 | 22.39 | 3.27 | 3.27 | 0.09 | 0.09 |
| gene-LOC105279437 | https://www.ncbi.nlm.nih.gov/gene/?term=LOC105279437 | 1069 | 41 | 383.54 | 1 | 9.35 | 2.44 | 22.95 | 20.6 | 2.35 | 2.35 | 0.09 | 0.09 |

...
... only first 25 lines shown ...

Directory: OGLall File: rnk-dmg-Ob_LI_BCphaseA2.vs.RphaseA2.txt

| gene_ID | gene_link | gwidth | #Sites | #per10Kb | #dmSites | #dmsp10kb | %dmSites | %pSite1 | %pSite2 | DMpSite | ADMpSite | DMpNucl | ADMpNucl |
|---|---|---|---|---|---|---|---|---|---|---|---|---|---|
| gene-Trnam-cau-2 | https://www.ncbi.nlm.nih.gov/gene/?term=Trnam-cau | 72 | 1 | 138.89 | 1 | 138.89 | 100 | 0 | 24 | -24 | 24 | -0.33 | 0.33 |
| gene-LOC113561930 | https://www.ncbi.nlm.nih.gov/gene/?term=LOC113561930 | 422 | 20 | 473.93 | 2 | 47.39 | 10 | 13.1 | 16.23 | -3.12 | 3.12 | -0.15 | 0.15 |
| gene-LOC109611472 | https://www.ncbi.nlm.nih.gov/gene/?term=LOC109611472 | 1351 | 27 | 199.85 | 2 | 14.8 | 7.41 | 32.9 | 39.85 | -6.95 | 6.95 | -0.14 | 0.14 |
| gene-LOC105286212 | https://www.ncbi.nlm.nih.gov/gene/?term=LOC105286212 | 1022 | 36 | 352.25 | 1 | 9.78 | 2.78 | 37.91 | 41.67 | -3.76 | 3.76 | -0.13 | 0.13 |
| gene-LOC105282572 | https://www.ncbi.nlm.nih.gov/gene/?term=LOC105282572 | 847 | 37 | 436.84 | 1 | 11.81 | 2.7 | 30.33 | 33.17 | -2.84 | 2.84 | -0.12 | 0.12 |
| gene-LOC105287692 | https://www.ncbi.nlm.nih.gov/gene/?term=LOC105287692 | 1505 | 100 | 664.45 | 3 | 19.93 | 3 | 27.86 | 29.51 | -1.66 | 1.66 | -0.11 | 0.11 |
| gene-LOC105278847 | https://www.ncbi.nlm.nih.gov/gene/?term=LOC105278847 | 1851 | 112 | 605.08 | 1 | 5.4 | 0.89 | 10.12 | 11.9 | -1.78 | 1.78 | -0.11 | 0.11 |
| gene-LOC105276920 | https://www.ncbi.nlm.nih.gov/gene/?term=LOC105276920 | 2623 | 95 | 362.18 | 1 | 3.81 | 1.05 | 19.9 | 22.94 | -3.04 | 3.04 | -0.11 | 0.11 |
| gene-LOC105274524 | https://www.ncbi.nlm.nih.gov/gene/?term=LOC105274524 | 1126 | 38 | 337.48 | 1 | 8.88 | 2.63 | 28.9 | 32.2 | -3.3 | 3.3 | -0.11 | 0.11 |
| gene-LOC105284411 | https://www.ncbi.nlm.nih.gov/gene/?term=LOC105284411 | 1795 | 63 | 350.97 | 1 | 5.57 | 1.59 | 35.7 | 32.95 | 2.74 | 2.74 | 0.1 | 0.1 |
| gene-LOC105276360 | https://www.ncbi.nlm.nih.gov/gene/?term=LOC105276360 | 1466 | 108 | 736.7 | 1 | 6.82 | 0.93 | 29.44 | 30.83 | -1.39 | 1.39 | -0.1 | 0.1 |
| gene-LOC105276423 | https://www.ncbi.nlm.nih.gov/gene/?term=LOC105276423 | 1689 | 77 | 455.89 | 1 | 5.92 | 1.3 | 16.63 | 18.76 | -2.13 | 2.13 | -0.1 | 0.1 |
| gene-LOC105287677 | https://www.ncbi.nlm.nih.gov/gene/?term=LOC105287677 | 1588 | 75 | 472.29 | 1 | 6.3 | 1.33 | 40.32 | 38.45 | 1.87 | 1.87 | 0.09 | 0.09 |
| gene-LOC105278824 | https://www.ncbi.nlm.nih.gov/gene/?term=LOC105278824 | 1230 | 61 | 495.93 | 1 | 8.13 | 1.64 | 27.78 | 29.67 | -1.89 | 1.89 | -0.09 | 0.09 |
| gene-LOC105275307 | https://www.ncbi.nlm.nih.gov/gene/?term=LOC105275307 | 1420 | 53 | 373.24 | 2 | 14.08 | 3.77 | 40.01 | 42.31 | -2.31 | 2.31 | -0.09 | 0.09 |
| gene-LOC105280327 | https://www.ncbi.nlm.nih.gov/gene/?term=LOC105280327 | 2670 | 100 | 374.53 | 1 | 3.75 | 1 | 29.14 | 31.51 | -2.37 | 2.37 | -0.09 | 0.09 |
| gene-LOC105281038 | https://www.ncbi.nlm.nih.gov/gene/?term=LOC105281038 | 2221 | 78 | 351.19 | 1 | 4.5 | 1.28 | 24.56 | 27.03 | -2.46 | 2.46 | -0.09 | 0.09 |
| gene-LOC105277012 | https://www.ncbi.nlm.nih.gov/gene/?term=LOC105277012 | 1494 | 52 | 348.06 | 3 | 20.08 | 5.77 | 24.71 | 27.23 | -2.52 | 2.52 | -0.09 | 0.09 |
| gene-LOC105281009 | https://www.ncbi.nlm.nih.gov/gene/?term=LOC105281009 | 1562 | 42 | 268.89 | 3 | 19.21 | 7.14 | 50.63 | 54.06 | -3.42 | 3.42 | -0.09 | 0.09 |
| gene-LOC105284400 | https://www.ncbi.nlm.nih.gov/gene/?term=LOC105284400 | 1906 | 44 | 230.85 | 3 | 15.74 | 6.82 | 27.71 | 31.42 | -3.71 | 3.71 | -0.09 | 0.09 |
| gene-LOC105278074 | https://www.ncbi.nlm.nih.gov/gene/?term=LOC105278074 | 623 | 11 | 176.57 | 2 | 32.1 | 18.18 | 11.57 | 16.88 | -5.31 | 5.31 | -0.09 | 0.09 |
| gene-LOC105281691 | https://www.ncbi.nlm.nih.gov/gene/?term=LOC105281691 | 2096 | 13 | 62.02 | 4 | 19.08 | 30.77 | 7.23 | 22.12 | -14.89 | 14.89 | -0.09 | 0.09 |
| gene-LOC105279538 | https://www.ncbi.nlm.nih.gov/gene/?term=LOC105279538 | 889 | 20 | 224.97 | 2 | 22.5 | 10 | 30.72 | 27.36 | 3.36 | 3.36 | 0.08 | 0.08 |
| gene-LOC113561393 | https://www.ncbi.nlm.nih.gov/gene/?term=LOC113561393 | 1872 | 63 | 336.54 | 2 | 10.68 | 3.17 | 21.1 | 18.81 | 2.28 | 2.28 | 0.08 | 0.08 |

...
... only first 25 lines shown ...

Directory: OGLall File: rnk-dmg-Ob_LI_BCphaseA2.vs.RphaseB1.txt

| gene_ID | gene_link | gwidth | #Sites | #per10Kb | #dmSites | #dmsp10kb | %dmSites | %pSite1 | %pSite2 | DMpSite | ADMpSite | DMpNucl | ADMpNucl |
|---|---|---|---|---|---|---|---|---|---|---|---|---|---|
| gene-LOC113562345 | https://www.ncbi.nlm.nih.gov/gene/?term=LOC113562345 | 2290 | 71 | 310.04 | 15 | 65.5 | 21.13 | 28.2 | 15.69 | 12.51 | 12.51 | 0.39 | 0.39 |
| gene-LOC105285695 | https://www.ncbi.nlm.nih.gov/gene/?term=LOC105285695 | 1371 | 74 | 539.75 | 1 | 7.29 | 1.35 | 27.97 | 24.61 | 3.36 | 3.36 | 0.18 | 0.18 |
| gene-LOC105275070 | https://www.ncbi.nlm.nih.gov/gene/?term=LOC105275070 | 5667 | 53 | 93.52 | 4 | 7.06 | 7.55 | 16.59 | 5.92 | 10.67 | 10.67 | 0.1 | 0.1 |
| gene-LOC105282156 | https://www.ncbi.nlm.nih.gov/gene/?term=LOC105282156 | 2577 | 128 | 496.7 | 1 | 3.88 | 0.78 | 28.45 | 26.37 | 2.08 | 2.08 | 0.1 | 0.1 |
| gene-LOC105282149 | https://www.ncbi.nlm.nih.gov/gene/?term=LOC105282149 | 1787 | 105 | 587.58 | 1 | 5.6 | 0.95 | 43.64 | 41.97 | 1.67 | 1.67 | 0.1 | 0.1 |
| gene-LOC105276704 | https://www.ncbi.nlm.nih.gov/gene/?term=LOC105276704 | 2063 | 89 | 431.41 | 1 | 4.85 | 1.12 | 14.86 | 13.07 | 1.79 | 1.79 | 0.08 | 0.08 |
| gene-LOC105276078 | https://www.ncbi.nlm.nih.gov/gene/?term=LOC105276078 | 1051 | 14 | 133.21 | 1 | 9.51 | 7.14 | 31.96 | 27.06 | 4.9 | 4.9 | 0.07 | 0.07 |
| gene-LOC105283552 | https://www.ncbi.nlm.nih.gov/gene/?term=LOC105283552 | 1562 | 36 | 230.47 | 1 | 6.4 | 2.78 | 56.28 | 53.31 | 2.97 | 2.97 | 0.07 | 0.07 |
| gene-LOC105287063 | https://www.ncbi.nlm.nih.gov/gene/?term=LOC105287063 | 1874 | 49 | 261.47 | 1 | 5.34 | 2.04 | 32.42 | 29.81 | 2.61 | 2.61 | 0.07 | 0.07 |
| gene-LOC105280774 | https://www.ncbi.nlm.nih.gov/gene/?term=LOC105280774 | 2793 | 101 | 361.62 | 1 | 3.58 | 0.99 | 32.38 | 30.39 | 1.99 | 1.99 | 0.07 | 0.07 |
| gene-LOC105274534 | https://www.ncbi.nlm.nih.gov/gene/?term=LOC105274534 | 1329 | 55 | 413.84 | 1 | 7.52 | 1.82 | 46.07 | 47.79 | -1.72 | 1.72 | -0.07 | 0.07 |
| gene-LOC105285060 | https://www.ncbi.nlm.nih.gov/gene/?term=LOC105285060 | 1285 | 13 | 101.17 | 1 | 7.78 | 7.69 | 56.31 | 50.43 | 5.87 | 5.87 | 0.06 | 0.06 |
| gene-LOC105284704 | https://www.ncbi.nlm.nih.gov/gene/?term=LOC105284704 | 2400 | 62 | 258.33 | 1 | 4.17 | 1.61 | 26.43 | 23.93 | 2.5 | 2.5 | 0.06 | 0.06 |
| gene-LOC105275708 | https://www.ncbi.nlm.nih.gov/gene/?term=LOC105275708 | 1772 | 10 | 56.43 | 1 | 5.64 | 10 | 50.31 | 40.78 | 9.53 | 9.53 | 0.05 | 0.05 |
| gene-LOC105285674 | https://www.ncbi.nlm.nih.gov/gene/?term=LOC105285674 | 3035 | 93 | 306.43 | 1 | 3.29 | 1.08 | 26.91 | 25.27 | 1.64 | 1.64 | 0.05 | 0.05 |
| gene-LOC105284184 | https://www.ncbi.nlm.nih.gov/gene/?term=LOC105284184 | 2535 | 87 | 343.2 | 1 | 3.94 | 1.15 | 34.29 | 32.68 | 1.6 | 1.6 | 0.05 | 0.05 |
| gene-LOC105283123 | https://www.ncbi.nlm.nih.gov/gene/?term=LOC105283123 | 1507 | 51 | 338.42 | 1 | 6.64 | 1.96 | 16.3 | 14.72 | 1.58 | 1.58 | 0.05 | 0.05 |
| gene-LOC105280786 | https://www.ncbi.nlm.nih.gov/gene/?term=LOC105280786 | 2573 | 122 | 474.15 | 1 | 3.89 | 0.82 | 30.88 | 29.81 | 1.07 | 1.07 | 0.05 | 0.05 |
| gene-LOC105285055 | https://www.ncbi.nlm.nih.gov/gene/?term=LOC105285055 | 4108 | 241 | 586.66 | 2 | 4.87 | 0.83 | 12.15 | 11.34 | 0.81 | 0.81 | 0.05 | 0.05 |
| gene-LOC105284213 | https://www.ncbi.nlm.nih.gov/gene/?term=LOC105284213 | 868 | 15 | 172.81 | 1 | 11.52 | 6.67 | 15.38 | 18.37 | -2.99 | 2.99 | -0.05 | 0.05 |
| gene-LOC105285514 | https://www.ncbi.nlm.nih.gov/gene/?term=LOC105285514 | 1277 | 11 | 86.14 | 1 | 7.83 | 9.09 | 90.26 | 85.12 | 5.14 | 5.14 | 0.04 | 0.04 |
| gene-LOC105283712 | https://www.ncbi.nlm.nih.gov/gene/?term=LOC105283712 | 1910 | 29 | 151.83 | 1 | 5.24 | 3.45 | 33.08 | 30.49 | 2.59 | 2.59 | 0.04 | 0.04 |
| gene-LOC105276272 | https://www.ncbi.nlm.nih.gov/gene/?term=LOC105276272 | 1965 | 38 | 193.38 | 1 | 5.09 | 2.63 | 30.65 | 28.5 | 2.15 | 2.15 | 0.04 | 0.04 |
| gene-LOC105281848 | https://www.ncbi.nlm.nih.gov/gene/?term=LOC105281848 | 2122 | 52 | 245.05 | 1 | 4.71 | 1.92 | 26.53 | 24.7 | 1.83 | 1.83 | 0.04 | 0.04 |
| gene-LOC105283697 | https://www.ncbi.nlm.nih.gov/gene/?term=LOC105283697 | 5301 | 137 | 258.44 | 1 | 1.89 | 0.73 | 17.96 | 16.39 | 1.57 | 1.57 | 0.04 | 0.04 |
| gene-LOC105288079 | https://www.ncbi.nlm.nih.gov/gene/?term=LOC105288079 | 3473 | 92 | 264.9 | 1 | 2.88 | 1.09 | 19.48 | 18 | 1.49 | 1.49 | 0.04 | 0.04 |
| gene-LOC105274449 | https://www.ncbi.nlm.nih.gov/gene/?term=LOC105274449 | 1549 | 53 | 342.16 | 1 | 6.46 | 1.89 | 50.38 | 49.29 | 1.09 | 1.09 | 0.04 | 0.04 |
| gene-LOC105287095 | https://www.ncbi.nlm.nih.gov/gene/?term=LOC105287095 | 1435 | 56 | 390.24 | 1 | 6.97 | 1.79 | 30.4 | 29.35 | 1.05 | 1.05 | 0.04 | 0.04 |
| gene-LOC105276449 | https://www.ncbi.nlm.nih.gov/gene/?term=LOC105276449 | 1231 | 43 | 349.31 | 1 | 8.12 | 2.33 | 30.8 | 29.78 | 1.02 | 1.02 | 0.04 | 0.04 |
| gene-LOC105279298 | https://www.ncbi.nlm.nih.gov/gene/?term=LOC105279298 | 2773 | 123 | 443.56 | 1 | 3.61 | 0.81 | 24.88 | 23.88 | 1 | 1 | 0.04 | 0.04 |
| gene-LOC105278497 | https://www.ncbi.nlm.nih.gov/gene/?term=LOC105278497 | 2072 | 89 | 429.54 | 1 | 4.83 | 1.12 | 21.38 | 20.51 | 0.87 | 0.87 | 0.04 | 0.04 |
| gene-LOC105278283 | https://www.ncbi.nlm.nih.gov/gene/?term=LOC105278283 | 2317 | 126 | 543.81 | 1 | 4.32 | 0.79 | 27.76 | 27.04 | 0.72 | 0.72 | 0.04 | 0.04 |
| gene-LOC105278282 | https://www.ncbi.nlm.nih.gov/gene/?term=LOC105278282 | 1775 | 56 | 315.49 | 1 | 5.63 | 1.79 | 33.68 | 34.81 | -1.13 | 1.13 | -0.04 | 0.04 |
| gene-LOC105274900 | https://www.ncbi.nlm.nih.gov/gene/?term=LOC105274900 | 5561 | 51 | 91.71 | 2 | 3.6 | 3.92 | 39.2 | 36.36 | 2.84 | 2.84 | 0.03 | 0.03 |
| gene-LOC105279898 | https://www.ncbi.nlm.nih.gov/gene/?term=LOC105279898 | 1343 | 13 | 96.8 | 1 | 7.45 | 7.69 | 58.71 | 56.01 | 2.69 | 2.69 | 0.03 | 0.03 |
| gene-LOC105288066 | https://www.ncbi.nlm.nih.gov/gene/?term=LOC105288066 | 1964 | 23 | 117.11 | 1 | 5.09 | 4.35 | 46.39 | 44.06 | 2.34 | 2.34 | 0.03 | 0.03 |
| gene-LOC105280348 | https://www.ncbi.nlm.nih.gov/gene/?term=LOC105280348 | 3797 | 65 | 171.19 | 1 | 2.63 | 1.54 | 21.64 | 19.63 | 2.01 | 2.01 | 0.03 | 0.03 |
| gene-LOC105280641 | https://www.ncbi.nlm.nih.gov/gene/?term=LOC105280641 | 3682 | 56 | 152.09 | 1 | 2.72 | 1.79 | 17.9 | 16.23 | 1.66 | 1.66 | 0.03 | 0.03 |
| gene-LOC105280972 | https://www.ncbi.nlm.nih.gov/gene/?term=LOC105280972 | 1532 | 32 | 208.88 | 1 | 6.53 | 3.12 | 19.4 | 18.05 | 1.35 | 1.35 | 0.03 | 0.03 |
| gene-LOC105282885 | https://www.ncbi.nlm.nih.gov/gene/?term=LOC105282885 | 1870 | 38 | 203.21 | 1 | 5.35 | 2.63 | 45.14 | 43.81 | 1.33 | 1.33 | 0.03 | 0.03 |
| gene-LOC105286265 | https://www.ncbi.nlm.nih.gov/gene/?term=LOC105286265 | 2507 | 64 | 255.29 | 1 | 3.99 | 1.56 | 30.69 | 29.49 | 1.21 | 1.21 | 0.03 | 0.03 |
| gene-LOC105278807 | https://www.ncbi.nlm.nih.gov/gene/?term=LOC105278807 | 2166 | 71 | 327.79 | 1 | 4.62 | 1.41 | 19.81 | 18.81 | 1 | 1 | 0.03 | 0.03 |
| gene-LOC105274823 | https://www.ncbi.nlm.nih.gov/gene/?term=LOC105274823 | 4620 | 152 | 329 | 1 | 2.16 | 0.66 | 19.69 | 18.71 | 0.99 | 0.99 | 0.03 | 0.03 |
| gene-LOC105283728 | https://www.ncbi.nlm.nih.gov/gene/?term=LOC105283728 | 3256 | 104 | 319.41 | 2 | 6.14 | 1.92 | 22.23 | 21.4 | 0.83 | 0.83 | 0.03 | 0.03 |
| gene-LOC105285179 | https://www.ncbi.nlm.nih.gov/gene/?term=LOC105285179 | 4191 | 178 | 424.72 | 1 | 2.39 | 0.56 | 17.79 | 17.08 | 0.71 | 0.71 | 0.03 | 0.03 |
| gene-LOC105278145 | https://www.ncbi.nlm.nih.gov/gene/?term=LOC105278145 | 4470 | 250 | 559.28 | 1 | 2.24 | 0.4 | 10.94 | 10.34 | 0.6 | 0.6 | 0.03 | 0.03 |
| gene-LOC105278459 | https://www.ncbi.nlm.nih.gov/gene/?term=LOC105278459 | 2936 | 97 | 330.38 | 1 | 3.41 | 1.03 | 20.46 | 21.23 | -0.77 | 0.77 | -0.03 | 0.03 |
| gene-LOC105281173 | https://www.ncbi.nlm.nih.gov/gene/?term=LOC105281173 | 1595 | 39 | 244.51 | 1 | 6.27 | 2.56 | 60.96 | 62.08 | -1.12 | 1.12 | -0.03 | 0.03 |
| gene-LOC105284507 | https://www.ncbi.nlm.nih.gov/gene/?term=LOC105284507 | 5207 | 86 | 165.16 | 1 | 1.92 | 1.16 | 21.7 | 23.52 | -1.83 | 1.83 | -0.03 | 0.03 |
| gene-LOC105275347 | https://www.ncbi.nlm.nih.gov/gene/?term=LOC105275347 | 2526 | 14 | 55.42 | 1 | 3.96 | 7.14 | 33.57 | 29.73 | 3.84 | 3.84 | 0.02 | 0.02 |
| gene-LOC105288010 | https://www.ncbi.nlm.nih.gov/gene/?term=LOC105288010 | 3189 | 51 | 159.92 | 1 | 3.14 | 1.96 | 34.39 | 32.85 | 1.54 | 1.54 | 0.02 | 0.02 |
| gene-LOC105277243 | https://www.ncbi.nlm.nih.gov/gene/?term=LOC105277243 | 2284 | 32 | 140.11 | 1 | 4.38 | 3.12 | 44.88 | 43.35 | 1.53 | 1.53 | 0.02 | 0.02 |
| gene-LOC105276538 | https://www.ncbi.nlm.nih.gov/gene/?term=LOC105276538 | 1511 | 19 | 125.74 | 1 | 6.62 | 5.26 | 70.28 | 68.86 | 1.42 | 1.42 | 0.02 | 0.02 |
| gene-LOC105278384 | https://www.ncbi.nlm.nih.gov/gene/?term=LOC105278384 | 1390 | 33 | 237.41 | 1 | 7.19 | 3.03 | 46.69 | 45.77 | 0.92 | 0.92 | 0.02 | 0.02 |
| gene-LOC113561805 | https://www.ncbi.nlm.nih.gov/gene/?term=LOC113561805 | 3337 | 69 | 206.77 | 1 | 3 | 1.45 | 38.18 | 37.26 | 0.92 | 0.92 | 0.02 | 0.02 |
| gene-LOC105282794 | https://www.ncbi.nlm.nih.gov/gene/?term=LOC105282794 | 4571 | 99 | 216.58 | 1 | 2.19 | 1.01 | 18.66 | 17.76 | 0.9 | 0.9 | 0.02 | 0.02 |
| gene-LOC105286644 | https://www.ncbi.nlm.nih.gov/gene/?term=LOC105286644 | 1235 | 22 | 178.14 | 1 | 8.1 | 4.55 | 42.04 | 41.14 | 0.89 | 0.89 | 0.02 | 0.02 |
| gene-LOC105283592 | https://www.ncbi.nlm.nih.gov/gene/?term=LOC105283592 | 3479 | 75 | 215.58 | 1 | 2.87 | 1.33 | 25.19 | 24.33 | 0.86 | 0.86 | 0.02 | 0.02 |
| gene-LOC105281312 | https://www.ncbi.nlm.nih.gov/gene/?term=LOC105281312 | 2551 | 68 | 266.56 | 1 | 3.92 | 1.47 | 31.46 | 30.8 | 0.65 | 0.65 | 0.02 | 0.02 |
| gene-LOC105284355 | https://www.ncbi.nlm.nih.gov/gene/?term=LOC105284355 | 3904 | 143 | 366.29 | 1 | 2.56 | 0.7 | 22.99 | 22.35 | 0.64 | 0.64 | 0.02 | 0.02 |

| | | | | | | | | | | | | | | | |
|---|---|---|---|---|---|---|---|---|---|---|---|---|---|---|---|
| gene-LOC105285671 | https://www.ncbi.nlm.nih.gov/gene/?term=LOC105285671 | 2263 | 99 | 437.47 | | 1 | 4.42 | 1.01 | 30.99 | 30.44 | 0.55 | 0.55 | 0.02 | 0.02 | |
| gene-LOC105275460 | https://www.ncbi.nlm.nih.gov/gene/?term=LOC105275460 | 2473 | 118 | 477.15 | | 1 | 4.04 | 0.85 | 13.46 | 12.95 | 0.51 | 0.51 | 0.02 | 0.02 | |
| gene-LOC105278632 | https://www.ncbi.nlm.nih.gov/gene/?term=LOC105278632 | 3624 | 131 | 361.48 | | 1 | 2.76 | 0.76 | 14.47 | 14.04 | 0.43 | 0.43 | 0.02 | 0.02 | |
| gene-LOC105276462 | https://www.ncbi.nlm.nih.gov/gene/?term=LOC105276462 | 6561 | 275 | 419.14 | | 1 | 1.52 | 0.36 | 8.32 | 7.94 | 0.37 | 0.37 | 0.02 | 0.02 | |
| gene-LOC105277435 | https://www.ncbi.nlm.nih.gov/gene/?term=LOC105277435 | 3643 | 175 | 480.37 | | 1 | 2.74 | 0.57 | 11.3 | 10.97 | 0.33 | 0.33 | 0.02 | 0.02 | |
| gene-LOC105286649 | https://www.ncbi.nlm.nih.gov/gene/?term=LOC105286649 | 14178 | 785 | 553.67 | | 1 | 0.71 | 0.13 | 0.94 | 1.3 | -0.36 | 0.36 | -0.02 | 0.02 | |
| gene-LOC105280956 | https://www.ncbi.nlm.nih.gov/gene/?term=LOC105280956 | 2639 | 77 | 291.78 | | 1 | 3.79 | 1.3 | 18.18 | 18.82 | -0.64 | 0.64 | -0.02 | 0.02 | |
| gene-LOC105286515 | https://www.ncbi.nlm.nih.gov/gene/?term=LOC105286515 | 3751 | 113 | 301.25 | | 1 | 2.67 | 0.88 | 10.68 | 11.31 | -0.64 | 0.64 | -0.02 | 0.02 | |
| gene-LOC105278454 | https://www.ncbi.nlm.nih.gov/gene/?term=LOC105278454 | 1972 | 41 | 207.91 | | 1 | 5.07 | 2.44 | 43.16 | 44.07 | -0.91 | 0.91 | -0.02 | 0.02 | |
| gene-LOC105287633 | https://www.ncbi.nlm.nih.gov/gene/?term=LOC105287633 | 1524 | 23 | 150.92 | | 1 | 6.56 | 4.35 | 68.63 | 69.73 | -1.11 | 1.11 | -0.02 | 0.02 | |
| gene-LOC105285252 | https://www.ncbi.nlm.nih.gov/gene/?term=LOC105285252 | 7427 | 124 | 166.96 | | 2 | 2.69 | 1.61 | 12.83 | 14.05 | -1.22 | 1.22 | -0.02 | 0.02 | |
| gene-LOC105285315 | https://www.ncbi.nlm.nih.gov/gene/?term=LOC105285315 | 5458 | 21 | 38.48 | 1 | 1.83 | 4.76 | 43.16 | 40.32 | 2.84 | 2.84 | 0.01 | 0.01 | | |
| gene-LOC105280408 | https://www.ncbi.nlm.nih.gov/gene/?term=LOC105280408 | 5983 | 50 | 83.57 | 1 | 1.67 | 2 | 19.04 | 17.37 | 1.67 | 1.67 | 0.01 | 0.01 | | |
| gene-LOC105281515 | https://www.ncbi.nlm.nih.gov/gene/?term=LOC105281515 | 8827 | 48 | 54.38 | 2 | 2.27 | 4.17 | 32.1 | 30.7 | 1.4 | 1.4 | 0.01 | 0.01 | | |
| gene-LOC105279500 | https://www.ncbi.nlm.nih.gov/gene/?term=LOC105279500 | 3587 | 33 | 92 | 1 | 2.79 | 3.03 | 51.88 | 50.67 | 1.21 | 1.21 | 0.01 | 0.01 | | |
| gene-LOC105275518 | https://www.ncbi.nlm.nih.gov/gene/?term=LOC105275518 | 2335 | 19 | 81.37 | 1 | 4.28 | 5.26 | 51.76 | 50.67 | 1.09 | 1.09 | 0.01 | 0.01 | | |
| gene-LOC105279664 | https://www.ncbi.nlm.nih.gov/gene/?term=LOC105279664 | 4023 | 57 | 141.69 | | 1 | 2.49 | 1.75 | 22.82 | 21.77 | 1.05 | 1.05 | 0.01 | 0.01 | |
| gene-LOC105277215 | https://www.ncbi.nlm.nih.gov/gene/?term=LOC105277215 | 2619 | 23 | 87.82 | 1 | 3.82 | 4.35 | 35.64 | 34.71 | 0.93 | 0.93 | 0.01 | 0.01 | | |
| gene-LOC105283838 | https://www.ncbi.nlm.nih.gov/gene/?term=LOC105283838 | 1748 | 21 | 120.14 | | 1 | 5.72 | 4.76 | 46.37 | 45.45 | 0.92 | 0.92 | 0.01 | 0.01 | |
| gene-LOC105278109 | https://www.ncbi.nlm.nih.gov/gene/?term=LOC105278109 | 4029 | 65 | 161.33 | | 1 | 2.48 | 1.54 | 46.32 | 45.45 | 0.88 | 0.88 | 0.01 | 0.01 | |
| gene-LOC105276560 | https://www.ncbi.nlm.nih.gov/gene/?term=LOC105276560 | 18607 | 319 | 171.44 | | 1 | 0.54 | 0.31 | 2.53 | 1.85 | 0.67 | 0.67 | 0.01 | 0.01 | |
| gene-LOC105286514 | https://www.ncbi.nlm.nih.gov/gene/?term=LOC105286514 | 2528 | 43 | 170.09 | | 1 | 3.96 | 2.33 | 34.72 | 34.07 | 0.65 | 0.65 | 0.01 | 0.01 | |
| gene-LOC105276059 | https://www.ncbi.nlm.nih.gov/gene/?term=LOC105276059 | 6256 | 123 | 196.61 | | 1 | 1.6 | 0.81 | 12.44 | 11.86 | 0.58 | 0.58 | 0.01 | 0.01 | |
| gene-LOC105280407 | https://www.ncbi.nlm.nih.gov/gene/?term=LOC105280407 | 10950 | 203 | 185.39 | | 1 | 0.91 | 0.49 | 15.56 | 14.99 | 0.56 | 0.56 | 0.01 | 0.01 | |
| gene-LOC105285395 | https://www.ncbi.nlm.nih.gov/gene/?term=LOC105285395 | 8085 | 207 | 256.03 | | 1 | 1.24 | 0.48 | 14.3 | 13.76 | 0.54 | 0.54 | 0.01 | 0.01 | |
| gene-LOC105280819 | https://www.ncbi.nlm.nih.gov/gene/?term=LOC105280819 | 1639 | 39 | 237.95 | | 1 | 6.1 | 2.56 | 35.99 | 35.46 | 0.53 | 0.53 | 0.01 | 0.01 | |
| gene-LOC105275451 | https://www.ncbi.nlm.nih.gov/gene/?term=LOC105275451 | 6035 | 133 | 220.38 | | 1 | 1.66 | 0.75 | 22.24 | 21.72 | 0.51 | 0.51 | 0.01 | 0.01 | |
| gene-LOC105278534 | https://www.ncbi.nlm.nih.gov/gene/?term=LOC105278534 | 7364 | 180 | 244.43 | | 1 | 1.36 | 0.56 | 18.01 | 17.6 | 0.41 | 0.41 | 0.01 | 0.01 | |
| gene-LOC105280890 | https://www.ncbi.nlm.nih.gov/gene/?term=LOC105280890 | 1139 | 29 | 254.61 | | 1 | 8.78 | 3.45 | 19.62 | 19.23 | 0.39 | 0.39 | 0.01 | 0.01 | |
| gene-LOC105286999 | https://www.ncbi.nlm.nih.gov/gene/?term=LOC105286999 | 2141 | 44 | 205.51 | | 1 | 4.67 | 2.27 | 35.31 | 34.92 | 0.39 | 0.39 | 0.01 | 0.01 | |
| gene-LOC105280410 | https://www.ncbi.nlm.nih.gov/gene/?term=LOC105280410 | 12580 | 265 | 210.65 | | 1 | 0.79 | 0.38 | 19.92 | 19.57 | 0.35 | 0.35 | 0.01 | 0.01 | |
| gene-LOC105283296 | https://www.ncbi.nlm.nih.gov/gene/?term=LOC105283296 | 2174 | 86 | 395.58 | | 1 | 4.6 | 1.16 | 16.05 | 15.7 | 0.35 | 0.35 | 0.01 | 0.01 | |
| gene-LOC105278666 | https://www.ncbi.nlm.nih.gov/gene/?term=LOC105278666 | 10341 | 456 | 440.96 | | 1 | 0.97 | 0.22 | 7.52 | 7.19 | 0.34 | 0.34 | 0.01 | 0.01 | |
| gene-LOC105279657 | https://www.ncbi.nlm.nih.gov/gene/?term=LOC105279657 | 5878 | 96 | 163.32 | | 1 | 1.7 | 1.04 | 18.1 | 17.76 | 0.34 | 0.34 | 0.01 | 0.01 | |
| gene-LOC105284036 | https://www.ncbi.nlm.nih.gov/gene/?term=LOC105284036 | 3254 | 87 | 267.36 | | 1 | 3.07 | 1.15 | 24.27 | 23.96 | 0.31 | 0.31 | 0.01 | 0.01 | |
| gene-LOC105281272 | https://www.ncbi.nlm.nih.gov/gene/?term=LOC105281272 | 5252 | 175 | 333.21 | | 1 | 1.9 | 0.57 | 16.26 | 15.96 | 0.3 | 0.3 | 0.01 | 0.01 | |
| gene-LOC105285807 | https://www.ncbi.nlm.nih.gov/gene/?term=LOC105285807 | 6063 | 207 | 341.42 | | 1 | 1.65 | 0.48 | 7.17 | 6.87 | 0.3 | 0.3 | 0.01 | 0.01 | |
| gene-LOC105282491 | https://www.ncbi.nlm.nih.gov/gene/?term=LOC105282491 | 4761 | 170 | 357.07 | | 1 | 2.1 | 0.59 | 7.92 | 7.67 | 0.25 | 0.25 | 0.01 | 0.01 | |
| gene-LOC105274706 | https://www.ncbi.nlm.nih.gov/gene/?term=LOC105274706 | 1446 | 50 | 345.78 | | 1 | 6.92 | 2 | 39.23 | 39 | 0.23 | 0.23 | 0.01 | 0.01 | |
| gene-LOC105283649 | https://www.ncbi.nlm.nih.gov/gene/?term=LOC105283649 | 1918 | 55 | 286.76 | | 1 | 5.21 | 1.82 | 38.85 | 38.63 | 0.21 | 0.21 | 0.01 | 0.01 | |
| gene-LOC105275374 | https://www.ncbi.nlm.nih.gov/gene/?term=LOC105275374 | 11376 | 525 | 461.5 | 1 | 0.88 | 0.19 | 10.23 | 10.06 | 0.17 | 0.17 | 0.01 | 0.01 | | |
| gene-LOC105282658 | https://www.ncbi.nlm.nih.gov/gene/?term=LOC105282658 | 2421 | 67 | 276.75 | | 1 | 4.13 | 1.49 | 20.83 | 21.03 | -0.21 | 0.21 | -0.01 | 0.01 | |
| gene-LOC105285981 | https://www.ncbi.nlm.nih.gov/gene/?term=LOC105285981 | 7666 | 183 | 238.72 | | 1 | 1.3 | 0.55 | 18.71 | 18.92 | -0.21 | 0.21 | -0.01 | 0.01 | |
| gene-LOC105278877 | https://www.ncbi.nlm.nih.gov/gene/?term=LOC105278877 | 1968 | 77 | 391.26 | | 1 | 5.08 | 1.3 | 34.55 | 34.76 | -0.22 | 0.22 | -0.01 | 0.01 | |
| gene-LOC105274851 | https://www.ncbi.nlm.nih.gov/gene/?term=LOC105274851 | 6616 | 127 | 191.96 | | 1 | 1.51 | 0.79 | 11.51 | 11.83 | -0.33 | 0.33 | -0.01 | 0.01 | |
| gene-LOC105285742 | https://www.ncbi.nlm.nih.gov/gene/?term=LOC105285742 | 3446 | 97 | 281.49 | | 1 | 2.9 | 1.03 | 20.21 | 20.62 | -0.41 | 0.41 | -0.01 | 0.01 | |
| gene-LOC105274590 | https://www.ncbi.nlm.nih.gov/gene/?term=LOC105274590 | 4516 | 64 | 141.72 | | 2 | 4.43 | 3.12 | 32.42 | 32.94 | -0.51 | 0.51 | -0.01 | 0.01 | |
| gene-LOC105284146 | https://www.ncbi.nlm.nih.gov/gene/?term=LOC105284146 | 2488 | 66 | 265.27 | | 1 | 4.02 | 1.52 | 23.09 | 23.62 | -0.54 | 0.54 | -0.01 | 0.01 | |
| gene-LOC105278228 | https://www.ncbi.nlm.nih.gov/gene/?term=LOC105278228 | 8610 | 169 | 196.28 | | 1 | 1.16 | 0.59 | 21.96 | 22.51 | -0.55 | 0.55 | -0.01 | 0.01 | |
| gene-LOC105283989 | https://www.ncbi.nlm.nih.gov/gene/?term=LOC105283989 | 4948 | 25 | 50.53 | 1 | 2.02 | 4 | 46.26 | 45.67 | 0.59 | 0.59 | 0 | 0 | | |
| gene-LOC105275187 | https://www.ncbi.nlm.nih.gov/gene/?term=LOC105275187 | 5534 | 69 | 124.68 | | 1 | 1.81 | 1.45 | 14.14 | 13.85 | 0.29 | 0.29 | 0 | 0 | |
| gene-LOC105281700 | https://www.ncbi.nlm.nih.gov/gene/?term=LOC105281700 | 2208 | 38 | 172.1 | 1 | 4.53 | 2.63 | 39.22 | 39.09 | 0.14 | 0.14 | 0 | 0 | | |
| gene-LOC105274531 | https://www.ncbi.nlm.nih.gov/gene/?term=LOC105274531 | 7774 | 283 | 364.03 | | 1 | 1.29 | 0.35 | 10.82 | 10.73 | 0.09 | 0.09 | 0 | 0 | |
| gene-LOC105282825 | https://www.ncbi.nlm.nih.gov/gene/?term=LOC105282825 | 3919 | 78 | 199.03 | | 1 | 2.55 | 1.28 | 12.99 | 12.9 | 0.09 | 0.09 | 0 | 0 | |
| gene-LOC105275179 | https://www.ncbi.nlm.nih.gov/gene/?term=LOC105275179 | 3271 | 71 | 217.06 | | 1 | 3.06 | 1.41 | 16.78 | 16.7 | 0.08 | 0.08 | 0 | 0 | |
| gene-LOC105279038 | https://www.ncbi.nlm.nih.gov/gene/?term=LOC105279038 | 9453 | 317 | 335.34 | | 1 | 1.06 | 0.32 | 9.53 | 9.46 | 0.07 | 0.07 | 0 | 0 | |
| gene-LOC105283987 | https://www.ncbi.nlm.nih.gov/gene/?term=LOC105283987 | 4841 | 71 | 146.66 | | 1 | 2.07 | 1.41 | 32.31 | 32.24 | 0.07 | 0.07 | 0 | 0 | |
| gene-LOC105284329 | https://www.ncbi.nlm.nih.gov/gene/?term=LOC105284329 | 4051 | 51 | 125.89 | | 1 | 2.47 | 1.96 | 19.38 | 19.32 | 0.06 | 0.06 | 0 | 0 | |
| gene-LOC105286194 | https://www.ncbi.nlm.nih.gov/gene/?term=LOC105286194 | 152619 | | 2202 | 144.28 | | 1 | 0.07 | 0.05 | 6.22 | 6.17 | 0.04 | 0.04 | 0 | 0 |
| gene-LOC105288078 | https://www.ncbi.nlm.nih.gov/gene/?term=LOC105288078 | 19126 | 307 | 160.51 | | 1 | 0.52 | 0.33 | 16.81 | 16.78 | 0.03 | 0.03 | 0 | 0 | |
| gene-LOC105276527 | https://www.ncbi.nlm.nih.gov/gene/?term=LOC105276527 | 7923 | 334 | 421.56 | | 1 | 1.26 | 0.3 | 8.56 | 8.54 | 0.02 | 0.02 | 0 | 0 | |
| gene-LOC105286452 | https://www.ncbi.nlm.nih.gov/gene/?term=LOC105286452 | 6888 | 68 | 98.72 | 1 | 1.45 | 1.47 | 28.88 | 28.89 | 0 | 0 | 0 | 0 | | |
| gene-LOC105274856 | https://www.ncbi.nlm.nih.gov/gene/?term=LOC105274856 | 5216 | 212 | 406.44 | | 1 | 1.92 | 0.47 | 18.3 | 18.31 | -0.01 | 0.01 | 0 | 0 | |
| gene-LOC105275327 | https://www.ncbi.nlm.nih.gov/gene/?term=LOC105275327 | 14245 | 472 | 331.34 | | 1 | 0.7 | 0.21 | 7.13 | 7.15 | -0.03 | 0.03 | 0 | 0 | |

| | | | | | | | | | | | | | |
|---|---|---|---|---|---|---|---|---|---|---|---|---|---|
| gene-LOC105283943 | https://www.ncbi.nlm.nih.gov/gene/?term=LOC105283943 | 3559 | 56 | 157.35 | 1 | 2.81 | 1.79 | 16.7 | 16.73 | -0.03 | 0.03 | 0 | 0 |
| gene-LOC105284043 | https://www.ncbi.nlm.nih.gov/gene/?term=LOC105284043 | 5661 | 278 | 491.08 | 1 | 1.77 | 0.36 | 10.84 | 10.87 | -0.03 | 0.03 | 0 | 0 |
| gene-LOC105275683 | https://www.ncbi.nlm.nih.gov/gene/?term=LOC105275683 | 4256 | 191 | 448.78 | 1 | 2.35 | 0.52 | 12.17 | 12.21 | -0.04 | 0.04 | 0 | 0 |
| gene-LOC105277866 | https://www.ncbi.nlm.nih.gov/gene/?term=LOC105277866 | 33045 | 409 | 123.77 | 1 | 0.3 | 0.24 | 3.12 | 3.19 | -0.07 | 0.07 | 0 | 0 |
| gene-LOC105277216 | https://www.ncbi.nlm.nih.gov/gene/?term=LOC105277216 | 7627 | 87 | 114.07 | 1 | 1.31 | 1.15 | 9.6 | 9.67 | -0.08 | 0.08 | 0 | 0 |
| gene-LOC105277125 | https://www.ncbi.nlm.nih.gov/gene/?term=LOC105277125 | 51079 | 1802 | 352.79 | 1 | 0.2 | 0.06 | 1.61 | 1.7 | -0.09 | 0.09 | 0 | 0 |
| gene-LOC105278455 | https://www.ncbi.nlm.nih.gov/gene/?term=LOC105278455 | 27181 | 915 | 336.63 | 1 | 0.37 | 0.11 | 7.11 | 7.23 | -0.12 | 0.12 | 0 | 0 |
| gene-LOC105280265 | https://www.ncbi.nlm.nih.gov/gene/?term=LOC105280265 | 85984 | 3009 | 349.95 | 1 | 0.12 | 0.03 | 1.05 | 1.18 | -0.12 | 0.12 | 0 | 0 |
| gene-LOC105287417 | https://www.ncbi.nlm.nih.gov/gene/?term=LOC105287417 | 1025 | 28 | 273.17 | 1 | 9.76 | 3.57 | 42.07 | 42.23 | -0.16 | 0.16 | 0 | 0 |
| gene-LOC105284147 | https://www.ncbi.nlm.nih.gov/gene/?term=LOC105284147 | 77820 | 534 | 68.62 1 | 0.13 | 0.19 | 9 | 9.26 | -0.26 | 0.26 | 0 | 0 | |
| gene-LOC105284774 | https://www.ncbi.nlm.nih.gov/gene/?term=LOC105284774 | 12089 | 81 | 67 1 | 0.83 | 1.23 | 21 | 21.27 | -0.27 | 0.27 | 0 | 0 | |
| gene-LOC105276196 | https://www.ncbi.nlm.nih.gov/gene/?term=LOC105276196 | 2953 | 45 | 152.39 | 1 | 3.39 | 2.22 | 48.59 | 48.91 | -0.31 | 0.31 | 0 | 0 |

Directory: OGLall File: rnk-dmg-Ob_LI_BCphaseA2.vs.RphaseB2.txt

| gene_ID | gene_link | gwidth | #Sites | #per10Kb | #dmSites | #dmsp10kb | %dmSites | %pSite1 | %pSite2 | DMpSite | ADMpSite | DMpNucl | ADMpNucl |
|---|---|---|---|---|---|---|---|---|---|---|---|---|---|
| gene-LOC113562345 | https://www.ncbi.nlm.nih.gov/gene/?term=LOC113562345 | 2290 | 67 | 292.58 | 11 | 48.03 | 16.42 | 29.88 | 17.32 | 12.57 | 12.57 | 0.37 | 0.37 |
| gene-LOC105284677 | https://www.ncbi.nlm.nih.gov/gene/?term=LOC105284677 | 1905 | 39 | 204.72 | 3 | 15.75 | 7.69 | 19.98 | 31.6 | -11.62 | 11.62 | -0.24 | 0.24 |
| gene-LOC113562901 | https://www.ncbi.nlm.nih.gov/gene/?term=LOC113562901 | 1531 | 22 | 143.7 | 1 | 6.53 | 4.55 | 60.1 | 44.57 | 15.53 | 15.53 | 0.22 | 0.22 |
| gene-LOC105282976 | https://www.ncbi.nlm.nih.gov/gene/?term=LOC105282976 | 1648 | 47 | 285.19 | 1 | 6.07 | 2.13 | 28.02 | 22.43 | 5.59 | 5.59 | 0.16 | 0.16 |
| gene-LOC105283552 | https://www.ncbi.nlm.nih.gov/gene/?term=LOC105283552 | 1562 | 36 | 230.47 | 1 | 6.4 | 2.78 | 56.28 | 51.7 | 4.58 | 4.58 | 0.11 | 0.11 |
| gene-LOC105282149 | https://www.ncbi.nlm.nih.gov/gene/?term=LOC105282149 | 1787 | 108 | 604.36 | 1 | 5.6 | 0.93 | 43.65 | 41.89 | 1.76 | 1.76 | 0.11 | 0.11 |
| gene-LOC105279898 | https://www.ncbi.nlm.nih.gov/gene/?term=LOC105279898 | 1343 | 10 | 74.46 | 1 | 7.45 | 10 | 84.43 | 71.53 | 12.89 | 12.89 | 0.1 | 0.1 |
| gene-LOC105275070 | https://www.ncbi.nlm.nih.gov/gene/?term=LOC105275070 | 5667 | 41 | 72.35 | 5 | 8.82 | 12.2 | 13.7 | 1.06 | 12.65 | 12.65 | 0.09 | 0.09 |
| gene-LOC105286644 | https://www.ncbi.nlm.nih.gov/gene/?term=LOC105286644 | 1235 | 22 | 178.14 | 1 | 8.1 | 4.55 | 42.35 | 37.26 | 5.09 | 5.09 | 0.09 | 0.09 |
| gene-LOC105285674 | https://www.ncbi.nlm.nih.gov/gene/?term=LOC105285674 | 3035 | 93 | 306.43 | 2 | 6.59 | 2.15 | 26.74 | 23.71 | 3.03 | 3.03 | 0.09 | 0.09 |
| gene-LOC105276704 | https://www.ncbi.nlm.nih.gov/gene/?term=LOC105276704 | 2063 | 88 | 426.56 | 1 | 4.85 | 1.14 | 15.96 | 13.92 | 2.05 | 2.05 | 0.09 | 0.09 |
| gene-LOC105285671 | https://www.ncbi.nlm.nih.gov/gene/?term=LOC105285671 | 2263 | 103 | 455.15 | 1 | 4.42 | 0.97 | 29.21 | 27.3 | 1.91 | 1.91 | 0.09 | 0.09 |
| gene-LOC105282156 | https://www.ncbi.nlm.nih.gov/gene/?term=LOC105282156 | 2577 | 129 | 500.58 | 1 | 3.88 | 0.78 | 28.68 | 26.84 | 1.84 | 1.84 | 0.09 | 0.09 |
| gene-LOC105277565 | https://www.ncbi.nlm.nih.gov/gene/?term=LOC105277565 | 3216 | 169 | 525.5 | 1 | 3.11 | 0.59 | 20.65 | 19.2 | 1.45 | 1.45 | 0.08 | 0.08 |
| gene-LOC105283649 | https://www.ncbi.nlm.nih.gov/gene/?term=LOC105283649 | 1918 | 54 | 281.54 | 1 | 5.21 | 1.85 | 37.63 | 35.03 | 2.6 | 2.6 | 0.07 | 0.07 |
| gene-LOC105280786 | https://www.ncbi.nlm.nih.gov/gene/?term=LOC105280786 | 2573 | 113 | 439.18 | 1 | 3.89 | 0.88 | 32.35 | 30.66 | 1.69 | 1.69 | 0.07 | 0.07 |
| gene-LOC105278215 | https://www.ncbi.nlm.nih.gov/gene/?term=LOC105278215 | 1700 | 29 | 170.59 | 1 | 5.88 | 3.45 | 57.89 | 54.66 | 3.23 | 3.23 | 0.06 | 0.06 |
| gene-LOC105283123 | https://www.ncbi.nlm.nih.gov/gene/?term=LOC105283123 | 1507 | 44 | 291.97 | 1 | 6.64 | 2.27 | 19.91 | 17.87 | 2.05 | 2.05 | 0.06 | 0.06 |
| gene-LOC105278497 | https://www.ncbi.nlm.nih.gov/gene/?term=LOC105278497 | 2072 | 92 | 444.02 | 1 | 4.83 | 1.09 | 21.61 | 20.36 | 1.26 | 1.26 | 0.06 | 0.06 |
| gene-LOC105279298 | https://www.ncbi.nlm.nih.gov/gene/?term=LOC105279298 | 2773 | 131 | 472.41 | 2 | 7.21 | 1.53 | 24.81 | 23.62 | 1.18 | 1.18 | 0.06 | 0.06 |
| gene-LOC105276078 | https://www.ncbi.nlm.nih.gov/gene/?term=LOC105276078 | 1051 | 14 | 133.21 | 1 | 9.51 | 7.14 | 36.98 | 32.93 | 4.05 | 4.05 | 0.05 | 0.05 |
| gene-LOC105288063 | https://www.ncbi.nlm.nih.gov/gene/?term=LOC105288063 | 3400 | 43 | 126.47 | 1 | 2.94 | 2.33 | 18.5 | 14.57 | 3.92 | 3.92 | 0.05 | 0.05 |
| gene-LOC105278109 | https://www.ncbi.nlm.nih.gov/gene/?term=LOC105278109 | 4029 | 59 | 146.44 | 1 | 2.48 | 1.69 | 50.99 | 47.24 | 3.75 | 3.75 | 0.05 | 0.05 |
| gene-LOC105281754 | https://www.ncbi.nlm.nih.gov/gene/?term=LOC105281754 | 4300 | 110 | 255.81 | 1 | 2.33 | 0.91 | 16.49 | 14.51 | 1.98 | 1.98 | 0.05 | 0.05 |
| gene-LOC105284184 | https://www.ncbi.nlm.nih.gov/gene/?term=LOC105284184 | 2535 | 79 | 311.64 | 1 | 3.94 | 1.27 | 37.02 | 35.48 | 1.54 | 1.54 | 0.05 | 0.05 |
| gene-LOC105277761 | https://www.ncbi.nlm.nih.gov/gene/?term=LOC105277761 | 4648 | 143 | 307.66 | 1 | 2.15 | 0.7 | 15.33 | 13.81 | 1.52 | 1.52 | 0.05 | 0.05 |
| gene-LOC105285055 | https://www.ncbi.nlm.nih.gov/gene/?term=LOC105285055 | 4108 | 255 | 620.74 | 2 | 4.87 | 0.78 | 11.66 | 10.91 | 0.76 | 0.76 | 0.05 | 0.05 |
| gene-LOC105274534 | https://www.ncbi.nlm.nih.gov/gene/?term=LOC105274534 | 1329 | 56 | 421.37 | 1 | 7.52 | 1.79 | 47.04 | 48.26 | -1.22 | 1.22 | -0.05 | 0.05 |
| gene-LOC105285060 | https://www.ncbi.nlm.nih.gov/gene/?term=LOC105285060 | 1285 | 17 | 132.3 | 1 | 7.78 | 5.88 | 51.2 | 48.29 | 2.91 | 2.91 | 0.04 | 0.04 |
| gene-LOC113561805 | https://www.ncbi.nlm.nih.gov/gene/?term=LOC113561805 | 3337 | 68 | 203.78 | 1 | 3 | 1.47 | 35.02 | 33.08 | 1.94 | 1.94 | 0.04 | 0.04 |
| gene-LOC105282970 | https://www.ncbi.nlm.nih.gov/gene/?term=LOC105282970 | 1529 | 39 | 255.07 | 1 | 6.54 | 2.56 | 51.08 | 49.46 | 1.62 | 1.62 | 0.04 | 0.04 |
| gene-LOC105278807 | https://www.ncbi.nlm.nih.gov/gene/?term=LOC105278807 | 2166 | 62 | 286.24 | 1 | 4.62 | 1.61 | 21.98 | 20.52 | 1.46 | 1.46 | 0.04 | 0.04 |
| gene-LOC105283728 | https://www.ncbi.nlm.nih.gov/gene/?term=LOC105283728 | 3256 | 107 | 328.62 | 2 | 6.14 | 1.87 | 20.96 | 19.59 | 1.37 | 1.37 | 0.04 | 0.04 |
| gene-LOC105276875 | https://www.ncbi.nlm.nih.gov/gene/?term=LOC105276875 | 3151 | 109 | 345.92 | 1 | 3.17 | 0.92 | 25.45 | 24.19 | 1.26 | 1.26 | 0.04 | 0.04 |
| gene-LOC105284355 | https://www.ncbi.nlm.nih.gov/gene/?term=LOC105284355 | 3904 | 143 | 366.29 | 1 | 2.56 | 0.7 | 22.85 | 21.77 | 1.08 | 1.08 | 0.04 | 0.04 |
| gene-LOC105285179 | https://www.ncbi.nlm.nih.gov/gene/?term=LOC105285179 | 4191 | 177 | 422.33 | 1 | 2.39 | 0.56 | 19.66 | 18.63 | 1.04 | 1.04 | 0.04 | 0.04 |
| gene-LOC105276196 | https://www.ncbi.nlm.nih.gov/gene/?term=LOC105276196 | 2953 | 48 | 162.55 | 1 | 3.39 | 2.08 | 47.64 | 49.83 | -2.19 | 2.19 | -0.04 | 0.04 |
| gene-LOC105275708 | https://www.ncbi.nlm.nih.gov/gene/?term=LOC105275708 | 1772 | 10 | 56.43 | 1 | 5.64 | 10 | 50.31 | 44.22 | 6.09 | 6.09 | 0.03 | 0.03 |
| gene-LOC105279664 | https://www.ncbi.nlm.nih.gov/gene/?term=LOC105279664 | 4023 | 48 | 119.31 | 1 | 2.49 | 2.08 | 23.51 | 21.34 | 2.18 | 2.18 | 0.03 | 0.03 |
| gene-LOC105276190 | https://www.ncbi.nlm.nih.gov/gene/?term=LOC105276190 | 4839 | 82 | 169.46 | 1 | 2.07 | 1.22 | 11.63 | 9.92 | 1.71 | 1.71 | 0.03 | 0.03 |
| gene-LOC105282794 | https://www.ncbi.nlm.nih.gov/gene/?term=LOC105282794 | 4571 | 92 | 201.27 | 1 | 2.19 | 1.09 | 18.21 | 16.51 | 1.7 | 1.7 | 0.03 | 0.03 |
| gene-LOC105276059 | https://www.ncbi.nlm.nih.gov/gene/?term=LOC105276059 | 6256 | 121 | 193.41 | 1 | 1.6 | 0.83 | 10.95 | 9.59 | 1.36 | 1.36 | 0.03 | 0.03 |
| gene-LOC105281173 | https://www.ncbi.nlm.nih.gov/gene/?term=LOC105281173 | 1595 | 38 | 238.24 | 1 | 6.27 | 2.63 | 60.05 | 58.85 | 1.2 | 1.2 | 0.03 | 0.03 |
| gene-LOC105287063 | https://www.ncbi.nlm.nih.gov/gene/?term=LOC105287063 | 1874 | 48 | 256.14 | 1 | 5.34 | 2.08 | 31.5 | 30.33 | 1.17 | 1.17 | 0.03 | 0.03 |
| gene-LOC105281272 | https://www.ncbi.nlm.nih.gov/gene/?term=LOC105281272 | 5252 | 156 | 297.03 | 1 | 1.9 | 0.64 | 18.91 | 17.94 | 0.97 | 0.97 | 0.03 | 0.03 |
| gene-LOC105280774 | https://www.ncbi.nlm.nih.gov/gene/?term=LOC105280774 | 2793 | 89 | 318.65 | 1 | 3.58 | 1.12 | 36.33 | 35.38 | 0.94 | 0.94 | 0.03 | 0.03 |
| gene-LOC105276449 | https://www.ncbi.nlm.nih.gov/gene/?term=LOC105276449 | 1231 | 42 | 341.19 | 1 | 8.12 | 2.38 | 35.95 | 35.13 | 0.82 | 0.82 | 0.03 | 0.03 |
| gene-LOC105285202 | https://www.ncbi.nlm.nih.gov/gene/?term=LOC105285202 | 1838 | 67 | 364.53 | 1 | 5.44 | 1.49 | 26.25 | 25.49 | 0.76 | 0.76 | 0.03 | 0.03 |
| gene-LOC105275363 | https://www.ncbi.nlm.nih.gov/gene/?term=LOC105275363 | 3798 | 150 | 394.94 | 1 | 2.63 | 0.67 | 18.12 | 17.37 | 0.75 | 0.75 | 0.03 | 0.03 |
| gene-LOC105287095 | https://www.ncbi.nlm.nih.gov/gene/?term=LOC105287095 | 1435 | 57 | 397.21 | 1 | 6.97 | 1.75 | 29.87 | 29.16 | 0.71 | 0.71 | 0.03 | 0.03 |
| gene-LOC105275374 | https://www.ncbi.nlm.nih.gov/gene/?term=LOC105275374 | 11376 | 489 | 429.85 | 1 | 0.88 | 0.2 | 11.7 | 11.06 | 0.64 | 0.64 | 0.03 | 0.03 |
| gene-LOC105278145 | https://www.ncbi.nlm.nih.gov/gene/?term=LOC105278145 | 4470 | 246 | 550.34 | 1 | 2.24 | 0.41 | 11.03 | 10.48 | 0.55 | 0.55 | 0.03 | 0.03 |
| gene-LOC105278283 | https://www.ncbi.nlm.nih.gov/gene/?term=LOC105278283 | 2317 | 125 | 539.49 | 1 | 4.32 | 0.8 | 28.1 | 27.55 | 0.55 | 0.55 | 0.03 | 0.03 |
| gene-LOC105276452 | https://www.ncbi.nlm.nih.gov/gene/?term=LOC105276452 | 2917 | 263 | 901.61 | 1 | 3.43 | 0.38 | 9.41 | 9.13 | 0.28 | 0.28 | 0.03 | 0.03 |
| gene-LOC109611283 | https://www.ncbi.nlm.nih.gov/gene/?term=LOC109611283 | 1451 | 36 | 248.1 | 1 | 6.89 | 2.78 | 35.62 | 36.97 | -1.35 | 1.35 | -0.03 | 0.03 |
| gene-LOC105280819 | https://www.ncbi.nlm.nih.gov/gene/?term=LOC105280819 | 1639 | 37 | 225.75 | 1 | 6.1 | 2.7 | 36.55 | 38.07 | -1.52 | 1.52 | -0.03 | 0.03 |
| gene-LOC105286694 | https://www.ncbi.nlm.nih.gov/gene/?term=LOC105286694 | 3409 | 9 | 26.4 | 1 | 2.93 | 11.11 | 71.7 | 62.28 | 9.42 | 9.42 | 0.02 | 0.02 |
| gene-LOC105285315 | https://www.ncbi.nlm.nih.gov/gene/?term=LOC105285315 | 5458 | 19 | 34.81 | 1 | 1.83 | 5.26 | 47.88 | 43.22 | 4.66 | 4.66 | 0.02 | 0.02 |
| gene-LOC105275518 | https://www.ncbi.nlm.nih.gov/gene/?term=LOC105275518 | 2335 | 18 | 77.09 | 1 | 4.28 | 5.56 | 54.44 | 51.41 | 3.03 | 3.03 | 0.02 | 0.02 |
| gene-LOC105275347 | https://www.ncbi.nlm.nih.gov/gene/?term=LOC105275347 | 2526 | 14 | 55.42 | 1 | 3.96 | 7.14 | 33.57 | 30.77 | 2.8 | 2.8 | 0.02 | 0.02 |

| | | | | | | | | | | | | | | | |
|---|---|---|---|---|---|---|---|---|---|---|---|---|---|---|---|
| gene-LOC105274900 | https://www.ncbi.nlm.nih.gov/gene/?term=LOC105274900 | 5561 | 47 | 84.52 | 2 | 3.6 | 4.26 | 37.9 | 35.14 | 2.76 | 2.76 | 0.02 | 0.02 | | |
| gene-LOC105288066 | https://www.ncbi.nlm.nih.gov/gene/?term=LOC105288066 | 1964 | 20 | 101.83 | | 1 | 5.09 | 5 | 44.28 | 42.38 | 1.9 | 1.9 | 0.02 | 0.02 | |
| gene-LOC105275966 | https://www.ncbi.nlm.nih.gov/gene/?term=LOC105275966 | 3299 | 53 | 160.65 | | 1 | 3.03 | 1.89 | 54.5 | 52.99 | 1.51 | 1.51 | 0.02 | 0.02 | |
| gene-LOC105283712 | https://www.ncbi.nlm.nih.gov/gene/?term=LOC105283712 | 1910 | 31 | 162.3 | 1 | 5.24 | 3.23 | 30.95 | 29.59 | 1.36 | 1.36 | 0.02 | 0.02 | | |
| gene-LOC105281848 | https://www.ncbi.nlm.nih.gov/gene/?term=LOC105281848 | 2122 | 47 | 221.49 | | 1 | 4.71 | 2.13 | 27.8 | 26.69 | 1.11 | 1.11 | 0.02 | 0.02 | |
| gene-LOC105275451 | https://www.ncbi.nlm.nih.gov/gene/?term=LOC105275451 | 6035 | 118 | 195.53 | | 1 | 1.66 | 0.85 | 23.45 | 22.41 | 1.04 | 1.04 | 0.02 | 0.02 | |
| gene-LOC105276272 | https://www.ncbi.nlm.nih.gov/gene/?term=LOC105276272 | 1965 | 34 | 173.03 | | 1 | 5.09 | 2.94 | 33.29 | 32.32 | 0.98 | 0.98 | 0.02 | 0.02 | |
| gene-LOC105287417 | https://www.ncbi.nlm.nih.gov/gene/?term=LOC105287417 | 1025 | 22 | 214.63 | | 1 | 9.76 | 4.55 | 50.13 | 49.23 | 0.89 | 0.89 | 0.02 | 0.02 | |
| gene-LOC105284036 | https://www.ncbi.nlm.nih.gov/gene/?term=LOC105284036 | 3254 | 96 | 295.02 | | 1 | 3.07 | 1.04 | 22.78 | 21.95 | 0.83 | 0.83 | 0.02 | 0.02 | |
| gene-LOC105283592 | https://www.ncbi.nlm.nih.gov/gene/?term=LOC105283592 | 3479 | 72 | 206.96 | | 1 | 2.87 | 1.39 | 23.84 | 23.1 | 0.74 | 0.74 | 0.02 | 0.02 | |
| gene-LOC105285395 | https://www.ncbi.nlm.nih.gov/gene/?term=LOC105285395 | 8085 | 185 | 228.82 | | 1 | 1.24 | 0.54 | 16.99 | 16.27 | 0.72 | 0.72 | 0.02 | 0.02 | |
| gene-LOC113561621 | https://www.ncbi.nlm.nih.gov/gene/?term=LOC113561621 | 2249 | 82 | 364.61 | | 1 | 4.45 | 1.22 | 20.63 | 19.95 | 0.68 | 0.68 | 0.02 | 0.02 | |
| gene-LOC105285639 | https://www.ncbi.nlm.nih.gov/gene/?term=LOC105285639 | 10325 | 319 | 308.96 | | 1 | 0.97 | 0.31 | 12.35 | 11.71 | 0.65 | 0.65 | 0.02 | 0.02 | |
| gene-LOC105274452 | https://www.ncbi.nlm.nih.gov/gene/?term=LOC105274452 | 6595 | 252 | 382.11 | | 1 | 1.52 | 0.4 | 13.74 | 13.13 | 0.61 | 0.61 | 0.02 | 0.02 | |
| gene-LOC105274706 | https://www.ncbi.nlm.nih.gov/gene/?term=LOC105274706 | 1446 | 55 | 380.36 | | 1 | 6.92 | 1.82 | 37.05 | 36.54 | 0.5 | 0.5 | 0.02 | 0.02 | |
| gene-LOC105274856 | https://www.ncbi.nlm.nih.gov/gene/?term=LOC105274856 | 5216 | 207 | 396.86 | | 1 | 1.92 | 0.48 | 19.99 | 19.54 | 0.45 | 0.45 | 0.02 | 0.02 | |
| gene-LOC105278666 | https://www.ncbi.nlm.nih.gov/gene/?term=LOC105278666 | 10341 | 429 | 414.85 | | 1 | 0.97 | 0.23 | 7.71 | 7.27 | 0.44 | 0.44 | 0.02 | 0.02 | |
| gene-LOC105285807 | https://www.ncbi.nlm.nih.gov/gene/?term=LOC105285807 | 6063 | 214 | 352.96 | | 1 | 1.65 | 0.47 | 7.55 | 7.11 | 0.44 | 0.44 | 0.02 | 0.02 | |
| gene-LOC105276612 | https://www.ncbi.nlm.nih.gov/gene/?term=LOC105276612 | 4157 | 181 | 435.41 | | 1 | 2.41 | 0.55 | 10.16 | 9.76 | 0.4 | 0.4 | 0.02 | 0.02 | |
| gene-LOC105277423 | https://www.ncbi.nlm.nih.gov/gene/?term=LOC105277423 | 2340 | 108 | 461.54 | | 1 | 4.27 | 0.93 | 30.29 | 29.91 | 0.38 | 0.38 | 0.02 | 0.02 | |
| gene-LOC105286649 | https://www.ncbi.nlm.nih.gov/gene/?term=LOC105286649 | 14178 | 778 | 548.74 | | 1 | 0.71 | 0.13 | 0.86 | 1.14 | -0.27 | 0.27 | -0.02 | 0.02 | |
| gene-LOC105285252 | https://www.ncbi.nlm.nih.gov/gene/?term=LOC105285252 | 7427 | 112 | 150.8 | 2 | 2.69 | 1.79 | 14.17 | 15.17 | -1 | 1 | -0.02 | 0.02 | | |
| gene-LOC105281700 | https://www.ncbi.nlm.nih.gov/gene/?term=LOC105281700 | 2208 | 34 | 153.99 | | 1 | 4.53 | 2.94 | 35.91 | 36.92 | -1.02 | 1.02 | -0.02 | 0.02 | |
| gene-LOC105285860 | https://www.ncbi.nlm.nih.gov/gene/?term=LOC105285860 | 3110 | 52 | 167.2 | 1 | 3.22 | 1.92 | 20.92 | 22.04 | -1.12 | 1.12 | -0.02 | 0.02 | | |
| gene-LOC105284507 | https://www.ncbi.nlm.nih.gov/gene/?term=LOC105284507 | 5207 | 77 | 147.88 | | 1 | 1.92 | 1.3 | 22.93 | 24.13 | -1.2 | 1.2 | -0.02 | 0.02 | |
| gene-LOC105281515 | https://www.ncbi.nlm.nih.gov/gene/?term=LOC105281515 | 8827 | 38 | 43.05 | 1 | 1.13 | 2.63 | 31.29 | 29.13 | 2.16 | 2.16 | 0.01 | 0.01 | | |
| gene-LOC105280408 | https://www.ncbi.nlm.nih.gov/gene/?term=LOC105280408 | 5983 | 47 | 78.56 | 1 | 1.67 | 2.13 | 19.12 | 17.33 | 1.78 | 1.78 | 0.01 | 0.01 | | |
| gene-LOC105274590 | https://www.ncbi.nlm.nih.gov/gene/?term=LOC105274590 | 4516 | 57 | 126.22 | | 2 | 4.43 | 3.51 | 34.78 | 33.73 | 1.06 | 1.06 | 0.01 | 0.01 | |
| gene-LOC105279500 | https://www.ncbi.nlm.nih.gov/gene/?term=LOC105279500 | 3587 | 34 | 94.79 | 1 | 2.79 | 2.94 | 52.36 | 51.39 | 0.97 | 0.97 | 0.01 | 0.01 | | |
| gene-LOC105283987 | https://www.ncbi.nlm.nih.gov/gene/?term=LOC105283987 | 4841 | 64 | 132.2 | 1 | 2.07 | 1.56 | 30.28 | 29.39 | 0.89 | 0.89 | 0.01 | 0.01 | | |
| gene-LOC105274851 | https://www.ncbi.nlm.nih.gov/gene/?term=LOC105274851 | 6616 | 123 | 185.91 | | 1 | 1.51 | 0.81 | 11.44 | 10.81 | 0.63 | 0.63 | 0.01 | 0.01 | |
| gene-LOC105282885 | https://www.ncbi.nlm.nih.gov/gene/?term=LOC105282885 | 1870 | 45 | 240.64 | | 1 | 5.35 | 2.22 | 42.36 | 41.77 | 0.58 | 0.58 | 0.01 | 0.01 | |
| gene-LOC105276692 | https://www.ncbi.nlm.nih.gov/gene/?term=LOC105276692 | 1745 | 29 | 166.19 | | 1 | 5.73 | 3.45 | 25.51 | 24.96 | 0.55 | 0.55 | 0.01 | 0.01 | |
| gene-LOC105284704 | https://www.ncbi.nlm.nih.gov/gene/?term=LOC105284704 | 2400 | 57 | 237.5 | 1 | 4.17 | 1.75 | 25.25 | 24.69 | 0.55 | 0.55 | 0.01 | 0.01 | | |
| gene-LOC105282151 | https://www.ncbi.nlm.nih.gov/gene/?term=LOC105282151 | 1601 | 35 | 218.61 | | 1 | 6.25 | 2.86 | 42.13 | 41.61 | 0.53 | 0.53 | 0.01 | 0.01 | |
| gene-LOC105280410 | https://www.ncbi.nlm.nih.gov/gene/?term=LOC105280410 | 12580 | 235 | 186.8 | 1 | 0.79 | 0.43 | 21.34 | 20.84 | 0.5 | 0.5 | 0.01 | 0.01 | | |
| gene-LOC105274823 | https://www.ncbi.nlm.nih.gov/gene/?term=LOC105274823 | 4620 | 132 | 285.71 | | 1 | 2.16 | 0.76 | 21.94 | 21.47 | 0.47 | 0.47 | 0.01 | 0.01 | |
| gene-LOC105279038 | https://www.ncbi.nlm.nih.gov/gene/?term=LOC105279038 | 9453 | 306 | 323.71 | | 1 | 1.06 | 0.33 | 10.13 | 9.67 | 0.46 | 0.46 | 0.01 | 0.01 | |
| gene-LOC105286514 | https://www.ncbi.nlm.nih.gov/gene/?term=LOC105286514 | 2528 | 44 | 174.05 | | 1 | 3.96 | 2.27 | 33.92 | 33.47 | 0.45 | 0.45 | 0.01 | 0.01 | |
| gene-LOC105278282 | https://www.ncbi.nlm.nih.gov/gene/?term=LOC105278282 | 1775 | 52 | 292.96 | | 1 | 5.63 | 1.92 | 38.05 | 37.63 | 0.42 | 0.42 | 0.01 | 0.01 | |
| gene-LOC105276941 | https://www.ncbi.nlm.nih.gov/gene/?term=LOC105276941 | 15956 | 494 | 309.6 | 1 | 0.63 | 0.2 | 5.44 | 5.07 | 0.38 | 0.38 | 0.01 | 0.01 | | |
| gene-LOC105287689 | https://www.ncbi.nlm.nih.gov/gene/?term=LOC105287689 | 10867 | 331 | 304.59 | | 1 | 0.92 | 0.3 | 9.53 | 9.17 | 0.36 | 0.36 | 0.01 | 0.01 | |
| gene-LOC105278454 | https://www.ncbi.nlm.nih.gov/gene/?term=LOC105278454 | 1972 | 44 | 223.12 | | 1 | 5.07 | 2.27 | 46.65 | 46.38 | 0.26 | 0.26 | 0.01 | 0.01 | |
| gene-LOC105278877 | https://www.ncbi.nlm.nih.gov/gene/?term=LOC105278877 | 1968 | 69 | 350.61 | | 1 | 5.08 | 1.45 | 35.47 | 35.23 | 0.25 | 0.25 | 0.01 | 0.01 | |
| gene-LOC105275460 | https://www.ncbi.nlm.nih.gov/gene/?term=LOC105275460 | 2473 | 116 | 469.07 | | 1 | 4.04 | 0.86 | 13.65 | 13.42 | 0.23 | 0.23 | 0.01 | 0.01 | |
| gene-LOC113562050 | https://www.ncbi.nlm.nih.gov/gene/?term=LOC113562050 | 17097 | 540 | 315.84 | | 1 | 0.58 | 0.19 | 5.38 | 5.15 | 0.23 | 0.23 | 0.01 | 0.01 | |
| gene-LOC105278534 | https://www.ncbi.nlm.nih.gov/gene/?term=LOC105278534 | 7364 | 175 | 237.64 | | 1 | 1.36 | 0.57 | 18.73 | 18.51 | 0.22 | 0.22 | 0.01 | 0.01 | |
| gene-LOC105286166 | https://www.ncbi.nlm.nih.gov/gene/?term=LOC105286166 | 3050 | 103 | 337.7 | 1 | 3.28 | 0.97 | 25.5 | 25.28 | 0.22 | 0.22 | 0.01 | 0.01 | | |
| gene-LOC105274449 | https://www.ncbi.nlm.nih.gov/gene/?term=LOC105274449 | 1549 | 48 | 309.88 | | 1 | 6.46 | 2.08 | 51.17 | 50.96 | 0.21 | 0.21 | 0.01 | 0.01 | |
| gene-LOC105280956 | https://www.ncbi.nlm.nih.gov/gene/?term=LOC105280956 | 2639 | 69 | 261.46 | | 1 | 3.79 | 1.45 | 18.51 | 18.31 | 0.21 | 0.21 | 0.01 | 0.01 | |
| gene-LOC105275683 | https://www.ncbi.nlm.nih.gov/gene/?term=LOC105275683 | 4256 | 183 | 429.98 | | 1 | 2.35 | 0.55 | 12.85 | 12.65 | 0.2 | 0.2 | 0.01 | 0.01 | |
| gene-LOC105276462 | https://www.ncbi.nlm.nih.gov/gene/?term=LOC105276462 | 6561 | 246 | 374.94 | | 1 | 1.52 | 0.41 | 8.14 | 7.99 | 0.15 | 0.15 | 0.01 | 0.01 | |
| gene-LOC105278649 | https://www.ncbi.nlm.nih.gov/gene/?term=LOC105278649 | 5616 | 161 | 286.68 | | 1 | 1.78 | 0.62 | 11.64 | 11.9 | -0.26 | 0.26 | -0.01 | 0.01 | |
| gene-LOC105276527 | https://www.ncbi.nlm.nih.gov/gene/?term=LOC105276527 | 7923 | 309 | 390 | 1 | 1.26 | 0.32 | 9.63 | 9.9 | -0.27 | 0.27 | -0.01 | 0.01 | | |
| gene-LOC105286274 | https://www.ncbi.nlm.nih.gov/gene/?term=LOC105286274 | 3817 | 193 | 505.63 | | 1 | 2.62 | 0.52 | 16.76 | 17.05 | -0.29 | 0.29 | -0.01 | 0.01 | |
| gene-LOC105286515 | https://www.ncbi.nlm.nih.gov/gene/?term=LOC105286515 | 3751 | 99 | 263.93 | | 1 | 2.67 | 1.01 | 13.14 | 13.49 | -0.36 | 0.36 | -0.01 | 0.01 | |
| gene-LOC105287633 | https://www.ncbi.nlm.nih.gov/gene/?term=LOC105287633 | 1524 | 27 | 177.17 | | 1 | 6.56 | 3.7 | 63.52 | 64 | -0.48 | 0.48 | -0.01 | 0.01 | |
| gene-LOC105286452 | https://www.ncbi.nlm.nih.gov/gene/?term=LOC105286452 | 6888 | 62 | 90.01 | 1 | 1.45 | 1.61 | 32.65 | 33.21 | -0.56 | 0.56 | -0.01 | 0.01 | | |
| gene-LOC105278108 | https://www.ncbi.nlm.nih.gov/gene/?term=LOC105278108 | 4392 | 44 | 100.18 | | 1 | 2.28 | 2.27 | 24.81 | 25.4 | -0.59 | 0.59 | -0.01 | 0.01 | |
| gene-LOC105275187 | https://www.ncbi.nlm.nih.gov/gene/?term=LOC105275187 | 5534 | 66 | 119.26 | | 1 | 1.81 | 1.52 | 11.92 | 12.68 | -0.77 | 0.77 | -0.01 | 0.01 | |
| gene-LOC105282067 | https://www.ncbi.nlm.nih.gov/gene/?term=LOC105282067 | 5941 | 51 | 85.84 | 1 | 1.68 | 1.96 | 26.09 | 27.84 | -1.75 | 1.75 | -0.01 | 0.01 | | |
| gene-LOC105283989 | https://www.ncbi.nlm.nih.gov/gene/?term=LOC105283989 | 4948 | 26 | 52.55 | 1 | 2.02 | 3.85 | 37.24 | 39.59 | -2.34 | 2.34 | -0.01 | 0.01 | | |
| gene-LOC105285406 | https://www.ncbi.nlm.nih.gov/gene/?term=LOC105285406 | 8015 | 60 | 74.86 | 1 | 1.25 | 1.67 | 25.96 | 25.52 | 0.44 | 0.44 | 0 | 0 | | |
| gene-LOC105286194 | https://www.ncbi.nlm.nih.gov/gene/?term=LOC105286194 | 152619 | | 2040 | 133.67 | | 1 | 0.07 | 0.05 | 6.36 | 6.07 | 0.3 | 0.3 | 0 | 0 |

| | | | | | | | | | | | | | | |
|---|---|---|---|---|---|---|---|---|---|---|---|---|---|---|
| gene-LOC105280407 | https://www.ncbi.nlm.nih.gov/gene/?term=LOC105280407 | 10950 | 189 | 172.6 | 1 | 0.91 | 0.53 | 16.21 | 15.97 | 0.24 | 0.24 | 0 | 0 | |
| gene-LOC105287764 | https://www.ncbi.nlm.nih.gov/gene/?term=LOC105287764 | 18359 | 375 | 204.26 | | 1 | 0.54 | 0.27 | 3.25 | 3.09 | 0.17 | 0.17 | 0 | 0 |
| gene-LOC105282658 | https://www.ncbi.nlm.nih.gov/gene/?term=LOC105282658 | 2421 | 54 | 223.05 | | 1 | 4.13 | 1.85 | 19.57 | 19.42 | 0.16 | 0.16 | 0 | 0 |
| gene-LOC105282825 | https://www.ncbi.nlm.nih.gov/gene/?term=LOC105282825 | 3919 | 69 | 176.07 | | 1 | 2.55 | 1.45 | 13.52 | 13.36 | 0.16 | 0.16 | 0 | 0 |
| gene-LOC105278455 | https://www.ncbi.nlm.nih.gov/gene/?term=LOC105278455 | 27181 | 882 | 324.49 | | 1 | 0.37 | 0.11 | 7.78 | 7.66 | 0.12 | 0.12 | 0 | 0 |
| gene-LOC105286999 | https://www.ncbi.nlm.nih.gov/gene/?term=LOC105286999 | 2141 | 42 | 196.17 | | 1 | 4.67 | 2.38 | 36.17 | 36.05 | 0.12 | 0.12 | 0 | 0 |
| gene-LOC105282811 | https://www.ncbi.nlm.nih.gov/gene/?term=LOC105282811 | 5765 | 198 | 343.45 | | 1 | 1.73 | 0.51 | 7.89 | 7.77 | 0.11 | 0.11 | 0 | 0 |
| gene-LOC105275327 | https://www.ncbi.nlm.nih.gov/gene/?term=LOC105275327 | 14245 | 460 | 322.92 | | 1 | 0.7 | 0.22 | 7.25 | 7.2 | 0.05 | 0.05 | 0 | 0 |
| gene-LOC105280265 | https://www.ncbi.nlm.nih.gov/gene/?term=LOC105280265 | 85984 | 2858 | 332.39 | | 1 | 0.12 | 0.03 | 1.09 | 1.08 | 0.01 | 0.01 | 0 | 0 |
| gene-LOC105277125 | https://www.ncbi.nlm.nih.gov/gene/?term=LOC105277125 | 51079 | 1693 | 331.45 | | 1 | 0.2 | 0.06 | 1.64 | 1.64 | 0 | 0 | 0 | 0 |
| gene-LOC105277866 | https://www.ncbi.nlm.nih.gov/gene/?term=LOC105277866 | 33045 | 352 | 106.52 | | 1 | 0.3 | 0.28 | 3.39 | 3.44 | -0.04 | 0.04 | 0 | 0 |
| gene-LOC105282646 | https://www.ncbi.nlm.nih.gov/gene/?term=LOC105282646 | 3317 | 99 | 298.46 | | 1 | 3.01 | 1.01 | 25.07 | 25.11 | -0.04 | 0.04 | 0 | 0 |
| gene-LOC105274531 | https://www.ncbi.nlm.nih.gov/gene/?term=LOC105274531 | 7774 | 285 | 366.61 | | 1 | 1.29 | 0.35 | 10.62 | 10.74 | -0.11 | 0.11 | 0 | 0 |
| gene-LOC105280377 | https://www.ncbi.nlm.nih.gov/gene/?term=LOC105280377 | 2024 | 58 | 286.56 | | 1 | 4.94 | 1.72 | 49.97 | 50.08 | -0.11 | 0.11 | 0 | 0 |
| gene-LOC105275848 | https://www.ncbi.nlm.nih.gov/gene/?term=LOC105275848 | 3718 | 133 | 357.72 | | 1 | 2.69 | 0.75 | 21.71 | 21.82 | -0.12 | 0.12 | 0 | 0 |
| gene-LOC105276538 | https://www.ncbi.nlm.nih.gov/gene/?term=LOC105276538 | 1511 | 19 | 125.74 | | 1 | 6.62 | 5.26 | 67.47 | 67.59 | -0.12 | 0.12 | 0 | 0 |
| gene-LOC105283296 | https://www.ncbi.nlm.nih.gov/gene/?term=LOC105283296 | 2174 | 84 | 386.38 | | 1 | 4.6 | 1.19 | 19.37 | 19.49 | -0.12 | 0.12 | 0 | 0 |
| gene-LOC105278632 | https://www.ncbi.nlm.nih.gov/gene/?term=LOC105278632 | 3624 | 113 | 311.81 | | 1 | 2.76 | 0.88 | 16.03 | 16.16 | -0.13 | 0.13 | 0 | 0 |
| gene-LOC105283943 | https://www.ncbi.nlm.nih.gov/gene/?term=LOC105283943 | 3559 | 42 | 118.01 | | 1 | 2.81 | 2.38 | 14.31 | 14.48 | -0.17 | 0.17 | 0 | 0 |
| gene-LOC105275179 | https://www.ncbi.nlm.nih.gov/gene/?term=LOC105275179 | 3271 | 66 | 201.77 | | 1 | 3.06 | 1.52 | 20.6 | 20.81 | -0.21 | 0.21 | 0 | 0 |
| gene-LOC105285514 | https://www.ncbi.nlm.nih.gov/gene/?term=LOC105285514 | 1277 | 14 | 109.63 | | 1 | 7.83 | 7.14 | 85.59 | 85.88 | -0.28 | 0.28 | 0 | 0 |
| gene-LOC109611234 | https://www.ncbi.nlm.nih.gov/gene/?term=LOC109611234 | 3663 | 28 | 76.44 | 1 | 2.73 | 3.57 | 16.2 | 16.56 | -0.36 | 0.36 | 0 | 0 | |

Directory: OGLall File: rnk-dmg-Ob_LI_BCphaseB1.vs.BCphaseB2.txt

| gene_ID | gene_link | gwidth | #Sites | #per10Kb | #dmSites | #dmsp10kb | %dmSites | %pSite1 | %pSite2 | DMpSite | ADMpSite | DMpNucl | ADMpNucl |
|---|---|---|---|---|---|---|---|---|---|---|---|---|---|
| gene-LOC113562345 | https://www.ncbi.nlm.nih.gov/gene/?term=LOC113562345 | 2290 | 64 | 279.48 | 15 | 65.5 | 23.44 | 32.83 | 17.93 | 14.9 | 14.9 | 0.42 | 0.42 |
| gene-LOC105284677 | https://www.ncbi.nlm.nih.gov/gene/?term=LOC105284677 | 1905 | 42 | 220.47 | 1 | 5.25 | 2.38 | 26.24 | 37.79 | -11.55 | 11.55 | -0.25 | 0.25 |
| gene-LOC105283649 | https://www.ncbi.nlm.nih.gov/gene/?term=LOC105283649 | 1918 | 56 | 291.97 | 1 | 5.21 | 1.79 | 39.64 | 36.17 | 3.47 | 3.47 | 0.1 | 0.1 |
| gene-LOC105276449 | https://www.ncbi.nlm.nih.gov/gene/?term=LOC105276449 | 1231 | 51 | 414.3 | 1 | 8.12 | 1.96 | 27.95 | 25.74 | 2.21 | 2.21 | 0.09 | 0.09 |
| gene-LOC105275070 | https://www.ncbi.nlm.nih.gov/gene/?term=LOC105275070 | 5667 | 49 | 86.47 | 6 | 10.59 | 12.24 | 10.93 | 1.84 | 9.09 | 9.09 | 0.08 | 0.08 |
| gene-LOC105275447 | https://www.ncbi.nlm.nih.gov/gene/?term=LOC105275447 | 1846 | 32 | 173.35 | 1 | 5.42 | 3.12 | 8.01 | 3.17 | 4.84 | 4.84 | 0.08 | 0.08 |
| gene-LOC105285674 | https://www.ncbi.nlm.nih.gov/gene/?term=LOC105285674 | 3035 | 94 | 309.72 | 2 | 6.59 | 2.13 | 27.11 | 24.65 | 2.46 | 2.46 | 0.08 | 0.08 |
| gene-LOC105278877 | https://www.ncbi.nlm.nih.gov/gene/?term=LOC105278877 | 1968 | 82 | 416.67 | 1 | 5.08 | 1.22 | 32.75 | 30.76 | 1.99 | 1.99 | 0.08 | 0.08 |
| gene-LOC105282156 | https://www.ncbi.nlm.nih.gov/gene/?term=LOC105282156 | 2577 | 137 | 531.63 | 1 | 3.88 | 0.73 | 24.98 | 23.48 | 1.5 | 1.5 | 0.08 | 0.08 |
| gene-LOC105282885 | https://www.ncbi.nlm.nih.gov/gene/?term=LOC105282885 | 1870 | 47 | 251.34 | 1 | 5.35 | 2.13 | 35.98 | 39.14 | -3.16 | 3.16 | -0.08 | 0.08 |
| gene-LOC105275708 | https://www.ncbi.nlm.nih.gov/gene/?term=LOC105275708 | 1772 | 15 | 84.65 | 1 | 5.64 | 6.67 | 46.38 | 38.5 | 7.88 | 7.88 | 0.07 | 0.07 |
| gene-LOC105281039 | https://www.ncbi.nlm.nih.gov/gene/?term=LOC105281039 | 1471 | 12 | 81.58 | 1 | 6.8 | 8.33 | 38.6 | 31.46 | 7.13 | 7.13 | 0.06 | 0.06 |
| gene-LOC105279900 | https://www.ncbi.nlm.nih.gov/gene/?term=LOC105279900 | 2191 | 56 | 255.59 | 1 | 4.56 | 1.79 | 33.98 | 31.69 | 2.29 | 2.29 | 0.06 | 0.06 |
| gene-LOC105279898 | https://www.ncbi.nlm.nih.gov/gene/?term=LOC105279898 | 1343 | 13 | 96.8 | 1 | 7.45 | 7.69 | 64.63 | 59.11 | 5.52 | 5.52 | 0.05 | 0.05 |
| gene-LOC105275518 | https://www.ncbi.nlm.nih.gov/gene/?term=LOC105275518 | 2335 | 23 | 98.5 | 1 | 4.28 | 4.35 | 43.86 | 38.74 | 5.13 | 5.13 | 0.05 | 0.05 |
| gene-LOC105287417 | https://www.ncbi.nlm.nih.gov/gene/?term=LOC105287417 | 1025 | 36 | 351.22 | 1 | 9.76 | 2.78 | 35.49 | 34.17 | 1.32 | 1.32 | 0.05 | 0.05 |
| gene-LOC105280786 | https://www.ncbi.nlm.nih.gov/gene/?term=LOC105280786 | 2573 | 119 | 462.5 | 1 | 3.89 | 0.84 | 30.85 | 29.73 | 1.12 | 1.12 | 0.05 | 0.05 |
| gene-LOC105274706 | https://www.ncbi.nlm.nih.gov/gene/?term=LOC105274706 | 1446 | 50 | 345.78 | 1 | 6.92 | 2 | 40.41 | 41.89 | -1.49 | 1.49 | -0.05 | 0.05 |
| gene-LOC105287633 | https://www.ncbi.nlm.nih.gov/gene/?term=LOC105287633 | 1524 | 31 | 203.41 | 1 | 6.56 | 3.23 | 53.47 | 55.78 | -2.31 | 2.31 | -0.05 | 0.05 |
| gene-LOC105274900 | https://www.ncbi.nlm.nih.gov/gene/?term=LOC105274900 | 5561 | 54 | 97.1 | 2 | 3.6 | 3.7 | 35.58 | 31 | 4.57 | 4.57 | 0.04 | 0.04 |
| gene-LOC105281420 | https://www.ncbi.nlm.nih.gov/gene/?term=LOC105281420 | 3882 | 67 | 172.59 | 1 | 2.58 | 1.49 | 15.32 | 12.95 | 2.37 | 2.37 | 0.04 | 0.04 |
| gene-LOC105283592 | https://www.ncbi.nlm.nih.gov/gene/?term=LOC105283592 | 3479 | 71 | 204.08 | 1 | 2.87 | 1.41 | 27.65 | 25.73 | 1.93 | 1.93 | 0.04 | 0.04 |
| gene-LOC105278454 | https://www.ncbi.nlm.nih.gov/gene/?term=LOC105278454 | 1972 | 48 | 243.41 | 1 | 5.07 | 2.08 | 42.9 | 41.15 | 1.75 | 1.75 | 0.04 | 0.04 |
| gene-LOC105285583 | https://www.ncbi.nlm.nih.gov/gene/?term=LOC105285583 | 1947 | 45 | 231.12 | 1 | 5.14 | 2.22 | 29.78 | 28.25 | 1.53 | 1.53 | 0.04 | 0.04 |
| gene-LOC105278632 | https://www.ncbi.nlm.nih.gov/gene/?term=LOC105278632 | 3624 | 126 | 347.68 | 1 | 2.76 | 0.79 | 16.08 | 14.93 | 1.15 | 1.15 | 0.04 | 0.04 |
| gene-LOC105285202 | https://www.ncbi.nlm.nih.gov/gene/?term=LOC105285202 | 1838 | 63 | 342.76 | 1 | 5.44 | 1.59 | 23.99 | 22.95 | 1.05 | 1.05 | 0.04 | 0.04 |
| gene-LOC105284355 | https://www.ncbi.nlm.nih.gov/gene/?term=LOC105284355 | 3904 | 158 | 404.71 | 1 | 2.56 | 0.63 | 20.65 | 19.62 | 1.02 | 1.02 | 0.04 | 0.04 |
| gene-LOC105288036 | https://www.ncbi.nlm.nih.gov/gene/?term=LOC105288036 | 5110 | 227 | 444.23 | 1 | 1.96 | 0.44 | 15.78 | 14.87 | 0.91 | 0.91 | 0.04 | 0.04 |
| gene-LOC105275384 | https://www.ncbi.nlm.nih.gov/gene/?term=LOC105275384 | 1987 | 92 | 463.01 | 1 | 5.03 | 1.09 | 32.77 | 31.88 | 0.89 | 0.89 | 0.04 | 0.04 |
| gene-LOC105285055 | https://www.ncbi.nlm.nih.gov/gene/?term=LOC105285055 | 4108 | 253 | 615.87 | 1 | 2.43 | 0.4 | 12.35 | 11.74 | 0.61 | 0.61 | 0.04 | 0.04 |
| gene-LOC105274534 | https://www.ncbi.nlm.nih.gov/gene/?term=LOC105274534 | 1329 | 60 | 451.47 | 1 | 7.52 | 1.67 | 40.57 | 41.38 | -0.82 | 0.82 | -0.04 | 0.04 |
| gene-LOC105276429 | https://www.ncbi.nlm.nih.gov/gene/?term=LOC105276429 | 2901 | 108 | 372.29 | 1 | 3.45 | 0.93 | 24.11 | 25.23 | -1.12 | 1.12 | -0.04 | 0.04 |
| gene-LOC105276538 | https://www.ncbi.nlm.nih.gov/gene/?term=LOC105276538 | 1511 | 17 | 112.51 | 1 | 6.62 | 5.88 | 72.11 | 69.24 | 2.87 | 2.87 | 0.03 | 0.03 |
| gene-LOC105285514 | https://www.ncbi.nlm.nih.gov/gene/?term=LOC105285514 | 1277 | 15 | 117.46 | 1 | 7.83 | 6.67 | 80.63 | 78.25 | 2.38 | 2.38 | 0.03 | 0.03 |
| gene-LOC105279664 | https://www.ncbi.nlm.nih.gov/gene/?term=LOC105279664 | 4023 | 65 | 161.57 | 1 | 2.49 | 1.54 | 19.54 | 17.85 | 1.7 | 1.7 | 0.03 | 0.03 |
| gene-LOC105286514 | https://www.ncbi.nlm.nih.gov/gene/?term=LOC105286514 | 2528 | 46 | 181.96 | 1 | 3.96 | 2.17 | 37.97 | 36.35 | 1.61 | 1.61 | 0.03 | 0.03 |
| gene-LOC105283943 | https://www.ncbi.nlm.nih.gov/gene/?term=LOC105283943 | 3559 | 62 | 174.21 | 1 | 2.81 | 1.61 | 15.22 | 13.63 | 1.6 | 1.6 | 0.03 | 0.03 |
| gene-LOC105277903 | https://www.ncbi.nlm.nih.gov/gene/?term=LOC105277903 | 1835 | 37 | 201.63 | 1 | 5.45 | 2.7 | 54.03 | 52.65 | 1.39 | 1.39 | 0.03 | 0.03 |
| gene-LOC105282794 | https://www.ncbi.nlm.nih.gov/gene/?term=LOC105282794 | 4571 | 98 | 214.4 | 1 | 2.19 | 1.02 | 19.72 | 18.47 | 1.25 | 1.25 | 0.03 | 0.03 |
| gene-LOC105280956 | https://www.ncbi.nlm.nih.gov/gene/?term=LOC105280956 | 2639 | 75 | 284.2 | 1 | 3.79 | 1.33 | 17.61 | 16.45 | 1.16 | 1.16 | 0.03 | 0.03 |
| gene-LOC105287063 | https://www.ncbi.nlm.nih.gov/gene/?term=LOC105287063 | 1874 | 51 | 272.15 | 1 | 5.34 | 1.96 | 30.2 | 29.07 | 1.12 | 1.12 | 0.03 | 0.03 |
| gene-LOC105275752 | https://www.ncbi.nlm.nih.gov/gene/?term=LOC105275752 | 3462 | 130 | 375.51 | 1 | 2.89 | 0.77 | 31.52 | 30.74 | 0.77 | 0.77 | 0.03 | 0.03 |
| gene-LOC105285639 | https://www.ncbi.nlm.nih.gov/gene/?term=LOC105285639 | 10325 | 378 | 366.1 | 1 | 0.97 | 0.26 | 11.07 | 10.35 | 0.72 | 0.72 | 0.03 | 0.03 |
| gene-LOC105278807 | https://www.ncbi.nlm.nih.gov/gene/?term=LOC105278807 | 2166 | 80 | 369.34 | 1 | 4.62 | 1.25 | 18.57 | 17.87 | 0.69 | 0.69 | 0.03 | 0.03 |
| gene-LOC105278497 | https://www.ncbi.nlm.nih.gov/gene/?term=LOC105278497 | 2072 | 95 | 458.49 | 1 | 4.83 | 1.05 | 19.7 | 19.09 | 0.61 | 0.61 | 0.03 | 0.03 |
| gene-LOC105282149 | https://www.ncbi.nlm.nih.gov/gene/?term=LOC105282149 | 1787 | 107 | 598.77 | 1 | 5.6 | 0.93 | 41.49 | 40.93 | 0.57 | 0.57 | 0.03 | 0.03 |
| gene-LOC105279298 | https://www.ncbi.nlm.nih.gov/gene/?term=LOC105279298 | 2773 | 132 | 476.02 | 2 | 7.21 | 1.52 | 23.66 | 23.13 | 0.53 | 0.53 | 0.03 | 0.03 |
| gene-LOC105276704 | https://www.ncbi.nlm.nih.gov/gene/?term=LOC105276704 | 2063 | 99 | 479.88 | 2 | 9.69 | 2.02 | 13.44 | 13.97 | -0.53 | 0.53 | -0.03 | 0.03 |
| gene-LOC105285533 | https://www.ncbi.nlm.nih.gov/gene/?term=LOC105285533 | 1640 | 70 | 426.83 | 1 | 6.1 | 1.43 | 22.75 | 23.49 | -0.75 | 0.75 | -0.03 | 0.03 |
| gene-LOC105274449 | https://www.ncbi.nlm.nih.gov/gene/?term=LOC105274449 | 1549 | 53 | 342.16 | 1 | 6.46 | 1.89 | 50.86 | 51.77 | -0.91 | 0.91 | -0.03 | 0.03 |
| gene-LOC105281312 | https://www.ncbi.nlm.nih.gov/gene/?term=LOC105281312 | 2551 | 65 | 254.8 | 1 | 3.92 | 1.54 | 31.99 | 33.22 | -1.23 | 1.23 | -0.03 | 0.03 |
| gene-LOC105281173 | https://www.ncbi.nlm.nih.gov/gene/?term=LOC105281173 | 1595 | 38 | 238.24 | 1 | 6.27 | 2.63 | 59.28 | 60.67 | -1.38 | 1.38 | -0.03 | 0.03 |
| gene-LOC105279500 | https://www.ncbi.nlm.nih.gov/gene/?term=LOC105279500 | 3587 | 33 | 92 | 1 | 2.79 | 3.03 | 43.74 | 47.01 | -3.27 | 3.27 | -0.03 | 0.03 |
| gene-LOC105281515 | https://www.ncbi.nlm.nih.gov/gene/?term=LOC105281515 | 8827 | 57 | 64.57 | 2 | 2.27 | 3.51 | 30.56 | 27.6 | 2.96 | 2.96 | 0.02 | 0.02 |
| gene-LOC105287648 | https://www.ncbi.nlm.nih.gov/gene/?term=LOC105287648 | 2205 | 30 | 136.05 | 1 | 4.54 | 3.33 | 43.37 | 41.93 | 1.44 | 1.44 | 0.02 | 0.02 |
| gene-LOC105287854 | https://www.ncbi.nlm.nih.gov/gene/?term=LOC105287854 | 3644 | 67 | 183.86 | 1 | 2.74 | 1.49 | 16.53 | 15.24 | 1.29 | 1.29 | 0.02 | 0.02 |
| gene-LOC105282991 | https://www.ncbi.nlm.nih.gov/gene/?term=LOC105282991 | 5617 | 105 | 186.93 | 1 | 1.78 | 0.95 | 19.71 | 18.51 | 1.19 | 1.19 | 0.02 | 0.02 |
| gene-LOC105288078 | https://www.ncbi.nlm.nih.gov/gene/?term=LOC105288078 | 19126 | 309 | 161.56 | 1 | 0.52 | 0.32 | 17.45 | 16.3 | 1.15 | 1.15 | 0.02 | 0.02 |
| gene-LOC105280641 | https://www.ncbi.nlm.nih.gov/gene/?term=LOC105280641 | 3682 | 55 | 149.38 | 1 | 2.72 | 1.82 | 19.12 | 18.02 | 1.1 | 1.1 | 0.02 | 0.02 |
| gene-LOC105280002 | https://www.ncbi.nlm.nih.gov/gene/?term=LOC105280002 | 1940 | 28 | 144.33 | 1 | 5.15 | 3.57 | 40.24 | 39.15 | 1.09 | 1.09 | 0.02 | 0.02 |

| | | | | | | | | | | | | | |
|---|---|---|---|---|---|---|---|---|---|---|---|---|---|
| gene-LOC105280348 | https://www.ncbi.nlm.nih.gov/gene/?term=LOC105280348 | 3797 | 76 | 200.16 | 1 | 2.63 | 1.32 | 20.16 | 19.19 | 0.97 | 0.97 | 0.02 | 0.02 |
| gene-LOC105288042 | https://www.ncbi.nlm.nih.gov/gene/?term=LOC105288042 | 3507 | 99 | 282.29 | 1 | 2.85 | 1.01 | 23.14 | 22.25 | 0.89 | 0.89 | 0.02 | 0.02 |
| gene-LOC105284117 | https://www.ncbi.nlm.nih.gov/gene/?term=LOC105284117 | 4340 | 94 | 216.59 | 1 | 2.3 | 1.06 | 11.08 | 10.26 | 0.83 | 0.83 | 0.02 | 0.02 |
| gene-LOC105284116 | https://www.ncbi.nlm.nih.gov/gene/?term=LOC105284116 | 3970 | 107 | 269.52 | 1 | 2.52 | 0.93 | 7.59 | 6.83 | 0.75 | 0.75 | 0.02 | 0.02 |
| gene-LOC105280405 | https://www.ncbi.nlm.nih.gov/gene/?term=LOC105280405 | 5422 | 161 | 296.94 | 1 | 1.84 | 0.62 | 10.76 | 10.05 | 0.71 | 0.71 | 0.02 | 0.02 |
| gene-LOC105281848 | https://www.ncbi.nlm.nih.gov/gene/?term=LOC105281848 | 2122 | 60 | 282.75 | 1 | 4.71 | 1.67 | 20.32 | 19.71 | 0.62 | 0.62 | 0.02 | 0.02 |
| gene-LOC105283728 | https://www.ncbi.nlm.nih.gov/gene/?term=LOC105283728 | 3256 | 122 | 374.69 | 2 | 6.14 | 1.64 | 19.35 | 18.81 | 0.54 | 0.54 | 0.02 | 0.02 |
| gene-LOC105275363 | https://www.ncbi.nlm.nih.gov/gene/?term=LOC105275363 | 3798 | 168 | 442.34 | 1 | 2.63 | 0.6 | 16.96 | 16.44 | 0.52 | 0.52 | 0.02 | 0.02 |
| gene-LOC105275683 | https://www.ncbi.nlm.nih.gov/gene/?term=LOC105275683 | 4256 | 205 | 481.67 | 1 | 2.35 | 0.49 | 13.27 | 12.75 | 0.52 | 0.52 | 0.02 | 0.02 |
| gene-LOC105282951 | https://www.ncbi.nlm.nih.gov/gene/?term=LOC105282951 | 3366 | 121 | 359.48 | 1 | 2.97 | 0.83 | 20.13 | 19.67 | 0.46 | 0.46 | 0.02 | 0.02 |
| gene-LOC105274856 | https://www.ncbi.nlm.nih.gov/gene/?term=LOC105274856 | 5216 | 222 | 425.61 | 1 | 1.92 | 0.45 | 17.35 | 16.9 | 0.45 | 0.45 | 0.02 | 0.02 |
| gene-LOC105274531 | https://www.ncbi.nlm.nih.gov/gene/?term=LOC105274531 | 7774 | 291 | 374.32 | 1 | 1.29 | 0.34 | 10.83 | 10.42 | 0.41 | 0.41 | 0.02 | 0.02 |
| gene-LOC105286274 | https://www.ncbi.nlm.nih.gov/gene/?term=LOC105286274 | 3817 | 200 | 523.97 | 1 | 2.62 | 0.5 | 16.54 | 16.86 | -0.32 | 0.32 | -0.02 | 0.02 |
| gene-LOC105278283 | https://www.ncbi.nlm.nih.gov/gene/?term=LOC105278283 | 2317 | 139 | 599.91 | 1 | 4.32 | 0.72 | 25.04 | 25.46 | -0.41 | 0.41 | -0.02 | 0.02 |
| gene-LOC105285409 | https://www.ncbi.nlm.nih.gov/gene/?term=LOC105285409 | 5099 | 58 | 113.75 | 1 | 1.96 | 1.72 | 1.58 | 3.25 | -1.67 | 1.67 | -0.02 | 0.02 |
| gene-LOC105282914 | https://www.ncbi.nlm.nih.gov/gene/?term=LOC105282914 | 5623 | 41 | 72.91 1 | 1.78 | 2.44 | 49.33 | 52.21 | -2.88 | 2.88 | -0.02 | 0.02 | |
| gene-LOC105283989 | https://www.ncbi.nlm.nih.gov/gene/?term=LOC105283989 | 4948 | 31 | 62.65 1 | 2.02 | 3.23 | 46.67 | 44.28 | 2.39 | 2.39 | 0.01 | 0.01 | |
| gene-LOC105285315 | https://www.ncbi.nlm.nih.gov/gene/?term=LOC105285315 | 5458 | 26 | 47.64 1 | 1.83 | 3.85 | 38.49 | 36.2 | 2.29 | 2.29 | 0.01 | 0.01 | |
| gene-LOC105276635 | https://www.ncbi.nlm.nih.gov/gene/?term=LOC105276635 | 4962 | 33 | 66.51 1 | 2.02 | 3.03 | 19.08 | 17.59 | 1.49 | 1.49 | 0.01 | 0.01 | |
| gene-LOC105276692 | https://www.ncbi.nlm.nih.gov/gene/?term=LOC105276692 | 1745 | 34 | 194.84 | 1 | 5.73 | 2.94 | 22.19 | 21.46 | 0.73 | 0.73 | 0.01 | 0.01 |
| gene-LOC105282600 | https://www.ncbi.nlm.nih.gov/gene/?term=LOC105282600 | 3715 | 50 | 134.59 | 1 | 2.69 | 2 | 33.68 | 32.96 | 0.73 | 0.73 | 0.01 | 0.01 |
| gene-LOC105288066 | https://www.ncbi.nlm.nih.gov/gene/?term=LOC105288066 | 1964 | 30 | 152.75 | 1 | 5.09 | 3.33 | 35.84 | 35.13 | 0.71 | 0.71 | 0.01 | 0.01 |
| gene-LOC105281266 | https://www.ncbi.nlm.nih.gov/gene/?term=LOC105281266 | 4102 | 79 | 192.59 | 1 | 2.44 | 1.27 | 15.99 | 15.33 | 0.66 | 0.66 | 0.01 | 0.01 |
| gene-LOC105278962 | https://www.ncbi.nlm.nih.gov/gene/?term=LOC105278962 | 4491 | 106 | 236.03 | 1 | 2.23 | 0.94 | 20.64 | 20.08 | 0.56 | 0.56 | 0.01 | 0.01 |
| gene-LOC105288010 | https://www.ncbi.nlm.nih.gov/gene/?term=LOC105288010 | 3189 | 45 | 141.11 | 1 | 3.14 | 2.22 | 33.55 | 32.98 | 0.56 | 0.56 | 0.01 | 0.01 |
| gene-LOC105275187 | https://www.ncbi.nlm.nih.gov/gene/?term=LOC105275187 | 5534 | 74 | 133.72 | 1 | 1.81 | 1.35 | 16.17 | 15.62 | 0.54 | 0.54 | 0.01 | 0.01 |
| gene-LOC105278228 | https://www.ncbi.nlm.nih.gov/gene/?term=LOC105278228 | 8610 | 172 | 199.77 | 1 | 1.16 | 0.58 | 23.43 | 22.95 | 0.48 | 0.48 | 0.01 | 0.01 |
| gene-LOC105275170 | https://www.ncbi.nlm.nih.gov/gene/?term=LOC105275170 | 12918 | 298 | 230.69 | 1 | 0.77 | 0.34 | 2.35 | 1.88 | 0.47 | 0.47 | 0.01 | 0.01 |
| gene-LOC105276059 | https://www.ncbi.nlm.nih.gov/gene/?term=LOC105276059 | 6256 | 138 | 220.59 | 1 | 1.6 | 0.72 | 9.13 | 8.71 | 0.42 | 0.42 | 0.01 | 0.01 |
| gene-LOC105282811 | https://www.ncbi.nlm.nih.gov/gene/?term=LOC105282811 | 5765 | 218 | 378.14 | 1 | 1.73 | 0.46 | 8.32 | 7.93 | 0.39 | 0.39 | 0.01 | 0.01 |
| gene-LOC105285395 | https://www.ncbi.nlm.nih.gov/gene/?term=LOC105285395 | 8085 | 208 | 257.27 | 1 | 1.24 | 0.48 | 14.72 | 14.34 | 0.39 | 0.39 | 0.01 | 0.01 |
| gene-LOC105274590 | https://www.ncbi.nlm.nih.gov/gene/?term=LOC105274590 | 4516 | 75 | 166.08 | 2 | 4.43 | 2.67 | 29.15 | 28.79 | 0.35 | 0.35 | 0.01 | 0.01 |
| gene-LOC105275451 | https://www.ncbi.nlm.nih.gov/gene/?term=LOC105275451 | 6035 | 136 | 225.35 | 1 | 1.66 | 0.74 | 21.08 | 20.75 | 0.33 | 0.33 | 0.01 | 0.01 |
| gene-LOC105277761 | https://www.ncbi.nlm.nih.gov/gene/?term=LOC105277761 | 4648 | 148 | 318.42 | 1 | 2.15 | 0.68 | 15.39 | 15.08 | 0.31 | 0.31 | 0.01 | 0.01 |
| gene-LOC105279038 | https://www.ncbi.nlm.nih.gov/gene/?term=LOC105279038 | 9453 | 363 | 384.01 | 1 | 1.06 | 0.28 | 8.55 | 8.25 | 0.3 | 0.3 | 0.01 | 0.01 |
| gene-LOC105285807 | https://www.ncbi.nlm.nih.gov/gene/?term=LOC105285807 | 6063 | 228 | 376.05 | 1 | 1.65 | 0.44 | 8.95 | 8.68 | 0.28 | 0.28 | 0.01 | 0.01 |
| gene-LOC105274452 | https://www.ncbi.nlm.nih.gov/gene/?term=LOC105274452 | 6595 | 260 | 394.24 | 1 | 1.52 | 0.38 | 13.07 | 12.82 | 0.25 | 0.25 | 0.01 | 0.01 |
| gene-LOC105278666 | https://www.ncbi.nlm.nih.gov/gene/?term=LOC105278666 | 10341 | 477 | 461.27 | 1 | 0.97 | 0.21 | 6.78 | 6.53 | 0.24 | 0.24 | 0.01 | 0.01 |
| gene-LOC105276527 | https://www.ncbi.nlm.nih.gov/gene/?term=LOC105276527 | 7923 | 332 | 419.03 | 1 | 1.26 | 0.3 | 9.67 | 9.44 | 0.23 | 0.23 | 0.01 | 0.01 |
| gene-LOC113562050 | https://www.ncbi.nlm.nih.gov/gene/?term=LOC113562050 | 17097 | 605 | 353.86 | 1 | 0.58 | 0.17 | 5.18 | 4.96 | 0.21 | 0.21 | 0.01 | 0.01 |
| gene-LOC105275327 | https://www.ncbi.nlm.nih.gov/gene/?term=LOC105275327 | 14245 | 543 | 381.19 | 1 | 0.7 | 0.18 | 6.62 | 6.43 | 0.19 | 0.19 | 0.01 | 0.01 |
| gene-LOC105282491 | https://www.ncbi.nlm.nih.gov/gene/?term=LOC105282491 | 4761 | 177 | 371.77 | 1 | 2.1 | 0.56 | 8.66 | 8.47 | 0.19 | 0.19 | 0.01 | 0.01 |
| gene-LOC105275460 | https://www.ncbi.nlm.nih.gov/gene/?term=LOC105275460 | 2473 | 123 | 497.37 | 1 | 4.04 | 0.81 | 13.15 | 12.97 | 0.18 | 0.18 | 0.01 | 0.01 |
| gene-LOC105284262 | https://www.ncbi.nlm.nih.gov/gene/?term=LOC105284262 | 6669 | 235 | 352.38 | 1 | 1.5 | 0.43 | 4.64 | 4.46 | 0.18 | 0.18 | 0.01 | 0.01 |
| gene-LOC105284043 | https://www.ncbi.nlm.nih.gov/gene/?term=LOC105284043 | 5661 | 311 | 549.37 | 1 | 1.77 | 0.32 | 9.86 | 9.71 | 0.15 | 0.15 | 0.01 | 0.01 |
| gene-LOC105278145 | https://www.ncbi.nlm.nih.gov/gene/?term=LOC105278145 | 4470 | 235 | 525.73 | 1 | 2.24 | 0.43 | 10.67 | 10.54 | 0.14 | 0.14 | 0.01 | 0.01 |
| gene-LOC105275374 | https://www.ncbi.nlm.nih.gov/gene/?term=LOC105275374 | 11376 | 539 | 473.8 1 | 0.88 | 0.19 | 9.85 | 10 | -0.15 | 0.15 | -0.01 | 0.01 | |
| gene-LOC105278282 | https://www.ncbi.nlm.nih.gov/gene/?term=LOC105278282 | 1775 | 62 | 349.3 1 | 5.63 | 1.61 | 32.74 | 32.89 | -0.15 | 0.15 | -0.01 | 0.01 | |
| gene-LOC105285742 | https://www.ncbi.nlm.nih.gov/gene/?term=LOC105285742 | 3446 | 123 | 356.94 | 1 | 2.9 | 0.81 | 19.52 | 19.69 | -0.16 | 0.16 | -0.01 | 0.01 |
| gene-LOC105280723 | https://www.ncbi.nlm.nih.gov/gene/?term=LOC105280723 | 3572 | 181 | 506.72 | 1 | 2.8 | 0.55 | 15.21 | 15.4 | -0.2 | 0.2 | -0.01 | 0.01 |
| gene-LOC105284184 | https://www.ncbi.nlm.nih.gov/gene/?term=LOC105284184 | 2535 | 99 | 390.53 | 1 | 3.94 | 1.01 | 29.28 | 29.51 | -0.23 | 0.23 | -0.01 | 0.01 |
| gene-LOC105275587 | https://www.ncbi.nlm.nih.gov/gene/?term=LOC105275587 | 2404 | 55 | 228.79 | 1 | 4.16 | 1.82 | 36.62 | 36.89 | -0.27 | 0.27 | -0.01 | 0.01 |
| gene-LOC105278691 | https://www.ncbi.nlm.nih.gov/gene/?term=LOC105278691 | 3871 | 168 | 434 1 | 2.58 | 0.6 | 14.64 | 14.98 | -0.34 | 0.34 | -0.01 | 0.01 | |
| gene-LOC105283967 | https://www.ncbi.nlm.nih.gov/gene/?term=LOC105283967 | 2393 | 53 | 221.48 | 1 | 4.18 | 1.89 | 45.17 | 45.77 | -0.59 | 0.59 | -0.01 | 0.01 |
| gene-LOC105286452 | https://www.ncbi.nlm.nih.gov/gene/?term=LOC105286452 | 6888 | 60 | 87.11 1 | 1.45 | 1.67 | 31.09 | 30.62 | 0.47 | 0.47 | 0 | 0 | |
| gene-LOC105280408 | https://www.ncbi.nlm.nih.gov/gene/?term=LOC105280408 | 5983 | 56 | 93.6 1 | 1.67 | 1.79 | 20.14 | 19.81 | 0.33 | 0.33 | 0 | 0 | |
| gene-LOC105276683 | https://www.ncbi.nlm.nih.gov/gene/?term=LOC105276683 | 17000 | 137 | 80.59 1 | 0.59 | 0.73 | 6.59 | 6.28 | 0.3 | 0.3 | 0 | 0 | |
| gene-LOC105276196 | https://www.ncbi.nlm.nih.gov/gene/?term=LOC105276196 | 2953 | 58 | 196.41 | 1 | 3.39 | 1.72 | 38.52 | 38.28 | 0.24 | 0.24 | 0 | 0 |
| gene-LOC105288161 | https://www.ncbi.nlm.nih.gov/gene/?term=LOC105288161 | 26065 | 321 | 123.15 | 1 | 0.38 | 0.31 | 11.9 | 11.67 | 0.23 | 0.23 | 0 | 0 |
| gene-LOC105279657 | https://www.ncbi.nlm.nih.gov/gene/?term=LOC105279657 | 5878 | 88 | 149.71 | 1 | 1.7 | 1.14 | 19.35 | 19.14 | 0.21 | 0.21 | 0 | 0 |
| gene-LOC105282151 | https://www.ncbi.nlm.nih.gov/gene/?term=LOC105282151 | 1601 | 44 | 274.83 | 1 | 6.25 | 2.27 | 35.59 | 35.42 | 0.18 | 0.18 | 0 | 0 |
| gene-LOC105287764 | https://www.ncbi.nlm.nih.gov/gene/?term=LOC105287764 | 18359 | 402 | 218.97 | 1 | 0.54 | 0.25 | 3.16 | 2.99 | 0.17 | 0.17 | 0 | 0 |
| gene-LOC105283712 | https://www.ncbi.nlm.nih.gov/gene/?term=LOC105283712 | 1910 | 36 | 188.48 | 1 | 5.24 | 2.78 | 29.2 | 29.04 | 0.16 | 0.16 | 0 | 0 |
| gene-LOC105284036 | https://www.ncbi.nlm.nih.gov/gene/?term=LOC105284036 | 3254 | 107 | 328.83 | 1 | 3.07 | 0.93 | 20.44 | 20.34 | 0.09 | 0.09 | 0 | 0 |

| | | | | | | | | | | | | | |
|---|---|---|---|---|---|---|---|---|---|---|---|---|---|
| gene-LOC105274823 | https://www.ncbi.nlm.nih.gov/gene/?term=LOC105274823 | 4620 | 146 | 316.02 | 1 | 2.16 | 0.68 | 20.43 | 20.37 | 0.06 | 0.06 | 0 | 0 |
| gene-LOC105280265 | https://www.ncbi.nlm.nih.gov/gene/?term=LOC105280265 | 85984 | 3274 | 380.77 | 1 | 0.12 | 0.03 | 1.1 | 1.05 | 0.06 | 0.06 | 0 | 0 |
| gene-LOC105280513 | https://www.ncbi.nlm.nih.gov/gene/?term=LOC105280513 | 69542 | 1377 | 198.01 | 1 | 0.14 | 0.07 | 0.72 | 0.66 | 0.06 | 0.06 | 0 | 0 |
| gene-LOC105286194 | https://www.ncbi.nlm.nih.gov/gene/?term=LOC105286194 | 152619 | 2206 | 144.54 | 1 | 0.07 | 0.05 | 6.36 | 6.3 | 0.06 | 0.06 | 0 | 0 |
| gene-LOC105281700 | https://www.ncbi.nlm.nih.gov/gene/?term=LOC105281700 | 2208 | 37 | 167.57 | 1 | 4.53 | 2.7 | 39.8 | 39.75 | 0.05 | 0.05 | 0 | 0 |
| gene-LOC105286515 | https://www.ncbi.nlm.nih.gov/gene/?term=LOC105286515 | 3751 | 106 | 282.59 | 1 | 2.67 | 0.94 | 10.25 | 10.22 | 0.04 | 0.04 | 0 | 0 |
| gene-LOC105278455 | https://www.ncbi.nlm.nih.gov/gene/?term=LOC105278455 | 27181 | 985 | 362.39 | 1 | 0.37 | 0.1 | 6.75 | 6.75 | 0 | 0 | 0 | 0 |
| gene-LOC105280407 | https://www.ncbi.nlm.nih.gov/gene/?term=LOC105280407 | 10950 | 210 | 191.78 | 1 | 0.91 | 0.48 | 16.39 | 16.4 | -0.01 | 0.01 | 0 | 0 |
| gene-LOC105283123 | https://www.ncbi.nlm.nih.gov/gene/?term=LOC105283123 | 1507 | 55 | 364.96 | 1 | 6.64 | 1.82 | 18.11 | 18.13 | -0.02 | 0.02 | 0 | 0 |
| gene-LOC105274669 | https://www.ncbi.nlm.nih.gov/gene/?term=LOC105274669 | 8555 | 270 | 315.6 | 1 | 1.17 | 0.37 | 5.54 | 5.56 | -0.03 | 0.03 | 0 | 0 |
| gene-LOC105278534 | https://www.ncbi.nlm.nih.gov/gene/?term=LOC105278534 | 7364 | 196 | 266.16 | 1 | 1.36 | 0.51 | 17.75 | 17.8 | -0.05 | 0.05 | 0 | 0 |
| gene-LOC105287550 | https://www.ncbi.nlm.nih.gov/gene/?term=LOC105287550 | 15659 | 616 | 393.38 | 1 | 0.64 | 0.16 | 5.82 | 5.86 | -0.05 | 0.05 | 0 | 0 |
| gene-LOC105285496 | https://www.ncbi.nlm.nih.gov/gene/?term=LOC105285496 | 4080 | 110 | 269.61 | 1 | 2.45 | 0.91 | 40.35 | 40.5 | -0.15 | 0.15 | 0 | 0 |
| gene-LOC105274851 | https://www.ncbi.nlm.nih.gov/gene/?term=LOC105274851 | 6616 | 147 | 222.19 | 1 | 1.51 | 0.68 | 9.12 | 9.33 | -0.21 | 0.21 | 0 | 0 |
| gene-LOC105277866 | https://www.ncbi.nlm.nih.gov/gene/?term=LOC105277866 | 33045 | 423 | 128.01 | 1 | 0.3 | 0.24 | 3 | 3.27 | -0.28 | 0.28 | 0 | 0 |
| gene-LOC105285406 | https://www.ncbi.nlm.nih.gov/gene/?term=LOC105285406 | 8015 | 55 | 68.62 | 1 | 1.25 | 1.82 | 29.11 | 29.59 | -0.47 | 0.47 | 0 | 0 |

Directory: OGLall File: rnk-dmg-Ob_LI_BCphaseB1.vs.RphaseA1.txt

| gene_ID | gene_link | gwidth | #Sites | #per10Kb | #dmSites | #dmsp10kb | %dmSites | %pSite1 | %pSite2 | DMpSite | ADMpSite | DMpNucl | ADMpNucl |
|---|---|---|---|---|---|---|---|---|---|---|---|---|---|
| gene-LOC105285392 | https://www.ncbi.nlm.nih.gov/gene/?term=LOC105285392 | 1601 | 91 | 568.39 | 6 | 37.48 | 6.59 | 35.68 | 31.54 | 4.15 | 4.15 | 0.24 | 0.24 |
| gene-LOC113561930 | https://www.ncbi.nlm.nih.gov/gene/?term=LOC113561930 | 422 | 22 | 521.33 | 1 | 23.7 | 4.55 | 10.08 | 13.72 | -3.64 | 3.64 | -0.19 | 0.19 |
| gene-LOC105275378 | https://www.ncbi.nlm.nih.gov/gene/?term=LOC105275378 | 1189 | 21 | 176.62 | 3 | 25.23 | 14.29 | 38.9 | 30.15 | 8.75 | 8.75 | 0.15 | 0.15 |
| gene-LOC105278153 | https://www.ncbi.nlm.nih.gov/gene/?term=LOC105278153 | 550 | 23 | 418.18 | 3 | 54.55 | 13.04 | 16.52 | 13.29 | 3.23 | 3.23 | 0.14 | 0.14 |
| gene-LOC109610897 | https://www.ncbi.nlm.nih.gov/gene/?term=LOC109610897 | 1561 | 124 | 794.36 | 2 | 12.81 | 1.61 | 14.34 | 16.15 | -1.8 | 1.8 | -0.14 | 0.14 |
| gene-LOC105286645 | https://www.ncbi.nlm.nih.gov/gene/?term=LOC105286645 | 1499 | 55 | 366.91 | 1 | 6.67 | 1.82 | 37.78 | 41.38 | -3.6 | 3.6 | -0.13 | 0.13 |
| gene-LOC105274528 | https://www.ncbi.nlm.nih.gov/gene/?term=LOC105274528 | 2564 | 155 | 604.52 | 3 | 11.7 | 1.94 | 18.68 | 16.63 | 2.05 | 2.05 | 0.12 | 0.12 |
| gene-LOC105278849 | https://www.ncbi.nlm.nih.gov/gene/?term=LOC105278849 | 776 | 75 | 966.49 | 1 | 12.89 | 1.33 | 24.51 | 23.23 | 1.28 | 1.28 | 0.12 | 0.12 |
| gene-LOC105284174 | https://www.ncbi.nlm.nih.gov/gene/?term=LOC105284174 | 1627 | 83 | 510.14 | 1 | 6.15 | 1.2 | 19.97 | 22.34 | -2.36 | 2.36 | -0.12 | 0.12 |
| gene-LOC105286269 | https://www.ncbi.nlm.nih.gov/gene/?term=LOC105286269 | 1401 | 51 | 364.03 | 4 | 28.55 | 7.84 | 50.68 | 53.95 | -3.27 | 3.27 | -0.12 | 0.12 |
| gene-LOC105283717 | https://www.ncbi.nlm.nih.gov/gene/?term=LOC105283717 | 1373 | 87 | 633.65 | 2 | 14.57 | 2.3 | 29.33 | 27.67 | 1.66 | 1.66 | 0.11 | 0.11 |
| gene-LOC105285635 | https://www.ncbi.nlm.nih.gov/gene/?term=LOC105285635 | 1175 | 92 | 782.98 | 1 | 8.51 | 1.09 | 28.39 | 26.97 | 1.42 | 1.42 | 0.11 | 0.11 |
| gene-LOC105288242 | https://www.ncbi.nlm.nih.gov/gene/?term=LOC105288242 | 1260 | 36 | 285.71 | 1 | 7.94 | 2.78 | 42.4 | 46.4 | -4 | 4 | -0.11 | 0.11 |
| gene-LOC113562457 | https://www.ncbi.nlm.nih.gov/gene/?term=LOC113562457 | 739 | 11 | 148.85 | 2 | 27.06 | 18.18 | 47.39 | 54.48 | -7.08 | 7.08 | -0.11 | 0.11 |
| gene-LOC105275214 | https://www.ncbi.nlm.nih.gov/gene/?term=LOC105275214 | 1299 | 44 | 338.72 | 1 | 7.7 | 2.27 | 33.78 | 30.91 | 2.87 | 2.87 | 0.1 | 0.1 |
| gene-LOC105276563 | https://www.ncbi.nlm.nih.gov/gene/?term=LOC105276563 | 1958 | 116 | 592.44 | 5 | 25.54 | 4.31 | 17.76 | 16.02 | 1.74 | 1.74 | 0.1 | 0.1 |
| gene-LOC105285716 | https://www.ncbi.nlm.nih.gov/gene/?term=LOC105285716 | 1138 | 36 | 316.34 | 1 | 8.79 | 2.78 | 36.22 | 39.41 | -3.19 | 3.19 | -0.1 | 0.1 |
| gene-LOC105280327 | https://www.ncbi.nlm.nih.gov/gene/?term=LOC105280327 | 2670 | 109 | 408.24 | 6 | 22.47 | 5.5 | 29.28 | 27.03 | 2.25 | 2.25 | 0.09 | 0.09 |
| gene-LOC105274711 | https://www.ncbi.nlm.nih.gov/gene/?term=LOC105274711 | 1655 | 86 | 519.64 | 2 | 12.08 | 2.33 | 46.42 | 44.73 | 1.69 | 1.69 | 0.09 | 0.09 |
| gene-LOC105278873 | https://www.ncbi.nlm.nih.gov/gene/?term=LOC105278873 | 1897 | 149 | 785.45 | 6 | 31.63 | 4.03 | 20.63 | 19.44 | 1.19 | 1.19 | 0.09 | 0.09 |
| gene-LOC105287690 | https://www.ncbi.nlm.nih.gov/gene/?term=LOC105287690 | 1378 | 108 | 783.74 | 3 | 21.77 | 2.78 | 18.67 | 17.5 | 1.16 | 1.16 | 0.09 | 0.09 |
| gene-LOC105287942 | https://www.ncbi.nlm.nih.gov/gene/?term=LOC105287942 | 2147 | 78 | 363.3 | 5 | 23.29 | 6.41 | 31.04 | 33.39 | -2.35 | 2.35 | -0.09 | 0.09 |
| gene-LOC105286212 | https://www.ncbi.nlm.nih.gov/gene/?term=LOC105286212 | 1022 | 32 | 313.11 | 1 | 9.78 | 3.12 | 35.65 | 38.39 | -2.74 | 2.74 | -0.09 | 0.09 |
| gene-LOC105286107 | https://www.ncbi.nlm.nih.gov/gene/?term=LOC105286107 | 1902 | 47 | 247.11 | 2 | 10.52 | 4.26 | 14.16 | 17.81 | -3.65 | 3.65 | -0.09 | 0.09 |

...

... only first 25 lines shown ...

Directory: OGLall File: rnk-dmg-Ob_LI_BCphaseB1.vs.RphaseA2.txt

| gene_ID | gene_link | gwidth | #Sites | #per10Kb | #dmSites | #dmsp10kb | %dmSites | %pSite1 | %pSite2 | DMpSite | ADMpSite | DMpNucl | ADMpNucl |
|---|---|---|---|---|---|---|---|---|---|---|---|---|---|
| gene-LOC109611487 | https://www.ncbi.nlm.nih.gov/gene/?term=LOC109611487 | 437 | 32 | 732.27 | 1 | 22.88 | 3.12 | 14.09 | 17.07 | -2.98 | 2.98 | -0.22 | 0.22 |
| gene-LOC105275355 | https://www.ncbi.nlm.nih.gov/gene/?term=LOC105275355 | 3197 | 169 | 528.62 | 10 | 31.28 | 5.92 | 27.68 | 24.07 | 3.61 | 3.61 | 0.19 | 0.19 |
| gene-LOC105286212 | https://www.ncbi.nlm.nih.gov/gene/?term=LOC105286212 | 1022 | 39 | 381.6 | 3 | 29.35 | 7.69 | 33.25 | 37.99 | -4.73 | 4.73 | -0.18 | 0.18 |
| gene-LOC105276451 | https://www.ncbi.nlm.nih.gov/gene/?term=LOC105276451 | 1360 | 40 | 294.12 | 3 | 22.06 | 7.5 | 35.85 | 41.18 | -5.34 | 5.34 | -0.16 | 0.16 |
| gene-LOC105279049 | https://www.ncbi.nlm.nih.gov/gene/?term=LOC105279049 | 1285 | 100 | 778.21 | 1 | 7.78 | 1 | 16.79 | 18.78 | -1.99 | 1.99 | -0.15 | 0.15 |
| gene-LOC105282220 | https://www.ncbi.nlm.nih.gov/gene/?term=LOC105282220 | 2298 | 236 | 1026.98 | 3 | 13.05 | 1.27 | 7.12 | 8.51 | -1.39 | 1.39 | -0.14 | 0.14 |
| gene-LOC105275307 | https://www.ncbi.nlm.nih.gov/gene/?term=LOC105275307 | 1420 | 54 | 380.28 | 2 | 14.08 | 3.7 | 38.11 | 41.77 | -3.67 | 3.67 | -0.14 | 0.14 |
| gene-LOC109610897 | https://www.ncbi.nlm.nih.gov/gene/?term=LOC109610897 | 1561 | 147 | 941.7 | 2 | 12.81 | 1.36 | 13.43 | 14.76 | -1.34 | 1.34 | -0.13 | 0.13 |
| gene-LOC105281172 | https://www.ncbi.nlm.nih.gov/gene/?term=LOC105281172 | 1310 | 30 | 229.01 | 3 | 22.9 | 10 | 42.61 | 48.18 | -5.57 | 5.57 | -0.13 | 0.13 |
| gene-LOC105285628 | https://www.ncbi.nlm.nih.gov/gene/?term=LOC105285628 | 1033 | 22 | 212.97 | 3 | 29.04 | 13.64 | 32.6 | 38.87 | -6.27 | 6.27 | -0.13 | 0.13 |
| gene-LOC105278849 | https://www.ncbi.nlm.nih.gov/gene/?term=LOC105278849 | 776 | 76 | 979.38 | 1 | 12.89 | 1.32 | 24.69 | 23.43 | 1.26 | 1.26 | 0.12 | 0.12 |
| gene-LOC105279035 | https://www.ncbi.nlm.nih.gov/gene/?term=LOC105279035 | 2108 | 166 | 787.48 | 4 | 18.98 | 2.41 | 18.4 | 19.91 | -1.5 | 1.5 | -0.12 | 0.12 |
| gene-LOC105278396 | https://www.ncbi.nlm.nih.gov/gene/?term=LOC105278396 | 3012 | 190 | 630.81 | 5 | 16.6 | 2.63 | 24.21 | 26.15 | -1.94 | 1.94 | -0.12 | 0.12 |
| gene-LOC105285716 | https://www.ncbi.nlm.nih.gov/gene/?term=LOC105285716 | 1138 | 42 | 369.07 | 3 | 26.36 | 7.14 | 36.53 | 39.66 | -3.13 | 3.13 | -0.12 | 0.12 |
| gene-LOC113562901 | https://www.ncbi.nlm.nih.gov/gene/?term=LOC113562901 | 1531 | 25 | 163.29 | 3 | 19.6 | 12 | 50.23 | 57.46 | -7.23 | 7.23 | -0.12 | 0.12 |
| gene-LOC105284174 | https://www.ncbi.nlm.nih.gov/gene/?term=LOC105284174 | 1627 | 92 | 565.46 | 1 | 6.15 | 1.09 | 19.2 | 21.11 | -1.92 | 1.92 | -0.11 | 0.11 |
| gene-LOC113561930 | https://www.ncbi.nlm.nih.gov/gene/?term=LOC113561930 | 422 | 23 | 545.02 | 1 | 23.7 | 4.35 | 12.86 | 14.92 | -2.06 | 2.06 | -0.11 | 0.11 |
| gene-LOC105274542 | https://www.ncbi.nlm.nih.gov/gene/?term=LOC105274542 | 1266 | 61 | 481.83 | 2 | 15.8 | 3.28 | 29.9 | 32.16 | -2.26 | 2.26 | -0.11 | 0.11 |
| gene-LOC105285056 | https://www.ncbi.nlm.nih.gov/gene/?term=LOC105285056 | 1849 | 88 | 475.93 | 1 | 5.41 | 1.14 | 26.08 | 28.38 | -2.3 | 2.3 | -0.11 | 0.11 |
| gene-LOC105283847 | https://www.ncbi.nlm.nih.gov/gene/?term=LOC105283847 | 1590 | 74 | 465.41 | 2 | 12.58 | 2.7 | 33.62 | 36.08 | -2.47 | 2.47 | -0.11 | 0.11 |
| gene-LOC105274524 | https://www.ncbi.nlm.nih.gov/gene/?term=LOC105274524 | 1126 | 43 | 381.88 | 2 | 17.76 | 4.65 | 27.35 | 30.21 | -2.85 | 2.85 | -0.11 | 0.11 |
| gene-LOC105278282 | https://www.ncbi.nlm.nih.gov/gene/?term=LOC105278282 | 1775 | 68 | 383.1 | 3 | 16.9 | 4.41 | 31.98 | 34.93 | -2.95 | 2.95 | -0.11 | 0.11 |
| gene-LOC105286269 | https://www.ncbi.nlm.nih.gov/gene/?term=LOC105286269 | 1401 | 51 | 364.03 | 1 | 7.14 | 1.96 | 50.68 | 53.75 | -3.07 | 3.07 | -0.11 | 0.11 |
| gene-LOC105280117 | https://www.ncbi.nlm.nih.gov/gene/?term=LOC105280117 | 1817 | 63 | 346.73 | 2 | 11.01 | 3.17 | 46.36 | 49.66 | -3.3 | 3.3 | -0.11 | 0.11 |

...
... only first 25 lines shown ...

Directory: OGLall File: rnk-dmg-Ob_LI_BCphaseB1.vs.RphaseB1.txt

| gene_ID | gene_link | gwidth | #Sites | #per10Kb | #dmSites | #dmsp10kb | %dmSites | %pSite1 | %pSite2 | DMpSite | ADMpSite | DMpNucl | ADMpNucl |
|---|---|---|---|---|---|---|---|---|---|---|---|---|---|
| gene-LOC113562345 | https://www.ncbi.nlm.nih.gov/gene/?term=LOC113562345 | 2290 | 77 | 336.24 | 12 | 52.4 | 15.58 | 26.12 | 14.52 | 11.59 | 11.59 | 0.39 | 0.39 |
| gene-LOC105275355 | https://www.ncbi.nlm.nih.gov/gene/?term=LOC105275355 | 3197 | 174 | 544.26 | 1 | 3.13 | 0.57 | 26.94 | 21.77 | 5.17 | 5.17 | 0.28 | 0.28 |
| gene-LOC105284677 | https://www.ncbi.nlm.nih.gov/gene/?term=LOC105284677 | 1905 | 42 | 220.47 | 1 | 5.25 | 2.38 | 23.34 | 35.09 | -11.75 | 11.75 | -0.26 | 0.26 |
| gene-LOC105275384 | https://www.ncbi.nlm.nih.gov/gene/?term=LOC105275384 | 1987 | 92 | 463.01 | 1 | 5.03 | 1.09 | 34.18 | 31.19 | 2.98 | 2.98 | 0.14 | 0.14 |
| gene-LOC105275070 | https://www.ncbi.nlm.nih.gov/gene/?term=LOC105275070 | 5667 | 61 | 107.64 | 2 | 3.53 | 3.28 | 14.18 | 5.29 | 8.9 | 8.9 | 0.1 | 0.1 |
| gene-LOC105274534 | https://www.ncbi.nlm.nih.gov/gene/?term=LOC105274534 | 1329 | 59 | 443.94 | 1 | 7.52 | 1.69 | 41.22 | 43.19 | -1.98 | 1.98 | -0.09 | 0.09 |
| gene-LOC105279900 | https://www.ncbi.nlm.nih.gov/gene/?term=LOC105279900 | 2191 | 56 | 255.59 | 1 | 4.56 | 1.79 | 31.89 | 28.91 | 2.98 | 2.98 | 0.08 | 0.08 |
| gene-LOC105275708 | https://www.ncbi.nlm.nih.gov/gene/?term=LOC105275708 | 1772 | 15 | 84.65 | 1 | 5.64 | 6.67 | 46.38 | 38.49 | 7.9 | 7.9 | 0.07 | 0.07 |
| gene-LOC105278282 | https://www.ncbi.nlm.nih.gov/gene/?term=LOC105278282 | 1775 | 65 | 366.2 | 1 | 5.63 | 1.54 | 30.53 | 32.37 | -1.84 | 1.84 | -0.07 | 0.07 |
| gene-LOC105285514 | https://www.ncbi.nlm.nih.gov/gene/?term=LOC105285514 | 1277 | 11 | 86.14 | 1 | 7.83 | 9.09 | 92.13 | 85.12 | 7.01 | 7.01 | 0.06 | 0.06 |
| gene-LOC105276272 | https://www.ncbi.nlm.nih.gov/gene/?term=LOC105276272 | 1965 | 40 | 203.56 | 1 | 5.09 | 2.5 | 30.29 | 27.31 | 2.98 | 2.98 | 0.06 | 0.06 |
| gene-LOC105276611 | https://www.ncbi.nlm.nih.gov/gene/?term=LOC105276611 | 1731 | 36 | 207.97 | 1 | 5.78 | 2.78 | 30.25 | 27.6 | 2.65 | 2.65 | 0.06 | 0.06 |
| gene-LOC105287063 | https://www.ncbi.nlm.nih.gov/gene/?term=LOC105287063 | 1874 | 50 | 266.81 | 1 | 5.34 | 2 | 31.85 | 29.62 | 2.23 | 2.23 | 0.06 | 0.06 |
| gene-LOC105283552 | https://www.ncbi.nlm.nih.gov/gene/?term=LOC105283552 | 1562 | 41 | 262.48 | 1 | 6.4 | 2.44 | 53.31 | 51.1 | 2.21 | 2.21 | 0.06 | 0.06 |
| gene-LOC105280704 | https://www.ncbi.nlm.nih.gov/gene/?term=LOC105280704 | 4266 | 147 | 344.59 | 1 | 2.34 | 0.68 | 18.45 | 16.63 | 1.81 | 1.81 | 0.06 | 0.06 |
| gene-LOC105287095 | https://www.ncbi.nlm.nih.gov/gene/?term=LOC105287095 | 1435 | 70 | 487.8 | 1 | 6.97 | 1.43 | 25.15 | 23.89 | 1.26 | 1.26 | 0.06 | 0.06 |
| gene-LOC105288036 | https://www.ncbi.nlm.nih.gov/gene/?term=LOC105288036 | 5110 | 232 | 454.01 | 1 | 1.96 | 0.43 | 15.89 | 14.67 | 1.22 | 1.22 | 0.06 | 0.06 |
| gene-LOC105283712 | https://www.ncbi.nlm.nih.gov/gene/?term=LOC105283712 | 1910 | 35 | 183.25 | 1 | 5.24 | 2.86 | 31.43 | 28.9 | 2.53 | 2.53 | 0.05 | 0.05 |
| gene-LOC113561805 | https://www.ncbi.nlm.nih.gov/gene/?term=LOC113561805 | 3337 | 76 | 227.75 | 1 | 3 | 1.32 | 32.51 | 30.53 | 1.98 | 1.98 | 0.05 | 0.05 |
| gene-LOC105274449 | https://www.ncbi.nlm.nih.gov/gene/?term=LOC105274449 | 1549 | 53 | 342.16 | 1 | 6.46 | 1.89 | 50.86 | 49.29 | 1.57 | 1.57 | 0.05 | 0.05 |
| gene-LOC105287417 | https://www.ncbi.nlm.nih.gov/gene/?term=LOC105287417 | 1025 | 40 | 390.24 | 1 | 9.76 | 2.5 | 32.13 | 30.76 | 1.38 | 1.38 | 0.05 | 0.05 |
| gene-LOC105282156 | https://www.ncbi.nlm.nih.gov/gene/?term=LOC105282156 | 2577 | 132 | 512.22 | 1 | 3.88 | 0.76 | 26.63 | 25.67 | 0.96 | 0.96 | 0.05 | 0.05 |
| gene-LOC105285055 | https://www.ncbi.nlm.nih.gov/gene/?term=LOC105285055 | 4108 | 248 | 603.7 | 1 | 2.43 | 0.4 | 12.58 | 11.79 | 0.79 | 0.79 | 0.05 | 0.05 |
| gene-LOC105282885 | https://www.ncbi.nlm.nih.gov/gene/?term=LOC105282885 | 1870 | 41 | 219.25 | 1 | 5.35 | 2.44 | 39.07 | 41.13 | -2.06 | 2.06 | -0.05 | 0.05 |
| gene-LOC105278454 | https://www.ncbi.nlm.nih.gov/gene/?term=LOC105278454 | 1972 | 41 | 207.91 | 1 | 5.07 | 2.44 | 41.77 | 44.02 | -2.26 | 2.26 | -0.05 | 0.05 |
| gene-LOC105281039 | https://www.ncbi.nlm.nih.gov/gene/?term=LOC105281039 | 1471 | 14 | 95.17 | 1 | 6.8 | 7.14 | 44.63 | 40.12 | 4.51 | 4.51 | 0.04 | 0.04 |
| gene-LOC105275518 | https://www.ncbi.nlm.nih.gov/gene/?term=LOC105275518 | 2335 | 25 | 107.07 | 1 | 4.28 | 4 | 44.99 | 41.35 | 3.64 | 3.64 | 0.04 | 0.04 |
| gene-LOC105285179 | https://www.ncbi.nlm.nih.gov/gene/?term=LOC105285179 | 4191 | 194 | 462.9 | 1 | 2.39 | 0.52 | 17.62 | 16.69 | 0.93 | 0.93 | 0.04 | 0.04 |
| gene-LOC105285695 | https://www.ncbi.nlm.nih.gov/gene/?term=LOC105285695 | 1371 | 75 | 547.05 | 1 | 7.29 | 1.33 | 27.5 | 26.71 | 0.79 | 0.79 | 0.04 | 0.04 |
| gene-LOC105275460 | https://www.ncbi.nlm.nih.gov/gene/?term=LOC105275460 | 2473 | 127 | 513.55 | 1 | 4.04 | 0.79 | 13.03 | 12.3 | 0.73 | 0.73 | 0.04 | 0.04 |
| gene-LOC105287633 | https://www.ncbi.nlm.nih.gov/gene/?term=LOC105287633 | 1524 | 25 | 164.04 | 1 | 6.56 | 4 | 61.97 | 64.15 | -2.18 | 2.18 | -0.04 | 0.04 |
| gene-LOC105283989 | https://www.ncbi.nlm.nih.gov/gene/?term=LOC105283989 | 4948 | 45 | 90.95 | 2 | 4.04 | 4.44 | 38.35 | 35.25 | 3.1 | 3.1 | 0.03 | 0.03 |
| gene-LOC105274900 | https://www.ncbi.nlm.nih.gov/gene/?term=LOC105274900 | 5561 | 63 | 113.29 | 2 | 3.6 | 3.17 | 32.28 | 30.06 | 2.22 | 2.22 | 0.03 | 0.03 |
| gene-LOC105279898 | https://www.ncbi.nlm.nih.gov/gene/?term=LOC105279898 | 1343 | 16 | 119.14 | 1 | 7.45 | 6.25 | 58.69 | 56.55 | 2.14 | 2.14 | 0.03 | 0.03 |
| gene-LOC105280348 | https://www.ncbi.nlm.nih.gov/gene/?term=LOC105280348 | 3797 | 79 | 208.06 | 1 | 2.63 | 1.27 | 18.26 | 16.64 | 1.61 | 1.61 | 0.03 | 0.03 |
| gene-LOC105276692 | https://www.ncbi.nlm.nih.gov/gene/?term=LOC105276692 | 1745 | 35 | 200.57 | 1 | 5.73 | 2.86 | 24.46 | 23.1 | 1.36 | 1.36 | 0.03 | 0.03 |
| gene-LOC105278384 | https://www.ncbi.nlm.nih.gov/gene/?term=LOC105278384 | 1390 | 33 | 237.41 | 1 | 7.19 | 3.03 | 46.84 | 45.77 | 1.07 | 1.07 | 0.03 | 0.03 |
| gene-LOC105288079 | https://www.ncbi.nlm.nih.gov/gene/?term=LOC105288079 | 3473 | 110 | 316.73 | 1 | 2.88 | 0.91 | 18.5 | 17.43 | 1.07 | 1.07 | 0.03 | 0.03 |
| gene-LOC105278632 | https://www.ncbi.nlm.nih.gov/gene/?term=LOC105278632 | 3624 | 139 | 383.55 | 1 | 2.76 | 0.72 | 14.57 | 13.67 | 0.9 | 0.9 | 0.03 | 0.03 |
| gene-LOC105283123 | https://www.ncbi.nlm.nih.gov/gene/?term=LOC105283123 | 1507 | 55 | 364.96 | 1 | 6.64 | 1.82 | 19.83 | 18.96 | 0.86 | 0.86 | 0.03 | 0.03 |
| gene-LOC105285674 | https://www.ncbi.nlm.nih.gov/gene/?term=LOC105285674 | 3035 | 109 | 359.14 | 1 | 3.29 | 0.92 | 24.18 | 23.43 | 0.75 | 0.75 | 0.03 | 0.03 |
| gene-LOC105285639 | https://www.ncbi.nlm.nih.gov/gene/?term=LOC105285639 | 10325 | 372 | 360.29 | 2 | 1.94 | 0.54 | 11.5 | 10.78 | 0.72 | 0.72 | 0.03 | 0.03 |
| gene-LOC105283728 | https://www.ncbi.nlm.nih.gov/gene/?term=LOC105283728 | 3256 | 146 | 448.4 | 1 | 3.07 | 0.68 | 16.37 | 15.66 | 0.71 | 0.71 | 0.03 | 0.03 |
| gene-LOC105277435 | https://www.ncbi.nlm.nih.gov/gene/?term=LOC105277435 | 3643 | 179 | 491.35 | 1 | 2.74 | 0.56 | 10.99 | 10.3 | 0.7 | 0.7 | 0.03 | 0.03 |
| gene-LOC105285671 | https://www.ncbi.nlm.nih.gov/gene/?term=LOC105285671 | 2263 | 103 | 455.15 | 1 | 4.42 | 0.97 | 30.33 | 29.64 | 0.69 | 0.69 | 0.03 | 0.03 |
| gene-LOC105276449 | https://www.ncbi.nlm.nih.gov/gene/?term=LOC105276449 | 1231 | 55 | 446.79 | 1 | 8.12 | 1.82 | 22.22 | 21.54 | 0.67 | 0.67 | 0.03 | 0.03 |
| gene-LOC105284355 | https://www.ncbi.nlm.nih.gov/gene/?term=LOC105284355 | 3904 | 154 | 394.47 | 1 | 2.56 | 0.65 | 22.13 | 21.47 | 0.66 | 0.66 | 0.03 | 0.03 |
| gene-LOC105282480 | https://www.ncbi.nlm.nih.gov/gene/?term=LOC105282480 | 4177 | 166 | 397.41 | 1 | 2.39 | 0.6 | 12.65 | 12.01 | 0.65 | 0.65 | 0.03 | 0.03 |
| gene-LOC105276704 | https://www.ncbi.nlm.nih.gov/gene/?term=LOC105276704 | 2063 | 87 | 421.72 | 2 | 9.69 | 2.3 | 14.01 | 13.37 | 0.64 | 0.64 | 0.03 | 0.03 |
| gene-LOC105286997 | https://www.ncbi.nlm.nih.gov/gene/?term=LOC105286997 | 2850 | 150 | 526.32 | 1 | 3.51 | 0.67 | 11.33 | 10.85 | 0.48 | 0.48 | 0.03 | 0.03 |
| gene-LOC105277631 | https://www.ncbi.nlm.nih.gov/gene/?term=LOC105277631 | 4387 | 313 | 713.47 | 1 | 2.28 | 0.32 | 10.74 | 10.28 | 0.46 | 0.46 | 0.03 | 0.03 |
| gene-LOC105274856 | https://www.ncbi.nlm.nih.gov/gene/?term=LOC105274856 | 5216 | 222 | 425.61 | 1 | 1.92 | 0.45 | 16.84 | 17.52 | -0.68 | 0.68 | -0.03 | 0.03 |
| gene-LOC105279778 | https://www.ncbi.nlm.nih.gov/gene/?term=LOC105279778 | 4347 | 163 | 374.97 | 1 | 2.3 | 0.61 | 8.62 | 9.47 | -0.85 | 0.85 | -0.03 | 0.03 |
| gene-LOC105283132 | https://www.ncbi.nlm.nih.gov/gene/?term=LOC105283132 | 1819 | 56 | 307.86 | 1 | 5.5 | 1.79 | 39.1 | 40.18 | -1.09 | 1.09 | -0.03 | 0.03 |
| gene-LOC105285409 | https://www.ncbi.nlm.nih.gov/gene/?term=LOC105285409 | 5099 | 66 | 129.44 | 1 | 1.96 | 1.52 | 2.12 | 4.53 | -2.41 | 2.41 | -0.03 | 0.03 |
| gene-LOC105280419 | https://www.ncbi.nlm.nih.gov/gene/?term=LOC105280419 | 1601 | 11 | 68.71 | 1 | 6.25 | 9.09 | 46.06 | 42.48 | 3.57 | 3.57 | 0.02 | 0.02 |
| gene-LOC105285315 | https://www.ncbi.nlm.nih.gov/gene/?term=LOC105285315 | 5458 | 32 | 58.63 | 1 | 1.83 | 3.12 | 31.91 | 29.18 | 2.72 | 2.72 | 0.02 | 0.02 |
| gene-LOC105276538 | https://www.ncbi.nlm.nih.gov/gene/?term=LOC105276538 | 1511 | 20 | 132.36 | 1 | 6.62 | 5 | 72.51 | 71.18 | 1.32 | 1.32 | 0.02 | 0.02 |
| gene-LOC105280972 | https://www.ncbi.nlm.nih.gov/gene/?term=LOC105280972 | 1532 | 35 | 228.46 | 1 | 6.53 | 2.86 | 17.63 | 16.59 | 1.04 | 1.04 | 0.02 | 0.02 |
| gene-LOC105282794 | https://www.ncbi.nlm.nih.gov/gene/?term=LOC105282794 | 4571 | 110 | 240.65 | 1 | 2.19 | 0.91 | 18.14 | 17.16 | 0.98 | 0.98 | 0.02 | 0.02 |

| | | | | | | | | | | | | | | |
|---|---|---|---|---|---|---|---|---|---|---|---|---|---|---|
| gene-LOC105281266 | https://www.ncbi.nlm.nih.gov/gene/?term=LOC105281266 | 4102 | 88 | 214.53 | | 1 | 2.44 | 1.14 | 15.7 | 14.74 | 0.96 | 0.96 | 0.02 | 0.02 |
| gene-LOC105281848 | https://www.ncbi.nlm.nih.gov/gene/?term=LOC105281848 | 2122 | 58 | 273.33 | | 1 | 4.71 | 1.72 | 22.33 | 21.51 | 0.82 | 0.82 | 0.02 | 0.02 |
| gene-LOC105276410 | https://www.ncbi.nlm.nih.gov/gene/?term=LOC105276410 | 11085 | 341 | 307.62 | | 1 | 0.9 | 0.29 | 9.89 | 9.31 | 0.58 | 0.58 | 0.02 | 0.02 |
| gene-LOC105274823 | https://www.ncbi.nlm.nih.gov/gene/?term=LOC105274823 | 4620 | 172 | 372.29 | | 1 | 2.16 | 0.58 | 17.73 | 17.2 | 0.53 | 0.53 | 0.02 | 0.02 |
| gene-LOC105278497 | https://www.ncbi.nlm.nih.gov/gene/?term=LOC105278497 | 2072 | 92 | 444.02 | | 1 | 4.83 | 1.09 | 20.29 | 19.78 | 0.51 | 0.51 | 0.02 | 0.02 |
| gene-LOC105279298 | https://www.ncbi.nlm.nih.gov/gene/?term=LOC105279298 | 2773 | 130 | 468.81 | | 1 | 3.61 | 0.77 | 23.65 | 23.17 | 0.49 | 0.49 | 0.02 | 0.02 |
| gene-LOC105282491 | https://www.ncbi.nlm.nih.gov/gene/?term=LOC105282491 | 4761 | 193 | 405.38 | | 1 | 2.1 | 0.52 | 8.12 | 7.73 | 0.38 | 0.38 | 0.02 | 0.02 |
| gene-LOC105276527 | https://www.ncbi.nlm.nih.gov/gene/?term=LOC105276527 | 7923 | 370 | 466.99 | | 1 | 1.26 | 0.27 | 8.27 | 7.9 | 0.37 | 0.37 | 0.02 | 0.02 |
| gene-LOC105276429 | https://www.ncbi.nlm.nih.gov/gene/?term=LOC105276429 | 2901 | 132 | 455.02 | | 1 | 3.45 | 0.76 | 22.03 | 22.48 | -0.45 | 0.45 | -0.02 | 0.02 |
| gene-LOC105282151 | https://www.ncbi.nlm.nih.gov/gene/?term=LOC105282151 | 1601 | 43 | 268.58 | | 1 | 6.25 | 2.33 | 38.6 | 39.18 | -0.58 | 0.58 | -0.02 | 0.02 |
| gene-LOC105284036 | https://www.ncbi.nlm.nih.gov/gene/?term=LOC105284036 | 3254 | 103 | 316.53 | | 1 | 3.07 | 0.97 | 23.72 | 24.37 | -0.65 | 0.65 | -0.02 | 0.02 |
| gene-LOC105286265 | https://www.ncbi.nlm.nih.gov/gene/?term=LOC105286265 | 2507 | 67 | 267.25 | | 1 | 3.99 | 1.49 | 29.04 | 29.68 | -0.65 | 0.65 | -0.02 | 0.02 |
| gene-LOC105286515 | https://www.ncbi.nlm.nih.gov/gene/?term=LOC105286515 | 3751 | 111 | 295.92 | | 1 | 2.67 | 0.9 | 10.2 | 10.99 | -0.8 | 0.8 | -0.02 | 0.02 |
| gene-LOC105277903 | https://www.ncbi.nlm.nih.gov/gene/?term=LOC105277903 | 1835 | 38 | 207.08 | | 1 | 5.45 | 2.63 | 52.47 | 53.46 | -0.99 | 0.99 | -0.02 | 0.02 |
| gene-LOC105286770 | https://www.ncbi.nlm.nih.gov/gene/?term=LOC105286770 | 3055 | 49 | 160.39 | | 1 | 3.27 | 2.04 | 11.46 | 12.59 | -1.13 | 1.13 | -0.02 | 0.02 |
| gene-LOC105276196 | https://www.ncbi.nlm.nih.gov/gene/?term=LOC105276196 | 2953 | 60 | 203.18 | | 1 | 3.39 | 1.67 | 37.66 | 38.86 | -1.2 | 1.2 | -0.02 | 0.02 |
| gene-LOC105279528 | https://www.ncbi.nlm.nih.gov/gene/?term=LOC105279528 | 5802 | 41 | 70.67 | 1 | 1.72 | 2.44 | 17.37 | 19.59 | -2.22 | 2.22 | -0.02 | 0.02 | |
| gene-LOC105280447 | https://www.ncbi.nlm.nih.gov/gene/?term=LOC105280447 | 2215 | 19 | 85.78 | 1 | 4.51 | 5.26 | 33.1 | 35.5 | -2.4 | 2.4 | -0.02 | 0.02 | |
| gene-LOC105281515 | https://www.ncbi.nlm.nih.gov/gene/?term=LOC105281515 | 8827 | 55 | 62.31 | 2 | 2.27 | 3.64 | 34.93 | 32.95 | 1.98 | 1.98 | 0.01 | 0.01 | |
| gene-LOC105285060 | https://www.ncbi.nlm.nih.gov/gene/?term=LOC105285060 | 1285 | 16 | 124.51 | | 1 | 7.78 | 6.25 | 42.41 | 41.27 | 1.13 | 1.13 | 0.01 | 0.01 |
| gene-LOC105277215 | https://www.ncbi.nlm.nih.gov/gene/?term=LOC105277215 | 2619 | 29 | 110.73 | | 1 | 3.82 | 3.45 | 25.46 | 24.64 | 0.82 | 0.82 | 0.01 | 0.01 |
| gene-LOC105283943 | https://www.ncbi.nlm.nih.gov/gene/?term=LOC105283943 | 3559 | 69 | 193.87 | | 1 | 2.81 | 1.45 | 20.61 | 20.02 | 0.59 | 0.59 | 0.01 | 0.01 |
| gene-LOC105279664 | https://www.ncbi.nlm.nih.gov/gene/?term=LOC105279664 | 4023 | 67 | 166.54 | | 1 | 2.49 | 1.49 | 18.98 | 18.43 | 0.55 | 0.55 | 0.01 | 0.01 |
| gene-LOC105280408 | https://www.ncbi.nlm.nih.gov/gene/?term=LOC105280408 | 5983 | 63 | 105.3 | 1 | 1.67 | 1.59 | 16.29 | 15.78 | 0.51 | 0.51 | 0.01 | 0.01 | |
| gene-LOC105278534 | https://www.ncbi.nlm.nih.gov/gene/?term=LOC105278534 | 7364 | 204 | 277.02 | | 1 | 1.36 | 0.49 | 17.28 | 16.79 | 0.5 | 0.5 | 0.01 | 0.01 |
| gene-LOC105283649 | https://www.ncbi.nlm.nih.gov/gene/?term=LOC105283649 | 1918 | 57 | 297.18 | | 1 | 5.21 | 1.75 | 40.29 | 39.8 | 0.48 | 0.48 | 0.01 | 0.01 |
| gene-LOC105288078 | https://www.ncbi.nlm.nih.gov/gene/?term=LOC105288078 | 19126 | 350 | 183 | 1 | 0.52 | 0.29 | 15.9 | 15.43 | 0.47 | 0.47 | 0.01 | 0.01 | |
| gene-LOC105280641 | https://www.ncbi.nlm.nih.gov/gene/?term=LOC105280641 | 3682 | 84 | 228.14 | | 1 | 2.72 | 1.19 | 13.99 | 13.53 | 0.46 | 0.46 | 0.01 | 0.01 |
| gene-LOC105275187 | https://www.ncbi.nlm.nih.gov/gene/?term=LOC105275187 | 5534 | 78 | 140.95 | | 1 | 1.81 | 1.28 | 16.44 | 16.02 | 0.42 | 0.42 | 0.01 | 0.01 |
| gene-LOC105278807 | https://www.ncbi.nlm.nih.gov/gene/?term=LOC105278807 | 2166 | 75 | 346.26 | | 1 | 4.62 | 1.33 | 20.48 | 20.06 | 0.41 | 0.41 | 0.01 | 0.01 |
| gene-LOC105286099 | https://www.ncbi.nlm.nih.gov/gene/?term=LOC105286099 | 5919 | 77 | 130.09 | | 1 | 1.69 | 1.3 | 12.19 | 11.79 | 0.4 | 0.4 | 0.01 | 0.01 |
| gene-LOC105286514 | https://www.ncbi.nlm.nih.gov/gene/?term=LOC105286514 | 2528 | 51 | 201.74 | | 1 | 3.96 | 1.96 | 35.82 | 35.46 | 0.36 | 0.36 | 0.01 | 0.01 |
| gene-LOC105275170 | https://www.ncbi.nlm.nih.gov/gene/?term=LOC105275170 | 12918 | 359 | 277.91 | | 1 | 0.77 | 0.28 | 2.02 | 1.81 | 0.21 | 0.21 | 0.01 | 0.01 |
| gene-LOC105284262 | https://www.ncbi.nlm.nih.gov/gene/?term=LOC105284262 | 6669 | 241 | 361.37 | | 1 | 1.5 | 0.41 | 5.35 | 5.16 | 0.19 | 0.19 | 0.01 | 0.01 |
| gene-LOC105285742 | https://www.ncbi.nlm.nih.gov/gene/?term=LOC105285742 | 3446 | 118 | 342.43 | | 1 | 2.9 | 0.85 | 17.32 | 17.14 | 0.18 | 0.18 | 0.01 | 0.01 |
| gene-LOC105274531 | https://www.ncbi.nlm.nih.gov/gene/?term=LOC105274531 | 7774 | 295 | 379.47 | | 1 | 1.29 | 0.34 | 10.65 | 10.48 | 0.16 | 0.16 | 0.01 | 0.01 |
| gene-LOC105279038 | https://www.ncbi.nlm.nih.gov/gene/?term=LOC105279038 | 9453 | 346 | 366.02 | | 1 | 1.06 | 0.29 | 8.85 | 9.03 | -0.18 | 0.18 | -0.01 | 0.01 |
| gene-LOC105288042 | https://www.ncbi.nlm.nih.gov/gene/?term=LOC105288042 | 3507 | 98 | 279.44 | | 1 | 2.85 | 1.02 | 22.85 | 23.09 | -0.24 | 0.24 | -0.01 | 0.01 |
| gene-LOC105280956 | https://www.ncbi.nlm.nih.gov/gene/?term=LOC105280956 | 2639 | 81 | 306.93 | | 1 | 3.79 | 1.23 | 18.56 | 18.86 | -0.29 | 0.29 | -0.01 | 0.01 |
| gene-LOC105281173 | https://www.ncbi.nlm.nih.gov/gene/?term=LOC105281173 | 1595 | 42 | 263.32 | | 1 | 6.27 | 2.38 | 62.5 | 62.82 | -0.31 | 0.31 | -0.01 | 0.01 |
| gene-LOC105285004 | https://www.ncbi.nlm.nih.gov/gene/?term=LOC105285004 | 1334 | 35 | 262.37 | | 1 | 7.5 | 2.86 | 19.44 | 19.75 | -0.31 | 0.31 | -0.01 | 0.01 |
| gene-LOC105278962 | https://www.ncbi.nlm.nih.gov/gene/?term=LOC105278962 | 4491 | 129 | 287.24 | | 1 | 2.23 | 0.78 | 17.72 | 18.05 | -0.34 | 0.34 | -0.01 | 0.01 |
| gene-LOC105278455 | https://www.ncbi.nlm.nih.gov/gene/?term=LOC105278455 | 27181 | 996 | 366.43 | | 1 | 0.37 | 0.1 | 6.32 | 6.68 | -0.36 | 0.36 | -0.01 | 0.01 |
| gene-LOC105282110 | https://www.ncbi.nlm.nih.gov/gene/?term=LOC105282110 | 7748 | 148 | 191.02 | | 1 | 1.29 | 0.68 | 11.12 | 11.53 | -0.42 | 0.42 | -0.01 | 0.01 |
| gene-LOC105282825 | https://www.ncbi.nlm.nih.gov/gene/?term=LOC105282825 | 3919 | 98 | 250.06 | | 1 | 2.55 | 1.02 | 13.43 | 13.91 | -0.48 | 0.48 | -0.01 | 0.01 |
| gene-LOC105274590 | https://www.ncbi.nlm.nih.gov/gene/?term=LOC105274590 | 4516 | 84 | 186.01 | | 2 | 4.43 | 2.38 | 24.43 | 25 | -0.57 | 0.57 | -0.01 | 0.01 |
| gene-LOC105276745 | https://www.ncbi.nlm.nih.gov/gene/?term=LOC105276745 | 3953 | 76 | 192.26 | | 1 | 2.53 | 1.32 | 18.95 | 19.72 | -0.76 | 0.76 | -0.01 | 0.01 |
| gene-LOC105282914 | https://www.ncbi.nlm.nih.gov/gene/?term=LOC105282914 | 5623 | 43 | 76.47 | 1 | 1.78 | 2.33 | 48 | 49.79 | -1.79 | 1.79 | -0.01 | 0.01 | |
| gene-LOC105280407 | https://www.ncbi.nlm.nih.gov/gene/?term=LOC105280407 | 10950 | 227 | 207.31 | | 1 | 0.91 | 0.44 | 15.19 | 14.98 | 0.21 | 0.21 | 0 | 0 |
| gene-LOC105279500 | https://www.ncbi.nlm.nih.gov/gene/?term=LOC105279500 | 3587 | 35 | 97.57 | 1 | 2.79 | 2.86 | 48.82 | 48.61 | 0.2 | 0.2 | 0 | 0 | |
| gene-LOC105288066 | https://www.ncbi.nlm.nih.gov/gene/?term=LOC105288066 | 1964 | 31 | 157.84 | | 1 | 5.09 | 3.23 | 36.75 | 36.57 | 0.19 | 0.19 | 0 | 0 |
| gene-LOC105285981 | https://www.ncbi.nlm.nih.gov/gene/?term=LOC105285981 | 7666 | 219 | 285.68 | | 1 | 1.3 | 0.46 | 18.34 | 18.17 | 0.17 | 0.17 | 0 | 0 |
| gene-LOC105278228 | https://www.ncbi.nlm.nih.gov/gene/?term=LOC105278228 | 8610 | 199 | 231.13 | | 1 | 1.16 | 0.5 | 21.48 | 21.33 | 0.15 | 0.15 | 0 | 0 |
| gene-LOC105275451 | https://www.ncbi.nlm.nih.gov/gene/?term=LOC105275451 | 6035 | 142 | 235.29 | | 1 | 1.66 | 0.7 | 20 | 19.9 | 0.1 | 0.1 | 0 | 0 |
| gene-LOC105283838 | https://www.ncbi.nlm.nih.gov/gene/?term=LOC105283838 | 1748 | 30 | 171.62 | | 1 | 5.72 | 3.33 | 35.36 | 35.26 | 0.1 | 0.1 | 0 | 0 |
| gene-LOC105277216 | https://www.ncbi.nlm.nih.gov/gene/?term=LOC105277216 | 7627 | 117 | 153.4 | 1 | 1.31 | 0.85 | 6.74 | 6.65 | 0.09 | 0.09 | 0 | 0 | |
| gene-LOC105278145 | https://www.ncbi.nlm.nih.gov/gene/?term=LOC105278145 | 4470 | 261 | 583.89 | | 1 | 2.24 | 0.38 | 10.03 | 9.95 | 0.08 | 0.08 | 0 | 0 |
| gene-LOC105280002 | https://www.ncbi.nlm.nih.gov/gene/?term=LOC105280002 | 1940 | 32 | 164.95 | | 1 | 5.15 | 3.12 | 34.33 | 34.27 | 0.06 | 0.06 | 0 | 0 |
| gene-LOC105284043 | https://www.ncbi.nlm.nih.gov/gene/?term=LOC105284043 | 5661 | 295 | 521.11 | | 1 | 1.77 | 0.34 | 10.33 | 10.27 | 0.06 | 0.06 | 0 | 0 |
| gene-LOC105284184 | https://www.ncbi.nlm.nih.gov/gene/?term=LOC105284184 | 2535 | 102 | 402.37 | | 1 | 3.94 | 0.98 | 30 | 29.94 | 0.06 | 0.06 | 0 | 0 |
| gene-LOC105282811 | https://www.ncbi.nlm.nih.gov/gene/?term=LOC105282811 | 5765 | 235 | 407.63 | | 1 | 1.73 | 0.43 | 8.46 | 8.42 | 0.04 | 0.04 | 0 | 0 |
| gene-LOC105278283 | https://www.ncbi.nlm.nih.gov/gene/?term=LOC105278283 | 2317 | 138 | 595.6 | 1 | 4.32 | 0.72 | 25.04 | 25.03 | 0.02 | 0.02 | 0 | 0 | |
| gene-LOC105278666 | https://www.ncbi.nlm.nih.gov/gene/?term=LOC105278666 | 10341 | 484 | 468.04 | | 1 | 0.97 | 0.21 | 6.48 | 6.46 | 0.02 | 0.02 | 0 | 0 |
| gene-LOC105274706 | https://www.ncbi.nlm.nih.gov/gene/?term=LOC105274706 | 1446 | 46 | 318.12 | | 1 | 6.92 | 2.17 | 42.4 | 42.39 | 0 | 0 | 0 | 0 |

| | | | | | | | | | | | | | | | |
|---|---|---|---|---|---|---|---|---|---|---|---|---|---|---|---|
| gene-LOC105276059 | https://www.ncbi.nlm.nih.gov/gene/?term=LOC105276059 | 6256 | 146 | 233.38 | | 1 | 1.6 | 0.68 | 11.41 | 11.42 | 0 | 0 | 0 | 0 | |
| gene-LOC105280265 | https://www.ncbi.nlm.nih.gov/gene/?term=LOC105280265 | 85984 | 3391 | 394.38 | | 1 | 0.12 | 0.03 | 1.11 | 1.13 | -0.03 | 0.03 | 0 | 0 | |
| gene-LOC105278877 | https://www.ncbi.nlm.nih.gov/gene/?term=LOC105278877 | 1968 | 88 | 447.15 | | 1 | 5.08 | 1.14 | 30.96 | 31.01 | -0.05 | 0.05 | 0 | 0 | |
| gene-LOC105287065 | https://www.ncbi.nlm.nih.gov/gene/?term=LOC105287065 | 4859 | 249 | 512.45 | | 1 | 2.06 | 0.4 | 12.25 | 12.3 | -0.05 | 0.05 | 0 | 0 | |
| gene-LOC105284774 | https://www.ncbi.nlm.nih.gov/gene/?term=LOC105284774 | 12089 | 100 | 82.72 | 1 | 0.83 | 1 | 19.53 | 19.59 | -0.06 | 0.06 | 0 | 0 | | |
| gene-LOC105275327 | https://www.ncbi.nlm.nih.gov/gene/?term=LOC105275327 | 14245 | 552 | 387.5 | 1 | 0.7 | 0.18 | 6.33 | 6.41 | -0.08 | 0.08 | 0 | 0 | | |
| gene-LOC105282600 | https://www.ncbi.nlm.nih.gov/gene/?term=LOC105282600 | 3715 | 60 | 161.51 | | 1 | 2.69 | 1.67 | 28.86 | 28.93 | -0.08 | 0.08 | 0 | 0 | |
| gene-LOC105286339 | https://www.ncbi.nlm.nih.gov/gene/?term=LOC105286339 | 5230 | 184 | 351.82 | | 1 | 1.91 | 0.54 | 16.14 | 16.23 | -0.09 | 0.09 | 0 | 0 | |
| gene-LOC105288010 | https://www.ncbi.nlm.nih.gov/gene/?term=LOC105288010 | 3189 | 52 | 163.06 | | 1 | 3.14 | 1.92 | 33.66 | 33.76 | -0.1 | 0.1 | 0 | 0 | |
| gene-LOC105285807 | https://www.ncbi.nlm.nih.gov/gene/?term=LOC105285807 | 6063 | 221 | 364.51 | | 1 | 1.65 | 0.45 | 9.8 | 9.91 | -0.11 | 0.11 | 0 | 0 | |
| gene-LOC105286194 | https://www.ncbi.nlm.nih.gov/gene/?term=LOC105286194 | 152619 | | 2446 | 160.27 | | 1 | 0.07 | 0.04 | 6.09 | 6.23 | -0.14 | 0.14 | 0 | 0 |
| gene-LOC105277866 | https://www.ncbi.nlm.nih.gov/gene/?term=LOC105277866 | 33045 | 544 | 164.62 | | 1 | 0.3 | 0.18 | 2.82 | 3.03 | -0.21 | 0.21 | 0 | 0 | |
| gene-LOC105285406 | https://www.ncbi.nlm.nih.gov/gene/?term=LOC105285406 | 8015 | 57 | 71.12 | 1 | 1.25 | 1.75 | 28.09 | 28.72 | -0.63 | 0.63 | 0 | 0 | | |

Directory: OGLall File: rnk-dmg-Ob_LI_BCphaseB1.vs.RphaseB2.txt

| gene_ID | gene_link | gwidth | #Sites | #per10Kb | #dmSites | #dmsp10kb | %dmSites | %pSite1 | %pSite2 | DMpSite | ADMpSite | DMpNucl | ADMpNucl |
|---|---|---|---|---|---|---|---|---|---|---|---|---|---|
| gene-LOC113562345 | https://www.ncbi.nlm.nih.gov/gene/?term=LOC113562345 | 2290 | 66 | 288.21 | 12 | 52.4 | 18.18 | 30.39 | 17.58 | 12.82 | 12.82 | 0.37 | 0.37 |
| gene-LOC105283552 | https://www.ncbi.nlm.nih.gov/gene/?term=LOC105283552 | 1562 | 41 | 262.48 | 1 | 6.4 | 2.44 | 53.31 | 48.83 | 4.48 | 4.48 | 0.12 | 0.12 |
| gene-LOC105286683 | https://www.ncbi.nlm.nih.gov/gene/?term=LOC105286683 | 1665 | 55 | 330.33 | 1 | 6.01 | 1.82 | 34.95 | 31.78 | 3.17 | 3.17 | 0.1 | 0.1 |
| gene-LOC105275070 | https://www.ncbi.nlm.nih.gov/gene/?term=LOC105275070 | 5667 | 41 | 72.35 | 6 | 10.59 | 14.63 | 12.96 | 1.06 | 11.9 | 11.9 | 0.09 | 0.09 |
| gene-LOC105279898 | https://www.ncbi.nlm.nih.gov/gene/?term=LOC105279898 | 1343 | 10 | 74.46 | 1 | 7.45 | 10 | 83.43 | 71.53 | 11.89 | 11.89 | 0.09 | 0.09 |
| gene-LOC105285671 | https://www.ncbi.nlm.nih.gov/gene/?term=LOC105285671 | 2263 | 105 | 463.99 | 1 | 4.42 | 0.95 | 28.66 | 26.78 | 1.88 | 1.88 | 0.09 | 0.09 |
| gene-LOC105280104 | https://www.ncbi.nlm.nih.gov/gene/?term=LOC105280104 | 3960 | 135 | 340.91 | 1 | 2.53 | 0.74 | 18.4 | 16.2 | 2.2 | 2.2 | 0.07 | 0.07 |
| gene-LOC105274534 | https://www.ncbi.nlm.nih.gov/gene/?term=LOC105274534 | 1329 | 55 | 413.84 | 1 | 7.52 | 1.82 | 44.22 | 45.89 | -1.68 | 1.68 | -0.07 | 0.07 |
| gene-LOC105279356 | https://www.ncbi.nlm.nih.gov/gene/?term=LOC105279356 | 1314 | 43 | 327.25 | 1 | 7.61 | 2.33 | 32.17 | 30.3 | 1.87 | 1.87 | 0.06 | 0.06 |
| gene-LOC105281419 | https://www.ncbi.nlm.nih.gov/gene/?term=LOC105281419 | 2366 | 80 | 338.12 | 1 | 4.23 | 1.25 | 22.59 | 20.79 | 1.81 | 1.81 | 0.06 | 0.06 |
| gene-LOC105280786 | https://www.ncbi.nlm.nih.gov/gene/?term=LOC105280786 | 2573 | 123 | 478.04 | 1 | 3.89 | 0.81 | 29.6 | 28.26 | 1.34 | 1.34 | 0.06 | 0.06 |
| gene-LOC105282156 | https://www.ncbi.nlm.nih.gov/gene/?term=LOC105282156 | 2577 | 127 | 492.82 | 1 | 3.88 | 0.79 | 27.5 | 26.37 | 1.13 | 1.13 | 0.06 | 0.06 |
| gene-LOC105275708 | https://www.ncbi.nlm.nih.gov/gene/?term=LOC105275708 | 1772 | 14 | 79.01 | 1 | 5.64 | 7.14 | 49.7 | 43.39 | 6.31 | 6.31 | 0.05 | 0.05 |
| gene-LOC105275518 | https://www.ncbi.nlm.nih.gov/gene/?term=LOC105275518 | 2335 | 20 | 85.65 | 1 | 4.28 | 5 | 58.2 | 52.79 | 5.42 | 5.42 | 0.05 | 0.05 |
| gene-LOC105282794 | https://www.ncbi.nlm.nih.gov/gene/?term=LOC105282794 | 4571 | 91 | 199.08 | 1 | 2.19 | 1.1 | 19.77 | 17.09 | 2.68 | 2.68 | 0.05 | 0.05 |
| gene-LOC105285674 | https://www.ncbi.nlm.nih.gov/gene/?term=LOC105285674 | 3035 | 98 | 322.9 | 2 | 6.59 | 2.04 | 25.36 | 23.72 | 1.64 | 1.64 | 0.05 | 0.05 |
| gene-LOC105283728 | https://www.ncbi.nlm.nih.gov/gene/?term=LOC105283728 | 3256 | 115 | 353.19 | 1 | 3.07 | 0.87 | 19.83 | 18.35 | 1.48 | 1.48 | 0.05 | 0.05 |
| gene-LOC105288036 | https://www.ncbi.nlm.nih.gov/gene/?term=LOC105288036 | 5110 | 200 | 391.39 | 1 | 1.96 | 0.5 | 17.99 | 16.61 | 1.37 | 1.37 | 0.05 | 0.05 |
| gene-LOC113561621 | https://www.ncbi.nlm.nih.gov/gene/?term=LOC113561621 | 2249 | 83 | 369.05 | 1 | 4.45 | 1.2 | 21.13 | 19.82 | 1.31 | 1.31 | 0.05 | 0.05 |
| gene-LOC105285179 | https://www.ncbi.nlm.nih.gov/gene/?term=LOC105285179 | 4191 | 175 | 417.56 | 1 | 2.39 | 0.57 | 18.73 | 17.48 | 1.25 | 1.25 | 0.05 | 0.05 |
| gene-LOC105275363 | https://www.ncbi.nlm.nih.gov/gene/?term=LOC105275363 | 3798 | 160 | 421.27 | 1 | 2.63 | 0.62 | 17.46 | 16.32 | 1.14 | 1.14 | 0.05 | 0.05 |
| gene-LOC105282885 | https://www.ncbi.nlm.nih.gov/gene/?term=LOC105282885 | 1870 | 47 | 251.34 | 1 | 5.35 | 2.13 | 35.98 | 38.05 | -2.07 | 2.07 | -0.05 | 0.05 |
| gene-LOC105276196 | https://www.ncbi.nlm.nih.gov/gene/?term=LOC105276196 | 2953 | 60 | 203.18 | 1 | 3.39 | 1.67 | 38.8 | 41.25 | -2.45 | 2.45 | -0.05 | 0.05 |
| gene-LOC105283492 | https://www.ncbi.nlm.nih.gov/gene/?term=LOC105283492 | 1535 | 10 | 65.15 | 1 | 6.51 | 10 | 34.7 | 42.8 | -8.1 | 8.1 | -0.05 | 0.05 |
| gene-LOC105287648 | https://www.ncbi.nlm.nih.gov/gene/?term=LOC105287648 | 2205 | 25 | 113.38 | 1 | 4.54 | 4 | 44.43 | 41.17 | 3.26 | 3.26 | 0.04 | 0.04 |
| gene-LOC105285514 | https://www.ncbi.nlm.nih.gov/gene/?term=LOC105285514 | 1277 | 17 | 133.12 | 1 | 7.83 | 5.88 | 73.52 | 70.72 | 2.8 | 2.8 | 0.04 | 0.04 |
| gene-LOC105279664 | https://www.ncbi.nlm.nih.gov/gene/?term=LOC105279664 | 4023 | 54 | 134.23 | 1 | 2.49 | 1.85 | 21.88 | 19.13 | 2.74 | 2.74 | 0.04 | 0.04 |
| gene-LOC105287417 | https://www.ncbi.nlm.nih.gov/gene/?term=LOC105287417 | 1025 | 23 | 224.39 | 1 | 9.76 | 4.35 | 51.1 | 49.27 | 1.83 | 1.83 | 0.04 | 0.04 |
| gene-LOC105284355 | https://www.ncbi.nlm.nih.gov/gene/?term=LOC105284355 | 3904 | 151 | 386.78 | 1 | 2.56 | 0.66 | 21.57 | 20.54 | 1.02 | 1.02 | 0.04 | 0.04 |
| gene-LOC105276704 | https://www.ncbi.nlm.nih.gov/gene/?term=LOC105276704 | 2063 | 87 | 421.72 | 1 | 4.85 | 1.15 | 14.98 | 14.08 | 0.9 | 0.9 | 0.04 | 0.04 |
| gene-LOC105284043 | https://www.ncbi.nlm.nih.gov/gene/?term=LOC105284043 | 5661 | 294 | 519.34 | 1 | 1.77 | 0.34 | 10.31 | 9.51 | 0.8 | 0.8 | 0.04 | 0.04 |
| gene-LOC105285055 | https://www.ncbi.nlm.nih.gov/gene/?term=LOC105285055 | 4108 | 256 | 623.17 | 1 | 2.43 | 0.39 | 12.1 | 11.41 | 0.69 | 0.69 | 0.04 | 0.04 |
| gene-LOC105280377 | https://www.ncbi.nlm.nih.gov/gene/?term=LOC105280377 | 2024 | 69 | 340.91 | 1 | 4.94 | 1.45 | 43.74 | 44.8 | -1.06 | 1.06 | -0.04 | 0.04 |
| gene-LOC105276429 | https://www.ncbi.nlm.nih.gov/gene/?term=LOC105276429 | 2901 | 114 | 392.97 | 1 | 3.45 | 0.88 | 24.72 | 25.87 | -1.14 | 1.14 | -0.04 | 0.04 |
| gene-LOC105287633 | https://www.ncbi.nlm.nih.gov/gene/?term=LOC105287633 | 1524 | 31 | 203.41 | 1 | 6.56 | 3.23 | 53.47 | 55.25 | -1.78 | 1.78 | -0.04 | 0.04 |
| gene-LOC105274900 | https://www.ncbi.nlm.nih.gov/gene/?term=LOC105274900 | 5561 | 55 | 98.9 | 2 | 3.6 | 3.64 | 33.24 | 30.24 | 3 | 3 | 0.03 | 0.03 |
| gene-LOC105280641 | https://www.ncbi.nlm.nih.gov/gene/?term=LOC105280641 | 3682 | 50 | 135.8 | 1 | 2.72 | 2 | 17.97 | 16.08 | 1.89 | 1.89 | 0.03 | 0.03 |
| gene-LOC105288042 | https://www.ncbi.nlm.nih.gov/gene/?term=LOC105288042 | 3507 | 89 | 253.78 | 1 | 2.85 | 1.12 | 22.75 | 21.38 | 1.37 | 1.37 | 0.03 | 0.03 |
| gene-LOC105276901 | https://www.ncbi.nlm.nih.gov/gene/?term=LOC105276901 | 3254 | 82 | 252 | 1 | 3.07 | 1.22 | 28 | 26.83 | 1.17 | 1.17 | 0.03 | 0.03 |
| gene-LOC105274449 | https://www.ncbi.nlm.nih.gov/gene/?term=LOC105274449 | 1549 | 48 | 309.88 | 1 | 6.46 | 2.08 | 51.97 | 50.96 | 1 | 1 | 0.03 | 0.03 |
| gene-LOC105285639 | https://www.ncbi.nlm.nih.gov/gene/?term=LOC105285639 | 10325 | 332 | 321.55 | 2 | 1.94 | 0.6 | 12.25 | 11.3 | 0.95 | 0.95 | 0.03 | 0.03 |
| gene-LOC105278877 | https://www.ncbi.nlm.nih.gov/gene/?term=LOC105278877 | 1968 | 69 | 350.61 | 1 | 5.08 | 1.45 | 36.22 | 35.3 | 0.91 | 0.91 | 0.03 | 0.03 |
| gene-LOC105278807 | https://www.ncbi.nlm.nih.gov/gene/?term=LOC105278807 | 2166 | 67 | 309.33 | 1 | 4.62 | 1.49 | 22.11 | 21.25 | 0.87 | 0.87 | 0.03 | 0.03 |
| gene-LOC105276875 | https://www.ncbi.nlm.nih.gov/gene/?term=LOC105276875 | 3151 | 116 | 368.14 | 1 | 3.17 | 0.86 | 22.78 | 21.93 | 0.84 | 0.84 | 0.03 | 0.03 |
| gene-LOC105283123 | https://www.ncbi.nlm.nih.gov/gene/?term=LOC105283123 | 1507 | 46 | 305.24 | 1 | 6.64 | 2.17 | 17.92 | 17.09 | 0.83 | 0.83 | 0.03 | 0.03 |
| gene-LOC105287095 | https://www.ncbi.nlm.nih.gov/gene/?term=LOC105287095 | 1435 | 58 | 404.18 | 1 | 6.97 | 1.72 | 29.38 | 28.65 | 0.73 | 0.73 | 0.03 | 0.03 |
| gene-LOC105278497 | https://www.ncbi.nlm.nih.gov/gene/?term=LOC105278497 | 2072 | 92 | 444.02 | 1 | 4.83 | 1.09 | 19.2 | 18.5 | 0.7 | 0.7 | 0.03 | 0.03 |
| gene-LOC105277423 | https://www.ncbi.nlm.nih.gov/gene/?term=LOC105277423 | 2340 | 106 | 452.99 | 1 | 4.27 | 0.94 | 29 | 28.34 | 0.66 | 0.66 | 0.03 | 0.03 |
| gene-LOC105286166 | https://www.ncbi.nlm.nih.gov/gene/?term=LOC105286166 | 3050 | 110 | 360.66 | 1 | 3.28 | 0.91 | 23.43 | 24.21 | -0.78 | 0.78 | -0.03 | 0.03 |
| gene-LOC105278282 | https://www.ncbi.nlm.nih.gov/gene/?term=LOC105278282 | 1775 | 56 | 315.49 | 1 | 5.63 | 1.79 | 37.29 | 38.34 | -1.05 | 1.05 | -0.03 | 0.03 |
| gene-LOC105275966 | https://www.ncbi.nlm.nih.gov/gene/?term=LOC105275966 | 3299 | 59 | 178.84 | 1 | 3.03 | 1.69 | 49.71 | 51.14 | -1.42 | 1.42 | -0.03 | 0.03 |
| gene-LOC105285409 | https://www.ncbi.nlm.nih.gov/gene/?term=LOC105285409 | 5099 | 49 | 96.1 | 2 | 3.92 | 4.08 | 2.16 | 4.82 | -2.66 | 2.66 | -0.03 | 0.03 |
| gene-LOC105278108 | https://www.ncbi.nlm.nih.gov/gene/?term=LOC105278108 | 4392 | 46 | 104.74 | 1 | 2.28 | 2.17 | 27.18 | 25.03 | 2.14 | 2.14 | 0.02 | 0.02 |
| gene-LOC113561805 | https://www.ncbi.nlm.nih.gov/gene/?term=LOC113561805 | 3337 | 65 | 194.79 | 1 | 3 | 1.54 | 35.44 | 34.26 | 1.18 | 1.18 | 0.02 | 0.02 |
| gene-LOC105281848 | https://www.ncbi.nlm.nih.gov/gene/?term=LOC105281848 | 2122 | 47 | 221.49 | 1 | 4.71 | 2.13 | 27.57 | 26.57 | 1 | 1 | 0.02 | 0.02 |
| gene-LOC105283592 | https://www.ncbi.nlm.nih.gov/gene/?term=LOC105283592 | 3479 | 79 | 227.08 | 1 | 2.87 | 1.27 | 22.25 | 21.39 | 0.85 | 0.85 | 0.02 | 0.02 |
| gene-LOC105276059 | https://www.ncbi.nlm.nih.gov/gene/?term=LOC105276059 | 6256 | 132 | 211 | 1 | 1.6 | 0.76 | 10.3 | 9.47 | 0.83 | 0.83 | 0.02 | 0.02 |
| gene-LOC105278632 | https://www.ncbi.nlm.nih.gov/gene/?term=LOC105278632 | 3624 | 108 | 298.01 | 1 | 2.76 | 0.93 | 17.43 | 16.66 | 0.77 | 0.77 | 0.02 | 0.02 |
| gene-LOC105285395 | https://www.ncbi.nlm.nih.gov/gene/?term=LOC105285395 | 8085 | 196 | 242.42 | 1 | 1.24 | 0.51 | 16.42 | 15.66 | 0.76 | 0.76 | 0.02 | 0.02 |
| gene-LOC105280956 | https://www.ncbi.nlm.nih.gov/gene/?term=LOC105280956 | 2639 | 71 | 269.04 | 1 | 3.79 | 1.41 | 18.55 | 17.79 | 0.75 | 0.75 | 0.02 | 0.02 |

| | | | | | | | | | | | | | |
|---|---|---|---|---|---|---|---|---|---|---|---|---|---|
| gene-LOC105278534 | https://www.ncbi.nlm.nih.gov/gene/?term=LOC105278534 | 7364 | 187 | 253.94 | 1 | 1.36 | 0.53 | 19.13 | 18.41 | 0.72 | 0.72 | 0.02 | 0.02 |
| gene-LOC105275584 | https://www.ncbi.nlm.nih.gov/gene/?term=LOC105275584 | 3070 | 89 | 289.9 | 1 | 3.26 | 1.12 | 19.48 | 18.76 | 0.71 | 0.71 | 0.02 | 0.02 |
| gene-LOC105284184 | https://www.ncbi.nlm.nih.gov/gene/?term=LOC105284184 | 2535 | 86 | 339.25 | 1 | 3.94 | 1.16 | 33.56 | 32.88 | 0.68 | 0.68 | 0.02 | 0.02 |
| gene-LOC105282811 | https://www.ncbi.nlm.nih.gov/gene/?term=LOC105282811 | 5765 | 215 | 372.94 | 1 | 1.73 | 0.47 | 8.61 | 8.16 | 0.45 | 0.45 | 0.02 | 0.02 |
| gene-LOC105275460 | https://www.ncbi.nlm.nih.gov/gene/?term=LOC105275460 | 2473 | 117 | 473.11 | 1 | 4.04 | 0.85 | 13.75 | 13.31 | 0.44 | 0.44 | 0.02 | 0.02 |
| gene-LOC105279298 | https://www.ncbi.nlm.nih.gov/gene/?term=LOC105279298 | 2773 | 133 | 479.62 | 2 | 7.21 | 1.5 | 23.5 | 23.06 | 0.44 | 0.44 | 0.02 | 0.02 |
| gene-LOC105278454 | https://www.ncbi.nlm.nih.gov/gene/?term=LOC105278454 | 1972 | 44 | 223.12 | 1 | 5.07 | 2.27 | 45.26 | 45.99 | -0.74 | 0.74 | -0.02 | 0.02 |
| gene-LOC105282151 | https://www.ncbi.nlm.nih.gov/gene/?term=LOC105282151 | 1601 | 39 | 243.6 | 1 | 6.25 | 2.56 | 40.09 | 41 | -0.91 | 0.91 | -0.02 | 0.02 |
| gene-LOC105286999 | https://www.ncbi.nlm.nih.gov/gene/?term=LOC105286999 | 2141 | 46 | 214.85 | 1 | 4.67 | 2.17 | 31.8 | 32.92 | -1.12 | 1.12 | -0.02 | 0.02 |
| gene-LOC105281700 | https://www.ncbi.nlm.nih.gov/gene/?term=LOC105281700 | 2208 | 32 | 144.93 | 1 | 4.53 | 3.12 | 37.39 | 39.09 | -1.7 | 1.7 | -0.02 | 0.02 |
| gene-LOC105286770 | https://www.ncbi.nlm.nih.gov/gene/?term=LOC105286770 | 3055 | 30 | 98.2 | 1 | 3.27 | 3.33 | 17.03 | 18.87 | -1.84 | 1.84 | -0.02 | 0.02 |
| gene-LOC105285315 | https://www.ncbi.nlm.nih.gov/gene/?term=LOC105285315 | 5458 | 19 | 34.81 | 1 | 1.83 | 5.26 | 47.14 | 43.22 | 3.93 | 3.93 | 0.01 | 0.01 |
| gene-LOC105281515 | https://www.ncbi.nlm.nih.gov/gene/?term=LOC105281515 | 8827 | 40 | 45.32 | 2 | 2.27 | 5 | 31.53 | 28.62 | 2.91 | 2.91 | 0.01 | 0.01 |
| gene-LOC105280408 | https://www.ncbi.nlm.nih.gov/gene/?term=LOC105280408 | 5983 | 56 | 93.6 | 1 | 1.67 | 1.79 | 17.1 | 16.06 | 1.03 | 1.03 | 0.01 | 0.01 |
| gene-LOC105286099 | https://www.ncbi.nlm.nih.gov/gene/?term=LOC105286099 | 5919 | 72 | 121.64 | 1 | 1.69 | 1.39 | 10.67 | 9.65 | 1.01 | 1.01 | 0.01 | 0.01 |
| gene-LOC105280447 | https://www.ncbi.nlm.nih.gov/gene/?term=LOC105280447 | 2215 | 21 | 94.81 | 1 | 4.51 | 4.76 | 30.62 | 29.74 | 0.88 | 0.88 | 0.01 | 0.01 |
| gene-LOC105275451 | https://www.ncbi.nlm.nih.gov/gene/?term=LOC105275451 | 6035 | 126 | 208.78 | 1 | 1.66 | 0.79 | 22.21 | 21.64 | 0.57 | 0.57 | 0.01 | 0.01 |
| gene-LOC105281599 | https://www.ncbi.nlm.nih.gov/gene/?term=LOC105281599 | 3859 | 49 | 126.98 | 1 | 2.59 | 2.04 | 4.62 | 4.08 | 0.54 | 0.54 | 0.01 | 0.01 |
| gene-LOC105285742 | https://www.ncbi.nlm.nih.gov/gene/?term=LOC105285742 | 3446 | 110 | 319.21 | 1 | 2.9 | 0.91 | 21.56 | 21.15 | 0.4 | 0.4 | 0.01 | 0.01 |
| gene-LOC105274706 | https://www.ncbi.nlm.nih.gov/gene/?term=LOC105274706 | 1446 | 50 | 345.78 | 1 | 6.92 | 2 | 40.48 | 40.09 | 0.39 | 0.39 | 0.01 | 0.01 |
| gene-LOC105280002 | https://www.ncbi.nlm.nih.gov/gene/?term=LOC105280002 | 1940 | 28 | 144.33 | 1 | 5.15 | 3.57 | 37.11 | 36.72 | 0.39 | 0.39 | 0.01 | 0.01 |
| gene-LOC105279038 | https://www.ncbi.nlm.nih.gov/gene/?term=LOC105279038 | 9453 | 314 | 332.17 | 1 | 1.06 | 0.32 | 9.5 | 9.12 | 0.38 | 0.38 | 0.01 | 0.01 |
| gene-LOC105280774 | https://www.ncbi.nlm.nih.gov/gene/?term=LOC105280774 | 2793 | 99 | 354.46 | 1 | 3.58 | 1.01 | 32.85 | 32.47 | 0.38 | 0.38 | 0.01 | 0.01 |
| gene-LOC105284036 | https://www.ncbi.nlm.nih.gov/gene/?term=LOC105284036 | 3254 | 102 | 313.46 | 1 | 3.07 | 0.98 | 22.68 | 22.3 | 0.38 | 0.38 | 0.01 | 0.01 |
| gene-LOC105274590 | https://www.ncbi.nlm.nih.gov/gene/?term=LOC105274590 | 4516 | 69 | 152.79 | 2 | 4.43 | 2.9 | 27.64 | 27.27 | 0.37 | 0.37 | 0.01 | 0.01 |
| gene-LOC105277079 | https://www.ncbi.nlm.nih.gov/gene/?term=LOC105277079 | 13009 | 273 | 209.85 | 1 | 0.77 | 0.37 | 9.7 | 9.33 | 0.37 | 0.37 | 0.01 | 0.01 |
| gene-LOC105278962 | https://www.ncbi.nlm.nih.gov/gene/?term=LOC105278962 | 4491 | 100 | 222.67 | 1 | 2.23 | 1 | 21.33 | 20.97 | 0.36 | 0.36 | 0.01 | 0.01 |
| gene-LOC105287764 | https://www.ncbi.nlm.nih.gov/gene/?term=LOC105287764 | 18359 | 394 | 214.61 | 1 | 0.54 | 0.25 | 3.28 | 2.97 | 0.31 | 0.31 | 0.01 | 0.01 |
| gene-LOC105281266 | https://www.ncbi.nlm.nih.gov/gene/?term=LOC105281266 | 4102 | 78 | 190.15 | 1 | 2.44 | 1.28 | 16.25 | 15.95 | 0.3 | 0.3 | 0.01 | 0.01 |
| gene-LOC105278145 | https://www.ncbi.nlm.nih.gov/gene/?term=LOC105278145 | 4470 | 235 | 525.73 | 1 | 2.24 | 0.43 | 10.78 | 10.55 | 0.23 | 0.23 | 0.01 | 0.01 |
| gene-LOC105284262 | https://www.ncbi.nlm.nih.gov/gene/?term=LOC105284262 | 6669 | 215 | 322.39 | 1 | 1.5 | 0.47 | 5.95 | 5.73 | 0.22 | 0.22 | 0.01 | 0.01 |
| gene-LOC105278666 | https://www.ncbi.nlm.nih.gov/gene/?term=LOC105278666 | 10341 | 444 | 429.36 | 1 | 0.97 | 0.23 | 6.98 | 6.79 | 0.19 | 0.19 | 0.01 | 0.01 |
| gene-LOC105282149 | https://www.ncbi.nlm.nih.gov/gene/?term=LOC105282149 | 1787 | 108 | 604.36 | 1 | 5.6 | 0.93 | 41.15 | 40.97 | 0.18 | 0.18 | 0.01 | 0.01 |
| gene-LOC105276527 | https://www.ncbi.nlm.nih.gov/gene/?term=LOC105276527 | 7923 | 325 | 410.2 | 1 | 1.26 | 0.31 | 9.54 | 9.36 | 0.17 | 0.17 | 0.01 | 0.01 |
| gene-LOC105287550 | https://www.ncbi.nlm.nih.gov/gene/?term=LOC105287550 | 15659 | 590 | 376.78 | 1 | 0.64 | 0.17 | 6.31 | 6.15 | 0.16 | 0.16 | 0.01 | 0.01 |
| gene-LOC105276612 | https://www.ncbi.nlm.nih.gov/gene/?term=LOC105276612 | 4157 | 183 | 440.22 | 1 | 2.41 | 0.55 | 9.73 | 9.61 | 0.11 | 0.11 | 0.01 | 0.01 |
| gene-LOC105286274 | https://www.ncbi.nlm.nih.gov/gene/?term=LOC105286274 | 3817 | 191 | 500.39 | 1 | 2.62 | 0.52 | 17.09 | 17.23 | -0.14 | 0.14 | -0.01 | 0.01 |
| gene-LOC105278455 | https://www.ncbi.nlm.nih.gov/gene/?term=LOC105278455 | 27181 | 921 | 338.84 | 1 | 0.37 | 0.11 | 7.15 | 7.32 | -0.18 | 0.18 | -0.01 | 0.01 |
| gene-LOC105274856 | https://www.ncbi.nlm.nih.gov/gene/?term=LOC105274856 | 5216 | 210 | 402.61 | 1 | 1.92 | 0.48 | 18.25 | 18.52 | -0.26 | 0.26 | -0.01 | 0.01 |
| gene-LOC105277903 | https://www.ncbi.nlm.nih.gov/gene/?term=LOC105277903 | 1835 | 36 | 196.19 | 1 | 5.45 | 2.78 | 49.51 | 49.79 | -0.27 | 0.27 | -0.01 | 0.01 |
| gene-LOC105282491 | https://www.ncbi.nlm.nih.gov/gene/?term=LOC105282491 | 4761 | 165 | 346.57 | 1 | 2.1 | 0.61 | 8.1 | 8.48 | -0.38 | 0.38 | -0.01 | 0.01 |
| gene-LOC105274851 | https://www.ncbi.nlm.nih.gov/gene/?term=LOC105274851 | 6616 | 136 | 205.56 | 1 | 1.51 | 0.74 | 10.78 | 11.18 | -0.4 | 0.4 | -0.01 | 0.01 |
| gene-LOC105286515 | https://www.ncbi.nlm.nih.gov/gene/?term=LOC105286515 | 3751 | 102 | 271.93 | 1 | 2.67 | 0.98 | 12.76 | 13.19 | -0.43 | 0.43 | -0.01 | 0.01 |
| gene-LOC105282399 | https://www.ncbi.nlm.nih.gov/gene/?term=LOC105282399 | 2682 | 23 | 85.76 | 1 | 3.73 | 4.35 | 34.71 | 35.34 | -0.63 | 0.63 | -0.01 | 0.01 |
| gene-LOC105288010 | https://www.ncbi.nlm.nih.gov/gene/?term=LOC105288010 | 3189 | 44 | 137.97 | 1 | 3.14 | 2.27 | 30.11 | 30.74 | -0.63 | 0.63 | -0.01 | 0.01 |
| gene-LOC105285060 | https://www.ncbi.nlm.nih.gov/gene/?term=LOC105285060 | 1285 | 18 | 140.08 | 1 | 7.78 | 5.56 | 44.89 | 45.6 | -0.72 | 0.72 | -0.01 | 0.01 |
| gene-LOC105283712 | https://www.ncbi.nlm.nih.gov/gene/?term=LOC105283712 | 1910 | 35 | 183.25 | 1 | 5.24 | 2.86 | 26.72 | 27.47 | -0.75 | 0.75 | -0.01 | 0.01 |
| gene-LOC105286452 | https://www.ncbi.nlm.nih.gov/gene/?term=LOC105286452 | 6888 | 59 | 85.66 | 1 | 1.45 | 1.69 | 32.42 | 31.88 | 0.55 | 0.55 | 0 | 0 |
| gene-LOC105285406 | https://www.ncbi.nlm.nih.gov/gene/?term=LOC105285406 | 8015 | 53 | 66.13 | 1 | 1.25 | 1.89 | 28.7 | 28.32 | 0.39 | 0.39 | 0 | 0 |
| gene-LOC105283989 | https://www.ncbi.nlm.nih.gov/gene/?term=LOC105283989 | 4948 | 31 | 62.65 | 1 | 2.02 | 3.23 | 36.84 | 36.46 | 0.37 | 0.37 | 0 | 0 |
| gene-LOC105288066 | https://www.ncbi.nlm.nih.gov/gene/?term=LOC105288066 | 1964 | 28 | 142.57 | 1 | 5.09 | 3.57 | 32.79 | 32.53 | 0.26 | 0.26 | 0 | 0 |
| gene-LOC105283943 | https://www.ncbi.nlm.nih.gov/gene/?term=LOC105283943 | 3559 | 47 | 132.06 | 1 | 2.81 | 2.13 | 17.56 | 17.32 | 0.24 | 0.24 | 0 | 0 |
| gene-LOC105286194 | https://www.ncbi.nlm.nih.gov/gene/?term=LOC105286194 | 152619 | 2124 | 139.17 | 1 | 0.07 | 0.05 | 6.36 | 6.23 | 0.14 | 0.14 | 0 | 0 |
| gene-LOC105274531 | https://www.ncbi.nlm.nih.gov/gene/?term=LOC105274531 | 7774 | 291 | 374.32 | 1 | 1.29 | 0.34 | 10.67 | 10.57 | 0.1 | 0.1 | 0 | 0 |
| gene-LOC105275327 | https://www.ncbi.nlm.nih.gov/gene/?term=LOC105275327 | 14245 | 475 | 333.45 | 1 | 0.7 | 0.21 | 7.23 | 7.14 | 0.1 | 0.1 | 0 | 0 |
| gene-LOC105280265 | https://www.ncbi.nlm.nih.gov/gene/?term=LOC105280265 | 85984 | 3003 | 349.25 | 1 | 0.12 | 0.03 | 1.15 | 1.07 | 0.08 | 0.08 | 0 | 0 |
| gene-LOC105282110 | https://www.ncbi.nlm.nih.gov/gene/?term=LOC105282110 | 7748 | 119 | 153.59 | 1 | 1.29 | 0.84 | 13.03 | 12.95 | 0.08 | 0.08 | 0 | 0 |
| gene-LOC105285807 | https://www.ncbi.nlm.nih.gov/gene/?term=LOC105285807 | 6063 | 223 | 367.8 | 1 | 1.65 | 0.45 | 9.47 | 9.4 | 0.06 | 0.06 | 0 | 0 |
| gene-LOC105278283 | https://www.ncbi.nlm.nih.gov/gene/?term=LOC105278283 | 2317 | 135 | 582.65 | 1 | 4.32 | 0.74 | 25.81 | 25.76 | 0.05 | 0.05 | 0 | 0 |
| gene-LOC105276538 | https://www.ncbi.nlm.nih.gov/gene/?term=LOC105276538 | 1511 | 17 | 112.51 | 1 | 6.62 | 5.88 | 69.95 | 69.91 | 0.04 | 0.04 | 0 | 0 |
| gene-LOC105287528 | https://www.ncbi.nlm.nih.gov/gene/?term=LOC105287528 | 14195 | 398 | 280.38 | 1 | 0.7 | 0.25 | 4.97 | 4.95 | 0.02 | 0.02 | 0 | 0 |
| gene-LOC105274823 | https://www.ncbi.nlm.nih.gov/gene/?term=LOC105274823 | 4620 | 138 | 298.7 | 1 | 2.16 | 0.72 | 21.27 | 21.26 | 0.01 | 0.01 | 0 | 0 |
| gene-LOC105275127 | https://www.ncbi.nlm.nih.gov/gene/?term=LOC105275127 | 9377 | 295 | 314.6 | 1 | 1.07 | 0.34 | 7.39 | 7.38 | 0.01 | 0.01 | 0 | 0 |
| gene-LOC105280407 | https://www.ncbi.nlm.nih.gov/gene/?term=LOC105280407 | 10950 | 207 | 189.04 | 1 | 0.91 | 0.48 | 16.49 | 16.48 | 0.01 | 0.01 | 0 | 0 |

| | | | | | | | | | | | | | | |
|---|---|---|---|---|---|---|---|---|---|---|---|---|---|---|
| gene-LOC105275187 | https://www.ncbi.nlm.nih.gov/gene/?term=LOC105275187 | 5534 | 70 | 126.49 | | 1 | 1.81 | 1.43 | 12.34 | 12.39 | -0.05 | 0.05 | 0 | 0 |
| gene-LOC105279500 | https://www.ncbi.nlm.nih.gov/gene/?term=LOC105279500 | 3587 | 36 | 100.36 | | 1 | 2.79 | 2.78 | 49.66 | 49.71 | -0.05 | 0.05 | 0 | 0 |
| gene-LOC105276692 | https://www.ncbi.nlm.nih.gov/gene/?term=LOC105276692 | 1745 | 29 | 166.19 | | 1 | 5.73 | 3.45 | 24.9 | 24.96 | -0.07 | 0.07 | 0 | 0 |
| gene-LOC105275374 | https://www.ncbi.nlm.nih.gov/gene/?term=LOC105275374 | 11376 | 502 | 441.28 | | 1 | 0.88 | 0.2 | 10.16 | 10.25 | -0.09 | 0.09 | 0 | 0 |
| gene-LOC105281173 | https://www.ncbi.nlm.nih.gov/gene/?term=LOC105281173 | 1595 | 39 | 244.51 | | 1 | 6.27 | 2.56 | 59.82 | 59.91 | -0.09 | 0.09 | 0 | 0 |
| gene-LOC105278245 | https://www.ncbi.nlm.nih.gov/gene/?term=LOC105278245 | 6445 | 117 | 181.54 | | 1 | 1.55 | 0.85 | 22.04 | 22.2 | -0.17 | 0.17 | 0 | 0 |
| gene-LOC105277866 | https://www.ncbi.nlm.nih.gov/gene/?term=LOC105277866 | 33045 | 390 | 118.02 | | 1 | 0.3 | 0.26 | 3.22 | 3.41 | -0.19 | 0.19 | 0 | 0 |
| gene-LOC105286514 | https://www.ncbi.nlm.nih.gov/gene/?term=LOC105286514 | 2528 | 48 | 189.87 | | 1 | 3.96 | 2.08 | 36.18 | 36.38 | -0.2 | 0.2 | 0 | 0 |
| gene-LOC105275500 | https://www.ncbi.nlm.nih.gov/gene/?term=LOC105275500 | 10943 | 242 | 221.15 | | 1 | 0.91 | 0.41 | 7.04 | 7.25 | -0.21 | 0.21 | 0 | 0 |
| gene-LOC105282067 | https://www.ncbi.nlm.nih.gov/gene/?term=LOC105282067 | 5941 | 49 | 82.48 | 1 | 1.68 | 2.04 | 26.22 | 26.65 | -0.43 | 0.43 | 0 | 0 | |

Directory: OGLall File: rnk-dmg-Ob_LI_BCphaseB2.vs.RphaseA1.txt

| gene_ID | gene_link | gwidth | #Sites | #per10Kb | #dmSites | #dmsp10kb | %dmSites | %pSite1 | %pSite2 | DMpSite | ADMpSite | DMpNucl | ADMpNucl |
|---|---|---|---|---|---|---|---|---|---|---|---|---|---|
| gene-LOC113562345 | https://www.ncbi.nlm.nih.gov/gene/?term=LOC113562345 | 2290 | 63 | 275.11 | 8 | 34.93 | 12.7 | 17.51 | 30.84 | -13.33 | 13.33 | -0.37 | 0.37 |
| gene-LOC105285674 | https://www.ncbi.nlm.nih.gov/gene/?term=LOC105285674 | 3035 | 84 | 276.77 | 2 | 6.59 | 2.38 | 24.69 | 27.77 | -3.08 | 3.08 | -0.09 | 0.09 |
| gene-LOC105275070 | https://www.ncbi.nlm.nih.gov/gene/?term=LOC105275070 | 5667 | 40 | 70.58 | 6 | 10.59 | 15 | 2.13 | 14.38 | -12.25 | 12.25 | -0.09 | 0.09 |
| gene-LOC105284396 | https://www.ncbi.nlm.nih.gov/gene/?term=LOC105284396 | 1976 | 66 | 334.01 | 1 | 5.06 | 1.52 | 34.48 | 32.1 | 2.38 | 2.38 | 0.08 | 0.08 |
| gene-LOC105276449 | https://www.ncbi.nlm.nih.gov/gene/?term=LOC105276449 | 1231 | 45 | 365.56 | 1 | 8.12 | 2.22 | 27.05 | 29.32 | -2.27 | 2.27 | -0.08 | 0.08 |
| gene-LOC105274706 | https://www.ncbi.nlm.nih.gov/gene/?term=LOC105274706 | 1446 | 50 | 345.78 | 1 | 6.92 | 2 | 39.84 | 37.9 | 1.94 | 1.94 | 0.07 | 0.07 |
| gene-LOC105282156 | https://www.ncbi.nlm.nih.gov/gene/?term=LOC105282156 | 2577 | 133 | 516.1 | 1 | 3.88 | 0.75 | 22.7 | 24.15 | -1.45 | 1.45 | -0.07 | 0.07 |
| gene-LOC105280786 | https://www.ncbi.nlm.nih.gov/gene/?term=LOC105280786 | 2573 | 109 | 423.63 | 1 | 3.89 | 0.92 | 28.63 | 30.24 | -1.61 | 1.61 | -0.07 | 0.07 |
| gene-LOC105275384 | https://www.ncbi.nlm.nih.gov/gene/?term=LOC105275384 | 1987 | 87 | 437.85 | 1 | 5.03 | 1.15 | 34.15 | 35.77 | -1.63 | 1.63 | -0.07 | 0.07 |
| gene-LOC105285202 | https://www.ncbi.nlm.nih.gov/gene/?term=LOC105285202 | 1838 | 60 | 326.44 | 1 | 5.44 | 1.67 | 23.97 | 26.11 | -2.15 | 2.15 | -0.07 | 0.07 |
| gene-LOC105283649 | https://www.ncbi.nlm.nih.gov/gene/?term=LOC105283649 | 1918 | 56 | 291.97 | 1 | 5.21 | 1.79 | 36.17 | 38.48 | -2.31 | 2.31 | -0.07 | 0.07 |
| gene-LOC105275708 | https://www.ncbi.nlm.nih.gov/gene/?term=LOC105275708 | 1772 | 15 | 84.65 | 1 | 5.64 | 6.67 | 45.9 | 54.1 | -8.2 | 8.2 | -0.07 | 0.07 |
| gene-LOC105274449 | https://www.ncbi.nlm.nih.gov/gene/?term=LOC105274449 | 1549 | 53 | 342.16 | 1 | 6.46 | 1.89 | 51.77 | 49.89 | 1.87 | 1.87 | 0.06 | 0.06 |
| gene-LOC105277903 | https://www.ncbi.nlm.nih.gov/gene/?term=LOC105277903 | 1835 | 36 | 196.19 | 1 | 5.45 | 2.78 | 53.1 | 56.25 | -3.14 | 3.14 | -0.06 | 0.06 |
| gene-LOC105274900 | https://www.ncbi.nlm.nih.gov/gene/?term=LOC105274900 | 5561 | 54 | 97.1 | 3 | 5.39 | 5.56 | 30 | 35.69 | -5.68 | 5.68 | -0.06 | 0.06 |
| gene-LOC105279898 | https://www.ncbi.nlm.nih.gov/gene/?term=LOC105279898 | 1343 | 11 | 81.91 | 1 | 7.45 | 9.09 | 61.61 | 68.61 | -7 | 7 | -0.06 | 0.06 |
| gene-LOC105278567 | https://www.ncbi.nlm.nih.gov/gene/?term=LOC105278567 | 3627 | 29 | 79.96 | 1 | 2.76 | 3.45 | 11.32 | 18.71 | -7.39 | 7.39 | -0.06 | 0.06 |
| gene-LOC105278454 | https://www.ncbi.nlm.nih.gov/gene/?term=LOC105278454 | 1972 | 43 | 218.05 | 1 | 5.07 | 2.33 | 44.38 | 46.55 | -2.17 | 2.17 | -0.05 | 0.05 |
| gene-LOC105281420 | https://www.ncbi.nlm.nih.gov/gene/?term=LOC105281420 | 3882 | 55 | 141.68 | 1 | 2.58 | 1.82 | 13.05 | 16.87 | -3.82 | 3.82 | -0.05 | 0.05 |
| gene-LOC105282885 | https://www.ncbi.nlm.nih.gov/gene/?term=LOC105282885 | 1870 | 46 | 245.99 | 1 | 5.35 | 2.17 | 37.06 | 35.54 | 1.52 | 1.52 | 0.04 | 0.04 |
| gene-LOC105285055 | https://www.ncbi.nlm.nih.gov/gene/?term=LOC105285055 | 4108 | 256 | 623.17 | 1 | 2.43 | 0.39 | 10.36 | 10.97 | -0.61 | 0.61 | -0.04 | 0.04 |
| gene-LOC105280723 | https://www.ncbi.nlm.nih.gov/gene/?term=LOC105280723 | 3572 | 182 | 509.52 | 1 | 2.8 | 0.55 | 15.32 | 16.11 | -0.79 | 0.79 | -0.04 | 0.04 |
| gene-LOC105278877 | https://www.ncbi.nlm.nih.gov/gene/?term=LOC105278877 | 1968 | 82 | 416.67 | 1 | 5.08 | 1.22 | 30.66 | 31.6 | -0.94 | 0.94 | -0.04 | 0.04 |
| gene-LOC105280956 | https://www.ncbi.nlm.nih.gov/gene/?term=LOC105280956 | 2639 | 72 | 272.83 | 1 | 3.79 | 1.39 | 17.08 | 18.51 | -1.43 | 1.43 | -0.04 | 0.04 |
| gene-LOC105278436 | https://www.ncbi.nlm.nih.gov/gene/?term=LOC105278436 | 2556 | 54 | 211.27 | 1 | 3.91 | 1.85 | 41.69 | 43.45 | -1.76 | 1.76 | -0.04 | 0.04 |
| gene-LOC105283592 | https://www.ncbi.nlm.nih.gov/gene/?term=LOC105283592 | 3479 | 64 | 183.96 | 1 | 2.87 | 1.56 | 24.19 | 26.35 | -2.16 | 2.16 | -0.04 | 0.04 |
| gene-LOC105281039 | https://www.ncbi.nlm.nih.gov/gene/?term=LOC105281039 | 1471 | 11 | 74.78 | 1 | 6.8 | 9.09 | 29.26 | 35.1 | -5.84 | 5.84 | -0.04 | 0.04 |
| gene-LOC105282914 | https://www.ncbi.nlm.nih.gov/gene/?term=LOC105282914 | 5623 | 41 | 72.91 | 1 | 1.78 | 2.44 | 52.21 | 48.43 | 3.78 | 3.78 | 0.03 | 0.03 |
| gene-LOC105276272 | https://www.ncbi.nlm.nih.gov/gene/?term=LOC105276272 | 1965 | 46 | 234.1 | 1 | 5.09 | 2.17 | 27.86 | 26.72 | 1.14 | 1.14 | 0.03 | 0.03 |
| gene-LOC105275587 | https://www.ncbi.nlm.nih.gov/gene/?term=LOC105275587 | 2404 | 57 | 237.1 | 1 | 4.16 | 1.75 | 37.35 | 36.26 | 1.09 | 1.09 | 0.03 | 0.03 |
| gene-LOC105278145 | https://www.ncbi.nlm.nih.gov/gene/?term=LOC105278145 | 4470 | 235 | 525.73 | 2 | 4.47 | 0.85 | 10.77 | 11.29 | -0.52 | 0.52 | -0.03 | 0.03 |
| gene-LOC105275460 | https://www.ncbi.nlm.nih.gov/gene/?term=LOC105275460 | 2473 | 120 | 485.24 | 1 | 4.04 | 0.83 | 13.18 | 13.75 | -0.57 | 0.57 | -0.03 | 0.03 |
| gene-LOC105279298 | https://www.ncbi.nlm.nih.gov/gene/?term=LOC105279298 | 2773 | 130 | 468.81 | 2 | 7.21 | 1.54 | 23.75 | 24.38 | -0.63 | 0.63 | -0.03 | 0.03 |
| gene-LOC105274856 | https://www.ncbi.nlm.nih.gov/gene/?term=LOC105274856 | 5216 | 225 | 431.37 | 1 | 1.92 | 0.44 | 17.14 | 17.89 | -0.76 | 0.76 | -0.03 | 0.03 |
| gene-LOC105282491 | https://www.ncbi.nlm.nih.gov/gene/?term=LOC105282491 | 4761 | 174 | 365.47 | 1 | 2.1 | 0.57 | 8.54 | 9.39 | -0.85 | 0.85 | -0.03 | 0.03 |
| gene-LOC105278807 | https://www.ncbi.nlm.nih.gov/gene/?term=LOC105278807 | 2166 | 78 | 360.11 | 1 | 4.62 | 1.28 | 18.54 | 19.5 | -0.97 | 0.97 | -0.03 | 0.03 |
| gene-LOC105282151 | https://www.ncbi.nlm.nih.gov/gene/?term=LOC105282151 | 1601 | 42 | 262.34 | 1 | 6.25 | 2.38 | 34.92 | 36.1 | -1.17 | 1.17 | -0.03 | 0.03 |
| gene-LOC105281266 | https://www.ncbi.nlm.nih.gov/gene/?term=LOC105281266 | 4102 | 81 | 197.46 | 1 | 2.44 | 1.23 | 16.07 | 17.51 | -1.44 | 1.44 | -0.03 | 0.03 |
| gene-LOC105288066 | https://www.ncbi.nlm.nih.gov/gene/?term=LOC105288066 | 1964 | 29 | 147.66 | 1 | 5.09 | 3.45 | 35.48 | 37.25 | -1.78 | 1.78 | -0.03 | 0.03 |
| gene-LOC105285514 | https://www.ncbi.nlm.nih.gov/gene/?term=LOC105285514 | 1277 | 11 | 86.14 | 1 | 7.83 | 9.09 | 70.96 | 74.98 | -4.02 | 4.02 | -0.03 | 0.03 |
| gene-LOC105279500 | https://www.ncbi.nlm.nih.gov/gene/?term=LOC105279500 | 3587 | 30 | 83.64 | 1 | 2.79 | 3.33 | 48.41 | 46.03 | 2.38 | 2.38 | 0.02 | 0.02 |
| gene-LOC105283967 | https://www.ncbi.nlm.nih.gov/gene/?term=LOC105283967 | 2393 | 50 | 208.94 | 1 | 4.18 | 2 | 45.88 | 44.9 | 0.98 | 0.98 | 0.02 | 0.02 |
| gene-LOC105282970 | https://www.ncbi.nlm.nih.gov/gene/?term=LOC105282970 | 1529 | 40 | 261.61 | 1 | 6.54 | 2.5 | 50.96 | 50.14 | 0.82 | 0.82 | 0.02 | 0.02 |
| gene-LOC105281173 | https://www.ncbi.nlm.nih.gov/gene/?term=LOC105281173 | 1595 | 38 | 238.24 | 1 | 6.27 | 2.63 | 60.67 | 59.87 | 0.8 | 0.8 | 0.02 | 0.02 |
| gene-LOC105281312 | https://www.ncbi.nlm.nih.gov/gene/?term=LOC105281312 | 2551 | 65 | 254.8 | 1 | 3.92 | 1.54 | 33.11 | 32.31 | 0.8 | 0.8 | 0.02 | 0.02 |
| gene-LOC105278962 | https://www.ncbi.nlm.nih.gov/gene/?term=LOC105278962 | 4491 | 93 | 207.08 | 1 | 2.23 | 1.08 | 22.79 | 22.03 | 0.76 | 0.76 | 0.02 | 0.02 |
| gene-LOC105280819 | https://www.ncbi.nlm.nih.gov/gene/?term=LOC105280819 | 1639 | 45 | 274.56 | 1 | 6.1 | 2.22 | 37.3 | 36.73 | 0.57 | 0.57 | 0.02 | 0.02 |
| gene-LOC105286274 | https://www.ncbi.nlm.nih.gov/gene/?term=LOC105286274 | 3817 | 187 | 489.91 | 1 | 2.62 | 0.53 | 16.7 | 16.2 | 0.51 | 0.51 | 0.02 | 0.02 |
| gene-LOC105284184 | https://www.ncbi.nlm.nih.gov/gene/?term=LOC105284184 | 2535 | 92 | 362.92 | 1 | 3.94 | 1.09 | 31.62 | 31.13 | 0.49 | 0.49 | 0.02 | 0.02 |
| gene-LOC105278283 | https://www.ncbi.nlm.nih.gov/gene/?term=LOC105278283 | 2317 | 139 | 599.91 | 1 | 4.32 | 0.72 | 25.21 | 25.46 | -0.25 | 0.25 | -0.02 | 0.02 |
| gene-LOC105282149 | https://www.ncbi.nlm.nih.gov/gene/?term=LOC105282149 | 1787 | 103 | 576.39 | 1 | 5.6 | 0.97 | 40.98 | 41.32 | -0.34 | 0.34 | -0.02 | 0.02 |
| gene-LOC105280774 | https://www.ncbi.nlm.nih.gov/gene/?term=LOC105280774 | 2793 | 117 | 418.9 | 1 | 3.58 | 0.85 | 27.14 | 27.51 | -0.36 | 0.36 | -0.02 | 0.02 |
| gene-LOC105274531 | https://www.ncbi.nlm.nih.gov/gene/?term=LOC105274531 | 7774 | 275 | 353.74 | 1 | 1.29 | 0.36 | 10.93 | 11.37 | -0.43 | 0.43 | -0.02 | 0.02 |
| gene-LOC105287095 | https://www.ncbi.nlm.nih.gov/gene/?term=LOC105287095 | 1435 | 64 | 445.99 | 1 | 6.97 | 1.56 | 25.57 | 26.08 | -0.51 | 0.51 | -0.02 | 0.02 |
| gene-LOC105275752 | https://www.ncbi.nlm.nih.gov/gene/?term=LOC105275752 | 3462 | 128 | 369.73 | 1 | 2.89 | 0.78 | 29.22 | 29.76 | -0.53 | 0.53 | -0.02 | 0.02 |
| gene-LOC105285742 | https://www.ncbi.nlm.nih.gov/gene/?term=LOC105285742 | 3446 | 105 | 304.7 | 1 | 2.9 | 0.95 | 17.29 | 17.85 | -0.55 | 0.55 | -0.02 | 0.02 |
| gene-LOC105288036 | https://www.ncbi.nlm.nih.gov/gene/?term=LOC105288036 | 5110 | 200 | 391.39 | 1 | 1.96 | 0.5 | 16.38 | 16.94 | -0.55 | 0.55 | -0.02 | 0.02 |
| gene-LOC105282658 | https://www.ncbi.nlm.nih.gov/gene/?term=LOC105282658 | 2421 | 59 | 243.7 | 1 | 4.13 | 1.69 | 24.61 | 25.23 | -0.62 | 0.62 | -0.02 | 0.02 |
| gene-LOC105287063 | https://www.ncbi.nlm.nih.gov/gene/?term=LOC105287063 | 1874 | 49 | 261.47 | 1 | 5.34 | 2.04 | 29.55 | 30.23 | -0.68 | 0.68 | -0.02 | 0.02 |
| gene-LOC105279900 | https://www.ncbi.nlm.nih.gov/gene/?term=LOC105279900 | 2191 | 54 | 246.46 | 1 | 4.56 | 1.85 | 30.89 | 31.61 | -0.72 | 0.72 | -0.02 | 0.02 |

| | | | | | | | | | | | | | | |
|---|---|---|---|---|---|---|---|---|---|---|---|---|---|---|
| gene-LOC105275451 | https://www.ncbi.nlm.nih.gov/gene/?term=LOC105275451 | 6035 | 123 | 203.81 | | 1 | 1.66 | 0.81 | 22.94 | 23.69 | -0.75 | 0.75 | -0.02 | 0.02 |
| gene-LOC105278384 | https://www.ncbi.nlm.nih.gov/gene/?term=LOC105278384 | 1390 | 35 | 251.8 | 1 | 7.19 | 2.86 | 45.32 | 46.11 | -0.79 | 0.79 | -0.02 | 0.02 | |
| gene-LOC105284116 | https://www.ncbi.nlm.nih.gov/gene/?term=LOC105284116 | 3970 | 107 | 269.52 | | 1 | 2.52 | 0.93 | 7.18 | 8 | -0.82 | 0.82 | -0.02 | 0.02 |
| gene-LOC105284117 | https://www.ncbi.nlm.nih.gov/gene/?term=LOC105284117 | 4340 | 94 | 216.59 | | 1 | 2.3 | 1.06 | 9.25 | 10.09 | -0.84 | 0.84 | -0.02 | 0.02 |
| gene-LOC105282991 | https://www.ncbi.nlm.nih.gov/gene/?term=LOC105282991 | 5617 | 101 | 179.81 | | 1 | 1.78 | 0.99 | 19.27 | 20.29 | -1.02 | 1.02 | -0.02 | 0.02 |
| gene-LOC105279664 | https://www.ncbi.nlm.nih.gov/gene/?term=LOC105279664 | 4023 | 60 | 149.14 | | 1 | 2.49 | 1.67 | 17.78 | 18.81 | -1.03 | 1.03 | -0.02 | 0.02 |
| gene-LOC105280348 | https://www.ncbi.nlm.nih.gov/gene/?term=LOC105280348 | 3797 | 71 | 186.99 | | 1 | 2.63 | 1.41 | 18.59 | 19.68 | -1.09 | 1.09 | -0.02 | 0.02 |
| gene-LOC105287854 | https://www.ncbi.nlm.nih.gov/gene/?term=LOC105287854 | 3644 | 64 | 175.63 | | 1 | 2.74 | 1.56 | 14.92 | 16.11 | -1.19 | 1.19 | -0.02 | 0.02 |
| gene-LOC105276692 | https://www.ncbi.nlm.nih.gov/gene/?term=LOC105276692 | 1745 | 33 | 189.11 | | 1 | 5.73 | 3.03 | 21.96 | 23.19 | -1.23 | 1.23 | -0.02 | 0.02 |
| gene-LOC105288078 | https://www.ncbi.nlm.nih.gov/gene/?term=LOC105288078 | 19126 | 298 | 155.81 | | 1 | 0.52 | 0.34 | 16.07 | 17.31 | -1.24 | 1.24 | -0.02 | 0.02 |
| gene-LOC105280641 | https://www.ncbi.nlm.nih.gov/gene/?term=LOC105280641 | 3682 | 55 | 149.38 | | 1 | 2.72 | 1.82 | 18.09 | 19.48 | -1.39 | 1.39 | -0.02 | 0.02 |
| gene-LOC105287648 | https://www.ncbi.nlm.nih.gov/gene/?term=LOC105287648 | 2205 | 35 | 158.73 | | 1 | 4.54 | 2.86 | 35.94 | 37.4 | -1.45 | 1.45 | -0.02 | 0.02 |
| gene-LOC105282600 | https://www.ncbi.nlm.nih.gov/gene/?term=LOC105282600 | 3715 | 52 | 139.97 | | 1 | 2.69 | 1.92 | 31.36 | 33.08 | -1.72 | 1.72 | -0.02 | 0.02 |
| gene-LOC105283943 | https://www.ncbi.nlm.nih.gov/gene/?term=LOC105283943 | 3559 | 50 | 140.49 | | 1 | 2.81 | 2 | 16.35 | 18.09 | -1.74 | 1.74 | -0.02 | 0.02 |
| gene-LOC105284329 | https://www.ncbi.nlm.nih.gov/gene/?term=LOC105284329 | 4051 | 50 | 123.43 | | 1 | 2.47 | 2 | 17.13 | 19.12 | -1.99 | 1.99 | -0.02 | 0.02 |
| gene-LOC105281515 | https://www.ncbi.nlm.nih.gov/gene/?term=LOC105281515 | 8827 | 60 | 67.97 | 2 | 2.27 | 3.33 | 26.32 | 29.67 | -3.35 | 3.35 | -0.02 | 0.02 | |
| gene-LOC105283712 | https://www.ncbi.nlm.nih.gov/gene/?term=LOC105283712 | 1910 | 36 | 188.48 | | 1 | 5.24 | 2.78 | 26.26 | 25.56 | 0.7 | 0.7 | 0.01 | 0.01 |
| gene-LOC105288042 | https://www.ncbi.nlm.nih.gov/gene/?term=LOC105288042 | 3507 | 98 | 279.44 | | 1 | 2.85 | 1.02 | 22.45 | 21.99 | 0.46 | 0.46 | 0.01 | 0.01 |
| gene-LOC105282646 | https://www.ncbi.nlm.nih.gov/gene/?term=LOC105282646 | 3317 | 99 | 298.46 | | 1 | 3.01 | 1.01 | 21.06 | 20.67 | 0.39 | 0.39 | 0.01 | 0.01 |
| gene-LOC105274590 | https://www.ncbi.nlm.nih.gov/gene/?term=LOC105274590 | 4516 | 74 | 163.86 | | 1 | 2.21 | 1.35 | 30.44 | 30.06 | 0.38 | 0.38 | 0.01 | 0.01 |
| gene-LOC105276527 | https://www.ncbi.nlm.nih.gov/gene/?term=LOC105276527 | 7923 | 317 | 400.1 | 1 | 1.26 | 0.32 | 9.8 | 9.43 | 0.37 | 0.37 | 0.01 | 0.01 | |
| gene-LOC105282951 | https://www.ncbi.nlm.nih.gov/gene/?term=LOC105282951 | 3366 | 120 | 356.51 | | 1 | 2.97 | 0.83 | 19.9 | 19.57 | 0.33 | 0.33 | 0.01 | 0.01 |
| gene-LOC105276196 | https://www.ncbi.nlm.nih.gov/gene/?term=LOC105276196 | 2953 | 56 | 189.64 | | 1 | 3.39 | 1.79 | 39.44 | 39.14 | 0.3 | 0.3 | 0.01 | 0.01 |
| gene-LOC105278534 | https://www.ncbi.nlm.nih.gov/gene/?term=LOC105278534 | 7364 | 185 | 251.22 | | 2 | 2.72 | 1.08 | 18.38 | 18.12 | 0.25 | 0.25 | 0.01 | 0.01 |
| gene-LOC105274534 | https://www.ncbi.nlm.nih.gov/gene/?term=LOC105274534 | 1329 | 59 | 443.94 | | 1 | 7.52 | 1.69 | 42.17 | 42.03 | 0.14 | 0.14 | 0.01 | 0.01 |
| gene-LOC105278632 | https://www.ncbi.nlm.nih.gov/gene/?term=LOC105278632 | 3624 | 133 | 367 | 1 | 2.76 | 0.75 | 14.21 | 14.06 | 0.14 | 0.14 | 0.01 | 0.01 | |
| gene-LOC105278691 | https://www.ncbi.nlm.nih.gov/gene/?term=LOC105278691 | 3871 | 151 | 390.08 | | 1 | 2.58 | 0.66 | 14.98 | 14.84 | 0.14 | 0.14 | 0.01 | 0.01 |
| gene-LOC105275683 | https://www.ncbi.nlm.nih.gov/gene/?term=LOC105275683 | 4256 | 187 | 439.38 | | 1 | 2.35 | 0.53 | 13.19 | 13.34 | -0.15 | 0.15 | -0.01 | 0.01 |
| gene-LOC105278666 | https://www.ncbi.nlm.nih.gov/gene/?term=LOC105278666 | 10341 | 471 | 455.47 | | 1 | 0.97 | 0.21 | 6.62 | 6.83 | -0.21 | 0.21 | -0.01 | 0.01 |
| gene-LOC105275179 | https://www.ncbi.nlm.nih.gov/gene/?term=LOC105275179 | 3271 | 77 | 235.4 | 1 | 3.06 | 1.3 | 17.93 | 18.15 | -0.22 | 0.22 | -0.01 | 0.01 | |
| gene-LOC105284355 | https://www.ncbi.nlm.nih.gov/gene/?term=LOC105284355 | 3904 | 162 | 414.96 | | 1 | 2.56 | 0.62 | 19.15 | 19.36 | -0.22 | 0.22 | -0.01 | 0.01 |
| gene-LOC105275363 | https://www.ncbi.nlm.nih.gov/gene/?term=LOC105275363 | 3798 | 161 | 423.91 | | 1 | 2.63 | 0.62 | 16.94 | 17.21 | -0.26 | 0.26 | -0.01 | 0.01 |
| gene-LOC105279038 | https://www.ncbi.nlm.nih.gov/gene/?term=LOC105279038 | 9453 | 340 | 359.67 | | 1 | 1.06 | 0.29 | 8.66 | 8.92 | -0.26 | 0.26 | -0.01 | 0.01 |
| gene-LOC105275374 | https://www.ncbi.nlm.nih.gov/gene/?term=LOC105275374 | 11376 | 488 | 428.97 | | 1 | 0.88 | 0.2 | 10.37 | 10.64 | -0.27 | 0.27 | -0.01 | 0.01 |
| gene-LOC105275327 | https://www.ncbi.nlm.nih.gov/gene/?term=LOC105275327 | 14245 | 511 | 358.72 | | 1 | 0.7 | 0.2 | 6.72 | 7.02 | -0.3 | 0.3 | -0.01 | 0.01 |
| gene-LOC105278497 | https://www.ncbi.nlm.nih.gov/gene/?term=LOC105278497 | 2072 | 96 | 463.32 | | 1 | 4.83 | 1.04 | 18.89 | 19.19 | -0.31 | 0.31 | -0.01 | 0.01 |
| gene-LOC105275170 | https://www.ncbi.nlm.nih.gov/gene/?term=LOC105275170 | 12918 | 279 | 215.98 | | 1 | 0.77 | 0.36 | 2.01 | 2.33 | -0.32 | 0.32 | -0.01 | 0.01 |
| gene-LOC105278282 | https://www.ncbi.nlm.nih.gov/gene/?term=LOC105278282 | 1775 | 51 | 287.32 | | 1 | 5.63 | 1.96 | 40.43 | 40.76 | -0.33 | 0.33 | -0.01 | 0.01 |
| gene-LOC105282794 | https://www.ncbi.nlm.nih.gov/gene/?term=LOC105282794 | 4571 | 93 | 203.46 | | 1 | 2.19 | 1.08 | 16.99 | 17.32 | -0.33 | 0.33 | -0.01 | 0.01 |
| gene-LOC105274669 | https://www.ncbi.nlm.nih.gov/gene/?term=LOC105274669 | 8555 | 261 | 305.08 | | 1 | 1.17 | 0.38 | 5.68 | 6.02 | -0.35 | 0.35 | -0.01 | 0.01 |
| gene-LOC105278455 | https://www.ncbi.nlm.nih.gov/gene/?term=LOC105278455 | 27181 | 922 | 339.21 | | 1 | 0.37 | 0.11 | 7.06 | 7.41 | -0.35 | 0.35 | -0.01 | 0.01 |
| gene-LOC105277761 | https://www.ncbi.nlm.nih.gov/gene/?term=LOC105277761 | 4648 | 144 | 309.81 | | 1 | 2.15 | 0.69 | 14.39 | 14.8 | -0.41 | 0.41 | -0.01 | 0.01 |
| gene-LOC105274823 | https://www.ncbi.nlm.nih.gov/gene/?term=LOC105274823 | 4620 | 145 | 313.85 | | 1 | 2.16 | 0.69 | 19.86 | 20.29 | -0.43 | 0.43 | -0.01 | 0.01 |
| gene-LOC105285639 | https://www.ncbi.nlm.nih.gov/gene/?term=LOC105285639 | 10325 | 344 | 333.17 | | 1 | 0.97 | 0.29 | 11.23 | 11.66 | -0.43 | 0.43 | -0.01 | 0.01 |
| gene-LOC105278649 | https://www.ncbi.nlm.nih.gov/gene/?term=LOC105278649 | 5616 | 137 | 243.95 | | 1 | 1.78 | 0.73 | 13.17 | 13.73 | -0.56 | 0.56 | -0.01 | 0.01 |
| gene-LOC105277856 | https://www.ncbi.nlm.nih.gov/gene/?term=LOC105277856 | 3442 | 31 | 90.06 | 1 | 2.91 | 3.23 | 38.58 | 39.29 | -0.71 | 0.71 | -0.01 | 0.01 | |
| gene-LOC105276059 | https://www.ncbi.nlm.nih.gov/gene/?term=LOC105276059 | 6256 | 129 | 206.2 | 1 | 1.6 | 0.78 | 9.24 | 9.96 | -0.72 | 0.72 | -0.01 | 0.01 | |
| gene-LOC105275187 | https://www.ncbi.nlm.nih.gov/gene/?term=LOC105275187 | 5534 | 69 | 124.68 | | 1 | 1.81 | 1.45 | 13.92 | 14.78 | -0.86 | 0.86 | -0.01 | 0.01 |
| gene-LOC105283989 | https://www.ncbi.nlm.nih.gov/gene/?term=LOC105283989 | 4948 | 27 | 54.57 | 1 | 2.02 | 3.7 | 44.98 | 46.16 | -1.18 | 1.18 | -0.01 | 0.01 | |
| gene-LOC105275347 | https://www.ncbi.nlm.nih.gov/gene/?term=LOC105275347 | 2526 | 16 | 63.34 | 1 | 3.96 | 6.25 | 40.9 | 42.25 | -1.34 | 1.34 | -0.01 | 0.01 | |
| gene-LOC105280002 | https://www.ncbi.nlm.nih.gov/gene/?term=LOC105280002 | 1940 | 26 | 134.02 | | 1 | 5.15 | 3.85 | 34.83 | 34.64 | 0.19 | 0.19 | 0 | 0 |
| gene-LOC105282811 | https://www.ncbi.nlm.nih.gov/gene/?term=LOC105282811 | 5765 | 199 | 345.19 | | 1 | 1.73 | 0.5 | 8.45 | 8.31 | 0.14 | 0.14 | 0 | 0 |
| gene-LOC105285807 | https://www.ncbi.nlm.nih.gov/gene/?term=LOC105285807 | 6063 | 217 | 357.91 | | 1 | 1.65 | 0.46 | 8.91 | 8.79 | 0.12 | 0.12 | 0 | 0 |
| gene-LOC105281700 | https://www.ncbi.nlm.nih.gov/gene/?term=LOC105281700 | 2208 | 37 | 167.57 | | 1 | 4.53 | 2.7 | 39.75 | 39.65 | 0.11 | 0.11 | 0 | 0 |
| gene-LOC105283728 | https://www.ncbi.nlm.nih.gov/gene/?term=LOC105283728 | 3256 | 122 | 374.69 | | 2 | 6.14 | 1.64 | 18.7 | 18.6 | 0.1 | 0.1 | 0 | 0 |
| gene-LOC105276538 | https://www.ncbi.nlm.nih.gov/gene/?term=LOC105276538 | 1511 | 18 | 119.13 | | 1 | 6.62 | 5.56 | 68.54 | 68.54 | 0 | 0 | 0 | 0 |
| gene-LOC105280265 | https://www.ncbi.nlm.nih.gov/gene/?term=LOC105280265 | 85984 | 2983 | 346.93 | | 1 | 0.12 | 0.03 | 1.1 | 1.12 | -0.02 | 0.02 | 0 | 0 |
| gene-LOC105283123 | https://www.ncbi.nlm.nih.gov/gene/?term=LOC105283123 | 1507 | 47 | 311.88 | | 1 | 6.64 | 2.13 | 17.1 | 17.12 | -0.02 | 0.02 | 0 | 0 |
| gene-LOC105286514 | https://www.ncbi.nlm.nih.gov/gene/?term=LOC105286514 | 2528 | 44 | 174.05 | | 1 | 3.96 | 2.27 | 37.14 | 37.18 | -0.04 | 0.04 | 0 | 0 |
| gene-LOC105280513 | https://www.ncbi.nlm.nih.gov/gene/?term=LOC105280513 | 69542 | 1333 | 191.68 | | 1 | 0.14 | 0.08 | 0.66 | 0.72 | -0.05 | 0.05 | 0 | 0 |
| gene-LOC105285179 | https://www.ncbi.nlm.nih.gov/gene/?term=LOC105285179 | 4191 | 176 | 419.95 | | 1 | 2.39 | 0.57 | 19.46 | 19.52 | -0.06 | 0.06 | 0 | 0 |
| gene-LOC105285406 | https://www.ncbi.nlm.nih.gov/gene/?term=LOC105285406 | 8015 | 60 | 74.86 | 1 | 1.25 | 1.67 | 28.89 | 28.94 | -0.06 | 0.06 | 0 | 0 | |
| gene-LOC105284043 | https://www.ncbi.nlm.nih.gov/gene/?term=LOC105284043 | 5661 | 295 | 521.11 | | 1 | 1.77 | 0.34 | 9.91 | 9.97 | -0.07 | 0.07 | 0 | 0 |
| gene-LOC105280421 | https://www.ncbi.nlm.nih.gov/gene/?term=LOC105280421 | 54824 | 1624 | 296.22 | | 1 | 0.18 | 0.06 | 2.3 | 2.37 | -0.08 | 0.08 | 0 | 0 |

| | | | | | | | | | | | | | |
|---|---|---|---|---|---|---|---|---|---|---|---|---|---|
| gene-LOC105277866 | https://www.ncbi.nlm.nih.gov/gene/?term=LOC105277866 | 33045 | 394 | 119.23 | 1 | 0.3 | 0.25 | 3.34 | 3.43 | -0.09 | 0.09 | 0 | 0 |
| gene-LOC105286194 | https://www.ncbi.nlm.nih.gov/gene/?term=LOC105286194 | 152619 | 2103 | 137.79 | 1 | 0.07 | 0.05 | 6.13 | 6.22 | -0.09 | 0.09 | 0 | 0 |
| gene-LOC105287417 | https://www.ncbi.nlm.nih.gov/gene/?term=LOC105287417 | 1025 | 32 | 312.2 | 1 | 9.76 | 3.12 | 37.56 | 37.69 | -0.12 | 0.12 | 0 | 0 |
| gene-LOC113562050 | https://www.ncbi.nlm.nih.gov/gene/?term=LOC113562050 | 17097 | 587 | 343.34 | 1 | 0.58 | 0.17 | 4.91 | 5.05 | -0.13 | 0.13 | 0 | 0 |
| gene-LOC105278228 | https://www.ncbi.nlm.nih.gov/gene/?term=LOC105278228 | 8610 | 154 | 178.86 | 1 | 1.16 | 0.65 | 23.95 | 24.14 | -0.2 | 0.2 | 0 | 0 |
| gene-LOC105274851 | https://www.ncbi.nlm.nih.gov/gene/?term=LOC105274851 | 6616 | 130 | 196.49 | 1 | 1.51 | 0.77 | 11.46 | 11.67 | -0.21 | 0.21 | 0 | 0 |
| gene-LOC105286452 | https://www.ncbi.nlm.nih.gov/gene/?term=LOC105286452 | 6888 | 61 | 88.56 | 1 | 1.45 | 1.64 | 30.12 | 30.34 | -0.22 | 0.22 | 0 | 0 |
| gene-LOC105288010 | https://www.ncbi.nlm.nih.gov/gene/?term=LOC105288010 | 3189 | 39 | 122.3 | 1 | 3.14 | 2.56 | 33.4 | 33.65 | -0.26 | 0.26 | 0 | 0 |
| gene-LOC105282067 | https://www.ncbi.nlm.nih.gov/gene/?term=LOC105282067 | 5941 | 41 | 69.01 | 1 | 1.68 | 2.44 | 26.24 | 26.89 | -0.64 | 0.64 | 0 | 0 |

Directory: OGLall File: rnk-dmg-Ob_LI_BCphaseB2.vs.RphaseA2.txt

| gene_ID | gene_link | gwidth | #Sites | #per10Kb | #dmSites | #dmsp10kb | %dmSites | %pSite1 | %pSite2 | DMpSite | ADMpSite | DMpNucl | ADMpNucl |
|---|---|---|---|---|---|---|---|---|---|---|---|---|---|
| gene-LOC113562345 | https://www.ncbi.nlm.nih.gov/gene/?term=LOC113562345 | 2290 | 69 | 301.31 | 19 | 82.97 | 27.54 | 16.9 | 33.89 | -17 | 17 | -0.51 | 0.51 |
| gene-LOC105284677 | https://www.ncbi.nlm.nih.gov/gene/?term=LOC105284677 | 1905 | 43 | 225.72 | 2 | 10.5 | 4.65 | 38.99 | 29.38 | 9.62 | 9.62 | 0.22 | 0.22 |
| gene-LOC113562901 | https://www.ncbi.nlm.nih.gov/gene/?term=LOC113562901 | 1531 | 22 | 143.7 | 3 | 19.6 | 13.64 | 45.03 | 59.16 | -14.13 | 14.13 | -0.2 | 0.2 |
| gene-LOC105280327 | https://www.ncbi.nlm.nih.gov/gene/?term=LOC105280327 | 2670 | 104 | 389.51 | 1 | 3.75 | 0.96 | 27.09 | 30.78 | -3.69 | 3.69 | -0.14 | 0.14 |
| gene-LOC105282220 | https://www.ncbi.nlm.nih.gov/gene/?term=LOC105282220 | 2298 | 224 | 974.76 | 1 | 4.35 | 0.45 | 7.52 | 8.84 | -1.32 | 1.32 | -0.13 | 0.13 |
| gene-LOC105282156 | https://www.ncbi.nlm.nih.gov/gene/?term=LOC105282156 | 2577 | 145 | 562.67 | 1 | 3.88 | 0.69 | 23.3 | 25.57 | -2.27 | 2.27 | -0.13 | 0.13 |
| gene-LOC105280786 | https://www.ncbi.nlm.nih.gov/gene/?term=LOC105280786 | 2573 | 119 | 462.5 | 1 | 3.89 | 0.84 | 29.73 | 32.5 | -2.77 | 2.77 | -0.13 | 0.13 |
| gene-LOC105276449 | https://www.ncbi.nlm.nih.gov/gene/?term=LOC105276449 | 1231 | 56 | 454.91 | 1 | 8.12 | 1.79 | 30.06 | 32.73 | -2.67 | 2.67 | -0.12 | 0.12 |
| gene-LOC105285674 | https://www.ncbi.nlm.nih.gov/gene/?term=LOC105285674 | 3035 | 97 | 319.6 | 2 | 6.59 | 2.06 | 24.1 | 27.79 | -3.69 | 3.69 | -0.12 | 0.12 |
| gene-LOC105287417 | https://www.ncbi.nlm.nih.gov/gene/?term=LOC105287417 | 1025 | 36 | 351.22 | 1 | 9.76 | 2.78 | 34.17 | 37.2 | -3.03 | 3.03 | -0.11 | 0.11 |
| gene-LOC105279900 | https://www.ncbi.nlm.nih.gov/gene/?term=LOC105279900 | 2191 | 62 | 282.98 | 1 | 4.56 | 1.61 | 30.24 | 34.06 | -3.83 | 3.83 | -0.11 | 0.11 |
| gene-LOC105277631 | https://www.ncbi.nlm.nih.gov/gene/?term=LOC105277631 | 4387 | 333 | 759.06 | 1 | 2.28 | 0.3 | 9.46 | 10.66 | -1.19 | 1.19 | -0.09 | 0.09 |
| gene-LOC105282149 | https://www.ncbi.nlm.nih.gov/gene/?term=LOC105282149 | 1787 | 109 | 609.96 | 1 | 5.6 | 0.92 | 41.83 | 43.33 | -1.5 | 1.5 | -0.09 | 0.09 |
| gene-LOC105283649 | https://www.ncbi.nlm.nih.gov/gene/?term=LOC105283649 | 1918 | 57 | 297.18 | 2 | 10.43 | 3.51 | 35.86 | 38.92 | -3.06 | 3.06 | -0.09 | 0.09 |
| gene-LOC113561805 | https://www.ncbi.nlm.nih.gov/gene/?term=LOC113561805 | 3337 | 84 | 251.72 | 1 | 3 | 1.19 | 34.18 | 37.85 | -3.66 | 3.66 | -0.09 | 0.09 |
| gene-LOC105274449 | https://www.ncbi.nlm.nih.gov/gene/?term=LOC105274449 | 1549 | 53 | 342.16 | 1 | 6.46 | 1.89 | 51.77 | 49.51 | 2.26 | 2.26 | 0.08 | 0.08 |
| gene-LOC105286799 | https://www.ncbi.nlm.nih.gov/gene/?term=LOC105286799 | 3557 | 188 | 528.54 | 1 | 2.81 | 0.53 | 15.27 | 16.76 | -1.49 | 1.49 | -0.08 | 0.08 |
| gene-LOC105284557 | https://www.ncbi.nlm.nih.gov/gene/?term=LOC105284557 | 1672 | 84 | 502.39 | 1 | 5.98 | 1.19 | 28.16 | 29.69 | -1.53 | 1.53 | -0.08 | 0.08 |
| gene-LOC105275070 | https://www.ncbi.nlm.nih.gov/gene/?term=LOC105275070 | 5667 | 53 | 93.52 | 6 | 10.59 | 11.32 | 1.77 | 10.27 | -8.5 | 8.5 | -0.08 | 0.08 |
| gene-LOC105279298 | https://www.ncbi.nlm.nih.gov/gene/?term=LOC105279298 | 2773 | 138 | 497.66 | 2 | 7.21 | 1.45 | 22.5 | 23.97 | -1.47 | 1.47 | -0.07 | 0.07 |
| gene-LOC105276704 | https://www.ncbi.nlm.nih.gov/gene/?term=LOC105276704 | 2063 | 101 | 489.58 | 2 | 9.69 | 1.98 | 13.1 | 14.6 | -1.5 | 1.5 | -0.07 | 0.07 |
| gene-LOC105286683 | https://www.ncbi.nlm.nih.gov/gene/?term=LOC105286683 | 1665 | 53 | 318.32 | 1 | 6.01 | 1.89 | 31.4 | 33.71 | -2.31 | 2.31 | -0.07 | 0.07 |
| gene-LOC105278454 | https://www.ncbi.nlm.nih.gov/gene/?term=LOC105278454 | 1972 | 51 | 258.62 | 1 | 5.07 | 1.96 | 39.78 | 42.63 | -2.85 | 2.85 | -0.07 | 0.07 |
| gene-LOC105286514 | https://www.ncbi.nlm.nih.gov/gene/?term=LOC105286514 | 2528 | 47 | 185.92 | 1 | 3.96 | 2.13 | 35.58 | 39.1 | -3.52 | 3.52 | -0.07 | 0.07 |
| gene-LOC105286644 | https://www.ncbi.nlm.nih.gov/gene/?term=LOC105286644 | 1235 | 20 | 161.94 | 1 | 8.1 | 5 | 37.53 | 41.72 | -4.19 | 4.19 | -0.07 | 0.07 |
| gene-LOC105275743 | https://www.ncbi.nlm.nih.gov/gene/?term=LOC105275743 | 1782 | 76 | 426.49 | 1 | 5.61 | 1.32 | 23.56 | 22.24 | 1.32 | 1.32 | 0.06 | 0.06 |
| gene-LOC105285055 | https://www.ncbi.nlm.nih.gov/gene/?term=LOC105285055 | 4108 | 274 | 666.99 | 2 | 4.87 | 0.73 | 10.9 | 11.79 | -0.89 | 0.89 | -0.06 | 0.06 |
| gene-LOC105274856 | https://www.ncbi.nlm.nih.gov/gene/?term=LOC105274856 | 5216 | 232 | 444.79 | 1 | 1.92 | 0.43 | 17.14 | 18.43 | -1.29 | 1.29 | -0.06 | 0.06 |
| gene-LOC105274534 | https://www.ncbi.nlm.nih.gov/gene/?term=LOC105274534 | 1329 | 61 | 458.99 | 1 | 7.52 | 1.64 | 40.79 | 42.14 | -1.35 | 1.35 | -0.06 | 0.06 |
| gene-LOC105278282 | https://www.ncbi.nlm.nih.gov/gene/?term=LOC105278282 | 1775 | 64 | 360.56 | 1 | 5.63 | 1.56 | 33.91 | 35.49 | -1.58 | 1.58 | -0.06 | 0.06 |
| gene-LOC105282151 | https://www.ncbi.nlm.nih.gov/gene/?term=LOC105282151 | 1601 | 46 | 287.32 | 2 | 12.49 | 4.35 | 33.88 | 35.82 | -1.94 | 1.94 | -0.06 | 0.06 |
| gene-LOC105287063 | https://www.ncbi.nlm.nih.gov/gene/?term=LOC105287063 | 1874 | 54 | 288.15 | 1 | 5.34 | 1.85 | 27.46 | 29.52 | -2.06 | 2.06 | -0.06 | 0.06 |
| gene-LOC105285583 | https://www.ncbi.nlm.nih.gov/gene/?term=LOC105285583 | 1947 | 47 | 241.4 | 1 | 5.14 | 2.13 | 27.05 | 29.68 | -2.63 | 2.63 | -0.06 | 0.06 |
| gene-LOC105276538 | https://www.ncbi.nlm.nih.gov/gene/?term=LOC105276538 | 1511 | 19 | 125.74 | 1 | 6.62 | 5.26 | 69.47 | 74.28 | -4.82 | 4.82 | -0.06 | 0.06 |
| gene-LOC105279898 | https://www.ncbi.nlm.nih.gov/gene/?term=LOC105279898 | 1343 | 14 | 104.24 | 1 | 7.45 | 7.14 | 54.89 | 60.18 | -5.29 | 5.29 | -0.06 | 0.06 |
| gene-LOC105275708 | https://www.ncbi.nlm.nih.gov/gene/?term=LOC105275708 | 1772 | 18 | 101.58 | 1 | 5.64 | 5.56 | 43.28 | 48.95 | -5.67 | 5.67 | -0.06 | 0.06 |
| gene-LOC105282885 | https://www.ncbi.nlm.nih.gov/gene/?term=LOC105282885 | 1870 | 49 | 262.03 | 1 | 5.35 | 2.04 | 39.47 | 37.7 | 1.77 | 1.77 | 0.05 | 0.05 |
| gene-LOC105282682 | https://www.ncbi.nlm.nih.gov/gene/?term=LOC105282682 | 4127 | 241 | 583.96 | 1 | 2.42 | 0.41 | 12.88 | 13.79 | -0.91 | 0.91 | -0.05 | 0.05 |
| gene-LOC105275683 | https://www.ncbi.nlm.nih.gov/gene/?term=LOC105275683 | 4256 | 210 | 493.42 | 1 | 2.35 | 0.48 | 13.47 | 14.47 | -1.01 | 1.01 | -0.05 | 0.05 |
| gene-LOC105278497 | https://www.ncbi.nlm.nih.gov/gene/?term=LOC105278497 | 2072 | 103 | 497.1 | 1 | 4.83 | 0.97 | 19.13 | 20.15 | -1.02 | 1.02 | -0.05 | 0.05 |
| gene-LOC105284355 | https://www.ncbi.nlm.nih.gov/gene/?term=LOC105284355 | 3904 | 165 | 422.64 | 1 | 2.56 | 0.61 | 19.1 | 20.26 | -1.16 | 1.16 | -0.05 | 0.05 |
| gene-LOC109611283 | https://www.ncbi.nlm.nih.gov/gene/?term=LOC109611283 | 1451 | 39 | 268.78 | 1 | 6.89 | 2.56 | 35.83 | 37.66 | -1.83 | 1.83 | -0.05 | 0.05 |
| gene-LOC105275966 | https://www.ncbi.nlm.nih.gov/gene/?term=LOC105275966 | 3299 | 73 | 221.28 | 1 | 3.03 | 1.37 | 43.49 | 45.58 | -2.08 | 2.08 | -0.05 | 0.05 |
| gene-LOC105288063 | https://www.ncbi.nlm.nih.gov/gene/?term=LOC105288063 | 3400 | 57 | 167.65 | 1 | 2.94 | 1.75 | 12.44 | 15.19 | -2.75 | 2.75 | -0.05 | 0.05 |
| gene-LOC105276680 | https://www.ncbi.nlm.nih.gov/gene/?term=LOC105276680 | 1977 | 35 | 177.04 | 1 | 5.06 | 2.86 | 58.22 | 61.02 | -2.8 | 2.8 | -0.05 | 0.05 |
| gene-LOC105288066 | https://www.ncbi.nlm.nih.gov/gene/?term=LOC105288066 | 1964 | 32 | 162.93 | 1 | 5.09 | 3.12 | 32.93 | 36 | -3.07 | 3.07 | -0.05 | 0.05 |
| gene-LOC105276078 | https://www.ncbi.nlm.nih.gov/gene/?term=LOC105276078 | 1051 | 17 | 161.75 | 1 | 9.51 | 5.88 | 25.17 | 28.55 | -3.38 | 3.38 | -0.05 | 0.05 |
| gene-LOC105281039 | https://www.ncbi.nlm.nih.gov/gene/?term=LOC105281039 | 1471 | 12 | 81.58 | 1 | 6.8 | 8.33 | 31.46 | 37.55 | -6.08 | 6.08 | -0.05 | 0.05 |
| gene-LOC105284396 | https://www.ncbi.nlm.nih.gov/gene/?term=LOC105284396 | 1976 | 70 | 354.25 | 1 | 5.06 | 1.43 | 32.98 | 31.96 | 1.02 | 1.02 | 0.04 | 0.04 |
| gene-LOC105282491 | https://www.ncbi.nlm.nih.gov/gene/?term=LOC105282491 | 4761 | 184 | 386.47 | 1 | 2.1 | 0.54 | 8.25 | 9.16 | -0.91 | 0.91 | -0.04 | 0.04 |
| gene-LOC105278877 | https://www.ncbi.nlm.nih.gov/gene/?term=LOC105278877 | 1968 | 86 | 436.99 | 1 | 5.08 | 1.16 | 31.6 | 32.52 | -0.92 | 0.92 | -0.04 | 0.04 |
| gene-LOC105280774 | https://www.ncbi.nlm.nih.gov/gene/?term=LOC105280774 | 2793 | 122 | 436.81 | 1 | 3.58 | 0.82 | 26.14 | 27.14 | -0.99 | 0.99 | -0.04 | 0.04 |
| gene-LOC105285639 | https://www.ncbi.nlm.nih.gov/gene/?term=LOC105285639 | 10325 | 408 | 395.16 | 1 | 0.97 | 0.25 | 10.25 | 11.26 | -1.02 | 1.02 | -0.04 | 0.04 |
| gene-LOC105277761 | https://www.ncbi.nlm.nih.gov/gene/?term=LOC105277761 | 4648 | 160 | 344.23 | 1 | 2.15 | 0.62 | 14.78 | 16.05 | -1.27 | 1.27 | -0.04 | 0.04 |
| gene-LOC105280956 | https://www.ncbi.nlm.nih.gov/gene/?term=LOC105280956 | 2639 | 77 | 291.78 | 1 | 3.79 | 1.3 | 16.02 | 17.39 | -1.37 | 1.37 | -0.04 | 0.04 |
| gene-LOC105282970 | https://www.ncbi.nlm.nih.gov/gene/?term=LOC105282970 | 1529 | 40 | 261.61 | 1 | 6.54 | 2.5 | 50.96 | 52.56 | -1.6 | 1.6 | -0.04 | 0.04 |
| gene-LOC105278384 | https://www.ncbi.nlm.nih.gov/gene/?term=LOC105278384 | 1390 | 35 | 251.8 | 1 | 7.19 | 2.86 | 45.32 | 46.94 | -1.62 | 1.62 | -0.04 | 0.04 |
| gene-LOC105283592 | https://www.ncbi.nlm.nih.gov/gene/?term=LOC105283592 | 3479 | 72 | 206.96 | 1 | 2.87 | 1.39 | 25.41 | 27.37 | -1.95 | 1.95 | -0.04 | 0.04 |
| gene-LOC105281420 | https://www.ncbi.nlm.nih.gov/gene/?term=LOC105281420 | 3882 | 67 | 172.59 | 1 | 2.58 | 1.49 | 12.82 | 15.24 | -2.42 | 2.42 | -0.04 | 0.04 |
| gene-LOC105274900 | https://www.ncbi.nlm.nih.gov/gene/?term=LOC105274900 | 5561 | 55 | 98.9 | 2 | 3.6 | 3.64 | 30.44 | 34.13 | -3.69 | 3.69 | -0.04 | 0.04 |

| | | | | | | | | | | | | | | |
|---|---|---|---|---|---|---|---|---|---|---|---|---|---|---|
| gene-LOC105281282 | https://www.ncbi.nlm.nih.gov/gene/?term=LOC105281282 | 3250 | 3 | 9.23 | 1 | 3.08 | 33.33 | 23.25 | 65.16 | -41.92 | | 41.92 | -0.04 | 0.04 |
| gene-LOC105276452 | https://www.ncbi.nlm.nih.gov/gene/?term=LOC105276452 | 2917 | 262 | 898.18 | | 1 | 3.43 | 0.38 | 9.36 | 9.65 | -0.29 | 0.29 | -0.03 | 0.03 |
| gene-LOC105278283 | https://www.ncbi.nlm.nih.gov/gene/?term=LOC105278283 | 2317 | 146 | 630.13 | | 1 | 4.32 | 0.68 | 24.36 | 24.8 | -0.45 | 0.45 | -0.03 | 0.03 |
| gene-LOC105275460 | https://www.ncbi.nlm.nih.gov/gene/?term=LOC105275460 | 2473 | 122 | 493.33 | | 1 | 4.04 | 0.82 | 12.96 | 13.51 | -0.55 | 0.55 | -0.03 | 0.03 |
| gene-LOC105285807 | https://www.ncbi.nlm.nih.gov/gene/?term=LOC105285807 | 6063 | 237 | 390.9 | 1 | 1.65 | 0.42 | 8.52 | 9.16 | -0.64 | 0.64 | -0.03 | 0.03 | |
| gene-LOC105274452 | https://www.ncbi.nlm.nih.gov/gene/?term=LOC105274452 | 6595 | 279 | 423.05 | | 1 | 1.52 | 0.36 | 12.34 | 12.98 | -0.65 | 0.65 | -0.03 | 0.03 |
| gene-LOC105280723 | https://www.ncbi.nlm.nih.gov/gene/?term=LOC105280723 | 3572 | 185 | 517.92 | | 1 | 2.8 | 0.54 | 15.07 | 15.74 | -0.67 | 0.67 | -0.03 | 0.03 |
| gene-LOC105275752 | https://www.ncbi.nlm.nih.gov/gene/?term=LOC105275752 | 3462 | 141 | 407.28 | | 1 | 2.89 | 0.71 | 30.44 | 31.2 | -0.75 | 0.75 | -0.03 | 0.03 |
| gene-LOC105279038 | https://www.ncbi.nlm.nih.gov/gene/?term=LOC105279038 | 9453 | 369 | 390.35 | | 1 | 1.06 | 0.27 | 8.12 | 8.94 | -0.82 | 0.82 | -0.03 | 0.03 |
| gene-LOC105278632 | https://www.ncbi.nlm.nih.gov/gene/?term=LOC105278632 | 3624 | 141 | 389.07 | | 1 | 2.76 | 0.71 | 13.56 | 14.44 | -0.87 | 0.87 | -0.03 | 0.03 |
| gene-LOC105285202 | https://www.ncbi.nlm.nih.gov/gene/?term=LOC105285202 | 1838 | 66 | 359.09 | | 1 | 5.44 | 1.52 | 24.58 | 25.52 | -0.94 | 0.94 | -0.03 | 0.03 |
| gene-LOC105281261 | https://www.ncbi.nlm.nih.gov/gene/?term=LOC105281261 | 5947 | 158 | 265.68 | | 1 | 1.68 | 0.63 | 14.18 | 15.18 | -1 | 1 | -0.03 | 0.03 |
| gene-LOC105281848 | https://www.ncbi.nlm.nih.gov/gene/?term=LOC105281848 | 2122 | 64 | 301.6 | 1 | 4.71 | 1.56 | 21.13 | 22.15 | -1.02 | 1.02 | -0.03 | 0.03 | |
| gene-LOC105284116 | https://www.ncbi.nlm.nih.gov/gene/?term=LOC105284116 | 3970 | 113 | 284.63 | | 1 | 2.52 | 0.88 | 7.21 | 8.25 | -1.03 | 1.03 | -0.03 | 0.03 |
| gene-LOC105282991 | https://www.ncbi.nlm.nih.gov/gene/?term=LOC105282991 | 5617 | 112 | 199.39 | | 1 | 1.78 | 0.89 | 19.26 | 20.58 | -1.32 | 1.32 | -0.03 | 0.03 |
| gene-LOC105278962 | https://www.ncbi.nlm.nih.gov/gene/?term=LOC105278962 | 4491 | 107 | 238.25 | | 1 | 2.23 | 0.93 | 20.7 | 22.05 | -1.35 | 1.35 | -0.03 | 0.03 |
| gene-LOC105279664 | https://www.ncbi.nlm.nih.gov/gene/?term=LOC105279664 | 4023 | 74 | 183.94 | | 1 | 2.49 | 1.35 | 16.49 | 17.94 | -1.45 | 1.45 | -0.03 | 0.03 |
| gene-LOC105278109 | https://www.ncbi.nlm.nih.gov/gene/?term=LOC105278109 | 4029 | 72 | 178.7 | 1 | 2.48 | 1.39 | 42.29 | 43.78 | -1.48 | 1.48 | -0.03 | 0.03 | |
| gene-LOC105282893 | https://www.ncbi.nlm.nih.gov/gene/?term=LOC105282893 | 2645 | 47 | 177.69 | | 1 | 3.78 | 2.13 | 28.96 | 30.53 | -1.56 | 1.56 | -0.03 | 0.03 |
| gene-LOC105287854 | https://www.ncbi.nlm.nih.gov/gene/?term=LOC105287854 | 3644 | 66 | 181.12 | | 1 | 2.74 | 1.52 | 15.87 | 17.63 | -1.76 | 1.76 | -0.03 | 0.03 |
| gene-LOC105282600 | https://www.ncbi.nlm.nih.gov/gene/?term=LOC105282600 | 3715 | 56 | 150.74 | | 1 | 2.69 | 1.79 | 29.97 | 31.8 | -1.83 | 1.83 | -0.03 | 0.03 |
| gene-LOC105288010 | https://www.ncbi.nlm.nih.gov/gene/?term=LOC105288010 | 3189 | 45 | 141.11 | | 1 | 3.14 | 2.22 | 34.76 | 36.59 | -1.83 | 1.83 | -0.03 | 0.03 |
| gene-LOC105276635 | https://www.ncbi.nlm.nih.gov/gene/?term=LOC105276635 | 4962 | 40 | 80.61 | 1 | 2.02 | 2.5 | 23.53 | 27.39 | -3.86 | 3.86 | -0.03 | 0.03 | |
| gene-LOC105282914 | https://www.ncbi.nlm.nih.gov/gene/?term=LOC105282914 | 5623 | 43 | 76.47 | 1 | 1.78 | 2.33 | 49.78 | 47.18 | 2.6 | 2.6 | 0.02 | 0.02 | |
| gene-LOC105285409 | https://www.ncbi.nlm.nih.gov/gene/?term=LOC105285409 | 5099 | 63 | 123.55 | | 1 | 1.96 | 1.59 | 3.13 | 1.16 | 1.97 | 1.97 | 0.02 | 0.02 |
| gene-LOC105281296 | https://www.ncbi.nlm.nih.gov/gene/?term=LOC105281296 | 2434 | 43 | 176.66 | | 1 | 4.11 | 2.33 | 29.93 | 28.76 | 1.17 | 1.17 | 0.02 | 0.02 |
| gene-LOC105285252 | https://www.ncbi.nlm.nih.gov/gene/?term=LOC105285252 | 7427 | 125 | 168.3 | 1 | 1.35 | 0.8 | 14.66 | 13.59 | 1.08 | 1.08 | 0.02 | 0.02 | |
| gene-LOC105280819 | https://www.ncbi.nlm.nih.gov/gene/?term=LOC105280819 | 1639 | 43 | 262.36 | | 1 | 6.1 | 2.33 | 39.04 | 38.17 | 0.87 | 0.87 | 0.02 | 0.02 |
| gene-LOC105288105 | https://www.ncbi.nlm.nih.gov/gene/?term=LOC105288105 | 1460 | 37 | 253.42 | | 1 | 6.85 | 2.7 | 53 | 52.16 | 0.84 | 0.84 | 0.02 | 0.02 |
| gene-LOC105275587 | https://www.ncbi.nlm.nih.gov/gene/?term=LOC105275587 | 2404 | 57 | 237.1 | 1 | 4.16 | 1.75 | 36.63 | 35.9 | 0.73 | 0.73 | 0.02 | 0.02 | |
| gene-LOC105276272 | https://www.ncbi.nlm.nih.gov/gene/?term=LOC105276272 | 1965 | 50 | 254.45 | | 1 | 5.09 | 2 | 27.69 | 27.01 | 0.68 | 0.68 | 0.02 | 0.02 |
| gene-LOC105287633 | https://www.ncbi.nlm.nih.gov/gene/?term=LOC105287633 | 1524 | 35 | 229.66 | | 1 | 6.56 | 2.86 | 49.8 | 49.12 | 0.68 | 0.68 | 0.02 | 0.02 |
| gene-LOC105278145 | https://www.ncbi.nlm.nih.gov/gene/?term=LOC105278145 | 4470 | 252 | 563.76 | | 1 | 2.24 | 0.4 | 10.24 | 10.56 | -0.32 | 0.32 | -0.02 | 0.02 |
| gene-LOC105288055 | https://www.ncbi.nlm.nih.gov/gene/?term=LOC105288055 | 9934 | 548 | 551.64 | | 1 | 1.01 | 0.18 | 7.05 | 7.45 | -0.4 | 0.4 | -0.02 | 0.02 |
| gene-LOC105275374 | https://www.ncbi.nlm.nih.gov/gene/?term=LOC105275374 | 11376 | 567 | 498.42 | | 1 | 0.88 | 0.18 | 10.6 | 11.02 | -0.41 | 0.41 | -0.02 | 0.02 |
| gene-LOC105285742 | https://www.ncbi.nlm.nih.gov/gene/?term=LOC105285742 | 3446 | 132 | 383.05 | | 1 | 2.9 | 0.76 | 18.9 | 19.31 | -0.41 | 0.41 | -0.02 | 0.02 |
| gene-LOC105284043 | https://www.ncbi.nlm.nih.gov/gene/?term=LOC105284043 | 5661 | 325 | 574.1 | 1 | 1.77 | 0.31 | 9.91 | 10.34 | -0.43 | 0.43 | -0.02 | 0.02 | |
| gene-LOC105278666 | https://www.ncbi.nlm.nih.gov/gene/?term=LOC105278666 | 10341 | 494 | 477.71 | | 1 | 0.97 | 0.2 | 6.74 | 7.18 | -0.44 | 0.44 | -0.02 | 0.02 |
| gene-LOC105281272 | https://www.ncbi.nlm.nih.gov/gene/?term=LOC105281272 | 5252 | 203 | 386.52 | | 1 | 1.9 | 0.49 | 14.55 | 14.99 | -0.44 | 0.44 | -0.02 | 0.02 |
| gene-LOC105274669 | https://www.ncbi.nlm.nih.gov/gene/?term=LOC105274669 | 8555 | 293 | 342.49 | | 1 | 1.17 | 0.34 | 5.34 | 5.82 | -0.48 | 0.48 | -0.02 | 0.02 |
| gene-LOC105275363 | https://www.ncbi.nlm.nih.gov/gene/?term=LOC105275363 | 3798 | 176 | 463.4 | 1 | 2.63 | 0.57 | 15.74 | 16.23 | -0.5 | 0.5 | -0.02 | 0.02 | |
| gene-LOC113562050 | https://www.ncbi.nlm.nih.gov/gene/?term=LOC113562050 | 17097 | 643 | 376.09 | | 1 | 0.58 | 0.16 | 4.96 | 5.49 | -0.53 | 0.53 | -0.02 | 0.02 |
| gene-LOC105284036 | https://www.ncbi.nlm.nih.gov/gene/?term=LOC105284036 | 3254 | 112 | 344.19 | | 1 | 3.07 | 0.89 | 21.95 | 22.5 | -0.55 | 0.55 | -0.02 | 0.02 |
| gene-LOC105283728 | https://www.ncbi.nlm.nih.gov/gene/?term=LOC105283728 | 3256 | 123 | 377.76 | | 2 | 6.14 | 1.63 | 18.65 | 19.22 | -0.57 | 0.57 | -0.02 | 0.02 |
| gene-LOC105276875 | https://www.ncbi.nlm.nih.gov/gene/?term=LOC105276875 | 3151 | 124 | 393.53 | | 1 | 3.17 | 0.81 | 21.3 | 21.9 | -0.6 | 0.6 | -0.02 | 0.02 |
| gene-LOC105274706 | https://www.ncbi.nlm.nih.gov/gene/?term=LOC105274706 | 1446 | 57 | 394.19 | | 1 | 6.92 | 1.75 | 36.89 | 37.51 | -0.61 | 0.61 | -0.02 | 0.02 |
| gene-LOC105275451 | https://www.ncbi.nlm.nih.gov/gene/?term=LOC105275451 | 6035 | 142 | 235.29 | | 1 | 1.66 | 0.7 | 20.66 | 21.45 | -0.78 | 0.78 | -0.02 | 0.02 |
| gene-LOC105280348 | https://www.ncbi.nlm.nih.gov/gene/?term=LOC105280348 | 3797 | 79 | 208.06 | | 1 | 2.63 | 1.27 | 19.69 | 20.49 | -0.8 | 0.8 | -0.02 | 0.02 |
| gene-LOC105283967 | https://www.ncbi.nlm.nih.gov/gene/?term=LOC105283967 | 2393 | 60 | 250.73 | | 1 | 4.18 | 1.67 | 41.56 | 42.36 | -0.8 | 0.8 | -0.02 | 0.02 |
| gene-LOC105285395 | https://www.ncbi.nlm.nih.gov/gene/?term=LOC105285395 | 8085 | 224 | 277.06 | | 1 | 1.24 | 0.45 | 14.28 | 15.08 | -0.8 | 0.8 | -0.02 | 0.02 |
| gene-LOC105286332 | https://www.ncbi.nlm.nih.gov/gene/?term=LOC105286332 | 27577 | 532 | 192.91 | | 1 | 0.36 | 0.19 | 7.3 | 8.12 | -0.82 | 0.82 | -0.02 | 0.02 |
| gene-LOC105280407 | https://www.ncbi.nlm.nih.gov/gene/?term=LOC105280407 | 10950 | 218 | 199.09 | | 1 | 0.91 | 0.46 | 16.66 | 17.49 | -0.83 | 0.83 | -0.02 | 0.02 |
| gene-LOC105278228 | https://www.ncbi.nlm.nih.gov/gene/?term=LOC105278228 | 8610 | 176 | 204.41 | | 1 | 1.16 | 0.57 | 22.43 | 23.28 | -0.84 | 0.84 | -0.02 | 0.02 |
| gene-LOC105274590 | https://www.ncbi.nlm.nih.gov/gene/?term=LOC105274590 | 4516 | 80 | 177.15 | | 2 | 4.43 | 2.5 | 29.75 | 30.69 | -0.95 | 0.95 | -0.02 | 0.02 |
| gene-LOC105282825 | https://www.ncbi.nlm.nih.gov/gene/?term=LOC105282825 | 3919 | 98 | 250.06 | | 1 | 2.55 | 1.02 | 14.18 | 15.14 | -0.96 | 0.96 | -0.02 | 0.02 |
| gene-LOC105284117 | https://www.ncbi.nlm.nih.gov/gene/?term=LOC105284117 | 4340 | 101 | 232.72 | | 1 | 2.3 | 0.99 | 10.38 | 11.36 | -0.98 | 0.98 | -0.02 | 0.02 |
| gene-LOC105276196 | https://www.ncbi.nlm.nih.gov/gene/?term=LOC105276196 | 2953 | 59 | 199.8 | 1 | 3.39 | 1.69 | 37.63 | 38.7 | -1.07 | 1.07 | -0.02 | 0.02 | |
| gene-LOC105283943 | https://www.ncbi.nlm.nih.gov/gene/?term=LOC105283943 | 3559 | 64 | 179.83 | | 1 | 2.81 | 1.56 | 14.69 | 15.79 | -1.1 | 1.1 | -0.02 | 0.02 |
| gene-LOC105276501 | https://www.ncbi.nlm.nih.gov/gene/?term=LOC105276501 | 6089 | 110 | 180.65 | | 1 | 1.64 | 0.91 | 15.38 | 16.49 | -1.11 | 1.11 | -0.02 | 0.02 |
| gene-LOC105288078 | https://www.ncbi.nlm.nih.gov/gene/?term=LOC105288078 | 19126 | 332 | 173.59 | | 1 | 0.52 | 0.3 | 16.16 | 17.36 | -1.2 | 1.2 | -0.02 | 0.02 |
| gene-LOC105283712 | https://www.ncbi.nlm.nih.gov/gene/?term=LOC105283712 | 1910 | 36 | 188.48 | | 1 | 5.24 | 2.78 | 28.48 | 29.79 | -1.31 | 1.31 | -0.02 | 0.02 |
| gene-LOC105280641 | https://www.ncbi.nlm.nih.gov/gene/?term=LOC105280641 | 3682 | 61 | 165.67 | | 1 | 2.72 | 1.64 | 16.75 | 18.07 | -1.32 | 1.32 | -0.02 | 0.02 |
| gene-LOC105280002 | https://www.ncbi.nlm.nih.gov/gene/?term=LOC105280002 | 1940 | 28 | 144.33 | | 1 | 5.15 | 3.57 | 39.15 | 40.87 | -1.73 | 1.73 | -0.02 | 0.02 |
| gene-LOC105275518 | https://www.ncbi.nlm.nih.gov/gene/?term=LOC105275518 | 2335 | 26 | 111.35 | | 1 | 4.28 | 3.85 | 36.28 | 38.04 | -1.76 | 1.76 | -0.02 | 0.02 |

| | | | | | | | | | | | | | | | |
|---|---|---|---|---|---|---|---|---|---|---|---|---|---|---|---|
| gene-LOC105277856 | https://www.ncbi.nlm.nih.gov/gene/?term=LOC105277856 | 3442 | 37 | 107.5 | 1 | 2.91 | 2.7 | 37.9 | 39.79 | -1.89 | 1.89 | -0.02 | 0.02 | | |
| gene-LOC105280738 | https://www.ncbi.nlm.nih.gov/gene/?term=LOC105280738 | 4287 | 41 | 95.64 | 1 | 2.33 | 2.44 | 24.39 | 26.44 | -2.04 | 2.04 | -0.02 | 0.02 | | |
| gene-LOC105284609 | https://www.ncbi.nlm.nih.gov/gene/?term=LOC105284609 | 5678 | 35 | 61.64 | 1 | 1.76 | 2.86 | 28.06 | 31.35 | -3.29 | 3.29 | -0.02 | 0.02 | | |
| gene-LOC105281515 | https://www.ncbi.nlm.nih.gov/gene/?term=LOC105281515 | 8827 | 54 | 61.18 | 2 | 2.27 | 3.7 | 27.93 | 31.25 | -3.31 | 3.31 | -0.02 | 0.02 | | |
| gene-LOC105286983 | https://www.ncbi.nlm.nih.gov/gene/?term=LOC105286983 | 16901 | 64 | 37.87 | 2 | 1.18 | 3.12 | 0.67 | 5.99 | -5.32 | 5.32 | -0.02 | 0.02 | | |
| gene-LOC105279500 | https://www.ncbi.nlm.nih.gov/gene/?term=LOC105279500 | 3587 | 33 | 92 | 1 | 2.79 | 3.03 | 47.01 | 45.48 | 1.53 | 1.53 | 0.01 | 0.01 | | |
| gene-LOC105281173 | https://www.ncbi.nlm.nih.gov/gene/?term=LOC105281173 | 1595 | 38 | 238.24 | | 1 | 6.27 | 2.63 | 60.67 | 60.19 | 0.48 | 0.48 | 0.01 | 0.01 | |
| gene-LOC105280304 | https://www.ncbi.nlm.nih.gov/gene/?term=LOC105280304 | 8919 | 115 | 128.94 | | 1 | 1.12 | 0.87 | 7.4 | 6.97 | 0.43 | 0.43 | 0.01 | 0.01 | |
| gene-LOC105284507 | https://www.ncbi.nlm.nih.gov/gene/?term=LOC105284507 | 5207 | 91 | 174.76 | | 1 | 1.92 | 1.1 | 18.91 | 18.57 | 0.35 | 0.35 | 0.01 | 0.01 | |
| gene-LOC105283296 | https://www.ncbi.nlm.nih.gov/gene/?term=LOC105283296 | 2174 | 96 | 441.58 | | 1 | 4.6 | 1.04 | 20.81 | 20.51 | 0.3 | 0.3 | 0.01 | 0.01 | |
| gene-LOC105278807 | https://www.ncbi.nlm.nih.gov/gene/?term=LOC105278807 | 2166 | 81 | 373.96 | | 1 | 4.62 | 1.23 | 19.2 | 19.33 | -0.14 | 0.14 | -0.01 | 0.01 | |
| gene-LOC105285179 | https://www.ncbi.nlm.nih.gov/gene/?term=LOC105285179 | 4191 | 183 | 436.65 | | 1 | 2.39 | 0.55 | 19.23 | 19.39 | -0.16 | 0.16 | -0.01 | 0.01 | |
| gene-LOC105282811 | https://www.ncbi.nlm.nih.gov/gene/?term=LOC105282811 | 5765 | 213 | 369.47 | | 1 | 1.73 | 0.47 | 8.07 | 8.24 | -0.17 | 0.17 | -0.01 | 0.01 | |
| gene-LOC105287550 | https://www.ncbi.nlm.nih.gov/gene/?term=LOC105287550 | 15659 | 656 | 418.93 | | 1 | 0.64 | 0.15 | 6.08 | 6.26 | -0.18 | 0.18 | -0.01 | 0.01 | |
| gene-LOC105287095 | https://www.ncbi.nlm.nih.gov/gene/?term=LOC105287095 | 1435 | 65 | 452.96 | | 1 | 6.97 | 1.54 | 25.24 | 25.44 | -0.2 | 0.2 | -0.01 | 0.01 | |
| gene-LOC105286274 | https://www.ncbi.nlm.nih.gov/gene/?term=LOC105286274 | 3817 | 206 | 539.69 | | 1 | 2.62 | 0.49 | 16.61 | 16.82 | -0.21 | 0.21 | -0.01 | 0.01 | |
| gene-LOC105278455 | https://www.ncbi.nlm.nih.gov/gene/?term=LOC105278455 | 27181 | 1027 | 377.84 | | 2 | 0.74 | 0.19 | 6.56 | 6.78 | -0.22 | 0.22 | -0.01 | 0.01 | |
| gene-LOC105274699 | https://www.ncbi.nlm.nih.gov/gene/?term=LOC105274699 | 2386 | 63 | 264.04 | | 1 | 4.19 | 1.59 | 22.17 | 22.41 | -0.24 | 0.24 | -0.01 | 0.01 | |
| gene-LOC105274531 | https://www.ncbi.nlm.nih.gov/gene/?term=LOC105274531 | 7774 | 298 | 383.33 | | 1 | 1.29 | 0.34 | 10.39 | 10.72 | -0.33 | 0.33 | -0.01 | 0.01 | |
| gene-LOC105275327 | https://www.ncbi.nlm.nih.gov/gene/?term=LOC105275327 | 14245 | 568 | 398.74 | | 1 | 0.7 | 0.18 | 6.19 | 6.55 | -0.36 | 0.36 | -0.01 | 0.01 | |
| gene-LOC105285163 | https://www.ncbi.nlm.nih.gov/gene/?term=LOC105285163 | 5061 | 85 | 167.95 | | 1 | 1.98 | 1.18 | 19.84 | 20.21 | -0.37 | 0.37 | -0.01 | 0.01 | |
| gene-LOC105278771 | https://www.ncbi.nlm.nih.gov/gene/?term=LOC105278771 | 10457 | 248 | 237.16 | | 1 | 0.96 | 0.4 | 8.97 | 9.36 | -0.39 | 0.39 | -0.01 | 0.01 | |
| gene-LOC105282794 | https://www.ncbi.nlm.nih.gov/gene/?term=LOC105282794 | 4571 | 100 | 218.77 | | 1 | 2.19 | 1 | 16.99 | 17.37 | -0.39 | 0.39 | -0.01 | 0.01 | |
| gene-LOC105283123 | https://www.ncbi.nlm.nih.gov/gene/?term=LOC105283123 | 1507 | 57 | 378.23 | | 1 | 6.64 | 1.75 | 17.49 | 17.88 | -0.39 | 0.39 | -0.01 | 0.01 | |
| gene-LOC105287309 | https://www.ncbi.nlm.nih.gov/gene/?term=LOC105287309 | 3553 | 96 | 270.19 | | 1 | 2.81 | 1.04 | 7.77 | 8.16 | -0.39 | 0.39 | -0.01 | 0.01 | |
| gene-LOC105287648 | https://www.ncbi.nlm.nih.gov/gene/?term=LOC105287648 | 2205 | 35 | 158.73 | | 1 | 4.54 | 2.86 | 35.94 | 36.45 | -0.5 | 0.5 | -0.01 | 0.01 | |
| gene-LOC105276059 | https://www.ncbi.nlm.nih.gov/gene/?term=LOC105276059 | 6256 | 145 | 231.78 | | 1 | 1.6 | 0.69 | 8.95 | 9.49 | -0.54 | 0.54 | -0.01 | 0.01 | |
| gene-LOC105285502 | https://www.ncbi.nlm.nih.gov/gene/?term=LOC105285502 | 9990 | 164 | 164.16 | | 1 | 1 | 0.61 | 8.55 | 9.16 | -0.62 | 0.62 | -0.01 | 0.01 | |
| gene-LOC105276683 | https://www.ncbi.nlm.nih.gov/gene/?term=LOC105276683 | 17000 | 149 | 87.65 | 1 | 0.59 | 0.67 | 8.22 | 8.88 | -0.65 | 0.65 | -0.01 | 0.01 | | |
| gene-LOC105286452 | https://www.ncbi.nlm.nih.gov/gene/?term=LOC105286452 | 6888 | 66 | 95.82 | 1 | 1.45 | 1.52 | 30.11 | 30.78 | -0.67 | 0.67 | -0.01 | 0.01 | | |
| gene-LOC105279657 | https://www.ncbi.nlm.nih.gov/gene/?term=LOC105279657 | 5878 | 91 | 154.81 | | 1 | 1.7 | 1.1 | 18.9 | 19.62 | -0.72 | 0.72 | -0.01 | 0.01 | |
| gene-LOC105288161 | https://www.ncbi.nlm.nih.gov/gene/?term=LOC105288161 | 26065 | 356 | 136.58 | | 1 | 0.38 | 0.28 | 10.97 | 11.71 | -0.74 | 0.74 | -0.01 | 0.01 | |
| gene-LOC105275187 | https://www.ncbi.nlm.nih.gov/gene/?term=LOC105275187 | 5534 | 77 | 139.14 | | 1 | 1.81 | 1.3 | 15.38 | 16.21 | -0.83 | 0.83 | -0.01 | 0.01 | |
| gene-LOC105276917 | https://www.ncbi.nlm.nih.gov/gene/?term=LOC105276917 | 2336 | 33 | 141.27 | | 1 | 4.28 | 3.03 | 30.13 | 31.08 | -0.95 | 0.95 | -0.01 | 0.01 | |
| gene-LOC105275347 | https://www.ncbi.nlm.nih.gov/gene/?term=LOC105275347 | 2526 | 20 | 79.18 | 1 | 3.96 | 5 | 37.1 | 38.21 | -1.12 | 1.12 | -0.01 | 0.01 | | |
| gene-LOC105285514 | https://www.ncbi.nlm.nih.gov/gene/?term=LOC105285514 | 1277 | 15 | 117.46 | | 1 | 7.83 | 6.67 | 78.25 | 79.38 | -1.12 | 1.12 | -0.01 | 0.01 | |
| gene-LOC105283989 | https://www.ncbi.nlm.nih.gov/gene/?term=LOC105283989 | 4948 | 32 | 64.67 | 2 | 4.04 | 6.25 | 43.84 | 45.43 | -1.59 | 1.59 | -0.01 | 0.01 | | |
| gene-LOC105285315 | https://www.ncbi.nlm.nih.gov/gene/?term=LOC105285315 | 5458 | 26 | 47.64 | 1 | 1.83 | 3.85 | 36.55 | 38.99 | -2.44 | 2.44 | -0.01 | 0.01 | | |
| gene-LOC105284329 | https://www.ncbi.nlm.nih.gov/gene/?term=LOC105284329 | 4051 | 53 | 130.83 | | 1 | 2.47 | 1.89 | 19.36 | 19.04 | 0.32 | 0.32 | 0 | 0 | |
| gene-LOC105286770 | https://www.ncbi.nlm.nih.gov/gene/?term=LOC105286770 | 3055 | 38 | 124.39 | | 1 | 3.27 | 2.63 | 15.65 | 15.48 | 0.17 | 0.17 | 0 | 0 | |
| gene-LOC105287842 | https://www.ncbi.nlm.nih.gov/gene/?term=LOC105287842 | 2301 | 57 | 247.72 | | 1 | 4.35 | 1.75 | 21.58 | 21.41 | 0.17 | 0.17 | 0 | 0 | |
| gene-LOC105285496 | https://www.ncbi.nlm.nih.gov/gene/?term=LOC105285496 | 4080 | 112 | 274.51 | | 1 | 2.45 | 0.89 | 40.26 | 40.12 | 0.14 | 0.14 | 0 | 0 | |
| gene-LOC105278534 | https://www.ncbi.nlm.nih.gov/gene/?term=LOC105278534 | 7364 | 200 | 271.59 | | 2 | 2.72 | 1 | 17.58 | 17.48 | 0.1 | 0.1 | 0 | 0 | |
| gene-LOC105281312 | https://www.ncbi.nlm.nih.gov/gene/?term=LOC105281312 | 2551 | 74 | 290.08 | | 1 | 3.92 | 1.35 | 29.31 | 29.21 | 0.1 | 0.1 | 0 | 0 | |
| gene-LOC105282951 | https://www.ncbi.nlm.nih.gov/gene/?term=LOC105282951 | 3366 | 131 | 389.19 | | 1 | 2.97 | 0.76 | 18.77 | 18.69 | 0.08 | 0.08 | 0 | 0 | |
| gene-LOC105277273 | https://www.ncbi.nlm.nih.gov/gene/?term=LOC105277273 | 3711 | 169 | 455.4 | 1 | 2.69 | 0.59 | 12.24 | 12.17 | 0.06 | 0.06 | 0 | 0 | | |
| gene-LOC105281700 | https://www.ncbi.nlm.nih.gov/gene/?term=LOC105281700 | 2208 | 39 | 176.63 | | 1 | 4.53 | 2.56 | 37.83 | 37.79 | 0.04 | 0.04 | 0 | 0 | |
| gene-LOC105284184 | https://www.ncbi.nlm.nih.gov/gene/?term=LOC105284184 | 2535 | 102 | 402.37 | | 1 | 3.94 | 0.98 | 28.97 | 28.93 | 0.03 | 0.03 | 0 | 0 | |
| gene-LOC105285533 | https://www.ncbi.nlm.nih.gov/gene/?term=LOC105285533 | 1640 | 71 | 432.93 | | 1 | 6.1 | 1.41 | 23.22 | 23.19 | 0.03 | 0.03 | 0 | 0 | |
| gene-LOC105281754 | https://www.ncbi.nlm.nih.gov/gene/?term=LOC105281754 | 4300 | 122 | 283.72 | | 1 | 2.33 | 0.82 | 16.75 | 16.74 | 0.01 | 0.01 | 0 | 0 | |
| gene-LOC105285838 | https://www.ncbi.nlm.nih.gov/gene/?term=LOC105285838 | 21225 | 249 | 117.31 | | 1 | 0.47 | 0.4 | 4.78 | 4.78 | 0 | 0 | 0 | 0 | |
| gene-LOC105274851 | https://www.ncbi.nlm.nih.gov/gene/?term=LOC105274851 | 6616 | 150 | 226.72 | | 1 | 1.51 | 0.67 | 10.51 | 10.53 | -0.02 | 0.02 | 0 | 0 | |
| gene-LOC105277866 | https://www.ncbi.nlm.nih.gov/gene/?term=LOC105277866 | 33045 | 432 | 130.73 | | 1 | 0.3 | 0.23 | 3.25 | 3.28 | -0.03 | 0.03 | 0 | 0 | |
| gene-LOC105280806 | https://www.ncbi.nlm.nih.gov/gene/?term=LOC105280806 | 357459 | | 15148 | 423.77 | | 1 | 0.03 | 0.01 | 0.61 | 0.64 | -0.03 | 0.03 | 0 | 0 |
| gene-LOC105278691 | https://www.ncbi.nlm.nih.gov/gene/?term=LOC105278691 | 3871 | 172 | 444.33 | | 1 | 2.58 | 0.58 | 14.5 | 14.55 | -0.05 | 0.05 | 0 | 0 | |
| gene-LOC105280265 | https://www.ncbi.nlm.nih.gov/gene/?term=LOC105280265 | 85984 | 3467 | 403.21 | | 1 | 0.12 | 0.03 | 1.03 | 1.08 | -0.05 | 0.05 | 0 | 0 | |
| gene-LOC105286649 | https://www.ncbi.nlm.nih.gov/gene/?term=LOC105286649 | 14178 | 892 | 629.14 | | 1 | 0.71 | 0.11 | 1.01 | 1.06 | -0.05 | 0.05 | 0 | 0 | |
| gene-LOC105275848 | https://www.ncbi.nlm.nih.gov/gene/?term=LOC105275848 | 3718 | 149 | 400.75 | | 1 | 2.69 | 0.67 | 21.41 | 21.47 | -0.06 | 0.06 | 0 | 0 | |
| gene-LOC105277125 | https://www.ncbi.nlm.nih.gov/gene/?term=LOC105277125 | 51079 | 2082 | 407.6 | 1 | 0.2 | 0.05 | 1.65 | 1.71 | -0.06 | 0.06 | 0 | 0 | | |
| gene-LOC105280513 | https://www.ncbi.nlm.nih.gov/gene/?term=LOC105280513 | 69542 | 1415 | 203.47 | | 1 | 0.14 | 0.07 | 0.66 | 0.73 | -0.06 | 0.06 | 0 | 0 | |
| gene-LOC105282646 | https://www.ncbi.nlm.nih.gov/gene/?term=LOC105282646 | 3317 | 110 | 331.62 | | 1 | 3.01 | 0.91 | 19.95 | 20.03 | -0.08 | 0.08 | 0 | 0 | |
| gene-LOC105276462 | https://www.ncbi.nlm.nih.gov/gene/?term=LOC105276462 | 6561 | 296 | 451.15 | | 1 | 1.52 | 0.34 | 7.73 | 7.83 | -0.1 | 0.1 | 0 | 0 | |
| gene-LOC105276527 | https://www.ncbi.nlm.nih.gov/gene/?term=LOC105276527 | 7923 | 340 | 429.13 | | 1 | 1.26 | 0.29 | 9.36 | 9.47 | -0.11 | 0.11 | 0 | 0 | |
| gene-LOC105280408 | https://www.ncbi.nlm.nih.gov/gene/?term=LOC105280408 | 5983 | 60 | 100.28 | | 1 | 1.67 | 1.67 | 21.93 | 22.03 | -0.11 | 0.11 | 0 | 0 | |

| | | | | | | | | | | | | | |
|---|---|---|---|---|---|---|---|---|---|---|---|---|---|
| gene-LOC105280421 | https://www.ncbi.nlm.nih.gov/gene/?term=LOC105280421 | 54824 | 1900 | 346.56 | 1 | 0.18 | 0.05 | 2.13 | 2.24 | -0.11 | 0.11 | 0 | 0 |
| gene-LOC105280722 | https://www.ncbi.nlm.nih.gov/gene/?term=LOC105280722 | 3237 | 98 | 302.75 | 1 | 3.09 | 1.02 | 29.35 | 29.46 | -0.11 | 0.11 | 0 | 0 |
| gene-LOC105275179 | https://www.ncbi.nlm.nih.gov/gene/?term=LOC105275179 | 3271 | 85 | 259.86 | 1 | 3.06 | 1.18 | 19.41 | 19.53 | -0.12 | 0.12 | 0 | 0 |
| gene-LOC105284262 | https://www.ncbi.nlm.nih.gov/gene/?term=LOC105284262 | 6669 | 247 | 370.37 | 1 | 1.5 | 0.4 | 4.68 | 4.81 | -0.13 | 0.13 | 0 | 0 |
| gene-LOC105274823 | https://www.ncbi.nlm.nih.gov/gene/?term=LOC105274823 | 4620 | 148 | 320.35 | 1 | 2.16 | 0.68 | 20.3 | 20.45 | -0.14 | 0.14 | 0 | 0 |
| gene-LOC105280410 | https://www.ncbi.nlm.nih.gov/gene/?term=LOC105280410 | 12580 | 263 | 209.06 | 1 | 0.79 | 0.38 | 19.71 | 19.86 | -0.15 | 0.15 | 0 | 0 |
| gene-LOC105286515 | https://www.ncbi.nlm.nih.gov/gene/?term=LOC105286515 | 3751 | 123 | 327.91 | 1 | 2.67 | 0.81 | 10.35 | 10.5 | -0.15 | 0.15 | 0 | 0 |
| gene-LOC105285406 | https://www.ncbi.nlm.nih.gov/gene/?term=LOC105285406 | 8015 | 59 | 73.61 | 1 | 1.25 | 1.69 | 29.55 | 29.82 | -0.27 | 0.27 | 0 | 0 |
| gene-LOC105286194 | https://www.ncbi.nlm.nih.gov/gene/?term=LOC105286194 | 152619 | 2371 | 155.35 | 1 | 0.07 | 0.04 | 6.01 | 6.33 | -0.32 | 0.32 | 0 | 0 |
| gene-LOC105282067 | https://www.ncbi.nlm.nih.gov/gene/?term=LOC105282067 | 5941 | 44 | 74.06 | 1 | 1.68 | 2.27 | 30.63 | 31 | -0.36 | 0.36 | 0 | 0 |

Directory: OGLall File: rnk-dmg-Ob_LI_BCphaseB2.vs.RphaseB1.txt

| gene_ID | gene_link | gwidth | #Sites | #per10Kb | #dmSites | #dmsp10kb | %dmSites | %pSite1 | %pSite2 | DMpSite | ADMpSite | DMpNucl | ADMpNucl |
|---|---|---|---|---|---|---|---|---|---|---|---|---|---|
| gene-LOC105283830 | https://www.ncbi.nlm.nih.gov/gene/?term=LOC105283830 | 252 | 14 | 555.56 | 1 | 39.68 | 7.14 | 34.01 | 40.39 | -6.38 | 6.38 | -0.35 | 0.35 |
| gene-LOC105288193 | https://www.ncbi.nlm.nih.gov/gene/?term=LOC105288193 | 1495 | 85 | 568.56 | 5 | 33.44 | 5.88 | 31.62 | 28.31 | 3.31 | 3.31 | 0.19 | 0.19 |
| gene-LOC105275833 | https://www.ncbi.nlm.nih.gov/gene/?term=LOC105275833 | 981 | 41 | 417.94 | 1 | 10.19 | 2.44 | 35.54 | 39.29 | -3.75 | 3.75 | -0.16 | 0.16 |
| gene-LOC105276202 | https://www.ncbi.nlm.nih.gov/gene/?term=LOC105276202 | 1540 | 43 | 279.22 | 5 | 32.47 | 11.63 | 24.19 | 18.86 | 5.33 | 5.33 | 0.15 | 0.15 |
| gene-LOC113562453 | https://www.ncbi.nlm.nih.gov/gene/?term=LOC113562453 | 763 | 32 | 419.4 | 2 | 26.21 | 6.25 | 26.93 | 30.48 | -3.55 | 3.55 | -0.15 | 0.15 |
| gene-LOC105287692 | https://www.ncbi.nlm.nih.gov/gene/?term=LOC105287692 | 1505 | 97 | 644.52 | 5 | 33.22 | 5.15 | 29.7 | 27.6 | 2.1 | 2.1 | 0.14 | 0.14 |
| gene-LOC105278849 | https://www.ncbi.nlm.nih.gov/gene/?term=LOC105278849 | 776 | 75 | 966.49 | 3 | 38.66 | 4 | 22.31 | 21.08 | 1.24 | 1.24 | 0.12 | 0.12 |
| gene-LOC105283267 | https://www.ncbi.nlm.nih.gov/gene/?term=LOC105283267 | 2571 | 73 | 283.94 | 5 | 19.45 | 6.85 | 39.09 | 43.39 | -4.29 | 4.29 | -0.12 | 0.12 |
| gene-LOC105276115 | https://www.ncbi.nlm.nih.gov/gene/?term=LOC105276115 | 1520 | 36 | 236.84 | 3 | 19.74 | 8.33 | 20.99 | 26.1 | -5.11 | 5.11 | -0.12 | 0.12 |
| gene-LOC105280891 | https://www.ncbi.nlm.nih.gov/gene/?term=LOC105280891 | 1844 | 36 | 195.23 | 2 | 10.85 | 5.56 | 18.81 | 24.77 | -5.96 | 5.96 | -0.12 | 0.12 |
| gene-LOC105285004 | https://www.ncbi.nlm.nih.gov/gene/?term=LOC105285004 | 1334 | 33 | 247.38 | 3 | 22.49 | 9.09 | 22.51 | 18.04 | 4.47 | 4.47 | 0.11 | 0.11 |
| gene-LOC105277371 | https://www.ncbi.nlm.nih.gov/gene/?term=LOC105277371 | 1816 | 74 | 407.49 | 2 | 11.01 | 2.7 | 20.36 | 17.6 | 2.76 | 2.76 | 0.11 | 0.11 |
| gene-LOC105286845 | https://www.ncbi.nlm.nih.gov/gene/?term=LOC105286845 | 1159 | 81 | 698.88 | 1 | 8.63 | 1.23 | 14.6 | 12.97 | 1.63 | 1.63 | 0.11 | 0.11 |
| gene-LOC105284093 | https://www.ncbi.nlm.nih.gov/gene/?term=LOC105284093 | 809 | 59 | 729.3 | 1 | 12.36 | 1.69 | 8.36 | 9.93 | -1.57 | 1.57 | -0.11 | 0.11 |
| gene-LOC105280710 | https://www.ncbi.nlm.nih.gov/gene/?term=LOC105280710 | 1215 | 68 | 559.67 | 2 | 16.46 | 2.94 | 20.85 | 22.76 | -1.91 | 1.91 | -0.11 | 0.11 |
| gene-LOC105285203 | https://www.ncbi.nlm.nih.gov/gene/?term=LOC105285203 | 1020 | 27 | 264.71 | 1 | 9.8 | 3.7 | 30.05 | 26.28 | 3.77 | 3.77 | 0.1 | 0.1 |
| gene-LOC105284168 | https://www.ncbi.nlm.nih.gov/gene/?term=LOC105284168 | 1400 | 76 | 542.86 | 2 | 14.29 | 2.63 | 24.73 | 22.83 | 1.9 | 1.9 | 0.1 | 0.1 |
| gene-LOC105283909 | https://www.ncbi.nlm.nih.gov/gene/?term=LOC105283909 | 2234 | 112 | 501.34 | 2 | 8.95 | 1.79 | 10.52 | 12.48 | -1.96 | 1.96 | -0.1 | 0.1 |
| gene-LOC105284189 | https://www.ncbi.nlm.nih.gov/gene/?term=LOC105284189 | 2601 | 130 | 499.81 | 3 | 11.53 | 2.31 | 14.87 | 16.94 | -2.07 | 2.07 | -0.1 | 0.1 |
| gene-LOC105275897 | https://www.ncbi.nlm.nih.gov/gene/?term=LOC105275897 | 1684 | 72 | 427.55 | 1 | 5.94 | 1.39 | 36.2 | 38.44 | -2.24 | 2.24 | -0.1 | 0.1 |
| gene-LOC105277345 | https://www.ncbi.nlm.nih.gov/gene/?term=LOC105277345 | 2151 | 82 | 381.22 | 1 | 4.65 | 1.22 | 16.59 | 19.1 | -2.51 | 2.51 | -0.1 | 0.1 |
| gene-LOC105278537 | https://www.ncbi.nlm.nih.gov/gene/?term=LOC105278537 | 1660 | 62 | 373.49 | 4 | 24.1 | 6.45 | 28.42 | 31.22 | -2.8 | 2.8 | -0.1 | 0.1 |
| gene-LOC105284046 | https://www.ncbi.nlm.nih.gov/gene/?term=LOC105284046 | 1027 | 28 | 272.64 | 4 | 38.95 | 14.29 | 19.04 | 15.74 | 3.3 | 3.3 | 0.09 | 0.09 |
| gene-LOC105282209 | https://www.ncbi.nlm.nih.gov/gene/?term=LOC105282209 | 1041 | 35 | 336.22 | 1 | 9.61 | 2.86 | 31.01 | 28.36 | 2.65 | 2.65 | 0.09 | 0.09 |

...
... only first 25 lines shown ...

Directory: OGLall File: rnk-dmg-Ob_LI_BCphaseB2.vs.RphaseB2.txt

| gene_ID | gene_link | gwidth | #Sites | #per10Kb | #dmSites | #dmsp10kb | %dmSites | %pSite1 | %pSite2 | DMpSite | ADMpSite | DMpNucl | ADMpNucl |
|---|---|---|---|---|---|---|---|---|---|---|---|---|---|
| gene-LOC105278458 | https://www.ncbi.nlm.nih.gov/gene/?term=LOC105278458 | 216 | 3 | 138.89 | 1 | 46.3 | 33.33 | 10.63 | 29.54 | -18.91 | 18.91 | -0.26 | 0.26 |
| gene-LOC105283830 | https://www.ncbi.nlm.nih.gov/gene/?term=LOC105283830 | 252 | 5 | 198.41 | 1 | 39.68 | 20 | 31.26 | 44.09 | -12.83 | 12.83 | -0.25 | 0.25 |
| gene-LOC105285203 | https://www.ncbi.nlm.nih.gov/gene/?term=LOC105285203 | 1020 | 27 | 264.71 | 2 | 19.61 | 7.41 | 30.05 | 24.28 | 5.76 | 5.76 | 0.15 | 0.15 |
| gene-LOC105275897 | https://www.ncbi.nlm.nih.gov/gene/?term=LOC105275897 | 1684 | 72 | 427.55 | 1 | 5.94 | 1.39 | 36.2 | 39.56 | -3.37 | 3.37 | -0.14 | 0.14 |
| gene-LOC105278938 | https://www.ncbi.nlm.nih.gov/gene/?term=LOC105278938 | 1818 | 110 | 605.06 | 1 | 5.5 | 0.91 | 20.68 | 22.88 | -2.2 | 2.2 | -0.13 | 0.13 |
| gene-LOC105276563 | https://www.ncbi.nlm.nih.gov/gene/?term=LOC105276563 | 1958 | 113 | 577.12 | 6 | 30.64 | 5.31 | 19.41 | 17.48 | 1.93 | 1.93 | 0.11 | 0.11 |
| gene-LOC105287692 | https://www.ncbi.nlm.nih.gov/gene/?term=LOC105287692 | 1505 | 89 | 591.36 | 5 | 33.22 | 5.62 | 31.34 | 29.41 | 1.93 | 1.93 | 0.11 | 0.11 |
| gene-LOC105278737 | https://www.ncbi.nlm.nih.gov/gene/?term=LOC105278737 | 1346 | 98 | 728.08 | 2 | 14.86 | 2.04 | 19.42 | 17.93 | 1.49 | 1.49 | 0.11 | 0.11 |
| gene-LOC105278873 | https://www.ncbi.nlm.nih.gov/gene/?term=LOC105278873 | 1897 | 147 | 774.91 | 4 | 21.09 | 2.72 | 19.42 | 20.81 | -1.39 | 1.39 | -0.11 | 0.11 |
| gene-LOC105279049 | https://www.ncbi.nlm.nih.gov/gene/?term=LOC105279049 | 1285 | 90 | 700.39 | 1 | 7.78 | 1.11 | 17.36 | 18.97 | -1.61 | 1.61 | -0.11 | 0.11 |
| gene-LOC105280710 | https://www.ncbi.nlm.nih.gov/gene/?term=LOC105280710 | 1215 | 64 | 526.75 | 2 | 16.46 | 3.12 | 22.16 | 24.28 | -2.12 | 2.12 | -0.11 | 0.11 |
| gene-LOC105274670 | https://www.ncbi.nlm.nih.gov/gene/?term=LOC105274670 | 1306 | 39 | 298.62 | 2 | 15.31 | 5.13 | 33.71 | 37.28 | -3.56 | 3.56 | -0.11 | 0.11 |
| gene-LOC105284320 | https://www.ncbi.nlm.nih.gov/gene/?term=LOC105284320 | 390 | 10 | 256.41 | 1 | 25.64 | 10 | 21.93 | 26.26 | -4.33 | 4.33 | -0.11 | 0.11 |
| gene-LOC105275176 | https://www.ncbi.nlm.nih.gov/gene/?term=LOC105275176 | 656 | 8 | 121.95 | 2 | 30.49 | 25 | 14.79 | 6.32 | 8.47 | 8.47 | 0.1 | 0.1 |
| gene-LOC105283552 | https://www.ncbi.nlm.nih.gov/gene/?term=LOC105283552 | 1562 | 37 | 236.88 | 1 | 6.4 | 2.7 | 55 | 50.64 | 4.36 | 4.36 | 0.1 | 0.1 |
| gene-LOC105285914 | https://www.ncbi.nlm.nih.gov/gene/?term=LOC105285914 | 867 | 28 | 322.95 | 1 | 11.53 | 3.57 | 16.16 | 12.99 | 3.17 | 3.17 | 0.1 | 0.1 |
| gene-LOC105284253 | https://www.ncbi.nlm.nih.gov/gene/?term=LOC105284253 | 1255 | 42 | 334.66 | 1 | 7.97 | 2.38 | 28.76 | 31.67 | -2.91 | 2.91 | -0.1 | 0.1 |
| gene-LOC105282481 | https://www.ncbi.nlm.nih.gov/gene/?term=LOC105282481 | 1900 | 59 | 310.53 | 2 | 10.53 | 3.39 | 33.33 | 36.46 | -3.13 | 3.13 | -0.1 | 0.1 |
| gene-LOC105274938 | https://www.ncbi.nlm.nih.gov/gene/?term=LOC105274938 | 1964 | 54 | 274.95 | 4 | 20.37 | 7.41 | 27.19 | 30.79 | -3.6 | 3.6 | -0.1 | 0.1 |
| gene-LOC105275219 | https://www.ncbi.nlm.nih.gov/gene/?term=LOC105275219 | 997 | 18 | 180.54 | 1 | 10.03 | 5.56 | 37.08 | 42.66 | -5.58 | 5.58 | -0.1 | 0.1 |
| gene-LOC105278083 | https://www.ncbi.nlm.nih.gov/gene/?term=LOC105278083 | 1605 | 38 | 236.76 | 1 | 6.23 | 2.63 | 32.5 | 28.84 | 3.66 | 3.66 | 0.09 | 0.09 |
| gene-LOC105281704 | https://www.ncbi.nlm.nih.gov/gene/?term=LOC105281704 | 2195 | 57 | 259.68 | 3 | 13.67 | 5.26 | 54.26 | 50.77 | 3.49 | 3.49 | 0.09 | 0.09 |
| gene-LOC105277371 | https://www.ncbi.nlm.nih.gov/gene/?term=LOC105277371 | 1816 | 71 | 390.97 | 1 | 5.51 | 1.41 | 20.94 | 18.75 | 2.2 | 2.2 | 0.09 | 0.09 |
| gene-LOC105277398 | https://www.ncbi.nlm.nih.gov/gene/?term=LOC105277398 | 1963 | 70 | 356.6 | 3 | 15.28 | 4.29 | 35.51 | 37.95 | -2.44 | 2.44 | -0.09 | 0.09 |

...

... only first 25 lines shown ...

Directory: OGLall File: rnk-dmg-Ob_LI_RphaseA1.vs.RphaseA2.txt

| gene_ID | gene_link | gwidth | #Sites | #per10Kb | #dmSites | #dmsp10kb | %dmSites | %pSite1 | %pSite2 | DMpSite | ADMpSite | DMpNucl | ADMpNucl |
|---|---|---|---|---|---|---|---|---|---|---|---|---|---|
| gene-LOC105288042 | https://www.ncbi.nlm.nih.gov/gene/?term=LOC105288042 | 3507 | 111 | 316.51 | 1 | 2.85 | 0.9 | 22.13 | 23.4 | -1.28 | 1.28 | -0.04 | 0.04 |
| gene-LOC105283264 | https://www.ncbi.nlm.nih.gov/gene/?term=LOC105283264 | 4005 | 96 | 239.7 | 1 | 2.5 | 1.04 | 23.23 | 22.15 | 1.09 | 1.09 | 0.03 | 0.03 |
| gene-LOC105283296 | https://www.ncbi.nlm.nih.gov/gene/?term=LOC105283296 | 2174 | 101 | 464.58 | 1 | 4.6 | 0.99 | 18.85 | 18.17 | 0.67 | 0.67 | 0.03 | 0.03 |
| gene-LOC105288036 | https://www.ncbi.nlm.nih.gov/gene/?term=LOC105288036 | 5110 | 209 | 409 | 1 | 1.96 | 0.48 | 17.07 | 17.79 | -0.71 | 0.71 | -0.03 | 0.03 |
| gene-LOC105277903 | https://www.ncbi.nlm.nih.gov/gene/?term=LOC105277903 | 1835 | 39 | 212.53 | 1 | 5.45 | 2.56 | 54.3 | 53.51 | 0.79 | 0.79 | 0.02 | 0.02 |
| gene-LOC105284557 | https://www.ncbi.nlm.nih.gov/gene/?term=LOC105284557 | 1672 | 84 | 502.39 | 1 | 5.98 | 1.19 | 28.63 | 28.17 | 0.46 | 0.46 | 0.02 | 0.02 |
| gene-LOC105275384 | https://www.ncbi.nlm.nih.gov/gene/?term=LOC105275384 | 1987 | 86 | 432.81 | 1 | 5.03 | 1.16 | 36.56 | 36.97 | -0.41 | 0.41 | -0.02 | 0.02 |
| gene-LOC105281272 | https://www.ncbi.nlm.nih.gov/gene/?term=LOC105281272 | 5252 | 199 | 378.9 | 1 | 1.9 | 0.5 | 14.62 | 15.07 | -0.45 | 0.45 | -0.02 | 0.02 |
| gene-LOC105284507 | https://www.ncbi.nlm.nih.gov/gene/?term=LOC105284507 | 5207 | 112 | 215.1 | 1 | 1.92 | 0.89 | 18.79 | 19.06 | -0.28 | 0.28 | -0.01 | 0.01 |
| gene-LOC105277125 | https://www.ncbi.nlm.nih.gov/gene/?term=LOC105277125 | 51079 | 2004 | 392.33 | 1 | 0.2 | 0.05 | 1.65 | 1.59 | 0.06 | 0.06 | 0 | 0 |
| gene-LOC105280304 | https://www.ncbi.nlm.nih.gov/gene/?term=LOC105280304 | 8919 | 150 | 168.18 | 1 | 1.12 | 0.67 | 6.83 | 6.96 | -0.12 | 0.12 | 0 | 0 |

Directory: OGLall File: rnk-dmg-Ob_LI_RphaseA1.vs.RphaseB1.txt

| gene_ID | gene_link | gwidth | #Sites | #per10Kb | #dmSites | #dmsp10kb | %dmSites | %pSite1 | %pSite2 | DMpSite | ADMpSite | DMpNucl | ADMpNucl |
|---|---|---|---|---|---|---|---|---|---|---|---|---|---|
| gene-LOC113562345 | https://www.ncbi.nlm.nih.gov/gene/?term=LOC113562345 | 2290 | 78 | 340.61 | 11 | 48.03 | 14.1 | 24.77 | 13.87 | 10.9 | 10.9 | 0.37 | 0.37 |
| gene-LOC105275355 | https://www.ncbi.nlm.nih.gov/gene/?term=LOC105275355 | 3197 | 174 | 544.26 | 1 | 3.13 | 0.57 | 25.86 | 21.8 | 4.06 | 4.06 | 0.22 | 0.22 |
| gene-LOC105275384 | https://www.ncbi.nlm.nih.gov/gene/?term=LOC105275384 | 1987 | 88 | 442.88 | 1 | 5.03 | 1.14 | 36.35 | 32.79 | 3.56 | 3.56 | 0.16 | 0.16 |
| gene-LOC105275070 | https://www.ncbi.nlm.nih.gov/gene/?term=LOC105275070 | 5667 | 56 | 98.82 | 3 | 5.29 | 5.36 | 17.83 | 5.85 | 11.98 | 11.98 | 0.12 | 0.12 |
| gene-LOC105285695 | https://www.ncbi.nlm.nih.gov/gene/?term=LOC105285695 | 1371 | 68 | 495.99 | 1 | 7.29 | 1.47 | 23.53 | 21.55 | 1.99 | 1.99 | 0.1 | 0.1 |
| gene-LOC105285671 | https://www.ncbi.nlm.nih.gov/gene/?term=LOC105285671 | 2263 | 100 | 441.89 | 1 | 4.42 | 1 | 32.24 | 30.53 | 1.71 | 1.71 | 0.08 | 0.08 |
| gene-LOC105275708 | https://www.ncbi.nlm.nih.gov/gene/?term=LOC105275708 | 1772 | 14 | 79.01 | 1 | 5.64 | 7.14 | 50.83 | 42.47 | 8.36 | 8.36 | 0.07 | 0.07 |
| gene-LOC105287063 | https://www.ncbi.nlm.nih.gov/gene/?term=LOC105287063 | 1874 | 49 | 261.47 | 1 | 5.34 | 2.04 | 32.77 | 30.12 | 2.65 | 2.65 | 0.07 | 0.07 |
| gene-LOC105285004 | https://www.ncbi.nlm.nih.gov/gene/?term=LOC105285004 | 1334 | 37 | 277.36 | 1 | 7.5 | 2.7 | 16.38 | 18.79 | -2.41 | 2.41 | -0.07 | 0.07 |
| gene-LOC105285514 | https://www.ncbi.nlm.nih.gov/gene/?term=LOC105285514 | 1277 | 7 | 54.82 | 1 | 7.83 | 14.29 | 90.55 | 80 | 10.54 | 10.54 | 0.06 | 0.06 |
| gene-LOC105274534 | https://www.ncbi.nlm.nih.gov/gene/?term=LOC105274534 | 1329 | 59 | 443.94 | 1 | 7.52 | 1.69 | 43.5 | 44.89 | -1.39 | 1.39 | -0.06 | 0.06 |
| gene-LOC105276272 | https://www.ncbi.nlm.nih.gov/gene/?term=LOC105276272 | 1965 | 44 | 223.92 | 1 | 5.09 | 2.27 | 27.22 | 24.96 | 2.26 | 2.26 | 0.05 | 0.05 |
| gene-LOC105278807 | https://www.ncbi.nlm.nih.gov/gene/?term=LOC105278807 | 2166 | 74 | 341.64 | 1 | 4.62 | 1.35 | 21.42 | 20.03 | 1.39 | 1.39 | 0.05 | 0.05 |
| gene-LOC105275460 | https://www.ncbi.nlm.nih.gov/gene/?term=LOC105275460 | 2473 | 126 | 509.5 | 1 | 4.04 | 0.79 | 13.41 | 12.4 | 1.01 | 1.01 | 0.05 | 0.05 |
| gene-LOC105278282 | https://www.ncbi.nlm.nih.gov/gene/?term=LOC105278282 | 1775 | 51 | 287.32 | 1 | 5.63 | 1.96 | 39.13 | 40.69 | -1.57 | 1.57 | -0.05 | 0.05 |
| gene-LOC113562102 | https://www.ncbi.nlm.nih.gov/gene/?term=LOC113562102 | 3433 | 63 | 183.51 | 1 | 2.91 | 1.59 | 17.89 | 15.59 | 2.3 | 2.3 | 0.04 | 0.04 |
| gene-LOC105282991 | https://www.ncbi.nlm.nih.gov/gene/?term=LOC105282991 | 5617 | 103 | 183.37 | 1 | 1.78 | 0.97 | 18.36 | 16.13 | 2.23 | 2.23 | 0.04 | 0.04 |
| gene-LOC105283712 | https://www.ncbi.nlm.nih.gov/gene/?term=LOC105283712 | 1910 | 35 | 183.25 | 1 | 5.24 | 2.86 | 28.45 | 26.47 | 1.98 | 1.98 | 0.04 | 0.04 |
| gene-LOC105286265 | https://www.ncbi.nlm.nih.gov/gene/?term=LOC105286265 | 2507 | 67 | 267.25 | 1 | 3.99 | 1.49 | 32.57 | 31.11 | 1.46 | 1.46 | 0.04 | 0.04 |
| gene-LOC105285742 | https://www.ncbi.nlm.nih.gov/gene/?term=LOC105285742 | 3446 | 109 | 316.31 | 1 | 2.9 | 0.92 | 17.79 | 16.59 | 1.2 | 1.2 | 0.04 | 0.04 |
| gene-LOC105285674 | https://www.ncbi.nlm.nih.gov/gene/?term=LOC105285674 | 3035 | 100 | 329.49 | 2 | 6.59 | 2 | 25.96 | 24.78 | 1.18 | 1.18 | 0.04 | 0.04 |
| gene-LOC105276449 | https://www.ncbi.nlm.nih.gov/gene/?term=LOC105276449 | 1231 | 48 | 389.93 | 1 | 8.12 | 2.08 | 25.67 | 24.69 | 0.99 | 0.99 | 0.04 | 0.04 |
| gene-LOC105274823 | https://www.ncbi.nlm.nih.gov/gene/?term=LOC105274823 | 4620 | 177 | 383.12 | 1 | 2.16 | 0.56 | 17.15 | 16.21 | 0.95 | 0.95 | 0.04 | 0.04 |
| gene-LOC105282156 | https://www.ncbi.nlm.nih.gov/gene/?term=LOC105282156 | 2577 | 127 | 492.82 | 1 | 3.88 | 0.79 | 25.13 | 24.3 | 0.82 | 0.82 | 0.04 | 0.04 |
| gene-LOC105278283 | https://www.ncbi.nlm.nih.gov/gene/?term=LOC105278283 | 2317 | 142 | 612.86 | 1 | 4.32 | 0.7 | 24.95 | 24.22 | 0.73 | 0.73 | 0.04 | 0.04 |
| gene-LOC105285055 | https://www.ncbi.nlm.nih.gov/gene/?term=LOC105285055 | 4108 | 253 | 615.87 | 1 | 2.43 | 0.4 | 10.93 | 10.3 | 0.63 | 0.63 | 0.04 | 0.04 |
| gene-LOC105283649 | https://www.ncbi.nlm.nih.gov/gene/?term=LOC105283649 | 1918 | 56 | 291.97 | 1 | 5.21 | 1.79 | 37.76 | 39.05 | -1.29 | 1.29 | -0.04 | 0.04 |
| gene-LOC105288042 | https://www.ncbi.nlm.nih.gov/gene/?term=LOC105288042 | 3507 | 100 | 285.14 | 1 | 2.85 | 1 | 20.77 | 22.23 | -1.46 | 1.46 | -0.04 | 0.04 |
| gene-LOC105278962 | https://www.ncbi.nlm.nih.gov/gene/?term=LOC105278962 | 4491 | 109 | 242.71 | 1 | 2.23 | 0.92 | 19.66 | 21.14 | -1.48 | 1.48 | -0.04 | 0.04 |
| gene-LOC105286514 | https://www.ncbi.nlm.nih.gov/gene/?term=LOC105286514 | 2528 | 50 | 197.78 | 1 | 3.96 | 2 | 35.29 | 37.3 | -2.01 | 2.01 | -0.04 | 0.04 |
| gene-LOC105281039 | https://www.ncbi.nlm.nih.gov/gene/?term=LOC105281039 | 1471 | 13 | 88.38 | 1 | 6.8 | 7.69 | 31.24 | 27.65 | 3.59 | 3.59 | 0.03 | 0.03 |
| gene-LOC105274900 | https://www.ncbi.nlm.nih.gov/gene/?term=LOC105274900 | 5561 | 61 | 109.69 | 3 | 5.39 | 4.92 | 33.5 | 30.47 | 3.02 | 3.02 | 0.03 | 0.03 |
| gene-LOC105279898 | https://www.ncbi.nlm.nih.gov/gene/?term=LOC105279898 | 1343 | 15 | 111.69 | 1 | 7.45 | 6.67 | 61.47 | 58.93 | 2.54 | 2.54 | 0.03 | 0.03 |
| gene-LOC105282652 | https://www.ncbi.nlm.nih.gov/gene/?term=LOC105282652 | 5065 | 85 | 167.82 | 1 | 1.97 | 1.18 | 16.28 | 14.6 | 1.68 | 1.68 | 0.03 | 0.03 |
| gene-LOC105276692 | https://www.ncbi.nlm.nih.gov/gene/?term=LOC105276692 | 1745 | 34 | 194.84 | 1 | 5.73 | 2.94 | 25.17 | 23.64 | 1.53 | 1.53 | 0.03 | 0.03 |
| gene-LOC113561805 | https://www.ncbi.nlm.nih.gov/gene/?term=LOC113561805 | 3337 | 75 | 224.75 | 1 | 3 | 1.33 | 34.35 | 32.91 | 1.44 | 1.44 | 0.03 | 0.03 |
| gene-LOC105280348 | https://www.ncbi.nlm.nih.gov/gene/?term=LOC105280348 | 3797 | 79 | 208.06 | 1 | 2.63 | 1.27 | 18.28 | 16.89 | 1.4 | 1.4 | 0.03 | 0.03 |
| gene-LOC105283697 | https://www.ncbi.nlm.nih.gov/gene/?term=LOC105283697 | 5301 | 153 | 288.62 | 1 | 1.89 | 0.65 | 16.46 | 15.43 | 1.02 | 1.02 | 0.03 | 0.03 |
| gene-LOC105283552 | https://www.ncbi.nlm.nih.gov/gene/?term=LOC105283552 | 1562 | 40 | 256.08 | 1 | 6.4 | 2.5 | 50.43 | 49.44 | 0.99 | 0.99 | 0.03 | 0.03 |
| gene-LOC105288079 | https://www.ncbi.nlm.nih.gov/gene/?term=LOC105288079 | 3473 | 113 | 325.37 | 1 | 2.88 | 0.88 | 17.44 | 16.56 | 0.89 | 0.89 | 0.03 | 0.03 |
| gene-LOC105282491 | https://www.ncbi.nlm.nih.gov/gene/?term=LOC105282491 | 4761 | 179 | 375.97 | 1 | 2.1 | 0.56 | 9.15 | 8.31 | 0.83 | 0.83 | 0.03 | 0.03 |
| gene-LOC105288036 | https://www.ncbi.nlm.nih.gov/gene/?term=LOC105288036 | 5110 | 200 | 391.39 | 1 | 1.96 | 0.5 | 17.43 | 16.6 | 0.83 | 0.83 | 0.03 | 0.03 |
| gene-LOC105278145 | https://www.ncbi.nlm.nih.gov/gene/?term=LOC105278145 | 4470 | 252 | 563.76 | 2 | 4.47 | 0.79 | 10.98 | 10.51 | 0.48 | 0.48 | 0.03 | 0.03 |
| gene-LOC105282658 | https://www.ncbi.nlm.nih.gov/gene/?term=LOC105282658 | 2421 | 64 | 264.35 | 1 | 4.13 | 1.56 | 23.38 | 24.34 | -0.95 | 0.95 | -0.03 | 0.03 |
| gene-LOC105284036 | https://www.ncbi.nlm.nih.gov/gene/?term=LOC105284036 | 3254 | 98 | 301.17 | 1 | 3.07 | 1.02 | 22.14 | 23.09 | -0.95 | 0.95 | -0.03 | 0.03 |
| gene-LOC105281173 | https://www.ncbi.nlm.nih.gov/gene/?term=LOC105281173 | 1595 | 38 | 238.24 | 1 | 6.27 | 2.63 | 59.87 | 61.09 | -1.22 | 1.22 | -0.03 | 0.03 |
| gene-LOC105276196 | https://www.ncbi.nlm.nih.gov/gene/?term=LOC105276196 | 2953 | 57 | 193.02 | 1 | 3.39 | 1.75 | 39.57 | 40.89 | -1.32 | 1.32 | -0.03 | 0.03 |
| gene-LOC105275571 | https://www.ncbi.nlm.nih.gov/gene/?term=LOC105275571 | 1926 | 30 | 155.76 | 1 | 5.19 | 3.33 | 42.71 | 44.75 | -2.04 | 2.04 | -0.03 | 0.03 |
| gene-LOC105276538 | https://www.ncbi.nlm.nih.gov/gene/?term=LOC105276538 | 1511 | 21 | 138.98 | 1 | 6.62 | 4.76 | 57.36 | 59.41 | -2.05 | 2.05 | -0.03 | 0.03 |
| gene-LOC105282914 | https://www.ncbi.nlm.nih.gov/gene/?term=LOC105282914 | 5623 | 43 | 76.47 | 1 | 1.78 | 2.33 | 46.2 | 49.79 | -3.59 | 3.59 | -0.03 | 0.03 |
| gene-LOC105275347 | https://www.ncbi.nlm.nih.gov/gene/?term=LOC105275347 | 2526 | 17 | 67.3 | 1 | 3.96 | 5.88 | 45.17 | 41.88 | 3.29 | 3.29 | 0.02 | 0.02 |
| gene-LOC105288066 | https://www.ncbi.nlm.nih.gov/gene/?term=LOC105288066 | 1964 | 30 | 152.75 | 1 | 5.09 | 3.33 | 38.49 | 37.01 | 1.48 | 1.48 | 0.02 | 0.02 |
| gene-LOC105282600 | https://www.ncbi.nlm.nih.gov/gene/?term=LOC105282600 | 3715 | 60 | 161.51 | 1 | 2.69 | 1.67 | 27.74 | 26.44 | 1.3 | 1.3 | 0.02 | 0.02 |
| gene-LOC105285060 | https://www.ncbi.nlm.nih.gov/gene/?term=LOC105285060 | 1285 | 16 | 124.51 | 1 | 7.78 | 6.25 | 42.55 | 41.27 | 1.27 | 1.27 | 0.02 | 0.02 |
| gene-LOC105279900 | https://www.ncbi.nlm.nih.gov/gene/?term=LOC105279900 | 2191 | 55 | 251.03 | 1 | 4.56 | 1.82 | 30.23 | 29.28 | 0.95 | 0.95 | 0.02 | 0.02 |
| gene-LOC105281848 | https://www.ncbi.nlm.nih.gov/gene/?term=LOC105281848 | 2122 | 57 | 268.61 | 1 | 4.71 | 1.75 | 23.82 | 22.98 | 0.84 | 0.84 | 0.02 | 0.02 |
| gene-LOC105285981 | https://www.ncbi.nlm.nih.gov/gene/?term=LOC105285981 | 7666 | 179 | 233.5 | 1 | 1.3 | 0.56 | 19.94 | 19.14 | 0.81 | 0.81 | 0.02 | 0.02 |
| gene-LOC105282794 | https://www.ncbi.nlm.nih.gov/gene/?term=LOC105282794 | 4571 | 104 | 227.52 | 1 | 2.19 | 0.96 | 16.85 | 16.16 | 0.69 | 0.69 | 0.02 | 0.02 |
| gene-LOC105274449 | https://www.ncbi.nlm.nih.gov/gene/?term=LOC105274449 | 1549 | 53 | 342.16 | 1 | 6.46 | 1.89 | 49.89 | 49.29 | 0.6 | 0.6 | 0.02 | 0.02 |
| gene-LOC105285179 | https://www.ncbi.nlm.nih.gov/gene/?term=LOC105285179 | 4191 | 189 | 450.97 | 1 | 2.39 | 0.53 | 18.34 | 17.82 | 0.52 | 0.52 | 0.02 | 0.02 |

| | | | | | | | | | | | | | |
|---|---|---|---|---|---|---|---|---|---|---|---|---|---|
| gene-LOC105274531 | https://www.ncbi.nlm.nih.gov/gene/?term=LOC105274531 | 7774 | 286 | 367.89 | 1 | 1.29 | 0.35 | 11.23 | 10.77 | 0.46 | 0.46 | 0.02 | 0.02 |
| gene-LOC105275363 | https://www.ncbi.nlm.nih.gov/gene/?term=LOC105275363 | 3798 | 179 | 471.3 | 1 | 2.63 | 0.56 | 15.94 | 15.54 | 0.4 | 0.4 | 0.02 | 0.02 |
| gene-LOC105286355 | https://www.ncbi.nlm.nih.gov/gene/?term=LOC105286355 | 3245 | 149 | 459.17 | 1 | 3.08 | 0.67 | 21.45 | 21.07 | 0.38 | 0.38 | 0.02 | 0.02 |
| gene-LOC105278497 | https://www.ncbi.nlm.nih.gov/gene/?term=LOC105278497 | 2072 | 100 | 482.63 | 1 | 4.83 | 1 | 18.63 | 18.28 | 0.35 | 0.35 | 0.02 | 0.02 |
| gene-LOC105278632 | https://www.ncbi.nlm.nih.gov/gene/?term=LOC105278632 | 3624 | 143 | 394.59 | 1 | 2.76 | 0.7 | 13.02 | 13.42 | -0.4 | 0.4 | -0.02 | 0.02 |
| gene-LOC105284355 | https://www.ncbi.nlm.nih.gov/gene/?term=LOC105284355 | 3904 | 155 | 397.03 | 1 | 2.56 | 0.65 | 20.33 | 20.86 | -0.53 | 0.53 | -0.02 | 0.02 |
| gene-LOC105274851 | https://www.ncbi.nlm.nih.gov/gene/?term=LOC105274851 | 6616 | 144 | 217.65 | 1 | 1.51 | 0.69 | 11.65 | 12.43 | -0.79 | 0.79 | -0.02 | 0.02 |
| gene-LOC105279596 | https://www.ncbi.nlm.nih.gov/gene/?term=LOC105279596 | 7695 | 115 | 149.45 | 1 | 1.3 | 0.87 | 16.77 | 17.78 | -1 | 1 | -0.02 | 0.02 |
| gene-LOC105274590 | https://www.ncbi.nlm.nih.gov/gene/?term=LOC105274590 | 4516 | 82 | 181.58 | 1 | 2.21 | 1.22 | 25.47 | 26.68 | -1.21 | 1.21 | -0.02 | 0.02 |
| gene-LOC105281515 | https://www.ncbi.nlm.nih.gov/gene/?term=LOC105281515 | 8827 | 57 | 64.57 | 2 | 2.27 | 3.51 | 32.7 | 30.37 | 2.32 | 2.32 | 0.01 | 0.01 |
| gene-LOC105279528 | https://www.ncbi.nlm.nih.gov/gene/?term=LOC105279528 | 5802 | 40 | 68.94 | 1 | 1.72 | 2.5 | 21.84 | 20.71 | 1.12 | 1.12 | 0.01 | 0.01 |
| gene-LOC105275187 | https://www.ncbi.nlm.nih.gov/gene/?term=LOC105275187 | 5534 | 73 | 131.91 | 1 | 1.81 | 1.37 | 14.7 | 13.62 | 1.08 | 1.08 | 0.01 | 0.01 |
| gene-LOC105283943 | https://www.ncbi.nlm.nih.gov/gene/?term=LOC105283943 | 3559 | 57 | 160.16 | 1 | 2.81 | 1.75 | 23.04 | 22.27 | 0.77 | 0.77 | 0.01 | 0.01 |
| gene-LOC105288237 | https://www.ncbi.nlm.nih.gov/gene/?term=LOC105288237 | 2226 | 49 | 220.13 | 1 | 4.49 | 2.04 | 43.01 | 42.34 | 0.67 | 0.67 | 0.01 | 0.01 |
| gene-LOC105275451 | https://www.ncbi.nlm.nih.gov/gene/?term=LOC105275451 | 6035 | 135 | 223.7 | 1 | 1.66 | 0.74 | 21.36 | 20.75 | 0.61 | 0.61 | 0.01 | 0.01 |
| gene-LOC105277215 | https://www.ncbi.nlm.nih.gov/gene/?term=LOC105277215 | 2619 | 29 | 110.73 | 1 | 3.82 | 3.45 | 25.25 | 24.64 | 0.61 | 0.61 | 0.01 | 0.01 |
| gene-LOC105288078 | https://www.ncbi.nlm.nih.gov/gene/?term=LOC105288078 | 19126 | 337 | 176.2 | 1 | 0.52 | 0.3 | 15.64 | 15.05 | 0.59 | 0.59 | 0.01 | 0.01 |
| gene-LOC105280819 | https://www.ncbi.nlm.nih.gov/gene/?term=LOC105280819 | 1639 | 42 | 256.25 | 1 | 6.1 | 2.38 | 39.26 | 38.69 | 0.58 | 0.58 | 0.01 | 0.01 |
| gene-LOC105280641 | https://www.ncbi.nlm.nih.gov/gene/?term=LOC105280641 | 3682 | 75 | 203.69 | 1 | 2.72 | 1.33 | 13.21 | 12.66 | 0.55 | 0.55 | 0.01 | 0.01 |
| gene-LOC105287417 | https://www.ncbi.nlm.nih.gov/gene/?term=LOC105287417 | 1025 | 32 | 312.2 | 1 | 9.76 | 3.12 | 37.69 | 37.23 | 0.46 | 0.46 | 0.01 | 0.01 |
| gene-LOC105277903 | https://www.ncbi.nlm.nih.gov/gene/?term=LOC105277903 | 1835 | 37 | 201.63 | 1 | 5.45 | 2.7 | 54.5 | 54.08 | 0.42 | 0.42 | 0.01 | 0.01 |
| gene-LOC105283592 | https://www.ncbi.nlm.nih.gov/gene/?term=LOC105283592 | 3479 | 80 | 229.95 | 1 | 2.87 | 1.25 | 21.51 | 21.14 | 0.37 | 0.37 | 0.01 | 0.01 |
| gene-LOC105283838 | https://www.ncbi.nlm.nih.gov/gene/?term=LOC105283838 | 1748 | 30 | 171.62 | 1 | 5.72 | 3.33 | 33.04 | 32.68 | 0.36 | 0.36 | 0.01 | 0.01 |
| gene-LOC105278384 | https://www.ncbi.nlm.nih.gov/gene/?term=LOC105278384 | 1390 | 33 | 237.41 | 1 | 7.19 | 3.03 | 46.11 | 45.77 | 0.34 | 0.34 | 0.01 | 0.01 |
| gene-LOC105285639 | https://www.ncbi.nlm.nih.gov/gene/?term=LOC105285639 | 10325 | 351 | 339.95 | 2 | 1.94 | 0.57 | 11.49 | 11.2 | 0.29 | 0.29 | 0.01 | 0.01 |
| gene-LOC105279298 | https://www.ncbi.nlm.nih.gov/gene/?term=LOC105279298 | 2773 | 127 | 457.99 | 1 | 3.61 | 0.79 | 24.14 | 23.89 | 0.26 | 0.26 | 0.01 | 0.01 |
| gene-LOC105283123 | https://www.ncbi.nlm.nih.gov/gene/?term=LOC105283123 | 1507 | 47 | 311.88 | 1 | 6.64 | 2.13 | 20.21 | 19.96 | 0.25 | 0.25 | 0.01 | 0.01 |
| gene-LOC105284184 | https://www.ncbi.nlm.nih.gov/gene/?term=LOC105284184 | 2535 | 92 | 362.92 | 1 | 3.94 | 1.09 | 31.61 | 31.41 | 0.2 | 0.2 | 0.01 | 0.01 |
| gene-LOC105286997 | https://www.ncbi.nlm.nih.gov/gene/?term=LOC105286997 | 2850 | 142 | 498.25 | 1 | 3.51 | 0.7 | 11.2 | 11.03 | 0.17 | 0.17 | 0.01 | 0.01 |
| gene-LOC105275374 | https://www.ncbi.nlm.nih.gov/gene/?term=LOC105275374 | 11376 | 514 | 451.83 | 1 | 0.88 | 0.19 | 9.9 | 9.75 | 0.15 | 0.15 | 0.01 | 0.01 |
| gene-LOC105283728 | https://www.ncbi.nlm.nih.gov/gene/?term=LOC105283728 | 3256 | 152 | 466.83 | 2 | 6.14 | 1.32 | 15.71 | 15.57 | 0.14 | 0.14 | 0.01 | 0.01 |
| gene-LOC105276527 | https://www.ncbi.nlm.nih.gov/gene/?term=LOC105276527 | 7923 | 338 | 426.61 | 1 | 1.26 | 0.3 | 8.29 | 8.43 | -0.14 | 0.14 | -0.01 | 0.01 |
| gene-LOC105279038 | https://www.ncbi.nlm.nih.gov/gene/?term=LOC105279038 | 9453 | 329 | 348.04 | 1 | 1.06 | 0.3 | 9.14 | 9.32 | -0.18 | 0.18 | -0.01 | 0.01 |
| gene-LOC105282811 | https://www.ncbi.nlm.nih.gov/gene/?term=LOC105282811 | 5765 | 203 | 352.12 | 1 | 1.73 | 0.49 | 8.59 | 8.78 | -0.19 | 0.19 | -0.01 | 0.01 |
| gene-LOC105274856 | https://www.ncbi.nlm.nih.gov/gene/?term=LOC105274856 | 5216 | 228 | 437.12 | 1 | 1.92 | 0.44 | 17.7 | 17.9 | -0.2 | 0.2 | -0.01 | 0.01 |
| gene-LOC105285807 | https://www.ncbi.nlm.nih.gov/gene/?term=LOC105285807 | 6063 | 215 | 354.61 | 1 | 1.65 | 0.47 | 8.65 | 8.94 | -0.28 | 0.28 | -0.01 | 0.01 |
| gene-LOC105274706 | https://www.ncbi.nlm.nih.gov/gene/?term=LOC105274706 | 1446 | 50 | 345.78 | 1 | 6.92 | 2 | 37.45 | 37.82 | -0.37 | 0.37 | -0.01 | 0.01 |
| gene-LOC105282825 | https://www.ncbi.nlm.nih.gov/gene/?term=LOC105282825 | 3919 | 96 | 244.96 | 1 | 2.55 | 1.04 | 14.53 | 14.91 | -0.38 | 0.38 | -0.01 | 0.01 |
| gene-LOC105280002 | https://www.ncbi.nlm.nih.gov/gene/?term=LOC105280002 | 1940 | 31 | 159.79 | 1 | 5.15 | 3.23 | 31.38 | 32.27 | -0.88 | 0.88 | -0.01 | 0.01 |
| gene-LOC105275518 | https://www.ncbi.nlm.nih.gov/gene/?term=LOC105275518 | 2335 | 23 | 98.5 | 1 | 4.28 | 4.35 | 45.78 | 46.9 | -1.12 | 1.12 | -0.01 | 0.01 |
| gene-LOC105283989 | https://www.ncbi.nlm.nih.gov/gene/?term=LOC105283989 | 4948 | 32 | 64.67 | 1 | 2.02 | 3.12 | 46.57 | 45.88 | 0.69 | 0.69 | 0 | 0 |
| gene-LOC105286452 | https://www.ncbi.nlm.nih.gov/gene/?term=LOC105286452 | 6888 | 68 | 98.72 | 1 | 1.45 | 1.47 | 28.16 | 27.75 | 0.4 | 0.4 | 0 | 0 |
| gene-LOC105288010 | https://www.ncbi.nlm.nih.gov/gene/?term=LOC105288010 | 3189 | 52 | 163.06 | 1 | 3.14 | 1.92 | 29.32 | 29.04 | 0.28 | 0.28 | 0 | 0 |
| gene-LOC105284329 | https://www.ncbi.nlm.nih.gov/gene/?term=LOC105284329 | 4051 | 71 | 175.27 | 1 | 2.47 | 1.41 | 19.25 | 18.99 | 0.25 | 0.25 | 0 | 0 |
| gene-LOC105275170 | https://www.ncbi.nlm.nih.gov/gene/?term=LOC105275170 | 12918 | 350 | 270.94 | 1 | 0.77 | 0.29 | 1.94 | 1.8 | 0.15 | 0.15 | 0 | 0 |
| gene-LOC105276059 | https://www.ncbi.nlm.nih.gov/gene/?term=LOC105276059 | 6256 | 134 | 214.19 | 1 | 1.6 | 0.75 | 11.07 | 10.91 | 0.15 | 0.15 | 0 | 0 |
| gene-LOC105286099 | https://www.ncbi.nlm.nih.gov/gene/?term=LOC105286099 | 5919 | 70 | 118.26 | 1 | 1.69 | 1.43 | 12.4 | 12.26 | 0.14 | 0.14 | 0 | 0 |
| gene-LOC105278534 | https://www.ncbi.nlm.nih.gov/gene/?term=LOC105278534 | 7364 | 198 | 268.88 | 1 | 1.36 | 0.51 | 17.97 | 17.87 | 0.1 | 0.1 | 0 | 0 |
| gene-LOC105280774 | https://www.ncbi.nlm.nih.gov/gene/?term=LOC105280774 | 2793 | 132 | 472.61 | 1 | 3.58 | 0.76 | 25.2 | 25.1 | 0.1 | 0.1 | 0 | 0 |
| gene-LOC105282149 | https://www.ncbi.nlm.nih.gov/gene/?term=LOC105282149 | 1787 | 103 | 576.39 | 1 | 5.6 | 0.97 | 41.42 | 41.34 | 0.08 | 0.08 | 0 | 0 |
| gene-LOC105278666 | https://www.ncbi.nlm.nih.gov/gene/?term=LOC105278666 | 10341 | 474 | 458.37 | 1 | 0.97 | 0.21 | 6.99 | 6.95 | 0.05 | 0.05 | 0 | 0 |
| gene-LOC105280956 | https://www.ncbi.nlm.nih.gov/gene/?term=LOC105280956 | 2639 | 80 | 303.15 | 1 | 3.79 | 1.25 | 18.94 | 18.89 | 0.05 | 0.05 | 0 | 0 |
| gene-LOC105275327 | https://www.ncbi.nlm.nih.gov/gene/?term=LOC105275327 | 14245 | 529 | 371.36 | 1 | 0.7 | 0.19 | 6.66 | 6.63 | 0.03 | 0.03 | 0 | 0 |
| gene-LOC105284043 | https://www.ncbi.nlm.nih.gov/gene/?term=LOC105284043 | 5661 | 284 | 501.68 | 1 | 1.77 | 0.35 | 10.32 | 10.3 | 0.03 | 0.03 | 0 | 0 |
| gene-LOC105277216 | https://www.ncbi.nlm.nih.gov/gene/?term=LOC105277216 | 7627 | 106 | 138.98 | 1 | 1.31 | 0.94 | 7.36 | 7.34 | 0.02 | 0.02 | 0 | 0 |
| gene-LOC105277866 | https://www.ncbi.nlm.nih.gov/gene/?term=LOC105277866 | 33045 | 482 | 145.86 | 1 | 0.3 | 0.21 | 3.05 | 3.04 | 0.02 | 0.02 | 0 | 0 |
| gene-LOC105279500 | https://www.ncbi.nlm.nih.gov/gene/?term=LOC105279500 | 3587 | 32 | 89.21 | 1 | 2.79 | 3.12 | 46.58 | 46.57 | 0.01 | 0.01 | 0 | 0 |
| gene-LOC105286515 | https://www.ncbi.nlm.nih.gov/gene/?term=LOC105286515 | 3751 | 117 | 311.92 | 1 | 2.67 | 0.85 | 10.68 | 10.68 | 0 | 0 | 0 | 0 |
| gene-LOC105278228 | https://www.ncbi.nlm.nih.gov/gene/?term=LOC105278228 | 8610 | 176 | 204.41 | 1 | 1.16 | 0.57 | 22.62 | 22.66 | -0.04 | 0.04 | 0 | 0 |
| gene-LOC105278877 | https://www.ncbi.nlm.nih.gov/gene/?term=LOC105278877 | 1968 | 87 | 442.07 | 1 | 5.08 | 1.15 | 29.15 | 29.21 | -0.06 | 0.06 | 0 | 0 |
| gene-LOC105287095 | https://www.ncbi.nlm.nih.gov/gene/?term=LOC105287095 | 1435 | 66 | 459.93 | 1 | 6.97 | 1.52 | 24.97 | 25.03 | -0.06 | 0.06 | 0 | 0 |
| gene-LOC105286194 | https://www.ncbi.nlm.nih.gov/gene/?term=LOC105286194 | 152619 | 2310 | 151.36 | 1 | 0.07 | 0.04 | 5.89 | 6 | -0.11 | 0.11 | 0 | 0 |
| gene-LOC105279664 | https://www.ncbi.nlm.nih.gov/gene/?term=LOC105279664 | 4023 | 64 | 159.09 | 1 | 2.49 | 1.56 | 17.75 | 17.94 | -0.19 | 0.19 | 0 | 0 |
| gene-LOC105277856 | https://www.ncbi.nlm.nih.gov/gene/?term=LOC105277856 | 3442 | 33 | 95.87 | 1 | 2.91 | 3.03 | 39.28 | 39.53 | -0.25 | 0.25 | 0 | 0 |

Directory: OGLall File: rnk-dmg-Ob_LI_RphaseA1.vs.RphaseB2.txt

| gene_ID | gene_link | gwidth | #Sites | #per10Kb | #dmSites | #dmsp10kb | %dmSites | %pSite1 | %pSite2 | DMpSite | ADMpSite | DMpNucl | ADMpNucl |
|---|---|---|---|---|---|---|---|---|---|---|---|---|---|
| gene-LOC113562345 | https://www.ncbi.nlm.nih.gov/gene/?term=LOC113562345 | 2290 | 67 | 292.58 | 6 | 26.2 | 8.96 | 28.84 | 16.72 | 12.12 | 12.12 | 0.35 | 0.35 |
| gene-LOC105285671 | https://www.ncbi.nlm.nih.gov/gene/?term=LOC105285671 | 2263 | 101 | 446.31 | 1 | 4.42 | 0.99 | 30.8 | 27.84 | 2.96 | 2.96 | 0.13 | 0.13 |
| gene-LOC105275070 | https://www.ncbi.nlm.nih.gov/gene/?term=LOC105275070 | 5667 | 37 | 65.29 | 7 | 12.35 | 18.92 | 16.3 | 1.04 | 15.26 | 15.26 | 0.1 | 0.1 |
| gene-LOC105283552 | https://www.ncbi.nlm.nih.gov/gene/?term=LOC105283552 | 1562 | 41 | 262.48 | 1 | 6.4 | 2.44 | 50.09 | 46.71 | 3.37 | 3.37 | 0.09 | 0.09 |
| gene-LOC105279356 | https://www.ncbi.nlm.nih.gov/gene/?term=LOC105279356 | 1314 | 43 | 327.25 | 1 | 7.61 | 2.33 | 32.62 | 30.3 | 2.31 | 2.31 | 0.08 | 0.08 |
| gene-LOC105278436 | https://www.ncbi.nlm.nih.gov/gene/?term=LOC105278436 | 2556 | 54 | 211.27 | 1 | 3.91 | 1.85 | 40.35 | 37.22 | 3.13 | 3.13 | 0.07 | 0.07 |
| gene-LOC105285202 | https://www.ncbi.nlm.nih.gov/gene/?term=LOC105285202 | 1838 | 61 | 331.88 | 1 | 5.44 | 1.64 | 26.9 | 24.8 | 2.1 | 2.1 | 0.07 | 0.07 |
| gene-LOC105279898 | https://www.ncbi.nlm.nih.gov/gene/?term=LOC105279898 | 1343 | 9 | 67.01 | 1 | 7.45 | 11.11 | 83.37 | 74.78 | 8.59 | 8.59 | 0.06 | 0.06 |
| gene-LOC105275708 | https://www.ncbi.nlm.nih.gov/gene/?term=LOC105275708 | 1772 | 14 | 79.01 | 1 | 5.64 | 7.14 | 50.83 | 43.74 | 7.08 | 7.08 | 0.06 | 0.06 |
| gene-LOC105285674 | https://www.ncbi.nlm.nih.gov/gene/?term=LOC105285674 | 3035 | 92 | 303.13 | 2 | 6.59 | 2.17 | 26.97 | 24.84 | 2.13 | 2.13 | 0.06 | 0.06 |
| gene-LOC105280786 | https://www.ncbi.nlm.nih.gov/gene/?term=LOC105280786 | 2573 | 115 | 446.95 | 1 | 3.89 | 0.87 | 28.39 | 27.06 | 1.33 | 1.33 | 0.06 | 0.06 |
| gene-LOC105280596 | https://www.ncbi.nlm.nih.gov/gene/?term=LOC105280596 | 3434 | 104 | 302.85 | 1 | 2.91 | 0.96 | 16.78 | 18.68 | -1.9 | 1.9 | -0.06 | 0.06 |
| gene-LOC105276196 | https://www.ncbi.nlm.nih.gov/gene/?term=LOC105276196 | 2953 | 59 | 199.8 | 1 | 3.39 | 1.69 | 38.71 | 41.49 | -2.77 | 2.77 | -0.06 | 0.06 |
| gene-LOC105285514 | https://www.ncbi.nlm.nih.gov/gene/?term=LOC105285514 | 1277 | 14 | 109.63 | 1 | 7.83 | 7.14 | 69.69 | 65.1 | 4.59 | 4.59 | 0.05 | 0.05 |
| gene-LOC105278807 | https://www.ncbi.nlm.nih.gov/gene/?term=LOC105278807 | 2166 | 65 | 300.09 | 1 | 4.62 | 1.54 | 22.17 | 20.43 | 1.74 | 1.74 | 0.05 | 0.05 |
| gene-LOC105280104 | https://www.ncbi.nlm.nih.gov/gene/?term=LOC105280104 | 3960 | 123 | 310.61 | 1 | 2.53 | 0.81 | 18.15 | 16.51 | 1.64 | 1.64 | 0.05 | 0.05 |
| gene-LOC105275363 | https://www.ncbi.nlm.nih.gov/gene/?term=LOC105275363 | 3798 | 157 | 413.38 | 1 | 2.63 | 0.64 | 17.94 | 16.87 | 1.07 | 1.07 | 0.04 | 0.04 |
| gene-LOC105284043 | https://www.ncbi.nlm.nih.gov/gene/?term=LOC105284043 | 5661 | 283 | 499.91 | 1 | 1.77 | 0.35 | 10.38 | 9.58 | 0.79 | 0.79 | 0.04 | 0.04 |
| gene-LOC105282156 | https://www.ncbi.nlm.nih.gov/gene/?term=LOC105282156 | 2577 | 122 | 473.42 | 1 | 3.88 | 0.82 | 25.73 | 24.95 | 0.78 | 0.78 | 0.04 | 0.04 |
| gene-LOC105285534 | https://www.ncbi.nlm.nih.gov/gene/?term=LOC105285534 | 3944 | 232 | 588.24 | 1 | 2.54 | 0.43 | 11.59 | 10.9 | 0.69 | 0.69 | 0.04 | 0.04 |
| gene-LOC105285055 | https://www.ncbi.nlm.nih.gov/gene/?term=LOC105285055 | 4108 | 261 | 635.35 | 1 | 2.43 | 0.38 | 10.75 | 10.11 | 0.64 | 0.64 | 0.04 | 0.04 |
| gene-LOC105278283 | https://www.ncbi.nlm.nih.gov/gene/?term=LOC105278283 | 2317 | 138 | 595.6 | 1 | 4.32 | 0.72 | 25.64 | 25.05 | 0.59 | 0.59 | 0.04 | 0.04 |
| gene-LOC105274534 | https://www.ncbi.nlm.nih.gov/gene/?term=LOC105274534 | 1329 | 57 | 428.89 | 1 | 7.52 | 1.75 | 46.71 | 47.65 | -0.94 | 0.94 | -0.04 | 0.04 |
| gene-LOC105286999 | https://www.ncbi.nlm.nih.gov/gene/?term=LOC105286999 | 2141 | 42 | 196.17 | 1 | 4.67 | 2.38 | 33.74 | 35.94 | -2.2 | 2.2 | -0.04 | 0.04 |
| gene-LOC105283967 | https://www.ncbi.nlm.nih.gov/gene/?term=LOC105283967 | 2393 | 40 | 167.15 | 1 | 4.18 | 2.5 | 52.37 | 54.65 | -2.28 | 2.28 | -0.04 | 0.04 |
| gene-LOC105283492 | https://www.ncbi.nlm.nih.gov/gene/?term=LOC105283492 | 1535 | 9 | 58.63 | 1 | 6.51 | 11.11 | 30.57 | 36.97 | -6.4 | 6.4 | -0.04 | 0.04 |
| gene-LOC105274900 | https://www.ncbi.nlm.nih.gov/gene/?term=LOC105274900 | 5561 | 55 | 98.9 | 3 | 5.39 | 5.45 | 32.79 | 29.47 | 3.31 | 3.31 | 0.03 | 0.03 |
| gene-LOC105281266 | https://www.ncbi.nlm.nih.gov/gene/?term=LOC105281266 | 4102 | 78 | 190.15 | 1 | 2.44 | 1.28 | 18.58 | 17.01 | 1.57 | 1.57 | 0.03 | 0.03 |
| gene-LOC105282794 | https://www.ncbi.nlm.nih.gov/gene/?term=LOC105282794 | 4571 | 90 | 196.89 | 1 | 2.19 | 1.11 | 17.25 | 15.76 | 1.49 | 1.49 | 0.03 | 0.03 |
| gene-LOC105277903 | https://www.ncbi.nlm.nih.gov/gene/?term=LOC105277903 | 1835 | 36 | 196.19 | 1 | 5.45 | 2.78 | 51.23 | 49.79 | 1.45 | 1.45 | 0.03 | 0.03 |
| gene-LOC105280956 | https://www.ncbi.nlm.nih.gov/gene/?term=LOC105280956 | 2639 | 71 | 269.04 | 1 | 3.79 | 1.41 | 18.77 | 17.79 | 0.98 | 0.98 | 0.03 | 0.03 |
| gene-LOC105288036 | https://www.ncbi.nlm.nih.gov/gene/?term=LOC105288036 | 5110 | 181 | 354.21 | 1 | 1.96 | 0.55 | 18.99 | 18.01 | 0.98 | 0.98 | 0.03 | 0.03 |
| gene-LOC105283123 | https://www.ncbi.nlm.nih.gov/gene/?term=LOC105283123 | 1507 | 40 | 265.43 | 1 | 6.64 | 2.5 | 16.19 | 15.23 | 0.96 | 0.96 | 0.03 | 0.03 |
| gene-LOC113561621 | https://www.ncbi.nlm.nih.gov/gene/?term=LOC113561621 | 2249 | 76 | 337.93 | 1 | 4.45 | 1.32 | 22.44 | 21.49 | 0.95 | 0.95 | 0.03 | 0.03 |
| gene-LOC105284184 | https://www.ncbi.nlm.nih.gov/gene/?term=LOC105284184 | 2535 | 80 | 315.58 | 1 | 3.94 | 1.25 | 35.73 | 34.87 | 0.86 | 0.86 | 0.03 | 0.03 |
| gene-LOC105276612 | https://www.ncbi.nlm.nih.gov/gene/?term=LOC105276612 | 4157 | 178 | 428.19 | 1 | 2.41 | 0.56 | 10.68 | 9.88 | 0.8 | 0.8 | 0.03 | 0.03 |
| gene-LOC105283728 | https://www.ncbi.nlm.nih.gov/gene/?term=LOC105283728 | 3256 | 117 | 359.34 | 2 | 6.14 | 1.71 | 18.77 | 18.07 | 0.7 | 0.7 | 0.03 | 0.03 |
| gene-LOC105275460 | https://www.ncbi.nlm.nih.gov/gene/?term=LOC105275460 | 2473 | 118 | 477.15 | 1 | 4.04 | 0.85 | 13.95 | 13.28 | 0.67 | 0.67 | 0.03 | 0.03 |
| gene-LOC105279298 | https://www.ncbi.nlm.nih.gov/gene/?term=LOC105279298 | 2773 | 133 | 479.62 | 2 | 7.21 | 1.5 | 23.97 | 23.38 | 0.59 | 0.59 | 0.03 | 0.03 |
| gene-LOC105278145 | https://www.ncbi.nlm.nih.gov/gene/?term=LOC105278145 | 4470 | 235 | 525.73 | 1 | 2.24 | 0.43 | 11.34 | 10.86 | 0.48 | 0.48 | 0.03 | 0.03 |
| gene-LOC105280819 | https://www.ncbi.nlm.nih.gov/gene/?term=LOC105280819 | 1639 | 37 | 225.75 | 1 | 6.1 | 2.7 | 36.94 | 38.07 | -1.12 | 1.12 | -0.03 | 0.03 |
| gene-LOC105278384 | https://www.ncbi.nlm.nih.gov/gene/?term=LOC105278384 | 1390 | 35 | 251.8 | 1 | 7.19 | 2.86 | 48.96 | 50.26 | -1.3 | 1.3 | -0.03 | 0.03 |
| gene-LOC105283712 | https://www.ncbi.nlm.nih.gov/gene/?term=LOC105283712 | 1910 | 36 | 188.48 | 1 | 5.24 | 2.78 | 22.92 | 24.43 | -1.51 | 1.51 | -0.03 | 0.03 |
| gene-LOC105286514 | https://www.ncbi.nlm.nih.gov/gene/?term=LOC105286514 | 2528 | 46 | 181.96 | 1 | 3.96 | 2.17 | 35.96 | 37.79 | -1.83 | 1.83 | -0.03 | 0.03 |
| gene-LOC105285409 | https://www.ncbi.nlm.nih.gov/gene/?term=LOC105285409 | 5099 | 49 | 96.1 | 1 | 1.96 | 2.04 | 1.72 | 4.82 | -3.09 | 3.09 | -0.03 | 0.03 |
| gene-LOC105286694 | https://www.ncbi.nlm.nih.gov/gene/?term=LOC105286694 | 3409 | 8 | 23.47 | 1 | 2.93 | 12.5 | 66.22 | 57.86 | 8.36 | 8.36 | 0.02 | 0.02 |
| gene-LOC105281515 | https://www.ncbi.nlm.nih.gov/gene/?term=LOC105281515 | 8827 | 41 | 46.45 | 1 | 1.13 | 2.44 | 31.51 | 28.2 | 3.31 | 3.31 | 0.02 | 0.02 |
| gene-LOC105286099 | https://www.ncbi.nlm.nih.gov/gene/?term=LOC105286099 | 5919 | 67 | 113.19 | 1 | 1.69 | 1.49 | 11.4 | 9.83 | 1.57 | 1.57 | 0.02 | 0.02 |
| gene-LOC105284329 | https://www.ncbi.nlm.nih.gov/gene/?term=LOC105284329 | 4051 | 52 | 128.36 | 1 | 2.47 | 1.92 | 19.88 | 18.38 | 1.5 | 1.5 | 0.02 | 0.02 |
| gene-LOC105288066 | https://www.ncbi.nlm.nih.gov/gene/?term=LOC105288066 | 1964 | 27 | 137.47 | 1 | 5.09 | 3.7 | 34.07 | 32.8 | 1.27 | 1.27 | 0.02 | 0.02 |
| gene-LOC105275451 | https://www.ncbi.nlm.nih.gov/gene/?term=LOC105275451 | 6035 | 113 | 187.24 | 1 | 1.66 | 0.88 | 24.46 | 23.26 | 1.2 | 1.2 | 0.02 | 0.02 |
| gene-LOC105279664 | https://www.ncbi.nlm.nih.gov/gene/?term=LOC105279664 | 4023 | 53 | 131.74 | 1 | 2.49 | 1.89 | 19.07 | 17.88 | 1.19 | 1.19 | 0.02 | 0.02 |
| gene-LOC113561805 | https://www.ncbi.nlm.nih.gov/gene/?term=LOC113561805 | 3337 | 67 | 200.78 | 1 | 3 | 1.49 | 36.13 | 35.02 | 1.11 | 1.11 | 0.02 | 0.02 |
| gene-LOC105281173 | https://www.ncbi.nlm.nih.gov/gene/?term=LOC105281173 | 1595 | 38 | 238.24 | 1 | 6.27 | 2.63 | 59.87 | 58.85 | 1.02 | 1.02 | 0.02 | 0.02 |
| gene-LOC105276692 | https://www.ncbi.nlm.nih.gov/gene/?term=LOC105276692 | 1745 | 29 | 166.19 | 1 | 5.73 | 3.45 | 25.96 | 24.96 | 1 | 1 | 0.02 | 0.02 |
| gene-LOC105276059 | https://www.ncbi.nlm.nih.gov/gene/?term=LOC105276059 | 6256 | 125 | 199.81 | 1 | 1.6 | 0.8 | 10.91 | 10 | 0.91 | 0.91 | 0.02 | 0.02 |
| gene-LOC105283592 | https://www.ncbi.nlm.nih.gov/gene/?term=LOC105283592 | 3479 | 71 | 204.08 | 1 | 2.87 | 1.41 | 21.54 | 20.67 | 0.88 | 0.88 | 0.02 | 0.02 |
| gene-LOC105276901 | https://www.ncbi.nlm.nih.gov/gene/?term=LOC105276901 | 3254 | 84 | 258.14 | 1 | 3.07 | 1.19 | 27.21 | 26.37 | 0.84 | 0.84 | 0.02 | 0.02 |
| gene-LOC105274823 | https://www.ncbi.nlm.nih.gov/gene/?term=LOC105274823 | 4620 | 138 | 298.7 | 1 | 2.16 | 0.72 | 21.4 | 20.58 | 0.82 | 0.82 | 0.02 | 0.02 |
| gene-LOC105285639 | https://www.ncbi.nlm.nih.gov/gene/?term=LOC105285639 | 10325 | 312 | 302.18 | 2 | 1.94 | 0.64 | 12.89 | 12.24 | 0.65 | 0.65 | 0.02 | 0.02 |

| | | | | | | | | | | | | | | | |
|---|---|---|---|---|---|---|---|---|---|---|---|---|---|---|---|
| gene-LOC105278534 | https://www.ncbi.nlm.nih.gov/gene/?term=LOC105278534 | 7364 | 176 | 239 | 2 | 2.72 | 1.14 | 19.64 | 19.01 | 0.63 | 0.63 | 0.02 | 0.02 | | |
| gene-LOC105285179 | https://www.ncbi.nlm.nih.gov/gene/?term=LOC105285179 | 4191 | 171 | 408.02 | | 1 | 2.39 | 0.58 | 19 | 18.45 | 0.55 | 0.55 | 0.02 | 0.02 | |
| gene-LOC105278497 | https://www.ncbi.nlm.nih.gov/gene/?term=LOC105278497 | 2072 | 95 | 458.49 | | 1 | 4.83 | 1.05 | 19.39 | 18.89 | 0.5 | 0.5 | 0.02 | 0.02 | |
| gene-LOC105286997 | https://www.ncbi.nlm.nih.gov/gene/?term=LOC105286997 | 2850 | 106 | 371.93 | | 1 | 3.51 | 0.94 | 12.07 | 11.62 | 0.46 | 0.46 | 0.02 | 0.02 | |
| gene-LOC105275374 | https://www.ncbi.nlm.nih.gov/gene/?term=LOC105275374 | 11376 | 454 | 399.09 | | 1 | 0.88 | 0.22 | 10.82 | 10.43 | 0.38 | 0.38 | 0.02 | 0.02 | |
| gene-LOC105274449 | https://www.ncbi.nlm.nih.gov/gene/?term=LOC105274449 | 1549 | 48 | 309.88 | | 1 | 6.46 | 2.08 | 50.45 | 50.96 | -0.51 | 0.51 | -0.02 | 0.02 | |
| gene-LOC105278877 | https://www.ncbi.nlm.nih.gov/gene/?term=LOC105278877 | 1968 | 70 | 355.69 | | 1 | 5.08 | 1.43 | 34.63 | 35.21 | -0.58 | 0.58 | -0.02 | 0.02 | |
| gene-LOC105280377 | https://www.ncbi.nlm.nih.gov/gene/?term=LOC105280377 | 2024 | 65 | 321.15 | | 1 | 4.94 | 1.54 | 47.38 | 47.97 | -0.59 | 0.59 | -0.02 | 0.02 | |
| gene-LOC105281700 | https://www.ncbi.nlm.nih.gov/gene/?term=LOC105281700 | 2208 | 32 | 144.93 | | 1 | 4.53 | 3.12 | 37.54 | 39.09 | -1.55 | 1.55 | -0.02 | 0.02 | |
| gene-LOC105276538 | https://www.ncbi.nlm.nih.gov/gene/?term=LOC105276538 | 1511 | 18 | 119.13 | | 1 | 6.62 | 5.56 | 64.69 | 66.35 | -1.65 | 1.65 | -0.02 | 0.02 | |
| gene-LOC105275347 | https://www.ncbi.nlm.nih.gov/gene/?term=LOC105275347 | 2526 | 16 | 63.34 | 1 | 3.96 | 6.25 | 41.65 | 39.67 | 1.98 | 1.98 | 0.01 | 0.01 | | |
| gene-LOC105283943 | https://www.ncbi.nlm.nih.gov/gene/?term=LOC105283943 | 3559 | 45 | 126.44 | | 1 | 2.81 | 2.22 | 16.9 | 15.77 | 1.14 | 1.14 | 0.01 | 0.01 | |
| gene-LOC105288078 | https://www.ncbi.nlm.nih.gov/gene/?term=LOC105288078 | 19126 | 286 | 149.53 | | 1 | 0.52 | 0.35 | 17.66 | 16.67 | 0.99 | 0.99 | 0.01 | 0.01 | |
| gene-LOC105285060 | https://www.ncbi.nlm.nih.gov/gene/?term=LOC105285060 | 1285 | 16 | 124.51 | | 1 | 7.78 | 6.25 | 47.94 | 47.04 | 0.9 | 0.9 | 0.01 | 0.01 | |
| gene-LOC105287854 | https://www.ncbi.nlm.nih.gov/gene/?term=LOC105287854 | 3644 | 49 | 134.47 | | 1 | 2.74 | 2.04 | 24.37 | 23.53 | 0.84 | 0.84 | 0.01 | 0.01 | |
| gene-LOC105287417 | https://www.ncbi.nlm.nih.gov/gene/?term=LOC105287417 | 1025 | 19 | 185.37 | | 1 | 9.76 | 5.26 | 59.44 | 58.65 | 0.79 | 0.79 | 0.01 | 0.01 | |
| gene-LOC105286452 | https://www.ncbi.nlm.nih.gov/gene/?term=LOC105286452 | 6888 | 64 | 92.92 | 1 | 1.45 | 1.56 | 30.17 | 29.46 | 0.72 | 0.72 | 0.01 | 0.01 | | |
| gene-LOC105281848 | https://www.ncbi.nlm.nih.gov/gene/?term=LOC105281848 | 2122 | 47 | 221.49 | | 1 | 4.71 | 2.13 | 27.19 | 26.57 | 0.63 | 0.63 | 0.01 | 0.01 | |
| gene-LOC105275179 | https://www.ncbi.nlm.nih.gov/gene/?term=LOC105275179 | 3271 | 69 | 210.94 | | 1 | 3.06 | 1.45 | 18.8 | 18.36 | 0.45 | 0.45 | 0.01 | 0.01 | |
| gene-LOC105287646 | https://www.ncbi.nlm.nih.gov/gene/?term=LOC105287646 | 3556 | 83 | 233.41 | | 1 | 2.81 | 1.2 | 17.22 | 16.85 | 0.38 | 0.38 | 0.01 | 0.01 | |
| gene-LOC105287550 | https://www.ncbi.nlm.nih.gov/gene/?term=LOC105287550 | 15659 | 549 | 350.6 | 1 | 0.64 | 0.18 | 6.33 | 5.97 | 0.37 | 0.37 | 0.01 | 0.01 | | |
| gene-LOC105274856 | https://www.ncbi.nlm.nih.gov/gene/?term=LOC105274856 | 5216 | 209 | 400.69 | | 1 | 1.92 | 0.48 | 18.76 | 18.44 | 0.31 | 0.31 | 0.01 | 0.01 | |
| gene-LOC105274699 | https://www.ncbi.nlm.nih.gov/gene/?term=LOC105274699 | 2386 | 49 | 205.36 | | 1 | 4.19 | 2.04 | 23.32 | 23.03 | 0.29 | 0.29 | 0.01 | 0.01 | |
| gene-LOC105274531 | https://www.ncbi.nlm.nih.gov/gene/?term=LOC105274531 | 7774 | 273 | 351.17 | | 1 | 1.29 | 0.37 | 11.43 | 11.16 | 0.27 | 0.27 | 0.01 | 0.01 | |
| gene-LOC105283649 | https://www.ncbi.nlm.nih.gov/gene/?term=LOC105283649 | 1918 | 53 | 276.33 | | 1 | 5.21 | 1.89 | 37.66 | 37.39 | 0.27 | 0.27 | 0.01 | 0.01 | |
| gene-LOC105279038 | https://www.ncbi.nlm.nih.gov/gene/?term=LOC105279038 | 9453 | 297 | 314.19 | | 1 | 1.06 | 0.34 | 9.64 | 9.43 | 0.21 | 0.21 | 0.01 | 0.01 | |
| gene-LOC105282149 | https://www.ncbi.nlm.nih.gov/gene/?term=LOC105282149 | 1787 | 103 | 576.39 | | 1 | 5.6 | 0.97 | 41.32 | 41.12 | 0.2 | 0.2 | 0.01 | 0.01 | |
| gene-LOC105275327 | https://www.ncbi.nlm.nih.gov/gene/?term=LOC105275327 | 14245 | 458 | 321.52 | | 1 | 0.7 | 0.22 | 7.5 | 7.33 | 0.17 | 0.17 | 0.01 | 0.01 | |
| gene-LOC105278666 | https://www.ncbi.nlm.nih.gov/gene/?term=LOC105278666 | 10341 | 439 | 424.52 | | 1 | 0.97 | 0.23 | 7.06 | 6.9 | 0.16 | 0.16 | 0.01 | 0.01 | |
| gene-LOC105286274 | https://www.ncbi.nlm.nih.gov/gene/?term=LOC105286274 | 3817 | 179 | 468.95 | | 1 | 2.62 | 0.56 | 16.95 | 17.06 | -0.11 | 0.11 | -0.01 | 0.01 | |
| gene-LOC105276527 | https://www.ncbi.nlm.nih.gov/gene/?term=LOC105276527 | 7923 | 305 | 384.96 | | 1 | 1.26 | 0.33 | 9.71 | 9.92 | -0.21 | 0.21 | -0.01 | 0.01 | |
| gene-LOC105287095 | https://www.ncbi.nlm.nih.gov/gene/?term=LOC105287095 | 1435 | 58 | 404.18 | | 1 | 6.97 | 1.72 | 28.41 | 28.65 | -0.24 | 0.24 | -0.01 | 0.01 | |
| gene-LOC105288042 | https://www.ncbi.nlm.nih.gov/gene/?term=LOC105288042 | 3507 | 88 | 250.93 | | 1 | 2.85 | 1.14 | 21.32 | 21.59 | -0.27 | 0.27 | -0.01 | 0.01 | |
| gene-LOC105285807 | https://www.ncbi.nlm.nih.gov/gene/?term=LOC105285807 | 6063 | 213 | 351.31 | | 1 | 1.65 | 0.47 | 9.29 | 9.56 | -0.28 | 0.28 | -0.01 | 0.01 | |
| gene-LOC105286515 | https://www.ncbi.nlm.nih.gov/gene/?term=LOC105286515 | 3751 | 107 | 285.26 | | 1 | 2.67 | 0.93 | 12.3 | 12.58 | -0.28 | 0.28 | -0.01 | 0.01 | |
| gene-LOC105280774 | https://www.ncbi.nlm.nih.gov/gene/?term=LOC105280774 | 2793 | 100 | 358.04 | | 1 | 3.58 | 1 | 31.9 | 32.19 | -0.29 | 0.29 | -0.01 | 0.01 | |
| gene-LOC105278228 | https://www.ncbi.nlm.nih.gov/gene/?term=LOC105278228 | 8610 | 156 | 181.18 | | 1 | 1.16 | 0.64 | 22.2 | 22.5 | -0.3 | 0.3 | -0.01 | 0.01 | |
| gene-LOC105284036 | https://www.ncbi.nlm.nih.gov/gene/?term=LOC105284036 | 3254 | 96 | 295.02 | | 1 | 3.07 | 1.04 | 23.11 | 23.46 | -0.35 | 0.35 | -0.01 | 0.01 | |
| gene-LOC105280348 | https://www.ncbi.nlm.nih.gov/gene/?term=LOC105280348 | 3797 | 71 | 186.99 | | 1 | 2.63 | 1.41 | 20.69 | 21.13 | -0.44 | 0.44 | -0.01 | 0.01 | |
| gene-LOC105278962 | https://www.ncbi.nlm.nih.gov/gene/?term=LOC105278962 | 4491 | 89 | 198.17 | | 1 | 2.23 | 1.12 | 22.04 | 22.67 | -0.63 | 0.63 | -0.01 | 0.01 | |
| gene-LOC105278454 | https://www.ncbi.nlm.nih.gov/gene/?term=LOC105278454 | 1972 | 40 | 202.84 | | 1 | 5.07 | 2.5 | 48.64 | 49.28 | -0.64 | 0.64 | -0.01 | 0.01 | |
| gene-LOC105280002 | https://www.ncbi.nlm.nih.gov/gene/?term=LOC105280002 | 1940 | 27 | 139.18 | | 1 | 5.15 | 3.7 | 33.73 | 34.37 | -0.64 | 0.64 | -0.01 | 0.01 | |
| gene-LOC105283989 | https://www.ncbi.nlm.nih.gov/gene/?term=LOC105283989 | 4948 | 25 | 50.53 | 1 | 2.02 | 4 | 35.77 | 37.79 | -2.02 | 2.02 | -0.01 | 0.01 | | |
| gene-LOC105278108 | https://www.ncbi.nlm.nih.gov/gene/?term=LOC105278108 | 4392 | 43 | 97.91 | 1 | 2.28 | 2.33 | 26.73 | 26.25 | 0.49 | 0.49 | 0 | 0 | | |
| gene-LOC105285406 | https://www.ncbi.nlm.nih.gov/gene/?term=LOC105285406 | 8015 | 60 | 74.86 | 1 | 1.25 | 1.67 | 25.86 | 25.52 | 0.34 | 0.34 | 0 | 0 | | |
| gene-LOC105274851 | https://www.ncbi.nlm.nih.gov/gene/?term=LOC105274851 | 6616 | 129 | 194.98 | | 1 | 1.51 | 0.78 | 11.16 | 10.94 | 0.22 | 0.22 | 0 | 0 | |
| gene-LOC105275187 | https://www.ncbi.nlm.nih.gov/gene/?term=LOC105275187 | 5534 | 65 | 117.46 | | 1 | 1.81 | 1.54 | 11.46 | 11.27 | 0.19 | 0.19 | 0 | 0 | |
| gene-LOC105286194 | https://www.ncbi.nlm.nih.gov/gene/?term=LOC105286194 | 152619 | | 1987 | 130.19 | | 1 | 0.07 | 0.05 | 6.27 | 6.13 | 0.14 | 0.14 | 0 | 0 |
| gene-LOC105278649 | https://www.ncbi.nlm.nih.gov/gene/?term=LOC105278649 | 5616 | 138 | 245.73 | | 1 | 1.78 | 0.72 | 13.54 | 13.42 | 0.12 | 0.12 | 0 | 0 | |
| gene-LOC105288079 | https://www.ncbi.nlm.nih.gov/gene/?term=LOC105288079 | 3473 | 96 | 276.42 | | 1 | 2.88 | 1.04 | 19.82 | 19.7 | 0.12 | 0.12 | 0 | 0 | |
| gene-LOC105278455 | https://www.ncbi.nlm.nih.gov/gene/?term=LOC105278455 | 27181 | 875 | 321.92 | | 1 | 0.37 | 0.11 | 7.69 | 7.59 | 0.1 | 0.1 | 0 | 0 | |
| gene-LOC105277856 | https://www.ncbi.nlm.nih.gov/gene/?term=LOC105277856 | 3442 | 29 | 84.25 | 1 | 2.91 | 3.45 | 32.81 | 32.75 | 0.06 | 0.06 | 0 | 0 | | |
| gene-LOC105284355 | https://www.ncbi.nlm.nih.gov/gene/?term=LOC105284355 | 3904 | 152 | 389.34 | | 1 | 2.56 | 0.66 | 20.02 | 19.96 | 0.06 | 0.06 | 0 | 0 | |
| gene-LOC105287528 | https://www.ncbi.nlm.nih.gov/gene/?term=LOC105287528 | 14195 | 375 | 264.18 | | 1 | 0.7 | 0.27 | 5.12 | 5.07 | 0.05 | 0.05 | 0 | 0 | |
| gene-LOC105277866 | https://www.ncbi.nlm.nih.gov/gene/?term=LOC105277866 | 33045 | 368 | 111.36 | | 1 | 0.3 | 0.27 | 3.48 | 3.54 | -0.06 | 0.06 | 0 | 0 | |
| gene-LOC105282811 | https://www.ncbi.nlm.nih.gov/gene/?term=LOC105282811 | 5765 | 196 | 339.98 | | 1 | 1.73 | 0.51 | 8.25 | 8.31 | -0.06 | 0.06 | 0 | 0 | |
| gene-LOC105284704 | https://www.ncbi.nlm.nih.gov/gene/?term=LOC105284704 | 2400 | 59 | 245.83 | | 1 | 4.17 | 1.69 | 22.63 | 22.7 | -0.07 | 0.07 | 0 | 0 | |
| gene-LOC105274706 | https://www.ncbi.nlm.nih.gov/gene/?term=LOC105274706 | 1446 | 49 | 338.87 | | 1 | 6.92 | 2.04 | 38.21 | 38.31 | -0.1 | 0.1 | 0 | 0 | |
| gene-LOC105278282 | https://www.ncbi.nlm.nih.gov/gene/?term=LOC105278282 | 1775 | 52 | 292.96 | | 1 | 5.63 | 1.92 | 41.01 | 41.14 | -0.14 | 0.14 | 0 | 0 | |
| gene-LOC105274590 | https://www.ncbi.nlm.nih.gov/gene/?term=LOC105274590 | 4516 | 67 | 148.36 | | 1 | 2.21 | 1.49 | 28.5 | 28.65 | -0.15 | 0.15 | 0 | 0 | |
| gene-LOC105282067 | https://www.ncbi.nlm.nih.gov/gene/?term=LOC105282067 | 5941 | 49 | 82.48 | 1 | 1.68 | 2.04 | 25.2 | 25.4 | -0.2 | 0.2 | 0 | 0 | | |
| gene-LOC105282658 | https://www.ncbi.nlm.nih.gov/gene/?term=LOC105282658 | 2421 | 52 | 214.79 | | 1 | 4.13 | 1.92 | 21.81 | 22.01 | -0.2 | 0.2 | 0 | 0 | |

Directory: OGLall File: rnk-dmg-Ob_LI_RphaseA2.vs.RphaseB1.txt

| gene_ID | gene_link | gwidth | #Sites | #per10Kb | #dmSites | #dmsp10kb | %dmSites | %pSite1 | %pSite2 | DMpSite | ADMpSite | DMpNucl | ADMpNucl |
|---|---|---|---|---|---|---|---|---|---|---|---|---|---|
| gene-LOC113562345 | https://www.ncbi.nlm.nih.gov/gene/?term=LOC113562345 | 2290 | 80 | 349.34 | 16 | 69.87 | 20 | 25.78 | 13.98 | 11.8 | 11.8 | 0.41 | 0.41 |
| gene-LOC105279900 | https://www.ncbi.nlm.nih.gov/gene/?term=LOC105279900 | 2191 | 56 | 255.59 | 1 | 4.56 | 1.79 | 33.63 | 28.91 | 4.72 | 4.72 | 0.12 | 0.12 |
| gene-LOC105285695 | https://www.ncbi.nlm.nih.gov/gene/?term=LOC105285695 | 1371 | 76 | 554.34 | 1 | 7.29 | 1.32 | 28.59 | 26.36 | 2.24 | 2.24 | 0.12 | 0.12 |
| gene-LOC105287417 | https://www.ncbi.nlm.nih.gov/gene/?term=LOC105287417 | 1025 | 38 | 370.73 | 1 | 9.76 | 2.63 | 35.24 | 32.37 | 2.87 | 2.87 | 0.11 | 0.11 |
| gene-LOC105282156 | https://www.ncbi.nlm.nih.gov/gene/?term=LOC105282156 | 2577 | 134 | 519.98 | 1 | 3.88 | 0.75 | 27.52 | 25.46 | 2.06 | 2.06 | 0.11 | 0.11 |
| gene-LOC105280786 | https://www.ncbi.nlm.nih.gov/gene/?term=LOC105280786 | 2573 | 143 | 555.77 | 1 | 3.89 | 0.7 | 27.63 | 25.63 | 2 | 2 | 0.11 | 0.11 |
| gene-LOC105275070 | https://www.ncbi.nlm.nih.gov/gene/?term=LOC105275070 | 5667 | 64 | 112.93 | 6 | 10.59 | 9.38 | 14.32 | 5.2 | 9.13 | 9.13 | 0.1 | 0.1 |
| gene-LOC105287063 | https://www.ncbi.nlm.nih.gov/gene/?term=LOC105287063 | 1874 | 55 | 293.49 | 1 | 5.34 | 1.82 | 30.47 | 27.11 | 3.37 | 3.37 | 0.1 | 0.1 |
| gene-LOC105276704 | https://www.ncbi.nlm.nih.gov/gene/?term=LOC105276704 | 2063 | 93 | 450.8 5 | 24.24 | 5.38 | 14.49 | 12.51 | 1.99 | 1.99 | 0.09 | 0.09 | |
| gene-LOC105283457 | https://www.ncbi.nlm.nih.gov/gene/?term=LOC105283457 | 1882 | 49 | 260.36 | 1 | 5.31 | 2.04 | 35.19 | 32.03 | 3.16 | 3.16 | 0.08 | 0.08 |
| gene-LOC105285583 | https://www.ncbi.nlm.nih.gov/gene/?term=LOC105285583 | 1947 | 62 | 318.44 | 1 | 5.14 | 1.61 | 27.07 | 24.61 | 2.46 | 2.46 | 0.08 | 0.08 |
| gene-LOC105274706 | https://www.ncbi.nlm.nih.gov/gene/?term=LOC105274706 | 1446 | 56 | 387.28 | 1 | 6.92 | 1.79 | 39.8 | 37.63 | 2.17 | 2.17 | 0.08 | 0.08 |
| gene-LOC105285674 | https://www.ncbi.nlm.nih.gov/gene/?term=LOC105285674 | 3035 | 111 | 365.73 | 2 | 6.59 | 1.8 | 25.15 | 23.09 | 2.06 | 2.06 | 0.08 | 0.08 |
| gene-LOC105282893 | https://www.ncbi.nlm.nih.gov/gene/?term=LOC105282893 | 2645 | 50 | 189.04 | 1 | 3.78 | 2 | 28.92 | 25.31 | 3.6 | 3.6 | 0.07 | 0.07 |
| gene-LOC105284704 | https://www.ncbi.nlm.nih.gov/gene/?term=LOC105284704 | 2400 | 66 | 275 1 | 4.17 | 1.52 | 25.49 | 22.92 | 2.57 | 2.57 | 0.07 | 0.07 | |
| gene-LOC105286683 | https://www.ncbi.nlm.nih.gov/gene/?term=LOC105286683 | 1665 | 57 | 342.34 | 1 | 6.01 | 1.75 | 35.23 | 33.27 | 1.97 | 1.97 | 0.07 | 0.07 |
| gene-LOC105276449 | https://www.ncbi.nlm.nih.gov/gene/?term=LOC105276449 | 1231 | 55 | 446.79 | 1 | 8.12 | 1.82 | 24.75 | 23.28 | 1.47 | 1.47 | 0.07 | 0.07 |
| gene-LOC105277435 | https://www.ncbi.nlm.nih.gov/gene/?term=LOC105277435 | 3643 | 184 | 505.08 | 1 | 2.74 | 0.54 | 12.13 | 10.74 | 1.39 | 1.39 | 0.07 | 0.07 |
| gene-LOC105279298 | https://www.ncbi.nlm.nih.gov/gene/?term=LOC105279298 | 2773 | 134 | 483.23 | 2 | 7.21 | 1.49 | 24.2 | 22.81 | 1.39 | 1.39 | 0.07 | 0.07 |
| gene-LOC105282682 | https://www.ncbi.nlm.nih.gov/gene/?term=LOC105282682 | 4127 | 240 | 581.54 | 1 | 2.42 | 0.42 | 13.64 | 12.51 | 1.14 | 1.14 | 0.07 | 0.07 |
| gene-LOC105284213 | https://www.ncbi.nlm.nih.gov/gene/?term=LOC105284213 | 868 | 23 | 264.98 | 1 | 11.52 | 4.35 | 9.72 | 12.3 | -2.58 | 2.58 | -0.07 | 0.07 |
| gene-LOC105275708 | https://www.ncbi.nlm.nih.gov/gene/?term=LOC105275708 | 1772 | 19 | 107.22 | 1 | 5.64 | 5.26 | 37.25 | 31.29 | 5.96 | 5.96 | 0.06 | 0.06 |
| gene-LOC105276538 | https://www.ncbi.nlm.nih.gov/gene/?term=LOC105276538 | 1511 | 23 | 152.22 | 1 | 6.62 | 4.35 | 69.63 | 65.38 | 4.25 | 4.25 | 0.06 | 0.06 |
| gene-LOC105283712 | https://www.ncbi.nlm.nih.gov/gene/?term=LOC105283712 | 1910 | 33 | 172.77 | 1 | 5.24 | 3.03 | 32.87 | 29.11 | 3.76 | 3.76 | 0.06 | 0.06 |
| gene-LOC105281848 | https://www.ncbi.nlm.nih.gov/gene/?term=LOC105281848 | 2122 | 60 | 282.75 | 1 | 4.71 | 1.67 | 24.48 | 22.53 | 1.95 | 1.95 | 0.06 | 0.06 |
| gene-LOC105280704 | https://www.ncbi.nlm.nih.gov/gene/?term=LOC105280704 | 4266 | 155 | 363.34 | 1 | 2.34 | 0.65 | 17.59 | 16.02 | 1.58 | 1.58 | 0.06 | 0.06 |
| gene-LOC105278282 | https://www.ncbi.nlm.nih.gov/gene/?term=LOC105278282 | 1775 | 67 | 377.46 | 1 | 5.63 | 1.49 | 34.11 | 32.61 | 1.51 | 1.51 | 0.06 | 0.06 |
| gene-LOC105285671 | https://www.ncbi.nlm.nih.gov/gene/?term=LOC105285671 | 2263 | 103 | 455.15 | 1 | 4.42 | 0.97 | 30.91 | 29.64 | 1.27 | 1.27 | 0.06 | 0.06 |
| gene-LOC105278283 | https://www.ncbi.nlm.nih.gov/gene/?term=LOC105278283 | 2317 | 145 | 625.81 | 1 | 4.32 | 0.69 | 24.77 | 23.82 | 0.95 | 0.95 | 0.06 | 0.06 |
| gene-LOC105285055 | https://www.ncbi.nlm.nih.gov/gene/?term=LOC105285055 | 4108 | 264 | 642.65 | 2 | 4.87 | 0.76 | 12 | 11.12 | 0.87 | 0.87 | 0.06 | 0.06 |
| gene-LOC105277631 | https://www.ncbi.nlm.nih.gov/gene/?term=LOC105277631 | 4387 | 327 | 745.38 | 1 | 2.28 | 0.31 | 11.07 | 10.23 | 0.85 | 0.85 | 0.06 | 0.06 |
| gene-LOC105275328 | https://www.ncbi.nlm.nih.gov/gene/?term=LOC105275328 | 2227 | 85 | 381.68 | 1 | 4.49 | 1.18 | 17.9 | 19.55 | -1.65 | 1.65 | -0.06 | 0.06 |
| gene-LOC113562974 | https://www.ncbi.nlm.nih.gov/gene/?term=LOC113562974 | 1832 | 2 | 10.92 1 | 5.46 | 50 | 57.94 | 9 | 48.94 | 48.94 | 0.05 | 0.05 | |
| gene-LOC105285514 | https://www.ncbi.nlm.nih.gov/gene/?term=LOC105285514 | 1277 | 11 | 86.14 1 | 7.83 | 9.09 | 90.74 | 85.12 | 5.62 | 5.62 | 0.05 | 0.05 | |
| gene-LOC105281039 | https://www.ncbi.nlm.nih.gov/gene/?term=LOC105281039 | 1471 | 15 | 101.97 | 1 | 6.8 | 6.67 | 36.89 | 31.93 | 4.96 | 4.96 | 0.05 | 0.05 |
| gene-LOC105275190 | https://www.ncbi.nlm.nih.gov/gene/?term=LOC105275190 | 3837 | 77 | 200.68 | 1 | 2.61 | 1.3 | 21.84 | 19.45 | 2.39 | 2.39 | 0.05 | 0.05 |
| gene-LOC113561805 | https://www.ncbi.nlm.nih.gov/gene/?term=LOC113561805 | 3337 | 73 | 218.76 | 1 | 3 | 1.37 | 38.02 | 35.67 | 2.34 | 2.34 | 0.05 | 0.05 |
| gene-LOC105276272 | https://www.ncbi.nlm.nih.gov/gene/?term=LOC105276272 | 1965 | 47 | 239.19 | 1 | 5.09 | 2.13 | 25.43 | 23.44 | 1.98 | 1.98 | 0.05 | 0.05 |
| gene-LOC105288079 | https://www.ncbi.nlm.nih.gov/gene/?term=LOC105288079 | 3473 | 112 | 322.49 | 1 | 2.88 | 0.89 | 18.94 | 17.43 | 1.51 | 1.51 | 0.05 | 0.05 |
| gene-LOC105275460 | https://www.ncbi.nlm.nih.gov/gene/?term=LOC105275460 | 2473 | 123 | 497.37 | 1 | 4.04 | 0.81 | 13.29 | 12.32 | 0.97 | 0.97 | 0.05 | 0.05 |
| gene-LOC105280774 | https://www.ncbi.nlm.nih.gov/gene/?term=LOC105280774 | 2793 | 135 | 483.35 | 1 | 3.58 | 0.74 | 25.37 | 24.41 | 0.96 | 0.96 | 0.05 | 0.05 |
| gene-LOC105288066 | https://www.ncbi.nlm.nih.gov/gene/?term=LOC105288066 | 1964 | 31 | 157.84 | 1 | 5.09 | 3.23 | 39.38 | 36.57 | 2.81 | 2.81 | 0.04 | 0.04 |
| gene-LOC105288010 | https://www.ncbi.nlm.nih.gov/gene/?term=LOC105288010 | 3189 | 61 | 191.28 | 1 | 3.14 | 1.64 | 34.85 | 32.7 | 2.15 | 2.15 | 0.04 | 0.04 |
| gene-LOC105286644 | https://www.ncbi.nlm.nih.gov/gene/?term=LOC105286644 | 1235 | 23 | 186.23 | 1 | 8.1 | 4.35 | 43.78 | 41.75 | 2.04 | 2.04 | 0.04 | 0.04 |
| gene-LOC105282151 | https://www.ncbi.nlm.nih.gov/gene/?term=LOC105282151 | 1601 | 45 | 281.07 | 2 | 12.49 | 4.44 | 42.82 | 41.43 | 1.39 | 1.39 | 0.04 | 0.04 |
| gene-LOC105283123 | https://www.ncbi.nlm.nih.gov/gene/?term=LOC105283123 | 1507 | 55 | 364.96 | 1 | 6.64 | 1.82 | 20.08 | 18.96 | 1.12 | 1.12 | 0.04 | 0.04 |
| gene-LOC105276408 | https://www.ncbi.nlm.nih.gov/gene/?term=LOC105276408 | 5298 | 201 | 379.39 | 1 | 1.89 | 0.5 | 21.53 | 20.53 | 1 | 1 | 0.04 | 0.04 |
| gene-LOC105284355 | https://www.ncbi.nlm.nih.gov/gene/?term=LOC105284355 | 3904 | 155 | 397.03 | 1 | 2.56 | 0.65 | 22.33 | 21.37 | 0.95 | 0.95 | 0.04 | 0.04 |
| gene-LOC105278497 | https://www.ncbi.nlm.nih.gov/gene/?term=LOC105278497 | 2072 | 103 | 497.1 1 | 4.83 | 0.97 | 19.51 | 18.76 | 0.75 | 0.75 | 0.04 | 0.04 | |
| gene-LOC105284557 | https://www.ncbi.nlm.nih.gov/gene/?term=LOC105284557 | 1672 | 91 | 544.26 | 1 | 5.98 | 1.1 | 27.82 | 27.15 | 0.67 | 0.67 | 0.04 | 0.04 |
| gene-LOC105285409 | https://www.ncbi.nlm.nih.gov/gene/?term=LOC105285409 | 5099 | 67 | 131.4 2 | 3.92 | 2.99 | 1.72 | 4.47 | -2.75 | 2.75 | -0.04 | 0.04 | |
| gene-LOC105286302 | https://www.ncbi.nlm.nih.gov/gene/?term=LOC105286302 | 4536 | 24 | 52.91 1 | 2.2 | 4.17 | 20.45 | 15.15 | 5.3 | 5.3 | 0.03 | 0.03 | |
| gene-LOC105284609 | https://www.ncbi.nlm.nih.gov/gene/?term=LOC105284609 | 5678 | 44 | 77.49 1 | 1.76 | 2.27 | 32.19 | 27.8 | 4.39 | 4.39 | 0.03 | 0.03 | |
| gene-LOC105279898 | https://www.ncbi.nlm.nih.gov/gene/?term=LOC105279898 | 1343 | 16 | 119.14 | 1 | 7.45 | 6.25 | 58.57 | 56.24 | 2.33 | 2.33 | 0.03 | 0.03 |
| gene-LOC105275518 | https://www.ncbi.nlm.nih.gov/gene/?term=LOC105275518 | 2335 | 28 | 119.91 | 1 | 4.28 | 3.57 | 41.74 | 39.46 | 2.28 | 2.28 | 0.03 | 0.03 |
| gene-LOC105285060 | https://www.ncbi.nlm.nih.gov/gene/?term=LOC105285060 | 1285 | 18 | 140.08 | 1 | 7.78 | 5.56 | 38.72 | 36.69 | 2.03 | 2.03 | 0.03 | 0.03 |
| gene-LOC105276078 | https://www.ncbi.nlm.nih.gov/gene/?term=LOC105276078 | 1051 | 18 | 171.27 | 1 | 9.51 | 5.56 | 28.41 | 26.59 | 1.83 | 1.83 | 0.03 | 0.03 |
| gene-LOC105280348 | https://www.ncbi.nlm.nih.gov/gene/?term=LOC105280348 | 3797 | 78 | 205.43 | 1 | 2.63 | 1.28 | 18.55 | 17.12 | 1.44 | 1.44 | 0.03 | 0.03 |
| gene-LOC105282794 | https://www.ncbi.nlm.nih.gov/gene/?term=LOC105282794 | 4571 | 109 | 238.46 | 1 | 2.19 | 0.92 | 17.81 | 16.42 | 1.38 | 1.38 | 0.03 | 0.03 |
| gene-LOC105286514 | https://www.ncbi.nlm.nih.gov/gene/?term=LOC105286514 | 2528 | 55 | 217.56 | 1 | 3.96 | 1.82 | 37.15 | 35.79 | 1.36 | 1.36 | 0.03 | 0.03 |

| | | | | | | | | | | | | | |
|---|---|---|---|---|---|---|---|---|---|---|---|---|---|
| gene-LOC105280722 | https://www.ncbi.nlm.nih.gov/gene/?term=LOC105280722 | 3237 | 93 | 287.3 | 1 | 3.09 | 1.08 | 27.81 | 26.63 | 1.18 | 1.18 | 0.03 | 0.03 |
| gene-LOC105276969 | https://www.ncbi.nlm.nih.gov/gene/?term=LOC105276969 | 2997 | 104 | 347.01 | 1 | 3.34 | 0.96 | 11.67 | 10.81 | 0.86 | 0.86 | 0.03 | 0.03 |
| gene-LOC105278807 | https://www.ncbi.nlm.nih.gov/gene/?term=LOC105278807 | 2166 | 77 | 355.49 | 1 | 4.62 | 1.3 | 21.89 | 21.08 | 0.81 | 0.81 | 0.03 | 0.03 |
| gene-LOC105274823 | https://www.ncbi.nlm.nih.gov/gene/?term=LOC105274823 | 4620 | 171 | 370.13 | 1 | 2.16 | 0.58 | 18.07 | 17.27 | 0.8 | 0.8 | 0.03 | 0.03 |
| gene-LOC105280104 | https://www.ncbi.nlm.nih.gov/gene/?term=LOC105280104 | 3960 | 132 | 333.33 | 1 | 2.53 | 0.76 | 17.97 | 17.22 | 0.76 | 0.76 | 0.03 | 0.03 |
| gene-LOC105285742 | https://www.ncbi.nlm.nih.gov/gene/?term=LOC105285742 | 3446 | 122 | 354.03 | 1 | 2.9 | 0.82 | 17.63 | 16.88 | 0.76 | 0.76 | 0.03 | 0.03 |
| gene-LOC105282491 | https://www.ncbi.nlm.nih.gov/gene/?term=LOC105282491 | 4761 | 191 | 401.18 | 1 | 2.1 | 0.52 | 9 | 8.31 | 0.69 | 0.69 | 0.03 | 0.03 |
| gene-LOC105283728 | https://www.ncbi.nlm.nih.gov/gene/?term=LOC105283728 | 3256 | 159 | 488.33 | 2 | 6.14 | 1.26 | 15.6 | 14.91 | 0.69 | 0.69 | 0.03 | 0.03 |
| gene-LOC105287704 | https://www.ncbi.nlm.nih.gov/gene/?term=LOC105287704 | 3019 | 142 | 470.35 | 1 | 3.31 | 0.7 | 19.98 | 19.34 | 0.63 | 0.63 | 0.03 | 0.03 |
| gene-LOC105275683 | https://www.ncbi.nlm.nih.gov/gene/?term=LOC105275683 | 4256 | 204 | 479.32 | 1 | 2.35 | 0.49 | 13.38 | 12.77 | 0.61 | 0.61 | 0.03 | 0.03 |
| gene-LOC105286279 | https://www.ncbi.nlm.nih.gov/gene/?term=LOC105286279 | 5864 | 277 | 472.37 | 1 | 1.71 | 0.36 | 2.15 | 2.72 | -0.57 | 0.57 | -0.03 | 0.03 |
| gene-LOC105284329 | https://www.ncbi.nlm.nih.gov/gene/?term=LOC105284329 | 4051 | 72 | 177.73 | 1 | 2.47 | 1.39 | 18.86 | 20.42 | -1.56 | 1.56 | -0.03 | 0.03 |
| gene-LOC105286983 | https://www.ncbi.nlm.nih.gov/gene/?term=LOC105286983 | 16901 | 70 | 41.42 | 1 | 0.59 | 1.43 | 6.21 | 1.14 | 5.06 | 5.06 | 0.02 | 0.02 |
| gene-LOC105285315 | https://www.ncbi.nlm.nih.gov/gene/?term=LOC105285315 | 5458 | 33 | 60.46 | 1 | 1.83 | 3.03 | 33.27 | 29.4 | 3.88 | 3.88 | 0.02 | 0.02 |
| gene-LOC105281515 | https://www.ncbi.nlm.nih.gov/gene/?term=LOC105281515 | 8827 | 56 | 63.44 | 2 | 2.27 | 3.57 | 36.05 | 32.61 | 3.44 | 3.44 | 0.02 | 0.02 |
| gene-LOC105274900 | https://www.ncbi.nlm.nih.gov/gene/?term=LOC105274900 | 5561 | 61 | 109.69 | 2 | 3.6 | 3.28 | 32.53 | 30.66 | 1.87 | 1.87 | 0.02 | 0.02 |
| gene-LOC105278108 | https://www.ncbi.nlm.nih.gov/gene/?term=LOC105278108 | 4392 | 57 | 129.78 | 1 | 2.28 | 1.75 | 25.91 | 24.39 | 1.52 | 1.52 | 0.02 | 0.02 |
| gene-LOC105280002 | https://www.ncbi.nlm.nih.gov/gene/?term=LOC105280002 | 1940 | 34 | 175.26 | 1 | 5.15 | 2.94 | 33.64 | 32.26 | 1.38 | 1.38 | 0.02 | 0.02 |
| gene-LOC105288063 | https://www.ncbi.nlm.nih.gov/gene/?term=LOC105288063 | 3400 | 71 | 208.82 | 1 | 2.94 | 1.41 | 12.55 | 11.41 | 1.14 | 1.14 | 0.02 | 0.02 |
| gene-LOC105282600 | https://www.ncbi.nlm.nih.gov/gene/?term=LOC105282600 | 3715 | 67 | 180.35 | 1 | 2.69 | 1.49 | 27.6 | 26.48 | 1.12 | 1.12 | 0.02 | 0.02 |
| gene-LOC105278071 | https://www.ncbi.nlm.nih.gov/gene/?term=LOC105278071 | 4307 | 61 | 141.63 | 1 | 2.32 | 1.64 | 11.12 | 10.02 | 1.1 | 1.1 | 0.02 | 0.02 |
| gene-LOC105285416 | https://www.ncbi.nlm.nih.gov/gene/?term=LOC105285416 | 3820 | 76 | 198.95 | 1 | 2.62 | 1.32 | 22.22 | 21.12 | 1.1 | 1.1 | 0.02 | 0.02 |
| gene-LOC105280407 | https://www.ncbi.nlm.nih.gov/gene/?term=LOC105280407 | 10950 | 231 | 210.96 | 1 | 0.91 | 0.43 | 15.59 | 14.54 | 1.05 | 1.05 | 0.02 | 0.02 |
| gene-LOC105283838 | https://www.ncbi.nlm.nih.gov/gene/?term=LOC105283838 | 1748 | 33 | 188.79 | 1 | 5.72 | 3.03 | 33.49 | 32.48 | 1.01 | 1.01 | 0.02 | 0.02 |
| gene-LOC105283592 | https://www.ncbi.nlm.nih.gov/gene/?term=LOC105283592 | 3479 | 87 | 250.07 | 1 | 2.87 | 1.15 | 22.28 | 21.36 | 0.92 | 0.92 | 0.02 | 0.02 |
| gene-LOC105275179 | https://www.ncbi.nlm.nih.gov/gene/?term=LOC105275179 | 3271 | 85 | 259.86 | 1 | 3.06 | 1.18 | 16.68 | 15.92 | 0.76 | 0.76 | 0.02 | 0.02 |
| gene-LOC105281312 | https://www.ncbi.nlm.nih.gov/gene/?term=LOC105281312 | 2551 | 69 | 270.48 | 1 | 3.92 | 1.45 | 31.17 | 30.43 | 0.74 | 0.74 | 0.02 | 0.02 |
| gene-LOC105282825 | https://www.ncbi.nlm.nih.gov/gene/?term=LOC105282825 | 3919 | 102 | 260.27 | 1 | 2.55 | 0.98 | 14.91 | 14.23 | 0.68 | 0.68 | 0.02 | 0.02 |
| gene-LOC105275451 | https://www.ncbi.nlm.nih.gov/gene/?term=LOC105275451 | 6035 | 149 | 246.89 | 1 | 1.66 | 0.67 | 20.25 | 19.59 | 0.66 | 0.66 | 0.02 | 0.02 |
| gene-LOC105278632 | https://www.ncbi.nlm.nih.gov/gene/?term=LOC105278632 | 3624 | 150 | 413.91 | 1 | 2.76 | 0.67 | 13.57 | 13.05 | 0.53 | 0.53 | 0.02 | 0.02 |
| gene-LOC105285179 | https://www.ncbi.nlm.nih.gov/gene/?term=LOC105285179 | 4191 | 204 | 486.76 | 1 | 2.39 | 0.49 | 16.93 | 16.42 | 0.51 | 0.51 | 0.02 | 0.02 |
| gene-LOC105275752 | https://www.ncbi.nlm.nih.gov/gene/?term=LOC105275752 | 3462 | 138 | 398.61 | 1 | 2.89 | 0.72 | 31.47 | 31.01 | 0.46 | 0.46 | 0.02 | 0.02 |
| gene-LOC105279038 | https://www.ncbi.nlm.nih.gov/gene/?term=LOC105279038 | 9453 | 345 | 364.96 | 1 | 1.06 | 0.29 | 9.49 | 9.03 | 0.45 | 0.45 | 0.02 | 0.02 |
| gene-LOC105275363 | https://www.ncbi.nlm.nih.gov/gene/?term=LOC105275363 | 3798 | 212 | 558.19 | 1 | 2.63 | 0.47 | 14.03 | 13.61 | 0.43 | 0.43 | 0.02 | 0.02 |
| gene-LOC105285807 | https://www.ncbi.nlm.nih.gov/gene/?term=LOC105285807 | 6063 | 231 | 381 | 1 | 1.65 | 0.43 | 10.41 | 9.98 | 0.43 | 0.43 | 0.02 | 0.02 |
| gene-LOC105274531 | https://www.ncbi.nlm.nih.gov/gene/?term=LOC105274531 | 7774 | 307 | 394.91 | 1 | 1.29 | 0.33 | 10.8 | 10.38 | 0.42 | 0.42 | 0.02 | 0.02 |
| gene-LOC105281272 | https://www.ncbi.nlm.nih.gov/gene/?term=LOC105281272 | 5252 | 201 | 382.71 | 1 | 1.9 | 0.5 | 15.32 | 14.89 | 0.42 | 0.42 | 0.02 | 0.02 |
| gene-LOC105275374 | https://www.ncbi.nlm.nih.gov/gene/?term=LOC105275374 | 11376 | 587 | 516 | 1 | 0.88 | 0.17 | 9.68 | 9.27 | 0.41 | 0.41 | 0.02 | 0.02 |
| gene-LOC105284184 | https://www.ncbi.nlm.nih.gov/gene/?term=LOC105284184 | 2535 | 105 | 414.2 | 1 | 3.94 | 0.95 | 28.89 | 28.5 | 0.39 | 0.39 | 0.02 | 0.02 |
| gene-LOC105276462 | https://www.ncbi.nlm.nih.gov/gene/?term=LOC105276462 | 6561 | 289 | 440.48 | 1 | 1.52 | 0.35 | 8.58 | 8.21 | 0.38 | 0.38 | 0.02 | 0.02 |
| gene-LOC105278145 | https://www.ncbi.nlm.nih.gov/gene/?term=LOC105278145 | 4470 | 274 | 612.98 | 1 | 2.24 | 0.36 | 10.24 | 9.87 | 0.37 | 0.37 | 0.02 | 0.02 |
| gene-LOC105284043 | https://www.ncbi.nlm.nih.gov/gene/?term=LOC105284043 | 5661 | 312 | 551.14 | 1 | 1.77 | 0.32 | 10.26 | 9.89 | 0.37 | 0.37 | 0.02 | 0.02 |
| gene-LOC105276527 | https://www.ncbi.nlm.nih.gov/gene/?term=LOC105276527 | 7923 | 381 | 480.88 | 1 | 1.26 | 0.26 | 8.09 | 7.73 | 0.36 | 0.36 | 0.02 | 0.02 |
| gene-LOC105278877 | https://www.ncbi.nlm.nih.gov/gene/?term=LOC105278877 | 1968 | 91 | 462.4 | 2 | 10.16 | 2.2 | 30.61 | 31.07 | -0.45 | 0.45 | -0.02 | 0.02 |
| gene-LOC105284036 | https://www.ncbi.nlm.nih.gov/gene/?term=LOC105284036 | 3254 | 102 | 313.46 | 1 | 3.07 | 0.98 | 24.26 | 24.78 | -0.52 | 0.52 | -0.02 | 0.02 |
| gene-LOC105274851 | https://www.ncbi.nlm.nih.gov/gene/?term=LOC105274851 | 6616 | 160 | 241.84 | 1 | 1.51 | 0.62 | 10.97 | 11.63 | -0.66 | 0.66 | -0.02 | 0.02 |
| gene-LOC105275966 | https://www.ncbi.nlm.nih.gov/gene/?term=LOC105275966 | 3299 | 75 | 227.34 | 1 | 3.03 | 1.33 | 43.86 | 44.6 | -0.74 | 0.74 | -0.02 | 0.02 |
| gene-LOC105284507 | https://www.ncbi.nlm.nih.gov/gene/?term=LOC105284507 | 5207 | 123 | 236.22 | 1 | 1.92 | 0.81 | 18.34 | 19.09 | -0.75 | 0.75 | -0.02 | 0.02 |
| gene-LOC105280890 | https://www.ncbi.nlm.nih.gov/gene/?term=LOC105280890 | 1139 | 31 | 272.17 | 1 | 8.78 | 3.23 | 17.12 | 17.99 | -0.87 | 0.87 | -0.02 | 0.02 |
| gene-LOC105281173 | https://www.ncbi.nlm.nih.gov/gene/?term=LOC105281173 | 1595 | 39 | 244.51 | 1 | 6.27 | 2.56 | 61.21 | 62.08 | -0.87 | 0.87 | -0.02 | 0.02 |
| gene-LOC105282885 | https://www.ncbi.nlm.nih.gov/gene/?term=LOC105282885 | 1870 | 41 | 219.25 | 1 | 5.35 | 2.44 | 40.22 | 41.13 | -0.91 | 0.91 | -0.02 | 0.02 |
| gene-LOC105285252 | https://www.ncbi.nlm.nih.gov/gene/?term=LOC105285252 | 7427 | 148 | 199.27 | 1 | 1.35 | 0.68 | 11.91 | 12.93 | -1.01 | 1.01 | -0.02 | 0.02 |
| gene-LOC105287633 | https://www.ncbi.nlm.nih.gov/gene/?term=LOC105287633 | 1524 | 26 | 170.6 | 1 | 6.56 | 3.85 | 60.3 | 61.69 | -1.38 | 1.38 | -0.02 | 0.02 |
| gene-LOC105282914 | https://www.ncbi.nlm.nih.gov/gene/?term=LOC105282914 | 5623 | 43 | 76.47 | 1 | 1.78 | 2.33 | 47.44 | 49.79 | -2.35 | 2.35 | -0.02 | 0.02 |
| gene-LOC105279500 | https://www.ncbi.nlm.nih.gov/gene/?term=LOC105279500 | 3587 | 36 | 100.36 | 1 | 2.79 | 2.78 | 48.81 | 47.64 | 1.17 | 1.17 | 0.01 | 0.01 |
| gene-LOC105277856 | https://www.ncbi.nlm.nih.gov/gene/?term=LOC105277856 | 3442 | 36 | 104.59 | 1 | 2.91 | 2.78 | 37.81 | 36.75 | 1.06 | 1.06 | 0.01 | 0.01 |
| gene-LOC105275347 | https://www.ncbi.nlm.nih.gov/gene/?term=LOC105275347 | 2526 | 21 | 83.14 | 1 | 3.96 | 4.76 | 35.82 | 34.96 | 0.86 | 0.86 | 0.01 | 0.01 |
| gene-LOC105283989 | https://www.ncbi.nlm.nih.gov/gene/?term=LOC105283989 | 4948 | 44 | 88.92 | 2 | 4.04 | 4.55 | 37.47 | 36.79 | 0.68 | 0.68 | 0.01 | 0.01 |
| gene-LOC105283943 | https://www.ncbi.nlm.nih.gov/gene/?term=LOC105283943 | 3559 | 68 | 191.06 | 1 | 2.81 | 1.47 | 20.98 | 20.31 | 0.67 | 0.67 | 0.01 | 0.01 |
| gene-LOC105276560 | https://www.ncbi.nlm.nih.gov/gene/?term=LOC105276560 | 18607 | 384 | 206.37 | 3 | 1.61 | 0.78 | 2.53 | 1.87 | 0.66 | 0.66 | 0.01 | 0.01 |
| gene-LOC105288078 | https://www.ncbi.nlm.nih.gov/gene/?term=LOC105288078 | 19126 | 359 | 187.7 | 1 | 0.52 | 0.28 | 15.58 | 14.92 | 0.66 | 0.66 | 0.01 | 0.01 |
| gene-LOC105277215 | https://www.ncbi.nlm.nih.gov/gene/?term=LOC105277215 | 2619 | 30 | 114.55 | 1 | 3.82 | 3.33 | 27.61 | 26.99 | 0.61 | 0.61 | 0.01 | 0.01 |
| gene-LOC105281296 | https://www.ncbi.nlm.nih.gov/gene/?term=LOC105281296 | 2434 | 48 | 197.21 | 1 | 4.11 | 2.08 | 28.14 | 27.55 | 0.59 | 0.59 | 0.01 | 0.01 |
| gene-LOC105278384 | https://www.ncbi.nlm.nih.gov/gene/?term=LOC105278384 | 1390 | 33 | 237.41 | 1 | 7.19 | 3.03 | 46.32 | 45.77 | 0.55 | 0.55 | 0.01 | 0.01 |

| | | | | | | | | | | | | | |
|---|---|---|---|---|---|---|---|---|---|---|---|---|---|
| gene-LOC105274699 | https://www.ncbi.nlm.nih.gov/gene/?term=LOC105274699 | 2386 | 67 | 280.8 | 1 | 4.19 | 1.49 | 18.3 | 17.79 | 0.51 | 0.51 | 0.01 | 0.01 |
| gene-LOC105280641 | https://www.ncbi.nlm.nih.gov/gene/?term=LOC105280641 | 3682 | 78 | 211.84 | 1 | 2.72 | 1.28 | 13.91 | 13.42 | 0.49 | 0.49 | 0.01 | 0.01 |
| gene-LOC105274773 | https://www.ncbi.nlm.nih.gov/gene/?term=LOC105274773 | 11034 | 187 | 169.48 | 1 | 0.91 | 0.53 | 16.91 | 16.52 | 0.39 | 0.39 | 0.01 | 0.01 |
| gene-LOC105281700 | https://www.ncbi.nlm.nih.gov/gene/?term=LOC105281700 | 2208 | 43 | 194.75 | 1 | 4.53 | 2.33 | 36.88 | 36.49 | 0.39 | 0.39 | 0.01 | 0.01 |
| gene-LOC105278962 | https://www.ncbi.nlm.nih.gov/gene/?term=LOC105278962 | 4491 | 129 | 287.24 | 1 | 2.23 | 0.78 | 19.14 | 18.79 | 0.35 | 0.35 | 0.01 | 0.01 |
| gene-LOC105274534 | https://www.ncbi.nlm.nih.gov/gene/?term=LOC105274534 | 1329 | 60 | 451.47 | 1 | 7.52 | 1.67 | 44.47 | 44.14 | 0.32 | 0.32 | 0.01 | 0.01 |
| gene-LOC105286332 | https://www.ncbi.nlm.nih.gov/gene/?term=LOC105286332 | 27577 | 573 | 207.78 | 1 | 0.36 | 0.17 | 7.89 | 7.57 | 0.32 | 0.32 | 0.01 | 0.01 |
| gene-LOC105285395 | https://www.ncbi.nlm.nih.gov/gene/?term=LOC105285395 | 8085 | 260 | 321.58 | 1 | 1.24 | 0.38 | 13.21 | 12.9 | 0.31 | 0.31 | 0.01 | 0.01 |
| gene-LOC105286265 | https://www.ncbi.nlm.nih.gov/gene/?term=LOC105286265 | 2507 | 68 | 271.24 | 1 | 3.99 | 1.47 | 31.01 | 30.72 | 0.29 | 0.29 | 0.01 | 0.01 |
| gene-LOC105278666 | https://www.ncbi.nlm.nih.gov/gene/?term=LOC105278666 | 10341 | 492 | 475.78 | 1 | 0.97 | 0.2 | 7.02 | 6.75 | 0.27 | 0.27 | 0.01 | 0.01 |
| gene-LOC105274449 | https://www.ncbi.nlm.nih.gov/gene/?term=LOC105274449 | 1549 | 53 | 342.16 | 1 | 6.46 | 1.89 | 49.51 | 49.29 | 0.22 | 0.22 | 0.01 | 0.01 |
| gene-LOC105275327 | https://www.ncbi.nlm.nih.gov/gene/?term=LOC105275327 | 14245 | 550 | 386.1 | 1 | 0.7 | 0.18 | 6.58 | 6.38 | 0.2 | 0.2 | 0.01 | 0.01 |
| gene-LOC105284262 | https://www.ncbi.nlm.nih.gov/gene/?term=LOC105284262 | 6669 | 248 | 371.87 | 1 | 1.5 | 0.4 | 5.25 | 5.05 | 0.2 | 0.2 | 0.01 | 0.01 |
| gene-LOC113562050 | https://www.ncbi.nlm.nih.gov/gene/?term=LOC113562050 | 17097 | 698 | 408.26 | 1 | 0.58 | 0.14 | 5.09 | 4.89 | 0.2 | 0.2 | 0.01 | 0.01 |
| gene-LOC105278534 | https://www.ncbi.nlm.nih.gov/gene/?term=LOC105278534 | 7364 | 214 | 290.6 | 1 | 1.36 | 0.47 | 16.92 | 16.72 | 0.19 | 0.19 | 0.01 | 0.01 |
| gene-LOC105274856 | https://www.ncbi.nlm.nih.gov/gene/?term=LOC105274856 | 5216 | 233 | 446.7 | 1 | 1.92 | 0.43 | 17.97 | 17.83 | 0.14 | 0.14 | 0.01 | 0.01 |
| gene-LOC105287065 | https://www.ncbi.nlm.nih.gov/gene/?term=LOC105287065 | 4859 | 259 | 533.03 | 1 | 2.06 | 0.39 | 12.11 | 12 | 0.11 | 0.11 | 0.01 | 0.01 |
| gene-LOC105286649 | https://www.ncbi.nlm.nih.gov/gene/?term=LOC105286649 | 14178 | 852 | 600.93 | 1 | 0.71 | 0.12 | 1.32 | 1.49 | -0.16 | 0.16 | -0.01 | 0.01 |
| gene-LOC105284116 | https://www.ncbi.nlm.nih.gov/gene/?term=LOC105284116 | 3970 | 134 | 337.53 | 1 | 2.52 | 0.75 | 7.83 | 8 | -0.17 | 0.17 | -0.01 | 0.01 |
| gene-LOC105278455 | https://www.ncbi.nlm.nih.gov/gene/?term=LOC105278455 | 27181 | 1036 | 381.15 | 1 | 0.37 | 0.1 | 6.32 | 6.5 | -0.18 | 0.18 | -0.01 | 0.01 |
| gene-LOC105287058 | https://www.ncbi.nlm.nih.gov/gene/?term=LOC105287058 | 4788 | 247 | 515.87 | 1 | 2.09 | 0.4 | 8.85 | 9.06 | -0.2 | 0.2 | -0.01 | 0.01 |
| gene-LOC105278771 | https://www.ncbi.nlm.nih.gov/gene/?term=LOC105278771 | 10457 | 267 | 255.33 | 1 | 0.96 | 0.37 | 9.06 | 9.27 | -0.21 | 0.21 | -0.01 | 0.01 |
| gene-LOC105286515 | https://www.ncbi.nlm.nih.gov/gene/?term=LOC105286515 | 3751 | 133 | 354.57 | 1 | 2.67 | 0.75 | 10.29 | 10.56 | -0.27 | 0.27 | -0.01 | 0.01 |
| gene-LOC105285981 | https://www.ncbi.nlm.nih.gov/gene/?term=LOC105285981 | 7666 | 224 | 292.2 | 1 | 1.3 | 0.45 | 17.35 | 17.63 | -0.28 | 0.28 | -0.01 | 0.01 |
| gene-LOC105280956 | https://www.ncbi.nlm.nih.gov/gene/?term=LOC105280956 | 2639 | 83 | 314.51 | 1 | 3.79 | 1.2 | 18.28 | 18.58 | -0.3 | 0.3 | -0.01 | 0.01 |
| gene-LOC105283649 | https://www.ncbi.nlm.nih.gov/gene/?term=LOC105283649 | 1918 | 59 | 307.61 | 2 | 10.43 | 3.39 | 41.12 | 41.42 | -0.31 | 0.31 | -0.01 | 0.01 |
| gene-LOC105285202 | https://www.ncbi.nlm.nih.gov/gene/?term=LOC105285202 | 1838 | 74 | 402.61 | 1 | 5.44 | 1.35 | 23.68 | 24 | -0.32 | 0.32 | -0.01 | 0.01 |
| gene-LOC105283132 | https://www.ncbi.nlm.nih.gov/gene/?term=LOC105283132 | 1819 | 56 | 307.86 | 1 | 5.5 | 1.79 | 39.81 | 40.18 | -0.38 | 0.38 | -0.01 | 0.01 |
| gene-LOC105280304 | https://www.ncbi.nlm.nih.gov/gene/?term=LOC105280304 | 8919 | 143 | 160.33 | 1 | 1.12 | 0.7 | 6.95 | 7.37 | -0.42 | 0.42 | -0.01 | 0.01 |
| gene-LOC105278454 | https://www.ncbi.nlm.nih.gov/gene/?term=LOC105278454 | 1972 | 43 | 218.05 | 1 | 5.07 | 2.33 | 42.19 | 42.86 | -0.67 | 0.67 | -0.01 | 0.01 |
| gene-LOC105280408 | https://www.ncbi.nlm.nih.gov/gene/?term=LOC105280408 | 5983 | 65 | 108.64 | 1 | 1.67 | 1.54 | 15.19 | 14.75 | 0.44 | 0.44 | 0 | 0 |
| gene-LOC105285406 | https://www.ncbi.nlm.nih.gov/gene/?term=LOC105285406 | 8015 | 63 | 78.6 | 1 | 1.25 | 1.59 | 28.98 | 28.57 | 0.41 | 0.41 | 0 | 0 |
| gene-LOC105275187 | https://www.ncbi.nlm.nih.gov/gene/?term=LOC105275187 | 5534 | 87 | 157.21 | 1 | 1.81 | 1.15 | 15.13 | 14.88 | 0.25 | 0.25 | 0 | 0 |
| gene-LOC105286194 | https://www.ncbi.nlm.nih.gov/gene/?term=LOC105286194 | 152619 | 2577 | 168.85 | 1 | 0.07 | 0.04 | 6.51 | 6.31 | 0.19 | 0.19 | 0 | 0 |
| gene-LOC105274590 | https://www.ncbi.nlm.nih.gov/gene/?term=LOC105274590 | 4516 | 88 | 194.86 | 2 | 4.43 | 2.27 | 24.95 | 24.81 | 0.14 | 0.14 | 0 | 0 |
| gene-LOC105274669 | https://www.ncbi.nlm.nih.gov/gene/?term=LOC105274669 | 8555 | 274 | 320.28 | 2 | 2.34 | 0.73 | 5.92 | 5.78 | 0.14 | 0.14 | 0 | 0 |
| gene-LOC105277866 | https://www.ncbi.nlm.nih.gov/gene/?term=LOC105277866 | 33045 | 539 | 163.11 | 1 | 0.3 | 0.19 | 3.2 | 3.07 | 0.14 | 0.14 | 0 | 0 |
| gene-LOC105286452 | https://www.ncbi.nlm.nih.gov/gene/?term=LOC105286452 | 6888 | 78 | 113.24 | 1 | 1.45 | 1.28 | 26.06 | 25.92 | 0.14 | 0.14 | 0 | 0 |
| gene-LOC105286770 | https://www.ncbi.nlm.nih.gov/gene/?term=LOC105286770 | 3055 | 50 | 163.67 | 1 | 3.27 | 2 | 14.75 | 14.61 | 0.14 | 0.14 | 0 | 0 |
| gene-LOC105280410 | https://www.ncbi.nlm.nih.gov/gene/?term=LOC105280410 | 12580 | 320 | 254.37 | 1 | 0.79 | 0.31 | 17.81 | 17.7 | 0.11 | 0.11 | 0 | 0 |
| gene-LOC105278109 | https://www.ncbi.nlm.nih.gov/gene/?term=LOC105278109 | 4029 | 89 | 220.9 | 1 | 2.48 | 1.12 | 34.5 | 34.41 | 0.09 | 0.09 | 0 | 0 |
| gene-LOC105282811 | https://www.ncbi.nlm.nih.gov/gene/?term=LOC105282811 | 5765 | 232 | 402.43 | 1 | 1.73 | 0.43 | 8.6 | 8.51 | 0.09 | 0.09 | 0 | 0 |
| gene-LOC105287095 | https://www.ncbi.nlm.nih.gov/gene/?term=LOC105287095 | 1435 | 71 | 494.77 | 1 | 6.97 | 1.41 | 23.64 | 23.55 | 0.09 | 0.09 | 0 | 0 |
| gene-LOC105276917 | https://www.ncbi.nlm.nih.gov/gene/?term=LOC105276917 | 2336 | 42 | 179.79 | 1 | 4.28 | 2.38 | 28.76 | 28.68 | 0.08 | 0.08 | 0 | 0 |
| gene-LOC105276059 | https://www.ncbi.nlm.nih.gov/gene/?term=LOC105276059 | 6256 | 148 | 236.57 | 1 | 1.6 | 0.68 | 11.97 | 11.9 | 0.07 | 0.07 | 0 | 0 |
| gene-LOC105283264 | https://www.ncbi.nlm.nih.gov/gene/?term=LOC105283264 | 4005 | 94 | 234.71 | 1 | 2.5 | 1.06 | 22.97 | 22.91 | 0.07 | 0.07 | 0 | 0 |
| gene-LOC105283296 | https://www.ncbi.nlm.nih.gov/gene/?term=LOC105283296 | 2174 | 96 | 441.58 | 1 | 4.6 | 1.04 | 14.24 | 14.2 | 0.04 | 0.04 | 0 | 0 |
| gene-LOC105280819 | https://www.ncbi.nlm.nih.gov/gene/?term=LOC105280819 | 1639 | 44 | 268.46 | 1 | 6.1 | 2.27 | 37.71 | 37.7 | 0.02 | 0.02 | 0 | 0 |
| gene-LOC105280513 | https://www.ncbi.nlm.nih.gov/gene/?term=LOC105280513 | 69542 | 1608 | 231.23 | 1 | 0.14 | 0.06 | 0.74 | 0.72 | 0.01 | 0.01 | 0 | 0 |
| gene-LOC105277216 | https://www.ncbi.nlm.nih.gov/gene/?term=LOC105277216 | 7627 | 101 | 132.42 | 1 | 1.31 | 0.99 | 8.48 | 8.48 | 0 | 0 | 0 | 0 |
| gene-LOC105279664 | https://www.ncbi.nlm.nih.gov/gene/?term=LOC105279664 | 4023 | 75 | 186.43 | 1 | 2.49 | 1.33 | 19.05 | 19.05 | 0 | 0 | 0 | 0 |
| gene-LOC105277125 | https://www.ncbi.nlm.nih.gov/gene/?term=LOC105277125 | 51079 | 2092 | 409.56 | 1 | 0.2 | 0.05 | 1.63 | 1.65 | -0.02 | 0.02 | 0 | 0 |
| gene-LOC105279528 | https://www.ncbi.nlm.nih.gov/gene/?term=LOC105279528 | 5802 | 40 | 68.94 | 1 | 1.72 | 2.5 | 19.68 | 19.72 | -0.04 | 0.04 | 0 | 0 |
| gene-LOC105280265 | https://www.ncbi.nlm.nih.gov/gene/?term=LOC105280265 | 85984 | 3486 | 405.42 | 1 | 0.12 | 0.03 | 1.07 | 1.12 | -0.04 | 0.04 | 0 | 0 |
| gene-LOC105277588 | https://www.ncbi.nlm.nih.gov/gene/?term=LOC105277588 | 3955 | 98 | 247.79 | 1 | 2.53 | 1.02 | 8.55 | 8.61 | -0.06 | 0.06 | 0 | 0 |
| gene-LOC105277761 | https://www.ncbi.nlm.nih.gov/gene/?term=LOC105277761 | 4648 | 174 | 374.35 | 1 | 2.15 | 0.57 | 14.23 | 14.3 | -0.07 | 0.07 | 0 | 0 |
| gene-LOC105279657 | https://www.ncbi.nlm.nih.gov/gene/?term=LOC105279657 | 5878 | 112 | 190.54 | 1 | 1.7 | 0.89 | 16.48 | 16.55 | -0.07 | 0.07 | 0 | 0 |
| gene-LOC105284774 | https://www.ncbi.nlm.nih.gov/gene/?term=LOC105284774 | 12089 | 106 | 87.68 | 1 | 0.83 | 0.94 | 19.59 | 19.72 | -0.12 | 0.12 | 0 | 0 |
| gene-LOC105276196 | https://www.ncbi.nlm.nih.gov/gene/?term=LOC105276196 | 2953 | 61 | 206.57 | 1 | 3.39 | 1.64 | 38.98 | 39.11 | -0.13 | 0.13 | 0 | 0 |
| gene-LOC105284117 | https://www.ncbi.nlm.nih.gov/gene/?term=LOC105284117 | 4340 | 116 | 267.28 | 1 | 2.3 | 0.86 | 11.19 | 11.33 | -0.14 | 0.14 | 0 | 0 |

Directory: OGLall File: rnk-dmg-Ob_LI_RphaseA2.vs.RphaseB2.txt

| gene_ID | gene_link | gwidth | #Sites | #per10Kb | #dmSites | #dmsp10kb | %dmSites | %pSite1 | %pSite2 | DMpSite | ADMpSite | DMpNucl | ADMpNucl |
|---|---|---|---|---|---|---|---|---|---|---|---|---|---|
| gene-LOC113562345 | https://www.ncbi.nlm.nih.gov/gene/?term=LOC113562345 | 2290 | 69 | 301.31 | 14 | 61.14 | 20.29 | 29.74 | 16.81 | 12.93 | 12.93 | 0.39 | 0.39 |
| gene-LOC105284677 | https://www.ncbi.nlm.nih.gov/gene/?term=LOC105284677 | 1905 | 42 | 220.47 | 1 | 5.25 | 2.38 | 23.81 | 34.99 | -11.18 | 11.18 | -0.25 | 0.25 |
| gene-LOC113562901 | https://www.ncbi.nlm.nih.gov/gene/?term=LOC113562901 | 1531 | 24 | 156.76 | 3 | 19.6 | 12.5 | 59.3 | 45.31 | 14 | 14 | 0.22 | 0.22 |
| gene-LOC105286644 | https://www.ncbi.nlm.nih.gov/gene/?term=LOC105286644 | 1235 | 22 | 178.14 | 1 | 8.1 | 4.55 | 44.4 | 37.26 | 7.14 | 7.14 | 0.13 | 0.13 |
| gene-LOC105280786 | https://www.ncbi.nlm.nih.gov/gene/?term=LOC105280786 | 2573 | 127 | 493.59 | 1 | 3.89 | 0.79 | 30.11 | 27.41 | 2.7 | 2.7 | 0.13 | 0.13 |
| gene-LOC105274706 | https://www.ncbi.nlm.nih.gov/gene/?term=LOC105274706 | 1446 | 56 | 387.28 | 1 | 6.92 | 1.79 | 39.74 | 36.7 | 3.05 | 3.05 | 0.12 | 0.12 |
| gene-LOC105275447 | https://www.ncbi.nlm.nih.gov/gene/?term=LOC105275447 | 1846 | 31 | 167.93 | 1 | 5.42 | 3.23 | 9.13 | 2.78 | 6.34 | 6.34 | 0.11 | 0.11 |
| gene-LOC105286683 | https://www.ncbi.nlm.nih.gov/gene/?term=LOC105286683 | 1665 | 57 | 342.34 | 2 | 12.01 | 3.51 | 34.89 | 31.76 | 3.13 | 3.13 | 0.11 | 0.11 |
| gene-LOC105285671 | https://www.ncbi.nlm.nih.gov/gene/?term=LOC105285671 | 2263 | 106 | 468.4 | 1 | 4.42 | 0.94 | 28.93 | 26.53 | 2.4 | 2.4 | 0.11 | 0.11 |
| gene-LOC105287417 | https://www.ncbi.nlm.nih.gov/gene/?term=LOC105287417 | 1025 | 23 | 224.39 | 1 | 9.76 | 4.35 | 53.58 | 49.27 | 4.31 | 4.31 | 0.1 | 0.1 |
| gene-LOC105285583 | https://www.ncbi.nlm.nih.gov/gene/?term=LOC105285583 | 1947 | 49 | 251.67 | 1 | 5.14 | 2.04 | 30.89 | 26.83 | 4.06 | 4.06 | 0.1 | 0.1 |
| gene-LOC105285674 | https://www.ncbi.nlm.nih.gov/gene/?term=LOC105285674 | 3035 | 102 | 336.08 | 2 | 6.59 | 1.96 | 26.23 | 23.21 | 3.03 | 3.03 | 0.1 | 0.1 |
| gene-LOC105276704 | https://www.ncbi.nlm.nih.gov/gene/?term=LOC105276704 | 2063 | 88 | 426.56 | 1 | 4.85 | 1.14 | 16.14 | 13.92 | 2.23 | 2.23 | 0.1 | 0.1 |
| gene-LOC105282156 | https://www.ncbi.nlm.nih.gov/gene/?term=LOC105282156 | 2577 | 130 | 504.46 | 1 | 3.88 | 0.77 | 28.69 | 26.74 | 1.95 | 1.95 | 0.1 | 0.1 |
| gene-LOC105275070 | https://www.ncbi.nlm.nih.gov/gene/?term=LOC105275070 | 5667 | 42 | 74.11 | 5 | 8.82 | 11.9 | 13.23 | 1.03 | 12.2 | 12.2 | 0.09 | 0.09 |
| gene-LOC105279898 | https://www.ncbi.nlm.nih.gov/gene/?term=LOC105279898 | 1343 | 10 | 74.46 | 1 | 7.45 | 10 | 83.16 | 71.53 | 11.63 | 11.63 | 0.09 | 0.09 |
| gene-LOC105282388 | https://www.ncbi.nlm.nih.gov/gene/?term=LOC105282388 | 1661 | 34 | 204.7 | 1 | 6.02 | 2.94 | 48.95 | 44.6 | 4.35 | 4.35 | 0.09 | 0.09 |
| gene-LOC105280555 | https://www.ncbi.nlm.nih.gov/gene/?term=LOC105280555 | 1496 | 31 | 207.22 | 1 | 6.68 | 3.23 | 61.05 | 57.2 | 3.85 | 3.85 | 0.08 | 0.08 |
| gene-LOC105277631 | https://www.ncbi.nlm.nih.gov/gene/?term=LOC105277631 | 4387 | 304 | 692.96 | 1 | 2.28 | 0.33 | 10.76 | 9.61 | 1.15 | 1.15 | 0.08 | 0.08 |
| gene-LOC105282893 | https://www.ncbi.nlm.nih.gov/gene/?term=LOC105282893 | 2645 | 47 | 177.69 | 1 | 3.78 | 2.13 | 30.08 | 25.9 | 4.18 | 4.18 | 0.07 | 0.07 |
| gene-LOC105280104 | https://www.ncbi.nlm.nih.gov/gene/?term=LOC105280104 | 3960 | 139 | 351.01 | 1 | 2.53 | 0.72 | 17.44 | 15.39 | 2.05 | 2.05 | 0.07 | 0.07 |
| gene-LOC105280377 | https://www.ncbi.nlm.nih.gov/gene/?term=LOC105280377 | 2024 | 70 | 345.85 | 1 | 4.94 | 1.43 | 48.1 | 46.12 | 1.98 | 1.98 | 0.07 | 0.07 |
| gene-LOC105282947 | https://www.ncbi.nlm.nih.gov/gene/?term=LOC105282947 | 1162 | 41 | 352.84 | 1 | 8.61 | 2.44 | 36.57 | 34.72 | 1.85 | 1.85 | 0.07 | 0.07 |
| gene-LOC105287063 | https://www.ncbi.nlm.nih.gov/gene/?term=LOC105287063 | 1874 | 48 | 256.14 | 1 | 5.34 | 2.08 | 32.76 | 30.33 | 2.44 | 2.44 | 0.06 | 0.06 |
| gene-LOC105283132 | https://www.ncbi.nlm.nih.gov/gene/?term=LOC105283132 | 1819 | 57 | 313.36 | 1 | 5.5 | 1.75 | 39.92 | 37.88 | 2.04 | 2.04 | 0.06 | 0.06 |
| gene-LOC105283182 | https://www.ncbi.nlm.nih.gov/gene/?term=LOC105283182 | 3585 | 131 | 365.41 | 1 | 2.79 | 0.76 | 14.16 | 12.63 | 1.52 | 1.52 | 0.06 | 0.06 |
| gene-LOC105276449 | https://www.ncbi.nlm.nih.gov/gene/?term=LOC105276449 | 1231 | 50 | 406.17 | 1 | 8.12 | 2 | 32.79 | 31.42 | 1.37 | 1.37 | 0.06 | 0.06 |
| gene-LOC105278497 | https://www.ncbi.nlm.nih.gov/gene/?term=LOC105278497 | 2072 | 100 | 482.63 | 1 | 4.83 | 1 | 19.97 | 18.78 | 1.2 | 1.2 | 0.06 | 0.06 |
| gene-LOC105284043 | https://www.ncbi.nlm.nih.gov/gene/?term=LOC105284043 | 5661 | 308 | 544.07 | 1 | 1.77 | 0.32 | 10.53 | 9.46 | 1.06 | 1.06 | 0.06 | 0.06 |
| gene-LOC105278283 | https://www.ncbi.nlm.nih.gov/gene/?term=LOC105278283 | 2317 | 142 | 612.86 | 1 | 4.32 | 0.7 | 25.61 | 24.56 | 1.05 | 1.05 | 0.06 | 0.06 |
| gene-LOC105282149 | https://www.ncbi.nlm.nih.gov/gene/?term=LOC105282149 | 1787 | 110 | 615.56 | 1 | 5.6 | 0.91 | 43.04 | 42 | 1.04 | 1.04 | 0.06 | 0.06 |
| gene-LOC105275708 | https://www.ncbi.nlm.nih.gov/gene/?term=LOC105275708 | 1772 | 15 | 84.65 | 1 | 5.64 | 6.67 | 47.19 | 40.83 | 6.36 | 6.36 | 0.05 | 0.05 |
| gene-LOC105276538 | https://www.ncbi.nlm.nih.gov/gene/?term=LOC105276538 | 1511 | 20 | 132.36 | 1 | 6.62 | 5 | 73.31 | 69.21 | 4.1 | 4.1 | 0.05 | 0.05 |
| gene-LOC105278215 | https://www.ncbi.nlm.nih.gov/gene/?term=LOC105278215 | 1700 | 29 | 170.59 | 1 | 5.88 | 3.45 | 57.59 | 54.66 | 2.93 | 2.93 | 0.05 | 0.05 |
| gene-LOC105278109 | https://www.ncbi.nlm.nih.gov/gene/?term=LOC105278109 | 4029 | 67 | 166.29 | 1 | 2.48 | 1.49 | 44.53 | 41.68 | 2.86 | 2.86 | 0.05 | 0.05 |
| gene-LOC105282794 | https://www.ncbi.nlm.nih.gov/gene/?term=LOC105282794 | 4571 | 95 | 207.83 | 1 | 2.19 | 1.05 | 18.64 | 16.23 | 2.41 | 2.41 | 0.05 | 0.05 |
| gene-LOC105287646 | https://www.ncbi.nlm.nih.gov/gene/?term=LOC105287646 | 3556 | 90 | 253.09 | 1 | 2.81 | 1.11 | 18.4 | 16.59 | 1.81 | 1.81 | 0.05 | 0.05 |
| gene-LOC105278282 | https://www.ncbi.nlm.nih.gov/gene/?term=LOC105278282 | 1775 | 56 | 315.49 | 1 | 5.63 | 1.79 | 39.77 | 38.34 | 1.43 | 1.43 | 0.05 | 0.05 |
| gene-LOC105276875 | https://www.ncbi.nlm.nih.gov/gene/?term=LOC105276875 | 3151 | 118 | 374.48 | 1 | 3.17 | 0.85 | 23.91 | 22.53 | 1.38 | 1.38 | 0.05 | 0.05 |
| gene-LOC105284355 | https://www.ncbi.nlm.nih.gov/gene/?term=LOC105284355 | 3904 | 153 | 391.91 | 1 | 2.56 | 0.65 | 21.82 | 20.47 | 1.35 | 1.35 | 0.05 | 0.05 |
| gene-LOC105283728 | https://www.ncbi.nlm.nih.gov/gene/?term=LOC105283728 | 3256 | 120 | 368.55 | 2 | 6.14 | 1.67 | 19.01 | 17.69 | 1.32 | 1.32 | 0.05 | 0.05 |
| gene-LOC105281419 | https://www.ncbi.nlm.nih.gov/gene/?term=LOC105281419 | 2366 | 83 | 350.8 | 1 | 4.23 | 1.2 | 23.42 | 22.13 | 1.3 | 1.3 | 0.05 | 0.05 |
| gene-LOC105275363 | https://www.ncbi.nlm.nih.gov/gene/?term=LOC105275363 | 3798 | 165 | 434.44 | 1 | 2.63 | 0.61 | 17.38 | 16.26 | 1.12 | 1.12 | 0.05 | 0.05 |
| gene-LOC105279298 | https://www.ncbi.nlm.nih.gov/gene/?term=LOC105279298 | 2773 | 136 | 490.44 | 2 | 7.21 | 1.47 | 23.98 | 22.86 | 1.11 | 1.11 | 0.05 | 0.05 |
| gene-LOC105282682 | https://www.ncbi.nlm.nih.gov/gene/?term=LOC105282682 | 4127 | 223 | 540.34 | 1 | 2.42 | 0.45 | 13.69 | 12.76 | 0.93 | 0.93 | 0.05 | 0.05 |
| gene-LOC105285055 | https://www.ncbi.nlm.nih.gov/gene/?term=LOC105285055 | 4108 | 276 | 671.86 | 2 | 4.87 | 0.72 | 11.41 | 10.6 | 0.8 | 0.8 | 0.05 | 0.05 |
| gene-LOC105285534 | https://www.ncbi.nlm.nih.gov/gene/?term=LOC105285534 | 3944 | 237 | 600.91 | 1 | 2.54 | 0.42 | 11.45 | 10.67 | 0.78 | 0.78 | 0.05 | 0.05 |
| gene-LOC105287854 | https://www.ncbi.nlm.nih.gov/gene/?term=LOC105287854 | 3644 | 54 | 148.19 | 1 | 2.74 | 1.85 | 25.58 | 23.15 | 2.43 | 2.43 | 0.04 | 0.04 |
| gene-LOC105276917 | https://www.ncbi.nlm.nih.gov/gene/?term=LOC105276917 | 2336 | 38 | 162.67 | 1 | 4.28 | 2.63 | 28.71 | 26.35 | 2.36 | 2.36 | 0.04 | 0.04 |
| gene-LOC113561805 | https://www.ncbi.nlm.nih.gov/gene/?term=LOC113561805 | 3337 | 71 | 212.77 | 1 | 3 | 1.41 | 36.21 | 34.33 | 1.88 | 1.88 | 0.04 | 0.04 |
| gene-LOC105283649 | https://www.ncbi.nlm.nih.gov/gene/?term=LOC105283649 | 1918 | 56 | 291.97 | 2 | 10.43 | 3.57 | 37.98 | 36.48 | 1.5 | 1.5 | 0.04 | 0.04 |
| gene-LOC105282970 | https://www.ncbi.nlm.nih.gov/gene/?term=LOC105282970 | 1529 | 40 | 261.61 | 1 | 6.54 | 2.5 | 49.63 | 48.22 | 1.41 | 1.41 | 0.04 | 0.04 |
| gene-LOC105281272 | https://www.ncbi.nlm.nih.gov/gene/?term=LOC105281272 | 5252 | 164 | 312.26 | 1 | 1.9 | 0.61 | 19.36 | 17.96 | 1.4 | 1.4 | 0.04 | 0.04 |
| gene-LOC105284704 | https://www.ncbi.nlm.nih.gov/gene/?term=LOC105284704 | 2400 | 62 | 258.33 | 1 | 4.17 | 1.61 | 24.3 | 22.92 | 1.38 | 1.38 | 0.04 | 0.04 |
| gene-LOC105283123 | https://www.ncbi.nlm.nih.gov/gene/?term=LOC105283123 | 1507 | 48 | 318.51 | 1 | 6.64 | 2.08 | 17.79 | 16.48 | 1.31 | 1.31 | 0.04 | 0.04 |
| gene-LOC105288079 | https://www.ncbi.nlm.nih.gov/gene/?term=LOC105288079 | 3473 | 101 | 290.81 | 1 | 2.88 | 0.99 | 20.37 | 19.15 | 1.21 | 1.21 | 0.04 | 0.04 |
| gene-LOC105277238 | https://www.ncbi.nlm.nih.gov/gene/?term=LOC105277238 | 2708 | 88 | 324.96 | 1 | 3.69 | 1.14 | 27.16 | 25.96 | 1.2 | 1.2 | 0.04 | 0.04 |
| gene-LOC105277761 | https://www.ncbi.nlm.nih.gov/gene/?term=LOC105277761 | 4648 | 150 | 322.72 | 1 | 2.15 | 0.67 | 14.38 | 13.19 | 1.19 | 1.19 | 0.04 | 0.04 |
| gene-LOC105284184 | https://www.ncbi.nlm.nih.gov/gene/?term=LOC105284184 | 2535 | 86 | 339.25 | 1 | 3.94 | 1.16 | 34.02 | 32.88 | 1.13 | 1.13 | 0.04 | 0.04 |
| gene-LOC113561621 | https://www.ncbi.nlm.nih.gov/gene/?term=LOC113561621 | 2249 | 86 | 382.39 | 1 | 4.45 | 1.16 | 20.44 | 19.32 | 1.11 | 1.11 | 0.04 | 0.04 |

| | | | | | | | | | | | | | | |
|---|---|---|---|---|---|---|---|---|---|---|---|---|---|---|
| gene-LOC105285639 | https://www.ncbi.nlm.nih.gov/gene/?term=LOC105285639 | 10325 | 349 | 338.01 | | 1 | 0.97 | 0.29 | 13.03 | 11.95 | 1.08 | 1.08 | 0.04 | 0.04 |
| gene-LOC105285742 | https://www.ncbi.nlm.nih.gov/gene/?term=LOC105285742 | 3446 | 123 | 356.94 | | 1 | 2.9 | 0.81 | 20.56 | 19.51 | 1.04 | 1.04 | 0.04 | 0.04 |
| gene-LOC105274856 | https://www.ncbi.nlm.nih.gov/gene/?term=LOC105274856 | 5216 | 216 | 414.11 | | 1 | 1.92 | 0.46 | 19.8 | 18.89 | 0.91 | 0.91 | 0.04 | 0.04 |
| gene-LOC105275460 | https://www.ncbi.nlm.nih.gov/gene/?term=LOC105275460 | 2473 | 117 | 473.11 | | 1 | 4.04 | 0.85 | 14.05 | 13.26 | 0.78 | 0.78 | 0.04 | 0.04 |
| gene-LOC105284047 | https://www.ncbi.nlm.nih.gov/gene/?term=LOC105284047 | 1212 | 24 | 198.02 | | 1 | 8.25 | 4.17 | 25.81 | 27.68 | -1.87 | 1.87 | -0.04 | 0.04 |
| gene-LOC105274778 | https://www.ncbi.nlm.nih.gov/gene/?term=LOC105274778 | 4662 | 40 | 85.8 | 1 | 2.15 | 2.5 | 47.99 | 44.37 | 3.62 | 3.62 | 0.03 | 0.03 | |
| gene-LOC105275518 | https://www.ncbi.nlm.nih.gov/gene/?term=LOC105275518 | 2335 | 24 | 102.78 | | 1 | 4.28 | 4.17 | 49.73 | 46.6 | 3.13 | 3.13 | 0.03 | 0.03 |
| gene-LOC105279664 | https://www.ncbi.nlm.nih.gov/gene/?term=LOC105279664 | 4023 | 56 | 139.2 | 1 | 2.49 | 1.79 | 21.11 | 18.99 | 2.12 | 2.12 | 0.03 | 0.03 | |
| gene-LOC105280641 | https://www.ncbi.nlm.nih.gov/gene/?term=LOC105280641 | 3682 | 53 | 143.94 | | 1 | 2.72 | 1.89 | 17.27 | 15.26 | 2.01 | 2.01 | 0.03 | 0.03 |
| gene-LOC105282600 | https://www.ncbi.nlm.nih.gov/gene/?term=LOC105282600 | 3715 | 56 | 150.74 | | 1 | 2.69 | 1.79 | 27.25 | 25.37 | 1.87 | 1.87 | 0.03 | 0.03 |
| gene-LOC105276190 | https://www.ncbi.nlm.nih.gov/gene/?term=LOC105276190 | 4839 | 93 | 192.19 | | 1 | 2.07 | 1.08 | 11.07 | 9.37 | 1.7 | 1.7 | 0.03 | 0.03 |
| gene-LOC105281420 | https://www.ncbi.nlm.nih.gov/gene/?term=LOC105281420 | 3882 | 62 | 159.71 | | 1 | 2.58 | 1.61 | 15.69 | 14 | 1.7 | 1.7 | 0.03 | 0.03 |
| gene-LOC105276692 | https://www.ncbi.nlm.nih.gov/gene/?term=LOC105276692 | 1745 | 35 | 200.57 | | 1 | 5.73 | 2.86 | 22.12 | 20.68 | 1.44 | 1.44 | 0.03 | 0.03 |
| gene-LOC105282151 | https://www.ncbi.nlm.nih.gov/gene/?term=LOC105282151 | 1601 | 39 | 243.6 | 2 | 12.49 | 5.13 | 42.25 | 41 | 1.25 | 1.25 | 0.03 | 0.03 | |
| gene-LOC105281848 | https://www.ncbi.nlm.nih.gov/gene/?term=LOC105281848 | 2122 | 48 | 226.2 | 1 | 4.71 | 2.08 | 27.66 | 26.43 | 1.23 | 1.23 | 0.03 | 0.03 | |
| gene-LOC105281173 | https://www.ncbi.nlm.nih.gov/gene/?term=LOC105281173 | 1595 | 39 | 244.51 | | 1 | 6.27 | 2.56 | 61.03 | 59.91 | 1.13 | 1.13 | 0.03 | 0.03 |
| gene-LOC109611283 | https://www.ncbi.nlm.nih.gov/gene/?term=LOC109611283 | 1451 | 37 | 255 | 1 | 6.89 | 2.7 | 37.11 | 36.1 | 1 | 1 | 0.03 | 0.03 | |
| gene-LOC105285395 | https://www.ncbi.nlm.nih.gov/gene/?term=LOC105285395 | 8085 | 218 | 269.64 | | 1 | 1.24 | 0.46 | 17.01 | 16.05 | 0.96 | 0.96 | 0.03 | 0.03 |
| gene-LOC105278807 | https://www.ncbi.nlm.nih.gov/gene/?term=LOC105278807 | 2166 | 67 | 309.33 | | 1 | 4.62 | 1.49 | 22.84 | 21.92 | 0.93 | 0.93 | 0.03 | 0.03 |
| gene-LOC105279038 | https://www.ncbi.nlm.nih.gov/gene/?term=LOC105279038 | 9453 | 313 | 331.11 | | 1 | 1.06 | 0.32 | 10.35 | 9.46 | 0.9 | 0.9 | 0.03 | 0.03 |
| gene-LOC105284036 | https://www.ncbi.nlm.nih.gov/gene/?term=LOC105284036 | 3254 | 108 | 331.9 | 1 | 3.07 | 0.93 | 22.88 | 22.09 | 0.79 | 0.79 | 0.03 | 0.03 | |
| gene-LOC105280774 | https://www.ncbi.nlm.nih.gov/gene/?term=LOC105280774 | 2793 | 100 | 358.04 | | 1 | 3.58 | 1 | 32.96 | 32.18 | 0.78 | 0.78 | 0.03 | 0.03 |
| gene-LOC105274534 | https://www.ncbi.nlm.nih.gov/gene/?term=LOC105274534 | 1329 | 57 | 428.89 | | 1 | 7.52 | 1.75 | 48.36 | 47.65 | 0.71 | 0.71 | 0.03 | 0.03 |
| gene-LOC105284557 | https://www.ncbi.nlm.nih.gov/gene/?term=LOC105284557 | 1672 | 84 | 502.39 | | 1 | 5.98 | 1.19 | 29.36 | 28.7 | 0.66 | 0.66 | 0.03 | 0.03 |
| gene-LOC105275683 | https://www.ncbi.nlm.nih.gov/gene/?term=LOC105275683 | 4256 | 193 | 453.48 | | 1 | 2.35 | 0.52 | 13.76 | 13.13 | 0.63 | 0.63 | 0.03 | 0.03 |
| gene-LOC105285179 | https://www.ncbi.nlm.nih.gov/gene/?term=LOC105285179 | 4191 | 185 | 441.42 | | 1 | 2.39 | 0.54 | 18.94 | 18.34 | 0.6 | 0.6 | 0.03 | 0.03 |
| gene-LOC105275374 | https://www.ncbi.nlm.nih.gov/gene/?term=LOC105275374 | 11376 | 523 | 459.74 | | 1 | 0.88 | 0.19 | 11.54 | 10.97 | 0.57 | 0.57 | 0.03 | 0.03 |
| gene-LOC105276612 | https://www.ncbi.nlm.nih.gov/gene/?term=LOC105276612 | 4157 | 187 | 449.84 | | 1 | 2.41 | 0.53 | 10.04 | 9.47 | 0.57 | 0.57 | 0.03 | 0.03 |
| gene-LOC105276452 | https://www.ncbi.nlm.nih.gov/gene/?term=LOC105276452 | 2917 | 269 | 922.18 | | 1 | 3.43 | 0.37 | 9.31 | 8.95 | 0.36 | 0.36 | 0.03 | 0.03 |
| gene-LOC105280596 | https://www.ncbi.nlm.nih.gov/gene/?term=LOC105280596 | 3434 | 113 | 329.06 | | 1 | 2.91 | 0.88 | 17.62 | 18.56 | -0.94 | 0.94 | -0.03 | 0.03 |
| gene-LOC105282885 | https://www.ncbi.nlm.nih.gov/gene/?term=LOC105282885 | 1870 | 48 | 256.68 | | 1 | 5.35 | 2.08 | 38.27 | 39.26 | -0.99 | 0.99 | -0.03 | 0.03 |
| gene-LOC105278384 | https://www.ncbi.nlm.nih.gov/gene/?term=LOC105278384 | 1390 | 36 | 258.99 | | 1 | 7.19 | 2.78 | 50.65 | 51.65 | -1 | 1 | -0.03 | 0.03 |
| gene-LOC105280819 | https://www.ncbi.nlm.nih.gov/gene/?term=LOC105280819 | 1639 | 38 | 231.85 | | 1 | 6.1 | 2.63 | 35.58 | 37.06 | -1.49 | 1.49 | -0.03 | 0.03 |
| gene-LOC105276196 | https://www.ncbi.nlm.nih.gov/gene/?term=LOC105276196 | 2953 | 62 | 209.96 | | 1 | 3.39 | 1.61 | 36.95 | 38.53 | -1.58 | 1.58 | -0.03 | 0.03 |
| gene-LOC105281700 | https://www.ncbi.nlm.nih.gov/gene/?term=LOC105281700 | 2208 | 34 | 153.99 | | 1 | 4.53 | 2.94 | 35.23 | 36.92 | -1.69 | 1.69 | -0.03 | 0.03 |
| gene-LOC105285409 | https://www.ncbi.nlm.nih.gov/gene/?term=LOC105285409 | 5099 | 49 | 96.1 | 2 | 3.92 | 4.08 | 1.55 | 4.82 | -3.27 | 3.27 | -0.03 | 0.03 | |
| gene-LOC105283492 | https://www.ncbi.nlm.nih.gov/gene/?term=LOC105283492 | 1535 | 10 | 65.15 | 1 | 6.51 | 10 | 37.74 | 42.8 | -5.06 | 5.06 | -0.03 | 0.03 | |
| gene-LOC105286694 | https://www.ncbi.nlm.nih.gov/gene/?term=LOC105286694 | 3409 | 12 | 35.2 | 1 | 2.93 | 8.33 | 52.59 | 46.71 | 5.88 | 5.88 | 0.02 | 0.02 | |
| gene-LOC105284609 | https://www.ncbi.nlm.nih.gov/gene/?term=LOC105284609 | 5678 | 33 | 58.12 | 1 | 1.76 | 3.03 | 34 | 30.46 | 3.54 | 3.54 | 0.02 | 0.02 | |
| gene-LOC105274900 | https://www.ncbi.nlm.nih.gov/gene/?term=LOC105274900 | 5561 | 56 | 100.7 | 2 | 3.6 | 3.57 | 32.09 | 29.76 | 2.33 | 2.33 | 0.02 | 0.02 | |
| gene-LOC105277856 | https://www.ncbi.nlm.nih.gov/gene/?term=LOC105277856 | 3442 | 35 | 101.69 | | 1 | 2.91 | 2.86 | 35 | 32.98 | 2.02 | 2.02 | 0.02 | 0.02 |
| gene-LOC105278108 | https://www.ncbi.nlm.nih.gov/gene/?term=LOC105278108 | 4392 | 51 | 116.12 | | 1 | 2.28 | 1.96 | 25.15 | 23.14 | 2.01 | 2.01 | 0.02 | 0.02 |
| gene-LOC105284117 | https://www.ncbi.nlm.nih.gov/gene/?term=LOC105284117 | 4340 | 65 | 149.77 | | 1 | 2.3 | 1.54 | 12.82 | 11.26 | 1.56 | 1.56 | 0.02 | 0.02 |
| gene-LOC105287648 | https://www.ncbi.nlm.nih.gov/gene/?term=LOC105287648 | 2205 | 26 | 117.91 | | 1 | 4.54 | 3.85 | 41.13 | 39.59 | 1.54 | 1.54 | 0.02 | 0.02 |
| gene-LOC105288063 | https://www.ncbi.nlm.nih.gov/gene/?term=LOC105288063 | 3400 | 47 | 138.24 | | 1 | 2.94 | 2.13 | 15.4 | 13.86 | 1.54 | 1.54 | 0.02 | 0.02 |
| gene-LOC105274590 | https://www.ncbi.nlm.nih.gov/gene/?term=LOC105274590 | 4516 | 71 | 157.22 | | 2 | 4.43 | 2.82 | 29.16 | 27.84 | 1.32 | 1.32 | 0.02 | 0.02 |
| gene-LOC105283712 | https://www.ncbi.nlm.nih.gov/gene/?term=LOC105283712 | 1910 | 36 | 188.48 | | 1 | 5.24 | 2.78 | 27.71 | 26.46 | 1.25 | 1.25 | 0.02 | 0.02 |
| gene-LOC105288066 | https://www.ncbi.nlm.nih.gov/gene/?term=LOC105288066 | 1964 | 30 | 152.75 | | 1 | 5.09 | 3.33 | 32.32 | 31.12 | 1.2 | 1.2 | 0.02 | 0.02 |
| gene-LOC105287720 | https://www.ncbi.nlm.nih.gov/gene/?term=LOC105287720 | 4843 | 99 | 204.42 | | 1 | 2.06 | 1.01 | 16.39 | 15.24 | 1.16 | 1.16 | 0.02 | 0.02 |
| gene-LOC105275451 | https://www.ncbi.nlm.nih.gov/gene/?term=LOC105275451 | 6035 | 130 | 215.41 | | 1 | 1.66 | 0.77 | 22.16 | 21.09 | 1.07 | 1.07 | 0.02 | 0.02 |
| gene-LOC105288010 | https://www.ncbi.nlm.nih.gov/gene/?term=LOC105288010 | 3189 | 51 | 159.92 | | 1 | 3.14 | 1.96 | 29.8 | 28.73 | 1.07 | 1.07 | 0.02 | 0.02 |
| gene-LOC105286514 | https://www.ncbi.nlm.nih.gov/gene/?term=LOC105286514 | 2528 | 49 | 193.83 | | 1 | 3.96 | 2.04 | 36.75 | 35.71 | 1.04 | 1.04 | 0.02 | 0.02 |
| gene-LOC105283592 | https://www.ncbi.nlm.nih.gov/gene/?term=LOC105283592 | 3479 | 80 | 229.95 | | 1 | 2.87 | 1.25 | 23.33 | 22.32 | 1.01 | 1.01 | 0.02 | 0.02 |
| gene-LOC105278962 | https://www.ncbi.nlm.nih.gov/gene/?term=LOC105278962 | 4491 | 104 | 231.57 | | 1 | 2.23 | 0.96 | 22.01 | 21.09 | 0.92 | 0.92 | 0.02 | 0.02 |
| gene-LOC105276059 | https://www.ncbi.nlm.nih.gov/gene/?term=LOC105276059 | 6256 | 133 | 212.6 | 1 | 1.6 | 0.75 | 10.26 | 9.4 | 0.86 | 0.86 | 0.02 | 0.02 | |
| gene-LOC105280956 | https://www.ncbi.nlm.nih.gov/gene/?term=LOC105280956 | 2639 | 71 | 269.04 | | 1 | 3.79 | 1.41 | 18.65 | 17.79 | 0.86 | 0.86 | 0.02 | 0.02 |
| gene-LOC105280407 | https://www.ncbi.nlm.nih.gov/gene/?term=LOC105280407 | 10950 | 211 | 192.69 | | 1 | 0.91 | 0.47 | 16.65 | 15.81 | 0.84 | 0.84 | 0.02 | 0.02 |
| gene-LOC105274823 | https://www.ncbi.nlm.nih.gov/gene/?term=LOC105274823 | 4620 | 141 | 305.19 | | 1 | 2.16 | 0.71 | 21.37 | 20.81 | 0.56 | 0.56 | 0.02 | 0.02 |
| gene-LOC105275127 | https://www.ncbi.nlm.nih.gov/gene/?term=LOC105275127 | 9377 | 301 | 321 | 1 | 1.07 | 0.33 | 7.86 | 7.34 | 0.52 | 0.52 | 0.02 | 0.02 | |
| gene-LOC105285202 | https://www.ncbi.nlm.nih.gov/gene/?term=LOC105285202 | 1838 | 70 | 380.85 | | 1 | 5.44 | 1.43 | 25.94 | 25.46 | 0.48 | 0.48 | 0.02 | 0.02 |
| gene-LOC105282491 | https://www.ncbi.nlm.nih.gov/gene/?term=LOC105282491 | 4761 | 170 | 357.07 | | 1 | 2.1 | 0.59 | 10.04 | 9.59 | 0.45 | 0.45 | 0.02 | 0.02 |
| gene-LOC105278666 | https://www.ncbi.nlm.nih.gov/gene/?term=LOC105278666 | 10341 | 451 | 436.13 | | 1 | 0.97 | 0.22 | 7.32 | 6.95 | 0.38 | 0.38 | 0.02 | 0.02 |
| gene-LOC105278145 | https://www.ncbi.nlm.nih.gov/gene/?term=LOC105278145 | 4470 | 251 | 561.52 | | 1 | 2.24 | 0.4 | 10.77 | 10.43 | 0.34 | 0.34 | 0.02 | 0.02 |
| gene-LOC105286274 | https://www.ncbi.nlm.nih.gov/gene/?term=LOC105286274 | 3817 | 193 | 505.63 | | 1 | 2.62 | 0.52 | 17.37 | 17.05 | 0.32 | 0.32 | 0.02 | 0.02 |

| | | | | | | | | | | | | | |
|---|---|---|---|---|---|---|---|---|---|---|---|---|---|
| gene-LOC105287095 | https://www.ncbi.nlm.nih.gov/gene/?term=LOC105287095 | 1435 | 58 | 404.18 | 1 | 6.97 | 1.72 | 28.05 | 28.65 | -0.6 | 0.6 | -0.02 | 0.02 |
| gene-LOC105274449 | https://www.ncbi.nlm.nih.gov/gene/?term=LOC105274449 | 1549 | 48 | 309.88 | 1 | 6.46 | 2.08 | 50.3 | 50.96 | -0.66 | 0.66 | -0.02 | 0.02 |
| gene-LOC105285315 | https://www.ncbi.nlm.nih.gov/gene/?term=LOC105285315 | 5458 | 19 | 34.81 | 1 | 1.83 | 5.26 | 46.71 | 43.22 | 3.49 | 3.49 | 0.01 | 0.01 |
| gene-LOC105281515 | https://www.ncbi.nlm.nih.gov/gene/?term=LOC105281515 | 8827 | 42 | 47.58 | 1 | 1.13 | 2.38 | 30.33 | 27.53 | 2.81 | 2.81 | 0.01 | 0.01 |
| gene-LOC105275347 | https://www.ncbi.nlm.nih.gov/gene/?term=LOC105275347 | 2526 | 19 | 75.22 | 1 | 3.96 | 5.26 | 38.97 | 37.49 | 1.48 | 1.48 | 0.01 | 0.01 |
| gene-LOC105281532 | https://www.ncbi.nlm.nih.gov/gene/?term=LOC105281532 | 7793 | 81 | 103.94 | 1 | 1.28 | 1.23 | 12.92 | 11.51 | 1.4 | 1.4 | 0.01 | 0.01 |
| gene-LOC105279500 | https://www.ncbi.nlm.nih.gov/gene/?term=LOC105279500 | 3587 | 37 | 103.15 | 1 | 2.79 | 2.7 | 50.42 | 49.31 | 1.1 | 1.1 | 0.01 | 0.01 |
| gene-LOC105283987 | https://www.ncbi.nlm.nih.gov/gene/?term=LOC105283987 | 4841 | 68 | 140.47 | 1 | 2.07 | 1.47 | 29.57 | 28.67 | 0.91 | 0.91 | 0.01 | 0.01 |
| gene-LOC105285406 | https://www.ncbi.nlm.nih.gov/gene/?term=LOC105285406 | 8015 | 60 | 74.86 | 1 | 1.25 | 1.67 | 26.39 | 25.52 | 0.87 | 0.87 | 0.01 | 0.01 |
| gene-LOC105280408 | https://www.ncbi.nlm.nih.gov/gene/?term=LOC105280408 | 5983 | 58 | 96.94 | 1 | 1.67 | 1.72 | 16.75 | 15.89 | 0.86 | 0.86 | 0.01 | 0.01 |
| gene-LOC105280002 | https://www.ncbi.nlm.nih.gov/gene/?term=LOC105280002 | 1940 | 28 | 144.33 | 1 | 5.15 | 3.57 | 37.5 | 36.72 | 0.79 | 0.79 | 0.01 | 0.01 |
| gene-LOC105285060 | https://www.ncbi.nlm.nih.gov/gene/?term=LOC105285060 | 1285 | 21 | 163.42 | 1 | 7.78 | 4.76 | 39.84 | 39.09 | 0.75 | 0.75 | 0.01 | 0.01 |
| gene-LOC105284116 | https://www.ncbi.nlm.nih.gov/gene/?term=LOC105284116 | 3970 | 78 | 196.47 | 1 | 2.52 | 1.28 | 7.09 | 6.36 | 0.73 | 0.73 | 0.01 | 0.01 |
| gene-LOC105282067 | https://www.ncbi.nlm.nih.gov/gene/?term=LOC105282067 | 5941 | 54 | 90.89 | 1 | 1.68 | 1.85 | 28.74 | 28.06 | 0.67 | 0.67 | 0.01 | 0.01 |
| gene-LOC105285514 | https://www.ncbi.nlm.nih.gov/gene/?term=LOC105285514 | 1277 | 17 | 133.12 | 1 | 7.83 | 5.88 | 71.39 | 70.72 | 0.67 | 0.67 | 0.01 | 0.01 |
| gene-LOC105274699 | https://www.ncbi.nlm.nih.gov/gene/?term=LOC105274699 | 2386 | 52 | 217.94 | 1 | 4.19 | 1.92 | 24.82 | 24.15 | 0.66 | 0.66 | 0.01 | 0.01 |
| gene-LOC105275966 | https://www.ncbi.nlm.nih.gov/gene/?term=LOC105275966 | 3299 | 64 | 194 | 1 | 3.03 | 1.56 | 48.15 | 47.49 | 0.65 | 0.65 | 0.01 | 0.01 |
| gene-LOC105282399 | https://www.ncbi.nlm.nih.gov/gene/?term=LOC105282399 | 2682 | 24 | 89.49 | 1 | 3.73 | 4.17 | 35.91 | 35.25 | 0.65 | 0.65 | 0.01 | 0.01 |
| gene-LOC105283943 | https://www.ncbi.nlm.nih.gov/gene/?term=LOC105283943 | 3559 | 48 | 134.87 | 1 | 2.81 | 2.08 | 17.55 | 16.96 | 0.59 | 0.59 | 0.01 | 0.01 |
| gene-LOC105282825 | https://www.ncbi.nlm.nih.gov/gene/?term=LOC105282825 | 3919 | 85 | 216.89 | 1 | 2.55 | 1.18 | 14.44 | 13.87 | 0.57 | 0.57 | 0.01 | 0.01 |
| gene-LOC105278534 | https://www.ncbi.nlm.nih.gov/gene/?term=LOC105278534 | 7364 | 197 | 267.52 | 2 | 2.72 | 1.02 | 18.17 | 17.62 | 0.55 | 0.55 | 0.01 | 0.01 |
| gene-LOC105281599 | https://www.ncbi.nlm.nih.gov/gene/?term=LOC105281599 | 3859 | 49 | 126.98 | 1 | 2.59 | 2.04 | 4.6 | 4.08 | 0.53 | 0.53 | 0.01 | 0.01 |
| gene-LOC105280410 | https://www.ncbi.nlm.nih.gov/gene/?term=LOC105280410 | 12580 | 264 | 209.86 | 1 | 0.79 | 0.38 | 19.99 | 19.51 | 0.48 | 0.48 | 0.01 | 0.01 |
| gene-LOC105278454 | https://www.ncbi.nlm.nih.gov/gene/?term=LOC105278454 | 1972 | 46 | 233.27 | 1 | 5.07 | 2.17 | 44.92 | 44.47 | 0.46 | 0.46 | 0.01 | 0.01 |
| gene-LOC105275179 | https://www.ncbi.nlm.nih.gov/gene/?term=LOC105275179 | 3271 | 72 | 220.12 | 1 | 3.06 | 1.39 | 21.5 | 21.05 | 0.45 | 0.45 | 0.01 | 0.01 |
| gene-LOC105276078 | https://www.ncbi.nlm.nih.gov/gene/?term=LOC105276078 | 1051 | 14 | 133.21 | 1 | 9.51 | 7.14 | 33.32 | 32.93 | 0.39 | 0.39 | 0.01 | 0.01 |
| gene-LOC105285860 | https://www.ncbi.nlm.nih.gov/gene/?term=LOC105285860 | 3110 | 55 | 176.85 | 1 | 3.22 | 1.82 | 24.17 | 23.77 | 0.39 | 0.39 | 0.01 | 0.01 |
| gene-LOC105275327 | https://www.ncbi.nlm.nih.gov/gene/?term=LOC105275327 | 14245 | 492 | 345.38 | 1 | 0.7 | 0.2 | 7.28 | 6.9 | 0.38 | 0.38 | 0.01 | 0.01 |
| gene-LOC105286194 | https://www.ncbi.nlm.nih.gov/gene/?term=LOC105286194 | 152619 | 2242 | 146.9 | 1 | 0.07 | 0.04 | 6.47 | 6.09 | 0.38 | 0.38 | 0.01 | 0.01 |
| gene-LOC105274531 | https://www.ncbi.nlm.nih.gov/gene/?term=LOC105274531 | 7774 | 301 | 387.19 | 1 | 1.29 | 0.33 | 10.99 | 10.62 | 0.37 | 0.37 | 0.01 | 0.01 |
| gene-LOC105282811 | https://www.ncbi.nlm.nih.gov/gene/?term=LOC105282811 | 5765 | 213 | 369.47 | 1 | 1.73 | 0.47 | 8.63 | 8.26 | 0.37 | 0.37 | 0.01 | 0.01 |
| gene-LOC105287371 | https://www.ncbi.nlm.nih.gov/gene/?term=LOC105287371 | 8224 | 351 | 426.8 | 1 | 1.22 | 0.28 | 7.67 | 7.34 | 0.33 | 0.33 | 0.01 | 0.01 |
| gene-LOC105287528 | https://www.ncbi.nlm.nih.gov/gene/?term=LOC105287528 | 14195 | 419 | 295.17 | 1 | 0.7 | 0.24 | 5.19 | 4.86 | 0.33 | 0.33 | 0.01 | 0.01 |
| gene-LOC105287550 | https://www.ncbi.nlm.nih.gov/gene/?term=LOC105287550 | 15659 | 613 | 391.47 | 1 | 0.64 | 0.16 | 6.43 | 6.12 | 0.31 | 0.31 | 0.01 | 0.01 |
| gene-LOC105286166 | https://www.ncbi.nlm.nih.gov/gene/?term=LOC105286166 | 3050 | 110 | 360.66 | 1 | 3.28 | 0.91 | 24.5 | 24.21 | 0.29 | 0.29 | 0.01 | 0.01 |
| gene-LOC113562050 | https://www.ncbi.nlm.nih.gov/gene/?term=LOC113562050 | 17097 | 576 | 336.9 | 1 | 0.58 | 0.17 | 5.24 | 4.95 | 0.29 | 0.29 | 0.01 | 0.01 |
| gene-LOC105278877 | https://www.ncbi.nlm.nih.gov/gene/?term=LOC105278877 | 1968 | 72 | 365.85 | 1 | 5.08 | 1.39 | 36.04 | 35.76 | 0.28 | 0.28 | 0.01 | 0.01 |
| gene-LOC105277079 | https://www.ncbi.nlm.nih.gov/gene/?term=LOC105277079 | 13009 | 288 | 221.39 | 1 | 0.77 | 0.35 | 9.86 | 9.59 | 0.27 | 0.27 | 0.01 | 0.01 |
| gene-LOC105284262 | https://www.ncbi.nlm.nih.gov/gene/?term=LOC105284262 | 6669 | 225 | 337.38 | 1 | 1.5 | 0.44 | 6.17 | 5.94 | 0.23 | 0.23 | 0.01 | 0.01 |
| gene-LOC105278632 | https://www.ncbi.nlm.nih.gov/gene/?term=LOC105278632 | 3624 | 120 | 331.13 | 1 | 2.76 | 0.83 | 15.73 | 15.53 | 0.2 | 0.2 | 0.01 | 0.01 |
| gene-LOC105285807 | https://www.ncbi.nlm.nih.gov/gene/?term=LOC105285807 | 6063 | 227 | 374.4 | 1 | 1.65 | 0.44 | 9.73 | 9.55 | 0.18 | 0.18 | 0.01 | 0.01 |
| gene-LOC105276527 | https://www.ncbi.nlm.nih.gov/gene/?term=LOC105276527 | 7923 | 329 | 415.25 | 1 | 1.26 | 0.3 | 9.51 | 9.36 | 0.15 | 0.15 | 0.01 | 0.01 |
| gene-LOC105286649 | https://www.ncbi.nlm.nih.gov/gene/?term=LOC105286649 | 14178 | 817 | 576.24 | 1 | 0.71 | 0.12 | 1 | 1.1 | -0.1 | 0.1 | -0.01 | 0.01 |
| gene-LOC105275848 | https://www.ncbi.nlm.nih.gov/gene/?term=LOC105275848 | 3718 | 144 | 387.31 | 1 | 2.69 | 0.69 | 20.8 | 20.95 | -0.15 | 0.15 | -0.01 | 0.01 |
| gene-LOC105287633 | https://www.ncbi.nlm.nih.gov/gene/?term=LOC105287633 | 1524 | 34 | 223.1 | 1 | 6.56 | 2.94 | 50.71 | 50.96 | -0.26 | 0.26 | -0.01 | 0.01 |
| gene-LOC105281312 | https://www.ncbi.nlm.nih.gov/gene/?term=LOC105281312 | 2551 | 63 | 246.96 | 1 | 3.92 | 1.59 | 34.27 | 34.56 | -0.29 | 0.29 | -0.01 | 0.01 |
| gene-LOC105286999 | https://www.ncbi.nlm.nih.gov/gene/?term=LOC105286999 | 2141 | 44 | 205.51 | 1 | 4.67 | 2.27 | 34.01 | 34.41 | -0.41 | 0.41 | -0.01 | 0.01 |
| gene-LOC105285252 | https://www.ncbi.nlm.nih.gov/gene/?term=LOC105285252 | 7427 | 117 | 157.53 | 1 | 1.35 | 0.85 | 14.46 | 15.16 | -0.7 | 0.7 | -0.01 | 0.01 |
| gene-LOC105281296 | https://www.ncbi.nlm.nih.gov/gene/?term=LOC105281296 | 2434 | 37 | 152.01 | 1 | 4.11 | 2.7 | 30.9 | 31.72 | -0.83 | 0.83 | -0.01 | 0.01 |
| gene-LOC105286770 | https://www.ncbi.nlm.nih.gov/gene/?term=LOC105286770 | 3055 | 33 | 108.02 | 1 | 3.27 | 3.03 | 17.27 | 18.34 | -1.07 | 1.07 | -0.01 | 0.01 |
| gene-LOC105283989 | https://www.ncbi.nlm.nih.gov/gene/?term=LOC105283989 | 4948 | 31 | 62.65 | 2 | 4.04 | 6.45 | 34.97 | 36.46 | -1.49 | 1.49 | -0.01 | 0.01 |
| gene-LOC105286452 | https://www.ncbi.nlm.nih.gov/gene/?term=LOC105286452 | 6888 | 67 | 97.27 | 1 | 1.45 | 1.49 | 31.04 | 30.73 | 0.31 | 0.31 | 0 | 0 |
| gene-LOC105278245 | https://www.ncbi.nlm.nih.gov/gene/?term=LOC105278245 | 6445 | 117 | 181.54 | 1 | 1.55 | 0.85 | 22.48 | 22.2 | 0.27 | 0.27 | 0 | 0 |
| gene-LOC105276272 | https://www.ncbi.nlm.nih.gov/gene/?term=LOC105276272 | 1965 | 48 | 244.27 | 1 | 5.09 | 2.08 | 27.74 | 27.58 | 0.17 | 0.17 | 0 | 0 |
| gene-LOC105277125 | https://www.ncbi.nlm.nih.gov/gene/?term=LOC105277125 | 51079 | 1858 | 363.75 | 1 | 0.2 | 0.05 | 1.73 | 1.6 | 0.13 | 0.13 | 0 | 0 |
| gene-LOC105277866 | https://www.ncbi.nlm.nih.gov/gene/?term=LOC105277866 | 33045 | 400 | 121.05 | 1 | 0.3 | 0.25 | 3.44 | 3.35 | 0.1 | 0.1 | 0 | 0 |
| gene-LOC105280265 | https://www.ncbi.nlm.nih.gov/gene/?term=LOC105280265 | 85984 | 3130 | 364.02 | 1 | 0.12 | 0.03 | 1.14 | 1.05 | 0.09 | 0.09 | 0 | 0 |
| gene-LOC105274669 | https://www.ncbi.nlm.nih.gov/gene/?term=LOC105274669 | 8555 | 256 | 299.24 | 1 | 1.17 | 0.39 | 6.43 | 6.35 | 0.08 | 0.08 | 0 | 0 |
| gene-LOC105275752 | https://www.ncbi.nlm.nih.gov/gene/?term=LOC105275752 | 3462 | 132 | 381.28 | 1 | 2.89 | 0.76 | 32.74 | 32.68 | 0.06 | 0.06 | 0 | 0 |
| gene-LOC105278455 | https://www.ncbi.nlm.nih.gov/gene/?term=LOC105278455 | 27181 | 961 | 353.56 | 1 | 0.37 | 0.1 | 7.16 | 7.14 | 0.02 | 0.02 | 0 | 0 |
| gene-LOC105286515 | https://www.ncbi.nlm.nih.gov/gene/?term=LOC105286515 | 3751 | 114 | 303.92 | 1 | 2.67 | 0.88 | 11.87 | 11.89 | -0.02 | 0.02 | 0 | 0 |
| gene-LOC105283296 | https://www.ncbi.nlm.nih.gov/gene/?term=LOC105283296 | 2174 | 94 | 432.38 | 1 | 4.6 | 1.06 | 18.21 | 18.24 | -0.03 | 0.03 | 0 | 0 |
| gene-LOC105287695 | https://www.ncbi.nlm.nih.gov/gene/?term=LOC105287695 | 6106 | 201 | 329.18 | 1 | 1.64 | 0.5 | 6.09 | 6.16 | -0.06 | 0.06 | 0 | 0 |
| gene-LOC105274851 | https://www.ncbi.nlm.nih.gov/gene/?term=LOC105274851 | 6616 | 139 | 210.1 | 1 | 1.51 | 0.72 | 11.55 | 11.65 | -0.1 | 0.1 | 0 | 0 |

| | | | | | | | | | | | | | | |
|---|---|---|---|---|---|---|---|---|---|---|---|---|---|---|
| gene-LOC105275187 | https://www.ncbi.nlm.nih.gov/gene/?term=LOC105275187 | 5534 | 71 | 128.3 | 1 | 1.81 | 1.41 | 12.33 | 12.43 | -0.1 | 0.1 | 0 | 0 | |
| gene-LOC105282646 | https://www.ncbi.nlm.nih.gov/gene/?term=LOC105282646 | 3317 | 116 | 349.71 | | 1 | 3.01 | 0.86 | 24.81 | 24.92 | -0.11 | 0.11 | 0 | 0 |
| gene-LOC105283806 | https://www.ncbi.nlm.nih.gov/gene/?term=LOC105283806 | 10501 | 74 | 70.47 | 1 | 0.95 | 1.35 | 16.11 | 16.23 | -0.12 | 0.12 | 0 | 0 | |
| gene-LOC105284507 | https://www.ncbi.nlm.nih.gov/gene/?term=LOC105284507 | 5207 | 87 | 167.08 | | 1 | 1.92 | 1.15 | 21.48 | 21.62 | -0.13 | 0.13 | 0 | 0 |
| gene-LOC105280704 | https://www.ncbi.nlm.nih.gov/gene/?term=LOC105280704 | 4266 | 134 | 314.11 | | 1 | 2.34 | 0.75 | 19.89 | 20.04 | -0.15 | 0.15 | 0 | 0 |
| gene-LOC105280348 | https://www.ncbi.nlm.nih.gov/gene/?term=LOC105280348 | 3797 | 70 | 184.36 | | 1 | 2.63 | 1.43 | 20.64 | 20.81 | -0.17 | 0.17 | 0 | 0 |
| gene-LOC105283967 | https://www.ncbi.nlm.nih.gov/gene/?term=LOC105283967 | 2393 | 47 | 196.41 | | 1 | 4.18 | 2.13 | 48.9 | 49.11 | -0.21 | 0.21 | 0 | 0 |
| gene-LOC105285163 | https://www.ncbi.nlm.nih.gov/gene/?term=LOC105285163 | 5061 | 87 | 171.9 | 1 | 1.98 | 1.15 | 21.7 | 21.95 | -0.26 | 0.26 | 0 | 0 | |

Directory: OGLall File: rnk-dmg-Ob_LI_RphaseB1.vs.RphaseB2.txt

| gene_ID | gene_link | gwidth | #Sites | #per10Kb | #dmSites | #dmsp10kb | %dmSites | %pSite1 | %pSite2 | DMpSite | ADMpSite | DMpNucl | ADMpNucl |
|---|---|---|---|---|---|---|---|---|---|---|---|---|---|
| gene-LOC105278849 | https://www.ncbi.nlm.nih.gov/gene/?term=LOC105278849 | 776 | 74 | 953.61 | 2 | 25.77 | 2.7 | 20.95 | 22.99 | -2.05 | 2.05 | -0.2 | 0.2 |
| gene-LOC113562453 | https://www.ncbi.nlm.nih.gov/gene/?term=LOC113562453 | 763 | 32 | 419.4 | 2 | 26.21 | 6.25 | 30.48 | 26.18 | 4.3 | 4.3 | 0.18 | 0.18 |
| gene-LOC105284093 | https://www.ncbi.nlm.nih.gov/gene/?term=LOC105284093 | 809 | 60 | 741.66 | 2 | 24.72 | 3.33 | 10.26 | 7.81 | 2.45 | 2.45 | 0.18 | 0.18 |
| gene-LOC105284253 | https://www.ncbi.nlm.nih.gov/gene/?term=LOC105284253 | 1255 | 41 | 326.69 | 5 | 39.84 | 12.2 | 26.29 | 31.86 | -5.58 | 5.58 | -0.18 | 0.18 |
| gene-LOC105284320 | https://www.ncbi.nlm.nih.gov/gene/?term=LOC105284320 | 390 | 11 | 282.05 | 1 | 25.64 | 9.09 | 22.16 | 27.96 | -5.81 | 5.81 | -0.16 | 0.16 |
| gene-LOC105288193 | https://www.ncbi.nlm.nih.gov/gene/?term=LOC105288193 | 1495 | 76 | 508.36 | 5 | 33.44 | 6.58 | 30.07 | 33 | -2.93 | 2.93 | -0.15 | 0.15 |
| gene-LOC113561900 | https://www.ncbi.nlm.nih.gov/gene/?term=LOC113561900 | 629 | 11 | 174.88 | 1 | 15.9 | 9.09 | 33.39 | 25.51 | 7.88 | 7.88 | 0.14 | 0.14 |
| gene-LOC105282481 | https://www.ncbi.nlm.nih.gov/gene/?term=LOC105282481 | 1900 | 60 | 315.79 | 4 | 21.05 | 6.67 | 31.49 | 36.02 | -4.52 | 4.52 | -0.14 | 0.14 |
| gene-LOC105284189 | https://www.ncbi.nlm.nih.gov/gene/?term=LOC105284189 | 2601 | 133 | 511.34 | 3 | 11.53 | 2.26 | 17.44 | 14.95 | 2.49 | 2.49 | 0.13 | 0.13 |
| gene-LOC105278160 | https://www.ncbi.nlm.nih.gov/gene/?term=LOC105278160 | 1872 | 91 | 486.11 | 1 | 5.34 | 1.1 | 34.58 | 37.24 | -2.67 | 2.67 | -0.13 | 0.13 |
| gene-LOC105277345 | https://www.ncbi.nlm.nih.gov/gene/?term=LOC105277345 | 2151 | 77 | 357.97 | 3 | 13.95 | 3.9 | 18.53 | 15.19 | 3.34 | 3.34 | 0.12 | 0.12 |
| gene-LOC105276563 | https://www.ncbi.nlm.nih.gov/gene/?term=LOC105276563 | 1958 | 114 | 582.23 | 5 | 25.54 | 4.39 | 18.98 | 16.96 | 2.02 | 2.02 | 0.12 | 0.12 |
| gene-LOC105279346 | https://www.ncbi.nlm.nih.gov/gene/?term=LOC105279346 | 4649 | 252 | 542.05 | 6 | 12.91 | 2.38 | 14.38 | 16.52 | -2.15 | 2.15 | -0.12 | 0.12 |
| gene-LOC105278458 | https://www.ncbi.nlm.nih.gov/gene/?term=LOC105278458 | 216 | 3 | 138.89 | 1 | 46.3 | 33.33 | 20.6 | 29.54 | -8.94 | 8.94 | -0.12 | 0.12 |
| gene-LOC105285203 | https://www.ncbi.nlm.nih.gov/gene/?term=LOC105285203 | 1020 | 32 | 313.73 | 2 | 19.61 | 6.25 | 29.55 | 25.89 | 3.66 | 3.66 | 0.11 | 0.11 |
| gene-LOC105278025 | https://www.ncbi.nlm.nih.gov/gene/?term=LOC105278025 | 1625 | 61 | 375.38 | 2 | 12.31 | 3.28 | 30.61 | 33.53 | -2.91 | 2.91 | -0.11 | 0.11 |
| gene-LOC105286845 | https://www.ncbi.nlm.nih.gov/gene/?term=LOC105286845 | 1159 | 78 | 672.99 | 2 | 17.26 | 2.56 | 11.01 | 12.57 | -1.56 | 1.56 | -0.1 | 0.1 |
| gene-LOC105278938 | https://www.ncbi.nlm.nih.gov/gene/?term=LOC105278938 | 1818 | 110 | 605.06 | 4 | 22 | 3.64 | 19.53 | 21.23 | -1.71 | 1.71 | -0.1 | 0.1 |
| gene-LOC105284047 | https://www.ncbi.nlm.nih.gov/gene/?term=LOC105284047 | 1212 | 20 | 165.02 | 4 | 33 | 20 | 20.52 | 26.47 | -5.95 | 5.95 | -0.1 | 0.1 |
| gene-LOC105285713 | https://www.ncbi.nlm.nih.gov/gene/?term=LOC105285713 | 1048 | 39 | 372.14 | 2 | 19.08 | 5.13 | 46.18 | 43.72 | 2.46 | 2.46 | 0.09 | 0.09 |
| gene-LOC105274542 | https://www.ncbi.nlm.nih.gov/gene/?term=LOC105274542 | 1266 | 52 | 410.74 | 2 | 15.8 | 3.85 | 30.12 | 27.91 | 2.21 | 2.21 | 0.09 | 0.09 |
| gene-LOC105279487 | https://www.ncbi.nlm.nih.gov/gene/?term=LOC105279487 | 2117 | 101 | 477.09 | 2 | 9.45 | 1.98 | 14.93 | 12.99 | 1.94 | 1.94 | 0.09 | 0.09 |
| gene-LOC105277091 | https://www.ncbi.nlm.nih.gov/gene/?term=LOC105277091 | 902 | 43 | 476.72 | 2 | 22.17 | 4.65 | 23.8 | 21.99 | 1.82 | 1.82 | 0.09 | 0.09 |
| gene-LOC105285392 | https://www.ncbi.nlm.nih.gov/gene/?term=LOC105285392 | 1601 | 82 | 512.18 | 9 | 56.21 | 10.98 | 37.61 | 35.85 | 1.76 | 1.76 | 0.09 | 0.09 |

...

... only first 25 lines shown ...

```
Directory: OGLall File: wrt-Ob_LI.txt

... comparing  BCphaseA1.vs.BCphaseA2 ...

Number of genes with #dmsites >=  1 :  124

sample1     sum of percent methylation:  3175.16     average:    25.61
sample2     sum of percent methylation:  3294.59     average:    26.57

sum difference:  -119.39     average difference:  -0.9628

      Wilcoxon signed rank test with continuity correction

data:  pw_summary$`%pSite1` and pw_summary$`%pSite2`
V = 2323, p-value = 1e-04
alternative hypothesis: true location shift is not equal to 0

... comparing  BCphaseA1.vs.BCphaseB1 ...

Number of genes with #dmsites >=  1 :  127

sample1     sum of percent methylation:  3016.97     average:    23.76
sample2     sum of percent methylation:  3105.42     average:    24.45

sum difference:   -88.42     average difference:  -0.6962
...
... only first 25 lines shown ...
```

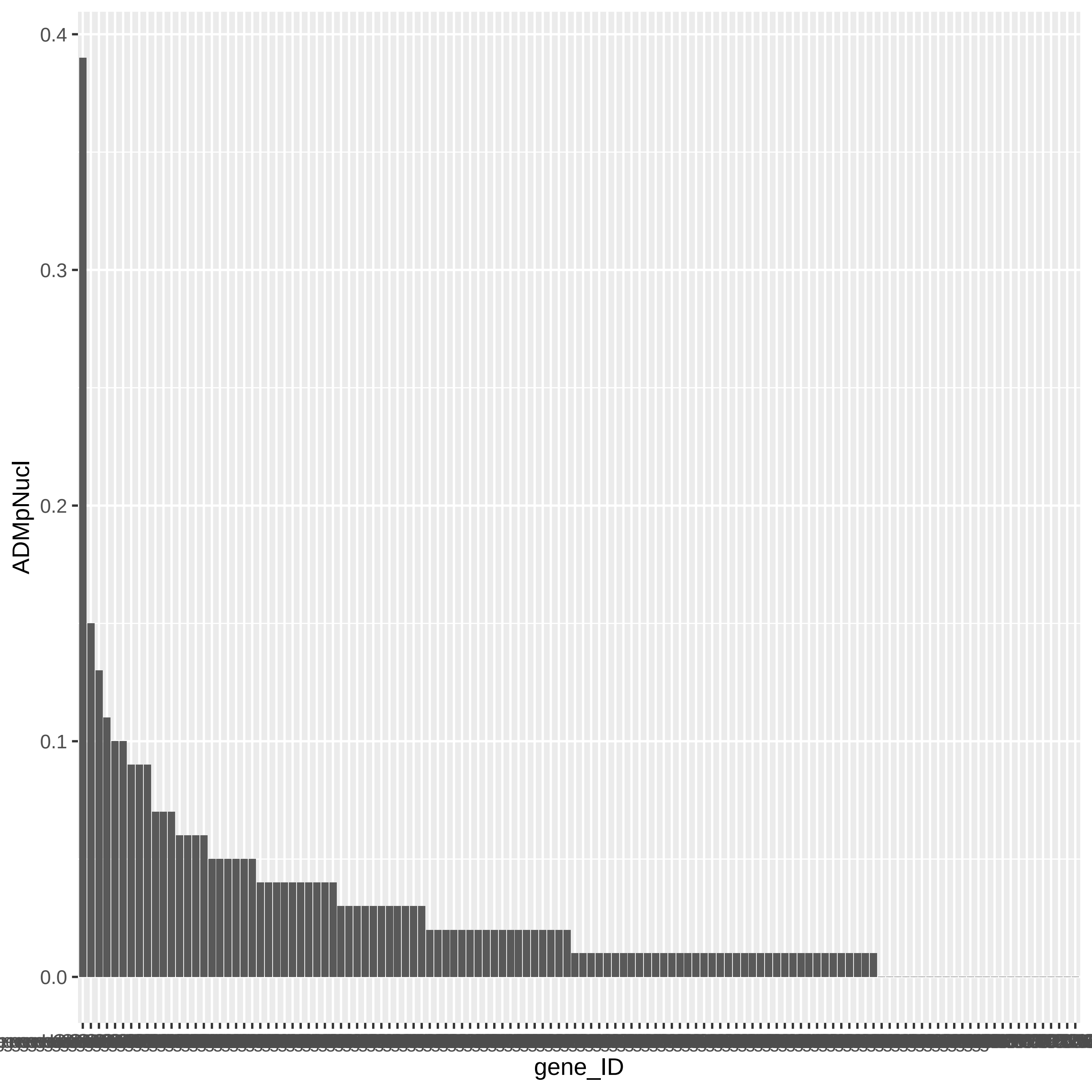

0.4
0.3
0.2
0.1
0.0
ADMpNucl
gene_ID

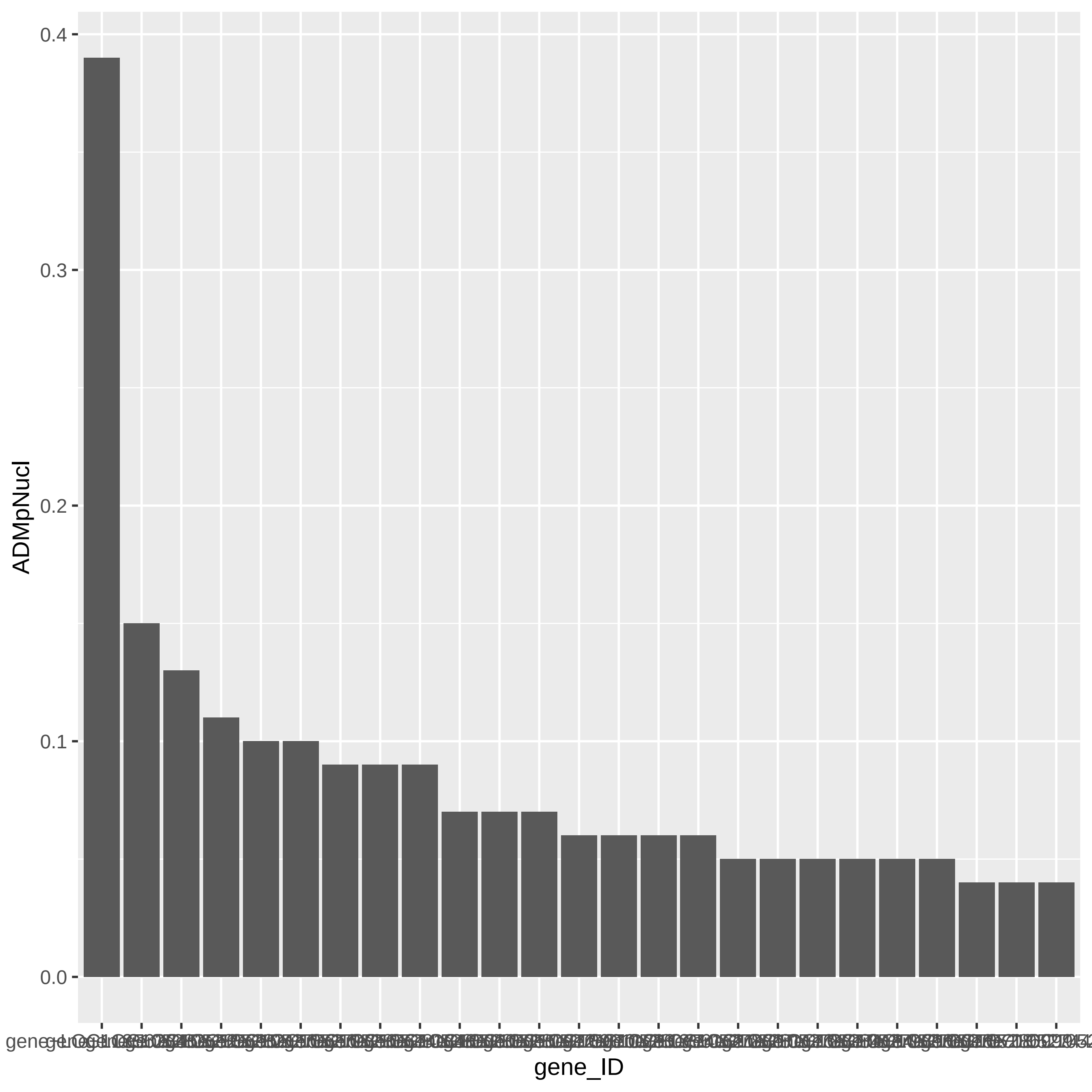
0.4
0.3
0.2
0.1
0.0
ADMpNucl
gene_ID

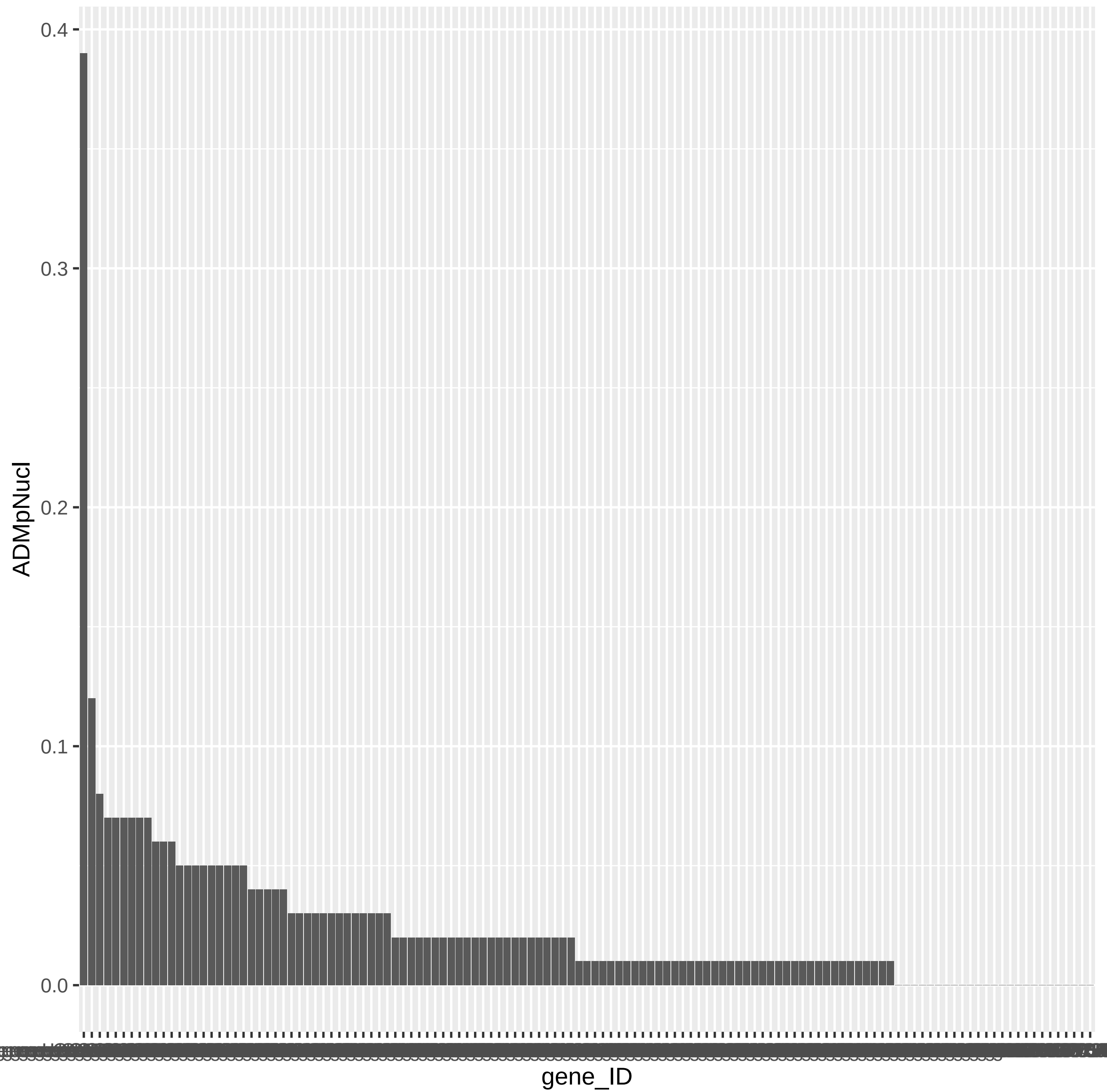

ADMpNucl
0.4
0.3
0.2
0.1
0.0
gene_ID

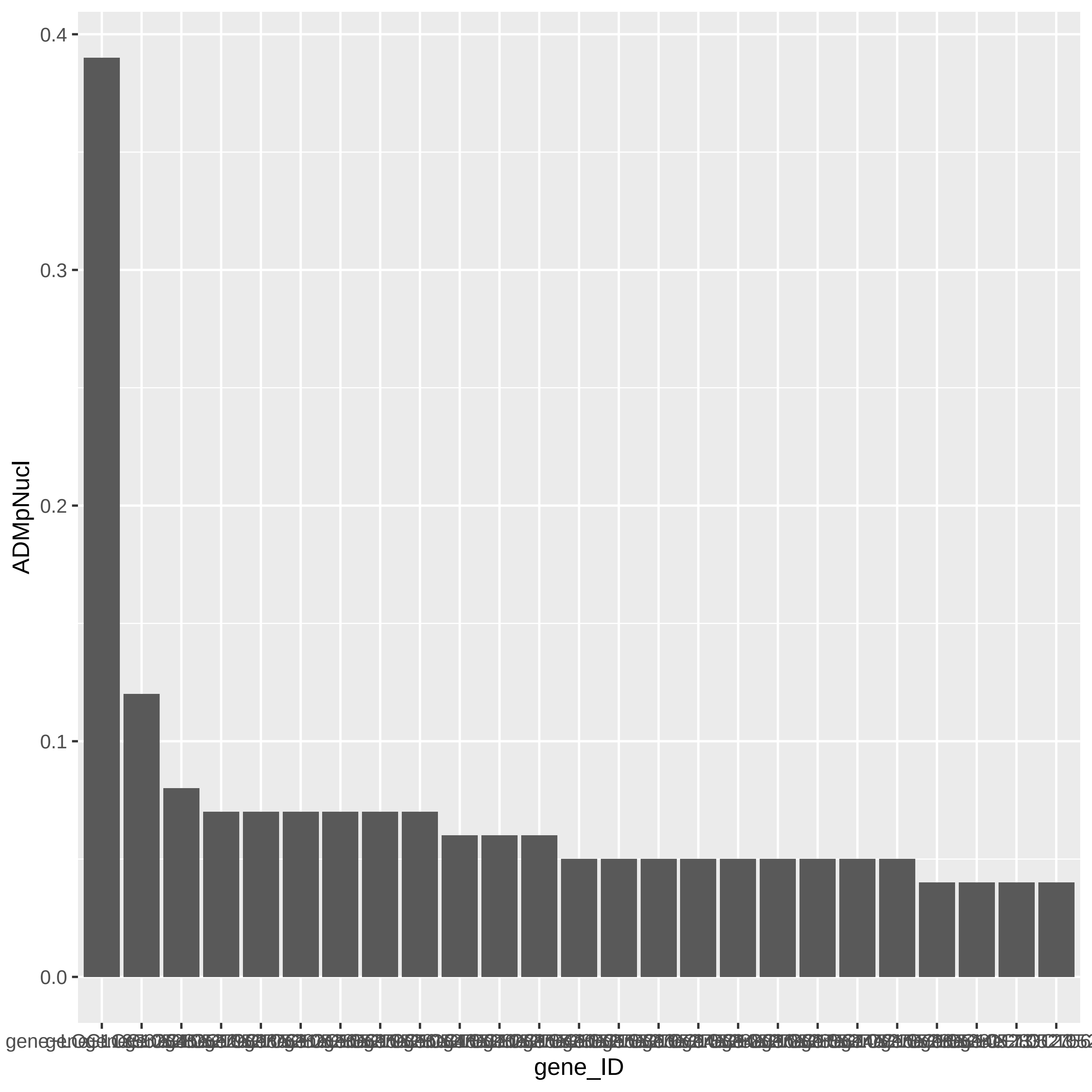

0.4
0.3
0.2
0.1
0.0
ADMpNucl
gene_ID

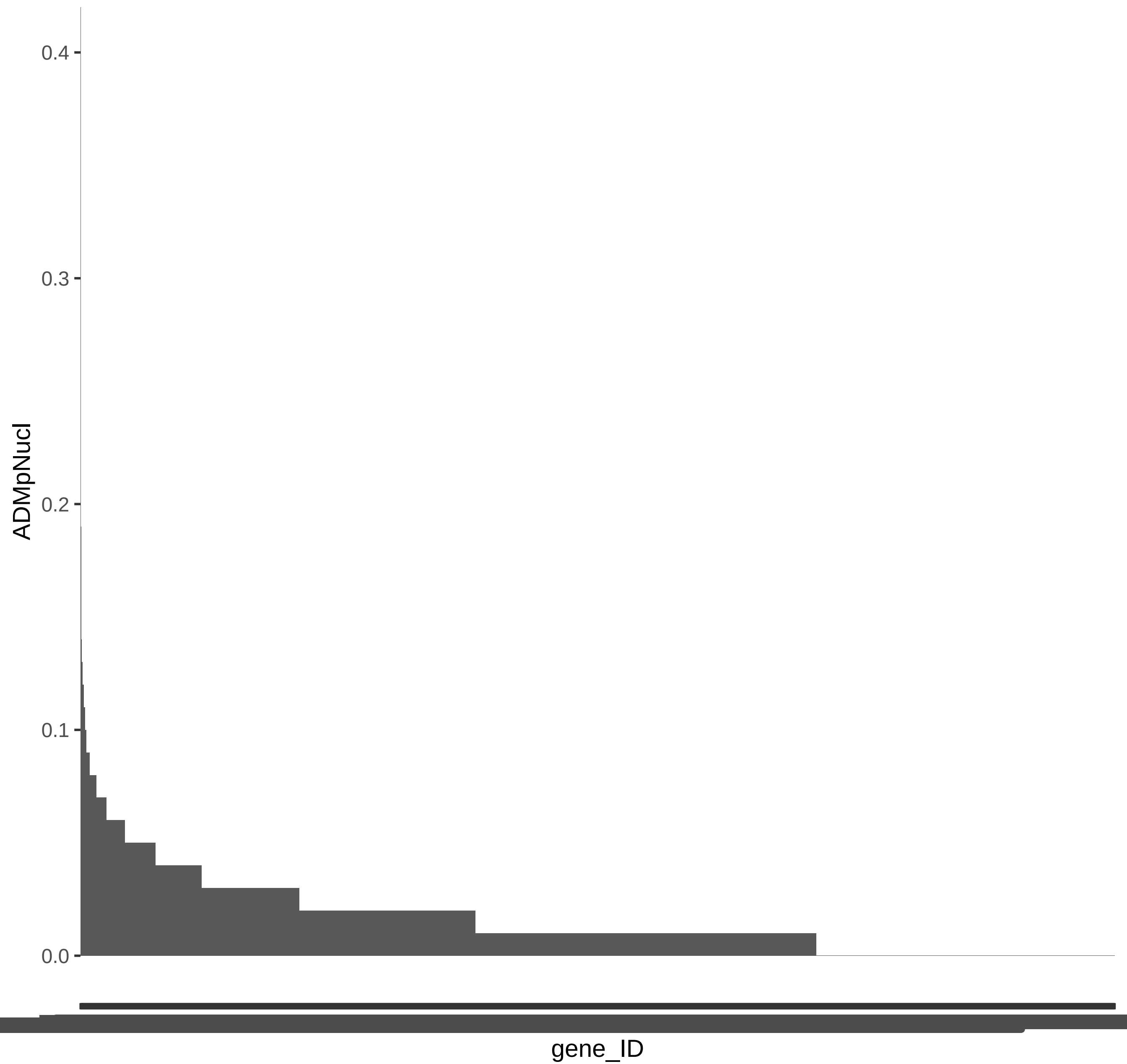

0.4
0.3
0.2
0.1
0.0
ADMpNucl
gene_ID

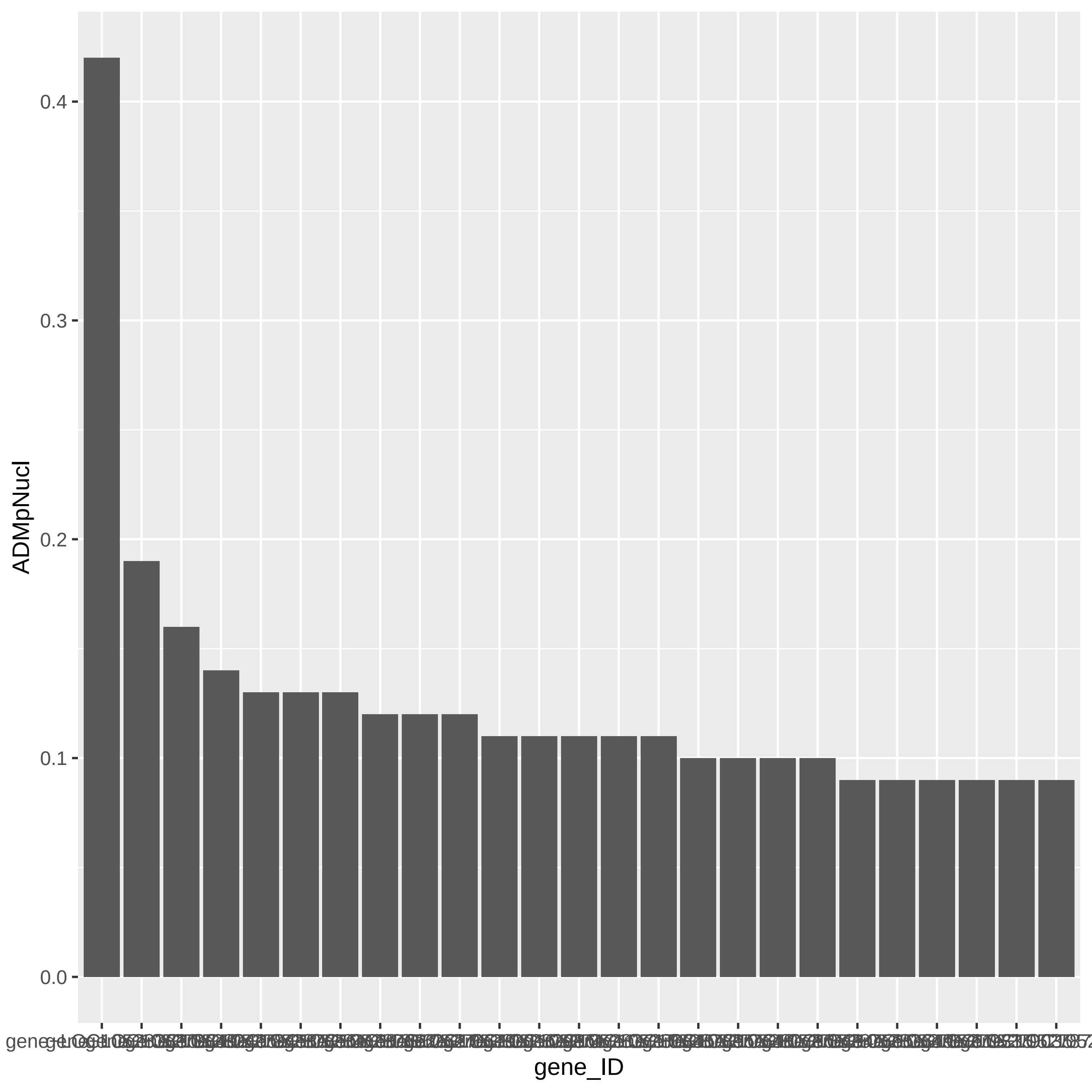
0.4
0.3
0.2
0.1
0.0
ADMpNucl
gene_ID

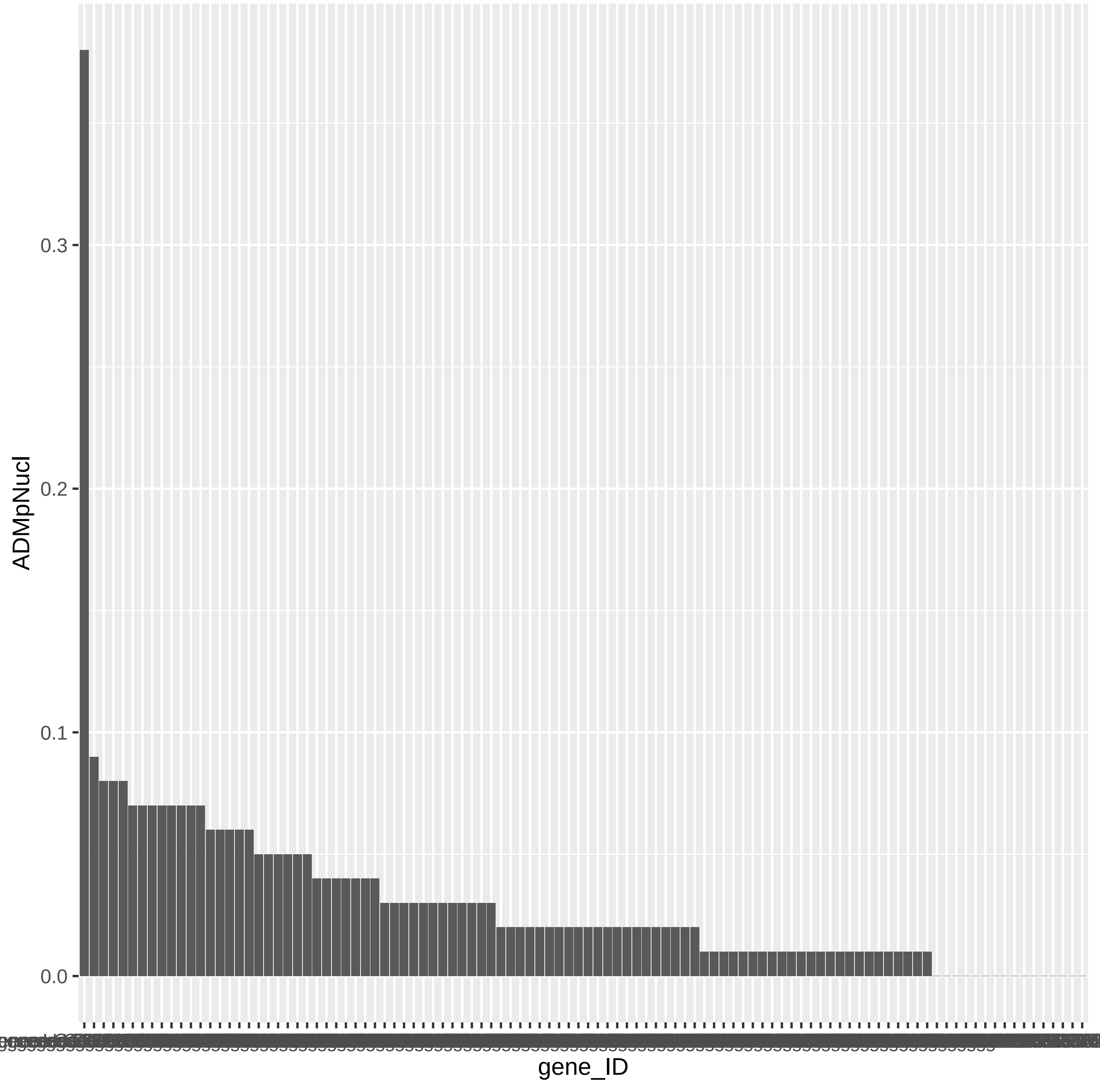

ADMpNucl
0.3
0.2
0.1
0.0
gene_ID

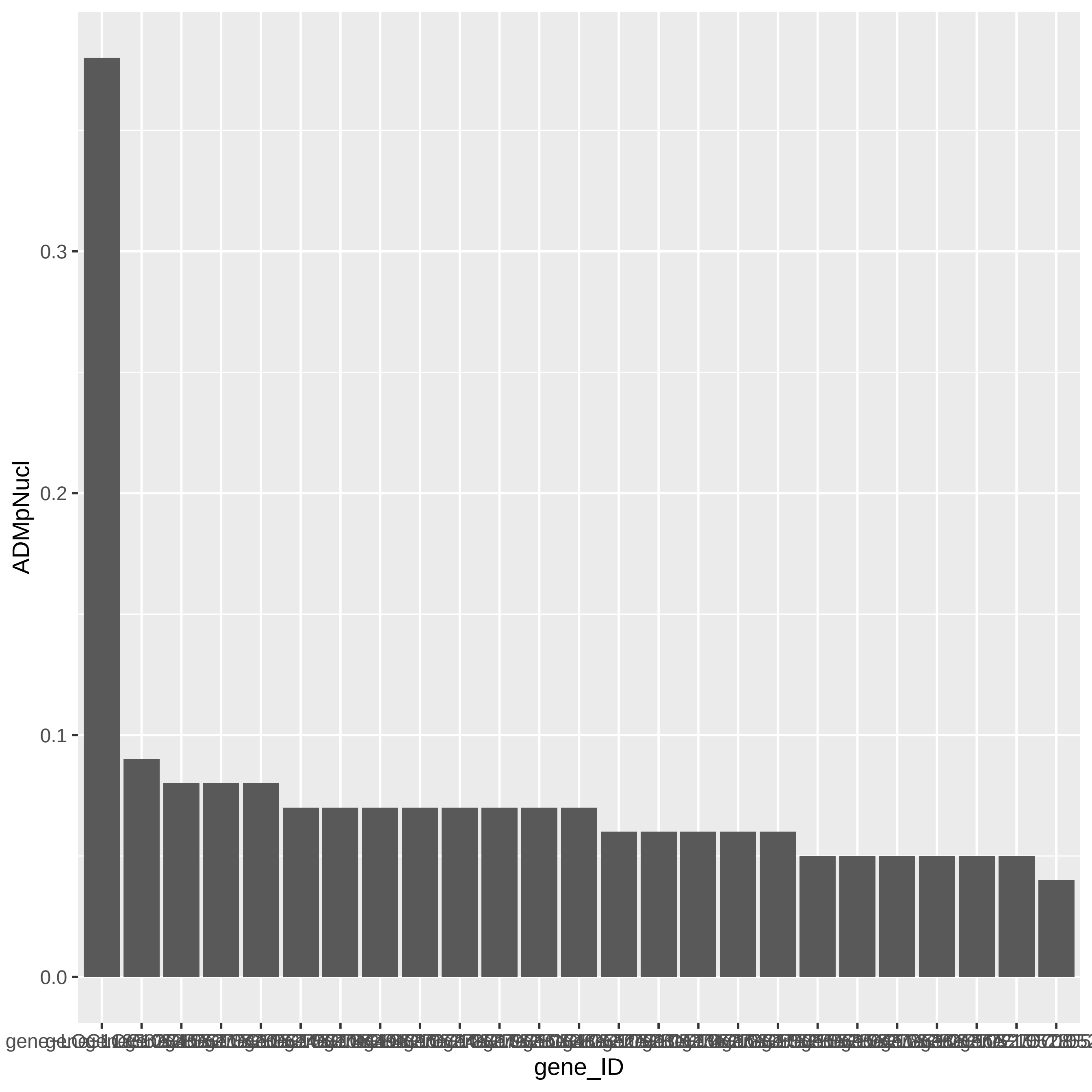
ADMpNucl
0.3
0.2
0.1
0.0
gene_ID

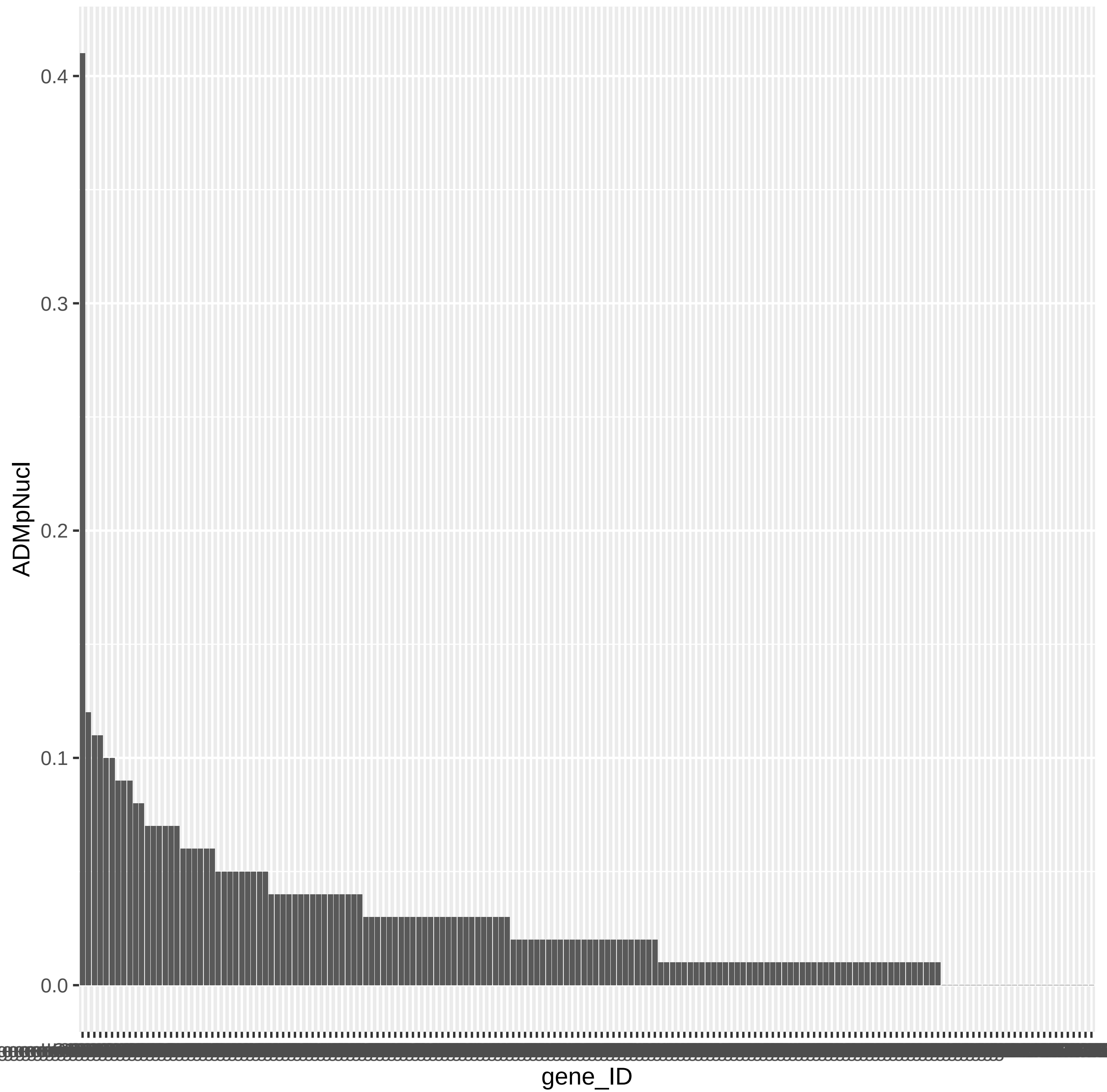

0.4
0.3
0.2
0.1
0.0
ADMpNucl
gene_ID

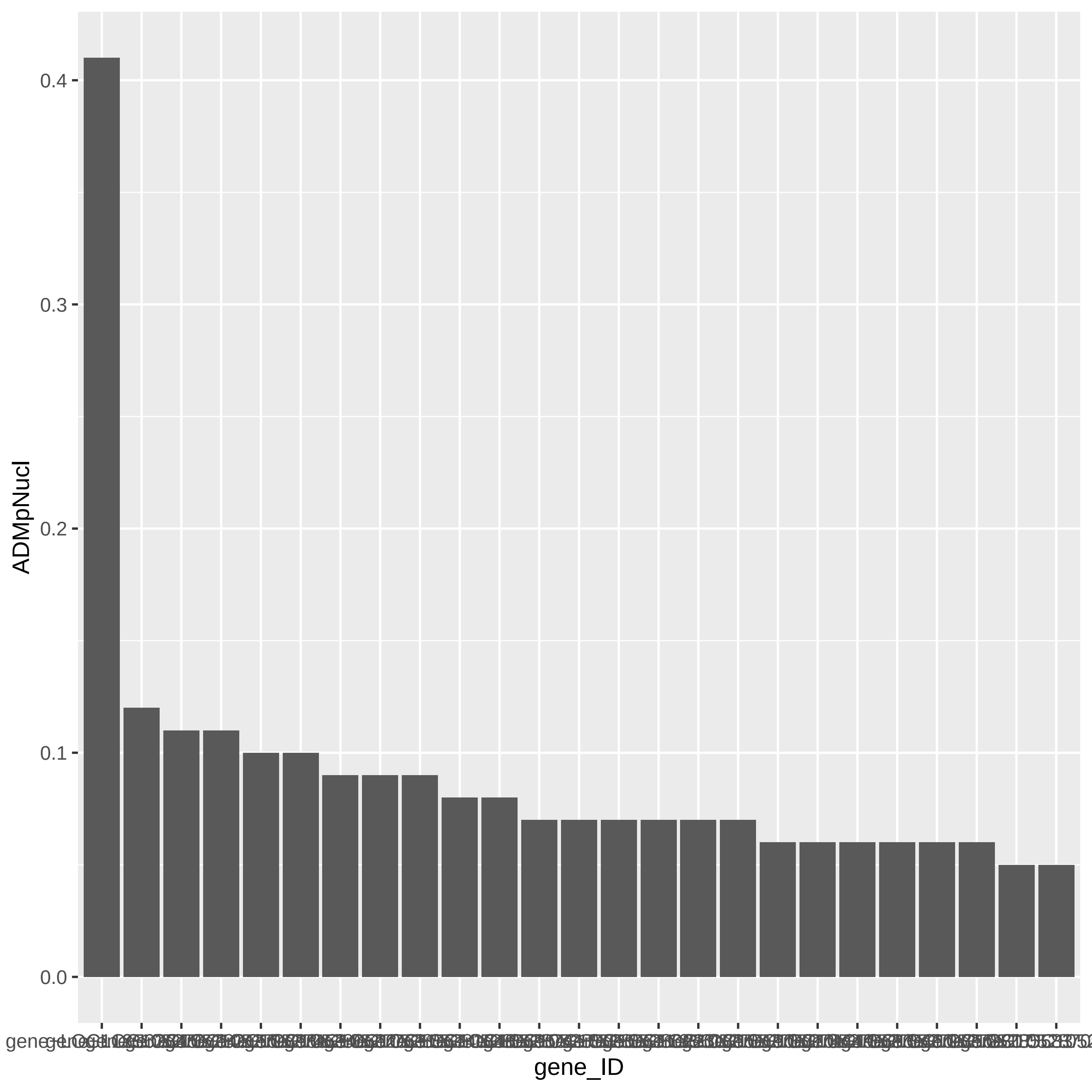
0.4
0.3
0.2
0.1
0.0
ADMpNucl
gene_ID

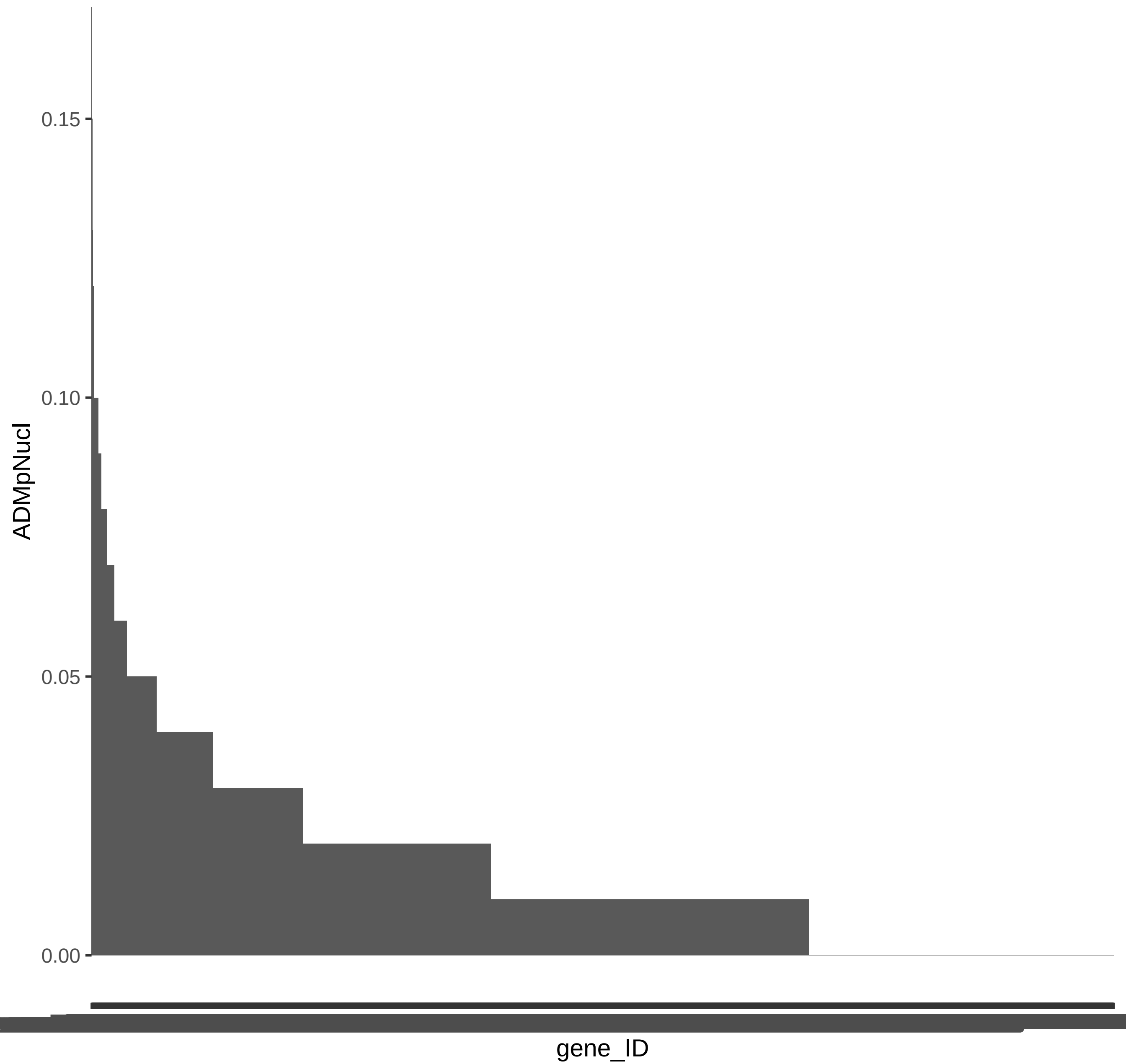

ADMpNucl
0.15
0.10
0.05
0.00
gene_ID

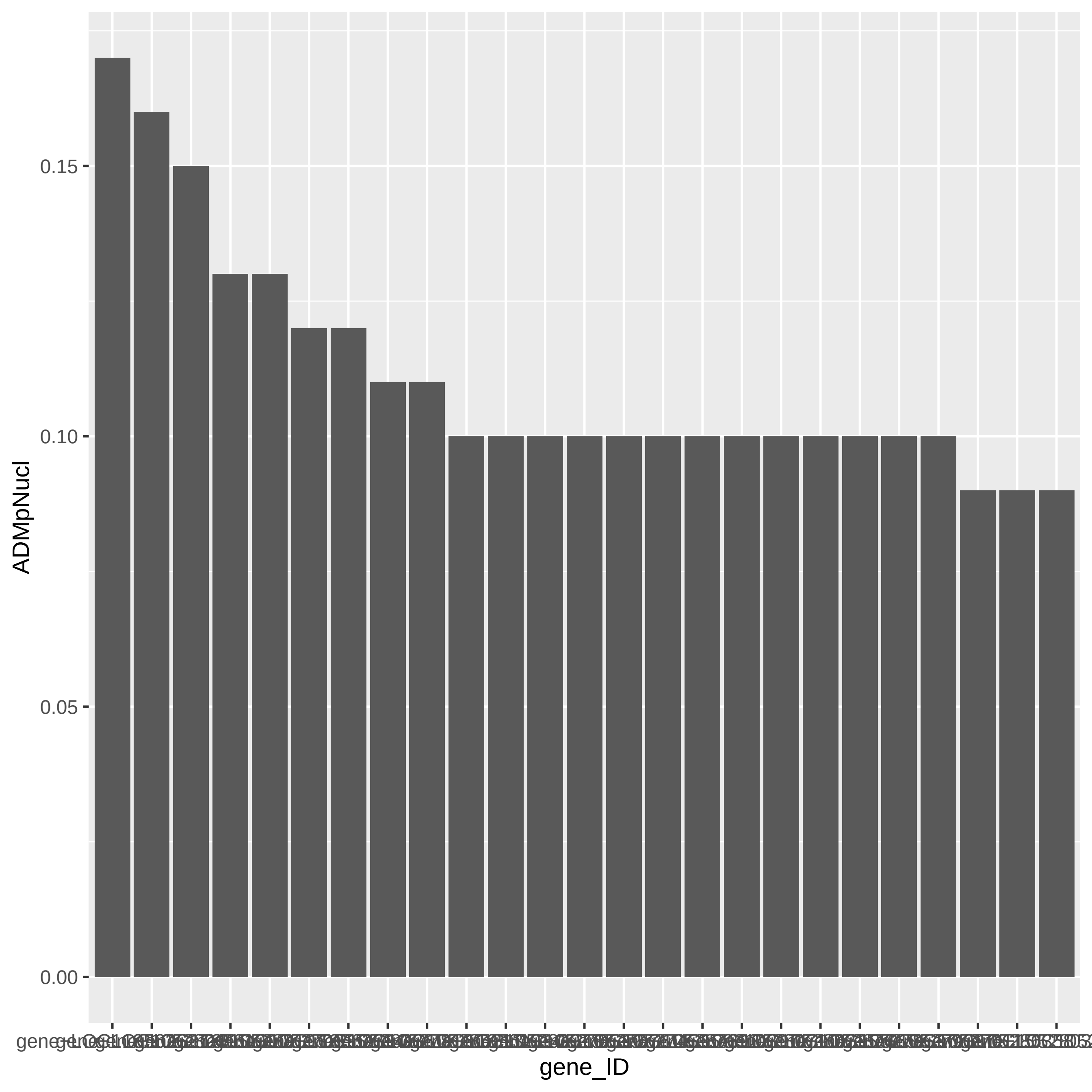

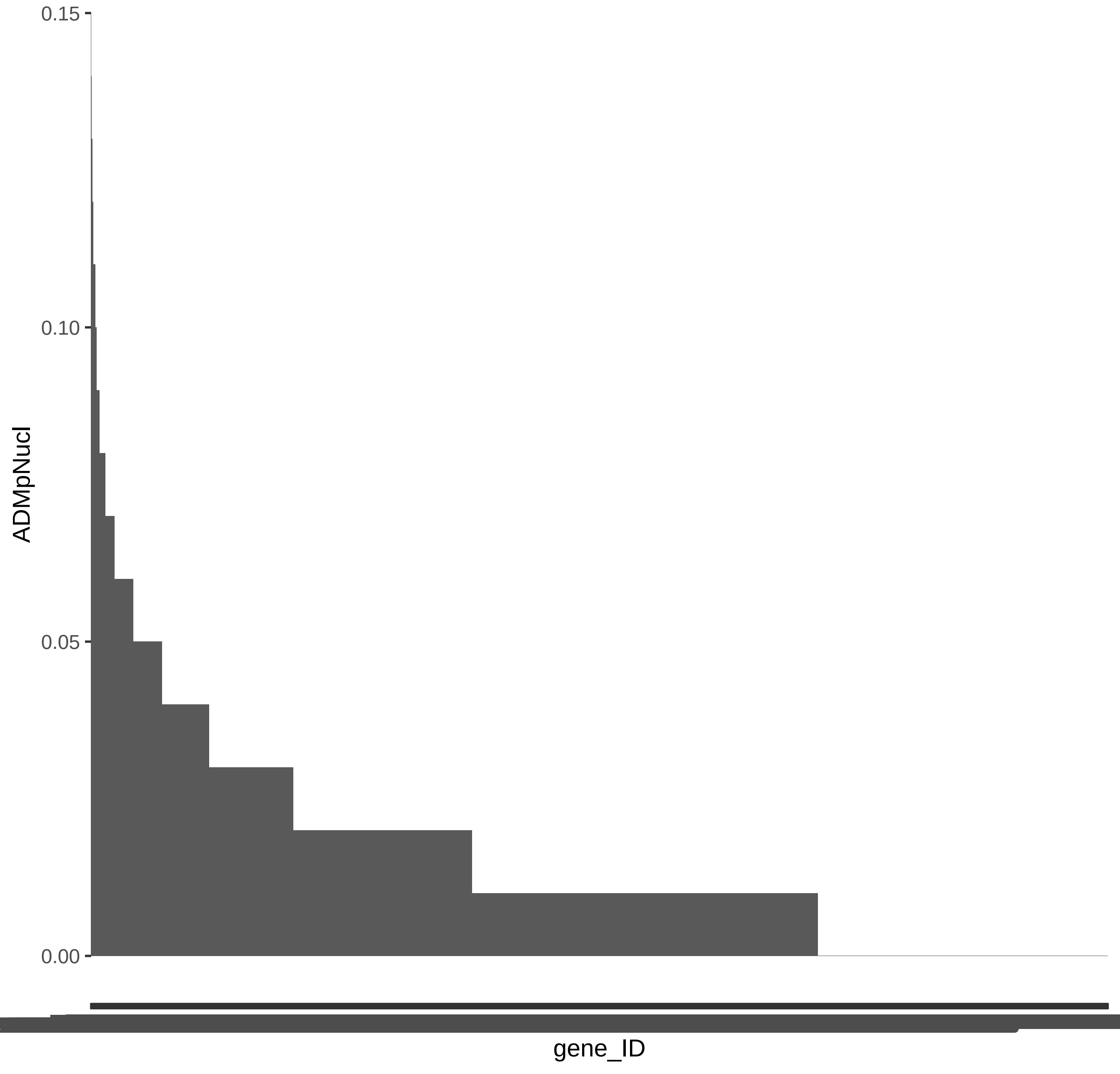

0.15
0.10
0.05
0.00
ADMpNucl
gene_ID

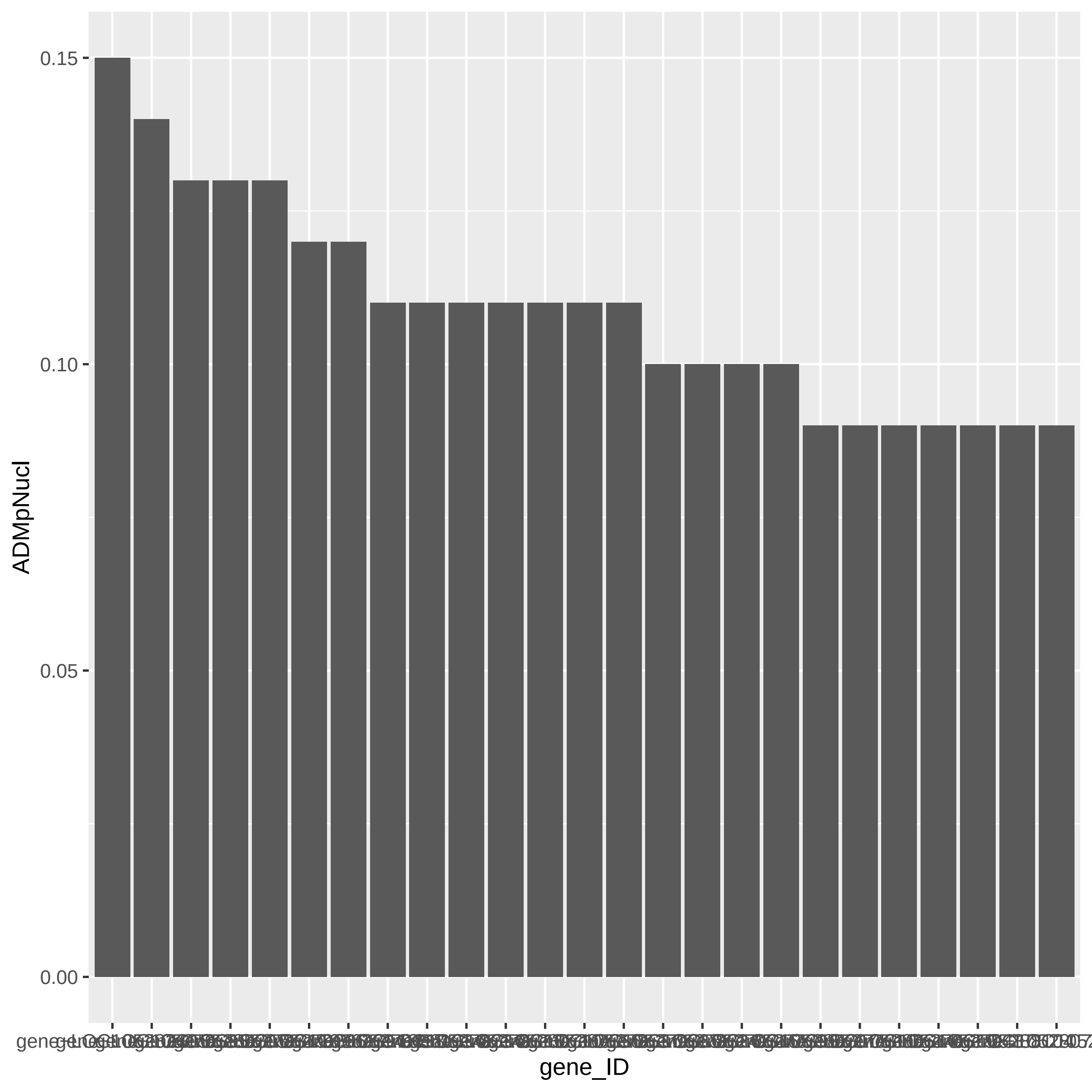

ADMpNucl
0.15
0.10
0.05
0.00
gene_ID

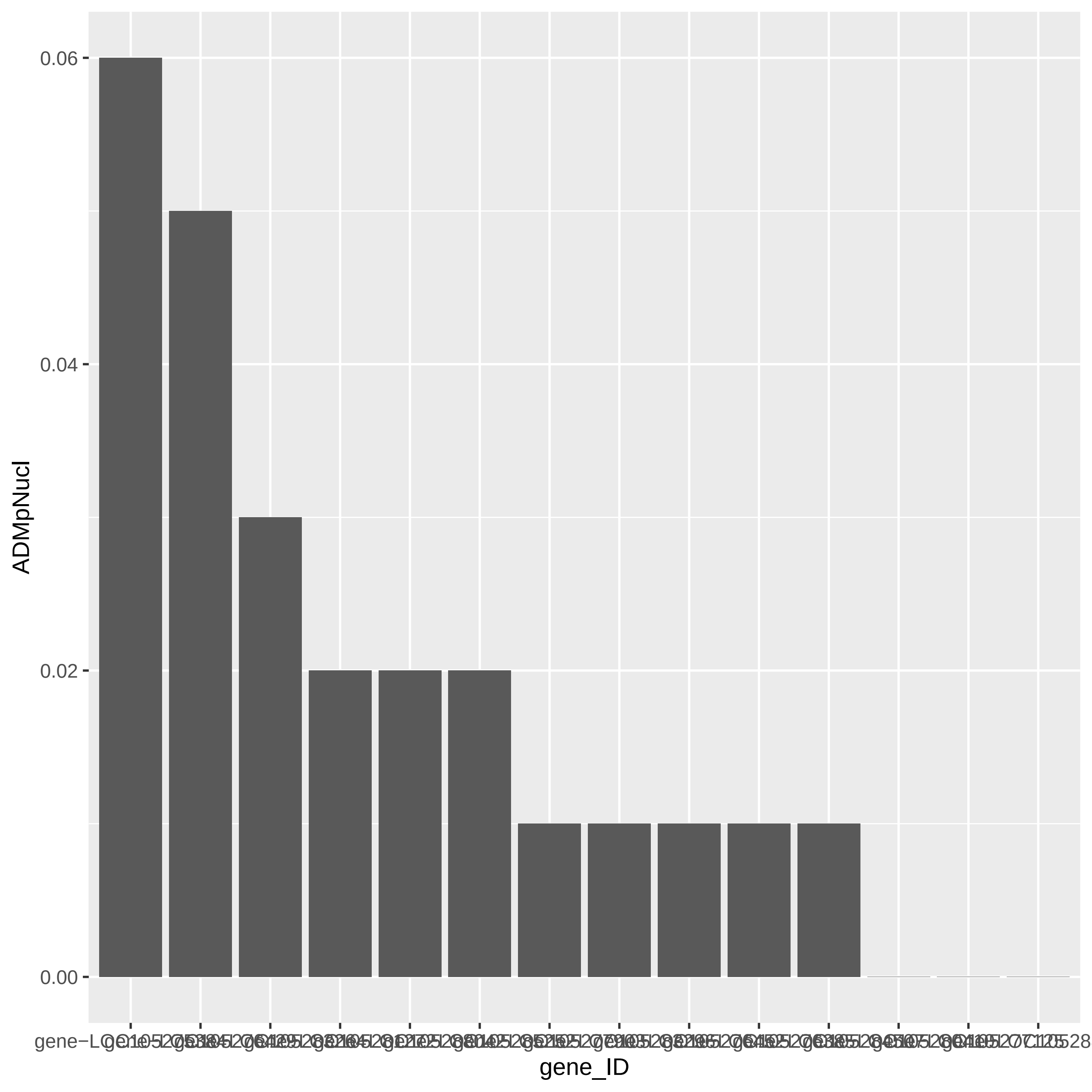

0.06
0.04
0.02
0.00
ADMpNucl
gene_ID

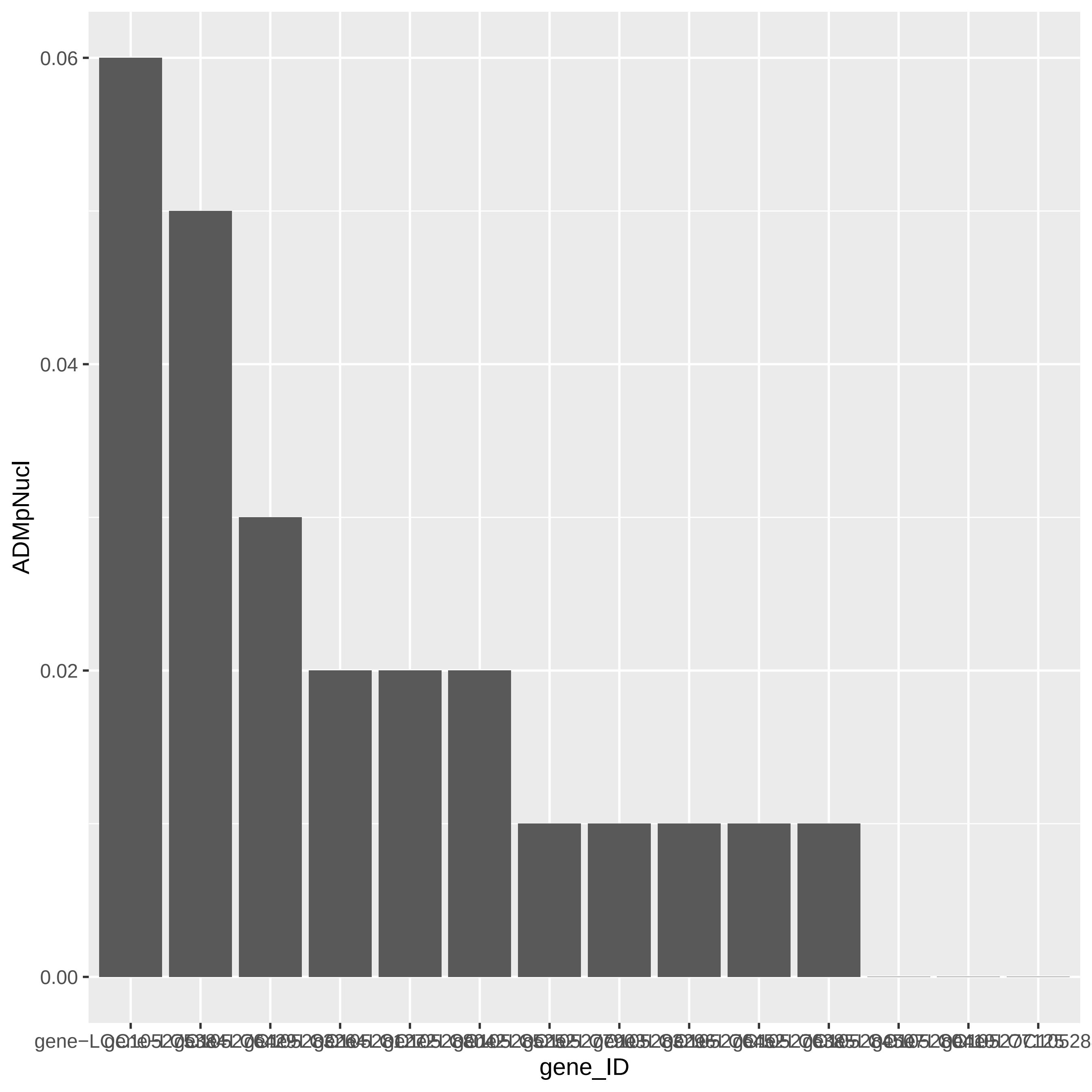

0.06
0.04
0.02
0.00
ADMpNucl
gene_ID

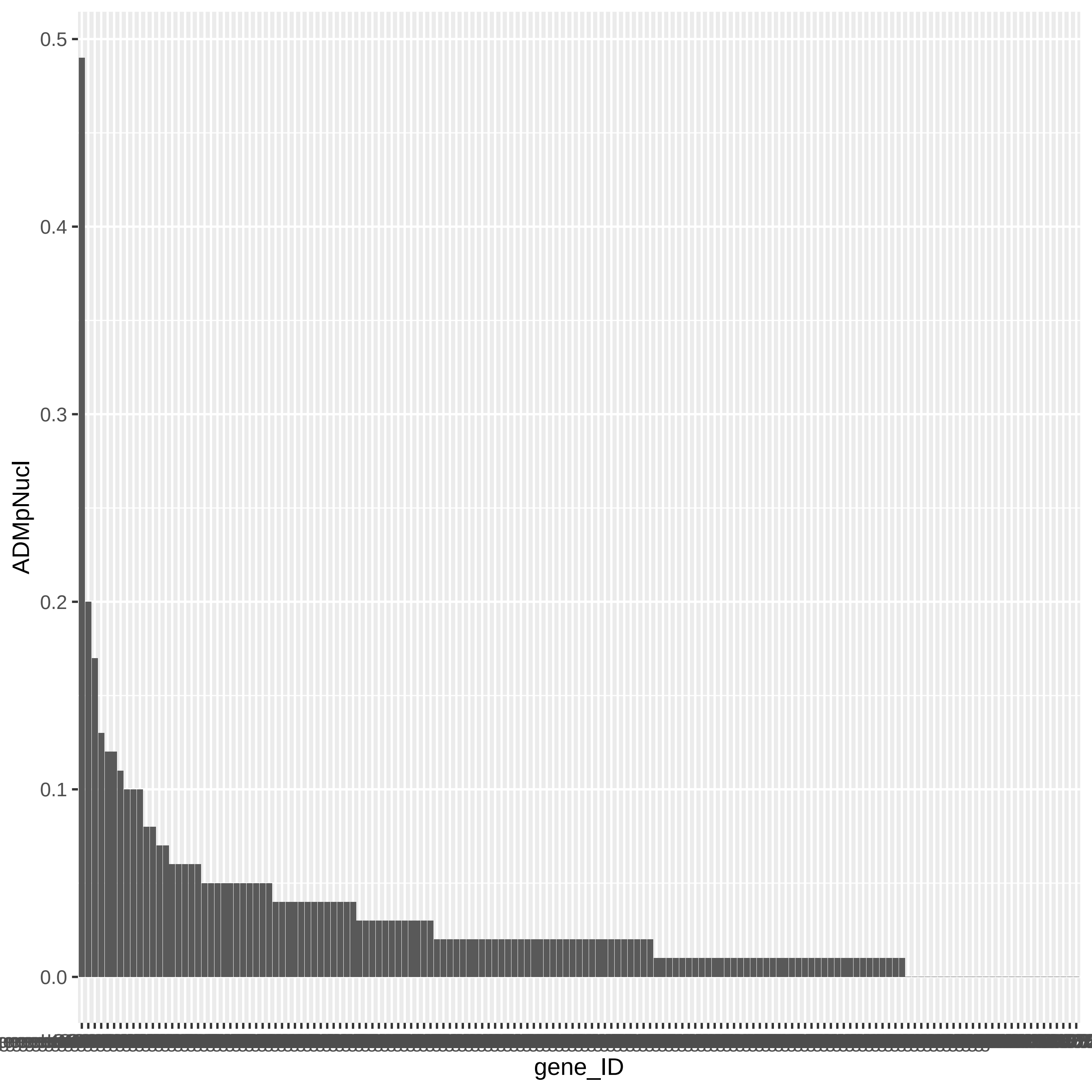

0.5
0.4
0.3
0.2
0.1
0.0
ADMpNucl
gene_ID

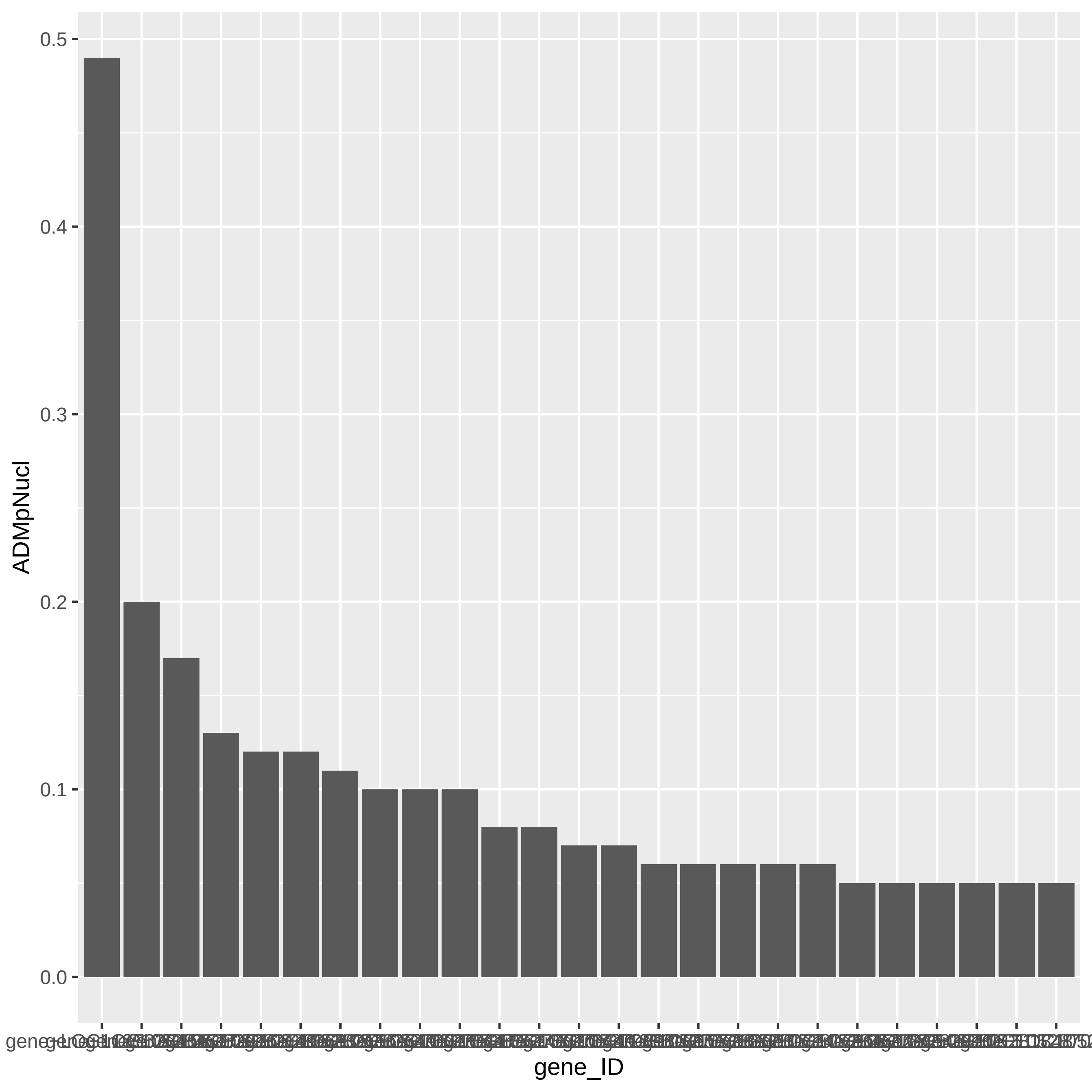

ADMpNucl
0.5
0.4
0.3
0.2
0.1
0.0
gene_ID

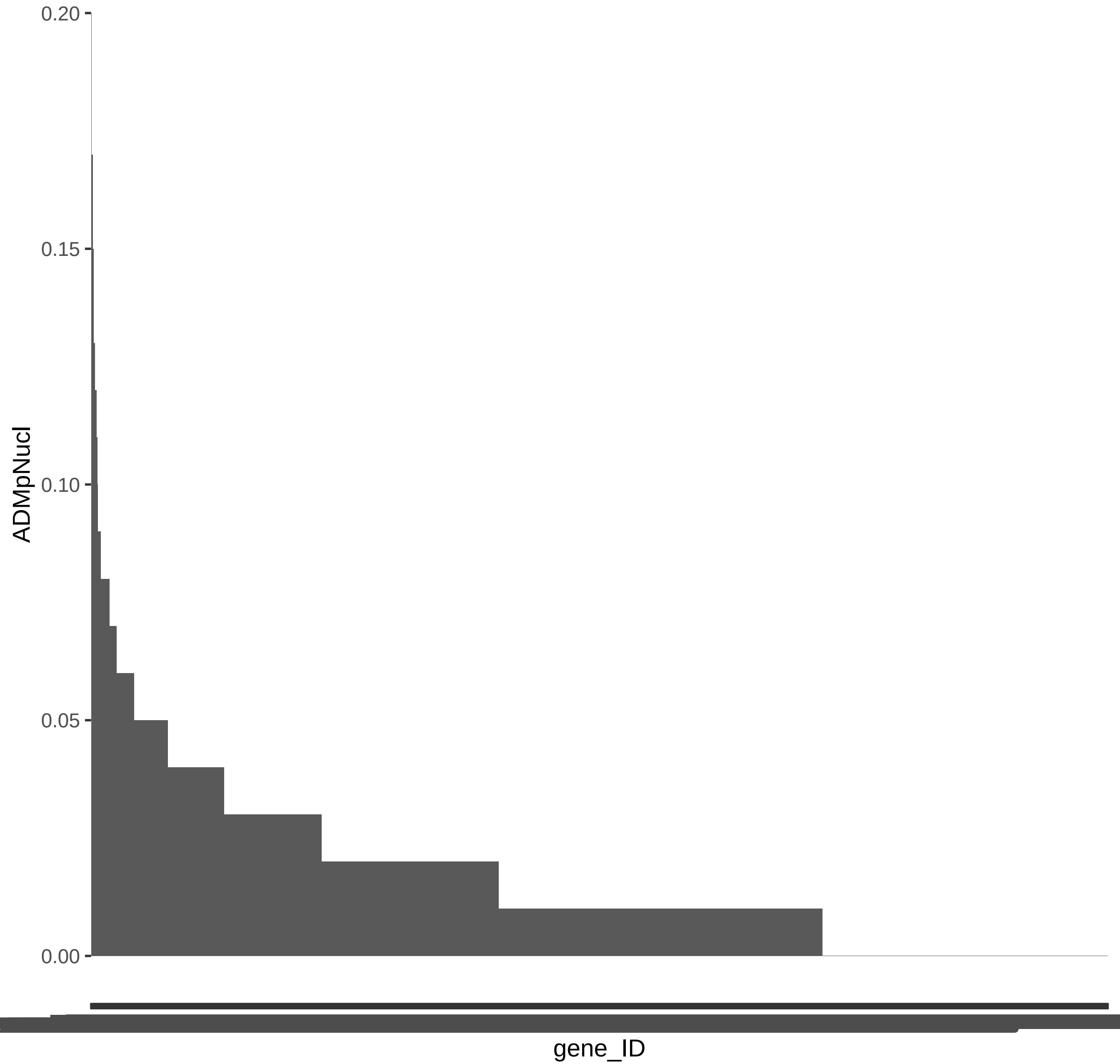
0.20
0.15
0.10
0.05
0.00
ADMpNucl
gene_ID

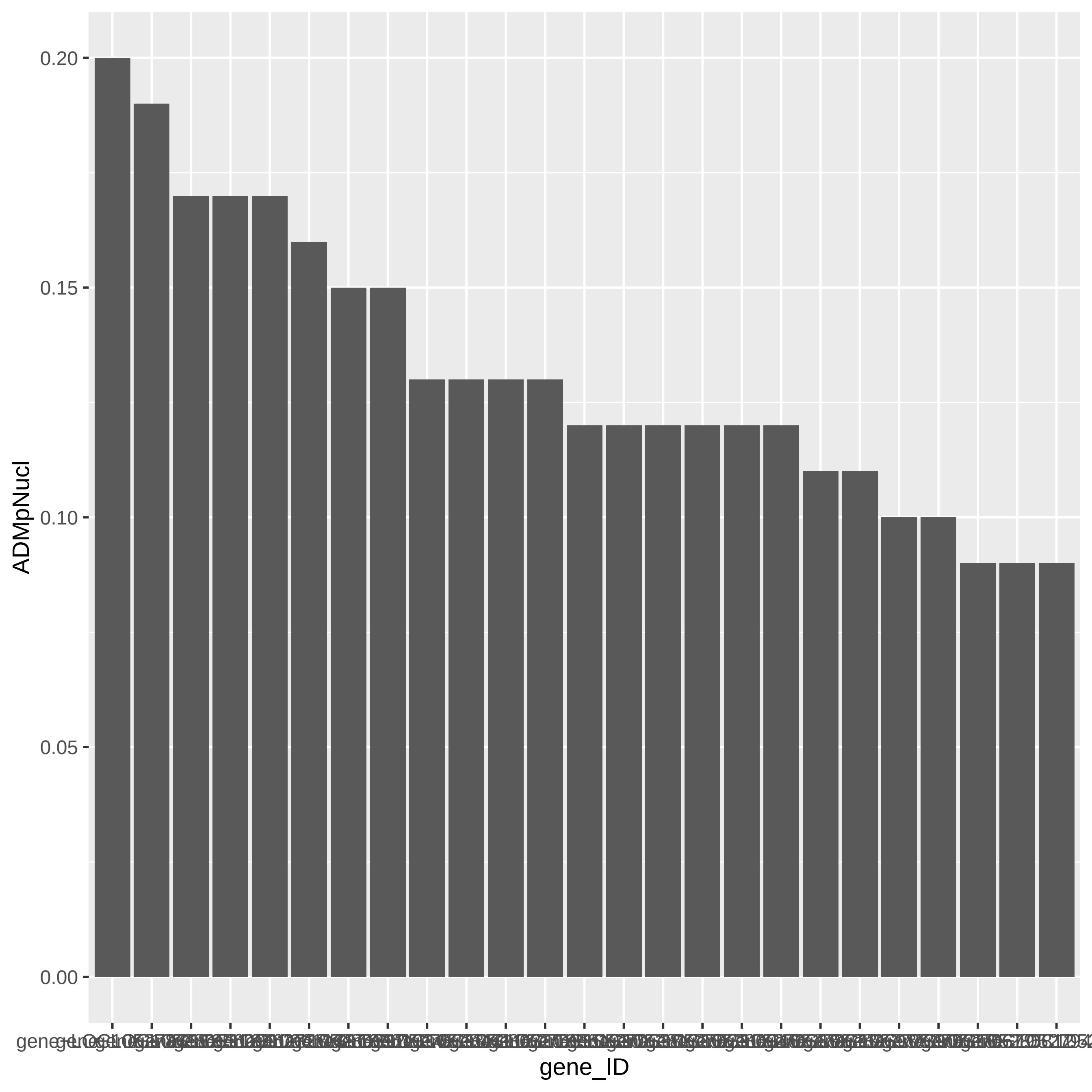

0.20
0.15
0.10
0.05
0.00
ADMpNucl
gene_ID

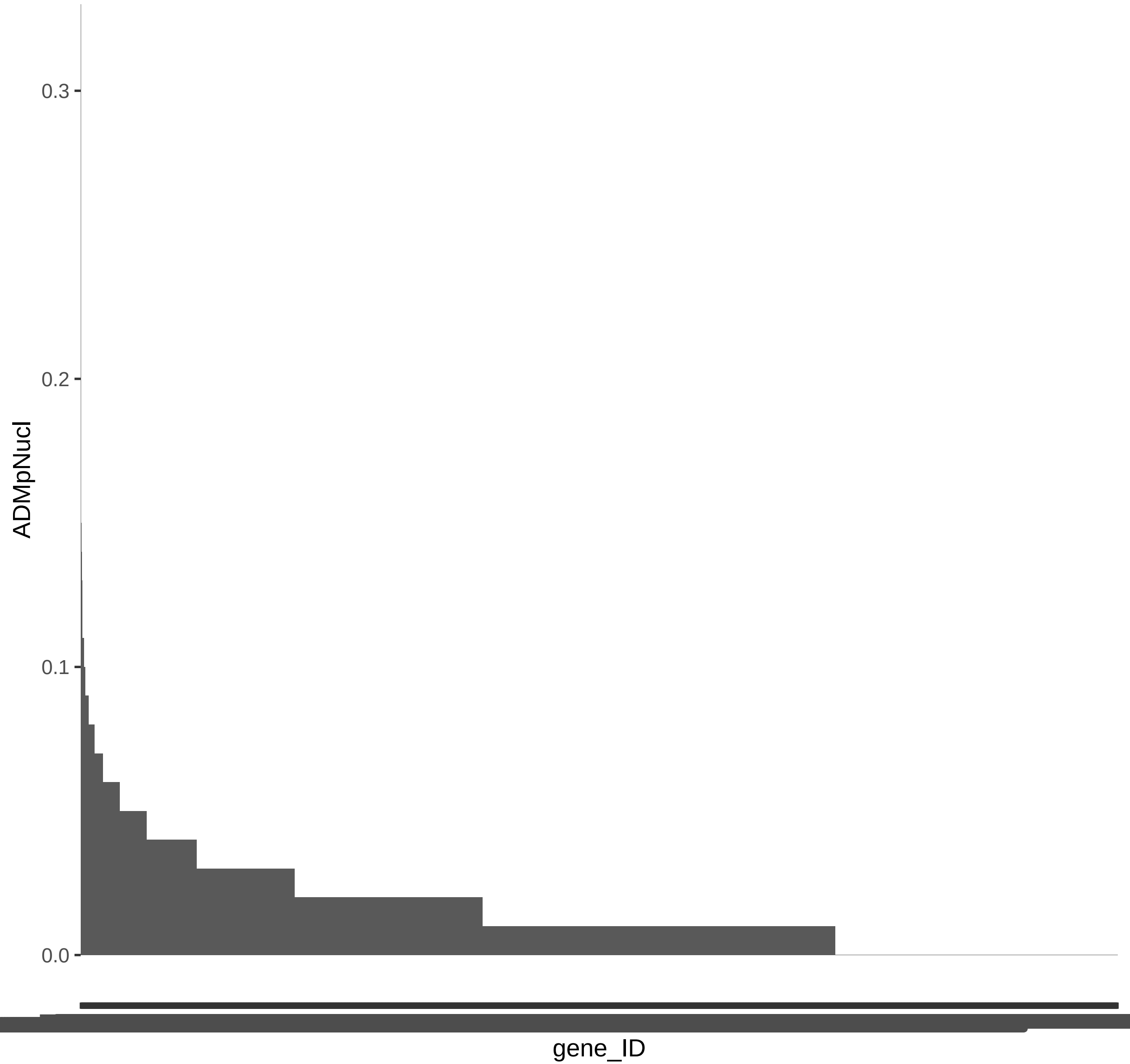
0.3
0.2
0.1
0.0
ADMpNucl
gene_ID

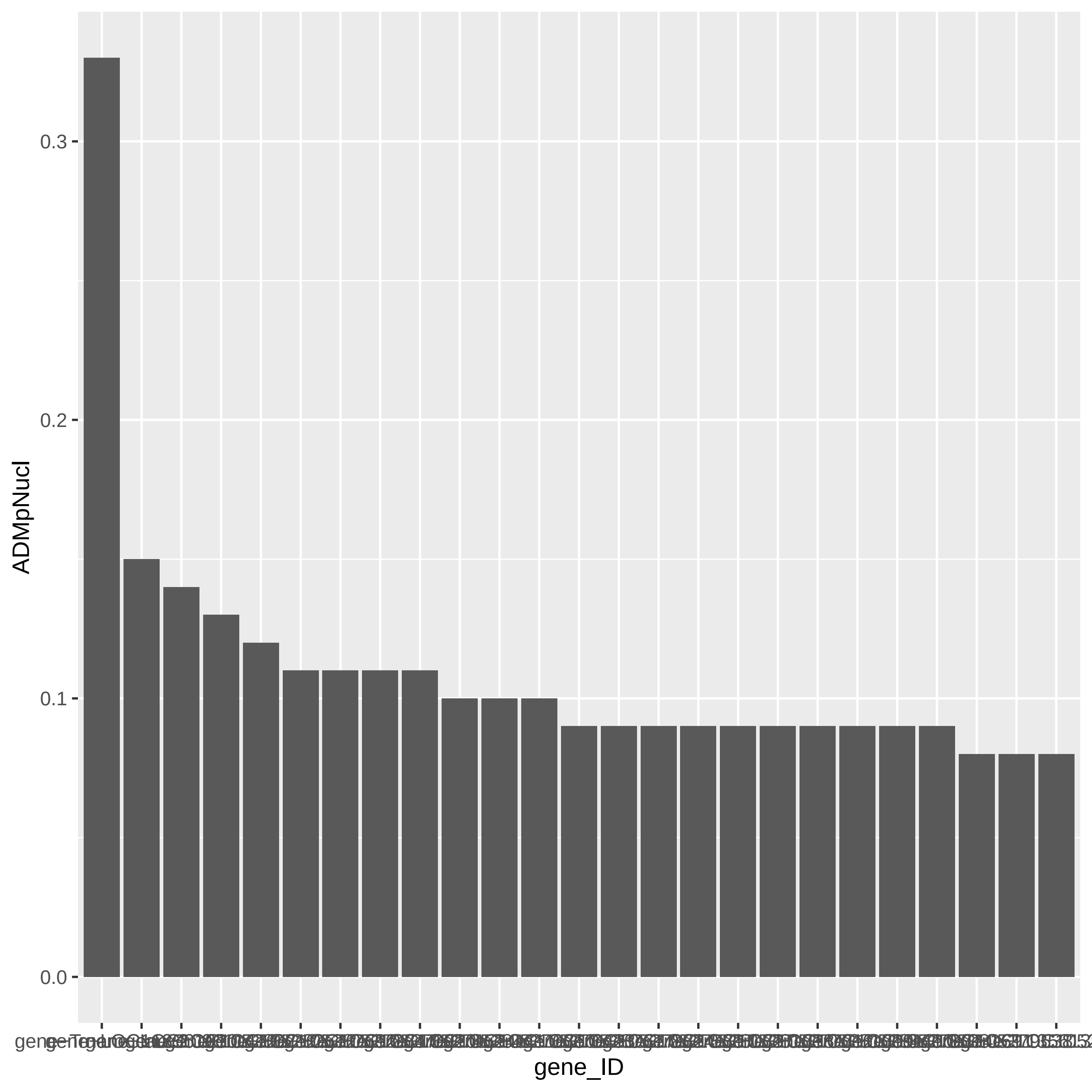

0.3
0.2
0.1
0.0
ADMpNucl
gene_ID

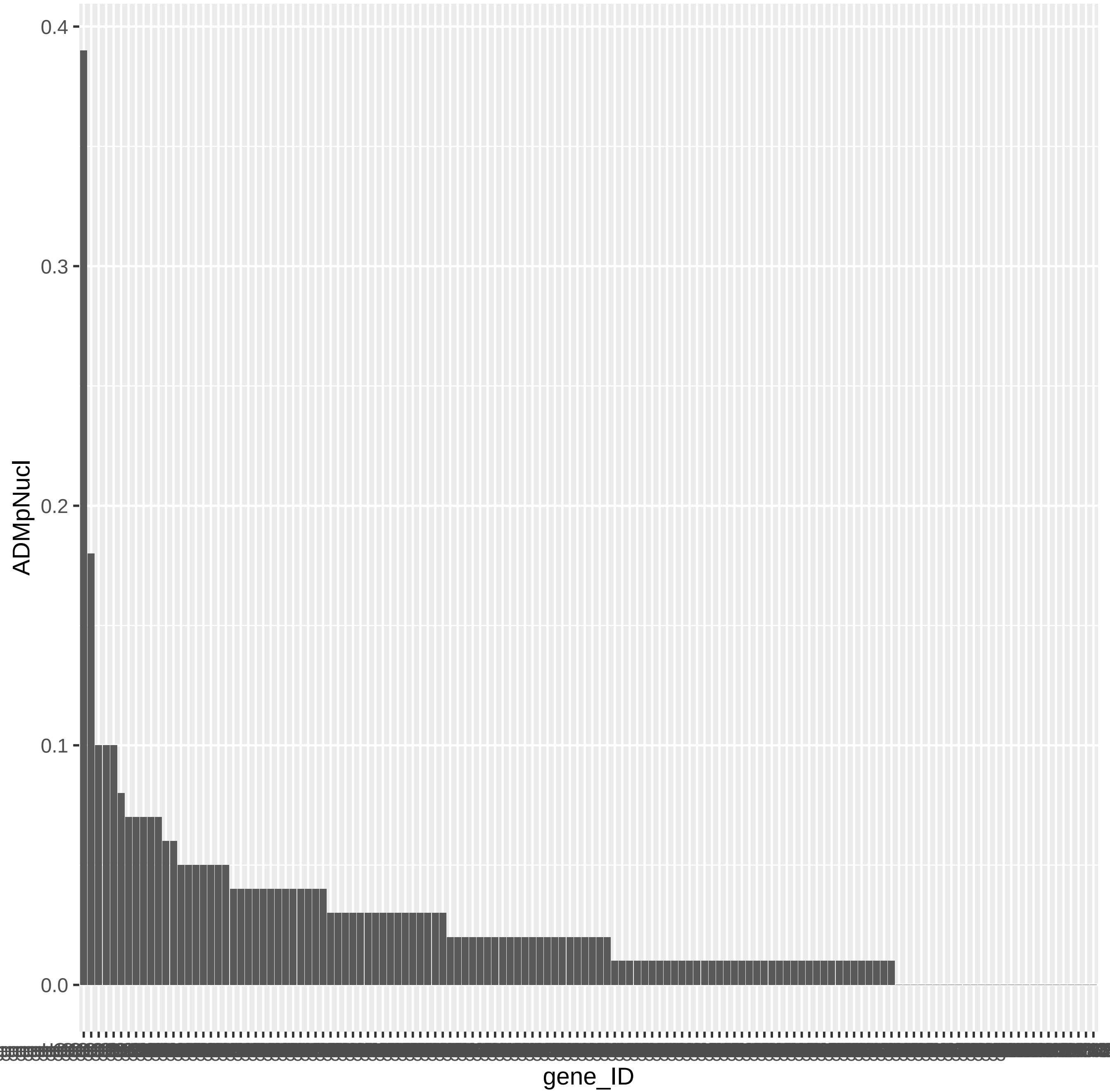

0.4
0.3
0.2
0.1
0.0
ADMpNucl
gene_ID

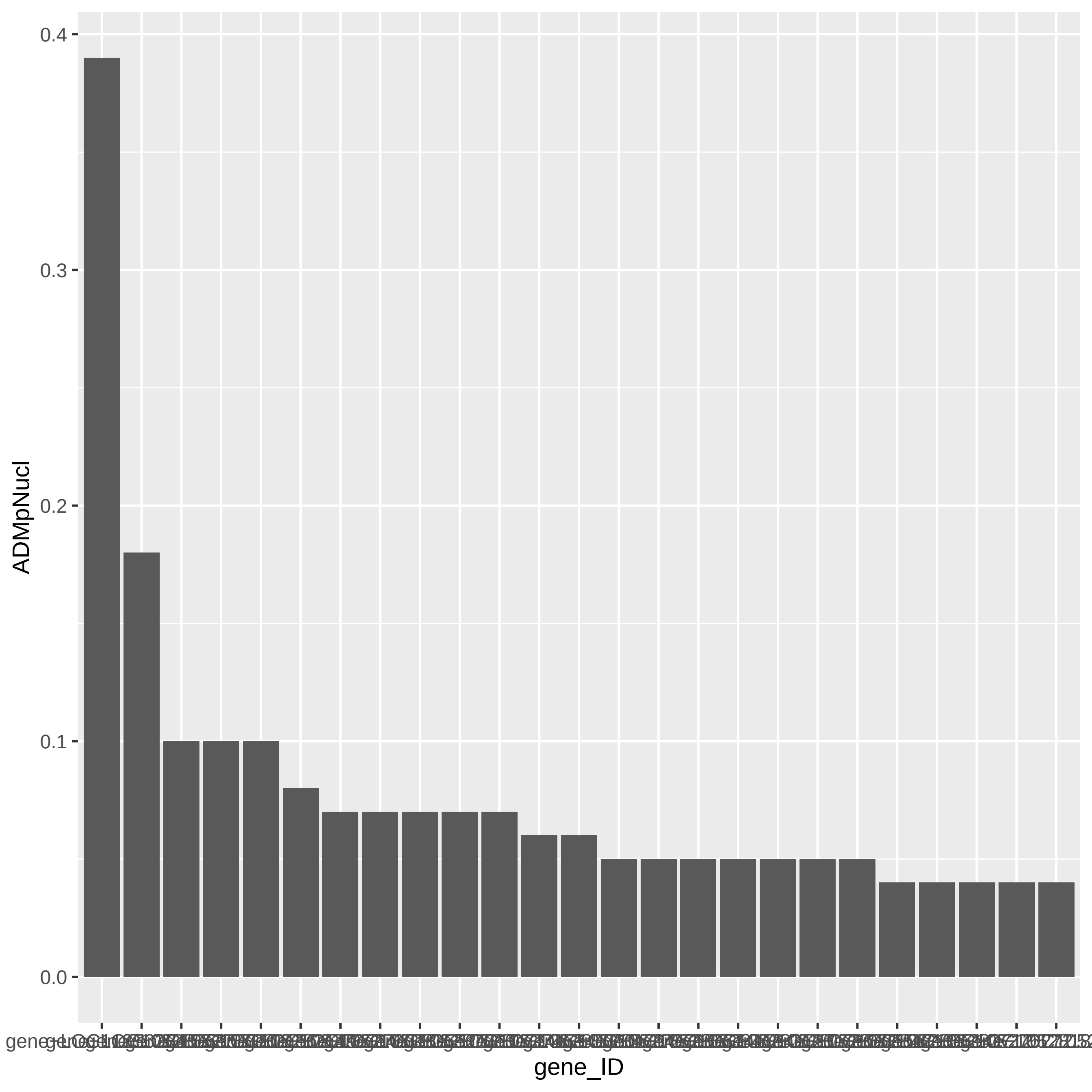

0.4
0.3
0.2
0.1
0.0
ADMpNucl
gene_ID

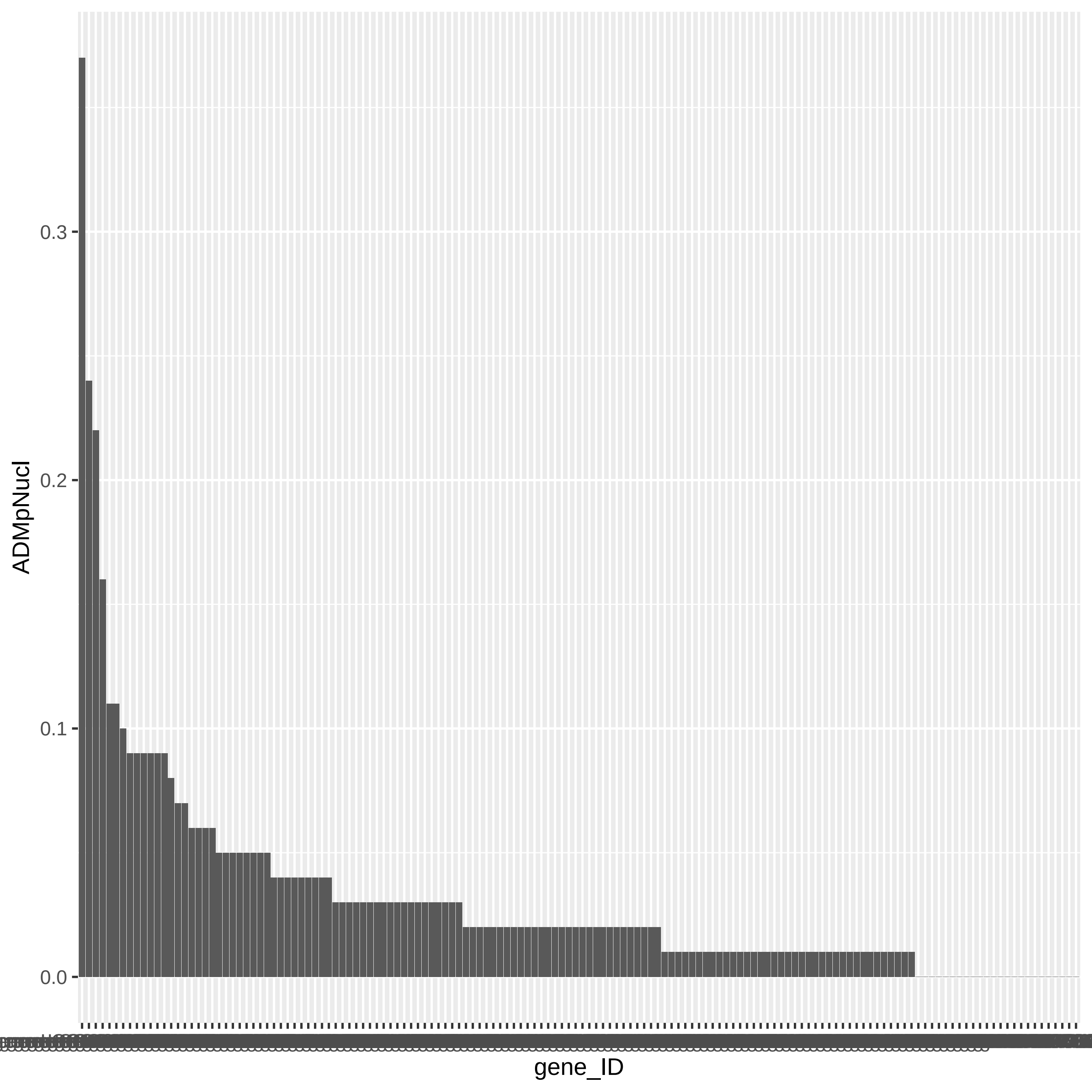

ADMpNucl
0.3
0.2
0.1
0.0
gene_ID

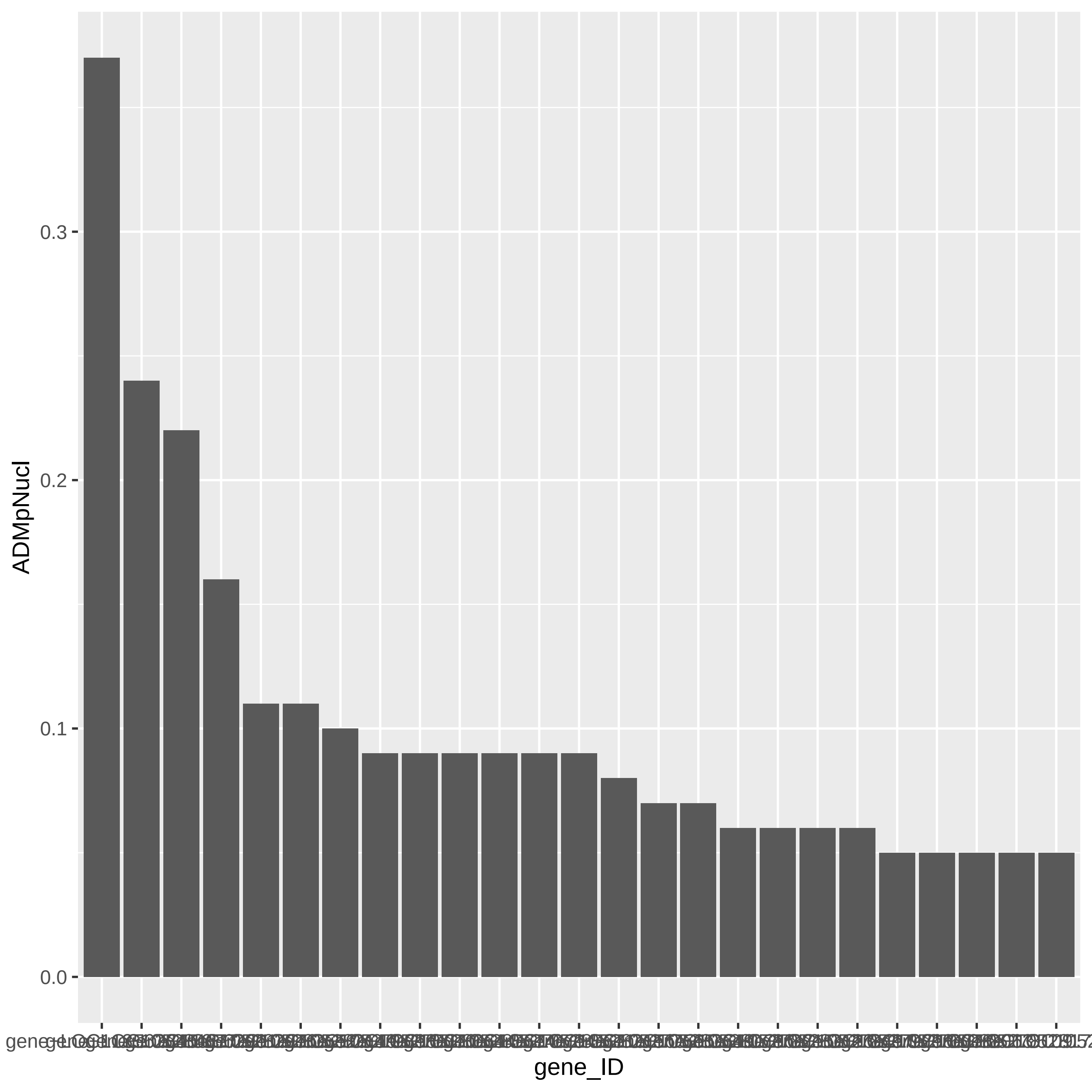

ADMpNucl
0.0
0.1
0.2
0.3
gene_ID

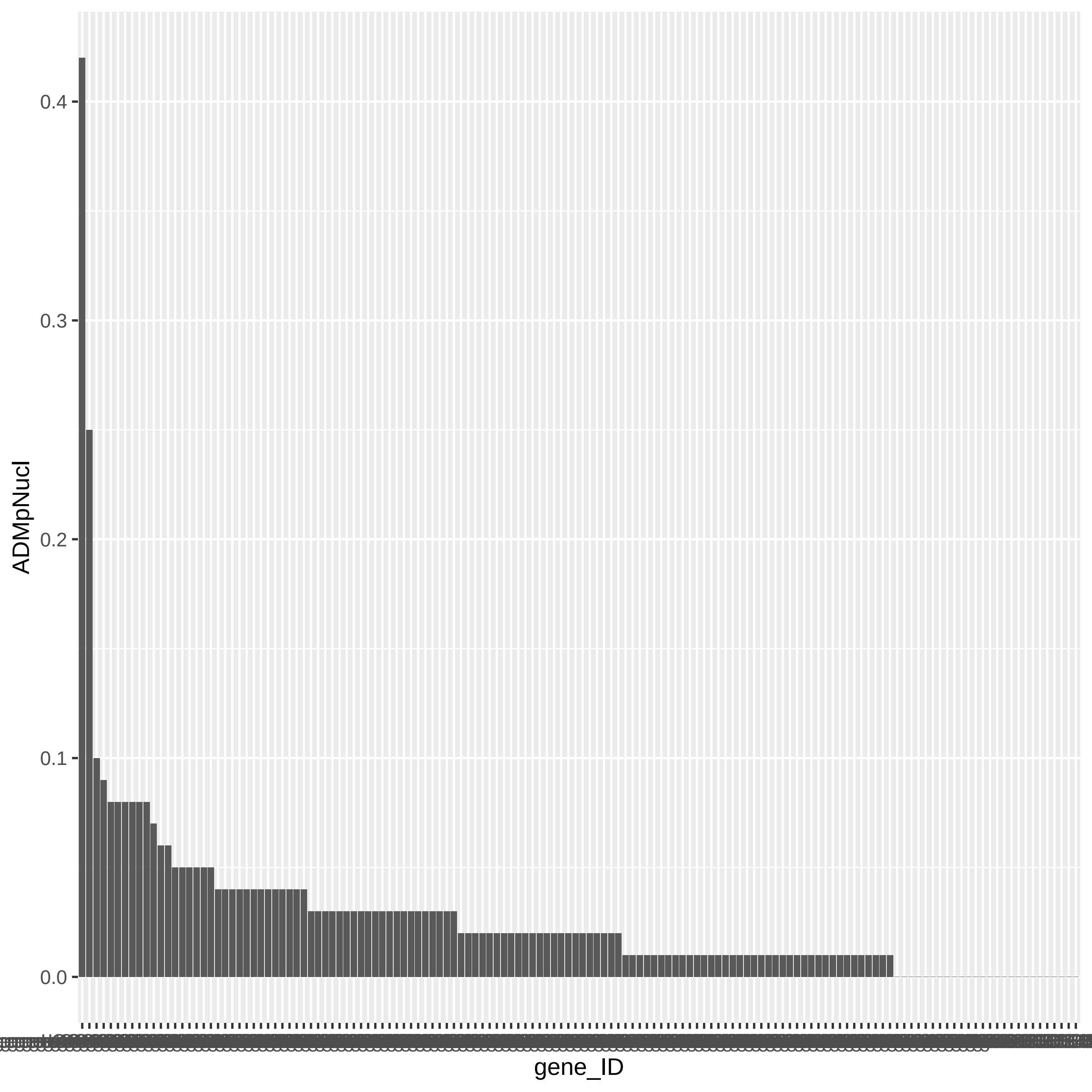
0.4
0.3
0.2
0.1
0.0
ADMpNucl
gene_ID

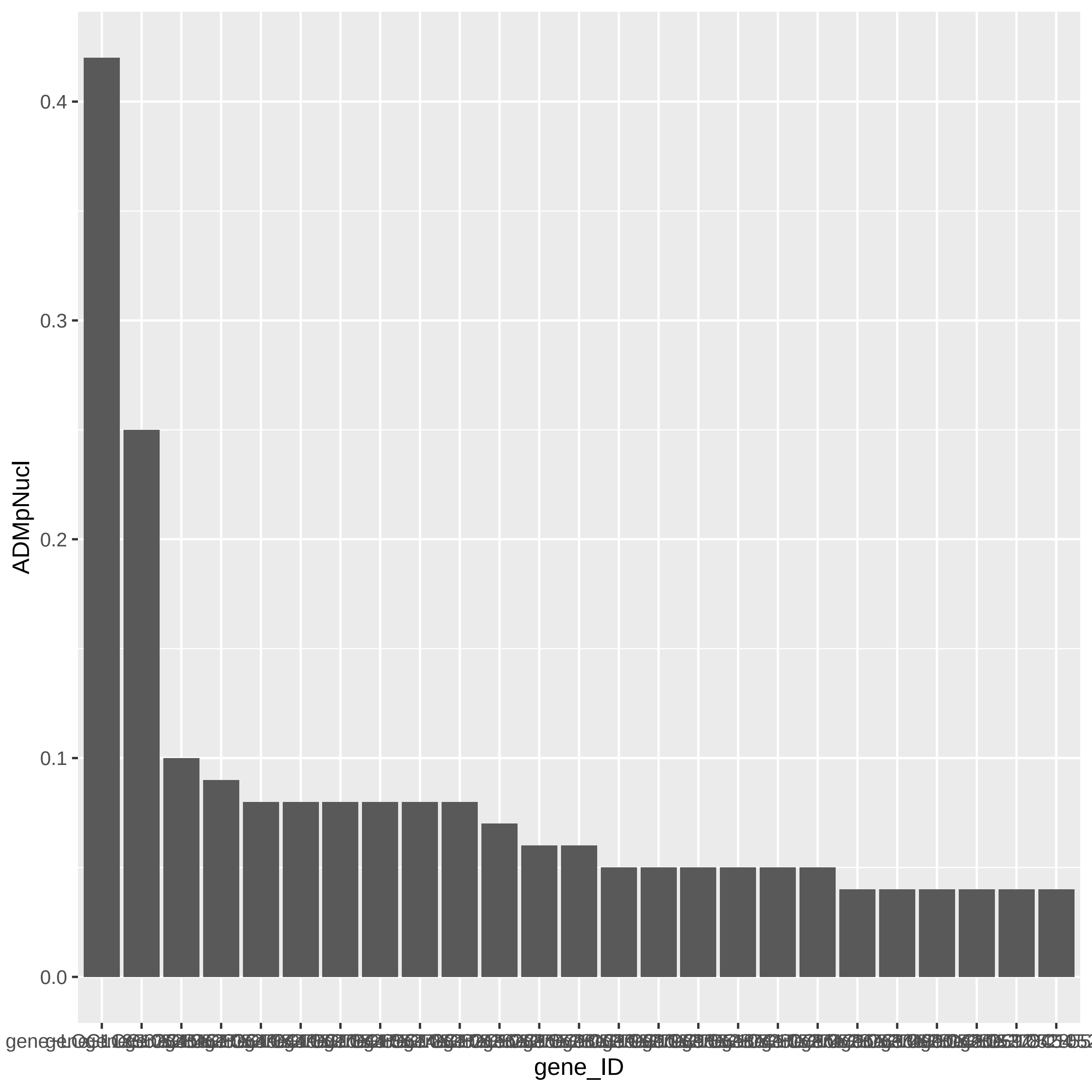
0.4
0.3
0.2
0.1
0.0
ADMpNucl
gene_ID

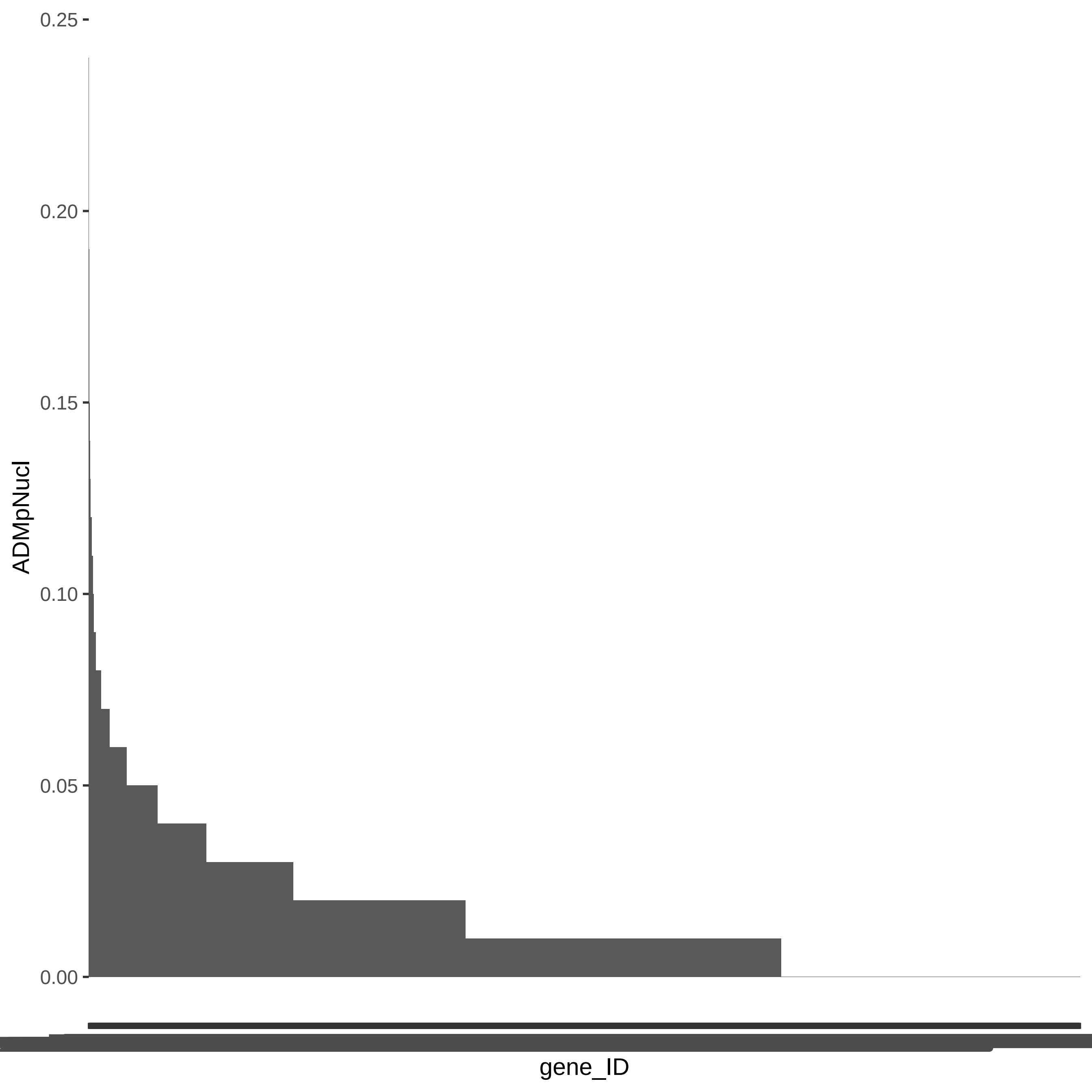

0.25
0.20
0.15
0.10
0.05
0.00
ADMpNucl
gene_ID

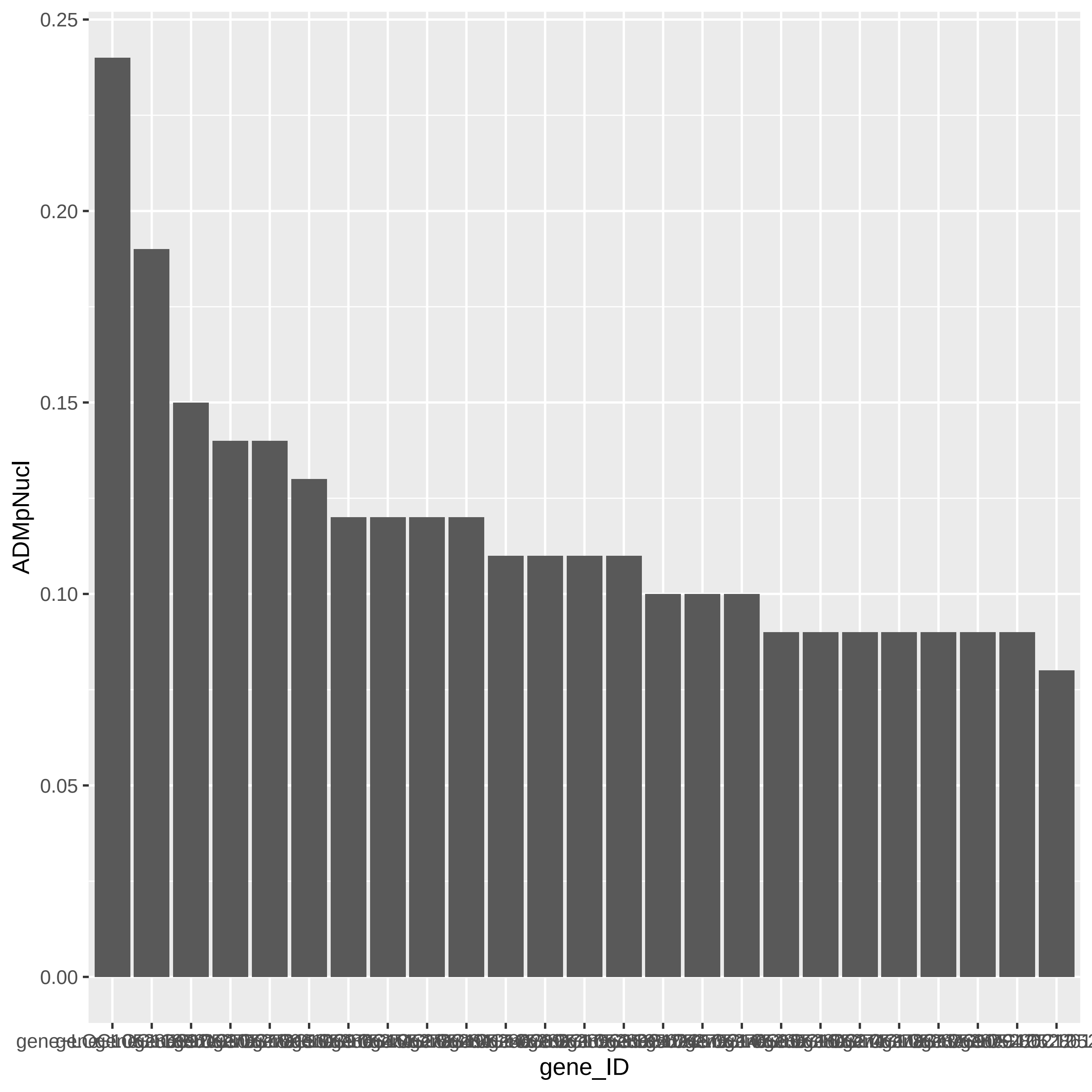

0.25
0.20
0.15
0.10
0.05
0.00
ADMpNucl
gene_ID

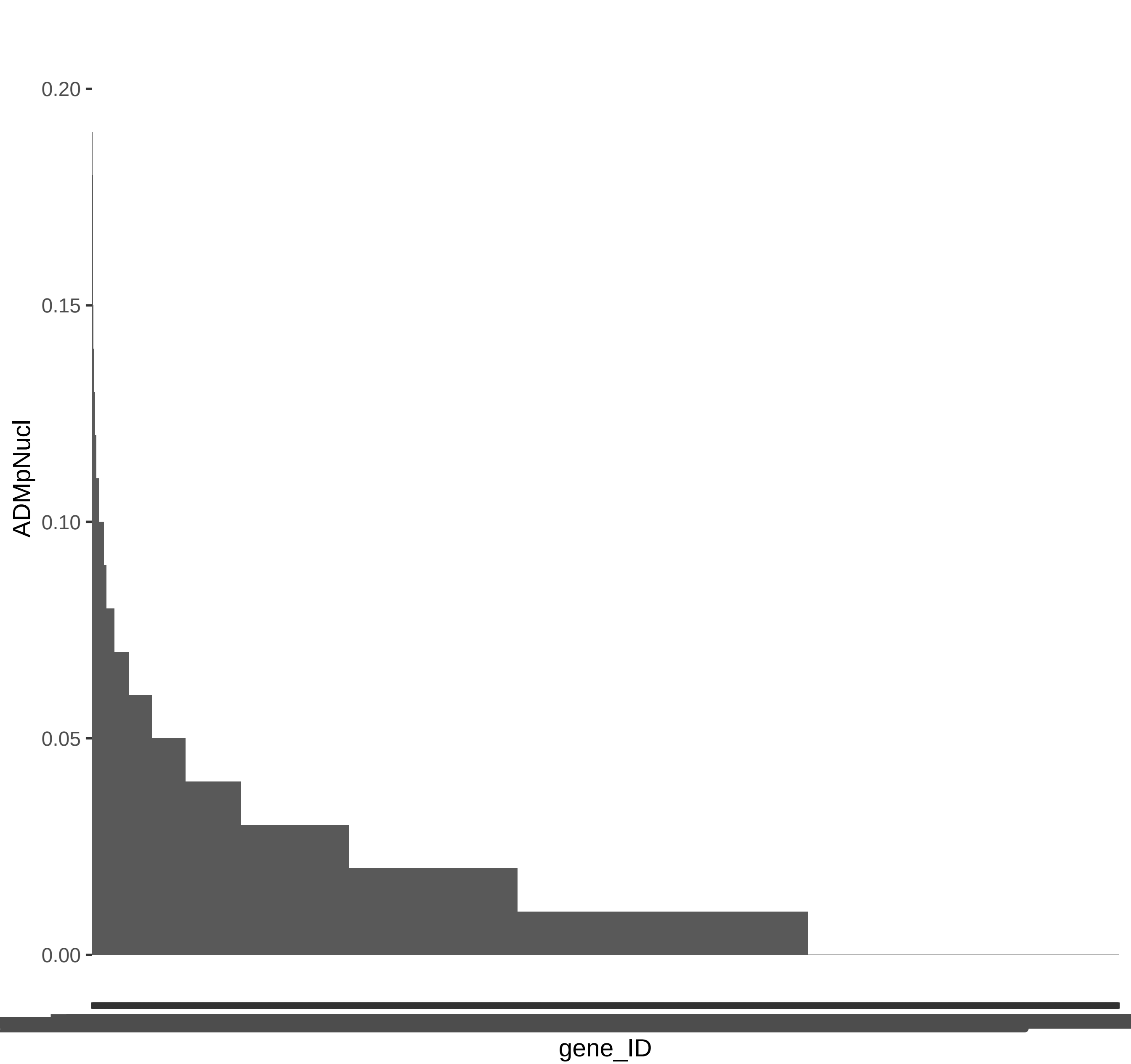

0.20
0.15
0.10
0.05
0.00
ADMpNucl
gene_ID

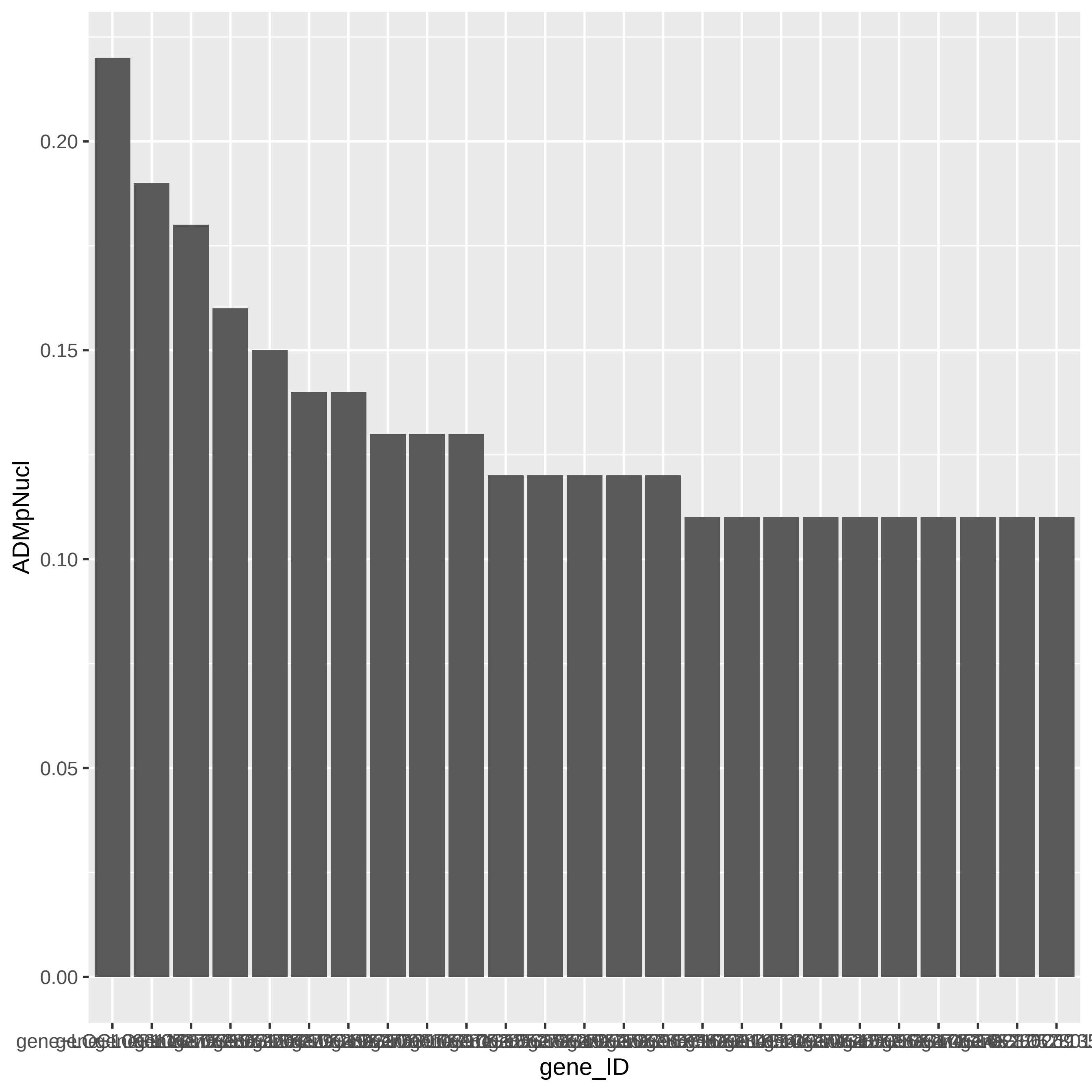
ADMpNucl
0.00
0.05
0.10
0.15
0.20
gene_ID

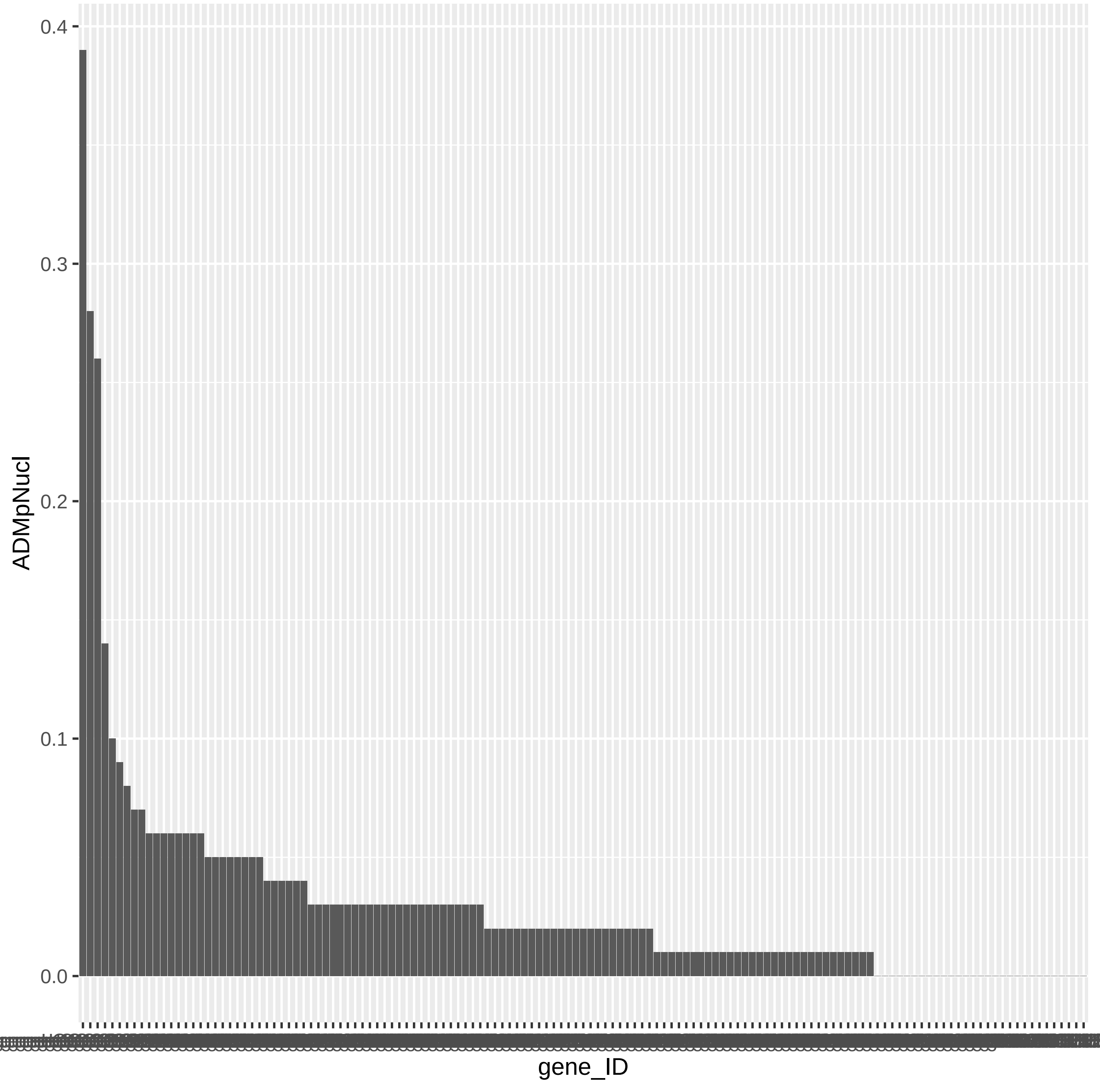
0.4
0.3
0.2
0.1
0.0
ADMpNucl
gene_ID

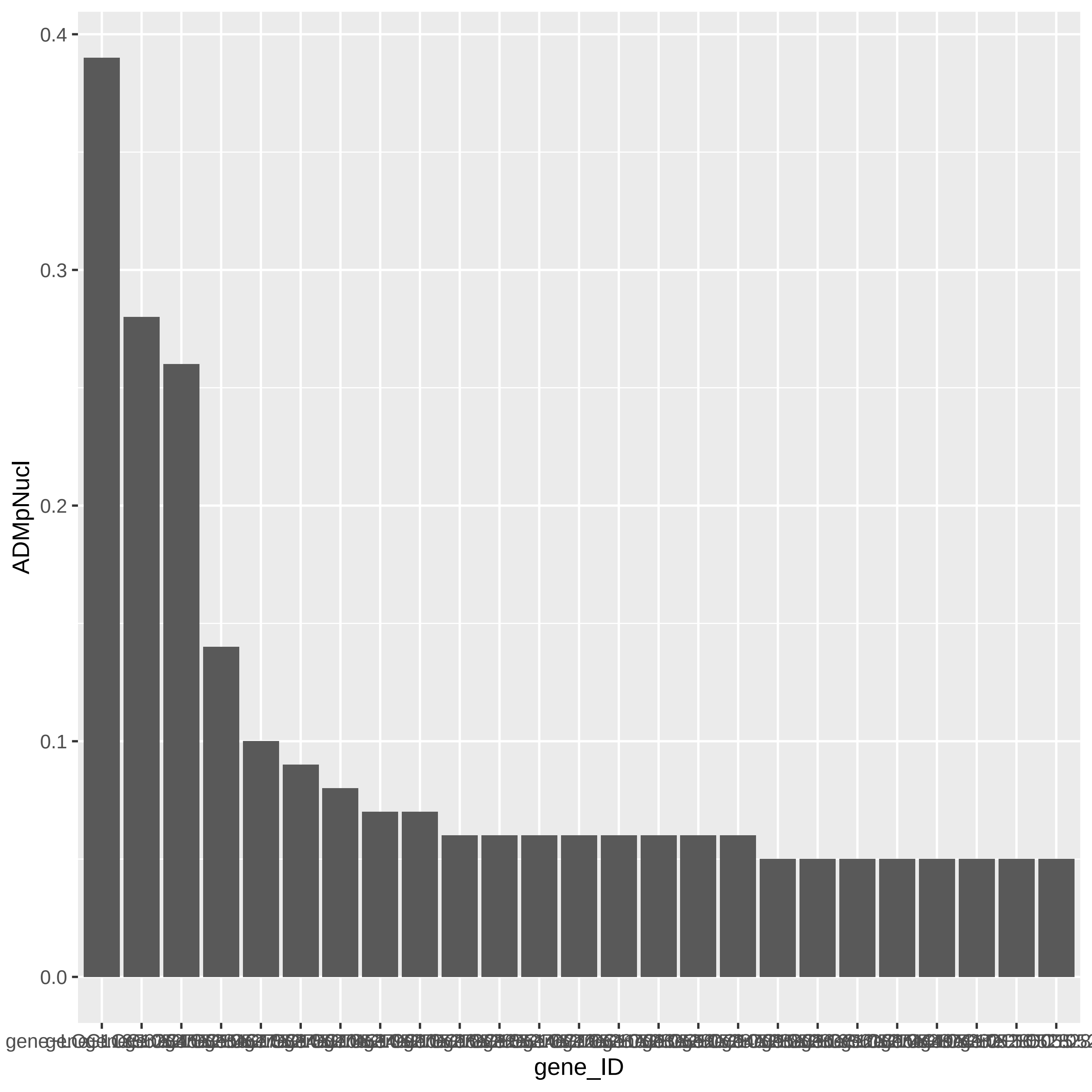
0.4
0.3
0.2
0.1
0.0
ADMpNucl
gene_ID

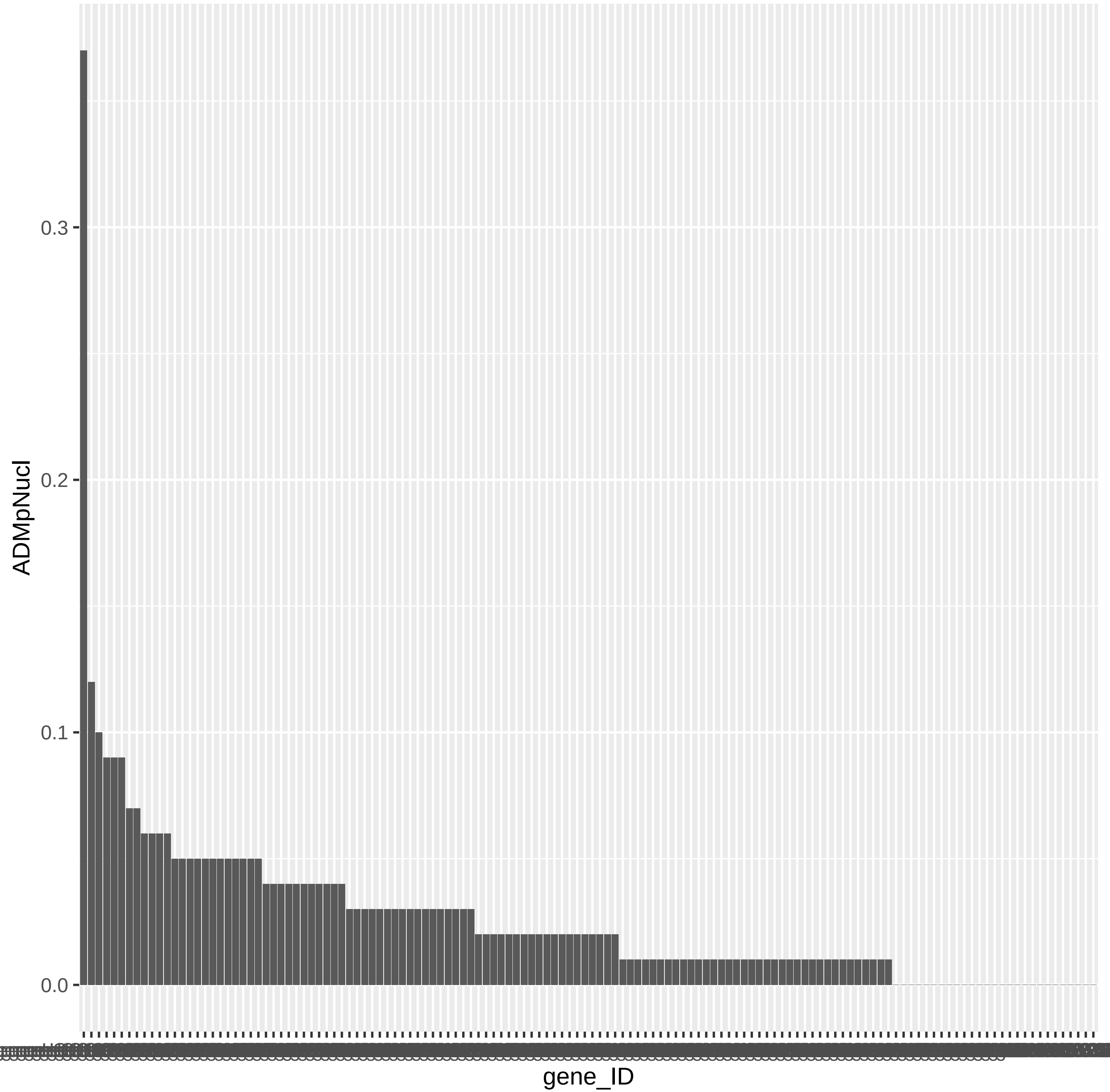

ADMpNucl
0.3
0.2
0.1
0.0
gene_ID

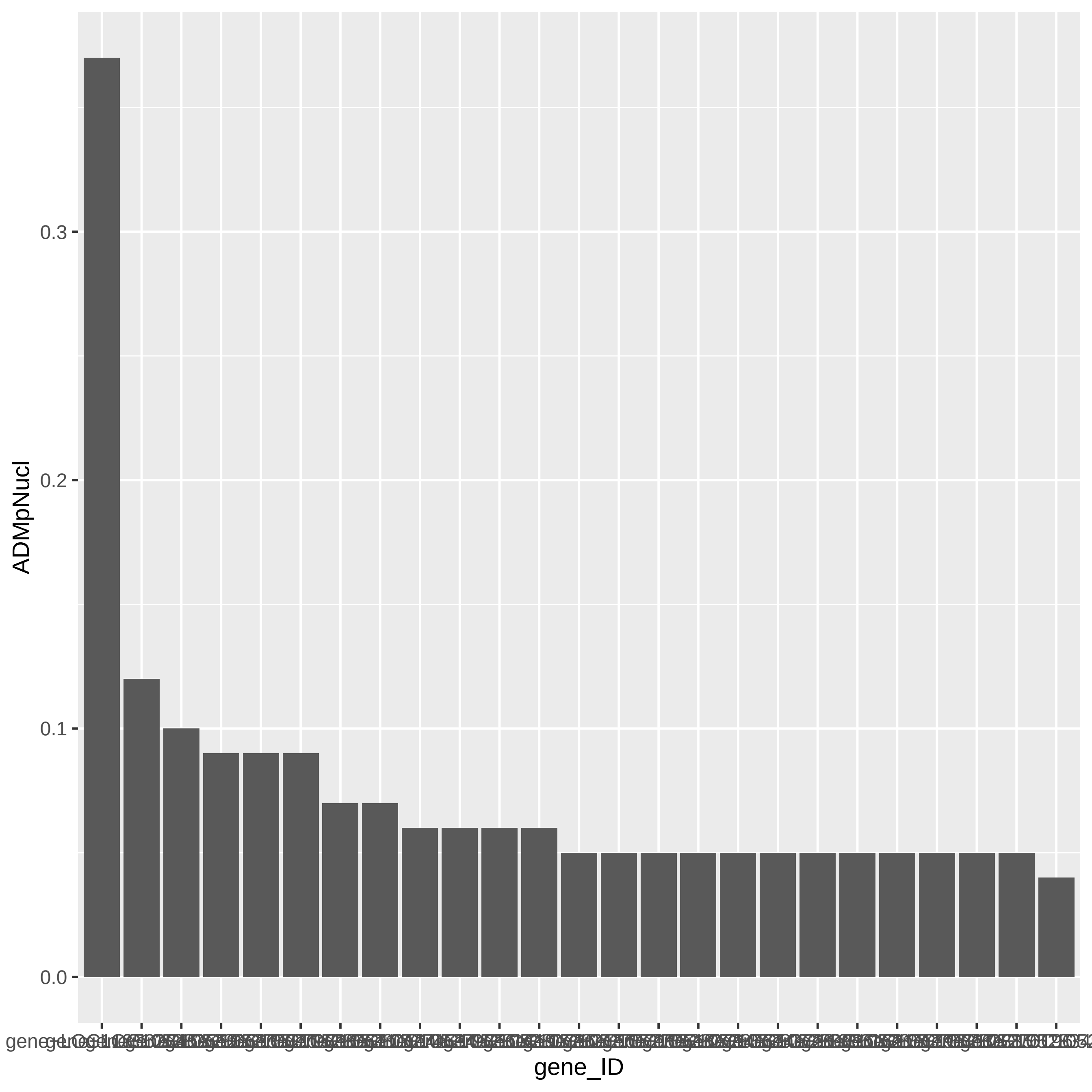

0.3
0.2
0.1
0.0
ADMpNucl
gene_ID

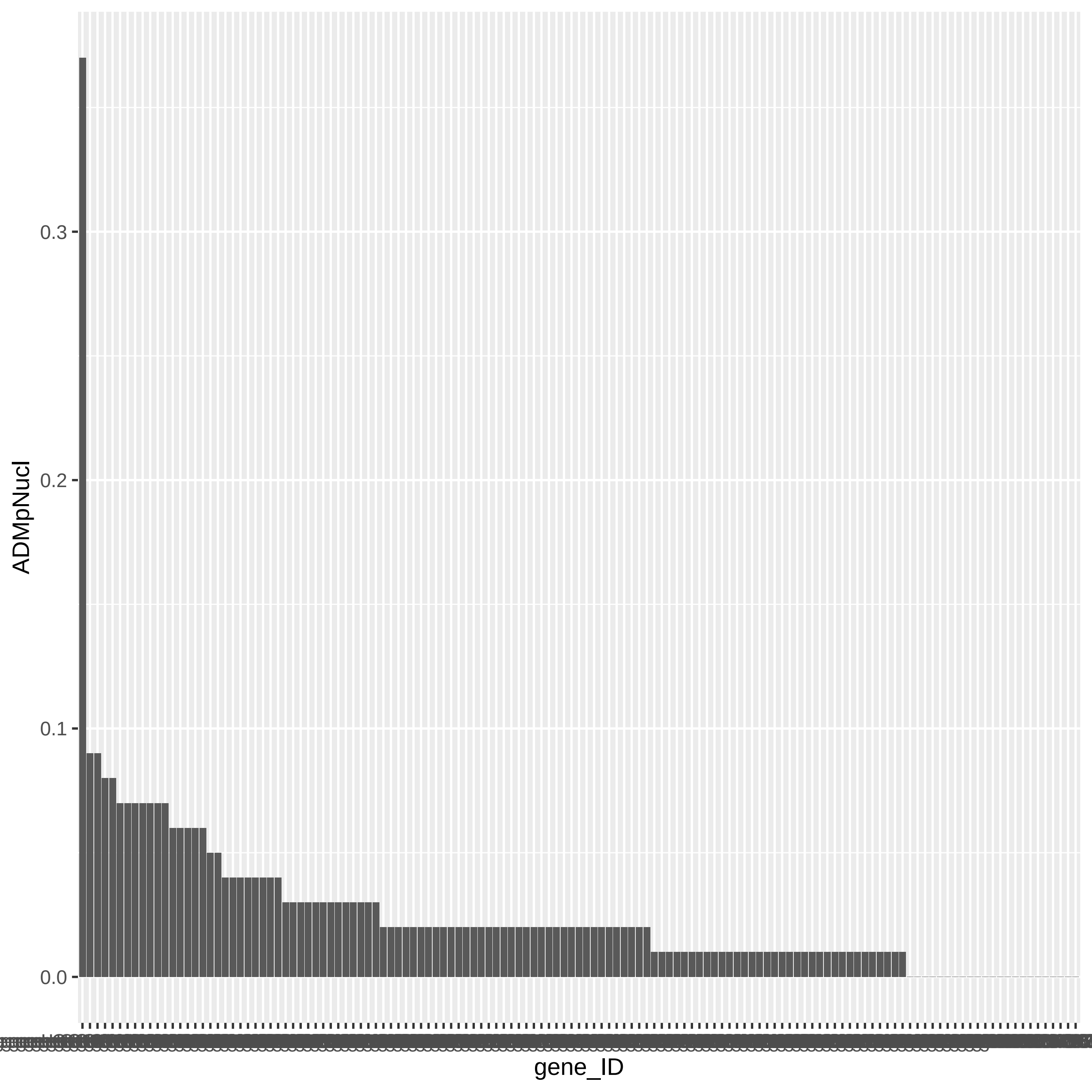

0.3
0.2
0.1
0.0
ADMpNucl
gene_ID

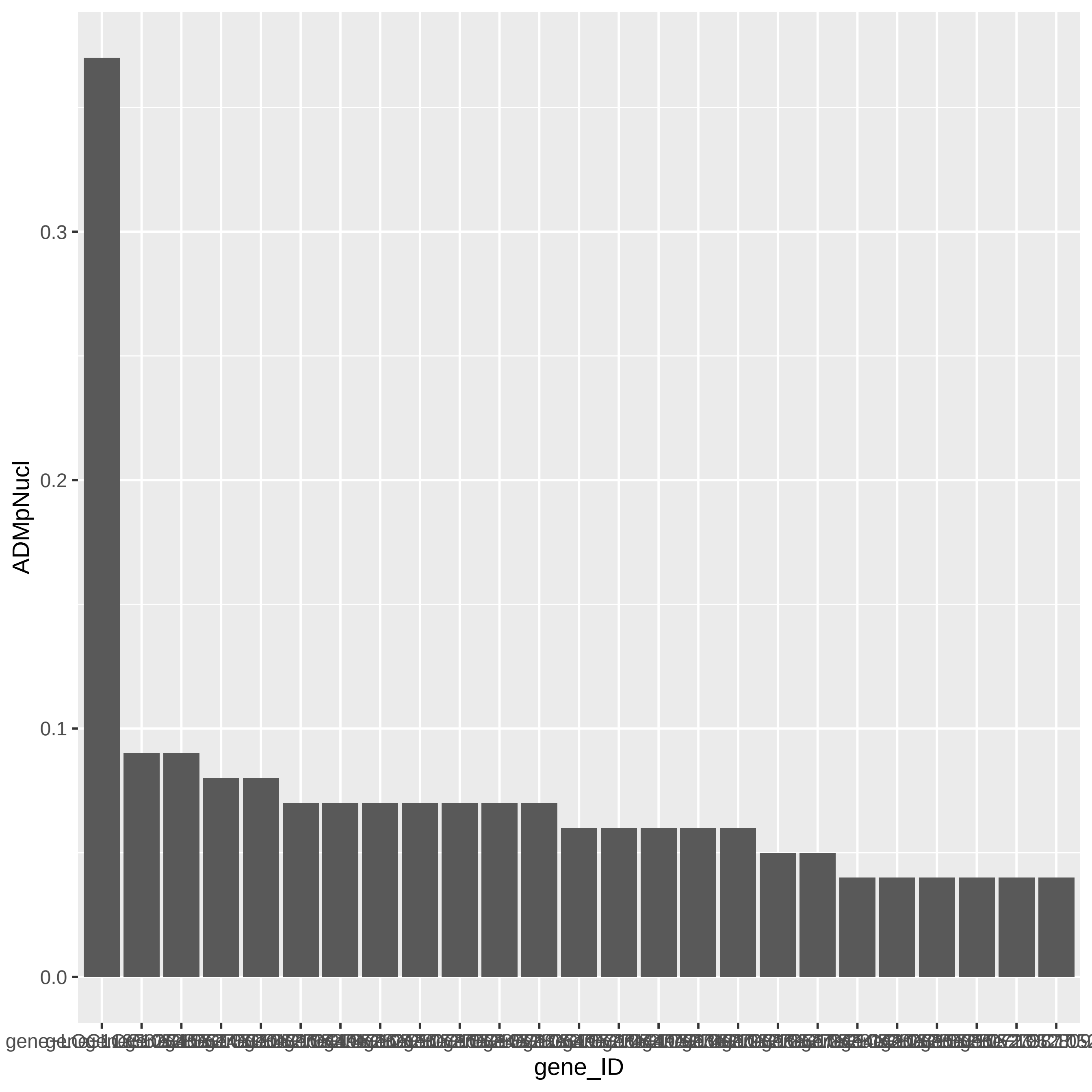
ADMpNucl
0.3
0.2
0.1
0.0
gene_ID

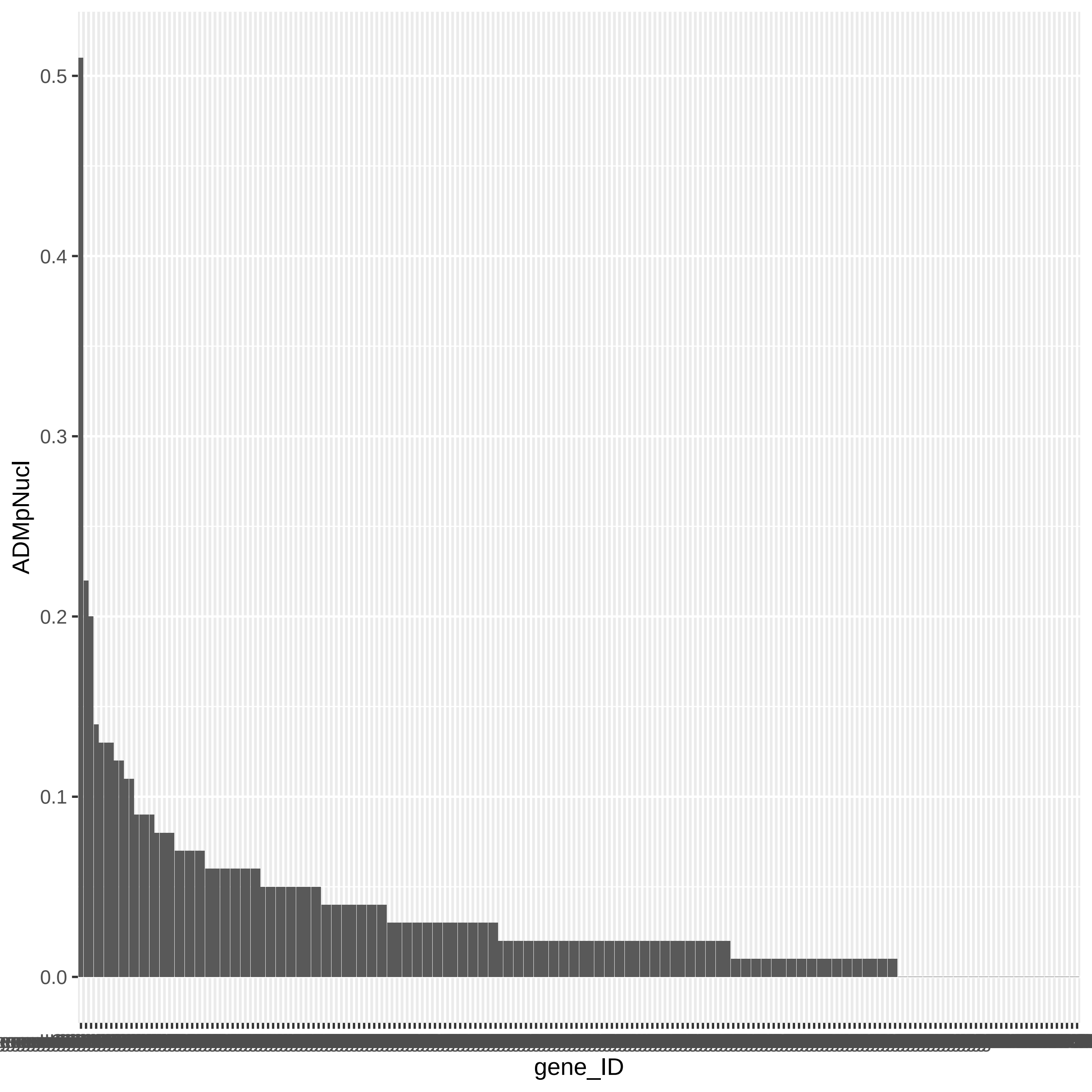

0.5
0.4
0.3
0.2
0.1
0.0
ADMpNucl
gene_ID

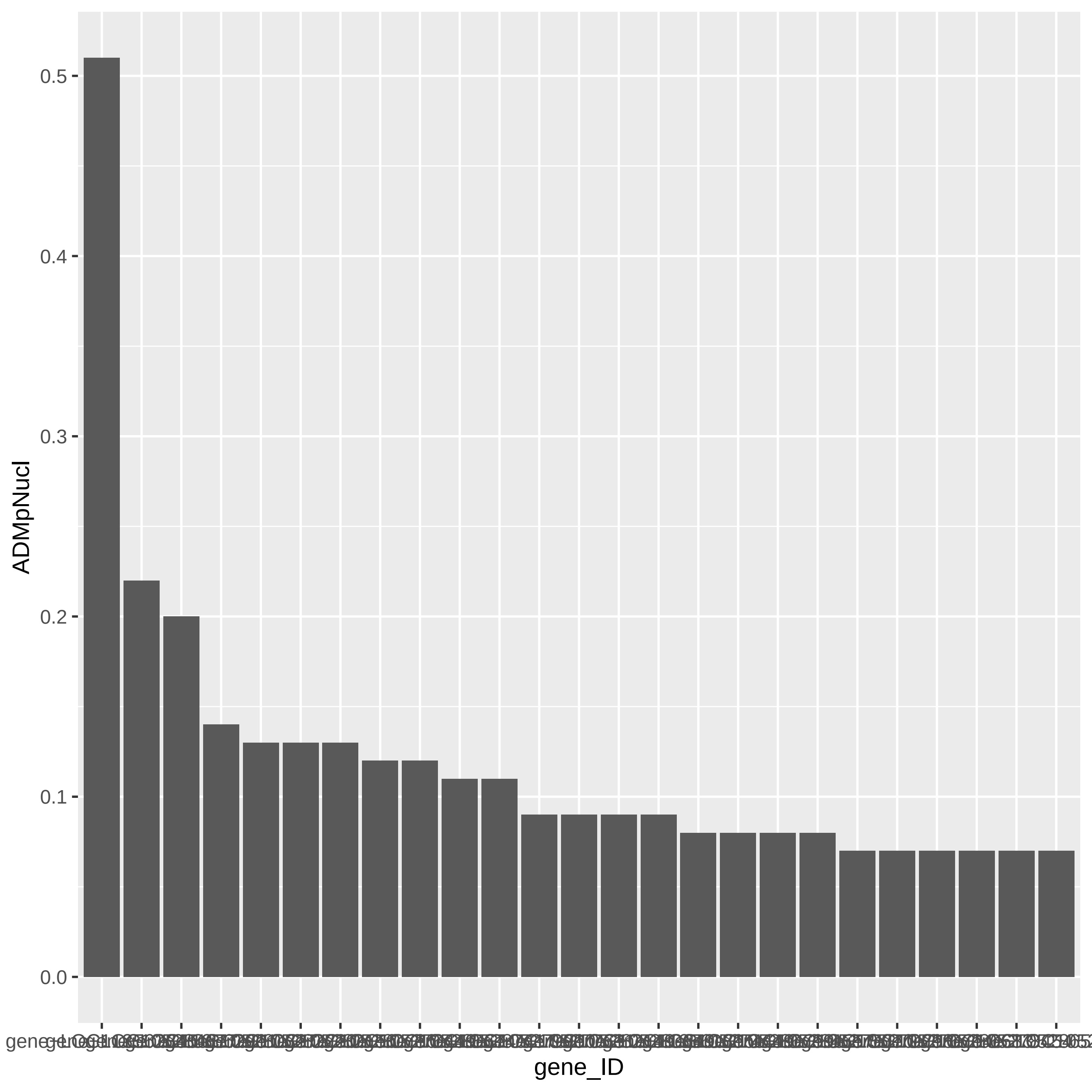
ADMpNucl
0.5
0.4
0.3
0.2
0.1
0.0
gene_ID

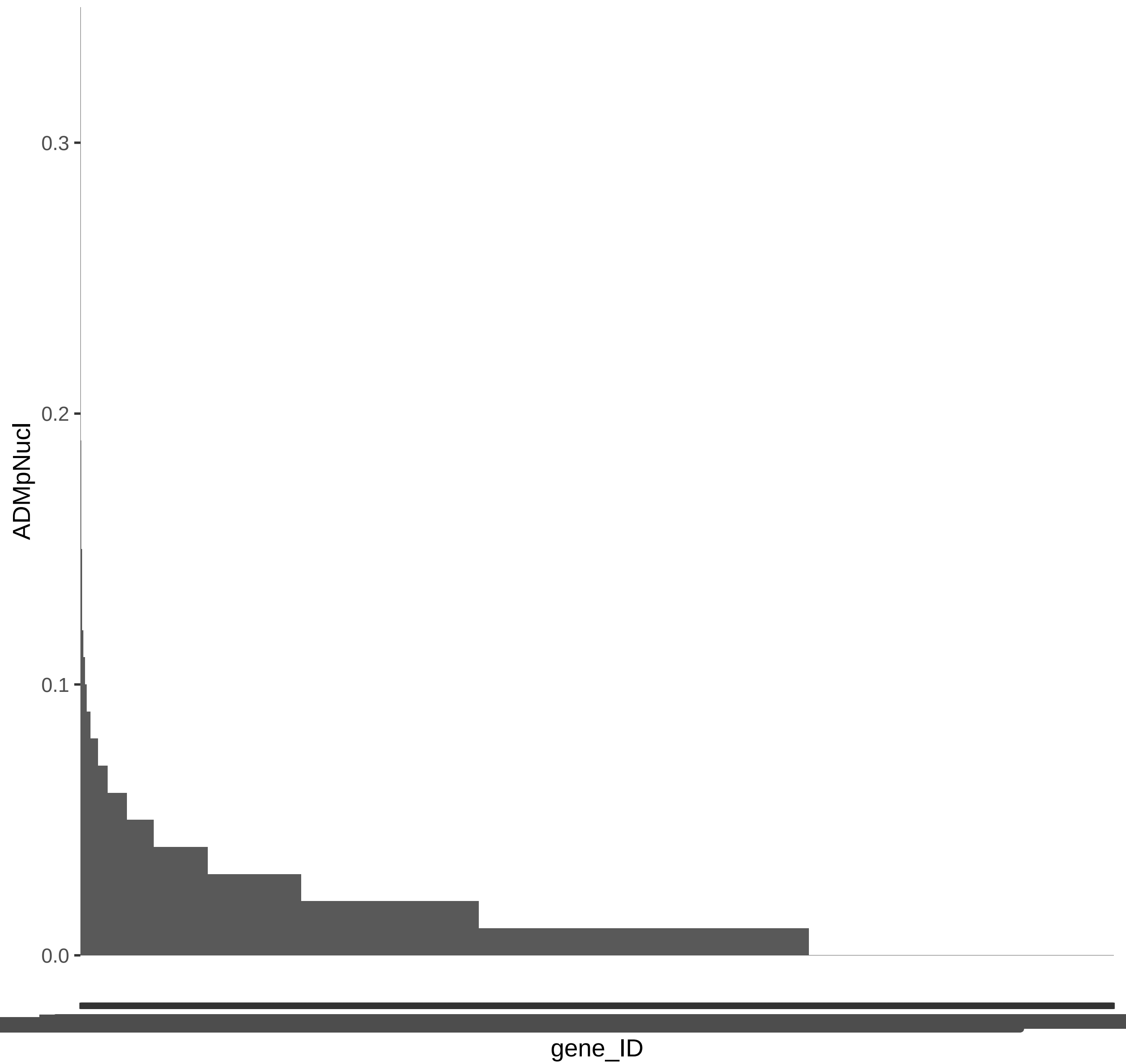

0.3
0.2
0.1
0.0
ADMpNucl
gene_ID

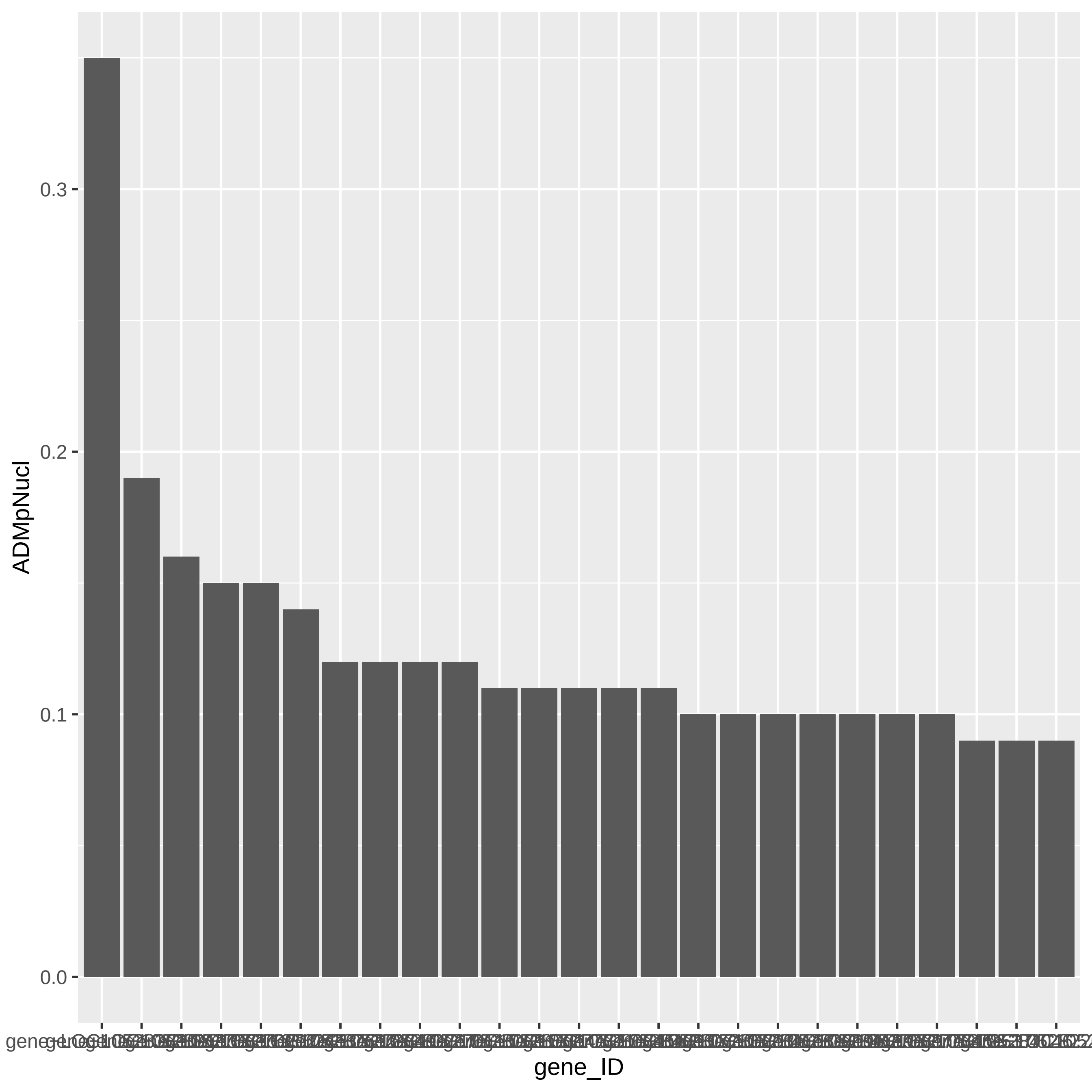

ADMpNucl
0.0
0.1
0.2
0.3
gene_ID

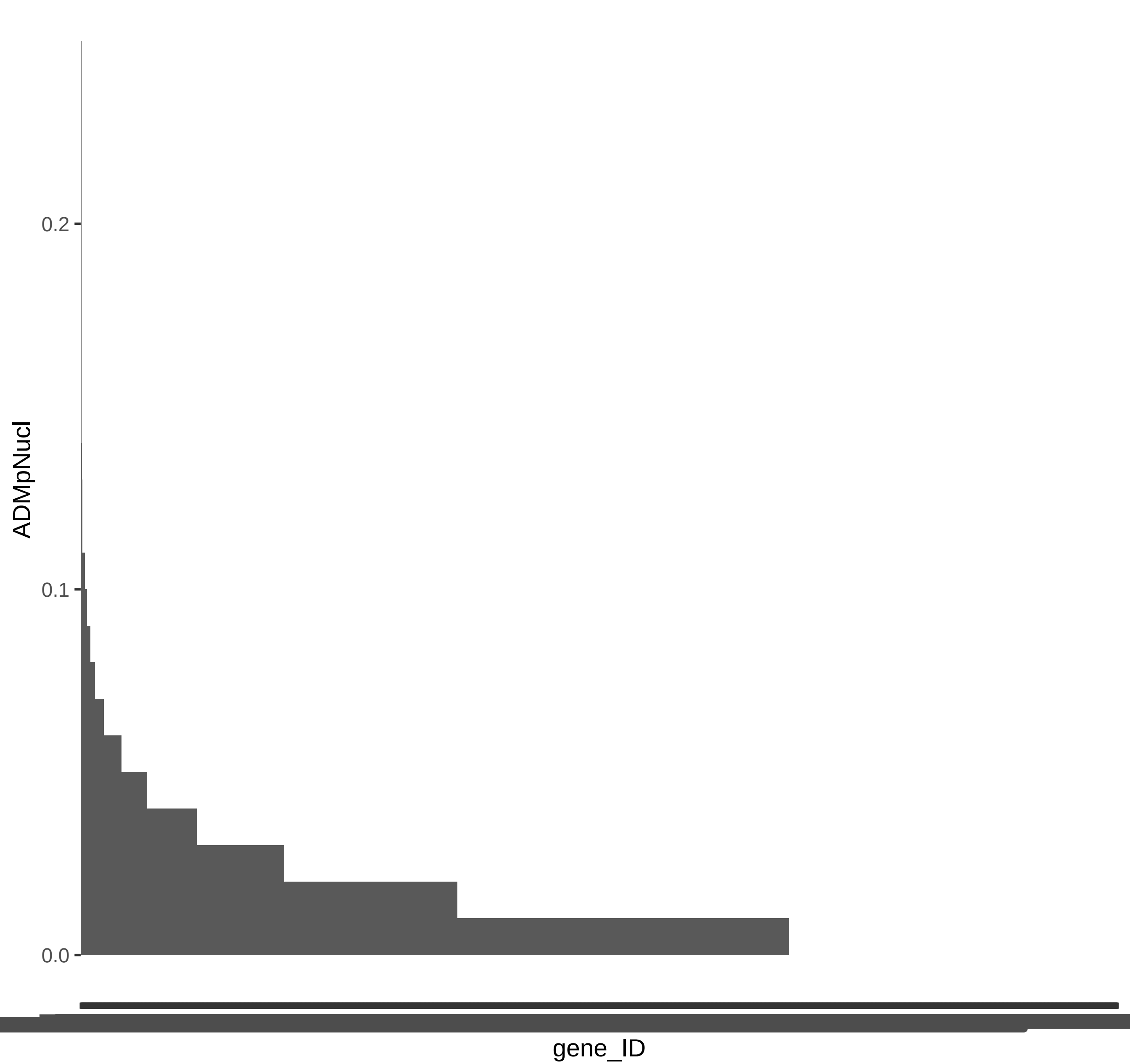

ADMpNucl
0.2
0.1
0.0
gene_ID

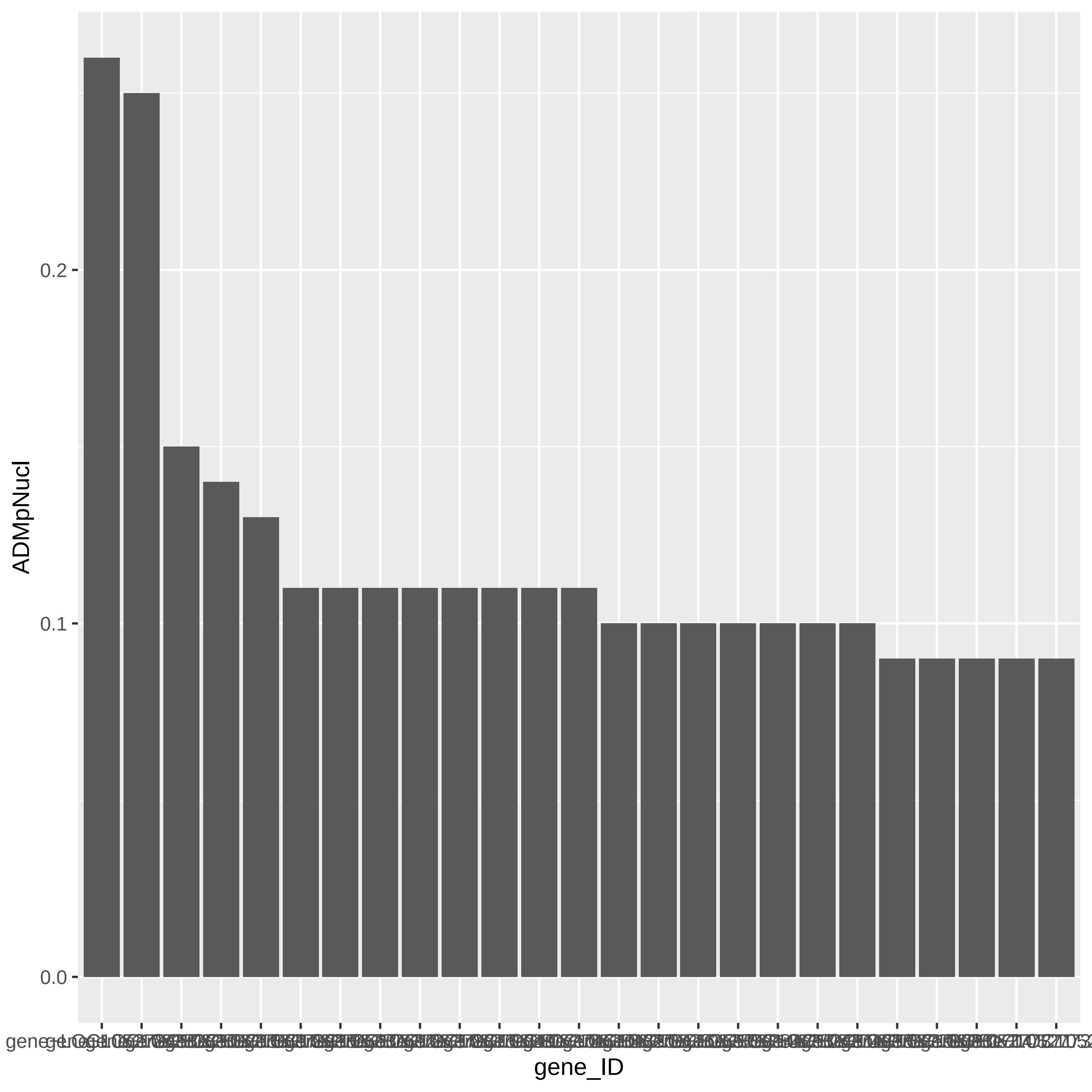

ADMpNucl
0.2
0.1
0.0
gene_ID

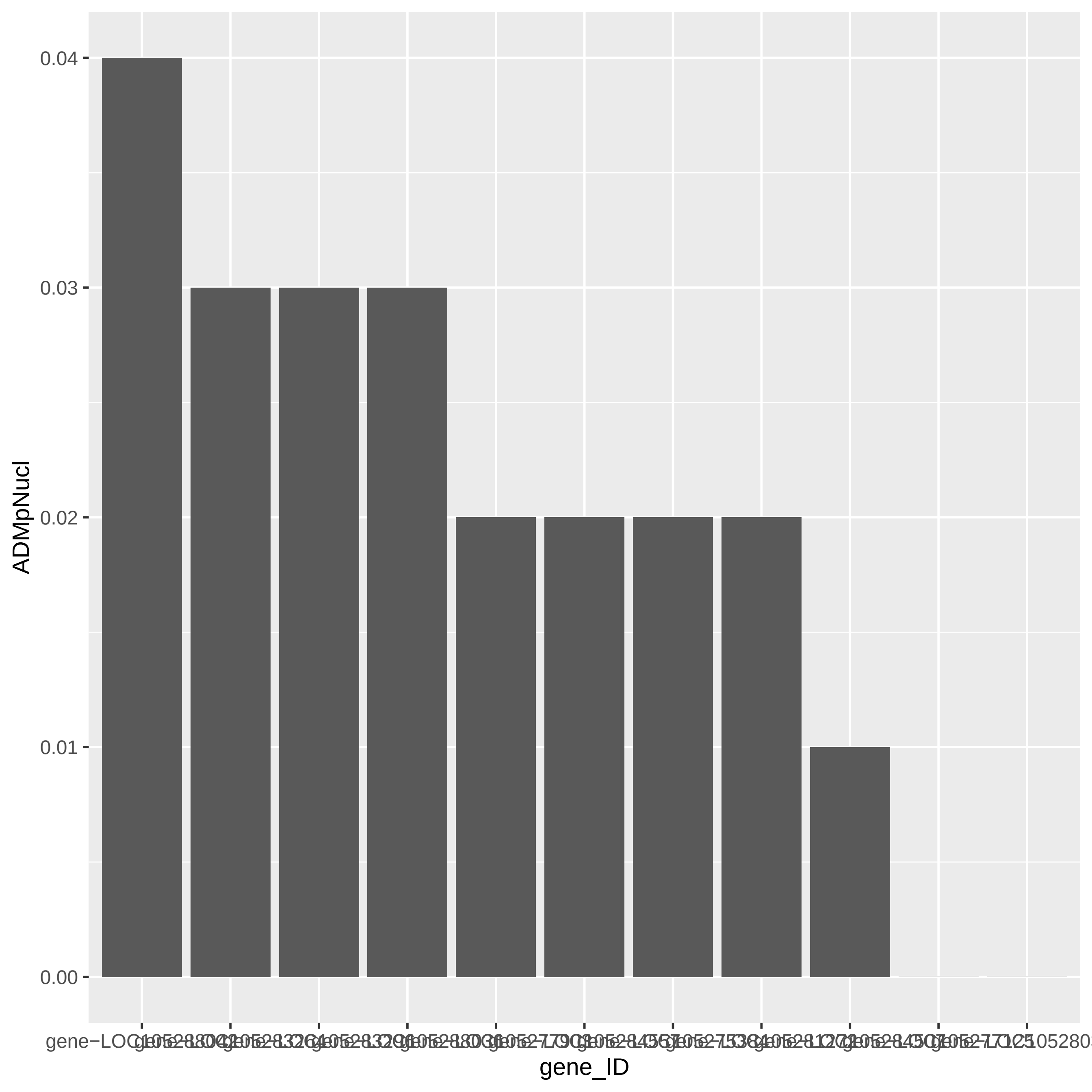

0.04
0.03
0.02
0.01
0.00
ADMpNucl
gene_ID

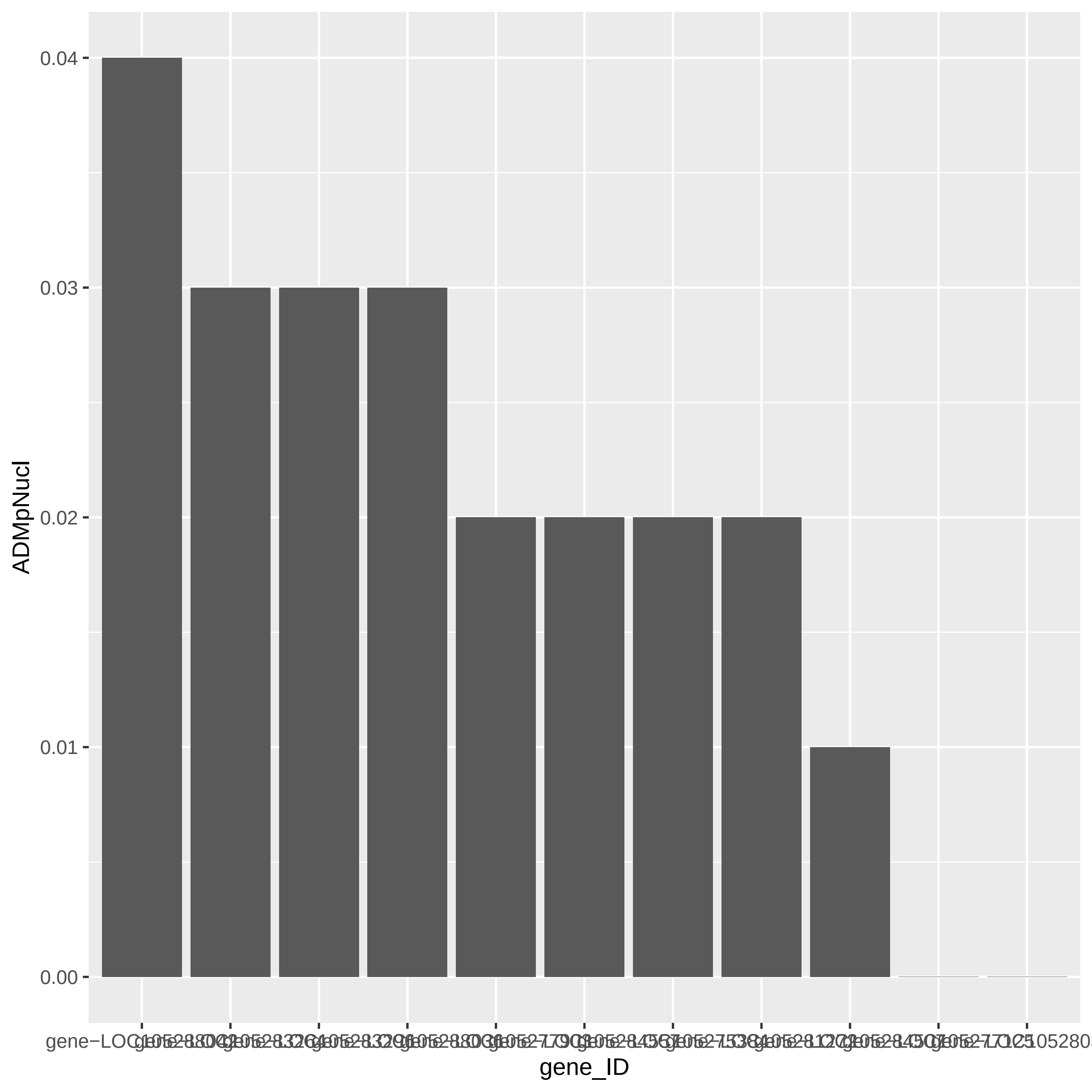
0.04
0.03
0.02
0.01
0.00
ADMpNucl
gene_ID

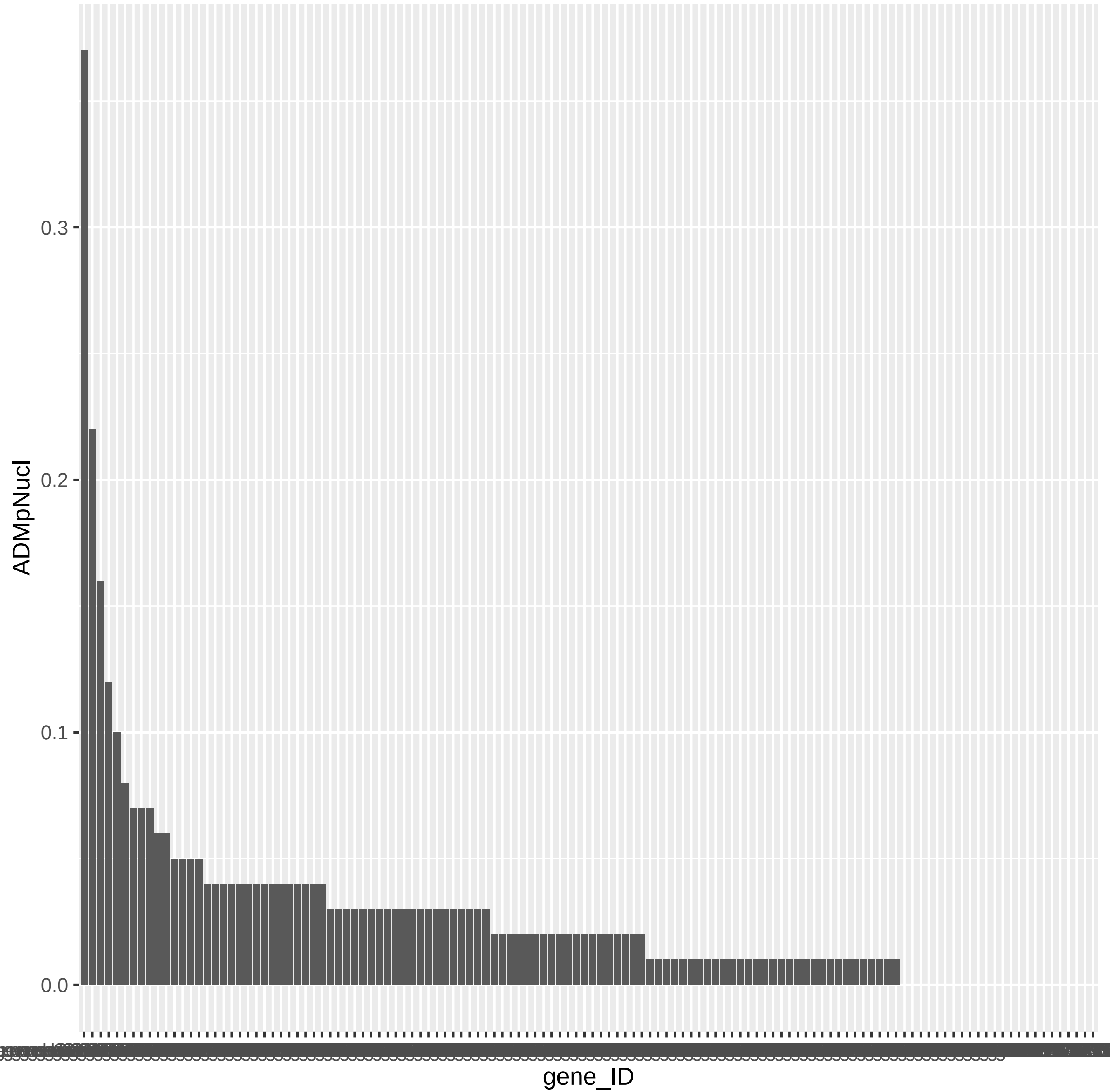

ADMpNucl
0.3
0.2
0.1
0.0
gene_ID

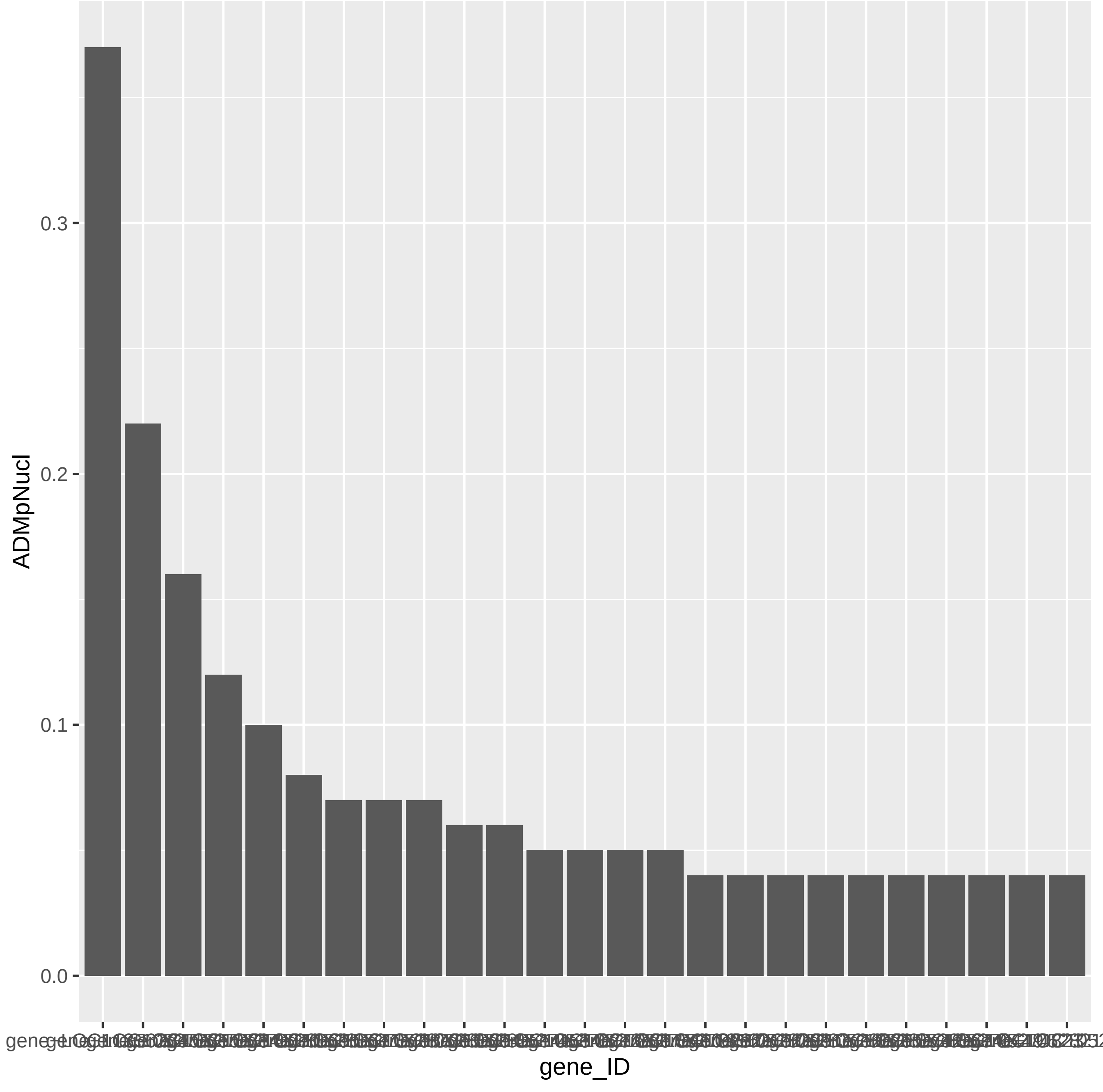

ADMpNucl
0.3
0.2
0.1
0.0
gene_ID

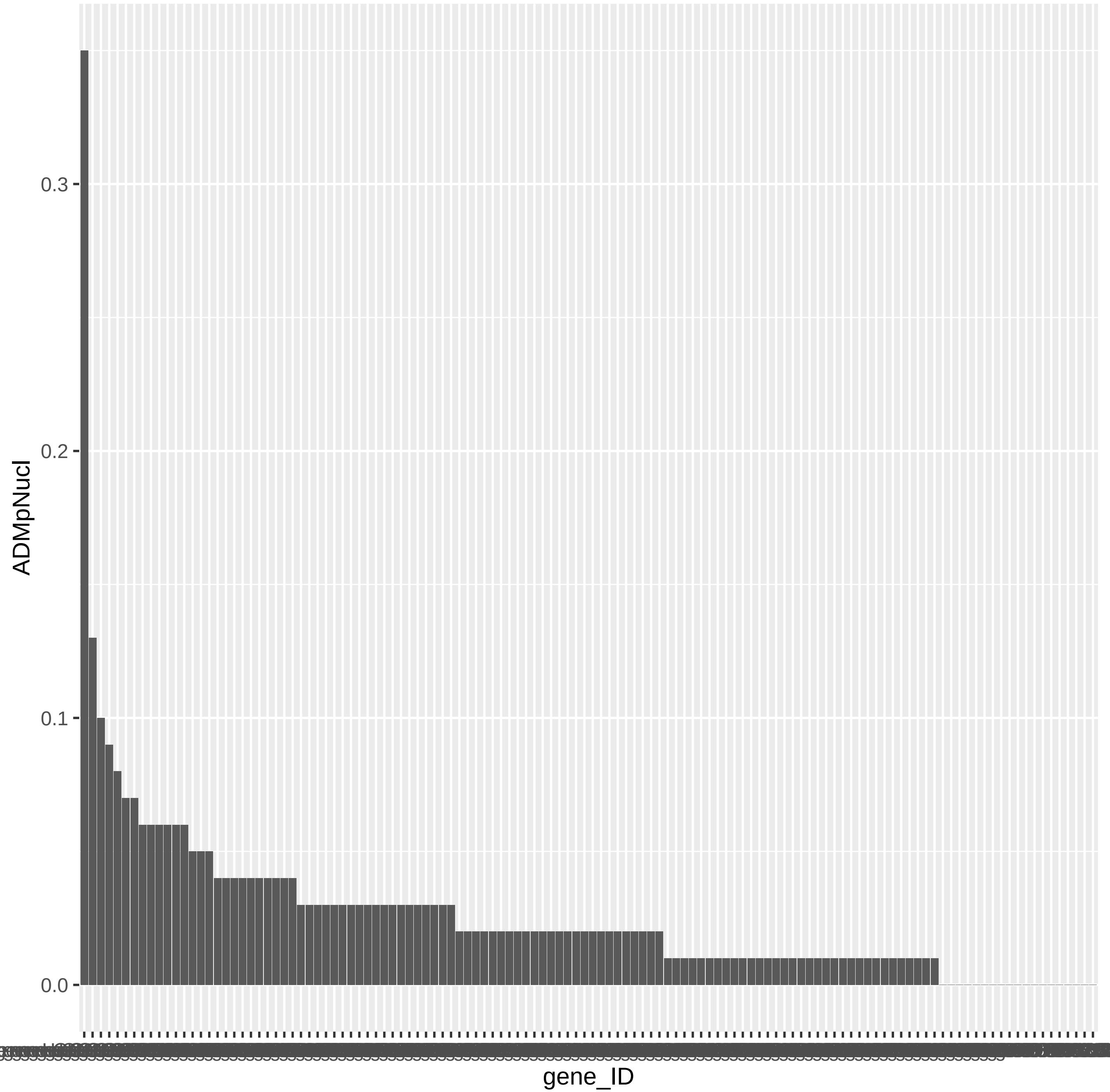
0.3
0.2
0.1
0.0
ADMpNucl
gene_ID

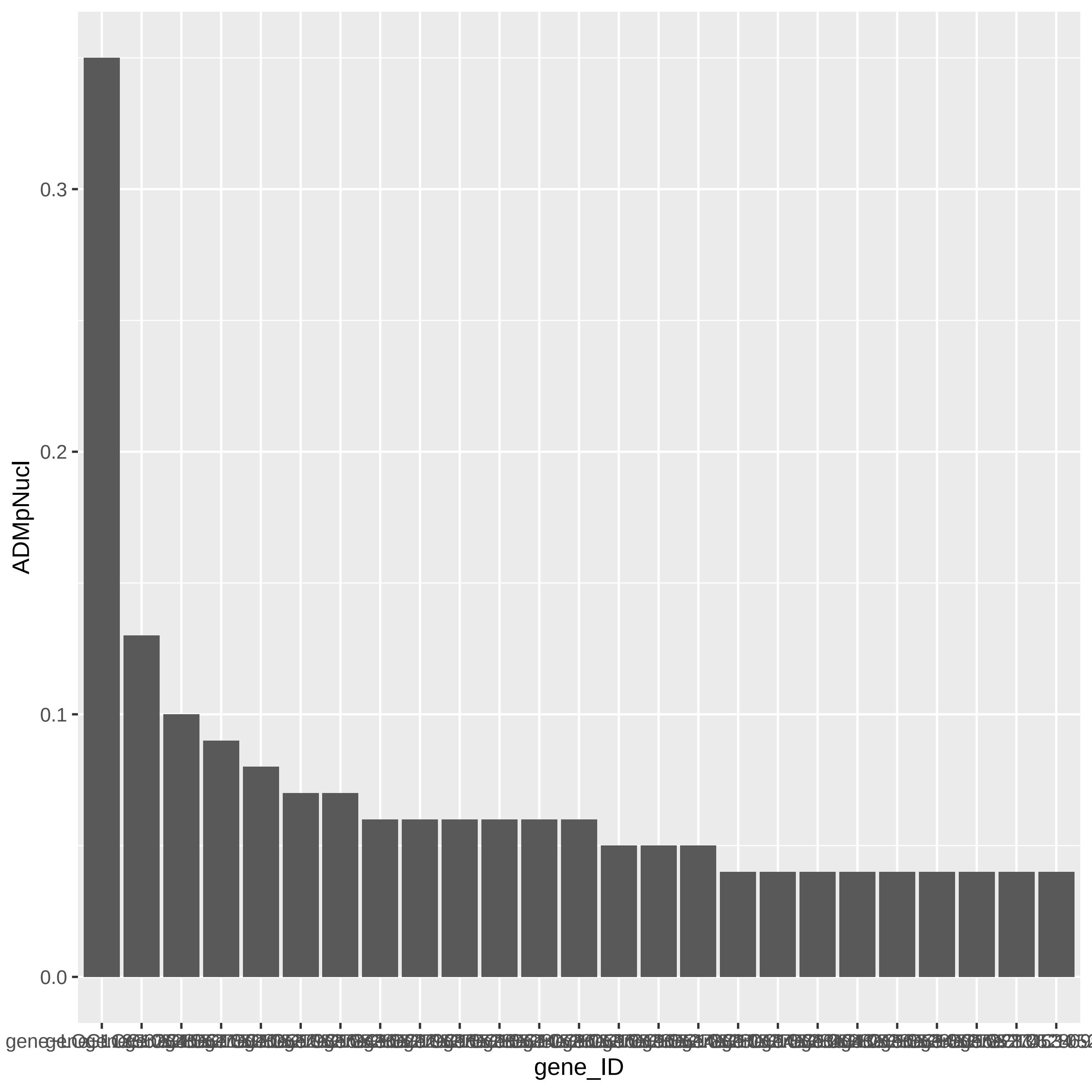

0.3
0.2
0.1
0.0
ADMpNucl
gene_ID

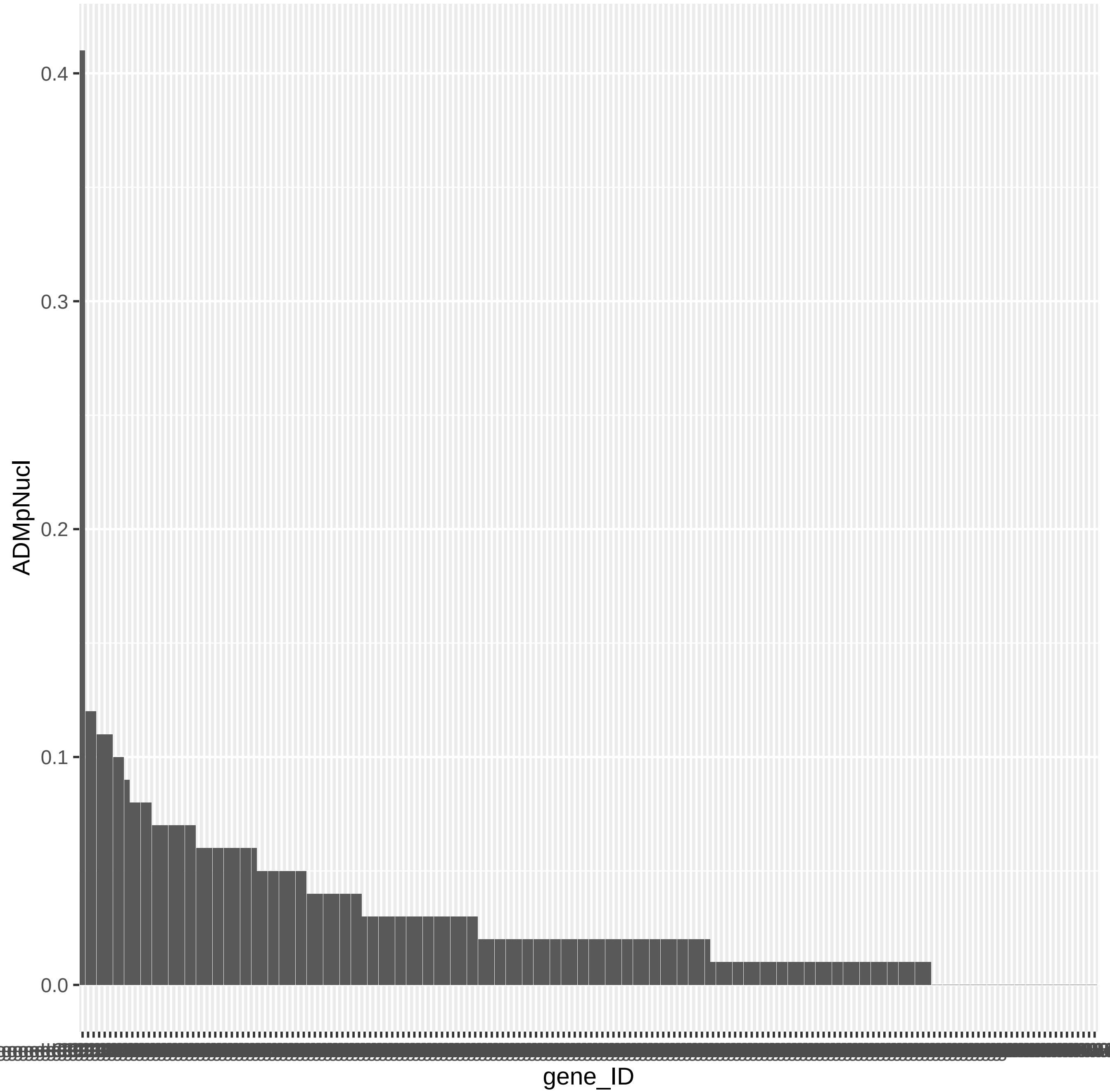
0.4
0.3
0.2
0.1
0.0
ADMpNucl
gene_ID

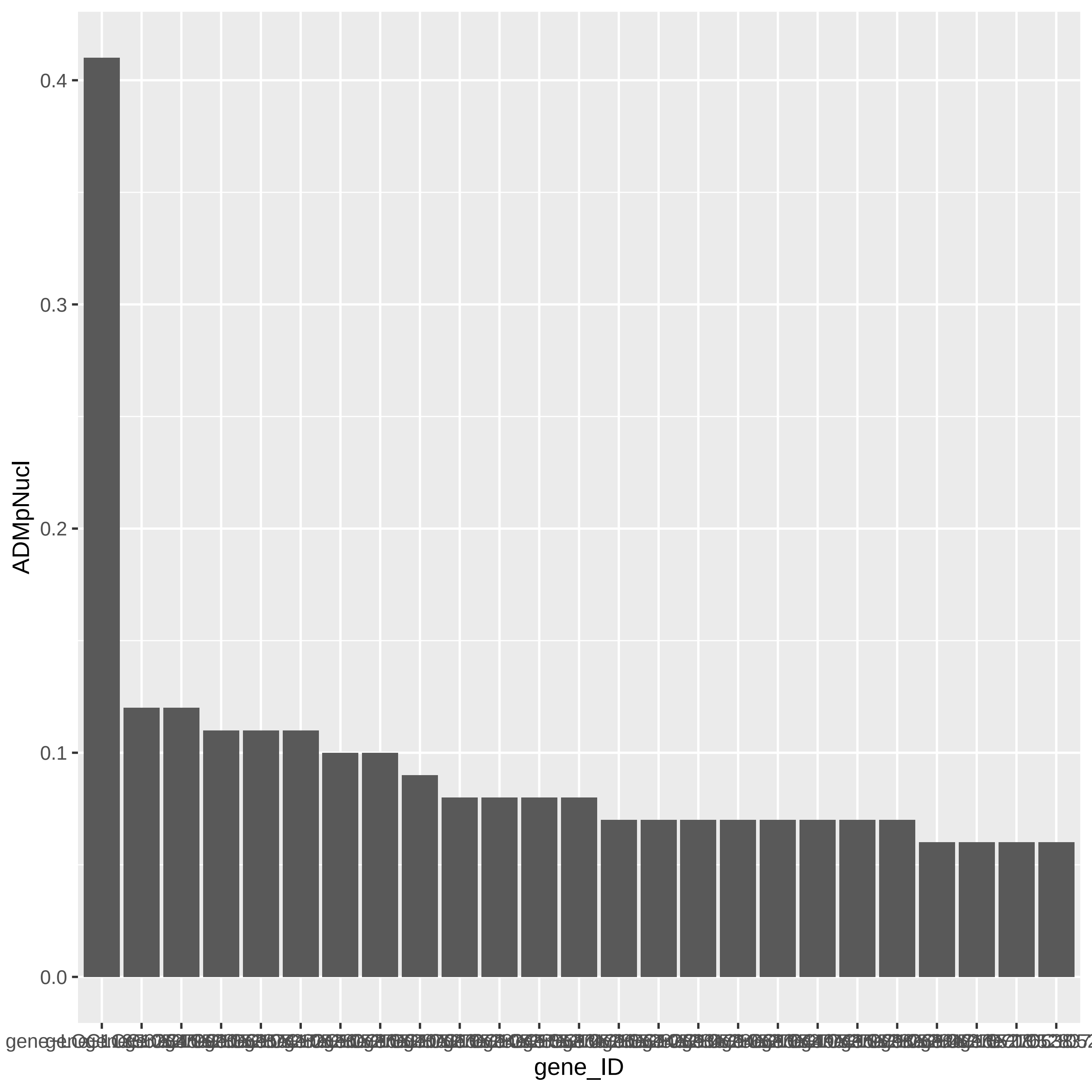

ADMpNucl
0.0
0.1
0.2
0.3
0.4
gene_ID

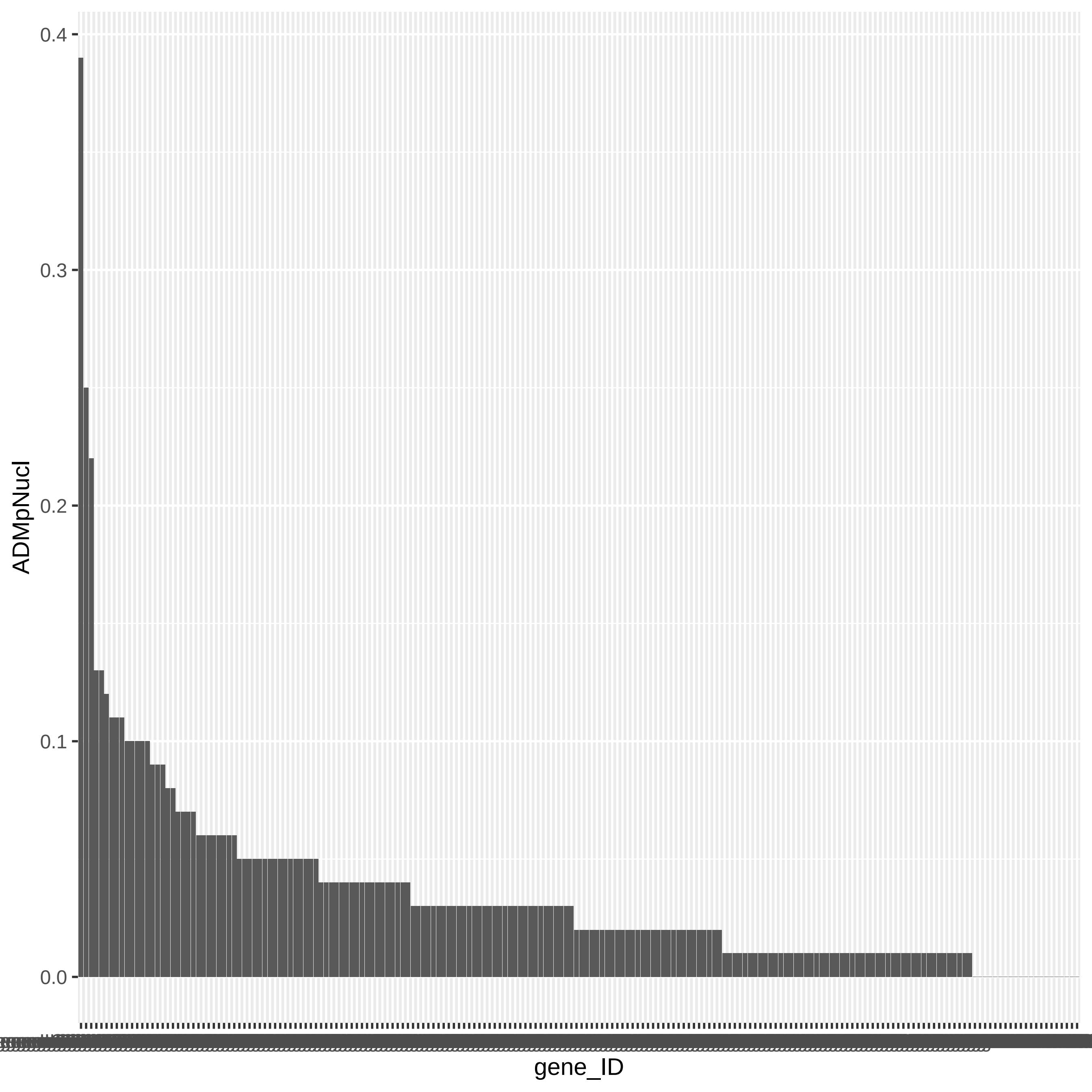

0.4
0.3
0.2
0.1
0.0
ADMpNucl
gene_ID

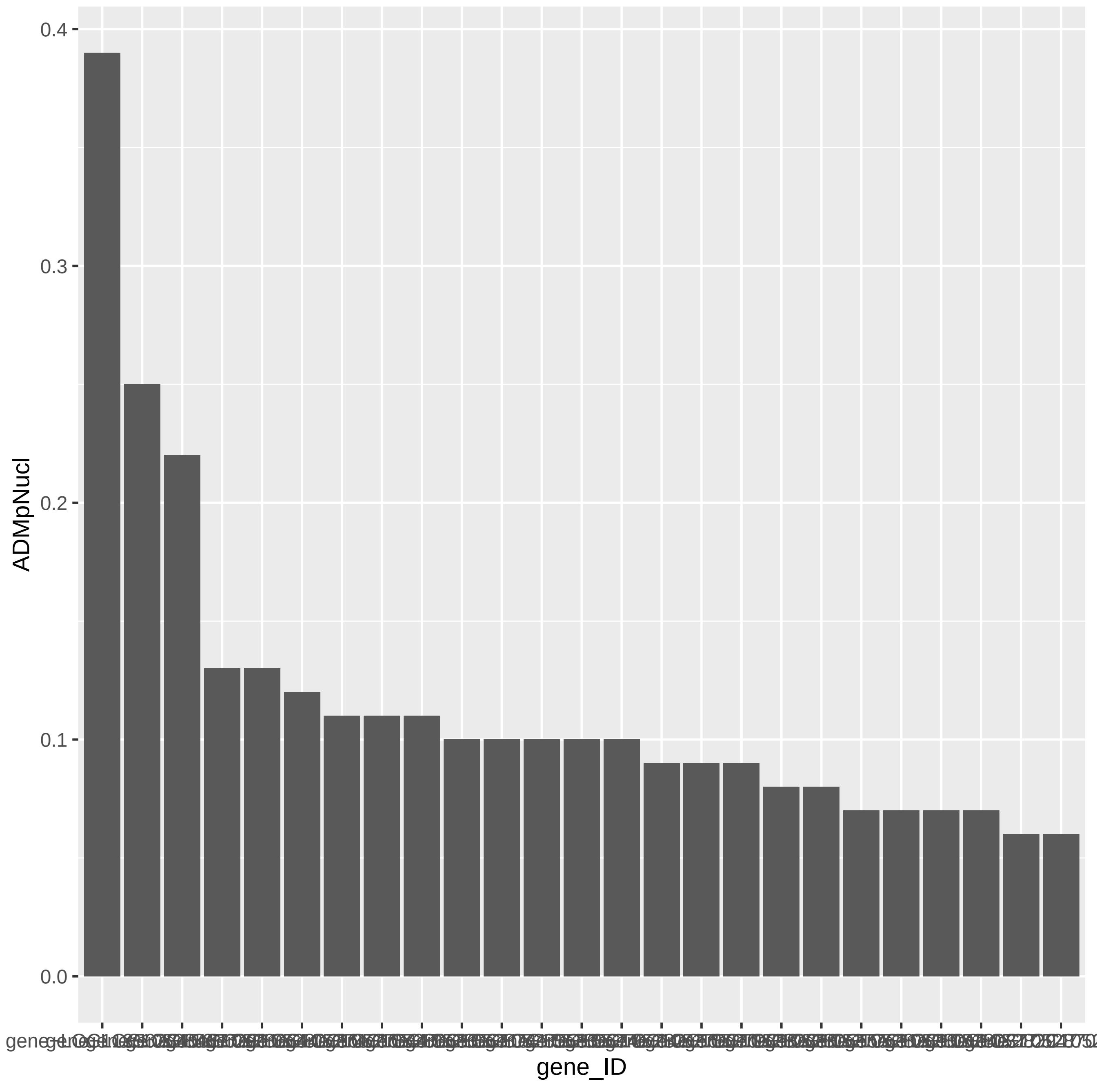
0.4
0.3
0.2
0.1
0.0
ADMpNucl
gene_ID

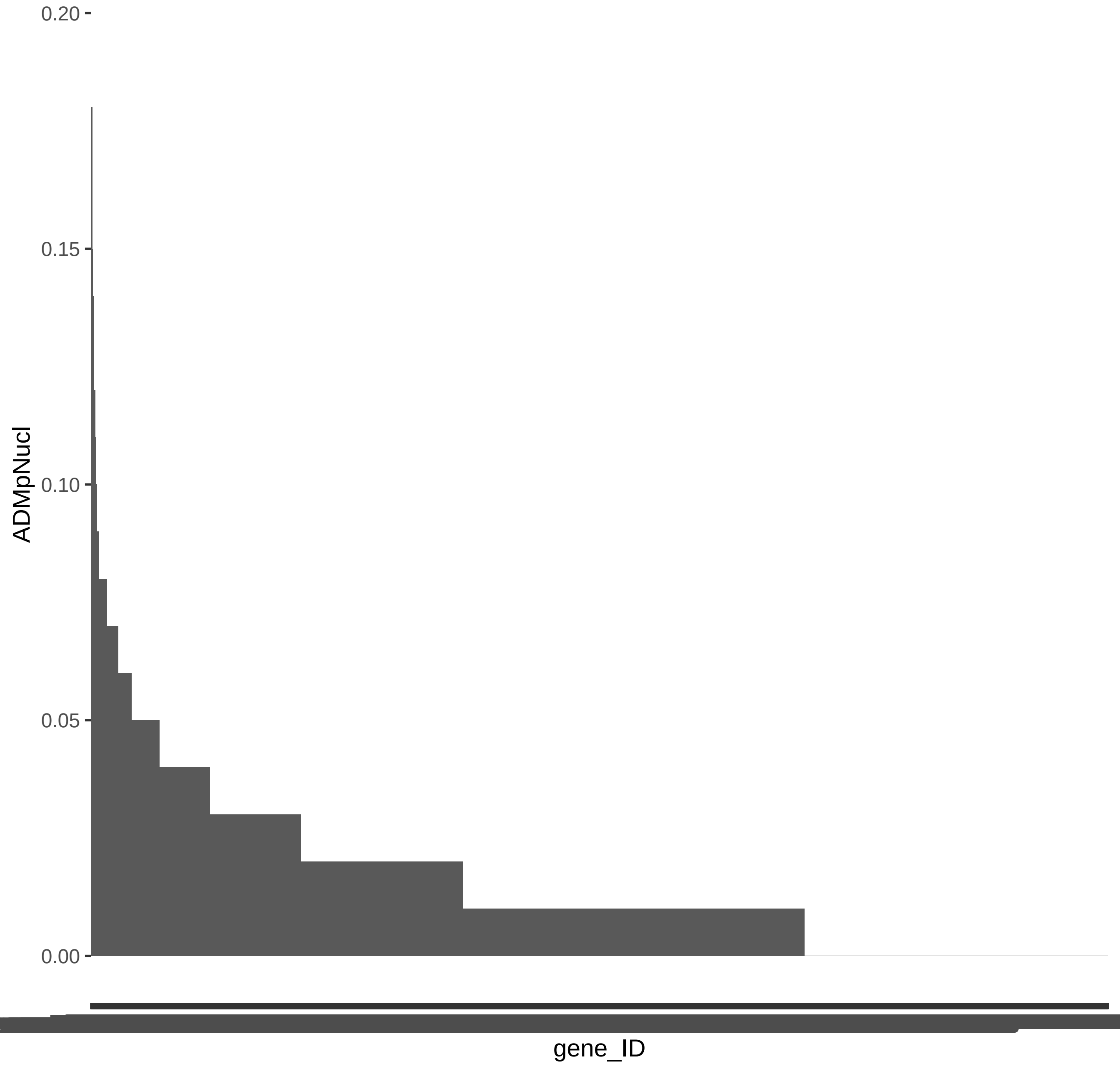

ADMpNucl
0.20
0.15
0.10
0.05
0.00
gene_ID

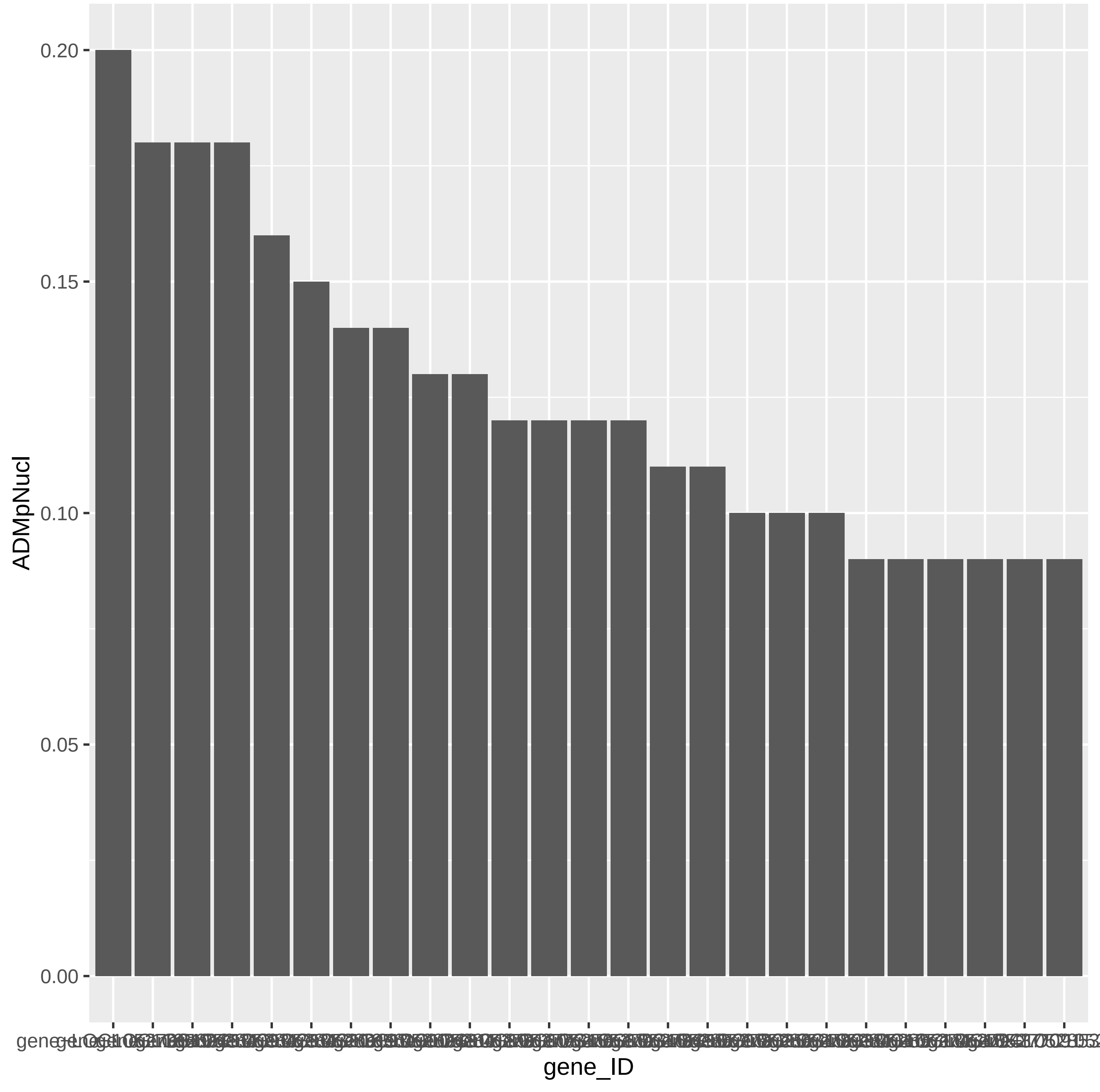

0.20
0.15
0.10
0.05
0.00
ADMpNucl
gene_ID
