## Supplementary material for "Tools and applications for integrative analysis of DNA methylation in social insects": Dataset S21

Ob_LIp.conf

```
#Customize variables here:######################################################
################################################################################

#Load the files:
#
infiles       <- setup_BWASPR(datafile="./Ob.dat",
                              parfile="./Ob.par")

#Set the study and samples in the study:
#
species       <- "Ob"
study         <- "LI"
samplelist    <- list("BCphase", "Rphase")
## The following two variables are used for output file labeling:
studyLabel    <- "LIp"
sampleLabels  <- list("Ob_LI_BCphase", "Ob_LI_Rphase")

hasreplicates <- TRUE
type          <- "CpG"
destrand      <- TRUE
covlist       <- c(6,20,50)
locount       <- 100
hicount       <- 1000
repcovlist    <- c(6,10,15)
replocount    <- 10
rephicount    <- 100
hheight       <- 0.035
nbrpnts       <- 5000

## The following four variables are sent to rank_rbm() and determine what data
#  points get plotted:
minnbrmsprm   <- 5      # minimum number of methylation sites per promoter
mingenewidth  <- 500    # minimum gene width
maxgenewidth  <- 5000   # maximum gene width
minnbrmsgene  <- 5      # minimum number of methylation sites per gene

## Other parameters (see Rscript.BWASPR for usage notes):
#
highcoverage  <- 20     # high read coverage threshold for studyhc methylRawList object
threshold     <- 20.0   # "difference" threshold for getMethylDiff(), called by det_dmsg()
qvalue        <- 0.05   # "qvalue" setting for getMethylDiff(), called by det_dmsg()
wsize         <- 500    # "win.size" parameter for tileMethylCounts() in det_dmt()
stepsize      <- 500    # "step.size" parameter for tileMethylCounts() in det_dmt()

minNsites     <-  1     # minimum number of hc sites in a gene to be heatmapped in show_dmsg()
maxNsites     <- 60     # maximum number of hc sites in a gene to be heatmapped in show_dmsg()
minPdmsites   <- 10     # minimum ratio of dm/hc (in %) in a gene to be heatmapped in show_dmsg()

maxgwidth     <- 20000  # maximal gene width for a gene to be considered by explore_dmsg()
minnbrdmsites <- 1      # minimum number of differentially methylated sites for a gene to be
                        #  considered in sample comparisons by explore_dmsg()
glink         <- "NCBIgene"   # URL to show in explore_dmsg(); options: "" or "NCBIgene"

#Set the number of processors to use:
#
numprc <- 6

#Determine what analyses to run:
#
RUNload    <- FALSE
```

```
Directory: CMS File: cms-Ob_LIp_BCphase.txt

Number of "BCphase" CpGhsm-sites with minimal and higher level coverage:

  number of "BCphase" CpGhsm-sites with coverage >=  5: 415425
  number of "BCphase" CpGhsm-sites with coverage >=  6: 414390
  number of "BCphase" CpGhsm-sites with coverage >= 20: 381770
  number of "BCphase" CpGhsm-sites with coverage >= 50: 291394

Coverage and methylation statistics for "BCphase" CpGhsm-sites at different levels of minimum coverage:

methylKit::getCoverageStats output for "BCphase" CpGhsm-sites (#: 415425) at minimum coverage 5      - read coverage statistics per base
summary:
   Min. 1st Qu.  Median    Mean 3rd Qu.    Max.
      5      43      79      93     131    2762
percentiles:
   0%   10%   20%   30%   40%   50%   60%   70%   80%   90%   95%   99% 99.5% 99.9%  100%
    5    22    36    50    64    79    98   119   146   185   218   283   309   373  2762

methylKit::getMethylationStats output for "BCphase" CpGhsm-sites (#: 415425) at minimum coverage 5   - methylation statistics per base
summary:
   Min. 1st Qu.  Median    Mean 3rd Qu.    Max.
    2.0    20.3    41.5    48.3    77.4   100.0
percentiles:
    0%    10%    20%    30%    40%    50%    60%    70%    80%    90%    95%    99%  99.5%  99.9%   100%
  1.96  12.77  17.50  23.26  31.09  41.51  54.94  70.00  84.00  94.12  97.40 100.00 100.00 100.00 100.00

methylKit::getCoverageStats output for "BCphase" CpGhsm-sites (#: 414390) at minimum coverage 6      - read coverage statistics per base
summary:
   Min. 1st Qu.  Median    Mean 3rd Qu.    Max.
      6      43      79      94     132    2762
percentiles:
   0%   10%   20%   30%   40%   50%   60%   70%   80%   90%   95%   99% 99.5% 99.9%  100%
    6    23    36    50    64    79    98   119   146   185   218   284   309   373  2762

methylKit::getMethylationStats output for "BCphase" CpGhsm-sites (#: 414390) at minimum coverage 6   - methylation statistics per base
summary:
   Min. 1st Qu.  Median    Mean 3rd Qu.    Max.
    2.0    20.3    41.4    48.2    77.0   100.0
percentiles:
    0%    10%    20%    30%    40%    50%    60%    70%    80%    90%    95%    99%  99.5%  99.9%   100%
  1.96  12.75  17.46  23.22  31.03  41.38  54.67  69.81  83.75  93.94  97.22 100.00 100.00 100.00 100.00

methylKit::getCoverageStats output for "BCphase" CpGhsm-sites (#: 381770) at minimum coverage 20     - read coverage statistics per base
summary:
   Min. 1st Qu.  Median    Mean 3rd Qu.    Max.
     20      51      86     101     137    2762
percentiles:
   0%   10%   20%   30%   40%   50%   60%   70%   80%   90%   95%   99% 99.5% 99.9%  100%
   20    33    45    57    71    86   104   125   151   189   222   287   313   377  2762

methylKit::getMethylationStats output for "BCphase" CpGhsm-sites (#: 381770) at minimum coverage 20  - methylation statistics per base
summary:
   Min. 1st Qu.  Median    Mean 3rd Qu.    Max.
    2.0    19.1    37.3    45.9    73.5   100.0
percentiles:
    0%    10%    20%    30%    40%    50%    60%    70%    80%    90%    95%    99%  99.5%  99.9%   100%
```

```
  1.96  12.36  16.67  21.74  28.24  37.27  50.00  65.38  81.25  93.02  96.47 100.00 100.00 100.00 100.00

methylKit::getCoverageStats output for "BCphase" CpGhsm-sites (#: 291394) at minimum coverage 50    - read coverage statistics per base
summary:
   Min. 1st Qu.  Median    Mean 3rd Qu.    Max.
     50      75     108     121     153    2762
percentiles:
   0%   10%   20%   30%   40%   50%   60%   70%   80%   90%   95%   99% 99.5% 99.9%  100%
   50    59    70    81    93   108   124   142   166   202   233   296   323   385  2762

methylKit::getMethylationStats output for "BCphase" CpGhsm-sites (#: 291394) at minimum coverage 50  - methylation statistics per base
summary:
   Min. 1st Qu.  Median    Mean 3rd Qu.    Max.
    2.0    16.3    32.9    43.2    70.3   100.0
percentiles:
    0%    10%    20%    30%    40%    50%    60%    70%    80%    90%    95%    99%  99.5%  99.9%   100%
  1.96  11.36  14.71  18.33  24.09  32.86  45.37  61.54  78.75  92.21  96.08  98.67  99.28 100.00 100.00

methylKit::getCoverageStats output for "BCphase" CpGhsm-sites (#: 162107) in coverage range [100-1000]    - read coverage statistics per base
summary:
   Min. 1st Qu.  Median    Mean 3rd Qu.    Max.
    100     121     147     160     186     956
percentiles:
   0%   10%   20%   30%   40%   50%   60%   70%   80%   90%   95%   99% 99.5% 99.9%  100%
  100   107   116   125   136   147   160   176   197   229   258   319   345   408   956

methylKit::getMethylationStats output for "BCphase" CpGhsm-sites (#: 162107) in coverage range [100-1000] - methylation statistics per base
summary:
   Min. 1st Qu.  Median    Mean 3rd Qu.    Max.
    2.6    14.5    32.8    42.7    70.6   100.0
percentiles:
    0%    10%    20%    30%    40%    50%    60%    70%    80%    90%    95%    99%  99.5%  99.9%   100%
  2.62   9.68  12.50  16.88  23.40  32.79  45.74  62.04  78.99  92.16  95.98  98.38  99.01  99.62 100.00
```

```
Directory: CMS File: cms-Ob_LIp_Rphase.txt

Number of "Rphase" CpGhsm-sites with minimal and higher level coverage:

  number of "Rphase" CpGhsm-sites with coverage >=  5: 445068
  number of "Rphase" CpGhsm-sites with coverage >=  6: 444146
  number of "Rphase" CpGhsm-sites with coverage >= 20: 413885
  number of "Rphase" CpGhsm-sites with coverage >= 50: 327310

Coverage and methylation statistics for "Rphase" CpGhsm-sites at different levels of minimum coverage:

methylKit::getCoverageStats output for "Rphase" CpGhsm-sites (#: 445068) at minimum coverage 5 - read coverage statistics per base
summary:
   Min. 1st Qu.  Median    Mean 3rd Qu.    Max.
      5      47      87     102     143    2877
percentiles:
   0%   10%   20%   30%   40%   50%   60%   70%   80%   90%   95%   99% 99.5% 99.9%  100%
    5    25    40    54    70    87   106   130   159   202   240   312   342   410  2877

methylKit::getMethylationStats output for "Rphase" CpGhsm-sites (#: 445068) at minimum coverage 5    - methylation statistics per base
summary:
   Min. 1st Qu.  Median    Mean 3rd Qu.    Max.
    1.8    18.5    38.5    46.5    75.0   100.0
percentiles:
    0%    10%    20%    30%    40%    50%    60%    70%    80%    90%    95%    99%  99.5%  99.9%   100%
  1.77  11.84  16.07  21.25  28.33  38.46  51.81  67.35  82.81  93.66  97.14 100.00 100.00 100.00 100.00

methylKit::getCoverageStats output for "Rphase" CpGhsm-sites (#: 444146) at minimum coverage 6 - read coverage statistics per base
summary:
   Min. 1st Qu.  Median    Mean 3rd Qu.    Max.
      6      48      87     103     144    2877
percentiles:
   0%   10%   20%   30%   40%   50%   60%   70%   80%   90%   95%   99% 99.5% 99.9%  100%
    6    25    40    55    70    87   106   130   159   203   240   312   342   410  2877

methylKit::getMethylationStats output for "Rphase" CpGhsm-sites (#: 444146) at minimum coverage 6    - methylation statistics per base
summary:
   Min. 1st Qu.  Median    Mean 3rd Qu.    Max.
    1.8    18.5    38.3    46.4    75.0   100.0
percentiles:
    0%    10%    20%    30%    40%    50%    60%    70%    80%    90%    95%    99%  99.5%  99.9%   100%
  1.77  11.83  16.07  21.21  28.26  38.27  51.61  67.05  82.56  93.48  97.00 100.00 100.00 100.00 100.00

methylKit::getCoverageStats output for "Rphase" CpGhsm-sites (#: 413885) at minimum coverage 20      - read coverage statistics per base
summary:
   Min. 1st Qu.  Median    Mean 3rd Qu.    Max.
     20      55      93     109     149    2877
percentiles:
   0%   10%   20%   30%   40%   50%   60%   70%   80%   90%   95%   99% 99.5% 99.9%  100%
   20    34    48    62    77    93   112   135   164   207   244   315   345   413  2877

methylKit::getMethylationStats output for "Rphase" CpGhsm-sites (#: 413885) at minimum coverage 20   - methylation statistics per base
summary:
   Min. 1st Qu.  Median    Mean 3rd Qu.    Max.
    1.8    17.6    34.7    44.3    71.4   100.0
percentiles:
    0%    10%    20%    30%    40%    50%    60%    70%    80%    90%    95%    99%  99.5%  99.9%   100%
```

```
  1.77  11.54  15.48  20.00  26.02  34.69  47.06  62.90  79.87  92.59  96.35 100.00 100.00 100.00 100.00

methylKit::getCoverageStats output for "Rphase" CpGhsm-sites (#: 327310) at minimum coverage 50     - read coverage statistics per base
summary:
   Min. 1st Qu.  Median    Mean 3rd Qu.    Max.
     50      78     113     129     165    2877
percentiles:
   0%   10%   20%   30%   40%   50%   60%   70%   80%   90%   95%   99% 99.5% 99.9%  100%
   50    60    72    84    98   113   131   152   179   220   255   326   355   423  2877

methylKit::getMethylationStats output for "Rphase" CpGhsm-sites (#: 327310) at minimum coverage 50  - methylation statistics per base
summary:
   Min. 1st Qu.  Median    Mean 3rd Qu.    Max.
    1.8    15.4    30.3    41.7    68.1   100.0
percentiles:
    0%    10%    20%    30%    40%    50%    60%    70%    80%    90%    95%    99%  99.5%  99.9%   100%
  1.77  10.72  13.85  17.14  22.16  30.26  42.55  58.90  77.11  91.76  95.98  98.57  99.16 100.00 100.00

methylKit::getCoverageStats output for "Rphase" CpGhsm-sites (#: 192756) in coverage range [100-1000]     - read coverage statistics per base
summary:
   Min. 1st Qu.  Median    Mean 3rd Qu.    Max.
    100     123     154     168     198     875
percentiles:
   0%   10%   20%   30%   40%   50%   60%   70%   80%   90%   95%   99% 99.5% 99.9%  100%
  100   108   118   129   141   154   169   187   211   247   279   348   377   446   875

methylKit::getMethylationStats output for "Rphase" CpGhsm-sites (#: 192756) in coverage range [100-1000]  - methylation statistics per base
summary:
   Min. 1st Qu.  Median    Mean 3rd Qu.    Max.
    2.9    13.5    30.6    41.4    68.9   100.0
percentiles:
    0%    10%    20%    30%    40%    50%    60%    70%    80%    90%    95%    99%  99.5%  99.9%   100%
  2.86   9.26  11.76  15.60  21.62  30.60  43.42  59.84  77.59  91.84  95.92  98.33  98.95  99.57 100.00
```

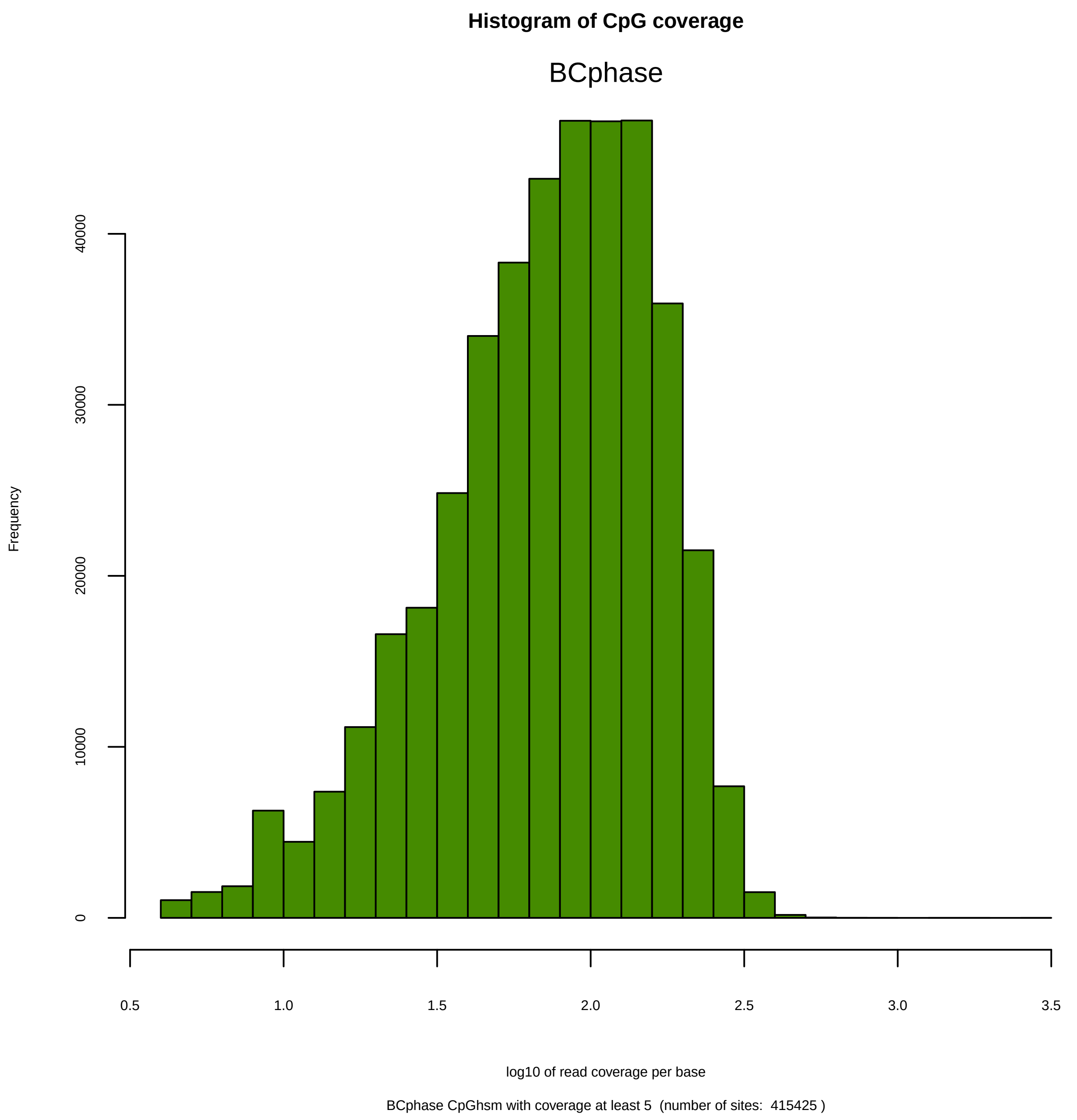

Histogram of CpG coverage
BCphase
Frequency
0
10000
20000
30000
40000
0.5
1.0
1.5
2.0
2.5
3.0
3.5
log10 of read coverage per base
BCphase CpGhsm with coverage at least 5 (number of sites: 415425 )

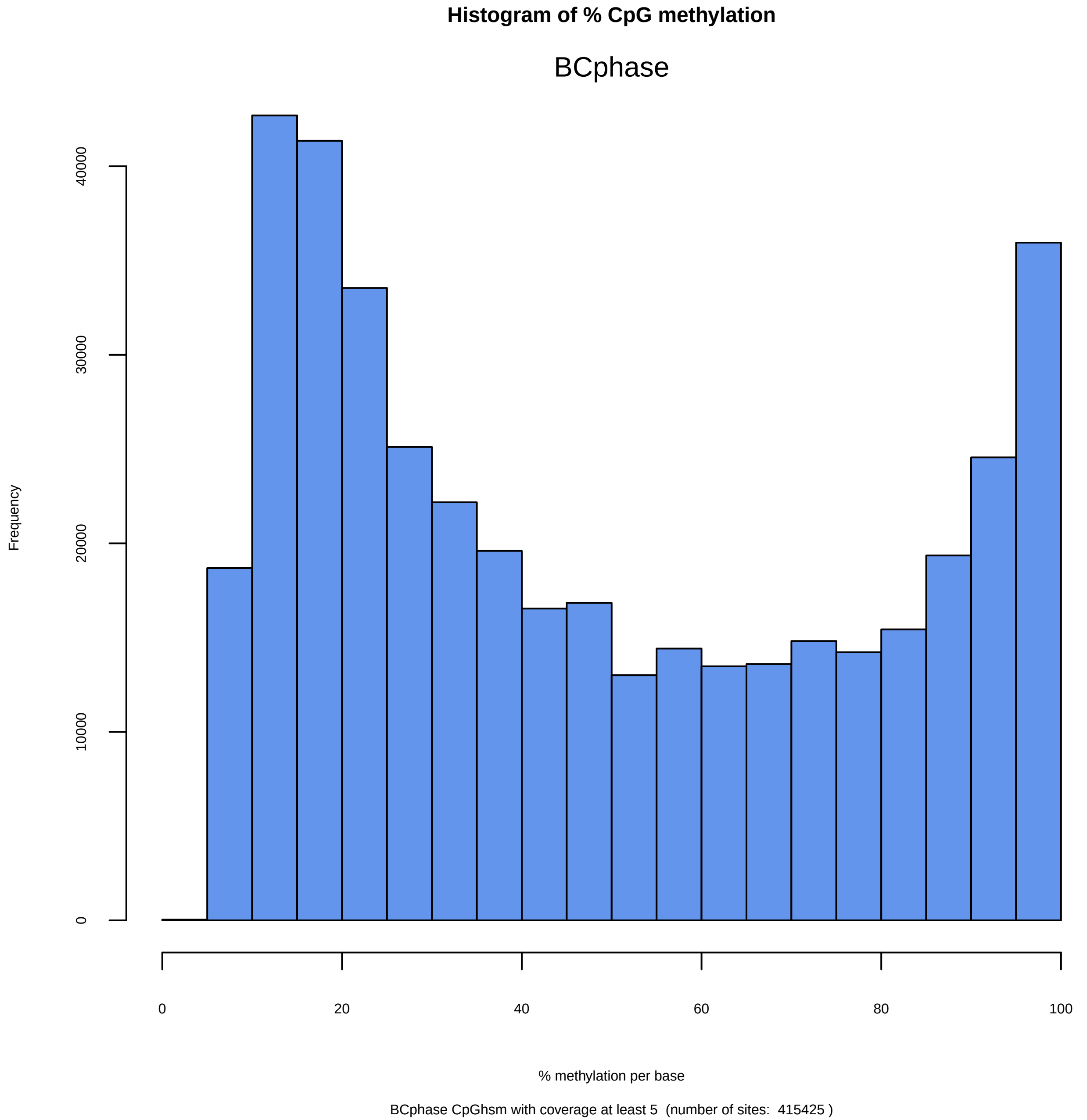
Histogram of % CpG methylation
BCphase
Frequency
0
10000
20000
30000
40000
0
20
40
60
80
100
% methylation per base
BCphase CpGhsm with coverage at least 5  (number of sites:  415425 )

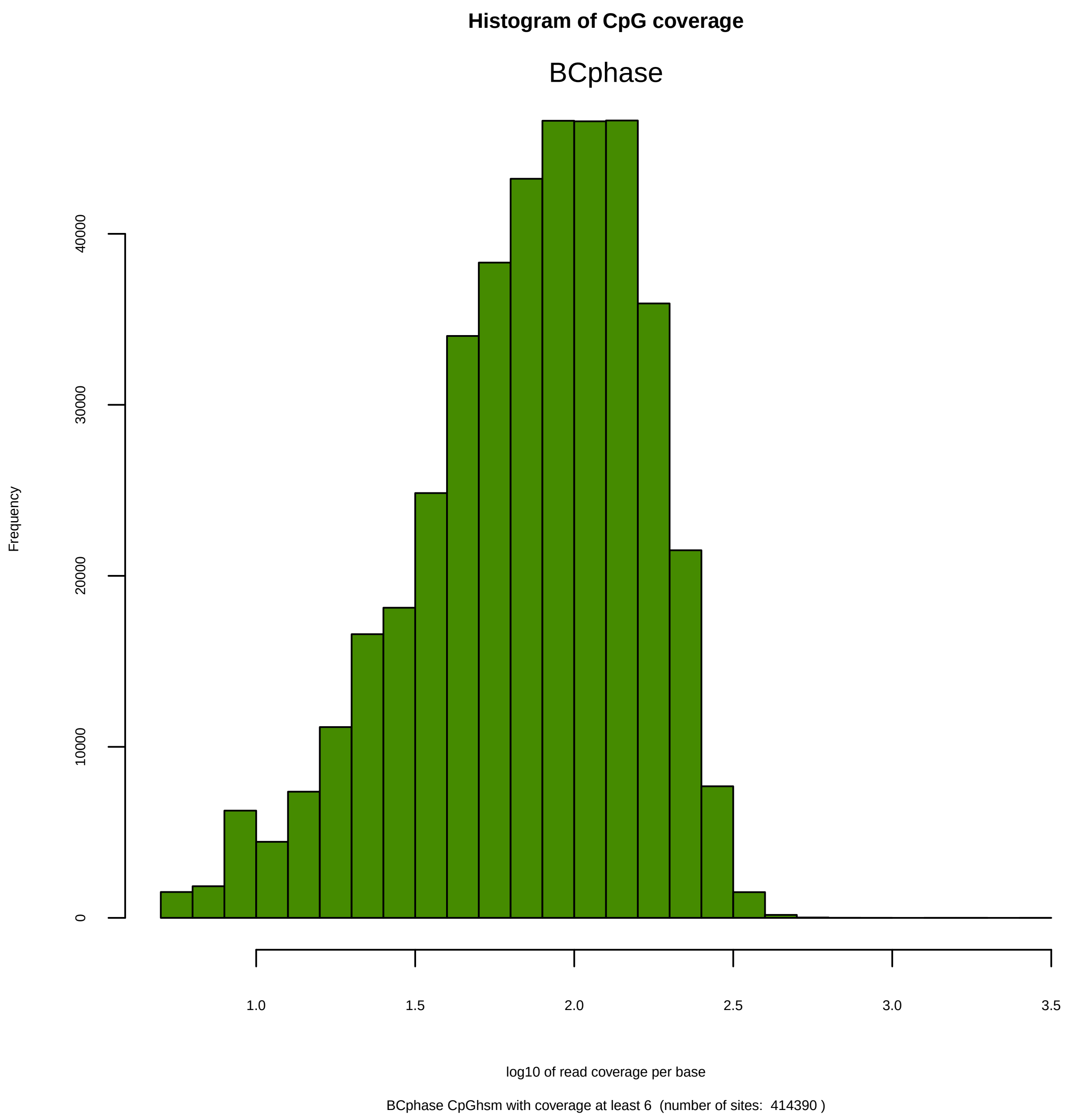

Histogram of CpG coverage
BCphase
Frequency
0
10000
20000
30000
40000
1.0
1.5
2.0
2.5
3.0
3.5
log10 of read coverage per base
BCphase CpGhsm with coverage at least 6  (number of sites:  414390 )

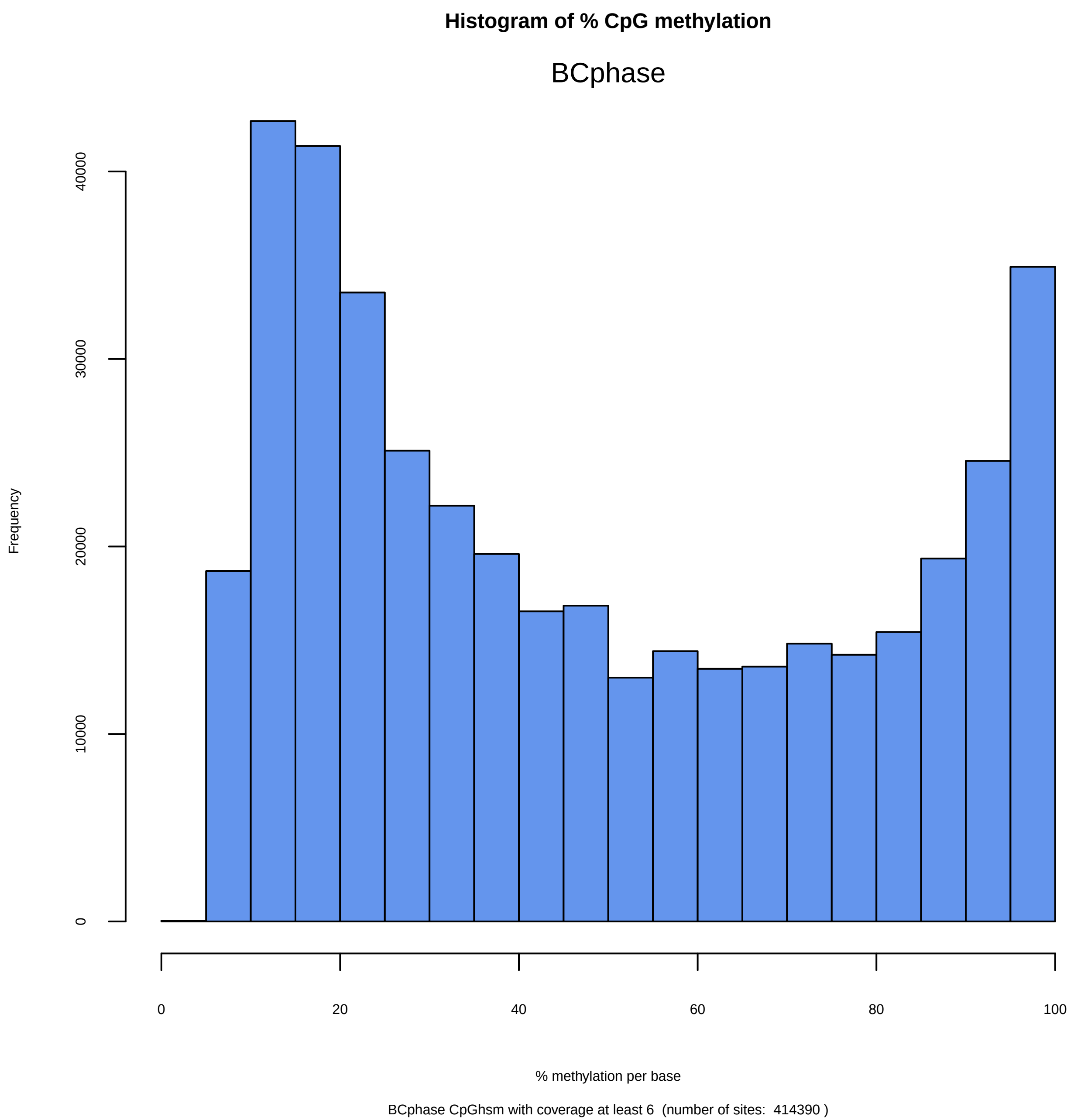
Histogram of % CpG methylation
BCphase
Frequency
0
10000
20000
30000
40000
0
20
40
60
80
100
% methylation per base
BCphase CpGhsm with coverage at least 6  (number of sites:  414390 )

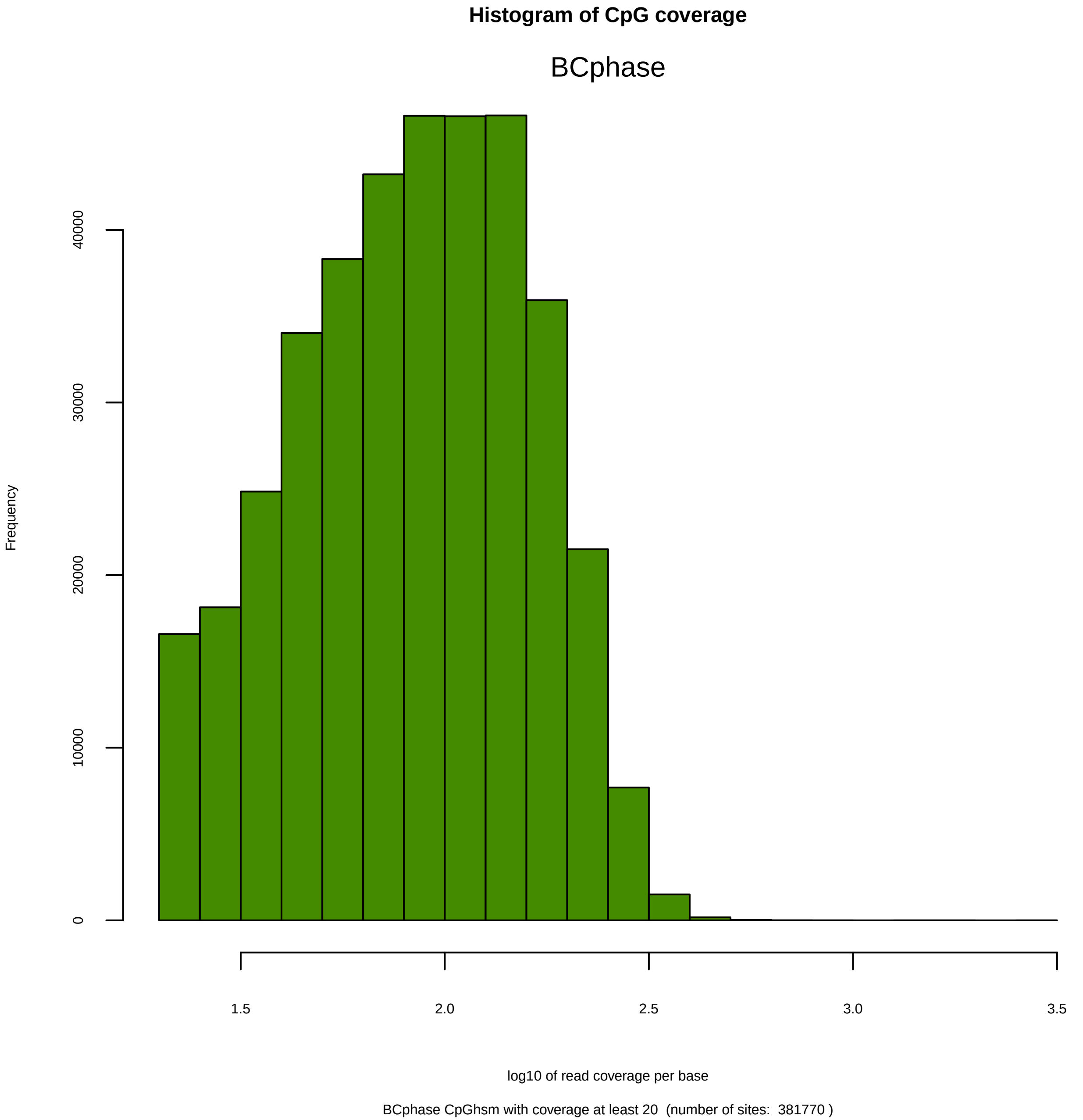

Histogram of CpG coverage
BCphase
Frequency
0
10000
20000
30000
40000
1.5
2.0
2.5
3.0
3.5
log10 of read coverage per base
BCphase CpGhsm with coverage at least 20 (number of sites: 381770 )

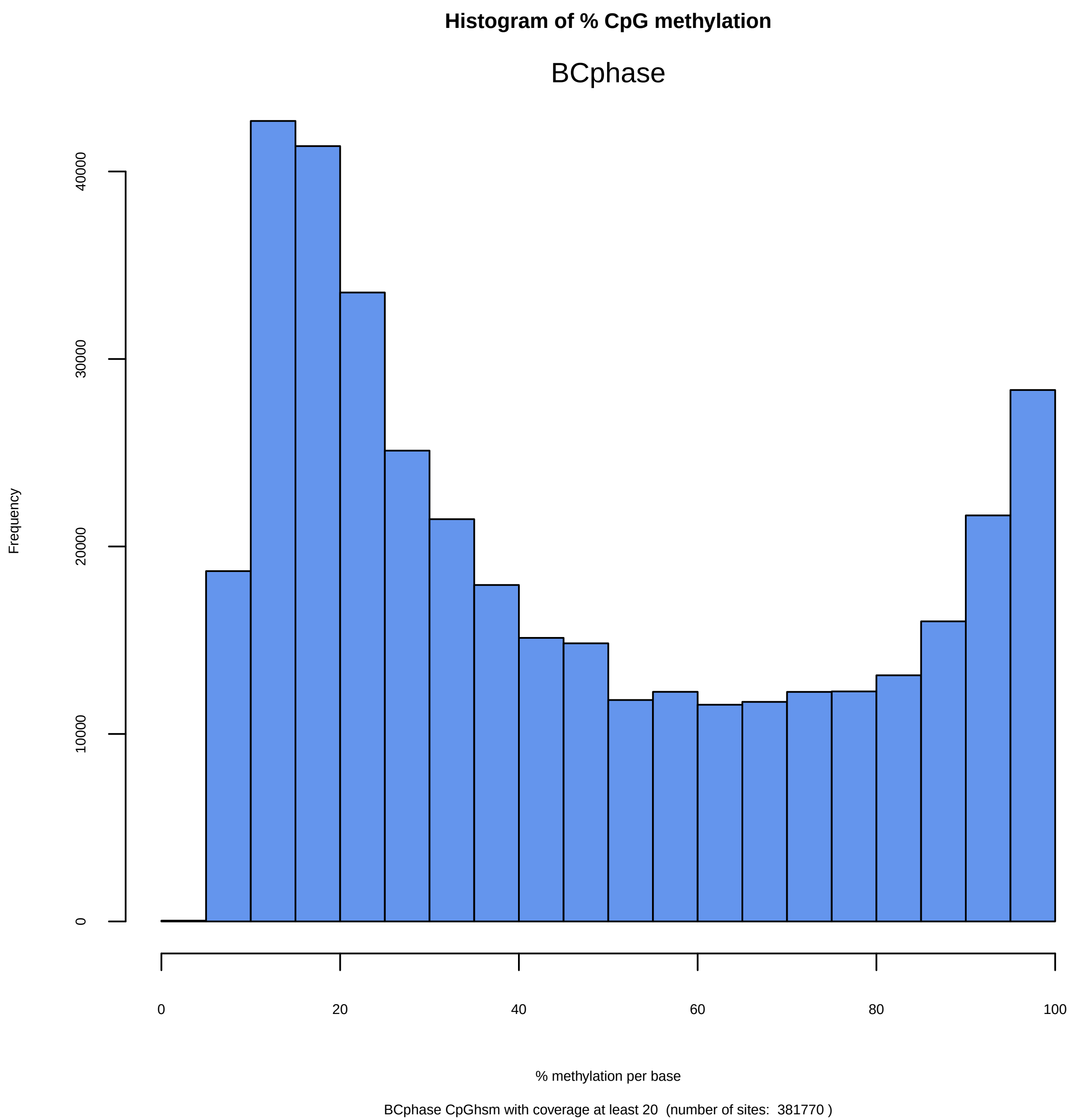
Histogram of % CpG methylation
BCphase
Frequency
0
10000
20000
30000
40000
0
20
40
60
80
100
% methylation per base
BCphase CpGhsm with coverage at least 20 (number of sites: 381770 )

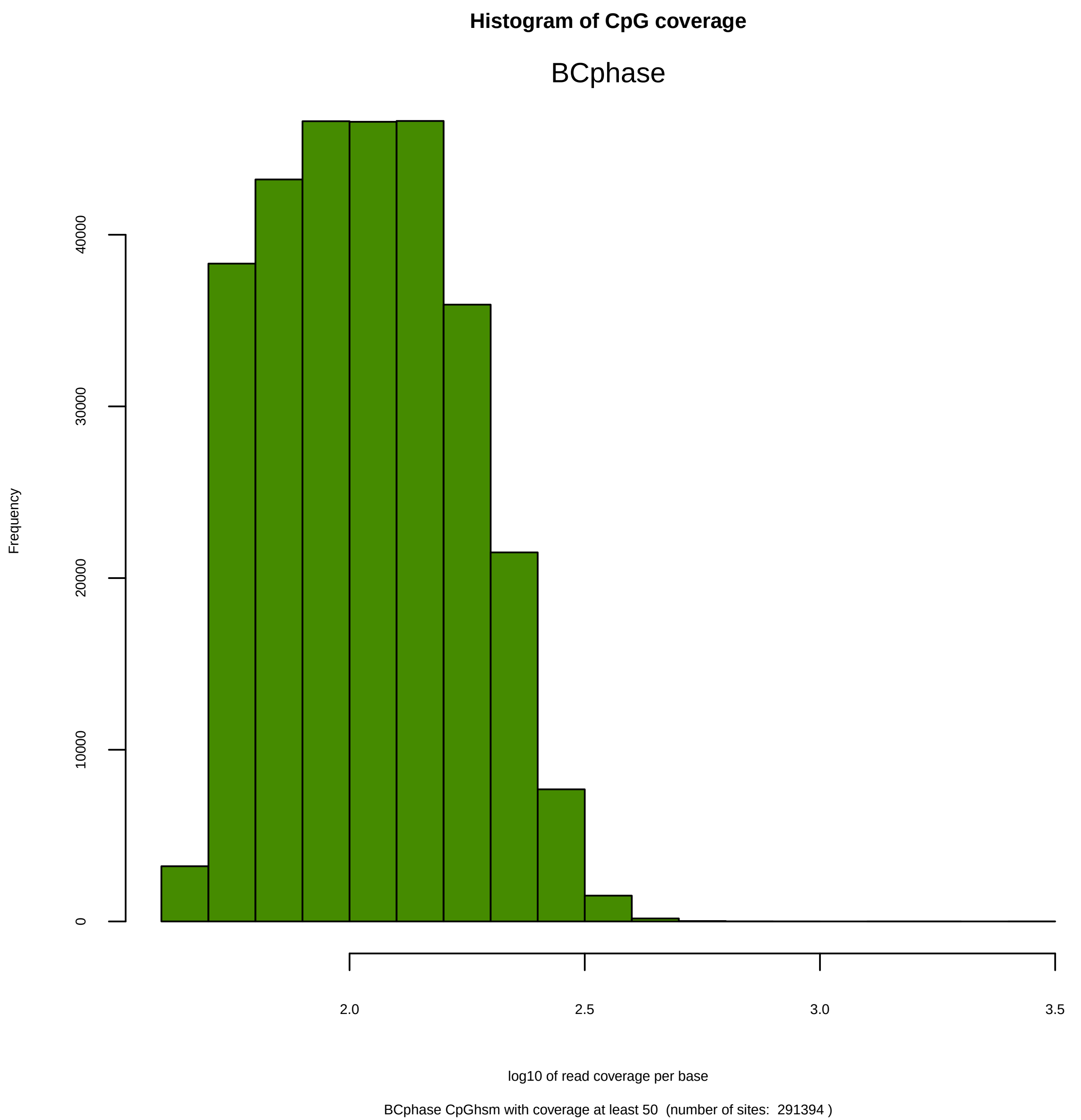
Histogram of CpG coverage
BCphase
Frequency
0
10000
20000
30000
40000
2.0
2.5
3.0
3.5
log10 of read coverage per base
BCphase CpGhsm with coverage at least 50 (number of sites: 291394 )

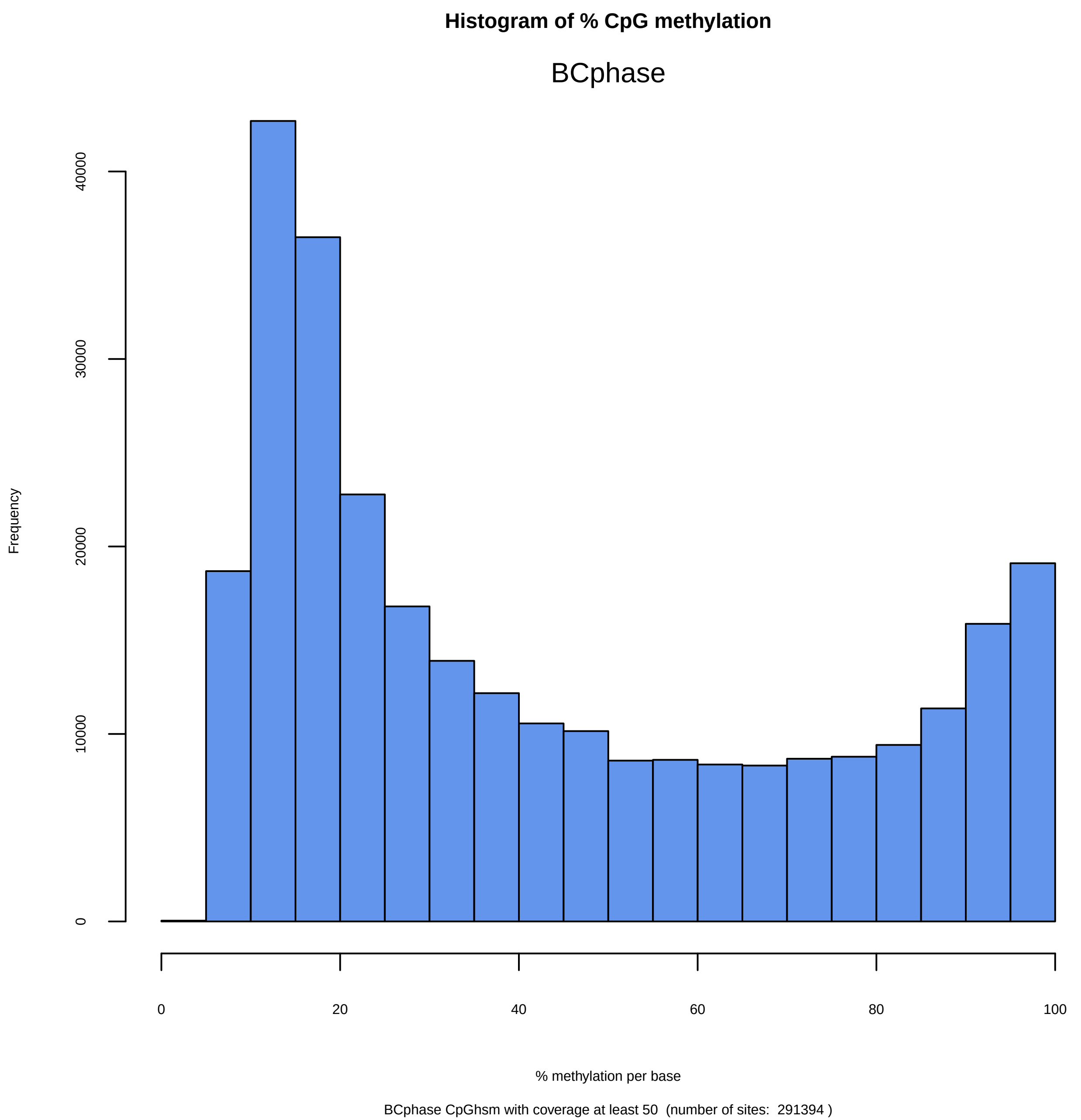
Histogram of % CpG methylation
BCphase
Frequency
0
10000
20000
30000
40000
0
20
40
60
80
100
% methylation per base
BCphase CpGhsm with coverage at least 50  (number of sites:  291394 )

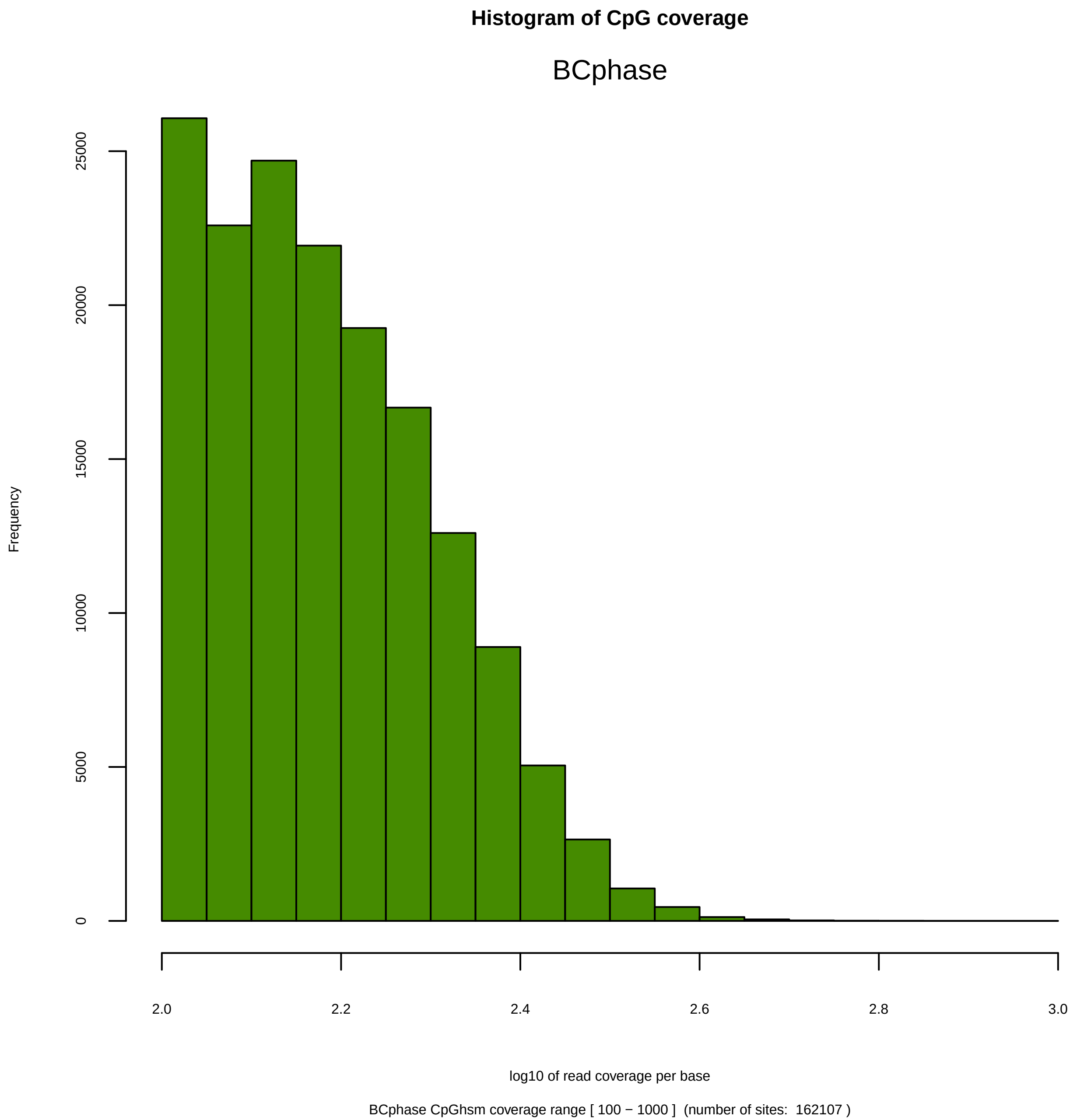
Histogram of CpG coverage
BCphase
Frequency
0
5000
10000
15000
20000
25000
2.0
2.2
2.4
2.6
2.8
3.0
log10 of read coverage per base
BCphase CpGhsm coverage range [ 100 − 1000 ]  (number of sites:  162107 )

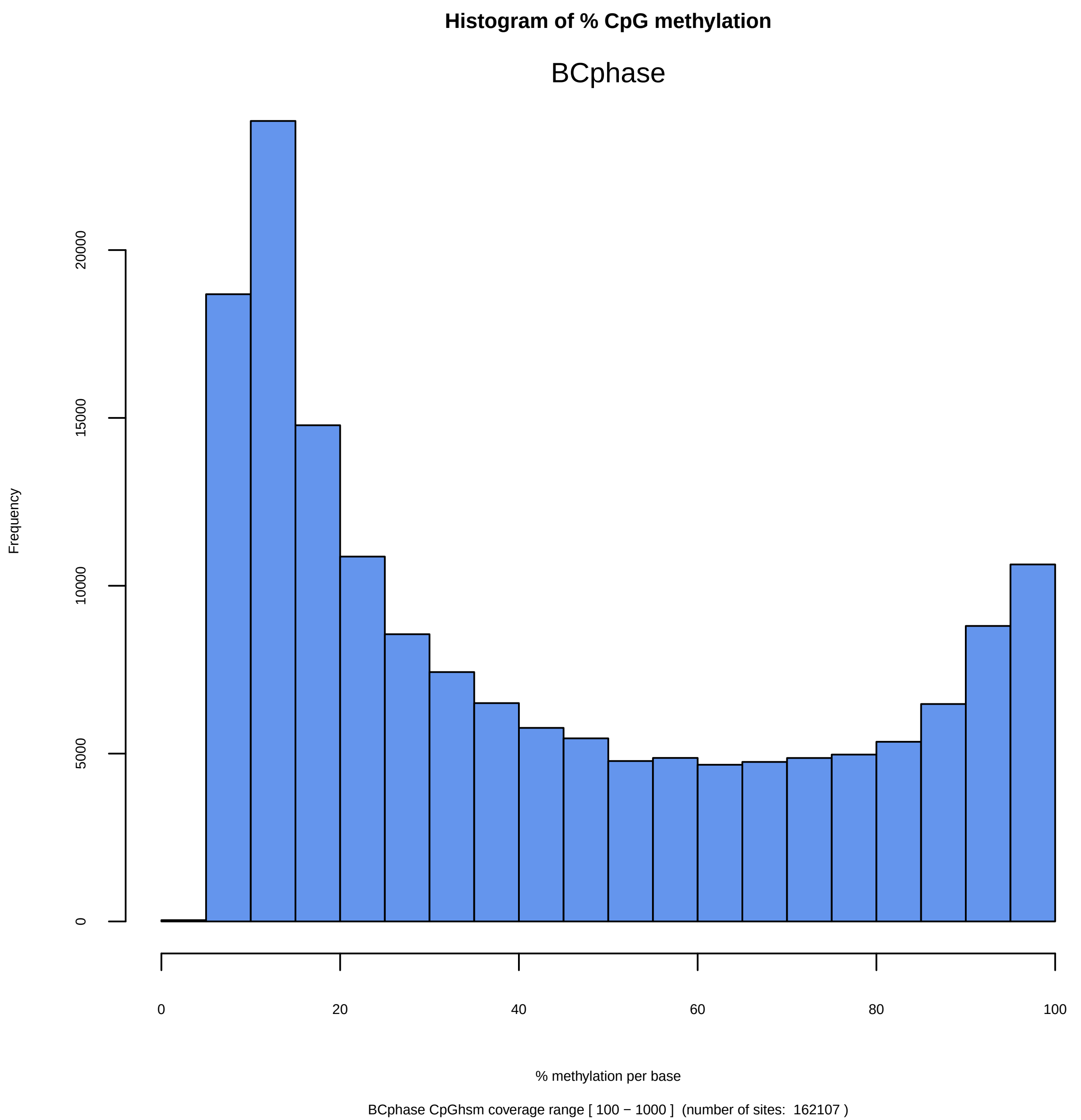

Histogram of % CpG methylation
BCphase
Frequency
0
5000
10000
15000
20000
0
20
40
60
80
100
% methylation per base
BCphase CpGhsm coverage range [ 100 − 1000 ]  (number of sites:  162107 )

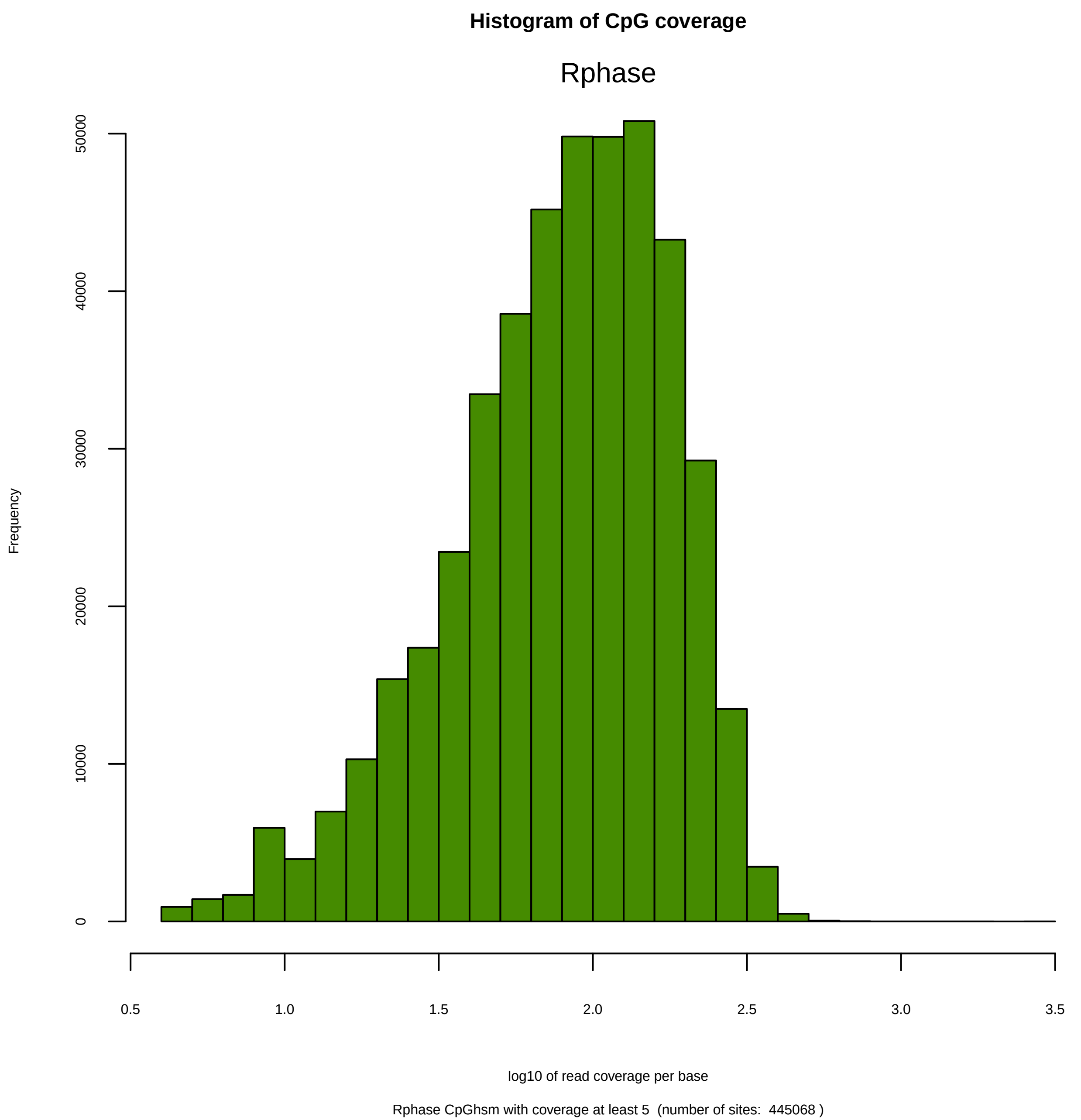

Histogram of CpG coverage
Rphase
Frequency
0
10000
20000
30000
40000
50000
0.5
1.0
1.5
2.0
2.5
3.0
3.5
log10 of read coverage per base
Rphase CpGhsm with coverage at least 5 (number of sites: 445068 )

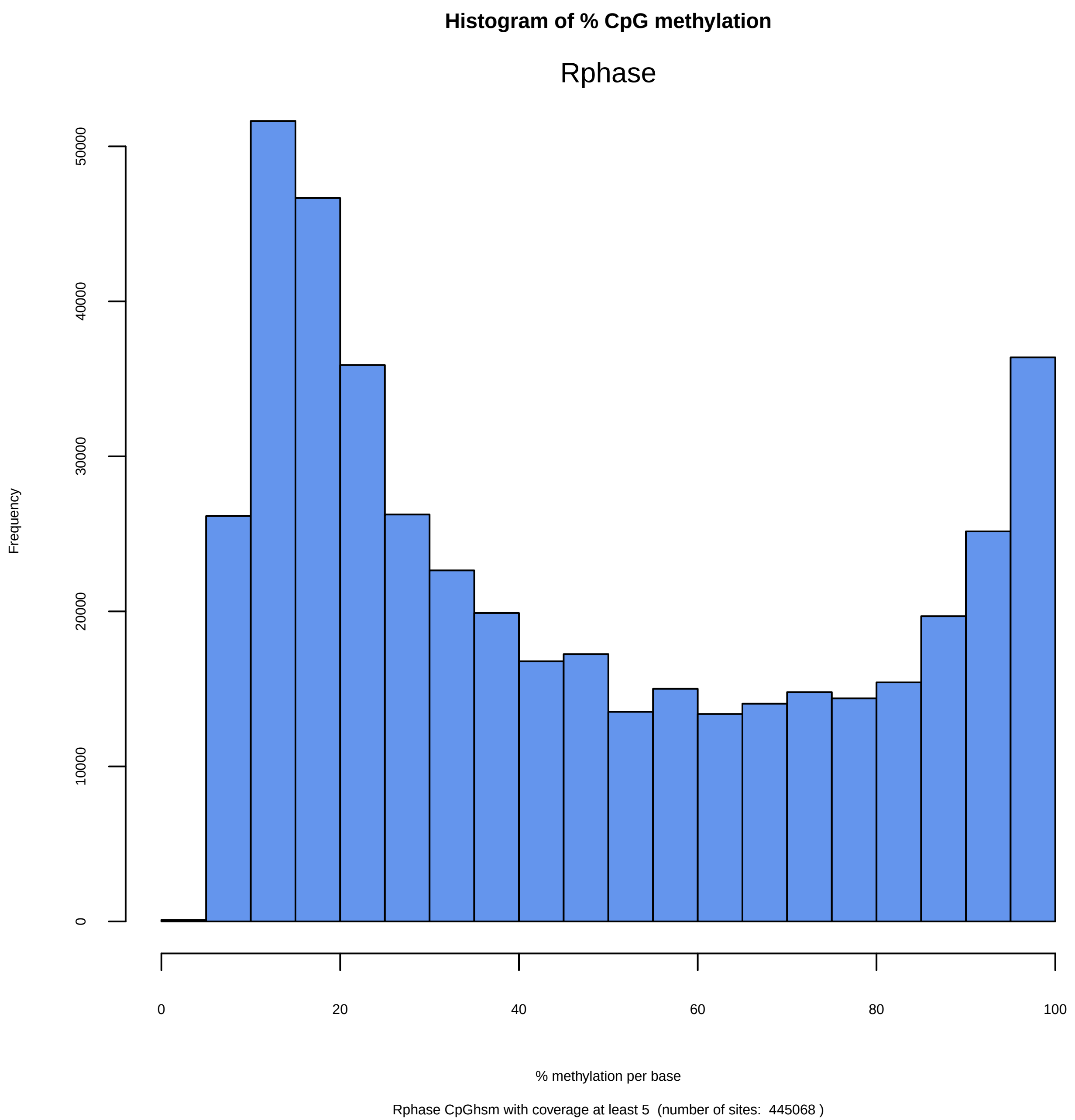

Histogram of % CpG methylation
Rphase
Frequency
0
10000
20000
30000
40000
50000
0
20
40
60
80
100
% methylation per base
Rphase CpGhsm with coverage at least 5 (number of sites: 445068 )

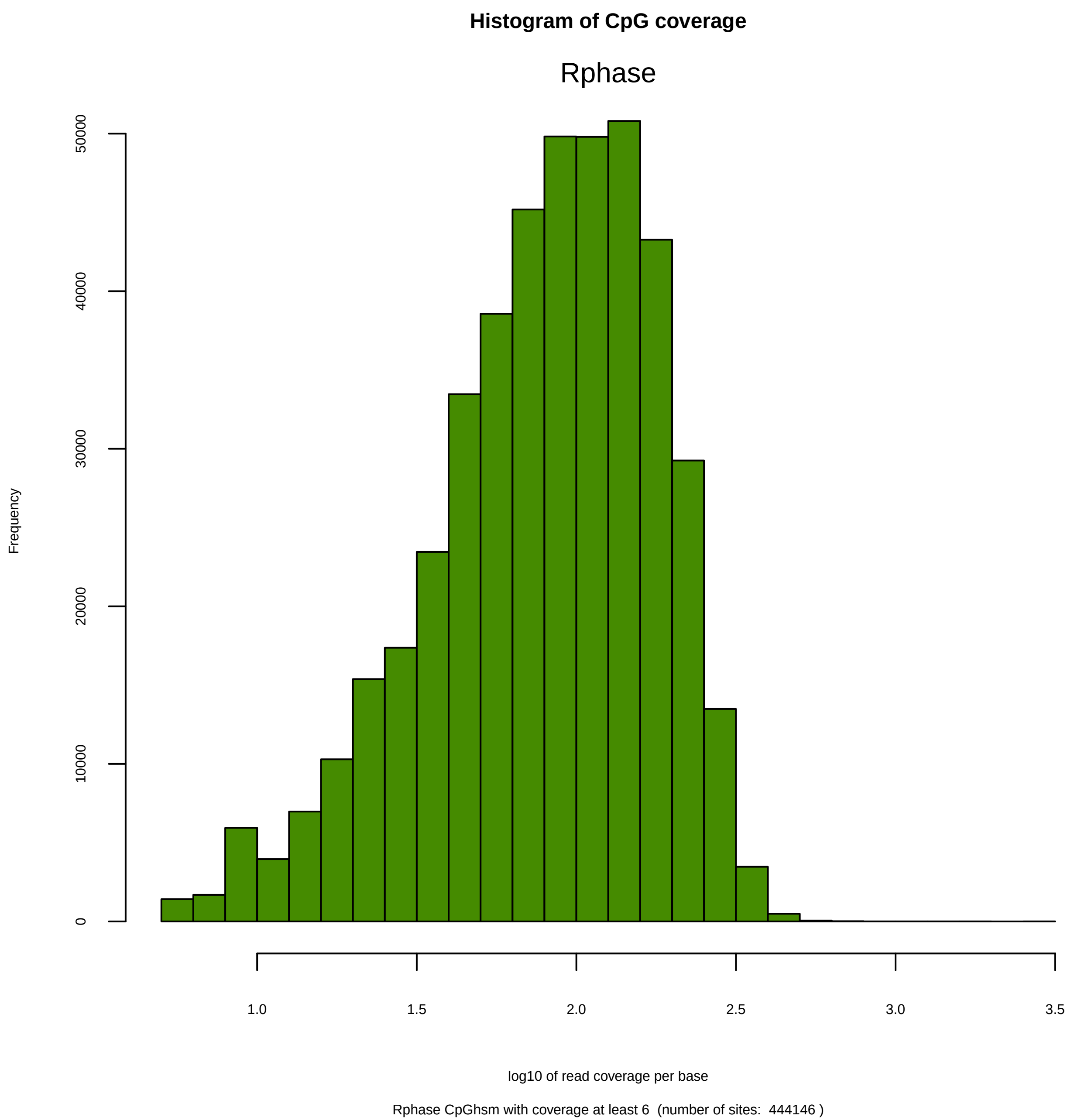

Histogram of CpG coverage
Rphase
Frequency
0
10000
20000
30000
40000
50000
1.0
1.5
2.0
2.5
3.0
3.5
log10 of read coverage per base
Rphase CpGhsm with coverage at least 6 (number of sites: 444146 )

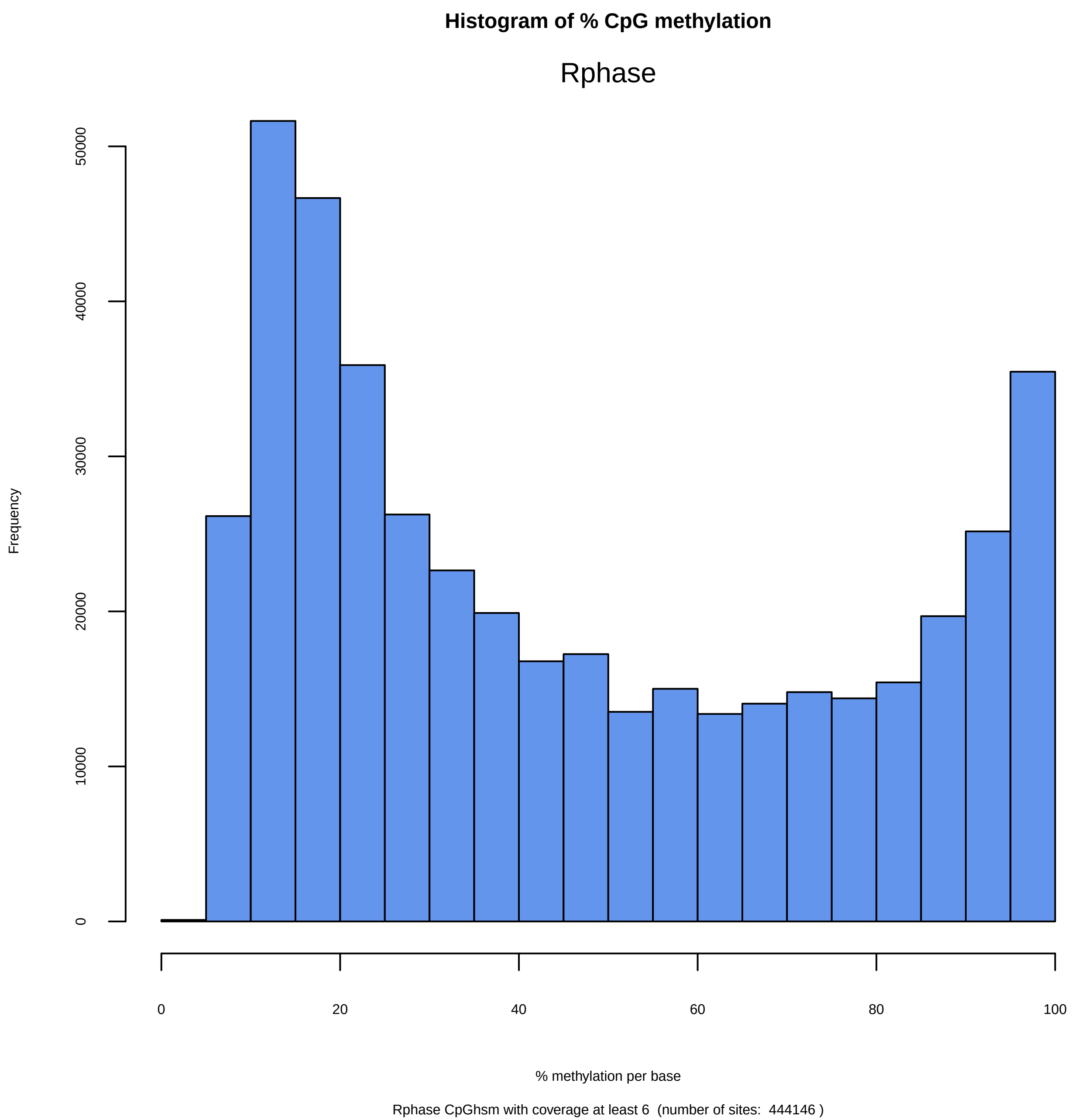
Histogram of % CpG methylation
Rphase
Frequency
0
10000
20000
30000
40000
50000
0
20
40
60
80
100
% methylation per base
Rphase CpGhsm with coverage at least 6 (number of sites: 444146 )

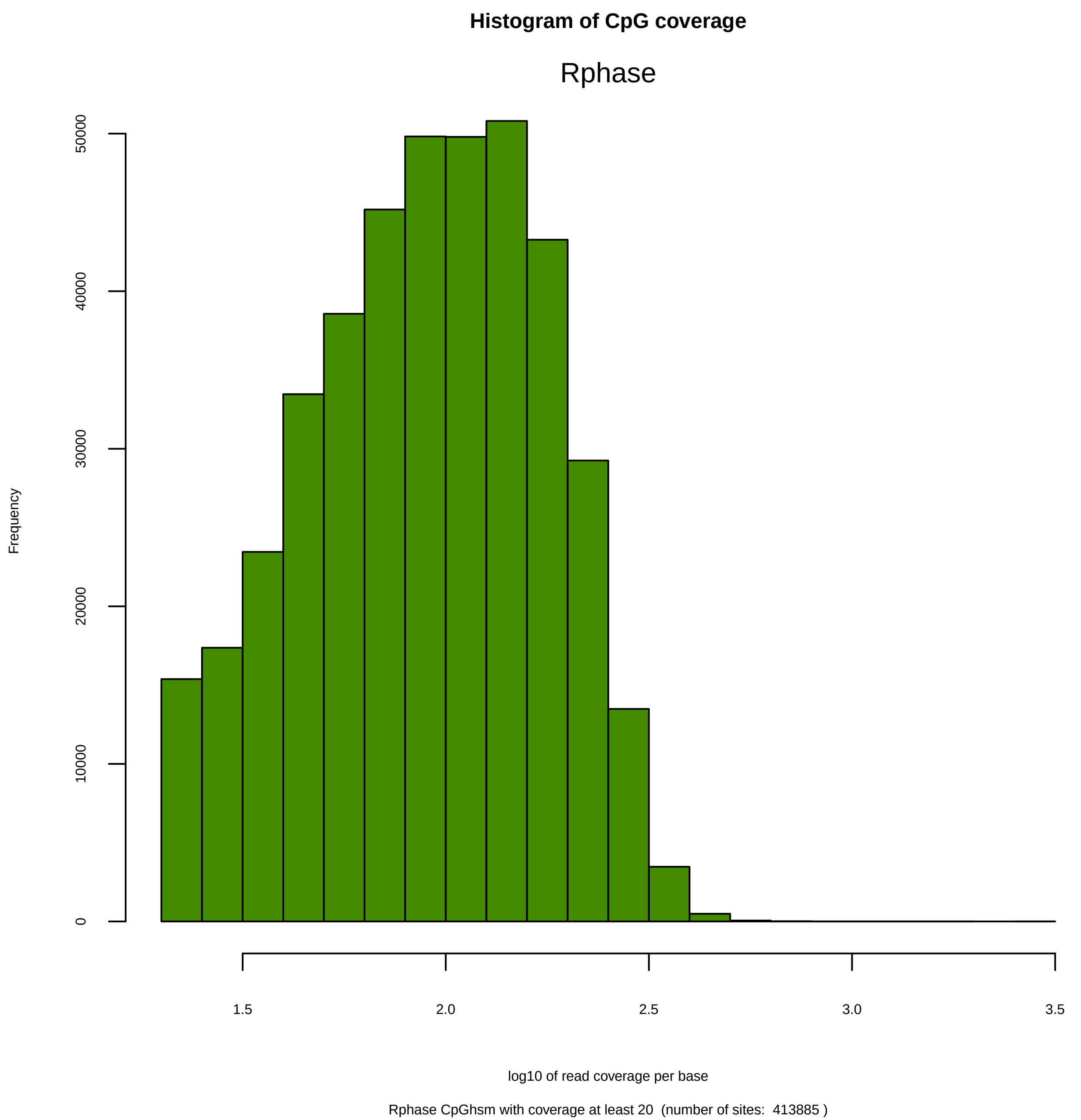
Histogram of CpG coverage
Rphase
Frequency
0
10000
20000
30000
40000
50000
1.5
2.0
2.5
3.0
3.5
log10 of read coverage per base
Rphase CpGhsm with coverage at least 20 (number of sites: 413885 )

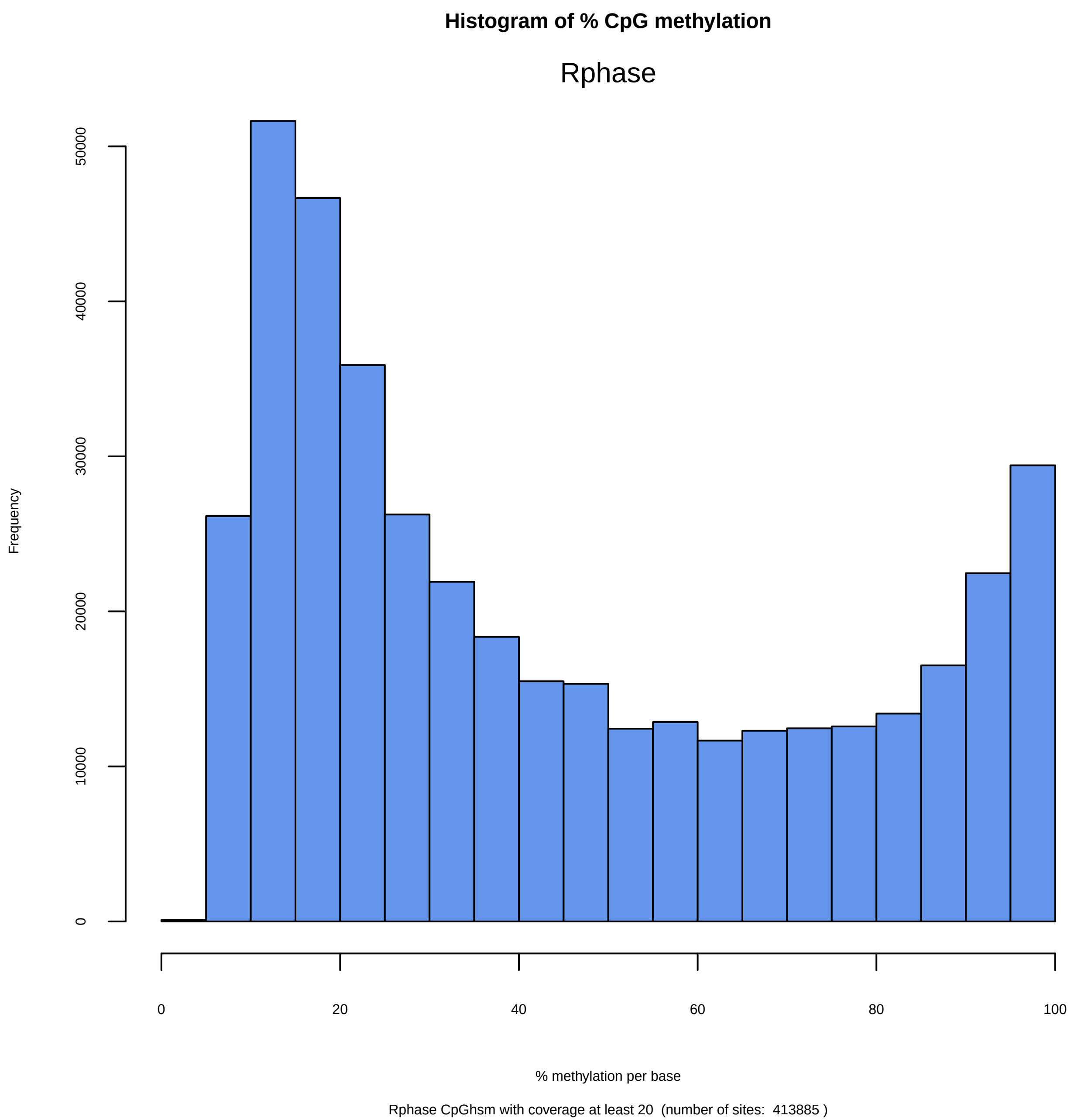

Histogram of % CpG methylation
Rphase
Frequency
0
10000
20000
30000
40000
50000
0
20
40
60
80
100
% methylation per base
Rphase CpGhsm with coverage at least 20  (number of sites:  413885 )

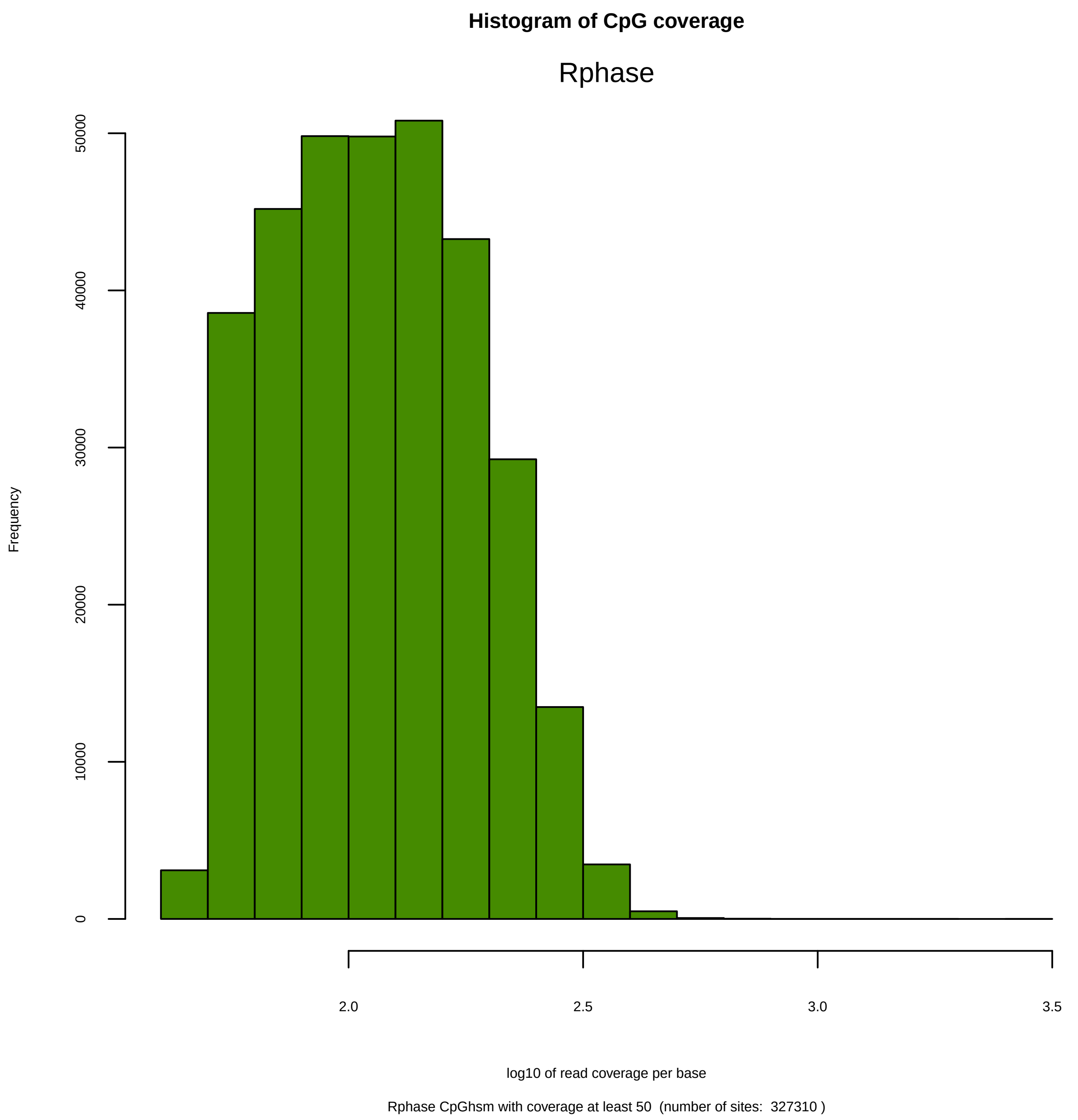

Histogram of CpG coverage
Rphase
Frequency
0
10000
20000
30000
40000
50000
2.0
2.5
3.0
3.5
log10 of read coverage per base
Rphase CpGhsm with coverage at least 50  (number of sites:  327310 )

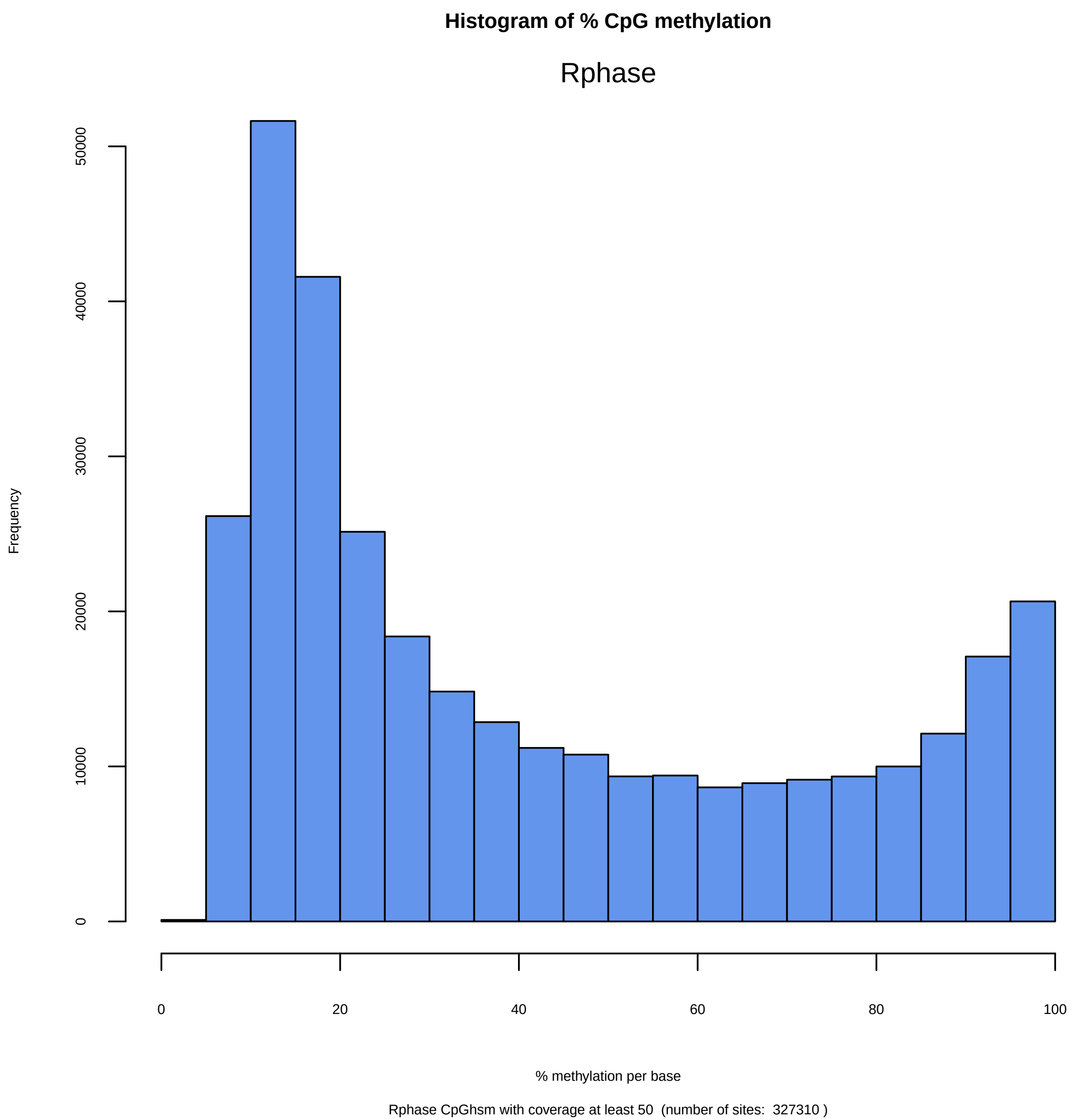
Histogram of % CpG methylation
Rphase
Frequency
0
10000
20000
30000
40000
50000
0
20
40
60
80
100
% methylation per base
Rphase CpGhsm with coverage at least 50 (number of sites: 327310 )

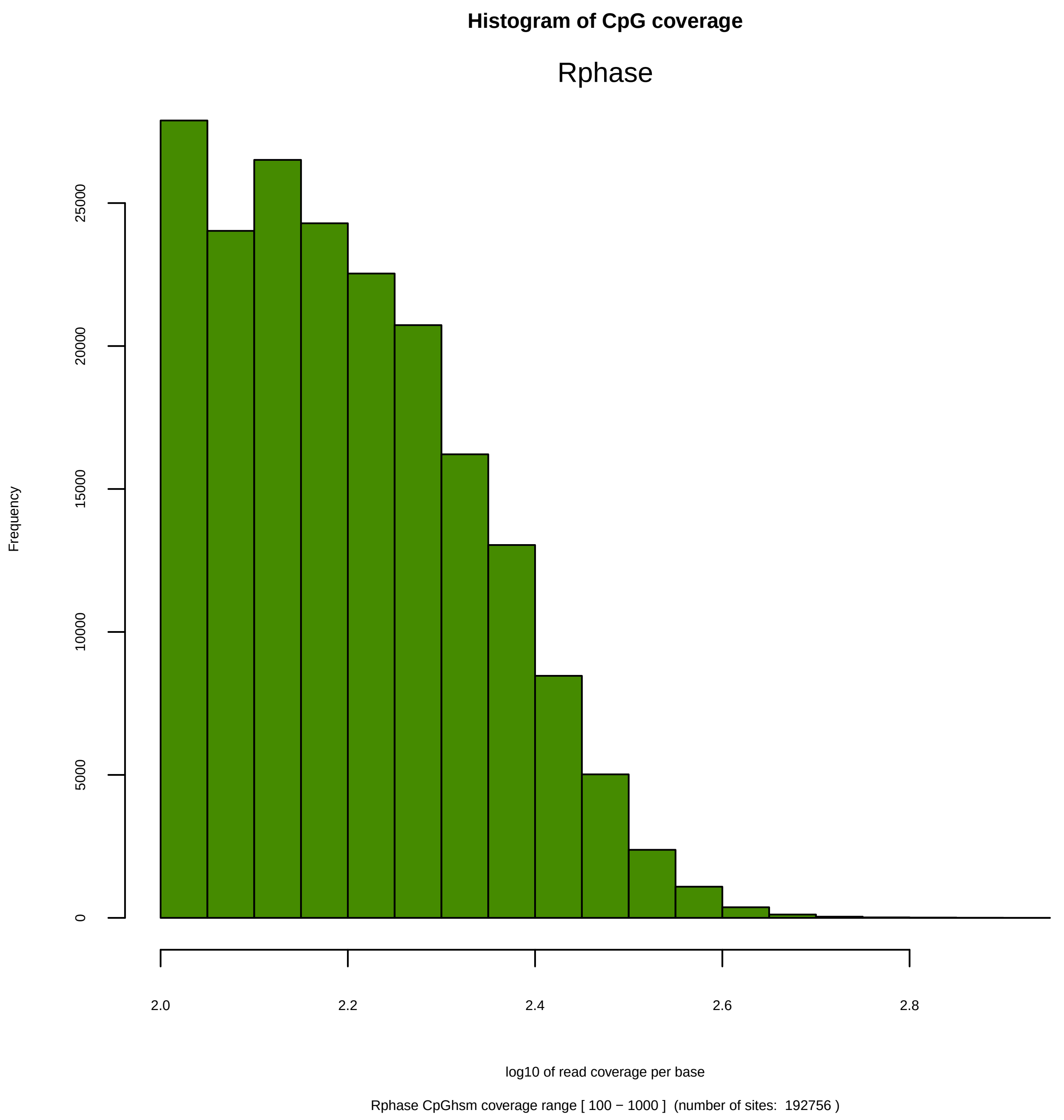
Histogram of CpG coverage
Rphase
Frequency
0
5000
10000
15000
20000
25000
2.0
2.2
2.4
2.6
2.8
log10 of read coverage per base
Rphase CpGhsm coverage range [ 100 − 1000 ]  (number of sites:  192756 )

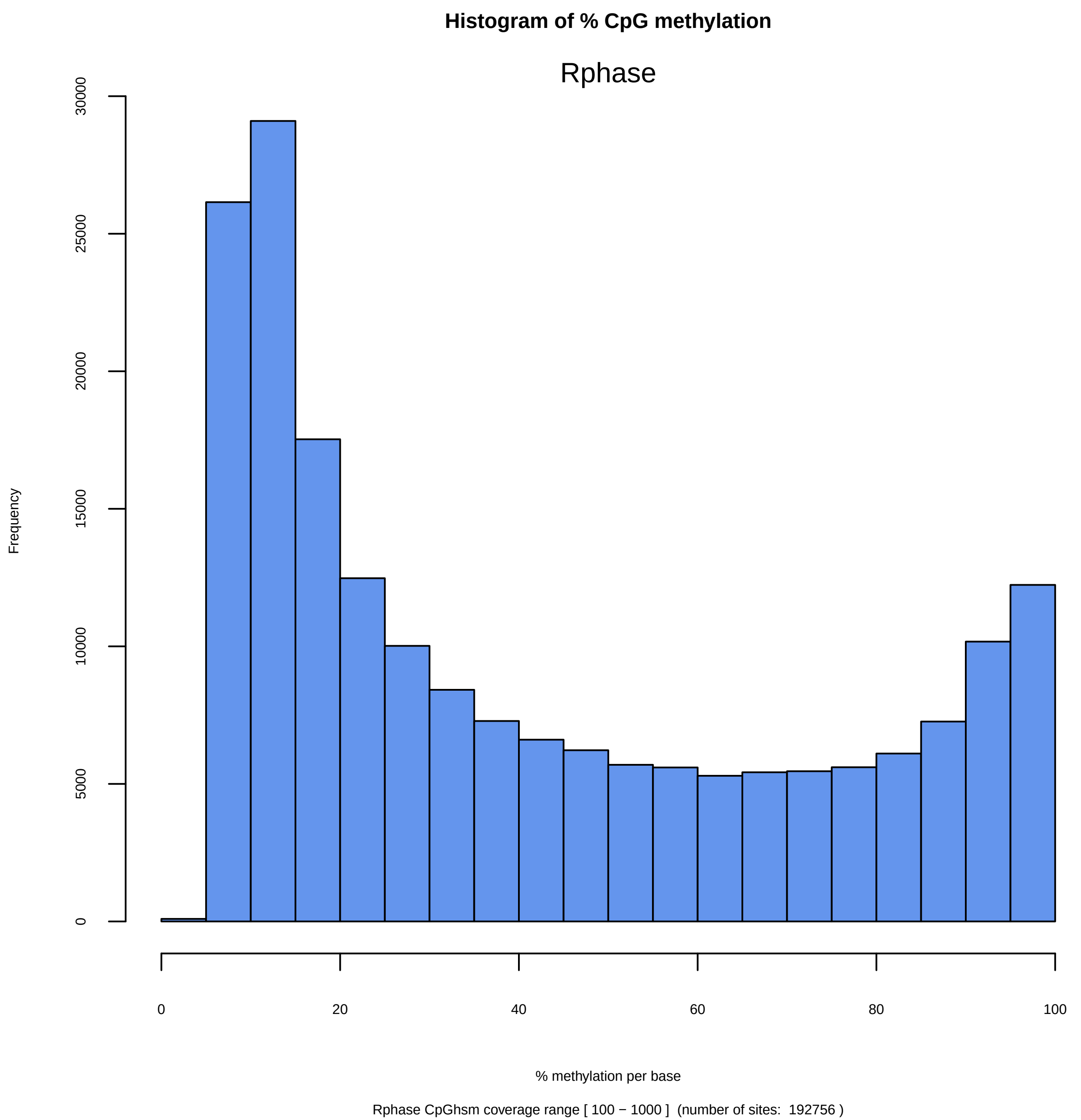

Histogram of % CpG methylation
Rphase
Frequency
0
5000
10000
15000
20000
25000
30000
0
20
40
60
80
100
% methylation per base
Rphase CpGhsm coverage range [ 100 − 1000 ] (number of sites: 192756 )

```
Directory: PWC File: 0READMEpwc

PWC - Pairwise comparisons between all samples.

Input: studymk, studymc, nbrpms, hheight, nbrpnts

Output: files pwc-*.txt pwc-.*pdf

Notes: The output is generated by BWASPR::cmpSites(), which determines for each pairwise
        comparison the numbers of common and distinct sites.  The function also calculates
        the overlap index between the samples and estimates the size of the common pool of
        potential methylation sites.

       The CpGscd data input (studymc) is necessary to determine sites that are detectable
        in both samples, and nbrpms (listed in the *.par file specified in the Ob_LIp.conf
        configuration file as argument to TOTALNBRPMSITES) provides the total number of
        potential methylation sites (typically, all CpG sites), a necessary parameter for
        the estimations.
       hheight (default: 0.10) specifies the y-axis limit in the histogram plots.
       nbrpnts (default: 5000) specifies the number of common sites to be plotted in the
        methylation levels scatter plot.
```

```
Directory: PWC File: pwc-Ob_LI_BCphase.vs.Ob_LI_Rphase.txt

Numbers of common and distinct sites comparing Ob_LI_BCphase versus Ob_LI_Rphase
================================================================================

  total number of potential sites:  28317694

  number of "Ob_LI_BCphase_hsm"         sites: 415425
  number of "Ob_LI_Rphase_hsm"         sites:  445068
    number of "Ob_LI_BCphase_hsm"-unique sites: 47418
    number of common sites:        368007
    number of "Ob_LI_Rphase_hsm"-unique sites:  77061
  total number of "Ob_LI_BCphase_hsm+Ob_LI_Rphase_hsm"-sites observed: 492486

  number of "Ob_LI_BCphase_scd"        sites:   24546430 ( 86.68% of total)
  number of "Ob_LI_Rphase_scd"        sites:    24689788 ( 87.19% of total)
    number of "Ob_LI_BCphase_scd"-unique sites:   418333
    number of sites in common:      24128097   (Expected:  21401677; O/E:       1.1)
    number of "Ob_LI_Rphase_scd"-unique sites:    561691
  total number of "Ob_LI_BCphase_scd+Ob_LI_Rphase_scd"-sites observed:  25108121

  number of sites in "Ob_LI_BCphase_hsm" that are not  detectable in "Ob_LI_Rphase_hsm":   1501
  number of sites in "Ob_LI_BCphase_hsm" that are also detectable in "Ob_LI_Rphase_hsm": 413924
    number of sites unique to "Ob_LI_BCphase_hsm" although detectable in "Ob_LI_Rphase_hsm":         45917
    number of "Ob_LI_BCphase_hsm" / "Ob_LI_Rphase_hsm" common sites                        :        368007     (Expected:   7588; O/E:      48.5)

  number of sites in "Ob_LI_Rphase_hsm" that are not  detectable in "Ob_LI_BCphase_hsm":   2744
  number of sites in "Ob_LI_Rphase_hsm" that are also detectable in "Ob_LI_BCphase_hsm": 442324
    number of sites unique to "Ob_LI_Rphase_hsm" although detectable in "Ob_LI_BCphase_hsm":         74317
    number of "Ob_LI_Rphase_hsm" / "Ob_LI_BCphase_hsm" common sites                        :        368007     (Expected:   7588; O/E:      48.5)

  Overlap index of "Ob_LI_BCphase_hsm" with "Ob_LI_Rphase_hsm":        0.805
  Estimated number of "Ob_LI_BCphase_hsm" = "Ob_LI_Rphase_hsm" sites (assuming sampling from one population):            497514
  Adjusted population size of "Ob_LI_BCphase_hsm" = "Ob_LI_Rphase_hsm" sites (assuming all sites detectable):            583902 (  1.19x of observed)

  number of "Ob_LI_BCphase_hsm" sites with coverage >=  5: 415425
  number of "Ob_LI_Rphase_hsm" sites with coverage >=  5: 445068
  number of common sites with coverage >=  5: 368007
  number of "Ob_LI_BCphase_hsm" sites with coverage >=  6: 414390
  number of "Ob_LI_Rphase_hsm" sites with coverage >=  6: 444146
  number of common sites with coverage >=  6: 367077
  number of "Ob_LI_BCphase_hsm" sites with coverage >= 20: 381770
  number of "Ob_LI_Rphase_hsm" sites with coverage >= 20: 413885
  number of common sites with coverage >= 20: 337795
  number of "Ob_LI_BCphase_hsm" sites with coverage >= 50: 291394
  number of "Ob_LI_Rphase_hsm" sites with coverage >= 50: 327310
  number of common sites with coverage >= 50: 259719
```

Methylation Levels in Common Sites (Coverage >=  5)

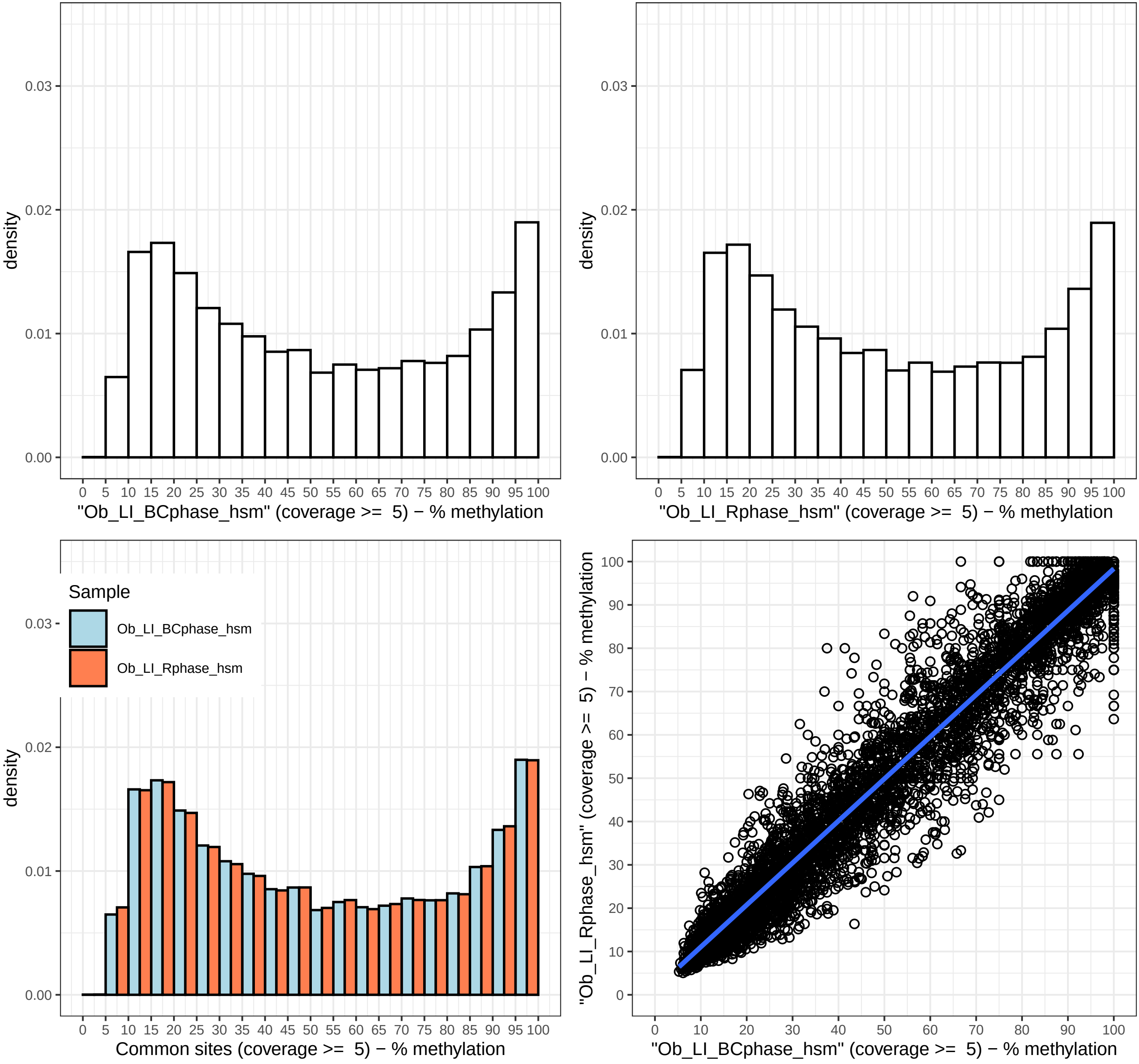

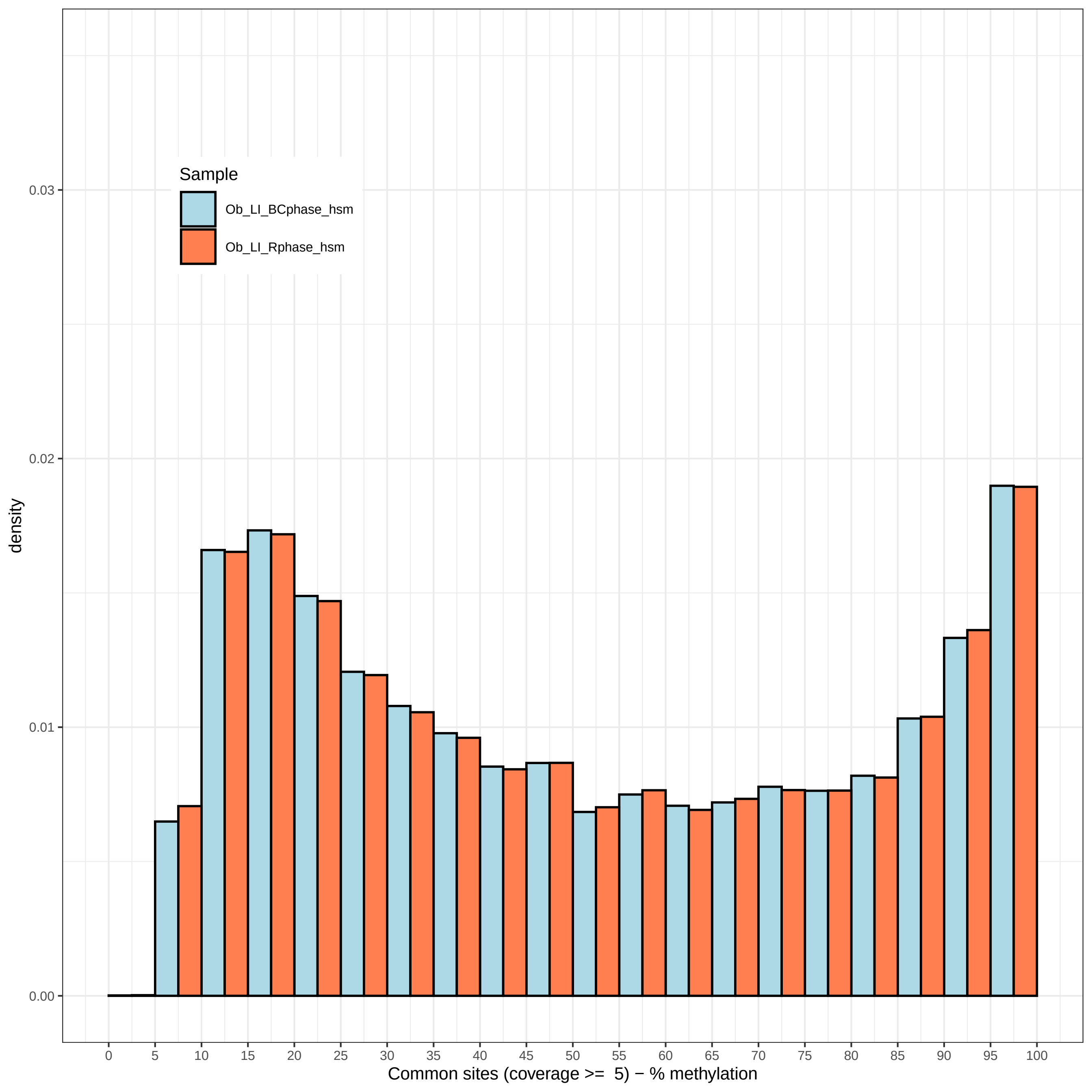

Sample
Ob_LI_BCphase_hsm
Ob_LI_Rphase_hsm
density
0.00
0.01
0.02
0.03
0
5
10
15
20
25
30
35
40
45
50
55
60
65
70
75
80
85
90
95
100
Common sites (coverage >= 5) − % methylation

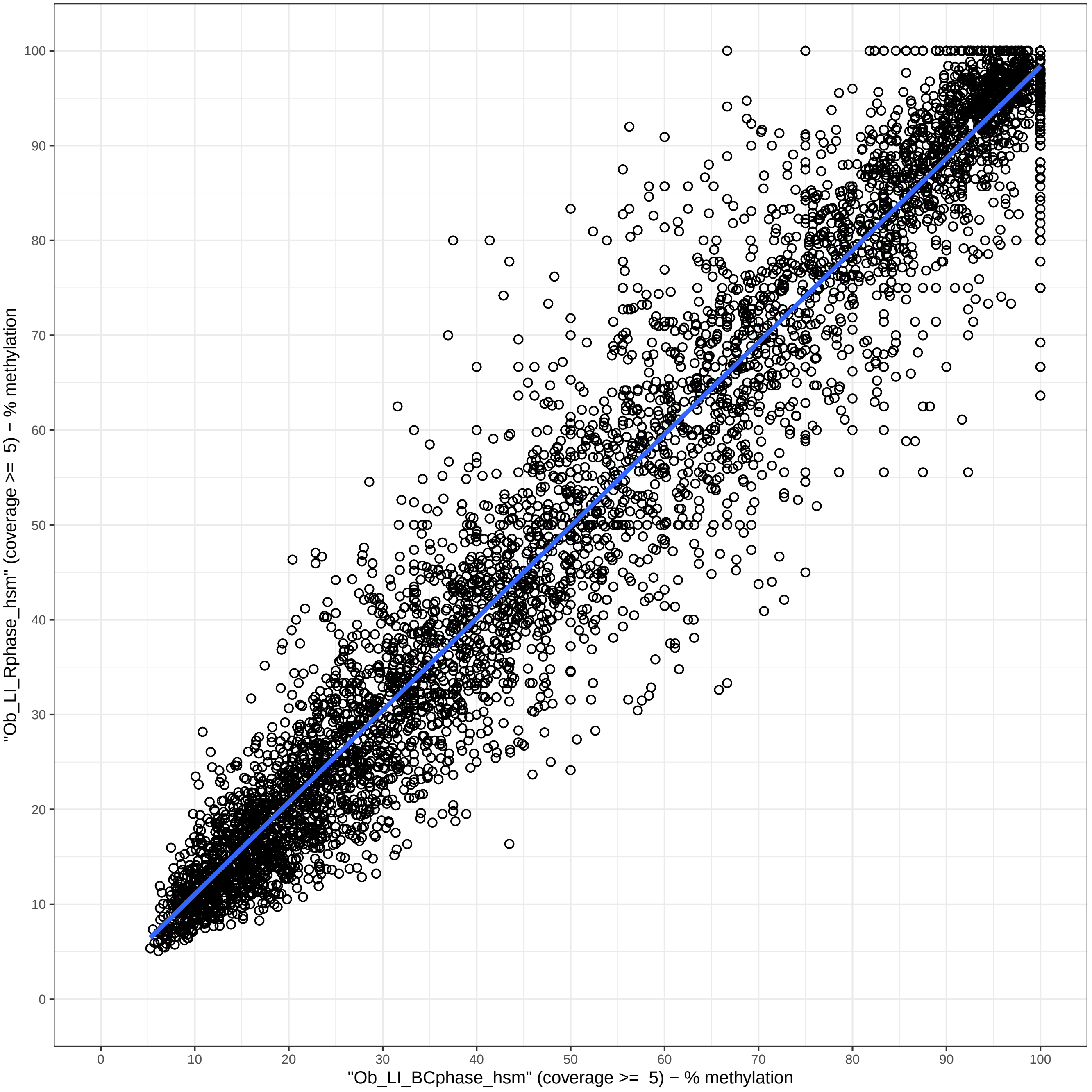

"Ob_LI_Rphase_hsm" (coverage >= 5) − % methylation
"Ob_LI_BCphase_hsm" (coverage >= 5) − % methylation
0
10
20
30
40
50
60
70
80
90
100

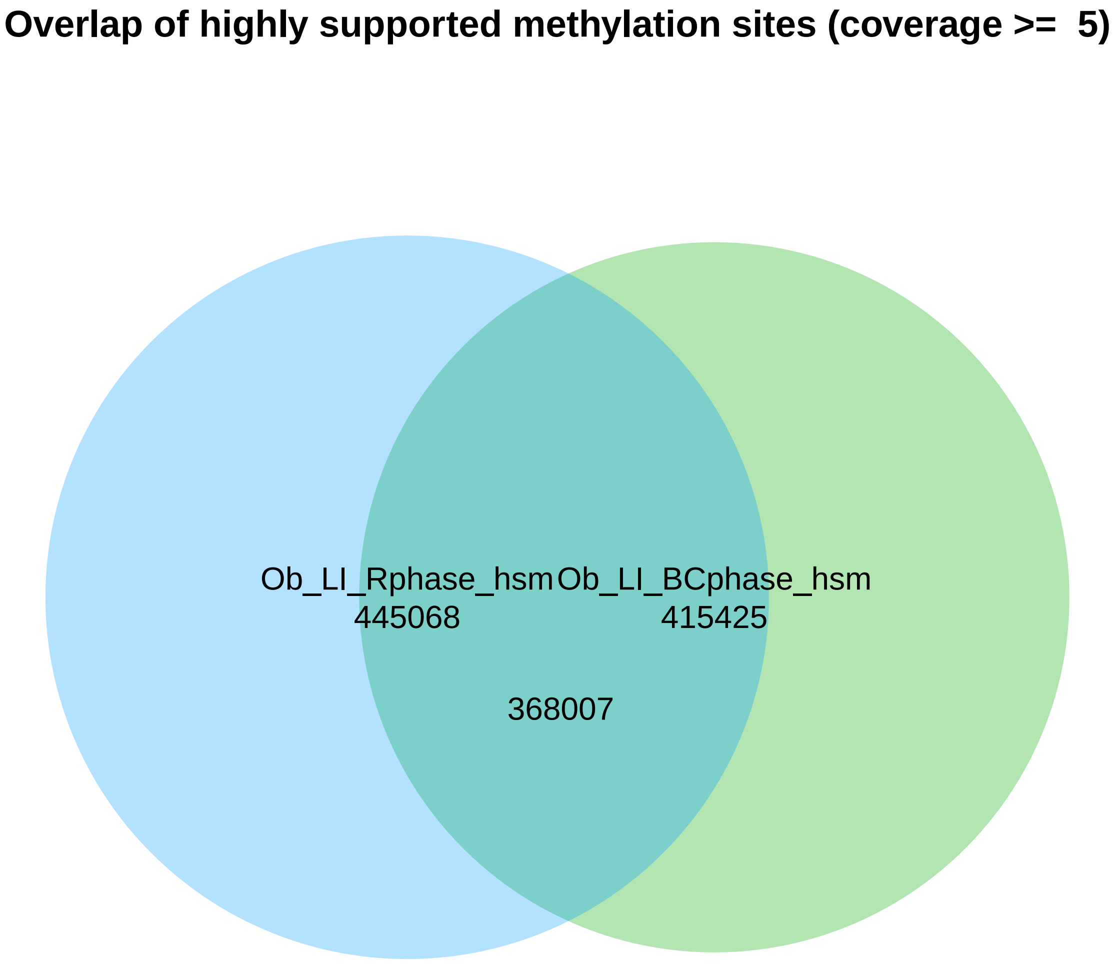
Overlap of highly supported methylation sites (coverage >= 5)
Ob_LI_Rphase_hsm
445068
Ob_LI_BCphase_hsm
415425
368007

### Methylation Levels in Common Sites (Coverage >=  6)

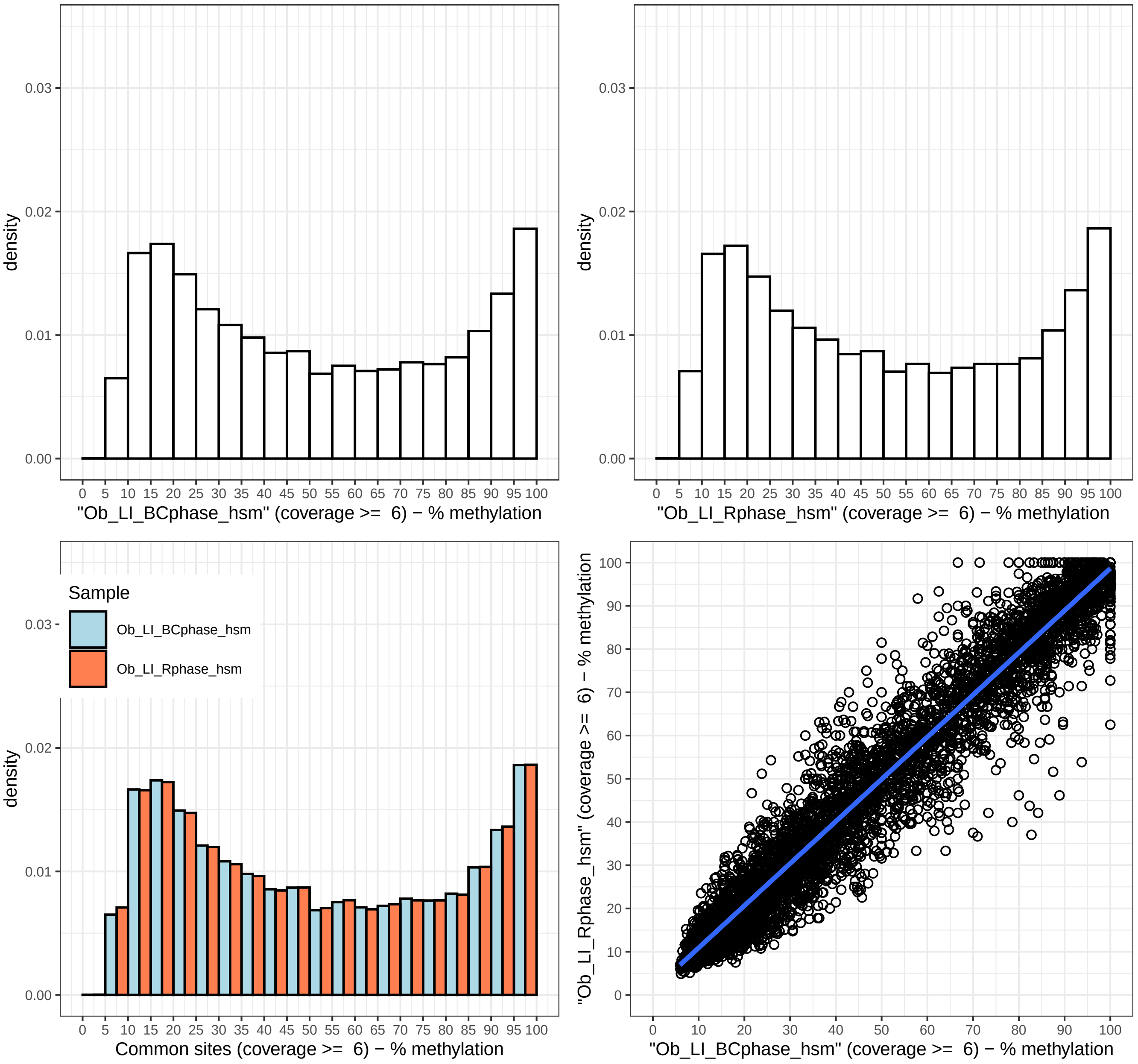

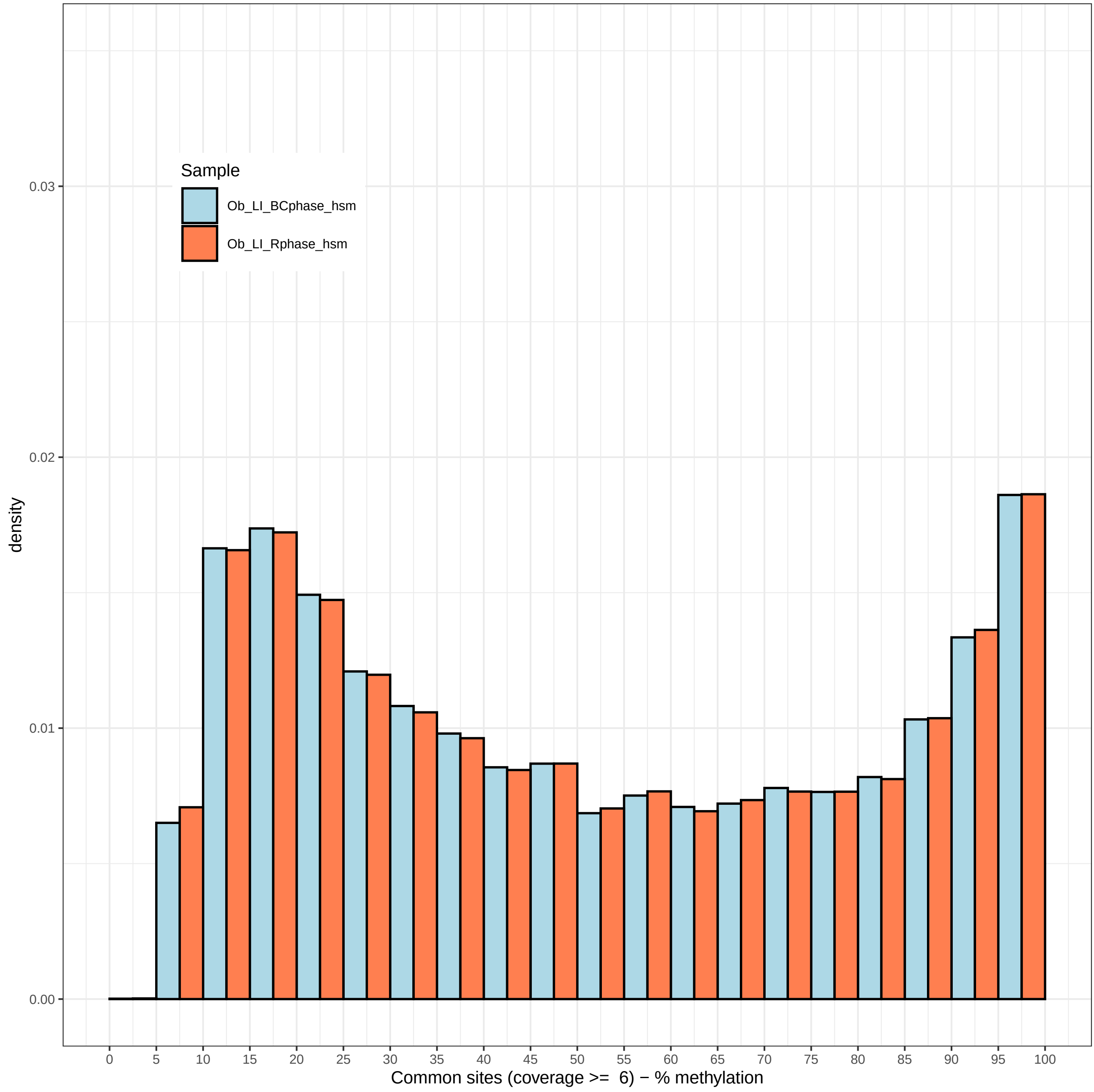
Sample
Ob_LI_BCphase_hsm
Ob_LI_Rphase_hsm
density
0.00
0.01
0.02
0.03
0
5
10
15
20
25
30
35
40
45
50
55
60
65
70
75
80
85
90
95
100
Common sites (coverage >= 6) − % methylation

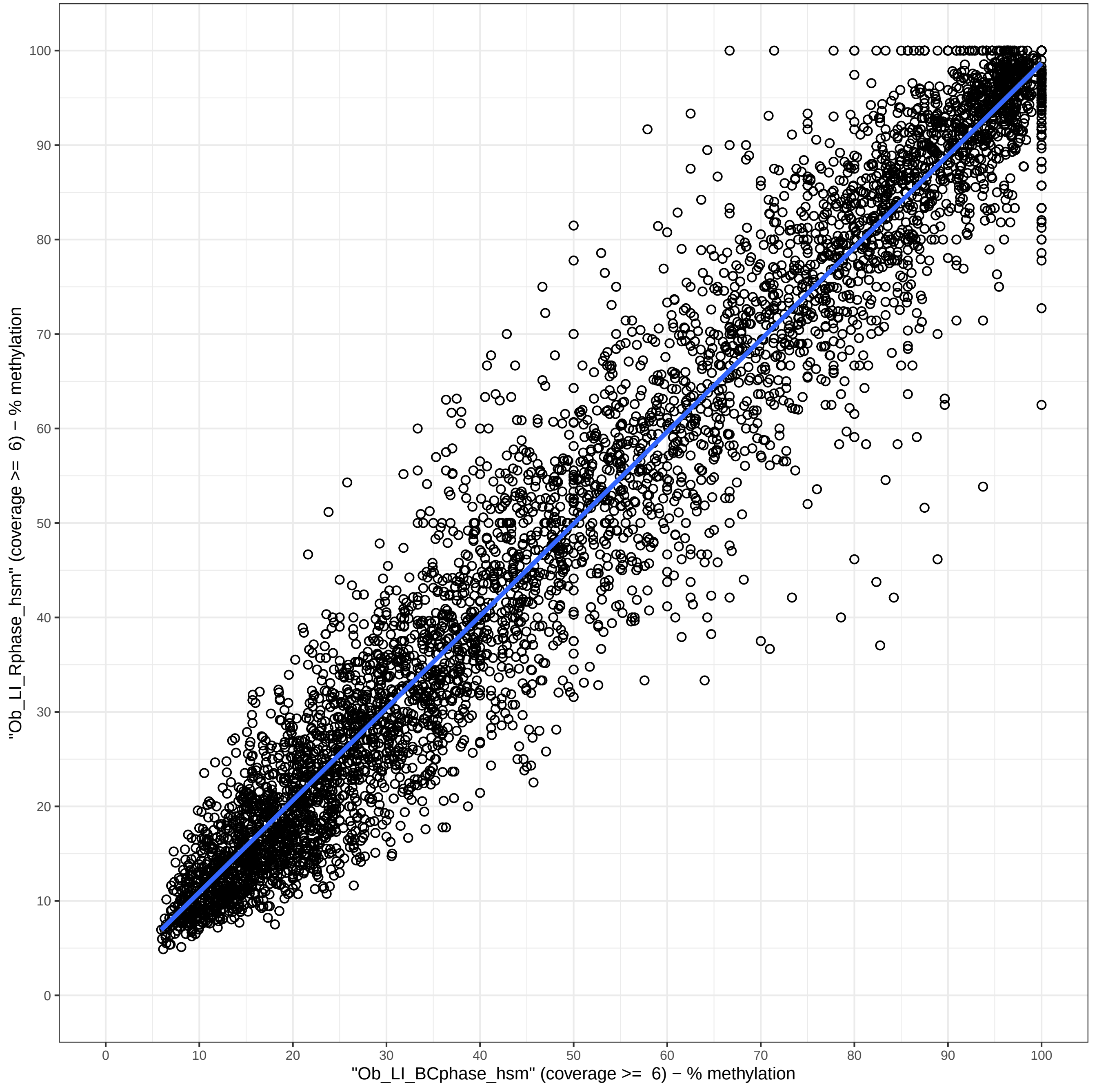

"Ob_LI_Rphase_hsm" (coverage >= 6) − % methylation
"Ob_LI_BCphase_hsm" (coverage >= 6) − % methylation
0
10
20
30
40
50
60
70
80
90
100

**Overlap of highly supported methylation sites (coverage >= 6)**

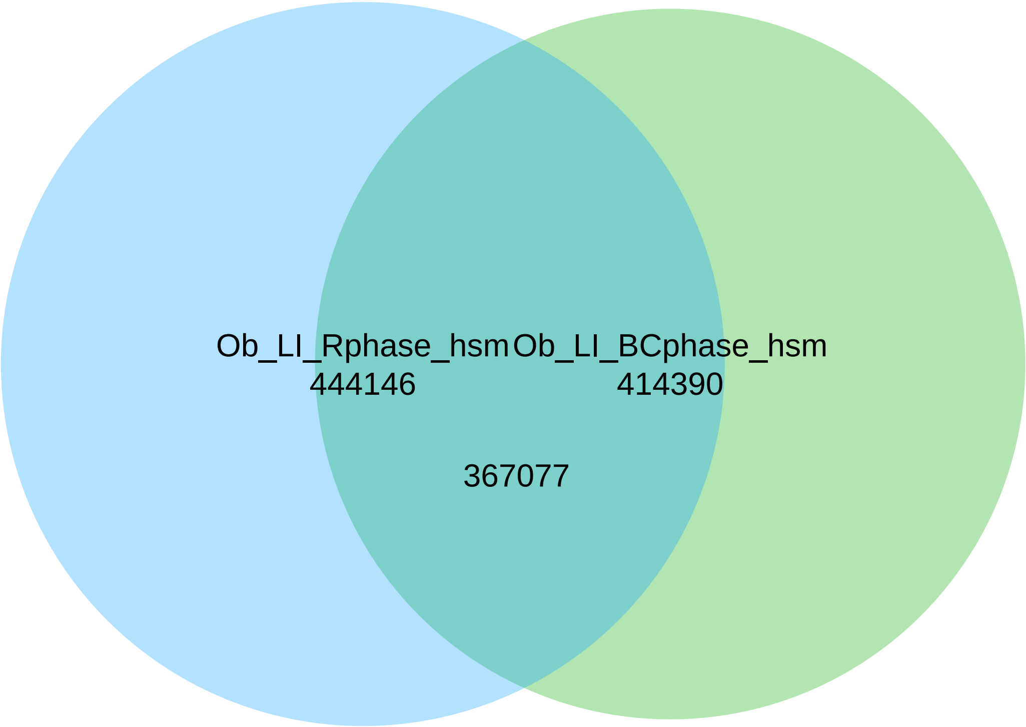

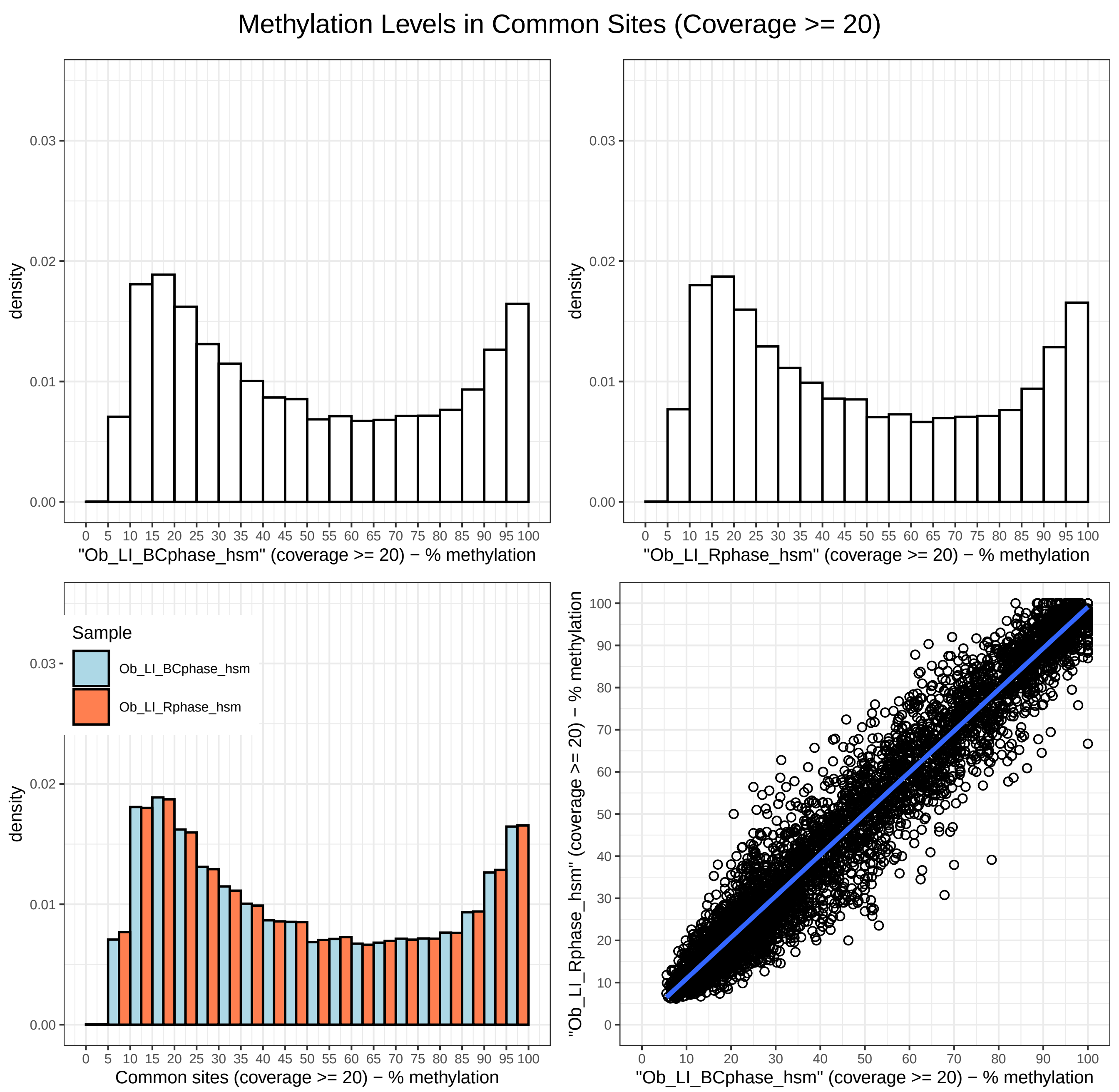

Methylation Levels in Common Sites (Coverage >= 20)
density
"Ob_LI_BCphase_hsm" (coverage >= 20) − % methylation
"Ob_LI_Rphase_hsm" (coverage >= 20) − % methylation
Sample
Ob_LI_BCphase_hsm
Ob_LI_Rphase_hsm
Common sites (coverage >= 20) − % methylation
"Ob_LI_Rphase_hsm" (coverage >= 20) − % methylation
"Ob_LI_BCphase_hsm" (coverage >= 20) − % methylation

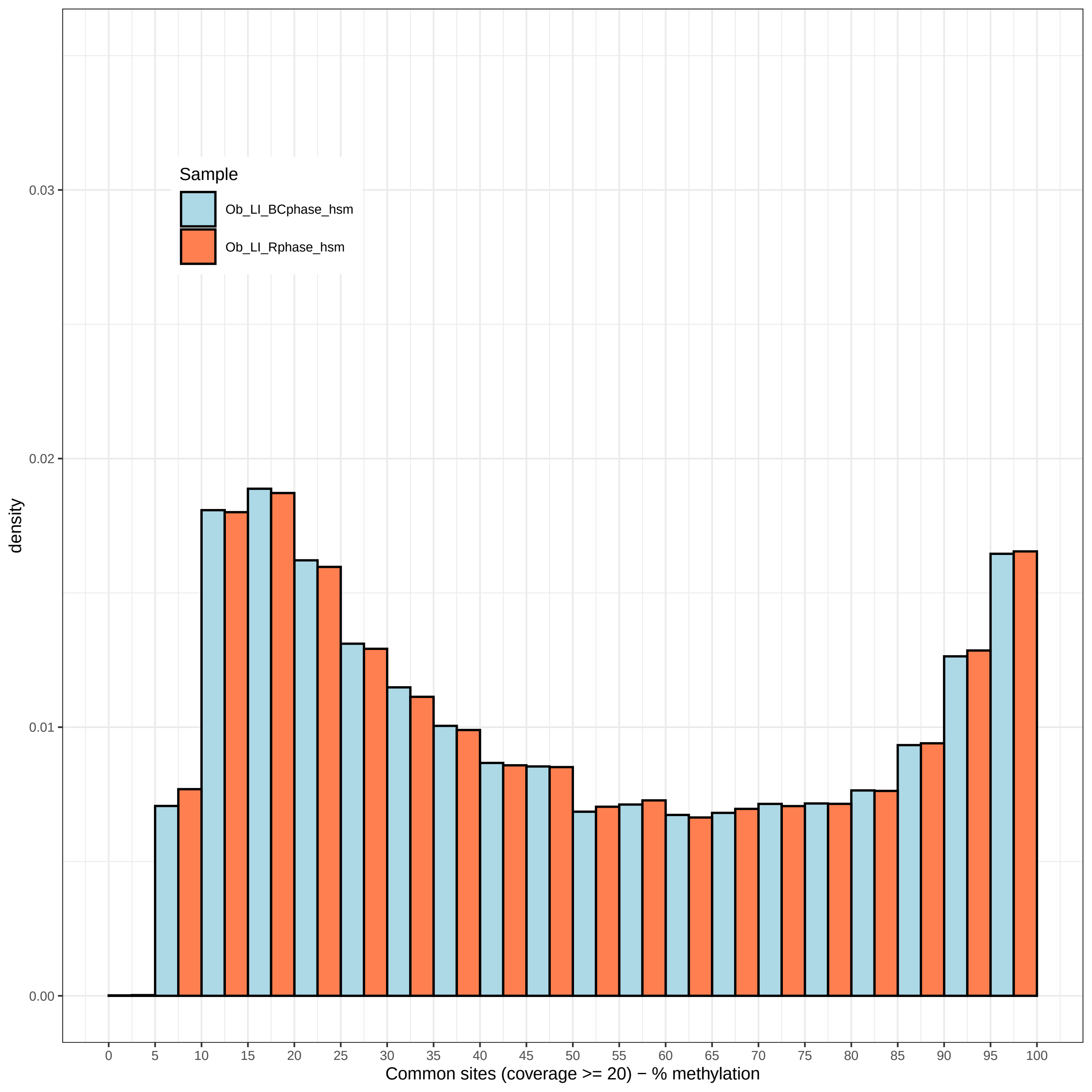

Sample
Ob_LI_BCphase_hsm
Ob_LI_Rphase_hsm
density
0.00
0.01
0.02
0.03
0
5
10
15
20
25
30
35
40
45
50
55
60
65
70
75
80
85
90
95
100
Common sites (coverage >= 20) − % methylation

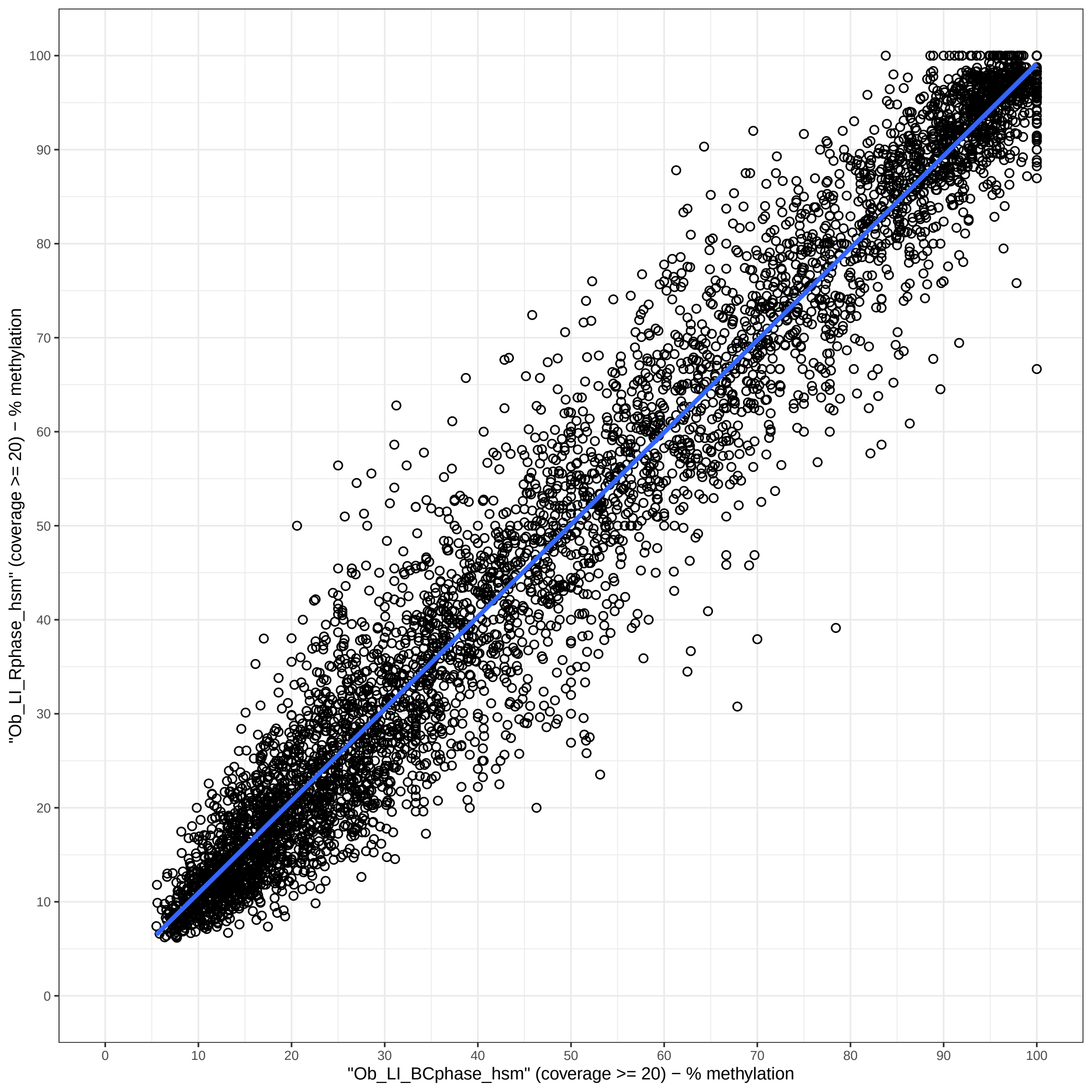
"Ob_LI_Rphase_hsm" (coverage >= 20) − % methylation
"Ob_LI_BCphase_hsm" (coverage >= 20) − % methylation
0
10
20
30
40
50
60
70
80
90
100

Overlap of highly supported methylation sites (coverage >= 20)

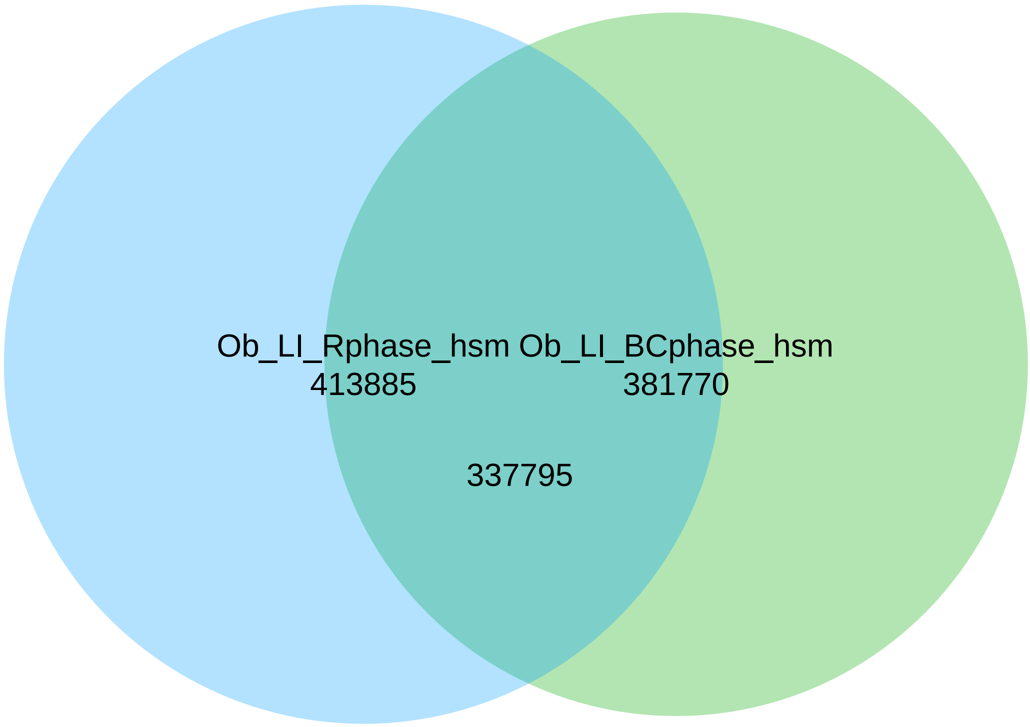

### Methylation Levels in Common Sites (Coverage >= 50)

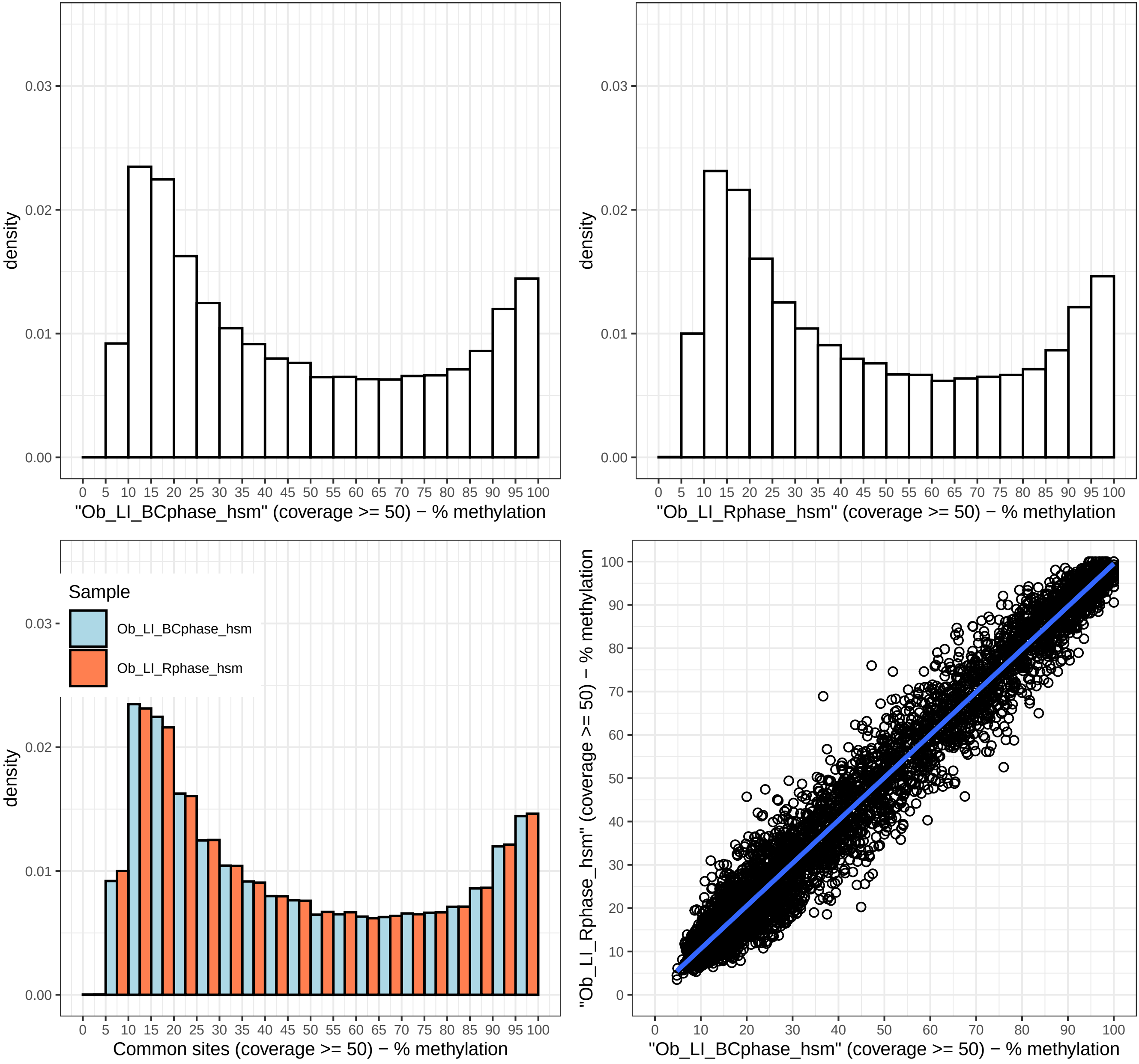

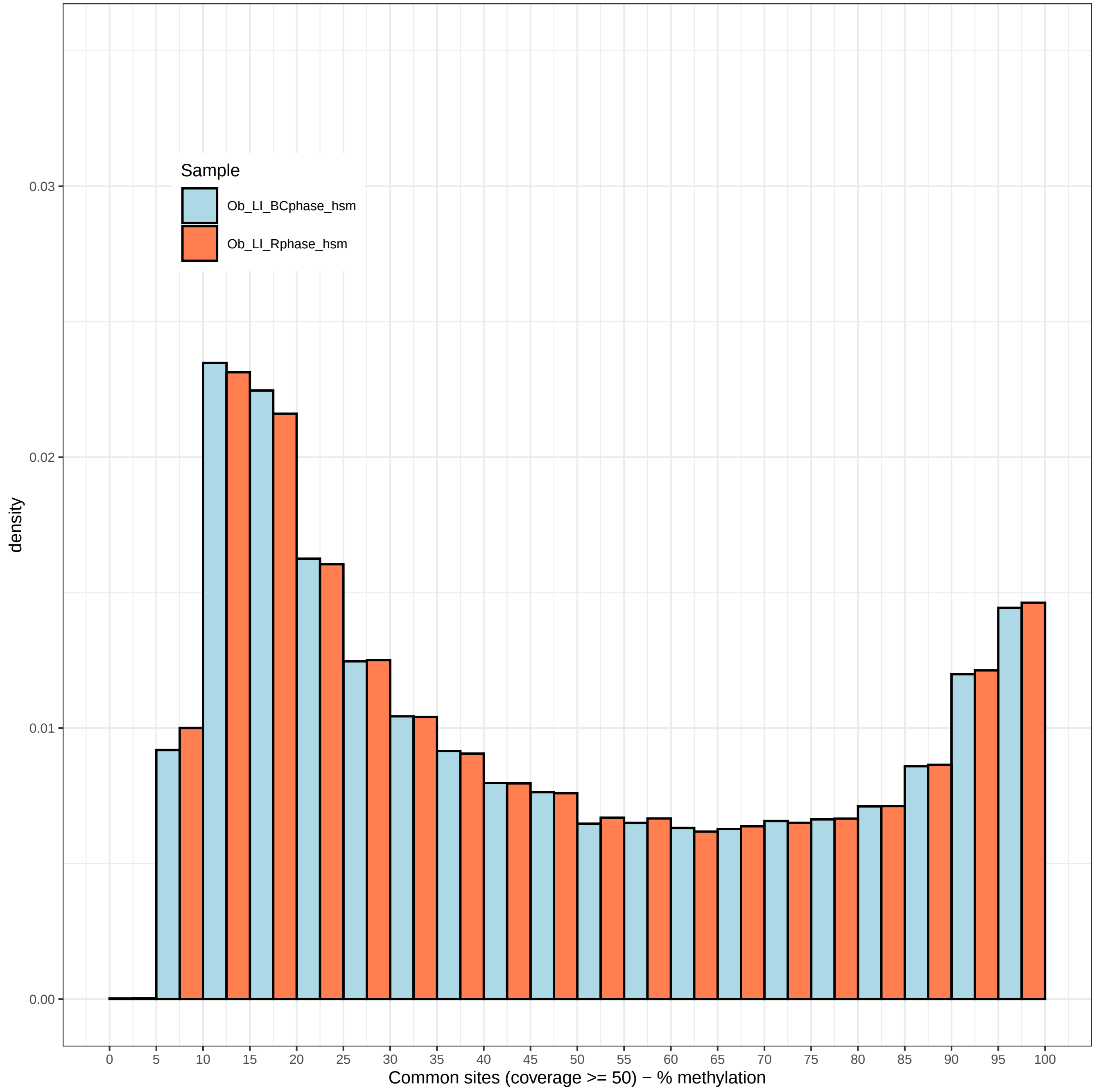
Sample
Ob_LI_BCphase_hsm
Ob_LI_Rphase_hsm
density
0.00
0.01
0.02
0.03
0
5
10
15
20
25
30
35
40
45
50
55
60
65
70
75
80
85
90
95
100
Common sites (coverage >= 50) − % methylation

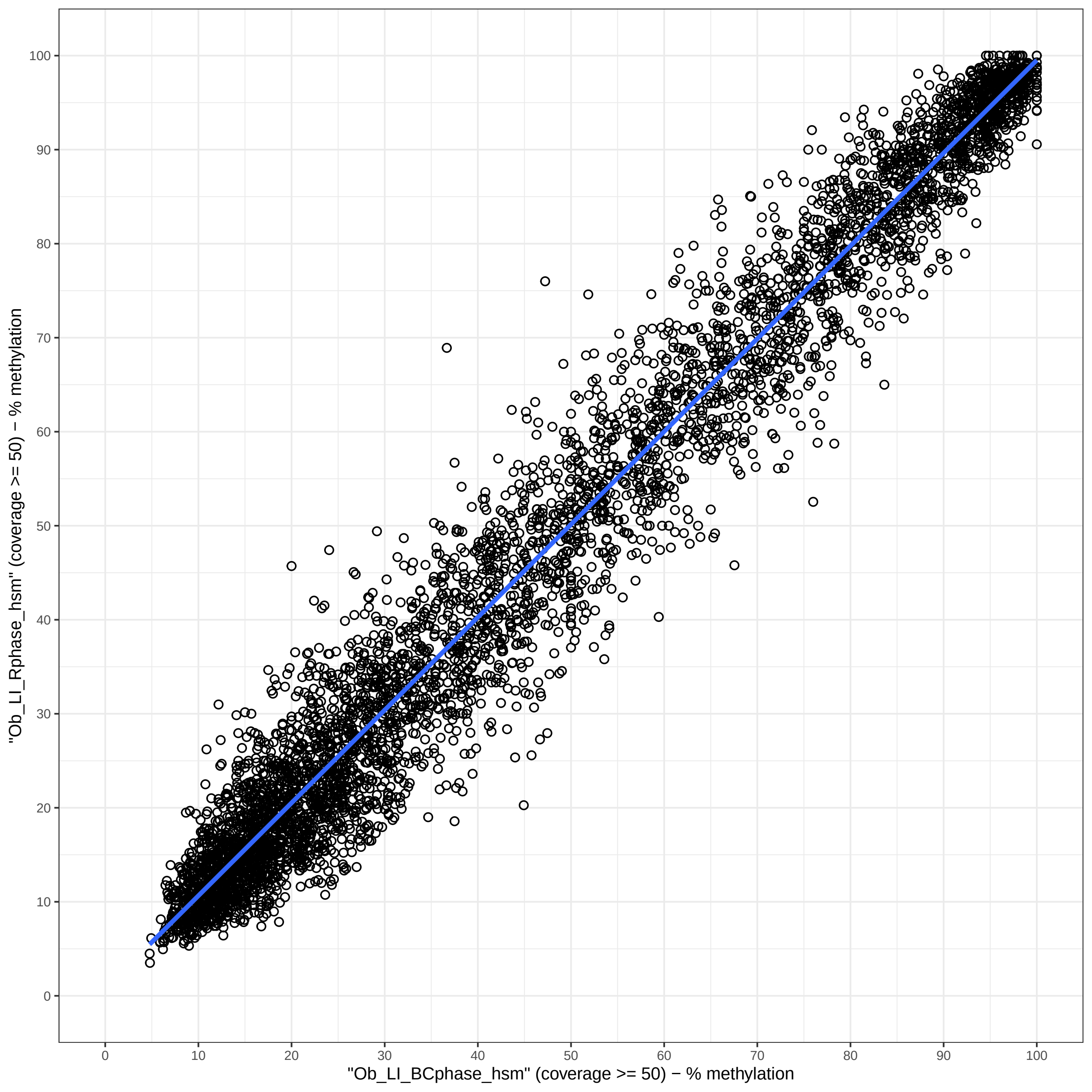

"Ob_LI_Rphase_hsm" (coverage >= 50) − % methylation
"Ob_LI_BCphase_hsm" (coverage >= 50) − % methylation
0
10
20
30
40
50
60
70
80
90
100

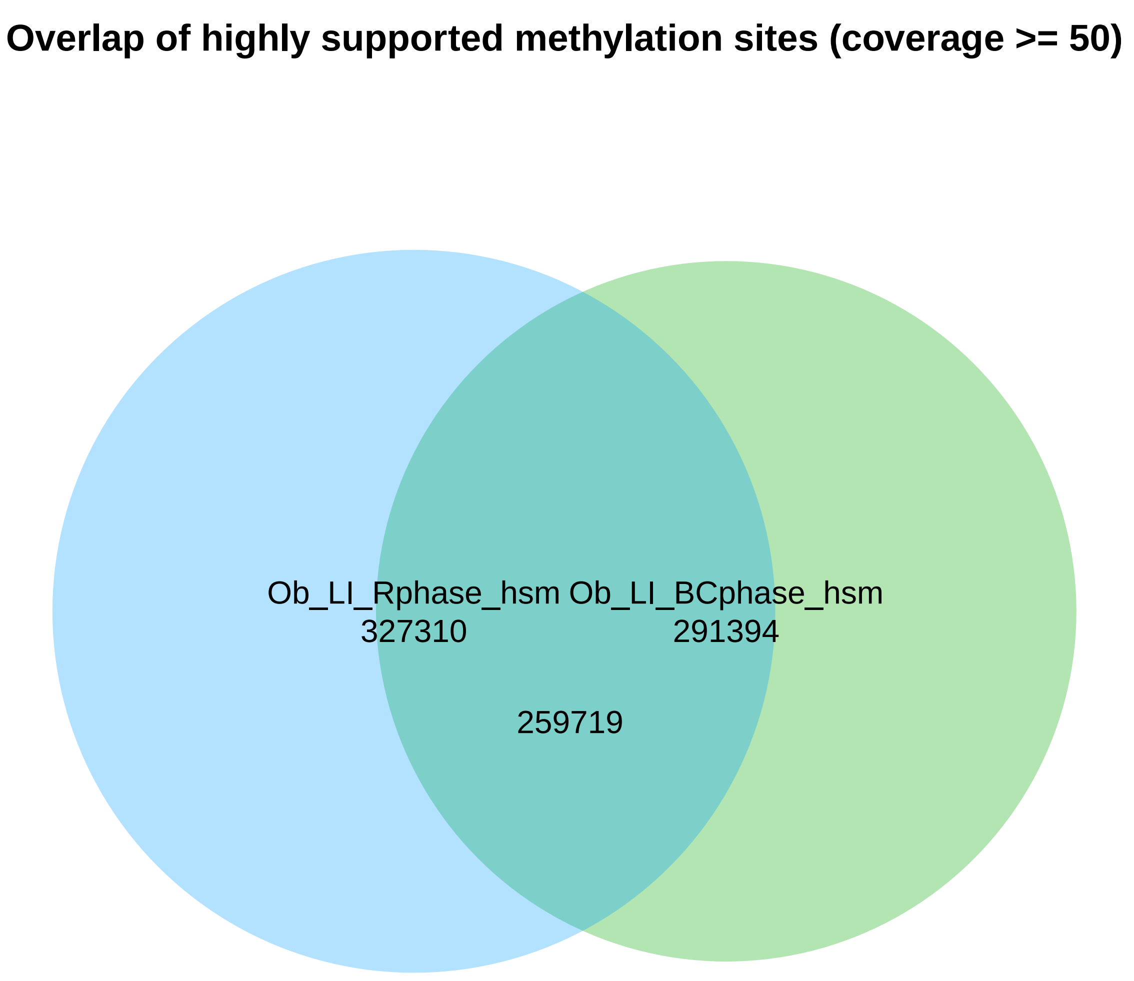
Overlap of highly supported methylation sites (coverage >= 50)
Ob_LI_Rphase_hsm
327310
Ob_LI_BCphase_hsm
291394
259719

```
Directory: CRL File: 0READMEcrl

CRL - Correlations between aggregate samples.

Input: studymk

Output: files crl-*.txt crl-*.pdf

Notes: The output is generated by BWASPR::cmpSamples(), which uses methylKit::unite()
        to determine the sites common to all samples and applies methylKit::getCorrelation()
        and methylKit::PCASamples().  The *.txt files show the correlations in a table, and
        the *.pdf files show graphics.

       Beware that the number of common sites (shown at the bottom of the *.txt files) may be
        small, which may make the correlations less informative.
```

Directory: CRL File: crl-Ob_LIp.txt

| | BCphase | Rphase |
|---|---|---|
| BCphase | 1.000 | 0.983 |
| Rphase | 0.983 | 1.000 |

The number of conserved sites is 237866.

CpG base pearson cor.

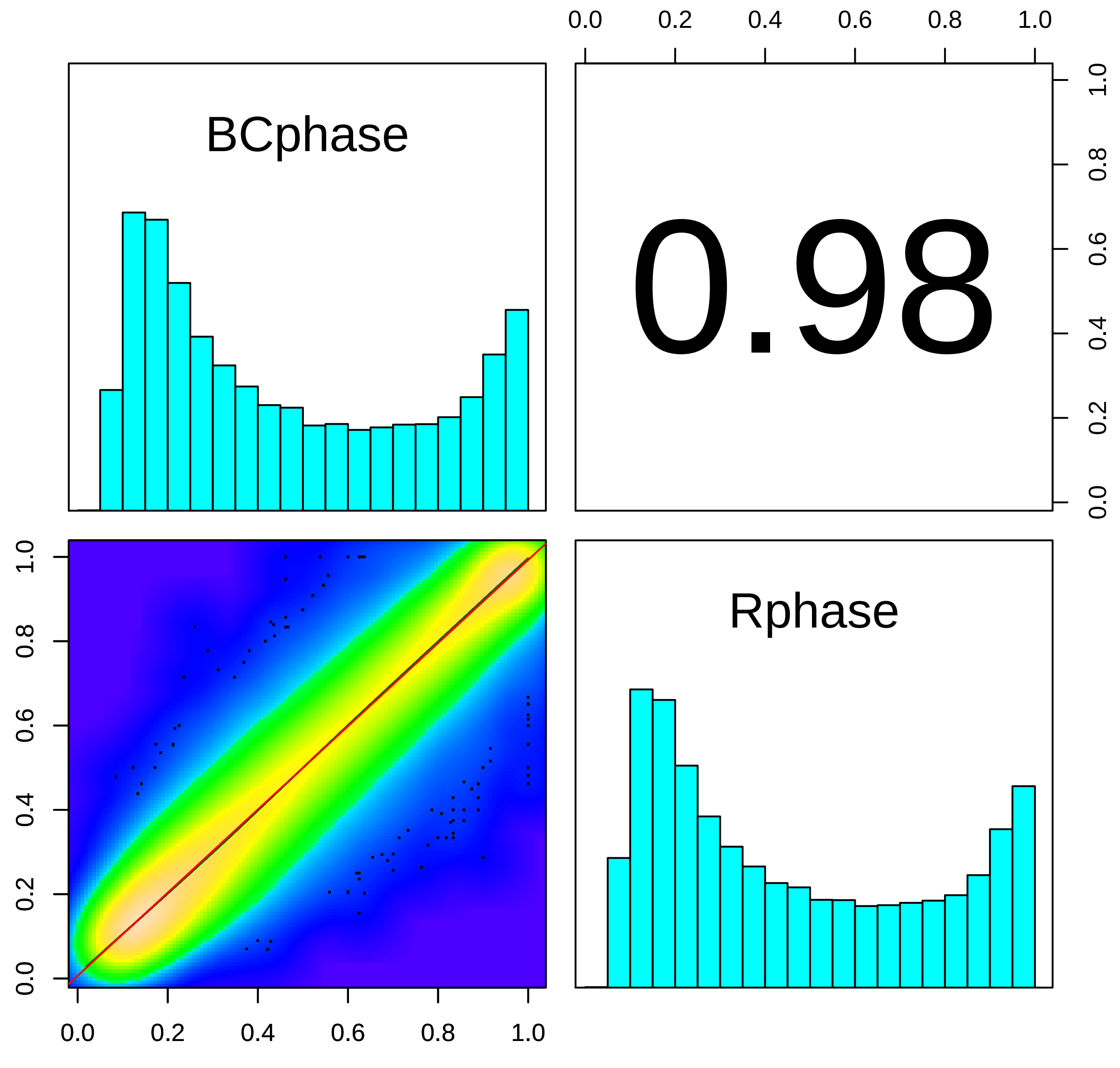

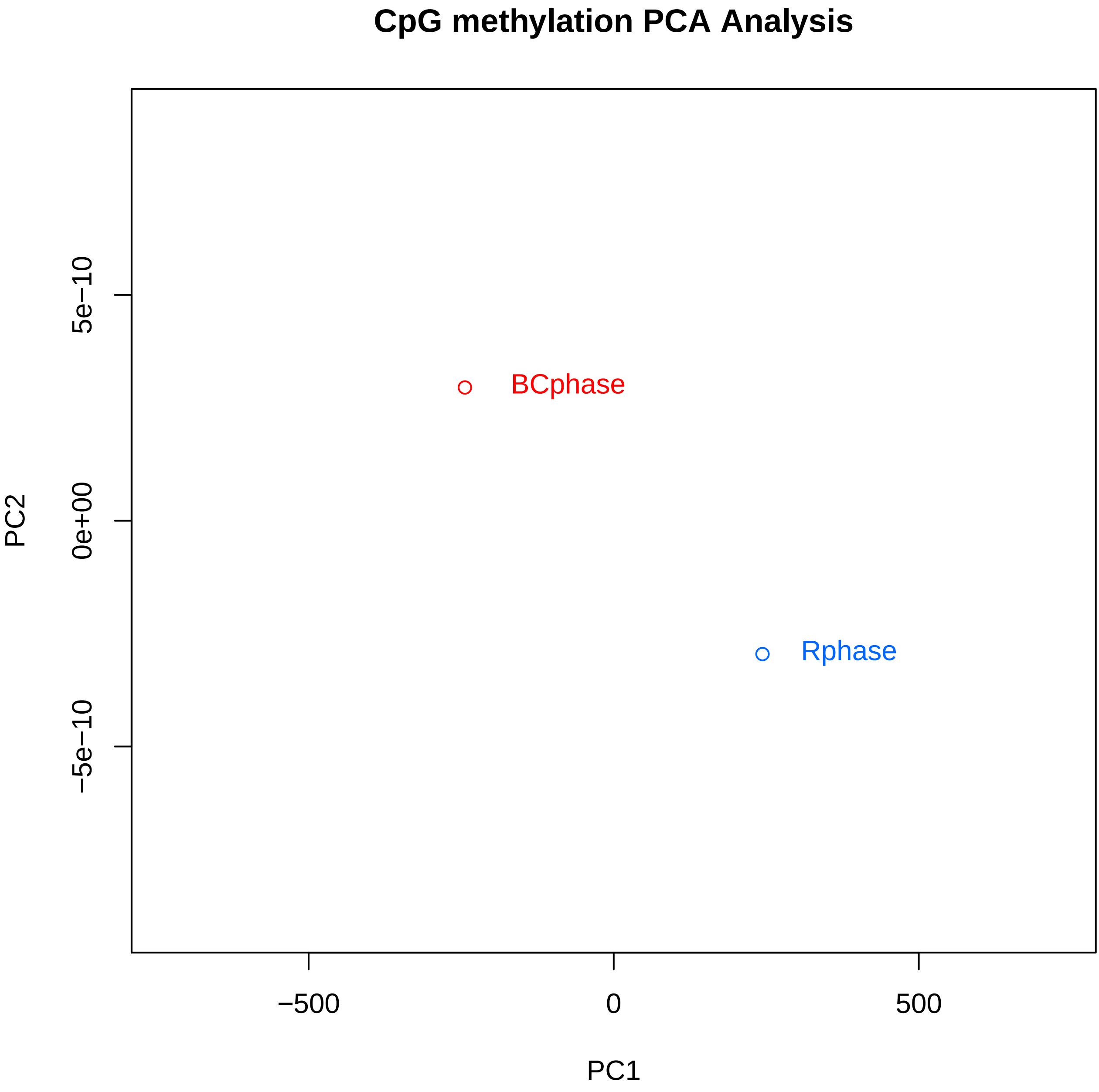
CpG methylation PCA Analysis
BCphase
Rphase
5e−10
0e+00
−5e−10
PC2
−500
0
500
PC1

Directory: REPCMS File: 0READMErepcms

```
REPCMS - Coverage and methylation statistics for replicate samples.

Input: studymk, repcovlist, replocount, rephicount

Output: files repcms-*txt and repcms-*pdf

Notes: The statistics are given for CpGhsm sites restricted to minimum or
        higher level coverage.  At the minimum level, the number of sites will be
        the number of lines of the corresponding *CpGhsm.mcalls file minus one
        (consistency check).  The distribution of methylation levels is really only
        interesting for high coverage because at low coverage there will be a strong
        bias towards high methylation levels (by definition of hsm sites).
        The same bias at high coverage would suggest that the sample includes a
        preponderance of consistently methylated genomic sites.

        replocount and rephicount set bounds on the coverage to exclude sites with too few
        or too many covering reads to provide statistics on a typical range.
```

```
Directory: REPCMS File: repcms-Ob_LIp_BCphase_1.txt

Number of "BCphase_1" CpGhsm-sites with minimal and higher level coverage:

  number of "BCphase_1" CpGhsm-sites with coverage >=  5: 197029
  number of "BCphase_1" CpGhsm-sites with coverage >=  6: 193888
  number of "BCphase_1" CpGhsm-sites with coverage >= 10: 170972
  number of "BCphase_1" CpGhsm-sites with coverage >= 15: 141998

Coverage and methylation statistics for "BCphase_1" CpGhsm-sites at different levels of minimum coverage:

methylKit::getCoverageStats output for "BCphase_1" CpGhsm-sites (#: 197029) at minimum coverage 5   - read coverage statistics per base
summary:
   Min. 1st Qu.  Median    Mean 3rd Qu.    Max.
      5      14      22      25      33     601
percentiles:
   0%   10%   20%   30%   40%   50%   60%   70%   80%   90%   95%   99% 99.5% 99.9%  100%
    5     9    12    15    18    22    26    31    36    44    50    63    68    80   601

methylKit::getMethylationStats output for "BCphase_1" CpGhsm-sites (#: 197029) at minimum coverage 5 - methylation statistics per base
summary:
   Min. 1st Qu.  Median    Mean 3rd Qu.    Max.
    5.3    46.7    71.4    68.0    90.6   100.0
percentiles:
    0%    10%    20%    30%    40%    50%    60%    70%    80%    90%    95%    99%  99.5%  99.9%   100%
  5.32  31.82  41.67  52.17  61.70  71.43  80.00  87.50  93.55 100.00 100.00 100.00 100.00 100.00 100.00

methylKit::getCoverageStats output for "BCphase_1" CpGhsm-sites (#: 193888) at minimum coverage 6   - read coverage statistics per base
summary:
   Min. 1st Qu.  Median    Mean 3rd Qu.    Max.
      6      14      23      25      33     601
percentiles:
   0%   10%   20%   30%   40%   50%   60%   70%   80%   90%   95%   99% 99.5% 99.9%  100%
    6     9    12    16    19    23    27    31    36    44    50    63    68    80   601

methylKit::getMethylationStats output for "BCphase_1" CpGhsm-sites (#: 193888) at minimum coverage 6 - methylation statistics per base
summary:
   Min. 1st Qu.  Median    Mean 3rd Qu.    Max.
    5.3    46.2    70.6    67.5    90.0   100.0
percentiles:
    0%    10%    20%    30%    40%    50%    60%    70%    80%    90%    95%    99%  99.5%  99.9%   100%
  5.32  31.63  41.18  51.52  61.11  70.59  79.41  86.67  92.86 100.00 100.00 100.00 100.00 100.00 100.00

methylKit::getCoverageStats output for "BCphase_1" CpGhsm-sites (#: 170972) at minimum coverage 10   - read coverage statistics per base
summary:
   Min. 1st Qu.  Median    Mean 3rd Qu.    Max.
     10      17      25      27      35     601
percentiles:
   0%   10%   20%   30%   40%   50%   60%   70%   80%   90%   95%   99% 99.5% 99.9%  100%
   10    12    15    18    21    25    29    33    38    45    52    64    69    82   601

methylKit::getMethylationStats output for "BCphase_1" CpGhsm-sites (#: 170972) at minimum coverage 10     - methylation statistics per base
summary:
   Min. 1st Qu.  Median    Mean 3rd Qu.    Max.
    5.3    43.3    66.7    65.1    88.9   100.0
percentiles:
    0%    10%    20%    30%    40%    50%    60%    70%    80%    90%    95%    99%  99.5%  99.9%   100%
```

```
  5.32  30.43  39.13  47.62  57.14  66.67  76.00  84.62  91.67  96.97 100.00 100.00 100.00 100.00 100.00

methylKit::getCoverageStats output for "BCphase_1" CpGhsm-sites (#: 141998) at minimum coverage 15  - read coverage statistics per base
summary:
   Min. 1st Qu.  Median    Mean 3rd Qu.    Max.
     15      21      28      30      37     601
percentiles:
   0%   10%   20%   30%   40%   50%   60%   70%   80%   90%   95%   99% 99.5% 99.9%  100%
   15    17    19    22    25    28    31    35    40    47    53    66    70    83   601

methylKit::getMethylationStats output for "BCphase_1" CpGhsm-sites (#: 141998) at minimum coverage 15     - methylation statistics per base
summary:
   Min. 1st Qu.  Median    Mean 3rd Qu.    Max.
    5.3    40.0    62.5    62.5    86.8   100.0
percentiles:
    0%    10%    20%    30%    40%    50%    60%    70%    80%    90%    95%    99%  99.5%  99.9%   100%
  5.32  28.57  36.21  43.75  52.63  62.50  72.73  82.35  90.48  95.92 100.00 100.00 100.00 100.00 100.00

methylKit::getCoverageStats output for "BCphase_1" CpGhsm-sites (#: 170947) in coverage range [10-100]    - read coverage statistics per base
summary:
   Min. 1st Qu.  Median    Mean 3rd Qu.    Max.
   10.0    17.0    25.0    27.2    35.0   100.0
percentiles:
   0%   10%   20%   30%   40%   50%   60%   70%   80%   90%   95%   99% 99.5% 99.9%  100%
   10    12    15    18    21    25    29    33    38    45    51    64    69    80   100

methylKit::getMethylationStats output for "BCphase_1" CpGhsm-sites (#: 170947) in coverage range [10-100] - methylation statistics per base
summary:
   Min. 1st Qu.  Median    Mean 3rd Qu.    Max.
   10.7    43.3    66.7    65.1    88.9   100.0
percentiles:
   0%   10%   20%   30%   40%   50%   60%   70%   80%   90%   95%   99% 99.5% 99.9%  100%
 10.7  30.4  39.1  47.6  57.1  66.7  76.0  84.6  91.7  97.0 100.0 100.0 100.0 100.0 100.0
```

```
Directory: REPCMS File: repcms-Ob_LIp_BCphase_2.txt

Number of "BCphase_2" CpGhsm-sites with minimal and higher level coverage:

  number of "BCphase_2" CpGhsm-sites with coverage >=  5: 213136
  number of "BCphase_2" CpGhsm-sites with coverage >=  6: 210280
  number of "BCphase_2" CpGhsm-sites with coverage >= 10: 189906
  number of "BCphase_2" CpGhsm-sites with coverage >= 15: 163705

Coverage and methylation statistics for "BCphase_2" CpGhsm-sites at different levels of minimum coverage:

methylKit::getCoverageStats output for "BCphase_2" CpGhsm-sites (#: 213136) at minimum coverage 5    - read coverage statistics per base
summary:
   Min. 1st Qu.  Median    Mean 3rd Qu.    Max.
    5.0    15.0    27.0    31.4    43.0   203.0
percentiles:
   0%   10%   20%   30%   40%   50%   60%   70%   80%   90%   95%   99% 99.5% 99.9%  100%
    5     9    13    17    22    27    32    39    48    61    72    95   105   126   203

methylKit::getMethylationStats output for "BCphase_2" CpGhsm-sites (#: 213136) at minimum coverage 5 - methylation statistics per base
summary:
   Min. 1st Qu.  Median    Mean 3rd Qu.    Max.
    7.7    42.9    69.0    65.7    90.0   100.0
percentiles:
    0%    10%    20%    30%    40%    50%    60%    70%    80%    90%    95%    99%  99.5%  99.9%   100%
  7.75  27.66  37.50  48.00  58.33  69.05  78.57  86.67  92.98 100.00 100.00 100.00 100.00 100.00 100.00

methylKit::getCoverageStats output for "BCphase_2" CpGhsm-sites (#: 210280) at minimum coverage 6    - read coverage statistics per base
summary:
   Min. 1st Qu.  Median    Mean 3rd Qu.    Max.
    6.0    16.0    27.0    31.8    43.0   203.0
percentiles:
   0%   10%   20%   30%   40%   50%   60%   70%   80%   90%   95%   99% 99.5% 99.9%  100%
    6    10    14    18    22    27    33    39    48    61    72    95   105   126   203

methylKit::getMethylationStats output for "BCphase_2" CpGhsm-sites (#: 210280) at minimum coverage 6 - methylation statistics per base
summary:
   Min. 1st Qu.  Median    Mean 3rd Qu.    Max.
    7.7    42.6    68.2    65.2    89.3   100.0
percentiles:
    0%    10%    20%    30%    40%    50%    60%    70%    80%    90%    95%    99%  99.5%  99.9%   100%
  7.75  27.59  37.50  47.62  57.92  68.18  77.78  85.71  92.31 100.00 100.00 100.00 100.00 100.00 100.00

methylKit::getCoverageStats output for "BCphase_2" CpGhsm-sites (#: 189906) at minimum coverage 10   - read coverage statistics per base
summary:
   Min. 1st Qu.  Median    Mean 3rd Qu.    Max.
   10.0    18.0    30.0    34.4    45.0   203.0
percentiles:
   0%   10%   20%   30%   40%   50%   60%   70%   80%   90%   95%   99% 99.5% 99.9%  100%
   10    13    17    20    25    30    35    41    50    63    74    97   106   128   203

methylKit::getMethylationStats output for "BCphase_2" CpGhsm-sites (#: 189906) at minimum coverage 10      - methylation statistics per base
summary:
   Min. 1st Qu.  Median    Mean 3rd Qu.    Max.
    7.7    40.0    64.2    63.0    88.1   100.0
percentiles:
    0%    10%    20%    30%    40%    50%    60%    70%    80%    90%    95%    99%  99.5%  99.9%   100%
```

```
  7.75  26.47  35.42  44.44  54.29  64.15  74.58  83.67  91.30  96.67 100.00 100.00 100.00 100.00 100.00

methylKit::getCoverageStats output for "BCphase_2" CpGhsm-sites (#: 163705) at minimum coverage 15  - read coverage statistics per base
summary:
   Min. 1st Qu.  Median    Mean 3rd Qu.    Max.
     15      23      33      38      49     203
percentiles:
   0%   10%   20%   30%   40%   50%   60%   70%   80%   90%   95%   99% 99.5% 99.9%  100%
   15    18    21    25    29    33    38    45    53    65    76    99   108   130   203

methylKit::getMethylationStats output for "BCphase_2" CpGhsm-sites (#: 163705) at minimum coverage 15     - methylation statistics per base
summary:
   Min. 1st Qu.  Median    Mean 3rd Qu.    Max.
    7.7    36.9    60.0    60.7    86.4   100.0
percentiles:
    0%    10%    20%    30%    40%    50%    60%    70%    80%    90%    95%    99%  99.5%  99.9%   100%
  7.75  25.00  33.33  40.91  50.00  60.00  71.43  81.58  90.00  95.83 100.00 100.00 100.00 100.00 100.00

methylKit::getCoverageStats output for "BCphase_2" CpGhsm-sites (#: 188483) in coverage range [10-100]    - read coverage statistics per base
summary:
   Min. 1st Qu.  Median    Mean 3rd Qu.    Max.
   10.0    18.0    29.0    33.8    45.0   100.0
percentiles:
   0%   10%   20%   30%   40%   50%   60%   70%   80%   90%   95%   99% 99.5% 99.9%  100%
   10    13    17    20    25    29    35    41    49    62    72    89    93    99   100

methylKit::getMethylationStats output for "BCphase_2" CpGhsm-sites (#: 188483) in coverage range [10-100] - methylation statistics per base
summary:
   Min. 1st Qu.  Median    Mean 3rd Qu.    Max.
   10.0    40.0    64.3    63.1    88.1   100.0
percentiles:
   0%   10%   20%   30%   40%   50%   60%   70%   80%   90%   95%   99% 99.5% 99.9%  100%
 10.0  26.5  35.5  44.4  54.3  64.3  74.7  83.7  91.3  96.7 100.0 100.0 100.0 100.0 100.0
```

```
Directory: REPCMS File: repcms-Ob_LIp_BCphase_3.txt

Number of "BCphase_3" CpGhsm-sites with minimal and higher level coverage:

  number of "BCphase_3" CpGhsm-sites with coverage >=  5: 226855
  number of "BCphase_3" CpGhsm-sites with coverage >=  6: 224253
  number of "BCphase_3" CpGhsm-sites with coverage >= 10: 204750
  number of "BCphase_3" CpGhsm-sites with coverage >= 15: 178265

Coverage and methylation statistics for "BCphase_3" CpGhsm-sites at different levels of minimum coverage:

methylKit::getCoverageStats output for "BCphase_3" CpGhsm-sites (#: 226855) at minimum coverage 5    - read coverage statistics per base
summary:
   Min. 1st Qu.  Median    Mean 3rd Qu.    Max.
    5.0    16.0    27.0    29.5    40.0   247.0
percentiles:
   0%   10%   20%   30%   40%   50%   60%   70%   80%   90%   95%   99% 99.5% 99.9%  100%
    5    10    14    18    22    27    32    37    44    53    60    76    82    96   247

methylKit::getMethylationStats output for "BCphase_3" CpGhsm-sites (#: 226855) at minimum coverage 5 - methylation statistics per base
summary:
   Min. 1st Qu.  Median    Mean 3rd Qu.    Max.
    6.9    42.1    67.4    65.1    89.5   100.0
percentiles:
    0%    10%    20%    30%    40%    50%    60%    70%    80%    90%    95%    99%  99.5%  99.9%   100%
  6.88  28.00  37.50  47.06  57.14  67.39  77.50  85.71  92.59 100.00 100.00 100.00 100.00 100.00 100.00

methylKit::getCoverageStats output for "BCphase_3" CpGhsm-sites (#: 224253) at minimum coverage 6    - read coverage statistics per base
summary:
   Min. 1st Qu.  Median    Mean 3rd Qu.    Max.
    6.0    16.0    27.0    29.7    40.0   247.0
percentiles:
   0%   10%   20%   30%   40%   50%   60%   70%   80%   90%   95%   99% 99.5% 99.9%  100%
    6    10    14    18    23    27    32    37    44    53    60    76    82    96   247

methylKit::getMethylationStats output for "BCphase_3" CpGhsm-sites (#: 224253) at minimum coverage 6 - methylation statistics per base
summary:
   Min. 1st Qu.  Median    Mean 3rd Qu.    Max.
    6.9    41.7    66.7    64.7    88.9   100.0
percentiles:
    0%    10%    20%    30%    40%    50%    60%    70%    80%    90%    95%    99%  99.5%  99.9%   100%
  6.88  27.78  37.04  46.67  56.67  66.67  76.81  85.71  92.00 100.00 100.00 100.00 100.00 100.00 100.00

methylKit::getCoverageStats output for "BCphase_3" CpGhsm-sites (#: 204750) at minimum coverage 10   - read coverage statistics per base
summary:
   Min. 1st Qu.  Median    Mean 3rd Qu.    Max.
   10.0    19.0    29.0    31.8    42.0   247.0
percentiles:
   0%   10%   20%   30%   40%   50%   60%   70%   80%   90%   95%   99% 99.5% 99.9%  100%
   10    13    17    21    25    29    34    39    45    54    61    77    82    97   247

methylKit::getMethylationStats output for "BCphase_3" CpGhsm-sites (#: 204750) at minimum coverage 10      - methylation statistics per base
summary:
   Min. 1st Qu.  Median    Mean 3rd Qu.    Max.
    6.9    40.0    63.6    62.8    87.5   100.0
percentiles:
    0%    10%    20%    30%    40%    50%    60%    70%    80%    90%    95%    99%  99.5%  99.9%   100%
```

```
  6.88  26.92  35.29  44.00  53.57  63.64  73.53  83.33  90.91  96.55 100.00 100.00 100.00 100.00 100.00
```

```
methylKit::getCoverageStats output for "BCphase_3" CpGhsm-sites (#: 178265) at minimum coverage 15  - read coverage statistics per base
summary:
   Min. 1st Qu.  Median    Mean 3rd Qu.    Max.
   15.0    23.0    32.0    34.8    44.0   247.0
percentiles:
   0%   10%   20%   30%   40%   50%   60%   70%   80%   90%   95%   99% 99.5% 99.9%  100%
   15    18    21    25    28    32    36    41    47    55    63    78    84    98   247

methylKit::getMethylationStats output for "BCphase_3" CpGhsm-sites (#: 178265) at minimum coverage 15     - methylation statistics per base
summary:
   Min. 1st Qu.  Median    Mean 3rd Qu.    Max.
    6.9    37.0    60.0    60.6    86.1   100.0
percentiles:
     0%    10%    20%    30%    40%    50%    60%    70%    80%    90%    95%    99%  99.5%  99.9%   100%
   6.88  25.71  33.33  40.62  50.00  60.00  70.59  81.25  90.00  95.74 100.00 100.00 100.00 100.00 100.00
```

```
methylKit::getCoverageStats output for "BCphase_3" CpGhsm-sites (#: 204626) in coverage range [10-100]    - read coverage statistics per base
summary:
   Min. 1st Qu.  Median    Mean 3rd Qu.    Max.
   10.0    19.0    29.0    31.8    42.0   100.0
percentiles:
   0%   10%   20%   30%   40%   50%   60%   70%   80%   90%   95%   99% 99.5% 99.9%  100%
   10    13    17    21    25    29    34    39    45    54    61    76    82    93   100

methylKit::getMethylationStats output for "BCphase_3" CpGhsm-sites (#: 204626) in coverage range [10-100] - methylation statistics per base
summary:
   Min. 1st Qu.  Median    Mean 3rd Qu.    Max.
   10.1    40.0    63.6    62.8    87.5   100.0
percentiles:
   0%   10%   20%   30%   40%   50%   60%   70%   80%   90%   95%   99% 99.5% 99.9%  100%
 10.1  26.9  35.3  44.0  53.6  63.6  73.5  83.3  90.9  96.6 100.0 100.0 100.0 100.0 100.0
```

```
Directory: REPCMS File: repcms-Ob_LIp_BCphase_4.txt

Number of "BCphase_4" CpGhsm-sites with minimal and higher level coverage:

  number of "BCphase_4" CpGhsm-sites with coverage >=  5: 209549
  number of "BCphase_4" CpGhsm-sites with coverage >=  6: 206680
  number of "BCphase_4" CpGhsm-sites with coverage >= 10: 186650
  number of "BCphase_4" CpGhsm-sites with coverage >= 15: 160680

Coverage and methylation statistics for "BCphase_4" CpGhsm-sites at different levels of minimum coverage:

methylKit::getCoverageStats output for "BCphase_4" CpGhsm-sites (#: 209549) at minimum coverage 5    - read coverage statistics per base
summary:
   Min. 1st Qu.  Median    Mean 3rd Qu.    Max.
      5      15      26      30      40     582
percentiles:
   0%   10%   20%   30%   40%   50%   60%   70%   80%   90%   95%   99% 99.5% 99.9%  100%
    5     9    13    17    21    26    31    37    44    56    65    85    93   111   582

methylKit::getMethylationStats output for "BCphase_4" CpGhsm-sites (#: 209549) at minimum coverage 5 - methylation statistics per base
summary:
   Min. 1st Qu.  Median    Mean 3rd Qu.    Max.
    4.0    42.9    68.8    65.8    89.7   100.0
percentiles:
    0%    10%    20%    30%    40%    50%    60%    70%    80%    90%    95%    99%  99.5%  99.9%   100%
  3.95  28.57  38.10  48.28  58.46  68.75  78.05  86.21  92.86 100.00 100.00 100.00 100.00 100.00 100.00

methylKit::getCoverageStats output for "BCphase_4" CpGhsm-sites (#: 206680) at minimum coverage 6    - read coverage statistics per base
summary:
   Min. 1st Qu.  Median    Mean 3rd Qu.    Max.
      6      16      26      30      41     582
percentiles:
   0%   10%   20%   30%   40%   50%   60%   70%   80%   90%   95%   99% 99.5% 99.9%  100%
    6    10    14    17    21    26    31    37    45    56    65    85    93   111   582

methylKit::getMethylationStats output for "BCphase_4" CpGhsm-sites (#: 206680) at minimum coverage 6 - methylation statistics per base
summary:
   Min. 1st Qu.  Median    Mean 3rd Qu.    Max.
    4.0    42.9    67.9    65.3    88.9   100.0
percentiles:
    0%    10%    20%    30%    40%    50%    60%    70%    80%    90%    95%    99%  99.5%  99.9%   100%
  3.95  28.21  37.84  47.83  58.06  67.86  77.55  85.71  92.31 100.00 100.00 100.00 100.00 100.00 100.00

methylKit::getCoverageStats output for "BCphase_4" CpGhsm-sites (#: 186650) at minimum coverage 10   - read coverage statistics per base
summary:
   Min. 1st Qu.  Median    Mean 3rd Qu.    Max.
     10      18      29      32      42     582
percentiles:
   0%   10%   20%   30%   40%   50%   60%   70%   80%   90%   95%   99% 99.5% 99.9%  100%
   10    13    17    20    24    29    33    39    46    57    67    86    94   112   582

methylKit::getMethylationStats output for "BCphase_4" CpGhsm-sites (#: 186650) at minimum coverage 10      - methylation statistics per base
summary:
   Min. 1st Qu.  Median    Mean 3rd Qu.    Max.
    4.0    40.0    64.0    63.1    87.5   100.0
percentiles:
    0%    10%    20%    30%    40%    50%    60%    70%    80%    90%    95%    99%  99.5%  99.9%   100%
```

```
  3.95  27.27  36.07  45.00  54.55  64.00  73.91  83.33  90.91  96.43 100.00 100.00 100.00 100.00 100.00
```

```
methylKit::getCoverageStats output for "BCphase_4" CpGhsm-sites (#: 160680) at minimum coverage 15  - read coverage statistics per base
summary:
   Min. 1st Qu.  Median    Mean 3rd Qu.    Max. 
     15      22      32      36      45     582 
percentiles:
   0%   10%   20%   30%   40%   50%   60%   70%   80%   90%   95%   99% 99.5% 99.9%  100% 
   15    17    21    24    28    32    36    42    49    59    69    88    96   114   582 

methylKit::getMethylationStats output for "BCphase_4" CpGhsm-sites (#: 160680) at minimum coverage 15      - methylation statistics per base
summary:
   Min. 1st Qu.  Median    Mean 3rd Qu.    Max. 
    4.0    37.5    60.0    60.8    86.2   100.0 
percentiles:
    0%    10%    20%    30%    40%    50%    60%    70%    80%    90%    95%    99%  99.5%  99.9%   100% 
  3.95  25.86  33.33  41.18  50.00  60.00  70.97  81.25  89.74  95.65 100.00 100.00 100.00 100.00 100.00
```

```
methylKit::getCoverageStats output for "BCphase_4" CpGhsm-sites (#: 186125) in coverage range [10-100]    - read coverage statistics per base
summary:
   Min. 1st Qu.  Median    Mean 3rd Qu.    Max. 
   10.0    18.0    28.0    32.2    42.0   100.0 
percentiles:
   0%   10%   20%   30%   40%   50%   60%   70%   80%   90%   95%   99% 99.5% 99.9%  100% 
   10    13    17    20    24    28    33    39    46    57    66    83    89    97   100 

methylKit::getMethylationStats output for "BCphase_4" CpGhsm-sites (#: 186125) in coverage range [10-100] - methylation statistics per base
summary:
   Min. 1st Qu.  Median    Mean 3rd Qu.    Max. 
   10.0    40.0    64.0    63.1    87.5   100.0 
percentiles:
   0%   10%   20%   30%   40%   50%   60%   70%   80%   90%   95%   99% 99.5% 99.9%  100% 
 10.0  27.3  36.1  45.0  54.5  64.0  73.9  83.3  90.9  96.4 100.0 100.0 100.0 100.0 100.0
```

```
Directory: REPCMS File: repcms-Ob_LIp_Rphase_1.txt

Number of "Rphase_1" CpGhsm-sites with minimal and higher level coverage:

  number of "Rphase_1" CpGhsm-sites with coverage >=  5: 213360
  number of "Rphase_1" CpGhsm-sites with coverage >=  6: 210617
  number of "Rphase_1" CpGhsm-sites with coverage >= 10: 189946
  number of "Rphase_1" CpGhsm-sites with coverage >= 15: 162121

Coverage and methylation statistics for "Rphase_1" CpGhsm-sites at different levels of minimum coverage:

methylKit::getCoverageStats output for "Rphase_1" CpGhsm-sites (#: 213360) at minimum coverage 5     - read coverage statistics per base
summary:
   Min. 1st Qu.  Median    Mean 3rd Qu.    Max.
      5      15      25      27      37     133
percentiles:
   0%   10%   20%   30%   40%   50%   60%   70%   80%   90%   95%   99% 99.5% 99.9%  100%
    5     9    13    17    21    25    29    34    40    48    54    68    73    86   133

methylKit::getMethylationStats output for "Rphase_1" CpGhsm-sites (#: 213360) at minimum coverage 5  - methylation statistics per base
summary:
   Min. 1st Qu.  Median    Mean 3rd Qu.    Max.
    8.3    44.4    70.0    66.7    90.0   100.0
percentiles:
    0%    10%    20%    30%    40%    50%    60%    70%    80%    90%    95%    99%  99.5%  99.9%   100%
  8.27  30.00  39.53  50.00  60.00  70.00  78.95  87.04  93.33 100.00 100.00 100.00 100.00 100.00 100.00

methylKit::getCoverageStats output for "Rphase_1" CpGhsm-sites (#: 210617) at minimum coverage 6     - read coverage statistics per base
summary:
   Min. 1st Qu.  Median    Mean 3rd Qu.    Max.
    6.0    15.0    25.0    27.3    37.0   133.0
percentiles:
   0%   10%   20%   30%   40%   50%   60%   70%   80%   90%   95%   99% 99.5% 99.9%  100%
    6    10    13    17    21    25    29    34    40    48    54    68    73    86   133

methylKit::getMethylationStats output for "Rphase_1" CpGhsm-sites (#: 210617) at minimum coverage 6  - methylation statistics per base
summary:
   Min. 1st Qu.  Median    Mean 3rd Qu.    Max.
    8.3    44.0    69.2    66.2    89.7   100.0
percentiles:
    0%    10%    20%    30%    40%    50%    60%    70%    80%    90%    95%    99%  99.5%  99.9%   100%
  8.27  29.69  39.29  50.00  59.09  69.23  78.26  86.27  92.59 100.00 100.00 100.00 100.00 100.00 100.00

methylKit::getCoverageStats output for "Rphase_1" CpGhsm-sites (#: 189946) at minimum coverage 10    - read coverage statistics per base
summary:
   Min. 1st Qu.  Median    Mean 3rd Qu.    Max.
   10.0    18.0    27.0    29.4    38.0   133.0
percentiles:
   0%   10%   20%   30%   40%   50%   60%   70%   80%   90%   95%   99% 99.5% 99.9%  100%
   10    13    16    20    23    27    31    36    41    49    55    69    74    87   133

methylKit::getMethylationStats output for "Rphase_1" CpGhsm-sites (#: 189946) at minimum coverage 10 - methylation statistics per base
summary:
   Min. 1st Qu.  Median    Mean 3rd Qu.    Max.
    8.3    41.7    65.0    64.1    88.5   100.0
percentiles:
    0%    10%    20%    30%    40%    50%    60%    70%    80%    90%    95%    99%  99.5%  99.9%   100%
```

```
  8.27  28.57  37.50  46.15  55.56  65.00  75.00  84.38  91.67  96.97 100.00 100.00 100.00 100.00 100.00

methylKit::getCoverageStats output for "Rphase_1" CpGhsm-sites (#: 162121) at minimum coverage 15    - read coverage statistics per base
summary:
   Min. 1st Qu.  Median    Mean 3rd Qu.    Max. 
   15.0    22.0    30.0    32.4    40.0   133.0 
percentiles:
   0%   10%   20%   30%   40%   50%   60%   70%   80%   90%   95%   99% 99.5% 99.9%  100% 
   15    17    20    23    27    30    34    38    43    51    57    70    75    88   133 

methylKit::getMethylationStats output for "Rphase_1" CpGhsm-sites (#: 162121) at minimum coverage 15 - methylation statistics per base
summary:
   Min. 1st Qu.  Median    Mean 3rd Qu.    Max. 
    8.3    38.5    61.3    61.8    86.7   100.0 
percentiles:
    0%    10%    20%    30%    40%    50%    60%    70%    80%    90%    95%    99%  99.5%  99.9%   100% 
  8.27  27.08  35.00  42.31  51.43  61.29  72.22  82.26  90.32  96.00 100.00 100.00 100.00 100.00 100.00

methylKit::getCoverageStats output for "Rphase_1" CpGhsm-sites (#: 189917) in coverage range [10-100]      - read coverage statistics per base
summary:
   Min. 1st Qu.  Median    Mean 3rd Qu.    Max. 
   10.0    18.0    27.0    29.4    38.0   100.0 
percentiles:
   0%   10%   20%   30%   40%   50%   60%   70%   80%   90%   95%   99% 99.5% 99.9%  100% 
   10    13    16    20    23    27    31    36    41    49    55    69    74    86   100 

methylKit::getMethylationStats output for "Rphase_1" CpGhsm-sites (#: 189917) in coverage range [10-100]   - methylation statistics per base
summary:
   Min. 1st Qu.  Median    Mean 3rd Qu.    Max. 
   10.5    41.7    65.0    64.1    88.5   100.0 
percentiles:
   0%   10%   20%   30%   40%   50%   60%   70%   80%   90%   95%   99% 99.5% 99.9%  100% 
 10.5  28.6  37.5  46.2  55.6  65.0  75.0  84.4  91.7  97.0 100.0 100.0 100.0 100.0 100.0 
```

```
Directory: REPCMS File: repcms-Ob_LIp_Rphase_2.txt

Number of "Rphase_2" CpGhsm-sites with minimal and higher level coverage:

  number of "Rphase_2" CpGhsm-sites with coverage >=  5: 265465
  number of "Rphase_2" CpGhsm-sites with coverage >=  6: 263149
  number of "Rphase_2" CpGhsm-sites with coverage >= 10: 246598
  number of "Rphase_2" CpGhsm-sites with coverage >= 15: 224377

Coverage and methylation statistics for "Rphase_2" CpGhsm-sites at different levels of minimum coverage:

methylKit::getCoverageStats output for "Rphase_2" CpGhsm-sites (#: 265465) at minimum coverage 5    - read coverage statistics per base
summary:
   Min. 1st Qu.  Median    Mean 3rd Qu.    Max.
    5.0    20.0    36.0    42.2    58.0   281.0
percentiles:
   0%   10%   20%   30%   40%   50%   60%   70%   80%   90%   95%   99% 99.5% 99.9%  100%
    5    11    17    23    29    36    43    53    65    83    98   129   141   167   281

methylKit::getMethylationStats output for "Rphase_2" CpGhsm-sites (#: 265465) at minimum coverage 5  - methylation statistics per base
summary:
   Min. 1st Qu.  Median    Mean 3rd Qu.    Max.
    5.0    35.6    62.5    61.1    87.5   100.0
percentiles:
    0%    10%    20%    30%    40%    50%    60%    70%    80%    90%    95%    99%  99.5%  99.9%   100%
  4.98  22.50  31.37  40.54  51.16  62.50  73.33  83.33  91.30  97.44 100.00 100.00 100.00 100.00 100.00

methylKit::getCoverageStats output for "Rphase_2" CpGhsm-sites (#: 263149) at minimum coverage 6    - read coverage statistics per base
summary:
   Min. 1st Qu.  Median    Mean 3rd Qu.    Max.
    6.0    20.0    36.0    42.5    58.0   281.0
percentiles:
   0%   10%   20%   30%   40%   50%   60%   70%   80%   90%   95%   99% 99.5% 99.9%  100%
    6    12    17    23    29    36    44    53    65    83    98   129   141   167   281

methylKit::getMethylationStats output for "Rphase_2" CpGhsm-sites (#: 263149) at minimum coverage 6  - methylation statistics per base
summary:
   Min. 1st Qu.  Median    Mean 3rd Qu.    Max.
    5.0    35.3    61.8    60.7    87.5   100.0
percentiles:
    0%    10%    20%    30%    40%    50%    60%    70%    80%    90%    95%    99%  99.5%  99.9%   100%
  4.98  22.45  31.17  40.00  50.00  61.76  72.73  83.33  90.91  96.88 100.00 100.00 100.00 100.00 100.00

methylKit::getCoverageStats output for "Rphase_2" CpGhsm-sites (#: 246598) at minimum coverage 10   - read coverage statistics per base
summary:
   Min. 1st Qu.  Median    Mean 3rd Qu.    Max.
   10.0    23.0    38.0    44.9    60.0   281.0
percentiles:
   0%   10%   20%   30%   40%   50%   60%   70%   80%   90%   95%   99% 99.5% 99.9%  100%
   10    15    20    26    32    38    46    55    67    84   100   130   143   168   281

methylKit::getMethylationStats output for "Rphase_2" CpGhsm-sites (#: 246598) at minimum coverage 10 - methylation statistics per base
summary:
   Min. 1st Qu.  Median    Mean 3rd Qu.    Max.
    5.0    34.1    58.7    59.1    85.7   100.0
percentiles:
    0%    10%    20%    30%    40%    50%    60%    70%    80%    90%    95%    99%  99.5%  99.9%   100%
```

```
  4.98  21.88  30.00  38.46  48.00  58.70  70.00  80.95  90.00  95.83 100.00 100.00 100.00 100.00 100.00

methylKit::getCoverageStats output for "Rphase_2" CpGhsm-sites (#: 224377) at minimum coverage 15    - read coverage statistics per base
summary:
   Min. 1st Qu.  Median    Mean 3rd Qu.    Max.
   15.0    27.0    42.0    48.1    63.0   281.0
percentiles:
   0%   10%   20%   30%   40%   50%   60%   70%   80%   90%   95%   99% 99.5% 99.9%  100%
   15    19    24    29    35    42    49    58    69    87   102   132   144   170   281

methylKit::getMethylationStats output for "Rphase_2" CpGhsm-sites (#: 224377) at minimum coverage 15 - methylation statistics per base
summary:
   Min. 1st Qu.  Median    Mean 3rd Qu.    Max.
    5.0    32.1    55.2    57.3    84.2   100.0
percentiles:
     0%    10%    20%    30%    40%    50%    60%    70%    80%    90%    95%    99%  99.5%  99.9%   100%
   4.98  21.21  28.57  36.00  44.68  55.17  66.67  78.95  88.89  95.35  98.00 100.00 100.00 100.00 100.00

methylKit::getCoverageStats output for "Rphase_2" CpGhsm-sites (#: 234744) in coverage range [10-100]      - read coverage statistics per base
summary:
   Min. 1st Qu.  Median    Mean 3rd Qu.    Max.
   10.0    22.0    36.0    41.1    56.0   100.0
percentiles:
   0%   10%   20%   30%   40%   50%   60%   70%   80%   90%   95%   99% 99.5% 99.9%  100%
   10    15    20    25    30    36    44    52    62    76    85    97    99   100   100

methylKit::getMethylationStats output for "Rphase_2" CpGhsm-sites (#: 234744) in coverage range [10-100]   - methylation statistics per base
summary:
   Min. 1st Qu.  Median    Mean 3rd Qu.    Max.
   10.0    34.5    58.8    59.3    85.7   100.0
percentiles:
   0%   10%   20%   30%   40%   50%   60%   70%   80%   90%   95%   99% 99.5% 99.9%  100%
 10.0  22.2  30.4  38.9  48.3  58.8  70.0  81.2  90.0  95.9 100.0 100.0 100.0 100.0 100.0
```

```
Directory: REPCMS File: repcms-Ob_LIp_Rphase_3.txt

Number of "Rphase_3" CpGhsm-sites with minimal and higher level coverage:

  number of "Rphase_3" CpGhsm-sites with coverage >=  5: 217571
  number of "Rphase_3" CpGhsm-sites with coverage >=  6: 214719
  number of "Rphase_3" CpGhsm-sites with coverage >= 10: 194269
  number of "Rphase_3" CpGhsm-sites with coverage >= 15: 167143

Coverage and methylation statistics for "Rphase_3" CpGhsm-sites at different levels of minimum coverage:

methylKit::getCoverageStats output for "Rphase_3" CpGhsm-sites (#: 217571) at minimum coverage 5    - read coverage statistics per base
summary:
   Min. 1st Qu.  Median    Mean 3rd Qu.    Max.
      5      15      25      28      38     692
percentiles:
   0%   10%   20%   30%   40%   50%   60%   70%   80%   90%   95%   99% 99.5% 99.9%  100%
    5     9    13    17    21    25    30    35    41    50    57    73    79    94   692

methylKit::getMethylationStats output for "Rphase_3" CpGhsm-sites (#: 217571) at minimum coverage 5  - methylation statistics per base
summary:
   Min. 1st Qu.  Median    Mean 3rd Qu.    Max.
    4.0    43.5    68.8    66.0    90.0   100.0
percentiles:
    0%    10%    20%    30%    40%    50%    60%    70%    80%    90%    95%    99%  99.5%  99.9%   100%
  4.05  29.17  38.71  48.65  58.33  68.75  77.78  86.36  92.86 100.00 100.00 100.00 100.00 100.00 100.00

methylKit::getCoverageStats output for "Rphase_3" CpGhsm-sites (#: 214719) at minimum coverage 6    - read coverage statistics per base
summary:
   Min. 1st Qu.  Median    Mean 3rd Qu.    Max.
      6      16      26      28      38     692
percentiles:
   0%   10%   20%   30%   40%   50%   60%   70%   80%   90%   95%   99% 99.5% 99.9%  100%
    6    10    14    17    21    26    30    35    42    50    58    73    79    94   692

methylKit::getMethylationStats output for "Rphase_3" CpGhsm-sites (#: 214719) at minimum coverage 6  - methylation statistics per base
summary:
   Min. 1st Qu.  Median    Mean 3rd Qu.    Max.
    4.0    43.1    67.9    65.5    88.9   100.0
percentiles:
    0%    10%    20%    30%    40%    50%    60%    70%    80%    90%    95%    99%  99.5%  99.9%   100%
  4.05  29.03  38.46  48.08  58.06  67.86  77.50  85.71  92.31 100.00 100.00 100.00 100.00 100.00 100.00

methylKit::getCoverageStats output for "Rphase_3" CpGhsm-sites (#: 194269) at minimum coverage 10   - read coverage statistics per base
summary:
   Min. 1st Qu.  Median    Mean 3rd Qu.    Max.
     10      18      28      30      40     692
percentiles:
   0%   10%   20%   30%   40%   50%   60%   70%   80%   90%   95%   99% 99.5% 99.9%  100%
   10    13    16    20    24    28    32    37    43    51    59    74    80    94   692

methylKit::getMethylationStats output for "Rphase_3" CpGhsm-sites (#: 194269) at minimum coverage 10 - methylation statistics per base
summary:
   Min. 1st Qu.  Median    Mean 3rd Qu.    Max.
    4.0    40.9    64.1    63.5    87.9   100.0
percentiles:
    0%    10%    20%    30%    40%    50%    60%    70%    80%    90%    95%    99%  99.5%  99.9%   100%
```

```
  4.05  28.00  36.67  45.45  54.55  64.10  74.19  83.33  91.30  96.67 100.00 100.00 100.00 100.00 100.00

methylKit::getCoverageStats output for "Rphase_3" CpGhsm-sites (#: 167143) at minimum coverage 15   - read coverage statistics per base
summary:
   Min. 1st Qu.  Median    Mean 3rd Qu.    Max. 
     15      22      31      33      42     692 
percentiles:
   0%   10%   20%   30%   40%   50%   60%   70%   80%   90%   95%   99% 99.5% 99.9%  100% 
 15.0  17.0  20.0  24.0  27.0  31.0  35.0  39.0  45.0  53.0  60.0  75.0  81.0  95.9 692.0 

methylKit::getMethylationStats output for "Rphase_3" CpGhsm-sites (#: 167143) at minimum coverage 15 - methylation statistics per base
summary:
   Min. 1st Qu.  Median    Mean 3rd Qu.    Max. 
    4.0    37.8    60.5    61.2    86.4   100.0 
percentiles:
    0%    10%    20%    30%    40%    50%    60%    70%    80%    90%    95%    99%  99.5%  99.9%   100% 
  4.05  26.67  34.38  41.67  50.00  60.47  71.43  81.48  90.00  95.83 100.00 100.00 100.00 100.00 100.00

methylKit::getCoverageStats output for "Rphase_3" CpGhsm-sites (#: 194168) in coverage range [10-100]     - read coverage statistics per base
summary:
   Min. 1st Qu.  Median    Mean 3rd Qu.    Max. 
   10.0    18.0    28.0    30.4    40.0   100.0 
percentiles:
   0%   10%   20%   30%   40%   50%   60%   70%   80%   90%   95%   99% 99.5% 99.9%  100% 
   10    13    16    20    24    28    32    37    43    51    58    73    79    91   100 

methylKit::getMethylationStats output for "Rphase_3" CpGhsm-sites (#: 194168) in coverage range [10-100]  - methylation statistics per base
summary:
   Min. 1st Qu.  Median    Mean 3rd Qu.    Max. 
   10.2    40.9    64.1    63.5    87.9   100.0 
percentiles:
   0%   10%   20%   30%   40%   50%   60%   70%   80%   90%   95%   99% 99.5% 99.9%  100% 
 10.2  28.0  36.7  45.5  54.5  64.1  74.2  83.3  91.3  96.7 100.0 100.0 100.0 100.0 100.0 
```

```
Directory: REPCMS File: repcms-Ob_LIp_Rphase_4.txt

Number of "Rphase_4" CpGhsm-sites with minimal and higher level coverage:

  number of "Rphase_4" CpGhsm-sites with coverage >=  5: 201205
  number of "Rphase_4" CpGhsm-sites with coverage >=  6: 198266
  number of "Rphase_4" CpGhsm-sites with coverage >= 10: 177006
  number of "Rphase_4" CpGhsm-sites with coverage >= 15: 149873

Coverage and methylation statistics for "Rphase_4" CpGhsm-sites at different levels of minimum coverage:

methylKit::getCoverageStats output for "Rphase_4" CpGhsm-sites (#: 201205) at minimum coverage 5     - read coverage statistics per base
summary:
   Min. 1st Qu.  Median    Mean 3rd Qu.    Max.
      5      14      24      28      38     541
percentiles:
   0%   10%   20%   30%   40%   50%   60%   70%   80%   90%   95%   99% 99.5% 99.9%  100%
    5     9    12    16    20    24    29    35    42    53    63    82    90   107   541

methylKit::getMethylationStats output for "Rphase_4" CpGhsm-sites (#: 201205) at minimum coverage 5  - methylation statistics per base
summary:
   Min. 1st Qu.  Median    Mean 3rd Qu.    Max.
    3.9    44.8    70.0    66.8    90.0   100.0
percentiles:
    0%    10%    20%    30%    40%    50%    60%    70%    80%    90%    95%    99%  99.5%  99.9%   100%
  3.88  29.73  40.00  50.00  60.00  70.00  79.31  86.96  93.33 100.00 100.00 100.00 100.00 100.00 100.00

methylKit::getCoverageStats output for "Rphase_4" CpGhsm-sites (#: 198266) at minimum coverage 6     - read coverage statistics per base
summary:
   Min. 1st Qu.  Median    Mean 3rd Qu.    Max.
      6      15      25      28      38     541
percentiles:
   0%   10%   20%   30%   40%   50%   60%   70%   80%   90%   95%   99% 99.5% 99.9%  100%
    6     9    13    16    20    25    29    35    42    53    63    82    90   107   541

methylKit::getMethylationStats output for "Rphase_4" CpGhsm-sites (#: 198266) at minimum coverage 6  - methylation statistics per base
summary:
   Min. 1st Qu.  Median    Mean 3rd Qu.    Max.
    3.9    44.4    69.4    66.3    89.5   100.0
percentiles:
    0%    10%    20%    30%    40%    50%    60%    70%    80%    90%    95%    99%  99.5%  99.9%   100%
  3.88  29.63  39.47  50.00  60.00  69.44  78.38  86.11  92.31 100.00 100.00 100.00 100.00 100.00 100.00

methylKit::getCoverageStats output for "Rphase_4" CpGhsm-sites (#: 177006) at minimum coverage 10    - read coverage statistics per base
summary:
   Min. 1st Qu.  Median    Mean 3rd Qu.    Max.
     10      18      27      31      40     541
percentiles:
   0%   10%   20%   30%   40%   50%   60%   70%   80%   90%   95%   99% 99.5% 99.9%  100%
   10    13    16    19    23    27    32    37    44    55    64    83    91   109   541

methylKit::getMethylationStats output for "Rphase_4" CpGhsm-sites (#: 177006) at minimum coverage 10 - methylation statistics per base
summary:
   Min. 1st Qu.  Median    Mean 3rd Qu.    Max.
    3.9    41.8    65.0    64.0    88.2   100.0
percentiles:
    0%    10%    20%    30%    40%    50%    60%    70%    80%    90%    95%    99%  99.5%  99.9%   100%
```

```
  3.88  28.57  37.50  46.15  55.56  65.00  75.00  84.09  91.38  96.77 100.00 100.00 100.00 100.00 100.00
```

```
methylKit::getCoverageStats output for "Rphase_4" CpGhsm-sites (#: 149873) at minimum coverage 15    - read coverage statistics per base
summary:
   Min. 1st Qu.  Median    Mean 3rd Qu.    Max.
     15      22      31      34      43     541
percentiles:
   0%   10%   20%   30%   40%   50%   60%   70%   80%   90%   95%   99% 99.5% 99.9%  100%
   15    17    20    23    27    31    35    40    47    57    66    85    93   111   541

methylKit::getMethylationStats output for "Rphase_4" CpGhsm-sites (#: 149873) at minimum coverage 15 - methylation statistics per base
summary:
   Min. 1st Qu.  Median    Mean 3rd Qu.    Max.
    3.9    38.3    61.1    61.6    86.7   100.0
percentiles:
    0%    10%    20%    30%    40%    50%    60%    70%    80%    90%    95%    99%  99.5%  99.9%   100%
  3.88  26.84  34.78  42.11  51.28  61.11  72.00  81.82  90.00  95.83 100.00 100.00 100.00 100.00 100.00
```

```
methylKit::getCoverageStats output for "Rphase_4" CpGhsm-sites (#: 176617) in coverage range [10-100]     - read coverage statistics per base
summary:
   Min. 1st Qu.  Median    Mean 3rd Qu.    Max.
   10.0    18.0    27.0    30.8    40.0   100.0
percentiles:
   0%   10%   20%   30%   40%   50%   60%   70%   80%   90%   95%   99% 99.5% 99.9%  100%
   10    13    16    19    23    27    32    37    44    55    64    81    87    96   100

methylKit::getMethylationStats output for "Rphase_4" CpGhsm-sites (#: 176617) in coverage range [10-100]  - methylation statistics per base
summary:
   Min. 1st Qu.  Median    Mean 3rd Qu.    Max.
   10.5    41.8    65.0    64.0    88.2   100.0
percentiles:
   0%   10%   20%   30%   40%   50%   60%   70%   80%   90%   95%   99% 99.5% 99.9%  100%
 10.5  28.6  37.5  46.2  55.6  65.0  75.0  84.1  91.4  96.8 100.0 100.0 100.0 100.0 100.0
```

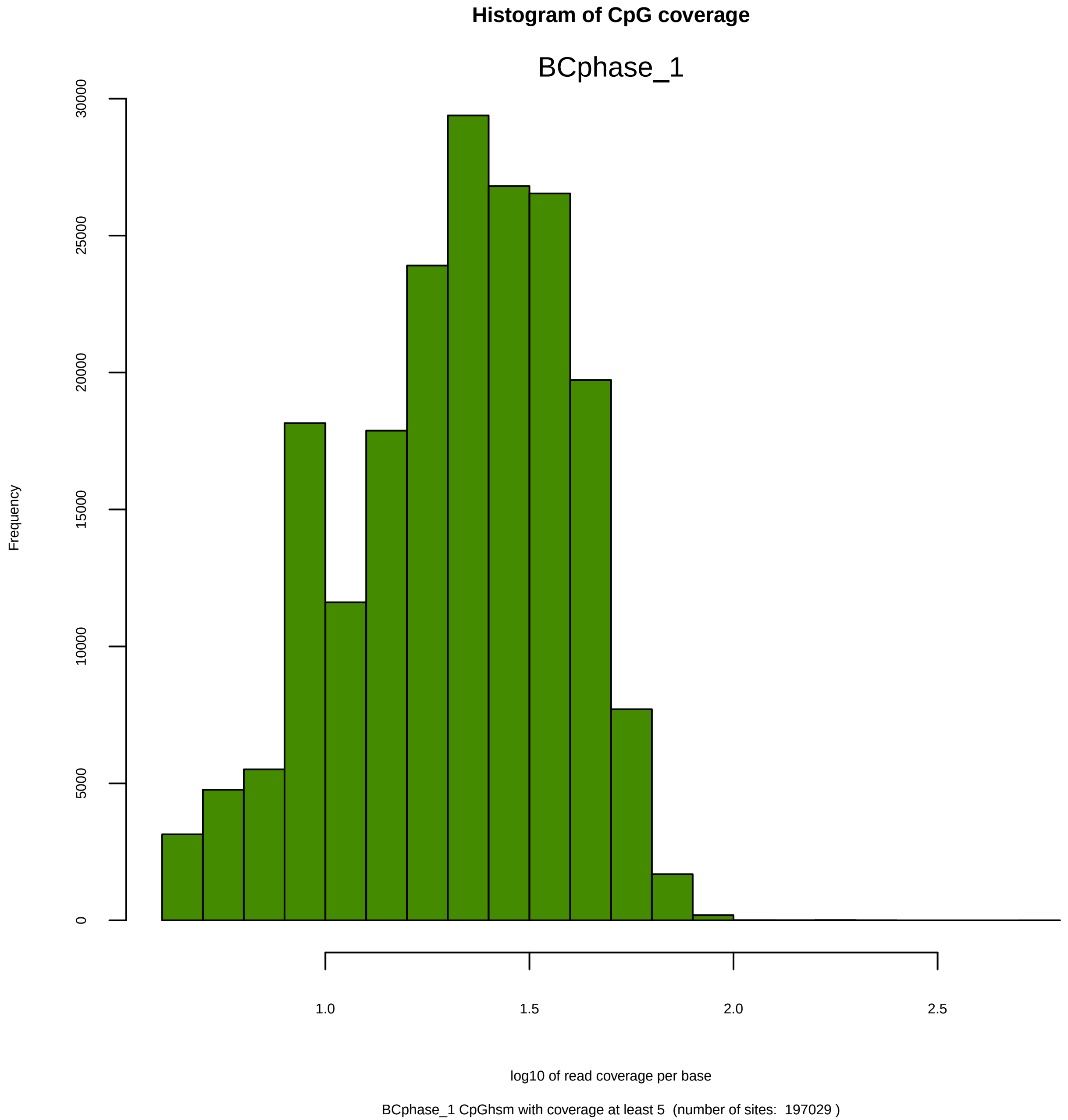

Histogram of CpG coverage
BCphase_1
Frequency
0
5000
10000
15000
20000
25000
30000
1.0
1.5
2.0
2.5
log10 of read coverage per base
BCphase_1 CpGhsm with coverage at least 5  (number of sites:  197029 )

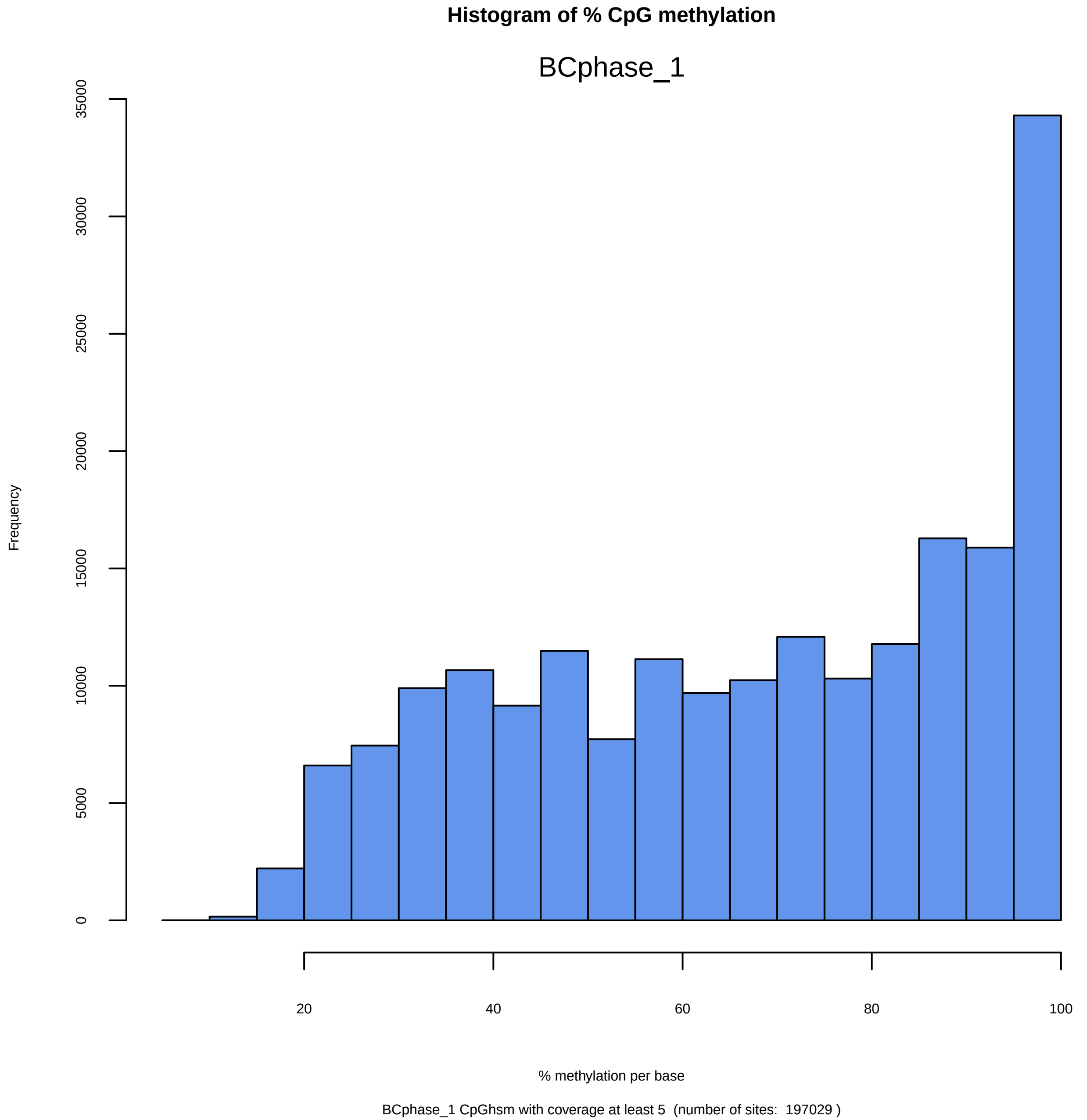
Histogram of % CpG methylation
BCphase_1
Frequency
0
5000
10000
15000
20000
25000
30000
35000
20
40
60
80
100
% methylation per base
BCphase_1 CpGhsm with coverage at least 5 (number of sites: 197029 )

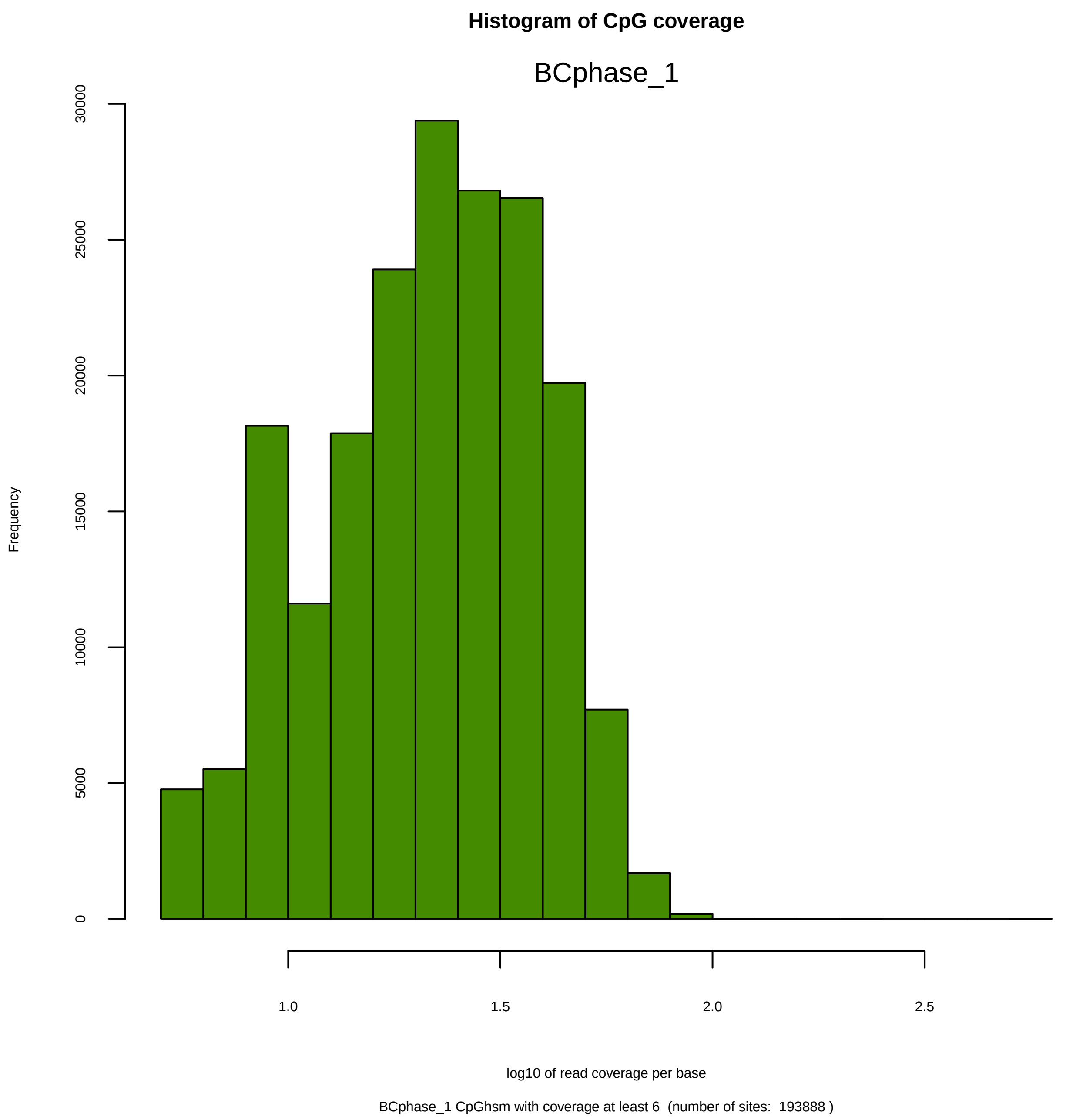
Histogram of CpG coverage
BCphase_1
Frequency
0
5000
10000
15000
20000
25000
30000
1.0
1.5
2.0
2.5
log10 of read coverage per base
BCphase_1 CpGhsm with coverage at least 6  (number of sites:  193888 )

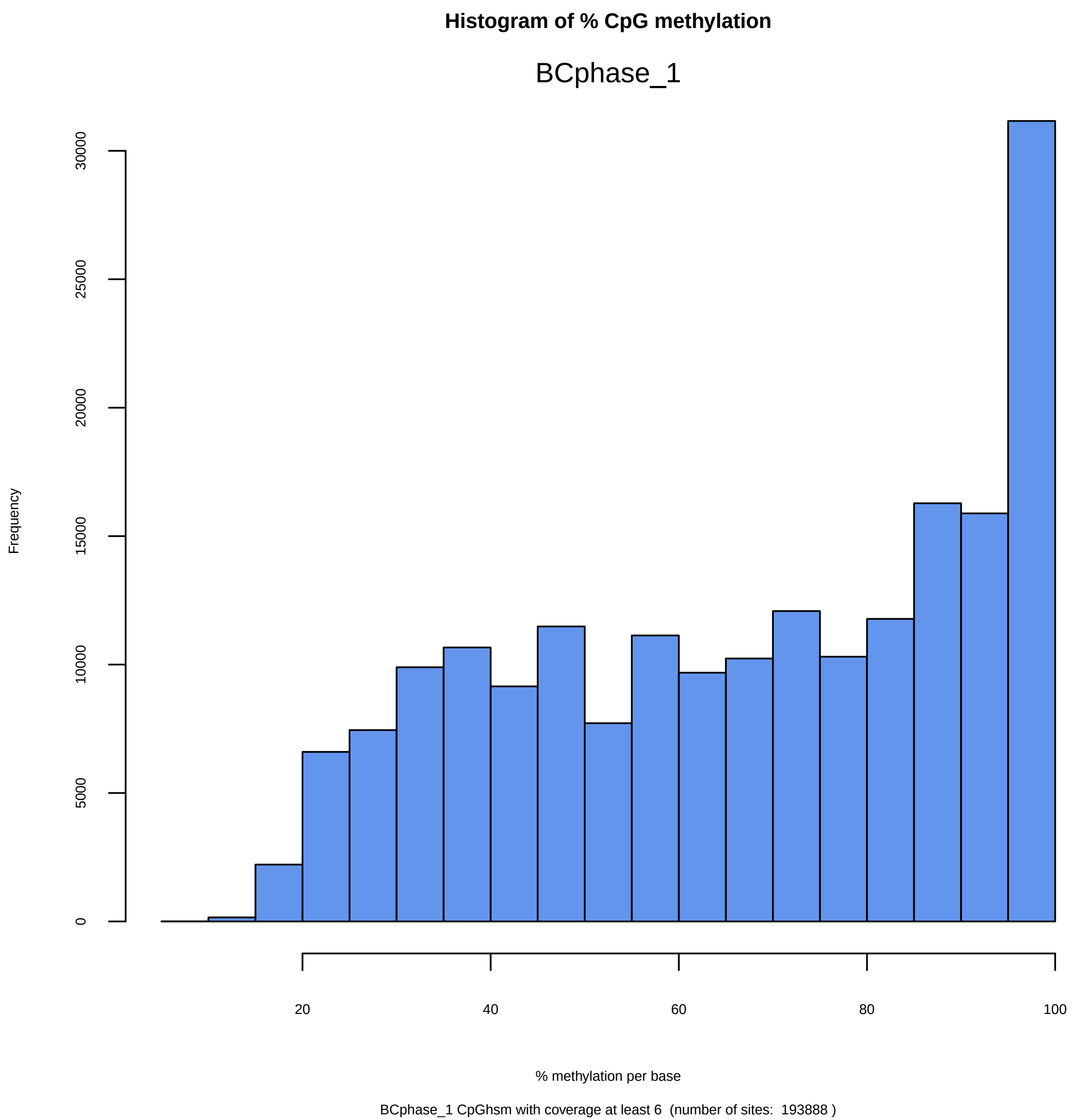
Histogram of % CpG methylation
BCphase_1
Frequency
0
5000
10000
15000
20000
25000
30000
20
40
60
80
100
% methylation per base
BCphase_1 CpGhsm with coverage at least 6 (number of sites: 193888 )

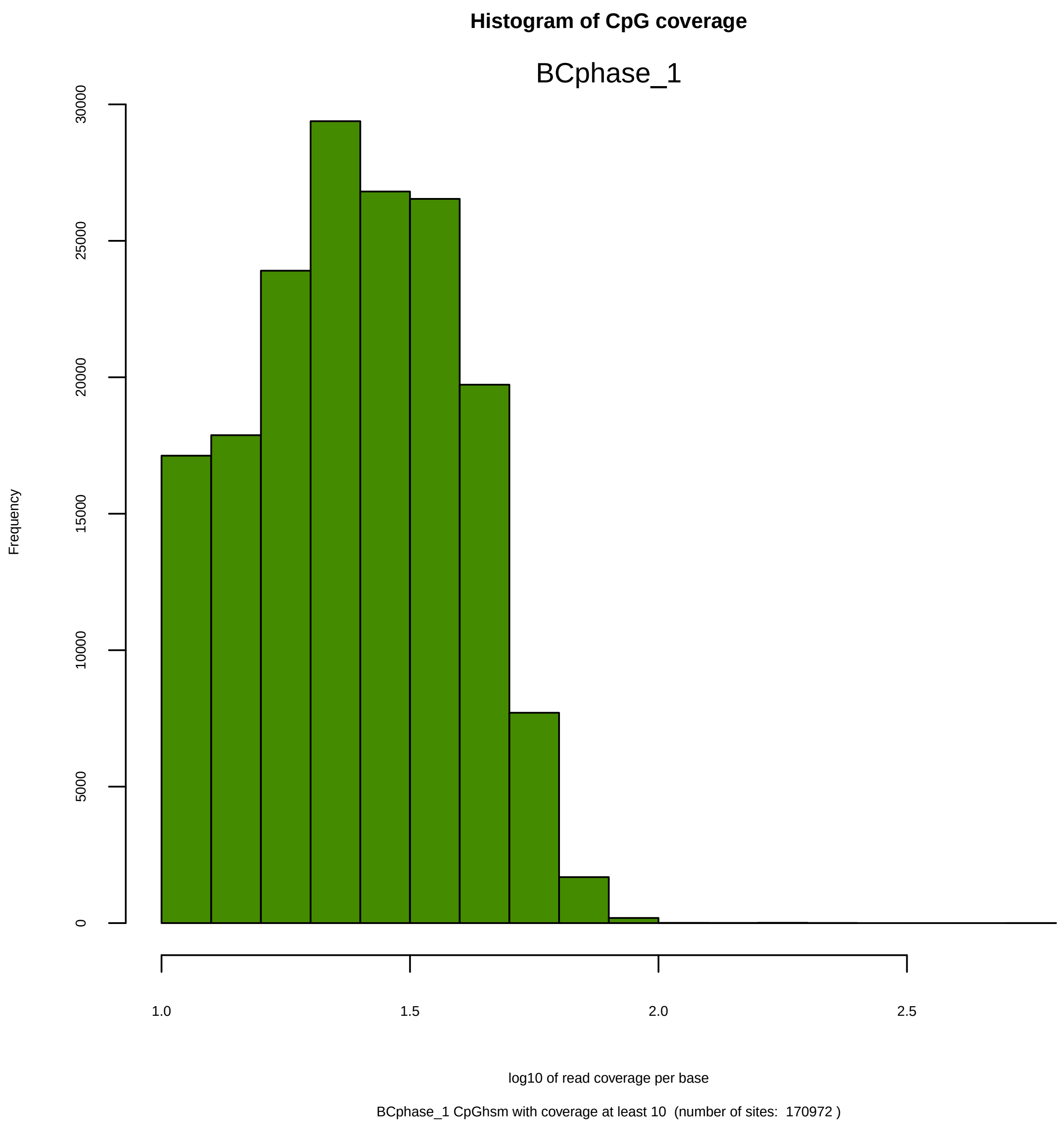

Histogram of CpG coverage
BCphase_1
Frequency
0
5000
10000
15000
20000
25000
30000
1.0
1.5
2.0
2.5
log10 of read coverage per base
BCphase_1 CpGhsm with coverage at least 10  (number of sites:  170972 )

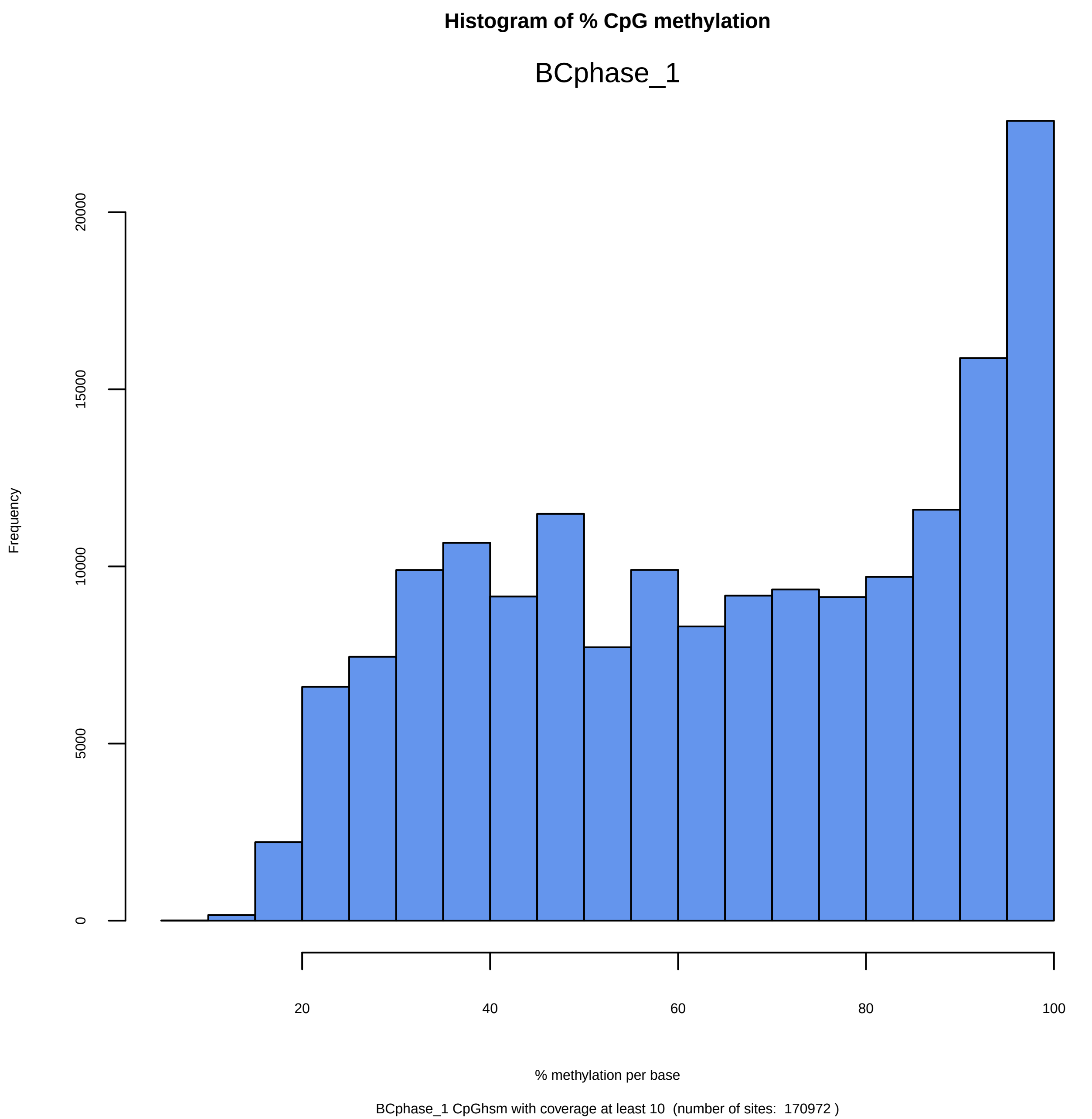

Histogram of % CpG methylation
BCphase_1
Frequency
0
5000
10000
15000
20000
20
40
60
80
100
% methylation per base
BCphase_1 CpGhsm with coverage at least 10 (number of sites: 170972 )

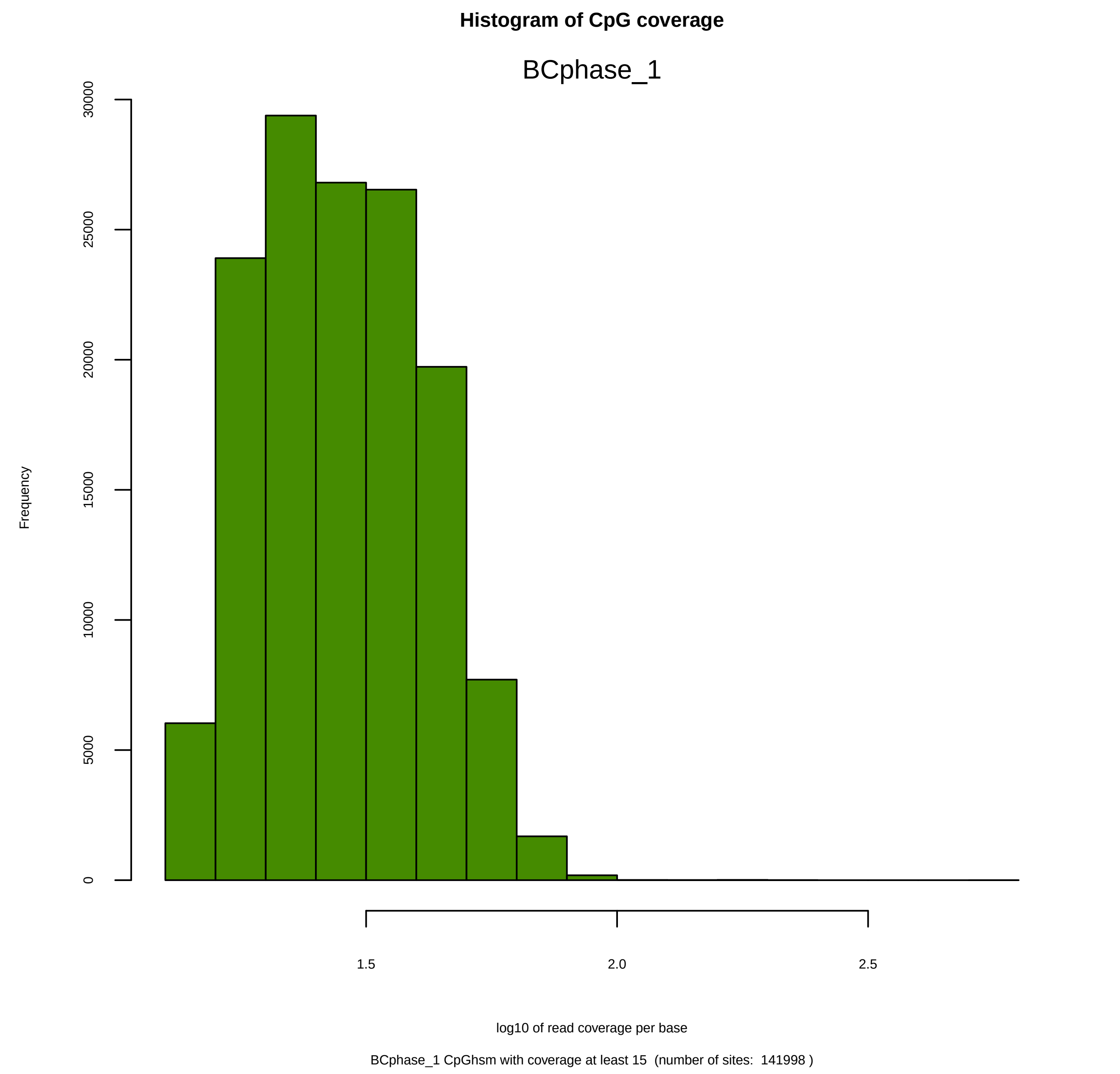

Histogram of CpG coverage
BCphase_1
Frequency
0
5000
10000
15000
20000
25000
30000
1.5
2.0
2.5
log10 of read coverage per base
BCphase_1 CpGhsm with coverage at least 15  (number of sites:  141998 )

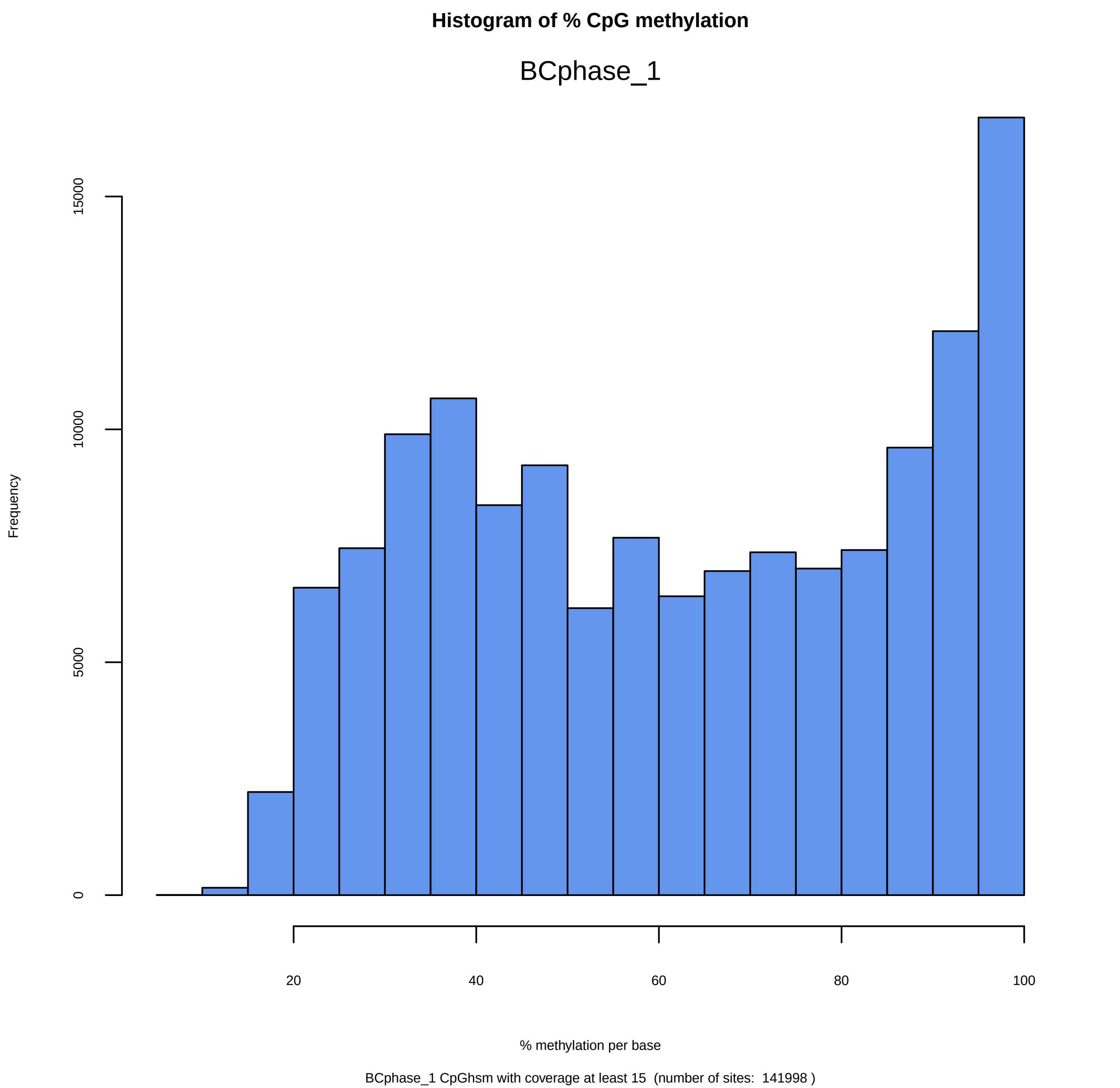
Histogram of % CpG methylation
BCphase_1
Frequency
0
5000
10000
15000
20
40
60
80
100
% methylation per base
BCphase_1 CpGhsm with coverage at least 15 (number of sites: 141998 )

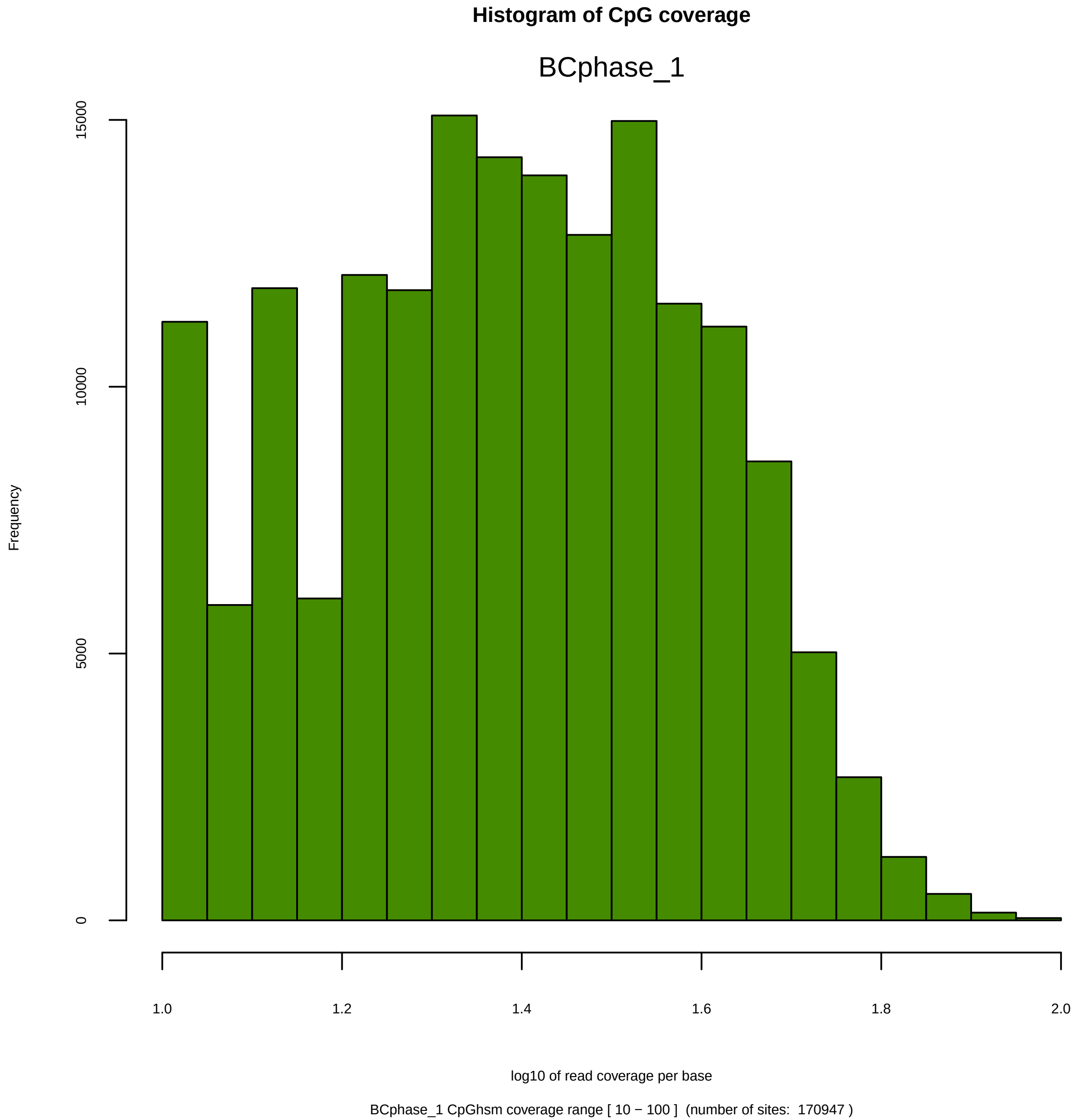
Histogram of CpG coverage
BCphase_1
Frequency
0
5000
10000
15000
1.0
1.2
1.4
1.6
1.8
2.0
log10 of read coverage per base
BCphase_1 CpGhsm coverage range [ 10 − 100 ] (number of sites: 170947 )

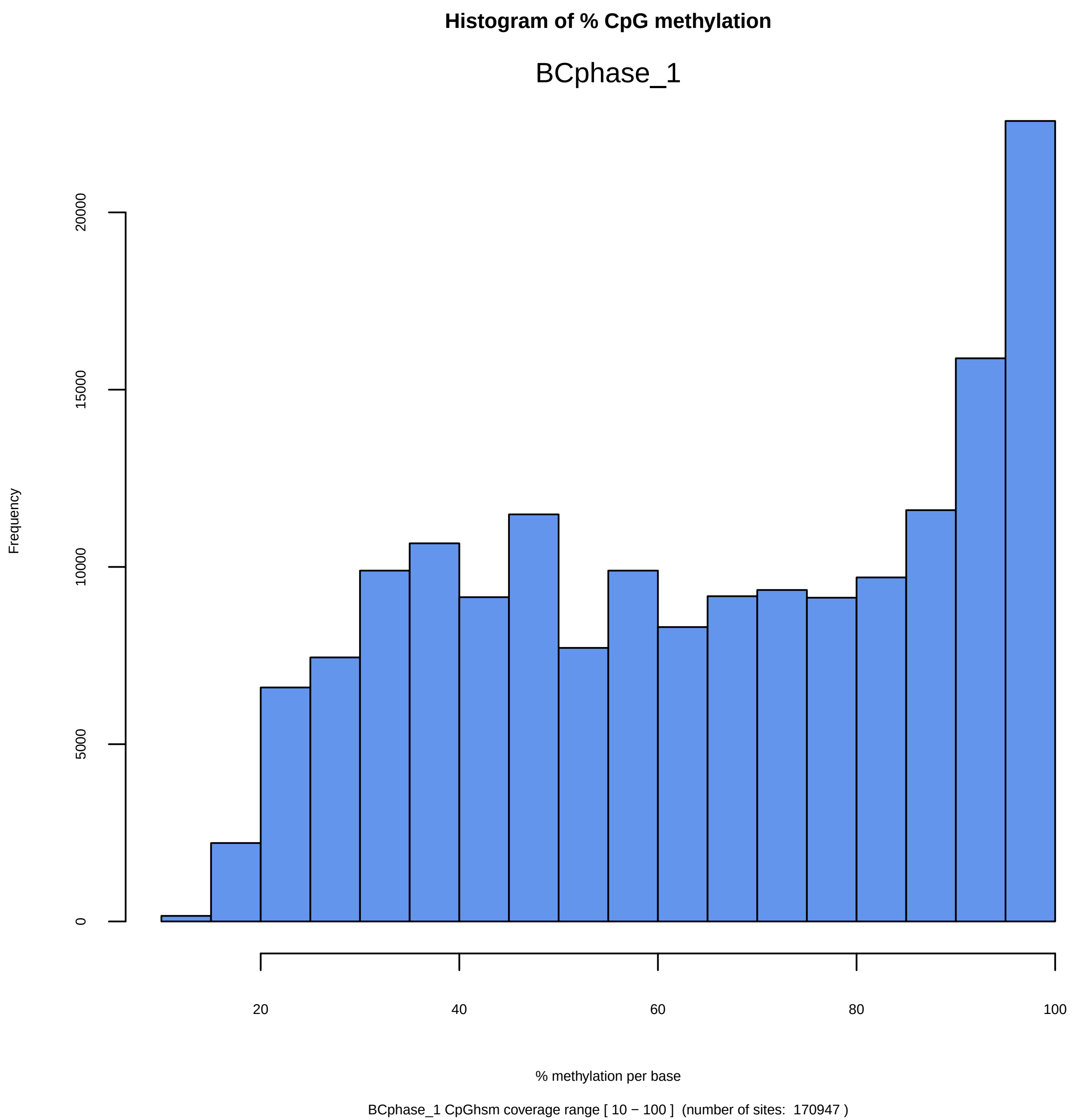
Histogram of % CpG methylation
BCphase_1
Frequency
0
5000
10000
15000
20000
20
40
60
80
100
% methylation per base
BCphase_1 CpGhsm coverage range [ 10 − 100 ]  (number of sites:  170947 )

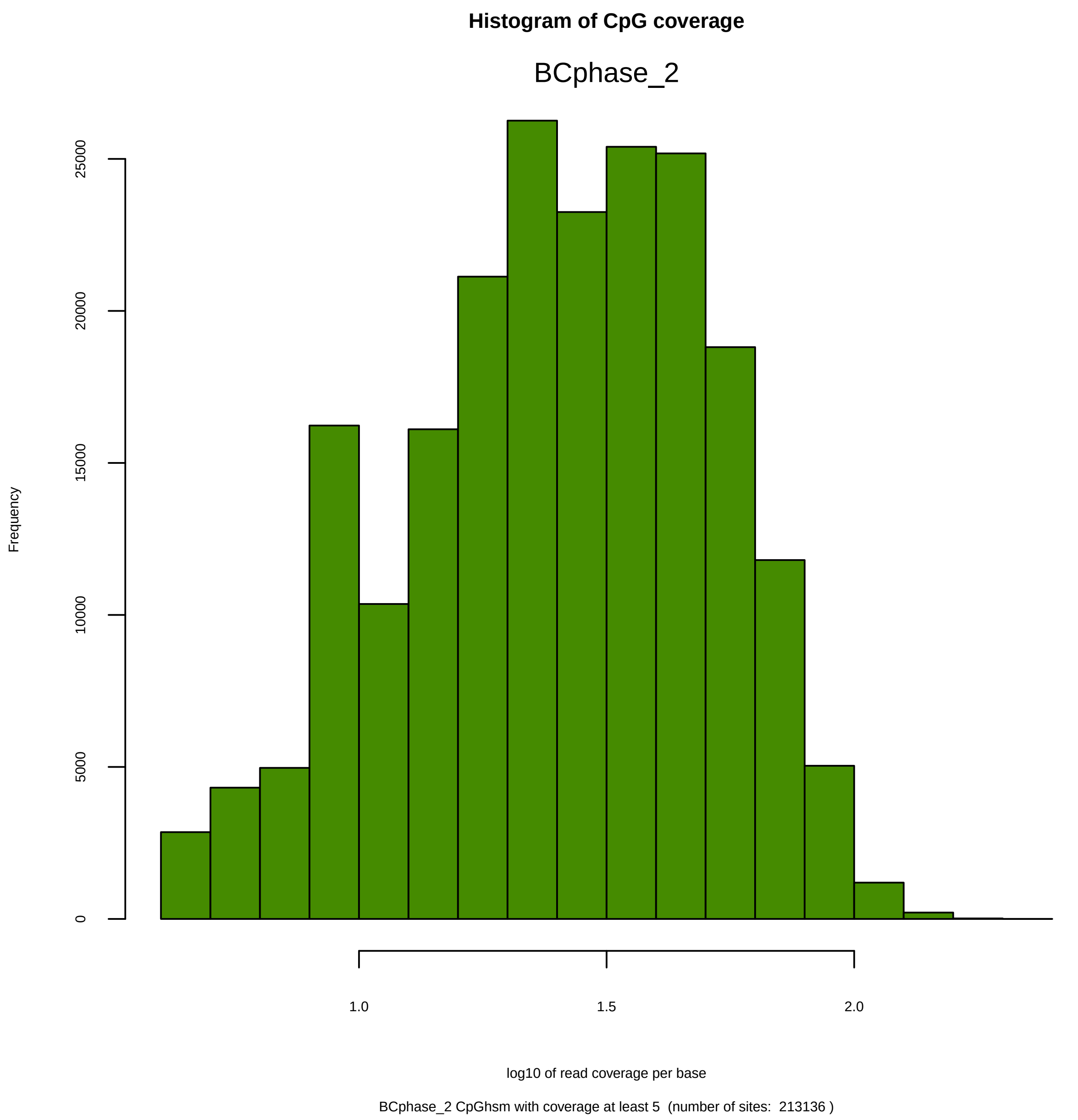
Histogram of CpG coverage
BCphase_2
Frequency
0
5000
10000
15000
20000
25000
1.0
1.5
2.0
log10 of read coverage per base
BCphase_2 CpGhsm with coverage at least 5  (number of sites:  213136 )

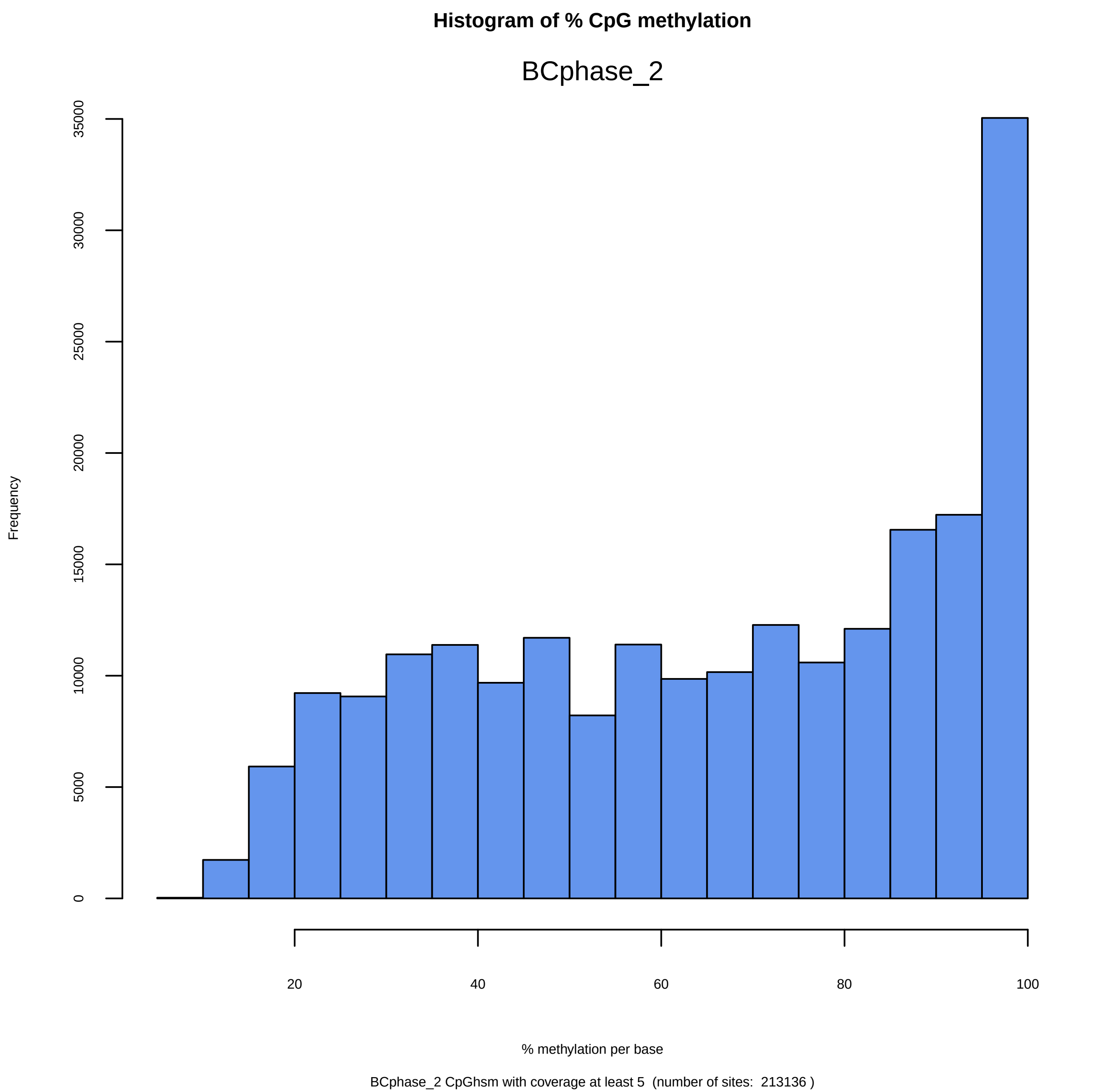

Histogram of % CpG methylation
BCphase_2
Frequency
0
5000
10000
15000
20000
25000
30000
35000
20
40
60
80
100
% methylation per base
BCphase_2 CpGhsm with coverage at least 5 (number of sites: 213136 )

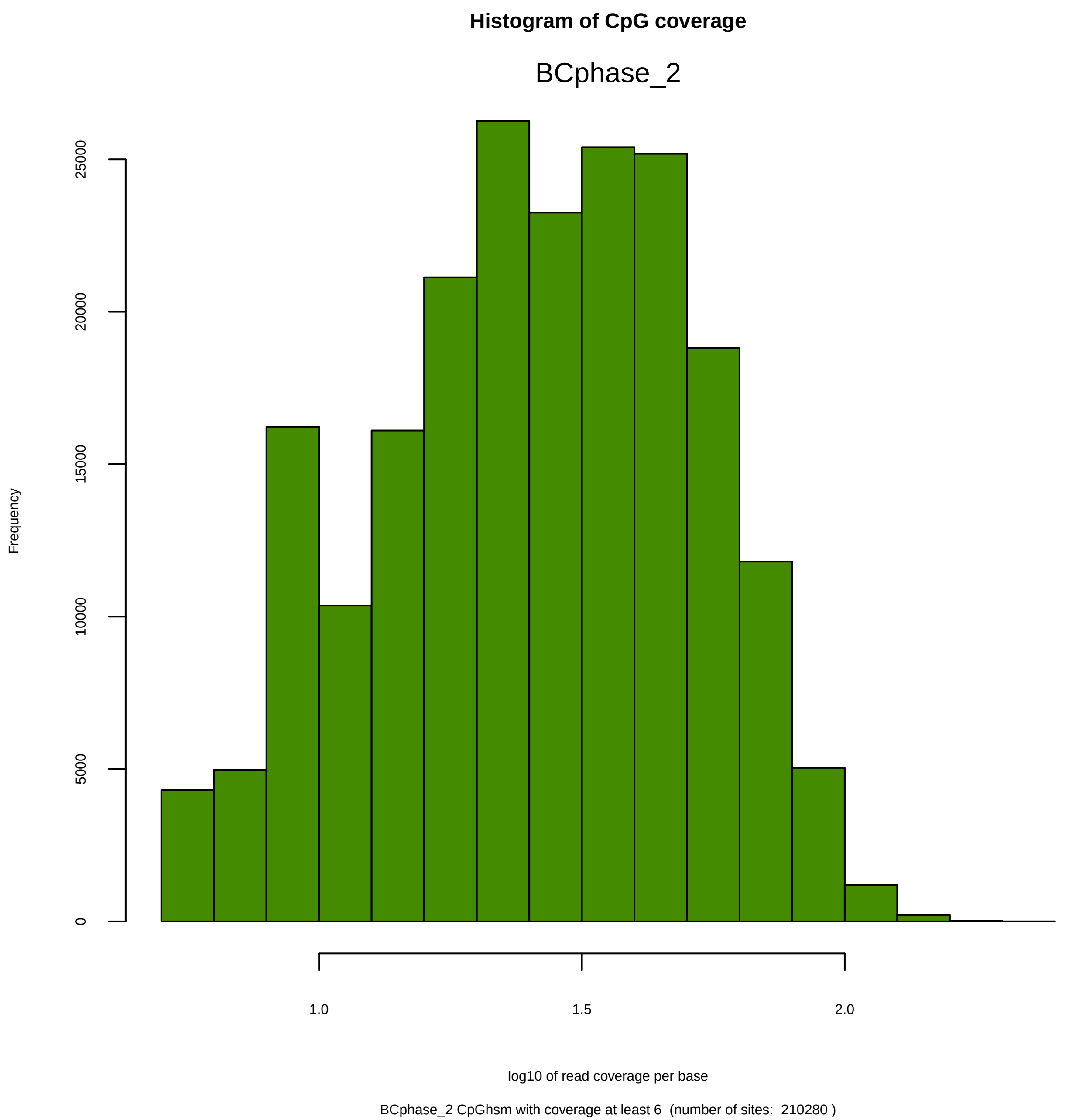
Histogram of CpG coverage
BCphase_2
Frequency
0
5000
10000
15000
20000
25000
1.0
1.5
2.0
log10 of read coverage per base
BCphase_2 CpGhsm with coverage at least 6 (number of sites: 210280 )

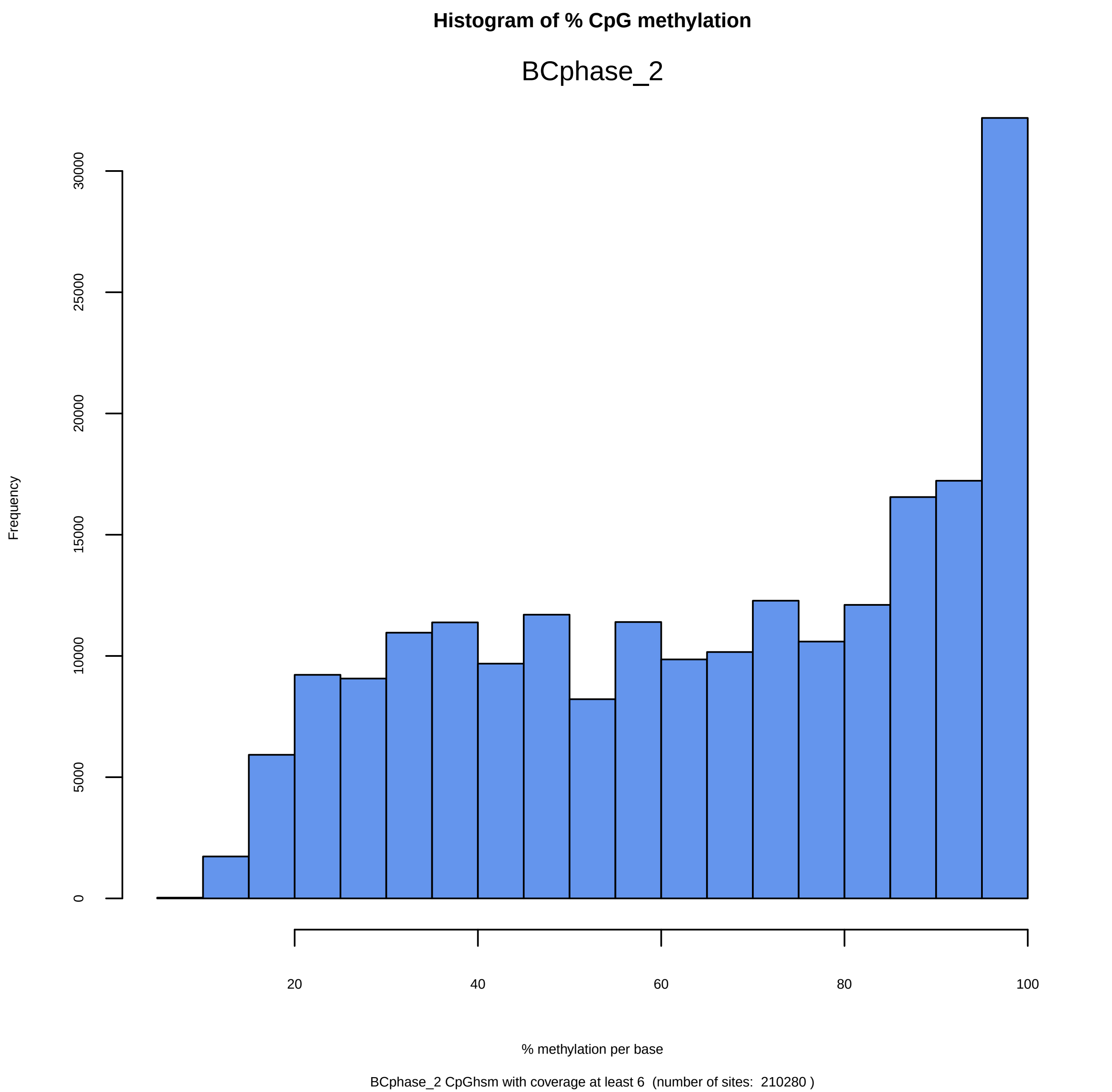

Histogram of % CpG methylation
BCphase_2
Frequency
0
5000
10000
15000
20000
25000
30000
20
40
60
80
100
% methylation per base
BCphase_2 CpGhsm with coverage at least 6 (number of sites: 210280 )

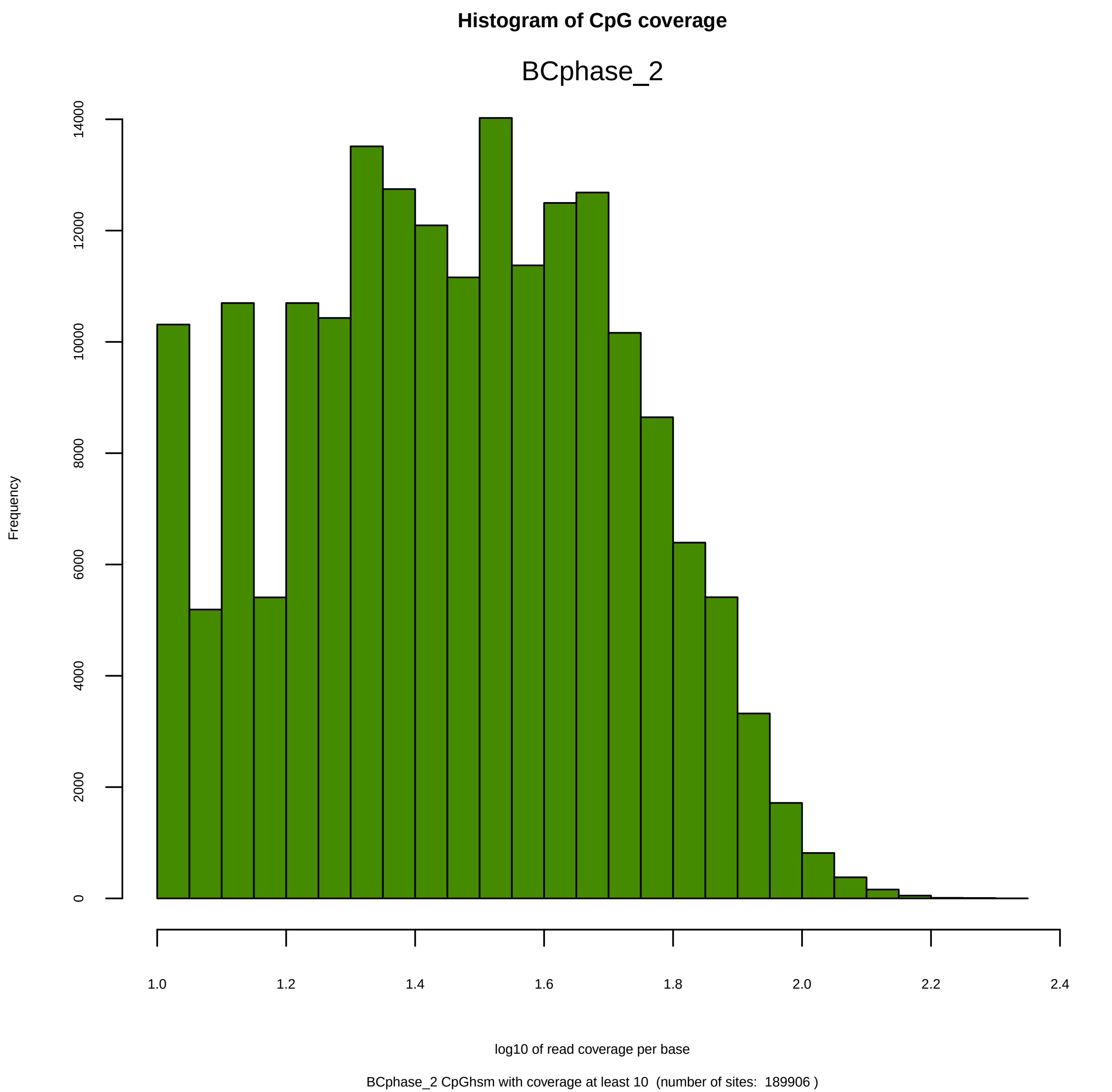

Histogram of CpG coverage
BCphase_2
Frequency
0
2000
4000
6000
8000
10000
12000
14000
1.0
1.2
1.4
1.6
1.8
2.0
2.2
2.4
log10 of read coverage per base
BCphase_2 CpGhsm with coverage at least 10 (number of sites: 189906 )

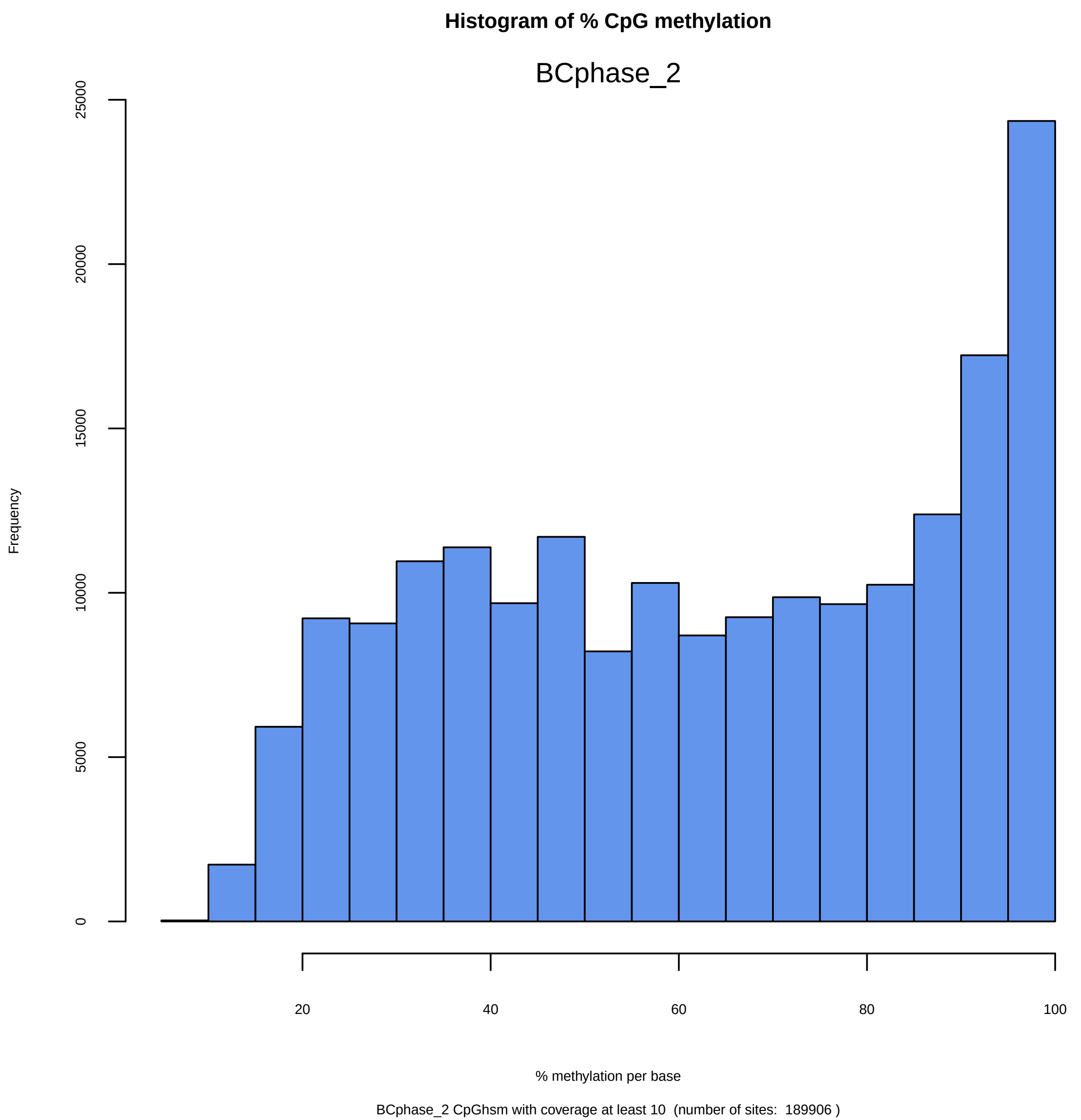
Histogram of % CpG methylation
BCphase_2
Frequency
0
5000
10000
15000
20000
25000
20
40
60
80
100
% methylation per base
BCphase_2 CpGhsm with coverage at least 10  (number of sites:  189906 )

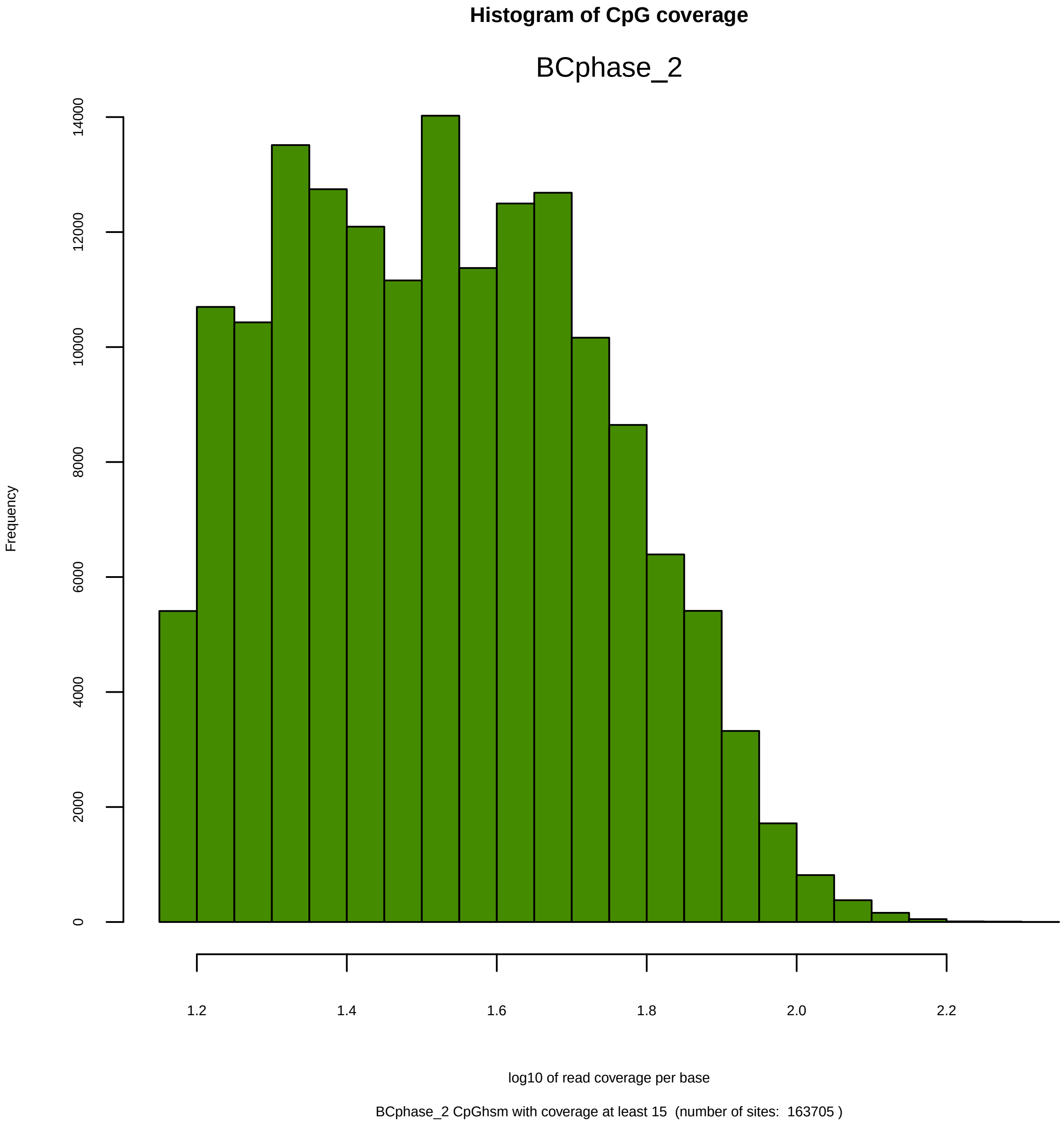

Histogram of CpG coverage
BCphase_2
Frequency
0
2000
4000
6000
8000
10000
12000
14000
1.2
1.4
1.6
1.8
2.0
2.2
log10 of read coverage per base
BCphase_2 CpGhsm with coverage at least 15 (number of sites: 163705 )

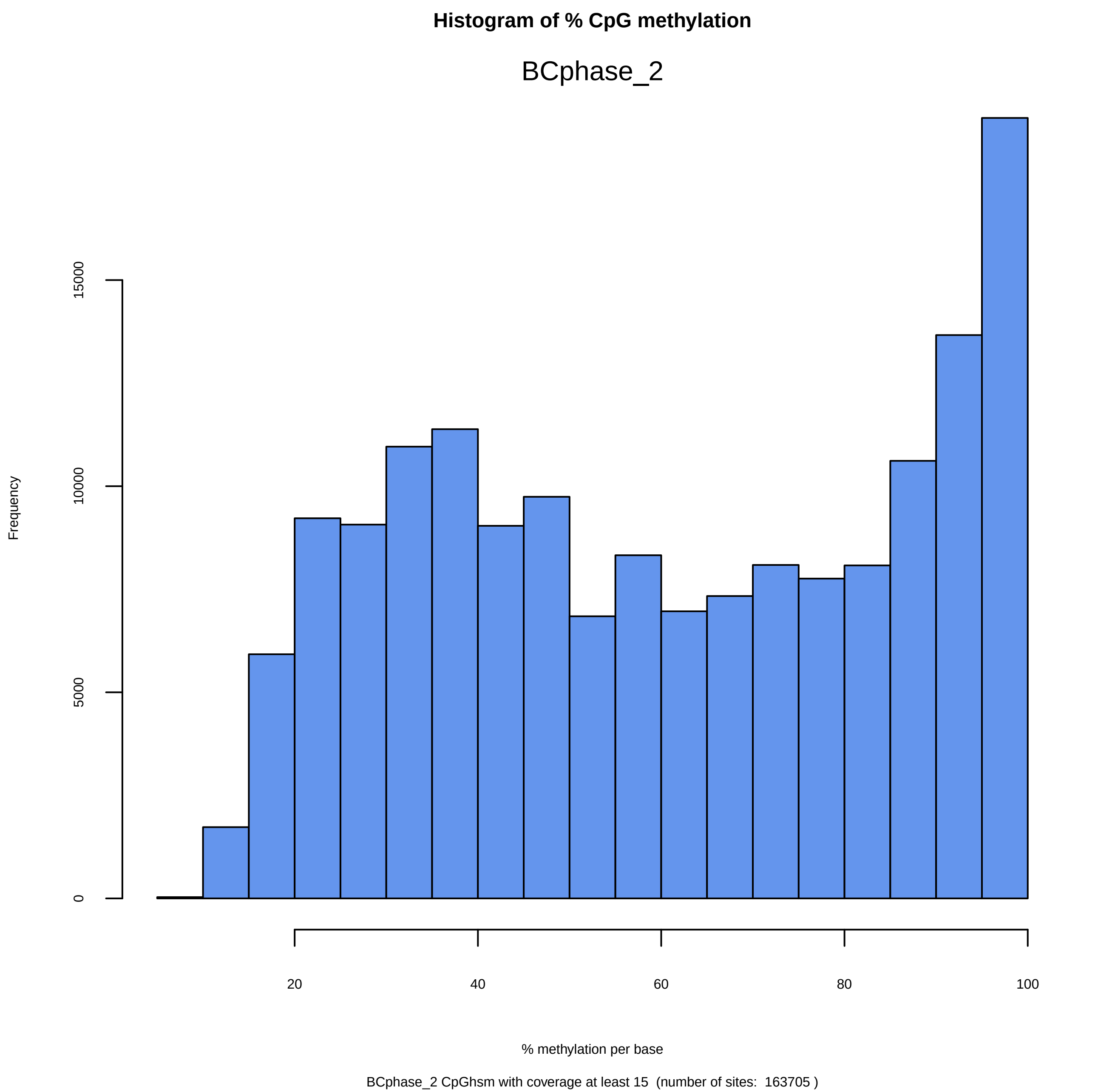
Histogram of % CpG methylation
BCphase_2
Frequency
0
5000
10000
15000
20
40
60
80
100
% methylation per base
BCphase_2 CpGhsm with coverage at least 15  (number of sites:  163705 )

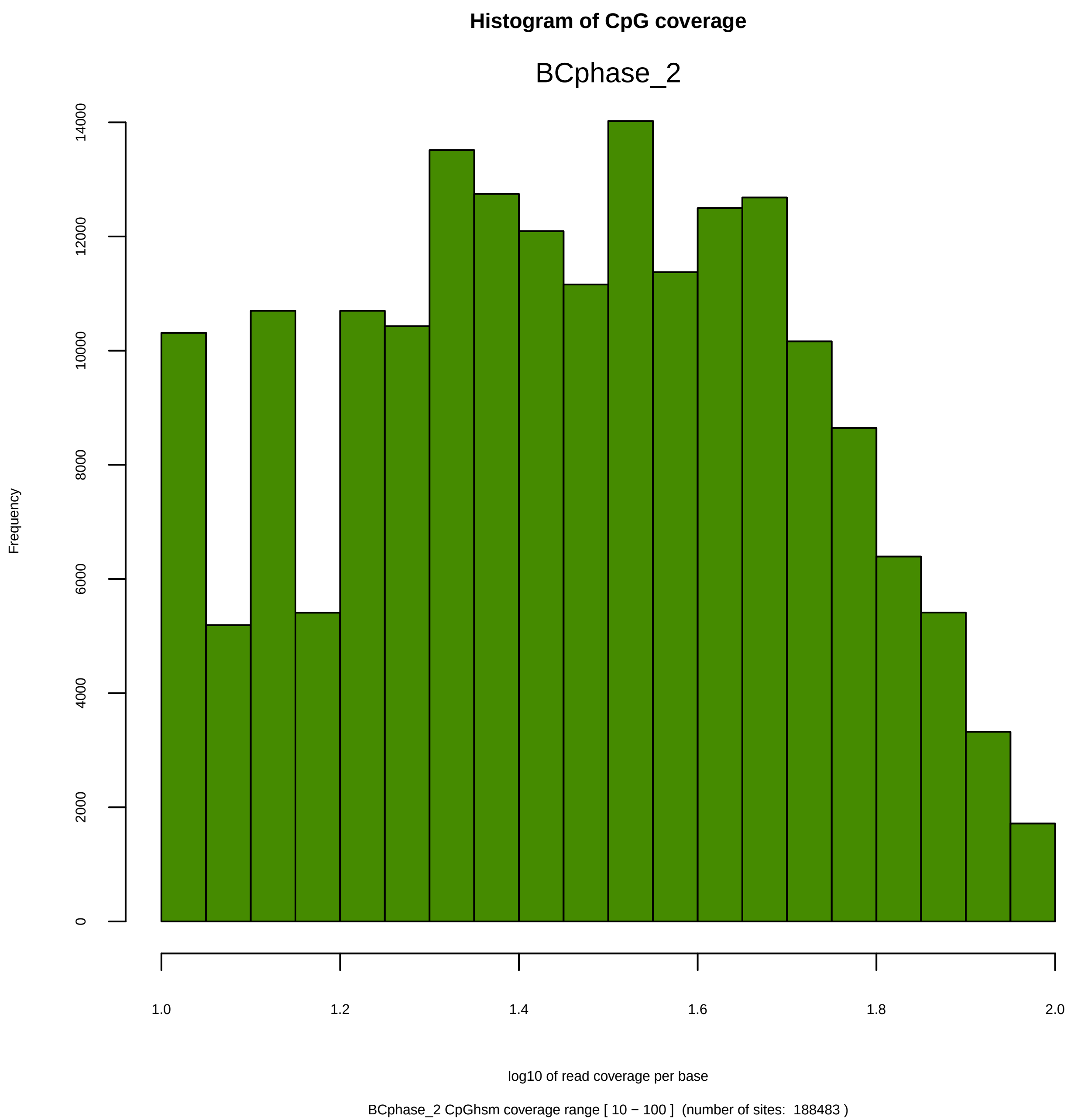
Histogram of CpG coverage
BCphase_2
Frequency
0
2000
4000
6000
8000
10000
12000
14000
1.0
1.2
1.4
1.6
1.8
2.0
log10 of read coverage per base
BCphase_2 CpGhsm coverage range [ 10 − 100 ] (number of sites: 188483 )

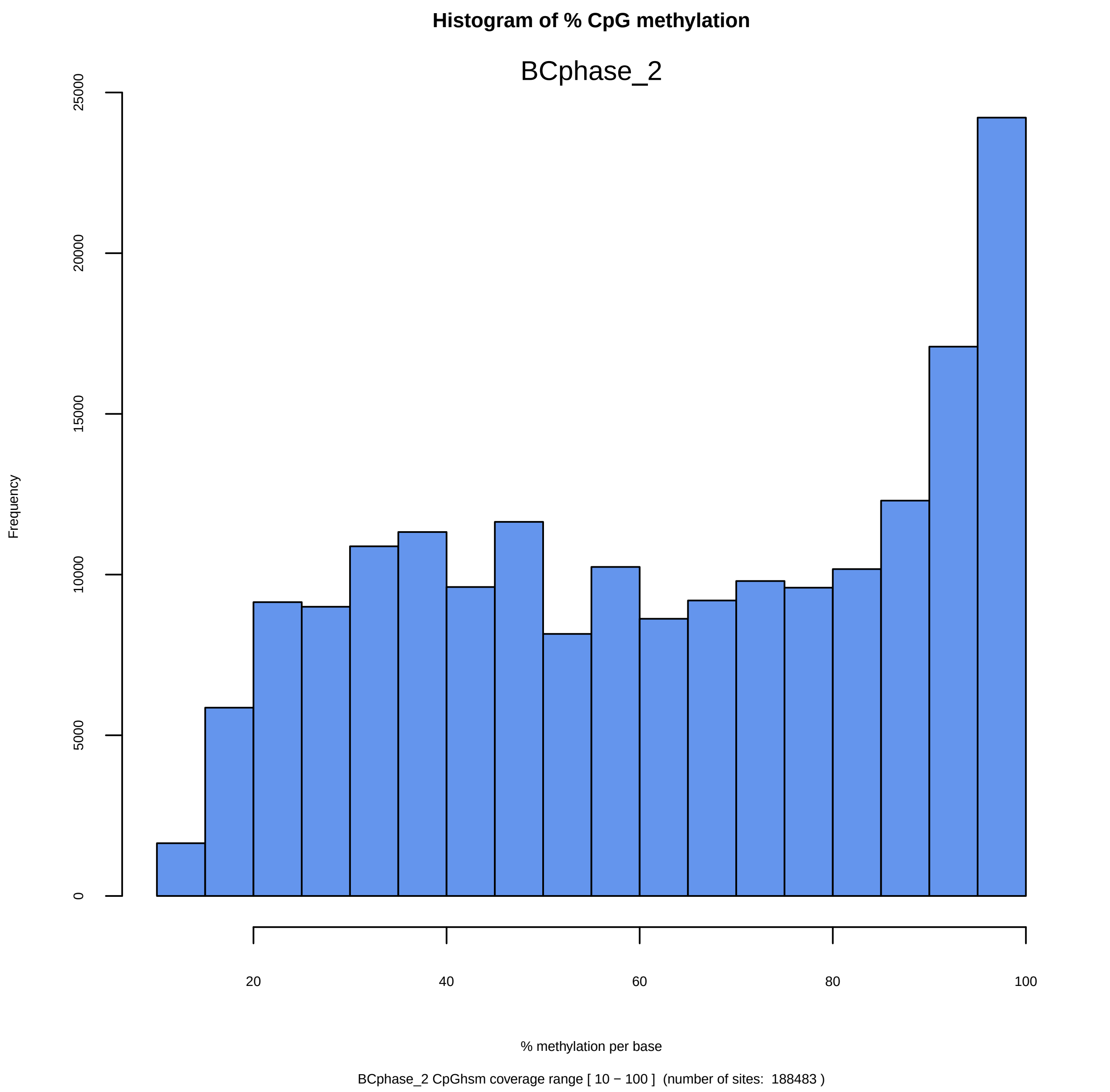

Histogram of % CpG methylation
BCphase_2
Frequency
0
5000
10000
15000
20000
25000
20
40
60
80
100
% methylation per base
BCphase_2 CpGhsm coverage range [ 10 − 100 ] (number of sites: 188483 )

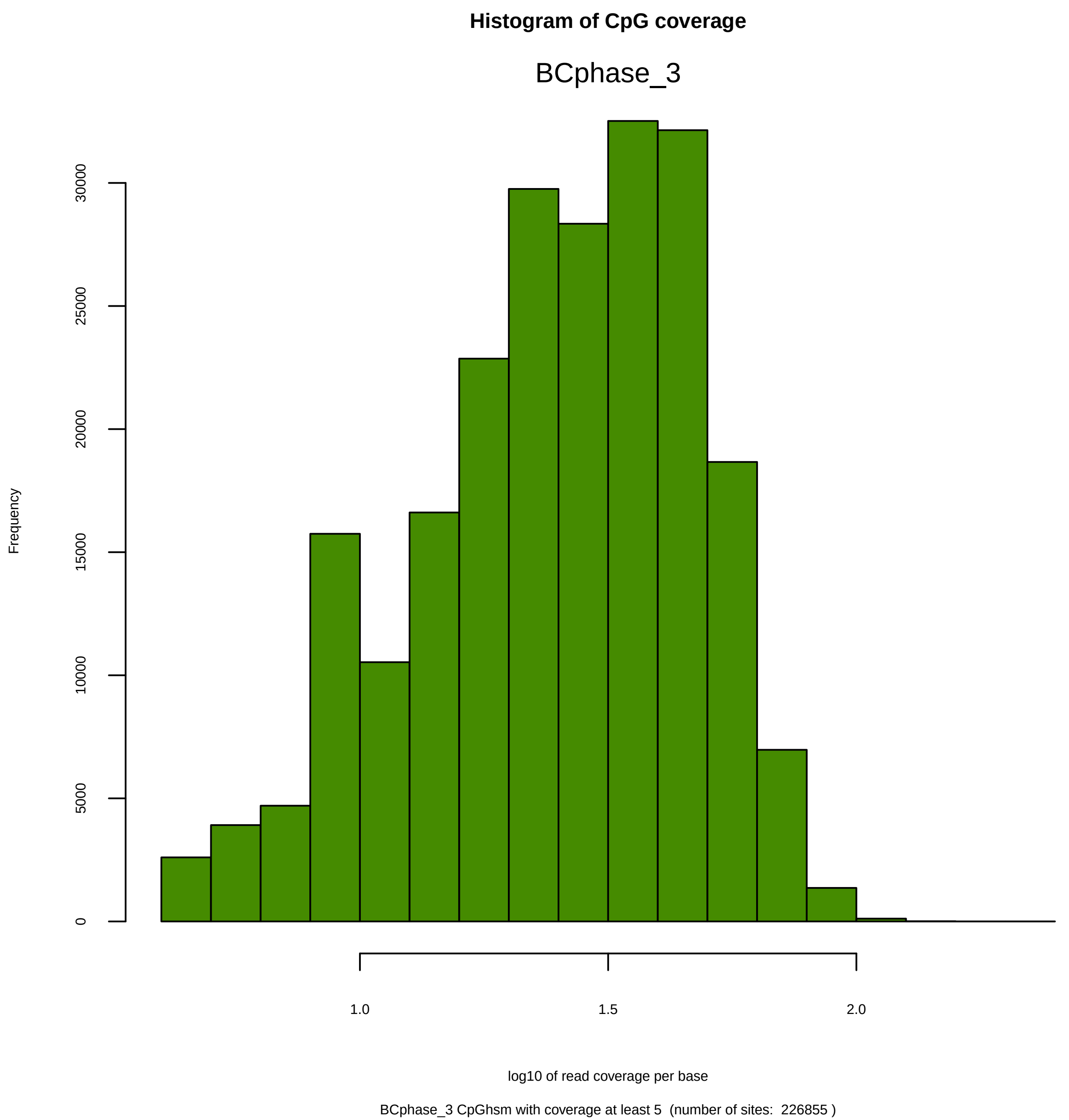

Histogram of CpG coverage
BCphase_3
Frequency
0
5000
10000
15000
20000
25000
30000
1.0
1.5
2.0
log10 of read coverage per base
BCphase_3 CpGhsm with coverage at least 5 (number of sites: 226855 )

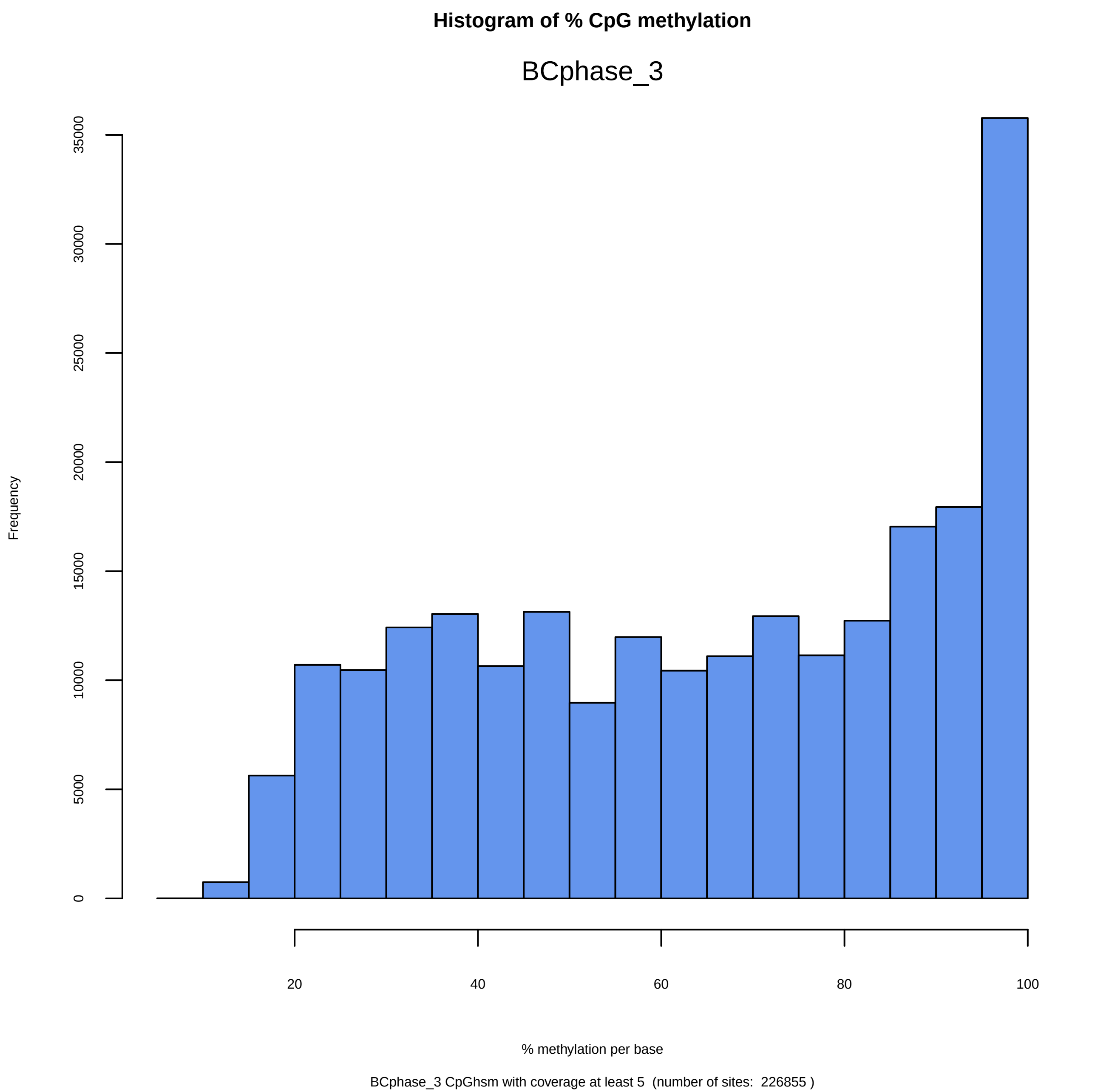
Histogram of % CpG methylation
BCphase_3
Frequency
0
5000
10000
15000
20000
25000
30000
35000
20
40
60
80
100
% methylation per base
BCphase_3 CpGhsm with coverage at least 5 (number of sites: 226855 )

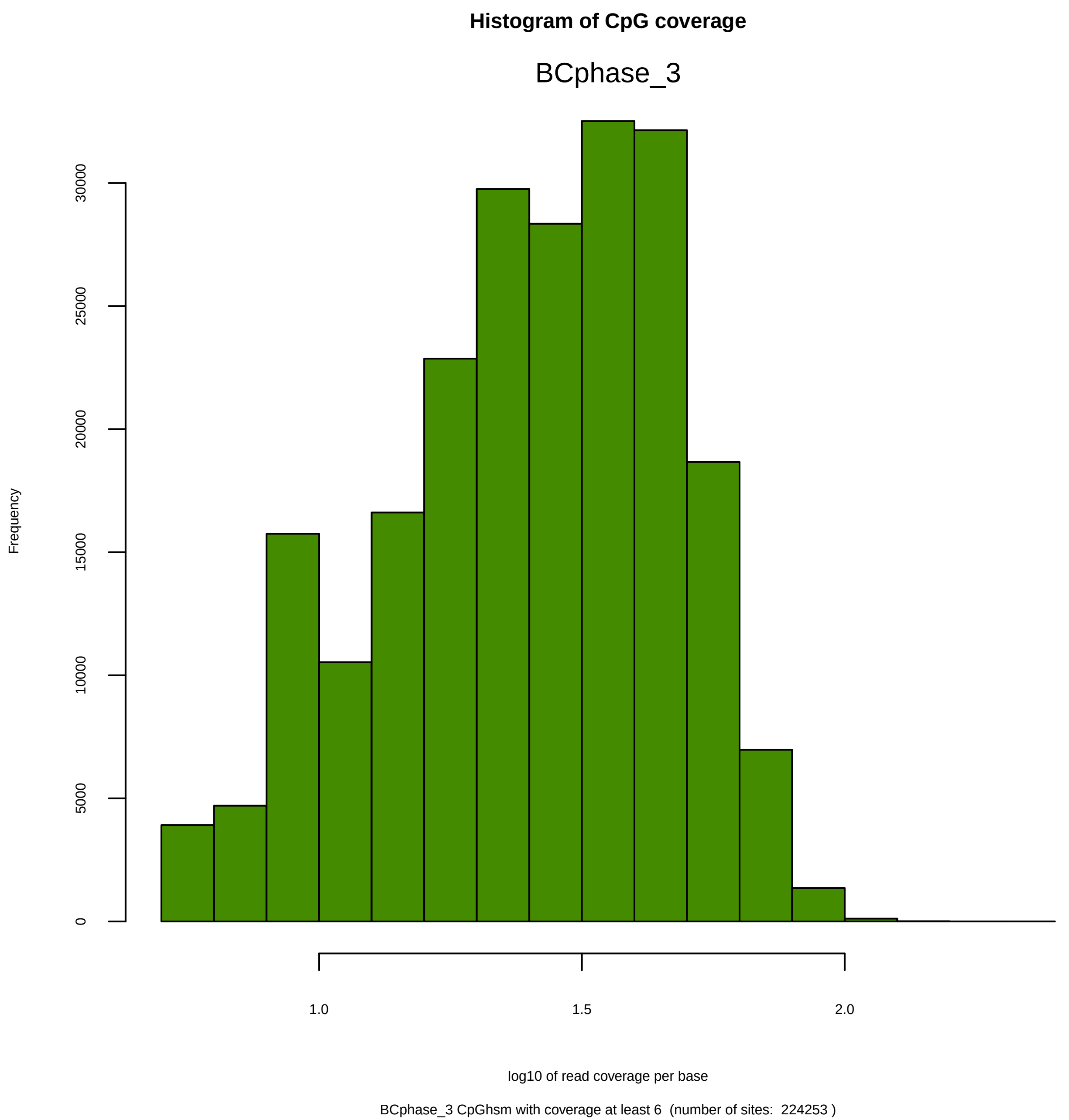

Histogram of CpG coverage
BCphase_3
Frequency
0
5000
10000
15000
20000
25000
30000
1.0
1.5
2.0
log10 of read coverage per base
BCphase_3 CpGhsm with coverage at least 6  (number of sites:  224253 )

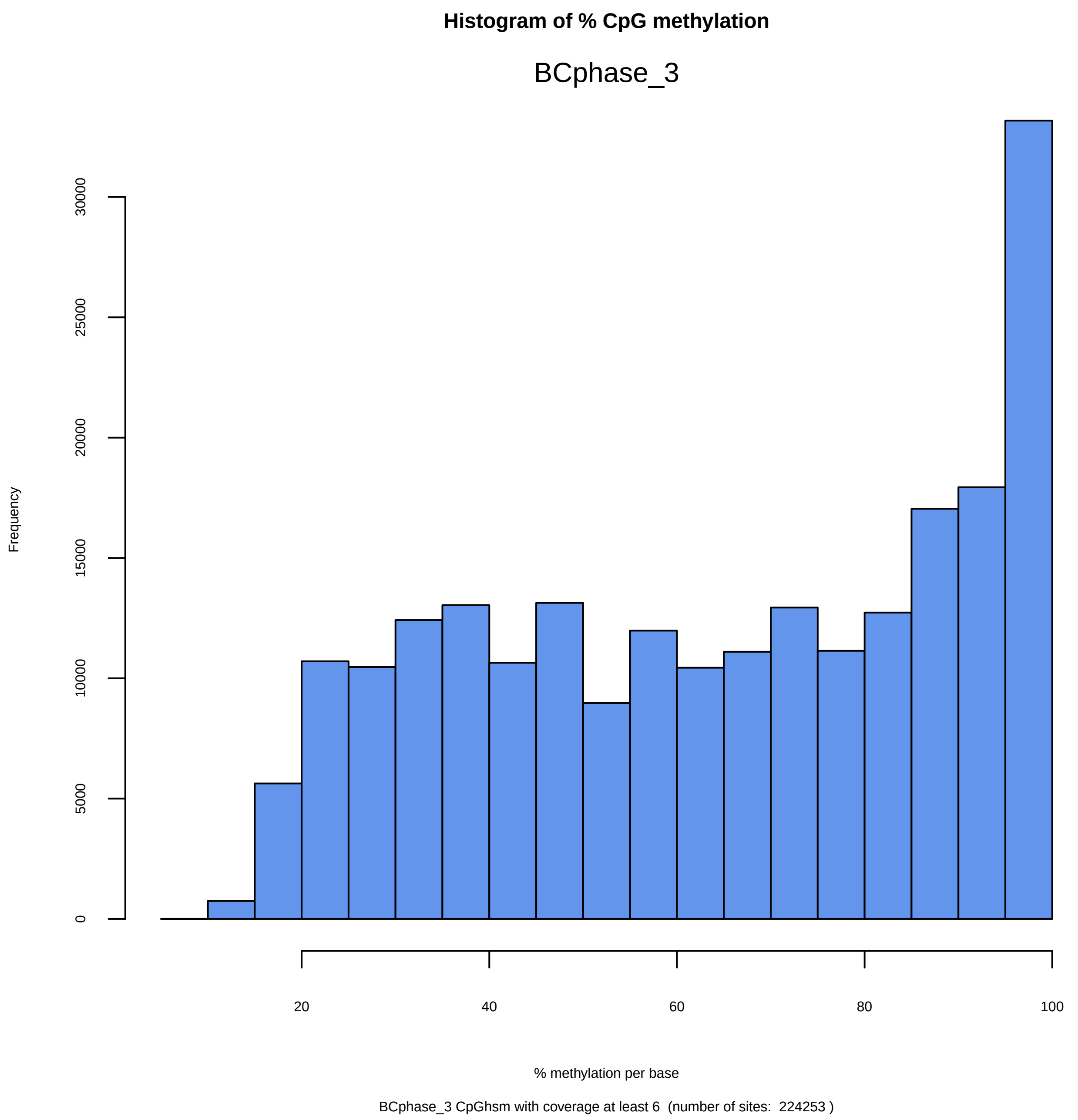

Histogram of % CpG methylation
BCphase_3
Frequency
0
5000
10000
15000
20000
25000
30000
20
40
60
80
100
% methylation per base
BCphase_3 CpGhsm with coverage at least 6  (number of sites:  224253 )

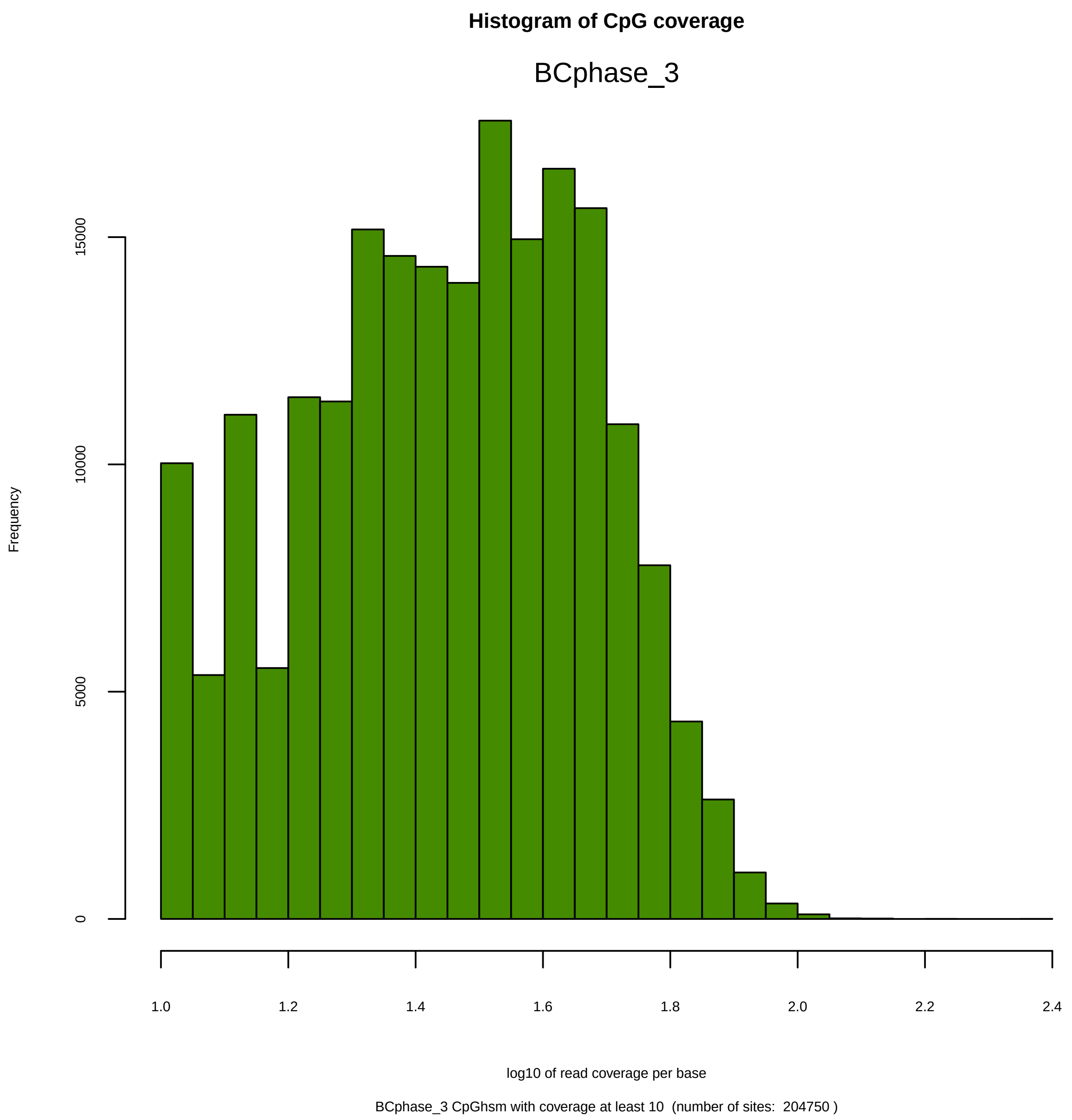
Histogram of CpG coverage
BCphase_3
Frequency
0
5000
10000
15000
1.0
1.2
1.4
1.6
1.8
2.0
2.2
2.4
log10 of read coverage per base
BCphase_3 CpGhsm with coverage at least 10  (number of sites:  204750 )

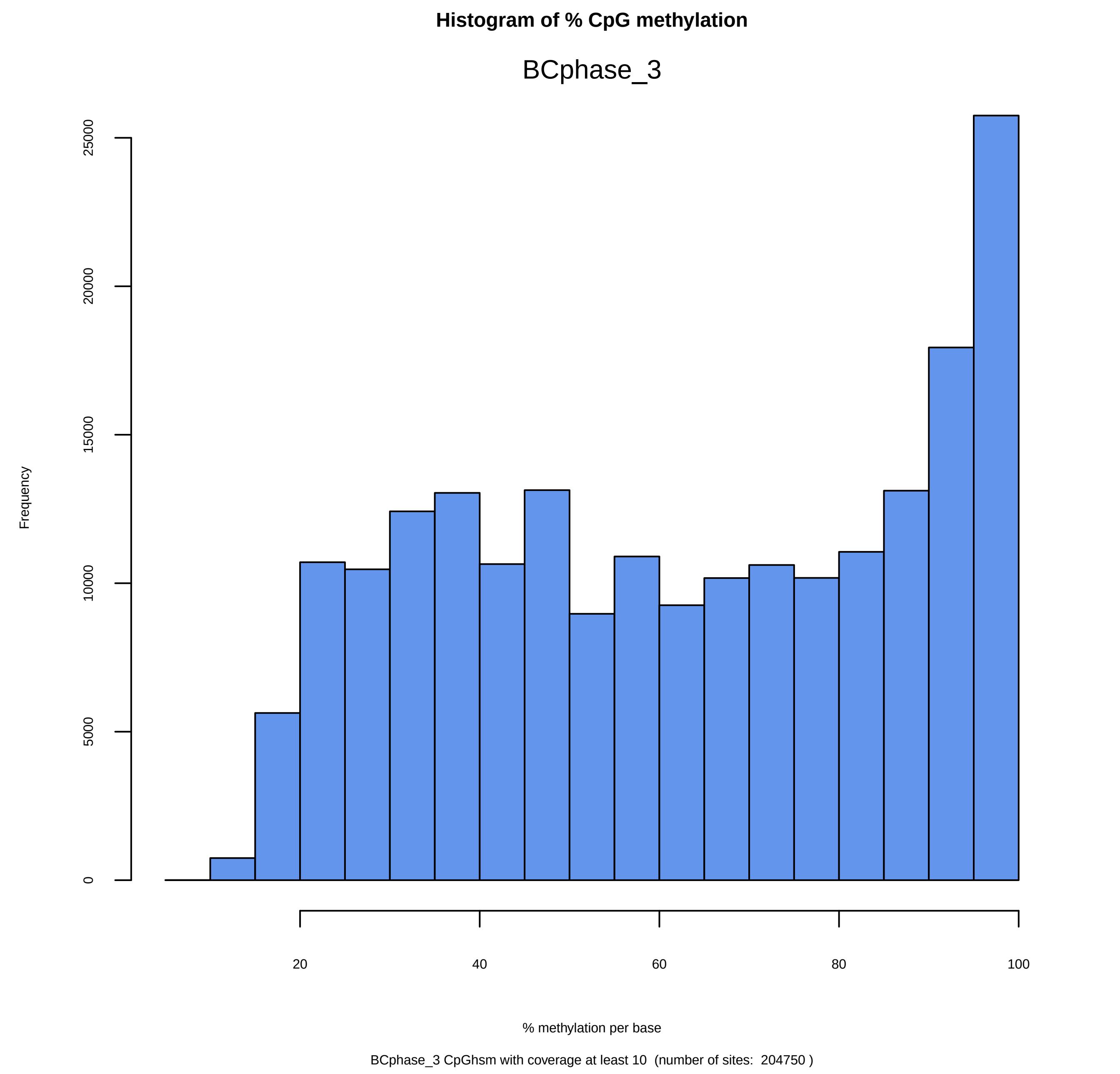
Histogram of % CpG methylation
BCphase_3
Frequency
0
5000
10000
15000
20000
25000
20
40
60
80
100
% methylation per base
BCphase_3 CpGhsm with coverage at least 10  (number of sites:  204750 )

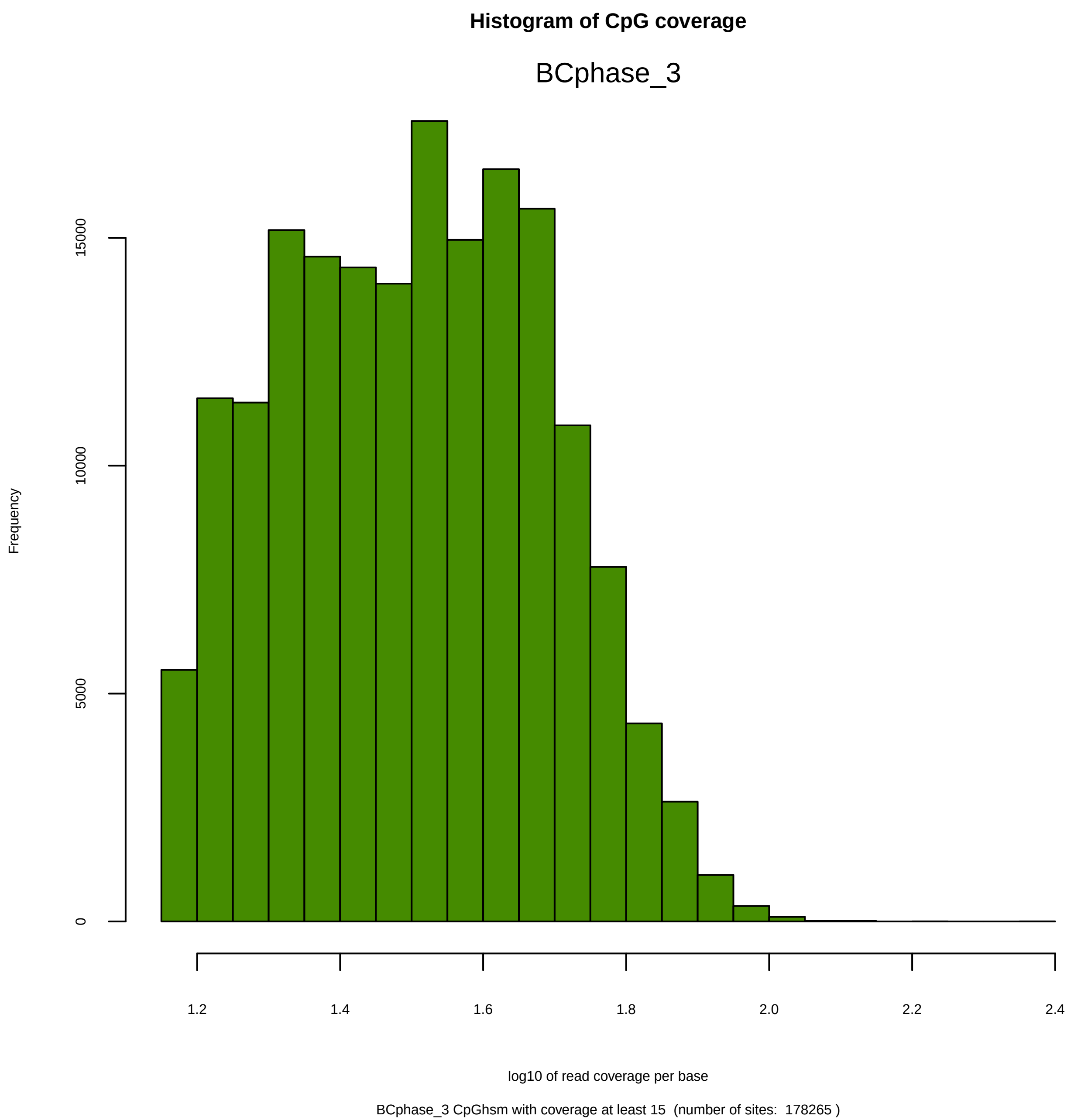

Histogram of CpG coverage
BCphase_3
Frequency
0
5000
10000
15000
1.2
1.4
1.6
1.8
2.0
2.2
2.4
log10 of read coverage per base
BCphase_3 CpGhsm with coverage at least 15 (number of sites: 178265 )

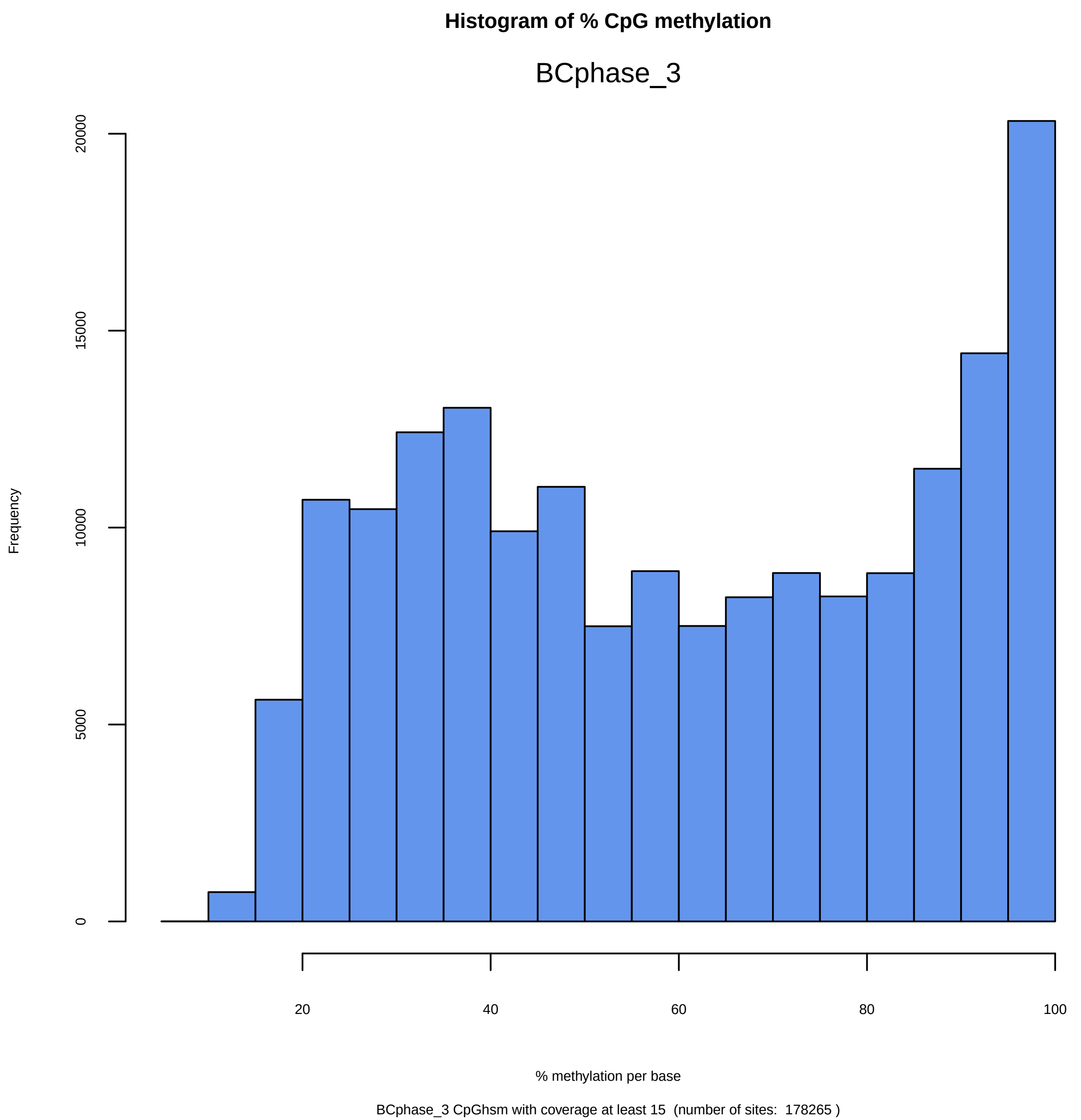

Histogram of % CpG methylation
BCphase_3
Frequency
0
5000
10000
15000
20000
20
40
60
80
100
% methylation per base
BCphase_3 CpGhsm with coverage at least 15 (number of sites: 178265 )

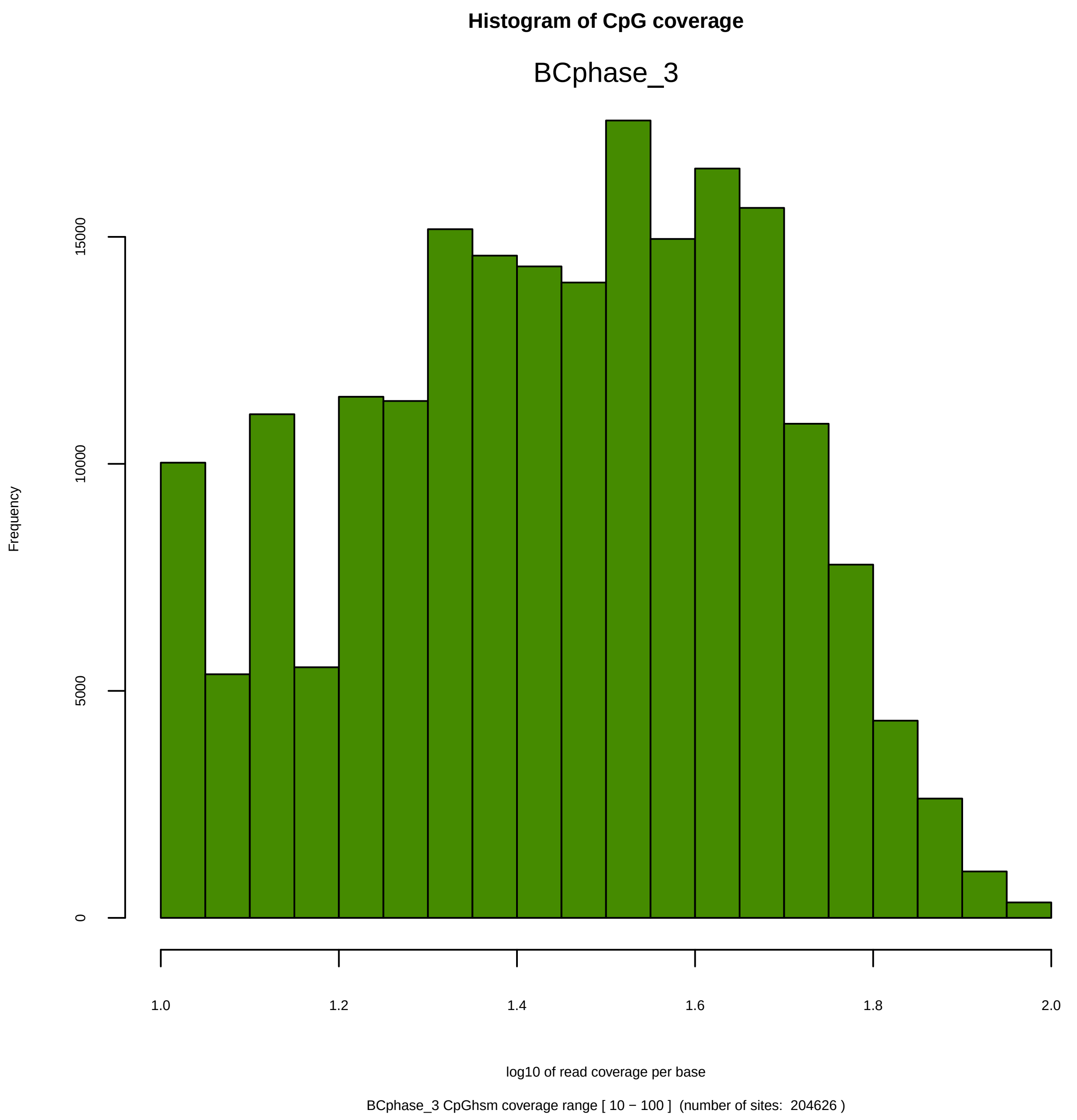
Histogram of CpG coverage
BCphase_3
Frequency
0
5000
10000
15000
1.0
1.2
1.4
1.6
1.8
2.0
log10 of read coverage per base
BCphase_3 CpGhsm coverage range [ 10 − 100 ]  (number of sites:  204626 )

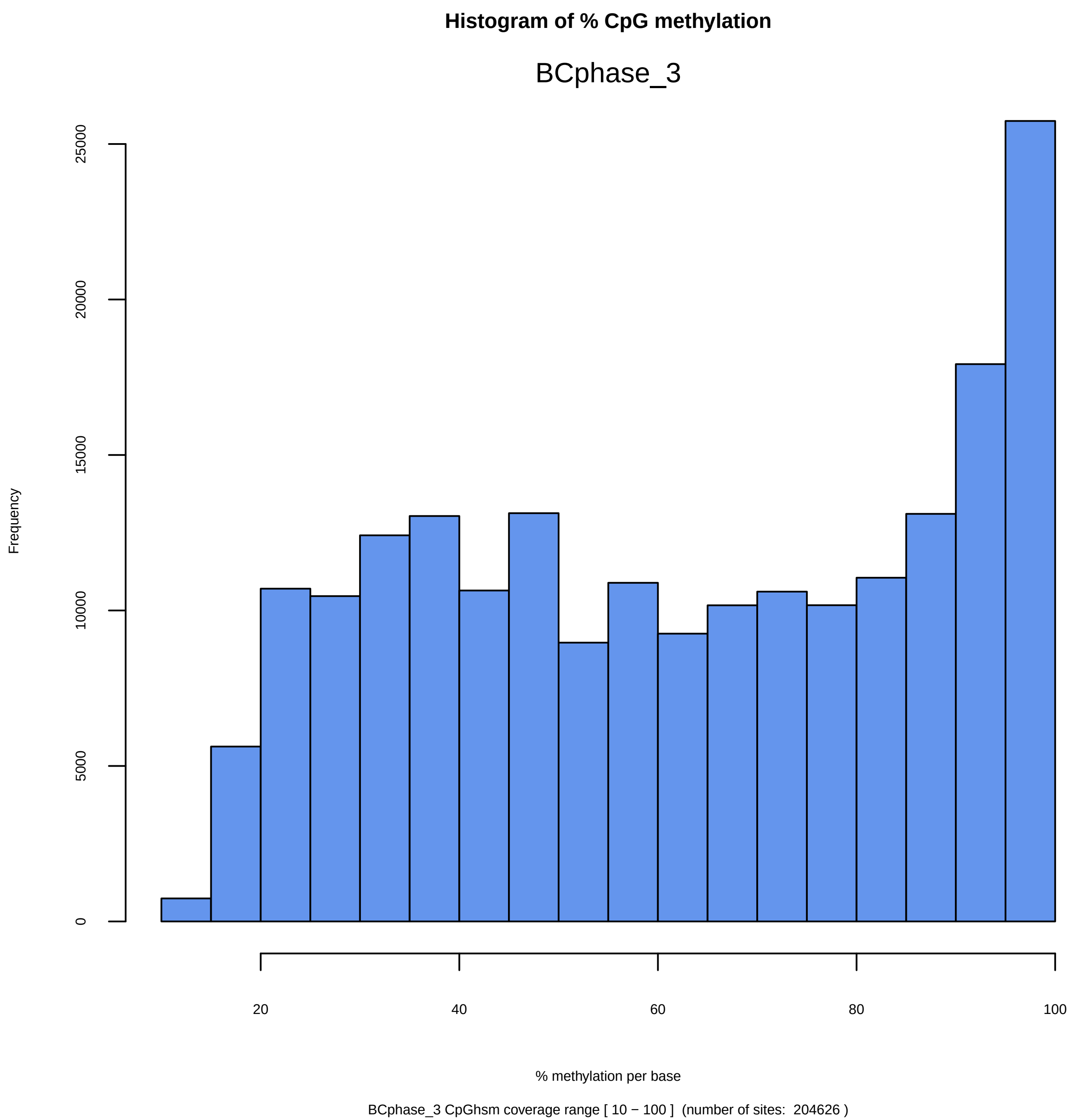

Histogram of % CpG methylation
BCphase_3
Frequency
0
5000
10000
15000
20000
25000
20
40
60
80
100
% methylation per base
BCphase_3 CpGhsm coverage range [ 10 − 100 ]  (number of sites:  204626 )

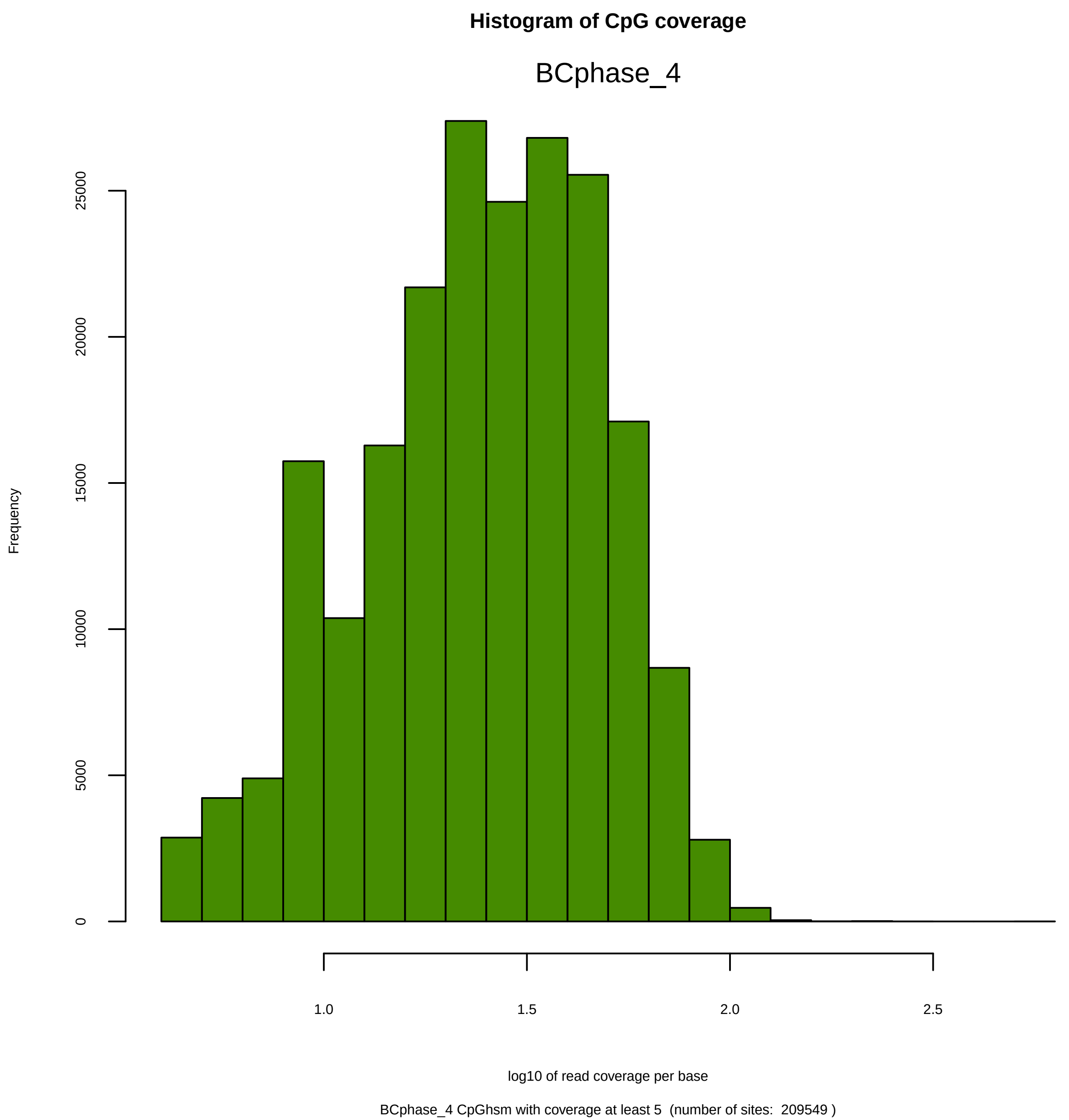
Histogram of CpG coverage
BCphase_4
Frequency
0
5000
10000
15000
20000
25000
1.0
1.5
2.0
2.5
log10 of read coverage per base
BCphase_4 CpGhsm with coverage at least 5  (number of sites:  209549 )

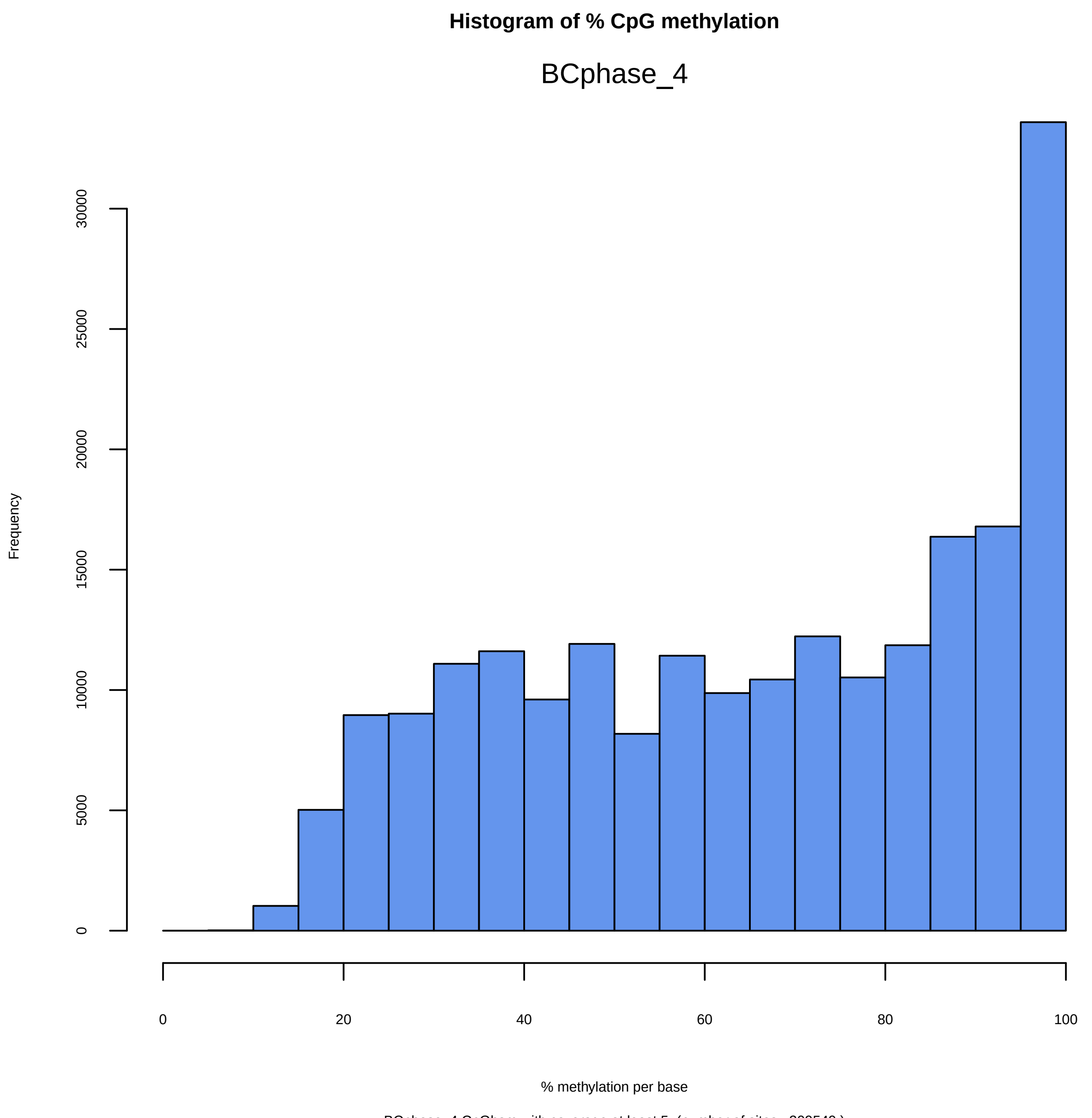
Histogram of % CpG methylation
BCphase_4
Frequency
0
5000
10000
15000
20000
25000
30000
0
20
40
60
80
100
% methylation per base
BCphase_4 CpGhsm with coverage at least 5  (number of sites:  209549 )

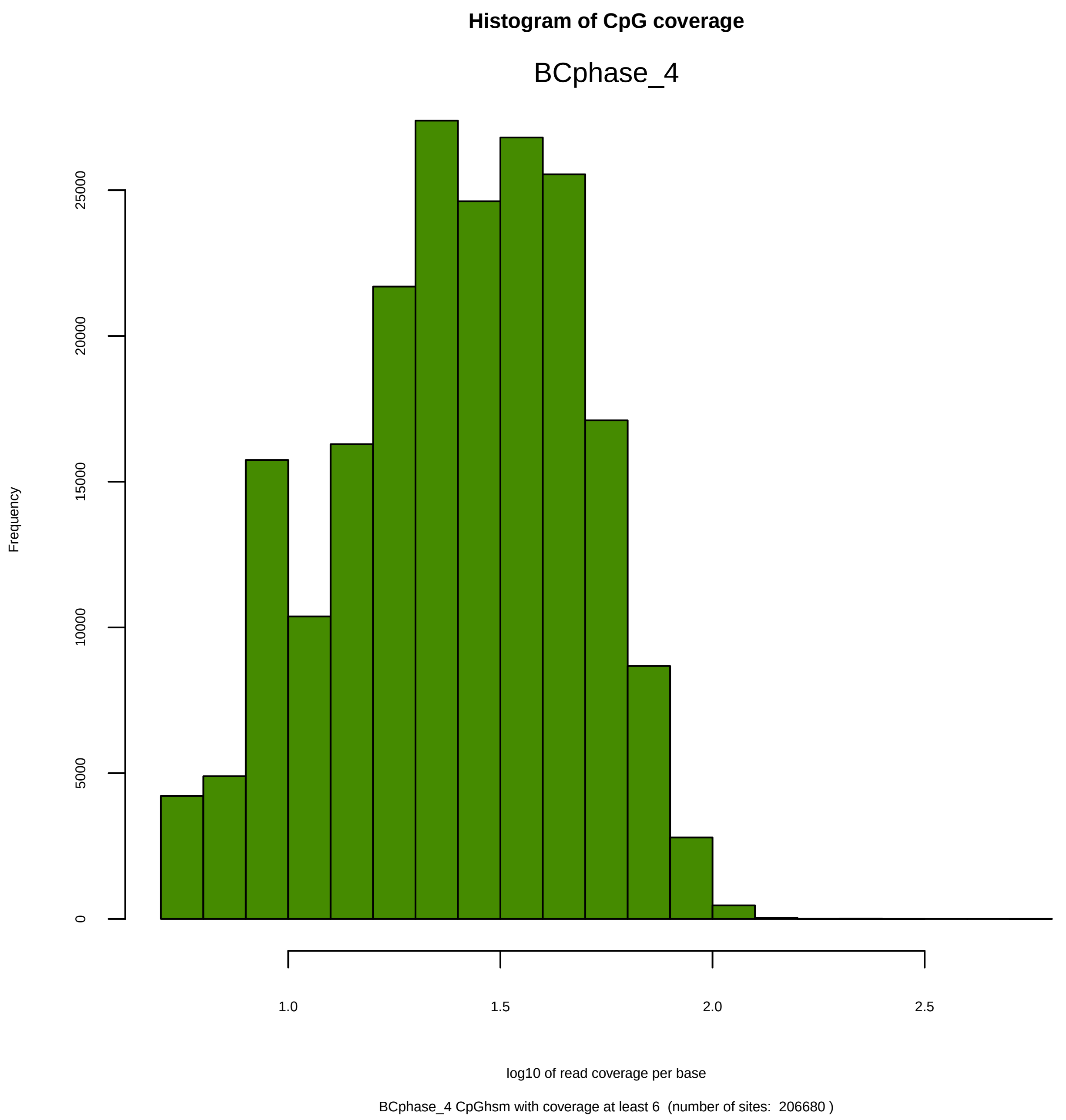

Histogram of CpG coverage
BCphase_4
Frequency
0
5000
10000
15000
20000
25000
1.0
1.5
2.0
2.5
log10 of read coverage per base
BCphase_4 CpGhsm with coverage at least 6  (number of sites:  206680 )

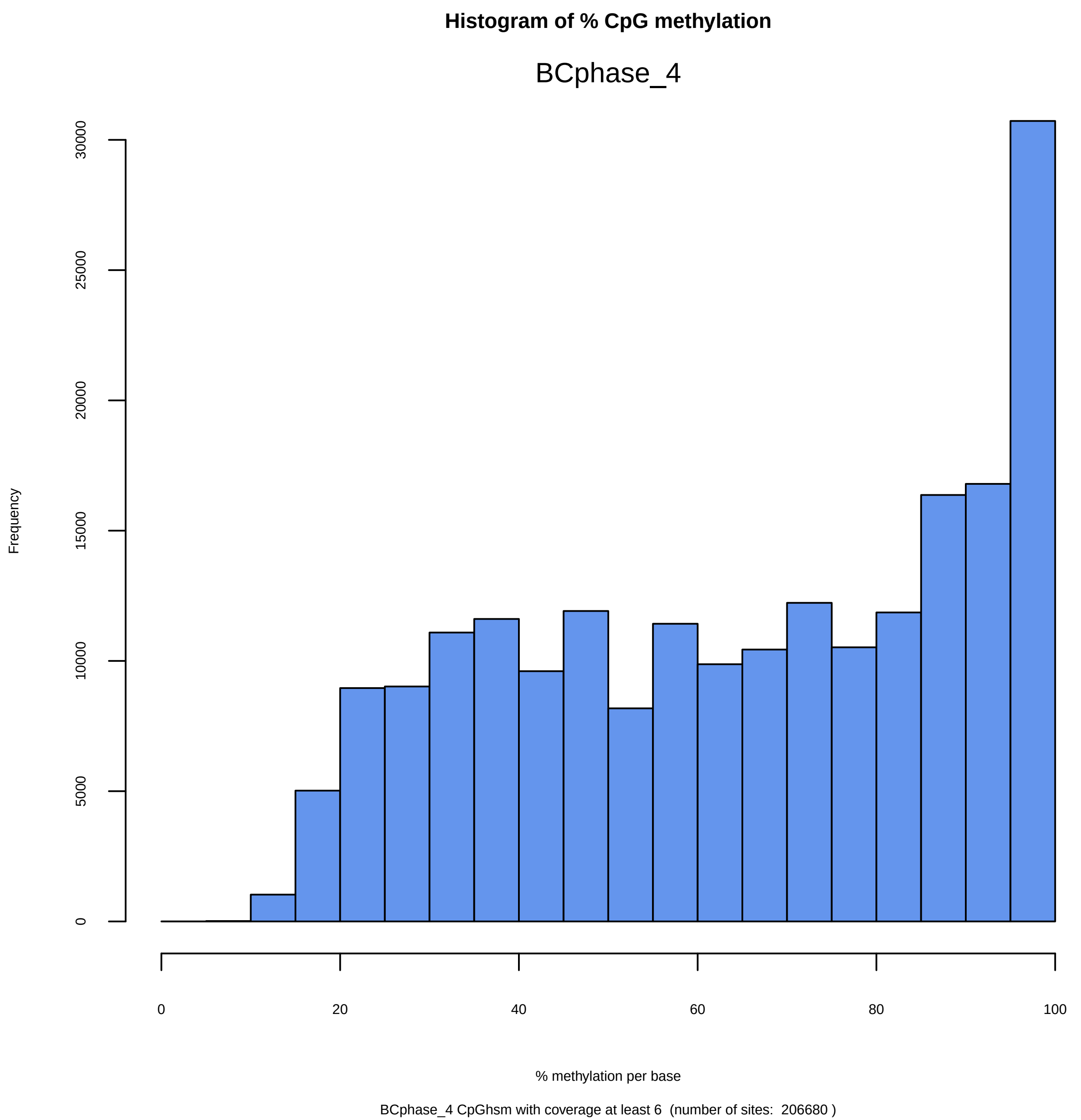
Histogram of % CpG methylation
BCphase_4
Frequency
0
5000
10000
15000
20000
25000
30000
0
20
40
60
80
100
% methylation per base
BCphase_4 CpGhsm with coverage at least 6 (number of sites: 206680 )

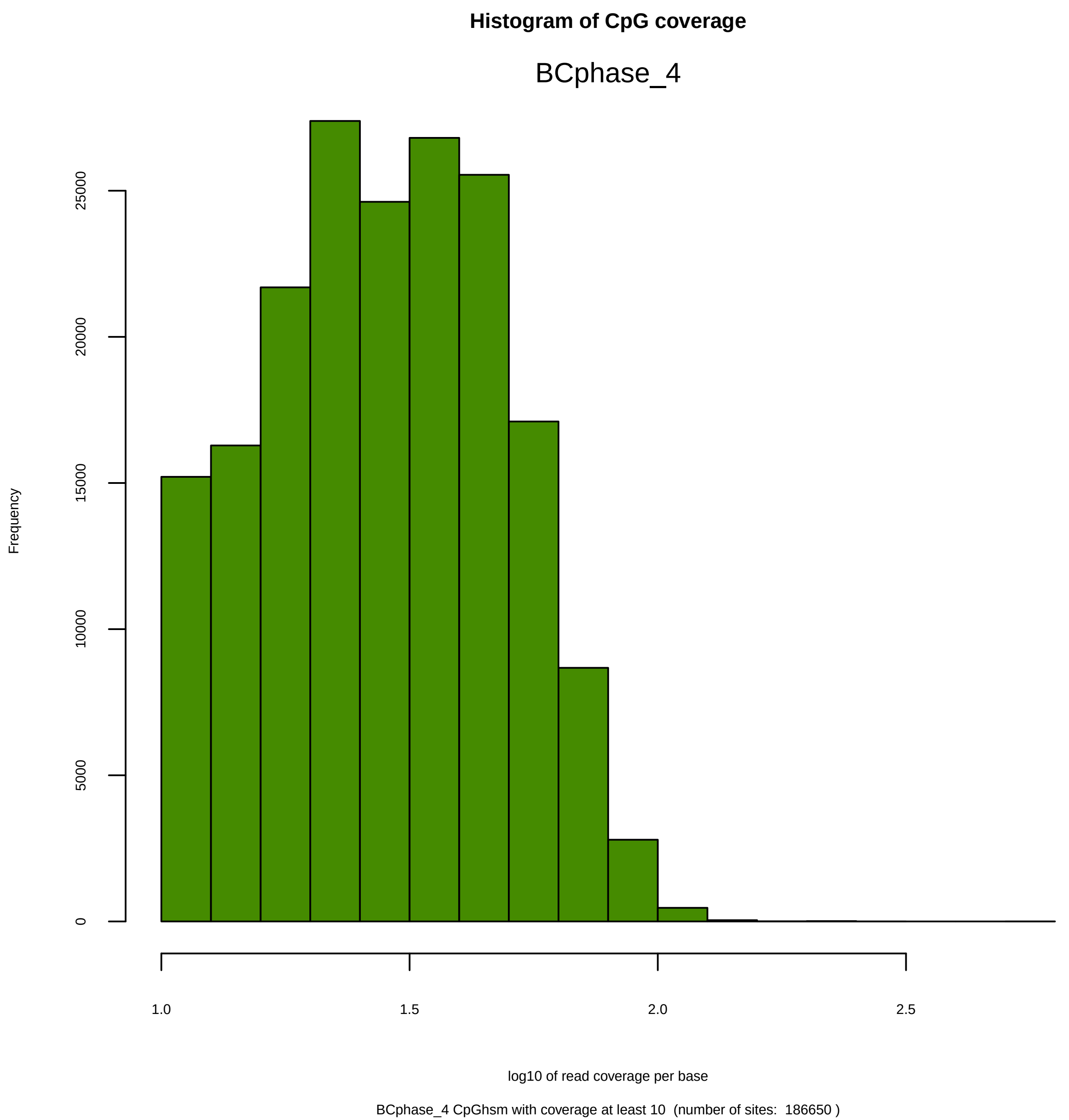
Histogram of CpG coverage
BCphase_4
Frequency
0
5000
10000
15000
20000
25000
1.0
1.5
2.0
2.5
log10 of read coverage per base
BCphase_4 CpGhsm with coverage at least 10 (number of sites: 186650 )

#### Histogram of % CpG methylation

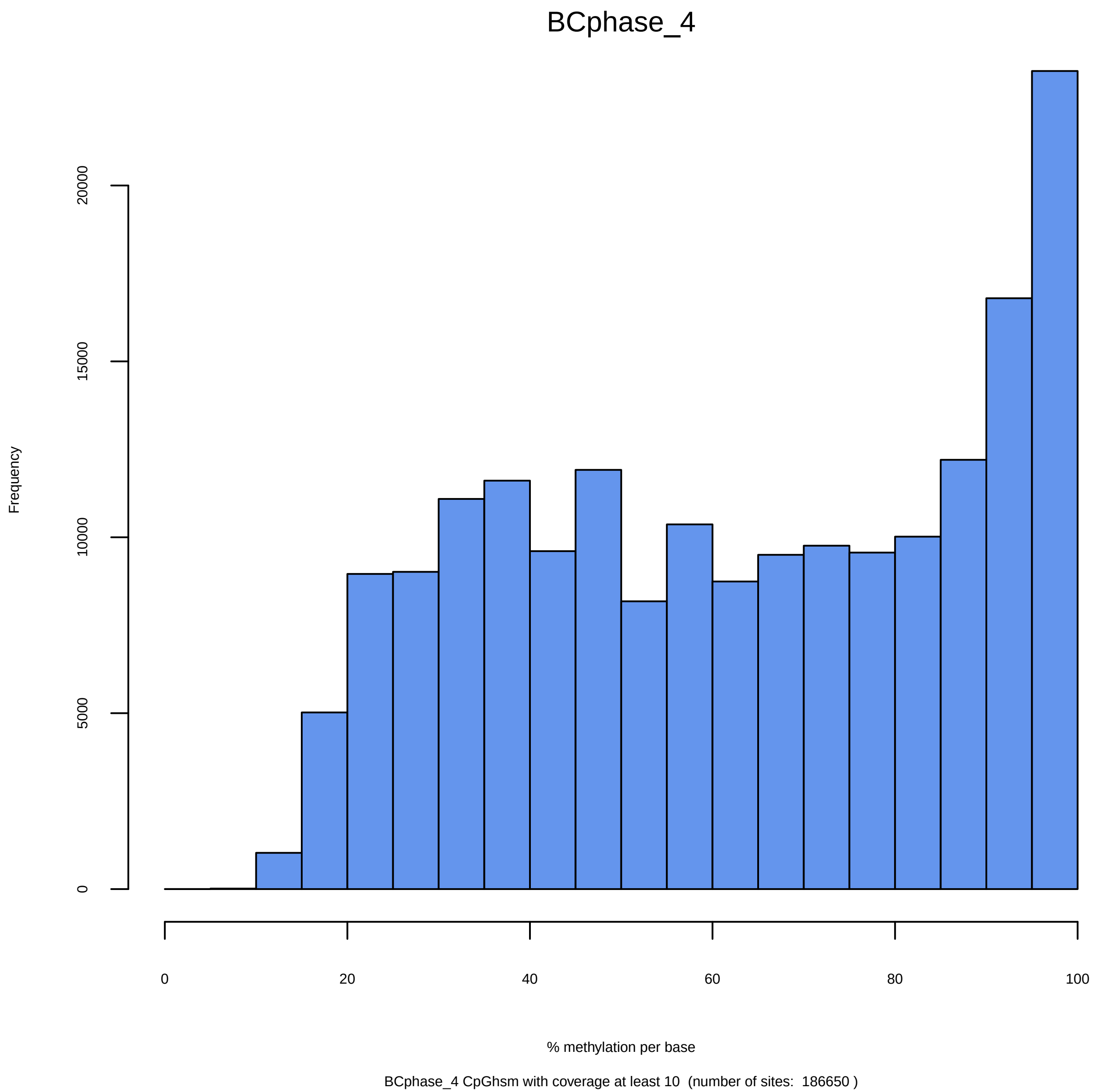

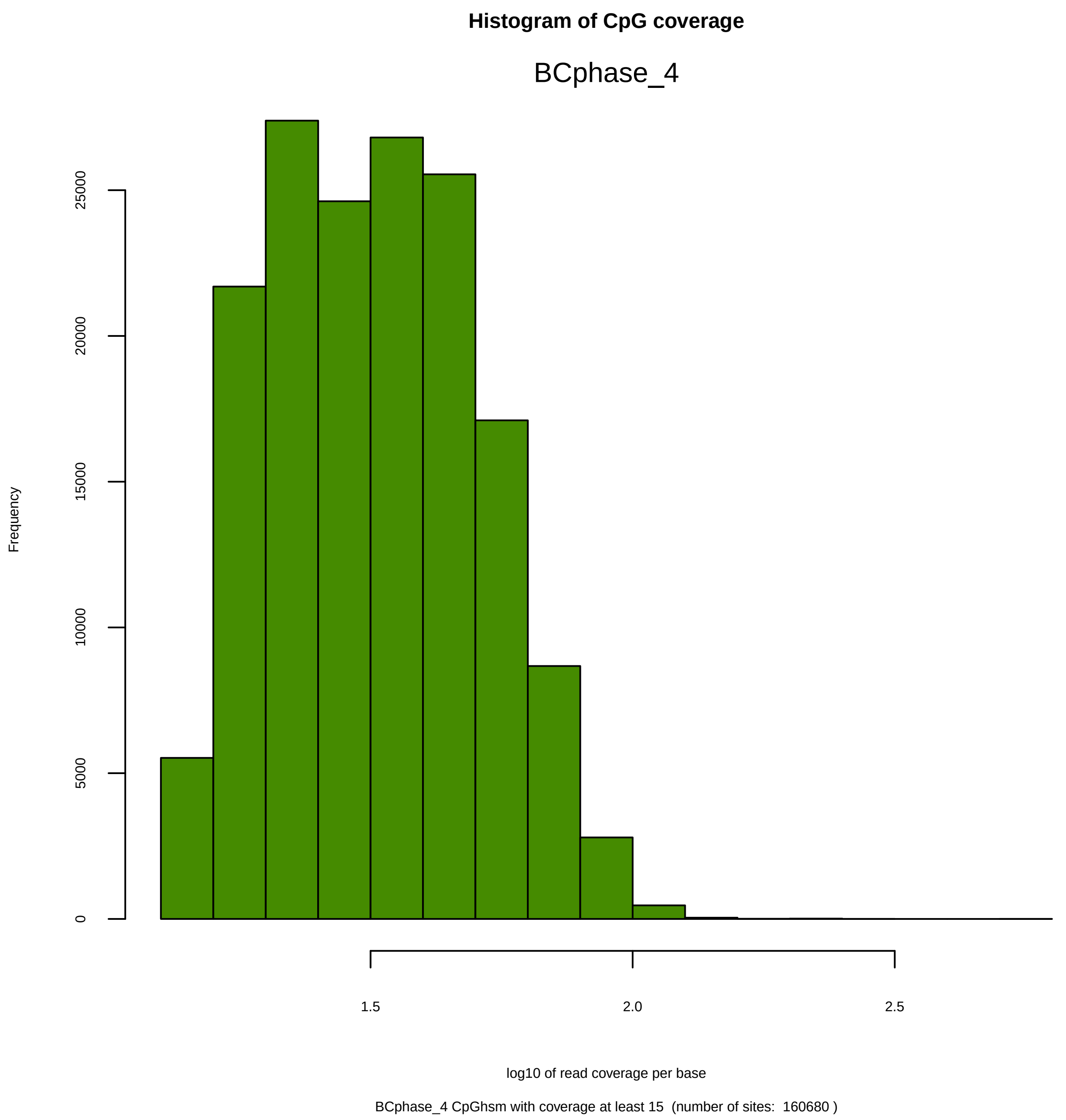

Histogram of CpG coverage
BCphase_4
Frequency
0
5000
10000
15000
20000
25000
1.5
2.0
2.5
log10 of read coverage per base
BCphase_4 CpGhsm with coverage at least 15  (number of sites:  160680 )

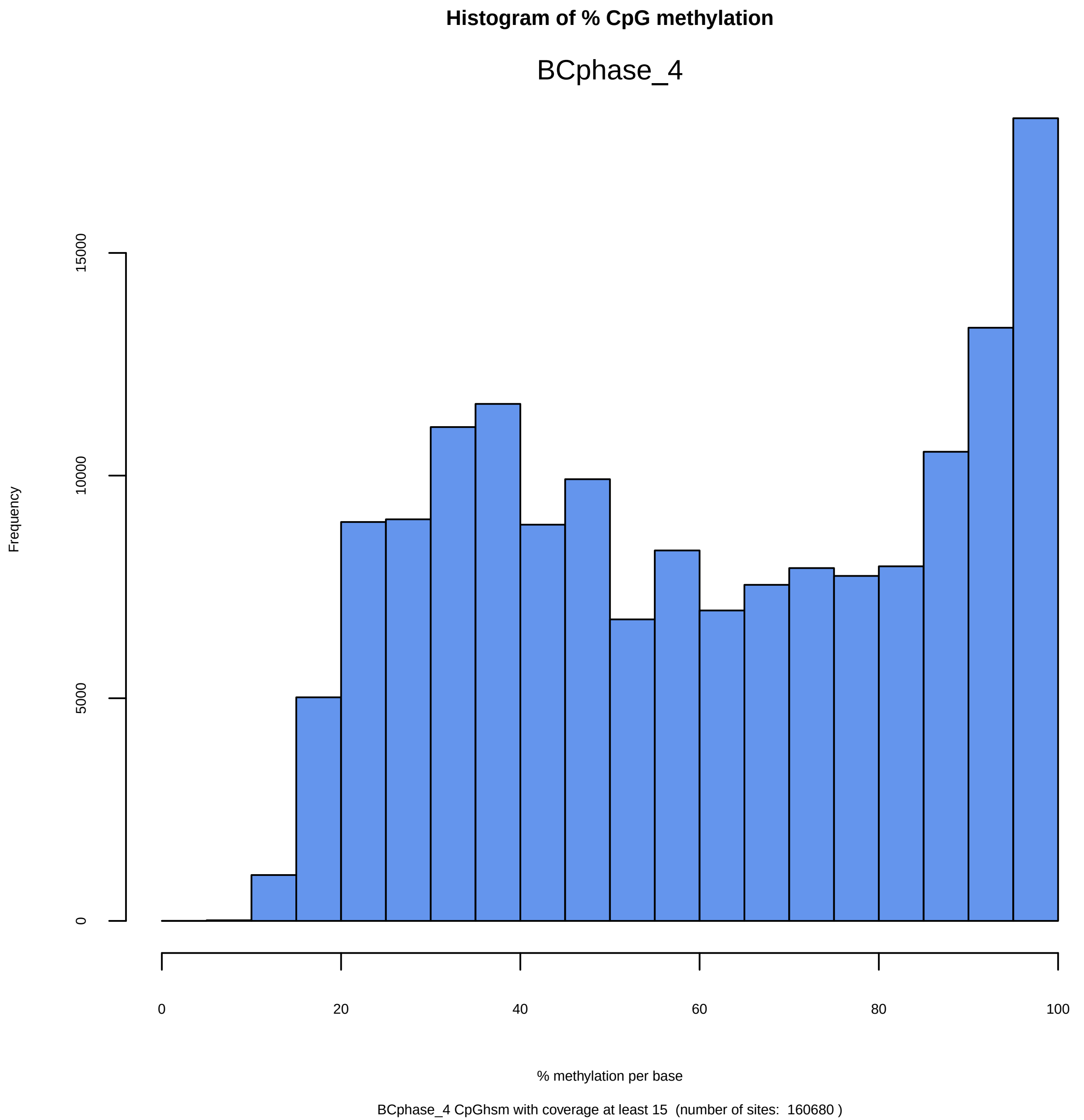

Histogram of % CpG methylation
BCphase_4
Frequency
0
5000
10000
15000
0
20
40
60
80
100
% methylation per base
BCphase_4 CpGhsm with coverage at least 15 (number of sites: 160680 )

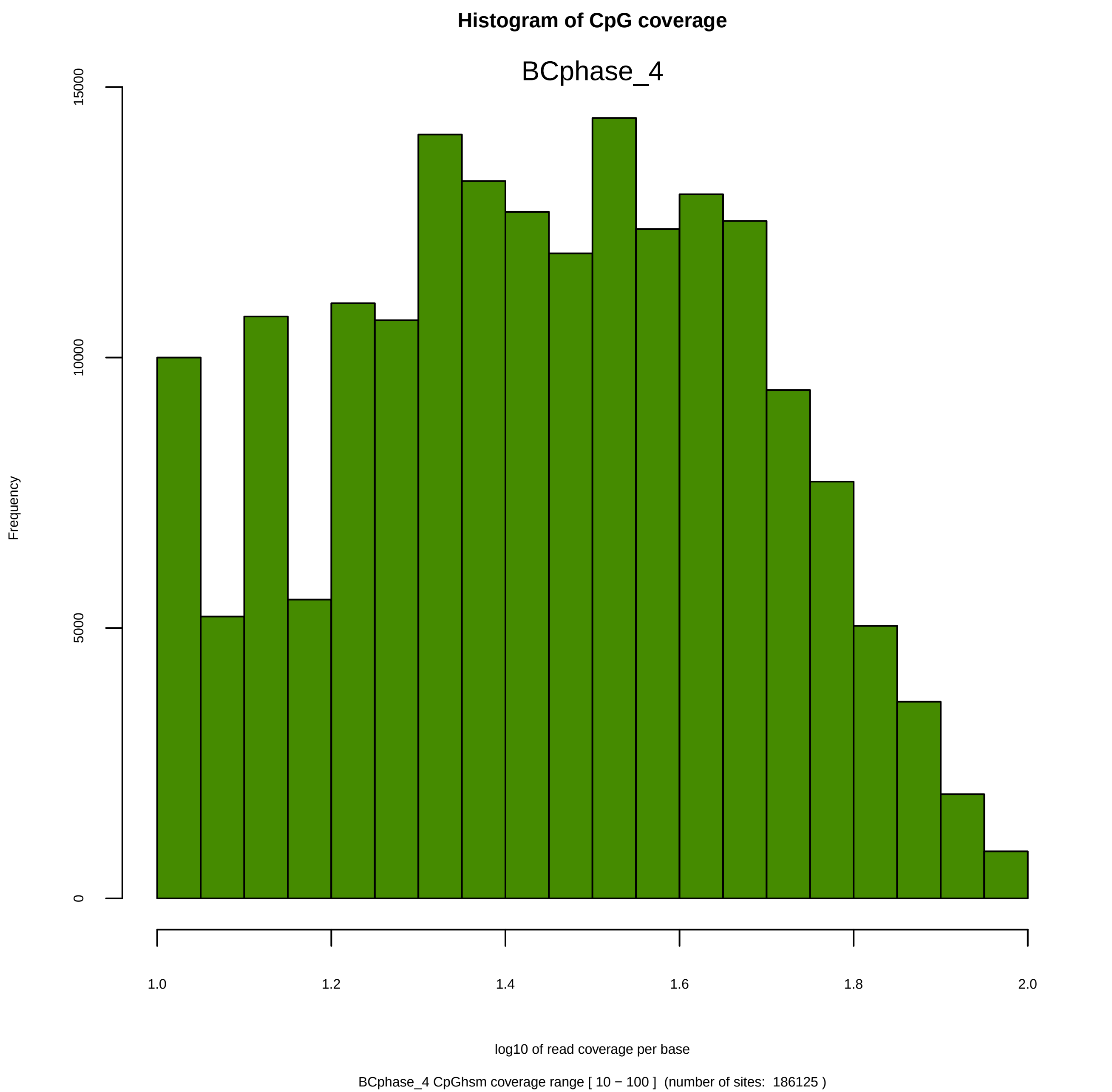

Histogram of CpG coverage
BCphase_4
Frequency
0
5000
10000
15000
1.0
1.2
1.4
1.6
1.8
2.0
log10 of read coverage per base
BCphase_4 CpGhsm coverage range [ 10 − 100 ] (number of sites: 186125 )

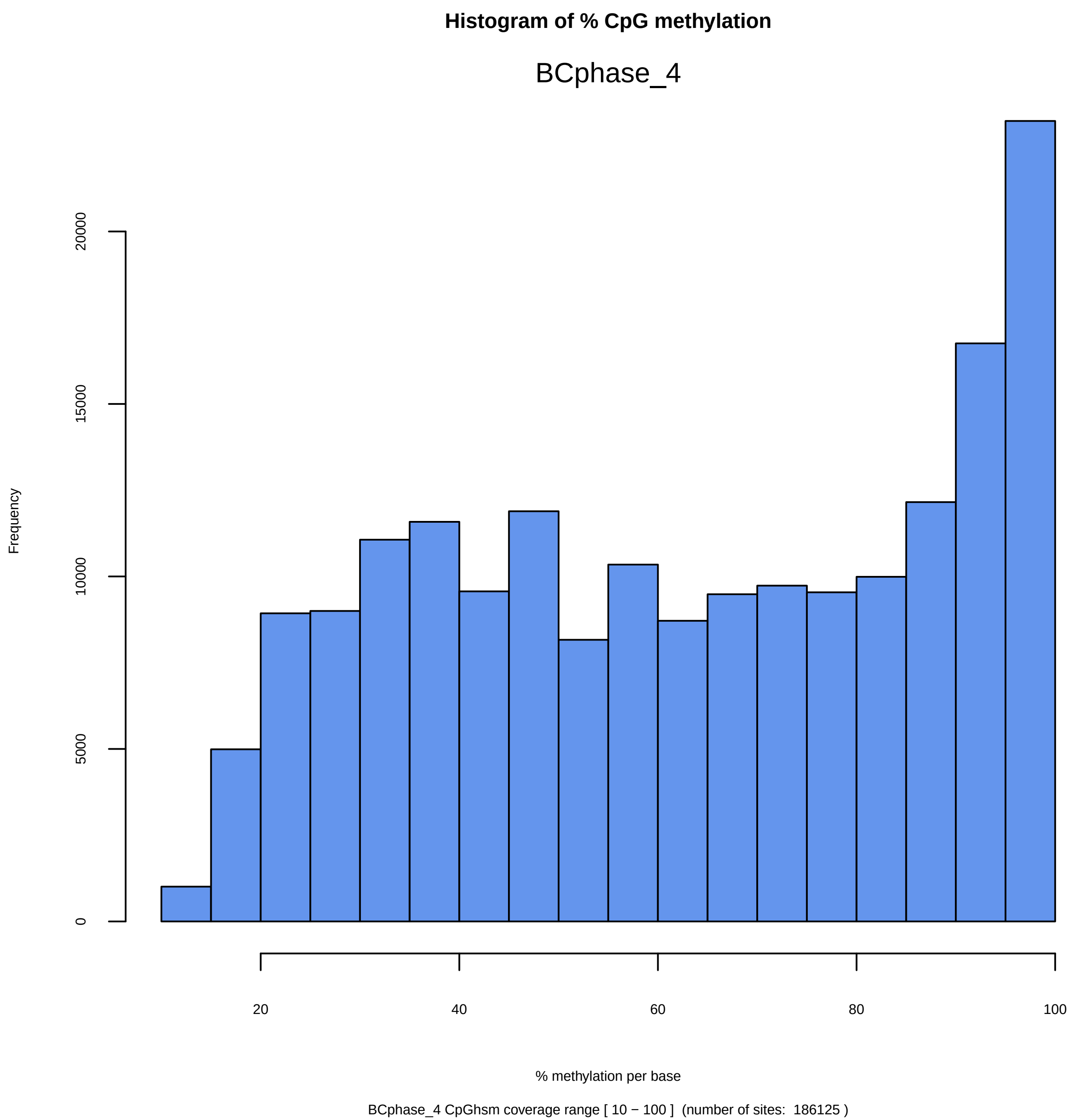

Histogram of % CpG methylation
BCphase_4
Frequency
0
5000
10000
15000
20000
20
40
60
80
100
% methylation per base
BCphase_4 CpGhsm coverage range [ 10 − 100 ] (number of sites: 186125 )

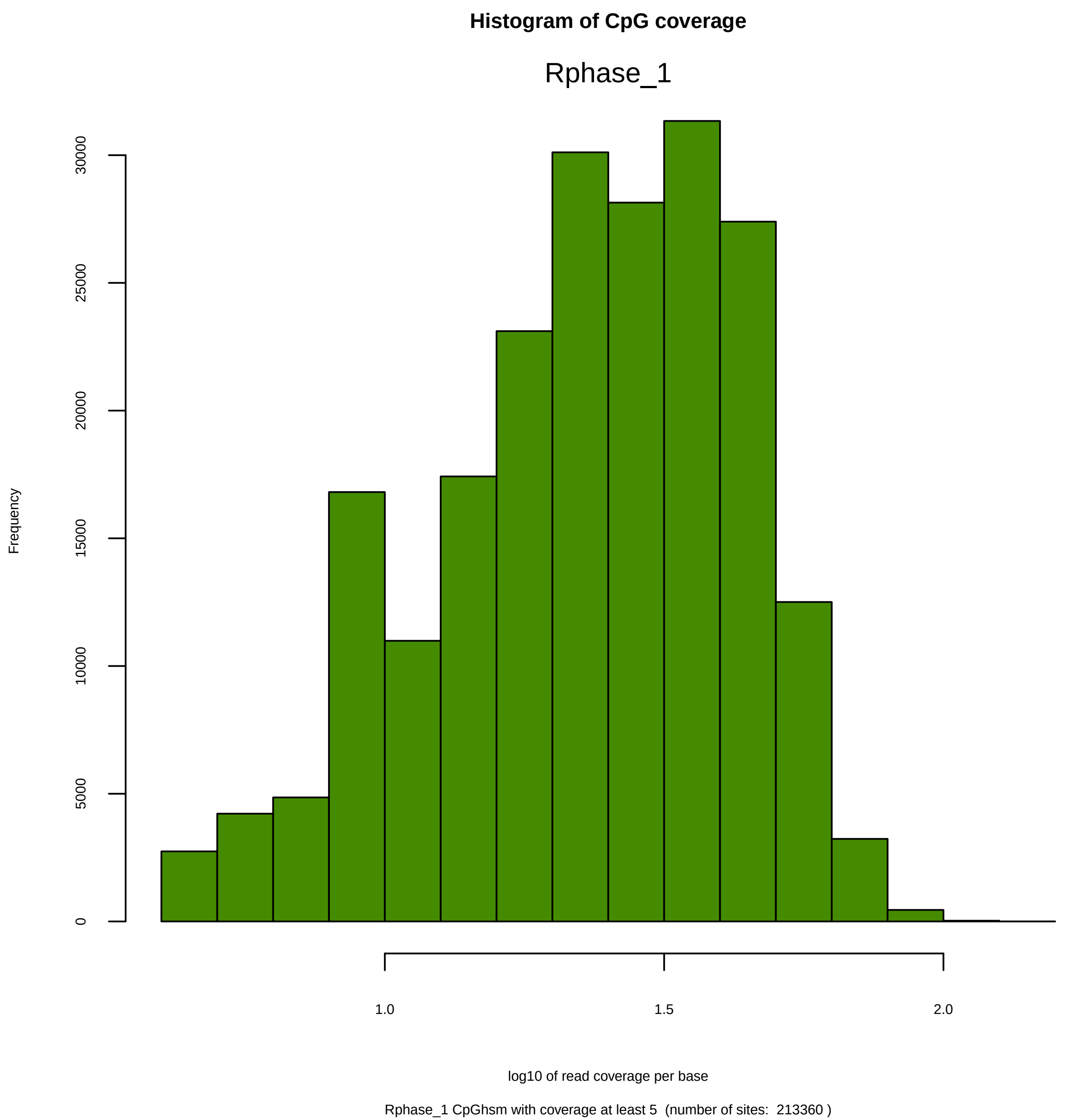

Histogram of CpG coverage
Rphase_1
Frequency
0
5000
10000
15000
20000
25000
30000
1.0
1.5
2.0
log10 of read coverage per base
Rphase_1 CpGhsm with coverage at least 5 (number of sites: 213360 )

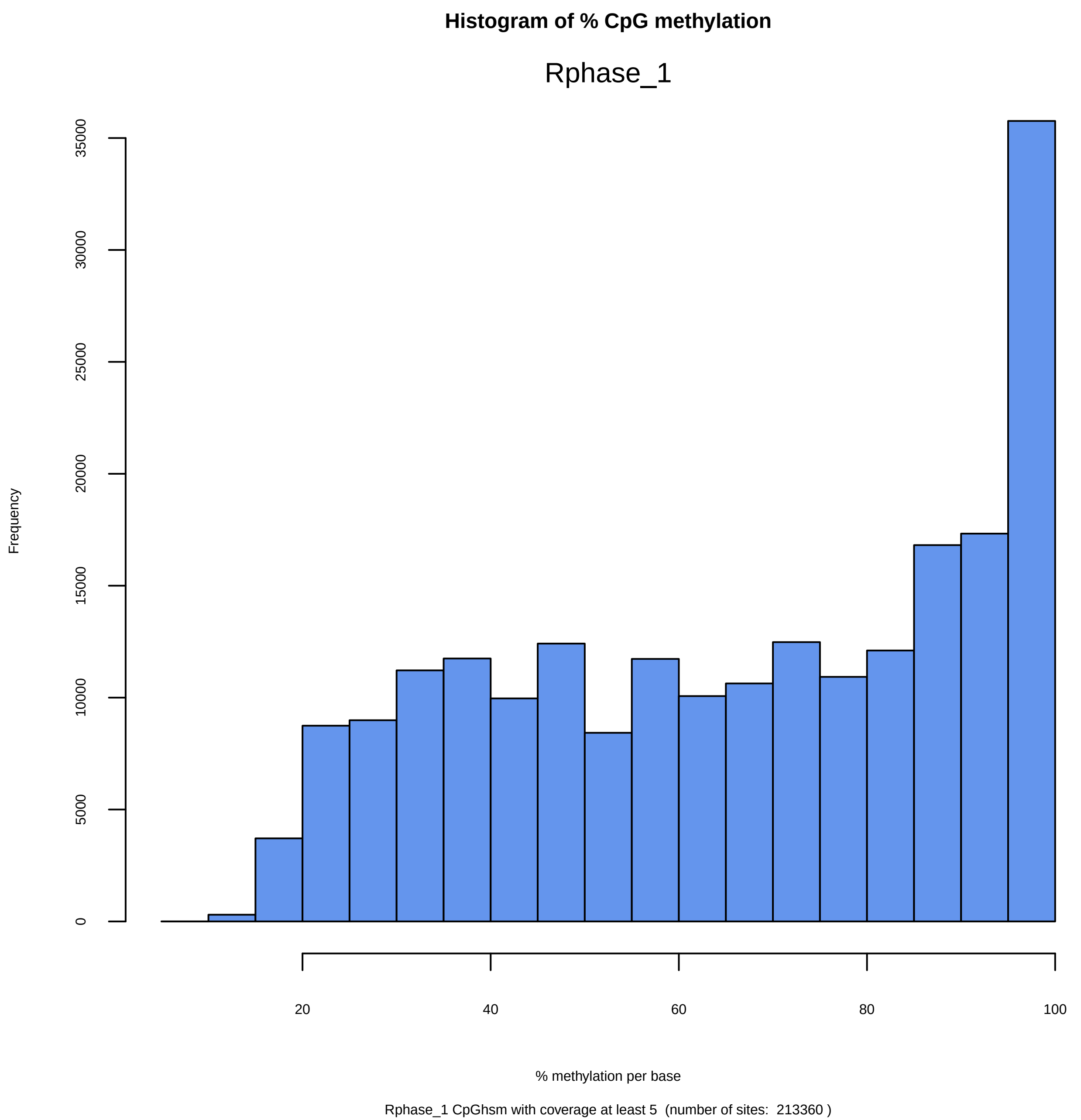
Histogram of % CpG methylation
Rphase_1
Frequency
0
5000
10000
15000
20000
25000
30000
35000
20
40
60
80
100
% methylation per base
Rphase_1 CpGhsm with coverage at least 5  (number of sites:  213360 )

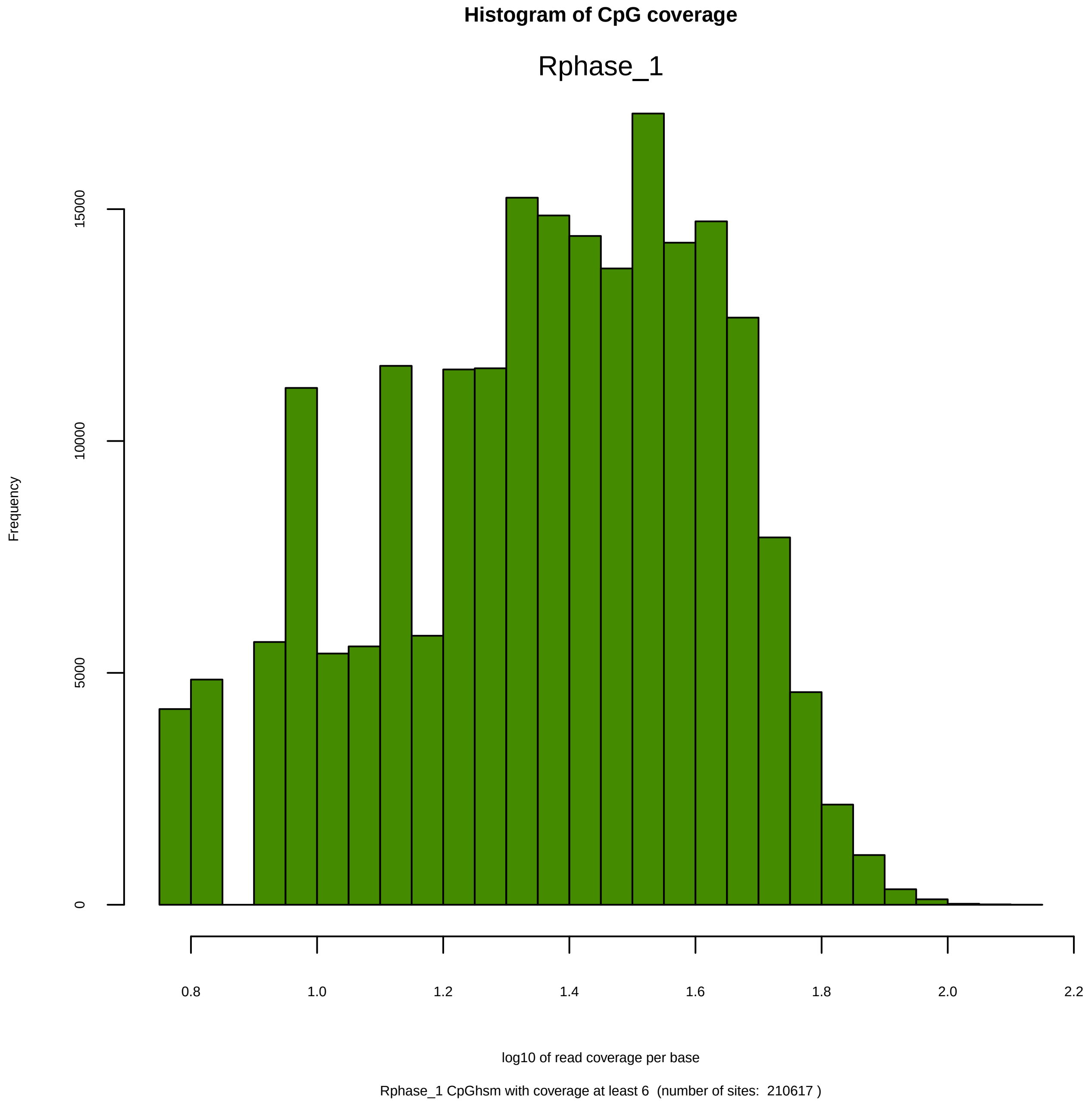
Histogram of CpG coverage
Rphase_1
Frequency
0
5000
10000
15000
0.8
1.0
1.2
1.4
1.6
1.8
2.0
2.2
log10 of read coverage per base
Rphase_1 CpGhsm with coverage at least 6 (number of sites: 210617 )

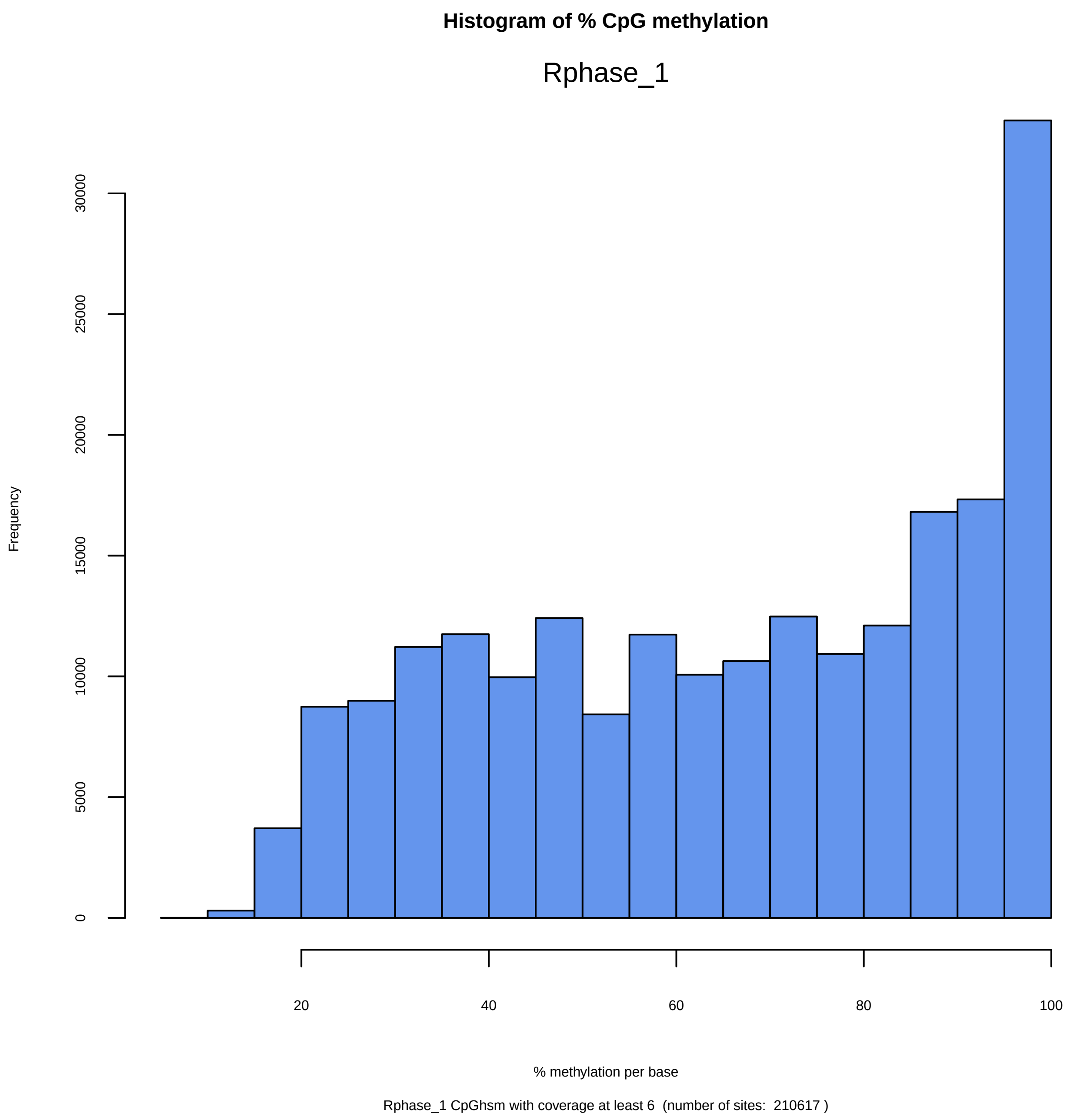

Histogram of % CpG methylation
Rphase_1
Frequency
0
5000
10000
15000
20000
25000
30000
20
40
60
80
100
% methylation per base
Rphase_1 CpGhsm with coverage at least 6  (number of sites:  210617 )

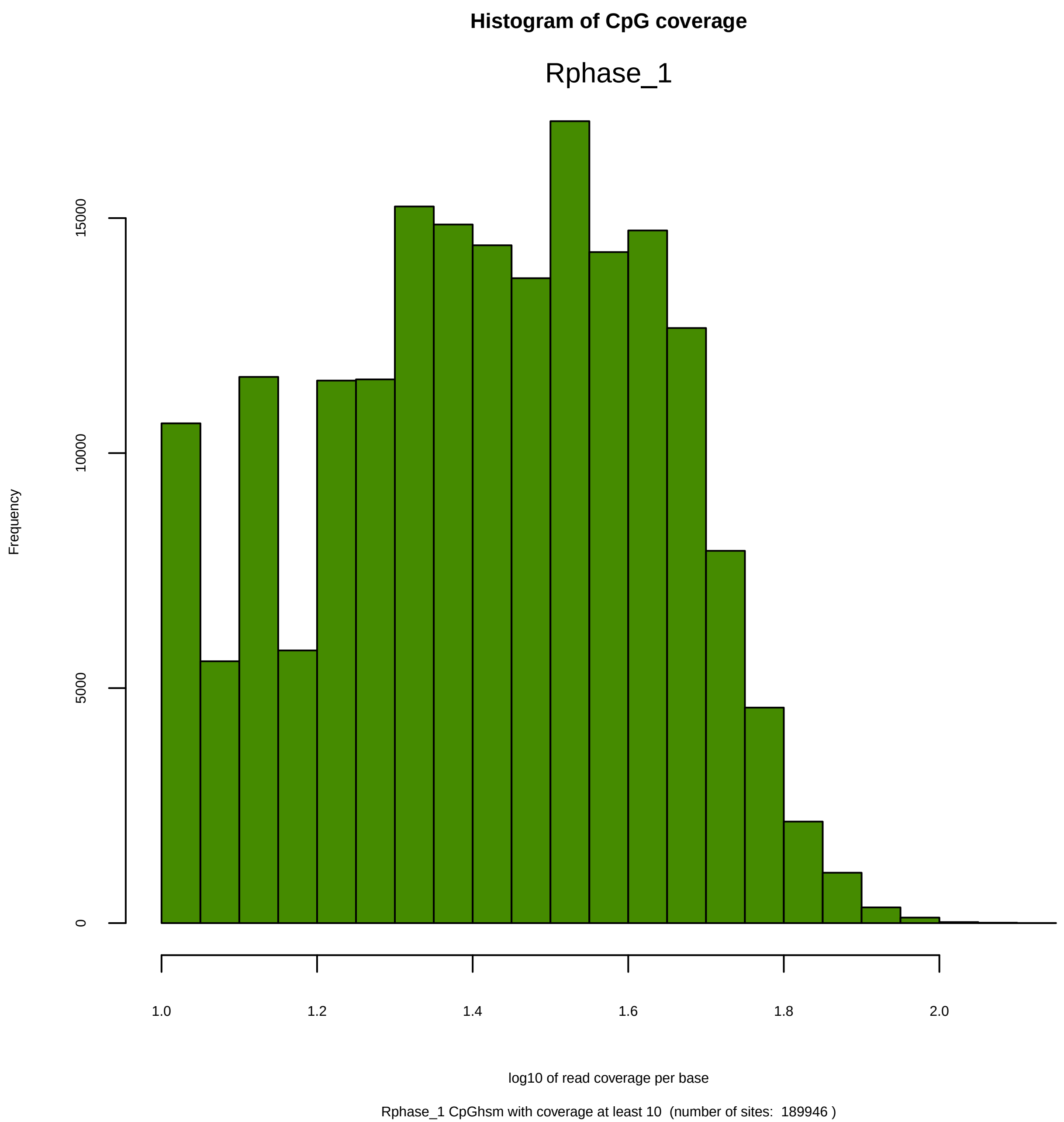
Histogram of CpG coverage
Rphase_1
Frequency
0
5000
10000
15000
1.0
1.2
1.4
1.6
1.8
2.0
log10 of read coverage per base
Rphase_1 CpGhsm with coverage at least 10 (number of sites: 189946 )

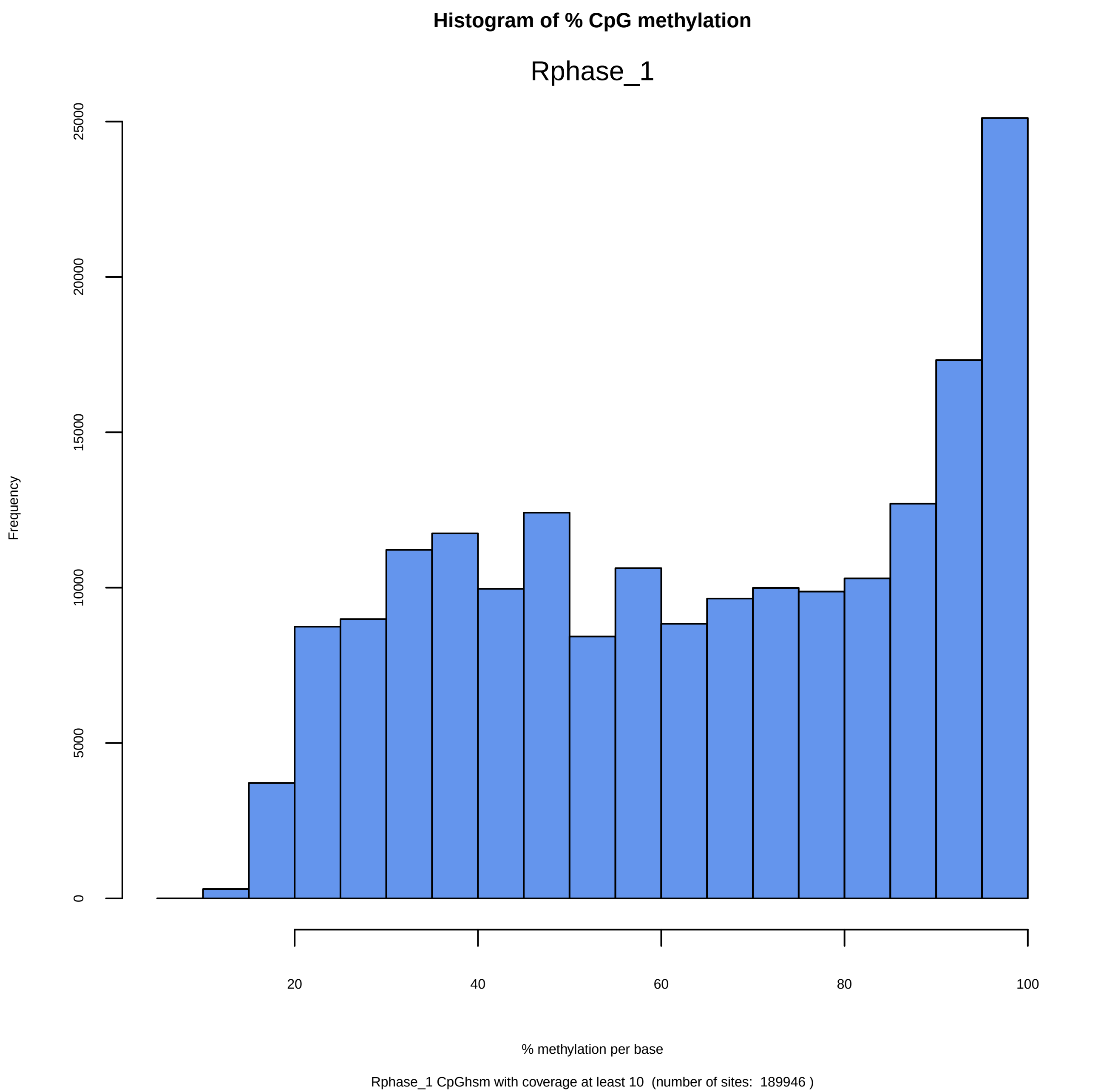

Histogram of % CpG methylation
Rphase_1
Frequency
0
5000
10000
15000
20000
25000
20
40
60
80
100
% methylation per base
Rphase_1 CpGhsm with coverage at least 10  (number of sites:  189946 )

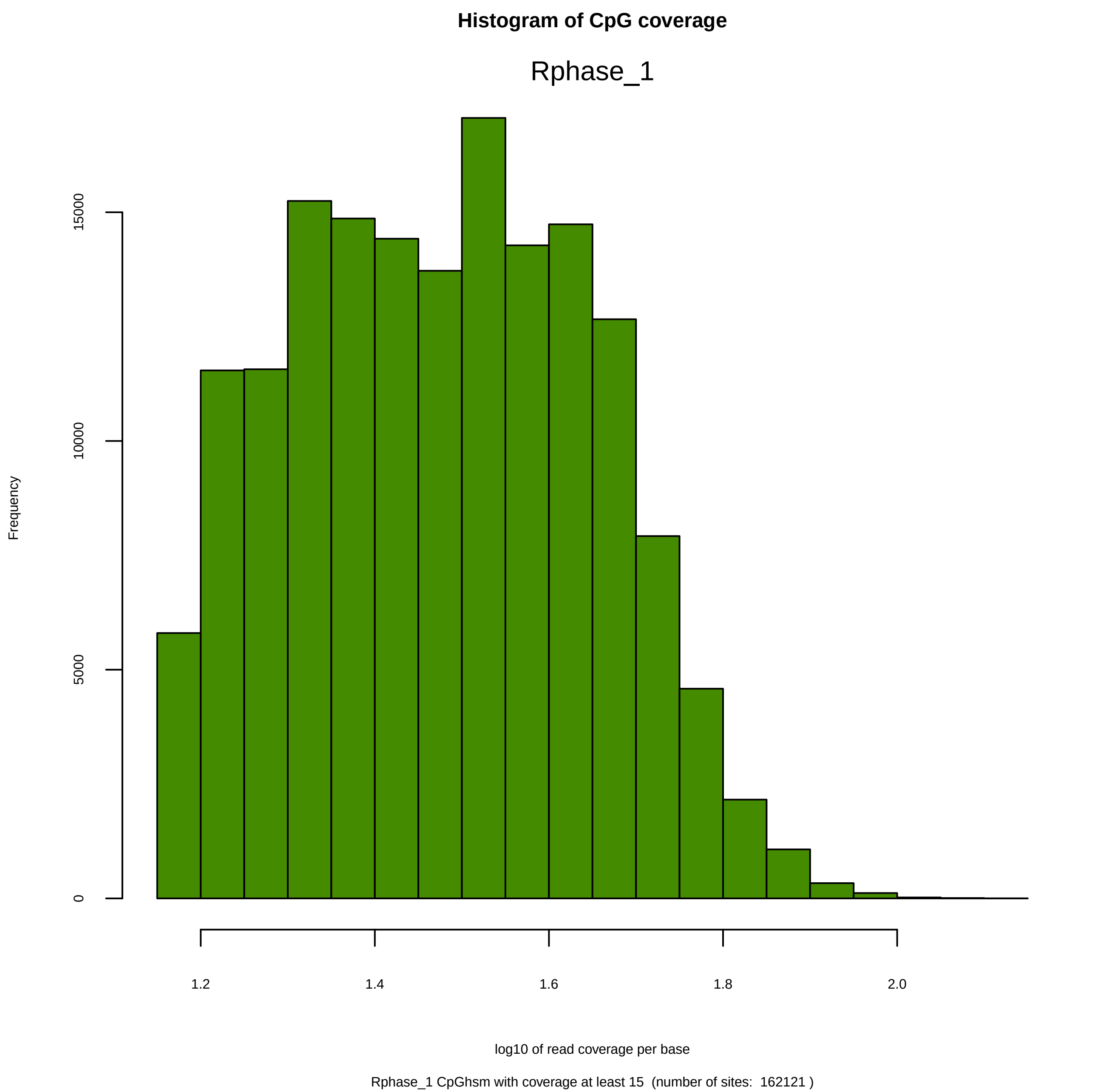
Histogram of CpG coverage
Rphase_1
Frequency
0
5000
10000
15000
1.2
1.4
1.6
1.8
2.0
log10 of read coverage per base
Rphase_1 CpGhsm with coverage at least 15  (number of sites:  162121 )

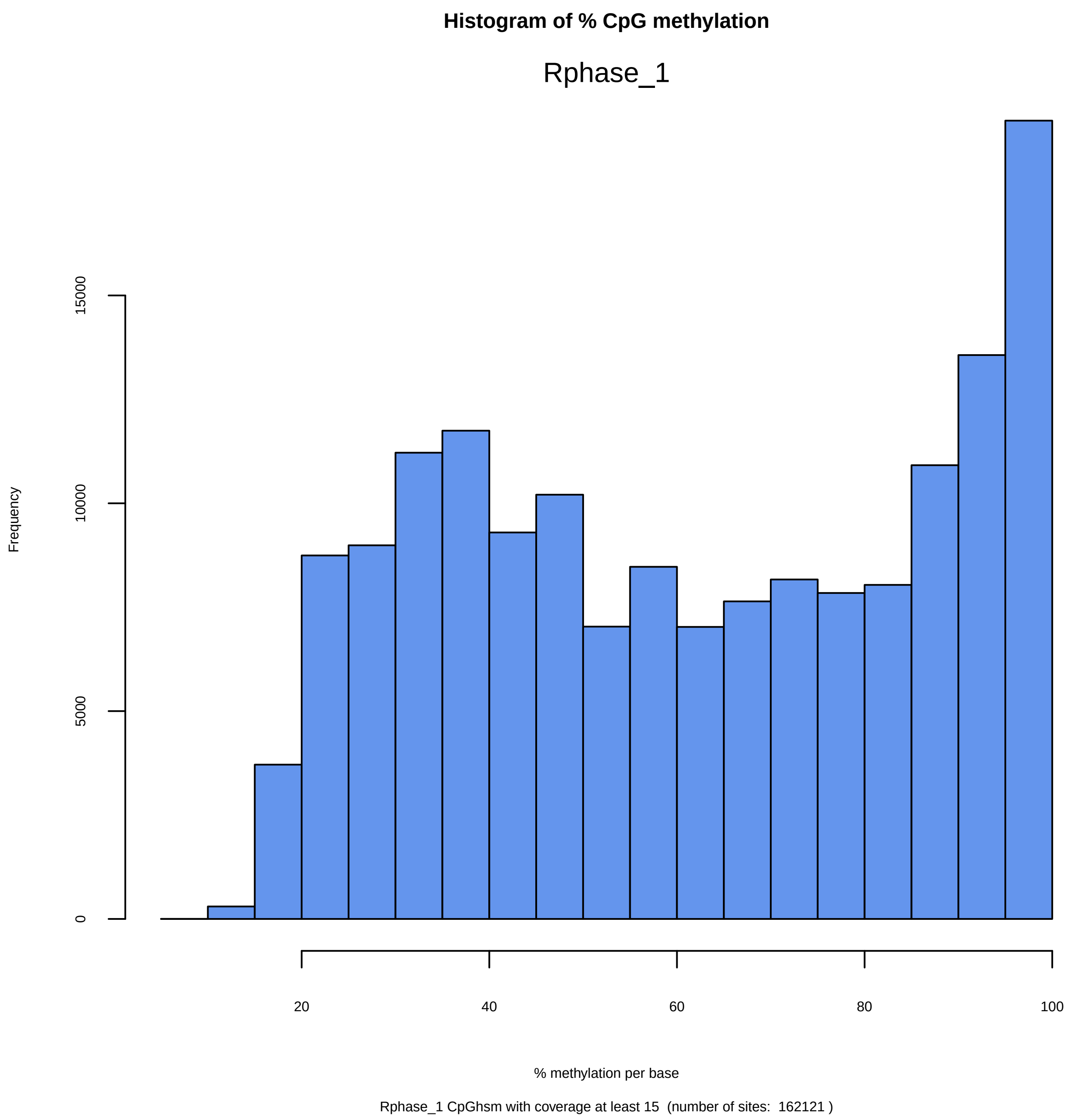

Histogram of % CpG methylation
Rphase_1
Frequency
0
5000
10000
15000
20
40
60
80
100
% methylation per base
Rphase_1 CpGhsm with coverage at least 15  (number of sites:  162121 )

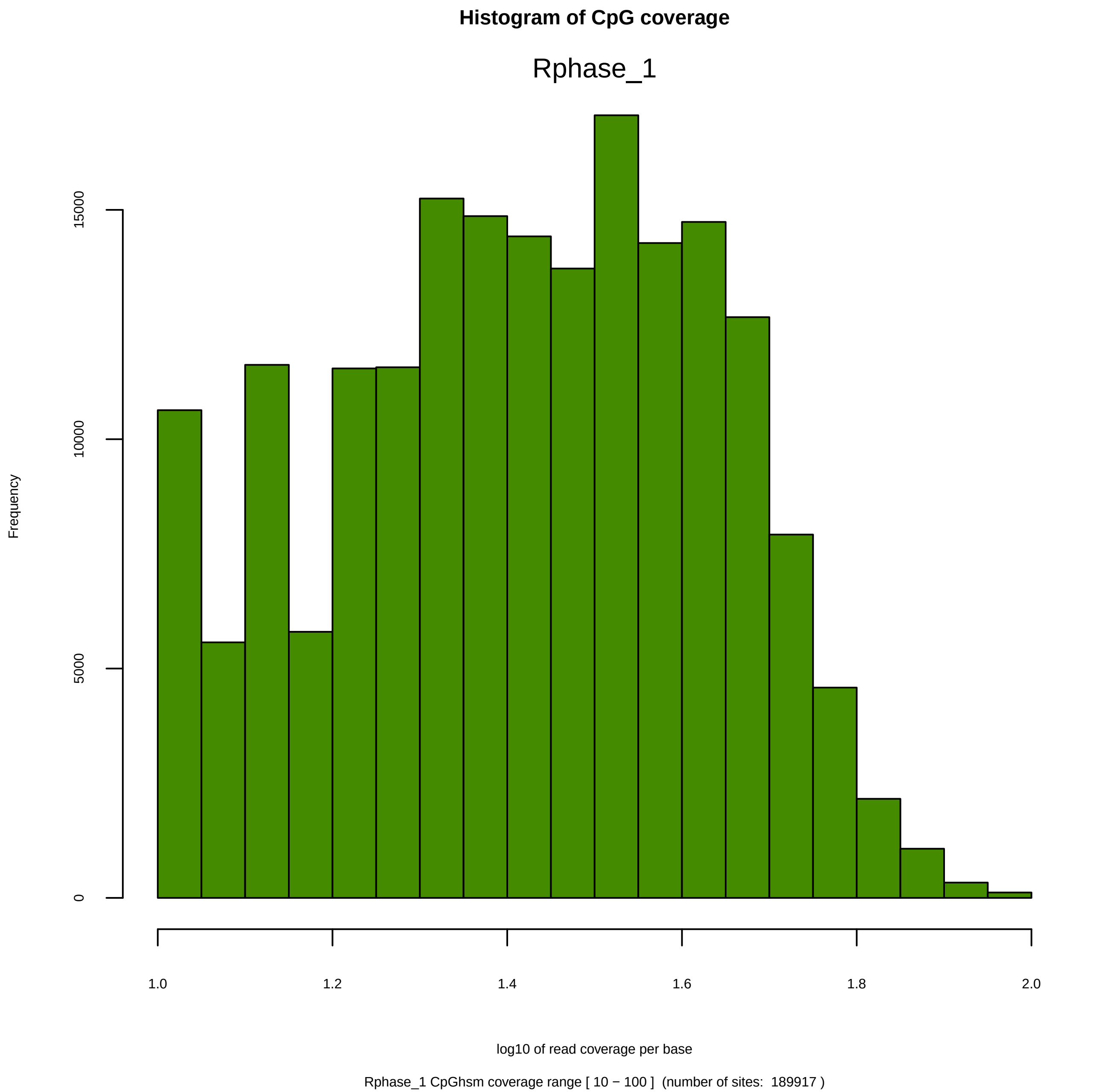

Histogram of CpG coverage
Rphase_1
Frequency
0
5000
10000
15000
1.0
1.2
1.4
1.6
1.8
2.0
log10 of read coverage per base
Rphase_1 CpGhsm coverage range [ 10 − 100 ]  (number of sites:  189917 )

### Histogram of % CpG methylation

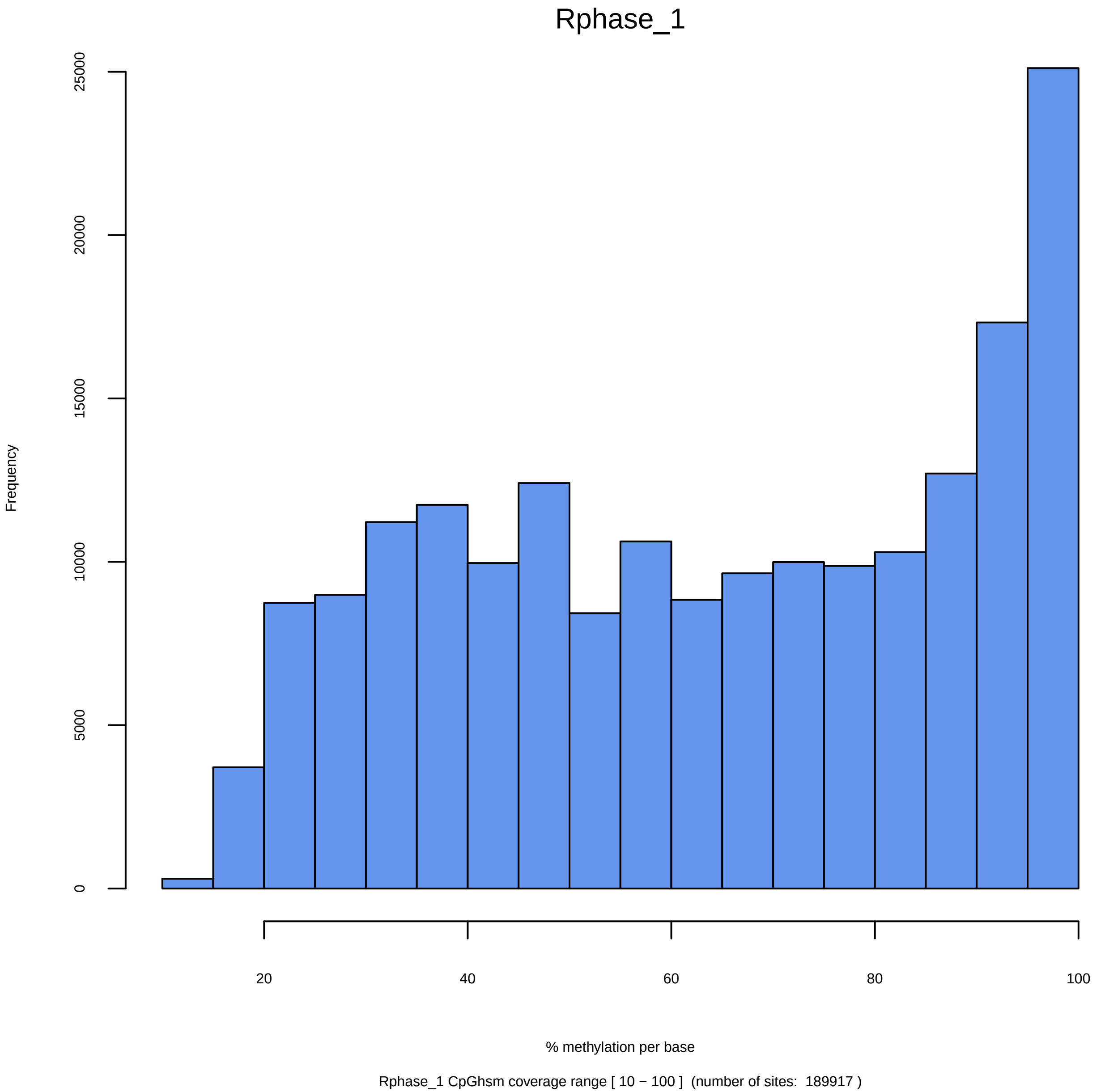

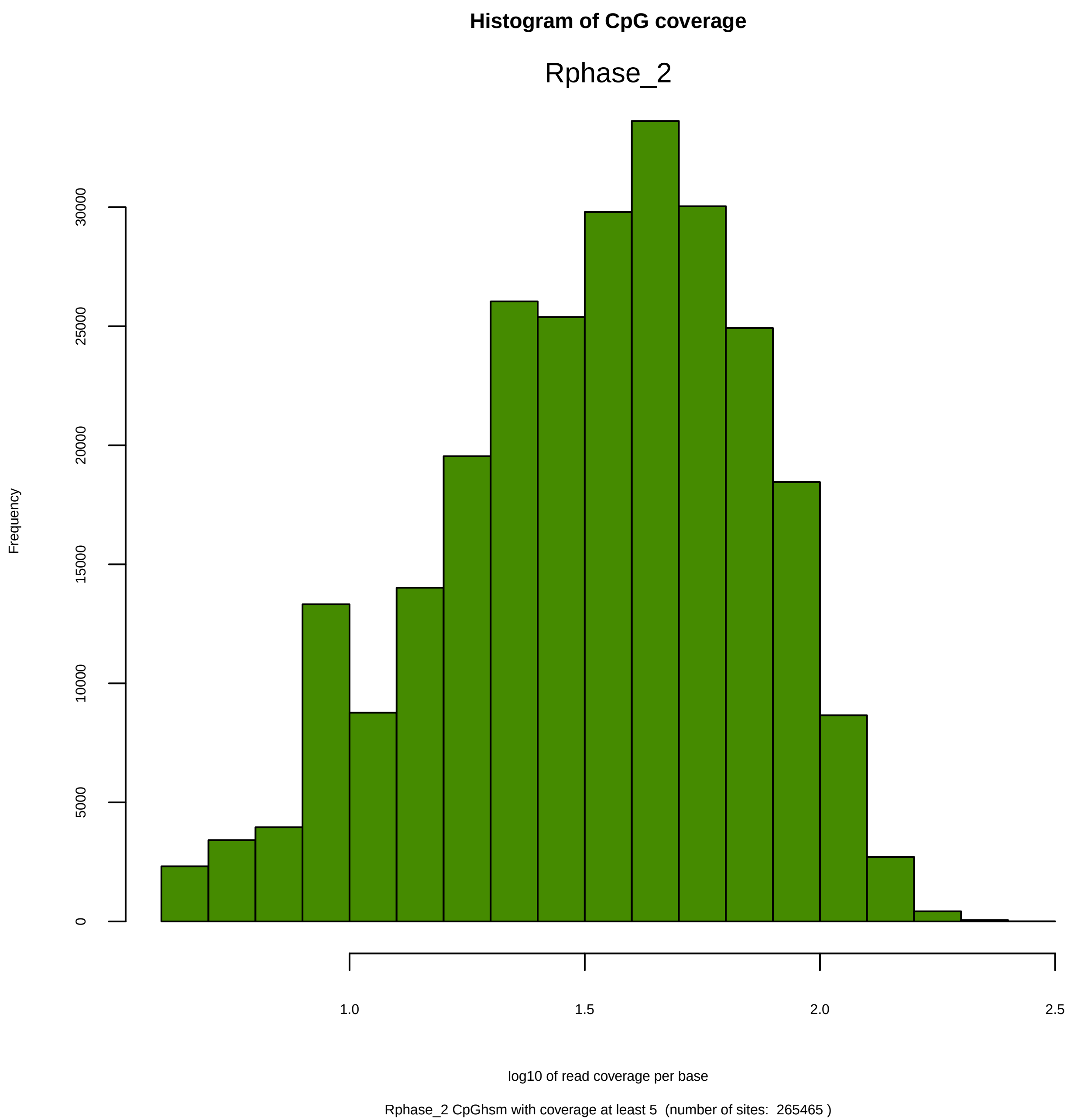

Histogram of CpG coverage
Rphase_2
Frequency
0
5000
10000
15000
20000
25000
30000
1.0
1.5
2.0
2.5
log10 of read coverage per base
Rphase_2 CpGhsm with coverage at least 5  (number of sites:  265465 )

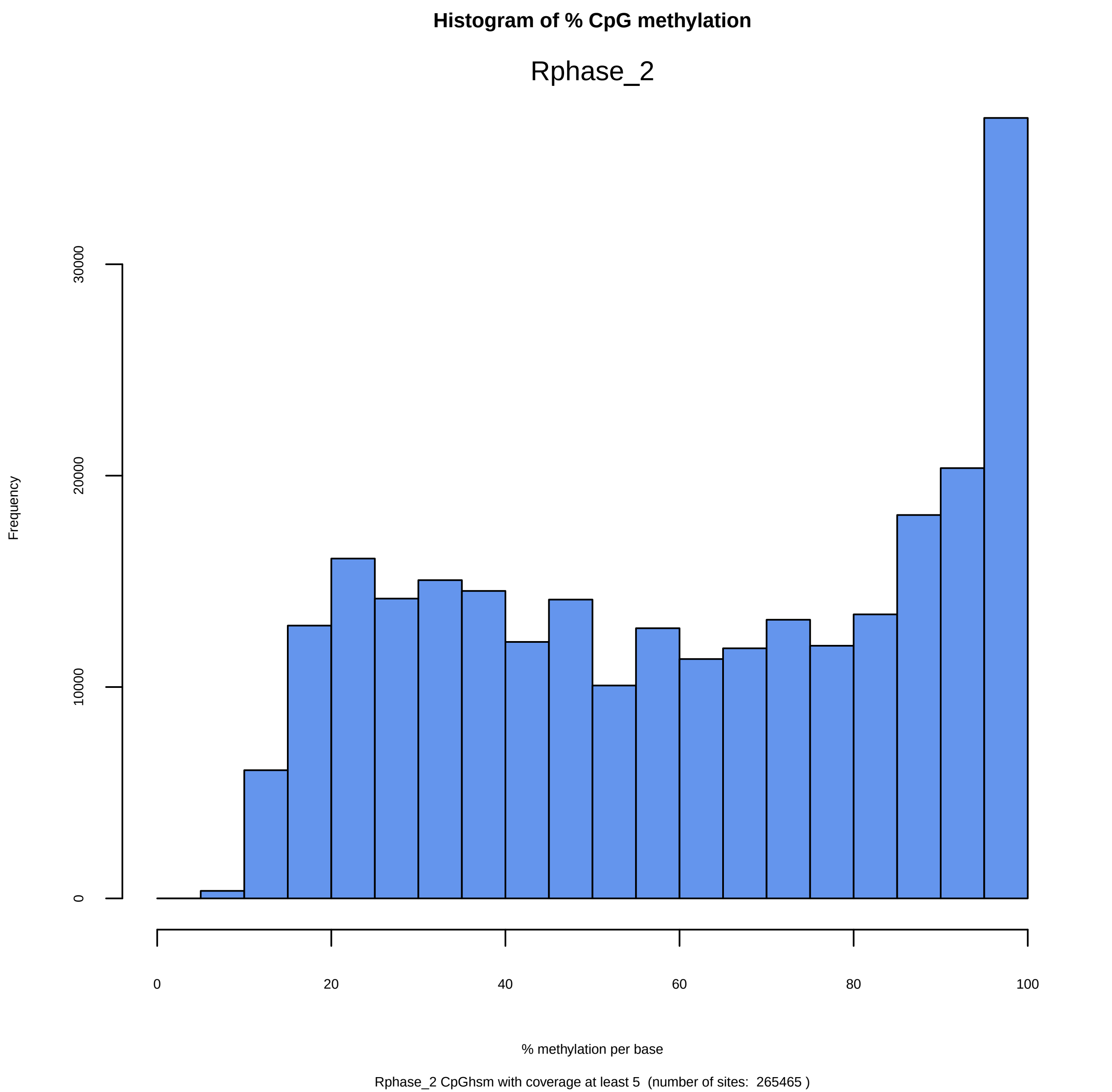
Histogram of % CpG methylation
Rphase_2
Frequency
0
10000
20000
30000
0
20
40
60
80
100
% methylation per base
Rphase_2 CpGhsm with coverage at least 5 (number of sites: 265465 )

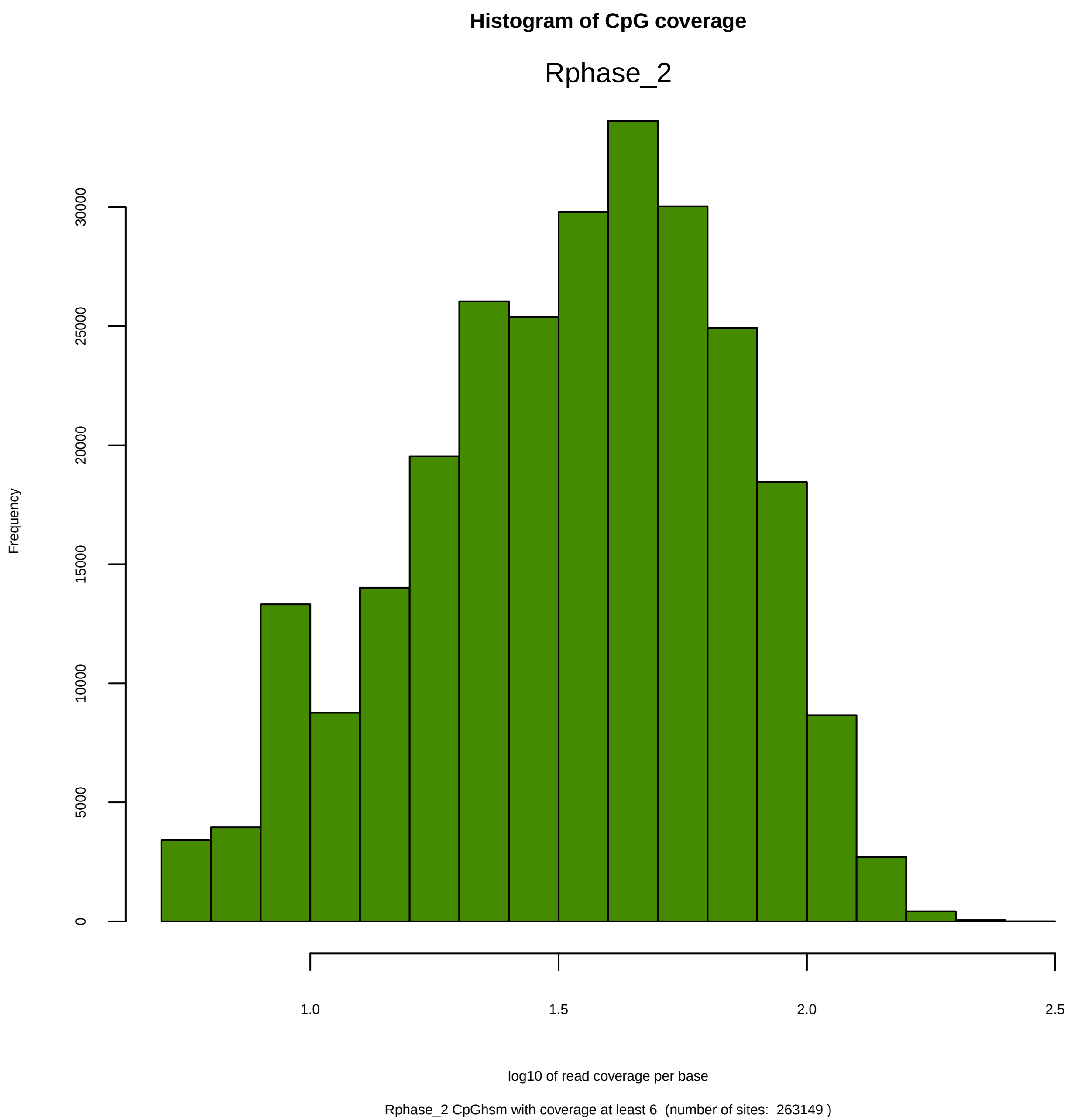
Histogram of CpG coverage
Rphase_2
Frequency
0
5000
10000
15000
20000
25000
30000
1.0
1.5
2.0
2.5
log10 of read coverage per base
Rphase_2 CpGhsm with coverage at least 6 (number of sites: 263149 )

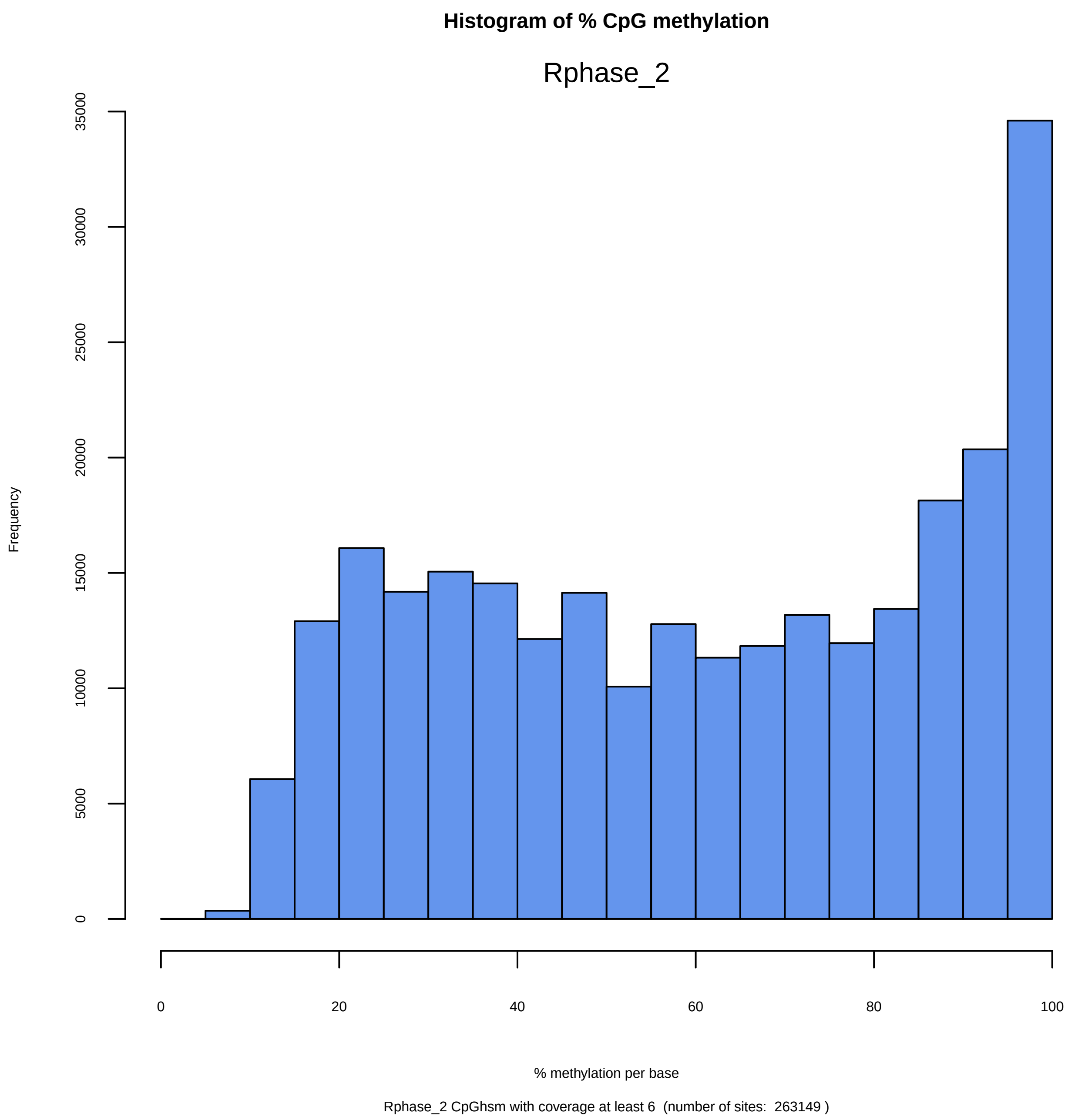
Histogram of % CpG methylation
Rphase_2
Frequency
0
5000
10000
15000
20000
25000
30000
35000
0
20
40
60
80
100
% methylation per base
Rphase_2 CpGhsm with coverage at least 6 (number of sites: 263149 )

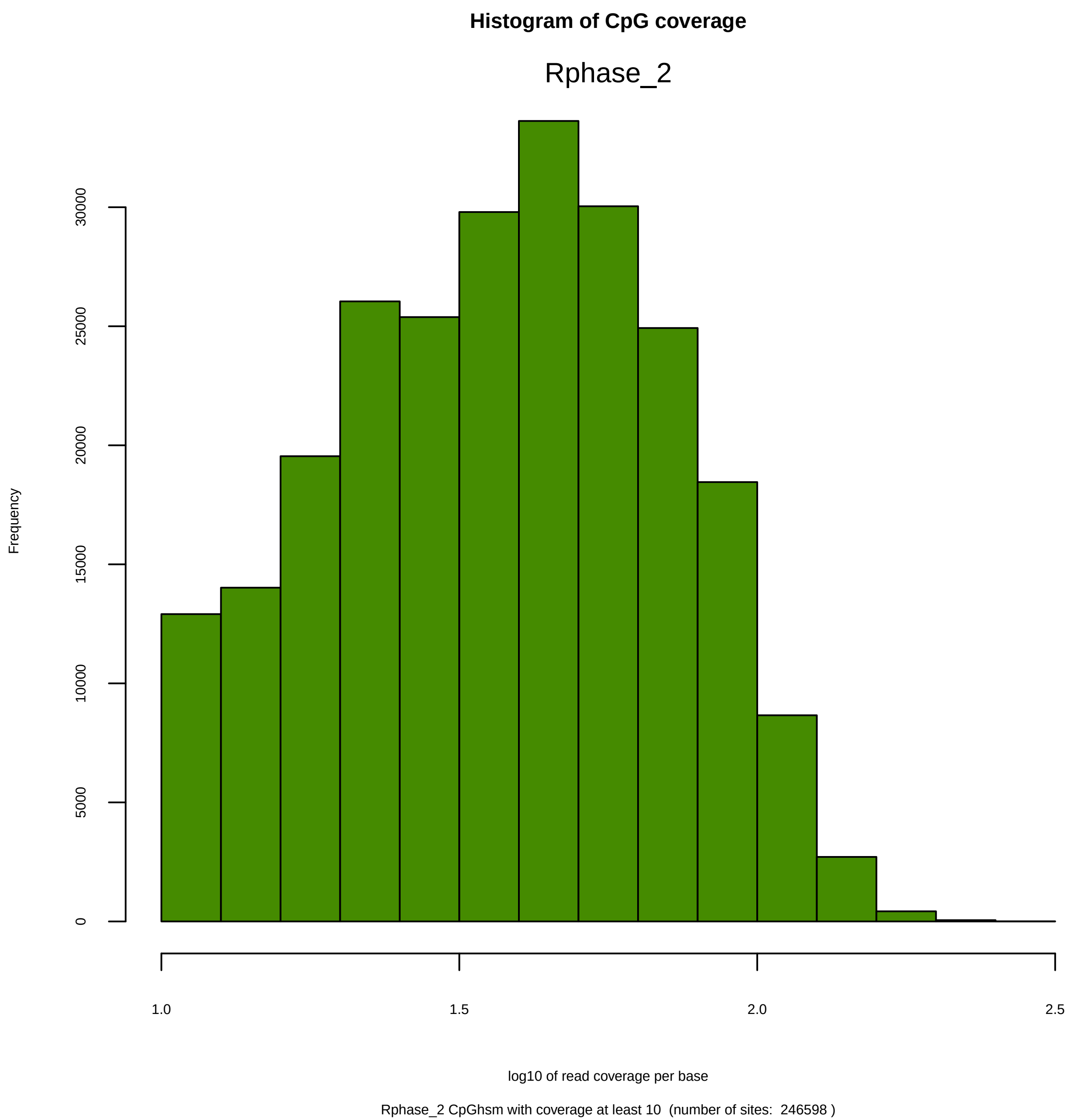
Histogram of CpG coverage
Rphase_2
Frequency
0
5000
10000
15000
20000
25000
30000
1.0
1.5
2.0
2.5
log10 of read coverage per base
Rphase_2 CpGhsm with coverage at least 10 (number of sites: 246598 )

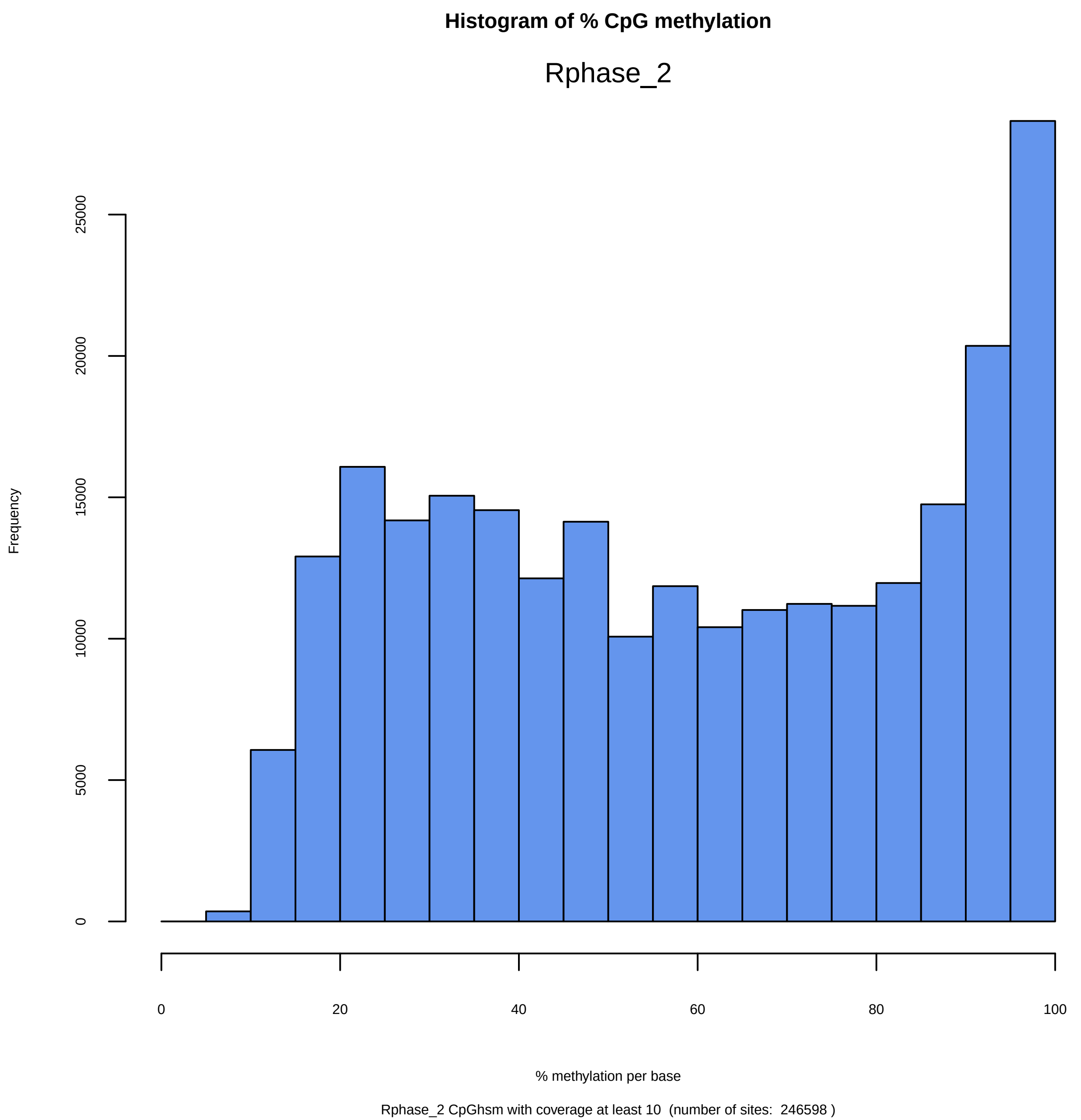

Histogram of % CpG methylation
Rphase_2
Frequency
0
5000
10000
15000
20000
25000
0
20
40
60
80
100
% methylation per base
Rphase_2 CpGhsm with coverage at least 10 (number of sites: 246598 )

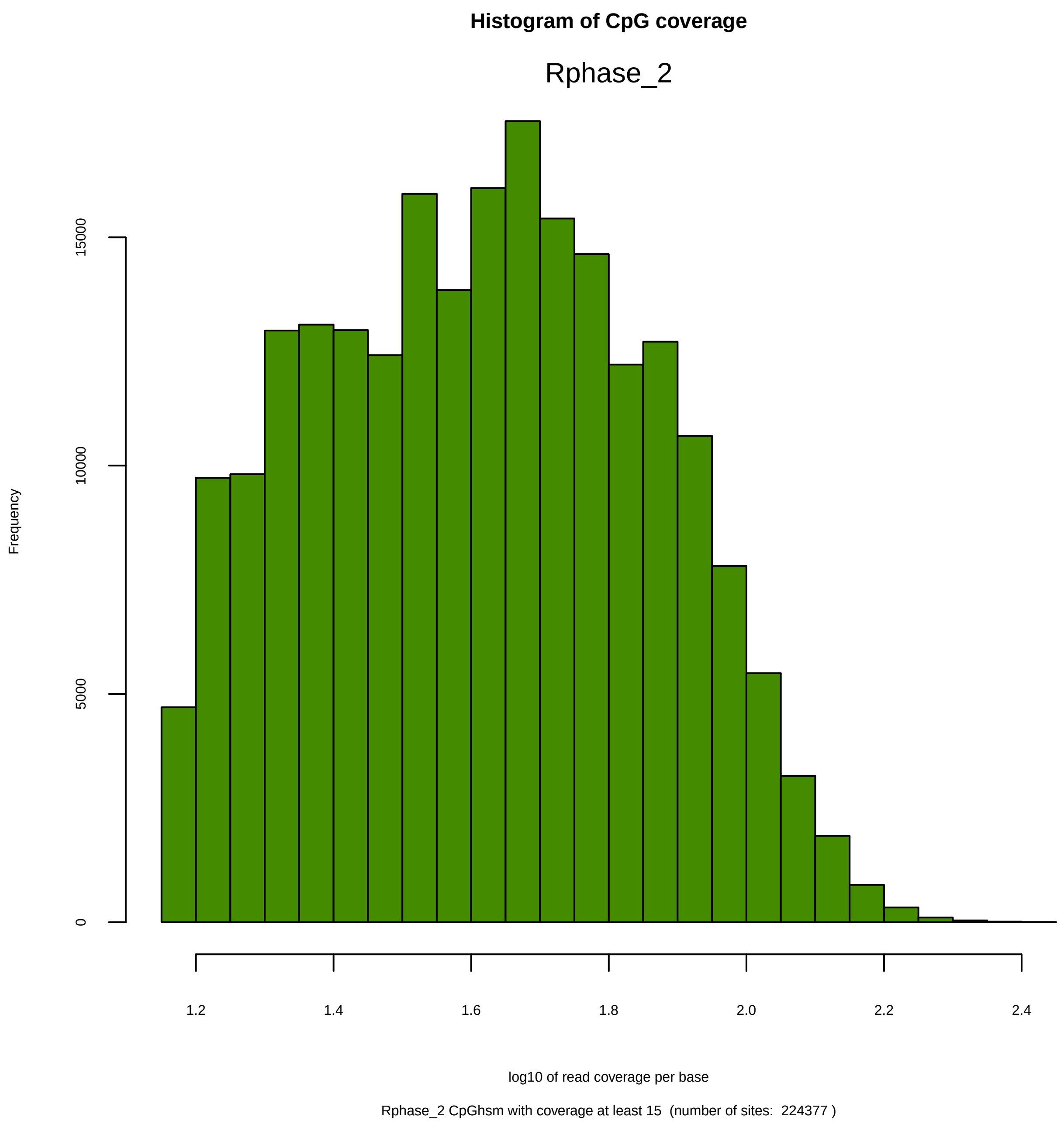
Histogram of CpG coverage
Rphase_2
Frequency
0
5000
10000
15000
1.2
1.4
1.6
1.8
2.0
2.2
2.4
log10 of read coverage per base
Rphase_2 CpGhsm with coverage at least 15 (number of sites: 224377 )

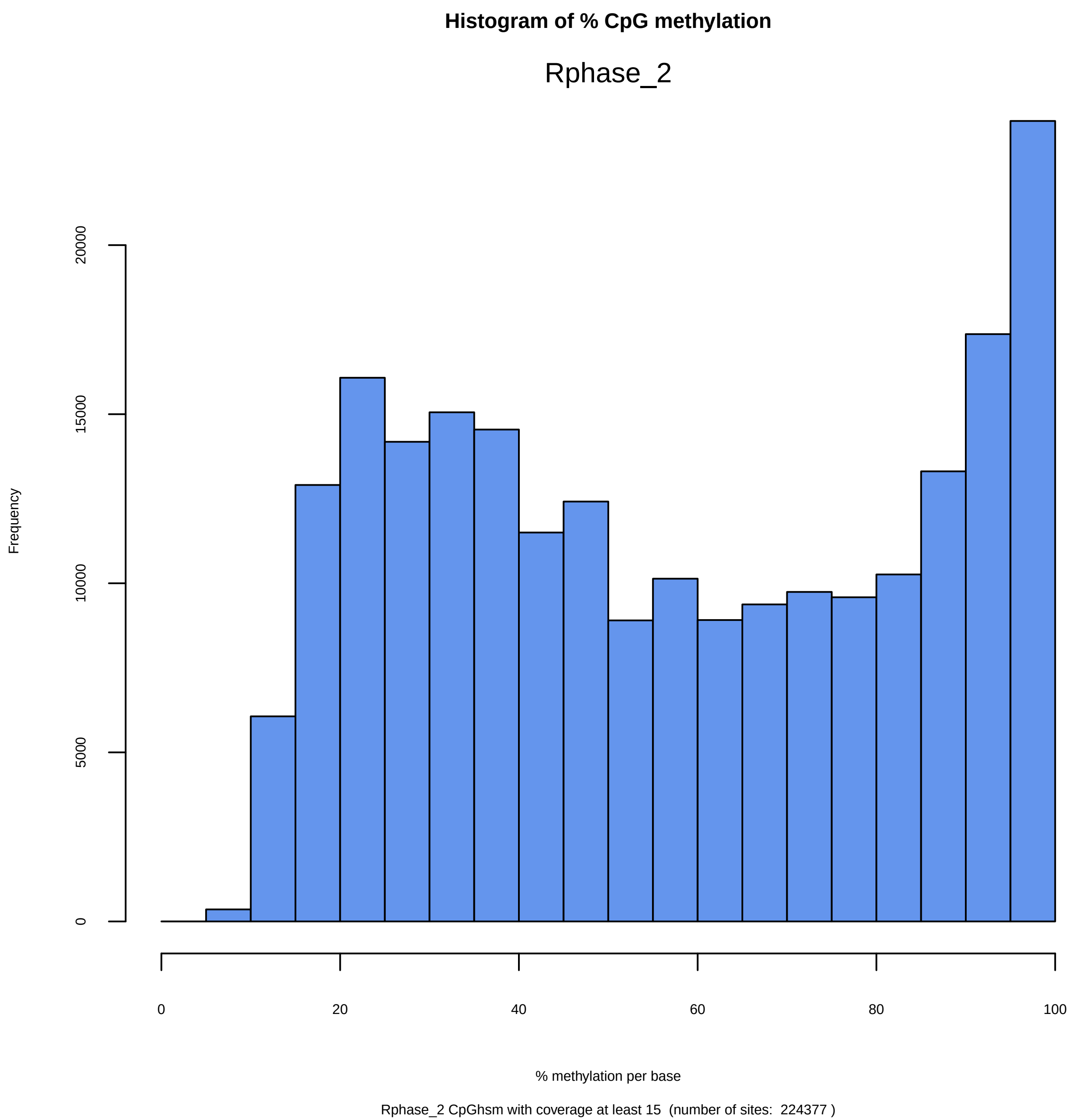
Histogram of % CpG methylation
Rphase_2
Frequency
0
5000
10000
15000
20000
0
20
40
60
80
100
% methylation per base
Rphase_2 CpGhsm with coverage at least 15 (number of sites: 224377 )

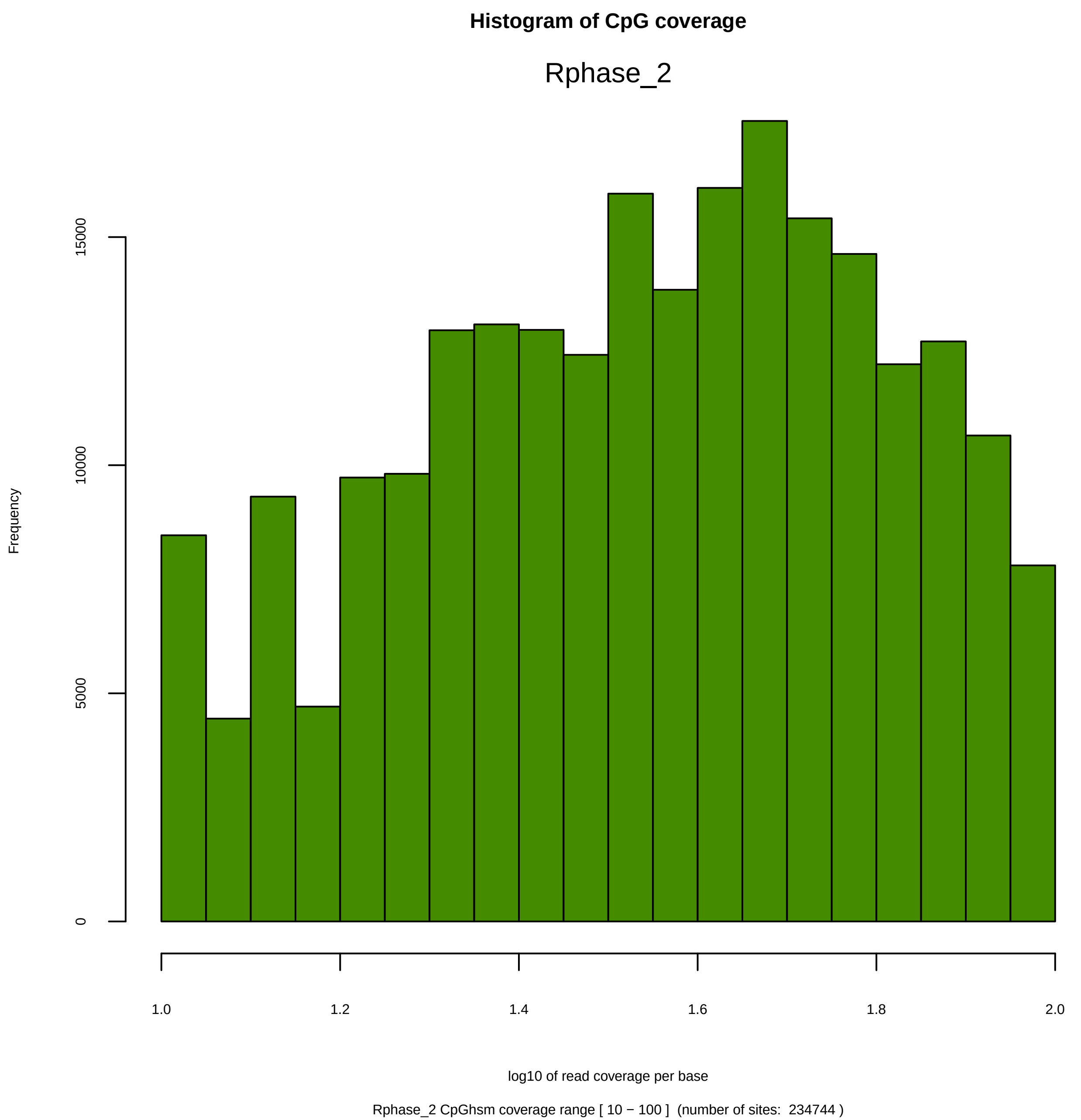
Histogram of CpG coverage
Rphase_2
Frequency
0
5000
10000
15000
1.0
1.2
1.4
1.6
1.8
2.0
log10 of read coverage per base
Rphase_2 CpGhsm coverage range [ 10 − 100 ]  (number of sites:  234744 )

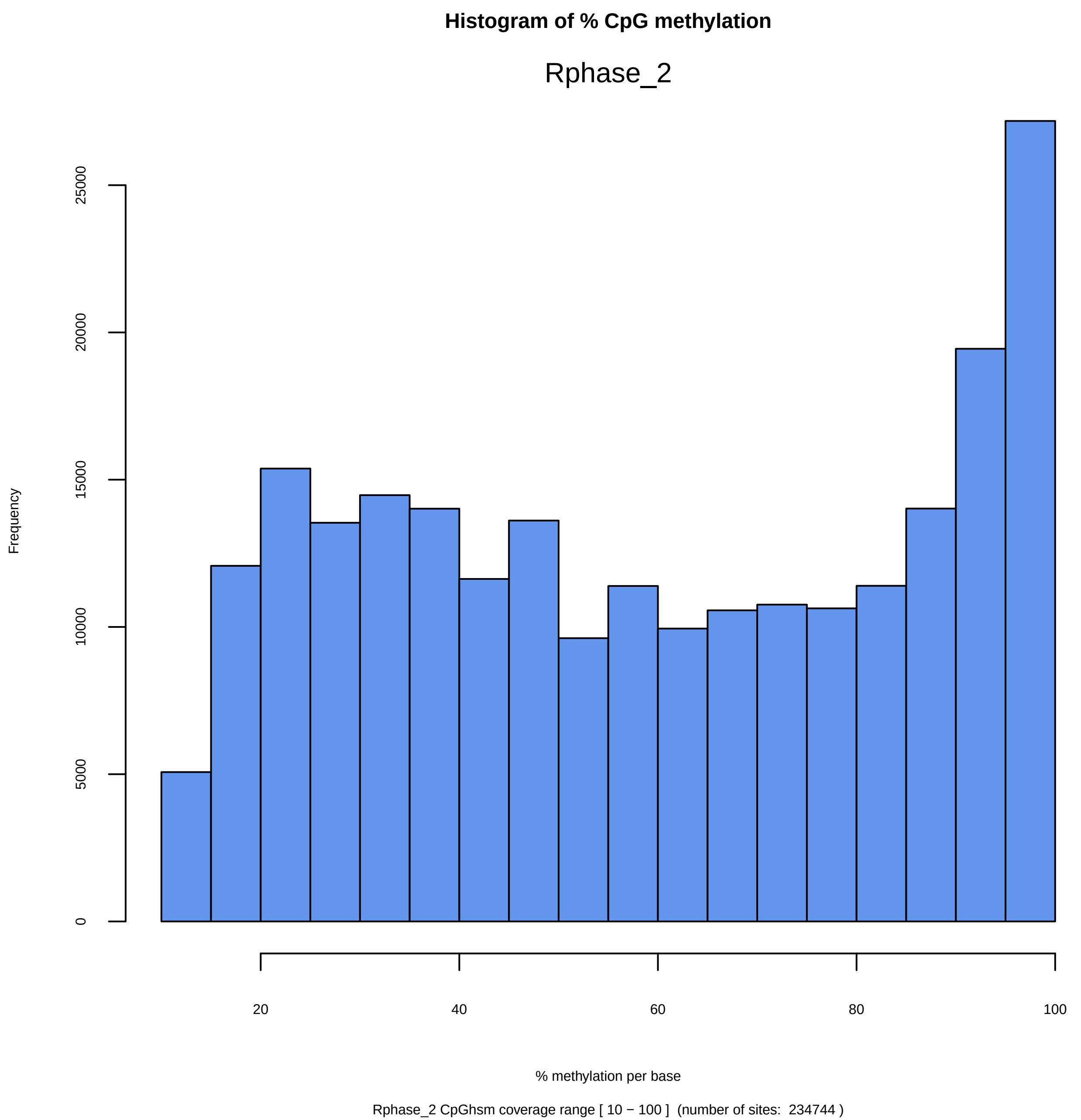
Histogram of % CpG methylation
Rphase_2
Frequency
0
5000
10000
15000
20000
25000
20
40
60
80
100
% methylation per base
Rphase_2 CpGhsm coverage range [ 10 − 100 ] (number of sites: 234744 )

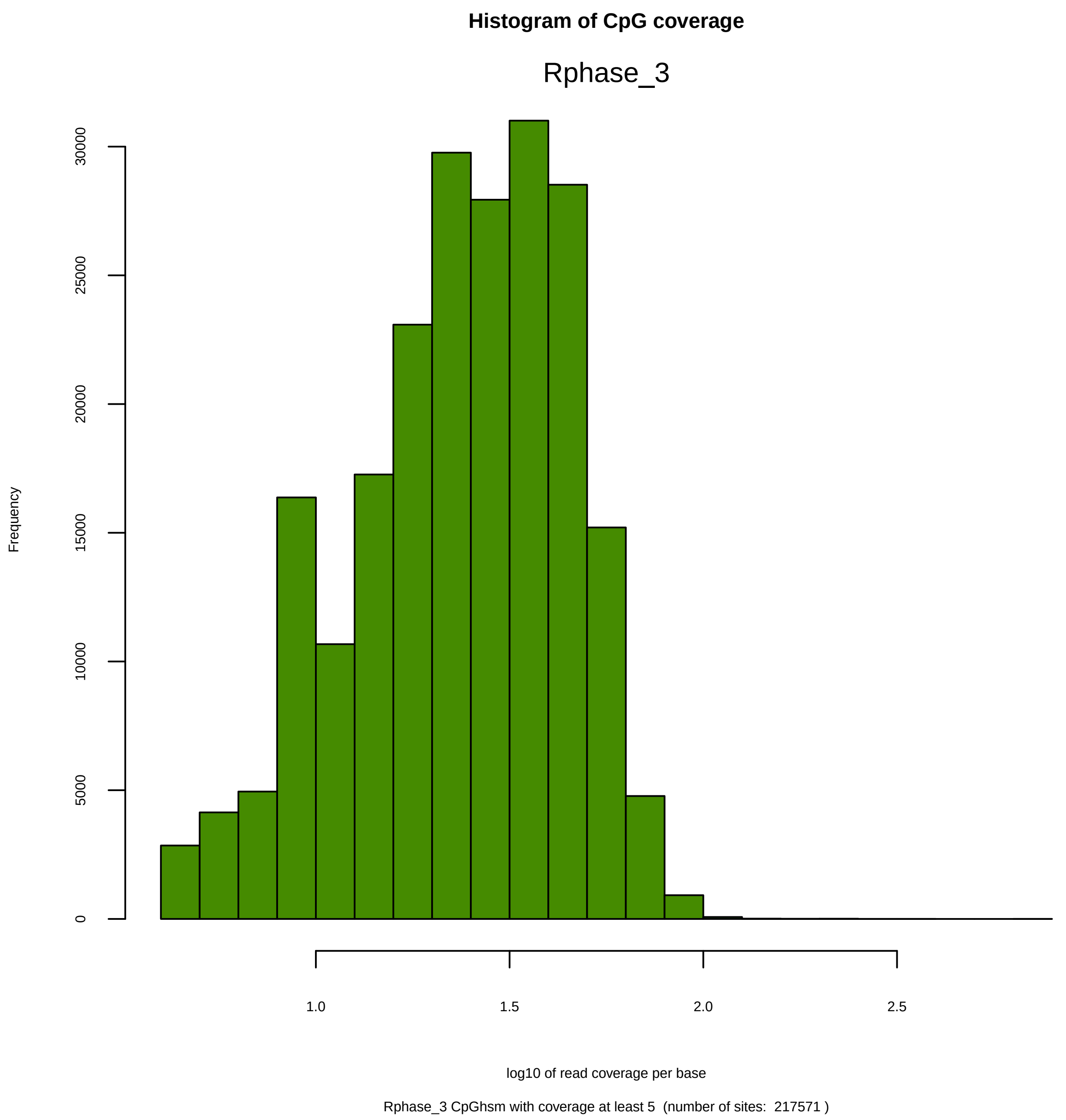
Histogram of CpG coverage
Rphase_3
Frequency
0
5000
10000
15000
20000
25000
30000
1.0
1.5
2.0
2.5
log10 of read coverage per base
Rphase_3 CpGhsm with coverage at least 5  (number of sites:  217571 )

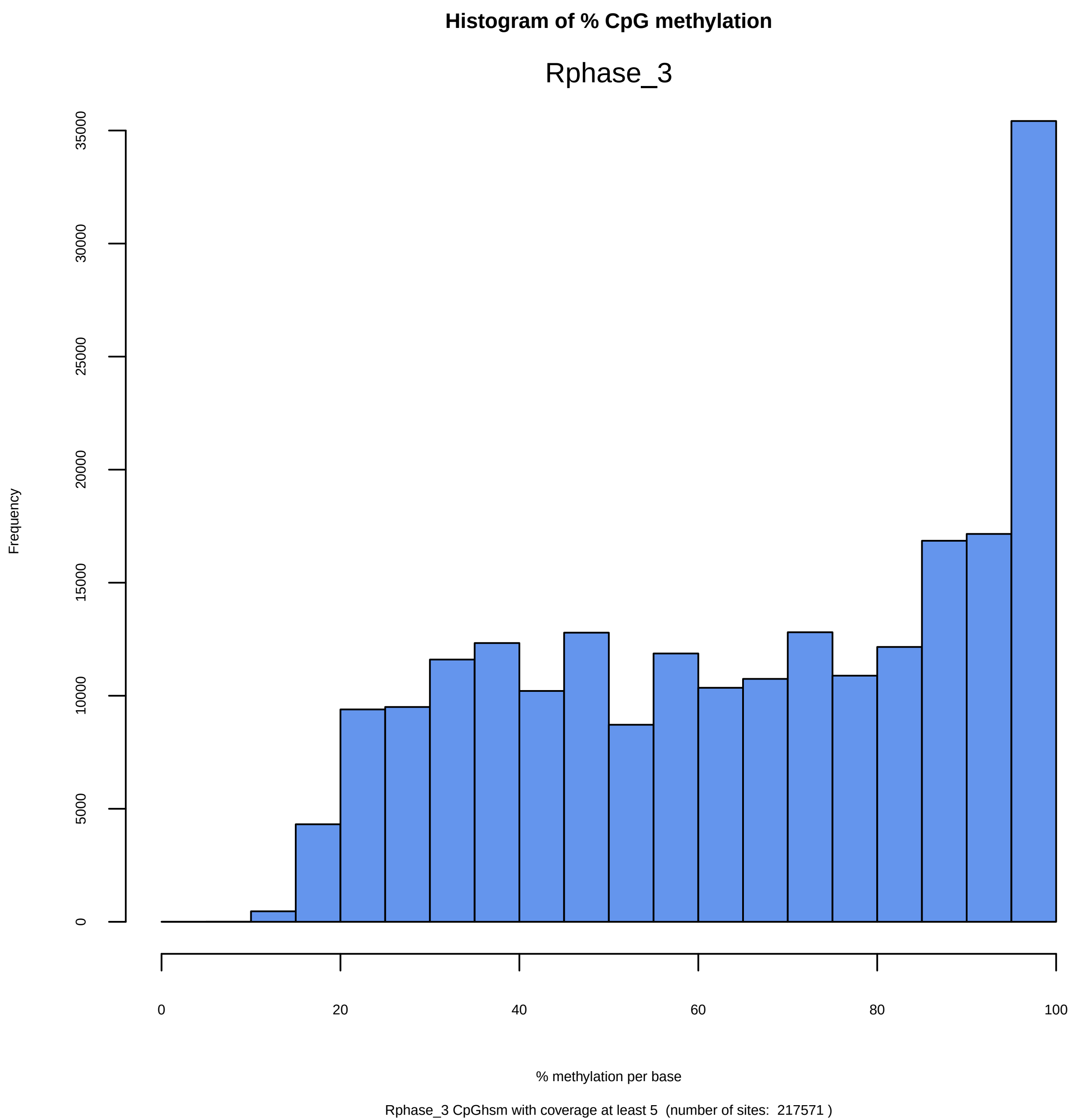

Histogram of % CpG methylation
Rphase_3
Frequency
0
5000
10000
15000
20000
25000
30000
35000
0
20
40
60
80
100
% methylation per base
Rphase_3 CpGhsm with coverage at least 5  (number of sites:  217571 )

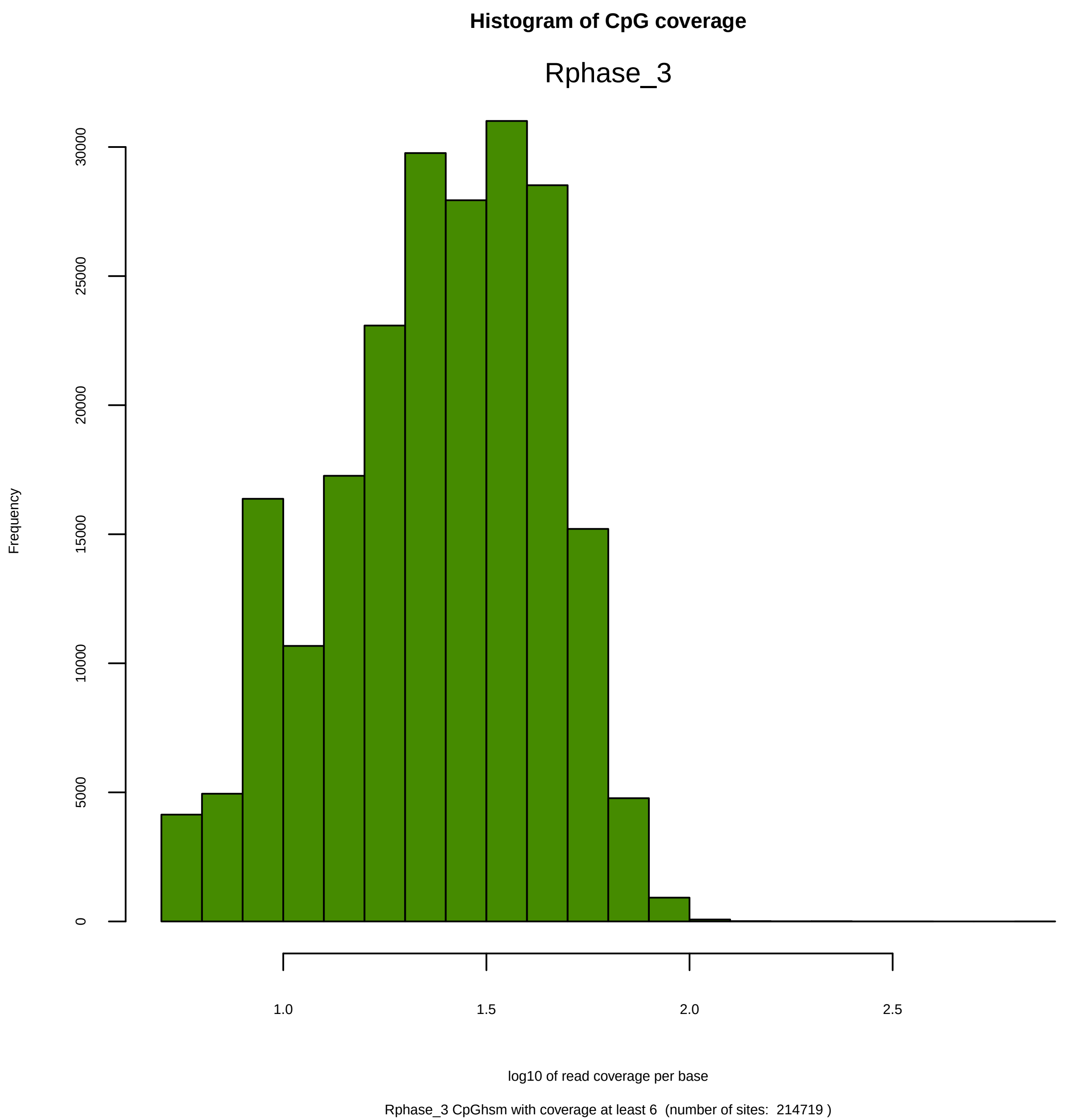

Histogram of CpG coverage
Rphase_3
Frequency
0
5000
10000
15000
20000
25000
30000
1.0
1.5
2.0
2.5
log10 of read coverage per base
Rphase_3 CpGhsm with coverage at least 6 (number of sites: 214719 )

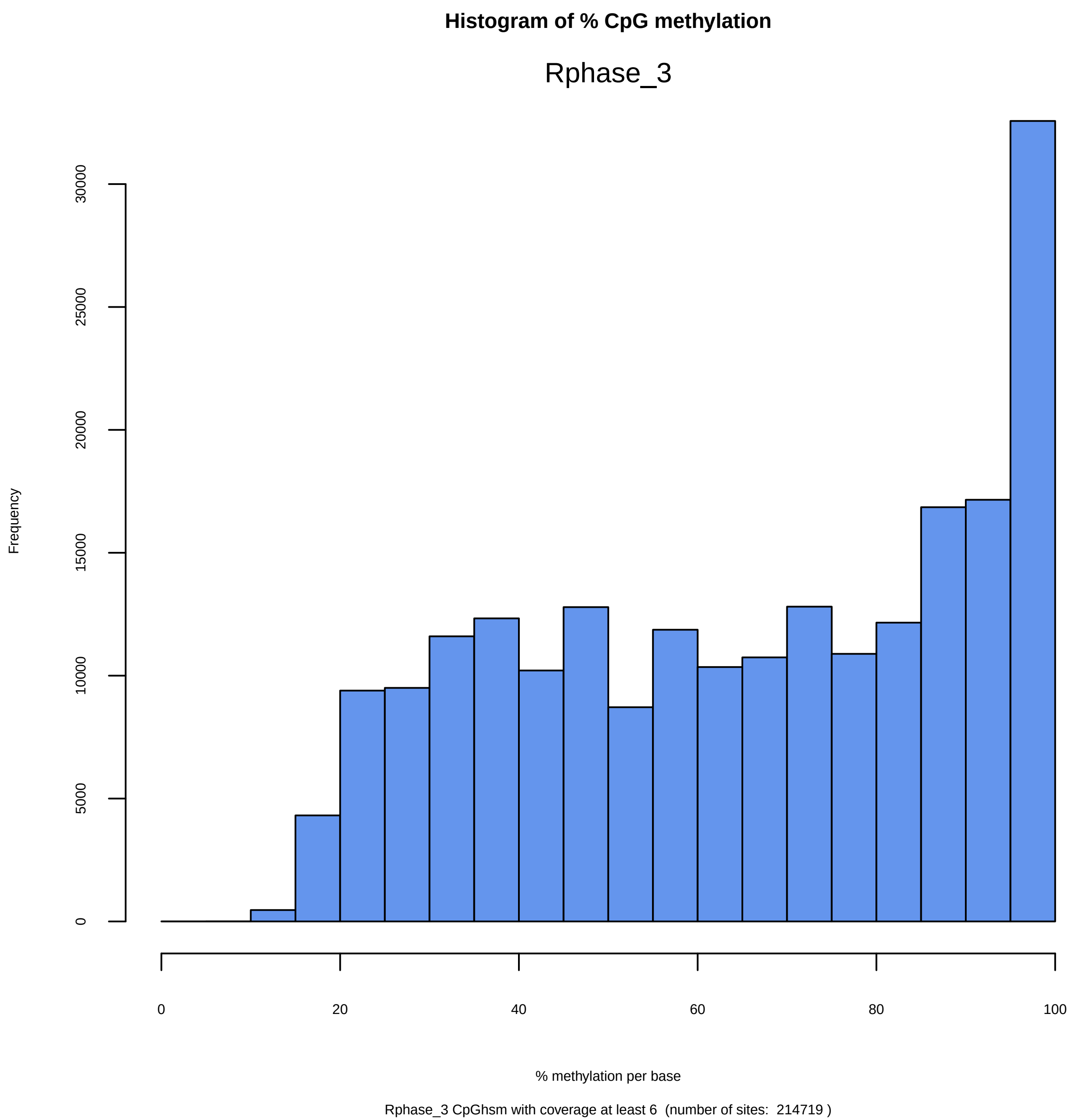

Histogram of % CpG methylation
Rphase_3
Frequency
0
5000
10000
15000
20000
25000
30000
0
20
40
60
80
100
% methylation per base
Rphase_3 CpGhsm with coverage at least 6  (number of sites:  214719 )

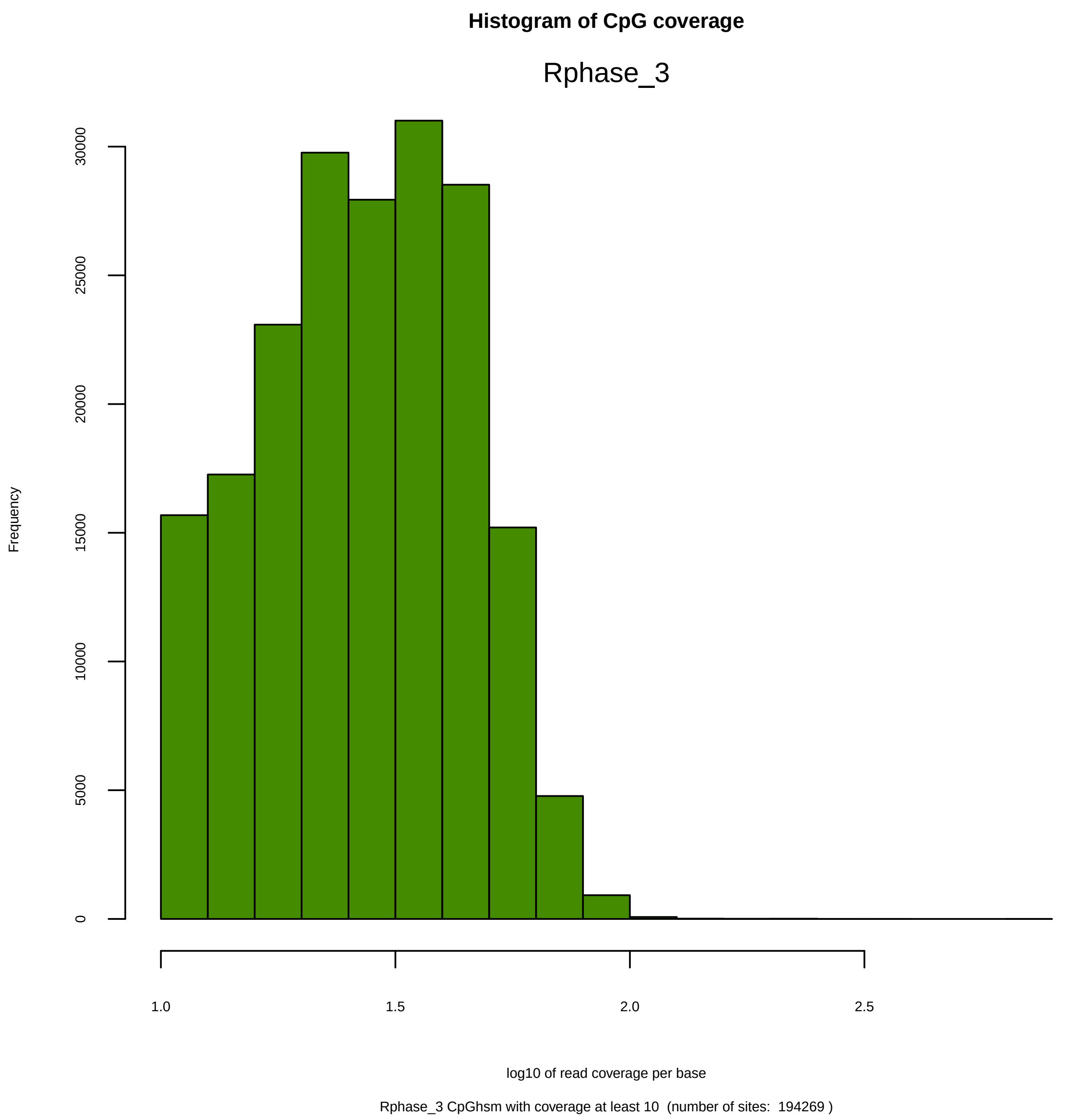

Histogram of CpG coverage
Rphase_3
Frequency
0
5000
10000
15000
20000
25000
30000
1.0
1.5
2.0
2.5
log10 of read coverage per base
Rphase_3 CpGhsm with coverage at least 10 (number of sites: 194269 )

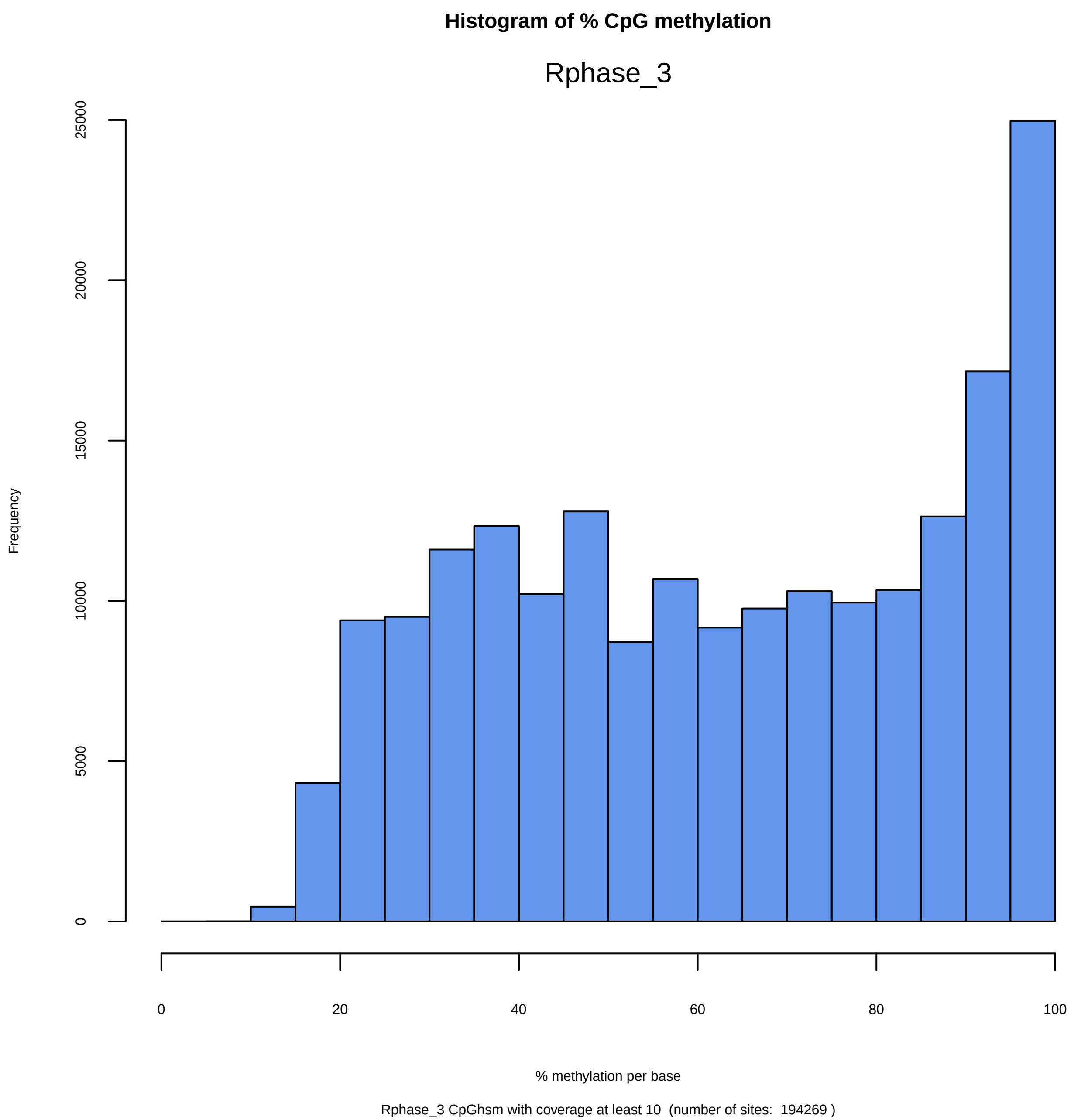

Histogram of % CpG methylation
Rphase_3
Frequency
0
5000
10000
15000
20000
25000
0
20
40
60
80
100
% methylation per base
Rphase_3 CpGhsm with coverage at least 10 (number of sites: 194269 )

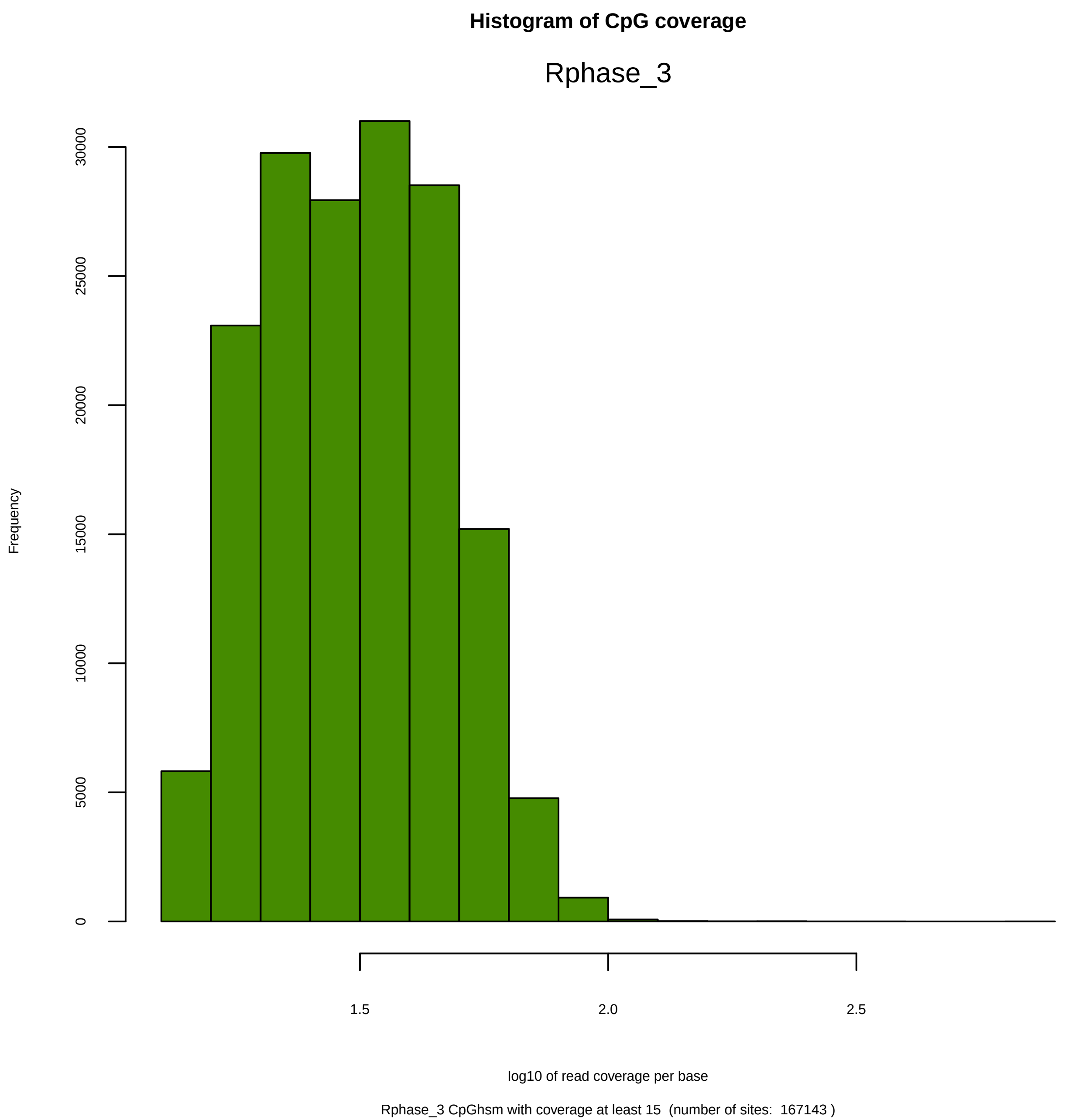
Histogram of CpG coverage
Rphase_3
Frequency
0
5000
10000
15000
20000
25000
30000
1.5
2.0
2.5
log10 of read coverage per base
Rphase_3 CpGhsm with coverage at least 15  (number of sites:  167143 )

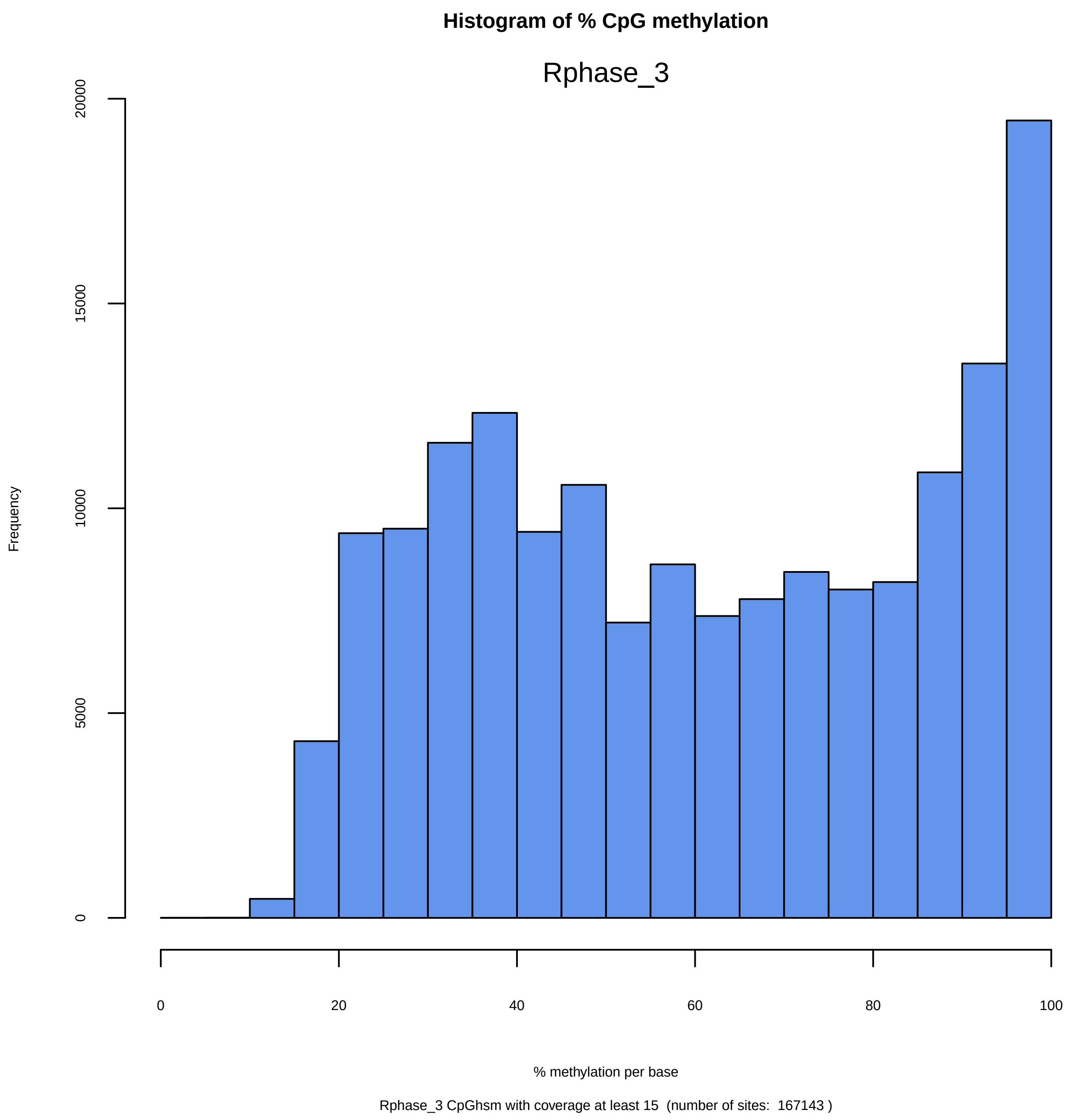

Histogram of % CpG methylation
Rphase_3
Frequency
0
5000
10000
15000
20000
0
20
40
60
80
100
% methylation per base
Rphase_3 CpGhsm with coverage at least 15 (number of sites: 167143 )

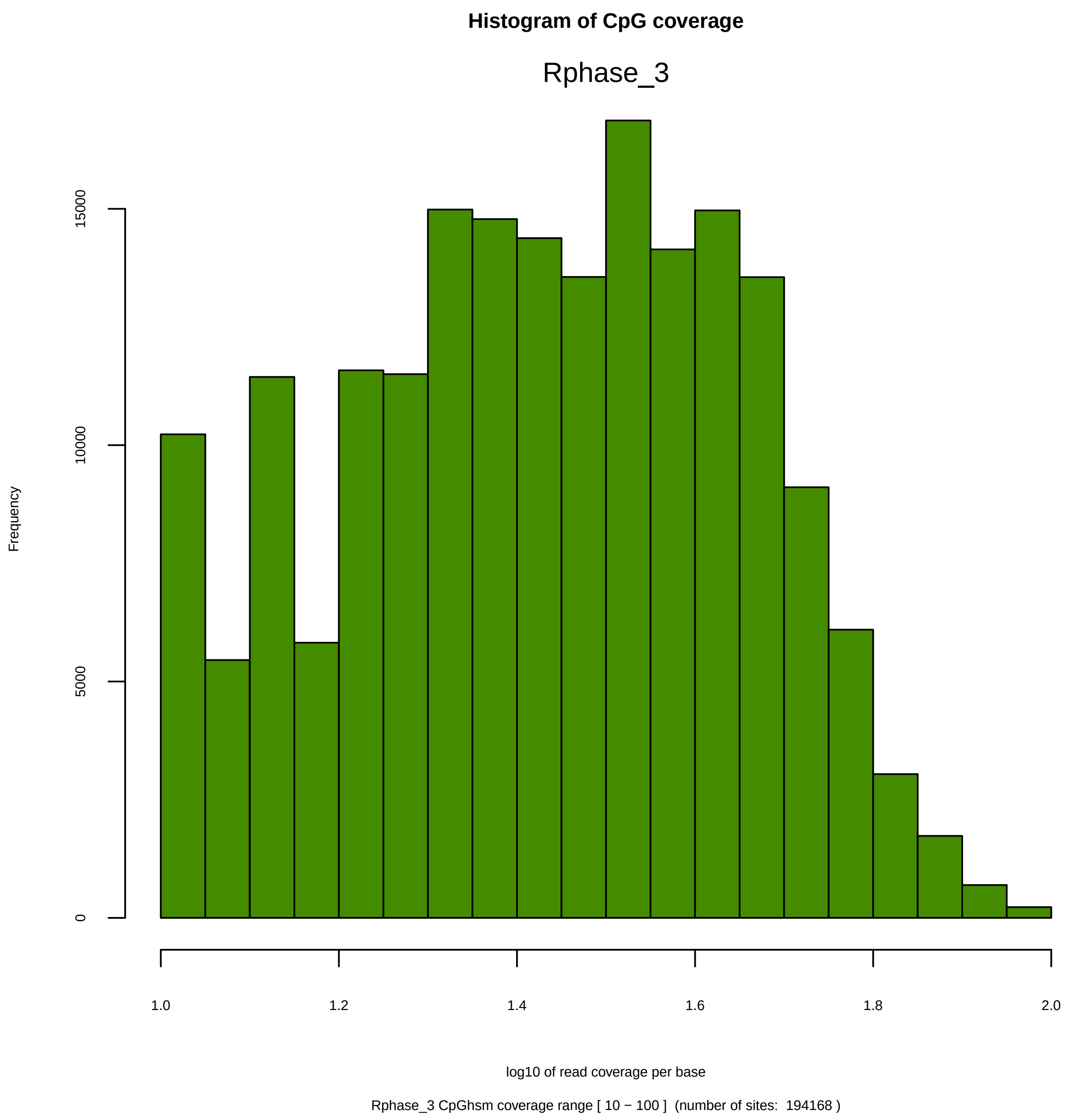
Histogram of CpG coverage
Rphase_3
Frequency
0
5000
10000
15000
1.0
1.2
1.4
1.6
1.8
2.0
log10 of read coverage per base
Rphase_3 CpGhsm coverage range [ 10 − 100 ]  (number of sites:  194168 )

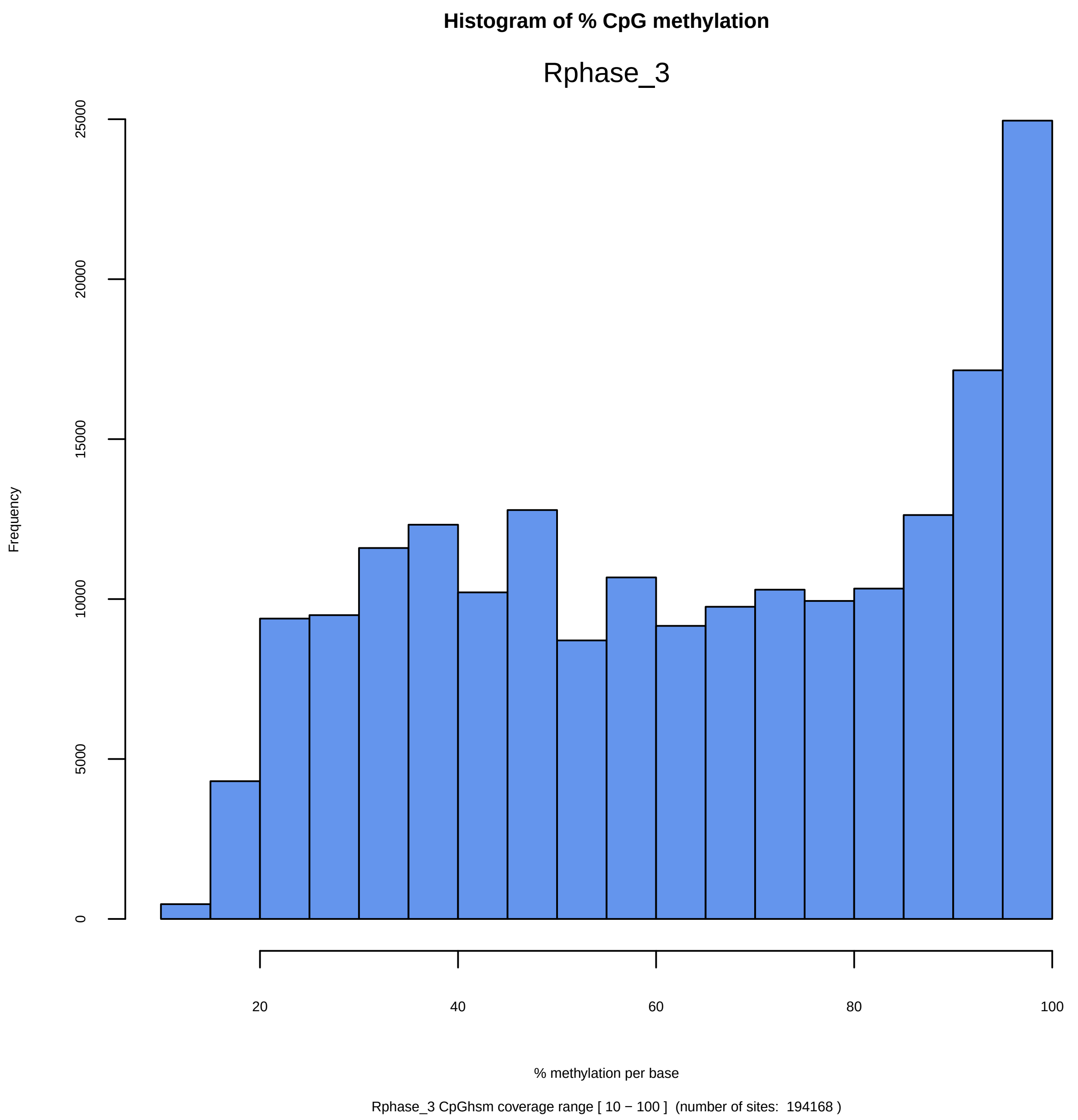

Histogram of % CpG methylation
Rphase_3
Frequency
0
5000
10000
15000
20000
25000
20
40
60
80
100
% methylation per base
Rphase_3 CpGhsm coverage range [ 10 − 100 ]  (number of sites:  194168 )

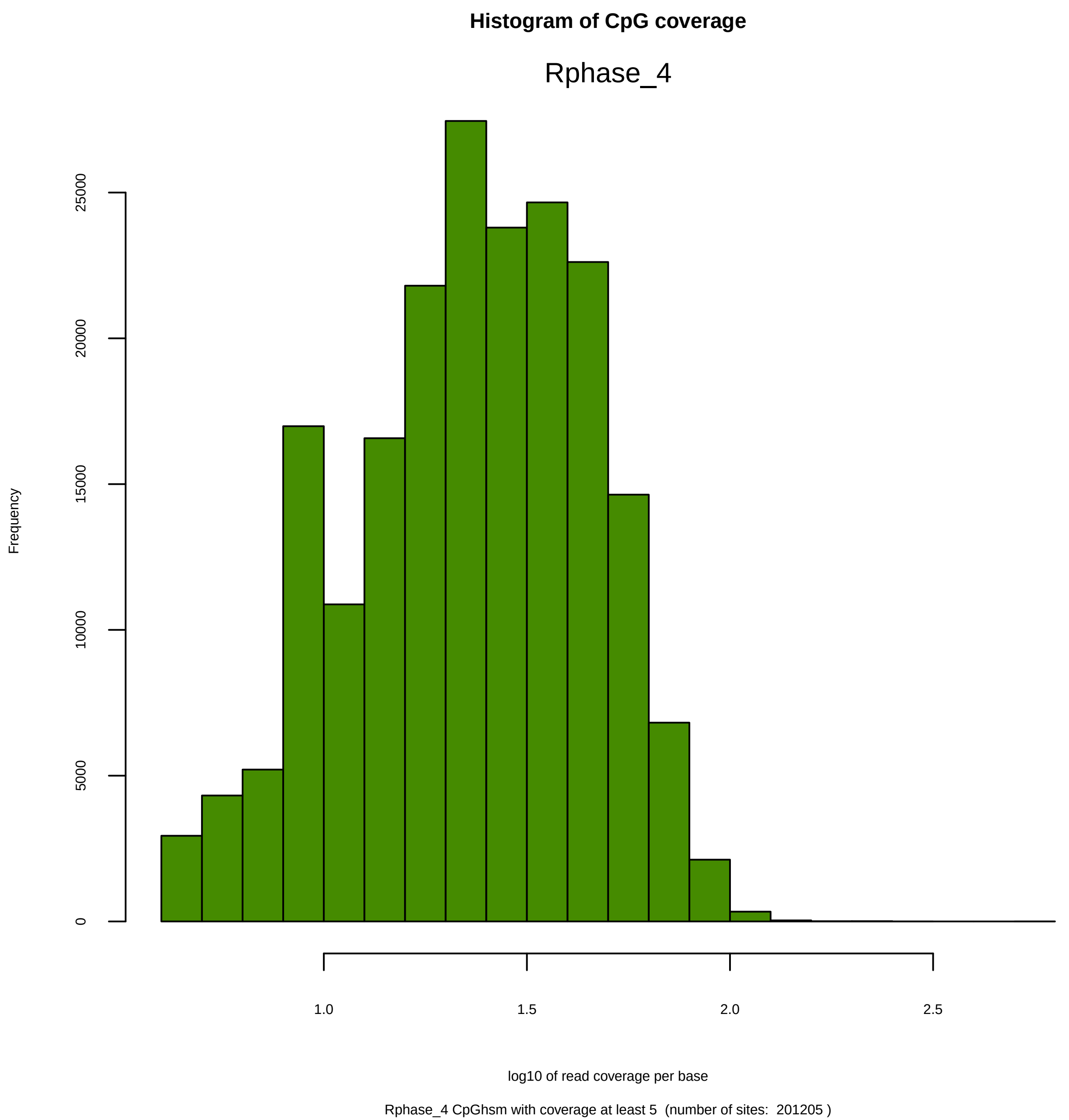
Histogram of CpG coverage
Rphase_4
Frequency
0
5000
10000
15000
20000
25000
1.0
1.5
2.0
2.5
log10 of read coverage per base
Rphase_4 CpGhsm with coverage at least 5 (number of sites: 201205 )

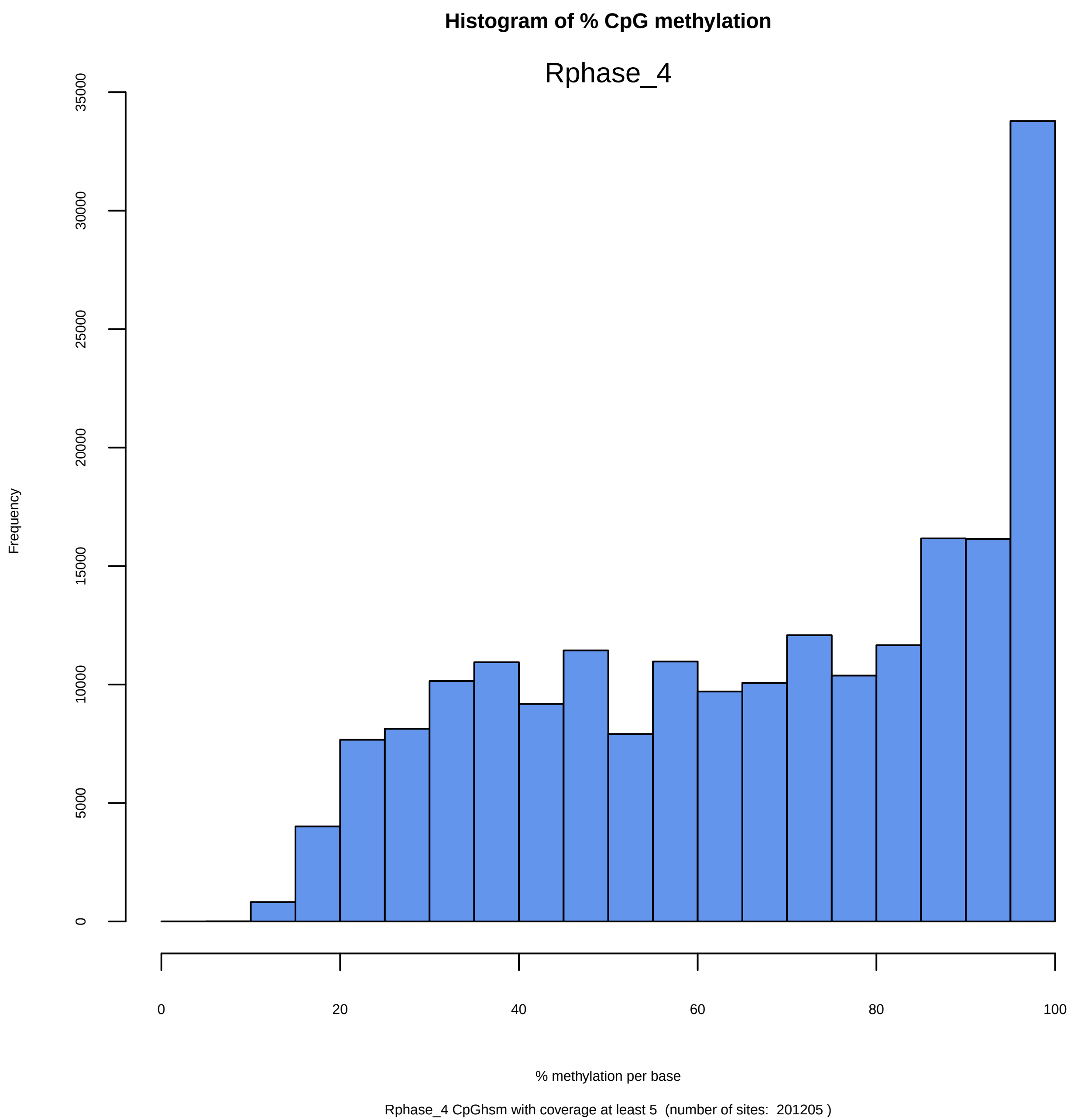

Histogram of % CpG methylation
Rphase_4
Frequency
0
5000
10000
15000
20000
25000
30000
35000
0
20
40
60
80
100
% methylation per base
Rphase_4 CpGhsm with coverage at least 5 (number of sites: 201205 )

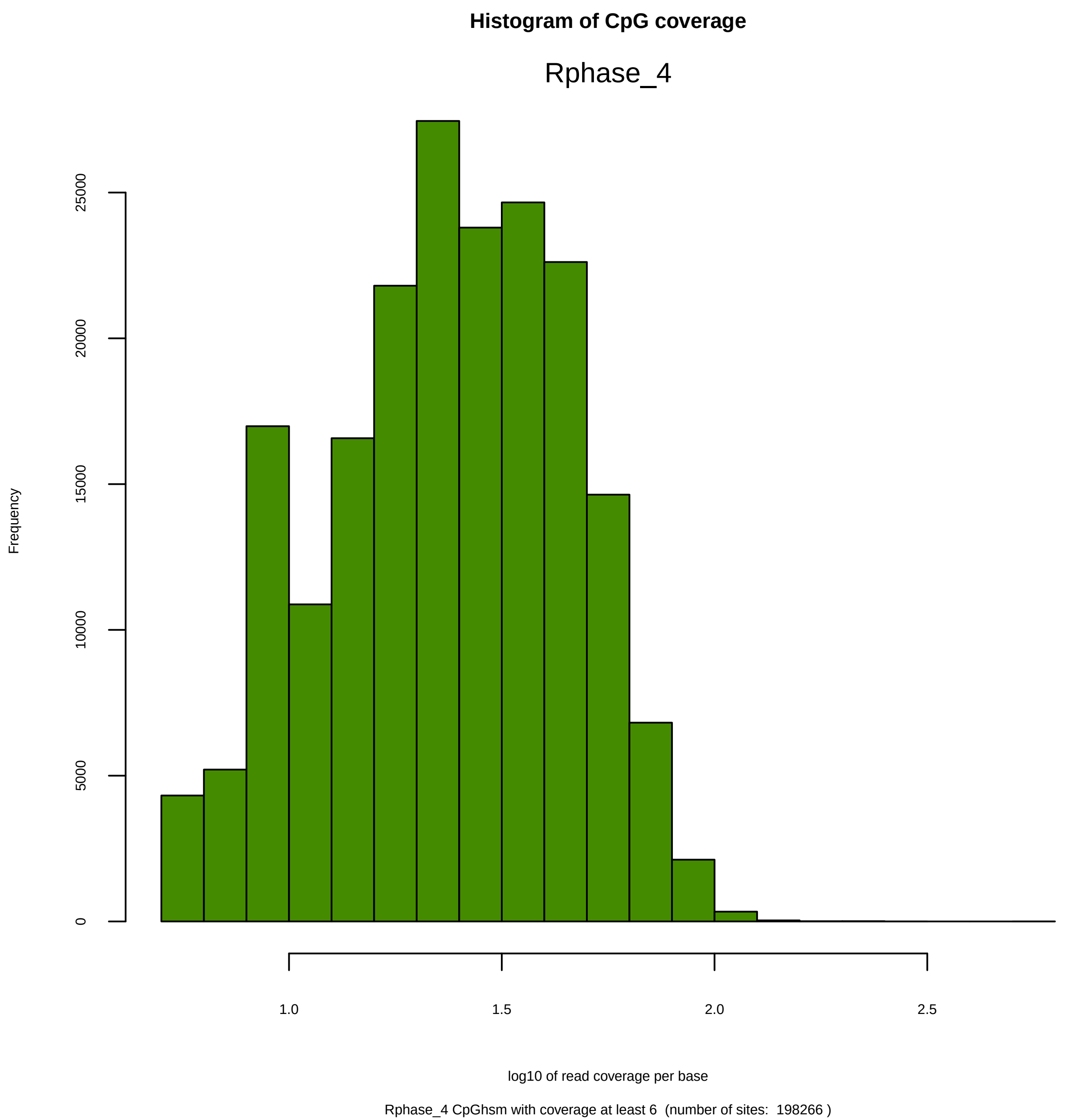
Histogram of CpG coverage
Rphase_4
Frequency
0
5000
10000
15000
20000
25000
1.0
1.5
2.0
2.5
log10 of read coverage per base
Rphase_4 CpGhsm with coverage at least 6 (number of sites: 198266 )

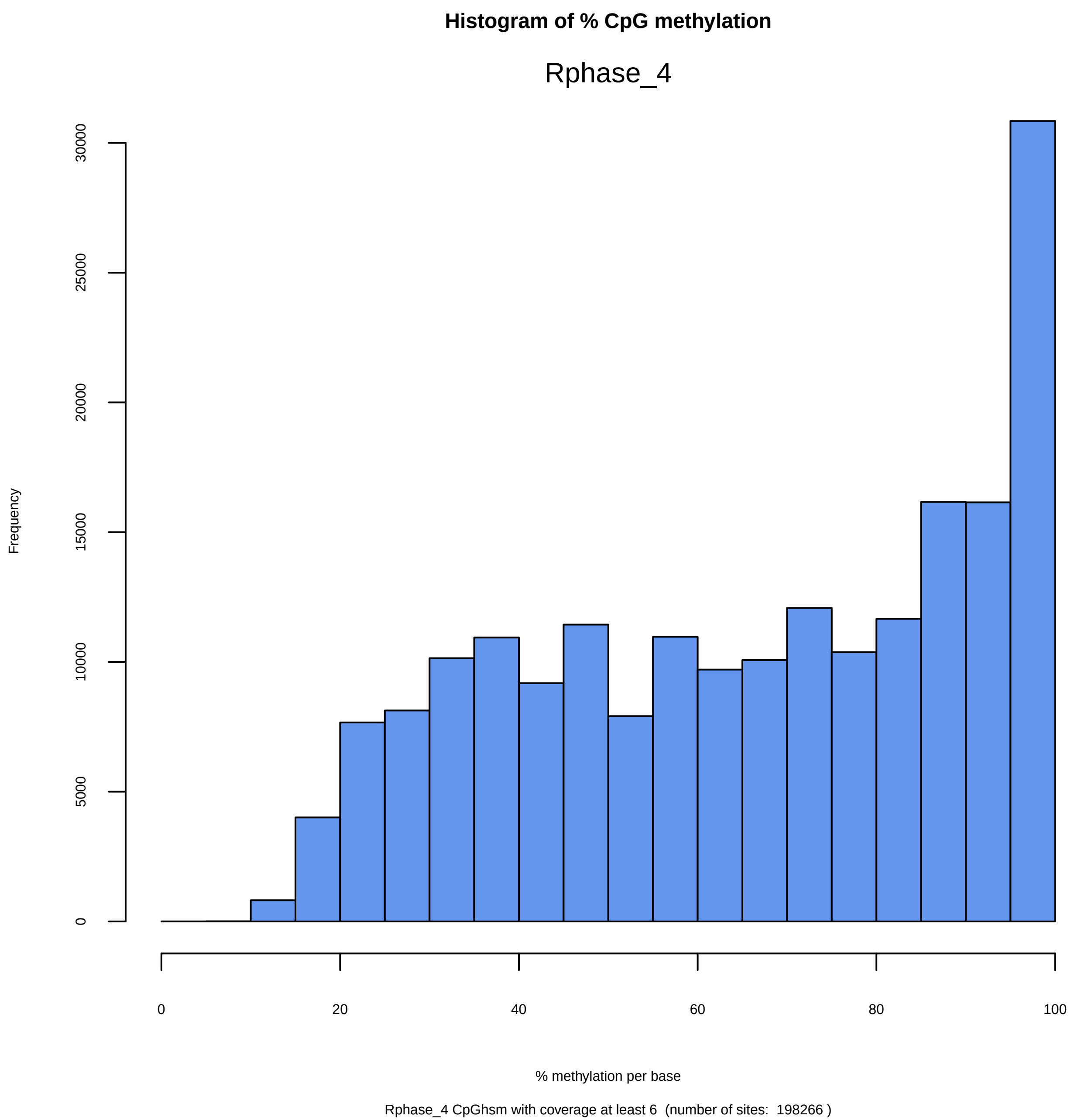

Histogram of % CpG methylation
Rphase_4
Frequency
0
5000
10000
15000
20000
25000
30000
0
20
40
60
80
100
% methylation per base
Rphase_4 CpGhsm with coverage at least 6  (number of sites:  198266 )

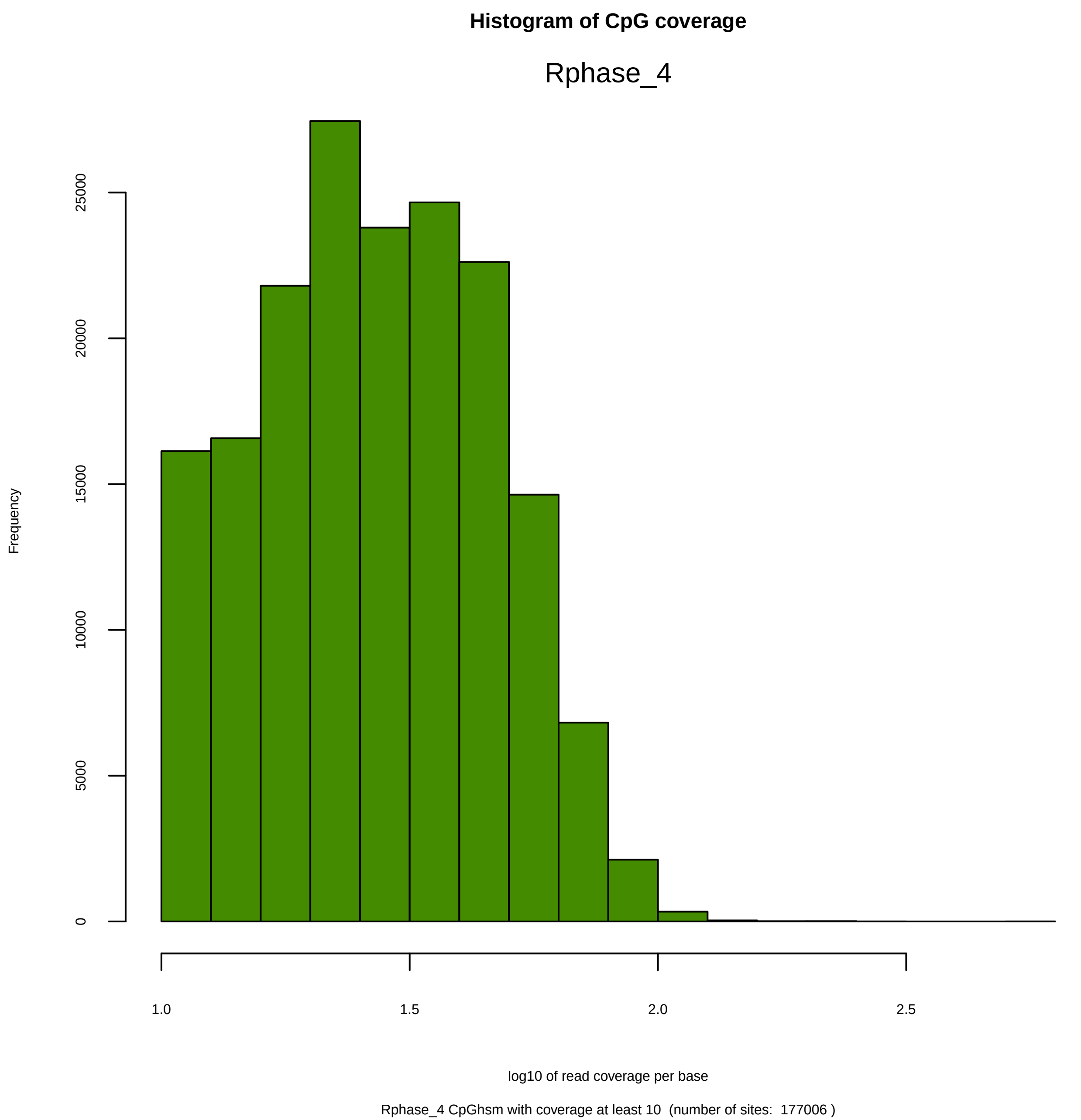
Histogram of CpG coverage
Rphase_4
Frequency
0
5000
10000
15000
20000
25000
1.0
1.5
2.0
2.5
log10 of read coverage per base
Rphase_4 CpGhsm with coverage at least 10  (number of sites:  177006 )

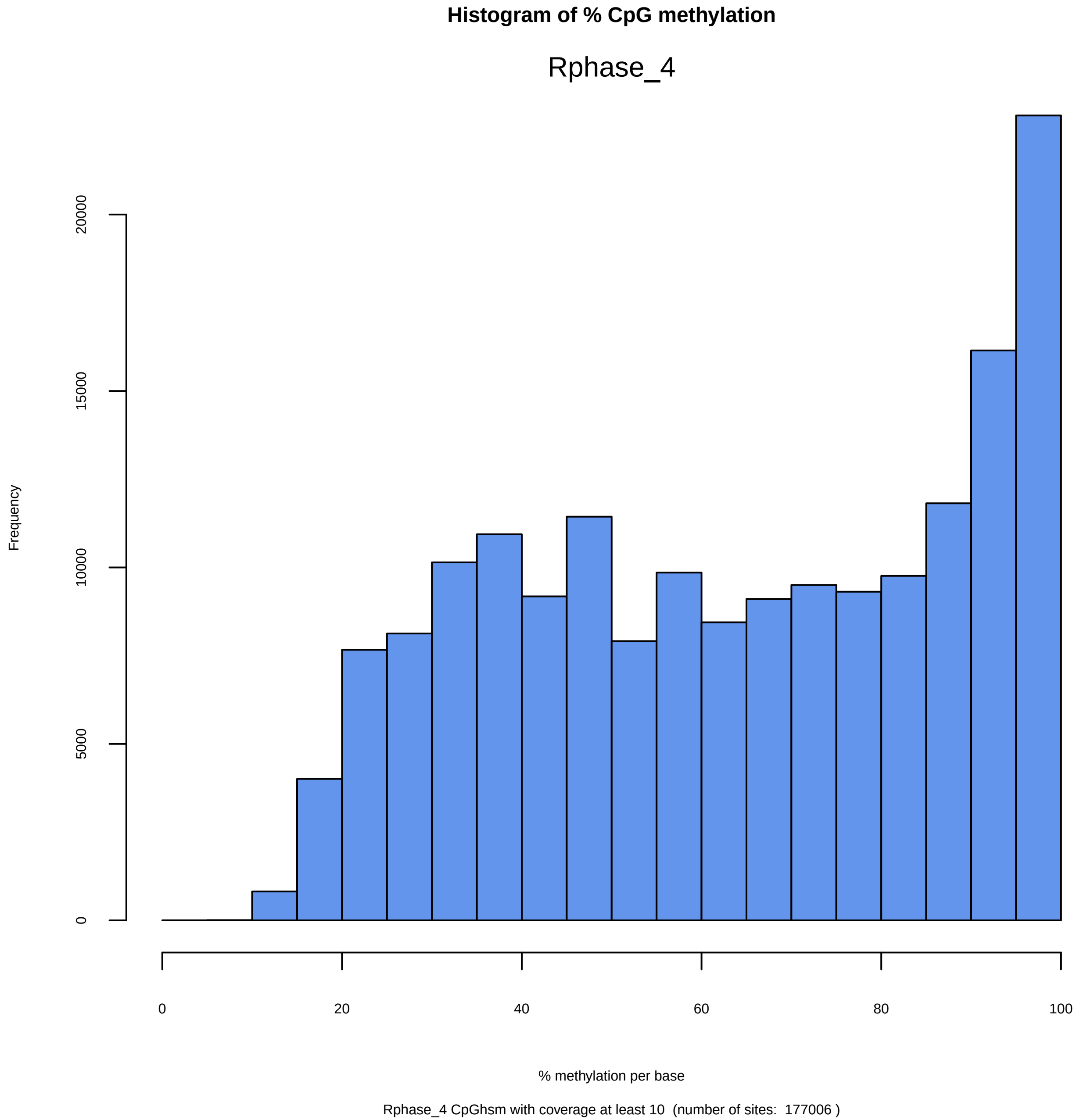
Histogram of % CpG methylation
Rphase_4
Frequency
0
5000
10000
15000
20000
0
20
40
60
80
100
% methylation per base
Rphase_4 CpGhsm with coverage at least 10 (number of sites: 177006 )

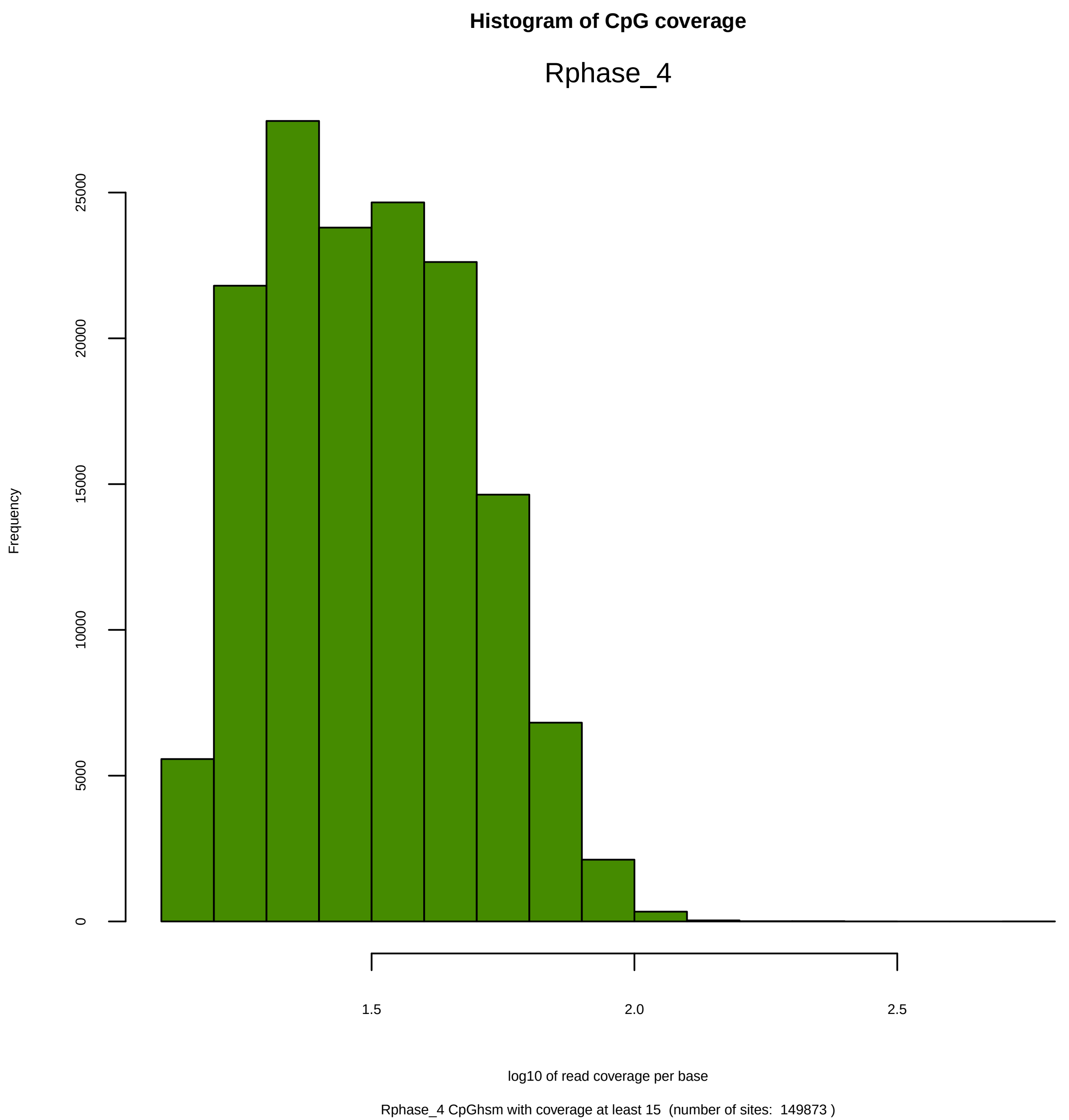

Histogram of CpG coverage
Rphase_4
Frequency
0
5000
10000
15000
20000
25000
1.5
2.0
2.5
log10 of read coverage per base
Rphase_4 CpGhsm with coverage at least 15  (number of sites:  149873 )

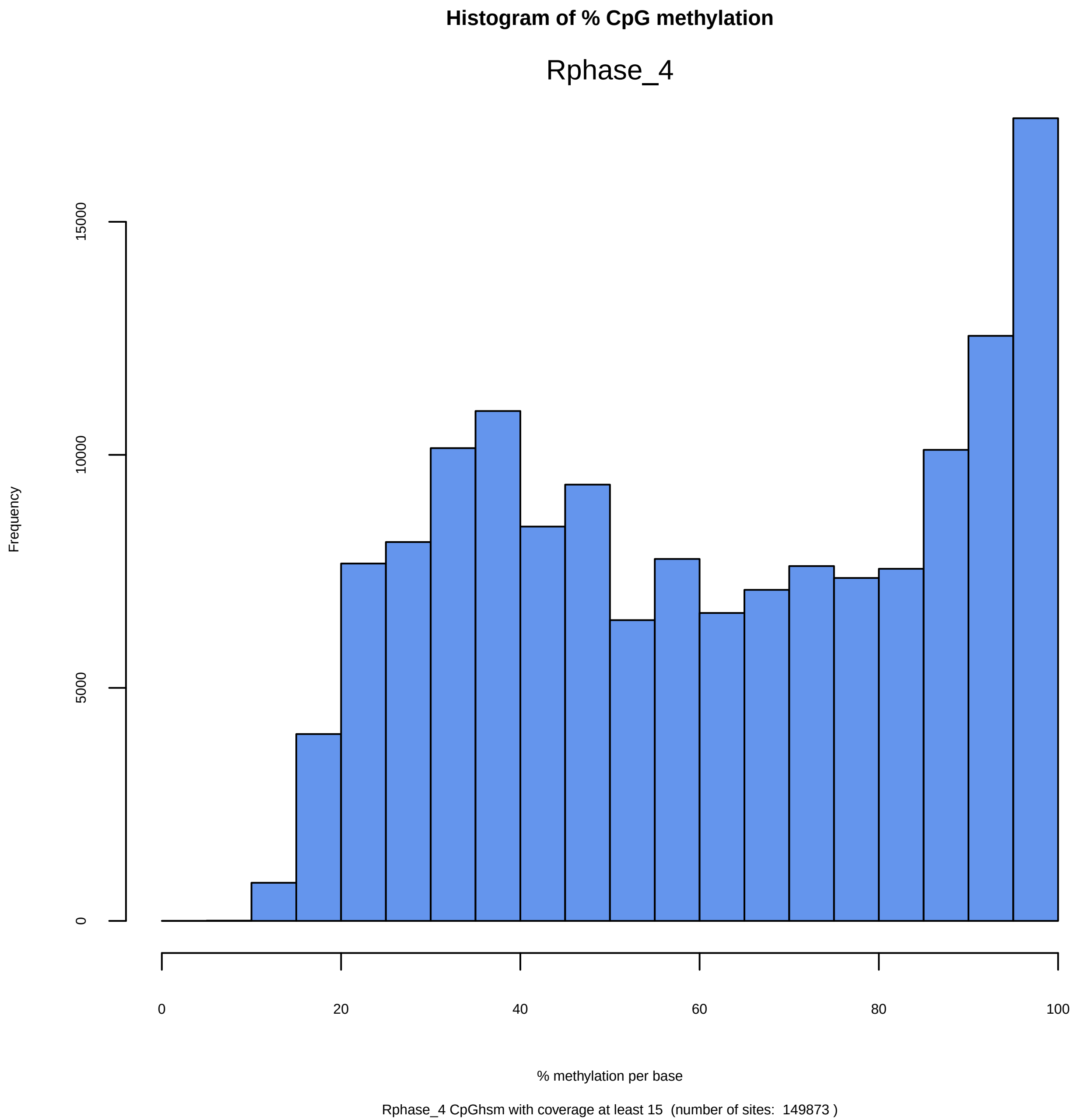

Histogram of % CpG methylation
Rphase_4
Frequency
0
5000
10000
15000
0
20
40
60
80
100
% methylation per base
Rphase_4 CpGhsm with coverage at least 15 (number of sites: 149873 )

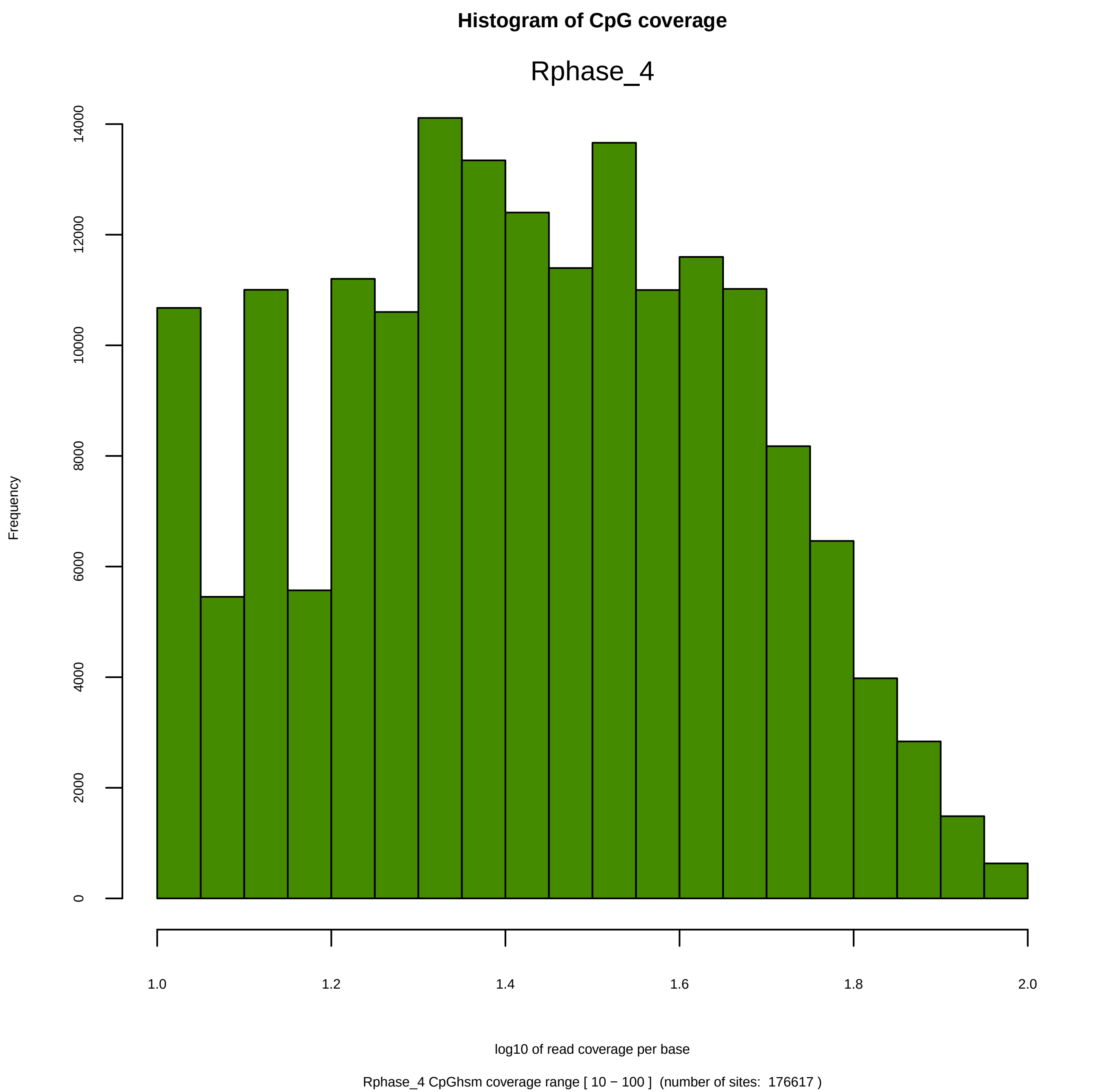

Histogram of CpG coverage
Rphase_4
Frequency
0
2000
4000
6000
8000
10000
12000
14000
1.0
1.2
1.4
1.6
1.8
2.0
log10 of read coverage per base
Rphase_4 CpGhsm coverage range [ 10 − 100 ]  (number of sites:  176617 )

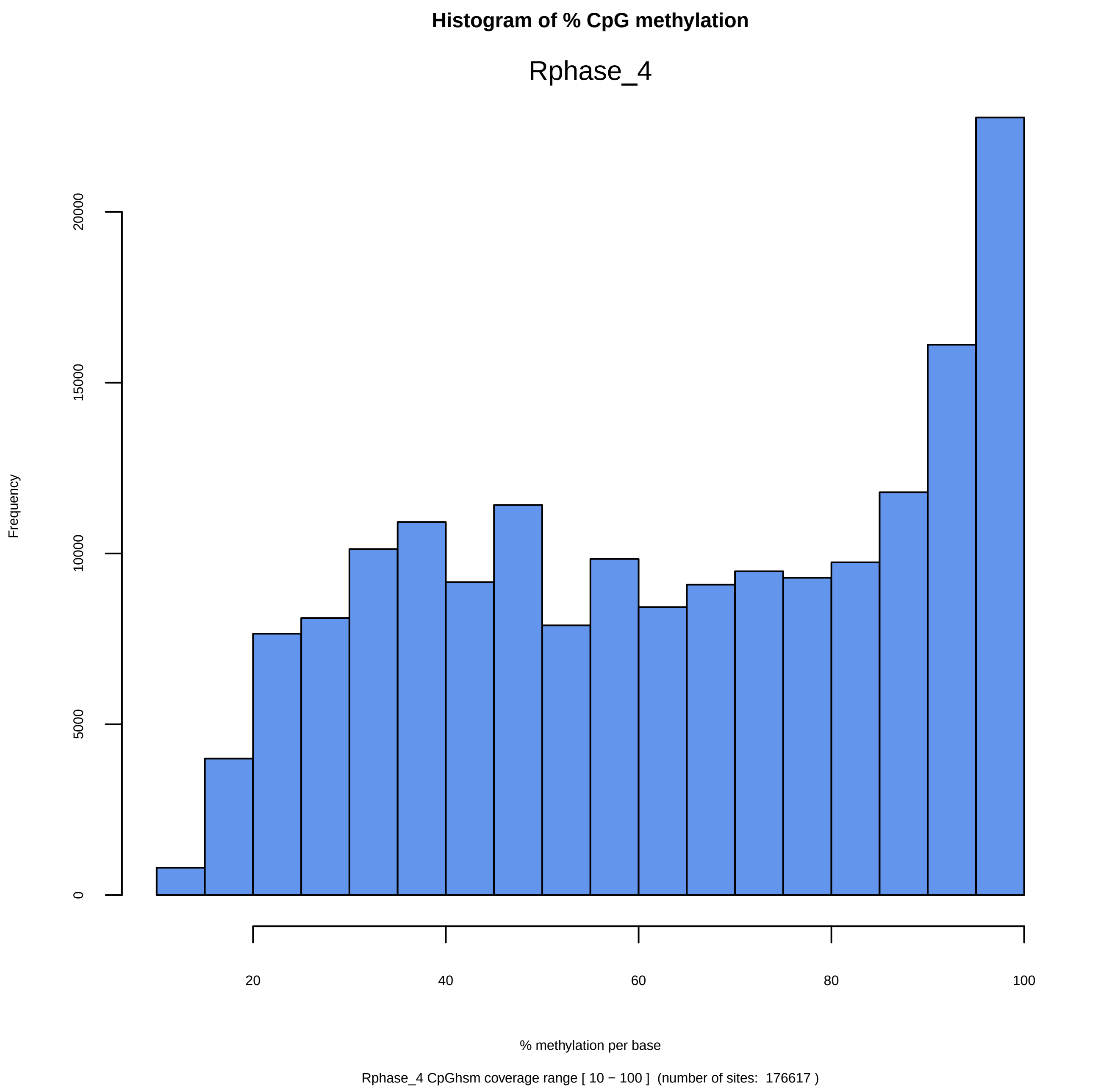
Histogram of % CpG methylation
Rphase_4
Frequency
0
5000
10000
15000
20000
20
40
60
80
100
% methylation per base
Rphase_4 CpGhsm coverage range [ 10 − 100 ]  (number of sites:  176617 )

```
Directory: REPCRL File: 0READMErepcrl

REPCRL - Correlations between replicates.

Input: mkrd (methylKit methylRawList object of replicate data)

Output: files repcrl-*.txt repcrl-*.pdf

Notes: The output is generated by BWASPR::cmpSamples(), which uses methylKit::unite()
        to determine the sites common to all replicates and applies methylKit::getCorrelation()
        and methylKit::PCASamples().  The *.txt files show the correlations in a table, and
        the *.pdf files show graphics.

       Beware that the number of common sites (shown at the bottom of the *.txt files) may be
        small, which may make the correlations less informative.
```

Directory: REPCRL File: repcrl-Ob_LIp_BCphase.txt

| | BCphase_1 | BCphase_2 | BCphase_3 | BCphase_4 |
|---|---|---|---|---|
| BCphase_1 | 1.000 | 0.906 | 0.909 | 0.915 |
| BCphase_2 | 0.906 | 1.000 | 0.924 | 0.915 |
| BCphase_3 | 0.909 | 0.924 | 1.000 | 0.916 |
| BCphase_4 | 0.915 | 0.915 | 0.916 | 1.000 |

The number of conserved sites is 112637.

Directory: REPCRL File: repcrl-Ob_LIp_Rphase.txt

| | Rphase_1 | Rphase_2 | Rphase_3 | Rphase_4 |
|---|---|---|---|---|
| Rphase_1 | 1.000 | 0.930 | 0.913 | 0.909 |
| Rphase_2 | 0.930 | 1.000 | 0.922 | 0.920 |
| Rphase_3 | 0.913 | 0.922 | 1.000 | 0.917 |
| Rphase_4 | 0.909 | 0.920 | 0.917 | 1.000 |

The number of conserved sites is 116407.

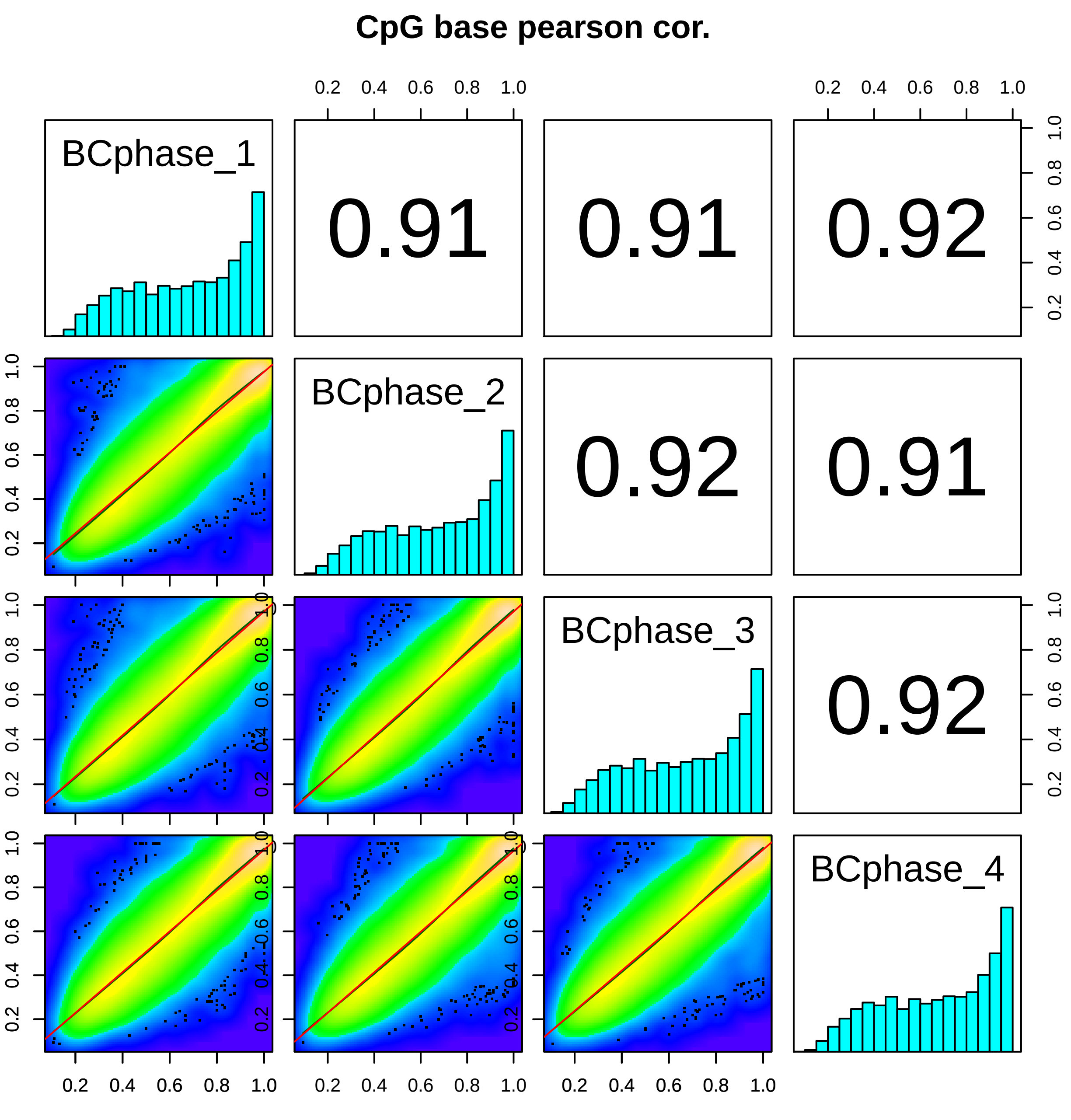
CpG base pearson cor.
BCphase_1
BCphase_2
BCphase_3
BCphase_4
0.91
0.91
0.92
0.92
0.91
0.92
0.2 0.4 0.6 0.8 1.0

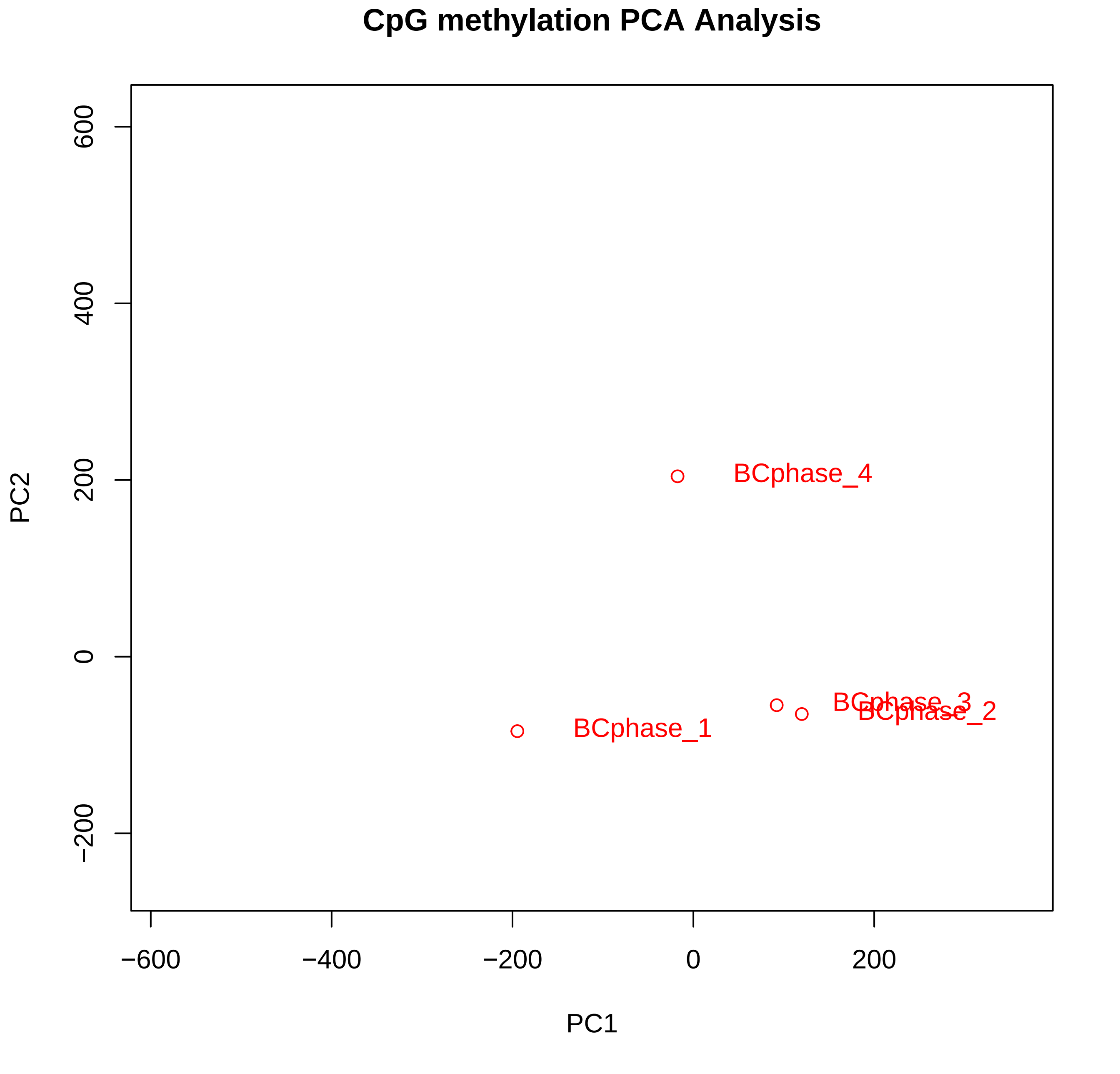
CpG methylation PCA Analysis
BCphase_4
BCphase_3
BCphase_2
BCphase_1
600
400
200
0
−200
−600
−400
−200
0
200
PC1
PC2

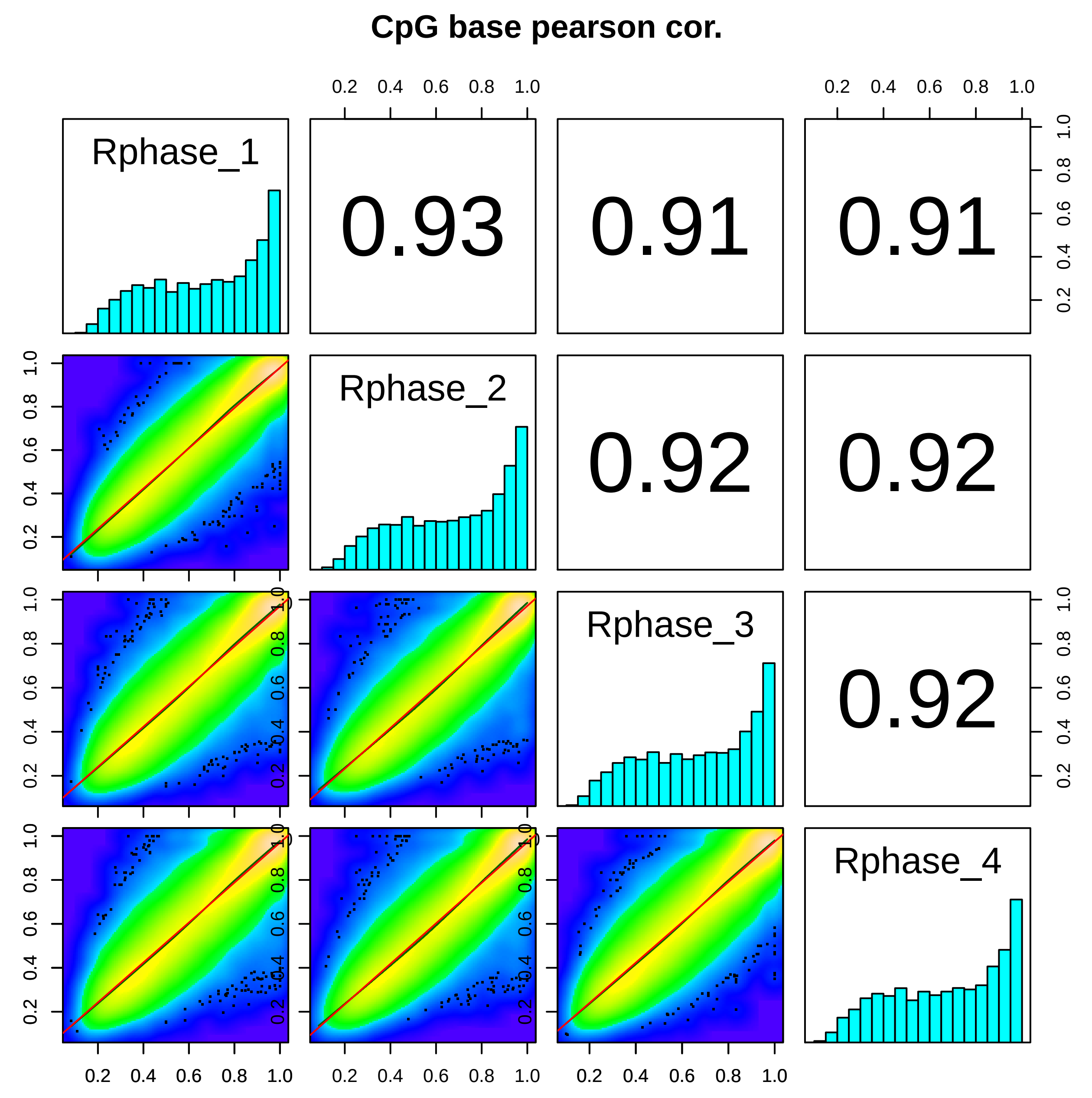
CpG base pearson cor.
Rphase_1
Rphase_2
Rphase_3
Rphase_4
0.93
0.91
0.91
0.92
0.92
0.92
0.2
0.4
0.6
0.8
1.0

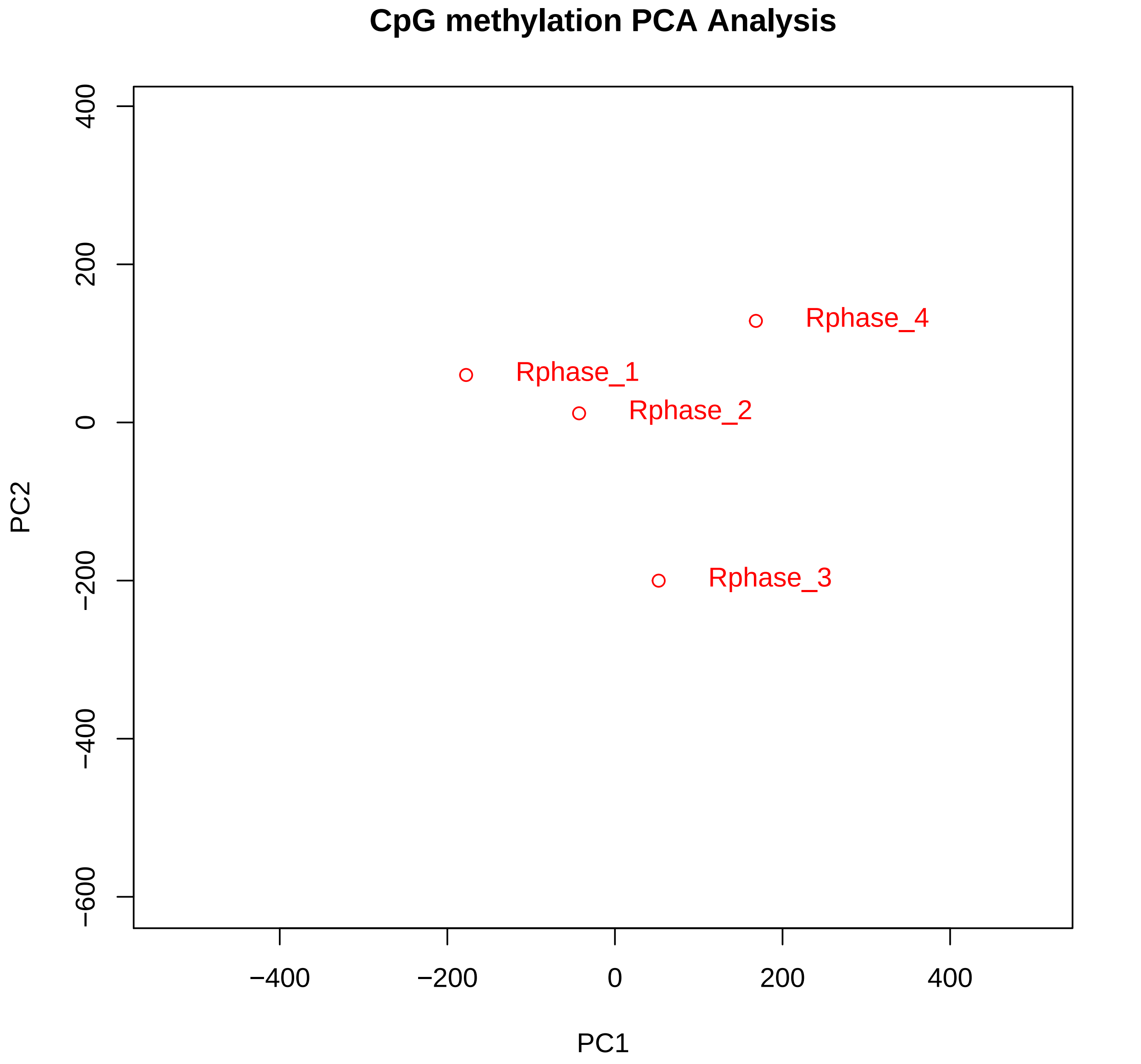

CpG methylation PCA Analysis
PC2
400
200
0
−200
−400
−600
PC1
−400
−200
0
200
400
Rphase_4
Rphase_1
Rphase_2
Rphase_3

```
Directory: MMP File: 0READMEmmp

MMP - Mapping of methylation sites on genome annotation.

Input: studymk, studymc, genome_ann (a list of GRanges objects providing annotated region labels
        and boundaries; output of BWASPR::get_genome_annotation())

Output: files mmp-*.txt

Notes: The output is generated by BWASPR::map_methylome() and gives an accounting for every sample
        as to where the CpGhsm and CpGscd (control) sites reside relative to the genome annotation.
       Abbreviations used: O/E, observed over expected
                           (expected percentages are based on the respective feature proportions
                            in the genome, as annotated).
```

```
Directory: MMP File: mmp-Ob_LI_BCphase.txt

================================================================================
Genomic composition in terms of feature regions
================================================================================

Ob genome size:                        223876465 bp
  Ob genic region size:                  166752117 bp ( 74.5%)
    Ob exon region size:                    44907373 bp ( 20.1%)
      Ob CDS region size:                     20651162 bp ( 46.0%)
      Ob five-prime UTR region size:           5434886 bp ( 12.1%)
      Ob three-prime UTR region size:         10723012 bp ( 23.9%)
      Ob other exon region size:               8098313 bp ( 18.0%)
    Ob intron region size:                 121844744 bp ( 54.4%)
  Ob intergenic region size:               57124348 bp ( 25.5%)
    Ob promoter region size:                 6874701 bp (  3.1%)
    Ob other intergenic region size:        50249647 bp ( 22.4%)

================================================================================
Methylation sites in genomic feature regions
================================================================================

Number and density (per 10kb) of sites in Ob_LI_BCphase_scd

Number of sites identified in Ob Ob_LI_BCphase_scd:                              24546430     ( 1096.43)
Number of sites identified in Ob Ob_LI_BCphase_scd genic regions:                17732952     ( 1063.43)                         72.24%     ( 0.97 O/E)
Number of sites identified in Ob Ob_LI_BCphase_scd   exon regions:                  4826112   ( 1074.68)                           19.66%   ( 0.98 O/E)
Number of sites identified in Ob Ob_LI_BCphase_scd     CDS regions:                   2636718 ( 1276.79)                             54.63% ( 1.19 O/E)
Number of sites identified in Ob Ob_LI_BCphase_scd     five-prime UTR regions:         749276 ( 1378.64)                             15.53% ( 1.28 O/E)
Number of sites identified in Ob Ob_LI_BCphase_scd     three-prime UTR regions:        946239 (  882.44)                             19.61% ( 0.82 O/E)
Number of sites identified in Ob Ob_LI_BCphase_scd     other exon regions:             493879 (  609.85)                             10.23% ( 0.57 O/E)
Number of sites identified in Ob Ob_LI_BCphase_scd   intron regions:               12906840   ( 1059.29)                           52.58%   ( 0.97 O/E)
Number of sites identified in Ob Ob_LI_BCphase_scd intergenic regions:            6813478     ( 1192.74)                         27.76%     ( 1.09 O/E)
Number of sites identified in Ob Ob_LI_BCphase_scd   promoter regions:               406408   (  591.16)                            1.66%   ( 0.54 O/E)
Number of sites identified in Ob Ob_LI_BCphase_scd   other intergenic regions:      6407070   ( 1275.05)                           26.10%   ( 1.16 O/E)

Number and density (per 10kb) of sites in Ob_LI_BCphase_hsm

Number of sites identified in Ob Ob_LI_BCphase_hsm:                                415425     (   18.56) (  1.69% of control)
Number of sites identified in Ob Ob_LI_BCphase_hsm genic regions:                  401349     (   24.07) (  2.26% of control)  96.61%     ( 1.30 O/E)
Number of sites identified in Ob Ob_LI_BCphase_hsm   exon regions:                   310975   (   69.25) (  6.44% of control)    74.86%   ( 3.73 O/E)
Number of sites identified in Ob Ob_LI_BCphase_hsm     CDS regions:                   287719 (  139.32) ( 10.91% of control)      92.52% ( 2.01 O/E)
Number of sites identified in Ob Ob_LI_BCphase_hsm     five-prime UTR regions:          5403 (    9.94) (  0.72% of control)       1.74% ( 0.14 O/E)
Number of sites identified in Ob Ob_LI_BCphase_hsm     three-prime UTR regions:        14589 (   13.61) (  1.54% of control)       4.69% ( 0.20 O/E)
Number of sites identified in Ob Ob_LI_BCphase_hsm     other exon regions:              3264 (    4.03) (  0.66% of control)       1.05% ( 0.06 O/E)
Number of sites identified in Ob Ob_LI_BCphase_hsm   intron regions:                  90374   (    7.42) (  0.70% of control)    21.75%   ( 0.40 O/E)
Number of sites identified in Ob Ob_LI_BCphase_hsm intergenic regions:              14076     (    2.46) (  0.21% of control)   3.39%     ( 0.13 O/E)
Number of sites identified in Ob Ob_LI_BCphase_hsm   promoter regions:                1344   (    1.95) (  0.33% of control)     0.32%   ( 0.11 O/E)
Number of sites identified in Ob Ob_LI_BCphase_hsm   other intergenic regions:       12732   (    2.53) (  0.20% of control)     3.06%   ( 0.14 O/E)
```

Directory: MMP File: mmp-Ob_LI_Rphase.txt

#### Genomic composition in terms of feature regions

| | | |
|---|---|---|
| Ob genome size: | 223876465 bp | |
| Ob genic region size: | 166752117 bp | ( 74.5%) |
| Ob exon region size: | 44907373 bp | ( 20.1%) |
| Ob CDS region size: | 20651162 bp | ( 46.0%) |
| Ob five-prime UTR region size: | 5434886 bp | ( 12.1%) |
| Ob three-prime UTR region size: | 10723012 bp | ( 23.9%) |
| Ob other exon region size: | 8098313 bp | ( 18.0%) |
| Ob intron region size: | 121844744 bp | ( 54.4%) |
| Ob intergenic region size: | 57124348 bp | ( 25.5%) |
| Ob promoter region size: | 6874701 bp | ( 3.1%) |
| Ob other intergenic region size: | 50249647 bp | ( 22.4%) |

#### Methylation sites in genomic feature regions

Number and density (per 10kb) of sites in Ob_LI_Rphase_scd

| | | | | |
|---|---|---|---|---|
| Number of sites identified in Ob Ob_LI_Rphase_scd: | 24689788 | ( 1102.83) | | |
| Number of sites identified in Ob Ob_LI_Rphase_scd genic regions: | 17834275 | ( 1069.51) | 72.23% | ( 0.97 O/E) |
| Number of sites identified in Ob Ob_LI_Rphase_scd exon regions: | 4853634 | ( 1080.81) | 19.66% | ( 0.98 O/E) |
| Number of sites identified in Ob Ob_LI_Rphase_scd CDS regions: | 2647636 | ( 1282.08) | 54.55% | ( 1.19 O/E) |
| Number of sites identified in Ob Ob_LI_Rphase_scd five-prime UTR regions: | 753239 | ( 1385.93) | 15.52% | ( 1.28 O/E) |
| Number of sites identified in Ob Ob_LI_Rphase_scd three-prime UTR regions: | 955702 | ( 891.26) | 19.69% | ( 0.82 O/E) |
| Number of sites identified in Ob Ob_LI_Rphase_scd other exon regions: | 497057 | ( 613.78) | 10.24% | ( 0.57 O/E) |
| Number of sites identified in Ob Ob_LI_Rphase_scd intron regions: | 12980641 | ( 1065.34) | 52.57% | ( 0.97 O/E) |
| Number of sites identified in Ob Ob_LI_Rphase_scd intergenic regions: | 6855513 | ( 1200.10) | 27.77% | ( 1.09 O/E) |
| Number of sites identified in Ob Ob_LI_Rphase_scd promoter regions: | 409346 | ( 595.44) | 1.66% | ( 0.54 O/E) |
| Number of sites identified in Ob Ob_LI_Rphase_scd other intergenic regions: | 6446167 | ( 1282.83) | 26.11% | ( 1.16 O/E) |

Number and density (per 10kb) of sites in Ob_LI_Rphase_hsm

| | | | | | |
|---|---|---|---|---|---|
| Number of sites identified in Ob Ob_LI_Rphase_hsm: | 445068 | ( 19.88) | ( 1.80% of control) | | |
| Number of sites identified in Ob Ob_LI_Rphase_hsm genic regions: | 427606 | ( 25.64) | ( 2.40% of control) | 96.08% | ( 1.29 O/E) |
| Number of sites identified in Ob Ob_LI_Rphase_hsm exon regions: | 327034 | ( 72.82) | ( 6.74% of control) | 73.48% | ( 3.66 O/E) |
| Number of sites identified in Ob Ob_LI_Rphase_hsm CDS regions: | 300947 | ( 145.73) | ( 11.37% of control) | 92.02% | ( 2.00 O/E) |
| Number of sites identified in Ob Ob_LI_Rphase_hsm five-prime UTR regions: | 6009 | ( 11.06) | ( 0.80% of control) | 1.84% | ( 0.15 O/E) |
| Number of sites identified in Ob Ob_LI_Rphase_hsm three-prime UTR regions: | 16375 | ( 15.27) | ( 1.71% of control) | 5.01% | ( 0.21 O/E) |
| Number of sites identified in Ob Ob_LI_Rphase_hsm other exon regions: | 3703 | ( 4.57) | ( 0.74% of control) | 1.13% | ( 0.06 O/E) |
| Number of sites identified in Ob Ob_LI_Rphase_hsm intron regions: | 100572 | ( 8.25) | ( 0.77% of control) | 22.60% | ( 0.42 O/E) |
| Number of sites identified in Ob Ob_LI_Rphase_hsm intergenic regions: | 17462 | ( 3.06) | ( 0.25% of control) | 3.92% | ( 0.15 O/E) |
| Number of sites identified in Ob Ob_LI_Rphase_hsm promoter regions: | 1648 | ( 2.40) | ( 0.40% of control) | 0.37% | ( 0.12 O/E) |
| Number of sites identified in Ob Ob_LI_Rphase_hsm other intergenic regions: | 15814 | ( 3.15) | ( 0.25% of control) | 3.55% | ( 0.16 O/E) |

```
Directory: ACS File: 0READMEacs

ACS - Annotation of conserved methylation sites.

Input: studymk, genome_ann (a list of GRanges objects providing annotated region labels and bounds;
                            output of BWASPR::get_genome_annotation())

Output: file acs-*.txt

Notes: The output is generated by BWASPR::annotate_methylome(), which uses methylKit::unite()
        to determine the sites common to all experiments and applies genomation::annotateWithFeature().
        The acs-*.txt files show the common sites with methylation levels in all samples
        and columns with booleans indicating whether the site fits a specific annotation feature.

       Abbreviations used: pc, protein coding; nc, non-coding; fp, five prime; tp, three prime.
                           UTR, untranslated region; unique, not overlapping with other categories.
```

Directory: ACS File: acs-Ob_LIp.txt

| BCphase | Rphase | chr | start | end | strand | gene | exon | pcexon | promoter | CDS | fpUTR | tpUTR | fpUTRnotCDS | tpUTRnotCDS | tpUTRunique | ncexon |
|---|---|---|---|---|---|---|---|---|---|---|---|---|---|---|---|---|
| 14.75 | 15.91 | NC_039506.1 | 55481 | 55481 | + | 0 | 0 | 0 | 0 | 0 | 0 | 0 | 0 | 0 | 0 | 0 |
| 9.62 | 10.38 | NC_039506.1 | 137597 | 137597 | + | 1 | 1 | 1 | 0 | 1 | 0 | 0 | 0 | 0 | 0 | 0 |
| 13.64 | 15.58 | NC_039506.1 | 138205 | 138205 | + | 1 | 1 | 1 | 0 | 1 | 0 | 0 | 0 | 0 | 0 | 0 |
| 10.99 | 11.46 | NC_039506.1 | 138592 | 138592 | + | 1 | 1 | 1 | 0 | 1 | 0 | 0 | 0 | 0 | 0 | 0 |
| 12.82 | 14.47 | NC_039506.1 | 138611 | 138611 | + | 1 | 1 | 1 | 0 | 1 | 0 | 0 | 0 | 0 | 0 | 0 |
| 29.17 | 17.46 | NC_039506.1 | 310073 | 310073 | + | 0 | 0 | 0 | 0 | 0 | 0 | 0 | 0 | 0 | 0 | 0 |
| 21.88 | 19.15 | NC_039506.1 | 376412 | 376412 | + | 1 | 1 | 1 | 0 | 0 | 0 | 1 | 0 | 1 | 1 | 0 |
| 40 | 37.93 | NC_039506.1 | 463931 | 463931 | + | 1 | 0 | 0 | 0 | 0 | 0 | 0 | 0 | 0 | 0 | 0 |
| 15.57 | 19.42 | NC_039506.1 | 633497 | 633497 | + | 0 | 0 | 0 | 0 | 0 | 0 | 0 | 0 | 0 | 0 | 0 |
| 20.45 | 23.33 | NC_039506.1 | 834762 | 834762 | + | 0 | 0 | 0 | 0 | 0 | 0 | 0 | 0 | 0 | 0 | 0 |
| 23.08 | 26.83 | NC_039506.1 | 926754 | 926754 | + | 1 | 1 | 1 | 0 | 0 | 0 | 1 | 0 | 1 | 1 | 0 |
| 17.65 | 19.2 | NC_039506.1 | 956202 | 956202 | + | 1 | 0 | 0 | 0 | 0 | 0 | 0 | 0 | 0 | 0 | 0 |
| 20.79 | 18.56 | NC_039506.1 | 1058205 | 1058205 | + | 1 | 1 | 0 | 0 | 0 | 0 | 0 | 0 | 0 | 0 | 1 |
| 14.55 | 15.38 | NC_039506.1 | 1097573 | 1097573 | + | 1 | 1 | 1 | 0 | 1 | 0 | 0 | 0 | 0 | 0 | 0 |
| 36 | 42 | NC_039506.1 | 1097588 | 1097588 | + | 1 | 1 | 1 | 0 | 1 | 0 | 0 | 0 | 0 | 0 | 0 |
| 52.3 | 59.54 | NC_039506.1 | 1097598 | 1097598 | + | 1 | 1 | 1 | 0 | 1 | 0 | 0 | 0 | 0 | 0 | 0 |
| 10.23 | 13.2 | NC_039506.1 | 1097647 | 1097647 | + | 1 | 1 | 1 | 0 | 1 | 0 | 0 | 0 | 0 | 0 | 0 |
| 18.85 | 23.38 | NC_039506.1 | 1097702 | 1097702 | + | 1 | 1 | 1 | 0 | 1 | 0 | 0 | 0 | 0 | 0 | 0 |
| 96 | 94.33 | NC_039506.1 | 1097773 | 1097773 | + | 1 | 1 | 1 | 0 | 1 | 0 | 0 | 0 | 0 | 0 | 0 |
| 80 | 86.96 | NC_039506.1 | 1097865 | 1097865 | + | 1 | 1 | 1 | 0 | 1 | 0 | 0 | 0 | 0 | 0 | 0 |
| 54.17 | 70.37 | NC_039506.1 | 1097884 | 1097884 | + | 1 | 1 | 1 | 0 | 1 | 0 | 0 | 0 | 0 | 0 | 0 |
| 93.55 | 96.77 | NC_039506.1 | 1097924 | 1097924 | + | 1 | 1 | 1 | 0 | 1 | 0 | 0 | 0 | 0 | 0 | 0 |
| 100 | 96.88 | NC_039506.1 | 1097943 | 1097943 | + | 1 | 1 | 1 | 0 | 1 | 0 | 0 | 0 | 0 | 0 | 0 |
| 86.96 | 95.45 | NC_039506.1 | 1097981 | 1097981 | + | 1 | 0 | 0 | 0 | 0 | 0 | 0 | 0 | 0 | 0 | 0 |

...
... only first 25 lines shown ...

```
Directory: RNK File: 0READMErnk

RNK - Genomic feature regions ranked by CpG methylation and hsm statistics.

Input: studymc, studymk, region.gr (GRanges object providing annotated region labels and bounds)

Output: files ranked-genes-*.txt sites-in-genes-*.txt
              ranked-promoters-*.txt sites-in-promoters-*.txt
              plot*.pdf

Notes: The output is generated by BWASPR::rank_rbm().
       The files ranked-genes-*.txt provide ranked lists of genes based on
        overall methylation percentage or the occurrence of CpGhms sites
        within the annotated gene bounds.  The output columns include
          region_ID    (= region/gene name)
          rwidth       (= region/gene width)
          nbrsites     (= number of CpGhsm sites in the region/gene)
          nbrper10kb   (= number of CpGhsm sites in the region/gene normalized to 10kb width)
          pmrpersite   (= average % methylation per CpGhsm site in the region/gene)
          pmrpernucl   (= average % CpGhsm methylation per nucleotide in the region/gene)
          prcntM       (= overall % CpGhsm methylation in the region/gene)

       The tables are sorted by prcntM (*byPrcntM* files) or by nbper10kb
         (*bySiteDensity* files) in descending order.

       The ranked-promoters-*.txt files provide analogous tables for promoter regions.
        If the promoter annotation was derived with BWASP, then these regions are simply
        defined here as 500 nucleotides upstream of the 5'-end of the gene (shorter if the
        scaffold ends before).

       The files sites-*.txt provide details for the CpGhsm sites in the respective
        regions.
       The files plot-*.pdf explore concordance of the two methylation measures.
       The optional parameters minrwidth, maxrwidth, and minnbrsites specify
       the length range and minimal number of methylation sites for regions
       to be included in the plot.
```

Directory: RNK File: ranked-genes-byPrcntM-Ob_LIp_BCphase.txt

| region_ID | seqnames | start | end | width | strand | coverage | numCs | numTs | prcntM |
|---|---|---|---|---|---|---|---|---|---|
| gene-LOC113562209 | NC_039511.1 | 14632766 | 14633743 | 978 | - | 55 | 54 | 1 | 98.18 |
| gene-LOC113561532 | NC_039507.1 | 7278071 | 7278142 | 72 | + | 512 | 491 | 21 | 95.9 |
| gene-LOC113563560 | NC_039506.1 | 6691659 | 6696603 | 4945 | + | 909 | 796 | 113 | 87.57 |
| gene-LOC113563324 | NC_039518.1 | 9465393 | 9469066 | 3674 | + | 1492 | 1177 | 315 | 78.89 |
| gene-LOC105286755 | NC_039506.1 | 6803777 | 6805333 | 1557 | - | 2372 | 1808 | 564 | 76.22 |
| gene-LOC105275557 | NC_039513.1 | 10333074 | 10333884 | 811 | + | 2768 | 2051 | 717 | 74.1 |
| gene-LOC105277804 | NC_039506.1 | 6089683 | 6090856 | 1174 | + | 1922 | 1397 | 525 | 72.68 |
| gene-LOC105281543 | NC_039517.1 | 3028824 | 3030314 | 1491 | - | 2194 | 1545 | 649 | 70.42 |
| gene-LOC105276706 | NC_039511.1 | 13248978 | 13250417 | 1440 | - | 4976 | 3503 | 1473 | 70.4 |
| gene-LOC105283535 | NC_039509.1 | 110345 | 111564 | 1220 | + | 2465 | 1723 | 742 | 69.9 |
| gene-LOC105283045 | NC_039519.1 | 7797834 | 7799515 | 1682 | + | 3733 | 2600 | 1133 | 69.65 |
| gene-LOC105277649 | NC_039512.1 | 2275901 | 2276899 | 999 | - | 2291 | 1591 | 700 | 69.45 |
| gene-LOC105281669 | NC_039507.1 | 5181562 | 5182434 | 873 | - | 2472 | 1711 | 761 | 69.22 |
| gene-LOC113562627 | NC_039514.1 | 15577696 | 15578092 | 397 | + | 3527 | 2432 | 1095 | 68.95 |
| gene-LOC109610680 | NC_039515.1 | 1495098 | 1496589 | 1492 | - | 8349 | 5607 | 2742 | 67.16 |
| gene-LOC105283854 | NC_039509.1 | 2130793 | 2133136 | 2344 | - | 1428 | 951 | 477 | 66.6 |
| gene-LOC113562396 | NC_039513.1 | 5906271 | 5907921 | 1651 | + | 239 | 158 | 81 | 66.11 |
| gene-LOC105281831 | NC_039511.1 | 2540409 | 2541476 | 1068 | - | 2562 | 1691 | 871 | 66 |
| gene-LOC105281226 | NC_039510.1 | 10003292 | 10004599 | 1308 | + | 2529 | 1646 | 883 | 65.09 |
| gene-LOC105286178 | NC_039509.1 | 7797927 | 7799516 | 1590 | + | 883 | 574 | 309 | 65.01 |
| gene-LOC105276680 | NC_039511.1 | 14307738 | 14309714 | 1977 | - | 9805 | 6359 | 3446 | 64.85 |
| gene-LOC105287955 | NC_039507.1 | 1428396 | 1430881 | 2486 | - | 4105 | 2645 | 1460 | 64.43 |
| gene-LOC105282390 | NC_039512.1 | 8801935 | 8803557 | 1623 | + | 8735 | 5616 | 3119 | 64.29 |
| gene-LOC105283997 | NC_039507.1 | 14517342 | 14519998 | 2657 | - | 162 | 104 | 58 | 64.2 |

...

... only first 25 lines shown ...

Directory: RNK File: ranked-genes-byPrcntM-Ob_LIp_Rphase.txt

| region_ID | seqnames | start | end | width | strand | coverage | numCs | numTs | prcntM |
|---|---|---|---|---|---|---|---|---|---|
| gene-LOC113561532 | NC_039507.1 | 7278071 | 7278142 | 72 | + | 643 | 617 | 26 | 95.96 |
| gene-LOC113563560 | NC_039506.1 | 6691659 | 6696603 | 4945 | + | 998 | 851 | 147 | 85.27 |
| gene-LOC113563324 | NC_039518.1 | 9465393 | 9469066 | 3674 | + | 2135 | 1686 | 449 | 78.97 |
| gene-LOC105286755 | NC_039506.1 | 6803777 | 6805333 | 1557 | - | 2427 | 1900 | 527 | 78.29 |
| gene-LOC105275557 | NC_039513.1 | 10333074 | 10333884 | 811 | + | 2945 | 2185 | 760 | 74.19 |
| gene-LOC105277649 | NC_039512.1 | 2275901 | 2276899 | 999 | - | 2541 | 1843 | 698 | 72.53 |
| gene-LOC113563370 | NC_039519.1 | 415372 | 417073 | 1702 | - | 21 | 15 | 6 | 71.43 |
| gene-LOC105277804 | NC_039506.1 | 6089683 | 6090856 | 1174 | + | 2025 | 1432 | 593 | 70.72 |
| gene-LOC105281543 | NC_039517.1 | 3028824 | 3030314 | 1491 | - | 2571 | 1808 | 763 | 70.32 |
| gene-LOC105276706 | NC_039511.1 | 13248978 | 13250417 | 1440 | - | 5762 | 4038 | 1724 | 70.08 |
| gene-LOC113562209 | NC_039511.1 | 14632766 | 14633743 | 978 | - | 70 | 49 | 21 | 70 |
| gene-LOC105283045 | NC_039519.1 | 7797834 | 7799515 | 1682 | + | 3810 | 2667 | 1143 | 70 |
| gene-LOC105283854 | NC_039509.1 | 2130793 | 2133136 | 2344 | - | 1599 | 1109 | 490 | 69.36 |
| gene-LOC105281669 | NC_039507.1 | 5181562 | 5182434 | 873 | - | 2877 | 1979 | 898 | 68.79 |
| gene-LOC105283535 | NC_039509.1 | 110345 | 111564 | 1220 | + | 2878 | 1958 | 920 | 68.03 |
| gene-LOC113562627 | NC_039514.1 | 15577696 | 15578092 | 397 | + | 3761 | 2547 | 1214 | 67.72 |
| gene-LOC109610680 | NC_039515.1 | 1495098 | 1496589 | 1492 | - | 9550 | 6463 | 3087 | 67.68 |
| gene-LOC113562396 | NC_039513.1 | 5906271 | 5907921 | 1651 | + | 261 | 176 | 85 | 67.43 |
| gene-LOC105281226 | NC_039510.1 | 10003292 | 10004599 | 1308 | + | 2763 | 1815 | 948 | 65.69 |
| gene-LOC105277325 | NC_039514.1 | 13671986 | 13674003 | 2018 | + | 3662 | 2390 | 1272 | 65.26 |
| gene-LOC105286178 | NC_039509.1 | 7797927 | 7799516 | 1590 | + | 1017 | 662 | 355 | 65.09 |
| gene-LOC105287955 | NC_039507.1 | 1428396 | 1430881 | 2486 | - | 4374 | 2846 | 1528 | 65.07 |
| gene-LOC105276680 | NC_039511.1 | 14307738 | 14309714 | 1977 | - | 11683 | 7583 | 4100 | 64.91 |
| gene-LOC105281831 | NC_039511.1 | 2540409 | 2541476 | 1068 | - | 2958 | 1918 | 1040 | 64.84 |

...

... only first 25 lines shown ...

Directory: RNK File: ranked-genes-bySiteDensity-Ob_LIp_BCphase.txt

| region_ID | rwidth | nbrsites | nbrper10kb | pmpersite | pmpernucl | pglink | seqnames | start | end | strand | coverage | numCs | numTs | prcntM |
|---|---|---|---|---|---|---|---|---|---|---|---|---|---|---|
| gene-LOC105278849 | 776 | 106 | 1365.98 | 31 | 4.23 | https://www.ncbi.nlm.nih.gov/gene/?term=LOC105278849 | NC_039517.1 | 644964 | 645739 | - | 21371 | 4972 | 16399 | 23.27 |
| gene-LOC105283830 | 252 | 26 | 1031.75 | 46.38 | 4.78 | https://www.ncbi.nlm.nih.gov/gene/?term=LOC105283830 | NC_039509.1 | 2003101 | 2003352 | - | 3256 | 1149 | 2107 | 35.29 |
| gene-LOC113561532 | 72 | 7 | 972.22 | 96.41 | 9.37 | https://www.ncbi.nlm.nih.gov/gene/?term=LOC113561532 | NC_039507.1 | 7278071 | 7278142 | + | 512 | 491 | 21 | 95.9 |
| gene-LOC105278873 | 1897 | 183 | 964.68 | 31.84 | 3.07 | https://www.ncbi.nlm.nih.gov/gene/?term=LOC105278873 | NC_039517.1 | 502319 | 504215 | + | 41100 | 8557 | 32543 | 20.82 |
| gene-LOC113562627 | 397 | 38 | 957.18 | 69.18 | 6.62 | https://www.ncbi.nlm.nih.gov/gene/?term=LOC113562627 | NC_039514.1 | 15577696 | 15578092 | + | 3527 | 2432 | 1095 | 68.95 |
| gene-LOC105285635 | 1175 | 106 | 902.13 | 45.45 | 4.1 | https://www.ncbi.nlm.nih.gov/gene/?term=LOC105285635 | NC_039513.1 | 15171298 | 15172472 | + | 22563 | 7285 | 15278 | 32.29 |
| gene-LOC105282149 | 1787 | 161 | 900.95 | 57.72 | 5.2 | https://www.ncbi.nlm.nih.gov/gene/?term=LOC105282149 | NC_039515.1 | 12586764 | 12588550 | + | 29466 | 12566 | 16900 | 42.65 |
| gene-LOC105286531 | 372 | 33 | 887.1 | 49.83 | 4.42 | https://www.ncbi.nlm.nih.gov/gene/?term=LOC105286531 | NC_039510.1 | 15744784 | 15745155 | - | 6618 | 2384 | 4234 | 36.02 |
| gene-LOC105278160 | 1872 | 162 | 865.38 | 45.23 | 3.91 | https://www.ncbi.nlm.nih.gov/gene/?term=LOC105278160 | NC_039507.1 | 17010609 | 17012480 | - | 29107 | 11295 | 17812 | 38.81 |
| gene-LOC105287677 | 1588 | 132 | 831.23 | 47.08 | 3.91 | https://www.ncbi.nlm.nih.gov/gene/?term=LOC105287677 | NC_039510.1 | 3210462 | 3212049 | - | 19610 | 8104 | 11506 | 41.33 |
| gene-LOC105285392 | 1601 | 133 | 830.73 | 46.81 | 3.89 | https://www.ncbi.nlm.nih.gov/gene/?term=LOC105285392 | NC_039518.1 | 2657034 | 2658634 | + | 22063 | 7877 | 14186 | 35.7 |
| gene-LOC105274711 | 1655 | 137 | 827.79 | 56.52 | 4.68 | https://www.ncbi.nlm.nih.gov/gene/?term=LOC105274711 | NC_039515.1 | 1527980 | 1529634 | + | 22014 | 10158 | 11856 | 46.14 |
| gene-LOC105285394 | 1307 | 108 | 826.32 | 72.6 | 6 | https://www.ncbi.nlm.nih.gov/gene/?term=LOC105285394 | NC_039518.1 | 2681489 | 2682795 | - | 13267 | 8083 | 5184 | 60.93 |
| gene-LOC105288193 | 1495 | 120 | 802.68 | 42.71 | 3.43 | https://www.ncbi.nlm.nih.gov/gene/?term=LOC105288193 | NC_039516.1 | 3100198 | 3101692 | + | 20589 | 5792 | 14797 | 28.13 |
| gene-LOC109611487 | 437 | 35 | 800.92 | 24.94 | 2 | https://www.ncbi.nlm.nih.gov/gene/?term=LOC109611487 | NC_039506.1 | 13096874 | 13097310 | + | 7475 | 1161 | 6314 | 15.53 |
| gene-LOC105279035 | 2108 | 167 | 792.22 | 43.99 | 3.49 | https://www.ncbi.nlm.nih.gov/gene/?term=LOC105279035 | NC_039514.1 | 7348206 | 7350313 | + | 40608 | 7780 | 32828 | 19.16 |
| gene-LOC109610897 | 1561 | 122 | 781.55 | 30.67 | 2.4 | https://www.ncbi.nlm.nih.gov/gene/?term=LOC109610897 | NC_039509.1 | 13081754 | 13083314 | - | 31416 | 4842 | 26574 | 15.41 |
| gene-LOC105276490 | 2575 | 201 | 780.58 | 33.93 | 2.65 | https://www.ncbi.nlm.nih.gov/gene/?term=LOC105276490 | NC_039511.1 | 697281 | 699855 | + | 53652 | 11377 | 42275 | 21.21 |
| gene-LOC105283847 | 1590 | 124 | 779.87 | 40.23 | 3.14 | https://www.ncbi.nlm.nih.gov/gene/?term=LOC105283847 | NC_039509.1 | 1750527 | 1752116 | - | 22395 | 7519 | 14876 | 33.57 |
| gene-LOC105276360 | 1466 | 114 | 777.63 | 57.53 | 4.47 | https://www.ncbi.nlm.nih.gov/gene/?term=LOC105276360 | NC_039513.1 | 12169421 | 12170886 | - | 26131 | 8305 | 17826 | 31.78 |
| gene-LOC105286269 | 1401 | 108 | 770.88 | 57.33 | 4.42 | https://www.ncbi.nlm.nih.gov/gene/?term=LOC105286269 | NC_039518.1 | 8735257 | 8736657 | - | 15972 | 7715 | 8257 | 48.3 |
| gene-LOC105287692 | 1505 | 115 | 764.12 | 49.3 | 3.77 | https://www.ncbi.nlm.nih.gov/gene/?term=LOC105287692 | NC_039510.1 | 3132542 | 3134046 | + | 23520 | 7507 | 16013 | 31.92 |
| gene-LOC105286135 | 2051 | 156 | 760.6 | 28.49 | 2.17 | https://www.ncbi.nlm.nih.gov/gene/?term=LOC105286135 | NC_039517.1 | 339965 | 342015 | + | 28786 | 5393 | 23393 | 18.73 |
| gene-LOC105278396 | 3012 | 228 | 756.97 | 41.6 | 3.15 | https://www.ncbi.nlm.nih.gov/gene/?term=LOC105278396 | NC_039515.1 | 2647513 | 2650524 | + | 48786 | 12462 | 36324 | 25.54 |

...

... only first 25 lines shown ...

Directory: RNK File: ranked-genes-bySiteDensity-Ob_LIp_Rphase.txt

| region_ID | rwidth | nbrsites | nbrper10kb | pmpersite | pmpernucl | pglink | seqnames | start | end | strand | coverage | numCs | numTs | prcntM |
|---|---|---|---|---|---|---|---|---|---|---|---|---|---|---|
| gene-LOC105278849 | 776 | 112 | 1443.3 | 29.54 | 4.26 | https://www.ncbi.nlm.nih.gov/gene/?term=LOC105278849 | NC_039517.1 | 644964 | 645739 | - | 23312 | 5249 | 18063 | 22.52 |
| gene-LOC105283830 | 252 | 25 | 992.06 | 49.15 | 4.88 | https://www.ncbi.nlm.nih.gov/gene/?term=LOC105283830 | NC_039509.1 | 2003101 | 2003352 | - | 3233 | 1218 | 2015 | 37.67 |
| gene-LOC105278873 | 1897 | 185 | 975.22 | 31.99 | 3.12 | https://www.ncbi.nlm.nih.gov/gene/?term=LOC105278873 | NC_039517.1 | 502319 | 504215 | + | 45260 | 9547 | 35713 | 21.09 |
| gene-LOC113561532 | 72 | 7 | 972.22 | 96.09 | 9.34 | https://www.ncbi.nlm.nih.gov/gene/?term=LOC113561532 | NC_039507.1 | 7278071 | 7278142 | + | 643 | 617 | 26 | 95.96 |
| gene-LOC113562627 | 397 | 38 | 957.18 | 69.14 | 6.62 | https://www.ncbi.nlm.nih.gov/gene/?term=LOC113562627 | NC_039514.1 | 15577696 | 15578092 | + | 3761 | 2547 | 1214 | 67.72 |
| gene-LOC105282149 | 1787 | 166 | 928.93 | 56.1 | 5.21 | https://www.ncbi.nlm.nih.gov/gene/?term=LOC105282149 | NC_039515.1 | 12586764 | 12588550 | + | 33188 | 14222 | 18966 | 42.85 |
| gene-LOC105285635 | 1175 | 104 | 885.11 | 46.08 | 4.08 | https://www.ncbi.nlm.nih.gov/gene/?term=LOC105285635 | NC_039513.1 | 15171298 | 15172472 | + | 25086 | 8092 | 16994 | 32.26 |
| gene-LOC105288193 | 1495 | 130 | 869.57 | 42.69 | 3.71 | https://www.ncbi.nlm.nih.gov/gene/?term=LOC105288193 | NC_039516.1 | 3100198 | 3101692 | + | 23357 | 6591 | 16766 | 28.22 |
| gene-LOC105278160 | 1872 | 162 | 865.38 | 43.7 | 3.78 | https://www.ncbi.nlm.nih.gov/gene/?term=LOC105278160 | NC_039507.1 | 17010609 | 17012480 | - | 33405 | 12631 | 20774 | 37.81 |
| gene-LOC105286531 | 372 | 32 | 860.22 | 50.66 | 4.36 | https://www.ncbi.nlm.nih.gov/gene/?term=LOC105286531 | NC_039510.1 | 15744784 | 15745155 | - | 7051 | 2474 | 4577 | 35.09 |
| gene-LOC105285392 | 1601 | 136 | 849.47 | 44.75 | 3.8 | https://www.ncbi.nlm.nih.gov/gene/?term=LOC105285392 | NC_039518.1 | 2657034 | 2658634 | + | 25449 | 9246 | 16203 | 36.33 |
| gene-LOC105274711 | 1655 | 140 | 845.92 | 54.15 | 4.58 | https://www.ncbi.nlm.nih.gov/gene/?term=LOC105274711 | NC_039515.1 | 1527980 | 1529634 | + | 24496 | 11222 | 13274 | 45.81 |
| gene-LOC105279035 | 2108 | 178 | 844.4 | 41.85 | 3.53 | https://www.ncbi.nlm.nih.gov/gene/?term=LOC105279035 | NC_039514.1 | 7348206 | 7350313 | + | 44448 | 9088 | 35360 | 20.45 |
| gene-LOC105285394 | 1307 | 110 | 841.62 | 71.57 | 6.02 | https://www.ncbi.nlm.nih.gov/gene/?term=LOC105285394 | NC_039518.1 | 2681489 | 2682795 | - | 14918 | 9290 | 5628 | 62.27 |
| gene-LOC109611487 | 437 | 36 | 823.8 | 26.29 | 2.17 | https://www.ncbi.nlm.nih.gov/gene/?term=LOC109611487 | NC_039506.1 | 13096874 | 13097310 | + | 8416 | 1387 | 7029 | 16.48 |
| gene-LOC105287677 | 1588 | 130 | 818.64 | 47.59 | 3.9 | https://www.ncbi.nlm.nih.gov/gene/?term=LOC105287677 | NC_039510.1 | 3210462 | 3212049 | - | 21898 | 9090 | 12808 | 41.51 |
| gene-LOC105276490 | 2575 | 210 | 815.53 | 33.79 | 2.76 | https://www.ncbi.nlm.nih.gov/gene/?term=LOC105276490 | NC_039511.1 | 697281 | 699855 | + | 60187 | 12583 | 47604 | 20.91 |
| gene-LOC109610897 | 1561 | 126 | 807.17 | 30.13 | 2.43 | https://www.ncbi.nlm.nih.gov/gene/?term=LOC109610897 | NC_039509.1 | 13081754 | 13083314 | - | 34155 | 5431 | 28724 | 15.9 |
| gene-LOC105274542 | 1266 | 102 | 805.69 | 41.11 | 3.31 | https://www.ncbi.nlm.nih.gov/gene/?term=LOC105274542 | NC_039516.1 | 1324720 | 1325985 | + | 15298 | 4362 | 10936 | 28.51 |
| gene-LOC105283847 | 1590 | 128 | 805.03 | 38.71 | 3.12 | https://www.ncbi.nlm.nih.gov/gene/?term=LOC105283847 | NC_039509.1 | 1750527 | 1752116 | - | 23914 | 8076 | 15838 | 33.77 |
| gene-LOC105286135 | 2051 | 163 | 794.73 | 28.15 | 2.24 | https://www.ncbi.nlm.nih.gov/gene/?term=LOC105286135 | NC_039517.1 | 339965 | 342015 | + | 31191 | 6130 | 25061 | 19.65 |
| gene-LOC105275231 | 1270 | 100 | 787.4 | 58.19 | 4.58 | https://www.ncbi.nlm.nih.gov/gene/?term=LOC105275231 | NC_039516.1 | 12121205 | 12122474 | - | 14689 | 6279 | 8410 | 42.75 |
| gene-LOC105276360 | 1466 | 115 | 784.45 | 57.19 | 4.49 | https://www.ncbi.nlm.nih.gov/gene/?term=LOC105276360 | NC_039513.1 | 12169421 | 12170886 | - | 29564 | 9528 | 20036 | 32.23 |
| gene-LOC105286737 | 2690 | 207 | 769.52 | 40.74 | 3.14 | https://www.ncbi.nlm.nih.gov/gene/?term=LOC105286737 | NC_039506.1 | 10795596 | 10798285 | + | 66851 | 18175 | 48676 | 27.19 |

...

... only first 25 lines shown ...

Directory: RNK File: ranked-promoters-byPrcntM-Ob_LIp_BCphase.txt

| region_ID | seqnames | start | end | width | strand | coverage | numCs | numTs | prcntM |
|---|---|---|---|---|---|---|---|---|---|
| gene-LOC105285355 | NC_039509.1 | 8035614 | 8036113 | 500 | - | 165 | 154 | 11 | 93.33 |
| gene-LOC105276139 | NC_039514.1 | 452530 | 453029 | 500 | - | 249 | 230 | 19 | 92.37 |
| gene-LOC105276685 | NC_039511.1 | 14299458 | 14299957 | 500 | - | 3437 | 3124 | 313 | 90.89 |
| gene-LOC105285429 | NC_039517.1 | 10183111 | 10183610 | 500 | + | 1358 | 1210 | 148 | 89.1 |
| gene-LOC113562329 | NC_039506.1 | 5740607 | 5741106 | 500 | + | 115 | 102 | 13 | 88.7 |
| gene-LOC113561391 | NC_039507.1 | 999503 | 1000002 | 500 | + | 508 | 442 | 66 | 87.01 |
| gene-LOC105276002 | NC_039511.1 | 12550061 | 12550560 | 500 | + | 308 | 264 | 44 | 85.71 |
| gene-LOC113561532 | NC_039507.1 | 7277571 | 7278070 | 500 | + | 4199 | 3597 | 602 | 85.66 |
| gene-LOC105277264 | NC_039518.1 | 2605800 | 2606299 | 500 | - | 19 | 16 | 3 | 84.21 |
| gene-LOC105281689 | NC_039507.1 | 5097604 | 5098103 | 500 | - | 742 | 623 | 119 | 83.96 |
| gene-LOC105280917 | NC_039506.1 | 11578183 | 11578682 | 500 | - | 377 | 310 | 67 | 82.23 |
| gene-LOC105277073 | NC_039508.1 | 12946948 | 12947447 | 500 | + | 206 | 168 | 38 | 81.55 |
| gene-LOC105282801 | NC_039512.1 | 16232132 | 16232631 | 500 | + | 807 | 650 | 157 | 80.55 |
| gene-LOC105279930 | NC_039512.1 | 11167002 | 11167501 | 500 | + | 424 | 337 | 87 | 79.48 |
| gene-LOC105282078 | NC_039512.1 | 6125187 | 6125686 | 500 | + | 355 | 282 | 73 | 79.44 |
| gene-LOC113561934 | NC_039506.1 | 15618494 | 15618993 | 500 | + | 4100 | 3253 | 847 | 79.34 |
| gene-LOC105281627 | NC_039514.1 | 11099400 | 11099899 | 500 | + | 1251 | 958 | 293 | 76.58 |
| gene-LOC105287833 | NC_039513.1 | 7297618 | 7298117 | 500 | + | 1276 | 962 | 314 | 75.39 |
| gene-LOC113562533 | NC_039514.1 | 1298855 | 1299354 | 500 | + | 1821 | 1338 | 483 | 73.48 |
| gene-LOC105279917 | NC_039512.1 | 13122631 | 13123130 | 500 | - | 861 | 631 | 230 | 73.29 |
| gene-LOC105282822 | NC_039512.1 | 16435014 | 16435513 | 500 | + | 1727 | 1263 | 464 | 73.13 |
| gene-LOC105288107 | NC_039515.1 | 1102618 | 1103117 | 500 | + | 3525 | 2568 | 957 | 72.85 |
| gene-LOC105281345 | NC_039508.1 | 2649490 | 2649989 | 500 | - | 375 | 273 | 102 | 72.8 |
| gene-LOC105281671 | NC_039507.1 | 5181504 | 5182003 | 500 | - | 2362 | 1708 | 654 | 72.31 |

...

... only first 25 lines shown ...

Directory: RNK File: ranked-promoters-byPrcntM-Ob_LIp_Rphase.txt

| region_ID | seqnames | start | end | width | strand | coverage | numCs | numTs | prcntM |
|---|---|---|---|---|---|---|---|---|---|
| gene-LOC105285355 | NC_039509.1 | 8035614 | 8036113 | 500 | - | 196 | 188 | 8 | 95.92 |
| gene-LOC105276685 | NC_039511.1 | 14299458 | 14299957 | 500 | - | 4071 | 3747 | 324 | 92.04 |
| gene-LOC105276139 | NC_039514.1 | 452530 | 453029 | 500 | - | 237 | 218 | 19 | 91.98 |
| gene-LOC105285429 | NC_039517.1 | 10183111 | 10183610 | 500 | + | 1436 | 1283 | 153 | 89.35 |
| gene-LOC113561391 | NC_039507.1 | 999503 | 1000002 | 500 | + | 507 | 448 | 59 | 88.36 |
| gene-LOC105277264 | NC_039518.1 | 2605800 | 2606299 | 500 | - | 23 | 20 | 3 | 86.96 |
| gene-LOC105280917 | NC_039506.1 | 11578183 | 11578682 | 500 | - | 364 | 315 | 49 | 86.54 |
| gene-LOC113562329 | NC_039506.1 | 5740607 | 5741106 | 500 | + | 140 | 119 | 21 | 85 |
| gene-LOC113561532 | NC_039507.1 | 7277571 | 7278070 | 500 | + | 4707 | 3969 | 738 | 84.32 |
| gene-LOC105282078 | NC_039512.1 | 6125187 | 6125686 | 500 | + | 381 | 311 | 70 | 81.63 |
| gene-LOC105281689 | NC_039507.1 | 5097604 | 5098103 | 500 | - | 909 | 734 | 175 | 80.75 |
| gene-LOC105282801 | NC_039512.1 | 16232132 | 16232631 | 500 | + | 994 | 799 | 195 | 80.38 |
| gene-LOC105276002 | NC_039511.1 | 12550061 | 12550560 | 500 | + | 408 | 326 | 82 | 79.9 |
| gene-LOC113561934 | NC_039506.1 | 15618494 | 15618993 | 500 | + | 3979 | 3174 | 805 | 79.77 |
| gene-LOC105279917 | NC_039512.1 | 13122631 | 13123130 | 500 | - | 965 | 756 | 209 | 78.34 |
| gene-LOC113562533 | NC_039514.1 | 1298855 | 1299354 | 500 | + | 2362 | 1815 | 547 | 76.84 |
| gene-LOC105277073 | NC_039508.1 | 12946948 | 12947447 | 500 | + | 241 | 185 | 56 | 76.76 |
| gene-LOC105279930 | NC_039512.1 | 11167002 | 11167501 | 500 | + | 394 | 300 | 94 | 76.14 |
| gene-LOC105282822 | NC_039512.1 | 16435014 | 16435513 | 500 | + | 2157 | 1641 | 516 | 76.08 |
| gene-LOC105276585 | NC_039517.1 | 4571850 | 4572349 | 500 | - | 3156 | 2390 | 766 | 75.73 |
| gene-LOC105287833 | NC_039513.1 | 7297618 | 7298117 | 500 | + | 1395 | 1056 | 339 | 75.7 |
| gene-LOC105281627 | NC_039514.1 | 11099400 | 11099899 | 500 | + | 1351 | 1020 | 331 | 75.5 |
| gene-LOC105288107 | NC_039515.1 | 1102618 | 1103117 | 500 | + | 3752 | 2764 | 988 | 73.67 |
| gene-LOC105281345 | NC_039508.1 | 2649490 | 2649989 | 500 | - | 422 | 308 | 114 | 72.99 |

...
... only first 25 lines shown ...

Directory: RNK File: ranked-promoters-bySiteDensity-Ob_LIp_BCphase.txt

| region_ID | rwidth | nbrsites | nbrper10kb | pmpersite | pmpernucl | pglink | seqnames | start | end | strand | coverage | numCs | numTs | prcntM |
|---|---|---|---|---|---|---|---|---|---|---|---|---|---|---|
| gene-LOC105274855 | 500 | 56 | 1120 | 30.12 | 3.37 | https://www.ncbi.nlm.nih.gov/gene/?term=LOC105274855 | NC_039513.1 | 3348900 | 3349399 | + | 7384 | 1667 | 5717 | 22.58 |
| gene-LOC105280861 | 500 | 53 | 1060 | 61.69 | 6.54 | https://www.ncbi.nlm.nih.gov/gene/?term=LOC105280861 | NC_039506.1 | 15371187 | 15371686 | + | 5271 | 2881 | 2390 | 54.66 |
| gene-LOC105277808 | 500 | 52 | 1040 | 45.48 | 4.73 | https://www.ncbi.nlm.nih.gov/gene/?term=LOC105277808 | NC_039516.1 | 13518519 | 13519018 | - | 10773 | 3340 | 7433 | 31 |
| gene-LOC105286234 | 500 | 51 | 1020 | 44.84 | 4.57 | https://www.ncbi.nlm.nih.gov/gene/?term=LOC105286234 | NC_039518.1 | 8582509 | 8583008 | + | 8651 | 3205 | 5446 | 37.05 |
| gene-LOC109611487 | 500 | 51 | 1020 | 39.04 | 3.98 | https://www.ncbi.nlm.nih.gov/gene/?term=LOC109611487 | NC_039506.1 | 13096374 | 13096873 | + | 9067 | 2366 | 6701 | 26.09 |
| gene-LOC105282404 | 500 | 49 | 980 | 47.44 | 4.65 | https://www.ncbi.nlm.nih.gov/gene/?term=LOC105282404 | NC_039512.1 | 8476210 | 8476709 | + | 6454 | 2650 | 3804 | 41.06 |
| gene-LOC105284094 | 500 | 47 | 940 | 28.34 | 2.66 | https://www.ncbi.nlm.nih.gov/gene/?term=LOC105284094 | NC_039511.1 | 3031639 | 3032138 | + | 8388 | 1682 | 6706 | 20.05 |
| gene-LOC113561934 | 500 | 47 | 940 | 80.03 | 7.52 | https://www.ncbi.nlm.nih.gov/gene/?term=LOC113561934 | NC_039506.1 | 15618494 | 15618993 | + | 4100 | 3253 | 847 | 79.34 |
| gene-LOC105284046 | 500 | 46 | 920 | 52.34 | 4.82 | https://www.ncbi.nlm.nih.gov/gene/?term=LOC105284046 | NC_039511.1 | 3657645 | 3658144 | - | 4364 | 1616 | 2748 | 37.03 |
| gene-LOC113563568 | 500 | 46 | 920 | 22.82 | 2.1 | https://www.ncbi.nlm.nih.gov/gene/?term=LOC113563568 | NC_039506.1 | 17097325 | 17097824 | + | 8408 | 1151 | 7257 | 13.69 |
| gene-LOC105277304 | 500 | 45 | 900 | 59.36 | 5.34 | https://www.ncbi.nlm.nih.gov/gene/?term=LOC105277304 | NC_039516.1 | 9356205 | 9356704 | - | 5264 | 2575 | 2689 | 48.92 |
| gene-LOC105280727 | 500 | 45 | 900 | 56.34 | 5.07 | https://www.ncbi.nlm.nih.gov/gene/?term=LOC105280727 | NC_039513.1 | 11520171 | 11520670 | + | 6885 | 3050 | 3835 | 44.3 |
| gene-LOC105279250 | 500 | 44 | 880 | 65.25 | 5.74 | https://www.ncbi.nlm.nih.gov/gene/?term=LOC105279250 | NC_039512.1 | 7901112 | 7901611 | - | 4307 | 2806 | 1501 | 65.15 |
| gene-LOC105280668 | 500 | 44 | 880 | 37.26 | 3.28 | https://www.ncbi.nlm.nih.gov/gene/?term=LOC105280668 | NC_039508.1 | 18012855 | 18013354 | - | 5489 | 1582 | 3907 | 28.82 |
| gene-LOC105283833 | 500 | 43 | 860 | 51.86 | 4.46 | https://www.ncbi.nlm.nih.gov/gene/?term=LOC105283833 | NC_039509.1 | 2003067 | 2003566 | - | 5169 | 2039 | 3130 | 39.45 |
| gene-LOC105286681 | 500 | 43 | 860 | 52.05 | 4.48 | https://www.ncbi.nlm.nih.gov/gene/?term=LOC105286681 | NC_039512.1 | 14397556 | 14398055 | + | 4366 | 1914 | 2452 | 43.84 |
| gene-LOC105286945 | 500 | 43 | 860 | 33.69 | 2.9 | https://www.ncbi.nlm.nih.gov/gene/?term=LOC105286945 | NC_039506.1 | 13096789 | 13097288 | + | 8185 | 1768 | 6417 | 21.6 |
| gene-LOC105288107 | 500 | 43 | 860 | 75.1 | 6.46 | https://www.ncbi.nlm.nih.gov/gene/?term=LOC105288107 | NC_039515.1 | 1102618 | 1103117 | + | 3525 | 2568 | 957 | 72.85 |
| gene-LOC105278673 | 500 | 42 | 840 | 74.36 | 6.25 | https://www.ncbi.nlm.nih.gov/gene/?term=LOC105278673 | NC_039516.1 | 11703885 | 11704384 | - | 4431 | 3051 | 1380 | 68.86 |
| gene-LOC105286944 | 500 | 42 | 840 | 73.15 | 6.14 | https://www.ncbi.nlm.nih.gov/gene/?term=LOC105286944 | NC_039506.1 | 13094778 | 13095277 | + | 2951 | 2069 | 882 | 70.11 |
| gene-LOC113561532 | 500 | 42 | 840 | 81.11 | 6.81 | https://www.ncbi.nlm.nih.gov/gene/?term=LOC113561532 | NC_039507.1 | 7277571 | 7278070 | + | 4199 | 3597 | 602 | 85.66 |
| gene-LOC113562689 | 500 | 40 | 800 | 56.67 | 4.53 | https://www.ncbi.nlm.nih.gov/gene/?term=LOC113562689 | NC_039515.1 | 5223138 | 5223637 | - | 4997 | 2052 | 2945 | 41.06 |
| gene-LOC105283684 | 500 | 38 | 760 | 45.81 | 3.48 | https://www.ncbi.nlm.nih.gov/gene/?term=LOC105283684 | NC_039515.1 | 9007620 | 9008119 | - | 4889 | 1879 | 3010 | 38.43 |
| gene-LOC105276585 | 500 | 37 | 740 | 75.66 | 5.6 | https://www.ncbi.nlm.nih.gov/gene/?term=LOC105276585 | NC_039517.1 | 4571850 | 4572349 | - | 2704 | 1929 | 775 | 71.34 |

...

... only first 25 lines shown ...

Directory: RNK File: ranked-promoters-bySiteDensity-Ob_LIp_Rphase.txt

| region_ID | rwidth | nbrsites | nbrper10kb | pmpersite | pmpernucl | pglink | seqnames | start | end | strand | coverage | numCs | numTs | prcntM |
|---|---|---|---|---|---|---|---|---|---|---|---|---|---|---|
| gene-LOC105277808 | 500 | 58 | 1160 | 43.81 | 5.08 | https://www.ncbi.nlm.nih.gov/gene/?term=LOC105277808 | NC_039516.1 | 13518519 | 13519018 | - | 12396 | 4025 | 8371 | 32.47 |
| gene-LOC109611487 | 500 | 55 | 1100 | 39 | 4.29 | https://www.ncbi.nlm.nih.gov/gene/?term=LOC109611487 | NC_039506.1 | 13096374 | 13096873 | + | 10805 | 2964 | 7841 | 27.43 |
| gene-LOC105280861 | 500 | 52 | 1040 | 62.14 | 6.46 | https://www.ncbi.nlm.nih.gov/gene/?term=LOC105280861 | NC_039506.1 | 15371187 | 15371686 | + | 5498 | 2891 | 2607 | 52.58 |
| gene-LOC105282404 | 500 | 52 | 1040 | 45.37 | 4.72 | https://www.ncbi.nlm.nih.gov/gene/?term=LOC105282404 | NC_039512.1 | 8476210 | 8476709 | + | 6774 | 2684 | 4090 | 39.62 |
| gene-LOC105286234 | 500 | 50 | 1000 | 44.06 | 4.41 | https://www.ncbi.nlm.nih.gov/gene/?term=LOC105286234 | NC_039518.1 | 8582509 | 8583008 | + | 9270 | 3473 | 5797 | 37.46 |
| gene-LOC105274855 | 500 | 49 | 980 | 32.42 | 3.18 | https://www.ncbi.nlm.nih.gov/gene/?term=LOC105274855 | NC_039513.1 | 3348900 | 3349399 | + | 8655 | 2059 | 6596 | 23.79 |
| gene-LOC113561934 | 500 | 49 | 980 | 79.71 | 7.81 | https://www.ncbi.nlm.nih.gov/gene/?term=LOC113561934 | NC_039506.1 | 15618494 | 15618993 | + | 3979 | 3174 | 805 | 79.77 |
| gene-LOC105280668 | 500 | 46 | 920 | 37.6 | 3.46 | https://www.ncbi.nlm.nih.gov/gene/?term=LOC105280668 | NC_039508.1 | 18012855 | 18013354 | - | 5389 | 1588 | 3801 | 29.47 |
| gene-LOC105280727 | 500 | 46 | 920 | 56.81 | 5.23 | https://www.ncbi.nlm.nih.gov/gene/?term=LOC105280727 | NC_039513.1 | 11520171 | 11520670 | + | 8210 | 3710 | 4500 | 45.19 |
| gene-LOC105284046 | 500 | 46 | 920 | 52.84 | 4.86 | https://www.ncbi.nlm.nih.gov/gene/?term=LOC105284046 | NC_039511.1 | 3657645 | 3658144 | - | 4309 | 1577 | 2732 | 36.6 |
| gene-LOC105275346 | 500 | 45 | 900 | 27.62 | 2.49 | https://www.ncbi.nlm.nih.gov/gene/?term=LOC105275346 | NC_039516.1 | 13041204 | 13041703 | + | 9749 | 1593 | 8156 | 16.34 |
| gene-LOC105277304 | 500 | 45 | 900 | 59.36 | 5.34 | https://www.ncbi.nlm.nih.gov/gene/?term=LOC105277304 | NC_039516.1 | 9356205 | 9356704 | - | 6051 | 3000 | 3051 | 49.58 |
| gene-LOC105279250 | 500 | 45 | 900 | 65.02 | 5.85 | https://www.ncbi.nlm.nih.gov/gene/?term=LOC105279250 | NC_039512.1 | 7901112 | 7901611 | - | 5279 | 3348 | 1931 | 63.42 |
| gene-LOC105286681 | 500 | 45 | 900 | 50.39 | 4.53 | https://www.ncbi.nlm.nih.gov/gene/?term=LOC105286681 | NC_039512.1 | 14397556 | 14398055 | + | 4776 | 2045 | 2731 | 42.82 |
| gene-LOC113562689 | 500 | 45 | 900 | 51.53 | 4.64 | https://www.ncbi.nlm.nih.gov/gene/?term=LOC113562689 | NC_039515.1 | 5223138 | 5223637 | - | 5977 | 2402 | 3575 | 40.19 |
| gene-LOC113563568 | 500 | 45 | 900 | 22 | 1.98 | https://www.ncbi.nlm.nih.gov/gene/?term=LOC113563568 | NC_039506.1 | 17097325 | 17097824 | + | 9597 | 1296 | 8301 | 13.5 |
| gene-LOC105286945 | 500 | 44 | 880 | 35.29 | 3.11 | https://www.ncbi.nlm.nih.gov/gene/?term=LOC105286945 | NC_039506.1 | 13096789 | 13097288 | + | 9270 | 2143 | 7127 | 23.12 |
| gene-LOC105283833 | 500 | 43 | 860 | 52.24 | 4.49 | https://www.ncbi.nlm.nih.gov/gene/?term=LOC105283833 | NC_039509.1 | 2003067 | 2003566 | - | 5356 | 2258 | 3098 | 42.16 |
| gene-LOC105284094 | 500 | 43 | 860 | 27.79 | 2.39 | https://www.ncbi.nlm.nih.gov/gene/?term=LOC105284094 | NC_039511.1 | 3031639 | 3032138 | + | 9420 | 1843 | 7577 | 19.56 |
| gene-LOC105288107 | 500 | 43 | 860 | 75.47 | 6.49 | https://www.ncbi.nlm.nih.gov/gene/?term=LOC105288107 | NC_039515.1 | 1102618 | 1103117 | + | 3752 | 2764 | 988 | 73.67 |
| gene-LOC105278673 | 500 | 42 | 840 | 72.82 | 6.12 | https://www.ncbi.nlm.nih.gov/gene/?term=LOC105278673 | NC_039516.1 | 11703885 | 11704384 | - | 4695 | 3149 | 1546 | 67.07 |
| gene-LOC105286944 | 500 | 42 | 840 | 72.91 | 6.12 | https://www.ncbi.nlm.nih.gov/gene/?term=LOC105286944 | NC_039506.1 | 13094778 | 13095277 | + | 3234 | 2348 | 886 | 72.6 |
| gene-LOC113561532 | 500 | 42 | 840 | 80.87 | 6.79 | https://www.ncbi.nlm.nih.gov/gene/?term=LOC113561532 | NC_039507.1 | 7277571 | 7278070 | + | 4707 | 3969 | 738 | 84.32 |
| gene-LOC105278632 | 500 | 39 | 780 | 65.42 | 5.1 | https://www.ncbi.nlm.nih.gov/gene/?term=LOC105278632 | NC_039516.1 | 11911628 | 11912127 | + | 3493 | 1844 | 1649 | 52.79 |

...

... only first 25 lines shown ...

Directory: RNK File: sites-in-genes-Ob_LIp_BCphase.txt

| seqnames | start | end | width | strand | coverage | numCs | numTs | perc_meth | region_seqnames | region_start | region_end | region_width | region_strand | region_source | region_type | region_score | region_phase | region_ID | region_Dbxref | region_Name | region_gbkey | region_gene | region_gene_biotype | region_end_range | region_partial | region_start_range |
|---|---|---|---|---|---|---|---|---|---|---|---|---|---|---|---|---|---|---|---|---|---|---|---|---|---|---|
| NC_039506.1 | 137598 | 137598 | 1 | - | 104 | 10 | 94 | 9.61538461538462 | NC_039506.1 | 134260 | 142204 | 7945 | - | Gnomon | gene | NA | NA | gene-LOC105288138 | GeneID:105288138 | LOC105288138 | Gene | LOC105288138 | protein_coding | character(0) | NA | character(0) |
| NC_039506.1 | 138205 | 138205 | 1 | + | 88 | 12 | 76 | 13.6363636363636 | NC_039506.1 | 134260 | 142204 | 7945 | - | Gnomon | gene | NA | NA | gene-LOC105288138 | GeneID:105288138 | LOC105288138 | Gene | LOC105288138 | protein_coding | character(0) | NA | character(0) |
| NC_039506.1 | 138593 | 138593 | 1 | - | 91 | 10 | 81 | 10.989010989011 | NC_039506.1 | 134260 | 142204 | 7945 | - | Gnomon | gene | NA | NA | gene-LOC105288138 | GeneID:105288138 | LOC105288138 | Gene | LOC105288138 | protein_coding | character(0) | NA | character(0) |
| NC_039506.1 | 138612 | 138612 | 1 | - | 78 | 10 | 68 | 12.8205128205128 | NC_039506.1 | 134260 | 142204 | 7945 | - | Gnomon | gene | NA | NA | gene-LOC105288138 | GeneID:105288138 | LOC105288138 | Gene | LOC105288138 | protein_coding | character(0) | NA | character(0) |
| NC_039506.1 | 376413 | 376413 | 1 | - | 32 | 7 | 25 | 21.875 | NC_039506.1 | 370197 | 377142 | 6946 | + | Gnomon | gene | NA | NA | gene-LOC105285439 | GeneID:105285439 | LOC105285439 | Gene | LOC105285439 | protein_coding | character(0) | NA | character(0) |
| NC_039506.1 | 458650 | 458650 | 1 | - | 56 | 9 | 47 | 16.0714285714286 | NC_039506.1 | 458174 | 503061 | 44888 | - | Gnomon | gene | NA | NA | gene-LOC105285445 | GeneID:105285445 | LOC105285445 | Gene | LOC105285445 | protein_coding | character(0) | NA | character(0) |
| NC_039506.1 | 463932 | 463932 | 1 | - | 30 | 12 | 18 | 40 | NC_039506.1 | 458174 | 503061 | 44888 | - | Gnomon | gene | NA | NA | gene-LOC105285445 | GeneID:105285445 | LOC105285445 | Gene | LOC105285445 | protein_coding | character(0) | NA | character(0) |
| NC_039506.1 | 572754 | 572754 | 1 | + | 46 | 9 | 37 | 19.5652173913043 | NC_039506.1 | 559004 | 580516 | 21513 | + | Gnomon | gene | NA | NA | gene-LOC105285447 | GeneID:105285447 | LOC105285447 | Gene | LOC105285447 | protein_coding | character(0) | NA | character(0) |
| NC_039506.1 | 623446 | 623446 | 1 | + | 43 | 8 | 35 | 18.6046511627907 | NC_039506.1 | 621875 | 626760 | 4886 | + | Gnomon | gene | NA | NA | gene-LOC105285450 | GeneID:105285450 | LOC105285450 | Gene | LOC105285450 | protein_coding | character(0) | NA | character(0) |
| NC_039506.1 | 735420 | 735420 | 1 | + | 29 | 7 | 22 | 24.1379310344828 | NC_039506.1 | 721010 | 827029 | 106020 | - | Gnomon | gene | NA | NA | gene-LOC113561370 | GeneID:113561370 | LOC113561370 | Gene | LOC113561370 | protein_coding | character(0) | NA | character(0) |
| NC_039506.1 | 926754 | 926754 | 1 | + | 39 | 9 | 30 | 23.0769230769231 | NC_039506.1 | 926117 | 968035 | 41919 | - | Gnomon | gene | NA | NA | gene-LOC113561445 | GeneID:113561445 | LOC113561445 | Gene | LOC113561445 | protein_coding | character(0) | NA | character(0) |
| NC_039506.1 | 928489 | 928489 | 1 | - | 80 | 9 | 71 | 11.25 | NC_039506.1 | 926117 | 968035 | 41919 | - | Gnomon | gene | NA | NA | gene-LOC113561445 | GeneID:113561445 | LOC113561445 | Gene | LOC113561445 | protein_coding | character(0) | NA | character(0) |
| NC_039506.1 | 955531 | 955531 | 1 | - | 27 | 7 | 20 | 25.9259259259259 | NC_039506.1 | 926117 | 968035 | 41919 | - | Gnomon | gene | NA | NA | gene-LOC113561445 | GeneID:113561445 | LOC113561445 | Gene | LOC113561445 | protein_coding | character(0) | NA | character(0) |
| NC_039506.1 | 956202 | 956202 | 1 | + | 102 | 18 | 84 | 17.6470588235294 | NC_039506.1 | 926117 | 968035 | 41919 | - | Gnomon | gene | NA | NA | gene-LOC113561445 | GeneID:113561445 | LOC113561445 | Gene | LOC113561445 | protein_coding | character(0) | NA | character(0) |
| NC_039506.1 | 1058205 | 1058205 | 1 | + | 101 | 21 | 80 | 20.7920792079208 | NC_039506.1 | 1056151 | 1064257 | 8107 | + | Gnomon | gene | NA | NA | gene-LOC105286747 | GeneID:105286747 | LOC105286747 | Gene | LOC105286747 | lncRNA | character(0) | NA | character(0) |
| NC_039506.1 | 1097561 | 1097561 | 1 | + | 67 | 9 | 58 | 13.4328358208955 | NC_039506.1 | 1095614 | 1100480 | 4867 | + | Gnomon | gene | NA | NA | gene-LOC105286750 | GeneID:105286750 | LOC105286750 | Gene | LOC105286750 | protein_coding | character(0) | NA | character(0) |
| NC_039506.1 | 1097573 | 1097573 | 1 | + | 55 | 8 | 47 | 14.5454545454545 | NC_039506.1 | 1095614 | 1100480 | 4867 | + | Gnomon | gene | NA | NA | gene-LOC105286750 | GeneID:105286750 | LOC105286750 | Gene | LOC105286750 | protein_coding | character(0) | NA | character(0) |
| NC_039506.1 | 1097588 | 1097588 | 1 | + | 121 | 42 | 79 | 34.7107438016529 | NC_039506.1 | 1095614 | 1100480 | 4867 | + | Gnomon | gene | NA | NA | gene-LOC105286750 | GeneID:105286750 | LOC105286750 | Gene | LOC105286750 | protein_coding | character(0) | NA | character(0) |
| NC_039506.1 | 1097589 | 1097589 | 1 | - | 29 | 12 | 17 | 41.3793103448276 | NC_039506.1 | 1095614 | 1100480 | 4867 | + | Gnomon | gene | NA | NA | gene-LOC105286750 | GeneID:105286750 | LOC105286750 | Gene | LOC105286750 | protein_coding | character(0) | NA | character(0) |
| NC_039506.1 | 1097598 | 1097598 | 1 | + | 134 | 71 | 63 | 52.9850746268657 | NC_039506.1 | 1095614 | 1100480 | 4867 | + | Gnomon | gene | NA | NA | gene-LOC105286750 | GeneID:105286750 | LOC105286750 | Gene | LOC105286750 | protein_coding | character(0) | NA | character(0) |
| NC_039506.1 | 1097599 | 1097599 | 1 | - | 40 | 20 | 20 | 50 | NC_039506.1 | 1095614 | 1100480 | 4867 | + | Gnomon | gene | NA | NA | gene-LOC105286750 | GeneID:105286750 | LOC105286750 | Gene | LOC105286750 | protein_coding | character(0) | NA | character(0) |
| NC_039506.1 | 1097647 | 1097647 | 1 | + | 176 | 18 | 158 | 10.2272727272727 | NC_039506.1 | 1095614 | 1100480 | 4867 | + | Gnomon | gene | NA | NA | gene-LOC105286750 | GeneID:105286750 | LOC105286750 | Gene | LOC105286750 | protein_coding | character(0) | NA | character(0) |
| NC_039506.1 | 1097702 | 1097702 | 1 | + | 180 | 30 | 150 | 16.6666666666667 | NC_039506.1 | 1095614 | 1100480 | 4867 | + | Gnomon | gene | NA | NA | gene-LOC105286750 | GeneID:105286750 | LOC105286750 | Gene | LOC105286750 | protein_coding | character(0) | NA | character(0) |
| NC_039506.1 | 1097703 | 1097703 | 1 | - | 80 | 19 | 61 | 23.75 | NC_039506.1 | 1095614 | 1100480 | 4867 | + | Gnomon | gene | NA | NA | gene-LOC105286750 | GeneID:105286750 | LOC105286750 | Gene | LOC105286750 | protein_coding | character(0) | NA | character(0) |

...
... only first 25 lines shown ...

Directory: RNK File: sites-in-genes-Ob_LIp_Rphase.txt

| seqnames | start | end | width | strand | coverage | numCs | numTs | perc_meth | region_seqnames | region_start | region_end | region_width | region_strand | region_source | region_type | region_score | region_phase | region_ID | region_Dbxref | region_Name | region_gbkey | region_gene | region_gene_biotype | region_end_range | region_partial | region_start_range |
|---|---|---|---|---|---|---|---|---|---|---|---|---|---|---|---|---|---|---|---|---|---|---|---|---|---|---|
| NC_039506.1 | 137159 | 137159 | 1 | - | 43 | 10 | 33 | 23.2558139534884 | NC_039506.1 | 134260 | 142204 | 7945 | - | Gnomon | gene | NA | NA | gene-LOC105288138 | GeneID:105288138 | LOC105288138 | Gene | LOC105288138 | protein_coding | character(0) | NA | character(0) |
| NC_039506.1 | 137598 | 137598 | 1 | - | 106 | 11 | 95 | 10.377358490566 | NC_039506.1 | 134260 | 142204 | 7945 | - | Gnomon | gene | NA | NA | gene-LOC105288138 | GeneID:105288138 | LOC105288138 | Gene | LOC105288138 | protein_coding | character(0) | NA | character(0) |
| NC_039506.1 | 137684 | 137684 | 1 | - | 139 | 17 | 122 | 12.2302158273381 | NC_039506.1 | 134260 | 142204 | 7945 | - | Gnomon | gene | NA | NA | gene-LOC105288138 | GeneID:105288138 | LOC105288138 | Gene | LOC105288138 | protein_coding | character(0) | NA | character(0) |
| NC_039506.1 | 138205 | 138205 | 1 | + | 78 | 14 | 64 | 17.9487179487179 | NC_039506.1 | 134260 | 142204 | 7945 | - | Gnomon | gene | NA | NA | gene-LOC105288138 | GeneID:105288138 | LOC105288138 | Gene | LOC105288138 | protein_coding | character(0) | NA | character(0) |
| NC_039506.1 | 138206 | 138206 | 1 | - | 76 | 10 | 66 | 13.1578947368421 | NC_039506.1 | 134260 | 142204 | 7945 | - | Gnomon | gene | NA | NA | gene-LOC105288138 | GeneID:105288138 | LOC105288138 | Gene | LOC105288138 | protein_coding | character(0) | NA | character(0) |
| NC_039506.1 | 138579 | 138579 | 1 | - | 116 | 14 | 102 | 12.0689655172414 | NC_039506.1 | 134260 | 142204 | 7945 | - | Gnomon | gene | NA | NA | gene-LOC105288138 | GeneID:105288138 | LOC105288138 | Gene | LOC105288138 | protein_coding | character(0) | NA | character(0) |
| NC_039506.1 | 138592 | 138592 | 1 | + | 80 | 9 | 71 | 11.25 | NC_039506.1 | 134260 | 142204 | 7945 | - | Gnomon | gene | NA | NA | gene-LOC105288138 | GeneID:105288138 | LOC105288138 | Gene | LOC105288138 | protein_coding | character(0) | NA | character(0) |
| NC_039506.1 | 138593 | 138593 | 1 | - | 112 | 13 | 99 | 11.6071428571429 | NC_039506.1 | 134260 | 142204 | 7945 | - | Gnomon | gene | NA | NA | gene-LOC105288138 | GeneID:105288138 | LOC105288138 | Gene | LOC105288138 | protein_coding | character(0) | NA | character(0) |
| NC_039506.1 | 138611 | 138611 | 1 | + | 76 | 11 | 65 | 14.4736842105263 | NC_039506.1 | 134260 | 142204 | 7945 | - | Gnomon | gene | NA | NA | gene-LOC105288138 | GeneID:105288138 | LOC105288138 | Gene | LOC105288138 | protein_coding | character(0) | NA | character(0) |
| NC_039506.1 | 138621 | 138621 | 1 | - | 88 | 10 | 78 | 11.3636363636364 | NC_039506.1 | 134260 | 142204 | 7945 | - | Gnomon | gene | NA | NA | gene-LOC105288138 | GeneID:105288138 | LOC105288138 | Gene | LOC105288138 | protein_coding | character(0) | NA | character(0) |
| NC_039506.1 | 224883 | 224883 | 1 | + | 45 | 8 | 37 | 17.7777777777778 | NC_039506.1 | 143775 | 267542 | 123768 | - | Gnomon | gene | NA | NA | gene-LOC105288140 | GeneID:105288140 | LOC105288140 | Gene | LOC105288140 | protein_coding | character(0) | NA | character(0) |
| NC_039506.1 | 242470 | 242470 | 1 | + | 59 | 9 | 50 | 15.2542372881356 | NC_039506.1 | 143775 | 267542 | 123768 | - | Gnomon | gene | NA | NA | gene-LOC105288140 | GeneID:105288140 | LOC105288140 | Gene | LOC105288140 | protein_coding | character(0) | NA | character(0) |
| NC_039506.1 | 376413 | 376413 | 1 | - | 47 | 9 | 38 | 19.1489361702128 | NC_039506.1 | 370197 | 377142 | 6946 | + | Gnomon | gene | NA | NA | gene-LOC105285439 | GeneID:105285439 | LOC105285439 | Gene | LOC105285439 | protein_coding | character(0) | NA | character(0) |
| NC_039506.1 | 424849 | 424849 | 1 | - | 55 | 9 | 46 | 16.3636363636364 | NC_039506.1 | 412793 | 450860 | 38068 | - | Gnomon | gene | NA | NA | gene-LOC105285468 | GeneID:105285468 | LOC105285468 | Gene | LOC105285468 | protein_coding | character(0) | NA | character(0) |
| NC_039506.1 | 440111 | 440111 | 1 | - | 92 | 10 | 82 | 10.8695652173913 | NC_039506.1 | 412793 | 450860 | 38068 | - | Gnomon | gene | NA | NA | gene-LOC105285468 | GeneID:105285468 | LOC105285468 | Gene | LOC105285468 | protein_coding | character(0) | NA | character(0) |
| NC_039506.1 | 462549 | 462549 | 1 | + | 50 | 8 | 42 | 16 | NC_039506.1 | 458174 | 503061 | 44888 | - | Gnomon | gene | NA | NA | gene-LOC105285445 | GeneID:105285445 | LOC105285445 | Gene | LOC105285445 | protein_coding | character(0) | NA | character(0) |
| NC_039506.1 | 463932 | 463932 | 1 | - | 29 | 11 | 18 | 37.9310344827586 | NC_039506.1 | 458174 | 503061 | 44888 | - | Gnomon | gene | NA | NA | gene-LOC105285445 | GeneID:105285445 | LOC105285445 | Gene | LOC105285445 | protein_coding | character(0) | NA | character(0) |
| NC_039506.1 | 926754 | 926754 | 1 | + | 41 | 11 | 30 | 26.8292682926829 | NC_039506.1 | 926117 | 968035 | 41919 | - | Gnomon | gene | NA | NA | gene-LOC113561445 | GeneID:113561445 | LOC113561445 | Gene | LOC113561445 | protein_coding | character(0) | NA | character(0) |
| NC_039506.1 | 930993 | 930993 | 1 | + | 108 | 10 | 98 | 9.25925925925926 | NC_039506.1 | 926117 | 968035 | 41919 | - | Gnomon | gene | NA | NA | gene-LOC113561445 | GeneID:113561445 | LOC113561445 | Gene | LOC113561445 | protein_coding | character(0) | NA | character(0) |
| NC_039506.1 | 956196 | 956196 | 1 | - | 162 | 15 | 147 | 9.25925925925926 | NC_039506.1 | 926117 | 968035 | 41919 | - | Gnomon | gene | NA | NA | gene-LOC113561445 | GeneID:113561445 | LOC113561445 | Gene | LOC113561445 | protein_coding | character(0) | NA | character(0) |
| NC_039506.1 | 956202 | 956202 | 1 | + | 125 | 24 | 101 | 19.2 | NC_039506.1 | 926117 | 968035 | 41919 | - | Gnomon | gene | NA | NA | gene-LOC113561445 | GeneID:113561445 | LOC113561445 | Gene | LOC113561445 | protein_coding | character(0) | NA | character(0) |
| NC_039506.1 | 1058205 | 1058205 | 1 | + | 97 | 18 | 79 | 18.5567010309278 | NC_039506.1 | 1056151 | 1064257 | 8107 | + | Gnomon | gene | NA | NA | gene-LOC105286747 | GeneID:105286747 | LOC105286747 | Gene | LOC105286747 | lncRNA | character(0) | NA | character(0) |
| NC_039506.1 | 1097547 | 1097547 | 1 | - | 27 | 7 | 20 | 25.9259259259259 | NC_039506.1 | 1095614 | 1100480 | 4867 | + | Gnomon | gene | NA | NA | gene-LOC105286750 | GeneID:105286750 | LOC105286750 | Gene | LOC105286750 | protein_coding | character(0) | NA | character(0) |
| NC_039506.1 | 1097573 | 1097573 | 1 | + | 65 | 10 | 55 | 15.3846153846154 | NC_039506.1 | 1095614 | 1100480 | 4867 | + | Gnomon | gene | NA | NA | gene-LOC105286750 | GeneID:105286750 | LOC105286750 | Gene | LOC105286750 | protein_coding | character(0) | NA | character(0) |

...

... only first 25 lines shown ...

Directory: RNK File: sites-in-promoters-Ob_LIp_BCphase.txt

| seqnames | start | end | width | strand | coverage | numCs | numTs | perc_meth | region_seqnames | region_start | region_end | region_width | region_strand | region_source | region_type | region_score | region_phase | region_ID | region_Dbxref | region_Name | region_gbkey | region_gene | region_gene_biotype | region_end_range | region_partial | region_start_range |
|---|---|---|---|---|---|---|---|---|---|---|---|---|---|---|---|---|---|---|---|---|---|---|---|---|---|---|
| NC_039506.1 | 1559274 | 1559274 | 1 | - | 27 | 10 | 17 | 37.037037037037 | NC_039506.1 | 1559129 | 1559628 | 500 | + | Gnomon | promoter | NA | NA | gene-LOC105280248 | GeneID:105280248 | LOC105280248 | Gene | LOC105280248 | protein_coding | character(0) | NA | character(0) |
| NC_039506.1 | 1627226 | 1627226 | 1 | - | 12 | 6 | 6 | 50 | NC_039506.1 | 1627051 | 1627550 | 500 | + | Gnomon | promoter | NA | NA | gene-LOC105280254 | GeneID:105280254 | LOC105280254 | Gene | LOC105280254 | protein_coding | character(0) | NA | character(0) |
| NC_039506.1 | 1627241 | 1627241 | 1 | - | 9 | 6 | 3 | 66.6666666666667 | NC_039506.1 | 1627051 | 1627550 | 500 | + | Gnomon | promoter | NA | NA | gene-LOC105280254 | GeneID:105280254 | LOC105280254 | Gene | LOC105280254 | protein_coding | character(0) | NA | character(0) |
| NC_039506.1 | 1664198 | 1664198 | 1 | + | 137 | 46 | 91 | 33.5766423357664 | NC_039506.1 | 1664107 | 1664606 | 500 | + | Gnomon | promoter | NA | NA | gene-LOC105280261 | GeneID:105280261 | LOC105280261 | Gene | LOC105280261 | protein_coding | character(0) | NA | character(0) |
| NC_039506.1 | 1664241 | 1664241 | 1 | + | 160 | 24 | 136 | 15 | NC_039506.1 | 1664107 | 1664606 | 500 | + | Gnomon | promoter | NA | NA | gene-LOC105280261 | GeneID:105280261 | LOC105280261 | Gene | LOC105280261 | protein_coding | character(0) | NA | character(0) |
| NC_039506.1 | 1664392 | 1664392 | 1 | + | 58 | 18 | 40 | 31.0344827586207 | NC_039506.1 | 1664107 | 1664606 | 500 | + | Gnomon | promoter | NA | NA | gene-LOC105280261 | GeneID:105280261 | LOC105280261 | Gene | LOC105280261 | protein_coding | character(0) | NA | character(0) |
| NC_039506.1 | 1664400 | 1664400 | 1 | + | 65 | 20 | 45 | 30.7692307692308 | NC_039506.1 | 1664107 | 1664606 | 500 | + | Gnomon | promoter | NA | NA | gene-LOC105280261 | GeneID:105280261 | LOC105280261 | Gene | LOC105280261 | protein_coding | character(0) | NA | character(0) |
| NC_039506.1 | 1673032 | 1673032 | 1 | + | 80 | 9 | 71 | 11.25 | NC_039506.1 | 1673027 | 1673526 | 500 | + | Gnomon | promoter | NA | NA | gene-LOC109611032 | GeneID:109611032 | LOC109611032 | Gene | LOC109611032 | lncRNA | character(0) | NA | character(0) |
| NC_039506.1 | 1673083 | 1673083 | 1 | + | 110 | 16 | 94 | 14.5454545454545 | NC_039506.1 | 1673027 | 1673526 | 500 | + | Gnomon | promoter | NA | NA | gene-LOC109611032 | GeneID:109611032 | LOC109611032 | Gene | LOC109611032 | lncRNA | character(0) | NA | character(0) |
| NC_039506.1 | 1673084 | 1673084 | 1 | - | 95 | 17 | 78 | 17.8947368421053 | NC_039506.1 | 1673027 | 1673526 | 500 | + | Gnomon | promoter | NA | NA | gene-LOC109611032 | GeneID:109611032 | LOC109611032 | Gene | LOC109611032 | lncRNA | character(0) | NA | character(0) |
| NC_039506.1 | 1673110 | 1673110 | 1 | + | 11 | 9 | 2 | 81.8181818181818 | NC_039506.1 | 1673027 | 1673526 | 500 | + | Gnomon | promoter | NA | NA | gene-LOC109611032 | GeneID:109611032 | LOC109611032 | Gene | LOC109611032 | lncRNA | character(0) | NA | character(0) |
| NC_039506.1 | 1673111 | 1673111 | 1 | - | 59 | 54 | 5 | 91.5254237288136 | NC_039506.1 | 1673027 | 1673526 | 500 | + | Gnomon | promoter | NA | NA | gene-LOC109611032 | GeneID:109611032 | LOC109611032 | Gene | LOC109611032 | lncRNA | character(0) | NA | character(0) |
| NC_039506.1 | 1673134 | 1673134 | 1 | + | 5 | 5 | 0 | 100 | NC_039506.1 | 1673027 | 1673526 | 500 | + | Gnomon | promoter | NA | NA | gene-LOC109611032 | GeneID:109611032 | LOC109611032 | Gene | LOC109611032 | lncRNA | character(0) | NA | character(0) |
| NC_039506.1 | 1673135 | 1673135 | 1 | - | 58 | 41 | 17 | 70.6896551724138 | NC_039506.1 | 1673027 | 1673526 | 500 | + | Gnomon | promoter | NA | NA | gene-LOC109611032 | GeneID:109611032 | LOC109611032 | Gene | LOC109611032 | lncRNA | character(0) | NA | character(0) |
| NC_039506.1 | 1673160 | 1673160 | 1 | - | 47 | 40 | 7 | 85.1063829787234 | NC_039506.1 | 1673027 | 1673526 | 500 | + | Gnomon | promoter | NA | NA | gene-LOC109611032 | GeneID:109611032 | LOC109611032 | Gene | LOC109611032 | lncRNA | character(0) | NA | character(0) |
| NC_039506.1 | 1673189 | 1673189 | 1 | - | 111 | 88 | 23 | 79.2792792792793 | NC_039506.1 | 1673027 | 1673526 | 500 | + | Gnomon | promoter | NA | NA | gene-LOC109611032 | GeneID:109611032 | LOC109611032 | Gene | LOC109611032 | lncRNA | character(0) | NA | character(0) |
| NC_039506.1 | 1673197 | 1673197 | 1 | - | 125 | 68 | 57 | 54.4 | NC_039506.1 | 1673027 | 1673526 | 500 | + | Gnomon | promoter | NA | NA | gene-LOC109611032 | GeneID:109611032 | LOC109611032 | Gene | LOC109611032 | lncRNA | character(0) | NA | character(0) |
| NC_039506.1 | 1673205 | 1673205 | 1 | - | 174 | 146 | 28 | 83.9080459770115 | NC_039506.1 | 1673027 | 1673526 | 500 | + | Gnomon | promoter | NA | NA | gene-LOC109611032 | GeneID:109611032 | LOC109611032 | Gene | LOC109611032 | lncRNA | character(0) | NA | character(0) |
| NC_039506.1 | 1673219 | 1673219 | 1 | - | 237 | 67 | 170 | 28.2700421940928 | NC_039506.1 | 1673027 | 1673526 | 500 | + | Gnomon | promoter | NA | NA | gene-LOC109611032 | GeneID:109611032 | LOC109611032 | Gene | LOC109611032 | lncRNA | character(0) | NA | character(0) |
| NC_039506.1 | 1673224 | 1673224 | 1 | + | 5 | 5 | 0 | 100 | NC_039506.1 | 1673027 | 1673526 | 500 | + | Gnomon | promoter | NA | NA | gene-LOC109611032 | GeneID:109611032 | LOC109611032 | Gene | LOC109611032 | lncRNA | character(0) | NA | character(0) |
| NC_039506.1 | 1673225 | 1673225 | 1 | - | 223 | 174 | 49 | 78.0269058295964 | NC_039506.1 | 1673027 | 1673526 | 500 | + | Gnomon | promoter | NA | NA | gene-LOC109611032 | GeneID:109611032 | LOC109611032 | Gene | LOC109611032 | lncRNA | character(0) | NA | character(0) |
| NC_039506.1 | 1673234 | 1673234 | 1 | - | 228 | 131 | 97 | 57.4561403508772 | NC_039506.1 | 1673027 | 1673526 | 500 | + | Gnomon | promoter | NA | NA | gene-LOC109611032 | GeneID:109611032 | LOC109611032 | Gene | LOC109611032 | lncRNA | character(0) | NA | character(0) |
| NC_039506.1 | 1673241 | 1673241 | 1 | - | 224 | 142 | 82 | 63.3928571428571 | NC_039506.1 | 1673027 | 1673526 | 500 | + | Gnomon | promoter | NA | NA | gene-LOC109611032 | GeneID:109611032 | LOC109611032 | Gene | LOC109611032 | lncRNA | character(0) | NA | character(0) |
| NC_039506.1 | 1673284 | 1673284 | 1 | - | 179 | 148 | 31 | 82.6815642458101 | NC_039506.1 | 1673027 | 1673526 | 500 | + | Gnomon | promoter | NA | NA | gene-LOC109611032 | GeneID:109611032 | LOC109611032 | Gene | LOC109611032 | lncRNA | character(0) | NA | character(0) |

...
... only first 25 lines shown ...

Directory: RNK File: sites-in-promoters-Ob_LIp_Rphase.txt

| seqnames | start | end | width | strand | coverage | numCs | numTs | perc_meth | region_seqnames | region_start | region_end | region_width | region_strand | region_source | region_type | region_score | region_phase | region_ID | region_Dbxref | region_Name | region_gbkey | region_gene | region_gene_biotype | region_end_range | region_partial | region_start_range |
|---|---|---|---|---|---|---|---|---|---|---|---|---|---|---|---|---|---|---|---|---|---|---|---|---|---|---|
| NC_039506.1 | 1034483 | 1034483 | 1 | - | 108 | 10 | 98 | 9.25925925925926 | NC_039506.1 | 1034377 | 1034876 | 500 | + | Gnomon | promoter | NA | NA | gene-LOC105286748 | GeneID:105286748 | LOC105286748 | Gene | LOC105286748 | protein_coding | character(0) | NA | character(0) |
| NC_039506.1 | 1034486 | 1034486 | 1 | - | 118 | 11 | 107 | 9.32203389830508 | NC_039506.1 | 1034377 | 1034876 | 500 | + | Gnomon | promoter | NA | NA | gene-LOC105286748 | GeneID:105286748 | LOC105286748 | Gene | LOC105286748 | protein_coding | character(0) | NA | character(0) |
| NC_039506.1 | 1487355 | 1487355 | 1 | + | 139 | 19 | 120 | 13.6690647482014 | NC_039506.1 | 1487279 | 1487778 | 500 | + | Gnomon | promoter | NA | NA | gene-LOC105280236 | GeneID:105280236 | LOC105280236 | Gene | LOC105280236 | protein_coding | character(0) | NA | character(0) |
| NC_039506.1 | 1548015 | 1548015 | 1 | - | 124 | 12 | 112 | 9.67741935483871 | NC_039506.1 | 1547999 | 1548498 | 500 | - | Gnomon | promoter | NA | NA | gene-LOC105280244 | GeneID:105280244 | LOC105280244 | Gene | LOC105280244 | protein_coding | character(0) | NA | character(0) |
| NC_039506.1 | 1559274 | 1559274 | 1 | - | 45 | 12 | 33 | 26.6666666666667 | NC_039506.1 | 1559129 | 1559628 | 500 | + | Gnomon | promoter | NA | NA | gene-LOC105280248 | GeneID:105280248 | LOC105280248 | Gene | LOC105280248 | protein_coding | character(0) | NA | character(0) |
| NC_039506.1 | 1627225 | 1627225 | 1 | + | 8 | 7 | 1 | 87.5 | NC_039506.1 | 1627051 | 1627550 | 500 | + | Gnomon | promoter | NA | NA | gene-LOC105280254 | GeneID:105280254 | LOC105280254 | Gene | LOC105280254 | protein_coding | character(0) | NA | character(0) |
| NC_039506.1 | 1627240 | 1627240 | 1 | + | 9 | 8 | 1 | 88.8888888888889 | NC_039506.1 | 1627051 | 1627550 | 500 | + | Gnomon | promoter | NA | NA | gene-LOC105280254 | GeneID:105280254 | LOC105280254 | Gene | LOC105280254 | protein_coding | character(0) | NA | character(0) |
| NC_039506.1 | 1627241 | 1627241 | 1 | - | 8 | 6 | 2 | 75 | NC_039506.1 | 1627051 | 1627550 | 500 | + | Gnomon | promoter | NA | NA | gene-LOC105280254 | GeneID:105280254 | LOC105280254 | Gene | LOC105280254 | protein_coding | character(0) | NA | character(0) |
| NC_039506.1 | 1630029 | 1630029 | 1 | - | 38 | 9 | 29 | 23.6842105263158 | NC_039506.1 | 1629671 | 1630170 | 500 | + | Gnomon | promoter | NA | NA | gene-LOC105280255 | GeneID:105280255 | LOC105280255 | Gene | LOC105280255 | protein_coding | character(0) | NA | character(0) |
| NC_039506.1 | 1664198 | 1664198 | 1 | + | 153 | 39 | 114 | 25.4901960784314 | NC_039506.1 | 1664107 | 1664606 | 500 | + | Gnomon | promoter | NA | NA | gene-LOC105280261 | GeneID:105280261 | LOC105280261 | Gene | LOC105280261 | protein_coding | character(0) | NA | character(0) |
| NC_039506.1 | 1664241 | 1664241 | 1 | + | 143 | 18 | 125 | 12.5874125874126 | NC_039506.1 | 1664107 | 1664606 | 500 | + | Gnomon | promoter | NA | NA | gene-LOC105280261 | GeneID:105280261 | LOC105280261 | Gene | LOC105280261 | protein_coding | character(0) | NA | character(0) |
| NC_039506.1 | 1664350 | 1664350 | 1 | + | 45 | 10 | 35 | 22.2222222222222 | NC_039506.1 | 1664107 | 1664606 | 500 | + | Gnomon | promoter | NA | NA | gene-LOC105280261 | GeneID:105280261 | LOC105280261 | Gene | LOC105280261 | protein_coding | character(0) | NA | character(0) |
| NC_039506.1 | 1664392 | 1664392 | 1 | + | 47 | 15 | 32 | 31.9148936170213 | NC_039506.1 | 1664107 | 1664606 | 500 | + | Gnomon | promoter | NA | NA | gene-LOC105280261 | GeneID:105280261 | LOC105280261 | Gene | LOC105280261 | protein_coding | character(0) | NA | character(0) |
| NC_039506.1 | 1664400 | 1664400 | 1 | + | 51 | 16 | 35 | 31.3725490196078 | NC_039506.1 | 1664107 | 1664606 | 500 | + | Gnomon | promoter | NA | NA | gene-LOC105280261 | GeneID:105280261 | LOC105280261 | Gene | LOC105280261 | protein_coding | character(0) | NA | character(0) |
| NC_039506.1 | 1673083 | 1673083 | 1 | + | 141 | 24 | 117 | 17.0212765957447 | NC_039506.1 | 1673027 | 1673526 | 500 | + | Gnomon | promoter | NA | NA | gene-LOC109611032 | GeneID:109611032 | LOC109611032 | Gene | LOC109611032 | lncRNA | character(0) | NA | character(0) |
| NC_039506.1 | 1673084 | 1673084 | 1 | - | 105 | 13 | 92 | 12.3809523809524 | NC_039506.1 | 1673027 | 1673526 | 500 | + | Gnomon | promoter | NA | NA | gene-LOC109611032 | GeneID:109611032 | LOC109611032 | Gene | LOC109611032 | lncRNA | character(0) | NA | character(0) |
| NC_039506.1 | 1673110 | 1673110 | 1 | + | 18 | 14 | 4 | 77.7777777777778 | NC_039506.1 | 1673027 | 1673526 | 500 | + | Gnomon | promoter | NA | NA | gene-LOC109611032 | GeneID:109611032 | LOC109611032 | Gene | LOC109611032 | lncRNA | character(0) | NA | character(0) |
| NC_039506.1 | 1673111 | 1673111 | 1 | - | 75 | 67 | 8 | 89.3333333333333 | NC_039506.1 | 1673027 | 1673526 | 500 | + | Gnomon | promoter | NA | NA | gene-LOC109611032 | GeneID:109611032 | LOC109611032 | Gene | LOC109611032 | lncRNA | character(0) | NA | character(0) |
| NC_039506.1 | 1673134 | 1673134 | 1 | + | 10 | 7 | 3 | 70 | NC_039506.1 | 1673027 | 1673526 | 500 | + | Gnomon | promoter | NA | NA | gene-LOC109611032 | GeneID:109611032 | LOC109611032 | Gene | LOC109611032 | lncRNA | character(0) | NA | character(0) |
| NC_039506.1 | 1673135 | 1673135 | 1 | - | 72 | 56 | 16 | 77.7777777777778 | NC_039506.1 | 1673027 | 1673526 | 500 | + | Gnomon | promoter | NA | NA | gene-LOC109611032 | GeneID:109611032 | LOC109611032 | Gene | LOC109611032 | lncRNA | character(0) | NA | character(0) |
| NC_039506.1 | 1673160 | 1673160 | 1 | - | 56 | 45 | 11 | 80.3571428571429 | NC_039506.1 | 1673027 | 1673526 | 500 | + | Gnomon | promoter | NA | NA | gene-LOC109611032 | GeneID:109611032 | LOC109611032 | Gene | LOC109611032 | lncRNA | character(0) | NA | character(0) |
| NC_039506.1 | 1673189 | 1673189 | 1 | - | 110 | 87 | 23 | 79.0909090909091 | NC_039506.1 | 1673027 | 1673526 | 500 | + | Gnomon | promoter | NA | NA | gene-LOC109611032 | GeneID:109611032 | LOC109611032 | Gene | LOC109611032 | lncRNA | character(0) | NA | character(0) |
| NC_039506.1 | 1673197 | 1673197 | 1 | - | 121 | 63 | 58 | 52.0661157024793 | NC_039506.1 | 1673027 | 1673526 | 500 | + | Gnomon | promoter | NA | NA | gene-LOC109611032 | GeneID:109611032 | LOC109611032 | Gene | LOC109611032 | lncRNA | character(0) | NA | character(0) |
| NC_039506.1 | 1673204 | 1673204 | 1 | + | 7 | 6 | 1 | 85.7142857142857 | NC_039506.1 | 1673027 | 1673526 | 500 | + | Gnomon | promoter | NA | NA | gene-LOC109611032 | GeneID:109611032 | LOC109611032 | Gene | LOC109611032 | lncRNA | character(0) | NA | character(0) |

...
... only first 25 lines shown ...

plot−genes−prcntM−vs−SiteDensity−Ob_LIp_BCphase

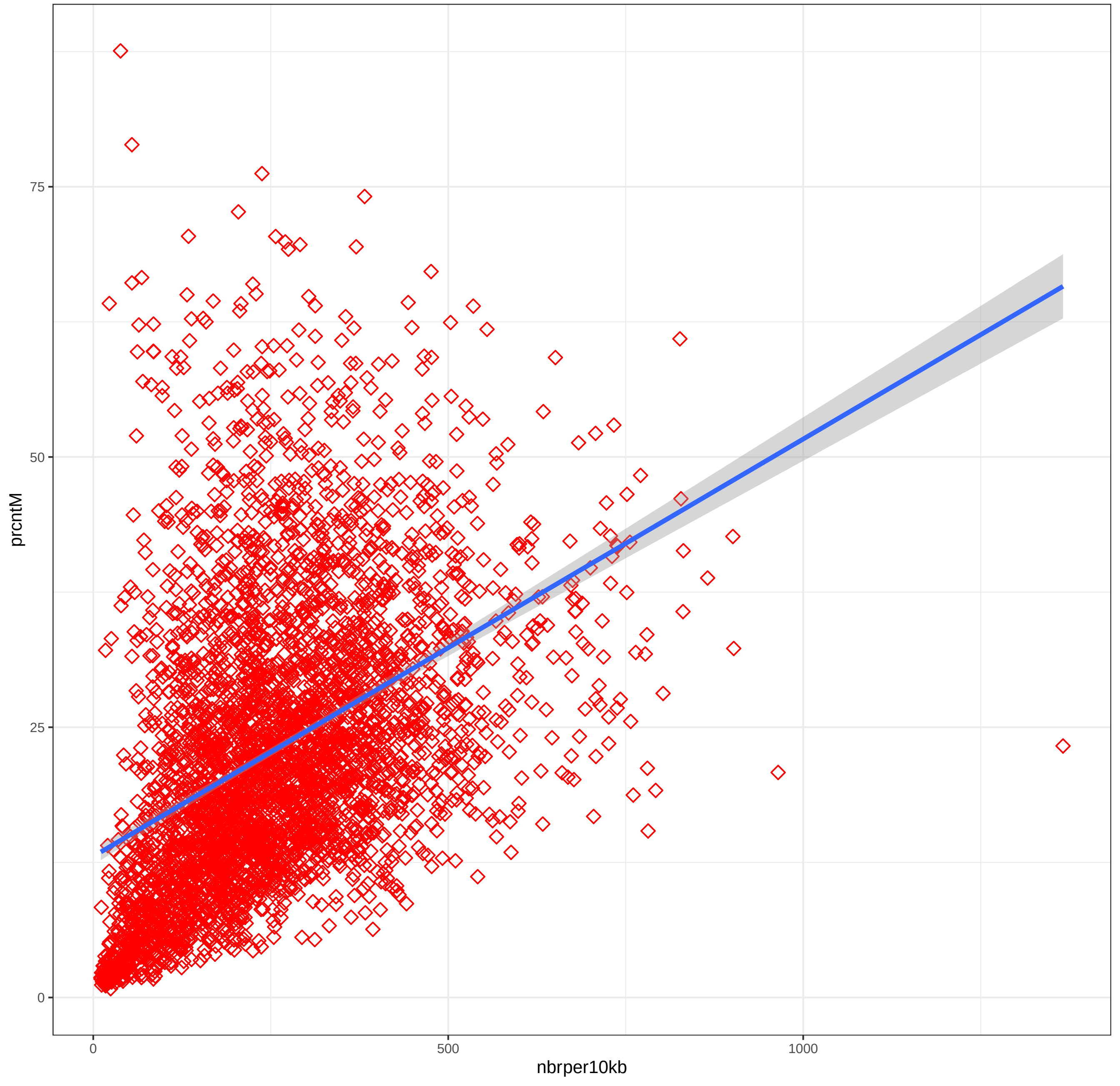

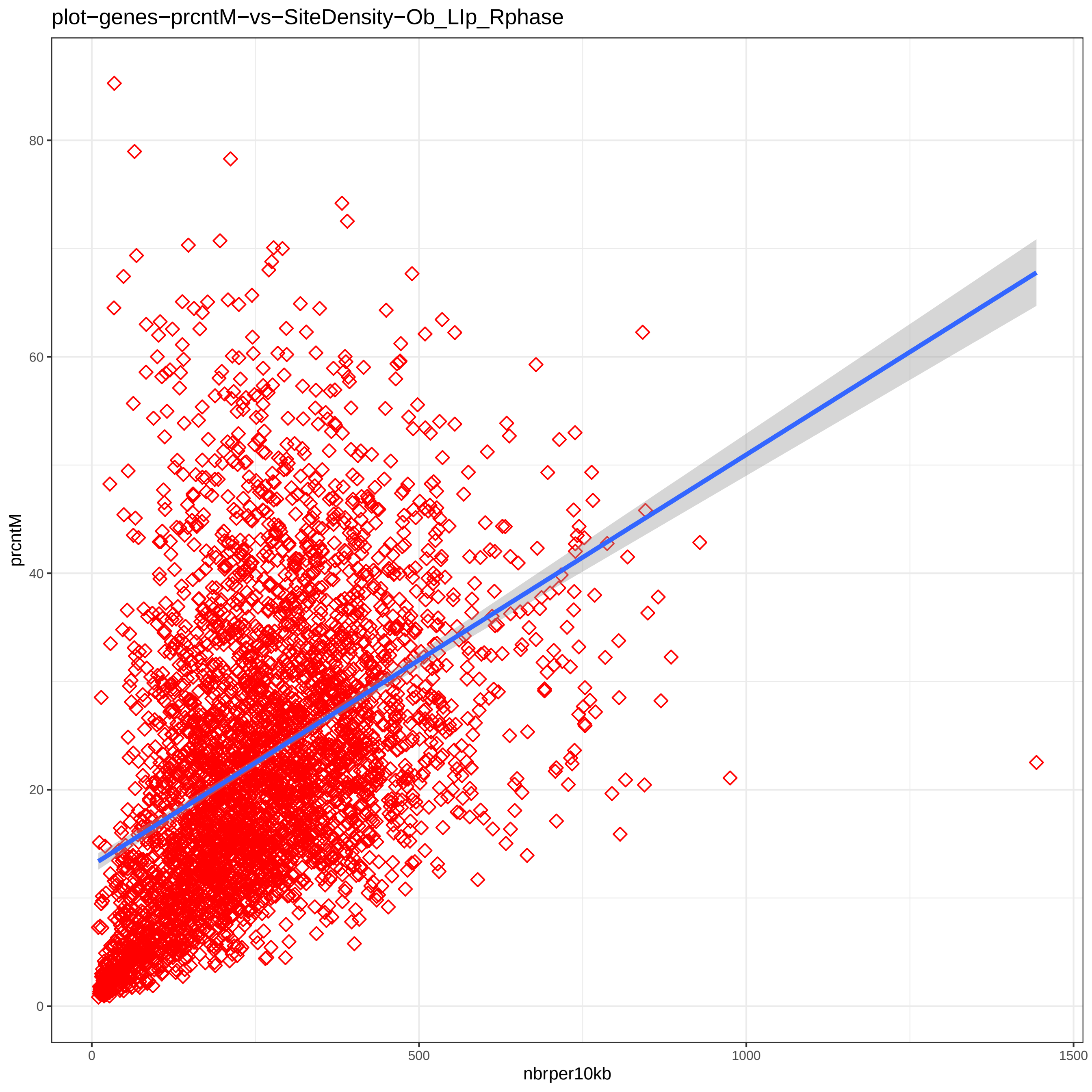
plot−genes−prcntM−vs−SiteDensity−Ob_LIp_Rphase
prcntM
0
20
40
60
80
0
500
1000
1500
nbrper10kb

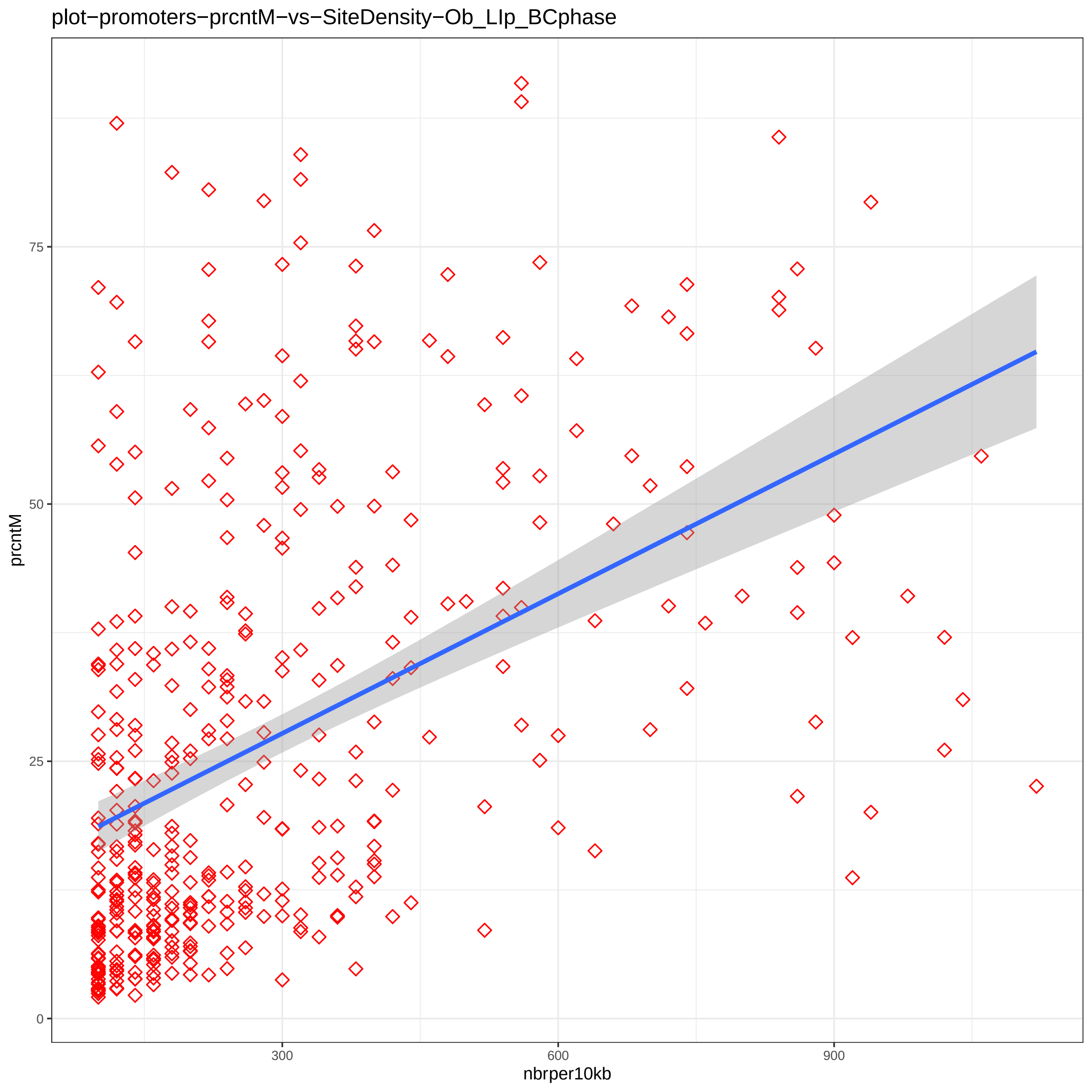

plot−promoters−prcntM−vs−SiteDensity−Ob_LIp_BCphase
prcntM
nbrper10kb
0
25
50
75
300
600
900

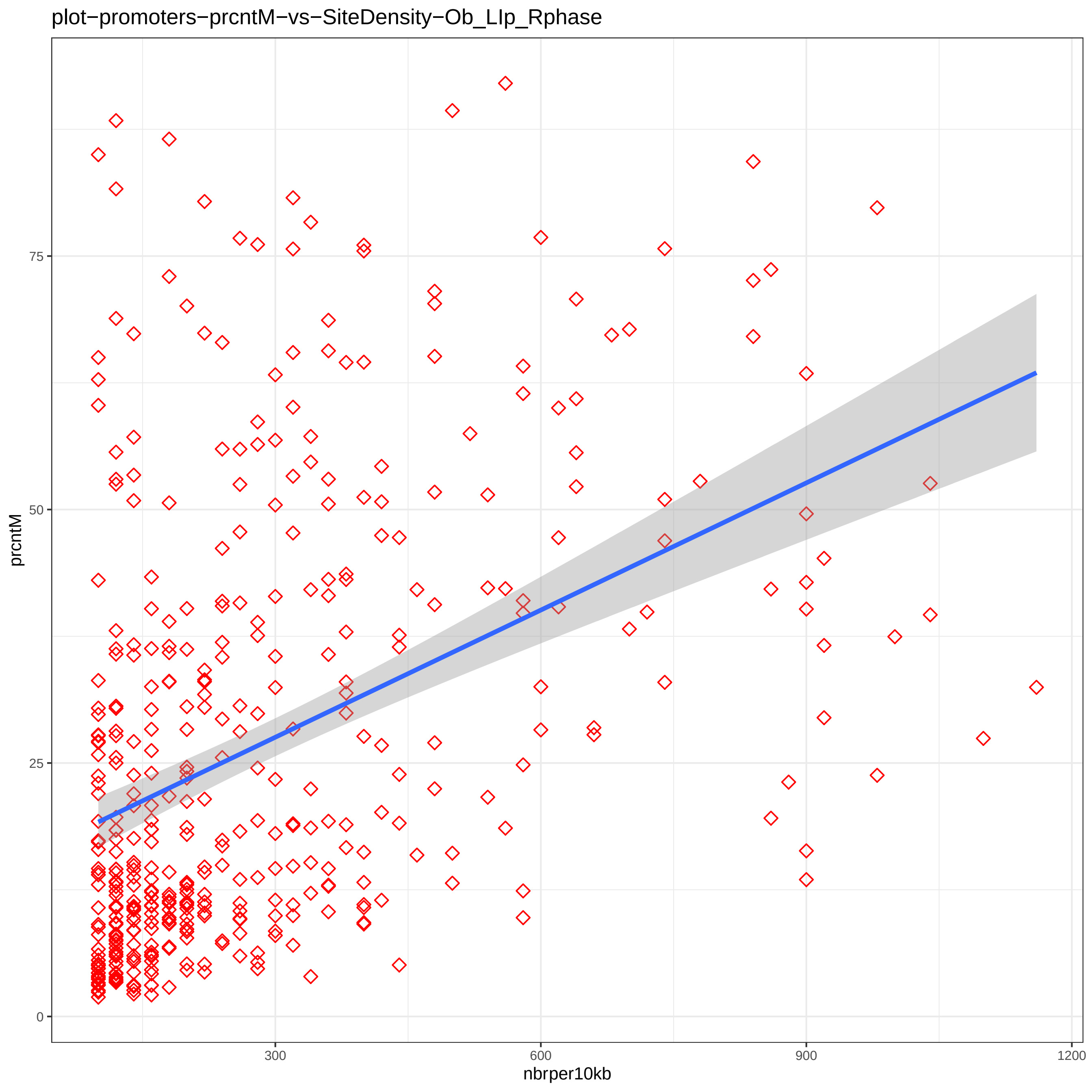
plot−promoters−prcntM−vs−SiteDensity−Ob_LIp_Rphase
prcntM
0
25
50
75
300
600
900
1200
nbrper10kb

```
Directory: MRPR File: 0READMEmrpr

MRPR - Methylation-rich and -poor regions.

Input: studymk, genome_ann (a list of GRanges objects providing annotated region labels and bounds;
                           output of BWASPR::get_genome_annotation())
       nbrxtrms (specifies the number of richest and poorest regions to be displayed; default: 100)

Output: files dst-*.txt 1ds-*.pdf 5ds-*.pdf    mdr-*.bed   rmp-* gwp-* gwr-*

Notes: The output is generated by BWASPR functions det_mrpr() and map_mrpr().
       Methylation-rich and -poor regions are determined based on the spacing between
        neighboring CpGhsm sites.  If sites occur at positions a, b, c, d, e, and f
        (and not in between), then b-a, c-b, ... are 1-distances, and f-a is a 5-distance.
        The file dst-*.txt shows the empirical distribution of d-distances for the sample.
        Methylation-poor regions are determined as long 1-distances (in the top <nbrxtrms>; merged
        if adjacent 1-distances are both in the top <nbrxtrms>).  Methylation-rich regions are determined
        as short 5-distances (in the low <nbrxtrms>; merged if adjacent 5-distances are both in the low
        <nbrxtrms>).  Parts of the distribution are plotted in the 1ds-*.pdf and 5ds-*.pdf files.

       The methylation-rich and -poor regions are listed in files dst-*.txt and mdr-*.bed,
        the latter in BED format for potential display in genome browsers.

       The overlap of methylation regions with genome features is summarized in files rmp-*.txt.
        Files gwr-*.txt show genes overlapping with methylation-rich regions, ordered by site
        density in the methylation-rich region.
        Files gwp-*.txt show genes overlapping with methylation-poor regions, ordered by site
        density in the methylation-poor region.
```

```
Directory: MRPR File: dst-Ob_LI_BCphase.txt

Analysis of  1-distances for sample "Ob_LI_BCphase" (415425 sites):

  Total number of  1-distances:     415382
  Median:     9.0 Mean:   532.4 Std:  2936.4
  Quantiles:
    0%     5%    10%    15%    20%    25%    30%    35%    40%    45%    50%    55%    60%    65%    70%    75%    80%    85%    90%    95%   100%
     1      1      1      1      1      1      1      1      4      6      9     13     17     22     29     39     57    102    286   2510 230101

  Low  density region distance cutoff (lowest  100):   61986
  High density region distance cutoff (highest 100):       1

Ordered lists of methylation-poor regions:

 Top 100 (out of    90) methylation-poor regions in "Ob_LI_BCphase":
 (Sdnsty = sites per 1kb)

   Rtype  Sample       SeqID     From       To  Rlgth NbrSites  Sdnsty
1   Poor BCphase NC_039506.1  7050654  7641130 590477        5 0.00847
2   Poor BCphase NC_039506.1  6804806  7034907 230102        2 0.00869
3   Poor BCphase NC_039514.1  3714651  3884987 170337        2 0.01174
4   Poor BCphase NC_039506.1 11732404 11983650 251247        3 0.01194
5   Poor BCphase NC_039506.1   138612   458650 320039        4 0.01250
6   Poor BCphase NC_039515.1  9763772  9912700 148929        2 0.01343
...
... only first 25 lines shown ...
```

Directory: MRPR File: dst-Ob_LI_Rphase.txt

Analysis of 1-distances for sample "Ob_LI_Rphase" (445068 sites):

Total number of 1-distances: 445012
Median: 10.0 Mean: 497.1 Std: 2536.7
Quantiles:

| 0% | 5% | 10% | 15% | 20% | 25% | 30% | 35% | 40% | 45% | 50% | 55% | 60% | 65% | 70% | 75% | 80% | 85% | 90% | 95% | 100% |
|---|---|---|---|---|---|---|---|---|---|---|---|---|---|---|---|---|---|---|---|---|
| 1 | 1 | 1 | 1 | 1 | 1 | 1 | 1 | 4 | 7 | 10 | 14 | 18 | 24 | 31 | 43 | 67 | 131 | 515 | 2759 | 248999 |

Low density region distance cutoff (lowest 100): 54759
High density region distance cutoff (highest 100): 1

Ordered lists of methylation-poor regions:

Top 100 (out of 92) methylation-poor regions in "Ob_LI_Rphase":
(Sdnsty = sites per 1kb)

| | Rtype | Sample | SeqID | From | To | Rlgth | NbrSites | Sdnsty |
|---|---|---|---|---|---|---|---|---|
| 1 | Poor | Rphase | NC_039506.1 | 7392270 | 7641269 | 249000 | 2 | 0.00803 |
| 2 | Poor | Rphase | NC_039506.1 | 463932 | 926754 | 462823 | 4 | 0.00864 |
| 3 | Poor | Rphase | NC_039506.1 | 6804755 | 7133639 | 328885 | 3 | 0.00912 |
| 4 | Poor | Rphase | NC_039506.1 | 7163033 | 7340882 | 177850 | 2 | 0.01125 |
| 5 | Poor | Rphase | NC_039515.1 | 9768933 | 9912700 | 143768 | 2 | 0.01391 |
| 6 | Poor | Rphase | NC_039514.1 | 3754274 | 3894633 | 140360 | 2 | 0.01425 |

...
... only first 25 lines shown ...

Directory: MRPR File: gwp-Ob_LI_BCphase.txt

| seqnames | start | end | width | strand | source | type | score | phase | ID | Dbxref | Name | gbkey | gene | gene_biotype | end_range | partial | start_range | seqnames | start | end | width | strand | NbrSites | Sdnsty |
|---|---|---|---|---|---|---|---|---|---|---|---|---|---|---|---|---|---|---|---|---|---|---|---|---|
| NC_039506.1 | 7068352 | 7072795 | 4444 | + | Gnomon | gene | NA | NA | gene-LOC105285461 | GeneID:105285461 | LOC105285461 | Gene | LOC105285461 | protein_coding | character(0) | NA | character(0) | NC_039506.1 | 7050654 | 7641130 | 590477 | * | 5 | 0.00846773032649875 |
| NC_039506.1 | 7087928 | 7195955 | 108028 | + | Gnomon | gene | NA | NA | gene-LOC105285459 | GeneID:105285459 | LOC105285459 | Gene | LOC105285459 | protein_coding | character(0) | NA | character(0) | NC_039506.1 | 7050654 | 7641130 | 590477 | * | 5 | 0.00846773032649875 |
| NC_039506.1 | 7200050 | 7204119 | 4070 | - | Gnomon | gene | NA | NA | gene-LOC105285458 | GeneID:105285458 | LOC105285458 | Gene | LOC105285458 | protein_coding | character(0) | NA | character(0) | NC_039506.1 | 7050654 | 7641130 | 590477 | * | 5 | 0.00846773032649875 |
| NC_039506.1 | 7218049 | 7225481 | 7433 | + | Gnomon | gene | NA | NA | gene-LOC105285457 | GeneID:105285457 | LOC105285457 | Gene | LOC105285457 | protein_coding | character(0) | NA | character(0) | NC_039506.1 | 7050654 | 7641130 | 590477 | * | 5 | 0.00846773032649875 |
| NC_039506.1 | 7233376 | 7276873 | 43498 | + | Gnomon | gene | NA | NA | gene-LOC113562292 | GeneID:113562292 | LOC113562292 | Gene | LOC113562292 | protein_coding | character(0) | NA | character(0) | NC_039506.1 | 7050654 | 7641130 | 590477 | * | 5 | 0.00846773032649875 |
| NC_039506.1 | 7277811 | 7279366 | 1556 | + | Gnomon | gene | NA | NA | gene-LOC113562297 | GeneID:113562297 | LOC113562297 | Gene | LOC113562297 | protein_coding | character(0) | NA | character(0) | NC_039506.1 | 7050654 | 7641130 | 590477 | * | 5 | 0.00846773032649875 |
| NC_039506.1 | 7279744 | 7280766 | 1023 | + | Gnomon | gene | NA | NA | gene-LOC105285452 | GeneID:105285452 | LOC105285452 | Gene | LOC105285452 | protein_coding | character(0) | NA | character(0) | NC_039506.1 | 7050654 | 7641130 | 590477 | * | 5 | 0.00846773032649875 |
| NC_039506.1 | 7288143 | 7290085 | 1943 | - | Gnomon | gene | NA | NA | gene-LOC113562235 | GeneID:113562235 | LOC113562235 | Gene | LOC113562235 | protein_coding | character(0) | NA | character(0) | NC_039506.1 | 7050654 | 7641130 | 590477 | * | 5 | 0.00846773032649875 |
| NC_039506.1 | 7303273 | 7305465 | 2193 | - | Gnomon | gene | NA | NA | gene-LOC105285451 | GeneID:105285451 | LOC105285451 | Gene | LOC105285451 | protein_coding | character(0) | NA | character(0) | NC_039506.1 | 7050654 | 7641130 | 590477 | * | 5 | 0.00846773032649875 |
| NC_039506.1 | 7307953 | 7323292 | 15340 | - | Gnomon | gene | NA | NA | gene-LOC105285449 | GeneID:105285449 | LOC105285449 | Gene | LOC105285449 | protein_coding | character(0) | NA | character(0) | NC_039506.1 | 7050654 | 7641130 | 590477 | * | 5 | 0.00846773032649875 |
| NC_039506.1 | 7361778 | 7368543 | 6766 | - | Gnomon | gene | NA | NA | gene-LOC105285446 | GeneID:105285446 | LOC105285446 | Gene | LOC105285446 | protein_coding | character(0) | NA | character(0) | NC_039506.1 | 7050654 | 7641130 | 590477 | * | 5 | 0.00846773032649875 |
| NC_039506.1 | 7371036 | 7371866 | 831 | - | Gnomon | gene | NA | NA | gene-LOC113562320 | GeneID:113562320 | LOC113562320 | Gene | LOC113562320 | protein_coding | character(0) | NA | character(0) | NC_039506.1 | 7050654 | 7641130 | 590477 | * | 5 | 0.00846773032649875 |
| NC_039506.1 | 7376295 | 7379683 | 3389 | + | Gnomon | gene | NA | NA | gene-LOC105285441 | GeneID:105285441 | LOC105285441 | Gene | LOC105285441 | protein_coding | character(0) | NA | character(0) | NC_039506.1 | 7050654 | 7641130 | 590477 | * | 5 | 0.00846773032649875 |
| NC_039506.1 | 7379406 | 7451038 | 71633 | - | Gnomon | gene | NA | NA | gene-LOC105285444 | GeneID:105285444 | LOC105285444 | Gene | LOC105285444 | protein_coding | character(0) | NA | character(0) | NC_039506.1 | 7050654 | 7641130 | 590477 | * | 5 | 0.00846773032649875 |
| NC_039506.1 | 7416229 | 7455130 | 38902 | + | Gnomon | gene | NA | NA | gene-LOC113561506 | GeneID:113561506 | LOC113561506 | Gene | LOC113561506 | protein_coding | character(0) | NA | character(0) | NC_039506.1 | 7050654 | 7641130 | 590477 | * | 5 | 0.00846773032649875 |
| NC_039506.1 | 7462673 | 7500744 | 38072 | + | Gnomon | gene | NA | NA | gene-LOC113563556 | GeneID:113563556 | LOC113563556 | Gene | LOC113563556 | protein_coding | character(0) | NA | character(0) | NC_039506.1 | 7050654 | 7641130 | 590477 | * | 5 | 0.00846773032649875 |
| NC_039506.1 | 7503462 | 7509165 | 5704 | + | Gnomon | gene | NA | NA | gene-LOC105285440 | GeneID:105285440 | LOC105285440 | Gene | LOC105285440 | protein_coding | character(0) | NA | character(0) | NC_039506.1 | 7050654 | 7641130 | 590477 | * | 5 | 0.00846773032649875 |
| NC_039506.1 | 7520890 | 7536243 | 15354 | + | Gnomon | gene | NA | NA | gene-LOC105285438 | GeneID:105285438 | LOC105285438 | Gene | LOC105285438 | protein_coding | character(0) | NA | character(0) | NC_039506.1 | 7050654 | 7641130 | 590477 | * | 5 | 0.00846773032649875 |
| NC_039506.1 | 7548941 | 7588987 | 40047 | + | Gnomon | gene | NA | NA | gene-LOC105286739 | GeneID:105286739 | LOC105286739 | Gene | LOC105286739 | protein_coding | character(0) | NA | character(0) | NC_039506.1 | 7050654 | 7641130 | 590477 | * | 5 | 0.00846773032649875 |
| NC_039506.1 | 7562407 | 7564340 | 1934 | + | Gnomon | gene | NA | NA | gene-LOC113562046 | GeneID:113562046 | LOC113562046 | Gene | LOC113562046 | protein_coding | character(0) | NA | character(0) | NC_039506.1 | 7050654 | 7641130 | 590477 | * | 5 | 0.00846773032649875 |
| NC_039506.1 | 7567762 | 7571445 | 3684 | + | Gnomon | gene | NA | NA | gene-LOC105285567 | GeneID:105285567 | LOC105285567 | Gene | LOC105285567 | protein_coding | character(0) | NA | character(0) | NC_039506.1 | 7050654 | 7641130 | 590477 | * | 5 | 0.00846773032649875 |
| NC_039506.1 | 7571564 | 7575323 | 3760 | + | Gnomon | gene | NA | NA | gene-LOC113562034 | GeneID:113562034 | LOC113562034 | Gene | LOC113562034 | protein_coding | character(0) | NA | character(0) | NC_039506.1 | 7050654 | 7641130 | 590477 | * | 5 | 0.00846773032649875 |
| NC_039506.1 | 7575125 | 7581202 | 6078 | + | Gnomon | gene | NA | NA | gene-LOC109611372 | GeneID:109611372 | LOC109611372 | Gene | LOC109611372 | protein_coding | character(0) | NA | character(0) | NC_039506.1 | 7050654 | 7641130 | 590477 | * | 5 | 0.00846773032649875 |
| NC_039506.1 | 7581490 | 7585137 | 3648 | + | Gnomon | gene | NA | NA | gene-LOC113562032 | GeneID:113562032 | LOC113562032 | Gene | LOC113562032 | protein_coding | character(0) | NA | character(0) | NC_039506.1 | 7050654 | 7641130 | 590477 | * | 5 | 0.00846773032649875 |

...

... only first 25 lines shown ...

Directory: MRPR File: gwp-Ob_LI_Rphase.txt

| seqnames | start | end | width | strand | source | type | score | phase | ID | Dbxref | Name | gbkey | gene | gene_biotype | end_range | partial | start_range | seqnames | start | end | width | strand | NbrSites | Sdnsty |
|---|---|---|---|---|---|---|---|---|---|---|---|---|---|---|---|---|---|---|---|---|---|---|---|---|
| NC_039506.1 | 7379406 | 7451038 | 71633 | - | Gnomon | gene | NA | NA | gene-LOC105285444 | GeneID:105285444 | LOC105285444 | Gene | LOC105285444 | protein_coding | character(0) | NA | character(0) | NC_039506.1 | 7392270 | 7641269 | 249000 | * | 2 | 0.00803212851405622 |
| NC_039506.1 | 7416229 | 7455130 | 38902 | + | Gnomon | gene | NA | NA | gene-LOC113561506 | GeneID:113561506 | LOC113561506 | Gene | LOC113561506 | protein_coding | character(0) | NA | character(0) | NC_039506.1 | 7392270 | 7641269 | 249000 | * | 2 | 0.00803212851405622 |
| NC_039506.1 | 7462673 | 7500744 | 38072 | + | Gnomon | gene | NA | NA | gene-LOC113563556 | GeneID:113563556 | LOC113563556 | Gene | LOC113563556 | protein_coding | character(0) | NA | character(0) | NC_039506.1 | 7392270 | 7641269 | 249000 | * | 2 | 0.00803212851405622 |
| NC_039506.1 | 7503462 | 7509165 | 5704 | + | Gnomon | gene | NA | NA | gene-LOC105285440 | GeneID:105285440 | LOC105285440 | Gene | LOC105285440 | protein_coding | character(0) | NA | character(0) | NC_039506.1 | 7392270 | 7641269 | 249000 | * | 2 | 0.00803212851405622 |
| NC_039506.1 | 7520890 | 7536243 | 15354 | + | Gnomon | gene | NA | NA | gene-LOC105285438 | GeneID:105285438 | LOC105285438 | Gene | LOC105285438 | protein_coding | character(0) | NA | character(0) | NC_039506.1 | 7392270 | 7641269 | 249000 | * | 2 | 0.00803212851405622 |
| NC_039506.1 | 7548941 | 7588987 | 40047 | + | Gnomon | gene | NA | NA | gene-LOC105286739 | GeneID:105286739 | LOC105286739 | Gene | LOC105286739 | protein_coding | character(0) | NA | character(0) | NC_039506.1 | 7392270 | 7641269 | 249000 | * | 2 | 0.00803212851405622 |
| NC_039506.1 | 7562407 | 7564340 | 1934 | + | Gnomon | gene | NA | NA | gene-LOC113562046 | GeneID:113562046 | LOC113562046 | Gene | LOC113562046 | protein_coding | character(0) | NA | character(0) | NC_039506.1 | 7392270 | 7641269 | 249000 | * | 2 | 0.00803212851405622 |
| NC_039506.1 | 7567762 | 7571445 | 3684 | + | Gnomon | gene | NA | NA | gene-LOC105285567 | GeneID:105285567 | LOC105285567 | Gene | LOC105285567 | protein_coding | character(0) | NA | character(0) | NC_039506.1 | 7392270 | 7641269 | 249000 | * | 2 | 0.00803212851405622 |
| NC_039506.1 | 7571564 | 7575323 | 3760 | + | Gnomon | gene | NA | NA | gene-LOC113562034 | GeneID:113562034 | LOC113562034 | Gene | LOC113562034 | protein_coding | character(0) | NA | character(0) | NC_039506.1 | 7392270 | 7641269 | 249000 | * | 2 | 0.00803212851405622 |
| NC_039506.1 | 7575125 | 7581202 | 6078 | + | Gnomon | gene | NA | NA | gene-LOC109611372 | GeneID:109611372 | LOC109611372 | Gene | LOC109611372 | protein_coding | character(0) | NA | character(0) | NC_039506.1 | 7392270 | 7641269 | 249000 | * | 2 | 0.00803212851405622 |
| NC_039506.1 | 7581490 | 7585137 | 3648 | + | Gnomon | gene | NA | NA | gene-LOC113562032 | GeneID:113562032 | LOC113562032 | Gene | LOC113562032 | protein_coding | character(0) | NA | character(0) | NC_039506.1 | 7392270 | 7641269 | 249000 | * | 2 | 0.00803212851405622 |
| NC_039506.1 | 7585265 | 7587090 | 1826 | + | Gnomon | gene | NA | NA | gene-LOC113562039 | GeneID:113562039 | LOC113562039 | Gene | LOC113562039 | protein_coding | character(0) | NA | character(0) | NC_039506.1 | 7392270 | 7641269 | 249000 | * | 2 | 0.00803212851405622 |
| NC_039506.1 | 7605350 | 7609877 | 4528 | + | Gnomon | gene | NA | NA | gene-LOC113563601 | GeneID:113563601 | LOC113563601 | Gene | LOC113563601 | protein_coding | character(0) | NA | character(0) | NC_039506.1 | 7392270 | 7641269 | 249000 | * | 2 | 0.00803212851405622 |
| NC_039506.1 | 7616669 | 7635522 | 18854 | + | Gnomon | gene | NA | NA | gene-LOC113563594 | GeneID:113563594 | LOC113563594 | Gene | LOC113563594 | protein_coding | character(0) | NA | character(0) | NC_039506.1 | 7392270 | 7641269 | 249000 | * | 2 | 0.00803212851405622 |
| NC_039506.1 | 7635628 | 7636926 | 1299 | - | Gnomon | gene | NA | NA | gene-LOC105288143 | GeneID:105288143 | LOC105288143 | Gene | LOC105288143 | protein_coding | character(0) | NA | character(0) | NC_039506.1 | 7392270 | 7641269 | 249000 | * | 2 | 0.00803212851405622 |
| NC_039506.1 | 7637029 | 7639864 | 2836 | - | Gnomon | gene | NA | NA | gene-LOC113563602 | GeneID:113563602 | LOC113563602 | Gene | LOC113563602 | protein_coding | character(0) | NA | character(0) | NC_039506.1 | 7392270 | 7641269 | 249000 | * | 2 | 0.00803212851405622 |
| NC_039506.1 | 7640376 | 7643220 | 2845 | + | Gnomon | gene | NA | NA | gene-LOC105288141 | GeneID:105288141 | LOC105288141 | Gene | LOC105288141 | protein_coding | character(0) | NA | character(0) | NC_039506.1 | 7392270 | 7641269 | 249000 | * | 2 | 0.00803212851405622 |
| NC_039506.1 | 458174 | 503061 | 44888 | - | Gnomon | gene | NA | NA | gene-LOC105285445 | GeneID:105285445 | LOC105285445 | Gene | LOC105285445 | protein_coding | character(0) | NA | character(0) | NC_039506.1 | 463932 | 926754 | 462823 | * | 4 | 0.00864261283471219 |
| NC_039506.1 | 534042 | 537304 | 3263 | - | Gnomon | gene | NA | NA | gene-LOC113562127 | GeneID:113562127 | LOC113562127 | Gene | LOC113562127 | protein_coding | character(0) | NA | character(0) | NC_039506.1 | 463932 | 926754 | 462823 | * | 4 | 0.00864261283471219 |
| NC_039506.1 | 541732 | 542562 | 831 | + | Gnomon | gene | NA | NA | gene-LOC113562288 | GeneID:113562288 | LOC113562288 | Gene | LOC113562288 | protein_coding | character(0) | NA | character(0) | NC_039506.1 | 463932 | 926754 | 462823 | * | 4 | 0.00864261283471219 |
| NC_039506.1 | 545053 | 551825 | 6773 | + | Gnomon | gene | NA | NA | gene-LOC113562287 | GeneID:113562287 | LOC113562287 | Gene | LOC113562287 | protein_coding | character(0) | NA | character(0) | NC_039506.1 | 463932 | 926754 | 462823 | * | 4 | 0.00864261283471219 |
| NC_039506.1 | 559004 | 580516 | 21513 | + | Gnomon | gene | NA | NA | gene-LOC105285447 | GeneID:105285447 | LOC105285447 | Gene | LOC105285447 | protein_coding | character(0) | NA | character(0) | NC_039506.1 | 463932 | 926754 | 462823 | * | 4 | 0.00864261283471219 |
| NC_039506.1 | 609439 | 611633 | 2195 | + | Gnomon | gene | NA | NA | gene-LOC113562164 | GeneID:113562164 | LOC113562164 | Gene | LOC113562164 | protein_coding | character(0) | NA | character(0) | NC_039506.1 | 463932 | 926754 | 462823 | * | 4 | 0.00864261283471219 |
| NC_039506.1 | 614991 | 620741 | 5751 | + | Gnomon | gene | NA | NA | gene-LOC113562147 | GeneID:113562147 | LOC113562147 | Gene | LOC113562147 | lncRNA | character(0) | NA | character(0) | NC_039506.1 | 463932 | 926754 | 462823 | * | 4 | 0.00864261283471219 |

...
... only first 25 lines shown ...

Directory: MRPR File: gwr-Ob_LI_BCphase.txt

| seqnames | start | end | width | strand | source | type | score | phase | ID | Dbxref | Name | gbkey | gene | gene_biotype | end_range | partial | start_range | seqnames | start | end | width | strand | NbrSites | Sdnsty |
|---|---|---|---|---|---|---|---|---|---|---|---|---|---|---|---|---|---|---|---|---|---|---|---|---|
| NC_039506.1 | 3208967 | 3211558 | 2592 | - | Gnomon | gene | NA | NA | gene-LOC105280373 | GeneID:105280373 | LOC105280373 | Gene | LOC105280373 | protein_coding | character(0) | NA | character(0) | NC_039506.1 | 3210393 | 3210398 | 6 | * | 6 | 1000 |
| NC_039506.1 | 3342881 | 3348391 | 5511 | + | Gnomon | gene | NA | NA | gene-LOC105280395 | GeneID:105280395 | LOC105280395 | Gene | LOC105280395 | protein_coding | character(0) | NA | character(0) | NC_039506.1 | 3344178 | 3344183 | 6 | * | 6 | 1000 |
| NC_039506.1 | 6089683 | 6090856 | 1174 | + | Gnomon | gene | NA | NA | gene-LOC105277804 | GeneID:105277804 | LOC105277804 | Gene | LOC105277804 | protein_coding | character(0) | NA | character(0) | NC_039506.1 | 6090624 | 6090629 | 6 | * | 6 | 1000 |
| NC_039506.1 | 13177256 | 13183827 | 6572 | + | Gnomon | gene | NA | NA | gene-LOC105284041 | GeneID:105284041 | LOC105284041 | Gene | LOC105284041 | protein_coding | character(0) | NA | character(0) | NC_039506.1 | 13181314 | 13181319 | 6 | * | 6 | 1000 |
| NC_039506.1 | 14188024 | 14192973 | 4950 | - | Gnomon | gene | NA | NA | gene-LOC105281111 | GeneID:105281111 | LOC105281111 | Gene | LOC105281111 | protein_coding | character(0) | NA | character(0) | NC_039506.1 | 14191640 | 14191645 | 6 | * | 6 | 1000 |
| NC_039506.1 | 15370515 | 15371538 | 1024 | + | Gnomon | gene | NA | NA | gene-LOC105280860 | GeneID:105280860 | LOC105280860 | Gene | LOC105280860 | protein_coding | character(0) | NA | character(0) | NC_039506.1 | 15371422 | 15371427 | 6 | * | 6 | 1000 |
| NC_039506.1 | 17130855 | 17136029 | 5175 | - | Gnomon | gene | NA | NA | gene-LOC105274952 | GeneID:105274952 | LOC105274952 | Gene | LOC105274952 | protein_coding | character(0) | NA | character(0) | NC_039506.1 | 17133800 | 17133805 | 6 | * | 6 | 1000 |
| NC_039506.1 | 23567682 | 23574351 | 6670 | + | Gnomon | gene | NA | NA | gene-LOC105279333 | GeneID:105279333 | LOC105279333 | Gene | LOC105279333 | protein_coding | character(0) | NA | character(0) | NC_039506.1 | 23568887 | 23568894 | 8 | * | 8 | 1000 |
| NC_039506.1 | 23689672 | 23694320 | 4649 | - | Gnomon | gene | NA | NA | gene-LOC105279346 | GeneID:105279346 | LOC105279346 | Gene | LOC105279346 | protein_coding | character(0) | NA | character(0) | NC_039506.1 | 23692892 | 23692897 | 6 | * | 6 | 1000 |
| NC_039506.1 | 24043149 | 24047806 | 4658 | + | Gnomon | gene | NA | NA | gene-LOC105279367 | GeneID:105279367 | LOC105279367 | Gene | LOC105279367 | protein_coding | character(0) | NA | character(0) | NC_039506.1 | 24044697 | 24044702 | 6 | * | 6 | 1000 |
| NC_039507.1 | 5093844 | 5095945 | 2102 | + | Gnomon | gene | NA | NA | gene-LOC105281690 | GeneID:105281690 | LOC105281690 | Gene | LOC105281690 | protein_coding | character(0) | NA | character(0) | NC_039507.1 | 5094466 | 5094471 | 6 | * | 6 | 1000 |
| NC_039507.1 | 5492373 | 5502473 | 10101 | + | Gnomon | gene | NA | NA | gene-LOC105281645 | GeneID:105281645 | LOC105281645 | Gene | LOC105281645 | protein_coding | character(0) | NA | character(0) | NC_039507.1 | 5493754 | 5493759 | 6 | * | 6 | 1000 |
| NC_039507.1 | 6546804 | 6552192 | 5389 | - | Gnomon | gene | NA | NA | gene-LOC105287207 | GeneID:105287207 | LOC105287207 | Gene | LOC105287207 | protein_coding | character(0) | NA | character(0) | NC_039507.1 | 6550682 | 6550687 | 6 | * | 6 | 1000 |
| NC_039507.1 | 7181199 | 7188522 | 7324 | + | Gnomon | gene | NA | NA | gene-LOC105283383 | GeneID:105283383 | LOC105283383 | Gene | LOC105283383 | protein_coding | character(0) | NA | character(0) | NC_039507.1 | 7182290 | 7182295 | 6 | * | 6 | 1000 |
| NC_039507.1 | 7276866 | 7278961 | 2096 | + | Gnomon | gene | NA | NA | gene-LOC105283373 | GeneID:105283373 | LOC105283373 | Gene | LOC105283373 | protein_coding | character(0) | NA | character(0) | NC_039507.1 | 7277947 | 7277952 | 6 | * | 6 | 1000 |
| NC_039507.1 | 9161485 | 9166239 | 4755 | + | Gnomon | gene | NA | NA | gene-LOC105282461 | GeneID:105282461 | LOC105282461 | Gene | LOC105282461 | protein_coding | character(0) | NA | character(0) | NC_039507.1 | 9162377 | 9162382 | 6 | * | 6 | 1000 |
| NC_039507.1 | 10633196 | 10639514 | 6319 | + | Gnomon | gene | NA | NA | gene-LOC105279155 | GeneID:105279155 | LOC105279155 | Gene | LOC105279155 | protein_coding | character(0) | NA | character(0) | NC_039507.1 | 10634249 | 10634254 | 6 | * | 6 | 1000 |
| NC_039507.1 | 13550180 | 13557055 | 6876 | - | Gnomon | gene | NA | NA | gene-LOC105285277 | GeneID:105285277 | LOC105285277 | Gene | LOC105285277 | protein_coding | character(0) | NA | character(0) | NC_039507.1 | 13555722 | 13555727 | 6 | * | 6 | 1000 |
| NC_039507.1 | 13552141 | 13556636 | 4496 | + | Gnomon | gene | NA | NA | gene-LOC105285278 | GeneID:105285278 | LOC105285278 | Gene | LOC105285278 | protein_coding | character(0) | NA | character(0) | NC_039507.1 | 13555722 | 13555727 | 6 | * | 6 | 1000 |
| NC_039507.1 | 15793673 | 15807525 | 13853 | - | Gnomon | gene | NA | NA | gene-LOC105280128 | GeneID:105280128 | LOC105280128 | Gene | LOC105280128 | protein_coding | character(0) | NA | character(0) | NC_039507.1 | 15805198 | 15805203 | 6 | * | 6 | 1000 |
| NC_039507.1 | 15793673 | 15807525 | 13853 | - | Gnomon | gene | NA | NA | gene-LOC105280128 | GeneID:105280128 | LOC105280128 | Gene | LOC105280128 | protein_coding | character(0) | NA | character(0) | NC_039507.1 | 15805657 | 15805662 | 6 | * | 6 | 1000 |
| NC_039507.1 | 18333035 | 18336377 | 3343 | - | Gnomon | gene | NA | NA | gene-LOC105281577 | GeneID:105281577 | LOC105281577 | Gene | LOC105281577 | protein_coding | character(0) | NA | character(0) | NC_039507.1 | 18335809 | 18335814 | 6 | * | 6 | 1000 |
| NC_039508.1 | 2590841 | 2594093 | 3253 | + | Gnomon | gene | NA | NA | gene-LOC105281341 | GeneID:105281341 | LOC105281341 | Gene | LOC105281341 | protein_coding | character(0) | NA | character(0) | NC_039508.1 | 2591612 | 2591617 | 6 | * | 6 | 1000 |
| NC_039508.1 | 10274002 | 10277440 | 3439 | + | Gnomon | gene | NA | NA | gene-LOC105276881 | GeneID:105276881 | LOC105276881 | Gene | LOC105276881 | protein_coding | character(0) | NA | character(0) | NC_039508.1 | 10275212 | 10275217 | 6 | * | 6 | 1000 |
| NC_039508.1 | 10716085 | 10732040 | 15956 | - | Gnomon | gene | NA | NA | gene-LOC105276941 | GeneID:105276941 | LOC105276941 | Gene | LOC105276941 | protein_coding | character(0) | NA | character(0) | NC_039508.1 | 10730559 | 10730564 | 6 | * | 6 | 1000 |
| NC_039508.1 | 10799899 | 10803333 | 3435 | + | Gnomon | gene | NA | NA | gene-LOC105276950 | GeneID:105276950 | LOC105276950 | Gene | LOC105276950 | protein_coding | character(0) | NA | character(0) | NC_039508.1 | 10801049 | 10801054 | 6 | * | 6 | 1000 |
| NC_039508.1 | 14022293 | 14024814 | 2522 | + | Gnomon | gene | NA | NA | gene-LOC105278683 | GeneID:105278683 | LOC105278683 | Gene | LOC105278683 | protein_coding | character(0) | NA | character(0) | NC_039508.1 | 14023382 | 14023387 | 6 | * | 6 | 1000 |
| NC_039508.1 | 15803095 | 15806139 | 3045 | + | Gnomon | gene | NA | NA | gene-LOC105277239 | GeneID:105277239 | LOC105277239 | Gene | LOC105277239 | protein_coding | character(0) | NA | character(0) | NC_039508.1 | 15804426 | 15804431 | 6 | * | 6 | 1000 |
| NC_039508.1 | 17548373 | 17550348 | 1976 | - | Gnomon | gene | NA | NA | gene-LOC105284396 | GeneID:105284396 | LOC105284396 | Gene | LOC105284396 | protein_coding | character(0) | NA | character(0) | NC_039508.1 | 17549290 | 17549295 | 6 | * | 6 | 1000 |
| NC_039509.1 | 1002697 | 1006107 | 3411 | - | Gnomon | gene | NA | NA | gene-LOC105283620 | GeneID:105283620 | LOC105283620 | Gene | LOC105283620 | protein_coding | | | | | | | | | | |

| | | | | | | | | | | | | | | |
|---|---|---|---|---|---|---|---|---|---|---|---|---|---|---|
| character(0) | NA | character(0) | NC_039509.1 | 1004827 | 1004832 | 6 | * | 6 | 1000 | | | | | |
| NC_039509.1 | 1135464 | 1137481 | 2018 | - | Gnomon | gene | NA | NA | gene-LOC105283634 | GeneID:105283634 | LOC105283634 | Gene | LOC105283634 | protein_coding |
| character(0) | NA | character(0) | NC_039509.1 | 1136266 | 1136271 | 6 | * | 6 | 1000 | | | | | |
| NC_039509.1 | 2825666 | 2828757 | 3092 | + | Gnomon | gene | NA | NA | gene-LOC105282589 | GeneID:105282589 | LOC105282589 | Gene | LOC105282589 | protein_coding |
| character(0) | NA | character(0) | NC_039509.1 | 2826537 | 2826542 | 6 | * | 6 | 1000 | | | | | |
| NC_039509.1 | 3925156 | 3928137 | 2982 | + | Gnomon | gene | NA | NA | gene-LOC105280619 | GeneID:105280619 | LOC105280619 | Gene | LOC105280619 | protein_coding |
| character(0) | NA | character(0) | NC_039509.1 | 3926254 | 3926259 | 6 | * | 6 | 1000 | | | | | |
| NC_039509.1 | 4882335 | 4886409 | 4075 | - | Gnomon | gene | NA | NA | gene-LOC105286160 | GeneID:105286160 | LOC105286160 | Gene | LOC105286160 | protein_coding |
| character(0) | NA | character(0) | NC_039509.1 | 4885053 | 4885058 | 6 | * | 6 | 1000 | | | | | |
| NC_039509.1 | 4882335 | 4886409 | 4075 | - | Gnomon | gene | NA | NA | gene-LOC105286160 | GeneID:105286160 | LOC105286160 | Gene | LOC105286160 | protein_coding |
| character(0) | NA | character(0) | NC_039509.1 | 4885072 | 4885077 | 6 | * | 6 | 1000 | | | | | |
| NC_039509.1 | 6318288 | 6320995 | 2708 | - | Gnomon | gene | NA | NA | gene-LOC105282525 | GeneID:105282525 | LOC105282525 | Gene | LOC105282525 | protein_coding |
| character(0) | NA | character(0) | NC_039509.1 | 6319895 | 6319900 | 6 | * | 6 | 1000 | | | | | |
| NC_039509.1 | 10209680 | 10213178 | 3499 | - | Gnomon | gene | NA | NA | gene-LOC105282662 | GeneID:105282662 | LOC105282662 | Gene | LOC105282662 | protein_coding |
| character(0) | NA | character(0) | NC_039509.1 | 10212587 | 10212592 | 6 | * | 6 | 1000 | | | | | |
| NC_039509.1 | 10209680 | 10213178 | 3499 | - | Gnomon | gene | NA | NA | gene-LOC105282662 | GeneID:105282662 | LOC105282662 | Gene | LOC105282662 | protein_coding |
| character(0) | NA | character(0) | NC_039509.1 | 10212613 | 10212618 | 6 | * | 6 | 1000 | | | | | |
| NC_039509.1 | 10745281 | 10747369 | 2089 | + | Gnomon | gene | NA | NA | gene-LOC105275200 | GeneID:105275200 | LOC105275200 | Gene | LOC105275200 | protein_coding |
| character(0) | NA | character(0) | NC_039509.1 | 10746212 | 10746217 | 6 | * | 6 | 1000 | | | | | |
| NC_039509.1 | 10756577 | 10757875 | 1299 | - | Gnomon | gene | NA | NA | gene-LOC105275214 | GeneID:105275214 | LOC105275214 | Gene | LOC105275214 | protein_coding |
| character(0) | NA | character(0) | NC_039509.1 | 10756803 | 10756808 | 6 | * | 6 | 1000 | | | | | |
| NC_039509.1 | 11047331 | 11049650 | 2320 | - | Gnomon | gene | NA | NA | gene-LOC105275731 | GeneID:105275731 | LOC105275731 | Gene | LOC105275731 | protein_coding |
| character(0) | NA | character(0) | NC_039509.1 | 11048986 | 11048991 | 6 | * | 6 | 1000 | | | | | |
| NC_039509.1 | 11068956 | 11086837 | 17882 | - | Gnomon | gene | NA | NA | gene-LOC105275770 | GeneID:105275770 | LOC105275770 | Gene | LOC105275770 | protein_coding |
| character(0) | NA | character(0) | NC_039509.1 | 11085457 | 11085462 | 6 | * | 6 | 1000 | | | | | |
| NC_039509.1 | 13033738 | 13035966 | 2229 | + | Gnomon | gene | NA | NA | gene-LOC105278527 | GeneID:105278527 | LOC105278527 | Gene | LOC105278527 | protein_coding |
| character(0) | NA | character(0) | NC_039509.1 | 13034669 | 13034674 | 6 | * | 6 | 1000 | | | | | |
| NC_039509.1 | 13067024 | 13068370 | 1347 | + | Gnomon | gene | NA | NA | gene-LOC105278478 | GeneID:105278478 | LOC105278478 | Gene | LOC105278478 | protein_coding |
| character(0) | NA | character(0) | NC_039509.1 | 13067776 | 13067781 | 6 | * | 6 | 1000 | | | | | |
| NC_039509.1 | 17207204 | 17209527 | 2324 | + | Gnomon | gene | NA | NA | gene-LOC105283016 | GeneID:105283016 | LOC105283016 | Gene | LOC105283016 | protein_coding |
| character(0) | NA | character(0) | NC_039509.1 | 17208074 | 17208079 | 6 | * | 6 | 1000 | | | | | |
| NC_039510.1 | 2607810 | 2610599 | 2790 | - | Gnomon | gene | NA | NA | gene-LOC105287754 | GeneID:105287754 | LOC105287754 | Gene | LOC105287754 | protein_coding |
| character(0) | NA | character(0) | NC_039510.1 | 2609179 | 2609184 | 6 | * | 6 | 1000 | | | | | |
| NC_039510.1 | 3032387 | 3034961 | 2575 | + | Gnomon | gene | NA | NA | gene-LOC105287715 | GeneID:105287715 | LOC105287715 | Gene | LOC105287715 | protein_coding |
| character(0) | NA | character(0) | NC_039510.1 | 3033855 | 3033860 | 6 | * | 6 | 1000 | | | | | |
| NC_039510.1 | 3126910 | 3147548 | 20639 | - | Gnomon | gene | NA | NA | gene-LOC105287694 | GeneID:105287694 | LOC105287694 | Gene | LOC105287694 | protein_coding |
| character(0) | NA | character(0) | NC_039510.1 | 3133558 | 3133563 | 6 | * | 6 | 1000 | | | | | |
| NC_039510.1 | 3132542 | 3134046 | 1505 | + | Gnomon | gene | NA | NA | gene-LOC105287692 | GeneID:105287692 | LOC105287692 | Gene | LOC105287692 | protein_coding |
| character(0) | NA | character(0) | NC_039510.1 | 3133558 | 3133563 | 6 | * | 6 | 1000 | | | | | |
| NC_039510.1 | 3210462 | 3212049 | 1588 | - | Gnomon | gene | NA | NA | gene-LOC105287677 | GeneID:105287677 | LOC105287677 | Gene | LOC105287677 | protein_coding |
| character(0) | NA | character(0) | NC_039510.1 | 3211153 | 3211158 | 6 | * | 6 | 1000 | | | | | |
| NC_039510.1 | 3319355 | 3322927 | 3573 | + | Gnomon | gene | NA | NA | gene-LOC105287663 | GeneID:105287663 | LOC105287663 | Gene | LOC105287663 | protein_coding |
| character(0) | NA | character(0) | NC_039510.1 | 3320255 | 3320260 | 6 | * | 6 | 1000 | | | | | |
| NC_039510.1 | 6135149 | 6137854 | 2706 | + | Gnomon | gene | NA | NA | gene-LOC105287333 | GeneID:105287333 | LOC105287333 | Gene | LOC105287333 | protein_coding |
| character(0) | NA | character(0) | NC_039510.1 | 6136456 | 6136461 | 6 | * | 6 | 1000 | | | | | |
| NC_039510.1 | 9658025 | 9667283 | 9259 | - | Gnomon | gene | NA | NA | gene-LOC105287015 | GeneID:105287015 | LOC105287015 | Gene | LOC105287015 | protein_coding |
| character(0) | NA | character(0) | NC_039510.1 | 9665819 | 9665824 | 6 | * | 6 | 1000 | | | | | |
| NC_039510.1 | 11515868 | 11522296 | 6429 | + | Gnomon | gene | NA | NA | gene-LOC105281115 | GeneID:105281115 | LOC105281115 | Gene | LOC105281115 | protein_coding |
| character(0) | NA | character(0) | NC_039510.1 | 11518799 | 11518804 | 6 | * | 6 | 1000 | | | | | |
| NC_039510.1 | 12000027 | 12025662 | 25636 | - | Gnomon | gene | NA | NA | gene-LOC105281066 | GeneID:105281066 | LOC105281066 | Gene | LOC105281066 | protein_coding |
| character(0) | NA | character(0) | NC_039510.1 | 12024411 | 12024416 | 6 | * | 6 | 1000 | | | | | |
| NC_039510.1 | 12023589 | 12029249 | 5661 | + | Gnomon | gene | NA | NA | gene-LOC105281067 | GeneID:105281067 | LOC105281067 | Gene | LOC105281067 | protein_coding |
| character(0) | NA | character(0) | NC_039510.1 | 12024411 | 12024416 | 6 | * | 6 | 1000 | | | | | |
| NC_039510.1 | 13595835 | 13598710 | 2876 | + | Gnomon | gene | NA | NA | gene-LOC105280955 | GeneID:105280955 | LOC105280955 | Gene | LOC105280955 | protein_coding |
| character(0) | NA | character(0) | NC_039510.1 | 13597157 | 13597162 | 6 | * | 6 | 1000 | | | | | |
| NC_039510.1 | 13947158 | 13949082 | 1925 | + | Gnomon | gene | NA | NA | gene-LOC105285205 | GeneID:105285205 | LOC105285205 | Gene | LOC105285205 | protein_coding |
| character(0) | NA | character(0) | NC_039510.1 | 13947830 | 13947835 | 6 | * | 6 | 1000 | | | | | |
| NC_039510.1 | 16287375 | 16291357 | 3983 | - | Gnomon | gene | NA | NA | gene-LOC105286509 | GeneID:105286509 | LOC105286509 | Gene | LOC105286509 | protein_coding |
| character(0) | NA | character(0) | NC_039510.1 | 16290598 | 16290603 | 6 | * | 6 | 1000 | | | | | |
| NC_039511.1 | 480353 | 488275 | 7923 | + | Gnomon | gene | NA | NA | gene-LOC105276527 | GeneID:105276527 | LOC105276527 | Gene | LOC105276527 | protein_coding |
| character(0) | NA | character(0) | NC_039511.1 | 481652 | 481657 | 6 | * | 6 | 1000 | | | | | |
| NC_039511.1 | 2234830 | 2236789 | 1960 | - | Gnomon | gene | NA | NA | gene-LOC105276382 | GeneID:105276382 | LOC105276382 | Gene | LOC105276382 | protein_coding |
| character(0) | NA | character(0) | NC_039511.1 | 2236016 | 2236021 | 6 | * | 6 | 1000 | | | | | |
| NC_039511.1 | 3614241 | 3616939 | 2699 | + | Gnomon | gene | NA | NA | gene-LOC105284057 | GeneID:105284057 | LOC105284057 | Gene | LOC105284057 | protein_coding |

character(0) NA character(0) NC_039511.1 3615462 3615467 6 * 6 1000
NC_039511.1 3625986 3631991 6006 - Gnomon gene NA NA gene-LOC105284053 GeneID:105284053 LOC105284053 Gene LOC105284053 protein_coding
character(0) NA character(0) NC_039511.1 3630771 3630776 6 * 6 1000
NC_039511.1 8407553 8425032 17480 + Gnomon gene NA NA gene-LOC105284845 GeneID:105284845 LOC105284845 Gene LOC105284845 protein_coding
character(0) NA character(0) NC_039511.1 8409706 8409711 6 * 6 1000
NC_039511.1 8695181 8698086 2906 + Gnomon gene NA NA gene-LOC105280856 GeneID:105280856 LOC105280856 Gene LOC105280856 protein_coding
character(0) NA character(0) NC_039511.1 8695844 8695849 6 * 6 1000
NC_039511.1 11533823 11536626 2804 + Gnomon gene NA NA gene-LOC105284137 GeneID:105284137 LOC105284137 Gene LOC105284137 protein_coding
character(0) NA character(0) NC_039511.1 11534787 11534792 6 * 6 1000
NC_039511.1 12778711 12781434 2724 + Gnomon gene NA NA gene-LOC105276248 GeneID:105276248 LOC105276248 Gene LOC105276248 protein_coding
character(0) NA character(0) NC_039511.1 12779347 12779354 8 * 8 1000
NC_039511.1 15681353 15682825 1473 + Gnomon gene NA NA gene-LOC105284582 GeneID:105284582 LOC105284582 Gene LOC105284582 protein_coding
character(0) NA character(0) NC_039511.1 15682121 15682126 6 * 6 1000
NC_039512.1 384950 392996 8047 - Gnomon gene NA NA gene-LOC105288044 GeneID:105288044 LOC105288044 Gene LOC105288044 protein_coding
character(0) NA character(0) NC_039512.1 391510 391515 6 * 6 1000
NC_039512.1 4385041 4390283 5243 - Gnomon gene NA NA gene-LOC105277552 GeneID:105277552 LOC105277552 Gene LOC105277552 protein_coding
character(0) NA character(0) NC_039512.1 4388975 4388980 6 * 6 1000
NC_039512.1 8874729 8882472 7744 - Gnomon gene NA NA gene-LOC105282356 GeneID:105282356 LOC105282356 Gene LOC105282356 protein_coding
character(0) NA character(0) NC_039512.1 8881441 8881446 6 * 6 1000
NC_039512.1 9800733 9802923 2191 - Gnomon gene NA NA gene-LOC105279900 GeneID:105279900 LOC105279900 Gene LOC105279900 protein_coding
character(0) NA character(0) NC_039512.1 9801464 9801469 6 * 6 1000
NC_039512.1 10275466 10280187 4722 + Gnomon gene NA NA gene-LOC105282847 GeneID:105282847 LOC105282847 Gene LOC105282847 protein_coding
character(0) NA character(0) NC_039512.1 10276481 10276486 6 * 6 1000
NC_039512.1 11140867 11151152 10286 - Gnomon gene NA NA gene-LOC105279854 GeneID:105279854 LOC105279854 Gene LOC105279854 protein_coding
character(0) NA character(0) NC_039512.1 11150038 11150045 8 * 8 1000
NC_039512.1 11239021 11242090 3070 + Gnomon gene NA NA gene-LOC105279838 GeneID:105279838 LOC105279838 Gene LOC105279838 protein_coding
character(0) NA character(0) NC_039512.1 11240529 11240534 6 * 6 1000
NC_039512.1 12092666 12097773 5108 + Gnomon gene NA NA gene-LOC105279779 GeneID:105279779 LOC105279779 Gene LOC105279779 protein_coding
character(0) NA character(0) NC_039512.1 12094200 12094205 6 * 6 1000
NC_039512.1 12752221 12767858 15638 - Gnomon gene NA NA gene-LOC105279717 GeneID:105279717 LOC105279717 Gene LOC105279717 protein_coding
character(0) NA character(0) NC_039512.1 12766252 12766257 6 * 6 1000
NC_039512.1 13230551 13243418 12868 - Gnomon gene NA NA gene-LOC113562252 GeneID:113562252 LOC113562252 Gene LOC113562252 lncRNA character(0)
NA character(0) NC_039512.1 13237133 13237138 6 * 6 1000
NC_039512.1 13235891 13243388 7498 + Gnomon gene NA NA gene-LOC105279649 GeneID:105279649 LOC105279649 Gene LOC105279649 protein_coding
character(0) NA character(0) NC_039512.1 13237133 13237138 6 * 6 1000
NC_039512.1 14248372 14257496 9125 + Gnomon gene NA NA gene-LOC105275078 GeneID:105275078 LOC105275078 Gene LOC105275078 misc_RNA character(0)
NA character(0) NC_039512.1 14249035 14249040 6 * 6 1000
NC_039512.1 15707994 15714445 6452 + Gnomon gene NA NA gene-LOC105274475 GeneID:105274475 LOC105274475 Gene LOC105274475 protein_coding
character(0) NA character(0) NC_039512.1 15711477 15711482 6 * 6 1000
NC_039512.1 15979275 15981529 2255 - Gnomon gene NA NA gene-LOC105274775 GeneID:105274775 LOC105274775 Gene LOC105274775 protein_coding
character(0) NA character(0) NC_039512.1 15980634 15980639 6 * 6 1000
NC_039512.1 16221715 16230076 8362 + Gnomon gene NA NA gene-LOC105282804 GeneID:105282804 LOC105282804 Gene LOC105282804 protein_coding
character(0) NA character(0) NC_039512.1 16225485 16225490 6 * 6 1000
NC_039513.1 3144818 3145709 892 + Gnomon gene NA NA gene-LOC105287814 GeneID:105287814 LOC105287814 Gene LOC105287814 protein_coding
character(0) NA character(0) NC_039513.1 3145174 3145179 6 * 6 1000
NC_039513.1 3298586 3302590 4005 - Gnomon gene NA NA gene-LOC105275760 GeneID:105275760 LOC105275760 Gene LOC105275760 protein_coding
character(0) NA character(0) NC_039513.1 3301859 3301864 6 * 6 1000
NC_039513.1 3303685 3306319 2635 + Gnomon gene NA NA gene-LOC105275761 GeneID:105275761 LOC105275761 Gene LOC105275761 protein_coding
character(0) NA character(0) NC_039513.1 3304809 3304814 6 * 6 1000
NC_039513.1 7416041 7419328 3288 - Gnomon gene NA NA gene-LOC105275746 GeneID:105275746 LOC105275746 Gene LOC105275746 protein_coding
character(0) NA character(0) NC_039513.1 7418089 7418094 6 * 6 1000
NC_039513.1 10401827 10404590 2764 - Gnomon gene NA NA gene-LOC105275550 GeneID:105275550 LOC105275550 Gene LOC105275550 protein_coding
character(0) NA character(0) NC_039513.1 10403469 10403474 6 * 6 1000
NC_039513.1 10540843 10548517 7675 - Gnomon gene NA NA gene-LOC105275526 GeneID:105275526 LOC105275526 Gene LOC105275526 protein_coding
character(0) NA character(0) NC_039513.1 10546428 10546433 6 * 6 1000
NC_039513.1 11029007 11033701 4695 - Gnomon gene NA NA gene-LOC105275763 GeneID:105275763 LOC105275763 Gene LOC105275763 protein_coding
character(0) NA character(0) NC_039513.1 11030126 11030131 6 * 6 1000
NC_039513.1 11993718 11996290 2573 - Gnomon gene NA NA gene-LOC105280786 GeneID:105280786 LOC105280786 Gene LOC105280786 protein_coding
character(0) NA character(0) NC_039513.1 11994675 11994680 6 * 6 1000
NC_039513.1 12028675 12030941 2267 - Gnomon gene NA NA gene-LOC105280791 GeneID:105280791 LOC105280791 Gene LOC105280791 protein_coding
character(0) NA character(0) NC_039513.1 12029791 12029796 6 * 6 1000
NC_039513.1 13361516 13366191 4676 - Gnomon gene NA NA gene-LOC105275047 GeneID:105275047 LOC105275047 Gene LOC105275047 protein_coding
character(0) NA character(0) NC_039513.1 13364779 13364784 6 * 6 1000
NC_039513.1 13434303 13438575 4273 - Gnomon gene NA NA gene-LOC105275036 GeneID:105275036 LOC105275036 Gene LOC105275036 protein_coding

| | | | | | | | | | | | | | |
|---|---|---|---|---|---|---|---|---|---|---|---|---|---|
| character(0) | NA | character(0) | NC_039513.1 | 13436991 | 13437000 | 10 | * | 10 | 1000 | | | | |
| NC_039513.1 | 14931340 | 14932816 | 1477 | + | Gnomon | gene | NA | NA | gene-LOC105285670 | GeneID:105285670 | LOC105285670 | Gene | LOC105285670 | protein_coding |
| character(0) | NA | character(0) | NC_039513.1 | 14932042 | 14932047 | 6 | * | 6 | 1000 | | | | |
| NC_039514.1 | 6622839 | 6626782 | 3944 | - | Gnomon | gene | NA | NA | gene-LOC105285534 | GeneID:105285534 | LOC105285534 | Gene | LOC105285534 | protein_coding |
| character(0) | NA | character(0) | NC_039514.1 | 6625853 | 6625858 | 6 | * | 6 | 1000 | | | | |
| NC_039514.1 | 8819238 | 8822718 | 3481 | - | Gnomon | gene | NA | NA | gene-LOC105279004 | GeneID:105279004 | LOC105279004 | Gene | LOC105279004 | protein_coding |
| character(0) | NA | character(0) | NC_039514.1 | 8821565 | 8821570 | 6 | * | 6 | 1000 | | | | |
| NC_039514.1 | 9286321 | 9287389 | 1069 | - | Gnomon | gene | NA | NA | gene-LOC105279437 | GeneID:105279437 | LOC105279437 | Gene | LOC105279437 | protein_coding |
| character(0) | NA | character(0) | NC_039514.1 | 9286345 | 9286350 | 6 | * | 6 | 1000 | | | | |
| NC_039514.1 | 9336154 | 9343762 | 7609 | + | Gnomon | gene | NA | NA | gene-LOC105284644 | GeneID:105284644 | LOC105284644 | Gene | LOC105284644 | protein_coding |
| character(0) | NA | character(0) | NC_039514.1 | 9337043 | 9337048 | 6 | * | 6 | 1000 | | | | |
| NC_039514.1 | 9564295 | 9566245 | 1951 | + | Gnomon | gene | NA | NA | gene-LOC105284676 | GeneID:105284676 | LOC105284676 | Gene | LOC105284676 | protein_coding |
| character(0) | NA | character(0) | NC_039514.1 | 9565156 | 9565161 | 6 | * | 6 | 1000 | | | | |
| NC_039514.1 | 11109536 | 11117116 | 7581 | - | Gnomon | gene | NA | NA | gene-LOC105281601 | GeneID:105281601 | LOC105281601 | Gene | LOC105281601 | protein_coding |
| character(0) | NA | character(0) | NC_039514.1 | 11112955 | 11112960 | 6 | * | 6 | 1000 | | | | |
| NC_039514.1 | 11111779 | 11115572 | 3794 | + | Gnomon | gene | NA | NA | gene-LOC105281603 | GeneID:105281603 | LOC105281603 | Gene | LOC105281603 | protein_coding |
| character(0) | NA | character(0) | NC_039514.1 | 11112955 | 11112960 | 6 | * | 6 | 1000 | | | | |
| NC_039514.1 | 11115729 | 11118339 | 2611 | + | Gnomon | gene | NA | NA | gene-LOC105281604 | GeneID:105281604 | LOC105281604 | Gene | LOC105281604 | protein_coding |
| character(0) | NA | character(0) | NC_039514.1 | 11117220 | 11117225 | 6 | * | 6 | 1000 | | | | |
| NC_039514.1 | 13115199 | 13117880 | 2682 | + | Gnomon | gene | NA | NA | gene-LOC105277357 | GeneID:105277357 | LOC105277357 | Gene | LOC105277357 | protein_coding |
| character(0) | NA | character(0) | NC_039514.1 | 13116759 | 13116766 | 8 | * | 8 | 1000 | | | | |
| NC_039515.1 | 1335326 | 1341505 | 6180 | + | Gnomon | gene | NA | NA | gene-LOC105288065 | GeneID:105288065 | LOC105288065 | Gene | LOC105288065 | protein_coding |
| character(0) | NA | character(0) | NC_039515.1 | 1337469 | 1337474 | 6 | * | 6 | 1000 | | | | |
| NC_039515.1 | 4503178 | 4505245 | 2068 | + | Gnomon | gene | NA | NA | gene-LOC105278234 | GeneID:105278234 | LOC105278234 | Gene | LOC105278234 | protein_coding |
| character(0) | NA | character(0) | NC_039515.1 | 4503560 | 4503565 | 6 | * | 6 | 1000 | | | | |
| NC_039515.1 | 6512460 | 6524444 | 11985 | + | Gnomon | gene | NA | NA | gene-LOC105282289 | GeneID:105282289 | LOC105282289 | Gene | LOC105282289 | protein_coding |
| character(0) | NA | character(0) | NC_039515.1 | 6513507 | 6513512 | 6 | * | 6 | 1000 | | | | |
| NC_039515.1 | 7098785 | 7103508 | 4724 | - | Gnomon | gene | NA | NA | gene-LOC105286003 | GeneID:105286003 | LOC105286003 | Gene | LOC105286003 | protein_coding |
| character(0) | NA | character(0) | NC_039515.1 | 7102685 | 7102690 | 6 | * | 6 | 1000 | | | | |
| NC_039515.1 | 7216666 | 7221298 | 4633 | + | Gnomon | gene | NA | NA | gene-LOC105283771 | GeneID:105283771 | LOC105283771 | Gene | LOC105283771 | protein_coding |
| character(0) | NA | character(0) | NC_039515.1 | 7217376 | 7217381 | 6 | * | 6 | 1000 | | | | |
| NC_039515.1 | 12582469 | 12585642 | 3174 | - | Gnomon | gene | NA | NA | gene-LOC105282150 | GeneID:105282150 | LOC105282150 | Gene | LOC105282150 | protein_coding |
| character(0) | NA | character(0) | NC_039515.1 | 12584748 | 12584753 | 6 | * | 6 | 1000 | | | | |
| NC_039515.1 | 12596735 | 12605489 | 8755 | - | Gnomon | gene | NA | NA | gene-LOC105282142 | GeneID:105282142 | LOC105282142 | Gene | LOC105282142 | protein_coding |
| character(0) | NA | character(0) | NC_039515.1 | 12604757 | 12604762 | 6 | * | 6 | 1000 | | | | |
| NC_039515.1 | 12601969 | 12605663 | 3695 | + | Gnomon | gene | NA | NA | gene-LOC105282144 | GeneID:105282144 | LOC105282144 | Gene | LOC105282144 | protein_coding |
| character(0) | NA | character(0) | NC_039515.1 | 12604757 | 12604762 | 6 | * | 6 | 1000 | | | | |
| NC_039516.1 | 1062627 | 1071280 | 8654 | + | Gnomon | gene | NA | NA | gene-LOC105274578 | GeneID:105274578 | LOC105274578 | Gene | LOC105274578 | protein_coding |
| character(0) | NA | character(0) | NC_039516.1 | 1064627 | 1064632 | 6 | * | 6 | 1000 | | | | |
| NC_039516.1 | 1359476 | 1362079 | 2604 | + | Gnomon | gene | NA | NA | gene-LOC113562925 | GeneID:113562925 | LOC113562925 | Gene | LOC113562925 | protein_coding |
| character(0) | NA | character(0) | NC_039516.1 | 1360969 | 1360974 | 6 | * | 6 | 1000 | | | | |
| NC_039516.1 | 2372548 | 2378885 | 6338 | + | Gnomon | gene | NA | NA | gene-LOC105274446 | GeneID:105274446 | LOC105274446 | Gene | LOC105274446 | protein_coding |
| character(0) | NA | character(0) | NC_039516.1 | 2377394 | 2377399 | 6 | * | 6 | 1000 | | | | |
| NC_039516.1 | 2464828 | 2468017 | 3190 | + | Gnomon | gene | NA | NA | gene-LOC105288248 | GeneID:105288248 | LOC105288248 | Gene | LOC105288248 | protein_coding |
| character(0) | NA | character(0) | NC_039516.1 | 2465766 | 2465771 | 6 | * | 6 | 1000 | | | | |
| NC_039516.1 | 3088715 | 3091076 | 2362 | + | Gnomon | gene | NA | NA | gene-LOC105288196 | GeneID:105288196 | LOC105288196 | Gene | LOC105288196 | protein_coding |
| character(0) | NA | character(0) | NC_039516.1 | 3089997 | 3090002 | 6 | * | 6 | 1000 | | | | |
| NC_039516.1 | 3100198 | 3101692 | 1495 | + | Gnomon | gene | NA | NA | gene-LOC105288193 | GeneID:105288193 | LOC105288193 | Gene | LOC105288193 | protein_coding |
| character(0) | NA | character(0) | NC_039516.1 | 3101072 | 3101077 | 6 | * | 6 | 1000 | | | | |
| NC_039516.1 | 4713084 | 4714767 | 1684 | - | Gnomon | gene | NA | NA | gene-LOC105275897 | GeneID:105275897 | LOC105275897 | Gene | LOC105275897 | protein_coding |
| character(0) | NA | character(0) | NC_039516.1 | 4713694 | 4713699 | 6 | * | 6 | 1000 | | | | |
| NC_039516.1 | 5472871 | 5475154 | 2284 | - | Gnomon | gene | NA | NA | gene-LOC105275878 | GeneID:105275878 | LOC105275878 | Gene | LOC105275878 | protein_coding |
| character(0) | NA | character(0) | NC_039516.1 | 5474292 | 5474297 | 6 | * | 6 | 1000 | | | | |
| NC_039516.1 | 5868176 | 5870154 | 1979 | - | Gnomon | gene | NA | NA | gene-LOC105275861 | GeneID:105275861 | LOC105275861 | Gene | LOC105275861 | protein_coding |
| character(0) | NA | character(0) | NC_039516.1 | 5869246 | 5869251 | 6 | * | 6 | 1000 | | | | |
| NC_039516.1 | 7444132 | 7447943 | 3812 | - | Gnomon | gene | NA | NA | gene-LOC105275781 | GeneID:105275781 | LOC105275781 | Gene | LOC105275781 | protein_coding |
| character(0) | NA | character(0) | NC_039516.1 | 7446427 | 7446432 | 6 | * | 6 | 1000 | | | | |
| NC_039516.1 | 9287279 | 9290515 | 3237 | - | Gnomon | gene | NA | NA | gene-LOC105277523 | GeneID:105277523 | LOC105277523 | Gene | LOC105277523 | protein_coding |
| character(0) | NA | character(0) | NC_039516.1 | 9289245 | 9289250 | 6 | * | 6 | 1000 | | | | |
| NC_039516.1 | 9389294 | 9390221 | 928 | + | Gnomon | gene | NA | NA | gene-LOC105277237 | GeneID:105277237 | LOC105277237 | Gene | LOC105277237 | protein_coding |
| character(0) | NA | character(0) | NC_039516.1 | 9390169 | 9390174 | 6 | * | 6 | 1000 | | | | |
| NC_039516.1 | 9555082 | 9557519 | 2438 | + | Gnomon | gene | NA | NA | gene-LOC105278789 | GeneID:105278789 | LOC105278789 | Gene | LOC105278789 | protein_coding |
| character(0) | NA | character(0) | NC_039516.1 | 9555938 | 9555943 | 6 | * | 6 | 1000 | | | | |
| NC_039516.1 | 10251143 | 10256122 | 4980 | - | Gnomon | gene | NA | NA | gene-LOC105287447 | GeneID:105287447 | LOC105287447 | Gene | LOC105287447 | protein_coding |

| | | | | | | | | | | | | | | | | | | | | | | | | |
|---|---|---|---|---|---|---|---|---|---|---|---|---|---|---|---|---|---|---|---|---|---|---|---|---|
| | | | | | | | | | | | | | | | character(0) | NA | character(0) | NC_039516.1 | 10254626 | 10254631 | 6 | * | 6 | 1000 |
| NC_039516.1 | 10251143 | 10256122 | 4980 | - | Gnomon | gene | NA | NA | gene-LOC105287447 | GeneID:105287447 | LOC105287447 | Gene | LOC105287447 | protein_coding | character(0) | NA | character(0) | NC_039516.1 | 10254677 | 10254682 | 6 | * | 6 | 1000 |
| NC_039516.1 | 12300144 | 12302800 | 2657 | - | Gnomon | gene | NA | NA | gene-LOC105275264 | GeneID:105275264 | LOC105275264 | Gene | LOC105275264 | protein_coding | character(0) | NA | character(0) | NC_039516.1 | 12301684 | 12301689 | 6 | * | 6 | 1000 |
| NC_039516.1 | 12307603 | 12308894 | 1292 | + | Gnomon | gene | NA | NA | gene-LOC105275270 | GeneID:105275270 | LOC105275270 | Gene | LOC105275270 | protein_coding | character(0) | NA | character(0) | NC_039516.1 | 12308308 | 12308313 | 6 | * | 6 | 1000 |
| NC_039516.1 | 12889538 | 12895772 | 6235 | + | Gnomon | gene | NA | NA | gene-LOC105275316 | GeneID:105275316 | LOC105275316 | Gene | LOC105275316 | protein_coding | character(0) | NA | character(0) | NC_039516.1 | 12890326 | 12890331 | 6 | * | 6 | 1000 |
| NC_039516.1 | 12996090 | 13006535 | 10446 | + | Gnomon | gene | NA | NA | gene-LOC105275342 | GeneID:105275342 | LOC105275342 | Gene | LOC105275342 | protein_coding | character(0) | NA | character(0) | NC_039516.1 | 12997443 | 12997448 | 6 | * | 6 | 1000 |
| NC_039516.1 | 13060943 | 13063893 | 2951 | + | Gnomon | gene | NA | NA | gene-LOC105275357 | GeneID:105275357 | LOC105275357 | Gene | LOC105275357 | protein_coding | character(0) | NA | character(0) | NC_039516.1 | 13062040 | 13062045 | 6 | * | 6 | 1000 |
| NC_039516.1 | 13065193 | 13068389 | 3197 | + | Gnomon | gene | NA | NA | gene-LOC105275355 | GeneID:105275355 | LOC105275355 | Gene | LOC105275355 | protein_coding | character(0) | NA | character(0) | NC_039516.1 | 13066190 | 13066195 | 6 | * | 6 | 1000 |
| NC_039516.1 | 13518014 | 13520692 | 2679 | + | Gnomon | gene | NA | NA | gene-LOC105277817 | GeneID:105277817 | LOC105277817 | Gene | LOC105277817 | protein_coding | character(0) | NA | character(0) | NC_039516.1 | 13519261 | 13519266 | 6 | * | 6 | 1000 |
| NC_039517.1 | 861480 | 867435 | 5956 | + | Gnomon | gene | NA | NA | gene-LOC105278828 | GeneID:105278828 | LOC105278828 | Gene | LOC105278828 | protein_coding | character(0) | NA | character(0) | NC_039517.1 | 862475 | 862480 | 6 | * | 6 | 1000 |
| NC_039517.1 | 861480 | 867435 | 5956 | + | Gnomon | gene | NA | NA | gene-LOC105278828 | GeneID:105278828 | LOC105278828 | Gene | LOC105278828 | protein_coding | character(0) | NA | character(0) | NC_039517.1 | 862920 | 862925 | 6 | * | 6 | 1000 |
| NC_039517.1 | 1360424 | 1367346 | 6923 | + | Gnomon | gene | NA | NA | gene-LOC105278812 | GeneID:105278812 | LOC105278812 | Gene | LOC105278812 | protein_coding | character(0) | NA | character(0) | NC_039517.1 | 1361411 | 1361416 | 6 | * | 6 | 1000 |
| NC_039517.1 | 2878959 | 2887374 | 8416 | - | Gnomon | gene | NA | NA | gene-LOC105281562 | GeneID:105281562 | LOC105281562 | Gene | LOC105281562 | protein_coding | character(0) | NA | character(0) | NC_039517.1 | 2885923 | 2885928 | 6 | * | 6 | 1000 |
| NC_039517.1 | 2989996 | 2992325 | 2330 | + | Gnomon | gene | NA | NA | gene-LOC105281549 | GeneID:105281549 | LOC105281549 | Gene | LOC105281549 | protein_coding | character(0) | NA | character(0) | NC_039517.1 | 2990626 | 2990631 | 6 | * | 6 | 1000 |
| NC_039517.1 | 7309114 | 7314427 | 5314 | - | Gnomon | gene | NA | NA | gene-LOC105284201 | GeneID:105284201 | LOC105284201 | Gene | LOC105284201 | protein_coding | character(0) | NA | character(0) | NC_039517.1 | 7312519 | 7312524 | 6 | * | 6 | 1000 |
| NC_039517.1 | 8175046 | 8181714 | 6669 | - | Gnomon | gene | NA | NA | gene-LOC105284262 | GeneID:105284262 | LOC105284262 | Gene | LOC105284262 | protein_coding | character(0) | NA | character(0) | NC_039517.1 | 8175507 | 8175512 | 6 | * | 6 | 1000 |
| NC_039517.1 | 9921720 | 9925663 | 3944 | + | Gnomon | gene | NA | NA | gene-LOC105287104 | GeneID:105287104 | LOC105287104 | Gene | LOC105287104 | protein_coding | character(0) | NA | character(0) | NC_039517.1 | 9922441 | 9922446 | 6 | * | 6 | 1000 |
| NC_039518.1 | 1893371 | 1896648 | 3278 | + | Gnomon | gene | NA | NA | gene-LOC105286395 | GeneID:105286395 | LOC105286395 | Gene | LOC105286395 | protein_coding | character(0) | NA | character(0) | NC_039518.1 | 1894511 | 1894516 | 6 | * | 6 | 1000 |
| NC_039518.1 | 2681489 | 2682795 | 1307 | - | Gnomon | gene | NA | NA | gene-LOC105285394 | GeneID:105285394 | LOC105285394 | Gene | LOC105285394 | protein_coding | character(0) | NA | character(0) | NC_039518.1 | 2681746 | 2681751 | 6 | * | 6 | 1000 |
| NC_039518.1 | 5496870 | 5510429 | 13560 | - | Gnomon | gene | NA | NA | gene-LOC105281906 | GeneID:105281906 | LOC105281906 | Gene | LOC105281906 | protein_coding | character(0) | NA | character(0) | NC_039518.1 | 5508860 | 5508865 | 6 | * | 6 | 1000 |
| NC_039518.1 | 8474988 | 8476009 | 1022 | + | Gnomon | gene | NA | NA | gene-LOC105286212 | GeneID:105286212 | LOC105286212 | Gene | LOC105286212 | protein_coding | character(0) | NA | character(0) | NC_039518.1 | 8475762 | 8475769 | 8 | * | 8 | 1000 |
| NC_039518.1 | 8721748 | 8727127 | 5380 | + | Gnomon | gene | NA | NA | gene-LOC105286364 | GeneID:105286364 | LOC105286364 | Gene | LOC105286364 | protein_coding | character(0) | NA | character(0) | NC_039518.1 | 8722922 | 8722927 | 6 | * | 6 | 1000 |
| NC_039518.1 | 9164450 | 9166108 | 1659 | - | Gnomon | gene | NA | NA | gene-LOC105286320 | GeneID:105286320 | LOC105286320 | Gene | LOC105286320 | protein_coding | character(0) | NA | character(0) | NC_039518.1 | 9164960 | 9164965 | 6 | * | 6 | 1000 |

Directory: MRPR File: gwr-Ob_LI_Rphase.txt

| seqnames | start | end | width | strand | source | type | score | phase | ID | Dbxref | Name | gbkey | gene | gene_biotype | end_range | partial | start_range | seqnames | start | end | width | strand | NbrSites | Sdnsty |
|---|---|---|---|---|---|---|---|---|---|---|---|---|---|---|---|---|---|---|---|---|---|---|---|---|
| NC_039506.1 | 3342881 | 3348391 | 5511 | + | Gnomon | gene | NA | NA | gene-LOC105280395 | GeneID:105280395 | LOC105280395 | Gene | LOC105280395 | protein_coding | character(0) | NA | character(0) | NC_039506.1 | 3344178 | 3344183 | 6 | * | 6 | 1000 |
| NC_039506.1 | 3620612 | 3622819 | 2208 | - | Gnomon | gene | NA | NA | gene-LOC105280424 | GeneID:105280424 | LOC105280424 | Gene | LOC105280424 | protein_coding | character(0) | NA | character(0) | NC_039506.1 | 3621090 | 3621095 | 6 | * | 6 | 1000 |
| NC_039506.1 | 6089683 | 6090856 | 1174 | + | Gnomon | gene | NA | NA | gene-LOC105277804 | GeneID:105277804 | LOC105277804 | Gene | LOC105277804 | protein_coding | character(0) | NA | character(0) | NC_039506.1 | 6090624 | 6090629 | 6 | * | 6 | 1000 |
| NC_039506.1 | 13091757 | 13093375 | 1619 | - | Gnomon | gene | NA | NA | gene-LOC105286948 | GeneID:105286948 | LOC105286948 | Gene | LOC105286948 | protein_coding | character(0) | NA | character(0) | NC_039506.1 | 13092491 | 13092496 | 6 | * | 6 | 1000 |
| NC_039506.1 | 13177256 | 13183827 | 6572 | + | Gnomon | gene | NA | NA | gene-LOC105284041 | GeneID:105284041 | LOC105284041 | Gene | LOC105284041 | protein_coding | character(0) | NA | character(0) | NC_039506.1 | 13181314 | 13181319 | 6 | * | 6 | 1000 |
| NC_039506.1 | 14188024 | 14192973 | 4950 | - | Gnomon | gene | NA | NA | gene-LOC105281111 | GeneID:105281111 | LOC105281111 | Gene | LOC105281111 | protein_coding | character(0) | NA | character(0) | NC_039506.1 | 14191640 | 14191645 | 6 | * | 6 | 1000 |
| NC_039506.1 | 15370515 | 15371538 | 1024 | + | Gnomon | gene | NA | NA | gene-LOC105280860 | GeneID:105280860 | LOC105280860 | Gene | LOC105280860 | protein_coding | character(0) | NA | character(0) | NC_039506.1 | 15371422 | 15371427 | 6 | * | 6 | 1000 |
| NC_039506.1 | 16242561 | 16257517 | 14957 | - | Gnomon | gene | NA | NA | gene-LOC105275005 | GeneID:105275005 | LOC105275005 | Gene | LOC105275005 | protein_coding | character(0) | NA | character(0) | NC_039506.1 | 16256384 | 16256389 | 6 | * | 6 | 1000 |
| NC_039506.1 | 23567682 | 23574351 | 6670 | + | Gnomon | gene | NA | NA | gene-LOC105279333 | GeneID:105279333 | LOC105279333 | Gene | LOC105279333 | protein_coding | character(0) | NA | character(0) | NC_039506.1 | 23568887 | 23568894 | 8 | * | 8 | 1000 |
| NC_039506.1 | 24043149 | 24047806 | 4658 | + | Gnomon | gene | NA | NA | gene-LOC105279367 | GeneID:105279367 | LOC105279367 | Gene | LOC105279367 | protein_coding | character(0) | NA | character(0) | NC_039506.1 | 24044697 | 24044702 | 6 | * | 6 | 1000 |
| NC_039507.1 | 5034413 | 5037096 | 2684 | - | Gnomon | gene | NA | NA | gene-LOC105281698 | GeneID:105281698 | LOC105281698 | Gene | LOC105281698 | protein_coding | character(0) | NA | character(0) | NC_039507.1 | 5035958 | 5035963 | 6 | * | 6 | 1000 |
| NC_039507.1 | 5093844 | 5095945 | 2102 | + | Gnomon | gene | NA | NA | gene-LOC105281690 | GeneID:105281690 | LOC105281690 | Gene | LOC105281690 | protein_coding | character(0) | NA | character(0) | NC_039507.1 | 5094466 | 5094471 | 6 | * | 6 | 1000 |
| NC_039507.1 | 5492373 | 5502473 | 10101 | + | Gnomon | gene | NA | NA | gene-LOC105281645 | GeneID:105281645 | LOC105281645 | Gene | LOC105281645 | protein_coding | character(0) | NA | character(0) | NC_039507.1 | 5493754 | 5493759 | 6 | * | 6 | 1000 |
| NC_039507.1 | 7181199 | 7188522 | 7324 | + | Gnomon | gene | NA | NA | gene-LOC105283383 | GeneID:105283383 | LOC105283383 | Gene | LOC105283383 | protein_coding | character(0) | NA | character(0) | NC_039507.1 | 7182290 | 7182295 | 6 | * | 6 | 1000 |
| NC_039507.1 | 7276866 | 7278961 | 2096 | + | Gnomon | gene | NA | NA | gene-LOC105283373 | GeneID:105283373 | LOC105283373 | Gene | LOC105283373 | protein_coding | character(0) | NA | character(0) | NC_039507.1 | 7277947 | 7277952 | 6 | * | 6 | 1000 |
| NC_039507.1 | 9161485 | 9166239 | 4755 | + | Gnomon | gene | NA | NA | gene-LOC105282461 | GeneID:105282461 | LOC105282461 | Gene | LOC105282461 | protein_coding | character(0) | NA | character(0) | NC_039507.1 | 9162377 | 9162382 | 6 | * | 6 | 1000 |
| NC_039507.1 | 10633196 | 10639514 | 6319 | + | Gnomon | gene | NA | NA | gene-LOC105279155 | GeneID:105279155 | LOC105279155 | Gene | LOC105279155 | protein_coding | character(0) | NA | character(0) | NC_039507.1 | 10634249 | 10634254 | 6 | * | 6 | 1000 |
| NC_039507.1 | 13550180 | 13557055 | 6876 | - | Gnomon | gene | NA | NA | gene-LOC105285277 | GeneID:105285277 | LOC105285277 | Gene | LOC105285277 | protein_coding | character(0) | NA | character(0) | NC_039507.1 | 13555722 | 13555727 | 6 | * | 6 | 1000 |
| NC_039507.1 | 13552141 | 13556636 | 4496 | + | Gnomon | gene | NA | NA | gene-LOC105285278 | GeneID:105285278 | LOC105285278 | Gene | LOC105285278 | protein_coding | character(0) | NA | character(0) | NC_039507.1 | 13555722 | 13555727 | 6 | * | 6 | 1000 |
| NC_039507.1 | 15793673 | 15807525 | 13853 | - | Gnomon | gene | NA | NA | gene-LOC105280128 | GeneID:105280128 | LOC105280128 | Gene | LOC105280128 | protein_coding | character(0) | NA | character(0) | NC_039507.1 | 15805657 | 15805662 | 6 | * | 6 | 1000 |
| NC_039507.1 | 18333035 | 18336377 | 3343 | - | Gnomon | gene | NA | NA | gene-LOC105281577 | GeneID:105281577 | LOC105281577 | Gene | LOC105281577 | protein_coding | character(0) | NA | character(0) | NC_039507.1 | 18335809 | 18335814 | 6 | * | 6 | 1000 |
| NC_039508.1 | 2590841 | 2594093 | 3253 | + | Gnomon | gene | NA | NA | gene-LOC105281341 | GeneID:105281341 | LOC105281341 | Gene | LOC105281341 | protein_coding | character(0) | NA | character(0) | NC_039508.1 | 2591612 | 2591617 | 6 | * | 6 | 1000 |
| NC_039508.1 | 10274002 | 10277440 | 3439 | + | Gnomon | gene | NA | NA | gene-LOC105276881 | GeneID:105276881 | LOC105276881 | Gene | LOC105276881 | protein_coding | character(0) | NA | character(0) | NC_039508.1 | 10275212 | 10275217 | 6 | * | 6 | 1000 |
| NC_039508.1 | 10716085 | 10732040 | 15956 | - | Gnomon | gene | NA | NA | gene-LOC105276941 | GeneID:105276941 | LOC105276941 | Gene | LOC105276941 | protein_coding | character(0) | NA | character(0) | NC_039508.1 | 10730559 | 10730564 | 6 | * | 6 | 1000 |
| NC_039508.1 | 10799899 | 10803333 | 3435 | + | Gnomon | gene | NA | NA | gene-LOC105276950 | GeneID:105276950 | LOC105276950 | Gene | LOC105276950 | protein_coding | character(0) | NA | character(0) | NC_039508.1 | 10801049 | 10801054 | 6 | * | 6 | 1000 |
| NC_039508.1 | 13125687 | 13133076 | 7390 | + | Gnomon | gene | NA | NA | gene-LOC105277094 | GeneID:105277094 | LOC105277094 | Gene | LOC105277094 | protein_coding | character(0) | NA | character(0) | NC_039508.1 | 13126612 | 13126617 | 6 | * | 6 | 1000 |
| NC_039508.1 | 14022293 | 14024814 | 2522 | + | Gnomon | gene | NA | NA | gene-LOC105278683 | GeneID:105278683 | LOC105278683 | Gene | LOC105278683 | protein_coding | character(0) | NA | character(0) | NC_039508.1 | 14023382 | 14023387 | 6 | * | 6 | 1000 |
| NC_039508.1 | 15803095 | 15806139 | 3045 | + | Gnomon | gene | NA | NA | gene-LOC105277239 | GeneID:105277239 | LOC105277239 | Gene | LOC105277239 | protein_coding | character(0) | NA | character(0) | NC_039508.1 | 15804426 | 15804431 | 6 | * | 6 | 1000 |
| NC_039508.1 | 17548373 | 17550348 | 1976 | - | Gnomon | gene | NA | NA | gene-LOC105284396 | GeneID:105284396 | LOC105284396 | Gene | LOC105284396 | protein_coding | character(0) | NA | character(0) | NC_039508.1 | 17549290 | 17549295 | 6 | * | 6 | 1000 |
| NC_039509.1 | 1002697 | 1006107 | 3411 | - | Gnomon | gene | NA | NA | gene-LOC105283620 | GeneID:105283620 | LOC105283620 | Gene | LOC105283620 | protein_coding | | | | | | | | | | |

| | | | | | | | | | | | | | | |
|---|---|---|---|---|---|---|---|---|---|---|---|---|---|---|
| character(0) | NA | character(0) | NC_039509.1 | 1004827 | 1004832 | 6 | * | 6 | 1000 | | | | | |
| NC_039509.1 | 1135464 | 1137481 | 2018 | - | Gnomon | gene | NA | NA | gene-LOC105283634 | GeneID:105283634 | LOC105283634 | Gene | LOC105283634 | protein_coding |
| character(0) | NA | character(0) | NC_039509.1 | 1136266 | 1136271 | 6 | * | 6 | 1000 | | | | | |
| NC_039509.1 | 2825666 | 2828757 | 3092 | + | Gnomon | gene | NA | NA | gene-LOC105282589 | GeneID:105282589 | LOC105282589 | Gene | LOC105282589 | protein_coding |
| character(0) | NA | character(0) | NC_039509.1 | 2826537 | 2826542 | 6 | * | 6 | 1000 | | | | | |
| NC_039509.1 | 3925156 | 3928137 | 2982 | + | Gnomon | gene | NA | NA | gene-LOC105280619 | GeneID:105280619 | LOC105280619 | Gene | LOC105280619 | protein_coding |
| character(0) | NA | character(0) | NC_039509.1 | 3926254 | 3926259 | 6 | * | 6 | 1000 | | | | | |
| NC_039509.1 | 4882335 | 4886409 | 4075 | - | Gnomon | gene | NA | NA | gene-LOC105286160 | GeneID:105286160 | LOC105286160 | Gene | LOC105286160 | protein_coding |
| character(0) | NA | character(0) | NC_039509.1 | 4885053 | 4885058 | 6 | * | 6 | 1000 | | | | | |
| NC_039509.1 | 4882335 | 4886409 | 4075 | - | Gnomon | gene | NA | NA | gene-LOC105286160 | GeneID:105286160 | LOC105286160 | Gene | LOC105286160 | protein_coding |
| character(0) | NA | character(0) | NC_039509.1 | 4885072 | 4885077 | 6 | * | 6 | 1000 | | | | | |
| NC_039509.1 | 6318288 | 6320995 | 2708 | - | Gnomon | gene | NA | NA | gene-LOC105282525 | GeneID:105282525 | LOC105282525 | Gene | LOC105282525 | protein_coding |
| character(0) | NA | character(0) | NC_039509.1 | 6319895 | 6319900 | 6 | * | 6 | 1000 | | | | | |
| NC_039509.1 | 10209680 | 10213178 | 3499 | - | Gnomon | gene | NA | NA | gene-LOC105282662 | GeneID:105282662 | LOC105282662 | Gene | LOC105282662 | protein_coding |
| character(0) | NA | character(0) | NC_039509.1 | 10212587 | 10212592 | 6 | * | 6 | 1000 | | | | | |
| NC_039509.1 | 10745281 | 10747369 | 2089 | + | Gnomon | gene | NA | NA | gene-LOC105275200 | GeneID:105275200 | LOC105275200 | Gene | LOC105275200 | protein_coding |
| character(0) | NA | character(0) | NC_039509.1 | 10746212 | 10746217 | 6 | * | 6 | 1000 | | | | | |
| NC_039509.1 | 10756577 | 10757875 | 1299 | - | Gnomon | gene | NA | NA | gene-LOC105275214 | GeneID:105275214 | LOC105275214 | Gene | LOC105275214 | protein_coding |
| character(0) | NA | character(0) | NC_039509.1 | 10756803 | 10756808 | 6 | * | 6 | 1000 | | | | | |
| NC_039509.1 | 11047331 | 11049650 | 2320 | - | Gnomon | gene | NA | NA | gene-LOC105275731 | GeneID:105275731 | LOC105275731 | Gene | LOC105275731 | protein_coding |
| character(0) | NA | character(0) | NC_039509.1 | 11048986 | 11048991 | 6 | * | 6 | 1000 | | | | | |
| NC_039509.1 | 11068956 | 11086837 | 17882 | - | Gnomon | gene | NA | NA | gene-LOC105275770 | GeneID:105275770 | LOC105275770 | Gene | LOC105275770 | protein_coding |
| character(0) | NA | character(0) | NC_039509.1 | 11085457 | 11085462 | 6 | * | 6 | 1000 | | | | | |
| NC_039509.1 | 13033738 | 13035966 | 2229 | + | Gnomon | gene | NA | NA | gene-LOC105278527 | GeneID:105278527 | LOC105278527 | Gene | LOC105278527 | protein_coding |
| character(0) | NA | character(0) | NC_039509.1 | 13034669 | 13034674 | 6 | * | 6 | 1000 | | | | | |
| NC_039509.1 | 13067024 | 13068370 | 1347 | + | Gnomon | gene | NA | NA | gene-LOC105278478 | GeneID:105278478 | LOC105278478 | Gene | LOC105278478 | protein_coding |
| character(0) | NA | character(0) | NC_039509.1 | 13067776 | 13067781 | 6 | * | 6 | 1000 | | | | | |
| NC_039509.1 | 17207204 | 17209527 | 2324 | + | Gnomon | gene | NA | NA | gene-LOC105283016 | GeneID:105283016 | LOC105283016 | Gene | LOC105283016 | protein_coding |
| character(0) | NA | character(0) | NC_039509.1 | 17208074 | 17208079 | 6 | * | 6 | 1000 | | | | | |
| NC_039510.1 | 2607810 | 2610599 | 2790 | - | Gnomon | gene | NA | NA | gene-LOC105287754 | GeneID:105287754 | LOC105287754 | Gene | LOC105287754 | protein_coding |
| character(0) | NA | character(0) | NC_039510.1 | 2609179 | 2609184 | 6 | * | 6 | 1000 | | | | | |
| NC_039510.1 | 3032387 | 3034961 | 2575 | + | Gnomon | gene | NA | NA | gene-LOC105287715 | GeneID:105287715 | LOC105287715 | Gene | LOC105287715 | protein_coding |
| character(0) | NA | character(0) | NC_039510.1 | 3033855 | 3033860 | 6 | * | 6 | 1000 | | | | | |
| NC_039510.1 | 3126910 | 3147548 | 20639 | - | Gnomon | gene | NA | NA | gene-LOC105287694 | GeneID:105287694 | LOC105287694 | Gene | LOC105287694 | protein_coding |
| character(0) | NA | character(0) | NC_039510.1 | 3133558 | 3133563 | 6 | * | 6 | 1000 | | | | | |
| NC_039510.1 | 3132542 | 3134046 | 1505 | + | Gnomon | gene | NA | NA | gene-LOC105287692 | GeneID:105287692 | LOC105287692 | Gene | LOC105287692 | protein_coding |
| character(0) | NA | character(0) | NC_039510.1 | 3133558 | 3133563 | 6 | * | 6 | 1000 | | | | | |
| NC_039510.1 | 3210462 | 3212049 | 1588 | - | Gnomon | gene | NA | NA | gene-LOC105287677 | GeneID:105287677 | LOC105287677 | Gene | LOC105287677 | protein_coding |
| character(0) | NA | character(0) | NC_039510.1 | 3211153 | 3211158 | 6 | * | 6 | 1000 | | | | | |
| NC_039510.1 | 3319355 | 3322927 | 3573 | + | Gnomon | gene | NA | NA | gene-LOC105287663 | GeneID:105287663 | LOC105287663 | Gene | LOC105287663 | protein_coding |
| character(0) | NA | character(0) | NC_039510.1 | 3320255 | 3320260 | 6 | * | 6 | 1000 | | | | | |
| NC_039510.1 | 3406804 | 3410359 | 3556 | - | Gnomon | gene | NA | NA | gene-LOC105287646 | GeneID:105287646 | LOC105287646 | Gene | LOC105287646 | protein_coding |
| character(0) | NA | character(0) | NC_039510.1 | 3409446 | 3409451 | 6 | * | 6 | 1000 | | | | | |
| NC_039510.1 | 3779009 | 3783006 | 3998 | + | Gnomon | gene | NA | NA | gene-LOC105287632 | GeneID:105287632 | LOC105287632 | Gene | LOC105287632 | protein_coding |
| character(0) | NA | character(0) | NC_039510.1 | 3780254 | 3780259 | 6 | * | 6 | 1000 | | | | | |
| NC_039510.1 | 4126542 | 4127912 | 1371 | - | Gnomon | gene | NA | NA | gene-LOC105287627 | GeneID:105287627 | LOC105287627 | Gene | LOC105287627 | protein_coding |
| character(0) | NA | character(0) | NC_039510.1 | 4126783 | 4126788 | 6 | * | 6 | 1000 | | | | | |
| NC_039510.1 | 6135149 | 6137854 | 2706 | + | Gnomon | gene | NA | NA | gene-LOC105287333 | GeneID:105287333 | LOC105287333 | Gene | LOC105287333 | protein_coding |
| character(0) | NA | character(0) | NC_039510.1 | 6136456 | 6136461 | 6 | * | 6 | 1000 | | | | | |
| NC_039510.1 | 9658025 | 9667283 | 9259 | - | Gnomon | gene | NA | NA | gene-LOC105287015 | GeneID:105287015 | LOC105287015 | Gene | LOC105287015 | protein_coding |
| character(0) | NA | character(0) | NC_039510.1 | 9665819 | 9665824 | 6 | * | 6 | 1000 | | | | | |
| NC_039510.1 | 11515868 | 11522296 | 6429 | + | Gnomon | gene | NA | NA | gene-LOC105281115 | GeneID:105281115 | LOC105281115 | Gene | LOC105281115 | protein_coding |
| character(0) | NA | character(0) | NC_039510.1 | 11518799 | 11518804 | 6 | * | 6 | 1000 | | | | | |
| NC_039510.1 | 11738075 | 11746888 | 8814 | - | Gnomon | gene | NA | NA | gene-LOC105281080 | GeneID:105281080 | LOC105281080 | Gene | LOC105281080 | protein_coding |
| character(0) | NA | character(0) | NC_039510.1 | 11745052 | 11745057 | 6 | * | 6 | 1000 | | | | | |
| NC_039510.1 | 12000027 | 12025662 | 25636 | - | Gnomon | gene | NA | NA | gene-LOC105281066 | GeneID:105281066 | LOC105281066 | Gene | LOC105281066 | protein_coding |
| character(0) | NA | character(0) | NC_039510.1 | 12024411 | 12024416 | 6 | * | 6 | 1000 | | | | | |
| NC_039510.1 | 12023589 | 12029249 | 5661 | + | Gnomon | gene | NA | NA | gene-LOC105281067 | GeneID:105281067 | LOC105281067 | Gene | LOC105281067 | protein_coding |
| character(0) | NA | character(0) | NC_039510.1 | 12024411 | 12024416 | 6 | * | 6 | 1000 | | | | | |
| NC_039510.1 | 13595835 | 13598710 | 2876 | + | Gnomon | gene | NA | NA | gene-LOC105280955 | GeneID:105280955 | LOC105280955 | Gene | LOC105280955 | protein_coding |
| character(0) | NA | character(0) | NC_039510.1 | 13597157 | 13597162 | 6 | * | 6 | 1000 | | | | | |
| NC_039510.1 | 13612884 | 13615835 | 2952 | - | Gnomon | gene | NA | NA | gene-LOC105280950 | GeneID:105280950 | LOC105280950 | Gene | LOC105280950 | protein_coding |
| character(0) | NA | character(0) | NC_039510.1 | 13614384 | 13614389 | 6 | * | 6 | 1000 | | | | | |
| NC_039510.1 | 13947158 | 13949082 | 1925 | + | Gnomon | gene | NA | NA | gene-LOC105285205 | GeneID:105285205 | LOC105285205 | Gene | LOC105285205 | protein_coding |

character(0) NA character(0) NC_039510.1 13947830 13947835 6 * 6 1000
NC_039510.1 16287375 16291357 3983 - Gnomon gene NA NA gene-LOC105286509 GeneID:105286509 LOC105286509 Gene LOC105286509 protein_coding
character(0) NA character(0) NC_039510.1 16290598 16290603 6 * 6 1000
NC_039511.1 480353 488275 7923 + Gnomon gene NA NA gene-LOC105276527 GeneID:105276527 LOC105276527 Gene LOC105276527 protein_coding
character(0) NA character(0) NC_039511.1 481652 481657 6 * 6 1000
NC_039511.1 2234830 2236789 1960 - Gnomon gene NA NA gene-LOC105276382 GeneID:105276382 LOC105276382 Gene LOC105276382 protein_coding
character(0) NA character(0) NC_039511.1 2236016 2236021 6 * 6 1000
NC_039511.1 3614241 3616939 2699 + Gnomon gene NA NA gene-LOC105284057 GeneID:105284057 LOC105284057 Gene LOC105284057 protein_coding
character(0) NA character(0) NC_039511.1 3615462 3615467 6 * 6 1000
NC_039511.1 3625986 3631991 6006 - Gnomon gene NA NA gene-LOC105284053 GeneID:105284053 LOC105284053 Gene LOC105284053 protein_coding
character(0) NA character(0) NC_039511.1 3630771 3630776 6 * 6 1000
NC_039511.1 4660409 4662530 2122 - Gnomon gene NA NA gene-LOC105281848 GeneID:105281848 LOC105281848 Gene LOC105281848 protein_coding
character(0) NA character(0) NC_039511.1 4661875 4661880 6 * 6 1000
NC_039511.1 8407553 8425032 17480 + Gnomon gene NA NA gene-LOC105284845 GeneID:105284845 LOC105284845 Gene LOC105284845 protein_coding
character(0) NA character(0) NC_039511.1 8409706 8409711 6 * 6 1000
NC_039511.1 8695181 8698086 2906 + Gnomon gene NA NA gene-LOC105280856 GeneID:105280856 LOC105280856 Gene LOC105280856 protein_coding
character(0) NA character(0) NC_039511.1 8695844 8695849 6 * 6 1000
NC_039511.1 11533823 11536626 2804 + Gnomon gene NA NA gene-LOC105284137 GeneID:105284137 LOC105284137 Gene LOC105284137 protein_coding
character(0) NA character(0) NC_039511.1 11534787 11534792 6 * 6 1000
NC_039511.1 12778711 12781434 2724 + Gnomon gene NA NA gene-LOC105276248 GeneID:105276248 LOC105276248 Gene LOC105276248 protein_coding
character(0) NA character(0) NC_039511.1 12779348 12779354 7 * 7 1000
NC_039511.1 15681353 15682825 1473 + Gnomon gene NA NA gene-LOC105284582 GeneID:105284582 LOC105284582 Gene LOC105284582 protein_coding
character(0) NA character(0) NC_039511.1 15682121 15682126 6 * 6 1000
NC_039511.1 16524025 16528441 4417 - Gnomon gene NA NA gene-LOC105284595 GeneID:105284595 LOC105284595 Gene LOC105284595 protein_coding
character(0) NA character(0) NC_039511.1 16526339 16526344 6 * 6 1000
NC_039512.1 384950 392996 8047 - Gnomon gene NA NA gene-LOC105288044 GeneID:105288044 LOC105288044 Gene LOC105288044 protein_coding
character(0) NA character(0) NC_039512.1 391510 391515 6 * 6 1000
NC_039512.1 1516399 1518988 2590 - Gnomon gene NA NA gene-LOC105277697 GeneID:105277697 LOC105277697 Gene LOC105277697 protein_coding
character(0) NA character(0) NC_039512.1 1518058 1518063 6 * 6 1000
NC_039512.1 4385041 4390283 5243 - Gnomon gene NA NA gene-LOC105277552 GeneID:105277552 LOC105277552 Gene LOC105277552 protein_coding
character(0) NA character(0) NC_039512.1 4388975 4388980 6 * 6 1000
NC_039512.1 8874729 8882472 7744 - Gnomon gene NA NA gene-LOC105282356 GeneID:105282356 LOC105282356 Gene LOC105282356 protein_coding
character(0) NA character(0) NC_039512.1 8881441 8881446 6 * 6 1000
NC_039512.1 9800733 9802923 2191 - Gnomon gene NA NA gene-LOC105279900 GeneID:105279900 LOC105279900 Gene LOC105279900 protein_coding
character(0) NA character(0) NC_039512.1 9801464 9801469 6 * 6 1000
NC_039512.1 9855608 9862550 6943 - Gnomon gene NA NA gene-LOC105279903 GeneID:105279903 LOC105279903 Gene LOC105279903 protein_coding
character(0) NA character(0) NC_039512.1 9860779 9860784 6 * 6 1000
NC_039512.1 10275466 10280187 4722 + Gnomon gene NA NA gene-LOC105282847 GeneID:105282847 LOC105282847 Gene LOC105282847 protein_coding
character(0) NA character(0) NC_039512.1 10276481 10276486 6 * 6 1000
NC_039512.1 11140867 11151152 10286 - Gnomon gene NA NA gene-LOC105279854 GeneID:105279854 LOC105279854 Gene LOC105279854 protein_coding
character(0) NA character(0) NC_039512.1 11149938 11149943 6 * 6 1000
NC_039512.1 11140867 11151152 10286 - Gnomon gene NA NA gene-LOC105279854 GeneID:105279854 LOC105279854 Gene LOC105279854 protein_coding
character(0) NA character(0) NC_039512.1 11150038 11150045 8 * 8 1000
NC_039512.1 11239021 11242090 3070 + Gnomon gene NA NA gene-LOC105279838 GeneID:105279838 LOC105279838 Gene LOC105279838 protein_coding
character(0) NA character(0) NC_039512.1 11240529 11240534 6 * 6 1000
NC_039512.1 12092666 12097773 5108 + Gnomon gene NA NA gene-LOC105279779 GeneID:105279779 LOC105279779 Gene LOC105279779 protein_coding
character(0) NA character(0) NC_039512.1 12094200 12094205 6 * 6 1000
NC_039512.1 12752221 12767858 15638 - Gnomon gene NA NA gene-LOC105279717 GeneID:105279717 LOC105279717 Gene LOC105279717 protein_coding
character(0) NA character(0) NC_039512.1 12766252 12766257 6 * 6 1000
NC_039512.1 14248372 14257496 9125 + Gnomon gene NA NA gene-LOC105275078 GeneID:105275078 LOC105275078 Gene LOC105275078 misc_RNA character(0)
NA character(0) NC_039512.1 14249035 14249040 6 * 6 1000
NC_039512.1 15707994 15714445 6452 + Gnomon gene NA NA gene-LOC105274475 GeneID:105274475 LOC105274475 Gene LOC105274475 protein_coding
character(0) NA character(0) NC_039512.1 15711477 15711482 6 * 6 1000
NC_039512.1 15979275 15981529 2255 - Gnomon gene NA NA gene-LOC105274775 GeneID:105274775 LOC105274775 Gene LOC105274775 protein_coding
character(0) NA character(0) NC_039512.1 15980634 15980639 6 * 6 1000
NC_039512.1 16221715 16230076 8362 + Gnomon gene NA NA gene-LOC105282804 GeneID:105282804 LOC105282804 Gene LOC105282804 protein_coding
character(0) NA character(0) NC_039512.1 16225485 16225490 6 * 6 1000
NC_039513.1 3144818 3145709 892 + Gnomon gene NA NA gene-LOC105287814 GeneID:105287814 LOC105287814 Gene LOC105287814 protein_coding
character(0) NA character(0) NC_039513.1 3145174 3145179 6 * 6 1000
NC_039513.1 3298586 3302590 4005 - Gnomon gene NA NA gene-LOC105275760 GeneID:105275760 LOC105275760 Gene LOC105275760 protein_coding
character(0) NA character(0) NC_039513.1 3301859 3301864 6 * 6 1000
NC_039513.1 3303685 3306319 2635 + Gnomon gene NA NA gene-LOC105275761 GeneID:105275761 LOC105275761 Gene LOC105275761 protein_coding
character(0) NA character(0) NC_039513.1 3304809 3304814 6 * 6 1000
NC_039513.1 9600901 9603687 2787 - Gnomon gene NA NA gene-LOC105275617 GeneID:105275617 LOC105275617 Gene LOC105275617 protein_coding

| | | | | | | | | | | | | | | |
|---|---|---|---|---|---|---|---|---|---|---|---|---|---|---|
| character(0) | NA | character(0) | NC_039513.1 | 9602265 | 9602270 | 6 | * | 6 | 1000 | | | | | |
| NC_039513.1 | 10401827 | 10404590 | 2764 | - | Gnomon | gene | NA | NA | gene-LOC105275550 | GeneID:105275550 | LOC105275550 | Gene | LOC105275550 | protein_coding |
| character(0) | NA | character(0) | NC_039513.1 | 10403469 | 10403474 | 6 | * | 6 | 1000 | | | | | |
| NC_039513.1 | 10540843 | 10548517 | 7675 | - | Gnomon | gene | NA | NA | gene-LOC105275526 | GeneID:105275526 | LOC105275526 | Gene | LOC105275526 | protein_coding |
| character(0) | NA | character(0) | NC_039513.1 | 10546428 | 10546433 | 6 | * | 6 | 1000 | | | | | |
| NC_039513.1 | 11029007 | 11033701 | 4695 | - | Gnomon | gene | NA | NA | gene-LOC105275763 | GeneID:105275763 | LOC105275763 | Gene | LOC105275763 | protein_coding |
| character(0) | NA | character(0) | NC_039513.1 | 11030126 | 11030131 | 6 | * | 6 | 1000 | | | | | |
| NC_039513.1 | 11993718 | 11996290 | 2573 | - | Gnomon | gene | NA | NA | gene-LOC105280786 | GeneID:105280786 | LOC105280786 | Gene | LOC105280786 | protein_coding |
| character(0) | NA | character(0) | NC_039513.1 | 11994675 | 11994680 | 6 | * | 6 | 1000 | | | | | |
| NC_039513.1 | 12028675 | 12030941 | 2267 | - | Gnomon | gene | NA | NA | gene-LOC105280791 | GeneID:105280791 | LOC105280791 | Gene | LOC105280791 | protein_coding |
| character(0) | NA | character(0) | NC_039513.1 | 12029791 | 12029796 | 6 | * | 6 | 1000 | | | | | |
| NC_039513.1 | 13361516 | 13366191 | 4676 | - | Gnomon | gene | NA | NA | gene-LOC105275047 | GeneID:105275047 | LOC105275047 | Gene | LOC105275047 | protein_coding |
| character(0) | NA | character(0) | NC_039513.1 | 13364779 | 13364784 | 6 | * | 6 | 1000 | | | | | |
| NC_039513.1 | 13434303 | 13438575 | 4273 | - | Gnomon | gene | NA | NA | gene-LOC105275036 | GeneID:105275036 | LOC105275036 | Gene | LOC105275036 | protein_coding |
| character(0) | NA | character(0) | NC_039513.1 | 13436991 | 13437000 | 10 | * | 10 | 1000 | | | | | |
| NC_039513.1 | 14931340 | 14932816 | 1477 | + | Gnomon | gene | NA | NA | gene-LOC105285670 | GeneID:105285670 | LOC105285670 | Gene | LOC105285670 | protein_coding |
| character(0) | NA | character(0) | NC_039513.1 | 14932042 | 14932047 | 6 | * | 6 | 1000 | | | | | |
| NC_039514.1 | 6622839 | 6626782 | 3944 | - | Gnomon | gene | NA | NA | gene-LOC105285534 | GeneID:105285534 | LOC105285534 | Gene | LOC105285534 | protein_coding |
| character(0) | NA | character(0) | NC_039514.1 | 6625853 | 6625858 | 6 | * | 6 | 1000 | | | | | |
| NC_039514.1 | 8004357 | 8009868 | 5512 | - | Gnomon | gene | NA | NA | gene-LOC105281166 | GeneID:105281166 | LOC105281166 | Gene | LOC105281166 | protein_coding |
| character(0) | NA | character(0) | NC_039514.1 | 8008572 | 8008577 | 6 | * | 6 | 1000 | | | | | |
| NC_039514.1 | 8819238 | 8822718 | 3481 | - | Gnomon | gene | NA | NA | gene-LOC105279004 | GeneID:105279004 | LOC105279004 | Gene | LOC105279004 | protein_coding |
| character(0) | NA | character(0) | NC_039514.1 | 8821565 | 8821570 | 6 | * | 6 | 1000 | | | | | |
| NC_039514.1 | 9286321 | 9287389 | 1069 | - | Gnomon | gene | NA | NA | gene-LOC105279437 | GeneID:105279437 | LOC105279437 | Gene | LOC105279437 | protein_coding |
| character(0) | NA | character(0) | NC_039514.1 | 9286345 | 9286350 | 6 | * | 6 | 1000 | | | | | |
| NC_039514.1 | 9336154 | 9343762 | 7609 | + | Gnomon | gene | NA | NA | gene-LOC105284644 | GeneID:105284644 | LOC105284644 | Gene | LOC105284644 | protein_coding |
| character(0) | NA | character(0) | NC_039514.1 | 9337043 | 9337048 | 6 | * | 6 | 1000 | | | | | |
| NC_039514.1 | 9564295 | 9566245 | 1951 | + | Gnomon | gene | NA | NA | gene-LOC105284676 | GeneID:105284676 | LOC105284676 | Gene | LOC105284676 | protein_coding |
| character(0) | NA | character(0) | NC_039514.1 | 9565156 | 9565161 | 6 | * | 6 | 1000 | | | | | |
| NC_039514.1 | 11109536 | 11117116 | 7581 | - | Gnomon | gene | NA | NA | gene-LOC105281601 | GeneID:105281601 | LOC105281601 | Gene | LOC105281601 | protein_coding |
| character(0) | NA | character(0) | NC_039514.1 | 11112955 | 11112960 | 6 | * | 6 | 1000 | | | | | |
| NC_039514.1 | 11111779 | 11115572 | 3794 | + | Gnomon | gene | NA | NA | gene-LOC105281603 | GeneID:105281603 | LOC105281603 | Gene | LOC105281603 | protein_coding |
| character(0) | NA | character(0) | NC_039514.1 | 11112955 | 11112960 | 6 | * | 6 | 1000 | | | | | |
| NC_039514.1 | 13115199 | 13117880 | 2682 | + | Gnomon | gene | NA | NA | gene-LOC105277357 | GeneID:105277357 | LOC105277357 | Gene | LOC105277357 | protein_coding |
| character(0) | NA | character(0) | NC_039514.1 | 13116759 | 13116766 | 8 | * | 8 | 1000 | | | | | |
| NC_039515.1 | 1335326 | 1341505 | 6180 | + | Gnomon | gene | NA | NA | gene-LOC105288065 | GeneID:105288065 | LOC105288065 | Gene | LOC105288065 | protein_coding |
| character(0) | NA | character(0) | NC_039515.1 | 1337469 | 1337474 | 6 | * | 6 | 1000 | | | | | |
| NC_039515.1 | 4503178 | 4505245 | 2068 | + | Gnomon | gene | NA | NA | gene-LOC105278234 | GeneID:105278234 | LOC105278234 | Gene | LOC105278234 | protein_coding |
| character(0) | NA | character(0) | NC_039515.1 | 4503560 | 4503565 | 6 | * | 6 | 1000 | | | | | |
| NC_039515.1 | 7098785 | 7103508 | 4724 | - | Gnomon | gene | NA | NA | gene-LOC105286003 | GeneID:105286003 | LOC105286003 | Gene | LOC105286003 | protein_coding |
| character(0) | NA | character(0) | NC_039515.1 | 7102685 | 7102690 | 6 | * | 6 | 1000 | | | | | |
| NC_039515.1 | 7216666 | 7221298 | 4633 | + | Gnomon | gene | NA | NA | gene-LOC105283771 | GeneID:105283771 | LOC105283771 | Gene | LOC105283771 | protein_coding |
| character(0) | NA | character(0) | NC_039515.1 | 7217376 | 7217381 | 6 | * | 6 | 1000 | | | | | |
| NC_039515.1 | 12582469 | 12585642 | 3174 | - | Gnomon | gene | NA | NA | gene-LOC105282150 | GeneID:105282150 | LOC105282150 | Gene | LOC105282150 | protein_coding |
| character(0) | NA | character(0) | NC_039515.1 | 12584748 | 12584753 | 6 | * | 6 | 1000 | | | | | |
| NC_039515.1 | 12596735 | 12605489 | 8755 | - | Gnomon | gene | NA | NA | gene-LOC105282142 | GeneID:105282142 | LOC105282142 | Gene | LOC105282142 | protein_coding |
| character(0) | NA | character(0) | NC_039515.1 | 12604757 | 12604762 | 6 | * | 6 | 1000 | | | | | |
| NC_039515.1 | 12601969 | 12605663 | 3695 | + | Gnomon | gene | NA | NA | gene-LOC105282144 | GeneID:105282144 | LOC105282144 | Gene | LOC105282144 | protein_coding |
| character(0) | NA | character(0) | NC_039515.1 | 12604757 | 12604762 | 6 | * | 6 | 1000 | | | | | |
| NC_039516.1 | 595169 | 598515 | 3347 | + | Gnomon | gene | NA | NA | gene-LOC105274673 | GeneID:105274673 | LOC105274673 | Gene | LOC105274673 | protein_coding |
| character(0) | NA | character(0) | NC_039516.1 | 597128 | 597133 | 6 | * | 6 | 1000 | | | | | |
| NC_039516.1 | 857783 | 860432 | 2650 | - | Gnomon | gene | NA | NA | gene-LOC105274613 | GeneID:105274613 | LOC105274613 | Gene | LOC105274613 | protein_coding |
| character(0) | NA | character(0) | NC_039516.1 | 859316 | 859321 | 6 | * | 6 | 1000 | | | | | |
| NC_039516.1 | 1062627 | 1071280 | 8654 | + | Gnomon | gene | NA | NA | gene-LOC105274578 | GeneID:105274578 | LOC105274578 | Gene | LOC105274578 | protein_coding |
| character(0) | NA | character(0) | NC_039516.1 | 1064627 | 1064632 | 6 | * | 6 | 1000 | | | | | |
| NC_039516.1 | 1359476 | 1362079 | 2604 | + | Gnomon | gene | NA | NA | gene-LOC113562925 | GeneID:113562925 | LOC113562925 | Gene | LOC113562925 | protein_coding |
| character(0) | NA | character(0) | NC_039516.1 | 1360969 | 1360974 | 6 | * | 6 | 1000 | | | | | |
| NC_039516.1 | 2372548 | 2378885 | 6338 | + | Gnomon | gene | NA | NA | gene-LOC105274446 | GeneID:105274446 | LOC105274446 | Gene | LOC105274446 | protein_coding |
| character(0) | NA | character(0) | NC_039516.1 | 2377394 | 2377399 | 6 | * | 6 | 1000 | | | | | |
| NC_039516.1 | 2464828 | 2468017 | 3190 | + | Gnomon | gene | NA | NA | gene-LOC105288248 | GeneID:105288248 | LOC105288248 | Gene | LOC105288248 | protein_coding |
| character(0) | NA | character(0) | NC_039516.1 | 2465766 | 2465771 | 6 | * | 6 | 1000 | | | | | |
| NC_039516.1 | 3088715 | 3091076 | 2362 | + | Gnomon | gene | NA | NA | gene-LOC105288196 | GeneID:105288196 | LOC105288196 | Gene | LOC105288196 | protein_coding |
| character(0) | NA | character(0) | NC_039516.1 | 3089997 | 3090002 | 6 | * | 6 | 1000 | | | | | |
| NC_039516.1 | 3100198 | 3101692 | 1495 | + | Gnomon | gene | NA | NA | gene-LOC105288193 | GeneID:105288193 | LOC105288193 | Gene | LOC105288193 | protein_coding |

character(0) NA character(0) NC_039516.1 3101072 3101077 6 * 6 1000
NC_039516.1 4713084 4714767 1684 - Gnomon gene NA NA gene-LOC105275897 GeneID:105275897 LOC105275897 Gene LOC105275897 protein_coding
character(0) NA character(0) NC_039516.1 4713694 4713699 6 * 6 1000
NC_039516.1 5472871 5475154 2284 - Gnomon gene NA NA gene-LOC105275878 GeneID:105275878 LOC105275878 Gene LOC105275878 protein_coding
character(0) NA character(0) NC_039516.1 5474292 5474297 6 * 6 1000
NC_039516.1 5868176 5870154 1979 - Gnomon gene NA NA gene-LOC105275861 GeneID:105275861 LOC105275861 Gene LOC105275861 protein_coding
character(0) NA character(0) NC_039516.1 5869246 5869251 6 * 6 1000
NC_039516.1 6141341 6145058 3718 + Gnomon gene NA NA gene-LOC105275848 GeneID:105275848 LOC105275848 Gene LOC105275848 protein_coding
character(0) NA character(0) NC_039516.1 6142911 6142916 6 * 6 1000
NC_039516.1 6142888 6144894 2007 - Gnomon gene NA NA gene-LOC113562897 GeneID:113562897 LOC113562897 Gene LOC113562897 lncRNA character(0)
NA character(0) NC_039516.1 6142911 6142916 6 * 6 1000
NC_039516.1 7444132 7447943 3812 - Gnomon gene NA NA gene-LOC105275781 GeneID:105275781 LOC105275781 Gene LOC105275781 protein_coding
character(0) NA character(0) NC_039516.1 7446427 7446432 6 * 6 1000
NC_039516.1 9287279 9290515 3237 - Gnomon gene NA NA gene-LOC105277523 GeneID:105277523 LOC105277523 Gene LOC105277523 protein_coding
character(0) NA character(0) NC_039516.1 9289245 9289250 6 * 6 1000
NC_039516.1 9389294 9390221 928 + Gnomon gene NA NA gene-LOC105277237 GeneID:105277237 LOC105277237 Gene LOC105277237 protein_coding
character(0) NA character(0) NC_039516.1 9390169 9390174 6 * 6 1000
NC_039516.1 9555082 9557519 2438 + Gnomon gene NA NA gene-LOC105278789 GeneID:105278789 LOC105278789 Gene LOC105278789 protein_coding
character(0) NA character(0) NC_039516.1 9555938 9555943 6 * 6 1000
NC_039516.1 10251143 10256122 4980 - Gnomon gene NA NA gene-LOC105287447 GeneID:105287447 LOC105287447 Gene LOC105287447 protein_coding
character(0) NA character(0) NC_039516.1 10254626 10254631 6 * 6 1000
NC_039516.1 10251143 10256122 4980 - Gnomon gene NA NA gene-LOC105287447 GeneID:105287447 LOC105287447 Gene LOC105287447 protein_coding
character(0) NA character(0) NC_039516.1 10254677 10254682 6 * 6 1000
NC_039516.1 11647758 11653686 5929 - Gnomon gene NA NA gene-LOC105278681 GeneID:105278681 LOC105278681 Gene LOC105278681 protein_coding
character(0) NA character(0) NC_039516.1 11652297 11652302 6 * 6 1000
NC_039516.1 12300144 12302800 2657 - Gnomon gene NA NA gene-LOC105275264 GeneID:105275264 LOC105275264 Gene LOC105275264 protein_coding
character(0) NA character(0) NC_039516.1 12301684 12301689 6 * 6 1000
NC_039516.1 12307603 12308894 1292 + Gnomon gene NA NA gene-LOC105275270 GeneID:105275270 LOC105275270 Gene LOC105275270 protein_coding
character(0) NA character(0) NC_039516.1 12308308 12308313 6 * 6 1000
NC_039516.1 12889538 12895772 6235 + Gnomon gene NA NA gene-LOC105275316 GeneID:105275316 LOC105275316 Gene LOC105275316 protein_coding
character(0) NA character(0) NC_039516.1 12890326 12890331 6 * 6 1000
NC_039516.1 12996090 13006535 10446 + Gnomon gene NA NA gene-LOC105275342 GeneID:105275342 LOC105275342 Gene LOC105275342 protein_coding
character(0) NA character(0) NC_039516.1 12997443 12997448 6 * 6 1000
NC_039516.1 13050216 13053802 3587 - Gnomon gene NA NA gene-LOC105275350 GeneID:105275350 LOC105275350 Gene LOC105275350 protein_coding
character(0) NA character(0) NC_039516.1 13052909 13052914 6 * 6 1000
NC_039516.1 13065193 13068389 3197 + Gnomon gene NA NA gene-LOC105275355 GeneID:105275355 LOC105275355 Gene LOC105275355 protein_coding
character(0) NA character(0) NC_039516.1 13066190 13066195 6 * 6 1000
NC_039517.1 1360424 1367346 6923 + Gnomon gene NA NA gene-LOC105278812 GeneID:105278812 LOC105278812 Gene LOC105278812 protein_coding
character(0) NA character(0) NC_039517.1 1361411 1361416 6 * 6 1000
NC_039517.1 1368615 1394804 26190 + Gnomon gene NA NA gene-LOC105278810 GeneID:105278810 LOC105278810 Gene LOC105278810 protein_coding
character(0) NA character(0) NC_039517.1 1369233 1369238 6 * 6 1000
NC_039517.1 2878959 2887374 8416 - Gnomon gene NA NA gene-LOC105281562 GeneID:105281562 LOC105281562 Gene LOC105281562 protein_coding
character(0) NA character(0) NC_039517.1 2885923 2885928 6 * 6 1000
NC_039517.1 2989996 2992325 2330 + Gnomon gene NA NA gene-LOC105281549 GeneID:105281549 LOC105281549 Gene LOC105281549 protein_coding
character(0) NA character(0) NC_039517.1 2990626 2990631 6 * 6 1000
NC_039517.1 4433765 4437688 3924 + Gnomon gene NA NA gene-LOC105276559 GeneID:105276559 LOC105276559 Gene LOC105276559 protein_coding
character(0) NA character(0) NC_039517.1 4434630 4434635 6 * 6 1000
NC_039517.1 7309114 7314427 5314 - Gnomon gene NA NA gene-LOC105284201 GeneID:105284201 LOC105284201 Gene LOC105284201 protein_coding
character(0) NA character(0) NC_039517.1 7312519 7312524 6 * 6 1000
NC_039517.1 8175046 8181714 6669 - Gnomon gene NA NA gene-LOC105284262 GeneID:105284262 LOC105284262 Gene LOC105284262 protein_coding
character(0) NA character(0) NC_039517.1 8175507 8175512 6 * 6 1000
NC_039517.1 9921720 9925663 3944 + Gnomon gene NA NA gene-LOC105287104 GeneID:105287104 LOC105287104 Gene LOC105287104 protein_coding
character(0) NA character(0) NC_039517.1 9922441 9922446 6 * 6 1000
NC_039518.1 2060545 2068664 8120 + Gnomon gene NA NA gene-LOC105280035 GeneID:105280035 LOC105280035 Gene LOC105280035 protein_coding
character(0) NA character(0) NC_039518.1 2061863 2061868 6 * 6 1000
NC_039518.1 2681489 2682795 1307 - Gnomon gene NA NA gene-LOC105285394 GeneID:105285394 LOC105285394 Gene LOC105285394 protein_coding
character(0) NA character(0) NC_039518.1 2681746 2681751 6 * 6 1000
NC_039518.1 5496870 5510429 13560 - Gnomon gene NA NA gene-LOC105281906 GeneID:105281906 LOC105281906 Gene LOC105281906 protein_coding
character(0) NA character(0) NC_039518.1 5508860 5508865 6 * 6 1000
NC_039518.1 8474988 8476009 1022 + Gnomon gene NA NA gene-LOC105286212 GeneID:105286212 LOC105286212 Gene LOC105286212 protein_coding
character(0) NA character(0) NC_039518.1 8475762 8475769 8 * 8 1000
NC_039518.1 8721748 8727127 5380 + Gnomon gene NA NA gene-LOC105286364 GeneID:105286364 LOC105286364 Gene LOC105286364 protein_coding
character(0) NA character(0) NC_039518.1 8722922 8722927 6 * 6 1000
NC_039518.1 9164450 9166108 1659 - Gnomon gene NA NA gene-LOC105286320 GeneID:105286320 LOC105286320 Gene LOC105286320 protein_coding

| | | | | | | | | | |
|---|---|---|---|---|---|---|---|---|---|
| character(0) | NA | character(0) | NC_039518.1 | 9164960 | 9164965 | 6 | * | 6 | 1000 |

Directory: MRPR File: rmp-Ob_LI_BCphase.txt

Overlap of methylation-rich regions with genome features for Ob sample Ob_LI_BCphase

```
Total size of the methylation-rich regions of Ob_LI_BCphase:                                       880 bp (  0.0% of genome)
Size of overlap of methylation-rich regions of Ob Ob_LI_BCphase with genic regions:                   866 bp ( 98.4% of total size)   ( 1.32 O/E)
Size of overlap of methylation-rich regions of Ob Ob_LI_BCphase with   exon regions:                    760 bp ( 86.4% of total size) ( 0.60 O/E)
Size of overlap of methylation-rich regions of Ob Ob_LI_BCphase with   intron regions:                  106 bp ( 12.1% of total size) ( 0.22 O/E)
Size of overlap of methylation-rich regions of Ob Ob_LI_BCphase with intergenic regions:               14 bp (  1.6% of total size)   ( 3.86 O/E)
Size of overlap of methylation-rich regions of Ob Ob_LI_BCphase with   promoter regions:                  0 bp (  0.0% of total size) ( 0.00 O/E)
Size of overlap of methylation-rich regions of Ob Ob_LI_BCphase with   other intergenic regions:         14 bp (  1.6% of total size) ( 0.07 O/E)
```

Overlap of methylation-poor regions with genome features for Ob sample Ob_LI_BCphase

```
Total size of the methylation-poor regions of Ob_LI_BCphase:                                   9338107 bp (  4.2% of genome)
Size of overlap of methylation-poor regions of Ob Ob_LI_BCphase with genic regions:               3665472 bp ( 39.2% of total size)   ( 0.53 O/E)
Size of overlap of methylation-poor regions of Ob Ob_LI_BCphase with   exon regions:                1147655 bp ( 12.3% of total size) ( 1.34 O/E)
Size of overlap of methylation-poor regions of Ob Ob_LI_BCphase with   intron regions:              2517817 bp ( 27.0% of total size) ( 0.50 O/E)
Size of overlap of methylation-poor regions of Ob Ob_LI_BCphase with intergenic regions:          5672635 bp ( 60.8% of total size)   ( 1.54 O/E)
Size of overlap of methylation-poor regions of Ob Ob_LI_BCphase with   promoter regions:             159844 bp (  1.7% of total size) ( 0.56 O/E)
Size of overlap of methylation-poor regions of Ob Ob_LI_BCphase with   other intergenic regions:    5512791 bp ( 59.0% of total size) ( 2.63 O/E)
```

Directory: MRPR File: rmp-Ob_LI_Rphase.txt

```
Overlap of methylation-rich regions with genome features for Ob sample Ob_LI_Rphase

Total size of the methylation-rich regions of Ob_LI_Rphase:                                       931 bp (  0.0% of genome)
Size of overlap of methylation-rich regions of Ob Ob_LI_Rphase with genic regions:                   925 bp ( 99.4% of total size)   ( 1.33 O/E)
Size of overlap of methylation-rich regions of Ob Ob_LI_Rphase with   exon regions:                    807 bp ( 86.7% of total size) ( 0.63 O/E)
Size of overlap of methylation-rich regions of Ob Ob_LI_Rphase with   intron regions:                  118 bp ( 12.7% of total size) ( 0.23 O/E)
Size of overlap of methylation-rich regions of Ob Ob_LI_Rphase with intergenic regions:                6 bp (  0.6% of total size)   ( 3.89 O/E)
Size of overlap of methylation-rich regions of Ob Ob_LI_Rphase with   promoter regions:                  0 bp (  0.0% of total size) ( 0.00 O/E)
Size of overlap of methylation-rich regions of Ob Ob_LI_Rphase with   other intergenic regions:          6 bp (  0.6% of total size) ( 0.03 O/E)

Overlap of methylation-poor regions with genome features for Ob sample Ob_LI_Rphase

Total size of the methylation-poor regions of Ob_LI_Rphase:                                   8226406 bp (  3.7% of genome)
Size of overlap of methylation-poor regions of Ob Ob_LI_Rphase with genic regions:               3227558 bp ( 39.2% of total size)   ( 0.53 O/E)
Size of overlap of methylation-poor regions of Ob Ob_LI_Rphase with   exon regions:                1008765 bp ( 12.3% of total size) ( 1.34 O/E)
Size of overlap of methylation-poor regions of Ob Ob_LI_Rphase with   intron regions:              2218793 bp ( 27.0% of total size) ( 0.50 O/E)
Size of overlap of methylation-poor regions of Ob Ob_LI_Rphase with intergenic regions:          4998848 bp ( 60.8% of total size)   ( 1.54 O/E)
Size of overlap of methylation-poor regions of Ob Ob_LI_Rphase with   promoter regions:             136640 bp (  1.7% of total size) ( 0.54 O/E)
Size of overlap of methylation-poor regions of Ob Ob_LI_Rphase with   other intergenic regions:    4862208 bp ( 59.1% of total size) ( 2.63 O/E)
```

Ob_LI_BCphase
1−distance distributions

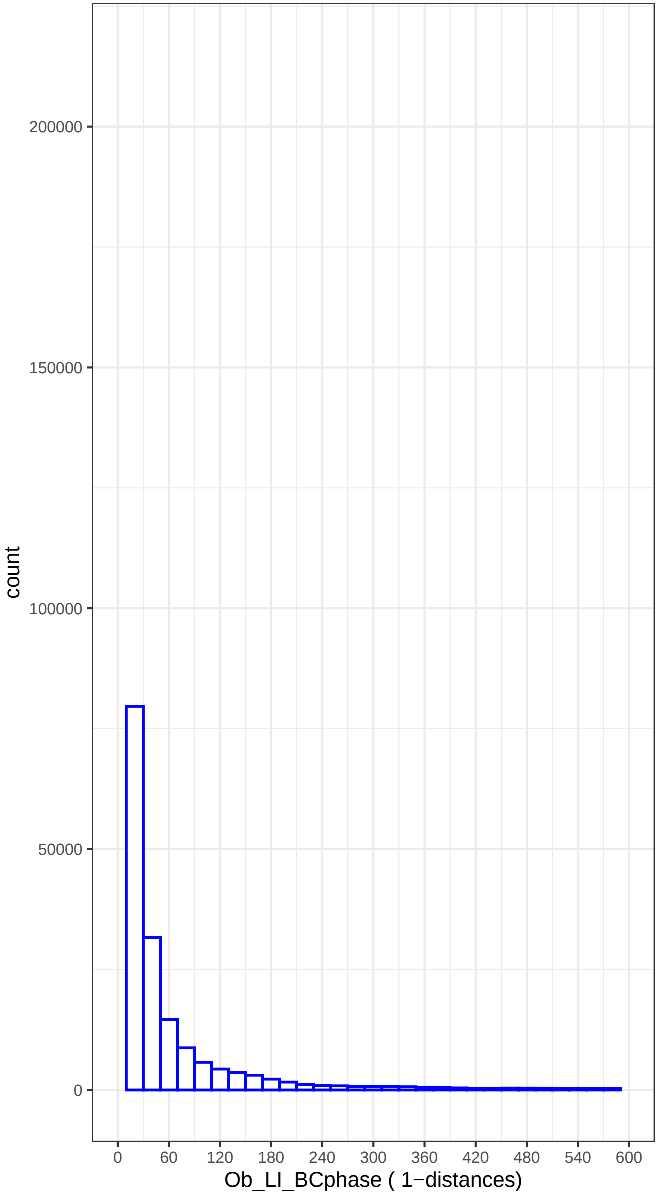

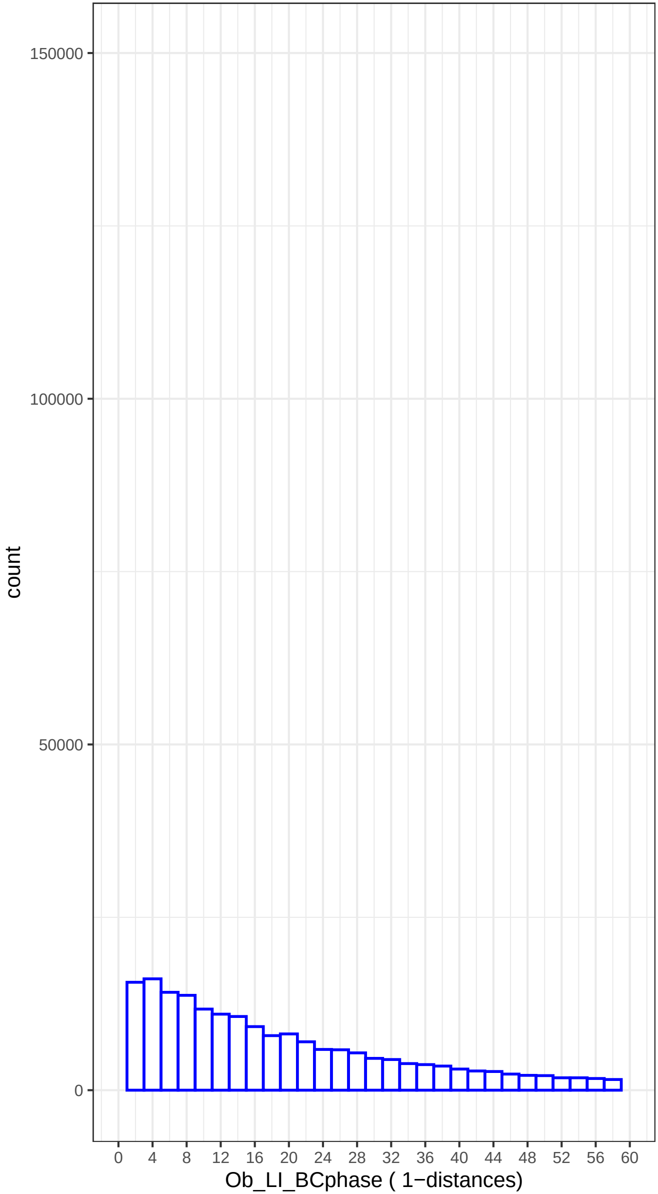

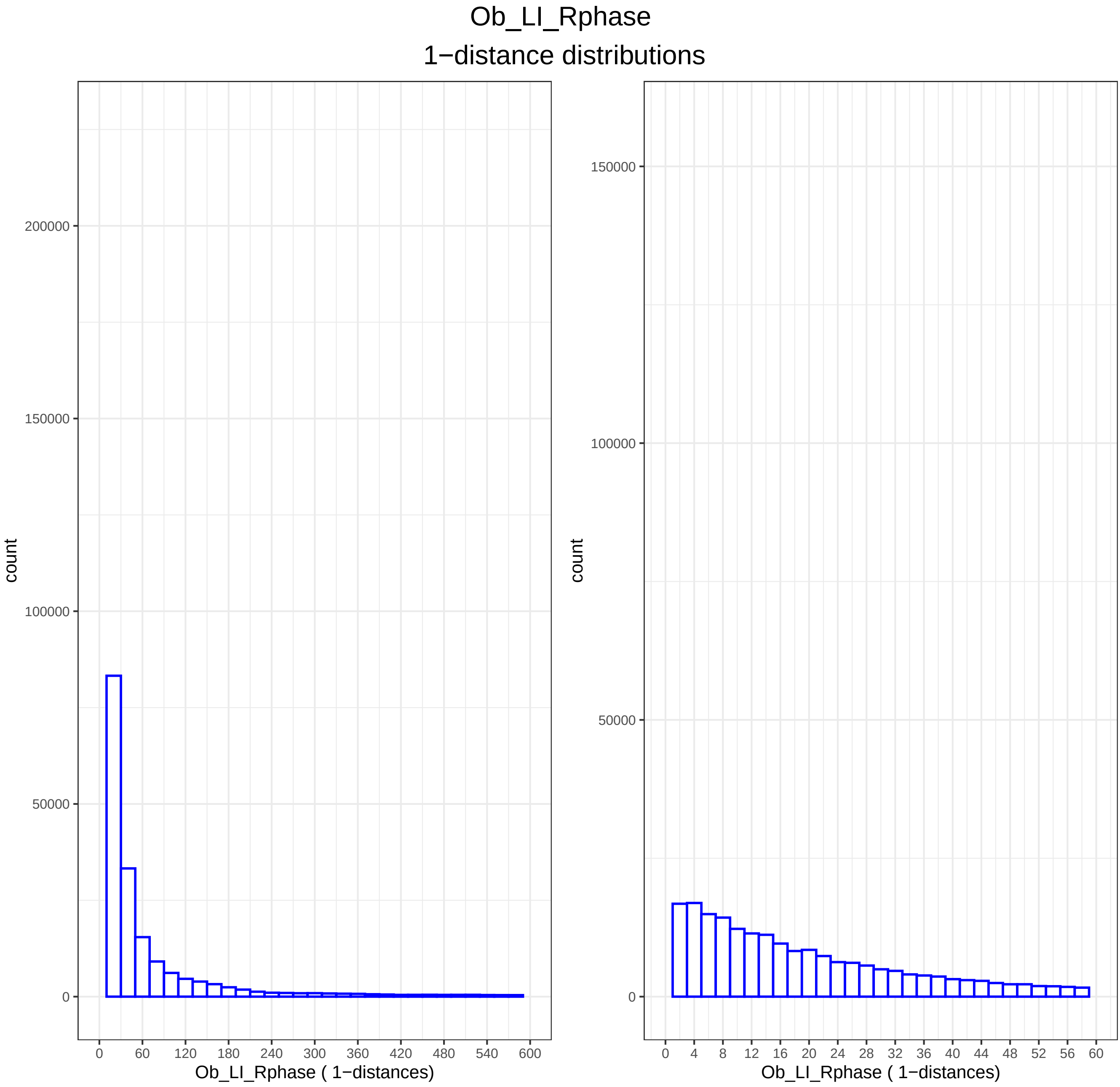

Ob_LI_Rphase
1−distance distributions
count
200000
150000
100000
50000
0
0 60 120 180 240 300 360 420 480 540 600
Ob_LI_Rphase ( 1−distances)
count
150000
100000
50000
0
0 4 8 12 16 20 24 28 32 36 40 44 48 52 56 60
Ob_LI_Rphase ( 1−distances)

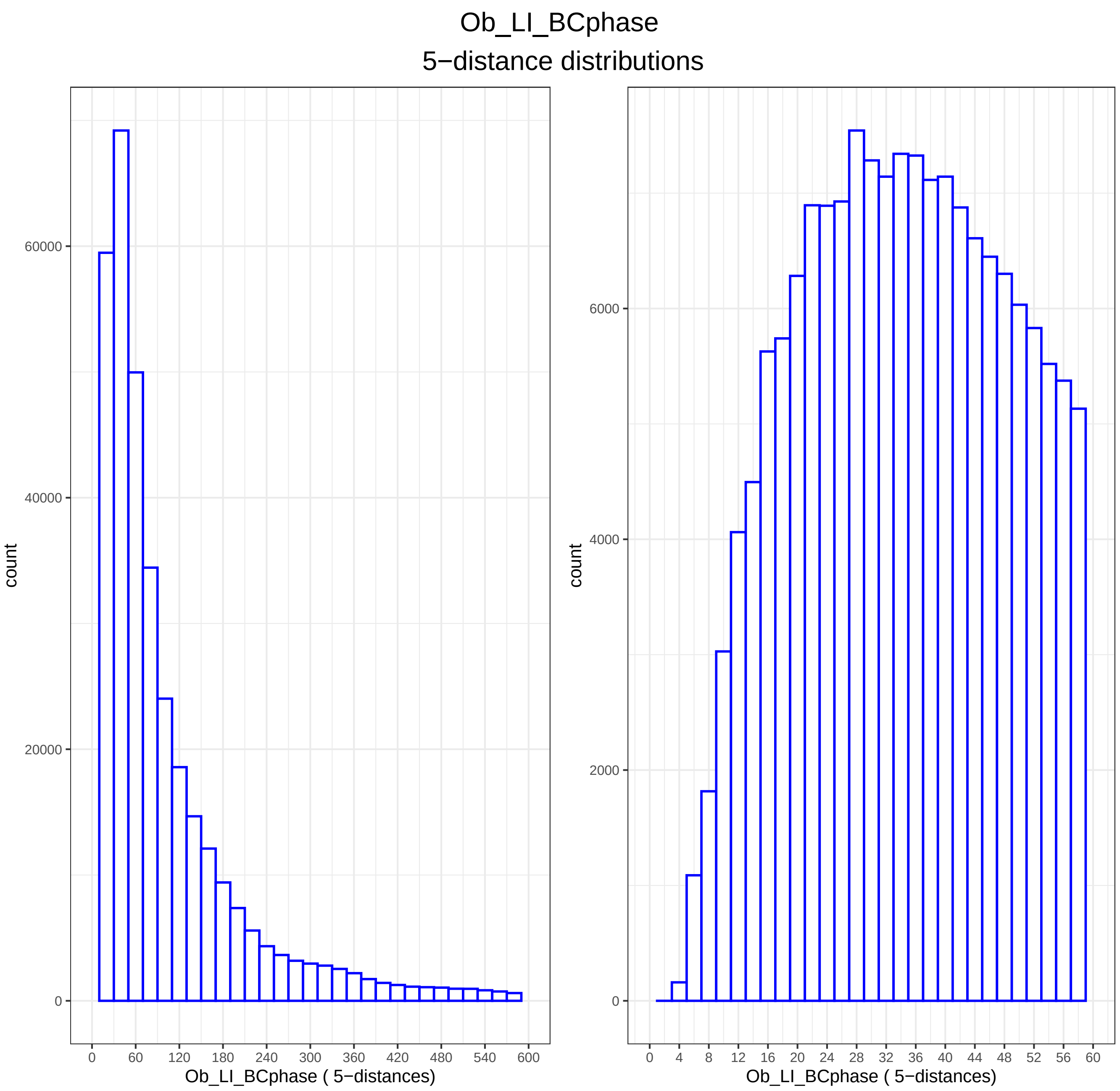

Ob_LI_BCphase
5−distance distributions
count
60000
40000
20000
0
0 60 120 180 240 300 360 420 480 540 600
Ob_LI_BCphase ( 5−distances)
count
6000
4000
2000
0
0 4 8 12 16 20 24 28 32 36 40 44 48 52 56 60
Ob_LI_BCphase ( 5−distances)

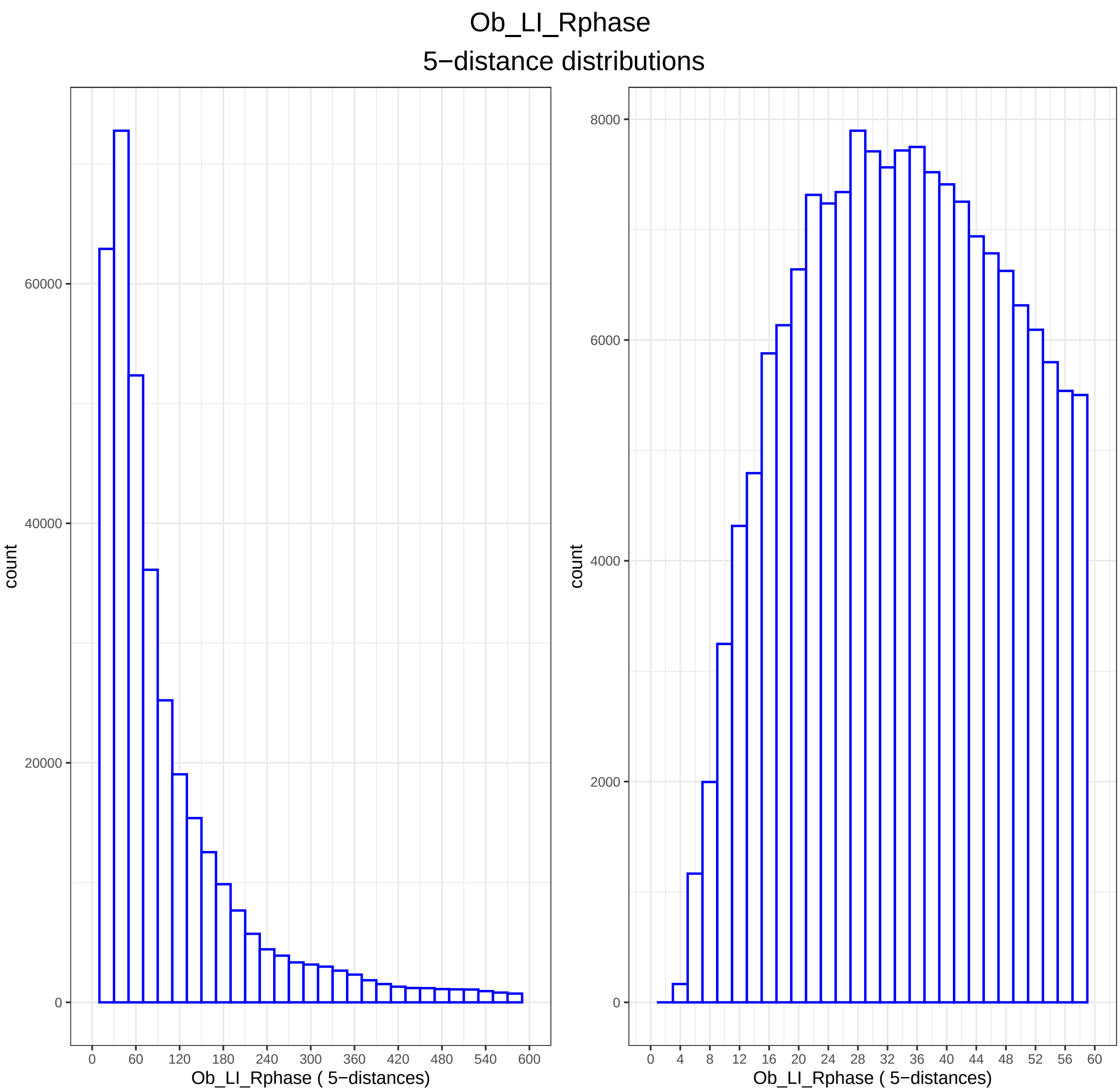

Ob_LI_Rphase
5−distance distributions
count
60000
40000
20000
0
0
60
120
180
240
300
360
420
480
540
600
Ob_LI_Rphase ( 5−distances)
count
8000
6000
4000
2000
0
0
4
8
12
16
20
24
28
32
36
40
44
48
52
56
60
Ob_LI_Rphase ( 5−distances)

```
Directory: DMT File: 0READMEdmt

DMT - Differentially methylated tiles.

Input: studymc wsize stepsize threshold qvalue
Output: files dmt-*.txt dmg-*.txt

Notes: Output files dmt-*.txt and dmg-*.txt are generated by BWASPR::det_dmt() and show the
        differentially methylated tiles and genes as determined by methylKit::getMethylDiff with
        parameters difference=threshold and qvalue, as provided in the Ob_LIp.conf
        configuration file.  Tiles refer to sliding windows along the genome within which
        methylation calls are cumulated by methylKit::tileMethylCounts.
        The dmg-*.txt files show all genes with at least one differentially methylated tile.

        A positive meth.diff value in comparison A.vs.B means that the B methylation percentage
        in that tile is higher than the A methylation percentage.
```

Directory: DMT File: dmg-Ob_LIp.txt

| seqnames | start | end | width | strand | source | type | score | phase | ID | Dbxref | Name | gbkey | gene | gene_biotype | end_range | partial | start_range |
|---|---|---|---|---|---|---|---|---|---|---|---|---|---|---|---|---|---|

Directory: DMT File: dmt-Ob_LIp.txt

| seqnames | start | end | width | strand | pvalue | qvalue | meth.diff |
| --- | --- | --- | --- | --- | --- | --- | --- |

```
Directory: DMSG File: 0READMEdmsg

DMSG - Differentially methylated sites and genes.

Input: studyhc threshold qvalue
       The studyhc methylKit raw object contains all CpGscd sites with coverage at least
        20 reads.
Output: files dms-*.txt dmg-*.txt  dmg-*details.txt dmg-*heatmaps.pdf

Notes: Output files dms-*.txt and dmg-*.txt are generated by BWASPR::det_dmsg() and show the
        differentially methylated sites and genes as determined by methylKit::getMethylDiff with
        parameters difference=threshold and qvalue, as provided in the Ob_LIp.conf
        configuration file.  The table of differentially methylated genes contains all genes
        with at least one differentially methylated site.

       The files dmg-*details.txt, generated by BWASPR::show_dmsg(), show all CpGscd sites in
        the differentially methylated genes, with coverage numbers and methylation percentages.

       The files dmg-*heatmaps.pdf, generated by BWASPR::show_dmsg(), give heatmap displays for
        genes meeting the following criteria: 1) there are between minNsites and maxNsites common
        CpGscd sites; 2) at least minPdmsites % of these sites are differentially methylated sites.
        The parameters are set in the Ob_LIp.conf configuration file.
```

Directory: DMSG File: dmg-Ob_LIp.txt

| seqnames | start | end | width | strand | source | type | score | phase | ID | Dbxref | Name | gbkey | gene | gene_biotype | end_range | partial | start_range | comparison |
|---|---|---|---|---|---|---|---|---|---|---|---|---|---|---|---|---|---|---|
| NC_039506.1 | 1279061 | 1289519 | 10459 | + | Gnomon | gene | NA | NA | gene-LOC105287325 | GeneID:105287325 | LOC105287325 | Gene | LOC105287325 | protein_coding | character(0) | NA | character(0) | BCphase.vs.Rphase |
| NC_039506.1 | 1279657 | 1286860 | 7204 | - | Gnomon | gene | NA | NA | gene-LOC109611476 | GeneID:109611476 | LOC109611476 | Gene | LOC109611476 | protein_coding | character(0) | NA | character(0) | BCphase.vs.Rphase |
| NC_039506.1 | 1483118 | 1487117 | 4000 | + | Gnomon | gene | NA | NA | gene-LOC105280235 | GeneID:105280235 | LOC105280235 | Gene | LOC105280235 | protein_coding | character(0) | NA | character(0) | BCphase.vs.Rphase |
| NC_039506.1 | 1726070 | 1728383 | 2314 | + | Gnomon | gene | NA | NA | gene-LOC105280262 | GeneID:105280262 | LOC105280262 | Gene | LOC105280262 | protein_coding | character(0) | NA | character(0) | BCphase.vs.Rphase |
| NC_039506.1 | 1728863 | 1739284 | 10422 | - | Gnomon | gene | NA | NA | gene-LOC105280269 | GeneID:105280269 | LOC105280269 | Gene | LOC105280269 | protein_coding | character(0) | NA | character(0) | BCphase.vs.Rphase |
| NC_039506.1 | 1740134 | 1743061 | 2928 | - | Gnomon | gene | NA | NA | gene-LOC105280270 | GeneID:105280270 | LOC105280270 | Gene | LOC105280270 | protein_coding | character(0) | NA | character(0) | BCphase.vs.Rphase |
| NC_039506.1 | 1812143 | 1818351 | 6209 | + | Gnomon | gene | NA | NA | gene-LOC105280272 | GeneID:105280272 | LOC105280272 | Gene | LOC105280272 | protein_coding | character(0) | NA | character(0) | BCphase.vs.Rphase |
| NC_039506.1 | 1831788 | 1843188 | 11401 | - | Gnomon | gene | NA | NA | gene-LOC105280275 | GeneID:105280275 | LOC105280275 | Gene | LOC105280275 | protein_coding | character(0) | NA | character(0) | BCphase.vs.Rphase |
| NC_039506.1 | 1844714 | 1846282 | 1569 | - | Gnomon | gene | NA | NA | gene-LOC105280279 | GeneID:105280279 | LOC105280279 | Gene | LOC105280279 | protein_coding | character(0) | NA | character(0) | BCphase.vs.Rphase |
| NC_039506.1 | 1846650 | 1853079 | 6430 | + | Gnomon | gene | NA | NA | gene-LOC105280280 | GeneID:105280280 | LOC105280280 | Gene | LOC105280280 | protein_coding | character(0) | NA | character(0) | BCphase.vs.Rphase |
| NC_039506.1 | 1912688 | 1914891 | 2204 | - | Gnomon | gene | NA | NA | gene-LOC105280289 | GeneID:105280289 | LOC105280289 | Gene | LOC105280289 | protein_coding | character(0) | NA | character(0) | BCphase.vs.Rphase |
| NC_039506.1 | 1919007 | 1920581 | 1575 | - | Gnomon | gene | NA | NA | gene-LOC105280287 | GeneID:105280287 | LOC105280287 | Gene | LOC105280287 | protein_coding | character(0) | NA | character(0) | BCphase.vs.Rphase |
| NC_039506.1 | 2106150 | 2112574 | 6425 | - | Gnomon | gene | NA | NA | gene-LOC105280296 | GeneID:105280296 | LOC105280296 | Gene | LOC105280296 | protein_coding | character(0) | NA | character(0) | BCphase.vs.Rphase |
| NC_039506.1 | 2169852 | 2172659 | 2808 | - | Gnomon | gene | NA | NA | gene-LOC105280305 | GeneID:105280305 | LOC105280305 | Gene | LOC105280305 | protein_coding | character(0) | NA | character(0) | BCphase.vs.Rphase |
| NC_039506.1 | 2201372 | 2203087 | 1716 | + | Gnomon | gene | NA | NA | gene-LOC105280312 | GeneID:105280312 | LOC105280312 | Gene | LOC105280312 | protein_coding | character(0) | NA | character(0) | BCphase.vs.Rphase |
| NC_039506.1 | 2203015 | 2205481 | 2467 | - | Gnomon | gene | NA | NA | gene-LOC105280311 | GeneID:105280311 | LOC105280311 | Gene | LOC105280311 | protein_coding | character(0) | NA | character(0) | BCphase.vs.Rphase |
| NC_039506.1 | 2274108 | 2276090 | 1983 | - | Gnomon | gene | NA | NA | gene-LOC105280319 | GeneID:105280319 | LOC105280319 | Gene | LOC105280319 | protein_coding | character(0) | NA | character(0) | BCphase.vs.Rphase |
| NC_039506.1 | 2445587 | 2449424 | 3838 | - | Gnomon | gene | NA | NA | gene-LOC105280479 | GeneID:105280479 | LOC105280479 | Gene | LOC105280479 | protein_coding | character(0) | NA | character(0) | BCphase.vs.Rphase |
| NC_039506.1 | 2980439 | 2984235 | 3797 | + | Gnomon | gene | NA | NA | gene-LOC105280348 | GeneID:105280348 | LOC105280348 | Gene | LOC105280348 | protein_coding | character(0) | NA | character(0) | BCphase.vs.Rphase |
| NC_039506.1 | 3253036 | 3254447 | 1412 | - | Gnomon | gene | NA | NA | gene-LOC105280380 | GeneID:105280380 | LOC105280380 | Gene | LOC105280380 | protein_coding | character(0) | NA | character(0) | BCphase.vs.Rphase |
| NC_039506.1 | 3254637 | 3259912 | 5276 | + | Gnomon | gene | NA | NA | gene-LOC105280381 | GeneID:105280381 | LOC105280381 | Gene | LOC105280381 | protein_coding | character(0) | NA | character(0) | BCphase.vs.Rphase |
| NC_039506.1 | 3267119 | 3278572 | 11454 | - | Gnomon | gene | NA | NA | gene-LOC105280383 | GeneID:105280383 | LOC105280383 | Gene | LOC105280383 | protein_coding | character(0) | NA | character(0) | BCphase.vs.Rphase |
| NC_039506.1 | 3347117 | 3353849 | 6733 | - | Gnomon | gene | NA | NA | gene-LOC105280394 | GeneID:105280394 | LOC105280394 | Gene | LOC105280394 | protein_coding | character(0) | NA | character(0) | BCphase.vs.Rphase |
| NC_039506.1 | 3353936 | 3370487 | 16552 | + | Gnomon | gene | NA | NA | gene-LOC105280397 | GeneID:105280397 | LOC105280397 | Gene | LOC105280397 | protein_coding | character(0) | NA | character(0) | BCphase.vs.Rphase |

...
... only first 25 lines shown ...

Directory: DMSG File: dmg-Ob_LIp_BCphase.vs.Rphase_details.txt

| seqnames | start | end | width | strand | coverage1 | numCs1 | numTs1 | coverage2 | numCs2 | numTs2 | BCphase | Rphase | gene_seqnames | gene_start | gene_end | gene_width | gene_strand | gene_source | gene_type | gene_ID | gene_Dbxref | gene_Name | gene_gbkey | gene_gene | gene_gene_biotype | gene_end_range | gene_partial | gene_start_range | gene_comparison | is.dm |
|---|---|---|---|---|---|---|---|---|---|---|---|---|---|---|---|---|---|---|---|---|---|---|---|---|---|---|---|---|---|---|
| NC_039506.1 | 1281995 | 1281995 | 1 | + | 33 | 0 | 33 | 61 | 0 | 61 | 0 | 0 | NC_039506.1 | 1279061 | 1289519 | 10459 | + | Gnomon | gene | gene-LOC105287325 | GeneID:105287325 | LOC105287325 | Gene | LOC105287325 | protein_coding | character(0) | NA | character(0) | BCphase.vs.Rphase | FALSE |
| NC_039506.1 | 1282025 | 1282025 | 1 | + | 26 | 1 | 25 | 32 | 1 | 31 | 3.85 | 3.12 | NC_039506.1 | 1279061 | 1289519 | 10459 | + | Gnomon | gene | gene-LOC105287325 | GeneID:105287325 | LOC105287325 | Gene | LOC105287325 | protein_coding | character(0) | NA | character(0) | BCphase.vs.Rphase | FALSE |
| NC_039506.1 | 1282033 | 1282033 | 1 | + | 25 | 1 | 24 | 30 | 2 | 28 | 4 | 6.67 | NC_039506.1 | 1279061 | 1289519 | 10459 | + | Gnomon | gene | gene-LOC105287325 | GeneID:105287325 | LOC105287325 | Gene | LOC105287325 | protein_coding | character(0) | NA | character(0) | BCphase.vs.Rphase | FALSE |
| NC_039506.1 | 1284692 | 1284692 | 1 | + | 36 | 18 | 18 | 68 | 34 | 34 | 50 | 50 | NC_039506.1 | 1279061 | 1289519 | 10459 | + | Gnomon | gene | gene-LOC105287325 | GeneID:105287325 | LOC105287325 | Gene | LOC105287325 | protein_coding | character(0) | NA | character(0) | BCphase.vs.Rphase | FALSE |
| NC_039506.1 | 1284710 | 1284710 | 1 | + | 84 | 23 | 61 | 116 | 35 | 81 | 27.38 | 30.17 | NC_039506.1 | 1279061 | 1289519 | 10459 | + | Gnomon | gene | gene-LOC105287325 | GeneID:105287325 | LOC105287325 | Gene | LOC105287325 | protein_coding | character(0) | NA | character(0) | BCphase.vs.Rphase | FALSE |
| NC_039506.1 | 1284713 | 1284713 | 1 | + | 85 | 3 | 82 | 116 | 8 | 108 | 3.53 | 6.9 | NC_039506.1 | 1279061 | 1289519 | 10459 | + | Gnomon | gene | gene-LOC105287325 | GeneID:105287325 | LOC105287325 | Gene | LOC105287325 | protein_coding | character(0) | NA | character(0) | BCphase.vs.Rphase | FALSE |
| NC_039506.1 | 1284718 | 1284718 | 1 | + | 84 | 1 | 83 | 117 | 6 | 111 | 1.19 | 5.13 | NC_039506.1 | 1279061 | 1289519 | 10459 | + | Gnomon | gene | gene-LOC105287325 | GeneID:105287325 | LOC105287325 | Gene | LOC105287325 | protein_coding | character(0) | NA | character(0) | BCphase.vs.Rphase | FALSE |
| NC_039506.1 | 1284731 | 1284731 | 1 | + | 86 | 14 | 72 | 117 | 11 | 106 | 16.28 | 9.4 | NC_039506.1 | 1279061 | 1289519 | 10459 | + | Gnomon | gene | gene-LOC105287325 | GeneID:105287325 | LOC105287325 | Gene | LOC105287325 | protein_coding | character(0) | NA | character(0) | BCphase.vs.Rphase | FALSE |
| NC_039506.1 | 1284758 | 1284758 | 1 | + | 73 | 51 | 22 | 86 | 58 | 28 | 69.86 | 67.44 | NC_039506.1 | 1279061 | 1289519 | 10459 | + | Gnomon | gene | gene-LOC105287325 | GeneID:105287325 | LOC105287325 | Gene | LOC105287325 | protein_coding | character(0) | NA | character(0) | BCphase.vs.Rphase | FALSE |
| NC_039506.1 | 1284791 | 1284791 | 1 | + | 35 | 22 | 13 | 42 | 16 | 26 | 62.86 | 38.1 | NC_039506.1 | 1279061 | 1289519 | 10459 | + | Gnomon | gene | gene-LOC105287325 | GeneID:105287325 | LOC105287325 | Gene | LOC105287325 | protein_coding | character(0) | NA | character(0) | BCphase.vs.Rphase | TRUE |
| NC_039506.1 | 1284799 | 1284799 | 1 | + | 32 | 19 | 13 | 46 | 20 | 26 | 59.38 | 43.48 | NC_039506.1 | 1279061 | 1289519 | 10459 | + | Gnomon | gene | gene-LOC105287325 | GeneID:105287325 | LOC105287325 | Gene | LOC105287325 | protein_coding | character(0) | NA | character(0) | BCphase.vs.Rphase | FALSE |
| NC_039506.1 | 1284839 | 1284839 | 1 | + | 30 | 7 | 23 | 33 | 11 | 22 | 23.33 | 33.33 | NC_039506.1 | 1279061 | 1289519 | 10459 | + | Gnomon | gene | gene-LOC105287325 | GeneID:105287325 | LOC105287325 | Gene | LOC105287325 | protein_coding | character(0) | NA | character(0) | BCphase.vs.Rphase | FALSE |
| NC_039506.1 | 1284851 | 1284851 | 1 | + | 27 | 1 | 26 | 28 | 2 | 26 | 3.7 | 7.14 | NC_039506.1 | 1279061 | 1289519 | 10459 | + | Gnomon | gene | gene-LOC105287325 | GeneID:105287325 | LOC105287325 | Gene | LOC105287325 | protein_coding | character(0) | NA | character(0) | BCphase.vs.Rphase | FALSE |
| NC_039506.1 | 1284859 | 1284859 | 1 | + | 26 | 0 | 26 | 31 | 5 | 26 | 0 | 16.13 | NC_039506.1 | 1279061 | 1289519 | 10459 | + | Gnomon | gene | gene-LOC105287325 | GeneID:105287325 | LOC105287325 | Gene | LOC105287325 | protein_coding | character(0) | NA | character(0) | BCphase.vs.Rphase | FALSE |
| NC_039506.1 | 1284863 | 1284863 | 1 | + | 25 | 1 | 24 | 32 | 7 | 25 | 4 | 21.88 | NC_039506.1 | 1279061 | 1289519 | 10459 | + | Gnomon | gene | gene-LOC105287325 | GeneID:105287325 | LOC105287325 | Gene | LOC105287325 | protein_coding | character(0) | NA | character(0) | BCphase.vs.Rphase | FALSE |
| NC_039506.1 | 1286932 | 1286932 | 1 | + | 23 | 0 | 23 | 22 | 0 | 22 | 0 | 0 | NC_039506.1 | 1279061 | 1289519 | 10459 | + | Gnomon | gene | gene-LOC105287325 | GeneID:105287325 | LOC105287325 | Gene | LOC105287325 | protein_coding | character(0) | NA | character(0) | BCphase.vs.Rphase | FALSE |
| NC_039506.1 | 1287139 | 1287139 | 1 | + | 22 | 0 | 22 | 59 | 0 | 59 | 0 | 0 | NC_039506.1 | 1279061 | 1289519 | 10459 | + | Gnomon | gene | gene-LOC105287325 | GeneID:105287325 | LOC105287325 | Gene | LOC105287325 | protein_coding | character(0) | NA | character(0) | BCphase.vs.Rphase | FALSE |
| NC_039506.1 | 1287335 | 1287335 | 1 | + | 25 | 1 | 24 | 38 | 0 | 38 | 4 | 0 | NC_039506.1 | 1279061 | 1289519 | 10459 | + | Gnomon | gene | gene-LOC105287325 | GeneID:105287325 | LOC105287325 | Gene | LOC105287325 | protein_coding | character(0) | NA | character(0) | BCphase.vs.Rphase | FALSE |
| NC_039506.1 | 1287350 | 1287350 | 1 | + | 23 | 0 | 23 | 38 | 0 | 38 | 0 | 0 | NC_039506.1 | 1279061 | 1289519 | 10459 | + | Gnomon | gene | gene-LOC105287325 | GeneID:105287325 | LOC105287325 | Gene | LOC105287325 | protein_coding | character(0) | NA | character(0) | BCphase.vs.Rphase | FALSE |
| NC_039506.1 | 1287900 | 1287900 | 1 | + | 48 | 1 | 47 | 26 | 0 | 26 | 2.08 | 0 | NC_039506.1 | 1279061 | 1289519 | 10459 | + | Gnomon | gene | gene-LOC105287325 | GeneID:105287325 | LOC105287325 | Gene | LOC105287325 | protein_coding | character(0) | NA | character(0) | BCphase.vs.Rphase | FALSE |
| NC_039506.1 | 1287915 | 1287915 | 1 | + | 51 | 0 | 51 | 26 | 0 | 26 | 0 | 0 | NC_039506.1 | 1279061 | 1289519 | 10459 | + | Gnomon | gene | gene-LOC105287325 | GeneID:105287325 | LOC105287325 | Gene | LOC105287325 | protein_coding | character(0) | NA | character(0) | BCphase.vs.Rphase | FALSE |
| NC_039506.1 | 1287925 | 1287925 | 1 | + | 59 | 0 | 59 | 26 | 0 | 26 | 0 | 0 | NC_039506.1 | 1279061 | 1289519 | 10459 | + | Gnomon | gene | gene-LOC105287325 | GeneID:105287325 | LOC105287325 | Gene | LOC105287325 | protein_coding | character(0) | NA | character(0) | BCphase.vs.Rphase | FALSE |
| NC_039506.1 | 1287938 | 1287938 | 1 | + | 35 | 0 | 35 | 29 | 0 | 29 | 0 | 0 | NC_039506.1 | 1279061 | 1289519 | 10459 | + | Gnomon | gene | gene-LOC105287325 | GeneID:105287325 | LOC105287325 | Gene | LOC105287325 | protein_coding | character(0) | NA | character(0) | BCphase.vs.Rphase | FALSE |
| NC_039506.1 | 1287962 | 1287962 | 1 | + | 35 | 0 | 35 | 25 | 0 | 25 | 0 | 0 | NC_039506.1 | 1279061 | 1289519 | 10459 | + | Gnomon | gene | gene-LOC105287325 | GeneID:105287325 | LOC105287325 | Gene | LOC105287325 | protein_coding | character(0) | NA | character(0) | BCphase.vs.Rphase | FALSE |

...
... only first 25 lines shown ...

Directory: DMSG File: dms-Ob_LIp.txt

| seqnames | start | end | width | strand | pvalue | qvalue | meth.diff | comparison |
|---|---|---|---|---|---|---|---|---|
| NC_039506.1 | 1260240 | 1260240 | 1 | + | 0.0561480851846645 | 0.031 | -26.36 | BCphase.vs.Rphase |
| NC_039506.1 | 1284791 | 1284791 | 1 | + | 0.0402471777976576 | 0.031 | -24.76 | BCphase.vs.Rphase |
| NC_039506.1 | 1486030 | 1486030 | 1 | + | 0.0936098551364574 | 0.031 | -22.82 | BCphase.vs.Rphase |
| NC_039506.1 | 1487054 | 1487054 | 1 | + | 0.208160926872775 | 0.031 | -22.38 | BCphase.vs.Rphase |
| NC_039506.1 | 1727582 | 1727582 | 1 | + | 0.0514197361297291 | 0.031 | 29.63 | BCphase.vs.Rphase |
| NC_039506.1 | 1727718 | 1727718 | 1 | + | 0.0967284392301921 | 0.031 | 23.28 | BCphase.vs.Rphase |
| NC_039506.1 | 1736219 | 1736219 | 1 | + | 0.0783930081734651 | 0.031 | 20.67 | BCphase.vs.Rphase |
| NC_039506.1 | 1741334 | 1741334 | 1 | + | 0.121937284673675 | 0.031 | 21.68 | BCphase.vs.Rphase |
| NC_039506.1 | 1812812 | 1812812 | 1 | + | 0.0336543144257268 | 0.031 | 26 | BCphase.vs.Rphase |
| NC_039506.1 | 1814180 | 1814180 | 1 | + | 0.0737549409645709 | 0.031 | 21.17 | BCphase.vs.Rphase |
| NC_039506.1 | 1814183 | 1814183 | 1 | + | 0.193019439588996 | 0.031 | 20.83 | BCphase.vs.Rphase |
| NC_039506.1 | 1841108 | 1841108 | 1 | + | 0.0500624361296581 | 0.031 | 22.76 | BCphase.vs.Rphase |
| NC_039506.1 | 1845378 | 1845378 | 1 | + | 0.0247978436657682 | 0.031 | 25 | BCphase.vs.Rphase |
| NC_039506.1 | 1849717 | 1849717 | 1 | + | 0.00951107075692951 | 0.031 | 20.5 | BCphase.vs.Rphase |
| NC_039506.1 | 1914415 | 1914415 | 1 | + | 0.042249871139523 | 0.031 | 20.3 | BCphase.vs.Rphase |
| NC_039506.1 | 1918935 | 1918935 | 1 | + | 0.00359498691690465 | 0.031 | -23.33 | BCphase.vs.Rphase |
| NC_039506.1 | 1919878 | 1919878 | 1 | + | 0.0111247804842326 | 0.031 | 34.52 | BCphase.vs.Rphase |
| NC_039506.1 | 2000674 | 2000674 | 1 | + | 4.89755265036887e-05 | 0.031 | 24 | BCphase.vs.Rphase |
| NC_039506.1 | 2108374 | 2108374 | 1 | + | 0.0655608753550934 | 0.031 | -25 | BCphase.vs.Rphase |
| NC_039506.1 | 2171100 | 2171100 | 1 | + | 0.0165211357440403 | 0.031 | 23.33 | BCphase.vs.Rphase |
| NC_039506.1 | 2202360 | 2202360 | 1 | + | 0.00345046333290056 | 0.031 | 23.13 | BCphase.vs.Rphase |
| NC_039506.1 | 2204653 | 2204653 | 1 | + | 0.0175207435401975 | 0.031 | 21.26 | BCphase.vs.Rphase |
| NC_039506.1 | 2205000 | 2205000 | 1 | + | 0.00230746887283885 | 0.031 | 33.33 | BCphase.vs.Rphase |
| NC_039506.1 | 2274652 | 2274652 | 1 | + | 0.00909245504095002 | 0.031 | 25.45 | BCphase.vs.Rphase |

...
... only first 25 lines shown ...

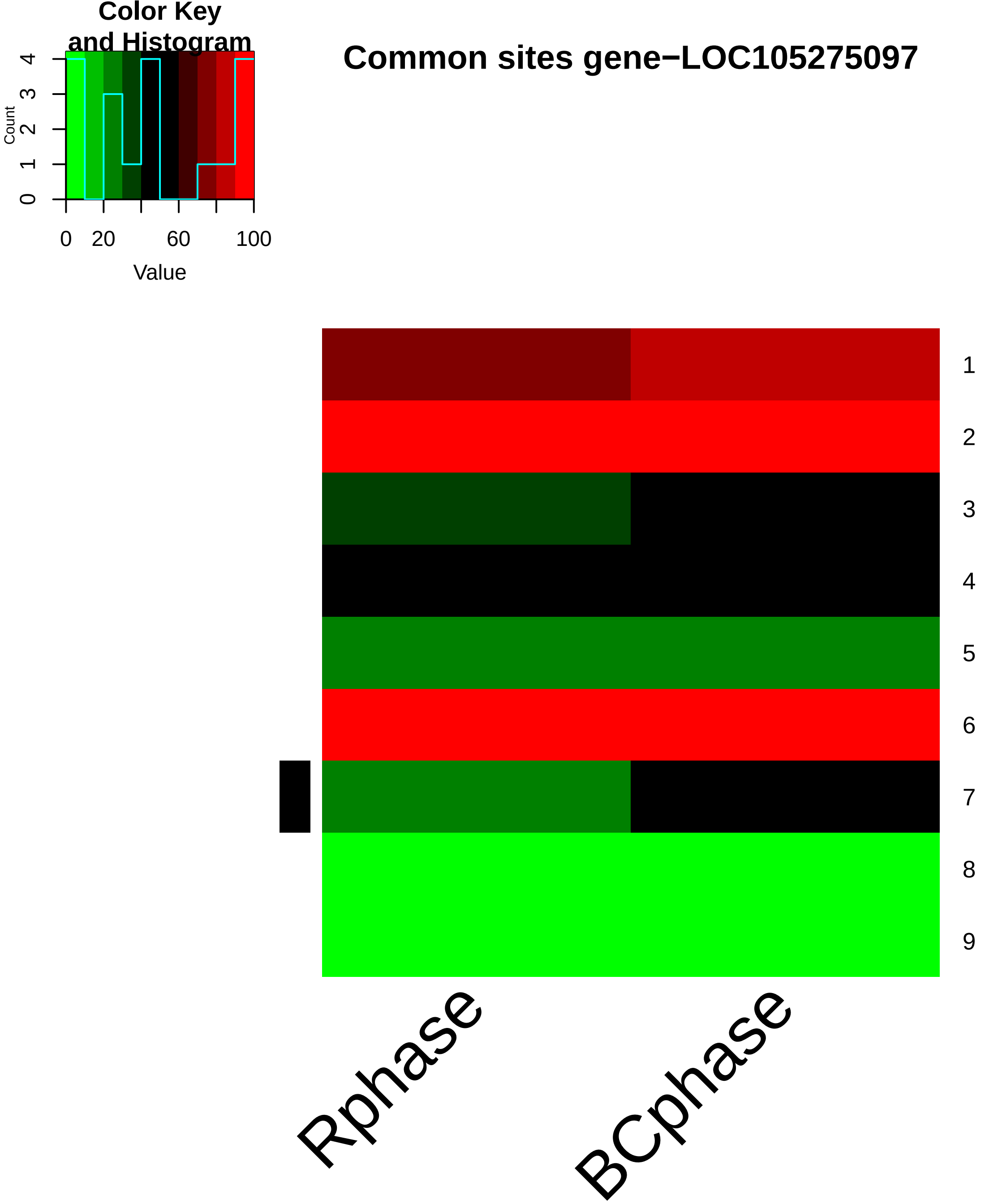
Color Key
and Histogram
Count
0
1
2
3
4
0
20
60
100
Value
Common sites gene−LOC105275097
1
2
3
4
5
6
7
8
9
Rphase
BCphase

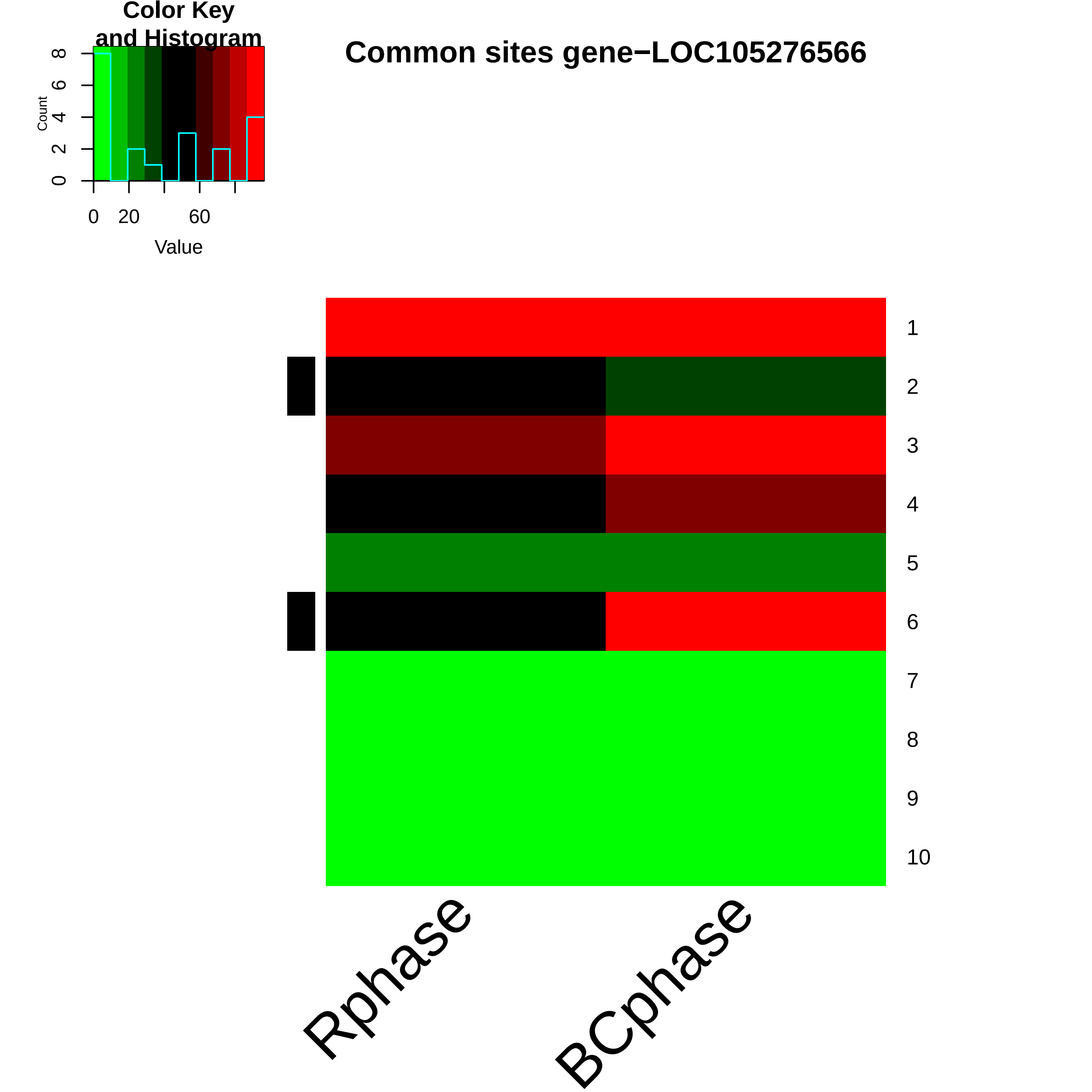
Color Key
and Histogram
Count
0
2
4
6
8
0
20
60
Value
Common sites gene−LOC105276566
1
2
3
4
5
6
7
8
9
10
Rphase
BCphase

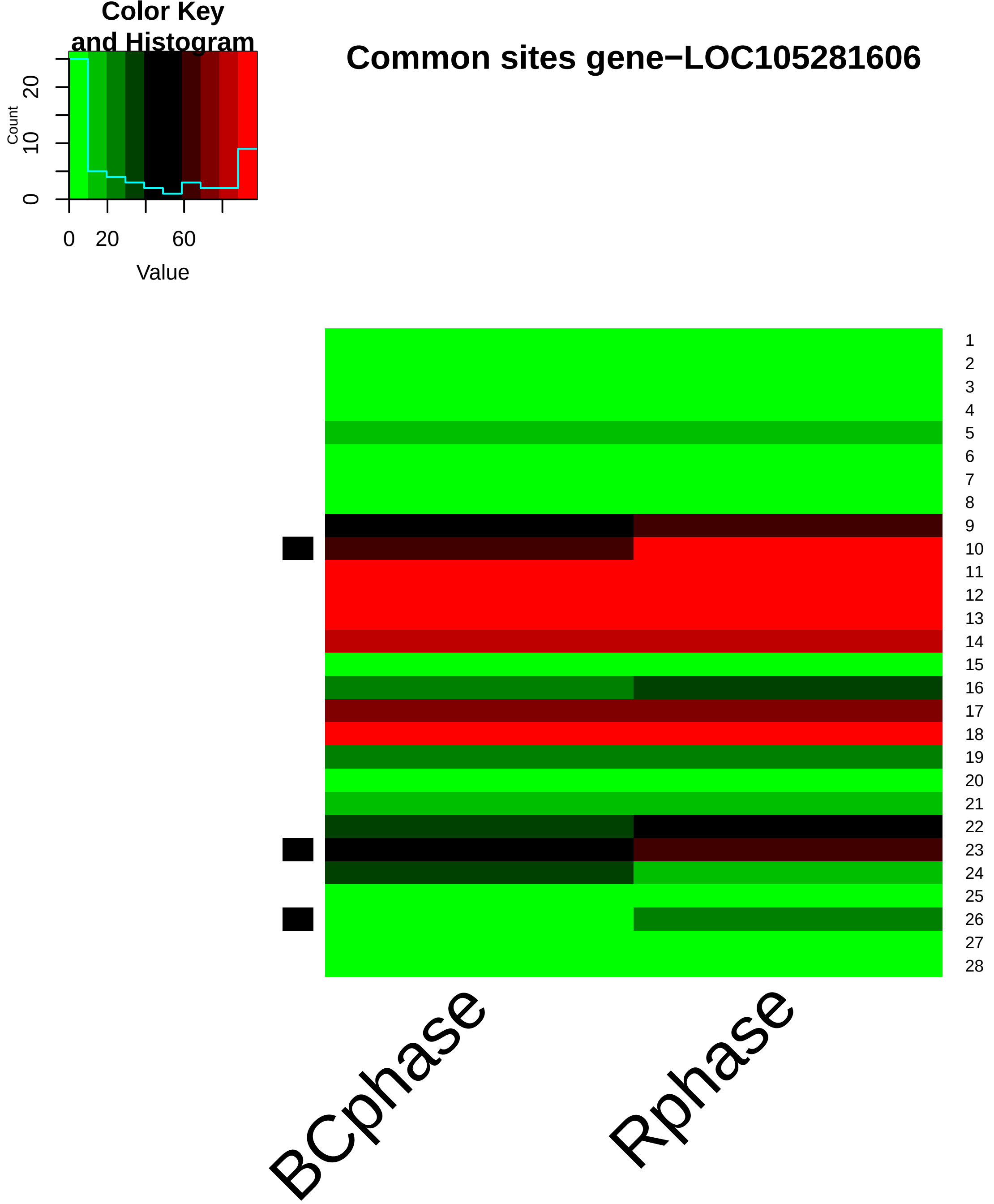

Color Key
and Histogram
Count
0
10
20
0
20
60
Value
Common sites gene−LOC105281606
BCphase
Rphase

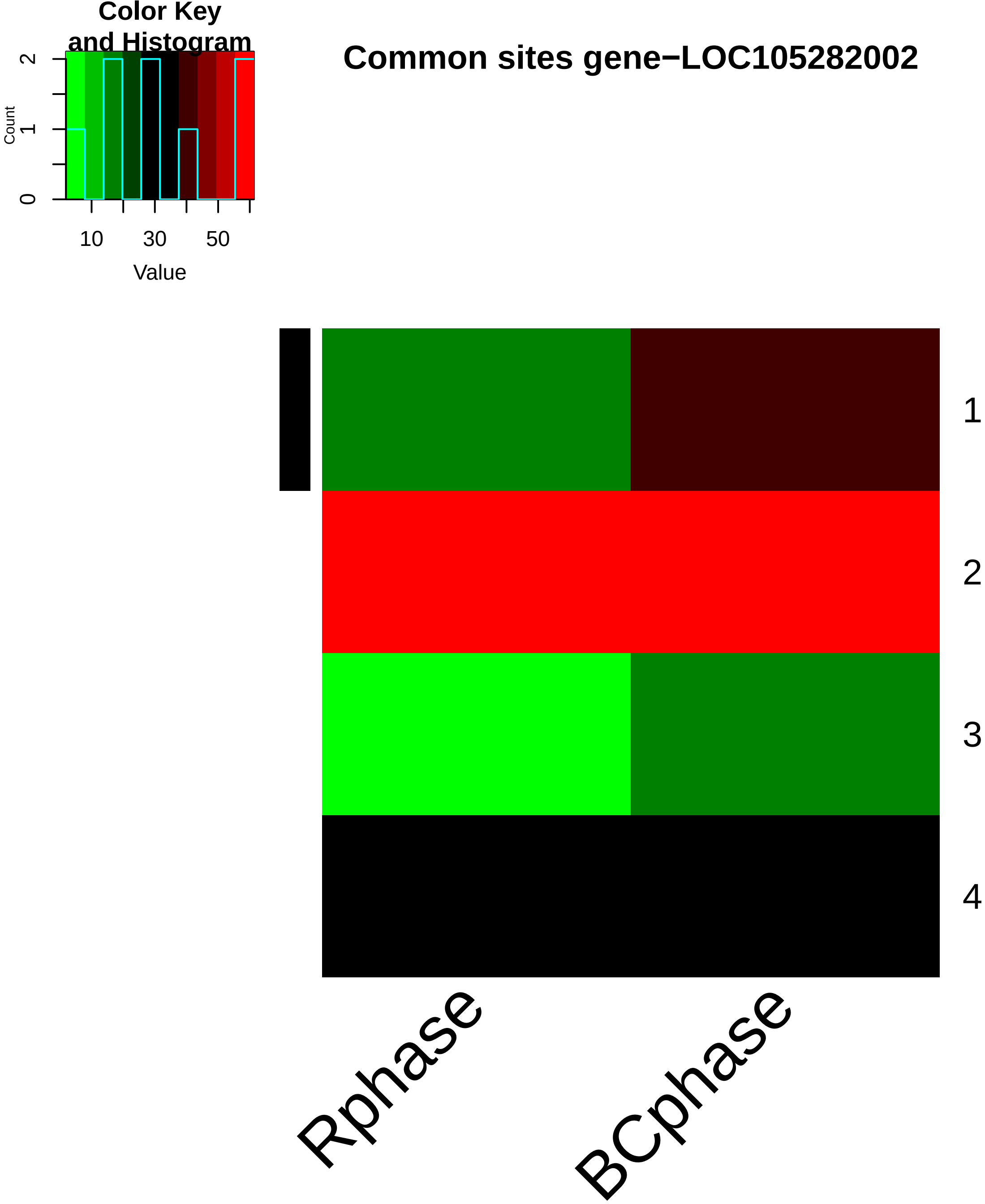

Color Key
and Histogram
Count
0
1
2
10
30
50
Value
Common sites gene−LOC105282002
1
2
3
4
Rphase
BCphase

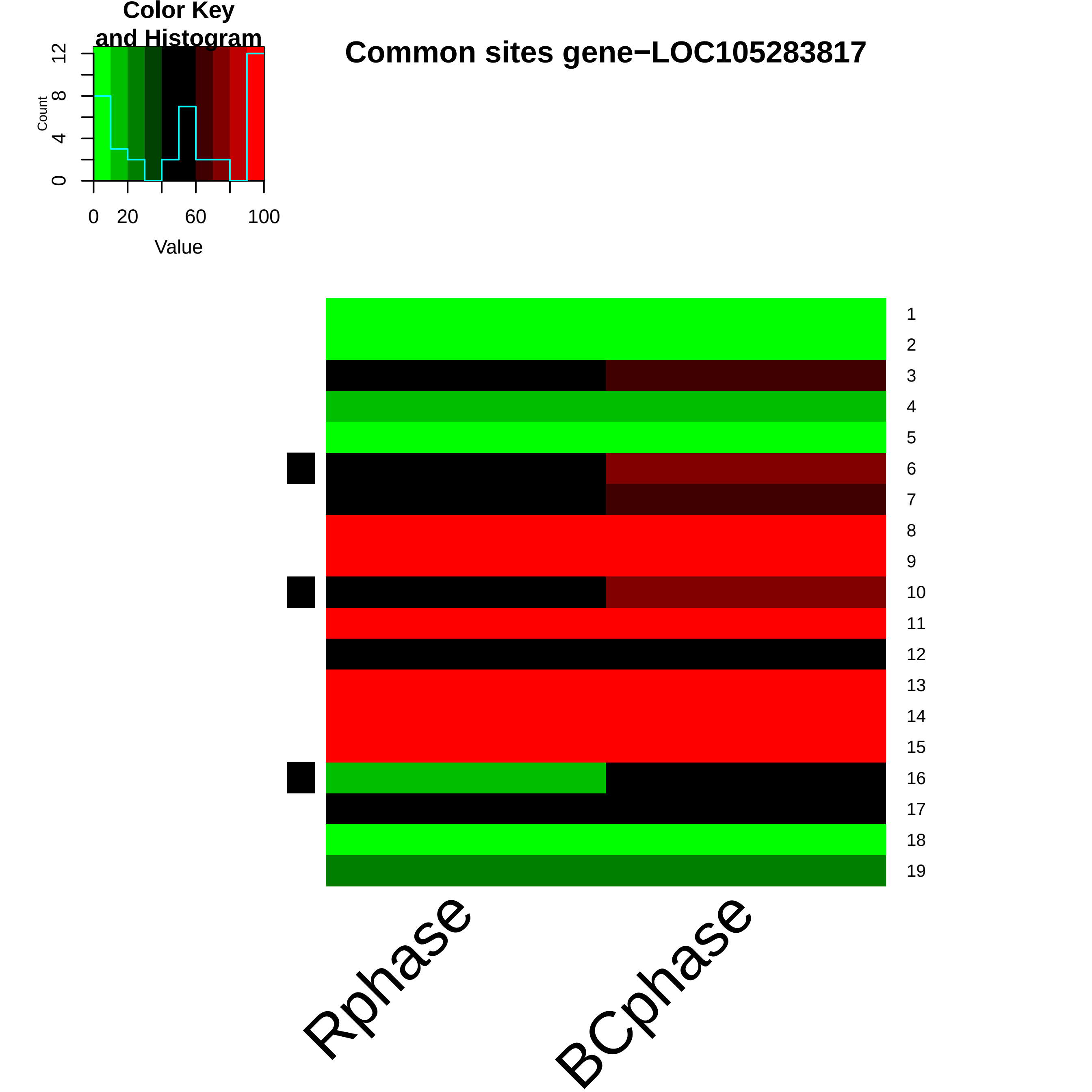

Color Key
and Histogram
Count
0
4
8
12
0
20
60
100
Value
Common sites gene−LOC105283817
Rphase
BCphase

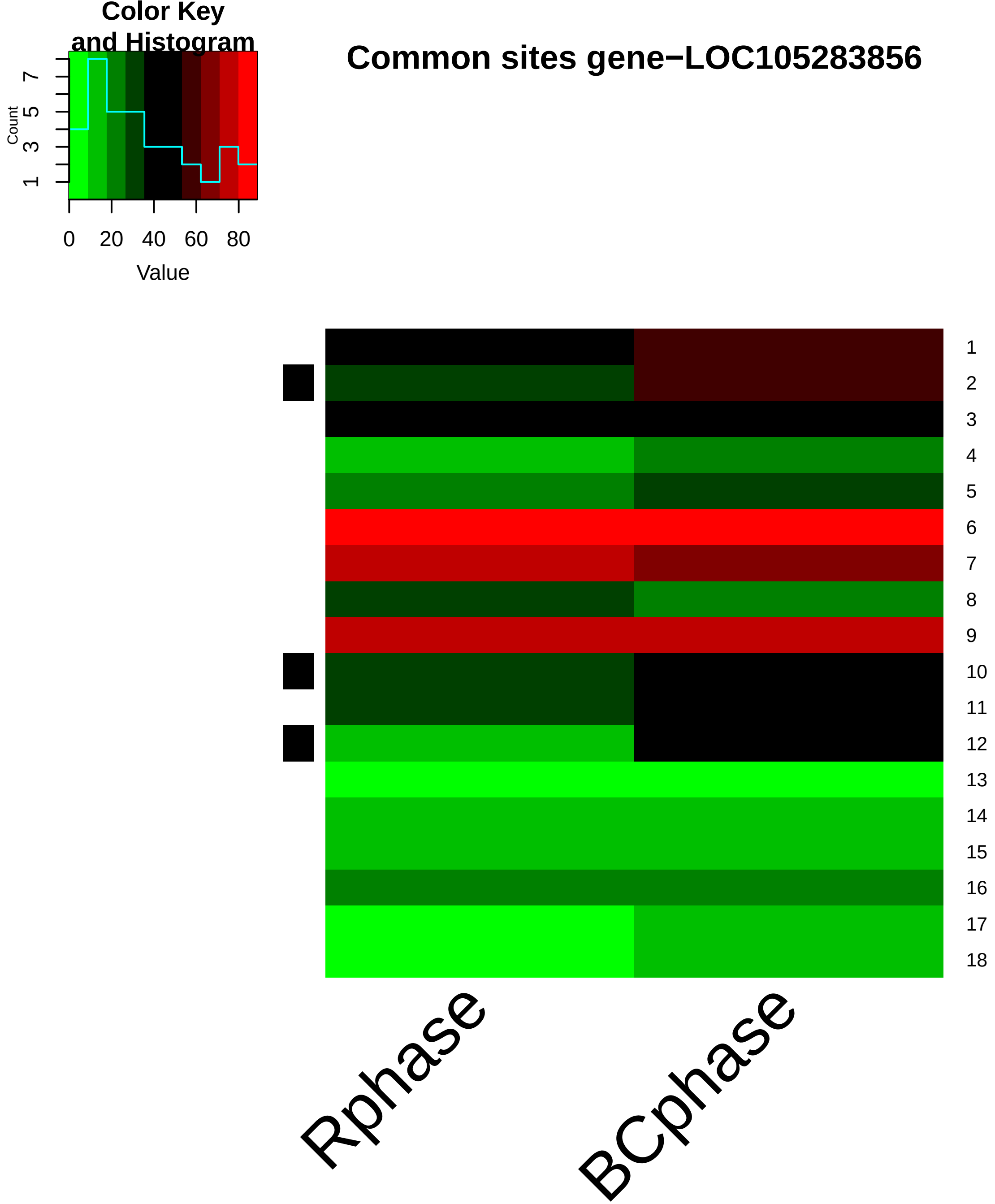

Color Key and Histogram
Count
1
3
5
7
0
20
40
60
80
Value
Common sites gene−LOC105283856
Rphase
BCphase

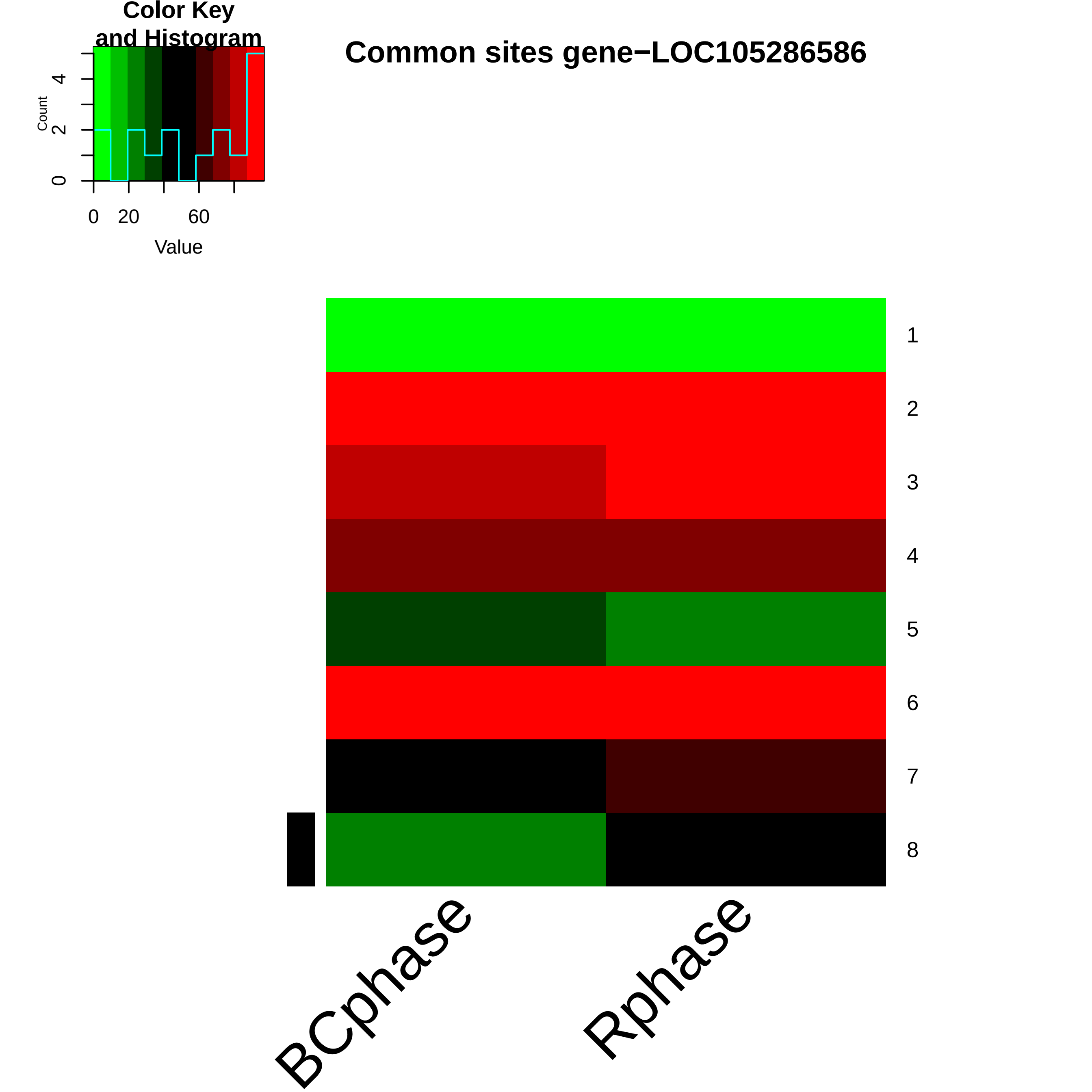

Color Key
and Histogram
Count
0
2
4
0
20
60
Value
Common sites gene−LOC105286586
1
2
3
4
5
6
7
8
BCphase
Rphase

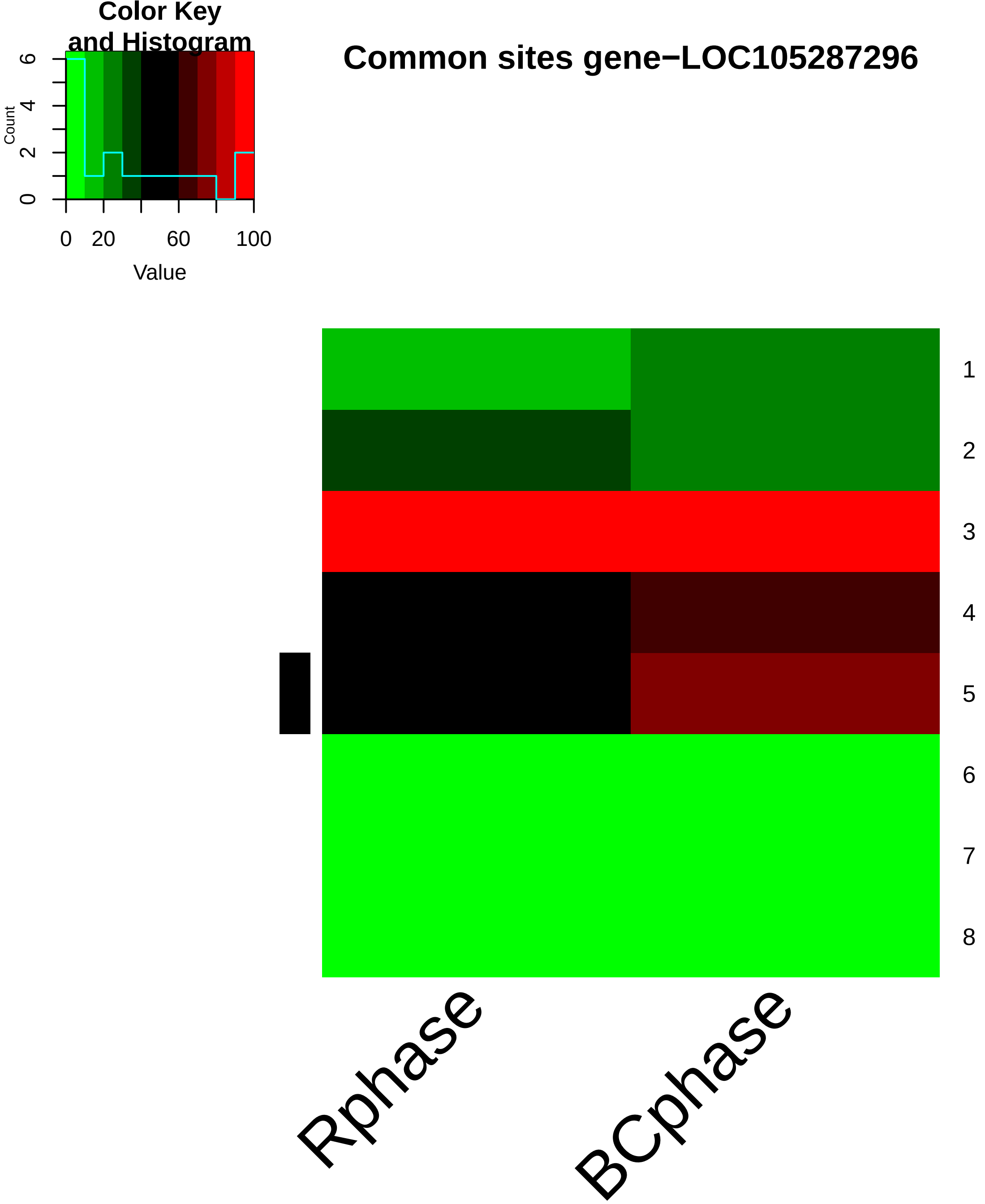

Color Key
and Histogram
Count
0
2
4
6
0
20
60
100
Value
Common sites gene−LOC105287296
1
2
3
4
5
6
7
8
Rphase
BCphase

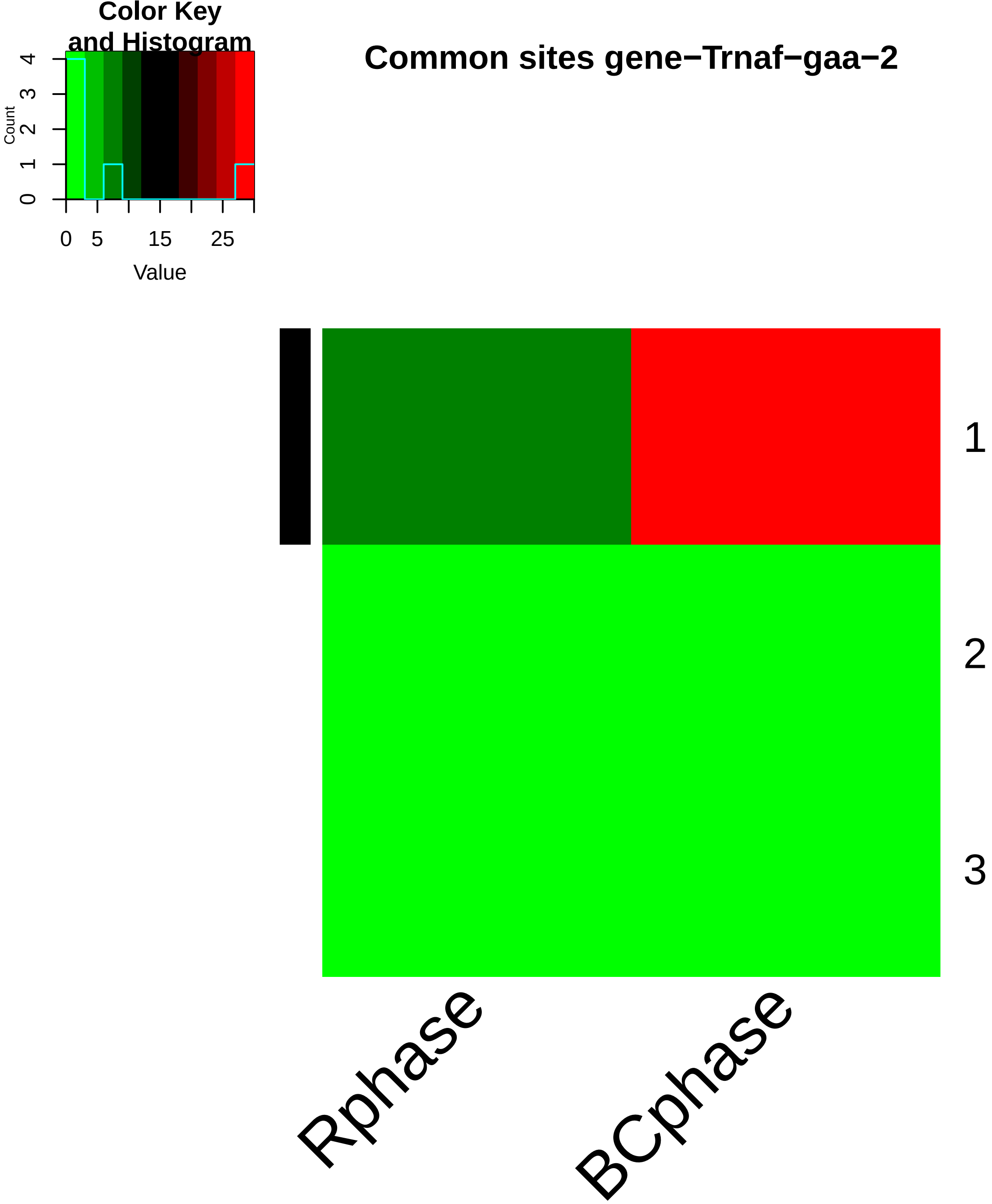

Color Key
and Histogram
Count
0
1
2
3
4
0
5
15
25
Value
Common sites gene−Trnaf−gaa−2
1
2
3
Rphase
BCphase

```
Directory: OGL File: 0READMEogl

OGL - Ordered gene lists.

Input: studyhc annotation   maxgwidth minnbrdmsites
       dmgprp (= output of BWASPR:show_dmsg())

Output: files ogl-*.txt rnk-dmg-*.txt
         ogl-<study>_<sample>.txt ogl-<study>_<sample1>.vs.<sample2>.txt
         rnk-dmg--<study>_<sample1>.vs.<sample2>.txt rnk-dmg--<study>_<sample1>.vs.<sample2>.pdf
         wrt-<study>.txt

Notes: Output files ogl-<study>_<sample>.txt give tables for each sample with columns

        gene_ID gwidth #Sites #per10Kb %perSite %pNucl

       ordered by %pNucl.  If available, a link to an NCBI entry of gene_ID is inserted as second column.
       Abbreviations used: %perSite, percent methylation per site
                           %perNucl, percent methylation per nucleotide of the gene

       Output files ogl-<study>_<sample1>.vs.<sample2>.txt give tables for each comparison with columns

        gene_ID gwidth #Sites #per10Kb #dmSites #dmsp10kb %dmSites %pSite1 %pSite2 DMpSite ADMpSite DMpNucl ADMpNucl

       ordered by DMpSite.  If available, a link to an NCBI entry of gene_ID is inserted as second column.
       Abbreviations used: %dmSites, percent sites that are differentially methylated
                           %pSite1, average per site % methylation for sample1
                           %pSite2, average per site % methylation for sample2
                           DMpSite, average per site difference in % methylation between sample1 and sample2
                           ADMpSite, absolute value of DMpSite
                           DMpNucl, average per nucleotide difference in % methylation between sample1 and sample2
                           ADMpNucl, absolute value of DMpNucl

       Output files rnk-dmg-<study>_<sample1>.vs.<sample2>.txt are equivalent to files
        ogl-<study>_<sample1>.vs.<sample2>.txt but ordered by ADMpNucl.
       Output files rnk-dmg-<study>_<sample1>.vs.<sample2>.pdf provide visualization of the distribution
        of ADMpNucl values.

       Output file wrt-<study>.txt gives results of the Wilcoxon signed rank test comparing
        the %pSite1 and %pSite2 vectors

       In output directory OGL, tables are restricted to genes with gwidth <= maxgwidth and, for
        pairwise comparisons, #dmSites >= minnbrdmsites

       If either maxgwidth or minnbrdmsites is set, then output directory OGLall will show the full tables
        with all genes (for reference).
```

Directory: OGL File: ogl-Ob_LIp_BCphase.txt

| gene_ID | gene_link | gwidth | #Sites | #per10Kb | %perSite | %pNucl |
|---|---|---|---|---|---|---|
| gene-LOC113561532 | https://www.ncbi.nlm.nih.gov/gene/?term=LOC113561532 | 72 | 6 | 833.33 | 95.81 | 7.98 |
| gene-LOC113562627 | https://www.ncbi.nlm.nih.gov/gene/?term=LOC113562627 | 397 | 37 | 931.99 | 69.15 | 6.45 |
| gene-LOC105285394 | https://www.ncbi.nlm.nih.gov/gene/?term=LOC105285394 | 1307 | 147 | 1124.71 | 50.61 | 5.69 |
| gene-LOC105282149 | https://www.ncbi.nlm.nih.gov/gene/?term=LOC105282149 | 1787 | 213 | 1191.94 | 42.72 | 5.09 |
| gene-LOC105283830 | https://www.ncbi.nlm.nih.gov/gene/?term=LOC105283830 | 252 | 34 | 1349.21 | 36.78 | 4.96 |
| gene-LOC105276360 | https://www.ncbi.nlm.nih.gov/gene/?term=LOC105276360 | 1466 | 237 | 1616.64 | 28.53 | 4.61 |
| gene-LOC105275231 | https://www.ncbi.nlm.nih.gov/gene/?term=LOC105275231 | 1270 | 149 | 1173.23 | 39.05 | 4.58 |
| gene-LOC105278849 | https://www.ncbi.nlm.nih.gov/gene/?term=LOC105278849 | 776 | 151 | 1945.88 | 23.52 | 4.58 |
| gene-LOC105277648 | https://www.ncbi.nlm.nih.gov/gene/?term=LOC105277648 | 722 | 53 | 734.07 | 62.32 | 4.57 |
| gene-LOC105286531 | https://www.ncbi.nlm.nih.gov/gene/?term=LOC105286531 | 372 | 46 | 1236.56 | 36.89 | 4.56 |
| gene-LOC105286269 | https://www.ncbi.nlm.nih.gov/gene/?term=LOC105286269 | 1401 | 156 | 1113.49 | 40.37 | 4.49 |
| gene-LOC105286562 | https://www.ncbi.nlm.nih.gov/gene/?term=LOC105286562 | 1484 | 129 | 869.27 | 51.38 | 4.47 |
| gene-LOC105274711 | https://www.ncbi.nlm.nih.gov/gene/?term=LOC105274711 | 1655 | 165 | 996.98 | 44.69 | 4.46 |
| gene-LOC105286801 | https://www.ncbi.nlm.nih.gov/gene/?term=LOC105286801 | 1102 | 120 | 1088.93 | 40.66 | 4.43 |
| gene-LOC105280739 | https://www.ncbi.nlm.nih.gov/gene/?term=LOC105280739 | 1991 | 182 | 914.11 | 47.53 | 4.34 |
| gene-LOC105287811 | https://www.ncbi.nlm.nih.gov/gene/?term=LOC105287811 | 2155 | 245 | 1136.89 | 38.05 | 4.33 |
| gene-LOC105285635 | https://www.ncbi.nlm.nih.gov/gene/?term=LOC105285635 | 1175 | 191 | 1625.53 | 26.28 | 4.27 |
| gene-LOC105275314 | https://www.ncbi.nlm.nih.gov/gene/?term=LOC105275314 | 1489 | 189 | 1269.31 | 33.05 | 4.19 |
| gene-LOC105279187 | https://www.ncbi.nlm.nih.gov/gene/?term=LOC105279187 | 2255 | 238 | 1055.43 | 39.14 | 4.13 |
| gene-LOC105286235 | https://www.ncbi.nlm.nih.gov/gene/?term=LOC105286235 | 1510 | 148 | 980.13 | 42.06 | 4.12 |
| gene-LOC105274534 | https://www.ncbi.nlm.nih.gov/gene/?term=LOC105274534 | 1329 | 148 | 1113.62 | 36.91 | 4.11 |
| gene-LOC105285392 | https://www.ncbi.nlm.nih.gov/gene/?term=LOC105285392 | 1601 | 230 | 1436.6 | 28.09 | 4.04 |
| gene-LOC105284659 | https://www.ncbi.nlm.nih.gov/gene/?term=LOC105284659 | 1988 | 217 | 1091.55 | 36.25 | 3.96 |
| gene-LOC105274863 | https://www.ncbi.nlm.nih.gov/gene/?term=LOC105274863 | 2035 | 178 | 874.69 | 44.78 | 3.92 |

...

... only first 25 lines shown ...

Directory: OGL File: ogl-Ob_LIp_BCphase.vs.Rphase.txt

| gene_ID | gene_link | gwidth | #Sites | #per10Kb | #dmSites | #dmsp10kb | %dmSites | %pSite1 | %pSite2 | DMpSite | ADMpSite | DMpNucl | ADMpNucl |
|---|---|---|---|---|---|---|---|---|---|---|---|---|---|
| gene-LOC105282002 | https://www.ncbi.nlm.nih.gov/gene/?term=LOC105282002 | 1311 | 4 | 30.51 | 1 | 7.63 | 25 | 37.08 | 25.87 | 11.2 | 11.2 | 0.03 | 0.03 |
| gene-Trnaf-gaa-2 | https://www.ncbi.nlm.nih.gov/gene/?term=Trnaf-gaa | 73 | 3 | 410.96 | 1 | 136.99 | 33.33 | 10 | 3.46 | 6.54 | 6.54 | 0.27 | 0.27 |
| gene-LOC105283856 | https://www.ncbi.nlm.nih.gov/gene/?term=LOC105283856 | 1988 | 18 | 90.54 | 3 | 15.09 | 16.67 | 37.55 | 31.12 | 6.44 | 6.44 | 0.06 | 0.06 |
| gene-LOC105276566 | https://www.ncbi.nlm.nih.gov/gene/?term=LOC105276566 | 1114 | 10 | 89.77 | 2 | 17.95 | 20 | 40.73 | 35.89 | 4.84 | 4.84 | 0.04 | 0.04 |
| gene-LOC105283817 | https://www.ncbi.nlm.nih.gov/gene/?term=LOC105283817 | 1309 | 19 | 145.15 | 3 | 22.92 | 15.79 | 55.23 | 50.62 | 4.61 | 4.61 | 0.07 | 0.07 |
| gene-LOC105287296 | https://www.ncbi.nlm.nih.gov/gene/?term=LOC105287296 | 1834 | 8 | 43.62 | 1 | 5.45 | 12.5 | 36.82 | 32.76 | 4.06 | 4.06 | 0.02 | 0.02 |
| gene-LOC105286178 | https://www.ncbi.nlm.nih.gov/gene/?term=LOC105286178 | 1590 | 13 | 81.76 | 1 | 6.29 | 7.69 | 65.07 | 61.17 | 3.91 | 3.91 | 0.03 | 0.03 |
| gene-LOC105275097 | https://www.ncbi.nlm.nih.gov/gene/?term=LOC105275097 | 1071 | 9 | 84.03 | 1 | 9.34 | 11.11 | 49.2 | 45.35 | 3.85 | 3.85 | 0.03 | 0.03 |
| gene-LOC105281507 | https://www.ncbi.nlm.nih.gov/gene/?term=LOC105281507 | 881 | 24 | 272.42 | 1 | 11.35 | 4.17 | 24.55 | 20.73 | 3.82 | 3.82 | 0.1 | 0.1 |
| gene-LOC105283800 | https://www.ncbi.nlm.nih.gov/gene/?term=LOC105283800 | 871 | 24 | 275.55 | 1 | 11.48 | 4.17 | 24.17 | 21.18 | 2.99 | 2.99 | 0.08 | 0.08 |
| gene-LOC113563090 | https://www.ncbi.nlm.nih.gov/gene/?term=LOC113563090 | 1729 | 22 | 127.24 | 2 | 11.57 | 9.09 | 57.11 | 54.13 | 2.98 | 2.98 | 0.04 | 0.04 |
| gene-LOC105287152 | https://www.ncbi.nlm.nih.gov/gene/?term=LOC105287152 | 1616 | 25 | 154.7 | 1 | 6.19 | 4 | 29.08 | 26.72 | 2.36 | 2.36 | 0.04 | 0.04 |
| gene-LOC105287372 | https://www.ncbi.nlm.nih.gov/gene/?term=LOC105287372 | 2659 | 50 | 188.04 | 1 | 3.76 | 2 | 10.93 | 8.75 | 2.18 | 2.18 | 0.04 | 0.04 |
| gene-LOC105285429 | https://www.ncbi.nlm.nih.gov/gene/?term=LOC105285429 | 1324 | 36 | 271.9 | 2 | 15.11 | 5.56 | 41.65 | 39.56 | 2.09 | 2.09 | 0.06 | 0.06 |
| gene-LOC105279501 | https://www.ncbi.nlm.nih.gov/gene/?term=LOC105279501 | 2114 | 40 | 189.21 | 1 | 4.73 | 2.5 | 51.95 | 49.94 | 2.01 | 2.01 | 0.04 | 0.04 |
| gene-LOC105275378 | https://www.ncbi.nlm.nih.gov/gene/?term=LOC105275378 | 1189 | 35 | 294.37 | 1 | 8.41 | 2.86 | 45.72 | 43.88 | 1.85 | 1.85 | 0.05 | 0.05 |
| gene-LOC113561875 | https://www.ncbi.nlm.nih.gov/gene/?term=LOC113561875 | 579 | 16 | 276.34 | 1 | 17.27 | 6.25 | 6.85 | 5.16 | 1.69 | 1.69 | 0.05 | 0.05 |
| gene-LOC105281162 | https://www.ncbi.nlm.nih.gov/gene/?term=LOC105281162 | 2174 | 27 | 124.2 | 1 | 4.6 | 3.7 | 32.48 | 30.86 | 1.62 | 1.62 | 0.02 | 0.02 |
| gene-LOC105287127 | https://www.ncbi.nlm.nih.gov/gene/?term=LOC105287127 | 2308 | 71 | 307.63 | 1 | 4.33 | 1.41 | 6.04 | 4.42 | 1.62 | 1.62 | 0.05 | 0.05 |
| gene-LOC105286157 | https://www.ncbi.nlm.nih.gov/gene/?term=LOC105286157 | 2004 | 73 | 364.27 | 2 | 9.98 | 2.74 | 25.32 | 23.75 | 1.57 | 1.57 | 0.06 | 0.06 |
| gene-LOC105276157 | https://www.ncbi.nlm.nih.gov/gene/?term=LOC105276157 | 1601 | 44 | 274.83 | 1 | 6.25 | 2.27 | 26.48 | 24.92 | 1.56 | 1.56 | 0.04 | 0.04 |
| gene-LOC105280380 | https://www.ncbi.nlm.nih.gov/gene/?term=LOC105280380 | 1412 | 38 | 269.12 | 1 | 7.08 | 2.63 | 6.6 | 5.05 | 1.55 | 1.55 | 0.04 | 0.04 |
| gene-LOC105285707 | https://www.ncbi.nlm.nih.gov/gene/?term=LOC105285707 | 2096 | 71 | 338.74 | 1 | 4.77 | 1.41 | 31.72 | 30.19 | 1.53 | 1.53 | 0.05 | 0.05 |
| gene-LOC105283991 | https://www.ncbi.nlm.nih.gov/gene/?term=LOC105283991 | 2852 | 59 | 206.87 | 1 | 3.51 | 1.69 | 33.92 | 32.4 | 1.52 | 1.52 | 0.03 | 0.03 |

...

... only first 25 lines shown ...

Directory: OGL File: ogl-Ob_LIp_Rphase.txt

| gene_ID | gene_link | gwidth | #Sites | #per10Kb | %perSite | %pNucl |
|---|---|---|---|---|---|---|
| gene-LOC113561532 | https://www.ncbi.nlm.nih.gov/gene/?term=LOC113561532 | 72 | 7 | 972.22 | 96.09 | 9.34 |
| gene-LOC113562627 | https://www.ncbi.nlm.nih.gov/gene/?term=LOC113562627 | 397 | 37 | 931.99 | 69.26 | 6.46 |
| gene-LOC105285394 | https://www.ncbi.nlm.nih.gov/gene/?term=LOC105285394 | 1307 | 148 | 1132.36 | 51.87 | 5.87 |
| gene-LOC105282149 | https://www.ncbi.nlm.nih.gov/gene/?term=LOC105282149 | 1787 | 217 | 1214.33 | 43.31 | 5.26 |
| gene-LOC105283830 | https://www.ncbi.nlm.nih.gov/gene/?term=LOC105283830 | 252 | 34 | 1349.21 | 37.52 | 5.06 |
| gene-LOC105286801 | https://www.ncbi.nlm.nih.gov/gene/?term=LOC105286801 | 1102 | 131 | 1188.75 | 38.91 | 4.62 |
| gene-LOC105276360 | https://www.ncbi.nlm.nih.gov/gene/?term=LOC105276360 | 1466 | 237 | 1616.64 | 28.5 | 4.61 |
| gene-LOC105278849 | https://www.ncbi.nlm.nih.gov/gene/?term=LOC105278849 | 776 | 151 | 1945.88 | 23.46 | 4.57 |
| gene-LOC105275231 | https://www.ncbi.nlm.nih.gov/gene/?term=LOC105275231 | 1270 | 151 | 1188.98 | 38.3 | 4.55 |
| gene-LOC105286269 | https://www.ncbi.nlm.nih.gov/gene/?term=LOC105286269 | 1401 | 155 | 1106.35 | 40.98 | 4.53 |
| gene-LOC105286531 | https://www.ncbi.nlm.nih.gov/gene/?term=LOC105286531 | 372 | 46 | 1236.56 | 36.39 | 4.5 |
| gene-LOC105286562 | https://www.ncbi.nlm.nih.gov/gene/?term=LOC105286562 | 1484 | 135 | 909.7 | 49.22 | 4.48 |
| gene-LOC105274711 | https://www.ncbi.nlm.nih.gov/gene/?term=LOC105274711 | 1655 | 169 | 1021.15 | 43.22 | 4.41 |
| gene-LOC105277648 | https://www.ncbi.nlm.nih.gov/gene/?term=LOC105277648 | 722 | 52 | 720.22 | 61.24 | 4.41 |
| gene-LOC105280739 | https://www.ncbi.nlm.nih.gov/gene/?term=LOC105280739 | 1991 | 182 | 914.11 | 47.75 | 4.36 |
| gene-LOC105287811 | https://www.ncbi.nlm.nih.gov/gene/?term=LOC105287811 | 2155 | 246 | 1141.53 | 38.2 | 4.36 |
| gene-LOC105275314 | https://www.ncbi.nlm.nih.gov/gene/?term=LOC105275314 | 1489 | 193 | 1296.17 | 33.12 | 4.29 |
| gene-LOC105274534 | https://www.ncbi.nlm.nih.gov/gene/?term=LOC105274534 | 1329 | 152 | 1143.72 | 37.41 | 4.28 |
| gene-LOC105285635 | https://www.ncbi.nlm.nih.gov/gene/?term=LOC105285635 | 1175 | 194 | 1651.06 | 25.86 | 4.27 |
| gene-LOC105284659 | https://www.ncbi.nlm.nih.gov/gene/?term=LOC105284659 | 1988 | 224 | 1126.76 | 35.93 | 4.05 |
| gene-LOC105286235 | https://www.ncbi.nlm.nih.gov/gene/?term=LOC105286235 | 1510 | 148 | 980.13 | 41.14 | 4.03 |
| gene-LOC105279187 | https://www.ncbi.nlm.nih.gov/gene/?term=LOC105279187 | 2255 | 236 | 1046.56 | 38.33 | 4.01 |
| gene-LOC105285392 | https://www.ncbi.nlm.nih.gov/gene/?term=LOC105285392 | 1601 | 230 | 1436.6 | 27.26 | 3.92 |
| gene-LOC105274863 | https://www.ncbi.nlm.nih.gov/gene/?term=LOC105274863 | 2035 | 182 | 894.35 | 43.72 | 3.91 |

...

... only first 25 lines shown ...

Directory: OGL File: rnk-dmg-Ob_LIp_BCphase.vs.Rphase.txt

| gene_ID | gene_link | gwidth | #Sites | #per10Kb | #dmSites | #dmsp10kb | %dmSites | %pSite1 | %pSite2 | DMpSite | ADMpSite | DMpNucl | ADMpNucl |
|---|---|---|---|---|---|---|---|---|---|---|---|---|---|
| gene-Trnaf-gaa-2 | https://www.ncbi.nlm.nih.gov/gene/?term=Trnaf-gaa | 73 | 3 | 410.96 | 1 | 136.99 | 33.33 | 10 | 3.46 | 6.54 | 6.54 | 0.27 | 0.27 |
| gene-LOC105279226 | https://www.ncbi.nlm.nih.gov/gene/?term=LOC105279226 | 782 | 28 | 358.06 | 1 | 12.79 | 3.57 | 19.64 | 23.14 | -3.5 | 3.5 | -0.13 | 0.13 |
| gene-LOC105281507 | https://www.ncbi.nlm.nih.gov/gene/?term=LOC105281507 | 881 | 24 | 272.42 | 1 | 11.35 | 4.17 | 24.55 | 20.73 | 3.82 | 3.82 | 0.1 | 0.1 |
| gene-LOC105284474 | https://www.ncbi.nlm.nih.gov/gene/?term=LOC105284474 | 479 | 14 | 292.28 | 1 | 20.88 | 7.14 | 19.98 | 23.06 | -3.08 | 3.08 | -0.09 | 0.09 |
| gene-LOC105275947 | https://www.ncbi.nlm.nih.gov/gene/?term=LOC105275947 | 940 | 24 | 255.32 | 1 | 10.64 | 4.17 | 10.71 | 14.08 | -3.37 | 3.37 | -0.09 | 0.09 |
| gene-LOC105283800 | https://www.ncbi.nlm.nih.gov/gene/?term=LOC105283800 | 871 | 24 | 275.55 | 1 | 11.48 | 4.17 | 24.17 | 21.18 | 2.99 | 2.99 | 0.08 | 0.08 |
| gene-LOC105275219 | https://www.ncbi.nlm.nih.gov/gene/?term=LOC105275219 | 997 | 36 | 361.08 | 1 | 10.03 | 2.78 | 41.71 | 43.84 | -2.13 | 2.13 | -0.08 | 0.08 |
| gene-LOC105283817 | https://www.ncbi.nlm.nih.gov/gene/?term=LOC105283817 | 1309 | 19 | 145.15 | 3 | 22.92 | 15.79 | 55.23 | 50.62 | 4.61 | 4.61 | 0.07 | 0.07 |
| gene-LOC105279725 | https://www.ncbi.nlm.nih.gov/gene/?term=LOC105279725 | 1608 | 101 | 628.11 | 2 | 12.44 | 1.98 | 19.93 | 18.85 | 1.09 | 1.09 | 0.07 | 0.07 |
| gene-LOC105278790 | https://www.ncbi.nlm.nih.gov/gene/?term=LOC105278790 | 1179 | 85 | 720.95 | 1 | 8.48 | 1.18 | 23.27 | 22.29 | 0.98 | 0.98 | 0.07 | 0.07 |
| gene-LOC105285817 | https://www.ncbi.nlm.nih.gov/gene/?term=LOC105285817 | 1417 | 69 | 486.94 | 1 | 7.06 | 1.45 | 16.72 | 18.19 | -1.47 | 1.47 | -0.07 | 0.07 |
| gene-LOC105277322 | https://www.ncbi.nlm.nih.gov/gene/?term=LOC105277322 | 1663 | 62 | 372.82 | 1 | 6.01 | 1.61 | 39.23 | 41.07 | -1.84 | 1.84 | -0.07 | 0.07 |
| gene-LOC105276706 | https://www.ncbi.nlm.nih.gov/gene/?term=LOC105276706 | 1440 | 35 | 243.06 | 1 | 6.94 | 2.86 | 50.39 | 53.13 | -2.74 | 2.74 | -0.07 | 0.07 |
| gene-LOC109611472 | https://www.ncbi.nlm.nih.gov/gene/?term=LOC109611472 | 1351 | 30 | 222.06 | 1 | 7.4 | 3.33 | 36.59 | 39.57 | -2.98 | 2.98 | -0.07 | 0.07 |
| gene-LOC105283856 | https://www.ncbi.nlm.nih.gov/gene/?term=LOC105283856 | 1988 | 18 | 90.54 | 3 | 15.09 | 16.67 | 37.55 | 31.12 | 6.44 | 6.44 | 0.06 | 0.06 |
| gene-LOC105285429 | https://www.ncbi.nlm.nih.gov/gene/?term=LOC105285429 | 1324 | 36 | 271.9 | 2 | 15.11 | 5.56 | 41.65 | 39.56 | 2.09 | 2.09 | 0.06 | 0.06 |
| gene-LOC105286157 | https://www.ncbi.nlm.nih.gov/gene/?term=LOC105286157 | 2004 | 73 | 364.27 | 2 | 9.98 | 2.74 | 25.32 | 23.75 | 1.57 | 1.57 | 0.06 | 0.06 |
| gene-LOC105281604 | https://www.ncbi.nlm.nih.gov/gene/?term=LOC105281604 | 2611 | 132 | 505.55 | 2 | 7.66 | 1.52 | 15 | 13.88 | 1.12 | 1.12 | 0.06 | 0.06 |
| gene-LOC105275539 | https://www.ncbi.nlm.nih.gov/gene/?term=LOC105275539 | 1272 | 67 | 526.73 | 1 | 7.86 | 1.49 | 29.6 | 28.55 | 1.06 | 1.06 | 0.06 | 0.06 |
| gene-LOC105283758 | https://www.ncbi.nlm.nih.gov/gene/?term=LOC105283758 | 1570 | 92 | 585.99 | 1 | 6.37 | 1.09 | 23.95 | 22.96 | 0.99 | 0.99 | 0.06 | 0.06 |
| gene-LOC105282952 | https://www.ncbi.nlm.nih.gov/gene/?term=LOC105282952 | 1699 | 112 | 659.21 | 1 | 5.89 | 0.89 | 15.29 | 16.19 | -0.89 | 0.89 | -0.06 | 0.06 |
| gene-LOC105274993 | https://www.ncbi.nlm.nih.gov/gene/?term=LOC105274993 | 2616 | 171 | 653.67 | 1 | 3.82 | 0.58 | 12.06 | 13 | -0.94 | 0.94 | -0.06 | 0.06 |
| gene-LOC105279437 | https://www.ncbi.nlm.nih.gov/gene/?term=LOC105279437 | 1069 | 70 | 654.82 | 1 | 9.35 | 1.43 | 21.76 | 22.71 | -0.95 | 0.95 | -0.06 | 0.06 |
| gene-LOC105276976 | https://www.ncbi.nlm.nih.gov/gene/?term=LOC105276976 | 1829 | 96 | 524.88 | 1 | 5.47 | 1.04 | 11.12 | 12.26 | -1.13 | 1.13 | -0.06 | 0.06 |

...

... only first 25 lines shown ...

```
Directory: OGL File: wrt-Ob_LIp.txt

... comparing  BCphase.vs.Rphase ...

Number of genes <=  20000  and with #dmsites >=  1 :  1337
sample1     sum of percent methylation: 20526.26     average:    15.35
sample2     sum of percent methylation: 20685.15     average:    15.47

sum difference:  -158.93      average difference:  -0.1189

      Wilcoxon signed rank test with continuity correction

data:  pw_summary$`%pSite1` and pw_summary$`%pSite2`
V = 3e+05, p-value <2e-16
alternative hypothesis: true location shift is not equal to 0
```

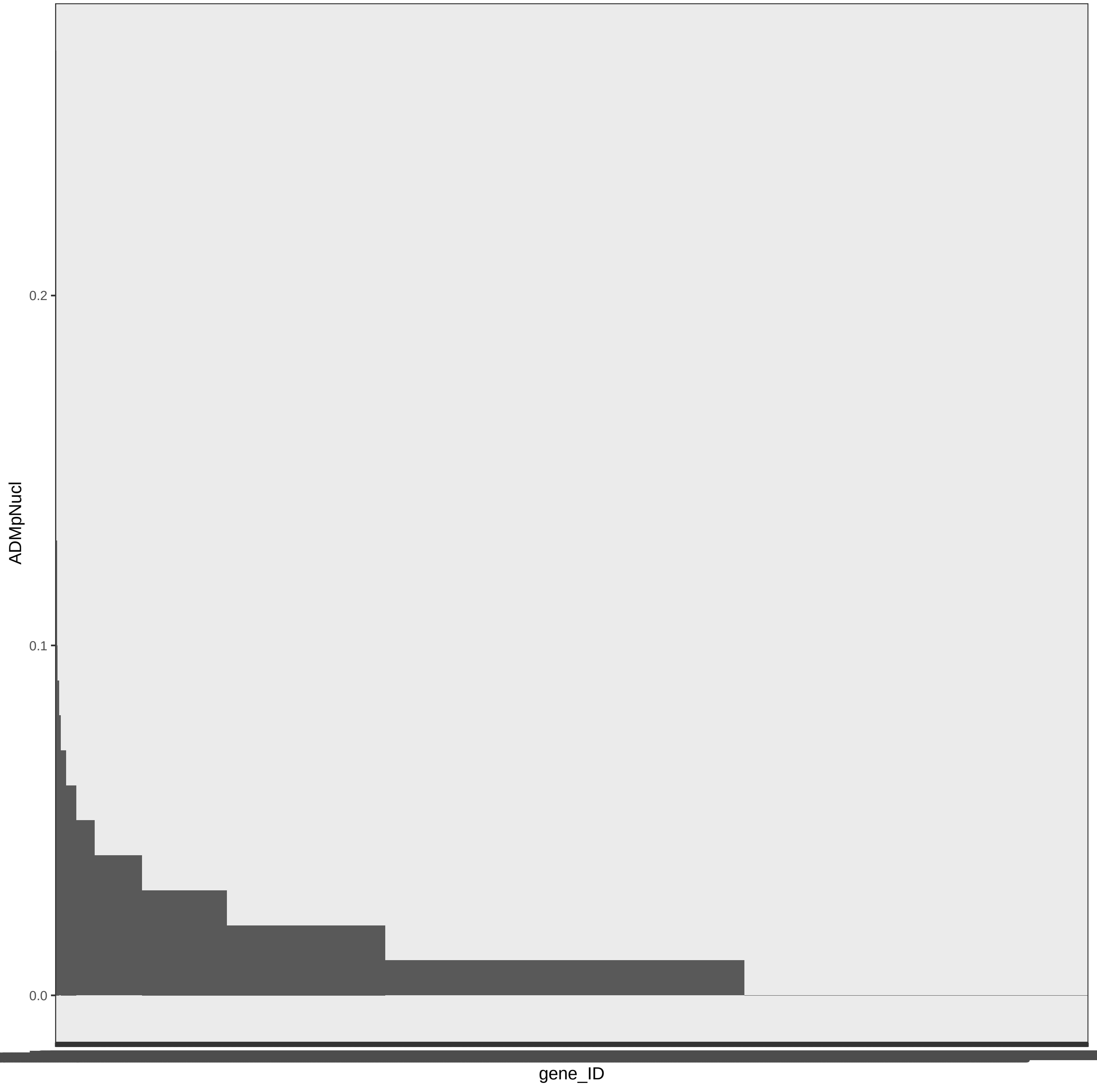

ADMpNucl
0.2
0.1
0.0
gene_ID

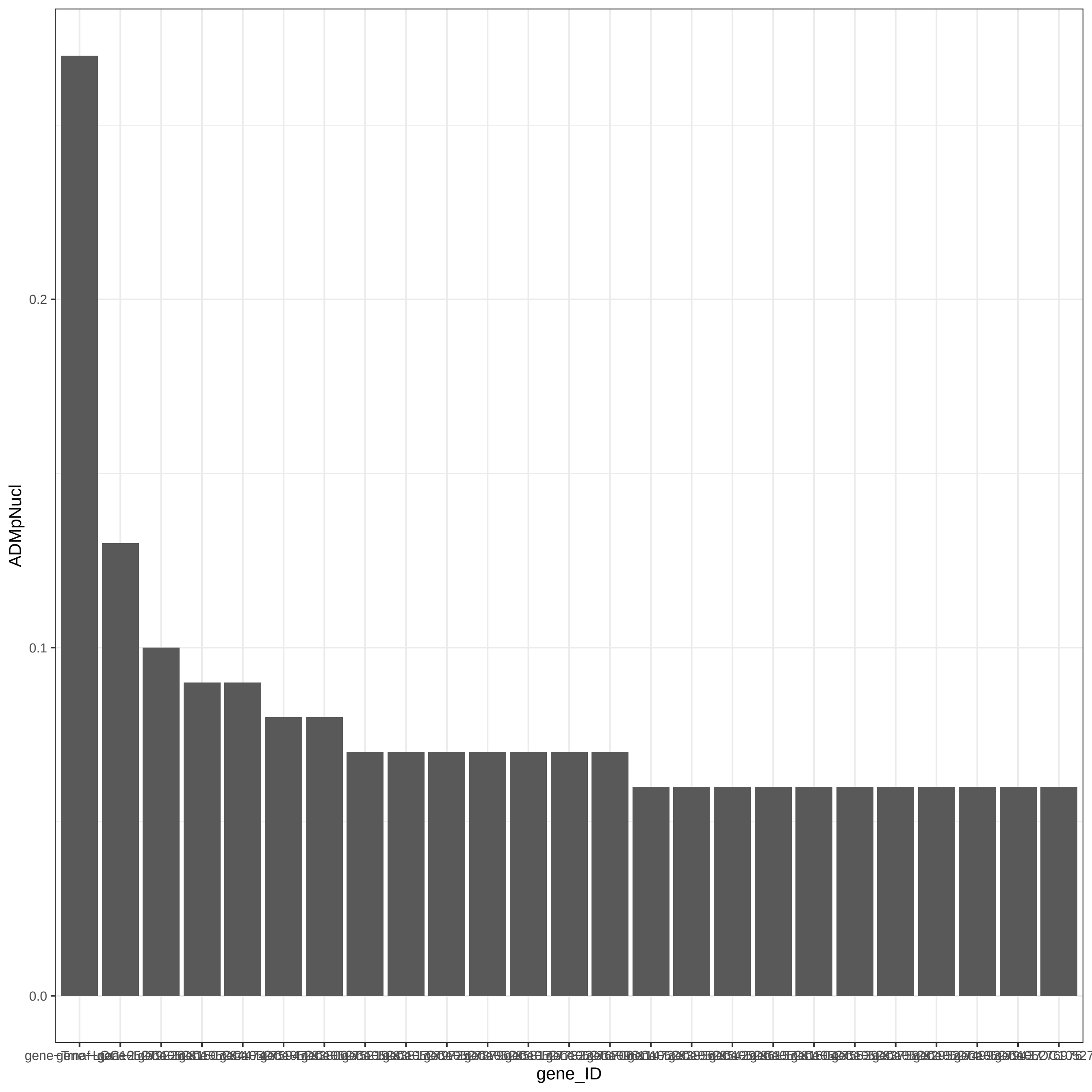

ADMpNucl
0.0
0.1
0.2
gene_ID

```
Directory: OGLall File: 0READMEogl

OGL - Ordered gene lists.

Input: studyhc annotation   maxgwidth minnbrdmsites
       dmgprp (= output of BWASPR:show_dmsg())

Output: files ogl-*.txt rnk-dmg-*.txt
         ogl-<study>_<sample>.txt ogl-<study>_<sample1>.vs.<sample2>.txt
         rnk-dmg--<study>_<sample1>.vs.<sample2>.txt rnk-dmg--<study>_<sample1>.vs.<sample2>.pdf
         wrt-<study>.txt

Notes: Output files ogl-<study>_<sample>.txt give tables for each sample with columns

        gene_ID gwidth #Sites #per10Kb %perSite %pNucl

       ordered by %pNucl.  If available, a link to an NCBI entry of gene_ID is inserted as second column.
       Abbreviations used: %perSite, percent methylation per site
                           %perNucl, percent methylation per nucleotide of the gene

       Output files ogl-<study>_<sample1>.vs.<sample2>.txt give tables for each comparison with columns

        gene_ID gwidth #Sites #per10Kb #dmSites #dmsp10kb %dmSites %pSite1 %pSite2 DMpSite ADMpSite DMpNucl ADMpNucl

       ordered by DMpSite.  If available, a link to an NCBI entry of gene_ID is inserted as second column.
       Abbreviations used: %dmSites, percent sites that are differentially methylated
                           %pSite1, average per site % methylation for sample1
                           %pSite2, average per site % methylation for sample2
                           DMpSite, average per site difference in % methylation between sample1 and sample2
                           ADMpSite, absolute value of DMpSite
                           DMpNucl, average per nucleotide difference in % methylation between sample1 and sample2
                           ADMpNucl, absolute value of DMpNucl

       Output files rnk-dmg-<study>_<sample1>.vs.<sample2>.txt are equivalent to files
        ogl-<study>_<sample1>.vs.<sample2>.txt but ordered by ADMpNucl.
       Output files rnk-dmg-<study>_<sample1>.vs.<sample2>.pdf provide visualization of the distribution
        of ADMpNucl values.

       Output file wrt-<study>.txt gives results of the Wilcoxon signed rank test comparing
        the %pSite1 and %pSite2 vectors

       In output directory OGL, tables are restricted to genes with gwidth <= maxgwidth and, for
        pairwise comparisons, #dmSites >= minnbrdmsites

       If either maxgwidth or minnbrdmsites is set, then output directory OGLall will show the full tables
        with all genes (for reference).
```

Directory: OGLall File: ogl-Ob_LIp_BCphase.txt

| gene_ID | gene_link | gwidth | #Sites | #per10Kb | %perSite | %pNucl |
|---|---|---|---|---|---|---|
| gene-LOC113561532 | https://www.ncbi.nlm.nih.gov/gene/?term=LOC113561532 | 72 | 6 | 833.33 | 95.81 | 7.98 |
| gene-LOC113562627 | https://www.ncbi.nlm.nih.gov/gene/?term=LOC113562627 | 397 | 37 | 931.99 | 69.15 | 6.45 |
| gene-LOC105285394 | https://www.ncbi.nlm.nih.gov/gene/?term=LOC105285394 | 1307 | 147 | 1124.71 | 50.61 | 5.69 |
| gene-LOC105282149 | https://www.ncbi.nlm.nih.gov/gene/?term=LOC105282149 | 1787 | 213 | 1191.94 | 42.72 | 5.09 |
| gene-LOC105283830 | https://www.ncbi.nlm.nih.gov/gene/?term=LOC105283830 | 252 | 34 | 1349.21 | 36.78 | 4.96 |
| gene-LOC105276360 | https://www.ncbi.nlm.nih.gov/gene/?term=LOC105276360 | 1466 | 237 | 1616.64 | 28.53 | 4.61 |
| gene-LOC105275231 | https://www.ncbi.nlm.nih.gov/gene/?term=LOC105275231 | 1270 | 149 | 1173.23 | 39.05 | 4.58 |
| gene-LOC105278849 | https://www.ncbi.nlm.nih.gov/gene/?term=LOC105278849 | 776 | 151 | 1945.88 | 23.52 | 4.58 |
| gene-LOC105277648 | https://www.ncbi.nlm.nih.gov/gene/?term=LOC105277648 | 722 | 53 | 734.07 | 62.32 | 4.57 |
| gene-LOC105286531 | https://www.ncbi.nlm.nih.gov/gene/?term=LOC105286531 | 372 | 46 | 1236.56 | 36.89 | 4.56 |
| gene-LOC105286269 | https://www.ncbi.nlm.nih.gov/gene/?term=LOC105286269 | 1401 | 156 | 1113.49 | 40.37 | 4.49 |
| gene-LOC105286562 | https://www.ncbi.nlm.nih.gov/gene/?term=LOC105286562 | 1484 | 129 | 869.27 | 51.38 | 4.47 |
| gene-LOC105274711 | https://www.ncbi.nlm.nih.gov/gene/?term=LOC105274711 | 1655 | 165 | 996.98 | 44.69 | 4.46 |
| gene-LOC105286801 | https://www.ncbi.nlm.nih.gov/gene/?term=LOC105286801 | 1102 | 120 | 1088.93 | 40.66 | 4.43 |
| gene-LOC105280739 | https://www.ncbi.nlm.nih.gov/gene/?term=LOC105280739 | 1991 | 182 | 914.11 | 47.53 | 4.34 |
| gene-LOC105287811 | https://www.ncbi.nlm.nih.gov/gene/?term=LOC105287811 | 2155 | 245 | 1136.89 | 38.05 | 4.33 |
| gene-LOC105285635 | https://www.ncbi.nlm.nih.gov/gene/?term=LOC105285635 | 1175 | 191 | 1625.53 | 26.28 | 4.27 |
| gene-LOC105275314 | https://www.ncbi.nlm.nih.gov/gene/?term=LOC105275314 | 1489 | 189 | 1269.31 | 33.05 | 4.19 |
| gene-LOC105279187 | https://www.ncbi.nlm.nih.gov/gene/?term=LOC105279187 | 2255 | 238 | 1055.43 | 39.14 | 4.13 |
| gene-LOC105286235 | https://www.ncbi.nlm.nih.gov/gene/?term=LOC105286235 | 1510 | 148 | 980.13 | 42.06 | 4.12 |
| gene-LOC105274534 | https://www.ncbi.nlm.nih.gov/gene/?term=LOC105274534 | 1329 | 148 | 1113.62 | 36.91 | 4.11 |
| gene-LOC105285392 | https://www.ncbi.nlm.nih.gov/gene/?term=LOC105285392 | 1601 | 230 | 1436.6 | 28.09 | 4.04 |
| gene-LOC105284659 | https://www.ncbi.nlm.nih.gov/gene/?term=LOC105284659 | 1988 | 217 | 1091.55 | 36.25 | 3.96 |
| gene-LOC105274863 | https://www.ncbi.nlm.nih.gov/gene/?term=LOC105274863 | 2035 | 178 | 874.69 | 44.78 | 3.92 |

...

... only first 25 lines shown ...

Directory: OGLall File: ogl-Ob_LIp_BCphase.vs.Rphase.txt

| gene_ID | gene_link | gwidth | #Sites | #per10Kb | #dmSites | #dmsp10kb | %dmSites | %pSite1 | %pSite2 | DMpSite | ADMpSite | DMpNucl | ADMpNucl |
|---|---|---|---|---|---|---|---|---|---|---|---|---|---|
| gene-LOC105282002 | https://www.ncbi.nlm.nih.gov/gene/?term=LOC105282002 | 1311 | 4 | 30.51 | 1 | 7.63 | 25 | 37.08 | 25.87 | 11.2 | 11.2 | 0.03 | 0.03 |
| gene-Trnaf-gaa-2 | https://www.ncbi.nlm.nih.gov/gene/?term=Trnaf-gaa | 73 | 3 | 410.96 | 1 | 136.99 | 33.33 | 10 | 3.46 | 6.54 | 6.54 | 0.27 | 0.27 |
| gene-LOC105283856 | https://www.ncbi.nlm.nih.gov/gene/?term=LOC105283856 | 1988 | 18 | 90.54 | 3 | 15.09 | 16.67 | 37.55 | 31.12 | 6.44 | 6.44 | 0.06 | 0.06 |
| gene-LOC105276566 | https://www.ncbi.nlm.nih.gov/gene/?term=LOC105276566 | 1114 | 10 | 89.77 | 2 | 17.95 | 20 | 40.73 | 35.89 | 4.84 | 4.84 | 0.04 | 0.04 |
| gene-LOC105283817 | https://www.ncbi.nlm.nih.gov/gene/?term=LOC105283817 | 1309 | 19 | 145.15 | 3 | 22.92 | 15.79 | 55.23 | 50.62 | 4.61 | 4.61 | 0.07 | 0.07 |
| gene-LOC105287296 | https://www.ncbi.nlm.nih.gov/gene/?term=LOC105287296 | 1834 | 8 | 43.62 | 1 | 5.45 | 12.5 | 36.82 | 32.76 | 4.06 | 4.06 | 0.02 | 0.02 |
| gene-LOC105286178 | https://www.ncbi.nlm.nih.gov/gene/?term=LOC105286178 | 1590 | 13 | 81.76 | 1 | 6.29 | 7.69 | 65.07 | 61.17 | 3.91 | 3.91 | 0.03 | 0.03 |
| gene-LOC105275097 | https://www.ncbi.nlm.nih.gov/gene/?term=LOC105275097 | 1071 | 9 | 84.03 | 1 | 9.34 | 11.11 | 49.2 | 45.35 | 3.85 | 3.85 | 0.03 | 0.03 |
| gene-LOC105281507 | https://www.ncbi.nlm.nih.gov/gene/?term=LOC105281507 | 881 | 24 | 272.42 | 1 | 11.35 | 4.17 | 24.55 | 20.73 | 3.82 | 3.82 | 0.1 | 0.1 |
| gene-LOC105283800 | https://www.ncbi.nlm.nih.gov/gene/?term=LOC105283800 | 871 | 24 | 275.55 | 1 | 11.48 | 4.17 | 24.17 | 21.18 | 2.99 | 2.99 | 0.08 | 0.08 |
| gene-LOC113563090 | https://www.ncbi.nlm.nih.gov/gene/?term=LOC113563090 | 1729 | 22 | 127.24 | 2 | 11.57 | 9.09 | 57.11 | 54.13 | 2.98 | 2.98 | 0.04 | 0.04 |
| gene-LOC105287152 | https://www.ncbi.nlm.nih.gov/gene/?term=LOC105287152 | 1616 | 25 | 154.7 | 1 | 6.19 | 4 | 29.08 | 26.72 | 2.36 | 2.36 | 0.04 | 0.04 |
| gene-LOC105287372 | https://www.ncbi.nlm.nih.gov/gene/?term=LOC105287372 | 2659 | 50 | 188.04 | 1 | 3.76 | 2 | 10.93 | 8.75 | 2.18 | 2.18 | 0.04 | 0.04 |
| gene-LOC105285429 | https://www.ncbi.nlm.nih.gov/gene/?term=LOC105285429 | 1324 | 36 | 271.9 | 2 | 15.11 | 5.56 | 41.65 | 39.56 | 2.09 | 2.09 | 0.06 | 0.06 |
| gene-LOC105279501 | https://www.ncbi.nlm.nih.gov/gene/?term=LOC105279501 | 2114 | 40 | 189.21 | 1 | 4.73 | 2.5 | 51.95 | 49.94 | 2.01 | 2.01 | 0.04 | 0.04 |
| gene-LOC105275378 | https://www.ncbi.nlm.nih.gov/gene/?term=LOC105275378 | 1189 | 35 | 294.37 | 1 | 8.41 | 2.86 | 45.72 | 43.88 | 1.85 | 1.85 | 0.05 | 0.05 |
| gene-LOC113561875 | https://www.ncbi.nlm.nih.gov/gene/?term=LOC113561875 | 579 | 16 | 276.34 | 1 | 17.27 | 6.25 | 6.85 | 5.16 | 1.69 | 1.69 | 0.05 | 0.05 |
| gene-LOC105281162 | https://www.ncbi.nlm.nih.gov/gene/?term=LOC105281162 | 2174 | 27 | 124.2 | 1 | 4.6 | 3.7 | 32.48 | 30.86 | 1.62 | 1.62 | 0.02 | 0.02 |
| gene-LOC105287127 | https://www.ncbi.nlm.nih.gov/gene/?term=LOC105287127 | 2308 | 71 | 307.63 | 1 | 4.33 | 1.41 | 6.04 | 4.42 | 1.62 | 1.62 | 0.05 | 0.05 |
| gene-LOC105286157 | https://www.ncbi.nlm.nih.gov/gene/?term=LOC105286157 | 2004 | 73 | 364.27 | 2 | 9.98 | 2.74 | 25.32 | 23.75 | 1.57 | 1.57 | 0.06 | 0.06 |
| gene-LOC105276157 | https://www.ncbi.nlm.nih.gov/gene/?term=LOC105276157 | 1601 | 44 | 274.83 | 1 | 6.25 | 2.27 | 26.48 | 24.92 | 1.56 | 1.56 | 0.04 | 0.04 |
| gene-LOC105280380 | https://www.ncbi.nlm.nih.gov/gene/?term=LOC105280380 | 1412 | 38 | 269.12 | 1 | 7.08 | 2.63 | 6.6 | 5.05 | 1.55 | 1.55 | 0.04 | 0.04 |
| gene-LOC105285707 | https://www.ncbi.nlm.nih.gov/gene/?term=LOC105285707 | 2096 | 71 | 338.74 | 1 | 4.77 | 1.41 | 31.72 | 30.19 | 1.53 | 1.53 | 0.05 | 0.05 |
| gene-LOC105283991 | https://www.ncbi.nlm.nih.gov/gene/?term=LOC105283991 | 2852 | 59 | 206.87 | 1 | 3.51 | 1.69 | 33.92 | 32.4 | 1.52 | 1.52 | 0.03 | 0.03 |

...

... only first 25 lines shown ...

Directory: OGLall File: ogl-Ob_LIp_Rphase.txt

| gene_ID | gene_link | gwidth | #Sites | #per10Kb | %perSite | %pNucl |
|---|---|---|---|---|---|---|
| gene-LOC113561532 | https://www.ncbi.nlm.nih.gov/gene/?term=LOC113561532 | 72 | 7 | 972.22 | 96.09 | 9.34 |
| gene-LOC113562627 | https://www.ncbi.nlm.nih.gov/gene/?term=LOC113562627 | 397 | 37 | 931.99 | 69.26 | 6.46 |
| gene-LOC105285394 | https://www.ncbi.nlm.nih.gov/gene/?term=LOC105285394 | 1307 | 148 | 1132.36 | 51.87 | 5.87 |
| gene-LOC105282149 | https://www.ncbi.nlm.nih.gov/gene/?term=LOC105282149 | 1787 | 217 | 1214.33 | 43.31 | 5.26 |
| gene-LOC105283830 | https://www.ncbi.nlm.nih.gov/gene/?term=LOC105283830 | 252 | 34 | 1349.21 | 37.52 | 5.06 |
| gene-LOC105286801 | https://www.ncbi.nlm.nih.gov/gene/?term=LOC105286801 | 1102 | 131 | 1188.75 | 38.91 | 4.62 |
| gene-LOC105276360 | https://www.ncbi.nlm.nih.gov/gene/?term=LOC105276360 | 1466 | 237 | 1616.64 | 28.5 | 4.61 |
| gene-LOC105278849 | https://www.ncbi.nlm.nih.gov/gene/?term=LOC105278849 | 776 | 151 | 1945.88 | 23.46 | 4.57 |
| gene-LOC105275231 | https://www.ncbi.nlm.nih.gov/gene/?term=LOC105275231 | 1270 | 151 | 1188.98 | 38.3 | 4.55 |
| gene-LOC105286269 | https://www.ncbi.nlm.nih.gov/gene/?term=LOC105286269 | 1401 | 155 | 1106.35 | 40.98 | 4.53 |
| gene-LOC105286531 | https://www.ncbi.nlm.nih.gov/gene/?term=LOC105286531 | 372 | 46 | 1236.56 | 36.39 | 4.5 |
| gene-LOC105286562 | https://www.ncbi.nlm.nih.gov/gene/?term=LOC105286562 | 1484 | 135 | 909.7 | 49.22 | 4.48 |
| gene-LOC105274711 | https://www.ncbi.nlm.nih.gov/gene/?term=LOC105274711 | 1655 | 169 | 1021.15 | 43.22 | 4.41 |
| gene-LOC105277648 | https://www.ncbi.nlm.nih.gov/gene/?term=LOC105277648 | 722 | 52 | 720.22 | 61.24 | 4.41 |
| gene-LOC105280739 | https://www.ncbi.nlm.nih.gov/gene/?term=LOC105280739 | 1991 | 182 | 914.11 | 47.75 | 4.36 |
| gene-LOC105287811 | https://www.ncbi.nlm.nih.gov/gene/?term=LOC105287811 | 2155 | 246 | 1141.53 | 38.2 | 4.36 |
| gene-LOC105275314 | https://www.ncbi.nlm.nih.gov/gene/?term=LOC105275314 | 1489 | 193 | 1296.17 | 33.12 | 4.29 |
| gene-LOC105274534 | https://www.ncbi.nlm.nih.gov/gene/?term=LOC105274534 | 1329 | 152 | 1143.72 | 37.41 | 4.28 |
| gene-LOC105285635 | https://www.ncbi.nlm.nih.gov/gene/?term=LOC105285635 | 1175 | 194 | 1651.06 | 25.86 | 4.27 |
| gene-LOC105284659 | https://www.ncbi.nlm.nih.gov/gene/?term=LOC105284659 | 1988 | 224 | 1126.76 | 35.93 | 4.05 |
| gene-LOC105286235 | https://www.ncbi.nlm.nih.gov/gene/?term=LOC105286235 | 1510 | 148 | 980.13 | 41.14 | 4.03 |
| gene-LOC105279187 | https://www.ncbi.nlm.nih.gov/gene/?term=LOC105279187 | 2255 | 236 | 1046.56 | 38.33 | 4.01 |
| gene-LOC105285392 | https://www.ncbi.nlm.nih.gov/gene/?term=LOC105285392 | 1601 | 230 | 1436.6 | 27.26 | 3.92 |
| gene-LOC105274863 | https://www.ncbi.nlm.nih.gov/gene/?term=LOC105274863 | 2035 | 182 | 894.35 | 43.72 | 3.91 |

...

... only first 25 lines shown ...

Directory: OGLall File: rnk-dmg-Ob_LIp_BCphase.vs.Rphase.txt

| gene_ID | gene_link | gwidth | #Sites | #per10Kb | #dmSites | #dmsp10kb | %dmSites | %pSite1 | %pSite2 | DMpSite | ADMpSite | DMpNucl | ADMpNucl |
|---|---|---|---|---|---|---|---|---|---|---|---|---|---|
| gene-Trnaf-gaa-2 | https://www.ncbi.nlm.nih.gov/gene/?term=Trnaf-gaa | 73 | 3 | 410.96 | 1 | 136.99 | 33.33 | 10 | 3.46 | 6.54 | 6.54 | 0.27 | 0.27 |
| gene-LOC105279226 | https://www.ncbi.nlm.nih.gov/gene/?term=LOC105279226 | 782 | 28 | 358.06 | 1 | 12.79 | 3.57 | 19.64 | 23.14 | -3.5 | 3.5 | -0.13 | 0.13 |
| gene-LOC105281507 | https://www.ncbi.nlm.nih.gov/gene/?term=LOC105281507 | 881 | 24 | 272.42 | 1 | 11.35 | 4.17 | 24.55 | 20.73 | 3.82 | 3.82 | 0.1 | 0.1 |
| gene-LOC105284474 | https://www.ncbi.nlm.nih.gov/gene/?term=LOC105284474 | 479 | 14 | 292.28 | 1 | 20.88 | 7.14 | 19.98 | 23.06 | -3.08 | 3.08 | -0.09 | 0.09 |
| gene-LOC105275947 | https://www.ncbi.nlm.nih.gov/gene/?term=LOC105275947 | 940 | 24 | 255.32 | 1 | 10.64 | 4.17 | 10.71 | 14.08 | -3.37 | 3.37 | -0.09 | 0.09 |
| gene-LOC105283800 | https://www.ncbi.nlm.nih.gov/gene/?term=LOC105283800 | 871 | 24 | 275.55 | 1 | 11.48 | 4.17 | 24.17 | 21.18 | 2.99 | 2.99 | 0.08 | 0.08 |
| gene-LOC105275219 | https://www.ncbi.nlm.nih.gov/gene/?term=LOC105275219 | 997 | 36 | 361.08 | 1 | 10.03 | 2.78 | 41.71 | 43.84 | -2.13 | 2.13 | -0.08 | 0.08 |
| gene-LOC105283817 | https://www.ncbi.nlm.nih.gov/gene/?term=LOC105283817 | 1309 | 19 | 145.15 | 3 | 22.92 | 15.79 | 55.23 | 50.62 | 4.61 | 4.61 | 0.07 | 0.07 |
| gene-LOC105279725 | https://www.ncbi.nlm.nih.gov/gene/?term=LOC105279725 | 1608 | 101 | 628.11 | 2 | 12.44 | 1.98 | 19.93 | 18.85 | 1.09 | 1.09 | 0.07 | 0.07 |
| gene-LOC105278790 | https://www.ncbi.nlm.nih.gov/gene/?term=LOC105278790 | 1179 | 85 | 720.95 | 1 | 8.48 | 1.18 | 23.27 | 22.29 | 0.98 | 0.98 | 0.07 | 0.07 |
| gene-LOC105285817 | https://www.ncbi.nlm.nih.gov/gene/?term=LOC105285817 | 1417 | 69 | 486.94 | 1 | 7.06 | 1.45 | 16.72 | 18.19 | -1.47 | 1.47 | -0.07 | 0.07 |
| gene-LOC105277322 | https://www.ncbi.nlm.nih.gov/gene/?term=LOC105277322 | 1663 | 62 | 372.82 | 1 | 6.01 | 1.61 | 39.23 | 41.07 | -1.84 | 1.84 | -0.07 | 0.07 |
| gene-LOC105276706 | https://www.ncbi.nlm.nih.gov/gene/?term=LOC105276706 | 1440 | 35 | 243.06 | 1 | 6.94 | 2.86 | 50.39 | 53.13 | -2.74 | 2.74 | -0.07 | 0.07 |
| gene-LOC109611472 | https://www.ncbi.nlm.nih.gov/gene/?term=LOC109611472 | 1351 | 30 | 222.06 | 1 | 7.4 | 3.33 | 36.59 | 39.57 | -2.98 | 2.98 | -0.07 | 0.07 |
| gene-LOC105283856 | https://www.ncbi.nlm.nih.gov/gene/?term=LOC105283856 | 1988 | 18 | 90.54 | 3 | 15.09 | 16.67 | 37.55 | 31.12 | 6.44 | 6.44 | 0.06 | 0.06 |
| gene-LOC105285429 | https://www.ncbi.nlm.nih.gov/gene/?term=LOC105285429 | 1324 | 36 | 271.9 | 2 | 15.11 | 5.56 | 41.65 | 39.56 | 2.09 | 2.09 | 0.06 | 0.06 |
| gene-LOC105286157 | https://www.ncbi.nlm.nih.gov/gene/?term=LOC105286157 | 2004 | 73 | 364.27 | 2 | 9.98 | 2.74 | 25.32 | 23.75 | 1.57 | 1.57 | 0.06 | 0.06 |
| gene-LOC105281604 | https://www.ncbi.nlm.nih.gov/gene/?term=LOC105281604 | 2611 | 132 | 505.55 | 2 | 7.66 | 1.52 | 15 | 13.88 | 1.12 | 1.12 | 0.06 | 0.06 |
| gene-LOC105275539 | https://www.ncbi.nlm.nih.gov/gene/?term=LOC105275539 | 1272 | 67 | 526.73 | 1 | 7.86 | 1.49 | 29.6 | 28.55 | 1.06 | 1.06 | 0.06 | 0.06 |
| gene-LOC105283758 | https://www.ncbi.nlm.nih.gov/gene/?term=LOC105283758 | 1570 | 92 | 585.99 | 1 | 6.37 | 1.09 | 23.95 | 22.96 | 0.99 | 0.99 | 0.06 | 0.06 |
| gene-LOC105282952 | https://www.ncbi.nlm.nih.gov/gene/?term=LOC105282952 | 1699 | 112 | 659.21 | 1 | 5.89 | 0.89 | 15.29 | 16.19 | -0.89 | 0.89 | -0.06 | 0.06 |
| gene-LOC105274993 | https://www.ncbi.nlm.nih.gov/gene/?term=LOC105274993 | 2616 | 171 | 653.67 | 1 | 3.82 | 0.58 | 12.06 | 13 | -0.94 | 0.94 | -0.06 | 0.06 |
| gene-LOC105279437 | https://www.ncbi.nlm.nih.gov/gene/?term=LOC105279437 | 1069 | 70 | 654.82 | 1 | 9.35 | 1.43 | 21.76 | 22.71 | -0.95 | 0.95 | -0.06 | 0.06 |
| gene-LOC105276976 | https://www.ncbi.nlm.nih.gov/gene/?term=LOC105276976 | 1829 | 96 | 524.88 | 1 | 5.47 | 1.04 | 11.12 | 12.26 | -1.13 | 1.13 | -0.06 | 0.06 |

...

... only first 25 lines shown ...

```
Directory: OGLall File: wrt-Ob_LIp.txt

... comparing  BCphase.vs.Rphase ...

Number of genes with #dmsites >=  1 :  1618

sample1     sum of percent methylation: 20788.57     average:    12.85
sample2     sum of percent methylation: 20952.70     average:    12.95

sum difference:  -164.20      average difference:  -0.1015

      Wilcoxon signed rank test with continuity correction

data:  pw_summary$`%pSite1` and pw_summary$`%pSite2`
V = 4e+05, p-value <2e-16
alternative hypothesis: true location shift is not equal to 0
```

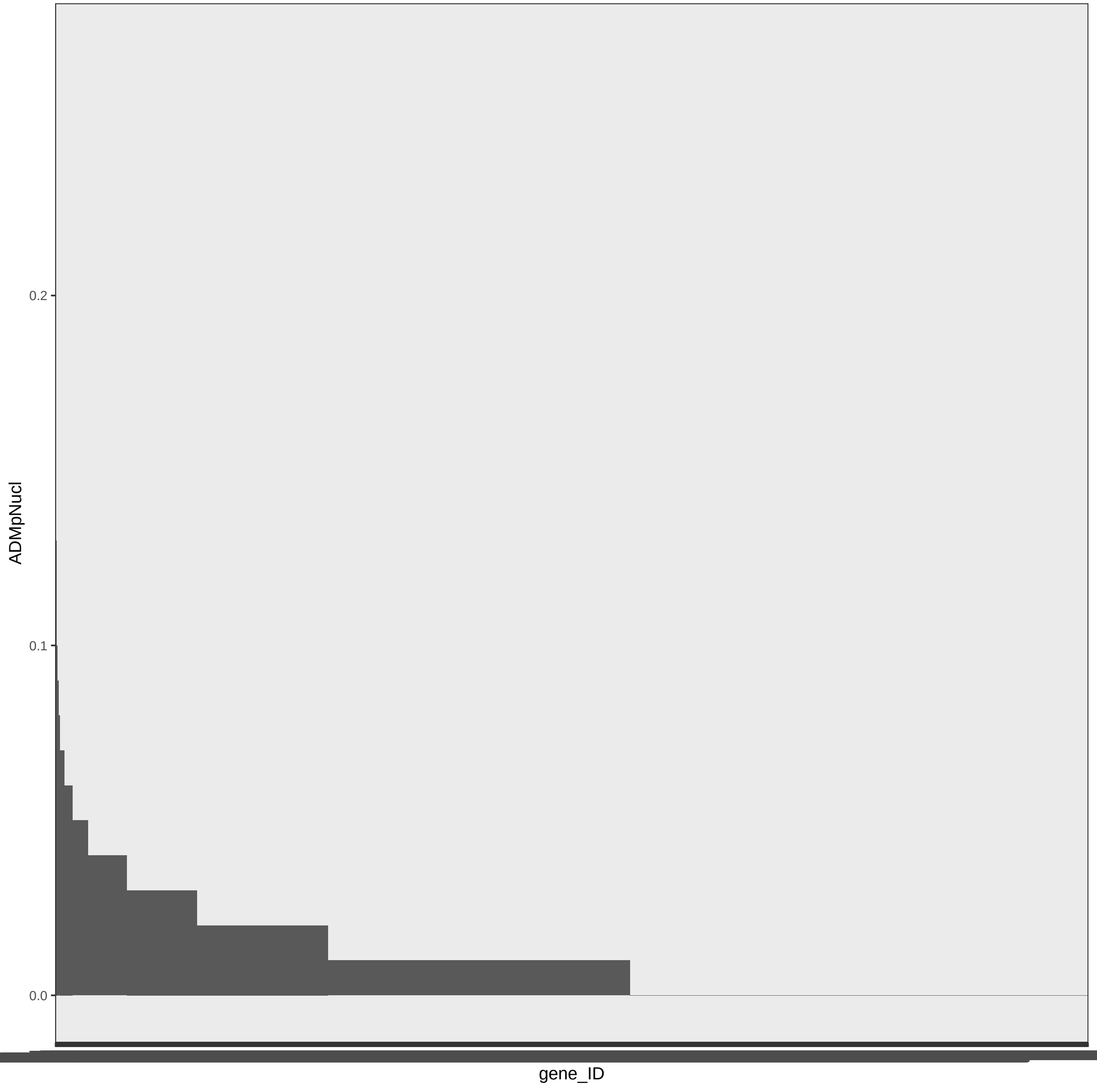

ADMpNucl
0.2
0.1
0.0
gene_ID

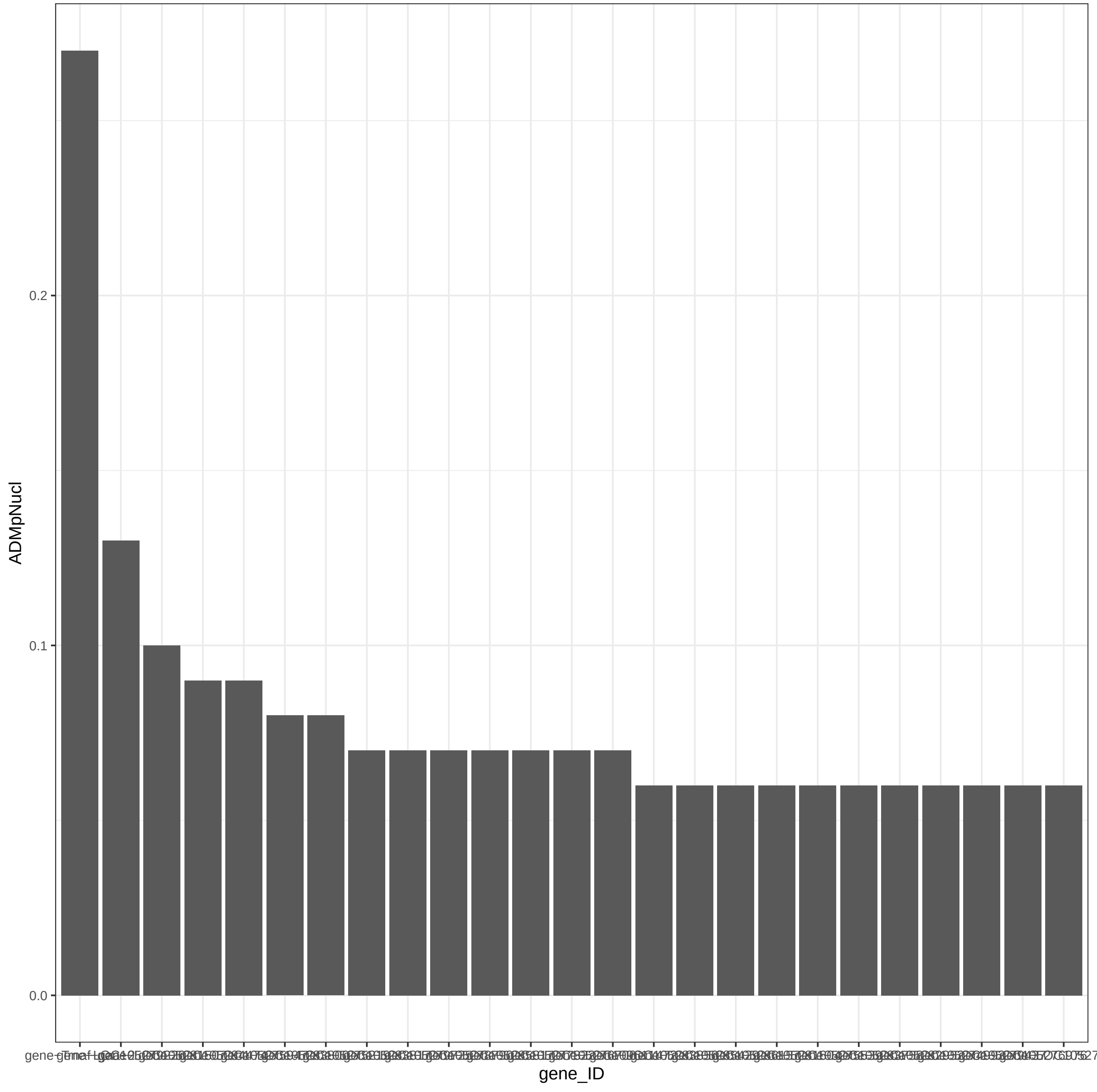
0.2
0.1
0.0
ADMpNucl
gene_ID
