## Supplementary material for "Tools and applications for integrative analysis of DNA methylation in social insects": Dataset S22

Sd_LI.conf

```
#Customize variables here:######################################################
################################################################################

#Load the files:
#
infiles       <- setup_BWASPR(datafile="./RWORK/Sdum/Sd.dat",
                              parfile="./RWORK/Sdum/Sd.par")

#Set the study and samples in the study:
#
species       <- "Sd"
study         <- "LI"
samplelist    <- list("be","ka")
## The following two variables are used for output file labeling:
studyLabel    <- "LI"
sampleLabels  <- list("Sd_LI_be","Sd_LI_ka")

hasreplicates <- TRUE
type          <- "CpG"
destrand      <- TRUE
covlist       <- c(6,20,50)
locount       <- 100
hicount       <- 1000
repcovlist    <- c(6,10,15)
replocount    <- 10
rephicount    <- 100
hheight       <- 0.120
nbrpnts       <- 5000

## The following four variables are sent to rank_rbm() and determine what data
#  points get plotted:
minnbrmsprm   <- 5      # minimum number of methylation sites per promoter
mingenewidth  <- 500    # minimum gene width
maxgenewidth  <- 5000   # maximum gene width
minnbrmsgene  <- 5      # minimum number of methylation sites per gene

## Other parameters (see Rscript.BWASPR for usage notes):
#
highcoverage  <- 20     # high read coverage threshold for studyhc methylRawList object
threshold     <- 25.0   # "difference" threshold for getMethylDiff(), called by det_dmsg()
qvalue        <- 0.01   # "qvalue" setting for getMethylDiff(), called by det_dmsg()
wsize         <- 1000   # "win.size" parameter for tileMethylCounts() in det_dmt()
stepsize      <- 1000   # "step.size" parameter for tileMethylCounts() in det_dmt()

minNsites     <- 10     # minimum number of hc sites in a gene to be heatmapped in show_dmsg()
maxNsites     <- 60     # maximum number of hc sites in a gene to be heatmapped in show_dmsg()
minPdmsites   <- 10     # minimum ratio of dm/hc (in %) in a gene to be heatmapped in show_dmsg()

maxgwidth     <- 20000  # maximal gene width for a gene to be considered by explore_dmsg()
minnbrdmsites <- 2      # minimum number of differentially methylated sites for a gene to be
                        #  considered in sample comparisons by explore_dmsg()
glink         <- "NCBIgene"   # URL to show in explore_dmsg(); options: "" or "NCBIgene"

#Set the number of processors to use:
#
numprc <- 6

#Determine what analyses to run:
#
RUNload    <- FALSE
```

```
RUNcms     <- TRUE
RUNpwc     <- TRUE
RUNcrl     <- TRUE

RUNrepcms  <- TRUE
RUNrepcrl  <- TRUE

RUNmmp     <- TRUE
RUNacs     <- TRUE
RUNrnk     <- TRUE
RUNmrpr    <- TRUE

RUNdmt     <- TRUE
RUNdmsg    <- TRUE
RUNdmgdtls <- TRUE
RUNogl     <- TRUE

RUNsave    <- TRUE

mymessage <- sprintf("\nAnalyzing %s study %s for type %s\n\n",species,studyLabel,type)
message(mymessage)

################################################################################
#End of typical customization.##################################################
```

Sd.par

```
SPECIESNAME	Stegodyphus dumicola
TOTALNBRPMSITES	123441397
ASSEMBLYVERSION	ASM1061486v1
GENOMESIZE	2551871595
SPECIESGFF3DIR	./MCALLS/Sdum/genome/GFF3DIR
GENELISTGFF3	Sdum.gene.gff3
EXONLISTGFF3	Sdum.exon.gff3
PCGEXNLISTGFF3	Sdum.pcg-exon.gff3
PROMOTRLISTGFF3	Sdum.promoter.gff3
CDSLISTGFF3	Sdum.pcg-CDS.gff3
UTRFLAGSET	1
5UTRLISTGFF3	Sdum.pcg-5pUTR.gff3
3UTRLISTGFF3	Sdum.pcg-3pUTR.gff3
```

Sd.dat

```
# Samples from Liu et al. (2019) Genes:
# be = Betta; ka = Karasburg
#
Sd    LI    be    0     CpGhsm      ./MCALLS/Sdum/Liu2019/Betta/Betta.CpGhsm.mcalls
Sd    LI    be    0     CpGscd      ./MCALLS/Sdum/Liu2019/Betta/Betta.CpGscd.mcalls
Sd    LI    ka    0     CpGhsm      ./MCALLS/Sdum/Liu2019/Karasburg/Karasburg.CpGhsm.mcalls
Sd    LI    ka    0     CpGscd      ./MCALLS/Sdum/Liu2019/Karasburg/Karasburg.CpGscd.mcalls

# ... continuing: replicates
#
Sd    LI    be    1     CpGhsm      ./MCALLS/Sdum/Liu2019/Betta/replicate1/Betta1.CpGhsm.mcalls
Sd    LI    be    1     CpGscd      ./MCALLS/Sdum/Liu2019/Betta/replicate1/Betta1.CpGscd.mcalls
Sd    LI    be    2     CpGhsm      ./MCALLS/Sdum/Liu2019/Betta/replicate2/Betta2.CpGhsm.mcalls
Sd    LI    be    2     CpGscd      ./MCALLS/Sdum/Liu2019/Betta/replicate2/Betta2.CpGscd.mcalls
Sd    LI    be    3     CpGhsm      ./MCALLS/Sdum/Liu2019/Betta/replicate3/Betta3.CpGhsm.mcalls
Sd    LI    be    3     CpGscd      ./MCALLS/Sdum/Liu2019/Betta/replicate3/Betta3.CpGscd.mcalls
Sd    LI    be    4     CpGhsm      ./MCALLS/Sdum/Liu2019/Betta/replicate4/Betta4.CpGhsm.mcalls
Sd    LI    be    4     CpGscd      ./MCALLS/Sdum/Liu2019/Betta/replicate4/Betta4.CpGscd.mcalls
Sd    LI    be    5     CpGhsm      ./MCALLS/Sdum/Liu2019/Betta/replicate5/Betta5.CpGhsm.mcalls
Sd    LI    be    5     CpGscd      ./MCALLS/Sdum/Liu2019/Betta/replicate5/Betta5.CpGscd.mcalls
Sd    LI    be    6     CpGhsm      ./MCALLS/Sdum/Liu2019/Betta/replicate6/Betta6.CpGhsm.mcalls
Sd    LI    be    6     CpGscd      ./MCALLS/Sdum/Liu2019/Betta/replicate6/Betta6.CpGscd.mcalls
Sd    LI    be    7     CpGhsm      ./MCALLS/Sdum/Liu2019/Betta/replicate7/Betta7.CpGhsm.mcalls
Sd    LI    be    7     CpGscd      ./MCALLS/Sdum/Liu2019/Betta/replicate7/Betta7.CpGscd.mcalls
Sd    LI    be    8     CpGhsm      ./MCALLS/Sdum/Liu2019/Betta/replicate8/Betta8.CpGhsm.mcalls
Sd    LI    be    8     CpGscd      ./MCALLS/Sdum/Liu2019/Betta/replicate8/Betta8.CpGscd.mcalls
Sd    LI    ka    1     CpGhsm      ./MCALLS/Sdum/Liu2019/Karasburg/replicate1/Karasburg1.CpGhsm.mcalls
Sd    LI    ka    1     CpGscd      ./MCALLS/Sdum/Liu2019/Karasburg/replicate1/Karasburg1.CpGscd.mcalls
Sd    LI    ka    2     CpGhsm      ./MCALLS/Sdum/Liu2019/Karasburg/replicate2/Karasburg2.CpGhsm.mcalls
Sd    LI    ka    2     CpGscd      ./MCALLS/Sdum/Liu2019/Karasburg/replicate2/Karasburg2.CpGscd.mcalls
Sd    LI    ka    3     CpGhsm      ./MCALLS/Sdum/Liu2019/Karasburg/replicate3/Karasburg3.CpGhsm.mcalls
Sd    LI    ka    3     CpGscd      ./MCALLS/Sdum/Liu2019/Karasburg/replicate3/Karasburg3.CpGscd.mcalls
Sd    LI    ka    4     CpGhsm      ./MCALLS/Sdum/Liu2019/Karasburg/replicate4/Karasburg4.CpGhsm.mcalls
Sd    LI    ka    4     CpGscd      ./MCALLS/Sdum/Liu2019/Karasburg/replicate4/Karasburg4.CpGscd.mcalls
Sd    LI    ka    5     CpGhsm      ./MCALLS/Sdum/Liu2019/Karasburg/replicate5/Karasburg5.CpGhsm.mcalls
Sd    LI    ka    5     CpGscd      ./MCALLS/Sdum/Liu2019/Karasburg/replicate5/Karasburg5.CpGscd.mcalls
Sd    LI    ka    6     CpGhsm      ./MCALLS/Sdum/Liu2019/Karasburg/replicate6/Karasburg6.CpGhsm.mcalls
Sd    LI    ka    6     CpGscd      ./MCALLS/Sdum/Liu2019/Karasburg/replicate6/Karasburg6.CpGscd.mcalls
Sd    LI    ka    7     CpGhsm      ./MCALLS/Sdum/Liu2019/Karasburg/replicate7/Karasburg7.CpGhsm.mcalls
Sd    LI    ka    7     CpGscd      ./MCALLS/Sdum/Liu2019/Karasburg/replicate7/Karasburg7.CpGscd.mcalls
Sd    LI    ka    8     CpGhsm      ./MCALLS/Sdum/Liu2019/Karasburg/replicate8/Karasburg8.CpGhsm.mcalls
Sd    LI    ka    8     CpGscd      ./MCALLS/Sdum/Liu2019/Karasburg/replicate8/Karasburg8.CpGscd.mcalls
```

```
0README

Rscript.BWASPR output for .conf file Sd_LI.conf.

Data input are the specified *.mcalls files and parameters set in Sd_LI.conf.
The *.mcalls input data are saved in the following data structures:

studymk - a methylKit methylRaw(List) object storing the CpGhsm site data for all samples
studymc - a methylKit methylRaw(List) object storing the CpGscd site data for all samples
studyhc - a methylKit methylRaw(List) object storing the CpGscd site data for all samples,
           restricted to sites with high coverage (here set to 20)
mkrd    - a methylKit methylRawList object storing the CpGhsm site data for all
           replicates of a given sample

Output of the analysis is stored in the following subdirectories (if the corresponding
 function calls were specified in Sd_LI.conf; please consult the 0README
 files in the subdirectories for details on the output files):

CMS - Coverage and methylation statistics for aggregate samples
PWC - Pairwise comparisons between all samples
CRL - Correlations between aggregate samples
REPCMS - Coverage and methylation statistics for replicate samples
REPCRL - Correlations between replicates
MMP - Mapping of methylation sites on genome annotation
ACS - Annotation of conserved methylation sites
RNK - Genomic feature regions ranked by CpGhsm statistics
MRPR - Methylation-rich and -poor regions
DMSG - Differentially methylateds sites and genes
OGL - Ordered gene lists
```

```
Directory: CMS File: 0READMEcms

CMS - Coverage and methylation statistics for aggregate samples.

Input: studymk, covlist, locount, hicount

Output: files cms-*.txt and cms-*.pdf

Notes: The statistics are given for CpGhsm sites restricted to minimum or
        higher level coverage.  At the minimum level, the number of sites will be
        the number of lines of the corresponding *CpGhsm.mcalls file minus one
        (consistency check).  The distribution of methylation levels is really only
        interesting for high coverage because at low coverage there will be a strong
        bias towards high methylation levels (by definition of hsm sites).
        The same bias at high coverage would suggest that the sample includes a
        preponderance of consistently methylated genomic sites.

        locount and hicount set bounds on the coverage to exclude sites with too few
        or too many covering reads to provide statistics on a typical range.
```

```
Directory: CMS File: cms-Sd_LI_be.txt

Number of "be" CpGhsm-sites with minimal and higher level coverage:

  number of "be" CpGhsm-sites with coverage >=  5: 6335325
  number of "be" CpGhsm-sites with coverage >=  6: 5822554
  number of "be" CpGhsm-sites with coverage >= 20: 774009
  number of "be" CpGhsm-sites with coverage >= 50: 108914

Coverage and methylation statistics for "be" CpGhsm-sites at different levels of minimum coverage:

methylKit::getCoverageStats output for "be" CpGhsm-sites (#: 6335325) at minimum coverage 5    - read coverage statistics per base
summary:
   Min. 1st Qu.  Median    Mean 3rd Qu.    Max.
      5       7      10      15      15   24537
percentiles:
   0%   10%   20%   30%   40%   50%   60%   70%   80%   90%   95%   99% 99.5% 99.9%  100%
    5     6     7     8     9    10    11    13    16    21    28    72   127   683 24537

methylKit::getMethylationStats output for "be" CpGhsm-sites (#: 6335325) at minimum coverage 5 - methylation statistics per base
summary:
   Min. 1st Qu.  Median    Mean 3rd Qu.    Max.
    1.0    81.8    90.9    87.4   100.0   100.0
percentiles:
     0%     10%     20%     30%     40%     50%     60%     70%     80%     90%     95%     99%   99.5%   99.9%    100%
  0.996  66.667  77.778  83.333  87.500  90.909 100.000 100.000 100.000 100.000 100.000 100.000 100.000 100.000 100.000

methylKit::getCoverageStats output for "be" CpGhsm-sites (#: 5822554) at minimum coverage 6    - read coverage statistics per base
summary:
   Min. 1st Qu.  Median    Mean 3rd Qu.    Max.
      6       8      11      16      15   24537
percentiles:
   0%   10%   20%   30%   40%   50%   60%   70%   80%   90%   95%   99% 99.5% 99.9%  100%
    6     6     7     8     9    11    12    14    17    22    29    77   136   747 24537

methylKit::getMethylationStats output for "be" CpGhsm-sites (#: 5822554) at minimum coverage 6 - methylation statistics per base
summary:
   Min. 1st Qu.  Median    Mean 3rd Qu.    Max.
    1.0    80.0    90.0    86.2   100.0   100.0
percentiles:
     0%     10%     20%     30%     40%     50%     60%     70%     80%     90%     95%     99%   99.5%   99.9%    100%
  0.996  66.667  76.190  83.333  85.714  90.000  93.333 100.000 100.000 100.000 100.000 100.000 100.000 100.000 100.000

methylKit::getCoverageStats output for "be" CpGhsm-sites (#: 774009) at minimum coverage 20    - read coverage statistics per base
summary:
   Min. 1st Qu.  Median    Mean 3rd Qu.    Max.
     20      22      26      55      35   24537
percentiles:
   0%   10%   20%   30%   40%   50%   60%   70%   80%   90%   95%   99% 99.5% 99.9%  100%
   20    20    21    22    24    26    28    32    40    63   107   544  1151  3522 24537

methylKit::getMethylationStats output for "be" CpGhsm-sites (#: 774009) at minimum coverage 20 - methylation statistics per base
summary:
   Min. 1st Qu.  Median    Mean 3rd Qu.    Max.
    1.0    53.2    80.0    71.0    91.3   100.0
percentiles:
     0%     10%     20%     30%     40%     50%     60%     70%     80%     90%     95%     99%   99.5%   99.9%    100%
```

```
  0.996  31.034  45.455  60.000  71.429  80.000  85.185  90.000  93.103  96.000 100.000 100.000 100.000 100.000 100.000

methylKit::getCoverageStats output for "be" CpGhsm-sites (#: 108914) at minimum coverage 50    - read coverage statistics per base
summary:
   Min. 1st Qu.  Median    Mean 3rd Qu.    Max. 
     50      60      81     228     145   24537 
percentiles:
   0%   10%   20%   30%   40%   50%   60%   70%   80%   90%   95%   99% 99.5% 99.9%  100% 
   50    53    58    63    71    81    97   124   177   365   808  2901  4208  8390 24537 

methylKit::getMethylationStats output for "be" CpGhsm-sites (#: 108914) at minimum coverage 50 - methylation statistics per base
summary:
   Min. 1st Qu.  Median    Mean 3rd Qu.    Max. 
    1.0    21.1    41.4    46.2    71.0   100.0 
percentiles:
     0%     10%     20%     30%     40%     50%     60%     70%     80%     90%     95%     99%   99.5%   99.9%    100% 
  0.996  13.483  18.367  24.074  31.646  41.379  53.247  65.385  76.548  86.858  91.480  96.226  97.260  99.099 100.000 

methylKit::getCoverageStats output for "be" CpGhsm-sites (#: 37905) in coverage range [100-1000]     - read coverage statistics per base
summary:
   Min. 1st Qu.  Median    Mean 3rd Qu.    Max. 
    100     123     165     241     277    1000 
percentiles:
   0%   10%   20%   30%   40%   50%   60%   70%   80%   90%   95%   99% 99.5% 99.9%  100% 
  100   107   117   129   145   165   195   241   326   504   679   911   957   991  1000 

methylKit::getMethylationStats output for "be" CpGhsm-sites (#: 37905) in coverage range [100-1000]  - methylation statistics per base
summary:
   Min. 1st Qu.  Median    Mean 3rd Qu.    Max. 
    2.5    16.2    39.1    44.9    74.1   100.0 
percentiles:
    0%    10%    20%    30%    40%    50%    60%    70%    80%    90%    95%    99%  99.5%  99.9%   100% 
  2.50   9.55  13.48  19.30  27.54  39.09  54.11  68.06  79.46  88.36  91.98  95.56  96.52  98.02 100.00 
```

```
Directory: CMS File: cms-Sd_LI_ka.txt

Number of "ka" CpGhsm-sites with minimal and higher level coverage:

  number of "ka" CpGhsm-sites with coverage >=  5: 6222608
  number of "ka" CpGhsm-sites with coverage >=  6: 5726881
  number of "ka" CpGhsm-sites with coverage >= 20: 842361
  number of "ka" CpGhsm-sites with coverage >= 50: 120244

Coverage and methylation statistics for "ka" CpGhsm-sites at different levels of minimum coverage:

methylKit::getCoverageStats output for "ka" CpGhsm-sites (#: 6222608) at minimum coverage 5    - read coverage statistics per base
summary:
   Min. 1st Qu.  Median    Mean 3rd Qu.    Max.
      5       7      10      17      15   30349
percentiles:
   0%   10%   20%   30%   40%   50%   60%   70%   80%   90%   95%   99% 99.5% 99.9%  100%
    5     6     7     8     9    10    12    14    17    22    29    80   149   966 30349

methylKit::getMethylationStats output for "ka" CpGhsm-sites (#: 6222608) at minimum coverage 5 - methylation statistics per base
summary:
   Min. 1st Qu.  Median    Mean 3rd Qu.    Max.
    0.9    81.2    90.9    86.9   100.0   100.0
percentiles:
     0%     10%     20%     30%     40%     50%     60%     70%     80%     90%     95%     99%   99.5%   99.9%    100%
  0.921  66.667  76.923  83.333  87.500  90.909  95.455 100.000 100.000 100.000 100.000 100.000 100.000 100.000 100.000

methylKit::getCoverageStats output for "ka" CpGhsm-sites (#: 5726881) at minimum coverage 6    - read coverage statistics per base
summary:
   Min. 1st Qu.  Median    Mean 3rd Qu.    Max.
      6       8      11      18      16   30349
percentiles:
   0%   10%   20%   30%   40%   50%   60%   70%   80%   90%   95%   99% 99.5% 99.9%  100%
    6     6     7     8     9    11    12    14    17    23    31    86   162  1067 30349

methylKit::getMethylationStats output for "ka" CpGhsm-sites (#: 5726881) at minimum coverage 6 - methylation statistics per base
summary:
   Min. 1st Qu.  Median    Mean 3rd Qu.    Max.
    0.9    80.0    88.9    85.8   100.0   100.0
percentiles:
     0%     10%     20%     30%     40%     50%     60%     70%     80%     90%     95%     99%   99.5%   99.9%    100%
  0.921  63.636  75.000  83.333  85.714  88.889  92.857 100.000 100.000 100.000 100.000 100.000 100.000 100.000 100.000

methylKit::getCoverageStats output for "ka" CpGhsm-sites (#: 842361) at minimum coverage 20    - read coverage statistics per base
summary:
   Min. 1st Qu.  Median    Mean 3rd Qu.    Max.
     20      22      26      63      36   30349
percentiles:
   0%   10%   20%   30%   40%   50%   60%   70%   80%   90%   95%   99% 99.5% 99.9%  100%
   20    20    21    23    24    26    28    32    40    64   112   690  1503  5057 30349

methylKit::getMethylationStats output for "ka" CpGhsm-sites (#: 842361) at minimum coverage 20 - methylation statistics per base
summary:
   Min. 1st Qu.  Median    Mean 3rd Qu.    Max.
    0.9    53.8    80.0    71.2    91.3   100.0
percentiles:
     0%     10%     20%     30%     40%     50%     60%     70%     80%     90%     95%     99%   99.5%   99.9%    100%
```

```
  0.921  31.481  46.154  60.377  71.429  80.000  85.294  90.000  93.182  96.000 100.000 100.000 100.000 100.000 100.000

methylKit::getCoverageStats output for "ka" CpGhsm-sites (#: 120244) at minimum coverage 50    - read coverage statistics per base
summary:
   Min. 1st Qu.  Median    Mean 3rd Qu.    Max.
     50      60      82     280     154   30349
percentiles:
   0%   10%   20%   30%   40%   50%   60%   70%   80%   90%   95%   99% 99.5% 99.9%  100%
   50    53    58    64    71    82    99   129   193   440  1008  4142  5923 11985 30349

methylKit::getMethylationStats output for "ka" CpGhsm-sites (#: 120244) at minimum coverage 50 - methylation statistics per base
summary:
   Min. 1st Qu.  Median    Mean 3rd Qu.    Max.
    0.9    21.4    42.3    46.7    71.8   100.0
percentiles:
     0%     10%     20%     30%     40%     50%     60%     70%     80%     90%     95%     99%   99.5%   99.9%    100%
  0.921  13.445  18.478  24.561  32.394  42.308  53.968  66.018  77.326  87.324  91.880  96.341  97.674 100.000 100.000

methylKit::getCoverageStats output for "ka" CpGhsm-sites (#: 41996) in coverage range [100-1000]    - read coverage statistics per base
summary:
   Min. 1st Qu.  Median    Mean 3rd Qu.    Max.
    100     124     171     252     297    1000
percentiles:
   0%   10%   20%   30%   40%   50%   60%   70%   80%   90%   95%   99% 99.5% 99.9%  100%
  100   108   118   131   148   171   205   257   350   542   721   926   959   993  1000

methylKit::getMethylationStats output for "ka" CpGhsm-sites (#: 41996) in coverage range [100-1000] - methylation statistics per base
summary:
   Min. 1st Qu.  Median    Mean 3rd Qu.    Max.
    2.6    15.7    38.1    44.5    73.9   100.0
percentiles:
    0%    10%    20%    30%    40%    50%    60%    70%    80%    90%    95%    99%  99.5%  99.9%   100%
  2.61   9.52  13.33  18.63  26.73  38.10  52.67  67.13  79.56  88.72  92.31  95.75  96.64  98.04 100.00
```

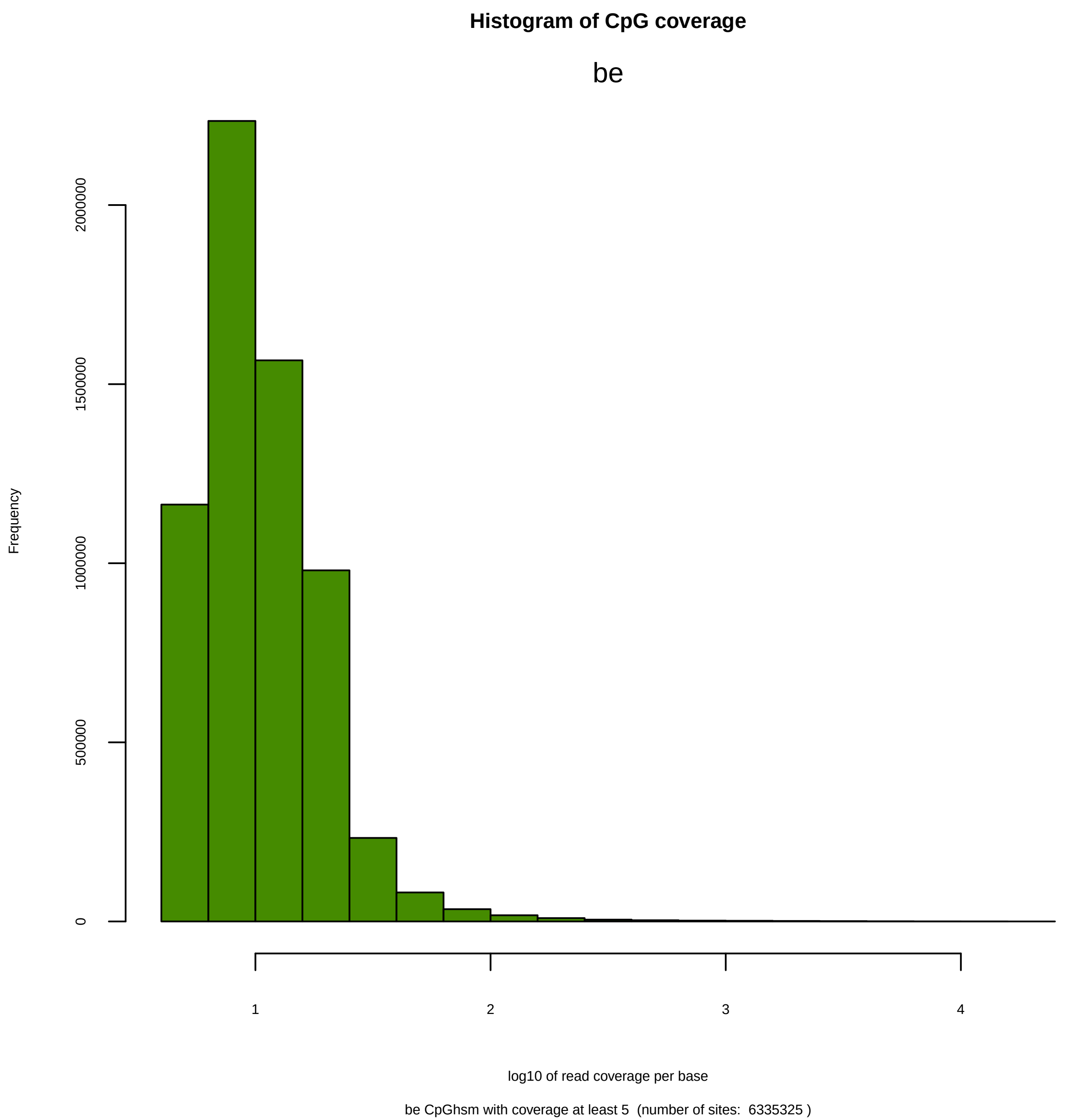

Histogram of CpG coverage
be
Frequency
0
500000
1000000
1500000
2000000
1
2
3
4
log10 of read coverage per base
be CpGhsm with coverage at least 5  (number of sites:  6335325 )

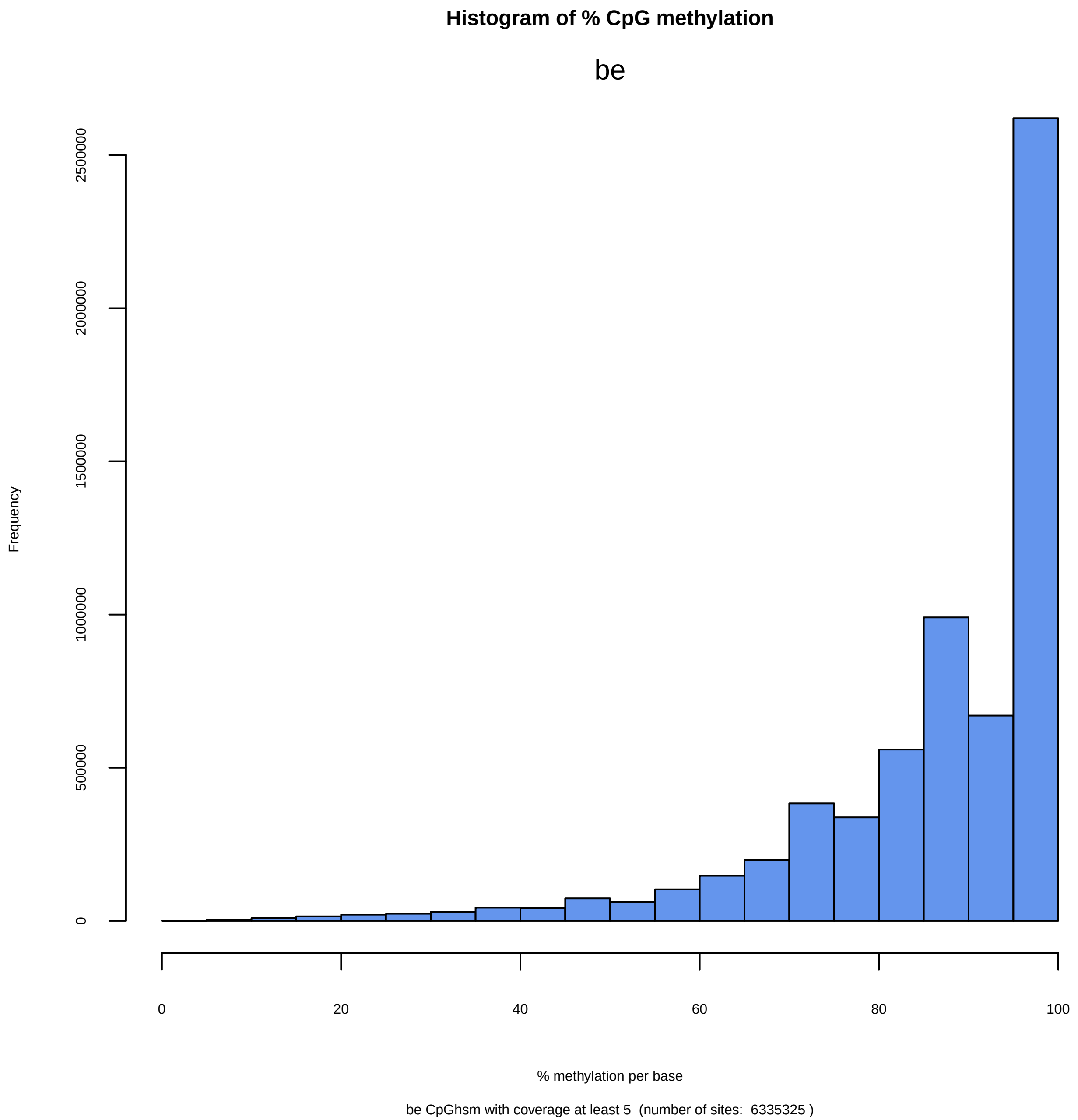

Histogram of % CpG methylation
be
Frequency
0
500000
1000000
1500000
2000000
2500000
0
20
40
60
80
100
% methylation per base
be CpGhsm with coverage at least 5  (number of sites:  6335325 )

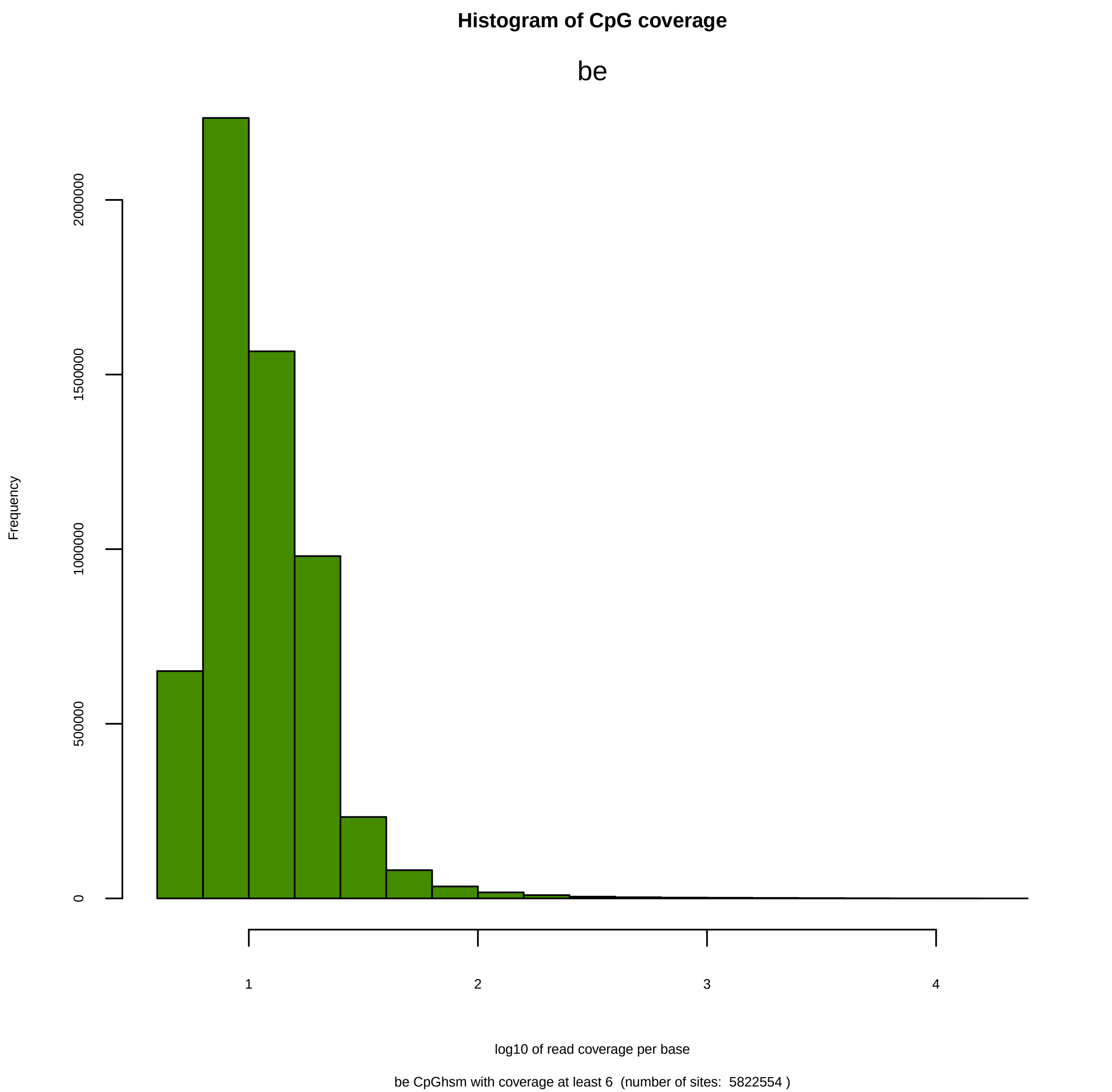
Histogram of CpG coverage
be
Frequency
0
500000
1000000
1500000
2000000
1
2
3
4
log10 of read coverage per base
be CpGhsm with coverage at least 6  (number of sites:  5822554 )

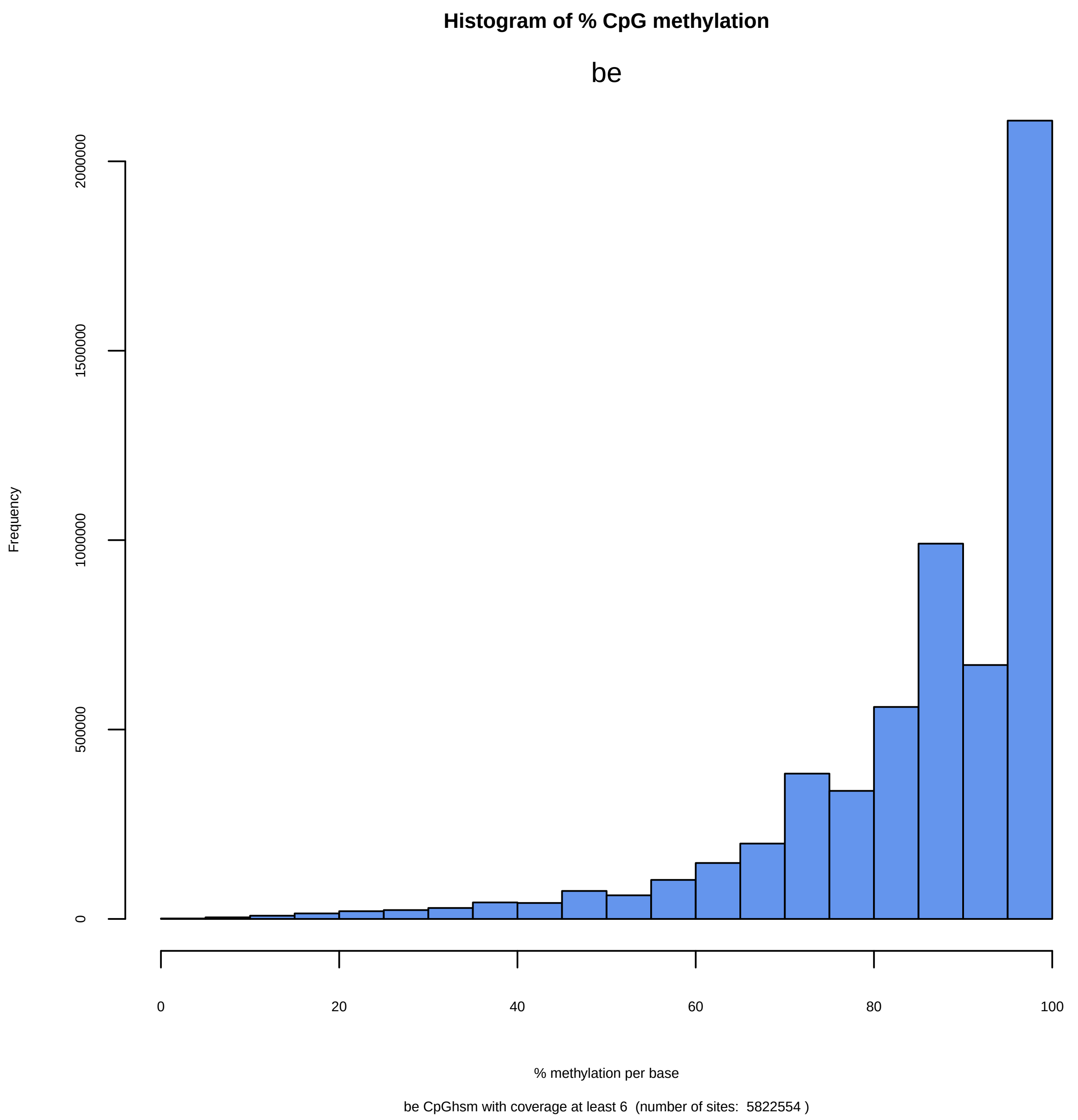

Histogram of % CpG methylation
be
Frequency
0
500000
1000000
1500000
2000000
0
20
40
60
80
100
% methylation per base
be CpGhsm with coverage at least 6 (number of sites: 5822554 )

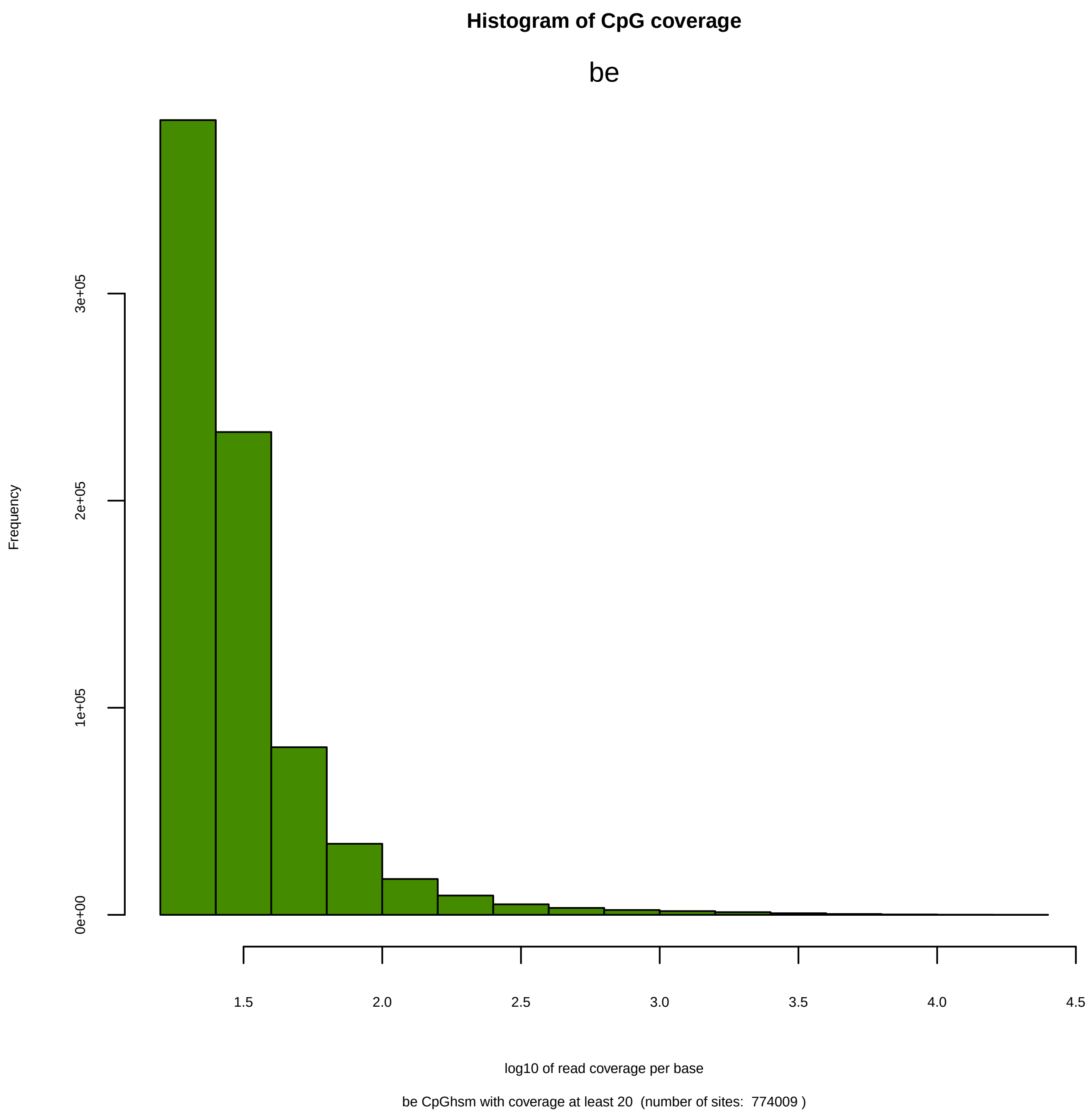

Histogram of CpG coverage
be
Frequency
0e+00
1e+05
2e+05
3e+05
1.5
2.0
2.5
3.0
3.5
4.0
4.5
log10 of read coverage per base
be CpGhsm with coverage at least 20 (number of sites: 774009 )

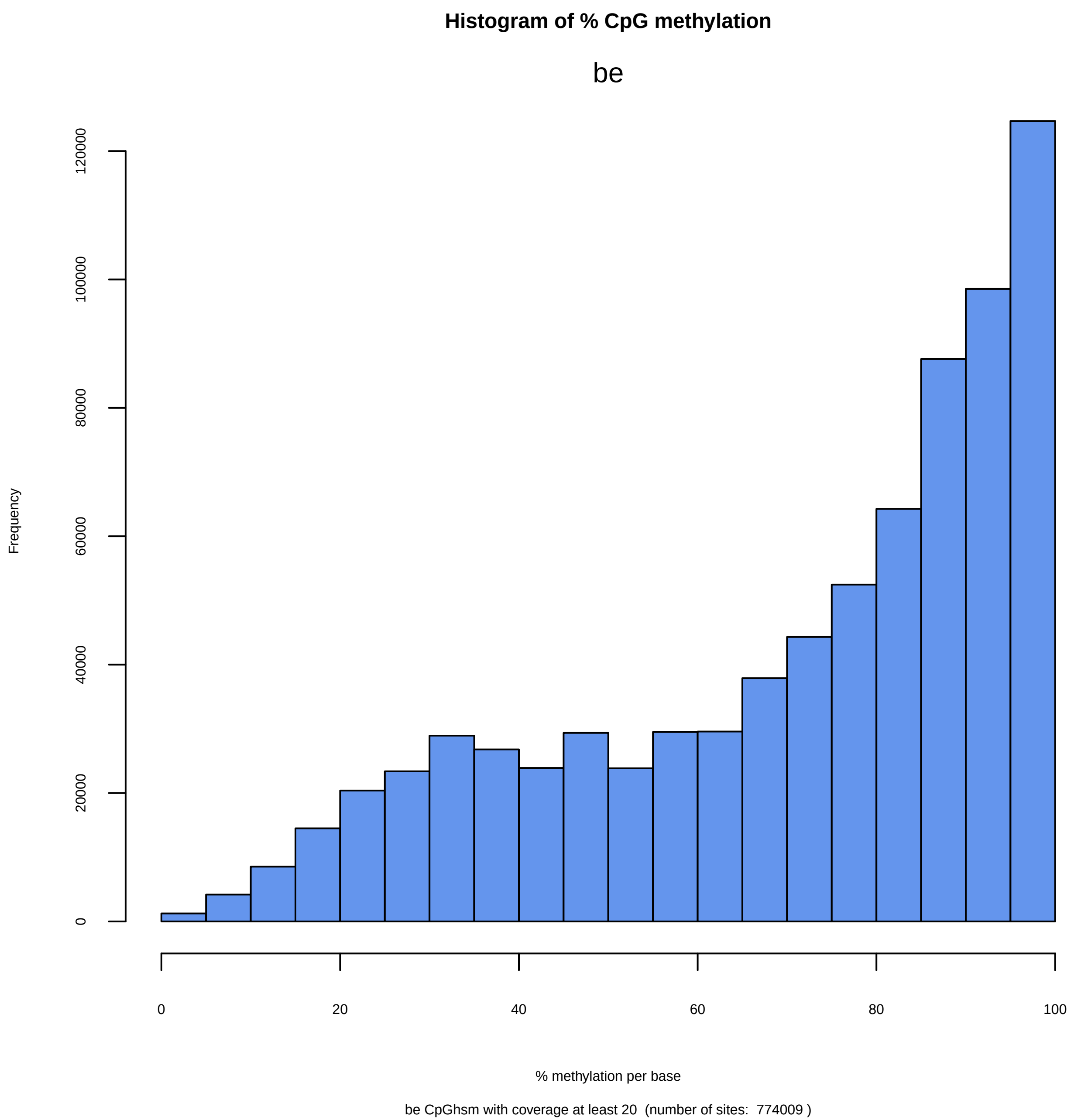
Histogram of % CpG methylation
be
Frequency
0
20000
40000
60000
80000
100000
120000
0
20
40
60
80
100
% methylation per base
be CpGhsm with coverage at least 20 (number of sites: 774009 )

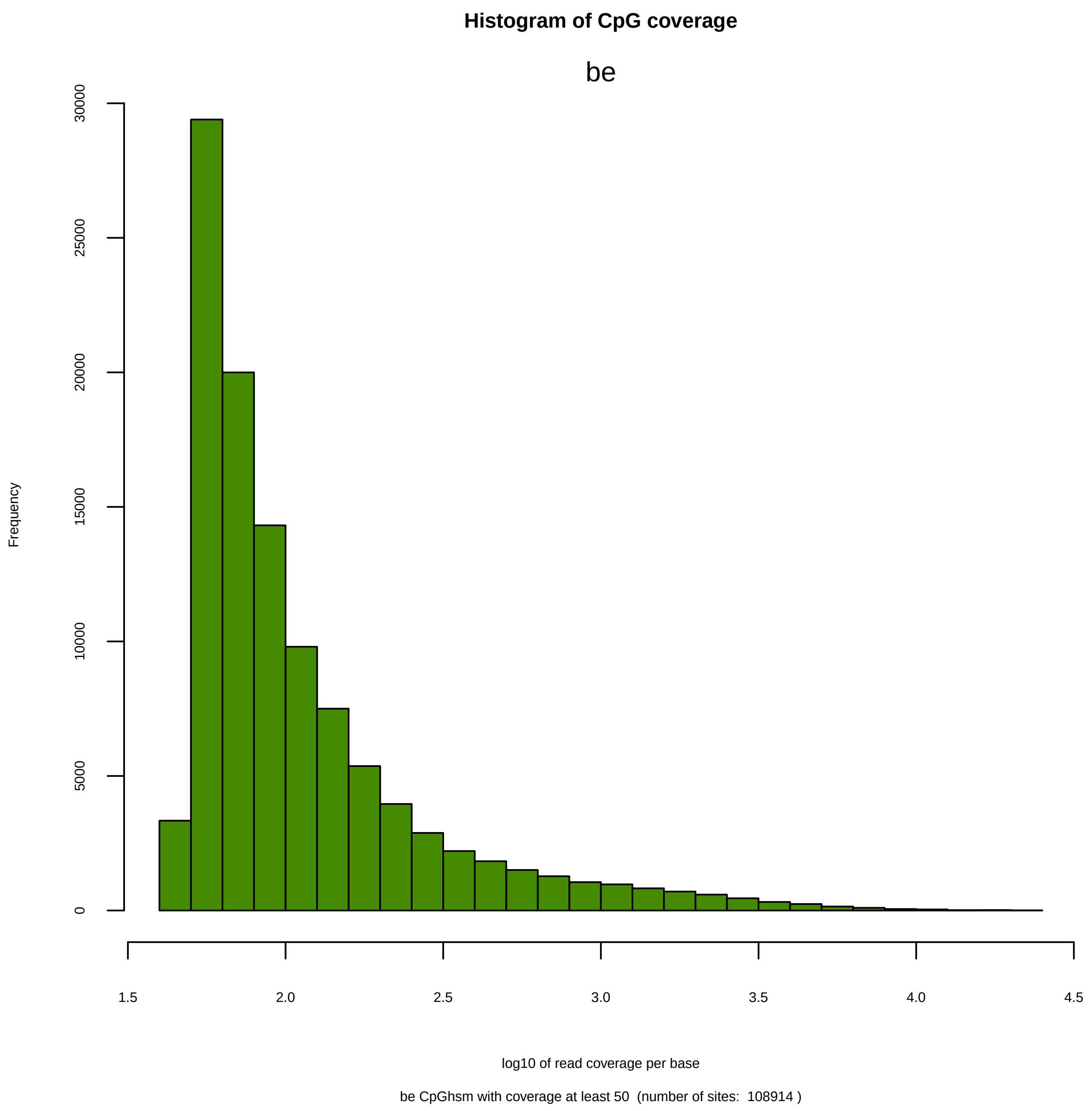

Histogram of CpG coverage
be
Frequency
0
5000
10000
15000
20000
25000
30000
1.5
2.0
2.5
3.0
3.5
4.0
4.5
log10 of read coverage per base
be CpGhsm with coverage at least 50  (number of sites:  108914 )

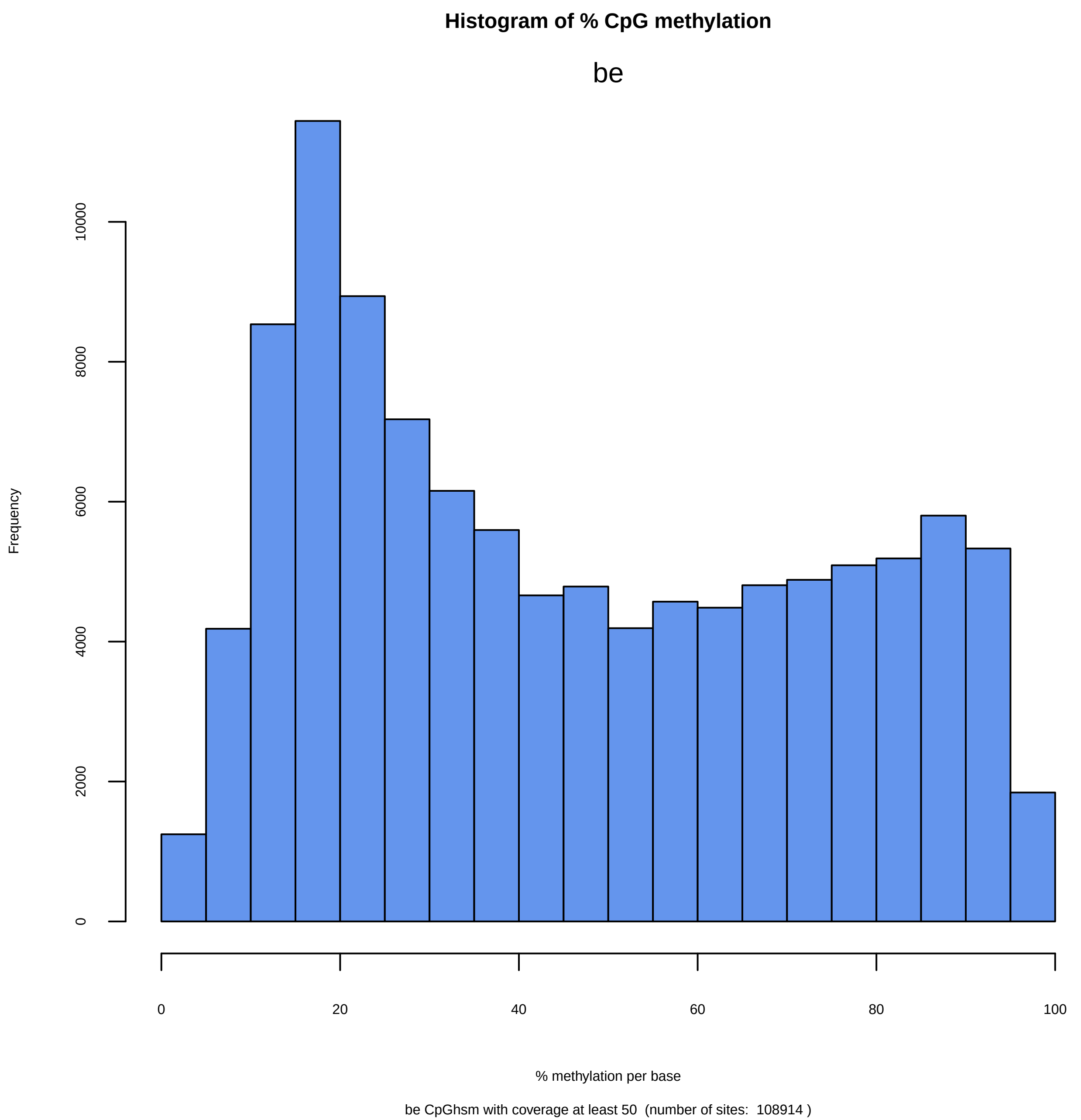

Histogram of % CpG methylation
be
Frequency
0
2000
4000
6000
8000
10000
0
20
40
60
80
100
% methylation per base
be CpGhsm with coverage at least 50 (number of sites: 108914 )

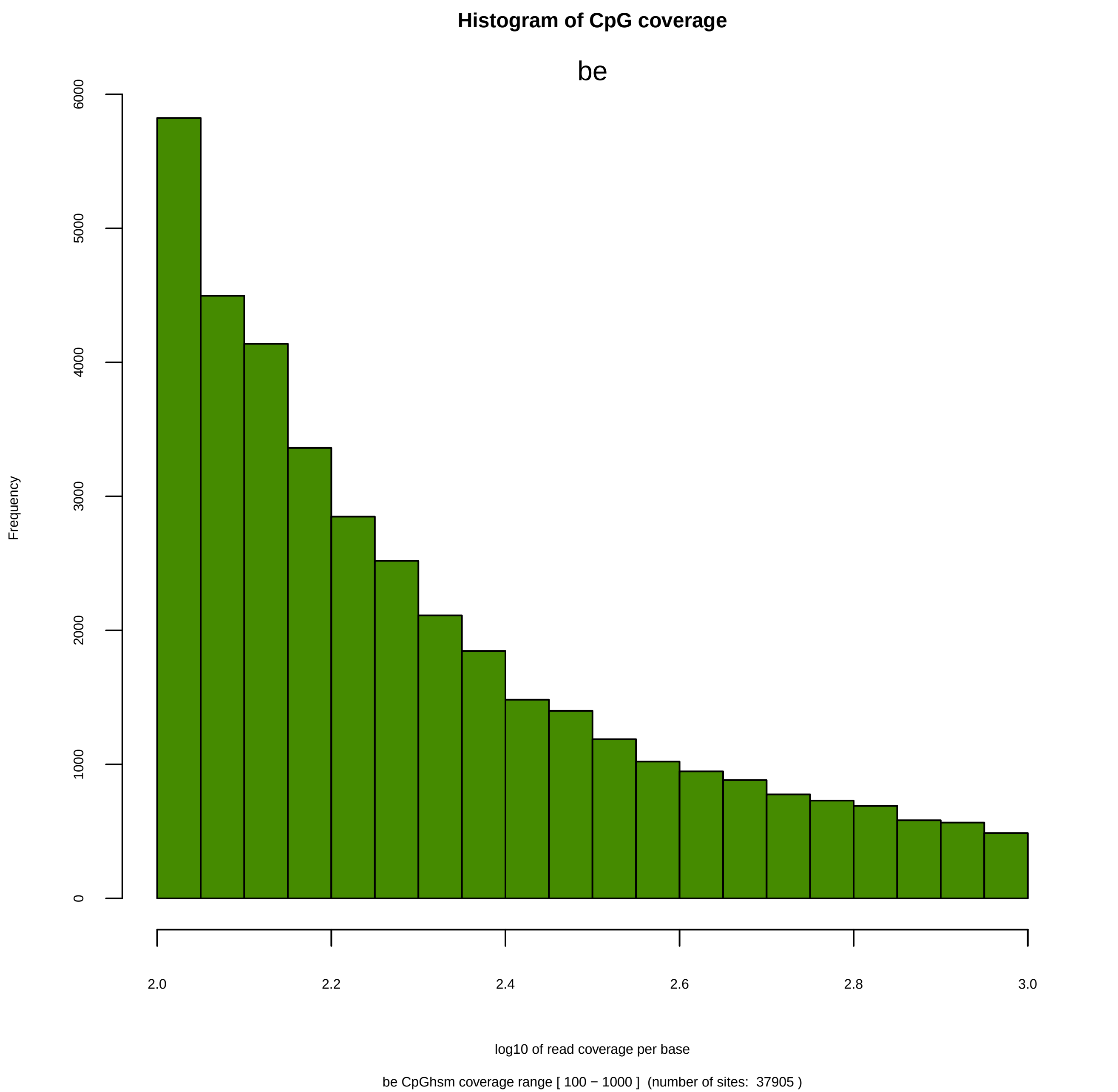

Histogram of CpG coverage
be
Frequency
0
1000
2000
3000
4000
5000
6000
2.0
2.2
2.4
2.6
2.8
3.0
log10 of read coverage per base
be CpGhsm coverage range [ 100 − 1000 ] (number of sites: 37905 )

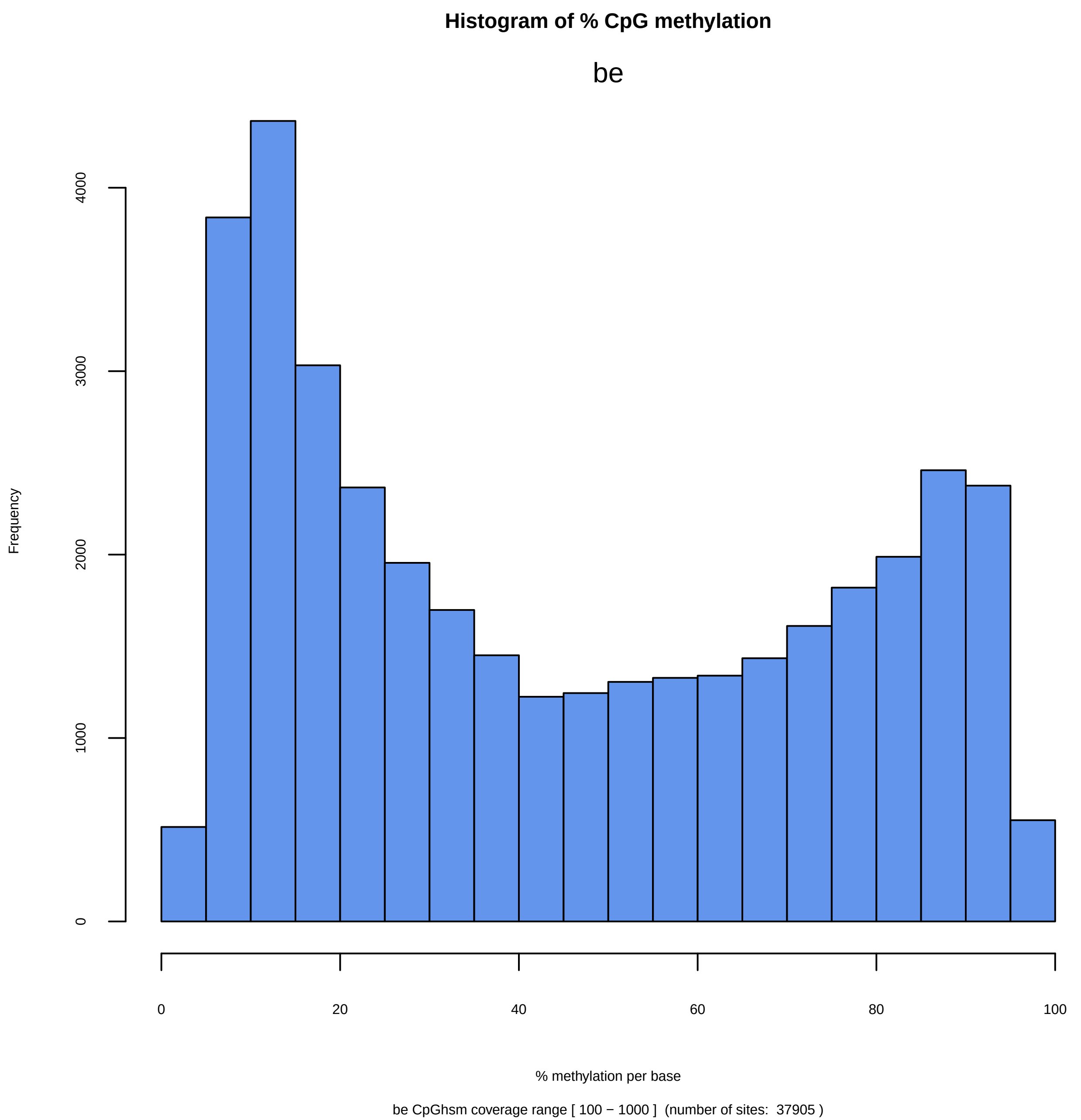

Histogram of % CpG methylation
be
Frequency
0
1000
2000
3000
4000
0
20
40
60
80
100
% methylation per base
be CpGhsm coverage range [ 100 − 1000 ]  (number of sites:  37905 )

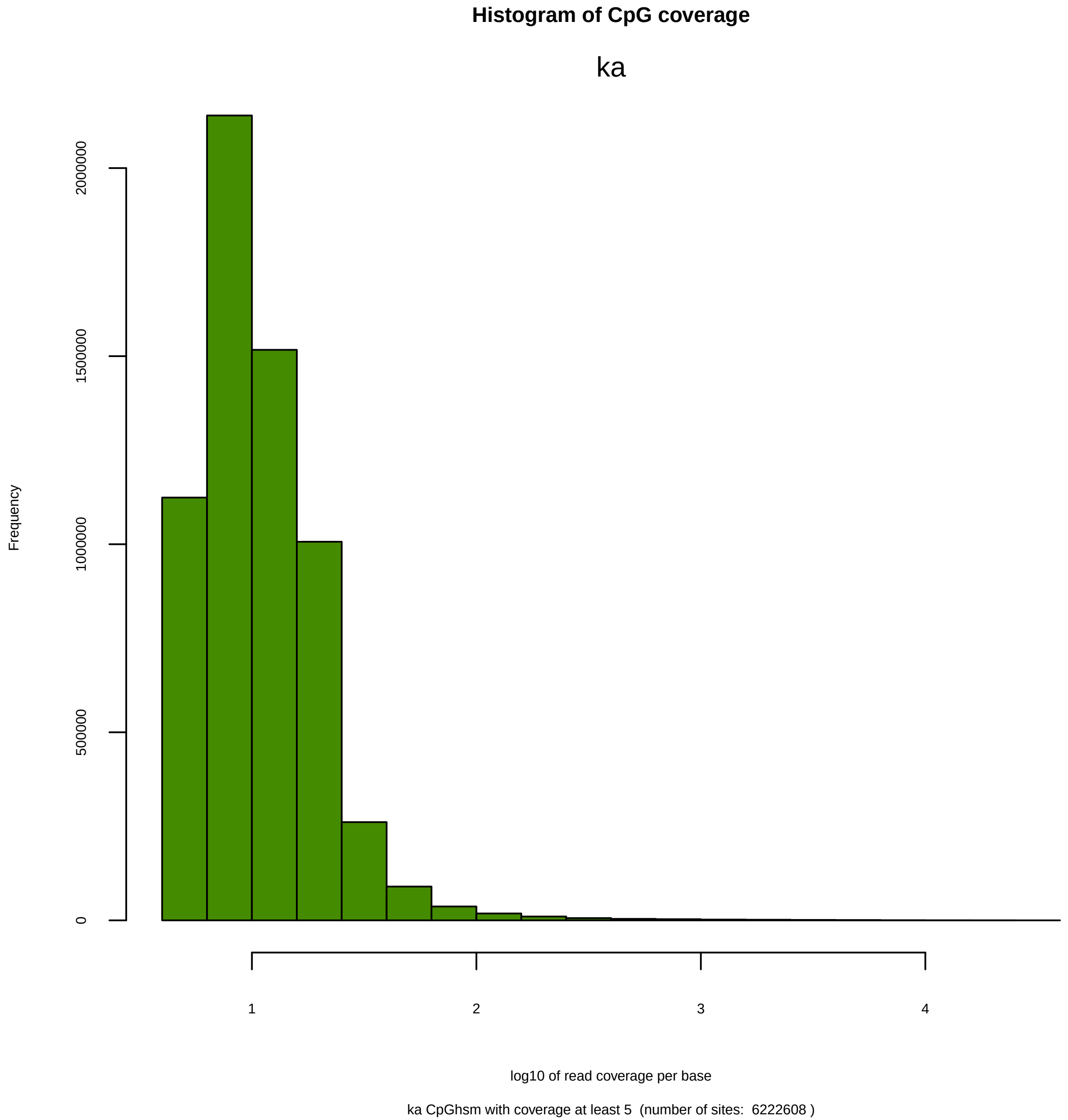

Histogram of CpG coverage
ka
Frequency
0
500000
1000000
1500000
2000000
1
2
3
4
log10 of read coverage per base
ka CpGhsm with coverage at least 5 (number of sites: 6222608 )

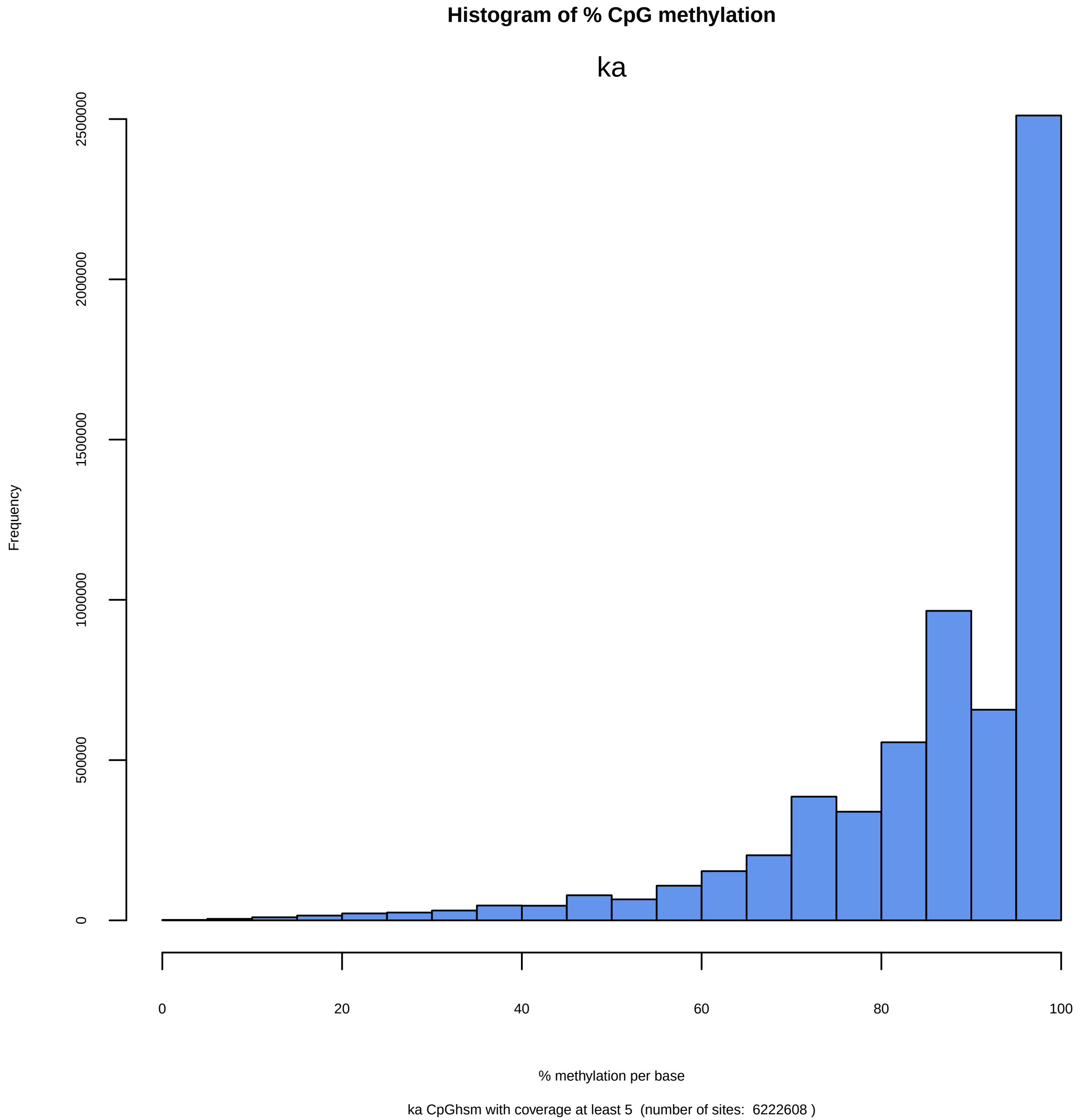

Histogram of % CpG methylation
ka
Frequency
0
500000
1000000
1500000
2000000
2500000
0
20
40
60
80
100
% methylation per base
ka CpGhsm with coverage at least 5 (number of sites: 6222608 )

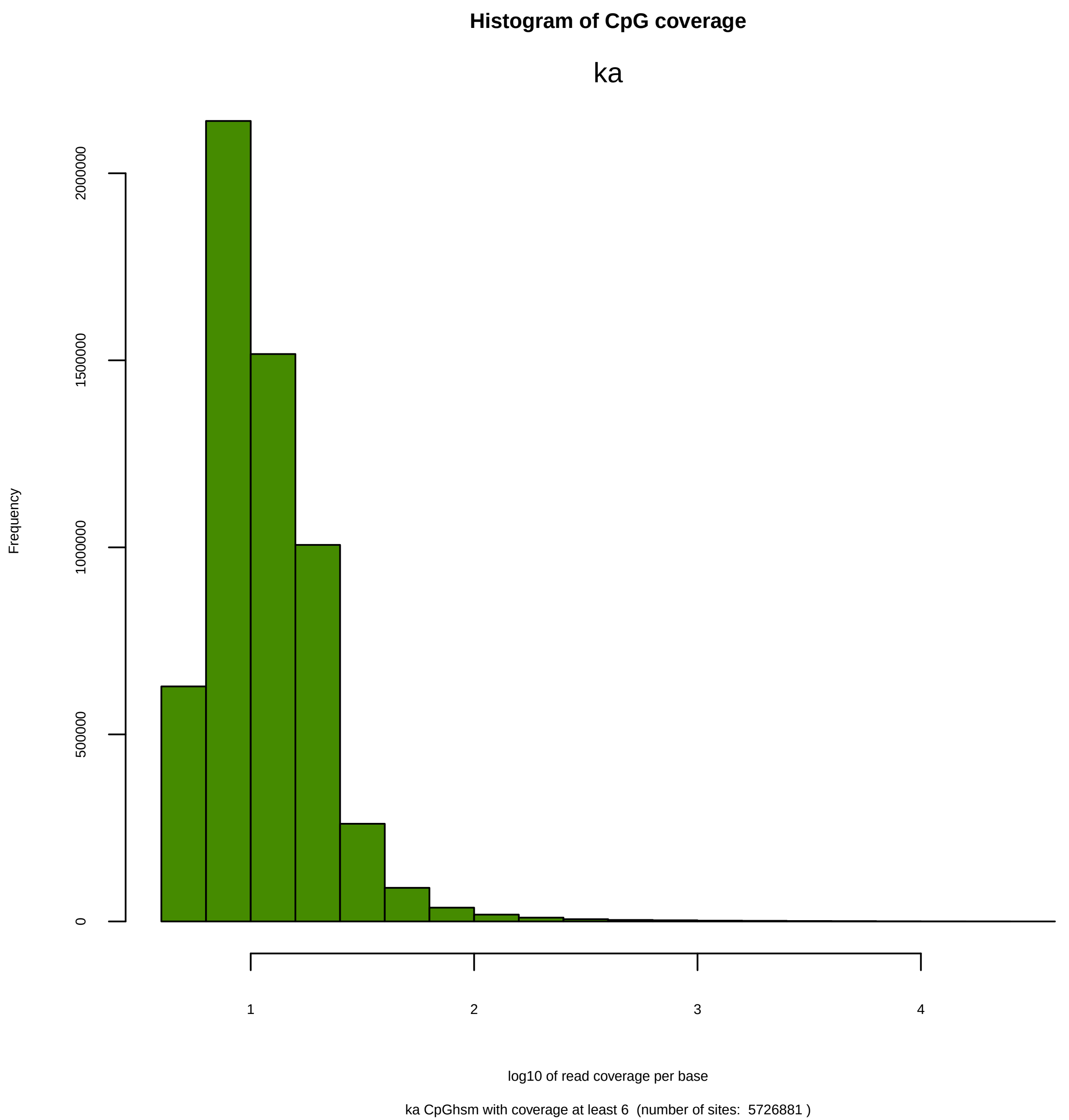

Histogram of CpG coverage
ka
Frequency
0
500000
1000000
1500000
2000000
1
2
3
4
log10 of read coverage per base
ka CpGhsm with coverage at least 6  (number of sites:  5726881 )

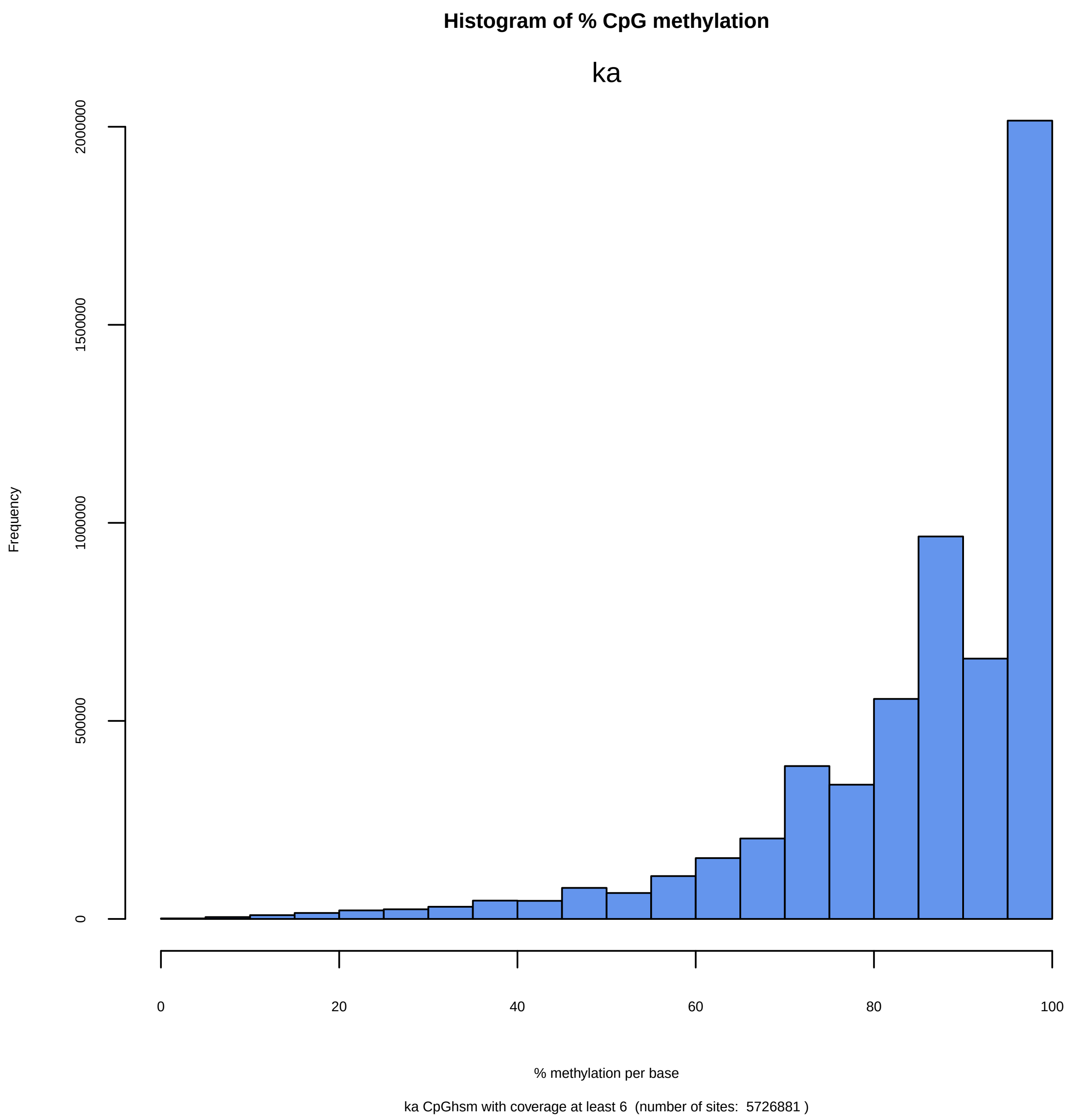

Histogram of % CpG methylation
ka
Frequency
0
500000
1000000
1500000
2000000
0
20
40
60
80
100
% methylation per base
ka CpGhsm with coverage at least 6 (number of sites: 5726881 )

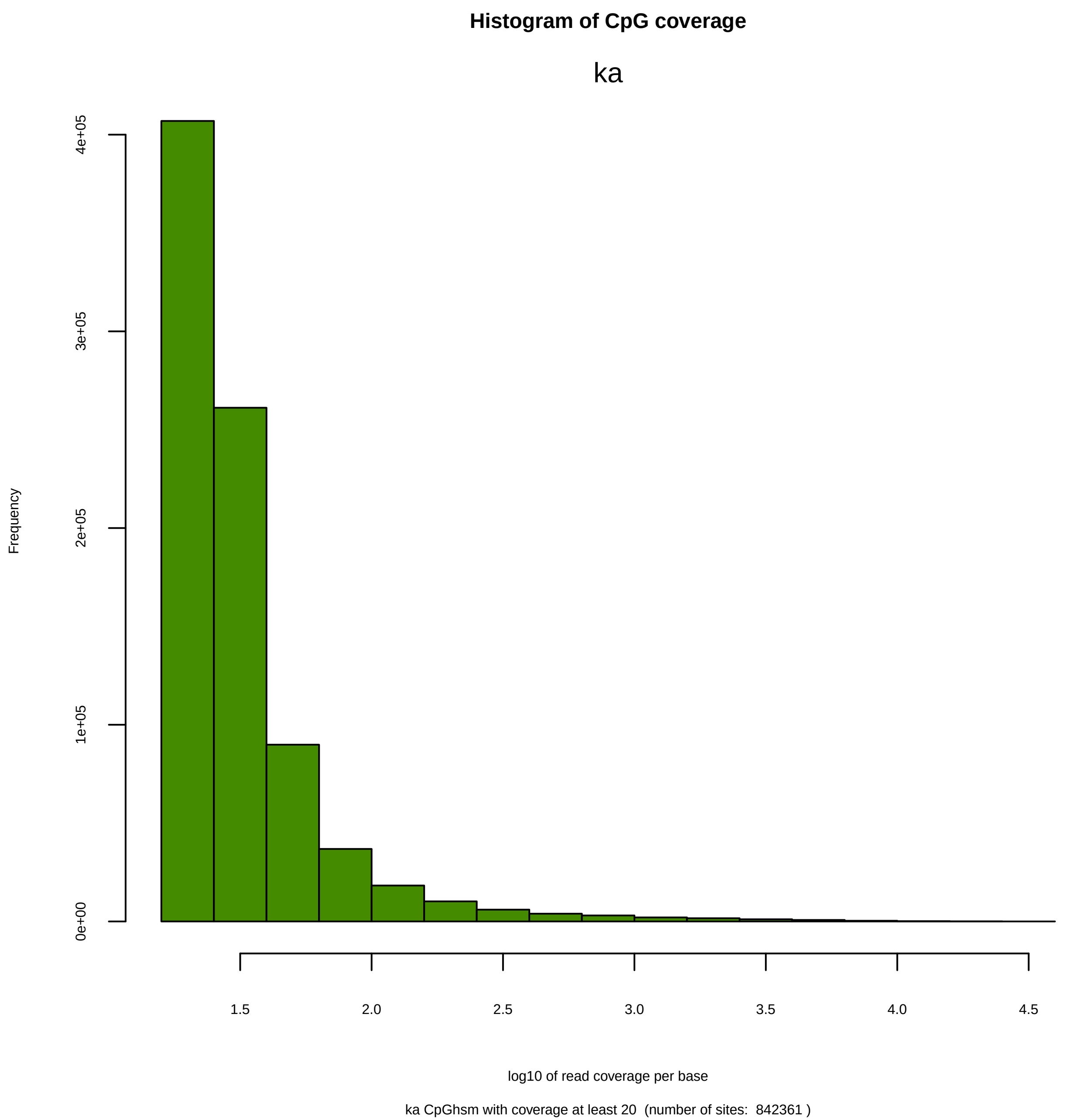

Histogram of CpG coverage
ka
Frequency
0e+00
1e+05
2e+05
3e+05
4e+05
1.5
2.0
2.5
3.0
3.5
4.0
4.5
log10 of read coverage per base
ka CpGhsm with coverage at least 20 (number of sites: 842361 )

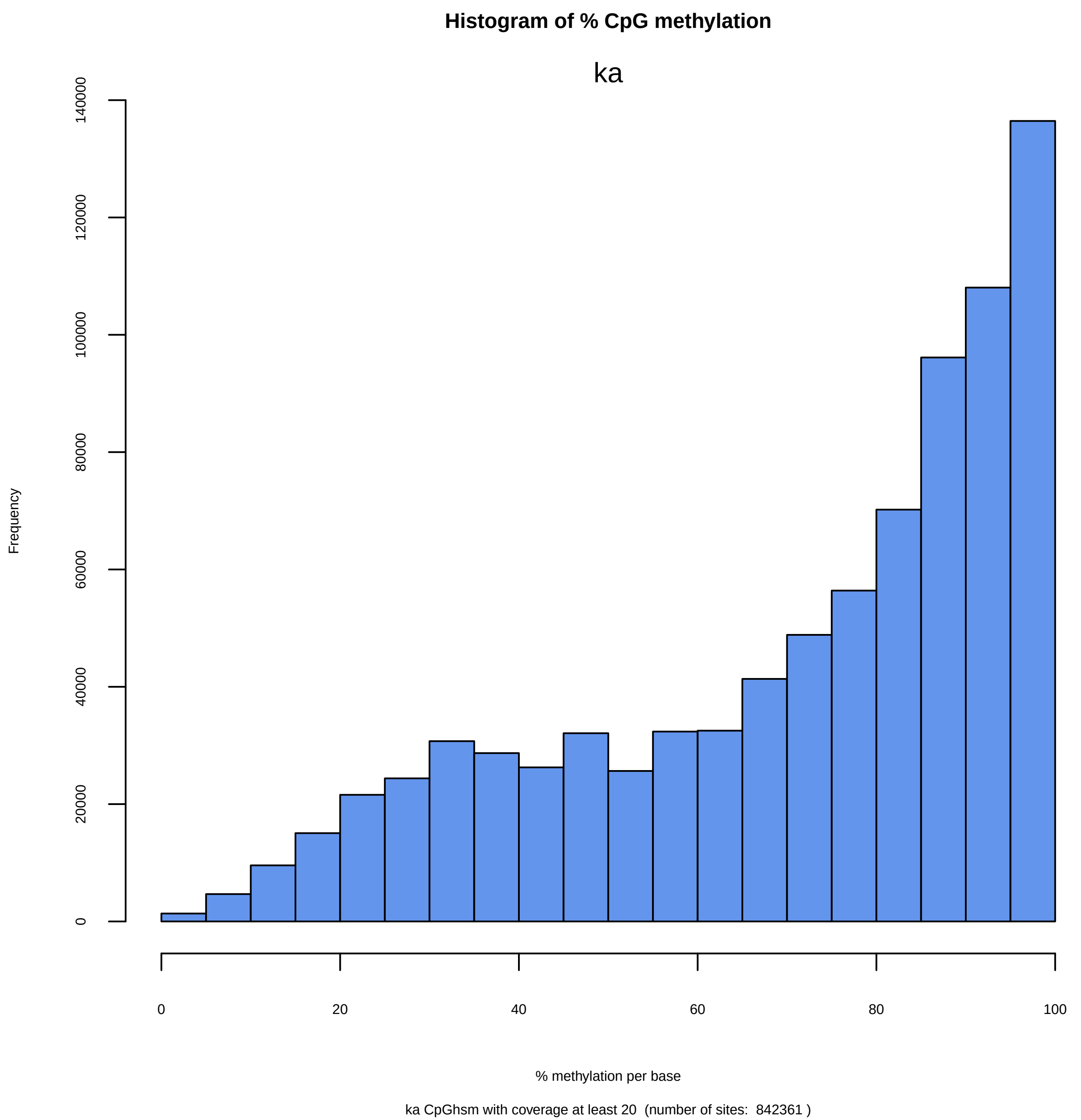
Histogram of % CpG methylation
ka
Frequency
0
20000
40000
60000
80000
100000
120000
140000
0
20
40
60
80
100
% methylation per base
ka CpGhsm with coverage at least 20 (number of sites: 842361 )

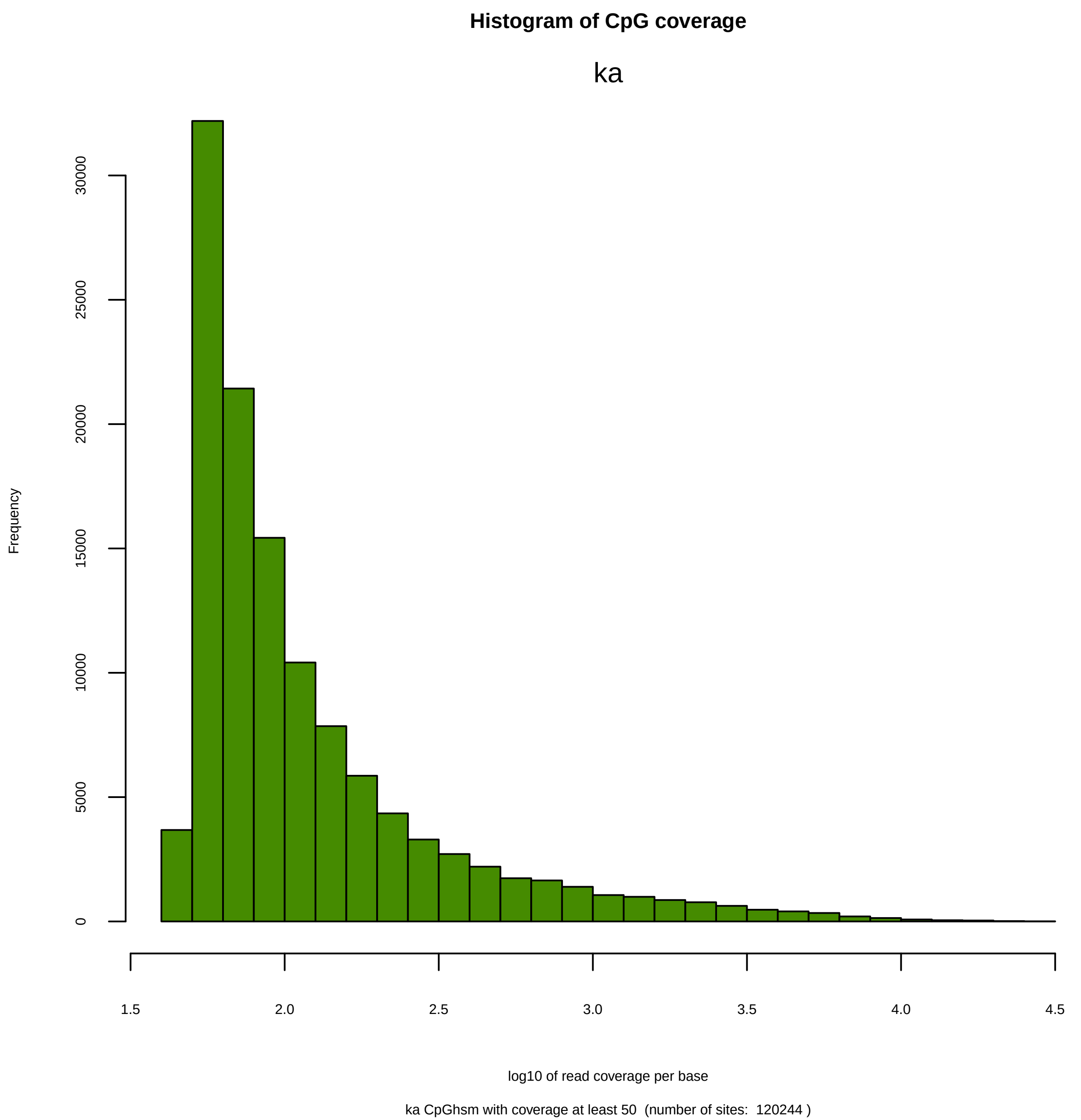
Histogram of CpG coverage
ka
Frequency
0
5000
10000
15000
20000
25000
30000
1.5
2.0
2.5
3.0
3.5
4.0
4.5
log10 of read coverage per base
ka CpGhsm with coverage at least 50 (number of sites: 120244 )

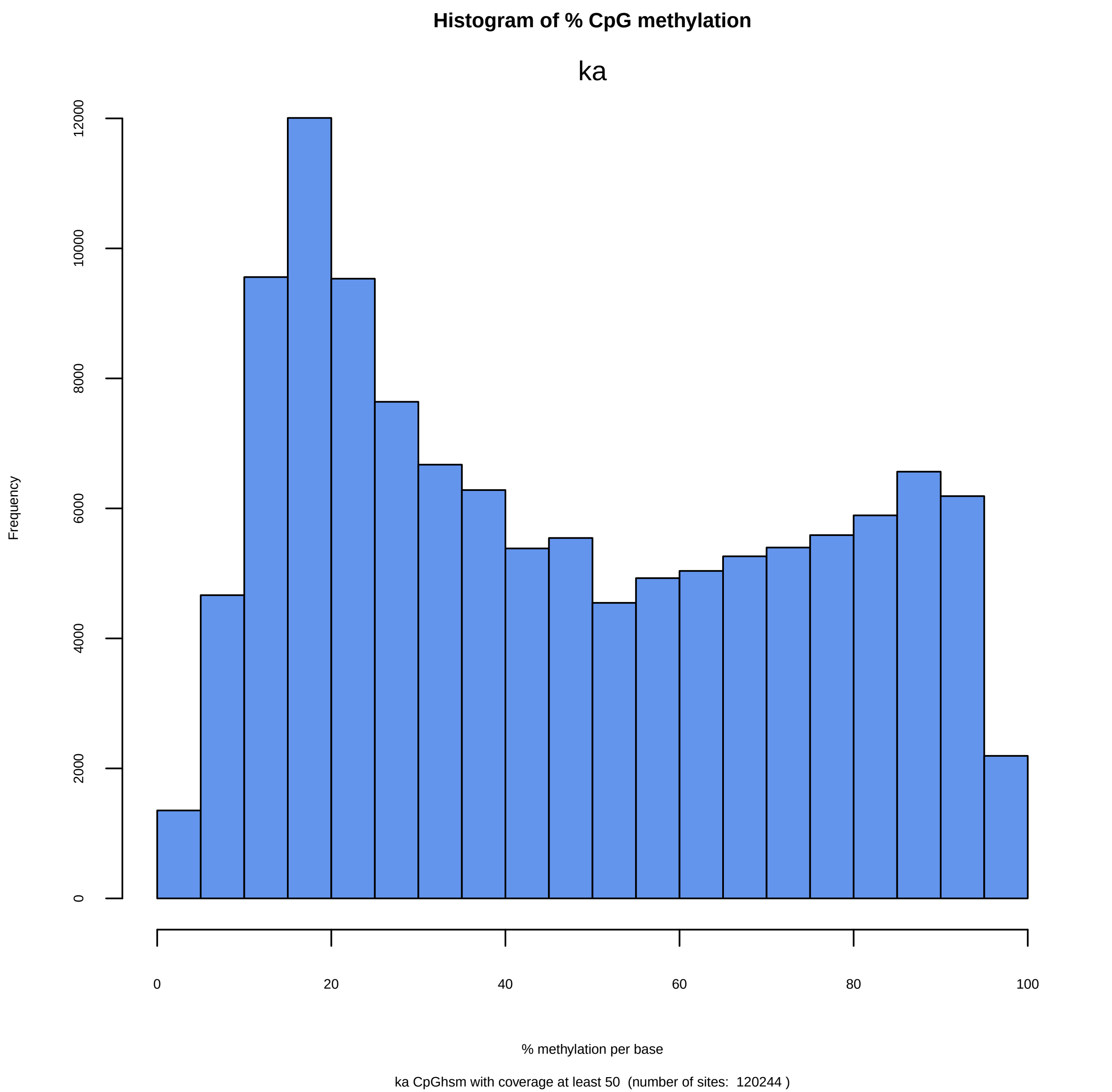

Histogram of % CpG methylation
ka
Frequency
0
2000
4000
6000
8000
10000
12000
0
20
40
60
80
100
% methylation per base
ka CpGhsm with coverage at least 50  (number of sites:  120244 )

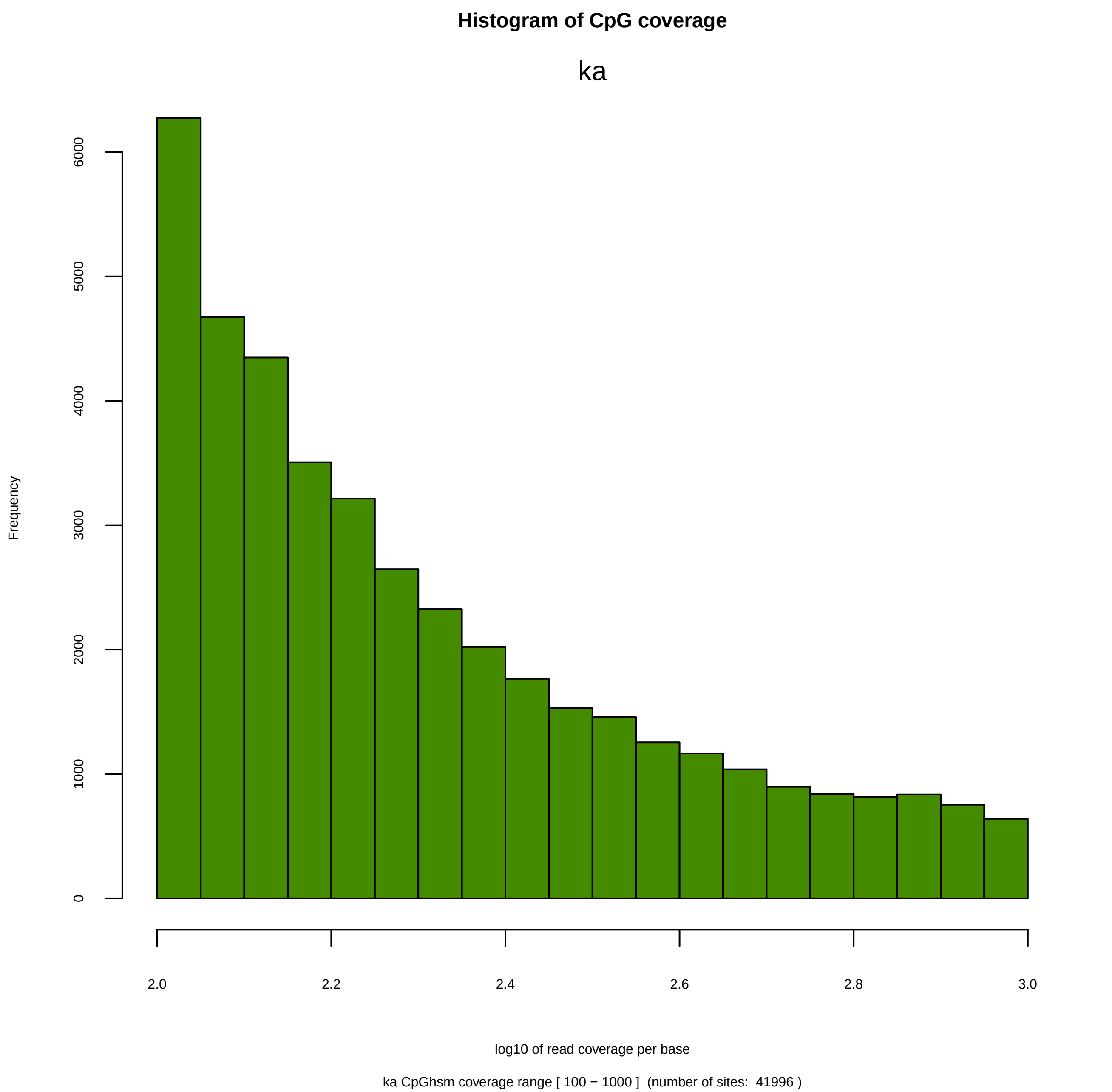

Histogram of CpG coverage
ka
Frequency
0
1000
2000
3000
4000
5000
6000
2.0
2.2
2.4
2.6
2.8
3.0
log10 of read coverage per base
ka CpGhsm coverage range [ 100 − 1000 ]  (number of sites:  41996 )

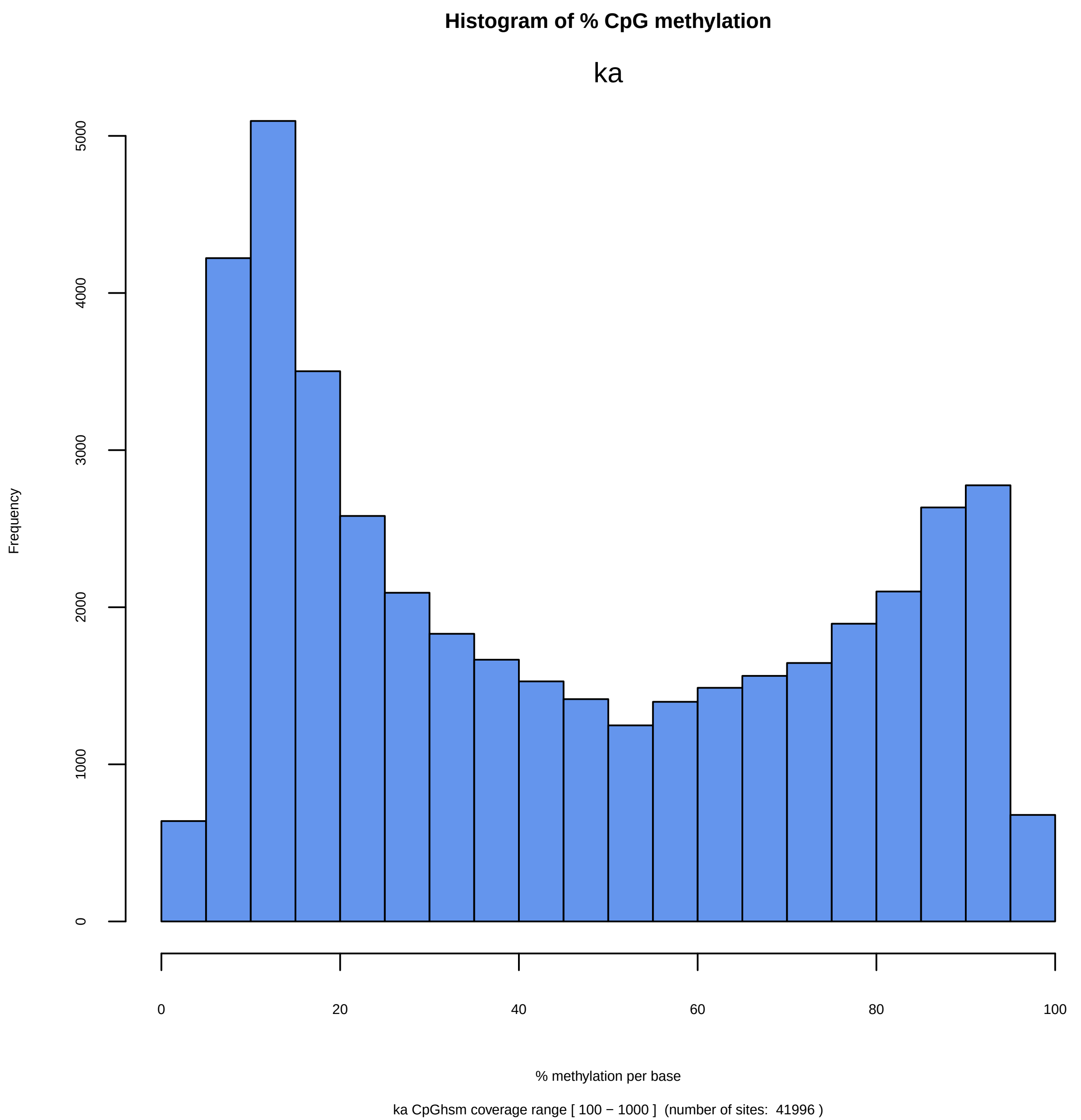

Histogram of % CpG methylation
ka
Frequency
0
1000
2000
3000
4000
5000
0
20
40
60
80
100
% methylation per base
ka CpGhsm coverage range [ 100 − 1000 ]  (number of sites:  41996 )

```
Directory: PWC File: 0READMEpwc

PWC - Pairwise comparisons between all samples.

Input: studymk, studymc, nbrpms, hheight, nbrpnts

Output: files pwc-*.txt pwc-.*pdf

Notes: The output is generated by BWASPR::cmpSites(), which determines for each pairwise
        comparison the numbers of common and distinct sites.  The function also calculates
        the overlap index between the samples and estimates the size of the common pool of
        potential methylation sites.

       The CpGscd data input (studymc) is necessary to determine sites that are detectable
        in both samples, and nbrpms (listed in the *.par file specified in the Sd_LI.conf
        configuration file as argument to TOTALNBRPMSITES) provides the total number of
        potential methylation sites (typically, all CpG sites), a necessary parameter for
        the estimations.
       hheight (default: 0.10) specifies the y-axis limit in the histogram plots.
       nbrpnts (default: 5000) specifies the number of common sites to be plotted in the
        methylation levels scatter plot.
```

```
Directory: PWC File: pwc-Sd_LI_be.vs.Sd_LI_ka.txt

Numbers of common and distinct sites comparing Sd_LI_be versus Sd_LI_ka
================================================================================

  total number of potential sites:  123441397

  number of "Sd_LI_be_hsm"         sites: 6335325
  number of "Sd_LI_ka_hsm"         sites: 6222608
    number of "Sd_LI_be_hsm"-unique sites:     1496854
    number of common sites:        4838471
    number of "Sd_LI_ka_hsm"-unique sites:     1384137
  total number of "Sd_LI_be_hsm+Sd_LI_ka_hsm"-sites observed:    7719462

  number of "Sd_LI_be_scd"        sites:  43378941 ( 35.14% of total)
  number of "Sd_LI_ka_scd"        sites:  42719577 ( 34.61% of total)
    number of "Sd_LI_be_scd"-unique sites:      10075542
    number of sites in common:      33303399   (Expected:  15012225; O/E:       2.2)
    number of "Sd_LI_ka_scd"-unique sites:       9416178
  total number of "Sd_LI_be_scd+Sd_LI_ka_scd"-sites observed:     52795119

  number of sites in "Sd_LI_be_hsm" that are not  detectable in "Sd_LI_ka_hsm":    1020533
  number of sites in "Sd_LI_be_hsm" that are also detectable in "Sd_LI_ka_hsm":    5314792
    number of sites unique to "Sd_LI_be_hsm" although detectable in "Sd_LI_ka_hsm":            476321
    number of "Sd_LI_be_hsm" / "Sd_LI_ka_hsm" common sites                        :            4838471     (Expected: 849184; O/E:       5.7)

  number of sites in "Sd_LI_ka_hsm" that are not  detectable in "Sd_LI_be_hsm":    901473
  number of sites in "Sd_LI_ka_hsm" that are also detectable in "Sd_LI_be_hsm":    5321135
    number of sites unique to "Sd_LI_ka_hsm" although detectable in "Sd_LI_be_hsm":            482664
    number of "Sd_LI_ka_hsm" / "Sd_LI_be_hsm" common sites                        :            4838471     (Expected: 849184; O/E:       5.7)

  Overlap index of "Sd_LI_be_hsm" with "Sd_LI_ka_hsm":           0.836
  Estimated number of "Sd_LI_be_hsm" = "Sd_LI_ka_hsm" sites (assuming sampling from one population):        5844972
  Adjusted population size of "Sd_LI_be_hsm" = "Sd_LI_ka_hsm" sites (assuming all sites detectable):       21664800 (  2.81x of observed)

  number of "Sd_LI_be_hsm" sites with coverage >=  5: 6335325
  number of "Sd_LI_ka_hsm" sites with coverage >=  5: 6222608
  number of common sites with coverage >=  5: 4838471
  number of "Sd_LI_be_hsm" sites with coverage >=  6: 5822554
  number of "Sd_LI_ka_hsm" sites with coverage >=  6: 5726881
  number of common sites with coverage >=  6: 4398087
  number of "Sd_LI_be_hsm" sites with coverage >= 20: 774009
  number of "Sd_LI_ka_hsm" sites with coverage >= 20: 842361
  number of common sites with coverage >= 20: 500559
  number of "Sd_LI_be_hsm" sites with coverage >= 50: 108914
  number of "Sd_LI_ka_hsm" sites with coverage >= 50: 120244
  number of common sites with coverage >= 50: 87098
```

### Methylation Levels in Common Sites (Coverage >=  5)

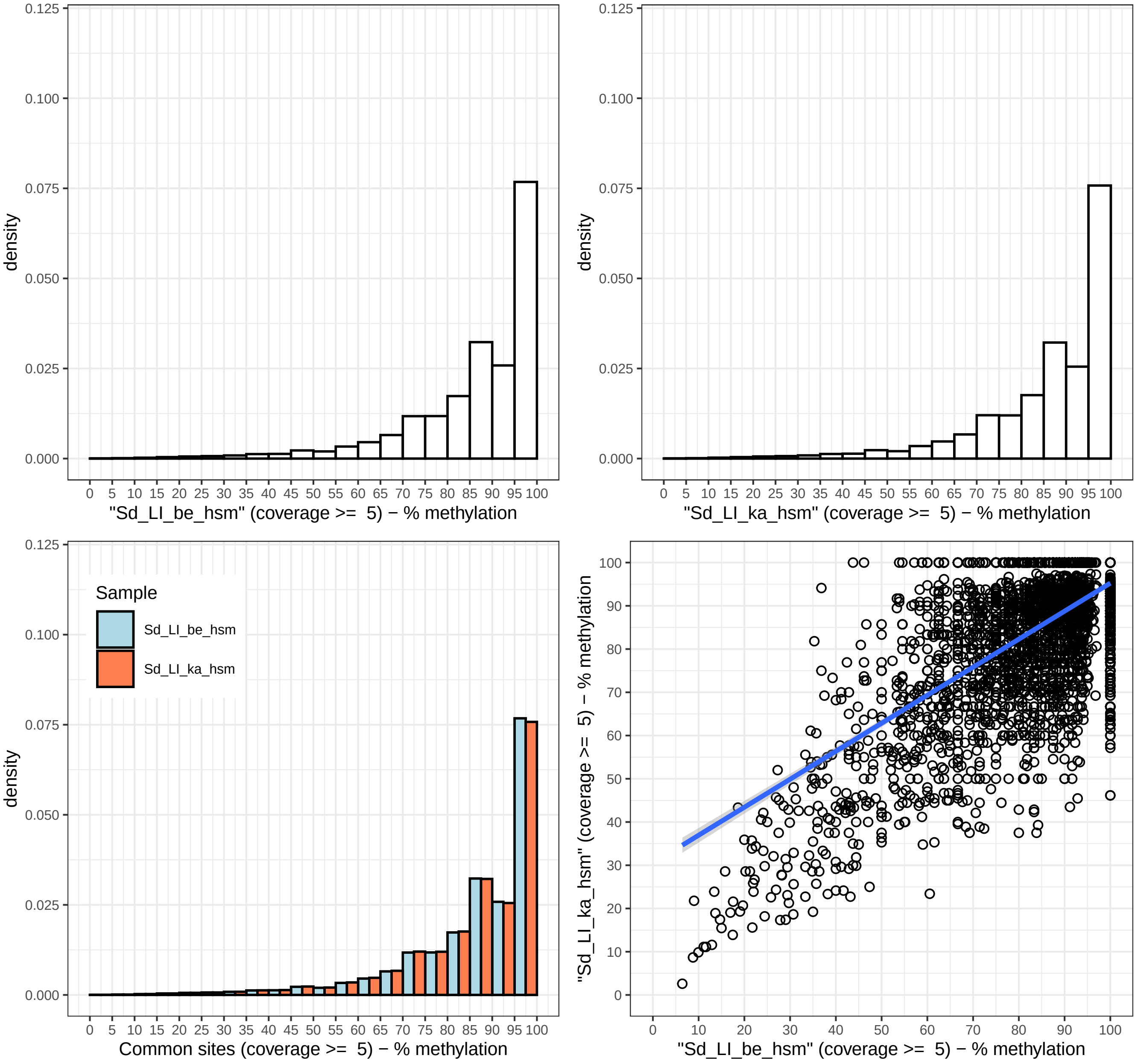

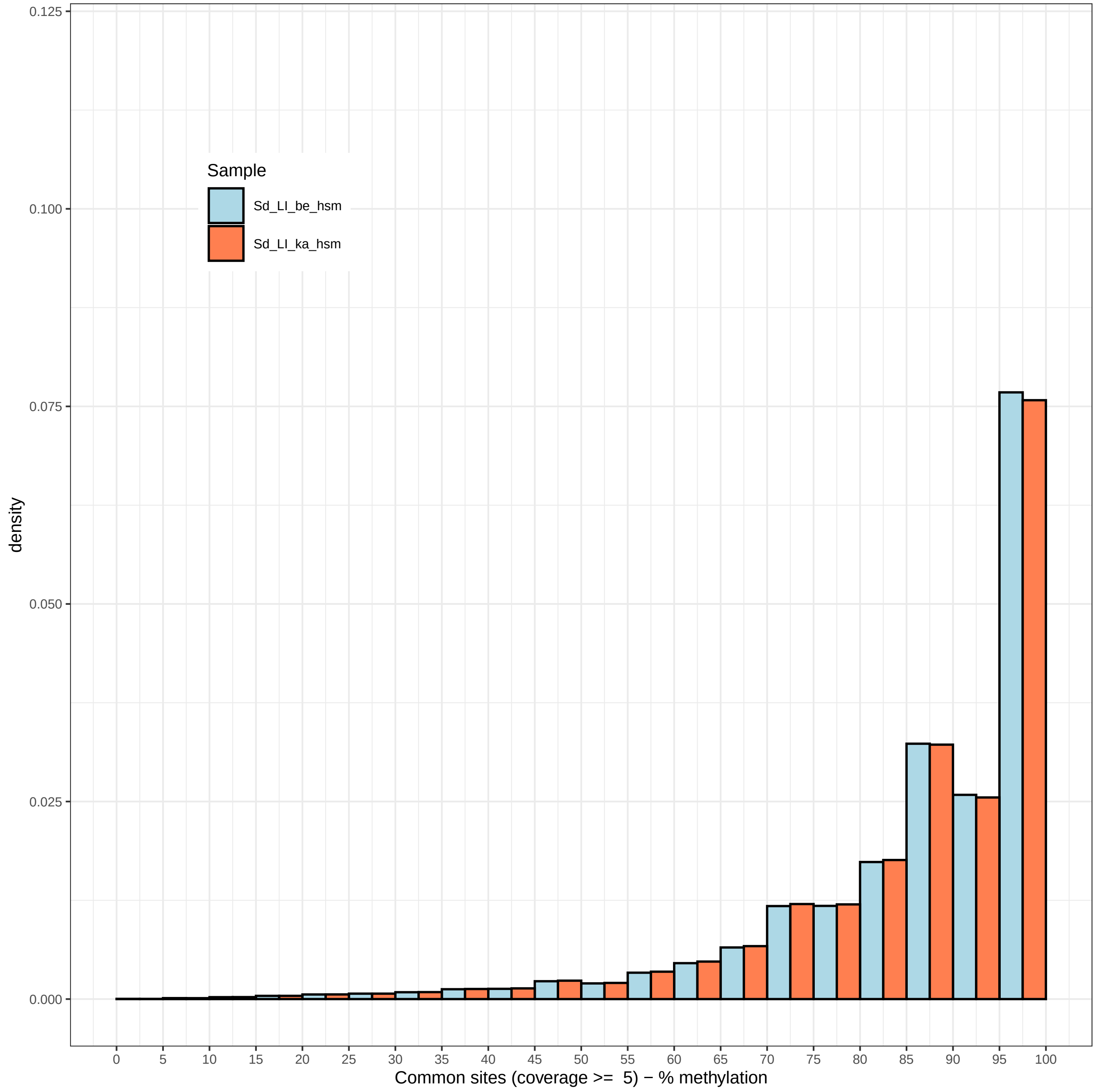

Sample
Sd_LI_be_hsm
Sd_LI_ka_hsm
density
0.000
0.025
0.050
0.075
0.100
0.125
0
5
10
15
20
25
30
35
40
45
50
55
60
65
70
75
80
85
90
95
100
Common sites (coverage >= 5) − % methylation

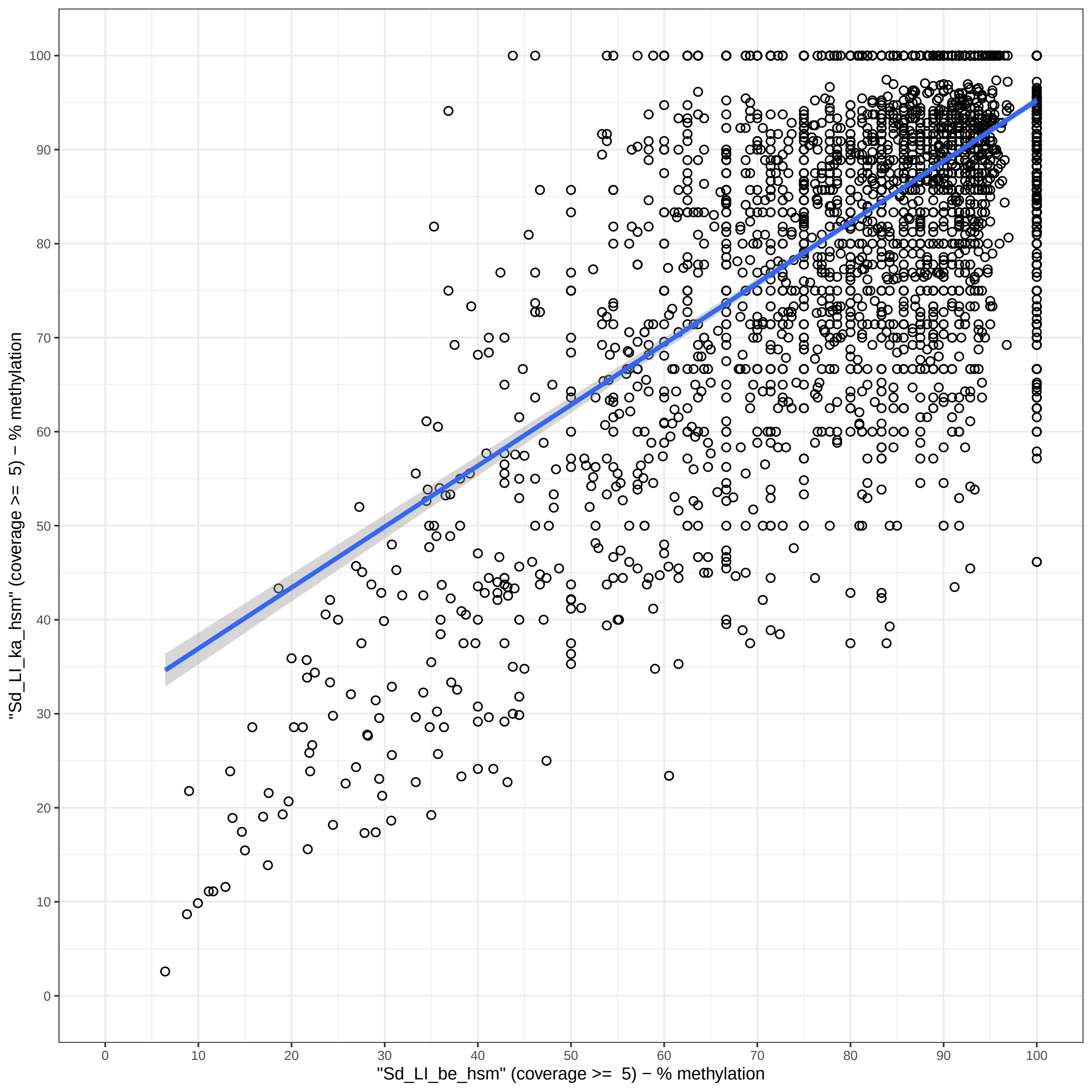

"Sd_LI_ka_hsm" (coverage >= 5) − % methylation
"Sd_LI_be_hsm" (coverage >= 5) − % methylation
0
10
20
30
40
50
60
70
80
90
100

#### Overlap of highly supported methylation sites (coverage >= 5)

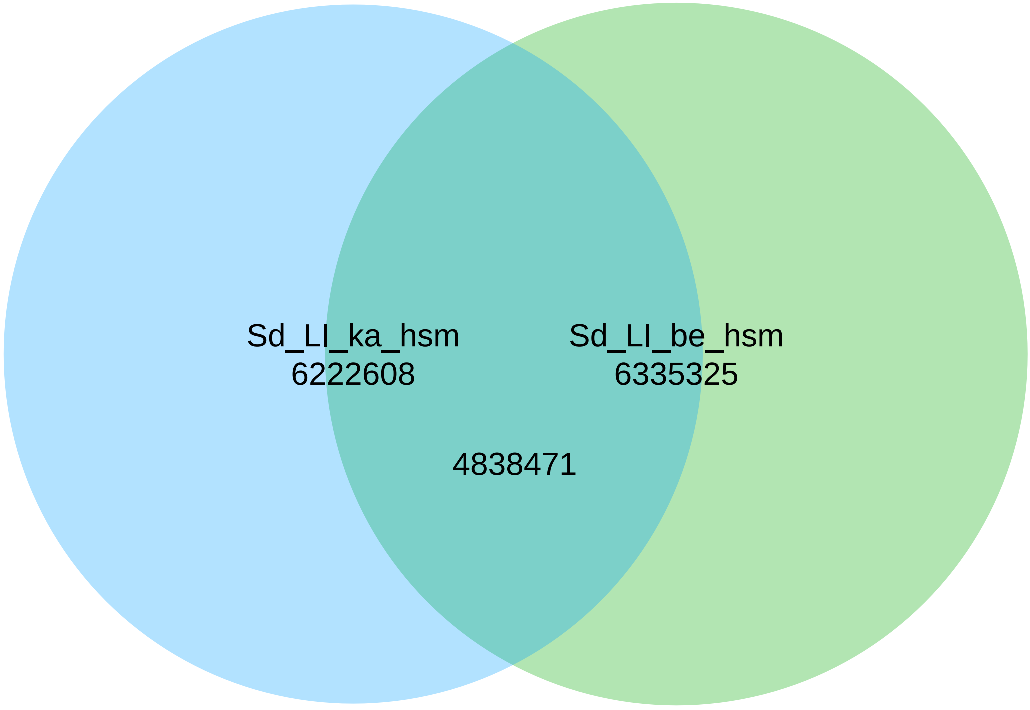

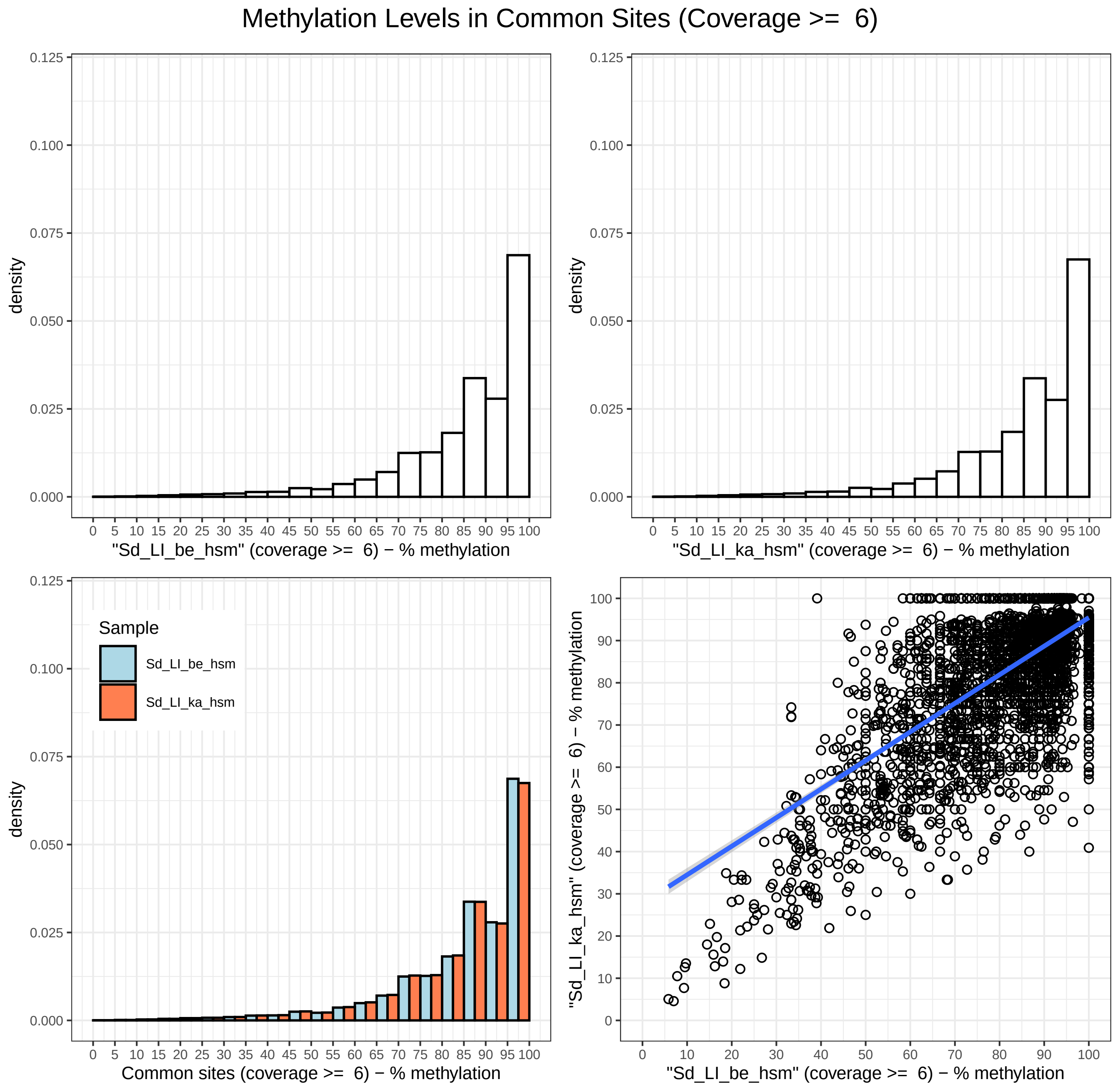

Methylation Levels in Common Sites (Coverage >= 6)
density
"Sd_LI_be_hsm" (coverage >= 6) − % methylation
"Sd_LI_ka_hsm" (coverage >= 6) − % methylation
Sample
Sd_LI_be_hsm
Sd_LI_ka_hsm
Common sites (coverage >= 6) − % methylation
"Sd_LI_ka_hsm" (coverage >= 6) − % methylation
"Sd_LI_be_hsm" (coverage >= 6) − % methylation

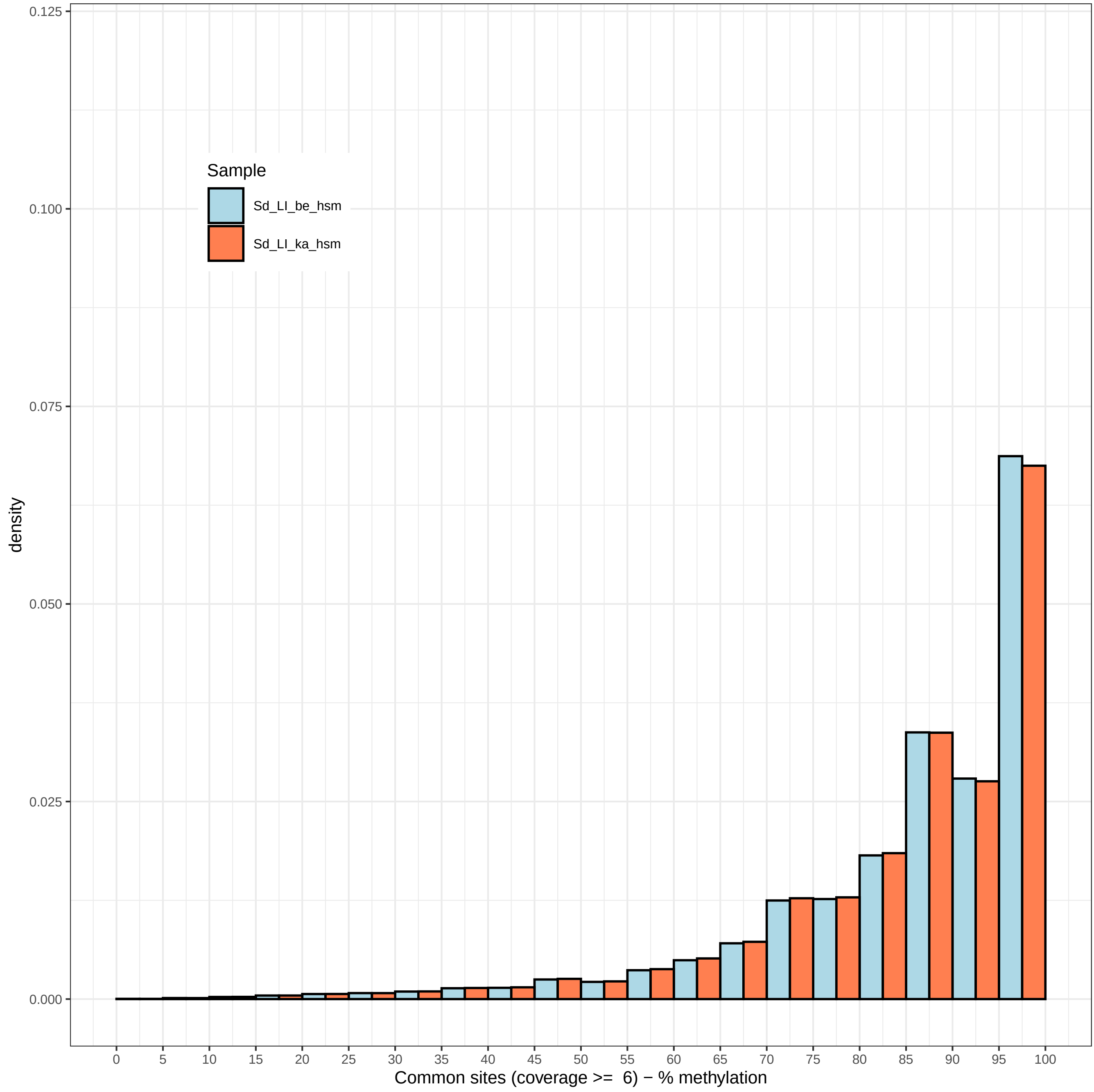

Sample
Sd_LI_be_hsm
Sd_LI_ka_hsm
density
0.000
0.025
0.050
0.075
0.100
0.125
0 5 10 15 20 25 30 35 40 45 50 55 60 65 70 75 80 85 90 95 100
Common sites (coverage >= 6) − % methylation

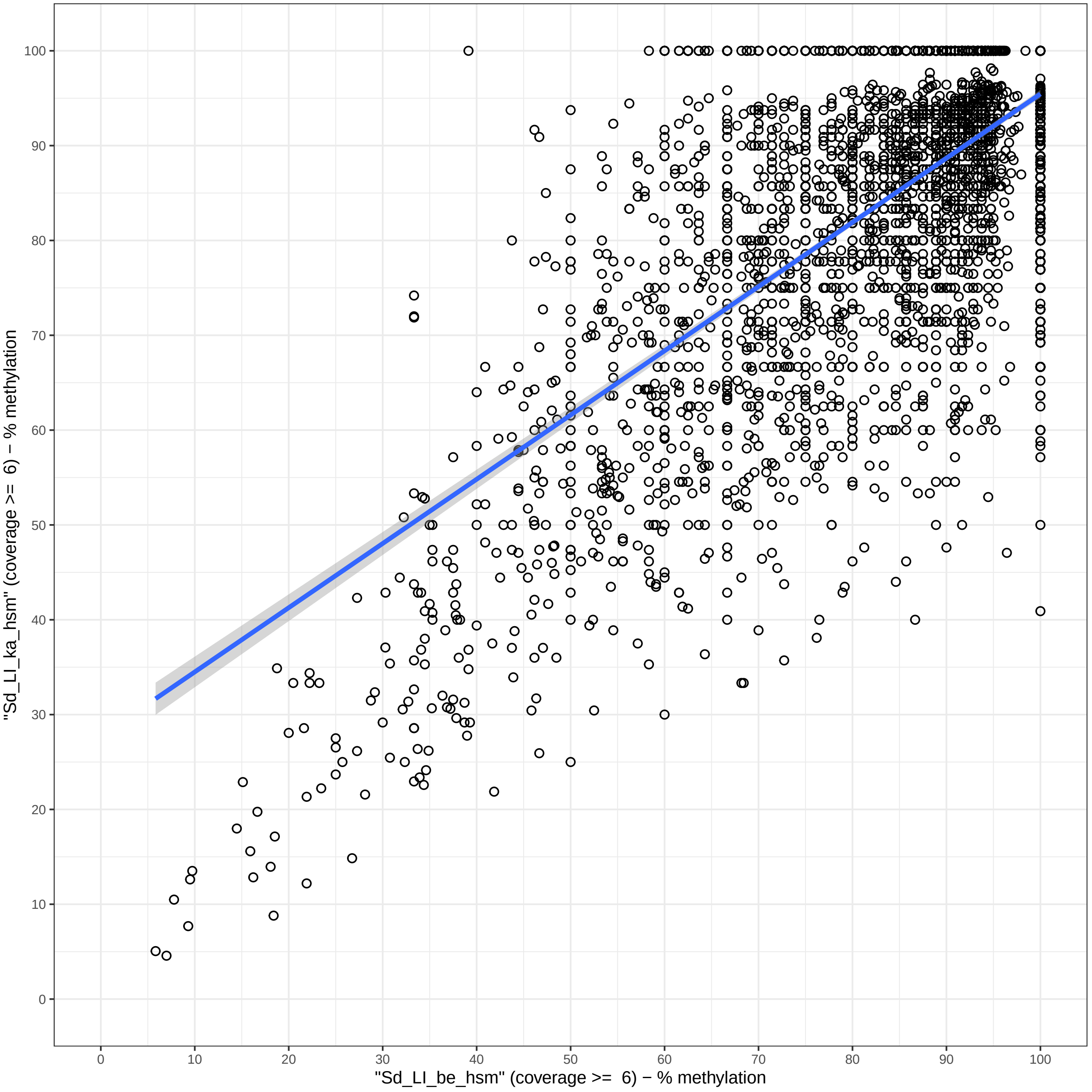
"Sd_LI_ka_hsm" (coverage >=  6) − % methylation
"Sd_LI_be_hsm" (coverage >=  6) − % methylation
0
10
20
30
40
50
60
70
80
90
100

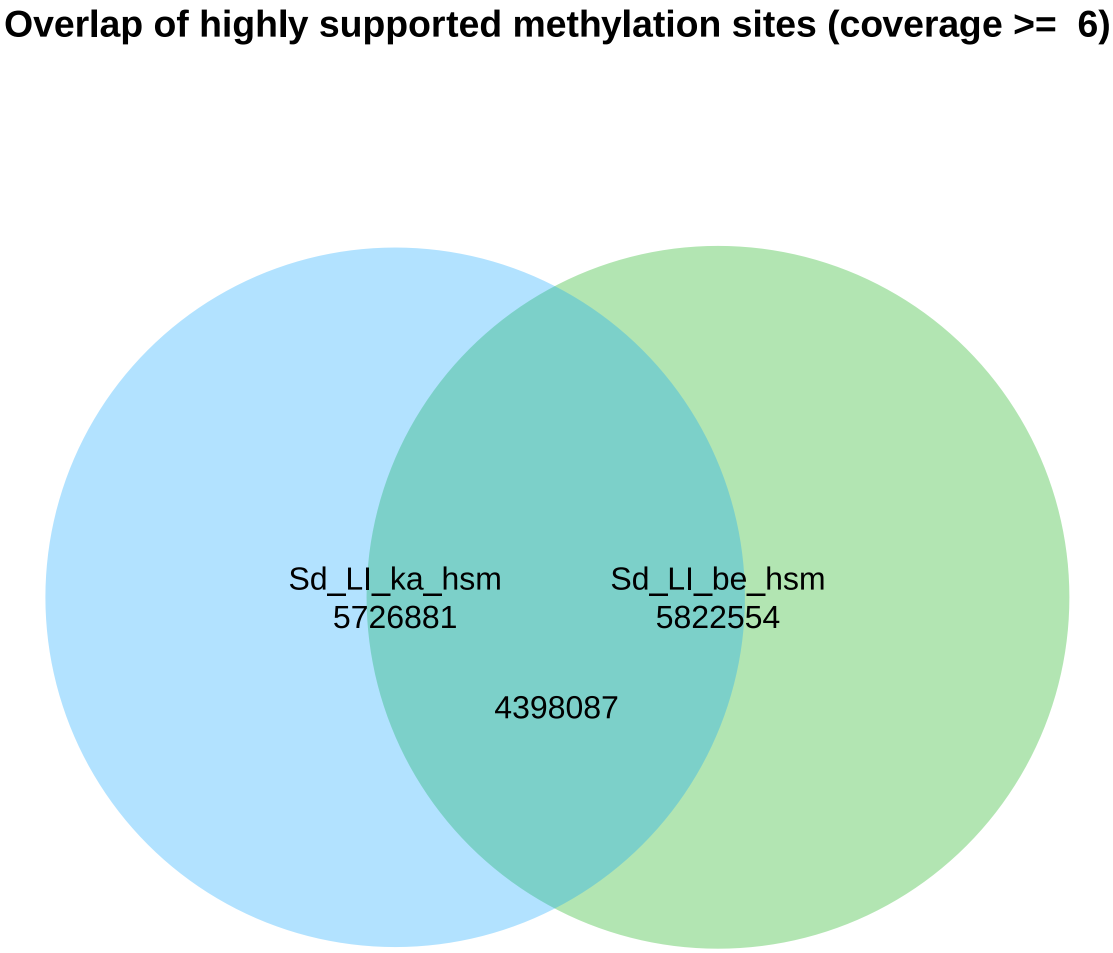
Overlap of highly supported methylation sites (coverage >= 6)
Sd_LI_ka_hsm
5726881
Sd_LI_be_hsm
5822554
4398087

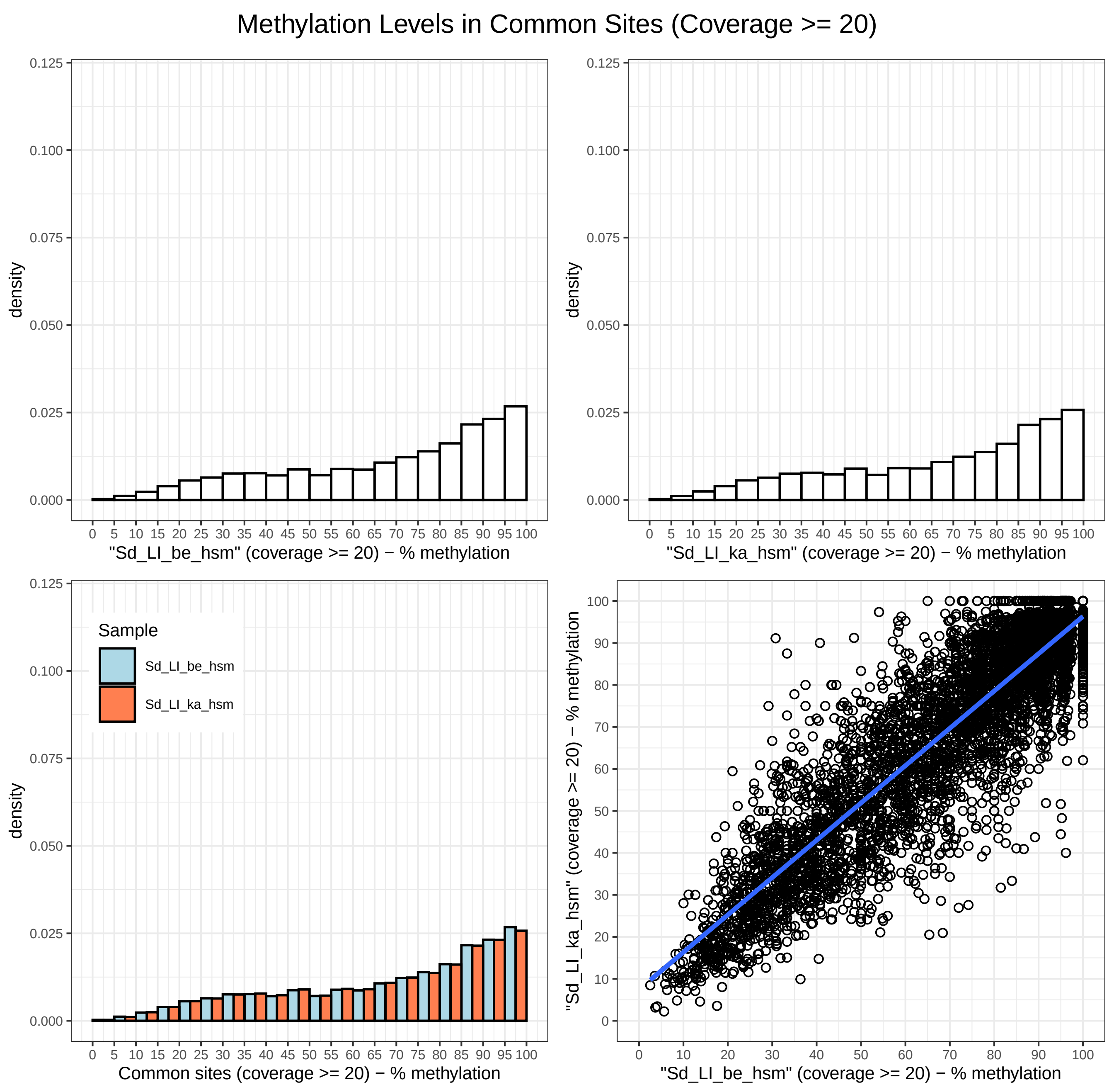

Methylation Levels in Common Sites (Coverage >= 20)
density
"Sd_LI_be_hsm" (coverage >= 20) − % methylation
density
"Sd_LI_ka_hsm" (coverage >= 20) − % methylation
Sample
Sd_LI_be_hsm
Sd_LI_ka_hsm
density
Common sites (coverage >= 20) − % methylation
"Sd_LI_ka_hsm" (coverage >= 20) − % methylation
"Sd_LI_be_hsm" (coverage >= 20) − % methylation

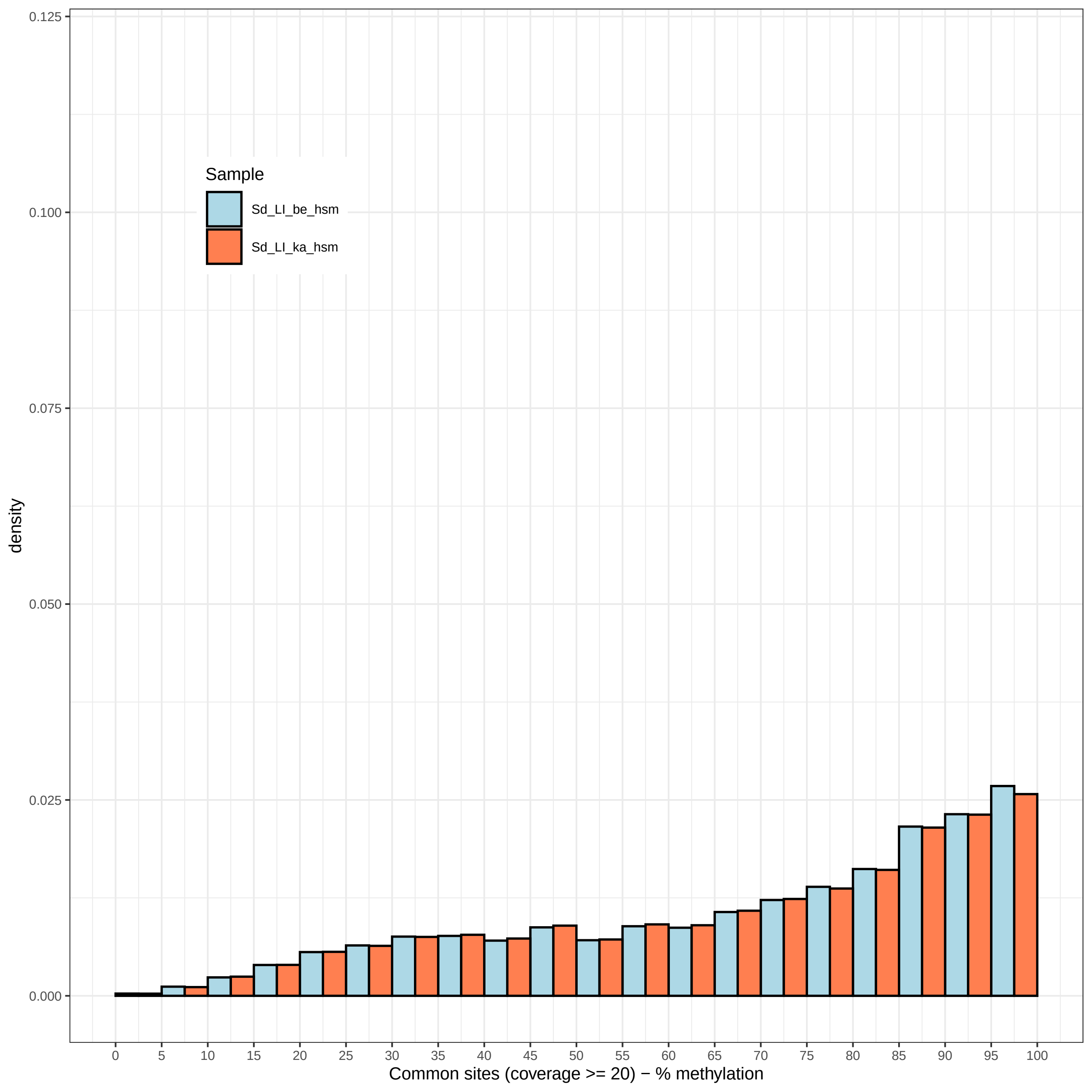
Sample
Sd_LI_be_hsm
Sd_LI_ka_hsm
density
0.125
0.100
0.075
0.050
0.025
0.000
0 5 10 15 20 25 30 35 40 45 50 55 60 65 70 75 80 85 90 95 100
Common sites (coverage >= 20) − % methylation

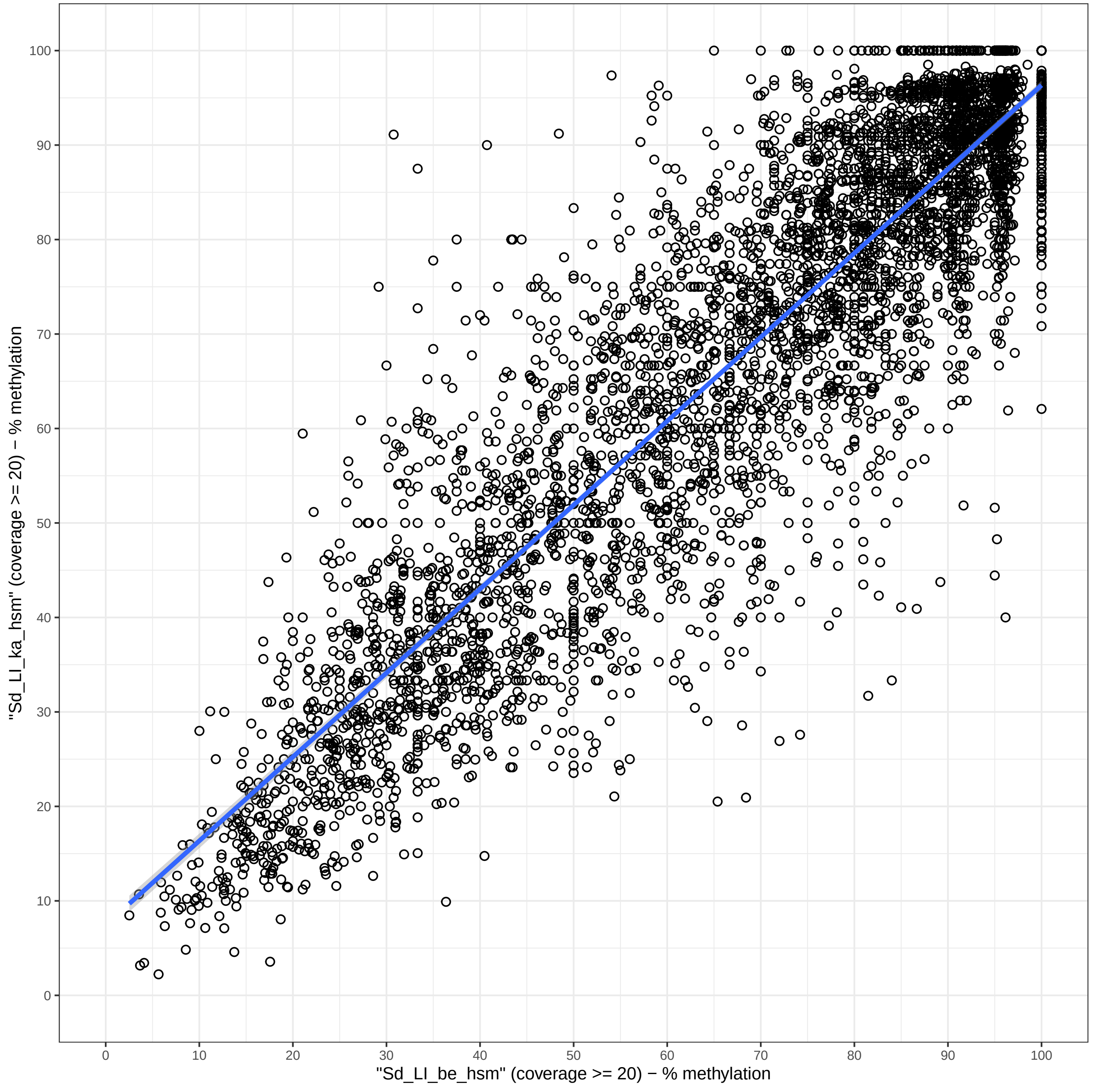

100
90
80
70
60
50
40
30
20
10
0
0
10
20
30
40
50
60
70
80
90
100
"Sd_LI_ka_hsm" (coverage >= 20) − % methylation
"Sd_LI_be_hsm" (coverage >= 20) − % methylation

#### Overlap of highly supported methylation sites (coverage >= 20)

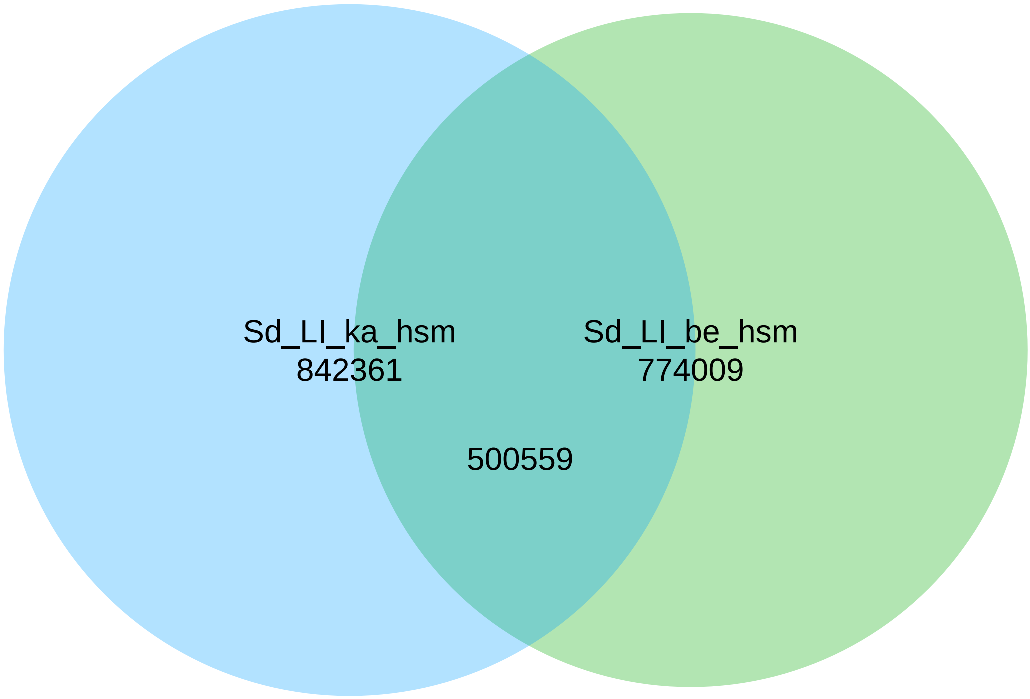

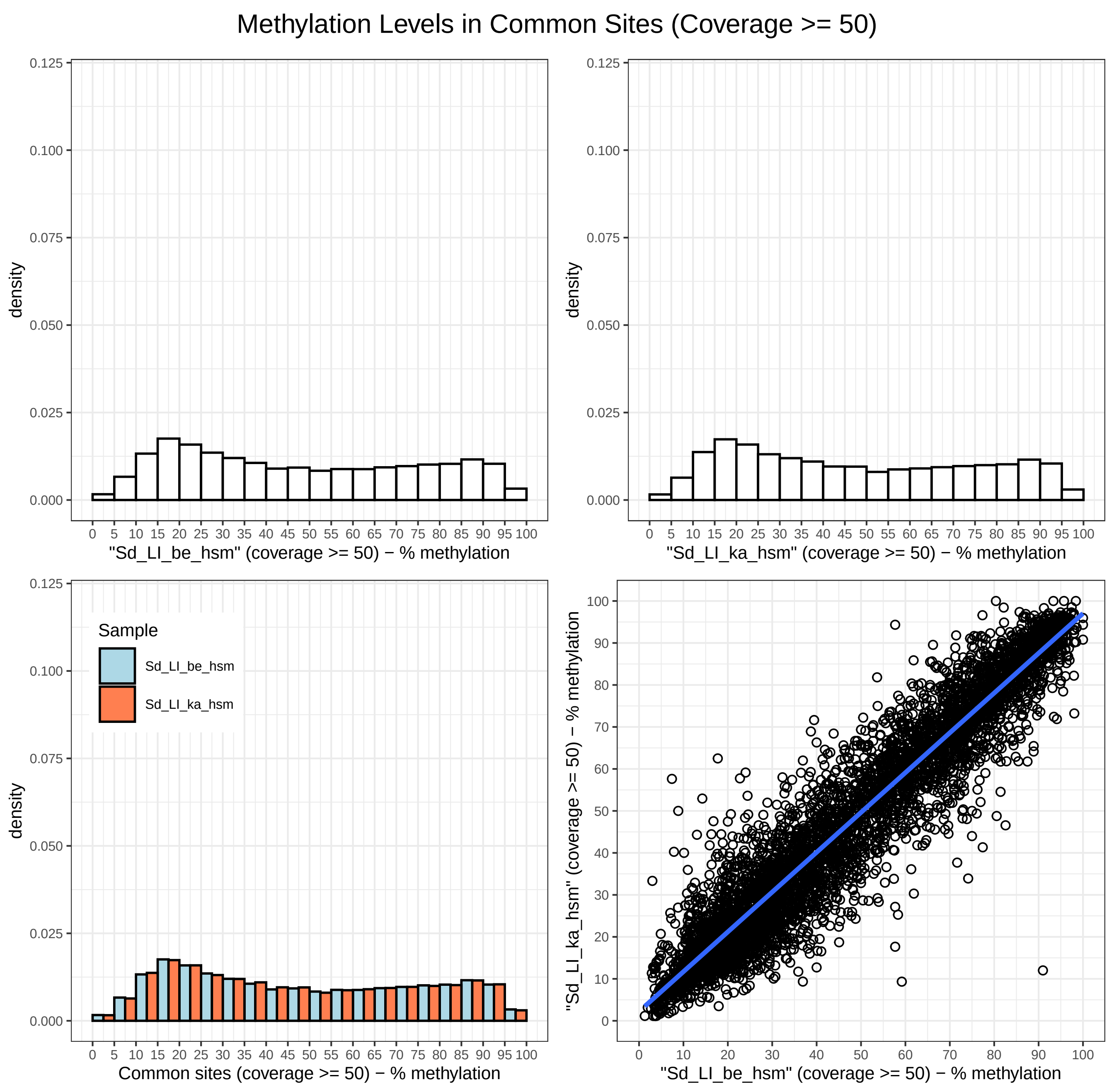

Methylation Levels in Common Sites (Coverage >= 50)
density
"Sd_LI_be_hsm" (coverage >= 50) − % methylation
"Sd_LI_ka_hsm" (coverage >= 50) − % methylation
Sample
Sd_LI_be_hsm
Sd_LI_ka_hsm
Common sites (coverage >= 50) − % methylation
"Sd_LI_ka_hsm" (coverage >= 50) − % methylation
"Sd_LI_be_hsm" (coverage >= 50) − % methylation

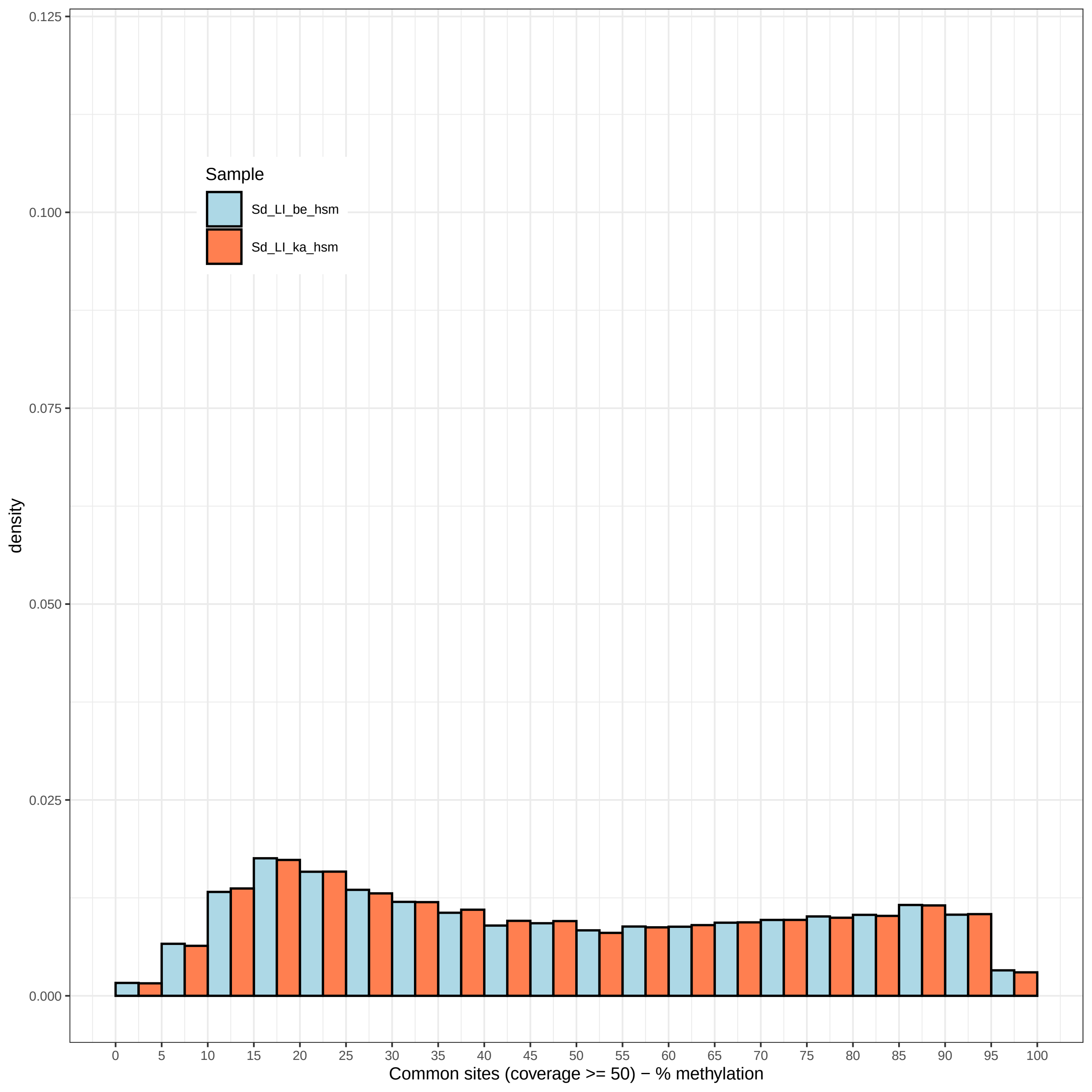

Sample
Sd_LI_be_hsm
Sd_LI_ka_hsm
density
0.125
0.100
0.075
0.050
0.025
0.000
0 5 10 15 20 25 30 35 40 45 50 55 60 65 70 75 80 85 90 95 100
Common sites (coverage >= 50) − % methylation

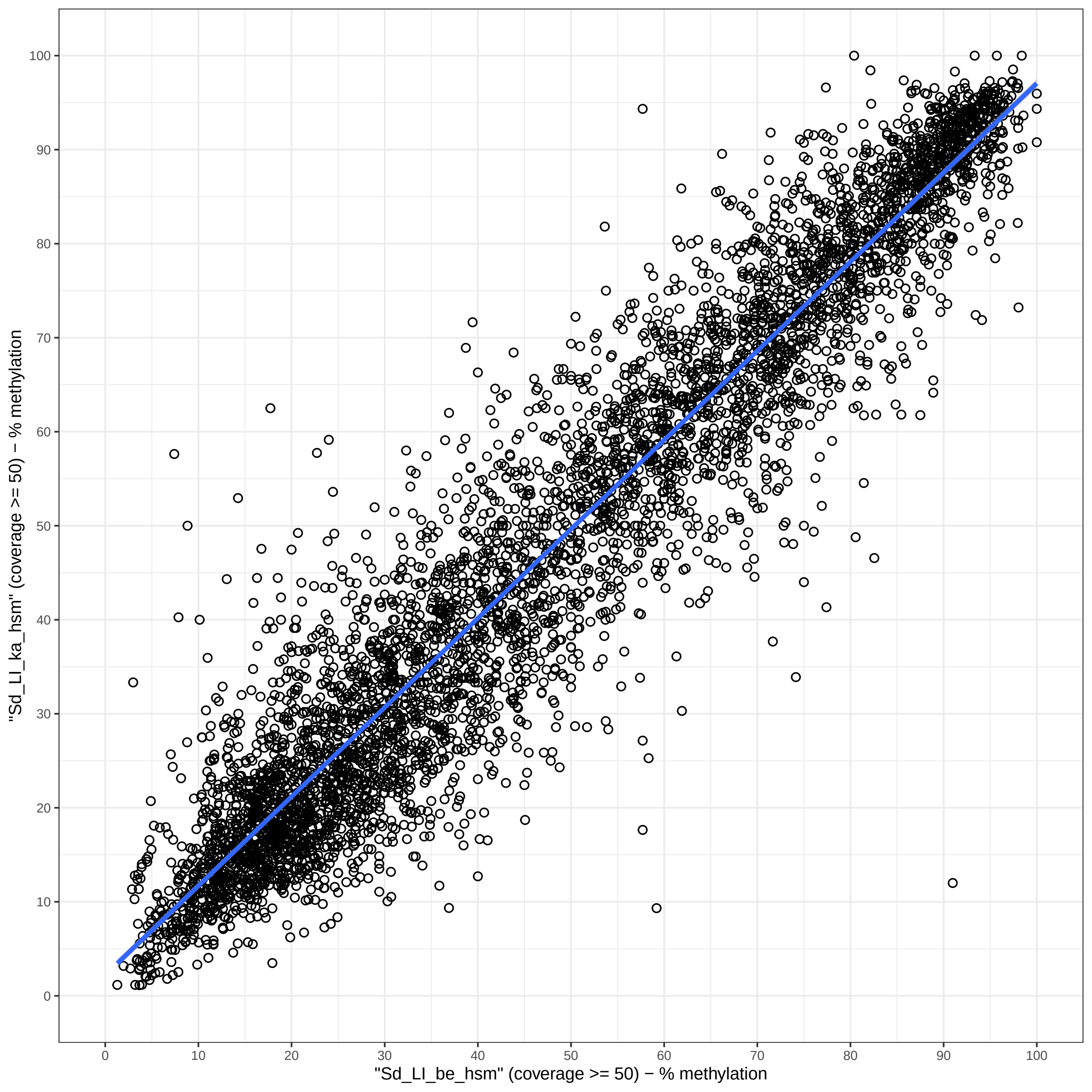
100
90
80
70
60
50
40
30
20
10
0
0
10
20
30
40
50
60
70
80
90
100
"Sd_LI_ka_hsm" (coverage >= 50) − % methylation
"Sd_LI_be_hsm" (coverage >= 50) − % methylation

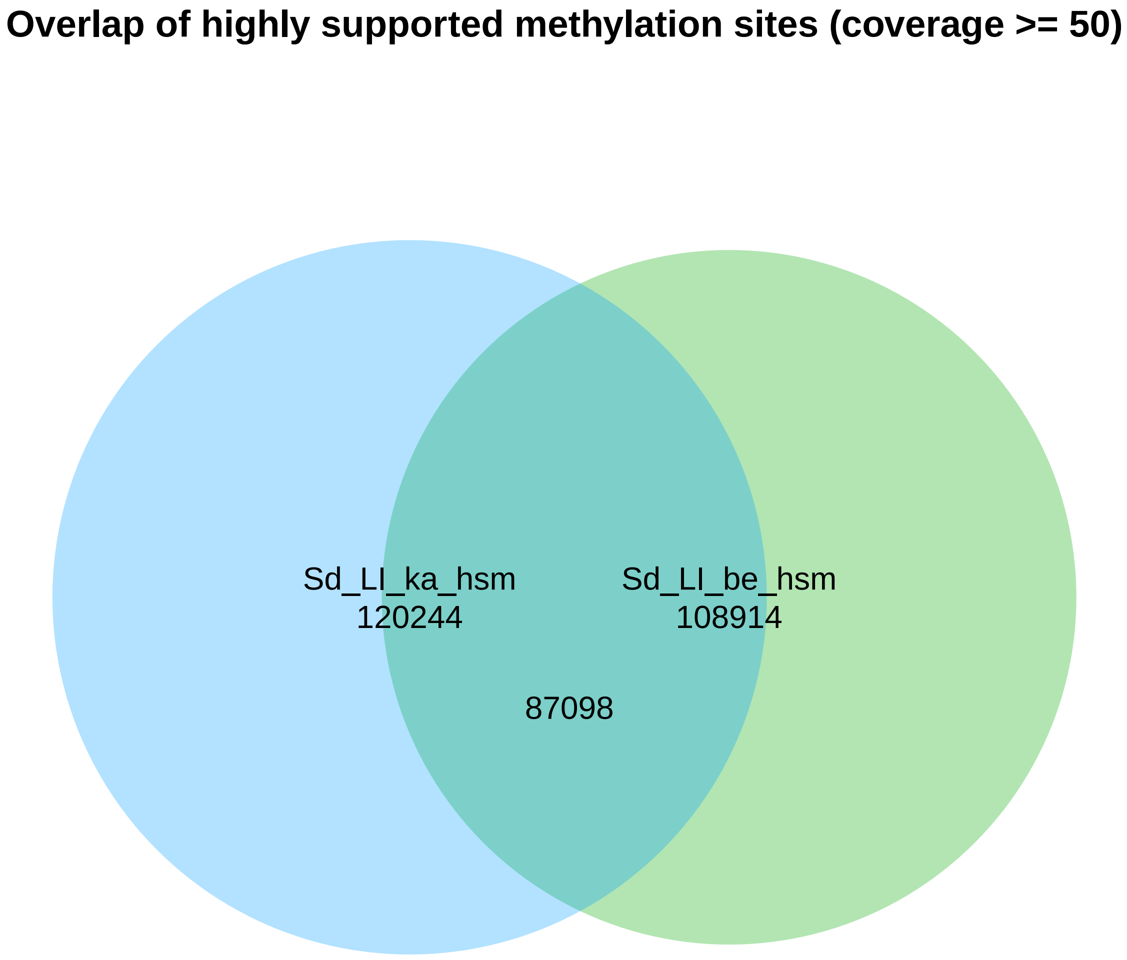

Overlap of highly supported methylation sites (coverage >= 50)
Sd_LI_ka_hsm
120244
Sd_LI_be_hsm
108914
87098

```
Directory: CRL File: 0READMEcrl

CRL - Correlations between aggregate samples.

Input: studymk

Output: files crl-*.txt crl-*.pdf

Notes: The output is generated by BWASPR::cmpSamples(), which uses methylKit::unite()
        to determine the sites common to all samples and applies methylKit::getCorrelation()
        and methylKit::PCASamples().  The *.txt files show the correlations in a table, and
        the *.pdf files show graphics.

       Beware that the number of common sites (shown at the bottom of the *.txt files) may be
        small, which may make the correlations less informative.
```

Directory: CRL File: crl-Sd_LI.txt

| | be | ka |
|---|---|---|
| be | 1.000 | 0.681 |
| ka | 0.681 | 1.000 |

The number of conserved sites is 4390509.

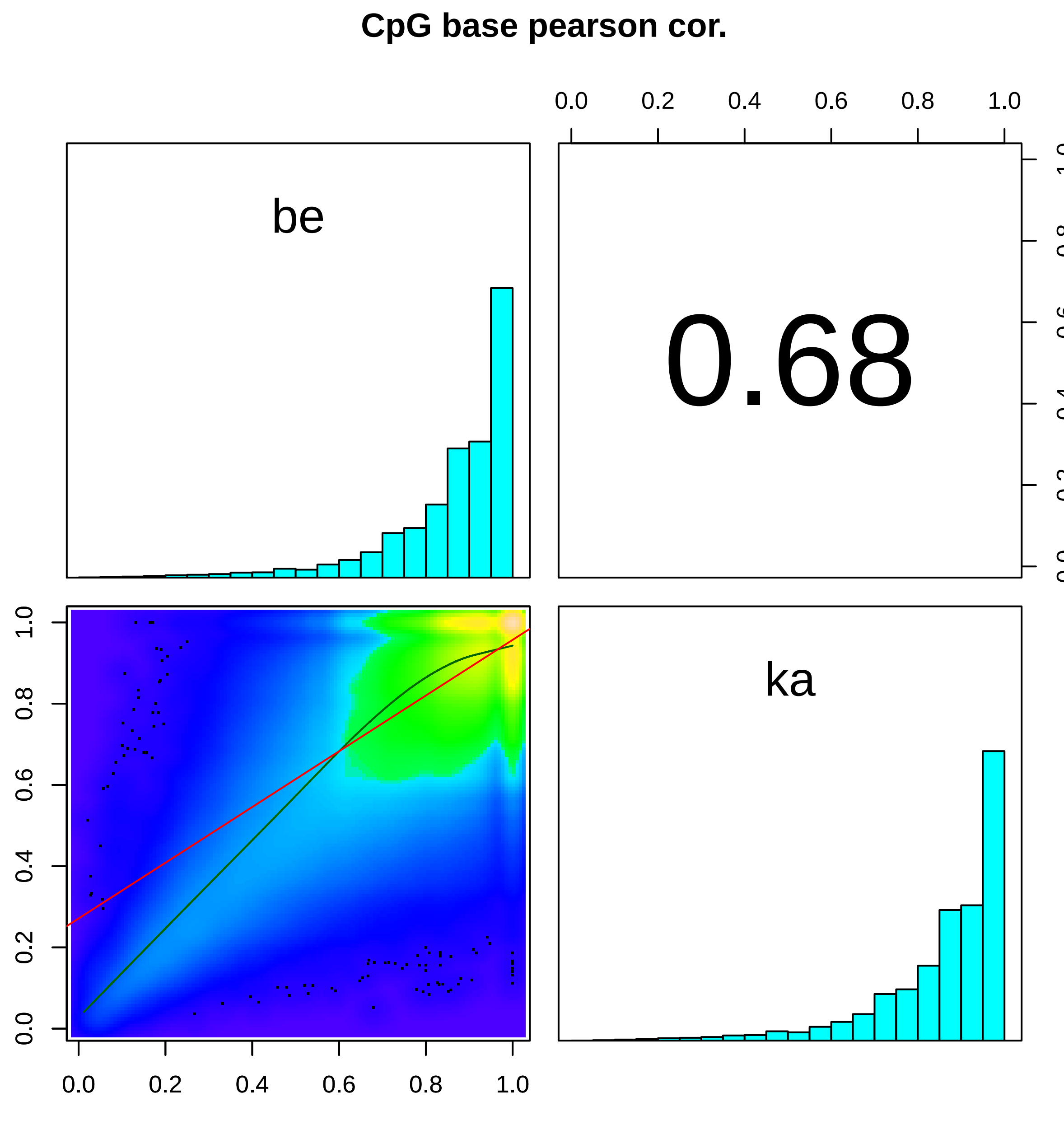

CpG base pearson cor.
be
0.68
ka
0.0
0.2
0.4
0.6
0.8
1.0

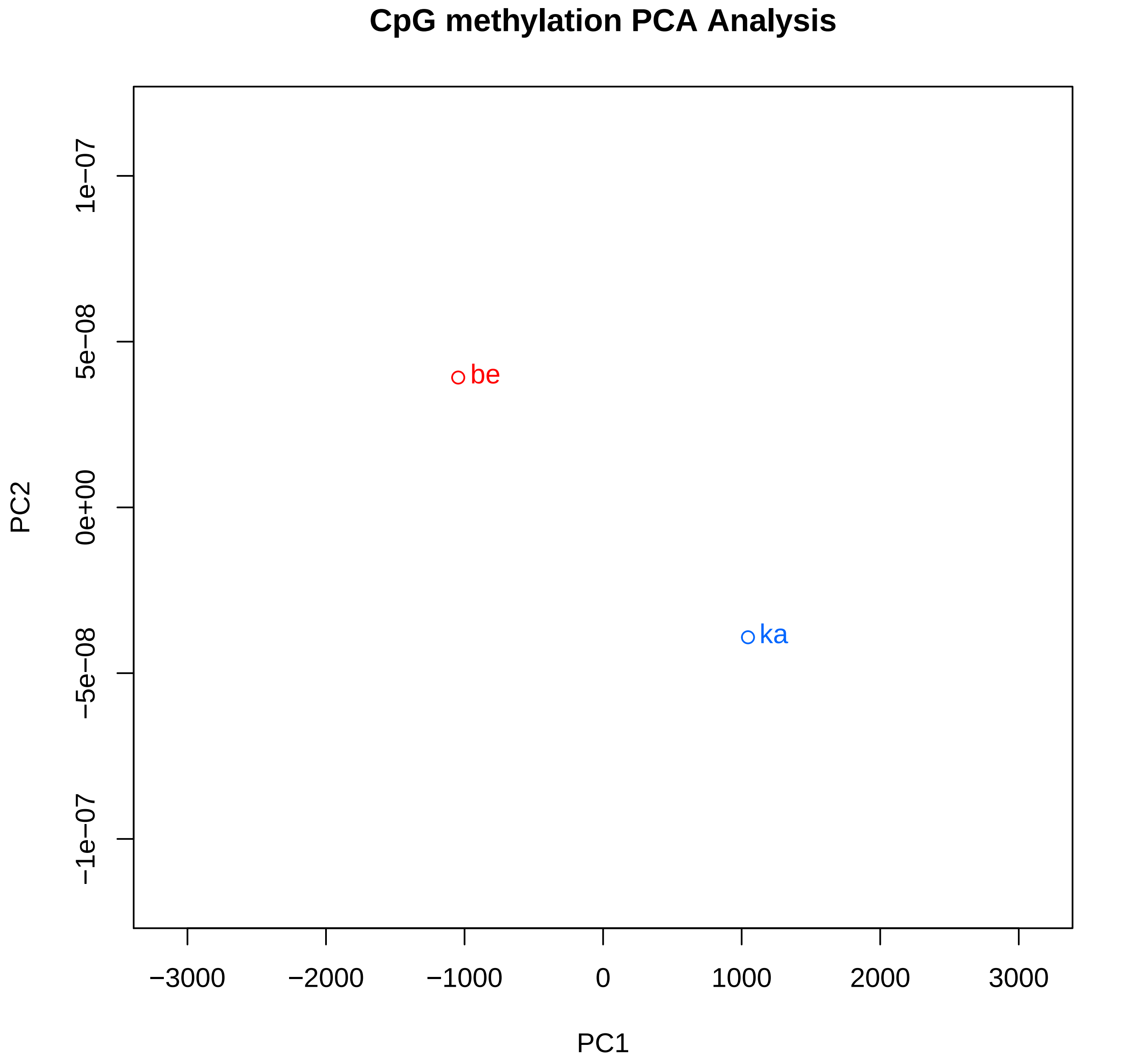

CpG methylation PCA Analysis
1e−07
5e−08
0e+00
−5e−08
−1e−07
PC2
be
ka
−3000
−2000
−1000
0
1000
2000
3000
PC1

Directory: REPCMS File: 0READMErepcms

REPCMS - Coverage and methylation statistics for replicate samples.

Input: studymk, repcovlist, replocount, rephicount

Output: files repcms-*txt and repcms-*pdf

```
Notes: The statistics are given for CpGhsm sites restricted to minimum or
        higher level coverage.  At the minimum level, the number of sites will be
        the number of lines of the corresponding *CpGhsm.mcalls file minus one
        (consistency check).  The distribution of methylation levels is really only
        interesting for high coverage because at low coverage there will be a strong
        bias towards high methylation levels (by definition of hsm sites).
        The same bias at high coverage would suggest that the sample includes a
        preponderance of consistently methylated genomic sites.

        replocount and rephicount set bounds on the coverage to exclude sites with too few
        or too many covering reads to provide statistics on a typical range.
```

```
Directory: REPCMS File: repcms-Sd_LI_be_1.txt

Number of "be_1" CpGhsm-sites with minimal and higher level coverage:

  number of "be_1" CpGhsm-sites with coverage >=  4: 306499
  number of "be_1" CpGhsm-sites with coverage >=  6:  94291
  number of "be_1" CpGhsm-sites with coverage >= 10:  36791
  number of "be_1" CpGhsm-sites with coverage >= 15:  23240

Coverage and methylation statistics for "be_1" CpGhsm-sites at different levels of minimum coverage:

methylKit::getCoverageStats output for "be_1" CpGhsm-sites (#: 306499) at minimum coverage 4   - read coverage statistics per base
summary:
   Min. 1st Qu.  Median    Mean 3rd Qu.    Max.
      4       4       5      11       6    2943
percentiles:
   0%   10%   20%   30%   40%   50%   60%   70%   80%   90%   95%   99% 99.5% 99.9%  100%
    4     4     4     4     4     5     5     6     7    11    23   145   258   625  2943

methylKit::getMethylationStats output for "be_1" CpGhsm-sites (#: 306499) at minimum coverage 4      - methylation statistics per base
summary:
   Min. 1st Qu.  Median    Mean 3rd Qu.    Max.
    1.7    83.3   100.0    91.6   100.0   100.0
percentiles:
   0%   10%   20%   30%   40%   50%   60%   70%   80%   90%   95%   99% 99.5% 99.9%  100%
  1.7  75.0  80.0  88.7 100.0 100.0 100.0 100.0 100.0 100.0 100.0 100.0 100.0 100.0 100.0

methylKit::getCoverageStats output for "be_1" CpGhsm-sites (#: 94291) at minimum coverage 6    - read coverage statistics per base
summary:
   Min. 1st Qu.  Median    Mean 3rd Qu.    Max.
      6       6       8      26      14    2943
percentiles:
   0%   10%   20%   30%   40%   50%   60%   70%   80%   90%   95%   99% 99.5% 99.9%  100%
    6     6     6     7     7     8     9    12    18    40    95   350   512  1035  2943

methylKit::getMethylationStats output for "be_1" CpGhsm-sites (#: 94291) at minimum coverage 6 - methylation statistics per base
summary:
   Min. 1st Qu.  Median    Mean 3rd Qu.    Max.
    1.7    70.0    83.3    79.2   100.0   100.0
percentiles:
   0%   10%   20%   30%   40%   50%   60%   70%   80%   90%   95%   99% 99.5% 99.9%  100%
  1.7  50.0  62.5  71.4  82.2  83.3  85.7  92.3 100.0 100.0 100.0 100.0 100.0 100.0 100.0

methylKit::getCoverageStats output for "be_1" CpGhsm-sites (#: 36791) at minimum coverage 10   - read coverage statistics per base
summary:
   Min. 1st Qu.  Median    Mean 3rd Qu.    Max.
     10      12      19      55      42    2943
percentiles:
   0%   10%   20%   30%   40%   50%   60%   70%   80%   90%   95%   99% 99.5% 99.9%  100%
   10    10    12    13    15    19    24    33    55   124   224   582   803  1401  2943

methylKit::getMethylationStats output for "be_1" CpGhsm-sites (#: 36791) at minimum coverage 10      - methylation statistics per base
summary:
   Min. 1st Qu.  Median    Mean 3rd Qu.    Max.
    1.7    50.0    68.8    65.2    84.6   100.0
percentiles:
   0%   10%   20%   30%   40%   50%   60%   70%   80%   90%   95%   99% 99.5% 99.9%  100%
```

```
  1.7  33.3  44.4  51.4  60.0  68.8  75.0  81.8  87.9  92.3  97.1 100.0 100.0 100.0 100.0

methylKit::getCoverageStats output for "be_1" CpGhsm-sites (#: 23240) at minimum coverage 15   - read coverage statistics per base
summary:
   Min. 1st Qu.  Median    Mean 3rd Qu.    Max. 
     15      20      31      80      74    2943 
percentiles:
   0%   10%   20%   30%   40%   50%   60%   70%   80%   90%   95%   99% 99.5% 99.9%  100% 
   15    16    18    21    25    31    41    59    96   186   306   705   948  1822  2943 

methylKit::getMethylationStats output for "be_1" CpGhsm-sites (#: 23240) at minimum coverage 15      - methylation statistics per base
summary:
   Min. 1st Qu.  Median    Mean 3rd Qu.    Max. 
    1.7    41.5    66.7    62.5    84.4   100.0 
percentiles:
   0%   10%   20%   30%   40%   50%   60%   70%   80%   90%   95%   99% 99.5% 99.9%  100% 
  1.7  26.9  37.0  46.9  57.1  66.7  75.0  81.4  87.0  92.2  94.7 100.0 100.0 100.0 100.0 

methylKit::getCoverageStats output for "be_1" CpGhsm-sites (#: 32336) in coverage range [10-100]     - read coverage statistics per base
summary:
   Min. 1st Qu.  Median    Mean 3rd Qu.    Max. 
   10.0    12.0    17.0    24.4    29.0   100.0 
percentiles:
   0%   10%   20%   30%   40%   50%   60%   70%   80%   90%   95%   99% 99.5% 99.9%  100% 
   10    10    11    12    14    17    20    25    34    52    70    93    97   100   100 

methylKit::getMethylationStats output for "be_1" CpGhsm-sites (#: 32336) in coverage range [10-100]  - methylation statistics per base
summary:
   Min. 1st Qu.  Median    Mean 3rd Qu.    Max. 
    9.0    50.0    69.2    66.6    84.6   100.0 
percentiles:
   0%   10%   20%   30%   40%   50%   60%   70%   80%   90%   95%   99% 99.5% 99.9%  100% 
  9.0  37.2  45.8  53.3  60.0  69.2  75.0  81.8  88.6  92.9 100.0 100.0 100.0 100.0 100.0 
```

```
Directory: REPCMS File: repcms-Sd_LI_be_2.txt

Number of "be_2" CpGhsm-sites with minimal and higher level coverage:

  number of "be_2" CpGhsm-sites with coverage >=  4: 394075
  number of "be_2" CpGhsm-sites with coverage >=  6: 121842
  number of "be_2" CpGhsm-sites with coverage >= 10:  42324
  number of "be_2" CpGhsm-sites with coverage >= 15:  26110

Coverage and methylation statistics for "be_2" CpGhsm-sites at different levels of minimum coverage:

methylKit::getCoverageStats output for "be_2" CpGhsm-sites (#: 394075) at minimum coverage 4   - read coverage statistics per base
summary:
   Min. 1st Qu.  Median    Mean 3rd Qu.    Max. 
      4       4       5      10       6    3220 
percentiles:
   0%   10%   20%   30%   40%   50%   60%   70%   80%   90%   95%   99% 99.5% 99.9%  100% 
    4     4     4     4     4     5     5     6     7    10    20   128   237   619  3220 

methylKit::getMethylationStats output for "be_2" CpGhsm-sites (#: 394075) at minimum coverage 4      - methylation statistics per base
summary:
   Min. 1st Qu.  Median    Mean 3rd Qu.    Max. 
    1.6    83.3   100.0    92.0   100.0   100.0 
percentiles:
    0%    10%    20%    30%    40%    50%    60%    70%    80%    90%    95%    99%  99.5%  99.9%   100% 
  1.63  75.36  80.00  88.89 100.00 100.00 100.00 100.00 100.00 100.00 100.00 100.00 100.00 100.00 100.00 

methylKit::getCoverageStats output for "be_2" CpGhsm-sites (#: 121842) at minimum coverage 6   - read coverage statistics per base
summary:
   Min. 1st Qu.  Median    Mean 3rd Qu.    Max. 
      6       6       8      24      12    3220 
percentiles:
   0%   10%   20%   30%   40%   50%   60%   70%   80%   90%   95%   99% 99.5% 99.9%  100% 
    6     6     6     6     7     8     9    11    16    34    80   331   500  1037  3220 

methylKit::getMethylationStats output for "be_2" CpGhsm-sites (#: 121842) at minimum coverage 6      - methylation statistics per base
summary:
   Min. 1st Qu.  Median    Mean 3rd Qu.    Max. 
    1.6    71.4    83.3    80.5   100.0   100.0 
percentiles:
    0%    10%    20%    30%    40%    50%    60%    70%    80%    90%    95%    99%  99.5%  99.9%   100% 
  1.63  50.00  66.67  75.00  83.33  83.33  87.50 100.00 100.00 100.00 100.00 100.00 100.00 100.00 100.00 

methylKit::getCoverageStats output for "be_2" CpGhsm-sites (#: 42324) at minimum coverage 10   - read coverage statistics per base
summary:
   Min. 1st Qu.  Median    Mean 3rd Qu.    Max. 
     10      12      18      55      40    3220 
percentiles:
   0%   10%   20%   30%   40%   50%   60%   70%   80%   90%   95%   99% 99.5% 99.9%  100% 
   10    10    11    13    15    18    23    32    53   119   225   593   823  1463  3220 

methylKit::getMethylationStats output for "be_2" CpGhsm-sites (#: 42324) at minimum coverage 10      - methylation statistics per base
summary:
   Min. 1st Qu.  Median    Mean 3rd Qu.    Max. 
    1.6    50.0    68.8    65.5    84.6   100.0 
percentiles:
    0%    10%    20%    30%    40%    50%    60%    70%    80%    90%    95%    99%  99.5%  99.9%   100% 
```

```
  1.63  33.33  45.45  52.63  60.00  68.75  75.00  81.82  87.76  92.31  97.96 100.00 100.00 100.00 100.00
```

```
methylKit::getCoverageStats output for "be_2" CpGhsm-sites (#: 26110) at minimum coverage 15   - read coverage statistics per base
summary:
   Min. 1st Qu.  Median    Mean 3rd Qu.    Max.
     15      20      31      82      73    3220
percentiles:
   0%   10%   20%   30%   40%   50%   60%   70%   80%   90%   95%   99% 99.5% 99.9%  100%
   15    16    18    21    25    31    41    59    94   188   318   733  1013  1917  3220

methylKit::getMethylationStats output for "be_2" CpGhsm-sites (#: 26110) at minimum coverage 15      - methylation statistics per base
summary:
   Min. 1st Qu.  Median    Mean 3rd Qu.    Max.
    1.6    41.2    65.9    62.0    83.9   100.0
percentiles:
    0%    10%    20%    30%    40%    50%    60%    70%    80%    90%    95%    99%  99.5%  99.9%   100%
  1.63  27.27  37.21  46.67  56.25  65.86  73.81  80.95  86.67  91.95  94.74 100.00 100.00 100.00 100.00
```

```
methylKit::getCoverageStats output for "be_2" CpGhsm-sites (#: 37385) in coverage range [10-100]    - read coverage statistics per base
summary:
   Min. 1st Qu.  Median    Mean 3rd Qu.    Max.
     10      12      16      24      28     100
percentiles:
   0%   10%   20%   30%   40%   50%   60%   70%   80%   90%   95%   99% 99.5% 99.9%  100%
   10    10    11    12    14    16    19    24    33    51    70    92    96   100   100

methylKit::getMethylationStats output for "be_2" CpGhsm-sites (#: 37385) in coverage range [10-100] - methylation statistics per base
summary:
   Min. 1st Qu.  Median    Mean 3rd Qu.    Max.
    9.0    50.0    69.2    66.8    84.6   100.0
percentiles:
   0%   10%   20%   30%   40%   50%   60%   70%   80%   90%   95%   99% 99.5% 99.9%  100%
  9.0  37.5  46.2  53.8  61.0  69.2  75.0  81.8  88.5  92.9 100.0 100.0 100.0 100.0 100.0
```

```
Directory: REPCMS File: repcms-Sd_LI_be_3.txt

Number of "be_3" CpGhsm-sites with minimal and higher level coverage:

  number of "be_3" CpGhsm-sites with coverage >=  4: 303128
  number of "be_3" CpGhsm-sites with coverage >=  6:  93096
  number of "be_3" CpGhsm-sites with coverage >= 10:  36325
  number of "be_3" CpGhsm-sites with coverage >= 15:  22983

Coverage and methylation statistics for "be_3" CpGhsm-sites at different levels of minimum coverage:

methylKit::getCoverageStats output for "be_3" CpGhsm-sites (#: 303128) at minimum coverage 4   - read coverage statistics per base
summary:
   Min. 1st Qu.  Median    Mean 3rd Qu.    Max.
      4       4       5      11       6    2917
percentiles:
   0%   10%   20%   30%   40%   50%   60%   70%   80%   90%   95%   99% 99.5% 99.9%  100%
    4     4     4     4     4     5     5     6     7    11    23   145   259   608  2917

methylKit::getMethylationStats output for "be_3" CpGhsm-sites (#: 303128) at minimum coverage 4      - methylation statistics per base
summary:
   Min. 1st Qu.  Median    Mean 3rd Qu.    Max.
    2.5    83.3   100.0    91.6   100.0   100.0
percentiles:
    0%    10%    20%    30%    40%    50%    60%    70%    80%    90%    95%    99%  99.5%  99.9%   100%
  2.45  75.00  80.00  87.88 100.00 100.00 100.00 100.00 100.00 100.00 100.00 100.00 100.00 100.00 100.00

methylKit::getCoverageStats output for "be_3" CpGhsm-sites (#: 93096) at minimum coverage 6    - read coverage statistics per base
summary:
   Min. 1st Qu.  Median    Mean 3rd Qu.    Max.
      6       6       8      26      14    2917
percentiles:
   0%   10%   20%   30%   40%   50%   60%   70%   80%   90%   95%   99% 99.5% 99.9%  100%
    6     6     6     7     7     8     9    12    18    41    94   354   511  1044  2917

methylKit::getMethylationStats output for "be_3" CpGhsm-sites (#: 93096) at minimum coverage 6 - methylation statistics per base
summary:
   Min. 1st Qu.  Median    Mean 3rd Qu.    Max.
    2.5    70.0    83.3    79.1   100.0   100.0
percentiles:
    0%    10%    20%    30%    40%    50%    60%    70%    80%    90%    95%    99%  99.5%  99.9%   100%
  2.45  50.00  62.50  71.43  81.82  83.33  85.71  92.16 100.00 100.00 100.00 100.00 100.00 100.00 100.00

methylKit::getCoverageStats output for "be_3" CpGhsm-sites (#: 36325) at minimum coverage 10   - read coverage statistics per base
summary:
   Min. 1st Qu.  Median    Mean 3rd Qu.    Max.
     10      12      19      55      42    2917
percentiles:
   0%   10%   20%   30%   40%   50%   60%   70%   80%   90%   95%   99% 99.5% 99.9%  100%
   10    10    12    13    15    19    24    34    56   122   225   566   796  1381  2917

methylKit::getMethylationStats output for "be_3" CpGhsm-sites (#: 36325) at minimum coverage 10      - methylation statistics per base
summary:
   Min. 1st Qu.  Median    Mean 3rd Qu.    Max.
    2.5    50.0    68.2    65.1    84.2   100.0
percentiles:
    0%    10%    20%    30%    40%    50%    60%    70%    80%    90%    95%    99%  99.5%  99.9%   100%
```

```
  2.45  33.33  45.00  51.85  60.00  68.16  75.00  81.45  87.30  92.31  97.14 100.00 100.00 100.00 100.00

methylKit::getCoverageStats output for "be_3" CpGhsm-sites (#: 22983) at minimum coverage 15   - read coverage statistics per base
summary:
   Min. 1st Qu.  Median    Mean 3rd Qu.    Max. 
     15      20      32      80      74    2917 
percentiles:
   0%   10%   20%   30%   40%   50%   60%   70%   80%   90%   95%   99% 99.5% 99.9%  100% 
   15    16    19    21    25    32    42    60    95   187   307   710   957  1824  2917 

methylKit::getMethylationStats output for "be_3" CpGhsm-sites (#: 22983) at minimum coverage 15      - methylation statistics per base
summary:
   Min. 1st Qu.  Median    Mean 3rd Qu.    Max. 
    2.5    42.1    66.7    62.4    84.2   100.0 
percentiles:
    0%    10%    20%    30%    40%    50%    60%    70%    80%    90%    95%    99%  99.5%  99.9%   100% 
  2.45  27.27  37.50  47.06  57.14  66.67  74.42  81.25  86.67  92.00  95.00 100.00 100.00 100.00 100.00 

methylKit::getCoverageStats output for "be_3" CpGhsm-sites (#: 31955) in coverage range [10-100]    - read coverage statistics per base
summary:
   Min. 1st Qu.  Median    Mean 3rd Qu.    Max. 
   10.0    12.0    17.0    24.6    29.0   100.0 
percentiles:
   0%   10%   20%   30%   40%   50%   60%   70%   80%   90%   95%   99% 99.5% 99.9%  100% 
   10    10    11    12    14    17    20    25    34    54    71    93    97   100   100 

methylKit::getMethylationStats output for "be_3" CpGhsm-sites (#: 31955) in coverage range [10-100] - methylation statistics per base
summary:
   Min. 1st Qu.  Median    Mean 3rd Qu.    Max. 
    9.0    50.0    68.8    66.4    84.5   100.0 
percentiles:
   0%   10%   20%   30%   40%   50%   60%   70%   80%   90%   95%   99% 99.5% 99.9%  100% 
  9.0  37.5  45.9  53.7  60.0  68.8  75.0  81.8  87.8  92.9 100.0 100.0 100.0 100.0 100.0 
```

```
Directory: REPCMS File: repcms-Sd_LI_be_4.txt

Number of "be_4" CpGhsm-sites with minimal and higher level coverage:

  number of "be_4" CpGhsm-sites with coverage >=  4: 316576
  number of "be_4" CpGhsm-sites with coverage >=  6:  97218
  number of "be_4" CpGhsm-sites with coverage >= 10:  37238
  number of "be_4" CpGhsm-sites with coverage >= 15:  23444

Coverage and methylation statistics for "be_4" CpGhsm-sites at different levels of minimum coverage:

methylKit::getCoverageStats output for "be_4" CpGhsm-sites (#: 316576) at minimum coverage 4   - read coverage statistics per base
summary:
   Min. 1st Qu.  Median    Mean 3rd Qu.    Max.
      4       4       5      11       6    2999
percentiles:
   0%   10%   20%   30%   40%   50%   60%   70%   80%   90%   95%   99% 99.5% 99.9%  100%
    4     4     4     4     4     5     5     6     7    11    22   144   254   623  2999

methylKit::getMethylationStats output for "be_4" CpGhsm-sites (#: 316576) at minimum coverage 4      - methylation statistics per base
summary:
   Min. 1st Qu.  Median    Mean 3rd Qu.    Max.
    2.5    83.3   100.0    91.7   100.0   100.0
percentiles:
   0%   10%   20%   30%   40%   50%   60%   70%   80%   90%   95%   99% 99.5% 99.9%  100%
  2.5  75.0  80.0  88.9 100.0 100.0 100.0 100.0 100.0 100.0 100.0 100.0 100.0 100.0 100.0

methylKit::getCoverageStats output for "be_4" CpGhsm-sites (#: 97218) at minimum coverage 6    - read coverage statistics per base
summary:
   Min. 1st Qu.  Median    Mean 3rd Qu.    Max.
      6       6       8      25      14    2999
percentiles:
   0%   10%   20%   30%   40%   50%   60%   70%   80%   90%   95%   99% 99.5% 99.9%  100%
    6     6     6     6     7     8     9    12    18    40    91   348   510  1020  2999

methylKit::getMethylationStats output for "be_4" CpGhsm-sites (#: 97218) at minimum coverage 6 - methylation statistics per base
summary:
   Min. 1st Qu.  Median    Mean 3rd Qu.    Max.
    2.5    70.4    83.3    79.4   100.0   100.0
percentiles:
   0%   10%   20%   30%   40%   50%   60%   70%   80%   90%   95%   99% 99.5% 99.9%  100%
  2.5  50.0  63.6  72.0  82.9  83.3  85.7  92.9 100.0 100.0 100.0 100.0 100.0 100.0 100.0

methylKit::getCoverageStats output for "be_4" CpGhsm-sites (#: 37238) at minimum coverage 10   - read coverage statistics per base
summary:
   Min. 1st Qu.  Median    Mean 3rd Qu.    Max.
     10      12      19      55      42    2999
percentiles:
   0%   10%   20%   30%   40%   50%   60%   70%   80%   90%   95%   99% 99.5% 99.9%  100%
   10    10    11    13    15    19    24    33    56   125   225   573   795  1412  2999

methylKit::getMethylationStats output for "be_4" CpGhsm-sites (#: 37238) at minimum coverage 10      - methylation statistics per base
summary:
   Min. 1st Qu.  Median    Mean 3rd Qu.    Max.
    2.5    48.4    68.8    65.3    84.6   100.0
percentiles:
   0%   10%   20%   30%   40%   50%   60%   70%   80%   90%   95%   99% 99.5% 99.9%  100%
```

```
  2.5  33.3  44.4  51.2  60.0  68.8  75.0  81.8  88.0  92.6  97.6 100.0 100.0 100.0 100.0

methylKit::getCoverageStats output for "be_4" CpGhsm-sites (#: 23444) at minimum coverage 15   - read coverage statistics per base
summary:
   Min. 1st Qu.  Median    Mean 3rd Qu.    Max.
     15      20      31      81      74    2999
percentiles:
   0%   10%   20%   30%   40%   50%   60%   70%   80%   90%   95%   99% 99.5% 99.9%  100%
   15    16    18    21    25    31    41    60    95   188   312   723   944  1783  2999

methylKit::getMethylationStats output for "be_4" CpGhsm-sites (#: 23444) at minimum coverage 15      - methylation statistics per base
summary:
   Min. 1st Qu.  Median    Mean 3rd Qu.    Max.
    2.5    41.9    66.7    62.4    84.2   100.0
percentiles:
   0%   10%   20%   30%   40%   50%   60%   70%   80%   90%   95%   99% 99.5% 99.9%  100%
  2.5  26.9  37.5  46.7  56.7  66.7  75.0  81.2  87.0  92.4  95.0 100.0 100.0 100.0 100.0

methylKit::getCoverageStats output for "be_4" CpGhsm-sites (#: 32758) in coverage range [10-100]    - read coverage statistics per base
summary:
   Min. 1st Qu.  Median    Mean 3rd Qu.    Max.
   10.0    12.0    16.0    24.4    29.0   100.0
percentiles:
   0%   10%   20%   30%   40%   50%   60%   70%   80%   90%   95%   99% 99.5% 99.9%  100%
   10    10    11    12    14    16    20    25    34    53    71    93    96   100   100

methylKit::getMethylationStats output for "be_4" CpGhsm-sites (#: 32758) in coverage range [10-100]  - methylation statistics per base
summary:
   Min. 1st Qu.  Median    Mean 3rd Qu.    Max.
    9.0    50.0    69.2    66.7    84.6   100.0
percentiles:
   0%   10%   20%   30%   40%   50%   60%   70%   80%   90%   95%   99% 99.5% 99.9%  100%
  9.0  37.5  45.8  53.4  60.2  69.2  75.0  81.8  88.9  93.2 100.0 100.0 100.0 100.0 100.0
```

```
Directory: REPCMS File: repcms-Sd_LI_be_5.txt

Number of "be_5" CpGhsm-sites with minimal and higher level coverage:

  number of "be_5" CpGhsm-sites with coverage >=  4: 411806
  number of "be_5" CpGhsm-sites with coverage >=  6: 127731
  number of "be_5" CpGhsm-sites with coverage >= 10:  43988
  number of "be_5" CpGhsm-sites with coverage >= 15:  26941

Coverage and methylation statistics for "be_5" CpGhsm-sites at different levels of minimum coverage:

methylKit::getCoverageStats output for "be_5" CpGhsm-sites (#: 411806) at minimum coverage 4   - read coverage statistics per base
summary:
   Min. 1st Qu.  Median    Mean 3rd Qu.    Max.
      4       4       5      10       6    3209
percentiles:
   0%   10%   20%   30%   40%   50%   60%   70%   80%   90%   95%   99% 99.5% 99.9%  100%
    4     4     4     4     4     5     5     6     7    10    19   124   235   615  3209

methylKit::getMethylationStats output for "be_5" CpGhsm-sites (#: 411806) at minimum coverage 4      - methylation statistics per base
summary:
   Min. 1st Qu.  Median    Mean 3rd Qu.    Max.
    2.3    83.3   100.0    91.9   100.0   100.0
percentiles:
    0%    10%    20%    30%    40%    50%    60%    70%    80%    90%    95%    99%  99.5%  99.9%   100%
  2.28  75.00  80.00  88.89 100.00 100.00 100.00 100.00 100.00 100.00 100.00 100.00 100.00 100.00 100.00

methylKit::getCoverageStats output for "be_5" CpGhsm-sites (#: 127731) at minimum coverage 6   - read coverage statistics per base
summary:
   Min. 1st Qu.  Median    Mean 3rd Qu.    Max.
      6       6       8      23      12    3209
percentiles:
   0%   10%   20%   30%   40%   50%   60%   70%   80%   90%   95%   99% 99.5% 99.9%  100%
    6     6     6     6     7     8     9    11    15    32    77   328   490  1021  3209

methylKit::getMethylationStats output for "be_5" CpGhsm-sites (#: 127731) at minimum coverage 6      - methylation statistics per base
summary:
   Min. 1st Qu.  Median    Mean 3rd Qu.    Max.
    2.3    71.4    83.3    80.5   100.0   100.0
percentiles:
    0%    10%    20%    30%    40%    50%    60%    70%    80%    90%    95%    99%  99.5%  99.9%   100%
  2.28  50.18  66.67  75.00  83.33  83.33  87.01 100.00 100.00 100.00 100.00 100.00 100.00 100.00 100.00

methylKit::getCoverageStats output for "be_5" CpGhsm-sites (#: 43988) at minimum coverage 10   - read coverage statistics per base
summary:
   Min. 1st Qu.  Median    Mean 3rd Qu.    Max.
     10      12      18      54      39    3209
percentiles:
   0%   10%   20%   30%   40%   50%   60%   70%   80%   90%   95%   99% 99.5% 99.9%  100%
   10    10    11    13    15    18    23    31    51   116   222   592   828  1472  3209

methylKit::getMethylationStats output for "be_5" CpGhsm-sites (#: 43988) at minimum coverage 10      - methylation statistics per base
summary:
   Min. 1st Qu.  Median    Mean 3rd Qu.    Max.
    2.3    50.0    68.8    65.6    84.6   100.0
percentiles:
    0%    10%    20%    30%    40%    50%    60%    70%    80%    90%    95%    99%  99.5%  99.9%   100%
```

```
  2.28  33.72  45.45  52.94  60.00  68.75  75.00  81.82  88.13  92.65 100.00 100.00 100.00 100.00 100.00

methylKit::getCoverageStats output for "be_5" CpGhsm-sites (#: 26941) at minimum coverage 15   - read coverage statistics per base
summary:
   Min. 1st Qu.  Median    Mean 3rd Qu.    Max.
     15      20      31      81      72    3209
percentiles:
   0%   10%   20%   30%   40%   50%   60%   70%   80%   90%   95%   99% 99.5% 99.9%  100%
   15    16    18    21    25    31    40    57    93   184   315   734  1004  1951  3209

methylKit::getMethylationStats output for "be_5" CpGhsm-sites (#: 26941) at minimum coverage 15      - methylation statistics per base
summary:
   Min. 1st Qu.  Median    Mean 3rd Qu.    Max.
    2.3    41.4    65.7    62.1    83.9   100.0
percentiles:
    0%    10%    20%    30%    40%    50%    60%    70%    80%    90%    95%    99%  99.5%  99.9%   100%
  2.28  27.43  37.50  46.67  56.25  65.71  73.68  80.95  86.67  92.05  95.00 100.00 100.00 100.00 100.00

methylKit::getCoverageStats output for "be_5" CpGhsm-sites (#: 38949) in coverage range [10-100]    - read coverage statistics per base
summary:
   Min. 1st Qu.  Median    Mean 3rd Qu.    Max.
   10.0    12.0    16.0    23.7    27.0   100.0
percentiles:
   0%   10%   20%   30%   40%   50%   60%   70%   80%   90%   95%   99% 99.5% 99.9%  100%
   10    10    11    12    14    16    19    24    32    50    69    93    96   100   100

methylKit::getMethylationStats output for "be_5" CpGhsm-sites (#: 38949) in coverage range [10-100] - methylation statistics per base
summary:
   Min. 1st Qu.  Median    Mean 3rd Qu.    Max.
    9.0    50.0    69.2    66.9    84.6   100.0
percentiles:
   0%   10%   20%   30%   40%   50%   60%   70%   80%   90%   95%   99% 99.5% 99.9%  100%
  9.0  37.5  46.2  53.8  60.9  69.2  75.0  81.8  88.9  93.3 100.0 100.0 100.0 100.0 100.0
```

```
Directory: REPCMS File: repcms-Sd_LI_be_6.txt

Number of "be_6" CpGhsm-sites with minimal and higher level coverage:

  number of "be_6" CpGhsm-sites with coverage >=  4: 294359
  number of "be_6" CpGhsm-sites with coverage >=  6:  90681
  number of "be_6" CpGhsm-sites with coverage >= 10:  35917
  number of "be_6" CpGhsm-sites with coverage >= 15:  22751

Coverage and methylation statistics for "be_6" CpGhsm-sites at different levels of minimum coverage:

methylKit::getCoverageStats output for "be_6" CpGhsm-sites (#: 294359) at minimum coverage 4   - read coverage statistics per base
summary:
   Min. 1st Qu.  Median    Mean 3rd Qu.    Max.
      4       4       5      11       6    3226
percentiles:
   0%   10%   20%   30%   40%   50%   60%   70%   80%   90%   95%   99% 99.5% 99.9%  100%
    4     4     4     4     4     5     5     6     7    11    23   146   253   635  3226

methylKit::getMethylationStats output for "be_6" CpGhsm-sites (#: 294359) at minimum coverage 4      - methylation statistics per base
summary:
   Min. 1st Qu.  Median    Mean 3rd Qu.    Max.
    1.5    83.3   100.0    91.5   100.0   100.0
percentiles:
    0%    10%    20%    30%    40%    50%    60%    70%    80%    90%    95%    99%  99.5%  99.9%   100%
  1.52  75.00  80.00  87.50 100.00 100.00 100.00 100.00 100.00 100.00 100.00 100.00 100.00 100.00 100.00

methylKit::getCoverageStats output for "be_6" CpGhsm-sites (#: 90681) at minimum coverage 6    - read coverage statistics per base
summary:
   Min. 1st Qu.  Median    Mean 3rd Qu.    Max.
      6       6       8      26      15    3226
percentiles:
   0%   10%   20%   30%   40%   50%   60%   70%   80%   90%   95%   99% 99.5% 99.9%  100%
    6     6     6     7     7     8     9    12    19    41    94   347   503  1048  3226

methylKit::getMethylationStats output for "be_6" CpGhsm-sites (#: 90681) at minimum coverage 6 - methylation statistics per base
summary:
   Min. 1st Qu.  Median    Mean 3rd Qu.    Max.
    1.5    70.0    83.3    79.0   100.0   100.0
percentiles:
    0%    10%    20%    30%    40%    50%    60%    70%    80%    90%    95%    99%  99.5%  99.9%   100%
  1.52  50.00  62.50  71.43  81.29  83.33  85.71  91.67 100.00 100.00 100.00 100.00 100.00 100.00 100.00

methylKit::getCoverageStats output for "be_6" CpGhsm-sites (#: 35917) at minimum coverage 10   - read coverage statistics per base
summary:
   Min. 1st Qu.  Median    Mean 3rd Qu.    Max.
     10      12      19      54      41    3226
percentiles:
   0%   10%   20%   30%   40%   50%   60%   70%   80%   90%   95%   99% 99.5% 99.9%  100%
   10    10    12    13    15    19    24    33    55   122   220   570   806  1409  3226

methylKit::getMethylationStats output for "be_6" CpGhsm-sites (#: 35917) at minimum coverage 10      - methylation statistics per base
summary:
   Min. 1st Qu.  Median    Mean 3rd Qu.    Max.
    1.5    50.0    68.7    65.3    84.2   100.0
percentiles:
    0%    10%    20%    30%    40%    50%    60%    70%    80%    90%    95%    99%  99.5%  99.9%   100%
```

```
  1.52  33.33  45.16  52.17  60.00  68.67  75.00  81.48  87.50  92.31  97.14 100.00 100.00 100.00 100.00

methylKit::getCoverageStats output for "be_6" CpGhsm-sites (#: 22751) at minimum coverage 15   - read coverage statistics per base
summary:
   Min. 1st Qu.  Median    Mean 3rd Qu.    Max.
     15      20      31      79      72    3226
percentiles:
   0%   10%   20%   30%   40%   50%   60%   70%   80%   90%   95%   99% 99.5% 99.9%  100%
   15    16    18    21    25    31    41    58    93   183   301   708   950  1779  3226

methylKit::getMethylationStats output for "be_6" CpGhsm-sites (#: 22751) at minimum coverage 15      - methylation statistics per base
summary:
   Min. 1st Qu.  Median    Mean 3rd Qu.    Max.
    1.5    42.1    66.7    62.5    84.2   100.0
percentiles:
    0%    10%    20%    30%    40%    50%    60%    70%    80%    90%    95%    99%  99.5%  99.9%   100%
  1.52  26.92  37.50  47.06  57.14  66.67  74.65  81.25  86.88  92.00  94.91 100.00 100.00 100.00 100.00

methylKit::getCoverageStats output for "be_6" CpGhsm-sites (#: 31626) in coverage range [10-100]    - read coverage statistics per base
summary:
   Min. 1st Qu.  Median    Mean 3rd Qu.    Max.
   10.0    12.0    17.0    24.4    29.0   100.0
percentiles:
   0%   10%   20%   30%   40%   50%   60%   70%   80%   90%   95%   99% 99.5% 99.9%  100%
   10    10    11    12    14    17    20    25    34    52    70    92    96    99   100

methylKit::getMethylationStats output for "be_6" CpGhsm-sites (#: 31626) in coverage range [10-100] - methylation statistics per base
summary:
   Min. 1st Qu.  Median    Mean 3rd Qu.    Max.
    9.0    50.0    69.2    66.6    84.6   100.0
percentiles:
   0%   10%   20%   30%   40%   50%   60%   70%   80%   90%   95%   99% 99.5% 99.9%  100%
  9.0  37.5  46.2  53.8  60.5  69.2  75.0  81.8  88.2  92.9 100.0 100.0 100.0 100.0 100.0
```

```
Directory: REPCMS File: repcms-Sd_LI_be_7.txt

Number of "be_7" CpGhsm-sites with minimal and higher level coverage:

  number of "be_7" CpGhsm-sites with coverage >=  4: 387816
  number of "be_7" CpGhsm-sites with coverage >=  6: 135235
  number of "be_7" CpGhsm-sites with coverage >= 10:  45510
  number of "be_7" CpGhsm-sites with coverage >= 15:  27343

Coverage and methylation statistics for "be_7" CpGhsm-sites at different levels of minimum coverage:

methylKit::getCoverageStats output for "be_7" CpGhsm-sites (#: 387816) at minimum coverage 4   - read coverage statistics per base
summary:
   Min. 1st Qu.  Median    Mean 3rd Qu.    Max.
      4       4       5      11       6    3416
percentiles:
   0%   10%   20%   30%   40%   50%   60%   70%   80%   90%   95%   99% 99.5% 99.9%  100%
    4     4     4     4     4     5     5     6     7    11    21   136   255   647  3416

methylKit::getMethylationStats output for "be_7" CpGhsm-sites (#: 387816) at minimum coverage 4      - methylation statistics per base
summary:
   Min. 1st Qu.  Median    Mean 3rd Qu.    Max.
    1.5   100.0   100.0    93.3   100.0   100.0
percentiles:
    0%    10%    20%    30%    40%    50%    60%    70%    80%    90%    95%    99%  99.5%  99.9%   100%
  1.47  75.00  85.71 100.00 100.00 100.00 100.00 100.00 100.00 100.00 100.00 100.00 100.00 100.00 100.00

methylKit::getCoverageStats output for "be_7" CpGhsm-sites (#: 135235) at minimum coverage 6   - read coverage statistics per base
summary:
   Min. 1st Qu.  Median    Mean 3rd Qu.    Max.
      6       6       7      23      12    3416
percentiles:
   0%   10%   20%   30%   40%   50%   60%   70%   80%   90%   95%   99% 99.5% 99.9%  100%
    6     6     6     6     7     7     8    10    15    31    74   328   501  1029  3416

methylKit::getMethylationStats output for "be_7" CpGhsm-sites (#: 135235) at minimum coverage 6      - methylation statistics per base
summary:
   Min. 1st Qu.  Median    Mean 3rd Qu.    Max.
    1.5    71.4    83.3    80.8   100.0   100.0
percentiles:
    0%    10%    20%    30%    40%    50%    60%    70%    80%    90%    95%    99%  99.5%  99.9%   100%
  1.47  52.63  66.67  75.00  83.33  83.33  87.50 100.00 100.00 100.00 100.00 100.00 100.00 100.00 100.00

methylKit::getCoverageStats output for "be_7" CpGhsm-sites (#: 45510) at minimum coverage 10   - read coverage statistics per base
summary:
   Min. 1st Qu.  Median    Mean 3rd Qu.    Max.
     10      12      18      54      38    3416
percentiles:
   0%   10%   20%   30%   40%   50%   60%   70%   80%   90%   95%   99% 99.5% 99.9%  100%
   10    10    11    12    15    18    22    31    50   114   223   602   824  1516  3416

methylKit::getMethylationStats output for "be_7" CpGhsm-sites (#: 45510) at minimum coverage 10      - methylation statistics per base
summary:
   Min. 1st Qu.  Median    Mean 3rd Qu.    Max.
    1.5    50.0    68.8    65.5    84.5   100.0
percentiles:
    0%    10%    20%    30%    40%    50%    60%    70%    80%    90%    95%    99%  99.5%  99.9%   100%
```

```
  1.47  33.57  45.45  52.94  60.00  68.75  75.00  81.58  87.78  92.31  97.94 100.00 100.00 100.00 100.00
```

```
methylKit::getCoverageStats output for "be_7" CpGhsm-sites (#: 27343) at minimum coverage 15   - read coverage statistics per base
summary:
   Min. 1st Qu.  Median    Mean 3rd Qu.    Max. 
     15      20      31      83      73    3416 
percentiles:
   0%   10%   20%   30%   40%   50%   60%   70%   80%   90%   95%   99% 99.5% 99.9%  100% 
   15    16    18    21    25    31    40    57    93   190   326   760  1017  2001  3416 

methylKit::getMethylationStats output for "be_7" CpGhsm-sites (#: 27343) at minimum coverage 15      - methylation statistics per base
summary:
   Min. 1st Qu.  Median    Mean 3rd Qu.    Max. 
    1.5    41.2    65.2    61.8    83.3   100.0 
percentiles:
    0%    10%    20%    30%    40%    50%    60%    70%    80%    90%    95%    99%  99.5%  99.9%   100% 
  1.47  27.08  37.21  46.67  55.89  65.22  73.33  80.44  86.67  91.67  94.74 100.00 100.00 100.00 100.00
```

```
methylKit::getCoverageStats output for "be_7" CpGhsm-sites (#: 40382) in coverage range [10-100]    - read coverage statistics per base
summary:
   Min. 1st Qu.  Median    Mean 3rd Qu.    Max. 
   10.0    11.0    16.0    23.5    27.0   100.0 
percentiles:
   0%   10%   20%   30%   40%   50%   60%   70%   80%   90%   95%   99% 99.5% 99.9%  100% 
   10    10    11    12    13    16    19    24    32    50    68    92    96   100   100 

methylKit::getMethylationStats output for "be_7" CpGhsm-sites (#: 40382) in coverage range [10-100] - methylation statistics per base
summary:
   Min. 1st Qu.  Median    Mean 3rd Qu.    Max. 
    9.2    50.0    69.2    66.8    84.6   100.0 
percentiles:
    0%    10%    20%    30%    40%    50%    60%    70%    80%    90%    95%    99%  99.5%  99.9%   100% 
  9.18  37.50  46.67  53.85  60.71  69.23  75.00  81.82  88.64  92.86 100.00 100.00 100.00 100.00 100.00
```

```
Directory: REPCMS File: repcms-Sd_LI_be_8.txt

Number of "be_8" CpGhsm-sites with minimal and higher level coverage:

  number of "be_8" CpGhsm-sites with coverage >=  4: 350436
  number of "be_8" CpGhsm-sites with coverage >=  6: 107453
  number of "be_8" CpGhsm-sites with coverage >= 10:  39496
  number of "be_8" CpGhsm-sites with coverage >= 15:  24727

Coverage and methylation statistics for "be_8" CpGhsm-sites at different levels of minimum coverage:

methylKit::getCoverageStats output for "be_8" CpGhsm-sites (#: 350436) at minimum coverage 4   - read coverage statistics per base
summary:
   Min. 1st Qu.  Median    Mean 3rd Qu.    Max. 
      4       4       5      11       6    2968 
percentiles:
   0%   10%   20%   30%   40%   50%   60%   70%   80%   90%   95%   99% 99.5% 99.9%  100% 
    4     4     4     4     4     5     5     6     7    10    21   138   247   607  2968 

methylKit::getMethylationStats output for "be_8" CpGhsm-sites (#: 350436) at minimum coverage 4      - methylation statistics per base
summary:
   Min. 1st Qu.  Median    Mean 3rd Qu.    Max. 
    2.2    83.3   100.0    91.9   100.0   100.0 
percentiles:
    0%    10%    20%    30%    40%    50%    60%    70%    80%    90%    95%    99%  99.5%  99.9%   100% 
  2.16  75.00  80.00  88.89 100.00 100.00 100.00 100.00 100.00 100.00 100.00 100.00 100.00 100.00 100.00 

methylKit::getCoverageStats output for "be_8" CpGhsm-sites (#: 107453) at minimum coverage 6   - read coverage statistics per base
summary:
   Min. 1st Qu.  Median    Mean 3rd Qu.    Max. 
      6       6       8      25      13    2968 
percentiles:
   0%   10%   20%   30%   40%   50%   60%   70%   80%   90%   95%   99% 99.5% 99.9%  100% 
    6     6     6     6     7     8     9    11    17    37    87   345   500  1007  2968 

methylKit::getMethylationStats output for "be_8" CpGhsm-sites (#: 107453) at minimum coverage 6      - methylation statistics per base
summary:
   Min. 1st Qu.  Median    Mean 3rd Qu.    Max. 
    2.2    71.4    83.3    80.0   100.0   100.0 
percentiles:
    0%    10%    20%    30%    40%    50%    60%    70%    80%    90%    95%    99%  99.5%  99.9%   100% 
  2.16  50.00  64.86  73.44  83.33  83.33  85.71  94.12 100.00 100.00 100.00 100.00 100.00 100.00 100.00 

methylKit::getCoverageStats output for "be_8" CpGhsm-sites (#: 39496) at minimum coverage 10   - read coverage statistics per base
summary:
   Min. 1st Qu.  Median    Mean 3rd Qu.    Max. 
     10      12      19      55      41    2968 
percentiles:
   0%   10%   20%   30%   40%   50%   60%   70%   80%   90%   95%   99% 99.5% 99.9%  100% 
   10    10    11    13    15    19    24    33    55   122   227   582   811  1465  2968 

methylKit::getMethylationStats output for "be_8" CpGhsm-sites (#: 39496) at minimum coverage 10      - methylation statistics per base
summary:
   Min. 1st Qu.  Median    Mean 3rd Qu.    Max. 
    2.2    50.0    69.2    65.6    84.6   100.0 
percentiles:
    0%    10%    20%    30%    40%    50%    60%    70%    80%    90%    95%    99%  99.5%  99.9%   100% 
```

```
  2.16  33.33  45.45  52.94  60.00  69.23  75.00  81.82  88.00  92.59  97.92 100.00 100.00 100.00 100.00

methylKit::getCoverageStats output for "be_8" CpGhsm-sites (#: 24727) at minimum coverage 15   - read coverage statistics per base
summary:
   Min. 1st Qu.  Median    Mean 3rd Qu.    Max.
     15      20      32      81      75    2968
percentiles:
   0%   10%   20%   30%   40%   50%   60%   70%   80%   90%   95%   99% 99.5% 99.9%  100%
   15    16    19    21    25    32    41    60    95   188   317   723   968  1896  2968

methylKit::getMethylationStats output for "be_8" CpGhsm-sites (#: 24727) at minimum coverage 15      - methylation statistics per base
summary:
   Min. 1st Qu.  Median    Mean 3rd Qu.    Max.
    2.2    41.7    66.7    62.3    84.2   100.0
percentiles:
    0%    10%    20%    30%    40%    50%    60%    70%    80%    90%    95%    99%  99.5%  99.9%   100%
  2.16  27.27  37.50  46.67  56.82  66.67  74.42  81.25  86.84  92.11  94.74 100.00 100.00 100.00 100.00

methylKit::getCoverageStats output for "be_8" CpGhsm-sites (#: 34789) in coverage range [10-100]     - read coverage statistics per base
summary:
   Min. 1st Qu.  Median    Mean 3rd Qu.    Max.
   10.0    12.0    16.0    24.4    29.0   100.0
percentiles:
   0%   10%   20%   30%   40%   50%   60%   70%   80%   90%   95%   99% 99.5% 99.9%  100%
   10    10    11    12    14    16    20    25    34    52    71    93    96   100   100

methylKit::getMethylationStats output for "be_8" CpGhsm-sites (#: 34789) in coverage range [10-100] - methylation statistics per base
summary:
   Min. 1st Qu.  Median    Mean 3rd Qu.    Max.
    9.1    50.0    70.0    67.1    84.6   100.0
percentiles:
    0%    10%    20%    30%    40%    50%    60%    70%    80%    90%    95%    99%  99.5%  99.9%   100%
  9.09  37.50  46.67  54.17  61.54  70.00  76.19  81.82  88.89  93.10 100.00 100.00 100.00 100.00 100.00
```

```
Directory: REPCMS File: repcms-Sd_LI_ka_1.txt

Number of "ka_1" CpGhsm-sites with minimal and higher level coverage:

  number of "ka_1" CpGhsm-sites with coverage >=  4: 381532
  number of "ka_1" CpGhsm-sites with coverage >=  6: 125360
  number of "ka_1" CpGhsm-sites with coverage >= 10:  46096
  number of "ka_1" CpGhsm-sites with coverage >= 15:  29235

Coverage and methylation statistics for "ka_1" CpGhsm-sites at different levels of minimum coverage:

methylKit::getCoverageStats output for "ka_1" CpGhsm-sites (#: 381532) at minimum coverage 4   - read coverage statistics per base
summary:
   Min. 1st Qu.  Median    Mean 3rd Qu.    Max.
      4       4       5      13       6    3854
percentiles:
   0%   10%   20%   30%   40%   50%   60%   70%   80%   90%   95%   99% 99.5% 99.9%  100%
    4     4     4     4     4     5     5     6     7    11    24   187   351   885  3854

methylKit::getMethylationStats output for "ka_1" CpGhsm-sites (#: 381532) at minimum coverage 4      - methylation statistics per base
summary:
   Min. 1st Qu.  Median    Mean 3rd Qu.    Max.
    1.9    83.3   100.0    91.3   100.0   100.0
percentiles:
    0%    10%    20%    30%    40%    50%    60%    70%    80%    90%    95%    99%  99.5%  99.9%   100%
  1.86  74.61  80.00  87.18 100.00 100.00 100.00 100.00 100.00 100.00 100.00 100.00 100.00 100.00 100.00

methylKit::getCoverageStats output for "ka_1" CpGhsm-sites (#: 125360) at minimum coverage 6   - read coverage statistics per base
summary:
   Min. 1st Qu.  Median    Mean 3rd Qu.    Max.
      6       6       8      30      13    3854
percentiles:
   0%   10%   20%   30%   40%   50%   60%   70%   80%   90%   95%   99% 99.5% 99.9%  100%
    6     6     6     6     7     8     9    11    17    42   107   475   714  1427  3854

methylKit::getMethylationStats output for "ka_1" CpGhsm-sites (#: 125360) at minimum coverage 6      - methylation statistics per base
summary:
   Min. 1st Qu.  Median    Mean 3rd Qu.    Max.
    1.9    71.4    83.3    79.7   100.0   100.0
percentiles:
    0%    10%    20%    30%    40%    50%    60%    70%    80%    90%    95%    99%  99.5%  99.9%   100%
  1.86  50.00  63.64  73.33  83.33  83.33  85.71  94.74 100.00 100.00 100.00 100.00 100.00 100.00 100.00

methylKit::getCoverageStats output for "ka_1" CpGhsm-sites (#: 46096) at minimum coverage 10   - read coverage statistics per base
summary:
   Min. 1st Qu.  Median    Mean 3rd Qu.    Max.
     10      12      19      69      47    3854
percentiles:
   0%   10%   20%   30%   40%   50%   60%   70%   80%   90%   95%   99% 99.5% 99.9%  100%
   10    10    11    13    15    19    25    37    64   153   299   804  1139  2065  3854

methylKit::getMethylationStats output for "ka_1" CpGhsm-sites (#: 46096) at minimum coverage 10      - methylation statistics per base
summary:
   Min. 1st Qu.  Median    Mean 3rd Qu.    Max.
    1.9    48.0    68.7    64.9    84.6   100.0
percentiles:
    0%    10%    20%    30%    40%    50%    60%    70%    80%    90%    95%    99%  99.5%  99.9%   100%
```

```
  1.86  31.58  43.75  50.00  60.00  68.65  75.00  81.82  87.88  92.42  97.50 100.00 100.00 100.00 100.00

methylKit::getCoverageStats output for "ka_1" CpGhsm-sites (#: 29235) at minimum coverage 15   - read coverage statistics per base
summary:
   Min. 1st Qu.  Median    Mean 3rd Qu.    Max.
     15      20      34     103      88    3854
percentiles:
   0%   10%   20%   30%   40%   50%   60%   70%   80%   90%   95%   99% 99.5% 99.9%  100%
   15    16    19    22    27    34    46    69   116   247   420  1008  1354  2255  3854

methylKit::getMethylationStats output for "ka_1" CpGhsm-sites (#: 29235) at minimum coverage 15      - methylation statistics per base
summary:
   Min. 1st Qu.  Median    Mean 3rd Qu.    Max.
    1.9    40.0    65.6    61.4    83.9   100.0
percentiles:
    0%    10%    20%    30%    40%    50%    60%    70%    80%    90%    95%    99%  99.5%  99.9%   100%
  1.86  25.00  35.56  44.83  55.00  65.62  73.98  81.08  86.67  92.00  94.74 100.00 100.00 100.00 100.00

methylKit::getCoverageStats output for "ka_1" CpGhsm-sites (#: 39517) in coverage range [10-100]    - read coverage statistics per base
summary:
   Min. 1st Qu.  Median    Mean 3rd Qu.    Max.
   10.0    12.0    16.0    24.6    29.0   100.0
percentiles:
   0%   10%   20%   30%   40%   50%   60%   70%   80%   90%   95%   99% 99.5% 99.9%  100%
   10    10    11    12    14    16    20    25    35    53    71    93    97   100   100

methylKit::getMethylationStats output for "ka_1" CpGhsm-sites (#: 39517) in coverage range [10-100] - methylation statistics per base
summary:
   Min. 1st Qu.  Median    Mean 3rd Qu.    Max.
    9.0    50.0    69.2    66.6    84.8   100.0
percentiles:
   0%   10%   20%   30%   40%   50%   60%   70%   80%   90%   95%   99% 99.5% 99.9%  100%
  9.0  36.8  46.2  53.5  60.0  69.2  75.0  81.8  88.9  93.1 100.0 100.0 100.0 100.0 100.0
```

```
Directory: REPCMS File: repcms-Sd_LI_ka_2.txt

Number of "ka_2" CpGhsm-sites with minimal and higher level coverage:

  number of "ka_2" CpGhsm-sites with coverage >=  4: 401846
  number of "ka_2" CpGhsm-sites with coverage >=  6: 131363
  number of "ka_2" CpGhsm-sites with coverage >= 10:  47658
  number of "ka_2" CpGhsm-sites with coverage >= 15:  29956

Coverage and methylation statistics for "ka_2" CpGhsm-sites at different levels of minimum coverage:

methylKit::getCoverageStats output for "ka_2" CpGhsm-sites (#: 401846) at minimum coverage 4   - read coverage statistics per base
summary:
   Min. 1st Qu.  Median    Mean 3rd Qu.    Max. 
      4       4       5      13       6    4023 
percentiles:
   0%   10%   20%   30%   40%   50%   60%   70%   80%   90%   95%   99% 99.5% 99.9%  100% 
    4     4     4     4     4     5     5     6     7    11    23   181   342   872  4023 

methylKit::getMethylationStats output for "ka_2" CpGhsm-sites (#: 401846) at minimum coverage 4      - methylation statistics per base
summary:
   Min. 1st Qu.  Median    Mean 3rd Qu.    Max. 
    2.1    83.3   100.0    91.4   100.0   100.0 
percentiles:
    0%    10%    20%    30%    40%    50%    60%    70%    80%    90%    95%    99%  99.5%  99.9%   100% 
  2.15  75.00  80.00  87.50 100.00 100.00 100.00 100.00 100.00 100.00 100.00 100.00 100.00 100.00 100.00 

methylKit::getCoverageStats output for "ka_2" CpGhsm-sites (#: 131363) at minimum coverage 6   - read coverage statistics per base
summary:
   Min. 1st Qu.  Median    Mean 3rd Qu.    Max. 
      6       6       8      29      13    4023 
percentiles:
   0%   10%   20%   30%   40%   50%   60%   70%   80%   90%   95%   99% 99.5% 99.9%  100% 
    6     6     6     6     7     8     9    11    17    41   103   467   704  1421  4023 

methylKit::getMethylationStats output for "ka_2" CpGhsm-sites (#: 131363) at minimum coverage 6      - methylation statistics per base
summary:
   Min. 1st Qu.  Median    Mean 3rd Qu.    Max. 
    2.1    71.4    83.3    79.8   100.0   100.0 
percentiles:
    0%    10%    20%    30%    40%    50%    60%    70%    80%    90%    95%    99%  99.5%  99.9%   100% 
  2.15  50.00  64.29  73.18  83.33  83.33  86.36  95.65 100.00 100.00 100.00 100.00 100.00 100.00 100.00 

methylKit::getCoverageStats output for "ka_2" CpGhsm-sites (#: 47658) at minimum coverage 10   - read coverage statistics per base
summary:
   Min. 1st Qu.  Median    Mean 3rd Qu.    Max. 
     10      12      19      69      46    4023 
percentiles:
   0%   10%   20%   30%   40%   50%   60%   70%   80%   90%   95%   99% 99.5% 99.9%  100% 
   10    10    11    13    15    19    25    36    63   151   297   808  1144  2108  4023 

methylKit::getMethylationStats output for "ka_2" CpGhsm-sites (#: 47658) at minimum coverage 10      - methylation statistics per base
summary:
   Min. 1st Qu.  Median    Mean 3rd Qu.    Max. 
    2.1    48.0    68.2    64.9    84.6   100.0 
percentiles:
    0%    10%    20%    30%    40%    50%    60%    70%    80%    90%    95%    99%  99.5%  99.9%   100% 
```

```
  2.15  31.71  43.75  50.00  60.00  68.18  75.00  81.82  88.04  92.80 100.00 100.00 100.00 100.00 100.00

methylKit::getCoverageStats output for "ka_2" CpGhsm-sites (#: 29956) at minimum coverage 15   - read coverage statistics per base
summary:
   Min. 1st Qu.  Median    Mean 3rd Qu.    Max.
     15      20      34     103      87    4023
percentiles:
   0%   10%   20%   30%   40%   50%   60%   70%   80%   90%   95%   99% 99.5% 99.9%  100%
   15    16    19    22    27    34    46    68   114   247   424  1010  1345  2332  4023

methylKit::getMethylationStats output for "ka_2" CpGhsm-sites (#: 29956) at minimum coverage 15      - methylation statistics per base
summary:
   Min. 1st Qu.  Median    Mean 3rd Qu.    Max.
    2.1    40.0    65.2    61.3    84.0   100.0
percentiles:
    0%    10%    20%    30%    40%    50%    60%    70%    80%    90%    95%    99%  99.5%  99.9%   100%
  2.15  25.49  35.35  44.76  55.32  65.22  73.68  80.89  86.67  92.15  95.00 100.00 100.00 100.00 100.00

methylKit::getCoverageStats output for "ka_2" CpGhsm-sites (#: 40962) in coverage range [10-100]     - read coverage statistics per base
summary:
   Min. 1st Qu.  Median    Mean 3rd Qu.    Max.
   10.0    12.0    16.0    24.4    29.0   100.0
percentiles:
   0%   10%   20%   30%   40%   50%   60%   70%   80%   90%   95%   99% 99.5% 99.9%  100%
   10    10    11    12    14    16    20    25    34    53    71    93    97   100   100

methylKit::getMethylationStats output for "ka_2" CpGhsm-sites (#: 40962) in coverage range [10-100]  - methylation statistics per base
summary:
   Min. 1st Qu.  Median    Mean 3rd Qu.    Max.
    9.0    50.0    69.2    66.5    85.0   100.0
percentiles:
   0%   10%   20%   30%   40%   50%   60%   70%   80%   90%   95%   99% 99.5% 99.9%  100%
  9.0  36.8  46.2  53.3  60.0  69.2  75.0  81.8  89.1  93.3 100.0 100.0 100.0 100.0 100.0
```

```
Directory: REPCMS File: repcms-Sd_LI_ka_3.txt

Number of "ka_3" CpGhsm-sites with minimal and higher level coverage:

  number of "ka_3" CpGhsm-sites with coverage >=  4: 346459
  number of "ka_3" CpGhsm-sites with coverage >=  6: 113203
  number of "ka_3" CpGhsm-sites with coverage >= 10:  43472
  number of "ka_3" CpGhsm-sites with coverage >= 15:  27998

Coverage and methylation statistics for "ka_3" CpGhsm-sites at different levels of minimum coverage:

methylKit::getCoverageStats output for "ka_3" CpGhsm-sites (#: 346459) at minimum coverage 4   - read coverage statistics per base
summary:
   Min. 1st Qu.  Median    Mean 3rd Qu.    Max. 
      4       4       5      13       6    3641 
percentiles:
   0%   10%   20%   30%   40%   50%   60%   70%   80%   90%   95%   99% 99.5% 99.9%  100% 
    4     4     4     4     4     5     5     6     7    12    26   198   358   884  3641 

methylKit::getMethylationStats output for "ka_3" CpGhsm-sites (#: 346459) at minimum coverage 4      - methylation statistics per base
summary:
   Min. 1st Qu.  Median    Mean 3rd Qu.    Max. 
    2.2    83.3   100.0    91.1   100.0   100.0 
percentiles:
    0%    10%    20%    30%    40%    50%    60%    70%    80%    90%    95%    99%  99.5%  99.9%   100% 
  2.21  72.22  80.00  85.71 100.00 100.00 100.00 100.00 100.00 100.00 100.00 100.00 100.00 100.00 100.00 

methylKit::getCoverageStats output for "ka_3" CpGhsm-sites (#: 113203) at minimum coverage 6   - read coverage statistics per base
summary:
   Min. 1st Qu.  Median    Mean 3rd Qu.    Max. 
      6       6       8      31      14    3641 
percentiles:
   0%   10%   20%   30%   40%   50%   60%   70%   80%   90%   95%   99% 99.5% 99.9%  100% 
    6     6     6     7     7     8     9    12    19    47   114   492   713  1456  3641 

methylKit::getMethylationStats output for "ka_3" CpGhsm-sites (#: 113203) at minimum coverage 6      - methylation statistics per base
summary:
   Min. 1st Qu.  Median    Mean 3rd Qu.    Max. 
    2.2    70.0    83.3    79.0   100.0   100.0 
percentiles:
    0%    10%    20%    30%    40%    50%    60%    70%    80%    90%    95%    99%  99.5%  99.9%   100% 
  2.21  48.72  62.50  71.43  82.05  83.33  85.71  92.31 100.00 100.00 100.00 100.00 100.00 100.00 100.00 

methylKit::getCoverageStats output for "ka_3" CpGhsm-sites (#: 43472) at minimum coverage 10   - read coverage statistics per base
summary:
   Min. 1st Qu.  Median    Mean 3rd Qu.    Max. 
     10      12      20      70      49    3641 
percentiles:
   0%   10%   20%   30%   40%   50%   60%   70%   80%   90%   95%   99% 99.5% 99.9%  100% 
   10    10    12    13    16    20    26    38    67   156   299   795  1117  1989  3641 

methylKit::getMethylationStats output for "ka_3" CpGhsm-sites (#: 43472) at minimum coverage 10      - methylation statistics per base
summary:
   Min. 1st Qu.  Median    Mean 3rd Qu.    Max. 
    2.2    47.1    66.9    64.5    84.4   100.0 
percentiles:
    0%    10%    20%    30%    40%    50%    60%    70%    80%    90%    95%    99%  99.5%  99.9%   100% 
```

```
  2.21  30.91  42.86  50.00  60.00  66.92  75.00  81.48  87.50  92.50  97.45 100.00 100.00 100.00 100.00

methylKit::getCoverageStats output for "ka_3" CpGhsm-sites (#: 27998) at minimum coverage 15   - read coverage statistics per base
summary:
   Min. 1st Qu.  Median    Mean 3rd Qu.    Max. 
     15      21      35     102      89    3641 
percentiles:
   0%   10%   20%   30%   40%   50%   60%   70%   80%   90%   95%   99% 99.5% 99.9%  100% 
   15    16    19    22    27    35    47    70   115   242   417   997  1309  2137  3641 

methylKit::getMethylationStats output for "ka_3" CpGhsm-sites (#: 27998) at minimum coverage 15      - methylation statistics per base
summary:
   Min. 1st Qu.  Median    Mean 3rd Qu.    Max. 
    2.2    40.0    64.7    61.1    83.9   100.0 
percentiles:
    0%    10%    20%    30%    40%    50%    60%    70%    80%    90%    95%    99%  99.5%  99.9%   100% 
  2.21  25.00  35.29  44.31  54.55  64.71  73.68  80.95  86.67  92.11  94.74 100.00 100.00 100.00 100.00

methylKit::getCoverageStats output for "ka_3" CpGhsm-sites (#: 37145) in coverage range [10-100]    - read coverage statistics per base
summary:
   Min. 1st Qu.  Median    Mean 3rd Qu.    Max. 
     10      12      17      25      30     100 
percentiles:
   0%   10%   20%   30%   40%   50%   60%   70%   80%   90%   95%   99% 99.5% 99.9%  100% 
   10    10    11    12    14    17    20    26    36    55    72    94    97   100   100 

methylKit::getMethylationStats output for "ka_3" CpGhsm-sites (#: 37145) in coverage range [10-100] - methylation statistics per base
summary:
   Min. 1st Qu.  Median    Mean 3rd Qu.    Max. 
    9.1    50.0    68.6    66.1    84.6   100.0 
percentiles:
    0%    10%    20%    30%    40%    50%    60%    70%    80%    90%    95%    99%  99.5%  99.9%   100% 
  9.09  35.62  45.45  52.63  60.00  68.57  75.00  81.82  88.71  93.33 100.00 100.00 100.00 100.00 100.00 
```

```
Directory: REPCMS File: repcms-Sd_LI_ka_4.txt

Number of "ka_4" CpGhsm-sites with minimal and higher level coverage:

  number of "ka_4" CpGhsm-sites with coverage >=  4: 403525
  number of "ka_4" CpGhsm-sites with coverage >=  6: 132384
  number of "ka_4" CpGhsm-sites with coverage >= 10:  47748
  number of "ka_4" CpGhsm-sites with coverage >= 15:  30166

Coverage and methylation statistics for "ka_4" CpGhsm-sites at different levels of minimum coverage:

methylKit::getCoverageStats output for "ka_4" CpGhsm-sites (#: 403525) at minimum coverage 4   - read coverage statistics per base
summary:
   Min. 1st Qu.  Median    Mean 3rd Qu.    Max.
      4       4       5      13       6    3921
percentiles:
   0%   10%   20%   30%   40%   50%   60%   70%   80%   90%   95%   99% 99.5% 99.9%  100%
    4     4     4     4     4     5     5     6     7    11    23   182   345   882  3921

methylKit::getMethylationStats output for "ka_4" CpGhsm-sites (#: 403525) at minimum coverage 4      - methylation statistics per base
summary:
   Min. 1st Qu.  Median    Mean 3rd Qu.    Max.
    2.3    83.3   100.0    91.4   100.0   100.0
percentiles:
    0%    10%    20%    30%    40%    50%    60%    70%    80%    90%    95%    99%  99.5%  99.9%   100%
  2.28  74.42  80.00  87.50 100.00 100.00 100.00 100.00 100.00 100.00 100.00 100.00 100.00 100.00 100.00

methylKit::getCoverageStats output for "ka_4" CpGhsm-sites (#: 132384) at minimum coverage 6   - read coverage statistics per base
summary:
   Min. 1st Qu.  Median    Mean 3rd Qu.    Max.
      6       6       8      29      13    3921
percentiles:
   0%   10%   20%   30%   40%   50%   60%   70%   80%   90%   95%   99% 99.5% 99.9%  100%
    6     6     6     6     7     8     9    11    17    41   103   464   711  1434  3921

methylKit::getMethylationStats output for "ka_4" CpGhsm-sites (#: 132384) at minimum coverage 6      - methylation statistics per base
summary:
   Min. 1st Qu.  Median    Mean 3rd Qu.    Max.
    2.3    71.4    83.3    79.8   100.0   100.0
percentiles:
    0%    10%    20%    30%    40%    50%    60%    70%    80%    90%    95%    99%  99.5%  99.9%   100%
  2.28  50.00  64.29  73.07  83.33  83.33  86.00  95.83 100.00 100.00 100.00 100.00 100.00 100.00 100.00

methylKit::getCoverageStats output for "ka_4" CpGhsm-sites (#: 47748) at minimum coverage 10   - read coverage statistics per base
summary:
   Min. 1st Qu.  Median    Mean 3rd Qu.    Max.
     10      12      19      69      47    3921
percentiles:
   0%   10%   20%   30%   40%   50%   60%   70%   80%   90%   95%   99% 99.5% 99.9%  100%
   10    10    11    13    15    19    25    36    64   152   298   827  1139  2110  3921

methylKit::getMethylationStats output for "ka_4" CpGhsm-sites (#: 47748) at minimum coverage 10      - methylation statistics per base
summary:
   Min. 1st Qu.  Median    Mean 3rd Qu.    Max.
    2.3    47.6    68.1    64.8    84.3   100.0
percentiles:
    0%    10%    20%    30%    40%    50%    60%    70%    80%    90%    95%    99%  99.5%  99.9%   100%
```

```
  2.28  31.82  43.75  50.00  60.00  68.09  75.00  81.48  87.50  92.54  98.74 100.00 100.00 100.00 100.00

methylKit::getCoverageStats output for "ka_4" CpGhsm-sites (#: 30166) at minimum coverage 15   - read coverage statistics per base
summary:
   Min. 1st Qu.  Median    Mean 3rd Qu.    Max.
     15      20      34     103      88    3921
percentiles:
   0%   10%   20%   30%   40%   50%   60%   70%   80%   90%   95%   99% 99.5% 99.9%  100%
   15    16    19    22    26    34    46    68   115   245   428  1003  1356  2299  3921

methylKit::getMethylationStats output for "ka_4" CpGhsm-sites (#: 30166) at minimum coverage 15      - methylation statistics per base
summary:
   Min. 1st Qu.  Median    Mean 3rd Qu.    Max.
    2.3    40.0    65.0    61.2    83.6   100.0
percentiles:
    0%    10%    20%    30%    40%    50%    60%    70%    80%    90%    95%    99%  99.5%  99.9%   100%
  2.28  25.00  35.56  44.44  54.84  65.00  73.33  80.68  86.67  92.00  94.85 100.00 100.00 100.00 100.00

methylKit::getCoverageStats output for "ka_4" CpGhsm-sites (#: 40985) in coverage range [10-100]     - read coverage statistics per base
summary:
   Min. 1st Qu.  Median    Mean 3rd Qu.    Max.
   10.0    12.0    16.0    24.5    29.0   100.0
percentiles:
   0%   10%   20%   30%   40%   50%   60%   70%   80%   90%   95%   99% 99.5% 99.9%  100%
   10    10    11    12    14    16    20    25    34    53    71    94    97   100   100

methylKit::getMethylationStats output for "ka_4" CpGhsm-sites (#: 40985) in coverage range [10-100]  - methylation statistics per base
summary:
   Min. 1st Qu.  Median    Mean 3rd Qu.    Max.
    9.4    50.0    69.2    66.5    84.6   100.0
percentiles:
    0%    10%    20%    30%    40%    50%    60%    70%    80%    90%    95%    99%  99.5%  99.9%   100%
  9.38  36.84  45.83  53.33  60.00  69.23  75.00  81.82  88.89  93.33 100.00 100.00 100.00 100.00 100.00
```

```
Directory: REPCMS File: repcms-Sd_LI_ka_5.txt

Number of "ka_5" CpGhsm-sites with minimal and higher level coverage:

  number of "ka_5" CpGhsm-sites with coverage >=  4: 327536
  number of "ka_5" CpGhsm-sites with coverage >=  6: 106346
  number of "ka_5" CpGhsm-sites with coverage >= 10:  41424
  number of "ka_5" CpGhsm-sites with coverage >= 15:  26706

Coverage and methylation statistics for "ka_5" CpGhsm-sites at different levels of minimum coverage:

methylKit::getCoverageStats output for "ka_5" CpGhsm-sites (#: 327536) at minimum coverage 4   - read coverage statistics per base
summary:
   Min. 1st Qu.  Median    Mean 3rd Qu.    Max.
      4       4       5      13       6    3713
percentiles:
   0%   10%   20%   30%   40%   50%   60%   70%   80%   90%   95%   99% 99.5% 99.9%  100%
    4     4     4     4     4     5     5     6     7    12    27   201   367   889  3713

methylKit::getMethylationStats output for "ka_5" CpGhsm-sites (#: 327536) at minimum coverage 4      - methylation statistics per base
summary:
   Min. 1st Qu.  Median    Mean 3rd Qu.    Max.
    3.0    83.3   100.0    91.1   100.0   100.0
percentiles:
    0%    10%    20%    30%    40%    50%    60%    70%    80%    90%    95%    99%  99.5%  99.9%   100%
  2.97  72.73  80.00  85.71 100.00 100.00 100.00 100.00 100.00 100.00 100.00 100.00 100.00 100.00 100.00

methylKit::getCoverageStats output for "ka_5" CpGhsm-sites (#: 106346) at minimum coverage 6   - read coverage statistics per base
summary:
   Min. 1st Qu.  Median    Mean 3rd Qu.    Max.
      6       6       8      32      15    3713
percentiles:
   0%   10%   20%   30%   40%   50%   60%   70%   80%   90%   95%   99% 99.5% 99.9%  100%
    6     6     6     7     7     8     9    12    19    48   119   496   719  1439  3713

methylKit::getMethylationStats output for "ka_5" CpGhsm-sites (#: 106346) at minimum coverage 6      - methylation statistics per base
summary:
   Min. 1st Qu.  Median    Mean 3rd Qu.    Max.
    3.0    70.0    83.3    78.9   100.0   100.0
percentiles:
    0%    10%    20%    30%    40%    50%    60%    70%    80%    90%    95%    99%  99.5%  99.9%   100%
  2.97  50.00  62.50  71.43  81.82  83.33  85.71  91.67 100.00 100.00 100.00 100.00 100.00 100.00 100.00

methylKit::getCoverageStats output for "ka_5" CpGhsm-sites (#: 41424) at minimum coverage 10   - read coverage statistics per base
summary:
   Min. 1st Qu.  Median    Mean 3rd Qu.    Max.
     10      12      20      70      50    3713
percentiles:
   0%   10%   20%   30%   40%   50%   60%   70%   80%   90%   95%   99% 99.5% 99.9%  100%
   10    10    12    13    16    20    26    39    68   160   306   795  1116  1960  3713

methylKit::getMethylationStats output for "ka_5" CpGhsm-sites (#: 41424) at minimum coverage 10      - methylation statistics per base
summary:
   Min. 1st Qu.  Median    Mean 3rd Qu.    Max.
    3.0    47.8    68.7    64.8    84.5   100.0
percentiles:
    0%    10%    20%    30%    40%    50%    60%    70%    80%    90%    95%    99%  99.5%  99.9%   100%
```

```
  2.97  31.25  43.75  50.00  60.00  68.66  75.00  81.82  87.50  92.31  96.97 100.00 100.00 100.00 100.00

methylKit::getCoverageStats output for "ka_5" CpGhsm-sites (#: 26706) at minimum coverage 15   - read coverage statistics per base
summary:
   Min. 1st Qu.  Median    Mean 3rd Qu.    Max. 
     15      21      35     103      91    3713 
percentiles:
   0%   10%   20%   30%   40%   50%   60%   70%   80%   90%   95%   99% 99.5% 99.9%  100% 
   15    17    19    23    28    35    48    72   118   245   420   990  1321  2197  3713 

methylKit::getMethylationStats output for "ka_5" CpGhsm-sites (#: 26706) at minimum coverage 15      - methylation statistics per base
summary:
   Min. 1st Qu.  Median    Mean 3rd Qu.    Max. 
    3.0    40.0    66.5    61.6    84.2   100.0 
percentiles:
     0%    10%    20%    30%    40%    50%    60%    70%    80%    90%    95%    99%  99.5%  99.9%   100% 
  2.97  25.00  35.62  45.20  55.74  66.53  74.58  81.25  86.96  92.16  94.92 100.00 100.00 100.00 100.00

methylKit::getCoverageStats output for "ka_5" CpGhsm-sites (#: 35307) in coverage range [10-100]    - read coverage statistics per base
summary:
   Min. 1st Qu.  Median    Mean 3rd Qu.    Max. 
   10.0    12.0    17.0    25.2    30.0   100.0 
percentiles:
   0%   10%   20%   30%   40%   50%   60%   70%   80%   90%   95%   99% 99.5% 99.9%  100% 
   10    10    11    12    14    17    20    26    36    55    73    94    97   100   100 

methylKit::getMethylationStats output for "ka_5" CpGhsm-sites (#: 35307) in coverage range [10-100] - methylation statistics per base
summary:
   Min. 1st Qu.  Median    Mean 3rd Qu.    Max. 
    9.0    50.0    69.2    66.6    84.8   100.0 
percentiles:
   0%   10%   20%   30%   40%   50%   60%   70%   80%   90%   95%   99% 99.5% 99.9%  100% 
  9.0  36.8  46.2  53.8  60.0  69.2  75.3  81.8  88.9  93.1 100.0 100.0 100.0 100.0 100.0 
```

```
Directory: REPCMS File: repcms-Sd_LI_ka_6.txt

Number of "ka_6" CpGhsm-sites with minimal and higher level coverage:

  number of "ka_6" CpGhsm-sites with coverage >=  4: 399083
  number of "ka_6" CpGhsm-sites with coverage >=  6: 146888
  number of "ka_6" CpGhsm-sites with coverage >= 10:  50613
  number of "ka_6" CpGhsm-sites with coverage >= 15:  31348

Coverage and methylation statistics for "ka_6" CpGhsm-sites at different levels of minimum coverage:

methylKit::getCoverageStats output for "ka_6" CpGhsm-sites (#: 399083) at minimum coverage 4   - read coverage statistics per base
summary:
   Min. 1st Qu.  Median    Mean 3rd Qu.    Max.
      4       4       5      13       7    4095
percentiles:
   0%   10%   20%   30%   40%   50%   60%   70%   80%   90%   95%   99% 99.5% 99.9%  100%
    4     4     4     4     4     5     5     6     7    11    25   192   362   919  4095

methylKit::getMethylationStats output for "ka_6" CpGhsm-sites (#: 399083) at minimum coverage 4      - methylation statistics per base
summary:
   Min. 1st Qu.  Median    Mean 3rd Qu.    Max.
    1.9    92.9   100.0    92.8   100.0   100.0
percentiles:
    0%    10%    20%    30%    40%    50%    60%    70%    80%    90%    95%    99%  99.5%  99.9%   100%
  1.93  71.43  85.71 100.00 100.00 100.00 100.00 100.00 100.00 100.00 100.00 100.00 100.00 100.00 100.00

methylKit::getCoverageStats output for "ka_6" CpGhsm-sites (#: 146888) at minimum coverage 6   - read coverage statistics per base
summary:
   Min. 1st Qu.  Median    Mean 3rd Qu.    Max.
      6       6       8      28      12    4095
percentiles:
   0%   10%   20%   30%   40%   50%   60%   70%   80%   90%   95%   99% 99.5% 99.9%  100%
    6     6     6     6     7     8     9    11    16    37    96   448   699  1438  4095

methylKit::getMethylationStats output for "ka_6" CpGhsm-sites (#: 146888) at minimum coverage 6      - methylation statistics per base
summary:
   Min. 1st Qu.  Median    Mean 3rd Qu.    Max.
    1.9    71.4    83.3    80.4   100.0   100.0
percentiles:
    0%    10%    20%    30%    40%    50%    60%    70%    80%    90%    95%    99%  99.5%  99.9%   100%
  1.93  50.00  66.67  75.00  83.33  83.33  87.50 100.00 100.00 100.00 100.00 100.00 100.00 100.00 100.00

methylKit::getCoverageStats output for "ka_6" CpGhsm-sites (#: 50613) at minimum coverage 10   - read coverage statistics per base
summary:
   Min. 1st Qu.  Median    Mean 3rd Qu.    Max.
     10      12      19      69      45    4095
percentiles:
   0%   10%   20%   30%   40%   50%   60%   70%   80%   90%   95%   99% 99.5% 99.9%  100%
   10    10    11    13    15    19    24    35    61   147   296   826  1159  2149  4095

methylKit::getMethylationStats output for "ka_6" CpGhsm-sites (#: 50613) at minimum coverage 10      - methylation statistics per base
summary:
   Min. 1st Qu.  Median    Mean 3rd Qu.    Max.
    1.9    48.1    68.1    64.9    84.2   100.0
percentiles:
    0%    10%    20%    30%    40%    50%    60%    70%    80%    90%    95%    99%  99.5%  99.9%   100%
```

```
  1.93  31.58  43.75  50.00  60.00  68.13  75.00  81.48  87.69  92.65 100.00 100.00 100.00 100.00 100.00

methylKit::getCoverageStats output for "ka_6" CpGhsm-sites (#: 31348) at minimum coverage 15   - read coverage statistics per base
summary:
   Min. 1st Qu.  Median    Mean 3rd Qu.    Max.
     15      20      34     104      88    4095
percentiles:
   0%   10%   20%   30%   40%   50%   60%   70%   80%   90%   95%   99% 99.5% 99.9%  100%
   15    16    19    22    27    34    46    68   116   246   429  1046  1391  2417  4095

methylKit::getMethylationStats output for "ka_6" CpGhsm-sites (#: 31348) at minimum coverage 15      - methylation statistics per base
summary:
   Min. 1st Qu.  Median    Mean 3rd Qu.    Max.
    1.9    40.0    64.7    60.9    83.3   100.0
percentiles:
    0%    10%    20%    30%    40%    50%    60%    70%    80%    90%    95%    99%  99.5%  99.9%   100%
  1.93  25.00  35.29  44.10  54.55  64.71  73.33  80.39  86.57  91.89  94.83 100.00 100.00 100.00 100.00

methylKit::getCoverageStats output for "ka_6" CpGhsm-sites (#: 43539) in coverage range [10-100]     - read coverage statistics per base
summary:
   Min. 1st Qu.  Median    Mean 3rd Qu.    Max.
   10.0    11.0    16.0    24.1    28.0   100.0
percentiles:
   0%   10%   20%   30%   40%   50%   60%   70%   80%   90%   95%   99% 99.5% 99.9%  100%
   10    10    11    12    14    16    19    24    34    52    70    93    96   100   100

methylKit::getMethylationStats output for "ka_6" CpGhsm-sites (#: 43539) in coverage range [10-100] - methylation statistics per base
summary:
   Min. 1st Qu.  Median    Mean 3rd Qu.    Max.
    9.0    50.0    69.2    66.5    84.6   100.0
percentiles:
   0%   10%   20%   30%   40%   50%   60%   70%   80%   90%   95%   99% 99.5% 99.9%  100%
  9.0  36.8  46.2  53.6  60.0  69.2  75.0  81.8  88.9  93.3 100.0 100.0 100.0 100.0 100.0
```

```
Directory: REPCMS File: repcms-Sd_LI_ka_7.txt

Number of "ka_7" CpGhsm-sites with minimal and higher level coverage:

  number of "ka_7" CpGhsm-sites with coverage >=  4: 393550
  number of "ka_7" CpGhsm-sites with coverage >=  6: 144066
  number of "ka_7" CpGhsm-sites with coverage >= 10:  50541
  number of "ka_7" CpGhsm-sites with coverage >= 15:  31342

Coverage and methylation statistics for "ka_7" CpGhsm-sites at different levels of minimum coverage:

methylKit::getCoverageStats output for "ka_7" CpGhsm-sites (#: 393550) at minimum coverage 4   - read coverage statistics per base
summary:
   Min. 1st Qu.  Median    Mean 3rd Qu.    Max.
      4       4       5      13       7    4011
percentiles:
   0%   10%   20%   30%   40%   50%   60%   70%   80%   90%   95%   99% 99.5% 99.9%  100%
    4     4     4     4     4     5     5     6     7    12    25   193   362   920  4011

methylKit::getMethylationStats output for "ka_7" CpGhsm-sites (#: 393550) at minimum coverage 4      - methylation statistics per base
summary:
   Min. 1st Qu.  Median    Mean 3rd Qu.    Max.
    2.1    93.3   100.0    92.8   100.0   100.0
percentiles:
    0%    10%    20%    30%    40%    50%    60%    70%    80%    90%    95%    99%  99.5%  99.9%   100%
  2.12  71.43  85.71 100.00 100.00 100.00 100.00 100.00 100.00 100.00 100.00 100.00 100.00 100.00 100.00

methylKit::getCoverageStats output for "ka_7" CpGhsm-sites (#: 144066) at minimum coverage 6   - read coverage statistics per base
summary:
   Min. 1st Qu.  Median    Mean 3rd Qu.    Max.
      6       6       8      28      12    4011
percentiles:
   0%   10%   20%   30%   40%   50%   60%   70%   80%   90%   95%   99% 99.5% 99.9%  100%
    6     6     6     6     7     8     9    11    16    37    97   452   698  1419  4011

methylKit::getMethylationStats output for "ka_7" CpGhsm-sites (#: 144066) at minimum coverage 6      - methylation statistics per base
summary:
   Min. 1st Qu.  Median    Mean 3rd Qu.    Max.
    2.1    71.4    83.3    80.3   100.0   100.0
percentiles:
    0%    10%    20%    30%    40%    50%    60%    70%    80%    90%    95%    99%  99.5%  99.9%   100%
  2.12  50.00  66.67  75.00  83.33  83.33  87.50 100.00 100.00 100.00 100.00 100.00 100.00 100.00 100.00

methylKit::getCoverageStats output for "ka_7" CpGhsm-sites (#: 50541) at minimum coverage 10   - read coverage statistics per base
summary:
   Min. 1st Qu.  Median    Mean 3rd Qu.    Max.
     10      12      19      68      44    4011
percentiles:
   0%   10%   20%   30%   40%   50%   60%   70%   80%   90%   95%   99% 99.5% 99.9%  100%
   10    10    11    13    15    19    24    35    61   145   293   814  1148  2155  4011

methylKit::getMethylationStats output for "ka_7" CpGhsm-sites (#: 50541) at minimum coverage 10      - methylation statistics per base
summary:
   Min. 1st Qu.  Median    Mean 3rd Qu.    Max.
    2.1    48.1    68.4    65.0    84.6   100.0
percentiles:
    0%    10%    20%    30%    40%    50%    60%    70%    80%    90%    95%    99%  99.5%  99.9%   100%
```

```
  2.12  32.00  43.75  50.00  60.00  68.42  75.00  81.82  88.10  92.70 100.00 100.00 100.00 100.00 100.00

methylKit::getCoverageStats output for "ka_7" CpGhsm-sites (#: 31342) at minimum coverage 15   - read coverage statistics per base
summary:
   Min. 1st Qu.  Median    Mean 3rd Qu.    Max.
     15      20      33     103      87    4011
percentiles:
   0%   10%   20%   30%   40%   50%   60%   70%   80%   90%   95%   99% 99.5% 99.9%  100%
   15    16    19    22    26    33    45    67   113   242   423  1041  1376  2377  4011

methylKit::getMethylationStats output for "ka_7" CpGhsm-sites (#: 31342) at minimum coverage 15      - methylation statistics per base
summary:
   Min. 1st Qu.  Median    Mean 3rd Qu.    Max.
    2.1    40.0    64.9    61.3    83.9   100.0
percentiles:
    0%    10%    20%    30%    40%    50%    60%    70%    80%    90%    95%    99%  99.5%  99.9%   100%
  2.12  25.63  35.71  44.44  55.00  64.86  73.33  80.69  86.67  92.00  94.92 100.00 100.00 100.00 100.00

methylKit::getCoverageStats output for "ka_7" CpGhsm-sites (#: 43588) in coverage range [10-100]     - read coverage statistics per base
summary:
   Min. 1st Qu.  Median    Mean 3rd Qu.    Max.
   10.0    11.0    16.0    24.1    28.0   100.0
percentiles:
   0%   10%   20%   30%   40%   50%   60%   70%   80%   90%   95%   99% 99.5% 99.9%  100%
   10    10    11    12    14    16    19    24    33    52    70    94    97   100   100

methylKit::getMethylationStats output for "ka_7" CpGhsm-sites (#: 43588) in coverage range [10-100]  - methylation statistics per base
summary:
   Min. 1st Qu.  Median    Mean 3rd Qu.    Max.
    9.0    50.0    69.2    66.6    85.0   100.0
percentiles:
   0%   10%   20%   30%   40%   50%   60%   70%   80%   90%   95%   99% 99.5% 99.9%  100%
  9.0  37.0  46.2  53.6  60.0  69.2  75.0  81.8  89.3  93.3 100.0 100.0 100.0 100.0 100.0
```

```
Directory: REPCMS File: repcms-Sd_LI_ka_8.txt

Number of "ka_8" CpGhsm-sites with minimal and higher level coverage:

  number of "ka_8" CpGhsm-sites with coverage >=  4: 222387
  number of "ka_8" CpGhsm-sites with coverage >=  6:  73393
  number of "ka_8" CpGhsm-sites with coverage >= 10:  33635
  number of "ka_8" CpGhsm-sites with coverage >= 15:  22822

Coverage and methylation statistics for "ka_8" CpGhsm-sites at different levels of minimum coverage:

methylKit::getCoverageStats output for "ka_8" CpGhsm-sites (#: 222387) at minimum coverage 4   - read coverage statistics per base
summary:
   Min. 1st Qu.  Median    Mean 3rd Qu.    Max.
      4       4       5      15       6    3091
percentiles:
   0%   10%   20%   30%   40%   50%   60%   70%   80%   90%   95%   99% 99.5% 99.9%  100%
    4     4     4     4     4     5     5     6     7    15    38   240   404   909  3091

methylKit::getMethylationStats output for "ka_8" CpGhsm-sites (#: 222387) at minimum coverage 4      - methylation statistics per base
summary:
   Min. 1st Qu.  Median    Mean 3rd Qu.    Max.
    2.8    83.3   100.0    90.5   100.0   100.0
percentiles:
    0%    10%    20%    30%    40%    50%    60%    70%    80%    90%    95%    99%  99.5%  99.9%   100%
  2.79  71.43  80.00  85.71 100.00 100.00 100.00 100.00 100.00 100.00 100.00 100.00 100.00 100.00 100.00

methylKit::getCoverageStats output for "ka_8" CpGhsm-sites (#: 73393) at minimum coverage 6    - read coverage statistics per base
summary:
   Min. 1st Qu.  Median    Mean 3rd Qu.    Max.
      6       6       9      36      19    3091
percentiles:
   0%   10%   20%   30%   40%   50%   60%   70%   80%   90%   95%   99% 99.5% 99.9%  100%
    6     6     6     7     8     9    11    15    26    68   151   514   707  1394  3091

methylKit::getMethylationStats output for "ka_8" CpGhsm-sites (#: 73393) at minimum coverage 6 - methylation statistics per base
summary:
   Min. 1st Qu.  Median    Mean 3rd Qu.    Max.
    2.8    66.7    83.3    77.1    94.1   100.0
percentiles:
    0%    10%    20%    30%    40%    50%    60%    70%    80%    90%    95%    99%  99.5%  99.9%   100%
  2.79  45.45  60.00  71.43  77.78  83.33  85.71  88.89 100.00 100.00 100.00 100.00 100.00 100.00 100.00

methylKit::getCoverageStats output for "ka_8" CpGhsm-sites (#: 33635) at minimum coverage 10   - read coverage statistics per base
summary:
   Min. 1st Qu.  Median    Mean 3rd Qu.    Max.
     10      13      22      71      56    3091
percentiles:
   0%   10%   20%   30%   40%   50%   60%   70%   80%   90%   95%   99% 99.5% 99.9%  100%
   10    11    12    14    17    22    29    43    75   165   300   740   998  1801  3091

methylKit::getMethylationStats output for "ka_8" CpGhsm-sites (#: 33635) at minimum coverage 10      - methylation statistics per base
summary:
   Min. 1st Qu.  Median    Mean 3rd Qu.    Max.
    2.8    47.4    69.0    65.0    84.9   100.0
percentiles:
    0%    10%    20%    30%    40%    50%    60%    70%    80%    90%    95%    99%  99.5%  99.9%   100%
```

```
  2.79  31.25  42.86  50.00  60.00  68.97  75.76  81.82  88.00  92.86  96.88 100.00 100.00 100.00 100.00

methylKit::getCoverageStats output for "ka_8" CpGhsm-sites (#: 22822) at minimum coverage 15   - read coverage statistics per base
summary:
   Min. 1st Qu.  Median    Mean 3rd Qu.    Max.
     15      21      36      99      92    3091
percentiles:
   0%   10%   20%   30%   40%   50%   60%   70%   80%   90%   95%   99% 99.5% 99.9%  100%
   15    17    19    23    28    36    50    74   119   233   398   902  1188  1938  3091

methylKit::getMethylationStats output for "ka_8" CpGhsm-sites (#: 22822) at minimum coverage 15      - methylation statistics per base
summary:
   Min. 1st Qu.  Median    Mean 3rd Qu.    Max.
    2.8    41.2    67.9    62.7    85.0   100.0
percentiles:
    0%    10%    20%    30%    40%    50%    60%    70%    80%    90%    95%    99%  99.5%  99.9%   100%
  2.79  25.42  36.74  46.83  58.11  67.86  75.85  82.27  87.50  92.55  95.00 100.00 100.00 100.00 100.00

methylKit::getCoverageStats output for "ka_8" CpGhsm-sites (#: 28345) in coverage range [10-100]    - read coverage statistics per base
summary:
   Min. 1st Qu.  Median    Mean 3rd Qu.    Max.
   10.0    12.0    18.0    26.5    32.0   100.0
percentiles:
   0%   10%   20%   30%   40%   50%   60%   70%   80%   90%   95%   99% 99.5% 99.9%  100%
   10    10    11    13    15    18    22    28    38    59    76    94    97   100   100

methylKit::getMethylationStats output for "ka_8" CpGhsm-sites (#: 28345) in coverage range [10-100] - methylation statistics per base
summary:
   Min. 1st Qu.  Median    Mean 3rd Qu.    Max.
    9.0    50.0    70.0    66.9    85.7   100.0
percentiles:
   0%   10%   20%   30%   40%   50%   60%   70%   80%   90%   95%   99% 99.5% 99.9%  100%
  9.0  36.8  45.5  53.8  61.4  70.0  76.8  82.6  89.2  93.3 100.0 100.0 100.0 100.0 100.0
```

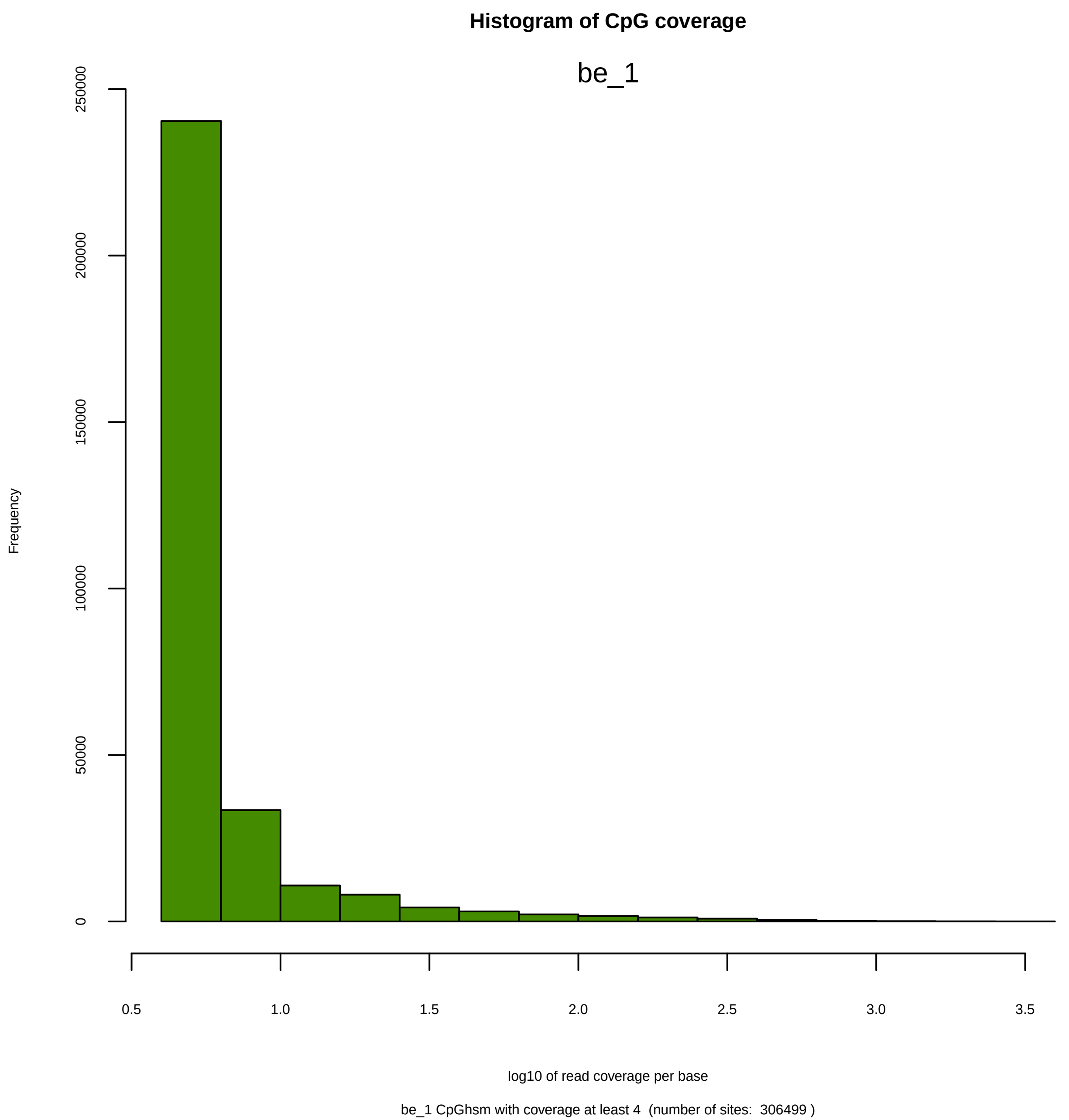

Histogram of CpG coverage
be_1
Frequency
0
50000
100000
150000
200000
250000
0.5
1.0
1.5
2.0
2.5
3.0
3.5
log10 of read coverage per base
be_1 CpGhsm with coverage at least 4 (number of sites: 306499 )

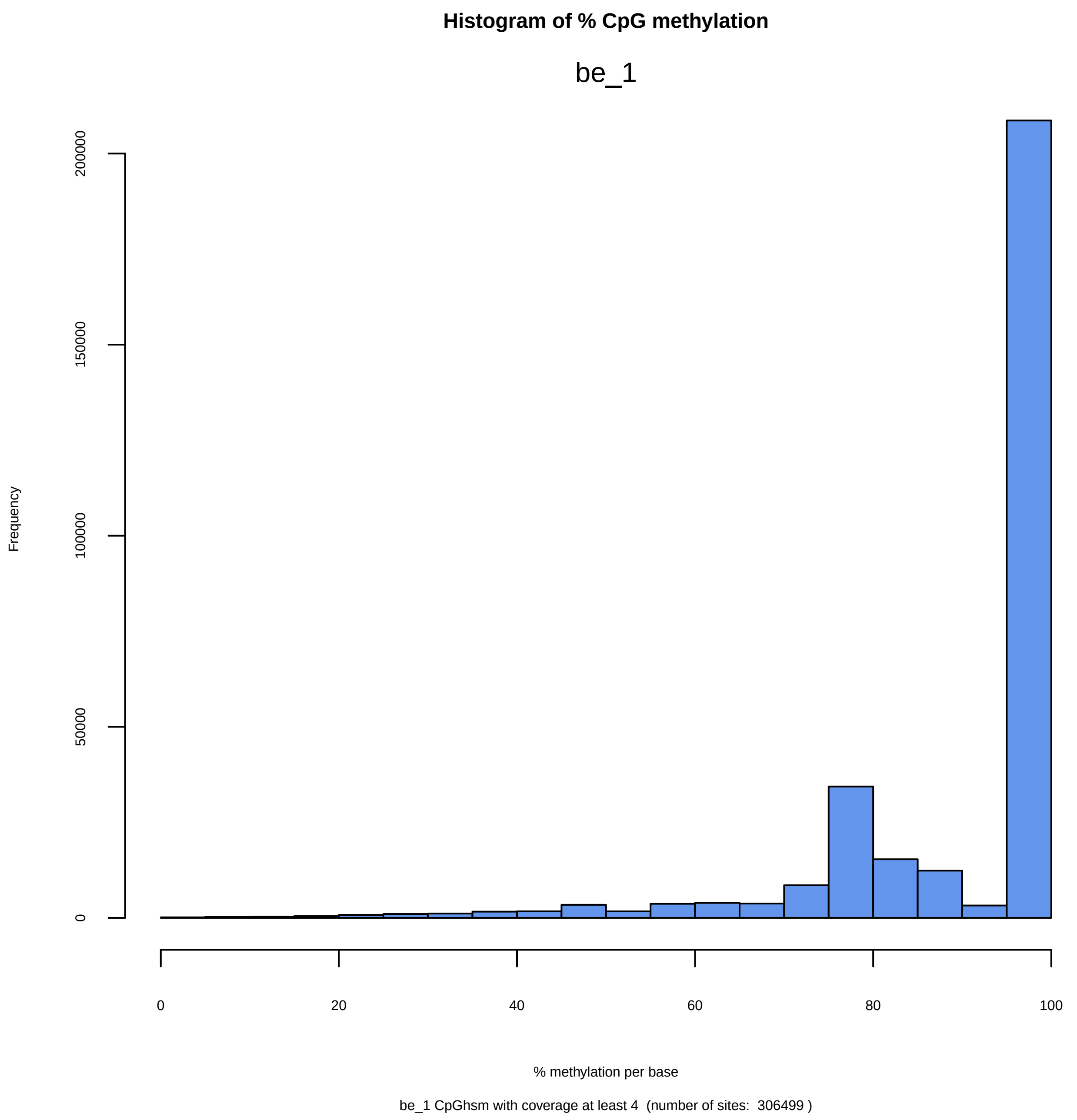

Histogram of % CpG methylation
be_1
Frequency
0
50000
100000
150000
200000
0
20
40
60
80
100
% methylation per base
be_1 CpGhsm with coverage at least 4 (number of sites: 306499 )

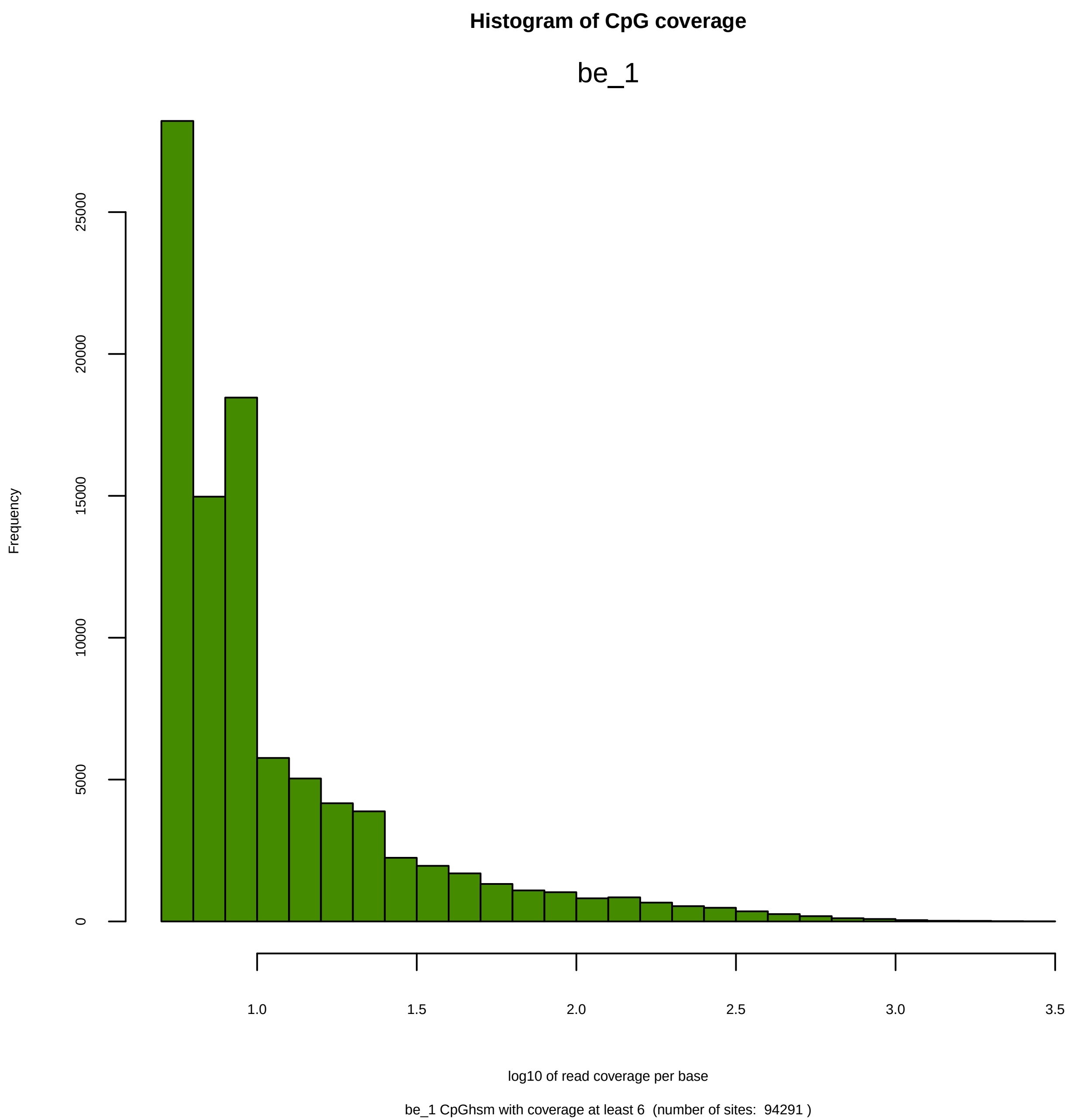

Histogram of CpG coverage
be_1
Frequency
0
5000
10000
15000
20000
25000
1.0
1.5
2.0
2.5
3.0
3.5
log10 of read coverage per base
be_1 CpGhsm with coverage at least 6  (number of sites:  94291 )

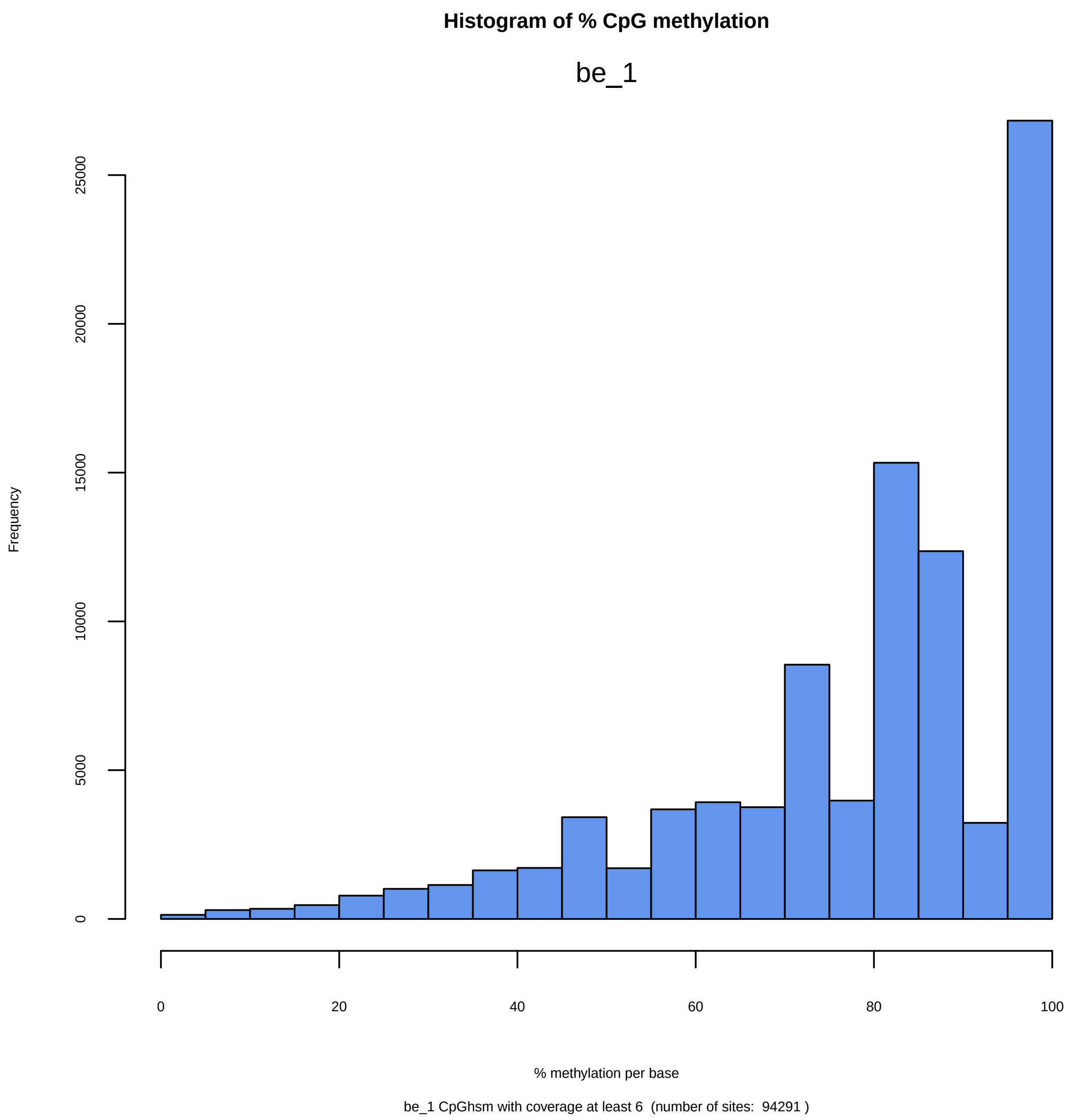

Histogram of % CpG methylation
be_1
Frequency
0
5000
10000
15000
20000
25000
0
20
40
60
80
100
% methylation per base
be_1 CpGhsm with coverage at least 6  (number of sites:  94291 )

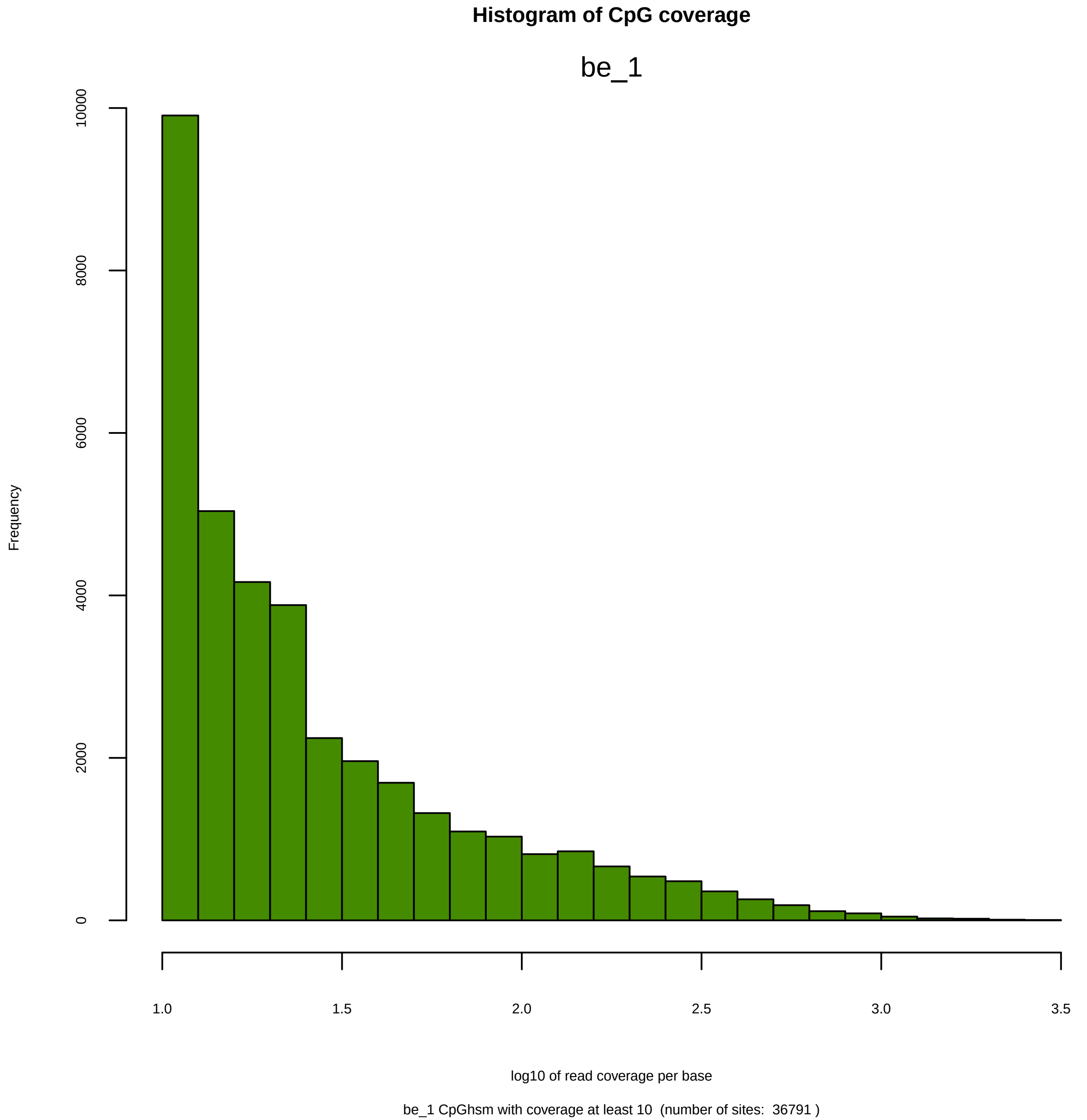
Histogram of CpG coverage
be_1
Frequency
0
2000
4000
6000
8000
10000
1.0
1.5
2.0
2.5
3.0
3.5
log10 of read coverage per base
be_1 CpGhsm with coverage at least 10 (number of sites: 36791 )

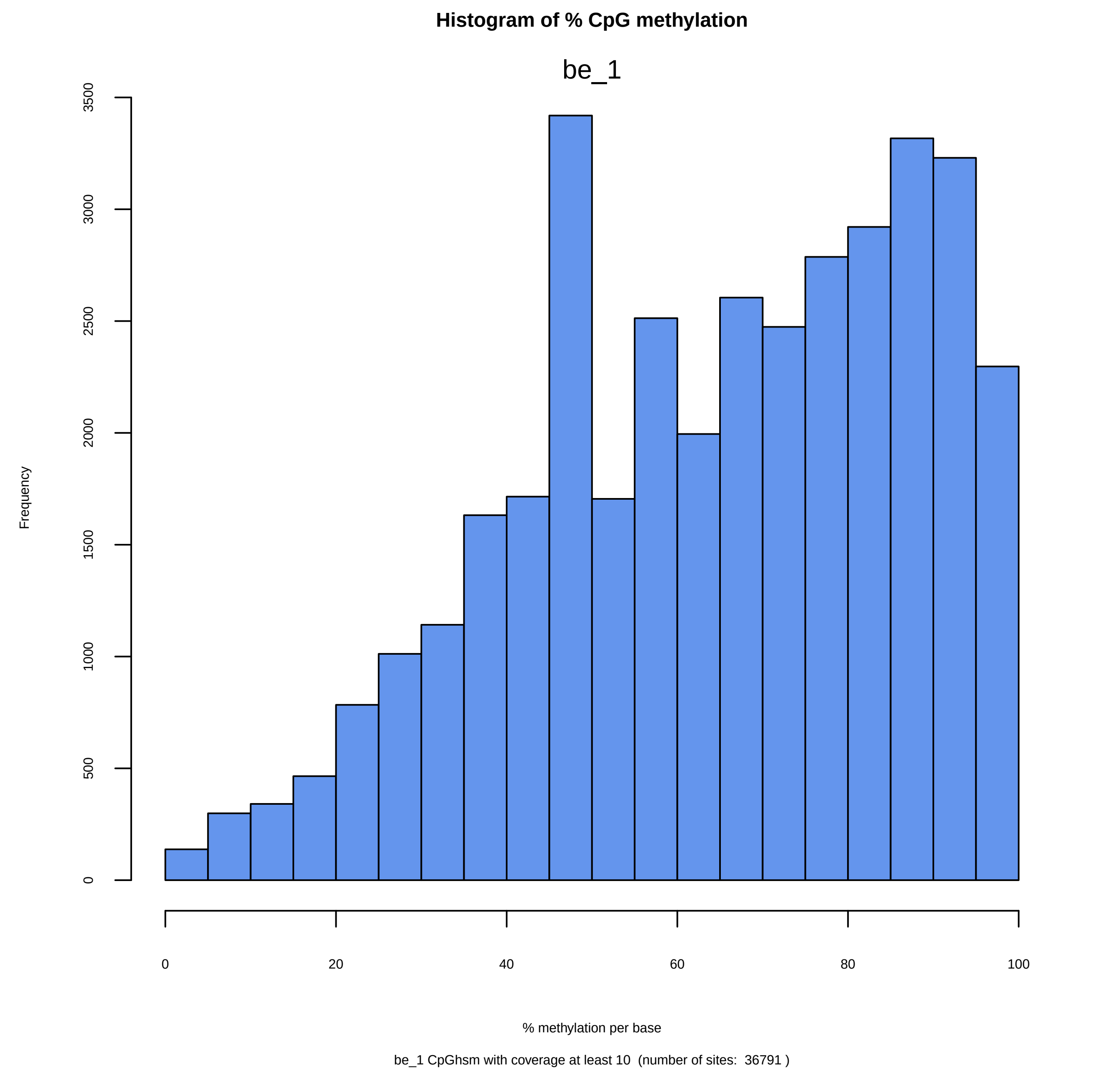

Histogram of % CpG methylation
be_1
Frequency
0
500
1000
1500
2000
2500
3000
3500
0
20
40
60
80
100
% methylation per base
be_1 CpGhsm with coverage at least 10 (number of sites: 36791 )

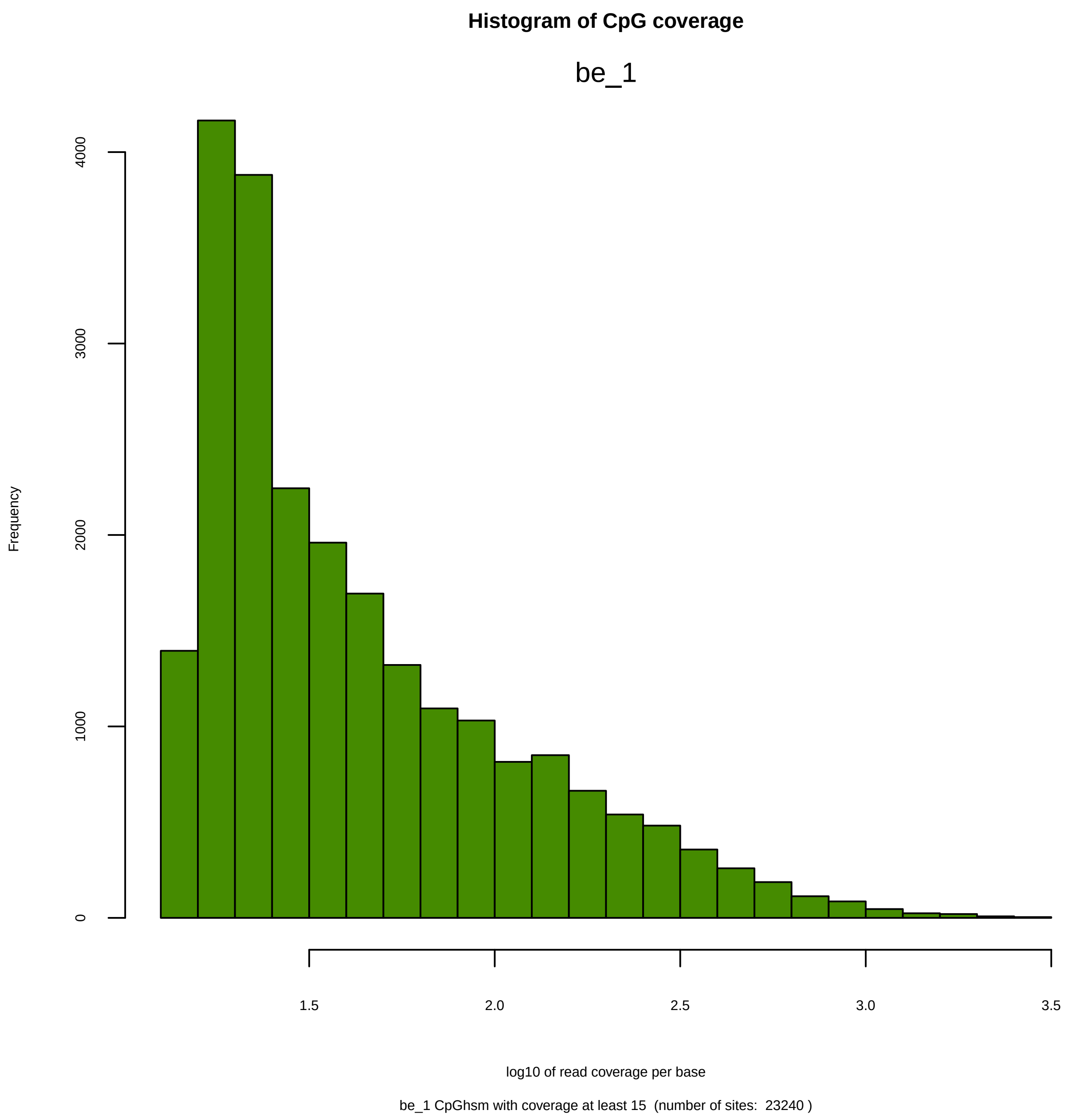

Histogram of CpG coverage
be_1
Frequency
0
1000
2000
3000
4000
1.5
2.0
2.5
3.0
3.5
log10 of read coverage per base
be_1 CpGhsm with coverage at least 15  (number of sites:  23240 )

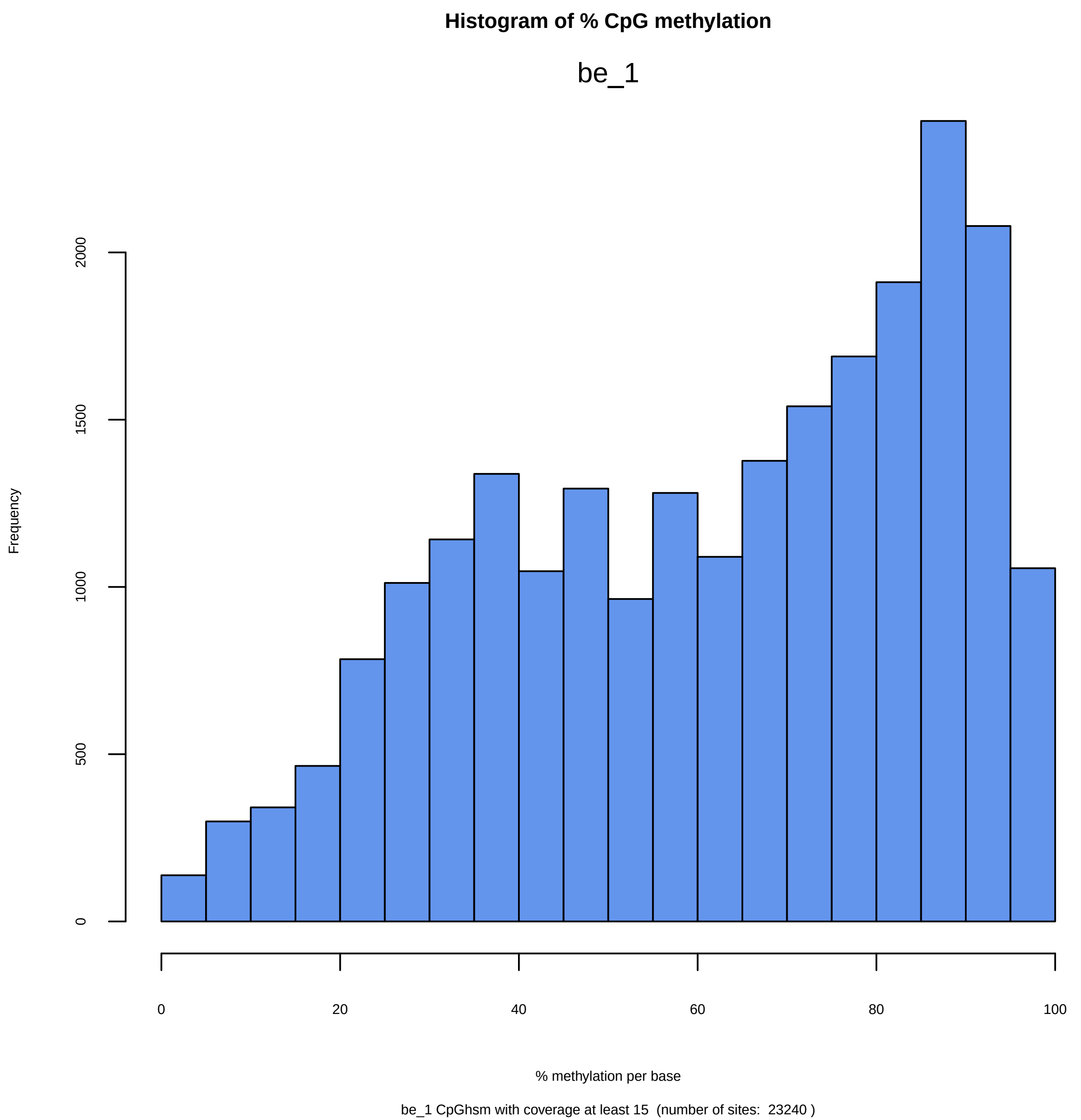
Histogram of % CpG methylation
be_1
Frequency
0
500
1000
1500
2000
0
20
40
60
80
100
% methylation per base
be_1 CpGhsm with coverage at least 15 (number of sites: 23240 )

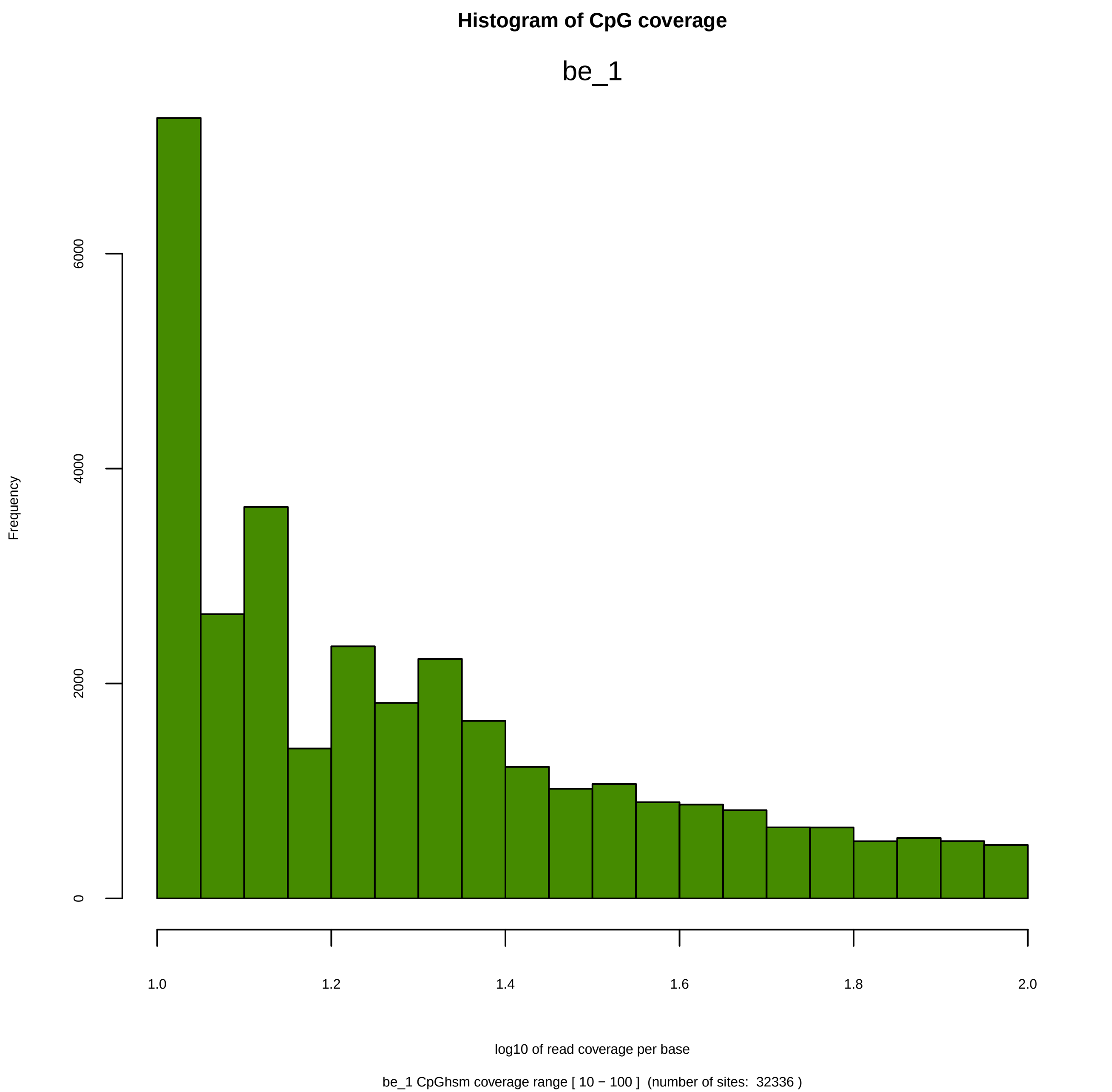
Histogram of CpG coverage
be_1
Frequency
0
2000
4000
6000
1.0
1.2
1.4
1.6
1.8
2.0
log10 of read coverage per base
be_1 CpGhsm coverage range [ 10 − 100 ] (number of sites: 32336 )

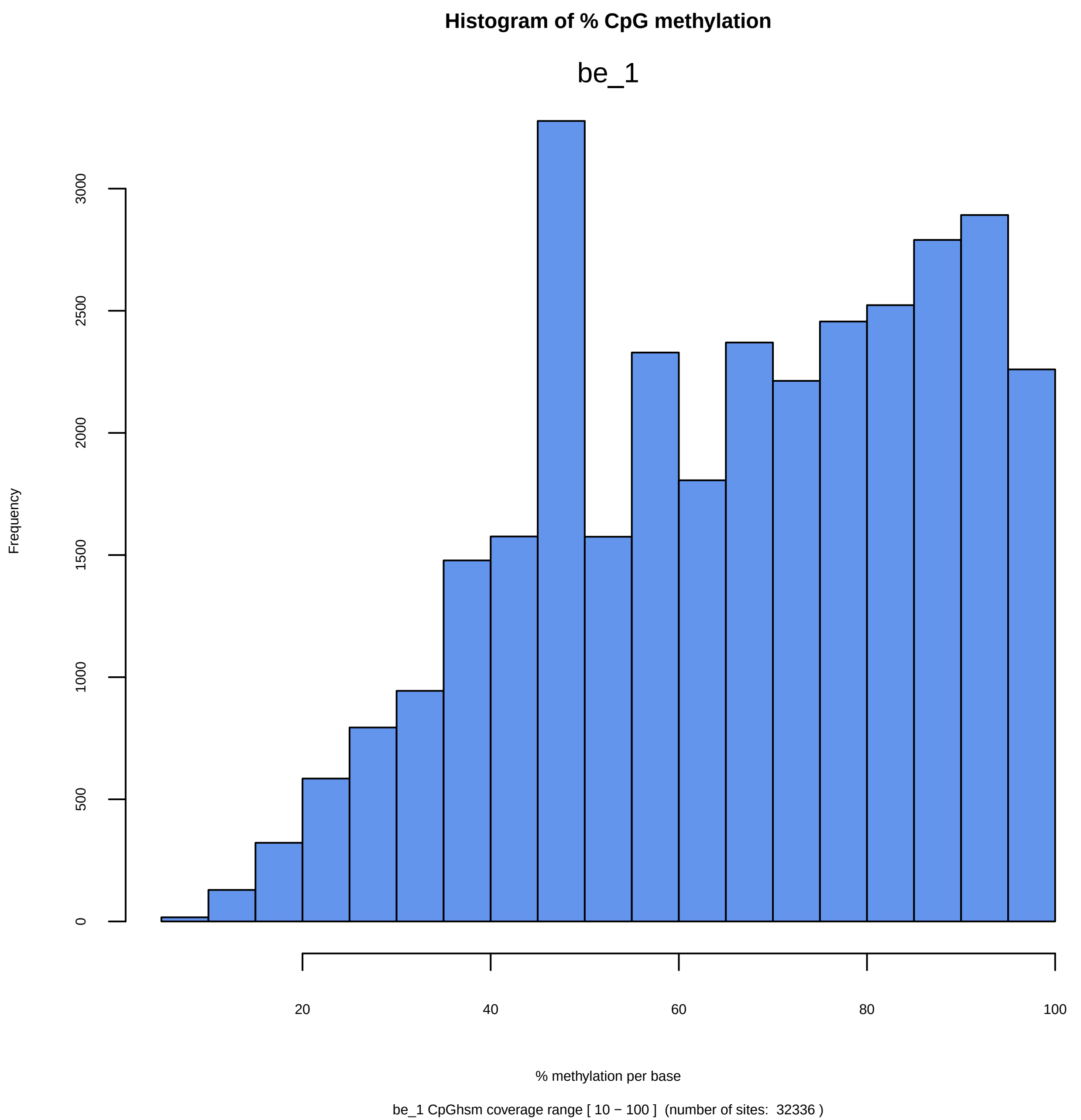
Histogram of % CpG methylation
be_1
Frequency
0
500
1000
1500
2000
2500
3000
20
40
60
80
100
% methylation per base
be_1 CpGhsm coverage range [ 10 − 100 ]  (number of sites:  32336 )

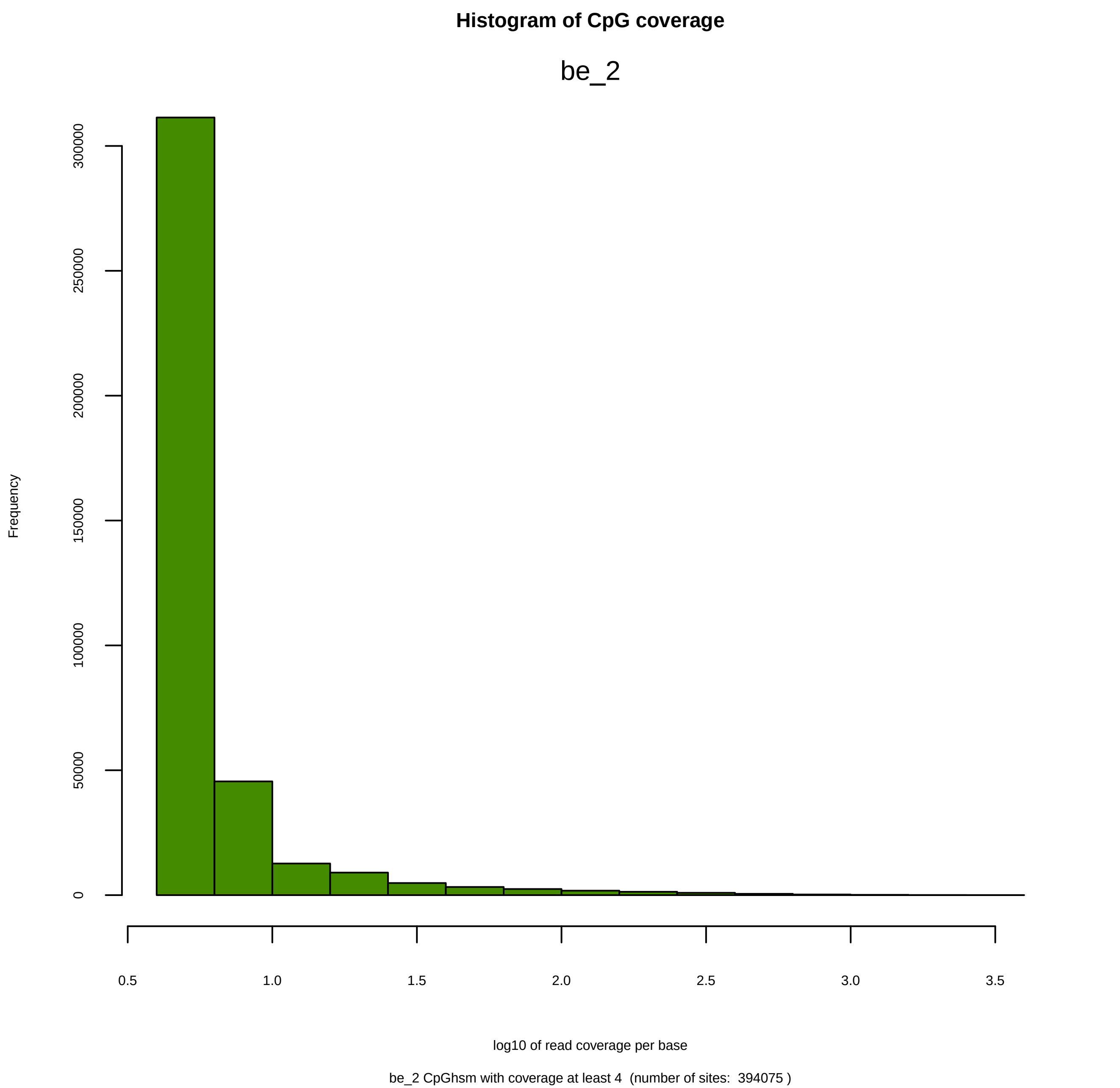
Histogram of CpG coverage
be_2
Frequency
0
50000
100000
150000
200000
250000
300000
0.5
1.0
1.5
2.0
2.5
3.0
3.5
log10 of read coverage per base
be_2 CpGhsm with coverage at least 4  (number of sites:  394075 )

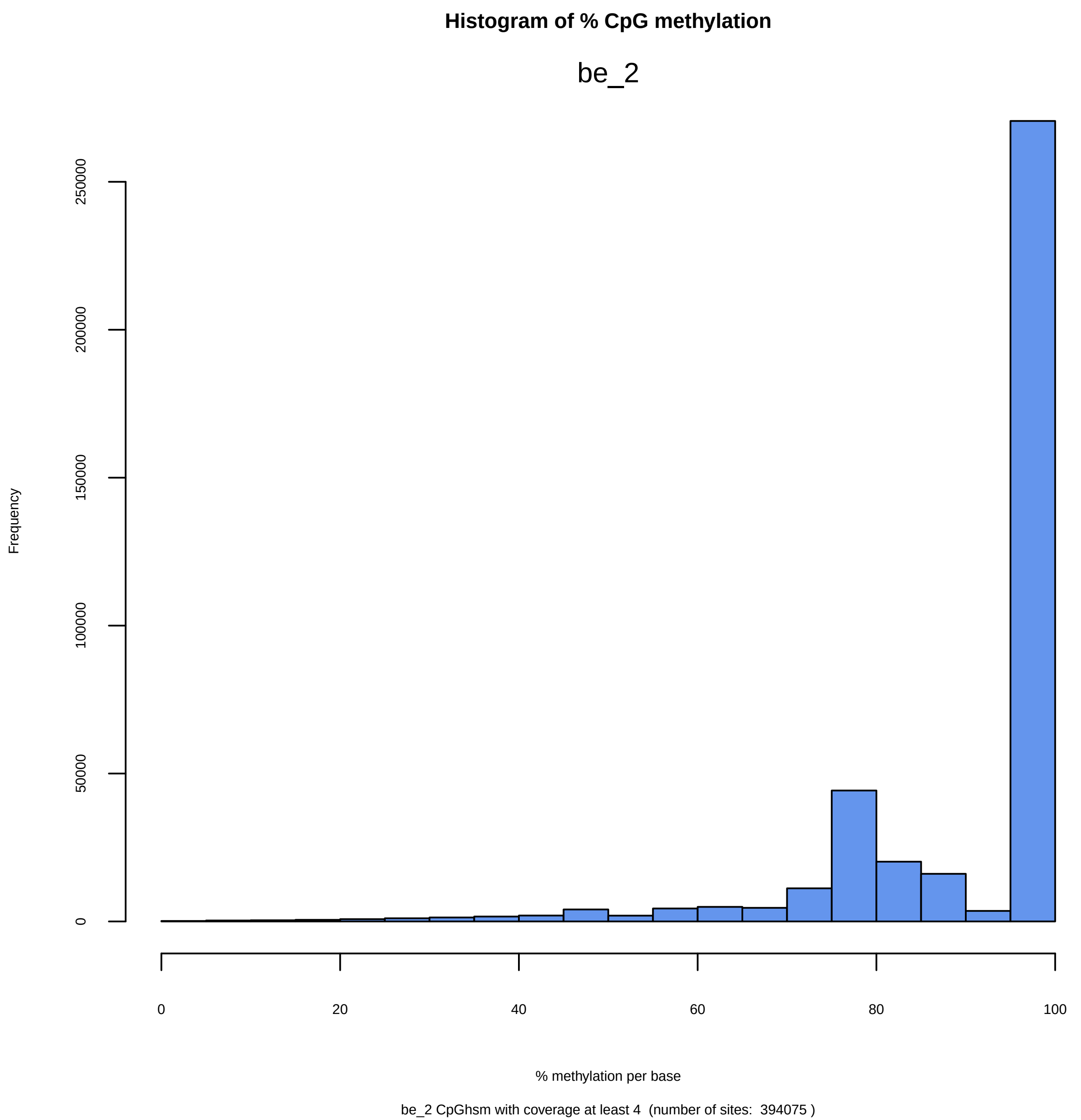

Histogram of % CpG methylation
be_2
Frequency
0
50000
100000
150000
200000
250000
0
20
40
60
80
100
% methylation per base
be_2 CpGhsm with coverage at least 4 (number of sites: 394075 )

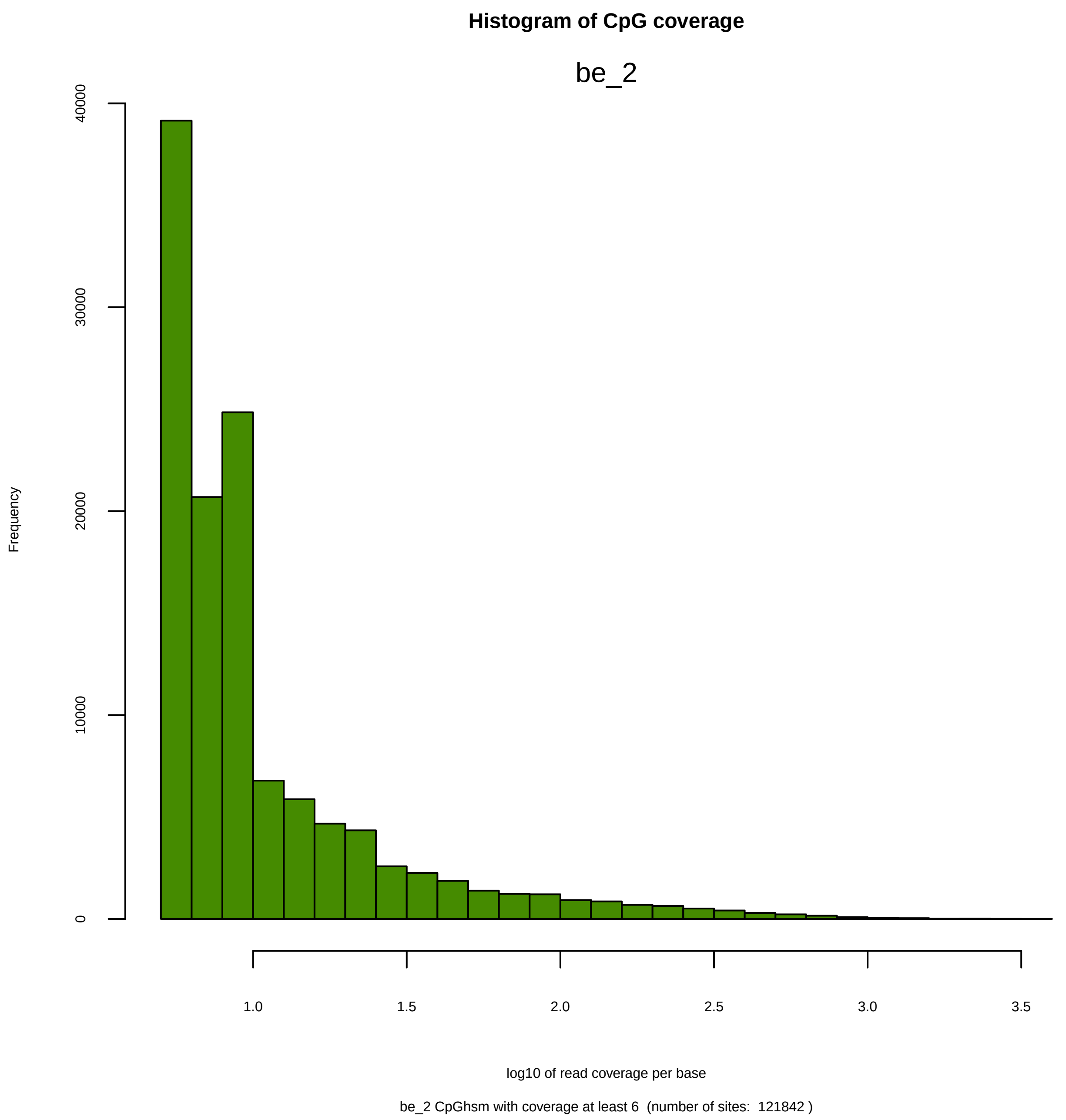

Histogram of CpG coverage
be_2
Frequency
0
10000
20000
30000
40000
1.0
1.5
2.0
2.5
3.0
3.5
log10 of read coverage per base
be_2 CpGhsm with coverage at least 6 (number of sites: 121842 )

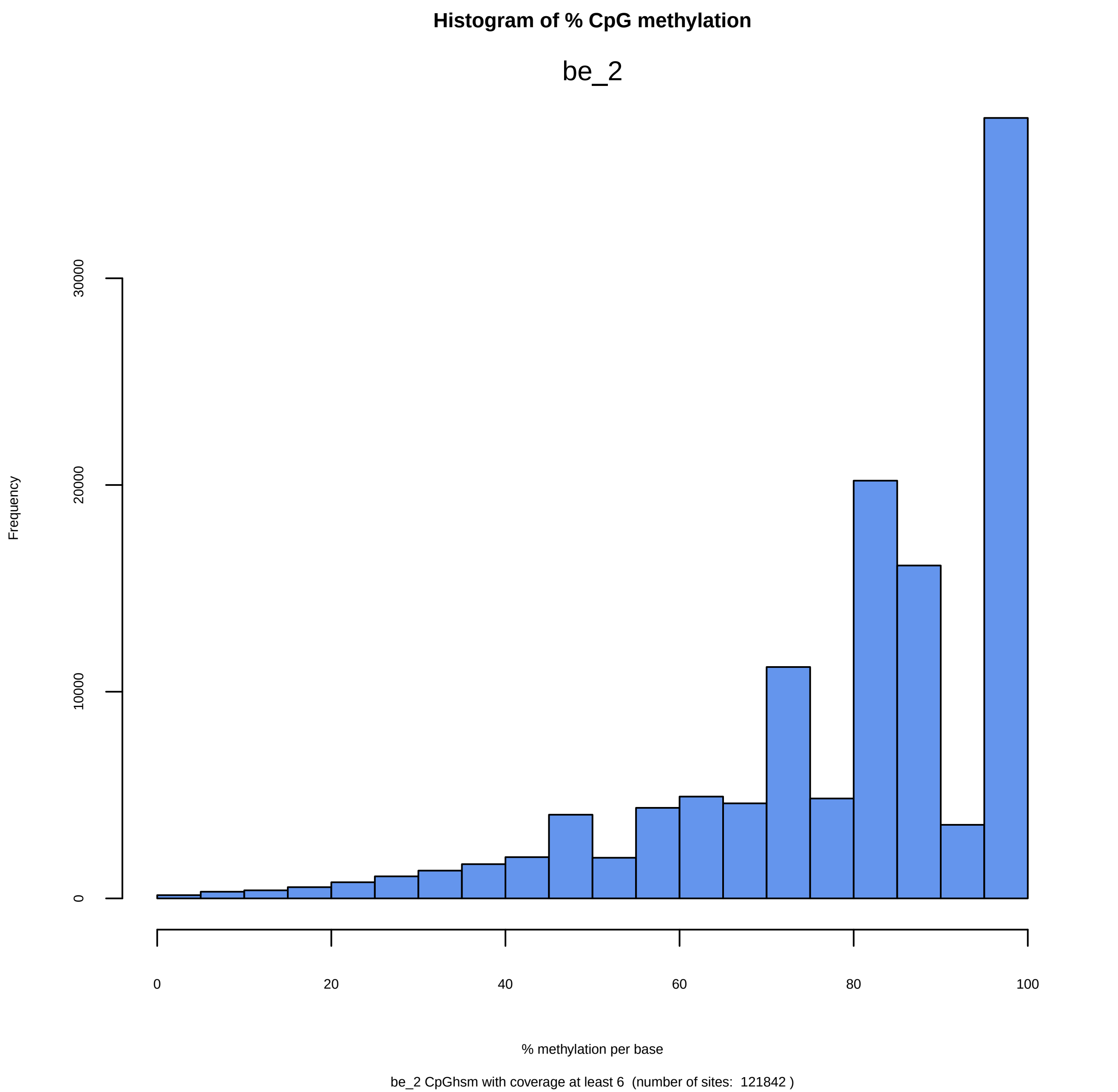

Histogram of % CpG methylation
be_2
Frequency
0
10000
20000
30000
0
20
40
60
80
100
% methylation per base
be_2 CpGhsm with coverage at least 6 (number of sites: 121842 )

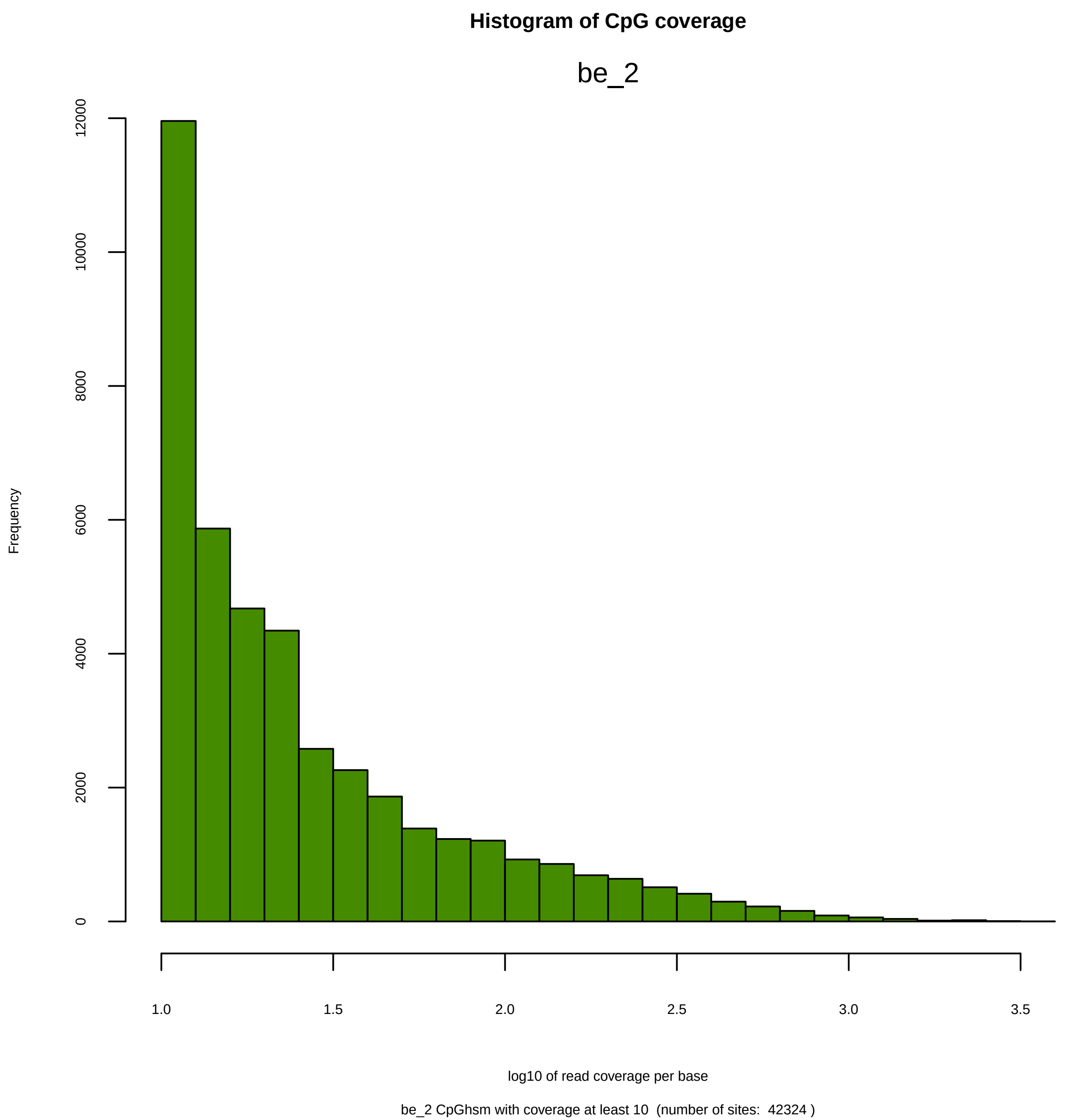

Histogram of CpG coverage
be_2
Frequency
0
2000
4000
6000
8000
10000
12000
1.0
1.5
2.0
2.5
3.0
3.5
log10 of read coverage per base
be_2 CpGhsm with coverage at least 10 (number of sites: 42324 )

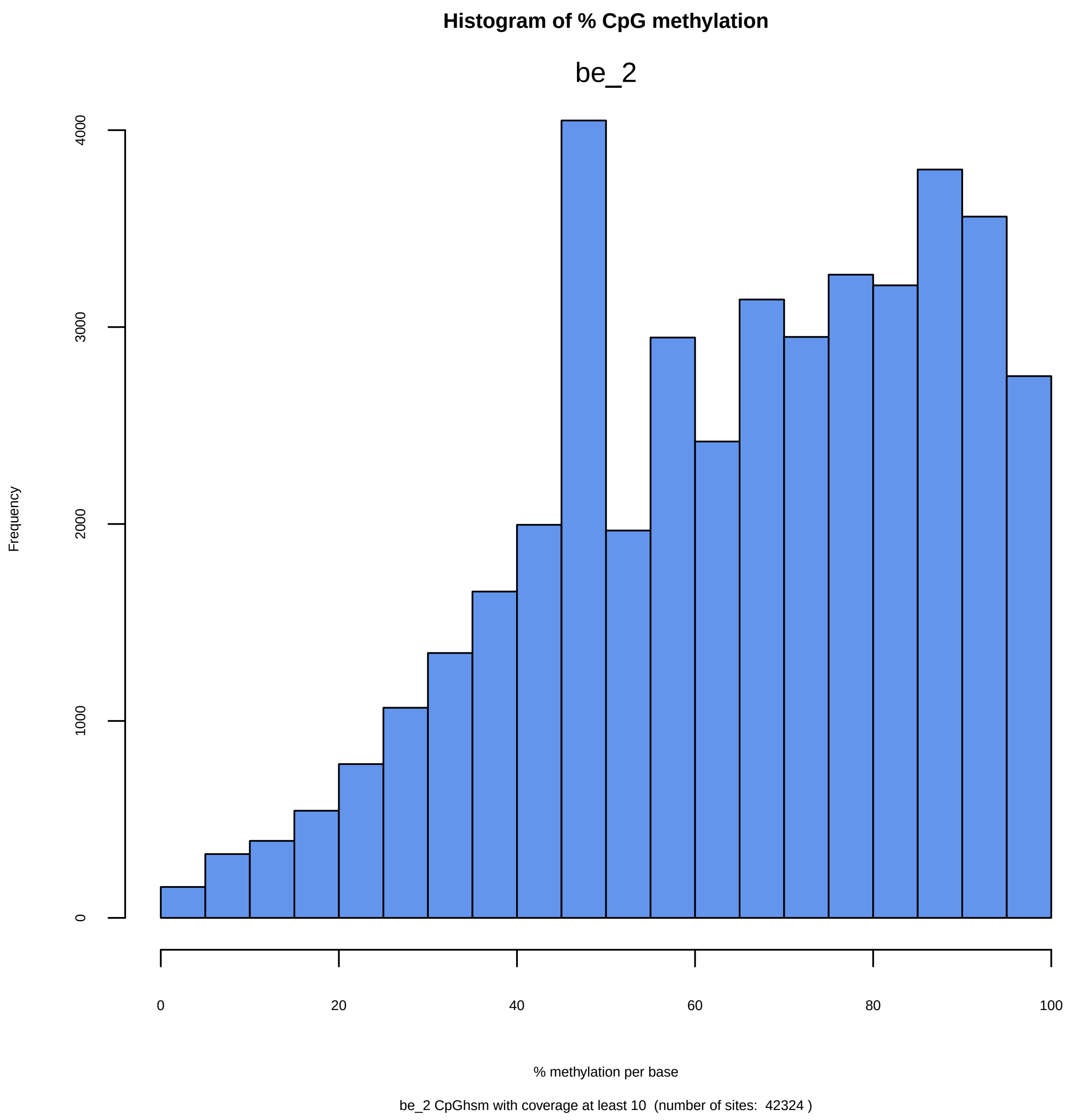

Histogram of % CpG methylation
be_2
Frequency
0
1000
2000
3000
4000
0
20
40
60
80
100
% methylation per base
be_2 CpGhsm with coverage at least 10 (number of sites: 42324 )

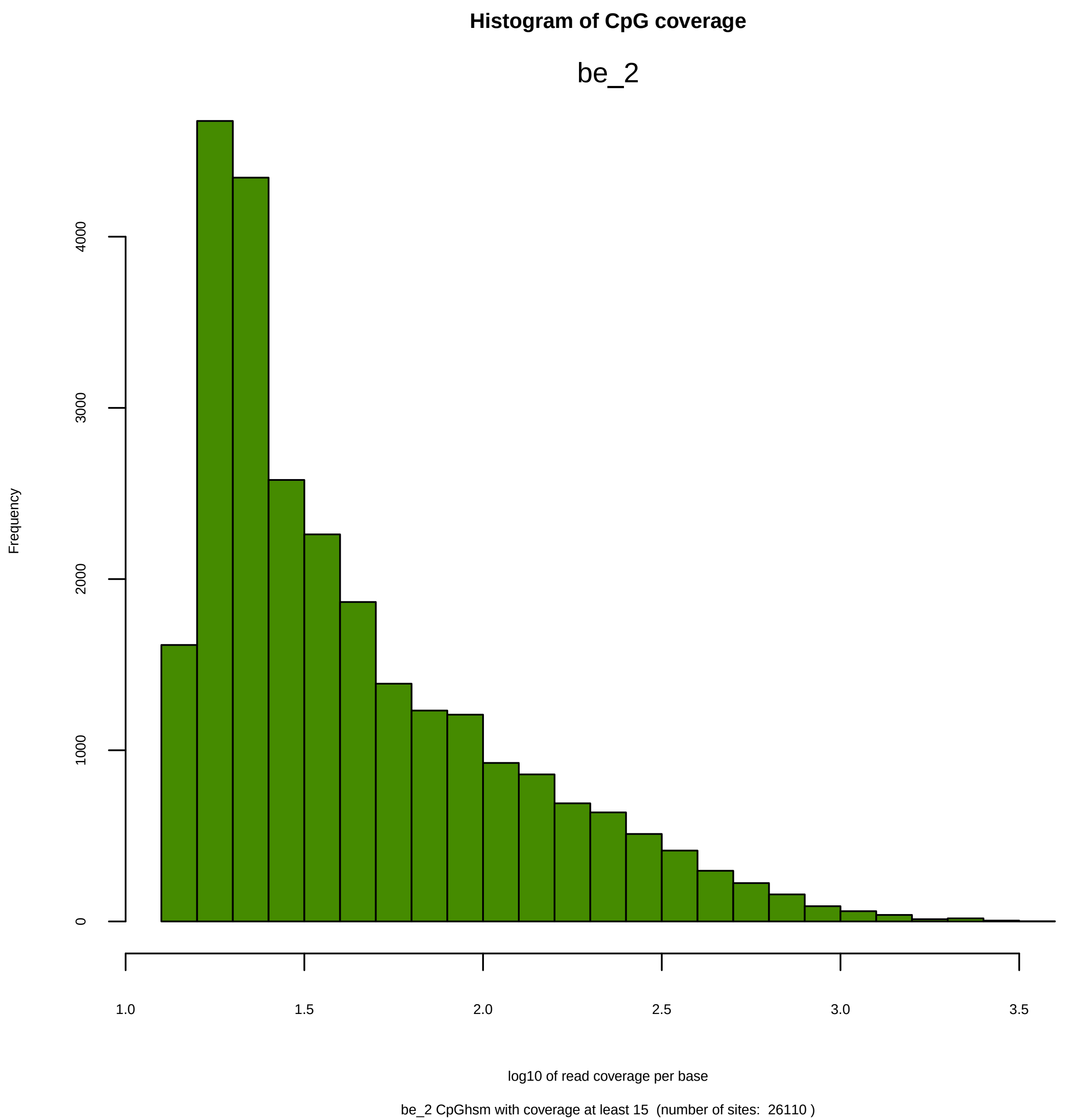
Histogram of CpG coverage
be_2
Frequency
0
1000
2000
3000
4000
1.0
1.5
2.0
2.5
3.0
3.5
log10 of read coverage per base
be_2 CpGhsm with coverage at least 15 (number of sites: 26110 )

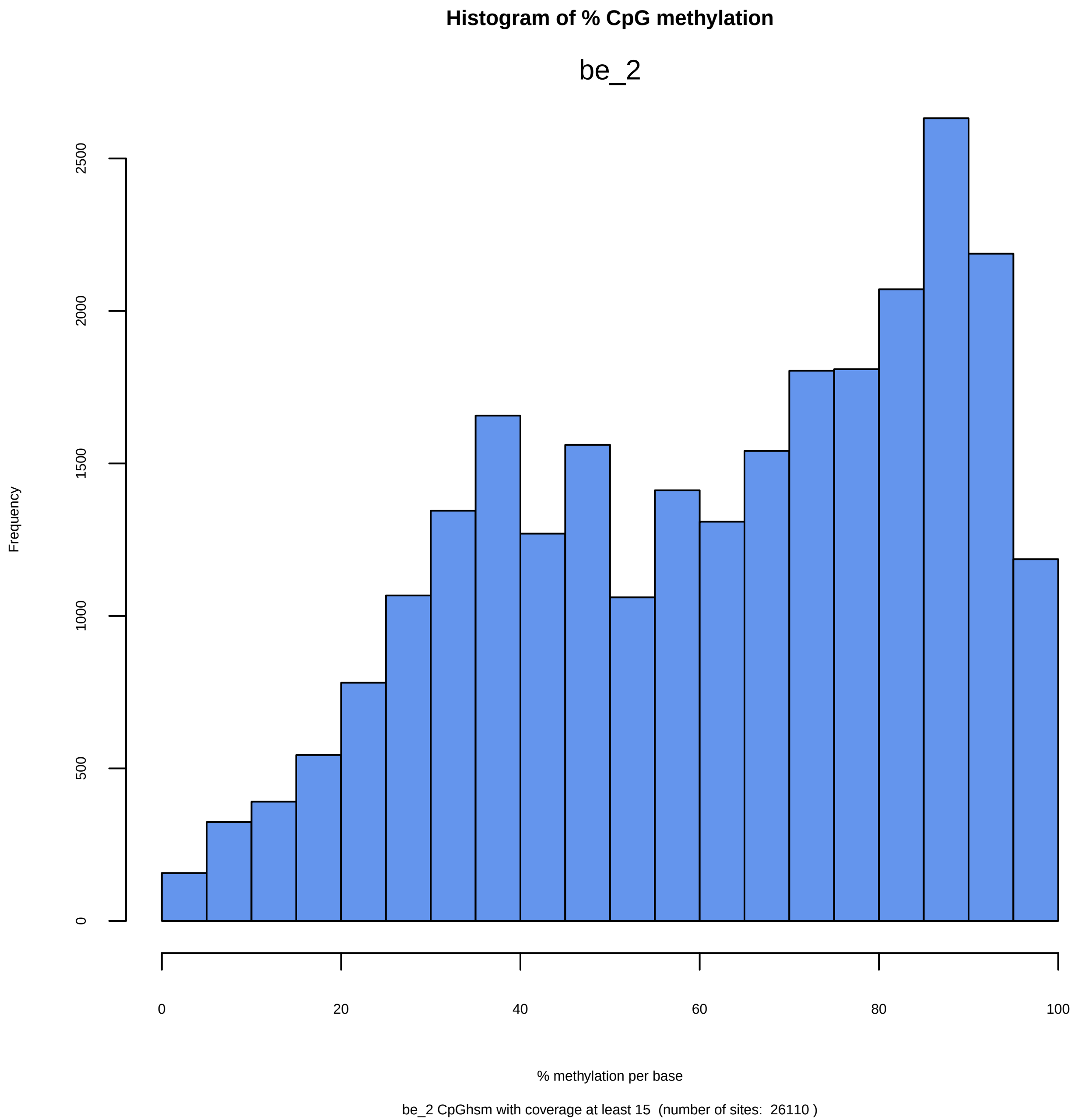
Histogram of % CpG methylation
be_2
Frequency
0
500
1000
1500
2000
2500
0
20
40
60
80
100
% methylation per base
be_2 CpGhsm with coverage at least 15  (number of sites:  26110 )

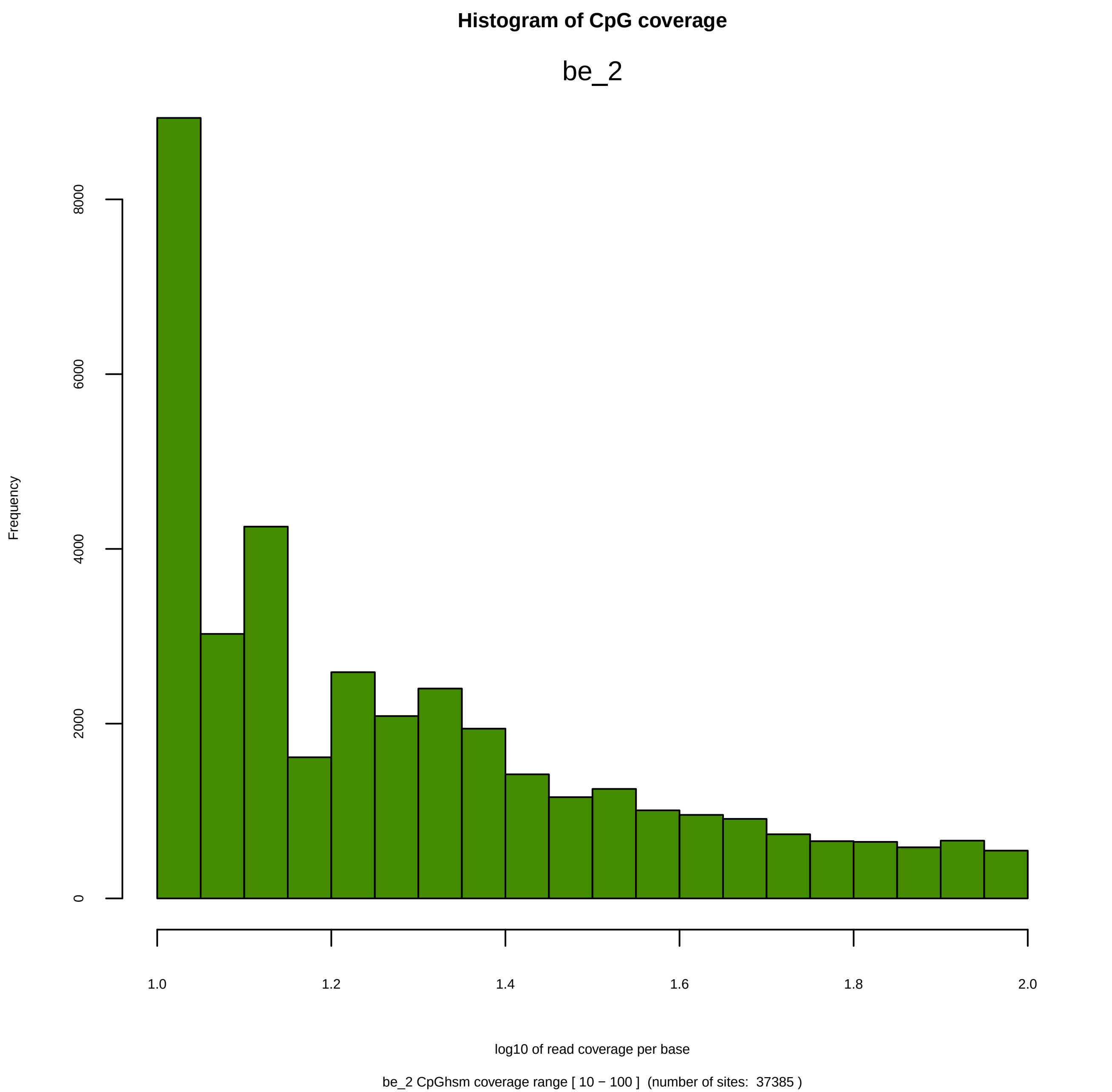

Histogram of CpG coverage
be_2
Frequency
0
2000
4000
6000
8000
1.0
1.2
1.4
1.6
1.8
2.0
log10 of read coverage per base
be_2 CpGhsm coverage range [ 10 − 100 ]  (number of sites:  37385 )

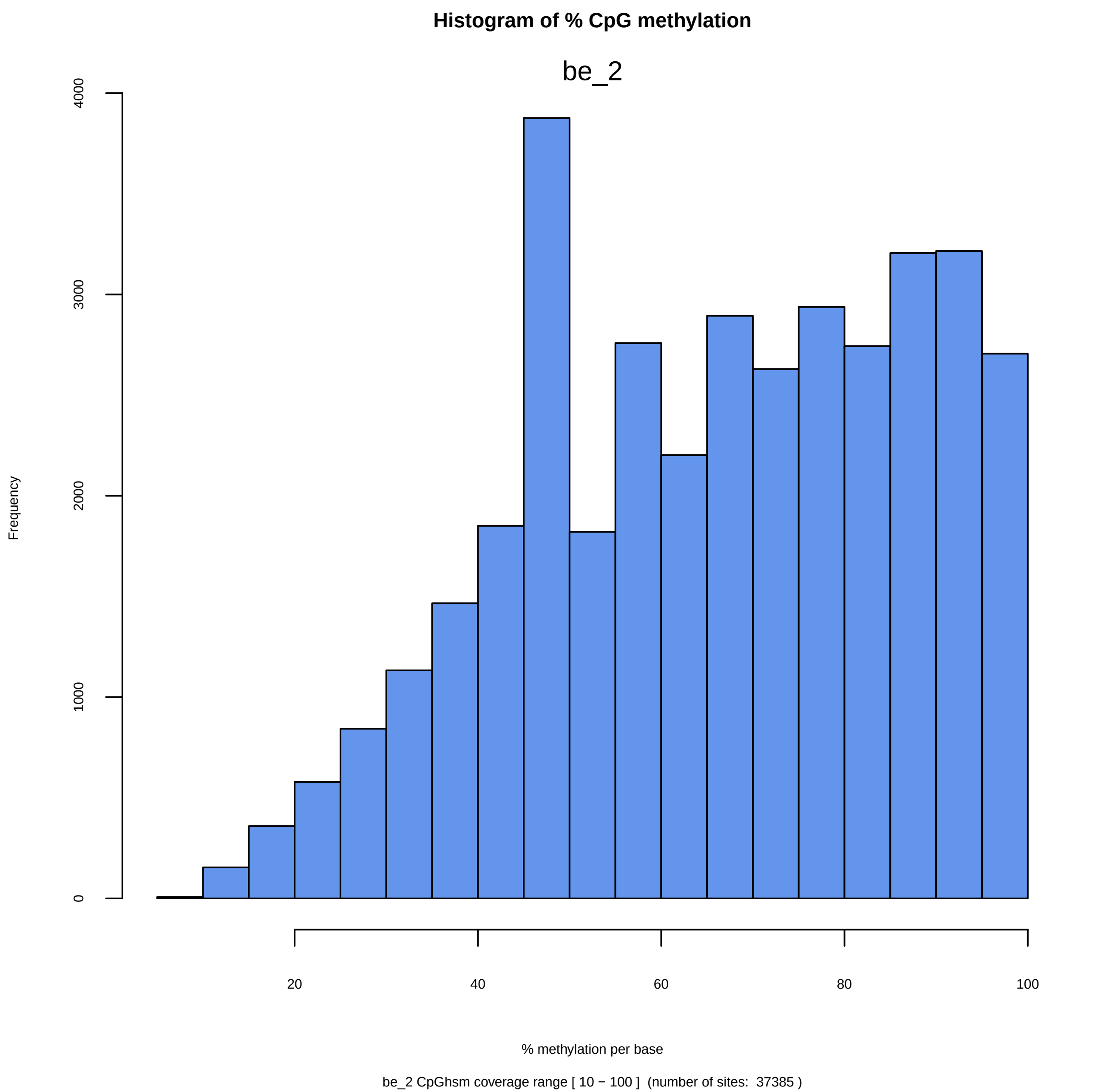

Histogram of % CpG methylation
be_2
Frequency
0
1000
2000
3000
4000
20
40
60
80
100
% methylation per base
be_2 CpGhsm coverage range [ 10 − 100 ]  (number of sites:  37385 )

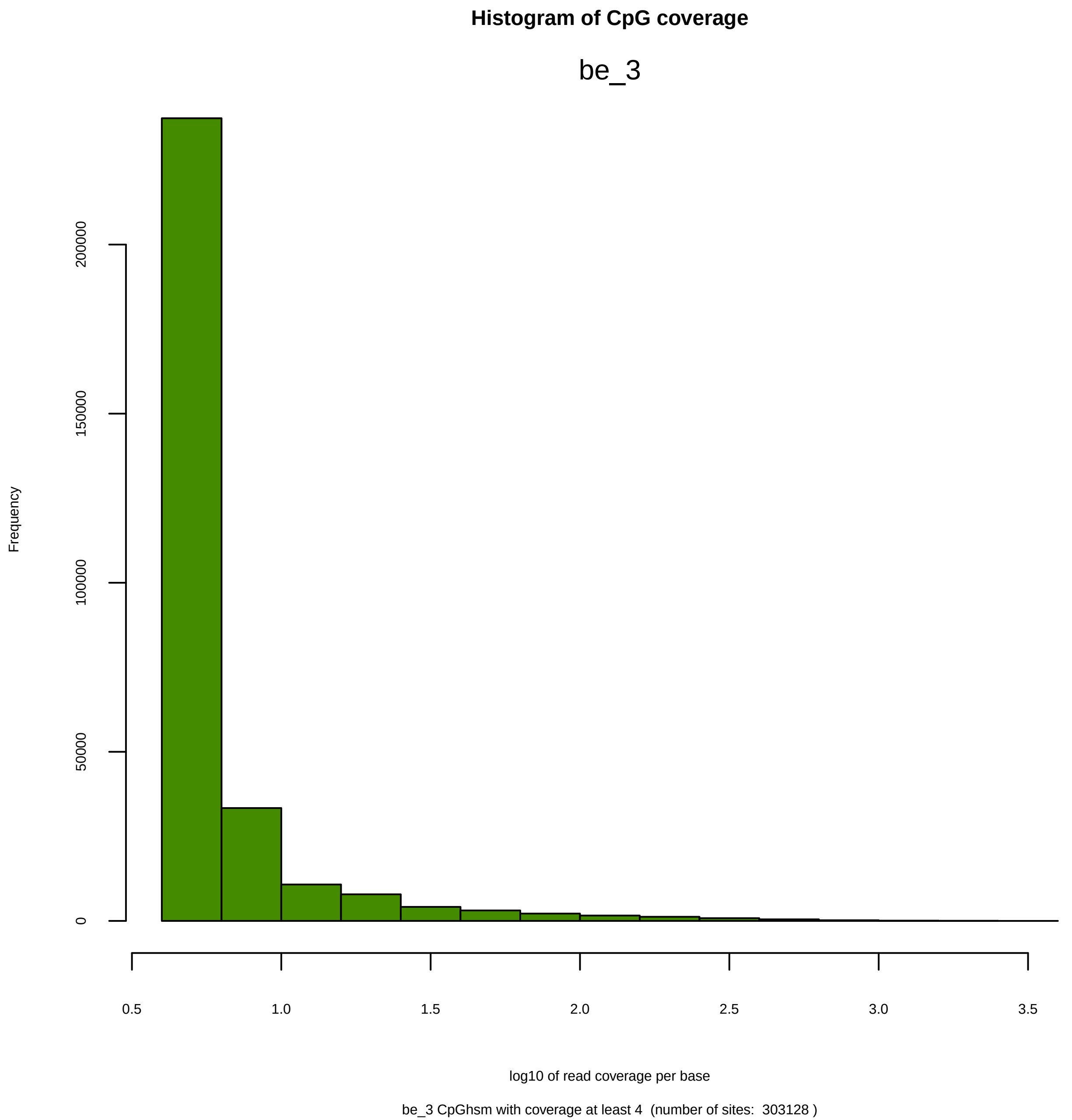
Histogram of CpG coverage
be_3
Frequency
0
50000
100000
150000
200000
0.5
1.0
1.5
2.0
2.5
3.0
3.5
log10 of read coverage per base
be_3 CpGhsm with coverage at least 4  (number of sites:  303128 )

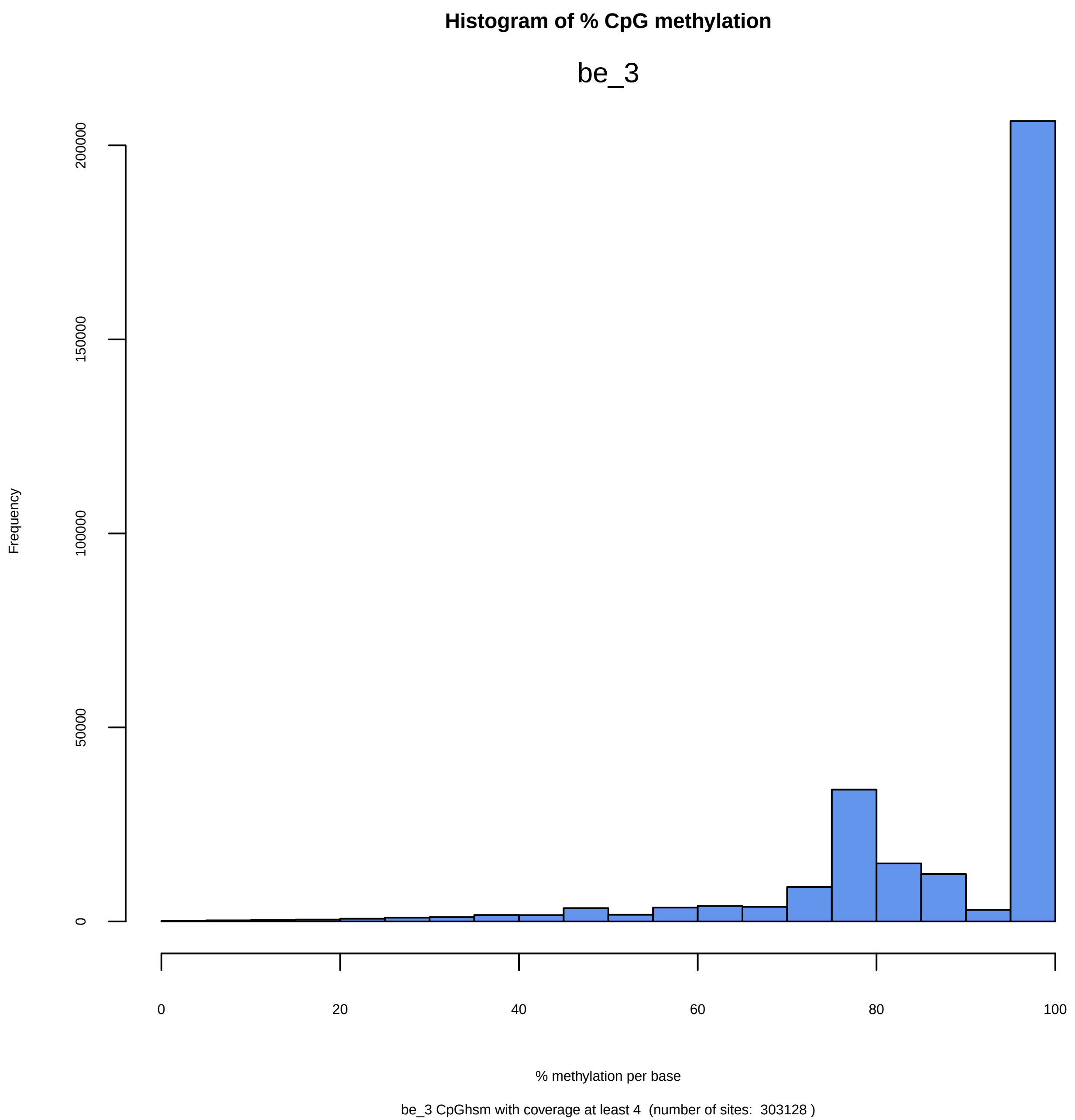

Histogram of % CpG methylation
be_3
Frequency
0
50000
100000
150000
200000
0
20
40
60
80
100
% methylation per base
be_3 CpGhsm with coverage at least 4 (number of sites: 303128 )

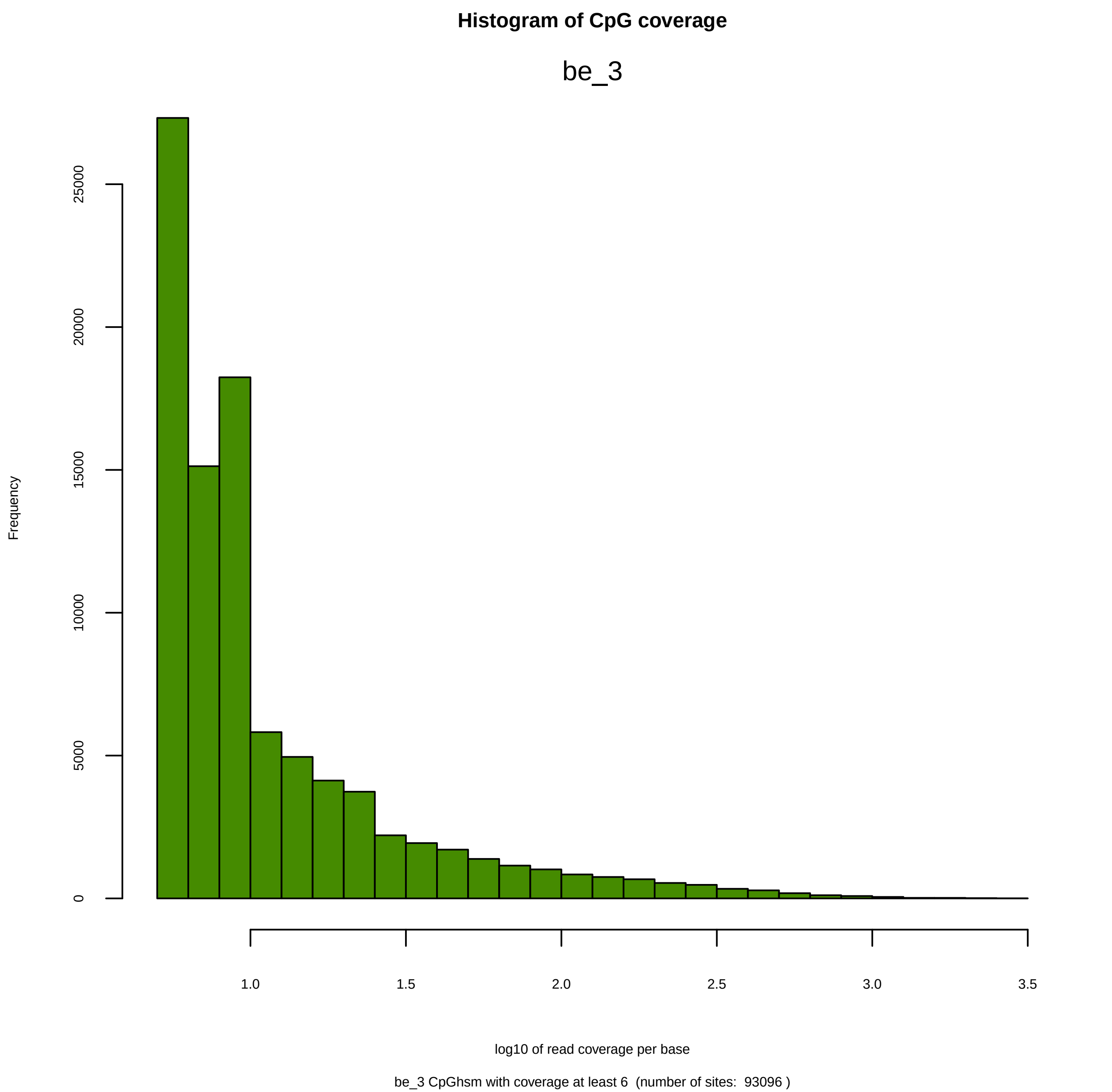

Histogram of CpG coverage
be_3
Frequency
0
5000
10000
15000
20000
25000
1.0
1.5
2.0
2.5
3.0
3.5
log10 of read coverage per base
be_3 CpGhsm with coverage at least 6  (number of sites:  93096 )

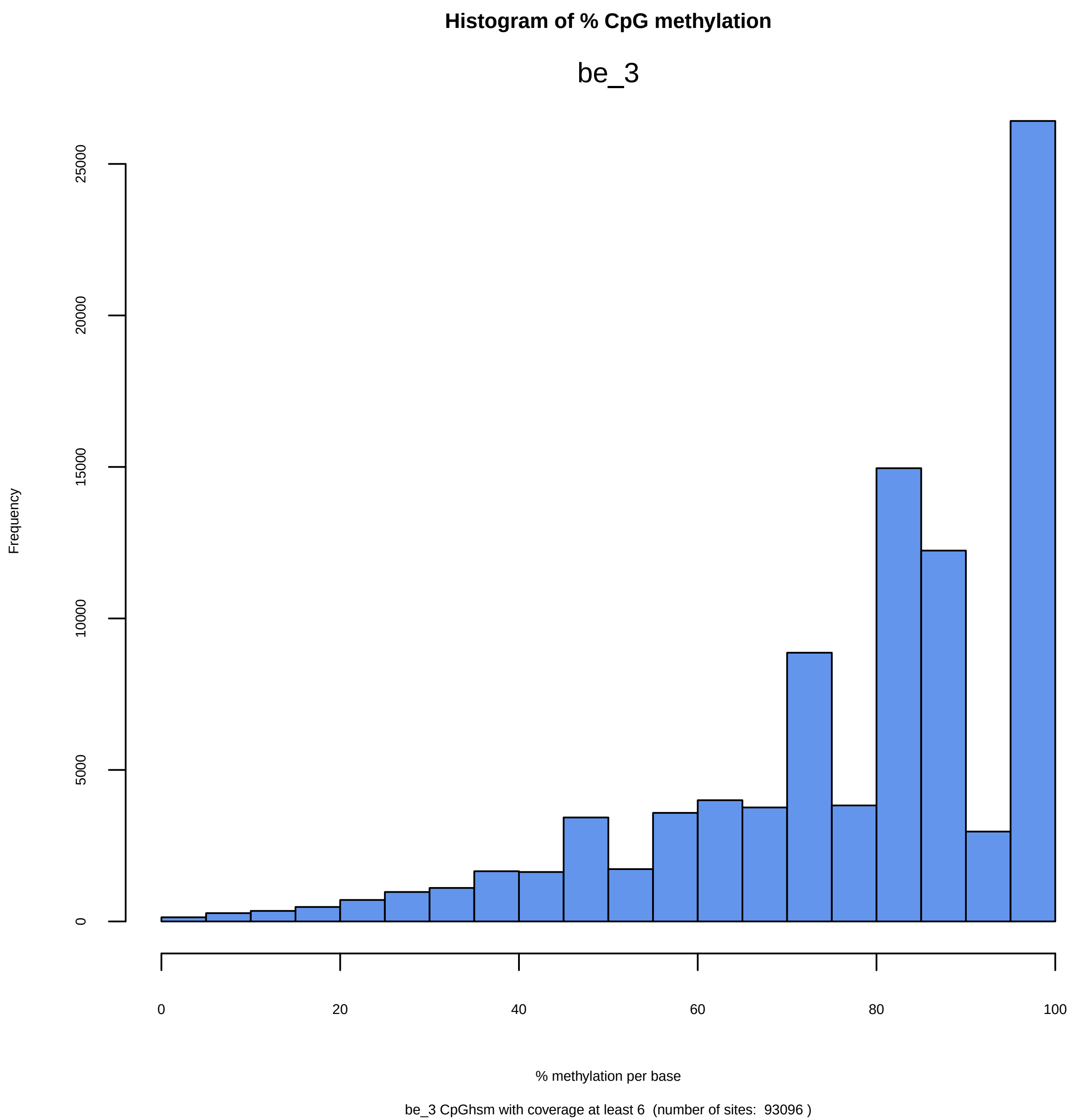
Histogram of % CpG methylation
be_3
Frequency
0
5000
10000
15000
20000
25000
0
20
40
60
80
100
% methylation per base
be_3 CpGhsm with coverage at least 6 (number of sites: 93096 )

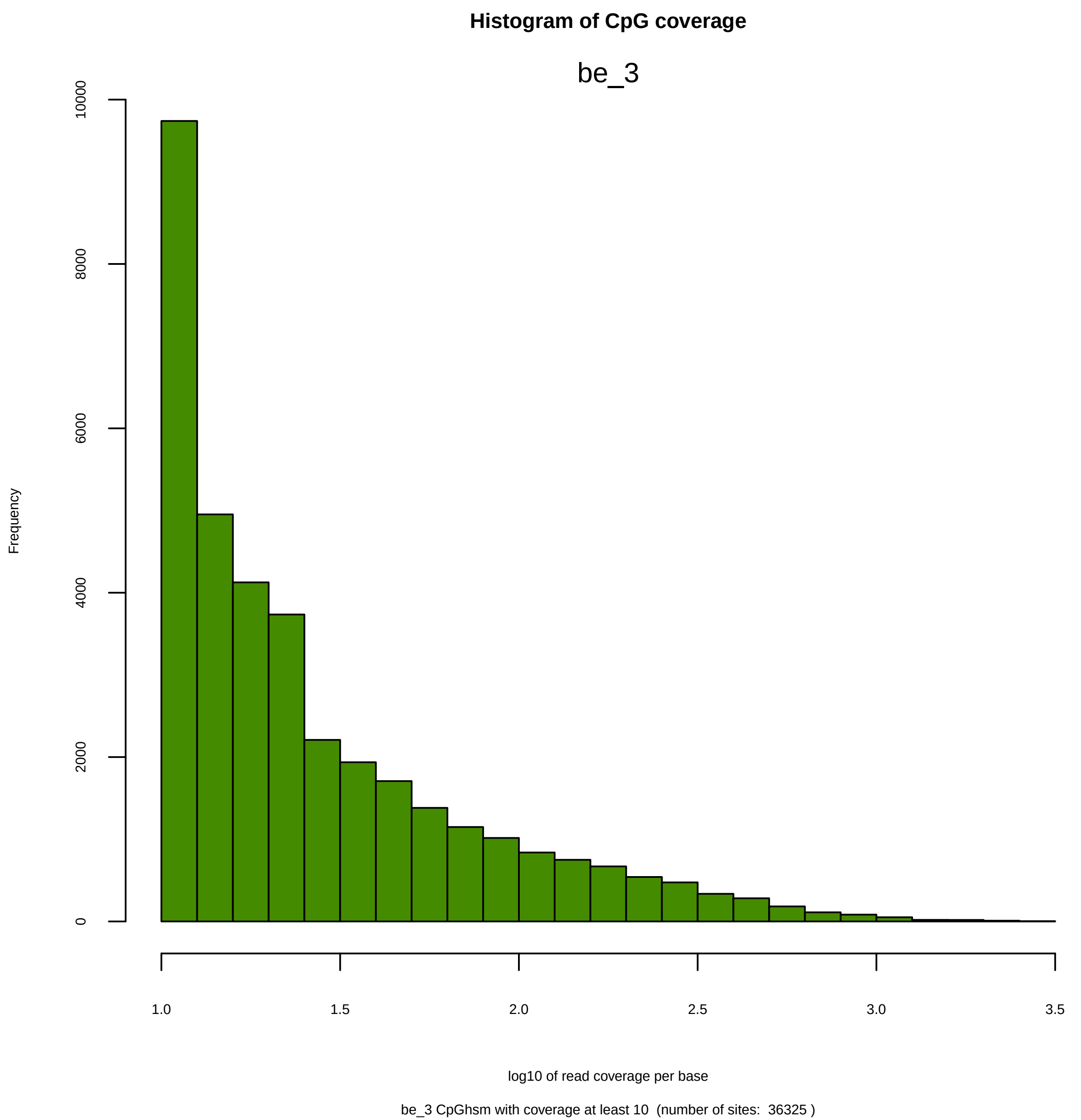
Histogram of CpG coverage
be_3
Frequency
0
2000
4000
6000
8000
10000
1.0
1.5
2.0
2.5
3.0
3.5
log10 of read coverage per base
be_3 CpGhsm with coverage at least 10  (number of sites:  36325 )

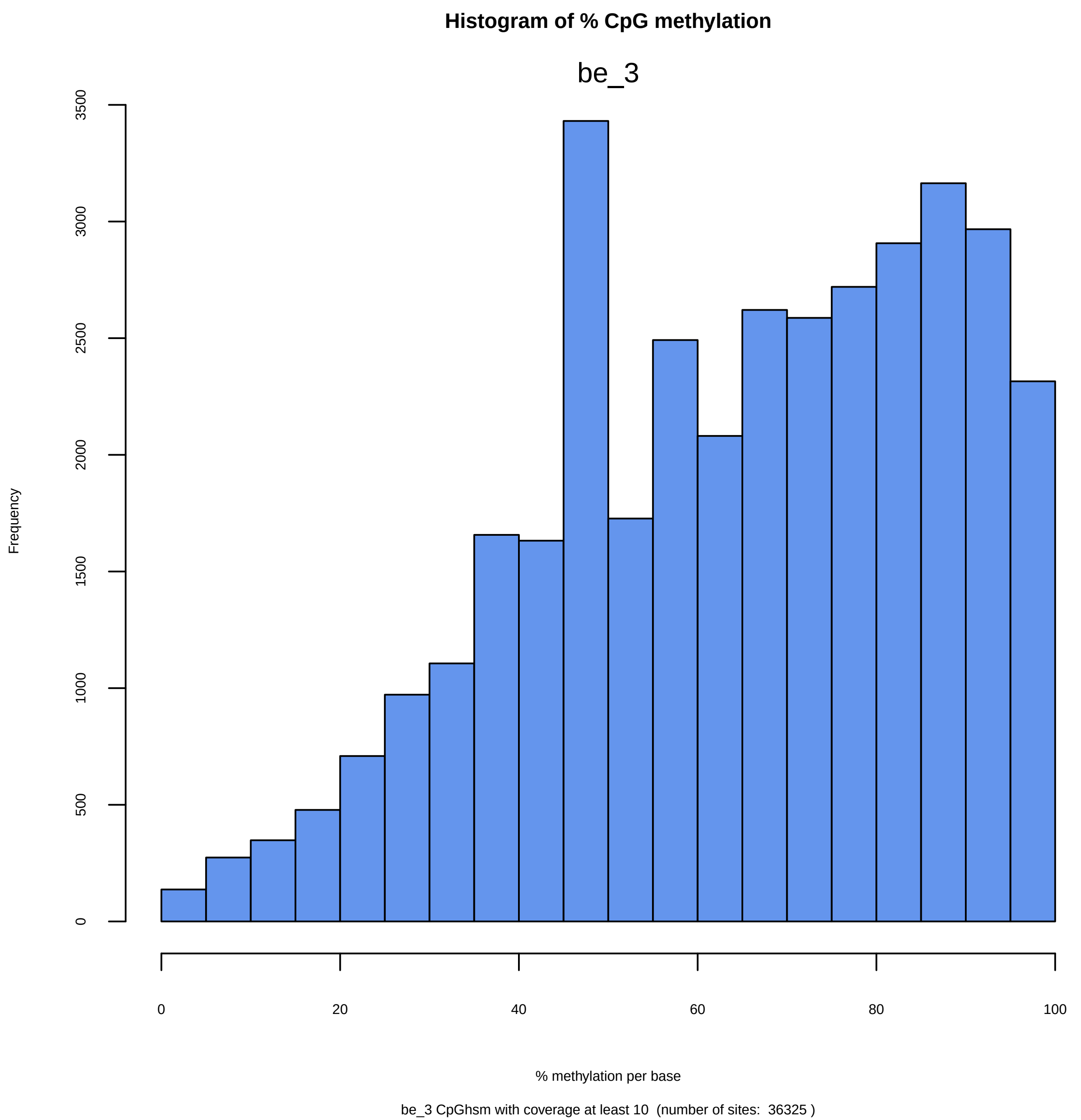

Histogram of % CpG methylation
be_3
Frequency
0
500
1000
1500
2000
2500
3000
3500
0
20
40
60
80
100
% methylation per base
be_3 CpGhsm with coverage at least 10 (number of sites: 36325 )

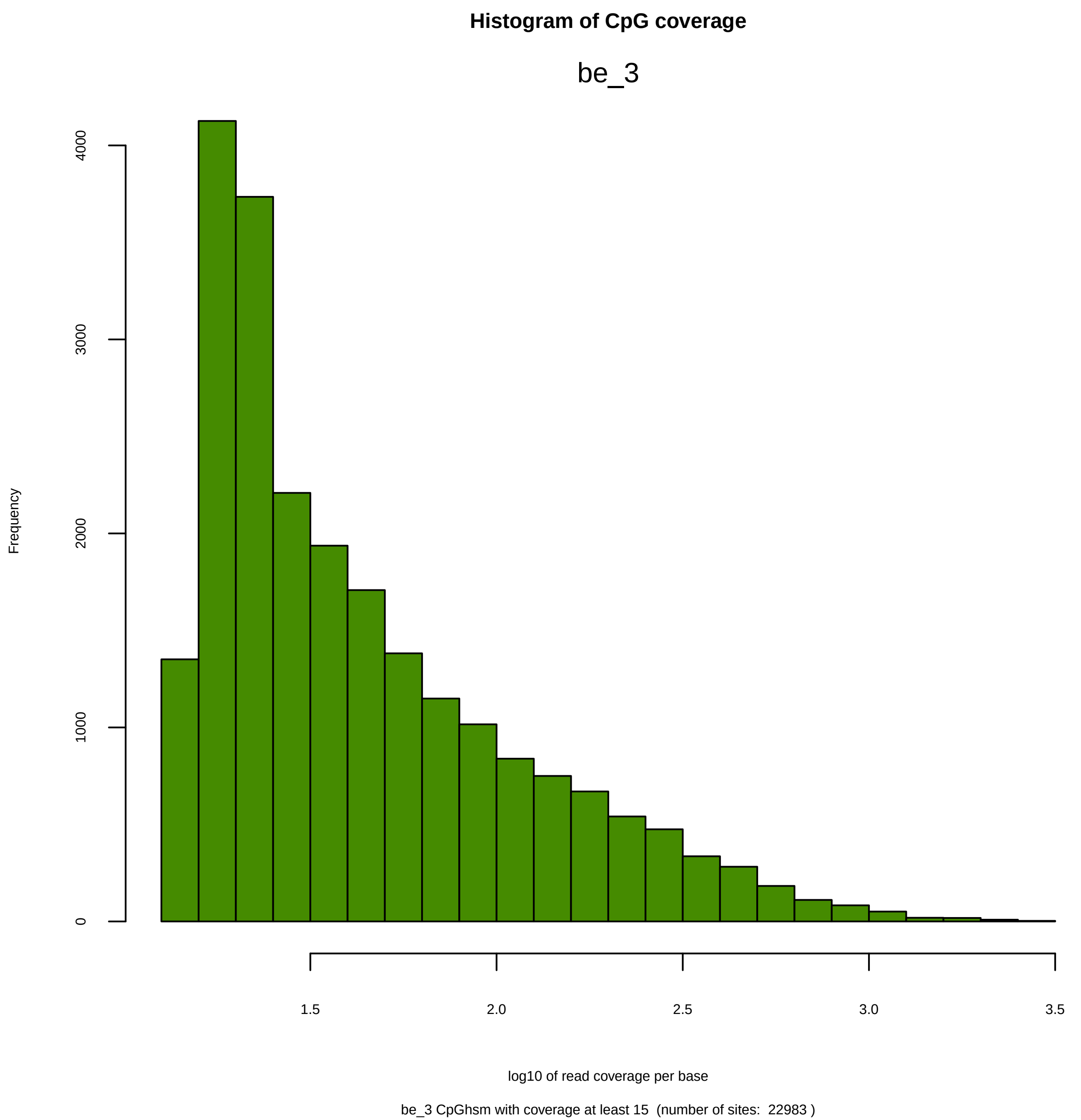

Histogram of CpG coverage
be_3
Frequency
0
1000
2000
3000
4000
1.5
2.0
2.5
3.0
3.5
log10 of read coverage per base
be_3 CpGhsm with coverage at least 15  (number of sites:  22983 )

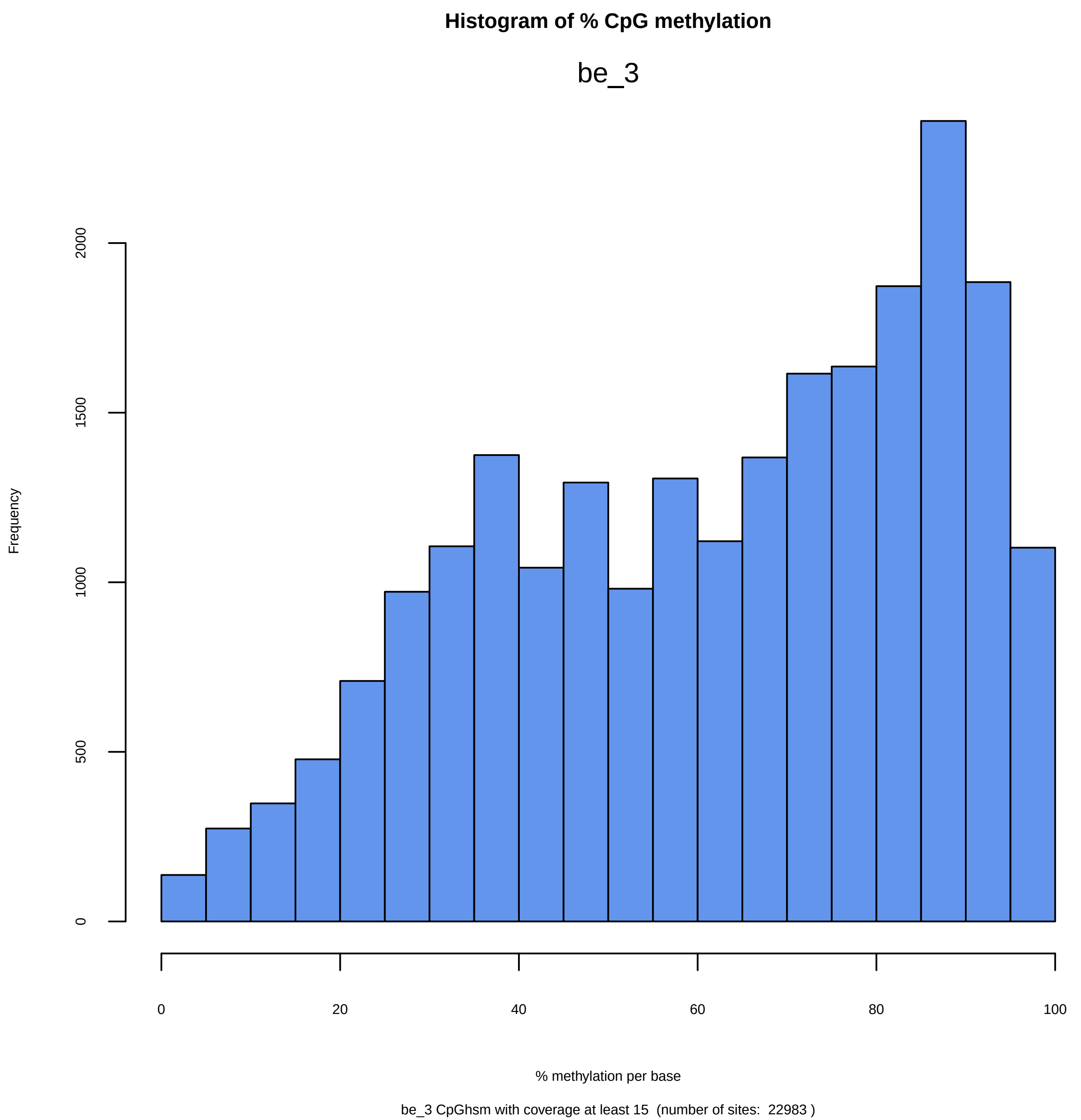
Histogram of % CpG methylation
be_3
Frequency
0
500
1000
1500
2000
0
20
40
60
80
100
% methylation per base
be_3 CpGhsm with coverage at least 15  (number of sites:  22983 )

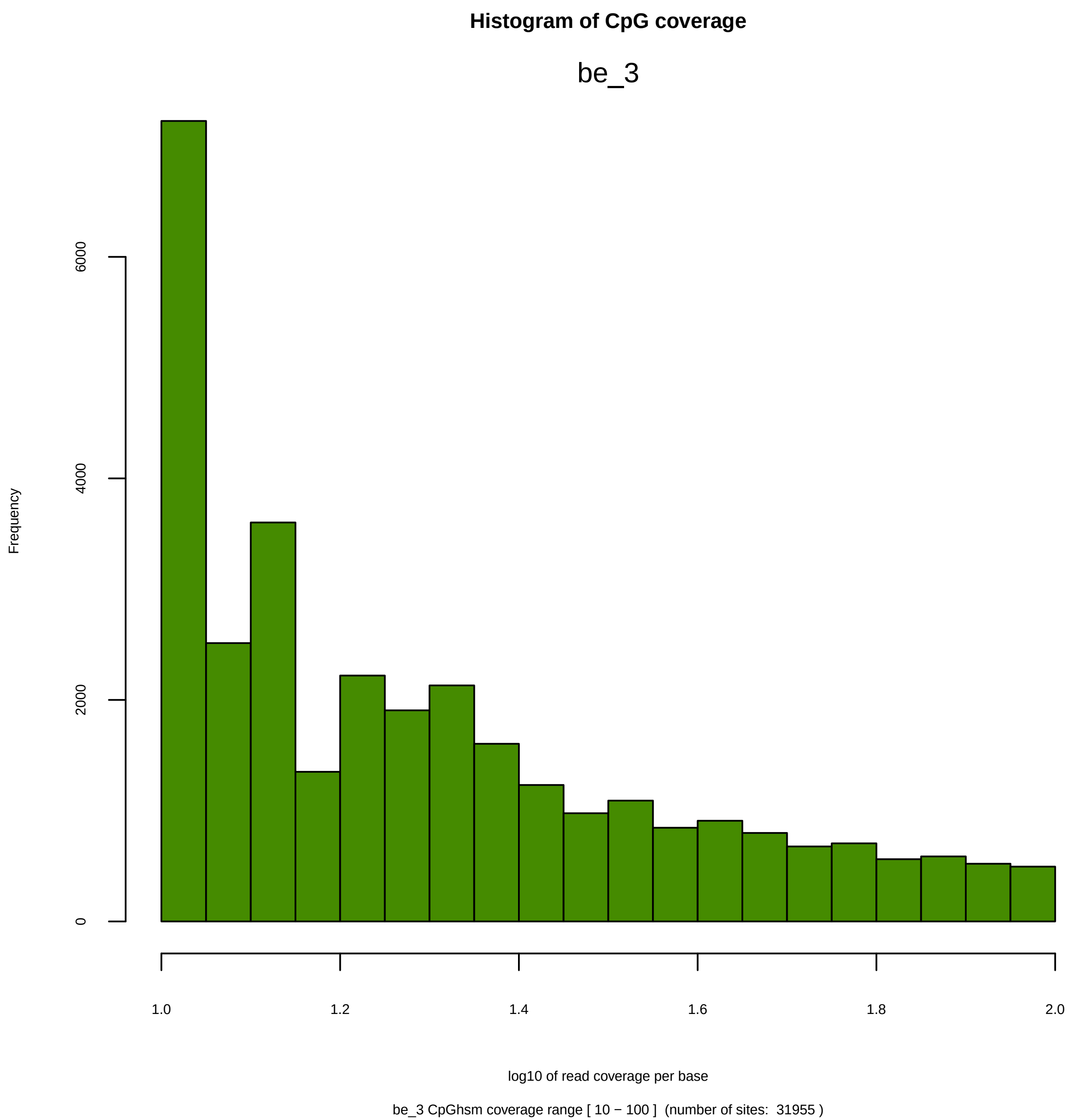

Histogram of CpG coverage
be_3
Frequency
0
2000
4000
6000
1.0
1.2
1.4
1.6
1.8
2.0
log10 of read coverage per base
be_3 CpGhsm coverage range [ 10 − 100 ]  (number of sites:  31955 )

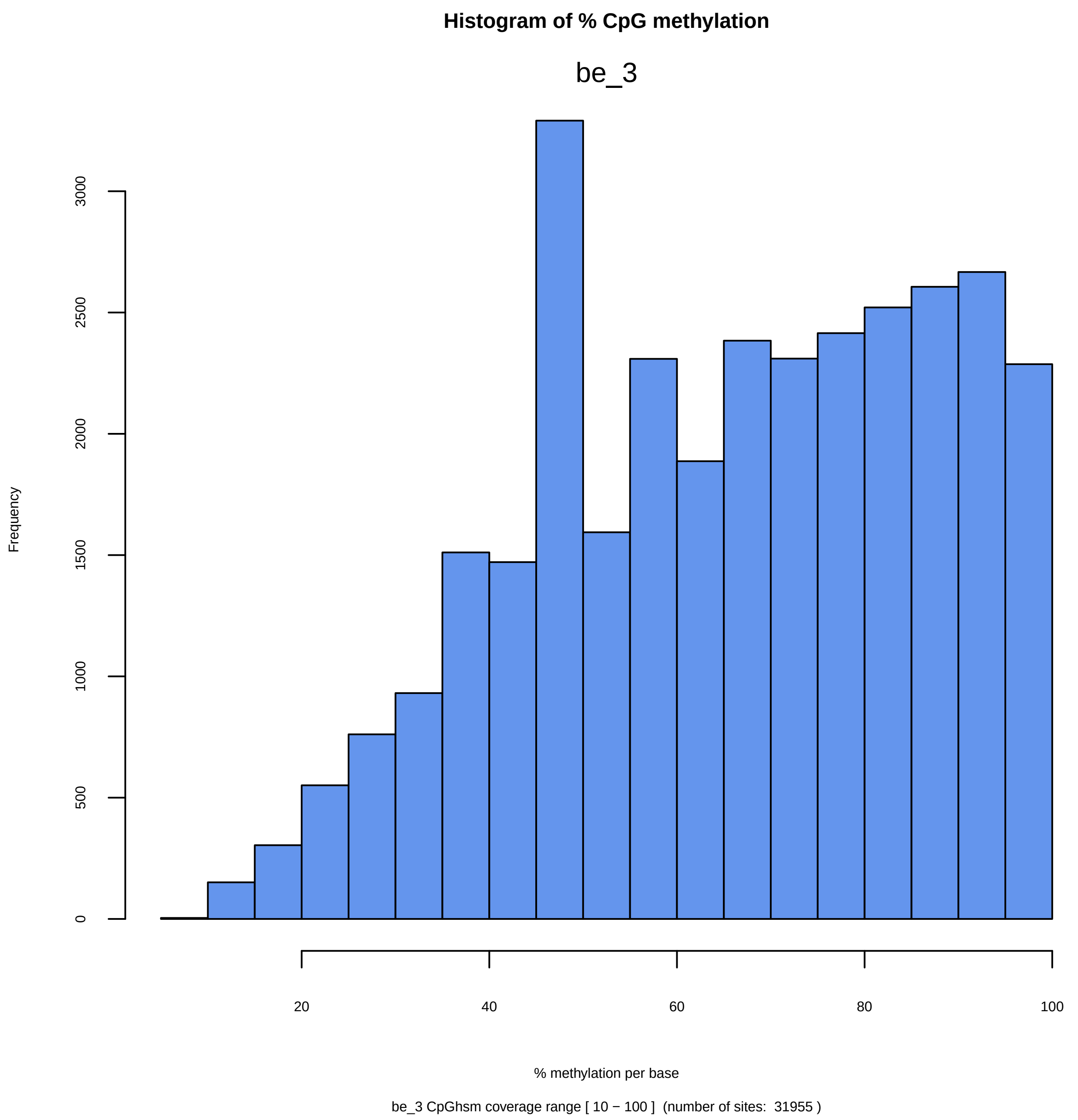
Histogram of % CpG methylation
be_3
Frequency
0
500
1000
1500
2000
2500
3000
20
40
60
80
100
% methylation per base
be_3 CpGhsm coverage range [ 10 − 100 ]  (number of sites:  31955 )

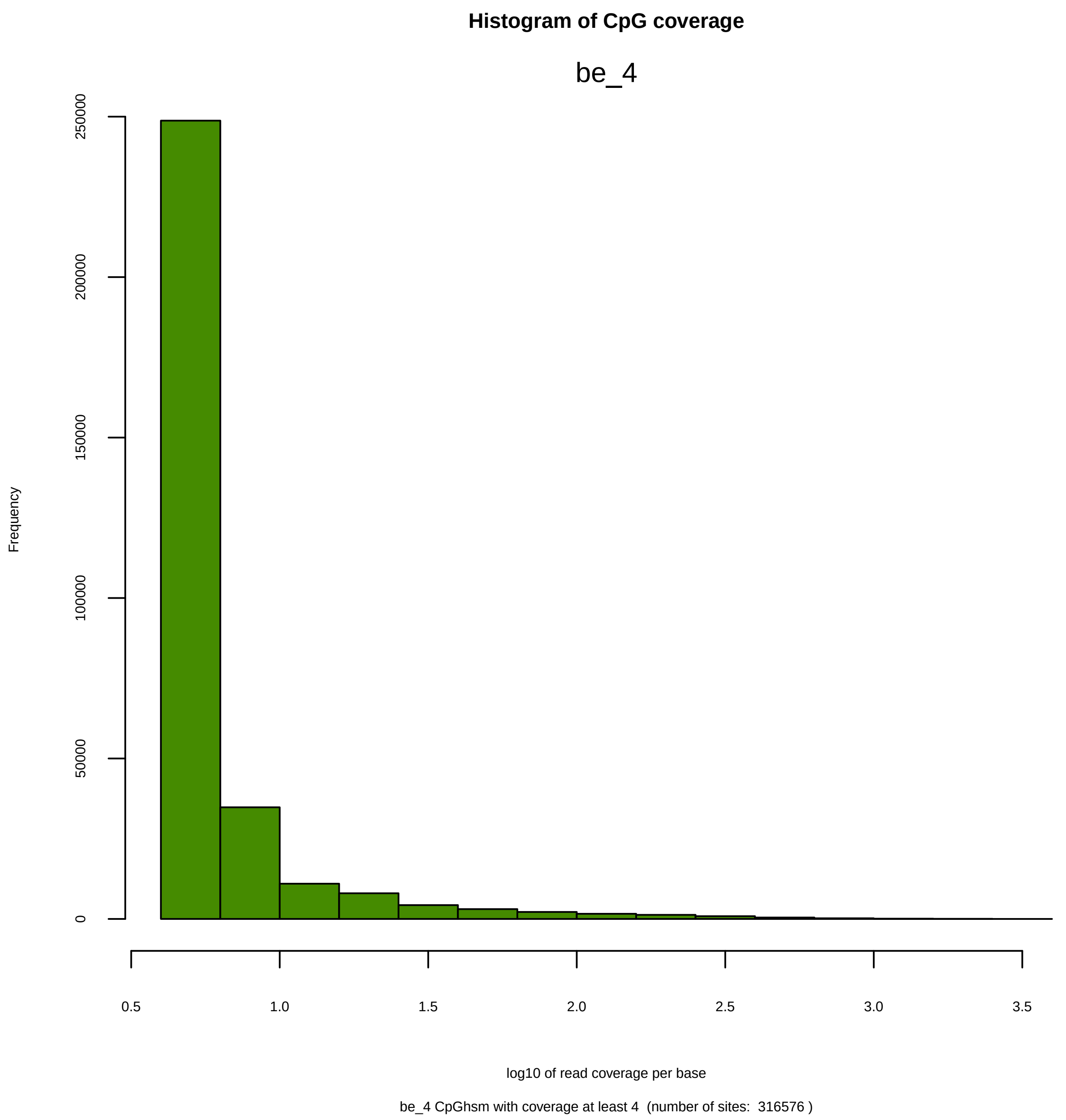

Histogram of CpG coverage
be_4
Frequency
0
50000
100000
150000
200000
250000
0.5
1.0
1.5
2.0
2.5
3.0
3.5
log10 of read coverage per base
be_4 CpGhsm with coverage at least 4 (number of sites: 316576 )

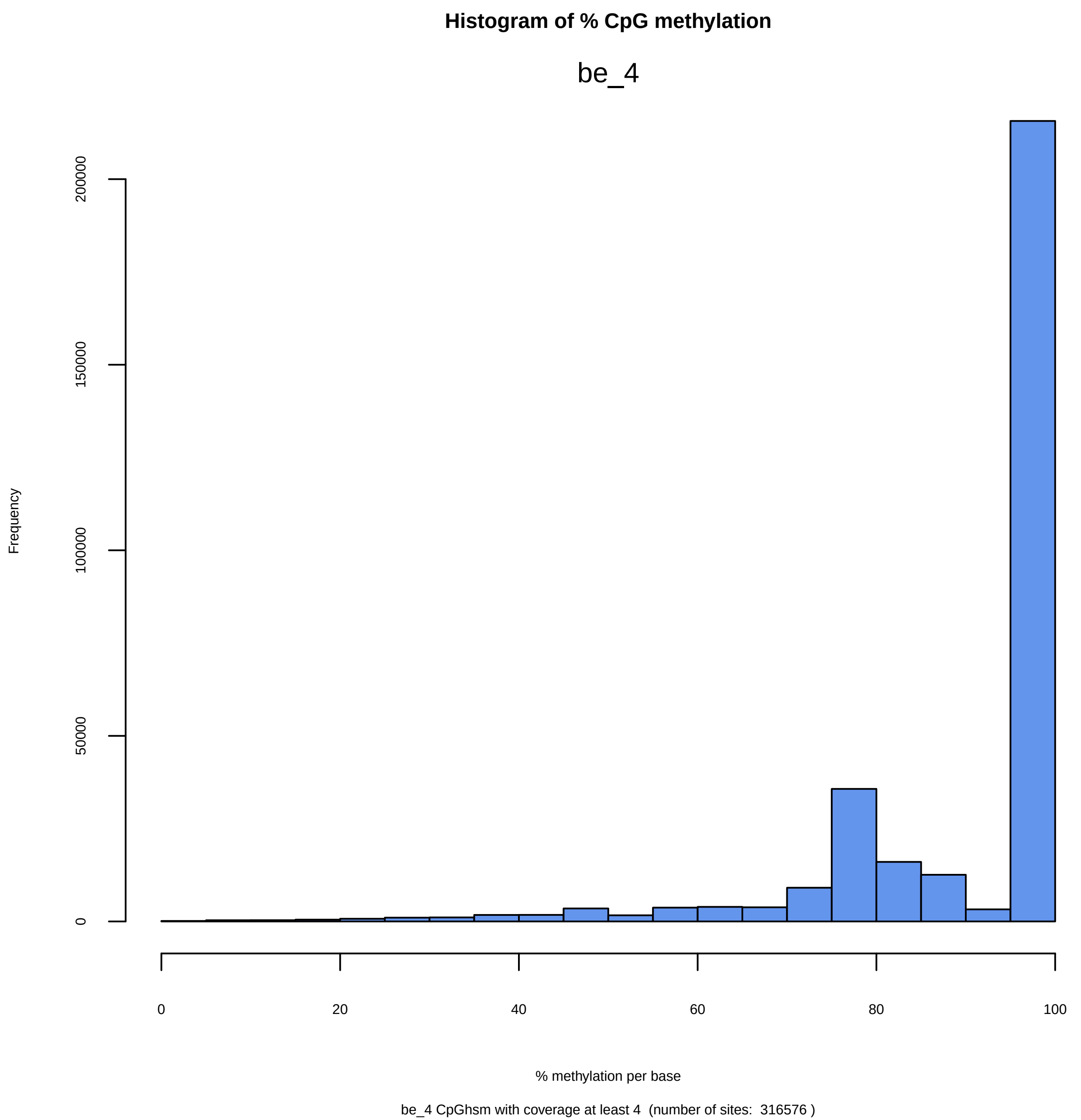
Histogram of % CpG methylation
be_4
Frequency
0
50000
100000
150000
200000
0
20
40
60
80
100
% methylation per base
be_4 CpGhsm with coverage at least 4 (number of sites: 316576 )

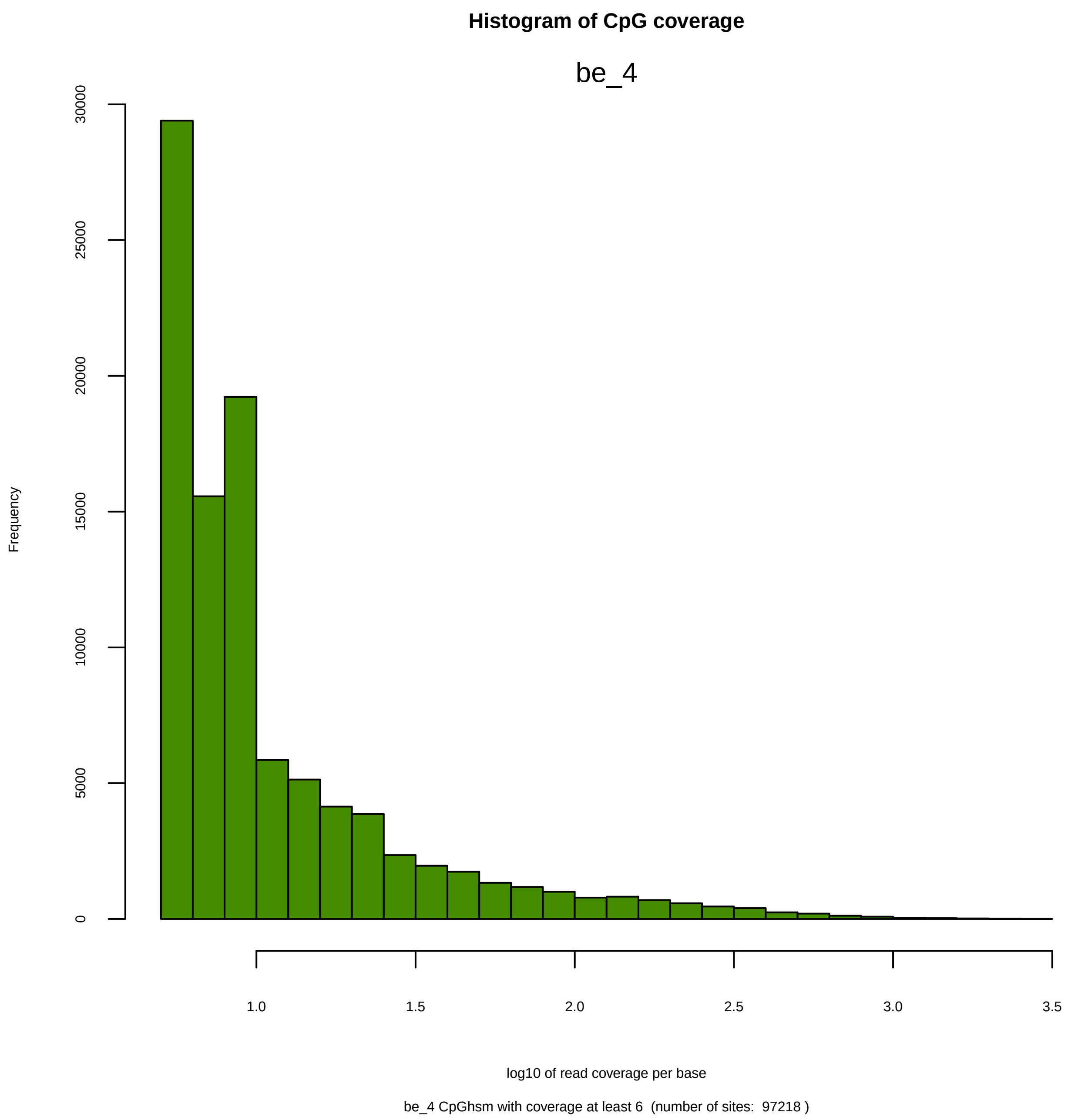

Histogram of CpG coverage
be_4
Frequency
0
5000
10000
15000
20000
25000
30000
1.0
1.5
2.0
2.5
3.0
3.5
log10 of read coverage per base
be_4 CpGhsm with coverage at least 6 (number of sites: 97218 )

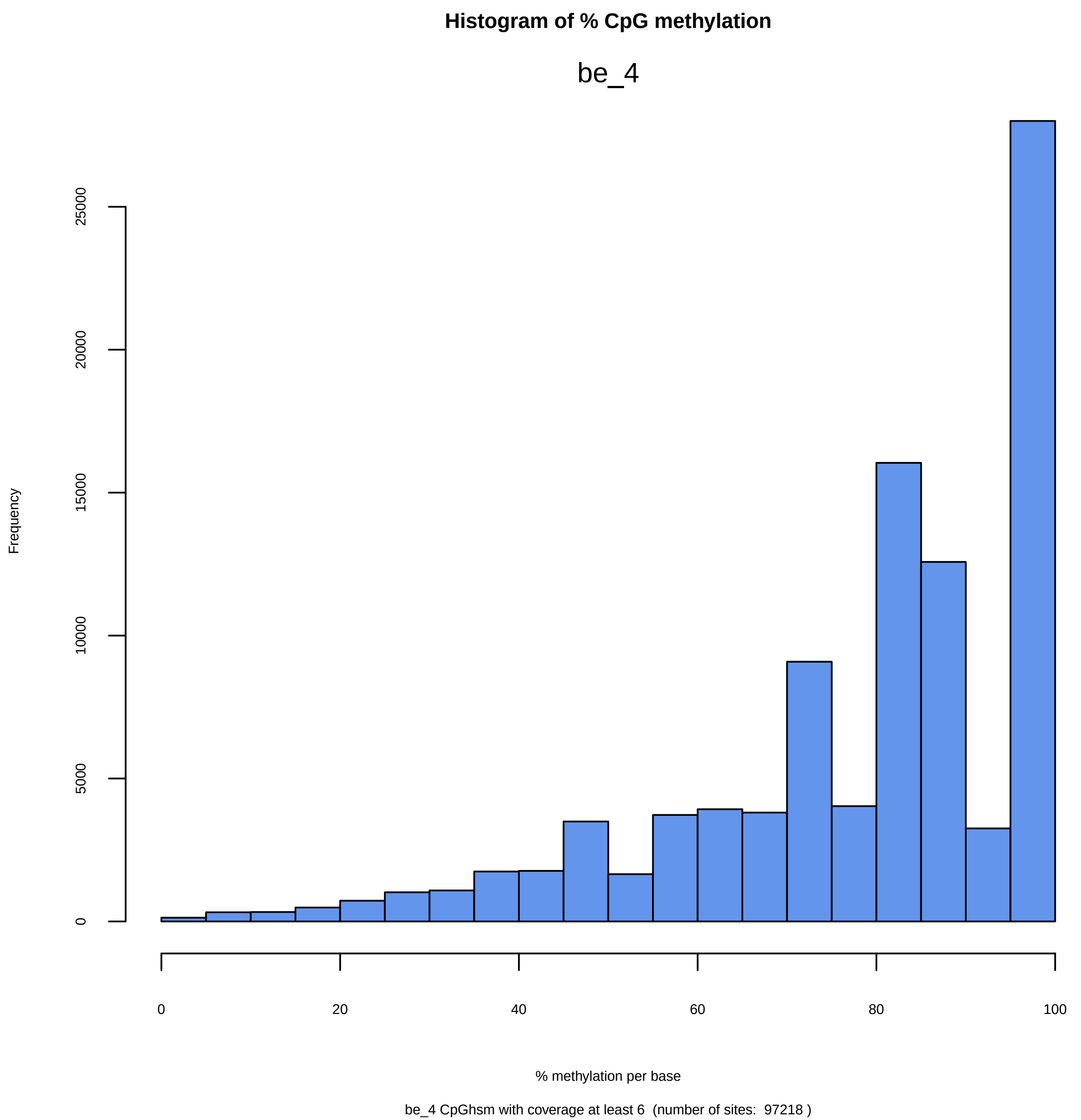
Histogram of % CpG methylation
be_4
Frequency
0
5000
10000
15000
20000
25000
0
20
40
60
80
100
% methylation per base
be_4 CpGhsm with coverage at least 6  (number of sites:  97218 )

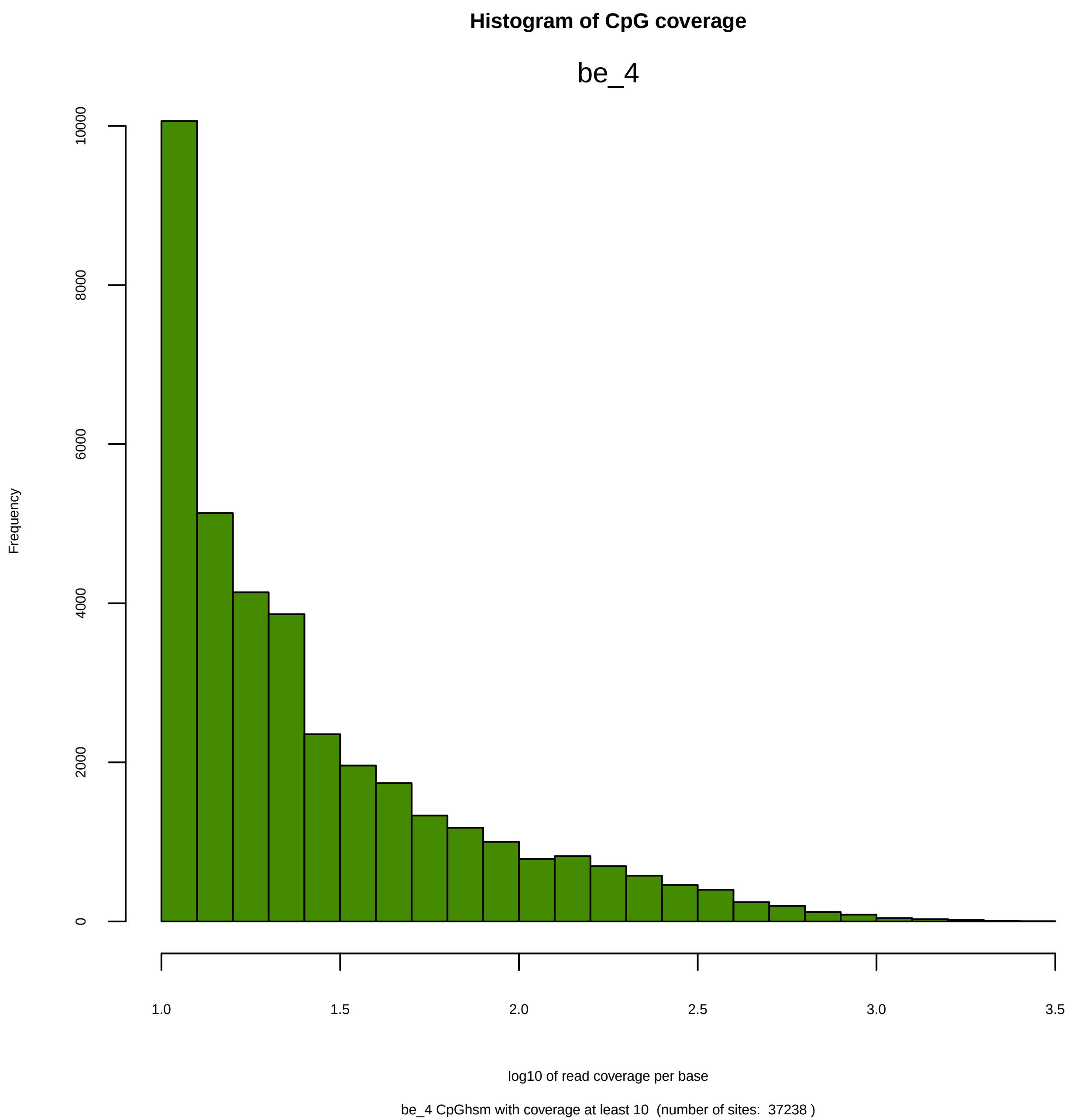

Histogram of CpG coverage
be_4
Frequency
0
2000
4000
6000
8000
10000
1.0
1.5
2.0
2.5
3.0
3.5
log10 of read coverage per base
be_4 CpGhsm with coverage at least 10  (number of sites:  37238 )

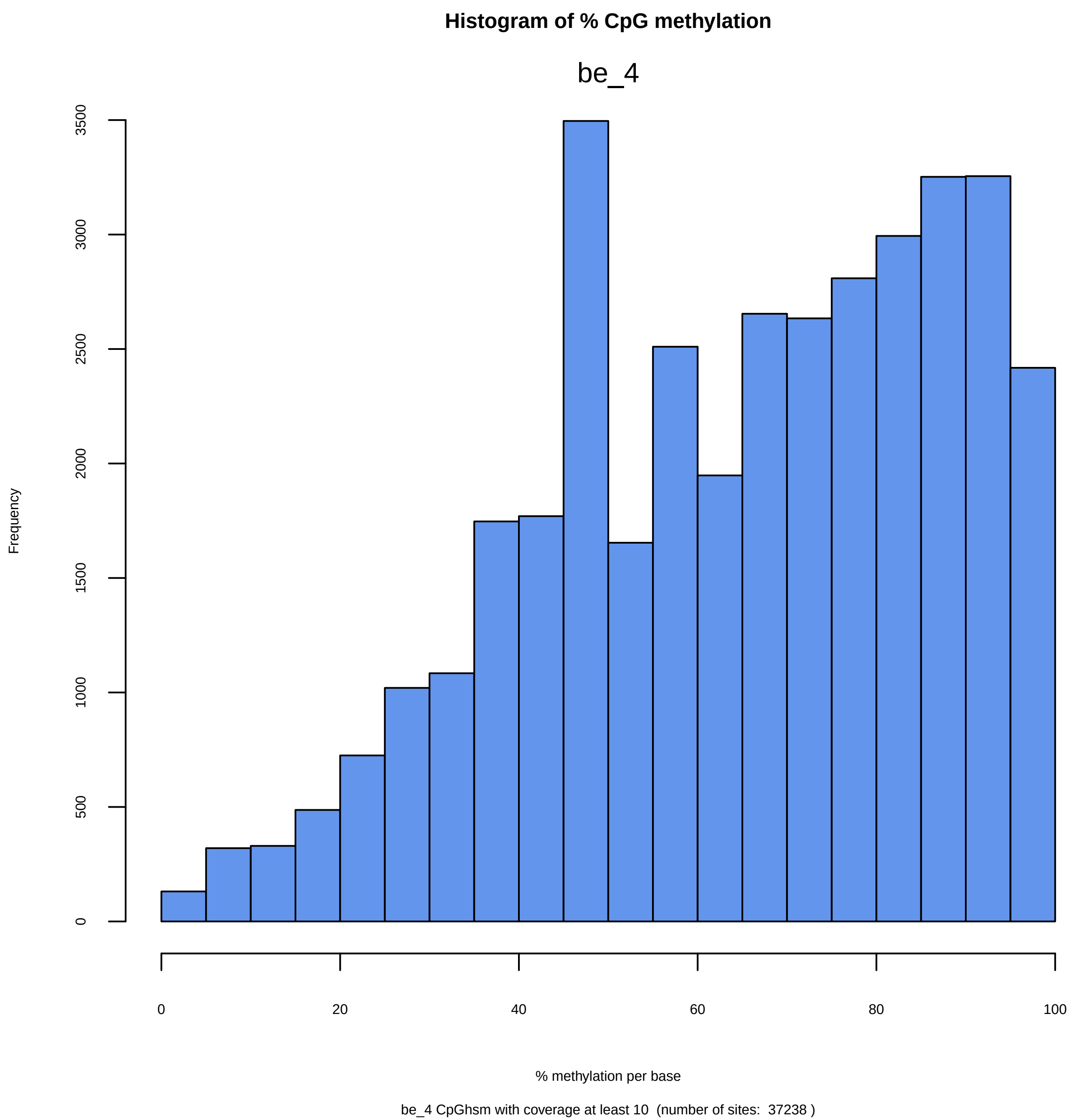
Histogram of % CpG methylation
be_4
Frequency
0
500
1000
1500
2000
2500
3000
3500
0
20
40
60
80
100
% methylation per base
be_4 CpGhsm with coverage at least 10 (number of sites: 37238 )

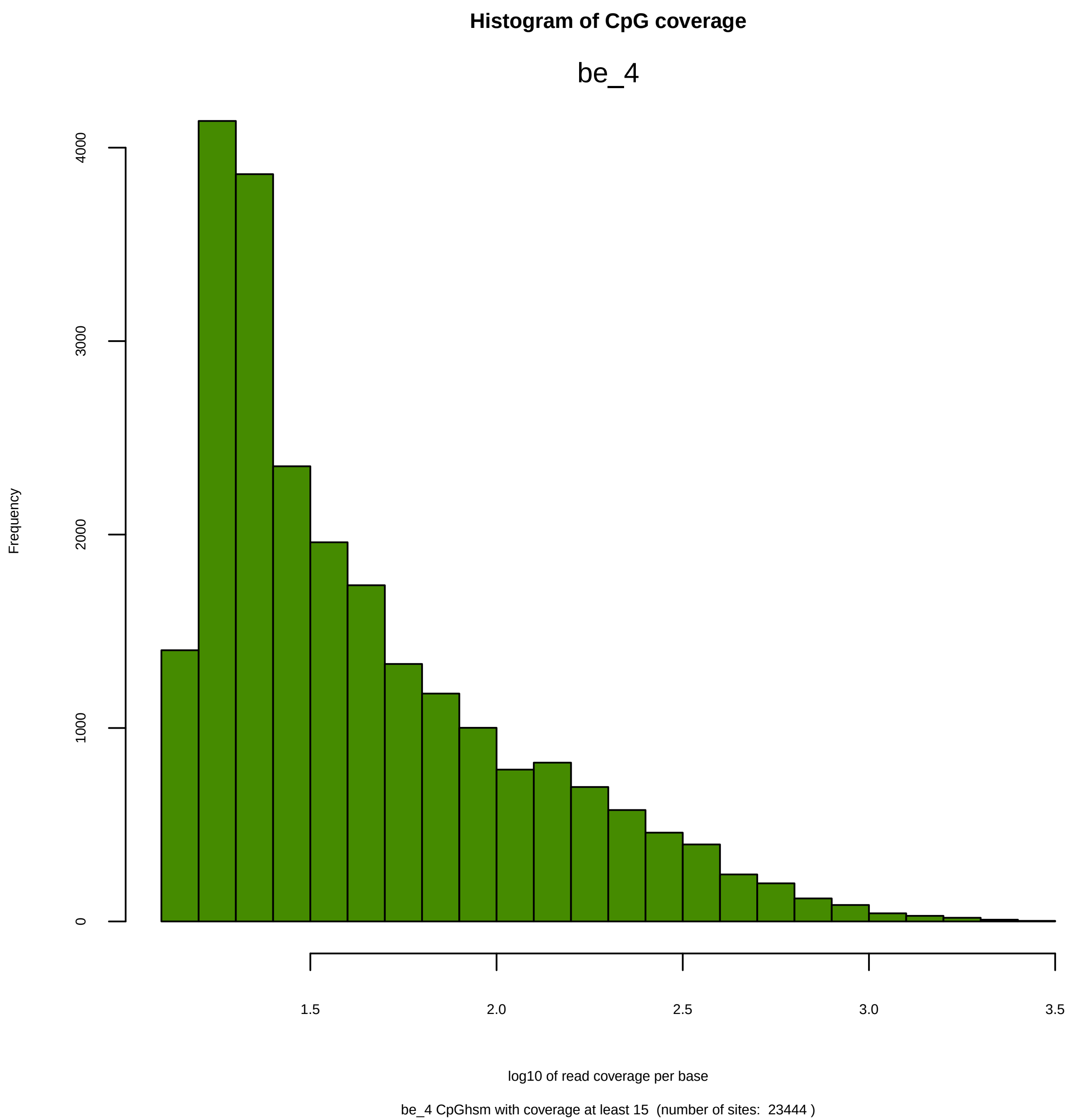
Histogram of CpG coverage
be_4
Frequency
0
1000
2000
3000
4000
1.5
2.0
2.5
3.0
3.5
log10 of read coverage per base
be_4 CpGhsm with coverage at least 15  (number of sites:  23444 )

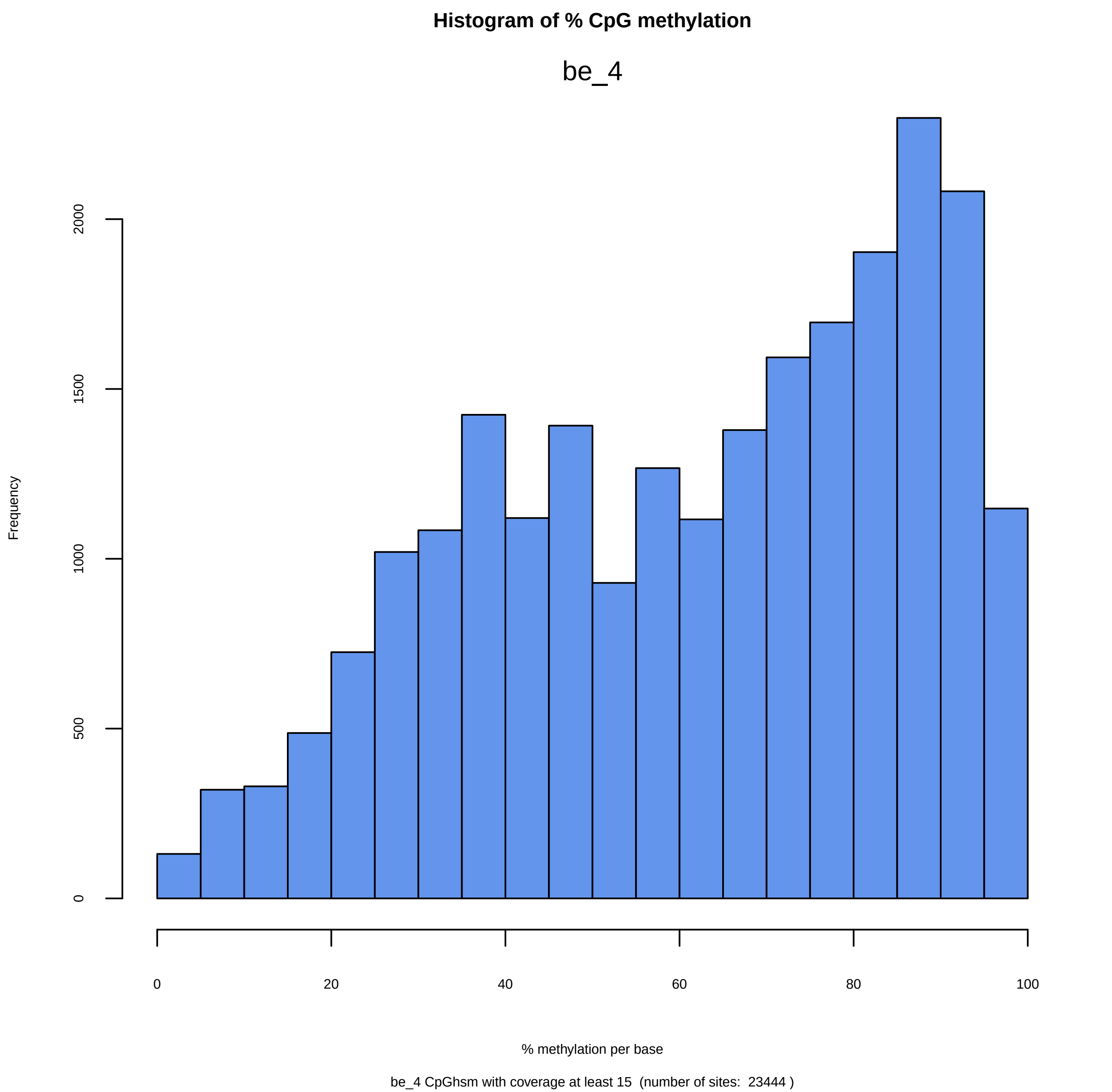

Histogram of % CpG methylation
be_4
Frequency
0
500
1000
1500
2000
0
20
40
60
80
100
% methylation per base
be_4 CpGhsm with coverage at least 15  (number of sites:  23444 )

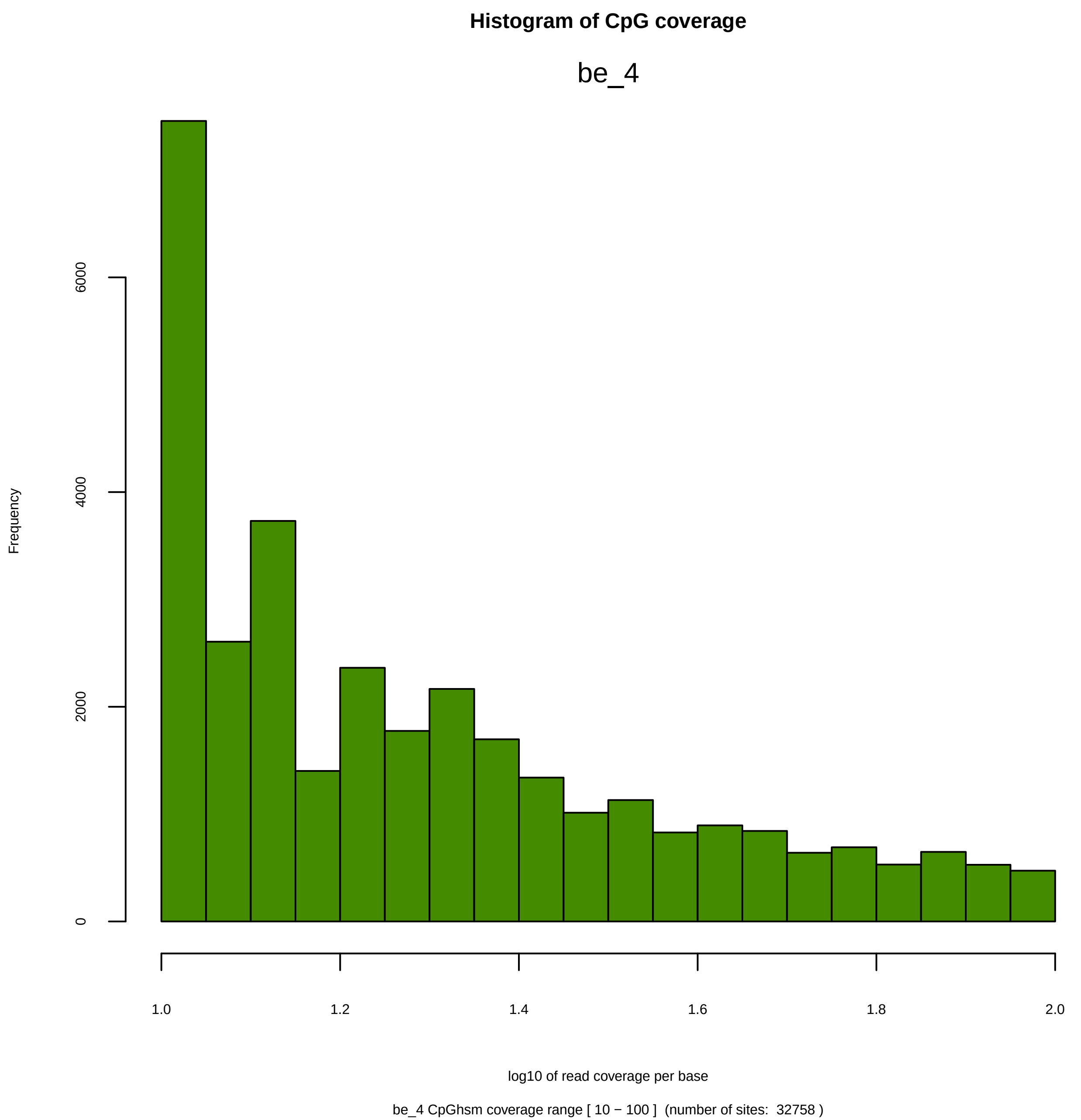
Histogram of CpG coverage
be_4
Frequency
0
2000
4000
6000
1.0
1.2
1.4
1.6
1.8
2.0
log10 of read coverage per base
be_4 CpGhsm coverage range [ 10 − 100 ]  (number of sites:  32758 )

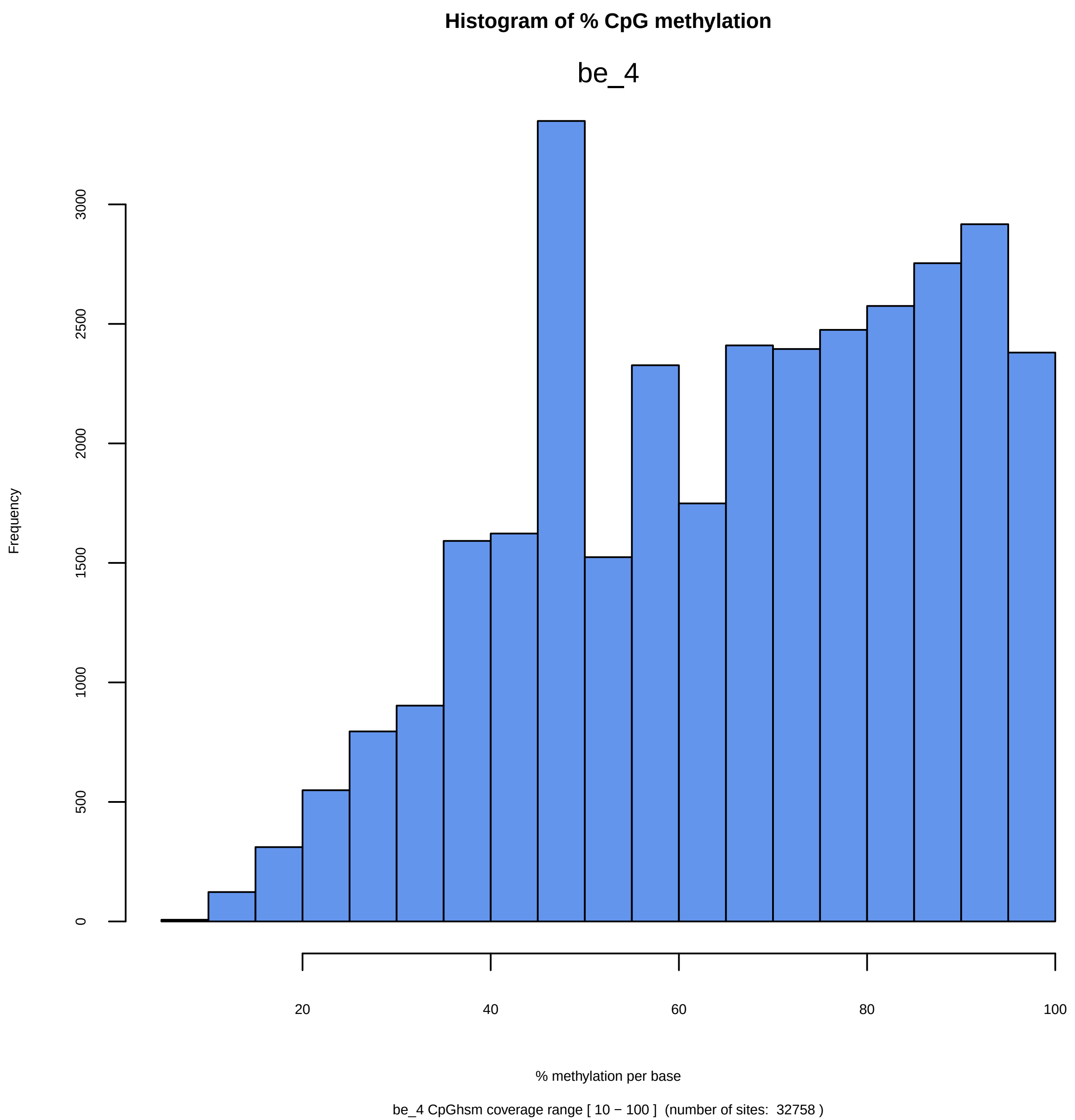

Histogram of % CpG methylation
be_4
Frequency
0
500
1000
1500
2000
2500
3000
20
40
60
80
100
% methylation per base
be_4 CpGhsm coverage range [ 10 − 100 ]  (number of sites:  32758 )

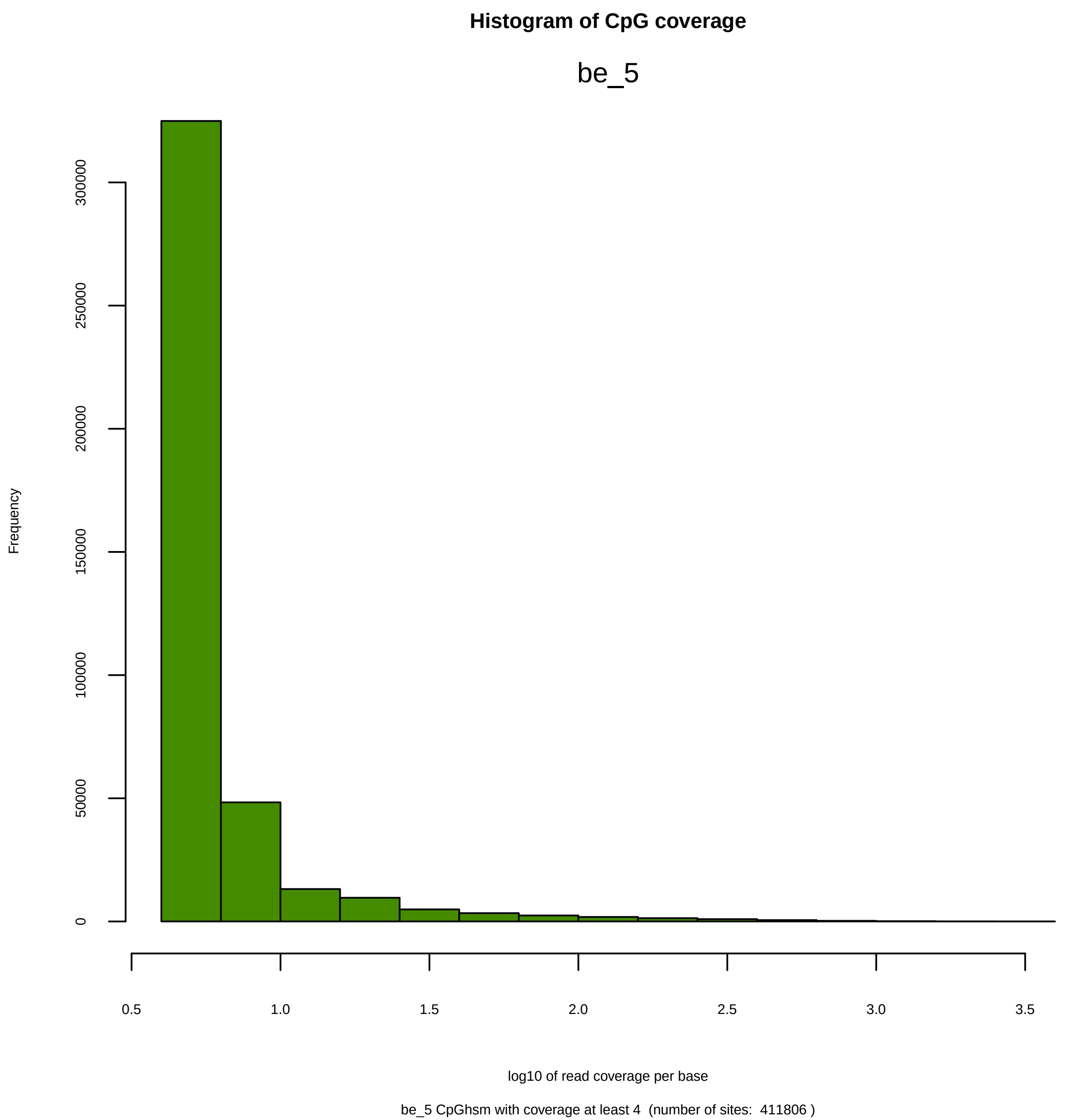

Histogram of CpG coverage
be_5
Frequency
0
50000
100000
150000
200000
250000
300000
0.5
1.0
1.5
2.0
2.5
3.0
3.5
log10 of read coverage per base
be_5 CpGhsm with coverage at least 4  (number of sites:  411806 )

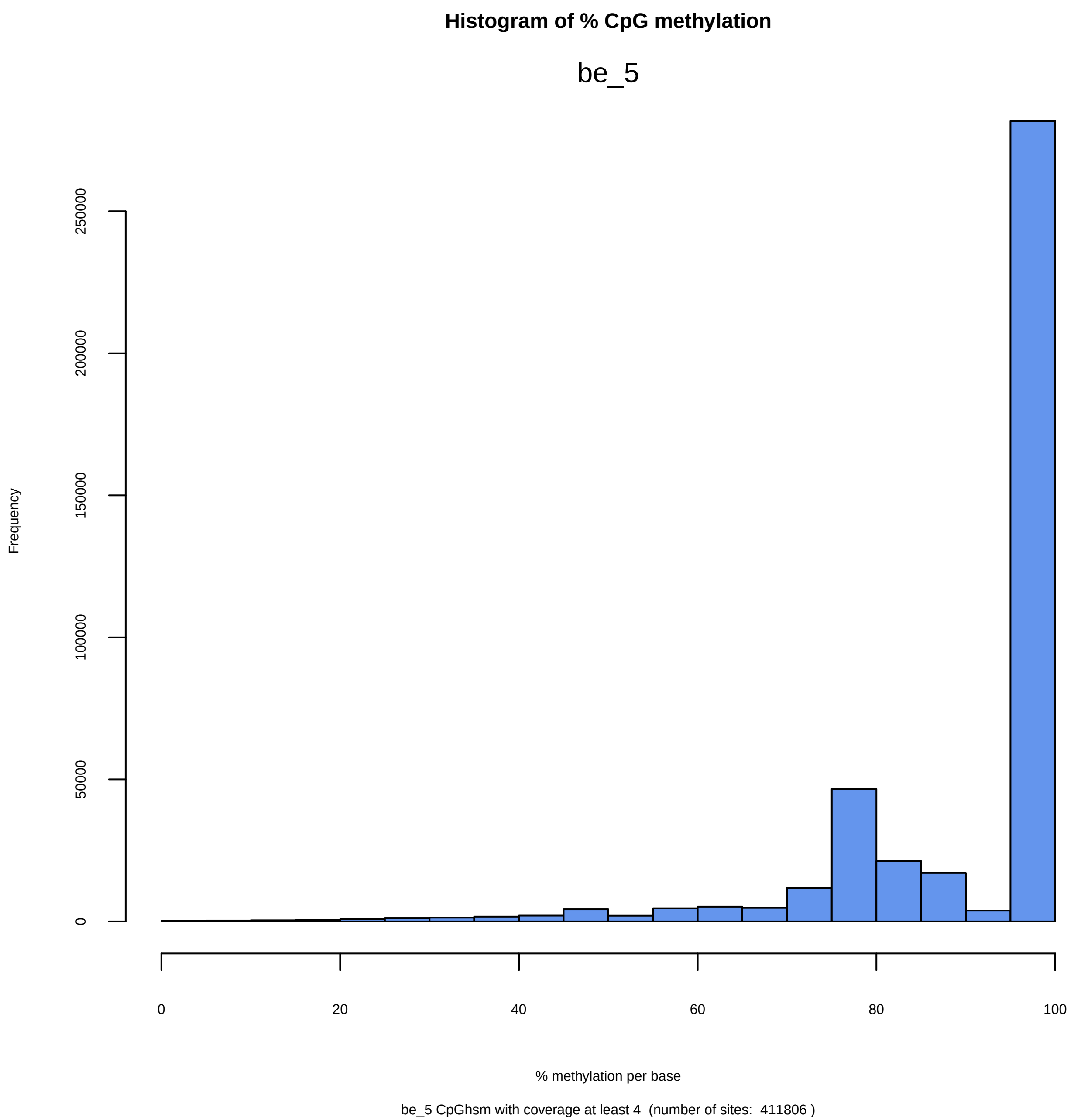
Histogram of % CpG methylation
be_5
Frequency
0
50000
100000
150000
200000
250000
0
20
40
60
80
100
% methylation per base
be_5 CpGhsm with coverage at least 4 (number of sites: 411806 )

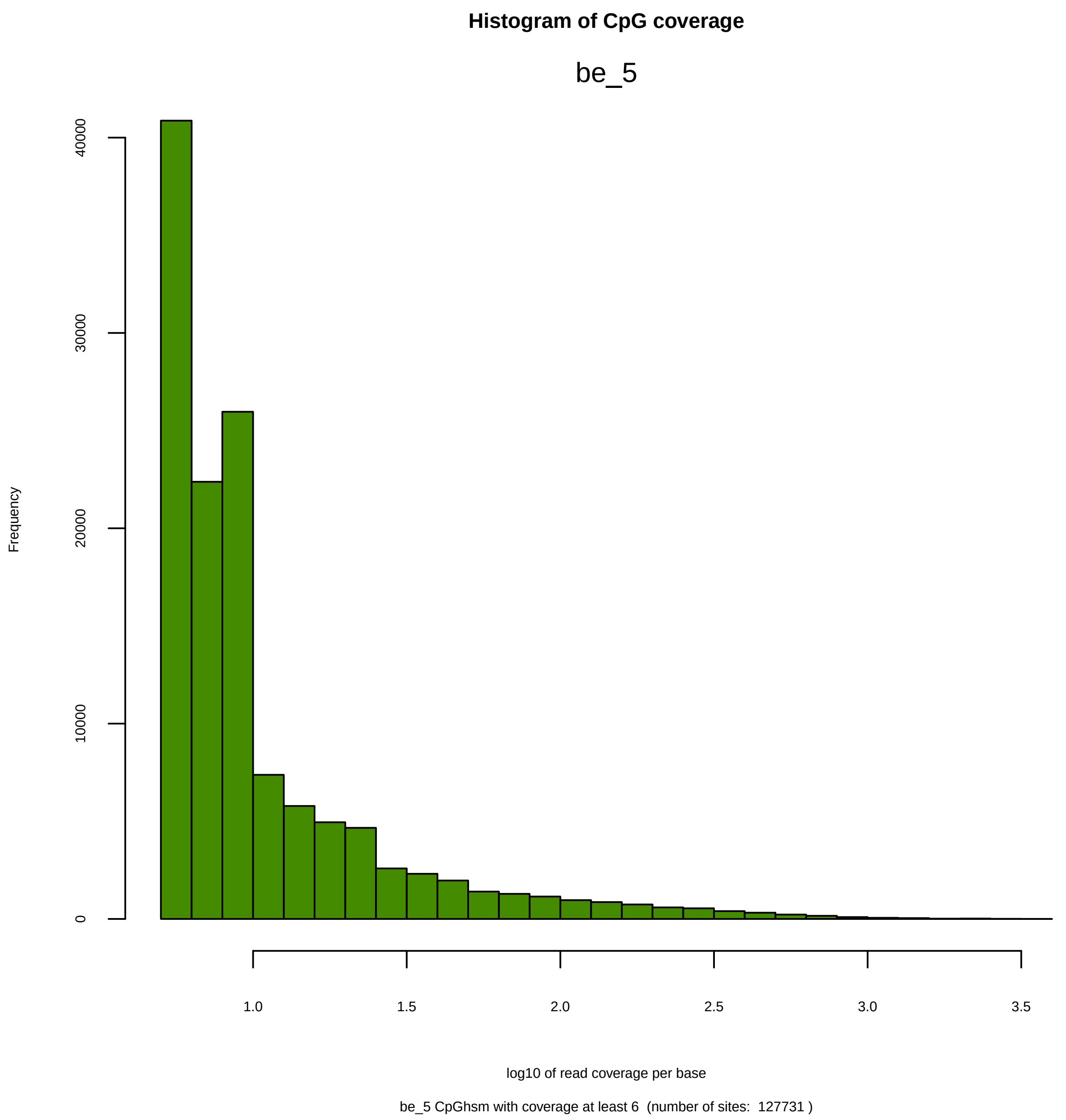

Histogram of CpG coverage
be_5
Frequency
0
10000
20000
30000
40000
1.0
1.5
2.0
2.5
3.0
3.5
log10 of read coverage per base
be_5 CpGhsm with coverage at least 6  (number of sites:  127731 )

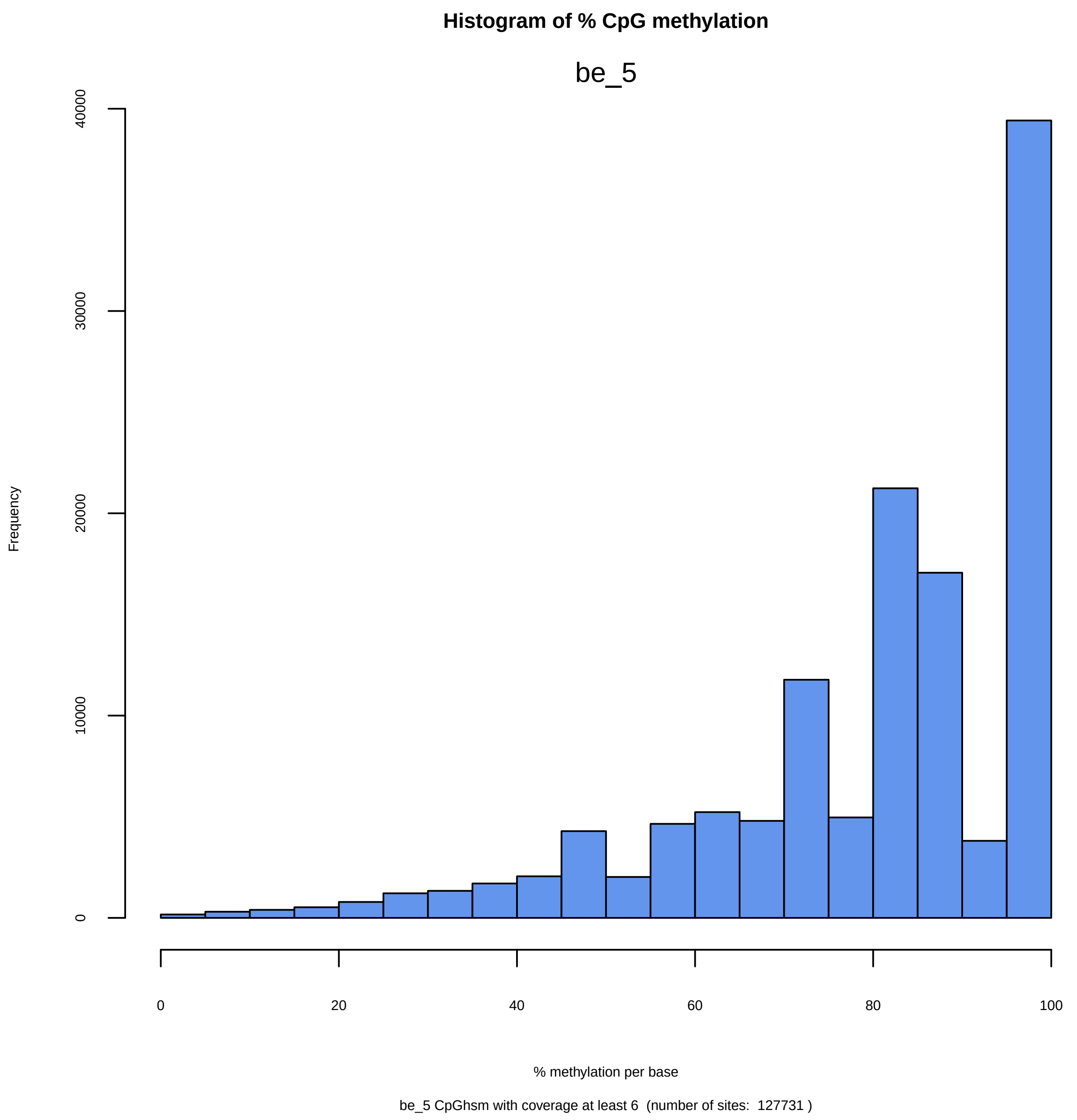

Histogram of % CpG methylation
be_5
Frequency
0
10000
20000
30000
40000
0
20
40
60
80
100
% methylation per base
be_5 CpGhsm with coverage at least 6  (number of sites:  127731 )

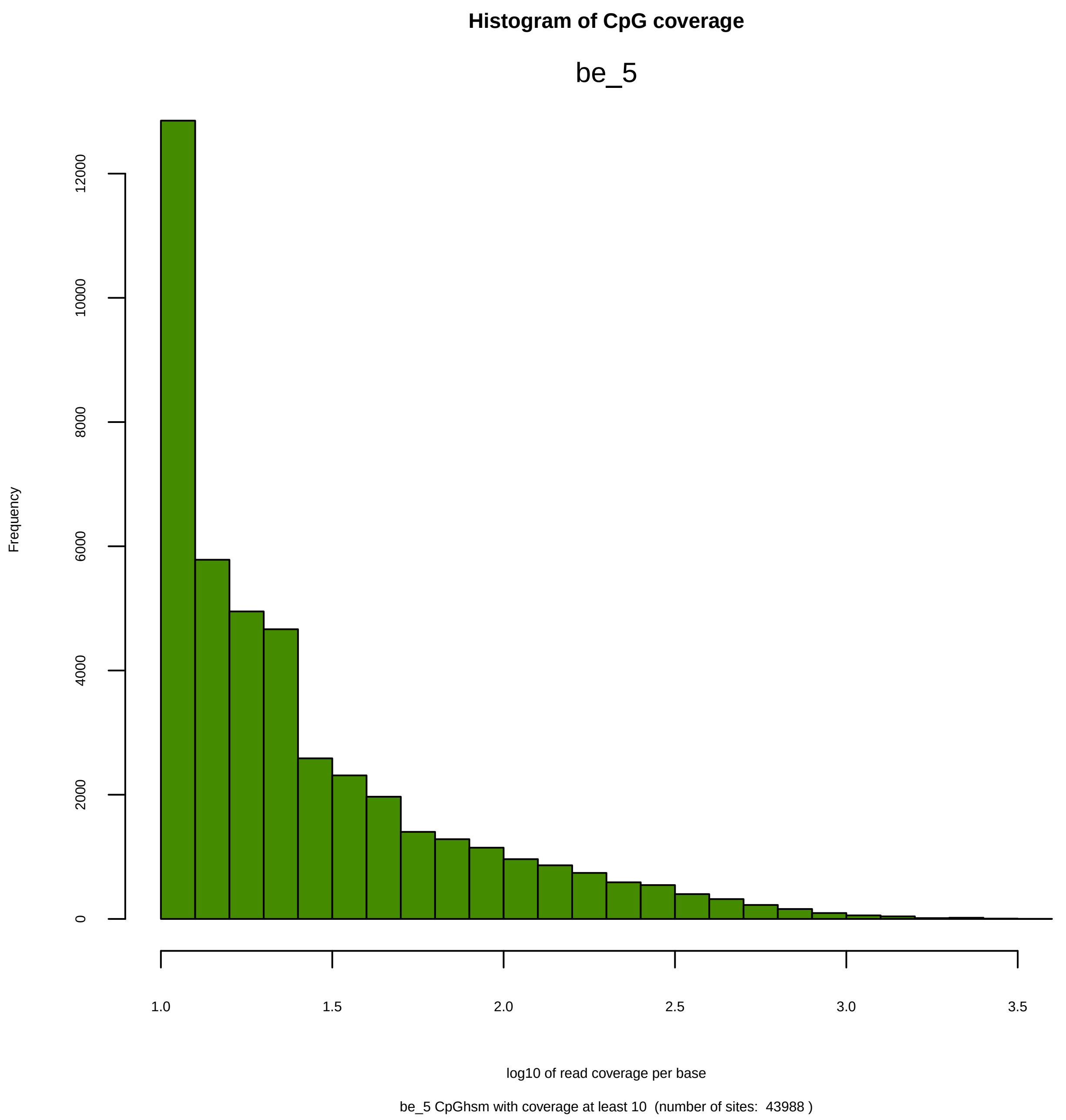

Histogram of CpG coverage
be_5
Frequency
0
2000
4000
6000
8000
10000
12000
1.0
1.5
2.0
2.5
3.0
3.5
log10 of read coverage per base
be_5 CpGhsm with coverage at least 10 (number of sites: 43988 )

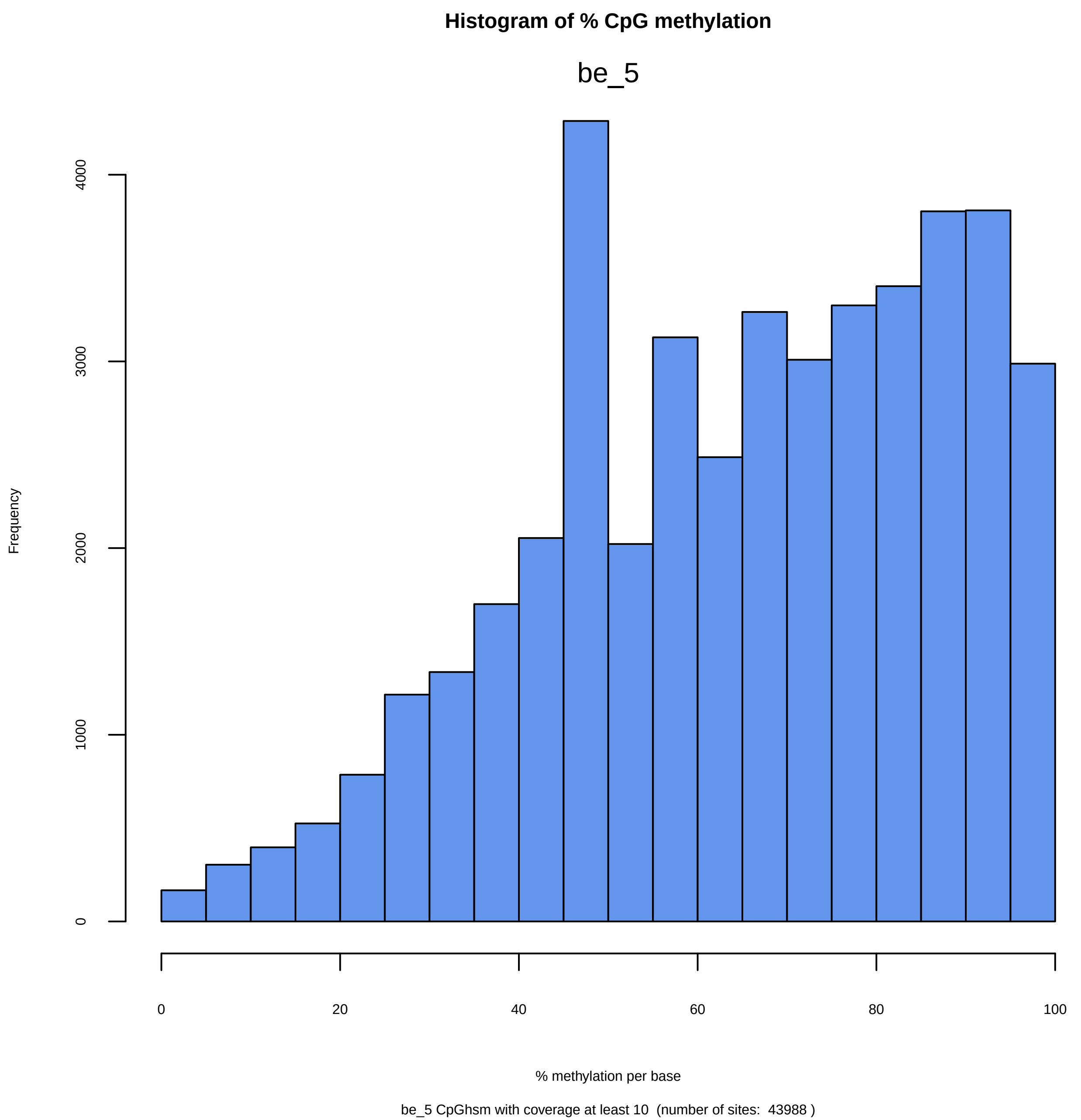

Histogram of % CpG methylation
be_5
Frequency
0
1000
2000
3000
4000
0
20
40
60
80
100
% methylation per base
be_5 CpGhsm with coverage at least 10  (number of sites:  43988 )

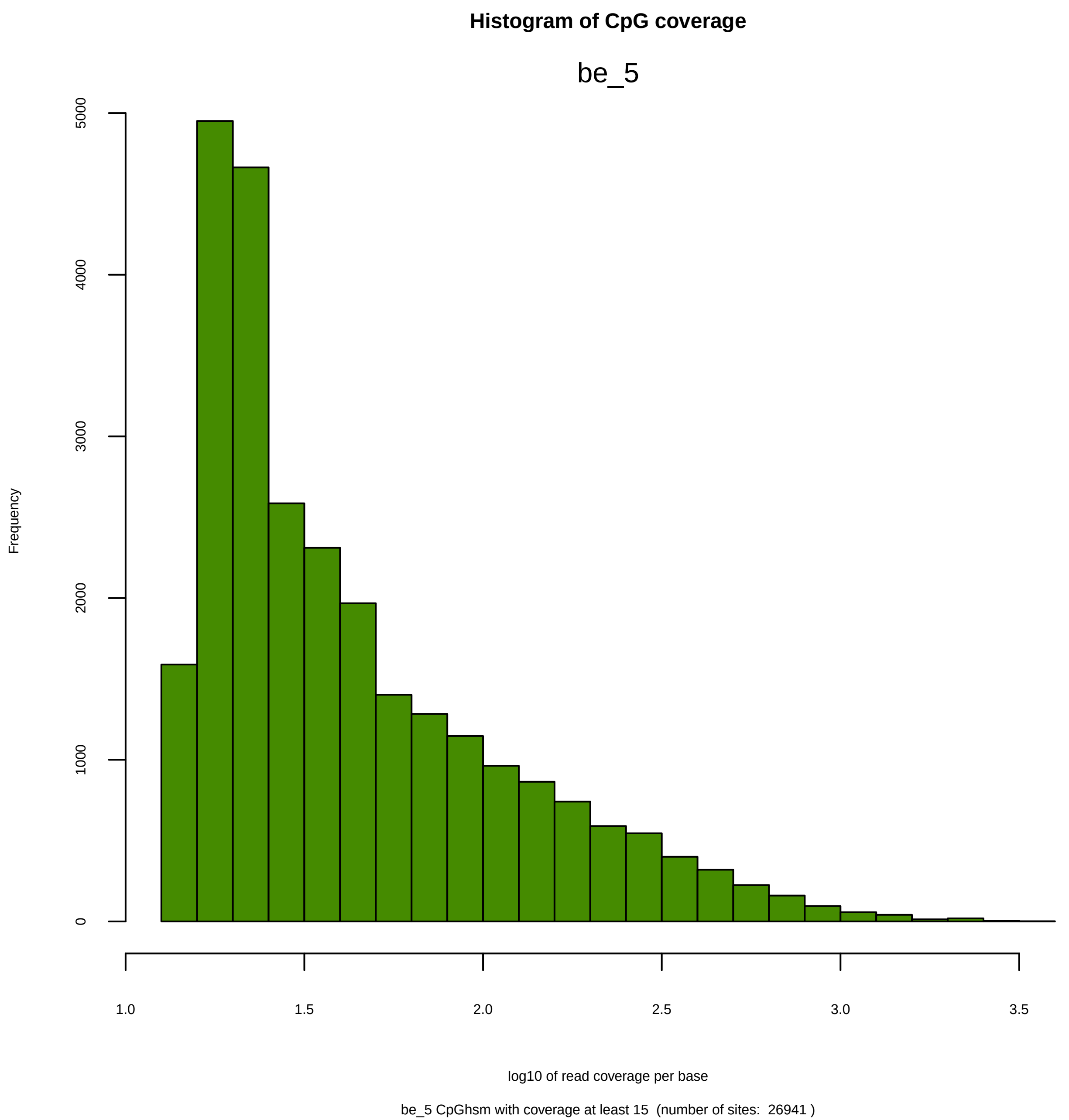

Histogram of CpG coverage
be_5
Frequency
0
1000
2000
3000
4000
5000
1.0
1.5
2.0
2.5
3.0
3.5
log10 of read coverage per base
be_5 CpGhsm with coverage at least 15 (number of sites: 26941 )

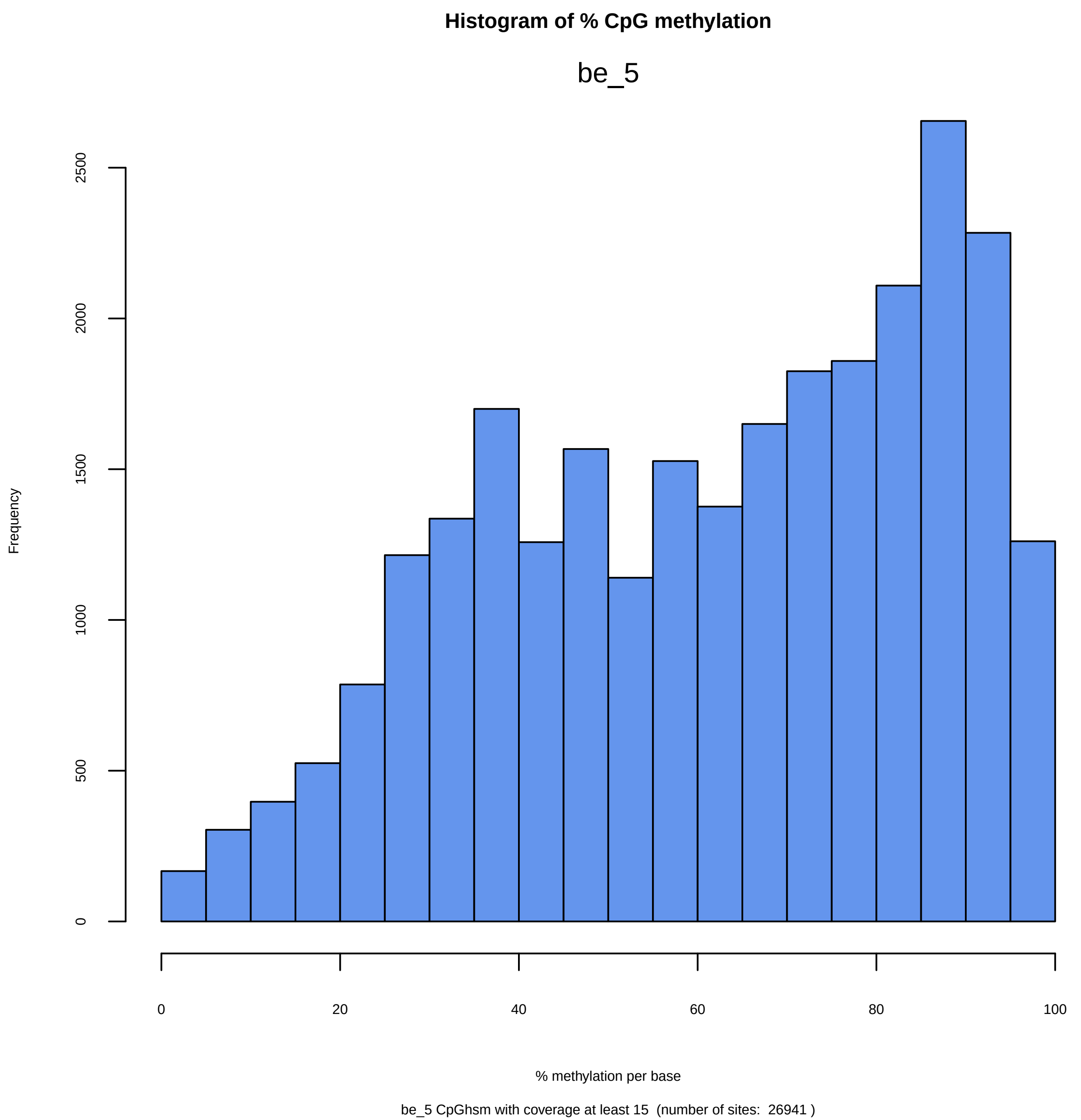
Histogram of % CpG methylation
be_5
Frequency
0
500
1000
1500
2000
2500
0
20
40
60
80
100
% methylation per base
be_5 CpGhsm with coverage at least 15 (number of sites: 26941 )

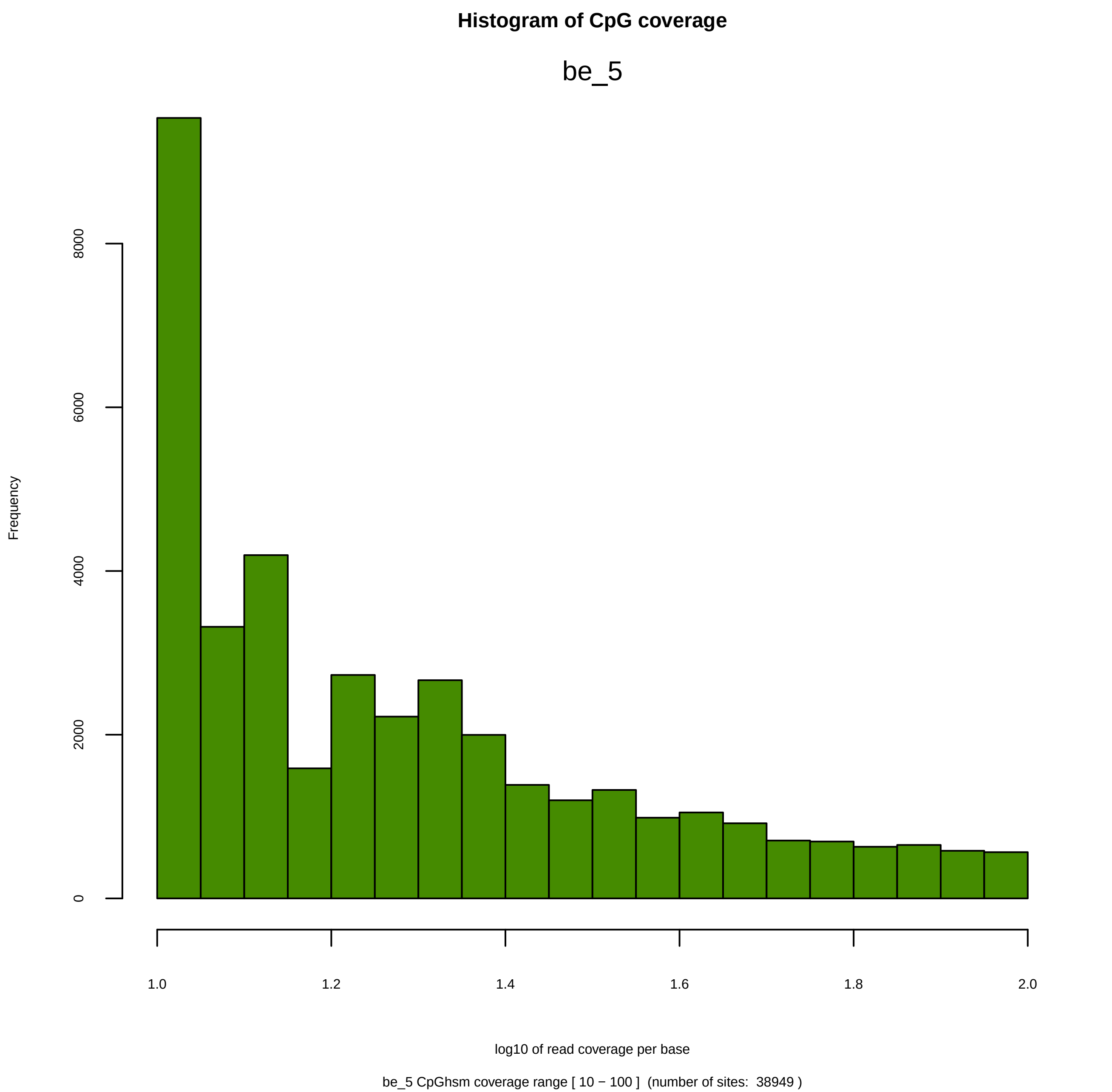

Histogram of CpG coverage
be_5
Frequency
0
2000
4000
6000
8000
1.0
1.2
1.4
1.6
1.8
2.0
log10 of read coverage per base
be_5 CpGhsm coverage range [ 10 − 100 ]  (number of sites:  38949 )

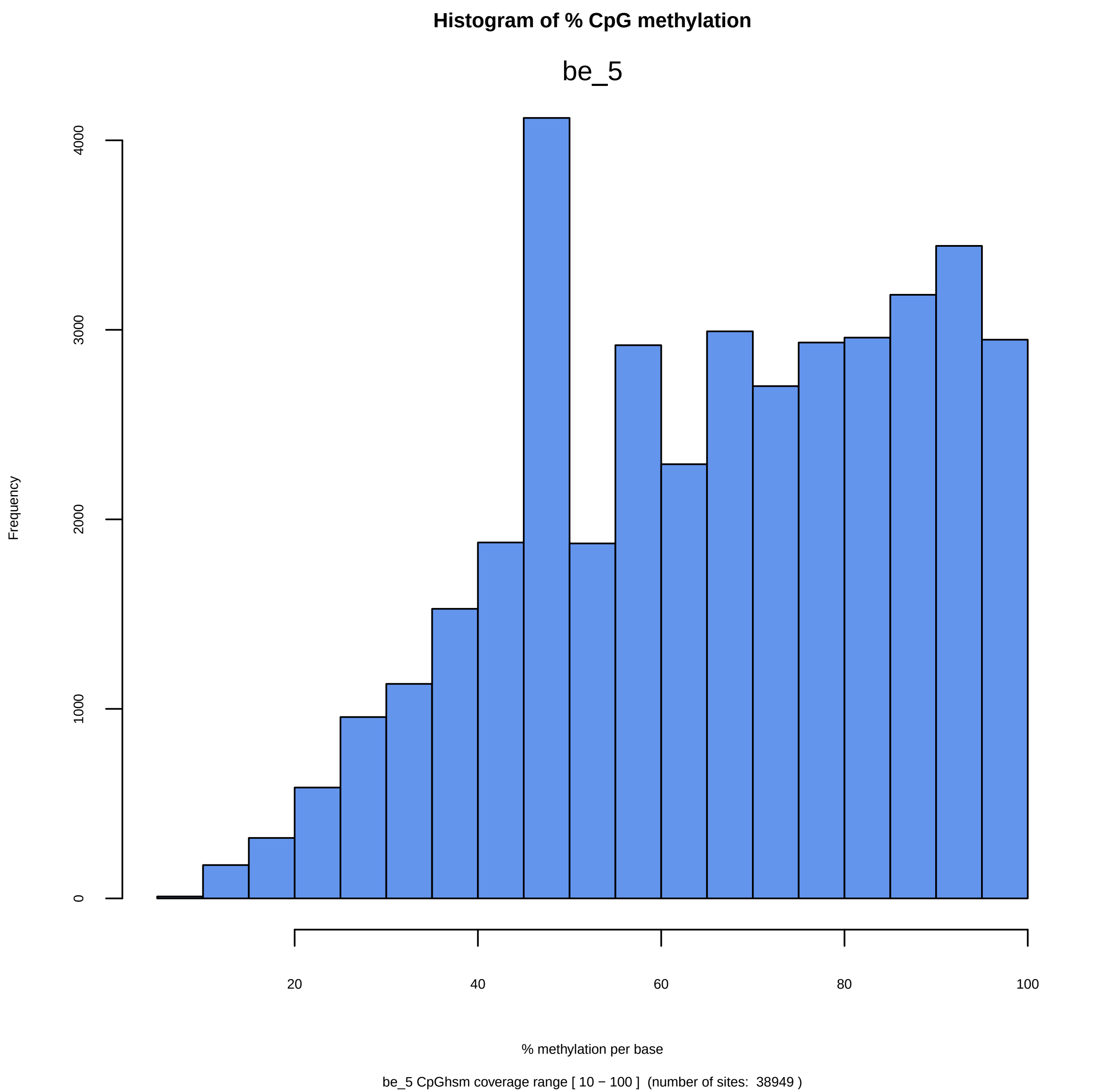

Histogram of % CpG methylation
be_5
Frequency
0
1000
2000
3000
4000
20
40
60
80
100
% methylation per base
be_5 CpGhsm coverage range [ 10 − 100 ]  (number of sites:  38949 )

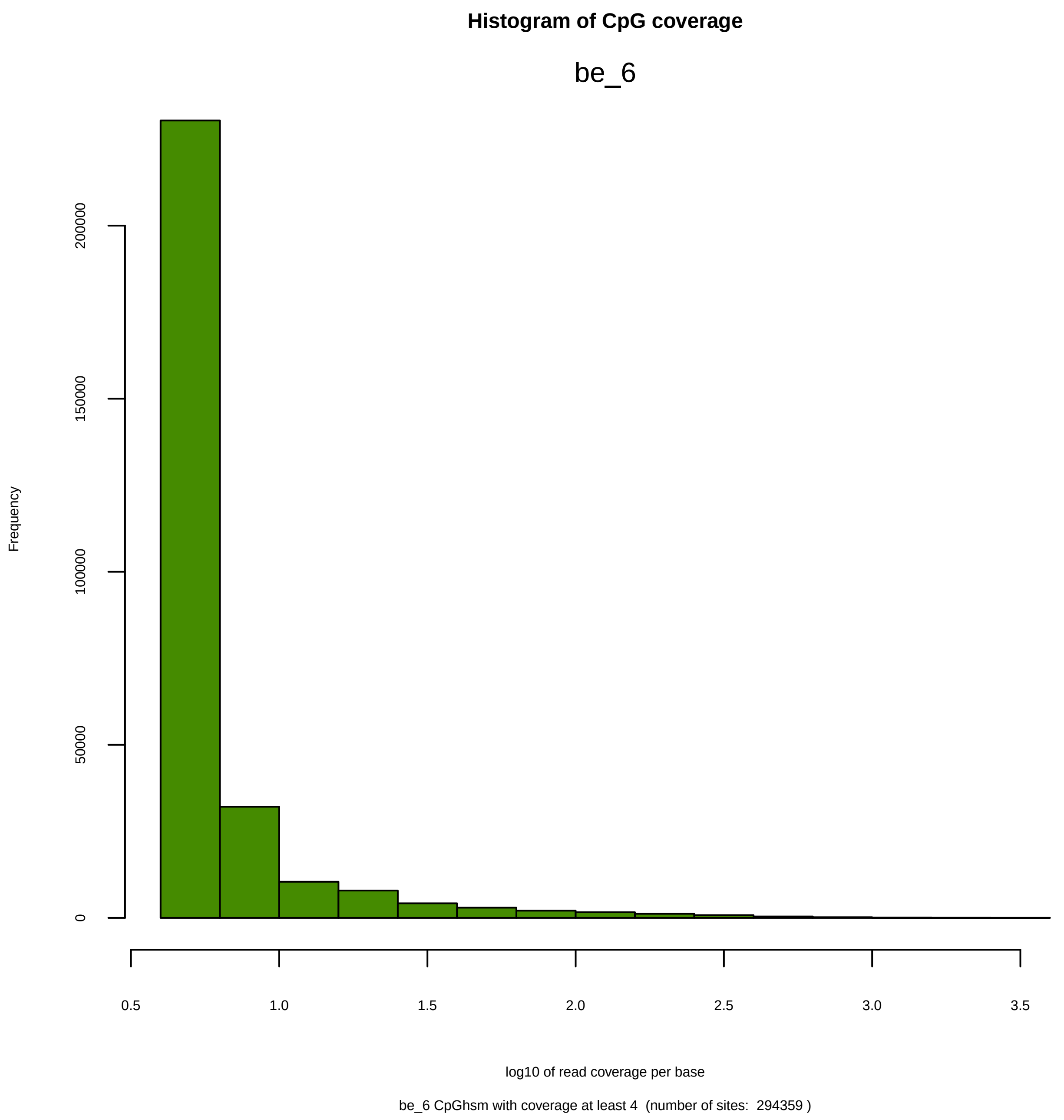
Histogram of CpG coverage
be_6
Frequency
0
50000
100000
150000
200000
0.5
1.0
1.5
2.0
2.5
3.0
3.5
log10 of read coverage per base
be_6 CpGhsm with coverage at least 4 (number of sites: 294359 )

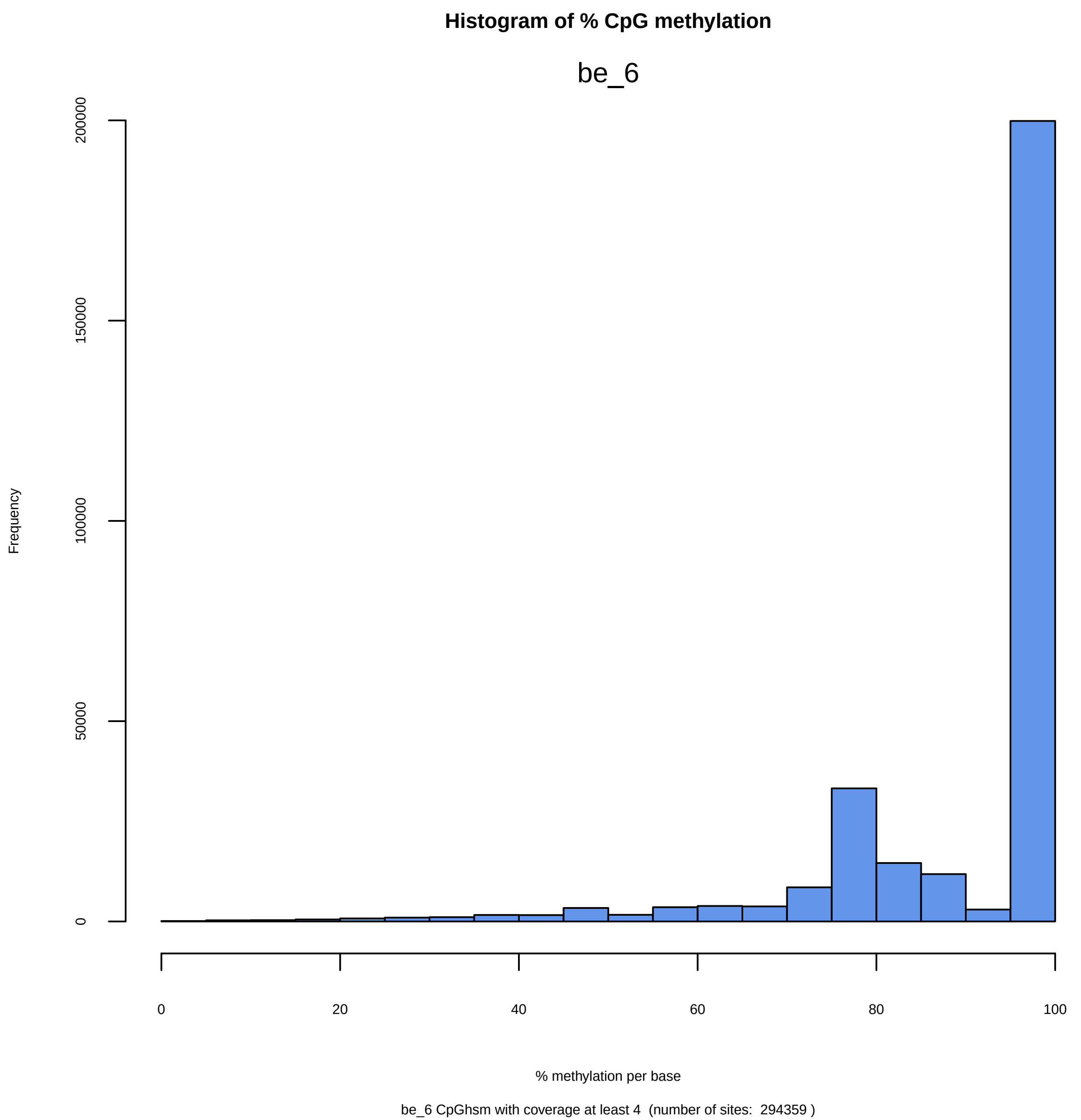

Histogram of % CpG methylation
be_6
Frequency
0
50000
100000
150000
200000
0
20
40
60
80
100
% methylation per base
be_6 CpGhsm with coverage at least 4 (number of sites: 294359 )

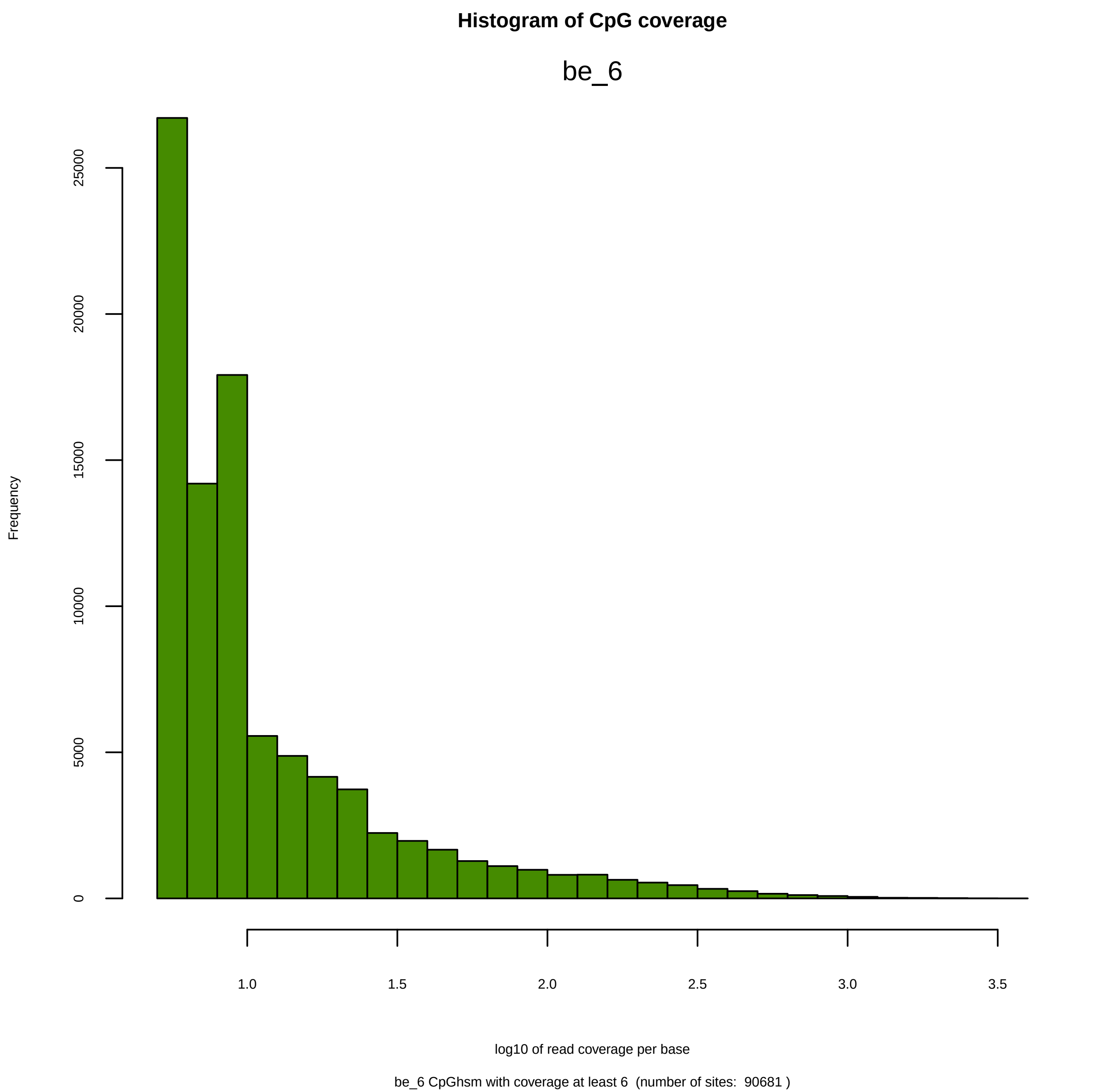

Histogram of CpG coverage
be_6
Frequency
0
5000
10000
15000
20000
25000
1.0
1.5
2.0
2.5
3.0
3.5
log10 of read coverage per base
be_6 CpGhsm with coverage at least 6 (number of sites: 90681 )

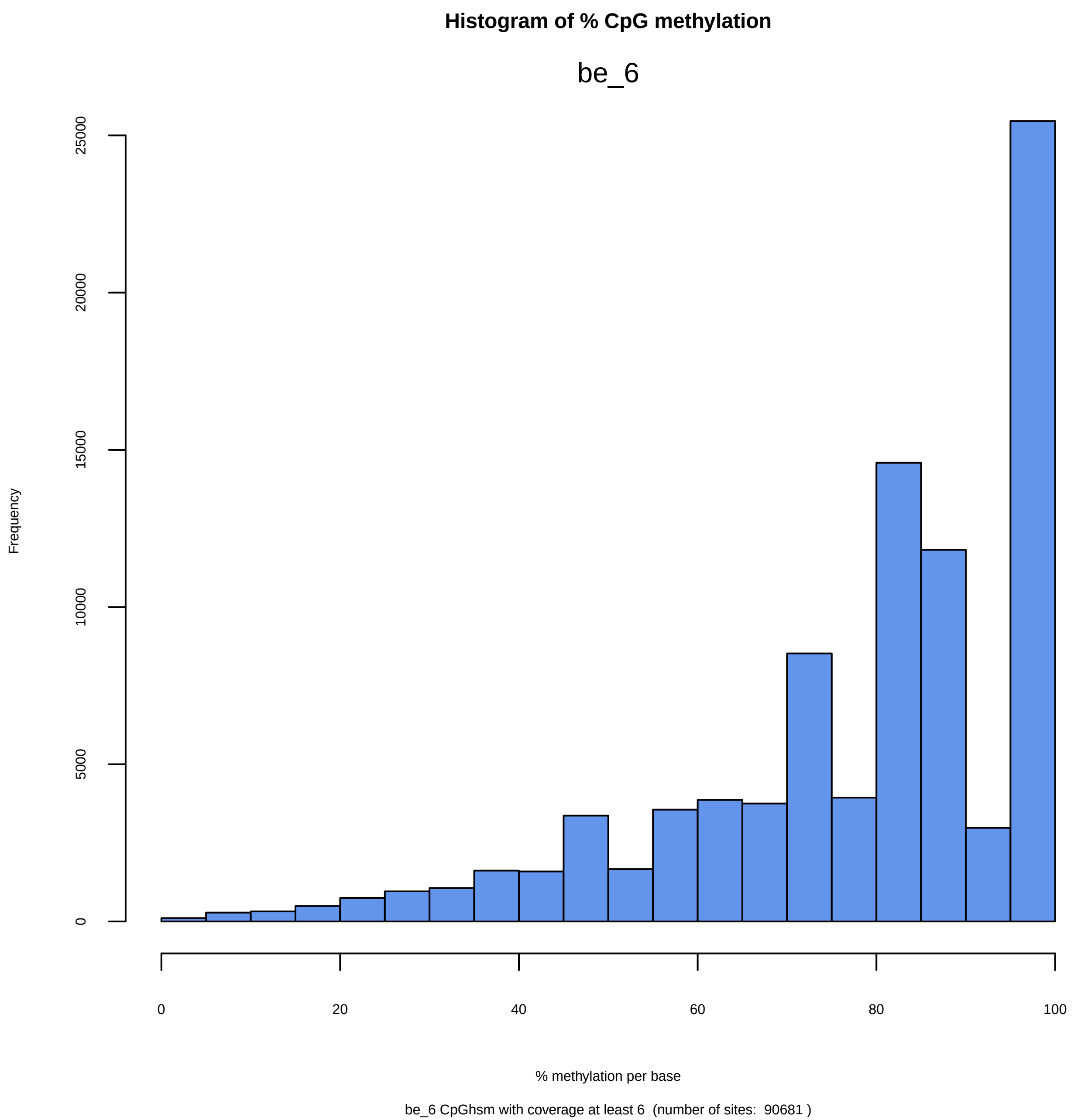

Histogram of % CpG methylation
be_6
Frequency
0
5000
10000
15000
20000
25000
0
20
40
60
80
100
% methylation per base
be_6 CpGhsm with coverage at least 6 (number of sites: 90681 )

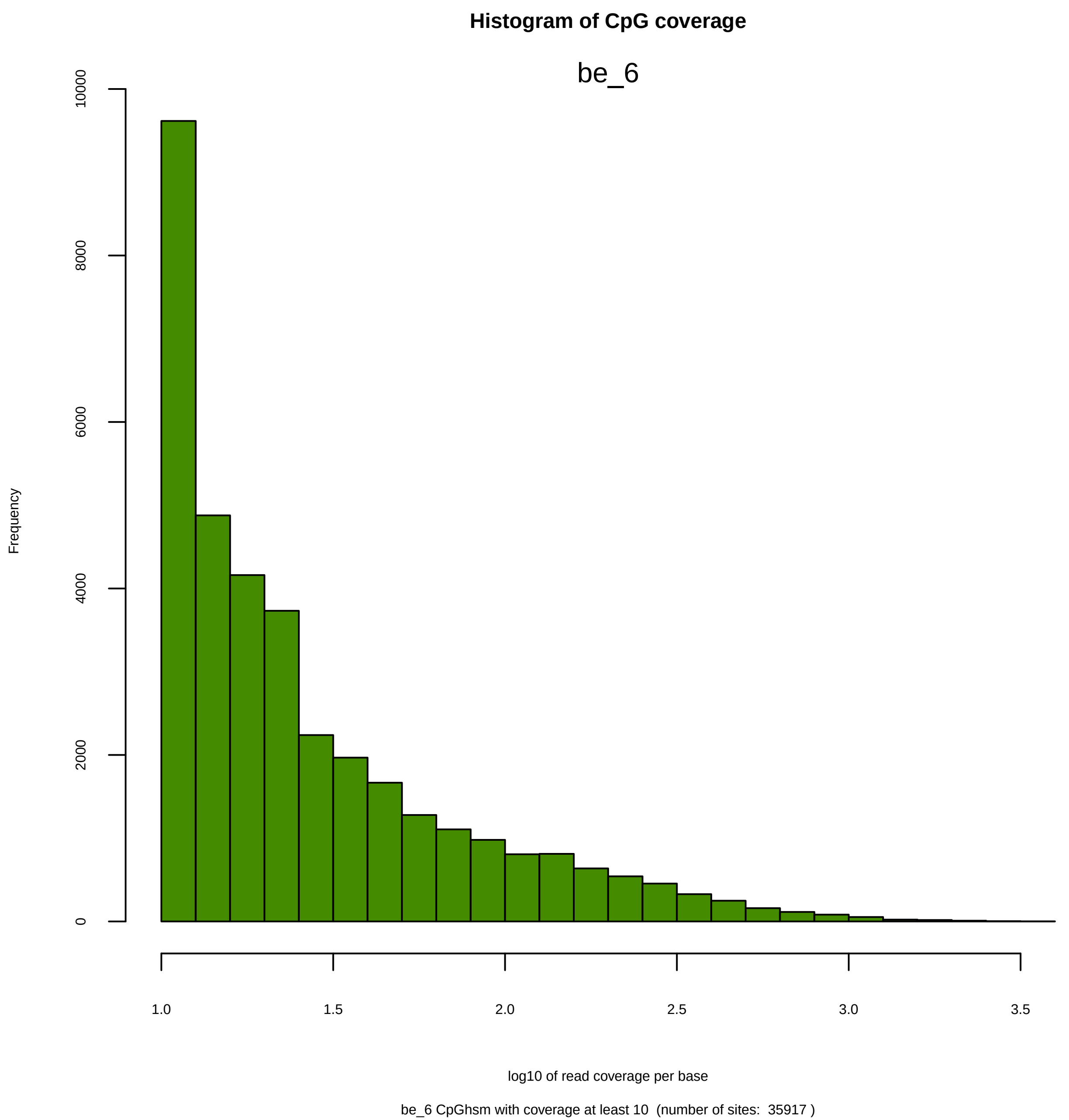

Histogram of CpG coverage
be_6
Frequency
0
2000
4000
6000
8000
10000
1.0
1.5
2.0
2.5
3.0
3.5
log10 of read coverage per base
be_6 CpGhsm with coverage at least 10  (number of sites:  35917 )

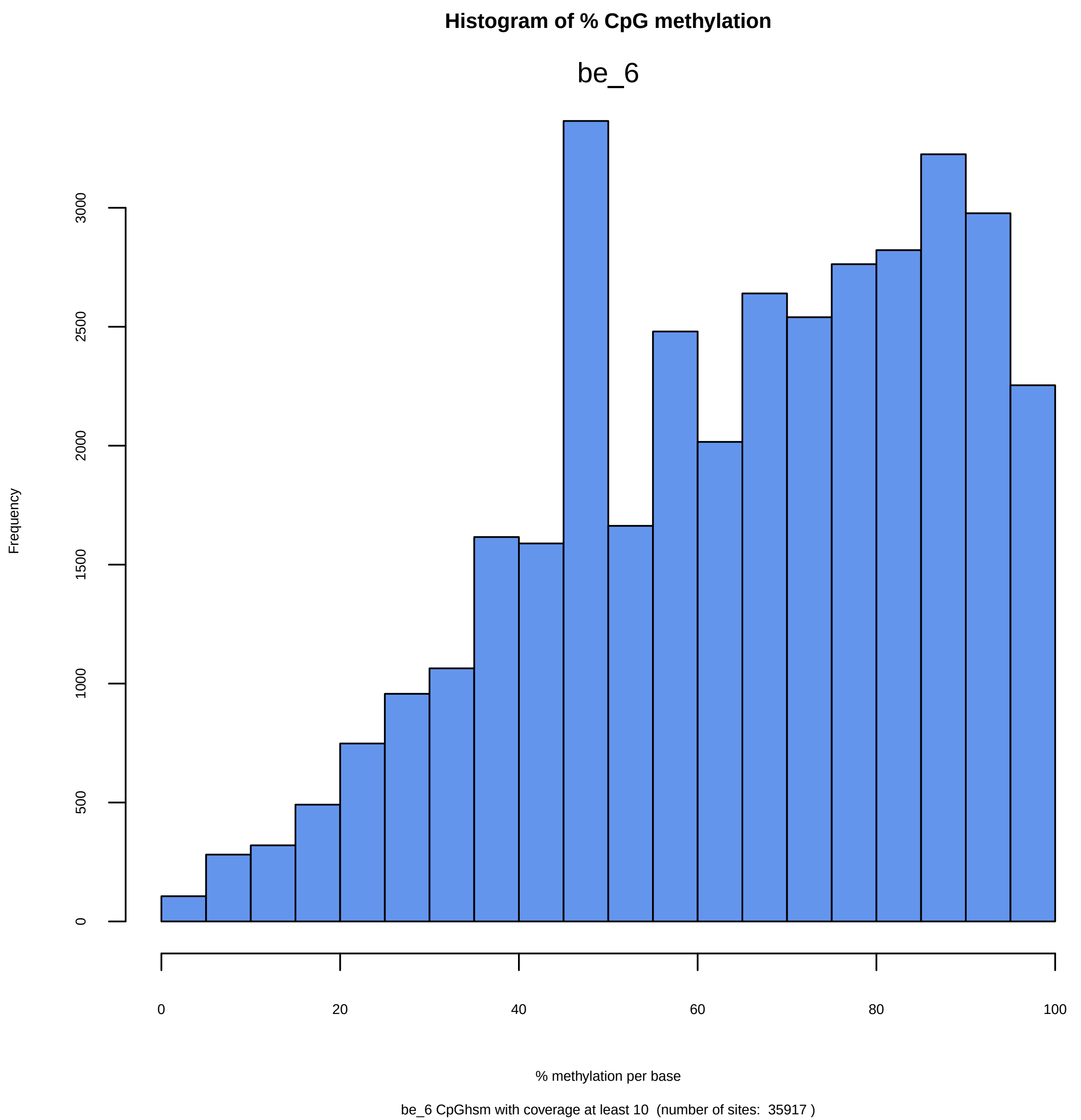
Histogram of % CpG methylation
be_6
Frequency
0
500
1000
1500
2000
2500
3000
0
20
40
60
80
100
% methylation per base
be_6 CpGhsm with coverage at least 10  (number of sites:  35917 )

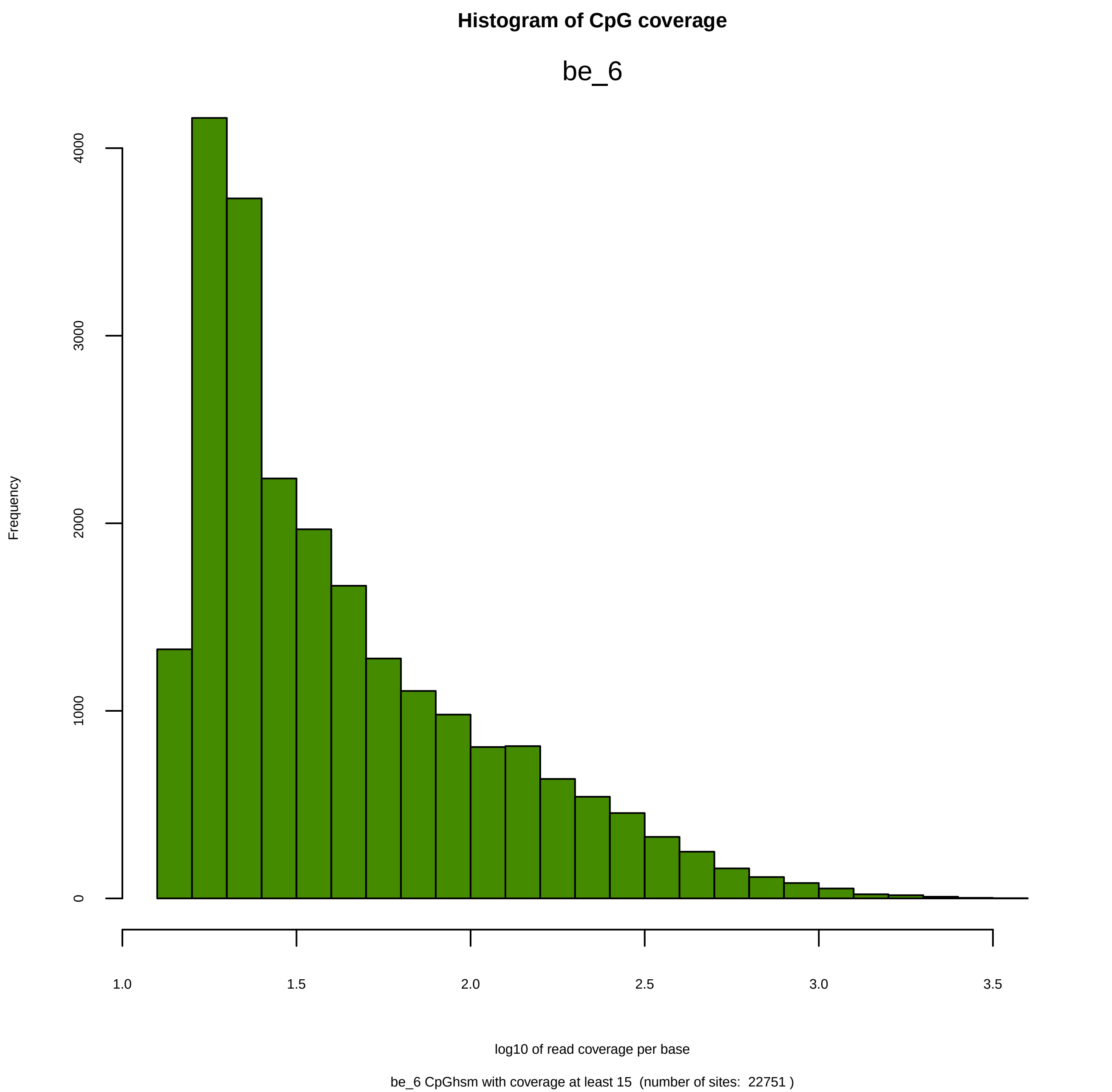

Histogram of CpG coverage
be_6
Frequency
0
1000
2000
3000
4000
1.0
1.5
2.0
2.5
3.0
3.5
log10 of read coverage per base
be_6 CpGhsm with coverage at least 15 (number of sites: 22751 )

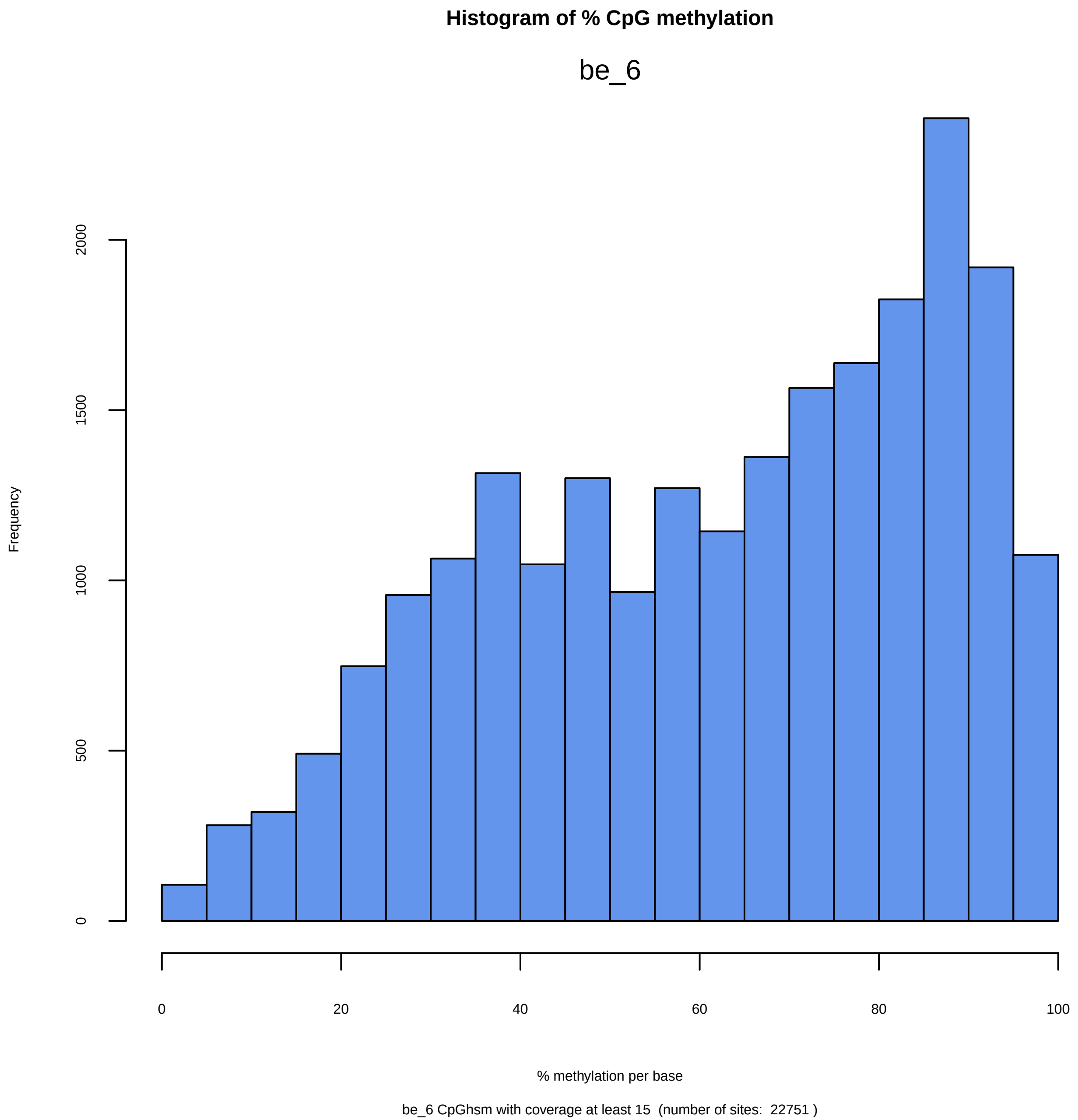
Histogram of % CpG methylation
be_6
Frequency
0
500
1000
1500
2000
0
20
40
60
80
100
% methylation per base
be_6 CpGhsm with coverage at least 15  (number of sites:  22751 )

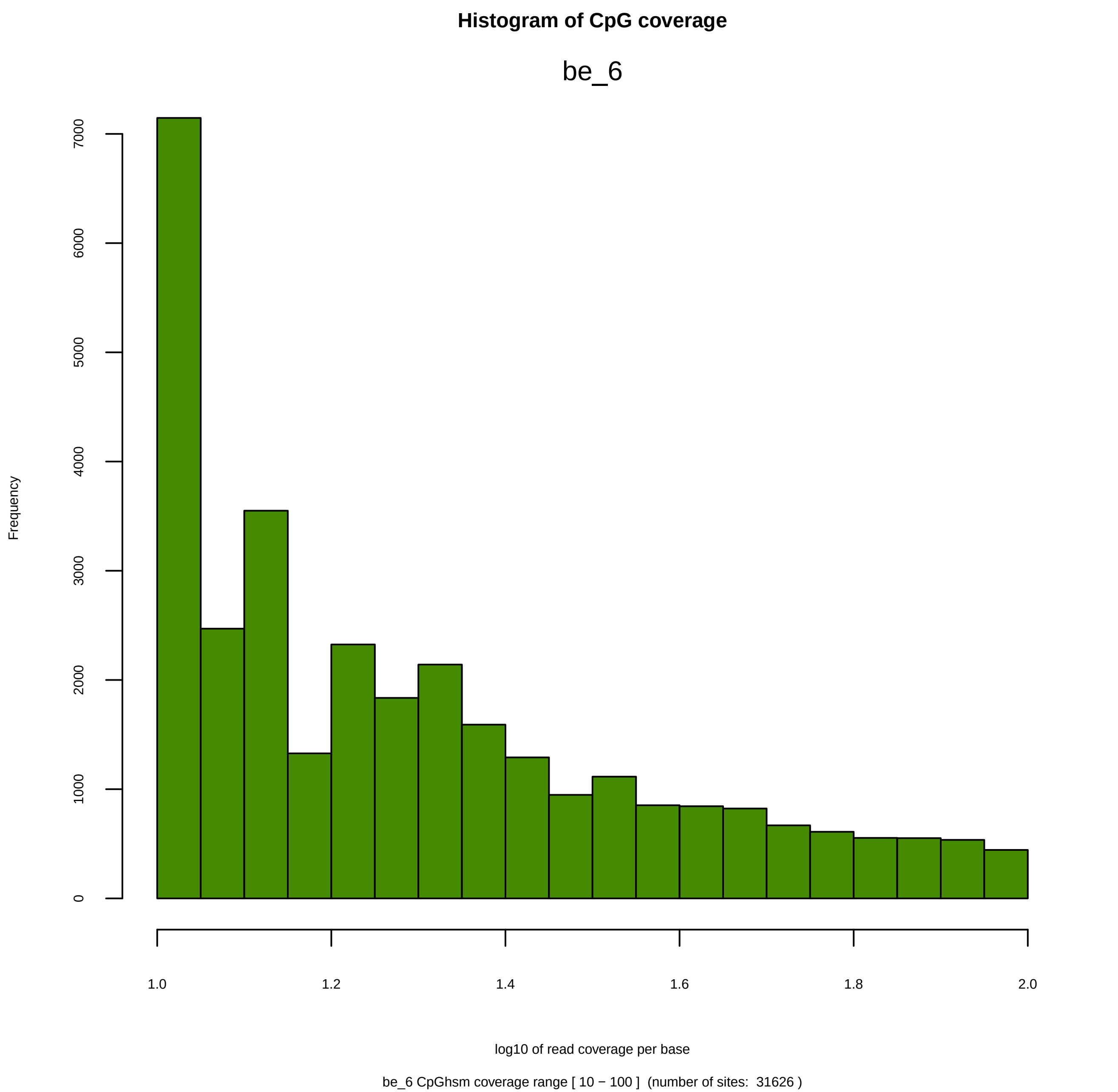

Histogram of CpG coverage
be_6
Frequency
0
1000
2000
3000
4000
5000
6000
7000
1.0
1.2
1.4
1.6
1.8
2.0
log10 of read coverage per base
be_6 CpGhsm coverage range [ 10 − 100 ]  (number of sites:  31626 )

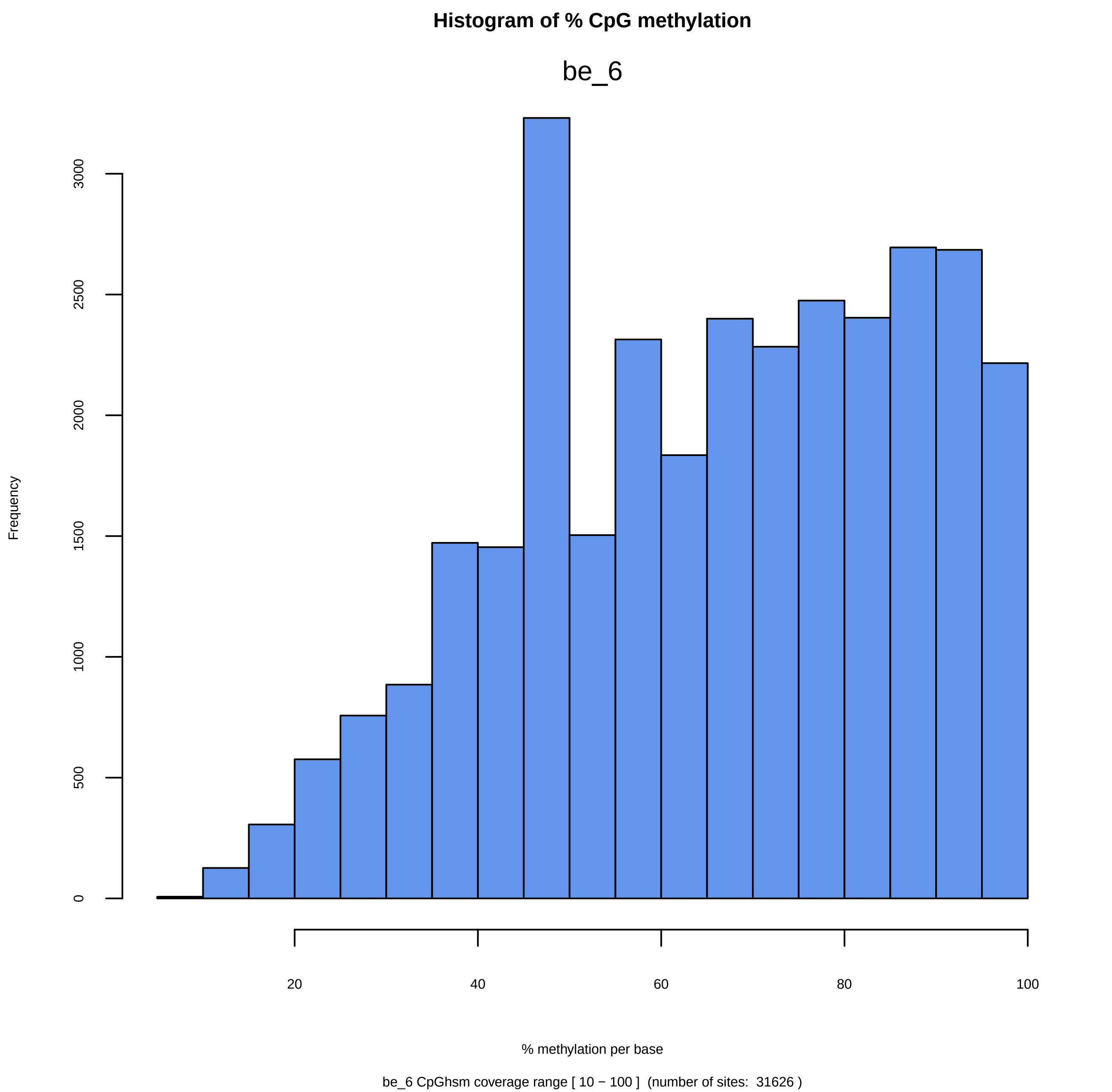

Histogram of % CpG methylation
be_6
Frequency
0
500
1000
1500
2000
2500
3000
20
40
60
80
100
% methylation per base
be_6 CpGhsm coverage range [ 10 − 100 ]  (number of sites:  31626 )

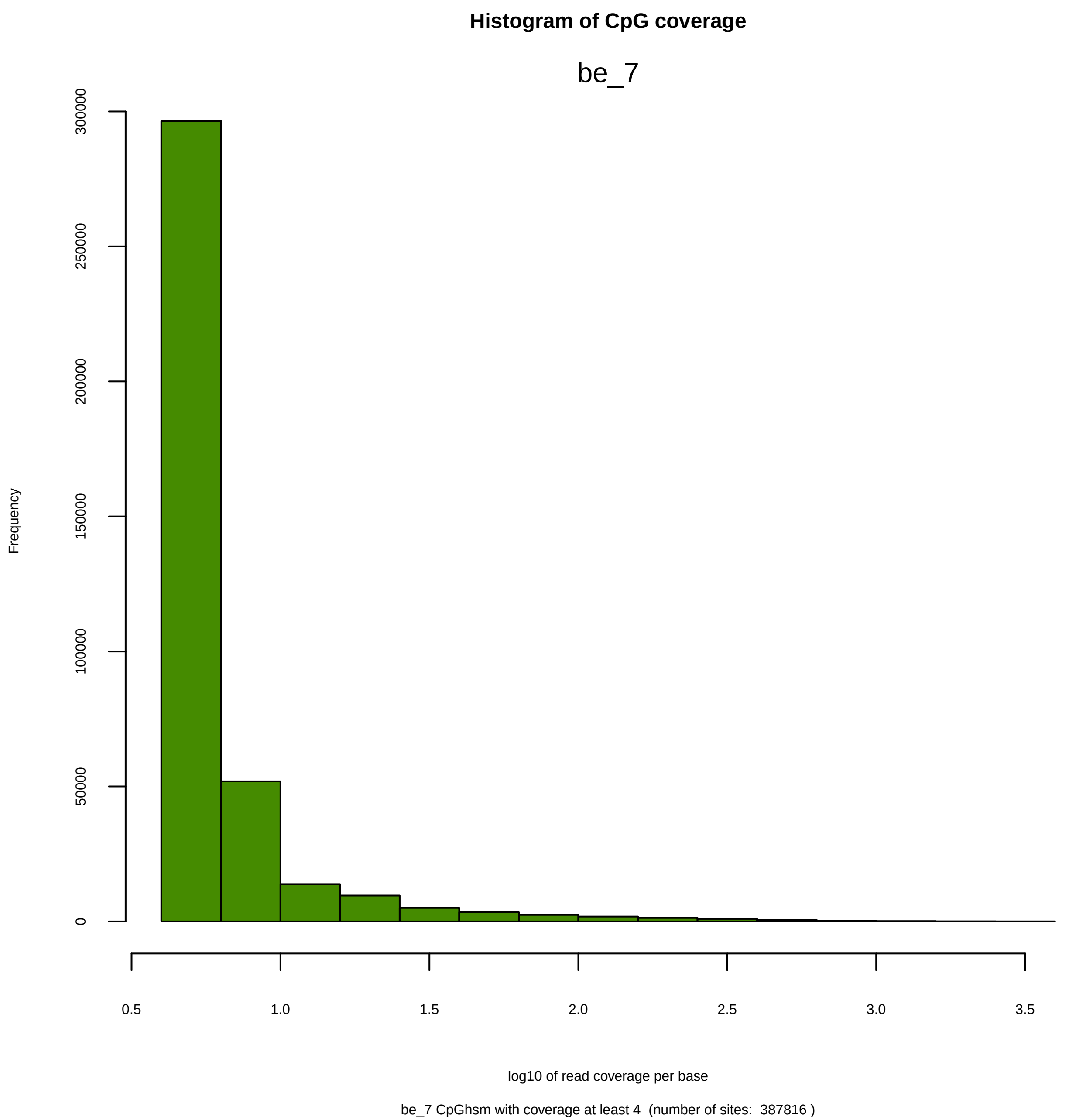

Histogram of CpG coverage
be_7
Frequency
0
50000
100000
150000
200000
250000
300000
0.5
1.0
1.5
2.0
2.5
3.0
3.5
log10 of read coverage per base
be_7 CpGhsm with coverage at least 4 (number of sites: 387816 )

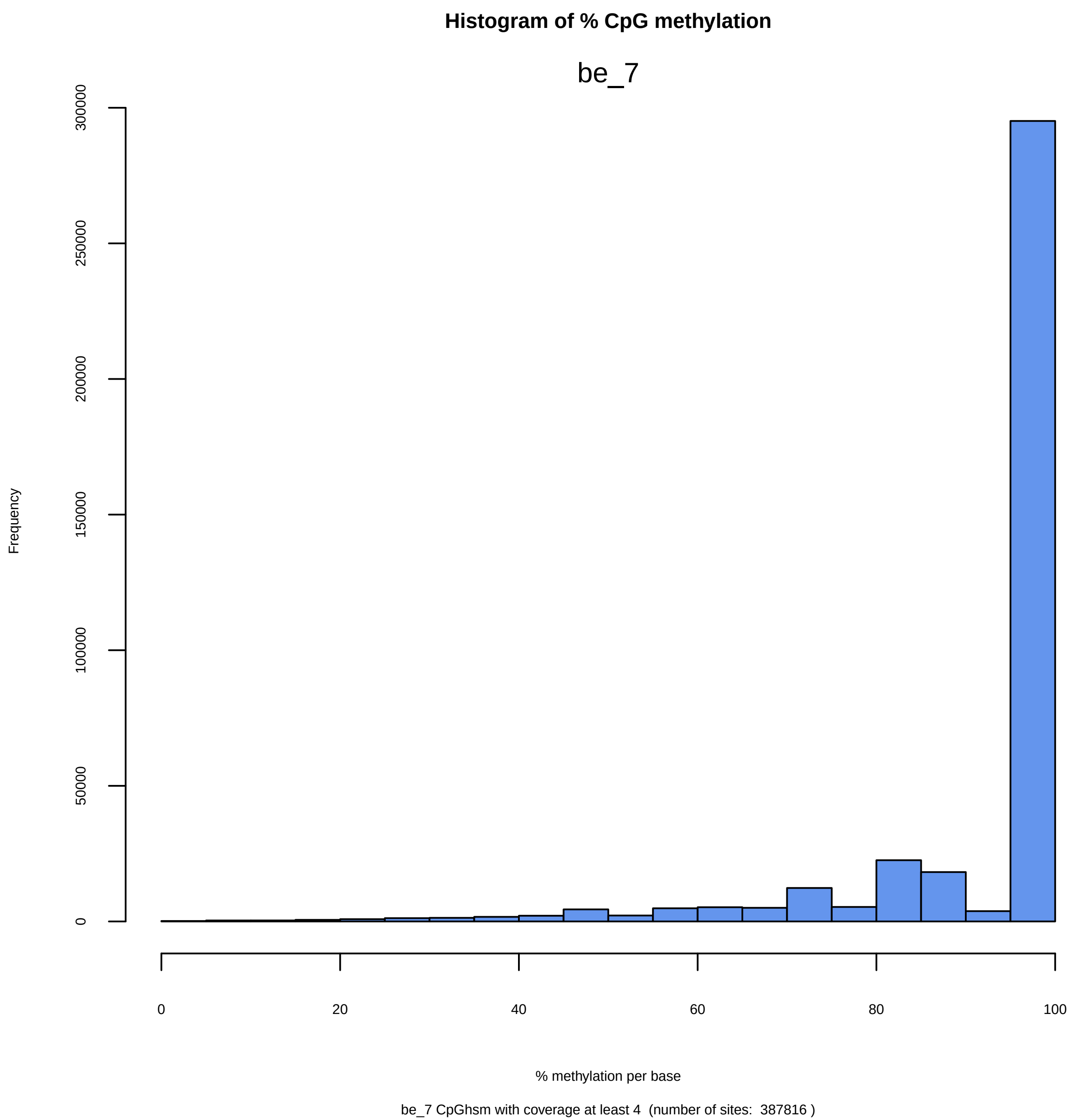
Histogram of % CpG methylation
be_7
Frequency
0
50000
100000
150000
200000
250000
300000
0
20
40
60
80
100
% methylation per base
be_7 CpGhsm with coverage at least 4 (number of sites: 387816 )

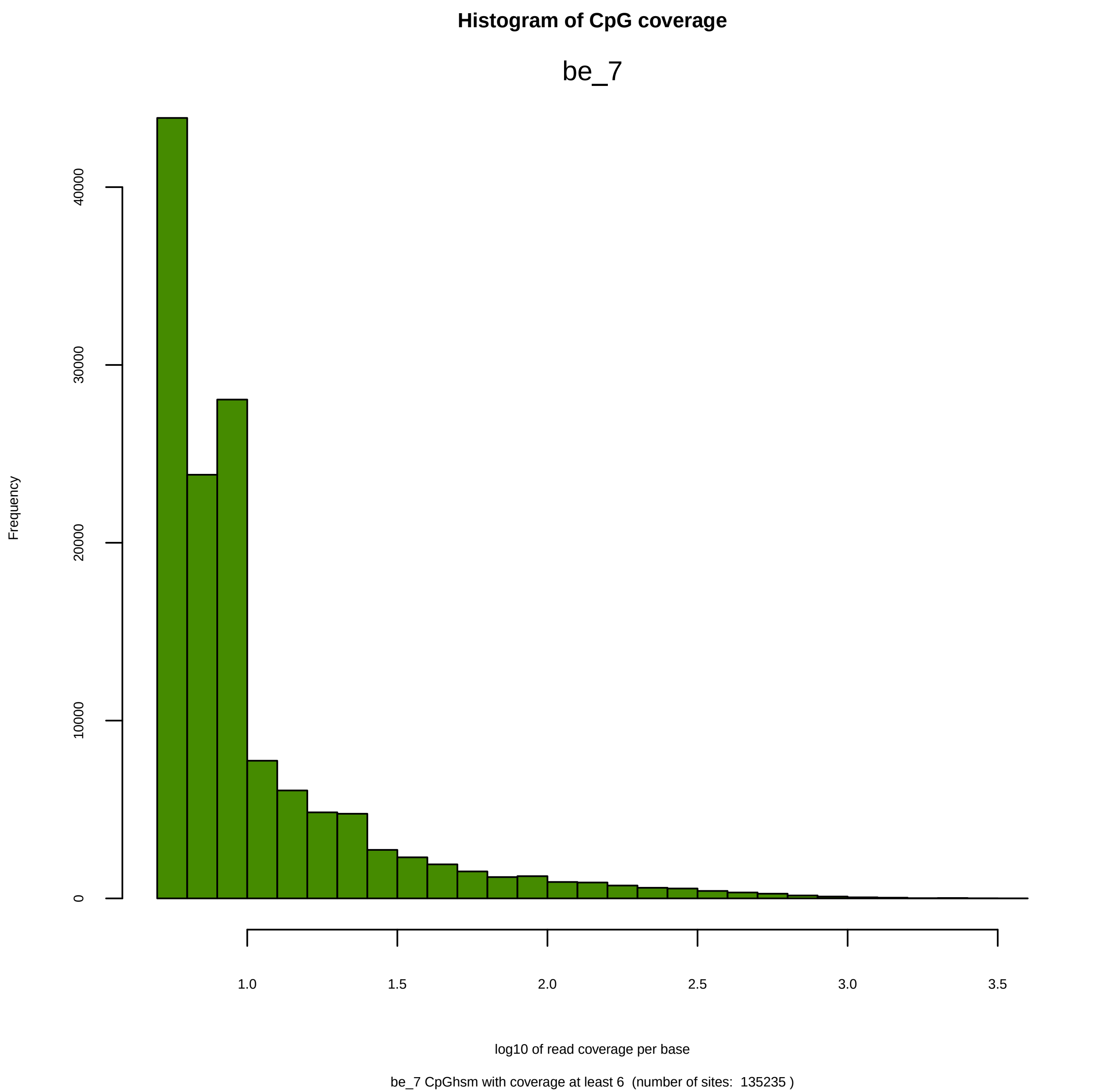

Histogram of CpG coverage
be_7
Frequency
0
10000
20000
30000
40000
1.0
1.5
2.0
2.5
3.0
3.5
log10 of read coverage per base
be_7 CpGhsm with coverage at least 6 (number of sites: 135235 )

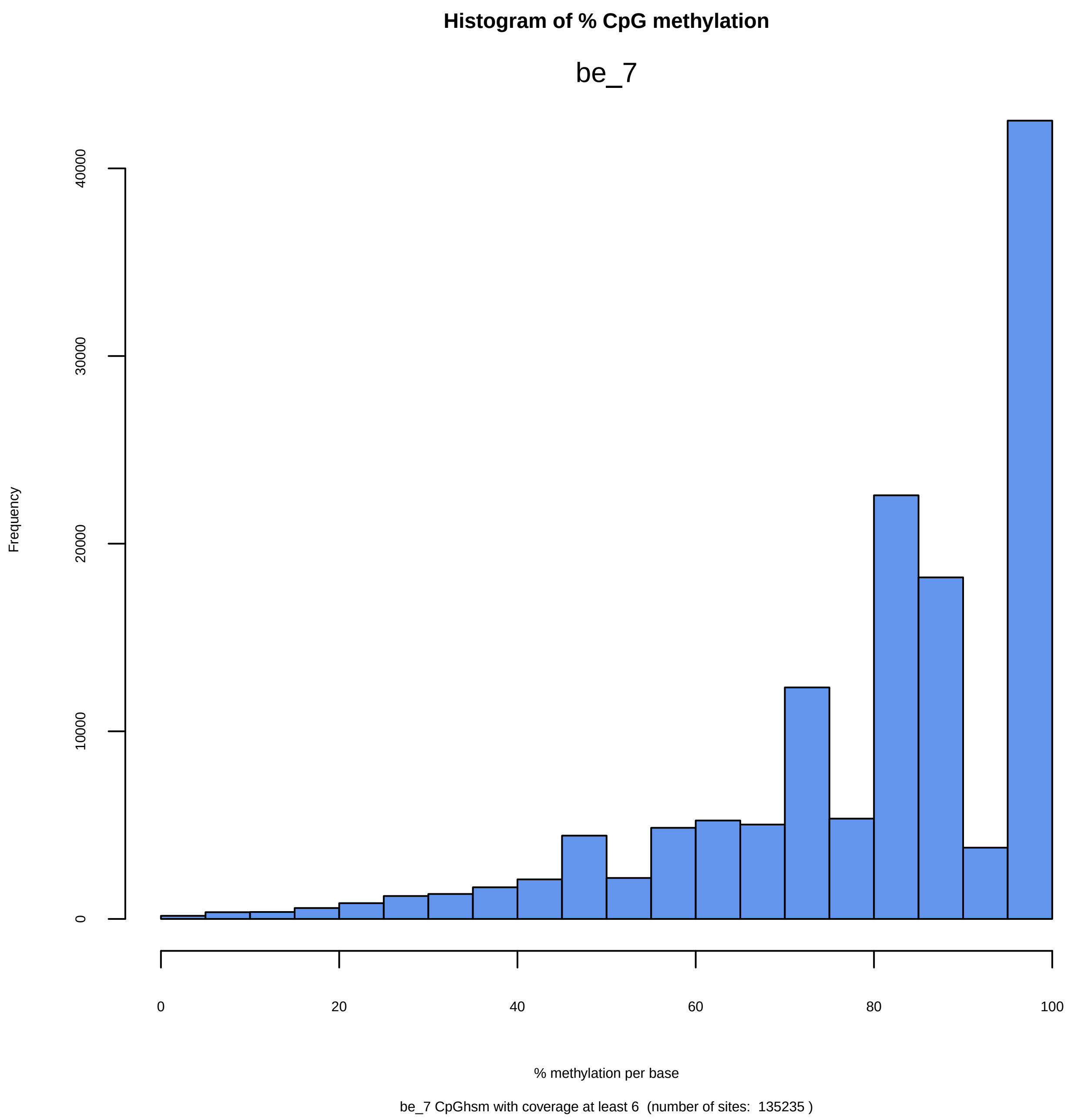
Histogram of % CpG methylation
be_7
Frequency
0
10000
20000
30000
40000
0
20
40
60
80
100
% methylation per base
be_7 CpGhsm with coverage at least 6 (number of sites: 135235 )

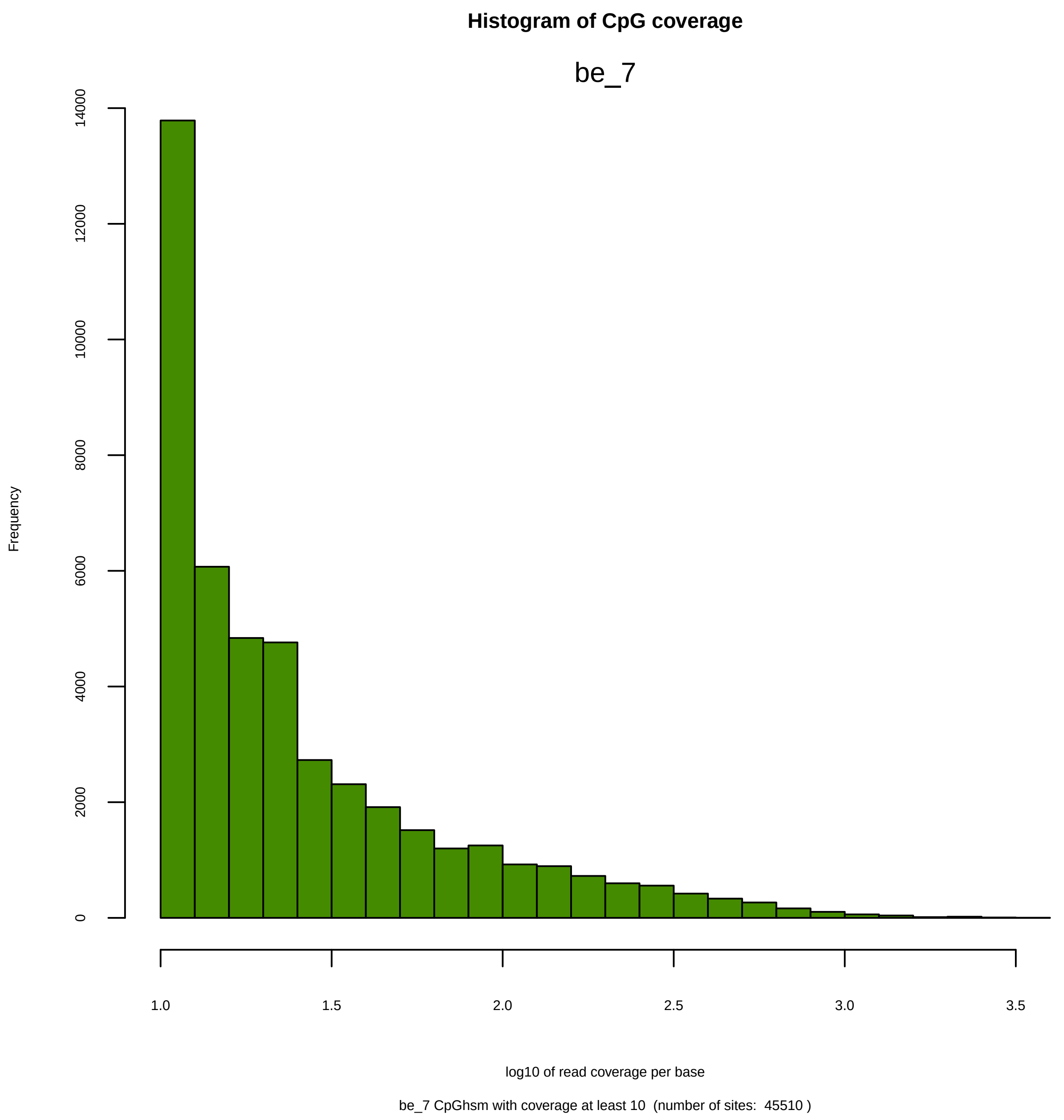
Histogram of CpG coverage
be_7
Frequency
0
2000
4000
6000
8000
10000
12000
14000
1.0
1.5
2.0
2.5
3.0
3.5
log10 of read coverage per base
be_7 CpGhsm with coverage at least 10  (number of sites:  45510 )

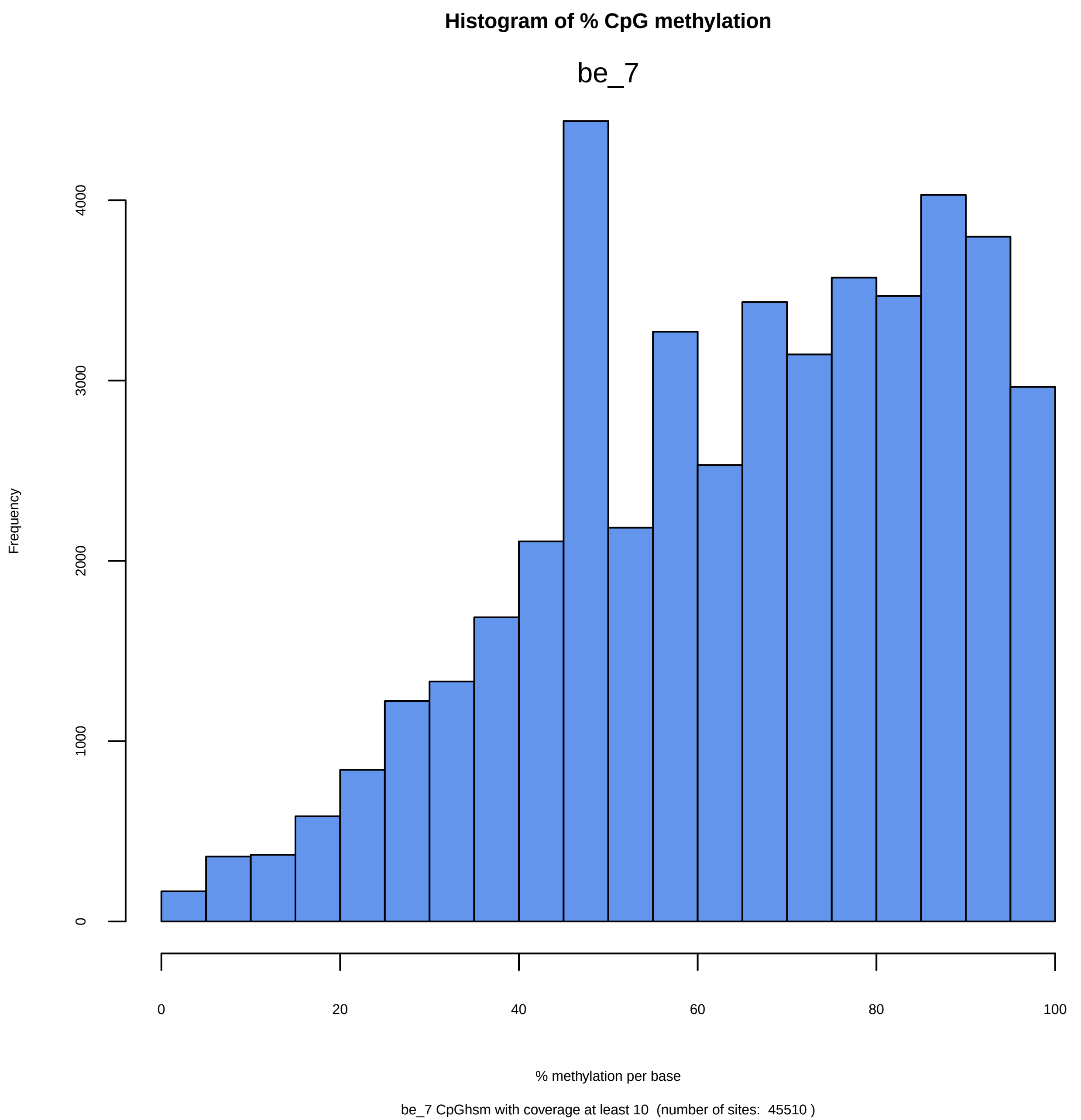
Histogram of % CpG methylation
be_7
Frequency
0
1000
2000
3000
4000
0
20
40
60
80
100
% methylation per base
be_7 CpGhsm with coverage at least 10 (number of sites: 45510 )

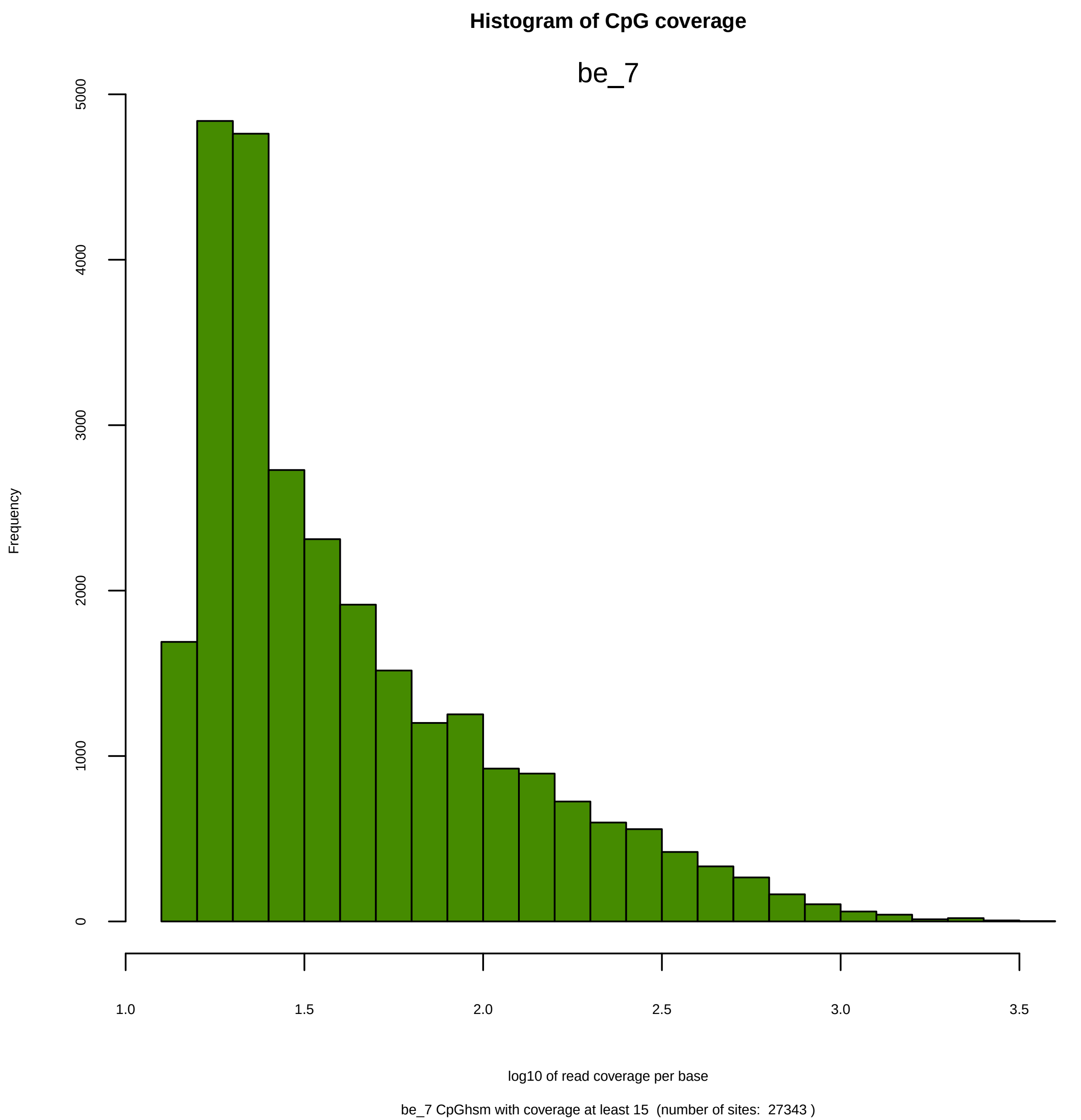
Histogram of CpG coverage
be_7
Frequency
0
1000
2000
3000
4000
5000
1.0
1.5
2.0
2.5
3.0
3.5
log10 of read coverage per base
be_7 CpGhsm with coverage at least 15  (number of sites:  27343 )

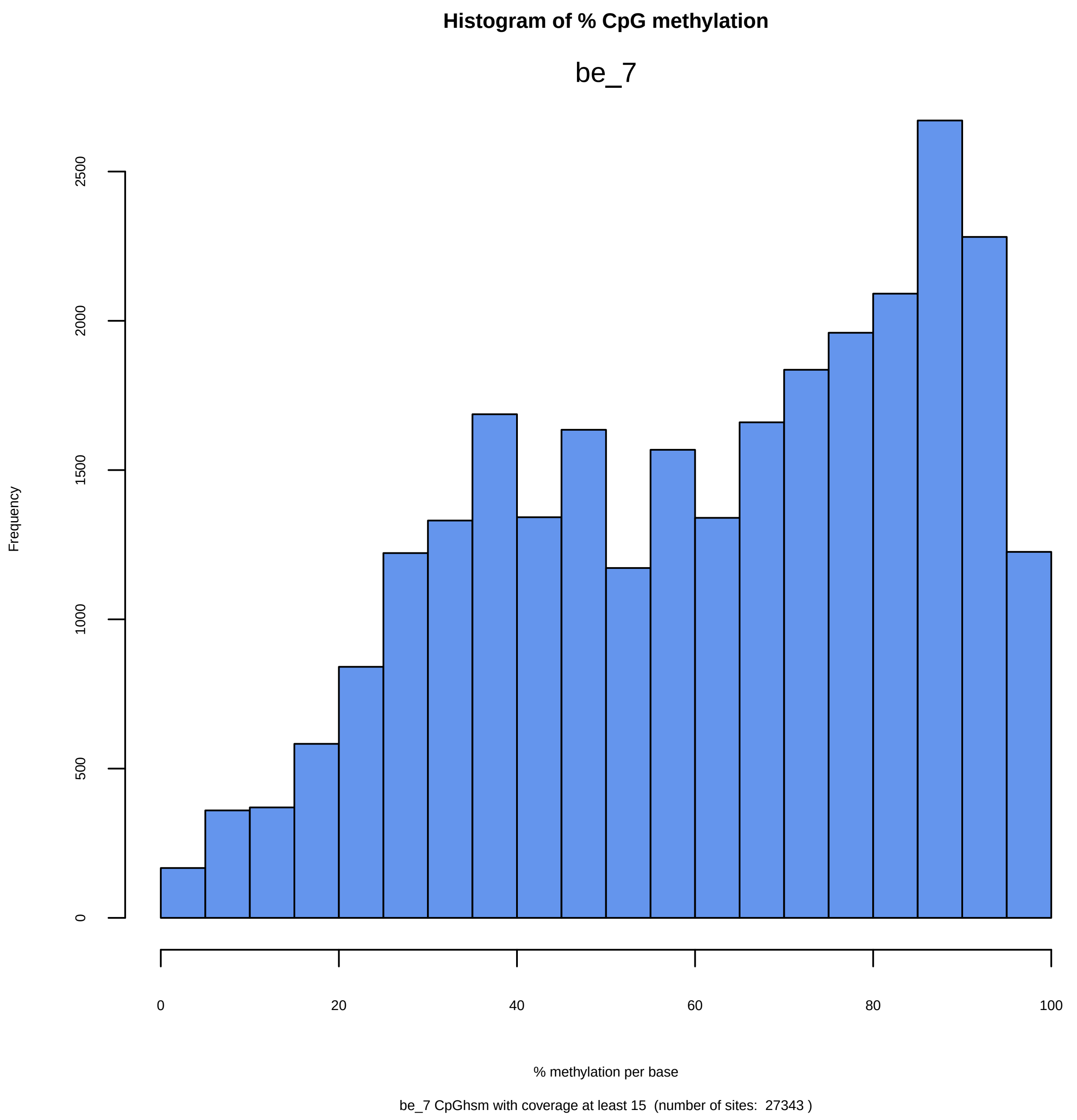
Histogram of % CpG methylation
be_7
Frequency
0
500
1000
1500
2000
2500
0
20
40
60
80
100
% methylation per base
be_7 CpGhsm with coverage at least 15  (number of sites:  27343 )

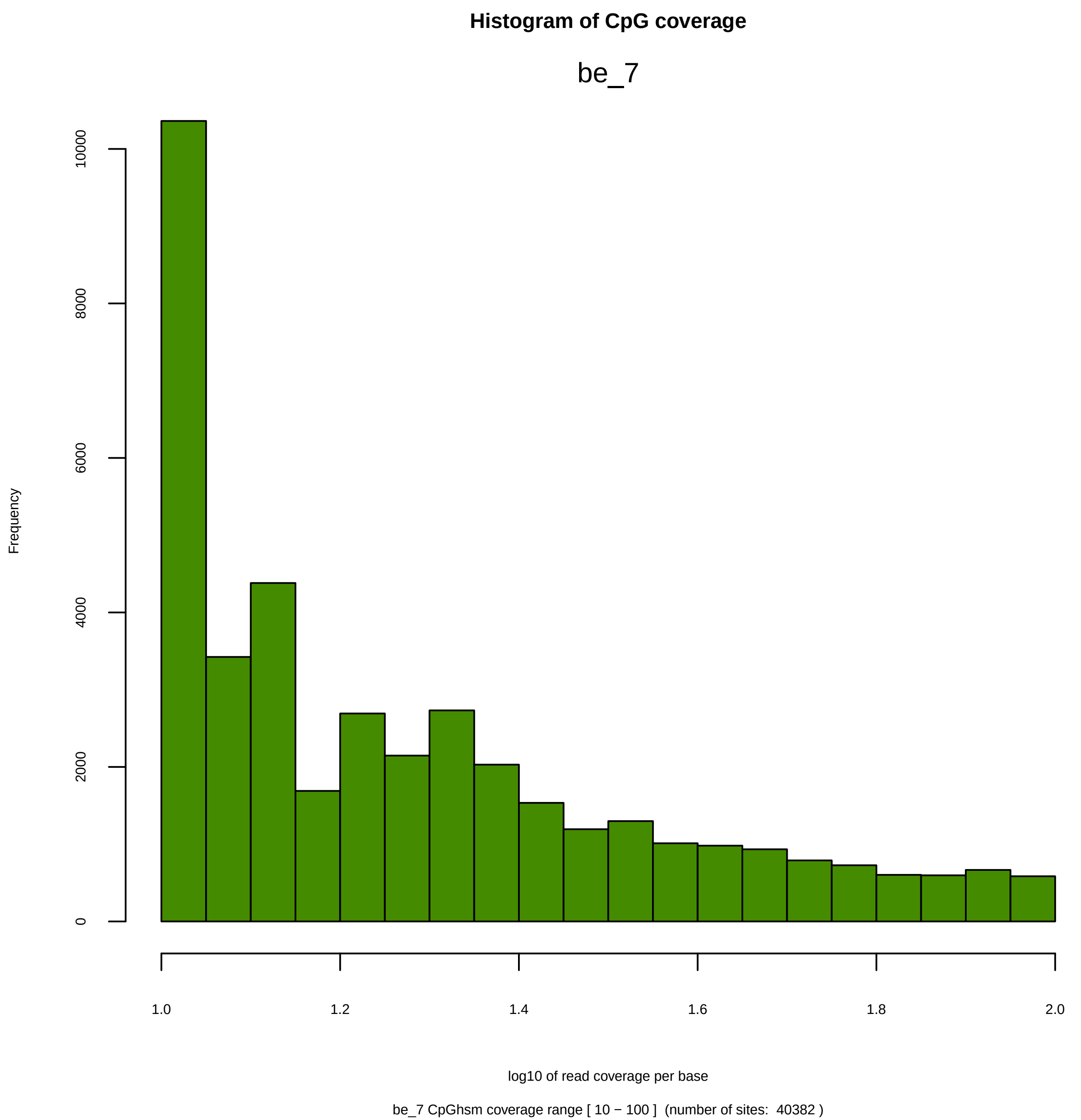

Histogram of CpG coverage
be_7
Frequency
0
2000
4000
6000
8000
10000
1.0
1.2
1.4
1.6
1.8
2.0
log10 of read coverage per base
be_7 CpGhsm coverage range [ 10 − 100 ] (number of sites: 40382 )

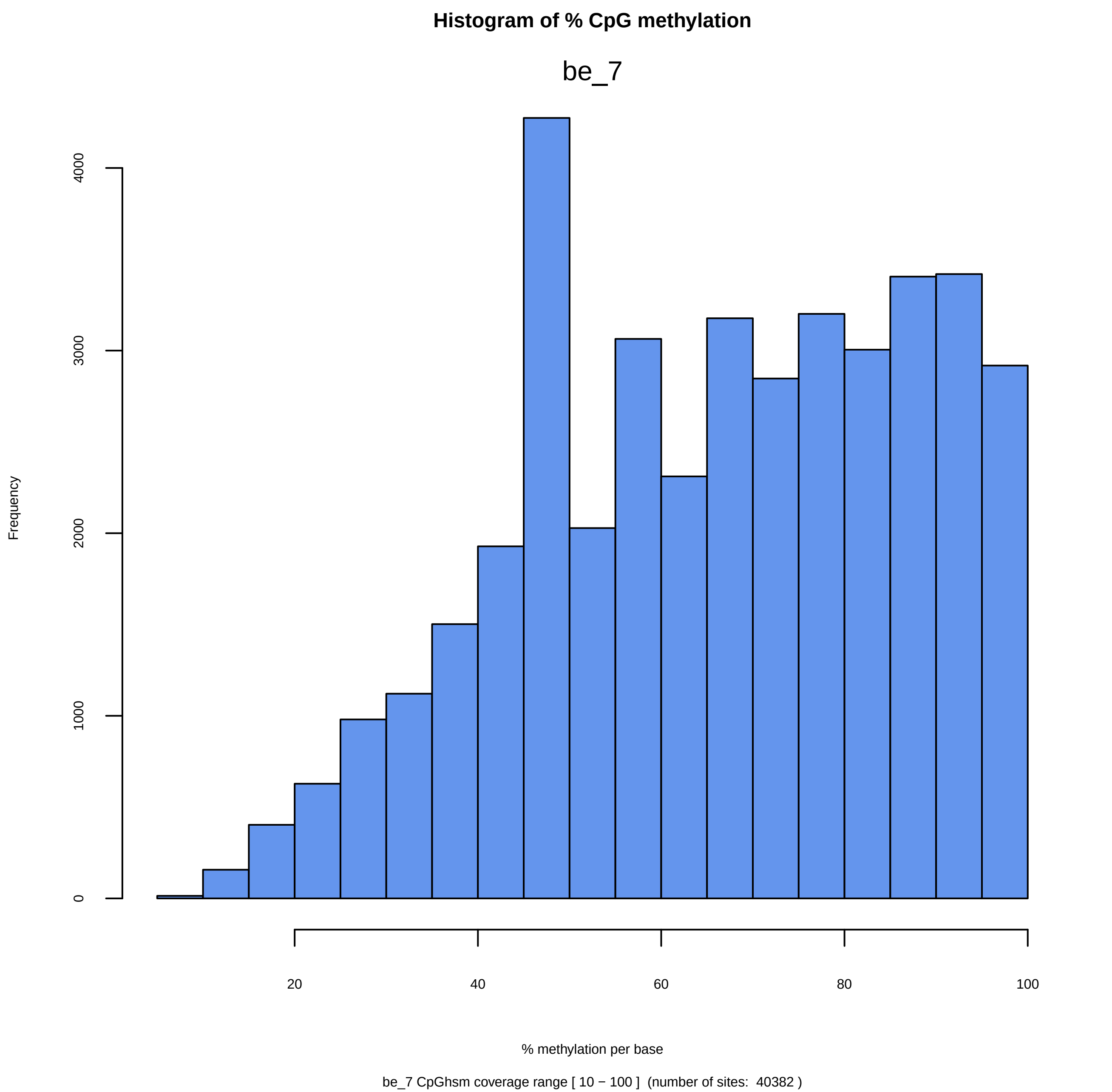

Histogram of % CpG methylation
be_7
Frequency
0
1000
2000
3000
4000
20
40
60
80
100
% methylation per base
be_7 CpGhsm coverage range [ 10 − 100 ]  (number of sites:  40382 )

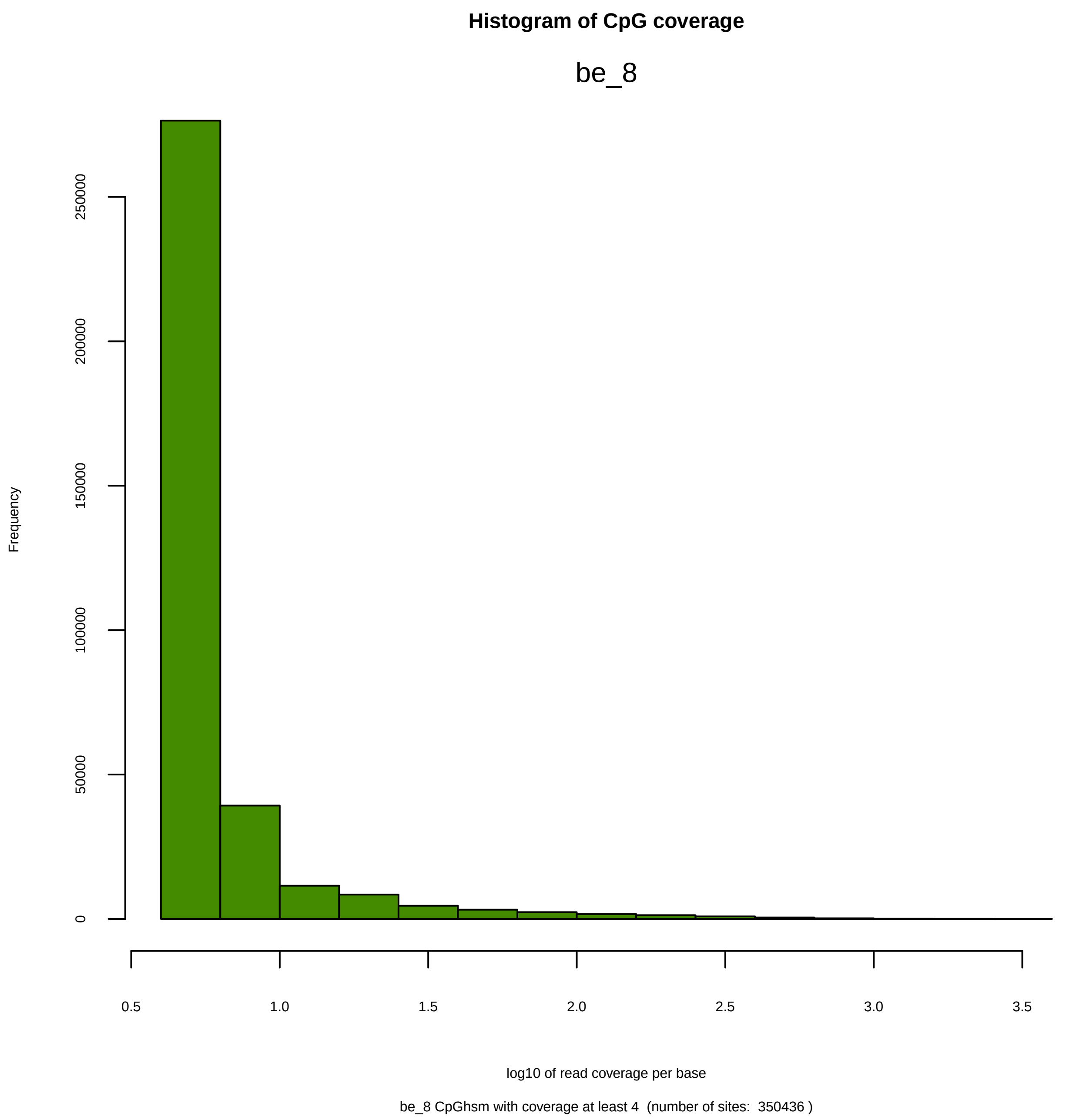
Histogram of CpG coverage
be_8
Frequency
0
50000
100000
150000
200000
250000
0.5
1.0
1.5
2.0
2.5
3.0
3.5
log10 of read coverage per base
be_8 CpGhsm with coverage at least 4  (number of sites:  350436 )

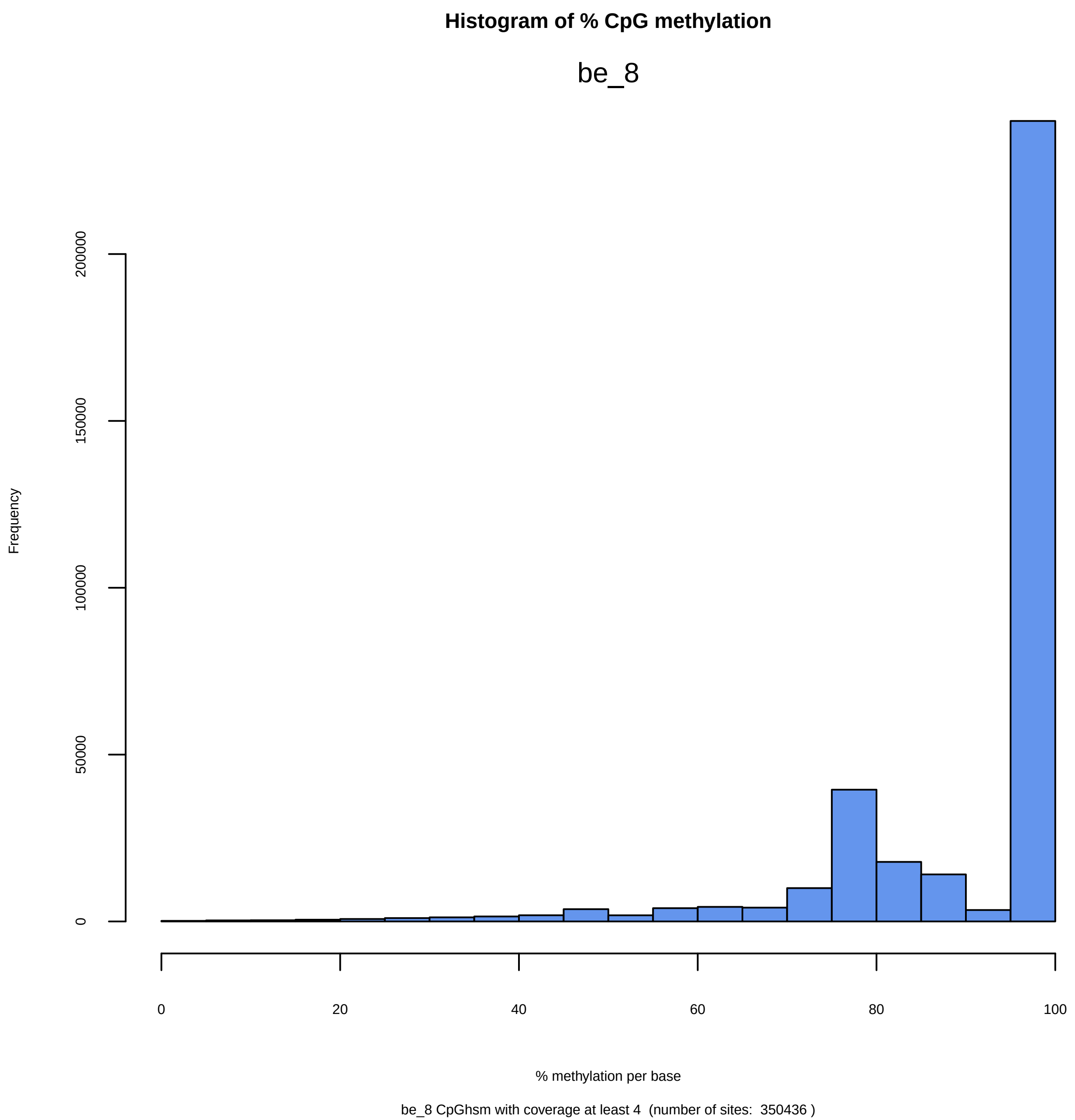
Histogram of % CpG methylation
be_8
Frequency
0
50000
100000
150000
200000
0
20
40
60
80
100
% methylation per base
be_8 CpGhsm with coverage at least 4  (number of sites:  350436 )

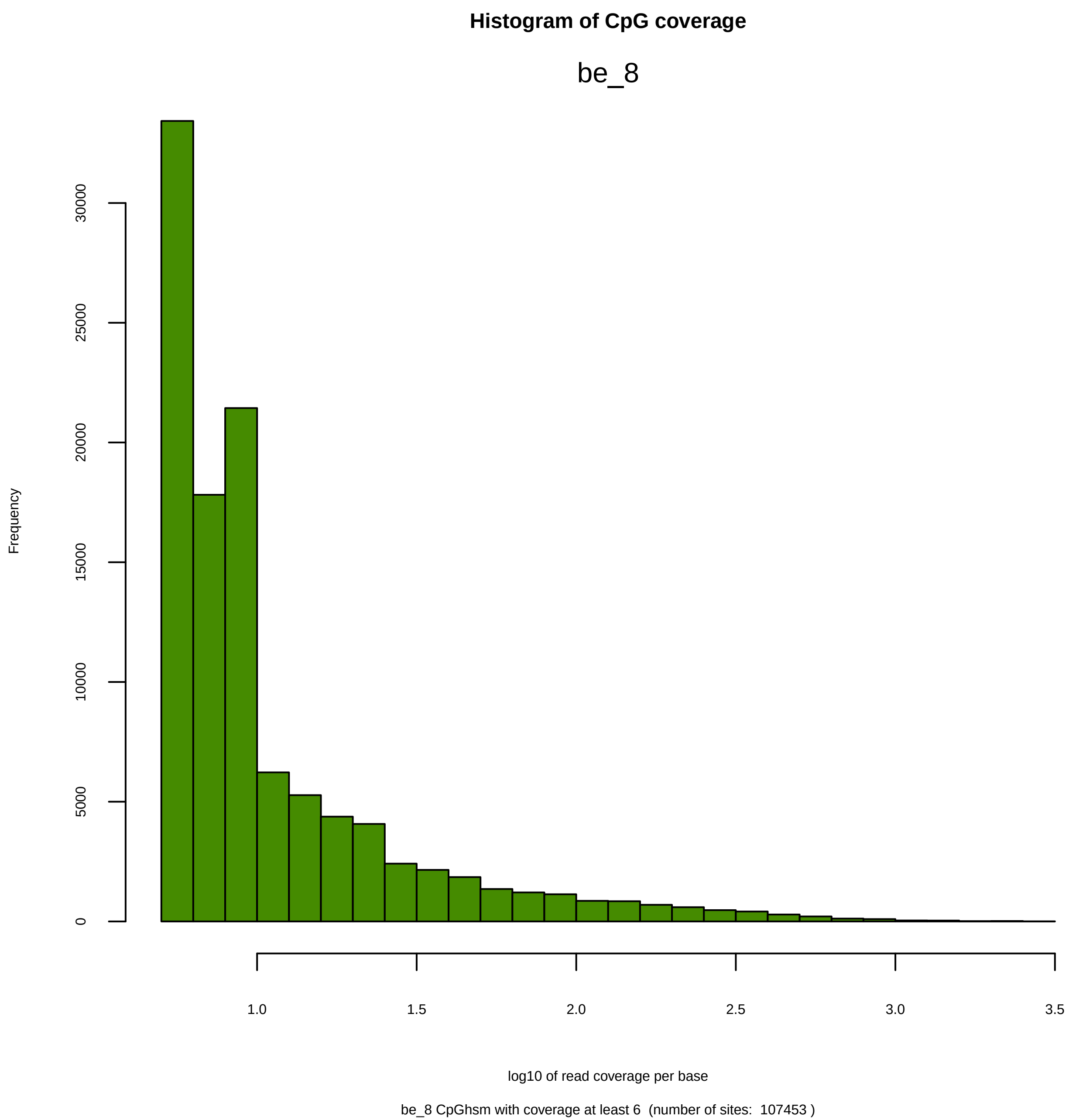

Histogram of CpG coverage
be_8
Frequency
0
5000
10000
15000
20000
25000
30000
1.0
1.5
2.0
2.5
3.0
3.5
log10 of read coverage per base
be_8 CpGhsm with coverage at least 6 (number of sites: 107453 )

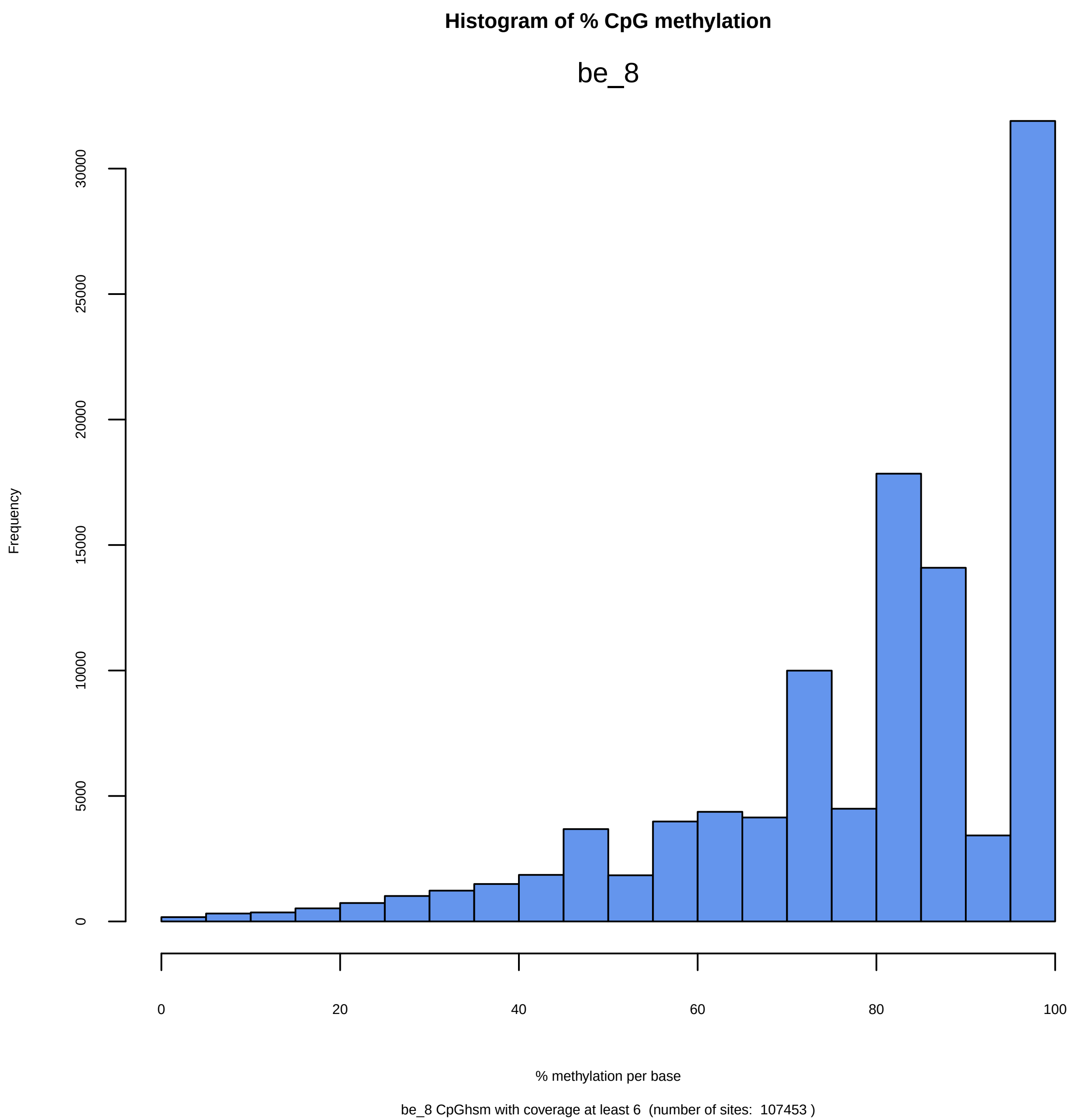

Histogram of % CpG methylation
be_8
Frequency
0
5000
10000
15000
20000
25000
30000
0
20
40
60
80
100
% methylation per base
be_8 CpGhsm with coverage at least 6  (number of sites:  107453 )

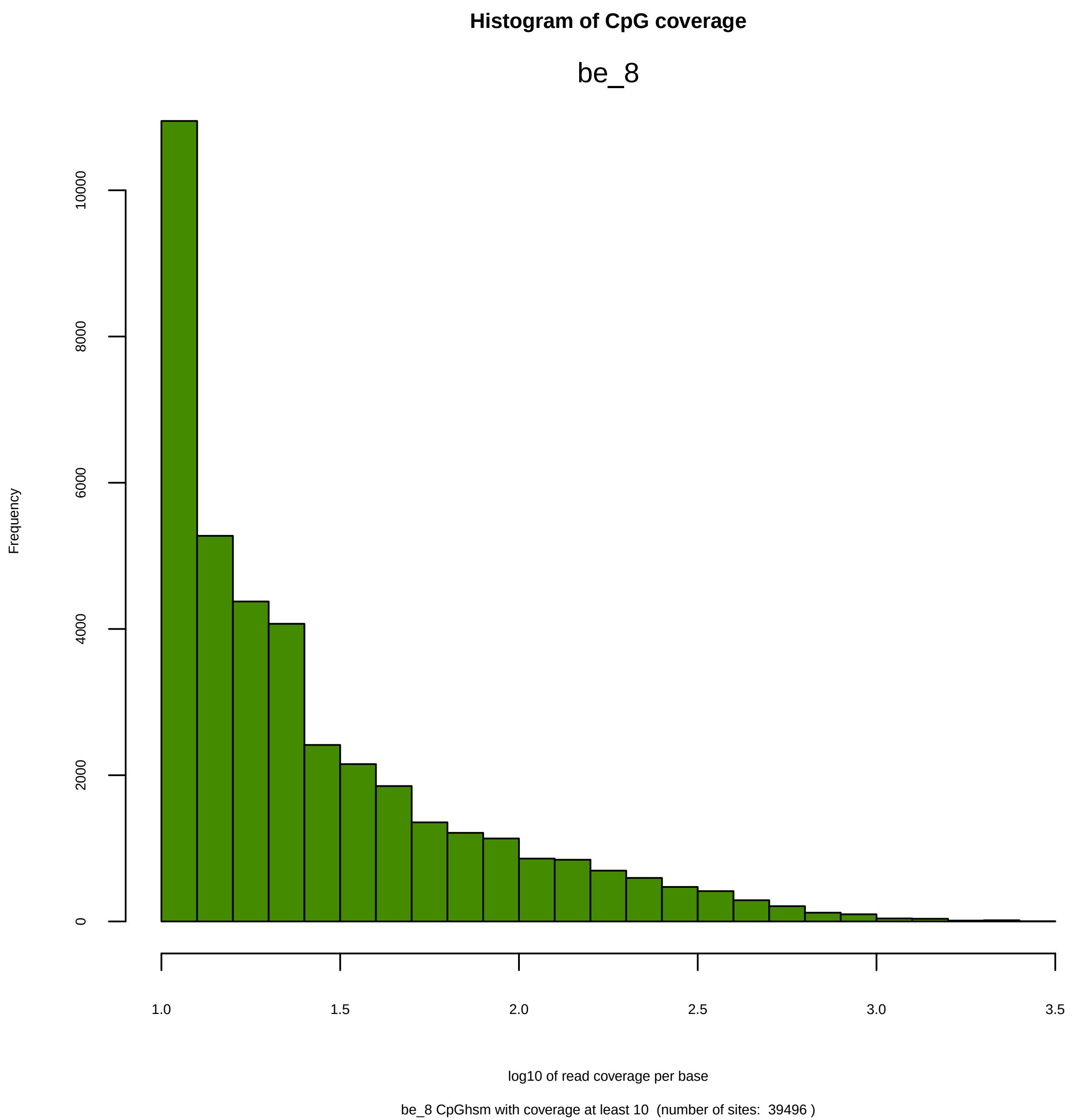
Histogram of CpG coverage
be_8
Frequency
0
2000
4000
6000
8000
10000
1.0
1.5
2.0
2.5
3.0
3.5
log10 of read coverage per base
be_8 CpGhsm with coverage at least 10  (number of sites:  39496 )

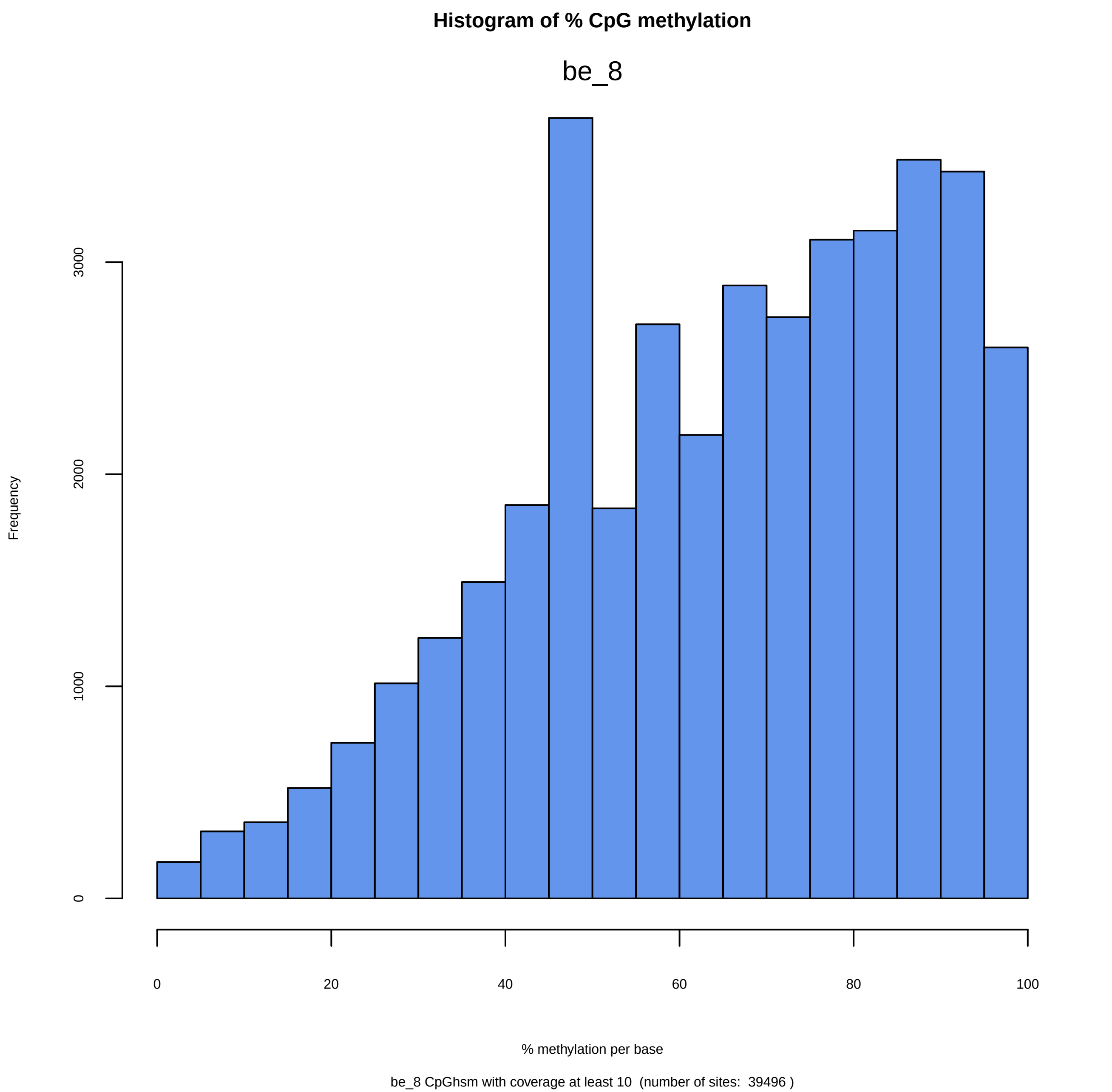
Histogram of % CpG methylation
be_8
Frequency
0
1000
2000
3000
0
20
40
60
80
100
% methylation per base
be_8 CpGhsm with coverage at least 10 (number of sites: 39496 )

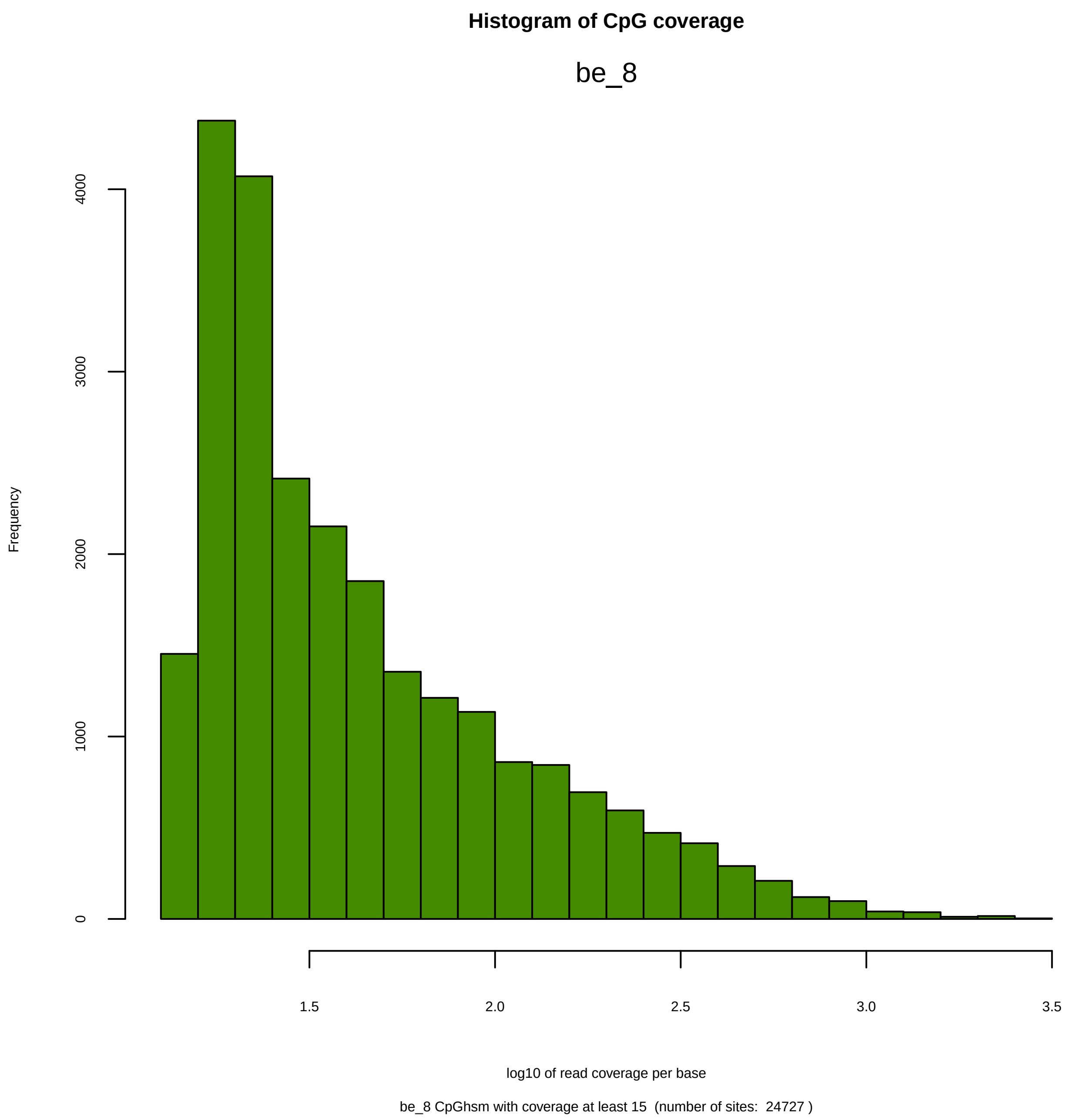

Histogram of CpG coverage
be_8
Frequency
0
1000
2000
3000
4000
1.5
2.0
2.5
3.0
3.5
log10 of read coverage per base
be_8 CpGhsm with coverage at least 15 (number of sites: 24727 )

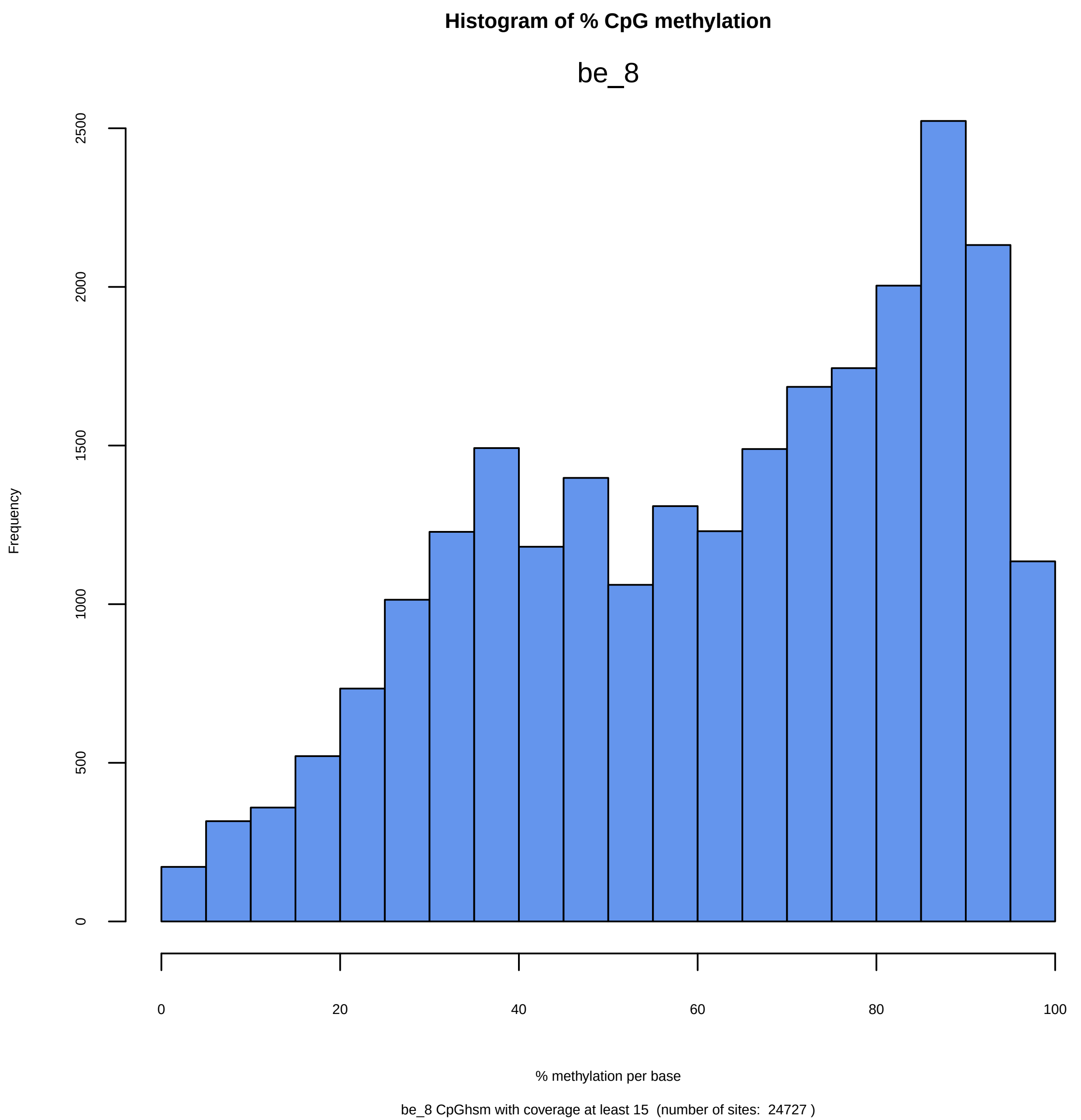
Histogram of % CpG methylation
be_8
Frequency
0
500
1000
1500
2000
2500
0
20
40
60
80
100
% methylation per base
be_8 CpGhsm with coverage at least 15  (number of sites:  24727 )

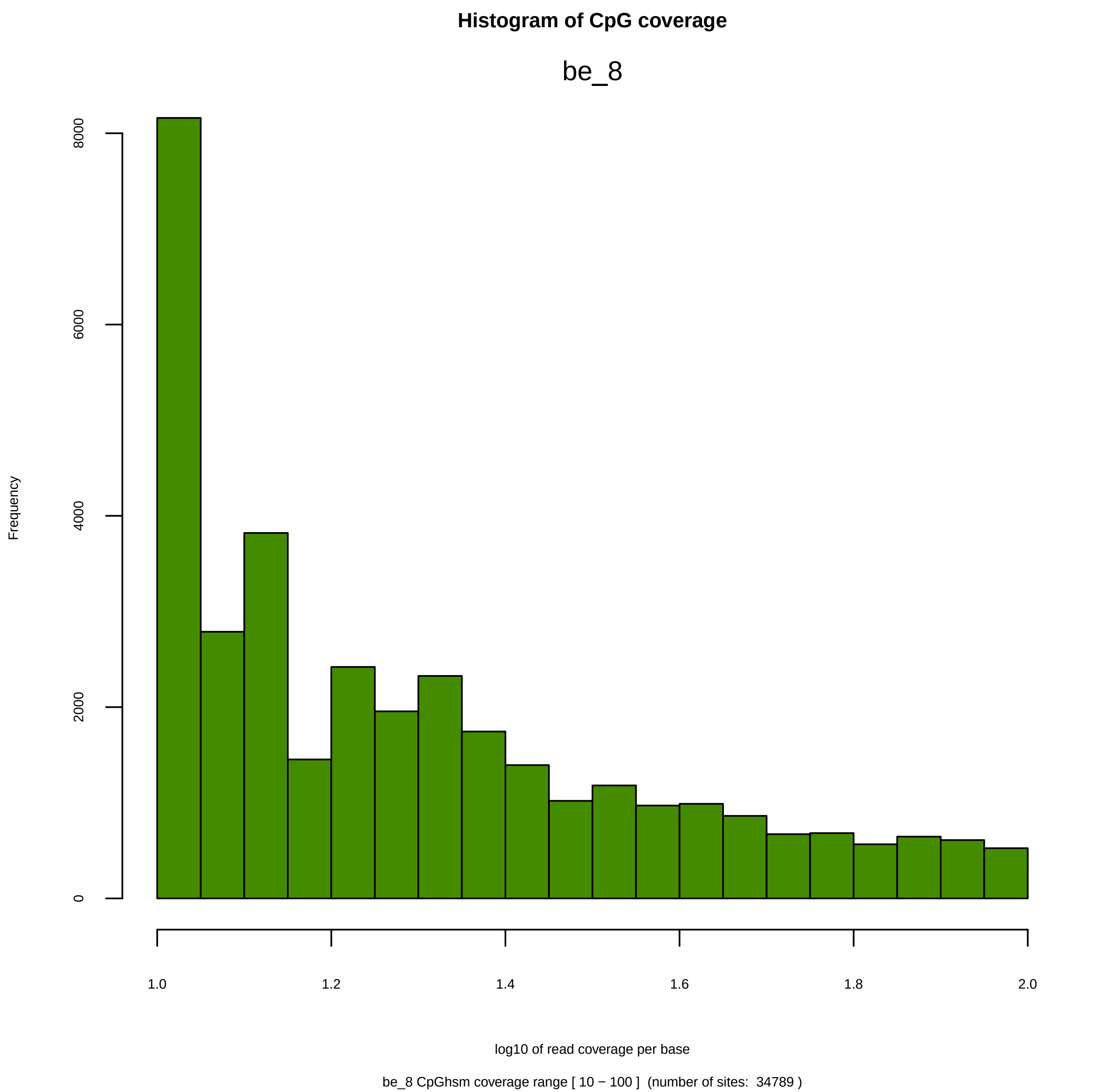

Histogram of CpG coverage
be_8
Frequency
0
2000
4000
6000
8000
1.0
1.2
1.4
1.6
1.8
2.0
log10 of read coverage per base
be_8 CpGhsm coverage range [ 10 − 100 ]  (number of sites:  34789 )

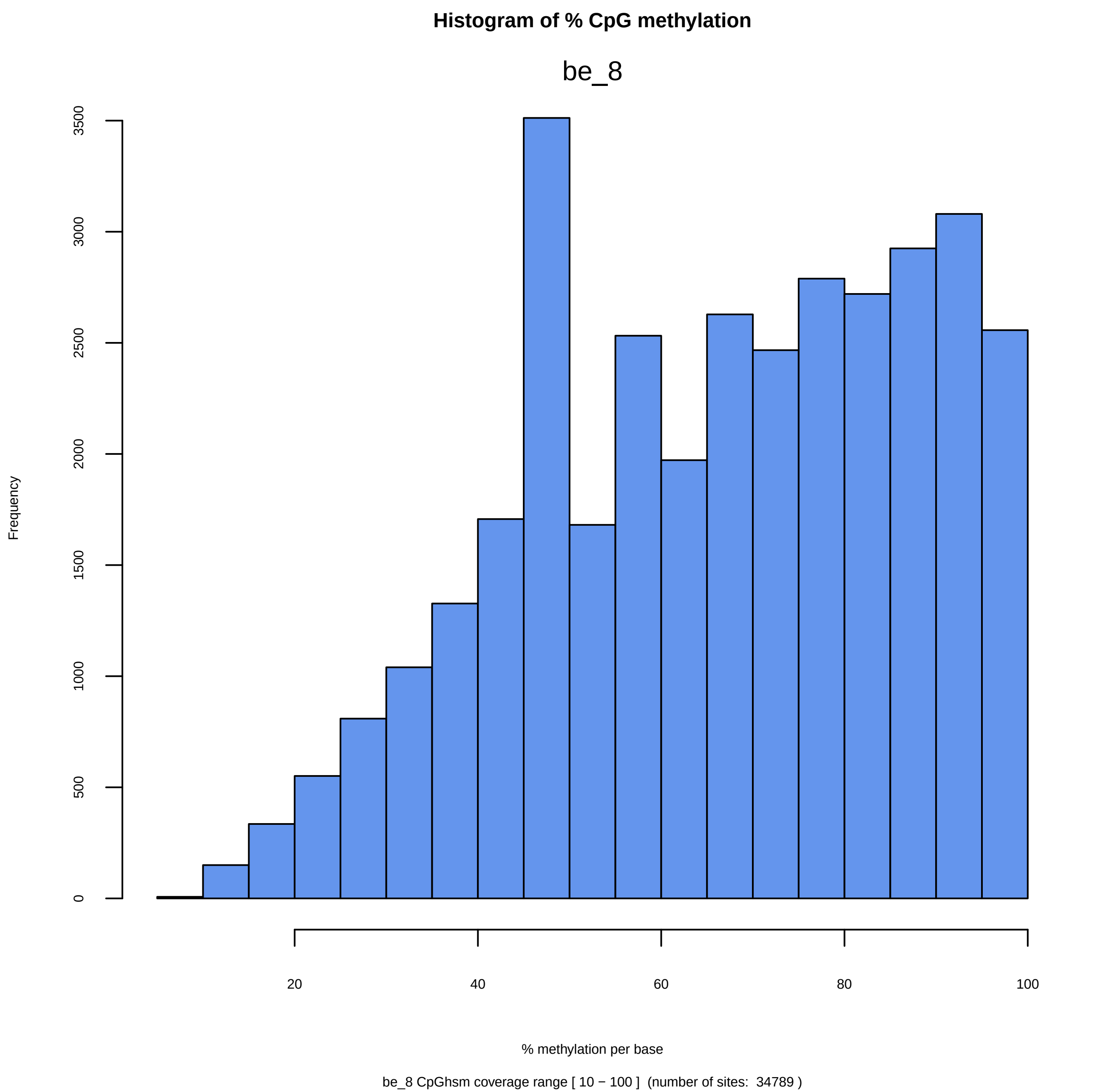

Histogram of % CpG methylation
be_8
Frequency
0
500
1000
1500
2000
2500
3000
3500
20
40
60
80
100
% methylation per base
be_8 CpGhsm coverage range [ 10 − 100 ] (number of sites: 34789 )

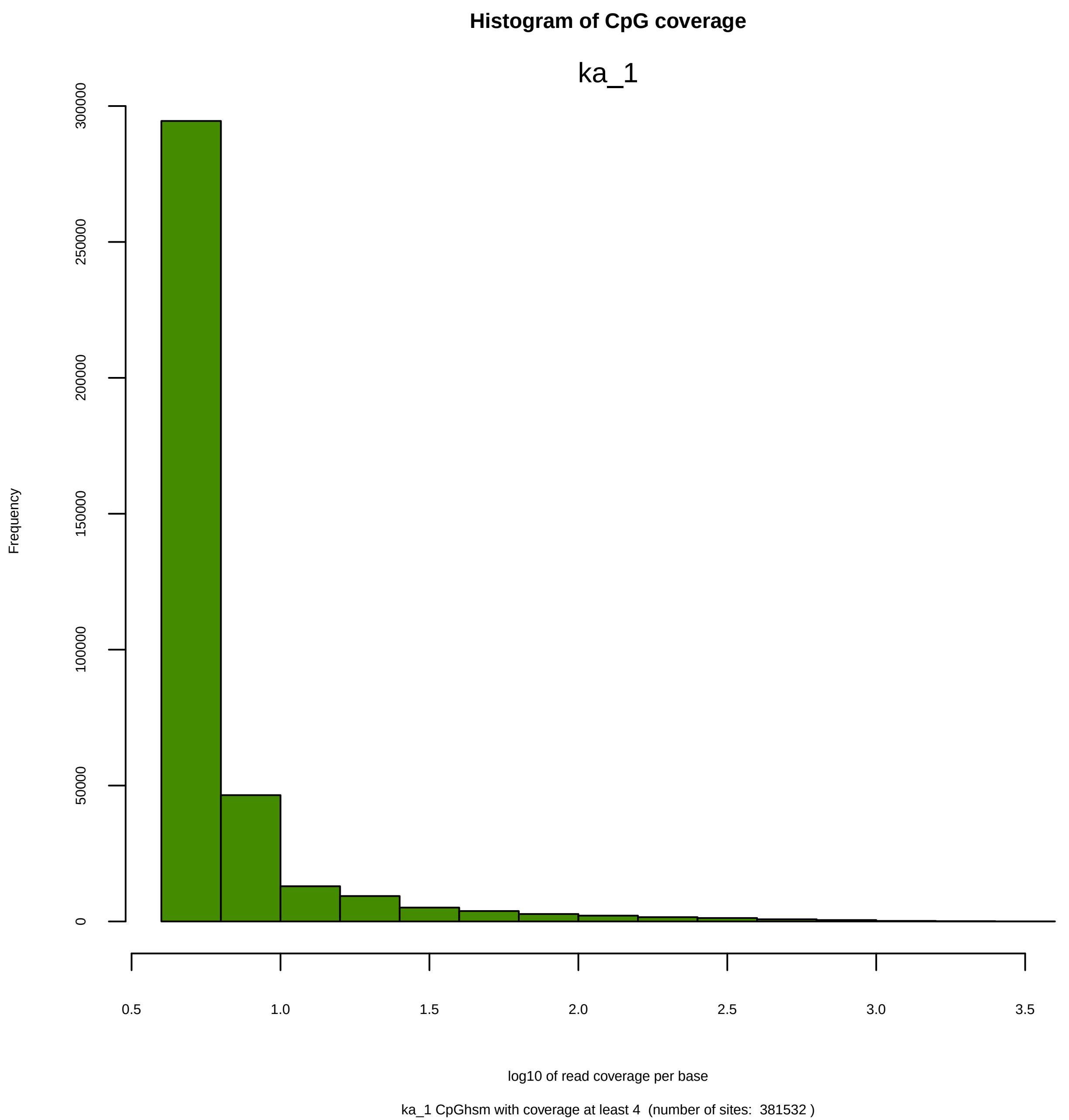

Histogram of CpG coverage
ka_1
Frequency
0
50000
100000
150000
200000
250000
300000
0.5
1.0
1.5
2.0
2.5
3.0
3.5
log10 of read coverage per base
ka_1 CpGhsm with coverage at least 4  (number of sites:  381532 )

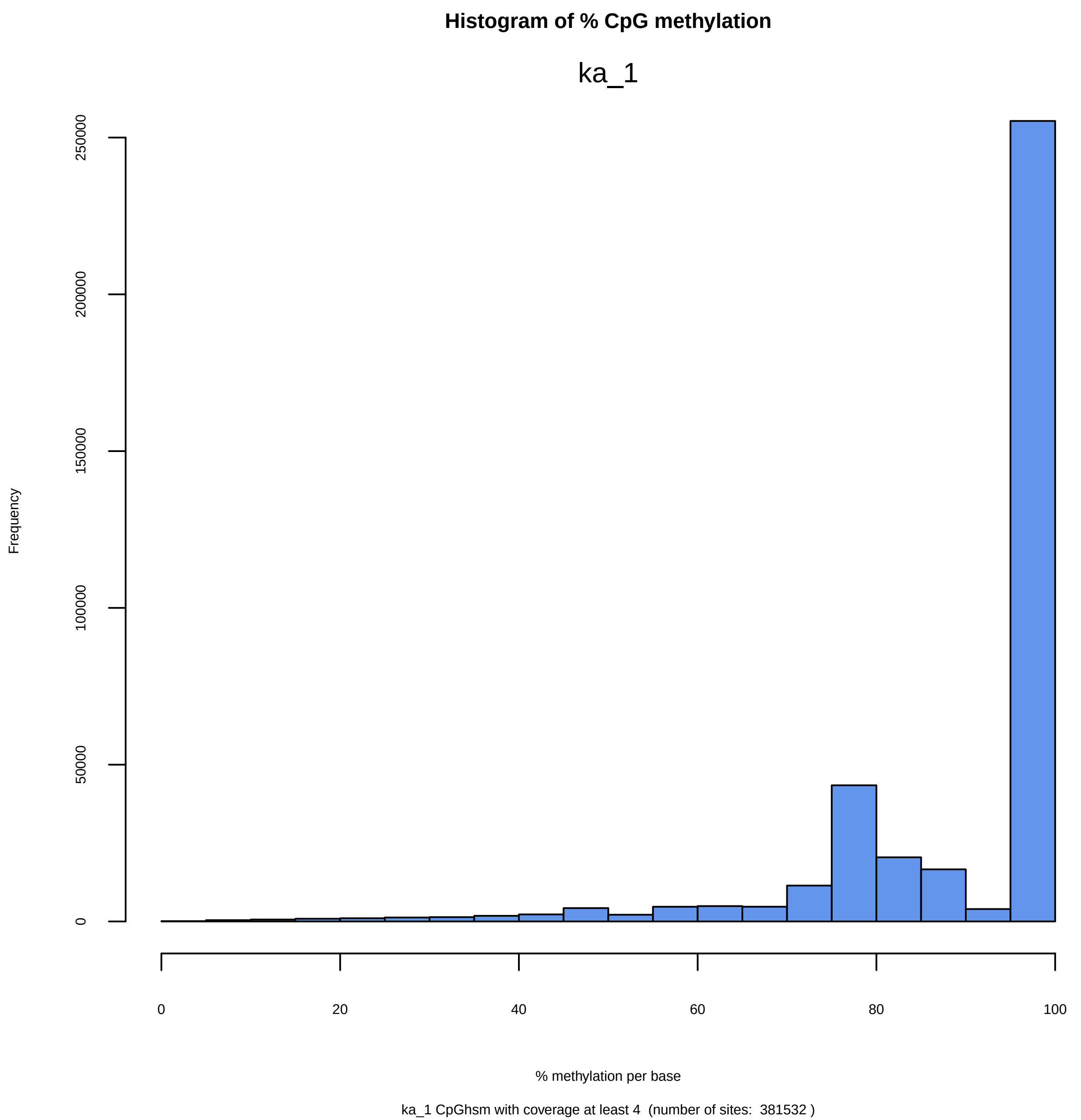

Histogram of % CpG methylation
ka_1
Frequency
0
50000
100000
150000
200000
250000
0
20
40
60
80
100
% methylation per base
ka_1 CpGhsm with coverage at least 4 (number of sites: 381532 )

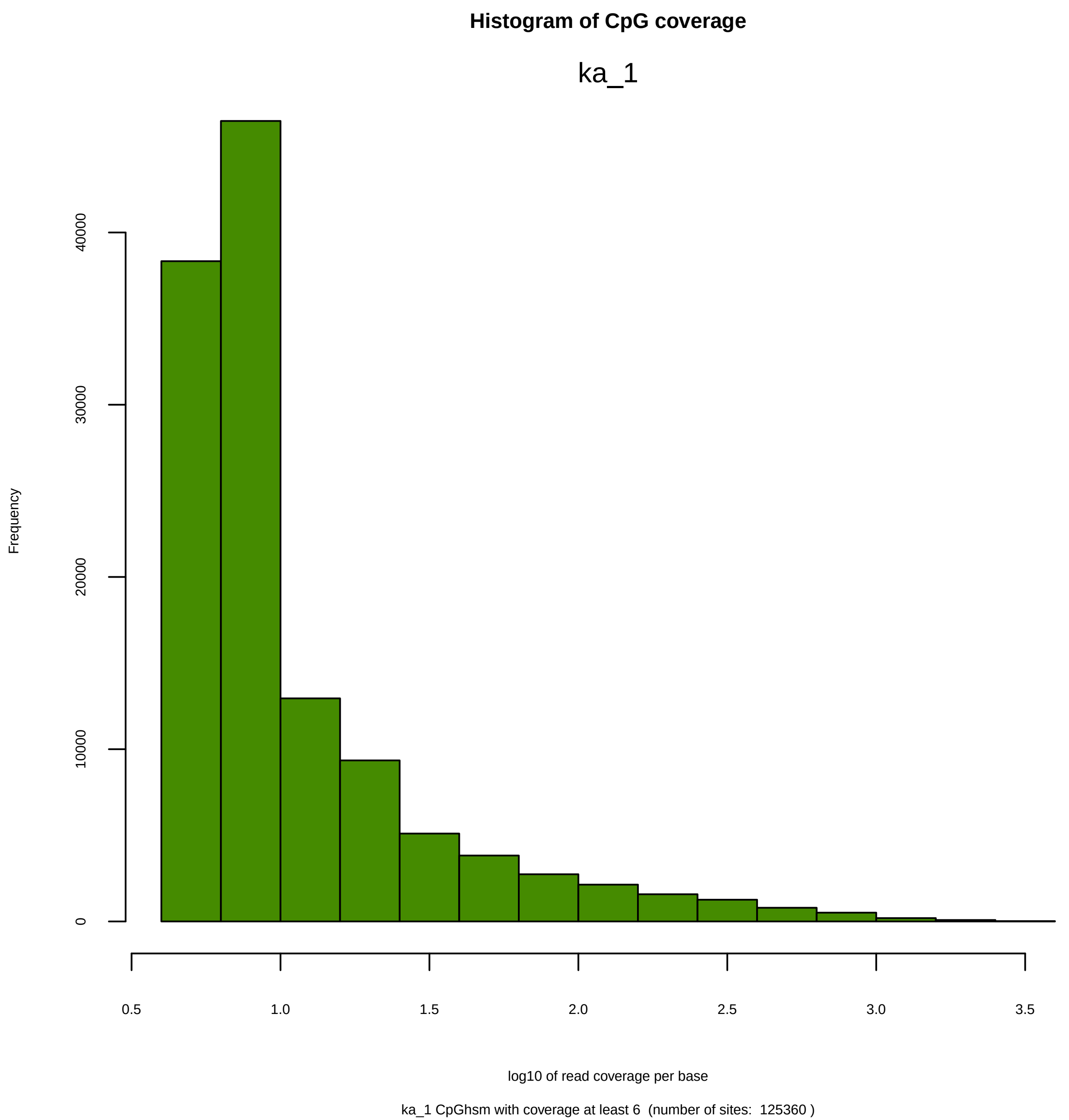
Histogram of CpG coverage
ka_1
Frequency
0
10000
20000
30000
40000
0.5
1.0
1.5
2.0
2.5
3.0
3.5
log10 of read coverage per base
ka_1 CpGhsm with coverage at least 6 (number of sites: 125360 )

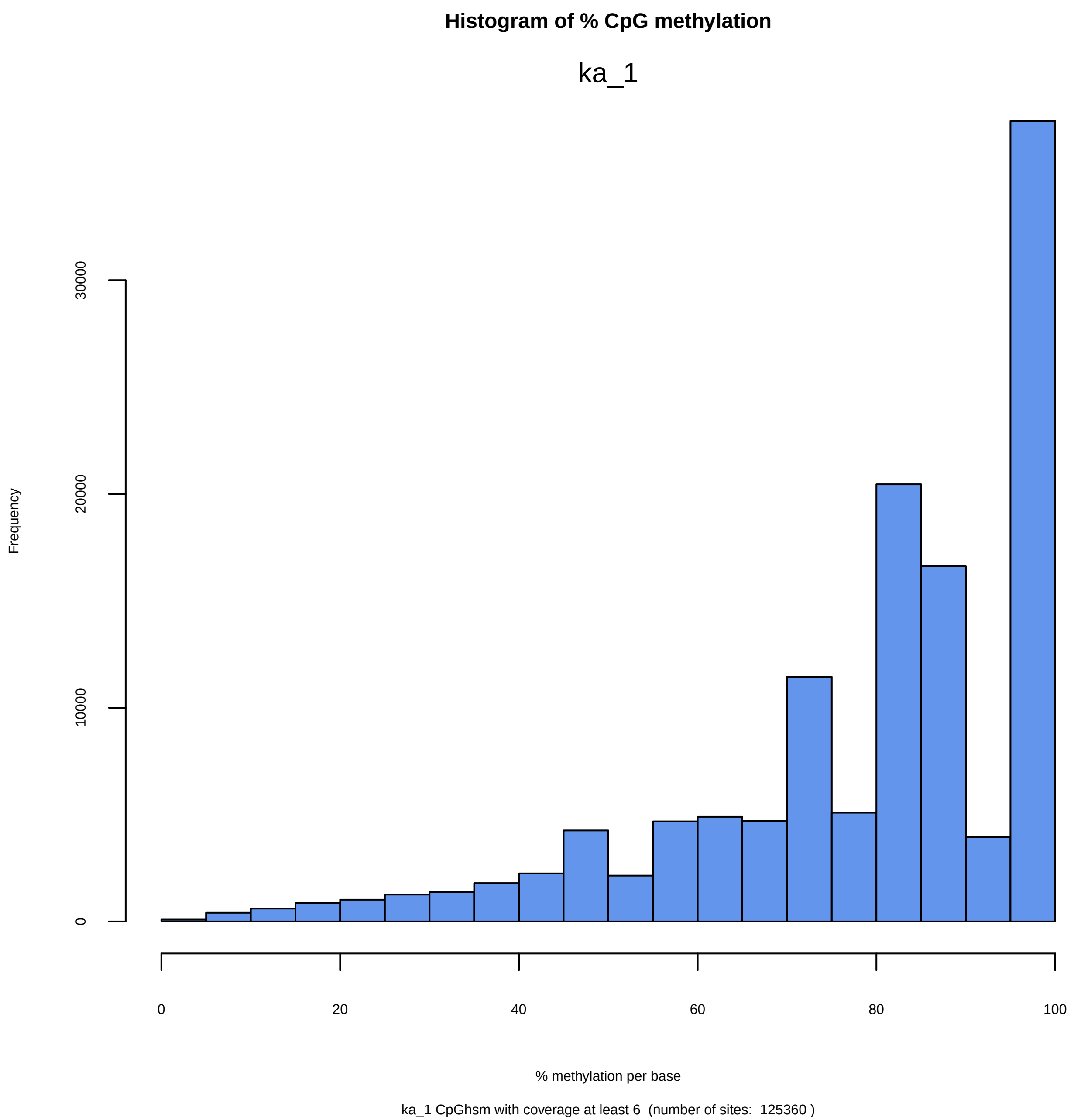
Histogram of % CpG methylation
ka_1
Frequency
0
10000
20000
30000
0
20
40
60
80
100
% methylation per base
ka_1 CpGhsm with coverage at least 6 (number of sites: 125360 )

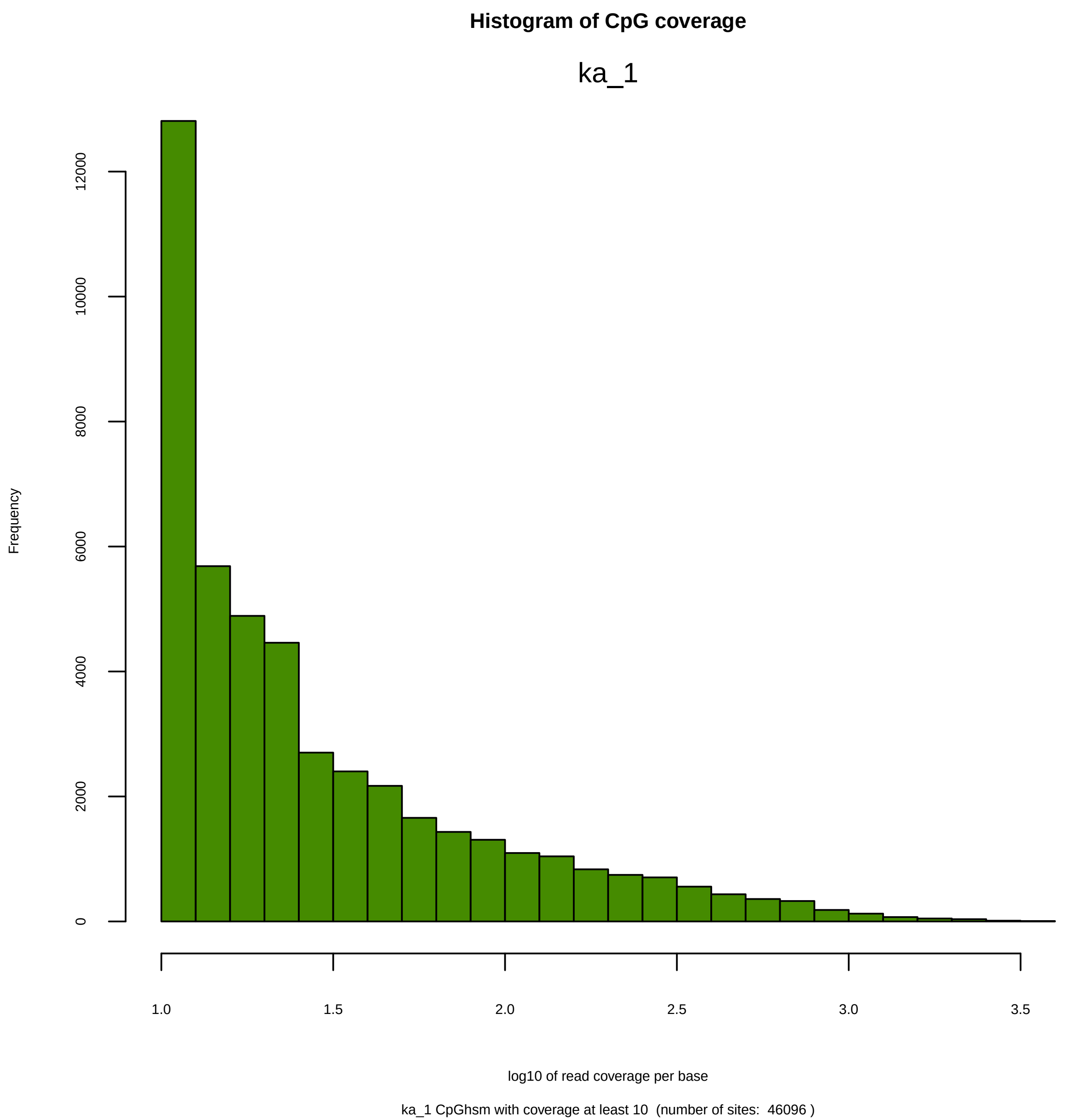

Histogram of CpG coverage
ka_1
Frequency
0
2000
4000
6000
8000
10000
12000
1.0
1.5
2.0
2.5
3.0
3.5
log10 of read coverage per base
ka_1 CpGhsm with coverage at least 10  (number of sites:  46096 )

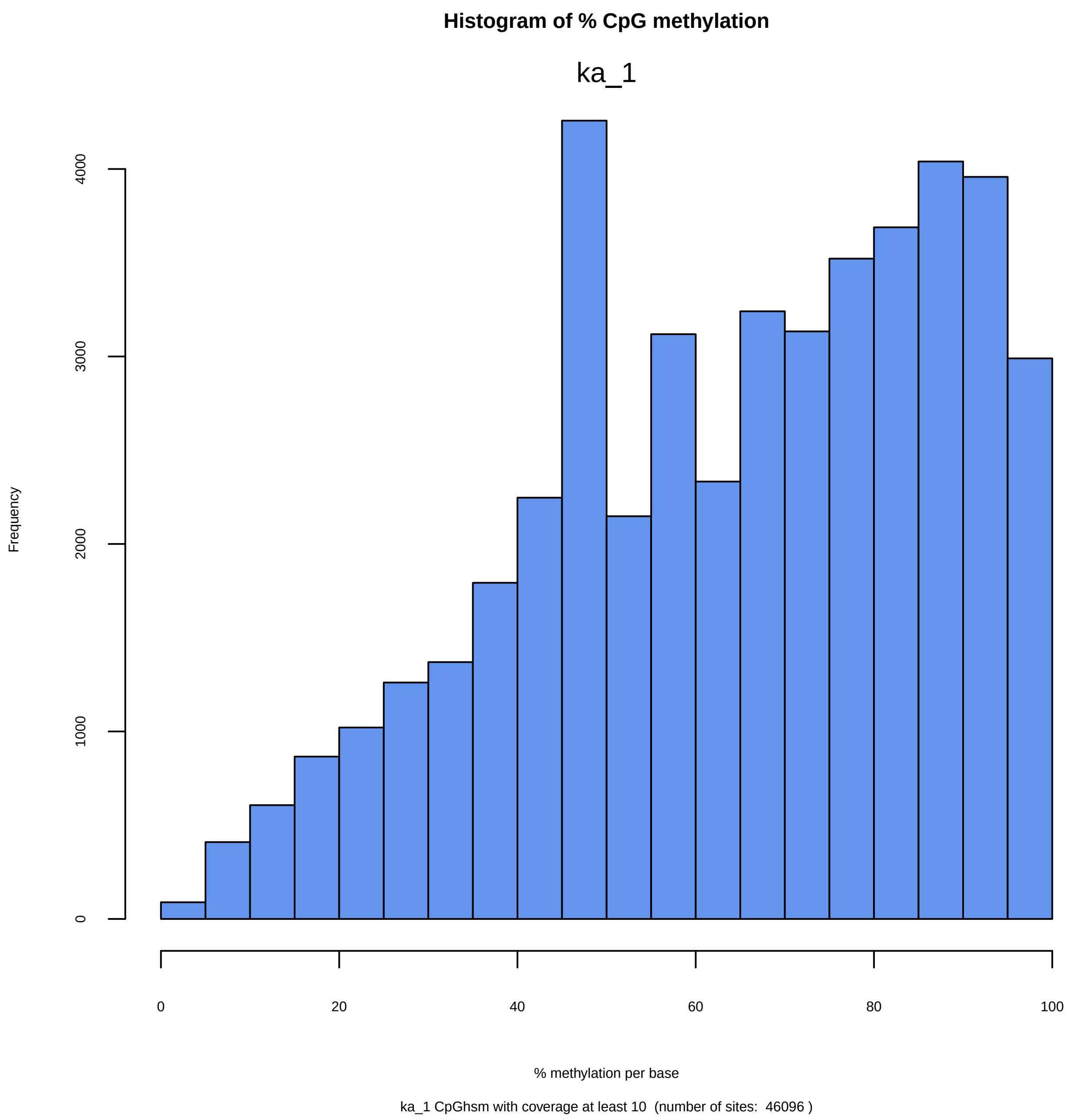
Histogram of % CpG methylation
ka_1
Frequency
0
1000
2000
3000
4000
0
20
40
60
80
100
% methylation per base
ka_1 CpGhsm with coverage at least 10  (number of sites:  46096 )

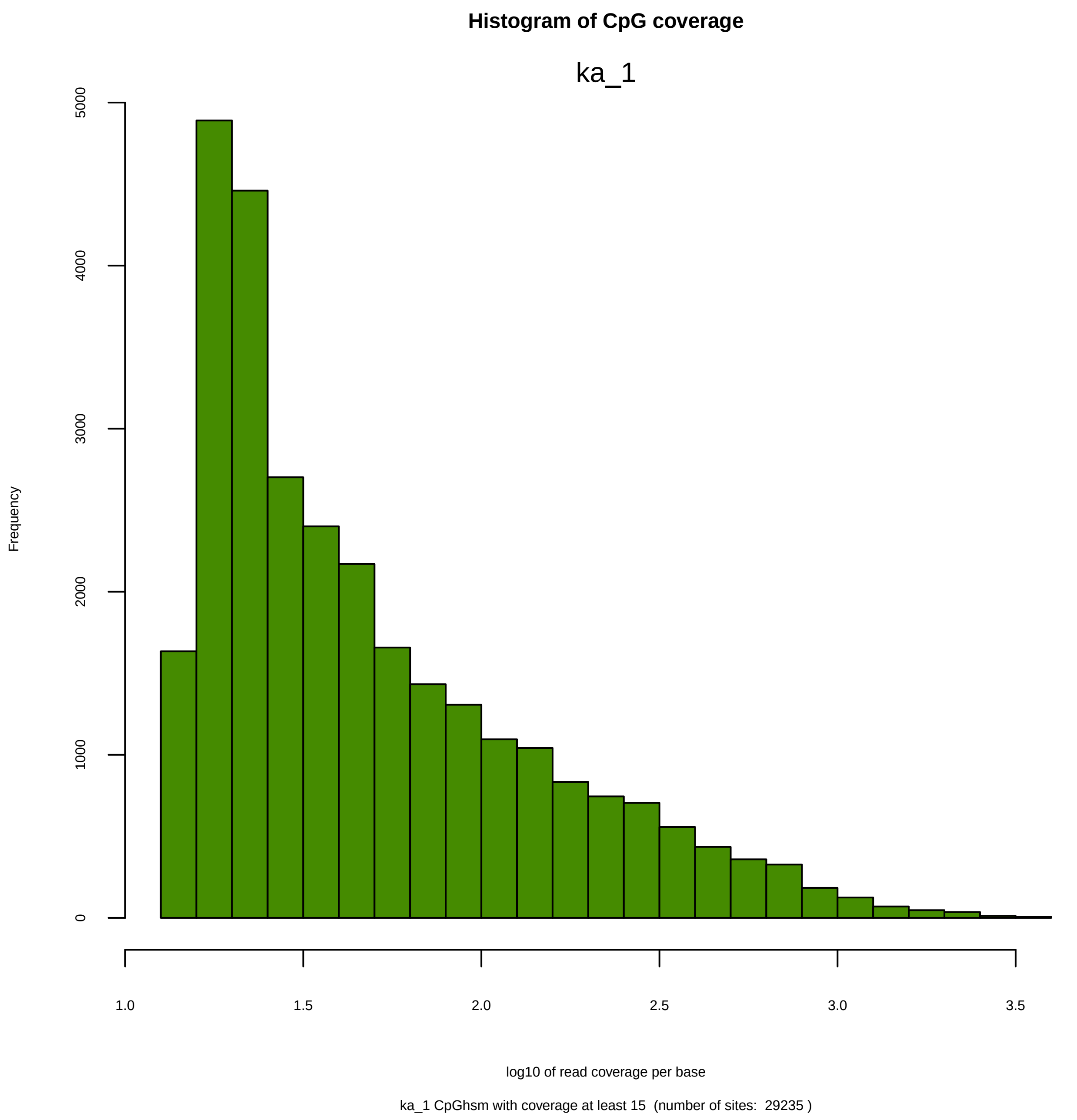
Histogram of CpG coverage
ka_1
Frequency
0
1000
2000
3000
4000
5000
1.0
1.5
2.0
2.5
3.0
3.5
log10 of read coverage per base
ka_1 CpGhsm with coverage at least 15  (number of sites:  29235 )

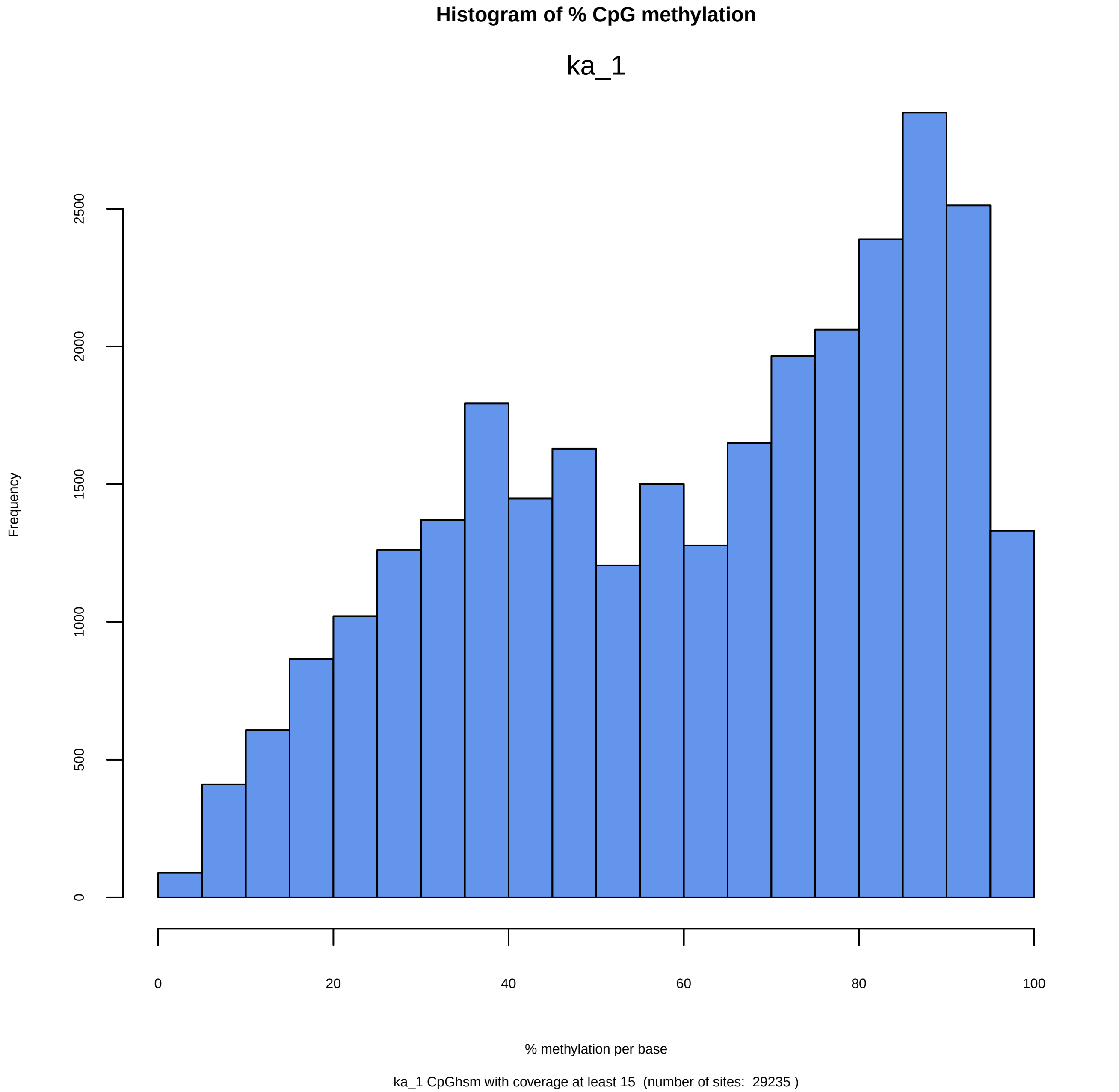

Histogram of % CpG methylation
ka_1
Frequency
0
500
1000
1500
2000
2500
0
20
40
60
80
100
% methylation per base
ka_1 CpGhsm with coverage at least 15 (number of sites: 29235 )

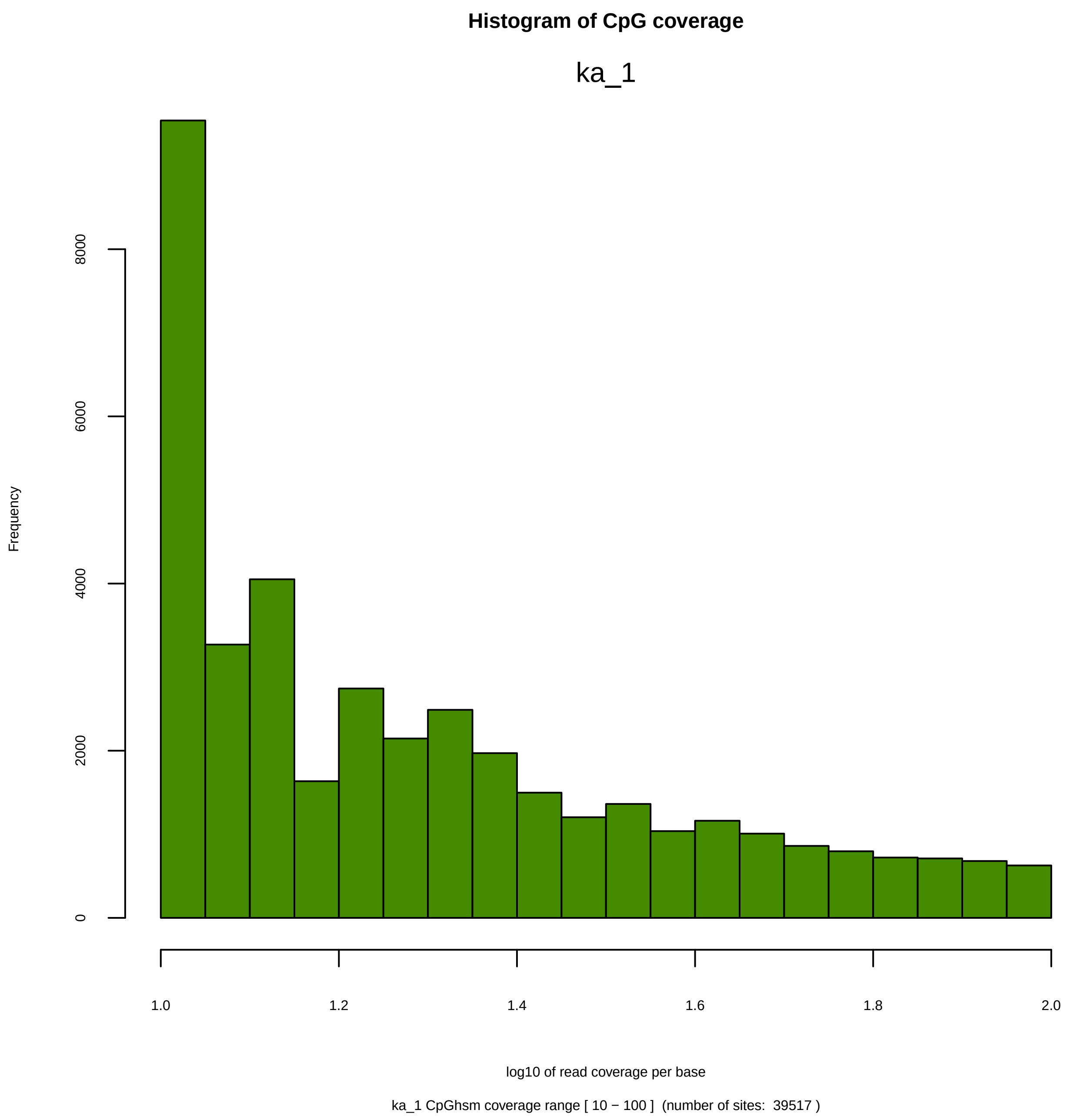
Histogram of CpG coverage
ka_1
Frequency
0
2000
4000
6000
8000
1.0
1.2
1.4
1.6
1.8
2.0
log10 of read coverage per base
ka_1 CpGhsm coverage range [ 10 − 100 ]  (number of sites:  39517 )

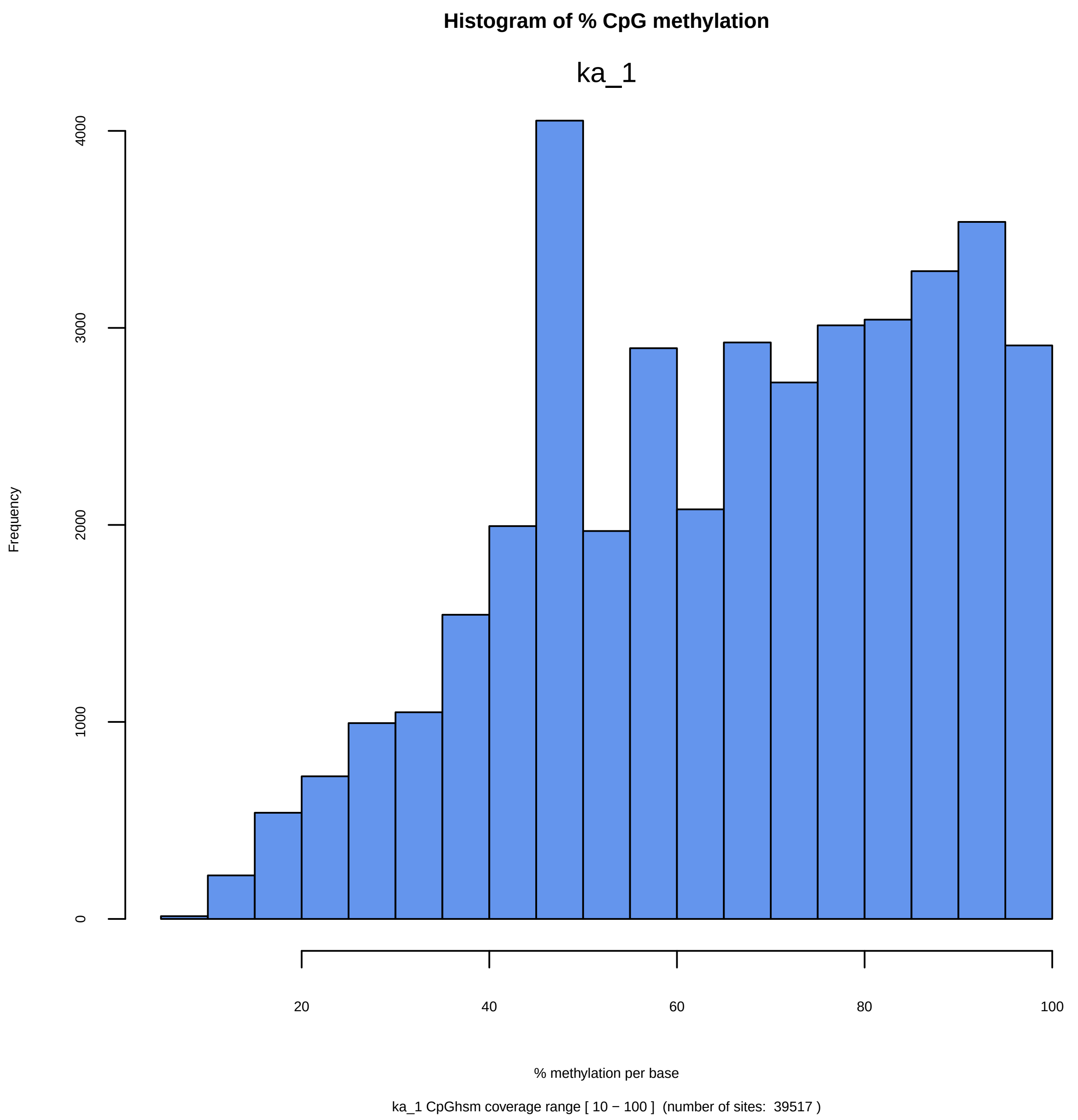

Histogram of % CpG methylation
ka_1
Frequency
0
1000
2000
3000
4000
20
40
60
80
100
% methylation per base
ka_1 CpGhsm coverage range [ 10 − 100 ] (number of sites: 39517 )

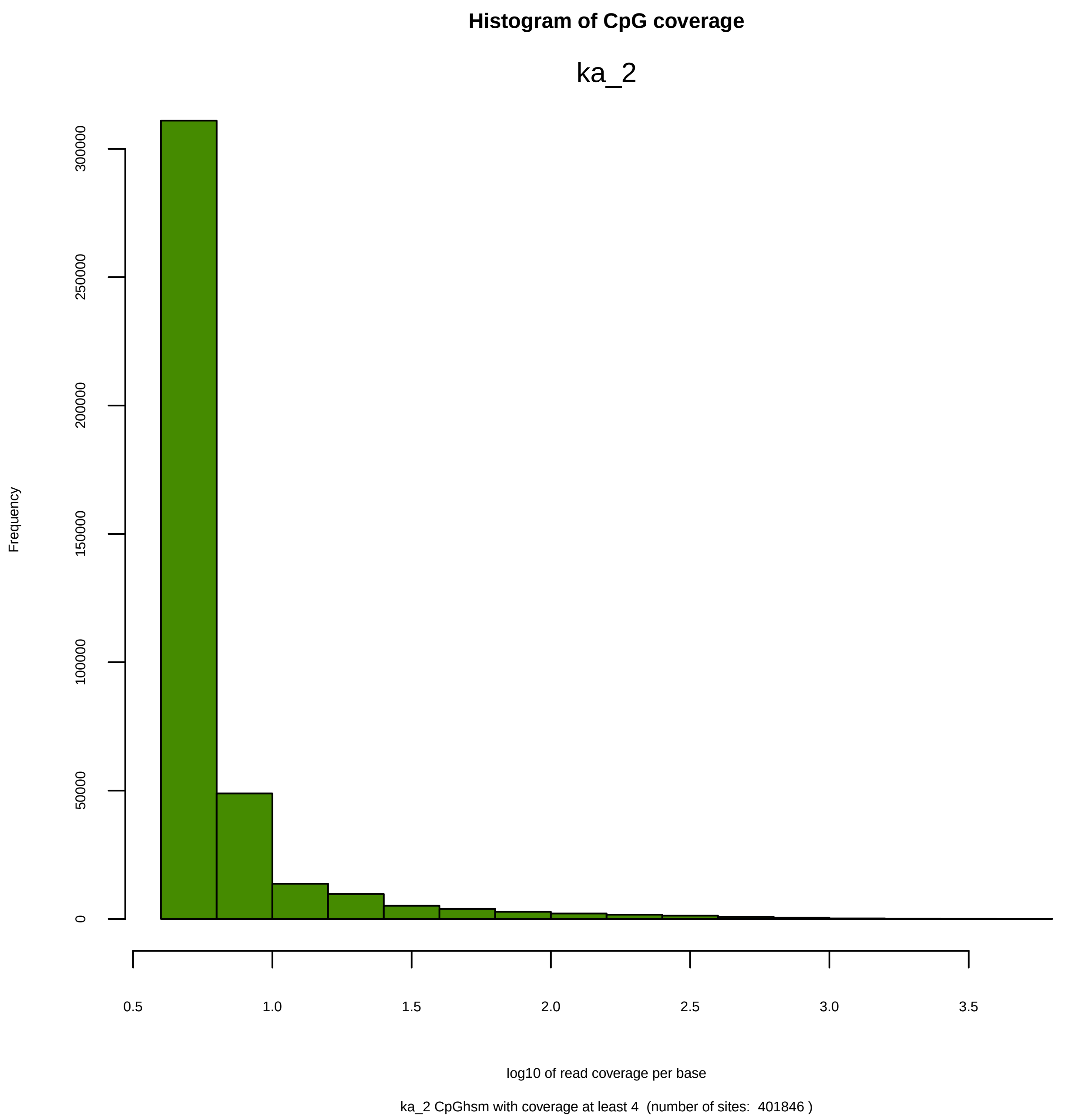
Histogram of CpG coverage
ka_2
Frequency
0
50000
100000
150000
200000
250000
300000
0.5
1.0
1.5
2.0
2.5
3.0
3.5
log10 of read coverage per base
ka_2 CpGhsm with coverage at least 4 (number of sites: 401846 )

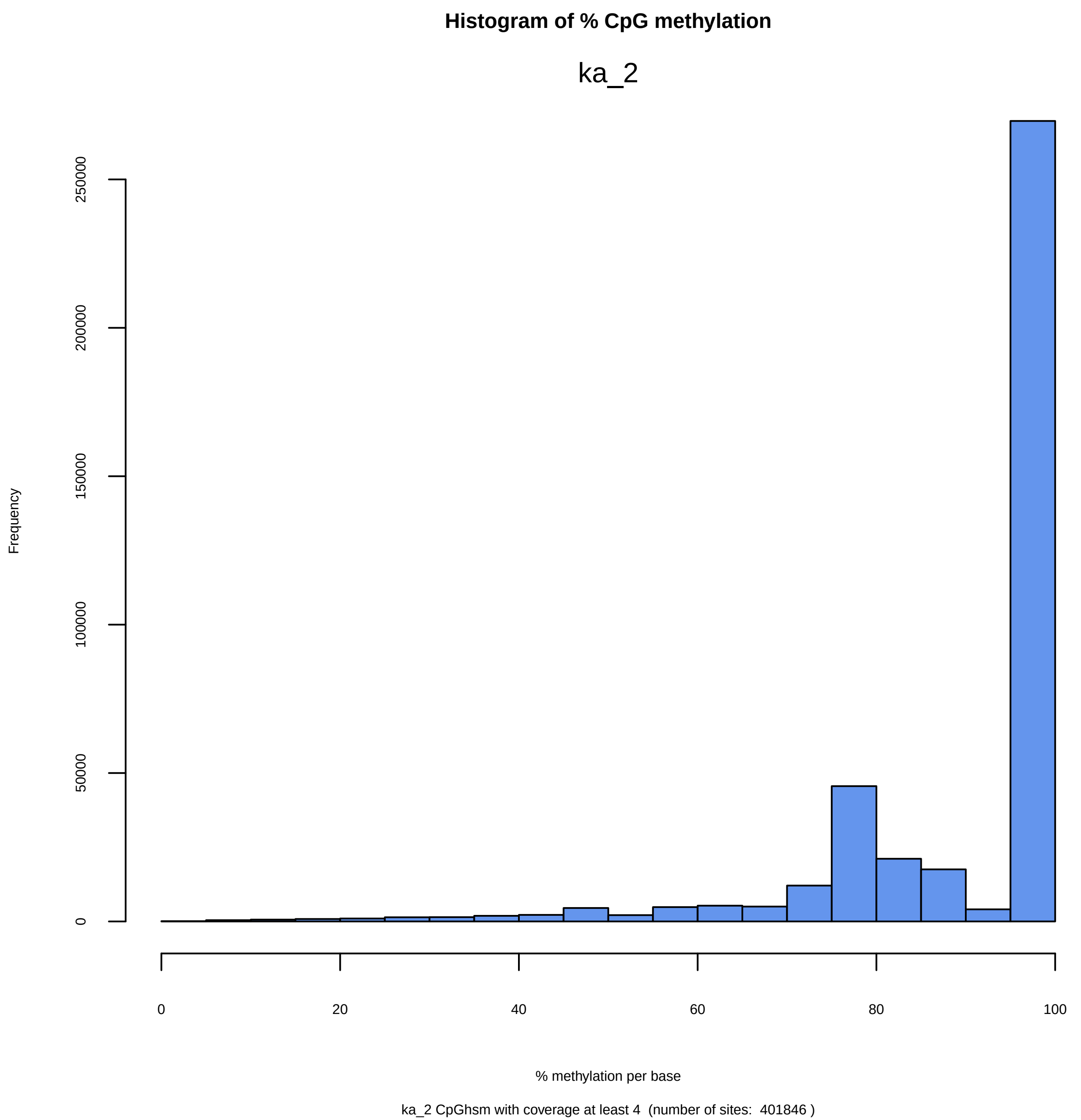

Histogram of % CpG methylation
ka_2
Frequency
0
50000
100000
150000
200000
250000
0
20
40
60
80
100
% methylation per base
ka_2 CpGhsm with coverage at least 4 (number of sites: 401846 )

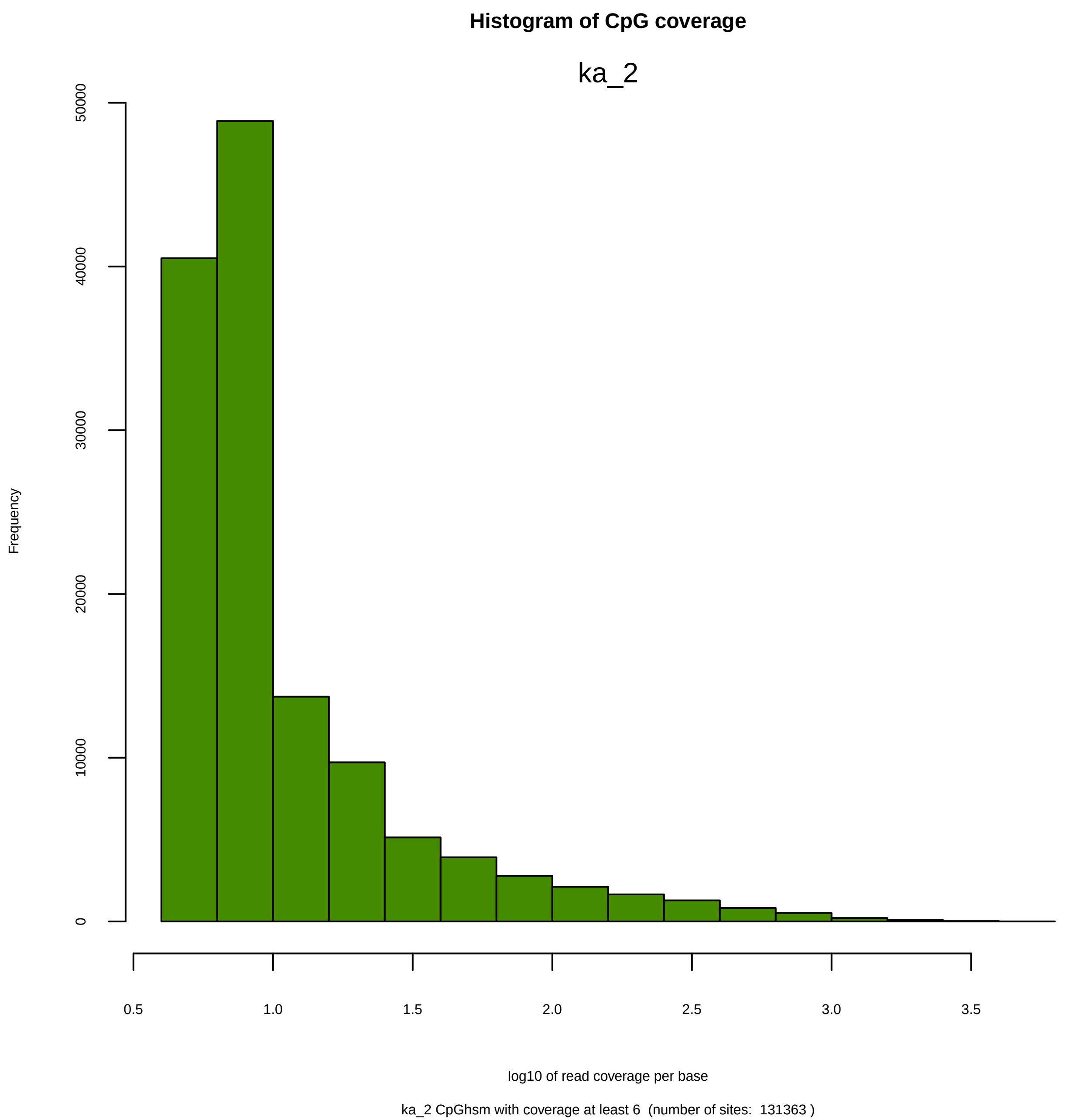
Histogram of CpG coverage
ka_2
Frequency
0
10000
20000
30000
40000
50000
0.5
1.0
1.5
2.0
2.5
3.0
3.5
log10 of read coverage per base
ka_2 CpGhsm with coverage at least 6 (number of sites: 131363 )

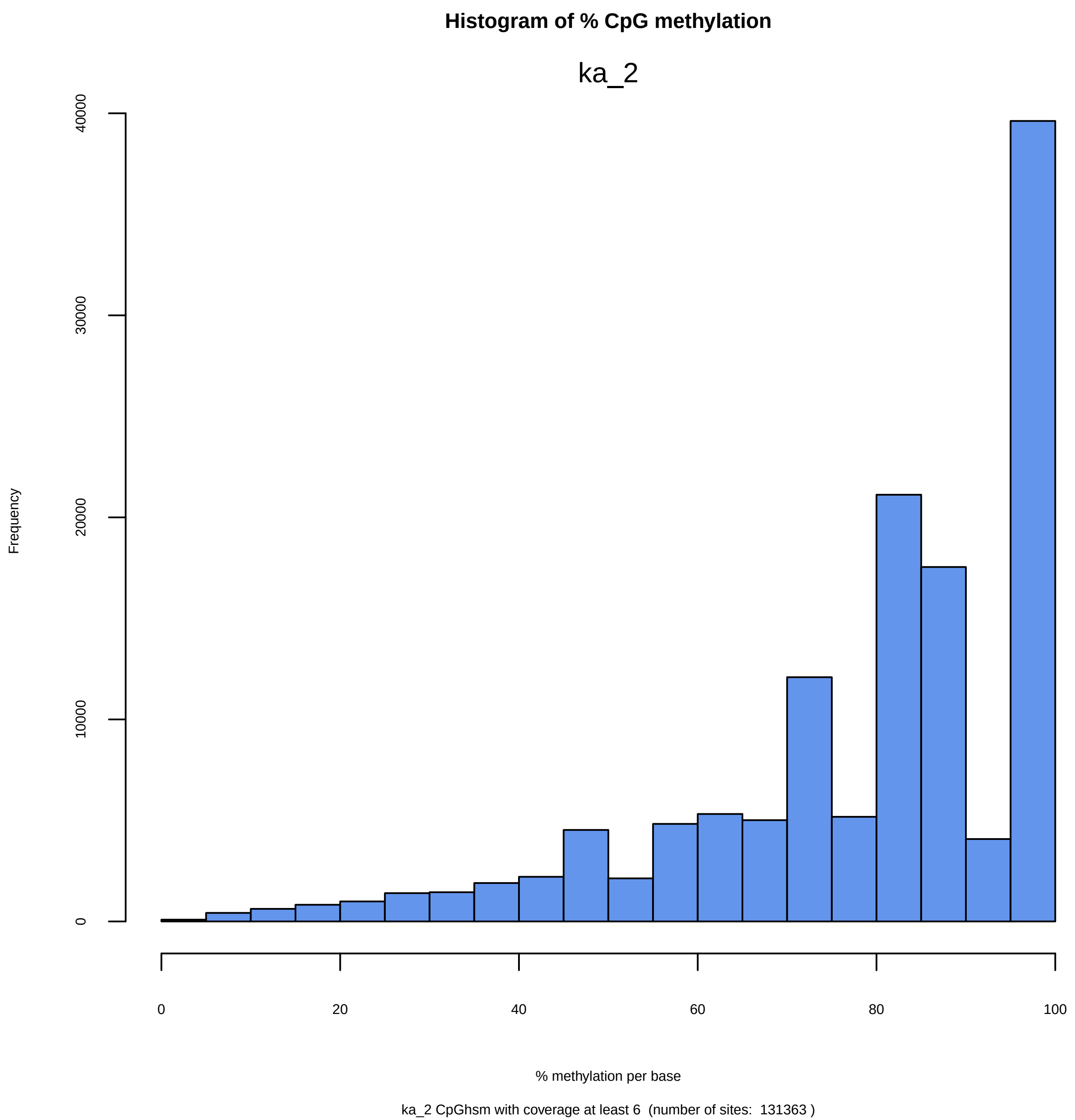
Histogram of % CpG methylation
ka_2
Frequency
0
10000
20000
30000
40000
0
20
40
60
80
100
% methylation per base
ka_2 CpGhsm with coverage at least 6  (number of sites:  131363 )

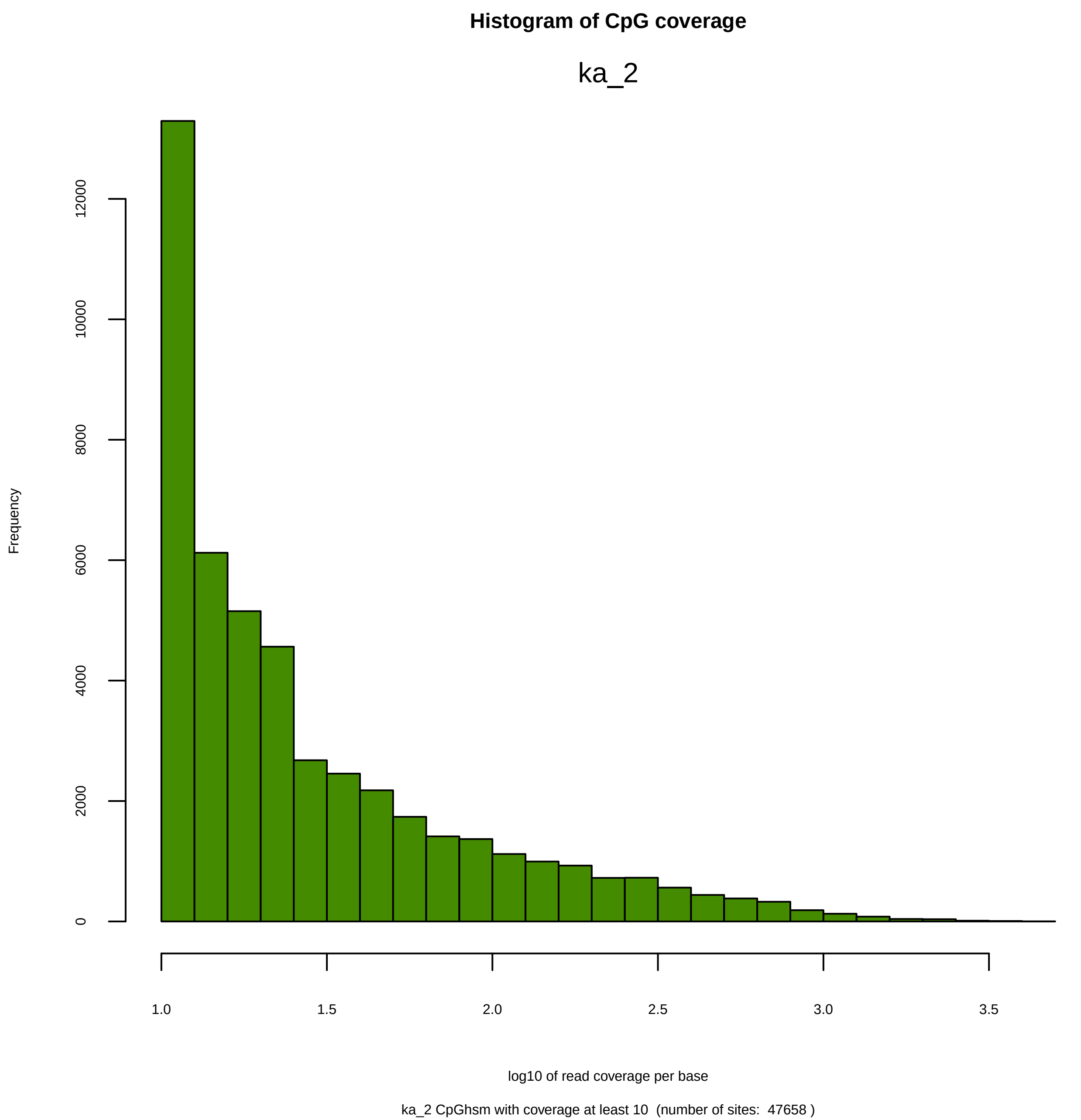

Histogram of CpG coverage
ka_2
Frequency
0
2000
4000
6000
8000
10000
12000
1.0
1.5
2.0
2.5
3.0
3.5
log10 of read coverage per base
ka_2 CpGhsm with coverage at least 10  (number of sites:  47658 )

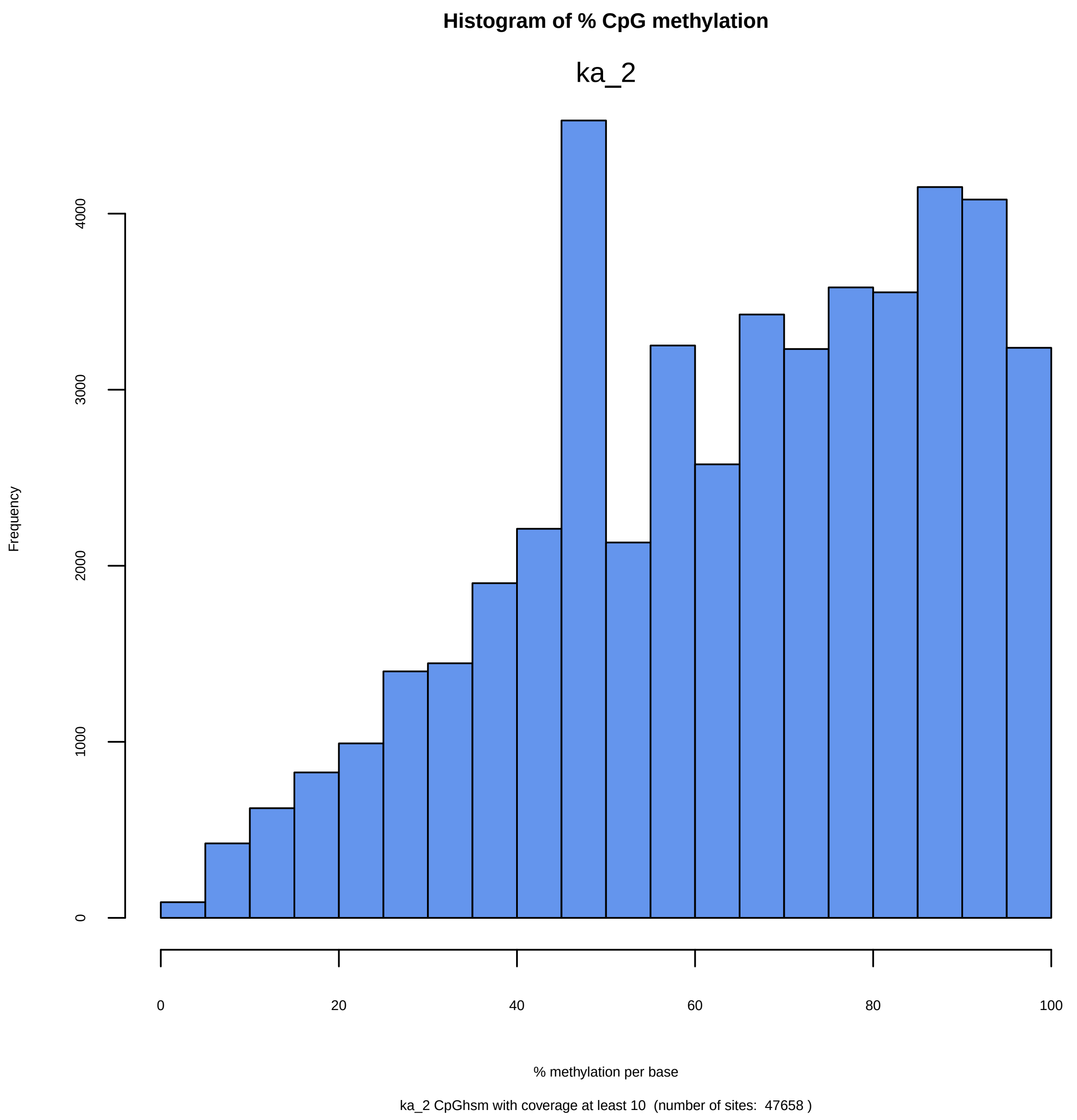
Histogram of % CpG methylation
ka_2
Frequency
0
1000
2000
3000
4000
0
20
40
60
80
100
% methylation per base
ka_2 CpGhsm with coverage at least 10 (number of sites: 47658 )

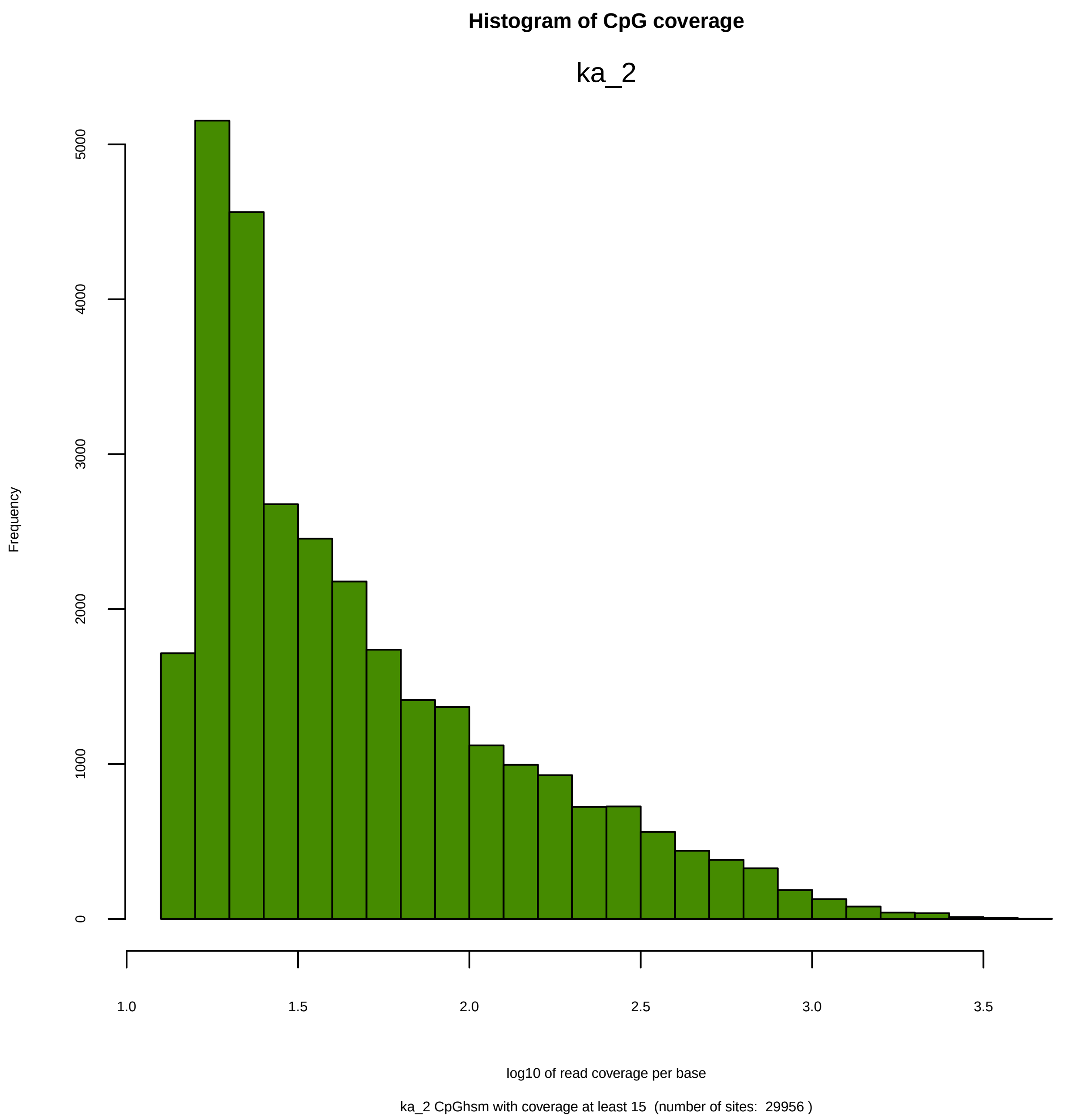

Histogram of CpG coverage
ka_2
Frequency
0
1000
2000
3000
4000
5000
1.0
1.5
2.0
2.5
3.0
3.5
log10 of read coverage per base
ka_2 CpGhsm with coverage at least 15  (number of sites:  29956 )

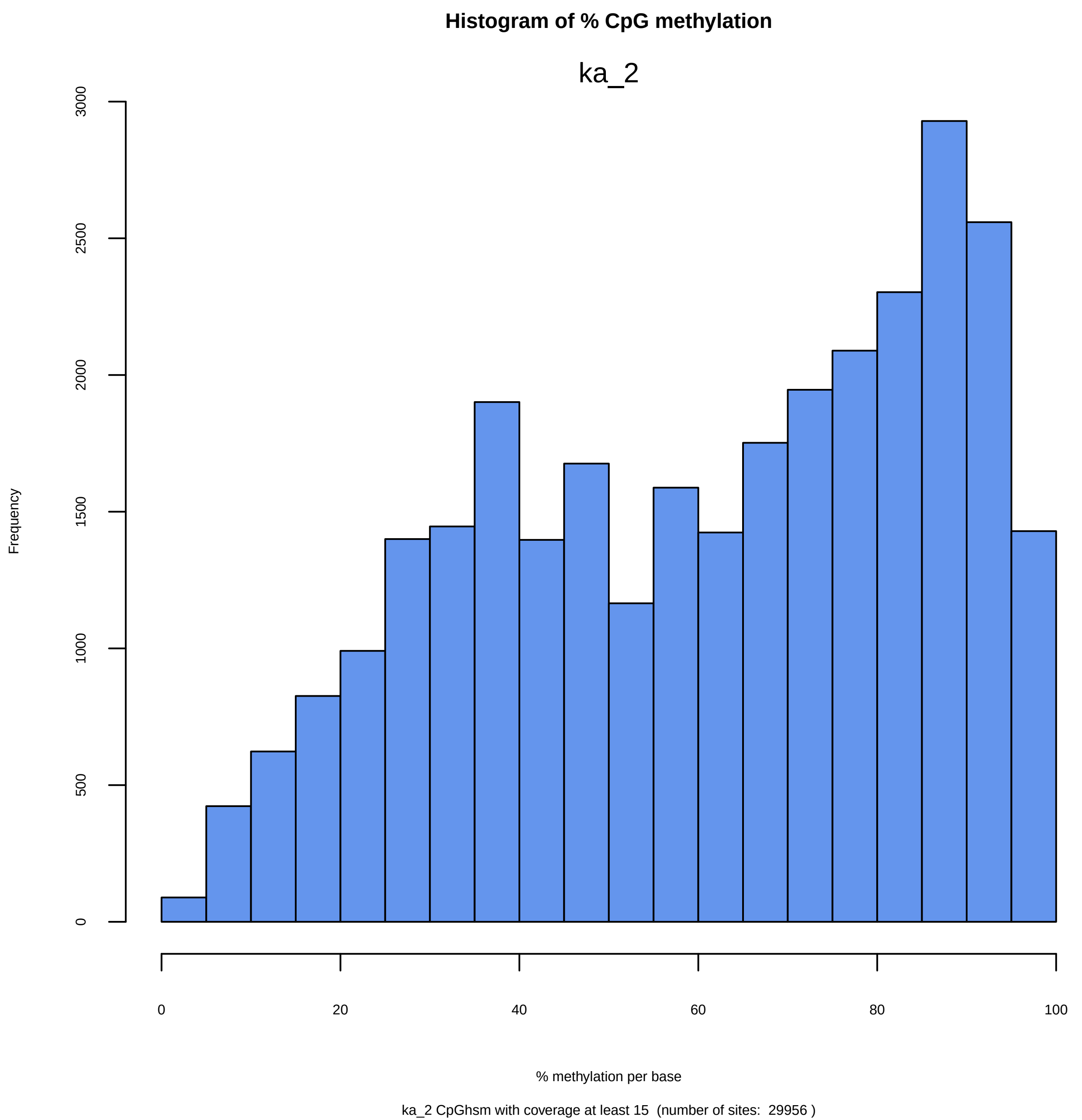

Histogram of % CpG methylation
ka_2
Frequency
0
500
1000
1500
2000
2500
3000
0
20
40
60
80
100
% methylation per base
ka_2 CpGhsm with coverage at least 15 (number of sites: 29956 )

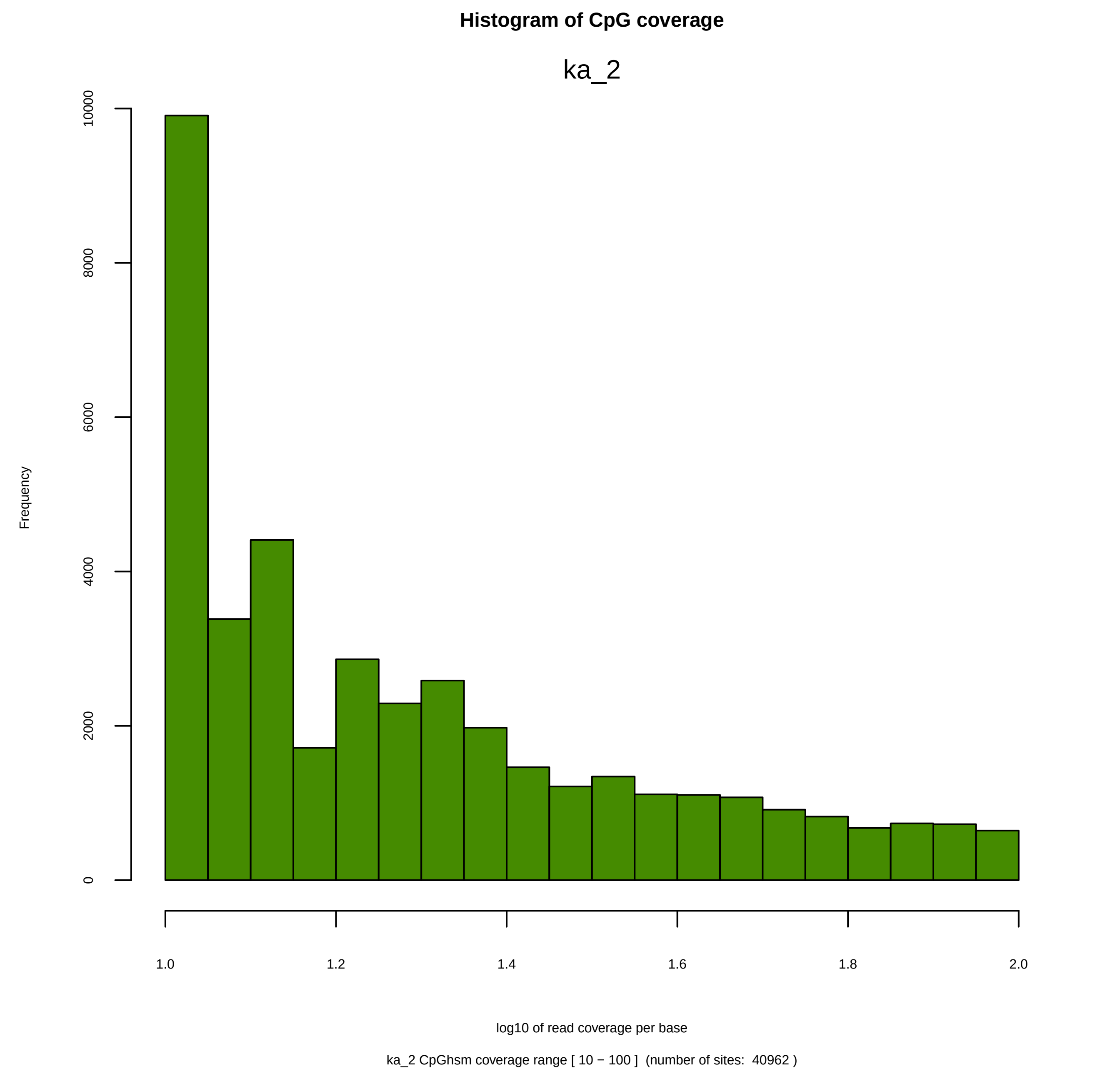
Histogram of CpG coverage
ka_2
Frequency
0
2000
4000
6000
8000
10000
1.0
1.2
1.4
1.6
1.8
2.0
log10 of read coverage per base
ka_2 CpGhsm coverage range [ 10 − 100 ] (number of sites: 40962 )

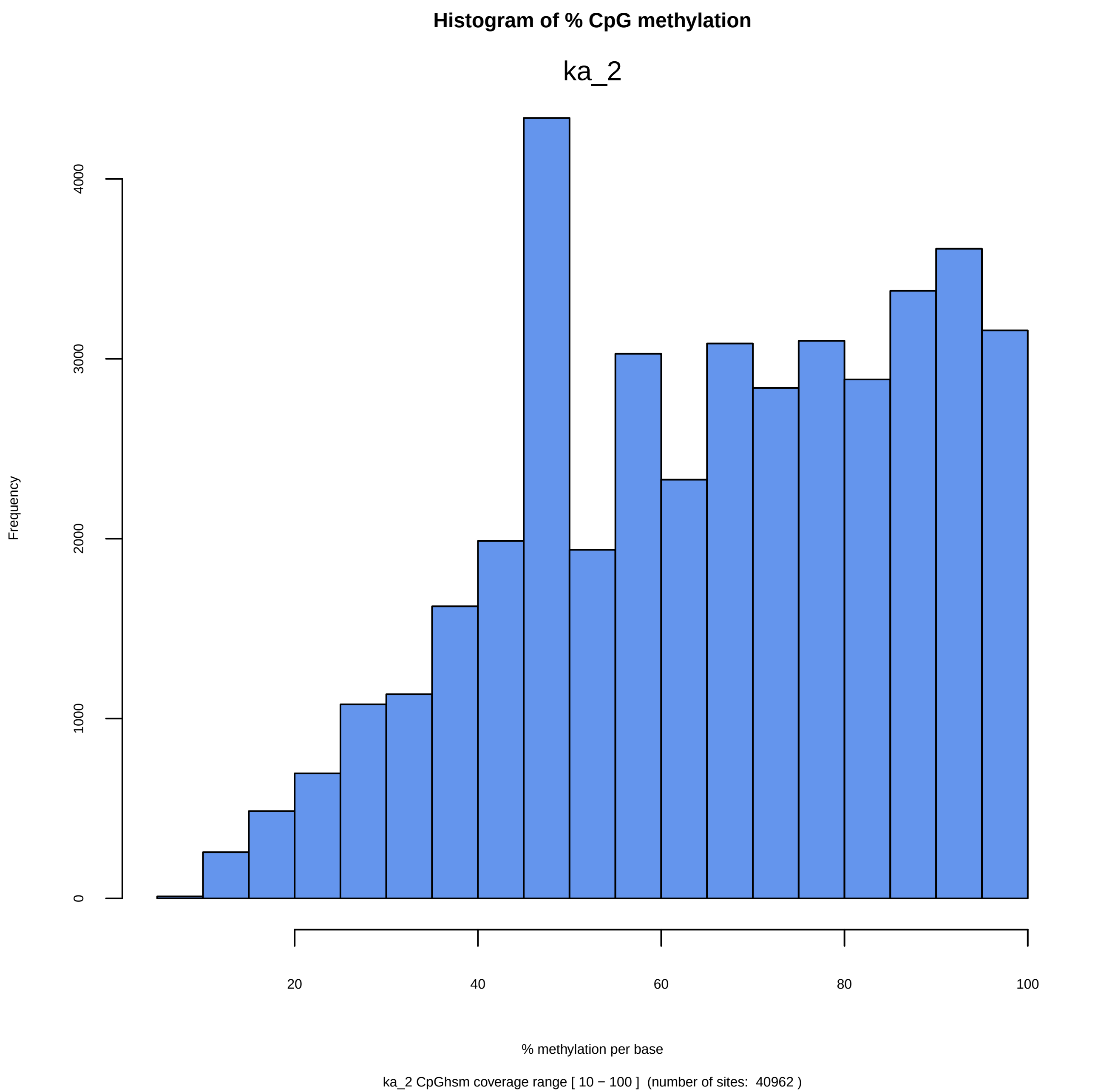

Histogram of % CpG methylation
ka_2
Frequency
0
1000
2000
3000
4000
20
40
60
80
100
% methylation per base
ka_2 CpGhsm coverage range [ 10 − 100 ] (number of sites: 40962 )

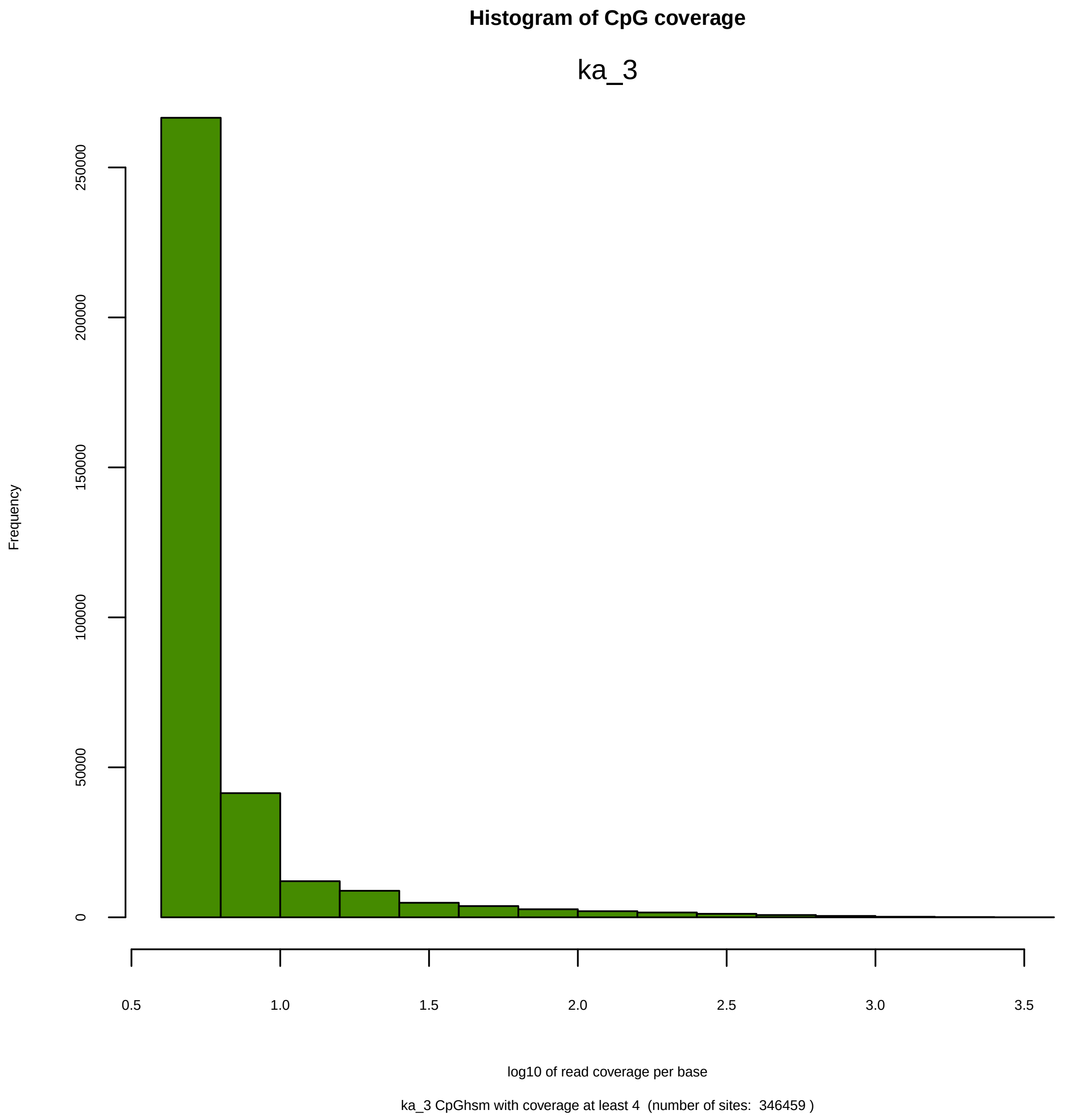

Histogram of CpG coverage
ka_3
Frequency
0
50000
100000
150000
200000
250000
0.5
1.0
1.5
2.0
2.5
3.0
3.5
log10 of read coverage per base
ka_3 CpGhsm with coverage at least 4 (number of sites: 346459 )

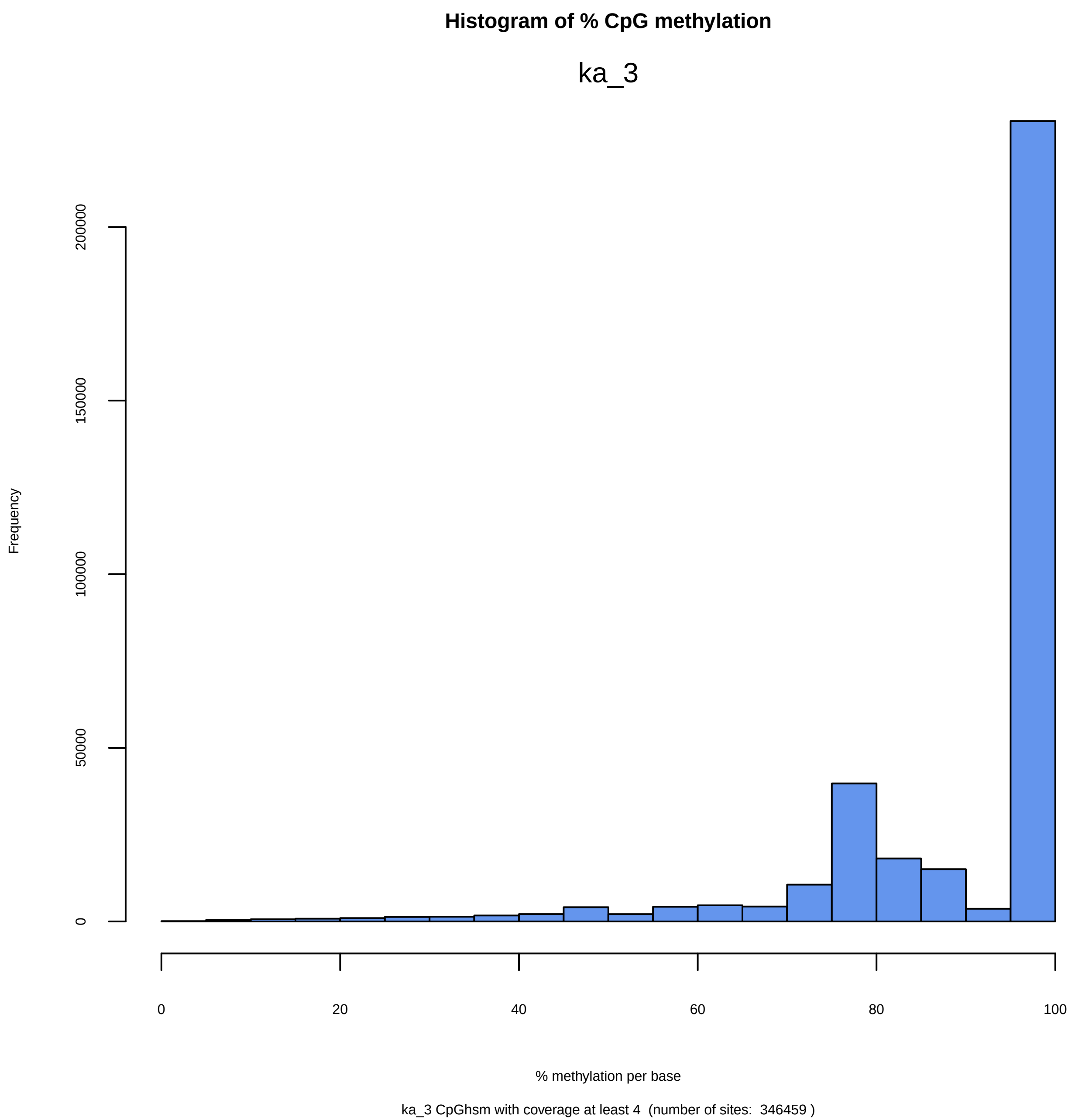

Histogram of % CpG methylation
ka_3
Frequency
0
50000
100000
150000
200000
0
20
40
60
80
100
% methylation per base
ka_3 CpGhsm with coverage at least 4  (number of sites:  346459 )

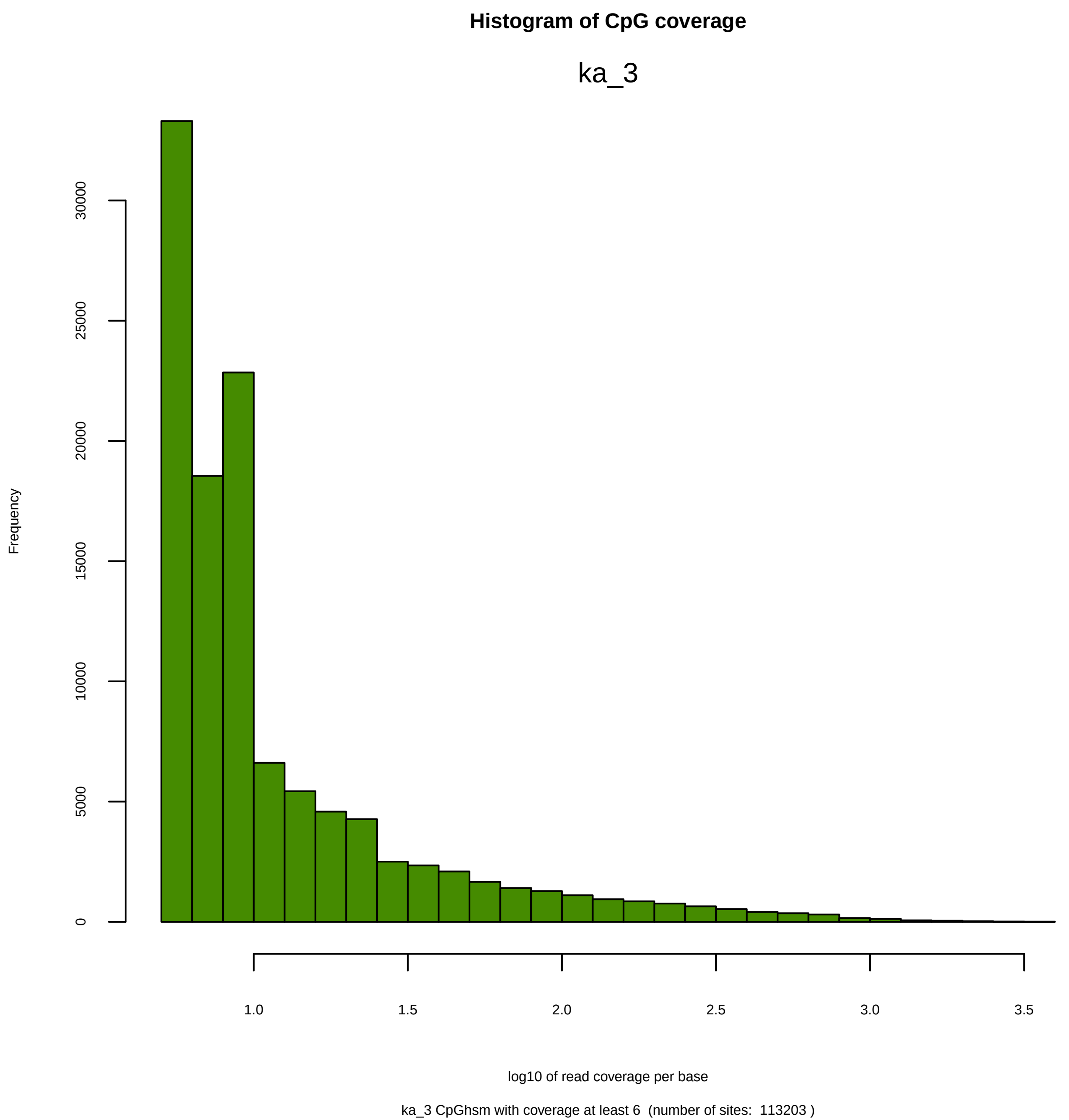

Histogram of CpG coverage
ka_3
Frequency
0
5000
10000
15000
20000
25000
30000
1.0
1.5
2.0
2.5
3.0
3.5
log10 of read coverage per base
ka_3 CpGhsm with coverage at least 6 (number of sites: 113203 )

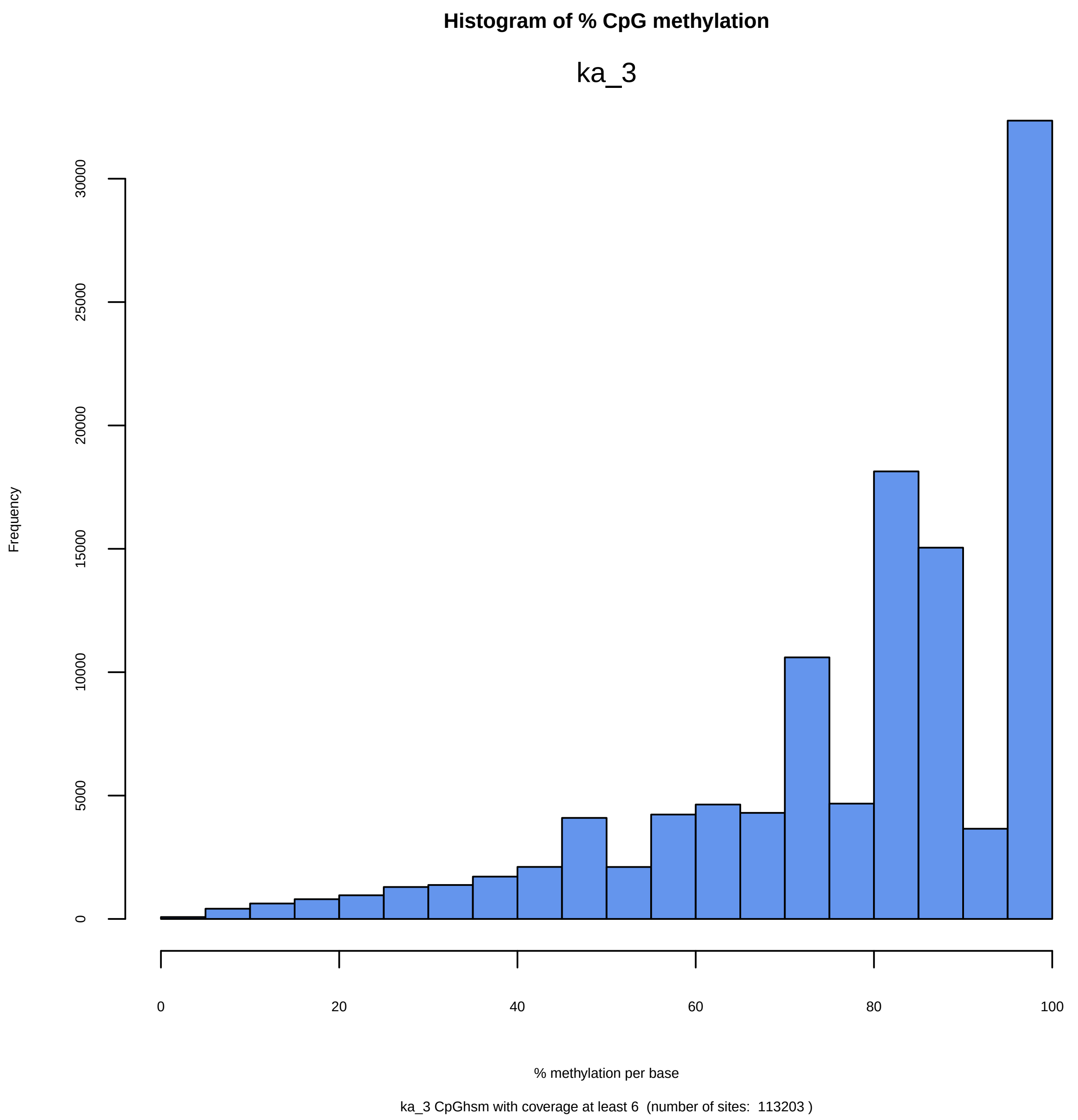

Histogram of % CpG methylation
ka_3
Frequency
0
5000
10000
15000
20000
25000
30000
0
20
40
60
80
100
% methylation per base
ka_3 CpGhsm with coverage at least 6 (number of sites: 113203 )

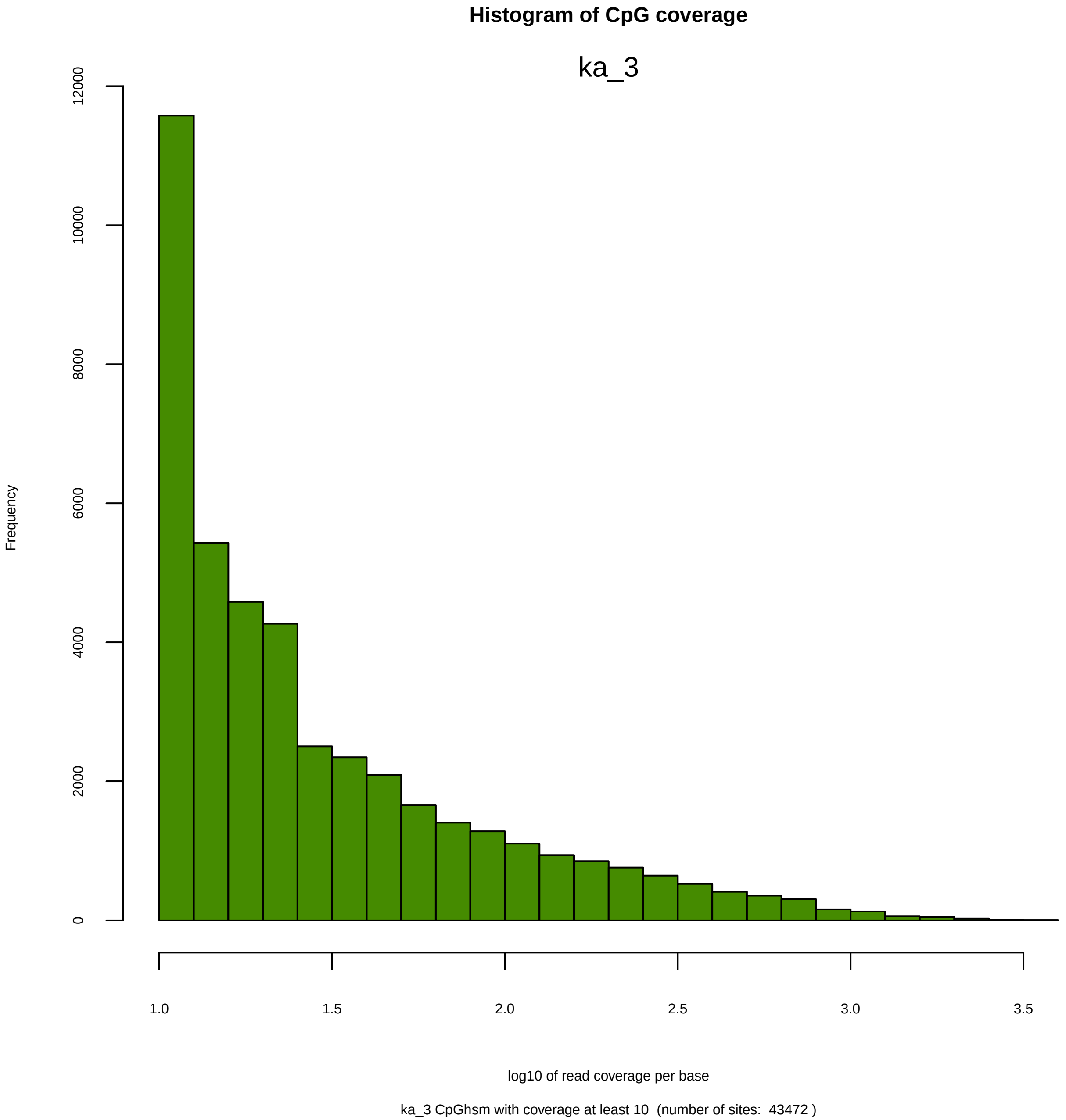
Histogram of CpG coverage
ka_3
Frequency
0
2000
4000
6000
8000
10000
12000
1.0
1.5
2.0
2.5
3.0
3.5
log10 of read coverage per base
ka_3 CpGhsm with coverage at least 10 (number of sites: 43472 )

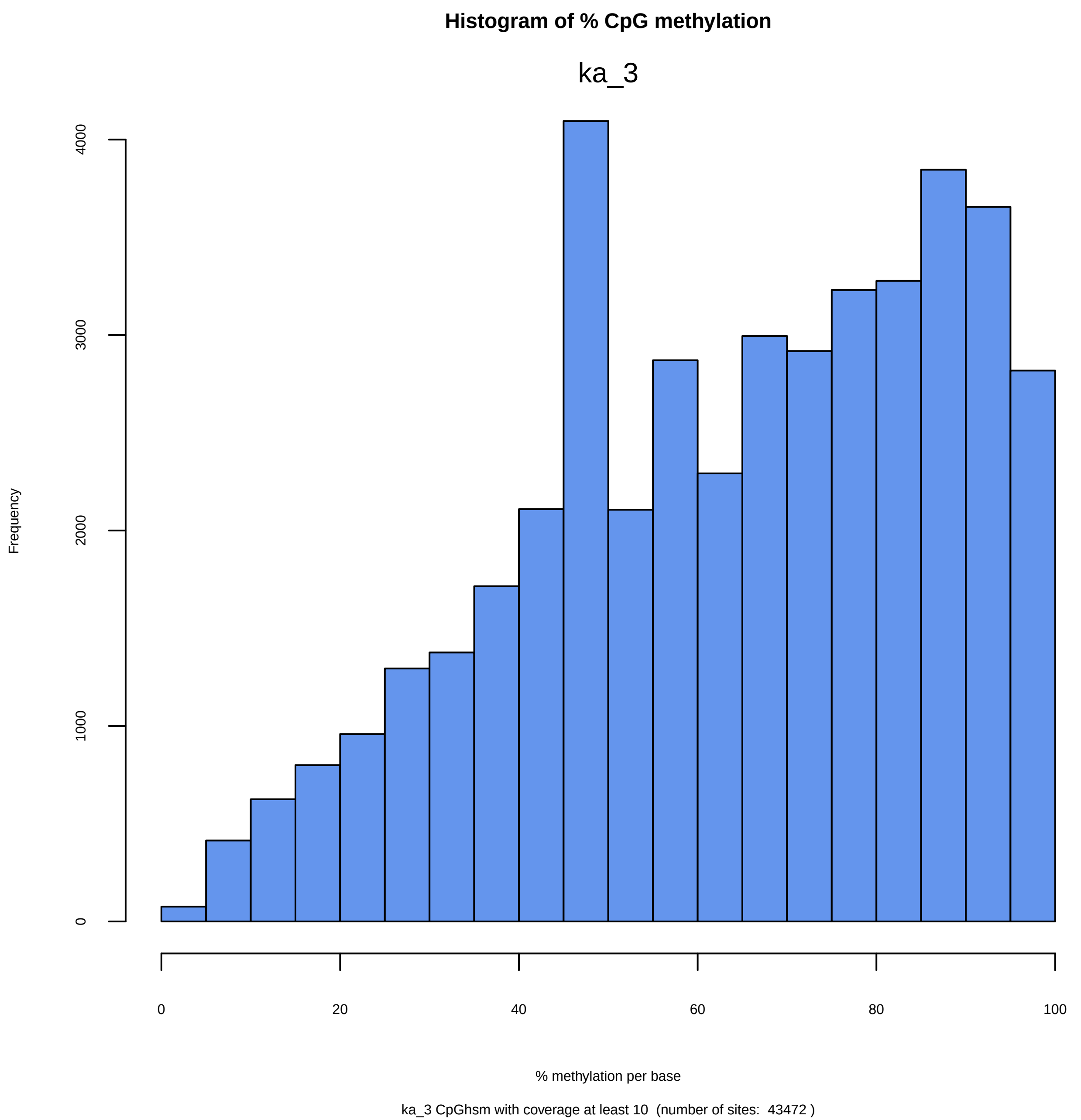
Histogram of % CpG methylation
ka_3
Frequency
0
1000
2000
3000
4000
0
20
40
60
80
100
% methylation per base
ka_3 CpGhsm with coverage at least 10  (number of sites:  43472 )

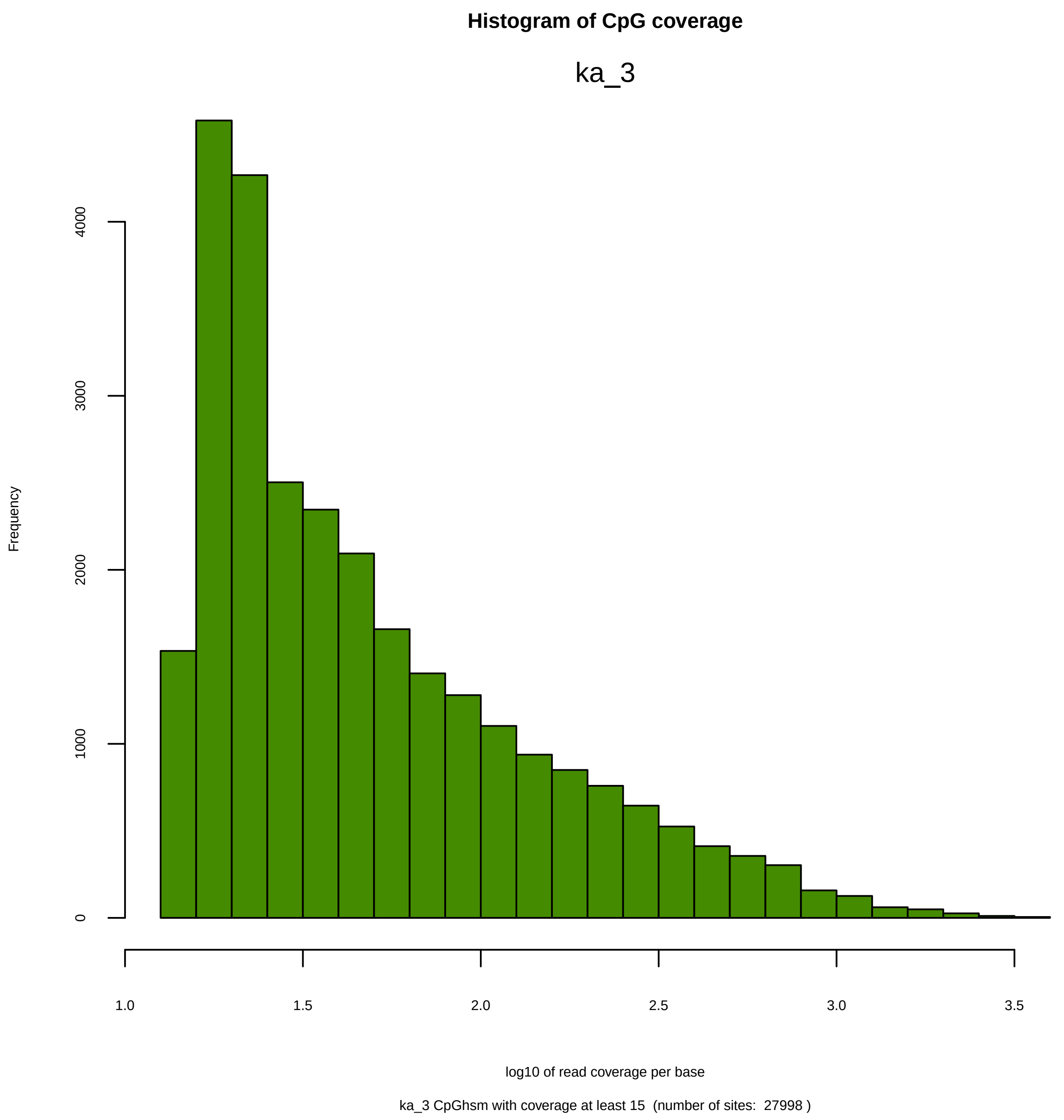

Histogram of CpG coverage
ka_3
Frequency
0
1000
2000
3000
4000
1.0
1.5
2.0
2.5
3.0
3.5
log10 of read coverage per base
ka_3 CpGhsm with coverage at least 15  (number of sites:  27998 )

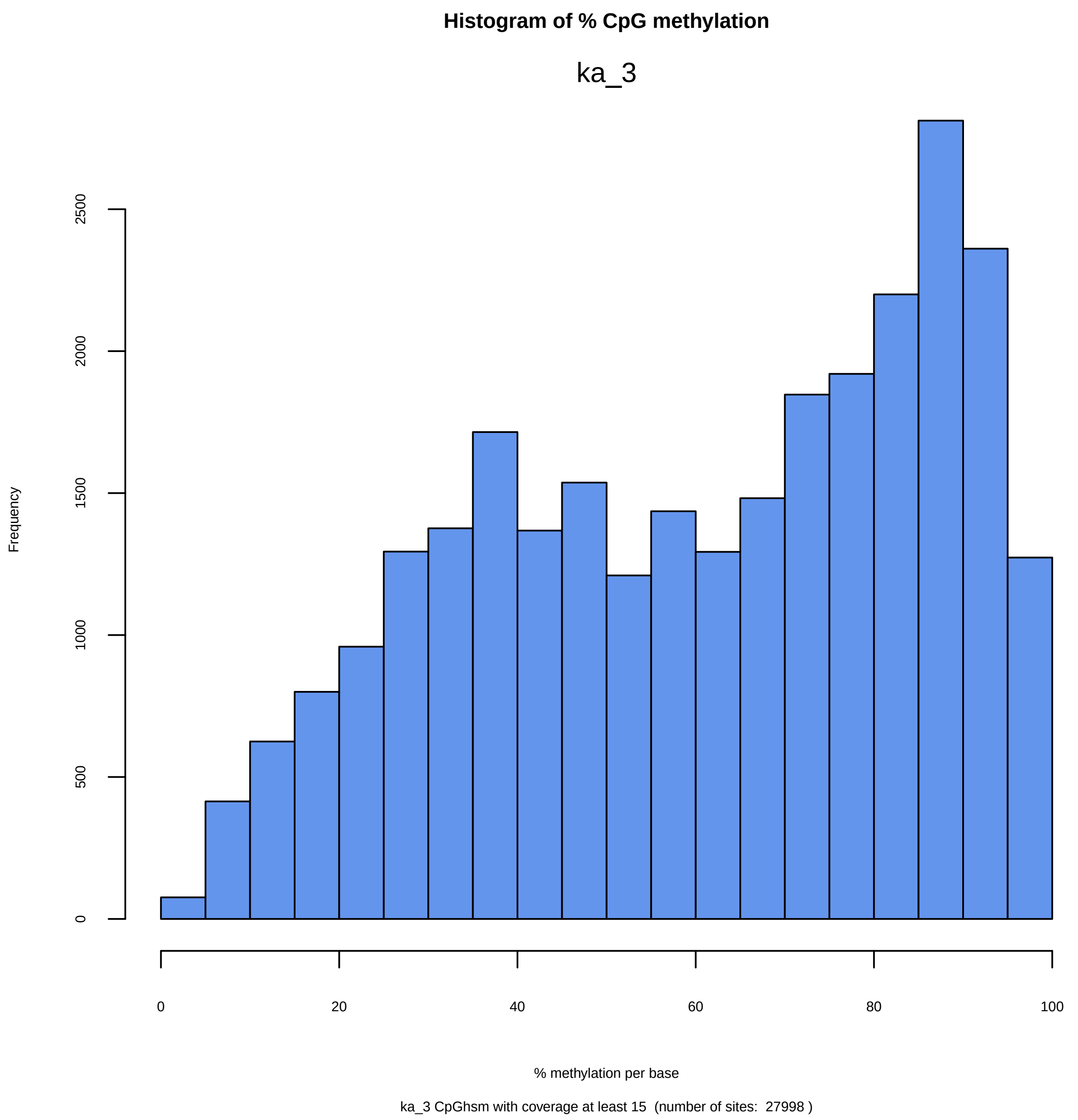
Histogram of % CpG methylation
ka_3
Frequency
0
500
1000
1500
2000
2500
0
20
40
60
80
100
% methylation per base
ka_3 CpGhsm with coverage at least 15  (number of sites:  27998 )

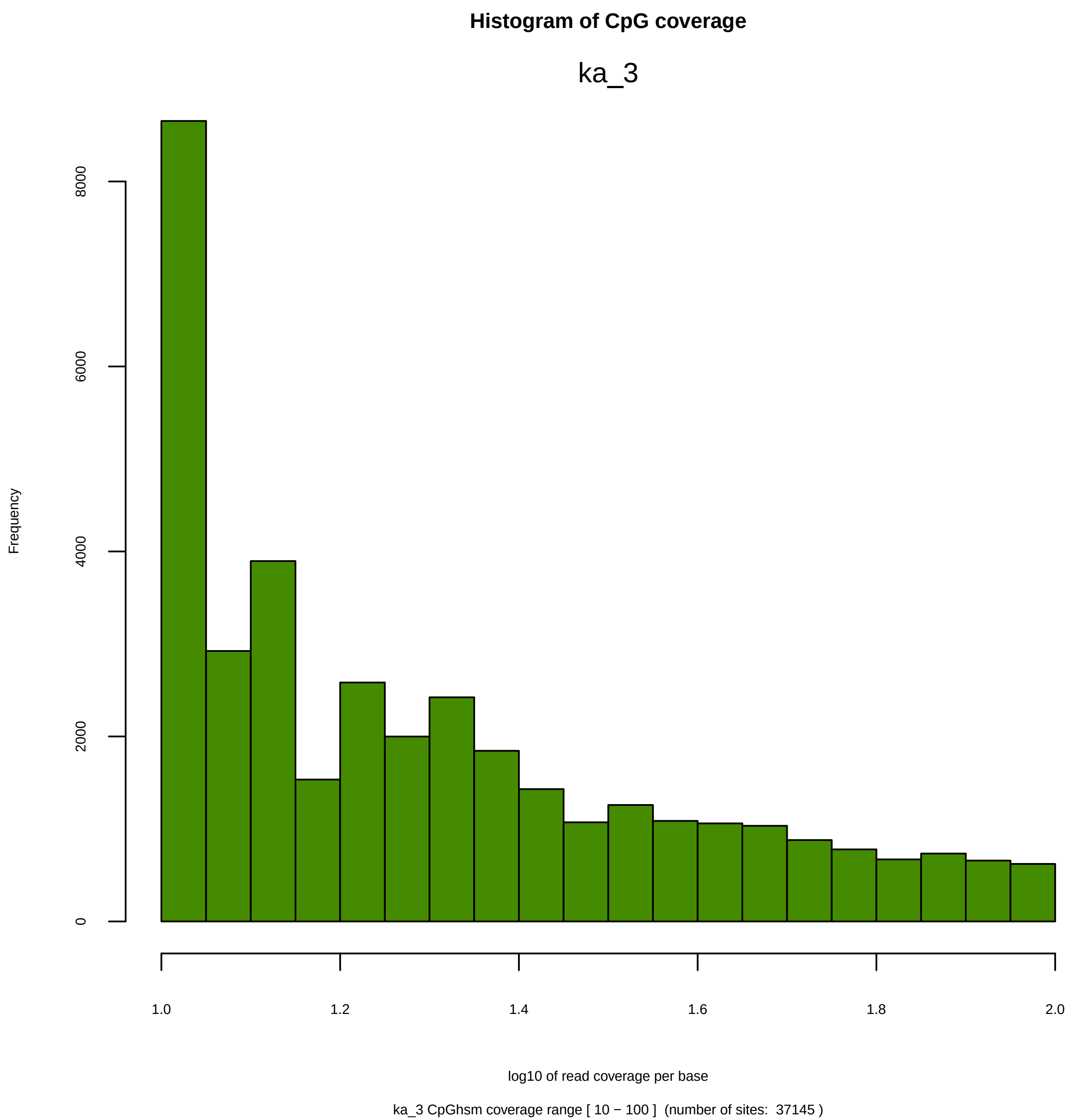
Histogram of CpG coverage
ka_3
Frequency
0
2000
4000
6000
8000
1.0
1.2
1.4
1.6
1.8
2.0
log10 of read coverage per base
ka_3 CpGhsm coverage range [ 10 − 100 ]  (number of sites:  37145 )

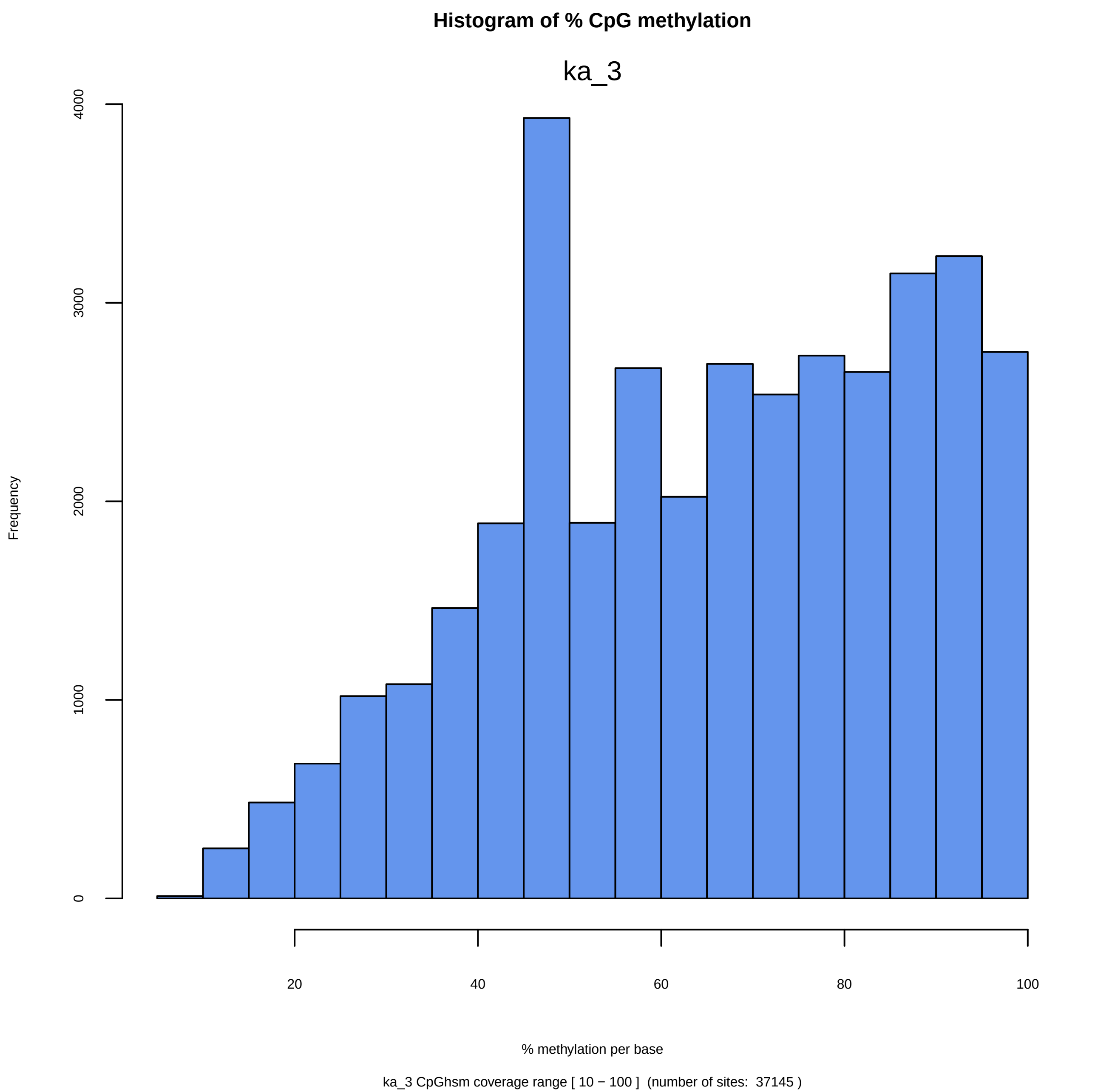

Histogram of % CpG methylation
ka_3
Frequency
0
1000
2000
3000
4000
20
40
60
80
100
% methylation per base
ka_3 CpGhsm coverage range [ 10 − 100 ] (number of sites: 37145 )

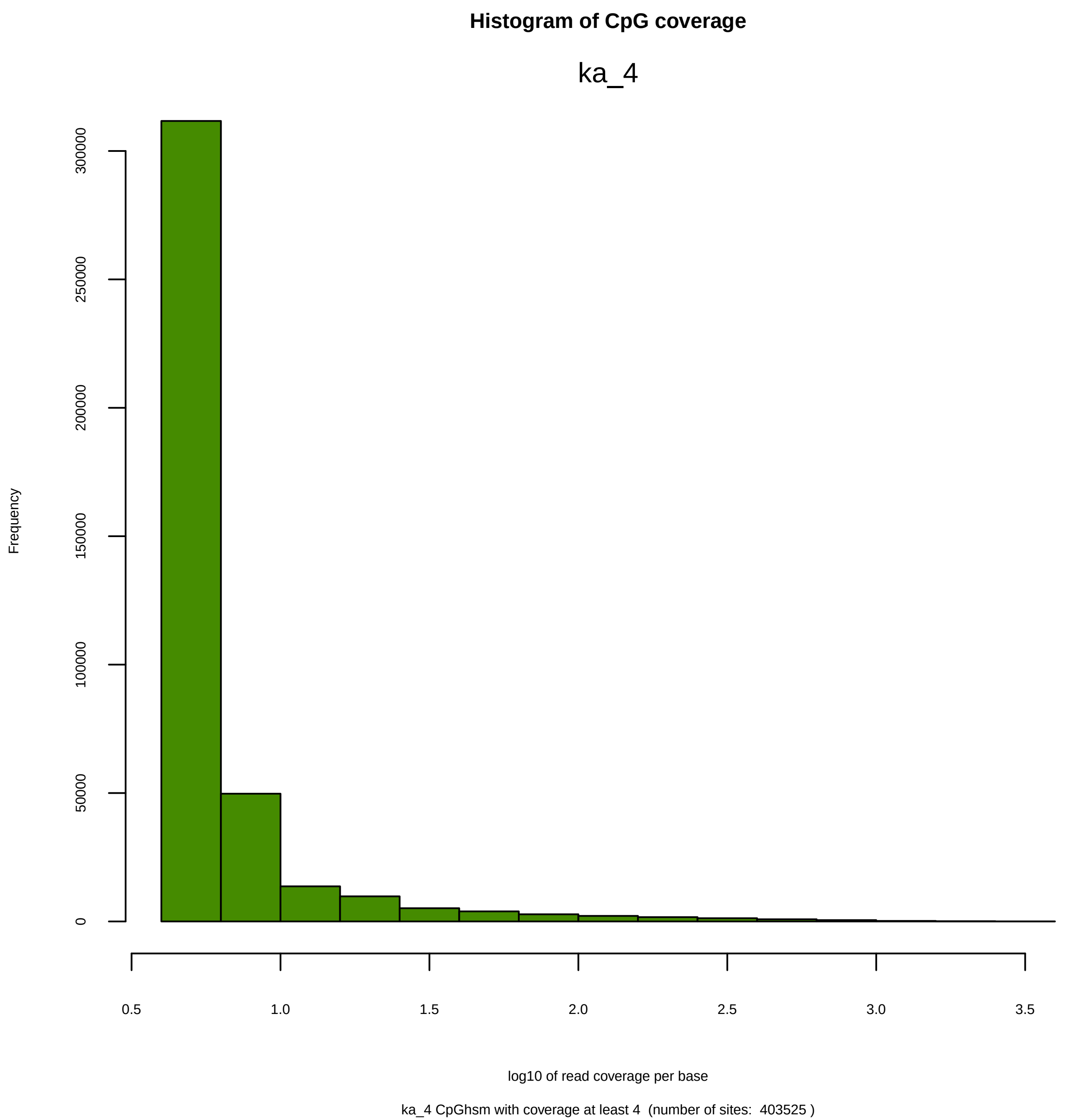

Histogram of CpG coverage
ka_4
Frequency
0
50000
100000
150000
200000
250000
300000
0.5
1.0
1.5
2.0
2.5
3.0
3.5
log10 of read coverage per base
ka_4 CpGhsm with coverage at least 4  (number of sites:  403525 )

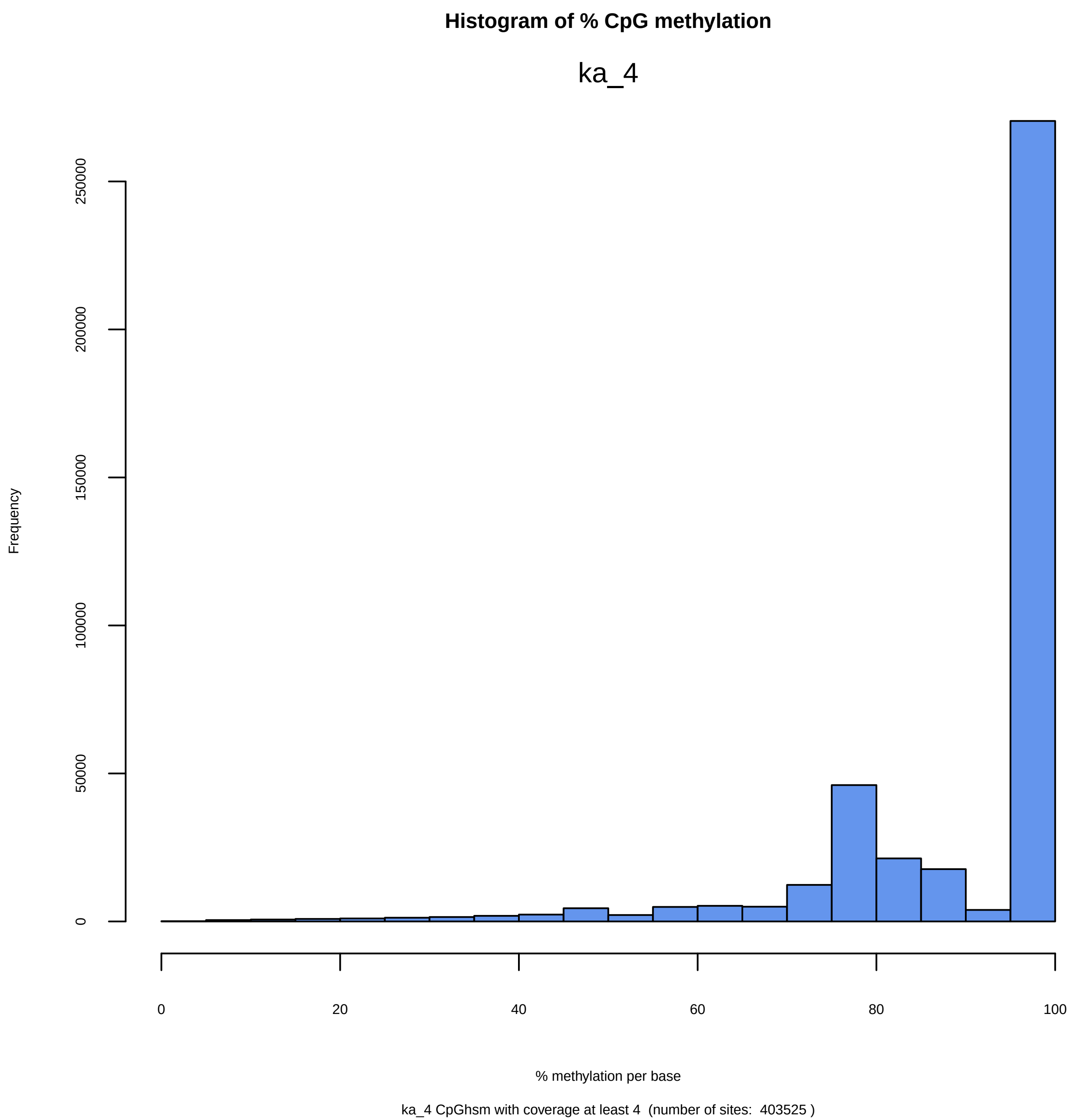

Histogram of % CpG methylation
ka_4
Frequency
0
50000
100000
150000
200000
250000
0
20
40
60
80
100
% methylation per base
ka_4 CpGhsm with coverage at least 4  (number of sites:  403525 )

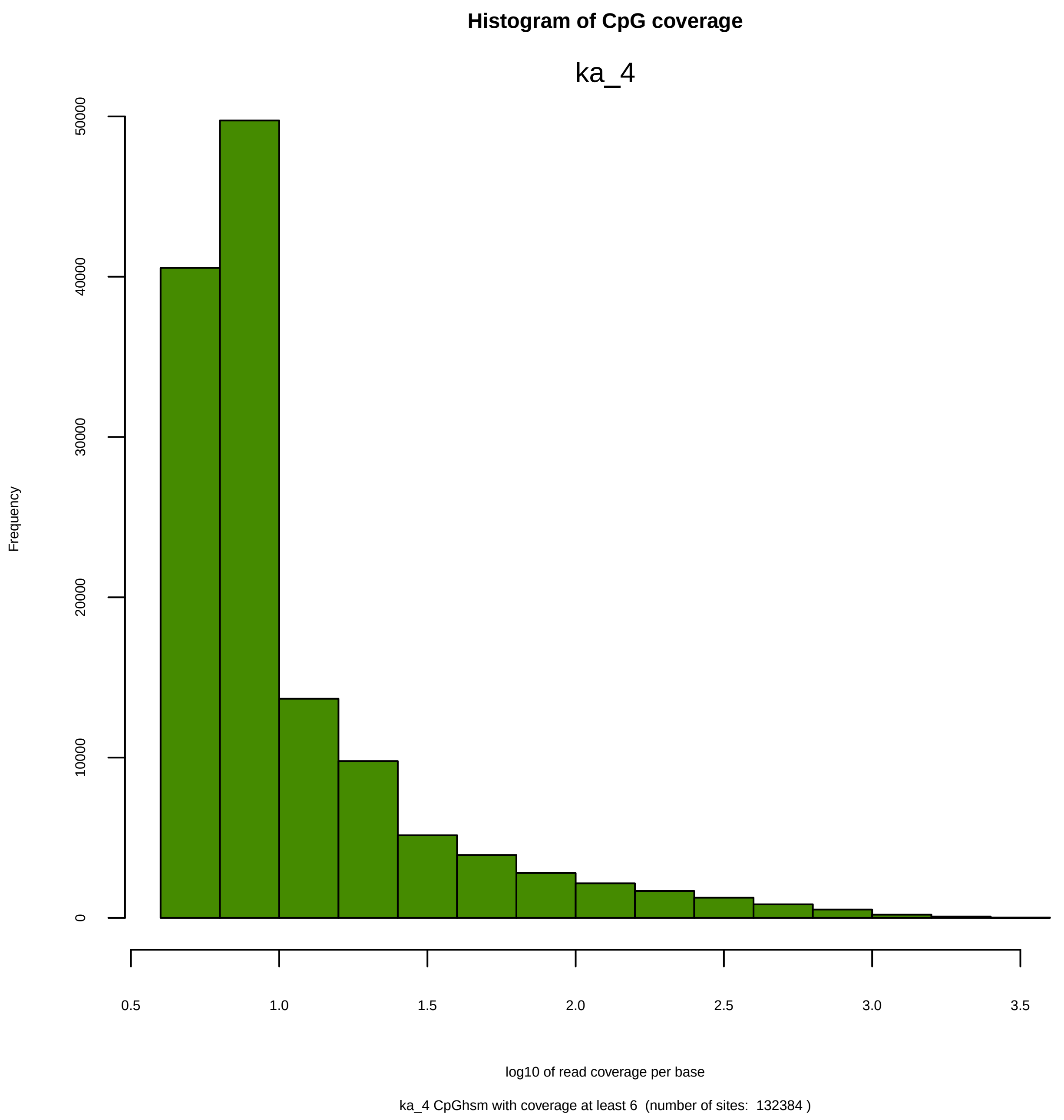
Histogram of CpG coverage
ka_4
Frequency
0
10000
20000
30000
40000
50000
0.5
1.0
1.5
2.0
2.5
3.0
3.5
log10 of read coverage per base
ka_4 CpGhsm with coverage at least 6 (number of sites: 132384 )

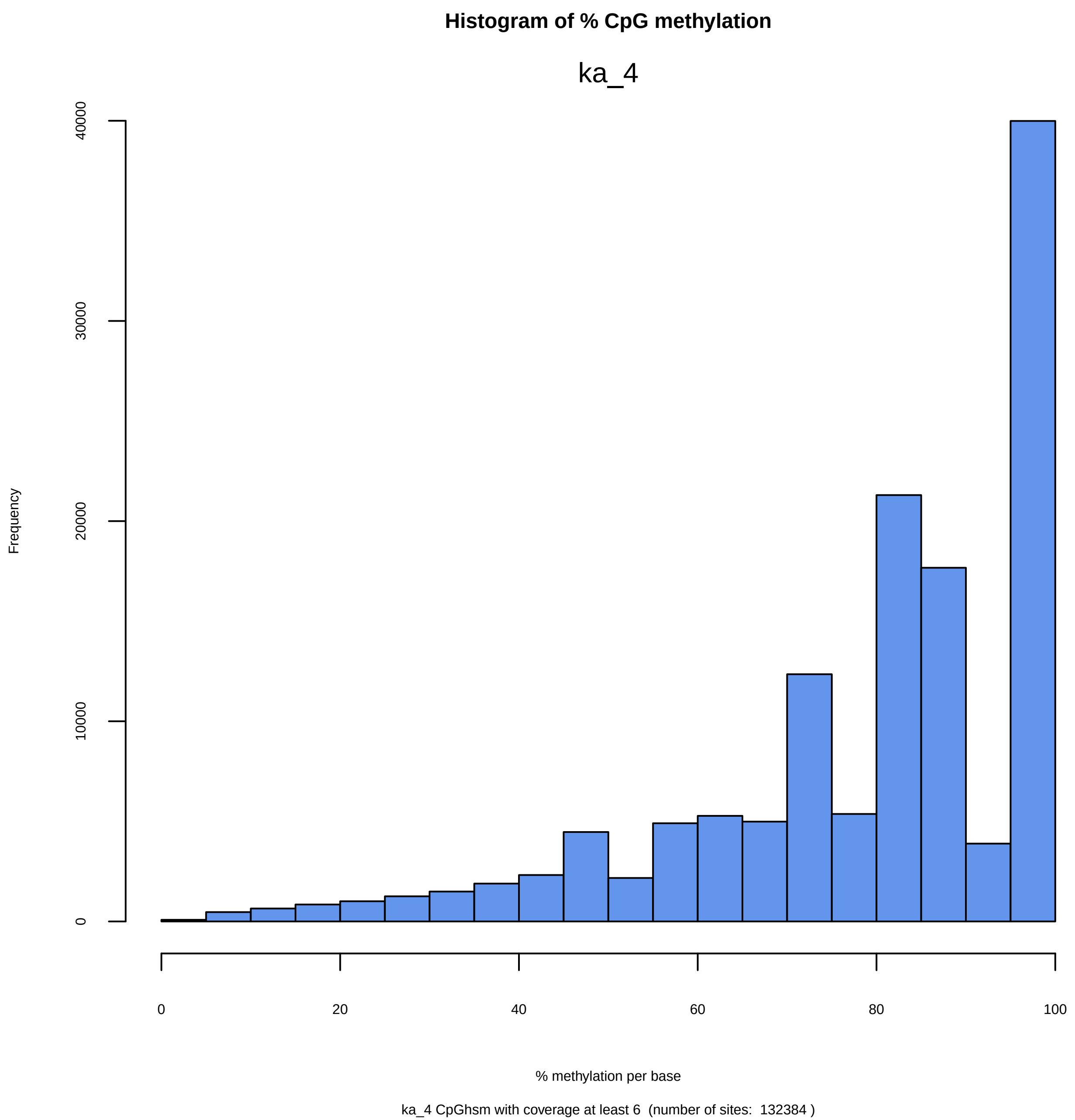

Histogram of % CpG methylation
ka_4
Frequency
0
10000
20000
30000
40000
0
20
40
60
80
100
% methylation per base
ka_4 CpGhsm with coverage at least 6  (number of sites:  132384 )

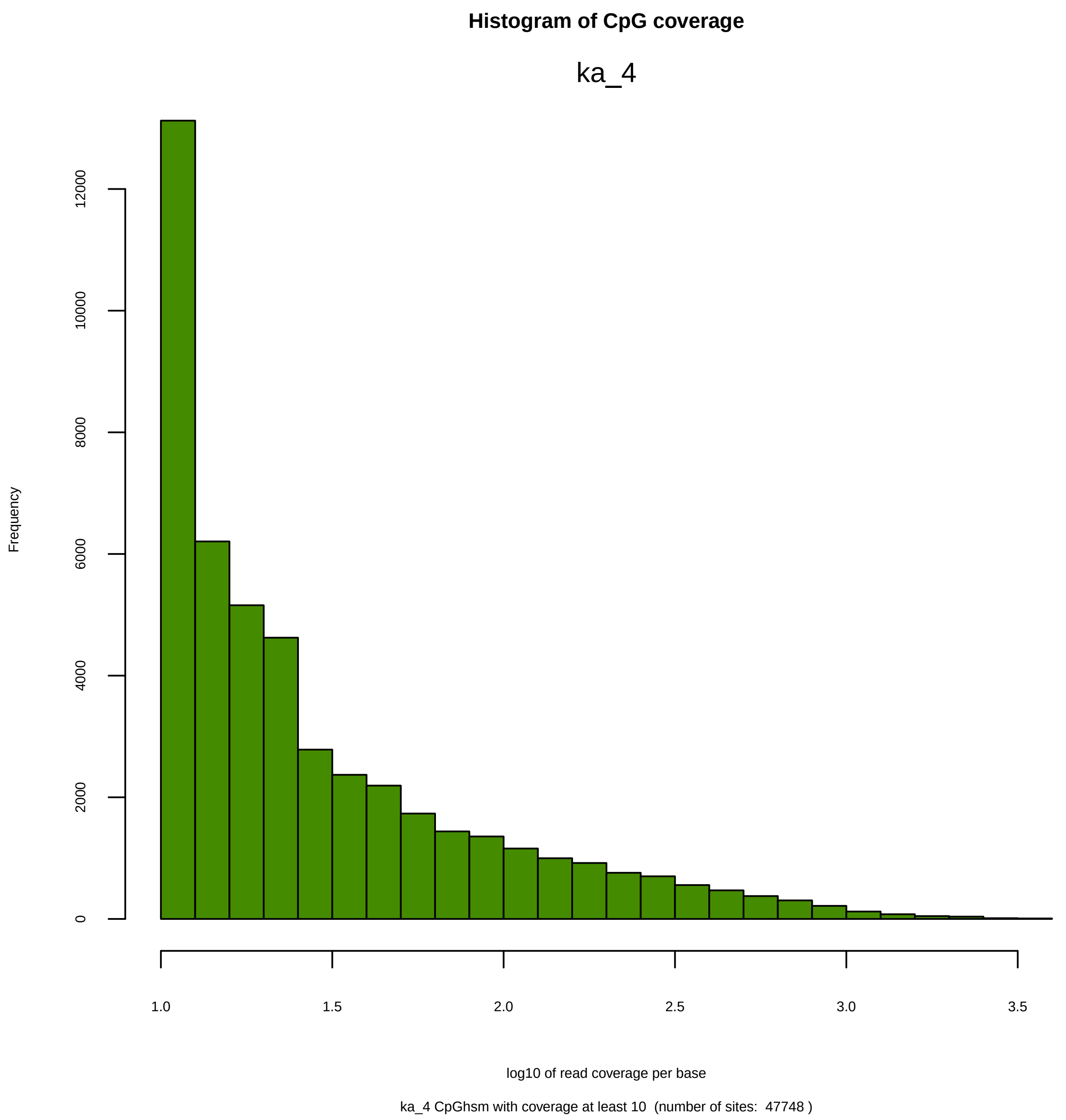

Histogram of CpG coverage
ka_4
Frequency
0
2000
4000
6000
8000
10000
12000
1.0
1.5
2.0
2.5
3.0
3.5
log10 of read coverage per base
ka_4 CpGhsm with coverage at least 10 (number of sites: 47748 )

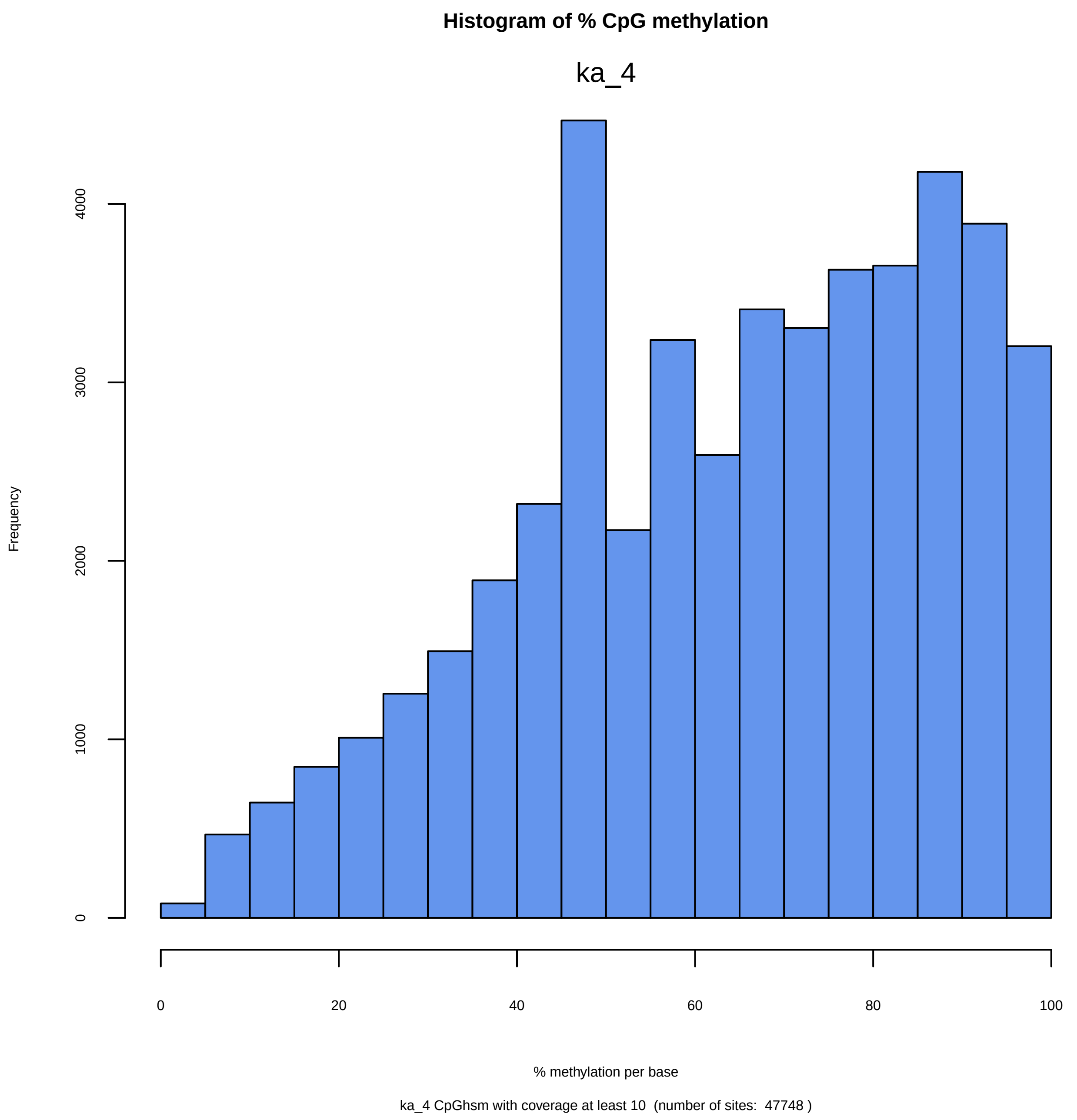

Histogram of % CpG methylation
ka_4
Frequency
0
1000
2000
3000
4000
0
20
40
60
80
100
% methylation per base
ka_4 CpGhsm with coverage at least 10  (number of sites:  47748 )

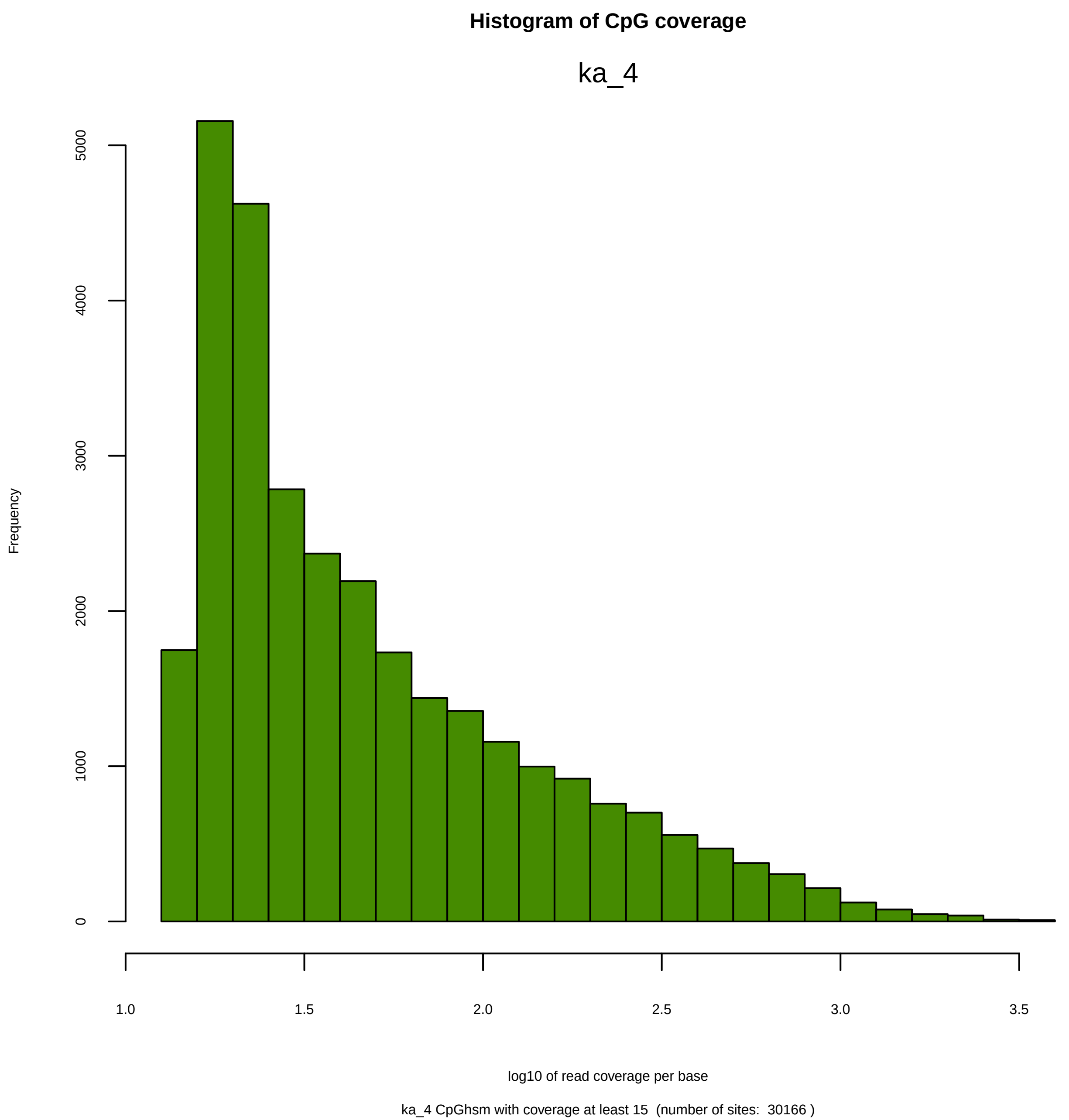

Histogram of CpG coverage
ka_4
Frequency
0
1000
2000
3000
4000
5000
1.0
1.5
2.0
2.5
3.0
3.5
log10 of read coverage per base
ka_4 CpGhsm with coverage at least 15  (number of sites:  30166 )

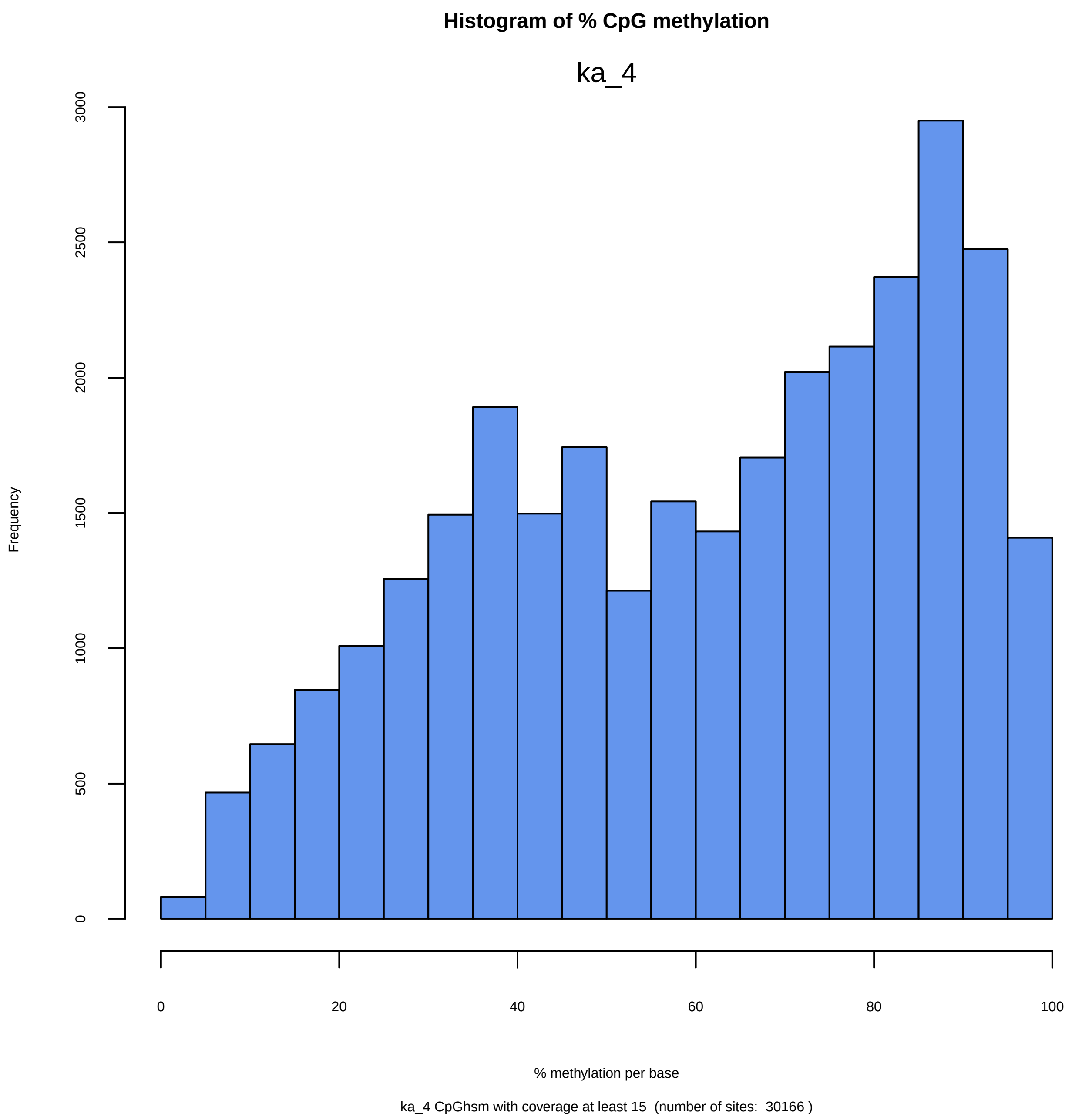

Histogram of % CpG methylation
ka_4
Frequency
0
500
1000
1500
2000
2500
3000
0
20
40
60
80
100
% methylation per base
ka_4 CpGhsm with coverage at least 15  (number of sites:  30166 )

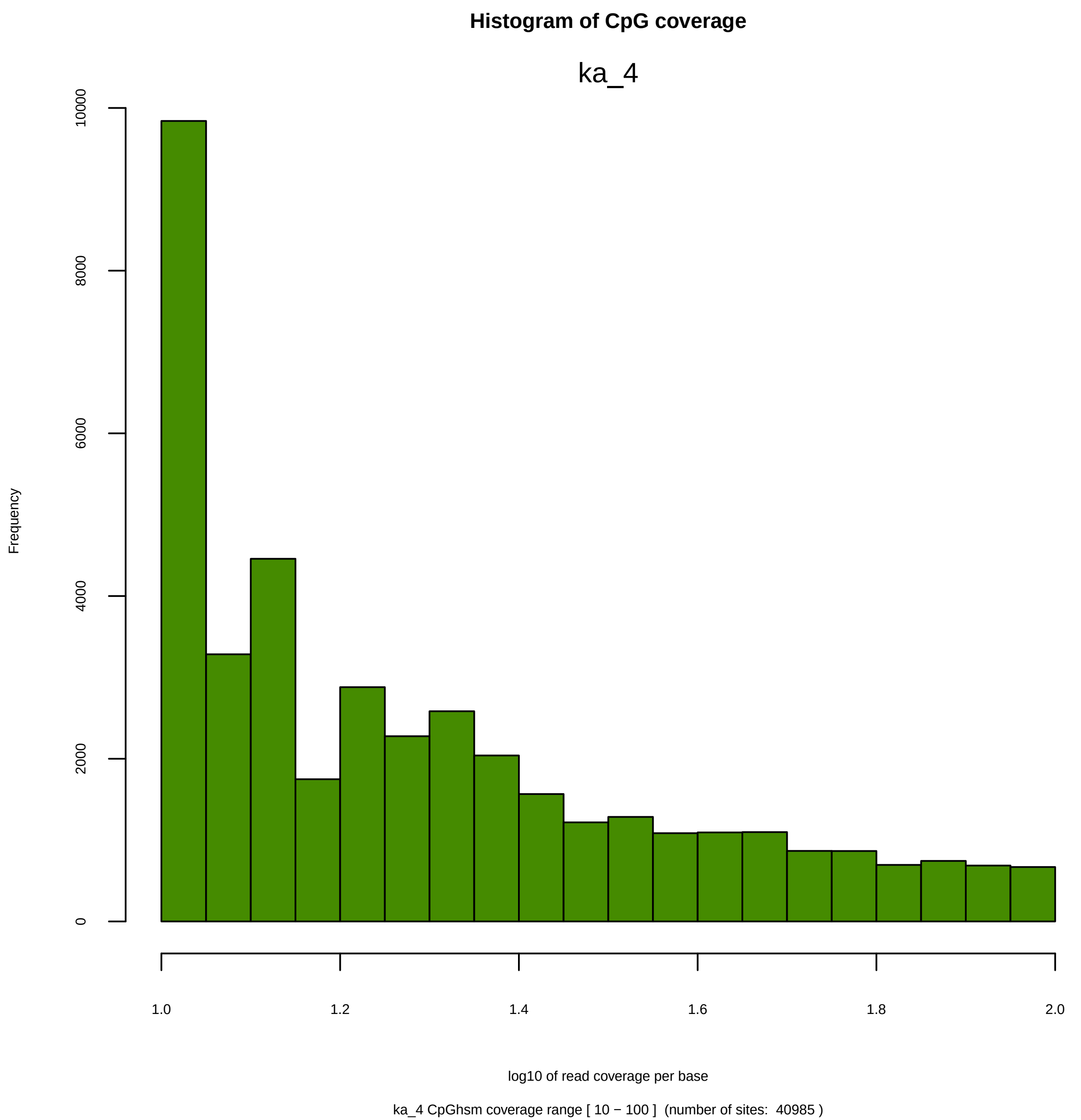

Histogram of CpG coverage
ka_4
Frequency
0
2000
4000
6000
8000
10000
1.0
1.2
1.4
1.6
1.8
2.0
log10 of read coverage per base
ka_4 CpGhsm coverage range [ 10 − 100 ]  (number of sites:  40985 )

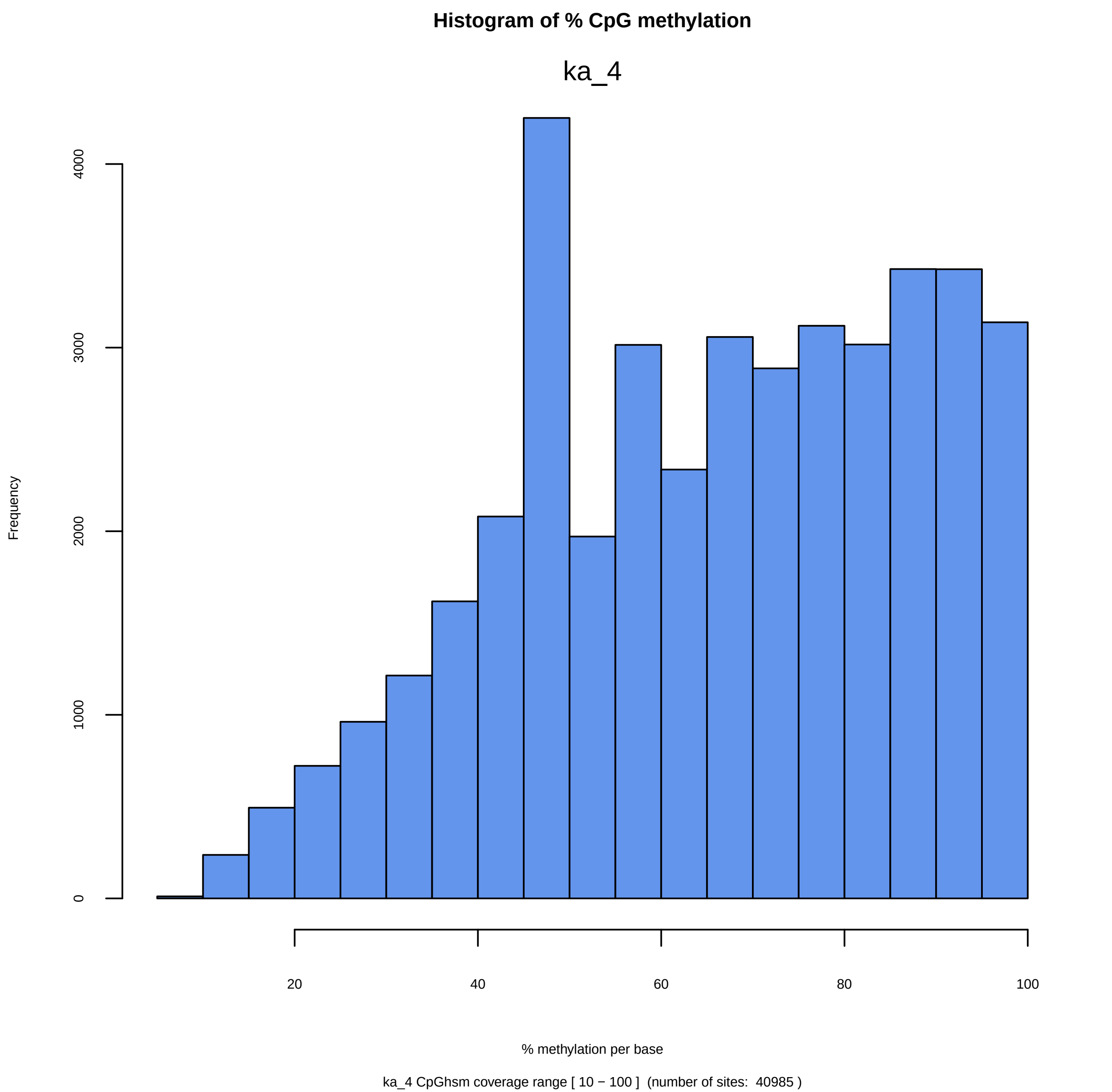
Histogram of % CpG methylation
ka_4
Frequency
0
1000
2000
3000
4000
20
40
60
80
100
% methylation per base
ka_4 CpGhsm coverage range [ 10 − 100 ]  (number of sites:  40985 )

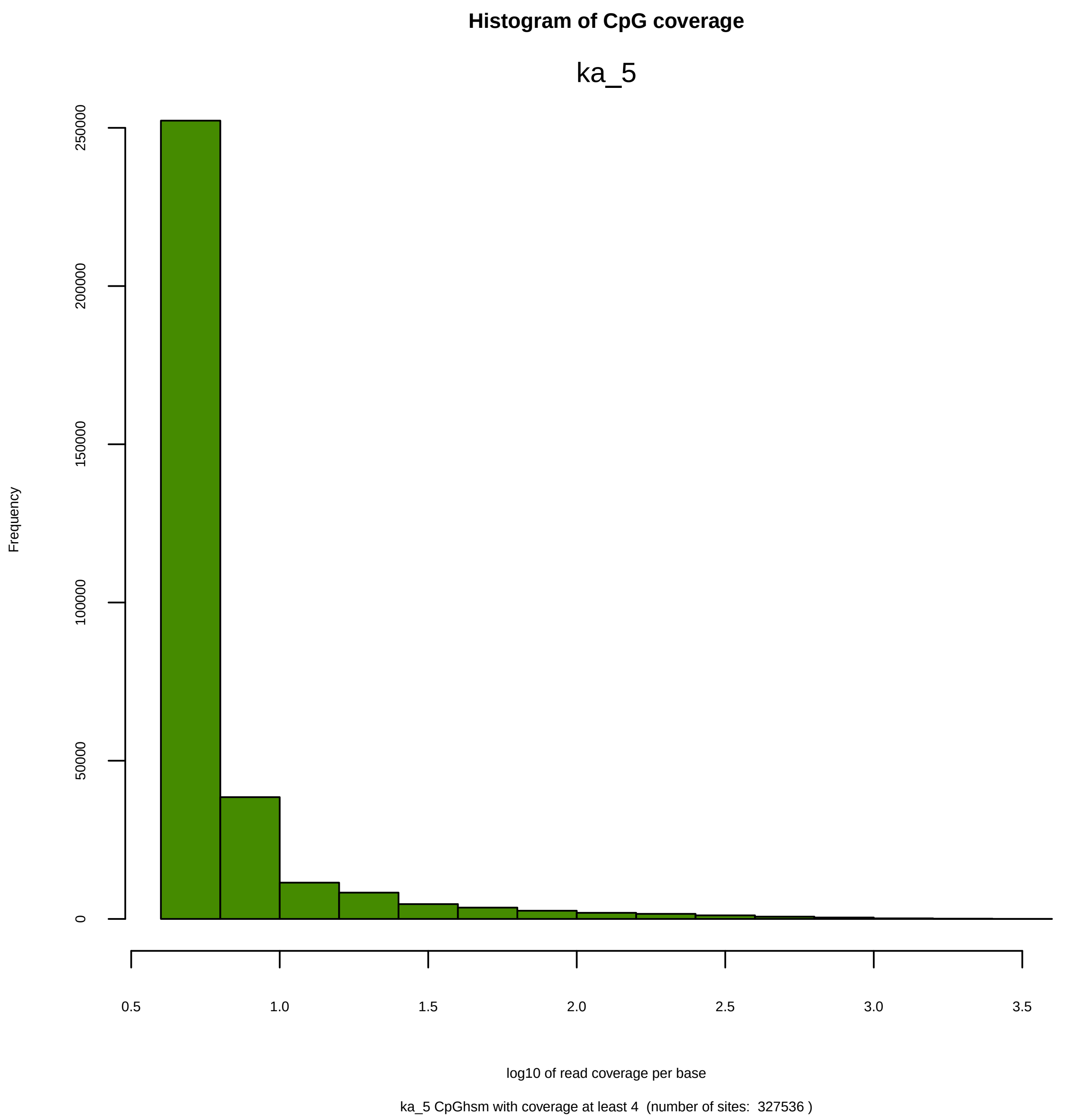

Histogram of CpG coverage
ka_5
Frequency
0
50000
100000
150000
200000
250000
0.5
1.0
1.5
2.0
2.5
3.0
3.5
log10 of read coverage per base
ka_5 CpGhsm with coverage at least 4 (number of sites: 327536 )

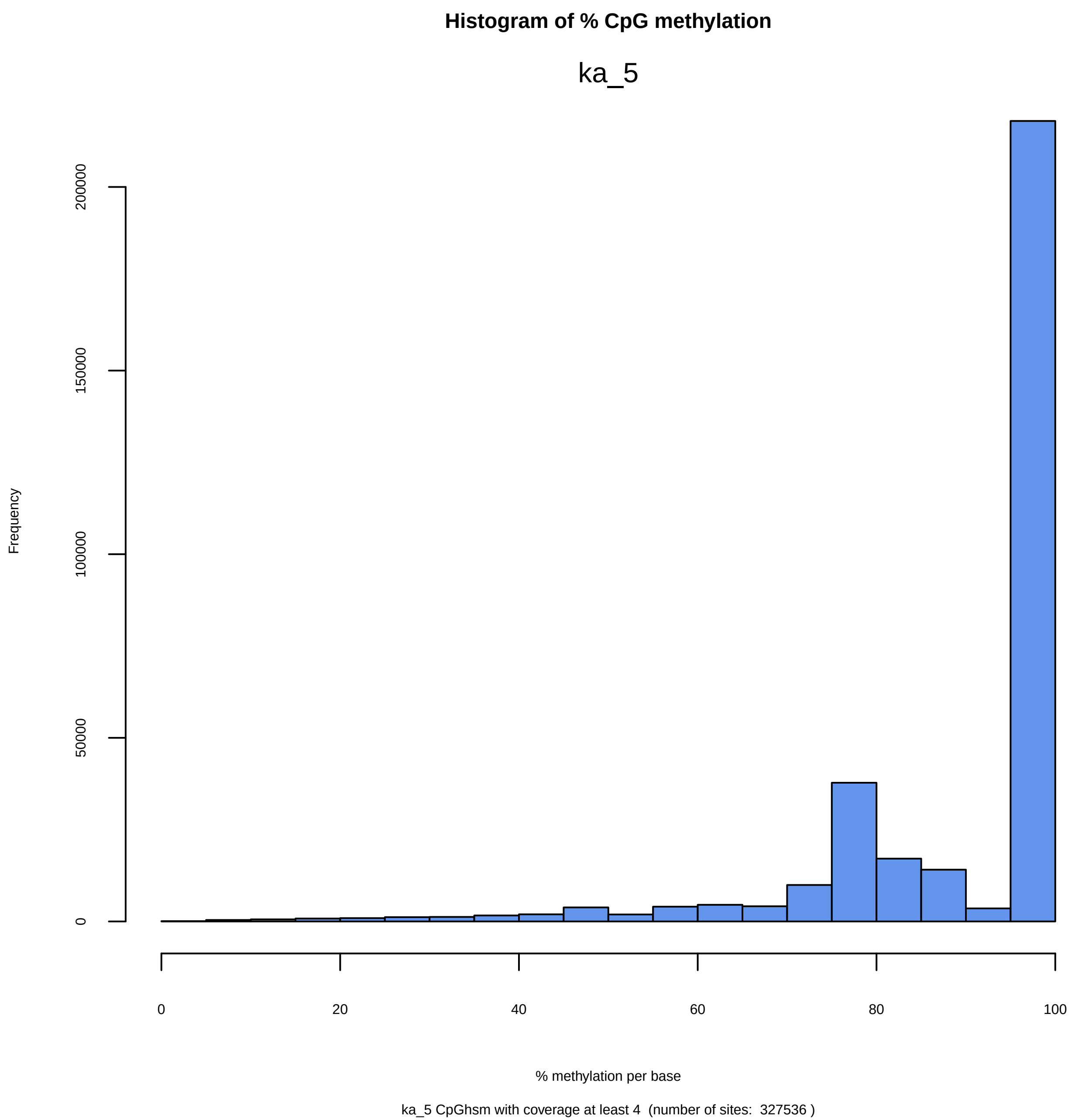

Histogram of % CpG methylation
ka_5
Frequency
0
50000
100000
150000
200000
0
20
40
60
80
100
% methylation per base
ka_5 CpGhsm with coverage at least 4 (number of sites: 327536 )

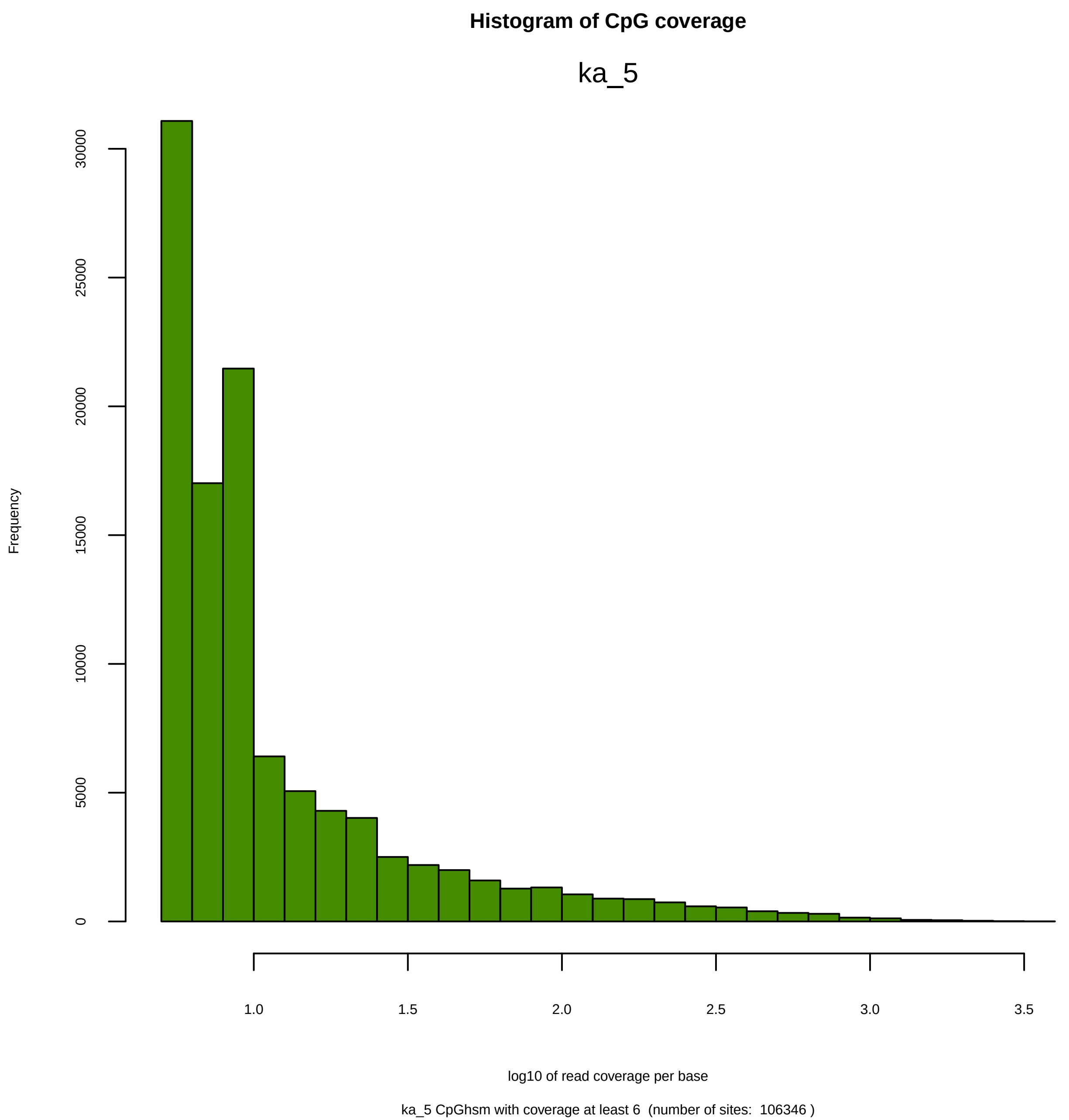

Histogram of CpG coverage
ka_5
Frequency
0
5000
10000
15000
20000
25000
30000
1.0
1.5
2.0
2.5
3.0
3.5
log10 of read coverage per base
ka_5 CpGhsm with coverage at least 6 (number of sites: 106346 )

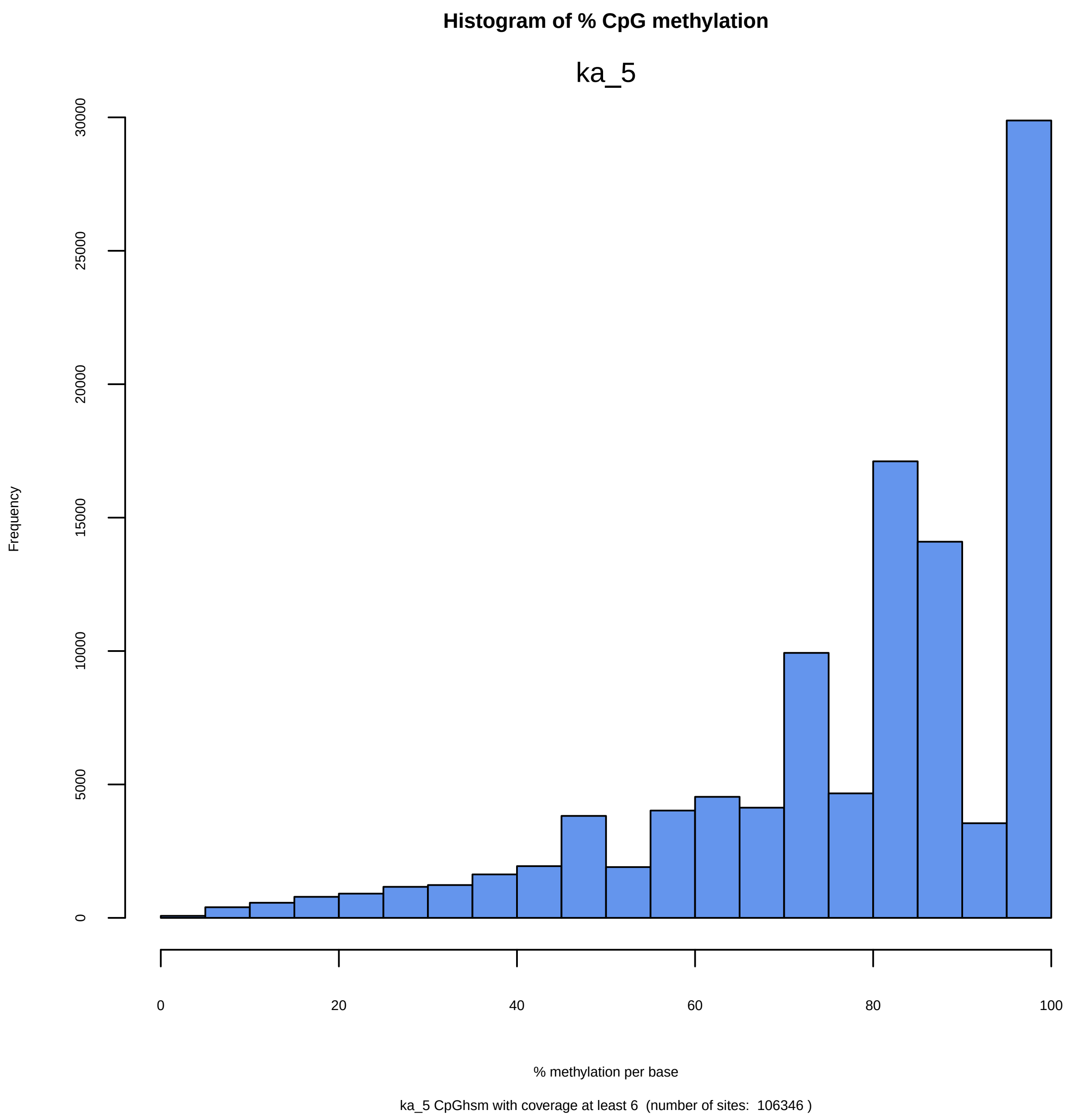

Histogram of % CpG methylation
ka_5
Frequency
0
5000
10000
15000
20000
25000
30000
0
20
40
60
80
100
% methylation per base
ka_5 CpGhsm with coverage at least 6 (number of sites: 106346 )

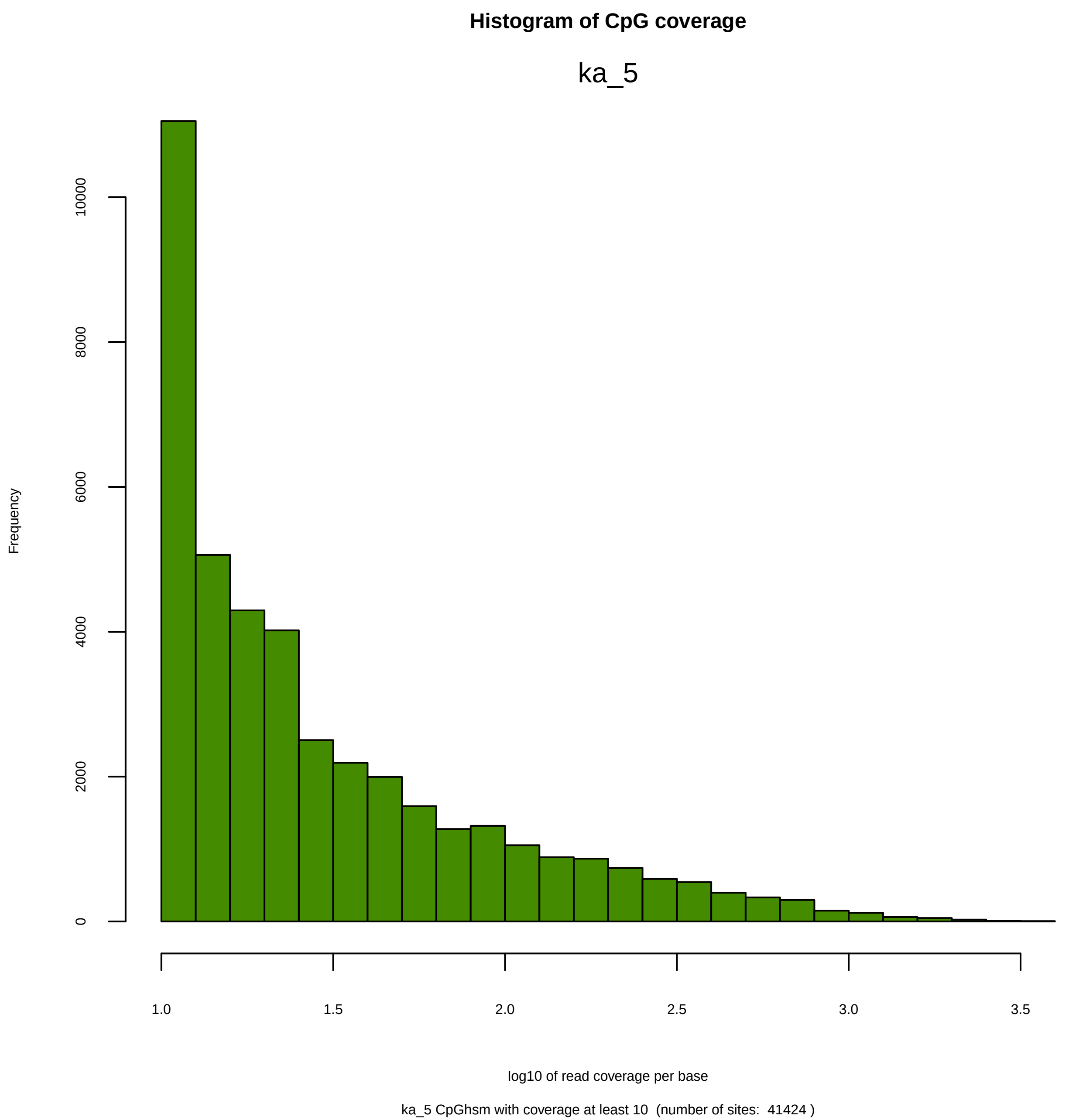
Histogram of CpG coverage
ka_5
Frequency
0
2000
4000
6000
8000
10000
1.0
1.5
2.0
2.5
3.0
3.5
log10 of read coverage per base
ka_5 CpGhsm with coverage at least 10  (number of sites:  41424 )

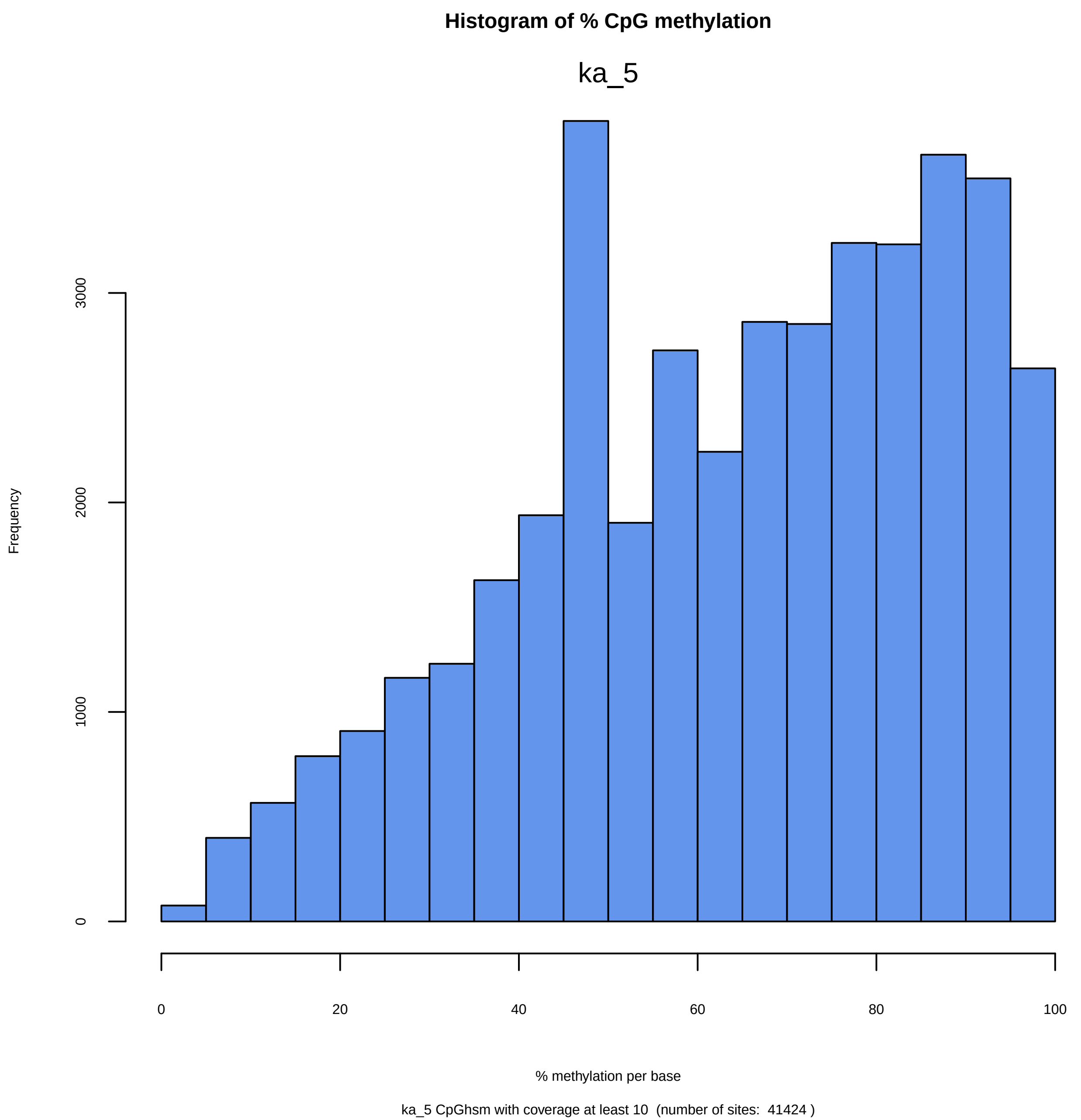
Histogram of % CpG methylation
ka_5
Frequency
0
1000
2000
3000
0
20
40
60
80
100
% methylation per base
ka_5 CpGhsm with coverage at least 10 (number of sites: 41424 )

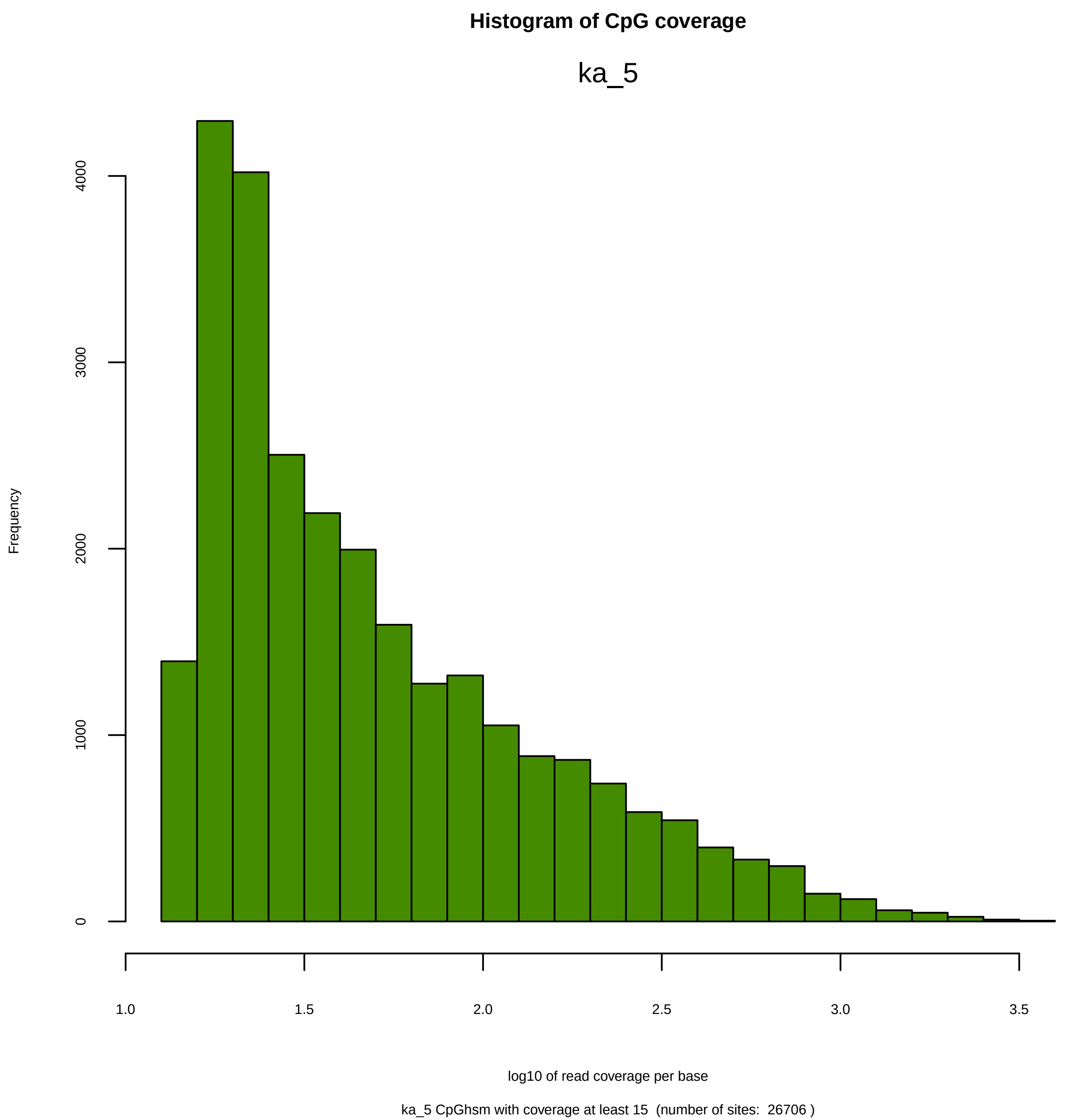
Histogram of CpG coverage
ka_5
Frequency
0
1000
2000
3000
4000
1.0
1.5
2.0
2.5
3.0
3.5
log10 of read coverage per base
ka_5 CpGhsm with coverage at least 15  (number of sites:  26706 )

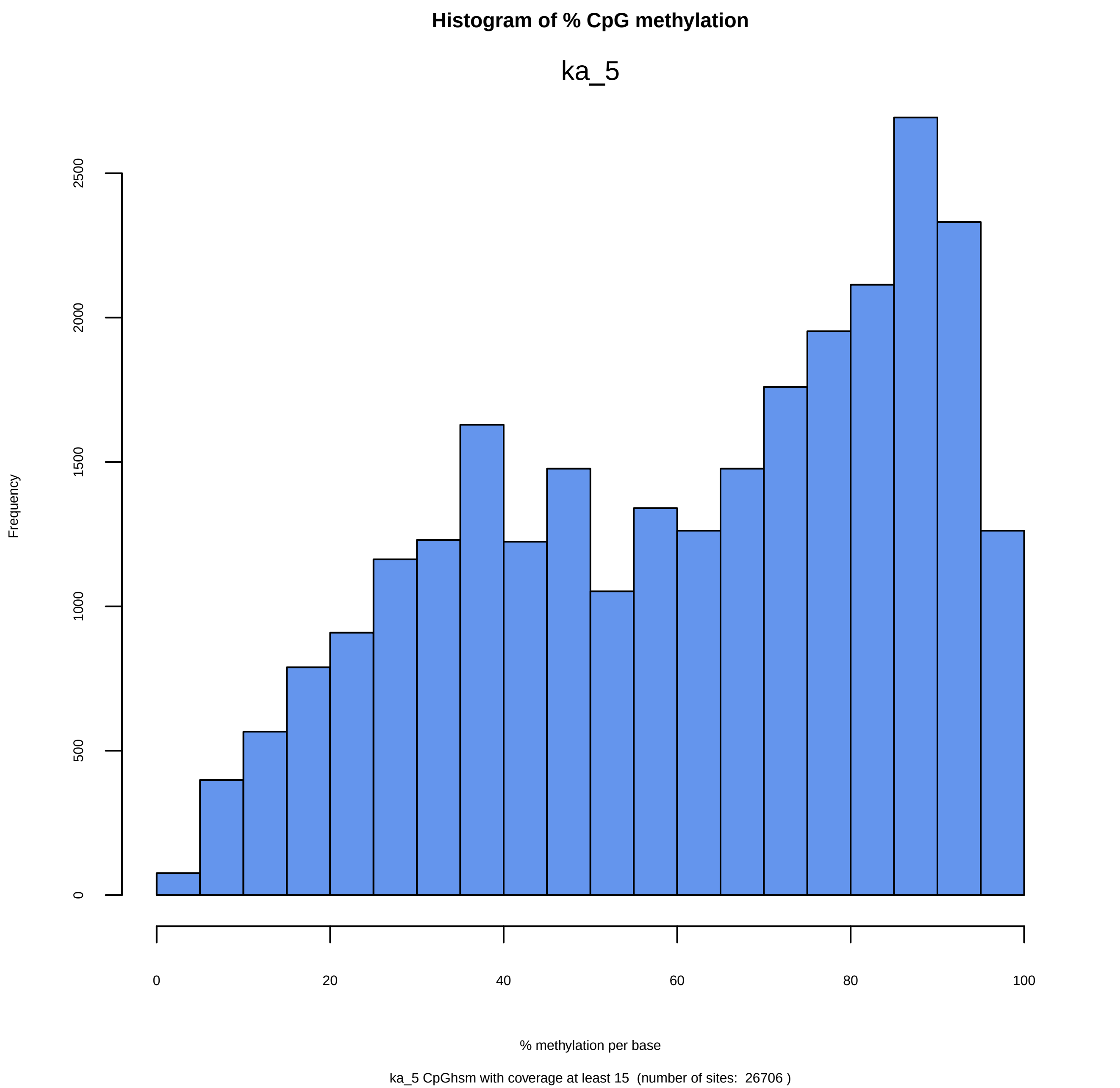
Histogram of % CpG methylation
ka_5
Frequency
0
500
1000
1500
2000
2500
0
20
40
60
80
100
% methylation per base
ka_5 CpGhsm with coverage at least 15 (number of sites: 26706 )

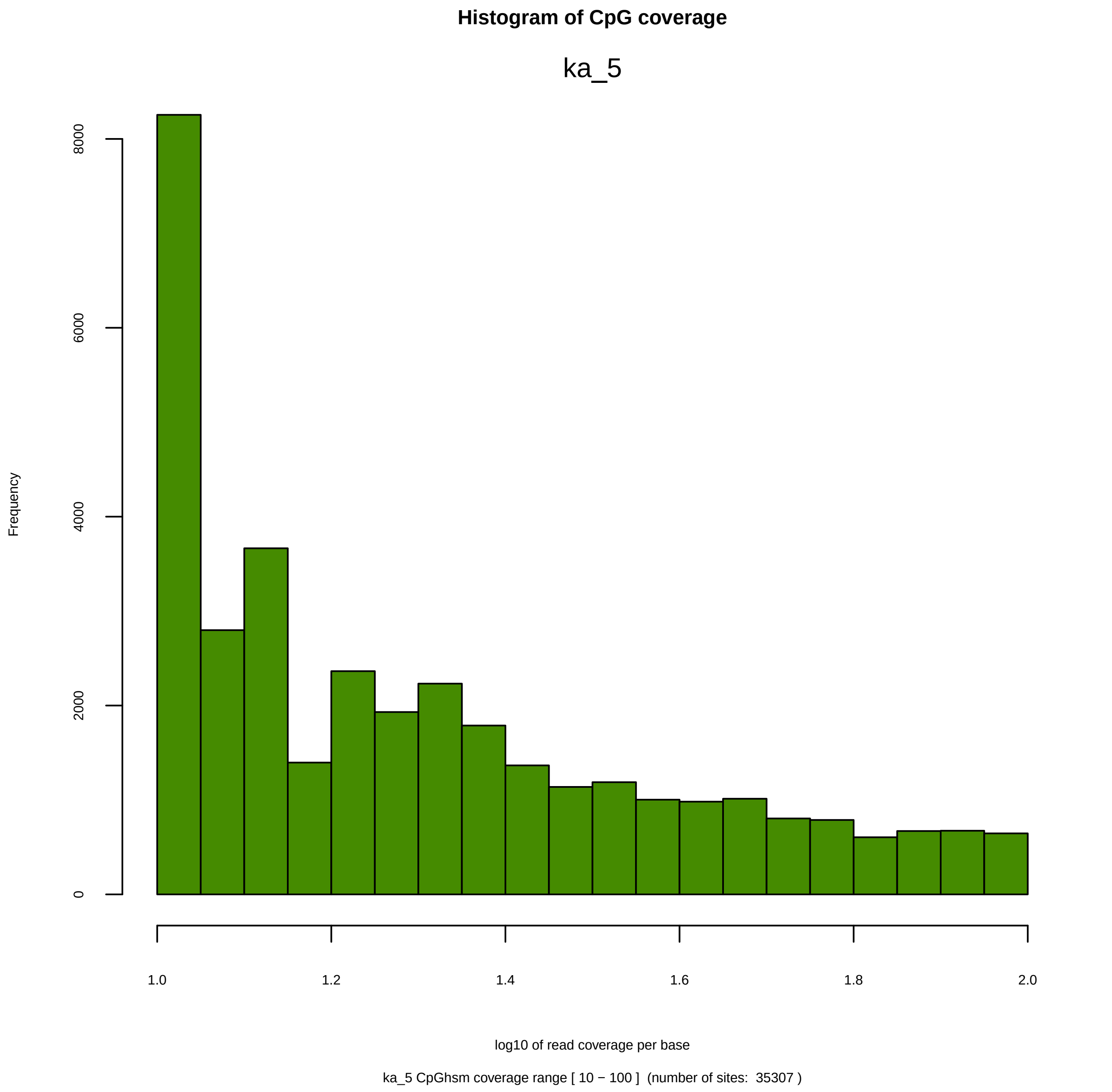

Histogram of CpG coverage
ka_5
Frequency
0
2000
4000
6000
8000
1.0
1.2
1.4
1.6
1.8
2.0
log10 of read coverage per base
ka_5 CpGhsm coverage range [ 10 − 100 ]  (number of sites:  35307 )

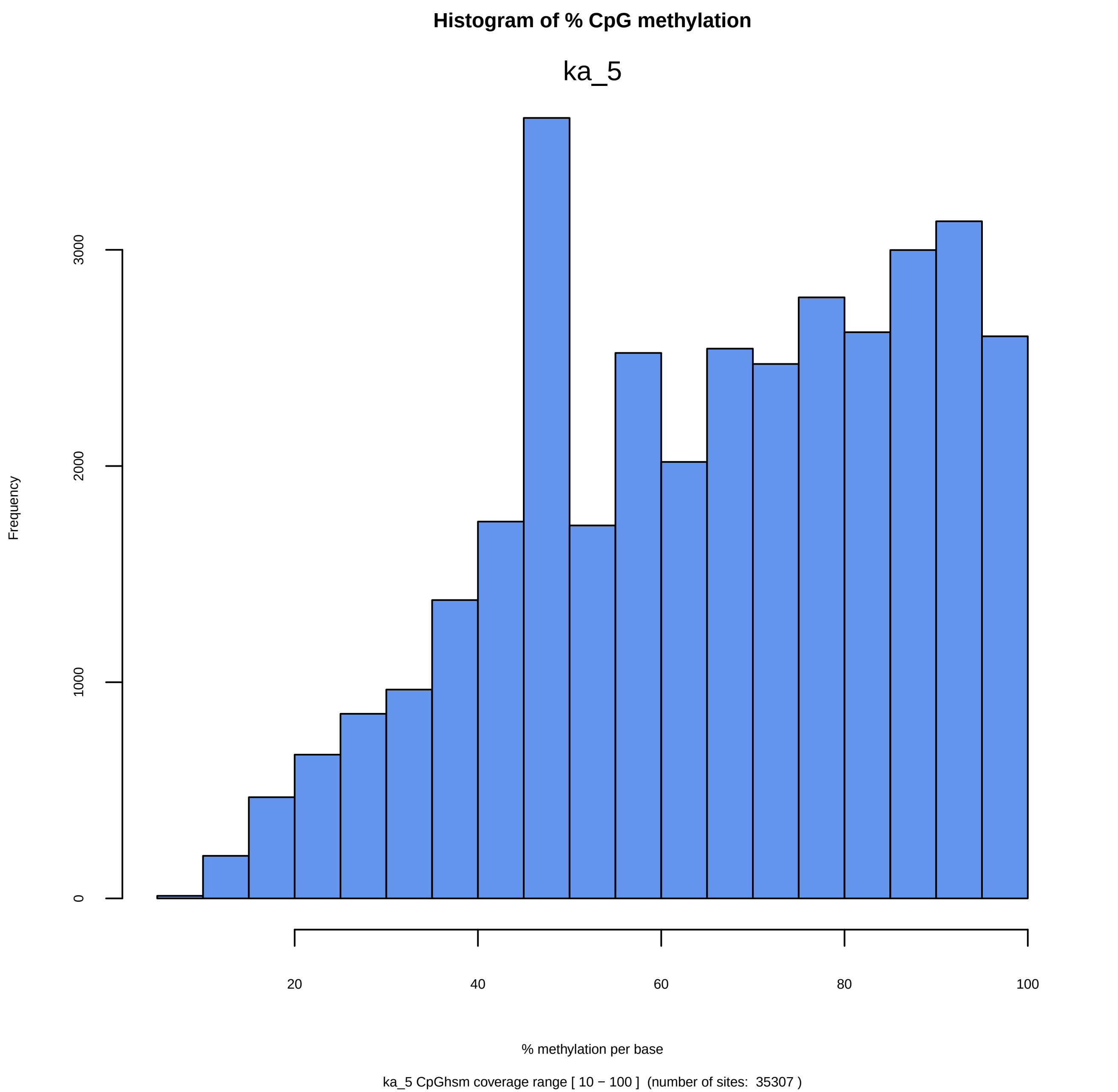
Histogram of % CpG methylation
ka_5
Frequency
0
1000
2000
3000
20
40
60
80
100
% methylation per base
ka_5 CpGhsm coverage range [ 10 − 100 ] (number of sites: 35307 )

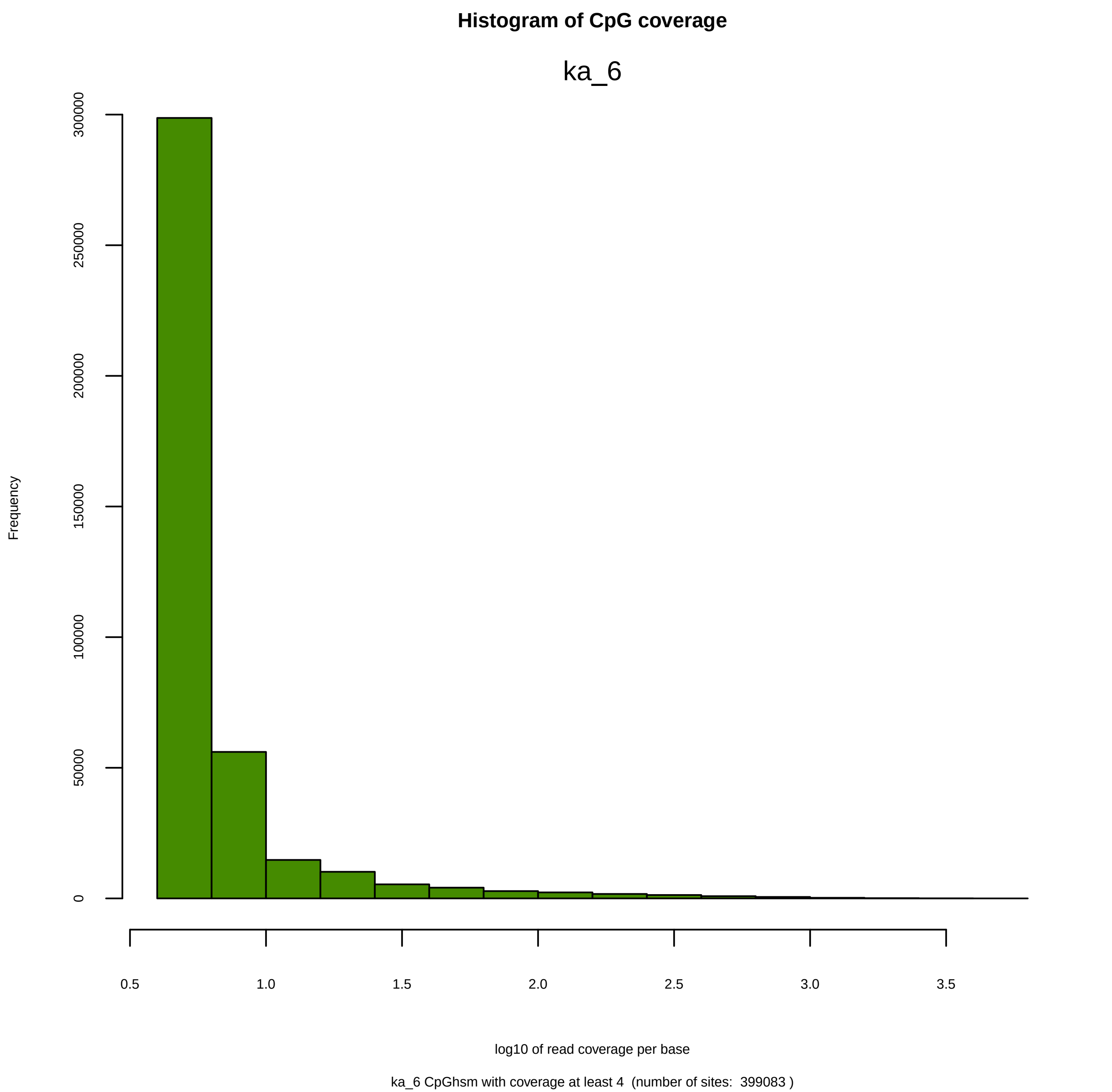
Histogram of CpG coverage
ka_6
Frequency
0
50000
100000
150000
200000
250000
300000
0.5
1.0
1.5
2.0
2.5
3.0
3.5
log10 of read coverage per base
ka_6 CpGhsm with coverage at least 4  (number of sites:  399083 )

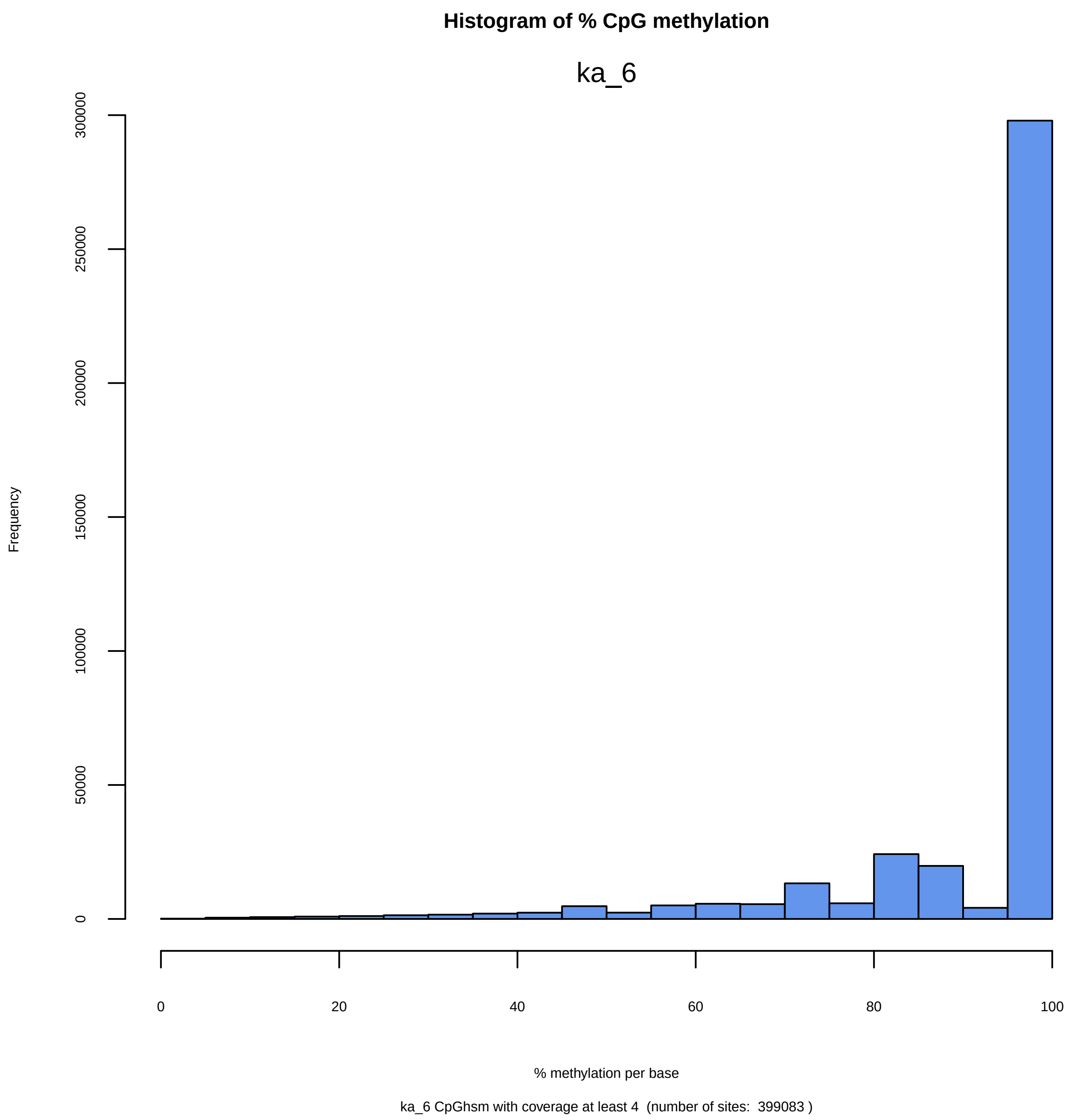

Histogram of % CpG methylation
ka_6
Frequency
0
50000
100000
150000
200000
250000
300000
0
20
40
60
80
100
% methylation per base
ka_6 CpGhsm with coverage at least 4 (number of sites: 399083 )

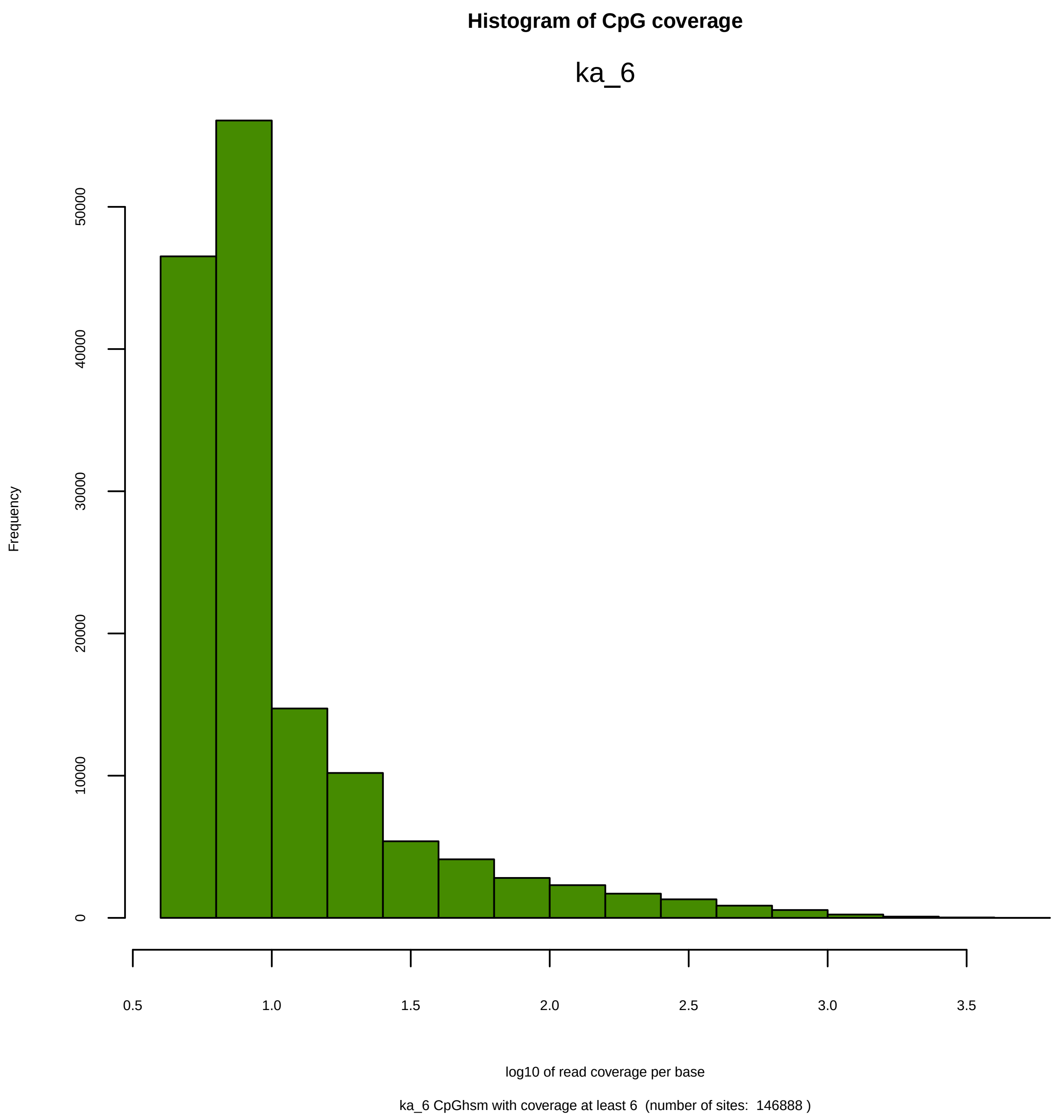
Histogram of CpG coverage
ka_6
Frequency
0
10000
20000
30000
40000
50000
0.5
1.0
1.5
2.0
2.5
3.0
3.5
log10 of read coverage per base
ka_6 CpGhsm with coverage at least 6  (number of sites:  146888 )

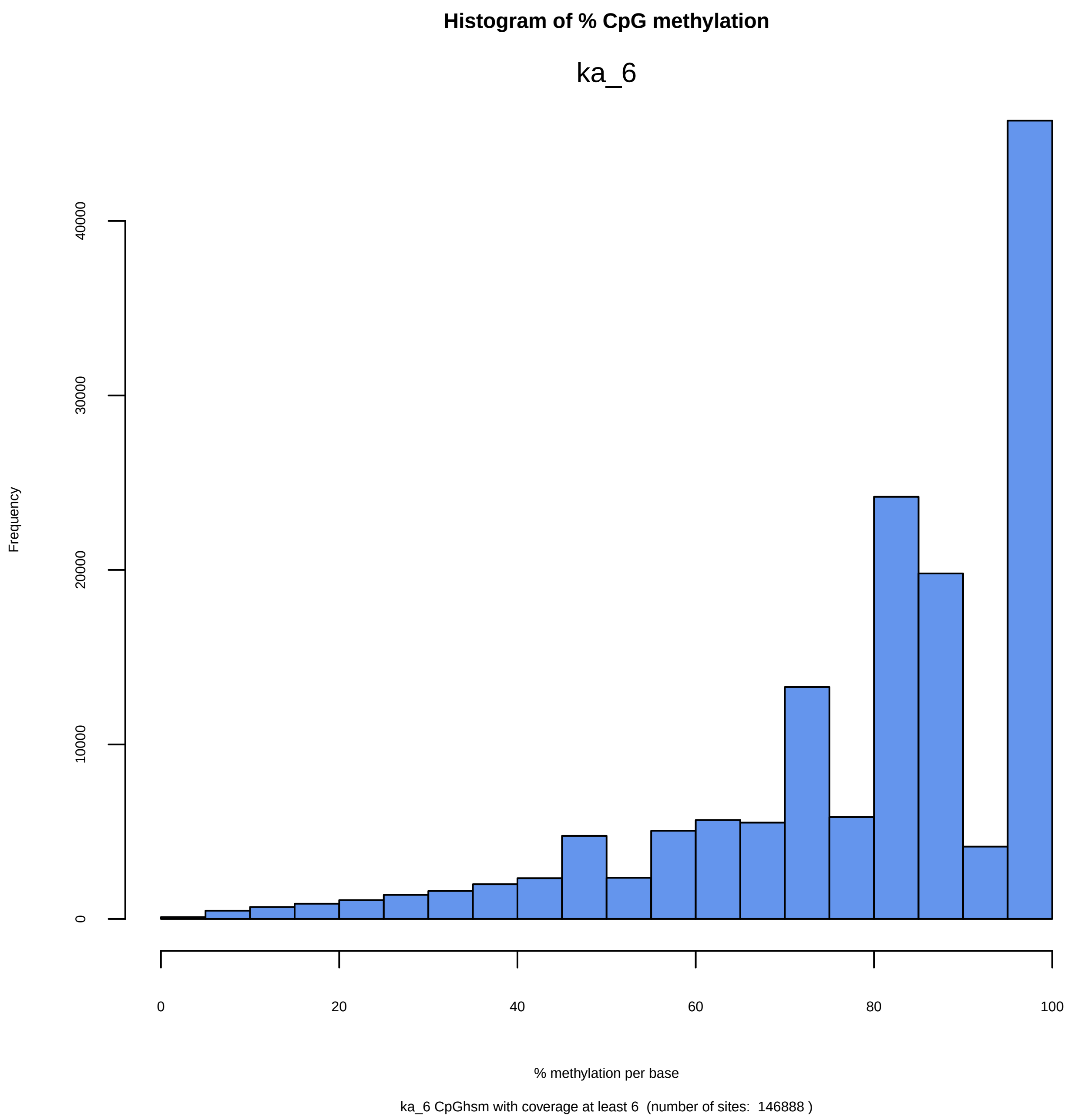

Histogram of % CpG methylation
ka_6
Frequency
0
10000
20000
30000
40000
0
20
40
60
80
100
% methylation per base
ka_6 CpGhsm with coverage at least 6 (number of sites: 146888 )

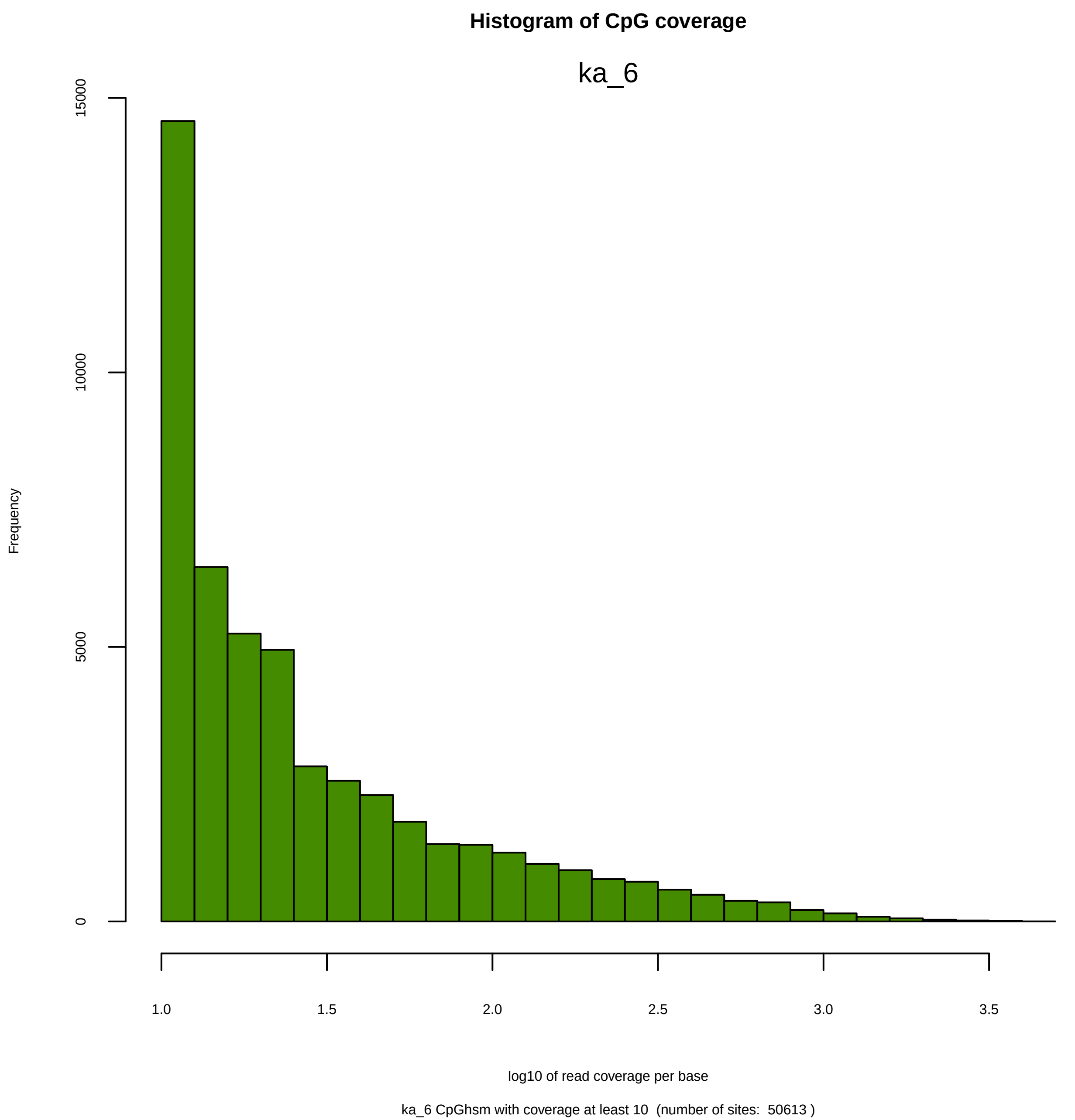
Histogram of CpG coverage
ka_6
Frequency
0
5000
10000
15000
1.0
1.5
2.0
2.5
3.0
3.5
log10 of read coverage per base
ka_6 CpGhsm with coverage at least 10 (number of sites: 50613 )

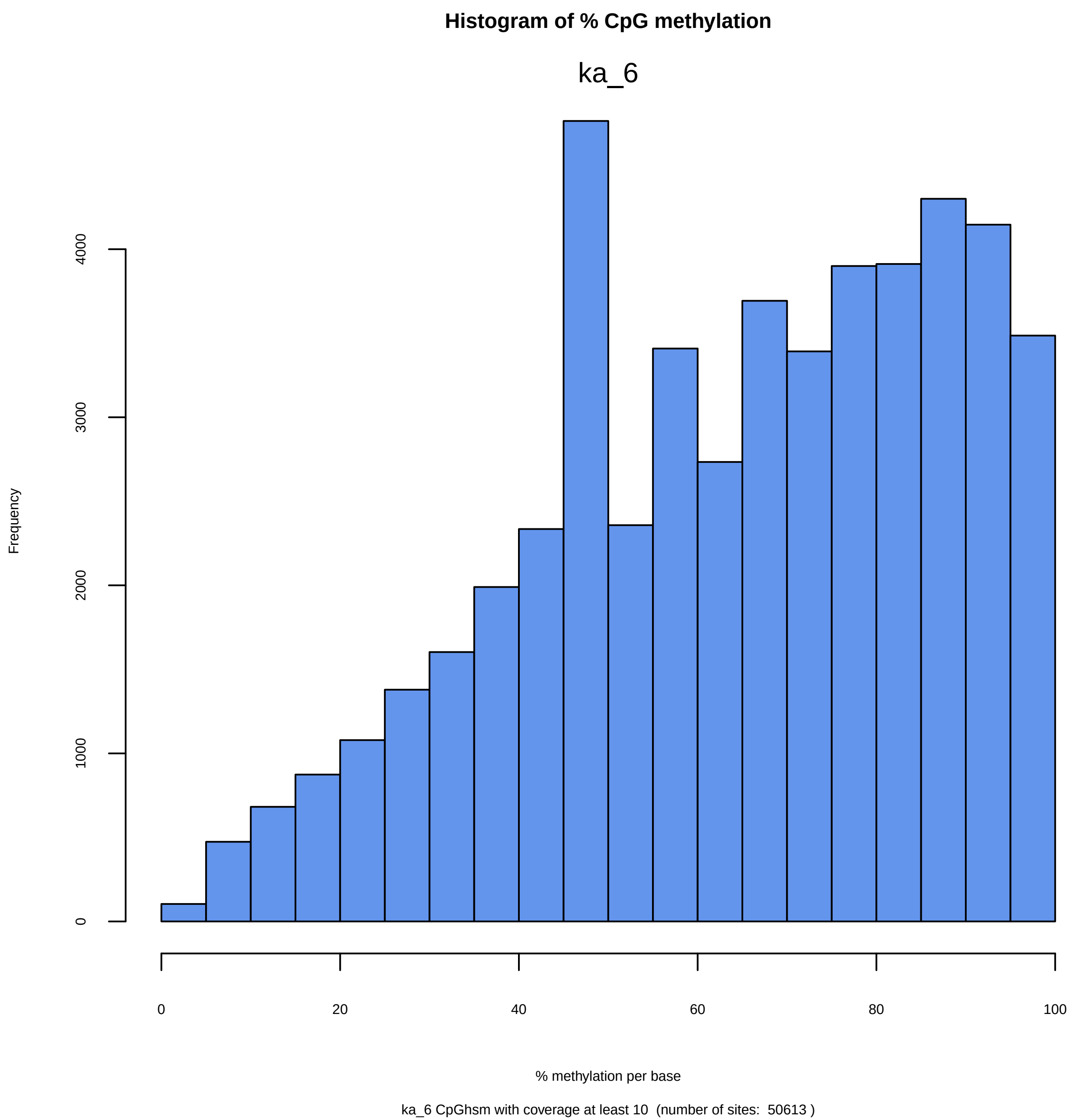
Histogram of % CpG methylation
ka_6
Frequency
0
1000
2000
3000
4000
0
20
40
60
80
100
% methylation per base
ka_6 CpGhsm with coverage at least 10  (number of sites:  50613 )

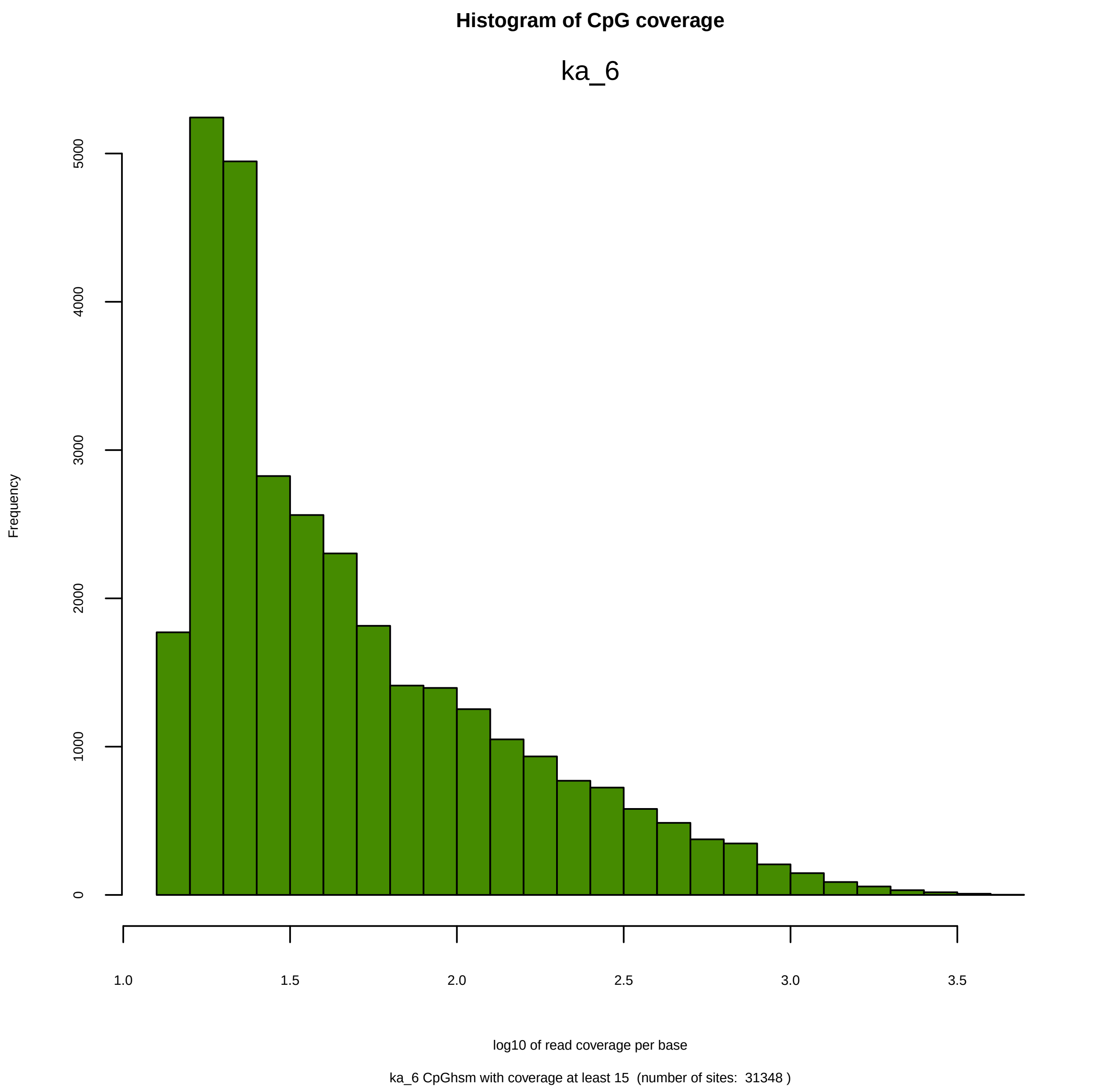
Histogram of CpG coverage
ka_6
Frequency
0
1000
2000
3000
4000
5000
1.0
1.5
2.0
2.5
3.0
3.5
log10 of read coverage per base
ka_6 CpGhsm with coverage at least 15  (number of sites:  31348 )

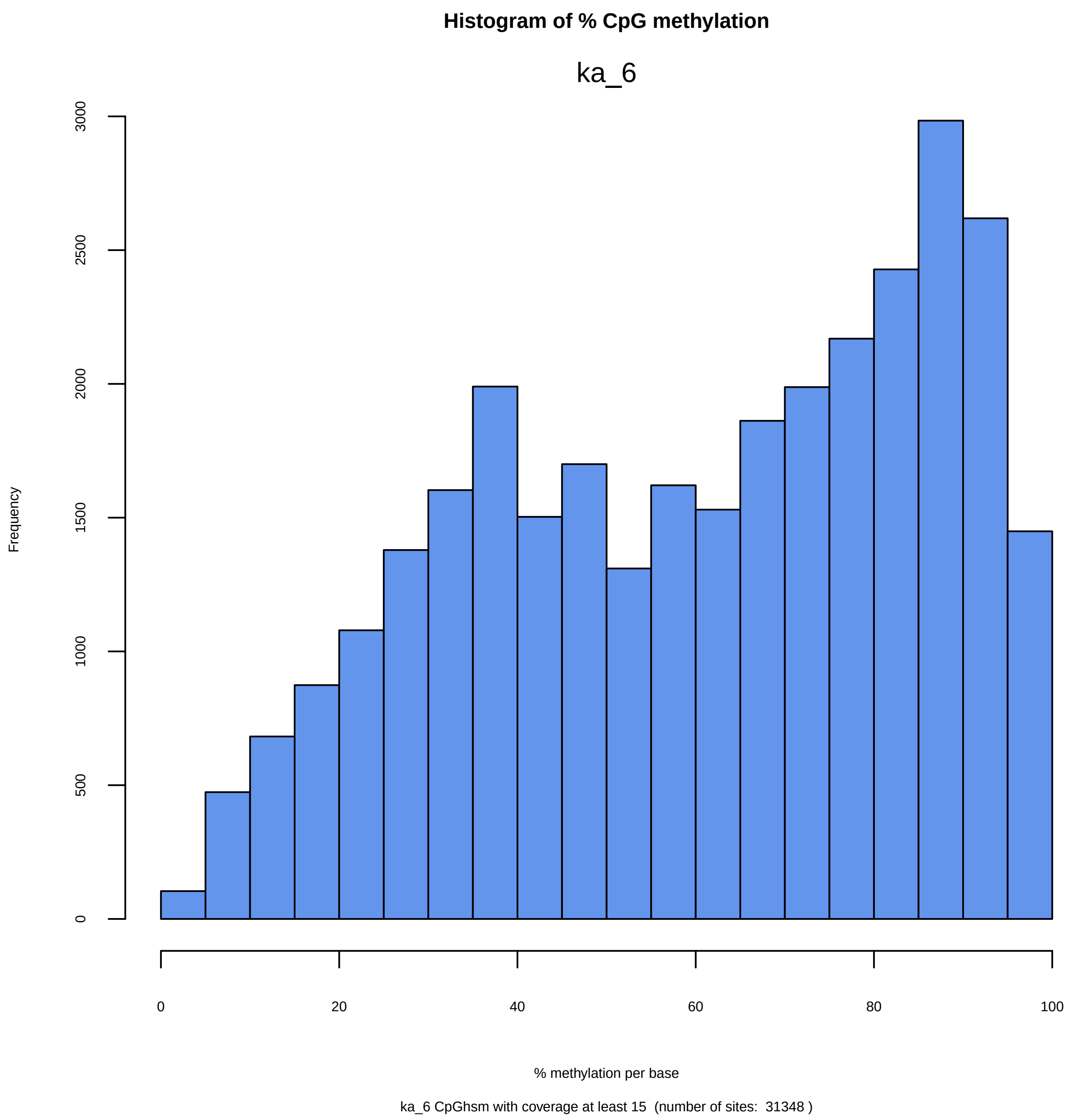

Histogram of % CpG methylation
ka_6
Frequency
0
500
1000
1500
2000
2500
3000
0
20
40
60
80
100
% methylation per base
ka_6 CpGhsm with coverage at least 15  (number of sites:  31348 )

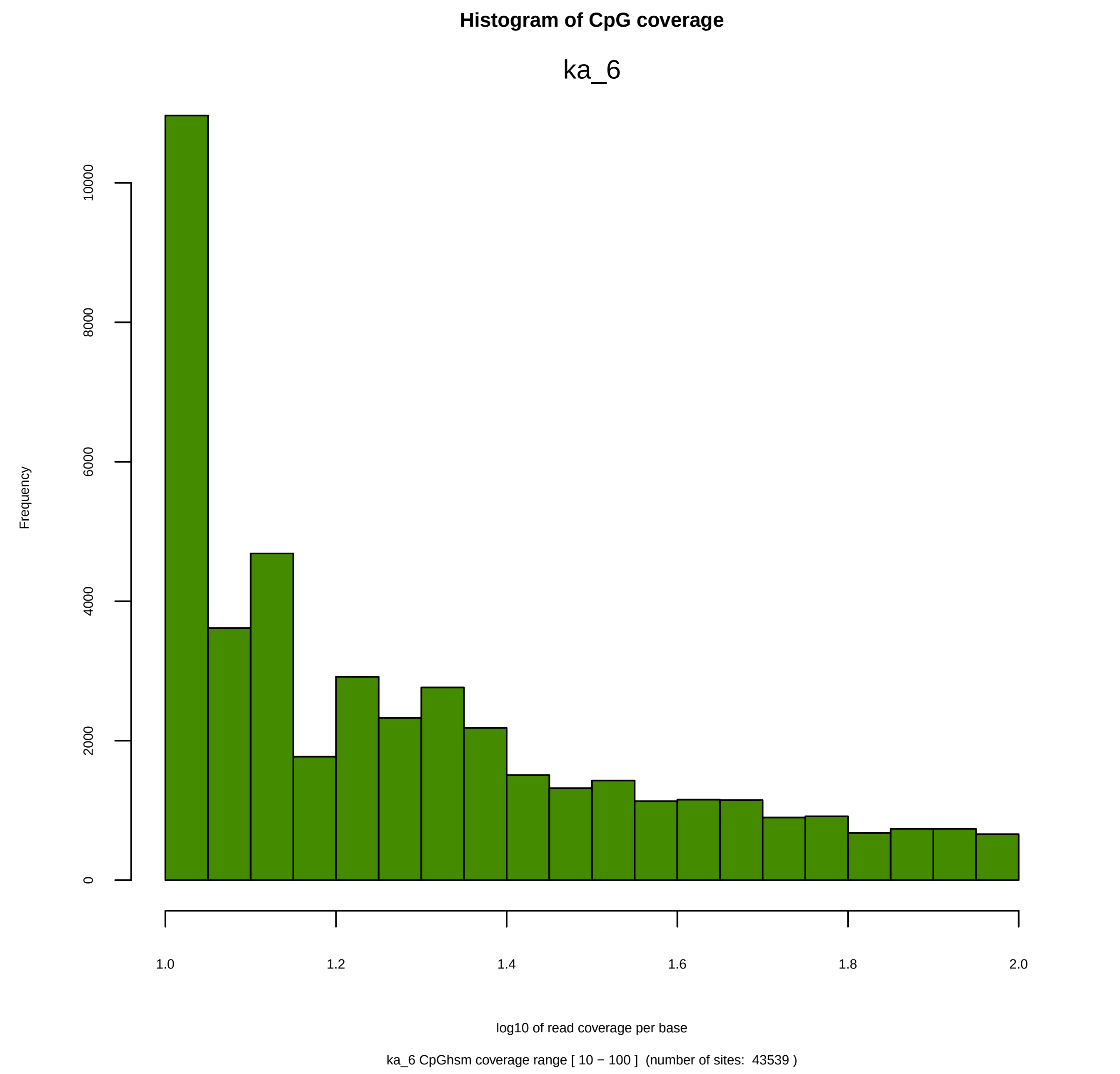
Histogram of CpG coverage
ka_6
Frequency
0
2000
4000
6000
8000
10000
1.0
1.2
1.4
1.6
1.8
2.0
log10 of read coverage per base
ka_6 CpGhsm coverage range [ 10 − 100 ]  (number of sites:  43539 )

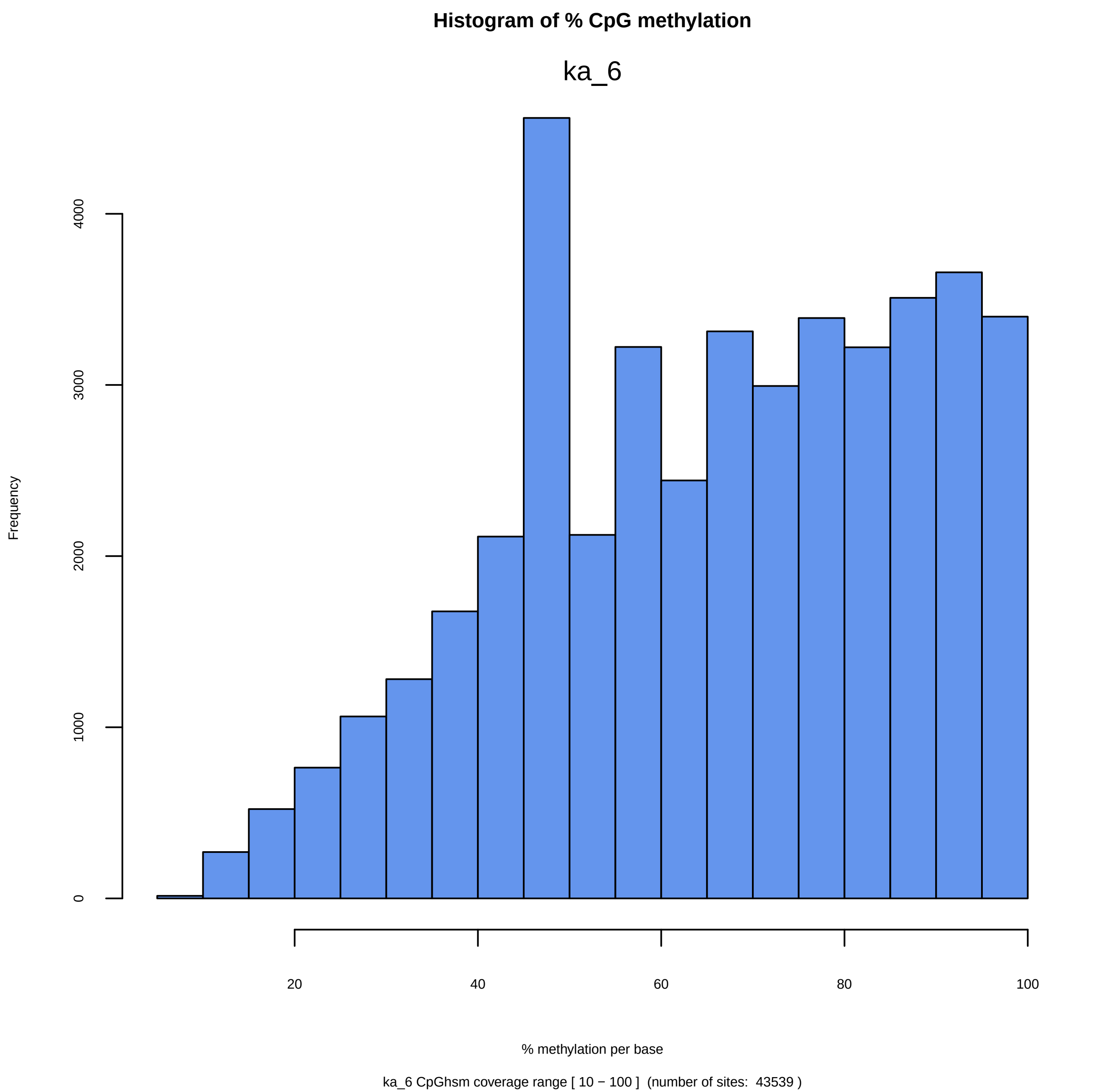

Histogram of % CpG methylation
ka_6
Frequency
0
1000
2000
3000
4000
20
40
60
80
100
% methylation per base
ka_6 CpGhsm coverage range [ 10 − 100 ]  (number of sites:  43539 )

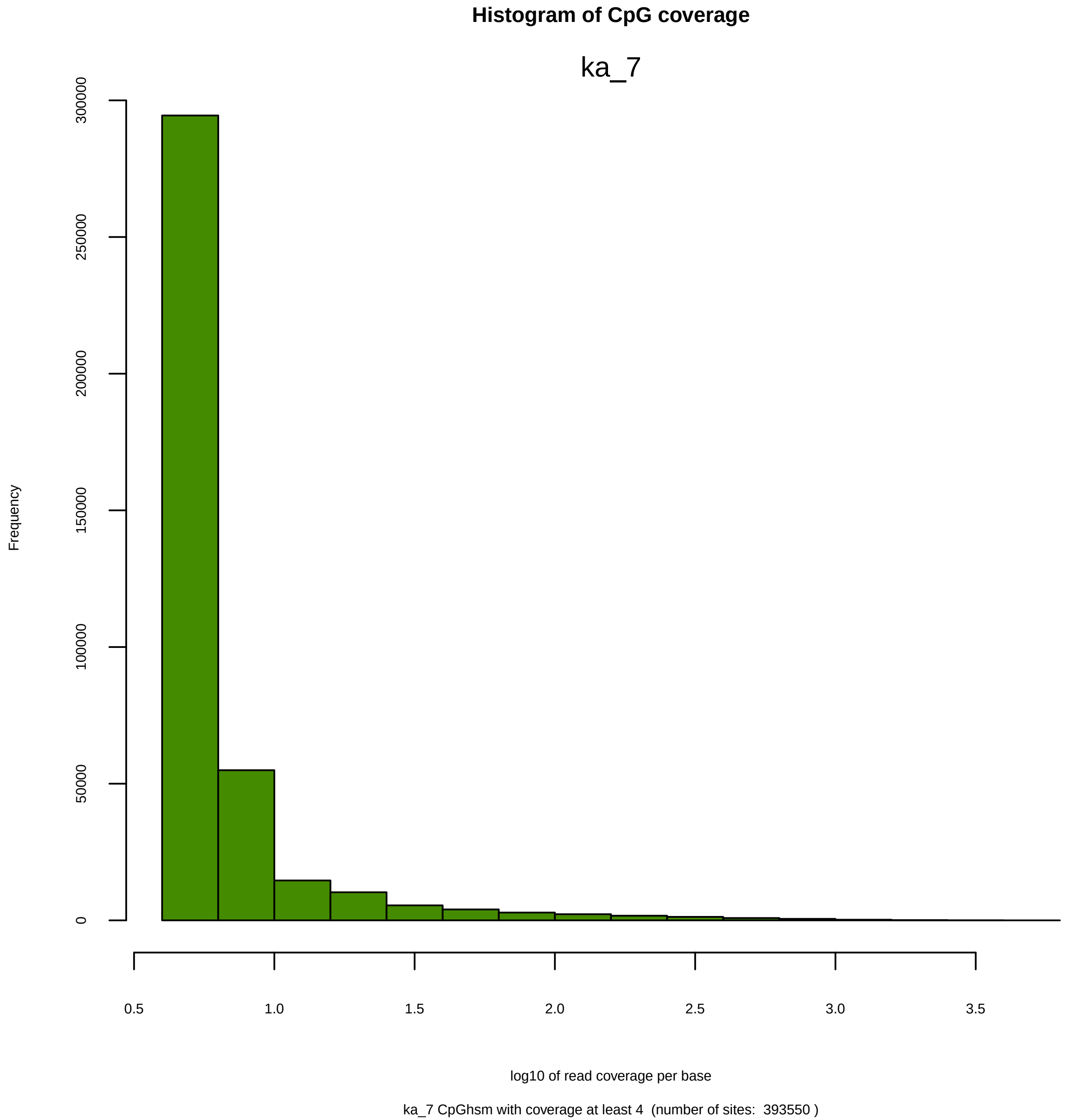
Histogram of CpG coverage
ka_7
Frequency
0
50000
100000
150000
200000
250000
300000
0.5
1.0
1.5
2.0
2.5
3.0
3.5
log10 of read coverage per base
ka_7 CpGhsm with coverage at least 4 (number of sites: 393550 )

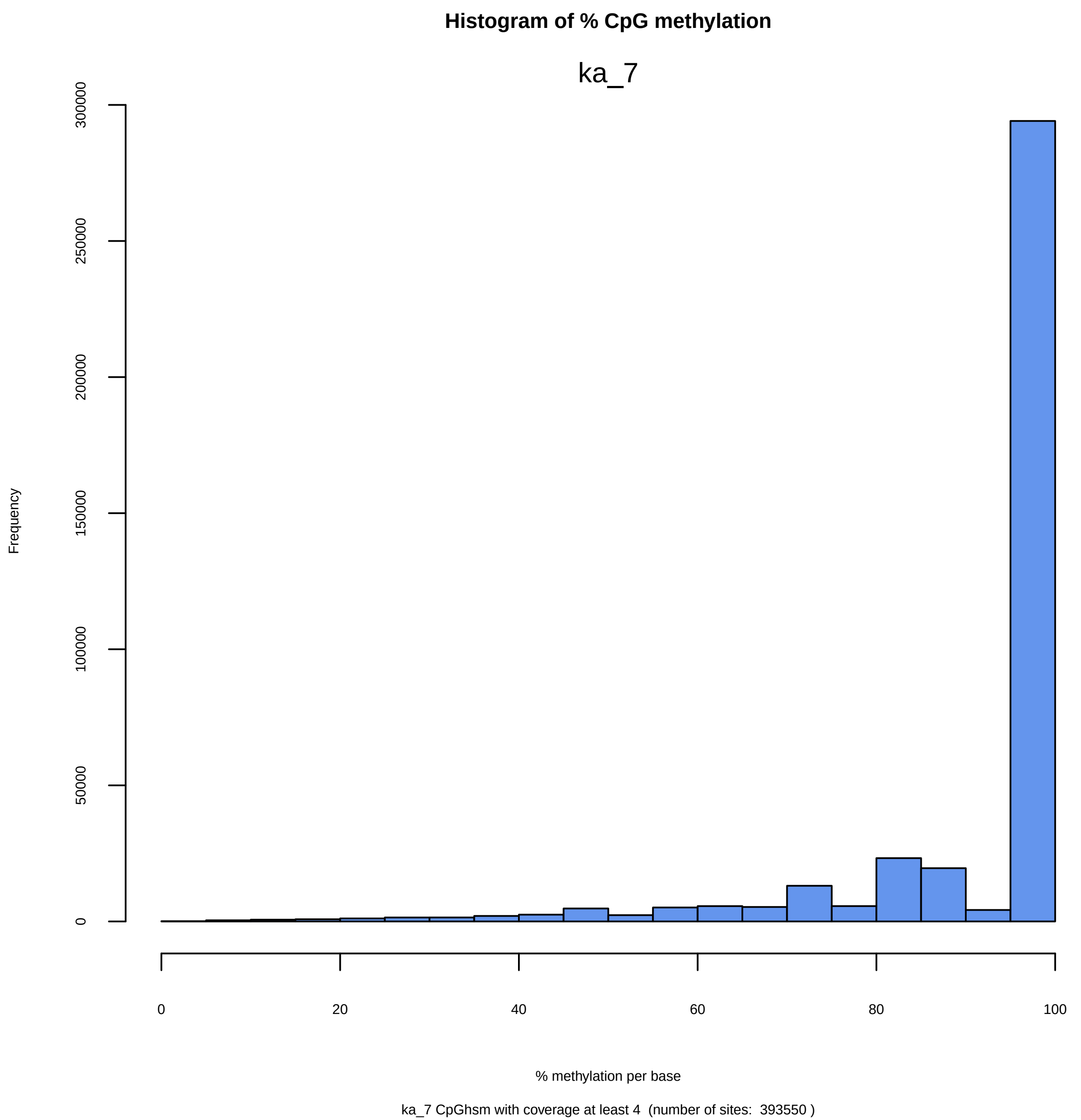

Histogram of % CpG methylation
ka_7
Frequency
0
50000
100000
150000
200000
250000
300000
0
20
40
60
80
100
% methylation per base
ka_7 CpGhsm with coverage at least 4  (number of sites:  393550 )

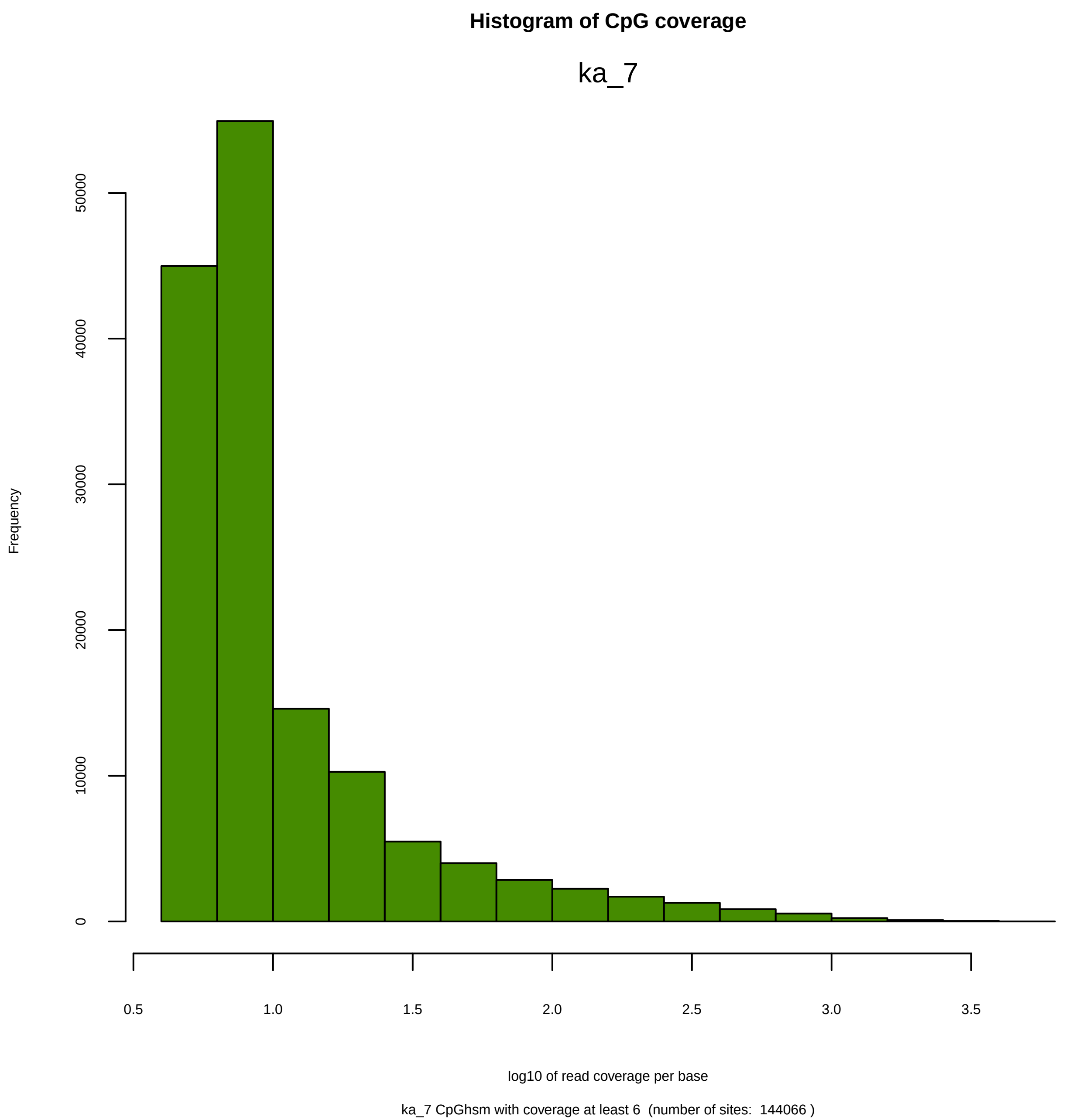

Histogram of CpG coverage
ka_7
Frequency
0
10000
20000
30000
40000
50000
0.5
1.0
1.5
2.0
2.5
3.0
3.5
log10 of read coverage per base
ka_7 CpGhsm with coverage at least 6 (number of sites: 144066 )

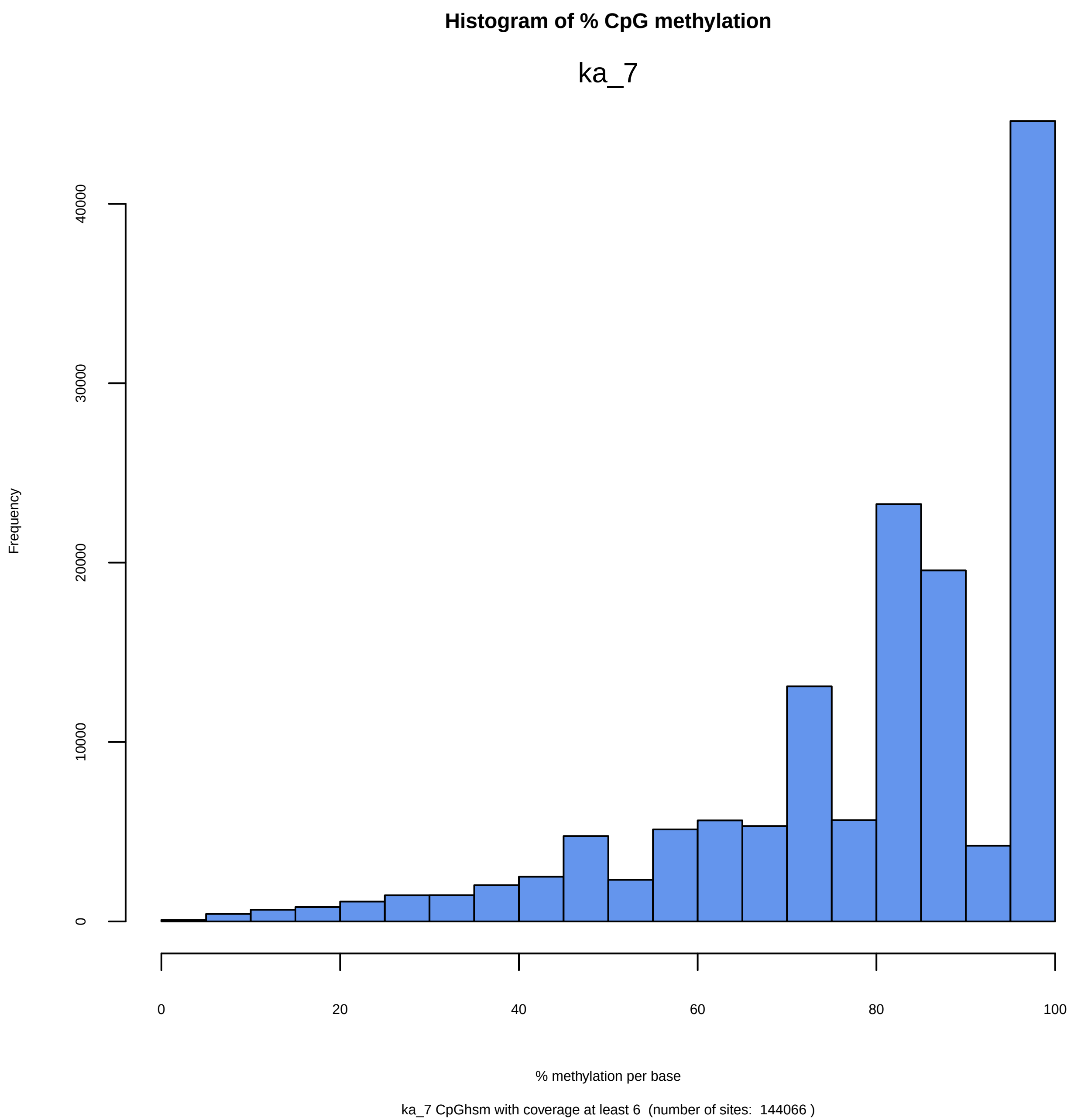

Histogram of % CpG methylation
ka_7
Frequency
0
10000
20000
30000
40000
0
20
40
60
80
100
% methylation per base
ka_7 CpGhsm with coverage at least 6 (number of sites: 144066 )

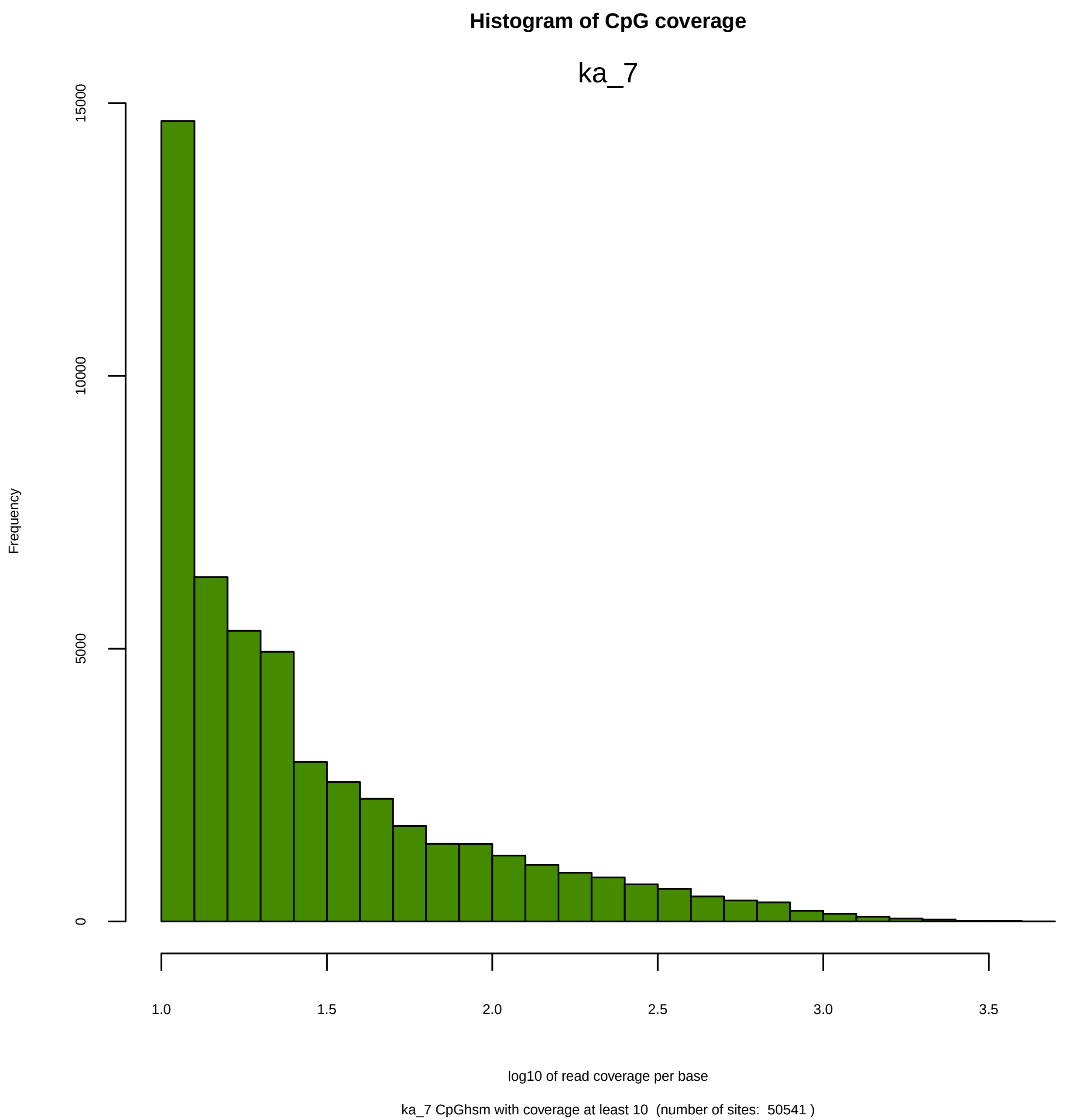
Histogram of CpG coverage
ka_7
Frequency
0
5000
10000
15000
1.0
1.5
2.0
2.5
3.0
3.5
log10 of read coverage per base
ka_7 CpGhsm with coverage at least 10  (number of sites:  50541 )

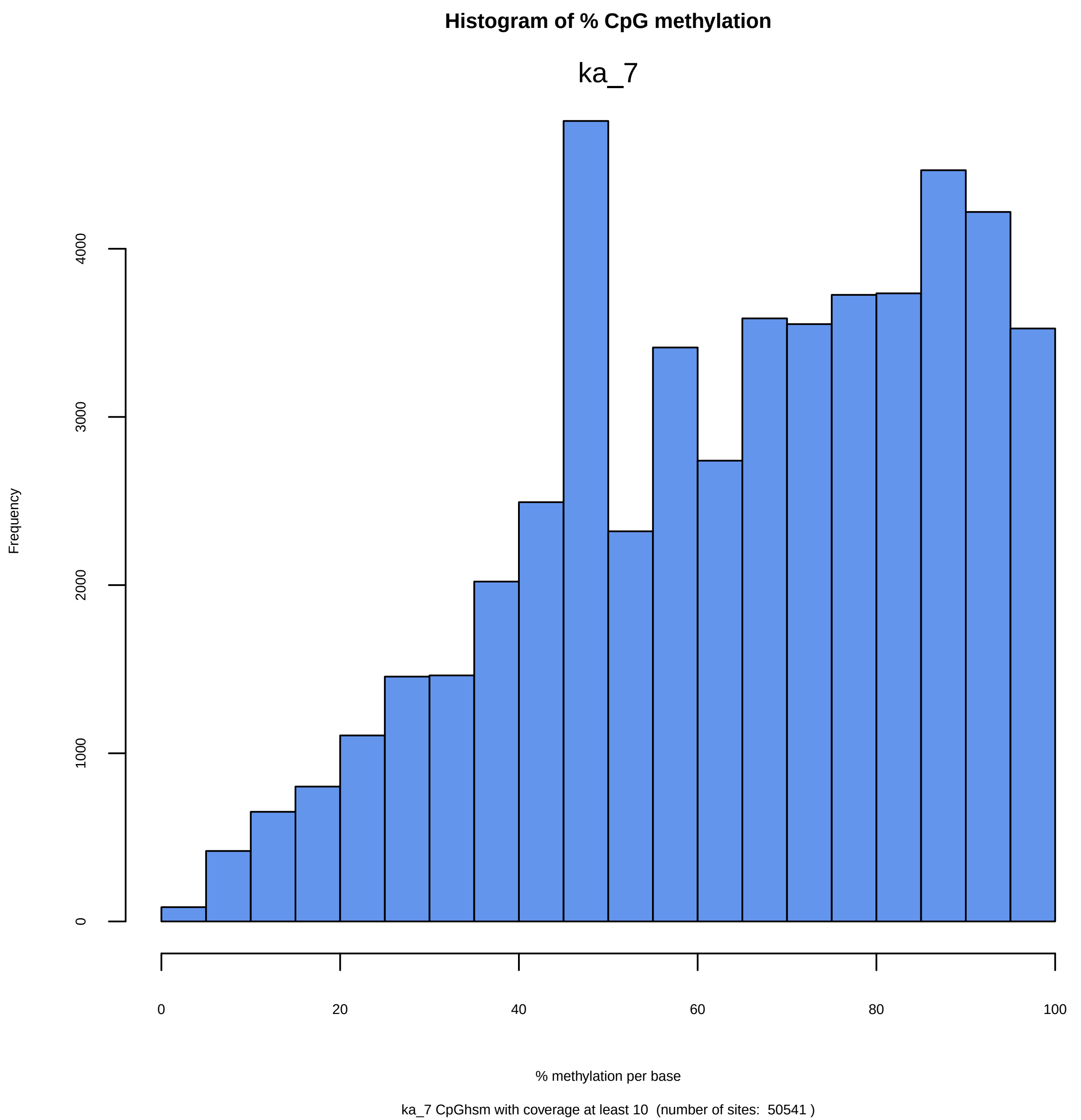

Histogram of % CpG methylation
ka_7
Frequency
0
1000
2000
3000
4000
0
20
40
60
80
100
% methylation per base
ka_7 CpGhsm with coverage at least 10 (number of sites: 50541 )

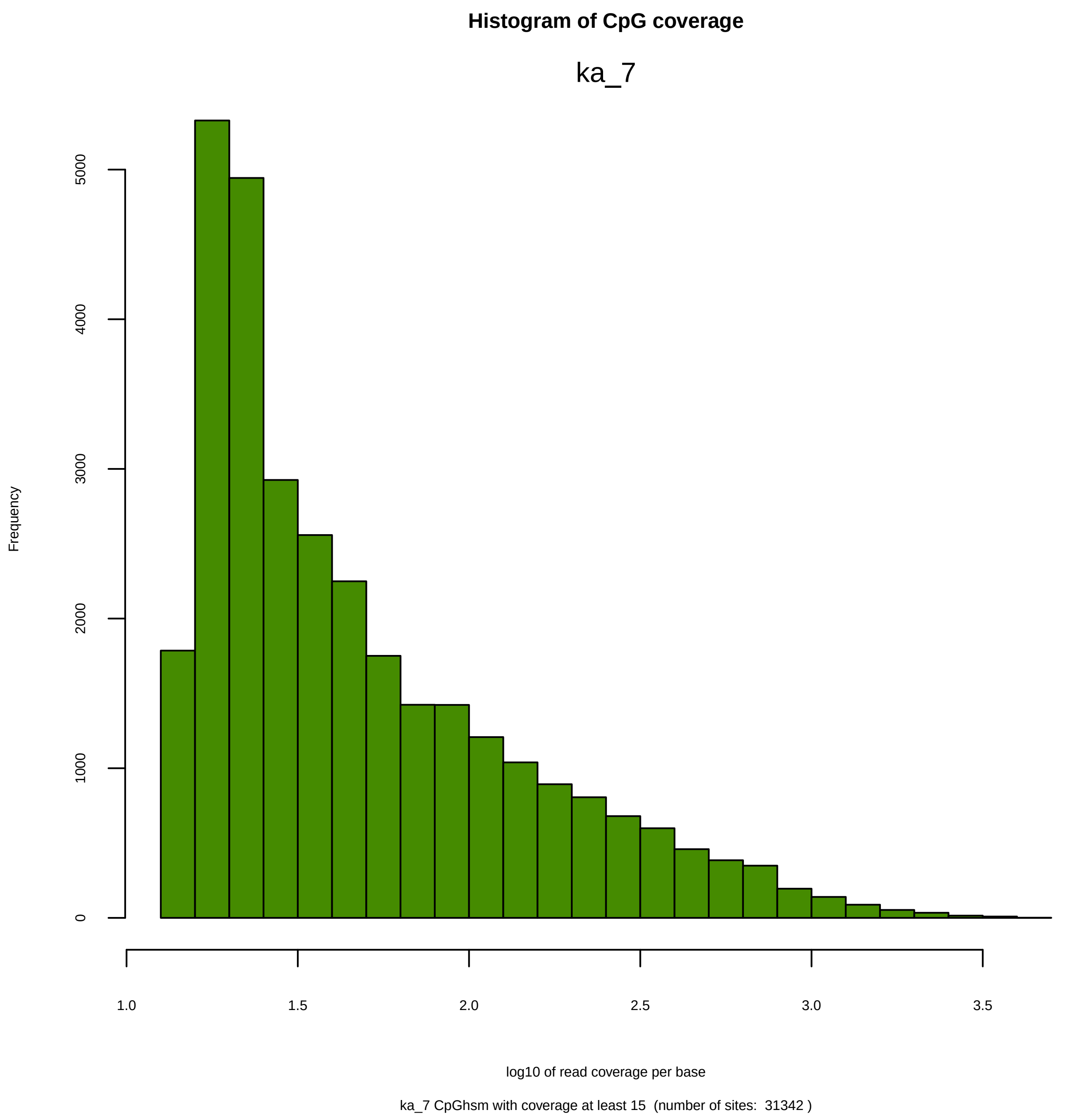
Histogram of CpG coverage
ka_7
Frequency
0
1000
2000
3000
4000
5000
1.0
1.5
2.0
2.5
3.0
3.5
log10 of read coverage per base
ka_7 CpGhsm with coverage at least 15  (number of sites:  31342 )

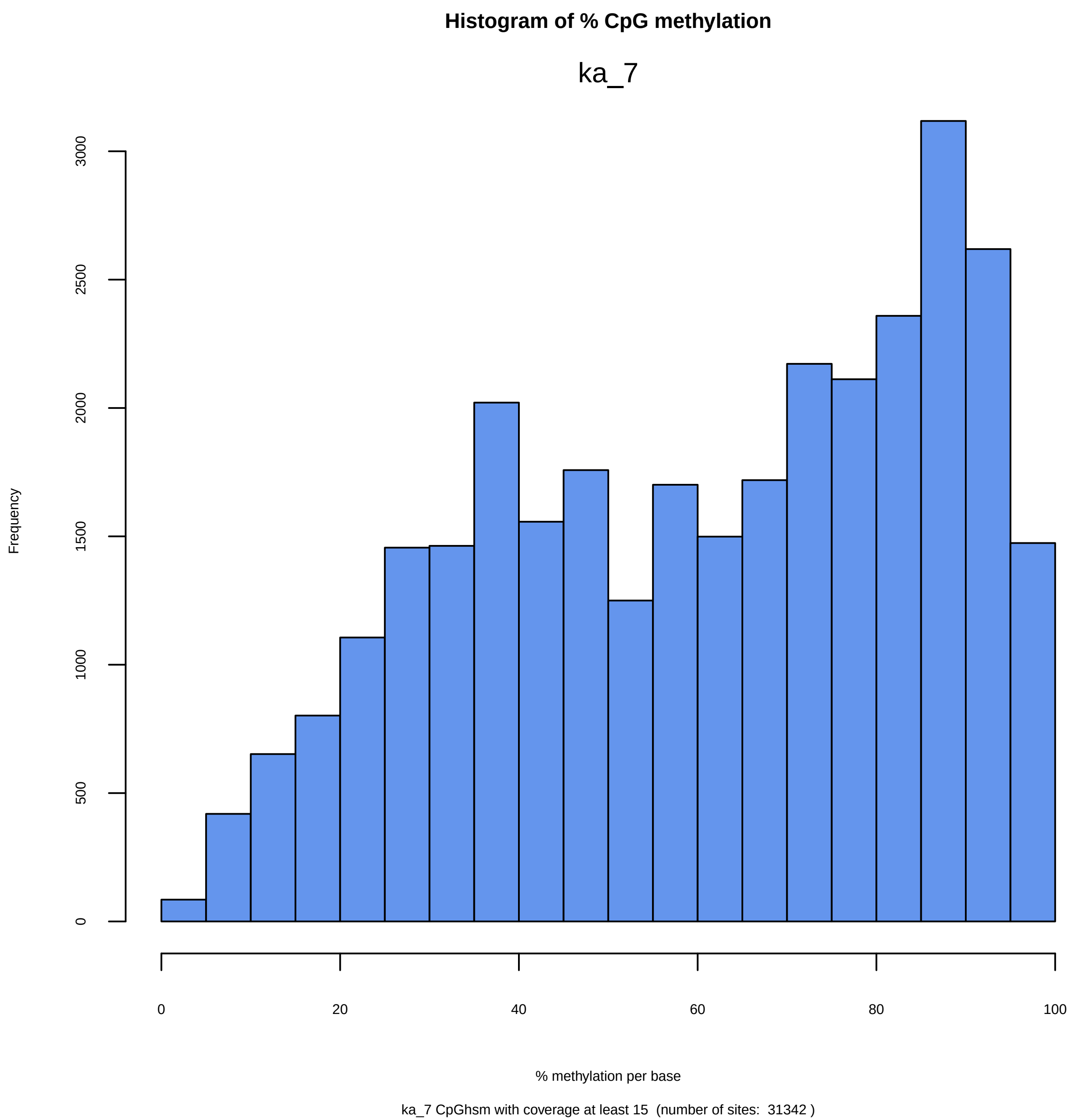
Histogram of % CpG methylation
ka_7
Frequency
0
500
1000
1500
2000
2500
3000
0
20
40
60
80
100
% methylation per base
ka_7 CpGhsm with coverage at least 15 (number of sites: 31342 )

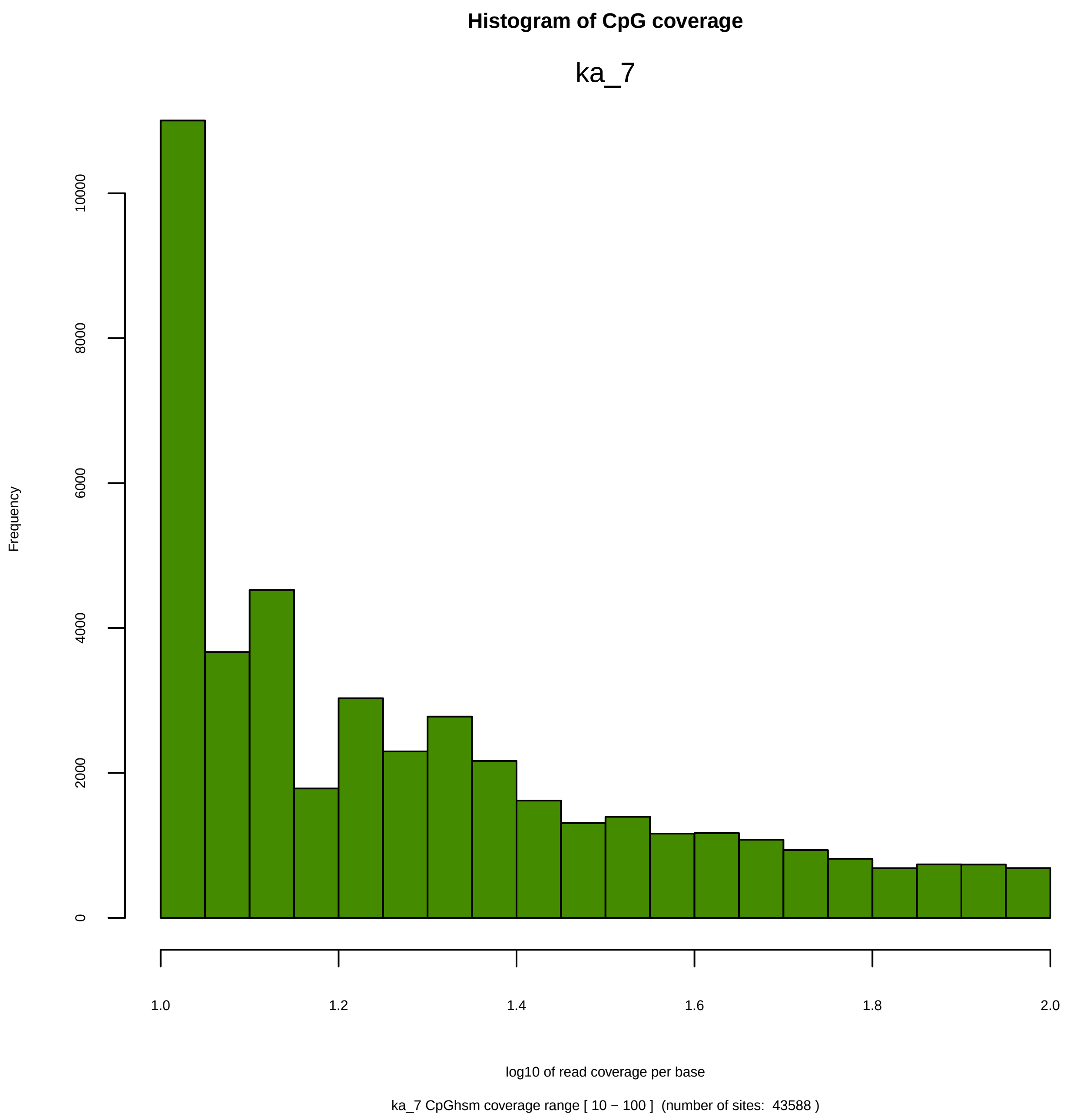

Histogram of CpG coverage
ka_7
Frequency
0
2000
4000
6000
8000
10000
1.0
1.2
1.4
1.6
1.8
2.0
log10 of read coverage per base
ka_7 CpGhsm coverage range [ 10 − 100 ]  (number of sites:  43588 )

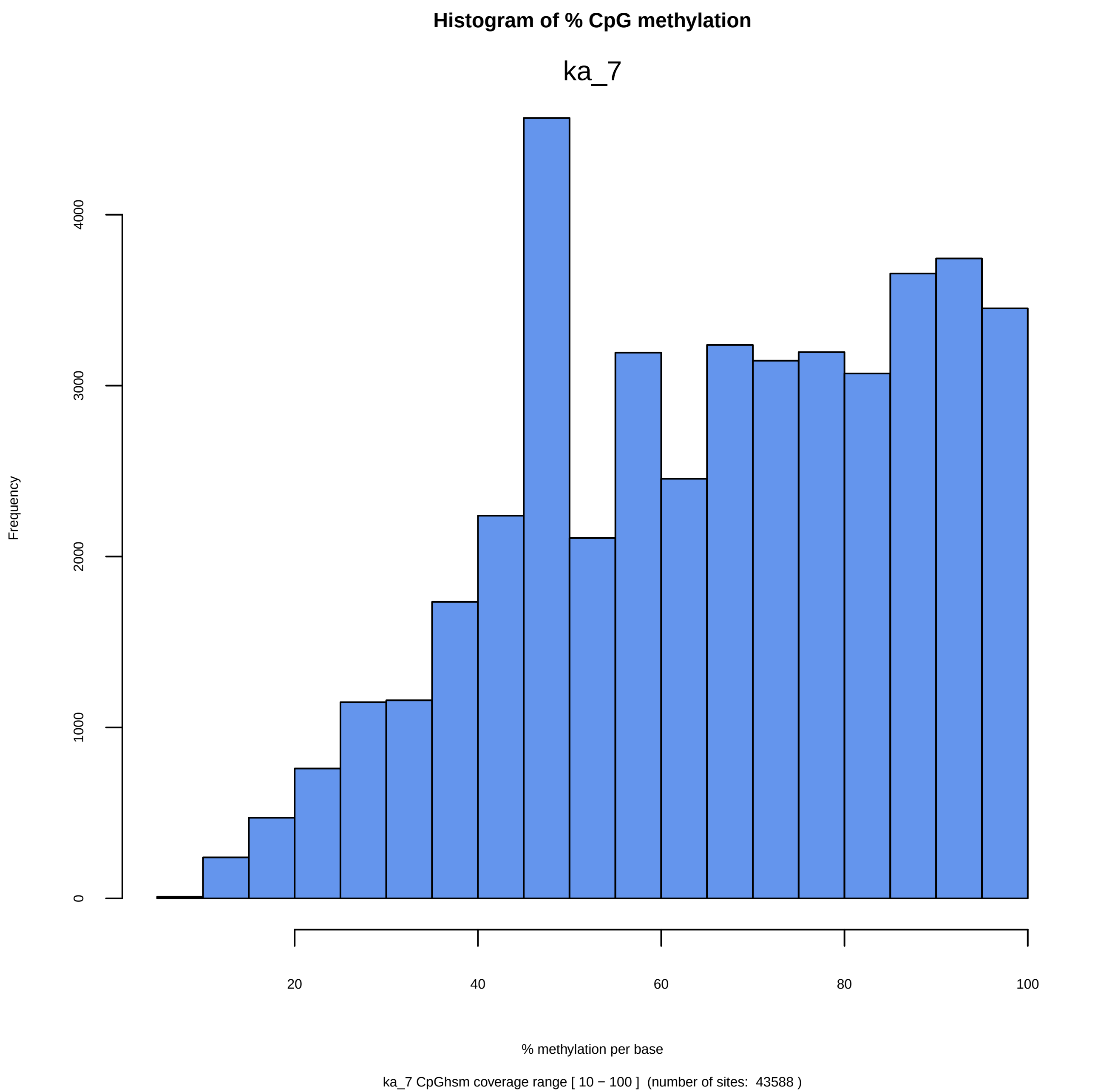

Histogram of % CpG methylation
ka_7
Frequency
0
1000
2000
3000
4000
20
40
60
80
100
% methylation per base
ka_7 CpGhsm coverage range [ 10 − 100 ]  (number of sites:  43588 )

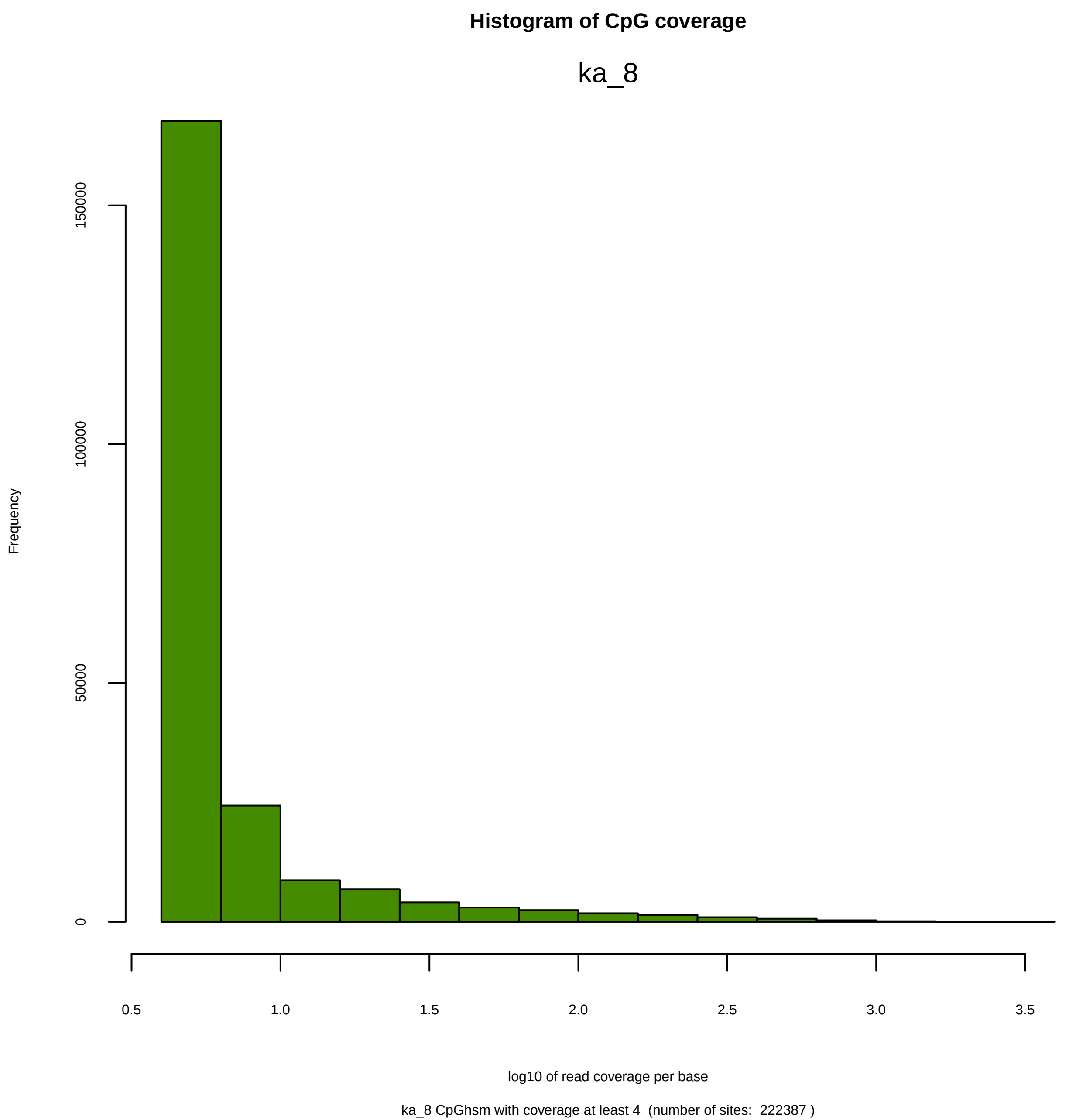

Histogram of CpG coverage
ka_8
Frequency
0
50000
100000
150000
0.5
1.0
1.5
2.0
2.5
3.0
3.5
log10 of read coverage per base
ka_8 CpGhsm with coverage at least 4 (number of sites: 222387 )

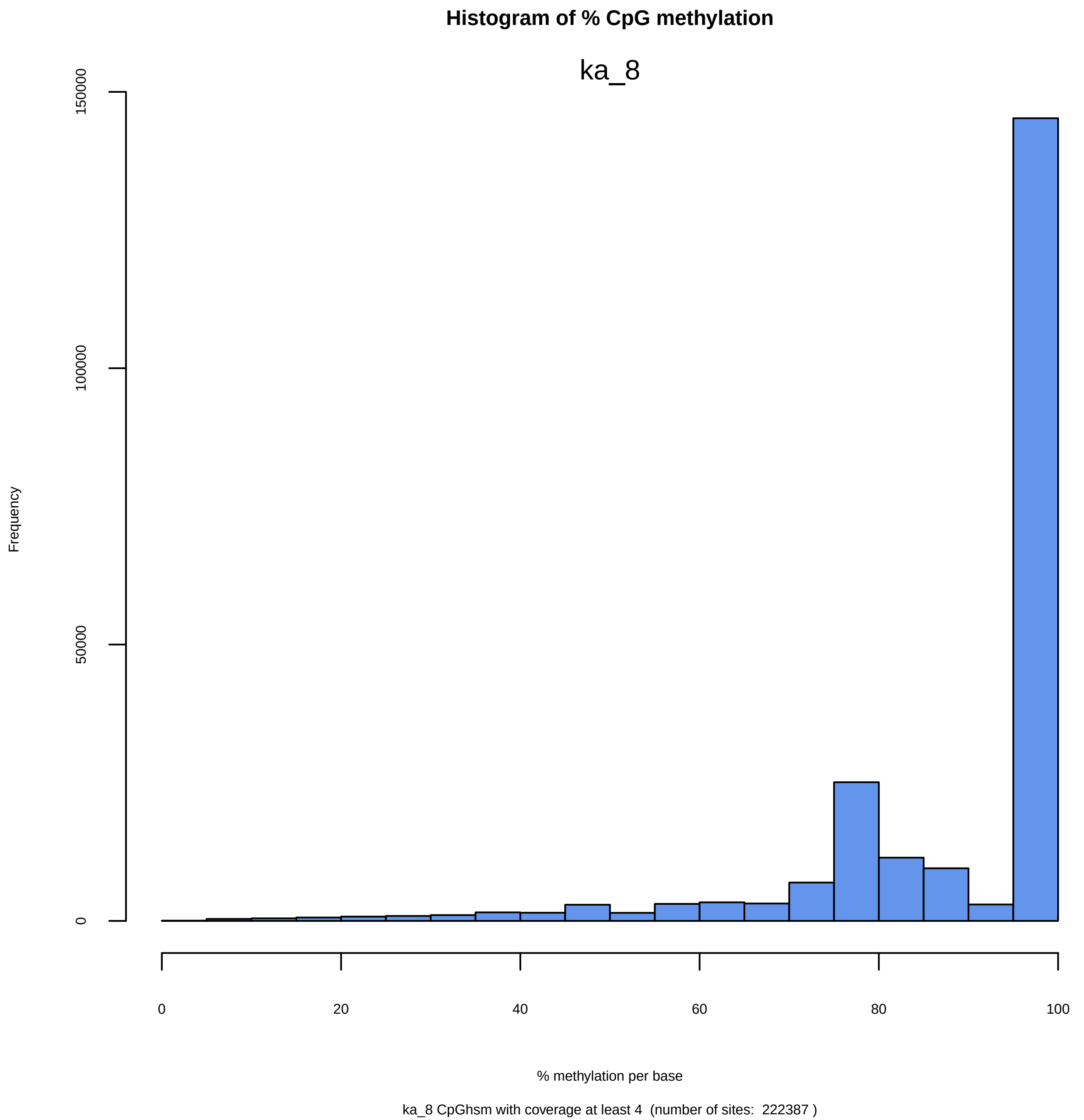

Histogram of % CpG methylation
ka_8
Frequency
0
50000
100000
150000
0
20
40
60
80
100
% methylation per base
ka_8 CpGhsm with coverage at least 4 (number of sites: 222387 )

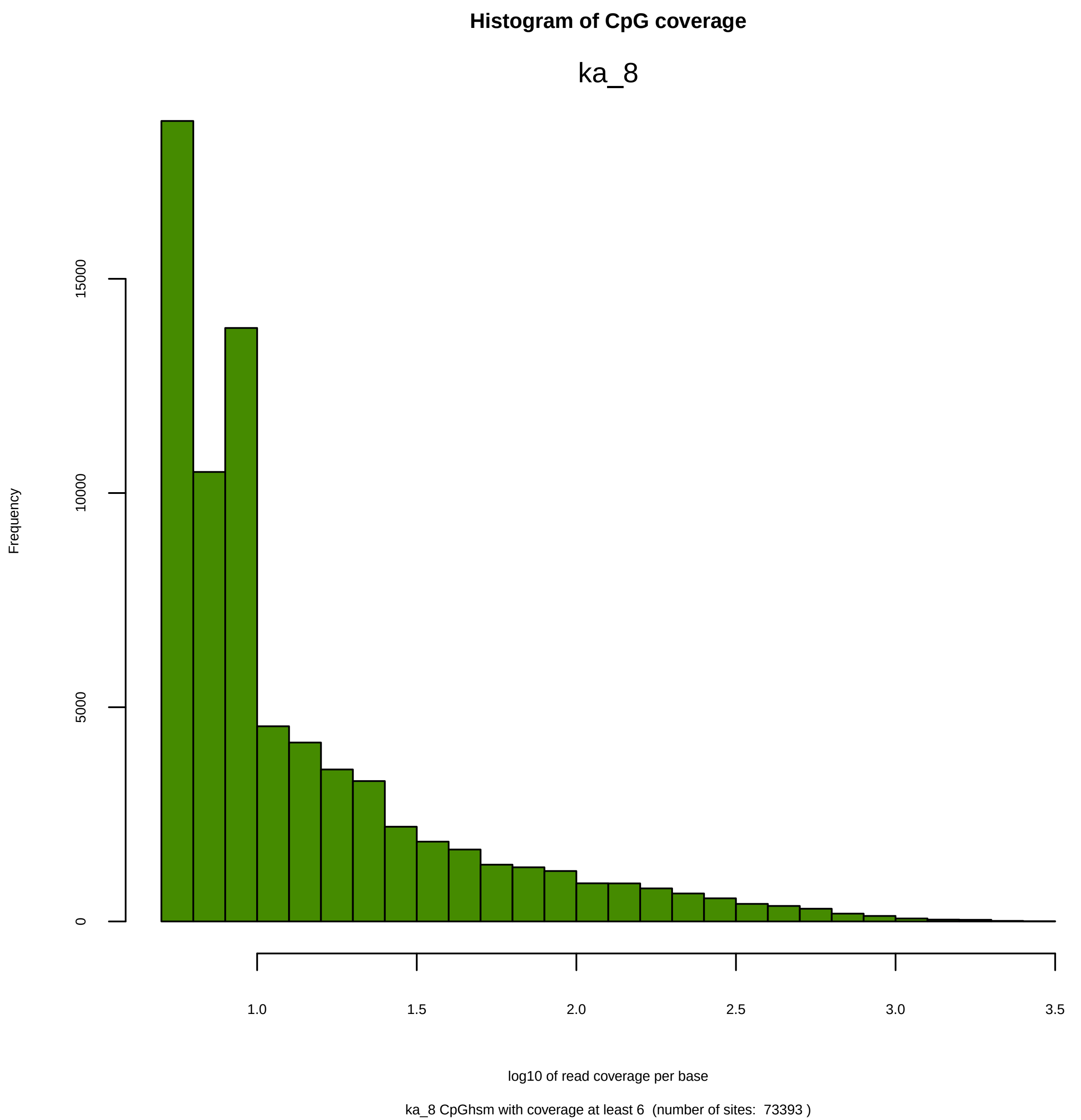

Histogram of CpG coverage
ka_8
Frequency
0
5000
10000
15000
1.0
1.5
2.0
2.5
3.0
3.5
log10 of read coverage per base
ka_8 CpGhsm with coverage at least 6 (number of sites: 73393 )

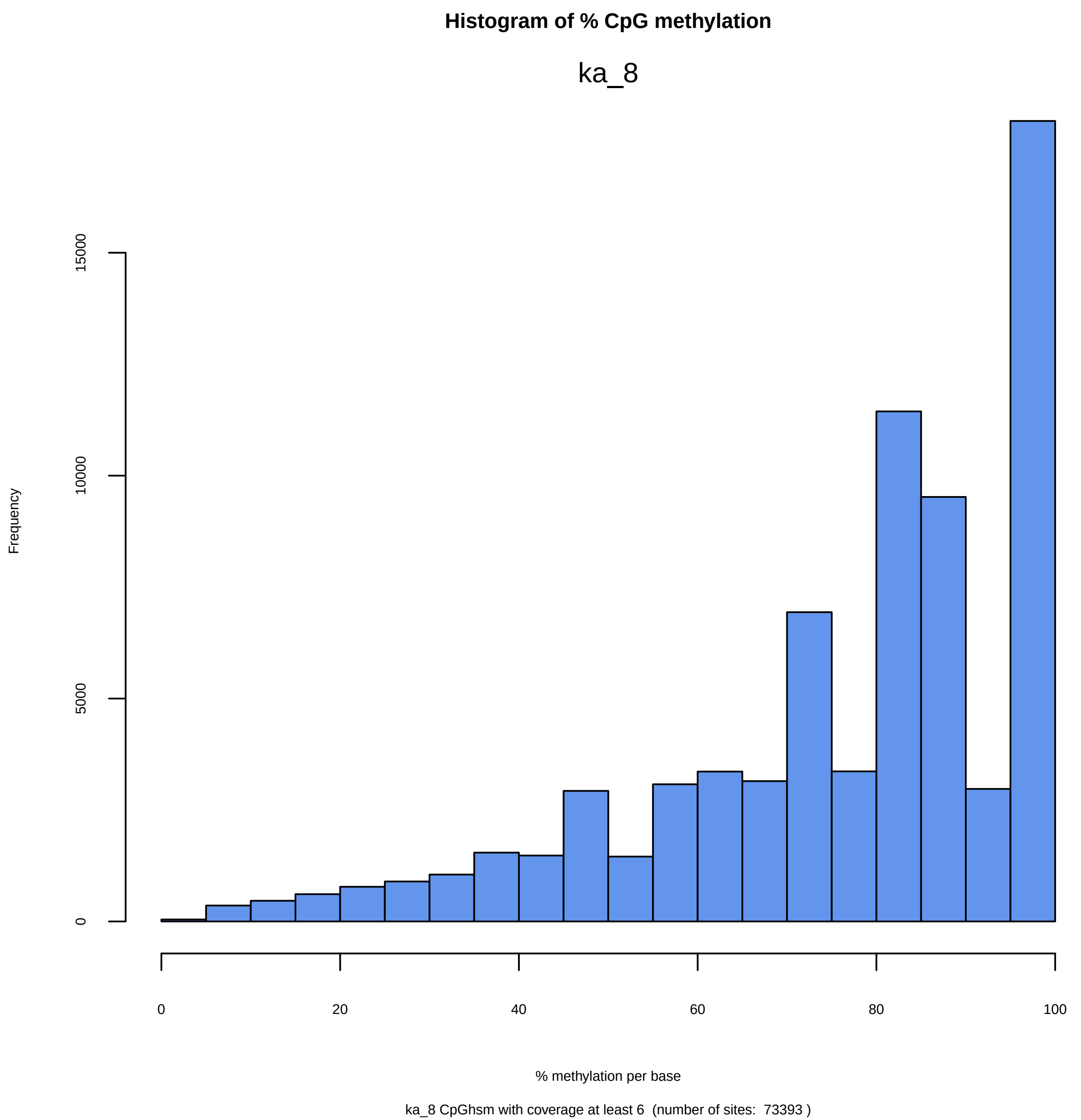
Histogram of % CpG methylation
ka_8
Frequency
0
5000
10000
15000
0
20
40
60
80
100
% methylation per base
ka_8 CpGhsm with coverage at least 6 (number of sites: 73393 )

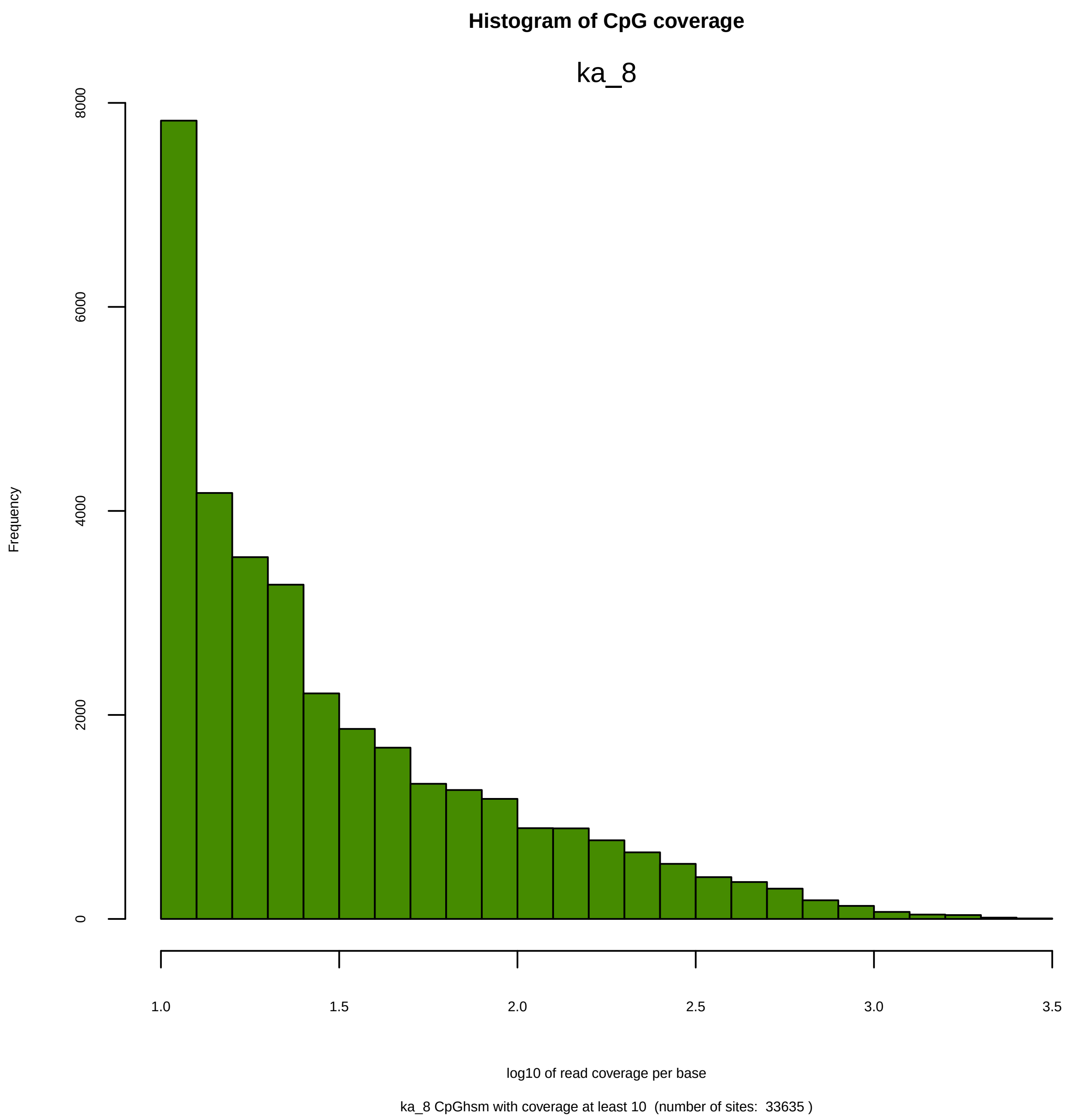

Histogram of CpG coverage
ka_8
Frequency
0
2000
4000
6000
8000
1.0
1.5
2.0
2.5
3.0
3.5
log10 of read coverage per base
ka_8 CpGhsm with coverage at least 10  (number of sites:  33635 )

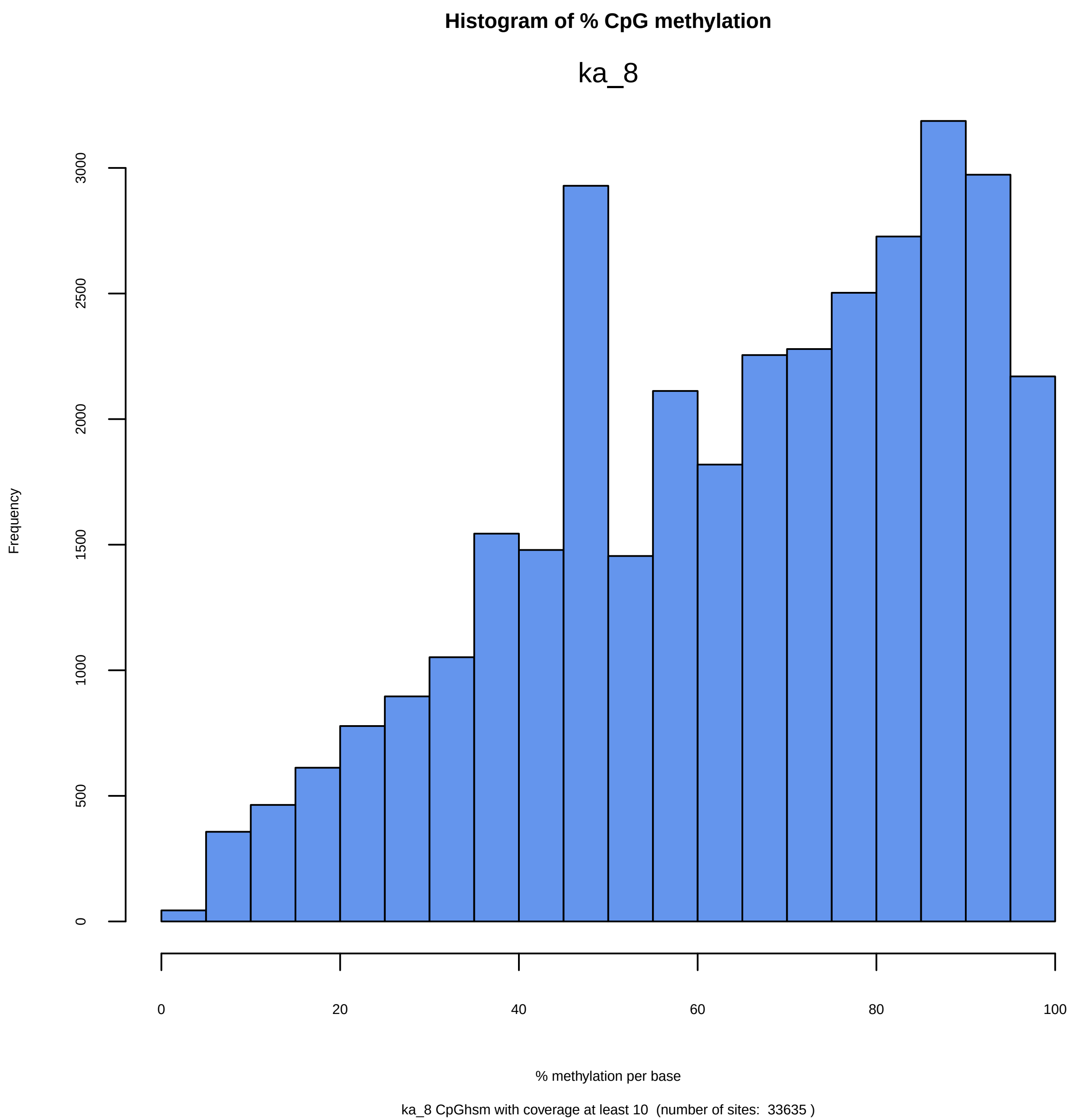
Histogram of % CpG methylation
ka_8
Frequency
0
500
1000
1500
2000
2500
3000
0
20
40
60
80
100
% methylation per base
ka_8 CpGhsm with coverage at least 10 (number of sites: 33635 )

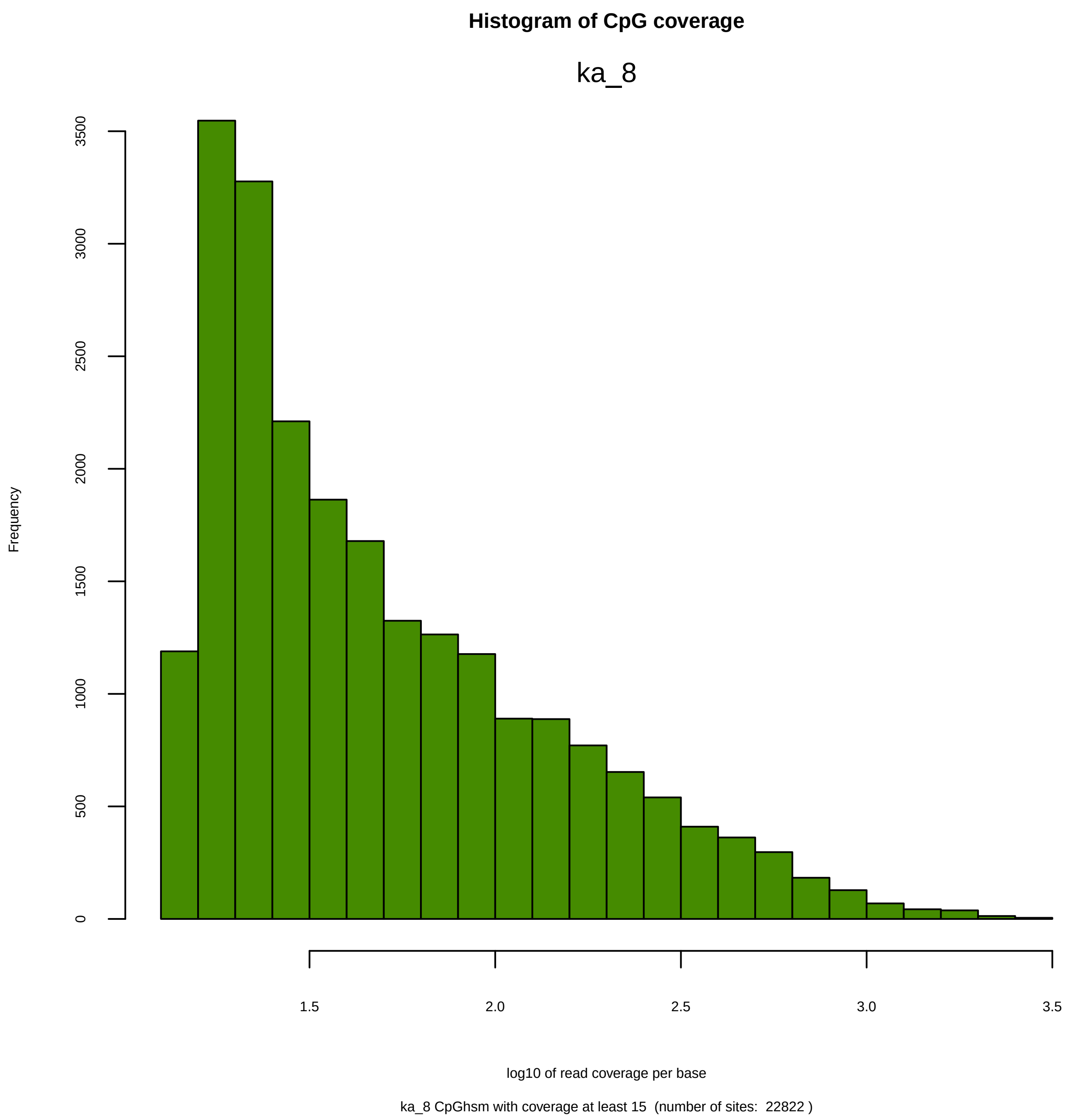

Histogram of CpG coverage
ka_8
Frequency
0
500
1000
1500
2000
2500
3000
3500
1.5
2.0
2.5
3.0
3.5
log10 of read coverage per base
ka_8 CpGhsm with coverage at least 15  (number of sites:  22822 )

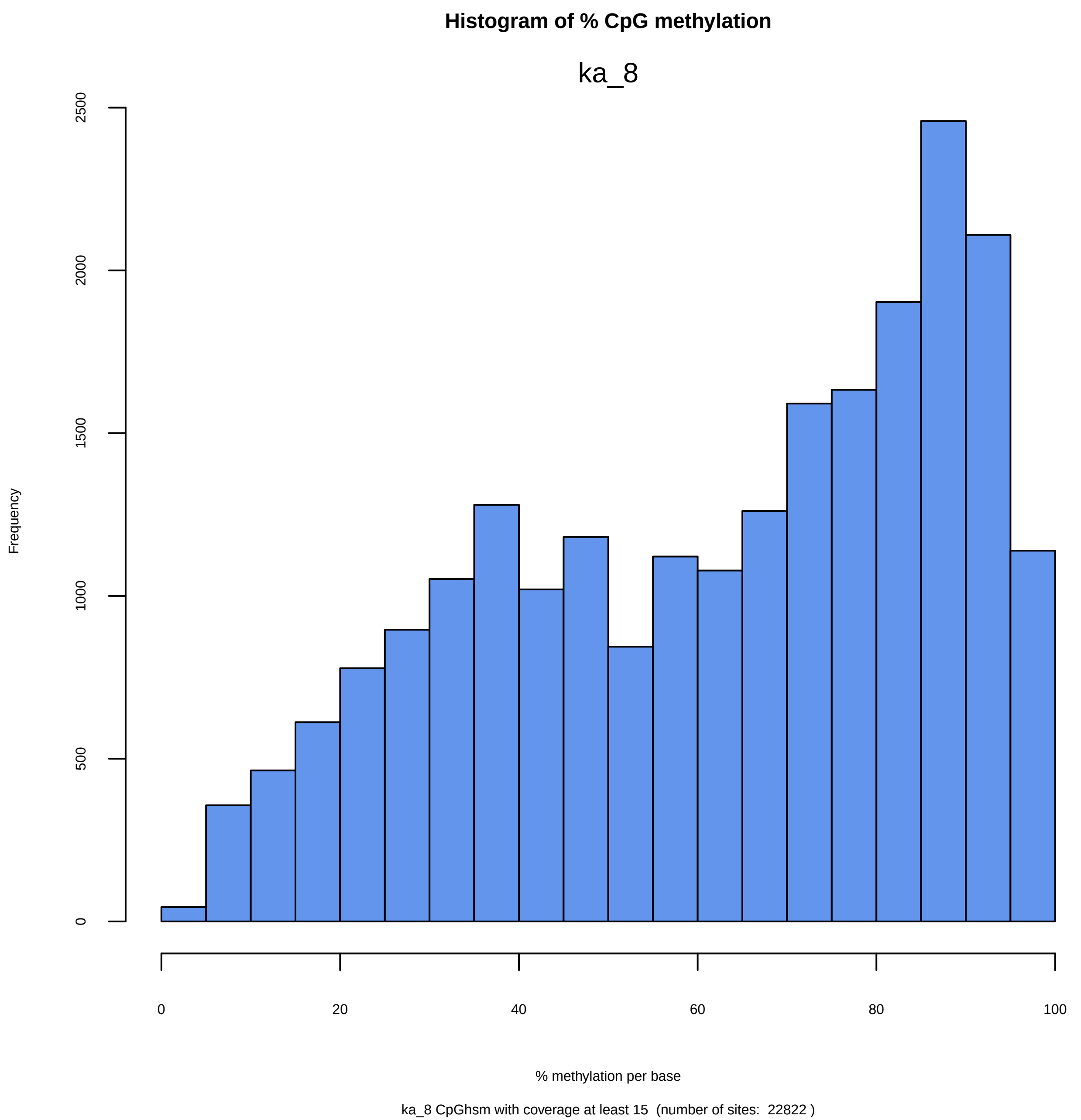
Histogram of % CpG methylation
ka_8
Frequency
0
500
1000
1500
2000
2500
0
20
40
60
80
100
% methylation per base
ka_8 CpGhsm with coverage at least 15  (number of sites:  22822 )

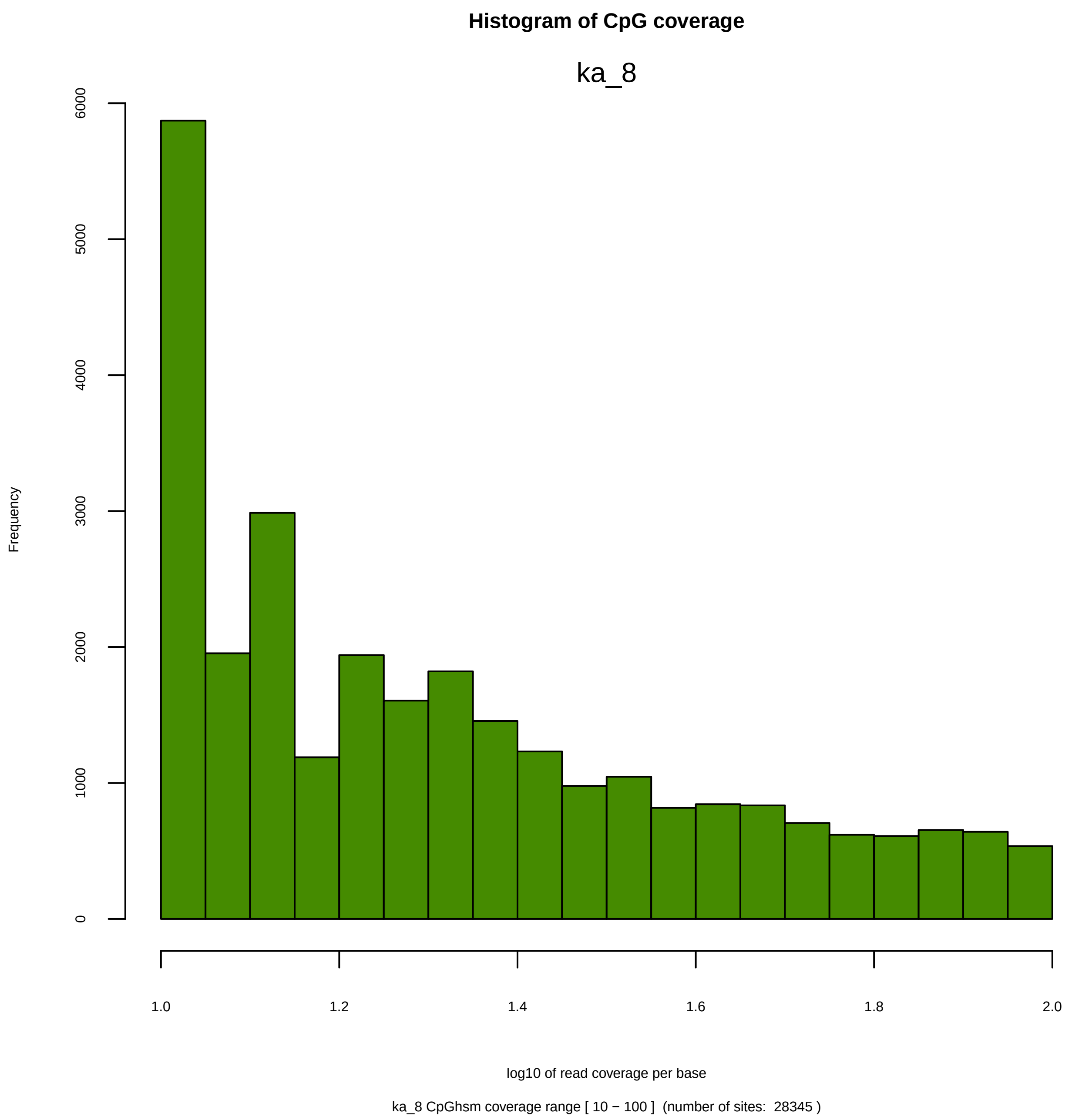

Histogram of CpG coverage
ka_8
Frequency
0
1000
2000
3000
4000
5000
6000
1.0
1.2
1.4
1.6
1.8
2.0
log10 of read coverage per base
ka_8 CpGhsm coverage range [ 10 − 100 ]  (number of sites:  28345 )

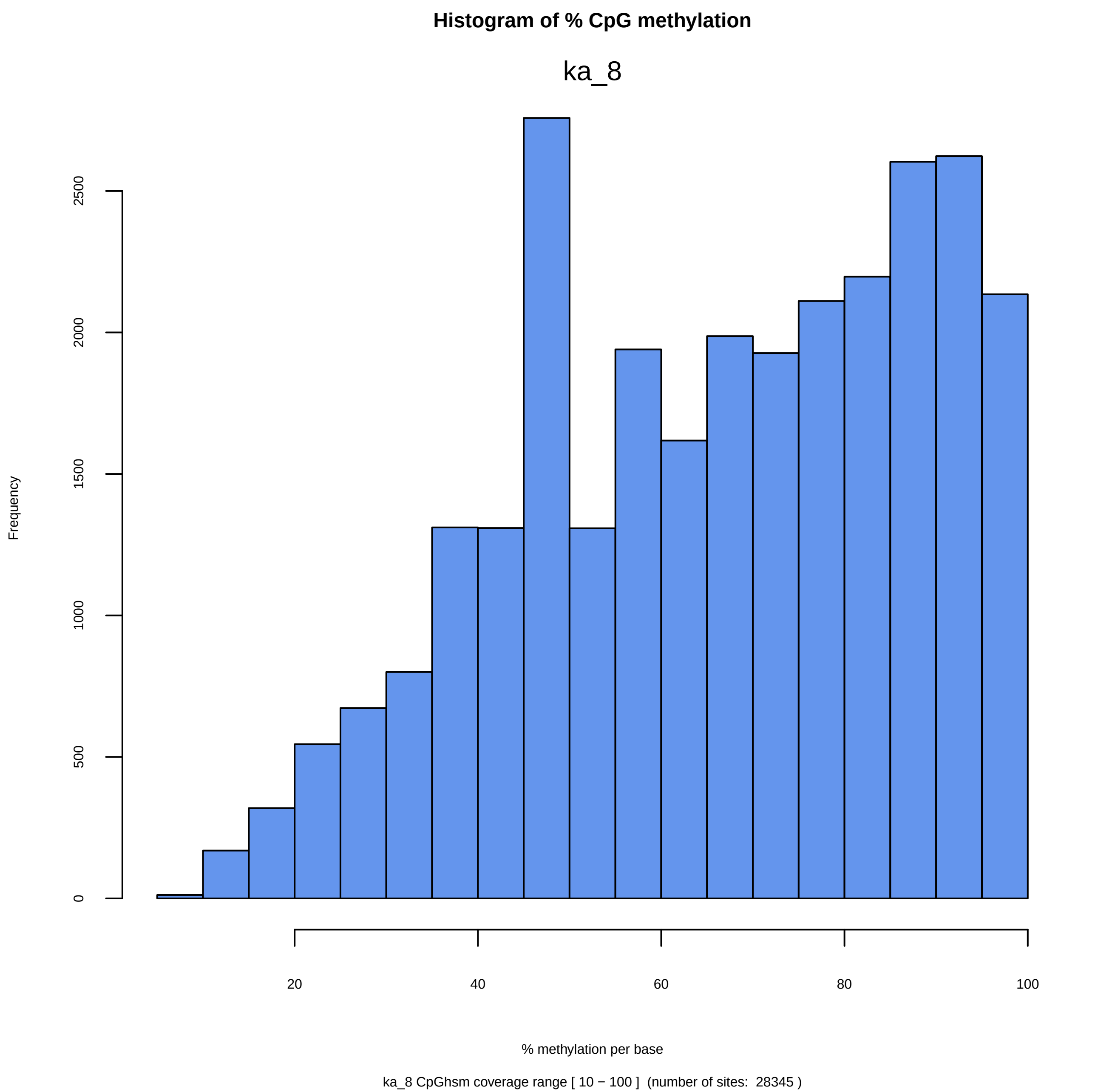

Histogram of % CpG methylation
ka_8
Frequency
0
500
1000
1500
2000
2500
20
40
60
80
100
% methylation per base
ka_8 CpGhsm coverage range [ 10 − 100 ] (number of sites: 28345 )

```
Directory: REPCRL File: 0READMErepcrl

REPCRL - Correlations between replicates.

Input: mkrd (methylKit methylRawList object of replicate data)

Output: files repcrl-*.txt repcrl-*.pdf

Notes: The output is generated by BWASPR::cmpSamples(), which uses methylKit::unite()
        to determine the sites common to all replicates and applies methylKit::getCorrelation()
        and methylKit::PCASamples().  The *.txt files show the correlations in a table, and
        the *.pdf files show graphics.

       Beware that the number of common sites (shown at the bottom of the *.txt files) may be
        small, which may make the correlations less informative.
```

Directory: REPCRL File: repcrl-Sd_LI_be.txt

| | be_1 | be_2 | be_3 | be_4 | be_5 | be_6 | be_7 | be_8 |
|---|---|---|---|---|---|---|---|---|
| be_1 | 1.000 | 0.871 | 0.867 | 0.868 | 0.870 | 0.868 | 0.871 | 0.868 |
| be_2 | 0.871 | 1.000 | 0.869 | 0.870 | 0.876 | 0.870 | 0.874 | 0.872 |
| be_3 | 0.867 | 0.869 | 1.000 | 0.867 | 0.870 | 0.866 | 0.869 | 0.865 |
| be_4 | 0.868 | 0.870 | 0.867 | 1.000 | 0.873 | 0.868 | 0.871 | 0.867 |
| be_5 | 0.870 | 0.876 | 0.870 | 0.873 | 1.000 | 0.871 | 0.877 | 0.871 |
| be_6 | 0.868 | 0.870 | 0.866 | 0.868 | 0.871 | 1.000 | 0.871 | 0.867 |
| be_7 | 0.871 | 0.874 | 0.869 | 0.871 | 0.877 | 0.871 | 1.000 | 0.871 |
| be_8 | 0.868 | 0.872 | 0.865 | 0.867 | 0.871 | 0.867 | 0.871 | 1.000 |

The number of conserved sites is 21749.

Directory: REPCRL File: repcrl-Sd_LI_ka.txt

| | ka_1 | ka_2 | ka_3 | ka_4 | ka_5 | ka_6 | ka_7 | ka_8 |
|---|---|---|---|---|---|---|---|---|
| ka_1 | 1.000 | 0.892 | 0.887 | 0.891 | 0.885 | 0.890 | 0.892 | 0.880 |
| ka_2 | 0.892 | 1.000 | 0.889 | 0.893 | 0.889 | 0.894 | 0.891 | 0.882 |
| ka_3 | 0.887 | 0.889 | 1.000 | 0.889 | 0.887 | 0.887 | 0.888 | 0.877 |
| ka_4 | 0.891 | 0.893 | 0.889 | 1.000 | 0.890 | 0.892 | 0.892 | 0.881 |
| ka_5 | 0.885 | 0.889 | 0.887 | 0.890 | 1.000 | 0.889 | 0.889 | 0.878 |
| ka_6 | 0.890 | 0.894 | 0.887 | 0.892 | 0.889 | 1.000 | 0.891 | 0.879 |
| ka_7 | 0.892 | 0.891 | 0.888 | 0.892 | 0.889 | 0.891 | 1.000 | 0.883 |
| ka_8 | 0.880 | 0.882 | 0.877 | 0.881 | 0.878 | 0.879 | 0.883 | 1.000 |

The number of conserved sites is 24193.

**CpG base pearson cor.**

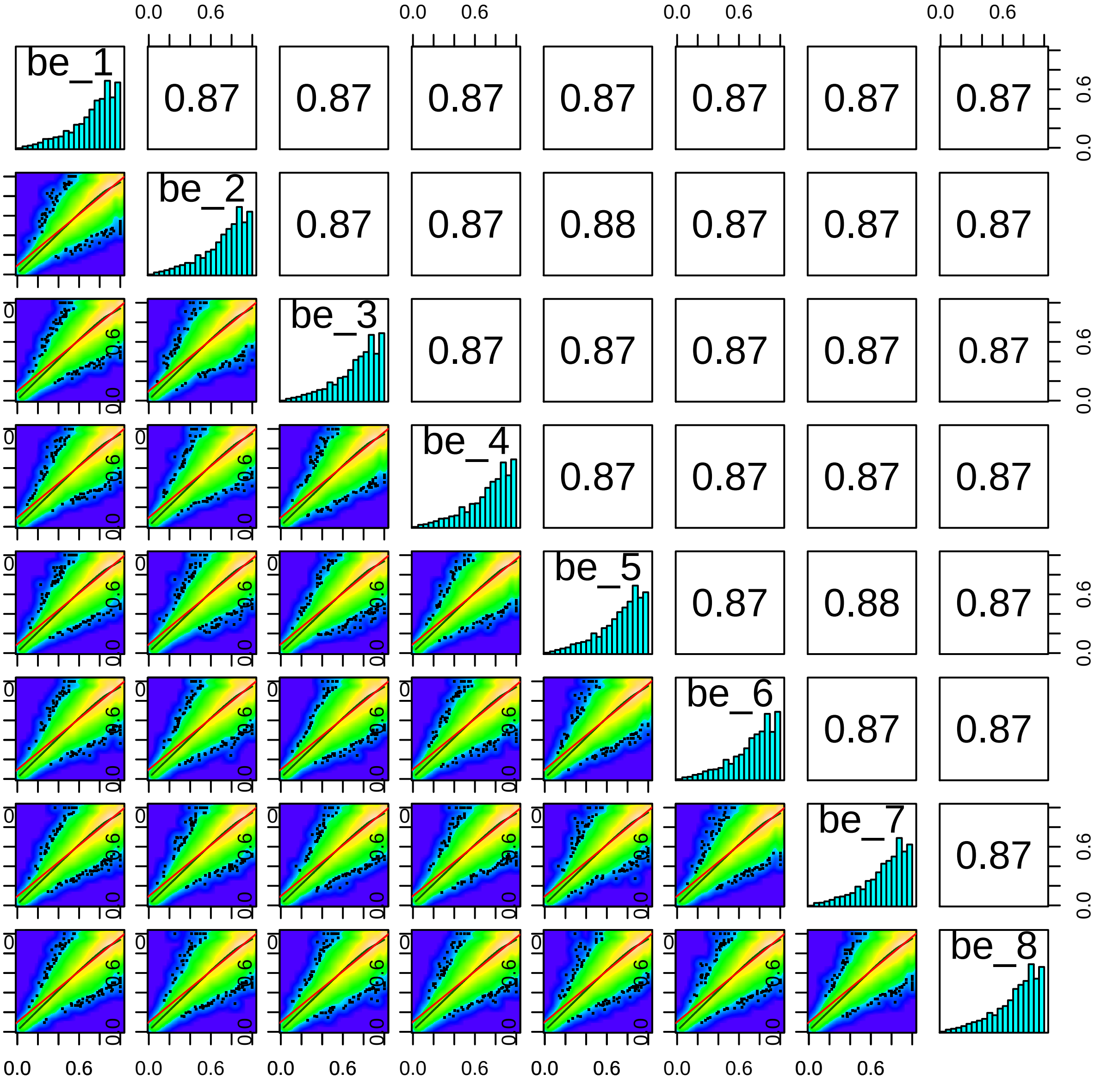

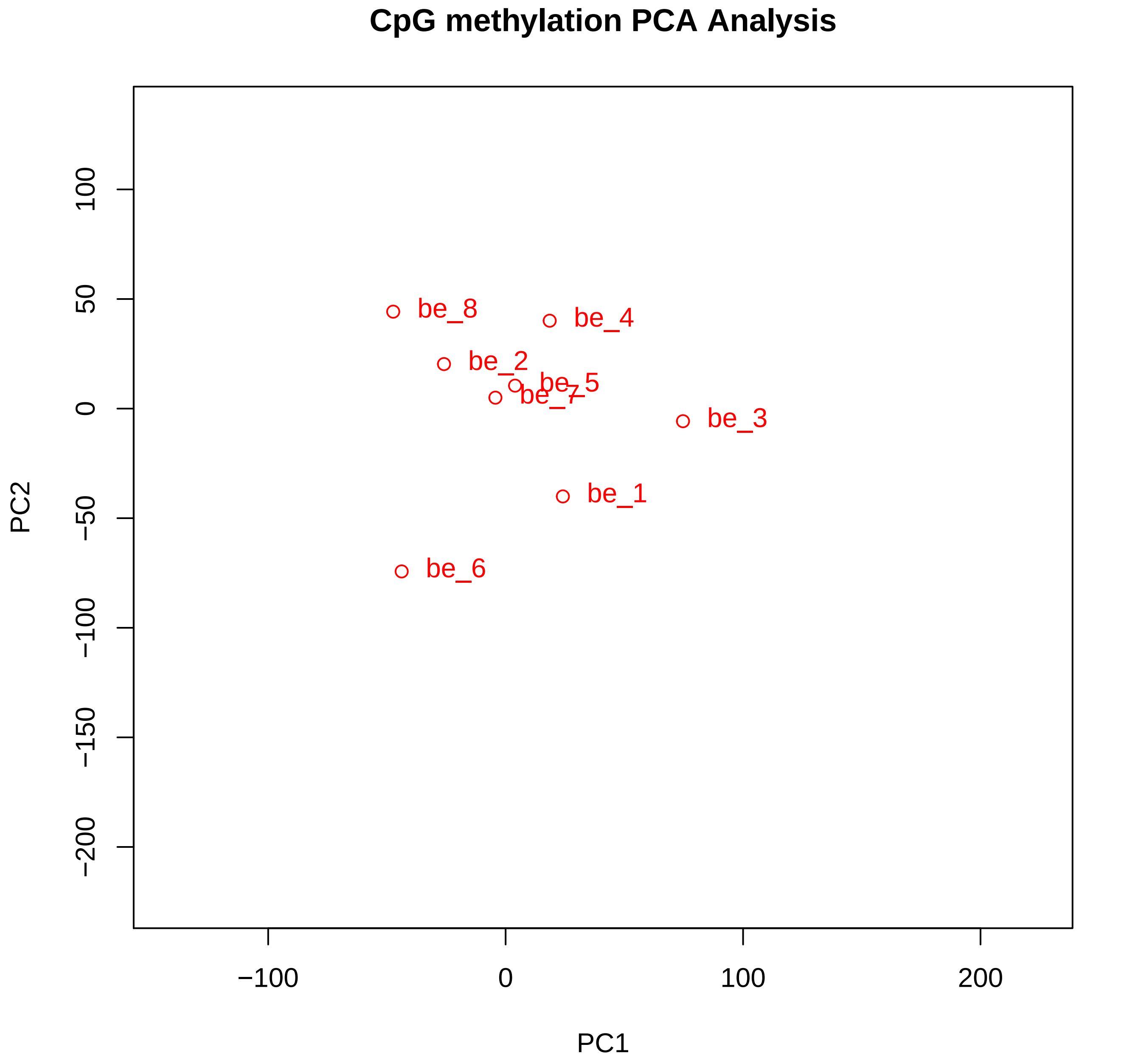

CpG methylation PCA Analysis
be_8
be_4
be_2
be_5
be_7
be_3
be_1
be_6
100
50
0
−50
−100
−150
−200
PC2
−100
0
100
200
PC1

### CpG base pearson cor.

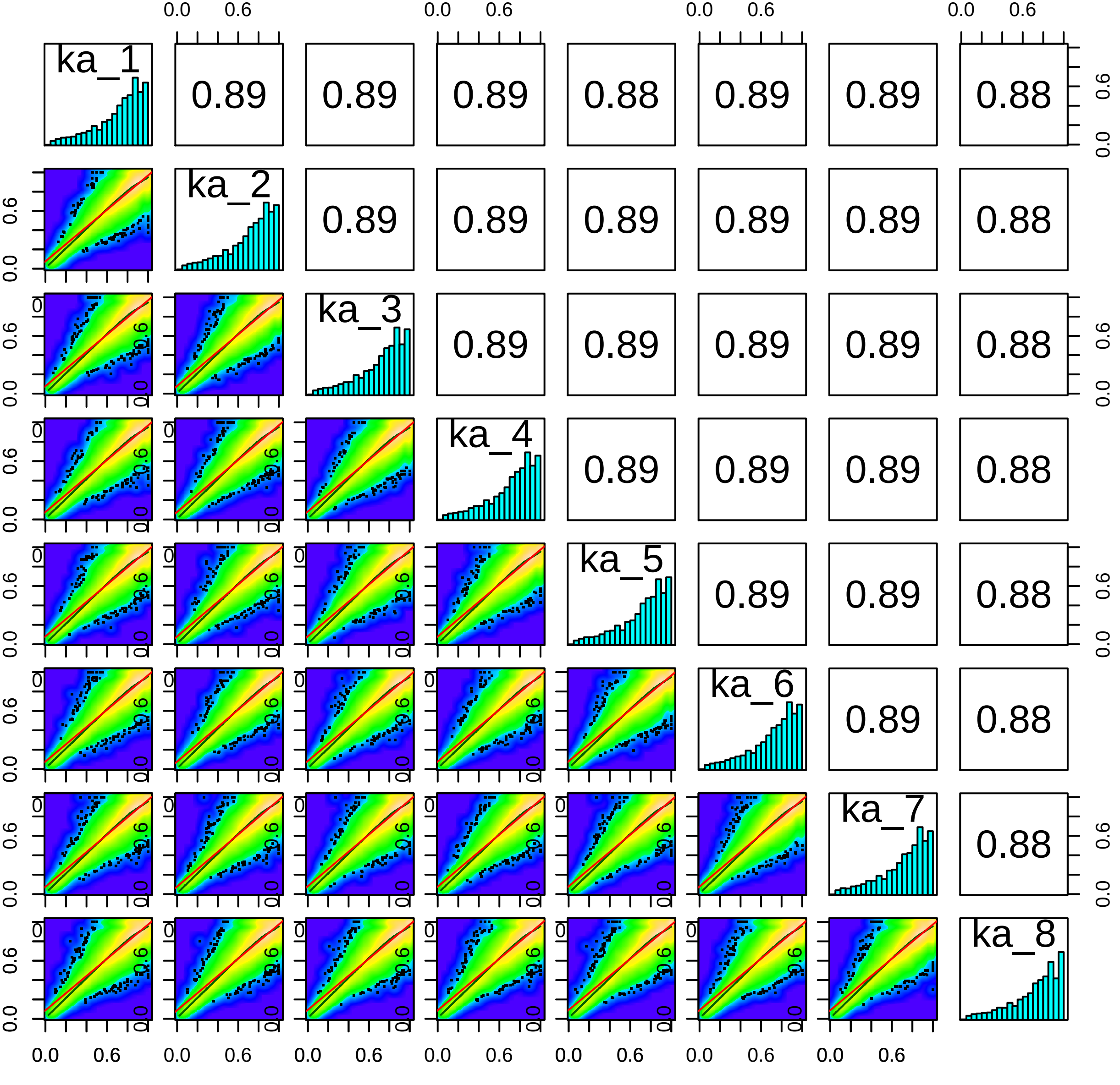

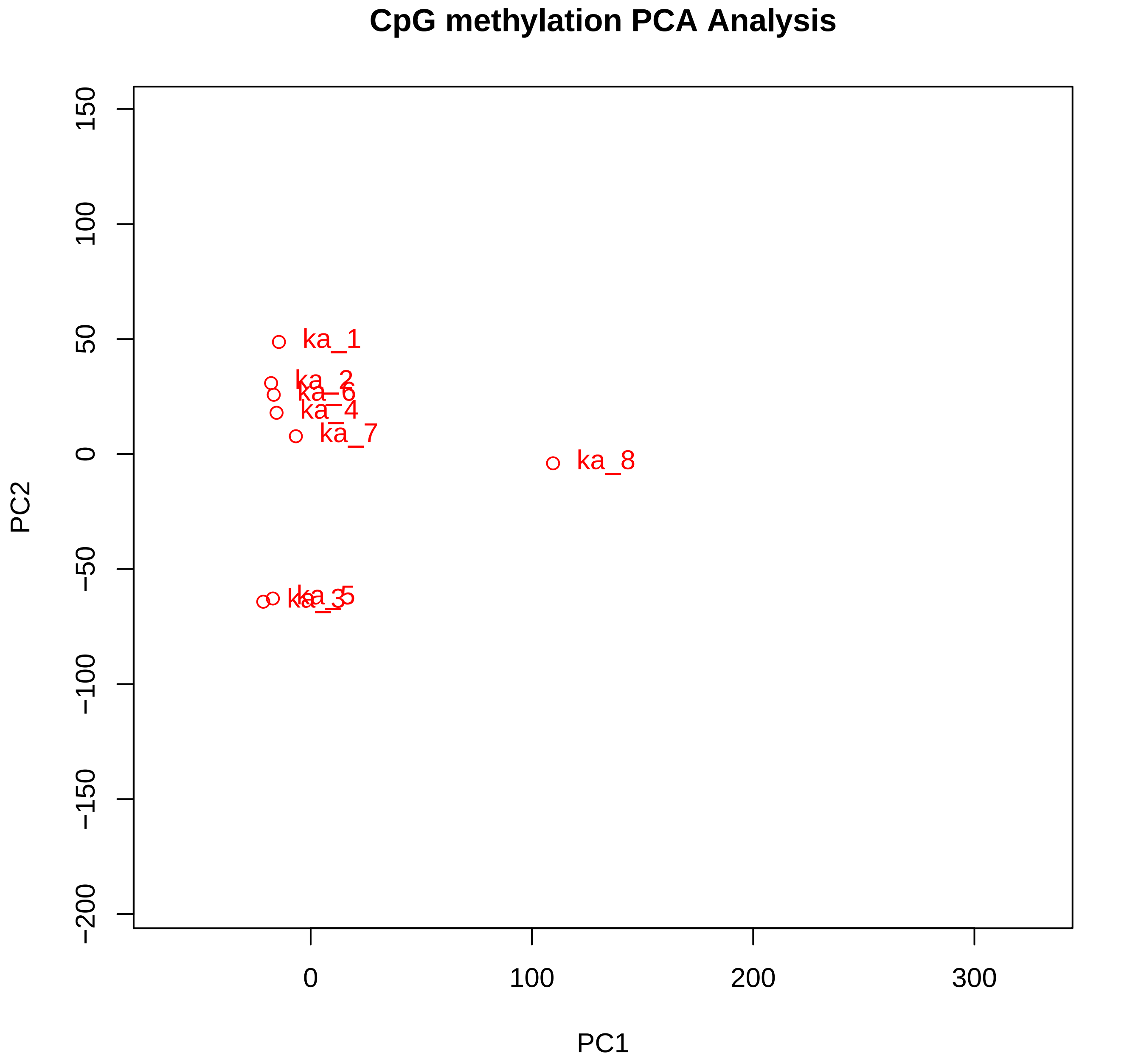

CpG methylation PCA Analysis
PC1
PC2
ka_1
ka_2
ka_6
ka_4
ka_7
ka_8
ka_3
ka_5
0
100
200
300
150
100
50
0
−50
−100
−150
−200

```
Directory: MMP File: 0READMEmmp

MMP - Mapping of methylation sites on genome annotation.

Input: studymk, studymc, genome_ann (a list of GRanges objects providing annotated region labels
        and boundaries; output of BWASPR::get_genome_annotation())

Output: files mmp-*.txt

Notes: The output is generated by BWASPR::map_methylome() and gives an accounting for every sample
        as to where the CpGhsm and CpGscd (control) sites reside relative to the genome annotation.
       Abbreviations used: O/E, observed over expected
                           (expected percentages are based on the respective feature proportions
                            in the genome, as annotated).
```

Directory: MMP File: mmp-Sd_LI_be.txt

#### Genomic composition in terms of feature regions

| | | |
|---|---|---|
| Sd genome size: | 2551871595 bp | |
| Sd genic region size: | 1056805054 bp | ( 41.4%) |
| Sd exon region size: | 53199269 bp | ( 2.1%) |
| Sd CDS region size: | 27472809 bp | ( 51.6%) |
| Sd five-prime UTR region size: | 3182761 bp | ( 6.0%) |
| Sd three-prime UTR region size: | 15900025 bp | ( 29.9%) |
| Sd other exon region size: | 6643674 bp | ( 12.5%) |
| Sd intron region size: | 1003605785 bp | ( 39.3%) |
| Sd intergenic region size: | 1495066541 bp | ( 58.6%) |
| Sd promoter region size: | 13041605 bp | ( 0.5%) |
| Sd other intergenic region size: | 1482024936 bp | ( 58.1%) |

#### Methylation sites in genomic feature regions

Number and density (per 10kb) of sites in Sd_LI_be_scd

| | | | | |
|---|---|---|---|---|
| Number of sites identified in Sd Sd_LI_be_scd: | 43378941 | ( 169.99) | | |
| Number of sites identified in Sd Sd_LI_be_scd genic regions: | 17168360 | ( 162.46) | 39.58% | ( 0.96 O/E) |
| Number of sites identified in Sd Sd_LI_be_scd exon regions: | 701817 | ( 131.92) | 1.62% | ( 0.78 O/E) |
| Number of sites identified in Sd Sd_LI_be_scd CDS regions: | 410412 | ( 149.39) | 58.48% | ( 1.13 O/E) |
| Number of sites identified in Sd Sd_LI_be_scd five-prime UTR regions: | 60414 | ( 189.82) | 8.61% | ( 1.44 O/E) |
| Number of sites identified in Sd Sd_LI_be_scd three-prime UTR regions: | 165041 | ( 103.80) | 23.52% | ( 0.79 O/E) |
| Number of sites identified in Sd Sd_LI_be_scd other exon regions: | 65950 | ( 99.27) | 9.40% | ( 0.75 O/E) |
| Number of sites identified in Sd Sd_LI_be_scd intron regions: | 16466543 | ( 164.07) | 37.96% | ( 0.97 O/E) |
| Number of sites identified in Sd Sd_LI_be_scd intergenic regions: | 26210581 | ( 175.31) | 60.42% | ( 1.03 O/E) |
| Number of sites identified in Sd Sd_LI_be_scd promoter regions: | 192935 | ( 147.94) | 0.44% | ( 0.87 O/E) |
| Number of sites identified in Sd Sd_LI_be_scd other intergenic regions: | 26017646 | ( 175.55) | 59.98% | ( 1.03 O/E) |

Number and density (per 10kb) of sites in Sd_LI_be_hsm

| | | | | | |
|---|---|---|---|---|---|
| Number of sites identified in Sd Sd_LI_be_hsm: | 6335325 | ( 24.83) | ( 14.60% of control) | | |
| Number of sites identified in Sd Sd_LI_be_hsm genic regions: | 4982483 | ( 47.15) | ( 29.02% of control) | 78.65% | ( 1.90 O/E) |
| Number of sites identified in Sd Sd_LI_be_hsm exon regions: | 219050 | ( 41.18) | ( 31.21% of control) | 3.46% | ( 1.66 O/E) |
| Number of sites identified in Sd Sd_LI_be_hsm CDS regions: | 146868 | ( 53.46) | ( 35.79% of control) | 67.05% | ( 1.30 O/E) |
| Number of sites identified in Sd Sd_LI_be_hsm five-prime UTR regions: | 4937 | ( 15.51) | ( 8.17% of control) | 2.25% | ( 0.38 O/E) |
| Number of sites identified in Sd Sd_LI_be_hsm three-prime UTR regions: | 53312 | ( 33.53) | ( 32.30% of control) | 24.34% | ( 0.81 O/E) |
| Number of sites identified in Sd Sd_LI_be_hsm other exon regions: | 13933 | ( 20.97) | ( 21.13% of control) | 6.36% | ( 0.51 O/E) |
| Number of sites identified in Sd Sd_LI_be_hsm intron regions: | 4763433 | ( 47.46) | ( 28.93% of control) | 75.19% | ( 1.91 O/E) |
| Number of sites identified in Sd Sd_LI_be_hsm intergenic regions: | 1352842 | ( 9.05) | ( 5.16% of control) | 21.35% | ( 0.36 O/E) |
| Number of sites identified in Sd Sd_LI_be_hsm promoter regions: | 15566 | ( 11.94) | ( 8.07% of control) | 0.25% | ( 0.48 O/E) |
| Number of sites identified in Sd Sd_LI_be_hsm other intergenic regions: | 1337276 | ( 9.02) | ( 5.14% of control) | 21.11% | ( 0.36 O/E) |

```
Directory: MMP File: mmp-Sd_LI_ka.txt

================================================================================
Genomic composition in terms of feature regions
================================================================================

Sd genome size:                       2551871595 bp
  Sd genic region size:                 1056805054 bp ( 41.4%)
    Sd exon region size:                    53199269 bp (  2.1%)
      Sd CDS region size:                     27472809 bp ( 51.6%)
      Sd five-prime UTR region size:           3182761 bp (  6.0%)
      Sd three-prime UTR region size:         15900025 bp ( 29.9%)
      Sd other exon region size:               6643674 bp ( 12.5%)
    Sd intron region size:                1003605785 bp ( 39.3%)
  Sd intergenic region size:            1495066541 bp ( 58.6%)
    Sd promoter region size:                13041605 bp (  0.5%)
    Sd other intergenic region size:      1482024936 bp ( 58.1%)

================================================================================
Methylation sites in genomic feature regions
================================================================================

Number and density (per 10kb) of sites in Sd_LI_ka_scd

Number of sites identified in Sd Sd_LI_ka_scd:                             42719577     (  167.40)                           
Number of sites identified in Sd Sd_LI_ka_scd genic regions:               16854990     (  159.49)                          39.45%     ( 0.95 O/E)
Number of sites identified in Sd Sd_LI_ka_scd   exon regions:                 687715    (  129.27)                             1.61%   ( 0.77 O/E)
Number of sites identified in Sd Sd_LI_ka_scd     CDS regions:                  402239 (  146.41)                              58.49% ( 1.13 O/E)
Number of sites identified in Sd Sd_LI_ka_scd     five-prime UTR regions:        58079 (  182.48)                               8.45% ( 1.41 O/E)
Number of sites identified in Sd Sd_LI_ka_scd     three-prime UTR regions:      162338 (  102.10)                              23.61% ( 0.79 O/E)
Number of sites identified in Sd Sd_LI_ka_scd     other exon regions:            65059 (   97.93)                               9.46% ( 0.76 O/E)
Number of sites identified in Sd Sd_LI_ka_scd   intron regions:             16167275    (  161.09)                            37.85%   ( 0.96 O/E)
Number of sites identified in Sd Sd_LI_ka_scd intergenic regions:          25864587     (  173.00)                          60.55%     ( 1.03 O/E)
Number of sites identified in Sd Sd_LI_ka_scd   promoter regions:             186297    (  142.85)                             0.44%   ( 0.85 O/E)
Number of sites identified in Sd Sd_LI_ka_scd   other intergenic regions:   25678290    (  173.26)                            60.11%   ( 1.04 O/E)

Number and density (per 10kb) of sites in Sd_LI_ka_hsm

Number of sites identified in Sd Sd_LI_ka_hsm:                              6222608     (   24.38) ( 14.57% of control)
Number of sites identified in Sd Sd_LI_ka_hsm genic regions:                4876145     (   46.14) ( 28.93% of control)  78.36%     ( 1.89 O/E)
Number of sites identified in Sd Sd_LI_ka_hsm   exon regions:                 217305    (   40.85) ( 31.60% of control)    3.49%   ( 1.68 O/E)
Number of sites identified in Sd Sd_LI_ka_hsm     CDS regions:                  145762 (   53.06) ( 36.24% of control)     67.08% ( 1.30 O/E)
Number of sites identified in Sd Sd_LI_ka_hsm     five-prime UTR regions:         4941 (   15.52) (  8.51% of control)      2.27% ( 0.38 O/E)
Number of sites identified in Sd Sd_LI_ka_hsm     three-prime UTR regions:       52787 (   33.20) ( 32.52% of control)     24.29% ( 0.81 O/E)
Number of sites identified in Sd Sd_LI_ka_hsm     other exon regions:            13815 (   20.79) ( 21.23% of control)      6.36% ( 0.51 O/E)
Number of sites identified in Sd Sd_LI_ka_hsm   intron regions:              4658840    (   46.42) ( 28.82% of control)   74.87%   ( 1.90 O/E)
Number of sites identified in Sd Sd_LI_ka_hsm intergenic regions:           1346463     (    9.01) (  5.21% of control)  21.64%     ( 0.37 O/E)
Number of sites identified in Sd Sd_LI_ka_hsm   promoter regions:              15470    (   11.86) (  8.30% of control)    0.25%   ( 0.49 O/E)
Number of sites identified in Sd Sd_LI_ka_hsm   other intergenic regions:    1330993    (    8.98) (  5.18% of control)   21.39%   ( 0.37 O/E)
```

```
Directory: ACS File: 0READMEacs

ACS - Annotation of conserved methylation sites.

Input: studymk, genome_ann (a list of GRanges objects providing annotated region labels and bounds;
                            output of BWASPR::get_genome_annotation())

Output: file acs-*.txt

Notes: The output is generated by BWASPR::annotate_methylome(), which uses methylKit::unite()
        to determine the sites common to all experiments and applies genomation::annotateWithFeature().
        The acs-*.txt files show the common sites with methylation levels in all samples
        and columns with booleans indicating whether the site fits a specific annotation feature.

       Abbreviations used: pc, protein coding; nc, non-coding; fp, five prime; tp, three prime.
                           UTR, untranslated region; unique, not overlapping with other categories.
```

Directory: ACS File: acs-Sd_LI.txt

| be | ka | chr | start | end | strand | gene | exon | pcexon | promoter | CDS | fpUTR | tpUTR | fpUTRnotCDS | tpUTRnotCDS | tpUTRunique | ncexon |
|---|---|---|---|---|---|---|---|---|---|---|---|---|---|---|---|---|
| 71.43 | 71.43 | NW_023312688.1 | 1111 | 1111 | + | 0 | 0 | 0 | 0 | 0 | 0 | 0 | 0 | 0 | 0 | 0 |
| 81.25 | 69.23 | NW_023312688.1 | 1158 | 1158 | + | 0 | 0 | 0 | 0 | 0 | 0 | 0 | 0 | 0 | 0 | 0 |
| 58.62 | 51.85 | NW_023312688.1 | 1175 | 1175 | + | 0 | 0 | 0 | 0 | 0 | 0 | 0 | 0 | 0 | 0 | 0 |
| 42.86 | 38.46 | NW_023312688.1 | 1282 | 1282 | + | 0 | 0 | 0 | 0 | 0 | 0 | 0 | 0 | 0 | 0 | 0 |
| 75 | 57.89 | NW_023312688.1 | 1345 | 1345 | + | 0 | 0 | 0 | 0 | 0 | 0 | 0 | 0 | 0 | 0 | 0 |
| 58.33 | 62.5 | NW_023312688.1 | 2569 | 2569 | + | 0 | 0 | 0 | 0 | 0 | 0 | 0 | 0 | 0 | 0 | 0 |
| 53.85 | 92.86 | NW_023312688.1 | 2834 | 2834 | + | 0 | 0 | 0 | 0 | 0 | 0 | 0 | 0 | 0 | 0 | 0 |
| 71.43 | 83.33 | NW_023312688.1 | 3536 | 3536 | + | 0 | 0 | 0 | 0 | 0 | 0 | 0 | 0 | 0 | 0 | 0 |
| 71.43 | 100 | NW_023312688.1 | 3929 | 3929 | + | 0 | 0 | 0 | 0 | 0 | 0 | 0 | 0 | 0 | 0 | 0 |
| 50 | 50 | NW_023312688.1 | 7630 | 7630 | + | 0 | 0 | 0 | 0 | 0 | 0 | 0 | 0 | 0 | 0 | 0 |
| 51.22 | 57.69 | NW_023312688.1 | 7650 | 7650 | + | 0 | 0 | 0 | 0 | 0 | 0 | 0 | 0 | 0 | 0 | 0 |
| 53.66 | 40 | NW_023312688.1 | 7659 | 7659 | + | 0 | 0 | 0 | 0 | 0 | 0 | 0 | 0 | 0 | 0 | 0 |
| 45.71 | 29.17 | NW_023312688.1 | 7676 | 7676 | + | 0 | 0 | 0 | 0 | 0 | 0 | 0 | 0 | 0 | 0 | 0 |
| 60.61 | 61.9 | NW_023312688.1 | 7681 | 7681 | + | 0 | 0 | 0 | 0 | 0 | 0 | 0 | 0 | 0 | 0 | 0 |
| 71.43 | 40 | NW_023312688.1 | 9664 | 9664 | + | 0 | 0 | 0 | 0 | 0 | 0 | 0 | 0 | 0 | 0 | 0 |
| 88.89 | 61.76 | NW_023312688.1 | 9721 | 9721 | + | 0 | 0 | 0 | 0 | 0 | 0 | 0 | 0 | 0 | 0 | 0 |
| 73.68 | 60 | NW_023312688.1 | 9773 | 9773 | + | 0 | 0 | 0 | 0 | 0 | 0 | 0 | 0 | 0 | 0 | 0 |
| 68.18 | 50 | NW_023312688.1 | 9776 | 9776 | + | 0 | 0 | 0 | 0 | 0 | 0 | 0 | 0 | 0 | 0 | 0 |
| 90 | 69.7 | NW_023312688.1 | 9794 | 9794 | + | 0 | 0 | 0 | 0 | 0 | 0 | 0 | 0 | 0 | 0 | 0 |
| 76.92 | 60 | NW_023312688.1 | 9817 | 9817 | + | 0 | 0 | 0 | 0 | 0 | 0 | 0 | 0 | 0 | 0 | 0 |
| 61.54 | 58.33 | NW_023312688.1 | 9819 | 9819 | + | 0 | 0 | 0 | 0 | 0 | 0 | 0 | 0 | 0 | 0 | 0 |
| 63.64 | 52 | NW_023312688.1 | 9833 | 9833 | + | 0 | 0 | 0 | 0 | 0 | 0 | 0 | 0 | 0 | 0 | 0 |
| 77.78 | 75 | NW_023312688.1 | 9857 | 9857 | + | 0 | 0 | 0 | 0 | 0 | 0 | 0 | 0 | 0 | 0 | 0 |
| 62.5 | 70 | NW_023312688.1 | 9870 | 9870 | + | 0 | 0 | 0 | 0 | 0 | 0 | 0 | 0 | 0 | 0 | 0 |

...
... only first 25 lines shown ...

```
Directory: RNK File: 0READMErnk

RNK - Genomic feature regions ranked by CpG methylation and hsm statistics.

Input: studymc, studymk, region.gr (GRanges object providing annotated region labels and bounds)

Output: files ranked-genes-*.txt sites-in-genes-*.txt
              ranked-promoters-*.txt sites-in-promoters-*.txt
              plot*.pdf

Notes: The output is generated by BWASPR::rank_rbm().
       The files ranked-genes-*.txt provide ranked lists of genes based on
        overall methylation percentage or the occurrence of CpGhms sites
        within the annotated gene bounds.  The output columns include
          region_ID    (= region/gene name)
          rwidth       (= region/gene width)
          nbrsites     (= number of CpGhsm sites in the region/gene)
          nbrper10kb   (= number of CpGhsm sites in the region/gene normalized to 10kb width)
          pmrpersite   (= average % methylation per CpGhsm site in the region/gene)
          pmrpernucl   (= average % CpGhsm methylation per nucleotide in the region/gene)
          prcntM       (= overall % CpGhsm methylation in the region/gene)

       The tables are sorted by prcntM (*byPrcntM* files) or by nbper10kb
         (*bySiteDensity* files) in descending order.

       The ranked-promoters-*.txt files provide analogous tables for promoter regions.
        If the promoter annotation was derived with BWASP, then these regions are simply
        defined here as 500 nucleotides upstream of the 5'-end of the gene (shorter if the
        scaffold ends before).

       The files sites-*.txt provide details for the CpGhsm sites in the respective
        regions.
       The files plot-*.pdf explore concordance of the two methylation measures.
       The optional parameters minrwidth, maxrwidth, and minnbrsites specify
       the length range and minimal number of methylation sites for regions
       to be included in the plot.
```

Directory: RNK File: ranked-genes-byPrcntM-Sd_LI_be.txt

| region_ID | seqnames | start | end | width | strand | coverage | numCs | numTs | prcntM |
|---|---|---|---|---|---|---|---|---|---|
| gene-LOC118184094 | NW_023312874.1 | 59741 | 63170 | 3430 | + | 18 | 18 | 0 | 100 |
| gene-Trnat-agu-24 | NW_023313939.1 | 23003 | 23075 | 73 | - | 22 | 22 | 0 | 100 |
| gene-Trnat-agu-41 | NW_023314769.1 | 146342 | 146413 | 72 | + | 5 | 5 | 0 | 100 |
| gene-LOC118184198 | NW_023314816.1 | 21196 | 21279 | 84 | - | 23 | 23 | 0 | 100 |
| gene-LOC118184196 | NW_023314816.1 | 30823 | 30893 | 71 | - | 7 | 7 | 0 | 100 |
| gene-Trnat-ggu-50 | NW_023314972.1 | 238786 | 238859 | 74 | + | 20 | 20 | 0 | 100 |
| gene-LOC118186326 | NW_023315624.1 | 379328 | 379900 | 573 | - | 7 | 7 | 0 | 100 |
| gene-LOC118187958 | NW_023316202.1 | 8595 | 9018 | 424 | + | 15 | 15 | 0 | 100 |
| gene-LOC118189665 | NW_023316887.1 | 5618 | 8290 | 2673 | - | 25 | 25 | 0 | 100 |
| gene-LOC118189893 | NW_023316988.1 | 134095 | 135039 | 945 | - | 7 | 7 | 0 | 100 |
| gene-LOC118189947 | NW_023317011.1 | 495525 | 495631 | 107 | + | 8 | 8 | 0 | 100 |
| gene-LOC118190335 | NW_023317152.1 | 25601 | 26537 | 937 | + | 6 | 6 | 0 | 100 |
| gene-Trnai-gau-89 | NW_023317293.1 | 231941 | 232013 | 73 | + | 22 | 22 | 0 | 100 |
| gene-LOC118195647 | NW_023319594.1 | 42477 | 43224 | 748 | - | 42 | 42 | 0 | 100 |
| gene-LOC118195884 | NW_023319718.1 | 38619 | 38726 | 108 | + | 8 | 8 | 0 | 100 |
| gene-LOC118196727 | NW_023320099.1 | 959 | 2159 | 1201 | - | 11 | 11 | 0 | 100 |
| gene-LOC118196923 | NW_023320200.1 | 192353 | 192910 | 558 | - | 10 | 10 | 0 | 100 |
| gene-LOC118197526 | NW_023320512.1 | 1461 | 4148 | 2688 | - | 7 | 7 | 0 | 100 |
| gene-LOC118198776 | NW_023321243.1 | 18153 | 21897 | 3745 | - | 8 | 8 | 0 | 100 |
| gene-LOC118198922 | NW_023321346.1 | 7 | 3002 | 2996 | + | 6 | 6 | 0 | 100 |
| gene-LOC118200580 | NW_023322419.1 | 149055 | 149381 | 327 | + | 25 | 25 | 0 | 100 |
| gene-LOC118201255 | NW_023322897.1 | 10627 | 15773 | 5147 | + | 10 | 10 | 0 | 100 |
| gene-LOC118201634 | NW_023323182.1 | 80377 | 81009 | 633 | + | 10 | 10 | 0 | 100 |
| gene-Trnat-ggu-152 | NW_023323893.1 | 23994 | 24065 | 72 | + | 35 | 35 | 0 | 100 |

...

... only first 25 lines shown ...

Directory: RNK File: ranked-genes-byPrcntM-Sd_LI_ka.txt

| region_ID | seqnames | start | end | width | strand | coverage | numCs | numTs | prcntM |
|---|---|---|---|---|---|---|---|---|---|
| gene-LOC118179585 | NW_023313611.1 | 136444 | 138913 | 2470 | - | 7 | 7 | 0 | 100 |
| gene-LOC118183224 | NW_023314471.1 | 93928 | 93995 | 68 | + | 7 | 7 | 0 | 100 |
| gene-LOC118184196 | NW_023314816.1 | 30823 | 30893 | 71 | - | 15 | 15 | 0 | 100 |
| gene-LOC118185454 | NW_023315292.1 | 75210 | 76118 | 909 | + | 6 | 6 | 0 | 100 |
| gene-LOC118186263 | NW_023315602.1 | 2487 | 2904 | 418 | - | 33 | 33 | 0 | 100 |
| gene-LOC118186326 | NW_023315624.1 | 379328 | 379900 | 573 | - | 11 | 11 | 0 | 100 |
| gene-LOC118186620 | NW_023315725.1 | 438470 | 442610 | 4141 | + | 14 | 14 | 0 | 100 |
| gene-LOC118189855 | NW_023316975.1 | 14098 | 14445 | 348 | + | 20 | 20 | 0 | 100 |
| gene-LOC118189947 | NW_023317011.1 | 495525 | 495631 | 107 | + | 10 | 10 | 0 | 100 |
| gene-LOC118190164 | NW_023317087.1 | 277829 | 278218 | 390 | + | 6 | 6 | 0 | 100 |
| gene-LOC118192831 | NW_023318290.1 | 4666 | 5147 | 482 | + | 14 | 14 | 0 | 100 |
| gene-LOC118193268 | NW_023318482.1 | 157807 | 158139 | 333 | - | 5 | 5 | 0 | 100 |
| gene-LOC118194236 | NW_023318942.1 | 242907 | 246670 | 3764 | + | 5 | 5 | 0 | 100 |
| gene-LOC118196923 | NW_023320200.1 | 192353 | 192910 | 558 | - | 5 | 5 | 0 | 100 |
| gene-LOC118197020 | NW_023320240.1 | 53580 | 53684 | 105 | - | 26 | 26 | 0 | 100 |
| gene-Trnat-agu-130 | NW_023320971.1 | 11220 | 11293 | 74 | - | 25 | 25 | 0 | 100 |
| gene-Trnas-cga-130 | NW_023321084.1 | 151042 | 151114 | 73 | - | 5 | 5 | 0 | 100 |
| gene-LOC118198776 | NW_023321243.1 | 18153 | 21897 | 3745 | - | 10 | 10 | 0 | 100 |
| gene-Trnat-ugu-138 | NW_023321978.1 | 133945 | 134016 | 72 | - | 68 | 68 | 0 | 100 |
| gene-LOC118200271 | NW_023322215.1 | 5844 | 6421 | 578 | - | 14 | 14 | 0 | 100 |
| gene-Trnam-cau-134 | NW_023322301.1 | 89447 | 89517 | 71 | - | 12 | 12 | 0 | 100 |
| gene-LOC118200580 | NW_023322419.1 | 149055 | 149381 | 327 | + | 25 | 25 | 0 | 100 |
| gene-Trnat-ugu-146 | NW_023322896.1 | 67966 | 68038 | 73 | - | 5 | 5 | 0 | 100 |
| gene-LOC118201582 | NW_023323149.1 | 11287 | 11736 | 450 | + | 62 | 62 | 0 | 100 |

...
... only first 25 lines shown ...

Directory: RNK File: ranked-genes-bySiteDensity-Sd_LI_be.txt

| region_ID | rwidth | nbrsites | nbrper10kb | pmpersite | pmpernucl | pglink | seqnames | start | end | strand | coverage | numCs | numTs | prcntM |
|---|---|---|---|---|---|---|---|---|---|---|---|---|---|---|
| gene-Trnas-cga-130 | 73 | 10 | 1369.86 | 94.82 | 12.99 | https://www.ncbi.nlm.nih.gov/gene/?term=Trnas-cga | NW_023321084.1 | 151042 | 151114 | - | 77 | 73 | 4 | 94.81 |
| gene-Trnam-cau-159 | 73 | 9 | 1232.88 | 97.22 | 11.99 | https://www.ncbi.nlm.nih.gov/gene/?term=Trnam-cau | NW_023328972.1 | 13638 | 13710 | + | 86 | 82 | 4 | 95.35 |
| gene-Trnas-gga-2 | 73 | 9 | 1232.88 | 74.16 | 9.14 | https://www.ncbi.nlm.nih.gov/gene/?term=Trnas-gga | NW_023312908.1 | 251357 | 251429 | + | 92 | 69 | 23 | 75 |
| gene-Trnag-gcc-15 | 71 | 8 | 1126.76 | 96.67 | 10.89 | https://www.ncbi.nlm.nih.gov/gene/?term=Trnag-gcc | NW_023316992.1 | 203136 | 203206 | + | 79 | 76 | 3 | 96.2 |
| gene-Trnai-gau-140 | 72 | 8 | 1111.11 | 84.38 | 9.38 | https://www.ncbi.nlm.nih.gov/gene/?term=Trnai-gau | NW_023323067.1 | 10341 | 10412 | - | 70 | 58 | 12 | 82.86 |
| gene-Trnat-agu-74 | 72 | 8 | 1111.11 | 90.06 | 10.01 | https://www.ncbi.nlm.nih.gov/gene/?term=Trnat-agu | NW_023316790.1 | 337422 | 337493 | + | 98 | 88 | 10 | 89.8 |
| gene-Trnat-ugu-106 | 73 | 8 | 1095.89 | 97.92 | 10.73 | https://www.ncbi.nlm.nih.gov/gene/?term=Trnat-ugu | NW_023318608.1 | 62412 | 62484 | - | 47 | 46 | 1 | 97.87 |
| gene-Trnat-ugu-149 | 73 | 8 | 1095.89 | 100 | 10.96 | https://www.ncbi.nlm.nih.gov/gene/?term=Trnat-ugu | NW_023324025.1 | 87554 | 87626 | + | 54 | 53 | 1 | 98.15 |
| gene-Trnai-gau-96 | 72 | 7 | 972.22 | 97.96 | 9.52 | https://www.ncbi.nlm.nih.gov/gene/?term=Trnai-gau | NW_023317572.1 | 88037 | 88108 | + | 98 | 96 | 2 | 97.96 |
| gene-Trnat-agu-138 | 72 | 7 | 972.22 | 87.9 | 8.55 | https://www.ncbi.nlm.nih.gov/gene/?term=Trnat-agu | NW_023322724.1 | 32438 | 32509 | - | 57 | 50 | 7 | 87.72 |
| gene-Trnat-ugu-138 | 72 | 7 | 972.22 | 98.81 | 9.61 | https://www.ncbi.nlm.nih.gov/gene/?term=Trnat-ugu | NW_023321978.1 | 133945 | 134016 | - | 69 | 68 | 1 | 98.55 |
| gene-Trnam-cau-137 | 73 | 7 | 958.9 | 77.21 | 7.4 | https://www.ncbi.nlm.nih.gov/gene/?term=Trnam-cau | NW_023322450.1 | 46243 | 46315 | + | 48 | 37 | 11 | 77.08 |
| gene-Trnas-gcu-9 | 74 | 7 | 945.95 | 94.15 | 8.91 | https://www.ncbi.nlm.nih.gov/gene/?term=Trnas-gcu | NW_023314130.1 | 367590 | 367663 | - | 50 | 47 | 3 | 94 |
| gene-LOC118185191 | 846 | 76 | 898.35 | 64.34 | 5.78 | https://www.ncbi.nlm.nih.gov/gene/?term=LOC118185191 | NW_023312912.1 | 746 | 1591 | - | 2547 | 1511 | 1036 | 59.32 |
| gene-Trnat-agu-145 | 72 | 6 | 833.33 | 94.44 | 7.87 | https://www.ncbi.nlm.nih.gov/gene/?term=Trnat-agu | NW_023325266.1 | 10451 | 10522 | - | 40 | 37 | 3 | 92.5 |
| gene-Trnat-agu-94 | 72 | 6 | 833.33 | 95 | 7.92 | https://www.ncbi.nlm.nih.gov/gene/?term=Trnat-agu | NW_023318075.1 | 145068 | 145139 | + | 61 | 58 | 3 | 95.08 |
| gene-Trnat-ggu-143 | 72 | 6 | 833.33 | 97.62 | 8.13 | https://www.ncbi.nlm.nih.gov/gene/?term=Trnat-ggu | NW_023322710.1 | 53923 | 53994 | + | 45 | 44 | 1 | 97.78 |
| gene-Trnal-caa-13 | 73 | 6 | 821.92 | 98.15 | 8.07 | https://www.ncbi.nlm.nih.gov/gene/?term=Trnal-caa | NW_023314314.1 | 128339 | 128411 | + | 52 | 51 | 1 | 98.08 |
| gene-Trnam-cau-2 | 73 | 6 | 821.92 | 92.25 | 7.58 | https://www.ncbi.nlm.nih.gov/gene/?term=Trnam-cau | NW_023312995.1 | 256484 | 256556 | - | 53 | 49 | 4 | 92.45 |
| gene-Trnat-ugu-146 | 73 | 6 | 821.92 | 91.54 | 7.52 | https://www.ncbi.nlm.nih.gov/gene/?term=Trnat-ugu | NW_023322896.1 | 67966 | 68038 | - | 44 | 40 | 4 | 90.91 |
| gene-Trnap-ggg-7 | 72 | 5 | 694.44 | 95 | 6.6 | https://www.ncbi.nlm.nih.gov/gene/?term=Trnap-ggg | NW_023316385.1 | 122186 | 122257 | - | 36 | 34 | 2 | 94.44 |
| gene-Trnat-ggu-152 | 72 | 5 | 694.44 | 100 | 6.94 | https://www.ncbi.nlm.nih.gov/gene/?term=Trnat-ggu | NW_023323893.1 | 23994 | 24065 | + | 35 | 35 | 0 | 100 |
| gene-Trnai-aau-41 | 74 | 5 | 675.68 | 87.86 | 5.94 | https://www.ncbi.nlm.nih.gov/gene/?term=Trnai-aau | NW_023325653.1 | 534 | 607 | - | 71 | 54 | 17 | 76.06 |
| gene-LOC118199038 | 1069 | 72 | 673.53 | 87.98 | 5.93 | https://www.ncbi.nlm.nih.gov/gene/?term=LOC118199038 | NW_023321425.1 | 10092 | 11160 | + | 2341 | 2062 | 279 | 88.08 |

...

... only first 25 lines shown ...

Directory: RNK File: ranked-genes-bySiteDensity-Sd_LI_ka.txt

| region_ID | rwidth | nbrsites | nbrper10kb | pmpersite | pmpernucl | pglink | seqnames | start | end | strand | coverage | numCs | numTs | prcntM |
|---|---|---|---|---|---|---|---|---|---|---|---|---|---|---|
| gene-Trnam-cau-159 | 73 | 9 | 1232.88 | 93.06 | 11.47 | https://www.ncbi.nlm.nih.gov/gene/?term=Trnam-cau | NW_023328972.1 | 13638 | 13710 | + | 69 | 64 | 5 | 92.75 |
| gene-Trnag-gcc-15 | 71 | 8 | 1126.76 | 94.06 | 10.6 | https://www.ncbi.nlm.nih.gov/gene/?term=Trnag-gcc | NW_023316992.1 | 203136 | 203206 | + | 106 | 100 | 6 | 94.34 |
| gene-Trnat-ugu-149 | 73 | 8 | 1095.89 | 94.79 | 10.39 | https://www.ncbi.nlm.nih.gov/gene/?term=Trnat-ugu | NW_023324025.1 | 87554 | 87626 | + | 51 | 48 | 3 | 94.12 |
| gene-Trnat-agu-47 | 74 | 8 | 1081.08 | 96.67 | 10.45 | https://www.ncbi.nlm.nih.gov/gene/?term=Trnat-agu | NW_023314948.1 | 149304 | 149377 | - | 67 | 65 | 2 | 97.01 |
| gene-Trnai-gau-140 | 72 | 7 | 972.22 | 97.62 | 9.49 | https://www.ncbi.nlm.nih.gov/gene/?term=Trnai-gau | NW_023323067.1 | 10341 | 10412 | - | 52 | 44 | 8 | 84.62 |
| gene-Trnai-gau-96 | 72 | 7 | 972.22 | 93.94 | 9.13 | https://www.ncbi.nlm.nih.gov/gene/?term=Trnai-gau | NW_023317572.1 | 88037 | 88108 | + | 59 | 55 | 4 | 93.22 |
| gene-Trnat-agu-138 | 72 | 7 | 972.22 | 91.16 | 8.86 | https://www.ncbi.nlm.nih.gov/gene/?term=Trnat-agu | NW_023322724.1 | 32438 | 32509 | - | 48 | 44 | 4 | 91.67 |
| gene-Trnat-ugu-138 | 72 | 7 | 972.22 | 100 | 9.72 | https://www.ncbi.nlm.nih.gov/gene/?term=Trnat-ugu | NW_023321978.1 | 133945 | 134016 | - | 68 | 68 | 0 | 100 |
| gene-Trnam-cau-137 | 73 | 7 | 958.9 | 96.1 | 9.22 | https://www.ncbi.nlm.nih.gov/gene/?term=Trnam-cau | NW_023322450.1 | 46243 | 46315 | + | 54 | 51 | 3 | 94.44 |
| gene-Trnam-cau-2 | 73 | 7 | 958.9 | 100 | 9.59 | https://www.ncbi.nlm.nih.gov/gene/?term=Trnam-cau | NW_023312995.1 | 256484 | 256556 | - | 51 | 49 | 2 | 96.08 |
| gene-Trnam-cau-33 | 73 | 7 | 958.9 | 66.33 | 6.36 | https://www.ncbi.nlm.nih.gov/gene/?term=Trnam-cau | NW_023314550.1 | 84579 | 84651 | + | 81 | 51 | 30 | 62.96 |
| gene-Trnas-gcu-9 | 74 | 7 | 945.95 | 98.9 | 9.36 | https://www.ncbi.nlm.nih.gov/gene/?term=Trnas-gcu | NW_023314130.1 | 367590 | 367663 | - | 89 | 88 | 1 | 98.88 |
| gene-Trnas-aga-10 | 82 | 7 | 853.66 | 93.69 | 8 | https://www.ncbi.nlm.nih.gov/gene/?term=Trnas-aga | NW_023315199.1 | 170704 | 170785 | - | 71 | 65 | 6 | 91.55 |
| gene-LOC118185830 | 72 | 6 | 833.33 | 92.36 | 7.7 | https://www.ncbi.nlm.nih.gov/gene/?term=LOC118185830 | NW_023315437.1 | 13994 | 14065 | + | 51 | 47 | 4 | 92.16 |
| gene-Trnat-agu-145 | 72 | 6 | 833.33 | 71.03 | 5.92 | https://www.ncbi.nlm.nih.gov/gene/?term=Trnat-agu | NW_023325266.1 | 10451 | 10522 | - | 52 | 35 | 17 | 67.31 |
| gene-Trnat-ggu-143 | 72 | 6 | 833.33 | 93.7 | 7.81 | https://www.ncbi.nlm.nih.gov/gene/?term=Trnat-ggu | NW_023322710.1 | 53923 | 53994 | + | 47 | 44 | 3 | 93.62 |
| gene-Trnat-ggu-6 | 72 | 6 | 833.33 | 34.01 | 2.83 | https://www.ncbi.nlm.nih.gov/gene/?term=Trnat-ggu | NW_023312877.1 | 112059 | 112130 | - | 380 | 102 | 278 | 26.84 |
| gene-Trnal-caa-13 | 73 | 6 | 821.92 | 95.77 | 7.87 | https://www.ncbi.nlm.nih.gov/gene/?term=Trnal-caa | NW_023314314.1 | 128339 | 128411 | + | 48 | 46 | 2 | 95.83 |
| gene-Trnat-ugu-106 | 73 | 6 | 821.92 | 94.44 | 7.76 | https://www.ncbi.nlm.nih.gov/gene/?term=Trnat-ugu | NW_023318608.1 | 62412 | 62484 | - | 42 | 37 | 5 | 88.1 |
| gene-LOC118185191 | 846 | 69 | 815.6 | 59.44 | 4.85 | https://www.ncbi.nlm.nih.gov/gene/?term=LOC118185191 | NW_023312912.1 | 746 | 1591 | - | 2374 | 1374 | 1000 | 57.88 |
| gene-Trnai-aau-41 | 74 | 6 | 810.81 | 96.65 | 7.84 | https://www.ncbi.nlm.nih.gov/gene/?term=Trnai-aau | NW_023325653.1 | 534 | 607 | - | 79 | 76 | 3 | 96.2 |
| gene-Trnap-ggg-7 | 72 | 5 | 694.44 | 90.95 | 6.32 | https://www.ncbi.nlm.nih.gov/gene/?term=Trnap-ggg | NW_023316385.1 | 122186 | 122257 | - | 40 | 35 | 5 | 87.5 |
| gene-Trnat-ggu-152 | 72 | 5 | 694.44 | 94 | 6.53 | https://www.ncbi.nlm.nih.gov/gene/?term=Trnat-ggu | NW_023323893.1 | 23994 | 24065 | + | 47 | 44 | 3 | 93.62 |
| gene-Trnat-cgu-144 | 73 | 5 | 684.93 | 100 | 6.85 | https://www.ncbi.nlm.nih.gov/gene/?term=Trnat-cgu | NW_023322570.1 | 1573 | 1645 | + | 30 | 29 | 1 | 96.67 |

...

... only first 25 lines shown ...

Directory: RNK File: ranked-promoters-byPrcntM-Sd_LI_be.txt

| region_ID | seqnames | start | end | width | strand | coverage | numCs | numTs | prcntM |
|---|---|---|---|---|---|---|---|---|---|
| gene-LOC118183835 | NW_023312867.1 | 15234 | 15733 | 500 | + | 15 | 15 | 0 | 100 |
| gene-LOC118186050 | NW_023312941.1 | 173625 | 174124 | 500 | - | 7 | 7 | 0 | 100 |
| gene-LOC118187247 | NW_023312980.1 | 1 | 365 | 365 | + | 36 | 36 | 0 | 100 |
| gene-LOC118189159 | NW_023313051.1 | 88274 | 88773 | 500 | - | 11 | 11 | 0 | 100 |
| gene-LOC118191111 | NW_023313119.1 | 1579 | 2078 | 500 | + | 29 | 29 | 0 | 100 |
| gene-LOC118193412 | NW_023313199.1 | 312854 | 313353 | 500 | - | 43 | 43 | 0 | 100 |
| gene-LOC118194168 | NW_023313221.1 | 48176 | 48675 | 500 | - | 20 | 20 | 0 | 100 |
| gene-LOC118200103 | NW_023313387.1 | 331142 | 331641 | 500 | + | 12 | 12 | 0 | 100 |
| gene-LOC118200907 | NW_023313421.1 | 404868 | 405367 | 500 | - | 59 | 59 | 0 | 100 |
| gene-LOC118204719 | NW_023313546.1 | 619 | 1118 | 500 | + | 21 | 21 | 0 | 100 |
| gene-LOC118179644 | NW_023313613.1 | 43853 | 44352 | 500 | + | 5 | 5 | 0 | 100 |
| gene-LOC118181001 | NW_023313703.1 | 2499 | 2998 | 500 | + | 13 | 13 | 0 | 100 |
| gene-LOC118181390 | NW_023313833.1 | 38947 | 39446 | 500 | + | 19 | 19 | 0 | 100 |
| gene-LOC118181680 | NW_023313919.1 | 5148 | 5647 | 500 | + | 17 | 17 | 0 | 100 |
| gene-LOC118181792 | NW_023313960.1 | 519078 | 519577 | 500 | - | 26 | 26 | 0 | 100 |
| gene-LOC118181955 | NW_023314023.1 | 40079 | 40578 | 500 | - | 7 | 7 | 0 | 100 |
| gene-LOC118182074 | NW_023314063.1 | 377840 | 378339 | 500 | - | 11 | 11 | 0 | 100 |
| gene-LOC118182222 | NW_023314100.1 | 464950 | 465449 | 500 | - | 5 | 5 | 0 | 100 |
| gene-LOC118182241 | NW_023314107.1 | 28456 | 28955 | 500 | - | 12 | 12 | 0 | 100 |
| gene-LOC118182384 | NW_023314147.1 | 2565 | 3064 | 500 | + | 5 | 5 | 0 | 100 |
| gene-LOC118182402 | NW_023314155.1 | 15992 | 16491 | 500 | + | 5 | 5 | 0 | 100 |
| gene-LOC118182704 | NW_023314281.1 | 284880 | 285224 | 345 | - | 10 | 10 | 0 | 100 |
| gene-LOC118183284 | NW_023314493.1 | 22314 | 22813 | 500 | + | 22 | 22 | 0 | 100 |
| gene-LOC118183523 | NW_023314587.1 | 91275 | 91774 | 500 | + | 6 | 6 | 0 | 100 |

...

... only first 25 lines shown ...

Directory: RNK File: ranked-promoters-byPrcntM-Sd_LI_ka.txt

| region_ID | seqnames | start | end | width | strand | coverage | numCs | numTs | prcntM |
|---|---|---|---|---|---|---|---|---|---|
| gene-LOC118183835 | NW_023312867.1 | 15234 | 15733 | 500 | + | 9 | 9 | 0 | 100 |
| gene-LOC118186050 | NW_023312941.1 | 173625 | 174124 | 500 | - | 8 | 8 | 0 | 100 |
| gene-LOC118194423 | NW_023313229.1 | 1299128 | 1299627 | 500 | - | 31 | 31 | 0 | 100 |
| gene-Trnaa-ggc-6 | NW_023313289.1 | 47843 | 48342 | 500 | + | 42 | 42 | 0 | 100 |
| gene-LOC118197224 | NW_023313308.1 | 137573 | 138072 | 500 | + | 10 | 10 | 0 | 100 |
| gene-LOC118200103 | NW_023313387.1 | 331142 | 331641 | 500 | + | 11 | 11 | 0 | 100 |
| gene-LOC118200799 | NW_023313415.1 | 84135 | 84634 | 500 | + | 8 | 8 | 0 | 100 |
| gene-LOC118202570 | NW_023313479.1 | 40513 | 41012 | 500 | + | 19 | 19 | 0 | 100 |
| gene-LOC118204705 | NW_023313546.1 | 102945 | 103444 | 500 | - | 40 | 40 | 0 | 100 |
| gene-LOC118179644 | NW_023313613.1 | 43853 | 44352 | 500 | + | 7 | 7 | 0 | 100 |
| gene-LOC118181001 | NW_023313703.1 | 2499 | 2998 | 500 | + | 15 | 15 | 0 | 100 |
| gene-LOC118181390 | NW_023313833.1 | 38947 | 39446 | 500 | + | 15 | 15 | 0 | 100 |
| gene-LOC118181462 | NW_023313852.1 | 105718 | 105939 | 222 | - | 6 | 6 | 0 | 100 |
| gene-LOC118181680 | NW_023313919.1 | 5148 | 5647 | 500 | + | 9 | 9 | 0 | 100 |
| gene-LOC118181687 | NW_023313921.1 | 5428 | 5927 | 500 | + | 13 | 13 | 0 | 100 |
| gene-LOC118181789 | NW_023313959.1 | 38309 | 38808 | 500 | - | 9 | 9 | 0 | 100 |
| gene-LOC118181944 | NW_023314020.1 | 12950 | 13449 | 500 | + | 21 | 21 | 0 | 100 |
| gene-LOC118181955 | NW_023314023.1 | 40079 | 40578 | 500 | - | 7 | 7 | 0 | 100 |
| gene-LOC118182241 | NW_023314107.1 | 28456 | 28955 | 500 | - | 9 | 9 | 0 | 100 |
| gene-LOC118182801 | NW_023314314.1 | 94107 | 94606 | 500 | - | 5 | 5 | 0 | 100 |
| gene-LOC118183234 | NW_023314473.1 | 319626 | 320125 | 500 | - | 6 | 6 | 0 | 100 |
| gene-LOC118183385 | NW_023314541.1 | 3854 | 4353 | 500 | + | 6 | 6 | 0 | 100 |
| gene-LOC118183525 | NW_023314588.1 | 2170 | 2669 | 500 | + | 5 | 5 | 0 | 100 |
| gene-LOC118183609 | NW_023314626.1 | 11482 | 11981 | 500 | + | 10 | 10 | 0 | 100 |

...

... only first 25 lines shown ...

Directory: RNK File: ranked-promoters-bySiteDensity-Sd_LI_be.txt

| region_ID | rwidth | nbrsites | nbrper10kb | pmpersite | pmpernucl | pglink | seqnames | start | end | strand | coverage | numCs | numTs | prcntM |
|---|---|---|---|---|---|---|---|---|---|---|---|---|---|---|
| gene-LOC118202462 | 500 | 49 | 980 | 89.47 | 8.77 | https://www.ncbi.nlm.nih.gov/gene/?term=LOC118202462 | NW_023323757.1 | 147005 | 147504 | + | 578 | 445 | 133 | 76.99 |
| gene-LOC118198320 | 500 | 43 | 860 | 94.47 | 8.12 | https://www.ncbi.nlm.nih.gov/gene/?term=LOC118198320 | NW_023320965.1 | 23272 | 23771 | - | 613 | 579 | 34 | 94.45 |
| gene-LOC118182676 | 500 | 40 | 800 | 80.48 | 6.44 | https://www.ncbi.nlm.nih.gov/gene/?term=LOC118182676 | NW_023312841.1 | 11701 | 12200 | - | 920 | 713 | 207 | 77.5 |
| gene-LOC118183077 | 500 | 40 | 800 | 92.44 | 7.39 | https://www.ncbi.nlm.nih.gov/gene/?term=LOC118183077 | NW_023314415.1 | 2785 | 3284 | + | 399 | 366 | 33 | 91.73 |
| gene-LOC118200883 | 500 | 38 | 760 | 98.12 | 7.46 | https://www.ncbi.nlm.nih.gov/gene/?term=LOC118200883 | NW_023322620.1 | 27247 | 27746 | - | 375 | 367 | 8 | 97.87 |
| gene-LOC118205510 | 500 | 38 | 760 | 92.24 | 7.01 | https://www.ncbi.nlm.nih.gov/gene/?term=LOC118205510 | NW_023326523.1 | 5370 | 5869 | - | 797 | 716 | 81 | 89.84 |
| gene-LOC118204740 | 500 | 37 | 740 | 85.74 | 6.34 | https://www.ncbi.nlm.nih.gov/gene/?term=LOC118204740 | NW_023325720.1 | 59895 | 60394 | - | 345 | 283 | 62 | 82.03 |
| gene-LOC118205209 | 500 | 37 | 740 | 76.43 | 5.66 | https://www.ncbi.nlm.nih.gov/gene/?term=LOC118205209 | NW_023326187.1 | 5974 | 6473 | + | 1124 | 805 | 319 | 71.62 |
| gene-LOC118179543 | 331 | 24 | 725.08 | 71.54 | 5.19 | https://www.ncbi.nlm.nih.gov/gene/?term=LOC118179543 | NW_023327590.1 | 1 | 331 | + | 468 | 322 | 146 | 68.8 |
| gene-LOC118192113 | 500 | 36 | 720 | 93.93 | 6.76 | https://www.ncbi.nlm.nih.gov/gene/?term=LOC118192113 | NW_023317940.1 | 13199 | 13698 | + | 440 | 411 | 29 | 93.41 |
| gene-LOC118202955 | 500 | 36 | 720 | 94.52 | 6.81 | https://www.ncbi.nlm.nih.gov/gene/?term=LOC118202955 | NW_023324125.1 | 2910 | 3409 | + | 326 | 307 | 19 | 94.17 |
| gene-LOC118192222 | 500 | 35 | 700 | 94.96 | 6.65 | https://www.ncbi.nlm.nih.gov/gene/?term=LOC118192222 | NW_023317979.1 | 50766 | 51265 | - | 365 | 345 | 20 | 94.52 |
| gene-LOC118195284 | 500 | 35 | 700 | 90.05 | 6.3 | https://www.ncbi.nlm.nih.gov/gene/?term=LOC118195284 | NW_023319403.1 | 91205 | 91704 | - | 305 | 274 | 31 | 89.84 |
| gene-LOC118204163 | 500 | 35 | 700 | 95.87 | 6.71 | https://www.ncbi.nlm.nih.gov/gene/?term=LOC118204163 | NW_023325170.1 | 77252 | 77751 | - | 289 | 275 | 14 | 95.16 |
| gene-LOC118179404 | 500 | 34 | 680 | 95.79 | 6.51 | https://www.ncbi.nlm.nih.gov/gene/?term=LOC118179404 | NW_023327404.1 | 19277 | 19776 | - | 254 | 233 | 21 | 91.73 |
| gene-LOC118180258 | 500 | 34 | 680 | 94.98 | 6.46 | https://www.ncbi.nlm.nih.gov/gene/?term=LOC118180258 | NW_023328434.1 | 1646 | 2145 | - | 303 | 277 | 26 | 91.42 |
| gene-LOC118183590 | 500 | 33 | 660 | 94.49 | 6.24 | https://www.ncbi.nlm.nih.gov/gene/?term=LOC118183590 | NW_023314619.1 | 62020 | 62519 | + | 340 | 322 | 18 | 94.71 |
| gene-LOC118204540 | 500 | 33 | 660 | 89.49 | 5.91 | https://www.ncbi.nlm.nih.gov/gene/?term=LOC118204540 | NW_023325538.1 | 31801 | 32300 | - | 539 | 459 | 80 | 85.16 |
| gene-LOC118185225 | 500 | 32 | 640 | 90.9 | 5.82 | https://www.ncbi.nlm.nih.gov/gene/?term=LOC118185225 | NW_023312914.1 | 746830 | 747329 | - | 304 | 268 | 36 | 88.16 |
| gene-LOC118186106 | 500 | 32 | 640 | 93.86 | 6.01 | https://www.ncbi.nlm.nih.gov/gene/?term=LOC118186106 | NW_023315545.1 | 82926 | 83425 | + | 434 | 402 | 32 | 92.63 |
| gene-LOC118191888 | 500 | 32 | 640 | 79.53 | 5.09 | https://www.ncbi.nlm.nih.gov/gene/?term=LOC118191888 | NW_023317814.1 | 10516 | 11015 | + | 2186 | 1691 | 495 | 77.36 |
| gene-LOC118197832 | 500 | 32 | 640 | 88.19 | 5.64 | https://www.ncbi.nlm.nih.gov/gene/?term=LOC118197832 | NW_023320679.1 | 178651 | 179150 | - | 332 | 281 | 51 | 84.64 |
| gene-LOC118204935 | 500 | 32 | 640 | 91.96 | 5.89 | https://www.ncbi.nlm.nih.gov/gene/?term=LOC118204935 | NW_023325921.1 | 8970 | 9469 | + | 407 | 369 | 38 | 90.66 |
| gene-LOC118182191 | 500 | 30 | 600 | 87.09 | 5.23 | https://www.ncbi.nlm.nih.gov/gene/?term=LOC118182191 | NW_023314092.1 | 190495 | 190994 | + | 376 | 321 | 55 | 85.37 |

...

... only first 25 lines shown ...

Directory: RNK File: ranked-promoters-bySiteDensity-Sd_LI_ka.txt

| region_ID | rwidth | nbrsites | nbrper10kb | pmpersite | pmpernucl | pglink | seqnames | start | end | strand | coverage | numCs | numTs | prcntM |
|---|---|---|---|---|---|---|---|---|---|---|---|---|---|---|
| gene-LOC118182676 | 500 | 57 | 1140 | 87.04 | 9.92 | https://www.ncbi.nlm.nih.gov/gene/?term=LOC118182676 | NW_023312841.1 | 11701 | 12200 | - | 1015 | 880 | 135 | 86.7 |
| gene-LOC118202462 | 500 | 54 | 1080 | 88.46 | 9.55 | https://www.ncbi.nlm.nih.gov/gene/?term=LOC118202462 | NW_023323757.1 | 147005 | 147504 | + | 846 | 642 | 204 | 75.89 |
| gene-LOC118198320 | 500 | 41 | 820 | 86.79 | 7.12 | https://www.ncbi.nlm.nih.gov/gene/?term=LOC118198320 | NW_023320965.1 | 23272 | 23771 | - | 356 | 299 | 57 | 83.99 |
| gene-LOC118183077 | 500 | 40 | 800 | 93.54 | 7.48 | https://www.ncbi.nlm.nih.gov/gene/?term=LOC118183077 | NW_023314415.1 | 2785 | 3284 | + | 443 | 411 | 32 | 92.78 |
| gene-LOC118204540 | 500 | 39 | 780 | 94.3 | 7.36 | https://www.ncbi.nlm.nih.gov/gene/?term=LOC118204540 | NW_023325538.1 | 31801 | 32300 | - | 505 | 474 | 31 | 93.86 |
| gene-LOC118205510 | 500 | 39 | 780 | 94.43 | 7.37 | https://www.ncbi.nlm.nih.gov/gene/?term=LOC118205510 | NW_023326523.1 | 5370 | 5869 | - | 901 | 842 | 59 | 93.45 |
| gene-LOC118185225 | 500 | 38 | 760 | 89.41 | 6.79 | https://www.ncbi.nlm.nih.gov/gene/?term=LOC118185225 | NW_023312914.1 | 746830 | 747329 | - | 448 | 398 | 50 | 88.84 |
| gene-LOC118192866 | 500 | 38 | 760 | 96.31 | 7.32 | https://www.ncbi.nlm.nih.gov/gene/?term=LOC118192866 | NW_023318309.1 | 238501 | 239000 | - | 275 | 262 | 13 | 95.27 |
| gene-LOC118179543 | 331 | 25 | 755.29 | 72.91 | 5.51 | https://www.ncbi.nlm.nih.gov/gene/?term=LOC118179543 | NW_023327590.1 | 1 | 331 | + | 567 | 398 | 169 | 70.19 |
| gene-LOC118204935 | 500 | 34 | 680 | 92.52 | 6.29 | https://www.ncbi.nlm.nih.gov/gene/?term=LOC118204935 | NW_023325921.1 | 8970 | 9469 | + | 325 | 292 | 33 | 89.85 |
| gene-LOC118180258 | 500 | 33 | 660 | 96.45 | 6.37 | https://www.ncbi.nlm.nih.gov/gene/?term=LOC118180258 | NW_023328434.1 | 1646 | 2145 | - | 300 | 285 | 15 | 95 |
| gene-LOC118185492 | 500 | 33 | 660 | 93.82 | 6.19 | https://www.ncbi.nlm.nih.gov/gene/?term=LOC118185492 | NW_023315307.1 | 349017 | 349516 | - | 354 | 329 | 25 | 92.94 |
| gene-LOC118191888 | 500 | 32 | 640 | 80.46 | 5.15 | https://www.ncbi.nlm.nih.gov/gene/?term=LOC118191888 | NW_023317814.1 | 10516 | 11015 | + | 2370 | 1866 | 504 | 78.73 |
| gene-LOC118192113 | 500 | 32 | 640 | 91.29 | 5.84 | https://www.ncbi.nlm.nih.gov/gene/?term=LOC118192113 | NW_023317940.1 | 13199 | 13698 | + | 412 | 359 | 53 | 87.14 |
| gene-LOC118198631 | 500 | 32 | 640 | 90.42 | 5.79 | https://www.ncbi.nlm.nih.gov/gene/?term=LOC118198631 | NW_023321150.1 | 85008 | 85507 | - | 300 | 256 | 44 | 85.33 |
| gene-LOC118200022 | 500 | 32 | 640 | 96.55 | 6.18 | https://www.ncbi.nlm.nih.gov/gene/?term=LOC118200022 | NW_023322048.1 | 11588 | 12087 | + | 322 | 309 | 13 | 95.96 |
| gene-LOC118180343 | 500 | 31 | 620 | 39.25 | 2.43 | https://www.ncbi.nlm.nih.gov/gene/?term=LOC118180343 | NW_023328529.1 | 7840 | 8339 | - | 1137 | 398 | 739 | 35 |
| gene-LOC118195284 | 500 | 31 | 620 | 94.41 | 5.85 | https://www.ncbi.nlm.nih.gov/gene/?term=LOC118195284 | NW_023319403.1 | 91205 | 91704 | - | 293 | 266 | 27 | 90.78 |
| gene-LOC118205247 | 500 | 31 | 620 | 91.6 | 5.68 | https://www.ncbi.nlm.nih.gov/gene/?term=LOC118205247 | NW_023326228.1 | 6853 | 7352 | + | 267 | 241 | 26 | 90.26 |
| gene-LOC118205456 | 500 | 31 | 620 | 94.72 | 5.87 | https://www.ncbi.nlm.nih.gov/gene/?term=LOC118205456 | NW_023326459.1 | 20553 | 21052 | + | 478 | 427 | 51 | 89.33 |
| gene-LOC118193842 | 500 | 30 | 600 | 74.24 | 4.45 | https://www.ncbi.nlm.nih.gov/gene/?term=LOC118193842 | NW_023318750.1 | 142791 | 143290 | + | 507 | 359 | 148 | 70.81 |
| gene-LOC118198350 | 500 | 30 | 600 | 75.1 | 4.51 | https://www.ncbi.nlm.nih.gov/gene/?term=LOC118198350 | NW_023320977.1 | 105789 | 106288 | + | 942 | 662 | 280 | 70.28 |
| gene-LOC118200883 | 500 | 30 | 600 | 96.85 | 5.81 | https://www.ncbi.nlm.nih.gov/gene/?term=LOC118200883 | NW_023322620.1 | 27247 | 27746 | - | 245 | 228 | 17 | 93.06 |
| gene-LOC118202955 | 500 | 30 | 600 | 99 | 5.94 | https://www.ncbi.nlm.nih.gov/gene/?term=LOC118202955 | NW_023324125.1 | 2910 | 3409 | + | 187 | 182 | 5 | 97.33 |

...

... only first 25 lines shown ...

Directory: RNK File: sites-in-genes-Sd_LI_be.txt

```
seqnames	start end	width strand	coverage	numCs numTs perc_meth	region_seqnames	region_start	region_end	region_width	region_strand	region_source
	region_type region_score	region_phase	region_ID	region_Dbxref	region_Name region_gbkey	region_gene region_gene_biotype	region_end_range region_partial
	region_start_range
NW_023312688.1	28083 28083 1	-	11	9	2	81.8181818181818	NW_023312688.1	27109 87988 60880 +	Gnomon	gene	NA	NA	gene-LOC118188962
	GeneID:118188962	LOC118188962	Gene	LOC118188962	protein_coding	character(0)	NA	character(0)
NW_023312688.1	28102 28102 1	-	10	7	3	70	NW_023312688.1	27109 87988 60880 +	Gnomon	gene	NA	NA	gene-LOC118188962 GeneID:118188962
	LOC118188962	Gene	LOC118188962	protein_coding	character(0)	NA	character(0)
NW_023312688.1	28127 28127 1	-	14	11	3	78.5714285714286	NW_023312688.1	27109 87988 60880 +	Gnomon	gene	NA	NA	gene-LOC118188962
	GeneID:118188962	LOC118188962	Gene	LOC118188962	protein_coding	character(0)	NA	character(0)
NW_023312688.1	28147 28147 1	-	14	12	2	85.7142857142857	NW_023312688.1	27109 87988 60880 +	Gnomon	gene	NA	NA	gene-LOC118188962
	GeneID:118188962	LOC118188962	Gene	LOC118188962	protein_coding	character(0)	NA	character(0)
NW_023312688.1	28156 28156 1	-	16	15	1	93.75 NW_023312688.1	27109 87988 60880 +	Gnomon	gene	NA	NA	gene-LOC118188962 GeneID:118188962
	LOC118188962	Gene	LOC118188962	protein_coding	character(0)	NA	character(0)
NW_023312688.1	28397 28397 1	-	7	5	2	71.4285714285714	NW_023312688.1	27109 87988 60880 +	Gnomon	gene	NA	NA	gene-LOC118188962
	GeneID:118188962	LOC118188962	Gene	LOC118188962	protein_coding	character(0)	NA	character(0)
NW_023312688.1	28499 28499 1	+	8	6	2	75	NW_023312688.1	27109 87988 60880 +	Gnomon	gene	NA	NA	gene-LOC118188962 GeneID:118188962
	LOC118188962	Gene	LOC118188962	protein_coding	character(0)	NA	character(0)
NW_023312688.1	28630 28630 1	+	16	16	0	100	NW_023312688.1	27109 87988 60880 +	Gnomon	gene	NA	NA	gene-LOC118188962 GeneID:118188962
	LOC118188962	Gene	LOC118188962	protein_coding	character(0)	NA	character(0)
NW_023312688.1	28761 28761 1	+	9	8	1	88.8888888888889	NW_023312688.1	27109 87988 60880 +	Gnomon	gene	NA	NA	gene-LOC118188962
	GeneID:118188962	LOC118188962	Gene	LOC118188962	protein_coding	character(0)	NA	character(0)
NW_023312688.1	28762 28762 1	-	9	9	0	100	NW_023312688.1	27109 87988 60880 +	Gnomon	gene	NA	NA	gene-LOC118188962 GeneID:118188962
	LOC118188962	Gene	LOC118188962	protein_coding	character(0)	NA	character(0)
NW_023312688.1	28853 28853 1	-	11	11	0	100	NW_023312688.1	27109 87988 60880 +	Gnomon	gene	NA	NA	gene-LOC118188962 GeneID:118188962
	LOC118188962	Gene	LOC118188962	protein_coding	character(0)	NA	character(0)
NW_023312688.1	28866 28866 1	-	11	11	0	100	NW_023312688.1	27109 87988 60880 +	Gnomon	gene	NA	NA	gene-LOC118188962 GeneID:118188962
	LOC118188962	Gene	LOC118188962	protein_coding	character(0)	NA	character(0)
NW_023312688.1	28902 28902 1	-	11	11	0	100	NW_023312688.1	27109 87988 60880 +	Gnomon	gene	NA	NA	gene-LOC118188962 GeneID:118188962
	LOC118188962	Gene	LOC118188962	protein_coding	character(0)	NA	character(0)
NW_023312688.1	29285 29285 1	-	6	6	0	100	NW_023312688.1	27109 87988 60880 +	Gnomon	gene	NA	NA	gene-LOC118188962 GeneID:118188962
	LOC118188962	Gene	LOC118188962	protein_coding	character(0)	NA	character(0)
NW_023312688.1	29406 29406 1	-	9	8	1	88.8888888888889	NW_023312688.1	27109 87988 60880 +	Gnomon	gene	NA	NA	gene-LOC118188962
	GeneID:118188962	LOC118188962	Gene	LOC118188962	protein_coding	character(0)	NA	character(0)
NW_023312688.1	29413 29413 1	-	11	10	1	90.9090909090909	NW_023312688.1	27109 87988 60880 +	Gnomon	gene	NA	NA	gene-LOC118188962
	GeneID:118188962	LOC118188962	Gene	LOC118188962	protein_coding	character(0)	NA	character(0)
NW_023312688.1	29425 29425 1	-	8	7	1	87.5	NW_023312688.1	27109 87988 60880 +	Gnomon	gene	NA	NA	gene-LOC118188962 GeneID:118188962
	LOC118188962	Gene	LOC118188962	protein_coding	character(0)	NA	character(0)
NW_023312688.1	29445 29445 1	-	8	8	0	100	NW_023312688.1	27109 87988 60880 +	Gnomon	gene	NA	NA	gene-LOC118188962 GeneID:118188962
	LOC118188962	Gene	LOC118188962	protein_coding	character(0)	NA	character(0)
NW_023312688.1	29653 29653 1	-	11	9	2	81.8181818181818	NW_023312688.1	27109 87988 60880 +	Gnomon	gene	NA	NA	gene-LOC118188962
	GeneID:118188962	LOC118188962	Gene	LOC118188962	protein_coding	character(0)	NA	character(0)
NW_023312688.1	29663 29663 1	-	11	9	2	81.8181818181818	NW_023312688.1	27109 87988 60880 +	Gnomon	gene	NA	NA	gene-LOC118188962
	GeneID:118188962	LOC118188962	Gene	LOC118188962	protein_coding	character(0)	NA	character(0)
NW_023312688.1	29677 29677 1	-	10	8	2	80	NW_023312688.1	27109 87988 60880 +	Gnomon	gene	NA	NA	gene-LOC118188962 GeneID:118188962
	LOC118188962	Gene	LOC118188962	protein_coding	character(0)	NA	character(0)
NW_023312688.1	29695 29695 1	-	11	8	3	72.7272727272727	NW_023312688.1	27109 87988 60880 +	Gnomon	gene	NA	NA	gene-LOC118188962
	GeneID:118188962	LOC118188962	Gene	LOC118188962	protein_coding	character(0)	NA	character(0)
NW_023312688.1	29700 29700 1	-	12	8	4	66.6666666666667	NW_023312688.1	27109 87988 60880 +	Gnomon	gene	NA	NA	gene-LOC118188962
	GeneID:118188962	LOC118188962	Gene	LOC118188962	protein_coding	character(0)	NA	character(0)
NW_023312688.1	29715 29715 1	-	10	7	3	70	NW_023312688.1	27109 87988 60880 +	Gnomon	gene	NA	NA	gene-LOC118188962 GeneID:118188962
	LOC118188962	Gene	LOC118188962	protein_coding	character(0)	NA	character(0)
...
... only first 25 lines shown ...
```

Directory: RNK File: sites-in-genes-Sd_LI_ka.txt

```
seqnames	start	end	width	strand	coverage	numCs	numTs	perc_meth	region_seqnames	region_start	region_end	region_width	region_strand	region_source
	region_type	region_score	region_phase	region_ID	region_Dbxref	region_Name	region_gbkey	region_gene	region_gene_biotype	region_end_range	region_partial
	region_start_range
NW_023312688.1	28083	28083	1	-	6	6	0	100	NW_023312688.1	27109	87988	60880	+	Gnomon	gene	NA	NA	gene-LOC118188962	GeneID:118188962
	LOC118188962	Gene	LOC118188962	protein_coding	character(0)	NA	character(0)
NW_023312688.1	28127	28127	1	-	6	6	0	100	NW_023312688.1	27109	87988	60880	+	Gnomon	gene	NA	NA	gene-LOC118188962	GeneID:118188962
	LOC118188962	Gene	LOC118188962	protein_coding	character(0)	NA	character(0)
NW_023312688.1	28397	28397	1	-	9	8	1	88.8888888888889	NW_023312688.1	27109	87988	60880	+	Gnomon	gene	NA	NA	gene-LOC118188962
	GeneID:118188962	LOC118188962	Gene	LOC118188962	protein_coding	character(0)	NA	character(0)
NW_023312688.1	28499	28499	1	+	7	7	0	100	NW_023312688.1	27109	87988	60880	+	Gnomon	gene	NA	NA	gene-LOC118188962	GeneID:118188962
	LOC118188962	Gene	LOC118188962	protein_coding	character(0)	NA	character(0)
NW_023312688.1	28630	28630	1	+	15	15	0	100	NW_023312688.1	27109	87988	60880	+	Gnomon	gene	NA	NA	gene-LOC118188962	GeneID:118188962
	LOC118188962	Gene	LOC118188962	protein_coding	character(0)	NA	character(0)
NW_023312688.1	28762	28762	1	-	14	13	1	92.8571428571429	NW_023312688.1	27109	87988	60880	+	Gnomon	gene	NA	NA	gene-LOC118188962
	GeneID:118188962	LOC118188962	Gene	LOC118188962	protein_coding	character(0)	NA	character(0)
NW_023312688.1	28853	28853	1	-	20	19	1	95	NW_023312688.1	27109	87988	60880	+	Gnomon	gene	NA	NA	gene-LOC118188962	GeneID:118188962
	LOC118188962	Gene	LOC118188962	protein_coding	character(0)	NA	character(0)
NW_023312688.1	28866	28866	1	-	23	23	0	100	NW_023312688.1	27109	87988	60880	+	Gnomon	gene	NA	NA	gene-LOC118188962	GeneID:118188962
	LOC118188962	Gene	LOC118188962	protein_coding	character(0)	NA	character(0)
NW_023312688.1	28902	28902	1	-	19	18	1	94.7368421052632	NW_023312688.1	27109	87988	60880	+	Gnomon	gene	NA	NA	gene-LOC118188962
	GeneID:118188962	LOC118188962	Gene	LOC118188962	protein_coding	character(0)	NA	character(0)
NW_023312688.1	29285	29285	1	-	10	10	0	100	NW_023312688.1	27109	87988	60880	+	Gnomon	gene	NA	NA	gene-LOC118188962	GeneID:118188962
	LOC118188962	Gene	LOC118188962	protein_coding	character(0)	NA	character(0)
NW_023312688.1	29445	29445	1	-	5	5	0	100	NW_023312688.1	27109	87988	60880	+	Gnomon	gene	NA	NA	gene-LOC118188962	GeneID:118188962
	LOC118188962	Gene	LOC118188962	protein_coding	character(0)	NA	character(0)
NW_023312688.1	29652	29652	1	+	5	5	0	100	NW_023312688.1	27109	87988	60880	+	Gnomon	gene	NA	NA	gene-LOC118188962	GeneID:118188962
	LOC118188962	Gene	LOC118188962	protein_coding	character(0)	NA	character(0)
NW_023312688.1	29653	29653	1	-	5	5	0	100	NW_023312688.1	27109	87988	60880	+	Gnomon	gene	NA	NA	gene-LOC118188962	GeneID:118188962
	LOC118188962	Gene	LOC118188962	protein_coding	character(0)	NA	character(0)
NW_023312688.1	29662	29662	1	+	5	5	0	100	NW_023312688.1	27109	87988	60880	+	Gnomon	gene	NA	NA	gene-LOC118188962	GeneID:118188962
	LOC118188962	Gene	LOC118188962	protein_coding	character(0)	NA	character(0)
NW_023312688.1	29663	29663	1	-	5	5	0	100	NW_023312688.1	27109	87988	60880	+	Gnomon	gene	NA	NA	gene-LOC118188962	GeneID:118188962
	LOC118188962	Gene	LOC118188962	protein_coding	character(0)	NA	character(0)
NW_023312688.1	29677	29677	1	-	5	5	0	100	NW_023312688.1	27109	87988	60880	+	Gnomon	gene	NA	NA	gene-LOC118188962	GeneID:118188962
	LOC118188962	Gene	LOC118188962	protein_coding	character(0)	NA	character(0)
NW_023312688.1	29694	29694	1	+	5	5	0	100	NW_023312688.1	27109	87988	60880	+	Gnomon	gene	NA	NA	gene-LOC118188962	GeneID:118188962
	LOC118188962	Gene	LOC118188962	protein_coding	character(0)	NA	character(0)
NW_023312688.1	29699	29699	1	+	5	5	0	100	NW_023312688.1	27109	87988	60880	+	Gnomon	gene	NA	NA	gene-LOC118188962	GeneID:118188962
	LOC118188962	Gene	LOC118188962	protein_coding	character(0)	NA	character(0)
NW_023312688.1	29700	29700	1	-	7	6	1	85.7142857142857	NW_023312688.1	27109	87988	60880	+	Gnomon	gene	NA	NA	gene-LOC118188962
	GeneID:118188962	LOC118188962	Gene	LOC118188962	protein_coding	character(0)	NA	character(0)
NW_023312688.1	29714	29714	1	+	6	6	0	100	NW_023312688.1	27109	87988	60880	+	Gnomon	gene	NA	NA	gene-LOC118188962	GeneID:118188962
	LOC118188962	Gene	LOC118188962	protein_coding	character(0)	NA	character(0)
NW_023312688.1	29715	29715	1	-	7	6	1	85.7142857142857	NW_023312688.1	27109	87988	60880	+	Gnomon	gene	NA	NA	gene-LOC118188962
	GeneID:118188962	LOC118188962	Gene	LOC118188962	protein_coding	character(0)	NA	character(0)
NW_023312688.1	29722	29722	1	+	6	5	1	83.3333333333333	NW_023312688.1	27109	87988	60880	+	Gnomon	gene	NA	NA	gene-LOC118188962
	GeneID:118188962	LOC118188962	Gene	LOC118188962	protein_coding	character(0)	NA	character(0)
NW_023312688.1	29723	29723	1	-	7	6	1	85.7142857142857	NW_023312688.1	27109	87988	60880	+	Gnomon	gene	NA	NA	gene-LOC118188962
	GeneID:118188962	LOC118188962	Gene	LOC118188962	protein_coding	character(0)	NA	character(0)
NW_023312688.1	29732	29732	1	+	6	6	0	100	NW_023312688.1	27109	87988	60880	+	Gnomon	gene	NA	NA	gene-LOC118188962	GeneID:118188962
	LOC118188962	Gene	LOC118188962	protein_coding	character(0)	NA	character(0)
...
... only first 25 lines shown ...
```

Directory: RNK File: sites-in-promoters-Sd_LI_be.txt

```
seqnames	start	end	width	strand	coverage	numCs	numTs	perc_meth	region_seqnames	region_start	region_end	region_width	region_strand	region_source
	region_type	region_score	region_phase	region_ID	region_Dbxref	region_Name	region_gbkey	region_gene	region_gene_biotype	region_end_range	region_partial
	region_start_range
NW_023312688.1	193860	193860	1	+	9	6	3	66.6666666666667	NW_023312688.1	193599	194098	500	-	Gnomon	promoter	NA	NA
	gene-LOC118179278	GeneID:118179278	LOC118179278	Gene	LOC118179278	protein_coding	character(0)	NA	character(0)
NW_023312688.1	193988	193988	1	-	16	14	2	87.5	NW_023312688.1	193599	194098	500	-	Gnomon	promoter	NA	NA	gene-
LOC118179278	GeneID:118179278	LOC118179278	Gene	LOC118179278	protein_coding	character(0)	NA	character(0)
NW_023312697.1	236521	236521	1	-	10	7	3	70	NW_023312697.1	236503	237002	500	-	Gnomon	promoter	NA	NA	gene-
LOC118200926	GeneID:118200926	LOC118200926	Gene	LOC118200926	protein_coding	character(0)	NA	character(0)
NW_023312697.1	236734	236734	1	-	12	9	3	75	NW_023312697.1	236503	237002	500	-	Gnomon	promoter	NA	NA	gene-
LOC118200926	GeneID:118200926	LOC118200926	Gene	LOC118200926	protein_coding	character(0)	NA	character(0)
NW_023312697.1	236741	236741	1	-	13	12	1	92.3076923076923	NW_023312697.1	236503	237002	500	-	Gnomon	promoter	NA	NA
	gene-LOC118200926	GeneID:118200926	LOC118200926	Gene	LOC118200926	protein_coding	character(0)	NA	character(0)
NW_023312702.1	259175	259175	1	+	9	7	2	77.7777777777778	NW_023312702.1	259059	259558	500	-	Gnomon	promoter	NA	NA
	gene-LOC118182411	GeneID:118182411	LOC118182411	Gene	LOC118182411	protein_coding	character(0)	NA	character(0)
NW_023312702.1	259176	259176	1	-	5	5	0	100	NW_023312702.1	259059	259558	500	-	Gnomon	promoter	NA	NA	gene-
LOC118182411	GeneID:118182411	LOC118182411	Gene	LOC118182411	protein_coding	character(0)	NA	character(0)
NW_023312713.1	3093	3093	1	-	11	8	3	72.7272727272727	NW_023312713.1	3050	3549	500	+	Gnomon	promoter	NA	NA	gene-LOC118184707
	GeneID:118184707	LOC118184707	Gene	LOC118184707	protein_coding	c("32666", ".")	true	c(".", "3550")
NW_023312713.1	3185	3185	1	-	11	7	4	63.6363636363636	NW_023312713.1	3050	3549	500	+	Gnomon	promoter	NA	NA	gene-LOC118184707
	GeneID:118184707	LOC118184707	Gene	LOC118184707	protein_coding	c("32666", ".")	true	c(".", "3550")
NW_023312713.1	3308	3308	1	-	19	17	2	89.4736842105263	NW_023312713.1	3050	3549	500	+	Gnomon	promoter	NA	NA	gene-LOC118184707
	GeneID:118184707	LOC118184707	Gene	LOC118184707	protein_coding	c("32666", ".")	true	c(".", "3550")
NW_023312713.1	3316	3316	1	-	18	17	1	94.4444444444444	NW_023312713.1	3050	3549	500	+	Gnomon	promoter	NA	NA	gene-LOC118184707
	GeneID:118184707	LOC118184707	Gene	LOC118184707	protein_coding	c("32666", ".")	true	c(".", "3550")
NW_023312713.1	3354	3354	1	-	23	22	1	95.6521739130435	NW_023312713.1	3050	3549	500	+	Gnomon	promoter	NA	NA	gene-LOC118184707
	GeneID:118184707	LOC118184707	Gene	LOC118184707	protein_coding	c("32666", ".")	true	c(".", "3550")
NW_023312713.1	3360	3360	1	-	23	22	1	95.6521739130435	NW_023312713.1	3050	3549	500	+	Gnomon	promoter	NA	NA	gene-LOC118184707
	GeneID:118184707	LOC118184707	Gene	LOC118184707	protein_coding	c("32666", ".")	true	c(".", "3550")
NW_023312717.1	3805	3805	1	+	16	15	1	93.75	NW_023312717.1	3700	4199	500	+	Gnomon	promoter	NA	NA	gene-LOC118185112	GeneID:118185112
	LOC118185112	Gene	LOC118185112	protein_coding	character(0)	true	c(".", "4200")
NW_023312717.1	3812	3812	1	+	16	15	1	93.75	NW_023312717.1	3700	4199	500	+	Gnomon	promoter	NA	NA	gene-LOC118185112	GeneID:118185112
	LOC118185112	Gene	LOC118185112	protein_coding	character(0)	true	c(".", "4200")
NW_023312717.1	3820	3820	1	+	15	12	3	80	NW_023312717.1	3700	4199	500	+	Gnomon	promoter	NA	NA	gene-LOC118185112	GeneID:118185112
	LOC118185112	Gene	LOC118185112	protein_coding	character(0)	true	c(".", "4200")
NW_023312719.1	13794	13794	1	-	7	7	0	100	NW_023312719.1	13740	14239	500	-	Gnomon	promoter	NA	NA	gene-LOC118185876	GeneID:118185876
	LOC118185876	Gene	LOC118185876	protein_coding	character(0)	NA	character(0)
NW_023312719.1	13809	13809	1	-	6	6	0	100	NW_023312719.1	13740	14239	500	-	Gnomon	promoter	NA	NA	gene-LOC118185876	GeneID:118185876
	LOC118185876	Gene	LOC118185876	protein_coding	character(0)	NA	character(0)
NW_023312719.1	13866	13866	1	-	8	7	1	87.5	NW_023312719.1	13740	14239	500	-	Gnomon	promoter	NA	NA	gene-LOC118185876	GeneID:118185876
	LOC118185876	Gene	LOC118185876	protein_coding	character(0)	NA	character(0)
NW_023312719.1	13871	13871	1	-	8	7	1	87.5	NW_023312719.1	13740	14239	500	-	Gnomon	promoter	NA	NA	gene-LOC118185876	GeneID:118185876
	LOC118185876	Gene	LOC118185876	protein_coding	character(0)	NA	character(0)
NW_023312719.1	13880	13880	1	-	10	10	0	100	NW_023312719.1	13740	14239	500	-	Gnomon	promoter	NA	NA	gene-LOC118185876	GeneID:118185876
	LOC118185876	Gene	LOC118185876	protein_coding	character(0)	NA	character(0)
NW_023312719.1	13930	13930	1	-	8	6	2	75	NW_023312719.1	13740	14239	500	-	Gnomon	promoter	NA	NA	gene-LOC118185876	GeneID:118185876
	LOC118185876	Gene	LOC118185876	protein_coding	character(0)	NA	character(0)
NW_023312719.1	13939	13939	1	-	8	8	0	100	NW_023312719.1	13740	14239	500	-	Gnomon	promoter	NA	NA	gene-LOC118185876	GeneID:118185876
	LOC118185876	Gene	LOC118185876	protein_coding	character(0)	NA	character(0)
NW_023312719.1	13970	13970	1	-	9	8	1	88.8888888888889	NW_023312719.1	13740	14239	500	-	Gnomon	promoter	NA	NA	gene-LOC118185876
	GeneID:118185876	LOC118185876	Gene	LOC118185876	protein_coding	character(0)	NA	character(0)
...
... only first 25 lines shown ...
```

Directory: RNK File: sites-in-promoters-Sd_LI_ka.txt

| seqnames | start | end | width | strand | coverage | numCs | numTs | perc_meth | region_seqnames | region_start | region_end | region_width | region_strand | region_source | region_type | region_score | region_phase | region_ID | region_Dbxref | region_Name | region_gbkey | region_gene | region_gene_biotype | region_end_range | region_partial | region_start_range |
|---|---|---|---|---|---|---|---|---|---|---|---|---|---|---|---|---|---|---|---|---|---|---|---|---|---|---|
| NW_023312688.1 | 193860 | 193860 | 1 | + | 12 | 9 | 3 | 75 | NW_023312688.1 | 193599 | 194098 | 500 | - | Gnomon | promoter | NA | NA | gene-LOC118179278 | GeneID:118179278 | LOC118179278 | Gene | LOC118179278 | protein_coding | character(0) | NA | character(0) |
| NW_023312688.1 | 193988 | 193988 | 1 | - | 10 | 9 | 1 | 90 | NW_023312688.1 | 193599 | 194098 | 500 | - | Gnomon | promoter | NA | NA | gene-LOC118179278 | GeneID:118179278 | LOC118179278 | Gene | LOC118179278 | protein_coding | character(0) | NA | character(0) |
| NW_023312697.1 | 236521 | 236521 | 1 | - | 10 | 7 | 3 | 70 | NW_023312697.1 | 236503 | 237002 | 500 | - | Gnomon | promoter | NA | NA | gene-LOC118200926 | GeneID:118200926 | LOC118200926 | Gene | LOC118200926 | protein_coding | character(0) | NA | character(0) |
| NW_023312702.1 | 259175 | 259175 | 1 | + | 13 | 9 | 4 | 69.2307692307692 | NW_023312702.1 | 259059 | 259558 | 500 | - | Gnomon | promoter | NA | NA | gene-LOC118182411 | GeneID:118182411 | LOC118182411 | Gene | LOC118182411 | protein_coding | character(0) | NA | character(0) |
| NW_023312707.1 | 10590 | 10590 | 1 | + | 10 | 8 | 2 | 80 | NW_023312707.1 | 10274 | 10773 | 500 | + | Gnomon | promoter | NA | NA | gene-LOC118183191 | GeneID:118183191 | LOC118183191 | Gene | LOC118183191 | protein_coding | character(0) | NA | character(0) |
| NW_023312707.1 | 10591 | 10591 | 1 | - | 5 | 5 | 0 | 100 | NW_023312707.1 | 10274 | 10773 | 500 | + | Gnomon | promoter | NA | NA | gene-LOC118183191 | GeneID:118183191 | LOC118183191 | Gene | LOC118183191 | protein_coding | character(0) | NA | character(0) |
| NW_023312713.1 | 3093 | 3093 | 1 | - | 11 | 11 | 0 | 100 | NW_023312713.1 | 3050 | 3549 | 500 | + | Gnomon | promoter | NA | NA | gene-LOC118184707 | GeneID:118184707 | LOC118184707 | Gene | LOC118184707 | protein_coding | c("32666", ".") | true | c(".", "3550") |
| NW_023312713.1 | 3185 | 3185 | 1 | - | 6 | 5 | 1 | 83.3333333333333 | NW_023312713.1 | 3050 | 3549 | 500 | + | Gnomon | promoter | NA | NA | gene-LOC118184707 | GeneID:118184707 | LOC118184707 | Gene | LOC118184707 | protein_coding | c("32666", ".") | true | c(".", "3550") |
| NW_023312713.1 | 3308 | 3308 | 1 | - | 9 | 8 | 1 | 88.8888888888889 | NW_023312713.1 | 3050 | 3549 | 500 | + | Gnomon | promoter | NA | NA | gene-LOC118184707 | GeneID:118184707 | LOC118184707 | Gene | LOC118184707 | protein_coding | c("32666", ".") | true | c(".", "3550") |
| NW_023312713.1 | 3316 | 3316 | 1 | - | 11 | 9 | 2 | 81.8181818181818 | NW_023312713.1 | 3050 | 3549 | 500 | + | Gnomon | promoter | NA | NA | gene-LOC118184707 | GeneID:118184707 | LOC118184707 | Gene | LOC118184707 | protein_coding | c("32666", ".") | true | c(".", "3550") |
| NW_023312713.1 | 3354 | 3354 | 1 | - | 12 | 12 | 0 | 100 | NW_023312713.1 | 3050 | 3549 | 500 | + | Gnomon | promoter | NA | NA | gene-LOC118184707 | GeneID:118184707 | LOC118184707 | Gene | LOC118184707 | protein_coding | c("32666", ".") | true | c(".", "3550") |
| NW_023312713.1 | 3360 | 3360 | 1 | - | 13 | 13 | 0 | 100 | NW_023312713.1 | 3050 | 3549 | 500 | + | Gnomon | promoter | NA | NA | gene-LOC118184707 | GeneID:118184707 | LOC118184707 | Gene | LOC118184707 | protein_coding | c("32666", ".") | true | c(".", "3550") |
| NW_023312716.1 | 279920 | 279920 | 1 | + | 7 | 5 | 2 | 71.4285714285714 | NW_023312716.1 | 279579 | 280078 | 500 | + | Gnomon | promoter | NA | NA | gene-LOC118185035 | GeneID:118185035 | LOC118185035 | Gene | LOC118185035 | protein_coding | character(0) | NA | character(0) |
| NW_023312716.1 | 279929 | 279929 | 1 | + | 7 | 6 | 1 | 85.7142857142857 | NW_023312716.1 | 279579 | 280078 | 500 | + | Gnomon | promoter | NA | NA | gene-LOC118185035 | GeneID:118185035 | LOC118185035 | Gene | LOC118185035 | protein_coding | character(0) | NA | character(0) |
| NW_023312716.1 | 279965 | 279965 | 1 | + | 6 | 5 | 1 | 83.3333333333333 | NW_023312716.1 | 279579 | 280078 | 500 | + | Gnomon | promoter | NA | NA | gene-LOC118185035 | GeneID:118185035 | LOC118185035 | Gene | LOC118185035 | protein_coding | character(0) | NA | character(0) |
| NW_023312716.1 | 280023 | 280023 | 1 | + | 7 | 6 | 1 | 85.7142857142857 | NW_023312716.1 | 279579 | 280078 | 500 | + | Gnomon | promoter | NA | NA | gene-LOC118185035 | GeneID:118185035 | LOC118185035 | Gene | LOC118185035 | protein_coding | character(0) | NA | character(0) |
| NW_023312716.1 | 280044 | 280044 | 1 | + | 5 | 5 | 0 | 100 | NW_023312716.1 | 279579 | 280078 | 500 | + | Gnomon | promoter | NA | NA | gene-LOC118185035 | GeneID:118185035 | LOC118185035 | Gene | LOC118185035 | protein_coding | character(0) | NA | character(0) |
| NW_023312717.1 | 3805 | 3805 | 1 | + | 26 | 26 | 0 | 100 | NW_023312717.1 | 3700 | 4199 | 500 | + | Gnomon | promoter | NA | NA | gene-LOC118185112 | GeneID:118185112 | LOC118185112 | Gene | LOC118185112 | protein_coding | character(0) | true | c(".", "4200") |
| NW_023312717.1 | 3812 | 3812 | 1 | + | 26 | 24 | 2 | 92.3076923076923 | NW_023312717.1 | 3700 | 4199 | 500 | + | Gnomon | promoter | NA | NA | gene-LOC118185112 | GeneID:118185112 | LOC118185112 | Gene | LOC118185112 | protein_coding | character(0) | true | c(".", "4200") |
| NW_023312717.1 | 3820 | 3820 | 1 | + | 27 | 22 | 5 | 81.4814814814815 | NW_023312717.1 | 3700 | 4199 | 500 | + | Gnomon | promoter | NA | NA | gene-LOC118185112 | GeneID:118185112 | LOC118185112 | Gene | LOC118185112 | protein_coding | character(0) | true | c(".", "4200") |
| NW_023312719.1 | 13866 | 13866 | 1 | - | 7 | 7 | 0 | 100 | NW_023312719.1 | 13740 | 14239 | 500 | - | Gnomon | promoter | NA | NA | gene-LOC118185876 | GeneID:118185876 | LOC118185876 | Gene | LOC118185876 | protein_coding | character(0) | NA | character(0) |
| NW_023312719.1 | 13871 | 13871 | 1 | - | 6 | 5 | 1 | 83.3333333333333 | NW_023312719.1 | 13740 | 14239 | 500 | - | Gnomon | promoter | NA | NA | gene-LOC118185876 | GeneID:118185876 | LOC118185876 | Gene | LOC118185876 | protein_coding | character(0) | NA | character(0) |
| NW_023312719.1 | 13880 | 13880 | 1 | - | 6 | 6 | 0 | 100 | NW_023312719.1 | 13740 | 14239 | 500 | - | Gnomon | promoter | NA | NA | gene-LOC118185876 | GeneID:118185876 | LOC118185876 | Gene | LOC118185876 | protein_coding | character(0) | NA | character(0) |
| NW_023312719.1 | 13930 | 13930 | 1 | - | 5 | 5 | 0 | 100 | NW_023312719.1 | 13740 | 14239 | 500 | - | Gnomon | promoter | NA | NA | gene-LOC118185876 | GeneID:118185876 | LOC118185876 | Gene | LOC118185876 | protein_coding | character(0) | NA | character(0) |

...

... only first 25 lines shown ...

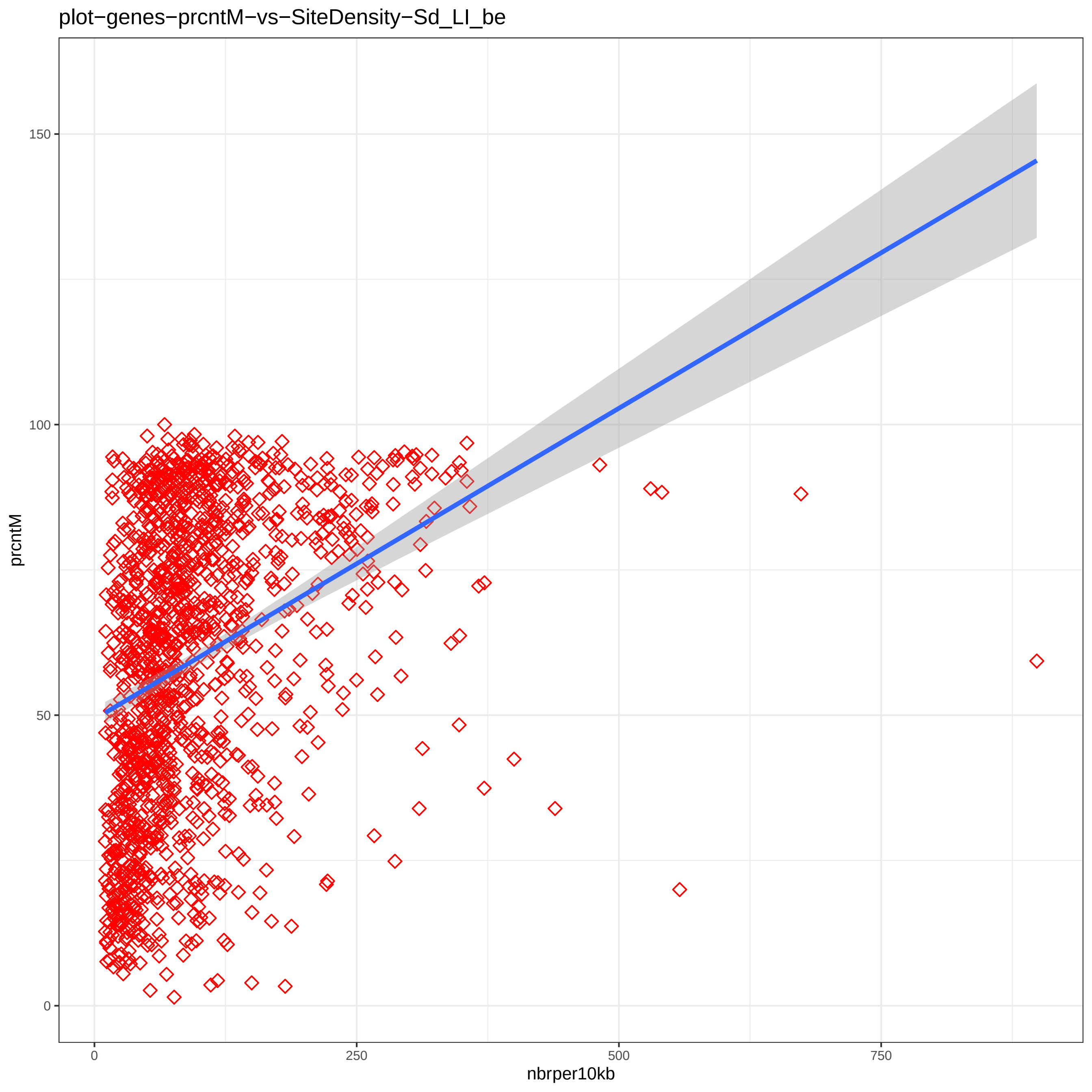
plot−genes−prcntM−vs−SiteDensity−Sd_LI_be
prcntM
nbrper10kb
0
50
100
150
0
250
500
750

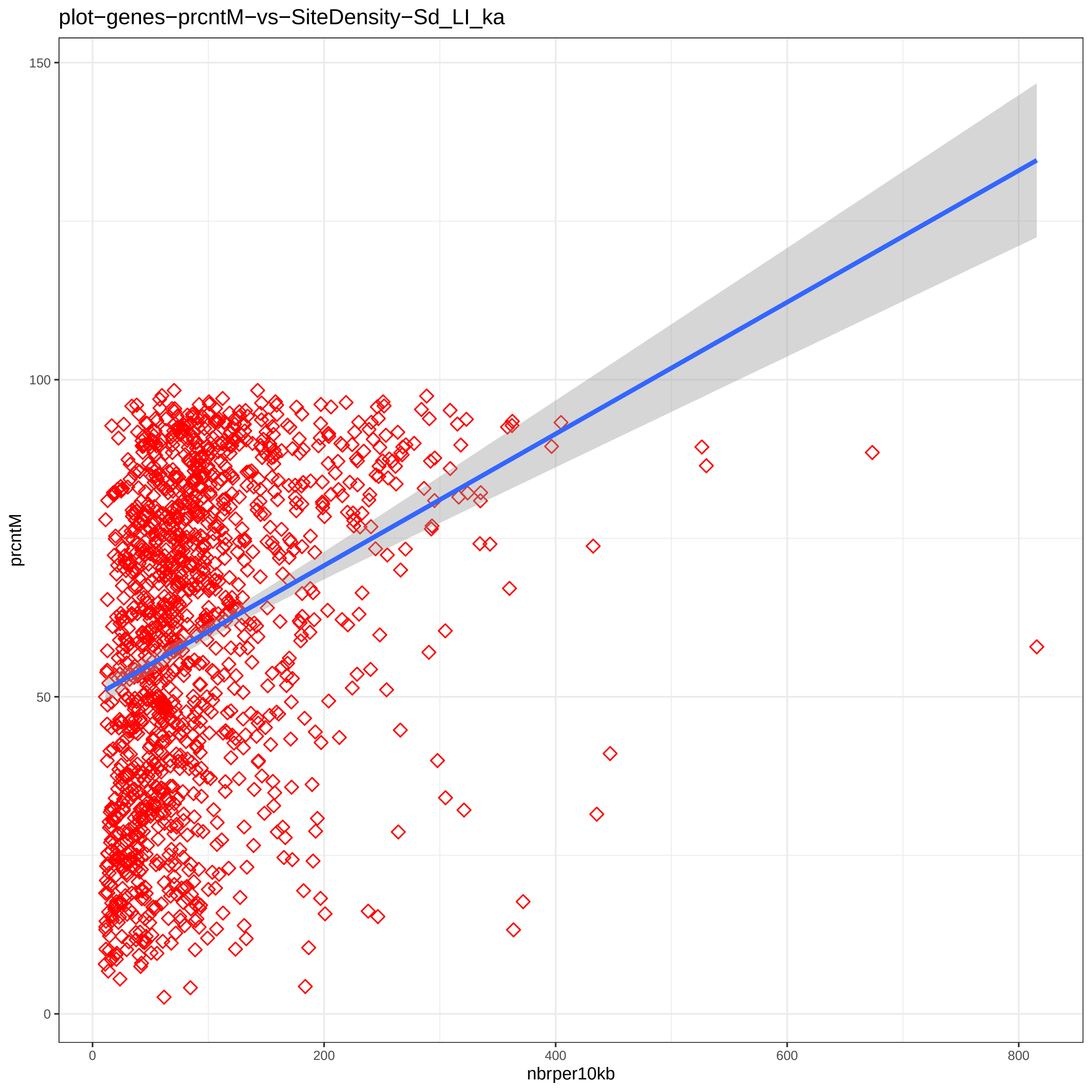

plot−genes−prcntM−vs−SiteDensity−Sd_LI_ka
prcntM
nbrper10kb
150
100
50
0
0
200
400
600
800

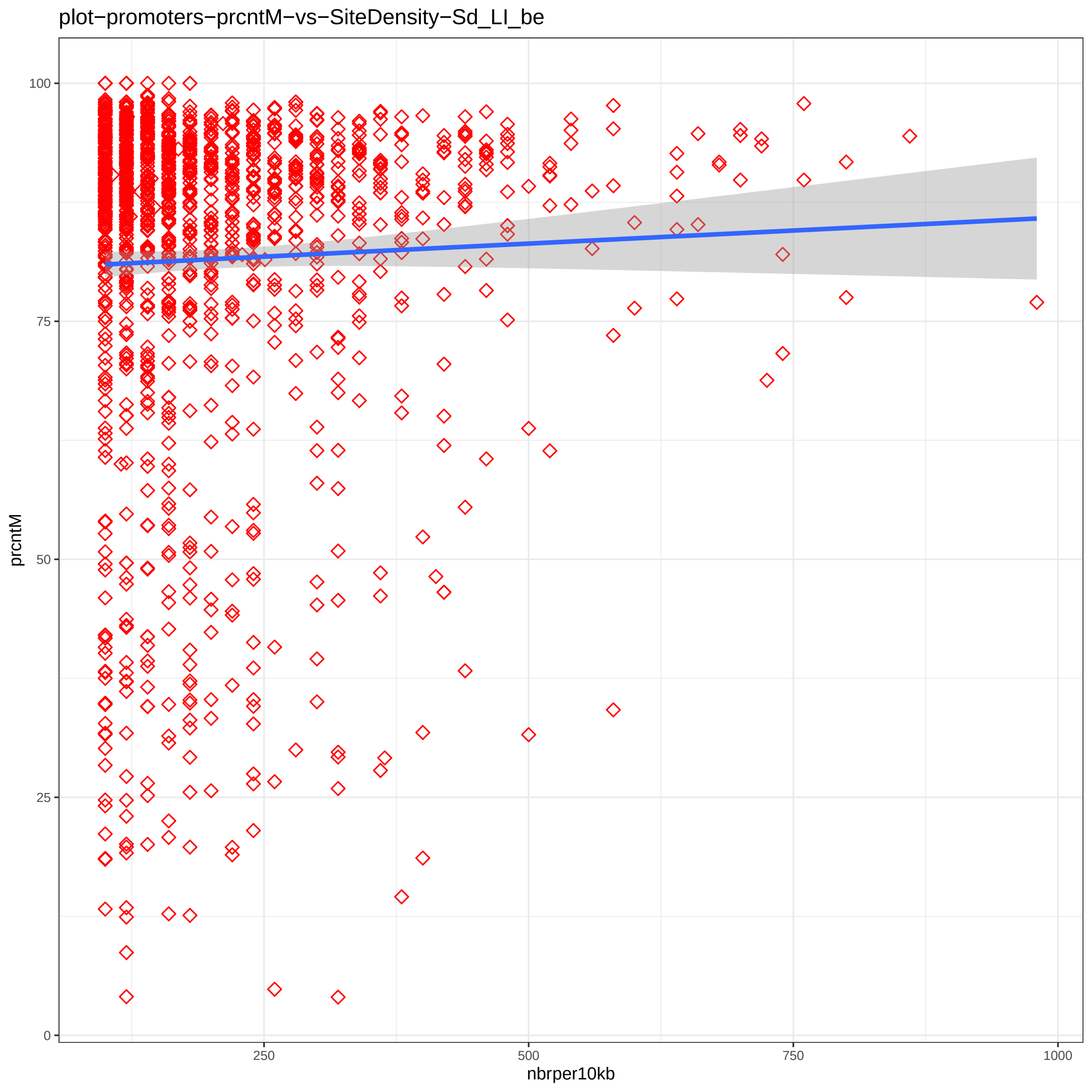
plot−promoters−prcntM−vs−SiteDensity−Sd_LI_be
prcntM
nbrper10kb
100
75
50
25
0
250
500
750
1000

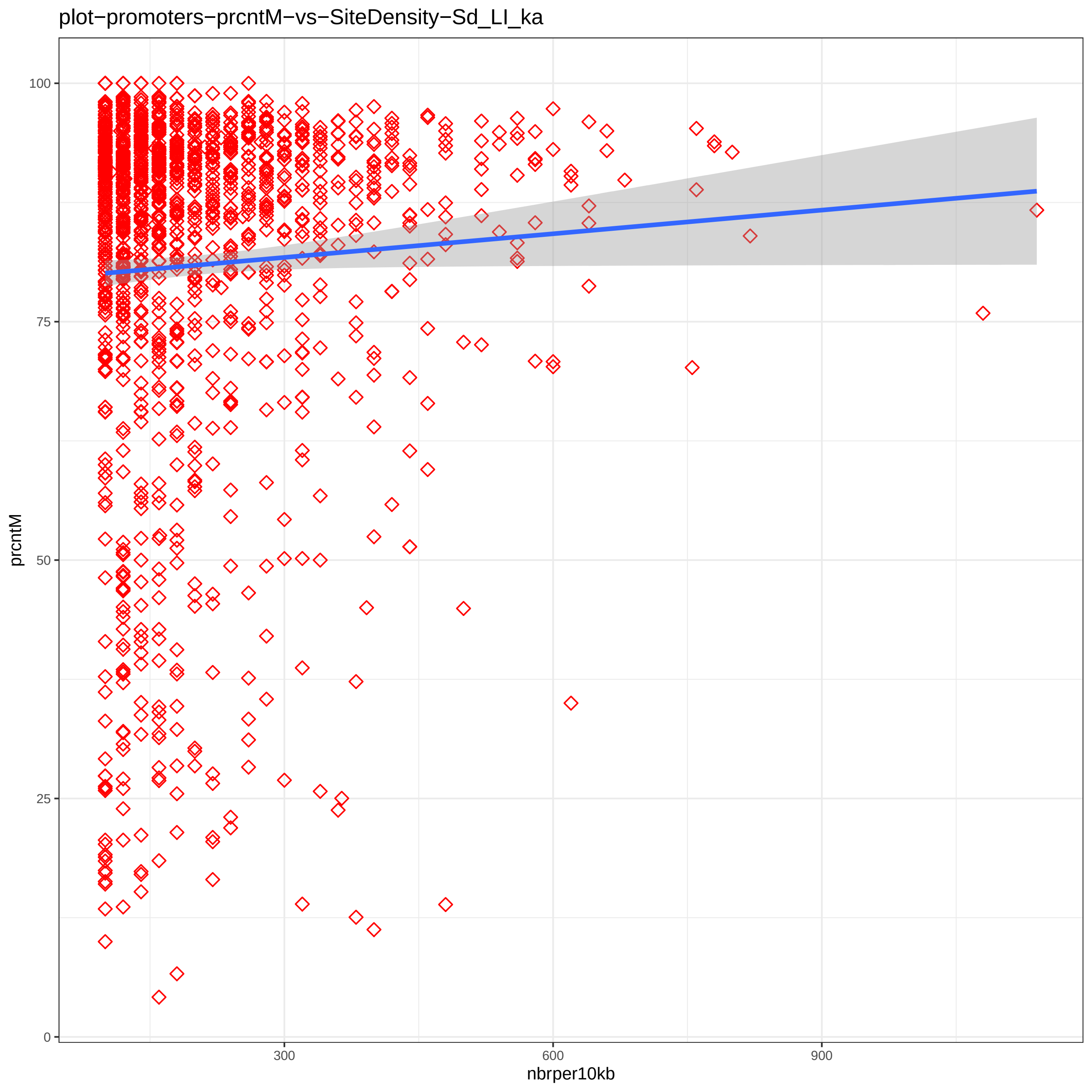
plot−promoters−prcntM−vs−SiteDensity−Sd_LI_ka
prcntM
100
75
50
25
0
300
600
900
nbrper10kb

Directory: MRPR File: 0READMEmrpr

MRPR - Methylation-rich and -poor regions.

Input: studymk, genome_ann (a list of GRanges objects providing annotated region labels and bounds;
output of BWASPR::get_genome_annotation())
nbrxtrms (specifies the number of richest and poorest regions to be displayed; default: 100)

Output: files dst-*.txt 1ds-*.pdf 5ds-*.pdf mdr-*.bed rmp-* gwp-* gwr-*

Notes: The output is generated by BWASPR functions det_mrpr() and map_mrpr().
Methylation-rich and -poor regions are determined based on the spacing between neighboring CpGhsm sites. If sites occur at positions a, b, c, d, e, and f (and not in between), then b-a, c-b, ... are 1-distances, and f-a is a 5-distance.
The file dst-*.txt shows the empirical distribution of d-distances for the sample.
Methylation-poor regions are determined as long 1-distances (in the top <nbrxtrms>; merged if adjacent 1-distances are both in the top <nbrxtrms>). Methylation-rich regions are determined as short 5-distances (in the low <nbrxtrms>; merged if adjacent 5-distances are both in the low <nbrxtrms>). Parts of the distribution are plotted in the 1ds-*.pdf and 5ds-*.pdf files.

The methylation-rich and -poor regions are listed in files dst-*.txt and mdr-*.bed, the latter in BED format for potential display in genome browsers.

The overlap of methylation regions with genome features is summarized in files rmp-*.txt.
Files gwr-*.txt show genes overlapping with methylation-rich regions, ordered by site density in the methylation-rich region.
Files gwp-*.txt show genes overlapping with methylation-poor regions, ordered by site density in the methylation-poor region.

Directory: MRPR File: dst-Sd_LI_be.txt

Analysis of 1-distances for sample "Sd_LI_be" (6335325 sites):

Total number of 1-distances: 6321826
Median: 21.0 Mean: 235.2 Std: 4049.6
Quantiles:

| 0% | 5% | 10% | 15% | 20% | 25% | 30% | 35% | 40% | 45% | 50% | 55% | 60% | 65% | 70% | 75% | 80% | 85% | 90% | 95% | 100% |
|---|---|---|---|---|---|---|---|---|---|---|---|---|---|---|---|---|---|---|---|---|
| 1 | 1 | 1 | 1 | 2 | 4 | 7 | 9 | 12 | 16 | 21 | 27 | 34 | 44 | 58 | 76 | 105 | 149 | 228 | 405 | 792419 |

Low density region distance cutoff (lowest 100): 310678
High density region distance cutoff (highest 100): 1

Ordered lists of methylation-poor regions:

Top 100 (out of 100) methylation-poor regions in "Sd_LI_be":
(Sdnsty = sites per 1kb)

| | Rtype | Sample | SeqID | From | To | Rlgth | NbrSites | Sdnsty |
|---|---|---|---|---|---|---|---|---|
| 1 | Poor | be | NW_023315573.1 | 595878 | 1388297 | 792420 | 2 | 0.00252 |
| 2 | Poor | be | NW_023314007.1 | 83019 | 763857 | 680839 | 2 | 0.00294 |
| 3 | Poor | be | NW_023313043.1 | 21721 | 558625 | 536905 | 2 | 0.00373 |
| 4 | Poor | be | NW_023321981.1 | 8499 | 528200 | 519702 | 2 | 0.00385 |
| 5 | Poor | be | NW_023314140.1 | 108837 | 611126 | 502290 | 2 | 0.00398 |
| 6 | Poor | be | NW_023316154.1 | 170209 | 671594 | 501386 | 2 | 0.00399 |

...
... only first 25 lines shown ...

```
Directory: MRPR File: dst-Sd_LI_ka.txt

Analysis of  1-distances for sample "Sd_LI_ka" (6222608 sites):

  Total number of  1-distances:     6208967
  Median:    22.0 Mean:   245.6 Std:  4056.4
  Quantiles:
    0%     5%    10%    15%    20%    25%    30%    35%    40%    45%    50%    55%    60%    65%    70%    75%    80%    85%    90%    95%   100%
     1      1      1      1      2      5      7     10     13     17     22     28     35     46     60     79    108    154    235    416 619206

  Low  density region distance cutoff (lowest  100):  312616
  High density region distance cutoff (highest 100):       1

Ordered lists of methylation-poor regions:

 Top 100 (out of   100) methylation-poor regions in "Sd_LI_ka":
 (Sdnsty = sites per 1kb)

    Rtype Sample          SeqID   From      To  Rlgth NbrSites  Sdnsty
1    Poor     ka NW_023315573.1 682240 1301446 619207        2 0.00323
2    Poor     ka NW_023313325.1 174338  751511 577174        2 0.00347
3    Poor     ka NW_023318051.1 140843  708761 567919        2 0.00352
4    Poor     ka NW_023314643.1 217844  748825 530982        2 0.00377
5    Poor     ka NW_023316154.1 170365  671594 501230        2 0.00399
6    Poor     ka NW_023313196.1 166973  637031 470059        2 0.00425
...
... only first 25 lines shown ...
```

Directory: MRPR File: gwp-Sd_LI_be.txt

```
seqnames	start	end	width	strand	source	type	score	phase	ID	Dbxref	Name	gbkey	gene	gene_biotype	end_range	partial	start_range	seqnames	start
	end	width	strand	NbrSites	Sdnsty
NW_023315573.1	444487	597831	153345	-	Gnomon	gene	NA	NA	gene-LOC118186187	GeneID:118186187	LOC118186187	Gene	LOC118186187	protein_coding
	character(0)	NA	character(0)	NW_023315573.1	595878	1388297	792420	*	2	0.00252391408596451
NW_023315573.1	685037	685109	73	+	tRNAscan-SE	gene	NA	NA	gene-Trnat-cgu-61	GeneID:118186190	Trnat-cgu	Gene	Trnat-cgu	tRNA	character(0)	NA
	character(0)	NW_023315573.1	595878	1388297	792420	*	2	0.00252391408596451
NW_023314007.1	487137	873595	386459	-	Gnomon	gene	NA	NA	gene-LOC118181919	GeneID:118181919	LOC118181919	Gene	LOC118181919	protein_coding
	character(0)	NA	character(0)	NW_023314007.1	83019	763857	680839	*	2	0.00293755204975038
NW_023313043.1	161239	600424	439186	+	Gnomon	gene	NA	NA	gene-LOC118188938	GeneID:118188938	LOC118188938	Gene	LOC118188938	protein_coding
	character(0)	NA	character(0)	NW_023313043.1	21721	558625	536905	*	2	0.00372505378046396
NW_023321981.1	11948	33913	21966	-	Gnomon	gene	NA	NA	gene-LOC118199930	GeneID:118199930	LOC118199930	Gene	LOC118199930	protein_coding	character(0)
	NA	character(0)	NW_023321981.1	8499	528200	519702	*	2	0.0038483592520329
NW_023321981.1	50053	75104	25052	-	Gnomon	gene	NA	NA	gene-LOC118199932	GeneID:118199932	LOC118199932	Gene	LOC118199932	protein_coding	character(0)
	NA	character(0)	NW_023321981.1	8499	528200	519702	*	2	0.0038483592520329
NW_023321981.1	82197	100173	17977	-	Gnomon	gene	NA	NA	gene-LOC118199931	GeneID:118199931	LOC118199931	Gene	LOC118199931	protein_coding
	character(0)	NA	character(0)	NW_023321981.1	8499	528200	519702	*	2	0.0038483592520329
NW_023321981.1	151428	201209	49782	+	Gnomon	gene	NA	NA	gene-LOC118199935	GeneID:118199935	LOC118199935	Gene	LOC118199935	protein_coding
	character(0)	NA	character(0)	NW_023321981.1	8499	528200	519702	*	2	0.0038483592520329
NW_023321981.1	203733	362228	158496	-	Gnomon	gene	NA	NA	gene-LOC118199928	GeneID:118199928	LOC118199928	Gene	LOC118199928	protein_coding
	character(0)	NA	character(0)	NW_023321981.1	8499	528200	519702	*	2	0.0038483592520329
NW_023321981.1	371388	418843	47456	+	Gnomon	gene	NA	NA	gene-LOC118199933	GeneID:118199933	LOC118199933	Gene	LOC118199933	lncRNA
	character(0)	NA	character(0)	NW_023321981.1	8499	528200	519702	*	2	0.0038483592520329
NW_023321981.1	454411	535254	80844	+	Gnomon	gene	NA	NA	gene-LOC118199929	GeneID:118199929	LOC118199929	Gene	LOC118199929	protein_coding
	character(0)	NA	character(0)	NW_023321981.1	8499	528200	519702	*	2	0.0038483592520329
NW_023314140.1	143423	708295	564873	+	Gnomon	gene	NA	NA	gene-LOC118182361	GeneID:118182361	LOC118182361	Gene	LOC118182361	protein_coding
	c("708295", ".")	true	c(".", "143423")	NW_023314140.1	108837	611126	502290	*	2	0.00398176352306437
NW_023316154.1	8179	859885	851707	-	Gnomon	gene	NA	NA	gene-LOC118187827	GeneID:118187827	LOC118187827	Gene	LOC118187827	protein_coding
	character(0)	true	c(".", "8179")	NW_023316154.1	170209	671594	501386	*	2	0.00398894265097151
NW_023316154.1	384852	384924	73	-	tRNAscan-SE	gene	NA	NA	gene-Trnat-agu-64	GeneID:118187830	Trnat-agu	Gene	Trnat-agu	tRNA	character(0)	NA
	character(0)	NW_023316154.1	170209	671594	501386	*	2	0.00398894265097151
NW_023312958.1	306023	657159	351137	-	Gnomon	gene	NA	NA	gene-LOC118186428	GeneID:118186428	LOC118186428	Gene	LOC118186428	protein_coding
	character(0)	NA	character(0)	NW_023312958.1	620974	1120610	499637	*	2	0.00400290610983574
NW_023312958.1	716464	717372	909	-	Gnomon	gene	NA	NA	gene-LOC118186435	GeneID:118186435	LOC118186435	Gene	LOC118186435	protein_coding
	character(0)	NA	character(0)	NW_023312958.1	620974	1120610	499637	*	2	0.00400290610983574
NW_023312958.1	717466	717906	441	-	Gnomon	gene	NA	NA	gene-LOC118186440	GeneID:118186440	LOC118186440	Gene	LOC118186440	protein_coding
	character(0)	NA	character(0)	NW_023312958.1	620974	1120610	499637	*	2	0.00400290610983574
NW_023312958.1	847009	853748	6740	-	Gnomon	gene	NA	NA	gene-LOC118186448	GeneID:118186448	LOC118186448	Gene	LOC118186448	lncRNA
	character(0)	NA	character(0)	NW_023312958.1	620974	1120610	499637	*	2	0.00400290610983574
NW_023316588.1	431551	530572	99022	-	Gnomon	gene	NA	NA	gene-LOC118188874	GeneID:118188874	LOC118188874	Gene	LOC118188874	protein_coding
	character(0)	NA	character(0)	NW_023316588.1	16337	515338	499002	*	2	0.004007999967936
NW_023313115.1	186997	626876	439880	+	Gnomon	gene	NA	NA	gene-LOC118191049	GeneID:118191049	LOC118191049	Gene	LOC118191049	protein_coding
	character(0)	NA	character(0)	NW_023313115.1	252045	750380	498336	*	2	0.00401335645026649
NW_023313115.1	279358	279430	73	-	tRNAscan-SE	gene	NA	NA	gene-Trnat-cgu-16	GeneID:118191055	Trnat-cgu	Gene	Trnat-cgu	tRNA	character(0)	NA
	character(0)	NW_023313115.1	252045	750380	498336	*	2	0.00401335645026649
NW_023313671.1	377148	452203	75056	+	Gnomon	gene	NA	NA	gene-LOC118180909	GeneID:118180909	LOC118180909	Gene	LOC118180909	protein_coding
	character(0)	NA	character(0)	NW_023313671.1	365749	860050	494302	*	2	0.00404610946344542
NW_023313671.1	416849	416921	73	-	tRNAscan-SE	gene	NA	NA	gene-Trnam-cau-18	GeneID:118180917	Trnam-cau	Gene	Trnam-cau	tRNA	character(0)	NA
	character(0)	NW_023313671.1	365749	860050	494302	*	2	0.00404610946344542
NW_023313671.1	456285	504712	48428	-	Gnomon	gene	NA	NA	gene-LOC118180910	GeneID:118180910	LOC118180910	Gene	LOC118180910	protein_coding
	character(0)	NA	character(0)	NW_023313671.1	365749	860050	494302	*	2	0.00404610946344542
...
... only first 25 lines shown ...
```

Directory: MRPR File: gwp-Sd_LI_ka.txt

| seqnames | start | end | width | strand | source | type | score | phase | ID | Dbxref | Name | gbkey | gene | gene_biotype | end_range | partial | start_range | seqnames | start | end | width | strand | NbrSites | Sdnsty |
|---|---|---|---|---|---|---|---|---|---|---|---|---|---|---|---|---|---|---|---|---|---|---|---|---|
| NW_023315573.1 | 685037 | 685109 | 73 | + | tRNAscan-SE | gene | NA | NA | gene-Trnat-cgu-61 | GeneID:118186190 | Trnat-cgu | Gene | Trnat-cgu | tRNA | character(0) | NA | character(0) | NW_023315573.1 | 682240 | 1301446 | 619207 | * | 2 | 0.00322993764605374 |
| NW_023313325.1 | 179715 | 192929 | 13215 | + | Gnomon | gene | NA | NA | gene-LOC118197645 | GeneID:118197645 | LOC118197645 | Gene | LOC118197645 | lncRNA | character(0) | NA | character(0) | NW_023313325.1 | 174338 | 751511 | 577174 | * | 2 | 0.00346515955327163 |
| NW_023313325.1 | 545579 | 545651 | 73 | + | tRNAscan-SE | gene | NA | NA | gene-Trnat-ugu-23 | GeneID:118197651 | Trnat-ugu | Gene | Trnat-ugu | tRNA | character(0) | NA | character(0) | NW_023313325.1 | 174338 | 751511 | 577174 | * | 2 | 0.00346515955327163 |
| NW_023318051.1 | 335946 | 336019 | 74 | + | tRNAscan-SE | gene | NA | NA | gene-Trnat-ggu-101 | GeneID:118192367 | Trnat-ggu | Gene | Trnat-ggu | tRNA | character(0) | NA | character(0) | NW_023318051.1 | 140843 | 708761 | 567919 | * | 2 | 0.00352162896469391 |
| NW_023318051.1 | 363247 | 749447 | 386201 | + | Gnomon | gene | NA | NA | gene-LOC118192366 | GeneID:118192366 | LOC118192366 | Gene | LOC118192366 | protein_coding | character(0) | NA | character(0) | NW_023318051.1 | 140843 | 708761 | 567919 | * | 2 | 0.00352162896469391 |
| NW_023318051.1 | 377786 | 377858 | 73 | - | tRNAscan-SE | gene | NA | NA | gene-Trnas-cga-98 | GeneID:118192368 | Trnas-cga | Gene | Trnas-cga | tRNA | character(0) | NA | character(0) | NW_023318051.1 | 140843 | 708761 | 567919 | * | 2 | 0.00352162896469391 |
| NW_023318051.1 | 435628 | 435700 | 73 | - | tRNAscan-SE | gene | NA | NA | gene-Trnat-cgu-107 | GeneID:118192369 | Trnat-cgu | Gene | Trnat-cgu | tRNA | character(0) | NA | character(0) | NW_023318051.1 | 140843 | 708761 | 567919 | * | 2 | 0.00352162896469391 |
| NW_023314643.1 | 202735 | 302831 | 100097 | + | Gnomon | gene | NA | NA | gene-LOC118183664 | GeneID:118183664 | LOC118183664 | Gene | LOC118183664 | protein_coding | character(0) | NA | character(0) | NW_023314643.1 | 217844 | 748825 | 530982 | * | 2 | 0.00376660602430968 |
| NW_023314643.1 | 563357 | 739928 | 176572 | + | Gnomon | gene | NA | NA | gene-LOC118183671 | GeneID:118183671 | LOC118183671 | Gene | LOC118183671 | protein_coding | character(0) | NA | character(0) | NW_023314643.1 | 217844 | 748825 | 530982 | * | 2 | 0.00376660602430968 |
| NW_023316154.1 | 8179 | 859885 | 851707 | - | Gnomon | gene | NA | NA | gene-LOC118187827 | GeneID:118187827 | LOC118187827 | Gene | LOC118187827 | protein_coding | character(0) | true | c(".", "8179") | NW_023316154.1 | 170365 | 671594 | 501230 | * | 2 | 0.00399018414699838 |
| NW_023316154.1 | 384852 | 384924 | 73 | - | tRNAscan-SE | gene | NA | NA | gene-Trnat-agu-64 | GeneID:118187830 | Trnat-agu | Gene | Trnat-agu | tRNA | character(0) | NA | character(0) | NW_023316154.1 | 170365 | 671594 | 501230 | * | 2 | 0.00399018414699838 |
| NW_023313196.1 | 242649 | 242720 | 72 | - | tRNAscan-SE | gene | NA | NA | gene-Trnaw-cca-2 | GeneID:118193366 | Trnaw-cca | Gene | Trnaw-cca | tRNA | character(0) | NA | character(0) | NW_023313196.1 | 166973 | 637031 | 470059 | * | 2 | 0.00425478503762294 |
| NW_023313196.1 | 290306 | 291118 | 813 | + | Gnomon | gene | NA | NA | gene-LOC118193335 | GeneID:118193335 | LOC118193335 | Gene | LOC118193335 | protein_coding | character(0) | NA | character(0) | NW_023313196.1 | 166973 | 637031 | 470059 | * | 2 | 0.00425478503762294 |
| NW_023313196.1 | 518187 | 518258 | 72 | + | tRNAscan-SE | gene | NA | NA | gene-Trnat-ugu-18 | GeneID:118193360 | Trnat-ugu | Gene | Trnat-ugu | tRNA | character(0) | NA | character(0) | NW_023313196.1 | 166973 | 637031 | 470059 | * | 2 | 0.00425478503762294 |
| NW_023313196.1 | 603005 | 604790 | 1786 | + | Gnomon | gene | NA | NA | gene-LOC118193347 | GeneID:118193347 | LOC118193347 | Gene | LOC118193347 | protein_coding | character(0) | NA | character(0) | NW_023313196.1 | 166973 | 637031 | 470059 | * | 2 | 0.00425478503762294 |
| NW_023313524.1 | 965 | 147607 | 146643 | - | Gnomon | gene | NA | NA | gene-LOC118204040 | GeneID:118204040 | LOC118204040 | Gene | LOC118204040 | protein_coding | character(0) | NA | character(0) | NW_023313524.1 | 146656 | 615794 | 469139 | * | 2 | 0.00426312883814818 |
| NW_023313524.1 | 126940 | 599845 | 472906 | + | Gnomon | gene | NA | NA | gene-LOC118203999 | GeneID:118203999 | LOC118203999 | Gene | LOC118203999 | protein_coding | character(0) | NA | character(0) | NW_023313524.1 | 146656 | 615794 | 469139 | * | 2 | 0.00426312883814818 |
| NW_023313524.1 | 614727 | 650207 | 35481 | + | Gnomon | gene | NA | NA | gene-LOC118204082 | GeneID:118204082 | LOC118204082 | Gene | LOC118204082 | protein_coding | character(0) | NA | character(0) | NW_023313524.1 | 146656 | 615794 | 469139 | * | 2 | 0.00426312883814818 |
| NW_023316098.1 | 728754 | 734237 | 5484 | + | Gnomon | gene | NA | NA | gene-LOC118187690 | GeneID:118187690 | LOC118187690 | Gene | LOC118187690 | protein_coding | character(0) | NA | character(0) | NW_023316098.1 | 377896 | 831866 | 453971 | * | 2 | 0.0044055677565307 |
| NW_023316098.1 | 761011 | 761436 | 426 | - | Gnomon | gene | NA | NA | gene-LOC118187692 | GeneID:118187692 | LOC118187692 | Gene | LOC118187692 | protein_coding | character(0) | NA | character(0) | NW_023316098.1 | 377896 | 831866 | 453971 | * | 2 | 0.0044055677565307 |
| NW_023316098.1 | 789380 | 789453 | 74 | - | tRNAscan-SE | gene | NA | NA | gene-Trnai-gau-68 | GeneID:118187694 | Trnai-gau | Gene | Trnai-gau | tRNA | character(0) | NA | character(0) | NW_023316098.1 | 377896 | 831866 | 453971 | * | 2 | 0.0044055677565307 |
| NW_023313561.1 | 246432 | 506093 | 259662 | - | Gnomon | gene | NA | NA | gene-LOC118205052 | GeneID:118205052 | LOC118205052 | Gene | LOC118205052 | protein_coding | character(0) | NA | character(0) | NW_023313561.1 | 505021 | 956724 | 451704 | * | 2 | 0.00442767830260525 |
| NW_023313561.1 | 527815 | 554656 | 26842 | + | Gnomon | gene | NA | NA | gene-LOC118205071 | GeneID:118205071 | LOC118205071 | Gene | LOC118205071 | protein_coding | character(0) | NA | character(0) | NW_023313561.1 | 505021 | 956724 | 451704 | * | 2 | 0.00442767830260525 |
| NW_023313561.1 | 557883 | 676173 | 118291 | - | Gnomon | gene | NA | NA | gene-LOC118205077 | GeneID:118205077 | LOC118205077 | Gene | LOC118205077 | protein_coding | character(0) | NA | character(0) | NW_023313561.1 | 505021 | 956724 | 451704 | * | 2 | 0.00442767830260525 |
| NW_023313561.1 | 712971 | 828626 | 115656 | - | Gnomon | gene | NA | NA | gene-LOC118205107 | GeneID:118205107 | LOC118205107 | Gene | LOC118205107 | protein_coding | character(0) | NA | character(0) | NW_023313561.1 | 505021 | 956724 | 451704 | * | 2 | 0.00442767830260525 |
| NW_023313561.1 | 845829 | 845901 | 73 | + | tRNAscan-SE | gene | NA | NA | gene-Trnas-cga-25 | GeneID:118205128 | Trnas-cga | Gene | Trnas-cga | tRNA | character(0) | NA | character(0) | NW_023313561.1 | 505021 | 956724 | 451704 | * | 2 | 0.00442767830260525 |
| NW_023313561.1 | 883235 | 884278 | 1044 | - | Gnomon | gene | NA | NA | gene-LOC118205117 | GeneID:118205117 | LOC118205117 | Gene | LOC118205117 | protein_coding | character(0) | NA | character(0) | NW_023313561.1 | 505021 | 956724 | 451704 | * | 2 | 0.00442767830260525 |
| NW_023313068.1 | 63 | 698055 | 697993 | - | Gnomon | gene | NA | NA | gene-LOC118189594 | GeneID:118189594 | LOC118189594 | Gene | LOC118189594 | protein_coding | character(0) | NA | character(0) | NW_023313068.1 | 336878 | 782435 | 445558 | * | 2 | 0.00448875342828543 |
| NW_023316428.1 | 3334 | 395850 | 392517 | - | Gnomon | gene | NA | NA | gene-LOC118188513 | GeneID:118188513 | LOC118188513 | Gene | LOC118188513 | protein_coding | character(0) | true | c(".", "3334") | NW_023316428.1 | 371528 | 811602 | 440075 | * | 2 | 0.00454467988411066 |
| NW_023316428.1 | 520332 | 520403 | 72 | + | tRNAscan-SE | gene | NA | NA | gene-Trnat-agu-70 | GeneID:118188515 | Trnat-agu | Gene | Trnat-agu | tRNA | character(0) | NA | | | | | | | | |

character(0) NW_023316428.1 371528 811602 440075 * 2 0.00454467988411066
NW_023316428.1 584950 730708 145759 - Gnomon gene NA NA gene-LOC118188514 GeneID:118188514 LOC118188514 Gene LOC118188514 protein_coding
character(0) NA character(0) NW_023316428.1 371528 811602 440075 * 2 0.00454467988411066
NW_023316428.1 752523 753128 606 + Gnomon gene NA NA gene-LOC118188512 GeneID:118188512 LOC118188512 Gene LOC118188512 protein_coding
character(0) NA character(0) NW_023316428.1 371528 811602 440075 * 2 0.00454467988411066
NW_023313086.1 132413 202576 70164 - Gnomon gene NA NA gene-LOC118190284 GeneID:118190284 LOC118190284 Gene LOC118190284 protein_coding
character(0) NA character(0) NW_023313086.1 201581 638556 436976 * 2 0.00457691040240196
NW_023313086.1 276458 614946 338489 + Gnomon gene NA NA gene-LOC118190295 GeneID:118190295 LOC118190295 Gene LOC118190295 protein_coding
character(0) NA character(0) NW_023313086.1 201581 638556 436976 * 2 0.00457691040240196
NW_023313086.1 512704 513420 717 + Gnomon gene NA NA gene-LOC118190318 GeneID:118190318 LOC118190318 Gene LOC118190318 protein_coding
character(0) NA character(0) NW_023313086.1 201581 638556 436976 * 2 0.00457691040240196
NW_023315156.1 30595 45185 14591 + Gnomon gene NA NA gene-LOC118185062 GeneID:118185062 LOC118185062 Gene LOC118185062 protein_coding character(0)
NA character(0) NW_023315156.1 37807 473719 435913 * 2 0.00458807147297741
NW_023315156.1 170339 626303 455965 - Gnomon gene NA NA gene-LOC118185063 GeneID:118185063 LOC118185063 Gene LOC118185063 protein_coding
character(0) NA character(0) NW_023315156.1 37807 473719 435913 * 2 0.00458807147297741
NW_023316797.1 340034 1174276 834243 - Gnomon gene NA NA gene-LOC118189460 GeneID:118189460 LOC118189460 Gene LOC118189460 protein_coding
character(0) NA character(0) NW_023316797.1 388926 824683 435758 * 2 0.0045897034592595
NW_023316797.1 429818 429890 73 + tRNAscan-SE gene NA NA gene-Trnav-cac-11 GeneID:118189461 Trnav-cac Gene Trnav-cac tRNA character(0) NA
character(0) NW_023316797.1 388926 824683 435758 * 2 0.0045897034592595
NW_023316797.1 779810 779882 73 + tRNAscan-SE gene NA NA gene-Trnay-gua-18 GeneID:118189462 Trnay-gua Gene Trnay-gua tRNA character(0) NA
character(0) NW_023316797.1 388926 824683 435758 * 2 0.0045897034592595
NW_023314828.1 405824 433831 28008 + Gnomon gene NA NA gene-LOC118184235 GeneID:118184235 LOC118184235 Gene LOC118184235 lncRNA
character(0) NA character(0) NW_023314828.1 6103 427366 421264 * 2 0.0047476166964184
NW_023315308.1 249135 281408 32274 + Gnomon gene NA NA gene-LOC118185497 GeneID:118185497 LOC118185497 Gene LOC118185497 protein_coding
character(0) NA character(0) NW_023315308.1 233892 653449 419558 * 2 0.00476692137916569
NW_023315308.1 372990 459957 86968 + Gnomon gene NA NA gene-LOC118185498 GeneID:118185498 LOC118185498 Gene LOC118185498 protein_coding
character(0) NA character(0) NW_023315308.1 233892 653449 419558 * 2 0.00476692137916569
NW_023315308.1 427798 427871 74 - tRNAscan-SE gene NA NA gene-Trnan-guu-21 GeneID:118185500 Trnan-guu Gene Trnan-guu tRNA character(0) NA
character(0) NW_023315308.1 233892 653449 419558 * 2 0.00476692137916569
NW_023315308.1 507975 508048 74 - tRNAscan-SE gene NA NA gene-Trnai-gau-52 GeneID:118185501 Trnai-gau Gene Trnai-gau tRNA character(0) NA
character(0) NW_023315308.1 233892 653449 419558 * 2 0.00476692137916569
NW_023315308.1 597603 638396 40794 - Gnomon gene NA NA gene-LOC118185499 GeneID:118185499 LOC118185499 Gene LOC118185499 protein_coding
character(0) NA character(0) NW_023315308.1 233892 653449 419558 * 2 0.00476692137916569
NW_023313292.1 241803 850679 608877 + Gnomon gene NA NA gene-LOC118196382 GeneID:118196382 LOC118196382 Gene LOC118196382 lncRNA
character(0) NA character(0) NW_023313292.1 225223 643683 418461 * 2 0.00477941791469217
NW_023312743.1 428425 428497 73 - tRNAscan-SE gene NA NA gene-Trnas-cga GeneID:118196092 Trnas-cga Gene Trnas-cga tRNA character(0) NA
character(0) NW_023312743.1 154321 572426 418106 * 2 0.00478347596064156
NW_023314236.1 803780 804524 745 - Gnomon gene NA NA gene-LOC118182605 GeneID:118182605 LOC118182605 Gene LOC118182605 protein_coding
character(0) NA character(0) NW_023314236.1 579485 996779 417295 * 2 0.0047927724990714
NW_023312795.1 252054 463806 211753 - Gnomon gene NA NA gene-LOC118181454 GeneID:118181454 LOC118181454 Gene LOC118181454 protein_coding
character(0) NA character(0) NW_023312795.1 263891 680713 416823 * 2 0.00479819971546676
NW_023315817.1 31815 65282 33468 - Gnomon gene NA NA gene-LOC118186878 GeneID:118186878 LOC118186878 Gene LOC118186878 protein_coding character(0)
NA character(0) NW_023315817.1 3388 418297 414910 * 2 0.00482032247957388
NW_023315817.1 211027 325421 114395 + Gnomon gene NA NA gene-LOC118186879 GeneID:118186879 LOC118186879 Gene LOC118186879 protein_coding
character(0) NA character(0) NW_023315817.1 3388 418297 414910 * 2 0.00482032247957388
NW_023315817.1 237584 237691 108 + cmsearch gene NA NA gene-LOC118186883 GeneID:118186883 LOC118186883 Gene LOC118186883 snRNA character(0)
NA character(0) NW_023315817.1 3388 418297 414910 * 2 0.00482032247957388
NW_023315817.1 320247 320355 109 + cmsearch gene NA NA gene-LOC118186884 GeneID:118186884 LOC118186884 Gene LOC118186884 snRNA character(0)
NA character(0) NW_023315817.1 3388 418297 414910 * 2 0.00482032247957388
NW_023315817.1 337164 429020 91857 - Gnomon gene NA NA gene-LOC118186880 GeneID:118186880 LOC118186880 Gene LOC118186880 protein_coding
character(0) NA character(0) NW_023315817.1 3388 418297 414910 * 2 0.00482032247957388
NW_023315817.1 372007 372079 73 - tRNAscan-SE gene NA NA gene-Trnat-cgu-68 GeneID:118186885 Trnat-cgu Gene Trnat-cgu tRNA character(0) NA
character(0) NW_023315817.1 3388 418297 414910 * 2 0.00482032247957388
NW_023314268.1 325075 325148 74 - tRNAscan-SE gene NA NA gene-Trnai-gau-30 GeneID:118182687 Trnai-gau Gene Trnai-gau tRNA character(0) NA
character(0) NW_023314268.1 167039 580160 413122 * 2 0.00484118492842308
NW_023313193.1 746121 825727 79607 - Gnomon gene NA NA gene-LOC118193212 GeneID:118193212 LOC118193212 Gene LOC118193212 protein_coding
character(0) NA character(0) NW_023313193.1 386504 793107 406604 * 2 0.00491879076447846
NW_023314649.1 113065 279476 166412 + Gnomon gene NA NA gene-LOC118183693 GeneID:118183693 LOC118183693 Gene LOC118183693 protein_coding
character(0) NA character(0) NW_023314649.1 183576 589423 405848 * 2 0.00492795332242613
NW_023314649.1 307427 307498 72 + tRNAscan-SE gene NA NA gene-Trnag-gcc-6 GeneID:118183696 Trnag-gcc Gene Trnag-gcc tRNA character(0) NA
character(0) NW_023314649.1 183576 589423 405848 * 2 0.00492795332242613
NW_023320080.1 306622 401935 95314 + Gnomon gene NA NA gene-LOC118196685 GeneID:118196685 LOC118196685 Gene LOC118196685 protein_coding
character(0) NA character(0) NW_023320080.1 32028 437676 405649 * 2 0.00493037083784257
NW_023316203.1 36711 175858 139148 - Gnomon gene NA NA gene-LOC118187962 GeneID:118187962 LOC118187962 Gene LOC118187962 protein_coding

character(0) NA character(0) NW_023316203.1 24991 425604 400614 * 2 0.00499233676306869
NW_023316203.1 212518 289386 76869 - Gnomon gene NA NA gene-LOC118187963 GeneID:118187963 LOC118187963 Gene LOC118187963 protein_coding
character(0) NA character(0) NW_023316203.1 24991 425604 400614 * 2 0.00499233676306869
NW_023315993.1 81916 379085 297170 - Gnomon gene NA NA gene-LOC118187447 GeneID:118187447 LOC118187447 Gene LOC118187447 protein_coding
character(0) NA character(0) NW_023315993.1 49697 445692 395996 * 2 0.00505055606622289
NW_023315993.1 431688 474550 42863 + Gnomon gene NA NA gene-LOC118187445 GeneID:118187445 LOC118187445 Gene LOC118187445 protein_coding
character(0) NA character(0) NW_023315993.1 49697 445692 395996 * 2 0.00505055606622289
NW_023321665.1 143708 574602 430895 + Gnomon gene NA NA gene-LOC118199411 GeneID:118199411 LOC118199411 Gene LOC118199411 lncRNA
character(0) NA character(0) NW_023321665.1 295372 688248 392877 * 2 0.00509065178160086
NW_023316199.1 307877 895571 587695 - Gnomon gene NA NA gene-LOC118187950 GeneID:118187950 LOC118187950 Gene LOC118187950 protein_coding
character(0) NA character(0) NW_023316199.1 683401 1075938 392538 * 2 0.00509504812272952
NW_023312866.1 77391 109909 32519 + Gnomon gene NA NA gene-LOC118183809 GeneID:118183809 LOC118183809 Gene LOC118183809 protein_coding
character(0) NA character(0) NW_023312866.1 61826 452478 390653 * 2 0.00511963302470479
NW_023312866.1 279487 300364 20878 + Gnomon gene NA NA gene-LOC118183818 GeneID:118183818 LOC118183818 Gene LOC118183818 protein_coding
character(0) NA character(0) NW_023312866.1 61826 452478 390653 * 2 0.00511963302470479
NW_023312866.1 351573 351644 72 + tRNAscan-SE gene NA NA gene-Trnap-ugg GeneID:118183831 Trnap-ugg Gene Trnap-ugg tRNA character(0) NA
character(0) NW_023312866.1 61826 452478 390653 * 2 0.00511963302470479
NW_023312866.1 372151 395080 22930 - Gnomon gene NA NA gene-LOC118183826 GeneID:118183826 LOC118183826 Gene LOC118183826 lncRNA
character(0) NA character(0) NW_023312866.1 61826 452478 390653 * 2 0.00511963302470479
NW_023312866.1 431035 431964 930 - Gnomon gene NA NA gene-LOC118183763 GeneID:118183763 LOC118183763 Gene LOC118183763 protein_coding
character(0) NA character(0) NW_023312866.1 61826 452478 390653 * 2 0.00511963302470479
NW_023312866.1 432004 432582 579 - Gnomon gene NA NA gene-LOC118183770 GeneID:118183770 LOC118183770 Gene LOC118183770 protein_coding
character(0) NA character(0) NW_023312866.1 61826 452478 390653 * 2 0.00511963302470479
NW_023314082.1 227427 511368 283942 - Gnomon gene NA NA gene-LOC118182139 GeneID:118182139 LOC118182139 Gene LOC118182139 protein_coding
character(0) NA character(0) NW_023314082.1 210551 599106 388556 * 2 0.00514726320015648
NW_023317533.1 67212 170253 103042 + Gnomon gene NA NA gene-LOC118191187 GeneID:118191187 LOC118191187 Gene LOC118191187 lncRNA
character(0) NA character(0) NW_023317533.1 110519 491304 380786 * 2 0.00525229393937802
NW_023317136.1 250269 424630 174362 + Gnomon gene NA NA gene-LOC118190301 GeneID:118190301 LOC118190301 Gene LOC118190301 lncRNA
character(0) NA character(0) NW_023317136.1 251172 631891 380720 * 2 0.00525320445471738
NW_023317136.1 569049 617081 48033 + Gnomon gene NA NA gene-LOC118190302 GeneID:118190302 LOC118190302 Gene LOC118190302 lncRNA
character(0) NA character(0) NW_023317136.1 251172 631891 380720 * 2 0.00525320445471738
NW_023315529.1 85805 529599 443795 + Gnomon gene NA NA gene-LOC118186067 GeneID:118186067 LOC118186067 Gene LOC118186067 protein_coding
character(0) NA character(0) NW_023315529.1 32934 412941 380008 * 2 0.00526304709374539
NW_023315529.1 401557 401629 73 - tRNAscan-SE gene NA NA gene-Trnas-uga-17 GeneID:118186068 Trnas-uga Gene Trnas-uga tRNA character(0) NA
character(0) NW_023315529.1 32934 412941 380008 * 2 0.00526304709374539
NW_023313671.1 377148 452203 75056 + Gnomon gene NA NA gene-LOC118180909 GeneID:118180909 LOC118180909 Gene LOC118180909 protein_coding
character(0) NA character(0) NW_023313671.1 365749 743473 377725 * 2 0.0052948573697796
NW_023313671.1 416849 416921 73 - tRNAscan-SE gene NA NA gene-Trnam-cau-18 GeneID:118180917 Trnam-cau Gene Trnam-cau tRNA character(0) NA
character(0) NW_023313671.1 365749 743473 377725 * 2 0.0052948573697796
NW_023313671.1 456285 504712 48428 - Gnomon gene NA NA gene-LOC118180910 GeneID:118180910 LOC118180910 Gene LOC118180910 protein_coding
character(0) NA character(0) NW_023313671.1 365749 743473 377725 * 2 0.0052948573697796
NW_023313671.1 718586 848995 130410 + Gnomon gene NA NA gene-LOC118180911 GeneID:118180911 LOC118180911 Gene LOC118180911 protein_coding
character(0) NA character(0) NW_023313671.1 365749 743473 377725 * 2 0.0052948573697796
NW_023315448.1 40331 518927 478597 - Gnomon gene NA NA gene-LOC118185853 GeneID:118185853 LOC118185853 Gene LOC118185853 protein_coding
character(0) NA character(0) NW_023315448.1 96685 473314 376630 * 2 0.0053102514404057
NW_023315448.1 458707 465452 6746 - Gnomon gene NA NA gene-LOC118185854 GeneID:118185854 LOC118185854 Gene LOC118185854 lncRNA
character(0) NA character(0) NW_023315448.1 96685 473314 376630 * 2 0.0053102514404057
NW_023313020.1 202483 535034 332552 + Gnomon gene NA NA gene-LOC118188465 GeneID:118188465 LOC118188465 Gene LOC118188465 protein_coding
character(0) NA character(0) NW_023313020.1 68175 441924 373750 * 2 0.00535117056856187
NW_023313523.1 120668 418344 297677 - Gnomon gene NA NA gene-LOC118203911 GeneID:118203911 LOC118203911 Gene LOC118203911 protein_coding
character(0) NA character(0) NW_023313523.1 224082 597145 373064 * 2 0.00536101044324834
NW_023313523.1 348728 348800 73 - tRNAscan-SE gene NA NA gene-Trnas-cga-23 GeneID:118203983 Trnas-cga Gene Trnas-cga tRNA character(0) NA
character(0) NW_023313523.1 224082 597145 373064 * 2 0.00536101044324834
NW_023313523.1 481066 481138 73 - tRNAscan-SE gene NA NA gene-Trnas-cga-24 GeneID:118203985 Trnas-cga Gene Trnas-cga tRNA character(0) NA
character(0) NW_023313523.1 224082 597145 373064 * 2 0.00536101044324834
NW_023313523.1 488044 586912 98869 - Gnomon gene NA NA gene-LOC118203919 GeneID:118203919 LOC118203919 Gene LOC118203919 protein_coding
character(0) NA character(0) NW_023313523.1 224082 597145 373064 * 2 0.00536101044324834
NW_023314412.1 168953 313042 144090 - Gnomon gene NA NA gene-LOC118183062 GeneID:118183062 LOC118183062 Gene LOC118183062 protein_coding
character(0) NA character(0) NW_023314412.1 48981 421626 372646 * 2 0.00536702393155971
NW_023317070.1 105958 199187 93230 - Gnomon gene NA NA gene-LOC118190127 GeneID:118190127 LOC118190127 Gene LOC118190127 lncRNA
character(0) NA character(0) NW_023317070.1 49388 421441 372054 * 2 0.00537556376224957
NW_023318742.1 9291 56230 46940 + Gnomon gene NA NA gene-LOC118193825 GeneID:118193825 LOC118193825 Gene LOC118193825 protein_coding character(0)
true c(".", "9291") NW_023318742.1 5293 374055 368763 * 2 0.00542353761087745
NW_023318742.1 139535 139608 74 + tRNAscan-SE gene NA NA gene-Trnan-guu-42 GeneID:118193828 Trnan-guu Gene Trnan-guu tRNA character(0) NA

character(0) NW_023318742.1 5293 374055 368763 * 2 0.00542353761087745
NW_023318742.1 156465 380789 224325 + Gnomon gene NA NA gene-LOC118193826 GeneID:118193826 LOC118193826 Gene LOC118193826 protein_coding
character(0) NA character(0) NW_023318742.1 5293 374055 368763 * 2 0.00542353761087745
NW_023322011.1 103306 600419 497114 - Gnomon gene NA NA gene-LOC118199969 GeneID:118199969 LOC118199969 Gene LOC118199969 protein_coding
character(0) NA character(0) NW_023322011.1 190266 558891 368626 * 2 0.00542555327079479
NW_023312806.1 252970 429347 176378 - Gnomon gene NA NA gene-LOC118181738 GeneID:118181738 LOC118181738 Gene LOC118181738 protein_coding
character(0) NA character(0) NW_023312806.1 42448 409580 367133 * 2 0.00544761707610049
NW_023314854.1 97225 341467 244243 + Gnomon gene NA NA gene-LOC118184292 GeneID:118184292 LOC118184292 Gene LOC118184292 protein_coding
character(0) NA character(0) NW_023314854.1 61441 426849 365409 * 2 0.00547331893850453
NW_023314854.1 370539 397595 27057 + Gnomon gene NA NA gene-LOC118184293 GeneID:118184293 LOC118184293 Gene LOC118184293 protein_coding
character(0) NA character(0) NW_023314854.1 61441 426849 365409 * 2 0.00547331893850453
NW_023314854.1 404473 421710 17238 + Gnomon gene NA NA gene-LOC118184294 GeneID:118184294 LOC118184294 Gene LOC118184294 lncRNA
character(0) NA character(0) NW_023314854.1 61441 426849 365409 * 2 0.00547331893850453
NW_023313387.1 944792 945664 873 - Gnomon gene NA NA gene-LOC118200166 GeneID:118200166 LOC118200166 Gene LOC118200166 protein_coding
character(0) NA character(0) NW_023313387.1 802198 1166740 364543 * 2 0.00548632122959431
NW_023313387.1 1038750 1631855 593106 + Gnomon gene NA NA gene-LOC118200115 GeneID:118200115 LOC118200115 Gene LOC118200115 protein_coding
character(0) NA character(0) NW_023313387.1 802198 1166740 364543 * 2 0.00548632122959431
NW_023315531.1 3990 328336 324347 + Gnomon gene NA NA gene-LOC118186070 GeneID:118186070 LOC118186070 Gene LOC118186070 protein_coding
character(0) NA character(0) NW_023315531.1 34165 398314 364150 * 2 0.00549224220788137
NW_023317100.1 150884 240449 89566 - Gnomon gene NA NA gene-LOC118190199 GeneID:118190199 LOC118190199 Gene LOC118190199 lncRNA
character(0) NA character(0) NW_023317100.1 44343 407527 363185 * 2 0.00550683535938984
NW_023312740.1 45699 111805 66107 - Gnomon gene NA NA gene-LOC118194736 GeneID:118194736 LOC118194736 Gene LOC118194736 protein_coding
character(0) NA character(0) NW_023312740.1 5551 368544 362994 * 2 0.00550973294324424
NW_023312740.1 117122 139734 22613 - Gnomon gene NA NA gene-LOC118194364 GeneID:118194364 LOC118194364 Gene LOC118194364 protein_coding
character(0) NA character(0) NW_023312740.1 5551 368544 362994 * 2 0.00550973294324424
NW_023315623.1 116861 629176 512316 + Gnomon gene NA NA gene-LOC118186316 GeneID:118186316 LOC118186316 Gene LOC118186316 protein_coding
character(0) NA character(0) NW_023315623.1 342498 703493 360996 * 2 0.0055402275925495
NW_023315623.1 427041 427113 73 - tRNAscan-SE gene NA NA gene-Trnas-cga-69 GeneID:118186320 Trnas-cga Gene Trnas-cga tRNA character(0) NA
character(0) NW_023315623.1 342498 703493 360996 * 2 0.0055402275925495
NW_023315623.1 553732 615175 61444 - Gnomon gene NA NA gene-LOC118186318 GeneID:118186318 LOC118186318 Gene LOC118186318 lncRNA
character(0) NA character(0) NW_023315623.1 342498 703493 360996 * 2 0.0055402275925495
NW_023319159.1 87200 161848 74649 - Gnomon gene NA NA gene-LOC118194745 GeneID:118194745 LOC118194745 Gene LOC118194745 protein_coding
character(0) NA character(0) NW_023319159.1 123590 483958 360369 * 2 0.00554986694194007
NW_023319159.1 164594 488760 324167 + Gnomon gene NA NA gene-LOC118194746 GeneID:118194746 LOC118194746 Gene LOC118194746 protein_coding
character(0) NA character(0) NW_023319159.1 123590 483958 360369 * 2 0.00554986694194007
NW_023315112.1 366657 366992 336 - Gnomon gene NA NA gene-LOC118184930 GeneID:118184930 LOC118184930 Gene LOC118184930 protein_coding
character(0) NA character(0) NW_023315112.1 242739 603087 360349 * 2 0.00555017496926591
NW_023315112.1 367821 368306 486 - Gnomon gene NA NA gene-LOC118184931 GeneID:118184931 LOC118184931 Gene LOC118184931 protein_coding
character(0) NA character(0) NW_023315112.1 242739 603087 360349 * 2 0.00555017496926591
NW_023313815.1 61070 456934 395865 + Gnomon gene NA NA gene-LOC118181355 GeneID:118181355 LOC118181355 Gene LOC118181355 protein_coding
character(0) NA character(0) NW_023313815.1 175735 534270 358536 * 2 0.00557824039984827
NW_023315038.1 77415 202990 125576 + Gnomon gene NA NA gene-LOC118184750 GeneID:118184750 LOC118184750 Gene LOC118184750 protein_coding
character(0) NA character(0) NW_023315038.1 117816 473262 355447 * 2 0.00562671790731108
NW_023315038.1 318799 350297 31499 - Gnomon gene NA NA gene-LOC118184751 GeneID:118184751 LOC118184751 Gene LOC118184751 lncRNA
character(0) NA character(0) NW_023315038.1 117816 473262 355447 * 2 0.00562671790731108
NW_023315038.1 382925 383491 567 + Gnomon gene NA NA gene-LOC118184752 GeneID:118184752 LOC118184752 Gene LOC118184752 protein_coding
character(0) NA character(0) NW_023315038.1 117816 473262 355447 * 2 0.00562671790731108
NW_023313412.1 243326 466348 223023 - Gnomon gene NA NA gene-LOC118200679 GeneID:118200679 LOC118200679 Gene LOC118200679 protein_coding
character(0) NA character(0) NW_023313412.1 216924 568310 351387 * 2 0.00569173020060503
NW_023317659.1 389751 473701 83951 + Gnomon gene NA NA gene-LOC118191501 GeneID:118191501 LOC118191501 Gene LOC118191501 protein_coding
character(0) NA character(0) NW_023317659.1 438127 786064 347938 * 2 0.00574815053256615
NW_023317659.1 518389 695873 177485 + Gnomon gene NA NA gene-LOC118191502 GeneID:118191502 LOC118191502 Gene LOC118191502 lncRNA
character(0) NA character(0) NW_023317659.1 438127 786064 347938 * 2 0.00574815053256615
NW_023317659.1 780085 833176 53092 - Gnomon gene NA NA gene-LOC118191505 GeneID:118191505 LOC118191505 Gene LOC118191505 protein_coding
character(0) NA character(0) NW_023317659.1 438127 786064 347938 * 2 0.00574815053256615
NW_023313787.1 379933 609163 229231 + Gnomon gene NA NA gene-LOC118181271 GeneID:118181271 LOC118181271 Gene LOC118181271 protein_coding
character(0) NA character(0) NW_023313787.1 94302 437103 342802 * 2 0.00583427167869499
NW_023319435.1 116341 545953 429613 + Gnomon gene NA NA gene-LOC118195362 GeneID:118195362 LOC118195362 Gene LOC118195362 protein_coding
character(0) NA character(0) NW_023319435.1 106914 449411 342498 * 2 0.00583945015737318
NW_023315478.1 197213 363987 166775 - Gnomon gene NA NA gene-LOC118185934 GeneID:118185934 LOC118185934 Gene LOC118185934 protein_coding
character(0) NA character(0) NW_023315478.1 202078 542257 340180 * 2 0.00587924040214004
NW_023315478.1 342170 425322 83153 + Gnomon gene NA NA gene-LOC118185933 GeneID:118185933 LOC118185933 Gene LOC118185933 protein_coding
character(0) NA character(0) NW_023315478.1 202078 542257 340180 * 2 0.00587924040214004
NW_023315478.1 486715 514309 27595 + Gnomon gene NA NA gene-LOC118185935 GeneID:118185935 LOC118185935 Gene LOC118185935 protein_coding

character(0) NA character(0) NW_023315478.1 202078 542257 340180 * 2 0.00587924040214004
NW_023313876.1 204076 225510 21435 - Gnomon gene NA NA gene-LOC118181519 GeneID:118181519 LOC118181519 Gene LOC118181519 protein_coding
character(0) NA character(0) NW_023313876.1 208418 548549 340132 * 2 0.00588007009043548
NW_023313876.1 243449 343187 99739 + Gnomon gene NA NA gene-LOC118181518 GeneID:118181518 LOC118181518 Gene LOC118181518 protein_coding
character(0) NA character(0) NW_023313876.1 208418 548549 340132 * 2 0.00588007009043548
NW_023313876.1 410828 522352 111525 + Gnomon gene NA NA gene-LOC118181520 GeneID:118181520 LOC118181520 Gene LOC118181520 protein_coding
character(0) NA character(0) NW_023313876.1 208418 548549 340132 * 2 0.00588007009043548
NW_023315008.1 255009 515701 260693 - Gnomon gene NA NA gene-LOC118184676 GeneID:118184676 LOC118184676 Gene LOC118184676 protein_coding
character(0) NA character(0) NW_023315008.1 215439 554234 338796 * 2 0.00590325741744295
NW_023315008.1 278998 320051 41054 + Gnomon gene NA NA gene-LOC118184678 GeneID:118184678 LOC118184678 Gene LOC118184678 lncRNA
character(0) NA character(0) NW_023315008.1 215439 554234 338796 * 2 0.00590325741744295
NW_023315008.1 515780 560275 44496 + Gnomon gene NA NA gene-LOC118184677 GeneID:118184677 LOC118184677 Gene LOC118184677 lncRNA
character(0) NA character(0) NW_023315008.1 215439 554234 338796 * 2 0.00590325741744295
NW_023313043.1 161239 600424 439186 + Gnomon gene NA NA gene-LOC118188938 GeneID:118188938 LOC118188938 Gene LOC118188938 protein_coding
character(0) NA character(0) NW_023313043.1 276196 612899 336704 * 2 0.00593993537350314
NW_023312810.1 30073 384342 354270 + Gnomon gene NA NA gene-LOC118181801 GeneID:118181801 LOC118181801 Gene LOC118181801 protein_coding
character(0) NA character(0) NW_023312810.1 217985 553922 335938 * 2 0.00595347951110026
NW_023318216.1 81357 198959 117603 + Gnomon gene NA NA gene-LOC118192675 GeneID:118192675 LOC118192675 Gene LOC118192675 lncRNA
character(0) NA character(0) NW_023318216.1 144161 478782 334622 * 2 0.00597689333038473
NW_023318216.1 291392 291464 73 - tRNAscan-SE gene NA NA gene-Trnaa-cgc-52 GeneID:118192676 Trnaa-cgc Gene Trnaa-cgc tRNA character(0) NA
character(0) NW_023318216.1 144161 478782 334622 * 2 0.00597689333038473
NW_023313804.1 33762 53469 19708 + Gnomon gene NA NA gene-LOC118181310 GeneID:118181310 LOC118181310 Gene LOC118181310 protein_coding character(0)
NA character(0) NW_023313804.1 28743 362612 333870 * 2 0.00599035552760056
NW_023313804.1 94669 367439 272771 + Gnomon gene NA NA gene-LOC118181311 GeneID:118181311 LOC118181311 Gene LOC118181311 protein_coding
character(0) NA character(0) NW_023313804.1 28743 362612 333870 * 2 0.00599035552760056
NW_023313804.1 176278 176350 73 - tRNAscan-SE gene NA NA gene-Trnat-cgu-30 GeneID:118181316 Trnat-cgu Gene Trnat-cgu tRNA character(0) NA
character(0) NW_023313804.1 28743 362612 333870 * 2 0.00599035552760056
NW_023313145.1 117491 312086 194596 - Gnomon gene NA NA gene-LOC118192025 GeneID:118192025 LOC118192025 Gene LOC118192025 lncRNA
character(0) NA character(0) NW_023313145.1 244724 577917 333194 * 2 0.00600250904878239
NW_023313145.1 300746 300817 72 - tRNAscan-SE gene NA NA gene-Trnat-ggu-10 GeneID:118192070 Trnat-ggu Gene Trnat-ggu tRNA character(0) NA
character(0) NW_023313145.1 244724 577917 333194 * 2 0.00600250904878239
NW_023313145.1 372095 598217 226123 - Gnomon gene NA NA gene-LOC118192030 GeneID:118192030 LOC118192030 Gene LOC118192030 protein_coding
character(0) NA character(0) NW_023313145.1 244724 577917 333194 * 2 0.00600250904878239
NW_023313145.1 553609 553681 73 + tRNAscan-SE gene NA NA gene-Trnak-uuu-2 GeneID:118192065 Trnak-uuu Gene Trnak-uuu tRNA character(0) NA
character(0) NW_023313145.1 244724 577917 333194 * 2 0.00600250904878239
NW_023312882.1 231226 605826 374601 - Gnomon gene NA NA gene-LOC118184280 GeneID:118184280 LOC118184280 Gene LOC118184280 protein_coding
character(0) NA character(0) NW_023312882.1 182837 515509 332673 * 2 0.00601190959290354
NW_023316215.1 496924 543731 46808 + Gnomon gene NA NA gene-LOC118187998 GeneID:118187998 LOC118187998 Gene LOC118187998 protein_coding
character(0) NA character(0) NW_023316215.1 469261 801583 332323 * 2 0.00601824128934801
NW_023316215.1 577507 885584 308078 - Gnomon gene NA NA gene-LOC118187999 GeneID:118187999 LOC118187999 Gene LOC118187999 protein_coding
character(0) NA character(0) NW_023316215.1 469261 801583 332323 * 2 0.00601824128934801
NW_023313927.1 262744 547960 285217 + Gnomon gene NA NA gene-LOC118181707 GeneID:118181707 LOC118181707 Gene LOC118181707 protein_coding
character(0) NA character(0) NW_023313927.1 204484 533097 328614 * 2 0.00608616796606353
NW_023317102.1 190244 313372 123129 + Gnomon gene NA NA gene-LOC118190204 GeneID:118190204 LOC118190204 Gene LOC118190204 protein_coding
character(0) NA character(0) NW_023317102.1 81584 409286 327703 * 2 0.00610308724668374
NW_023317102.1 382278 382350 73 + tRNAscan-SE gene NA NA gene-Trnak-cuu-44 GeneID:118190205 Trnak-cuu Gene Trnak-cuu tRNA character(0) NA
character(0) NW_023317102.1 81584 409286 327703 * 2 0.00610308724668374
NW_023312973.1 202062 709113 507052 + Gnomon gene NA NA gene-LOC118186875 GeneID:118186875 LOC118186875 Gene LOC118186875 protein_coding
character(0) NA character(0) NW_023312973.1 262946 588326 325381 * 2 0.00614664040002336
NW_023312973.1 416529 416599 71 + tRNAscan-SE gene NA NA gene-Trnaa-cgc-6 GeneID:118186895 Trnaa-cgc Gene Trnaa-cgc tRNA character(0) NA
character(0) NW_023312973.1 262946 588326 325381 * 2 0.00614664040002336
NW_023315623.1 116861 629176 512316 + Gnomon gene NA NA gene-LOC118186316 GeneID:118186316 LOC118186316 Gene LOC118186316 protein_coding
character(0) NA character(0) NW_023315623.1 13904 338969 325066 * 2 0.00615259670343869
NW_023315623.1 186733 186805 73 + tRNAscan-SE gene NA NA gene-Trnaa-cgc-26 GeneID:118186319 Trnaa-cgc Gene Trnaa-cgc tRNA character(0) NA
character(0) NW_023315623.1 13904 338969 325066 * 2 0.00615259670343869
NW_023316214.1 9383 818560 809178 - Gnomon gene NA NA gene-LOC118187993 GeneID:118187993 LOC118187993 Gene LOC118187993 protein_coding
character(0) NA character(0) NW_023316214.1 93902 416488 322587 * 2 0.00619987786240611
NW_023313628.1 330047 331169 1123 - Gnomon gene NA NA gene-LOC118180177 GeneID:118180177 LOC118180177 Gene LOC118180177 protein_coding
character(0) NA character(0) NW_023313628.1 149393 471574 322182 * 2 0.006207671440366
NW_023313628.1 338559 425590 87032 + Gnomon gene NA NA gene-LOC118180187 GeneID:118180187 LOC118180187 Gene LOC118180187 protein_coding
character(0) NA character(0) NW_023313628.1 149393 471574 322182 * 2 0.006207671440366
NW_023313169.1 447739 1065750 618012 - Gnomon gene NA NA gene-LOC118192559 GeneID:118192559 LOC118192559 Gene LOC118192559 protein_coding
character(0) NA character(0) NW_023313169.1 493685 814775 321091 * 2 0.00622876380839077
NW_023313295.1 212739 216080 3342 - Gnomon gene NA NA gene-LOC118196526 GeneID:118196526 LOC118196526 Gene LOC118196526 protein_coding

character(0) NA character(0) NW_023313295.1 212842 533724 320883 * 2 0.00623280136373694
NW_023316692.1 143331 366170 222840 - Gnomon gene NA NA gene-LOC118189141 GeneID:118189141 LOC118189141 Gene LOC118189141 protein_coding
character(0) NA character(0) NW_023316692.1 214201 531911 317711 * 2 0.0062950291302473
NW_023314546.1 395890 395962 73 + tRNAscan-SE gene NA NA gene-Trnaa-cgc-20 GeneID:118183410 Trnaa-cgc Gene Trnaa-cgc tRNA character(0) NA
character(0) NW_023314546.1 295546 613163 317618 * 2 0.00629687234350698
NW_023315600.1 26240 501250 475011 - Gnomon gene NA NA gene-LOC118186257 GeneID:118186257 LOC118186257 Gene LOC118186257 protein_coding
c("501250", ".") true character(0) NW_023315600.1 45680 363139 317460 * 2 0.0063000063000063
NW_023315600.1 135019 135091 73 - tRNAscan-SE gene NA NA gene-Trnak-cuu-24 GeneID:118186259 Trnak-cuu Gene Trnak-cuu tRNA character(0) NA
character(0) NW_023315600.1 45680 363139 317460 * 2 0.0063000063000063
NW_023316749.1 335013 335795 783 + Gnomon gene NA NA gene-LOC118189314 GeneID:118189314 LOC118189314 Gene LOC118189314 protein_coding
character(0) NA character(0) NW_023316749.1 277706 593808 316103 * 2 0.00632705162557774
NW_023316749.1 515604 694518 178915 - Gnomon gene NA NA gene-LOC118189316 GeneID:118189316 LOC118189316 Gene LOC118189316 protein_coding
character(0) NA character(0) NW_023316749.1 277706 593808 316103 * 2 0.00632705162557774
NW_023315711.1 124252 198891 74640 + Gnomon gene NA NA gene-LOC118186600 GeneID:118186600 LOC118186600 Gene LOC118186600 lncRNA
character(0) NA character(0) NW_023315711.1 64192 379966 315775 * 2 0.00633362362441612
NW_023315711.1 216771 217178 408 + Gnomon gene NA NA gene-LOC118186599 GeneID:118186599 LOC118186599 Gene LOC118186599 protein_coding
character(0) NA character(0) NW_023315711.1 64192 379966 315775 * 2 0.00633362362441612
NW_023315711.1 370068 428069 58002 - Gnomon gene NA NA gene-LOC118186601 GeneID:118186601 LOC118186601 Gene LOC118186601 lncRNA
character(0) NA character(0) NW_023315711.1 64192 379966 315775 * 2 0.00633362362441612
NW_023314414.1 539360 567313 27954 + Gnomon gene NA NA gene-LOC118183068 GeneID:118183068 LOC118183068 Gene LOC118183068 protein_coding
character(0) NA character(0) NW_023314414.1 501530 817145 315616 * 2 0.00633681435668661
NW_023312980.1 41359 132369 91011 - Gnomon gene NA NA gene-LOC118187202 GeneID:118187202 LOC118187202 Gene LOC118187202 protein_coding
character(0) NA character(0) NW_023312980.1 27359 342875 315517 * 2 0.00633880266356488
NW_023312980.1 97535 320226 222692 + Gnomon gene NA NA gene-LOC118187188 GeneID:118187188 LOC118187188 Gene LOC118187188 protein_coding
character(0) NA character(0) NW_023312980.1 27359 342875 315517 * 2 0.00633880266356488
NW_023316051.1 409329 485971 76643 - Gnomon gene NA NA gene-LOC118187589 GeneID:118187589 LOC118187589 Gene LOC118187589 protein_coding
character(0) NA character(0) NW_023316051.1 188458 503880 315423 * 2 0.00634069170605821
NW_023317215.1 284554 284955 402 - Gnomon gene NA NA gene-LOC118190464 GeneID:118190464 LOC118190464 Gene LOC118190464 protein_coding
character(0) NA character(0) NW_023317215.1 24514 338662 314149 * 2 0.00636640575013767
NW_023318544.1 112603 145817 33215 - Gnomon gene NA NA gene-LOC118193358 GeneID:118193358 LOC118193358 Gene LOC118193358 lncRNA
character(0) NA character(0) NW_023318544.1 53160 366858 313699 * 2 0.00637553833451812
NW_023321981.1 203733 362228 158496 - Gnomon gene NA NA gene-LOC118199928 GeneID:118199928 LOC118199928 Gene LOC118199928 protein_coding
character(0) NA character(0) NW_023321981.1 214675 528200 313526 * 2 0.00637905628241358
NW_023321981.1 371388 418843 47456 + Gnomon gene NA NA gene-LOC118199933 GeneID:118199933 LOC118199933 Gene LOC118199933 lncRNA
character(0) NA character(0) NW_023321981.1 214675 528200 313526 * 2 0.00637905628241358
NW_023321981.1 454411 535254 80844 + Gnomon gene NA NA gene-LOC118199929 GeneID:118199929 LOC118199929 Gene LOC118199929 protein_coding
character(0) NA character(0) NW_023321981.1 214675 528200 313526 * 2 0.00637905628241358
NW_023313617.1 188456 188812 357 + Gnomon gene NA NA gene-LOC118179809 GeneID:118179809 LOC118179809 Gene LOC118179809 protein_coding
character(0) NA character(0) NW_023313617.1 7007 320337 313331 * 2 0.00638302625657851
NW_023313549.1 153686 153759 74 + tRNAscan-SE gene NA NA gene-Trnai-gau-17 GeneID:118204796 Trnai-gau Gene Trnai-gau tRNA character(0) NA
character(0) NW_023313549.1 67128 380160 313033 * 2 0.00638910274635582
NW_023313549.1 292054 606570 314517 + Gnomon gene NA NA gene-LOC118204780 GeneID:118204780 LOC118204780 Gene LOC118204780 lncRNA
character(0) NA character(0) NW_023313549.1 67128 380160 313033 * 2 0.00638910274635582
NW_023316663.1 131717 282938 151222 - Gnomon gene NA NA gene-LOC118189076 GeneID:118189076 LOC118189076 Gene LOC118189076 protein_coding
character(0) NA character(0) NW_023316663.1 196 313050 312855 * 2 0.00639273784980262
NW_023314135.1 392516 729446 336931 - Gnomon gene NA NA gene-LOC118182347 GeneID:118182347 LOC118182347 Gene LOC118182347 protein_coding
character(0) NA character(0) NW_023314135.1 438651 751267 312617 * 2 0.00639760473678655
NW_023314135.1 560320 560392 73 - tRNAscan-SE gene NA NA gene-Trnam-cau-25 GeneID:118182349 Trnam-cau Gene Trnam-cau tRNA character(0) NA
character(0) NW_023314135.1 438651 751267 312617 * 2 0.00639760473678655

Directory: MRPR File: gwr-Sd_LI_be.txt

| seqnames | start | end | width | strand | source | type | score | phase | ID | Dbxref | Name | gbkey | gene | gene_biotype | end_range | partial | start_range | seqnames | start | end | width | strand | NbrSites | Sdnsty |
|---|---|---|---|---|---|---|---|---|---|---|---|---|---|---|---|---|---|---|---|---|---|---|---|---|
| NW_023312732.1 | 273883 | 345899 | 72017 | + | Gnomon | gene | NA | NA | gene-LOC118191020 | GeneID:118191020 | LOC118191020 | Gene | LOC118191020 | protein_coding | character(0) | NA | character(0) | NW_023312732.1 | 286760 | 286767 | 8 | * | 8 | 1000 |
| NW_023312732.1 | 451412 | 559259 | 107848 | + | Gnomon | gene | NA | NA | gene-LOC118191196 | GeneID:118191196 | LOC118191196 | Gene | LOC118191196 | protein_coding | character(0) | NA | character(0) | NW_023312732.1 | 510338 | 510343 | 6 | * | 6 | 1000 |
| NW_023312746.1 | 467813 | 535855 | 68043 | + | Gnomon | gene | NA | NA | gene-LOC118196644 | GeneID:118196644 | LOC118196644 | Gene | LOC118196644 | protein_coding | character(0) | NA | character(0) | NW_023312746.1 | 488111 | 488116 | 6 | * | 6 | 1000 |
| NW_023312753.1 | 105051 | 162160 | 57110 | - | Gnomon | gene | NA | NA | gene-LOC118199607 | GeneID:118199607 | LOC118199607 | Gene | LOC118199607 | protein_coding | character(0) | NA | character(0) | NW_023312753.1 | 128868 | 128873 | 6 | * | 6 | 1000 |
| NW_023312755.1 | 30608 | 136449 | 105842 | + | Gnomon | gene | NA | NA | gene-LOC118200099 | GeneID:118200099 | LOC118200099 | Gene | LOC118200099 | protein_coding | character(0) | NA | character(0) | NW_023312755.1 | 110038 | 110045 | 8 | * | 8 | 1000 |
| NW_023312755.1 | 30608 | 136449 | 105842 | + | Gnomon | gene | NA | NA | gene-LOC118200099 | GeneID:118200099 | LOC118200099 | Gene | LOC118200099 | protein_coding | character(0) | NA | character(0) | NW_023312755.1 | 114240 | 114245 | 6 | * | 6 | 1000 |
| NW_023312758.1 | 2535 | 177946 | 175412 | - | Gnomon | gene | NA | NA | gene-LOC118200694 | GeneID:118200694 | LOC118200694 | Gene | LOC118200694 | protein_coding | character(0) | NA | character(0) | NW_023312758.1 | 151906 | 151911 | 6 | * | 6 | 1000 |
| NW_023312782.1 | 88823 | 135338 | 46516 | + | Gnomon | gene | NA | NA | gene-LOC118180879 | GeneID:118180879 | LOC118180879 | Gene | LOC118180879 | protein_coding | character(0) | NA | character(0) | NW_023312782.1 | 106868 | 106873 | 6 | * | 6 | 1000 |
| NW_023312817.1 | 13538 | 81112 | 67575 | + | Gnomon | gene | NA | NA | gene-LOC118181999 | GeneID:118181999 | LOC118181999 | Gene | LOC118181999 | protein_coding | character(0) | NA | character(0) | NW_023312817.1 | 48211 | 48216 | 6 | * | 6 | 1000 |
| NW_023312819.1 | 222905 | 284616 | 61712 | + | Gnomon | gene | NA | NA | gene-LOC118182088 | GeneID:118182088 | LOC118182088 | Gene | LOC118182088 | protein_coding | character(0) | NA | character(0) | NW_023312819.1 | 240550 | 240555 | 6 | * | 6 | 1000 |
| NW_023312819.1 | 222905 | 284616 | 61712 | + | Gnomon | gene | NA | NA | gene-LOC118182088 | GeneID:118182088 | LOC118182088 | Gene | LOC118182088 | protein_coding | character(0) | NA | character(0) | NW_023312819.1 | 260689 | 260694 | 6 | * | 6 | 1000 |
| NW_023312824.1 | 152873 | 193878 | 41006 | - | Gnomon | gene | NA | NA | gene-LOC118182297 | GeneID:118182297 | LOC118182297 | Gene | LOC118182297 | protein_coding | character(0) | NA | character(0) | NW_023312824.1 | 162831 | 162836 | 6 | * | 6 | 1000 |
| NW_023312829.1 | 16054 | 211342 | 195289 | + | Gnomon | gene | NA | NA | gene-LOC118182373 | GeneID:118182373 | LOC118182373 | Gene | LOC118182373 | protein_coding | character(0) | NA | character(0) | NW_023312829.1 | 113486 | 113491 | 6 | * | 6 | 1000 |
| NW_023312837.1 | 5721 | 93064 | 87344 | - | Gnomon | gene | NA | NA | gene-LOC118182581 | GeneID:118182581 | LOC118182581 | Gene | LOC118182581 | protein_coding | character(0) | true | c(".", "5721") | NW_023312837.1 | 62620 | 62625 | 6 | * | 6 | 1000 |
| NW_023312841.1 | 3105 | 11700 | 8596 | - | Gnomon | gene | NA | NA | gene-LOC118182676 | GeneID:118182676 | LOC118182676 | Gene | LOC118182676 | lncRNA | character(0) | NA | character(0) | NW_023312841.1 | 9567 | 9574 | 8 | * | 8 | 1000 |
| NW_023312846.1 | 121760 | 204155 | 82396 | + | Gnomon | gene | NA | NA | gene-LOC118182870 | GeneID:118182870 | LOC118182870 | Gene | LOC118182870 | protein_coding | character(0) | NA | character(0) | NW_023312846.1 | 125814 | 125819 | 6 | * | 6 | 1000 |
| NW_023312846.1 | 121760 | 204155 | 82396 | + | Gnomon | gene | NA | NA | gene-LOC118182870 | GeneID:118182870 | LOC118182870 | Gene | LOC118182870 | protein_coding | character(0) | NA | character(0) | NW_023312846.1 | 140309 | 140314 | 6 | * | 6 | 1000 |
| NW_023312858.1 | 110039 | 221517 | 111479 | + | Gnomon | gene | NA | NA | gene-LOC118183482 | GeneID:118183482 | LOC118183482 | Gene | LOC118183482 | protein_coding | character(0) | NA | character(0) | NW_023312858.1 | 201097 | 201102 | 6 | * | 6 | 1000 |
| NW_023312859.1 | 15224 | 169817 | 154594 | - | Gnomon | gene | NA | NA | gene-LOC118183553 | GeneID:118183553 | LOC118183553 | Gene | LOC118183553 | protein_coding | character(0) | NA | character(0) | NW_023312859.1 | 82925 | 82930 | 6 | * | 6 | 1000 |
| NW_023312862.1 | 305971 | 329060 | 23090 | - | Gnomon | gene | NA | NA | gene-LOC118183641 | GeneID:118183641 | LOC118183641 | Gene | LOC118183641 | protein_coding | character(0) | NA | character(0) | NW_023312862.1 | 308115 | 308120 | 6 | * | 6 | 1000 |
| NW_023312863.1 | 59464 | 282910 | 223447 | + | Gnomon | gene | NA | NA | gene-LOC118183667 | GeneID:118183667 | LOC118183667 | Gene | LOC118183667 | protein_coding | c("282910", ".") | true | character(0) | NW_023312863.1 | 82853 | 82858 | 6 | * | 6 | 1000 |
| NW_023312863.1 | 59464 | 282910 | 223447 | + | Gnomon | gene | NA | NA | gene-LOC118183667 | GeneID:118183667 | LOC118183667 | Gene | LOC118183667 | protein_coding | c("282910", ".") | true | character(0) | NW_023312863.1 | 123837 | 123842 | 6 | * | 6 | 1000 |
| NW_023312863.1 | 59464 | 282910 | 223447 | + | Gnomon | gene | NA | NA | gene-LOC118183667 | GeneID:118183667 | LOC118183667 | Gene | LOC118183667 | protein_coding | c("282910", ".") | true | character(0) | NW_023312863.1 | 207646 | 207651 | 6 | * | 6 | 1000 |
| NW_023312888.1 | 141697 | 219678 | 77982 | - | Gnomon | gene | NA | NA | gene-LOC118184390 | GeneID:118184390 | LOC118184390 | Gene | LOC118184390 | protein_coding | character(0) | NA | character(0) | NW_023312888.1 | 193747 | 193752 | 6 | * | 6 | 1000 |

...

... only first 25 lines shown ...

Directory: MRPR File: gwr-Sd_LI_ka.txt

| seqnames | start | end | width | strand | source | type | score | phase | ID | Dbxref | Name | gbkey | gene | gene_biotype | end_range | partial | start_range | seqnames | start | end | width | strand | NbrSites | Sdnsty |
|---|---|---|---|---|---|---|---|---|---|---|---|---|---|---|---|---|---|---|---|---|---|---|---|---|
| NW_023312732.1 | 273883 | 345899 | 72017 | + | Gnomon | gene | NA | NA | gene-LOC118191020 | GeneID:118191020 | LOC118191020 | Gene | LOC118191020 | protein_coding | character(0) | NA | character(0) | NW_023312732.1 | 286760 | 286767 | 8 | * | 8 | 1000 |
| NW_023312741.1 | 313849 | 340725 | 26877 | - | Gnomon | gene | NA | NA | gene-LOC118195552 | GeneID:118195552 | LOC118195552 | Gene | LOC118195552 | protein_coding | character(0) | NA | character(0) | NW_023312741.1 | 315111 | 315116 | 6 | * | 6 | 1000 |
| NW_023312747.1 | 93615 | 138237 | 44623 | + | Gnomon | gene | NA | NA | gene-LOC118196990 | GeneID:118196990 | LOC118196990 | Gene | LOC118196990 | protein_coding | c("138237", ".") | true | character(0) | NW_023312747.1 | 124156 | 124161 | 6 | * | 6 | 1000 |
| NW_023312755.1 | 30608 | 136449 | 105842 | + | Gnomon | gene | NA | NA | gene-LOC118200099 | GeneID:118200099 | LOC118200099 | Gene | LOC118200099 | protein_coding | character(0) | NA | character(0) | NW_023312755.1 | 110038 | 110045 | 8 | * | 8 | 1000 |
| NW_023312755.1 | 30608 | 136449 | 105842 | + | Gnomon | gene | NA | NA | gene-LOC118200099 | GeneID:118200099 | LOC118200099 | Gene | LOC118200099 | protein_coding | character(0) | NA | character(0) | NW_023312755.1 | 114240 | 114245 | 6 | * | 6 | 1000 |
| NW_023312762.1 | 7519 | 174140 | 166622 | - | Gnomon | gene | NA | NA | gene-LOC118202388 | GeneID:118202388 | LOC118202388 | Gene | LOC118202388 | protein_coding | character(0) | NA | character(0) | NW_023312762.1 | 10182 | 10187 | 6 | * | 6 | 1000 |
| NW_023312795.1 | 107476 | 149877 | 42402 | - | Gnomon | gene | NA | NA | gene-LOC118181447 | GeneID:118181447 | LOC118181447 | Gene | LOC118181447 | protein_coding | character(0) | NA | character(0) | NW_023312795.1 | 146755 | 146760 | 6 | * | 6 | 1000 |
| NW_023312819.1 | 222905 | 284616 | 61712 | + | Gnomon | gene | NA | NA | gene-LOC118182088 | GeneID:118182088 | LOC118182088 | Gene | LOC118182088 | protein_coding | character(0) | NA | character(0) | NW_023312819.1 | 240550 | 240555 | 6 | * | 6 | 1000 |
| NW_023312824.1 | 152873 | 193878 | 41006 | - | Gnomon | gene | NA | NA | gene-LOC118182297 | GeneID:118182297 | LOC118182297 | Gene | LOC118182297 | protein_coding | character(0) | NA | character(0) | NW_023312824.1 | 162831 | 162836 | 6 | * | 6 | 1000 |
| NW_023312829.1 | 16054 | 211342 | 195289 | + | Gnomon | gene | NA | NA | gene-LOC118182373 | GeneID:118182373 | LOC118182373 | Gene | LOC118182373 | protein_coding | character(0) | NA | character(0) | NW_023312829.1 | 113486 | 113491 | 6 | * | 6 | 1000 |
| NW_023312841.1 | 3105 | 11700 | 8596 | - | Gnomon | gene | NA | NA | gene-LOC118182676 | GeneID:118182676 | LOC118182676 | Gene | LOC118182676 | lncRNA | character(0) | NA | character(0) | NW_023312841.1 | 9567 | 9574 | 8 | * | 8 | 1000 |
| NW_023312856.1 | 122962 | 162310 | 39349 | - | Gnomon | gene | NA | NA | gene-LOC118183394 | GeneID:118183394 | LOC118183394 | Gene | LOC118183394 | protein_coding | character(0) | NA | character(0) | NW_023312856.1 | 160522 | 160527 | 6 | * | 6 | 1000 |
| NW_023312859.1 | 15224 | 169817 | 154594 | - | Gnomon | gene | NA | NA | gene-LOC118183553 | GeneID:118183553 | LOC118183553 | Gene | LOC118183553 | protein_coding | character(0) | NA | character(0) | NW_023312859.1 | 82925 | 82930 | 6 | * | 6 | 1000 |
| NW_023312862.1 | 305971 | 329060 | 23090 | - | Gnomon | gene | NA | NA | gene-LOC118183641 | GeneID:118183641 | LOC118183641 | Gene | LOC118183641 | protein_coding | character(0) | NA | character(0) | NW_023312862.1 | 308115 | 308120 | 6 | * | 6 | 1000 |
| NW_023312887.1 | 16659 | 164623 | 147965 | + | Gnomon | gene | NA | NA | gene-LOC118184322 | GeneID:118184322 | LOC118184322 | Gene | LOC118184322 | protein_coding | character(0) | NA | character(0) | NW_023312887.1 | 38331 | 38336 | 6 | * | 6 | 1000 |
| NW_023312891.1 | 42448 | 174795 | 132348 | - | Gnomon | gene | NA | NA | gene-LOC118184543 | GeneID:118184543 | LOC118184543 | Gene | LOC118184543 | protein_coding | character(0) | NA | character(0) | NW_023312891.1 | 172238 | 172243 | 6 | * | 6 | 1000 |
| NW_023312896.1 | 184752 | 487505 | 302754 | - | Gnomon | gene | NA | NA | gene-LOC118184736 | GeneID:118184736 | LOC118184736 | Gene | LOC118184736 | protein_coding | character(0) | NA | character(0) | NW_023312896.1 | 232183 | 232188 | 6 | * | 6 | 1000 |
| NW_023312896.1 | 517802 | 560473 | 42672 | - | Gnomon | gene | NA | NA | gene-LOC118184778 | GeneID:118184778 | LOC118184778 | Gene | LOC118184778 | protein_coding | character(0) | NA | character(0) | NW_023312896.1 | 538217 | 538222 | 6 | * | 6 | 1000 |
| NW_023312902.1 | 2474 | 121088 | 118615 | + | Gnomon | gene | NA | NA | gene-LOC118184903 | GeneID:118184903 | LOC118184903 | Gene | LOC118184903 | protein_coding | character(0) | NA | character(0) | NW_023312902.1 | 24304 | 24309 | 6 | * | 6 | 1000 |
| NW_023312902.1 | 161458 | 227226 | 65769 | - | Gnomon | gene | NA | NA | gene-LOC118184910 | GeneID:118184910 | LOC118184910 | Gene | LOC118184910 | protein_coding | character(0) | NA | character(0) | NW_023312902.1 | 181563 | 181568 | 6 | * | 6 | 1000 |
| NW_023312919.1 | 49202 | 199856 | 150655 | - | Gnomon | gene | NA | NA | gene-LOC118185381 | GeneID:118185381 | LOC118185381 | Gene | LOC118185381 | protein_coding | character(0) | NA | character(0) | NW_023312919.1 | 58672 | 58677 | 6 | * | 6 | 1000 |
| NW_023312932.1 | 18207 | 103997 | 85791 | - | Gnomon | gene | NA | NA | gene-LOC118185808 | GeneID:118185808 | LOC118185808 | Gene | LOC118185808 | protein_coding | character(0) | NA | character(0) | NW_023312932.1 | 102399 | 102404 | 6 | * | 6 | 1000 |
| NW_023312955.1 | 59385 | 122306 | 62922 | - | Gnomon | gene | NA | NA | gene-LOC118186368 | GeneID:118186368 | LOC118186368 | Gene | LOC118186368 | protein_coding | character(0) | NA | character(0) | NW_023312955.1 | 99293 | 99298 | 6 | * | 6 | 1000 |
| NW_023312970.1 | 715 | 266769 | 266055 | + | Gnomon | gene | NA | NA | gene-LOC118186693 | GeneID:118186693 | LOC118186693 | Gene | LOC118186693 | protein_coding | character(0) | NA | character(0) | NW_023312970.1 | 250870 | 250875 | 6 | * | 6 | 1000 |

...

... only first 25 lines shown ...

```
Directory: MRPR File: rmp-Sd_LI_be.txt

Overlap of methylation-rich regions with genome features for Sd sample Sd_LI_be

Total size of the methylation-rich regions of Sd_LI_be:                                      11730 bp (  0.0% of genome)
Size of overlap of methylation-rich regions of Sd Sd_LI_be with genic regions:                   9276 bp ( 79.1% of total size)   ( 1.91 O/E)
Size of overlap of methylation-rich regions of Sd Sd_LI_be with   exon regions:                     22 bp (  0.2% of total size) (37.84 O/E)
Size of overlap of methylation-rich regions of Sd Sd_LI_be with   intron regions:                 9254 bp ( 78.9% of total size) ( 2.01 O/E)
Size of overlap of methylation-rich regions of Sd Sd_LI_be with intergenic regions:              2454 bp ( 20.9% of total size)   ( 1.35 O/E)
Size of overlap of methylation-rich regions of Sd Sd_LI_be with   promoter regions:                 18 bp (  0.1% of total size) ( 0.29 O/E)
Size of overlap of methylation-rich regions of Sd Sd_LI_be with   other intergenic regions:       2436 bp ( 20.8% of total size) ( 0.36 O/E)

Overlap of methylation-poor regions with genome features for Sd sample Sd_LI_be

Total size of the methylation-poor regions of Sd_LI_be:                                   38484235 bp (  1.5% of genome)
Size of overlap of methylation-poor regions of Sd Sd_LI_be with genic regions:               19069623 bp ( 49.5% of total size)   ( 1.20 O/E)
Size of overlap of methylation-poor regions of Sd Sd_LI_be with   exon regions:                 349072 bp (  0.9% of total size) (23.33 O/E)
Size of overlap of methylation-poor regions of Sd Sd_LI_be with   intron regions:             18720551 bp ( 48.6% of total size) ( 1.24 O/E)
Size of overlap of methylation-poor regions of Sd Sd_LI_be with intergenic regions:          19414612 bp ( 50.5% of total size)   ( 0.85 O/E)
Size of overlap of methylation-poor regions of Sd Sd_LI_be with   promoter regions:              63488 bp (  0.2% of total size) ( 0.31 O/E)
Size of overlap of methylation-poor regions of Sd Sd_LI_be with   other intergenic regions:   19351124 bp ( 50.3% of total size) ( 0.87 O/E)
```

```
Directory: MRPR File: rmp-Sd_LI_ka.txt

Overlap of methylation-rich regions with genome features for Sd sample Sd_LI_ka

Total size of the methylation-rich regions of Sd_LI_ka:                                      10743 bp (  0.0% of genome)
Size of overlap of methylation-rich regions of Sd Sd_LI_ka with genic regions:                   8562 bp ( 79.7% of total size)   ( 1.92 O/E)
Size of overlap of methylation-rich regions of Sd Sd_LI_ka with   exon regions:                    18 bp (  0.2% of total size) (38.15 O/E)
Size of overlap of methylation-rich regions of Sd Sd_LI_ka with   intron regions:                8544 bp ( 79.5% of total size) ( 2.02 O/E)
Size of overlap of methylation-rich regions of Sd Sd_LI_ka with intergenic regions:              2181 bp ( 20.3% of total size)   ( 1.36 O/E)
Size of overlap of methylation-rich regions of Sd Sd_LI_ka with   promoter regions:                36 bp (  0.3% of total size) ( 0.67 O/E)
Size of overlap of methylation-rich regions of Sd Sd_LI_ka with   other intergenic regions:      2145 bp ( 20.0% of total size) ( 0.34 O/E)

Overlap of methylation-poor regions with genome features for Sd sample Sd_LI_ka

Total size of the methylation-poor regions of Sd_LI_ka:                                   37582154 bp (  1.5% of genome)
Size of overlap of methylation-poor regions of Sd Sd_LI_ka with genic regions:               17522413 bp ( 46.6% of total size)   ( 1.13 O/E)
Size of overlap of methylation-poor regions of Sd Sd_LI_ka with   exon regions:                321708 bp (  0.9% of total size) (21.96 O/E)
Size of overlap of methylation-poor regions of Sd Sd_LI_ka with   intron regions:            17200705 bp ( 45.8% of total size) ( 1.16 O/E)
Size of overlap of methylation-poor regions of Sd Sd_LI_ka with intergenic regions:          20059741 bp ( 53.4% of total size)   ( 0.80 O/E)
Size of overlap of methylation-poor regions of Sd Sd_LI_ka with   promoter regions:             58617 bp (  0.2% of total size) ( 0.31 O/E)
Size of overlap of methylation-poor regions of Sd Sd_LI_ka with   other intergenic regions:  20001124 bp ( 53.2% of total size) ( 0.92 O/E)
```

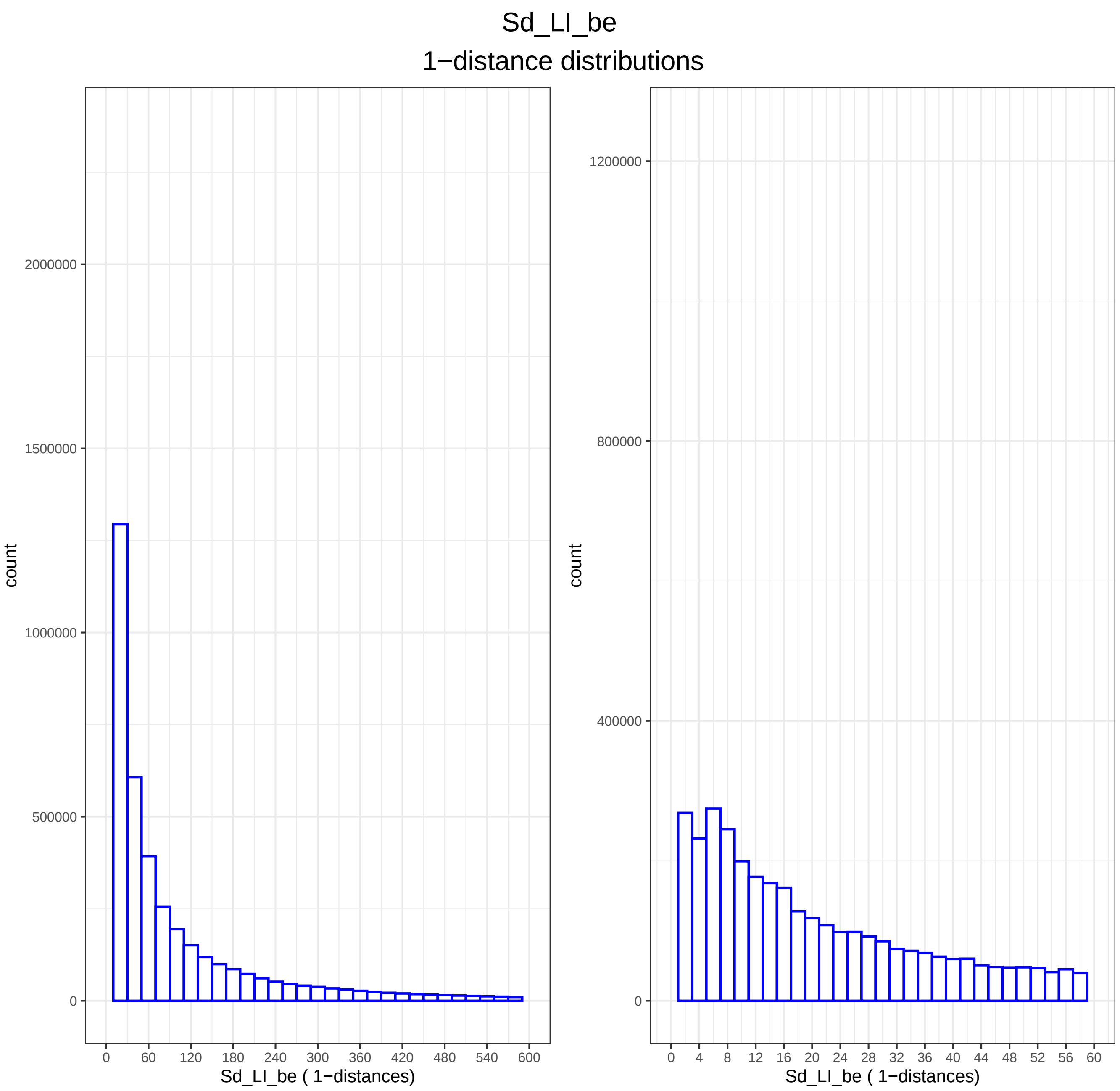

Sd_LI_be
1−distance distributions
count
2000000
1500000
1000000
500000
0
0
60
120
180
240
300
360
420
480
540
600
Sd_LI_be ( 1−distances)
count
1200000
800000
400000
0
0
4
8
12
16
20
24
28
32
36
40
44
48
52
56
60
Sd_LI_be ( 1−distances)

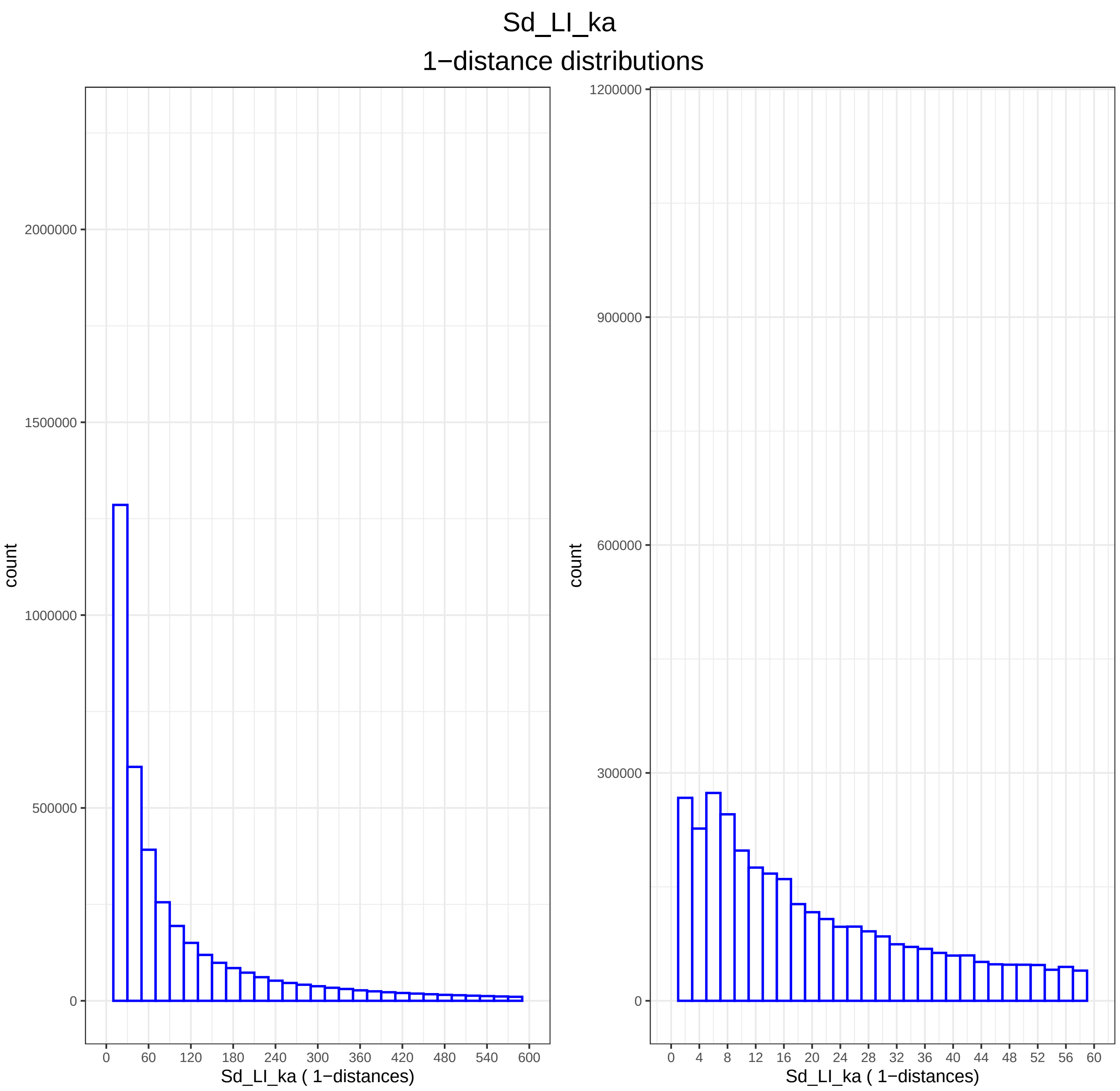

Sd_LI_ka
1−distance distributions
count
2000000
1500000
1000000
500000
0
0
60
120
180
240
300
360
420
480
540
600
Sd_LI_ka ( 1−distances)
count
1200000
900000
600000
300000
0
0
4
8
12
16
20
24
28
32
36
40
44
48
52
56
60
Sd_LI_ka ( 1−distances)

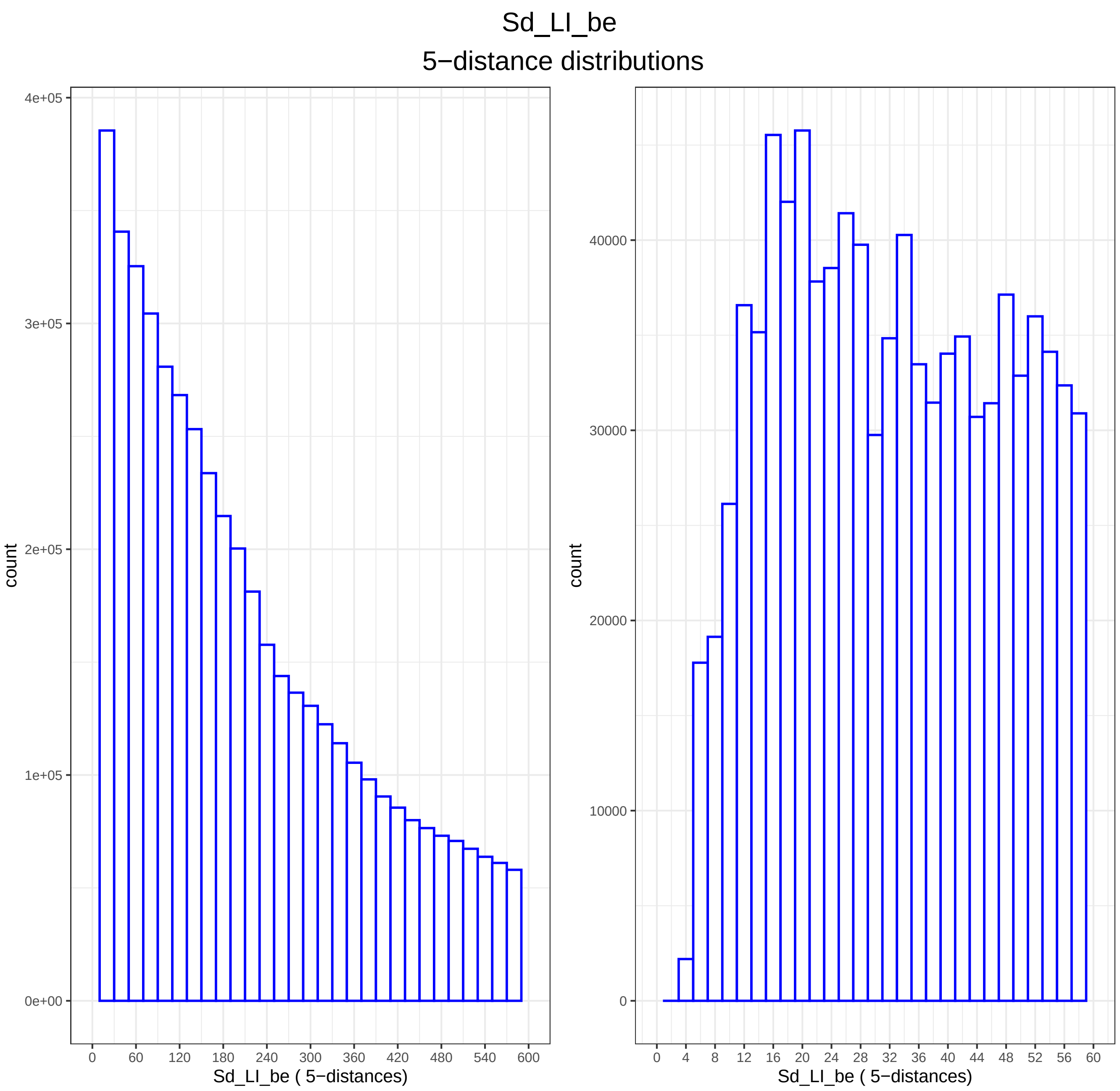

Sd_LI_be
5−distance distributions
count
4e+05
3e+05
2e+05
1e+05
0e+00
0
60
120
180
240
300
360
420
480
540
600
Sd_LI_be ( 5−distances)
count
40000
30000
20000
10000
0
0
4
8
12
16
20
24
28
32
36
40
44
48
52
56
60
Sd_LI_be ( 5−distances)

# Sd_LI_ka
#### 5−distance distributions

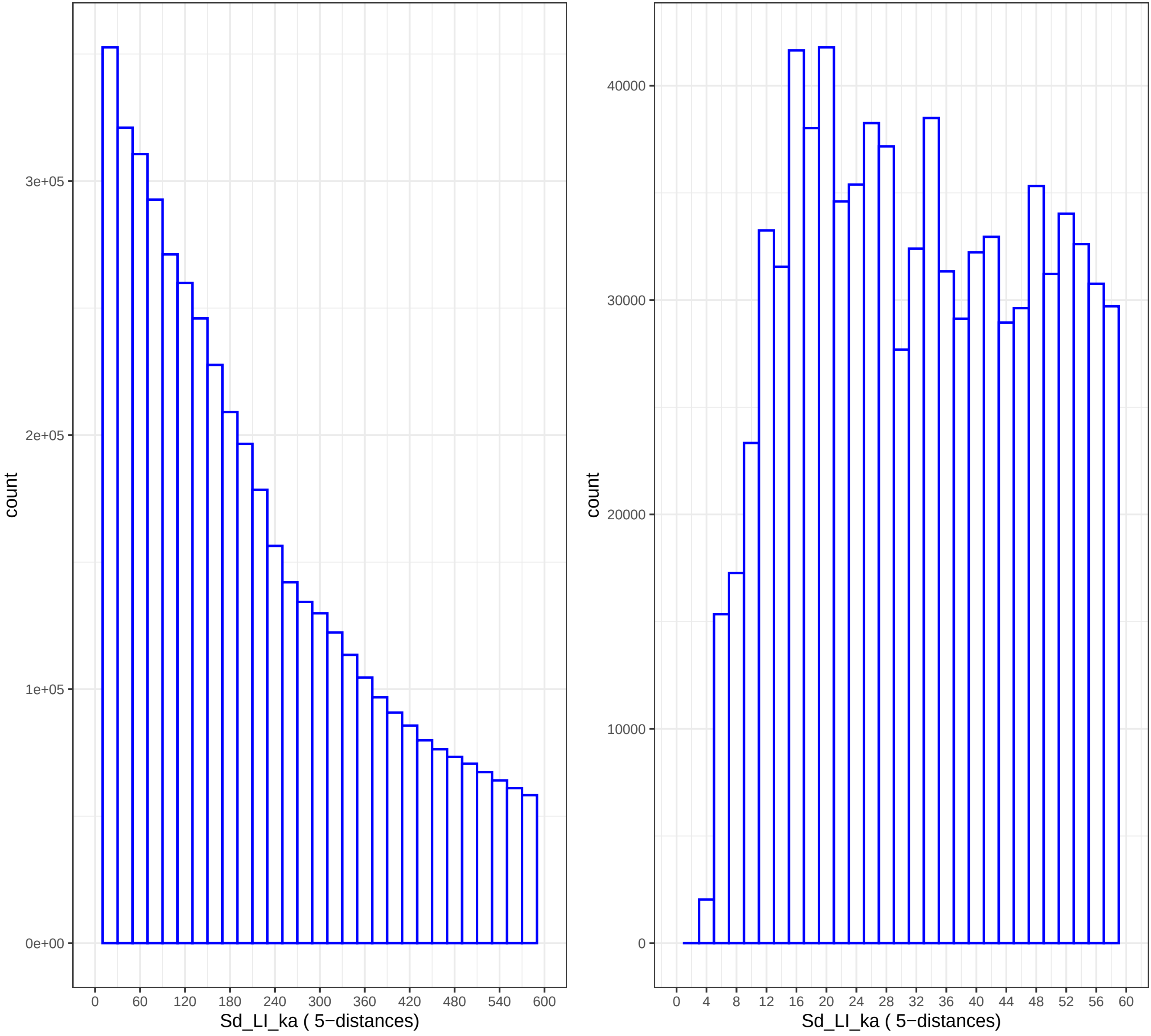

```
Directory: DMT File: 0READMEdmt

DMT - Differentially methylated tiles.

Input: studymc wsize stepsize threshold qvalue
Output: files dmt-*.txt dmg-*.txt

Notes: Output files dmt-*.txt and dmg-*.txt are generated by BWASPR::det_dmt() and show the
        differentially methylated tiles and genes as determined by methylKit::getMethylDiff with
        parameters difference=threshold and qvalue, as provided in the Sd_LI.conf
        configuration file.  Tiles refer to sliding windows along the genome within which
        methylation calls are cumulated by methylKit::tileMethylCounts.
        The dmg-*.txt files show all genes with at least one differentially methylated tile.

        A positive meth.diff value in comparison A.vs.B means that the B methylation percentage
        in that tile is higher than the A methylation percentage.
```

Directory: DMT File: dmg-Sd_LI.txt

| seqnames | start | end | width | strand | source | type | score | phase | ID | Dbxref | Name | gbkey | gene | gene_biotype | end_range | partial | start_range | pvalue | qvalue | meth.diff | comparison |
|---|---|---|---|---|---|---|---|---|---|---|---|---|---|---|---|---|---|---|---|---|---|
| NW_023312695.1 | 26038 | 120945 | 94908 | + | Gnomon | gene | NA | NA | gene-LOC118192109 | GeneID:118192109 | LOC118192109 | Gene | LOC118192109 | protein_coding | character(0) | NA | character(0) | 2.58638071673744e-34 | 2.36325704856201e-31 | -25.1711430798668 | be.vs.ka |
| NW_023312697.1 | 212878 | 236502 | 23625 | - | Gnomon | gene | NA | NA | gene-LOC118200926 | GeneID:118200926 | LOC118200926 | Gene | LOC118200926 | protein_coding | character(0) | NA | character(0) | 5.56681453621249e-05 | 0.00365896297965675 | -45.2275249722531 | be.vs.ka |
| NW_023312702.1 | 55813 | 105158 | 49346 | - | Gnomon | gene | NA | NA | gene-LOC118181825 | GeneID:118181825 | LOC118181825 | Gene | LOC118181825 | protein_coding | character(0) | NA | character(0) | 1.25182571392672e-05 | 0.00102956762943983 | -28.844169246646 | be.vs.ka |
| NW_023312706.1 | 36659 | 169459 | 132801 | - | Gnomon | gene | NA | NA | gene-LOC118182927 | GeneID:118182927 | LOC118182927 | Gene | LOC118182927 | protein_coding | character(0) | NA | character(0) | 3.39865743961762e-07 | 4.27414868516216e-05 | -26.176552492342 | be.vs.ka |
| NW_023312711.1 | 52293 | 291451 | 239159 | - | Gnomon | gene | NA | NA | gene-LOC118184206 | GeneID:118184206 | LOC118184206 | Gene | LOC118184206 | protein_coding | character(0) | NA | character(0) | 4.10908461913198e-06 | 0.000392279991422274 | 43.9325113562622 | be.vs.ka |
| NW_023312715.1 | 80286 | 109142 | 28857 | - | Gnomon | gene | NA | NA | gene-LOC118184790 | GeneID:118184790 | LOC118184790 | Gene | LOC118184790 | protein_coding | character(0) | NA | character(0) | 2.32445282522517e-05 | 0.00174855260122226 | 44.2857142857143 | be.vs.ka |
| NW_023312716.1 | 14006 | 270689 | 256684 | - | Gnomon | gene | NA | NA | gene-LOC118184954 | GeneID:118184954 | LOC118184954 | Gene | LOC118184954 | protein_coding | character(0) | NA | character(0) | 1.39074376493336e-55 | 2.82166919476171e-52 | 62.3662467831505 | be.vs.ka |
| NW_023312717.1 | 4200 | 221539 | 217340 | + | Gnomon | gene | NA | NA | gene-LOC118185112 | GeneID:118185112 | LOC118185112 | Gene | LOC118185112 | protein_coding | character(0) | true | c(".", "4200") | 4.21079873293984e-05 | 0.0028917912464893 | -39.1812865497076 | be.vs.ka |
| NW_023312729.1 | 74346 | 86709 | 12364 | + | Gnomon | gene | NA | NA | gene-LOC118189244 | GeneID:118189244 | LOC118189244 | Gene | LOC118189244 | protein_coding | character(0) | NA | character(0) | 0.000117205813408345 | 0.00677759198844193 | 60 | be.vs.ka |
| NW_023312731.1 | 35124 | 211977 | 176854 | - | Gnomon | gene | NA | NA | gene-LOC118189879 | GeneID:118189879 | LOC118189879 | Gene | LOC118189879 | protein_coding | character(0) | NA | character(0) | 1.0791322511233e-17 | 4.3695791640291e-15 | 29.2091648189209 | be.vs.ka |
| NW_023312731.1 | 258434 | 297820 | 39387 | - | Gnomon | gene | NA | NA | gene-LOC118190087 | GeneID:118190087 | LOC118190087 | Gene | LOC118190087 | protein_coding | character(0) | NA | character(0) | 2.46814631646951e-05 | 0.00184130440864212 | 35.5409050044951 | be.vs.ka |
| NW_023312731.1 | 298073 | 401198 | 103126 | + | Gnomon | gene | NA | NA | gene-LOC118190182 | GeneID:118190182 | LOC118190182 | Gene | LOC118190182 | protein_coding | character(0) | NA | character(0) | 0.000123293574997937 | 0.00706858891078224 | 30.7100859339665 | be.vs.ka |
| NW_023312737.1 | 106024 | 312214 | 206191 | - | Gnomon | gene | NA | NA | gene-LOC118193316 | GeneID:118193316 | LOC118193316 | Gene | LOC118193316 | protein_coding | character(0) | NA | character(0) | 4.00255007582718e-17 | 1.56289759724019e-14 | -34.9665425427667 | be.vs.ka |
| NW_023312738.1 | 32075 | 193979 | 161905 | - | Gnomon | gene | NA | NA | gene-LOC118193802 | GeneID:118193802 | LOC118193802 | Gene | LOC118193802 | protein_coding | character(0) | NA | character(0) | 2.82940285912634e-06 | 0.000283018989227712 | 53.7829912023461 | be.vs.ka |
| NW_023312740.1 | 570747 | 591326 | 20580 | + | Gnomon | gene | NA | NA | gene-LOC118194662 | GeneID:118194662 | LOC118194662 | Gene | LOC118194662 | protein_coding | character(0) | NA | character(0) | 6.75470888399472e-13 | 1.89439085872926e-10 | 27.1832884097035 | be.vs.ka |
| NW_023312741.1 | 288098 | 308610 | 20513 | + | Gnomon | gene | NA | NA | gene-LOC118195469 | GeneID:118195469 | LOC118195469 | Gene | LOC118195469 | protein_coding | character(0) | NA | character(0) | 4.95795584678161e-18 | 2.05293400797467e-15 | 47.50452079566 | be.vs.ka |
| NW_023312741.1 | 313849 | 340725 | 26877 | - | Gnomon | gene | NA | NA | gene-LOC118195552 | GeneID:118195552 | LOC118195552 | Gene | LOC118195552 | protein_coding | character(0) | NA | character(0) | 8.93270080748674e-07 | 0.000101751595782074 | -29.7076023391813 | be.vs.ka |
| NW_023312743.1 | 34674 | 75652 | 40979 | + | Gnomon | gene | NA | NA | gene-LOC118195734 | GeneID:118195734 | LOC118195734 | Gene | LOC118195734 | protein_coding | character(0) | NA | character(0) | 8.779439747514e-14 | 2.65323162815292e-11 | 73.5632183908046 | be.vs.ka |
| NW_023312746.1 | 159915 | 234902 | 74988 | + | Gnomon | gene | NA | NA | gene-LOC118196460 | GeneID:118196460 | LOC118196460 | Gene | LOC118196460 | protein_coding | character(0) | NA | character(0) | 2.15749632648472e-07 | 2.82748195557894e-05 | -28.5828877005348 | be.vs.ka |
| NW_023312748.1 | 218434 | 275732 | 57299 | - | Gnomon | gene | NA | NA | gene-LOC118197128 | GeneID:118197128 | LOC118197128 | Gene | LOC118197128 | protein_coding | character(0) | NA | character(0) | 3.03838713419943e-05 | 0.00219603241263743 | -30.7353150334234 | be.vs.ka |
| NW_023312751.1 | 646 | 91614 | 90969 | - | Gnomon | gene | NA | NA | gene-LOC118198557 | GeneID:118198557 | LOC118198557 | Gene | LOC118198557 | protein_coding | character(0) | NA | character(0) | 1.13297636006132e-10 | 2.50170029266958e-08 | -25.187969924812 | be.vs.ka |
| NW_023312754.1 | 23651 | 82082 | 58432 | - | Gnomon | gene | NA | NA | gene-LOC118199805 | GeneID:118199805 | LOC118199805 | Gene | LOC118199805 | lncRNA | character(0) | NA | character(0) | 4.38259247840242e-06 | 0.00041500111728903 | 32.7931172468988 | be.vs.ka |
| NW_023312758.1 | 2535 | 177946 | 175412 | - | Gnomon | gene | NA | NA | gene-LOC118200694 | GeneID:118200694 | LOC118200694 | Gene | LOC118200694 | protein_coding | character(0) | NA | character(0) | 1.74288699123387e-21 | 8.80516023311083e-19 | -43.5780321373542 | be.vs.ka |
| NW_023312758.1 | 197896 | 320783 | 122888 | - | Gnomon | gene | NA | NA | gene-LOC118200784 | GeneID:118200784 | LOC118200784 | Gene | LOC118200784 | protein_coding | character(0) | NA | character(0) | 2.74692627172902e-07 | 3.52621188961511e-05 | 60 | be.vs.ka |

...

... only first 25 lines shown ...

Directory: DMT File: dmt-Sd_LI.txt

| seqnames | start | end | width | strand | pvalue | qvalue | meth.diff | comparison |
|---|---|---|---|---|---|---|---|---|
| NW_023312690.1 | 341001 | 342000 | 1000 | * | 1.40374091451862e-13 | 0 | 29.84 | be.vs.ka |
| NW_023312690.1 | 484001 | 485000 | 1000 | * | 5.54744602524609e-05 | 0.004 | 37.72 | be.vs.ka |
| NW_023312691.1 | 47001 | 48000 | 1000 | * | 7.26545752836094e-16 | 0 | 46.46 | be.vs.ka |
| NW_023312692.1 | 35001 | 36000 | 1000 | * | 2.5765308254238e-13 | 0 | -34.12 | be.vs.ka |
| NW_023312692.1 | 37001 | 38000 | 1000 | * | 8.04945416335198e-09 | 0 | -38.63 | be.vs.ka |
| NW_023312692.1 | 42001 | 43000 | 1000 | * | 2.64574452312348e-07 | 0 | -35.65 | be.vs.ka |
| NW_023312694.1 | 355001 | 356000 | 1000 | * | 4.57118880640084e-05 | 0.003 | 27.14 | be.vs.ka |
| NW_023312695.1 | 27001 | 28000 | 1000 | * | 1.28927986458431e-05 | 0.001 | 32.59 | be.vs.ka |
| NW_023312695.1 | 42001 | 43000 | 1000 | * | 2.29616850246962e-12 | 0 | 33.67 | be.vs.ka |
| NW_023312695.1 | 43001 | 44000 | 1000 | * | 6.62009099697782e-10 | 0 | 36.25 | be.vs.ka |
| NW_023312695.1 | 71001 | 72000 | 1000 | * | 2.03151033227891e-45 | 0 | -58.38 | be.vs.ka |
| NW_023312695.1 | 72001 | 73000 | 1000 | * | 9.31337677836636e-33 | 0 | -62.7 | be.vs.ka |
| NW_023312695.1 | 73001 | 74000 | 1000 | * | 5.88390493568282e-36 | 0 | -49.92 | be.vs.ka |
| NW_023312695.1 | 78001 | 79000 | 1000 | * | 2.58638071673744e-34 | 0 | -25.17 | be.vs.ka |
| NW_023312696.1 | 55001 | 56000 | 1000 | * | 2.62861475386782e-05 | 0.002 | 40.62 | be.vs.ka |
| NW_023312697.1 | 210001 | 211000 | 1000 | * | 1.7325033588975e-27 | 0 | -69.38 | be.vs.ka |
| NW_023312697.1 | 211001 | 212000 | 1000 | * | 4.21046365036816e-07 | 0 | -26.21 | be.vs.ka |
| NW_023312697.1 | 218001 | 219000 | 1000 | * | 5.56681453621249e-05 | 0.004 | -45.23 | be.vs.ka |
| NW_023312697.1 | 227001 | 228000 | 1000 | * | 5.47856013369668e-08 | 0 | -46.6 | be.vs.ka |
| NW_023312698.1 | 46001 | 47000 | 1000 | * | 3.95711753578223e-08 | 0 | -26.66 | be.vs.ka |
| NW_023312702.1 | 100001 | 101000 | 1000 | * | 1.25182571392672e-05 | 0.001 | -28.84 | be.vs.ka |
| NW_023312706.1 | 152001 | 153000 | 1000 | * | 3.39865743961762e-07 | 0 | -26.18 | be.vs.ka |
| NW_023312707.1 | 190001 | 191000 | 1000 | * | 9.01799682401047e-19 | 0 | 45.02 | be.vs.ka |
| NW_023312707.1 | 375001 | 376000 | 1000 | * | 9.0005750174007e-07 | 0 | -51.83 | be.vs.ka |

...

... only first 25 lines shown ...

```
Directory: DMSG File: 0READMEdmsg

DMSG - Differentially methylated sites and genes.

Input: studyhc threshold qvalue
       The studyhc methylKit raw object contains all CpGscd sites with coverage at least
        20 reads.
Output: files dms-*.txt dmg-*.txt  dmg-*details.txt dmg-*heatmaps.pdf

Notes: Output files dms-*.txt and dmg-*.txt are generated by BWASPR::det_dmsg() and show the
        differentially methylated sites and genes as determined by methylKit::getMethylDiff with
        parameters difference=threshold and qvalue, as provided in the Sd_LI.conf
        configuration file.  The table of differentially methylated genes contains all genes
        with at least one differentially methylated site.

       The files dmg-*details.txt, generated by BWASPR::show_dmsg(), show all CpGscd sites in
        the differentially methylated genes, with coverage numbers and methylation percentages.

       The files dmg-*heatmaps.pdf, generated by BWASPR::show_dmsg(), give heatmap displays for
        genes meeting the following criteria: 1) there are between minNsites and maxNsites common
        CpGscd sites; 2) at least minPdmsites % of these sites are differentially methylated sites.
        The parameters are set in the Sd_LI.conf configuration file.
```

Directory: DMSG File: dmg-Sd_LI.txt

| seqnames | start | end | width | strand | source | type | score | phase | ID | Dbxref | Name | gbkey | gene | gene_biotype | end_range | partial | start_range | comparison |
|---|---|---|---|---|---|---|---|---|---|---|---|---|---|---|---|---|---|---|
| NW_023312695.1 | 26038 | 120945 | 94908 | + | Gnomon | gene | NA | NA | gene-LOC118192109 | GeneID:118192109 | LOC118192109 | Gene | LOC118192109 | protein_coding | character(0) | NA | character(0) | be.vs.ka |
| NW_023312704.1 | 57274 | 267424 | 210151 | + | Gnomon | gene | NA | NA | gene-LOC118182836 | GeneID:118182836 | LOC118182836 | Gene | LOC118182836 | protein_coding | character(0) | NA | character(0) | be.vs.ka |
| NW_023312716.1 | 14006 | 270689 | 256684 | - | Gnomon | gene | NA | NA | gene-LOC118184954 | GeneID:118184954 | LOC118184954 | Gene | LOC118184954 | protein_coding | character(0) | NA | character(0) | be.vs.ka |
| NW_023312726.1 | 201949 | 220759 | 18811 | - | Gnomon | gene | NA | NA | gene-LOC118187715 | GeneID:118187715 | LOC118187715 | Gene | LOC118187715 | protein_coding | character(0) | NA | character(0) | be.vs.ka |
| NW_023312731.1 | 298073 | 401198 | 103126 | + | Gnomon | gene | NA | NA | gene-LOC118190182 | GeneID:118190182 | LOC118190182 | Gene | LOC118190182 | protein_coding | character(0) | NA | character(0) | be.vs.ka |
| NW_023312735.1 | 22348 | 53922 | 31575 | - | Gnomon | gene | NA | NA | gene-LOC118191751 | GeneID:118191751 | LOC118191751 | Gene | LOC118191751 | protein_coding | character(0) | NA | character(0) | be.vs.ka |
| NW_023312735.1 | 53920 | 74196 | 20277 | + | Gnomon | gene | NA | NA | gene-LOC118191868 | GeneID:118191868 | LOC118191868 | Gene | LOC118191868 | protein_coding | character(0) | NA | character(0) | be.vs.ka |
| NW_023312741.1 | 288098 | 308610 | 20513 | + | Gnomon | gene | NA | NA | gene-LOC118195469 | GeneID:118195469 | LOC118195469 | Gene | LOC118195469 | protein_coding | character(0) | NA | character(0) | be.vs.ka |
| NW_023312765.1 | 208442 | 325427 | 116986 | - | Gnomon | gene | NA | NA | gene-LOC118202919 | GeneID:118202919 | LOC118202919 | Gene | LOC118202919 | protein_coding | character(0) | NA | character(0) | be.vs.ka |
| NW_023312767.1 | 33640 | 110297 | 76658 | - | Gnomon | gene | NA | NA | gene-LOC118203309 | GeneID:118203309 | LOC118203309 | Gene | LOC118203309 | protein_coding | character(0) | NA | character(0) | be.vs.ka |
| NW_023312768.1 | 44538 | 60365 | 15828 | + | Gnomon | gene | NA | NA | gene-LOC118203894 | GeneID:118203894 | LOC118203894 | Gene | LOC118203894 | lncRNA | character(0) | NA | character(0) | be.vs.ka |
| NW_023312776.1 | 18093 | 148391 | 130299 | + | Gnomon | gene | NA | NA | gene-LOC118205972 | GeneID:118205972 | LOC118205972 | Gene | LOC118205972 | protein_coding | character(0) | NA | character(0) | be.vs.ka |
| NW_023312782.1 | 31970 | 88461 | 56492 | - | Gnomon | gene | NA | NA | gene-LOC118180870 | GeneID:118180870 | LOC118180870 | Gene | LOC118180870 | protein_coding | character(0) | NA | character(0) | be.vs.ka |
| NW_023312813.1 | 38309 | 115925 | 77617 | - | Gnomon | gene | NA | NA | gene-LOC118181861 | GeneID:118181861 | LOC118181861 | Gene | LOC118181861 | protein_coding | character(0) | NA | character(0) | be.vs.ka |
| NW_023312816.1 | 115848 | 659441 | 543594 | - | Gnomon | gene | NA | NA | gene-LOC118181985 | GeneID:118181985 | LOC118181985 | Gene | LOC118181985 | protein_coding | character(0) | NA | character(0) | be.vs.ka |
| NW_023312817.1 | 13538 | 81112 | 67575 | + | Gnomon | gene | NA | NA | gene-LOC118181999 | GeneID:118181999 | LOC118181999 | Gene | LOC118181999 | protein_coding | character(0) | NA | character(0) | be.vs.ka |
| NW_023312842.1 | 46810 | 130248 | 83439 | - | Gnomon | gene | NA | NA | gene-LOC118182709 | GeneID:118182709 | LOC118182709 | Gene | LOC118182709 | protein_coding | character(0) | NA | character(0) | be.vs.ka |
| NW_023312846.1 | 59888 | 94725 | 34838 | + | Gnomon | gene | NA | NA | gene-LOC118182862 | GeneID:118182862 | LOC118182862 | Gene | LOC118182862 | protein_coding | character(0) | NA | character(0) | be.vs.ka |
| NW_023312846.1 | 121760 | 204155 | 82396 | + | Gnomon | gene | NA | NA | gene-LOC118182870 | GeneID:118182870 | LOC118182870 | Gene | LOC118182870 | protein_coding | character(0) | NA | character(0) | be.vs.ka |
| NW_023312858.1 | 110039 | 221517 | 111479 | + | Gnomon | gene | NA | NA | gene-LOC118183482 | GeneID:118183482 | LOC118183482 | Gene | LOC118183482 | protein_coding | character(0) | NA | character(0) | be.vs.ka |
| NW_023312858.1 | 267070 | 363467 | 96398 | + | Gnomon | gene | NA | NA | gene-LOC118183504 | GeneID:118183504 | LOC118183504 | Gene | LOC118183504 | protein_coding | character(0) | NA | character(0) | be.vs.ka |
| NW_023312863.1 | 59464 | 282910 | 223447 | + | Gnomon | gene | NA | NA | gene-LOC118183667 | GeneID:118183667 | LOC118183667 | Gene | LOC118183667 | protein_coding | c("282910", ".") | true | character(0) | be.vs.ka |
| NW_023312867.1 | 155630 | 180473 | 24844 | + | Gnomon | gene | NA | NA | gene-LOC118183868 | GeneID:118183868 | LOC118183868 | Gene | LOC118183868 | protein_coding | character(0) | NA | character(0) | be.vs.ka |
| NW_023312872.1 | 6786 | 118481 | 111696 | + | Gnomon | gene | NA | NA | gene-LOC118184060 | GeneID:118184060 | LOC118184060 | Gene | LOC118184060 | protein_coding | character(0) | true | c(".", "6786") | be.vs.ka |

...
... only first 25 lines shown ...

Directory: DMSG File: dmg-Sd_LI_be.vs.ka_details.txt

```
seqnames    start end   width strand      coverage1   numCs1      numTs1      coverage2   numCs2      numTs2      be    ka    gene_seqnames     gene_start  gene_end    gene_width
      gene_strand gene_source gene_type   gene_ID     gene_Dbxref gene_Name   gene_gbkey  gene_gene   gene_gene_biotype gene_end_range    gene_partial      gene_start_range
      gene_comparison   is.dm
NW_023312695.1    26415 26415 1     +     27    0     27    21    1     20    0     4.76  NW_023312695.1    26038 120945      94908 +     Gnomon      gene  gene-LOC118192109
      GeneID:118192109  LOC118192109      Gene  LOC118192109      protein_coding    character(0)      NA    character(0)      be.vs.ka    FALSE
NW_023312695.1    26426 26426 1     +     28    0     28    25    0     25    0     0     NW_023312695.1    26038 120945      94908 +     Gnomon      gene  gene-LOC118192109
      GeneID:118192109  LOC118192109      Gene  LOC118192109      protein_coding    character(0)      NA    character(0)      be.vs.ka    FALSE
NW_023312695.1    26442 26442 1     +     34    0     34    27    1     26    0     3.7   NW_023312695.1    26038 120945      94908 +     Gnomon      gene  gene-LOC118192109
      GeneID:118192109  LOC118192109      Gene  LOC118192109      protein_coding    character(0)      NA    character(0)      be.vs.ka    FALSE
NW_023312695.1    26474 26474 1     +     38    3     35    34    2     32    7.89  5.88  NW_023312695.1    26038 120945      94908 +     Gnomon      gene  gene-LOC118192109
      GeneID:118192109  LOC118192109      Gene  LOC118192109      protein_coding    character(0)      NA    character(0)      be.vs.ka    FALSE
NW_023312695.1    26511 26511 1     +     35    2     33    35    0     35    5.71  0     NW_023312695.1    26038 120945      94908 +     Gnomon      gene  gene-LOC118192109
      GeneID:118192109  LOC118192109      Gene  LOC118192109      protein_coding    character(0)      NA    character(0)      be.vs.ka    FALSE
NW_023312695.1    31133 31133 1     +     27    15    12    31    15    16    55.56 48.39 NW_023312695.1    26038 120945      94908 +     Gnomon      gene  gene-LOC118192109
      GeneID:118192109  LOC118192109      Gene  LOC118192109      protein_coding    character(0)      NA    character(0)      be.vs.ka    FALSE
NW_023312695.1    31151 31151 1     +     23    14    9     31    16    15    60.87 51.61 NW_023312695.1    26038 120945      94908 +     Gnomon      gene  gene-LOC118192109
      GeneID:118192109  LOC118192109      Gene  LOC118192109      protein_coding    character(0)      NA    character(0)      be.vs.ka    FALSE
NW_023312695.1    31180 31180 1     +     20    10    10    30    18    12    50    60    NW_023312695.1    26038 120945      94908 +     Gnomon      gene  gene-LOC118192109
      GeneID:118192109  LOC118192109      Gene  LOC118192109      protein_coding    character(0)      NA    character(0)      be.vs.ka    FALSE
NW_023312695.1    31182 31182 1     +     21    13    8     30    15    15    61.9  50    NW_023312695.1    26038 120945      94908 +     Gnomon      gene  gene-LOC118192109
      GeneID:118192109  LOC118192109      Gene  LOC118192109      protein_coding    character(0)      NA    character(0)      be.vs.ka    FALSE
NW_023312695.1    31192 31192 1     +     20    15    5     26    16    10    75    61.54 NW_023312695.1    26038 120945      94908 +     Gnomon      gene  gene-LOC118192109
      GeneID:118192109  LOC118192109      Gene  LOC118192109      protein_coding    character(0)      NA    character(0)      be.vs.ka    FALSE
NW_023312695.1    33933 33933 1     +     25    24    1     20    20    0     96    100   NW_023312695.1    26038 120945      94908 +     Gnomon      gene  gene-LOC118192109
      GeneID:118192109  LOC118192109      Gene  LOC118192109      protein_coding    character(0)      NA    character(0)      be.vs.ka    FALSE
NW_023312695.1    39544 39544 1     +     20    8     12    56    11    45    40    19.64 NW_023312695.1    26038 120945      94908 +     Gnomon      gene  gene-LOC118192109
      GeneID:118192109  LOC118192109      Gene  LOC118192109      protein_coding    character(0)      NA    character(0)      be.vs.ka    FALSE
NW_023312695.1    39561 39561 1     +     22    8     14    63    12    51    36.36 19.05 NW_023312695.1    26038 120945      94908 +     Gnomon      gene  gene-LOC118192109
      GeneID:118192109  LOC118192109      Gene  LOC118192109      protein_coding    character(0)      NA    character(0)      be.vs.ka    FALSE
NW_023312695.1    39576 39576 1     +     28    10    18    70    12    58    35.71 17.14 NW_023312695.1    26038 120945      94908 +     Gnomon      gene  gene-LOC118192109
      GeneID:118192109  LOC118192109      Gene  LOC118192109      protein_coding    character(0)      NA    character(0)      be.vs.ka    FALSE
NW_023312695.1    39579 39579 1     +     30    10    20    69    9     60    33.33 13.04 NW_023312695.1    26038 120945      94908 +     Gnomon      gene  gene-LOC118192109
      GeneID:118192109  LOC118192109      Gene  LOC118192109      protein_coding    character(0)      NA    character(0)      be.vs.ka    FALSE
NW_023312695.1    39583 39583 1     +     32    7     25    70    10    60    21.88 14.29 NW_023312695.1    26038 120945      94908 +     Gnomon      gene  gene-LOC118192109
      GeneID:118192109  LOC118192109      Gene  LOC118192109      protein_coding    character(0)      NA    character(0)      be.vs.ka    FALSE
NW_023312695.1    39594 39594 1     +     36    8     28    82    5     77    22.22 6.1   NW_023312695.1    26038 120945      94908 +     Gnomon      gene  gene-LOC118192109
      GeneID:118192109  LOC118192109      Gene  LOC118192109      protein_coding    character(0)      NA    character(0)      be.vs.ka    FALSE
NW_023312695.1    39711 39711 1     +     75    1     74    169   5     164   1.33  2.96  NW_023312695.1    26038 120945      94908 +     Gnomon      gene  gene-LOC118192109
      GeneID:118192109  LOC118192109      Gene  LOC118192109      protein_coding    character(0)      NA    character(0)      be.vs.ka    FALSE
NW_023312695.1    39727 39727 1     +     74    1     73    175   4     171   1.35  2.29  NW_023312695.1    26038 120945      94908 +     Gnomon      gene  gene-LOC118192109
      GeneID:118192109  LOC118192109      Gene  LOC118192109      protein_coding    character(0)      NA    character(0)      be.vs.ka    FALSE
NW_023312695.1    39741 39741 1     +     77    1     76    170   3     167   1.3   1.76  NW_023312695.1    26038 120945      94908 +     Gnomon      gene  gene-LOC118192109
      GeneID:118192109  LOC118192109      Gene  LOC118192109      protein_coding    character(0)      NA    character(0)      be.vs.ka    FALSE
NW_023312695.1    39754 39754 1     +     78    1     77    173   5     168   1.28  2.89  NW_023312695.1    26038 120945      94908 +     Gnomon      gene  gene-LOC118192109
      GeneID:118192109  LOC118192109      Gene  LOC118192109      protein_coding    character(0)      NA    character(0)      be.vs.ka    FALSE
NW_023312695.1    39800 39800 1     +     54    1     53    120   0     120   1.85  0     NW_023312695.1    26038 120945      94908 +     Gnomon      gene  gene-LOC118192109
      GeneID:118192109  LOC118192109      Gene  LOC118192109      protein_coding    character(0)      NA    character(0)      be.vs.ka    FALSE
NW_023312695.1    40053 40053 1     +     23    10    13    21    5     16    43.48 23.81 NW_023312695.1    26038 120945      94908 +     Gnomon      gene  gene-LOC118192109
      GeneID:118192109  LOC118192109      Gene  LOC118192109      protein_coding    character(0)      NA    character(0)      be.vs.ka    FALSE
NW_023312695.1    40065 40065 1     +     25    11    14    23    8     15    44    34.78 NW_023312695.1    26038 120945      94908 +     Gnomon      gene  gene-LOC118192109
      GeneID:118192109  LOC118192109      Gene  LOC118192109      protein_coding    character(0)      NA    character(0)      be.vs.ka    FALSE
...
... only first 25 lines shown ...
```

Directory: DMSG File: dms-Sd_LI.txt

| seqnames | start | end | width | strand | pvalue | qvalue | meth.diff | comparison |
|---|---|---|---|---|---|---|---|---|
| NW_023312695.1 | 72587 | 72587 | 1 | + | 2.61325069204052e-07 | 0 | -67.46 | be.vs.ka |
| NW_023312695.1 | 73542 | 73542 | 1 | + | 1.00211446151379e-05 | 0.002 | -70 | be.vs.ka |
| NW_023312695.1 | 73580 | 73580 | 1 | + | 7.67188819458404e-06 | 0.001 | -66.07 | be.vs.ka |
| NW_023312695.1 | 78288 | 78288 | 1 | + | 6.30449151534586e-05 | 0.009 | -50.66 | be.vs.ka |
| NW_023312695.1 | 78323 | 78323 | 1 | + | 4.19775293189531e-05 | 0.006 | -48.63 | be.vs.ka |
| NW_023312699.1 | 128230 | 128230 | 1 | + | 2.25395039523947e-05 | 0.004 | -42.47 | be.vs.ka |
| NW_023312704.1 | 258528 | 258528 | 1 | + | 7.18795680606445e-05 | 0.01 | -37.14 | be.vs.ka |
| NW_023312716.1 | 120406 | 120406 | 1 | + | 2.47059647673857e-05 | 0.004 | 38.1 | be.vs.ka |
| NW_023312716.1 | 120486 | 120486 | 1 | + | 9.12072863812676e-09 | 0 | 57.69 | be.vs.ka |
| NW_023312716.1 | 120488 | 120488 | 1 | + | 3.28478811295812e-08 | 0 | 53.85 | be.vs.ka |
| NW_023312726.1 | 216077 | 216077 | 1 | + | 2.01403473130494e-07 | 0 | -43.81 | be.vs.ka |
| NW_023312731.1 | 332557 | 332557 | 1 | + | 1.10445945750167e-06 | 0 | 75.71 | be.vs.ka |
| NW_023312732.1 | 349722 | 349722 | 1 | + | 4.0818704301528e-06 | 0.001 | 61.48 | be.vs.ka |
| NW_023312732.1 | 349745 | 349745 | 1 | + | 2.36222706197392e-10 | 0 | 79.15 | be.vs.ka |
| NW_023312732.1 | 349819 | 349819 | 1 | + | 5.91817685485486e-09 | 0 | 64.19 | be.vs.ka |
| NW_023312732.1 | 349887 | 349887 | 1 | + | 3.18092225080658e-05 | 0.005 | 38.74 | be.vs.ka |
| NW_023312732.1 | 349906 | 349906 | 1 | + | 8.66054463755933e-06 | 0.002 | 47.5 | be.vs.ka |
| NW_023312732.1 | 349994 | 349994 | 1 | + | 3.16052542850735e-10 | 0 | 73.33 | be.vs.ka |
| NW_023312732.1 | 350045 | 350045 | 1 | + | 1.78219754139806e-05 | 0.003 | 64.88 | be.vs.ka |
| NW_023312735.1 | 44911 | 44911 | 1 | + | 1.54231476990678e-05 | 0.003 | 63.46 | be.vs.ka |
| NW_023312735.1 | 69999 | 69999 | 1 | + | 1.03629389968165e-05 | 0.002 | 25.1 | be.vs.ka |
| NW_023312737.1 | 20955 | 20955 | 1 | + | 3.39756746478276e-08 | 0 | 78.26 | be.vs.ka |
| NW_023312741.1 | 307877 | 307877 | 1 | + | 6.36451199173005e-08 | 0 | 75.32 | be.vs.ka |
| NW_023312741.1 | 307918 | 307918 | 1 | + | 4.9217780222849e-10 | 0 | 76.3 | be.vs.ka |

...
... only first 25 lines shown ...

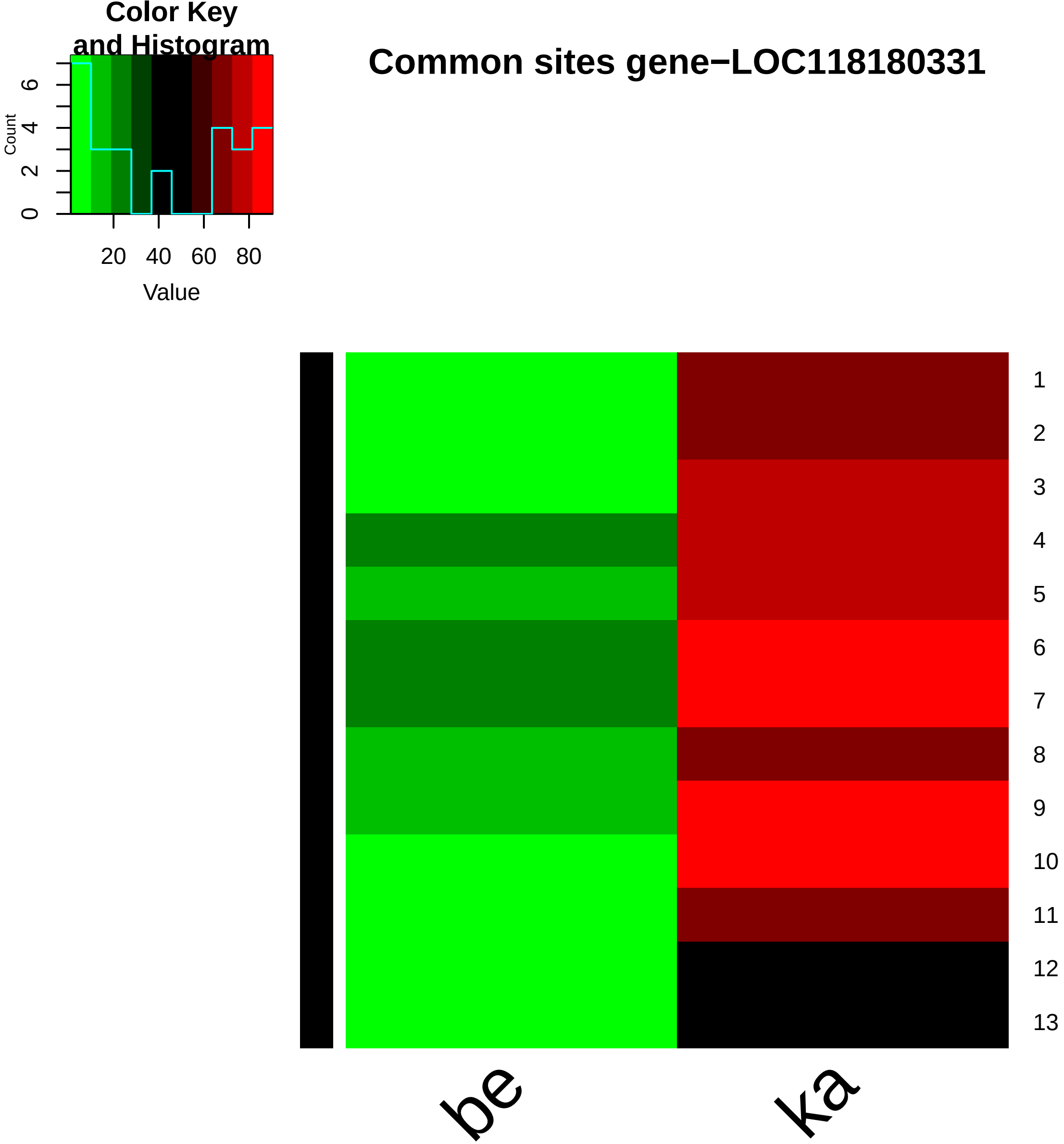

Color Key
and Histogram
Count
0
2
4
6
20
40
60
80
Value
Common sites gene−LOC118180331
1
2
3
4
5
6
7
8
9
10
11
12
13
be
ka

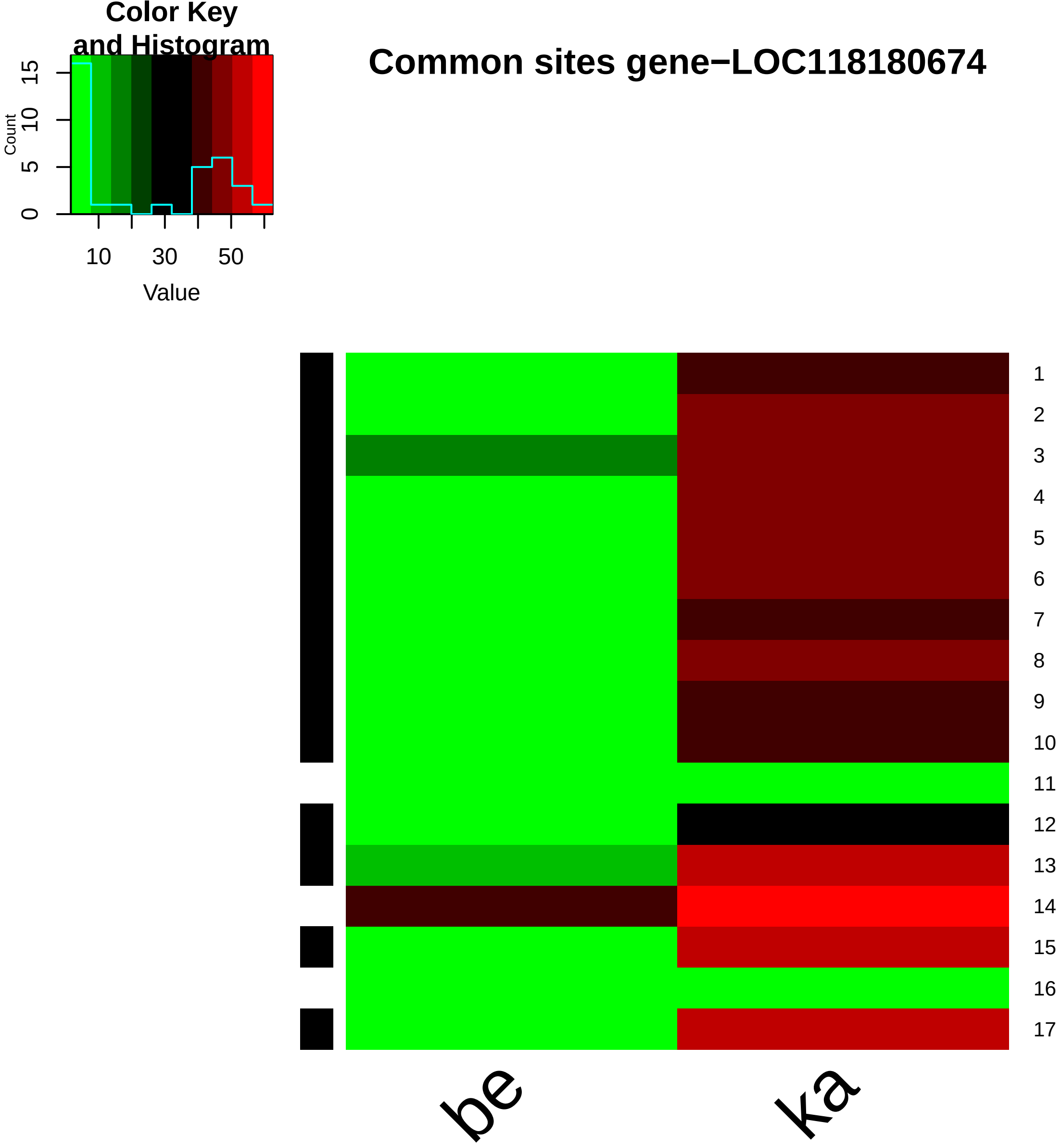
Color Key
and Histogram
Count
0
5
10
15
10
30
50
Value
Common sites gene−LOC118180674
be
ka

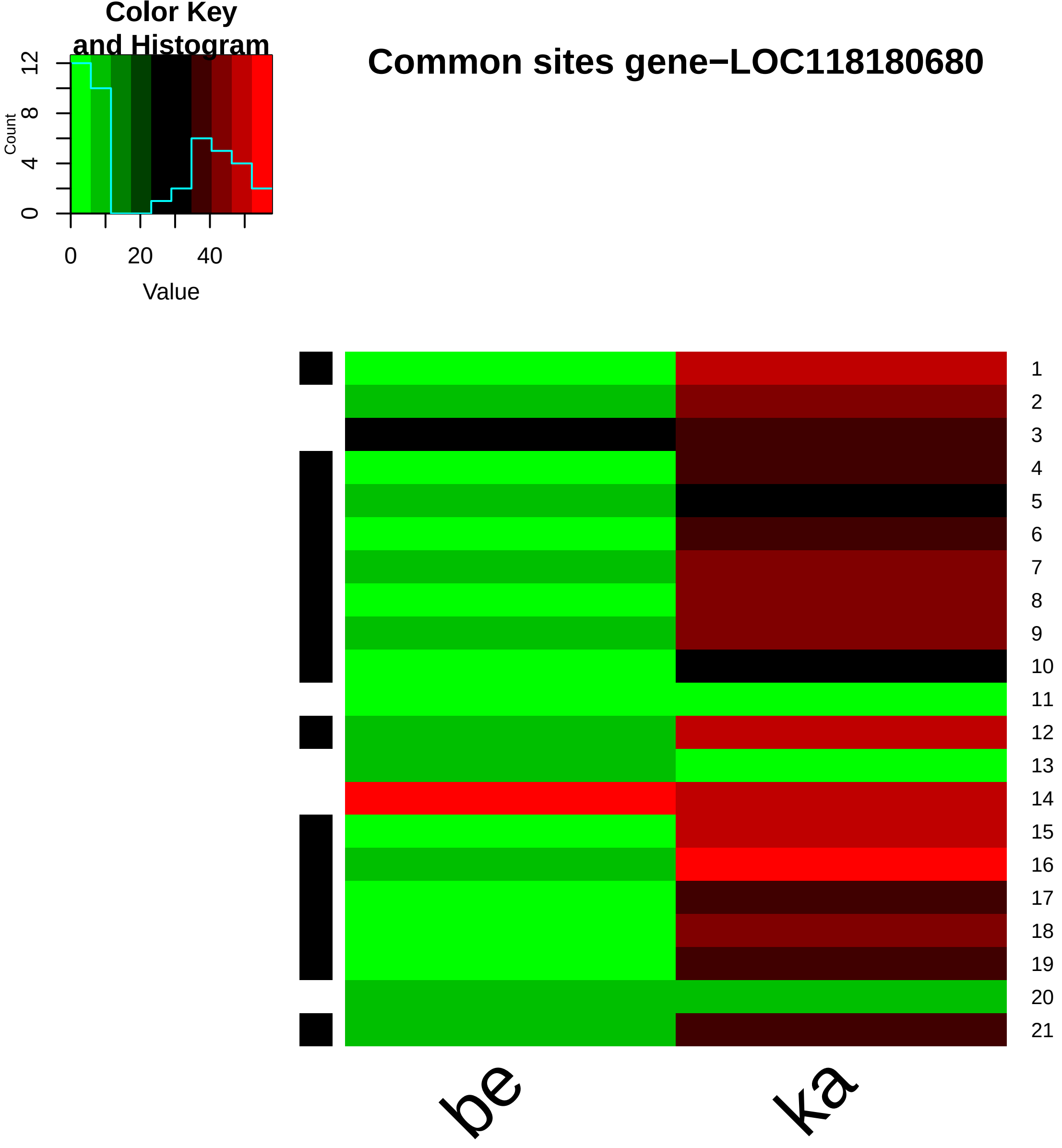

Color Key
and Histogram
Count
0
4
8
12
0
20
40
Value
Common sites gene−LOC118180680
be
ka

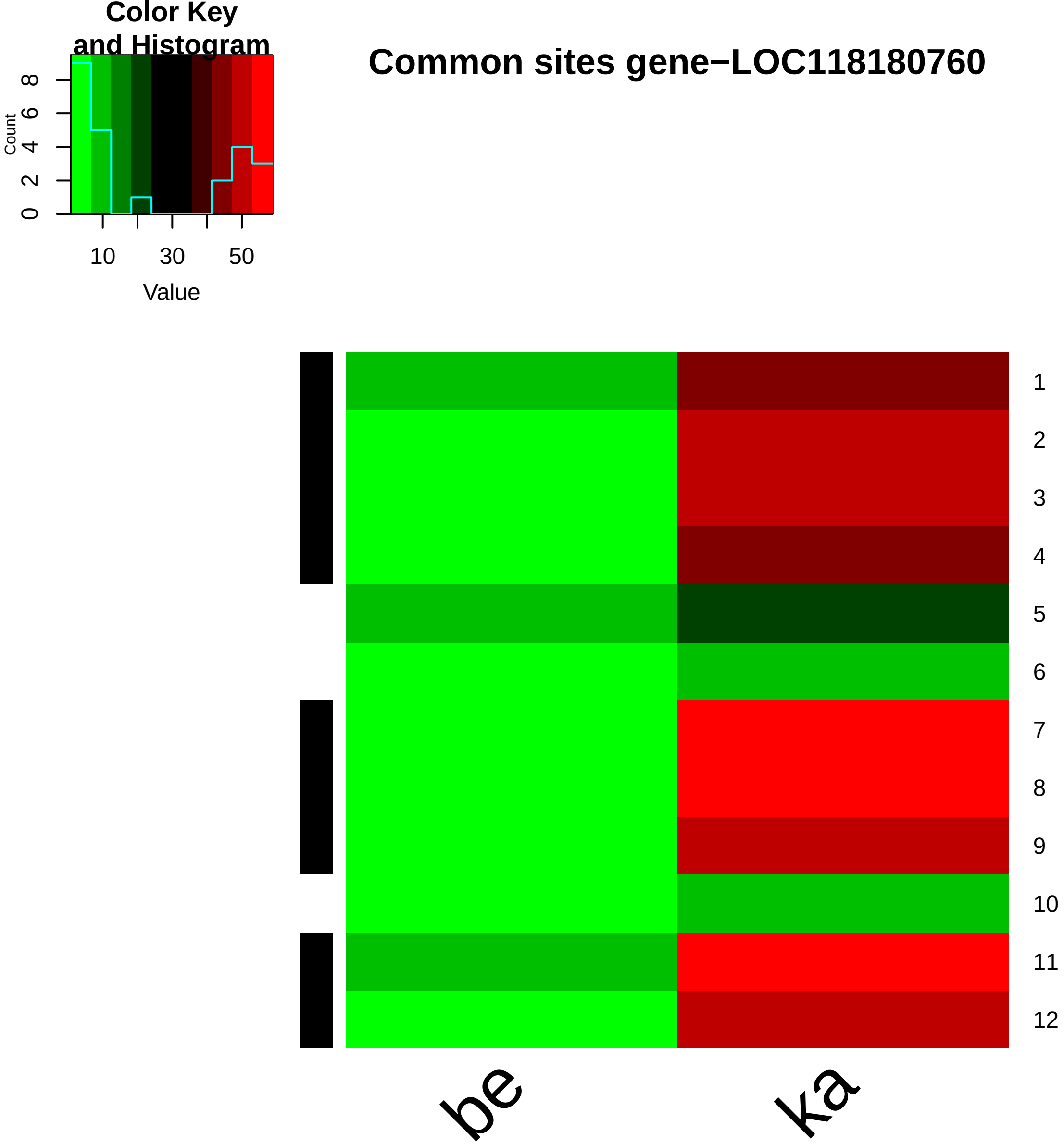
Color Key
and Histogram
Count
0 2 4 6 8
10 30 50
Value
Common sites gene−LOC118180760
1
2
3
4
5
6
7
8
9
10
11
12
be
ka

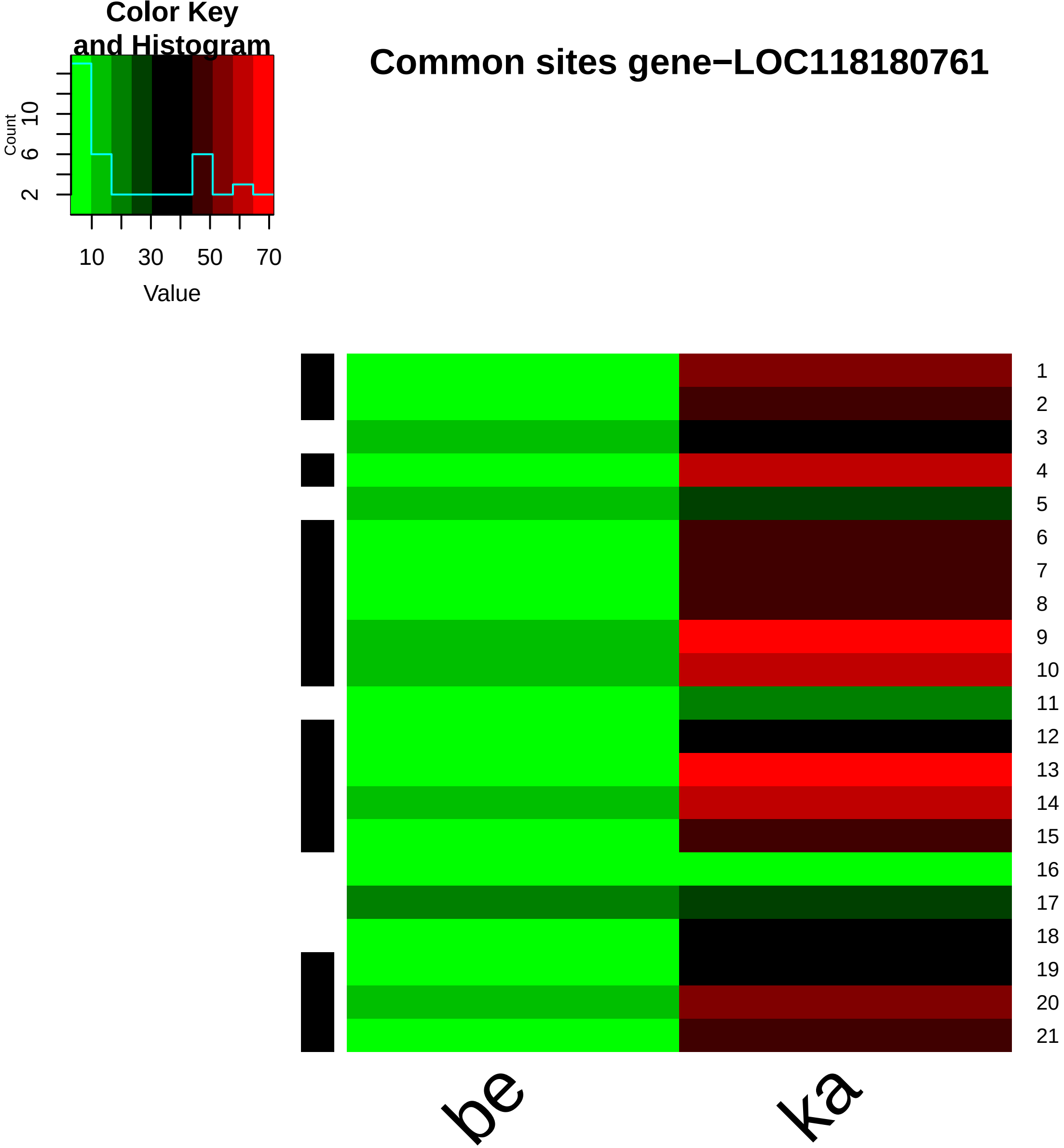

Color Key and Histogram
Count
2
6
10
10
30
50
70
Value
Common sites gene−LOC118180761
be
ka

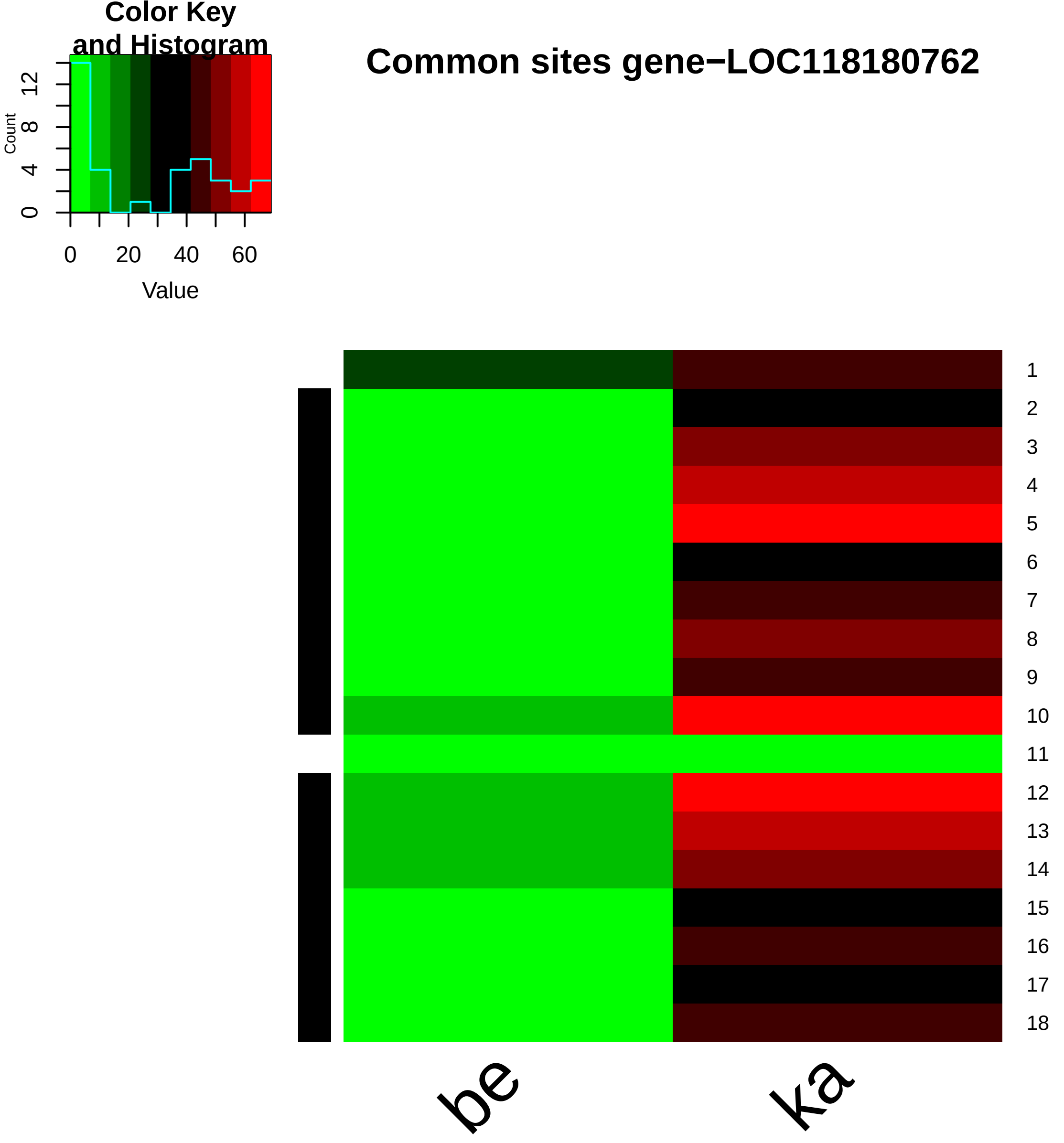

Color Key
and Histogram
Count
0
4
8
12
0
20
40
60
Value
Common sites gene−LOC118180762
be
ka

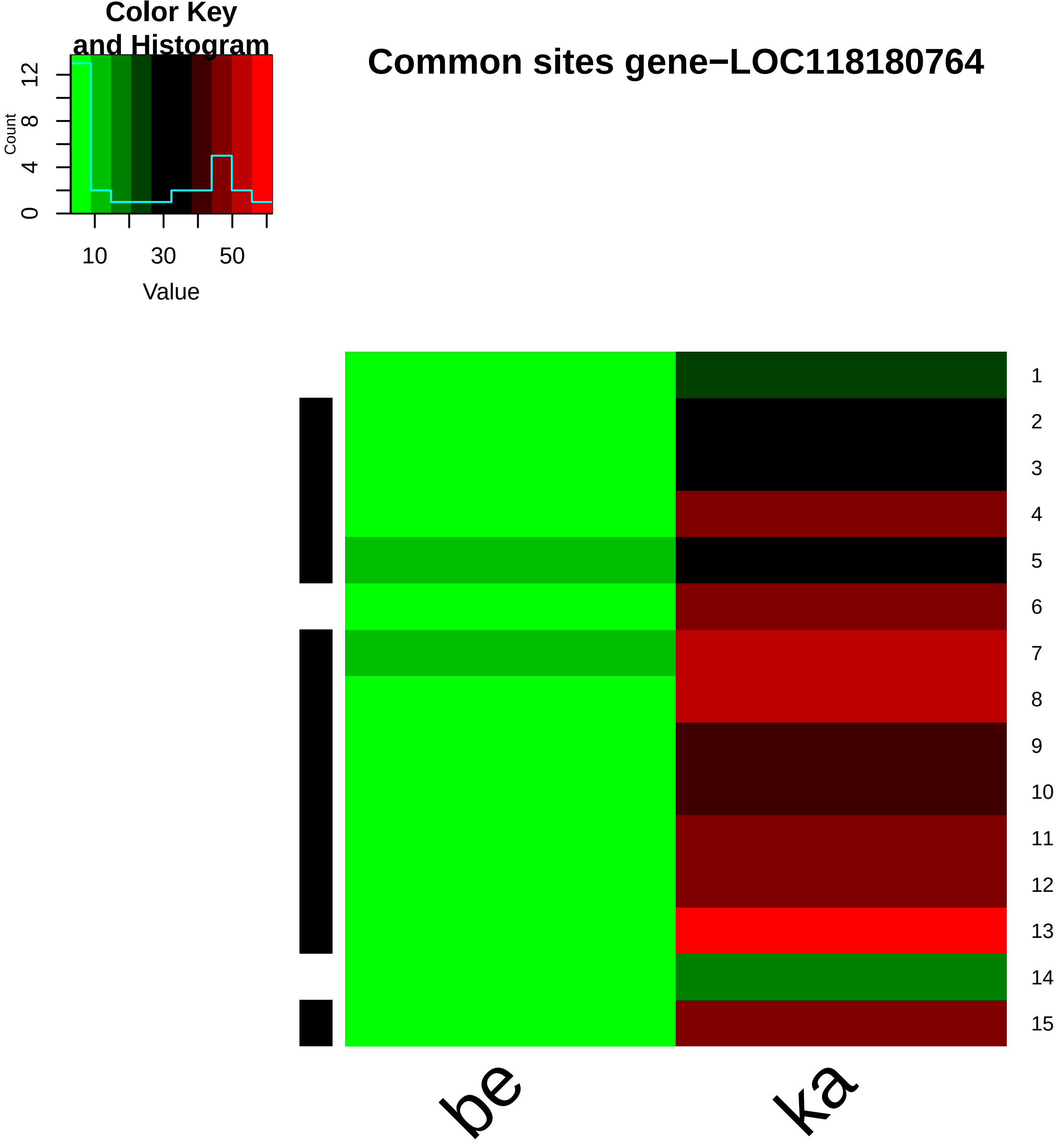
Color Key
and Histogram
Count
Value
0
4
8
12
10
30
50
Common sites gene−LOC118180764
be
ka

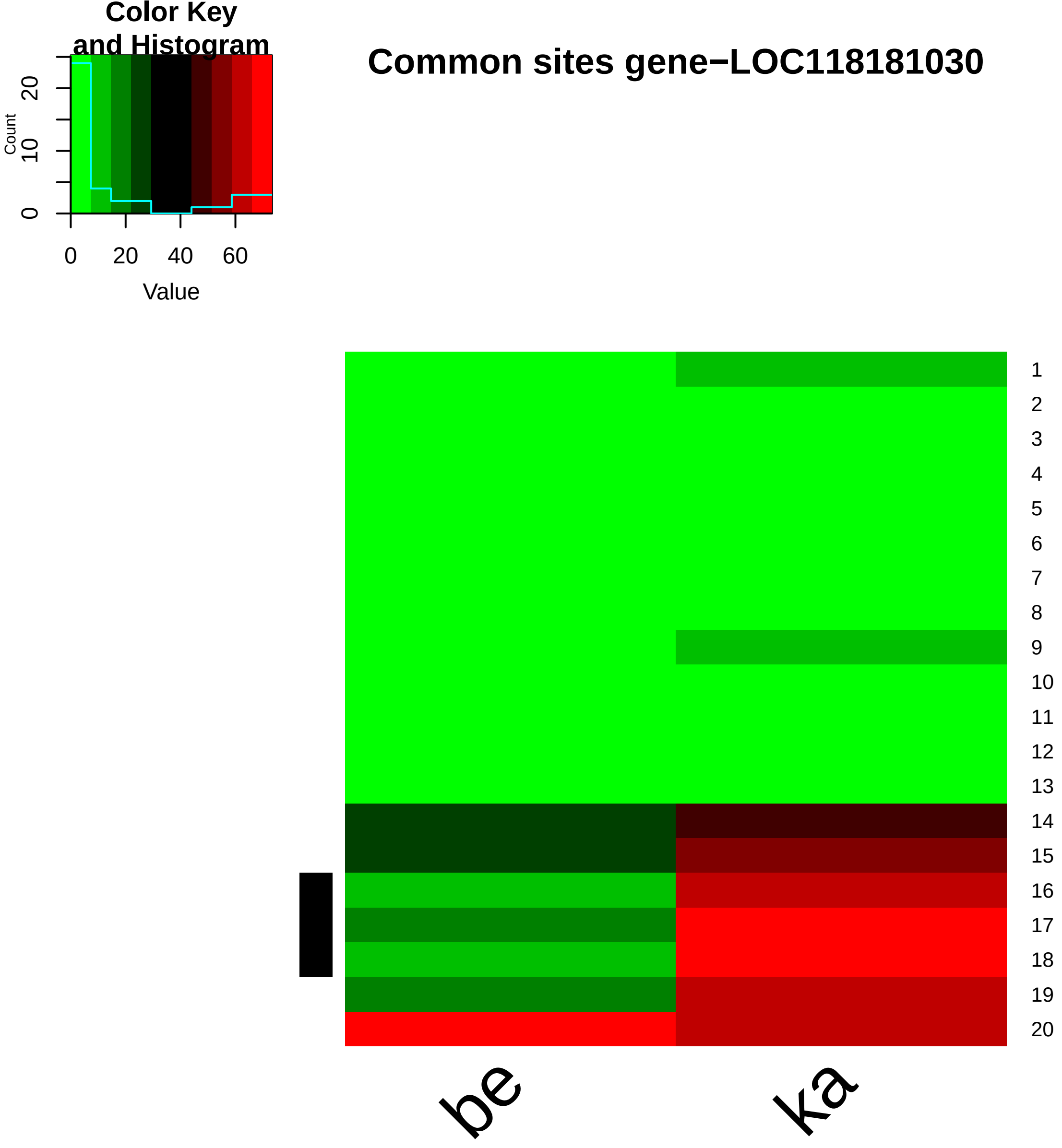

Color Key and Histogram
Count
Value
0
20
40
60
0
10
20
Common sites gene−LOC118181030
be
ka

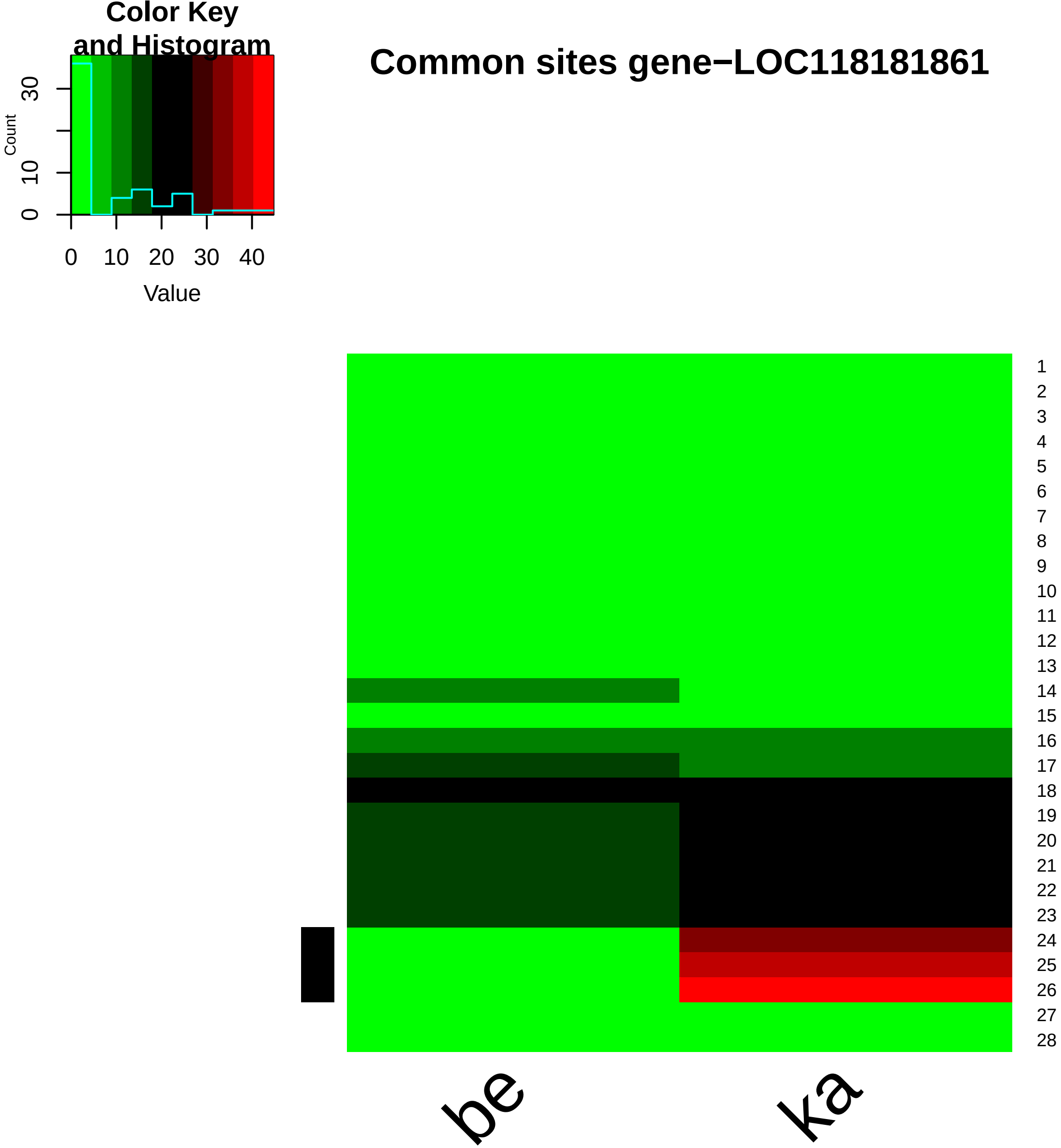

Color Key
and Histogram
Count
0
10
30
0
10
20
30
40
Value
Common sites gene−LOC118181861
be
ka

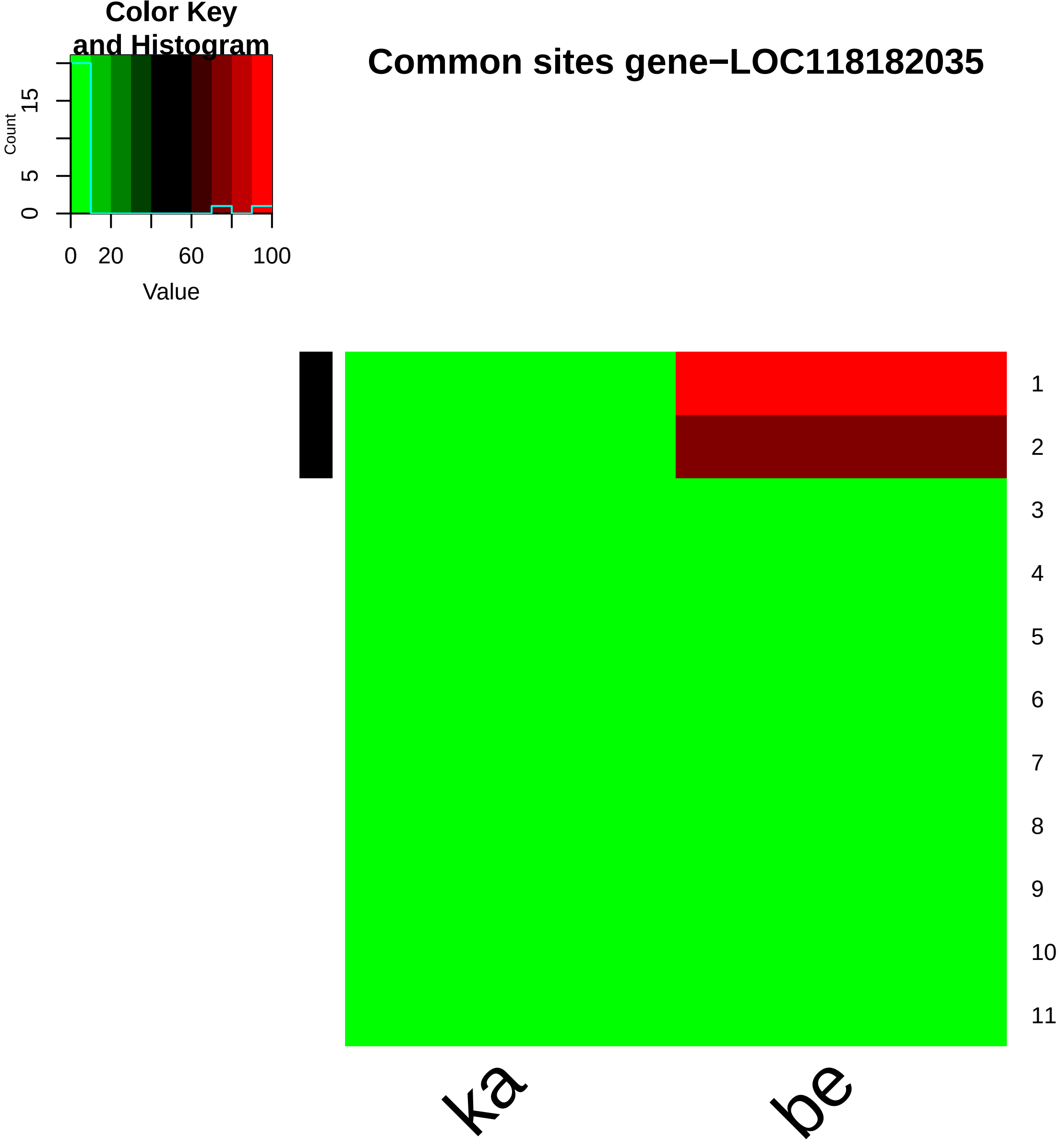

Color Key
and Histogram
Count
0
5
15
0
20
60
100
Value
Common sites gene−LOC118182035
1
2
3
4
5
6
7
8
9
10
11
ka
be

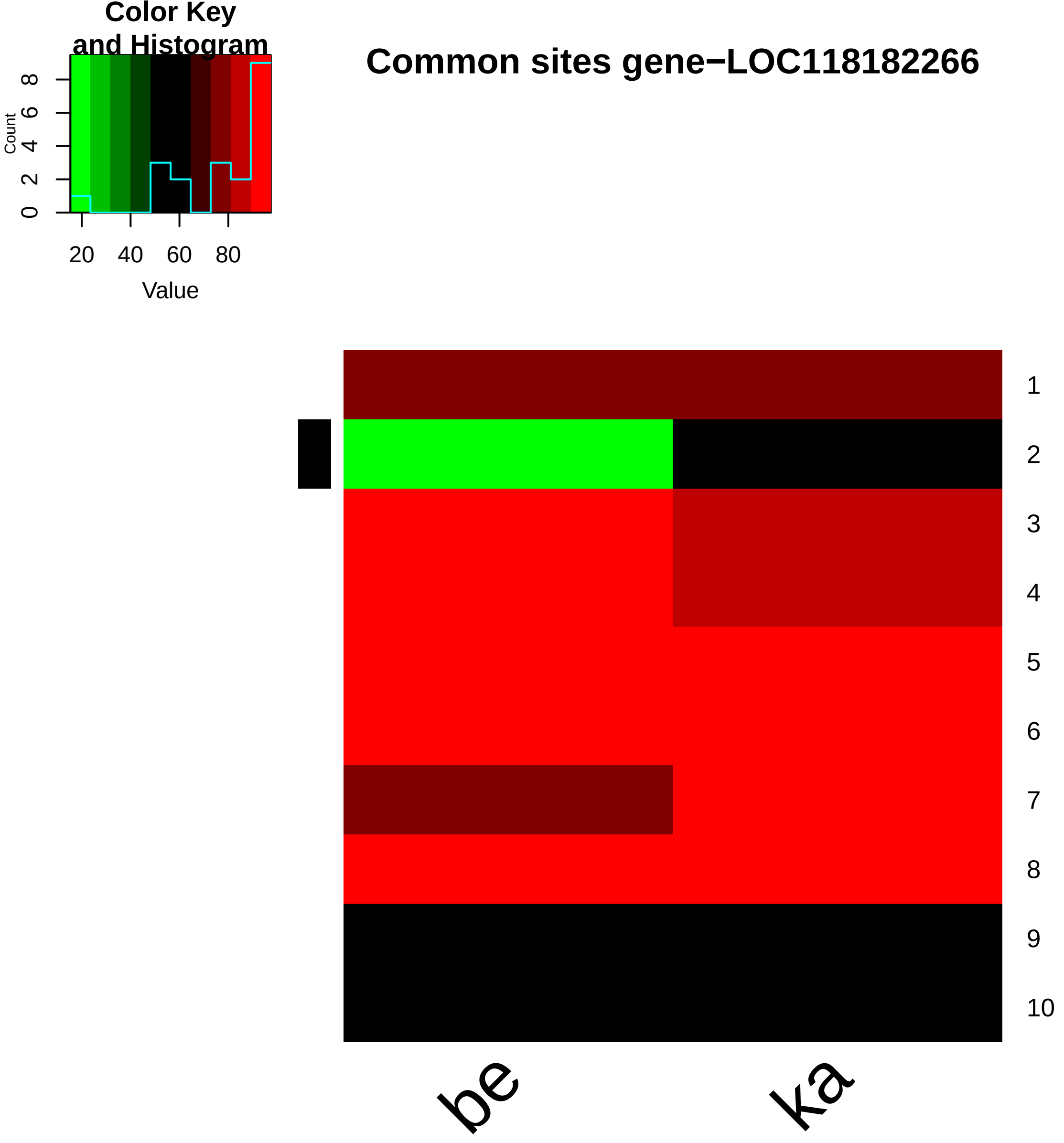
Color Key
and Histogram
Count
0
2
4
6
8
20
40
60
80
Value
Common sites gene−LOC118182266
1
2
3
4
5
6
7
8
9
10
be
ka

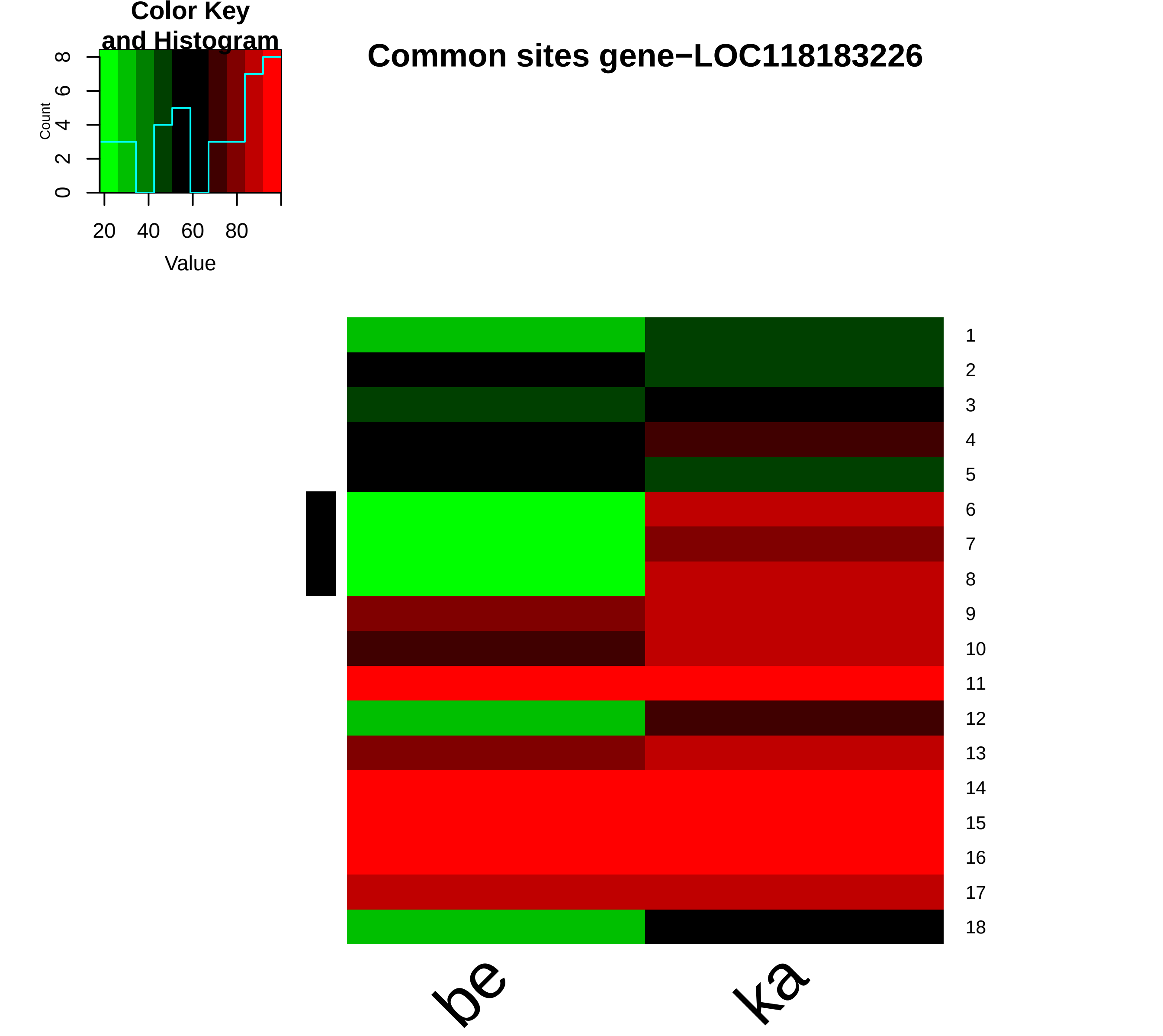
Color Key
and Histogram
Count
0
2
4
6
8
20
40
60
80
Value
Common sites gene−LOC118183226
be
ka

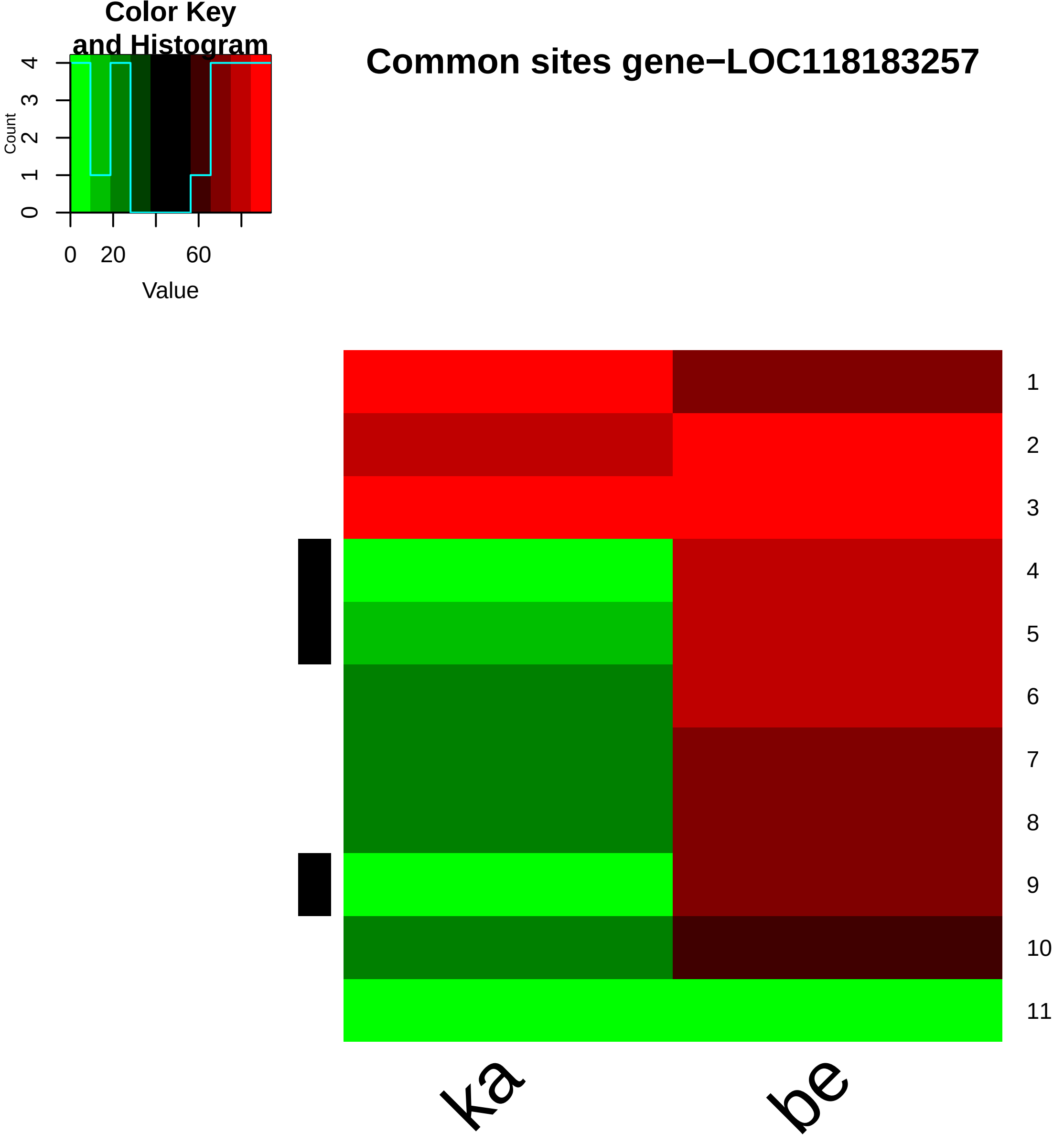
Color Key
and Histogram
Count
0
1
2
3
4
0
20
60
Value
Common sites gene−LOC118183257
1
2
3
4
5
6
7
8
9
10
11
ka
be

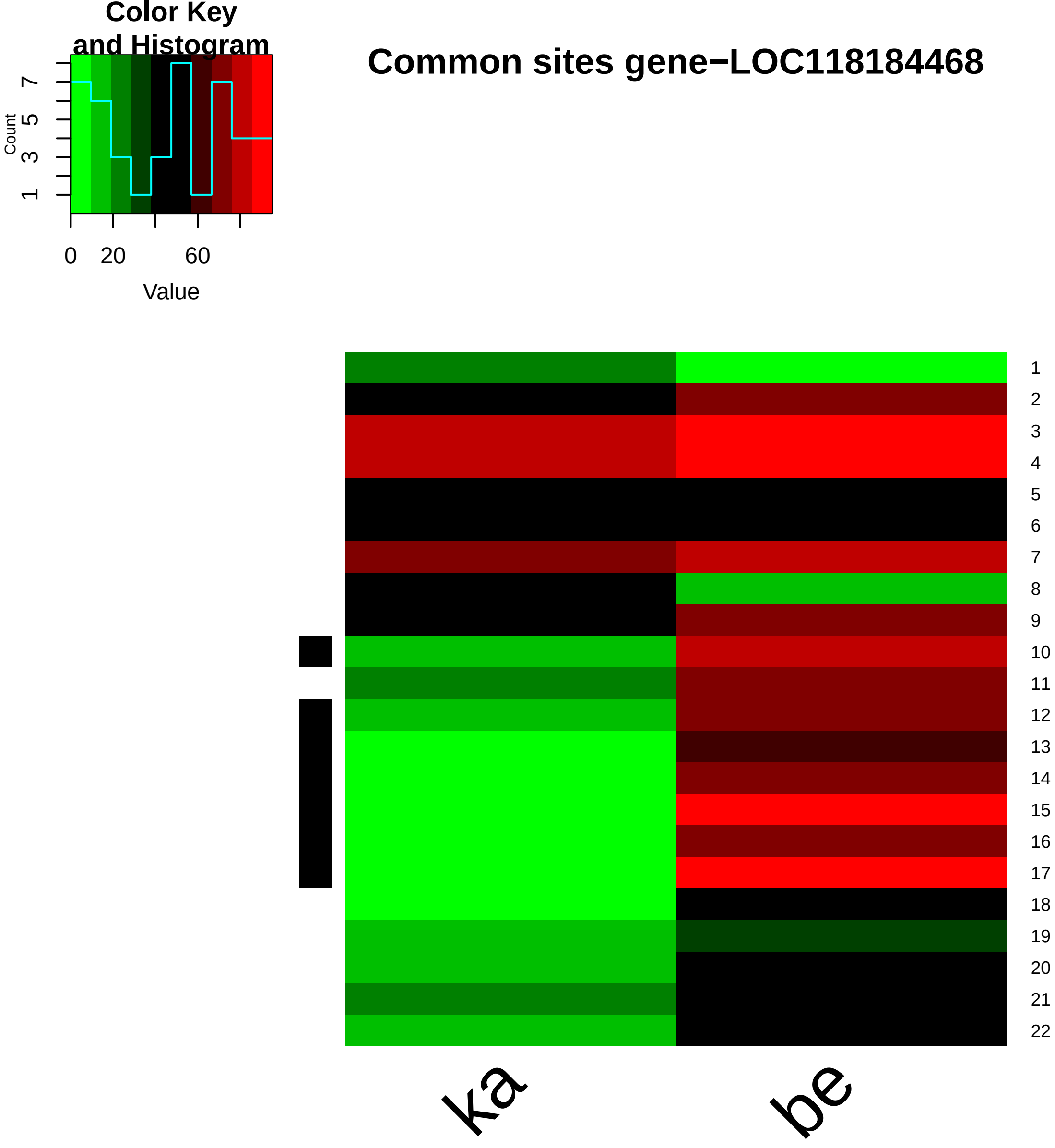

Color Key
and Histogram
Count
1
3
5
7
0
20
60
Value
Common sites gene−LOC118184468
ka
be

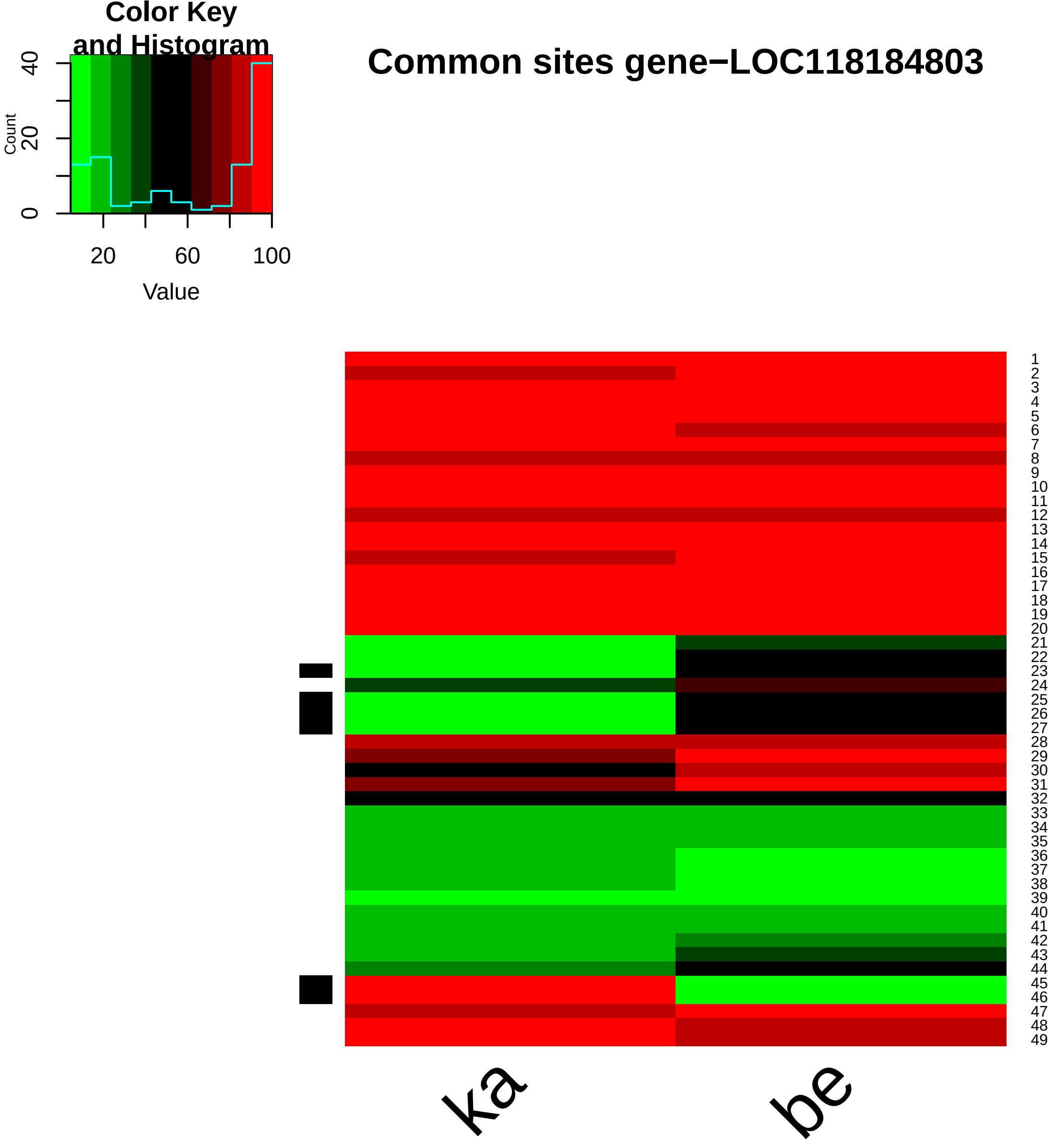

Color Key
and Histogram
Count
0
20
40
20
60
100
Value
Common sites gene−LOC118184803
ka
be

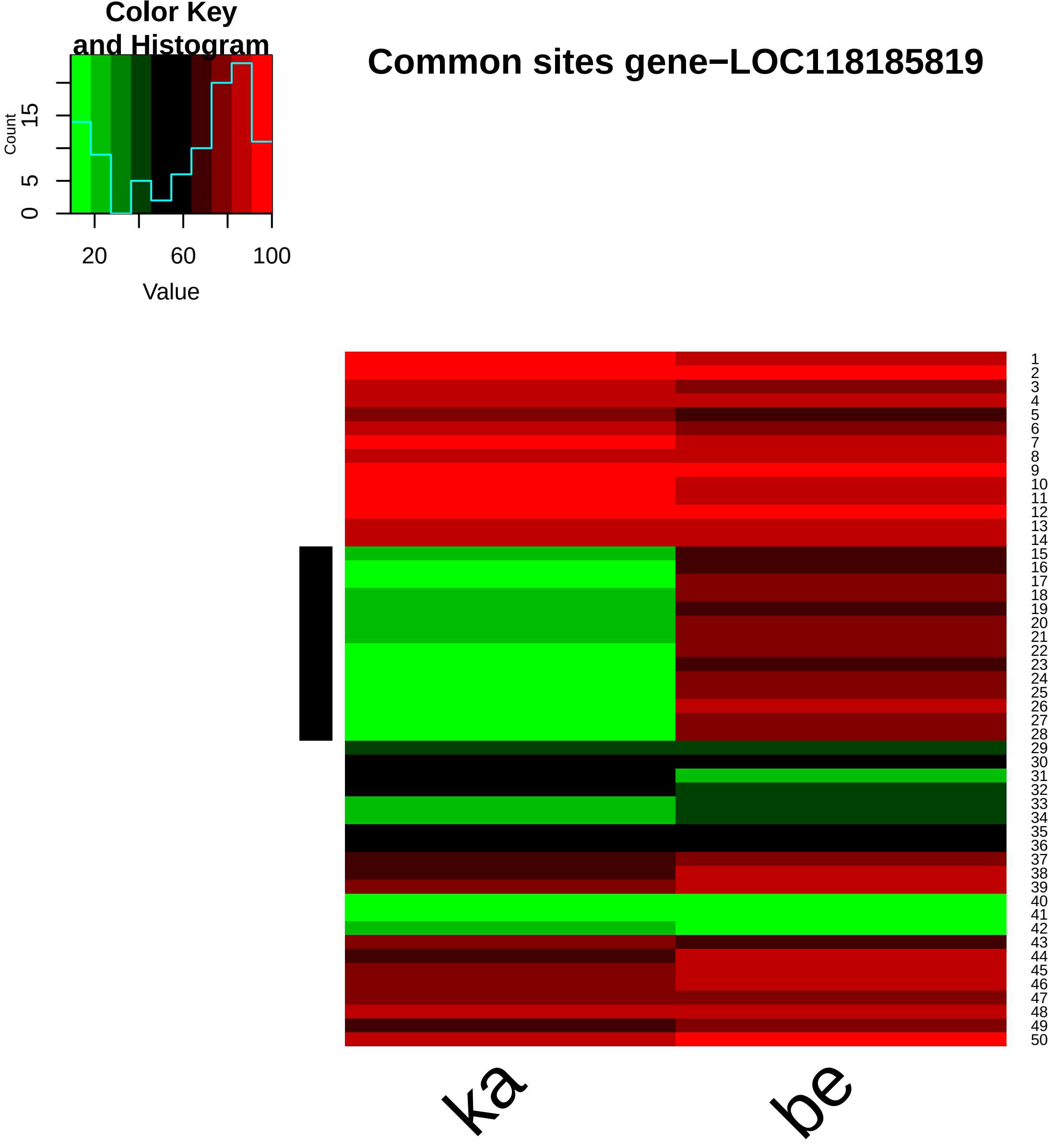

Color Key
and Histogram
Count
0
5
15
20
60
100
Value
Common sites gene−LOC118185819
ka
be

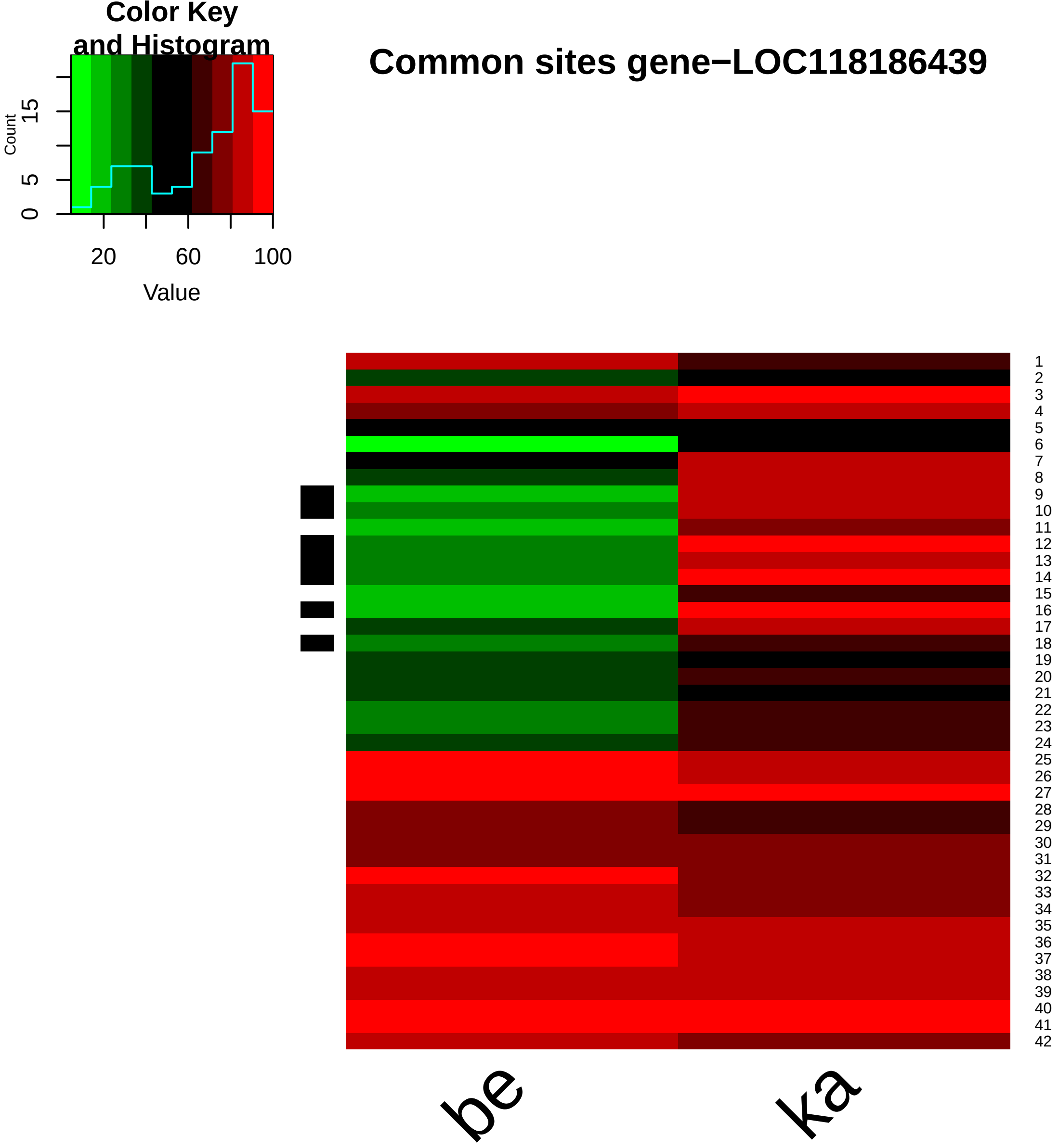
Color Key
and Histogram
Count
0
5
15
20
60
100
Value
Common sites gene−LOC118186439
be
ka

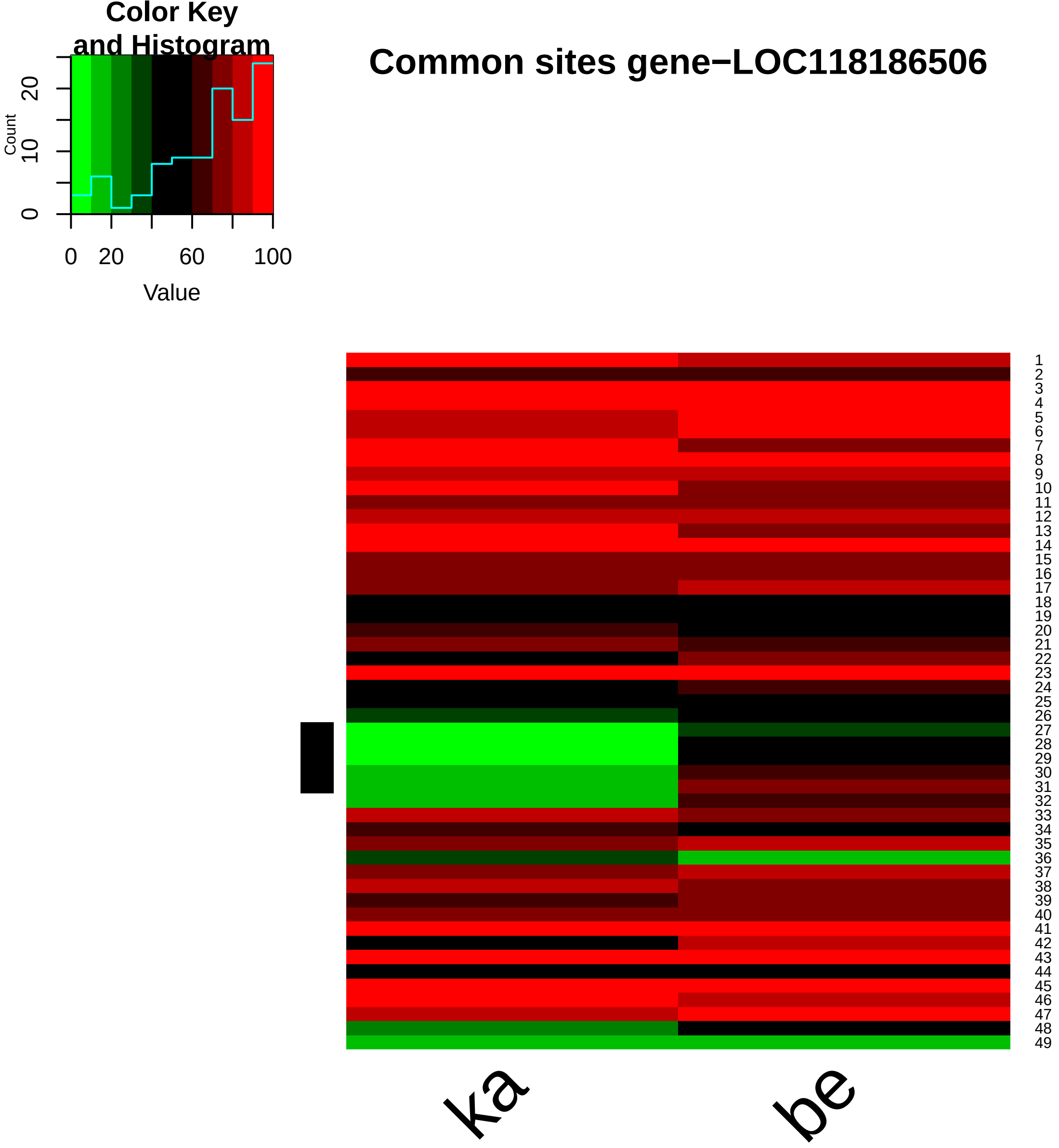
Color Key and Histogram
Count
0
10
20
0
20
60
100
Value
Common sites gene−LOC118186506
ka
be

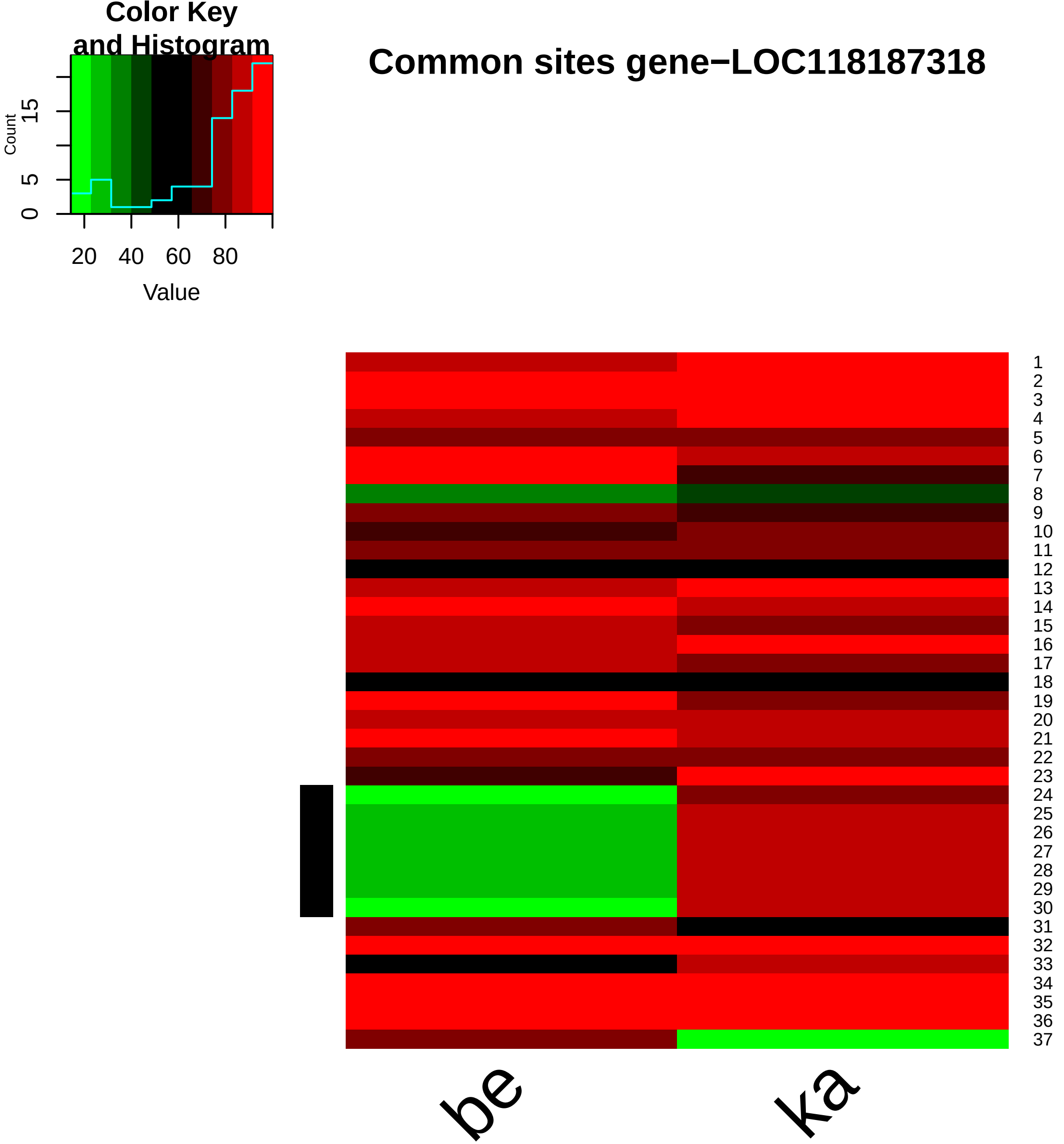

Color Key
and Histogram
Count
0
5
15
20
40
60
80
Value
Common sites gene−LOC118187318
be
ka

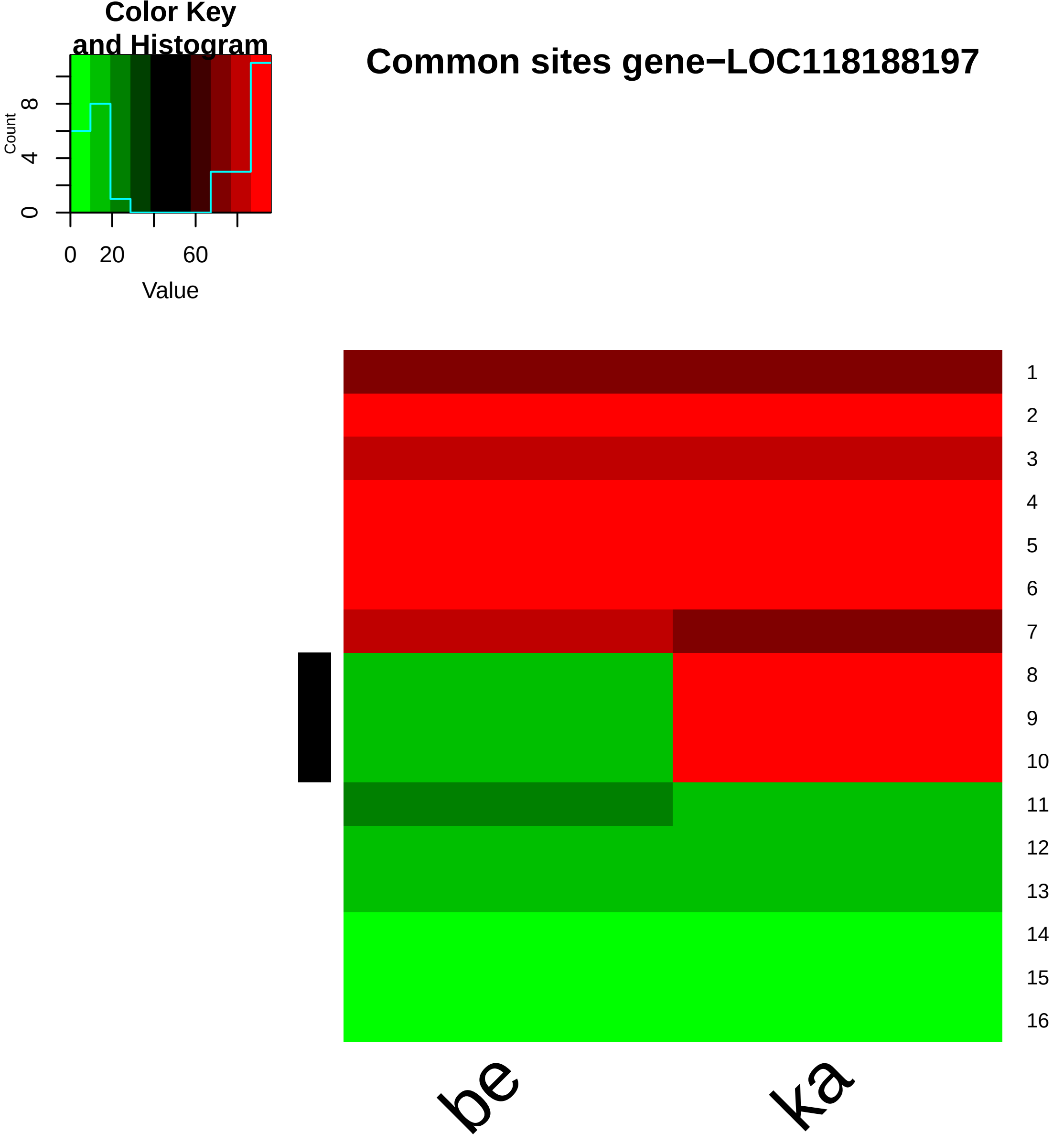

Color Key and Histogram
Count
0
4
8
0
20
60
Value
Common sites gene−LOC118188197
be
ka

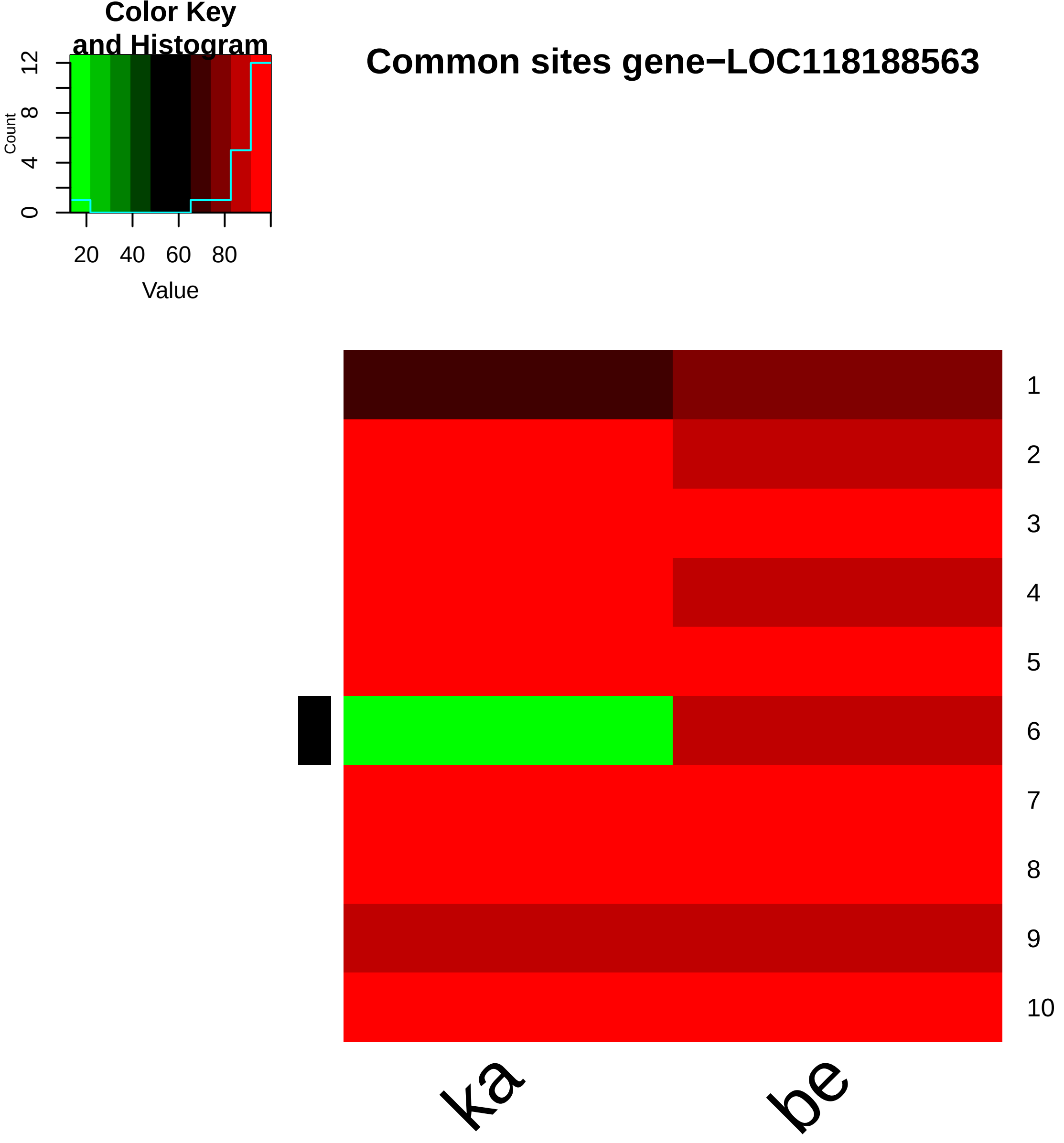

Color Key
and Histogram
Count
0
4
8
12
20
40
60
80
Value
Common sites gene−LOC118188563
1
2
3
4
5
6
7
8
9
10
ka
be

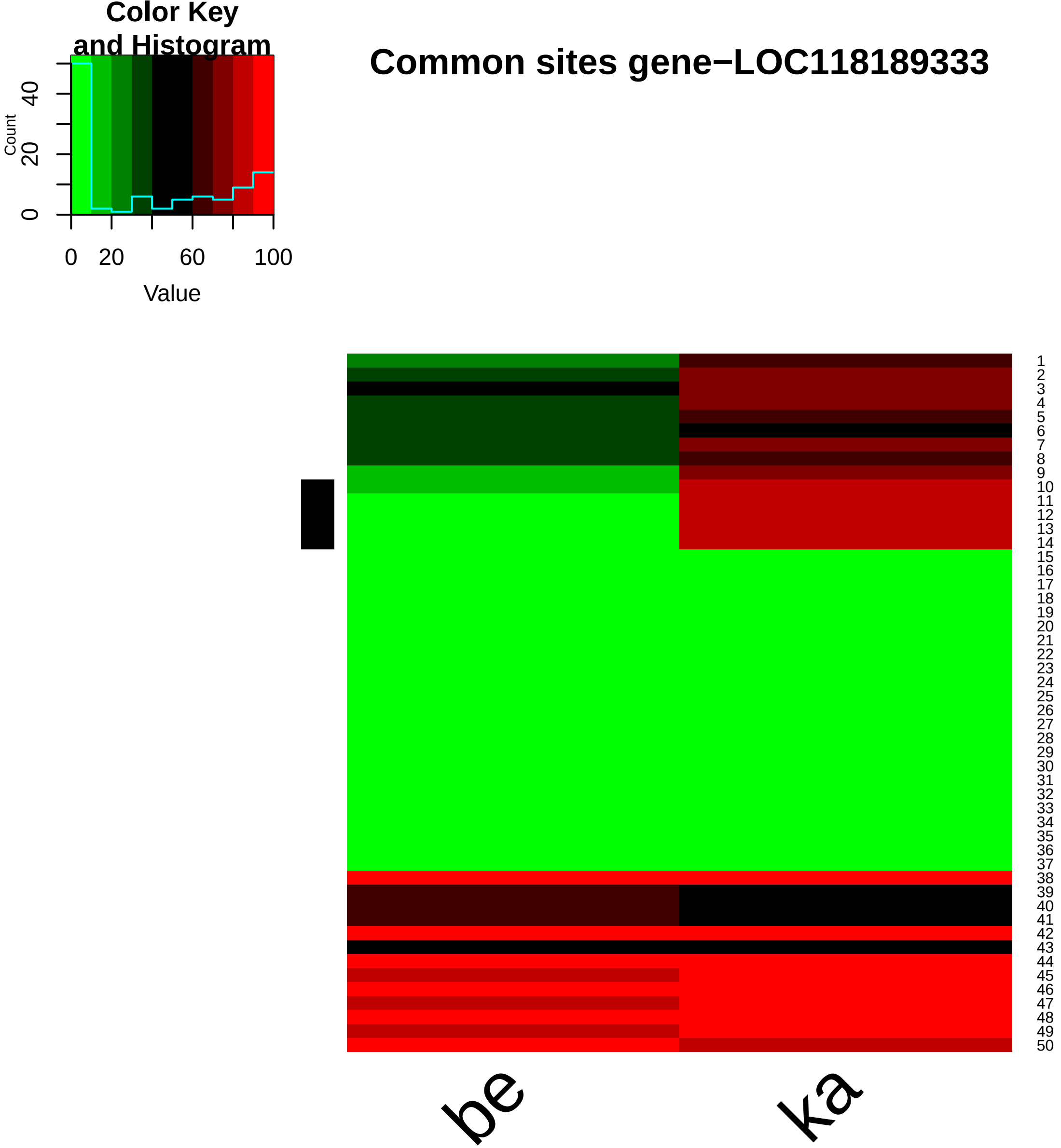
Color Key
and Histogram
Count
0
20
40
0
20
60
100
Value
Common sites gene−LOC118189333
be
ka

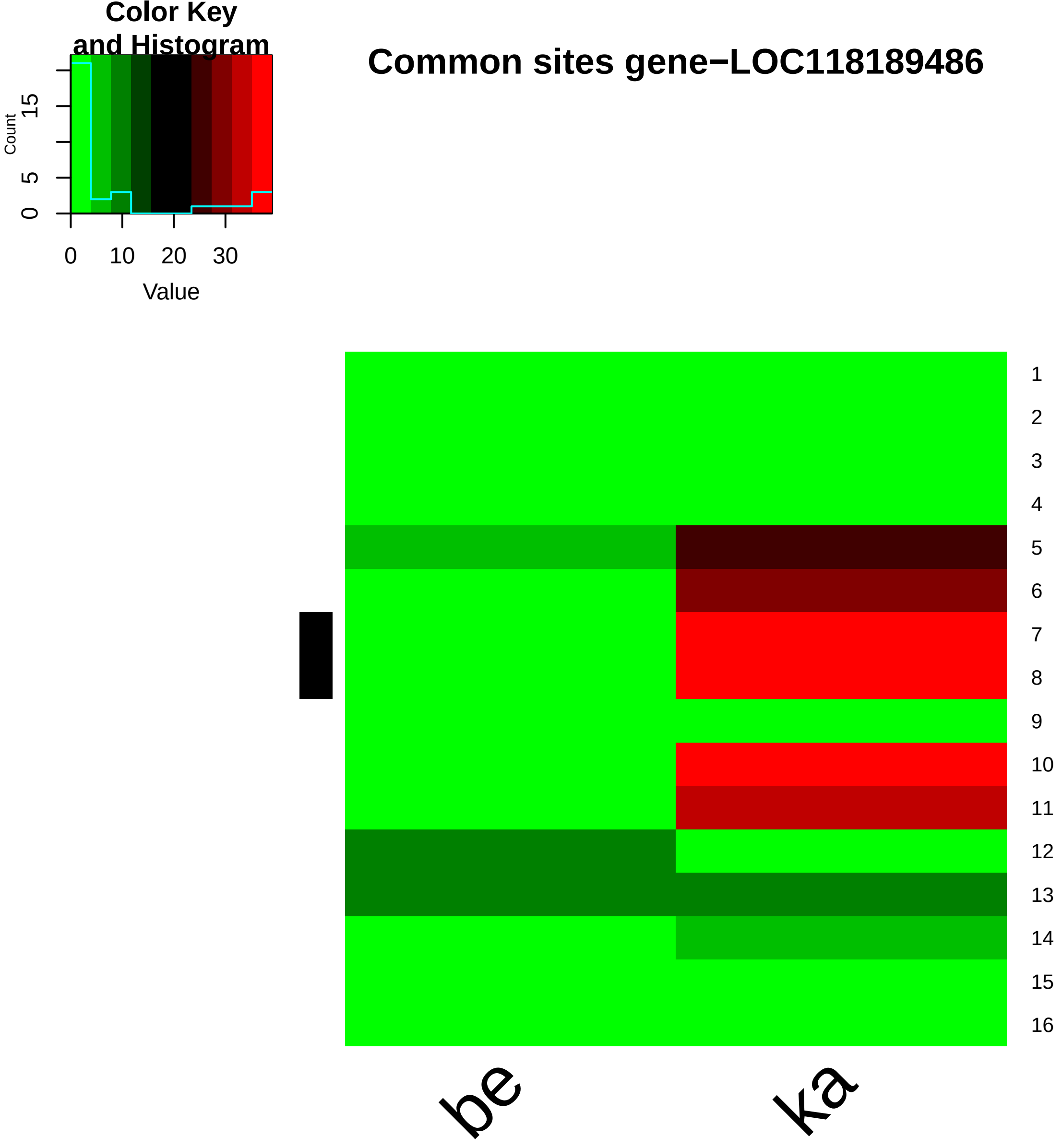

Color Key
and Histogram
Count
0
5
15
0
10
20
30
Value
Common sites gene−LOC118189486
be
ka

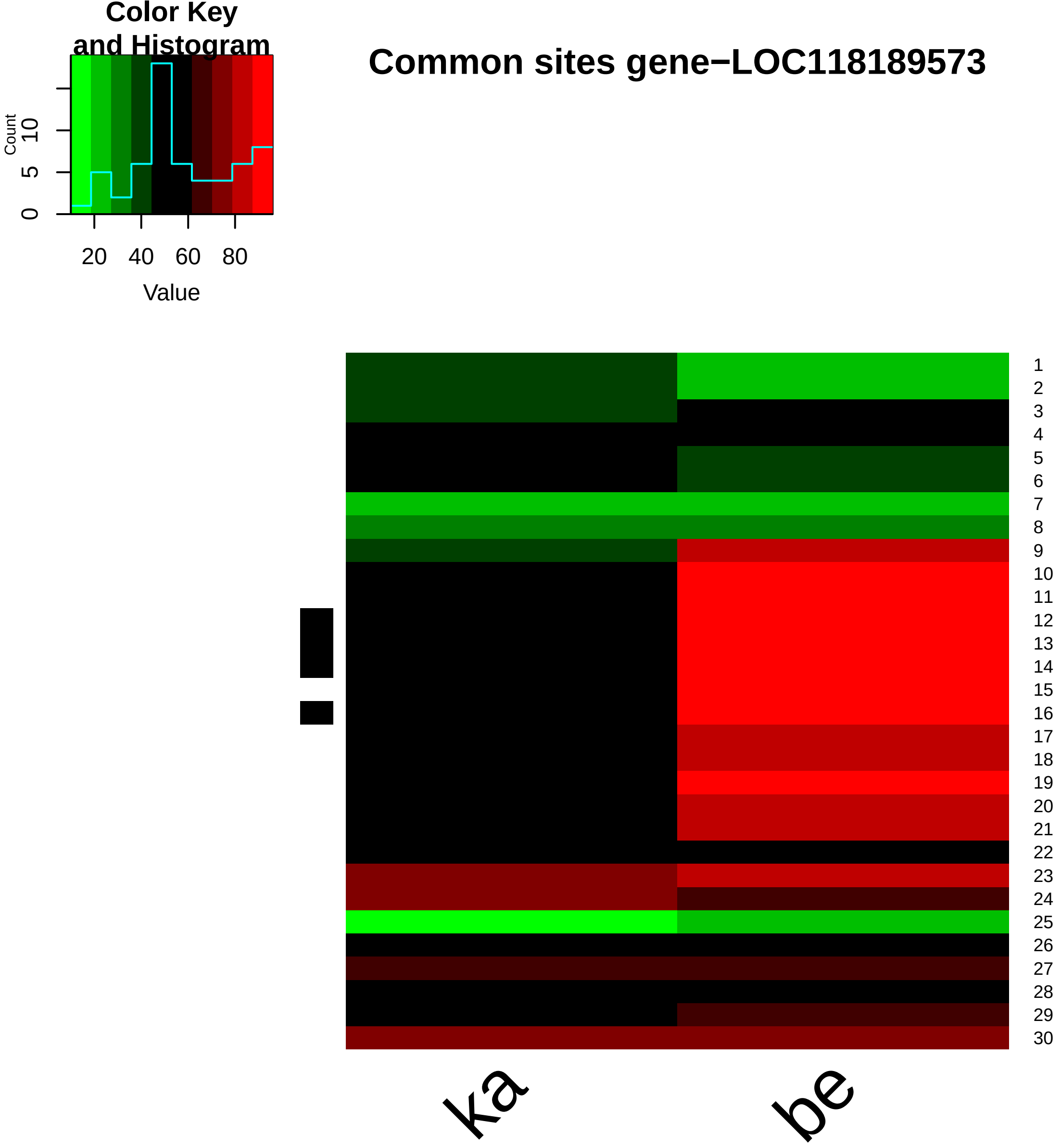
Color Key
and Histogram
Count
0
5
10
20
40
60
80
Value
Common sites gene−LOC118189573
ka
be

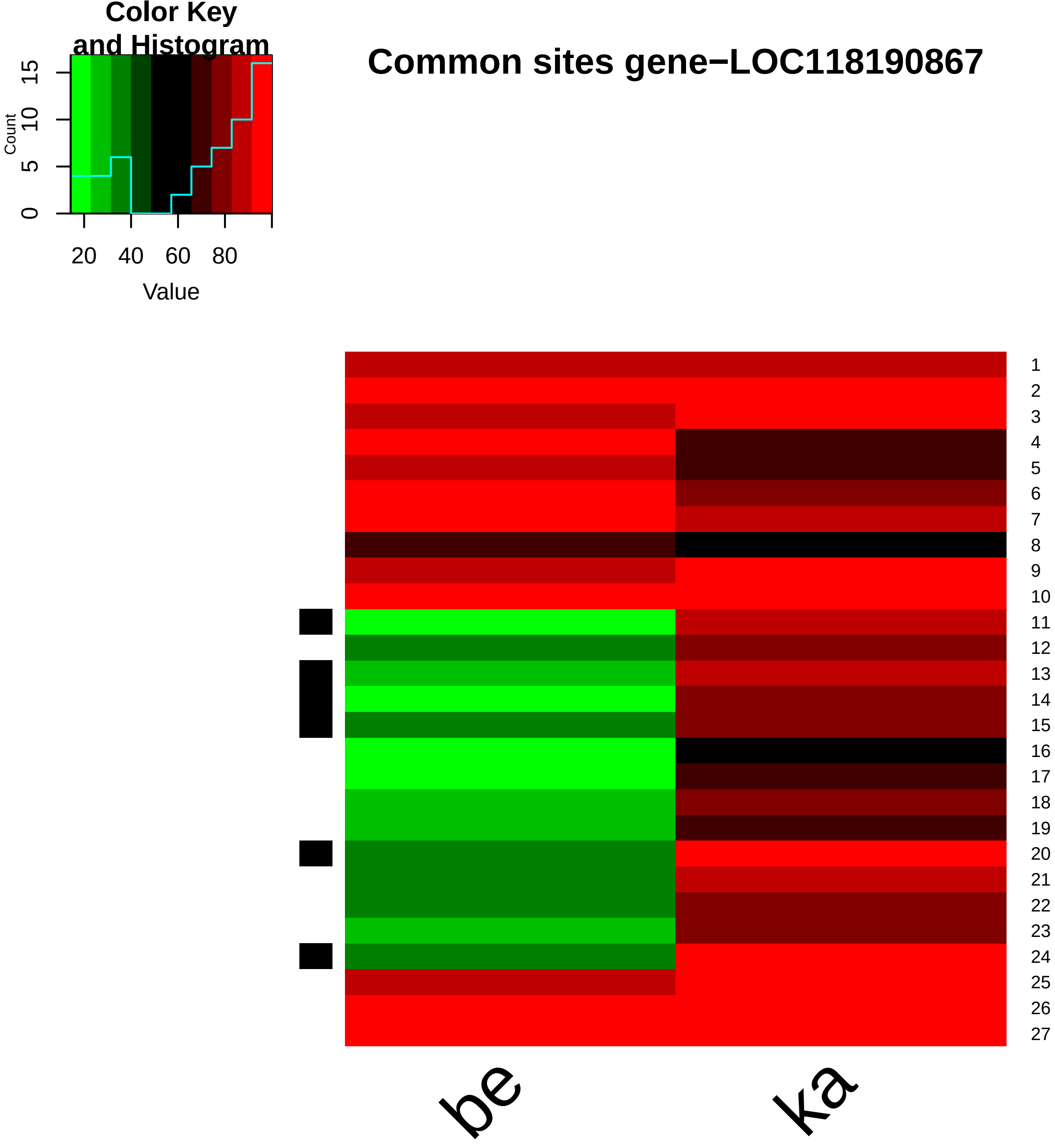

Color Key and Histogram
Count
0
5
10
15
20
40
60
80
Value
Common sites gene−LOC118190867
be
ka

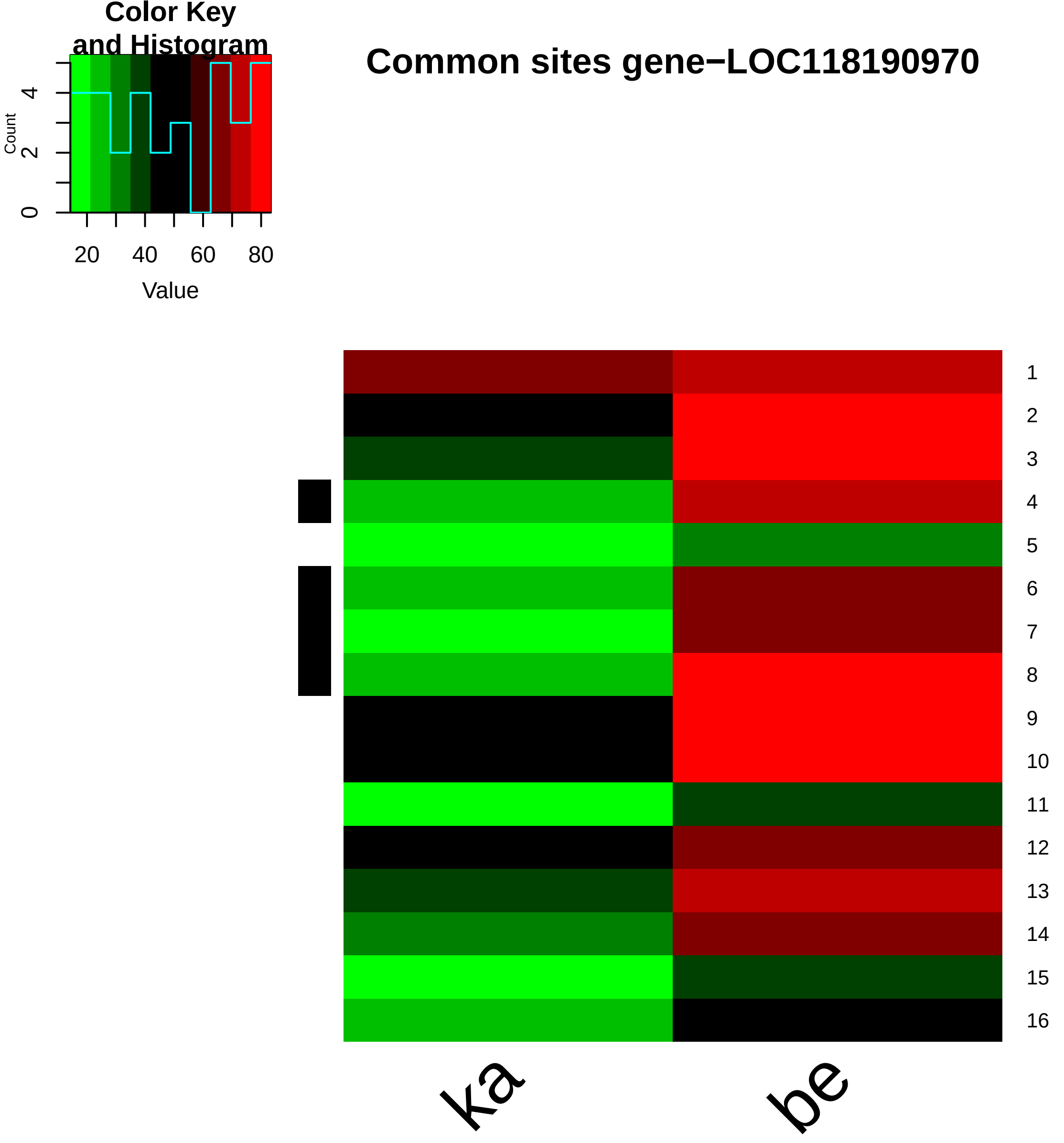

Color Key
and Histogram
Count
0
2
4
20
40
60
80
Value
Common sites gene−LOC118190970
ka
be

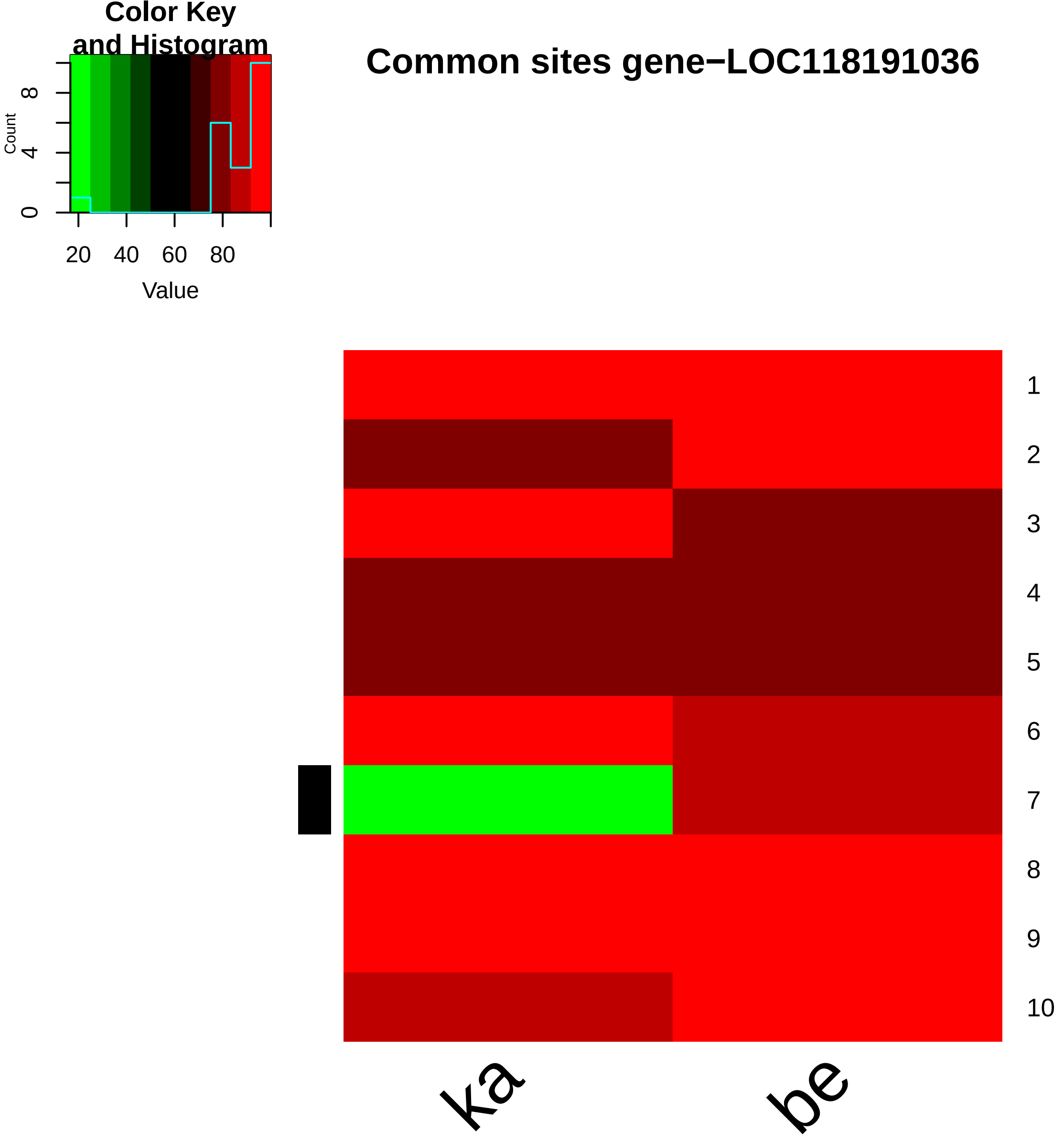
Color Key
and Histogram
Count
0
4
8
20
40
60
80
Value
Common sites gene−LOC118191036
1
2
3
4
5
6
7
8
9
10
ka
be

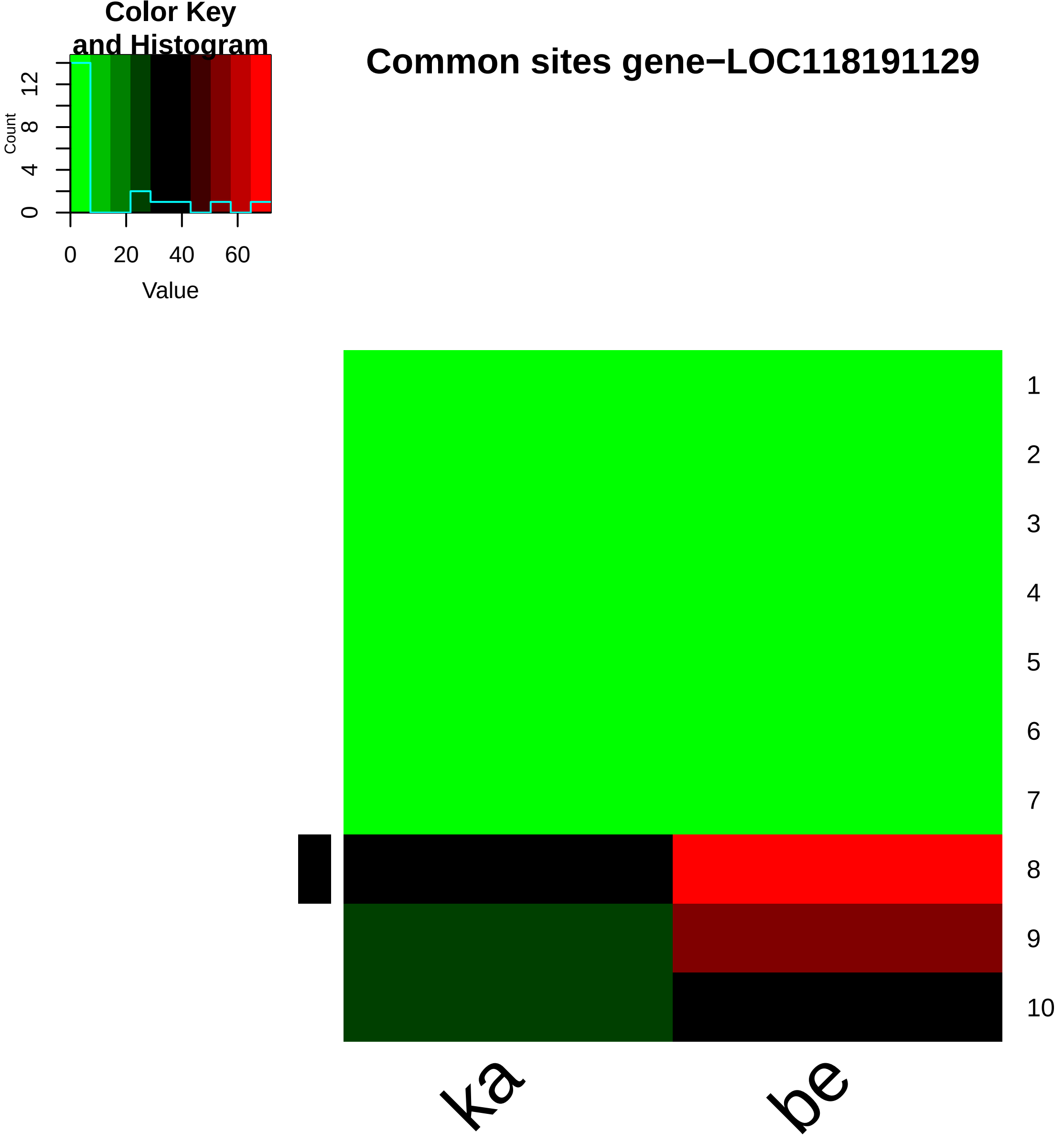
Color Key
and Histogram
Count
0
4
8
12
0
20
40
60
Value
Common sites gene−LOC118191129
1
2
3
4
5
6
7
8
9
10
ka
be

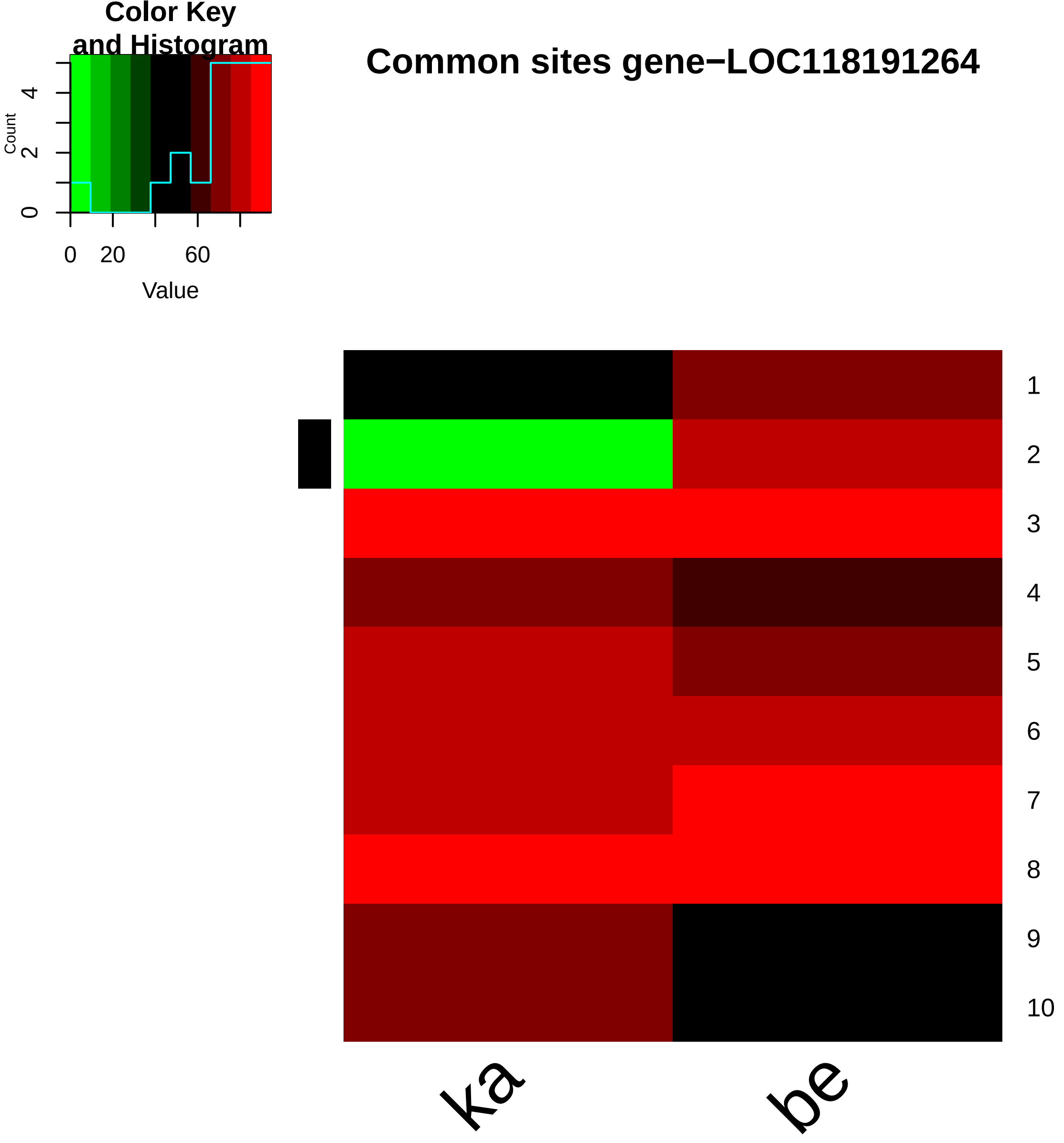

Color Key
and Histogram
Count
0
2
4
0
20
60
Value
Common sites gene−LOC118191264
1
2
3
4
5
6
7
8
9
10
ka
be

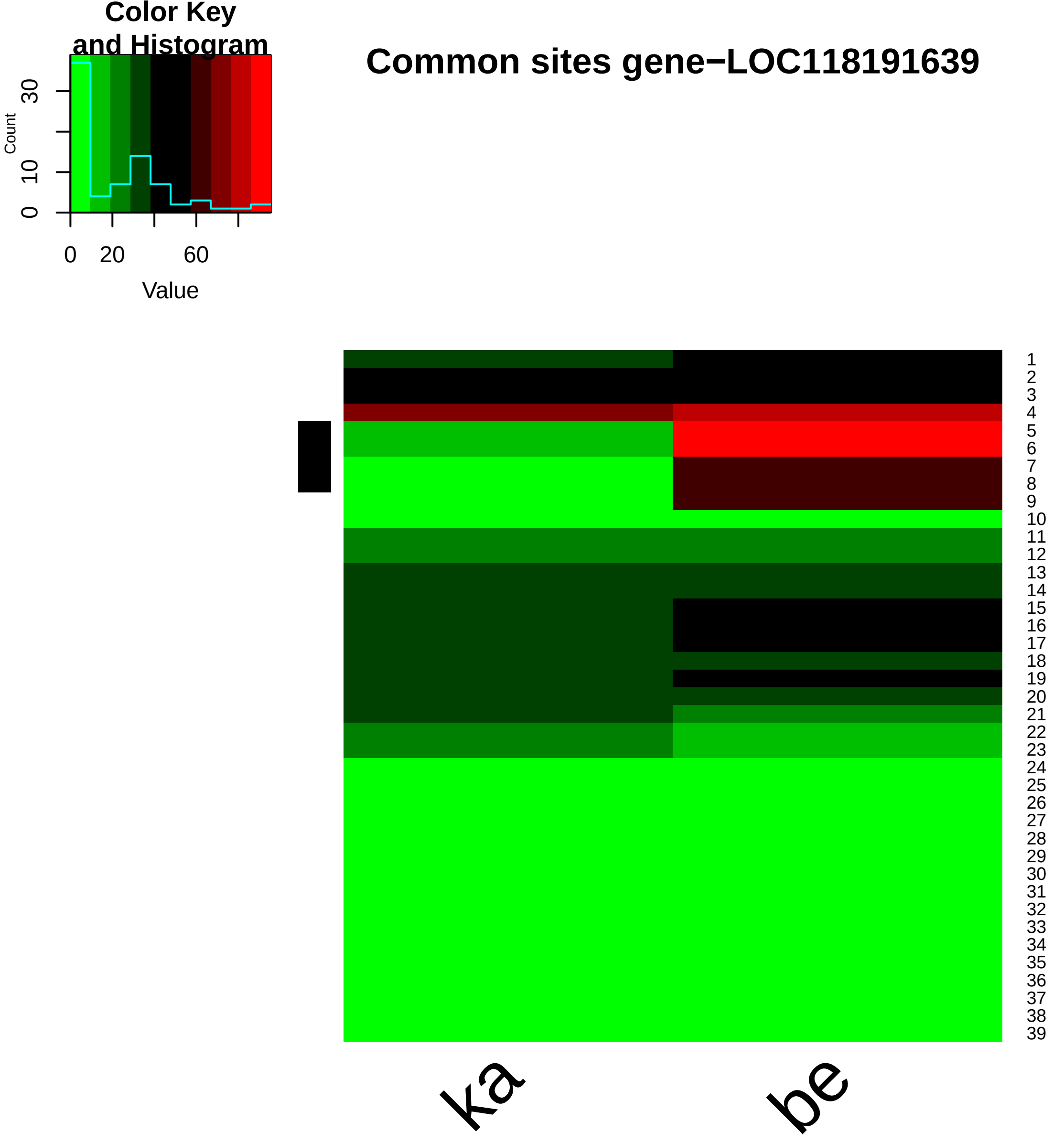

Color Key
and Histogram
Count
0
10
30
0
20
60
Value
Common sites gene−LOC118191639
ka
be

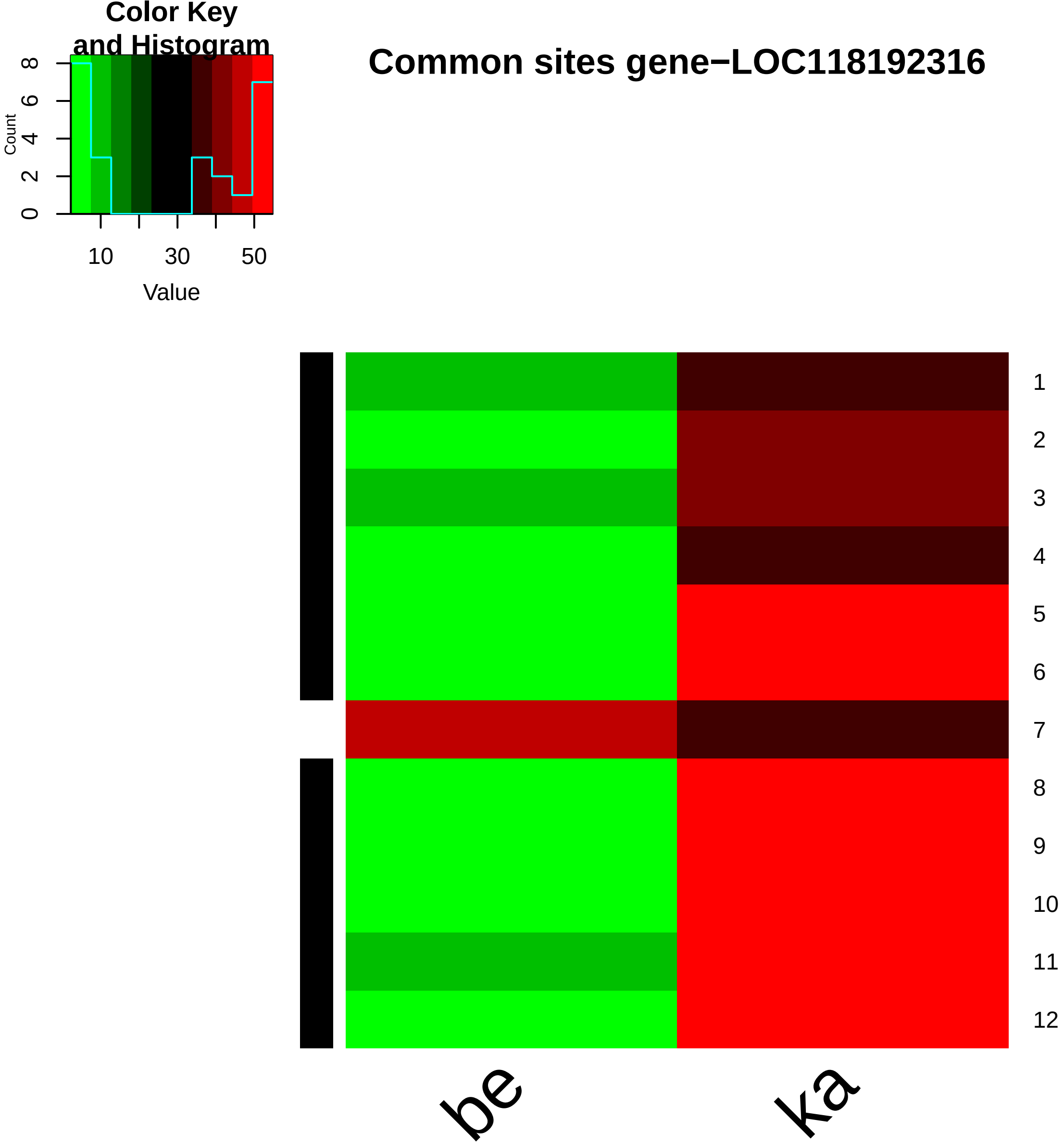

Color Key
and Histogram
Count
0
2
4
6
8
10
30
50
Value
Common sites gene−LOC118192316
1
2
3
4
5
6
7
8
9
10
11
12
be
ka

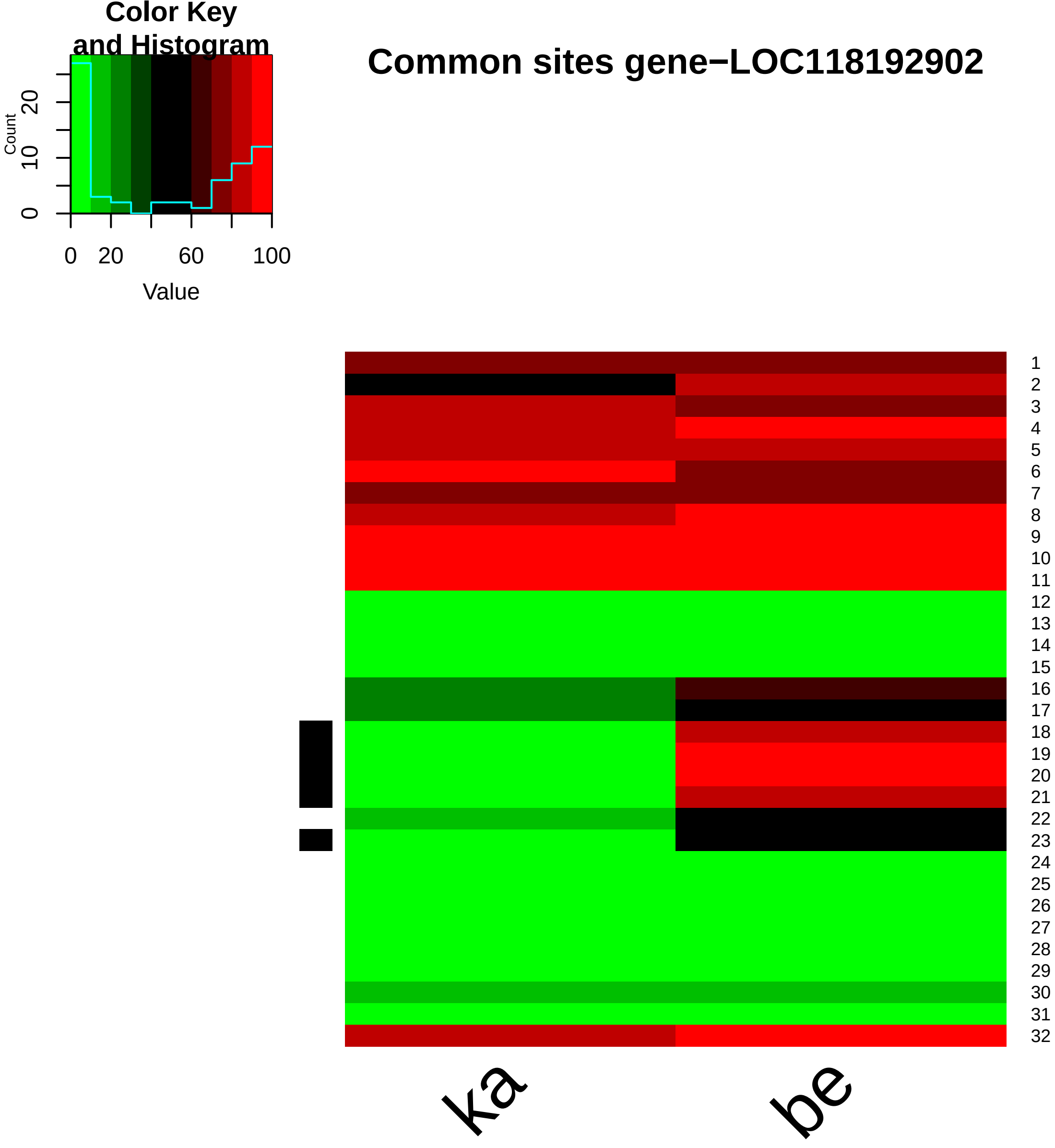
Color Key
and Histogram
Count
0
10
20
0
20
60
100
Value
Common sites gene−LOC118192902
ka
be

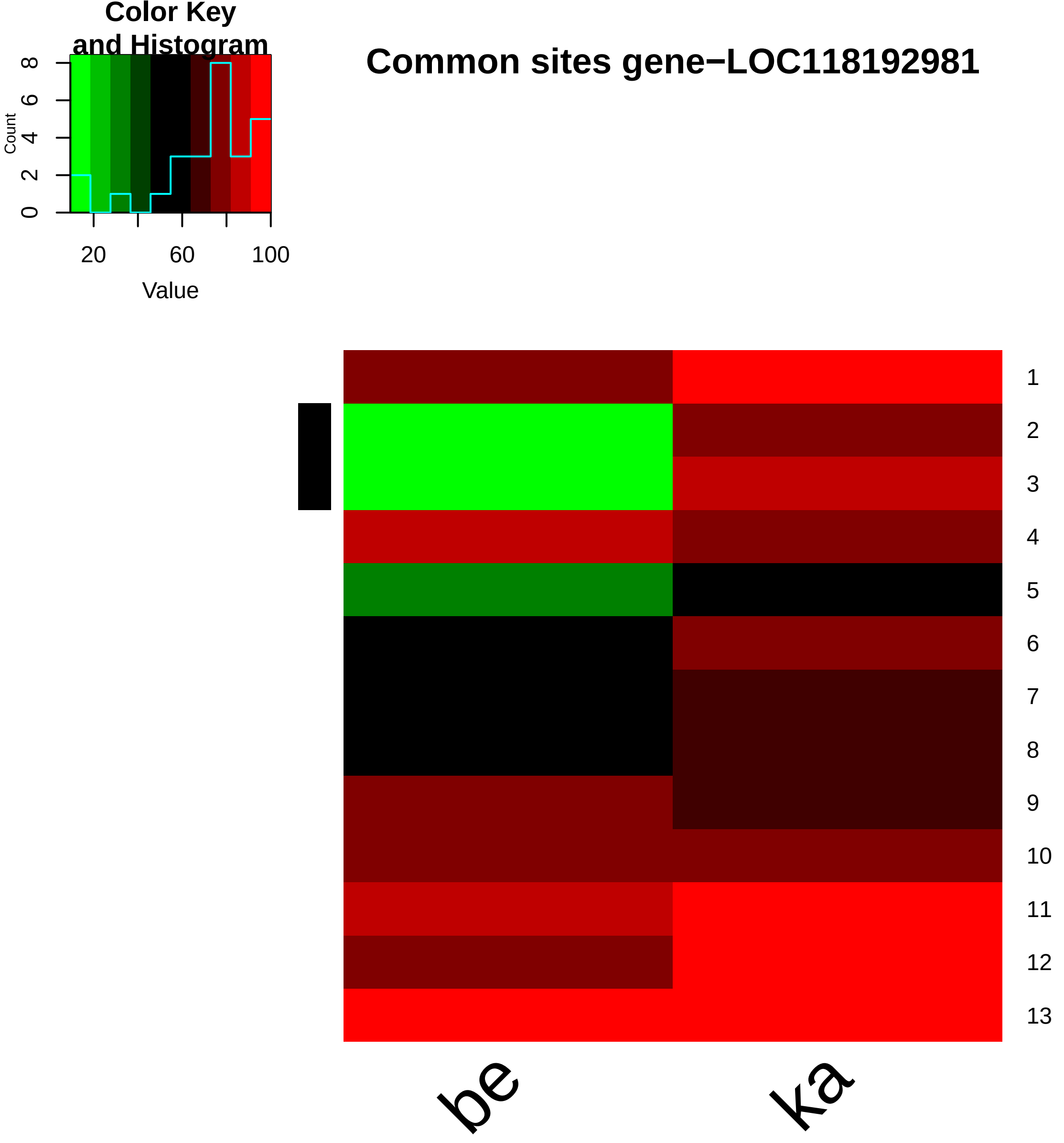
Color Key
and Histogram
Count
0
2
4
6
8
20
60
100
Value
Common sites gene−LOC118192981
1
2
3
4
5
6
7
8
9
10
11
12
13
be
ka

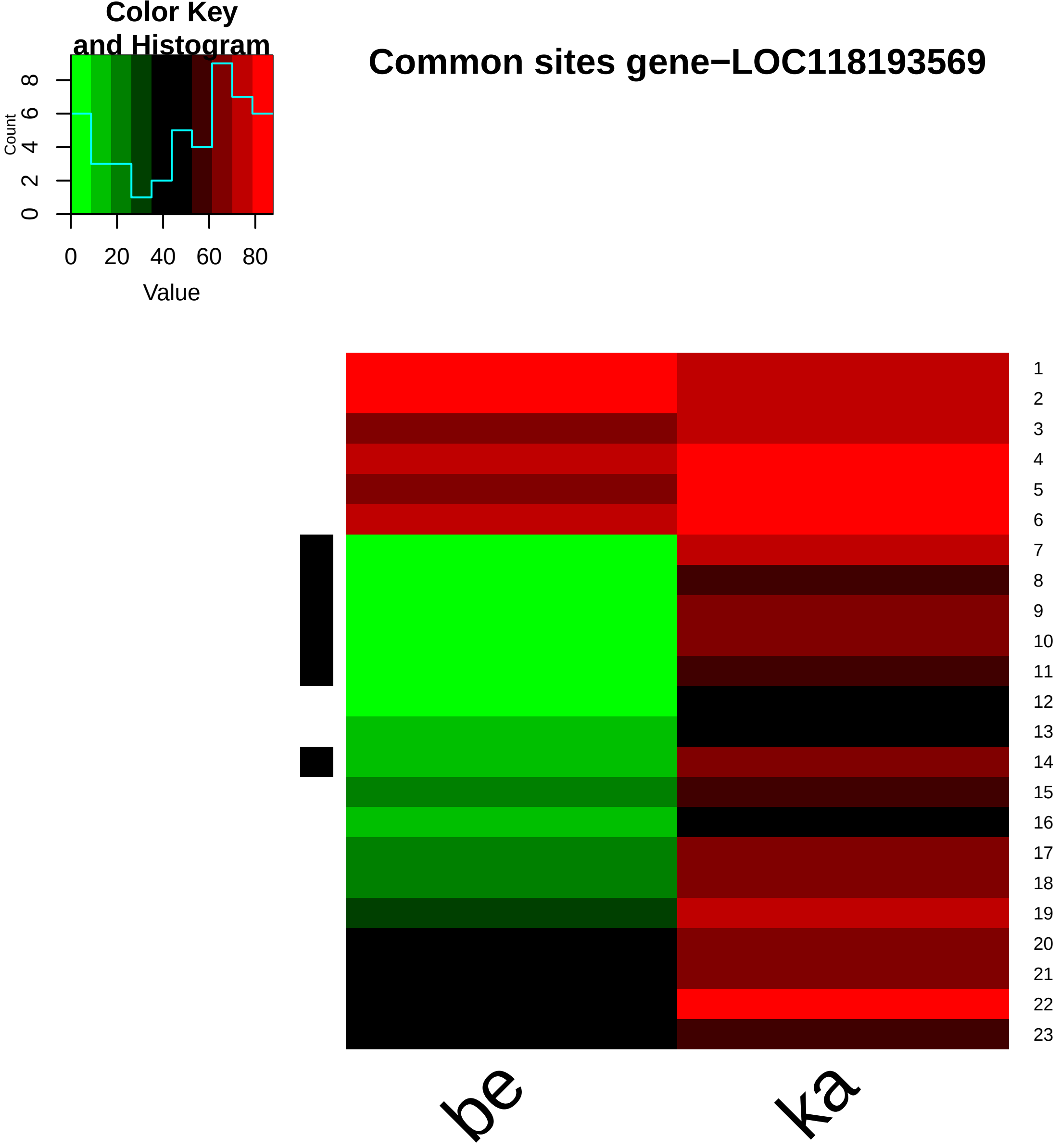
Color Key
and Histogram
Count
0 2 4 6 8
0 20 40 60 80
Value
Common sites gene−LOC118193569
be
ka

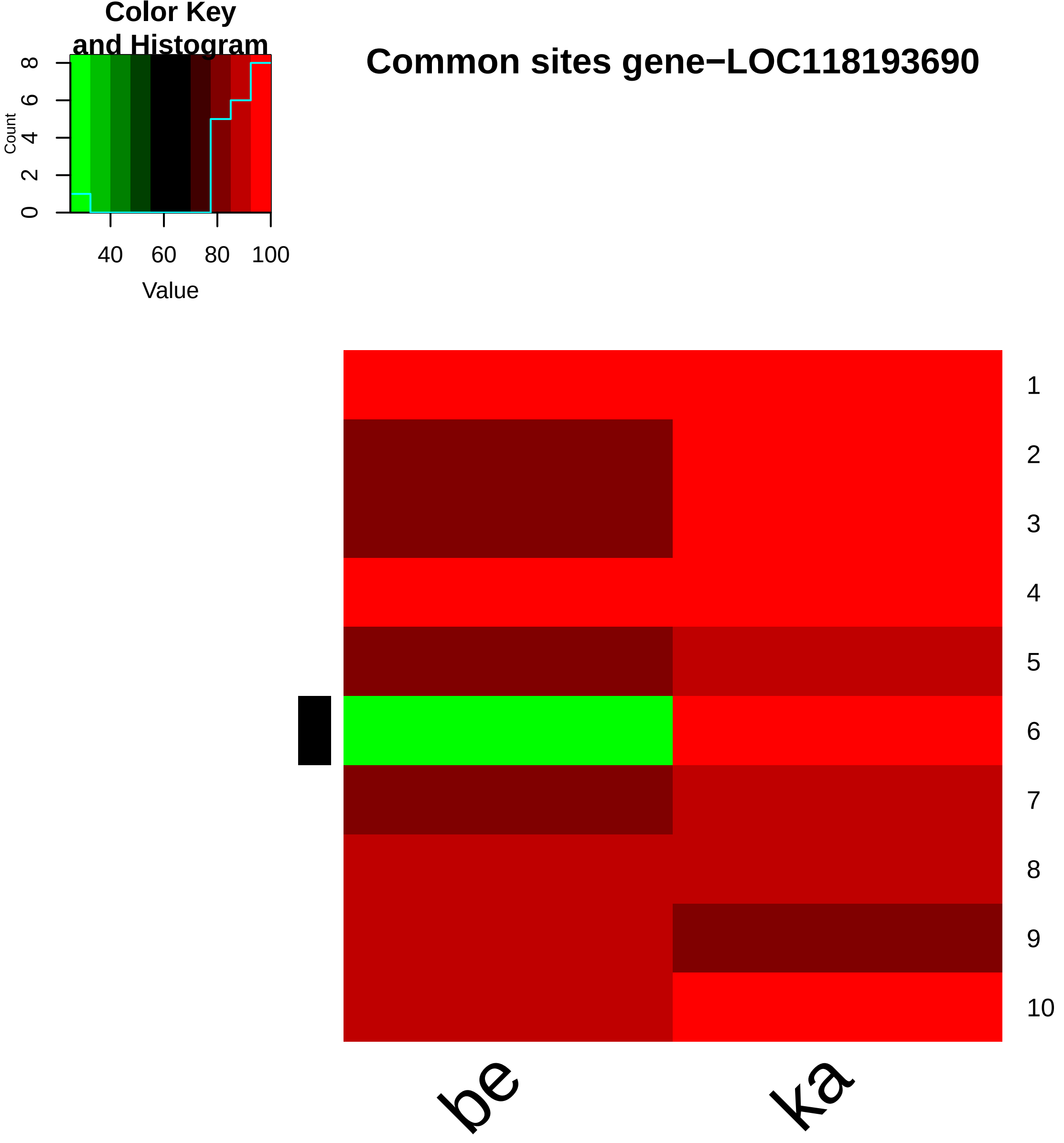
Color Key
and Histogram
Count
0
2
4
6
8
40
60
80
100
Value
Common sites gene−LOC118193690
1
2
3
4
5
6
7
8
9
10
be
ka

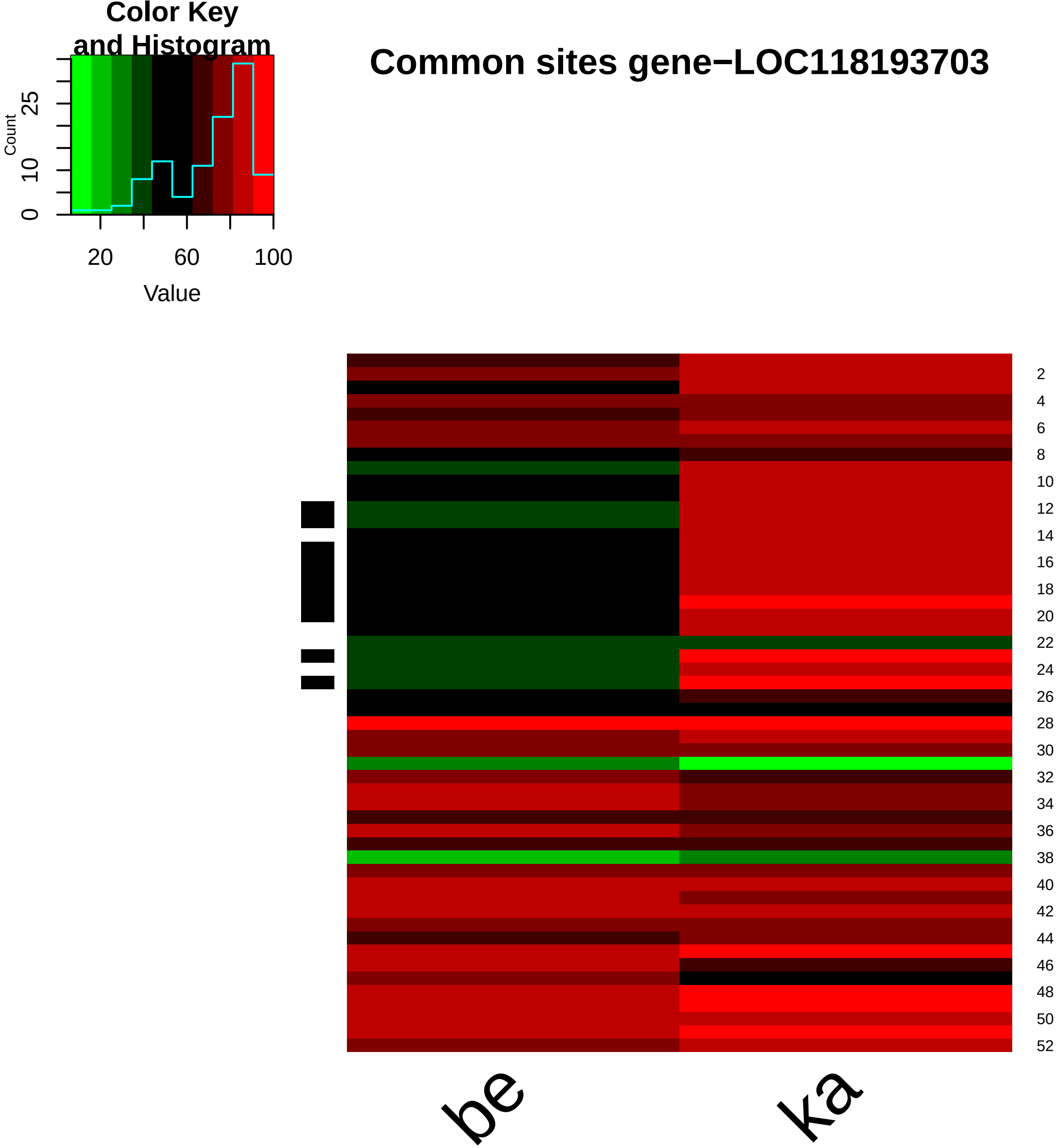

Color Key
and Histogram
Count
0
10
25
20
60
100
Value
Common sites gene−LOC118193703
2
4
6
8
10
12
14
16
18
20
22
24
26
28
30
32
34
36
38
40
42
44
46
48
50
52
be
ka

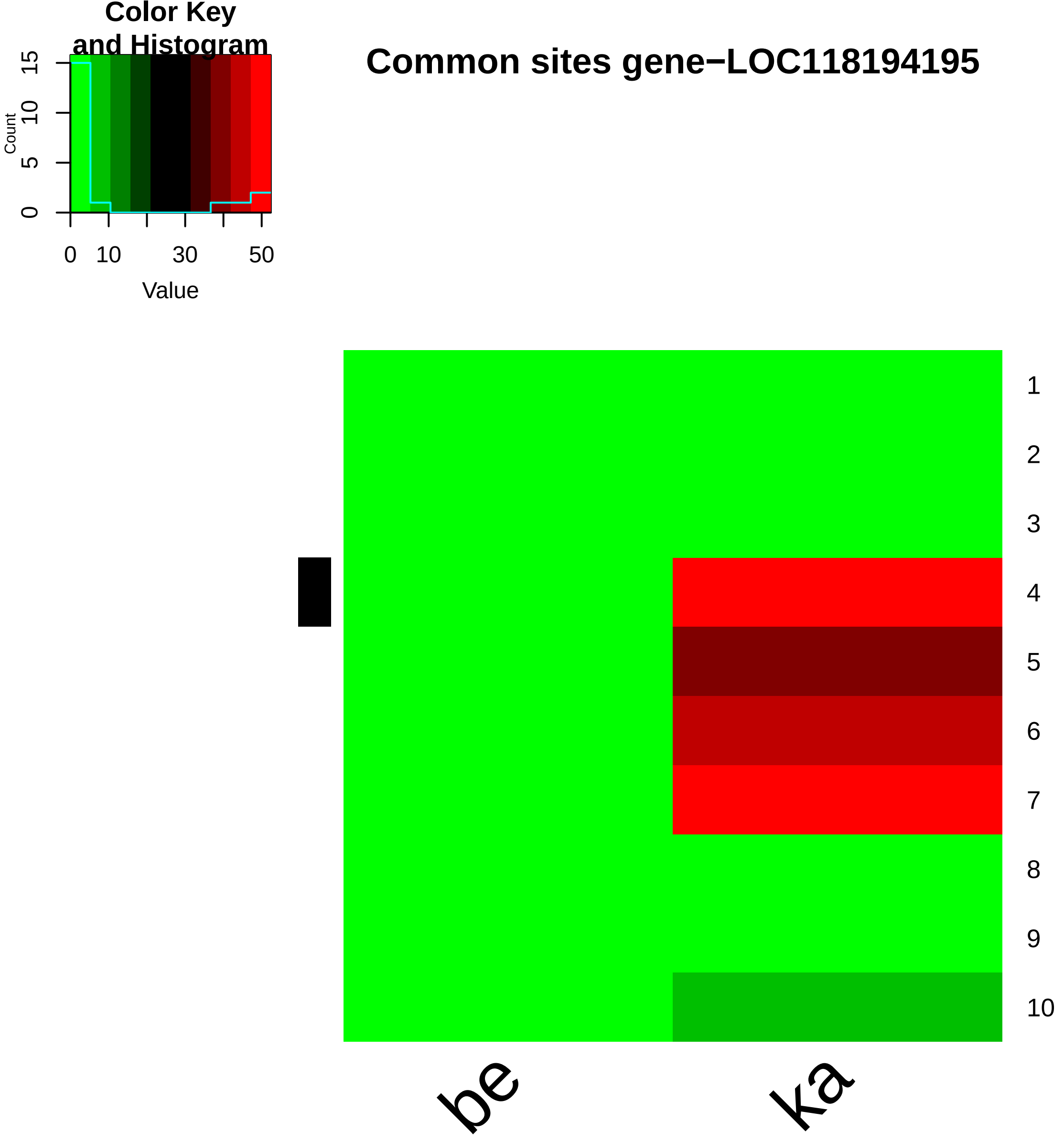
Color Key and Histogram
Count
0
5
10
15
0
10
30
50
Value
Common sites gene−LOC118194195
1
2
3
4
5
6
7
8
9
10
be
ka

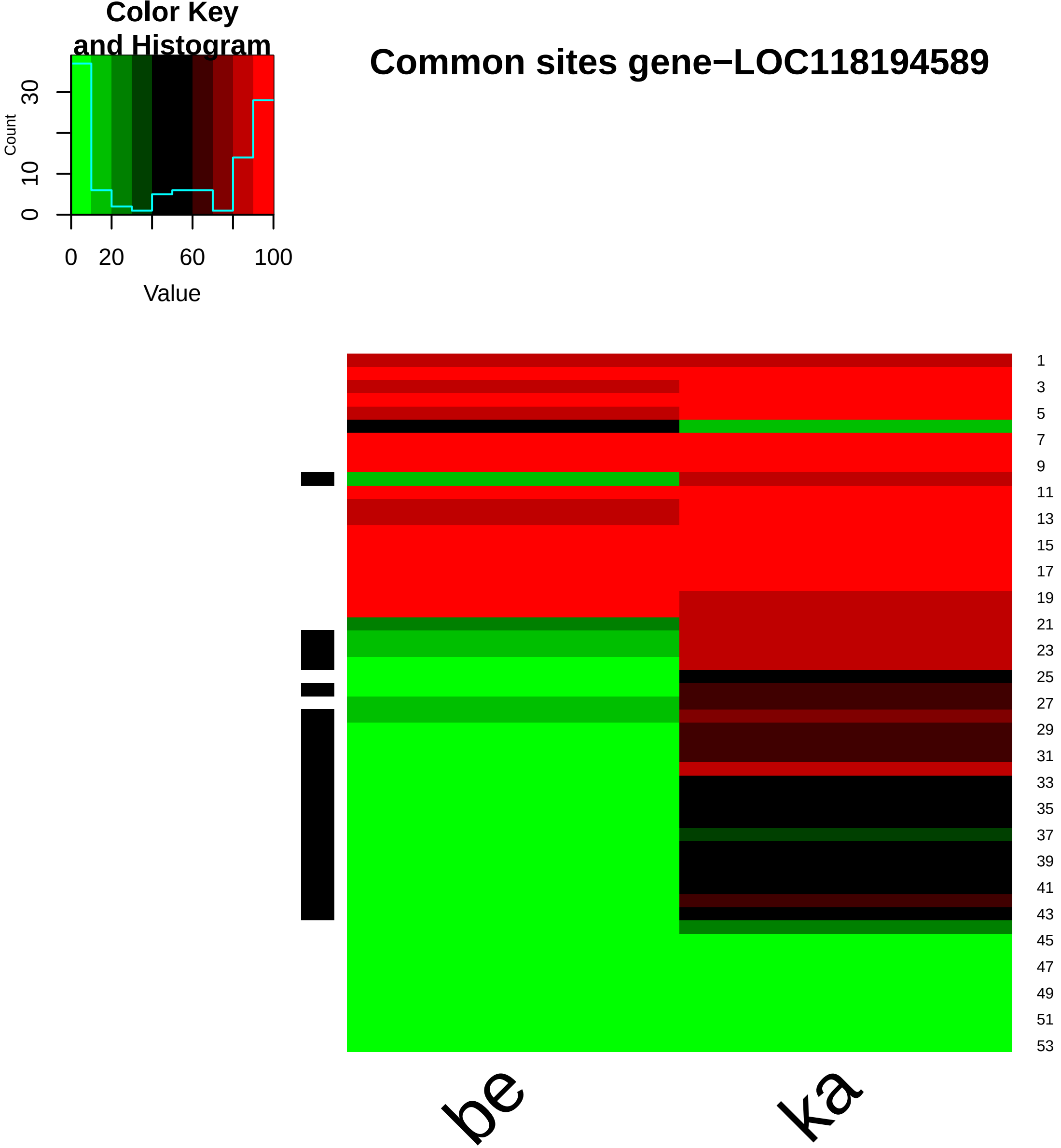

Color Key
and Histogram
Count
0
10
30
0
20
60
100
Value
Common sites gene−LOC118194589
1
3
5
7
9
11
13
15
17
19
21
23
25
27
29
31
33
35
37
39
41
43
45
47
49
51
53
be
ka

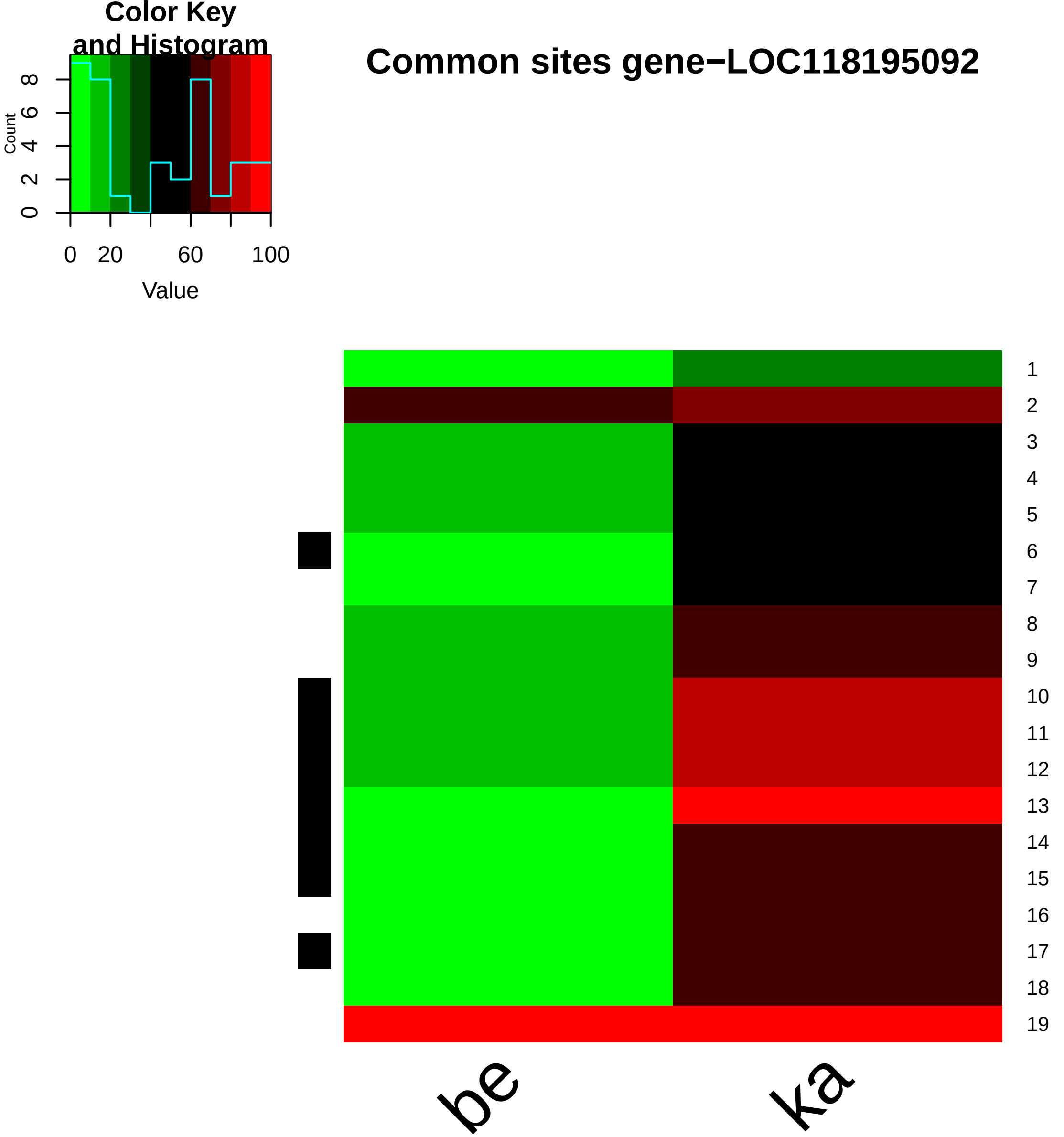
Color Key
and Histogram
Count
0
2
4
6
8
0
20
60
100
Value
Common sites gene−LOC118195092
be
ka

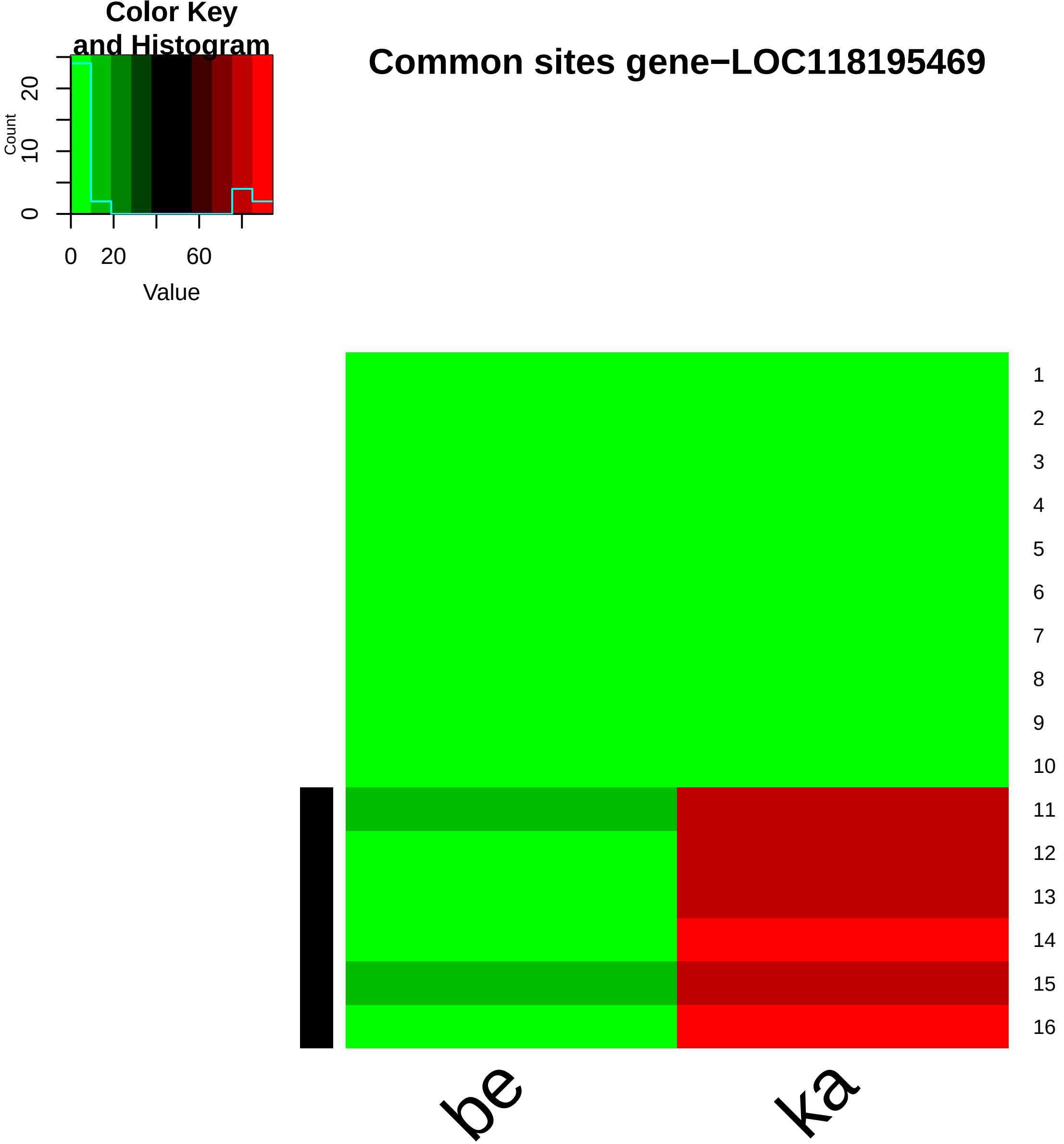
Color Key
and Histogram
Count
0
10
20
0
20
60
Value
Common sites gene−LOC118195469
be
ka

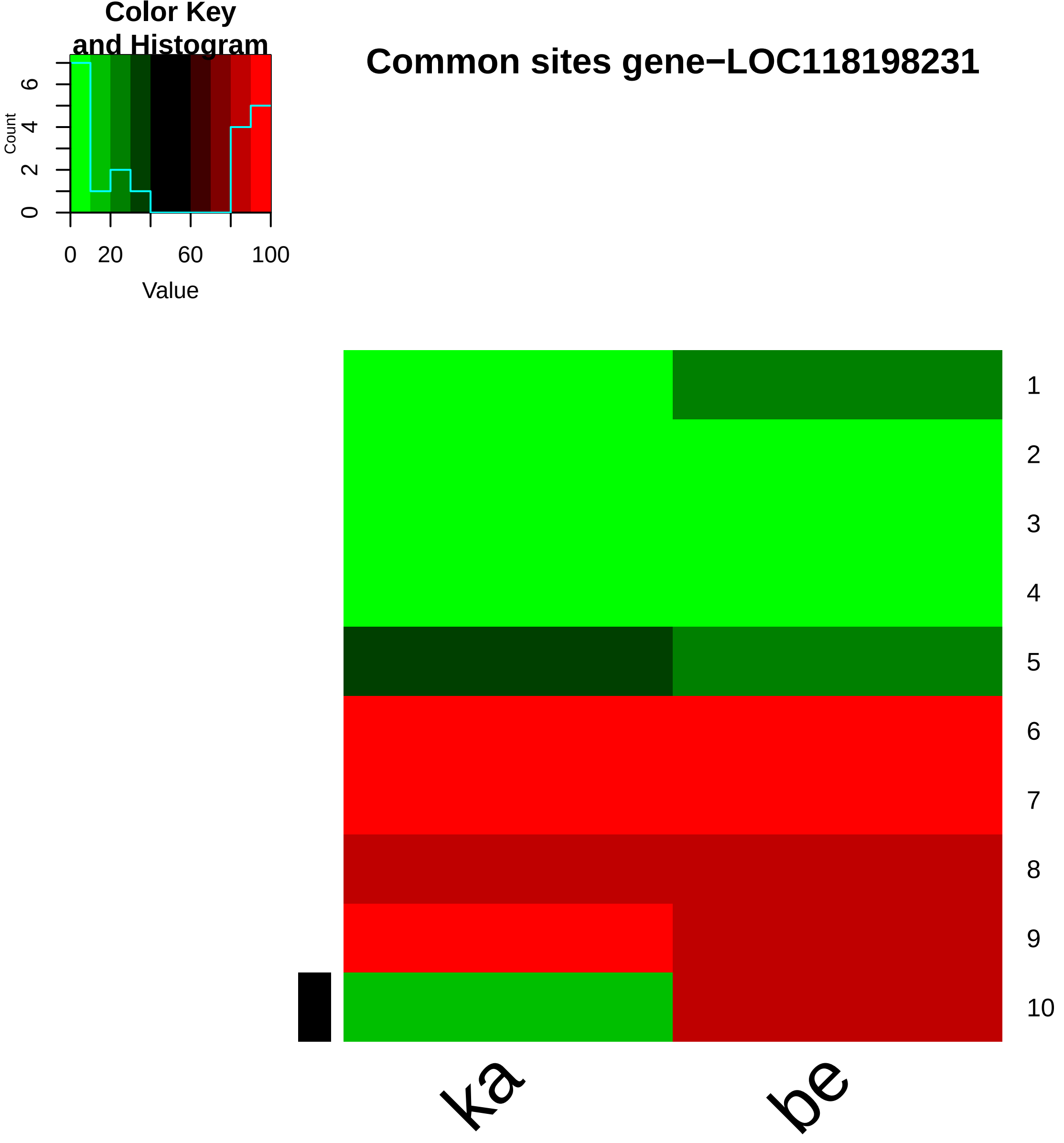
Color Key
and Histogram
Count
0
2
4
6
0
20
60
100
Value
Common sites gene−LOC118198231
1
2
3
4
5
6
7
8
9
10
ka
be

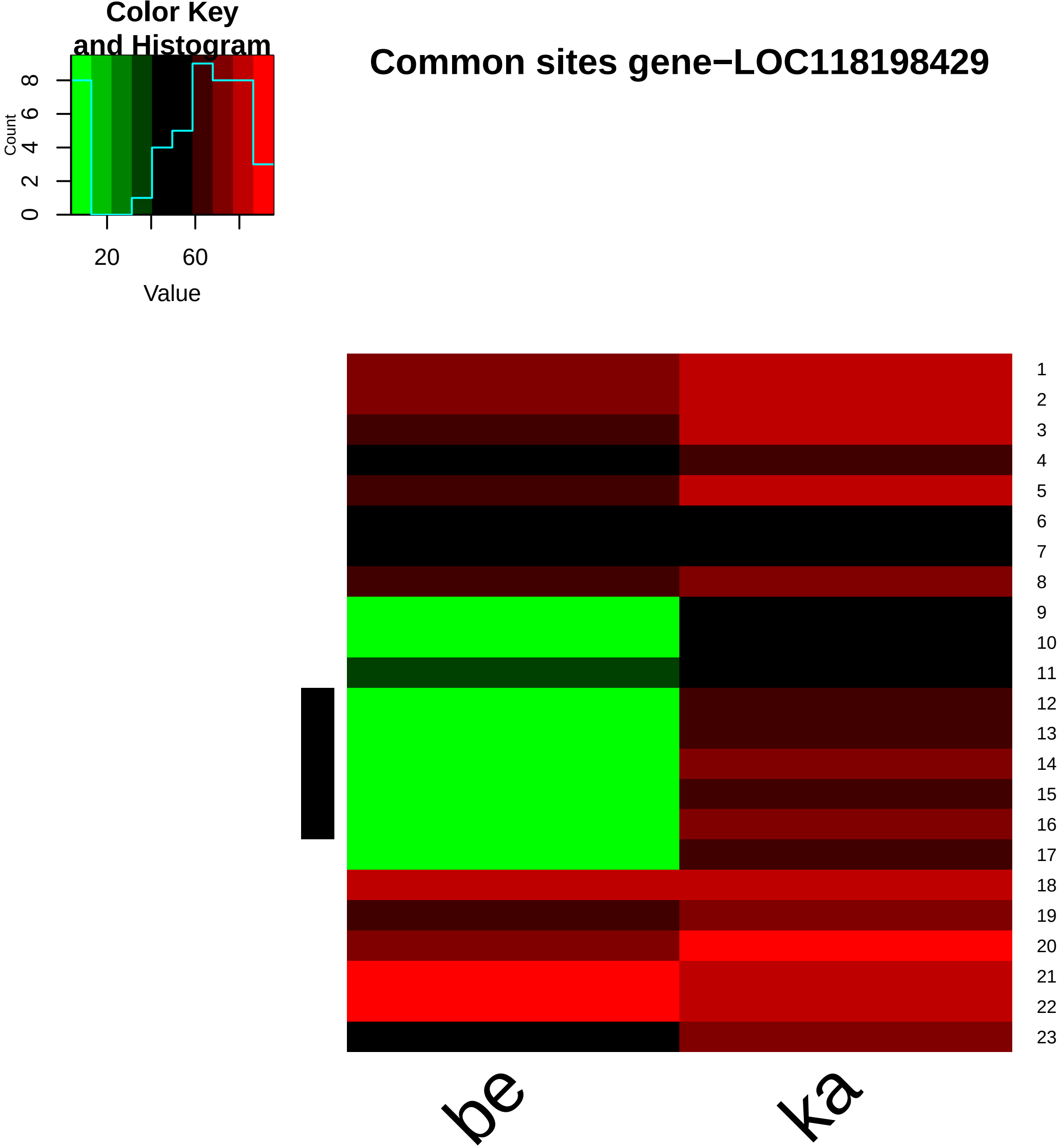

Color Key
and Histogram
Count
0
2
4
6
8
20
60
Value
Common sites gene−LOC118198429
be
ka

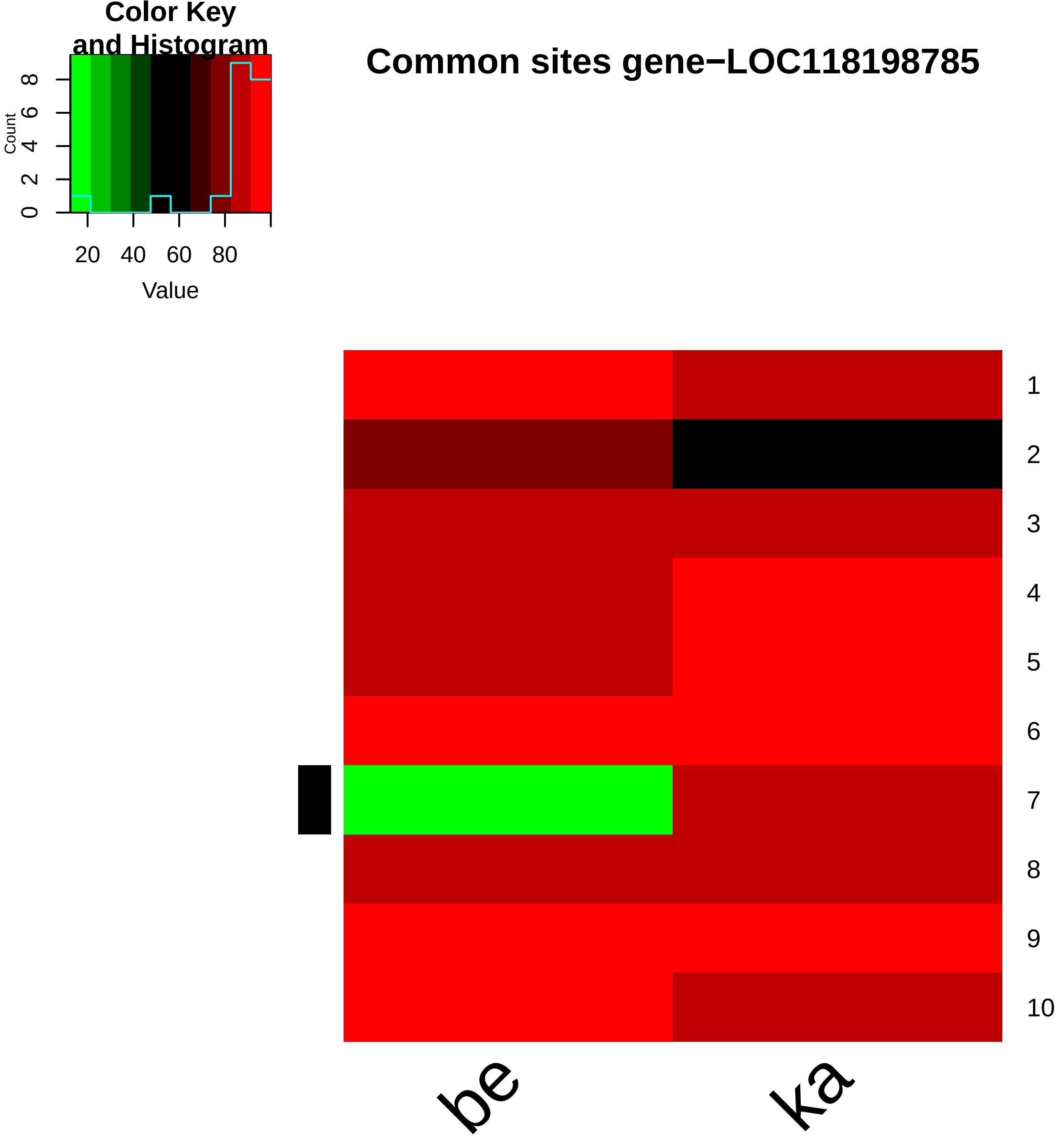
Color Key and Histogram
Count
0 2 4 6 8
20 40 60 80
Value
Common sites gene−LOC118198785
1
2
3
4
5
6
7
8
9
10
be
ka

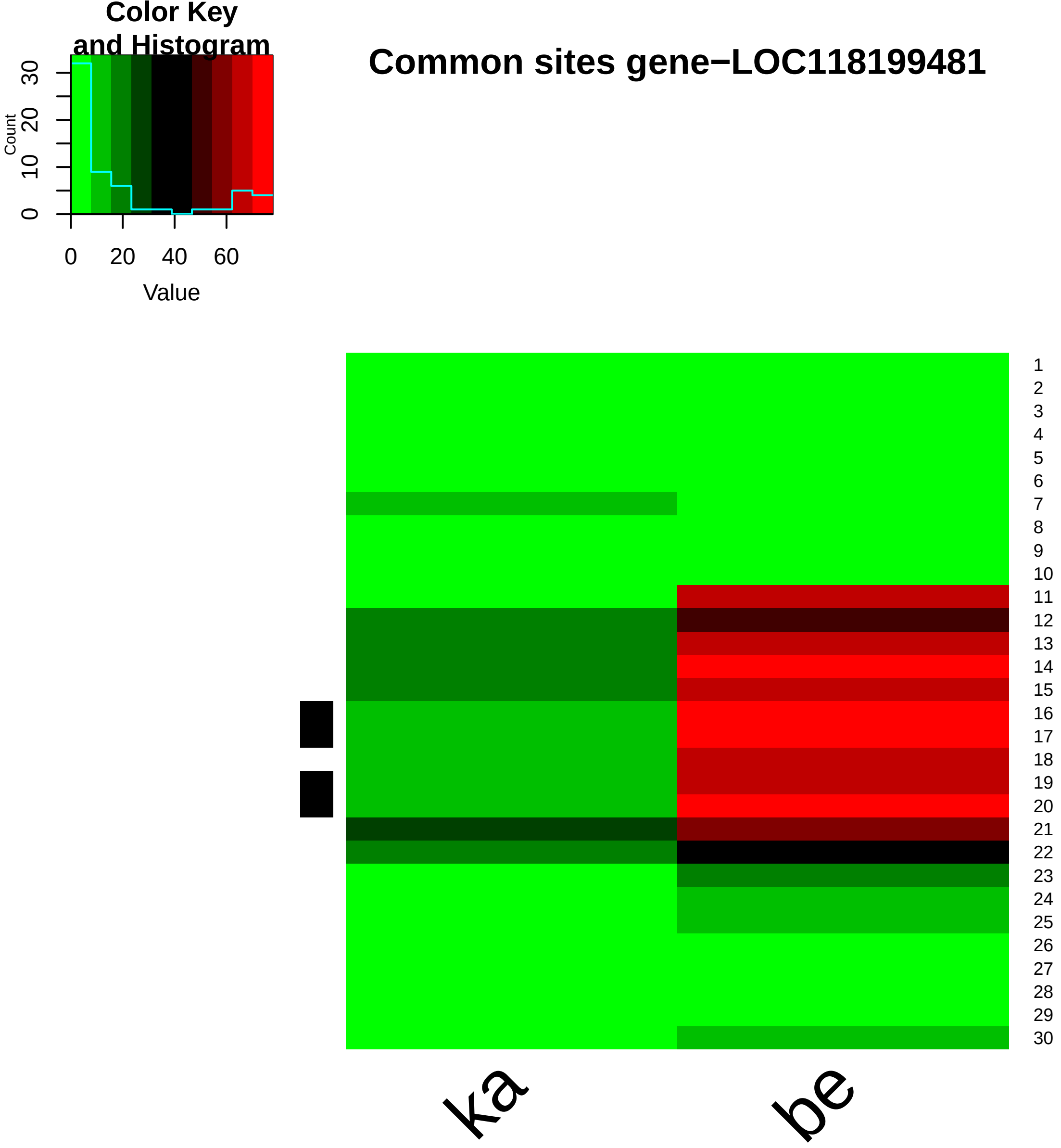
Color Key
and Histogram
Count
0
10
20
30
0
20
40
60
Value
Common sites gene−LOC118199481
ka
be

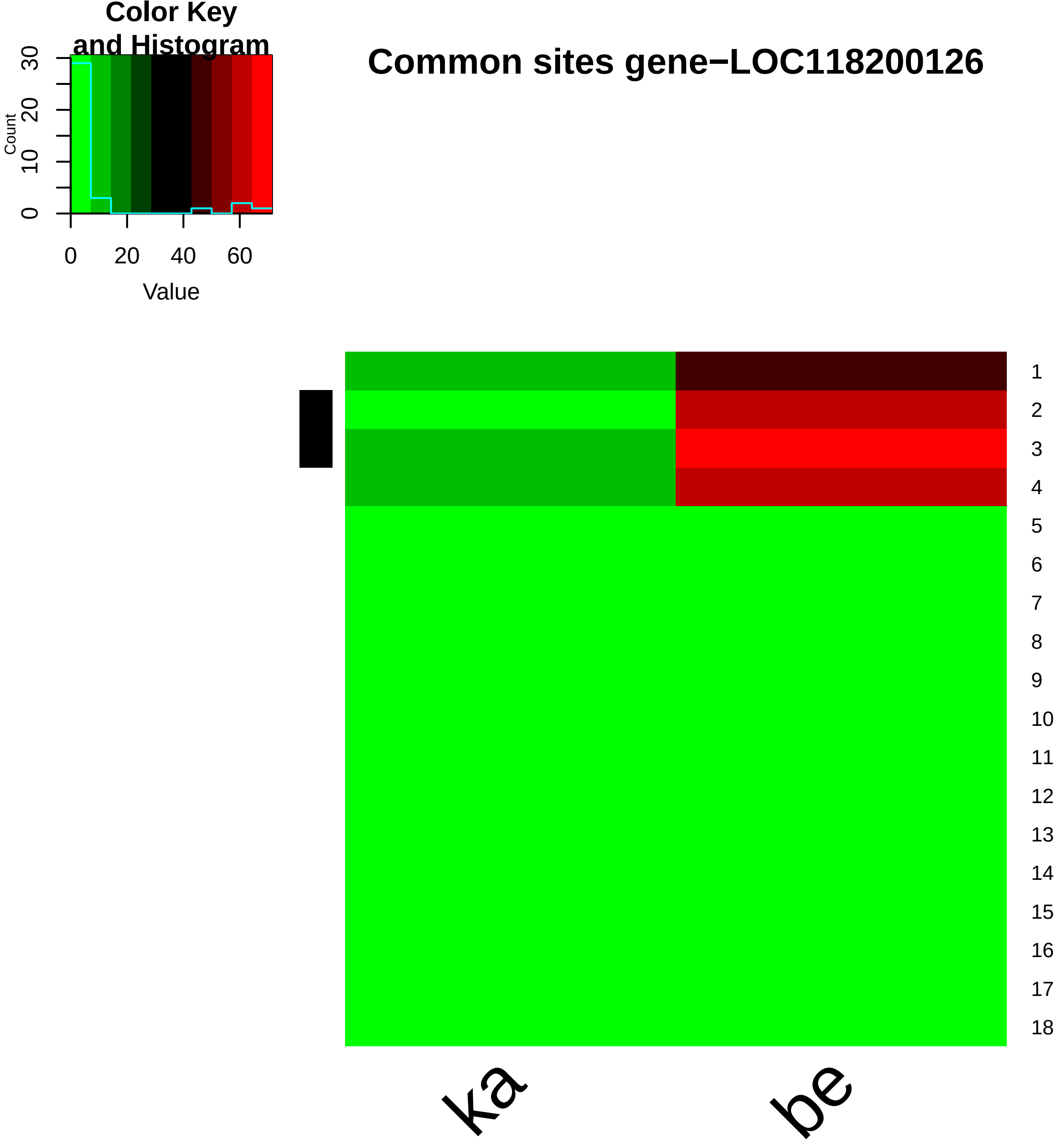

Color Key
and Histogram
Count
0
10
20
30
0
20
40
60
Value
Common sites gene−LOC118200126
ka
be

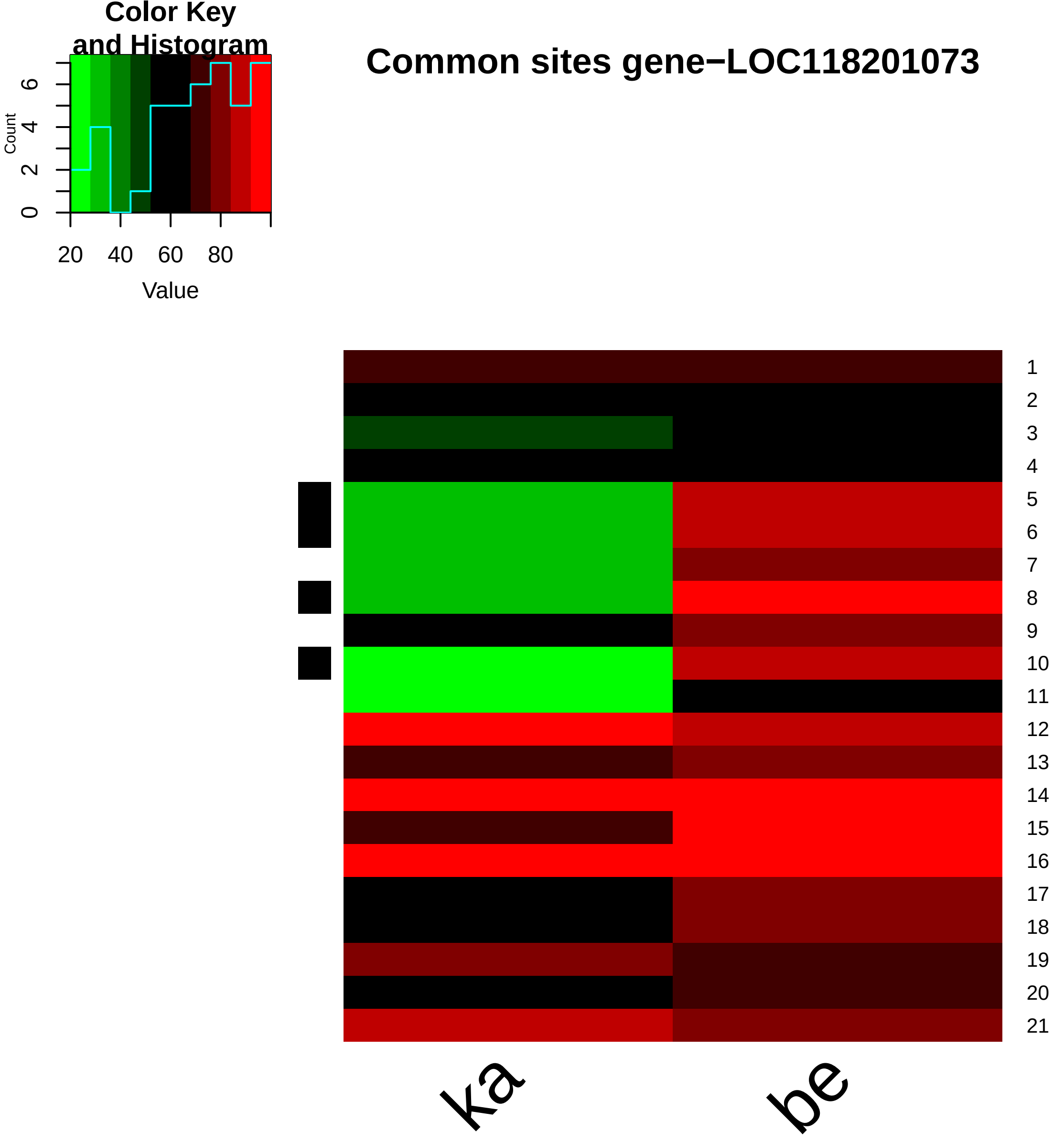
Color Key
and Histogram
Count
0
2
4
6
20
40
60
80
Value
Common sites gene−LOC118201073
ka
be

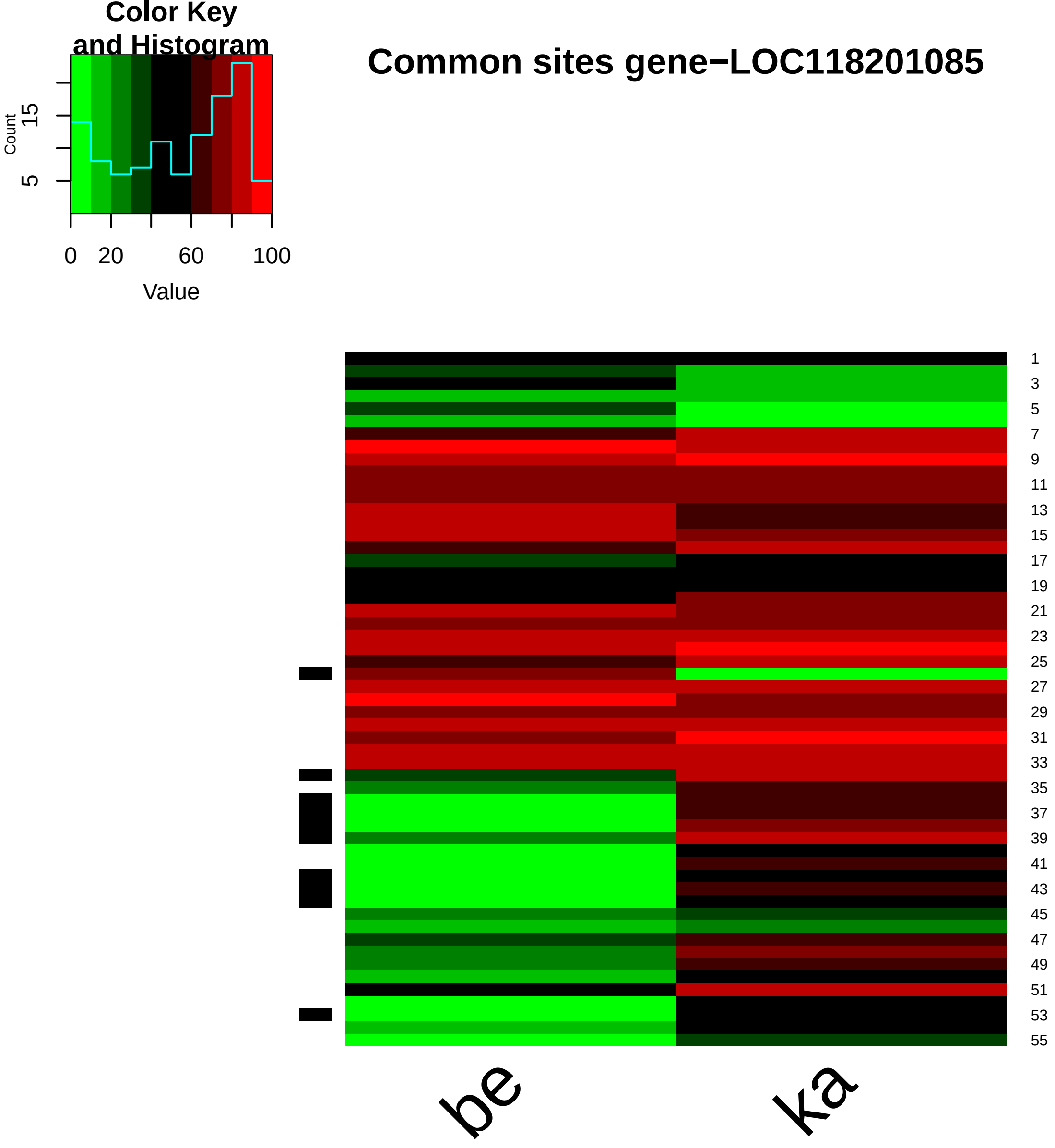
Color Key
and Histogram
Count
5
15
0
20
60
100
Value
Common sites gene−LOC118201085
1
3
5
7
9
11
13
15
17
19
21
23
25
27
29
31
33
35
37
39
41
43
45
47
49
51
53
55
be
ka

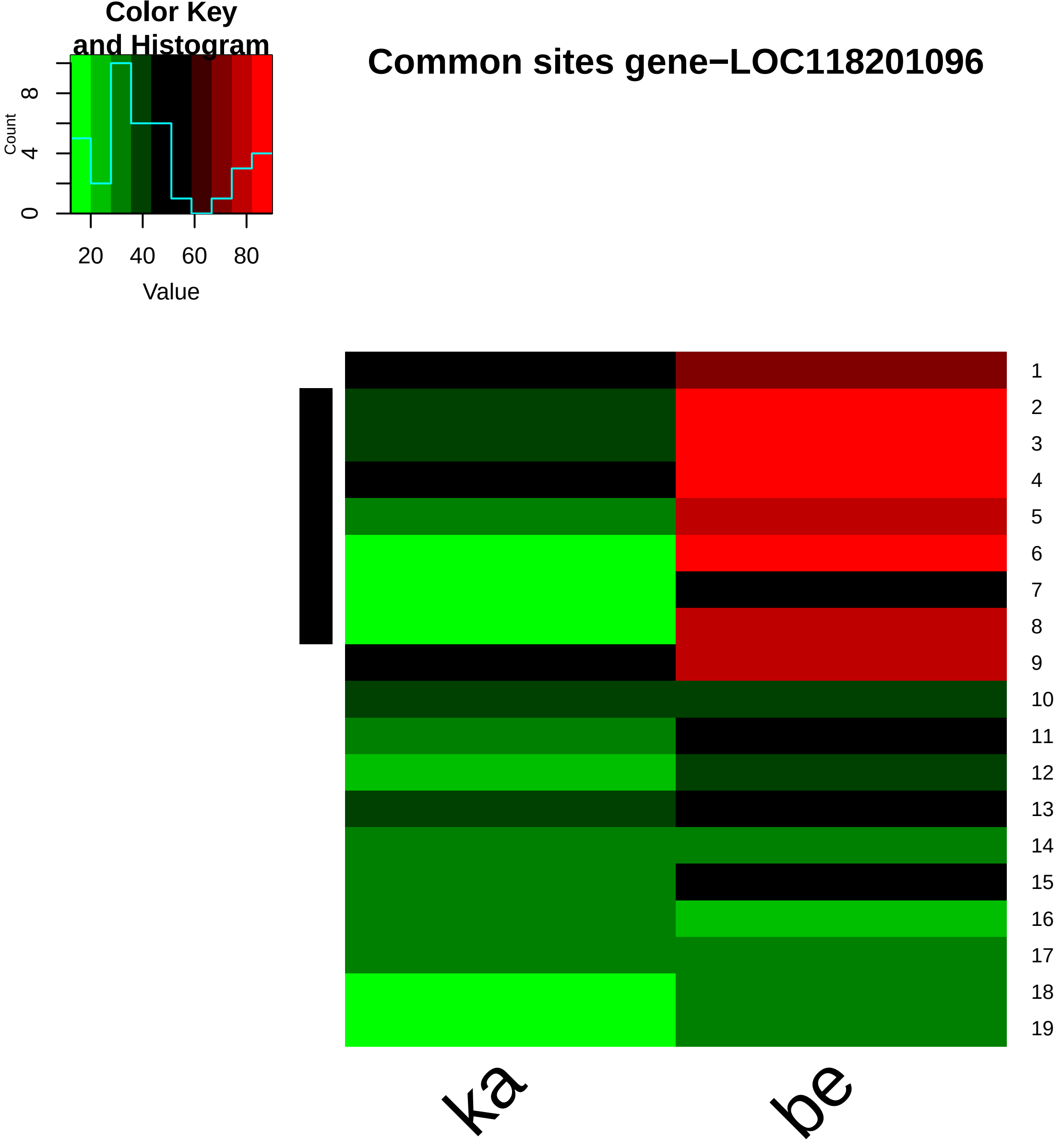

Color Key
and Histogram
Count
0
4
8
20
40
60
80
Value
Common sites gene−LOC118201096
ka
be

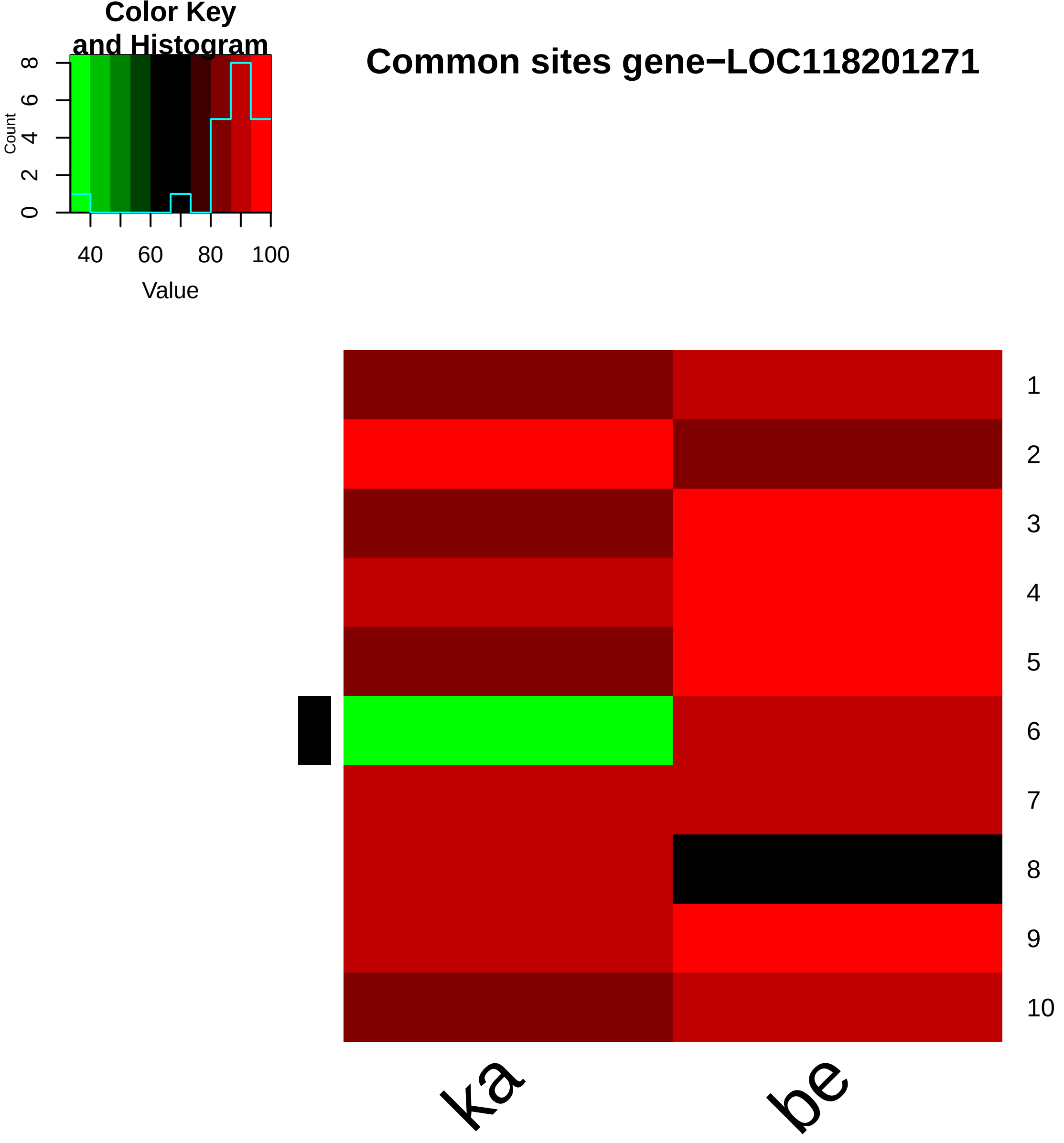

Color Key
and Histogram
Count
0
2
4
6
8
40
60
80
100
Value
Common sites gene−LOC118201271
1
2
3
4
5
6
7
8
9
10
ka
be

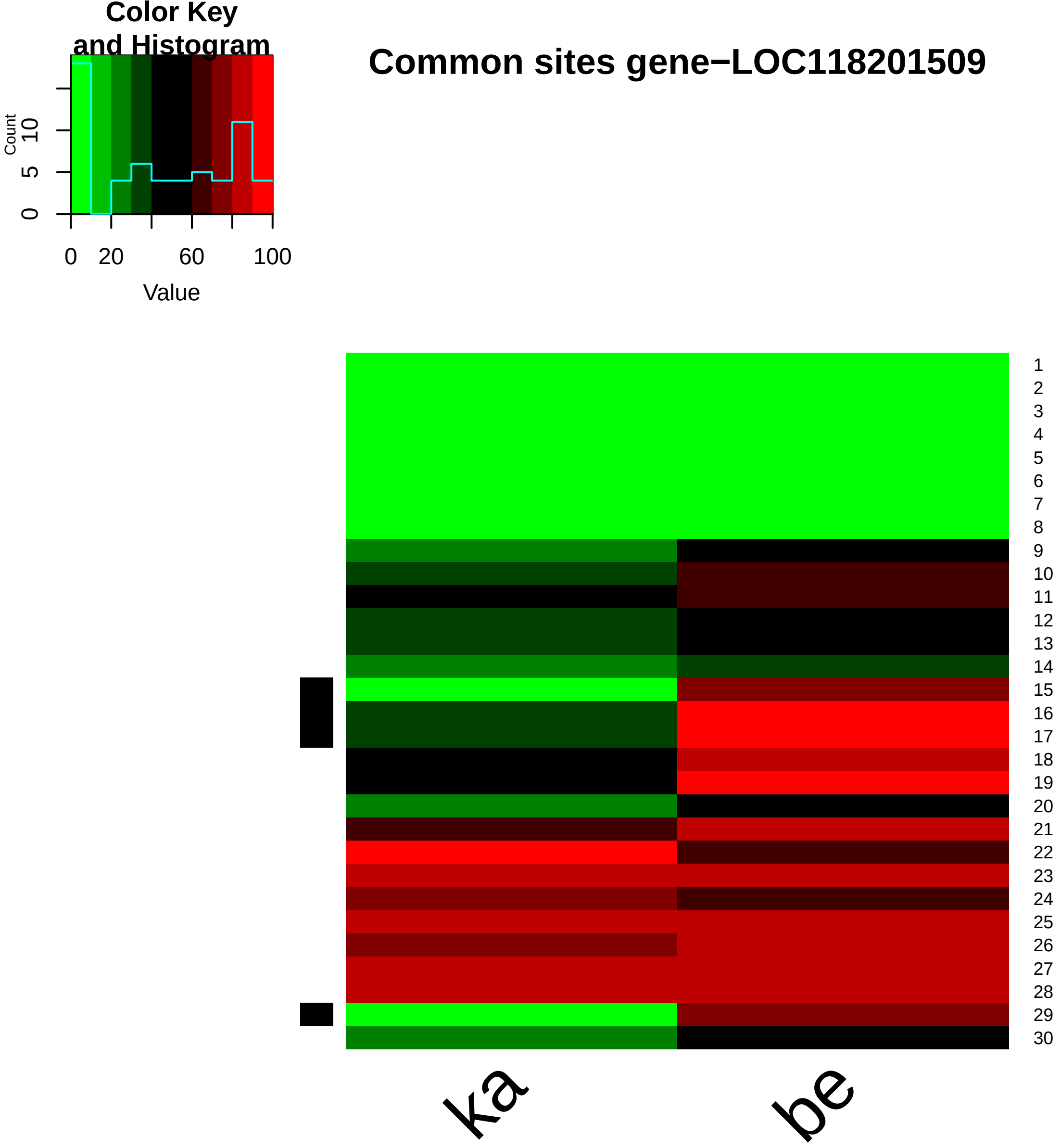
Color Key
and Histogram
Count
0
5
10
0
20
60
100
Value
Common sites gene−LOC118201509
ka
be

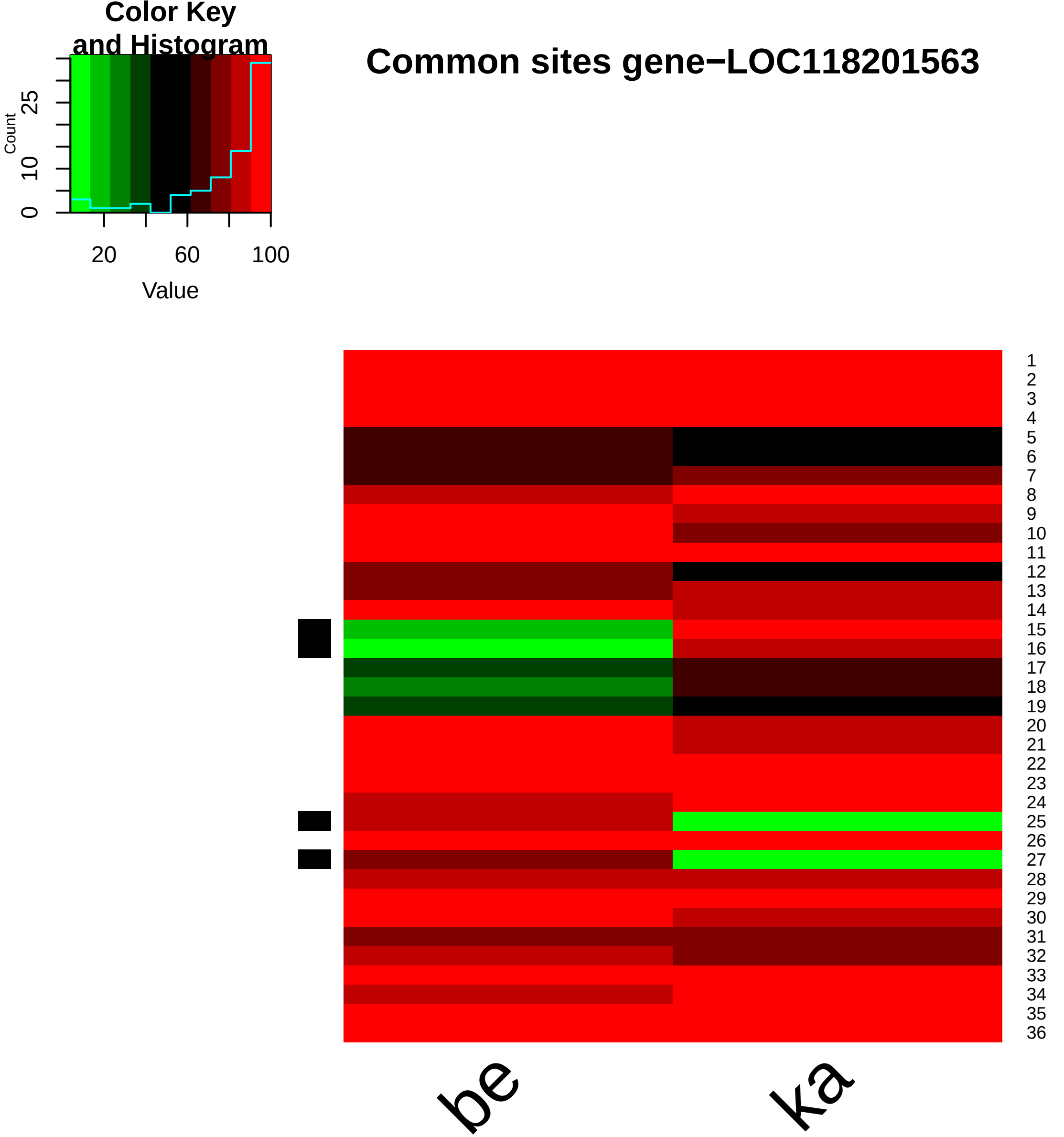
Color Key
and Histogram
Count
0
10
25
20
60
100
Value
Common sites gene−LOC118201563
be
ka

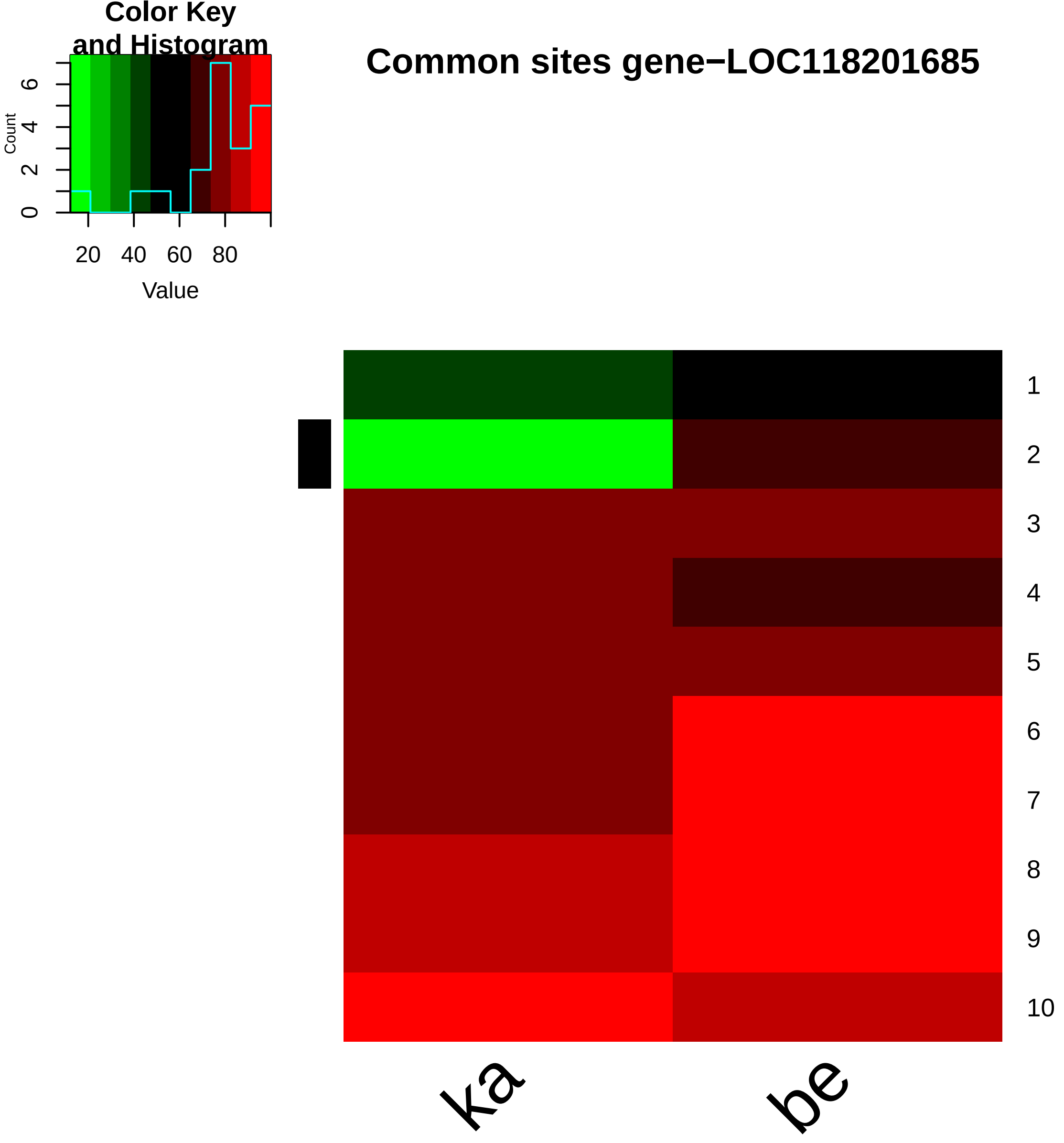

Color Key
and Histogram
Count
Value
Common sites gene−LOC118201685
1
2
3
4
5
6
7
8
9
10
ka
be

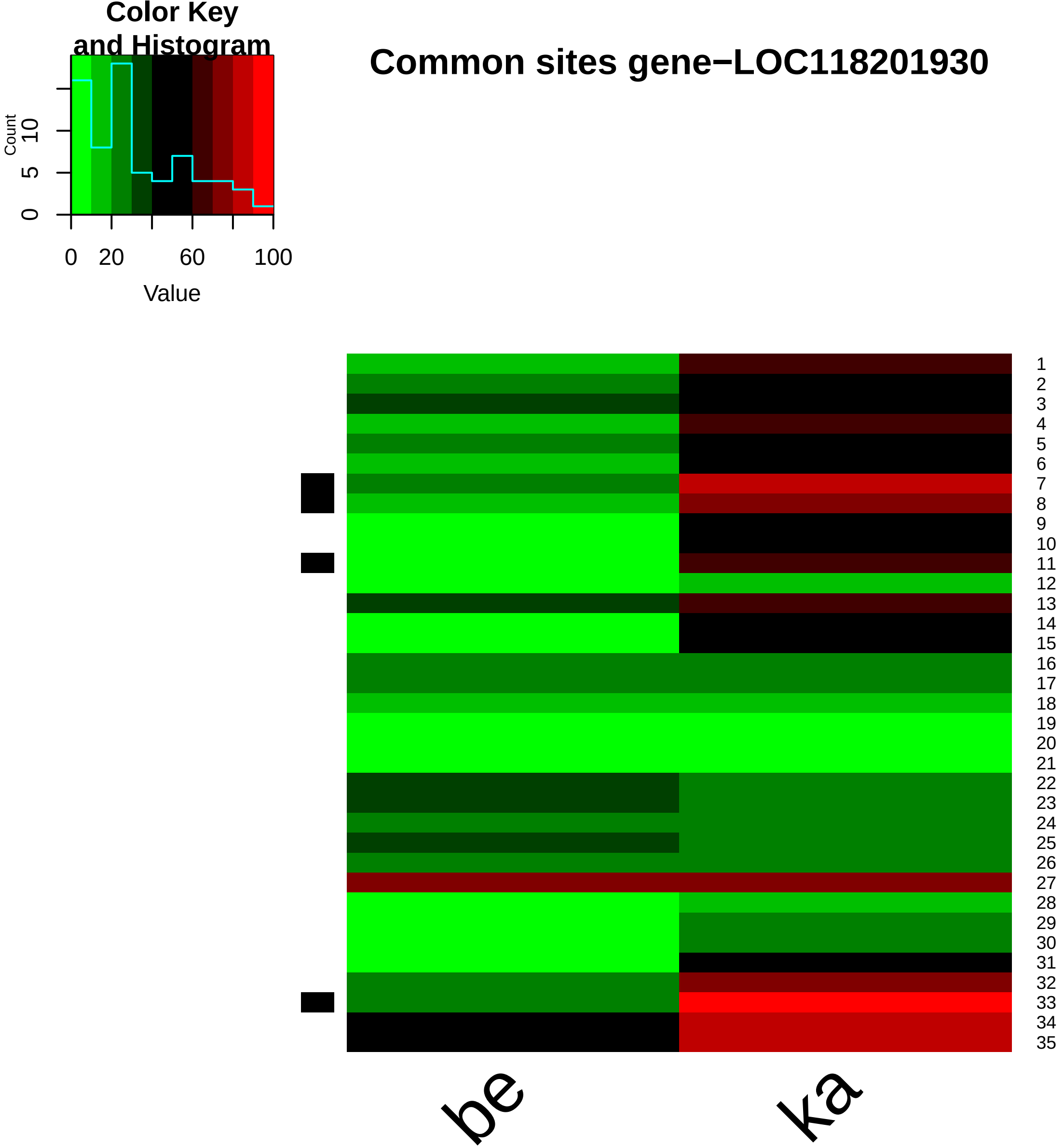
Color Key
and Histogram
Count
0
5
10
0
20
60
100
Value
Common sites gene−LOC118201930
be
ka

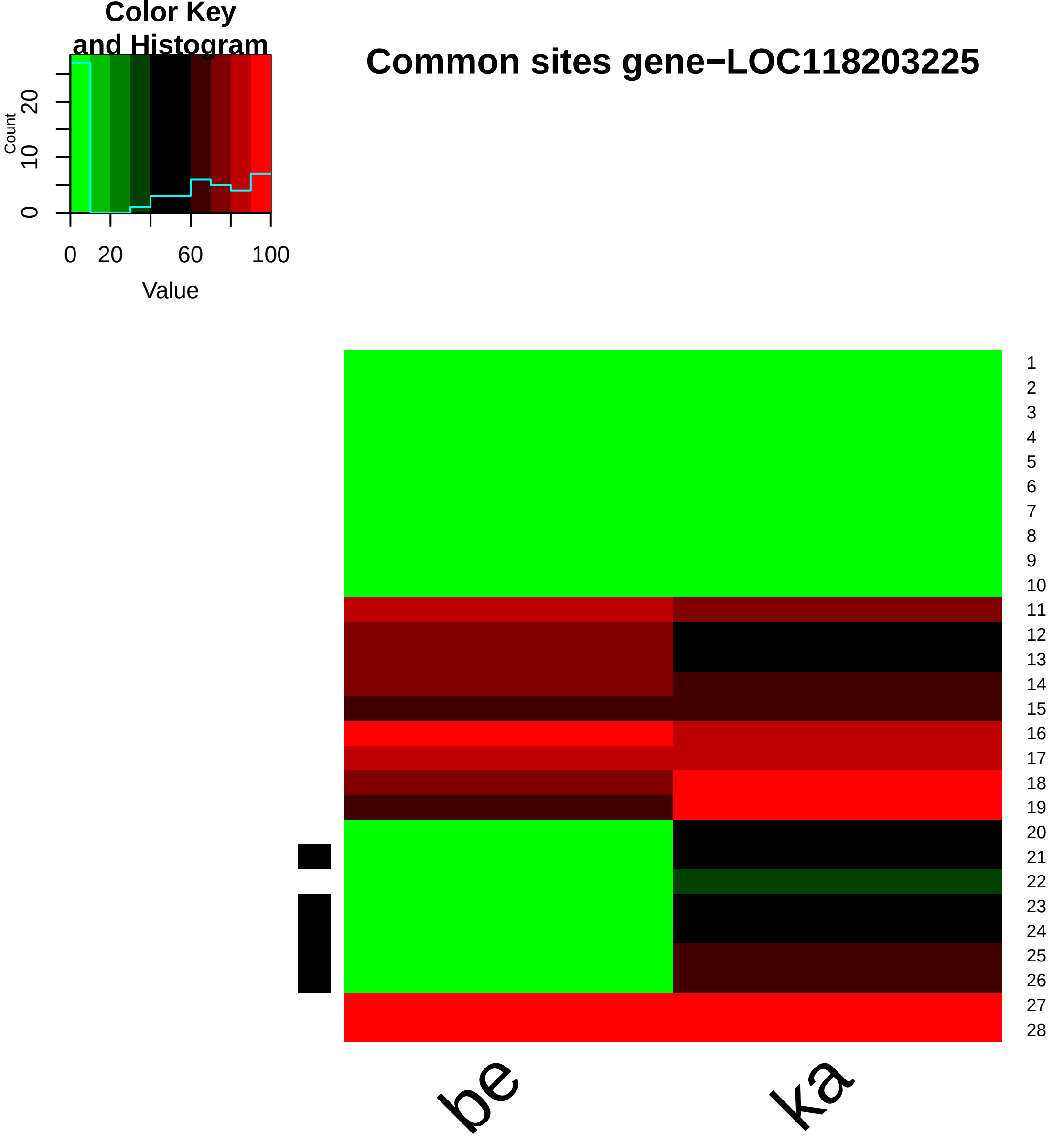
Color Key and Histogram
Count
0
10
20
0
20
60
100
Value
Common sites gene−LOC118203225
be
ka

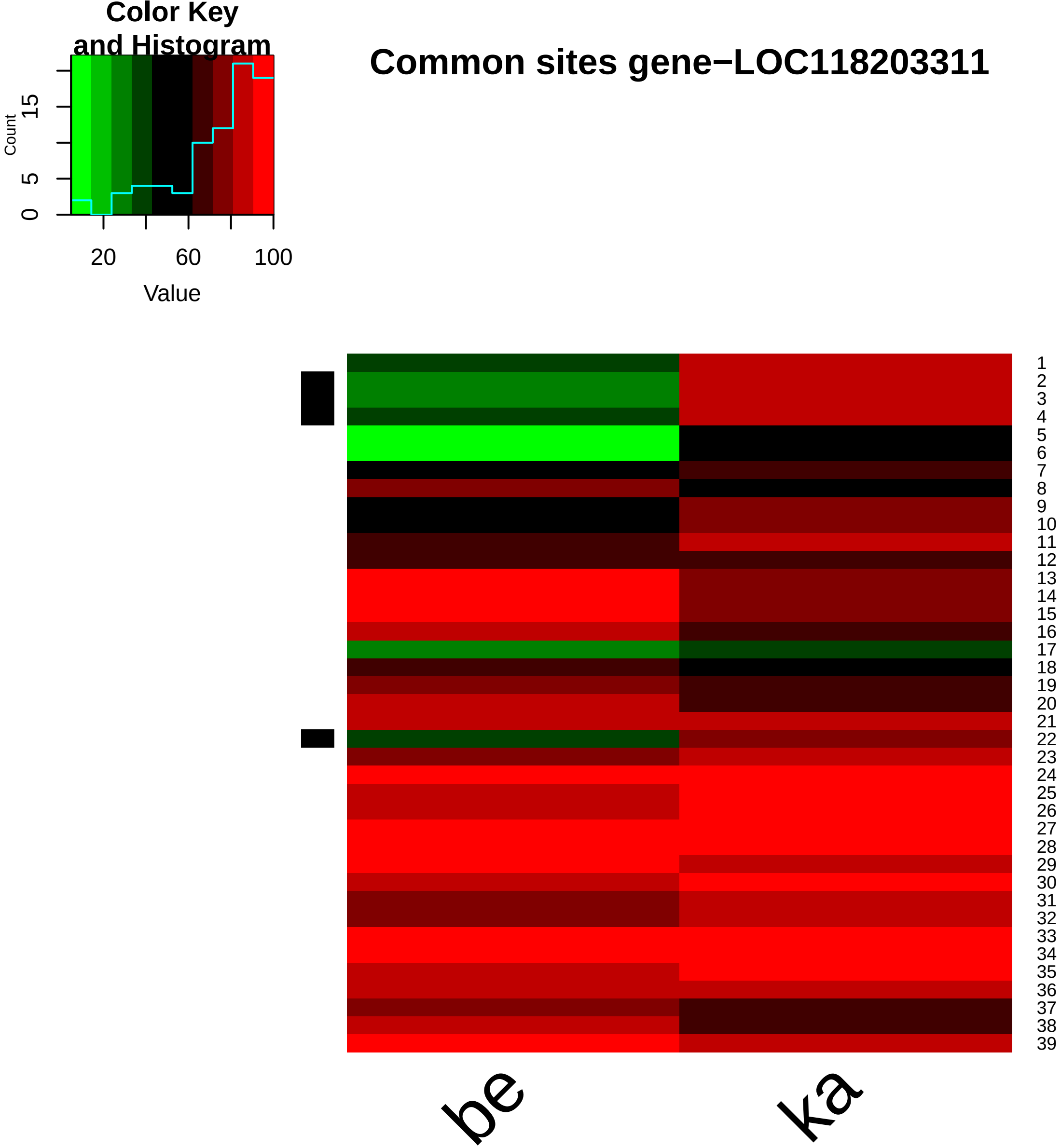

Color Key and Histogram
Count
0
5
15
20
60
100
Value
Common sites gene−LOC118203311
be
ka

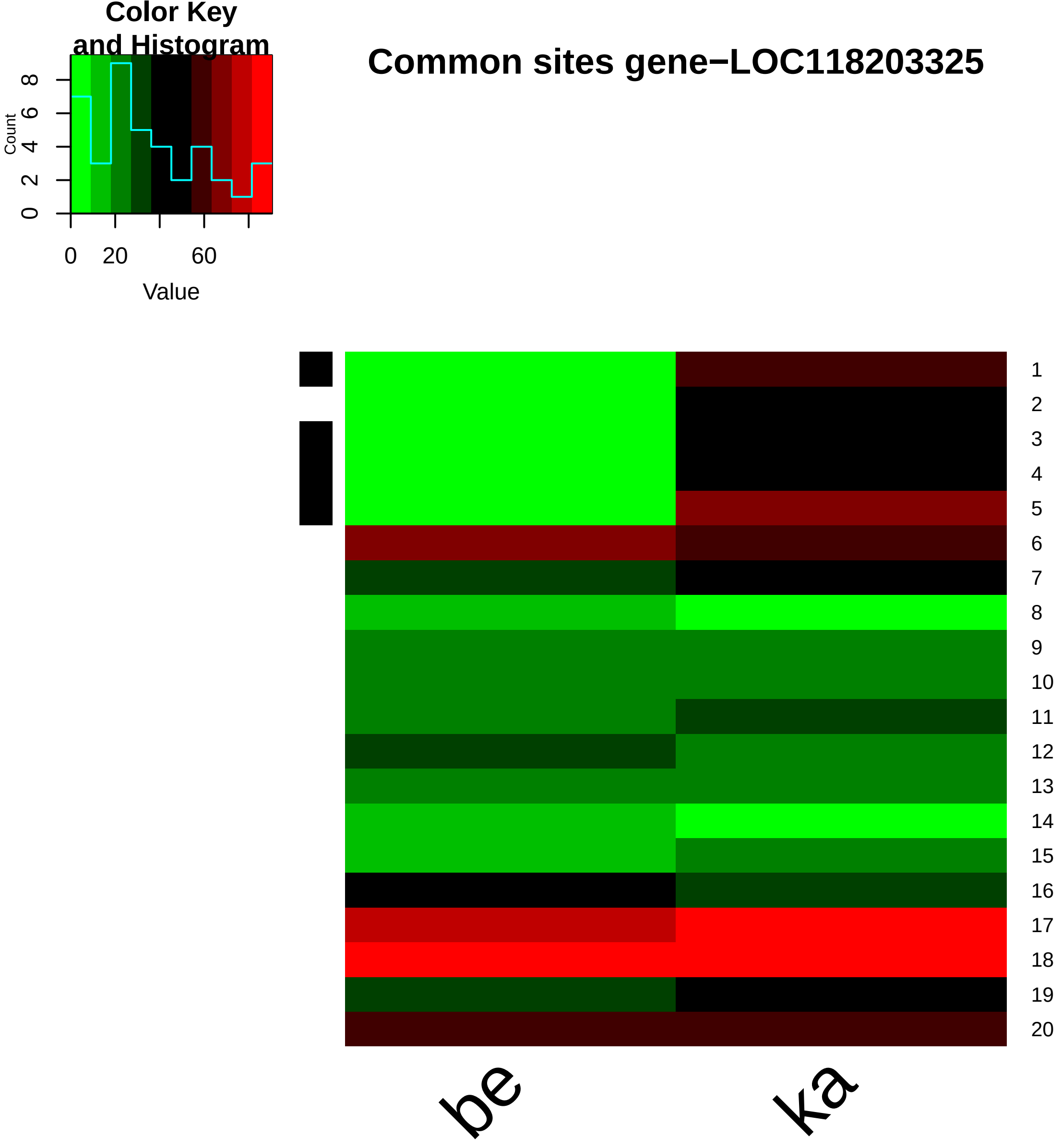
Color Key
and Histogram
Count
0
2
4
6
8
0
20
60
Value
Common sites gene−LOC118203325
be
ka

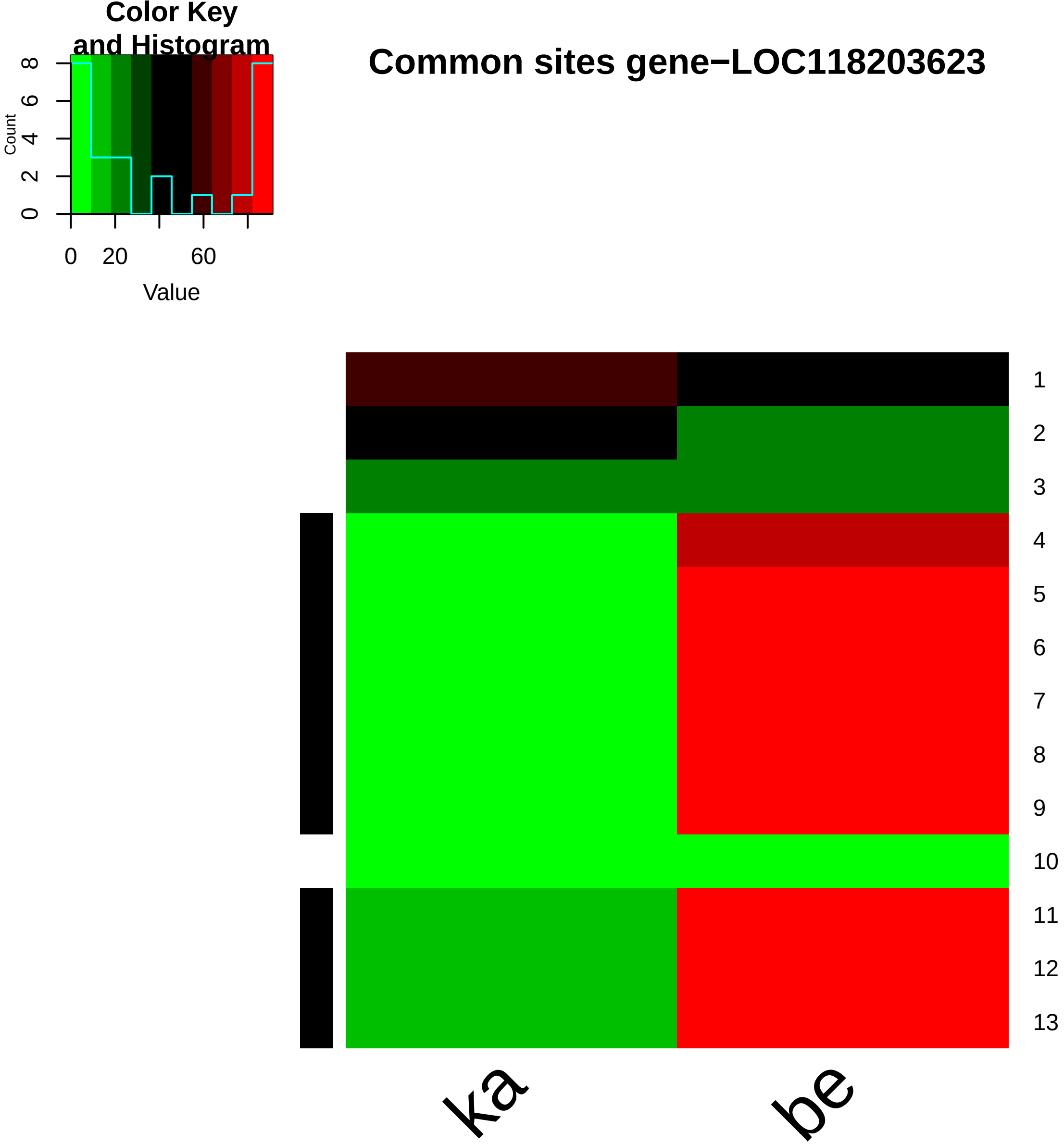
Color Key
and Histogram
Count
0
2
4
6
8
0
20
60
Value
Common sites gene−LOC118203623
1
2
3
4
5
6
7
8
9
10
11
12
13
ka
be

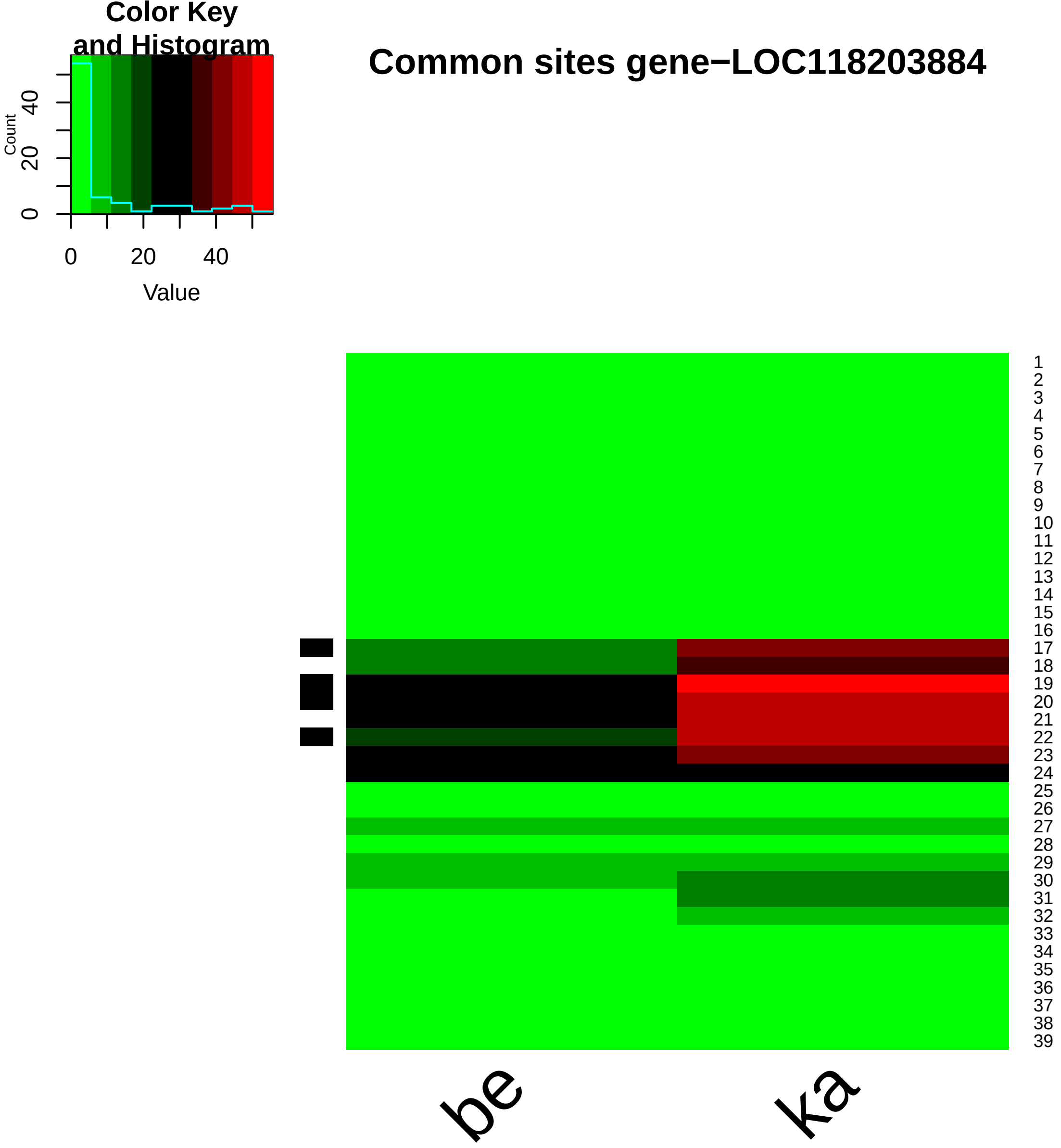
Color Key
and Histogram
Count
0
20
40
0
20
40
Value
Common sites gene−LOC118203884
be
ka

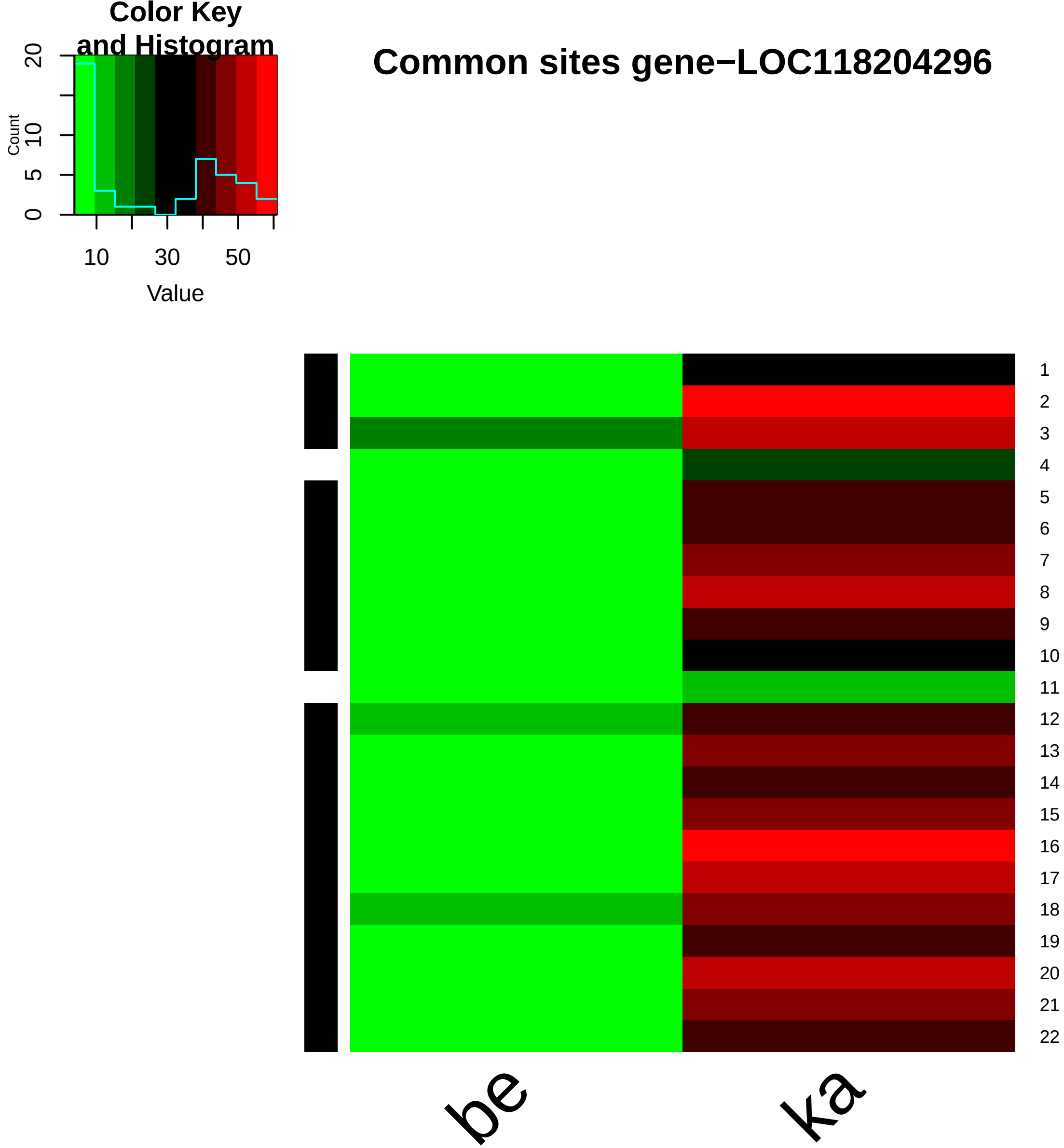

Color Key
and Histogram
Count
Value
0
5
10
20
10
30
50
Common sites gene−LOC118204296
be
ka

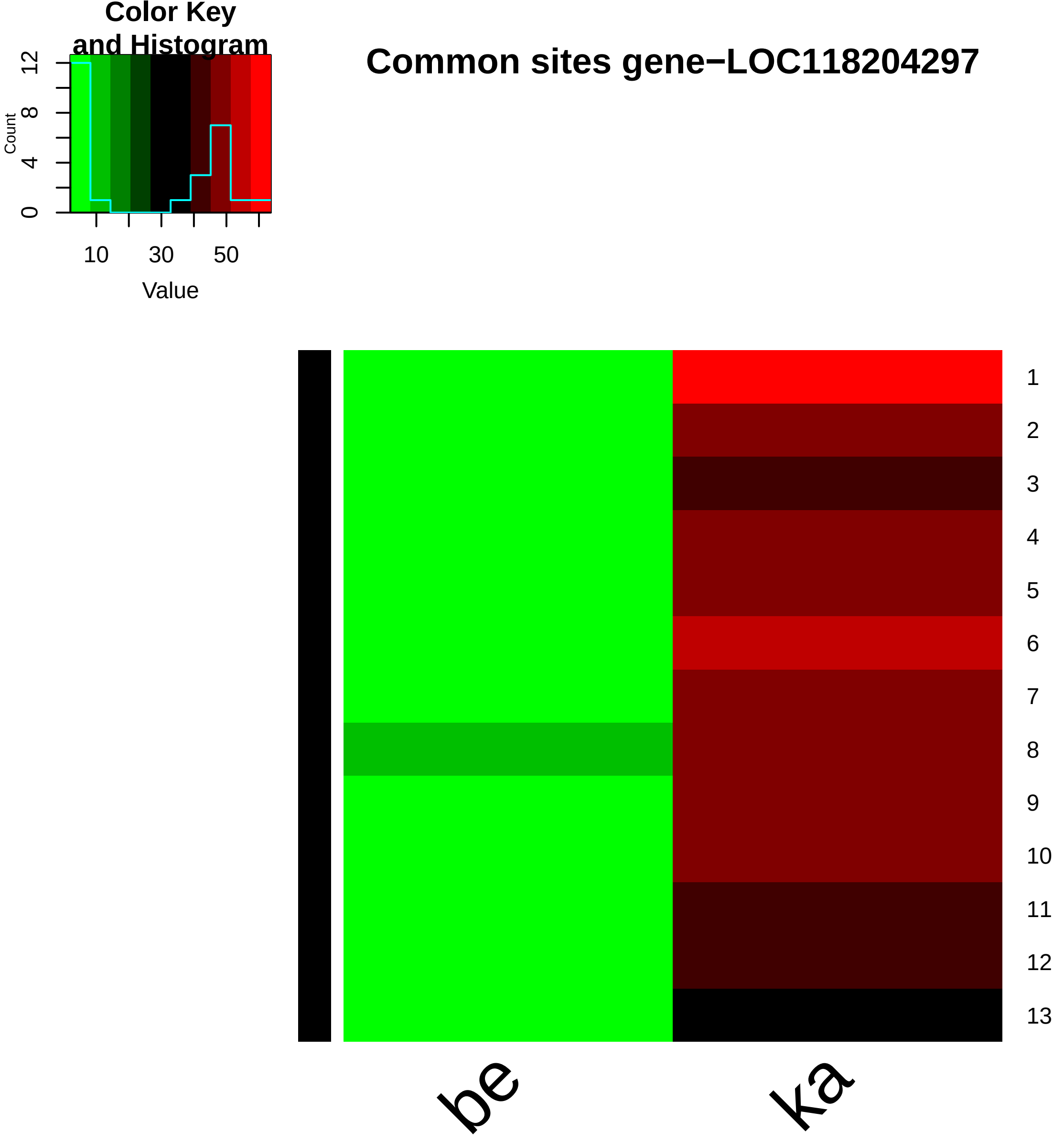

Color Key and Histogram
Count
0
4
8
12
10
30
50
Value
Common sites gene−LOC118204297
1
2
3
4
5
6
7
8
9
10
11
12
13
be
ka

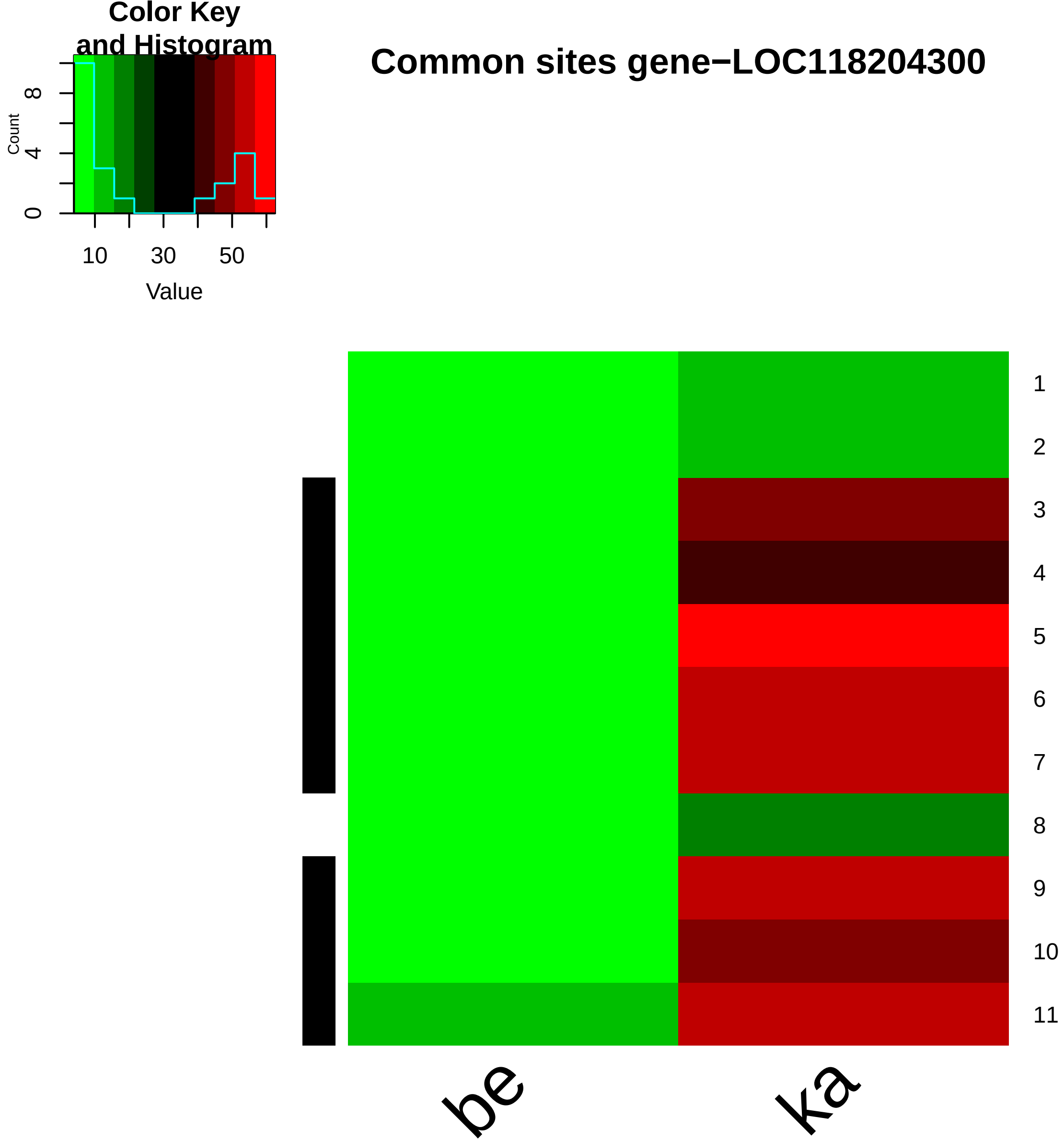

Color Key
and Histogram
Count
0
4
8
10
30
50
Value
Common sites gene−LOC118204300
1
2
3
4
5
6
7
8
9
10
11
be
ka

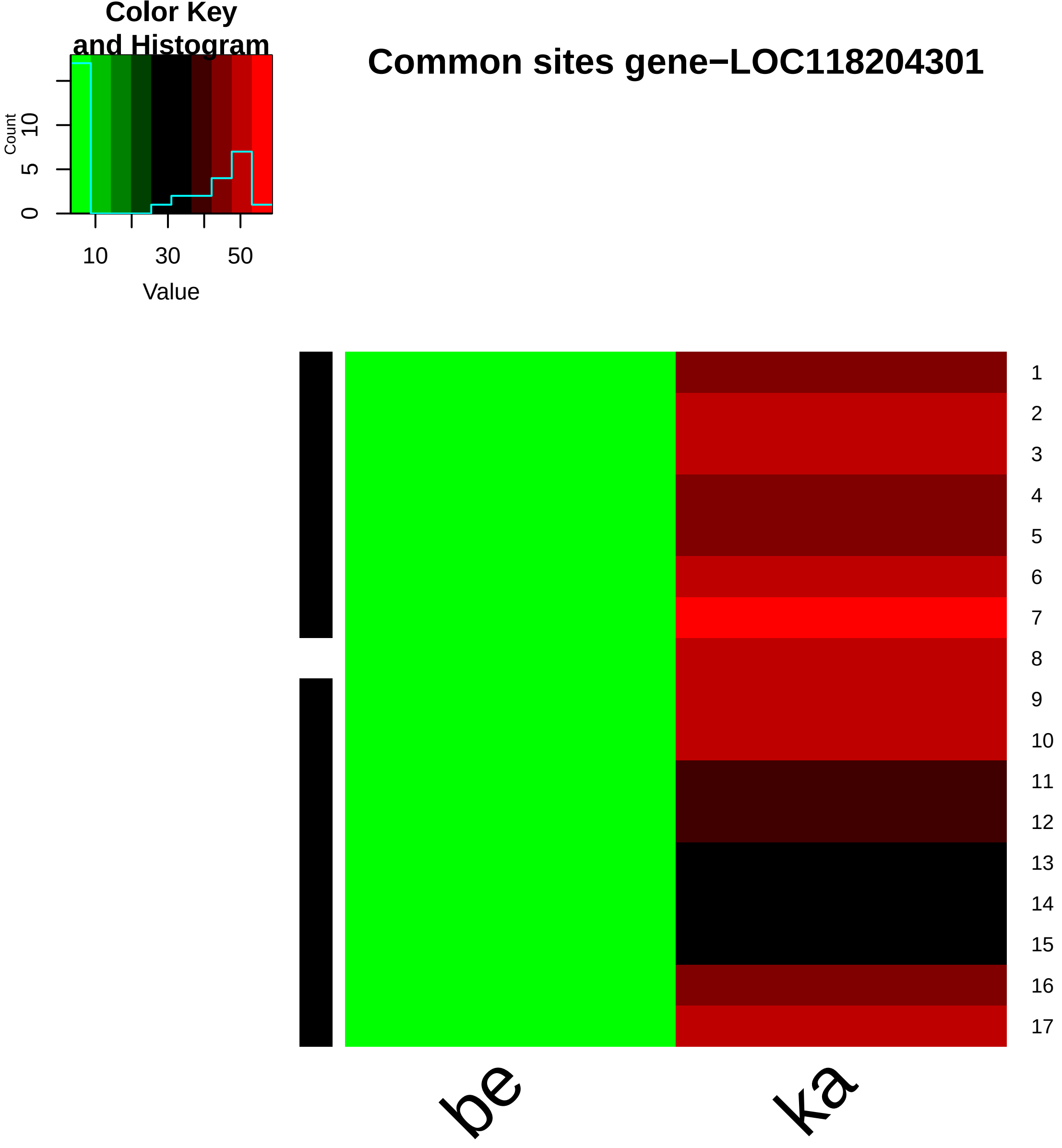

Color Key
and Histogram
Count
0
5
10
10
30
50
Value
Common sites gene−LOC118204301
be
ka

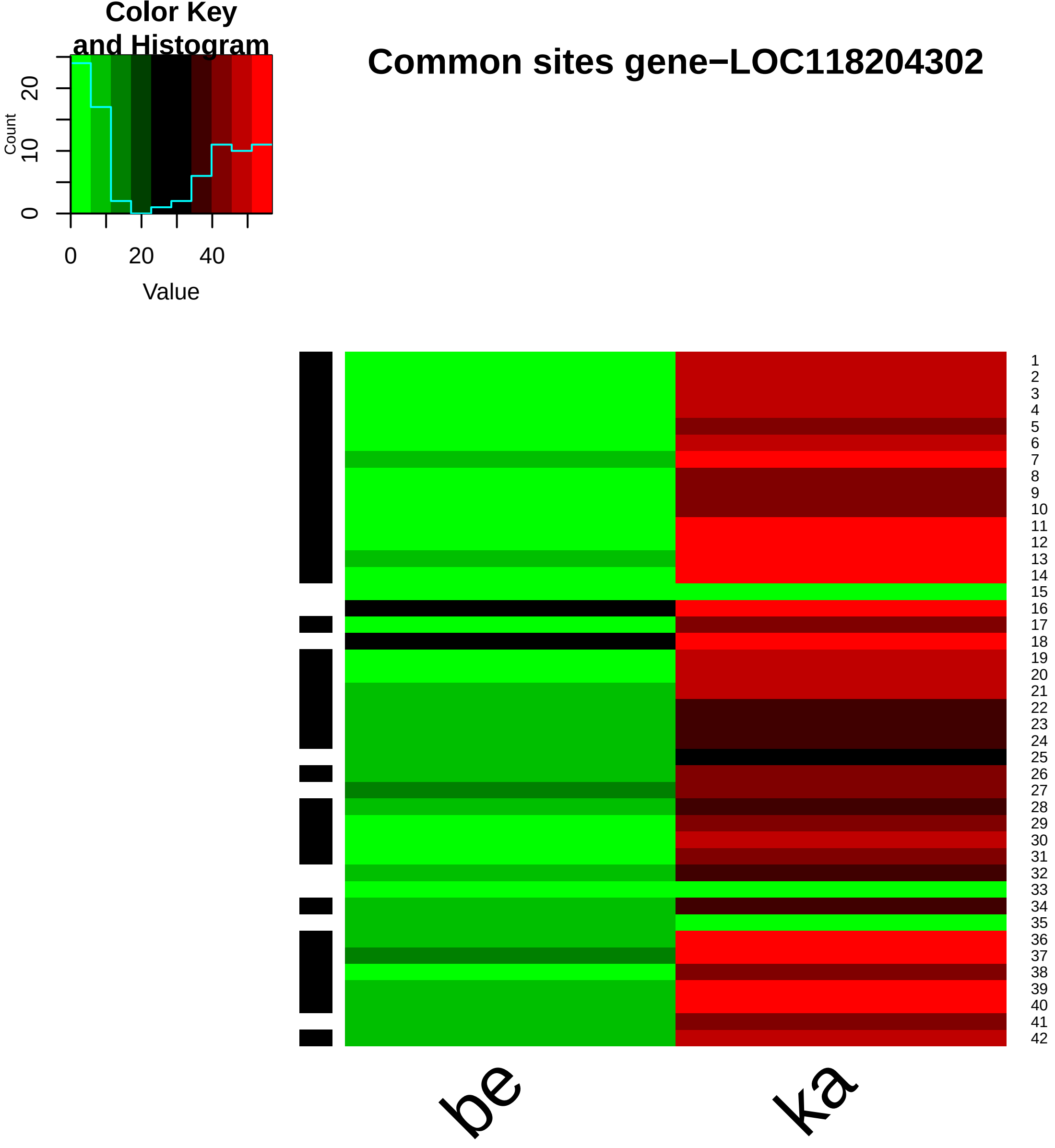
Color Key
and Histogram
Count
Value
0
10
20
0
20
40
Common sites gene−LOC118204302
be
ka

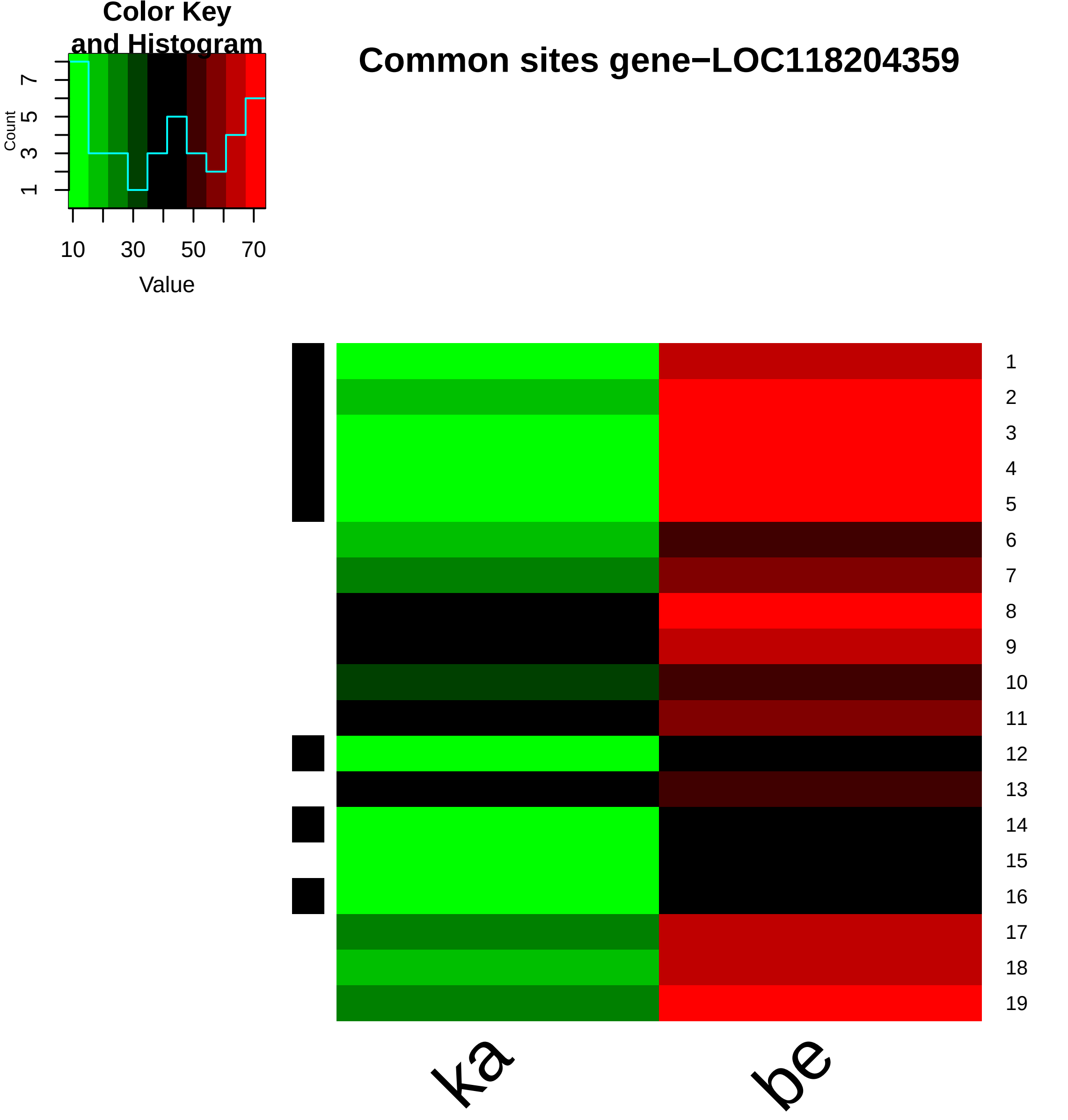
Color Key
and Histogram
Count
1
3
5
7
10
30
50
70
Value
Common sites gene−LOC118204359
ka
be

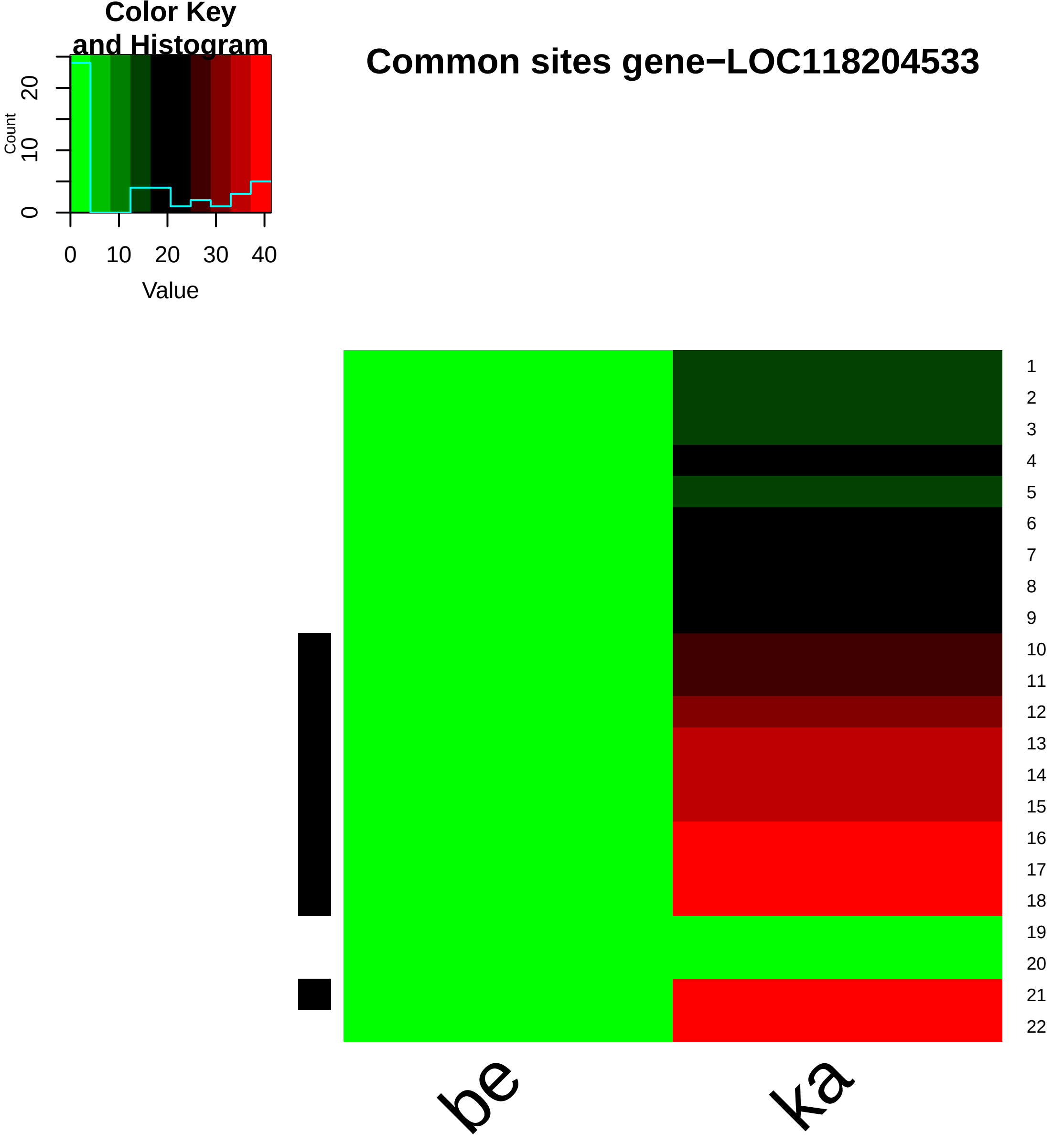
Color Key
and Histogram
Count
0
10
20
0
10
20
30
40
Value
Common sites gene−LOC118204533
be
ka

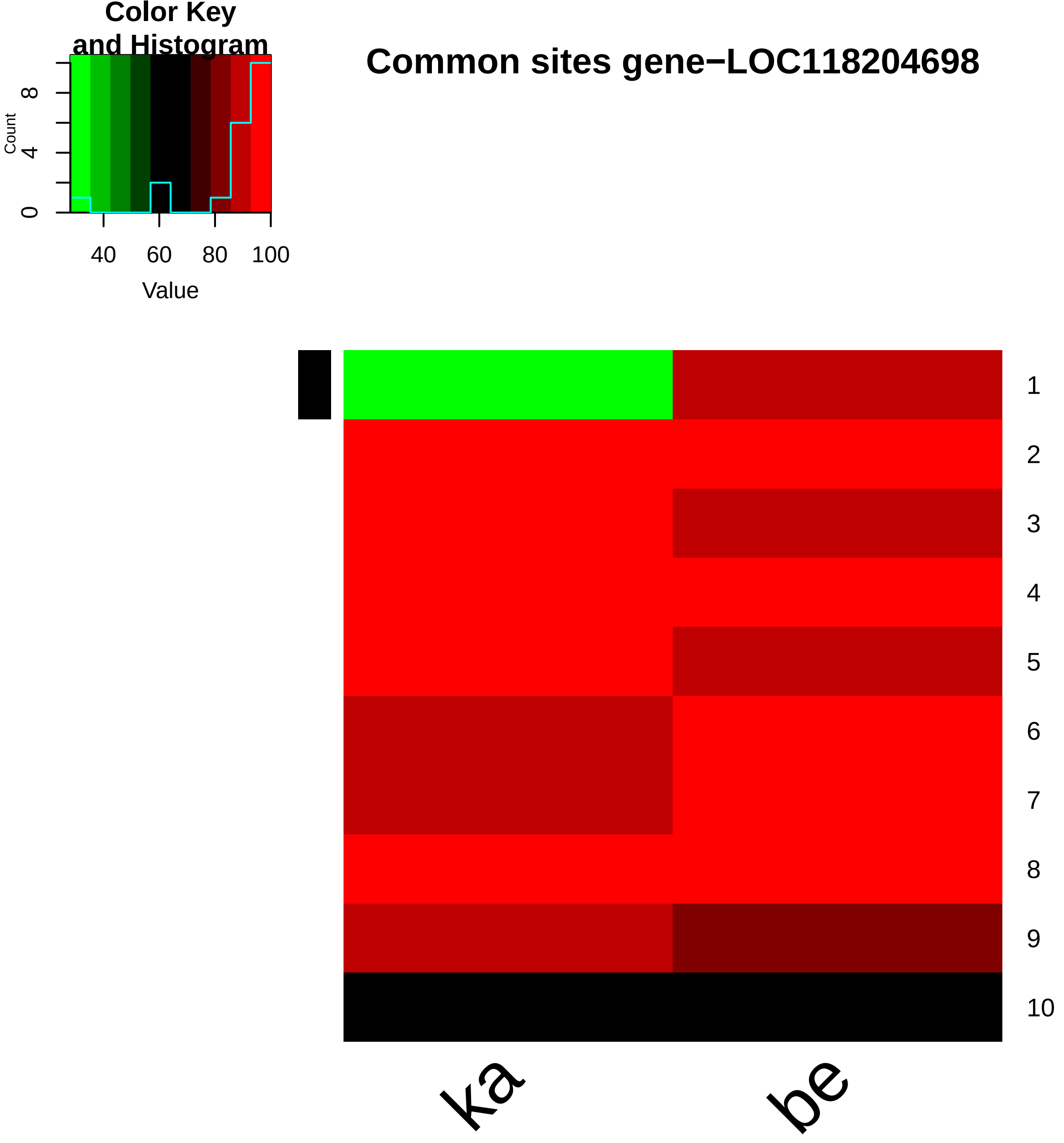
Color Key
and Histogram
Count
0
4
8
40
60
80
100
Value
Common sites gene−LOC118204698
1
2
3
4
5
6
7
8
9
10
ka
be

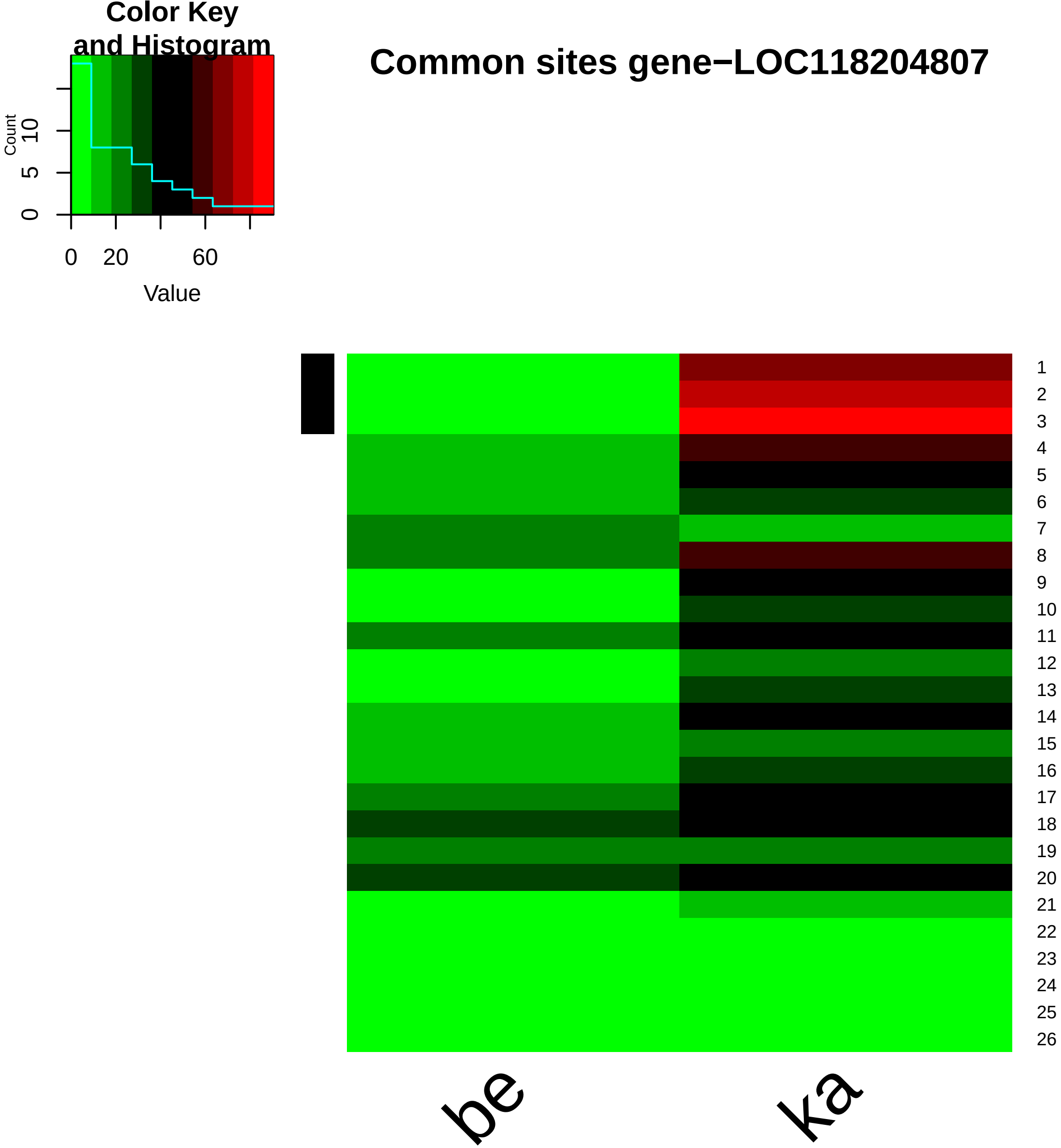

Color Key
and Histogram
Count
0
5
10
0
20
60
Value
Common sites gene−LOC118204807
be
ka

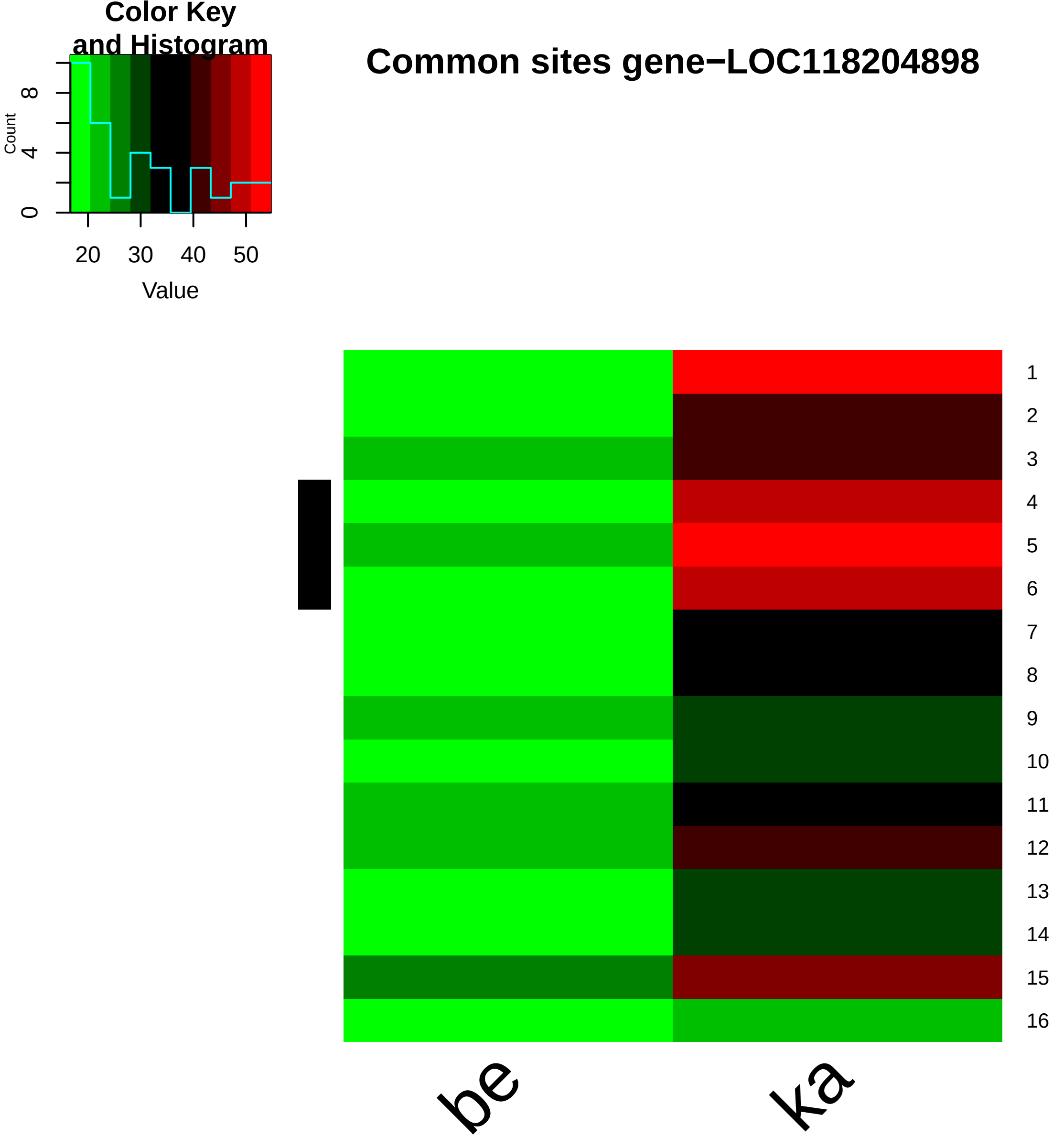

Color Key and Histogram
Count
0
4
8
20
30
40
50
Value
Common sites gene−LOC118204898
be
ka

#### Common sites gene−LOC118204950

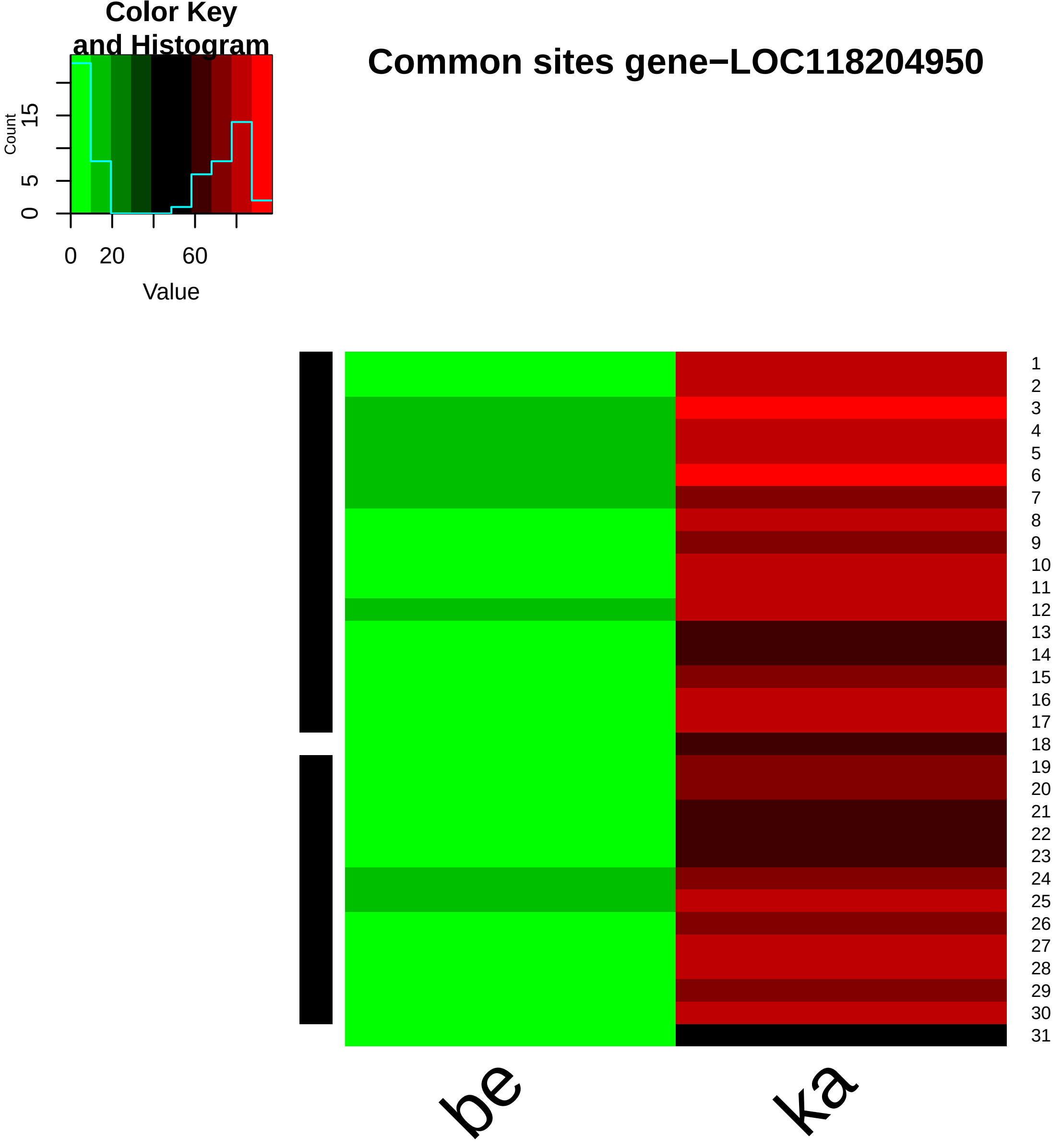

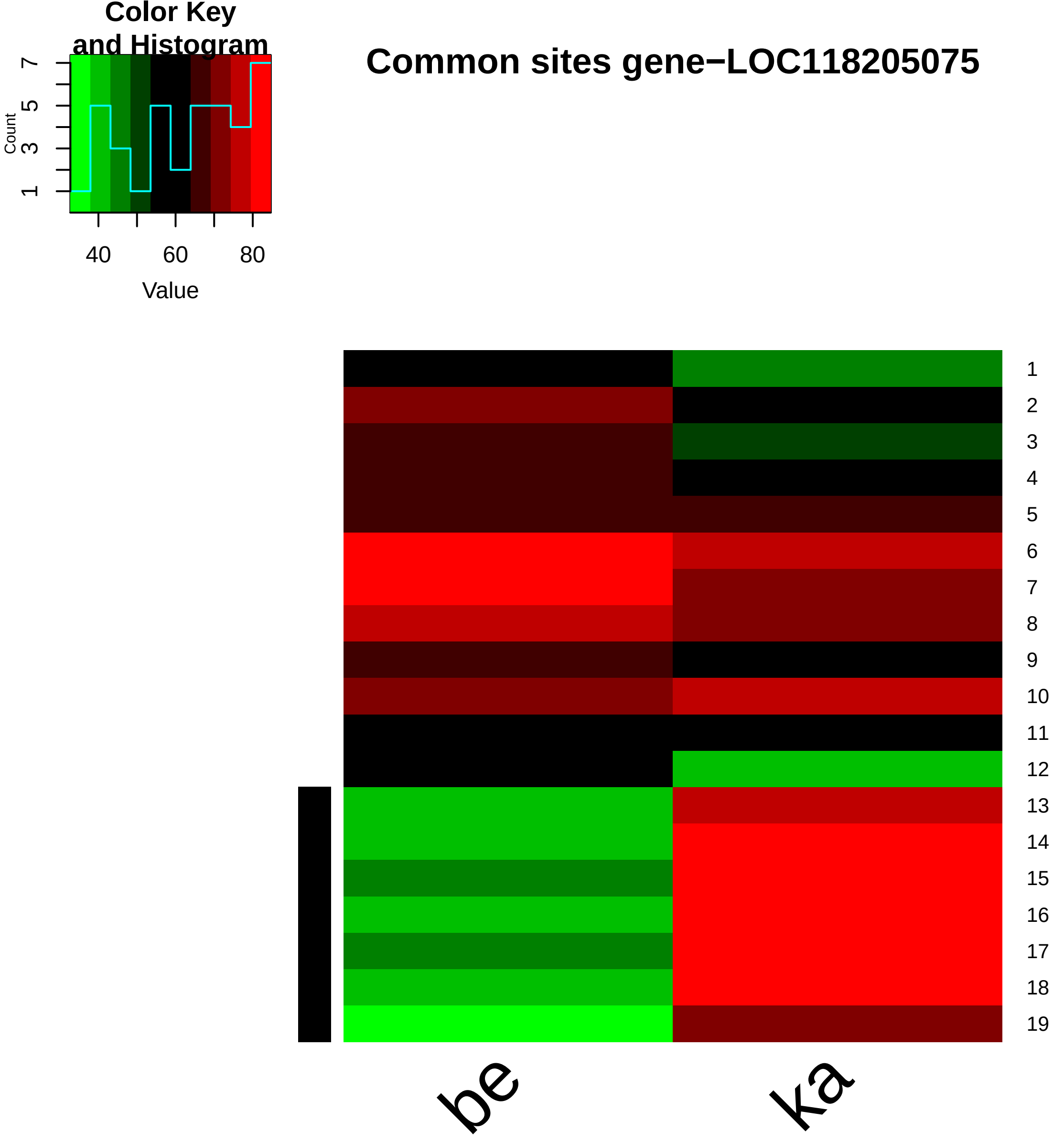

Color Key
and Histogram
Count
1
3
5
7
40
60
80
Value
Common sites gene−LOC118205075
be
ka

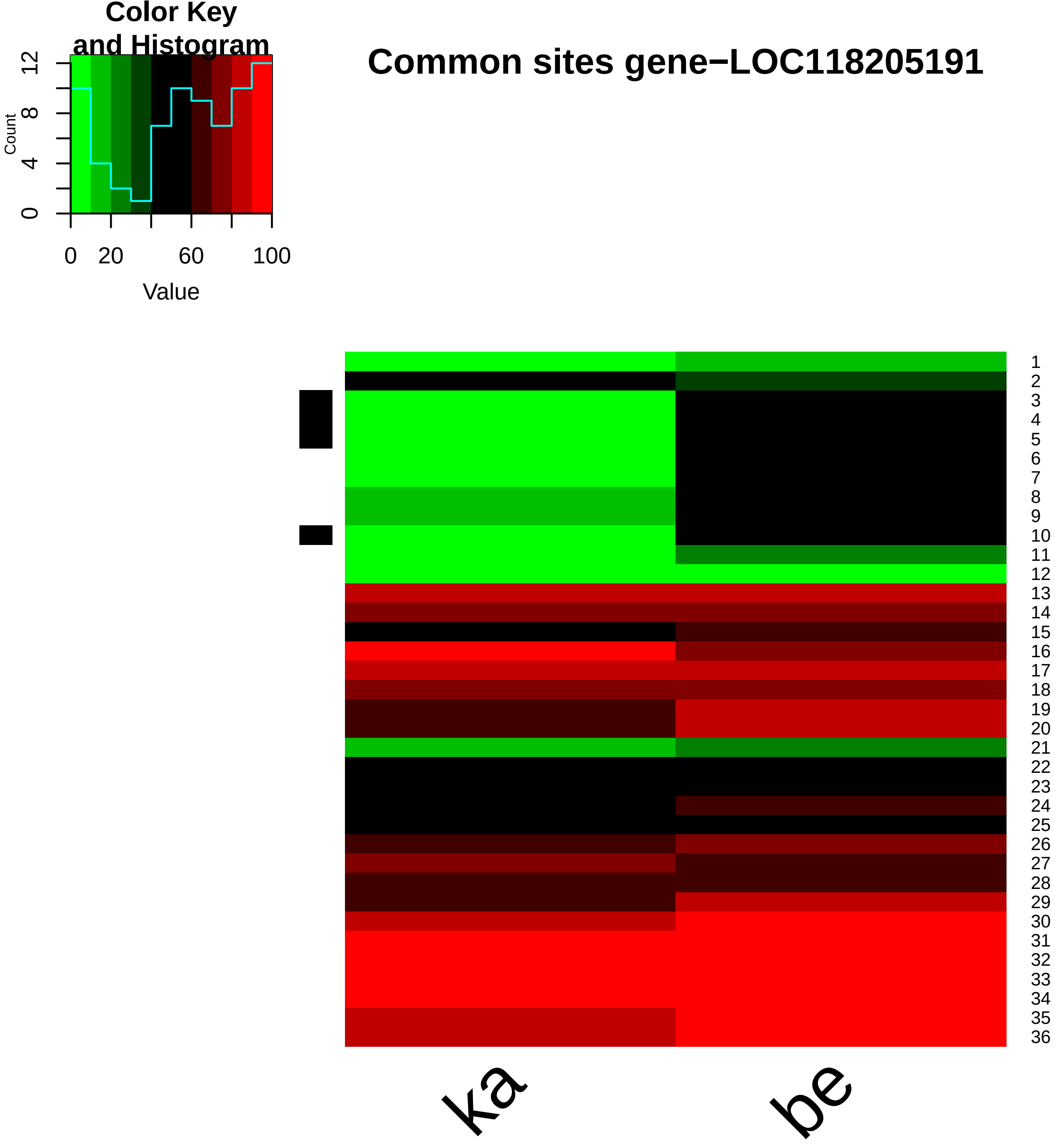
Color Key
and Histogram
Count
0
4
8
12
0
20
60
100
Value
Common sites gene−LOC118205191
ka
be

```
Directory: OGL File: 0READMEogl

OGL - Ordered gene lists.

Input: studyhc annotation   maxgwidth minnbrdmsites
       dmgprp (= output of BWASPR:show_dmsg())

Output: files ogl-*.txt rnk-dmg-*.txt
         ogl-<study>_<sample>.txt ogl-<study>_<sample1>.vs.<sample2>.txt
         rnk-dmg--<study>_<sample1>.vs.<sample2>.txt rnk-dmg--<study>_<sample1>.vs.<sample2>.pdf
         wrt-<study>.txt

Notes: Output files ogl-<study>_<sample>.txt give tables for each sample with columns

        gene_ID gwidth #Sites #per10Kb %perSite %pNucl

       ordered by %pNucl.  If available, a link to an NCBI entry of gene_ID is inserted as second column.
       Abbreviations used: %perSite, percent methylation per site
                           %perNucl, percent methylation per nucleotide of the gene

       Output files ogl-<study>_<sample1>.vs.<sample2>.txt give tables for each comparison with columns

        gene_ID gwidth #Sites #per10Kb #dmSites #dmsp10kb %dmSites %pSite1 %pSite2 DMpSite ADMpSite DMpNucl ADMpNucl

       ordered by DMpSite.  If available, a link to an NCBI entry of gene_ID is inserted as second column.
       Abbreviations used: %dmSites, percent sites that are differentially methylated
                           %pSite1, average per site % methylation for sample1
                           %pSite2, average per site % methylation for sample2
                           DMpSite, average per site difference in % methylation between sample1 and sample2
                           ADMpSite, absolute value of DMpSite
                           DMpNucl, average per nucleotide difference in % methylation between sample1 and sample2
                           ADMpNucl, absolute value of DMpNucl

       Output files rnk-dmg-<study>_<sample1>.vs.<sample2>.txt are equivalent to files
        ogl-<study>_<sample1>.vs.<sample2>.txt but ordered by ADMpNucl.
       Output files rnk-dmg-<study>_<sample1>.vs.<sample2>.pdf provide visualization of the distribution
        of ADMpNucl values.

       Output file wrt-<study>.txt gives results of the Wilcoxon signed rank test comparing
        the %pSite1 and %pSite2 vectors

       In output directory OGL, tables are restricted to genes with gwidth <= maxgwidth and, for
        pairwise comparisons, #dmSites >= minnbrdmsites

       If either maxgwidth or minnbrdmsites is set, then output directory OGLall will show the full tables
        with all genes (for reference).
```

Directory: OGL File: ogl-Sd_LI_be.txt

| gene_ID | gene_link | gwidth | #Sites | #per10Kb | %perSite | %pNucl |
|---|---|---|---|---|---|---|
| gene-LOC118199038 | https://www.ncbi.nlm.nih.gov/gene/?term=LOC118199038 | 1069 | 60 | 561.27 | 88.44 | 4.96 |
| gene-LOC118200794 | https://www.ncbi.nlm.nih.gov/gene/?term=LOC118200794 | 679 | 33 | 486.01 | 89.19 | 4.33 |
| gene-LOC118185191 | https://www.ncbi.nlm.nih.gov/gene/?term=LOC118185191 | 846 | 52 | 614.66 | 62 | 3.81 |
| gene-LOC118180258 | https://www.ncbi.nlm.nih.gov/gene/?term=LOC118180258 | 363 | 17 | 468.32 | 81 | 3.79 |
| gene-LOC118190272 | https://www.ncbi.nlm.nih.gov/gene/?term=LOC118190272 | 1960 | 62 | 316.33 | 82.96 | 2.62 |
| gene-Trnat-ggu-6 | https://www.ncbi.nlm.nih.gov/gene/?term=Trnat-ggu | 72 | 8 | 1111.11 | 21.07 | 2.34 |
| gene-LOC118193385 | https://www.ncbi.nlm.nih.gov/gene/?term=LOC118193385 | 363 | 9 | 247.93 | 90.23 | 2.24 |
| gene-LOC118194215 | https://www.ncbi.nlm.nih.gov/gene/?term=LOC118194215 | 6061 | 145 | 239.23 | 84.24 | 2.02 |
| gene-LOC118202299 | https://www.ncbi.nlm.nih.gov/gene/?term=LOC118202299 | 7958 | 207 | 260.12 | 77.6 | 2.02 |
| gene-LOC118191889 | https://www.ncbi.nlm.nih.gov/gene/?term=LOC118191889 | 13681 | 399 | 291.65 | 64.02 | 1.87 |
| gene-LOC118180687 | https://www.ncbi.nlm.nih.gov/gene/?term=LOC118180687 | 12663 | 289 | 228.22 | 81.06 | 1.85 |
| gene-LOC118205209 | https://www.ncbi.nlm.nih.gov/gene/?term=LOC118205209 | 7915 | 238 | 300.69 | 60.92 | 1.83 |
| gene-LOC118206125 | https://www.ncbi.nlm.nih.gov/gene/?term=LOC118206125 | 8096 | 171 | 211.22 | 82.62 | 1.75 |
| gene-LOC118205293 | https://www.ncbi.nlm.nih.gov/gene/?term=LOC118205293 | 850 | 35 | 411.76 | 42.14 | 1.74 |
| gene-LOC118195989 | https://www.ncbi.nlm.nih.gov/gene/?term=LOC118195989 | 756 | 15 | 198.41 | 86.25 | 1.71 |
| gene-LOC118187599 | https://www.ncbi.nlm.nih.gov/gene/?term=LOC118187599 | 447 | 9 | 201.34 | 84.23 | 1.7 |
| gene-LOC118204580 | https://www.ncbi.nlm.nih.gov/gene/?term=LOC118204580 | 13571 | 333 | 245.38 | 69.17 | 1.7 |
| gene-LOC118189203 | https://www.ncbi.nlm.nih.gov/gene/?term=LOC118189203 | 1497 | 32 | 213.76 | 78.15 | 1.67 |
| gene-LOC118205924 | https://www.ncbi.nlm.nih.gov/gene/?term=LOC118205924 | 9047 | 217 | 239.86 | 69.06 | 1.66 |
| gene-LOC118205294 | https://www.ncbi.nlm.nih.gov/gene/?term=LOC118205294 | 1667 | 123 | 737.85 | 22.23 | 1.64 |
| gene-LOC118188252 | https://www.ncbi.nlm.nih.gov/gene/?term=LOC118188252 | 531 | 10 | 188.32 | 86.41 | 1.63 |
| gene-LOC118204359 | https://www.ncbi.nlm.nih.gov/gene/?term=LOC118204359 | 1095 | 30 | 273.97 | 58.44 | 1.6 |
| gene-LOC118180265 | https://www.ncbi.nlm.nih.gov/gene/?term=LOC118180265 | 825 | 14 | 169.7 | 93.43 | 1.59 |
| gene-LOC118200043 | https://www.ncbi.nlm.nih.gov/gene/?term=LOC118200043 | 405 | 8 | 197.53 | 80.69 | 1.59 |

...

... only first 25 lines shown ...

Directory: OGL File: ogl-Sd_LI_be.vs.ka.txt

| gene_ID | gene_link | gwidth | #Sites | #per10Kb | #dmSites | #dmsp10kb | %dmSites | %pSite1 | %pSite2 | DMpSite | ADMpSite | DMpNucl | ADMpNucl |
|---|---|---|---|---|---|---|---|---|---|---|---|---|---|
| gene-LOC118196111 | https://www.ncbi.nlm.nih.gov/gene/?term=LOC118196111 | 17112 | 3 | 1.75 | 3 | 1.75 | 100 | 82.54 | 4.7 | 77.84 | 77.84 | 0.01 | 0.01 |
| gene-LOC118202498 | https://www.ncbi.nlm.nih.gov/gene/?term=LOC118202498 | 1721 | 6 | 34.86 | 5 | 29.05 | 83.33 | 65.1 | 4.6 | 60.5 | 60.5 | 0.21 | 0.21 |
| gene-LOC118204359 | https://www.ncbi.nlm.nih.gov/gene/?term=LOC118204359 | 1095 | 19 | 173.52 | 8 | 73.06 | 42.11 | 59.62 | 21.69 | 37.93 | 37.93 | 0.66 | 0.66 |
| gene-LOC118199292 | https://www.ncbi.nlm.nih.gov/gene/?term=LOC118199292 | 846 | 6 | 70.92 | 2 | 23.64 | 33.33 | 39.08 | 3.61 | 35.46 | 35.46 | 0.25 | 0.25 |
| gene-LOC118183257 | https://www.ncbi.nlm.nih.gov/gene/?term=LOC118183257 | 17126 | 11 | 6.42 | 3 | 1.75 | 27.27 | 69.5 | 34.75 | 34.75 | 34.75 | 0.02 | 0.02 |
| gene-LOC118190970 | https://www.ncbi.nlm.nih.gov/gene/?term=LOC118190970 | 1773 | 16 | 90.24 | 4 | 22.56 | 25 | 65.08 | 33.41 | 31.67 | 31.67 | 0.29 | 0.29 |
| gene-LOC118184468 | https://www.ncbi.nlm.nih.gov/gene/?term=LOC118184468 | 17739 | 22 | 12.4 | 7 | 3.95 | 31.82 | 61.08 | 30.15 | 30.93 | 30.93 | 0.04 | 0.04 |
| gene-LOC118201096 | https://www.ncbi.nlm.nih.gov/gene/?term=LOC118201096 | 4437 | 19 | 42.82 | 7 | 15.78 | 36.84 | 56.84 | 31.02 | 25.81 | 25.81 | 0.11 | 0.11 |
| gene-LOC118180489 | https://www.ncbi.nlm.nih.gov/gene/?term=LOC118180489 | 3793 | 4 | 10.55 | 2 | 5.27 | 50 | 33.84 | 16.14 | 17.7 | 17.7 | 0.02 | 0.02 |
| gene-LOC118182035 | https://www.ncbi.nlm.nih.gov/gene/?term=LOC118182035 | 11879 | 11 | 9.26 | 2 | 1.68 | 18.18 | 16.32 | 1.77 | 14.55 | 14.55 | 0.01 | 0.01 |
| gene-LOC118206129 | https://www.ncbi.nlm.nih.gov/gene/?term=LOC118206129 | 2062 | 32 | 155.19 | 2 | 9.7 | 6.25 | 40.75 | 29.4 | 11.35 | 11.35 | 0.18 | 0.18 |
| gene-LOC118192973 | https://www.ncbi.nlm.nih.gov/gene/?term=LOC118192973 | 17908 | 36 | 20.1 | 3 | 1.68 | 8.33 | 75.26 | 66.25 | 9.01 | 9.01 | 0.02 | 0.02 |
| gene-LOC118191639 | https://www.ncbi.nlm.nih.gov/gene/?term=LOC118191639 | 14462 | 39 | 26.97 | 4 | 2.77 | 10.26 | 25.59 | 16.63 | 8.96 | 8.96 | 0.02 | 0.02 |
| gene-LOC118199290 | https://www.ncbi.nlm.nih.gov/gene/?term=LOC118199290 | 15093 | 26 | 17.23 | 2 | 1.33 | 7.69 | 68.99 | 62.52 | 6.47 | 6.47 | 0.01 | 0.01 |
| gene-LOC118203894 | https://www.ncbi.nlm.nih.gov/gene/?term=LOC118203894 | 15828 | 34 | 21.48 | 3 | 1.9 | 8.82 | 27.49 | 22.85 | 4.64 | 4.64 | 0.01 | 0.01 |
| gene-LOC118200390 | https://www.ncbi.nlm.nih.gov/gene/?term=LOC118200390 | 13922 | 39 | 28.01 | 2 | 1.44 | 5.13 | 65.32 | 65.29 | 0.04 | 0.04 | 0 | 0 |
| gene-LOC118188906 | https://www.ncbi.nlm.nih.gov/gene/?term=LOC118188906 | 11213 | 40 | 35.67 | 2 | 1.78 | 5 | 82.47 | 83.49 | -1.02 | 1.02 | 0 | 0 |
| gene-LOC118205924 | https://www.ncbi.nlm.nih.gov/gene/?term=LOC118205924 | 9047 | 119 | 131.54 | 3 | 3.32 | 2.52 | 67.93 | 69.22 | -1.3 | 1.3 | -0.02 | 0.02 |
| gene-LOC118182290 | https://www.ncbi.nlm.nih.gov/gene/?term=LOC118182290 | 15134 | 27 | 17.84 | 2 | 1.32 | 7.41 | 38.35 | 41.25 | -2.9 | 2.9 | -0.01 | 0.01 |
| gene-LOC118205075 | https://www.ncbi.nlm.nih.gov/gene/?term=LOC118205075 | 6473 | 19 | 29.35 | 7 | 10.81 | 36.84 | 58.72 | 67.66 | -8.94 | 8.94 | -0.03 | 0.03 |
| gene-LOC118201196 | https://www.ncbi.nlm.nih.gov/gene/?term=LOC118201196 | 10621 | 84 | 79.09 | 3 | 2.82 | 3.57 | 29.17 | 39.29 | -10.12 | 10.12 | -0.08 | 0.08 |
| gene-LOC118189486 | https://www.ncbi.nlm.nih.gov/gene/?term=LOC118189486 | 14274 | 16 | 11.21 | 2 | 1.4 | 12.5 | 1.59 | 13.77 | -12.18 | 12.18 | -0.01 | 0.01 |
| gene-LOC118192981 | https://www.ncbi.nlm.nih.gov/gene/?term=LOC118192981 | 9333 | 13 | 13.93 | 2 | 2.14 | 15.38 | 63.22 | 81.48 | -18.26 | 18.26 | -0.03 | 0.03 |
| gene-LOC118204898 | https://www.ncbi.nlm.nih.gov/gene/?term=LOC118204898 | 2378 | 16 | 67.28 | 3 | 12.62 | 18.75 | 20.34 | 38.79 | -18.45 | 18.45 | -0.12 | 0.12 |
| gene-LOC118180675 | https://www.ncbi.nlm.nih.gov/gene/?term=LOC118180675 | 624 | 5 | 80.13 | 2 | 32.05 | 40 | 0.16 | 22.39 | -22.23 | 22.23 | -0.18 | 0.18 |
| gene-LOC118198429 | https://www.ncbi.nlm.nih.gov/gene/?term=LOC118198429 | 17137 | 23 | 13.42 | 5 | 2.92 | 21.74 | 45.23 | 69.23 | -24 | 24 | -0.03 | 0.03 |
| gene-LOC118204533 | https://www.ncbi.nlm.nih.gov/gene/?term=LOC118204533 | 446 | 22 | 493.27 | 10 | 224.22 | 45.45 | 0.38 | 24.93 | -24.55 | 24.55 | -1.21 | 1.21 |
| gene-LOC118180680 | https://www.ncbi.nlm.nih.gov/gene/?term=LOC118180680 | 3131 | 21 | 67.07 | 15 | 47.91 | 71.43 | 9.44 | 36.41 | -26.96 | 26.96 | -0.18 | 0.18 |
| gene-LOC118205893 | https://www.ncbi.nlm.nih.gov/gene/?term=LOC118205893 | 930 | 2 | 21.51 | 2 | 21.51 | 100 | 6.28 | 33.85 | -27.58 | 27.58 | -0.06 | 0.06 |
| gene-LOC118204298 | https://www.ncbi.nlm.nih.gov/gene/?term=LOC118204298 | 474 | 4 | 84.39 | 4 | 84.39 | 100 | 7.12 | 40.43 | -33.31 | 33.31 | -0.28 | 0.28 |
| gene-LOC118180674 | https://www.ncbi.nlm.nih.gov/gene/?term=LOC118180674 | 1344 | 17 | 126.49 | 14 | 104.17 | 82.35 | 7.39 | 42.16 | -34.78 | 34.78 | -0.44 | 0.44 |
| gene-LOC118204300 | https://www.ncbi.nlm.nih.gov/gene/?term=LOC118204300 | 1116 | 11 | 98.57 | 8 | 71.68 | 72.73 | 6.44 | 41.39 | -34.95 | 34.95 | -0.34 | 0.34 |
| gene-LOC118180764 | https://www.ncbi.nlm.nih.gov/gene/?term=LOC118180764 | 2040 | 15 | 73.53 | 12 | 58.82 | 80 | 6.29 | 41.3 | -35 | 35 | -0.26 | 0.26 |
| gene-LOC118180760 | https://www.ncbi.nlm.nih.gov/gene/?term=LOC118180760 | 978 | 12 | 122.7 | 9 | 92.02 | 75 | 5.55 | 41.25 | -35.7 | 35.7 | -0.44 | 0.44 |
| gene-LOC118204302 | https://www.ncbi.nlm.nih.gov/gene/?term=LOC118204302 | 4419 | 42 | 95.04 | 33 | 74.68 | 78.57 | 6.84 | 43 | -36.16 | 36.16 | -0.34 | 0.34 |
| gene-LOC118204296 | https://www.ncbi.nlm.nih.gov/gene/?term=LOC118204296 | 2118 | 22 | 103.87 | 20 | 94.43 | 90.91 | 7.23 | 43.59 | -36.36 | 36.36 | -0.38 | 0.38 |
| gene-LOC118180761 | https://www.ncbi.nlm.nih.gov/gene/?term=LOC118180761 | 3044 | 21 | 68.99 | 15 | 49.28 | 71.43 | 8.65 | 45.12 | -36.47 | 36.47 | -0.25 | 0.25 |
| gene-LOC118192316 | https://www.ncbi.nlm.nih.gov/gene/?term=LOC118192316 | 1188 | 12 | 101.01 | 11 | 92.59 | 91.67 | 9.13 | 46.82 | -37.68 | 37.68 | -0.38 | 0.38 |
| gene-LOC118180679 | https://www.ncbi.nlm.nih.gov/gene/?term=LOC118180679 | 1640 | 3 | 18.29 | 3 | 18.29 | 100 | 7.58 | 46.43 | -38.85 | 38.85 | -0.07 | 0.07 |
| gene-LOC118192320 | https://www.ncbi.nlm.nih.gov/gene/?term=LOC118192320 | 3613 | 2 | 5.54 | 2 | 5.54 | 100 | 3.32 | 43.12 | -39.8 | 39.8 | -0.02 | 0.02 |
| gene-LOC118204301 | https://www.ncbi.nlm.nih.gov/gene/?term=LOC118204301 | 1823 | 17 | 93.25 | 16 | 87.77 | 94.12 | 4.98 | 44.95 | -39.98 | 39.98 | -0.37 | 0.37 |
| gene-LOC118179631 | https://www.ncbi.nlm.nih.gov/gene/?term=LOC118179631 | 1049 | 7 | 66.73 | 6 | 57.2 | 85.71 | 4.77 | 45.73 | -40.96 | 40.96 | -0.27 | 0.27 |
| gene-LOC118180762 | https://www.ncbi.nlm.nih.gov/gene/?term=LOC118180762 | 1281 | 18 | 140.52 | 16 | 124.9 | 88.89 | 6.13 | 47.34 | -41.22 | 41.22 | -0.58 | 0.58 |
| gene-LOC118204297 | https://www.ncbi.nlm.nih.gov/gene/?term=LOC118204297 | 951 | 13 | 136.7 | 13 | 136.7 | 100 | 4.88 | 48.27 | -43.39 | 43.39 | -0.59 | 0.59 |
| gene-LOC118180763 | https://www.ncbi.nlm.nih.gov/gene/?term=LOC118180763 | 1104 | 4 | 36.23 | 4 | 36.23 | 100 | 7.51 | 51.27 | -43.75 | 43.75 | -0.16 | 0.16 |
| gene-LOC118180331 | https://www.ncbi.nlm.nih.gov/gene/?term=LOC118180331 | 1035 | 13 | 125.6 | 13 | 125.6 | 100 | 12.27 | 72.44 | -60.16 | 60.16 | -0.76 | 0.76 |

Directory: OGL File: ogl-Sd_LI_ka.txt

| gene_ID | gene_link | gwidth | #Sites | #per10Kb | %perSite | %pNucl |
|---|---|---|---|---|---|---|
| gene-LOC118200794 | https://www.ncbi.nlm.nih.gov/gene/?term=LOC118200794 | 679 | 36 | 530.19 | 86.07 | 4.56 |
| gene-LOC118199038 | https://www.ncbi.nlm.nih.gov/gene/?term=LOC118199038 | 1069 | 54 | 505.14 | 87.36 | 4.41 |
| gene-LOC118185191 | https://www.ncbi.nlm.nih.gov/gene/?term=LOC118185191 | 846 | 57 | 673.76 | 59.98 | 4.04 |
| gene-LOC118180258 | https://www.ncbi.nlm.nih.gov/gene/?term=LOC118180258 | 363 | 17 | 468.32 | 77.69 | 3.64 |
| gene-Trnat-ggu-6 | https://www.ncbi.nlm.nih.gov/gene/?term=Trnat-ggu | 72 | 8 | 1111.11 | 28.86 | 3.21 |
| gene-LOC118180265 | https://www.ncbi.nlm.nih.gov/gene/?term=LOC118180265 | 825 | 23 | 278.79 | 94.02 | 2.62 |
| gene-LOC118190272 | https://www.ncbi.nlm.nih.gov/gene/?term=LOC118190272 | 1960 | 62 | 316.33 | 81.07 | 2.56 |
| gene-LOC118198533 | https://www.ncbi.nlm.nih.gov/gene/?term=LOC118198533 | 333 | 8 | 240.24 | 96.08 | 2.31 |
| gene-LOC118193385 | https://www.ncbi.nlm.nih.gov/gene/?term=LOC118193385 | 363 | 9 | 247.93 | 83.77 | 2.08 |
| gene-LOC118202299 | https://www.ncbi.nlm.nih.gov/gene/?term=LOC118202299 | 7958 | 212 | 266.4 | 77.33 | 2.06 |
| gene-LOC118194215 | https://www.ncbi.nlm.nih.gov/gene/?term=LOC118194215 | 6061 | 147 | 242.53 | 83.85 | 2.03 |
| gene-LOC118191889 | https://www.ncbi.nlm.nih.gov/gene/?term=LOC118191889 | 13681 | 406 | 296.76 | 64.9 | 1.93 |
| gene-LOC118180687 | https://www.ncbi.nlm.nih.gov/gene/?term=LOC118180687 | 12663 | 296 | 233.75 | 81.41 | 1.9 |
| gene-LOC118205209 | https://www.ncbi.nlm.nih.gov/gene/?term=LOC118205209 | 7915 | 243 | 307.01 | 61.89 | 1.9 |
| gene-LOC118205293 | https://www.ncbi.nlm.nih.gov/gene/?term=LOC118205293 | 850 | 36 | 423.53 | 42.81 | 1.81 |
| gene-LOC118206125 | https://www.ncbi.nlm.nih.gov/gene/?term=LOC118206125 | 8096 | 170 | 209.98 | 81.25 | 1.71 |
| gene-LOC118205924 | https://www.ncbi.nlm.nih.gov/gene/?term=LOC118205924 | 9047 | 220 | 243.17 | 69.82 | 1.7 |
| gene-LOC118201471 | https://www.ncbi.nlm.nih.gov/gene/?term=LOC118201471 | 2104 | 42 | 199.62 | 84.85 | 1.69 |
| gene-LOC118204580 | https://www.ncbi.nlm.nih.gov/gene/?term=LOC118204580 | 13571 | 332 | 244.64 | 68.78 | 1.68 |
| gene-LOC118195989 | https://www.ncbi.nlm.nih.gov/gene/?term=LOC118195989 | 756 | 15 | 198.41 | 84.2 | 1.67 |
| gene-LOC118199153 | https://www.ncbi.nlm.nih.gov/gene/?term=LOC118199153 | 880 | 15 | 170.45 | 94.78 | 1.62 |
| gene-LOC118192985 | https://www.ncbi.nlm.nih.gov/gene/?term=LOC118192985 | 467 | 8 | 171.31 | 92.64 | 1.59 |
| gene-LOC118189203 | https://www.ncbi.nlm.nih.gov/gene/?term=LOC118189203 | 1497 | 30 | 200.4 | 78.04 | 1.56 |
| gene-LOC118205218 | https://www.ncbi.nlm.nih.gov/gene/?term=LOC118205218 | 684 | 12 | 175.44 | 87.51 | 1.54 |

...

... only first 25 lines shown ...

Directory: OGL File: rnk-dmg-Sd_LI_be.vs.ka.txt

| gene_ID | gene_link | gwidth | #Sites | #per10Kb | #dmSites | #dmsp10kb | %dmSites | %pSite1 | %pSite2 | DMpSite | ADMpSite | DMpNucl | ADMpNucl |
|---|---|---|---|---|---|---|---|---|---|---|---|---|---|
| gene-LOC118204533 | https://www.ncbi.nlm.nih.gov/gene/?term=LOC118204533 | 446 | 22 | 493.27 | 10 | 224.22 | 45.45 | 0.38 | 24.93 | -24.55 | 24.55 | -1.21 | 1.21 |
| gene-LOC118180331 | https://www.ncbi.nlm.nih.gov/gene/?term=LOC118180331 | 1035 | 13 | 125.6 | 13 | 125.6 | 100 | 12.27 | 72.44 | -60.16 | 60.16 | -0.76 | 0.76 |
| gene-LOC118204359 | https://www.ncbi.nlm.nih.gov/gene/?term=LOC118204359 | 1095 | 19 | 173.52 | 8 | 73.06 | 42.11 | 59.62 | 21.69 | 37.93 | 37.93 | 0.66 | 0.66 |
| gene-LOC118204297 | https://www.ncbi.nlm.nih.gov/gene/?term=LOC118204297 | 951 | 13 | 136.7 | 13 | 136.7 | 100 | 4.88 | 48.27 | -43.39 | 43.39 | -0.59 | 0.59 |
| gene-LOC118180762 | https://www.ncbi.nlm.nih.gov/gene/?term=LOC118180762 | 1281 | 18 | 140.52 | 16 | 124.9 | 88.89 | 6.13 | 47.34 | -41.22 | 41.22 | -0.58 | 0.58 |
| gene-LOC118180674 | https://www.ncbi.nlm.nih.gov/gene/?term=LOC118180674 | 1344 | 17 | 126.49 | 14 | 104.17 | 82.35 | 7.39 | 42.16 | -34.78 | 34.78 | -0.44 | 0.44 |
| gene-LOC118180760 | https://www.ncbi.nlm.nih.gov/gene/?term=LOC118180760 | 978 | 12 | 122.7 | 9 | 92.02 | 75 | 5.55 | 41.25 | -35.7 | 35.7 | -0.44 | 0.44 |
| gene-LOC118204296 | https://www.ncbi.nlm.nih.gov/gene/?term=LOC118204296 | 2118 | 22 | 103.87 | 20 | 94.43 | 90.91 | 7.23 | 43.59 | -36.36 | 36.36 | -0.38 | 0.38 |
| gene-LOC118192316 | https://www.ncbi.nlm.nih.gov/gene/?term=LOC118192316 | 1188 | 12 | 101.01 | 11 | 92.59 | 91.67 | 9.13 | 46.82 | -37.68 | 37.68 | -0.38 | 0.38 |
| gene-LOC118204301 | https://www.ncbi.nlm.nih.gov/gene/?term=LOC118204301 | 1823 | 17 | 93.25 | 16 | 87.77 | 94.12 | 4.98 | 44.95 | -39.98 | 39.98 | -0.37 | 0.37 |
| gene-LOC118204300 | https://www.ncbi.nlm.nih.gov/gene/?term=LOC118204300 | 1116 | 11 | 98.57 | 8 | 71.68 | 72.73 | 6.44 | 41.39 | -34.95 | 34.95 | -0.34 | 0.34 |
| gene-LOC118204302 | https://www.ncbi.nlm.nih.gov/gene/?term=LOC118204302 | 4419 | 42 | 95.04 | 33 | 74.68 | 78.57 | 6.84 | 43 | -36.16 | 36.16 | -0.34 | 0.34 |
| gene-LOC118190970 | https://www.ncbi.nlm.nih.gov/gene/?term=LOC118190970 | 1773 | 16 | 90.24 | 4 | 22.56 | 25 | 65.08 | 33.41 | 31.67 | 31.67 | 0.29 | 0.29 |
| gene-LOC118204298 | https://www.ncbi.nlm.nih.gov/gene/?term=LOC118204298 | 474 | 4 | 84.39 | 4 | 84.39 | 100 | 7.12 | 40.43 | -33.31 | 33.31 | -0.28 | 0.28 |
| gene-LOC118179631 | https://www.ncbi.nlm.nih.gov/gene/?term=LOC118179631 | 1049 | 7 | 66.73 | 6 | 57.2 | 85.71 | 4.77 | 45.73 | -40.96 | 40.96 | -0.27 | 0.27 |
| gene-LOC118180764 | https://www.ncbi.nlm.nih.gov/gene/?term=LOC118180764 | 2040 | 15 | 73.53 | 12 | 58.82 | 80 | 6.29 | 41.3 | -35 | 35 | -0.26 | 0.26 |
| gene-LOC118199292 | https://www.ncbi.nlm.nih.gov/gene/?term=LOC118199292 | 846 | 6 | 70.92 | 2 | 23.64 | 33.33 | 39.08 | 3.61 | 35.46 | 35.46 | 0.25 | 0.25 |
| gene-LOC118180761 | https://www.ncbi.nlm.nih.gov/gene/?term=LOC118180761 | 3044 | 21 | 68.99 | 15 | 49.28 | 71.43 | 8.65 | 45.12 | -36.47 | 36.47 | -0.25 | 0.25 |
| gene-LOC118202498 | https://www.ncbi.nlm.nih.gov/gene/?term=LOC118202498 | 1721 | 6 | 34.86 | 5 | 29.05 | 83.33 | 65.1 | 4.6 | 60.5 | 60.5 | 0.21 | 0.21 |
| gene-LOC118206129 | https://www.ncbi.nlm.nih.gov/gene/?term=LOC118206129 | 2062 | 32 | 155.19 | 2 | 9.7 | 6.25 | 40.75 | 29.4 | 11.35 | 11.35 | 0.18 | 0.18 |
| gene-LOC118180675 | https://www.ncbi.nlm.nih.gov/gene/?term=LOC118180675 | 624 | 5 | 80.13 | 2 | 32.05 | 40 | 0.16 | 22.39 | -22.23 | 22.23 | -0.18 | 0.18 |
| gene-LOC118180680 | https://www.ncbi.nlm.nih.gov/gene/?term=LOC118180680 | 3131 | 21 | 67.07 | 15 | 47.91 | 71.43 | 9.44 | 36.41 | -26.96 | 26.96 | -0.18 | 0.18 |
| gene-LOC118180763 | https://www.ncbi.nlm.nih.gov/gene/?term=LOC118180763 | 1104 | 4 | 36.23 | 4 | 36.23 | 100 | 7.51 | 51.27 | -43.75 | 43.75 | -0.16 | 0.16 |
| gene-LOC118204898 | https://www.ncbi.nlm.nih.gov/gene/?term=LOC118204898 | 2378 | 16 | 67.28 | 3 | 12.62 | 18.75 | 20.34 | 38.79 | -18.45 | 18.45 | -0.12 | 0.12 |
| gene-LOC118201096 | https://www.ncbi.nlm.nih.gov/gene/?term=LOC118201096 | 4437 | 19 | 42.82 | 7 | 15.78 | 36.84 | 56.84 | 31.02 | 25.81 | 25.81 | 0.11 | 0.11 |
| gene-LOC118201196 | https://www.ncbi.nlm.nih.gov/gene/?term=LOC118201196 | 10621 | 84 | 79.09 | 3 | 2.82 | 3.57 | 29.17 | 39.29 | -10.12 | 10.12 | -0.08 | 0.08 |
| gene-LOC118180679 | https://www.ncbi.nlm.nih.gov/gene/?term=LOC118180679 | 1640 | 3 | 18.29 | 3 | 18.29 | 100 | 7.58 | 46.43 | -38.85 | 38.85 | -0.07 | 0.07 |
| gene-LOC118205893 | https://www.ncbi.nlm.nih.gov/gene/?term=LOC118205893 | 930 | 2 | 21.51 | 2 | 21.51 | 100 | 6.28 | 33.85 | -27.58 | 27.58 | -0.06 | 0.06 |
| gene-LOC118184468 | https://www.ncbi.nlm.nih.gov/gene/?term=LOC118184468 | 17739 | 22 | 12.4 | 7 | 3.95 | 31.82 | 61.08 | 30.15 | 30.93 | 30.93 | 0.04 | 0.04 |
| gene-LOC118205075 | https://www.ncbi.nlm.nih.gov/gene/?term=LOC118205075 | 6473 | 19 | 29.35 | 7 | 10.81 | 36.84 | 58.72 | 67.66 | -8.94 | 8.94 | -0.03 | 0.03 |
| gene-LOC118192981 | https://www.ncbi.nlm.nih.gov/gene/?term=LOC118192981 | 9333 | 13 | 13.93 | 2 | 2.14 | 15.38 | 63.22 | 81.48 | -18.26 | 18.26 | -0.03 | 0.03 |
| gene-LOC118198429 | https://www.ncbi.nlm.nih.gov/gene/?term=LOC118198429 | 17137 | 23 | 13.42 | 5 | 2.92 | 21.74 | 45.23 | 69.23 | -24 | 24 | -0.03 | 0.03 |
| gene-LOC118183257 | https://www.ncbi.nlm.nih.gov/gene/?term=LOC118183257 | 17126 | 11 | 6.42 | 3 | 1.75 | 27.27 | 69.5 | 34.75 | 34.75 | 34.75 | 0.02 | 0.02 |
| gene-LOC118180489 | https://www.ncbi.nlm.nih.gov/gene/?term=LOC118180489 | 3793 | 4 | 10.55 | 2 | 5.27 | 50 | 33.84 | 16.14 | 17.7 | 17.7 | 0.02 | 0.02 |
| gene-LOC118192973 | https://www.ncbi.nlm.nih.gov/gene/?term=LOC118192973 | 17908 | 36 | 20.1 | 3 | 1.68 | 8.33 | 75.26 | 66.25 | 9.01 | 9.01 | 0.02 | 0.02 |
| gene-LOC118191639 | https://www.ncbi.nlm.nih.gov/gene/?term=LOC118191639 | 14462 | 39 | 26.97 | 4 | 2.77 | 10.26 | 25.59 | 16.63 | 8.96 | 8.96 | 0.02 | 0.02 |
| gene-LOC118205924 | https://www.ncbi.nlm.nih.gov/gene/?term=LOC118205924 | 9047 | 119 | 131.54 | 3 | 3.32 | 2.52 | 67.93 | 69.22 | -1.3 | 1.3 | -0.02 | 0.02 |
| gene-LOC118192320 | https://www.ncbi.nlm.nih.gov/gene/?term=LOC118192320 | 3613 | 2 | 5.54 | 2 | 5.54 | 100 | 3.32 | 43.12 | -39.8 | 39.8 | -0.02 | 0.02 |
| gene-LOC118196111 | https://www.ncbi.nlm.nih.gov/gene/?term=LOC118196111 | 17112 | 3 | 1.75 | 3 | 1.75 | 100 | 82.54 | 4.7 | 77.84 | 77.84 | 0.01 | 0.01 |
| gene-LOC118182035 | https://www.ncbi.nlm.nih.gov/gene/?term=LOC118182035 | 11879 | 11 | 9.26 | 2 | 1.68 | 18.18 | 16.32 | 1.77 | 14.55 | 14.55 | 0.01 | 0.01 |
| gene-LOC118199290 | https://www.ncbi.nlm.nih.gov/gene/?term=LOC118199290 | 15093 | 26 | 17.23 | 2 | 1.33 | 7.69 | 68.99 | 62.52 | 6.47 | 6.47 | 0.01 | 0.01 |
| gene-LOC118203894 | https://www.ncbi.nlm.nih.gov/gene/?term=LOC118203894 | 15828 | 34 | 21.48 | 3 | 1.9 | 8.82 | 27.49 | 22.85 | 4.64 | 4.64 | 0.01 | 0.01 |
| gene-LOC118182290 | https://www.ncbi.nlm.nih.gov/gene/?term=LOC118182290 | 15134 | 27 | 17.84 | 2 | 1.32 | 7.41 | 38.35 | 41.25 | -2.9 | 2.9 | -0.01 | 0.01 |
| gene-LOC118189486 | https://www.ncbi.nlm.nih.gov/gene/?term=LOC118189486 | 14274 | 16 | 11.21 | 2 | 1.4 | 12.5 | 1.59 | 13.77 | -12.18 | 12.18 | -0.01 | 0.01 |
| gene-LOC118200390 | https://www.ncbi.nlm.nih.gov/gene/?term=LOC118200390 | 13922 | 39 | 28.01 | 2 | 1.44 | 5.13 | 65.32 | 65.29 | 0.04 | 0.04 | 0 | 0 |
| gene-LOC118188906 | https://www.ncbi.nlm.nih.gov/gene/?term=LOC118188906 | 11213 | 40 | 35.67 | 2 | 1.78 | 5 | 82.47 | 83.49 | -1.02 | 1.02 | 0 | 0 |

```
Directory: OGL File: wrt-Sd_LI.txt

... comparing  be.vs.ka ...

Number of genes <=  20000  and with #dmsites >=  2 :  46
sample1     sum of percent methylation:  1391.76     average:    30.26
sample2     sum of percent methylation:  1851.15     average:    40.24

sum difference:  -459.40     average difference:  -9.9870

      Wilcoxon signed rank test with continuity correction

data:  pw_summary$`%pSite1` and pw_summary$`%pSite2`
V = 317, p-value = 0.01
alternative hypothesis: true location shift is not equal to 0
```

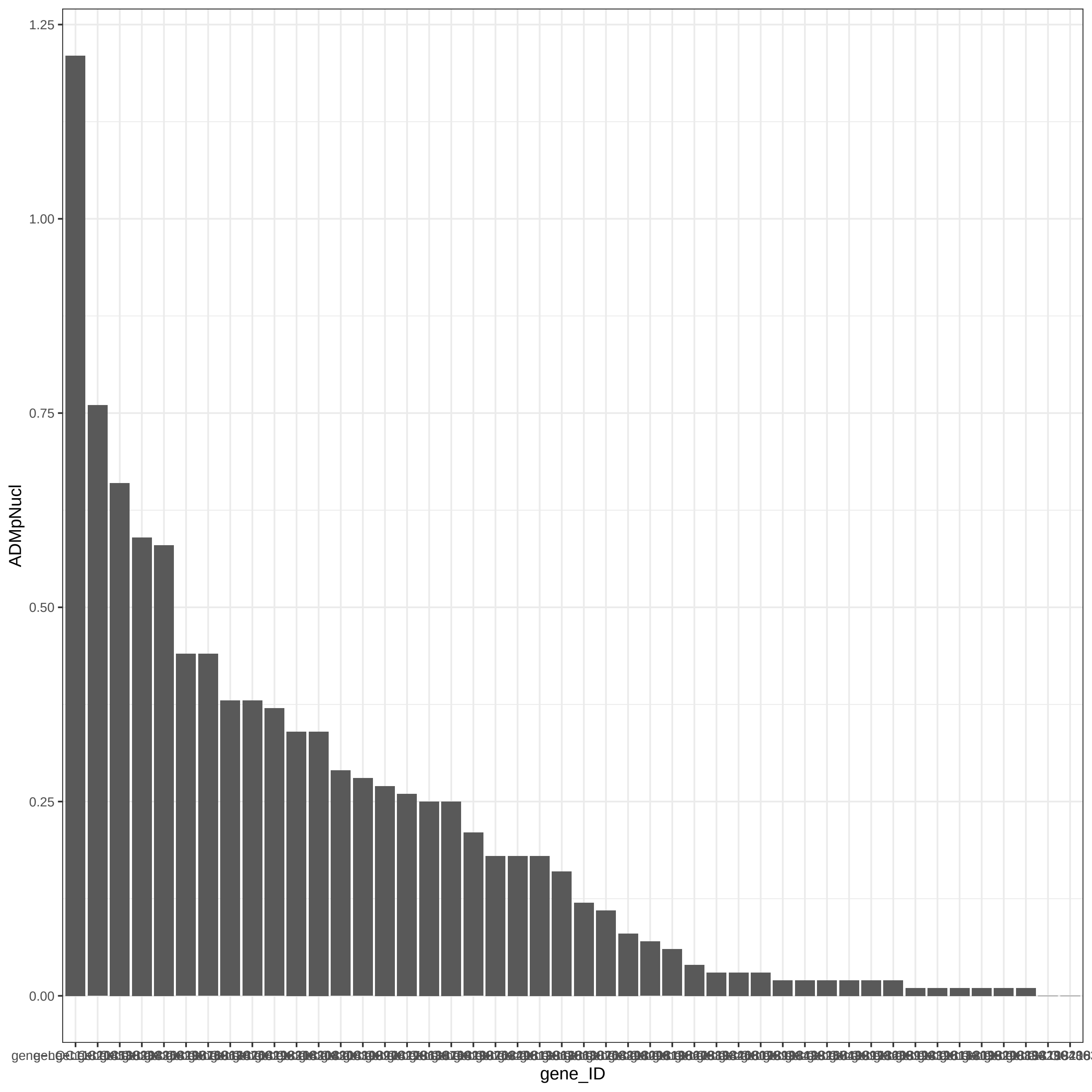

1.25
1.00
0.75
0.50
0.25
0.00
ADMpNucl
gene_ID

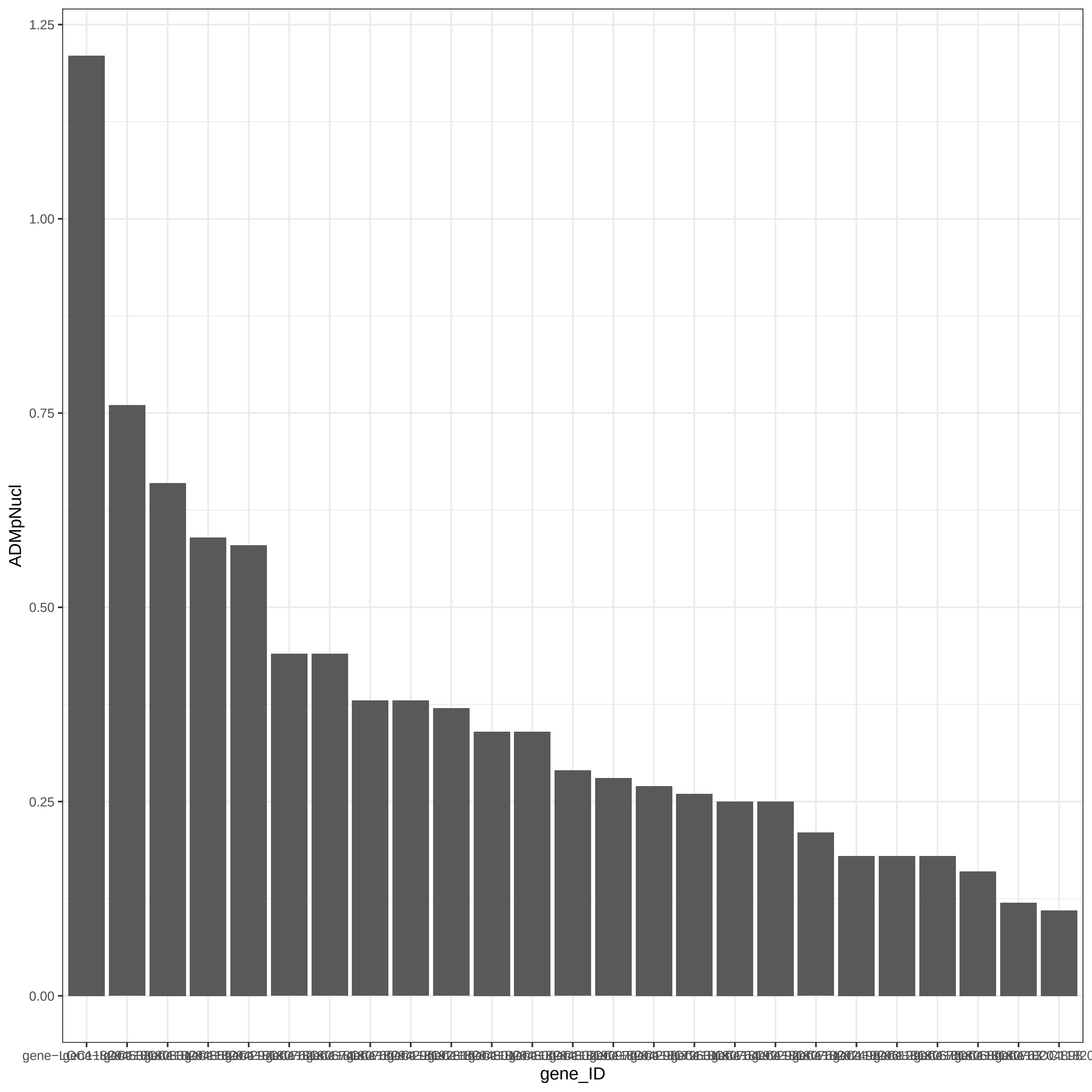

1.25
1.00
0.75
0.50
0.25
0.00
ADMpNucl
gene_ID

```
Directory: OGLall File: 0READMEogl

OGL - Ordered gene lists.

Input: studyhc annotation   maxgwidth minnbrdmsites
       dmgprp (= output of BWASPR:show_dmsg())

Output: files ogl-*.txt rnk-dmg-*.txt
         ogl-<study>_<sample>.txt ogl-<study>_<sample1>.vs.<sample2>.txt
         rnk-dmg--<study>_<sample1>.vs.<sample2>.txt rnk-dmg--<study>_<sample1>.vs.<sample2>.pdf
         wrt-<study>.txt

Notes: Output files ogl-<study>_<sample>.txt give tables for each sample with columns

        gene_ID gwidth #Sites #per10Kb %perSite %pNucl

       ordered by %pNucl.  If available, a link to an NCBI entry of gene_ID is inserted as second column.
       Abbreviations used: %perSite, percent methylation per site
                           %perNucl, percent methylation per nucleotide of the gene

       Output files ogl-<study>_<sample1>.vs.<sample2>.txt give tables for each comparison with columns

        gene_ID gwidth #Sites #per10Kb #dmSites #dmsp10kb %dmSites %pSite1 %pSite2 DMpSite ADMpSite DMpNucl ADMpNucl

       ordered by DMpSite.  If available, a link to an NCBI entry of gene_ID is inserted as second column.
       Abbreviations used: %dmSites, percent sites that are differentially methylated
                           %pSite1, average per site % methylation for sample1
                           %pSite2, average per site % methylation for sample2
                           DMpSite, average per site difference in % methylation between sample1 and sample2
                           ADMpSite, absolute value of DMpSite
                           DMpNucl, average per nucleotide difference in % methylation between sample1 and sample2
                           ADMpNucl, absolute value of DMpNucl

       Output files rnk-dmg-<study>_<sample1>.vs.<sample2>.txt are equivalent to files
        ogl-<study>_<sample1>.vs.<sample2>.txt but ordered by ADMpNucl.
       Output files rnk-dmg-<study>_<sample1>.vs.<sample2>.pdf provide visualization of the distribution
        of ADMpNucl values.

       Output file wrt-<study>.txt gives results of the Wilcoxon signed rank test comparing
        the %pSite1 and %pSite2 vectors

       In output directory OGL, tables are restricted to genes with gwidth <= maxgwidth and, for
        pairwise comparisons, #dmSites >= minnbrdmsites

       If either maxgwidth or minnbrdmsites is set, then output directory OGLall will show the full tables
        with all genes (for reference).
```

Directory: OGLall File: ogl-Sd_LI_be.txt

| gene_ID | gene_link | gwidth | #Sites | #per10Kb | %perSite | %pNucl |
|---|---|---|---|---|---|---|
| gene-LOC118199038 | https://www.ncbi.nlm.nih.gov/gene/?term=LOC118199038 | 1069 | 60 | 561.27 | 88.44 | 4.96 |
| gene-LOC118200794 | https://www.ncbi.nlm.nih.gov/gene/?term=LOC118200794 | 679 | 33 | 486.01 | 89.19 | 4.33 |
| gene-LOC118185191 | https://www.ncbi.nlm.nih.gov/gene/?term=LOC118185191 | 846 | 52 | 614.66 | 62 | 3.81 |
| gene-LOC118180258 | https://www.ncbi.nlm.nih.gov/gene/?term=LOC118180258 | 363 | 17 | 468.32 | 81 | 3.79 |
| gene-LOC118190272 | https://www.ncbi.nlm.nih.gov/gene/?term=LOC118190272 | 1960 | 62 | 316.33 | 82.96 | 2.62 |
| gene-Trnat-ggu-6 | https://www.ncbi.nlm.nih.gov/gene/?term=Trnat-ggu | 72 | 8 | 1111.11 | 21.07 | 2.34 |
| gene-LOC118193385 | https://www.ncbi.nlm.nih.gov/gene/?term=LOC118193385 | 363 | 9 | 247.93 | 90.23 | 2.24 |
| gene-LOC118194215 | https://www.ncbi.nlm.nih.gov/gene/?term=LOC118194215 | 6061 | 145 | 239.23 | 84.24 | 2.02 |
| gene-LOC118202299 | https://www.ncbi.nlm.nih.gov/gene/?term=LOC118202299 | 7958 | 207 | 260.12 | 77.6 | 2.02 |
| gene-LOC118191889 | https://www.ncbi.nlm.nih.gov/gene/?term=LOC118191889 | 13681 | 399 | 291.65 | 64.02 | 1.87 |
| gene-LOC118203587 | https://www.ncbi.nlm.nih.gov/gene/?term=LOC118203587 | 27117 | 695 | 256.3 | 72.74 | 1.86 |
| gene-LOC118180687 | https://www.ncbi.nlm.nih.gov/gene/?term=LOC118180687 | 12663 | 289 | 228.22 | 81.06 | 1.85 |
| gene-LOC118205209 | https://www.ncbi.nlm.nih.gov/gene/?term=LOC118205209 | 7915 | 238 | 300.69 | 60.92 | 1.83 |
| gene-LOC118195270 | https://www.ncbi.nlm.nih.gov/gene/?term=LOC118195270 | 39867 | 933 | 234.03 | 75.93 | 1.78 |
| gene-LOC118206125 | https://www.ncbi.nlm.nih.gov/gene/?term=LOC118206125 | 8096 | 171 | 211.22 | 82.62 | 1.75 |
| gene-LOC118205293 | https://www.ncbi.nlm.nih.gov/gene/?term=LOC118205293 | 850 | 35 | 411.76 | 42.14 | 1.74 |
| gene-LOC118195989 | https://www.ncbi.nlm.nih.gov/gene/?term=LOC118195989 | 756 | 15 | 198.41 | 86.25 | 1.71 |
| gene-LOC118187599 | https://www.ncbi.nlm.nih.gov/gene/?term=LOC118187599 | 447 | 9 | 201.34 | 84.23 | 1.7 |
| gene-LOC118204580 | https://www.ncbi.nlm.nih.gov/gene/?term=LOC118204580 | 13571 | 333 | 245.38 | 69.17 | 1.7 |
| gene-LOC118189203 | https://www.ncbi.nlm.nih.gov/gene/?term=LOC118189203 | 1497 | 32 | 213.76 | 78.15 | 1.67 |
| gene-LOC118205924 | https://www.ncbi.nlm.nih.gov/gene/?term=LOC118205924 | 9047 | 217 | 239.86 | 69.06 | 1.66 |
| gene-LOC118205294 | https://www.ncbi.nlm.nih.gov/gene/?term=LOC118205294 | 1667 | 123 | 737.85 | 22.23 | 1.64 |
| gene-LOC118188252 | https://www.ncbi.nlm.nih.gov/gene/?term=LOC118188252 | 531 | 10 | 188.32 | 86.41 | 1.63 |
| gene-LOC118204359 | https://www.ncbi.nlm.nih.gov/gene/?term=LOC118204359 | 1095 | 30 | 273.97 | 58.44 | 1.6 |

...

... only first 25 lines shown ...

Directory: OGLall File: ogl-Sd_LI_be.vs.ka.txt

| gene_ID | gene_link | gwidth | #Sites | #per10Kb | #dmSites | #dmsp10kb | %dmSites | %pSite1 | %pSite2 | DMpSite | ADMpSite | DMpNucl | ADMpNucl |
|---|---|---|---|---|---|---|---|---|---|---|---|---|---|
| gene-LOC118196111 | https://www.ncbi.nlm.nih.gov/gene/?term=LOC118196111 | 17112 | 3 | 1.75 | 3 | 1.75 | 100 | 82.54 | 4.7 | 77.84 | 77.84 | 0.01 | 0.01 |
| gene-LOC118204699 | https://www.ncbi.nlm.nih.gov/gene/?term=LOC118204699 | 21443 | 1 | 0.47 | 1 | 0.47 | 100 | 90 | 28.12 | 61.88 | 61.88 | 0 | 0 |
| gene-LOC118202498 | https://www.ncbi.nlm.nih.gov/gene/?term=LOC118202498 | 1721 | 6 | 34.86 | 5 | 29.05 | 83.33 | 65.1 | 4.6 | 60.5 | 60.5 | 0.21 | 0.21 |
| gene-LOC118203623 | https://www.ncbi.nlm.nih.gov/gene/?term=LOC118203623 | 22413 | 13 | 5.8 | 9 | 4.02 | 69.23 | 65.65 | 14.64 | 51.02 | 51.02 | 0.03 | 0.03 |
| gene-LOC118199636 | https://www.ncbi.nlm.nih.gov/gene/?term=LOC118199636 | 1885 | 1 | 5.31 | 1 | 5.31 | 100 | 48.57 | 6.67 | 41.9 | 41.9 | 0.02 | 0.02 |
| gene-LOC118204359 | https://www.ncbi.nlm.nih.gov/gene/?term=LOC118204359 | 1095 | 19 | 173.52 | 8 | 73.06 | 42.11 | 59.62 | 21.69 | 37.93 | 37.93 | 0.66 | 0.66 |
| gene-LOC118199292 | https://www.ncbi.nlm.nih.gov/gene/?term=LOC118199292 | 846 | 6 | 70.92 | 2 | 23.64 | 33.33 | 39.08 | 3.61 | 35.46 | 35.46 | 0.25 | 0.25 |
| gene-LOC118183257 | https://www.ncbi.nlm.nih.gov/gene/?term=LOC118183257 | 17126 | 11 | 6.42 | 3 | 1.75 | 27.27 | 69.5 | 34.75 | 34.75 | 34.75 | 0.02 | 0.02 |
| gene-LOC118190970 | https://www.ncbi.nlm.nih.gov/gene/?term=LOC118190970 | 1773 | 16 | 90.24 | 4 | 22.56 | 25 | 65.08 | 33.41 | 31.67 | 31.67 | 0.29 | 0.29 |
| gene-LOC118189387 | https://www.ncbi.nlm.nih.gov/gene/?term=LOC118189387 | 8258 | 2 | 2.42 | 1 | 1.21 | 50 | 91.66 | 60.42 | 31.25 | 31.25 | 0.01 | 0.01 |
| gene-LOC118184468 | https://www.ncbi.nlm.nih.gov/gene/?term=LOC118184468 | 17739 | 22 | 12.4 | 7 | 3.95 | 31.82 | 61.08 | 30.15 | 30.93 | 30.93 | 0.04 | 0.04 |
| gene-LOC118191662 | https://www.ncbi.nlm.nih.gov/gene/?term=LOC118191662 | 18543 | 13 | 7.01 | 1 | 0.54 | 7.69 | 40.22 | 10.93 | 29.29 | 29.29 | 0.02 | 0.02 |
| gene-LOC118204169 | https://www.ncbi.nlm.nih.gov/gene/?term=LOC118204169 | 6093 | 3 | 4.92 | 1 | 1.64 | 33.33 | 78 | 50.26 | 27.74 | 27.74 | 0.01 | 0.01 |
| gene-LOC118187297 | https://www.ncbi.nlm.nih.gov/gene/?term=LOC118187297 | 10135 | 9 | 8.88 | 1 | 0.99 | 11.11 | 41.41 | 14.63 | 26.78 | 26.78 | 0.02 | 0.02 |
| gene-LOC118201936 | https://www.ncbi.nlm.nih.gov/gene/?term=LOC118201936 | 9487 | 3 | 3.16 | 1 | 1.05 | 33.33 | 91.8 | 65.69 | 26.11 | 26.11 | 0.01 | 0.01 |
| gene-LOC118201096 | https://www.ncbi.nlm.nih.gov/gene/?term=LOC118201096 | 4437 | 19 | 42.82 | 7 | 15.78 | 36.84 | 56.84 | 31.02 | 25.81 | 25.81 | 0.11 | 0.11 |
| gene-LOC118183353 | https://www.ncbi.nlm.nih.gov/gene/?term=LOC118183353 | 1394 | 3 | 21.52 | 1 | 7.17 | 33.33 | 23.08 | 1.28 | 21.79 | 21.79 | 0.05 | 0.05 |
| gene-LOC118199481 | https://www.ncbi.nlm.nih.gov/gene/?term=LOC118199481 | 54320 | 30 | 5.52 | 4 | 0.74 | 13.33 | 28.06 | 7.89 | 20.17 | 20.17 | 0.01 | 0.01 |
| gene-LOC118201073 | https://www.ncbi.nlm.nih.gov/gene/?term=LOC118201073 | 60441 | 21 | 3.47 | 4 | 0.66 | 19.05 | 79.94 | 59.77 | 20.17 | 20.17 | 0.01 | 0.01 |
| gene-LOC118201509 | https://www.ncbi.nlm.nih.gov/gene/?term=LOC118201509 | 78877 | 30 | 3.8 | 4 | 0.51 | 13.33 | 54.77 | 36.16 | 18.61 | 18.61 | 0.01 | 0.01 |
| gene-LOC118204774 | https://www.ncbi.nlm.nih.gov/gene/?term=LOC118204774 | 25927 | 8 | 3.09 | 1 | 0.39 | 12.5 | 87.53 | 69.02 | 18.51 | 18.51 | 0.01 | 0.01 |
| gene-LOC118180489 | https://www.ncbi.nlm.nih.gov/gene/?term=LOC118180489 | 3793 | 4 | 10.55 | 2 | 5.27 | 50 | 33.84 | 16.14 | 17.7 | 17.7 | 0.02 | 0.02 |
| gene-LOC118189573 | https://www.ncbi.nlm.nih.gov/gene/?term=LOC118189573 | 78720 | 30 | 3.81 | 4 | 0.51 | 13.33 | 66.23 | 49.32 | 16.9 | 16.9 | 0.01 | 0.01 |
| gene-LOC118189564 | https://www.ncbi.nlm.nih.gov/gene/?term=LOC118189564 | 52694 | 53 | 10.06 | 4 | 0.76 | 7.55 | 63.46 | 47.12 | 16.34 | 16.34 | 0.02 | 0.02 |

...

... only first 25 lines shown ...

Directory: OGLall File: ogl-Sd_LI_ka.txt

| gene_ID | gene_link | gwidth | #Sites | #per10Kb | %perSite | %pNucl |
|---|---|---|---|---|---|---|
| gene-LOC118200794 | https://www.ncbi.nlm.nih.gov/gene/?term=LOC118200794 | 679 | 36 | 530.19 | 86.07 | 4.56 |
| gene-LOC118199038 | https://www.ncbi.nlm.nih.gov/gene/?term=LOC118199038 | 1069 | 54 | 505.14 | 87.36 | 4.41 |
| gene-LOC118185191 | https://www.ncbi.nlm.nih.gov/gene/?term=LOC118185191 | 846 | 57 | 673.76 | 59.98 | 4.04 |
| gene-LOC118180258 | https://www.ncbi.nlm.nih.gov/gene/?term=LOC118180258 | 363 | 17 | 468.32 | 77.69 | 3.64 |
| gene-Trnat-ggu-6 | https://www.ncbi.nlm.nih.gov/gene/?term=Trnat-ggu | 72 | 8 | 1111.11 | 28.86 | 3.21 |
| gene-LOC118180265 | https://www.ncbi.nlm.nih.gov/gene/?term=LOC118180265 | 825 | 23 | 278.79 | 94.02 | 2.62 |
| gene-LOC118190272 | https://www.ncbi.nlm.nih.gov/gene/?term=LOC118190272 | 1960 | 62 | 316.33 | 81.07 | 2.56 |
| gene-LOC118198533 | https://www.ncbi.nlm.nih.gov/gene/?term=LOC118198533 | 333 | 8 | 240.24 | 96.08 | 2.31 |
| gene-LOC118193385 | https://www.ncbi.nlm.nih.gov/gene/?term=LOC118193385 | 363 | 9 | 247.93 | 83.77 | 2.08 |
| gene-LOC118202299 | https://www.ncbi.nlm.nih.gov/gene/?term=LOC118202299 | 7958 | 212 | 266.4 | 77.33 | 2.06 |
| gene-LOC118194215 | https://www.ncbi.nlm.nih.gov/gene/?term=LOC118194215 | 6061 | 147 | 242.53 | 83.85 | 2.03 |
| gene-LOC118191889 | https://www.ncbi.nlm.nih.gov/gene/?term=LOC118191889 | 13681 | 406 | 296.76 | 64.9 | 1.93 |
| gene-LOC118180687 | https://www.ncbi.nlm.nih.gov/gene/?term=LOC118180687 | 12663 | 296 | 233.75 | 81.41 | 1.9 |
| gene-LOC118205209 | https://www.ncbi.nlm.nih.gov/gene/?term=LOC118205209 | 7915 | 243 | 307.01 | 61.89 | 1.9 |
| gene-LOC118195270 | https://www.ncbi.nlm.nih.gov/gene/?term=LOC118195270 | 39867 | 988 | 247.82 | 75.31 | 1.87 |
| gene-LOC118203587 | https://www.ncbi.nlm.nih.gov/gene/?term=LOC118203587 | 27117 | 692 | 255.19 | 73.17 | 1.87 |
| gene-LOC118205293 | https://www.ncbi.nlm.nih.gov/gene/?term=LOC118205293 | 850 | 36 | 423.53 | 42.81 | 1.81 |
| gene-LOC118206125 | https://www.ncbi.nlm.nih.gov/gene/?term=LOC118206125 | 8096 | 170 | 209.98 | 81.25 | 1.71 |
| gene-LOC118205924 | https://www.ncbi.nlm.nih.gov/gene/?term=LOC118205924 | 9047 | 220 | 243.17 | 69.82 | 1.7 |
| gene-LOC118201471 | https://www.ncbi.nlm.nih.gov/gene/?term=LOC118201471 | 2104 | 42 | 199.62 | 84.85 | 1.69 |
| gene-LOC118204580 | https://www.ncbi.nlm.nih.gov/gene/?term=LOC118204580 | 13571 | 332 | 244.64 | 68.78 | 1.68 |
| gene-LOC118195989 | https://www.ncbi.nlm.nih.gov/gene/?term=LOC118195989 | 756 | 15 | 198.41 | 84.2 | 1.67 |
| gene-LOC118199153 | https://www.ncbi.nlm.nih.gov/gene/?term=LOC118199153 | 880 | 15 | 170.45 | 94.78 | 1.62 |
| gene-LOC118192985 | https://www.ncbi.nlm.nih.gov/gene/?term=LOC118192985 | 467 | 8 | 171.31 | 92.64 | 1.59 |

...
... only first 25 lines shown ...

Directory: OGLall File: rnk-dmg-Sd_LI_be.vs.ka.txt

| gene_ID | gene_link | gwidth | #Sites | #per10Kb | #dmSites | #dmsp10kb | %dmSites | %pSite1 | %pSite2 | DMpSite | ADMpSite | DMpNucl | ADMpNucl |
|---|---|---|---|---|---|---|---|---|---|---|---|---|---|
| gene-Trnar-ucu-16 | https://www.ncbi.nlm.nih.gov/gene/?term=Trnar-ucu | 74 | 3 | 405.41 | 1 | 135.14 | 33.33 | 5.18 | 36.53 | -31.35 | 31.35 | -1.27 | 1.27 |
| gene-LOC118204533 | https://www.ncbi.nlm.nih.gov/gene/?term=LOC118204533 | 446 | 22 | 493.27 | 10 | 224.22 | 45.45 | 0.38 | 24.93 | -24.55 | 24.55 | -1.21 | 1.21 |
| gene-LOC118180331 | https://www.ncbi.nlm.nih.gov/gene/?term=LOC118180331 | 1035 | 13 | 125.6 | 13 | 125.6 | 100 | 12.27 | 72.44 | -60.16 | 60.16 | -0.76 | 0.76 |
| gene-LOC118204359 | https://www.ncbi.nlm.nih.gov/gene/?term=LOC118204359 | 1095 | 19 | 173.52 | 8 | 73.06 | 42.11 | 59.62 | 21.69 | 37.93 | 37.93 | 0.66 | 0.66 |
| gene-LOC118204297 | https://www.ncbi.nlm.nih.gov/gene/?term=LOC118204297 | 951 | 13 | 136.7 | 13 | 136.7 | 100 | 4.88 | 48.27 | -43.39 | 43.39 | -0.59 | 0.59 |
| gene-LOC118180762 | https://www.ncbi.nlm.nih.gov/gene/?term=LOC118180762 | 1281 | 18 | 140.52 | 16 | 124.9 | 88.89 | 6.13 | 47.34 | -41.22 | 41.22 | -0.58 | 0.58 |
| gene-LOC118180674 | https://www.ncbi.nlm.nih.gov/gene/?term=LOC118180674 | 1344 | 17 | 126.49 | 14 | 104.17 | 82.35 | 7.39 | 42.16 | -34.78 | 34.78 | -0.44 | 0.44 |
| gene-LOC118180760 | https://www.ncbi.nlm.nih.gov/gene/?term=LOC118180760 | 978 | 12 | 122.7 | 9 | 92.02 | 75 | 5.55 | 41.25 | -35.7 | 35.7 | -0.44 | 0.44 |
| gene-LOC118204296 | https://www.ncbi.nlm.nih.gov/gene/?term=LOC118204296 | 2118 | 22 | 103.87 | 20 | 94.43 | 90.91 | 7.23 | 43.59 | -36.36 | 36.36 | -0.38 | 0.38 |
| gene-LOC118192316 | https://www.ncbi.nlm.nih.gov/gene/?term=LOC118192316 | 1188 | 12 | 101.01 | 11 | 92.59 | 91.67 | 9.13 | 46.82 | -37.68 | 37.68 | -0.38 | 0.38 |
| gene-LOC118204301 | https://www.ncbi.nlm.nih.gov/gene/?term=LOC118204301 | 1823 | 17 | 93.25 | 16 | 87.77 | 94.12 | 4.98 | 44.95 | -39.98 | 39.98 | -0.37 | 0.37 |
| gene-LOC118204300 | https://www.ncbi.nlm.nih.gov/gene/?term=LOC118204300 | 1116 | 11 | 98.57 | 8 | 71.68 | 72.73 | 6.44 | 41.39 | -34.95 | 34.95 | -0.34 | 0.34 |
| gene-LOC118204302 | https://www.ncbi.nlm.nih.gov/gene/?term=LOC118204302 | 4419 | 42 | 95.04 | 33 | 74.68 | 78.57 | 6.84 | 43 | -36.16 | 36.16 | -0.34 | 0.34 |
| gene-LOC118190970 | https://www.ncbi.nlm.nih.gov/gene/?term=LOC118190970 | 1773 | 16 | 90.24 | 4 | 22.56 | 25 | 65.08 | 33.41 | 31.67 | 31.67 | 0.29 | 0.29 |
| gene-LOC118204298 | https://www.ncbi.nlm.nih.gov/gene/?term=LOC118204298 | 474 | 4 | 84.39 | 4 | 84.39 | 100 | 7.12 | 40.43 | -33.31 | 33.31 | -0.28 | 0.28 |
| gene-LOC118179631 | https://www.ncbi.nlm.nih.gov/gene/?term=LOC118179631 | 1049 | 7 | 66.73 | 6 | 57.2 | 85.71 | 4.77 | 45.73 | -40.96 | 40.96 | -0.27 | 0.27 |
| gene-LOC118180764 | https://www.ncbi.nlm.nih.gov/gene/?term=LOC118180764 | 2040 | 15 | 73.53 | 12 | 58.82 | 80 | 6.29 | 41.3 | -35 | 35 | -0.26 | 0.26 |
| gene-LOC118199292 | https://www.ncbi.nlm.nih.gov/gene/?term=LOC118199292 | 846 | 6 | 70.92 | 2 | 23.64 | 33.33 | 39.08 | 3.61 | 35.46 | 35.46 | 0.25 | 0.25 |
| gene-LOC118180761 | https://www.ncbi.nlm.nih.gov/gene/?term=LOC118180761 | 3044 | 21 | 68.99 | 15 | 49.28 | 71.43 | 8.65 | 45.12 | -36.47 | 36.47 | -0.25 | 0.25 |
| gene-LOC118202498 | https://www.ncbi.nlm.nih.gov/gene/?term=LOC118202498 | 1721 | 6 | 34.86 | 5 | 29.05 | 83.33 | 65.1 | 4.6 | 60.5 | 60.5 | 0.21 | 0.21 |
| gene-LOC118206129 | https://www.ncbi.nlm.nih.gov/gene/?term=LOC118206129 | 2062 | 32 | 155.19 | 2 | 9.7 | 6.25 | 40.75 | 29.4 | 11.35 | 11.35 | 0.18 | 0.18 |
| gene-LOC118180675 | https://www.ncbi.nlm.nih.gov/gene/?term=LOC118180675 | 624 | 5 | 80.13 | 2 | 32.05 | 40 | 0.16 | 22.39 | -22.23 | 22.23 | -0.18 | 0.18 |
| gene-LOC118180680 | https://www.ncbi.nlm.nih.gov/gene/?term=LOC118180680 | 3131 | 21 | 67.07 | 15 | 47.91 | 71.43 | 9.44 | 36.41 | -26.96 | 26.96 | -0.18 | 0.18 |
| gene-LOC118180763 | https://www.ncbi.nlm.nih.gov/gene/?term=LOC118180763 | 1104 | 4 | 36.23 | 4 | 36.23 | 100 | 7.51 | 51.27 | -43.75 | 43.75 | -0.16 | 0.16 |

...

... only first 25 lines shown ...

```
Directory: OGLall File: wrt-Sd_LI.txt

... comparing  be.vs.ka ...

Number of genes with #dmsites >=  1 :  1353

sample1     sum of percent methylation: 85487.53     average:    63.18
sample2     sum of percent methylation: 85949.53     average:    63.53

sum difference:  -462.10     average difference:  -0.3415

      Wilcoxon signed rank test with continuity correction

data:  pw_summary$`%pSite1` and pw_summary$`%pSite2`
V = 5e+05, p-value = 1e-04
alternative hypothesis: true location shift is not equal to 0
```

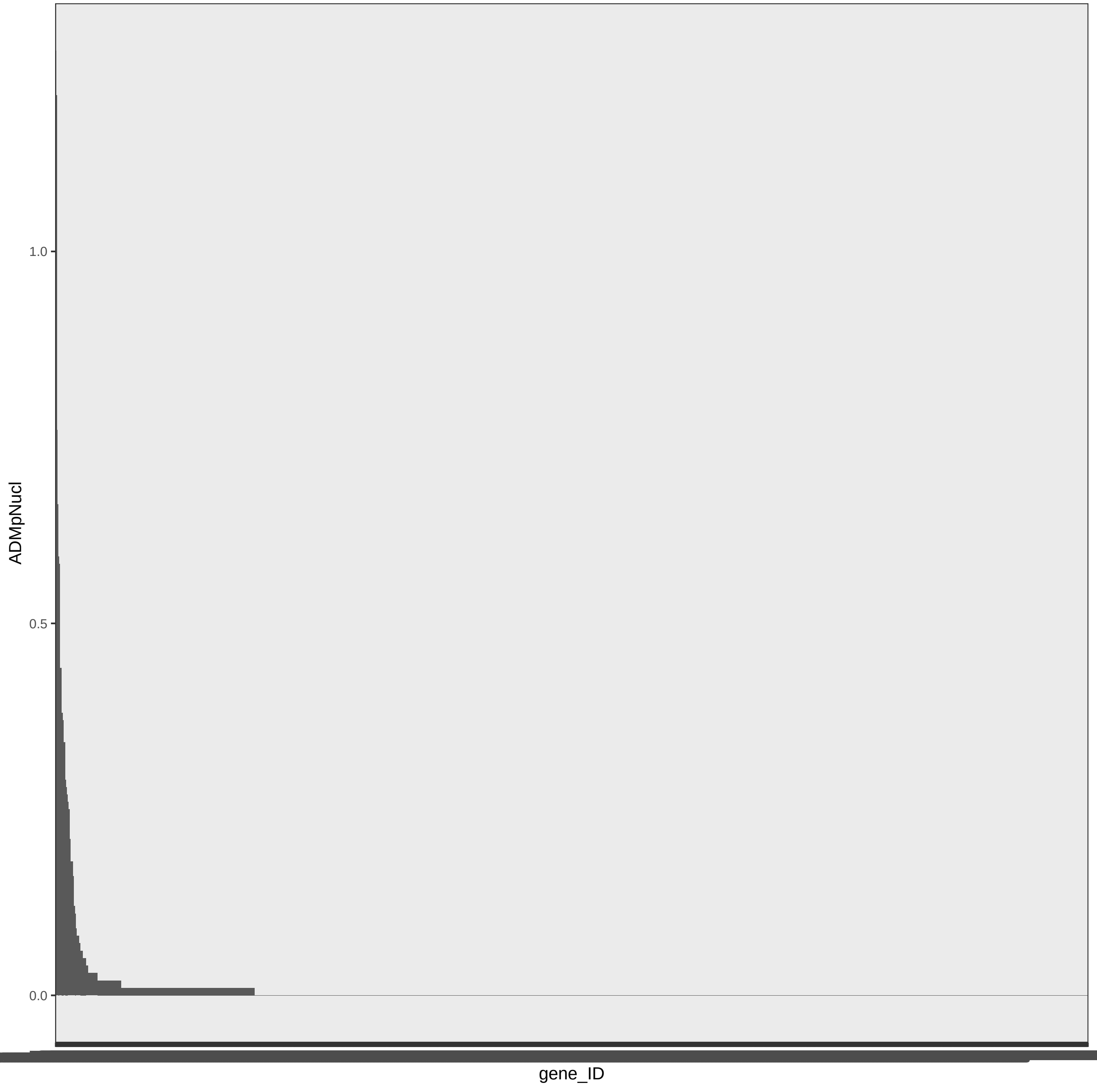

ADMpNucl
1.0
0.5
0.0
gene_ID

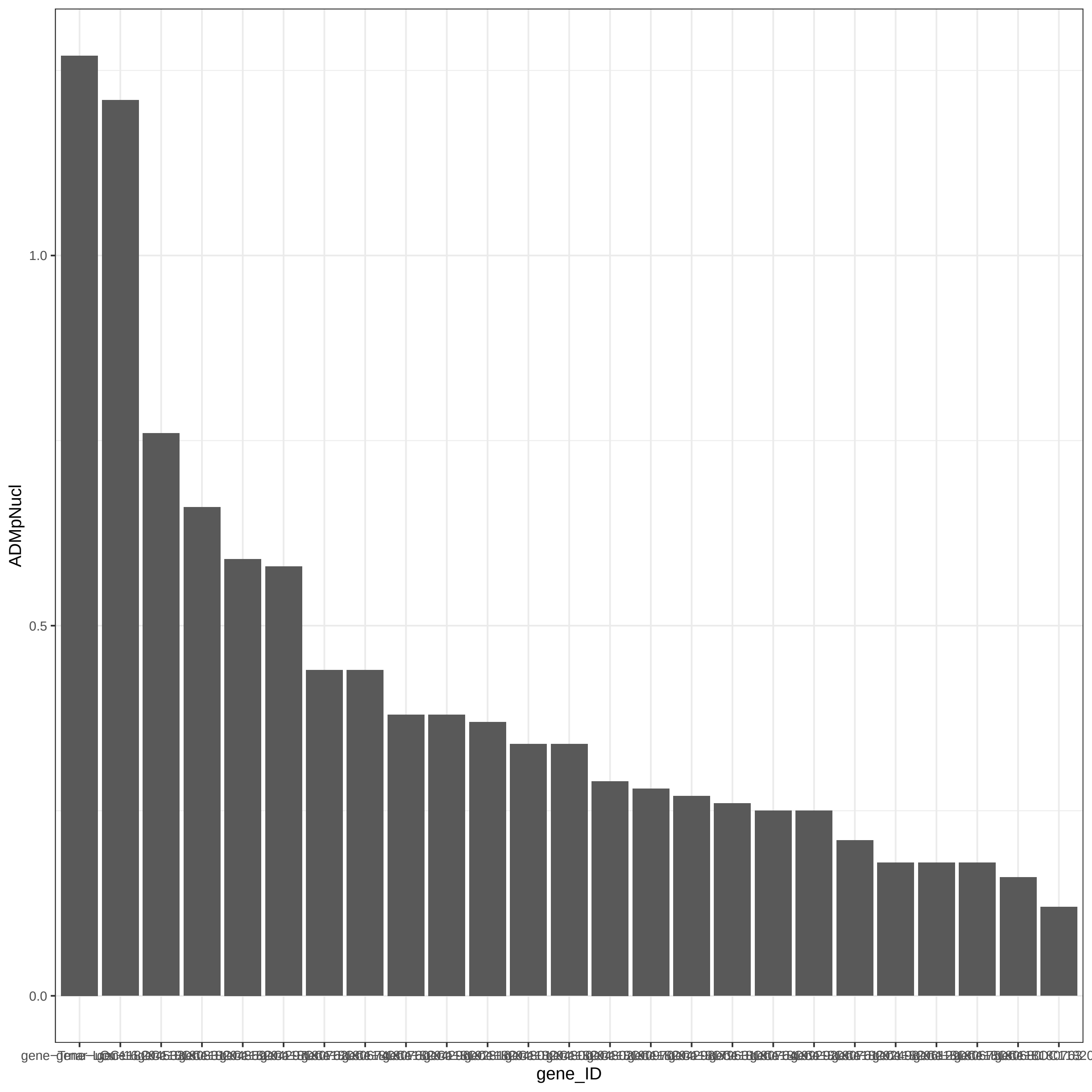

1.0
0.5
0.0
ADMpNucl
gene_ID
